# Retrospective cell lineage reconstruction in Humans using short tandem repeats

**DOI:** 10.1101/191296

**Authors:** Liming Tao, Ofir Raz, Zipora Marx, Manjusha Gosh, Sandra Huber, Julia Greindl-Junghans, Tamir Biezuner, Shiran Amir, Lilach Milo, Rivka Adar, Ron Levy, Amos Onn, Noa Chapal-Ilani, Veronika Berman, Asaf Ben Arie, Guy Rom, Barak Oron, Ruth Halaban, Zbigniew T. Czyz, Melanie Werner-Klein, Christoph A. Klein, Ehud Shapiro

## Abstract

Cell lineage analysis aims to uncover the developmental history of an organism back to its cell of origin^1^. Recently, novel *in vivo* methods and technologies utilizing genome editing enabled important insights into the cell lineages of animals^2–8^. In contrast, human cell lineage remains restricted to retrospective approaches, which still lack in resolution and cost-efficient solutions. Here we demonstrate a scalable platform for human cell lineage tracing based on Short Tandem Repeats (STRs) targeted by duplex Molecular Inversion Probes (MIPs). With this platform we accurately reproduced a known lineage of DU145 cell lines cells^9^ and reconstructed lineages of healthy and metastatic single cells from a melanoma patient. The reconstructed trees matched the anatomical and SNV references while adding further refinements. Our platform allowed to faithfully recapitulate lineages of developmental tissue formation in cells from healthy donors. In summary, our lineage discovery platform can profile informative STR somatic mutations efficiently and we provide a solid, high-resolution lineage reconstruction even in challenging low-mutation-rate healthy single cells.

---

Many fundamental open questions in human biology and medicine can be answered if the structure and dynamics of the human cell lineage tree are uncovered^1,8,10^. Yet *Caenorhabditis elegans* remains the only organism with a known cell lineage tree^11,12^, whereas more recently, high throughput molecular-recording cell lineage methods have been applied to model cell lineages in model organisms such as zebrafish and mouse^2–8^. All these studies demonstrate how valuable cell lineage information can be when done at scale and with tunable mutation rates. In contrast, the reconstruction of human cell lineages has to rely on somatic mutations occurring naturally during cell divisions, such as L1 retro transposition event, Copy Number Variants (CNV), Single Nucleotide Variant (SNV) and Short Tandem Repeats (STRs)^13–21^. Among those, STRs are the largest contributors of *de novo* mutations and are highly abundant^22^. Moreover, STRs’ mutation rates are predictable as they correspond to the type of the repeating unit and number of repeats^23^. Therefore, targeting a selection of STR loci is an effective method for tapping into that rich source of somatic mutations. However, existing methods for accurate genotyping of STRs such as the AccessArray™-based platform^24^ remain limited in their throughput or single cell (SC) compatibility (Supplemental Table1). To enable human cell lineage tracing we strived for (i) improving STR profiling of single cells; (ii) validating our lineage tree algorithms using an experimental set up with known ground truth; (iii) comparing generated lineage trees with current state-of-the-art results from SNV profiling of melanoma cells and (iv) exploring the power of the approach for healthy single cells from different human cell lineages.

We tested Molecular Inversion Probes (MIPs, also termed Padlock Probes) for high throughput (HT) and cost-effective cell lineage tracing. MIPs are single strand DNA molecules composed of two distal targeting arms connected with an internal linker region, which have the potential for much higher multiplexing capability and better specificity compared to multiplex PCR^25^. Several studies have optimized this approach for use with various informative markers^26,27,28,29^, MIPs have been shown to allow successful capture tri- and hexa-nucleotide STRs at a low scale, 102 loci per reaction^30^. We developed our STRs targeting platform with duplex MIPs produced from precursors synthesized on high-resolution microarray resulting in the capture of ~12K STR targets. Briefly, MIP precursors for selected STR loci (Supplementary “Hyper-mutable STRs”) and optional patient-specific areas of interest were designed, synthesized on a microarray (Figure 1a, Supplementary “Duplex MIPs structure”), and processed to duplex MIPs (Figure 1b, Supplementary “Duplex MIPs Preparation”). Target enrichment of each sample is performed in a series of reactions: hybridization between the duplex MIPs and the template DNA, gap filling, ligation, followed by nuclease digestion (Figure 1c, Supplementary “Duplex MIPs-based targeted enrichment pipeline”). Sequencing libraries are prepared by PCR, with primers that contain a unique Illumina barcode combination for each sample, pooled, and sequenced prior to analysis and cell lineage reconstruction (Figure 1c). Pipeline details are in Supplementary Methods.

**Figure 1.**
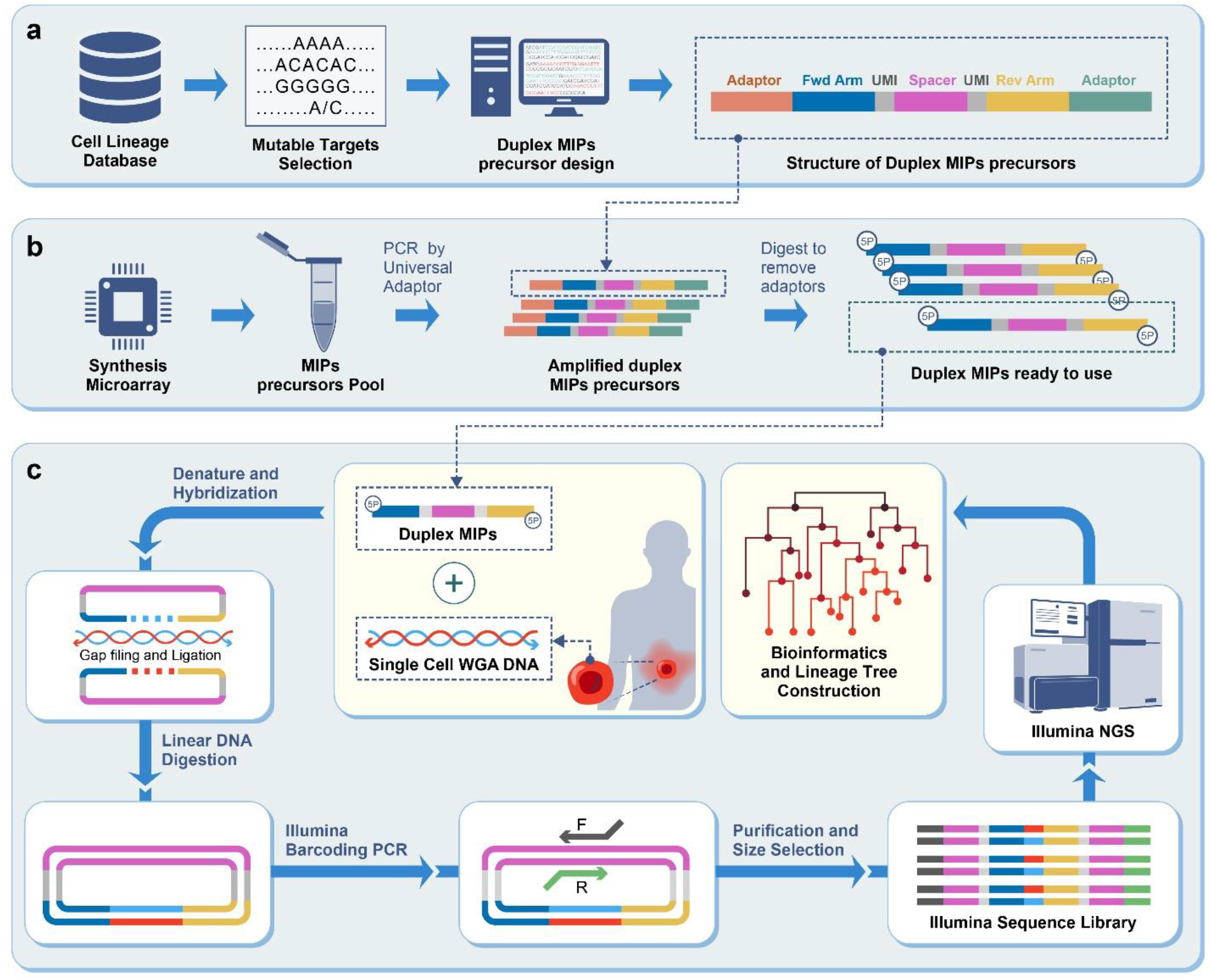
Duplex MIPs based cell lineage workflow. (a) Design of duplex MIPs precursor: desired targets are selected from our cell lineage database and precursors are designed; (b) Duplex MIPs preparation: duplex MIPs precursors are synthesized on a microarray, pooled and amplified by PCR. The product is digested by MlyI to remove the universal adaptors (orange and green), purified and diluted to obtain active duplex MIPs; (c) Duplex MIPs and template DNA are mixed together to allow annealing of the targeting arms (blue and yellow) to the regions flanking the targeted sequence in the template DNA, the products are then circularized by gap filling, facilitated by DNA polymerase and ligase. Linear DNA, including excess MIPs and template DNA, is eliminated by exonucleases digestion. The Illumina sequencing library is generated individually for each sample in a PCR step, facilitating addition of adaptors and barcodes. Libraries are pooled and sequenced using Illumina NGS platform, followed by analysis of the raw reads to detect mutations. The latter are then used to infer the cell lineage tree.

As first proof-of-concept for the duplex MIPs lineage system, we reproduced a known lineage history of single cells from a previously developed DU145 *in vitro* benchmark tree^9^ (Figure 2, Supplementary “DU145 *in vitro* tree”). Briefly, a SC from the DU145 human male prostate cancer cell line has been clonally cultured; iteratively single cells where picked from the previous culture and clonally cultured to generate an *in vitro* tree covering nine generations, each cultured for 12 to 15 cell divisions. In each iteration, SCs were picked using CellCelector, annotated with their plate/well coordinates in the lineage tree, WGA products were prepared and subjected to analysis using a duplex MIPs panel (“OM6”, Supplemental File 1), encompassing ~12K STR targets. Following the library’s sequencing by Illumina NextSeq, the raw data was subjected to tailored analysis in our computational pipeline (Supplementary “Integrated bioinformatics Database Management System (DBMS)” and “Reconstruction parameters”). Here and in all other experiments presented, the same tools and parameters were used: mapping a custom reference of STR variations using bowtie2, genotyping stutter profiles using R&B method^31^, coarse assignment of allelic identities and finally parsimony based phylogenetic reconstruction using FastTree2^32^. The results demonstrate imperfect but accurate reconstruction of the expected reference topology. Most of the major reference splits in this validation experiment are significantly supported in bootstrap analysis (Figure2, significance marked at transfer bootstrap expectation (TBE)^33^>70%).

**Figure 2.**
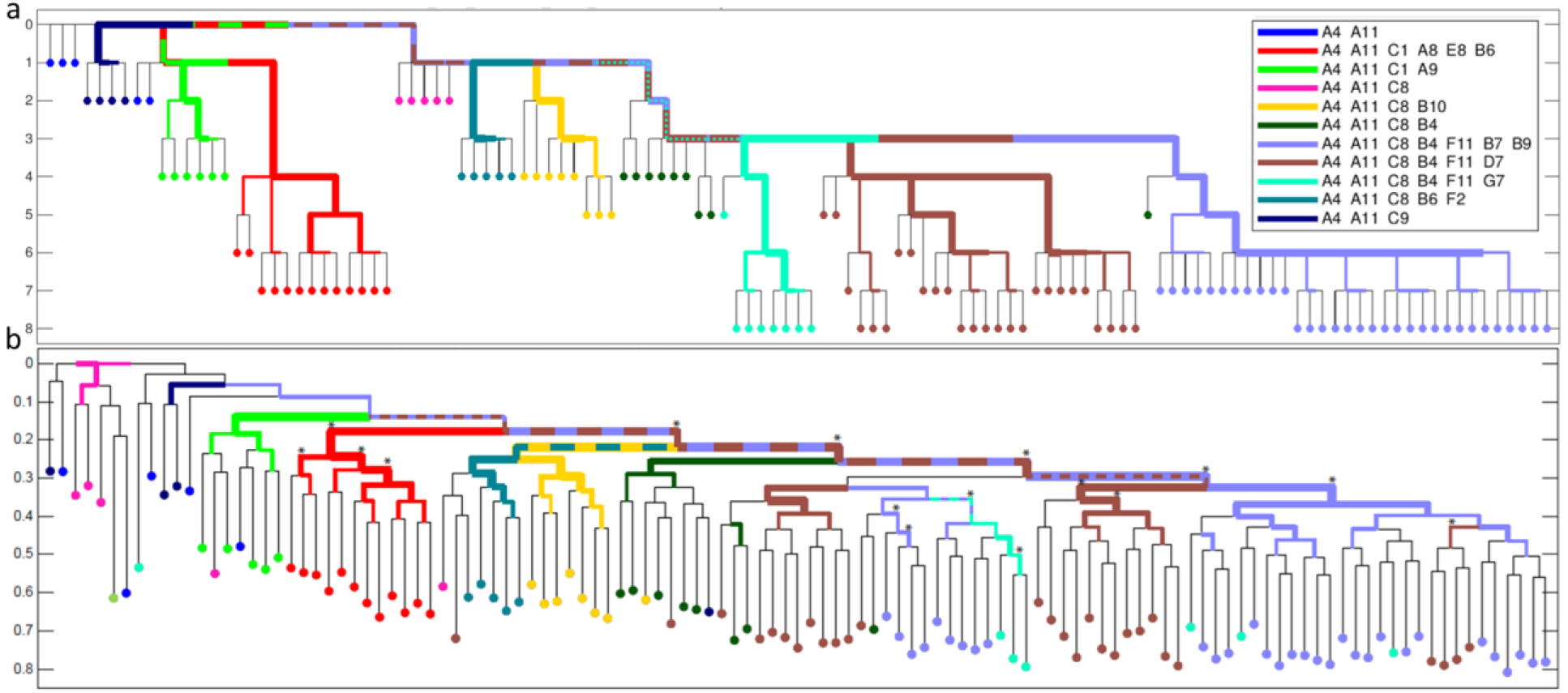
DU145 ex vivo tree comparison as reconstructed by the Duplex MIPS pipeline. (a). The in vitro reference tree, colors assigned by shared ancestors labeled by sampling wells prefix, Y axis indicated sampling generations. (b) The ex vivo lineage tree as reconstructed by duplex MIPs data using the FastTree2 algorithm and uniform transition probabilities. Hypergeometric tests are carried out for each internal branch to assess whether subtree leaves are enriched for a given cell population, the results are reflected in the highlighting and width of edges. Edges reproduced with significant support (TBE>70%) in bootstrap analysis of the duplex MIPs data are annotated with asterisks. Y-axis indicates relative depth when the tree is rooted at the median of the A4_A11 group, which is closest to the real root and therefore used for its approximation.

Consequently, in order to show the potential of duplex MIPs for *ex vivo* SC lineage reconstruction of real human clinical cases, we opted to reconstruct the lineage of SCs sampled from a melanoma patient termed YUCLAT^34^. Five sample groups were obtained from the patient: normal peripheral blood lymphocytes (PBL), skin metastases (Met 1, Met 2 and Met 3 recurring at the patient’s right scapular area excised several months apart, and a more distant metastasis 4, from an axillary lymph node. Primary tumor samples were not available. The cells were processed by our duplex MIPs platform using a panel that covers a similar set of ~12K STR targets as earlier, with the addition of cancer related SNV targets and patient-specific SNVs based on previous bulk exome sequencing data of the patient YUCLAT (Figure 3, Supplementary “WGA DNA of single cell from a Melanoma patient”, panel “OM7” described in Supplemental File 2). The reconstructed STR lineage tree demonstrated an effective *in vivo* separation, validating the expected groups by near-perfect separation of the healthy samples (PBL) from the metastatic ones (Met 1-4). The same separation can also be validated utilizing known melanoma and patient specific SNV mutations that are also sequenced by the same MIP panel (Figure 3 bottom table). Significantly, the STRs derived information for those cells goes further than the healthy/metastatic separation, refining the clusters and showing that Met4, sampled from an axillary lymph node, is clustered separately from the three other metastatic groups sampled from the patient’s back.

**Figure 3.**
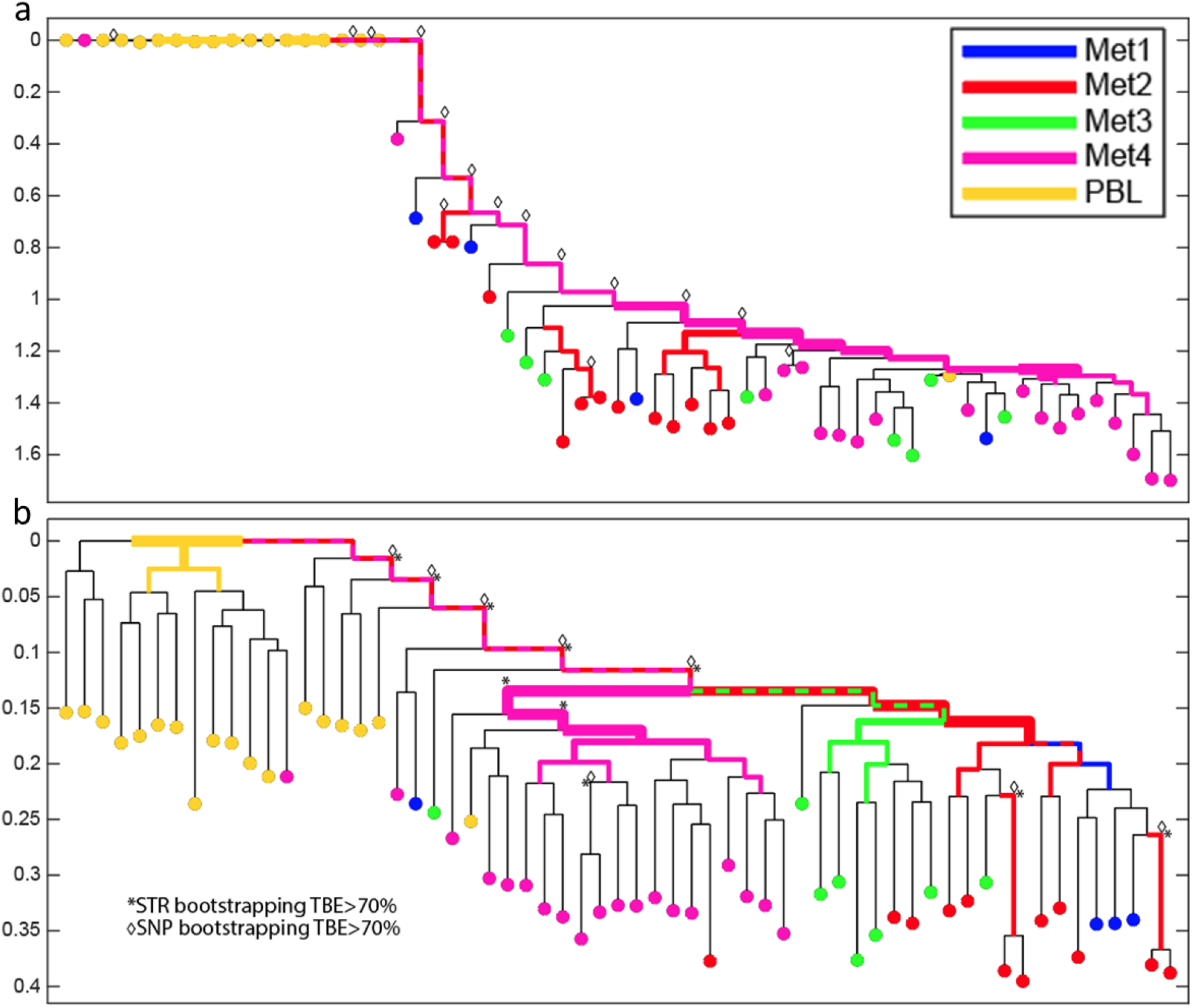
Cell lineage reconstruction of single cells from melanoma metastases and peripheral blood lymphocytes (PBL) from the patient YUCLAT. (a) SNV based reconstruction dendrogram of YUCLAT, presenting significant separation (TBE>70%, annotated by diamond symbols) of the expected separation of cancer cells from the PBLs. (b) STR based reconstruction dendrogram of YUCLAT, presenting significant separation (TBE>70%, annotated by asterisks) of the Met4 samples on top of the expected separation of cancer cells from the PBLs. In addition to the bootstrap analysis performed independently for the two trees, annotated by diamonds in (a) and asterisks in (b) when significant values were found, the SNV based support is also annotated on top of the STR based reconstruction in (b), emphasizing the refinement provided by STRs. Hypergeometric tests are carried out for each internal branch to assess whether subtree leaves are enriched for a given cell population, the results are reflected by the highlighting and width of the edges. Y-axis indicates relative depth when the tree is rooted at the median genotype of the PBL group, under the assumption that it would best approximate the true zygote root.

Finally, whereas cancer cell lineages can be tackled by assessing various DNA sequence alterations including SNVs, CNVs and STRs due to genetic and genomic instability, healthy cells display normal mutation rates, rendering their lineage reconstruction significantly more challenging. In order to test if we can recover the lineage history of healthy cell populations and even trace embryonic cell lineage commitment, we subjected cells of hematopoietic (CD3+ CD19+, CD68+), endothelial (CD31+) and epithelial (oral epithelial cells, OECs) origin (Supplementary “Healthy Human samples”) to whole genome amplification and MIP-analysis using the patient-generic duplex MIPs panel (“OM6”, Supplemental File 1). For WGA, we used the Ampli1-protocol^35,36^ that we found superior for SC lineage tree reconstruction compared to other methods^37^. Indeed, we demonstrate that this panel can identify mutations supporting the separation of embryonic germ layers in multiple patients (Figure 4).

**Figure 4.**
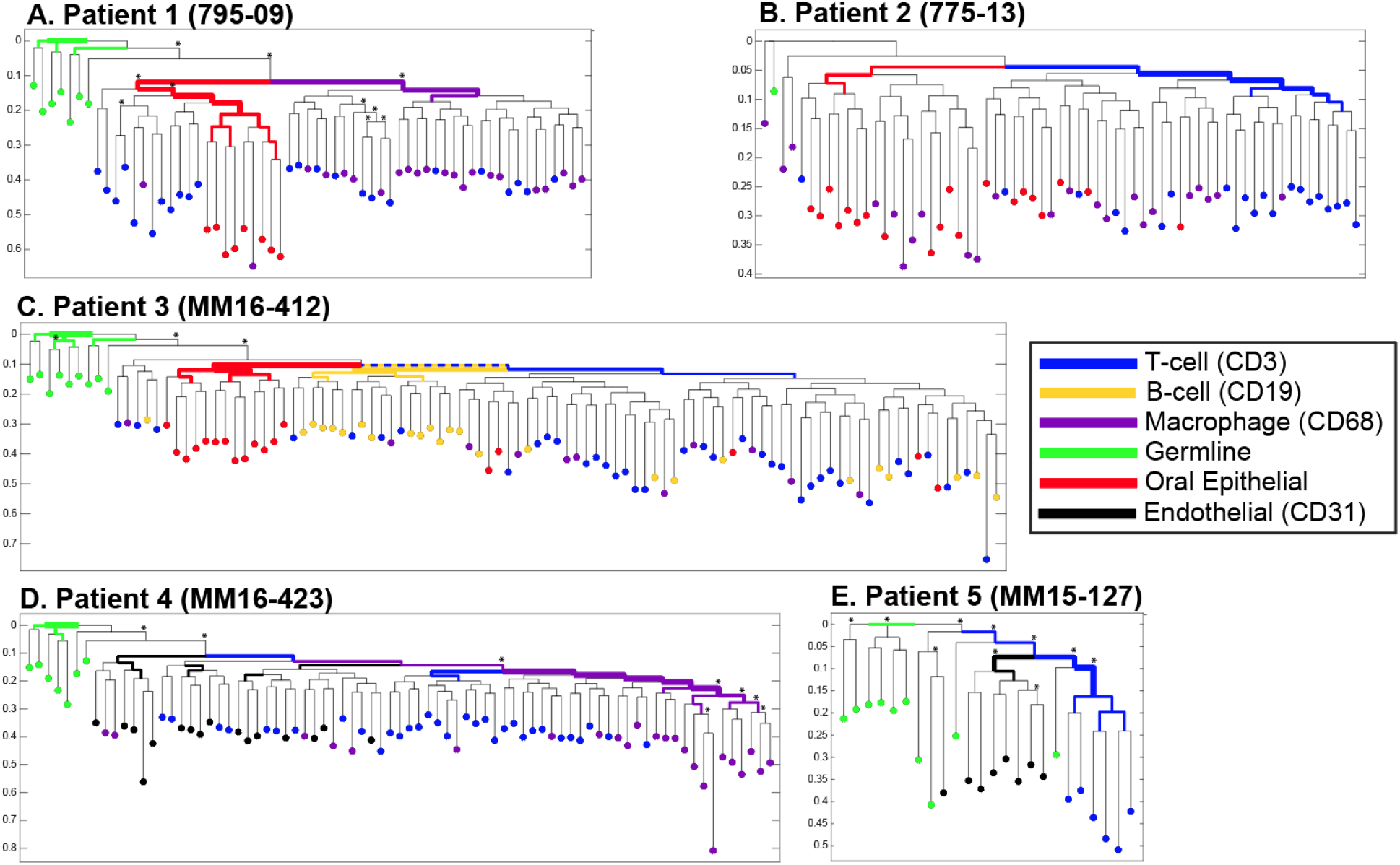
Cell lineage reconstruction of healthy cell populations showing significant separation (TBE>70%, annotated by asterisks) of oral epithelial cells in patient 1 and of endothelial cells in patients 4 and 5 from the hematopoietic populations. This is expected by our current understanding of embryonic development, marking the separation of the embryonic germ layers. As this separation is not necessarily clonal (a single cell gives rise to a germ layer), mixed clusters can also be observed. Hypergeometric tests are carried out for each internal branch to assess whether subtree leaves are enriched for a given cell population, the results are reflected by highlighting and width of the edges. Y-axis indicates relative depth in all the patient-trees and the trees are rooted at the median genotype of the Germline group (bulk samples), under the assumption that it would best approximate the true zygote root.

In complex organisms such as zebrafish, mouse and human, coalescence of cell lineage trees is highly challenging. We have developed a scalable and cost-effective cell lineage discovery platform with low initial costs (Supplementary “Comparison with other STR lineage platforms”) supported by a tailored bioinformatics analysis and Database Management System (Supplementary “Integrated bioinformatics DBMS”). By applying this platform to various types of human cells, we demonstrate that tens-of-thousands hyper-mutable STR targets in single-cell WGA DNA could be efficiently acquired, enabling lineage relations among these cells to be discovered.

After generating a proof-of-concept using single cancer cells cultured over several generations, we refined a SNV-based cell lineage tree of a melanoma patient by applying our STR-based approach. In this experiment, the distant anatomical location of a lymph node metastasis was reflected by a distinct cluster in the tree structure obtained by STR but not by SNV analysis. Pushing the envelope further, we successfully applied the approach to single non-cancer cells from different developmental cell lineages. In all five cases (Supplemental Table 6), we obtained a non-random clustering of hematopoietic either vs. oral epithelial cells or vs. endothelial cells, which reflect the early developmental separation of these lineages. Ectoderm-derived OECs and mesoderm-derived cells separate at gastrulation, whereas mesoderm-derived endothelial cells and hematopoietic cells are derived from a common ancestor, the hemangioblast during embryonic vasculogenesis^38^. Vascular regeneration, tissue maintenance and cancer angiogenesis are currently thought to be driven by vascular endothelial stem cells from pre-existing organ vessels and not from bone marrow-derived endothelial precursor cells^39,40^. In contrast, only in two patients (Pat. 1 and 4), we noted a separation of T cells (CD3) and macrophages (CD68). Since CD68-positive cells from patient 1 and 2 were isolated from peripheral blood and cells from patient 3 and 4 from lymph nodes, we could not establish a relation to the isolation source, i.e. between bone-marrow derived macrophages vs. putative tissue resident macrophages^41^. We conclude that our current ability to separate cells of different lineages depends on the time point of separation from common ancestors, the number of cell divisions over life-time and the mutation rate of each cell type. In addition, accurate reconstruction of hematopoietic lineage may be confounded by somatic mosaicism^42^ and, especially, clonal hematopoiesis resulting from accumulation of somatic mutations in hematopoietic stem cells and their clonal expansion through the lifetime of patients^43–46^.

Evidently, technical improvements will also have an impact on faithful cell lineage reconstruction, including (i) reduction of the artificial noise introduced during WGA. This can be achieved by reaction volume reduction^47^, by improvement of SC WGA increasing the uniformity of amplification across the genome^48^, utilization of UMIs or ideally the development of a direct MIP pipeline bypassing the WGA step. (ii) Acquisition of higher mutability STRs, such as mono-repeats, would improve performance in cases of fewer cell divisions and lower mutation rate. (iii) Duplex MIPs targeting joint SNV-STRs regions within one amplicon can be designed to aid bi-allelic STRs genotyping. Improved haplotyping strategies can be implemented to harness this additional information^49^. (iv) Selective targeting of heterozygous STRs with significant difference in repeat numbers, filtering out close bi-allelic loci like (AC)X14, (AC)X15. To do this, a more accurate STRs annotated human reference genome is needed^23^. (v) An improved loci selection during the panel planning might also allow higher success rates and uniformity, thus reducing the minimal sequencing coverage required per sample (Supplementary “Sample sequencing coverage”). Uniformity can be also improved by optimization of the MIP protocol (e.g. by either empirical optimization of the capture reaction or optimized design of capture probes. Finally, cell states, SC spatial information and phenotypes can be integrated into the cell lineage tree, which may be of significant interest for questions related to immune diseases, cancer, and tissue regeneration among others^50^. Thus, our platform will help study human development and cellular processes in both health and disease.

## CODE AVAILABILITY

All the codes of cell lineage analysis tools mentioned in the text are freely available under the GNU General Public License v2.0 at https://github.com/shapirolab/clineage

## DATA ACCESSION NUMBERS

Sequencing data has been deposited with ArrayExpress access number E-MTAB-6411.

## SUPPLEMENTARY DATA

Supplementary Data are available online.

## ACKNOWLEDGMENTS

Liming Tao was partially supported by VATAT postdoctoral fellowship from Israel’s Council for Higher Education Planning and Budgeting Committee. Ehud Shapiro is the incumbent of The Harry Weinrebe Professorial Chair of Computer Science and Biology. Sandra Huber was partially supported by a PhD scholarship of the Studienstiftung des Deutschen Volkes.

## Methods

### DU145 *in vitro* samples

Single cells (hereby SCs) from DU145 human male prostate cancer cell line was separated via an automated cell picking device (CellCelector, ALS) and cultured to generate a nine-level *ex vivo* tree. SCs were seeded in separate microwells and underwent clonal expansion. Then, repeatedly, SCs were picked from the clonal microwells, seeded separately in new microwells, and expanded. Overall, SCs were sampled nine times, every 12-15 cell divisions. WGA DNA of these single cells was prepared using a modified RepliG Mini protocol as described previously^6^.

### WGA DNA of single cell from a Melanoma patient

YUCLAT metastatic melanoma cells and Peripheral Blood Lymphocytes (PBL) were collected from a 64-yr-old male patient by the Tissue Resource Core of the Yale SPORE in Skin Cancer, with the participant’s signed informed consent according to Health Insurance Portability and Accountability Act (HIPAA) regulations with a Human Investigative Committee protocol as described^8^. For Single cell DNA WGA preparation, the RepliG protocol was applied as described previously^6^.

### Healthy Human samples

Cell of hematopoietic origin (T cells, B cells, macrophages) and endothelial cells (Supplemental Table 6), were isolated from various fresh, non-fixed tissues (blood, lymph nodes) sent for analysis to the Institute of Experimental Medicine and Therapy Research, University of Regensburg. From some patients, buccal swabs and peripheral blood were also obtained. The study was approved by the ethics committees of the University Regensburg and Tübingen (07-079, 18-948-101, 535/2016B02). Written informed consent was obtained from all patients.

### Preparation of adhesion slides for single cell isolation from human patient samples

#### Lymph nodes

Single cell suspension from lymph nodes were prepared as previously described^9,10^. Briefly, lymphatic tissue was cut into 1-mm pieces with a sterile scalpel and disaggregated mechanically into a single-cell suspension using the Medimachine (DAKO) equipped with sterile cartridges holding rotating knives (Medicons, DAKO), washed with HBSS (Life Technologies) and centrifuged for 20 min at 1000 g and room temperature (RT) on a density gradient consisting of a 60% Percoll solution (Amersham). Mononuclear cells were isolated from the interphase, washed with PBS (300g, 10 min, RT), resuspended in PBS and counted. Subsequently, cell density was set to 106 cell/ml in PBS and the resulting cell suspension was transferred onto adhesion slides (Thermo Fisher Scientific) at a density of 106 cells/side (0,33 x 106cells/spot). After sedimentation of cells for 1 hour at RT, residual PBS was discarded and slides were air-dried overnight at RT and stored at −20°C.

#### Metastasis

The metastatic tissue was put in Basal medium consisting of DMEM/Ham’s F12 Medium with 10mM Hepes, 1% penicillin-streptomycin (all from Pan Biotech GmbH) and 1% BSA (Sigma Aldrich) and cut into smaller pieces. To obtain single cell suspension, tissue pieces were enzymatically digested with collagenase (Sigma Aldrich), hyaluronidase (Sigma Aldrich) and DNaseI (Roche) at a final concentration of 0.33 mg/ml, 100 μg/ml and 100 μg/ml, respectively, for 20-40 minutes at 37°C. Next, samples were further disaggregated by pipetting and transferred into a 50 ml tube. Following a washing step with PBS (300g, 10 min, RT), the cell pellet was resuspended in pre-warmed (37°C) Trypsin 0.05%/EDTA 0.2% in PBS (Pan Biotech) and treated for five minutes at RT. Digestion was stopped by adding 5 ml of Basal medium supplemented with 10% FCS (Pan Biotech). Next, cells were filtered using a 40μm cell strainer (Becton Dickinson GmbH) and washed with 14 ml PBS (300g, 10 min, RT). The cell pellet was re-suspended in PBS (106cells/ml) and the cell suspension was transferred onto adhesion slides at a density of 106cells/side (0,33 x 106cells/spot). Following sedimentation of cells on the slide (incubation 1 hour at RT), residual PBS was discarded, slides were air-dried overnight at RT and stored at −20°C.

#### Buccal swabs

Oral epithelial cells (OECs) were isolated by rubbing a sterile cotton swab on the patient’s buccal mucosa. The cotton swab was stirred in 1ml PBS and the obtained cell solution was transferred onto adhesion slides without prior counting. Following sedimentation of cells for 1 hour at RT, residual PBS was discarded and slides were air-dried overnight at RT and stored at −20°C.

#### Peripheral blood

Peripheral blood mononuclear cells (PBMCs) were washed with PBS (CellSave tubes) or Hanks solution (Biochrom AG, EDTA vacutainers) and centrifuged (200 g, 10 min, 4°C). The cell pellet was diluted in Hanks solution and centrifuged (1000 g, 20 min, 4°C) on a 60% Percoll (Amersham) density gradient. The interphase containing PBMCs were collected, washed in PB S (200 g, 10 min, 4°C), resuspended and counted. PBMCs isolated from CellSave tubes were transferred onto adhesion slides at a density of 0.5×10^6^ cells/slide (0.17 x 10^6^ cells/spot). Following sedimentation of cells on the slide (incubation 1 hour at RT), residual PBS was discarded, slides were air-dried overnight at RT and stored at −20°C. PBMCs isolated from EDTA-vacutainers were cryo-conserved in 90% FCS and 10% DMSO (Applichem) in liquid nitrogen.

### Isolation of single cells from adhesion slides

#### T cells, macrophages, endothelial cells

After thawing of adhesion slides (lymph nodes, PBMCs from Cellsave tubes, metastatic tissue) for 30 min at RT, unspecific binding was blocked using PBS/10% AB-serum (Bio-Rad) for 30 min at RT. To isolate CD3+ T cells, CD68+ macrophages and CD31+ endothelial cells, slides were stained with an anti-CD3 (cat. No. C7930, Sigma Aldrich), anti-CD68 (clone KP1, Agilent Dako) or anti-CD31 antibody (clone JC70A, Agilent Dako), respectively, for 60 minutes at RT. Rabbit IgG (cat. No. 0111-01, Southern Biotech) was used as isotype control for CD3 and CD31 immunostaining, IgG1k (clone MOPC 21, Sigma-Aldrich) for CD68 immunostaining. Secondary staining was performed for CD3 and CD31 with an AP-polymer anti-rabbit solution (ZUC031-006, Zytomed Systems) or for CD68 with an AP-polymer anti-mouse solution (ZUC077-100) for 30 minutes at RT. Slides were washed with PBS (3 times, 3 min) and developed using the BCIP-NBT detection system (Bio-Rad Laboratories) for 12 min followed by washing with PBS (3 times, 3 min). If slide availability was limited, slides were subjected to a CD3/CD68 double staining: following the incubation with primary antibodies (see above), cells were stained with goat anti-rabbit IgG-Alexa Fluor 555 (Thermo Scientific) and goat anti-mouse IgG-Alexa Fluor 488 (Invitrogen) for 30 minutes at RT. Positively stained cells were isolated from slides with a micromanipulator (PatchMan NP2, Eppendorf) and subjected to whole genome amplification (see below).

#### Oral mucosal epithelial cells

After thawing adhesion slides for 30 min at RT, OECs were identified based on morphologic criteria – diameter of 60-80μm with clearly smaller nucleus and an irregular shape- and isolated using a micromanipulator (PatchMan NP2, Eppendorf). Isolated cells were subjected to whole genome amplification (see below).

### Isolation of single cells from frozen PBMCs

For isolation of single cells, cryo-conserved PBMCs were thawed in RPMI 1640 Medium (Pan Biotech) and 10% FCS, washed with PBS and counted. To identify single B and T cells, the single cell suspension was stained with anti-CD19 PE (clone HIB19, Biolegend) and anti-CD3 FITC-conjugated antibodies (clone HIT3a, Biolegend) for 15 min at 4°C, respectively. Unspecific antibody binding was blocked with PBS/10% AB-serum (Bio-Rad) for 5 min at 4°C prior to staining with CD19 or CD3. Cells were washed once with PBS/2% FCS/0.01% NaN3. Viability dye eFlour 780 (ebioscience) was used to enable discrimination of live and dead cells. CD19- and CD3-positive cells were isolated with a FACSAria cell sorter (BD Bioscience). Single cells were isolated from sorted populations using the micromanipulator (PatchMan NP2, Eppendorf) and subjected to whole genome amplification (see below).

#### Preparation of germline samples

Germline samples were prepared from bulk genomic DNA (gDNA) lymph node cells or PBMCs. gDNA was isolated using the DNeasy Blood & Tissue Kit (Qiagen, Hilden, Germany) following the manufacturer’s protocol. 200ng of unamplified bulk gDNA was used as template for library preparation with MIP-based probes and lineage analysis. In addition, 1ng and 10ng of genomic DNA were amplified (in three technical replicates for gDNA from the breast cancer patient and five technical replicates for gDNA from the melanoma patients) according to the whole genome amplification protocol (see below). The resulting samples were purified using the AMPure XP beads (bead to DNA ratio of 1.8X) following the manufacturer’s protocol.

#### Whole genome amplification

Whole genome amplification (WGA) from single cells was performed as previously described^11,12^. Following WGA, quality of the resulting products was controlled using a dedicated endpoint PCR assay (WGA-QC)^13^. Only DNA products with a genome integrity index (GII) of 3 and 4 were included in downstream analyses. As CD3+, CD19+, CD68+, CD31+ and OECs were isolated from tissues specimens of cancer patients, all cells were subjected to Ampli1TM LowPass Kit according to the manufacturer’s instructions (Menarini Silicon Biosystems). Only cells with a high quality balanced CNV-profile were considered for MIP-based analyses. Prior to MIP-based analyses, all WGA samples were re-amplified as described previously^14^. Every WGA product was re-amplified in two technical replicates. The temperature setting of the elongation steps included in re-amplification reaction was varied between the two technical replicates. One of the two re-amplification reaction was conducted with the temperature of the elongation step set to 65°C, while the other reaction was run with a corresponding setting set to 68°C. Once prepared both technical replicates were mixed in 1:1 ratio prior to further use. In addition, to eliminate residual oligonucleotides and reagents, pooled re-amplified samples were purified with AMPure XP beads with bead to DNA ratio of 1.8X (Beckman Coulter) following the protocol of the manufacturer.

### Target specific duplex MIPs design and preparation

Hyper-mutable STRs were chosen based on the hg19 reference human genome annotation. The selection criteria were that the AC- and AG-type STRs should be longer than 10 repeats; the A- and G-type STRs should be longer than six repeats. SNVs targets were chosen from cancer related highly mutable regions or known cancer associated regions. Furthermore, additional criteria were applied during selection of the target regions: amplicons containing TTAA sequences were ruled out to comply with the design of the Ampli1 WGA kit and amplicons were selected to be ~150bp in size.

Primer-3 based python script was used to design the targeting primers. Only top-scored primers were chosen as candidates for duplex MIPs precursor design. Additional filters were applied to the selected candidates: precursors harboring MlyI restriction site were ruled out; the total length of the precursors should not be longer than 150bp, which was the limit of the oligonucleotide synthesis provider.

Each duplex MIP comprises multiple functional elements (see below) ordered in the following sequence: 5’-[Mly1_F] + [FW_PRIMER_SEQ] + [UMI] + [Backbone] + [UMI] + [(RV_PRIMER_SEQ) in reverse-complement orientation] + [(Mly1_R) in reverse-complement orientation]-3’. The mentioned elements were designed as follows: Mly1_F:5’GTCTATGAGTGTGGAGTCGTTGC3’;

Mly1_R:5’CTAGCTTCCTGATGAGTCCGATG3’; FW_PRIMER_SEQ, RV_PRIMER_SEQ: variable primers designed to chosen targets; UMI: 5’-NNN-3’ (N stands for randomized base over A, C, G, T); Backbone 5’ -AGATCGGAAGAGCACACGTCTGAACTCTTTCCCTACACGACGCTCTTCCGATCT-3’ Ready-to-use duplex MIPS were generated from precursors by removing Mly1_F and Mly1_R with MlyI overnight digestion.

### Duplex MIPs Preparation

OM6 or OM7 pools (lists of the oligonucleotides comprised in each of the duplex MIP designs is listed in the Supplemental File 1 and 2, respectively) were synthesized by Custom Array (GenScript), Inc. The pools were diluted to a final concentration of 1ng/μl and used as templates. PCR primers (OM4_Mly_F:5’GTCTATGAGTGTGGAGTCGTTGC3’;

OM4_Mly_R:5’CTAGCTTCCTGATGAGTCCGATG3’) were designed to fit the universal adaptors.

#### (A). PreAmp PCR

Amplification of OM6 and OM7 precursor oligonucleotide pools was conducted in a 96 well plate by PCR reactions composed of the following reagents:

0.2ng/μl template, 0.3pmol/μl of each OM4_Mly_F and OM4_Mly_R primers, 1X KOD Polymerase MIX (Novagen, Toyobo) prepared according to manufacturers’ instructions, adding 0.5X SYBR green I (Lonza). Final volume of each reaction was 45ul, typically 4-16 reactions were done per panel. The PCR was executed in the LightCycler 480 (Roche) using the following program: 95°C for 2 min (denaturation), 18 cycles of 95°C for 20 sec, 60°C for 10 sec, 70°C for 5 sec followed by elongation at 70°C for 5 min and cool down. Following the PCR, the reactions were pooled in sets of four and purified on one column each (MinElute PCR purification kit, Qiagen) according to the manufacturer’s protocol, eluting in 45ul DDW. Concentration of the purified samples was measured using the Qubit dsDNA HS Assay Kit (Life Technologies). The purified PreAmp PCR product was diluted to 1ng/μl and served as a template for the next step (i.e. Production PCR).

#### (B). Production PCR

Production PCR reactions (final volume 45ul) were done in a 96 well plate using the same recipe and the same program as in the PreAmp PCR, but running 12 cycles, typically 32-48 reactions were done per panel. The same purification procedure was executed: 4 reactions per purification column eluting in 45ul DDW. The purified products were merged and the concentration was measured using the NanoDrop spectrophotometer (Thermo Scientific). The pool of purified PCR products was diluted to ~30ng/μl and subsequently processed in the next step.

#### (C). MlyI digestion of the duplex MIP precursors

In order to cleave the adapter sequences and generate the active form of duplex MIPs, MlyI digestion was carried out by mixing 84ul of the diluted DNA (30ng/μl), 10μl of 10x NEB CutSmart Buffer (NEB) and 6U MlyI (final concentration of 0.6U/μl MlyI) in a 100ul reaction. Several reactions were performed, keeping 20μl of the undigested sample for quality control. The reactions were incubated in Biometra T3 thermal cycler (Biometra) at 37°C for 12 hours, inactivated at 80°C for 20 min and finally kept at 4°C. Subsequently, each reaction was purified using the MinElute PCR Purification Kit (Qiagen) and eluted in 25ul DDW.

#### (D). Quality control of the duplex MIPs

All digested and purified products were merged and concentration measured using the Qubit dsDNA HS assay kit according to the manufacturer’s protocol. The size of the digested product (~105bp) and the undigested sample (~150bp) was verified by running them on a TapeStation (Agilent) screentape (Supplemental Figure 1a). If unwanted byproducts (represented by minor peaks in the TapeStation output plots) were detected in samples after digestion, BluePippin size selection method (Sage Science; 3% Gel Cassettes, 105bp tight mode) was used for further refinement of the MIPs with desired size (~105bp).

#### (E). Preparation of duplex MIPs working solution

Based on the average size of the duplex MIPs (105bp) and measured concentration, the purified digestion product was diluted to a final concentration of 80nM (80fmol/μl) (equivalent to 5.8ng/μl for OM6 and OM7 panels). This solution was used as a storage stock. The stock solution was further diluted 10 times to prepare a working solution (8nM) needed for the sequence capture experiments. Working solutions were stored at −20 °C. The list of all major reagents mentioned above appears in Supplemental Table 4.

### Duplex MIPs-based targeted enrichment pipeline

#### (A). Hybridization

Single cell WGA product concentration is generally 100-200ng/μl. To 2ul of single cell WGA DNA (200~500ng), 8ul of reaction mix was added with final concentration of 0.8fmol/μl duplex MIPs, 1X Ampligase Buffer and 0.9M betaine, in total volume of 10ul. The Hybridization was done using the following program:

100°C lid temperature, 98°C for 3 minutes, followed by a gradual decrease in temperature of 0.01°C per second to 56°C and incubated at 56°C for 17 hours.

Optionally, if the PCR machine cannot decrease the temperature as slowly as 0.01°C/second, an alternative strategy can be applied: 100°C lid temperature; denature at 98°C for 3 minutes; decreased by 0.1°C in a step-wise way, hold for 15 seconds for each step, until 56°C then incubated at 56°C for 17 hours.

#### (B). Gap filling

The gap filling mix was prepared before the end of the hybridization, with final concentration of: 0.3mM each dNTP, 2mM NAD (freshly thawed from −80°C), 1.1M betaine, 1X Ampligase buffer, 0.5U/μl Ampligase and 0.8U/μl Phusion^®^ High-Fidelity DNA Polymerase, 10ul/reaction and kept at 56°C in a heat block.

At the end of the hybridization, the plate was removed from the PCR machine to a 56°C heat block. 10μl of the gap filling mix were added to each well, carefully mixed by pipette, sealed tightly and quickly returned to the PCR machine at 56°C for 4 hours, followed by 68°C for 20 minutes and 4°C until next step. Optionally, after the gap filling, the reaction plate can be stored at 4°C for up to two days.

#### (C). Digestion of linear DNA

Digestion mix was prepared and 2ul/reaction were added with final concentration of: 3.5U/μl Exonuclease I, 18U/μl Exonuclease III, 4U/μl T7 Exonuclease, 0.4U/μl Exonuclease T, 3U/μl RecJf, and 0.2U/μl Lambda Exonuclease. Mix well. The reaction plate was sealed, spun down and incubated at 37°C for 60 minutes, 80°C for 10 minutes and 95°C for 5 minutes. The reactions can be stored at −20°C for months after the digestion step.

### Optimization of the duplex MIP target enrichment workflow

#### (A). Selection of optimal ratio of template DNA and duplex MIPs in the target enrichment step

Whole Genome Amplification (WGA) is a required step for most single cell genomics studies. However, the stochastic nature of WGA protocols can pose challenges for downstream analyses^7^. Assuming that a limiting concentration of the probe could make up for underlying WGA biases, we sought to calibrate the ratio between duplex MIPs concentration and template DNA amount. For this purpose, variable amounts of template Hela DNA (range 0.01-2000ng) were hybridized with various concentrations of MIPs (ranging from 0.08 to 80nM). Performance of sequence capture was evaluated based on the outcome of the subsequent sequencing analysis. Samples with less than 8000 reads in total or under 4000 detected unique loci were regarded as outliers due to low coverage and were excluded from further evaluation. Conditions with 8-80nM of duplex MIPs probes and 250-500ng input DNA produced the most robust target capture efficiency (Supplemental Figure 1b). Considering the yield of single-cell WGA reaction, we decided to use 8nM MIPs and 250-500ng single cell WGA DNA as the template DNA for our standard protocol.

#### (B). Optimization of reaction times of the critical steps of the duplex MIP-based target enrichment protocol

Next, we tested the impact of duration of hybridization, gap filling and nuclease digestion (the three key steps of the duplex MIP-based workflow) on the performance of the target capture. To this end, hybridization was run for 2, 4 and 18 hours, gap filling was conducted for 1, 2 and 4 hours and nuclease digestion was executed for 1 and 2 hours. Variations of the protocol were compared in all by all combinations, amounting to 18 combinations, using HeLa genomic DNA (NEB) at a concentration of 200ng as template and 80nM duplex MIPs (OM6, Supplemental File 1). The best performing protocol was selected following sequencing and assessment of target capture efficiency. Among all tested conditions, the protocol with hybridization for 18 hours, gap filling for 4 hours and nuclease digestion for 1 hour proved to be the best and was used in all succeeding experiments. Following the sequencing of the enriched libraries at a sequencing depth of 10~15 X, we found that under the optimized conditions ~83% of the resulting reads successfully mapped to the target region and a similar percentage of targets represented in the panel could be successfully detected (Supplemental Table 2, an average of 76% of the loci at an average coverage of 226K reads/sample).

#### (C). Library size selection

In order to choose the proper library size, several ranges were chosen for comparison in the BluePippin (Sage Science) size selection step: broad range 240-340bp, narrow range 270-310bp and tight range 300bp, based on the designed amplicon size distribution. The sequencing results of 7 SCs WGA samples and one DNA bulk sample, revealed that both, the 300 and 270-310bp selection ranges had slightly better success rate (2~3%) compared to the 240-340bp range; however, the 240-340bp size selection range stood out with more loci captured (8~15%). Therefore, 240-340bp size selection range was chosen for future experiments (Supplemental Table 3).

### The combination of two independent panel OM6 and OM8

To test the feasibility of combine two independent panel of duplex MIPs, an independent panel of duplex MIPs OM8 (Supplemental File3, Supplemental Figure 1c,d), no shared targets between OM8 and OM6, was prepared and tested with OM6. 8nM OM6 and 8nM OM8 were pooled by 1:1 and used to capture in a single reaction with a modified protocol where 59.7°C was used as Hyb and Gap steps instead of 56°C.

### Sequencing library preparation

#### (A). Sample specific barcoding PCR

Sample specific barcoding PCR was performed using dual-index barcoding primers and NEBNext Ultra II Q5 DNA Polymerase. The structure of the dual-index Illumina barcoding primers was previously reported Biezuner *et al*.^6^.

I5-index-primer: AATGATACGGCGACCACCGAGATCTACAC [i5-8bp-index] ACACTCTTTCCCTACACGACGCTCTTCCG;

I7-index-primer: CAAGCAGAAGACGGCATACGAGAT[i7-8bp-index] GTGACTGGAGTTCAGACGTGTGCTCTTCCG;

2μl of the target enriched product was used as template, 0.5pmol/μl (final concentration) of each unique barcoding pair of dual-index Illumina primers for each sample, 1X NEBNext Ultra II Q5 Master Mix; 0.5X SYBER Green in a 20μl PCR reaction. The PCR was performed in a Roche 480 Light Cycler using the following program: 98°C, 30 sec (denaturation), 5 cycles of 98°C for 10 sec, 56°C for 30 sec and 65°C for 45 sec; 17 cycles of 98°C for 10 sec, 65°C for 75 sec; followed by 65°C for 5 min (elongation); cool down.

#### (B). Purification and pooling of barcoding PCR Samples

The barcoding PCR product was cleaned by adding 0.8X (by volume) AMPure XP SPRI magnetic beads (Beckman Coulter) according to manufacturer’s manual and eluted in 38μl DDW. The cleaned PCR products were transferred by robot (Bravo, Agilent) to a 384 well plate and an equal volume of each of them pooled by Echo (Echo550, Labcyte). The pool was concentrated by MinElute column (Quiagen) to 35ul.

#### (C). Size Selection for Diagnostic Sequencing

The concentrated pool (30μl) was loaded on a 2% V1 cassette for BluePippin (Sage Science) size selection with setting range 240-340bp according to manufacturer’s protocol while 3μl of the concentrated pool was kept for later quality control. The size-selected elution was collected, cleaned by MinElute and eluted with 15μl DDW. The concentration was measured by Qubit dsDNA HS assay kit. The size distribution of the concentrated pool before and after BluePippin was checked by Tape Station dsDNA screen tape. The size-selected pool with a single peak around 300bp was used to prepare 12μl of 4nM (4fmol/μl) ready-to-run library for Illumina NGS.

#### (D). Diagnostic Sequencing

The ready-to-run library was sequenced on MiSeq Nano (MS-103-1001, Illumina) or Micro flow cell (MS-103-1002, Illumina) with 151×2 paired end run parameters according to manufacturer’s manual, the default sequencing primers were used.

#### (E). Production sequencing

Based on the diagnostic sequencing results, a new pool was produced by Echo550 with a volume adjustment for each sample in order to equalize the reads. The pool was concentrated by MiniElute to 35ul and used to prepare a sequencing library as in step (C).

The normalized library was sequenced on NextSeq 500) FC-404-2004-Illumina) flow cell with 151×2 paired end, run parameters according to the manufacturer’s manual, the default sequencing primers were used.

### Sequencing coverage

For the healthy human samples of Figure 4, a target coverage of 5M reads per cell was set in order to saturate coverage question and allow for accurate estimation of the required coverage. Considering the three main patients that had a substantial number of successful samples, we repeated the bootstrapping experiment with a range of computationally subsampled coverage values. The cases are compared against the tree reconstructed with full data, considering normalized triples distance as tree distance (TreeCmp^20^).

### Duplex MIPs workflow timeline

The whole workflow of duplex MIPs pipeline took 5days from hybridization to data analysis, with roughly 3-hour hands on time (Supplemental Figure 1e).

### Comparison with Access Array based STR lineage platforms

The cost between duplex MIPs pipeline and AA pipeline differs in two main aspects: the initial, one-time, synthesis of MIP precursors or primers and the per-reaction consumables such as reagents or microfluidics chip. While the initial synthesis cost remains affordable for both systems at the 2K loci scale, when scaling up to 100K loci, the cost of primers synthesis for AA increases to roughly $1.8M, while the cost of duplex MIPs synthesis remains reasonable, at around $12K (Supplemental Table 4). Once synthesized, sufficient duplex MIPs can be produced for millions of cells using PCR. The cost of reagents and consumables in targeting reactions does not change much as the duplex MIPs pipeline’s scales; but for the AA pipeline, several chips are required to capture more loci, increasing the cost significantly (Supplemental Figure 2d)

The synthesis cost of duplex MIPs pipeline is ~2,200$ for 12,000 loci panel compared with over 30,000$ for 2000 AA primer pairs. Duplex MIPs capture reaction per cell is ~2.33$, while the AA reaction cost per cell is ~19.69$. In total, the cost per locus per cell in duplex MIPs platform is reduced as much as ~8 times not mentioning the time saved.

The duplex MIPs pipeline uses a single PCR phase rather than two in the AA pipeline, resulting in a reduction of 15~20 cycles as inferred by the noise estimation of the STR genotyping algorithm (Supplemental Figure 2a). This translates to less in-vitro noise introduced into the STR repeats during the target enrichment process, which in turn allows a reduction in sequencing costs by enabling confident genotyping with lower sequencing coverage (Supplemental Figure 2c). Thus, STR genotyping was performed using 10X as the minimal coverage threshold in duplex MIPs pipeline, compared to 30X for AA pipeline. The STR genotyping results of the same single-cell WGA DNA sample produced by our MIPs pipeline were compared with the genotypes reported by Biezuner *et al*^6^. We find that the genotypes from both pipelines correlate with 0.994 (Supplemental Figure 2b).

Several protocols for STR genotyping or targeted sequencing of bulk DNA samples have been previously developed (Supplemental Table1). However, a scalable and affordable high throughput method for single-cell lineage reconstruction from WGA DNA is still lacking. During the past decade our lab developed two STR target enrichment protocols for single-cell whole genome amplified DNA. The first was targeting 128 STRs in 4X multiplex PCR; genotyping was analyzed by fragment length distribution using capillary electrophoresis (CE) ^15^. To overcome the low throughput of CE, a second protocol was developed with up to 50X multiplex PCR and subsequent genotyping using NGS sequencing ^6^. This protocol could target over 2000 STRs across 48 samples on a single Access Array chip. This protocol was however too expensive to scale-up further due to the accumulating cost of primers synthesis and AA chips ^6^. Stemming from a published MIPs protocol designed for complex targets ^16^, we developed a cost effective pipeline for STRs targeted sequencing based on duplex MIPs and demonstrated its performance across 12K and 50K targets (Supplemental Table 5). Since the duplex MIPs can be generated by microarray-based synthesis for as little as $11K for 100K probes, while maintaining a fixed reaction cost with the increment of targeting panel, the cost and scalability of STRs target enrichment is significantly improved (Supplemental Figure 2d).

Furthermore, the amplification during library construction was reduced by 15-20 cycles compared to the AA pipeline. This allows for accurate STR genotyping using as little as 5X reads, further reducing the sequencing costs. Currently, with a 12K panel, 150-200 cells can be deep sequenced in one NextSeq (400M paired-end reads) run as demonstrated with the *ex vivo* lineage reconstruction benchmark (Figure 2). We also demonstrated the feasibility to combine two independent duplex MIPs panels working in a single reaction (Supplemental Figure 1d). This allows us to design, purchase, and test our panels incrementally and flexibly. Thus, customized panels could be created and optimized by various combinations. Six base pairs of Unique Molecular Identifier (UMI) are implemented in the duplex MIPs structure, creating 4096 different UMI combinations that can be used to detect capture events for current and future applications. We note that due to the low coverage requirements of our pipeline (minimum of 5 reads), collapsing reads by their UMI content can be neglected. Running the whole workflow, from hybridization to data analysis, takes approximately 5 days, with roughly 3-5 hours of hands-on time. With more refinement in the duplex MIP arm design process, biochemical calibrations and improvements in the bioinformatic design and analysis, more single cell samples could fit in a single sequencing run, scaling up to 100K targets and allowing cell lineage reconstruction of more challenging cell sets, such as those with low mutation rate ^6^.

The DU145 cell line used in the *ex vivo* experiment carries various chromosomal aberrations, while the X chromosome remains mostly haploid ^6^, other chromosomes were found to be entirely or partially polyploidic. The AA platform was designed mostly for targeting loci on the X chromosome and the duplex MIPs platform is distributed more evenly in terms of loci selection across chromosomes. As an outcome, the scale advantage of duplex MIPs is mostly lost on the *ex vivo* benchmark as recently duplicated genomic regions are often highly similar and impossible to haplotype following WGA. Despite the significant loss of genomic information for this reason, similar reconstruction resolution was demonstrated using the duplex MIPs system comparing to AA with only 65% of the haplotype AA loci available via the duplex MIPs. This asserts the reproducibility of STR based lineage reconstruction and supports the relaxation of the strict male-patient X-chromosome policy of the AA system without loss of accuracy.

Another case for comparison is the YUCLAT melanoma cancer patient (Figure 3). Previously, only a separation of Met 4 from healthy PBL was demonstrated ^6^. Here we show a finer separation among the different metastases groups on top of the perfect separation of the PBL group. Met 1, 2 3 and 4 were collected from the scapula and from the axilla, respectively. This topology is supported by the resulting cell lineage tree, reproducing clonal features of the melanoma metastases and the spatial progression of the disease. Part of the duplex MIPs panel applied on the YUCLAT samples targets cancer specific mutations and hotspots, providing an independent source of variability that supported the STR based results.

### Integrated bioinformatics Database Management System (DBMS)

We designed and implemented a scalable architecture of Cell Lineage Discovery Workflow DBMS for collaborative cell lineage studies. The system supports (i) Data storage and labeling that allows access to workflow data, including each donor’s data (anonymously) as well as samples’ details like origin, cell type, physical location, downstream handling, sequencing results and analysis steps; (ii) Jupyter integration for customized data mining and analysis; (iii) Tracking of workflow protocols, algorithms, sequencing and data analysis steps.

The design, as outlined in Supplemental Figure 11, dissects both the biological and the computational workflow into atomic objects that are documented, referenced and stored for every experiment. The full Entity Relation Diagram (ERD) of this cell-lineage database structure is shown in Supplemental Figure 12. Samples are documented using the web based graphical user interface, from individual to SC DNA resolution (Supplemental Figure 3a). Processes such as targeted enrichment protocols, library preparation (Supplemental Figure 3b) and NGS outputs are documented for all the cells. The NGS raw data (BCL files) is uploaded to the system and demultiplexed according to the documented sample information. After merging paired-end reads, each sample can be processed by either of the two STR-aware alignment pipelines developed in-house. The alignment is implemented for parallel execution on a computing cluster using the Dask distributed package. This results in the creation of millions of histograms depicting the STR length distribution for each cell-locus combination. The analysis is designed with fail-safe points at every step so that any underlying malfunction can be recovered, if needed, from prior computational steps. Those histograms are then processed by the genotyping module employing our STRs amplification stutter model^17^. Individual genotyping results are aggregated for haplotyping, harnessing a population wide perspective to discern the two underlying alleles for each locus (Supplemental Figure 3c).

Following haplotyping, bi-allelic cases are treated as two distinct mono-allelic loci, allowing for integration with unaware phylogenetic reconstruction tools. The system accommodates multiple distance based phylogenetic reconstruction algorithms^18,19^ as well as other hierarchical clustering approaches. Reconstructed phylogenetic trees can be further processed by in-house adaptations of tree plotting tools as well as multiple methods for clustering assessment.

### Tree Reconstruction parameters

The same computational pipeline and set of parameters was used across all the experiments mentioned in the manuscript. A new mapping strategy was used for duplex MIPs sequencing data that improved computational efficiency. Reads were aligned against a custom reference genome of all possible STRs variations in the panel. Reference sequences for an STR locus are shown as an example (Supplemental Figure 4). Sequenced data with a minimal coverage of 10X reads was genotyped and a confidence threshold of 0.05 (correlation above 0.95) between the measured histogram and the reported model was set. The reconstruction based on these resulting genotypes was performed using the FastTree2^19^ algorithm with the mutation count distance matrix.

---

**Sup Fig1:**
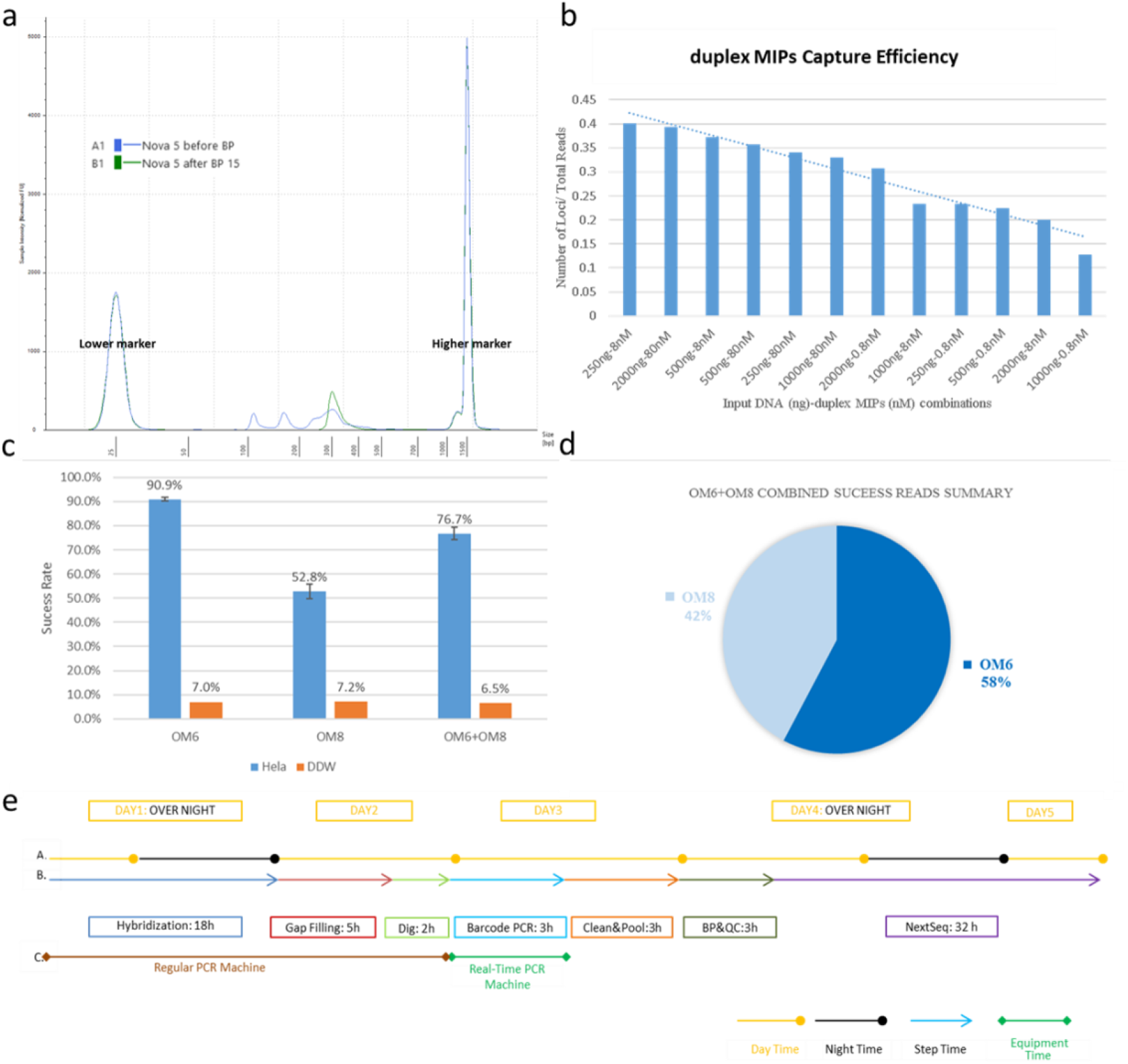
a. Library Quality Control: Tape Station Electropherogram before and after Blue Pippin size selection run b. Comparison of target capture efficiency between different experimental conditions utilizing various MIP to template ratio. Efficiency of capture was evaluated based on the ratio of the number of loci detected to the number of total reads. Average value of two replicates for each test was used in the figure. c,d. The combination of two independent panel OM6 and OM8 | Hela bar (blue) for OM6 is average value of two replicates. Hela bar for OM8 is average value of three replicates. OM6+OM8 bar is average value of two replicates. DDW (orange) is negative control, no replicates. Pie chart is average value of two OM6+OM8 replicates value, *e*. Duplex MIPs workflow timeline (A). Day counts (B). Reaction step time count (C). Machine time count

**Sup.Fig2:**
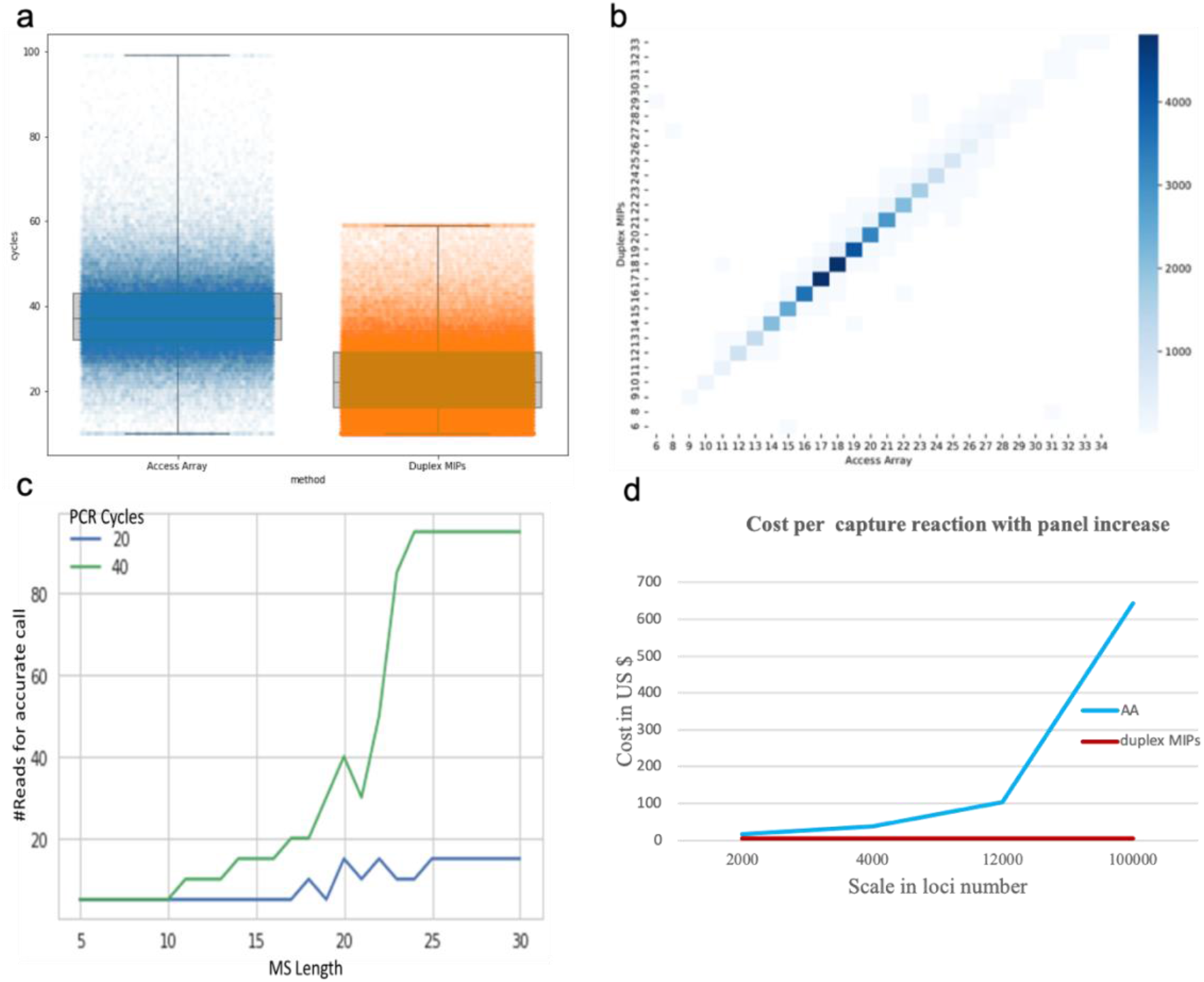
Comparison between Access Array and duplex MIPs pipeline: a, The amplification cycles comparison; b,c, Simulation analysis for the minimal number of reads required for accurate (less than one mistake in 1000 attempts) genotyping of AC microsatellite using AA protocol (estimated 40 amplification cycles) and using duplex MIPs protocol (estimated 20 amplification cycles). d, Genotyping correlation, AA vs Duplex MIPs. Correlation=0.994

**Sup.Fig3:**
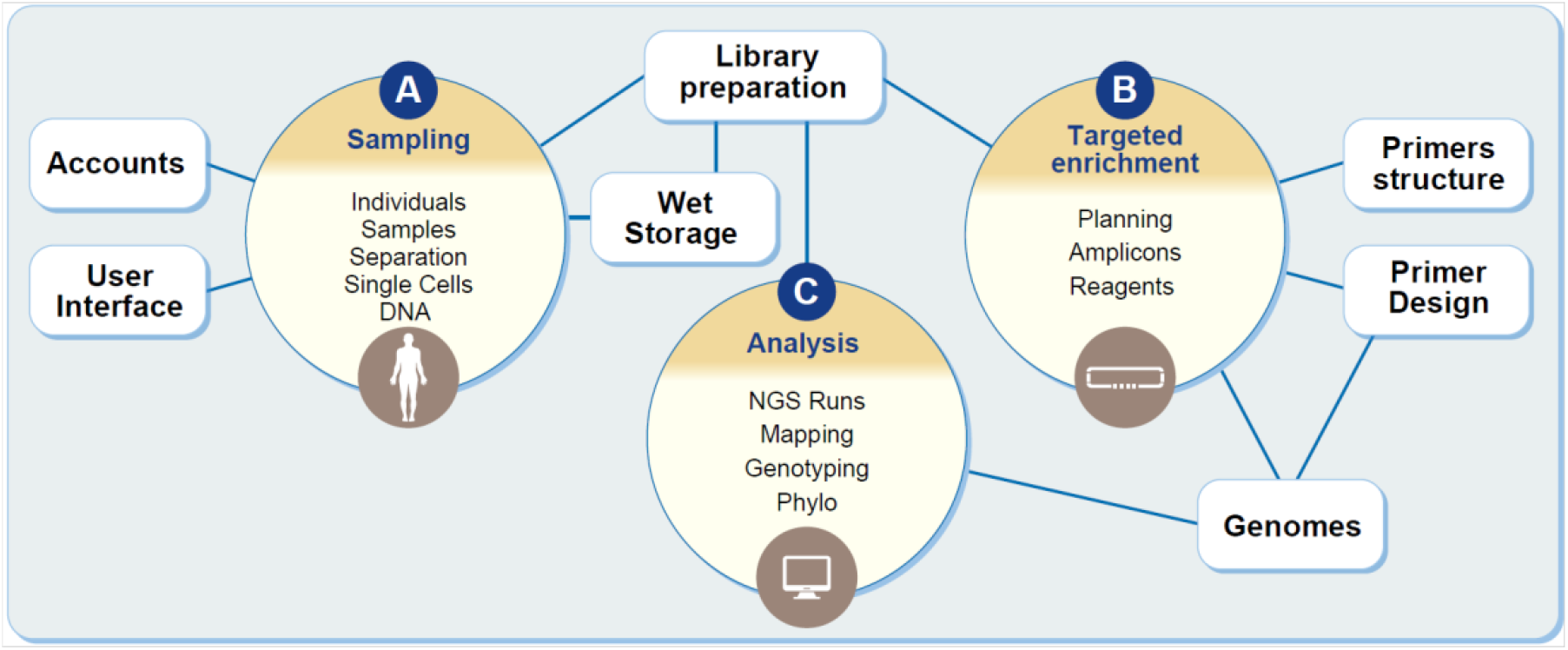
Outline of the integrated bioinformatics Database Management System. (a) Sampling: sampling documentation from patient to DNA, paired with User Interface for viewing, searching and documenting sampling components; (b) Targeted Enrichment: documenting target selection and probe design; (c) Analysis: steps from NGS raw data to Tree across multiple tools, versions and parameters. Paired with the Dask Distributed package for computing clusters. Entity Relation Diagram (ERD) of the cell lineage database structure is in supplemental file4

**Sup Fig4:**
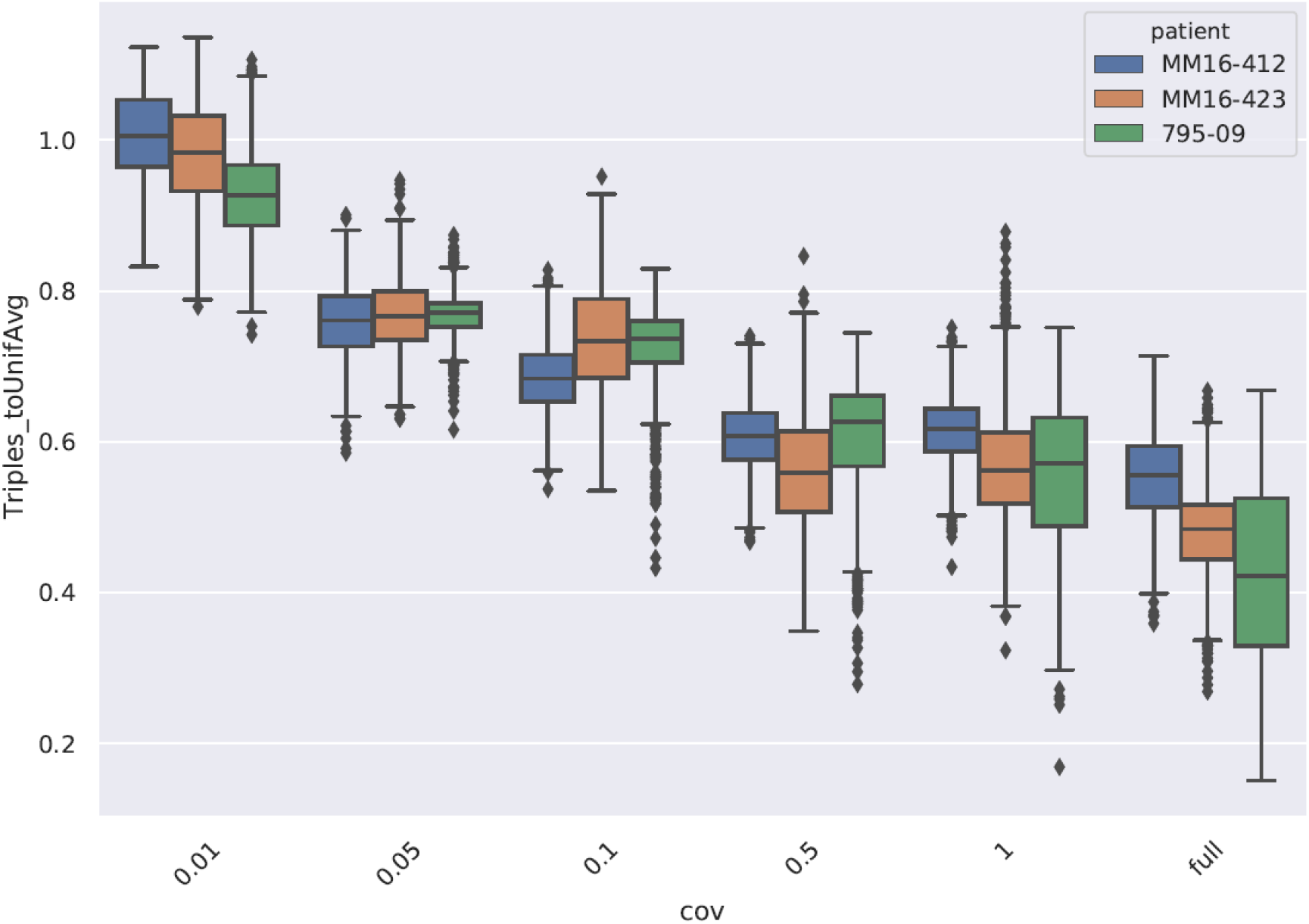
Repeated bootstrapping experiment with a range of computationally subsampled coverage values (x-axis, in million reads). The cases were compared against the tree reconstructed with full data, measuring normalized triples distance (y-axis).

