## Supplementary material for "Retrospective cell lineage reconstruction in Humans using short tandem repeats": sup tab 1

| Tissue | Template | Target Enrichment | Calling Method | Majority STR Type | Targets | Informative Reads | Purpose | Refs |
|---|---|---|---|---|---|---|---|---|
| human blood | Bulk | multiplex PCR | Capillary Electrophoresis | hexa- | ~20 | -- | Forensic | 1 |
| Human | Bulk | Array capture | Next Generation Sequencing | all types | 7851 | 7% | Mutation Discovery | 2 |
| human blood | Bulk | RNA Probes | Next Generation Sequencing | tri- and longer | 10764 | 40% | Mutation Discovery | 3 |
| A.thaliana | Bulk | MIPs | Next Generation Sequencing | tri- and hexa- | 102 | 55-64% | Evolution phylogeny | 4 |
| human leukemia | scWGA | multiplex PCR | Capillary Electrophoresis | di- | 128 | -- | Lineage Reconstruction | 5 |
| human cancer | scWGA | Access Array | Next Generation Sequencing | di- | ~2000 | 90% | Lineage Reconstruction | 6 |
| human cancer/normal | scWGA | duplex MIPs | Next Generation Sequencing | di-, mono- | 12473 | 96% | Lineage Reconstruction | |

**STR capture methods summary**
