## Supplementary material for "Retrospective cell lineage reconstruction in Humans using short tandem repeats": sup tab 2

| Probe Type | DNA | Hyb(hr) | Gap(hr) | Dig(hr) | Total Reads | Total Success | Success Rate | Loci>0 | Loci>4 | Loci>9 |
|---|---|---|---|---|---|---|---|---|---|---|
| OM6 | Hela | 2 | 1 | 1 | 91805 | 18568 | 20% | 6568 | 854 | 121 |
| OM6 | Hela | 2 | 1 | 1 | 115167 | 23632 | 21% | 7293 | 1322 | 243 |
| OM6 | Hela | 2 | 1 | 2 | 121728 | 71250 | 59% | 8892 | 4125 | 1770 |
| OM6 | Hela | 2 | 1 | 2 | 114540 | 71036 | 62% | 9229 | 4508 | 1960 |
| OM6 | Hela | 2 | 2 | 1 | 199923 | 39214 | 20% | 8365 | 2536 | 694 |
| OM6 | Hela | 2 | 2 | 1 | 195185 | 79740 | 41% | 9451 | 4911 | 2267 |
| OM6 | Hela | 2 | 2 | 2 | 100563 | 56274 | 56% | 8787 | 3641 | 1337 |
| OM6 | Hela | 2 | 2 | 2 | 88212 | 51594 | 58% | 8605 | 3247 | 1098 |
| OM6 | Hela | 2 | 4 | 1 | 151143 | 48412 | 32% | 8854 | 3198 | 997 |
| OM6 | Hela | 2 | 4 | 1 | 141481 | 45520 | 32% | 8390 | 3000 | 902 |
| OM6 | Hela | 2 | 4 | 2 | 157111 | 84307 | 54% | 9506 | 5147 | 2480 |
| OM6 | Hela | 2 | 4 | 2 | 129168 | 88406 | 68% | 9498 | 5333 | 2611 |
| OM6 | Hela | 4 | 1 | 1 | 212479 | 111956 | 53% | 10162 | 6138 | 3348 |
| OM6 | Hela | 4 | 1 | 1 | 234372 | 133546 | 57% | 10269 | 6808 | 4101 |
| OM6 | Hela | 4 | 1 | 2 | 129933 | 52523 | 40% | 8995 | 3295 | 1127 |
| OM6 | Hela | 4 | 1 | 2 | 141878 | 62774 | 44% | 9369 | 4097 | 1566 |
| OM6 | Hela | 4 | 2 | 1 | 291192 | 151906 | 52% | 10468 | 7360 | 4635 |
| OM6 | Hela | 4 | 2 | 1 | 261932 | 154769 | 59% | 10503 | 7442 | 4729 |
| OM6 | Hela | 4 | 2 | 2 | 2279390 | 960410 | 42% | 8474 | 8086 | 7674 |
| OM6 | Hela | 4 | 2 | 2 | 158861 | 119662 | 75% | 10064 | 6275 | 3624 |
| OM6 | Hela | 4 | 4 | 1 | 258732 | 93063 | 36% | 10062 | 5689 | 2785 |
| OM6 | Hela | 4 | 4 | 1 | 175854 | 107480 | 61% | 10156 | 6287 | 3512 |
| OM6 | Hela | 4 | 4 | 2 | 207550 | 156801 | 76% | 10395 | 7339 | 4781 |
| OM6 | Hela | 4 | 4 | 2 | 146975 | 112963 | 77% | 10028 | 6267 | 3519 |
| OM6 | Hela | 18 | 1 | 1 | 108935 | 75979 | 70% | 9946 | 5124 | 2297 |
| OM6 | Hela | 18 | 1 | 1 | 281556 | 218901 | 78% | 10831 | 8540 | 6092 |
| OM6 | Hela | 18 | 1 | 2 | 229945 | 82983 | 36% | 9935 | 5247 | 2571 |
| OM6 | Hela | 18 | 1 | 2 | 161878 | 80571 | 50% | 9948 | 5148 | 2376 |
| OM6 | Hela | 18 | 2 | 1 | 112089 | 80908 | 72% | 10092 | 5458 | 2587 |
| OM6 | Hela | 18 | 2 | 1 | 191178 | 154354 | 81% | 10649 | 7833 | 5016 |
| OM6 | Hela | 18 | 2 | 2 | 97018 | 39422 | 41% | 8628 | 2692 | 893 |
| OM6 | Hela | 18 | 2 | 2 | 111756 | 57099 | 51% | 9508 | 4006 | 1576 |
| OM6 | Hela | 18 | 4 | 1 | 105243 | 87278 | 83% | 10100 | 5780 | 2814 |
| OM6 | Hela | 18 | 4 | 1 | 240644 | 200976 | 84% | 10795 | 8679 | 6224 |
| OM6 | Hela | 18 | 4 | 2 | 223929 | 95769 | 43% | 10204 | 6009 | 3099 |
| OM6 | Hela | 18 | 4 | 2 | 183300 | 145216 | 79% | 10607 | 7781 | 4816 |

Calibration of duplex MIPs process: Hyb -Gap-Dig | Hyb means hybridization, the first step in duplex MIPs capture protocol. Gap means gap filing, the second step. Dig is the third step, linear DNA digestion. Green highlighted the protocol we chosen as standard. The success rate calculated as mapped reads/total reads. The loci captured defined as loci that has at least one mapped read.
