## Supplementary material for "Retrospective cell lineage reconstruction in Humans using short tandem repeats": sup tab 3

| BluePippin Size (bp) | Name | Total Reads | Total Success | Success Rate | Loci >0 |
|---|---|---|---|---|---|
| 300 | W151020 p2-C9 | 61816 | 57226 | 92.6% | 7944 |
| 240-340 | W151020 p2-C9 | 144518 | 130517 | 90.3% | 9783 |
| 270-310 | W151020 p2-C9 | 87924 | 82158 | 93.4% | 8791 |
| 300 | H1- 090215-B3 | 87665 | 83768 | 95.6% | 3075 |
| 240-340 | H1- 090215-B3 | 164359 | 155680 | 94.7% | 4046 |
| 270-310 | H1- 090215-B3 | 122252 | 117106 | 95.8% | 3574 |
| 300 | H1- 090215-B3 | 85631 | 81251 | 94.9% | 2891 |
| 240-340 | H1- 090215-B3 | 178585 | 168311 | 94.2% | 3985 |
| 270-310 | H1- 090215-B3 | 123557 | 117945 | 95.5% | 3411 |
| 300 | H1- 090215-B6 | 129546 | 123914 | 95.7% | 5568 |
| 240-340 | H1- 090215-B6 | 387493 | 368533 | 95.1% | 6850 |
| 270-310 | H1- 090215-B6 | 213020 | 203982 | 95.8% | 6209 |
| 300 | H1- 090215-E9 | 114190 | 109002 | 95.5% | 5194 |
| 240-340 | H1- 090215-E9 | 460648 | 436100 | 94.7% | 6728 |
| 270-310 | H1- 090215-E9 | 137124 | 131168 | 95.7% | 5569 |
| 300 | H1- 090215-A1 | 77196 | 73307 | 95.0% | 5230 |
| 240-340 | H1- 090215-A1 | 154987 | 146026 | 94.2% | 6327 |
| 270-310 | H1- 090215-A1 | 120505 | 114812 | 95.3% | 5882 |
| 300 | H1- 090215-F5 | 14620 | 13535 | 92.6% | 3706 |
| 240-340 | H1- 090215-F5 | 184932 | 170304 | 92.1% | 7744 |
| 270-310 | H1- 090215-F5 | 22392 | 20930 | 93.5% | 4533 |
| 300 | PC2 | 12488 | 11149 | 89.3% | 5004 |
| 240-340 | PC2 | 95192 | 81596 | 85.7% | 10347 |
| 270-310 | PC2 | 9078 | 8208 | 90.4% | 4454 |

Calibration of Sequencing Library Size Selection | PC2 was bulk DNA; all the other samples were single cell WGA DNA
