## Supplementary material for "Retrospective cell lineage reconstruction in Humans using short tandem repeats": sup tab4

| Reagents | Cat.No | Cost($) | Total Volume(ul or reactions) | Volume per Reaction (ul) | Cost per Reaction ($) |
|---|---|---|---|---|---|
| duplex MIPs | Home made | 2200 | 9400000 | 1 | 0.000234043 |
| Betaine solution 5M | B0306 1VL Sigma | 49 | 1500 | 4 | 0.13 |
| Phusion High-Fidelity DNA Polymerase | NEB-M0530L | 424 | 250 | 0.4 | 0.68 |
| Ampligase 10X Reaction Buffer | A1905B EPICENTRE | 66 | 5000 | 2 | 0.03 |
| Ampligase DNA Ligase W/O Buffer | A3210K EPICENTRE | 693 | 2000 | 1 | 0.35 |
| Exonuclease I (E.coli) | NEB-M0293L | 268 | 750 | 0.175 | 0.06 |
| Exonuclease III (E.coli) | NEB-M0206L | 236 | 250 | 0.18 | 0.17 |
| RecJf | NEB-M0264L | 272 | 167 | 0.1 | 0.16 |
| Exonuclease T - 1,250 units, | NEB-M0265L | 280 | 250 | 0.08 | 0.09 |
| T7 Exonuclease | NEB-M0263L | 248 | 500 | 0.4 | 0.20 |
| Lambda Exonuclease | M0262L | 268 | 1000 | 0.02 | 0.01 |
| NEBNext Ultra II Q5 MasterMix | NEB-M0544L | 395 | 12500 | 10 | 0.32 |
| MinElute PCR Purification Kit | QIAGEN 28006 | 594 | 250reactions | 2reaction/Run | 0.02 |
| Qubit® dsDNA HS Assay Kit, | Q32854 | 269 | 500reactions | 2reaction/Run | 0.01 |
| Agencourt Ampure XP Beads | BeckmanCoulter A63881 | 1485 | 600000 | 16 | 0.04 |
| 2% Agarose, dye-free, BluePippin, 100 - 600, | BDF2010 | 475 | 50reactions | 1reaction/Run | 0.05 |
| TapeStation Screen Tap | 5067-5582 | 211 | 112 reactions | 2reaction/Run | 0.02 |
| TapeStation Reagents | 5067-5583 | 90.33 | 112 reactions | 2reaction/Run | 0.01 |
| | | | | **Cost per Cell** | **2.33** |

The cost of duplex MIPs capture pipeline. The cost was calculated by 200 cells/run, WGA cost and sequencing run cost were not included.
