## Supplementary material for "Retrospective cell lineage reconstruction in Humans using short tandem repeats": sup tab5

| Barcodecontent ID | Tota lReads | Success Read | Success Rate | # Loci ( >=1X) | Panel Name | Template | Panel Size |
|---|---|---|---|---|---|---|---|
| 11094 | 83946 | 64784 | 77.2% | 15425 | OM9 | Hela | 50K |
| 11098 | 86039 | 66589 | 77.4% | 15531 | OM9 | Hela | 50K |
| 11102 | 46 | 9 | 19.6% | 9 | OM9 | DDW | 50K |
| 11730 | 205862 | 163105 | 79.2% | 13784 | OM6+OM8 | Hela | 25K |
| 11731 | 189982 | 140744 | 74.1% | 13312 | OM6+OM8 | Hela | 25K |
| 11732 | 184 | 12 | 6.5% | 12 | OM6+OM8 | DDW | 25K |

25K and 50K panels performance (Data from Miseq 70 and 71)
