## Supplementary material for "Retrospective cell lineage reconstruction in Humans using short tandem repeats": sup tab6

| | Patient 1 | Patient 2 | Patient 3 | Patient 4 | Patient 5 |
|---|---|---|---|---|---|
| **Germline (bulk)** | PB | PB | PB | PB | LN |
| **CD68+ Macrophages** | PB | PB | LN<br>Met | LN | LN |
| **CD3+ T cells** | PB | PB | LN<br>Met | LN | LN |
| **CD19 + B cells** | -- | -- | PB | -- | -- |
| **CD31+ endothelial cells** | | | | LN | LN |
| **Oral epithelial cells** | OM | OM | OM | | |

LN: lymph node; PB: peripheral blood, Met: metastases; bulk: bulk genomic DNA; OM: oral mucosa
