## Supplementary figures and images for "Retrospective cell lineage reconstruction in Humans using short tandem repeats"

### sup file5

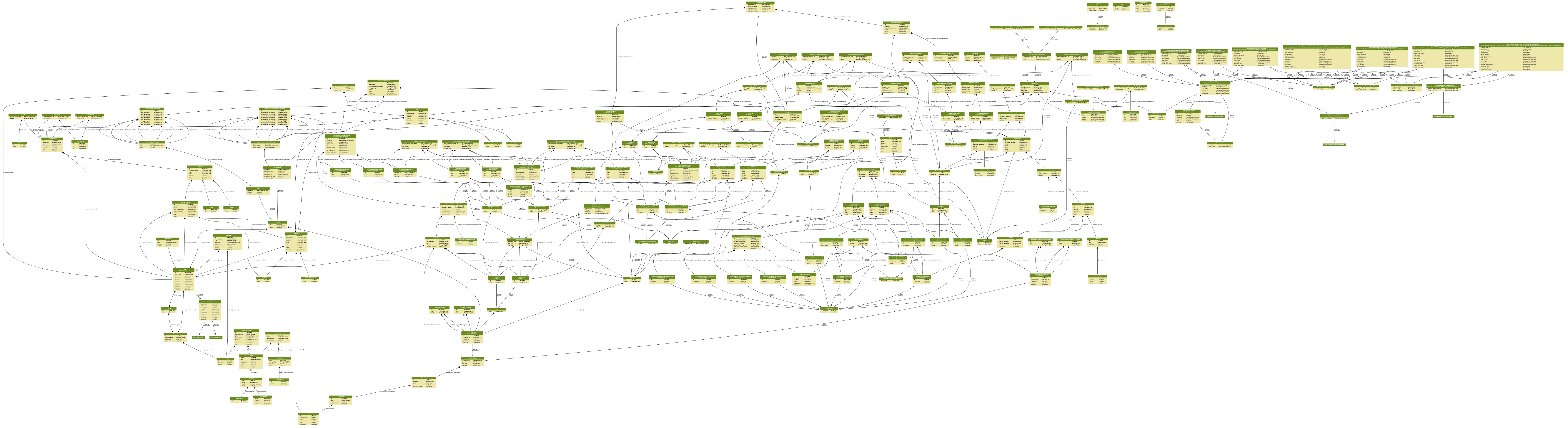
