## Supplementary material for "Retrospective cell lineage reconstruction in Humans using short tandem repeats": sup file1

| TEID | Sequence |
|---|---|
| 121394 | TATGAGTGTGGAGTCGTTGCAGTTATGCTCTTTCTCCCAGGCTANNNAGATCGGAAG |
| 120023 | TATGAGTGTGGAGTCGTTGCAGATACAGTCTCACTTTGTGGCCANNNAGATCGGAAG |
| 68760 | TATGAGTGTGGAGTCGTTGCAGAAGGGGCAAAAGGTTTTTGTTNNNAGATCGGAA |
| 121343 | TATGAGTGTGGAGTCGTTGCGGAGTTTTGGTGTTTCACCCCGGNNNAGATCGGAAG |
| 121104 | TATGAGTGTGGAGTCGTTGCATGGGCTCTCACTTTGATATGCANNNAGATCGGAAG |
| 121071 | TATGAGTGTGGAGTCGTTGCCATTGGTTTCCTAGGCCTGTTAGAANNNAGATCGGAA |
| 121158 | TATGAGTGTGGAGTCGTTGCGGTTTGTTTTTGTATTTGTCTCGCTNNNAGATCGGAA |
| 121359 | TATGAGTGTGGAGTCGTTGCGAGTGGAGACTGGGTTTCATTATGNNNAGATCGGAA |
| 36853 | TATGAGTGTGGAGTCGTTGCTTCTGGGAAGTAAACCACAAGTCANNNAGATCGGAA |
| 99042 | TATGAGTGTGGAGTCGTTGCCCCAGAATGTGTGAGCAGTTCATNNNAGATCGGAAG |
| 61402 | TATGAGTGTGGAGTCGTTGCTGACCCCTTGATTTACCATCTCCTNNNAGATCGGAAGA |
| 48447 | TATGAGTGTGGAGTCGTTGCTTGAGCAGCAGAAGTGACAGATCNNNAGATCGGAAG |
| 69560 | TATGAGTGTGGAGTCGTTGCGTTGTGTGTGTCTGCAAGGTTTTNNNAGATCGGAAG |
| 96115 | TATGAGTGTGGAGTCGTTGCGAAACACAGCCAAATACCAGTCGNNNAGATCGGAAG |
| 39350 | TATGAGTGTGGAGTCGTTGCTCAGCACCATCTACTTTCCTGGANNNAGATCGGAAGA |
| 39895 | TATGAGTGTGGAGTCGTTGCGATTCCCTGGTCTTACTCCCTGANNNAGATCGGAAGA |
| 58044 | TATGAGTGTGGAGTCGTTGCACTCCTGTACCTCTTGGCAGAATNNNAGATCGGAAGA |
| 64935 | TATGAGTGTGGAGTCGTTGCTGCTCCATGAATAACAGTGAGGTNNNAGATCGGAAG |
| 67681 | TATGAGTGTGGAGTCGTTGCCTCCAACCACATTTTTCCCCCTTNNNAGATCGGAAGAG |
| 121398 | TATGAGTGTGGAGTCGTTGCCCCCATAAAAGGAGTTAGGGCANNNAGATCGGAAGA |
| 34313 | TATGAGTGTGGAGTCGTTGCACTGGAAAAATATTGTGACCCCCANNNAGATCGGAA |
| 60882 | TATGAGTGTGGAGTCGTTGCAACATGCACATACACATACCCCTNNNAGATCGGAAGA |
| 58721 | TATGAGTGTGGAGTCGTTGCGCCCTATGTAATGAAGCCTGCATNNNAGATCGGAAG |
| 121388 | TATGAGTGTGGAGTCGTTGCGATAACCTCTGCTCACCGCAANNNAGATCGGAAGAG |
| 37069 | TATGAGTGTGGAGTCGTTGCAAGTCACCACAAAGACCTTGTCANNNAGATCGGAAG |
| 121362 | TATGAGTGTGGAGTCGTTGCTTTTATTTTGTTTCCCTCGCTACANNNAGATCGGAAGA |
| 120787 | TATGAGTGTGGAGTCGTTGCAGTATTGCTCTGTTGCCCTACGNNNAGATCGGAAGAG |
| 121321 | TATGAGTGTGGAGTCGTTGCCTGTTGGGAGCTTCTGCTCTGNNNAGATCGGAAGAG |
| 119705 | TATGAGTGTGGAGTCGTTGCGCTTGTCTGGAGCTCTTGGATNNNAGATCGGAAGAG |
| 121229 | TATGAGTGTGGAGTCGTTGCAGTACCTCTTTGACATTCCACTTTNNNAGATCGGAAG |
| 121406 | TATGAGTGTGGAGTCGTTGCCTGGGTTCAAGAATTTCTCCAGTNNNAGATCGGAAG |
| 121347 | TATGAGTGTGGAGTCGTTGCCTCCTGGTTCATCACAACCTCTNNNAGATCGGAAGAG |
| 40649 | TATGAGTGTGGAGTCGTTGCGCTTCTTTTTCGTTCCGCGAATANNNAGATCGGAAGA |
| 121422 | TATGAGTGTGGAGTCGTTGCTTTGTGTTGATTGCTTTCCAGACANNNAGATCGGAAG |
| 121738 | TATGAGTGTGGAGTCGTTGCGTCTGGTCTCACACTTCTGGATACNNNAGATCGGAAG |
| 35680 | TATGAGTGTGGAGTCGTTGCCAGAACACCATGTGATGTAGGCANNNAGATCGGAAG |
| 39727 | TATGAGTGTGGAGTCGTTGCTGGACCTGAAATATGGCTGATGANNNAGATCGGAAG |
| 121431 | TATGAGTGTGGAGTCGTTGCATTGATTCTCCTTCCTCTGCCTTCNNNAGATCGGAAGA |
| 99045 | TATGAGTGTGGAGTCGTTGCCGGTTTGTAGGTTTCTGCAGAGAGNNNAGATCGGAA |
| 45885 | TATGAGTGTGGAGTCGTTGCAGGTGGGATACACTTCTCCTTCCNNNAGATCGGAAGA |
| 58344 | TATGAGTGTGGAGTCGTTGCGGTACTCACAGCCAAACTCACATNNNAGATCGGAAGA |
| 121233 | TATGAGTGTGGAGTCGTTGCTGCTGTGGCTAGAACTTCCAAAAANNNAGATCGGAA |

| | |
|---|---|
| 33588 | TATGAGTGTGGAGTCGTTGCGCACACATGCAAAAATACAGGACANNNAGATCGGAA |
| 121209 | TATGAGTGTGGAGTCGTTGCTGTGATTGGTAACAGGTCAGGAAAGNNNAGATCGG |
| 121316 | TATGAGTGTGGAGTCGTTGCGACTGGAGTGGAGTGATGTAGTGATNNNAGATCGG |
| 121890 | TATGAGTGTGGAGTCGTTGCAAACACAATGCCACAGAAGCACNNNAGATCGGAAGA |
| 121003 | TATGAGTGTGGAGTCGTTGCCCTTGTATGCTGTGTACTGCCTGANNNAGATCGGAA |
| 121272 | TATGAGTGTGGAGTCGTTGCCACCCCTCAGCCCAACTACTTGNNNAGATCGGAAGAG |
| 34124 | TATGAGTGTGGAGTCGTTGCTGTTGTCTTCCAGTATGGGACCANNNAGATCGGAAG |
| 121379 | TATGAGTGTGGAGTCGTTGCGACAACCAGATCCCTTGACATCTGNNNAGATCGGAA |
| 121423 | TATGAGTGTGGAGTCGTTGCACAAAATGGATGGACTTCAAGCCAGNNNAGATCGGA |
| 121368 | TATGAGTGTGGAGTCGTTGCAGTAGGTGAGGTGCAGGCACNNNAGATCGGAAGA |
| 56111 | TATGAGTGTGGAGTCGTTGCACAGACTTCAAATCAAGAACAGCTGNNNAGATCGGA |
| 120191 | TATGAGTGTGGAGTCGTTGCCAGCCTACCTCCCAATTCACAGTNNNAGATCGGAAGA |
| 62688 | TATGAGTGTGGAGTCGTTGCCGAGATTTCAAGCCAGTGGATCTNNNAGATCGGAAG |
| 121376 | TATGAGTGTGGAGTCGTTGCTCAGAGCCGTAACAGCTTCCAGNNNAGATCGGAAGA |
| 31119 | TATGAGTGTGGAGTCGTTGCACCACACACAGGGAGATATCCAANNNAGATCGGAAG |
| 121281 | TATGAGTGTGGAGTCGTTGCTCTTGTTCAGCAGATTCCACCCTNNNAGATCGGAAGA |
| 121232 | TATGAGTGTGGAGTCGTTGCAACACCTGTGGGACAGAACTATTCANNNAGATCGGA |
| 121332 | TATGAGTGTGGAGTCGTTGCATGCCCACCCCAGAAATACATATNNNAGATCGGAAGA |
| 121367 | TATGAGTGTGGAGTCGTTGCACAACTATGTCCCTTCCCCTAGGNNNAGATCGGAAGA |
| 120049 | TATGAGTGTGGAGTCGTTGCGGGGAAGCAGCATGATCTCTGCNNNAGATCGGAAG |
| 52651 | TATGAGTGTGGAGTCGTTGCTTGATCCCTTTTGGACCAGTTCGNNNAGATCGGAAGA |
| 121231 | TATGAGTGTGGAGTCGTTGCTCCAAATGTTTCCAAAACCGCCTTTNNNAGATCGGAA |
| 121418 | TATGAGTGTGGAGTCGTTGCTCAGTGGTCCTCCCACAATCANNNAGATCGGAAGAG |
| 100467 | TATGAGTGTGGAGTCGTTGCATGTGTGTATGTACAGTGGCTTGTNNNAGATCGGAA |
| 121292 | TATGAGTGTGGAGTCGTTGCAGACAATGCAGGTGTGATCCCTNNNAGATCGGAAGA |
| 62174 | TATGAGTGTGGAGTCGTTGCAGTCTGCTTACAGGAATTTGGCTNNNAGATCGGAAG |
| 119992 | TATGAGTGTGGAGTCGTTGCAGAGCCTCACTCTGTTACCTAGAGNNNAGATCGGAA |
| 121220 | TATGAGTGTGGAGTCGTTGCGTGGGTTCAAGCACTTCTCAANNNAGATCGGAAGAG |
| 30185 | TATGAGTGTGGAGTCGTTGCGGGGAAGCTCAACCTCACTAAAANNNAGATCGGAAG |
| 49706 | TATGAGTGTGGAGTCGTTGCTGGAAGTCTAAGCAGCATGCTTCNNNAGATCGGAAG |
| 55721 | TATGAGTGTGGAGTCGTTGCTCTGCTCTCTCCTGGTCTGTATGNNNAGATCGGAAGA |
| 53070 | TATGAGTGTGGAGTCGTTGCAGATGTAAGATGTACATGGTGCAGGNNNAGATCGG |
| 99046 | TATGAGTGTGGAGTCGTTGCACTACGTCCAGTTAGGTTTCCAAGNNNAGATCGGAA |
| 56603 | TATGAGTGTGGAGTCGTTGCTGGTGTCTCTGATTTGCTCAGTGNNNAGATCGGAAG |
| 98724 | TATGAGTGTGGAGTCGTTGCCACCCACCATAGCTGAGGATTGAGNNNAGATCGGAA |
| 95606 | TATGAGTGTGGAGTCGTTGCGAAGTTTACAGTCTGGGCCACTTNNNAGATCGGAAG |
| 100459 | TATGAGTGTGGAGTCGTTGCATCAAAGAACACTGAGCACCTCNNNAGATCGGAAGA |
| 100470 | TATGAGTGTGGAGTCGTTGCTCCGTTTGGCAGGATATTTACTANNNAGATCGGAAG |
| 100495 | TATGAGTGTGGAGTCGTTGCGTTACCATGCCAGGTCTCCTATNNNAGATCGGAAGA |
| 98636 | TATGAGTGTGGAGTCGTTGCCCCCACTAGTAGATTATAAGTTCCTGNNNAGATCGGA |
| 95384 | TATGAGTGTGGAGTCGTTGCGCTTCAGATTCCGGAGTTCCTTGNNNAGATCGGAAG |
| 98548 | TATGAGTGTGGAGTCGTTGCTGTGATTGAATGTGATTTGAAGGCAGNNNAGATCG |
| 93674 | TATGAGTGTGGAGTCGTTGCCACACACACGACTTTTGCTCTTANNNAGATCGGAAGA |

| | |
|---|---|
| 100302 | TATGAGTGTGGAGTCGTTGCCTCCACTAGACCTATTCTCTGTGGTNNNAGATCGGAA |
| 99220 | TATGAGTGTGGAGTCGTTGCCGCTTGAGCAGGAGATCGAGNNNAGATCGGAAGAG |
| 99321 | TATGAGTGTGGAGTCGTTGCAATCCCTCATCTTGCTCCACCTANNNAGATCGGAAGA |
| 120205 | TATGAGTGTGGAGTCGTTGCCAGACTTTCGTCCAGGCTGGNNNAGATCGGAAGAG |
| 121164 | TATGAGTGTGGAGTCGTTGCAGACGCACTTTCACTGTGTCNNNAGATCGGAAGAGC |
| 70981 | TATGAGTGTGGAGTCGTTGCTTGGGGAGGGGGATACAATGAATNNNAGATCGGA |
| 35347 | TATGAGTGTGGAGTCGTTGCCACACATTTACCCCTAAATATTCAGCANNNAGATCGG |
| 70132 | TATGAGTGTGGAGTCGTTGCGGGGGTGAACTCGCCATTTTATANNNAGATCGGAA |
| 93750 | TATGAGTGTGGAGTCGTTGCTGTAATCTCAGCCCCAGCATCTANNNAGATCGGAAG |
| 120437 | TATGAGTGTGGAGTCGTTGCGGGGAGATGAAATGATATTCTAAACCNNNAGATCG |
| 97810 | TATGAGTGTGGAGTCGTTGCACCAATCTGCATTCTCTGTGGNNNAGATCGGAAGAG |
| 120138 | TATGAGTGTGGAGTCGTTGCAGGTCTCACTGTCACTCCAGCTANNNAGATCGGAAG |
| 70672 | TATGAGTGTGGAGTCGTTGCCGGCCCATCATAGCCAAAGATAGNNNAGATCGGAAG |
| 99705 | TATGAGTGTGGAGTCGTTGCGGAGAGAGAGAAAATGGGAGCAGNNNAGATCGGA |
| 69934 | TATGAGTGTGGAGTCGTTGCGGTAAATTTGGTCCCCCAACTCANNNAGATCGGAAG |
| 71345 | TATGAGTGTGGAGTCGTTGCTGAAGGAGTAACCAGAAGACAGGTNNNAGATCGGA |
| 91108 | TATGAGTGTGGAGTCGTTGCCACAATGAAGTGAAGCAGGTCCTNNNAGATCGGAA |
| 70537 | TATGAGTGTGGAGTCGTTGCCTTCCTCCCTCACCATCTCTCTCNNNAGATCGGAAGAG |
| 70579 | TATGAGTGTGGAGTCGTTGCCACCCCCGCTCTTTTTCTTTTTCNNNAGATCGGAAGAG |
| 121175 | TATGAGTGTGGAGTCGTTGCGGAGAGAGAATGAGTGGTTTCTANNNAGATCGGAA |
| 70844 | TATGAGTGTGGAGTCGTTGCGCACCTGTGTATGAATGCACATGNNNAGATCGGAAG |
| 97972 | TATGAGTGTGGAGTCGTTGCCAGAGCGGACACCTTTGTCCTNNNAGATCGGAAGAG |
| 100376 | TATGAGTGTGGAGTCGTTGCAGTTCTCTCTCTCCAGAATGCACNNNAGATCGGAAGA |
| 92544 | TATGAGTGTGGAGTCGTTGCACACACACTGTTCAGGGTGAAGNNNAGATCGGAAG |
| 90935 | TATGAGTGTGGAGTCGTTGCTCTCTCCTCTCCTCTTTTGATCTNNNAGATCGGAAGAG |
| 93536 | TATGAGTGTGGAGTCGTTGCGGGGAGGAGAGAGAATTTTCCACNNNAGATCGGAA |
| 120062 | TATGAGTGTGGAGTCGTTGCAGTAATGACCACAGATATTTTTGTTTGNNNAGATCG |
| 120461 | TATGAGTGTGGAGTCGTTGCACGGTCTTGCTCTGTCGCCATANNNAGATCGGAAGA |
| 69886 | TATGAGTGTGGAGTCGTTGCTTGTCTTCCCTTCTCTACCAGCANNNAGATCGGAAGA |
| 90147 | TATGAGTGTGGAGTCGTTGCGTTAGGGGCTGTGGTGAGAATTTNNNAGATCGGAA |
| 92295 | TATGAGTGTGGAGTCGTTGCGTGGAATTCTGTTTGTAATCAGCCGNNNAGATCGGA |
| 121013 | TATGAGTGTGGAGTCGTTGCGGGGTTTTCACTCTTGTCACCANNNAGATCGGAAGA |
| 70405 | TATGAGTGTGGAGTCGTTGCCCCAGAAGGAAACATAGTCCAGCNNNAGATCGGAA |
| 90252 | TATGAGTGTGGAGTCGTTGCTGTCTGTGTAATTTGGCAAGCTTNNNAGATCGGAAG |
| 91279 | TATGAGTGTGGAGTCGTTGCGTGTGTGTGTCTGTACTAAGCACTNNNAGATCGGAA |
| 119594 | TATGAGTGTGGAGTCGTTGCCCTCGCTGAGCATCTTTCTATATCAANNNAGATCGGA |
| 99108 | TATGAGTGTGGAGTCGTTGCGCCAGGTTCTCTTTGTTATGGCNNNAGATCGGAAGA |
| 118929 | TATGAGTGTGGAGTCGTTGCTCAGTGGTTATGGCCAATAGCTNNNAGATCGGAAG |
| 92826 | TATGAGTGTGGAGTCGTTGCGAGAGAGAAAGAGAGCGCACATCNNNAGATCGGA |
| 90879 | TATGAGTGTGGAGTCGTTGCGCATAGTCCTTTATTTAGCTGGTCTNNNAGATCGGA |
| 70489 | TATGAGTGTGGAGTCGTTGCCGCTGAGGGTACATAAGATGTGCNNNAGATCGGAA |
| 70930 | TATGAGTGTGGAGTCGTTGCTCCCCCAGGCTCAGATAAGATTGNNNAGATCGGAAG |
| 119640 | TATGAGTGTGGAGTCGTTGCGATGAAATCCTGCTGCTCTGTNNNAGATCGGAAGAG |

70526 TATGAGTGTGGAGTCGTTGCCCTGAATGTGGCTATGTGTCCTCNNNAGATCGGAAG
91805 TATGAGTGTGGAGTCGTTGCCTCTCACTCACTCACTGTTCCTGNNNAGATCGGAAGA
120777 TATGAGTGTGGAGTCGTTGCGTAGCACATACACACCTGTGGTNNNAGATCGGAAG
94078 TATGAGTGTGGAGTCGTTGCCGTGTGTGTTTGGTGATGAAGGANNNAGATCGGAA
90376 TATGAGTGTGGAGTCGTTGCAAGAGAGGAAGCCAGAAAGAAATTNNNAGATCGG
45510 TATGAGTGTGGAGTCGTTGCTAGGAGGGAAGGGTTCAAACTCCNNNAGATCGGAA
69673 TATGAGTGTGGAGTCGTTGCGCGGAGGGGAGATAGTCAAAAANNNAGATCGGAA
93102 TATGAGTGTGGAGTCGTTGCTAATAGCATCTGTTCCCACCAGCNNNAGATCGGAAG
69794 TATGAGTGTGGAGTCGTTGCGTGTGTGTGTGACTAAAGCTAACANNNAGATCGGA
92589 TATGAGTGTGGAGTCGTTGCACACACACATCCCCAACAGAAAGNNNAGATCGGAAG
71139 TATGAGTGTGGAGTCGTTGCGTCTTGCCACGTTATCTTCACCTNNNAGATCGGAAGA
96013 TATGAGTGTGGAGTCGTTGCAGACTTACCAGCTTGGGCCAATGNNNAGATCGGAA
91776 TATGAGTGTGGAGTCGTTGCCCCACAAAAGGCAAAGGAATCTGNNNAGATCGGAA
29500 TATGAGTGTGGAGTCGTTGCGAGGTCCAACTGAACAGCTCAATNNNAGATCGGAA
94482 TATGAGTGTGGAGTCGTTGCAGAGAAGCTATGGGAAAACAGCANNNAGATCGGA
98010 TATGAGTGTGGAGTCGTTGCCCGTATGCTTTTGTGTCCTCACNNNAGATCGGAAGA
99743 TATGAGTGTGGAGTCGTTGCTGTCAAATATCTCCTGTGTGTGTGTNNNAGATCGGA
94467 TATGAGTGTGGAGTCGTTGCAACACATTCACACAGTCAGAGCANNNAGATCGGAAG
119769 TATGAGTGTGGAGTCGTTGCTAGTGAGAGGAAGTTTCTAAAAAGTTTNNNAGATC
120183 TATGAGTGTGGAGTCGTTGCACGACAGAGTTTCACTTTCTTGNNNAGATCGGAAGA
29135 TATGAGTGTGGAGTCGTTGCGCAGCACGTTAGGTTTCATTTTCTNNNAGATCGGAA
29291 TATGAGTGTGGAGTCGTTGCCCCCAGCACACATTTTGTAACACNNNAGATCGGAAG
70459 TATGAGTGTGGAGTCGTTGCCCTGGCATATAGACACATAGTGGCNNNAGATCGGA
71319 TATGAGTGTGGAGTCGTTGCGGGATAGTAGGTGCTCAGTGAGTNNNAGATCGGA
94403 TATGAGTGTGGAGTCGTTGCACACACCCCTACCTGTATATGCANNNAGATCGGAAG
96066 TATGAGTGTGGAGTCGTTGCATTTGATGGCTGAGGAAACTGGGANNNAGATCGGA
91969 TATGAGTGTGGAGTCGTTGCCAGAGTGAAAGAGCGAGTTCTGGNNNAGATCGGA
29659 TATGAGTGTGGAGTCGTTGCCACACACAACAGACATAAGCTGGNNNAGATCGGAA
69830 TATGAGTGTGGAGTCGTTGCTGGAGCAAAGGAATTGTCTTGACANNNAGATCGGA
70188 TATGAGTGTGGAGTCGTTGCGTTCAGCACCACAGGTGTAGTTANNNAGATCGGAA
70241 TATGAGTGTGGAGTCGTTGCCCAACAGACAACTTGAACAGCACNNNAGATCGGAAG
92135 TATGAGTGTGGAGTCGTTGCGAGAGAGAGCAAACAACCTACGGNNNAGATCGGAA
92355 TATGAGTGTGGAGTCGTTGCTTTGGGAGGAGAAGGGTCCATAGNNNAGATCGGA
94696 TATGAGTGTGGAGTCGTTGCTTGGCAGAGATAAGAGCTAGACANNNAGATCGGAA
98785 TATGAGTGTGGAGTCGTTGCAGTAGTGGTTACTGGAATGCTCACANNNAGATCGG
90315 TATGAGTGTGGAGTCGTTGCTGGGAGGAGATGAAGTGGTGATTNNNAGATCGGA
117227 TATGAGTGTGGAGTCGTTGCTCTCCTCTTTTGATCTACAATTTTCGGNNNAGATCGG
93388 TATGAGTGTGGAGTCGTTGCTGTTGGTGAATAGTCACTGGACCNNNAGATCGGAA
29678 TATGAGTGTGGAGTCGTTGCCTCTTCTGGATGGCCCTGTTTTCNNNAGATCGGAAG
98285 TATGAGTGTGGAGTCGTTGCTTCACTTTCTCCTCCCACCCTGNNNAGATCGGAAGAG
98312 TATGAGTGTGGAGTCGTTGCGGTTTTCACAGTACCAGAGACATNNNAGATCGGAAG
98385 TATGAGTGTGGAGTCGTTGCGGAGCCTGCCAAAAAGATATGAANNNAGATCGGAA
92384 TATGAGTGTGGAGTCGTTGCTTCTGCATTGTAACAGGGTGGAGNNNAGATCGGAA

| | |
|---|---|
| 92304 | TATGAGTGTGGAGTCGTTGCTGAGGAGACTTAGGAAGCCTACGNNNAGATCGGAA |
| 94060 | TATGAGTGTGGAGTCGTTGCTCTCTCTCTTCAGTTGAGCAGGANNNAGATCGGAAG |
| 120389 | TATGAGTGTGGAGTCGTTGCGGAGCCTTGCTCTTATCGCTCNNNAGATCGGAAGAG |
| 29046 | TATGAGTGTGGAGTCGTTGCTATTTCTCCCCAGCTTCTGACCTNNNAGATCGGAAGA |
| 120763 | TATGAGTGTGGAGTCGTTGCGGAGACAAGAAGACAAGCTTTTGCNNNAGATCGGA |
| 121056 | TATGAGTGTGGAGTCGTTGCTGAAGTTTTGTTCTTGTTGCTGAGGNNNAGATCGGA |
| 29605 | TATGAGTGTGGAGTCGTTGCTGGTTATTGCTGGCCTTGTAGAANNNAGATCGGAA |
| 69837 | TATGAGTGTGGAGTCGTTGCTCATACCCATGTACCCACGTACANNNAGATCGGAAG |
| 70853 | TATGAGTGTGGAGTCGTTGCTTAGTTCAGTGCCTGGTGTGATGNNNAGATCGGAA |
| 93603 | TATGAGTGTGGAGTCGTTGCCTGCACTACCCAAAGGTAGGAACNNNAGATCGGAA |
| 91522 | TATGAGTGTGGAGTCGTTGCGCAATCAAGAATGAACAAGGAACATNNNAGATCGG |
| 29314 | TATGAGTGTGGAGTCGTTGCAATCACAAAACCAGCTATGGGCANNNAGATCGGAA |
| 29355 | TATGAGTGTGGAGTCGTTGCGAAGAAGTCCCACTATCCCAGGTNNNAGATCGGAAG |
| 29602 | TATGAGTGTGGAGTCGTTGCCAAAATCAGGTATCACAAGCTGAANNNAGATCGGA |
| 29762 | TATGAGTGTGGAGTCGTTGCAGGAAGTGTGTAGGAGGTGACAANNNAGATCGGA |
| 70721 | TATGAGTGTGGAGTCGTTGCCTGCTAACATGATCACCTCCAGGNNNAGATCGGAAG |
| 95340 | TATGAGTGTGGAGTCGTTGCAACATTATCTGTAGCAGTGAGGAAAANNNAGATCG |
| 28941 | TATGAGTGTGGAGTCGTTGCATTCTGTGAGGTGGTAGTGATGCNNNAGATCGGAA |
| 29233 | TATGAGTGTGGAGTCGTTGCGTGCAAACATACACACAGAGCCTNNNAGATCGGAAG |
| 29613 | TATGAGTGTGGAGTCGTTGCCCTCCAGAGCACTGAAAAGTCTTNNNAGATCGGAAG |
| 29623 | TATGAGTGTGGAGTCGTTGCGGAACTATGCGAGGTAGTATGTATGTNNNAGATCG |
| 94230 | TATGAGTGTGGAGTCGTTGCTTGGGTGATGAGGGATCCAAAGANNNAGATCGGA |
| 94928 | TATGAGTGTGGAGTCGTTGCTTGCACAGAACAACGCATTTCAANNNAGATCGGAAG |
| 95101 | TATGAGTGTGGAGTCGTTGCTACTGAATGACAGCCATCTGGTCNNNAGATCGGAAG |
| 98672 | TATGAGTGTGGAGTCGTTGCCTGTAGATGTTGAGCAGTGTTCCTNNNAGATCGGAA |
| 119954 | TATGAGTGTGGAGTCGTTGCTAGGCTGAAATGCAGTGTGGCAANNNAGATCGGAA |
| 28768 | TATGAGTGTGGAGTCGTTGCCGAAGTTTCTCTCTCATCACCCANNNAGATCGGAAGA |
| 29658 | TATGAGTGTGGAGTCGTTGCCTGGAAACAGTCTCACTCTCTGGNNNAGATCGGAAG |
| 29727 | TATGAGTGTGGAGTCGTTGCAGAGGCTGACTTTGCAGAGACTANNNAGATCGGAA |
| 71230 | TATGAGTGTGGAGTCGTTGCTTGGCATGATTGTTATGCTTGCTNNNAGATCGGAAG |
| 92206 | TATGAGTGTGGAGTCGTTGCGAGACCCCAGATAAATGTGCAGGNNNAGATCGGAA |
| 95733 | TATGAGTGTGGAGTCGTTGCATGTTCTTACACTGGCTTTCTGGTNNNAGATCGGAA |
| 46731 | TATGAGTGTGGAGTCGTTGCAGGGAGAAAGAGAGAGATCACGCNNNAGATCGGA |
| 69747 | TATGAGTGTGGAGTCGTTGCAGCAGCAAATTCACACAGGAGAANNNAGATCGGAA |
| 70918 | TATGAGTGTGGAGTCGTTGCGCTTGCGCACATGTGTATATGTGNNNAGATCGGAA |
| 28828 | TATGAGTGTGGAGTCGTTGCCAGGGGCCAGATAGAGGAAGATANNNAGATCGGA |
| 28840 | TATGAGTGTGGAGTCGTTGCCTCTGTTACCCAGATTGAAGGGCNNNAGATCGGAAG |
| 29054 | TATGAGTGTGGAGTCGTTGCTTGAGTTCAGTGAGGGCACAAATNNNAGATCGGAA |
| 29298 | TATGAGTGTGGAGTCGTTGCATGGCGGGGTGGAGTTATAATGANNNAGATCGGA |
| 29540 | TATGAGTGTGGAGTCGTTGCTACAGGTGCCAAGTAACAGCTTCNNNAGATCGGAAG |
| 29560 | TATGAGTGTGGAGTCGTTGCATACTGCTTGATGCTCTCTGCCNNNAGATCGGAAGA |
| 29563 | TATGAGTGTGGAGTCGTTGCTCAGGGATTTCAAAGGACAAGCCNNNAGATCGGAA |
| 29673 | TATGAGTGTGGAGTCGTTGCCACACACACTCCTATTGTTGCAGNNNAGATCGGAAG |

69783 TATGAGTGTGGAGTCGTTGCAGCATTCCATTTCACAGGAAACANNNAGATCGGAAG
90194 TATGAGTGTGGAGTCGTTGCGGTGTGTGTGGGTGTGTTTAGTTNNNAGATCGGAA
95441 TATGAGTGTGGAGTCGTTGCTCTGGACTACACTGGCGTGATNNNAGATCGGAAGA
97411 TATGAGTGTGGAGTCGTTGCCCCTGATTTCCCTTTCTGCTCTTNNNAGATCGGAAGA
92312 TATGAGTGTGGAGTCGTTGCTGTGTGTGAATGTTTGTATGCACGNNNAGATCGGA
28906 TATGAGTGTGGAGTCGTTGCCTCCAAATCCTAGCACCATTCATNNNAGATCGGAAGA
29011 TATGAGTGTGGAGTCGTTGCCTGAAGGTGATTCTGTCTGGTGANNNAGATCGGAA
29207 TATGAGTGTGGAGTCGTTGCGCAGGGGCTCAACATACGATAAANNNAGATCGGAA
29306 TATGAGTGTGGAGTCGTTGCTTGGCATTCTCTAGGCTTATTCANNNAGATCGGAAG
29503 TATGAGTGTGGAGTCGTTGCTACAGCCAGACTTCACACGATTGNNNAGATCGGAAG
69963 TATGAGTGTGGAGTCGTTGCGGTGTTTGGGGCATGGATATTCANNNAGATCGGAA
70005 TATGAGTGTGGAGTCGTTGCTGGCAAAAGCAAGAGATGAGAGANNNAGATCGGA
70259 TATGAGTGTGGAGTCGTTGCTGCTGTTGGAACGTGTGATAGACNNNAGATCGGAA
70646 TATGAGTGTGGAGTCGTTGCGAAAGGGAAGGTGGGTTTGAGAGNNNAGATCGGA
71581 TATGAGTGTGGAGTCGTTGCACCATGTCTTCTGCCTCACATTTNNNAGATCGGAAGA
92676 TATGAGTGTGGAGTCGTTGCCCCCCTGCTCCATTCATACATTCNNNAGATCGGAAGA
97520 TATGAGTGTGGAGTCGTTGCCTGCATCTACACTGCCTGCAGNNNAGATCGGAAGAG
91918 TATGAGTGTGGAGTCGTTGCTTTGGATTACAGCCAGGTGTTGGNNNAGATCGGAA
93055 TATGAGTGTGGAGTCGTTGCTTCTCTCTCTCTTCCCTGCTAGCNNNAGATCGGAAGA
120550 TATGAGTGTGGAGTCGTTGCTCTAATGAGGGTAGGCTGTTGAANNNAGATCGGAA
120945 TATGAGTGTGGAGTCGTTGCCGGTGGCAAGAACTTACTTCNNNAGATCGGAAGAG
92222 TATGAGTGTGGAGTCGTTGCATACAATACACTCCACCCCCAGGNNNAGATCGGAAG
94669 TATGAGTGTGGAGTCGTTGCAACTCGCCTGTGTAAAGCATACANNNAGATCGGAAG
28907 TATGAGTGTGGAGTCGTTGCTCCCCTGATTTCCATCTGGACATNNNAGATCGGAAG
29760 TATGAGTGTGGAGTCGTTGCACACATTGCTACTATCTGGTACAANNNAGATCGGAA
70789 TATGAGTGTGGAGTCGTTGCAAAAGTCTATGGTGTTGGCTGGGNNNAGATCGGAA
91155 TATGAGTGTGGAGTCGTTGCTCCTATATTTGTCCCAAGGGCCTNNNAGATCGGAAG
91920 TATGAGTGTGGAGTCGTTGCATCAATTTGGAGTTGGGGGTTGGNNNAGATCGGAA
93467 TATGAGTGTGGAGTCGTTGCGCACAGGTCATTTTGGGAATGACNNNAGATCGGAA
95402 TATGAGTGTGGAGTCGTTGCTCACAGTGGTCACAGCTACTTGNNNAGATCGGAAGA
120258 TATGAGTGTGGAGTCGTTGCTTTGCGGAGATATTGAGTGAGTGNNNAGATCGGAA
120590 TATGAGTGTGGAGTCGTTGCCTCCTGGTGAAACCTTATTCATCCAGNNNAGATCGGA
29122 TATGAGTGTGGAGTCGTTGCTCATGGACATTCCTCTCTACCTTNNNAGATCGGAAGA
29220 TATGAGTGTGGAGTCGTTGCACAGGCAGATCTAAAGTAAATGAGGTNNNAGATCG
29303 TATGAGTGTGGAGTCGTTGCATTGAAGGAAACCCGGTAGGAGANNNAGATCGGA
53719 TATGAGTGTGGAGTCGTTGCCCCTTACCTGATTTTGGTATCAGGGNNNAGATCGGA
69820 TATGAGTGTGGAGTCGTTGCTGACCACTTTCAAAGCAAAGACANNNAGATCGGAAG
70065 TATGAGTGTGGAGTCGTTGCAAGTTCCCTTGTTAGCCGAATGANNNAGATCGGAAG
94783 TATGAGTGTGGAGTCGTTGCCTGCCTATCTGGCTGCATAAACANNNAGATCGGAAG
93365 TATGAGTGTGGAGTCGTTGCAAAACTCTAGCAGGACCAAACCCNNNAGATCGGAAG
95072 TATGAGTGTGGAGTCGTTGCTAGAGTGTAGGGTTGGGCACAAANNNAGATCGGA
93078 TATGAGTGTGGAGTCGTTGCTTTCTTGACCTCAGATGCAGAGCNNNAGATCGGAAG
90369 TATGAGTGTGGAGTCGTTGCTGTAAAGTGTTAGCGGGGCAATTNNNAGATCGGAA

91511 TATGAGTGTGGAGTCGTTGCTCCTTTAGACTGGAACGTCACATNNNAGATCGGAAG
119839 TATGAGTGTGGAGTCGTTGCGCGTCTTCTCTGTCCAACTACTGNNNAGATCGGAAG
120653 TATGAGTGTGGAGTCGTTGCACTTGTTTCTGGACTTTGCCTTGNNNAGATCGGAAG
28820 TATGAGTGTGGAGTCGTTGCAGAGGCCTAGTGTAACTGAAGGGNNNAGATCGGA
28951 TATGAGTGTGGAGTCGTTGCACTTTCCCCTTTTGGATTTGACCANNNAGATCGGAAG
29082 TATGAGTGTGGAGTCGTTGCGGTATCAGGGTAATGCTCCTGACNNNAGATCGGAA
29256 TATGAGTGTGGAGTCGTTGCTGGTAAGTGGGTAGATTCCAGGGNNNAGATCGGA
69681 TATGAGTGTGGAGTCGTTGCGGCTTTGCTAGGGCATTTCAAAANNNAGATCGGAA
70412 TATGAGTGTGGAGTCGTTGCACTTTTCCCACACAACATTCAGCNNNAGATCGGAAGA
71544 TATGAGTGTGGAGTCGTTGCCACACACACCTTGGGGTATTCTTNNNAGATCGGAAG
91907 TATGAGTGTGGAGTCGTTGCGCAGTAGGTGCTCCACAAAAATGNNNAGATCGGAA
94002 TATGAGTGTGGAGTCGTTGCAAGCTAGGATAGACACACAATGGANNNAGATCGGA
103048 TATGAGTGTGGAGTCGTTGCTGGCTTTATGGGCATGTTCCAATNNNAGATCGGAAG
71208 TATGAGTGTGGAGTCGTTGCGCAATAAGAAGTCACAGCAGGCTNNNAGATCGGAA
91580 TATGAGTGTGGAGTCGTTGCAGGCAGCAGATGTGAACTGAAATNNNAGATCGGAA
120413 TATGAGTGTGGAGTCGTTGCCGGAGTTTTCCCTCTTGTCGACCNNNAGATCGGAAG
28736 TATGAGTGTGGAGTCGTTGCTTGCTTTGTTTTCAGTTGGAATAAAAGNNNAGATCG
29124 TATGAGTGTGGAGTCGTTGCGGAGACAACCTCATGTGCTCATTNNNAGATCGGAAG
29271 TATGAGTGTGGAGTCGTTGCCTCTTTCTTGATTTGGCTCTGAGCNNNAGATCGGAA
29382 TATGAGTGTGGAGTCGTTGCAGAGAGAGAGAGTTACCTCCTTTNNNAGATCGGAA
29413 TATGAGTGTGGAGTCGTTGCGACACACCCACCAGATTTTCTTTNNNAGATCGGAAG
29578 TATGAGTGTGGAGTCGTTGCTGCTGTGGCATTTCTGACACATANNNAGATCGGAAG
29764 TATGAGTGTGGAGTCGTTGCGCCTGTTGTCTGGGAAGGATAAANNNAGATCGGAA
29771 TATGAGTGTGGAGTCGTTGCTCACCCCACTCTGTAACCTAGTTNNNAGATCGGAAGA
69997 TATGAGTGTGGAGTCGTTGCTCTACACACAGAAGCCTTCCAGANNNAGATCGGAAG
70141 TATGAGTGTGGAGTCGTTGCAAGCCATTTGCAACTTGTCCCTANNNAGATCGGAAG
71099 TATGAGTGTGGAGTCGTTGCCTACTCACTCTGCCTTTTGCACTNNNAGATCGGAAGA
93856 TATGAGTGTGGAGTCGTTGCACACCTCTTGCAATACAGATCTTGANNNAGATCGGA
95974 TATGAGTGTGGAGTCGTTGCTCCTGTTCTGACTTTGGGGATAGANNNAGATCGGAA
96702 TATGAGTGTGGAGTCGTTGCGATTCTCTAGGTATTTTGGCCGATGATNNNAGATCG
93261 TATGAGTGTGGAGTCGTTGCTGTTATGGATCAGCAAGAAGGCCNNNAGATCGGAA
29026 TATGAGTGTGGAGTCGTTGCAGAGGATGAGAGGTTGTGGGAAANNNAGATCGGA
29584 TATGAGTGTGGAGTCGTTGCCACCACATACACACACACCTTCANNNAGATCGGAAGA
69916 TATGAGTGTGGAGTCGTTGCACACAGTCTGAGGTACTACTGCANNNAGATCGGAAG
70053 TATGAGTGTGGAGTCGTTGCACAGGAAATTGACAGATAGTTGTGGANNNAGATCG
71019 TATGAGTGTGGAGTCGTTGCGCCCATGTATGTGTGATTGGCATNNNAGATCGGAA
71065 TATGAGTGTGGAGTCGTTGCTGCCCATGCCTGTCTTTTACTATNNNAGATCGGAAGA
71083 TATGAGTGTGGAGTCGTTGCAGAAACAGAACTACACAGCAACTNNNAGATCGGAA
94831 TATGAGTGTGGAGTCGTTGCTGGGGTTTCTATGGAAGGGCTAANNNAGATCGGAA
95280 TATGAGTGTGGAGTCGTTGCAGGTCTAACCAGCTCTCTAGGTGNNNAGATCGGAAG
95442 TATGAGTGTGGAGTCGTTGCAAGATTGGGTTATTTCACGCATCNNNAGATCGGAAG
29273 TATGAGTGTGGAGTCGTTGCGGGGCATGCTTTTTATACTCAGCNNNAGATCGGAAG
69828 TATGAGTGTGGAGTCGTTGCAGTCTCACAACAGCAGAATGACANNNAGATCGGAA

| | |
|---|---|
| 70929 | TATGAGTGTGGAGTCGTTGCGCTGCATGTTCATTGATGCATTGNNNAGATCGGAAG |
| 70980 | TATGAGTGTGGAGTCGTTGCAACCAAATGACCTGGCTTGGAATNNNAGATCGGAA |
| 71094 | TATGAGTGTGGAGTCGTTGCTTCTCTCTCTCAAACTCACCACTNNNAGATCGGAAGA |
| 95462 | TATGAGTGTGGAGTCGTTGCAGTCTGTTTTGCCAGGCTGGAGNNNAGATCGGAAG |
| 98998 | TATGAGTGTGGAGTCGTTGCAGGGAGATGTAACAGGCGTCACTNNNAGATCGGAA |
| 93297 | TATGAGTGTGGAGTCGTTGCCACACTCATTTTCCCTCTCTCCCNNNAGATCGGAAGA |
| 91719 | TATGAGTGTGGAGTCGTTGCGACAGAGACAGAGAGCAAAGGTGNNNAGATCGGA |
| 93390 | TATGAGTGTGGAGTCGTTGCTTCAACTCTGTGTGTGTGAGACCNNNAGATCGGAAG |
| 28960 | TATGAGTGTGGAGTCGTTGCCTCTCACCCACTCCCTATTCCTTNNNAGATCGGAAGA |
| 29140 | TATGAGTGTGGAGTCGTTGCGGGCTAGGGGGTTGAGTATGTATNNNAGATCGGA |
| 69838 | TATGAGTGTGGAGTCGTTGCCTGTGGAACCTGAATGTGGTACANNNAGATCGGAA |
| 70193 | TATGAGTGTGGAGTCGTTGCCCTGGCACATAGCAGCTGATTTANNNAGATCGGAAG |
| 90649 | TATGAGTGTGGAGTCGTTGCTCAGTCCTAAAACAAACTTATGAGGTNNNAGATCGG |
| 96787 | TATGAGTGTGGAGTCGTTGCTTTTTGGGCACACATGCTCTGGNNNAGATCGGAAGA |
| 98100 | TATGAGTGTGGAGTCGTTGCGCTTCACAATCCTAGGCAGTTATNNNAGATCGGAAG |
| 28748 | TATGAGTGTGGAGTCGTTGCTTCTGCAGTTTTTGAACTTGGCTGNNNAGATCGGAA |
| 28841 | TATGAGTGTGGAGTCGTTGCCACCACTACTACCGCCAATGAAGNNNAGATCGGAAG |
| 29160 | TATGAGTGTGGAGTCGTTGCACAGGCGTTTATGATAGATTTCTAAAGNNNAGATCG |
| 29250 | TATGAGTGTGGAGTCGTTGCTAGGGATAAGTGATCAGGTTCCTGNNNAGATCGGA |
| 29496 | TATGAGTGTGGAGTCGTTGCCAACTGAACCAGAGCACTGTCATNNNAGATCGGAAG |
| 29691 | TATGAGTGTGGAGTCGTTGCCCACCCTGGGAAATTAGATGCTCNNNAGATCGGAAG |
| 70885 | TATGAGTGTGGAGTCGTTGCTGTGCTGCTTTTTCCGATTATCTGNNNAGATCGGAA |
| 71491 | TATGAGTGTGGAGTCGTTGCCGTCTATGGTGTGTGTTGGGATTNNNAGATCGGAA |
| 90477 | TATGAGTGTGGAGTCGTTGCTACTGTGGAAGAGATATGCCTGTNNNAGATCGGAA |
| 121014 | TATGAGTGTGGAGTCGTTGCGAGATGGCAGCCAATTGCAGTTGNNNAGATCGGAA |
| 28886 | TATGAGTGTGGAGTCGTTGCATTACGGTTTCATCTGCTGAAGCNNNAGATCGGAAG |
| 28888 | TATGAGTGTGGAGTCGTTGCAGGACAAGACAAATAGGGGACCANNNAGATCGGA |
| 28895 | TATGAGTGTGGAGTCGTTGCTGGAAAAGCACTCTGTAGCTCAGNNNAGATCGGAA |
| 29064 | TATGAGTGTGGAGTCGTTGCTTCATCACATGTGCAGGTGTAGGNNNAGATCGGAA |
| 29441 | TATGAGTGTGGAGTCGTTGCTCAGGGAGATTCATTCGGCTTTGNNNAGATCGGAA |
| 69734 | TATGAGTGTGGAGTCGTTGCTCAACCAGAAACATGTGGCTCAANNNAGATCGGAAG |
| 70125 | TATGAGTGTGGAGTCGTTGCCACTCCCCACAACGACTTTCATANNNAGATCGGAAGA |
| 70340 | TATGAGTGTGGAGTCGTTGCGGGCATTTGAAACATTTGAAGCCNNNAGATCGGAA |
| 70647 | TATGAGTGTGGAGTCGTTGCGTCAACAGAGCAGGCAGATAGAGNNNAGATCGGA |
| 70815 | TATGAGTGTGGAGTCGTTGCTAAAAGGCTTACTCCAGTGCTGGNNNAGATCGGAA |
| 93220 | TATGAGTGTGGAGTCGTTGCGTTCATTGGTACAATGCTTAGTGCCNNNAGATCGGA |
| 93957 | TATGAGTGTGGAGTCGTTGCTCTGTCCTCTATAACACCTGAATGANNNAGATCGGA |
| 98219 | TATGAGTGTGGAGTCGTTGCAGCCAATGCTAAAGATGCCTCAANNNAGATCGGAAG |
| 91777 | TATGAGTGTGGAGTCGTTGCACCTAAAGCAAAGGCTTTTTGCTGNNNAGATCGGAA |
| 120924 | TATGAGTGTGGAGTCGTTGCTGTCCACGATCACTTCTTTTGTTNNNAGATCGGAAGA |
| 28814 | TATGAGTGTGGAGTCGTTGCTGAAGTTGGTGGGGAATGTTTATNNNAGATCGGAA |
| 28953 | TATGAGTGTGGAGTCGTTGCGGCTGGGATGGCATAATTTGGAANNNAGATCGGAA |
| 29619 | TATGAGTGTGGAGTCGTTGCAGATTTGCGGGGAGTTCATTTGTNNNAGATCGGAA |

| | |
|---|---|
| 70249 | TATGAGTGTGGAGTCGTTGCAACATGCACACATCCCTACTCACNNNAGATCGGAAGA |
| 70267 | TATGAGTGTGGAGTCGTTGCGCAAGCAATTTGGTAAGTTTGGACNNNAGATCGGAA |
| 70499 | TATGAGTGTGGAGTCGTTGCTGGTGGTTTCCGCTATAATATTTGCNNNAGATCGGAA |
| 70587 | TATGAGTGTGGAGTCGTTGCGAAGCATGTGAGAACACAGCAAGNNNAGATCGGAA |
| 70771 | TATGAGTGTGGAGTCGTTGCCCTCTAAGTTCTCCTCTCAGGGGNNNAGATCGGAAGA |
| 70951 | TATGAGTGTGGAGTCGTTGCTCAGCTTCTCCACCATCTTGTTGNNNAGATCGGAAGA |
| 93600 | TATGAGTGTGGAGTCGTTGCTTCAGCTCCTGTTTCCTTGGAACNNNAGATCGGAAGA |
| 94149 | TATGAGTGTGGAGTCGTTGCACTGCTCTCTCAAAGCTCAGAGANNNAGATCGGAAG |
| 97708 | TATGAGTGTGGAGTCGTTGCCTGGAGCTCATTACTGGTACAACNNNAGATCGGAAG |
| 120673 | TATGAGTGTGGAGTCGTTGCCGAAGTTCACTCTGTCGCCCNNNAGATCGGAAGAGC |
| 28819 | TATGAGTGTGGAGTCGTTGCCTCTCTCCAGTTCTTGCAGCAATNNNAGATCGGAAGA |
| 28827 | TATGAGTGTGGAGTCGTTGCCAGTTCCTACTTCTGCGATCCAGNNNAGATCGGAAGA |
| 29332 | TATGAGTGTGGAGTCGTTGCACAGTTTGTGGAGGAGTATGCAANNNAGATCGGAA |
| 70302 | TATGAGTGTGGAGTCGTTGCTGCCCTGTTTCTCTAAAGAACCCNNNAGATCGGAAGA |
| 92612 | TATGAGTGTGGAGTCGTTGCCTCTCTAGCTTCCGCTTCCTTTCNNNAGATCGGAAGA |
| 94443 | TATGAGTGTGGAGTCGTTGCTCAATAAGGTCCCAGAAGAGGCANNNAGATCGGAA |
| 98360 | TATGAGTGTGGAGTCGTTGCGGAGATGGAGGGGAAAAAGGAGANNNAGATCGGA |
| 90162 | TATGAGTGTGGAGTCGTTGCCCTTGGGGAGAAGTTTCAGTGTTNNNAGATCGGAA |
| 119556 | TATGAGTGTGGAGTCGTTGCCGGGGAGGAAAATATGGGTGTTGNNNAGATCGGAA |
| 119980 | TATGAGTGTGGAGTCGTTGCTTGCAGAGGGGAGATGTCACCTTNNNAGATCGGAA |
| 120863 | TATGAGTGTGGAGTCGTTGCAAGCTGGAGTGCAGAGGTATAATTNNNAGATCGG |
| 29471 | TATGAGTGTGGAGTCGTTGCCCTTCCTCTCTGTCCGCTATTTTTNNNAGATCGGAAGA |
| 29576 | TATGAGTGTGGAGTCGTTGCCAACGCCAACACCACTACAGATANNNAGATCGGAAG |
| 29599 | TATGAGTGTGGAGTCGTTGCGCTCTGAAAACTCCCTTGCTACANNNAGATCGGAAGA |
| 29601 | TATGAGTGTGGAGTCGTTGCAGGCAGCCAGAAAGAAATAACACANNNAGATCGGA |
| 70261 | TATGAGTGTGGAGTCGTTGCAGTTGTAGAGATGGTTGCACGACNNNAGATCGGAA |
| 70371 | TATGAGTGTGGAGTCGTTGCCTCAGATAATGGTGCCACTGTCCNNNAGATCGGAAG |
| 70959 | TATGAGTGTGGAGTCGTTGCACTCGCCTTTGCATGACATTTTGNNNAGATCGGAAGA |
| 95080 | TATGAGTGTGGAGTCGTTGCCCCCAGGCTTGAGAAGACTAAAANNNAGATCGGAA |
| 95374 | TATGAGTGTGGAGTCGTTGCTTGTACTCCCAATACTTTGGGGGANNNAGATCGGAA |
| 95406 | TATGAGTGTGGAGTCGTTGCAGAACCTACGGATGCAGAGAAGAGNNNAGATCGGA |
| 98905 | TATGAGTGTGGAGTCGTTGCCAGTGTCAGCATTTTCCTGCTCNNNAGATCGGAAGA |
| 99320 | TATGAGTGTGGAGTCGTTGCCCCAAAAGTAGGGCTCTAAAGAAGCNNNAGATCGGA |
| 100122 | TATGAGTGTGGAGTCGTTGCTCTCAACAAACCAGTCCCGAAGNNNAGATCGGAAGA |
| 119630 | TATGAGTGTGGAGTCGTTGCTTCAGCGTCCTGTCCAAAGTCTANNNAGATCGGAAGA |
| 93030 | TATGAGTGTGGAGTCGTTGCAAGGAGCATAATGCCTAACAGGCNNNAGATCGGAA |
| 29211 | TATGAGTGTGGAGTCGTTGCACCAGTTTGGGCTTGAATACGTTNNNAGATCGGAAG |
| 29469 | TATGAGTGTGGAGTCGTTGCTGTGTAGGAGGATGGGTAGCTTTNNNAGATCGGAA |
| 61679 | TATGAGTGTGGAGTCGTTGCATACAGTGAAGGATTTCACAGCTNNNAGATCGGAA |
| 70595 | TATGAGTGTGGAGTCGTTGCTGGAGAGAGGAATGTGGAGGAAGNNNAGATCGGA |
| 92178 | TATGAGTGTGGAGTCGTTGCAGAGTAGAGACAAAGCAGGGAGGNNNAGATCGGA |
| 96900 | TATGAGTGTGGAGTCGTTGCAGTGGCTCCTTCTCTATCATCTCTTTNNNAGATCGGAA |
| 100370 | TATGAGTGTGGAGTCGTTGCTATTATCGGCAAGGCCTGGTAGCNNNAGATCGGAA |

| | |
|---|---|
| 102310 | TATGAGTGTGGAGTCGTTGCAGGCATCAAAGTCTGGAAGAGAGNNNAGATCGGAA |
| 90462 | TATGAGTGTGGAGTCGTTGCCAAGTAGGCCAATTACGTGCTGTNNNAGATCGGAAG |
| 119670 | TATGAGTGTGGAGTCGTTGCCTATCTGGGCTCATTGCTGGTTNNNAGATCGGAAGA |
| 119832 | TATGAGTGTGGAGTCGTTGCACGCATGAGAGTAAAGGCTCTGNNNAGATCGGAAG |
| 29004 | TATGAGTGTGGAGTCGTTGCACTTGCACACAAATGTTCAACCCNNNAGATCGGAAGA |
| 29098 | TATGAGTGTGGAGTCGTTGCTGGGTCCTTGTTTGTTTACATTCANNNAGATCGGAAG |
| 29202 | TATGAGTGTGGAGTCGTTGCCACCTACAGGCCAAATTGGGTAANNNAGATCGGAAG |
| 29354 | TATGAGTGTGGAGTCGTTGCAAGTATTGCTCTTCAGCAAGGGTNNNAGATCGGAAG |
| 29404 | TATGAGTGTGGAGTCGTTGCTGGCTACAGTGGAAGTTTCTTTTNNNAGATCGGAAG |
| 29408 | TATGAGTGTGGAGTCGTTGCGTGTGTGTGTGTGCTAAGCTTTTNNNAGATCGGAAG |
| 69922 | TATGAGTGTGGAGTCGTTGCCTGGGGTTACTATAAGGCTTGCANNNAGATCGGAAG |
| 70041 | TATGAGTGTGGAGTCGTTGCCTTGCCCATGAGATGAAATGGGANNNAGATCGGAAG |
| 71201 | TATGAGTGTGGAGTCGTTGCTTCTCATTGGTACTGCCCTAGCTNNNAGATCGGAAGA |
| 90763 | TATGAGTGTGGAGTCGTTGCTGCTTGAAAAGACACAAAGCAGTNNNAGATCGGAAG |
| 91632 | TATGAGTGTGGAGTCGTTGCTTGTGCATCTGTATACCGGGTTGNNNAGATCGGAAG |
| 92115 | TATGAGTGTGGAGTCGTTGCTACTCTTTGGCTCTGAGGAAGGGNNNAGATCGGAAG |
| 95805 | TATGAGTGTGGAGTCGTTGCCCCAATGCTGCCTATTGTCAGANNNAGATCGGAAGAG |
| 71270 | TATGAGTGTGGAGTCGTTGCAAATAGAGAGCTACCTGAAATAGTTCTNNNAGATCG |
| 95013 | TATGAGTGTGGAGTCGTTGCTGCTTCACTAACACCCCAAACAANNNAGATCGGAAGA |
| 28794 | TATGAGTGTGGAGTCGTTGCATAAAAGAGAAACCTTGGGCCGGNNNAGATCGGAA |
| 29192 | TATGAGTGTGGAGTCGTTGCAGAGCACATGGCATATTATTGGCANNNAGATCGGAA |
| 29327 | TATGAGTGTGGAGTCGTTGCAAACCTTCATGGGTACTGGGGAANNNAGATCGGAAG |
| 70220 | TATGAGTGTGGAGTCGTTGCTGAAGTGGGCAAAGAGAAGGAACNNNAGATCGGAA |
| 70775 | TATGAGTGTGGAGTCGTTGCTTAGTCTCTGAGAAACCTGGGGGNNNAGATCGGAAG |
| 71534 | TATGAGTGTGGAGTCGTTGCATTGGCTTGGAAAATAAGTGCTTNNNAGATCGGAAG |
| 90974 | TATGAGTGTGGAGTCGTTGCTGCTAGGTCATGTACTAAGTGCTNNNAGATCGGAAG |
| 95685 | TATGAGTGTGGAGTCGTTGCCAGGGCTATTGCAGTAGGTAAAAAGNNNAGATCGG |
| 120347 | TATGAGTGTGGAGTCGTTGCCAAGCAATAACTGATAAAGGCCAAANNNAGATCGGA |
| 120635 | TATGAGTGTGGAGTCGTTGCAACACACGCCTTCCTTGTACTTGNNNAGATCGGAAGA |
| 90212 | TATGAGTGTGGAGTCGTTGCCCTGGAGCCTGTGAATGTTTCTTNNNAGATCGGAAG |
| 91638 | TATGAGTGTGGAGTCGTTGCTACCCTGTACCCTGGTTCTCTTGNNNAGATCGGAAGA |
| 29583 | TATGAGTGTGGAGTCGTTGCCGAAGATGGGTGAAGAAGCAAGANNNAGATCGGAA |
| 70086 | TATGAGTGTGGAGTCGTTGCTCTTGCTTGTTCCTACCTCCTGANNNAGATCGGAAGA |
| 90115 | TATGAGTGTGGAGTCGTTGCATTTCCTCTTGCTTGACCCCTTTNNNAGATCGGAAGAG |
| 93640 | TATGAGTGTGGAGTCGTTGCCGTGTGTGTGTGTATGCCTTTTANNNAGATCGGAAG |
| 94206 | TATGAGTGTGGAGTCGTTGCACCAGTTCTAGCGGGAACTAAGANNNAGATCGGAAG |
| 97687 | TATGAGTGTGGAGTCGTTGCTCTGACATTGGAGAGAAGATGGAAATNNNAGATCGG |
| 93965 | TATGAGTGTGGAGTCGTTGCTTATTGGCAACTTCCAAATGTTGGANNNAGATCGGA |
| 91732 | TATGAGTGTGGAGTCGTTGCATGAGAGCACAAGAACGTGAGTGNNNAGATCGGAA |
| 119627 | TATGAGTGTGGAGTCGTTGCGGAATGTGGAAGGCTATTGGCNNNAGATCGGAAGA |
| 119790 | TATGAGTGTGGAGTCGTTGCGCAGCATAACAACTCACACCAAANNNAGATCGGAAG |
| 119665 | TATGAGTGTGGAGTCGTTGCCCGTGACAAACTAAACGGGAATNNNAGATCGGAAGA |
| 119878 | TATGAGTGTGGAGTCGTTGCGTGGGGGTGGTTGTGTGTTTTNNNAGATCGGAAGA |

| | |
|---|---|
| 120670 | TATGAGTGTGGAGTCGTTGCCTGTGATGCCCCTGTGCATAGANNNAGATCGGAAG |
| 93637 | TATGAGTGTGGAGTCGTTGCCTTTCAGCAGCCCATCCAAAAACNNNAGATCGGAAG |
| 29044 | TATGAGTGTGGAGTCGTTGCCTGCGGGAAATACATTTAGCCCTNNNAGATCGGAAG |
| 29203 | TATGAGTGTGGAGTCGTTGCCAGTGTTGAGTGGACTTGGTGAANNNAGATCGGAA |
| 29765 | TATGAGTGTGGAGTCGTTGCTGCTCAACACACAGAAGGATAAANNNAGATCGGAA |
| 70341 | TATGAGTGTGGAGTCGTTGCTGACCAAGTTCCTTATACCAGCCNNNAGATCGGAAG |
| 70353 | TATGAGTGTGGAGTCGTTGCCGTTTTTGGTAGTGAGTGTTGCCNNNAGATCGGAA |
| 70801 | TATGAGTGTGGAGTCGTTGCTGGCTCTCTGGCACTAAATATGGNNNAGATCGGAAG |
| 71158 | TATGAGTGTGGAGTCGTTGCTCTTCACTTCCATAAGGTTAGCCTNNNAGATCGGAAG |
| 71292 | TATGAGTGTGGAGTCGTTGCAGAACAGGATGGGATAATCCTAAGTNNNAGATCGG |
| 71642 | TATGAGTGTGGAGTCGTTGCAGCAGTGGCCAGAGGAGATGAANNNAGATCGGAA |
| 91149 | TATGAGTGTGGAGTCGTTGCACACACAGACACAATTCATGCCTNNNAGATCGGAAG |
| 100382 | TATGAGTGTGGAGTCGTTGCGCCCTCTTCTTCCACTGATTCGNNNAGATCGGAAGAG |
| 28882 | TATGAGTGTGGAGTCGTTGCTTGCACACAGTGGTACAGTAGTGNNNAGATCGGAA |
| 28949 | TATGAGTGTGGAGTCGTTGCCAGTGACTTGGATTGCAGTGAGANNNAGATCGGAA |
| 29269 | TATGAGTGTGGAGTCGTTGCGGTAGCACTTGCTGTAGGGAATCNNNAGATCGGAA |
| 29434 | TATGAGTGTGGAGTCGTTGCGCCATACCTCTCTGAGTATTTAGCTNNNAGATCGGA |
| 69872 | TATGAGTGTGGAGTCGTTGCAGCATAGTGAGTACACTGTTCCANNNAGATCGGAAG |
| 69909 | TATGAGTGTGGAGTCGTTGCACAGTAATAAACACTCATGGGCANNNAGATCGGAA |
| 70645 | TATGAGTGTGGAGTCGTTGCGGTCTGTTTCAGGTGGTTGAGAGNNNAGATCGGAA |
| 70793 | TATGAGTGTGGAGTCGTTGCGAAAAGAGAGAAAGGGGGTTGGGNNNAGATCGG |
| 70996 | TATGAGTGTGGAGTCGTTGCTGCGAAAATGTTCATTGCACCATNNNAGATCGGAAG |
| 71520 | TATGAGTGTGGAGTCGTTGCATCAATGCATAGGCTTAGCCCTTNNNAGATCGGAAG |
| 99963 | TATGAGTGTGGAGTCGTTGCGACAGGTGATTTCTGCATTGTGAACNNNAGATCGG |
| 29125 | TATGAGTGTGGAGTCGTTGCCGCATTGCCTTCCACTTCAAATTNNNAGATCGGAAGA |
| 69729 | TATGAGTGTGGAGTCGTTGCCACGCACAAAGAATCAACTGCAANNNAGATCGGAAG |
| 69950 | TATGAGTGTGGAGTCGTTGCAAGAACATGCATGCTGAGAAGTCANNNAGATCGG |
| 93502 | TATGAGTGTGGAGTCGTTGCAGAGAGAGAGGCAGAGATGTCACNNNAGATCGG |
| 119709 | TATGAGTGTGGAGTCGTTGCACAAAATCCAAAAAGCTAGAAGAAGANNNAGATCG |
| 121099 | TATGAGTGTGGAGTCGTTGCACTATTCTCTGCAGCCACAAGCCNNNAGATCGGAAG |
| 28931 | TATGAGTGTGGAGTCGTTGCGCCCAGAGGTGTATAGCCAAAATNNNAGATCGGAA |
| 29108 | TATGAGTGTGGAGTCGTTGCAGTAATGGGAAGGATGACCCACANNNAGATCGGAA |
| 29110 | TATGAGTGTGGAGTCGTTGCGGCAGGTTTCCTCTCTAAAGCAANNNAGATCGGAAG |
| 29168 | TATGAGTGTGGAGTCGTTGCATCCCCAGGAAGTTTTCTAGTGCNNNAGATCGGAAG |
| 29185 | TATGAGTGTGGAGTCGTTGCGAGAACGAAGGGAGACAGAGAGANNNAGATCGGA |
| 29369 | TATGAGTGTGGAGTCGTTGCCTTTGGTCATTGTGTCACTAGGCTNNNAGATCGGAA |
| 29464 | TATGAGTGTGGAGTCGTTGCGAATCTATTGCCTTCCCGGGTTTNNNAGATCGGAAG |
| 70465 | TATGAGTGTGGAGTCGTTGCCCATCTCTCTCCTGCTGTAATGCNNNAGATCGGAAGA |
| 70509 | TATGAGTGTGGAGTCGTTGCACCATGTACTCAACACTCGCATCNNNAGATCGGAAG |
| 70598 | TATGAGTGTGGAGTCGTTGCCACTCCGTCACTCTGGTTGAAGNNNAGATCGGAAGA |
| 70805 | TATGAGTGTGGAGTCGTTGCAGTGGAGGAGTAGAAAGCACTGGNNNAGATCGG |
| 71063 | TATGAGTGTGGAGTCGTTGCAAGCAGTTAGGGGCCTCAGATATNNNAGATCGGAA |
| 71115 | TATGAGTGTGGAGTCGTTGCACACATGAACACCTATATACACATACTNNNAGATCGG |

| | |
|---|---|
| 71391 | TATGAGTGTGGAGTCGTTGCCAATGTGTACCTCTCCTGACATGTNNNAGATCGGAA |
| 93620 | TATGAGTGTGGAGTCGTTGCCCATGCCTTTCAGGAAAACCAACNNNAGATCGGAAG |
| 93707 | TATGAGTGTGGAGTCGTTGCACACCACTCCTGTTGGTTCTGTANNNAGATCGGAAG |
| 94646 | TATGAGTGTGGAGTCGTTGCAAAGGCCTGAAAGAATACAAACCANNNAGATCGGA |
| 98194 | TATGAGTGTGGAGTCGTTGCCAACCAGACTTACATGCAGTTTCCNNNAGATCGGAA |
| 90829 | TATGAGTGTGGAGTCGTTGCACACACACACTCAAAAAGGTTTCTNNNAGATCGGAA |
| 28879 | TATGAGTGTGGAGTCGTTGCACCGACATTTTACCACCTCTAGTNNNAGATCGGAAG |
| 28881 | TATGAGTGTGGAGTCGTTGCTGGTTCCTGGCTCATAACTCTTGNNNAGATCGGAAG |
| 29008 | TATGAGTGTGGAGTCGTTGCGGGGAAAAGTCAGGAGAGAGAACNNNAGATCGGA |
| 29164 | TATGAGTGTGGAGTCGTTGCGAGCCACAGCCTATTTCCATCTCNNNAGATCGGAAG |
| 29343 | TATGAGTGTGGAGTCGTTGCAATCTTCAGGCTTCCCTGCATTTNNNAGATCGGAAG |
| 29625 | TATGAGTGTGGAGTCGTTGCAGGAGGTGAGTTTCATATTTTTGGTNNNAGATCGG |
| 70510 | TATGAGTGTGGAGTCGTTGCCAGTACACACAGGCAGATGCATCNNNAGATCGGAA |
| 94159 | TATGAGTGTGGAGTCGTTGCAATGAGTGTAGGAGGTGCTCAGANNNAGATCGGA |
| 95360 | TATGAGTGTGGAGTCGTTGCGGTCAGGATTTGAACCCATAGAGTNNNAGATCGGA |
| 90249 | TATGAGTGTGGAGTCGTTGCTCTGAGGAGAATGTCCAGAGCTTNNNAGATCGGAA |
| 91115 | TATGAGTGTGGAGTCGTTGCACACTGGGTTTGTGGTATCTCCTNNNAGATCGGAAG |
| 115017 | TATGAGTGTGGAGTCGTTGCTACACATATGCTCCCATCCAGGCNNNAGATCGGAAG |
| 28800 | TATGAGTGTGGAGTCGTTGCACCCATATTTCAAAGTGATGTCATGTNNNAGATCGG |
| 28901 | TATGAGTGTGGAGTCGTTGCCCTTTCTTTCTTGGTCGTGGGTANNNAGATCGGAAG |
| 28905 | TATGAGTGTGGAGTCGTTGCGTCACATGAAGGCTGCTATAGCTNNNAGATCGGAA |
| 29476 | TATGAGTGTGGAGTCGTTGCGAGAGAGAGACGCTGTGTGTTTTNNNAGATCGGAA |
| 29582 | TATGAGTGTGGAGTCGTTGCTTGCCAGAAGGTATAACTGAAGANNNAGATCGGAA |
| 29630 | TATGAGTGTGGAGTCGTTGCTGGAAATTCTGGAGTTGAAAAGGTNNNAGATCGGA |
| 70707 | TATGAGTGTGGAGTCGTTGCCAAGAGTAGGAGGGAGAGGAAGGNNNAGATCGG |
| 71186 | TATGAGTGTGGAGTCGTTGCGGACACACACACTGACATAAGCTNNNAGATCGGAAG |
| 71256 | TATGAGTGTGGAGTCGTTGCAAAGTCTGAAGTTTGGCCAGTCTNNNAGATCGGAA |
| 94857 | TATGAGTGTGGAGTCGTTGCGGAAGCCACATGAAGAAGGTGAANNNAGATCGGA |
| 98239 | TATGAGTGTGGAGTCGTTGCGGGTGAAAATCATACAGTGTGTCCNNNAGATCGGA |
| 98339 | TATGAGTGTGGAGTCGTTGCGCCCCTAGCCTGTAGTATTTCTTNNNAGATCGGAAG |
| 93931 | TATGAGTGTGGAGTCGTTGCGTGGCAGCAGTGGATATTGATGANNNAGATCGGAA |
| 90229 | TATGAGTGTGGAGTCGTTGCCCATAAGCCAAGACTGCCATCTTNNNAGATCGGAAG |
| 92925 | TATGAGTGTGGAGTCGTTGCATCACCTCCAAAATCCCTCATGCNNNAGATCGGAAGA |
| 28763 | TATGAGTGTGGAGTCGTTGCTGATACTGTCGCAGGTGCTAAACNNNAGATCGGAA |
| 28855 | TATGAGTGTGGAGTCGTTGCCTTTCCTTTCCTCCCCACTCTCTNNNAGATCGGAAGAG |
| 28996 | TATGAGTGTGGAGTCGTTGCCAAGGCCACAGCATTTTTGGAAGNNNAGATCGGAA |
| 29178 | TATGAGTGTGGAGTCGTTGCGGGAGGGGGTGGTGATATAAGTANNNAGATCGGA |
| 29200 | TATGAGTGTGGAGTCGTTGCCAGTTAGGGGGAAGGAGAAGACANNNAGATCGGA |
| 29392 | TATGAGTGTGGAGTCGTTGCGGCACAAGCACACACATTCTTTTNNNAGATCGGAAG |
| 69677 | TATGAGTGTGGAGTCGTTGCCAGGTGGAACAGTTTGCACAAAANNNAGATCGGAA |
| 69767 | TATGAGTGTGGAGTCGTTGCTGACTACCCCATTGCCAGGATAANNNAGATCGGAAG |
| 70165 | TATGAGTGTGGAGTCGTTGCATACATCTTGGCTAGGCTCGGTANNNAGATCGGAAG |
| 70692 | TATGAGTGTGGAGTCGTTGCAGAACATGAAGGGACACACATCGNNNAGATCGGAA |

71157 TATGAGTGTGGAGTCGTTGCGGCATGTTTTATGTTCTGAGCCTNNNAGATCGGAAG
71227 TATGAGTGTGGAGTCGTTGCCAGCCCAGAAAAACTAGAGTGCTNNNAGATCGGAA
96337 TATGAGTGTGGAGTCGTTGCGGGTTTATGTTGATGGGGAAAAGNNNAGATCGGA
96387 TATGAGTGTGGAGTCGTTGCTTCTTCACTGCCCCATTTCATCTAGNNNAGATCGGAA
96845 TATGAGTGTGGAGTCGTTGCGCCCTGAACTCCTGTACTCATGCNNNAGATCGGAAG
98007 TATGAGTGTGGAGTCGTTGCAAGAAAGACAACTCAGAATGGGAGNNNAGATCGG
103054 TATGAGTGTGGAGTCGTTGCTTTCTCATCGATTCATCCATCAGTGNNNAGATCGGAA
119968 TATGAGTGTGGAGTCGTTGCTCTCTGATAAAACTGGCTTTACACCANNNAGATCGG
120162 TATGAGTGTGGAGTCGTTGCGCTCATGGACACTATCCTCCATCTNNNAGATCGGAAG
120037 TATGAGTGTGGAGTCGTTGCTTGTATTGGGCCATGAGGTAGTTGNNNAGATCGGA
120290 TATGAGTGTGGAGTCGTTGCCCCTCGTCCGTTTAGAAAGCTCNNNAGATCGGAAGA
28825 TATGAGTGTGGAGTCGTTGCGGGAGCTTCAGTTCATAGCTTGANNNAGATCGGAA
28893 TATGAGTGTGGAGTCGTTGCGCCCAGCCAGGAAATAAACCTAGNNNAGATCGGAA
28950 TATGAGTGTGGAGTCGTTGCAGGCTTGGGGAGCTATGATAAAGANNNAGATCGG
28991 TATGAGTGTGGAGTCGTTGCGAATTGCAGGTAGCAGGAGAAGGNNNAGATCGGA
29253 TATGAGTGTGGAGTCGTTGCTAGCATTGTGCTGAGGAATGTGGNNNAGATCGGAA
29255 TATGAGTGTGGAGTCGTTGCATGAGTATGTGGGGTGTAGGGGNNNAGATCGGAA
29725 TATGAGTGTGGAGTCGTTGCCCAGGGATCCTCATCTGTTGGTANNNAGATCGGAAG
29731 TATGAGTGTGGAGTCGTTGCTCTGACATAACCTGGAATTTGTTGANNNAGATCGGA
69691 TATGAGTGTGGAGTCGTTGCGGCTGATGAGAGCGTCTTACAAANNNAGATCGGAA
70138 TATGAGTGTGGAGTCGTTGCCCAGGAAACCTTCTCTTCCCCTANNNAGATCGGAAGA
70998 TATGAGTGTGGAGTCGTTGCATGATCCAGGAGTTGGAACCCATNNNAGATCGGAA
71607 TATGAGTGTGGAGTCGTTGCATAAACTGCATTGAGCTGCGTTTNNNAGATCGGAAG
93170 TATGAGTGTGGAGTCGTTGCGAAGTTTGCCTTTTCAGCTGTCCNNNAGATCGGAAG
93811 TATGAGTGTGGAGTCGTTGCCCCTCTCCCTCTCTCTGTTCATANNNAGATCGGAAGA
96981 TATGAGTGTGGAGTCGTTGCATCCCCCTGCATCCTTTTCTTTNNNAGATCGGAAGAG
98303 TATGAGTGTGGAGTCGTTGCGCAGTGTTTCTAATTTCAGGGAGCNNNAGATCGGA
121987 TATGAGTGTGGAGTCGTTGCTGTAATTGACCTAGATGCCTGTCAACNNNAGATCGG
121994 TATGAGTGTGGAGTCGTTGCGCTCTGTCTTTCAGATTGGAGTGGNNNAGATCGGA
94312 TATGAGTGTGGAGTCGTTGCGGTGAGAAACATGAGTTGGCTCANNNAGATCGGAA
119935 TATGAGTGTGGAGTCGTTGCGTGGAAATGAGGCCTCGGTATGTNNNAGATCGGAA
120158 TATGAGTGTGGAGTCGTTGCTCTCCACTCACAGCCACTTCTANNNAGATCGGAAGAG
28737 TATGAGTGTGGAGTCGTTGCTCGGTCAGGCTACACATTTTCCANNNAGATCGGAAG
28762 TATGAGTGTGGAGTCGTTGCTACATCCCCCACAAAAAGAGCACNNNAGATCGGAAG
28914 TATGAGTGTGGAGTCGTTGCTGAAGCATGATTTTTAGTGGCTGCNNNAGATCGGA
28961 TATGAGTGTGGAGTCGTTGCATTTGATCTGTTGGGGGTGCATTNNNAGATCGGAA
29013 TATGAGTGTGGAGTCGTTGCGGTCATGGAGCTCCTATTCAGGANNNAGATCGGAA
29022 TATGAGTGTGGAGTCGTTGCCCCTCCCACGTGACATCTAGTAANNNAGATCGGAAG
29495 TATGAGTGTGGAGTCGTTGCTGGCAGTGATGTTCCAAGTTCATNNNAGATCGGAAG
29569 TATGAGTGTGGAGTCGTTGCCTCTTGTGCATAAATTGCCCCACNNNAGATCGGAAG
29651 TATGAGTGTGGAGTCGTTGCAGAGACCTGCTATGAGATGGCATNNNAGATCGGAA
29758 TATGAGTGTGGAGTCGTTGCTTATCAGTTCTGTTCTCTGGAGAANNNAGATCGGAA
69884 TATGAGTGTGGAGTCGTTGCTACTTACAGTTCCTTCTTTACAGCANNNAGATCGGAA

70050 TATGAGTGTGGAGTCGTTGCTCTTTGGGCATTGAAACTACTGGANNNAGATCGGAA
70271 TATGAGTGTGGAGTCGTTGCTGGTCTGTTTTCCGGGTCAATACNNNAGATCGGAAG
70298 TATGAGTGTGGAGTCGTTGCCCCATTCCTTGGCTGATTCTACCNNNAGATCGGAAGA
70983 TATGAGTGTGGAGTCGTTGCAAGGCATATCAGCCCCTTTGAATNNNAGATCGGAAG
71078 TATGAGTGTGGAGTCGTTGCAAGAAGGGGCAGATTTGCAAACTNNNAGATCGGAA
71081 TATGAGTGTGGAGTCGTTGCAGGTGCTCTGTTCTATCCCAACTNNNAGATCGGAAG
71161 TATGAGTGTGGAGTCGTTGCAGGAAGACAATGTACAGAGGCCTNNNAGATCGGAA
91184 TATGAGTGTGGAGTCGTTGCGCACCACATAATGACATTCCCCTNNNAGATCGGAAG
95797 TATGAGTGTGGAGTCGTTGCAAAATAATGCGTGAAAAGTGCTTGGNNNAGATCGG
90159 TATGAGTGTGGAGTCGTTGCCAGCATGACAAGAGCTGATTGTTNNNAGATCGGAA
93644 TATGAGTGTGGAGTCGTTGCTCATGGCCTCTCCCTTCTGTTTANNNAGATCGGAAGA
92009 TATGAGTGTGGAGTCGTTGCTGGAGACTGAGAGAGAGAGATGGNNNAGATCGGA
90285 TATGAGTGTGGAGTCGTTGCCCCCACCCTTCCTTTCTCTACTTNNNAGATCGGAAGAG
92425 TATGAGTGTGGAGTCGTTGCATTTAGGAGGCACATGCACAGAGNNNAGATCGGAA
29331 TATGAGTGTGGAGTCGTTGCGGTTTGTCTGTGTGTGAGCTCAANNNAGATCGGAA
29634 TATGAGTGTGGAGTCGTTGCAAACAAATTCTGCAAAGTTGCAAGTNNNAGATCGGA
35459 TATGAGTGTGGAGTCGTTGCGTCCACAGACCAATAGCATAGCANNNAGATCGGAAG
70194 TATGAGTGTGGAGTCGTTGCGGAGGTGGGGAGATGAAGCTTTANNNAGATCGGA
70463 TATGAGTGTGGAGTCGTTGCGAGGCATGAATGCTGAGAAATGCNNNAGATCGGAA
70799 TATGAGTGTGGAGTCGTTGCGAGTTTGAAGACCTAGCCCATGGNNNAGATCGGAA
94438 TATGAGTGTGGAGTCGTTGCTTACTTGGTCTGTGAGGTAGGCANNNAGATCGGAA
95880 TATGAGTGTGGAGTCGTTGCGCTTCTTTTTGGGGTGATATGTTCTANNNAGATCGG
96714 TATGAGTGTGGAGTCGTTGCTCACAGAAAATGAAAAGTTACCCTGANNNAGATCGG
96899 TATGAGTGTGGAGTCGTTGCTTCCAGTATATATCCCACTCCCTTTGNNNAGATCGGA
97764 TATGAGTGTGGAGTCGTTGCTTCTTGCAATCTCCCCTAAACGGNNNAGATCGGAAG
92566 TATGAGTGTGGAGTCGTTGCGGAGTAGAGCAACCAAGGTCAAGNNNAGATCGGAA
119683 TATGAGTGTGGAGTCGTTGCACGGATTTTCACTCTCTCAACCCCNNNAGATCGGAAG
29017 TATGAGTGTGGAGTCGTTGCCCCACAAGAAATACTCAGCAGCANNNAGATCGGAAG
29152 TATGAGTGTGGAGTCGTTGCTAAGAGCATGTCATCTACGAAATGAGGNNNAGATC
29258 TATGAGTGTGGAGTCGTTGCCAGAGGGACAGAGACAGAGGTAGNNNAGATCGGA
29511 TATGAGTGTGGAGTCGTTGCTCTTACTGGGTGCTTTTCCACTGNNNAGATCGGAAG
29562 TATGAGTGTGGAGTCGTTGCTACTTTTCCTGCAGTGTACAGCCNNNAGATCGGAAG
69679 TATGAGTGTGGAGTCGTTGCCCTGGATGAACACAGTTGCAAAANNNAGATCGGAA
69723 TATGAGTGTGGAGTCGTTGCTGTGCCTCTTTCTTTCTGAGCAANNNAGATCGGAAG
69908 TATGAGTGTGGAGTCGTTGCAGGAGACACAAATTTCTAGGGCANNNAGATCGGAA
69915 TATGAGTGTGGAGTCGTTGCCTGCAAGGTTTGGAAATACTGCANNNAGATCGGAA
69938 TATGAGTGTGGAGTCGTTGCATAATCCAGGAAGAGCACCCTCANNNAGATCGGAAG
70760 TATGAGTGTGGAGTCGTTGCACACATCTTTTCACCACACAGGGNNNAGATCGGAAG
90493 TATGAGTGTGGAGTCGTTGCATTGTGGGTGGCAAAGAAACTGTNNNAGATCGGAA
93612 TATGAGTGTGGAGTCGTTGCTGCTTACCCTTTCTACATGCAACNNNAGATCGGAAGA
95807 TATGAGTGTGGAGTCGTTGCGCAGTTACCCAGAGTTTTTGAAGGTANNNAGATCGG
106766 TATGAGTGTGGAGTCGTTGCTGTCTTTATTCATCCACTGATGGGANNNAGATCGGA
110170 TATGAGTGTGGAGTCGTTGCAGAGAGAAGGAGAGAAAGGCAGTGNNNAGATCGG

| | |
|---|---|
| 90921 | TATGAGTGTGGAGTCGTTGCTTCAACGGTCTGATGTACCCTCTNNNAGATCGGAAG |
| 94871 | TATGAGTGTGGAGTCGTTGCACACACACACGGAGACAAATGAANNNAGATCGGAA |
| 93756 | TATGAGTGTGGAGTCGTTGCCAAGCAAGCAAATAACTGGGCTANNNAGATCGGAA |
| 92138 | TATGAGTGTGGAGTCGTTGCCTCCTTGGGCCAATCAAGTTAGGNNNAGATCGGAAG |
| 120332 | TATGAGTGTGGAGTCGTTGCAGCGATTTGTGGGTTCACATCACNNNAGATCGGAAG |
| 120968 | TATGAGTGTGGAGTCGTTGCGTTGGAGGGCATACTTGTTGAANNNAGATCGGAAG |
| 121159 | TATGAGTGTGGAGTCGTTGCGGACAACCGTCATCTCTAGCTCNNNAGATCGGAAGA |
| 121022 | TATGAGTGTGGAGTCGTTGCGGTGGTGGGGACAGGATATTGNNNAGATCGGAAG |
| 28739 | TATGAGTGTGGAGTCGTTGCTCCACCTGACAAAAACGTCAGTTNNNAGATCGGAAG |
| 28742 | TATGAGTGTGGAGTCGTTGCACCAACTTTTTCAGAGCTTCGGTNNNAGATCGGAAG |
| 28948 | TATGAGTGTGGAGTCGTTGCCACCACCAACAAACACCTTAGGANNNAGATCGGAAG |
| 28992 | TATGAGTGTGGAGTCGTTGCGGCATCTTCTAAGCCACAAAAGGNNNAGATCGGAA |
| 29318 | TATGAGTGTGGAGTCGTTGCGCTCTACATTTCAGATGTGCCCANNNAGATCGGAAG |
| 29697 | TATGAGTGTGGAGTCGTTGCAGAGGAGAGAGAGACTTTGGTGCNNNAGATCGGA |
| 70017 | TATGAGTGTGGAGTCGTTGCGACACATTTCTGCAGTGGGTAGANNNAGATCGGAA |
| 71173 | TATGAGTGTGGAGTCGTTGCAATCCACTCATCCTACCCATCCTNNNAGATCGGAAGA |
| 119708 | TATGAGTGTGGAGTCGTTGCGGCAGACACAATAAAGACACAAANNNAGATCGGAA |
| 120451 | TATGAGTGTGGAGTCGTTGCTACAAGAGGTGCAAAGAAGAACAGGNNNAGATCG |
| 91675 | TATGAGTGTGGAGTCGTTGCGGTCCTCTATCTAGGCTGTTGTGNNNAGATCGGAAG |
| 28861 | TATGAGTGTGGAGTCGTTGCAAAGTTACAGCAATCTGGGTGGANNNAGATCGGAA |
| 29066 | TATGAGTGTGGAGTCGTTGCCTTTGCTTGCACATGCTGTCAGNNNAGATCGGAAGA |
| 29501 | TATGAGTGTGGAGTCGTTGCCAGGGGCACACTCACACATAAATNNNAGATCGGAAG |
| 69749 | TATGAGTGTGGAGTCGTTGCTTATGTGCTCCCCTTCCTGAGAANNNAGATCGGAAG |
| 70190 | TATGAGTGTGGAGTCGTTGCTCACCACACCCATGCTTTTGTTANNNAGATCGGAAGA |
| 70439 | TATGAGTGTGGAGTCGTTGCGCATAGACAAAGGTGAAGAGGGCNNNAGATCGGA |
| 70561 | TATGAGTGTGGAGTCGTTGCGGCACACTCTGGTCTTTCTCTTCNNNAGATCGGAAGA |
| 98622 | TATGAGTGTGGAGTCGTTGCCCGACTTTTCCACCATTCCTTACANNNAGATCGGAAG |
| 100422 | TATGAGTGTGGAGTCGTTGCGGTACTACGGTCCTTGGTAGAAGNNNAGATCGGAA |
| 70389 | TATGAGTGTGGAGTCGTTGCGGCATGAAAGGAGATAAGCAAAGCNNNAGATCGG |
| 119782 | TATGAGTGTGGAGTCGTTGCCTGCTACTGTTTTTGCATGTTGATTNNNAGATCGGA |
| 120771 | TATGAGTGTGGAGTCGTTGCTACTGAAATGGGCACTCTGATACATNNNAGATCGGA |
| 28765 | TATGAGTGTGGAGTCGTTGCGGTGAGAGAGCCCAAATCTCCTANNNAGATCGGAA |
| 28940 | TATGAGTGTGGAGTCGTTGCAGAAGTTTGCTTTCCCTAGCTGCNNNAGATCGGAAG |
| 28976 | TATGAGTGTGGAGTCGTTGCTTTCCTGGTCTTGTCAATGCACTNNNAGATCGGAAG |
| 29084 | TATGAGTGTGGAGTCGTTGCTGCTTGGCAATGGATGGAATAACNNNAGATCGGAA |
| 29123 | TATGAGTGTGGAGTCGTTGCAGAAGAAACAGGGAGTGGGGATTNNNAGATCGGA |
| 29159 | TATGAGTGTGGAGTCGTTGCGACGGGGAAAGAGAGAACAGAAGNNNAGATCGGA |
| 29574 | TATGAGTGTGGAGTCGTTGCTGGGAAGGGGAAAACAAAGACTANNNAGATCGGA |
| 29577 | TATGAGTGTGGAGTCGTTGCGGGCAGGAAGAGTGGTGTACATANNNAGATCGGA |
| 29763 | TATGAGTGTGGAGTCGTTGCTCAGGTCCCCATCCCAAACTAAANNNAGATCGGAAG |
| 69826 | TATGAGTGTGGAGTCGTTGCTGGAAGTGTGGGTAAAATGGACANNNAGATCGGA |
| 70159 | TATGAGTGTGGAGTCGTTGCACCAAAACATTGGAGCCACAGTANNNAGATCGGAA |
| 70169 | TATGAGTGTGGAGTCGTTGCGAGTGGATGTGCGTGCATATGTANNNAGATCGGAA |

| | |
|---|---|
| 71285 | TATGAGTGTGGAGTCGTTGCAACAGTGCATCCCAAGTGAAAGTNNNAGATCGGAA |
| 95382 | TATGAGTGTGGAGTCGTTGCGGAAACAATGAACCTGACTTGCAGNNNAGATCGGA |
| 98480 | TATGAGTGTGGAGTCGTTGCTGCCACACACACTTCCACATAANNNAGATCGGAAGA |
| 121936 | TATGAGTGTGGAGTCGTTGCATGTGAAAGCCAGAACTCAGCCNNNAGATCGGAAG |
| 91979 | TATGAGTGTGGAGTCGTTGCCCTAATACAACTGTGGGAGCTGGNNNAGATCGGAA |
| 119986 | TATGAGTGTGGAGTCGTTGCCCTCATTATTTTGGAGTGCTGTGTNNNAGATCGGA |
| 121114 | TATGAGTGTGGAGTCGTTGCGAGTTTCATTCACTCTGTCGCCAANNNAGATCGGAA |
| 28808 | TATGAGTGTGGAGTCGTTGCATGAGGTCTTTCCCAGTGGTCTTNNNAGATCGGAAG |
| 29047 | TATGAGTGTGGAGTCGTTGCATTCAACAAGGAGGGGTCTGACTNNNAGATCGGAA |
| 29208 | TATGAGTGTGGAGTCGTTGCCACACACACACTGAGCTCTCTTTNNNAGATCGGAAGA |
| 29515 | TATGAGTGTGGAGTCGTTGCGGAGCACACTTTTGATAATGCTGGNNNAGATCGGA |
| 29522 | TATGAGTGTGGAGTCGTTGCGGGCACCTTGACTAATACGAAGGNNNAGATCGGAA |
| 69680 | TATGAGTGTGGAGTCGTTGCCTGTTCCTGCTCTCAGGTCAAAANNNAGATCGGAAG |
| 69933 | TATGAGTGTGGAGTCGTTGCAGCACATAGCTTGTAGGGTTATCANNNAGATCGGAA |
| 70637 | TATGAGTGTGGAGTCGTTGCTCGCTCTTCGGTATCACAAAGAGNNNAGATCGGAAG |
| 71035 | TATGAGTGTGGAGTCGTTGCAACAACCAATTCCTGACCCAGATNNNAGATCGGAAG |
| 71438 | TATGAGTGTGGAGTCGTTGCGGGAGAGAGACTTATGCATGTGTNNNAGATCGGAA |
| 71559 | TATGAGTGTGGAGTCGTTGCGCATACCTTCTCTGGAAAGGGTTNNNAGATCGGAAG |
| 91729 | TATGAGTGTGGAGTCGTTGCGCATGGTTTCTGCCTTCATAGTGNNNAGATCGGAAG |
| 94527 | TATGAGTGTGGAGTCGTTGCATCTAATTGCCTCCTGTGCTCCANNNAGATCGGAAGA |
| 97681 | TATGAGTGTGGAGTCGTTGCTCAACCCCCACTCTTGAGAACATGNNNAGATCGGAA |
| 100283 | TATGAGTGTGGAGTCGTTGCGGGAAGCTAGGTGAAAGAACCCANNNAGATCGGAA |
| 120294 | TATGAGTGTGGAGTCGTTGCGGACCTGGGAAATACTTCCAAACTNNNAGATCGGAA |
| 94185 | TATGAGTGTGGAGTCGTTGCGATTTCCAAGCATGGCCTACAGANNNAGATCGGAAG |
| 28983 | TATGAGTGTGGAGTCGTTGCCTGTGGATAAGAGCTCACTGCTGNNNAGATCGGAA |
| 29317 | TATGAGTGTGGAGTCGTTGCAGTGGTTTGAATGGTTGTTCCCANNNAGATCGGAA |
| 71557 | TATGAGTGTGGAGTCGTTGCAAGACTGAGTTGTGGTCTTCGTTNNNAGATCGGAA |
| 90864 | TATGAGTGTGGAGTCGTTGCTGTGCTTGTACTTCTTCCTCATTCTNNNAGATCGGAA |
| 92350 | TATGAGTGTGGAGTCGTTGCTCCCACAGGTCGGCCTTATATAGNNNAGATCGGAAG |
| 96311 | TATGAGTGTGGAGTCGTTGCGAATGCCTCCTTCTCTCTGATGTTCNNNAGATCGGAA |
| 119662 | TATGAGTGTGGAGTCGTTGCAAGAAAGGAGGGGAAACCTTATTNNNAGATCGGAA |
| 91201 | TATGAGTGTGGAGTCGTTGCGGTGTGTGAAGGACAGAGTACCTNNNAGATCGGAA |
| 93880 | TATGAGTGTGGAGTCGTTGCGAAACCGGGAGAAATGATGGTGANNNAGATCGGA |
| 120260 | TATGAGTGTGGAGTCGTTGCGACTGGCATAAGATCCAGCAATCNNNAGATCGGAA |
| 121155 | TATGAGTGTGGAGTCGTTGCGACACTTTGGAAGACAATTGGTTNNNAGATCGGAA |
| 28746 | TATGAGTGTGGAGTCGTTGCAAAGGGGTAACCCAGCAAAGAAGNNNAGATCGGA |
| 28776 | TATGAGTGTGGAGTCGTTGCGTGTGCCAATTGCTGTTTCAAGTNNNAGATCGGAAG |
| 28947 | TATGAGTGTGGAGTCGTTGCCTCACCGTTCGCAGTACTGAATANNNAGATCGGAAG |
| 29076 | TATGAGTGTGGAGTCGTTGCTTGATCTTTGGTGCTATGAGGCCNNNAGATCGGAAG |
| 29254 | TATGAGTGTGGAGTCGTTGCGGAAGAATCCATCTGGAGCATGGNNNAGATCGGAA |
| 29502 | TATGAGTGTGGAGTCGTTGCGAACTGTTGAAGCACCTGCTTTGNNNAGATCGGAAG |
| 29768 | TATGAGTGTGGAGTCGTTGCAGGGGCTGGTGATAATCTCTGTTNNNAGATCGGAA |
| 69879 | TATGAGTGTGGAGTCGTTGCTAGGCAAAATGATGCGGTAAGCANNNAGATCGGAA |

| | |
|---|---|
| 69955 | TATGAGTGTGGAGTCGTTGCTCACGTTCCATATTTTCCTGTCANNNAGATCGGAAGA |
| 71565 | TATGAGTGTGGAGTCGTTGCTGCTTGTGTCCCCCTAAAATGTTNNNAGATCGGAAG |
| 93854 | TATGAGTGTGGAGTCGTTGCATTTCGGGGGACTAAGGAGTTGANNNAGATCGGAA |
| 99655 | TATGAGTGTGGAGTCGTTGCACTTGTGACCTGTAGTGGCTGTNNNAGATCGGAAG |
| 109481 | TATGAGTGTGGAGTCGTTGCGAGACAGAGACTGAATTGCTTGAGANNNAGATCGG |
| 90861 | TATGAGTGTGGAGTCGTTGCTACCCACACACCTCATGGATTCTNNNAGATCGGAAG |
| 119971 | TATGAGTGTGGAGTCGTTGCTGAACATGCAGTCTCTCTTGGAGATGNNNAGATCGG |
| 121072 | TATGAGTGTGGAGTCGTTGCGAACAGGGGCAAGAAGACGGACTNNNAGATCGGA |
| 28908 | TATGAGTGTGGAGTCGTTGCATTGAGAAGAGAGACTGGGGGAGNNNAGATCGGA |
| 28954 | TATGAGTGTGGAGTCGTTGCAAGCATGTGTGTATGCCGAGAAANNNAGATCGGAA |
| 29099 | TATGAGTGTGGAGTCGTTGCGAGTGGACCTGGTAAGAAGGTCANNNAGATCGGA |
| 29259 | TATGAGTGTGGAGTCGTTGCCACACACCTATCTGACAGCTGAGNNNAGATCGGAAG |
| 29660 | TATGAGTGTGGAGTCGTTGCACCTTCTGACCTATTTTGTCCTGGNNNAGATCGGAAG |
| 29709 | TATGAGTGTGGAGTCGTTGCGTAGCATTTCCACTTCTGAGGCCNNNAGATCGGAAG |
| 29739 | TATGAGTGTGGAGTCGTTGCGCATCACTCCATTGCACTACTCANNNAGATCGGAAG |
| 70567 | TATGAGTGTGGAGTCGTTGCGGGTGGGTTGTATTAGTCCGTTCNNNAGATCGGAA |
| 70677 | TATGAGTGTGGAGTCGTTGCTGCCTCTCTCTGATCACAGGTAGNNNAGATCGGAAG |
| 71266 | TATGAGTGTGGAGTCGTTGCAGTAGCCTCCACACATTCCTTCTNNNAGATCGGAAGA |
| 92607 | TATGAGTGTGGAGTCGTTGCCTCTCTGGGGAACTTCGGGTTTCNNNAGATCGGAAG |
| 95600 | TATGAGTGTGGAGTCGTTGCGGGTTTTACTTTGTTGCCACCAANNNAGATCGGAAG |
| 100308 | TATGAGTGTGGAGTCGTTGCTTAGCATCACGCAATATGCCCATANNNAGATCGGAA |
| 93204 | TATGAGTGTGGAGTCGTTGCATCCAAAGGCAGTTTCTCCATCCNNNAGATCGGAAG |
| 109108 | TATGAGTGTGGAGTCGTTGCTCGTTCGTGTGTTTGGATAGATANNNAGATCGGAA |
| 117322 | TATGAGTGTGGAGTCGTTGCGGAATCAAAGTCTGACAACAGCCANNNAGATCGGA |
| 120400 | TATGAGTGTGGAGTCGTTGCACTGCCTTCATGATTTAGGATGGACNNNAGATCGGA |
| 92803 | TATGAGTGTGGAGTCGTTGCCCCCTGGTTTTCTACCATGGATCNNNAGATCGGAAG |
| 28804 | TATGAGTGTGGAGTCGTTGCAGCATGAGGAATAGCAGAATGGCNNNAGATCGGA |
| 28868 | TATGAGTGTGGAGTCGTTGCAGGAGCAGGAGTTGAAGTGGATTNNNAGATCGGA |
| 29000 | TATGAGTGTGGAGTCGTTGCCACCAATCCCTTTTATCTGCTGCNNNAGATCGGAAGA |
| 29059 | TATGAGTGTGGAGTCGTTGCACTCTTTCCTCTATATGCACGTGGNNNAGATCGGAA |
| 29086 | TATGAGTGTGGAGTCGTTGCAAGTGTGGCTCCTGGAATTTGTANNNAGATCGGAA |
| 29320 | TATGAGTGTGGAGTCGTTGCTGCTGTTTCTTCAGCTATCACCANNNAGATCGGAAG |
| 29393 | TATGAGTGTGGAGTCGTTGCTTCAAGCACAGTAGCTTTGGTTTNNNAGATCGGAAG |
| 29458 | TATGAGTGTGGAGTCGTTGCCCAACCCCCGTTATCTGTGATTTNNNAGATCGGAAG |
| 29615 | TATGAGTGTGGAGTCGTTGCGAGCCTTTGGAGCAGCTTTACTTNNNAGATCGGAAG |
| 29736 | TATGAGTGTGGAGTCGTTGCGCACACACAAGCATTTTCTATTTCANNNAGATCGGAA |
| 70687 | TATGAGTGTGGAGTCGTTGCTTACCACTCTGTCTCTCTTCCCGNNNAGATCGGAAGA |
| 71445 | TATGAGTGTGGAGTCGTTGCAACTGGAGGAACAATGTGTGTGTNNNAGATCGGAA |
| 97077 | TATGAGTGTGGAGTCGTTGCCGGTGGAAATAGCCAGGAAATATNNNAGATCGGAA |
| 97690 | TATGAGTGTGGAGTCGTTGCTATTCCTCCTTTTGCATCTGGCTTTNNNAGATCGGAA |
| 93659 | TATGAGTGTGGAGTCGTTGCTGCCCAGAAAGCCAAACCATTTANNNAGATCGGAAG |
| 70923 | TATGAGTGTGGAGTCGTTGCAGGAATACGAAGTTGACGGTGTGNNNAGATCGGA |
| 120039 | TATGAGTGTGGAGTCGTTGCAAAATGCTGTTGGAGACTGGCAAAGNNNAGATCGG |

120013 TATGAGTGTGGAGTCGTTGCCTGGAGTTTCCCAAACTAAATGGCNNNAGATCGGAA
120796 TATGAGTGTGGAGTCGTTGCTGTTCCACTCTCTTCTGGTCTTTNNNAGATCGGAAGA
93309 TATGAGTGTGGAGTCGTTGCACGGTAGTGCTCTACTACATCCCNNNAGATCGGAAG
28756 TATGAGTGTGGAGTCGTTGCATTCTACCCTGGATTCCCTTCCCNNNAGATCGGAAGA
28852 TATGAGTGTGGAGTCGTTGCCCGCCAGGGTTTGTTTGTTTTAGNNNAGATCGGAAG
29240 TATGAGTGTGGAGTCGTTGCAGCCTCATGTTGTGGACCTCTATNNNAGATCGGAAG
29521 TATGAGTGTGGAGTCGTTGCTCCGCCTCCTGGTGTATATAAGGNNNAGATCGGAAG
29730 TATGAGTGTGGAGTCGTTGCAGCAACCTGAATATCAGCTTTGANNNAGATCGGAAG
69703 TATGAGTGTGGAGTCGTTGCCTGACCCCCTCCCCAGTTATAAANNNAGATCGGAAGA
70887 TATGAGTGTGGAGTCGTTGCCTGCCTACTGTGTGCTTACTCTGNNNAGATCGGAAG
71528 TATGAGTGTGGAGTCGTTGCACATAATTTGAGGCCCACAGCTTNNNAGATCGGAAG
71708 TATGAGTGTGGAGTCGTTGCCCACATCAGCCTAGTTTCGCTCNNNAGATCGGAAGA
95258 TATGAGTGTGGAGTCGTTGCTTCTTAGGCAGTAGAGGACTGCGNNNAGATCGGAA
95458 TATGAGTGTGGAGTCGTTGCAGAAATAACACAAAGGCAAAGTACCANNNAGATCG
97639 TATGAGTGTGGAGTCGTTGCCCATTCAACTCACTACAGCCTTNNNAGATCGGAAGAG
91827 TATGAGTGTGGAGTCGTTGCCAAGGCTGAGAAACCCTAACCTGNNNAGATCGGAA
91020 TATGAGTGTGGAGTCGTTGCAGGAAGCAATGCAGAATATGGCTNNNAGATCGGAA
119988 TATGAGTGTGGAGTCGTTGCAAACTCAGAGGGCCAGGATGGAANNNAGATCGGA
120978 TATGAGTGTGGAGTCGTTGCATTAGTGACAGACATTTTAGATGTGATNNNAGATCG
28740 TATGAGTGTGGAGTCGTTGCATTGATCGTAGGACAGGACATGCNNNAGATCGGAA
28894 TATGAGTGTGGAGTCGTTGCCACACGGGAACCCTAACCAATAGNNNAGATCGGAAG
29261 TATGAGTGTGGAGTCGTTGCACAGGGGAGATACATTCCTGGAGNNNAGATCGGAA
29288 TATGAGTGTGGAGTCGTTGCGGGGGAAGAGAAAGAGACTTATCACNNNAGATCG
29300 TATGAGTGTGGAGTCGTTGCGCCCCATTATACCTGGAAGAGGANNNAGATCGGAA
29326 TATGAGTGTGGAGTCGTTGCCAGACAGAGTGAAAGGCCTTGAANNNAGATCGGAA
29361 TATGAGTGTGGAGTCGTTGCAAAAACAACCTTCTTCCCCCAGTNNNAGATCGGAAG
29498 TATGAGTGTGGAGTCGTTGCGAGGCTACTTGCTGCTTCTGAATNNNAGATCGGAAG
29586 TATGAGTGTGGAGTCGTTGCAGGGGTAAGGTATTATTTCTCCTCANNNAGATCGGA
29681 TATGAGTGTGGAGTCGTTGCCACAGATGAGATTGAGGCAGGAGNNNAGATCGGA
70254 TATGAGTGTGGAGTCGTTGCAAGAACAGGCTGTCAACCAAGACNNNAGATCGGAA
70355 TATGAGTGTGGAGTCGTTGCACACTTCTTCCACAGCATCATCCNNNAGATCGGAAGA
95418 TATGAGTGTGGAGTCGTTGCTTGTGGCCCCTTTTCAGAGAACNNNAGATCGGAAGA
97118 TATGAGTGTGGAGTCGTTGCCTGGAAAGGAGGGGCTTGAATTNNNAGATCGGAA
99993 TATGAGTGTGGAGTCGTTGCAAGAAACTTCCGGATTATGGGCANNNAGATCGGAA
121992 TATGAGTGTGGAGTCGTTGCAACTGGCTCAGGCATGAAAACAANNNAGATCGGAA
92981 TATGAGTGTGGAGTCGTTGCTAATCCCATCATGAAGCTTGGGCNNNAGATCGGAAG
93824 TATGAGTGTGGAGTCGTTGCTCCAAGACCATCACTCAGCCATANNNAGATCGGAAG
117867 TATGAGTGTGGAGTCGTTGCCTTCCTTTTCTGCTCTCCATGTCNNNAGATCGGAAGA
119650 TATGAGTGTGGAGTCGTTGCTCAATTTCAGTCTTGGGAGGGTGTANNNAGATCGG
120377 TATGAGTGTGGAGTCGTTGCTATCTTCAAAGCCAGCAATCTGANNNAGATCGGAAG
28885 TATGAGTGTGGAGTCGTTGCTCATAACCTAACTGGGCATGTGCNNNAGATCGGAAG
28909 TATGAGTGTGGAGTCGTTGCGGAAGAGAAGAAAAGGGCTCGAGNNNAGATCGGA
29648 TATGAGTGTGGAGTCGTTGCTCCTTGTGTTTGGATAGCAAACTNNNAGATCGGAAG

| | |
|---|---|
| 70116 | TATGAGTGTGGAGTCGTTGCGGAGGTGGGAAGAGGGAGAAAATANNNAGATCG |
| 70897 | TATGAGTGTGGAGTCGTTGCGGGGCACAAGGAAATGTTTTCTGNNNAGATCGGAA |
| 71452 | TATGAGTGTGGAGTCGTTGCCTCAAGATTTCATGTCATTTCATTGTNNNAGATCGGA |
| 71482 | TATGAGTGTGGAGTCGTTGCCTTCCTGTCTTTGGCACCTCATTNNNAGATCGGAAGA |
| 90184 | TATGAGTGTGGAGTCGTTGCTCACATCTTCACCTTTCAGGGTTNNNAGATCGGAAG |
| 95438 | TATGAGTGTGGAGTCGTTGCAGTCCATGACAACGTAGAATGCTGANNNAGATCGG |
| 99587 | TATGAGTGTGGAGTCGTTGCAAAAAGGAATGTCAGGGTTGTGGTAANNNAGATCG |
| 120034 | TATGAGTGTGGAGTCGTTGCAGGCTGTCTCTCTCTCCCAGTNNNAGATCGGAAGAG |
| 92479 | TATGAGTGTGGAGTCGTTGCGATTCTCACCTGACTGAAGGCAGNNNAGATCGGAA |
| 28773 | TATGAGTGTGGAGTCGTTGCGGATTTTACCCAAGGTCACAGAACNNNAGATCGGAA |
| 28811 | TATGAGTGTGGAGTCGTTGCTGTCCAGGATGCCTTCAATGTTCNNNAGATCGGAAG |
| 29007 | TATGAGTGTGGAGTCGTTGCAAAGGACACGCTGAAGTCCTAACNNNAGATCGGAA |
| 29031 | TATGAGTGTGGAGTCGTTGCACAGGAAAACTGACATGCCACTTNNNAGATCGGAAG |
| 29102 | TATGAGTGTGGAGTCGTTGCTGGCTACAATGAATGCTTTTCCANNNAGATCGGAAG |
| 29191 | TATGAGTGTGGAGTCGTTGCTGACACCACTGAACTTTTATGCANNNAGATCGGAAG |
| 29230 | TATGAGTGTGGAGTCGTTGCTTGATTTGAAACACACCTCAAGCTNNNAGATCGGAA |
| 29396 | TATGAGTGTGGAGTCGTTGCCCTCTGGGCTCATGTTGAGTTTTNNNAGATCGGAAG |
| 29416 | TATGAGTGTGGAGTCGTTGCTAGCTCCAACCCTGGTTTTGTTTNNNAGATCGGAAG |
| 29419 | TATGAGTGTGGAGTCGTTGCATTCTGCTTCCAAGCCCTGTTTTNNNAGATCGGAAGA |
| 29566 | TATGAGTGTGGAGTCGTTGCACACTCCTGACATGTTTTTCCCCNNNAGATCGGAAGA |
| 29590 | TATGAGTGTGGAGTCGTTGCACTCGTATTCTCCTTCCTCTGCANNNAGATCGGAAGA |
| 29772 | TATGAGTGTGGAGTCGTTGCACGGGCATGTGTATGTGTTTCTTNNNAGATCGGAAG |
| 69725 | TATGAGTGTGGAGTCGTTGCGCACACATAAAACCTTGTGGCAANNNAGATCGGAAG |
| 70202 | TATGAGTGTGGAGTCGTTGCCGAGCAACAGACCAAAGGAAAACNNNAGATCGGAA |
| 70544 | TATGAGTGTGGAGTCGTTGCAAATGTTCCCAGTTTCCCAGGTCNNNAGATCGGAAG |
| 70580 | TATGAGTGTGGAGTCGTTGCGCTGCCACTATCATGCCAAAAAGNNNAGATCGGAAG |
| 70658 | TATGAGTGTGGAGTCGTTGCTGGAAAGTACCCAGAGTGAGGAGNNNAGATCGGA |
| 71027 | TATGAGTGTGGAGTCGTTGCCTCCCCTTGCCATTCTTCTTCATNNNAGATCGGAAGA |
| 95610 | TATGAGTGTGGAGTCGTTGCAGGACCCCTGGTATCAAACAACCNNNAGATCGGAAG |
| 90321 | TATGAGTGTGGAGTCGTTGCCCAGGCAGTCTGTGATGTATGATTNNNAGATCGGA |
| 121063 | TATGAGTGTGGAGTCGTTGCATTTCACTTTTGTTGCCCAAGAGTGNNNAGATCGGA |
| 28790 | TATGAGTGTGGAGTCGTTGCCCTCTTTCCATTCTAAGCACTCTCANNNAGATCGGAA |
| 28821 | TATGAGTGTGGAGTCGTTGCTGTCCTCTTTGACCTTGAGAGCTNNNAGATCGGAAG |
| 28932 | TATGAGTGTGGAGTCGTTGCCAAACCAAAGTAGGTGGGTCCTGNNNAGATCGGAA |
| 29090 | TATGAGTGTGGAGTCGTTGCAATATCTGTGGCAGCAACACTGANNNAGATCGGAA |
| 29096 | TATGAGTGTGGAGTCGTTGCATTCCCAGTCCACCAATACCAGANNNAGATCGGAAG |
| 29163 | TATGAGTGTGGAGTCGTTGCTGCTGTGAGGAAACTCAACAGTCNNNAGATCGGAA |
| 29449 | TATGAGTGTGGAGTCGTTGCGGAAGTTTTCTTTAGGGGCTGCTNNNAGATCGGAA |
| 29461 | TATGAGTGTGGAGTCGTTGCTTTTGCATTGCTCCCAAGTAGGTNNNAGATCGGAAG |
| 29537 | TATGAGTGTGGAGTCGTTGCTAGGATATGGGCCCAAGGAGAAGNNNAGATCGGA |
| 29614 | TATGAGTGTGGAGTCGTTGCGAAGGGAGTGTCCCTAGAACCTTNNNAGATCGGAA |
| 29644 | TATGAGTGTGGAGTCGTTGCCACATCCTGTTGCCATTGTACCTNNNAGATCGGAAGA |
| 70085 | TATGAGTGTGGAGTCGTTGCATTACCATTTGCTCTCAGCCTGANNNAGATCGGAAG |

93661 TATGAGTGTGGAGTCGTTGCTGTTGCTTGGCTCAGTTCTGTTANNNAGATCGGAAG
98794 TATGAGTGTGGAGTCGTTGCACGATCAGAAAGCAGCATTTCCGNNNAGATCGGAA
121957 TATGAGTGTGGAGTCGTTGCATGTACTCAAAGTGGAATTGATGGATNNNAGATCG
93445 TATGAGTGTGGAGTCGTTGCACCCAATGACCTTTGGCAAGTACNNNAGATCGGAAG
28902 TATGAGTGTGGAGTCGTTGCATGCCAGCTGATGGTAAGTACCANNNAGATCGGAA
28916 TATGAGTGTGGAGTCGTTGCCCACTCCTAACTTCGATGCTTCCNNNAGATCGGAAGA
29080 TATGAGTGTGGAGTCGTTGCCTCATTGCTTCTGCTCAACAACCNNNAGATCGGAAGA
29279 TATGAGTGTGGAGTCGTTGCTACCCCAATCTCCAAAGTTTGCCNNNAGATCGGAAG
29376 TATGAGTGTGGAGTCGTTGCTTTTGAGTGGGGTTTCAAGGTTTNNNAGATCGGAA
29705 TATGAGTGTGGAGTCGTTGCTGGGAAAGTGCATACATGTGTCCNNNAGATCGGAA
49454 TATGAGTGTGGAGTCGTTGCGTGAGGTGATGAAGACTGGTGTCNNNAGATCGGA
92189 TATGAGTGTGGAGTCGTTGCGGATCATTCAAAGGCTACCGAGGNNNAGATCGGAA
93819 TATGAGTGTGGAGTCGTTGCTAATCTGCTGAGTGGAGGGCATANNNAGATCGGAA
95042 TATGAGTGTGGAGTCGTTGCGGAGATGCACGTATGCTGAGAAANNNAGATCGGAA
95861 TATGAGTGTGGAGTCGTTGCTTTTACAGAACCAATACCTGGGCTCNNNAGATCGGA
96540 TATGAGTGTGGAGTCGTTGCCGTCCACAAGTTGTATGTCTTCTGNNNAGATCGGAA
99120 TATGAGTGTGGAGTCGTTGCCCCACATCAAATCAAACCCCCTNNNAGATCGGAAGAG
100130 TATGAGTGTGGAGTCGTTGCCGGTGCAAGGTGATGGGATCATNNNAGATCGGAAG
119890 TATGAGTGTGGAGTCGTTGCAGATAGAGTGGTGGGGCAAGAAGNNNAGATCGGA
119925 TATGAGTGTGGAGTCGTTGCTAACAATTTGTGCTCATAGGCAAGNNNAGATCGGAA
120979 TATGAGTGTGGAGTCGTTGCTTCACCTGGAACAGCATGTCAGGNNNAGATCGGAA
28780 TATGAGTGTGGAGTCGTTGCGCACACTGGAACAGAGAACAAGANNNAGATCGGAA
29241 TATGAGTGTGGAGTCGTTGCGCCACCTTTCATTGTGGACCTATNNNAGATCGGAAG
29335 TATGAGTGTGGAGTCGTTGCAAAACCTATGCCTGGCACTGAAANNNAGATCGGAAG
29497 TATGAGTGTGGAGTCGTTGCGGGTGTTTGTTACCCCAAGACATNNNAGATCGGAAG
29665 TATGAGTGTGGAGTCGTTGCAACTATTGTCACCATGCTCTCCGNNNAGATCGGAAG
29752 TATGAGTGTGGAGTCGTTGCATTGGCCCCTGAAGGTTCACTAANNNAGATCGGAAG
69818 TATGAGTGTGGAGTCGTTGCGGCTCTCACAACAGAGATTCACANNNAGATCGGAAG
70106 TATGAGTGTGGAGTCGTTGCTAACATGAGCTCAACGCTCTTGANNNAGATCGGAAG
95295 TATGAGTGTGGAGTCGTTGCCAGGAAACAGTATTTGTAGTTGACATGNNNAGATCG
96711 TATGAGTGTGGAGTCGTTGCCCTCCTGAATTTGTGCAATTTTTGAANNNAGATCGG
91318 TATGAGTGTGGAGTCGTTGCAAGCTCCAACTCAAACGACAACTNNNAGATCGGAAG
120365 TATGAGTGTGGAGTCGTTGCGCCACAGGAGTTTGGTGTTCAGNNNAGATCGGAAG
28777 TATGAGTGTGGAGTCGTTGCTGAACCTTATTTATTGTGCTTCTACTGNNNAGATCGG
28830 TATGAGTGTGGAGTCGTTGCCTGAGTGATATGGCAGCCAACTTNNNAGATCGGAA
28890 TATGAGTGTGGAGTCGTTGCGGAGTACGAAAGCATGAAGGTGTNNNAGATCGGA
29217 TATGAGTGTGGAGTCGTTGCGCTGGAGTGCAGTAGAATTCTGTNNNAGATCGGAA
29340 TATGAGTGTGGAGTCGTTGCGATGAGGATAGCTGCTGTGCTTTNNNAGATCGGAA
29363 TATGAGTGTGGAGTCGTTGCAATCAGCTGTCAAGTTCCACAGTNNNAGATCGGAAG
29456 TATGAGTGTGGAGTCGTTGCTCCCCACTCTCACGTTGTAATCTNNNAGATCGGAAGA
29487 TATGAGTGTGGAGTCGTTGCTCATAGCGCTCTTTTCCTTTCCTNNNAGATCGGAAGA
29617 TATGAGTGTGGAGTCGTTGCTGACCACAAATTGCCCAAGACTTNNNAGATCGGAAG
92520 TATGAGTGTGGAGTCGTTGCTGACTACCACCAAATGAGCACAGNNNAGATCGGAAG

93532 TATGAGTGTGGAGTCGTTGCGACCTTGAAGGAGTACTCAGCACNNNAGATCGGAA
93685 TATGAGTGTGGAGTCGTTGCTTTGGGCATTTGGGGAACACTTANNNAGATCGGAA
100072 TATGAGTGTGGAGTCGTTGCCATTTCACACTCTACATCCTTGGGTANNNAGATCGGA
121991 TATGAGTGTGGAGTCGTTGCAGTCATGGACGGTAAAAACTAGGNNNAGATCGGAA
92410 TATGAGTGTGGAGTCGTTGCCCAGCCAGAACATGAACCTAGAGNNNAGATCGGAA
90340 TATGAGTGTGGAGTCGTTGCCTTTGGAAGGGTAGTGTGGCATTNNNAGATCGGAA
120839 TATGAGTGTGGAGTCGTTGCTCACCATCACCTAGGTTGGAGTGNNNAGATCGGAAG
28982 TATGAGTGTGGAGTCGTTGCCTCACTCACTGCAACCTGTGTGNNNAGATCGGAAGA
29033 TATGAGTGTGGAGTCGTTGCCTCTCCCTGCAGCTCAGACTATTNNNAGATCGGAAGA
29040 TATGAGTGTGGAGTCGTTGCAAGAAGGCACGTGGAAAGTTTCTNNNAGATCGGAA
29451 TATGAGTGTGGAGTCGTTGCGATGACACGTGTAGGTTCACATCTNNNAGATCGGAA
29520 TATGAGTGTGGAGTCGTTGCTATGGGTGTGCTTGTGTGTAAGGNNNAGATCGGAA
29552 TATGAGTGTGGAGTCGTTGCCTTAGCTACTTTGGCATTGCTGCNNNAGATCGGAAG
29680 TATGAGTGTGGAGTCGTTGCTTGTTTCCTTGTCTTCACCCTGGNNNAGATCGGAAGA
67976 TATGAGTGTGGAGTCGTTGCGTTTCCAGGACTACTTGGTGCTTNNNAGATCGGAAG
69815 TATGAGTGTGGAGTCGTTGCTTTTCTGTAAGCTATCCCTCACANNNAGATCGGAAGA
69926 TATGAGTGTGGAGTCGTTGCGGGACACAGTGAGAAAGCAATCANNNAGATCGGAA
70313 TATGAGTGTGGAGTCGTTGCCCTTCATCTCTGTCCTTTTACCCCNNNAGATCGGAAG
90200 TATGAGTGTGGAGTCGTTGCACAGTCCCCTTGAGTGAAGAGTTNNNAGATCGGAA
90739 TATGAGTGTGGAGTCGTTGCACATTTGGTTCCTGTGCAAGAGTNNNAGATCGGAAG
92314 TATGAGTGTGGAGTCGTTGCTAAACATTGTTGCCAGTAGCACGNNNAGATCGGAAG
95356 TATGAGTGTGGAGTCGTTGCACCTTTTACACGGCTCTGTACTCNNNAGATCGGAAG
95450 TATGAGTGTGGAGTCGTTGCGTCATCAGTCATTTCCTCCAGTTTNNNAGATCGGAAG
96103 TATGAGTGTGGAGTCGTTGCTTGGGTCCTTATGACAGTTTGCTTANNNAGATCGGA
96139 TATGAGTGTGGAGTCGTTGCGCTTTTGTGTCTGGCTTACTCTTNNNAGATCGGAAG
97423 TATGAGTGTGGAGTCGTTGCGAATACCTGCTTTGCAAATGTCTTTNNNAGATCGGA
97597 TATGAGTGTGGAGTCGTTGCAGAGAAGTAGCCAGATGGCATGAGNNNAGATCGG
97760 TATGAGTGTGGAGTCGTTGCGAATGGGCCCTCACTTGACATGANNNAGATCGGAA
105584 TATGAGTGTGGAGTCGTTGCCCAACTCATTTCTGCTGCCCANNNAGATCGGAAGAG
93531 TATGAGTGTGGAGTCGTTGCATTGTCTATCTGCCCTTCAGCACNNNAGATCGGAAGA
93566 TATGAGTGTGGAGTCGTTGCGGGGGAGTGTTTTAGATGGACACNNNAGATCGGA
28787 TATGAGTGTGGAGTCGTTGCGCAACAGCCGCATACTTACTATGNNNAGATCGGAAG
29006 TATGAGTGTGGAGTCGTTGCTGCCCCAAGAAAACTCTTCTCACNNNAGATCGGAAG
29111 TATGAGTGTGGAGTCGTTGCCCAATGAGCTTCCATTCTCCCAANNNAGATCGGAAGA
29162 TATGAGTGTGGAGTCGTTGCCTTACTATGTGCCAGGCGTTGTCNNNAGATCGGAAG
29198 TATGAGTGTGGAGTCGTTGCATCTTGGTGCAGCACACTTTACANNNAGATCGGAAG
29260 TATGAGTGTGGAGTCGTTGCCCTCATTCTCCCCCAAACATGAGNNNAGATCGGAAGA
29285 TATGAGTGTGGAGTCGTTGCATACTCCCTTGGGTAACTCCACCNNNAGATCGGAAG
29527 TATGAGTGTGGAGTCGTTGCCACTGTAGGGAGACAGAGGAGAGNNNAGATCGGA
29565 TATGAGTGTGGAGTCGTTGCTAGTCCTTTCAATACTGCAGCCCNNNAGATCGGAAG
29573 TATGAGTGTGGAGTCGTTGCTTTCCCAGAACATGGCCTACCTANNNAGATCGGAAG
29640 TATGAGTGTGGAGTCGTTGCCATCTTTGTTGACATACGCAGCTNNNAGATCGGAAG
29724 TATGAGTGTGGAGTCGTTGCGGGTATAGAGGAGTGCACCTGTANNNAGATCGGAA

| | |
|---|---|
| 70835 | TATGAGTGTGGAGTCGTTGCGCTGAGGTATTTGTTTGCTCTTGGNNNAGATCGGAA |
| 70839 | TATGAGTGTGGAGTCGTTGCATCTATCTGCACCAGCAGTTTGGNNNAGATCGGAAG |
| 71290 | TATGAGTGTGGAGTCGTTGCGATGAACCTGCATGACCTGAAGTNNNAGATCGGAA |
| 92368 | TATGAGTGTGGAGTCGTTGCGCAACTCTGGCTTAGACTGTGAGNNNAGATCGGAA |
| 96090 | TATGAGTGTGGAGTCGTTGCGAGGGAGAGTTTGGAATTTCCCAGNNNAGATCGGA |
| 98985 | TATGAGTGTGGAGTCGTTGCCATCCGTGCAACAAACAACAACTNNNAGATCGGAAG |
| 98995 | TATGAGTGTGGAGTCGTTGCGGACTTATCCCTGGCCTTAGGANNNAGATCGGAAG |
| 100136 | TATGAGTGTGGAGTCGTTGCATCTGTTTCTCAATCCCCGAGTTGNNNAGATCGGAA |
| 94188 | TATGAGTGTGGAGTCGTTGCTCCCCATCAATTTGCATGACAGANNNAGATCGGAAG |
| 92810 | TATGAGTGTGGAGTCGTTGCGGAGGTGATGGGTCTTTGTCATCNNNAGATCGGAA |
| 119653 | TATGAGTGTGGAGTCGTTGCACTTCCTCTGGACCATGTTCATTNNNAGATCGGAAG |
| 120227 | TATGAGTGTGGAGTCGTTGCATTTCCTGCCTCCAGAGTAGCTNNNAGATCGGAAGA |
| 120810 | TATGAGTGTGGAGTCGTTGCTCTTGACTGTGCTTCTTTACTCCNNNAGATCGGAAGA |
| 121184 | TATGAGTGTGGAGTCGTTGCCATCATACCAATTGCGTCTTTCTGANNNAGATCGGAA |
| 121095 | TATGAGTGTGGAGTCGTTGCAACTGCAATCTCTTACTTCTGGGTTCNNNAGATCGGA |
| 28990 | TATGAGTGTGGAGTCGTTGCATAGTCCAAAGATGAGCTGCAGGNNNAGATCGGAA |
| 29112 | TATGAGTGTGGAGTCGTTGCCCGACCATATCTGCACAGAACAANNNAGATCGGAAG |
| 29585 | TATGAGTGTGGAGTCGTTGCTCAACCCTCCAATATGGCTCTCANNNAGATCGGAAG |
| 29685 | TATGAGTGTGGAGTCGTTGCCACACAGTGCAGCATAATGCAATNNNAGATCGGAAG |
| 69983 | TATGAGTGTGGAGTCGTTGCTGTGAGAGAGAGCCACTTGAAGANNNAGATCGGAA |
| 70111 | TATGAGTGTGGAGTCGTTGCATTGCCTCTGTTCTTGCCTTTGANNNAGATCGGAAG |
| 70143 | TATGAGTGTGGAGTCGTTGCAGAAAAGGGCTTGGTCAGTCCTANNNAGATCGGAA |
| 70276 | TATGAGTGTGGAGTCGTTGCTGGCCTCCATGAAACAGTTCTACNNNAGATCGGAAG |
| 70424 | TATGAGTGTGGAGTCGTTGCTTTACCTACTCCGATATGCACGCNNNAGATCGGAAG |
| 70478 | TATGAGTGTGGAGTCGTTGCTCAGGCAGAATAAATTTTCCATCTGCNNNAGATCGG |
| 71168 | TATGAGTGTGGAGTCGTTGCGTTTTGTTTCTGGCATGTGCCTNNNAGATCGGAAGA |
| 71232 | TATGAGTGTGGAGTCGTTGCCCAAAAGGCGCTATCTCCAATCTNNNAGATCGGAAG |
| 90684 | TATGAGTGTGGAGTCGTTGCAAGGAGTATTATAGGCCAGGCGTNNNAGATCGGAA |
| 91814 | TATGAGTGTGGAGTCGTTGCTTCTGTGGCTATTTCTCAGCCTGNNNAGATCGGAAG |
| 92759 | TATGAGTGTGGAGTCGTTGCTCGTCAACTTCAACAGTTCCCTCNNNAGATCGGAAGA |
| 95538 | TATGAGTGTGGAGTCGTTGCCAGAGGGATAGTGGAGGCCATANNNAGATCGGAA |
| 96699 | TATGAGTGTGGAGTCGTTGCGGGCTATCAGAGAAAATGTGCTTTCNNNAGATCGG |
| 96792 | TATGAGTGTGGAGTCGTTGCAAACAAAGCAGCAAAAGGCCCCNNNAGATCGGAAG |
| 100904 | TATGAGTGTGGAGTCGTTGCTGTGTTTTCATGTGTCTGCCAGATTTNNNAGATCGG |
| 94163 | TATGAGTGTGGAGTCGTTGCAAACGAAGGGAAGGGAGATCAGANNNAGATCGGA |
| 117766 | TATGAGTGTGGAGTCGTTGCAGGATTCGTACTTTACCTTTGGAGGNNNAGATCGGA |
| 90134 | TATGAGTGTGGAGTCGTTGCTTGGCTTCAGTAAGGTGGCATTTNNNAGATCGGAA |
| 120263 | TATGAGTGTGGAGTCGTTGCTGCTTGTGGTGTGTACTTCTAATTANNNAGATCGGA |
| 121100 | TATGAGTGTGGAGTCGTTGCGGGTCTCCCGTTCAAAGTGTTGNNNAGATCGGAAG |
| 90953 | TATGAGTGTGGAGTCGTTGCTTCTGCTTTGGTGGTGGAAATCTNNNAGATCGGAAG |
| 28896 | TATGAGTGTGGAGTCGTTGCCTGGATGAGGTTTCCCTTTGGTCNNNAGATCGGAAG |
| 29010 | TATGAGTGTGGAGTCGTTGCTGGACTGCAGTTGCTCTAGAGTANNNAGATCGGAA |
| 29042 | TATGAGTGTGGAGTCGTTGCGGCTGTCCTCAGATGCCTATTCTNNNAGATCGGAAG |

29068 TATGAGTGTGGAGTCGTTGCCCCCATTCTGTGGAAGTAACCAGNNNAGATCGGAAG
29147 TATGAGTGTGGAGTCGTTGCTGAATGTAGCATATTTGCAACTCTGNNNAGATCGGA
29290 TATGAGTGTGGAGTCGTTGCCTGAGTTTCTCCTGGGTGTACACNNNAGATCGGAAG
29426 TATGAGTGTGGAGTCGTTGCGGCTTCTTGGTAAAAGAAATGGTTNNNAGATCGGA
29589 TATGAGTGTGGAGTCGTTGCAGCATACCAAAAGATTGTAACATCANNNAGATCGGA
70720 TATGAGTGTGGAGTCGTTGCTATGCCTCTGTATGTTTGCCAGGNNNAGATCGGAAG
70725 TATGAGTGTGGAGTCGTTGCGTTTCTGATCTCATCCTGGCAGGNNNAGATCGGAAG
70758 TATGAGTGTGGAGTCGTTGCGCAAGGAATGGAGTATGGAAGGGNNNAGATCGGA
90895 TATGAGTGTGGAGTCGTTGCTGTCCCTGCTCCACTATTTCTCTNNNAGATCGGAAGA
93372 TATGAGTGTGGAGTCGTTGCTGTGGTTGGTAGGAAGAGGTACCNNNAGATCGGA
95383 TATGAGTGTGGAGTCGTTGCACGATCTCTCGGCTCATTGTAACNNNAGATCGGAAG
95390 TATGAGTGTGGAGTCGTTGCAGAAGGAAGTGCTCAATTTCAGTTNNNAGATCGGA
95473 TATGAGTGTGGAGTCGTTGCGTTTTCTCCCCCACCACCTTGNNNAGATCGGAAGAGC
97048 TATGAGTGTGGAGTCGTTGCTCCTGTCATTTTGTTATTTTGTTGTTNNNAGATCGGA
98930 TATGAGTGTGGAGTCGTTGCTCCTCCCACCCCAGACATTTTTCNNNAGATCGGAAGA
93984 TATGAGTGTGGAGTCGTTGCTGTATGCTACTGCTCTAGCTGGANNNAGATCGGAAG
70522 TATGAGTGTGGAGTCGTTGCTACCTGGCTGACTACAGGAACTCNNNAGATCGGAAG
91405 TATGAGTGTGGAGTCGTTGCCCAGGAGTTCCACATCTAGGGATNNNAGATCGGAA
119565 TATGAGTGTGGAGTCGTTGCGACTACAAGTGGGCCCTAAAGATTNNNAGATCGGA
94100 TATGAGTGTGGAGTCGTTGCCAAACTCTGACCTGATTCAGCGANNNAGATCGGAAG
29060 TATGAGTGTGGAGTCGTTGCAGAGATTCCTAAGGGAGTGTGGGNNNAGATCGGA
29087 TATGAGTGTGGAGTCGTTGCACTGGTCACAGGTCTGGATTGTANNNAGATCGGAA
29346 TATGAGTGTGGAGTCGTTGCTGAACCAAACAGCTGCTAGCATTNNNAGATCGGAAG
29473 TATGAGTGTGGAGTCGTTGCATCCCTCTCCTGTCTGCTTCTTTNNNAGATCGGAAGA
29544 TATGAGTGTGGAGTCGTTGCAAAAACCATGCCCAGGAGTACTCNNNAGATCGGAAG
29593 TATGAGTGTGGAGTCGTTGCACCAGCACCATTTATTGAAGGCANNNAGATCGGAAG
29755 TATGAGTGTGGAGTCGTTGCATTTGCAGCAACCTAGGTGGGAANNNAGATCGGAA
42966 TATGAGTGTGGAGTCGTTGCTTGAAACCTTATGGCCCCATCACNNNAGATCGGAAG
69731 TATGAGTGTGGAGTCGTTGCTACTAGCCAAGCCACTTTTGCAANNNAGATCGGAAG
70827 TATGAGTGTGGAGTCGTTGCTGGGAAGTTTTGAATAAGTTTGGTGGNNNAGATCG
70829 TATGAGTGTGGAGTCGTTGCAGGAGGGGGAAGGAAATATGTGGNNNAGATCGGA
71300 TATGAGTGTGGAGTCGTTGCACTTTCTCACAGAGCTCTCCAGTNNNAGATCGGAAG
71304 TATGAGTGTGGAGTCGTTGCTGTAAAGAACTCTGAAAGCTCAGTNNNAGATCGGA
92856 TATGAGTGTGGAGTCGTTGCTGGTCCTAAGGTTTTCTGGTTGCNNNAGATCGGAAG
93398 TATGAGTGTGGAGTCGTTGCCTGAGGTGGCAAGGAATATCACCNNNAGATCGGAA
94549 TATGAGTGTGGAGTCGTTGCGCCACTATATACACACACTGCCANNNAGATCGGAAG
95764 TATGAGTGTGGAGTCGTTGCTGGACTGGCTTGTAACTTGCTTTNNNAGATCGGAAG
94340 TATGAGTGTGGAGTCGTTGCATCAGTGGAGTGGCATCCTATCANNNAGATCGGAA
92999 TATGAGTGTGGAGTCGTTGCTTCGTACTTTACCTTTGGAGGGCNNNAGATCGGAAG
120363 TATGAGTGTGGAGTCGTTGCACGAGACCTGAATTATTCTGCTACCNNNAGATCGGA
95033 TATGAGTGTGGAGTCGTTGCACCCCAATAAAAATGGGTGGAAANNNAGATCGGAA
28810 TATGAGTGTGGAGTCGTTGCTGGATGCCCCAACATTATGATCGNNNAGATCGGAAG
28924 TATGAGTGTGGAGTCGTTGCACCCTGTGCATTTGAGAACACTTNNNAGATCGGAAG

28993 TATGAGTGTGGAGTCGTTGCCCTGTGCCTGCCTTGTACTATAGNNNAGATCGGAAG
29161 TATGAGTGTGGAGTCGTTGCCCTGGCTAGAGGTTTGTCAGTTTCNNNAGATCGGAA
29373 TATGAGTGTGGAGTCGTTGCCAGCCCCATCTCAATTGGTCTTTNNNAGATCGGAAG
69949 TATGAGTGTGGAGTCGTTGCGCTTTGAGCACAGTGAGTTCTCANNNAGATCGGAAG
71351 TATGAGTGTGGAGTCGTTGCTACCATTCTGGGCTTGATGAGGTNNNAGATCGGAAG
90946 TATGAGTGTGGAGTCGTTGCTCATATCCGTGGTTTCCCCATCTNNNAGATCGGAAGA
93211 TATGAGTGTGGAGTCGTTGCTAAGGGAAATCCTGCCATTTGCCNNNAGATCGGAAG
93894 TATGAGTGTGGAGTCGTTGCCCCATACATAAGGAACAGTGAAGTGANNNAGATCG
95391 TATGAGTGTGGAGTCGTTGCGAAACACCAGCTAGATTTATGCATGACNNNAGATCG
96710 TATGAGTGTGGAGTCGTTGCGCATGTGTTGGTGTTAGTGTCTGTNNNAGATCGGA
99895 TATGAGTGTGGAGTCGTTGCCACTCATGTCTCAGTCTCACAGANNNAGATCGGAAG
91664 TATGAGTGTGGAGTCGTTGCTTCAGAGAGCCATCCAACCATTGNNNAGATCGGAAG
90445 TATGAGTGTGGAGTCGTTGCTCTCCTCTCCTTGAGCTTTCTGTNNNAGATCGGAAGA
119568 TATGAGTGTGGAGTCGTTGCATAGAAATAACAGAGTGGAGGCCNNNAGATCGGAA
121062 TATGAGTGTGGAGTCGTTGCTCTCAGCTCACTTCAGCTCACTCNNNAGATCGGAAGA
28927 TATGAGTGTGGAGTCGTTGCATGGGAGTTCCTTGTACTGTGCTNNNAGATCGGAAG
28969 TATGAGTGTGGAGTCGTTGCTGCTTGTTGCCAATTGAATTCCTNNNAGATCGGAAG
29350 TATGAGTGTGGAGTCGTTGCGTATGTTGTGCGGCTGTGTGTNNNAGATCGGAAGA
29375 TATGAGTGTGGAGTCGTTGCTCAGTTGCCATGTGCTATGGTTTNNNAGATCGGAAG
29588 TATGAGTGTGGAGTCGTTGCATGCTCACTTCCTCCCATCATCANNNAGATCGGAAGA
29689 TATGAGTGTGGAGTCGTTGCTACCTCCCCTGACCTCTTTCTTCNNNAGATCGGAAGA
29707 TATGAGTGTGGAGTCGTTGCCTAAGGTGTTTTGTTCCCCCTCCNNNAGATCGGAAGA
69801 TATGAGTGTGGAGTCGTTGCATCCACCAAATTTCCACCACACANNNAGATCGGAAGA
69928 TATGAGTGTGGAGTCGTTGCCACTGTTCACAGCCAGAACATCANNNAGATCGGAAG
69947 TATGAGTGTGGAGTCGTTGCTGGGTTCAATAACATCATACTTCTCANNNAGATCGGA
70402 TATGAGTGTGGAGTCGTTGCGCCTCAGAGAAGTGTAAGACAGCNNNAGATCGGAA
70744 TATGAGTGTGGAGTCGTTGCTGTCCAACTCAAATTCAGGAATATAGGNNNAGATCG
70831 TATGAGTGTGGAGTCGTTGCAGAGATGATAGTGGGTCCTGTGGNNNAGATCGGA
91347 TATGAGTGTGGAGTCGTTGCCGTGTAGGTATATTTTGGGGGTATNNNAGATCGGA
93604 TATGAGTGTGGAGTCGTTGCCTGGGACTTCACTGGGTTAGAACNNNAGATCGGAAG
95410 TATGAGTGTGGAGTCGTTGCGATTGTTTACCTCAGCCTCCTGAGTNNNAGATCGGA
98262 TATGAGTGTGGAGTCGTTGCTATGTGCTAATTTCCATGGCCAGTNNNAGATCGGAA
121980 TATGAGTGTGGAGTCGTTGCTCCAGTGTAACTTATTTGGTTGAACCNNNAGATCGG
70777 TATGAGTGTGGAGTCGTTGCAACGAAGACGATTAGATTGGGGGNNNAGATCGGA
91702 TATGAGTGTGGAGTCGTTGCTCTTTGTCACTCAGGCAGATGTGNNNAGATCGGAAG
120387 TATGAGTGTGGAGTCGTTGCTCAGCTAGGCTCTGGACTACAGTNNNAGATCGGAAG
120750 TATGAGTGTGGAGTCGTTGCAATCCTCCAGTCCTAAGGGTGGNNNAGATCGGAAG
28913 TATGAGTGTGGAGTCGTTGCCAGCTAGGCAGCTTCACTACATCNNNAGATCGGAAG
29177 TATGAGTGTGGAGTCGTTGCGCAGGCTGGACAACCAAATCTTANNNAGATCGGAAG
29195 TATGAGTGTGGAGTCGTTGCTGGAGGGAGTAGAGAACAAAGCANNNAGATCGGA
29199 TATGAGTGTGGAGTCGTTGCTGGTTTTCAATGAGAGGTGGACANNNAGATCGGAA
29295 TATGAGTGTGGAGTCGTTGCGGCTACAGCTCTGAGGTACACTANNNAGATCGGAAG
29422 TATGAGTGTGGAGTCGTTGCTAGCCAGCTTCCACTGTTCAAACNNNAGATCGGAAG

29472 TATGAGTGTGGAGTCGTTGCGCCAGTTCAAGCTGTGTTGTTTTNNNAGATCGGAAG
29538 TATGAGTGTGGAGTCGTTGCTTTTTGCCAGCCAATTGGACAAGNNNAGATCGGAAG
29595 TATGAGTGTGGAGTCGTTGCTGTCTGTGTTGGTTGACAATAGCANNNAGATCGGAA
29726 TATGAGTGTGGAGTCGTTGCTCAGGTCCATCTTTCAAGCCCTANNNAGATCGGAAGA
69787 TATGAGTGTGGAGTCGTTGCTGCCAGGAAATTGAAATCAACAACANNNAGATCGGA
69869 TATGAGTGTGGAGTCGTTGCCCATGAGTTCATACTGTTGTCTCCANNNAGATCGGAA
70397 TATGAGTGTGGAGTCGTTGCAAACTGACGAGGTCGCTATAAGCNNNAGATCGGAA
70447 TATGAGTGTGGAGTCGTTGCCCATCCCATCTGCCTATTATGGCNNNAGATCGGAAGA
70665 TATGAGTGTGGAGTCGTTGCACTTCTGGGCTTCTACTCCTGAGNNNAGATCGGAAGA
71055 TATGAGTGTGGAGTCGTTGCGAGCAACTCTTGTGCTAGCTGATNNNAGATCGGAAG
71152 TATGAGTGTGGAGTCGTTGCAGCCAAGAAGTTACCAATTTCCCTNNNAGATCGGAA
92882 TATGAGTGTGGAGTCGTTGCAGGATGAAACCACCTAATGGTGCNNNAGATCGGAA
95044 TATGAGTGTGGAGTCGTTGCGGGGGTTGTCTGTGAATGAGAAANNNAGATCGGAA
95562 TATGAGTGTGGAGTCGTTGCCCCTGAGCTTCAGTTTCTGTGATTGNNNAGATCGGAA
96837 TATGAGTGTGGAGTCGTTGCCGGCAGCCAGCAAAGGAAATNNNAGATCGGAAGAG
97417 TATGAGTGTGGAGTCGTTGCTAAAAGACACATGCACCTGCACGNNNAGATCGGAAG
99025 TATGAGTGTGGAGTCGTTGCCCTTGCTATTTCATCAGTAGGCANNNAGATCGGAAGA
91845 TATGAGTGTGGAGTCGTTGCAACACCTGAAAGCCAGAGAACTGNNNAGATCGGAA
117247 TATGAGTGTGGAGTCGTTGCTCTCCTCTCCTTGAGCTTTCTGTACNNNAGATCGGAA
120480 TATGAGTGTGGAGTCGTTGCGTCAGGAGTTTGACACCATCGCNNNAGATCGGAAGA
121055 TATGAGTGTGGAGTCGTTGCAGTGGTGCATTCTAGGCTCACCANNNAGATCGGAAG
28734 TATGAGTGTGGAGTCGTTGCCTTTGGAATGGGGAGAAGCCTTTNNNAGATCGGAA
28761 TATGAGTGTGGAGTCGTTGCTACAGTGGAGCAATCTGGTTCACNNNAGATCGGAAG
28900 TATGAGTGTGGAGTCGTTGCCAGGAGAGGAGAGTTGCATGAACNNNAGATCGGAA
28966 TATGAGTGTGGAGTCGTTGCTCAGGATGCCAAAAACCACATCTNNNAGATCGGAAG
28977 TATGAGTGTGGAGTCGTTGCATGGTTAGACAGCAGGGAGTGATNNNAGATCGGAA
28981 TATGAGTGTGGAGTCGTTGCGTTCAAGAGCAGTGGTAGCCAATNNNAGATCGGAA
29053 TATGAGTGTGGAGTCGTTGCCCACTTTCCCACCAAGCAGAAATNNNAGATCGGAAGA
29089 TATGAGTGTGGAGTCGTTGCCCTGTAGCCACCAGAATTGTTGANNNAGATCGGAAG
69832 TATGAGTGTGGAGTCGTTGCTGGGTTTAGGTATATGGGCATACANNNAGATCGGA
70055 TATGAGTGTGGAGTCGTTGCGGGTCCAAATATGTGGCATTGGANNNAGATCGGAA
70071 TATGAGTGTGGAGTCGTTGCACACCAGAAAGGCTTCAAGATGANNNAGATCGGAA
70153 TATGAGTGTGGAGTCGTTGCCAAGTCCCAAGATCTGCAGTCTANNNAGATCGGAAG
70178 TATGAGTGTGGAGTCGTTGCGATGGAGCCTGAAACTGCTTGTANNNAGATCGGAA
70212 TATGAGTGTGGAGTCGTTGCTGTTGAATAAACTGTTGTCAGATCAACNNNAGATCG
70477 TATGAGTGTGGAGTCGTTGCAAGAGAAATCAAGGTCCACCTGCNNNAGATCGGAAG
71182 TATGAGTGTGGAGTCGTTGCCTATTCTTGCACCTTCGCTTCCTNNNAGATCGGAAGA
71192 TATGAGTGTGGAGTCGTTGCTGTCTGTTTTGTCTACTGCAGCTNNNAGATCGGAAGA
71250 TATGAGTGTGGAGTCGTTGCGCTGACATCACAGTCTTCTCTCTNNNAGATCGGAAGA
71282 TATGAGTGTGGAGTCGTTGCCCTGTCCTTGAGCAAGTCAAAGTNNNAGATCGGAAG
71457 TATGAGTGTGGAGTCGTTGCACACTGATAGACACTTGGGTTGTNNNAGATCGGAAG
71511 TATGAGTGTGGAGTCGTTGCATTGGGAAGCTGAGGTAGGACTTNNNAGATCGGAA
91082 TATGAGTGTGGAGTCGTTGCCAGTGACTATCCAAAGGCAAGCTNNNAGATCGGAAG

95412 TATGAGTGTGGAGTCGTTGCTCAAGAGAAAGCAGAGCCCCAANNNAGATCGGAAG
95455 TATGAGTGTGGAGTCGTTGCGCCCTGCAAAGGATTCCATGTTNNNAGATCGGAAGA
96508 TATGAGTGTGGAGTCGTTGCTCCACAGACAGTGACTTCTCCCNNNAGATCGGAAGA
91811 TATGAGTGTGGAGTCGTTGCTTGTGACCCTCCTAAGGATCCTGNNNAGATCGGAAG
90479 TATGAGTGTGGAGTCGTTGCCAACTGAGAGCAATAACCCCTGTNNNAGATCGGAAG
70906 TATGAGTGTGGAGTCGTTGCTGAGGTTGAGTTGGAATACCGTGNNNAGATCGGAA
70394 TATGAGTGTGGAGTCGTTGCCCCTTTGTGTCACTTTGTAGAAGCNNNAGATCGGAA
91964 TATGAGTGTGGAGTCGTTGCAACTGGAGAAAGAACTGCAGTGGNNNAGATCGGAA
120552 TATGAGTGTGGAGTCGTTGCAGTTCTTCAATGGGTCTGGCTTTNNNAGATCGGAAG
120571 TATGAGTGTGGAGTCGTTGCGGGATTACAGGTACGGGTCAACATNNNAGATCGGA
28729 TATGAGTGTGGAGTCGTTGCTCTGATCATTCACCATGGCAGTGNNNAGATCGGAAG
28793 TATGAGTGTGGAGTCGTTGCTGAACTACTTGTGCGTGTTAGGTNNNAGATCGGAAG
28919 TATGAGTGTGGAGTCGTTGCGCTCCTTTGGCATAATGACGGTANNNAGATCGGAAG
28922 TATGAGTGTGGAGTCGTTGCAAGGGGAAATGGTAATAGTAATAGCANNNAGATCG
28935 TATGAGTGTGGAGTCGTTGCTTTCCTGATTACCAGGCTGCAAGNNNAGATCGGAAG
29038 TATGAGTGTGGAGTCGTTGCGGAAAAGTAAAAGCGCTTGGAGTNNNAGATCGGAA
29121 TATGAGTGTGGAGTCGTTGCCCCATACAATCTTGGCTCTGCTTNNNAGATCGGAAGA
29218 TATGAGTGTGGAGTCGTTGCGTGTGCCTTGTACCAAGTGATGTNNNAGATCGGAAG
29308 TATGAGTGTGGAGTCGTTGCCAAGCCCACACATGTCTTAGTCANNNAGATCGGAAG
29360 TATGAGTGTGGAGTCGTTGCTGTTTAGGGAGTTCTCATATGGAGTNNNAGATCGGA
29397 TATGAGTGTGGAGTCGTTGCGCATGGGTGTCTGTCTCTCTTTTNNNAGATCGGAAG
29432 TATGAGTGTGGAGTCGTTGCCACTTAGGATCGCAGGAATCAGGNNNAGATCGGAA
29488 TATGAGTGTGGAGTCGTTGCTCAGCAAATGAGATGGTTTGCCTNNNAGATCGGAAG
45858 TATGAGTGTGGAGTCGTTGCTTGGAAATTGGCTAGCTGATTCCNNNAGATCGGAAG
70120 TATGAGTGTGGAGTCGTTGCAAAGAGATGCACAGGCACACATANNNAGATCGGAA
71320 TATGAGTGTGGAGTCGTTGCTGGTGTATGTCCAGGATGTGAGTNNNAGATCGGAA
71440 TATGAGTGTGGAGTCGTTGCTACAAGTCTGCACTTACGTGTGTNNNAGATCGGAAG
90576 TATGAGTGTGGAGTCGTTGCTGGGATAGAAGTGGTTTTCTGGTNNNAGATCGGAA
91862 TATGAGTGTGGAGTCGTTGCTGGCATGTTTGCAACAACTGATGNNNAGATCGGAAG
92733 TATGAGTGTGGAGTCGTTGCGAAGGTTGAGAAGTCCCATGCTCNNNAGATCGGAA
94998 TATGAGTGTGGAGTCGTTGCTCAAAATGTGGGGGATGAGACAANNNAGATCGGAA
95321 TATGAGTGTGGAGTCGTTGCCGCTTCGGGATTGTTTGGGGTNNNAGATCGGAAGA
96902 TATGAGTGTGGAGTCGTTGCTCAGAGTGAAGGACCTTTCAGCTNNNAGATCGGAAG
97758 TATGAGTGTGGAGTCGTTGCGGTGGGAAGAGTAATTTCAGAGGTAGNNNAGATCG
99989 TATGAGTGTGGAGTCGTTGCTGTCCTGTTTTGCTCACCCTCNNNAGATCGGAAGAGC
119679 TATGAGTGTGGAGTCGTTGCATGAATGAAAACAAACAGAGCTTCAGANNNAGATCG
119858 TATGAGTGTGGAGTCGTTGCTGGGAAGGCAGTACAAAAACTGTNNNAGATCGGAA
28875 TATGAGTGTGGAGTCGTTGCCCTGTTCCATATCTCTGGAGAACCNNNAGATCGGAA
28903 TATGAGTGTGGAGTCGTTGCGCGGTTCTTCCCAGTGATTGTAANNNAGATCGGAAG
28988 TATGAGTGTGGAGTCGTTGCAGAGAGGTTAGGAATTGGGGAGGNNNAGATCGGA
29074 TATGAGTGTGGAGTCGTTGCTTATAGCAGGCATGGATTCTGGCNNNAGATCGGAA
29083 TATGAGTGTGGAGTCGTTGCTCCAGCGTAAGAAATTGAGGGACNNNAGATCGGAA
29113 TATGAGTGTGGAGTCGTTGCGTTGGCAGAGATTAGCCCTGAAANNNAGATCGGAA

29194 TATGAGTGTGGAGTCGTTGCATCACTTCATAGTAGTGCAAGCANNNAGATCGGAAG
29245 TATGAGTGTGGAGTCGTTGCGTGTGTGTCAGGGGAGATTGTTGNNNAGATCGGA
29301 TATGAGTGTGGAGTCGTTGCAGGCTGTGTGGTATACCTAAGGANNNAGATCGGAA
29342 TATGAGTGTGGAGTCGTTGCCAAACACACGCACTCACTCATTTNNNAGATCGGAAG
29353 TATGAGTGTGGAGTCGTTGCTCCTATGTGTGATGAGCTCTGGTNNNAGATCGGAAG
29430 TATGAGTGTGGAGTCGTTGCATTTTCAGCATACCCAAGGGTGGNNNAGATCGGAA
29554 TATGAGTGTGGAGTCGTTGCTGTGTGTCATTCTAGGAGCATGCNNNAGATCGGAA
69835 TATGAGTGTGGAGTCGTTGCCTCTGCATCTGTATTTATCTCTCTACANNNAGATCGG
71217 TATGAGTGTGGAGTCGTTGCAAGAATCCTCAAATCACAGTGGCTNNNAGATCGGAA
90582 TATGAGTGTGGAGTCGTTGCGGCTATGAGATACTTCAGCTGGTNNNAGATCGGAA
91243 TATGAGTGTGGAGTCGTTGCTGCTGAAGATAGCAAGAGTGACTNNNAGATCGGAA
97358 TATGAGTGTGGAGTCGTTGCAGGACAAAGCCTGGGTTATGTTTCNNNAGATCGGA
101731 TATGAGTGTGGAGTCGTTGCCCTGGAGTGCCTAGCATTCTTNNNAGATCGGAAGAG
94376 TATGAGTGTGGAGTCGTTGCAGTTCAAAAGCAGTGACATTGCANNNAGATCGGAA
119855 TATGAGTGTGGAGTCGTTGCCTGACAAATTGGGATTCACCACANNNAGATCGGAAG
29009 TATGAGTGTGGAGTCGTTGCAGGAGCCAAATTGAAACTCAAGAACNNNAGATCGG
29239 TATGAGTGTGGAGTCGTTGCGTGGGTACGTGGAGGTTCATTATNNNAGATCGGAA
29270 TATGAGTGTGGAGTCGTTGCGAAAAATGCAATACAGGCTGGGCNNNAGATCGGAA
29366 TATGAGTGTGGAGTCGTTGCGGGGTAGGGAGGCTGATTATTCTNNNAGATCGGAA
29443 TATGAGTGTGGAGTCGTTGCAAAGAGGCCAGTCTCAAGCAAATNNNAGATCGGAA
29454 TATGAGTGTGGAGTCGTTGCTCCTCCCTCCCTTCTTCTCAATTNNNAGATCGGAAGAG
37856 TATGAGTGTGGAGTCGTTGCAAATGCCAATAGATGTGCCCAGANNNAGATCGGAA
66030 TATGAGTGTGGAGTCGTTGCTTATCGAGGCTGAGGATTCCTGTNNNAGATCGGAA
69942 TATGAGTGTGGAGTCGTTGCGTGACCCTTATGAAAGCAGCTCANNNAGATCGGAAG
70356 TATGAGTGTGGAGTCGTTGCTCATCTAATCCCCACAGACCTCCNNNAGATCGGAAGA
70379 TATGAGTGTGGAGTCGTTGCGAGAGGGAGAGAAATCTGCTTCCNNNAGATCGGAA
70934 TATGAGTGTGGAGTCGTTGCTCTTCTACTCCCACACCTCCTTGNNNAGATCGGAAGA
71028 TATGAGTGTGGAGTCGTTGCAAGTCTCCTGAAGGCTGGTTCATNNNAGATCGGAAG
71183 TATGAGTGTGGAGTCGTTGCTTTCCATTTCACATTGGTTTTCCTNNNAGATCGGAAG
71590 TATGAGTGTGGAGTCGTTGCGTTGGTGGACACTCGGGTTATTTNNNAGATCGGAA
93181 TATGAGTGTGGAGTCGTTGCTCTCTTCCCCTCCTCAATTCTCCNNNAGATCGGAAGAG
95454 TATGAGTGTGGAGTCGTTGCACAGAGTTTCGCTATTGCCCACNNNAGATCGGAAGA
99018 TATGAGTGTGGAGTCGTTGCCTGTTGGTCTCAGGTGCCTATNNNAGATCGGAAGAG
99122 TATGAGTGTGGAGTCGTTGCGAGTTTTGATTTGCTGATTGCCAGANNNAGATCGGA
99513 TATGAGTGTGGAGTCGTTGCGAATACAACAATTCCTGCCATCATGNNNAGATCGGA
100131 TATGAGTGTGGAGTCGTTGCTAGCGAAAATCGGCCCGGCCNNNAGATCGGAAGAG
28752 TATGAGTGTGGAGTCGTTGCTCCGAGTGTTCCAGTGTCTTTTTNNNAGATCGGAAG
28859 TATGAGTGTGGAGTCGTTGCAGTAGGTGCAGACTAGAAAGCCANNNAGATCGGAA
28926 TATGAGTGTGGAGTCGTTGCTGGTTTCCTGAATTCTGTCACAGTNNNAGATCGGAA
28975 TATGAGTGTGGAGTCGTTGCCCACTCATCCATCAACAGTCACTNNNAGATCGGAAGA
29325 TATGAGTGTGGAGTCGTTGCGCAATCGTGTTCAACTTGACAACANNNAGATCGGAA
29627 TATGAGTGTGGAGTCGTTGCTCCATTGTCTTTTTGTTTGCATGGTNNNAGATCGGAA
29649 TATGAGTGTGGAGTCGTTGCGAGCAGGAACAAGAGCTGCTTATNNNAGATCGGAA

| | |
|---|---|
| 29688 | TATGAGTGTGGAGTCGTTGCTGGGTTTGTAAGCATGCTGTTTCNNNAGATCGGAAG |
| 29728 | TATGAGTGTGGAGTCGTTGCGAGACTGTGTGTGTCTGTGCATANNNAGATCGGAA |
| 35146 | TATGAGTGTGGAGTCGTTGCACAGAGACACACTTTTCAAAGCANNNAGATCGGAAG |
| 69980 | TATGAGTGTGGAGTCGTTGCACTTCCAAAATCCAGAAGGAAGANNNAGATCGGAA |
| 71051 | TATGAGTGTGGAGTCGTTGCCCCAAATTGCTCCAGATGTGGATNNNAGATCGGAAG |
| 71465 | TATGAGTGTGGAGTCGTTGCACATGCTTCCCTCACCATTTTGTNNNAGATCGGAAGA |
| 71540 | TATGAGTGTGGAGTCGTTGCACACTGCTGGAGAGGTAAGTCTTNNNAGATCGGAA |
| 90278 | TATGAGTGTGGAGTCGTTGCTCTCAGTCCCTCACATTCACCTTNNNAGATCGGAAGA |
| 91514 | TATGAGTGTGGAGTCGTTGCTGTGTGAGAGAGATCAAGCACATNNNAGATCGGAA |
| 94032 | TATGAGTGTGGAGTCGTTGCTTTCATGTGTTTCCTGCAAGGGANNNAGATCGGAAG |
| 94257 | TATGAGTGTGGAGTCGTTGCTCTCTGGCCTTTCCATCTCTTCANNNAGATCGGAAGA |
| 100419 | TATGAGTGTGGAGTCGTTGCGGATCTGGGCGCTATGAAAAGTGTNNNAGATCGGA |
| 91880 | TATGAGTGTGGAGTCGTTGCACCGTGTAGTGAACAATTGCATGNNNAGATCGGAA |
| 120702 | TATGAGTGTGGAGTCGTTGCTCGGATTACAGGTGTGAGGAGTTACNNNAGATCGG |
| 28759 | TATGAGTGTGGAGTCGTTGCAACTGTGCTCGCTTCAGAGAAATNNNAGATCGGAAG |
| 28883 | TATGAGTGTGGAGTCGTTGCCCCTAAAGCATGGGGAGCTTTAGNNNAGATCGGAA |
| 28957 | TATGAGTGTGGAGTCGTTGCGCTCTGAGTGTCTTGTAGCCTTTNNNAGATCGGAAG |
| 28997 | TATGAGTGTGGAGTCGTTGCGATTCAACTGTGAGGTCTGGGTCNNNAGATCGGAA |
| 29045 | TATGAGTGTGGAGTCGTTGCGGGATAGACAGAAGGACAACCCTNNNAGATCGGAA |
| 29166 | TATGAGTGTGGAGTCGTTGCAAATTCCTGCAGCTAGATGGCTCNNNAGATCGGAAG |
| 29534 | TATGAGTGTGGAGTCGTTGCCCCAGAATCACTGGTCCTAACAGNNNAGATCGGAAG |
| 92022 | TATGAGTGTGGAGTCGTTGCATCCTTGTGATTTGGGGTTTGGGNNNAGATCGGAA |
| 95607 | TATGAGTGTGGAGTCGTTGCGAAATATGGGCTGCACTGAGCANNNAGATCGGAAG |
| 98394 | TATGAGTGTGGAGTCGTTGCACCAGGGGTGGGATGATACAGGNNNAGATCGGAA |
| 90855 | TATGAGTGTGGAGTCGTTGCGCCATGCTACAAGCCCTTATTCTNNNAGATCGGAAG |
| 90887 | TATGAGTGTGGAGTCGTTGCCTGCCCACAGTAGAACCTAGTCTNNNAGATCGGAAG |
| 120456 | TATGAGTGTGGAGTCGTTGCTCCAACTCTCATGATCCTCCCANNNAGATCGGAAGAG |
| 120903 | TATGAGTGTGGAGTCGTTGCTGCTCAATGGTGCGATCATAGCTNNNAGATCGGAAG |
| 120957 | TATGAGTGTGGAGTCGTTGCGCTCAAGTAATTATCCTGTCTCAGCANNNAGATCGG |
| 28939 | TATGAGTGTGGAGTCGTTGCTCTCCATACCCATTGTTGCCATCNNNAGATCGGAAGA |
| 29127 | TATGAGTGTGGAGTCGTTGCAAAAAGTGCTTGGAGGAAAATGTNNNAGATCGGAA |
| 29210 | TATGAGTGTGGAGTCGTTGCACAAAAATCACCTGGAGAGCTGTTNNNAGATCGGA |
| 29384 | TATGAGTGTGGAGTCGTTGCCCTGGGGAAAGGTTTTGGAGTTTNNNAGATCGGAA |
| 29401 | TATGAGTGTGGAGTCGTTGCCGGGCAGTTTCTAGAACAAGTTTNNNAGATCGGAA |
| 29661 | TATGAGTGTGGAGTCGTTGCGGTTGATTTTCTGCTAATATCAGGGGNNNAGATCGG |
| 29664 | TATGAGTGTGGAGTCGTTGCAATGCAAAGCCAAAGACACAAGGNNNAGATCGGAA |
| 69711 | TATGAGTGTGGAGTCGTTGCCCAAAGTATGCCAGGACAGACAANNNAGATCGGAA |
| 70927 | TATGAGTGTGGAGTCGTTGCTGTCCCCGTGGATCTATTTTGTGNNNAGATCGGAAG |
| 71149 | TATGAGTGTGGAGTCGTTGCAGAACCCTGCACTGAATATCCCTNNNAGATCGGAAG |
| 71490 | TATGAGTGTGGAGTCGTTGCTATTCCCTGTTGCTCAAGCGATTNNNAGATCGGAAG |
| 90677 | TATGAGTGTGGAGTCGTTGCTCTCCATTCTCCACTTTCCTCGTNNNAGATCGGAAGA |
| 91676 | TATGAGTGTGGAGTCGTTGCTAGTTGAGGGTTTGAGGGTTGTGNNNAGATCGGA |
| 94632 | TATGAGTGTGGAGTCGTTGCAAGATAGAATAACCTAGAATACCACCANNNAGATCG |

| | |
|---|---|
| 95470 | TATGAGTGTGGAGTCGTTGCTGGTATTCTCCCCTTGTGTCTGTNNNAGATCGGAAG |
| 97561 | TATGAGTGTGGAGTCGTTGCTGATGAGCAGCGTCAAGGACAACNNNAGATCGGAA |
| 98012 | TATGAGTGTGGAGTCGTTGCCAAACTGAACTTGGGAACTTGCANNNAGATCGGAA |
| 90531 | TATGAGTGTGGAGTCGTTGCCCAAACAGTGGGTGAGTGAATGTNNNAGATCGGAA |
| 119928 | TATGAGTGTGGAGTCGTTGCGCAGTTGACGGACAGATCATCATNNNAGATCGGAA |
| 120012 | TATGAGTGTGGAGTCGTTGCAGTGGGTCATGGGAGCATCTAGTANNNAGATCGGA |
| 120159 | TATGAGTGTGGAGTCGTTGCCAACAGGGCCACTGCCTTTGANNNAGATCGGAAGA |
| 120856 | TATGAGTGTGGAGTCGTTGCAAGTGGCATGATCTTTGCTCACGANNNAGATCGGAA |
| 90697 | TATGAGTGTGGAGTCGTTGCGGGTCCTCTCTCATCTTGGTTAGTNNNAGATCGGAA |
| 28834 | TATGAGTGTGGAGTCGTTGCGACTTTCCTTTGGAGCATTATTGGGNNNAGATCGGA |
| 28936 | TATGAGTGTGGAGTCGTTGCCAGACAGGAGTGAGTGCTACTTCNNNAGATCGGAA |
| 28937 | TATGAGTGTGGAGTCGTTGCACAGCAGAGGGCTCTTTGAATTCNNNAGATCGGAA |
| 28938 | TATGAGTGTGGAGTCGTTGCCATTAGGATGTTGGGGGTTGGTCNNNAGATCGGAA |
| 28963 | TATGAGTGTGGAGTCGTTGCCTCATTTAGGCCAGAGGCTTTGTNNNAGATCGGAAG |
| 29120 | TATGAGTGTGGAGTCGTTGCGTAATGGGCCTCCTTGTCTTCTTNNNAGATCGGAAG |
| 29134 | TATGAGTGTGGAGTCGTTGCAAAATGCAGGGGACAGTCAAAGTNNNAGATCGGAA |
| 29137 | TATGAGTGTGGAGTCGTTGCTATGCCCAGTGGTGAAAAATGCTNNNAGATCGGAA |
| 29555 | TATGAGTGTGGAGTCGTTGCGCAAATATCAACGACCTGTCAGCNNNAGATCGGAAG |
| 52161 | TATGAGTGTGGAGTCGTTGCCTGGTTGTGACAGTGTCCTGTAGNNNAGATCGGAA |
| 62182 | TATGAGTGTGGAGTCGTTGCTGTCATTGCATTCTTGTTTGGCTNNNAGATCGGAAG |
| 70122 | TATGAGTGTGGAGTCGTTGCGCCCAATCCCCTGTTTTCACATANNNAGATCGGAAGA |
| 70933 | TATGAGTGTGGAGTCGTTGCTCATGGAGCAGAAGACAACCTTGNNNAGATCGGAA |
| 71464 | TATGAGTGTGGAGTCGTTGCGTTTCTAGGAGGCAGCTGTTTGTNNNAGATCGGAA |
| 90733 | TATGAGTGTGGAGTCGTTGCTGTGGGAGTACAAGTACATAGAGTNNNAGATCGGA |
| 91016 | TATGAGTGTGGAGTCGTTGCCTTTCTGTCCCTACTTCCTGGCTNNNAGATCGGAAGA |
| 95680 | TATGAGTGTGGAGTCGTTGCATTCGGTTTTGCTGTTTGCTCANNNAGATCGGAAGA |
| 96148 | TATGAGTGTGGAGTCGTTGCCTCAGTAGTCTTCCACGTCATCANNNAGATCGGAAG |
| 96784 | TATGAGTGTGGAGTCGTTGCCACTTTTCCCTCTCTCCTTCACTTNNNAGATCGGAAGA |
| 98266 | TATGAGTGTGGAGTCGTTGCGTCCATTGCCCTGTCTCCTCTTTNNNAGATCGGAAGA |
| 98817 | TATGAGTGTGGAGTCGTTGCGGTAAAATAGGGGTGGAGGTCAAANNNAGATCGG |
| 104527 | TATGAGTGTGGAGTCGTTGCTGTGTAGCTTGAACCCAGGAGGTNNNAGATCGGAA |
| 94882 | TATGAGTGTGGAGTCGTTGCTGAGTTGTCTGGCTTTTTGGGAANNNAGATCGGAA |
| 93282 | TATGAGTGTGGAGTCGTTGCAAAGCACTATGGCACTAGAAGCCNNNAGATCGGAA |
| 92083 | TATGAGTGTGGAGTCGTTGCTGTGGGTAGGAAAGACCTTAGGGNNNAGATCGGA |
| 119728 | TATGAGTGTGGAGTCGTTGCCTGGCTGGAGAGCACGATTTTNNNAGATCGGAAGA |
| 119876 | TATGAGTGTGGAGTCGTTGCAACAAAGGAAACACAGAAGACGGNNNAGATCGGA |
| 28747 | TATGAGTGTGGAGTCGTTGCACCCAGAATCAACAGAAACAAACANNNAGATCGGAA |
| 28823 | TATGAGTGTGGAGTCGTTGCAAAGAAGAGGGTACCTGAGCCTTNNNAGATCGGAA |
| 28863 | TATGAGTGTGGAGTCGTTGCATCCAGTATGCAAGGGTCCTTATNNNAGATCGGAAG |
| 28871 | TATGAGTGTGGAGTCGTTGCGAAAAAGCATGAGTTGGTGGGTGNNNAGATCGGA |
| 28892 | TATGAGTGTGGAGTCGTTGCGCAGTTGGGAGAAAAGGAGTAGGNNNAGATCGGA |
| 28974 | TATGAGTGTGGAGTCGTTGCAAAACGTAAGGAGCGCACATACTNNNAGATCGGAA |
| 29232 | TATGAGTGTGGAGTCGTTGCATTTGAAGAGGACATGTGCCTNNNAGATCGGAAGA |

29251 TATGAGTGTGGAGTCGTTGCGAGGGCTATGAGCAGAAATCCTGNNNAGATCGGAA
29347 TATGAGTGTGGAGTCGTTGCTTGTCCCCAGAAAACAACCCATTNNNAGATCGGAAG
29348 TATGAGTGTGGAGTCGTTGCTTGCTGTGTCTAAGCATCTTTGTNNNAGATCGGAAG
29421 TATGAGTGTGGAGTCGTTGCTGCTTTACTGATGACCTTCTCAGCNNNAGATCGGAA
29423 TATGAGTGTGGAGTCGTTGCTGAGTAGATGTGGATGCAGCAAGNNNAGATCGGA
29750 TATGAGTGTGGAGTCGTTGCTCACCAAAAGAGCTCCTGATTTACANNNAGATCGGA
70662 TATGAGTGTGGAGTCGTTGCCGTGTGTGTAATAGGGAGTGGAGNNNAGATCGGA
71143 TATGAGTGTGGAGTCGTTGCACAACAGATTTGACTGAATACCCTNNNAGATCGGAA
91850 TATGAGTGTGGAGTCGTTGCGGAGACCACAGCACAGTCTTATGNNNAGATCGGAA
91889 TATGAGTGTGGAGTCGTTGCGTCTCTTTGTCTCTCTGGCCATGNNNAGATCGGAAG
93405 TATGAGTGTGGAGTCGTTGCTTCTCCCTCTGTGATTAGCCACCNNNAGATCGGAAGA
94995 TATGAGTGTGGAGTCGTTGCGTTGCTGTCTCTGTCCATGACAANNNAGATCGGAAG
109189 TATGAGTGTGGAGTCGTTGCATAATGTTAGGTGTGCTAGGCCTNNNAGATCGGAA
92272 TATGAGTGTGGAGTCGTTGCTCTAGTGGTGAGGAAAGACCTCGNNNAGATCGGAA
91910 TATGAGTGTGGAGTCGTTGCACAGGGTTTCACTCTCACTTTTGGNNNAGATCGGAA
119669 TATGAGTGTGGAGTCGTTGCACAACACATCAAAAAGATAGCATCCCNNNAGATCGG
119321 TATGAGTGTGGAGTCGTTGCCAGGCTACAGTGGTGCAATCTCAANNNAGATCGGA
90805 TATGAGTGTGGAGTCGTTGCTGAGGTTCAAATGGCTAGCAAGTNNNAGATCGGAA
28854 TATGAGTGTGGAGTCGTTGCGCATGGGCTTGGGGCTCAGTNNNAGATCGGAAGAG
28943 TATGAGTGTGGAGTCGTTGCTGCTGCCTTTTTCCTTTATTTTCTCCNNNAGATCGGAA
28979 TATGAGTGTGGAGTCGTTGCGAAGCCCCTCTCTCTAACTCCATNNNAGATCGGAAGA
29106 TATGAGTGTGGAGTCGTTGCGGTAACAAGGTCAAATCTGCACANNNAGATCGGAA
29299 TATGAGTGTGGAGTCGTTGCAAGCACGATTGCTAATTGTGGGANNNAGATCGGAA
29533 TATGAGTGTGGAGTCGTTGCACCTTTGAAAGCCACTACCACAGNNNAGATCGGAAG
29600 TATGAGTGTGGAGTCGTTGCGGAAACCATGTAGTTTGGGAGACANNNAGATCGGA
29610 TATGAGTGTGGAGTCGTTGCGGTCTTGGGGAACAAGTGATGTTNNNAGATCGGAA
69699 TATGAGTGTGGAGTCGTTGCGTGGGTAGAATGCCTTTGGGAAANNNAGATCGGAA
69755 TATGAGTGTGGAGTCGTTGCCTGAGAAGATATGCGCTGTGGAANNNAGATCGGAA
69761 TATGAGTGTGGAGTCGTTGCTTTCCCTGACCTGGAAGAGTGAANNNAGATCGGAA
69905 TATGAGTGTGGAGTCGTTGCATCATTTCCATCAGCATGAGGCANNNAGATCGGAAG
69998 TATGAGTGTGGAGTCGTTGCCAGAAGGGAGAATAAGGGGCAGANNNAGATCGGA
70824 TATGAGTGTGGAGTCGTTGCAGCCAAGGTTCTTGAAAGAGTGGNNNAGATCGGAA
71589 TATGAGTGTGGAGTCGTTGCTCACATGGTTGTGGCTGCTATTTNNNAGATCGGAAG
91468 TATGAGTGTGGAGTCGTTGCGCATACACACACCTAGAAGGCATNNNAGATCGGAAG
91476 TATGAGTGTGGAGTCGTTGCCTCCGAACCAATAGCATCAGCATNNNAGATCGGAAG
95619 TATGAGTGTGGAGTCGTTGCGAAGCAACCACATGCATGGATTTCNNNAGATCGGAA
97051 TATGAGTGTGGAGTCGTTGCTGCTCTTTTCATTCCACCTCCTCNNNAGATCGGAAGA
97120 TATGAGTGTGGAGTCGTTGCAAATTGAGAGAAACCAGCTGATGATNNNAGATCGG
98044 TATGAGTGTGGAGTCGTTGCTGTAGCAATAATAGAAAGCACCCAANNNAGATCGG
98141 TATGAGTGTGGAGTCGTTGCAGACTGAAACAACCACTAAAACAGANNNAGATCGG
98984 TATGAGTGTGGAGTCGTTGCTGATAAAAGCCAAGGACATGTGGNNNAGATCGGAA
105625 TATGAGTGTGGAGTCGTTGCGAAGCAGAAGTGTTTGGGATTTGGNNNAGATCGG
92337 TATGAGTGTGGAGTCGTTGCGCTCTTCTCCAGACCCTCTCTAGNNNAGATCGGAAGA

| | |
|---|---|
| 119727 | TATGAGTGTGGAGTCGTTGCAAGCAATGCAGAAAAGTTTCACTNNNAGATCGGAA |
| 119914 | TATGAGTGTGGAGTCGTTGCGACATCAAGTGTTTGACTGAAGGACNNNAGATCGG |
| 120545 | TATGAGTGTGGAGTCGTTGCATAGAACTTAGACAAGTGGCTCCTGANNNAGATCG |
| 120525 | TATGAGTGTGGAGTCGTTGCTTCAAGCCAACTACCCACCTCNNNAGATCGGAAGAG |
| 120567 | TATGAGTGTGGAGTCGTTGCAATGTCTGTCAAAACGGGAAAGGNNNAGATCGGAA |
| 48188 | TATGAGTGTGGAGTCGTTGCGCCACATTTATTGCTGCATTTGCNNNAGATCGGAAG |
| 28750 | TATGAGTGTGGAGTCGTTGCAGGGGGTAAAAGGGAGGAGTTTTNNNAGATCGGA |
| 28757 | TATGAGTGTGGAGTCGTTGCTTGAACAGGGGATCCTTACACCTNNNAGATCGGAAG |
| 28770 | TATGAGTGTGGAGTCGTTGCATTTCAGACACAGTCCCACTCTGNNNAGATCGGAAG |
| 28818 | TATGAGTGTGGAGTCGTTGCAGAGGGAATTCGCAAACTTACCCNNNAGATCGGAA |
| 28917 | TATGAGTGTGGAGTCGTTGCCTTCCACTGTTTCTCTCCCCAACNNNAGATCGGAAGA |
| 28964 | TATGAGTGTGGAGTCGTTGCCGCTTCTGTGTGTCTAGAAATGTNNNAGATCGGAAG |
| 28970 | TATGAGTGTGGAGTCGTTGCATACTCCCAAAAGTGTGGGTCCTNNNAGATCGGAAG |
| 29016 | TATGAGTGTGGAGTCGTTGCAGTATTAGTGGATGGGAGCAGCANNNAGATCGGA |
| 29035 | TATGAGTGTGGAGTCGTTGCTTATTTTGATCACCGTGGGGTGTNNNAGATCGGAAG |
| 29146 | TATGAGTGTGGAGTCGTTGCTCCTAGTGGTATCTGTGCATTTGTGNNNAGATCGGA |
| 29171 | TATGAGTGTGGAGTCGTTGCGACCCAAGGTTTGGAAAAACAGCNNNAGATCGGAA |
| 29307 | TATGAGTGTGGAGTCGTTGCCACACAGGCACTGGGATAATTCANNNAGATCGGAA |
| 29329 | TATGAGTGTGGAGTCGTTGCTTTGAGGAGAGGGAGCAAGAGAANNNAGATCGGA |
| 29349 | TATGAGTGTGGAGTCGTTGCTGCTTGGATTTGCCGATAGTTGTNNNAGATCGGAAG |
| 29389 | TATGAGTGTGGAGTCGTTGCTGGCAGAAGTATCAGCAGCATTTNNNAGATCGGAA |
| 29425 | TATGAGTGTGGAGTCGTTGCGCGAGGCAACATAGGGTACATAANNNAGATCGGAA |
| 29463 | TATGAGTGTGGAGTCGTTGCAGGTCCAGGAAATACTCCAGCTTNNNAGATCGGAAG |
| 29494 | TATGAGTGTGGAGTCGTTGCGGTCCTCAACCAGACATTCTGATNNNAGATCGGAAG |
| 29628 | TATGAGTGTGGAGTCGTTGCCAACATGGATAACCCTCAAGGGTNNNAGATCGGAA |
| 29733 | TATGAGTGTGGAGTCGTTGCCCATAGAAATTTGTACCTGTGGAGGANNNAGATCG |
| 69769 | TATGAGTGTGGAGTCGTTGCCTCCAGCTCTCATCTGGGTCTAANNNAGATCGGAAG |
| 69829 | TATGAGTGTGGAGTCGTTGCTGAATGTGTCAGCTTGCTTGACANNNAGATCGGAAG |
| 70443 | TATGAGTGTGGAGTCGTTGCTGCTTTCTTATTCAGGTCTGGGCNNNAGATCGGAAG |
| 70589 | TATGAGTGTGGAGTCGTTGCGAACTGGGGAGTGGTGATGCAAGNNNAGATCGGA |
| 70703 | TATGAGTGTGGAGTCGTTGCACTGACTTACTGCTGCATCAAGGNNNAGATCGGAAG |
| 71064 | TATGAGTGTGGAGTCGTTGCGGGCAGGAGGATGGTGTAGATATNNNAGATCGGA |
| 71472 | TATGAGTGTGGAGTCGTTGCTCGGCCTTGTGCATTGTAGAATTNNNAGATCGGAAG |
| 90213 | TATGAGTGTGGAGTCGTTGCTCCTGCAATCCTGCAAGTTTCTTNNNAGATCGGAAG |
| 91319 | TATGAGTGTGGAGTCGTTGCAGCCAGGTAGGTATATAGACAACTNNNAGATCGGA |
| 92599 | TATGAGTGTGGAGTCGTTGCTGTGTCTGCTCCCCTTTCTTTTCNNNAGATCGGAAGA |
| 92936 | TATGAGTGTGGAGTCGTTGCGCAGTGTGAACCTGAAGTAATGCNNNAGATCGGAA |
| 94775 | TATGAGTGTGGAGTCGTTGCGACACCATAAAGAGCAAGCAACANNNAGATCGGAA |
| 95567 | TATGAGTGTGGAGTCGTTGCAGTGCAGAGACATGATCTAGGTTCANNNAGATCGG |
| 97130 | TATGAGTGTGGAGTCGTTGCCCATTTTTCCCTAAAAGGCCTCTNNNAGATCGGAAG |
| 97754 | TATGAGTGTGGAGTCGTTGCAAGAAAGGGGCGAGACCCGGTNNNAGATCGGAAG |
| 98483 | TATGAGTGTGGAGTCGTTGCCCCTGCAATCCAAAAACAGGAAAGNNNAGATCGGA |
| 93571 | TATGAGTGTGGAGTCGTTGCCACACCTGATCACATCCACACACNNNAGATCGGAAG |

69948 TATGAGTGTGGAGTCGTTGCCCTTCTCCACACCTCTCTTCTCANNNAGATCGGAAGA
120681 TATGAGTGTGGAGTCGTTGCGGTTCGTTGCAGCCTCGATTTNNNAGATCGGAAGAG
121142 TATGAGTGTGGAGTCGTTGCGAATGTGAACAGGTCCAAACGACANNNAGATCGGA
120974 TATGAGTGTGGAGTCGTTGCTAGGGTCTACTCTGTCACCCAGGTNNNAGATCGGAA
28782 TATGAGTGTGGAGTCGTTGCGGAAAGCCTCCTCTGGAACATTNNNAGATCGGAAG
28805 TATGAGTGTGGAGTCGTTGCGGACAAAGAGAGCAGAGGTGATCNNNAGATCGGA
28812 TATGAGTGTGGAGTCGTTGCATCTGTTCCACCACTTTGCTGTTNNNAGATCGGAAGA
28862 TATGAGTGTGGAGTCGTTGCATGCCTGAGAATCAGTGAGGACTNNNAGATCGGAA
29003 TATGAGTGTGGAGTCGTTGCTGATGACCACTTCTACTTCTGCCNNNAGATCGGAAG
29062 TATGAGTGTGGAGTCGTTGCTACCAGAACCCAGAGATTGAGGGNNNAGATCGGAA
29316 TATGAGTGTGGAGTCGTTGCGTTTCCCCATACTGTTCTTGCCANNNAGATCGGAAGA
29372 TATGAGTGTGGAGTCGTTGCTTTTCTAGCCTTGGGCTTGGTTTNNNAGATCGGAAG
29379 TATGAGTGTGGAGTCGTTGCAGCCAAACGATTCAGTGCATTTTNNNAGATCGGAAG
29723 TATGAGTGTGGAGTCGTTGCTGGGGCAAAGGACAGAGAGATTANNNAGATCGGA
29748 TATGAGTGTGGAGTCGTTGCCCTGTCAAATCCTGATTCTGCCANNNAGATCGGAAG
65408 TATGAGTGTGGAGTCGTTGCAGCCTGAAAAAGAGGTAATTTCTGGTNNNAGATCG
69887 TATGAGTGTGGAGTCGTTGCACATGATTCTTTCTGTGCCAGCANNNAGATCGGAAG
70094 TATGAGTGTGGAGTCGTTGCGCAACTTGGATTGAGCTATAGGTGANNNAGATCGG
70232 TATGAGTGTGGAGTCGTTGCCTGGGCTACTAAAACCTTCCCACNNNAGATCGGAAG
71347 TATGAGTGTGGAGTCGTTGCTGAGTGTCTACTACGTGTCAGGTNNNAGATCGGAA
71420 TATGAGTGTGGAGTCGTTGCTGTAACTATGAAACCAGCTGTCTGTNNNAGATCGGA
93099 TATGAGTGTGGAGTCGTTGCTTGTTTTCAAAGTGCTAGCCAGCNNNAGATCGGAAG
93926 TATGAGTGTGGAGTCGTTGCACAGAACAAAACAAGCAAAACACTGANNNAGATCG
95477 TATGAGTGTGGAGTCGTTGCTATTTGCACACTTCAGATGGGTAANNNAGATCGGAA
96008 TATGAGTGTGGAGTCGTTGCCTGAGAGATGGAGAGAGAAGCCCNNNAGATCGGA
99188 TATGAGTGTGGAGTCGTTGCGACCACACAACTTTGCAACTTGNNNAGATCGGAAGA
106763 TATGAGTGTGGAGTCGTTGCCAATACTCATCCTCCTCCTACCCTCNNNAGATCGGAAG
90961 TATGAGTGTGGAGTCGTTGCCAACCCAGTAAGATTGCCTTGCTNNNAGATCGGAAG
120033 TATGAGTGTGGAGTCGTTGCGCGGTTGAGTTTTGCATATGAACCNNNAGATCGGA
94742 TATGAGTGTGGAGTCGTTGCACAGTTCTGGCAATTAGTGACACANNNAGATCGGAA
28833 TATGAGTGTGGAGTCGTTGCGATATTACCCTGATCCCCTCGCGNNNAGATCGGAAG
28845 TATGAGTGTGGAGTCGTTGCAGTAGCCCCATTATCTTGGGCTTNNNAGATCGGAAG
28884 TATGAGTGTGGAGTCGTTGCCCATCCTTTTCTCCCATCTGCTCNNNAGATCGGAAGA
29175 TATGAGTGTGGAGTCGTTGCGGAGGAATGAAAGCCATGCAAACNNNAGATCGGAA
29275 TATGAGTGTGGAGTCGTTGCTGGGATCCTCTTCAATGGAAAGCNNNAGATCGGAA
29591 TATGAGTGTGGAGTCGTTGCTAGTGCCTGGTAATTGGTAGGCANNNAGATCGGAA
70778 TATGAGTGTGGAGTCGTTGCATGTAGTGCTGGATTTGATGGGGNNNAGATCGGAA
94308 TATGAGTGTGGAGTCGTTGCCTCAGCATCCTTGAAAGCTCTCANNNAGATCGGAAG
95286 TATGAGTGTGGAGTCGTTGCGGGCGGTAAGGCTCAGGCATGNNNAGATCGGAAG
95400 TATGAGTGTGGAGTCGTTGCCGGCCACTACAAGCTGCTTTTNNNAGATCGGAAGAG
97846 TATGAGTGTGGAGTCGTTGCAACTAAGGTAGCAACAGAGCTTTNNNAGATCGGAA
107880 TATGAGTGTGGAGTCGTTGCGCAAGGTCAACAGAAACTTGTAAAGNNNAGATCGG
108642 TATGAGTGTGGAGTCGTTGCCAAAAGGTAGCATAGTATACACACAGCNNNAGATCG

| | |
|---|---|
| 90508 | TATGAGTGTGGAGTCGTTGCGGTTTCTGTGGCTAACTGGATGTNNNAGATCGGAA |
| 119779 | TATGAGTGTGGAGTCGTTGCACTGCAAGCTTTGCCAAGCGANNNAGATCGGAAGA |
| 119978 | TATGAGTGTGGAGTCGTTGCCCGATAACCAATGATTCACAAAGCCTNNNAGATCGG |
| 119907 | TATGAGTGTGGAGTCGTTGCCTGCAAGCTCTGCCTCCTATGANNNAGATCGGAAGA |
| 120353 | TATGAGTGTGGAGTCGTTGCAAACTAACTCTTCGATGCCCAGANNNAGATCGGAAG |
| 120981 | TATGAGTGTGGAGTCGTTGCATGTGGAGCTAGTACTTGTGCAGNNNAGATCGGAA |
| 120829 | TATGAGTGTGGAGTCGTTGCTTCTGCCATTCCATCTCCTTCTACNNNAGATCGGAAG |
| 28764 | TATGAGTGTGGAGTCGTTGCATCAAGCAAACAAAAAGGCCGTCNNNAGATCGGAA |
| 28809 | TATGAGTGTGGAGTCGTTGCACCACCACCACTACCATGACTTANNNAGATCGGAAGA |
| 28980 | TATGAGTGTGGAGTCGTTGCTGTCACTCCAAATTATGTTGCACATNNNAGATCGGAA |
| 29097 | TATGAGTGTGGAGTCGTTGCGGGCCTCAAAGACAGTTTCAAGANNNAGATCGGAA |
| 29118 | TATGAGTGTGGAGTCGTTGCTGTGTGTGTGTGAAAAGGAGGTTNNNAGATCGGAA |
| 29131 | TATGAGTGTGGAGTCGTTGCAGGGAAAAATAAACATGCAGAAGGTNNNAGATCGG |
| 29156 | TATGAGTGTGGAGTCGTTGCTTGTCTCCTTCCTTCCAGAGGAGNNNAGATCGGAAG |
| 29293 | TATGAGTGTGGAGTCGTTGCGTTGCTGAAATCCATGAGGGCTANNNAGATCGGAA |
| 29399 | TATGAGTGTGGAGTCGTTGCACACACAGTATGCTCCCGATTTTNNNAGATCGGAAG |
| 29462 | TATGAGTGTGGAGTCGTTGCCGGTACAATTAGCCCCTTTCCATNNNAGATCGGAAG |
| 29470 | TATGAGTGTGGAGTCGTTGCGTCTGTGTGTGTGTTTGTTGTTTTNNNAGATCGGAA |
| 29579 | TATGAGTGTGGAGTCGTTGCTCCTCCAAAGACATGAGGACTGANNNAGATCGGAA |
| 29636 | TATGAGTGTGGAGTCGTTGCGTGTAACCCTCGAAAACATATGCTNNNAGATCGGAA |
| 29686 | TATGAGTGTGGAGTCGTTGCAAAACACACCACCAAAATGCTGGNNNAGATCGGAAG |
| 29712 | TATGAGTGTGGAGTCGTTGCCCCCGTAGTAATGGAATAGAGCCNNNAGATCGGAA |
| 70941 | TATGAGTGTGGAGTCGTTGCCAAAAAGGAAAGGGGGCAAGTTGNNNAGATCGGA |
| 71005 | TATGAGTGTGGAGTCGTTGCGTTCCAGCCTCATAATCAGCCATNNNAGATCGGAAG |
| 71294 | TATGAGTGTGGAGTCGTTGCCTCTTGCCCATTCTTCCAACAGTNNNAGATCGGAAGA |
| 71458 | TATGAGTGTGGAGTCGTTGCGAAACCATCTTTGCAATGGTTGTNNNAGATCGGAAG |
| 93307 | TATGAGTGTGGAGTCGTTGCTTGTGACCTAGCATTTGCATCCCNNNAGATCGGAAG |
| 97379 | TATGAGTGTGGAGTCGTTGCTCTTCTTTGAGGTGGTAAGGCCNNNAGATCGGAAG |
| 98990 | TATGAGTGTGGAGTCGTTGCGATTCCTAATTGCCAGCCAGTANNNAGATCGGAAGA |
| 100322 | TATGAGTGTGGAGTCGTTGCCAAAGTTTCAGCTCAGATTGATCACCNNNAGATCGG |
| 100386 | TATGAGTGTGGAGTCGTTGCGGAAAGGCTGCTGGAGGGGCNNNAGATCGGAAGA |
| 94349 | TATGAGTGTGGAGTCGTTGCATGCCGCAAAAACATTTGCATCANNNAGATCGGAAG |
| 119550 | TATGAGTGTGGAGTCGTTGCGCATGAGAATCTTGAATCCCGGANNNAGATCGGAA |
| 28775 | TATGAGTGTGGAGTCGTTGCCCAAGGCATGTGTCTTACTGACANNNAGATCGGAAG |
| 28836 | TATGAGTGTGGAGTCGTTGCCGAGCTAAAATCATTTAGGGTGTTNNNAGATCGGAA |
| 28856 | TATGAGTGTGGAGTCGTTGCTCATGCATTGTCTGTCCTTTCCCNNNAGATCGGAAGA |
| 29071 | TATGAGTGTGGAGTCGTTGCGGGAGCAGGAGAGGAGAATTTTCNNNAGATCGGA |
| 29130 | TATGAGTGTGGAGTCGTTGCAGACAGTGTACCATACTGAGGGTNNNAGATCGGAA |
| 29142 | TATGAGTGTGGAGTCGTTGCCCCAAATAAATGGTGTATACCTACATNNNAGATCGG |
| 29157 | TATGAGTGTGGAGTCGTTGCTCCTGTGACTTCCTCAGACAGAGNNNAGATCGGAAG |
| 29172 | TATGAGTGTGGAGTCGTTGCTCCAAGCTGTTCTGGCTTATTCCNNNAGATCGGAAG |
| 29336 | TATGAGTGTGGAGTCGTTGCTGGGGGAGATTGATGTCCAGAAANNNAGATCGGAA |
| 29407 | TATGAGTGTGGAGTCGTTGCCTGAGGCTGGGTTACAGGTTTTTNNNAGATCGGAA |

29663 TATGAGTGTGGAGTCGTTGCATGAATAGCCCCGCATTTAGAGGNNNAGATCGGAA
29700 TATGAGTGTGGAGTCGTTGCGTGTGTGTGTTTTCTTAGCAGGCNNNAGATCGGAAG
69859 TATGAGTGTGGAGTCGTTGCATAGTTGTAATTCACCTGAGGCCANNNAGATCGGAA
69988 TATGAGTGTGGAGTCGTTGCAGATGAAATGGGTTATGCCACAGANNNAGATCGGA
70730 TATGAGTGTGGAGTCGTTGCTACAGCAAGGGCAGAGATAGAGGNNNAGATCGGA
70889 TATGAGTGTGGAGTCGTTGCTCTTTTTCTGAACCTGGCCTCTGNNNAGATCGGAAGA
71399 TATGAGTGTGGAGTCGTTGCATGAAACACACACAGCCAGATGTNNNAGATCGGAAG
95275 TATGAGTGTGGAGTCGTTGCCGCCCTTGGCTAAGTAGTTGGTTNNNAGATCGGAAG
97467 TATGAGTGTGGAGTCGTTGCAGAAAGATGGATGAGTTGAAGCTTCNNNAGATCGG
97733 TATGAGTGTGGAGTCGTTGCAGTGGGATGGATGTCTCGATTCCNNNAGATCGGAA
92525 TATGAGTGTGGAGTCGTTGCCCTACTTTGCTGTACCACCACAGNNNAGATCGGAAG
90781 TATGAGTGTGGAGTCGTTGCTGACTTCCATTCAGTGCAACAGTNNNAGATCGGAAG
90456 TATGAGTGTGGAGTCGTTGCCTCCTAGCTCAGATGCTCTCTGTNNNAGATCGGAAGA
119692 TATGAGTGTGGAGTCGTTGCTCTACCTCAGCCTGTCAAGTAGGTNNNAGATCGGAA
120937 TATGAGTGTGGAGTCGTTGCCTCTCCCCTCATGGTTACGAATCNNNAGATCGGAAGA
93670 TATGAGTGTGGAGTCGTTGCTTTCAAGTGTACGTGGAGCGTTANNNAGATCGGAA
28816 TATGAGTGTGGAGTCGTTGCTCATCACCTCCGGAAAGTGTAGTNNNAGATCGGAAG
28843 TATGAGTGTGGAGTCGTTGCTGATGGGAGCTGTATTAGGGGAGNNNAGATCGGA
28864 TATGAGTGTGGAGTCGTTGCAGGTGCTCAAACGTGAGTTGAATNNNAGATCGGAA
29014 TATGAGTGTGGAGTCGTTGCTTCCAGGATAGTGGCCTTTCAGANNNAGATCGGAAG
29151 TATGAGTGTGGAGTCGTTGCGTTTATCTGGCCCACCAGTTAGGNNNAGATCGGAAG
29247 TATGAGTGTGGAGTCGTTGCTATGCGGGATGTGTGTGTATGTGNNNAGATCGGAA
29268 TATGAGTGTGGAGTCGTTGCAAGAGACACAGCCAACACTCATCNNNAGATCGGAAG
29321 TATGAGTGTGGAGTCGTTGCATGCAAGAGAAAGACAAGCACCANNNAGATCGGAA
29442 TATGAGTGTGGAGTCGTTGCTTTCCTTTCTTCTCCACCTCCCTNNNAGATCGGAAGAG
29722 TATGAGTGTGGAGTCGTTGCAATACACACACTGCAATGGCAACNNNAGATCGGAAG
29735 TATGAGTGTGGAGTCGTTGCTGATAGTTCTTTGCCCCTCCAGANNNAGATCGGAAG
70571 TATGAGTGTGGAGTCGTTGCGCTCCCTTTGTGTTGCCAATTTCNNNAGATCGGAAGA
70697 TATGAGTGTGGAGTCGTTGCTTTCTCCTGTTCTTGCTCTGTCGNNNAGATCGGAAGA
70769 TATGAGTGTGGAGTCGTTGCGATAAAAACAAGTTTGAGGCCGGGNNNAGATCGGA
71343 TATGAGTGTGGAGTCGTTGCGTATCCACAACCTCAGTCAAGGTNNNAGATCGGAAG
93305 TATGAGTGTGGAGTCGTTGCCGTTCTGTTTACCTGGAGATCCCNNNAGATCGGAAG
93103 TATGAGTGTGGAGTCGTTGCTGCTGCTAGATGGAAATCACAGCNNNAGATCGGAA
120587 TATGAGTGTGGAGTCGTTGCTAGTGGCTAATTGACATTCCCAACNNNAGATCGGAA
120843 TATGAGTGTGGAGTCGTTGCCTGGGGTAAGTTTTCAAGGCTGTCNNNAGATCGGA
92365 TATGAGTGTGGAGTCGTTGCTTCCTGGGCATGAGTTTCTTGAGNNNAGATCGGAAG
28727 TATGAGTGTGGAGTCGTTGCAGCAGCAGCATGGCATAGTAAATNNNAGATCGGAA
28829 TATGAGTGTGGAGTCGTTGCCGTGACACCCATTGCTCTAGATTNNNAGATCGGAAG
28846 TATGAGTGTGGAGTCGTTGCTGCGAGACGTACCTGTATTGTTCNNNAGATCGGAAG
28869 TATGAGTGTGGAGTCGTTGCGGGCTGGTTTAGAGATCCTCTGTNNNAGATCGGAA
29001 TATGAGTGTGGAGTCGTTGCCCCACTCCAGTTACCATGTTAGCNNNAGATCGGAAG
29030 TATGAGTGTGGAGTCGTTGCATCTACAGAATACTCCACCAACCTTNNNAGATCGGAA
29114 TATGAGTGTGGAGTCGTTGCTCAGCAATCCCTGTTGACCTTTTNNNAGATCGGAAGA

29128 TATGAGTGTGGAGTCGTTGCAAAAACTGATGCTTTGGGGTGGTNNNAGATCGGAA
29170 TATGAGTGTGGAGTCGTTGCTAGGGCACCACCAAATGAATAGCNNNAGATCGGAA
29330 TATGAGTGTGGAGTCGTTGCGGTAGGAACCAGGGACTGAAGAANNNAGATCGGA
29568 TATGAGTGTGGAGTCGTTGCTTGCTGGGAGAAAGGAAAGTGACNNNAGATCGGA
29629 TATGAGTGTGGAGTCGTTGCTGTCATCGTCAAATAGGGAAAGGTNNNAGATCGGA
69497 TATGAGTGTGGAGTCGTTGCCAAGAAAGGGAGATGGGGCTTTTNNNAGATCGGA
69712 TATGAGTGTGGAGTCGTTGCACCTGACGATCCATGAGAGACAANNNAGATCGGAA
70148 TATGAGTGTGGAGTCGTTGCGTGAGGCAATGGATGTGTTAGCTANNNAGATCGGA
70785 TATGAGTGTGGAGTCGTTGCTCTATTCCCAGGCATCCTATGGGNNNAGATCGGAAG
95262 TATGAGTGTGGAGTCGTTGCCTTACCTGCCTCTATACAACCCGANNNAGATCGGAAG
95433 TATGAGTGTGGAGTCGTTGCAGGCAAACTGCTAGTCTCCAATTCNNNAGATCGGAA
97086 TATGAGTGTGGAGTCGTTGCCCCAGCCATTGTCCCCGTCGNNNAGATCGGAAGAGC
97380 TATGAGTGTGGAGTCGTTGCATCTCTCCCGCCTCACTCTTGTCNNNAGATCGGAAGA
97590 TATGAGTGTGGAGTCGTTGCATGTCTGAGGGCACAGGAGCCNNNAGATCGGAAGA
103094 TATGAGTGTGGAGTCGTTGCTCAAGTTGTTGGGTAGGGCTTACANNNAGATCGGA
94960 TATGAGTGTGGAGTCGTTGCTGAACTGATGGTATTCGCAGCAANNNAGATCGGAA
94949 TATGAGTGTGGAGTCGTTGCTCAGTTATGAGGTAGGGTGGCAANNNAGATCGGAA
93648 TATGAGTGTGGAGTCGTTGCAAGGGAGGATCACAGGCAGTTTANNNAGATCGGAA
28877 TATGAGTGTGGAGTCGTTGCTCCCAGGTCAACTTAGTCATGGTNNNAGATCGGAAG
28898 TATGAGTGTGGAGTCGTTGCAGTTGCCAGATAAATACAGGGAGCNNNAGATCGGA
28918 TATGAGTGTGGAGTCGTTGCAGGTATGGCAGCTTACCACTGTANNNAGATCGGAA
28955 TATGAGTGTGGAGTCGTTGCTCAAGAGGGAGAATGCTGACAAANNNAGATCGGAA
28984 TATGAGTGTGGAGTCGTTGCGACTTCTGGAAGGGTGGCAATGNNNAGATCGGAAG
29333 TATGAGTGTGGAGTCGTTGCATCCATGCTCTGTGTTCTAACAANNNAGATCGGAAG
29656 TATGAGTGTGGAGTCGTTGCCAGGAACCTTCAACTAGAGACCTGNNNAGATCGGAA
29741 TATGAGTGTGGAGTCGTTGCTTTTGAGGTGAATTGAATGCATGCANNNAGATCGG
29767 TATGAGTGTGGAGTCGTTGCTCAGTGTTGGTTTCCAGTGTGTTNNNAGATCGGAAG
32770 TATGAGTGTGGAGTCGTTGCAGCAGGCATGAAACACCTAAACANNNAGATCGGAA
70484 TATGAGTGTGGAGTCGTTGCTTCCTAGGCCAATTGACAAGTGCNNNAGATCGGAAG
70832 TATGAGTGTGGAGTCGTTGCCTCTCCTTCCTTGGTTTCTGTGGNNNAGATCGGAAGA
70979 TATGAGTGTGGAGTCGTTGCAAAGGGCGAAATGTGTGGAGAATNNNAGATCGGA
90361 TATGAGTGTGGAGTCGTTGCGGGCCAAATGTGTAGGGCTAATTNNNAGATCGGAA
92454 TATGAGTGTGGAGTCGTTGCAACACCAAAGACAGAAGGCTCAGNNNAGATCGGAA
95242 TATGAGTGTGGAGTCGTTGCCCCTCCTGGCTATTATCCTCAGGNNNAGATCGGAAG
95403 TATGAGTGTGGAGTCGTTGCATGGGATGATCTTGTCTCACTGCTANNNAGATCGGA
95446 TATGAGTGTGGAGTCGTTGCCAAGTCCGCTCTGGAGATAACTNNNAGATCGGAAGA
120169 TATGAGTGTGGAGTCGTTGCGCTAGGTGGATCACTTGAGTGCNNNAGATCGGAAG
120618 TATGAGTGTGGAGTCGTTGCAGAAATGAAACATATCCCATGCTCGTNNNAGATCGG
120906 TATGAGTGTGGAGTCGTTGCAGTGTAGTGTAGTGGTGCAATCACNNNAGATCGGA
90233 TATGAGTGTGGAGTCGTTGCTGTGTAGGCATGTGAATGTGCTTNNNAGATCGGAA
28891 TATGAGTGTGGAGTCGTTGCTCTGGAGCGGTTTATCACAAGTGNNNAGATCGGAA
29049 TATGAGTGTGGAGTCGTTGCACCCAAACATTACACCGTAGACTNNNAGATCGGAAG
29050 TATGAGTGTGGAGTCGTTGCTCTTCTGAATGCCGGGTTCCTATNNNAGATCGGAAG

29181 TATGAGTGTGGAGTCGTTGCTCTTATTCAATCATCTCTGCAGTGANNNAGATCGGAA
29238 TATGAGTGTGGAGTCGTTGCCTGAACTTCCACAAGGAAGCACTNNNAGATCGGAAG
29542 TATGAGTGTGGAGTCGTTGCTCAATGCGATGCAGCAATCATTCNNNAGATCGGAAG
29654 TATGAGTGTGGAGTCGTTGCTTGTCTTTAGGCTGGGTGTAGTGNNNAGATCGGAA
29676 TATGAGTGTGGAGTCGTTGCAACAGCTGGCCATTCTCTGAAAGNNNAGATCGGAAG
29699 TATGAGTGTGGAGTCGTTGCAAACAGAAATGAAAAACCCTGGGCNNNAGATCGGA
69764 TATGAGTGTGGAGTCGTTGCGGAAGGGTGAGAAAAGGCTTGAANNNAGATCGGA
70258 TATGAGTGTGGAGTCGTTGCAGGGACACACCCTTACCATAGACNNNAGATCGGAAG
70466 TATGAGTGTGGAGTCGTTGCTTTAGAGCCGAATACCCACATGCNNNAGATCGGAAG
70615 TATGAGTGTGGAGTCGTTGCTTATTCAAACGATCCCTGCCCAGNNNAGATCGGAAG
70724 TATGAGTGTGGAGTCGTTGCTCTGCCATTTACAAGCTAGCAGGNNNAGATCGGAAG
92187 TATGAGTGTGGAGTCGTTGCACAGTAGTCCCCTTTATCCGAGGNNNAGATCGGAAG
95921 TATGAGTGTGGAGTCGTTGCGTGGAGTATTCATTCCAGCACAGNNNAGATCGGAA
96084 TATGAGTGTGGAGTCGTTGCTGCGATGTGTGTTCCAATGACNNNAGATCGGAAGA
96643 TATGAGTGTGGAGTCGTTGCCTGGCTTCACTGGGATATCCGANNNAGATCGGAAGA
97682 TATGAGTGTGGAGTCGTTGCGCCAATTACTAGAGAGAGCTGAAANNNAGATCGGA
70794 TATGAGTGTGGAGTCGTTGCTTCTAATCTGCACACTGGTTGGGNNNAGATCGGAAG
120241 TATGAGTGTGGAGTCGTTGCGTCTACCCTATCCACTTGGTACTNNNAGATCGGAAGA
120584 TATGAGTGTGGAGTCGTTGCCCTTACCCAGCCTAGCATTGGTTNNNAGATCGGAAG
28866 TATGAGTGTGGAGTCGTTGCCCTCTCACCCCCTTATCTCTCTGNNNAGATCGGAAGA
28956 TATGAGTGTGGAGTCGTTGCGGCTCGTACACATGGAACTCTTTNNNAGATCGGAAG
29036 TATGAGTGTGGAGTCGTTGCAGAGAAAGAAACCTTGGATTCATGGTNNNAGATCG
29179 TATGAGTGTGGAGTCGTTGCCCCATGAAGGTAAACGCTGACTANNNAGATCGGAA
29440 TATGAGTGTGGAGTCGTTGCCAGCTAAGGAACTGAAGCCTCCTNNNAGATCGGAAG
29510 TATGAGTGTGGAGTCGTTGCTTGGTGTTTTCTTCAGGGACCTGNNNAGATCGGAAG
29518 TATGAGTGTGGAGTCGTTGCATAAGATGCTGAAAGGAGGGGGNNNAGATCGGAA
29581 TATGAGTGTGGAGTCGTTGCGAGATGCTGAGAAAGCACACAGANNNAGATCGGAA
29774 TATGAGTGTGGAGTCGTTGCCCCTCGAATGTTCTCCATTGCTTNNNAGATCGGAAGA
37960 TATGAGTGTGGAGTCGTTGCTGGTTGTCAATTATGATGGCAGANNNAGATCGGAA
49202 TATGAGTGTGGAGTCGTTGCCAGACCAAGATGTCACACCAGTCNNNAGATCGGAAG
70659 TATGAGTGTGGAGTCGTTGCAGATTGCTTCTTGAGCTTGGGAGNNNAGATCGGAA
71219 TATGAGTGTGGAGTCGTTGCCCATATTTTCTGGCCAGGATGCTNNNAGATCGGAAG
71484 TATGAGTGTGGAGTCGTTGCACTGGGGTGTATGTGAGGTCATTNNNAGATCGGAA
94309 TATGAGTGTGGAGTCGTTGCTCTTGTGTGTCTGTTCTTGCTCANNNAGATCGGAAG
97589 TATGAGTGTGGAGTCGTTGCGTTGGCTCAGATGGGAGAGGAGNNNAGATCGGAA
99703 TATGAGTGTGGAGTCGTTGCCTCTTCCAGCCATCATTGCATCANNNAGATCGGAAGA
116422 TATGAGTGTGGAGTCGTTGCTCTGTTTCCTCTCACTTGCTCAAATNNNAGATCGGAA
94841 TATGAGTGTGGAGTCGTTGCAGGCCATGCTAGCCAGAGTATAANNNAGATCGGAA
92141 TATGAGTGTGGAGTCGTTGCGGGGCCTTCGAAAGGTTATTAGGNNNAGATCGGAA
92512 TATGAGTGTGGAGTCGTTGCAGGTCAGTCAGACAATCAGACAGNNNAGATCGGAA
120193 TATGAGTGTGGAGTCGTTGCTGGGAAGCTCACTTTTGGCATGNNNAGATCGGAAG
28797 TATGAGTGTGGAGTCGTTGCTGAAGCAGCCACAACAATGAAAGNNNAGATCGGAA
28803 TATGAGTGTGGAGTCGTTGCCTGTTTGTCAGGATTACCCTCGGNNNAGATCGGAAG

| | |
|---|---|
| 28865 | TATGAGTGTGGAGTCGTTGCGCAGCCATGTACAAACAAGTCTGNNNAGATCGGAA |
| 29061 | TATGAGTGTGGAGTCGTTGCAATGCCTCTTAGAAGCTAGAGGGGNNNAGATCGGA |
| 29143 | TATGAGTGTGGAGTCGTTGCAGAGCCTGTGACAATGGTGAAATNNNAGATCGGAA |
| 29339 | TATGAGTGTGGAGTCGTTGCGCTTCAAGGCCAAACTTCAGTTTNNNAGATCGGAAG |
| 29459 | TATGAGTGTGGAGTCGTTGCGAGAAGAGAGAGAGGGTCTCTGTNNNAGATCGGA |
| 29556 | TATGAGTGTGGAGTCGTTGCCTCCCTCATGCCTCTTGATTTCCNNNAGATCGGAAGA |
| 70040 | TATGAGTGTGGAGTCGTTGCATTCTAGCCTGGGTATTTGGGGANNNAGATCGGAA |
| 70127 | TATGAGTGTGGAGTCGTTGCTCCACAAAGAGTGATGGGGGATANNNAGATCGGAA |
| 70129 | TATGAGTGTGGAGTCGTTGCCGTGTCATCAATGGGCACACTATANNNAGATCGGAA |
| 70312 | TATGAGTGTGGAGTCGTTGCACACTTCCTACCTTCCTTACCCCNNNAGATCGGAAGA |
| 71003 | TATGAGTGTGGAGTCGTTGCACACTCCTCCACTATCATCCCATNNNAGATCGGAAGA |
| 93908 | TATGAGTGTGGAGTCGTTGCTGTCTACTAGTTCAATCTATTGCTGANNNAGATCGGA |
| 96151 | TATGAGTGTGGAGTCGTTGCAGGACACATAATTACAAACACTGGCTGNNNAGATCG |
| 98451 | TATGAGTGTGGAGTCGTTGCGCATCTATTCACAGCTAAGCAGCTCNNNAGATCGGA |
| 106785 | TATGAGTGTGGAGTCGTTGCGGTTCTCTACTTCTTTTCTCTGTGGNNNAGATCGGAA |
| 91374 | TATGAGTGTGGAGTCGTTGCGGTGAATGGTGTGGAAGGTTGATNNNAGATCGGA |
| 90116 | TATGAGTGTGGAGTCGTTGCGCTCAGAGGTACCAAAAACCTTTNNNAGATCGGAAG |
| 93994 | TATGAGTGTGGAGTCGTTGCTTGGTTACTTTTGGGAGGATGGANNNAGATCGGAA |
| 94336 | TATGAGTGTGGAGTCGTTGCGGGAGAGGAAAGGAGAGGTATCANNNAGATCGGA |
| 120403 | TATGAGTGTGGAGTCGTTGCTAGGCAAGTCATACGGAAAACATTTTCNNNAGATCG |
| 120922 | TATGAGTGTGGAGTCGTTGCGCACCCAGCCTCAATATTGTTTCNNNAGATCGGAAG |
| 119458 | TATGAGTGTGGAGTCGTTGCGGGTGCACTGCAGTAGTGTAATNNNAGATCGGAAG |
| 92921 | TATGAGTGTGGAGTCGTTGCATTGGTCACTGGCCTTACTATGCNNNAGATCGGAAG |
| 28837 | TATGAGTGTGGAGTCGTTGCTTCAATGGATTCTGGTTGGGGACNNNAGATCGGAA |
| 28958 | TATGAGTGTGGAGTCGTTGCCTCACCTTGCATGGCACTATGTTNNNAGATCGGAAG |
| 29296 | TATGAGTGTGGAGTCGTTGCTAATTGAGGGTGCTCTCCCTTGANNNAGATCGGAAG |
| 29310 | TATGAGTGTGGAGTCGTTGCATGGGCTGTGCTTTCAATTCTCANNNAGATCGGAAG |
| 29398 | TATGAGTGTGGAGTCGTTGCGCATGTTCACACACACACTGTTTNNNAGATCGGAAG |
| 29749 | TATGAGTGTGGAGTCGTTGCATGTGTTTGGAACGTTTGAACCANNNAGATCGGAA |
| 28741 | TATGAGTGTGGAGTCGTTGCACTTCTCAAGGAAGAAACAAGGANNNAGATCGGAA |
| 28945 | TATGAGTGTGGAGTCGTTGCTGGGCAAAGGTTGGGAGAATTACNNNAGATCGGA |
| 29015 | TATGAGTGTGGAGTCGTTGCAGGACACCCTAGCAAATAGCTCANNNAGATCGGAAG |
| 29095 | TATGAGTGTGGAGTCGTTGCATCCACTGGGAGACACTTTGAGANNNAGATCGGAA |
| 29148 | TATGAGTGTGGAGTCGTTGCGGGGCTGGCAAATCCAAATTATGNNNAGATCGGAA |
| 29248 | TATGAGTGTGGAGTCGTTGCACAAAATTCAGGTCAGTTGTGGTGNNNAGATCGGA |
| 29357 | TATGAGTGTGGAGTCGTTGCTAGAGATTGTCTGGCATGTCCGTNNNAGATCGGAA |
| 29754 | TATGAGTGTGGAGTCGTTGCGTGTGGAGACAGAAGGAATGGAANNNAGATCGGA |
| 38543 | TATGAGTGTGGAGTCGTTGCTGGTTTGGCCTTCTATCTACCGANNNAGATCGGAAG |
| 28754 | TATGAGTGTGGAGTCGTTGCGCCTTTCTCTGCCTCCATTTCTTNNNAGATCGGAAGA |
| 28789 | TATGAGTGTGGAGTCGTTGCTCCCAAAGAGGCATTTCCTGAACNNNAGATCGGAAG |
| 28921 | TATGAGTGTGGAGTCGTTGCTGTGATTCAGGAGAGTGCATTCANNNAGATCGGAA |
| 29081 | TATGAGTGTGGAGTCGTTGCAAATTGGGTTGGAAAGAACTGATACNNNAGATCGG |
| 29188 | TATGAGTGTGGAGTCGTTGCAGCCAGGGTTAGTTTTGCAATCANNNAGATCGGAA |

29435 TATGAGTGTGGAGTCGTTGCGACCAACAGAGTGGCATGAAGATNNNAGATCGGAA
29516 TATGAGTGTGGAGTCGTTGCGGTTCCTGATGGTACTAACTGGGNNNAGATCGGAA
29575 TATGAGTGTGGAGTCGTTGCAGGCTACAGTGATCATGCCACTANNNAGATCGGAAG
29655 TATGAGTGTGGAGTCGTTGCACATACTGAGTTGGAGTTGCCTGNNNAGATCGGAA
29738 TATGAGTGTGGAGTCGTTGCAAAGAAAACAGTATGTGCTCCTCANNNAGATCGGAA
29115 TATGAGTGTGGAGTCGTTGCAGGTGGAGGATAACCAAGTGTTTNNNAGATCGGAA
29368 TATGAGTGTGGAGTCGTTGCGCTTCTTTCTTTACCCTTGGGCTNNNAGATCGGAAGA
29558 TATGAGTGTGGAGTCGTTGCATGACTTGGAGAACAGCATGTCCNNNAGATCGGAA
29635 TATGAGTGTGGAGTCGTTGCGTTATGGAATTCAGCTGGGCTCTNNNAGATCGGAA
29662 TATGAGTGTGGAGTCGTTGCGGCTCAGTAACTTTTCCTGGAGGNNNAGATCGGAA
29675 TATGAGTGTGGAGTCGTTGCTACATTGTTGCTGCTGTGGGAAGNNNAGATCGGAA
29753 TATGAGTGTGGAGTCGTTGCTGGTTGATTCTAGGCCCCACTAANNNAGATCGGAAG
70248 TATGAGTGTGGAGTCGTTGCTCAGTGGCACAAAGTACACTCACNNNAGATCGGAAG
70681 TATGAGTGTGGAGTCGTTGCATGGGAAACTAGGGGAGGCTTAGNNNAGATCGGA
71059 TATGAGTGTGGAGTCGTTGCGGGCAACCAGGTAGATCCCATATNNNAGATCGGAA
71080 TATGAGTGTGGAGTCGTTGCAGAATGAGCAGTGATGGACAACTNNNAGATCGGAA
28857 TATGAGTGTGGAGTCGTTGCTGACTAAAAACTGCATTATGTTGTGTNNNAGATCGG
29092 TATGAGTGTGGAGTCGTTGCATAAATGCAGGACTGGGCTAGGANNNAGATCGGAA
29132 TATGAGTGTGGAGTCGTTGCTGTGTTACCGAGGACATAGGAGTNNNAGATCGGAA
29223 TATGAGTGTGGAGTCGTTGCGGAGGGAGTAAAGCCAAGAGAGTNNNAGATCGGA
29277 TATGAGTGTGGAGTCGTTGCGCCTTTGTGGTCCTCTAGATTCCNNNAGATCGGAAG
29289 TATGAGTGTGGAGTCGTTGCACTCCTGACAAATACACTGCCACNNNAGATCGGAAG
29390 TATGAGTGTGGAGTCGTTGCACACAGTGGTAGTTGCTATTGTTNNNAGATCGGAAG
29468 TATGAGTGTGGAGTCGTTGCTTGGCCTCAGGATACTCAGGTTTNNNAGATCGGAAG
29742 TATGAGTGTGGAGTCGTTGCGTGGGGTTTTCTAGAGTTTGGCANNNAGATCGGAA
70144 TATGAGTGTGGAGTCGTTGCAGAGTGGAGGCTTGAAGTTCCTANNNAGATCGGAA
71225 TATGAGTGTGGAGTCGTTGCACAATCATCTGATCCAATTCTGCTNNNAGATCGGAAG
71505 TATGAGTGTGGAGTCGTTGCGCATTCCGGACTAAATCCCACTTNNNAGATCGGAAG
28744 TATGAGTGTGGAGTCGTTGCGCCTTTGGAAATGTGGTCATGTGNNNAGATCGGAA
28872 TATGAGTGTGGAGTCGTTGCTCTTCTGTTCTTCCTTGTGTGGGNNNAGATCGGAAG
28928 TATGAGTGTGGAGTCGTTGCCTTCTAAAACCTGCTCCAGTGCTNNNAGATCGGAAG
28952 TATGAGTGTGGAGTCGTTGCTTGTGAATGTGGCTTTTTGTACANNNAGATCGGAAG
28978 TATGAGTGTGGAGTCGTTGCCCCATCCACTCTCCATACCAGATNNNAGATCGGAAGA
29055 TATGAGTGTGGAGTCGTTGCCCCAACGAGATAGGTAACAGGTGNNNAGATCGGAA
29228 TATGAGTGTGGAGTCGTTGCAATAACTAGAACAGGGCCAGGCTNNNAGATCGGAA
29415 TATGAGTGTGGAGTCGTTGCAGCGCATTGTTTTGCATCCTTTTNNNAGATCGGAAG
29519 TATGAGTGTGGAGTCGTTGCCTCAGCCGGGACAAGGAGGGNNNAGATCGGAAGA
28753 TATGAGTGTGGAGTCGTTGCTTCTCCTGGTTGAAAGGGTTTGGNNNAGATCGGAA
28772 TATGAGTGTGGAGTCGTTGCGCTAGAAGCCAAGGGAAGAGAACNNNAGATCGGA
28784 TATGAGTGTGGAGTCGTTGCTTCTGTGCTCAAGCAATCAGGTTNNNAGATCGGAAG
28847 TATGAGTGTGGAGTCGTTGCGGTCCATACAACTGGCCTTCTTCNNNAGATCGGAAG
28946 TATGAGTGTGGAGTCGTTGCGAGTGTGCCTATAGCCCCAGATANNNAGATCGGAA
29186 TATGAGTGTGGAGTCGTTGCGGGCATGGCTTCTAAACAACAGANNNAGATCGGAA

29528 TATGAGTGTGGAGTCGTTGCTCCTGGTCTAAAATGTGGAGCAGNNNAGATCGGAA

29603 TATGAGTGTGGAGTCGTTGCGTGCAAACACATGCATTCTGGAANNNAGATCGGAA

28730 TATGAGTGTGGAGTCGTTGCTCCCACCCCTAGCCAAGATTTTTNNNAGATCGGAAGA

29073 TATGAGTGTGGAGTCGTTGCATTTTGCCTTTGCCTGCTTCATCNNNAGATCGGAAGA

29205 TATGAGTGTGGAGTCGTTGCGCCAGTTTTGCTTCTACTTGCAANNNAGATCGGAAG

29242 TATGAGTGTGGAGTCGTTGCTACCTTGAGTGTGGTGATGGGATNNNAGATCGGAA

29244 TATGAGTGTGGAGTCGTTGCAAGCCTGAGTGAGGTATGGGAATNNNAGATCGGA

29433 TATGAGTGTGGAGTCGTTGCACACACACACACTAAACAGAGCCNNNAGATCGGAAG

29530 TATGAGTGTGGAGTCGTTGCATTCCCTCAGTTCAACACAGCAGNNNAGATCGGAAG

29706 TATGAGTGTGGAGTCGTTGCTGGTTATGCAGGTATGCAAGTCCNNNAGATCGGAA

29032 TATGAGTGTGGAGTCGTTGCAGGTTCTCAAATGCAGGACTATTNNNAGATCGGAAG

29647 TATGAGTGTGGAGTCGTTGCTCAAATTACAGGTCAAACACCAACTNNNAGATCGGA

29708 TATGAGTGTGGAGTCGTTGCGCAGAAGACCAAAAATGGATGCCNNNAGATCGGAA

69957 TATGAGTGTGGAGTCGTTGCAGCCATAGCCTAATCCAATTGTCANNNAGATCGGAA

70623 TATGAGTGTGGAGTCGTTGCTAATTTCCCCAGTCTTGCTCCAGNNNAGATCGGAAG

70674 TATGAGTGTGGAGTCGTTGCGAGCTGGATTGAGTTGAGGCTAGNNNAGATCGGA

70883 TATGAGTGTGGAGTCGTTGCGTTTACATGAGCAGAGCACATCTGNNNAGATCGGA

28989 TATGAGTGTGGAGTCGTTGCAAAATGAAGGGGGAAGCAAGAGGNNNAGATCGGA

29069 TATGAGTGTGGAGTCGTTGCTTCTGTTCAAGAGCCTGTGGAAGNNNAGATCGGAA

29138 TATGAGTGTGGAGTCGTTGCCATTAGTCACATGAAGCTGGCCTNNNAGATCGGAAG

29197 TATGAGTGTGGAGTCGTTGCCACTGAAGTCCCACTATTACACCANNNAGATCGGAA

29209 TATGAGTGTGGAGTCGTTGCCCTGCTGTTCTGGTATCTGCTTTNNNAGATCGGAAG

29570 TATGAGTGTGGAGTCGTTGCGGCAAAGGCCCCAAATTATGAACNNNAGATCGGAA

69823 TATGAGTGTGGAGTCGTTGCCGAATGCCAGAAGGTAAAGGACANNNAGATCGGAA

69952 TATGAGTGTGGAGTCGTTGCTACAATGTTACCAGTCCTGGTCANNNAGATCGGAAG

70084 TATGAGTGTGGAGTCGTTGCAAATGACTGTGCTAACAGCCTGANNNAGATCGGAA

28738 TATGAGTGTGGAGTCGTTGCCCAGGCTGAGAGTACAGTGACTANNNAGATCGGAA

28849 TATGAGTGTGGAGTCGTTGCCGAGACCCTGAGCTCTCCCGNNNAGATCGGAAGAG

28867 TATGAGTGTGGAGTCGTTGCATCACCGTGCCATGAATCAATCANNNAGATCGGAAG

29094 TATGAGTGTGGAGTCGTTGCGTCTGAGTTCTAGGGAGCAAGGANNNAGATCGGAA

29474 TATGAGTGTGGAGTCGTTGCTGAATTGCTATGCTGCTGGCTTTNNNAGATCGGAAG

29529 TATGAGTGTGGAGTCGTTGCCATAATAGGCTCTCAGGCAGCAGNNNAGATCGGAA

29592 TATGAGTGTGGAGTCGTTGCAACTGGAAATGTTGCACTAGGCANNNAGATCGGAA

28801 TATGAGTGTGGAGTCGTTGCTCCTCCTCTCCCACTAAAGGGTANNNAGATCGGAAG

28851 TATGAGTGTGGAGTCGTTGCCCAAACGGAGAGAAGAGAAGCACNNNAGATCGGA

28899 TATGAGTGTGGAGTCGTTGCTTCTCAACATCCCTTTCCTGCACNNNAGATCGGAAGA

29387 TATGAGTGTGGAGTCGTTGCATGGACACCAACACAGGTGTTTTNNNAGATCGGAAG

29428 TATGAGTGTGGAGTCGTTGCTGGGGATAGGTGTGGAGAATGTTNNNAGATCGGA

29541 TATGAGTGTGGAGTCGTTGCACGAAGCCAAGATCCTGAGATTCNNNAGATCGGAA

29652 TATGAGTGTGGAGTCGTTGCTTGCAGTTTCTCTGGAGAACCATNNNAGATCGGAAG

56081 TATGAGTGTGGAGTCGTTGCCGTGTCTTAGGGATTGTCTTCCTGNNNAGATCGGAA

70114 TATGAGTGTGGAGTCGTTGCTGTGACGTTTTGGTGTAAATGTTTGANNNAGATCGG

70532 TATGAGTGTGGAGTCGTTGCGGGCAGTCTGGCAGAATTATCTCNNNAGATCGGAA

28876 TATGAGTGTGGAGTCGTTGCTACCTCTTCTCTTGGCACTCACANNNAGATCGGAAGA
29154 TATGAGTGTGGAGTCGTTGCCATAGGCCAAGTCATCAGGAAGGNNNAGATCGGAA
29287 TATGAGTGTGGAGTCGTTGCGCTCACACAGATATTGAAGGTTGACNNNAGATCGGA
29358 TATGAGTGTGGAGTCGTTGCACACCTTTTCACACTGATACACGTNNNAGATCGGAAG
29377 TATGAGTGTGGAGTCGTTGCTGTGGTTTATGGGATCAGAGTTTNNNAGATCGGAA
70221 TATGAGTGTGGAGTCGTTGCTTTCCCACATTTCCTGGAGGAACNNNAGATCGGAAGA
70325 TATGAGTGTGGAGTCGTTGCCCTTCATCCTTGCATTTTTGCCCNNNAGATCGGAAGA
70360 TATGAGTGTGGAGTCGTTGCAAACGATCAACCTCTTTGCCTCCNNNAGATCGGAAGA
70501 TATGAGTGTGGAGTCGTTGCACCAATGTCATTCTCTTGCTTTTGCNNNAGATCGGAA
70878 TATGAGTGTGGAGTCGTTGCCCATCTTTCCTCCAATCAGGCTGNNNAGATCGGAAGA
70982 TATGAGTGTGGAGTCGTTGCGGTGGTTTCAGAGCTGACTGAATNNNAGATCGGAA
28910 TATGAGTGTGGAGTCGTTGCACAGAAACATGCCAGCTACAGAGNNNAGATCGGAA
28962 TATGAGTGTGGAGTCGTTGCGCTCTGCCCTAAGAACTTCCATTNNNAGATCGGAAGA
28995 TATGAGTGTGGAGTCGTTGCTTTTCCCAGTGGGTTTGGATCAGNNNAGATCGGAAG
29388 TATGAGTGTGGAGTCGTTGCCCCTCAATGGACTACAGCTGTTTNNNAGATCGGAAGA
29639 TATGAGTGTGGAGTCGTTGCTTCAACATTTCCTTCCTGGAGCTNNNAGATCGGAAGA
69695 TATGAGTGTGGAGTCGTTGCGGCAGGAGAGACAAGACAAGAAANNNAGATCGGAA
28796 TATGAGTGTGGAGTCGTTGCAGTGAGACCCAGTCACCACATTANNNAGATCGGAAG
28933 TATGAGTGTGGAGTCGTTGCGCCAAGAGGAGATAACAAGGTGGNNNAGATCGGAA
29043 TATGAGTGTGGAGTCGTTGCTTCGCATTTGTTTAGTCCTGCCTNNNAGATCGGAAGA
29149 TATGAGTGTGGAGTCGTTGCGGATTTATGCATGCAGTGTTGGGNNNAGATCGGAA
29221 TATGAGTGTGGAGTCGTTGCAGTTTCTCATAAGGTGGTTGTCGTNNNAGATCGGAA
29309 TATGAGTGTGGAGTCGTTGCCTGCTGAACCTGATGCATAGTCANNNAGATCGGAAG
29514 TATGAGTGTGGAGTCGTTGCAAAGTGGCATACTACTAGCGTGGNNNAGATCGGAA
29598 TATGAGTGTGGAGTCGTTGCTAAGAGTTGCCGAGAAAATGCCANNNAGATCGGAA
29641 TATGAGTGTGGAGTCGTTGCTCTGATGAAGGGTGAAAGAAGCTNNNAGATCGGAA
29645 TATGAGTGTGGAGTCGTTGCTCGTGAGCCAATATCTCATCACTNNNAGATCGGAAGA
29720 TATGAGTGTGGAGTCGTTGCCCCAGTCTGCCTTAGGTAAACACNNNAGATCGGAAGA
69743 TATGAGTGTGGAGTCGTTGCCCGAAAAGTGAACTCACCCAGAANNNAGATCGGAAG
28838 TATGAGTGTGGAGTCGTTGCTGAACCGTTACTGTATCCCAAGANNNAGATCGGAAG
29136 TATGAGTGTGGAGTCGTTGCAGTGGTGCTTTATTTTTCAGCTCTNNNAGATCGGAAG
29196 TATGAGTGTGGAGTCGTTGCCTCCTATCCTCTACTTGGCACCANNNAGATCGGAAGA
29367 TATGAGTGTGGAGTCGTTGCACTGGAATCTGATGGGACATTCTNNNAGATCGGAAG
29535 TATGAGTGTGGAGTCGTTGCTGCTATTGGCTTTCAAGCAACAGNNNAGATCGGAAG
69839 TATGAGTGTGGAGTCGTTGCTTCCAAGACCCTCAGTTGGTACANNNAGATCGGAAGA
70619 TATGAGTGTGGAGTCGTTGCTCCATTACACAGCCAATAGCCAGNNNAGATCGGAAGA
71042 TATGAGTGTGGAGTCGTTGCCTTGTGGCCATCACCATTGAGATNNNAGATCGGAAG
91730 TATGAGTGTGGAGTCGTTGCGAGAGAGCGACAGAGATTGAGTGNNNAGATCGGAA
29029 TATGAGTGTGGAGTCGTTGCACTCTCAGGTTAGATGAGGGCTTNNNAGATCGGAAG
29048 TATGAGTGTGGAGTCGTTGCACATGCTATCAGGGTTACTGACTNNNAGATCGGAAG
29212 TATGAGTGTGGAGTCGTTGCGTGGGAGACTTTAGCATGCTCTTNNNAGATCGGAAG
29453 TATGAGTGTGGAGTCGTTGCAATAAAATCCACCCTGCCAGCATNNNAGATCGGAAGA
29612 TATGAGTGTGGAGTCGTTGCACTCATTCTTTCATTGCCCCGTTNNNAGATCGGAAGA

29622 TATGAGTGTGGAGTCGTTGCAGCACTTTGGAGAAACTACCTGTNNNAGATCGGAAG
70800 TATGAGTGTGGAGTCGTTGCTCTGCCATTTTCTAGGCTCATGGNNNAGATCGGAAG
28766 TATGAGTGTGGAGTCGTTGCGCCTCCCGAGTAGAGAAGTATGANNNAGATCGGAA
28771 TATGAGTGTGGAGTCGTTGCCTAATTGCGAACTCCTGAGGTCCNNNAGATCGGAAG
28915 TATGAGTGTGGAGTCGTTGCGCCCAAGATGTTCTGAAAATGGCNNNAGATCGGAA
29020 TATGAGTGTGGAGTCGTTGCGCCATGAAAGAGCAGATGGTACANNNAGATCGGAA
29119 TATGAGTGTGGAGTCGTTGCCGGGTTGCATATATGTCCGAGTTNNNAGATCGGAA
29174 TATGAGTGTGGAGTCGTTGCGCAAGGAAAACCTCAACTTGCACNNNAGATCGGAAG
29216 TATGAGTGTGGAGTCGTTGCTCACTTGGTCCTTTCTTGTGTGTNNNAGATCGGAAG
29302 TATGAGTGTGGAGTCGTTGCACTGTCTAAACGGATGGATGTAGANNNAGATCGGA
59959 TATGAGTGTGGAGTCGTTGCACCTGTCAAAACTCATCAAACACTNNNAGATCGGAA
69689 TATGAGTGTGGAGTCGTTGCTCATGGAGGAGACAGATGACAAANNNAGATCGGAA
69893 TATGAGTGTGGAGTCGTTGCATTGACAGCATCACAGCATAGCANNNAGATCGGAAG
70180 TATGAGTGTGGAGTCGTTGCAGCATCAAACGGGGCAAACATTANNNAGATCGGAA
28792 TATGAGTGTGGAGTCGTTGCGTGCCAACTCATACAGCACCTAANNNAGATCGGAAG
29323 TATGAGTGTGGAGTCGTTGCATTTAGTGCCCAGCAAGCATACANNNAGATCGGAAG
29418 TATGAGTGTGGAGTCGTTGCTGGTAGGTGTTTTTCTTTTGTATTTTTNNNAGATCGG
29572 TATGAGTGTGGAGTCGTTGCTCCCTATCCAGGCTTTTTGGGTANNNAGATCGGAAG
70007 TATGAGTGTGGAGTCGTTGCCCAATGACCCAGGAAAAACGAGANNNAGATCGGAA
70319 TATGAGTGTGGAGTCGTTGCAGTATGAACACAGCGATTTCCCCNNNAGATCGGAAG
71401 TATGAGTGTGGAGTCGTTGCTGTAAACGAAGCCCTCAAACTGTNNNAGATCGGAAG
29344 TATGAGTGTGGAGTCGTTGCATCTGAGTTTGGGGCCAACATTTNNNAGATCGGAAG
29405 TATGAGTGTGGAGTCGTTGCCACTTGGGACTTGGCAGTAGTTTNNNAGATCGGAA
29670 TATGAGTGTGGAGTCGTTGCGAAACAGTTCTTGGCCAGAGGAGNNNAGATCGGAA
29704 TATGAGTGTGGAGTCGTTGCTATTTGTGGTTGTCGGGGTTTCCNNNAGATCGGAAG
71089 TATGAGTGTGGAGTCGTTGCTGAGCTACTCTGTGTCAGGAACTNNNAGATCGGAAG
94677 TATGAGTGTGGAGTCGTTGCAGGATGTGGGAGATGTTCTGACANNNAGATCGGAA
95259 TATGAGTGTGGAGTCGTTGCTTTTGATGGTGGATGAACAGAGAGGNNNAGATCGG
98053 TATGAGTGTGGAGTCGTTGCTCCGCGTCAAGACGACAGCGNNNAGATCGGAAGAG
28839 TATGAGTGTGGAGTCGTTGCCATCCAACCCCTGTCTGTTTTCANNNAGATCGGAAGA
28942 TATGAGTGTGGAGTCGTTGCTAAAGATAGTCAGGCTGCTGTCCNNNAGATCGGAA
29532 TATGAGTGTGGAGTCGTTGCGGCTTTCCCTGATCTGAAGACAGNNNAGATCGGAAG
29690 TATGAGTGTGGAGTCGTTGCCCCAGCATCTCCCTAACCAATTCNNNAGATCGGAAGA
31093 TATGAGTGTGGAGTCGTTGCCTCAACAATTATCAACTTGTGGCCAANNNAGATCGG
70010 TATGAGTGTGGAGTCGTTGCACGGATGTGGAACTTAGGTGAGANNNAGATCGGA
70118 TATGAGTGTGGAGTCGTTGCCACTGCAAACCCCCAGGTAAATANNNAGATCGGAAG
70229 TATGAGTGTGGAGTCGTTGCAGGGAAGGAAATTTGACACCCACNNNAGATCGGAA
70612 TATGAGTGTGGAGTCGTTGCTTCCTTGTGCTGGAAAGTACCAGNNNAGATCGGAAG
28887 TATGAGTGTGGAGTCGTTGCACCTTTGACCGAGATCTGAAGGANNNAGATCGGAA
29075 TATGAGTGTGGAGTCGTTGCGACCAGTTAGAAGGCAAGAGAGCNNNAGATCGGA
29088 TATGAGTGTGGAGTCGTTGCAGAGAACAGCCCATCTGAGTGTANNNAGATCGGAA
29091 TATGAGTGTGGAGTCGTTGCTCTGGTAATGGCGGACTAGATGANNNAGATCGGAA
29183 TATGAGTGTGGAGTCGTTGCATCCCTGCCCATAACCGATATGANNNAGATCGGAAG

29262 TATGAGTGTGGAGTCGTTGCGGAACTCGAACCAGGAGCTAGAGNNNAGATCGGAA
29539 TATGAGTGTGGAGTCGTTGCTACTATGTGCCTCCTGCAGAAAGNNNAGATCGGAAG
29624 TATGAGTGTGGAGTCGTTGCGCATGAGAAAAACTCTAAATTCAATGTNNNAGATCG
29646 TATGAGTGTGGAGTCGTTGCAGGATGTTTGTACCTGTCCCACTNNNAGATCGGAAG
41848 TATGAGTGTGGAGTCGTTGCGGGGAAAACGTCATAGTGGCTTANNNAGATCGGAA
28835 TATGAGTGTGGAGTCGTTGCCACAAGACAAGAAACTGAGGCCTNNNAGATCGGAA
28853 TATGAGTGTGGAGTCGTTGCCTGCGGTGAGCACTTACAGTATTNNNAGATCGGAAG
28911 TATGAGTGTGGAGTCGTTGCTTGTAGGAGGGAGTGTCTCACTCNNNAGATCGGAA
28925 TATGAGTGTGGAGTCGTTGCACATGAACCCAAGCACAGAGATTNNNAGATCGGAAG
28985 TATGAGTGTGGAGTCGTTGCGGTCCTGGAGATGCATTCTAATGGNNNAGATCGGA
29746 TATGAGTGTGGAGTCGTTGCCTAAAATAGGCTGCAGCGATCCANNNAGATCGGAAG
71274 TATGAGTGTGGAGTCGTTGCTCTCTCCCACTCAGTCTCTTTCTNNNAGATCGGAAGA
93635 TATGAGTGTGGAGTCGTTGCGTAAGCCCTGAAACCAGCAAAACNNNAGATCGGAAG
95362 TATGAGTGTGGAGTCGTTGCTACCTGCTATAACTCCTGGCGNNNAGATCGGAAGAG
95506 TATGAGTGTGGAGTCGTTGCGCCACCAGGCACCAATATTTTTNNNAGATCGGAAGA
95908 TATGAGTGTGGAGTCGTTGCGTTCTGCTTTCAGGCTCTTTCAGNNNAGATCGGAAG
28802 TATGAGTGTGGAGTCGTTGCGGTCAAGAGCATCACTGAGCTTTNNNAGATCGGAAG
29034 TATGAGTGTGGAGTCGTTGCAAGGGTGTAAGAGTGTTCTTTGTNNNAGATCGGAA
29284 TATGAGTGTGGAGTCGTTGCATGCTTTGAACCTAGACGTGACCNNNAGATCGGAAG
70075 TATGAGTGTGGAGTCGTTGCTGCTACCTTGTTTGATGCTGATGANNNAGATCGGAA
71106 TATGAGTGTGGAGTCGTTGCGTCAAAGCATGTCGACTGAGACTNNNAGATCGGAAG
71130 TATGAGTGTGGAGTCGTTGCATGCTCACATCTTGGGAGTTACTNNNAGATCGGAAG
97584 TATGAGTGTGGAGTCGTTGCAGGCTCACATACGTGGAAAGGAAGNNNAGATCGGA
100398 TATGAGTGTGGAGTCGTTGCATAGCAGAATGTCACTAGACCCATNNNAGATCGGAA
105612 TATGAGTGTGGAGTCGTTGCTTCTGTTTTCTTCTGGGTTGGAANNNAGATCGGAAG
116249 TATGAGTGTGGAGTCGTTGCGTGTGTTGCACCTGGCAATTTTNNNAGATCGGAAGA
28769 TATGAGTGTGGAGTCGTTGCTGTTGTATGTTCTGTGGGTTTGACANNNAGATCGGA
28813 TATGAGTGTGGAGTCGTTGCGGACATTACAGGCCCCAAGAATTNNNAGATCGGAAG
29362 TATGAGTGTGGAGTCGTTGCATGCAGCACTTGGTAAAACCAGTNNNAGATCGGAAG
29417 TATGAGTGTGGAGTCGTTGCGTGGGTGGACATGATCGGTTTTTNNNAGATCGGAA
29512 TATGAGTGTGGAGTCGTTGCCCGAGGGAATCCTGTATGTTTGGNNNAGATCGGAA
69803 TATGAGTGTGGAGTCGTTGCTGTCCTCATATACGAACCTACACANNNAGATCGGAAG
69971 TATGAGTGTGGAGTCGTTGCTGGCTGTGTTCGTTTCTCTTTCANNNAGATCGGAAGA
71047 TATGAGTGTGGAGTCGTTGCTCATCGTGCAGTGTTTTGAGGATNNNAGATCGGAAG
94607 TATGAGTGTGGAGTCGTTGCGTGATTGCCACAAAATTCACCCANNNAGATCGGAAG
28798 TATGAGTGTGGAGTCGTTGCAGTTGTATGGCAGGTTGAATTCTNNNAGATCGGAAG
28971 TATGAGTGTGGAGTCGTTGCACCTTTTCCCTTGGATCTCTCCTNNNAGATCGGAAGA
29063 TATGAGTGTGGAGTCGTTGCGCATACATCAAGAAAGTGCAACGGNNNAGATCGGA
29093 TATGAGTGTGGAGTCGTTGCAGGAAGACAAGGAATAAAGGAAGGANNNAGATCG
29252 TATGAGTGTGGAGTCGTTGCTTCACCAGTTGGTTTCTCCACTGNNNAGATCGGAAGA
29334 TATGAGTGTGGAGTCGTTGCGTGTGATAGCTGCAGGAGTGAAANNNAGATCGGAA
29457 TATGAGTGTGGAGTCGTTGCCCATCTTTCCCACCTTGAGCTTTNNNAGATCGGAAGA
29467 TATGAGTGTGGAGTCGTTGCAGATCCTGTTGTGTGAAGGTGTTNNNAGATCGGAA

69850 TATGAGTGTGGAGTCGTTGCCCTTTTTGGGTAACCACTACCCANNNAGATCGGAAG
28751 TATGAGTGTGGAGTCGTTGCCTGGAAAAGTGGTGGCTAGGTTTNNNAGATCGGAA
28860 TATGAGTGTGGAGTCGTTGCATACCAAACTCCTGTCCTCAAGCNNNAGATCGGAAG
29378 TATGAGTGTGGAGTCGTTGCGTGTGGCCAATGCATCAGGATTTNNNAGATCGGAA
29383 TATGAGTGTGGAGTCGTTGCGCTTTCTCTCTTCCATGGGGTTTNNNAGATCGGAAG
29536 TATGAGTGTGGAGTCGTTGCGCTGGCTTCTTCTGTCCTGTAAGNNNAGATCGGAAG
29604 TATGAGTGTGGAGTCGTTGCCAACCCATAAGCCTGAACTGGAANNNAGATCGGAAG
70592 TATGAGTGTGGAGTCGTTGCTGTGTGCGTGTGTATGAAAGAAGNNNAGATCGGAA
70649 TATGAGTGTGGAGTCGTTGCTGGCTTGTGAGACAGAAAAGGAGNNNAGATCGGA
70911 TATGAGTGTGGAGTCGTTGCTTTCATCAGAAGTGAGCCAGGTGNNNAGATCGGAA
29028 TATGAGTGTGGAGTCGTTGCATGGAAGCCCTAGTGACCTCTTTNNNAGATCGGAAG
29085 TATGAGTGTGGAGTCGTTGCACACTTTCCTCCCCAAAGGAAACNNNAGATCGGAAG
29213 TATGAGTGTGGAGTCGTTGCATCCGACAATCTAGCAGACACTTNNNAGATCGGAAG
66332 TATGAGTGTGGAGTCGTTGCTCCCAGTTCCACCTTACTAGTGTNNNAGATCGGAAGA
69932 TATGAGTGTGGAGTCGTTGCCTGGCTGGTTGGTATTTGGATCANNNAGATCGGAA
70660 TATGAGTGTGGAGTCGTTGCGAGGCTGAGAAGTTCTGATGGAGNNNAGATCGGA
90941 TATGAGTGTGGAGTCGTTGCTTCGGCTATTTCCTACCCGATCTNNNAGATCGGAAGA
28760 TATGAGTGTGGAGTCGTTGCGGCTTACCATACCAATCAGCTCCNNNAGATCGGAAG
28973 TATGAGTGTGGAGTCGTTGCACAGGATGTACACTAGGAGACCTNNNAGATCGGAA
29687 TATGAGTGTGGAGTCGTTGCTAGGCATCTCCCAACTGCTTTTCNNNAGATCGGAAGA
69717 TATGAGTGTGGAGTCGTTGCCACAGTCCAGGTTGAGAAACCAANNNAGATCGGAA
69974 TATGAGTGTGGAGTCGTTGCAGGAGGGCAGCAATTTGAAAAGANNNAGATCGGA
95417 TATGAGTGTGGAGTCGTTGCCTCCAGGACAGAAATTAGCAAGCTANNNAGATCGGA
95521 TATGAGTGTGGAGTCGTTGCGATATGGTGGCACCCAGCTACTCNNNAGATCGGAAG
96568 TATGAGTGTGGAGTCGTTGCTGGGAAATTATAAAGTGTGTGTGTGTNNNAGATCG
98095 TATGAGTGTGGAGTCGTTGCGGATATATTGAGTACGGACCTCCCCNNNAGATCGGA
28815 TATGAGTGTGGAGTCGTTGCAGAACCTACTTTGTCCCTGAGCTNNNAGATCGGAAG
28870 TATGAGTGTGGAGTCGTTGCGGGGATAGGATTTCCCAAGGAGTNNNAGATCGGAA
28998 TATGAGTGTGGAGTCGTTGCTGGAGGGAGCAGAGGTGTTTATCNNNAGATCGGA
28999 TATGAGTGTGGAGTCGTTGCGGGTTCAGGTGAGTGTTGAGATCNNNAGATCGGAA
29371 TATGAGTGTGGAGTCGTTGCGGACATTTGGGGAACGCATTTTTNNNAGATCGGAA
29491 TATGAGTGTGGAGTCGTTGCCAGTGTCGTGTTCATAGCTGACTNNNAGATCGGAAG
29543 TATGAGTGTGGAGTCGTTGCATAGCATCCAAGGAGTGGTCCTCNNNAGATCGGAA
29548 TATGAGTGTGGAGTCGTTGCGCAGGTTGACTAATCTCTTTGTGCNNNAGATCGGAA
32295 TATGAGTGTGGAGTCGTTGCCCCTCATCCCCCTTCCATGATAANNNAGATCGGAAGA
69965 TATGAGTGTGGAGTCGTTGCCCATTCACCACTGACTGTCTTCANNNAGATCGGAAGA
70898 TATGAGTGTGGAGTCGTTGCGTGTGCTCCTTGTCCTCTAAGTGNNNAGATCGGAAG
29609 TATGAGTGTGGAGTCGTTGCCGGGTGGGCCGAACATATTATTTNNNAGATCGGAA
69728 TATGAGTGTGGAGTCGTTGCGCCCAAAATGACAGCTTATGCAANNNAGATCGGAAG
71125 TATGAGTGTGGAGTCGTTGCTCTCCCTCATTCATATCAGGTACTNNNAGATCGGAAG
97081 TATGAGTGTGGAGTCGTTGCCTGCCCTGTAGACTTATGACCCNNNAGATCGGAAGA
98725 TATGAGTGTGGAGTCGTTGCCTCGGGAAAGAACTGTAAAAACCCCNNNAGATCGGA
98856 TATGAGTGTGGAGTCGTTGCTGATTTTTGTATGGGCCCAGGGTNNNAGATCGGAA

99228 TATGAGTGTGGAGTCGTTGCGCAGTTAGAAAAGTGGCAGACACNNNAGATCGGAA

28824 TATGAGTGTGGAGTCGTTGCCTCTCAGTCCATCTTGTCATGGCNNNAGATCGGAAG

29190 TATGAGTGTGGAGTCGTTGCGACTTATCCATGCTTCCACTGCANNNAGATCGGAAG

29410 TATGAGTGTGGAGTCGTTGCAGGAATTTTGGGGAGTGGCTTTTNNNAGATCGGAA

29606 TATGAGTGTGGAGTCGTTGCTGTAAATAAAAGACAGTTGGCCAANNNAGATCGGA

70894 TATGAGTGTGGAGTCGTTGCTGACCCCTACAGACTGACTTCTGNNNAGATCGGAAG

91904 TATGAGTGTGGAGTCGTTGCTGGAGTTCACAGCAGGAGAAATGNNNAGATCGGAA

95291 TATGAGTGTGGAGTCGTTGCTTACATCATGATCAGCCATGGTCACNNNAGATCGGA

29356 TATGAGTGTGGAGTCGTTGCCATTTGGGGAATCAAGTGAAAGGTNNNAGATCGGA

29674 TATGAGTGTGGAGTCGTTGCATGAAGTTCGGGAGCTATTGCAGNNNAGATCGGAA

32785 TATGAGTGTGGAGTCGTTGCGCAAACACCAAGGGAAAACAACANNNAGATCGGAA

69757 TATGAGTGTGGAGTCGTTGCTGACTGCACATTCTAGCAATGAANNNAGATCGGAAG

70195 TATGAGTGTGGAGTCGTTGCCATCACGGCTCACTGAAGCTTTANNNAGATCGGAAG

91899 TATGAGTGTGGAGTCGTTGCACAGTAGCAGTGGCAAGAGAATGNNNAGATCGGAA

28923 TATGAGTGTGGAGTCGTTGCGGTAAGTCCCTGAAACTAGGTAANNNAGATCGGAA

29067 TATGAGTGTGGAGTCGTTGCCTGATAACCCTTGGTGCAGTCAGNNNAGATCGGAAG

29105 TATGAGTGTGGAGTCGTTGCTTGCCATGTATCTCCAGTGTACANNNAGATCGGAAG

29304 TATGAGTGTGGAGTCGTTGCACTCTTTTGAGCTTCTTTCATTCAAGANNNAGATCGG

29319 TATGAGTGTGGAGTCGTTGCTTGCCTTCTTTAGTCCTTCACCANNNAGATCGGAAGA

29508 TATGAGTGTGGAGTCGTTGCGTTTTCTCGTTTCCCTTTGCCTGNNNAGATCGGAAGA

29594 TATGAGTGTGGAGTCGTTGCTGAGTGCAGACGTCATTTTAGCANNNAGATCGGAA

29773 TATGAGTGTGGAGTCGTTGCTGTTCCAGGATCCTTCTGGTCTTNNNAGATCGGAAG

71245 TATGAGTGTGGAGTCGTTGCAGTACAGCTCACATGGTTCCTCTNNNAGATCGGAAG

91264 TATGAGTGTGGAGTCGTTGCGTAAGCTTTTTGTGAGCTGTCACTNNNAGATCGGAA

95476 TATGAGTGTGGAGTCGTTGCCAGAGACAATCAGAGGATCACGANNNAGATCGGAA

29445 TATGAGTGTGGAGTCGTTGCCTACACCATTCGCATCCCTTCTTNNNAGATCGGAAGA

29523 TATGAGTGTGGAGTCGTTGCAATGAACACTTGGTTCTCCAGCGNNNAGATCGGAAG

70196 TATGAGTGTGGAGTCGTTGCGCCTACACAACTGGGGCATTTTANNNAGATCGGAAG

70740 TATGAGTGTGGAGTCGTTGCTCAAGTTTCTGCCAGATTGGAGGNNNAGATCGGAA

71050 TATGAGTGTGGAGTCGTTGCACTTTGCTCCATAACATCTGGATNNNAGATCGGAAG

92954 TATGAGTGTGGAGTCGTTGCACAGCAAACATAGTAAATGGTGGCNNNAGATCGGA

93860 TATGAGTGTGGAGTCGTTGCGCCAGCTTTGAGAACTTCCTTGANNNAGATCGGAAG

95431 TATGAGTGTGGAGTCGTTGCACGGCCAAAAGTGACCTAAAGAGNNNAGATCGGAA

28850 TATGAGTGTGGAGTCGTTGCCTCCTGTTTGAGTGCTCCTGAAGNNNAGATCGGAAG

29201 TATGAGTGTGGAGTCGTTGCTAGTTGGCAGTAGTTCCCTCACANNNAGATCGGAAG

29225 TATGAGTGTGGAGTCGTTGCTGTTTTGGAAACCGGAGTGAAGTNNNAGATCGGAA

29436 TATGAGTGTGGAGTCGTTGCACGATTCCTTTGATTTGGAAACANNNAGATCGGAAG

29531 TATGAGTGTGGAGTCGTTGCTACTTGGCATGCTGCTATGACAGNNNAGATCGGAAG

29621 TATGAGTGTGGAGTCGTTGCAATAAAGCCTAGATATTGCCCTGTNNNAGATCGGAA

29719 TATGAGTGTGGAGTCGTTGCATGACAGAGGAGGGAGAGATCACNNNAGATCGGA

71228 TATGAGTGTGGAGTCGTTGCTTACTGCCACAAAATGGAGTGCTNNNAGATCGGAAG

71329 TATGAGTGTGGAGTCGTTGCACATTTCAGACAAAACCCTGTTAGTNNNAGATCGGA

29133 TATGAGTGTGGAGTCGTTGCAACTCTGGTTACCTTGGGGAAGTNNNAGATCGGAA

29145 TATGAGTGTGGAGTCGTTGCATAGGCCACTGGGGTCAAAATTGNNNAGATCGGAA
29231 TATGAGTGTGGAGTCGTTGCATTTCTCCCCACTGTCCTTTCCTNNNAGATCGGAAGA
29337 TATGAGTGTGGAGTCGTTGCGCTCCTCCCGTAAGTATGCAAAANNNAGATCGGAAG
29499 TATGAGTGTGGAGTCGTTGCCTGAGAAGCAGAGGGAGAGGAATNNNAGATCGGA
29557 TATGAGTGTGGAGTCGTTGCTGAACGGAATAGCCTGTACTTCCNNNAGATCGGAAG
29718 TATGAGTGTGGAGTCGTTGCCCCAGTACTCTTTCACAGCTGACNNNAGATCGGAAG
29761 TATGAGTGTGGAGTCGTTGCATGAGCACAGGTGTTGTTGACAANNNAGATCGGAA
70610 TATGAGTGTGGAGTCGTTGCTAGGGTTCTTGTCAGGAGACCAGNNNAGATCGGAA
71179 TATGAGTGTGGAGTCGTTGCTGTTCCCAGTGTTTTGTAGTCCTNNNAGATCGGAAG
28822 TATGAGTGTGGAGTCGTTGCGTGACCCTGGATACTTGAGACCTNNNAGATCGGAAG
29324 TATGAGTGTGGAGTCGTTGCACAACAATCTGCAGCTCAAGACANNNAGATCGGAAG
29427 TATGAGTGTGGAGTCGTTGCTACTTCTCTGACAATCCCAGCCANNNAGATCGGAAGA
29667 TATGAGTGTGGAGTCGTTGCCACCCCTTTTCACACCATCCTAGNNNAGATCGGAAGA
67216 TATGAGTGTGGAGTCGTTGCTTCACCTCTCCATGGGTCTCATTNNNAGATCGGAAGA
29206 TATGAGTGTGGAGTCGTTGCGTCAAACTTCCCCGTTATCCCAANNNAGATCGGAAGA
29713 TATGAGTGTGGAGTCGTTGCTTGAACGCATCCTACTCATTCCCNNNAGATCGGAAGA
70244 TATGAGTGTGGAGTCGTTGCAGTTGGTCTTATGTCGTCTGCACNNNAGATCGGAAG
71856 TATGAGTGTGGAGTCGTTGCTTCATTTGTTCAGCTTCTCCCCCNNNAGATCGGAAGA
91250 TATGAGTGTGGAGTCGTTGCTTTCCTACTCCAAGGTCATGACTNNNAGATCGGAAG
95421 TATGAGTGTGGAGTCGTTGCGGGTCCAATCTAGACTTTTACAAGCNNNAGATCGGA
116295 TATGAGTGTGGAGTCGTTGCGATGGCAAATGGGATTACAGATGTNNNAGATCGGA
28878 TATGAGTGTGGAGTCGTTGCTGTGTTGGGTTTAGGTTTTGGGTNNNAGATCGGAA
29024 TATGAGTGTGGAGTCGTTGCCTGAAGATTCCTGGGGTAGGGAANNNAGATCGGAA
29027 TATGAGTGTGGAGTCGTTGCAGGGGTCATGGGTGATACAGAAANNNAGATCGGA
29224 TATGAGTGTGGAGTCGTTGCCACCCATCCAATCAGTGTTCAGTNNNAGATCGGAAG
29381 TATGAGTGTGGAGTCGTTGCGTATGGATTCTGCGGTGAGCTTTNNNAGATCGGAAG
29452 TATGAGTGTGGAGTCGTTGCTCAAAAAGGCTCCTGTCTCTCCTNNNAGATCGGAAG
29682 TATGAGTGTGGAGTCGTTGCCTCTCTCCTCCTACTGTGTGAGCNNNAGATCGGAAGA
29683 TATGAGTGTGGAGTCGTTGCTTTGTAGCTTCACCAACTGCCTTNNNAGATCGGAAGA
69686 TATGAGTGTGGAGTCGTTGCAACAGCTGCAGACAGAGAACAAANNNAGATCGGAA
91269 TATGAGTGTGGAGTCGTTGCTTGTCTCATGTGTCCCTCTCACTNNNAGATCGGAAGA
28779 TATGAGTGTGGAGTCGTTGCGACCATTTGCCAATAGGAATGCCNNNAGATCGGAAG
29437 TATGAGTGTGGAGTCGTTGCTTGGAAAGGAAAAGCCAGAATCTNNNAGATCGGAA
29571 TATGAGTGTGGAGTCGTTGCTGGCAGCAACTCTGATTTTGGTANNNAGATCGGAAG
70709 TATGAGTGTGGAGTCGTTGCGACAGCTAGGAGCAAGATGAAGGNNNAGATCGGA
70727 TATGAGTGTGGAGTCGTTGCGAAGGGAAAAGGAAGAGCTCAGGNNNAGATCGGA
70944 TATGAGTGTGGAGTCGTTGCTCCCAAGATCTGTACTGCAGTTGNNNAGATCGGAAG
71041 TATGAGTGTGGAGTCGTTGCTGAAGAGACGAGGAAGCTGAGATNNNAGATCGGA
93782 TATGAGTGTGGAGTCGTTGCAGGGCAACAAGGTGGCTATACTANNNAGATCGGAA
96495 TATGAGTGTGGAGTCGTTGCCGTCAGAAAGAGGAGGAAGCCNNNAGATCGGAAG
99358 TATGAGTGTGGAGTCGTTGCAAGTAGCTGTCTCCACCCCTTTNNNAGATCGGAAGA
28965 TATGAGTGTGGAGTCGTTGCTAAGGCAGCACACTTCTACCTCTNNNAGATCGGAAG
29101 TATGAGTGTGGAGTCGTTGCCCTGGGTTCAAAGCCTAATAGCANNNAGATCGGAAG

29144 TATGAGTGTGGAGTCGTTGCACCTCTCTGAATTTATGTCTTCATTTGNNNAGATCGG

29394 TATGAGTGTGGAGTCGTTGCGGGTTCTGTGAAGTTCAGGGTTTNNNAGATCGGAA

70367 TATGAGTGTGGAGTCGTTGCTGGGTTATAAGGGTGCTTTCTCCNNNAGATCGGAAG

71002 TATGAGTGTGGAGTCGTTGCTTTTCTTCACTTGCTCTGCCCATNNNAGATCGGAAGA

71034 TATGAGTGTGGAGTCGTTGCATGTAGGTCCACAGACACCAGATNNNAGATCGGAAG

93118 TATGAGTGTGGAGTCGTTGCCCTTTTGTCAGAAATCATGCAAGCNNNAGATCGGAA

95474 TATGAGTGTGGAGTCGTTGCTCATTTGTGTAAAGGGGATGGGGAANNNAGATCGG

99124 TATGAGTGTGGAGTCGTTGCTTACTGGTTTCTACTTCCCAGGGNNNAGATCGGAAG

28987 TATGAGTGTGGAGTCGTTGCTTTGAAGGCATGGTTAGCTGAGGNNNAGATCGGAA

29351 TATGAGTGTGGAGTCGTTGCCTGGTCCATGAGAAATGACAGTGTNNNAGATCGGA

29365 TATGAGTGTGGAGTCGTTGCGCGTCTTTTGCTCTTGGTTTTCTNNNAGATCGGAAGA

29386 TATGAGTGTGGAGTCGTTGCTCTTTCCTAGGGATCCTTGCTTTNNNAGATCGGAAGA

71133 TATGAGTGTGGAGTCGTTGCTGAGGCCAAAGAAAGAACAACCTNNNAGATCGGAA

94686 TATGAGTGTGGAGTCGTTGCAAACTCCTCCAAGAAACTGGACANNNAGATCGGAAG

95359 TATGAGTGTGGAGTCGTTGCGACTAAACTCGACGAAGGCGAANNNAGATCGGAAG

96368 TATGAGTGTGGAGTCGTTGCGAGTGACTACCTCAACATTATCACAGNNNAGATCGG

97242 TATGAGTGTGGAGTCGTTGCGACAGAGCCGTTCAGGAGAAANNNAGATCGGAAGA

97352 TATGAGTGTGGAGTCGTTGCCTTGGTCTCTTCGCACTGGACANNNAGATCGGAAGA

98498 TATGAGTGTGGAGTCGTTGCTCTCTCAGCTCTCGGAAGGACCNNNAGATCGGAAGA

28783 TATGAGTGTGGAGTCGTTGCGGCAAGAAAAATGCCCCTTTTGTNNNAGATCGGAAG

29237 TATGAGTGTGGAGTCGTTGCCTTACAACCCAGCAAGTGTCACTNNNAGATCGGAAG

29341 TATGAGTGTGGAGTCGTTGCGGGTTCTTGCCCACCTTTACTTTNNNAGATCGGAAGA

29395 TATGAGTGTGGAGTCGTTGCACCTCTCACACGGTTTCTGTTTTNNNAGATCGGAAGA

29506 TATGAGTGTGGAGTCGTTGCAGGACACTTCATCCATGTTGCTGNNNAGATCGGAAG

70715 TATGAGTGTGGAGTCGTTGCAAGCACGCATATAACAGGACAGGNNNAGATCGGAA

70817 TATGAGTGTGGAGTCGTTGCAACCTTCACTGCAATTTTGCTGGNNNAGATCGGAAG

91165 TATGAGTGTGGAGTCGTTGCTTATCAGAAGACAGAGGCAGCCTNNNAGATCGGAA

95261 TATGAGTGTGGAGTCGTTGCTACAGGTGTGAGACACTGCGAGCNNNAGATCGGAA

28968 TATGAGTGTGGAGTCGTTGCTCCCCAATTTATGGTAGCTTGCTNNNAGATCGGAAG

29315 TATGAGTGTGGAGTCGTTGCCTTTCATTCCAAGTTGGTTGCCANNNAGATCGGAAG

100550 TATGAGTGTGGAGTCGTTGCCCTGGCCATAGCCTATTTGAAAGNNNAGATCGGAA

103042 TATGAGTGTGGAGTCGTTGCCCTGGCCTCATTTTACGTTTTGANNNAGATCGGAAG

116757 TATGAGTGTGGAGTCGTTGCCACCACCACTCGTATTACACTGTGNNNAGATCGGAA

28733 TATGAGTGTGGAGTCGTTGCTCGTGACCATGTGTTAGGTAGGTNNNAGATCGGAA

29078 TATGAGTGTGGAGTCGTTGCCAGTTGCAGAGGAATCCACTACCNNNAGATCGGAAG

29117 TATGAGTGTGGAGTCGTTGCGCTCAGACCTTCAGAACAATGTTNNNAGATCGGAAG

29139 TATGAGTGTGGAGTCGTTGCACACCATGTGCTTGCAAGATACTNNNAGATCGGAAG

29155 TATGAGTGTGGAGTCGTTGCTACATTCCTACACCCTCCACTCGNNNAGATCGGAAGA

35116 TATGAGTGTGGAGTCGTTGCAGGTTCTTTCCAATGCTTTTCCANNNAGATCGGAAGA

54863 TATGAGTGTGGAGTCGTTGCATAATCTCCAGGCCAGTGAGTGGNNNAGATCGGAA

69784 TATGAGTGTGGAGTCGTTGCAGAGAAAGCAGAAGCTTGGAAACANNNAGATCGGA

70635 TATGAGTGTGGAGTCGTTGCCGCAGACCCTGCATATAGTCAGNNNAGATCGGAAGA

29380 TATGAGTGTGGAGTCGTTGCTGTCAGTGTATTGCTTCTGCTTTNNNAGATCGGAAG

29431 TATGAGTGTGGAGTCGTTGCGACAGACAAAGAGCCCAAGAGGNNNAGATCGGAAG
93574 TATGAGTGTGGAGTCGTTGCCCAGCCCAAGAAATACAACACACNNNAGATCGGAAG
104526 TATGAGTGTGGAGTCGTTGCTGACTGGGAAGAGACTATGATGGAANNNAGATCGG
121950 TATGAGTGTGGAGTCGTTGCGGAGTGGCTCTTGCTATCTCAGGGNNNAGATCGGA
28774 TATGAGTGTGGAGTCGTTGCTGGGTTAGCTATCTGTGGAACCANNNAGATCGGAA
28972 TATGAGTGTGGAGTCGTTGCTCTGAAGCATGCAAAGTTCATCCTNNNAGATCGGAA
29517 TATGAGTGTGGAGTCGTTGCGGATGTTCAGAGCAGTGTATGGGNNNAGATCGGAA
29525 TATGAGTGTGGAGTCGTTGCAACCCTCGGTAACTGTGAATGAGNNNAGATCGGAA
29740 TATGAGTGTGGAGTCGTTGCGAAATCTTGATGGCCTCATTGCANNNAGATCGGAAG
70251 TATGAGTGTGGAGTCGTTGCAAAGACATGCATCTGTCCCTCACNNNAGATCGGAAG
90264 TATGAGTGTGGAGTCGTTGCAGACAAACAAGTGTAGGTGCCTTNNNAGATCGGAA
95338 TATGAGTGTGGAGTCGTTGCCGCTCTGTAGCCCTGTAGCCNNNAGATCGGAAGAGC
96137 TATGAGTGTGGAGTCGTTGCGGCTGTGGTTGTAGACATCTGAANNNAGATCGGAA
96639 TATGAGTGTGGAGTCGTTGCCAGCACATCTGGCAGAAAAATTCTTNNNAGATCGGA
28785 TATGAGTGTGGAGTCGTTGCTGAAGTTACGACGTAGAGCCTTTNNNAGATCGGAA
28848 TATGAGTGTGGAGTCGTTGCAGGAGGATAAGATGACATCCGGGNNNAGATCGGA
29126 TATGAGTGTGGAGTCGTTGCGCCCTAGAGAAGTCTGCATTTGTNNNAGATCGGAAG
29385 TATGAGTGTGGAGTCGTTGCACTGCTCTTTCTCACAACTCTTTNNNAGATCGGAAGA
29450 TATGAGTGTGGAGTCGTTGCAGACACTAGGCAGTCATACGGATNNNAGATCGGAA
70039 TATGAGTGTGGAGTCGTTGCAGATGTTACTGTGTTTTGGGGGANNNAGATCGGAA
70536 TATGAGTGTGGAGTCGTTGCTCTTCCAGAACCTTCCCTCTCTCNNNAGATCGGAAGA
71144 TATGAGTGTGGAGTCGTTGCTAAGTTTCTGACTGTCCAGCCCTNNNAGATCGGAAG
91317 TATGAGTGTGGAGTCGTTGCTCCTTCAAACAAATTTCAAAGCAACTNNNAGATCGGA
95432 TATGAGTGTGGAGTCGTTGCTTTATTGCTGGTGGGGACTTTGCNNNAGATCGGAAG
98633 TATGAGTGTGGAGTCGTTGCGTTTGTTGCTGTTGACTTTGAGGTNNNAGATCGGAA
29187 TATGAGTGTGGAGTCGTTGCCTGTTGATAGCAATGTCAATGTCANNNAGATCGGAA
29204 TATGAGTGTGGAGTCGTTGCTCTGGCCTAACACCAAAAGAGAANNNAGATCGGAAG
29757 TATGAGTGTGGAGTCGTTGCTGGAGGGAACAACAAAGAAGGAANNNAGATCGGA
69692 TATGAGTGTGGAGTCGTTGCTCTCTGGTTATCAGACTGGCAAANNNAGATCGGAAG
71859 TATGAGTGTGGAGTCGTTGCCACATTCCTGGCCTAGGTTTCTCNNNAGATCGGAAGA
71860 TATGAGTGTGGAGTCGTTGCTTACTTCCAGGCCTGTTCTTGTGNNNAGATCGGAAGA
116576 TATGAGTGTGGAGTCGTTGCATGGATTGGTTCTGTTTCTCTGAAANNNAGATCGGA
29051 TATGAGTGTGGAGTCGTTGCCTGTAATGAACGTGAGAGTGCATNNNAGATCGGAA
29402 TATGAGTGTGGAGTCGTTGCTGCGTGTTTGTTTTGCCTTCTTTNNNAGATCGGAAGA
90597 TATGAGTGTGGAGTCGTTGCTAAGGACATTGCAGCCAAATGGTNNNAGATCGGAA
95387 TATGAGTGTGGAGTCGTTGCAGGAGTAGGAGGTAGGGAGGAANNNAGATCGGA
95424 TATGAGTGTGGAGTCGTTGCAGTCAGATTGTCCCTCTTAGTTGNNNAGATCGGAAG
95630 TATGAGTGTGGAGTCGTTGCTCAGCCTAGGAGTTCATGGTTGCNNNAGATCGGAAG
96384 TATGAGTGTGGAGTCGTTGCGCCATCTTCATTTCTGGCTGCAGGAATNNNAGATCG
96996 TATGAGTGTGGAGTCGTTGCCCCCAGGTTGTGTAGATTCCCANNNAGATCGGAAGA
97317 TATGAGTGTGGAGTCGTTGCACAGGTGAATCAGTTGGACCTGNNNAGATCGGAAG
97689 TATGAGTGTGGAGTCGTTGCCTTGTCAACTGCATTTCCTCTTGCNNNAGATCGGAAG
97761 TATGAGTGTGGAGTCGTTGCTCCCCTTTTCCTTGGCATTTCTCNNNAGATCGGAAGA

| | |
|---|---|
| 97931 | TATGAGTGTGGAGTCGTTGCTGGAGTGTAGTGATTTGATCCCTTCANNNAGATCGG |
| 102297 | TATGAGTGTGGAGTCGTTGCCACATCTCCCTTTTCTTCATTAGGGNNNAGATCGGAA |
| 29292 | TATGAGTGTGGAGTCGTTGCCATACCTCACCTTGCCTCCTCTANNNAGATCGGAAGA |
| 29561 | TATGAGTGTGGAGTCGTTGCTAAGCACACTTTGTTGCTAGGCCNNNAGATCGGAAG |
| 29616 | TATGAGTGTGGAGTCGTTGCTCGCATCCCCACACTATTTACTTNNNAGATCGGAAGA |
| 29717 | TATGAGTGTGGAGTCGTTGCGAATGTGTTTTGCCAAGGCTGACNNNAGATCGGAA |
| 70034 | TATGAGTGTGGAGTCGTTGCGAAAACATGTGTGTGGTCAGGGANNNAGATCGGA |
| 71863 | TATGAGTGTGGAGTCGTTGCGGGAGGTGGCTAGACTAGTTGATNNNAGATCGGA |
| 96109 | TATGAGTGTGGAGTCGTTGCGTGCTAAAAGTGGAACAGTCCAANNNAGATCGGAA |
| 99121 | TATGAGTGTGGAGTCGTTGCATAGCCAGTTGGAAAGGGTGGAAANNNAGATCGG |
| 99219 | TATGAGTGTGGAGTCGTTGCAAACAAAAGATTGGCTAGGTGTGGNNNAGATCGGA |
| 100379 | TATGAGTGTGGAGTCGTTGCGTAAAATGTGCATTCCCTTTGCAGNNNAGATCGGAA |
| 28781 | TATGAGTGTGGAGTCGTTGCTTTGATGTTGAGGTGACCAAGCANNNAGATCGGAA |
| 29769 | TATGAGTGTGGAGTCGTTGCACACAATCAAAGGACTGCCTGTTNNNAGATCGGAAG |
| 70063 | TATGAGTGTGGAGTCGTTGCTGACCAAAATTAGCTGTGGTTGGANNNAGATCGGA |
| 70100 | TATGAGTGTGGAGTCGTTGCTGCTGTTTGCCTAAGAACTGTGANNNAGATCGGAAG |
| 70745 | TATGAGTGTGGAGTCGTTGCCACGCAAGCACATGAAATGTAGGNNNAGATCGGAA |
| 70937 | TATGAGTGTGGAGTCGTTGCATCTTACGCAGTTATGGGGCTTGNNNAGATCGGAA |
| 95429 | TATGAGTGTGGAGTCGTTGCACCATGTCACTGGGCTTCCATTTNNNAGATCGGAAG |
| 96927 | TATGAGTGTGGAGTCGTTGCATCGGGGCATAGGGATAAGAGGNNNAGATCGGAA |
| 29465 | TATGAGTGTGGAGTCGTTGCTAGGATCATGGGGTGGTCACTTTNNNAGATCGGAA |
| 29551 | TATGAGTGTGGAGTCGTTGCGGACAATAATTGGGTTACTTGCTGCNNNAGATCGG |
| 71396 | TATGAGTGTGGAGTCGTTGCTCGCATACTGGAATCTTGCATGTNNNAGATCGGAAG |
| 101761 | TATGAGTGTGGAGTCGTTGCGCTGATACTTGCTACTGCAGTTNNNAGATCGGAAGA |
| 29219 | TATGAGTGTGGAGTCGTTGCCACGTATCTAGCATGATTTTAGGGTNNNAGATCGGA |
| 29446 | TATGAGTGTGGAGTCGTTGCATCTGGAAGTGAAAGCCATCGTGNNNAGATCGGAA |
| 29597 | TATGAGTGTGGAGTCGTTGCATCCCCAAATACAGATGGGTCCANNNAGATCGGAAG |
| 70018 | TATGAGTGTGGAGTCGTTGCATGGCTCTGGTGCATCTCTTAGANNNAGATCGGAAG |
| 71029 | TATGAGTGTGGAGTCGTTGCCTGGTACGTGTGAACCTGTTCATNNNAGATCGGAAG |
| 93913 | TATGAGTGTGGAGTCGTTGCGGAGAATGACAAAAGACAGCTGANNNAGATCGGA |
| 95311 | TATGAGTGTGGAGTCGTTGCAAGTTCCTGTAGCGCCGGAAAACNNNAGATCGGAA |
| 95448 | TATGAGTGTGGAGTCGTTGCTAAAGCCTCTGGGAGGTAATGCGNNNAGATCGGAA |
| 97524 | TATGAGTGTGGAGTCGTTGCTGCTCCAAACCTTTCTGCTCACTNNNAGATCGGAAGA |
| 98814 | TATGAGTGTGGAGTCGTTGCCAAGATGGCAGTGCGATGACCGNNNAGATCGGAAG |
| 100305 | TATGAGTGTGGAGTCGTTGCGCTAGGGGGAATCCTAGGGTAGANNNAGATCGGA |
| 121947 | TATGAGTGTGGAGTCGTTGCTTCTATTGTGAGTGATCTCTGGTGANNNAGATCGGA |
| 91157 | TATGAGTGTGGAGTCGTTGCAGTCCAGTGTAAGTACTTCGCCTNNNAGATCGGAAG |
| 95334 | TATGAGTGTGGAGTCGTTGCGTGCCTGGAGACAGAACCATTTNNNAGATCGGAAG |
| 95407 | TATGAGTGTGGAGTCGTTGCGGTCCAGCCCCAATCCTTTGAGNNNAGATCGGAAGA |
| 95449 | TATGAGTGTGGAGTCGTTGCCTCCACCCCAGGTTATGACAGACNNNAGATCGGAAG |
| 96110 | TATGAGTGTGGAGTCGTTGCAGATTCCAATAGACCTTGAGGCCCNNNAGATCGGAA |
| 97742 | TATGAGTGTGGAGTCGTTGCCTTTTTGCATCTCCTGACCACAGNNNAGATCGGAAGA |
| 102288 | TATGAGTGTGGAGTCGTTGCTAGTGATGAGATTGCTGGGTCATATNNNAGATCGG |

28731 TATGAGTGTGGAGTCGTTGCCACGCACATGAATATTGGCCTTCNNNAGATCGGAAG
29312 TATGAGTGTGGAGTCGTTGCTCAGAGTTTCAAGTCACAGAAATCANNNAGATCGGA
29546 TATGAGTGTGGAGTCGTTGCATGACAACAAAAACCGTGCCATCNNNAGATCGGAAG
70115 TATGAGTGTGGAGTCGTTGCTGCCCTTACAATATTTTTCCTTTTGANNNAGATCGGA
95420 TATGAGTGTGGAGTCGTTGCCCTAACTAACAGACTGTACTCCCACNNNAGATCGGAA
97803 TATGAGTGTGGAGTCGTTGCGACTGACATTGGACACGGAACAAGNNNAGATCGGA
99503 TATGAGTGTGGAGTCGTTGCTCCCAGCCCTGAGCAAGACATNNNAGATCGGAAGAG
121952 TATGAGTGTGGAGTCGTTGCGCCATTATGTGTAGGGACCATTCTNNNAGATCGGAA
28788 TATGAGTGTGGAGTCGTTGCTGAGTGTTTACTAGGCACTGTGCNNNAGATCGGAAG
29264 TATGAGTGTGGAGTCGTTGCGGTAGGCGCCATTTCCTAGAAGNNNAGATCGGAAG
29524 TATGAGTGTGGAGTCGTTGCTACATTCATAGTGAGGACCAGCGNNNAGATCGGAA
29669 TATGAGTGTGGAGTCGTTGCCCTGCTGATAACCCAAAAGGGAGNNNAGATCGGAA
29696 TATGAGTGTGGAGTCGTTGCTGTTCCAGCTTGCTCTCAGAATCNNNAGATCGGAAG
71876 TATGAGTGTGGAGTCGTTGCTCAAGAGATGCCAAGTACTTCAGANNNAGATCGGA
91856 TATGAGTGTGGAGTCGTTGCAAGACTGTGTGTATGTGCATATATGNNNAGATCGG
95395 TATGAGTGTGGAGTCGTTGCATTGTTGTATCCTAAGTGTCCAGTAGNNNAGATCGG
95747 TATGAGTGTGGAGTCGTTGCCACTCCCATTAGCCTGTGGTCCCNNNAGATCGGAAG
95809 TATGAGTGTGGAGTCGTTGCTTTTCCCCAACCCCACTAATTCCNNNAGATCGGAAGA
99674 TATGAGTGTGGAGTCGTTGCATTCCTAGACAGTTGTAGCAGGGTNNNAGATCGGA
99886 TATGAGTGTGGAGTCGTTGCGGTGAACATCCACAGGGAAACAGNNNAGATCGGAA
100138 TATGAGTGTGGAGTCGTTGCAAAAGGGAGCCCAAACATCCCNNNAGATCGGAAGA
116329 TATGAGTGTGGAGTCGTTGCCTTTTTGGAGCTCCCTTGTATGATNNNAGATCGGAA
28758 TATGAGTGTGGAGTCGTTGCTTTGAGTTTGAGGATAGCCAACTNNNAGATCGGAAG
29002 TATGAGTGTGGAGTCGTTGCAGTTTCTCCATCCTCCTTTTCAGCNNNAGATCGGAAG
29286 TATGAGTGTGGAGTCGTTGCTGGTACATCAAAAGAATTGCACACCNNNAGATCGGA
29513 TATGAGTGTGGAGTCGTTGCTTGAGACATTACCACCACCTTGGNNNAGATCGGAAG
29564 TATGAGTGTGGAGTCGTTGCAGCAGATCAGTGCATTCTTTCCCNNNAGATCGGAAG
69827 TATGAGTGTGGAGTCGTTGCTGCTCTTCACAATGCAAAATGACANNNAGATCGGAA
71132 TATGAGTGTGGAGTCGTTGCGACAGATGCATAAGCTTTTGAAACCTNNNAGATCGG
95269 TATGAGTGTGGAGTCGTTGCGGAAAATAATAGCAGGCATGGTGGNNNAGATCGGA
95376 TATGAGTGTGGAGTCGTTGCAGCCTCCCGAGTATTAGTAGTTGNNNAGATCGGAAG
95710 TATGAGTGTGGAGTCGTTGCGCTACCTGGAACTATTGCAGCCNNNAGATCGGAAGA
96012 TATGAGTGTGGAGTCGTTGCGGCTCTCTTTAGGCTGGGTCTCNNNAGATCGGAAGA
97626 TATGAGTGTGGAGTCGTTGCCCTCTCAGTTCTCTCATGCTGCNNNAGATCGGAAGAG
97821 TATGAGTGTGGAGTCGTTGCAGTCATTGGGGAGAATCTGAGAGTNNNAGATCGGA
100281 TATGAGTGTGGAGTCGTTGCCCACTCCAGTTTCTCTGAGCATTANNNAGATCGGAAG
29587 TATGAGTGTGGAGTCGTTGCTTTTCCCTTTGTTTCCTGCCTCANNNAGATCGGAAGA
92142 TATGAGTGTGGAGTCGTTGCCCTGAAGCCAGCACCATATTAGGNNNAGATCGGAAG
94725 TATGAGTGTGGAGTCGTTGCGCTGTAAGGTACTAATGGGCACANNNAGATCGGAA
95378 TATGAGTGTGGAGTCGTTGCCTCTGAATGGTCTTTCCCCCACNNNAGATCGGAAGAG
95460 TATGAGTGTGGAGTCGTTGCGACAGGGTTTCAGTTTGTCACTCATNNNAGATCGGA
99748 TATGAGTGTGGAGTCGTTGCCAAGAGGAAGAGGGGGCTTTAGAANNNAGATCGG
99987 TATGAGTGTGGAGTCGTTGCACTCTCATTGCTGGCCTTGAAGANNNAGATCGGAAG

| | |
|---|---|
| 121990 | TATGAGTGTGGAGTCGTTGCGGAAAATCACAGGCTGAGTGAAAGNNNAGATCGGA |
| 28745 | TATGAGTGTGGAGTCGTTGCTAAGCACAGGAGGGAATCTCTCGNNNAGATCGGAA |
| 29489 | TATGAGTGTGGAGTCGTTGCTGTCCTTCAAAGTCTAGTCTCCCTNNNAGATCGGAAG |
| 29559 | TATGAGTGTGGAGTCGTTGCCGTGGTAAGCACAATTTCTTGCCNNNAGATCGGAAG |
| 70252 | TATGAGTGTGGAGTCGTTGCGATAGTGTTCCCTTGCCAGTCACNNNAGATCGGAAG |
| 71207 | TATGAGTGTGGAGTCGTTGCAAAGGAAAGATCACAAGCAGGCTNNNAGATCGGAA |
| 71858 | TATGAGTGTGGAGTCGTTGCCTGTTCTCCTTCCCTGTCCCTAANNNAGATCGGAAGA |
| 93649 | TATGAGTGTGGAGTCGTTGCGTGTGGAGAAAGAGGCCAGTTTANNNAGATCGGAA |
| 98895 | TATGAGTGTGGAGTCGTTGCCTTGAACTTCTCCCAGAAAATGGTTNNNAGATCGGA |
| 103086 | TATGAGTGTGGAGTCGTTGCAATGCAGGTTGTTGAGGATTTGANNNAGATCGGAA |
| 29729 | TATGAGTGTGGAGTCGTTGCTAGTGCTTTTGCTGCCCAACATANNNAGATCGGAAG |
| 95369 | TATGAGTGTGGAGTCGTTGCCATTTTAGACACATGACTTCCGCNNNAGATCGGAAG |
| 29504 | TATGAGTGTGGAGTCGTTGCGTTATGCTCTGTGTGGATCTGTGNNNAGATCGGAA |
| 90384 | TATGAGTGTGGAGTCGTTGCGCCTAGTTCTCCTCTGTTCTTTGTNNNAGATCGGAAG |
| 105600 | TATGAGTGTGGAGTCGTTGCCCTCCCAGATTCTCTCCCACCTTNNNAGATCGGAAGA |
| 29666 | TATGAGTGTGGAGTCGTTGCACCACACACAATAAGCAACTAACGNNNAGATCGGAA |
| 95399 | TATGAGTGTGGAGTCGTTGCTAACACAGGACACAGCACCCAAANNNAGATCGGAAG |
| 95451 | TATGAGTGTGGAGTCGTTGCCTCAGCCCAACCTTGCTCCAAGANNNAGATCGGAAG |
| 98311 | TATGAGTGTGGAGTCGTTGCAGGTAAACGTGCTATTCTGCCTGNNNAGATCGGAAG |
| 102915 | TATGAGTGTGGAGTCGTTGCGATGGTGGGGCTATGGCTAATTTTTANNNAGATCG |
| 29338 | TATGAGTGTGGAGTCGTTGCATAGCCACTGTGCATGGAGTTTTNNNAGATCGGAAG |
| 29715 | TATGAGTGTGGAGTCGTTGCTGCTAAACAGACACTTGCAGACCNNNAGATCGGAAG |
| 98098 | TATGAGTGTGGAGTCGTTGCGGATCTCAGGTTTAGAACTCCTAGTCNNNAGATCGG |
| 99191 | TATGAGTGTGGAGTCGTTGCTAGCCTTTCTGATAGCTCTGTGGTAANNNAGATCGG |
| 28832 | TATGAGTGTGGAGTCGTTGCCGAGCAATTTCCATGCAACAGTTNNNAGATCGGAAG |
| 29107 | TATGAGTGTGGAGTCGTTGCAGGAATAAGATGTTTTAGGAGGCACANNNAGATCG |
| 29403 | TATGAGTGTGGAGTCGTTGCATTTGCCCAGGACTGATGGTTTTNNNAGATCGGAAG |
| 69966 | TATGAGTGTGGAGTCGTTGCTAGATCATTTCTGTTACATGTTCTTCANNNAGATCGG |
| 70054 | TATGAGTGTGGAGTCGTTGCCTTTGCATGCCTACTCCATTGGANNNAGATCGGAAG |
| 93362 | TATGAGTGTGGAGTCGTTGCTACCCGTTTCATTTACCCAACCCNNNAGATCGGAAGA |
| 94153 | TATGAGTGTGGAGTCGTTGCACTTACCTAGCAACCCACAGAGANNNAGATCGGAAG |
| 98316 | TATGAGTGTGGAGTCGTTGCGACCAGAATGGATCACAGATGATCGNNNAGATCGG |
| 116419 | TATGAGTGTGGAGTCGTTGCAGCCATCTGAAGCTGGGGTTGNNNAGATCGGAAGA |
| 29439 | TATGAGTGTGGAGTCGTTGCGGTGTAATCATGGGTGCATGGTTNNNAGATCGGAA |
| 94218 | TATGAGTGTGGAGTCGTTGCTTTCAGTTGTGTTCTGGGCAAGANNNAGATCGGAA |
| 95478 | TATGAGTGTGGAGTCGTTGCTAAGGATGAGTGGGGTCACTCANNNAGATCGGAAG |
| 98681 | TATGAGTGTGGAGTCGTTGCCTCTGCAGCCTATGGAGCTGANNNAGATCGGAAGA |
| 116047 | TATGAGTGTGGAGTCGTTGCTTTTGACAGCCTCCTTCTGCTGNNNAGATCGGAAGA |
| 119628 | TATGAGTGTGGAGTCGTTGCCAAGCGATACTCCTATCCCAGGNNNAGATCGGAAGA |
| 28826 | TATGAGTGTGGAGTCGTTGCGCAGTAGGGCCCATTAGTCTAGANNNAGATCGGAA |
| 94465 | TATGAGTGTGGAGTCGTTGCGCTGCAATGAATACAAGAGAGCANNNAGATCGGAA |
| 95416 | TATGAGTGTGGAGTCGTTGCGGGTGGGAGAGGCTCAGGAATGNNNAGATCGGAA |
| 99232 | TATGAGTGTGGAGTCGTTGCTATCCATTAGCCAATTCTGTGAGCTNNNAGATCGGA |

| | |
|---|---|
| 116050 | TATGAGTGTGGAGTCGTTGCATTCACCGAGTGTTTTGTCAGCTCNNNAGATCGGAA |
| 118763 | TATGAGTGTGGAGTCGTTGCTCTCCTGTGTTGGTCCGGATGATATCANNNAGATCG |
| 71862 | TATGAGTGTGGAGTCGTTGCCCTGAGCACTTACACTCACTGTCNNNAGATCGGAAG |
| 95445 | TATGAGTGTGGAGTCGTTGCTCGGACAGTAACCCTAAGGCTAANNNAGATCGGAA |
| 98481 | TATGAGTGTGGAGTCGTTGCCATAACCTCAATTGCCAGCCATANNNAGATCGGAAG |
| 28889 | TATGAGTGTGGAGTCGTTGCAGCACAGAAGCATCTCCCAAAAANNNAGATCGGAAG |
| 95434 | TATGAGTGTGGAGTCGTTGCTTCTTCTGGCAATAGCATCATCATTGNNNAGATCGG |
| 118795 | TATGAGTGTGGAGTCGTTGCTTTCAGAGTGTGTTGTGCCCCTNNNAGATCGGAAGA |
| 121946 | TATGAGTGTGGAGTCGTTGCGGTCTCACTCTGTCCATGCTGTACNNNAGATCGGAA |
| 28817 | TATGAGTGTGGAGTCGTTGCGCGGGAGGAGATGCTGTATATTTNNNAGATCGGAA |
| 29448 | TATGAGTGTGGAGTCGTTGCGTCGAAACGTGATGCGATGAATTNNNAGATCGGAA |
| 70631 | TATGAGTGTGGAGTCGTTGCCCAACTTCTGCTTTTGACTGCAGNNNAGATCGGAAG |
| 71857 | TATGAGTGTGGAGTCGTTGCAACCAGATGCTGTCATAGGCAAGNNNAGATCGGAA |
| 96582 | TATGAGTGTGGAGTCGTTGCGGGTCCTACAGAACAAAGACAAGGNNNAGATCGGA |
| 97592 | TATGAGTGTGGAGTCGTTGCGTCAAGTAGCGGAAAATGTCTTCANNNAGATCGGA |
| 98015 | TATGAGTGTGGAGTCGTTGCCAAAATAAAAGCCAAAGTGCACTACCNNNAGATCGG |
| 29370 | TATGAGTGTGGAGTCGTTGCTCAGCATATTGCCAGAAAAGGCTNNNAGATCGGAA |
| 71024 | TATGAGTGTGGAGTCGTTGCGCTCAGGTGGTCGAGTATCTCATNNNAGATCGGAA |
| 94623 | TATGAGTGTGGAGTCGTTGCTCCAGCATACAATGTGTTTCACCANNNAGATCGGAA |
| 98047 | TATGAGTGTGGAGTCGTTGCCCCTCCTCAGGACACACCTAAACNNNAGATCGGAAG |
| 98058 | TATGAGTGTGGAGTCGTTGCACGGACCAGGGATTAGGAAAAACNNNAGATCGGAA |
| 99178 | TATGAGTGTGGAGTCGTTGCCAGAAGATAGTGTAGTGAGGCTACNNNAGATCGGA |
| 29226 | TATGAGTGTGGAGTCGTTGCTCAGTAGGAACGTTATCGGCTCTNNNAGATCGGAAG |
| 69902 | TATGAGTGTGGAGTCGTTGCTGATTATGCGTGATATGCAGGCANNNAGATCGGAA |
| 71313 | TATGAGTGTGGAGTCGTTGCGAGAAAGCCACTTATCAGCGAGTNNNAGATCGGAA |
| 95422 | TATGAGTGTGGAGTCGTTGCTCTGTCTTCCAGTAGCTGGGATTNNNAGATCGGAAG |
| 98192 | TATGAGTGTGGAGTCGTTGCTTGGTTATTGCTTGTATCCCCAANNNAGATCGGAAG |
| 98810 | TATGAGTGTGGAGTCGTTGCCAGTTGCTCGAAGGGGCTTCCCNNNAGATCGGAAG |
| 99523 | TATGAGTGTGGAGTCGTTGCAGTTATTTCACAGGACTGTTGCTTNNNAGATCGGAA |
| 105672 | TATGAGTGTGGAGTCGTTGCGTGATAGAACAGCTCTGCATCTTNNNAGATCGGAAG |
| 118730 | TATGAGTGTGGAGTCGTTGCATACAAATGTTAGCAGGCTATAAACANNNAGATCGG |
| 29643 | TATGAGTGTGGAGTCGTTGCAAACATCAGGAGTTGATGTCCCTNNNAGATCGGAAG |
| 121984 | TATGAGTGTGGAGTCGTTGCTTCCCCTCAGGATGTGTTTGTATAGNNNAGATCGGA |
| 29547 | TATGAGTGTGGAGTCGTTGCGCCAGGAAGGAAAAAGTTCTTGCNNNAGATCGGAA |
| 71875 | TATGAGTGTGGAGTCGTTGCTCTGTCGCAATAAGGCATCCAAANNNAGATCGGAAG |
| 97587 | TATGAGTGTGGAGTCGTTGCGGTGGTCCTGCTTGTGTCATNNNAGATCGGAAGAG |
| 105629 | TATGAGTGTGGAGTCGTTGCTGATGGGAAGCTCAACTAGCATGGNNNAGATCGGA |
| 121949 | TATGAGTGTGGAGTCGTTGCTGGCTATGGGGAAATAACTGAGCNNNAGATCGGAA |
| 29770 | TATGAGTGTGGAGTCGTTGCTCAGGGTAGGTCAAGGAACTGTTNNNAGATCGGAA |
| 94975 | TATGAGTGTGGAGTCGTTGCAACTGCAGAATGACAGAAGCCAANNNAGATCGGAA |
| 95601 | TATGAGTGTGGAGTCGTTGCATTTGTAACAGTGATTGGCCCATACNNNAGATCGGA |
| 98281 | TATGAGTGTGGAGTCGTTGCGCAAGATGTTCTCGGGCTTGATNNNAGATCGGAAG |
| 98631 | TATGAGTGTGGAGTCGTTGCCACAAACTAAGTGGCTCCAACAANNNAGATCGGAAG |

100311 TATGAGTGTGGAGTCGTTGCGTAACCACACAACTCACAGATCCANNNAGATCGGAA
116132 TATGAGTGTGGAGTCGTTGCTCTCTAAAAAGGATTTGTGAGGCTAGTNNNAGATCG
28786 TATGAGTGTGGAGTCGTTGCACGCCTCAAGTCTTTGTCCTTTTNNNAGATCGGAAGA
28807 TATGAGTGTGGAGTCGTTGCACTTGATCCCTCTCCTCCATCTTNNNAGATCGGAAGA
71312 TATGAGTGTGGAGTCGTTGCAGGCTGGGAAAGATATACCGAGTNNNAGATCGGAA
95753 TATGAGTGTGGAGTCGTTGCATAGGCAGCAGGTATTAGTGACACNNNAGATCGGA
100011 TATGAGTGTGGAGTCGTTGCATCGTACCTGAGTTCCTCTCTGGNNNAGATCGGAAG
29703 TATGAGTGTGGAGTCGTTGCAGGATTTGGTGCTATGGCAAAGCNNNAGATCGGAA
95377 TATGAGTGTGGAGTCGTTGCTCCCCCATTTTGATATAGCATAGCCTNNNAGATCGGA
95758 TATGAGTGTGGAGTCGTTGCGGGACAACAGTTGGACACCATGNNNAGATCGGAAG
99710 TATGAGTGTGGAGTCGTTGCATTTTAGAAACCACTGCTCTAAAGAACNNNAGATCG
119644 TATGAGTGTGGAGTCGTTGCCAGTATAAGCCAGTGTGCCTAGCNNNAGATCGGAA
71384 TATGAGTGTGGAGTCGTTGCTGTCTAGGTCTTGTAGGTGTGGTNNNAGATCGGAA
95322 TATGAGTGTGGAGTCGTTGCAGGCAAATGGGATTGCATCAATCANNNAGATCGGA
90607 TATGAGTGTGGAGTCGTTGCAAAGTACAGAGCAGTTCCTGGGTNNNAGATCGGAA
95398 TATGAGTGTGGAGTCGTTGCGGACAATAGCTGGCTGGGGAAGTNNNAGATCGGA
100383 TATGAGTGTGGAGTCGTTGCGTCAGGAGTTCTCCATATTGGTCNNNAGATCGGAAG
100384 TATGAGTGTGGAGTCGTTGCACAGATATCACCCACACACACCTNNNAGATCGGAAG
101749 TATGAGTGTGGAGTCGTTGCCACCTGACCTGTCCTGTCCTTTTNNNAGATCGGAAGA
117380 TATGAGTGTGGAGTCGTTGCCTCGGACCAGGAGCCAGAGGANNNAGATCGGAAGA
95894 TATGAGTGTGGAGTCGTTGCGGTCTCAGAGGTGGCTAGCTTATCNNNAGATCGGA
97368 TATGAGTGTGGAGTCGTTGCTCAACAATGTATAACCACACCAGCATNNNAGATCGG
98635 TATGAGTGTGGAGTCGTTGCTTTCATTCTTCTCCGGCTCCCGNNNAGATCGGAAGAG
29475 TATGAGTGTGGAGTCGTTGCTCATGGGGCAGATTTCCCTTTTTNNNAGATCGGAAG
97588 TATGAGTGTGGAGTCGTTGCAACTAGGCTGGGCCTGGGCTANNNAGATCGGAAGA
29391 TATGAGTGTGGAGTCGTTGCGCATATTGTATTCTGTACCTTGTTTTTNNNAGATCGG
95773 TATGAGTGTGGAGTCGTTGCCCTCCCTAGCTGGAATTACAGATGNNNAGATCGGAA
100339 TATGAGTGTGGAGTCGTTGCGGAGCACATGTATAGCAGGACAGNNNAGATCGGAA
28831 TATGAGTGTGGAGTCGTTGCCTTCGCTCGGGATAGAGGGAGNNNAGATCGGAAGA
70634 TATGAGTGTGGAGTCGTTGCTCTGGCAATGGGTTCATTCTCAGNNNAGATCGGAAG
71877 TATGAGTGTGGAGTCGTTGCAGCAGAATTCCCCAGTATTTTGGGNNNAGATCGGAA
98495 TATGAGTGTGGAGTCGTTGCCAAGTGGGTCTCTGTGATGGATTCNNNAGATCGGA
29058 TATGAGTGTGGAGTCGTTGCGAGGTTCCACTATTGGCTCCATGNNNAGATCGGAAG
97268 TATGAGTGTGGAGTCGTTGCACTTACCTGCTACCTGGAGCGTTNNNAGATCGGAAG
98790 TATGAGTGTGGAGTCGTTGCACCCAATCTCAGGTATTAGCAGACNNNAGATCGGAA
99020 TATGAGTGTGGAGTCGTTGCGCTTGTGGGGACAGAATCAAGANNNAGATCGGAAG
99968 TATGAGTGTGGAGTCGTTGCCTGGGAATCAGTGTGATGATTGGNNNAGATCGGAA
104477 TATGAGTGTGGAGTCGTTGCGCTGTAACCTCCACCACTCAGNNNAGATCGGAAGAG
97124 TATGAGTGTGGAGTCGTTGCAGACAGGATTTTACCGTGTTAGACTNNNAGATCGGA
97254 TATGAGTGTGGAGTCGTTGCACCTCTGGCTAGCCCTTGAAATCNNNAGATCGGAAG
98276 TATGAGTGTGGAGTCGTTGCCTCCACGGCTCTGTCTTGATCNNNAGATCGGAAGAG
99706 TATGAGTGTGGAGTCGTTGCCCCTAAAGTCACCCATGGACATCNNNAGATCGGAAG
107869 TATGAGTGTGGAGTCGTTGCTCAGGATTTCTGGTTTCTGGACAGNNNAGATCGGAA

| | |
|---|---|
| 119019 | TATGAGTGTGGAGTCGTTGCAAGGCACTGGAGTTTCTGAGNNNAGATCGGAAGAG |
| 119062 | TATGAGTGTGGAGTCGTTGCGGGATTATAGGTGATTCTCCTGCCTNNNAGATCGGA |
| 29052 | TATGAGTGTGGAGTCGTTGCGGGTTCAAGTGATTCAGTGGCATNNNAGATCGGAA |
| 90298 | TATGAGTGTGGAGTCGTTGCACTACAGCACTACTTCCCCACTTNNNAGATCGGAAGA |
| 98479 | TATGAGTGTGGAGTCGTTGCGACTACATCTCCCGTCCTGCNNNAGATCGGAAGAGC |
| 105665 | TATGAGTGTGGAGTCGTTGCAGACACATCAAGTCTCAGTTTCTTCATNNNAGATCGG |
| 121985 | TATGAGTGTGGAGTCGTTGCTTACAGGTGCACAACATCATGTCCNNNAGATCGGAA |
| 97599 | TATGAGTGTGGAGTCGTTGCTGAAACAACAACCTGATGAGGTAGNNNAGATCGGA |
| 106769 | TATGAGTGTGGAGTCGTTGCGAAAAACTGGGAGGTGGTCATAGGANNNAGATCG |
| 95357 | TATGAGTGTGGAGTCGTTGCAAGTCTGCCACAAGGACAAAGAGNNNAGATCGGAA |
| 95457 | TATGAGTGTGGAGTCGTTGCTGGAAATCAAGACTGCTGTGAGCTNNNAGATCGGA |
| 95469 | TATGAGTGTGGAGTCGTTGCCCAACACTCCTGGCTAAAACTTTTNNNAGATCGGAA |
| 100389 | TATGAGTGTGGAGTCGTTGCAAGCCATCCCTGACACTCTGTCNNNAGATCGGAAGA |
| 101762 | TATGAGTGTGGAGTCGTTGCGTGTGTGTGTATAATTGCCTTTATTCNNNAGATCGG |
| 106737 | TATGAGTGTGGAGTCGTTGCCAAGCAATCGGTCCATCTCGTCNNNAGATCGGAAGA |
| 95392 | TATGAGTGTGGAGTCGTTGCGTGCATGAGTGTGAAAAGGGAGNNNAGATCGGAA |
| 95414 | TATGAGTGTGGAGTCGTTGCTGGCTGCTTTTGCACTGTAGTTGNNNAGATCGGAAG |
| 95803 | TATGAGTGTGGAGTCGTTGCGTTACCAAGGGCAGAAACCACNNNAGATCGGAAGA |
| 96017 | TATGAGTGTGGAGTCGTTGCCTCCCAGAGCCAGATTATAGGGGGNNNAGATCGGA |
| 99698 | TATGAGTGTGGAGTCGTTGCTAGATTGTTGATAGCGTCCCTCTNNNAGATCGGAAG |
| 95638 | TATGAGTGTGGAGTCGTTGCTCAGCTCCCAAAGCAGTAGAATTANNNAGATCGGAA |
| 95663 | TATGAGTGTGGAGTCGTTGCATAGGTGTGTACCACCGCAGACNNNAGATCGGAAGA |
| 121996 | TATGAGTGTGGAGTCGTTGCGTGCTCTGATTTAGGTGATCCAAGGNNNAGATCGGA |
| 95569 | TATGAGTGTGGAGTCGTTGCTTTTAGAGACAGAGCTTTGCCAGNNNAGATCGGAAG |
| 98168 | TATGAGTGTGGAGTCGTTGCCACCAGGAAAGTGGAAAATGGGTANNNAGATCGGA |
| 98259 | TATGAGTGTGGAGTCGTTGCCAACGTGCTACTCATTCTCTGCTANNNAGATCGGAAG |
| 98453 | TATGAGTGTGGAGTCGTTGCCCTCCCTTATACTGGGCCCCTNNNAGATCGGAAGAG |
| 98590 | TATGAGTGTGGAGTCGTTGCTTTCTGAGCCTTCTTGCCCCAGNNNAGATCGGAAGA |
| 101733 | TATGAGTGTGGAGTCGTTGCGGGAGATTCTTCAAGTAAAGGGCANNNAGATCGGA |
| 116575 | TATGAGTGTGGAGTCGTTGCAATGAGGCTCAAAGCAGGTCACANNNAGATCGGAA |
| 118784 | TATGAGTGTGGAGTCGTTGCCTCTGCTTTCAGTTCTGTTCACTATNNNAGATCGGAA |
| 107881 | TATGAGTGTGGAGTCGTTGCATACTGAAAACCATGATATCGTCTGTTNNNAGATCG |
| 97076 | TATGAGTGTGGAGTCGTTGCTGATGTTTGCCTTCCCTGAGACTNNNAGATCGGAAG |
| 97812 | TATGAGTGTGGAGTCGTTGCACTTCCATCCCCATGAGGCAANNNAGATCGGAAGAG |
| 102947 | TATGAGTGTGGAGTCGTTGCTCATAATTTCAAGCATTTCCCCTGCNNNAGATCGGAA |
| 107912 | TATGAGTGTGGAGTCGTTGCATACCAACACCACAATCAAGATATGGNNNAGATCGG |
| 94901 | TATGAGTGTGGAGTCGTTGCACAGTGAATCAGGCCAAAAGGAANNNAGATCGGAA |
| 98682 | TATGAGTGTGGAGTCGTTGCTTAGTCCAGCCAGTCTCTTCCTCNNNAGATCGGAAGA |
| 95372 | TATGAGTGTGGAGTCGTTGCAGTGCCTTTGGAGGAGAGTATGATGTGNNNAGATC |
| 96842 | TATGAGTGTGGAGTCGTTGCATTATGGGCATCTACAACCATGCNNNAGATCGGAAG |
| 91217 | TATGAGTGTGGAGTCGTTGCAGTGGATTTCTTACTTGTAGTCACCTNNNAGATCGGA |
| 28858 | TATGAGTGTGGAGTCGTTGCACACACACACCAGGGAACAAAATNNNAGATCGGAAG |
| 95243 | TATGAGTGTGGAGTCGTTGCTTCCCACACTCCTCTGTCTGTGNNNAGATCGGAAGAG |

95452 TATGAGTGTGGAGTCGTTGCTTTATTTTTGCACCCCACTCGCAGNNNAGATCGGAAG

95364 TATGAGTGTGGAGTCGTTGCTGAACACCACCACAAAAATCAGANNNAGATCGGAAG

119642 TATGAGTGTGGAGTCGTTGCTTTGGTAGAGATGGAGTTTTGCATGNNNAGATCGGA

121993 TATGAGTGTGGAGTCGTTGCCACCTCAAGGAAGCAATCATCTAANNNAGATCGGAA

98172 TATGAGTGTGGAGTCGTTGCCATCCTTCCTCCTCTTTCCTCCGCANNNAGATCGGAAG

96034 TATGAGTGTGGAGTCGTTGCCCGCATGTAAGATGTGTCTTTCTCNNNAGATCGGAA

96244 TATGAGTGTGGAGTCGTTGCGGAAGGTGGTTGAACTATGTGTTNNNAGATCGGAA

96343 TATGAGTGTGGAGTCGTTGCACTGCTGCTAAAATTCCTACAGCNNNAGATCGGAAGA

97636 TATGAGTGTGGAGTCGTTGCCTCCATGTCTCCCTCCTCTTGTANNNAGATCGGAAGA

98986 TATGAGTGTGGAGTCGTTGCGAAGAAAGAGGCGAGGGCTAGGNNNAGATCGGAA

103081 TATGAGTGTGGAGTCGTTGCCCAAATACTCCCGACCCTTCCTNNNAGATCGGAAGAG

95254 TATGAGTGTGGAGTCGTTGCGCCTCCCAAGCTCTAGTGATACTNNNAGATCGGAAGA

95428 TATGAGTGTGGAGTCGTTGCTAGCAGAATGGAAGGAAAAGAGGNNNAGATCGGA

101732 TATGAGTGTGGAGTCGTTGCCAAAACACTGATGAAACAATTGAAGATNNNAGATCG

95467 TATGAGTGTGGAGTCGTTGCCCGCAGACCATGAAACCTAGAANNNAGATCGGAAGA

95668 TATGAGTGTGGAGTCGTTGCCAGCATCATGCCCGGATAATTTTCNNNAGATCGGAA

95408 TATGAGTGTGGAGTCGTTGCATAACAGGCCCCCACCACTACATNNNAGATCGGAAGA

95411 TATGAGTGTGGAGTCGTTGCCTTTTGTCAGCTGCATCCTGTGNNNAGATCGGAAGA

95444 TATGAGTGTGGAGTCGTTGCGCTGTCCTGTAAAATTGGTGACAANNNAGATCGGAA

97407 TATGAGTGTGGAGTCGTTGCAAGCCTGGAACGGGGACTTTTGAAAATNNNAGATC

98950 TATGAGTGTGGAGTCGTTGCCTGTCCTCCTTGTACTCAACTTGATTNNNAGATCGGA

99347 TATGAGTGTGGAGTCGTTGCAGTGTTTCAGACTTCGGCTCTGTNNNAGATCGGAAG

107914 TATGAGTGTGGAGTCGTTGCGGAGACCATGTCTATGCAAGAGAAANNNAGATCGGA

95296 TATGAGTGTGGAGTCGTTGCTCGTGTCTCAGCTACTCAGTGAGNNNAGATCGGAAG

95307 TATGAGTGTGGAGTCGTTGCGCCAAACCATCTGGAAAAAGCCNNNAGATCGGAAGA

95456 TATGAGTGTGGAGTCGTTGCTGTAGCCATTCATTTTCCACCCTNNNAGATCGGAAGA

97123 TATGAGTGTGGAGTCGTTGCCACAGAGAAGGGACAGAAGCAGNNNAGATCGGAA

97652 TATGAGTGTGGAGTCGTTGCGGATGGAGACTGGTTAGGAGATTNNNAGATCGGAA

97765 TATGAGTGTGGAGTCGTTGCAGGCCAGTAAACTTCCATCTTAGACNNNAGATCGGA

121944 TATGAGTGTGGAGTCGTTGCATGTAATAACTCACTGTGGCCTGGANNNAGATCGGA

95346 TATGAGTGTGGAGTCGTTGCGAGGAAGAAGCCGGCCCAGCNNNAGATCGGAAGA

95393 TATGAGTGTGGAGTCGTTGCGGAAGTTAGTGCCACATGTAAGGNNNAGATCGGAA

95423 TATGAGTGTGGAGTCGTTGCGCCAGCATCATCCTAATACCCCCNNNAGATCGGAAGA

97028 TATGAGTGTGGAGTCGTTGCCGACCCCCTGGCCAAGGTCTANNNAGATCGGAAGAG

97209 TATGAGTGTGGAGTCGTTGCCTGTGCTTGGCTGGAAATGTTGTAANNNAGATCGGA

98280 TATGAGTGTGGAGTCGTTGCACTCTGGTAACAGCCCAAAAAGTGNNNAGATCGGAA

98475 TATGAGTGTGGAGTCGTTGCAGGAAGTGTGACCTGGGAGTTCNNNAGATCGGAAG

100449 TATGAGTGTGGAGTCGTTGCCTGTCTTCTGCGCTTGCTCGTAANNNAGATCGGAAGA

95440 TATGAGTGTGGAGTCGTTGCTCAGAGCATGCAAATACCCCTCTANNNAGATCGGAA

96390 TATGAGTGTGGAGTCGTTGCGTTACTCTCCAGATCTGCCTTTCCNNNAGATCGGAAG

117276 TATGAGTGTGGAGTCGTTGCGAGAGAGAGACAGAAACAGTCTGTNNNAGATCGGA

98632 TATGAGTGTGGAGTCGTTGCCAATCAATTCTGCCACCTCAGTTTNNNAGATCGGAAG

100124 TATGAGTGTGGAGTCGTTGCTTTTTCTTCCCCCAGTCCCTCANNNAGATCGGAAGAG

| | |
|---|---|
| 97970 | TATGAGTGTGGAGTCGTTGCCTCGGGTGGAGCATCTCCATTNNNAGATCGGAAGAG |
| 100874 | TATGAGTGTGGAGTCGTTGCAACTATCCTCCCAACCTCAGCTTNNNAGATCGGAAGA |
| 95349 | TATGAGTGTGGAGTCGTTGCGCACCAGTAGAGACTGGGTTTCACNNNAGATCGGAA |
| 95468 | TATGAGTGTGGAGTCGTTGCATTTAGGCCCCCAGAACTCTTTANNNAGATCGGAAGA |
| 102913 | TATGAGTGTGGAGTCGTTGCCGCCCAGCCTTGATTTCTATTTTTATNNNAGATCGGAA |
| 95260 | TATGAGTGTGGAGTCGTTGCGGGTTCCACACTTGGTCAACANNNAGATCGGAAGAG |
| 97807 | TATGAGTGTGGAGTCGTTGCTTCTCTCCGGCGGATACGGCANNNAGATCGGAAGAG |
| 121934 | TATGAGTGTGGAGTCGTTGCTAGCTTACTCTGTTCTCTACTGCNNNAGATCGGAAGA |
| 95347 | TATGAGTGTGGAGTCGTTGCATTCCTCCACATCGTTCTGACAGNNNAGATCGGAAGA |
| 96188 | TATGAGTGTGGAGTCGTTGCCAGATACGTTACACGTTGGCGTNNNAGATCGGAAGA |
| 97651 | TATGAGTGTGGAGTCGTTGCTTCTTCCACCTCACCAAGGTAGCNNNAGATCGGAAGA |
| 95375 | TATGAGTGTGGAGTCGTTGCACTCTGGCCTGGGAAATACAGCNNNAGATCGGAAGA |
| 96498 | TATGAGTGTGGAGTCGTTGCGTTGGAAGGAGGGCTGAGGGGANNNAGATCGGAA |
| 118781 | TATGAGTGTGGAGTCGTTGCCTTGAGCCTGGGTGGGAGACNNNAGATCGGAAGAG |
| 95426 | TATGAGTGTGGAGTCGTTGCCCAGAGGCTATTCCAGTGATTGANNNAGATCGGAAG |
| 100300 | TATGAGTGTGGAGTCGTTGCGGGTGGTTTTAGGGCTCCATCTNNNAGATCGGAAGA |
| 120101 | TATGAGTGTGGAGTCGTTGCTGCTAGGCTCAGCTCTTGTTTCNNNAGATCGGAAGA |
| 95425 | TATGAGTGTGGAGTCGTTGCGACTGGGAGAGGGTCCATGTCNNNAGATCGGAAGA |
| 96111 | TATGAGTGTGGAGTCGTTGCGCTCAGTTAGGACGAACCCTGAGNNNAGATCGGAA |
| 98855 | TATGAGTGTGGAGTCGTTGCCGGTCTGTCTCAGTCTGCTAAANNNAGATCGGAAGA |
| 96839 | TATGAGTGTGGAGTCGTTGCACTTGGGTTGATTGCGTTTGTGNNNAGATCGGAAGA |
| 105637 | TATGAGTGTGGAGTCGTTGCGTTGTTGTTGTAGAGATGAGGTCTAACNNNAGATCG |
| 95401 | TATGAGTGTGGAGTCGTTGCTATCAGTAAAACCCACTTGGAAGGCNNNAGATCGGA |
| 96867 | TATGAGTGTGGAGTCGTTGCTTGTGAAGGTGATTCCCCTCCCNNNAGATCGGAAGA |
| 98050 | TATGAGTGTGGAGTCGTTGCACTGCCAGATGTCACCTCTCTGNNNAGATCGGAAGA |
| 98591 | TATGAGTGTGGAGTCGTTGCCGGTGGCTGTGGCAGGTGAGNNNAGATCGGAAGA |
| 106754 | TATGAGTGTGGAGTCGTTGCCAGAGAACTTATAGGACCAGAGGGTNNNAGATCGG |
| 122000 | TATGAGTGTGGAGTCGTTGCAAGTCCCTTCCCTTCCATCTATGAGNNNAGATCGGAA |
| 95461 | TATGAGTGTGGAGTCGTTGCTTCTCTACGCAGGCCCCCGTNNNAGATCGGAAGAGCA |
| 97855 | TATGAGTGTGGAGTCGTTGCCCAACACAAGCCTGGAAATGCNNNAGATCGGAAGAG |
| 98037 | TATGAGTGTGGAGTCGTTGCAGGTCACTTCCAACAGGGAAAATGNNNAGATCGGAA |
| 71844 | TATGAGTGTGGAGTCGTTGCAGTCAATACTTGGGTGGTACCCANNNAGATCGGAAG |
| 71843 | TATGAGTGTGGAGTCGTTGCCTTTGGGTGCTAATGGTGGAGTTNNNAGATCGGAA |
| 71895 | TATGAGTGTGGAGTCGTTGCTATGATGTCTGTGTGGAAAGCGGNNNAGATCGGAA |
| 71842 | TATGAGTGTGGAGTCGTTGCCTACAAGGACAGGCCCATTTGAGNNNAGATCGGAAG |
| 71898 | TATGAGTGTGGAGTCGTTGCTGAGGCAGGAATCAAAGACAGATNNNAGATCGGAA |
| 71897 | TATGAGTGTGGAGTCGTTGCCCCCAGACGAAAATACCAAATGCNNNAGATCGGAAG |
| 71846 | TATGAGTGTGGAGTCGTTGCTTGGAGTTGCAGTTGATGTGTGANNNAGATCGGAA |
| 71845 | TATGAGTGTGGAGTCGTTGCACAATATTCATGGTGGCTGGCAGNNNAGATCGGAA |
| 71735 | TATGAGTGTGGAGTCGTTGCCTCGTAGGTTCAGTACCATTGGTGNNNAGATCGGAA |
| 97722 | TATGAGTGTGGAGTCGTTGCCACTGTTTTCCAGGGTGGAGCTTNNNAGATCGGAAG |
| 95999 | TATGAGTGTGGAGTCGTTGCATTTCCAATTGCTGAAGCTGGAGANNNAGATCGGAA |
| 98945 | TATGAGTGTGGAGTCGTTGCACCCCAGCCACCATGAGTAGNNNAGATCGGAAGAG |

61452 TATGAGTGTGGAGTCGTTGCAAATCTTCCACCAGGAAGGTCCTNNNAGATCGGAAG
40599 TATGAGTGTGGAGTCGTTGCCGCATACCCCGAAAAAGCAAATANNNAGATCGGAAG
68988 TATGAGTGTGGAGTCGTTGCAGTGTTTCCAGCCCTTTGACTTTNNNAGATCGGAAG
46706 TATGAGTGTGGAGTCGTTGCTCTGCTGTGAATCTTGGGTTAGCNNNAGATCGGAAG
46818 TATGAGTGTGGAGTCGTTGCAAGAAAGAAAGGTGGGGAAAGGCNNNAGATCGGA
68459 TATGAGTGTGGAGTCGTTGCCAACAAGAGCCAGAGAATGGGTTNNNAGATCGGAA
48046 TATGAGTGTGGAGTCGTTGCCACCCTGTTCACTTCAGAATTGCNNNAGATCGGAAG
53835 TATGAGTGTGGAGTCGTTGCCCTGGCACTGTTGATTTTTAGGGNNNAGATCGGAAG
106290 TATGAGTGTGGAGTCGTTGCTAGGACCTGACCCTGTAGCTTNNNAGATCGGAAGAG
109074 TATGAGTGTGGAGTCGTTGCCACACACCCCCAAACACTTGTGNNNAGATCGGAAGA
40188 TATGAGTGTGGAGTCGTTGCAACGTAAACACCGCATAAGGTGANNNAGATCGGAA
30316 TATGAGTGTGGAGTCGTTGCCACCAACATCTCTGTTCCCCAAANNNAGATCGGAAGA
52135 TATGAGTGTGGAGTCGTTGCAGAAAGCACAAAGGAGCAGGTAGNNNAGATCGGA
36296 TATGAGTGTGGAGTCGTTGCGGATCATCAGTCGATGTATTTGCANNNAGATCGGAA
52067 TATGAGTGTGGAGTCGTTGCCCTGCTCACTTCAGAACTGCTAGNNNAGATCGGAAG
69227 TATGAGTGTGGAGTCGTTGCCACACCAGCATTTGCCTTTCTTTNNNAGATCGGAAGA
61505 TATGAGTGTGGAGTCGTTGCTCTGTCTGTCTGCCTGTATTCCTNNNAGATCGGAAGA
67062 TATGAGTGTGGAGTCGTTGCTGGGAGACTATTGGAGGCACATTNNNAGATCGGAA
117319 TATGAGTGTGGAGTCGTTGCACCAGCATTTGCCTTTCTTTCANNNAGATCGGAAGAG
58583 TATGAGTGTGGAGTCGTTGCTTTGGCGAGAACAGTGATTCCATNNNAGATCGGAA
32105 TATGAGTGTGGAGTCGTTGCACTTGTGGCTAGTGAAAGCTGAANNNAGATCGGAA
56542 TATGAGTGTGGAGTCGTTGCCCGTGAAAGTAGGGTACACAGTGNNNAGATCGGAA
99310 TATGAGTGTGGAGTCGTTGCCGATCTGGTCCTCTAAAGACAGGAACNNNAGATCGG
54501 TATGAGTGTGGAGTCGTTGCTCTCTAGGTGCCATCAGAACTGGNNNAGATCGGAAG
52298 TATGAGTGTGGAGTCGTTGCACAATGACACACTGATCTGCACGNNNAGATCGGAAG
45166 TATGAGTGTGGAGTCGTTGCAGGAGGTAGAGATACCCAATGCCNNNAGATCGGAA
99311 TATGAGTGTGGAGTCGTTGCGGGTCTCATTCGGCGTCTCCCNNNAGATCGGAAGAG
51836 TATGAGTGTGGAGTCGTTGCCTGTTCCAGTGGGATCTCCTGAGNNNAGATCGGAAG
58641 TATGAGTGTGGAGTCGTTGCGCACAGAGCTAACAAGCTAGCATNNNAGATCGGAA
49155 TATGAGTGTGGAGTCGTTGCTCTCAGGTCTTGGGGGTTTCTCNNNAGATCGGAAGA
106639 TATGAGTGTGGAGTCGTTGCAGTTTCTCATGTTTGTGTGGGTGAGNNNAGATCGG
69513 TATGAGTGTGGAGTCGTTGCCAGGTACGAGAGATGGTGCTTTTNNNAGATCGGAA
58491 TATGAGTGTGGAGTCGTTGCTGGTGGGGAGTTCAATATCCCATNNNAGATCGGAA
53303 TATGAGTGTGGAGTCGTTGCTTCTCATGTTTGTGTGGGTGAGGNNNAGATCGGAA
56339 TATGAGTGTGGAGTCGTTGCATTTGATGCTCTCGACCACTCTGNNNAGATCGGAAG
64330 TATGAGTGTGGAGTCGTTGCGAAGGGTCCTGGACAGTATGAGTNNNAGATCGGAA
37807 TATGAGTGTGGAGTCGTTGCGGCAAGGTCAAAAAGAATGACAGANNNAGATCGGA
104267 TATGAGTGTGGAGTCGTTGCAGACAGAAAGAAAGAAGGTGAAGANNNAGATCGG
107100 TATGAGTGTGGAGTCGTTGCACCAACCTGACAGTGGCTATCTCNNNAGATCGGAAG
106309 TATGAGTGTGGAGTCGTTGCAAGTGCCAACAGATAGAAGAGGGNNNAGATCGGA
53152 TATGAGTGTGGAGTCGTTGCGAGCACTGAACTTGGAAGAGAGGNNNAGATCGGA
44944 TATGAGTGTGGAGTCGTTGCTGTTTCCAATAGGGTCAAGAGCCNNNAGATCGGAA
48220 TATGAGTGTGGAGTCGTTGCCTCCTTCAAACCTTGGTGTTTGCNNNAGATCGGAAG

| | |
|---|---|
| 51391 | TATGAGTGTGGAGTCGTTGCATTAGGTGCTGACACACACAGAGNNNAGATCGGAA |
| 39614 | TATGAGTGTGGAGTCGTTGCACACCATCATTATTCAGAGGACATGANNNAGATCGG |
| 44599 | TATGAGTGTGGAGTCGTTGCTTTTTGTATTTGTCGTGCTGCCCNNNAGATCGGAAG |
| 96531 | TATGAGTGTGGAGTCGTTGCTACCACCCCAATGCAACCTATGNNNAGATCGGAAGA |
| 53397 | TATGAGTGTGGAGTCGTTGCCTTGGGTAAAGGTGTGCAGTAGGNNNAGATCGGA |
| 100752 | TATGAGTGTGGAGTCGTTGCAAAACCGATCTCTCTCTCTCTTNNNAGATCGGAAG |
| 52570 | TATGAGTGTGGAGTCGTTGCATTCTGATACACAGGCTGGATCGNNNAGATCGGAA |
| 112530 | TATGAGTGTGGAGTCGTTGCGTGTGTGTTGAGGGGATACACAGNNNAGATCGGA |
| 42163 | TATGAGTGTGGAGTCGTTGCCCCTTCCCCAGGTGTTCTAAAACNNNAGATCGGAAG |
| 46364 | TATGAGTGTGGAGTCGTTGCGAAACTGAGCAAAGAAAGGCAGCNNNAGATCGGA |
| 117205 | TATGAGTGTGGAGTCGTTGCGTTACAGGGTCTTGCTCTTTCCANNNAGATCGGAAG |
| 108625 | TATGAGTGTGGAGTCGTTGCCAAGACTGTGAATCCATAGGGCAGNNNAGATCGGA |
| 106327 | TATGAGTGTGGAGTCGTTGCAAACTGTTCCCTTTGCCTCCAATNNNAGATCGGAAGA |
| 99641 | TATGAGTGTGGAGTCGTTGCTCAAGCCACATTCTGGGTTTTNNNAGATCGGAAGAG |
| 38770 | TATGAGTGTGGAGTCGTTGCTGCTTCCACACAATTTCTCAGGANNNAGATCGGAAG |
| 99643 | TATGAGTGTGGAGTCGTTGCTTGTGTGTAGCGGTTTGAATGTTGNNNAGATCGGA |
| 45988 | TATGAGTGTGGAGTCGTTGCGGGGAACTAGCAAGTCACTTTCCNNNAGATCGGAA |
| 63013 | TATGAGTGTGGAGTCGTTGCTAGGAGTACCCAGAACCCTCTCTNNNAGATCGGAAG |
| 69414 | TATGAGTGTGGAGTCGTTGCGAAGAAACCCCCAGGGACATTTTNNNAGATCGGAA |
| 41473 | TATGAGTGTGGAGTCGTTGCGCCTGTGTATCCTTCTTTGCGTANNNAGATCGGAAG |
| 64403 | TATGAGTGTGGAGTCGTTGCATTCGCAGTTGATGGTGCATAGTNNNAGATCGGAA |
| 51115 | TATGAGTGTGGAGTCGTTGCCAGAGCCTAAAGTGAGTGGCAGNNNAGATCGGAAG |
| 56191 | TATGAGTGTGGAGTCGTTGCGGTCCATGTGATTGTAGAGGCTGNNNAGATCGGAA |
| 97719 | TATGAGTGTGGAGTCGTTGCATGGACCCACCCACATGCTCNNNAGATCGGAAGAGC |
| 98939 | TATGAGTGTGGAGTCGTTGCGCATCCTTTAGCACTTGGGAAAAGNNNAGATCGGA |
| 64232 | TATGAGTGTGGAGTCGTTGCCCGACCCAAGTGTGTTTTAGAGTNNNAGATCGGAAG |
| 32517 | TATGAGTGTGGAGTCGTTGCATTCTTGTTGGCCAAGCCTGTAANNNAGATCGGAAG |
| 58699 | TATGAGTGTGGAGTCGTTGCTCTCTCTCTCTCTTCGTTGGCATNNNAGATCGGAAGA |
| 54424 | TATGAGTGTGGAGTCGTTGCAGAGGCTTGGGGTGTATAGATGGNNNAGATCGGA |
| 65873 | TATGAGTGTGGAGTCGTTGCAGCCCACATTTCCTGAAAACTGTNNNAGATCGGAAG |
| 55722 | TATGAGTGTGGAGTCGTTGCGGTGGGAGATGACAGTGTGTATGNNNAGATCGGA |
| 62995 | TATGAGTGTGGAGTCGTTGCCTGCCTCCCGGTATTCTATCTCTNNNAGATCGGAAGA |
| 62716 | TATGAGTGTGGAGTCGTTGCATCCTGGGTTCACGCTGATATCTNNNAGATCGGAAG |
| 57529 | TATGAGTGTGGAGTCGTTGCTTGGGGCCAGAGAAAAACATTTGNNNAGATCGGAA |
| 101900 | TATGAGTGTGGAGTCGTTGCGAGGGGGCGGTTAGAATTCTCNNNAGATCGGAAGA |
| 66843 | TATGAGTGTGGAGTCGTTGCAGTGTCTTTGGACTGCTGTTTGTNNNAGATCGGAAG |
| 40962 | TATGAGTGTGGAGTCGTTGCCTTGCTCCCAAAGTTCACCACTANNNAGATCGGAAGA |
| 100518 | TATGAGTGTGGAGTCGTTGCGAGGTCGTGAGAGTGGTCTTCNNNAGATCGGAAGA |
| 67290 | TATGAGTGTGGAGTCGTTGCGGAGGTGCTAGTCTGGGTAGATTNNNAGATCGGAA |
| 54901 | TATGAGTGTGGAGTCGTTGCAAGTAGTGAAGGGTGAAGGGTGGNNNAGATCGGA |
| 69524 | TATGAGTGTGGAGTCGTTGCGGGGGCGGTTAGAATTCTCTTTTNNNAGATCGGAA |
| 110029 | TATGAGTGTGGAGTCGTTGCCCCTTTGGATTTCTCGTGGATAGTNNNAGATCGGAA |
| 50692 | TATGAGTGTGGAGTCGTTGCCTGGTCCCATTTTCCTCTCACAGNNNAGATCGGAAGA |

| | |
|---|---|
| 44768 | TATGAGTGTGGAGTCGTTGCGAGTTACGAAGCACAGATGTCCCNNNAGATCGGAA |
| 53714 | TATGAGTGTGGAGTCGTTGCACGCCAGTCCTATTAGATCAGGGNNNAGATCGGAA |
| 54544 | TATGAGTGTGGAGTCGTTGCCAGGGACTGTCATTTCAGACTGGNNNAGATCGGAA |
| 58412 | TATGAGTGTGGAGTCGTTGCGACACATGCAGACCCATGTACATNNNAGATCGGAAG |
| 97723 | TATGAGTGTGGAGTCGTTGCTACACTCACACACAATCACATGCANNNAGATCGGAA |
| 58913 | TATGAGTGTGGAGTCGTTGCCACAACTAACCCGTGAGGTTCATNNNAGATCGGAAG |
| 57907 | TATGAGTGTGGAGTCGTTGCGGCAAAGAGAACTTGGTCCCAATNNNAGATCGGAA |
| 47506 | TATGAGTGTGGAGTCGTTGCGCTCACTACCAGCTAGTACATGCNNNAGATCGGAAG |
| 33212 | TATGAGTGTGGAGTCGTTGCCTTCCTGATGATTCTTGGCCACANNNAGATCGGAAG |
| 48382 | TATGAGTGTGGAGTCGTTGCCAACAATTCCACTGCTCAGCATCNNNAGATCGGAAG |
| 60631 | TATGAGTGTGGAGTCGTTGCTGAGAGAGCAGAGAAAGAATAACCTNNNAGATCGG |
| 103427 | TATGAGTGTGGAGTCGTTGCTTCATCATTGGTGGGCACTTAGGTNNNAGATCGGAA |
| 59013 | TATGAGTGTGGAGTCGTTGCAAAGCTGGAGGAGTGTCTCAGATNNNAGATCGGAA |
| 34715 | TATGAGTGTGGAGTCGTTGCCAGGTTCTCTAGTGTGTATGCCANNNAGATCGGAAG |
| 67994 | TATGAGTGTGGAGTCGTTGCATGGTAAGCAGAAGCACTTGCTTNNNAGATCGGAA |
| 117131 | TATGAGTGTGGAGTCGTTGCTCTCTCTCTCTCTTGATCGACAGNNNAGATCGGAAGA |
| 115865 | TATGAGTGTGGAGTCGTTGCCTCTCTCTCTCTTGATCGACAGGNNNAGATCGGAAG |
| 52865 | TATGAGTGTGGAGTCGTTGCCACAGAAAAGCCCTGAGGTAAGGNNNAGATCGGAA |
| 48889 | TATGAGTGTGGAGTCGTTGCCACTATTCCAGGCACATAGGCTCNNNAGATCGGAAG |
| 33927 | TATGAGTGTGGAGTCGTTGCACATGGCTGCTGTTTCCTTTACANNNAGATCGGAAG |
| 62312 | TATGAGTGTGGAGTCGTTGCCTCCCCTAATCTCTACAGCTGCTNNNAGATCGGAAGA |
| 66531 | TATGAGTGTGGAGTCGTTGCCCAGGACAGGGATAGAGTTGTGTNNNAGATCGGAA |
| 37440 | TATGAGTGTGGAGTCGTTGCCAGACCTTGCAGACACCAAAAGANNNAGATCGGAA |
| 48758 | TATGAGTGTGGAGTCGTTGCCCAATGACACTATGTGAGGCCTCNNNAGATCGGAAG |
| 41826 | TATGAGTGTGGAGTCGTTGCCTCCTTGGCTTCAGGTGTCCTTANNNAGATCGGAAG |
| 46713 | TATGAGTGTGGAGTCGTTGCAGATGGCCTGGTAATGCTTTAGCNNNAGATCGGAA |
| 68942 | TATGAGTGTGGAGTCGTTGCGAAGGTGGTTGGTGGTGGTATTTNNNAGATCGGA |
| 68756 | TATGAGTGTGGAGTCGTTGCAGCATACAGAGCTGCCTTTTGTTNNNAGATCGGAAG |
| 69610 | TATGAGTGTGGAGTCGTTGCCGCACACAGGCTTAGCAATTTTTNNNAGATCGGAAG |
| 37149 | TATGAGTGTGGAGTCGTTGCTTAGAGCCCATCCTGGTGATTCANNNAGATCGGAAG |
| 56688 | TATGAGTGTGGAGTCGTTGCATAACATGACACTGGGAACCGTGNNNAGATCGGAA |
| 70661 | TATGAGTGTGGAGTCGTTGCGACCAAAAGCTACCACTCTGGAGNNNAGATCGGAA |
| 55060 | TATGAGTGTGGAGTCGTTGCTATGTCTGTGCAATCCCACTTGGNNNAGATCGGAAG |
| 71738 | TATGAGTGTGGAGTCGTTGCAAGAGGCCAAATGAGTGTAAGGCNNNAGATCGGA |
| 102702 | TATGAGTGTGGAGTCGTTGCGCATATTTTCCAGAACTGTAGAGCTNNNAGATCGGA |
| 68214 | TATGAGTGTGGAGTCGTTGCTGGAACAAAACCTCTGCCAAGTTNNNAGATCGGAAG |
| 71775 | TATGAGTGTGGAGTCGTTGCAATGGTGGGAGAAAAGGCTTTGTNNNAGATCGGA |
| 100998 | TATGAGTGTGGAGTCGTTGCTCAGCATCCTCCTTAGAGAAGTANNNAGATCGGAAG |
| 103369 | TATGAGTGTGGAGTCGTTGCAGCCAGTTGTAAAGTACAGTGTGNNNAGATCGGAA |
| 104897 | TATGAGTGTGGAGTCGTTGCTGCTCCCAAAGTTCACCACTATTNNNAGATCGGAAG |
| 100946 | TATGAGTGTGGAGTCGTTGCTCTATTTTGCACAATTTGCCCCAANNNAGATCGGAAG |
| 103458 | TATGAGTGTGGAGTCGTTGCGAATCACTAACAGCCCAGACTTCNNNAGATCGGAAG |
| 100843 | TATGAGTGTGGAGTCGTTGCTATTACTGCCAGATGCTGTGTGTNNNAGATCGGAAG |

| | |
|---|---|
| 71771 | TATGAGTGTGGAGTCGTTGCAGGGAGTGACAGTAGGCTCTTTGNNNAGATCGGAA |
| 101003 | TATGAGTGTGGAGTCGTTGCCTCTGTTGCCAGTGTGTGTGTNNNAGATCGGAAGAG |
| 103443 | TATGAGTGTGGAGTCGTTGCGTCTTGCTGGTCTAGAACTGGCCNNNAGATCGGAAG |
| 108660 | TATGAGTGTGGAGTCGTTGCACACACAAATTCATACAACAGGCCAGNNNAGATCGGA |
| 101934 | TATGAGTGTGGAGTCGTTGCCTCCATTCCCGTTTCTTTTCCTCNNNAGATCGGAAGAG |
| 108628 | TATGAGTGTGGAGTCGTTGCTGTGTGTGTGTCAGAATGAGTAGNNNAGATCGGAA |
| 100846 | TATGAGTGTGGAGTCGTTGCTGTGTGTGTGTGTCTCCAATAAATNNNAGATCGGAA |
| 116647 | TATGAGTGTGGAGTCGTTGCGCACATCACCACGACTGGTAATTNNNAGATCGGAAG |
| 97414 | TATGAGTGTGGAGTCGTTGCCCAAGGCCCTTCTGTCCTGGTNNNAGATCGGAAGAG |
| 116033 | TATGAGTGTGGAGTCGTTGCACATGATAGGACCTGGCTACCTGNNNAGATCGGAAG |
| 101954 | TATGAGTGTGGAGTCGTTGCTGCCAGAAAAATGGGACAAGGAANNNAGATCGGAA |
| 65600 | TATGAGTGTGGAGTCGTTGCAGAGACAGAGAGGGAGCAAATGTNNNAGATCGGA |
| 42469 | TATGAGTGTGGAGTCGTTGCGTGGGTGCAGTTCTCTCTCTAACNNNAGATCGGAAG |
| 39839 | TATGAGTGTGGAGTCGTTGCTCTCTTTTTCCCATTATGTGACTGANNNAGATCGGAAG |
| 51623 | TATGAGTGTGGAGTCGTTGCGTGAAAGAGTTCAGCTGGAGGAGNNNAGATCGGAA |
| 40222 | TATGAGTGTGGAGTCGTTGCAGAGAGAGAAATGGGAATGGTGANNNAGATCGGA |
| 52695 | TATGAGTGTGGAGTCGTTGCCCAACCTGGTCTATAGGCAAAGGNNNAGATCGGAAG |
| 107224 | TATGAGTGTGGAGTCGTTGCTAGATGTGTGTATGTGGGTATATGTGTNNNAGATCG |
| 42421 | TATGAGTGTGGAGTCGTTGCCTCTGGGTATCATGGCACTGAACNNNAGATCGGAAG |
| 32946 | TATGAGTGTGGAGTCGTTGCACATGATGGGGAGGAGGAATAACANNNAGATCGGA |
| 69381 | TATGAGTGTGGAGTCGTTGCGCAGAATACAGTGGCCTGTGTTTNNNAGATCGGAAG |
| 55235 | TATGAGTGTGGAGTCGTTGCAGAATCAGTTTGCTGGCCAAATGNNNAGATCGGAAG |
| 110005 | TATGAGTGTGGAGTCGTTGCTGTATGTGGAGAGAGAGAATCCGTNNNAGATCGGA |
| 63818 | TATGAGTGTGGAGTCGTTGCAGTCCACCACTAGGGTTTGAAGTNNNAGATCGGAAG |
| 40041 | TATGAGTGTGGAGTCGTTGCGAGTGTCTCAGCCTTAGCTCTGANNNAGATCGGAAG |
| 43565 | TATGAGTGTGGAGTCGTTGCAAACTCCCATTGTGACCAGCTACNNNAGATCGGAAGA |
| 59366 | TATGAGTGTGGAGTCGTTGCCCTCAGCTCACAACAGAGTTGATNNNAGATCGGAAG |
| 53454 | TATGAGTGTGGAGTCGTTGCAGCCAGAGCTTCATTTATCACGGNNNAGATCGGAAG |
| 103335 | TATGAGTGTGGAGTCGTTGCTGATGGGGAGGAGGAATAACACAAANNNAGATCGG |
| 68985 | TATGAGTGTGGAGTCGTTGCAGGGGGTGGTTATGAGTGACTTTNNNAGATCGGAA |
| 98822 | TATGAGTGTGGAGTCGTTGCTTCAACTCAAACCCTTTGTCTGGNNNAGATCGGAAGA |
| 53904 | TATGAGTGTGGAGTCGTTGCGGATGTGGGGTATGGTATAGGGGNNNAGATCGGA |
| 49138 | TATGAGTGTGGAGTCGTTGCAGACTGCTCTGTGGTTTGTTCTCNNNAGATCGGAAGA |
| 60907 | TATGAGTGTGGAGTCGTTGCTAAACACAATCTTCAAGGCCCCTNNNAGATCGGAAGA |
| 69597 | TATGAGTGTGGAGTCGTTGCGGTGGTTCTTTGCAGTGTGTTTTNNNAGATCGGAAG |
| 66210 | TATGAGTGTGGAGTCGTTGCGTCTATCCCCAGGTTCTGTCTGTNNNAGATCGGAAGA |
| 65742 | TATGAGTGTGGAGTCGTTGCCATCTGTGCCTGTGTACTCATGTNNNAGATCGGAAGA |
| 67593 | TATGAGTGTGGAGTCGTTGCCCTTCCTTGCTTTCTGGTGACTTNNNAGATCGGAAGA |
| 104261 | TATGAGTGTGGAGTCGTTGCCAATGGTTTGGTCTATTTGGGGANNNAGATCGGAAG |
| 96532 | TATGAGTGTGGAGTCGTTGCTCTCTACTGTGTGGTCTCCTCTTTNNNAGATCGGAAG |
| 63751 | TATGAGTGTGGAGTCGTTGCCTCCAATTCCTCCGTGTTCAAGTNNNAGATCGGAAGA |
| 31952 | TATGAGTGTGGAGTCGTTGCTCCTAACCCCCAGTTTCATGGAANNNAGATCGGAAGA |
| 57648 | TATGAGTGTGGAGTCGTTGCATGCCTCTCCTTTCCCAGTTTTGNNNAGATCGGAAGA |

32716 TATGAGTGTGGAGTCGTTGCGAAAGAGGATGTCCAGGCAAACANNNAGATCGGAA

67970 TATGAGTGTGGAGTCGTTGCTCTCACAGAAACATACAGTGCTTNNNAGATCGGAAG

69070 TATGAGTGTGGAGTCGTTGCTCTGGCCTTTCCTCCTTTCCTTTNNNAGATCGGAAGA

61332 TATGAGTGTGGAGTCGTTGCGGACCAACCTCTGTTCTAGATCCTNNNAGATCGGAA

38263 TATGAGTGTGGAGTCGTTGCATTGGCTCTGTCTTTCCTGGAGANNNAGATCGGAAG

62588 TATGAGTGTGGAGTCGTTGCCTGTTGGGTGAGACAAAACATCTNNNAGATCGGAA

51401 TATGAGTGTGGAGTCGTTGCATCAAGGTGGCTGCTTTACAGAGNNNAGATCGGAA

107080 TATGAGTGTGGAGTCGTTGCATTGGCTCTGTCTTTCCTGGAGAANNNAGATCGGAA

57160 TATGAGTGTGGAGTCGTTGCACACAAGACGTTGACTTGACATTGNNNAGATCGGAA

104224 TATGAGTGTGGAGTCGTTGCCTCTGCATGTCTGAATTTGGTGGGNNNAGATCGGA

41418 TATGAGTGTGGAGTCGTTGCCTCGGCATTACCTTCAAGGAGTANNNAGATCGGAAG

47202 TATGAGTGTGGAGTCGTTGCAGGTGTGTATCACAGTGAATGGCNNNAGATCGGAA

38107 TATGAGTGTGGAGTCGTTGCTGACGTGTCTTCTGTAAACAGAGANNNAGATCGGA

32493 TATGAGTGTGGAGTCGTTGCGGAGGGGTGGAGAATTTGGGTAANNNAGATCGGA

35706 TATGAGTGTGGAGTCGTTGCTTGCTTCCCTAGATTACAGGGCANNNAGATCGGAAG

60140 TATGAGTGTGGAGTCGTTGCGTCTCTTCTAATATGCCAGGCACTNNNAGATCGGAA

105077 TATGAGTGTGGAGTCGTTGCACCAAGTAAAGGCCATGCCAGTNNNAGATCGGAAG

66878 TATGAGTGTGGAGTCGTTGCTGAGAATCAAGCTGTTTTGTTTTGTNNNAGATCGGA

34171 TATGAGTGTGGAGTCGTTGCGCATCAGAAGACATGCACTACCANNNAGATCGGAAG

33956 TATGAGTGTGGAGTCGTTGCACAATGATTAGCCTGCCAAACCANNNAGATCGGAAG

31669 TATGAGTGTGGAGTCGTTGCAAAAAGGCCCAGCTTCTTCAGAANNNAGATCGGAAG

60038 TATGAGTGTGGAGTCGTTGCCTTTGCTGGATAGAAAATAACCACTNNNAGATCGGA

54594 TATGAGTGTGGAGTCGTTGCAACTAGAAAAAGGCAACCCCTGGNNNAGATCGGAA

107090 TATGAGTGTGGAGTCGTTGCCTCTGTTCTAGATCCTGAACTATCAATNNNAGATCGG

55977 TATGAGTGTGGAGTCGTTGCTCTTAGTCACGGATGGATCCCTGNNNAGATCGGAAG

61146 TATGAGTGTGGAGTCGTTGCTTTCATTCACCTCTTTTTCGCCTNNNAGATCGGAAGA

95922 TATGAGTGTGGAGTCGTTGCAAGTGGATCTTGAGCCTGGTGCNNNAGATCGGAAG

54515 TATGAGTGTGGAGTCGTTGCTCATTCCTCATTTGACCCACTGGNNNAGATCGGAAG

108663 TATGAGTGTGGAGTCGTTGCACACACACACACCAGTTGATAGCACNNNAGATCGGA

43756 TATGAGTGTGGAGTCGTTGCAGGAGAGAAGAGATGGGGAAACCNNNAGATCGGA

96044 TATGAGTGTGGAGTCGTTGCACTCCTCCCCACCACCGTGGNNNAGATCGGAAGAGC

45685 TATGAGTGTGGAGTCGTTGCCACCCAGCACTCCTAAATTCTCCNNNAGATCGGAAGA

60306 TATGAGTGTGGAGTCGTTGCGACCTGAACCTCCAGCTTAGACTNNNAGATCGGAAG

53990 TATGAGTGTGGAGTCGTTGCAGGAAGATCTATTGGAGGTGGGGNNNAGATCGGA

52277 TATGAGTGTGGAGTCGTTGCTTGAGAGTTCAGCCATTACCACGNNNAGATCGGAAG

49883 TATGAGTGTGGAGTCGTTGCAACGCTCTCCACTTCCTGATTTCNNNAGATCGGAAGA

71645 TATGAGTGTGGAGTCGTTGCAACATACAATAAGAACAGCATCAAACNNNAGATCGG

37142 TATGAGTGTGGAGTCGTTGCGAGAAACAGGTGACACGGATTCANNNAGATCGGAA

32196 TATGAGTGTGGAGTCGTTGCACTCCTGCTGAGAACAACTTGAANNNAGATCGGAAG

63465 TATGAGTGTGGAGTCGTTGCAGTGTTTTGTTGTGGGTGGTTCTNNNAGATCGGAA

51362 TATGAGTGTGGAGTCGTTGCACAGCTCCAACATACTGCAAGAGNNNAGATCGGAAG

67012 TATGAGTGTGGAGTCGTTGCGGACGGTGGGCATGAGAATAATTNNNAGATCGGA

30854 TATGAGTGTGGAGTCGTTGCCTGAGGAGAACACACACACACAANNNAGATCGGAA

| | |
|---|---|
| 51859 | TATGAGTGTGGAGTCGTTGCTTTCTTGGGGCAACTCATCTGAGNNNAGATCGGAAG |
| 58694 | TATGAGTGTGGAGTCGTTGCCCTACTTGAGAGCGTTATGGCATNNNAGATCGGAAG |
| 51780 | TATGAGTGTGGAGTCGTTGCGAAGACAGGCCAAGGAACATGAGNNNAGATCGGA |
| 35566 | TATGAGTGTGGAGTCGTTGCTCCCTTTGTTTTTCCAAAAGGCANNNAGATCGGAAG |
| 66077 | TATGAGTGTGGAGTCGTTGCTTTGAAGCTGGTAAATGGGCTGTNNNAGATCGGAA |
| 51899 | TATGAGTGTGGAGTCGTTGCTACCTGAACTCTCCAAGGGTGAGNNNAGATCGGAA |
| 102645 | TATGAGTGTGGAGTCGTTGCCCCAAGGTCACCTGAAAATCTCCNNNAGATCGGAAG |
| 103429 | TATGAGTGTGGAGTCGTTGCAATCCAACAGGCTTAGGACCCTANNNAGATCGGAAG |
| 67869 | TATGAGTGTGGAGTCGTTGCTGGAACCCATTTATCTGGAGCTTNNNAGATCGGAAG |
| 56198 | TATGAGTGTGGAGTCGTTGCACTTTACGTGGCTCATTAGGCTGNNNAGATCGGAAG |
| 68072 | TATGAGTGTGGAGTCGTTGCGCCTATCCTAACAGCCTCCTCTTNNNAGATCGGAAGA |
| 44298 | TATGAGTGTGGAGTCGTTGCGCATATTGTGGGTCAAGAGACCCNNNAGATCGGAA |
| 53992 | TATGAGTGTGGAGTCGTTGCAATACTACCCACTGTAGGTGGGGNNNAGATCGGAA |
| 68816 | TATGAGTGTGGAGTCGTTGCAGGGAGGGAGAATGTCACCATTTNNNAGATCGGAA |
| 37013 | TATGAGTGTGGAGTCGTTGCACCTTTTCACCCTCTTACTGTCANNNAGATCGGAAGA |
| 66109 | TATGAGTGTGGAGTCGTTGCTGATTTGTATGCGTAGCTGCTGTNNNAGATCGGAAG |
| 96041 | TATGAGTGTGGAGTCGTTGCCCTCCCACATCGGCCTATCAAATNNNAGATCGGAAG |
| 46863 | TATGAGTGTGGAGTCGTTGCAGAAACCATGTCTACGAACAGGCNNNAGATCGGAA |
| 34239 | TATGAGTGTGGAGTCGTTGCAAAGGATTGGAAGAAAATACACCCANNNAGATCGG |
| 51592 | TATGAGTGTGGAGTCGTTGCGTGCAGGGTATTCTCAACAGGAGNNNAGATCGGAA |
| 39632 | TATGAGTGTGGAGTCGTTGCGGATCTGAGGATGGAGACCATGANNNAGATCGGA |
| 101432 | TATGAGTGTGGAGTCGTTGCCTGCATGCCTTTTTCTTTTGTTTNNNAGATCGGAAGA |
| 47784 | TATGAGTGTGGAGTCGTTGCCTGGGCCATATTGTATATTGCTGCNNNAGATCGGAA |
| 60148 | TATGAGTGTGGAGTCGTTGCCTTGGTGATGAAGCTATGGCACTNNNAGATCGGAA |
| 101745 | TATGAGTGTGGAGTCGTTGCAAGTAGAAGCTCAGTTGCTCTAANNNAGATCGGAA |
| 45657 | TATGAGTGTGGAGTCGTTGCAACGGAAAGCACTGAAGATCTCCNNNAGATCGGAA |
| 115938 | TATGAGTGTGGAGTCGTTGCAGGGGGTCGTGGATGTTTACAATGNNNAGATCGGA |
| 49153 | TATGAGTGTGGAGTCGTTGCGCCATTCCATTTGCAGGTTTCTCNNNAGATCGGAAG |
| 61197 | TATGAGTGTGGAGTCGTTGCCTCTGTGTGTTGGTTATTGGCCTNNNAGATCGGAAG |
| 71386 | TATGAGTGTGGAGTCGTTGCGGGGTCGTGGATGTTTACAATGTNNNAGATCGGAA |
| 42519 | TATGAGTGTGGAGTCGTTGCCTGGAGGAGTATGCTGACAACACNNNAGATCGGAA |
| 59222 | TATGAGTGTGGAGTCGTTGCTTGCTGATACACACAAGCTGGATNNNAGATCGGAAG |
| 62218 | TATGAGTGTGGAGTCGTTGCACAGAAGAGGAGAAGTCCATGCTNNNAGATCGGAA |
| 29862 | TATGAGTGTGGAGTCGTTGCTGGAAGAGATCACCTGTGTGTCANNNAGATCGGAA |
| 63540 | TATGAGTGTGGAGTCGTTGCTCTGCTCACTTACATTCAGTTTCTNNNAGATCGGAAG |
| 42515 | TATGAGTGTGGAGTCGTTGCCTTCCTCCTCCCATCTGAAACACNNNAGATCGGAAGA |
| 56306 | TATGAGTGTGGAGTCGTTGCATACAGCCAGCCCTTTGAATCTGNNNAGATCGGAAG |
| 65757 | TATGAGTGTGGAGTCGTTGCACACAGTACACATGCTTTCATGTNNNAGATCGGAAG |
| 55870 | TATGAGTGTGGAGTCGTTGCTTCACAACTGTCTCTTGGTGACTGNNNAGATCGGAA |
| 69106 | TATGAGTGTGGAGTCGTTGCGCCTAAGAAAGACAGTCGGCTTTNNNAGATCGGAA |
| 107204 | TATGAGTGTGGAGTCGTTGCTACAAGTCATGGCTCACTACAGCCNNNAGATCGGAA |
| 42238 | TATGAGTGTGGAGTCGTTGCCCAGACCATCAGACAGCCTAAACNNNAGATCGGAAG |
| 52263 | TATGAGTGTGGAGTCGTTGCAGCAAAAAGCACAAGCCATAACGNNNAGATCGGAA |

| | |
|---|---|
| 110060 | TATGAGTGTGGAGTCGTTGCCTGGAGAATATACATACACACACACANNNAGATCGG |
| 70939 | TATGAGTGTGGAGTCGTTGCCTGAGGGGTTGTGTGTTCTCTTGNNNAGATCGGAA |
| 64934 | TATGAGTGTGGAGTCGTTGCTCTCATCTTCATCACAAGTGAGGTNNNAGATCGGAA |
| 109943 | TATGAGTGTGGAGTCGTTGCTCAGCAAATACAGCCAGCCCTTTNNNAGATCGGAAG |
| 71429 | TATGAGTGTGGAGTCGTTGCTGTCTAGAGTGTGAAGGCAGTGTNNNAGATCGGAA |
| 45971 | TATGAGTGTGGAGTCGTTGCGTCTAGTGGCTTCTCCCATTTCCNNNAGATCGGAAG |
| 50620 | TATGAGTGTGGAGTCGTTGCAACCCAGTACTTCCAACCAACAGNNNAGATCGGAAG |
| 61797 | TATGAGTGTGGAGTCGTTGCACCTGAAACTACTGAAGAATGAGCTNNNAGATCGG |
| 95927 | TATGAGTGTGGAGTCGTTGCTCCAGCCATTTTTGCACCCATTNNNAGATCGGAAGAG |
| 48371 | TATGAGTGTGGAGTCGTTGCTTTCCTCCTTCCCCTTTTCCATCNNNAGATCGGAAGAG |
| 37027 | TATGAGTGTGGAGTCGTTGCTCTACCATTCTTTCCTTGCTGTCANNNAGATCGGAAG |
| 63652 | TATGAGTGTGGAGTCGTTGCACTGCTTTTCCTCTGCAGAAAGTNNNAGATCGGAAG |
| 38819 | TATGAGTGTGGAGTCGTTGCGAAAAAGTTCCTGCTCTCGAGGANNNAGATCGGAA |
| 30012 | TATGAGTGTGGAGTCGTTGCCCAGAGAGGGTGCAACAGAAAAANNNAGATCGGA |
| 66111 | TATGAGTGTGGAGTCGTTGCGCTCTAAGAATGCTGTCTGCTGTNNNAGATCGGAAG |
| 65965 | TATGAGTGTGGAGTCGTTGCACGGAAACAAAACCTCAACCTGTNNNAGATCGGAAG |
| 57532 | TATGAGTGTGGAGTCGTTGCGCCGGAGGGATTTTTCACATTTGNNNAGATCGGAA |
| 71854 | TATGAGTGTGGAGTCGTTGCTGGAATCCCTAGAAACTCGGTCTNNNAGATCGGAAG |
| 57904 | TATGAGTGTGGAGTCGTTGCTCTGTTCTGTGTCTCCTCCCAATNNNAGATCGGAAGA |
| 101000 | TATGAGTGTGGAGTCGTTGCAGATAAGCGAAAACCCATTGTGGNNNAGATCGGAA |
| 71852 | TATGAGTGTGGAGTCGTTGCCGGATTCTGGAAAGGGGCCCNNNAGATCGGAAGAG |
| 71851 | TATGAGTGTGGAGTCGTTGCATTCCTTGGGTTCAACCTTTCCCNNNAGATCGGAAGA |
| 105081 | TATGAGTGTGGAGTCGTTGCAGAGATAGCGACCAGAAAAAGACNNNAGATCGGAA |
| 71855 | TATGAGTGTGGAGTCGTTGCTTCCTCTGCTCCGTGCTGCCTANNNAGATCGGAAGA |
| 106644 | TATGAGTGTGGAGTCGTTGCCTCCATTTCGTGTACTAACACATGTGNNNAGATCGG |
| 108678 | TATGAGTGTGGAGTCGTTGCAAGAGAGAAGGAAGAGGGAACTCCANNNAGATCG |
| 104983 | TATGAGTGTGGAGTCGTTGCTGACTATGGAGATGTGGGAGGGNNNAGATCGGAA |
| 106310 | TATGAGTGTGGAGTCGTTGCACTCTGTACTGCCTGCTTGATTNNNAGATCGGAAGA |
| 100719 | TATGAGTGTGGAGTCGTTGCTTGGAAACCTCCAGAGTTGCANNNAGATCGGAAGA |
| 95923 | TATGAGTGTGGAGTCGTTGCTTACGGCTTACACAGTCCCCTTGNNNAGATCGGAAG |
| 101861 | TATGAGTGTGGAGTCGTTGCCAGTTTGTAATCACATTCACGCANNNAGATCGGAAG |
| 109092 | TATGAGTGTGGAGTCGTTGCGTCACCTCCCTGAGATAGGAGGGANNNAGATCGGA |
| 102007 | TATGAGTGTGGAGTCGTTGCTGCTTCTCTAATTCTTTCACTGTGANNNAGATCGGAA |
| 110068 | TATGAGTGTGGAGTCGTTGCATGTAACAAGATGTGCCCAGATTCTTTNNNAGATCG |
| 45371 | TATGAGTGTGGAGTCGTTGCGCCTCTGTTGTTACCATTTTGCCNNNAGATCGGAAGA |
| 98866 | TATGAGTGTGGAGTCGTTGCTGAAGCAGTTTCCTCTTATCCCATANNNAGATCGGAA |
| 47882 | TATGAGTGTGGAGTCGTTGCCCAACCTTACACATAACCAGTGCNNNAGATCGGAAG |
| 63325 | TATGAGTGTGGAGTCGTTGCTTGTCTGAGTGCATTCCCATTCTNNNAGATCGGAAG |
| 50994 | TATGAGTGTGGAGTCGTTGCCTCGGGTCATACTGGTTTTCCAGNNNAGATCGGAAG |
| 71723 | TATGAGTGTGGAGTCGTTGCCAACAGCTACCTGGACAAGGTGNNNAGATCGGAAG |
| 99135 | TATGAGTGTGGAGTCGTTGCCTGTAGGAGCACAGCTTCAACCNNNAGATCGGAAGA |
| 37939 | TATGAGTGTGGAGTCGTTGCTCAGAGGGACAGAATTTAGCAGANNNAGATCGGAA |
| 60520 | TATGAGTGTGGAGTCGTTGCTGATGCAAAGCAAACCTGGTACTNNNAGATCGGAA |

32335 TATGAGTGTGGAGTCGTTGCAGCAAACAGGACTGAATCACTAANNNAGATCGGAAG
64538 TATGAGTGTGGAGTCGTTGCAGGAAATCATCTTGCAAAATACACGTNNNAGATCGG
59280 TATGAGTGTGGAGTCGTTGCCGTGCGCCACAGAGTAATATGATNNNAGATCGGAAG
58832 TATGAGTGTGGAGTCGTTGCGACCTCTTCTCCAAACATCTCATNNNAGATCGGAAGA
43227 TATGAGTGTGGAGTCGTTGCAATCGGCTTGATGGAACTGAGACNNNAGATCGGAAG
41078 TATGAGTGTGGAGTCGTTGCTAGGGTCATGTGCTTTCAGCCTANNNAGATCGGAAG
63951 TATGAGTGTGGAGTCGTTGCTACCTATTGGGTTTCAGCCCAGTNNNAGATCGGAAGA
48676 TATGAGTGTGGAGTCGTTGCGTTGGGATGAGTTTCCAGACCTCNNNAGATCGGAAG
65771 TATGAGTGTGGAGTCGTTGCGGGCTTTTCCAAGACACAGATGTNNNAGATCGGAAG
59397 TATGAGTGTGGAGTCGTTGCCACCCACGGAAACCCTGAAATATNNNAGATCGGAAG
35932 TATGAGTGTGGAGTCGTTGCCAACCCATTTATCTGCACATGCANNNAGATCGGAAGA
99544 TATGAGTGTGGAGTCGTTGCGGAGTTTCGCTCTTATAAACGGGNNNAGATCGGAAG
57009 TATGAGTGTGGAGTCGTTGCTGATCCATCCAGCACTTTCTGTGNNNAGATCGGAAGA
65884 TATGAGTGTGGAGTCGTTGCTCTGTCGAAACTCATGGAACTGTNNNAGATCGGAAG
116750 TATGAGTGTGGAGTCGTTGCTGCTGCTCACAGACAAGTGTTATTAGNNNAGATCGG
60371 TATGAGTGTGGAGTCGTTGCACCTTGACCTAAACTTCATGACTNNNAGATCGGAAGA
57169 TATGAGTGTGGAGTCGTTGCTCCTTACCAAGCGTGATCCATTGNNNAGATCGGAAGA
54702 TATGAGTGTGGAGTCGTTGCAACTGAAGAACCTTTCTGGCTGGNNNAGATCGGAAG
44260 TATGAGTGTGGAGTCGTTGCCAAGGTTTTGATTGCTGACACCCNNNAGATCGGAAG
53132 TATGAGTGTGGAGTCGTTGCTACACTGGATGGGTACACAGAGGNNNAGATCGGAA
106261 TATGAGTGTGGAGTCGTTGCCACACACACATATACACCACATCTCNNNAGATCGGAAG
51459 TATGAGTGTGGAGTCGTTGCGGGCCTGGAAACAGTAAAGAGAGNNNAGATCGGAA
51511 TATGAGTGTGGAGTCGTTGCGGAAGAAGTACAGCCACTGAGAGNNNAGATCGGAA
104213 TATGAGTGTGGAGTCGTTGCTTGGGGACAGGATAGGGTAGCATNNNAGATCGGAA
65314 TATGAGTGTGGAGTCGTTGCAGTAGATGGGGCCTTGATATGGTNNNAGATCGGAA
37342 TATGAGTGTGGAGTCGTTGCCCTGGGCTGATTGTCAGATTTCANNNAGATCGGAAG
32461 TATGAGTGTGGAGTCGTTGCGGAGGGAAGAAGCTGGAGAGTAANNNAGATCGGA
45488 TATGAGTGTGGAGTCGTTGCCATCTTGCTGTGTGGGGATATCCNNNAGATCGGAAG
117493 TATGAGTGTGGAGTCGTTGCTCTGCTGGAAAAGCTTGATTCCNNNAGATCGGAAGA
54964 TATGAGTGTGGAGTCGTTGCCCCTGATTTTGATTTCCCTGTGGNNNAGATCGGAAGA
35637 TATGAGTGTGGAGTCGTTGCACCAGAGACCTAATTTTCAGGCANNNAGATCGGAAG
57394 TATGAGTGTGGAGTCGTTGCCCCTGCAGATTTGCTTGTTCTTGNNNAGATCGGAAGA
68759 TATGAGTGTGGAGTCGTTGCTGGGGTACATATGAGGTTTTGTTNNNAGATCGGAAG
38582 TATGAGTGTGGAGTCGTTGCTCTGAAAGGTCACTGGAACTCGANNNAGATCGGAAG
46347 TATGAGTGTGGAGTCGTTGCTGAATCTAGGCTGTGTTTCCAGCNNNAGATCGGAAG
45303 TATGAGTGTGGAGTCGTTGCAGCACTGTCAGATTTTTAGTGCCNNNAGATCGGAAG
35764 TATGAGTGTGGAGTCGTTGCAGGAGAGAGAAGATGGAATGGCANNNAGATCGGA
30503 TATGAGTGTGGAGTCGTTGCGTGGGAGGCTTCTTTTCCAGAAANNNAGATCGGAAG
40336 TATGAGTGTGGAGTCGTTGCTGTTGGTAGTCTCTGGGTTGTGANNNAGATCGGAAG
71417 TATGAGTGTGGAGTCGTTGCTAATAAGGGTGAGGCTGCTCTGTNNNAGATCGGAAG
51470 TATGAGTGTGGAGTCGTTGCTGAGCTGGGAGAGAAAGAGAGAGNNNAGATCGGA
44572 TATGAGTGTGGAGTCGTTGCGGAGAATGAGGTAGTTTTGGCCCNNNAGATCGGAA
47923 TATGAGTGTGGAGTCGTTGCAGAGAGAGAAGGGGTAAAGGTGCNNNAGATCGGA

34899 TATGAGTGTGGAGTCGTTGCCTTTGCTTCTGCCATTTCCTCCANNNAGATCGGAGA
101508 TATGAGTGTGGAGTCGTTGCAATGCAGATTGAGATCTTTGAGGAANNNAGATCGG
59334 TATGAGTGTGGAGTCGTTGCAGGGATGCTATACAGAACGTGATNNNAGATCGGAA
35649 TATGAGTGTGGAGTCGTTGCAGAACTGTTACAAAATAGGAGGCANNNAGATCGGA
36784 TATGAGTGTGGAGTCGTTGCCATCCAAATCACACTGGCTCTCANNNAGATCGGAAGA
56999 TATGAGTGTGGAGTCGTTGCGGGGGAGCTCTAAATTGTCTGTGNNNAGATCGGAA
41940 TATGAGTGTGGAGTCGTTGCTACCAGGATCACAAGGCTGGTTANNNAGATCGGAAG
59632 TATGAGTGTGGAGTCGTTGCCATTCACAGTGGGCAACCTGTATNNNAGATCGGAAG
49878 TATGAGTGTGGAGTCGTTGCTTGTGGTCGGTCTAACGGATTTCNNNAGATCGGAAG
47752 TATGAGTGTGGAGTCGTTGCCAGTGGCTAACTAGAAGAGCTGCNNNAGATCGGAA
70566 TATGAGTGTGGAGTCGTTGCCTCTCTCTACCTCCGGGAAGTTCNNNAGATCGGAAGA
61432 TATGAGTGTGGAGTCGTTGCTTTCAACTTCTCTTTTGTTTCTCCTNNNAGATCGGAAG
107061 TATGAGTGTGGAGTCGTTGCTTTATGTATGTCTTCCCACGTGTNNNAGATCGGAAGA
63997 TATGAGTGTGGAGTCGTTGCGCCACAGAAACCTTAGTTCCAGTNNNAGATCGGAAG
55810 TATGAGTGTGGAGTCGTTGCCAGACCCTTGTGTGTATCCACTGNNNAGATCGGAAG
33688 TATGAGTGTGGAGTCGTTGCGAACAGTGGCAGAATAGCTGACANNNAGATCGGAA
57417 TATGAGTGTGGAGTCGTTGCGGTTTCACTTTGTCACCCAGTTGNNNAGATCGGAAG
57347 TATGAGTGTGGAGTCGTTGCACCCAGAGGTTTGCTAACTCTTGNNNAGATCGGAAG
47909 TATGAGTGTGGAGTCGTTGCTCCTGTCTTTCCTTAGTGAGTGCNNNAGATCGGAAG
42275 TATGAGTGTGGAGTCGTTGCGGGAAGACTGTGCTCTAGACAACNNNAGATCGGAA
41187 TATGAGTGTGGAGTCGTTGCAATGACAATGACCTCCAGGGCTANNNAGATCGGAAG
67195 TATGAGTGTGGAGTCGTTGCCCAGCACCATGTCAGTGATCATTNNNAGATCGGAAG
59325 TATGAGTGTGGAGTCGTTGCACTTGGGTGGGACTTACAGTGATNNNAGATCGGAA
43805 TATGAGTGTGGAGTCGTTGCTTTTGTTCACCACCCTGAGAACCNNNAGATCGGAAG
66065 TATGAGTGTGGAGTCGTTGCTTTTCATGACTTTGAACTAGCTGTNNNAGATCGGAA
53244 TATGAGTGTGGAGTCGTTGCTGTCTTTTGGATGGAACTGGAGGNNNAGATCGGAA
61869 TATGAGTGTGGAGTCGTTGCACATTAGGTCGGGTATGGTAGCTNNNAGATCGGAA
30549 TATGAGTGTGGAGTCGTTGCGGCGTAGAAGGCACAATTAGAAANNNAGATCGGAA
68268 TATGAGTGTGGAGTCGTTGCCCTTTTCAGTTGCCTCTCCAGTTNNNAGATCGGAAGA
34420 TATGAGTGTGGAGTCGTTGCAGGTTTCATCTCTGGACATCCCANNNAGATCGGAAG
30534 TATGAGTGTGGAGTCGTTGCTGTGTATGGCTCGCAATGAGAAANNNAGATCGGAA
62709 TATGAGTGTGGAGTCGTTGCTGTCTTCCCACGTGTTTTGATCTNNNAGATCGGAAGA
63612 TATGAGTGTGGAGTCGTTGCGCATCGGGTGTCCTATCAAAAGTNNNAGATCGGAA
104807 TATGAGTGTGGAGTCGTTGCGTCAGGGTCTGGAAAGTTTTGGGNNNAGATCGGAA
51162 TATGAGTGTGGAGTCGTTGCTAGGCGGAATTCGATAAGTGCAGNNNAGATCGGAA
104805 TATGAGTGTGGAGTCGTTGCAGGCGGAATTCGATAAGTGCAGNNNAGATCGGAAG
53087 TATGAGTGTGGAGTCGTTGCTAGAGACCACCCTATTGCTCAGGNNNAGATCGGAAG
60177 TATGAGTGTGGAGTCGTTGCTTCACGGTATCCAGGCTATCACTNNNAGATCGGAAG
53194 TATGAGTGTGGAGTCGTTGCGTATTGAGTGCTTTCCCAGGAGGNNNAGATCGGAA
31943 TATGAGTGTGGAGTCGTTGCCAGGGTCTGGAAAGTTTTGGGAANNNAGATCGGAA
106672 TATGAGTGTGGAGTCGTTGCACCACCCTATTGCTCAGGCTNNNAGATCGGAAGAGC
33909 TATGAGTGTGGAGTCGTTGCTGCTGAACTCTTTCTTGCCTTACANNNAGATCGGAAG
99549 TATGAGTGTGGAGTCGTTGCAATGCCACACAATGTTTCCTTCTGNNNAGATCGGAA

| | |
|---|---|
| 41217 | TATGAGTGTGGAGTCGTTGCCTCCTTTGTCTTCCAAGCTGCTANNNAGATCGGAAGA |
| 59591 | TATGAGTGTGGAGTCGTTGCCTGCTGGAGGCAAAACAGAGTATNNNAGATCGGAA |
| 40117 | TATGAGTGTGGAGTCGTTGCACAGTGAGAGGGAAAGACAGTGANNNAGATCGGA |
| 43310 | TATGAGTGTGGAGTCGTTGCTTAGGAAAGAGAGGACTGGGGACNNNAGATCGGA |
| 33868 | TATGAGTGTGGAGTCGTTGCATCCCAGGAGGGAACTGAGTACANNNAGATCGGAA |
| 52567 | TATGAGTGTGGAGTCGTTGCACTGTGGCATGACATGAGATCGNNNAGATCGGAAG |
| 53654 | TATGAGTGTGGAGTCGTTGCAACAACCAGAATCTCTCACAGGGNNNAGATCGGAAG |
| 71377 | TATGAGTGTGGAGTCGTTGCCAGATGTCTCTTTGCCAAGTGGTNNNAGATCGGAAG |
| 110075 | TATGAGTGTGGAGTCGTTGCTGGTTCTCTGTGTCATCCCCATAGNNNAGATCGGAA |
| 42863 | TATGAGTGTGGAGTCGTTGCGATACACTGCAAGAATGACGCACNNNAGATCGGAA |
| 33049 | TATGAGTGTGGAGTCGTTGCCCCAACAGACATATGGACACACANNNAGATCGGAAG |
| 103398 | TATGAGTGTGGAGTCGTTGCGTGCTGAGAGAGGGCTGGAAATGNNNAGATCGGA |
| 49860 | TATGAGTGTGGAGTCGTTGCGCCACATCACAATCCCCAATTTCNNNAGATCGGAAGA |
| 66577 | TATGAGTGTGGAGTCGTTGCAATGGCAAGGGTAATGGCATTGTNNNAGATCGGAA |
| 48442 | TATGAGTGTGGAGTCGTTGCAGTCTCAGCAACCGAACAAGATCNNNAGATCGGAAG |
| 45318 | TATGAGTGTGGAGTCGTTGCTTTTAGGATGATCGGTGTGTGCCNNNAGATCGGAA |
| 48876 | TATGAGTGTGGAGTCGTTGCTGCTGTACGCCTAAGTTTAGCTCNNNAGATCGGAAG |
| 33475 | TATGAGTGTGGAGTCGTTGCCGGCTTCAAAATACACAAAGAAGACANNNAGATCGG |
| 55409 | TATGAGTGTGGAGTCGTTGCAAAGCAGAAGGAAAGGACCCATGNNNAGATCGGAA |
| 50451 | TATGAGTGTGGAGTCGTTGCTTAGGGGAACATGAGAGGGGAAGNNNAGATCGGA |
| 30796 | TATGAGTGTGGAGTCGTTGCGCAGCATGGGTTTCAGAAAACAANNNAGATCGGAA |
| 29860 | TATGAGTGTGGAGTCGTTGCTGGGATCTAAACTCTATGCCTGCNNNAGATCGGAAG |
| 54175 | TATGAGTGTGGAGTCGTTGCAGAGAAAGAGAAGTCGGAGTGGGNNNAGATCGGA |
| 37307 | TATGAGTGTGGAGTCGTTGCTATCTGCCAGGTTTTGCTGTTCANNNAGATCGGAAG |
| 106685 | TATGAGTGTGGAGTCGTTGCTCCCCCTCCATGTATATGCATAANNNAGATCGGAAGA |
| 48291 | TATGAGTGTGGAGTCGTTGCGGACAAAGGTCAGGACAGGAATCNNNAGATCGGAA |
| 57957 | TATGAGTGTGGAGTCGTTGCAAGAATGAGGGAGAGGAGGCAATNNNAGATCGGA |
| 51937 | TATGAGTGTGGAGTCGTTGCAGAAAACACACGGTTTCGTTGAGNNNAGATCGGAA |
| 98561 | TATGAGTGTGGAGTCGTTGCAGCCATCTGTGTGATGGGACCNNNAGATCGGAAGA |
| 58140 | TATGAGTGTGGAGTCGTTGCGAGTTCAGGCAGAGAACGTGAATNNNAGATCGGAA |
| 56889 | TATGAGTGTGGAGTCGTTGCGTTCACAGCTCCACCCATGTGNNNAGATCGGAAGAG |
| 36675 | TATGAGTGTGGAGTCGTTGCCACAAGGTAGCTTTCCAAGCTCANNNAGATCGGAAG |
| 38480 | TATGAGTGTGGAGTCGTTGCAATGGCTCCACGCTACTCTTAGANNNAGATCGGAAG |
| 36926 | TATGAGTGTGGAGTCGTTGCAAGAAGAGCAAGTGCAAAGGTCANNNAGATCGGAA |
| 69505 | TATGAGTGTGGAGTCGTTGCAACTGCTTCAATGCCATGCTTTTNNNAGATCGGAAG |
| 66107 | TATGAGTGTGGAGTCGTTGCCCTTAGCCTTCATTTCCTGCTGTNNNAGATCGGAAGA |
| 106272 | TATGAGTGTGGAGTCGTTGCTTGTGTGTTTGTGTTTCTGTGAGTGNNNAGATCGGA |
| 47116 | TATGAGTGTGGAGTCGTTGCCCCAAAGGATAAATGTAGGGGGCNNNAGATCGGAA |
| 68325 | TATGAGTGTGGAGTCGTTGCAGCCAAAATGCCACAGATGAGTTNNNAGATCGGAA |
| 38053 | TATGAGTGTGGAGTCGTTGCGCCAAGTGTCAGAAACCTTCAGANNNAGATCGGAA |
| 67225 | TATGAGTGTGGAGTCGTTGCTCTGCCCACCCAGTATAGTCATTNNNAGATCGGAAG |
| 63927 | TATGAGTGTGGAGTCGTTGCGAAACTGCACCCATCAGTACAGTNNNAGATCGGAAG |
| 36467 | TATGAGTGTGGAGTCGTTGCCAGACCAAAGACAGGCAAGATCANNNAGATCGGAA |

| | |
|---|---|
| 101037 | TATGAGTGTGGAGTCGTTGCTACACAAAGAAGACAGCAAACTAAAANNNAGATCGG |
| 59184 | TATGAGTGTGGAGTCGTTGCGTATGAGGAAAAGGGGAGGGGATNNNAGATCGGA |
| 66567 | TATGAGTGTGGAGTCGTTGCTGTATCTGGCTTCTTCCACATTGTNNNAGATCGGAAG |
| 40035 | TATGAGTGTGGAGTCGTTGCTTTCACAACTATGTTCTTCCTCTGANNNAGATCGGAA |
| 53060 | TATGAGTGTGGAGTCGTTGCGCGTATATTTTTCTGCCTGCAGGNNNAGATCGGAAG |
| 40091 | TATGAGTGTGGAGTCGTTGCATTGAGCCAACTGTGGAAAGTGANNNAGATCGGAA |
| 34557 | TATGAGTGTGGAGTCGTTGCAGTCAAATGGTCTGTCAATAAGCCANNNAGATCGGA |
| 34461 | TATGAGTGTGGAGTCGTTGCCAAGAAAGTTCATCCTGCTCCCANNNAGATCGGAAG |
| 102002 | TATGAGTGTGGAGTCGTTGCTCTTTCCCTCTCTCAGTGGGTTGTNNNAGATCGGAAG |
| 65176 | TATGAGTGTGGAGTCGTTGCTGATAGGCAGTTTACCACTGGGTNNNAGATCGGAAG |
| 42621 | TATGAGTGTGGAGTCGTTGCAGGGAGAGAGAGAAACAGGACACNNNAGATCGGA |
| 59825 | TATGAGTGTGGAGTCGTTGCTTATCCTCAGAACCCACCCAACTNNNAGATCGGAAGA |
| 96681 | TATGAGTGTGGAGTCGTTGCATCAAATGGGAGAGTGTAAAAAGGCNNNAGATCGG |
| 71759 | TATGAGTGTGGAGTCGTTGCGGAAGGCAAAAACGTGAAATCGTNNNAGATCGGAA |
| 100190 | TATGAGTGTGGAGTCGTTGCGCACTTGGAAGAAAGCGCTATTCNNNAGATCGGAAG |
| 55812 | TATGAGTGTGGAGTCGTTGCGACAAGCTTCCTTACCTCCACTGNNNAGATCGGAAG |
| 39302 | TATGAGTGTGGAGTCGTTGCGTGGAGGTGCTTGTGTTTATGGANNNAGATCGGAA |
| 104969 | TATGAGTGTGGAGTCGTTGCTGTATCTGGCTTCTTCCACATTGTAATNNNAGATCGG |
| 104932 | TATGAGTGTGGAGTCGTTGCGATAGGCAGTTTACCACTGGGTAANNNAGATCGGA |
| 56439 | TATGAGTGTGGAGTCGTTGCACTACGTGACTTCTGGGTGTCTGNNNAGATCGGAAG |
| 68799 | TATGAGTGTGGAGTCGTTGCAGAGCCAGTGTTCGGTAACATTTNNNAGATCGGAAG |
| 30903 | TATGAGTGTGGAGTCGTTGCCCAAGTGAGATGCTGCAAGACAANNNAGATCGGAA |
| 42546 | TATGAGTGTGGAGTCGTTGCCCCAAAGCAAAACAAAACCTAACACNNNAGATCGGA |
| 63914 | TATGAGTGTGGAGTCGTTGCCACACTGTTATTCCACTGACAGTNNNAGATCGGAAG |
| 49961 | TATGAGTGTGGAGTCGTTGCCGTAGAAGCTGGAGGGATGTTTCNNNAGATCGGAA |
| 71633 | TATGAGTGTGGAGTCGTTGCCAGAGTTTTCAGCCAGACTTTGTCNNNAGATCGGAA |
| 35737 | TATGAGTGTGGAGTCGTTGCAACTTTGTGTTGTATCCTGGGCANNNAGATCGGAAG |
| 33296 | TATGAGTGTGGAGTCGTTGCAGAGAGAGAGAATTCAGGGCACANNNAGATCGGAA |
| 33143 | TATGAGTGTGGAGTCGTTGCGGCACAGGAAGATCATGTACACANNNAGATCGGAA |
| 69486 | TATGAGTGTGGAGTCGTTGCGCTAGTGGCAGTCATGTCCTTTTNNNAGATCGGAAG |
| 110021 | TATGAGTGTGGAGTCGTTGCCTGAGACAGAATCTCACTGTGTGTNNNAGATCGGAA |
| 45020 | TATGAGTGTGGAGTCGTTGCAAAGACATGAAAAACCAACCGCCNNNAGATCGGAAG |
| 43790 | TATGAGTGTGGAGTCGTTGCAGGGGTATGAGGAGACTTCAACCNNNAGATCGGAA |
| 116964 | TATGAGTGTGGAGTCGTTGCGAGATGGCTTCTTCTACAGGCCNNNAGATCGGAAGA |
| 99076 | TATGAGTGTGGAGTCGTTGCTTCAGGAGCTGGTGGGACACATNNNAGATCGGAAG |
| 59566 | TATGAGTGTGGAGTCGTTGCTCCCTCCCTGTCACTCTCTCTATNNNAGATCGGAAGA |
| 36196 | TATGAGTGTGGAGTCGTTGCGCATCTGTAGTGTCTCTTGTGCANNNAGATCGGAAG |
| 104259 | TATGAGTGTGGAGTCGTTGCAGATTGAGTTTCTTCCCAGGTCGNNNAGATCGGAAG |
| 51586 | TATGAGTGTGGAGTCGTTGCGGCAATGCTCACAGCTAAGGAGNNNAGATCGGAAG |
| 43955 | TATGAGTGTGGAGTCGTTGCTATGTCTCATGGTGTGATGCACCNNNAGATCGGAAG |
| 45897 | TATGAGTGTGGAGTCGTTGCCTTGATGTTGGAAGGTGGCTTCCNNNAGATCGGAAG |
| 104215 | TATGAGTGTGGAGTCGTTGCGGGCAGGATCACCCAGAGAAAGNNNAGATCGGAAG |
| 37547 | TATGAGTGTGGAGTCGTTGCAAAATCTGGAAGTCACCCCAAGANNNAGATCGGAAG |

| | |
|---|---|
| 29985 | TATGAGTGTGGAGTCGTTGCACACCATGCATGCCAAACAAAAANNNAGATCGGAAG |
| 53159 | TATGAGTGTGGAGTCGTTGCCTGGTGCAGAGTTAGGGAGAGGNNNAGATCGGAA |
| 43471 | TATGAGTGTGGAGTCGTTGCCTCCTCCTTCACCTGGCAAATACNNNAGATCGGAAGA |
| 47689 | TATGAGTGTGGAGTCGTTGCTCAGAGCCTTGTTCCATTACTGCNNNAGATCGGAAG |
| 44118 | TATGAGTGTGGAGTCGTTGCGTGTGTGAGAGAGAGACTTGACCNNNAGATCGGAA |
| 108620 | TATGAGTGTGGAGTCGTTGCGTCTGTGTGTATGCCTATGTGTCTGNNNAGATCGGA |
| 65318 | TATGAGTGTGGAGTCGTTGCAGAATGGGGCTTGGTACTATGGTNNNAGATCGGAA |
| 99860 | TATGAGTGTGGAGTCGTTGCTTTTATTCACTGGAGTGGCACTGTNNNAGATCGGAA |
| 69318 | TATGAGTGTGGAGTCGTTGCGGGAGAGGAATTGTTTGGGGTTTNNNAGATCGGA |
| 57167 | TATGAGTGTGGAGTCGTTGCTGGAGTTGAGGAATTTGCCATTGNNNAGATCGGAA |
| 42714 | TATGAGTGTGGAGTCGTTGCGGTGGGGAGAGAAAATTACCCACNNNAGATCGGAA |
| 58785 | TATGAGTGTGGAGTCGTTGCTAGGTCTGGAGCATGTGGATCATNNNAGATCGGAA |
| 60906 | TATGAGTGTGGAGTCGTTGCTGAACTAGAAAGACAAGGCCCCTNNNAGATCGGAA |
| 68346 | TATGAGTGTGGAGTCGTTGCTGTGTGTGTGTGTATAGTAACTAGTTNNNAGATCGG |
| 116854 | TATGAGTGTGGAGTCGTTGCTTCTCACTAGGCTTTTCCCCCANNNAGATCGGAAGAG |
| 31772 | TATGAGTGTGGAGTCGTTGCACAAAGGAGGAGGAGAGAACGAANNNAGATCGGA |
| 61179 | TATGAGTGTGGAGTCGTTGCATGGAAGAAACAAAGTATGGGCCTNNNAGATCGGA |
| 99546 | TATGAGTGTGGAGTCGTTGCCTCCTCCCTATGGCTTTCTGAGCNNNAGATCGGAAG |
| 54160 | TATGAGTGTGGAGTCGTTGCTTAGCAGGAATGCAACTTCTGGGNNNAGATCGGAA |
| 65241 | TATGAGTGTGGAGTCGTTGCATGAATGCAGGTTTAGAGAAATGGTNNNAGATCGG |
| 51699 | TATGAGTGTGGAGTCGTTGCTGTTGGGAACTGGTTCAATGGAGNNNAGATCGGAA |
| 29859 | TATGAGTGTGGAGTCGTTGCGTGGGTGTGGGAGAAGTAGATCANNNAGATCGGA |
| 101472 | TATGAGTGTGGAGTCGTTGCTGATTTCCATGCTGACCAACATTGNNNAGATCGGAA |
| 105005 | TATGAGTGTGGAGTCGTTGCCTGCTCCTGCACATGTTTTGAANNNAGATCGGAAGA |
| 30285 | TATGAGTGTGGAGTCGTTGCACCAGAACAAAGGACCAACCAAANNNAGATCGGAA |
| 50315 | TATGAGTGTGGAGTCGTTGCCCATTTGTCAAGCTTCCTGCAAGNNNAGATCGGAAG |
| 43942 | TATGAGTGTGGAGTCGTTGCTAGCTTAGTGTTTGCAGAGCACCNNNAGATCGGAAG |
| 41877 | TATGAGTGTGGAGTCGTTGCTGGTTCTGGAAGCTCCAGTCTTANNNAGATCGGAAG |
| 104879 | TATGAGTGTGGAGTCGTTGCAAAATCTGGAAGTCACCCCAAGNNNAGATCGGAAG |
| 106228 | TATGAGTGTGGAGTCGTTGCGAAAAGGCTGTTGTGCCAGAAGTNNNAGATCGGAA |
| 49868 | TATGAGTGTGGAGTCGTTGCCATCTCCTCCAGGAAGCCATTTCNNNAGATCGGAAG |
| 66685 | TATGAGTGTGGAGTCGTTGCCAAAGGAGACCCGAGAAAGTTGTNNNAGATCGGAA |
| 39049 | TATGAGTGTGGAGTCGTTGCTGGAGGTCAGTCCCATATAGGGANNNAGATCGGAA |
| 116453 | TATGAGTGTGGAGTCGTTGCGGGGCCAATAATATCAAAGACATTGNNNAGATCGG |
| 55311 | TATGAGTGTGGAGTCGTTGCGCTGTACCACTTGAGAGGGAATGNNNAGATCGGAA |
| 48470 | TATGAGTGTGGAGTCGTTGCGTCGAGTAAGCACACAGAGGATCNNNAGATCGGAA |
| 58263 | TATGAGTGTGGAGTCGTTGCAACTTTTGGATCCAGGGCAACATNNNAGATCGGAAG |
| 116305 | TATGAGTGTGGAGTCGTTGCGTTTGCACCTGTGTCTCTCAGTATNNNAGATCGGAA |
| 106227 | TATGAGTGTGGAGTCGTTGCTAGTATCCTCCTCAGTTCTCCCCCTNNNAGATCGGAA |
| 44324 | TATGAGTGTGGAGTCGTTGCCTGTCACACCCATGGTCTACCCNNNAGATCGGAAGA |
| 54461 | TATGAGTGTGGAGTCGTTGCGCAATCTACCACGACCCATATGGNNNAGATCGGAAG |
| 64217 | TATGAGTGTGGAGTCGTTGCTTTTTGGCCACGTACATGAGAGTNNNAGATCGGAAG |
| 102597 | TATGAGTGTGGAGTCGTTGCAGAGAGTAAGAGATGGGCAGACTGNNNAGATCGG |

| | |
|---|---|
| 52850 | TATGAGTGTGGAGTCGTTGCCAGAGTTGGACCTGCTCATAAGGNNNAGATCGGAA |
| 67284 | TATGAGTGTGGAGTCGTTGCTGCTCTTACCTCTGCCTGAGATTNNNAGATCGGAAG |
| 58321 | TATGAGTGTGGAGTCGTTGCACATACAAGGGTTGTGTCCACATNNNAGATCGGAAG |
| 35811 | TATGAGTGTGGAGTCGTTGCTGAGGAAATAGCAAAAGCTGGCANNNAGATCGGAA |
| 31257 | TATGAGTGTGGAGTCGTTGCAAGAGGATCAGTATGGAGGGCAANNNAGATCGGA |
| 115902 | TATGAGTGTGGAGTCGTTGCACCCTTCTTGCTGTATTTCTCTTCCNNNAGATCGGAA |
| 33923 | TATGAGTGTGGAGTCGTTGCGGACCAAGCAGCTGGTATTTACANNNAGATCGGAA |
| 37643 | TATGAGTGTGGAGTCGTTGCCTTTTAGTTGAGGCCTGGGAAGANNNAGATCGGAA |
| 71792 | TATGAGTGTGGAGTCGTTGCAGCAAAACCCTGTGGGTCTAGAANNNAGATCGGAA |
| 49667 | TATGAGTGTGGAGTCGTTGCTTTGAAGGTGACCAAGCTCCTTCNNNAGATCGGAAG |
| 33245 | TATGAGTGTGGAGTCGTTGCAGGGGTGAATTTTTGGTTCCACANNNAGATCGGAA |
| 59186 | TATGAGTGTGGAGTCGTTGCCTATGGCATCCTAAGAGGGGGATNNNAGATCGGAA |
| 61376 | TATGAGTGTGGAGTCGTTGCAAATAGTGGCTGTACACCCTCCTNNNAGATCGGAAG |
| 100919 | TATGAGTGTGGAGTCGTTGCAGTTATTGGGTGAGCTATGCCCNNNAGATCGGAAG |
| 101945 | TATGAGTGTGGAGTCGTTGCGGACAACAAACATGCACCAACGNNNAGATCGGAAG |
| 100899 | TATGAGTGTGGAGTCGTTGCTGACTTGGCCTTTTATTCTCCACCNNNAGATCGGAAG |
| 102016 | TATGAGTGTGGAGTCGTTGCTTTTAGATCTAGCCATGTTGACATTTANNNAGATCGG |
| 105060 | TATGAGTGTGGAGTCGTTGCCAGAGTTCTAAAGAGCAGGGCTCNNNAGATCGGAA |
| 102012 | TATGAGTGTGGAGTCGTTGCTCATCTAGCAATTCCATTCCTGGANNNAGATCGGAA |
| 106727 | TATGAGTGTGGAGTCGTTGCACACTACGTGACTTCTGGGTGNNNAGATCGGAAGA |
| 116397 | TATGAGTGTGGAGTCGTTGCGAAACCCCAGACCAGGTCCANNNAGATCGGAAGAG |
| 100748 | TATGAGTGTGGAGTCGTTGCCAAAGGGAACAAATCACTCGACGNNNAGATCGGAA |
| 108644 | TATGAGTGTGGAGTCGTTGCATAGCAGCCCTTTGCAAACAAANNNAGATCGGAAGA |
| 71689 | TATGAGTGTGGAGTCGTTGCCTTCATGCTTTCCTGTGGGAGAANNNAGATCGGAAG |
| 102807 | TATGAGTGTGGAGTCGTTGCATCCCTCCAAAAGCAATTGGAAGNNNAGATCGGAAG |
| 104277 | TATGAGTGTGGAGTCGTTGCGGGATGTTTGCAAGATGGTGCTNNNAGATCGGAAG |
| 116768 | TATGAGTGTGGAGTCGTTGCGCTTGTTTGACCTTCAGTAGGCAANNNAGATCGGAA |
| 117712 | TATGAGTGTGGAGTCGTTGCAGGCCCACAGTGAAATACAGTTANNNAGATCGGAA |
| 100567 | TATGAGTGTGGAGTCGTTGCAACTGGGAGTAAAAGTTAGCAGGCNNNAGATCGGA |
| 122019 | TATGAGTGTGGAGTCGTTGCTGAAAGTGAAATTGCTGGGTATCANNNAGATCGGA |
| 117009 | TATGAGTGTGGAGTCGTTGCCACCTGCCTATTCTCAGAAGCAGNNNAGATCGGAAG |
| 106270 | TATGAGTGTGGAGTCGTTGCTACATTGTCTTCCCAGCAGTGTTNNNAGATCGGAAG |
| 101963 | TATGAGTGTGGAGTCGTTGCGCAGGAGGGTCTCTCAGAACTNNNAGATCGGAAGA |
| 102563 | TATGAGTGTGGAGTCGTTGCAAGTTCAACCAATTCTTCCGCCTNNNAGATCGGAAGA |
| 103483 | TATGAGTGTGGAGTCGTTGCATTATTGTGCCTCAGTCTCCCCGNNNAGATCGGAAG |
| 117654 | TATGAGTGTGGAGTCGTTGCTTTCAGTTTTGTCAGTTTCTGCTCCNNNAGATCGGAA |
| 101507 | TATGAGTGTGGAGTCGTTGCCACCCAACAATGGTTACAACTGGTNNNAGATCGGAA |
| 99136 | TATGAGTGTGGAGTCGTTGCCCAAGAGGGGTGACAAGATCTCNNNAGATCGGAAG |
| 110026 | TATGAGTGTGGAGTCGTTGCATTCCCTCTTCCTCAAAGGACGNNNAGATCGGAAGA |
| 95485 | TATGAGTGTGGAGTCGTTGCTTTTTCCAGTGTGGTCTGAGAGTTTNNNAGATCGGA |
| 110057 | TATGAGTGTGGAGTCGTTGCAGTGAAGTGCAGCCAGTCTAGANNNAGATCGGAAG |
| 108659 | TATGAGTGTGGAGTCGTTGCTACAGACATGGCCGGCTAATTTNNNAGATCGGAAGA |
| 99817 | TATGAGTGTGGAGTCGTTGCTTGTTCCTGCCTATCTCCCTTGGNNNAGATCGGAAGA |

95102 TATGAGTGTGGAGTCGTTGCCAGAGAGGAAAAACACCCAGGACNNNAGATCGGAA
62620 TATGAGTGTGGAGTCGTTGCTGGCAACACAAACAGATCCATCTNNNAGATCGGAAG
95884 TATGAGTGTGGAGTCGTTGCTCAAACCCCAACTCTGGGCTCNNNAGATCGGAAGAG
71782 TATGAGTGTGGAGTCGTTGCACCATCAGGACACCCTTGTAGGNNNAGATCGGAAG
95120 TATGAGTGTGGAGTCGTTGCATCAACGCCAGGACCACTTTCATNNNAGATCGGAAG
54094 TATGAGTGTGGAGTCGTTGCTCTCTCTCACCAAACTACCTGGGNNNAGATCGGAAG
53810 TATGAGTGTGGAGTCGTTGCCCAGCCTGTTAGCATTTCTAGGGNNNAGATCGGAAG
55390 TATGAGTGTGGAGTCGTTGCGCAGAGTGGTTGTGCATTACATGNNNAGATCGGAA
95838 TATGAGTGTGGAGTCGTTGCGCCCCGCCCCAATAAGTCTGTCNNNAGATCGGAAGA
47345 TATGAGTGTGGAGTCGTTGCTGGAAAATAGGCTTGTAGGTGGCNNNAGATCGGAA
36299 TATGAGTGTGGAGTCGTTGCTATTCGCAGCCTAACACTTTGCANNNAGATCGGAAG
53264 TATGAGTGTGGAGTCGTTGCCTGGTGCTTAGATGGGAATGAGGNNNAGATCGGA
58727 TATGAGTGTGGAGTCGTTGCTATTACCCCCTCTTCTGCTGCATNNNAGATCGGAAGA
60833 TATGAGTGTGGAGTCGTTGCAGCACACTGAGTGAGAACCCTNNNAGATCGGAAGA
96249 TATGAGTGTGGAGTCGTTGCCTGTCTCTGTGGATCGGAGGACNNNAGATCGGAAG
45461 TATGAGTGTGGAGTCGTTGCAAGGTCAGATGTCTGGGAGATCCNNNAGATCGGAA
54909 TATGAGTGTGGAGTCGTTGCAGGAGAAGAAAGATTGGGGGTGGNNNAGATCGGA
62150 TATGAGTGTGGAGTCGTTGCAAAATCAACCAAGTGTCGTGGCTNNNAGATCGGAAG
110049 TATGAGTGTGGAGTCGTTGCTGAAAAGGTATTTGTGGCCGGGNNNAGATCGGAAG
96952 TATGAGTGTGGAGTCGTTGCGGTCACTCAGGGGATCACAGGTANNNAGATCGGAA
62130 TATGAGTGTGGAGTCGTTGCTTTATTGTGCTTTCCTGCTGGCTNNNAGATCGGAAG
39096 TATGAGTGTGGAGTCGTTGCTGAAAAGAGATGGGGAATGGGGANNNAGATCGGA
42549 TATGAGTGTGGAGTCGTTGCACAAGTGGGTTGGCTAGTAACACNNNAGATCGGAA
56221 TATGAGTGTGGAGTCGTTGCTTCTTTCTGACAGTGTCTGGCTGNNNAGATCGGAAG
106724 TATGAGTGTGGAGTCGTTGCGAGCGTGAATGAAAATACAGCTGGNNNAGATCGGA
54113 TATGAGTGTGGAGTCGTTGCGCGTGAATGAAAATACAGCTGGGNNNAGATCGGAA
54350 TATGAGTGTGGAGTCGTTGCGCTAGTCTGCTACAGTCACATGGNNNAGATCGGAAG
116924 TATGAGTGTGGAGTCGTTGCTCTGTCTTTCTTTGTCTGGCTCTNNNAGATCGGAAGA
117297 TATGAGTGTGGAGTCGTTGCCTGTGAATGCATGTACCCCTCTGNNNAGATCGGAAG
53331 TATGAGTGTGGAGTCGTTGCAGAGAGAGAGCTGGTCACATAGGNNNAGATCGGA
51068 TATGAGTGTGGAGTCGTTGCTGTAGTGTCCATCATTTCCGCAGNNNAGATCGGAAG
99471 TATGAGTGTGGAGTCGTTGCTCCTTCCTTTTCAAGGCCCCGCNNNAGATCGGAAGAG
40897 TATGAGTGTGGAGTCGTTGCCCCCTTCCTCAGCCGTTACTATANNNAGATCGGAAGA
58275 TATGAGTGTGGAGTCGTTGCGCTGGAGAGCATGGTATGAACATNNNAGATCGGAA
50759 TATGAGTGTGGAGTCGTTGCTGGCAGGGAAGAGGATACTACAGNNNAGATCGGA
96609 TATGAGTGTGGAGTCGTTGCGGCTCCACTGCTTCTCTTTCTCNNNAGATCGGAAGAG
29844 TATGAGTGTGGAGTCGTTGCGATACCACTCAGTACAGTGGGGANNNAGATCGGAA
44248 TATGAGTGTGGAGTCGTTGCTCTCTTGCCTTCTCCTCTAACCCNNNAGATCGGAAGA
58084 TATGAGTGTGGAGTCGTTGCCTATTCCCTCGGCATCCAGGAATNNNAGATCGGAAG
53697 TATGAGTGTGGAGTCGTTGCGTTCATCCTAGGTTACAGCAGGGNNNAGATCGGAA
105011 TATGAGTGTGGAGTCGTTGCAGGTAAAAGAGAAAGGCCAATGTNNNAGATCGGA
98223 TATGAGTGTGGAGTCGTTGCAGGTAGAAGCTAAGGGTGCTGCANNNAGATCGGA
54010 TATGAGTGTGGAGTCGTTGCTTGACTGGATCTCTCTCTTGGGGNNNAGATCGGAAG

| | |
|---|---|
| 52469 | TATGAGTGTGGAGTCGTTGCACAAAACACAAGAGACCAGTCCGNNNAGATCGGAA |
| 96257 | TATGAGTGTGGAGTCGTTGCACATCTTTCTCCTCCCTGGGACNNNAGATCGGAAGA |
| 52062 | TATGAGTGTGGAGTCGTTGCCAGCTTTCTCTGGTTCTGGCTAGNNNAGATCGGAAG |
| 31171 | TATGAGTGTGGAGTCGTTGCTGGAAGAAACACACAGAAAGCAANNNAGATCGGAA |
| 99463 | TATGAGTGTGGAGTCGTTGCCTCACCCTCCAATTACCGGGTNNNAGATCGGAAGAG |
| 38524 | TATGAGTGTGGAGTCGTTGCGAGTGAAGAGTGGAGAGAGACGANNNAGATCGGA |
| 43715 | TATGAGTGTGGAGTCGTTGCAGTGCAACCTGCTTTCAAAAACCNNNAGATCGGAAG |
| 108655 | TATGAGTGTGGAGTCGTTGCATAAAGTCCATACCAGGCCAGGNNNAGATCGGAAG |
| 32762 | TATGAGTGTGGAGTCGTTGCTCCTTCCCACTTTGCAAATAAACANNNAGATCGGAAG |
| 54694 | TATGAGTGTGGAGTCGTTGCTACATTCACAAGAATGGGGCTGGNNNAGATCGGAA |
| 38404 | TATGAGTGTGGAGTCGTTGCCCATTCCCCTTCATTCCCCTAGANNNAGATCGGAAGA |
| 96612 | TATGAGTGTGGAGTCGTTGCCACTCACTCCCCACCCCATACTANNNAGATCGGAAGA |
| 61488 | TATGAGTGTGGAGTCGTTGCTCCAGTCAACCACTTCAAATTCCTNNNAGATCGGAAG |
| 60087 | TATGAGTGTGGAGTCGTTGCACCACATGACTTAGTAACTCCACTNNNAGATCGGAA |
| 69099 | TATGAGTGTGGAGTCGTTGCCATAAGGCCACTGGAAAGGCTTTNNNAGATCGGAA |
| 116801 | TATGAGTGTGGAGTCGTTGCTTTCTTCTCCTGGGAATTTCAGAGNNNAGATCGGAA |
| 55502 | TATGAGTGTGGAGTCGTTGCTCTCTCTCTCTCCAAGGCTCATGNNNAGATCGGAAGA |
| 61010 | TATGAGTGTGGAGTCGTTGCTTGTCTCTTTTTCCTCCCTCCCTNNNAGATCGGAAGAG |
| 39453 | TATGAGTGTGGAGTCGTTGCGCATGTATGACTGTGAGTGTGGANNNAGATCGGAA |
| 49821 | TATGAGTGTGGAGTCGTTGCCGAGACAGGATCTCACCATGTTCNNNAGATCGGAAG |
| 71447 | TATGAGTGTGGAGTCGTTGCTTGAGGAAGGTGTTTGTGTGTGTNNNAGATCGGAA |
| 59529 | TATGAGTGTGGAGTCGTTGCAAGGTCAGCAGTGTGTCTCCTATNNNAGATCGGAAG |
| 36033 | TATGAGTGTGGAGTCGTTGCGTGCGGCCATGTATTTTTATGCANNNAGATCGGAAG |
| 38171 | TATGAGTGTGGAGTCGTTGCTGTGAAAGGAGGTAAGCTGAGAGANNNAGATCGG |
| 42420 | TATGAGTGTGGAGTCGTTGCTGGCTGGAACAGACTTGATGAACNNNAGATCGGAA |
| 46172 | TATGAGTGTGGAGTCGTTGCACATATAGCAGTGGCACAGAAGCNNNAGATCGGAA |
| 44296 | TATGAGTGTGGAGTCGTTGCATTGCAGAAACATCCTCAGACCCNNNAGATCGGAAG |
| 48348 | TATGAGTGTGGAGTCGTTGCCAGTGCTAAACCCAGGGTACATCNNNAGATCGGAAG |
| 32381 | TATGAGTGTGGAGTCGTTGCTCAGCTTGGGCAAAGAAAGCTAANNNAGATCGGAA |
| 30261 | TATGAGTGTGGAGTCGTTGCTACGTGTGTGTGCGTTAGACAAANNNAGATCGGAA |
| 45367 | TATGAGTGTGGAGTCGTTGCTCTCTCTCTCTCTCCAGTTTGCCNNNAGATCGGAAGA |
| 99113 | TATGAGTGTGGAGTCGTTGCAGAAGGTCAGGAAGGTCAGGAAAANNNAGATCGG |
| 50483 | TATGAGTGTGGAGTCGTTGCGGGCCATGGATTTGTGAATGAAGNNNAGATCGGAA |
| 39380 | TATGAGTGTGGAGTCGTTGCCTGAAGGAGTGTCATTGTCTGGANNNAGATCGGAA |
| 97262 | TATGAGTGTGGAGTCGTTGCAACTGCAGGGGTAATGGTAGGCTANNNAGATCGGA |
| 34838 | TATGAGTGTGGAGTCGTTGCTTTGCTGCATGGAATGTGATCCANNNAGATCGGAAG |
| 59750 | TATGAGTGTGGAGTCGTTGCCCCTCAGCAAAGAGTGGAAAACTNNNAGATCGGAA |
| 62127 | TATGAGTGTGGAGTCGTTGCTTCAAATGACTGAATGGCTGGCTNNNAGATCGGAA |
| 58760 | TATGAGTGTGGAGTCGTTGCACCACAAGGCTTACCTAAATCATNNNAGATCGGAAG |
| 101505 | TATGAGTGTGGAGTCGTTGCCTGCCAAGGAACCTCATCTCCGNNNAGATCGGAAGA |
| 63636 | TATGAGTGTGGAGTCGTTGCGAGTGCACATAAACCAGCAAAGTNNNAGATCGGAA |
| 58168 | TATGAGTGTGGAGTCGTTGCGAGCAACAGCCACCAGCTATAATNNNAGATCGGAAG |
| 106681 | TATGAGTGTGGAGTCGTTGCCTGGGCCTATGCAGCTACTGTTTNNNAGATCGGAAG |

96917 TATGAGTGTGGAGTCGTTGCGTCCATTCACAAACTCAGGATCCTNNNAGATCGGAAC
99392 TATGAGTGTGGAGTCGTTGCCCCTCCTTGACCATTCAGGTTCAANNNAGATCGGAAG
40575 TATGAGTGTGGAGTCGTTGCATGGACAAAACAACTGCTTTGANNNAGATCGGAAG
95103 TATGAGTGTGGAGTCGTTGCCAGTGTGGGAAAGCATTCAGTTTNNNAGATCGGAAC
54536 TATGAGTGTGGAGTCGTTGCGAGAGCACACAGTCATTCACTGGNNNAGATCGGAAC
43168 TATGAGTGTGGAGTCGTTGCGCAGAGAGATCAAGAAGCCAGACNNNAGATCGGAA
71813 TATGAGTGTGGAGTCGTTGCCTCTCTCCAGTGTGAATCCTCCTNNNAGATCGGAAGA
56013 TATGAGTGTGGAGTCGTTGCTCCATCTATCTCAGGATGGCCTGNNNAGATCGGAAGA
54023 TATGAGTGTGGAGTCGTTGCCCACATACCCCAGTAACAATGGGNNNAGATCGGAAG
37793 TATGAGTGTGGAGTCGTTGCGCTTCTGTTTCTCTGCAGACAGANNNAGATCGGAAGA
97259 TATGAGTGTGGAGTCGTTGCCCTCCTATTGGCTTCTTGTGGTNNNAGATCGGAAGAC
71653 TATGAGTGTGGAGTCGTTGCGGAAGCTGTGGGGAGAAGCAGNNNAGATCGGAAG
40153 TATGAGTGTGGAGTCGTTGCACATGGAGAAAGGACTGGAGTGANNNAGATCGGAA
58714 TATGAGTGTGGAGTCGTTGCGCAGAGAGATTGGAGTGATGCATNNNAGATCGGAA
62072 TATGAGTGTGGAGTCGTTGCCTATGCAGCTACTGTTTTGGGCTNNNAGATCGGAAG
100149 TATGAGTGTGGAGTCGTTGCGTGCCAGGCTGCTCTCTGAAATNNNAGATCGGAAGA
100847 TATGAGTGTGGAGTCGTTGCCCAACACCTAGGCTGAAGTATAGTNNNAGATCGGAA
56809 TATGAGTGTGGAGTCGTTGCGAAAGACCAGAGAGACTTGGGTGNNNAGATCGGAA
67407 TATGAGTGTGGAGTCGTTGCAATGCCTCCCTGGTCAGTCTATTNNNAGATCGGAAGA
51093 TATGAGTGTGGAGTCGTTGCCGAGGAAAAAGAGATGGAGGCAGNNNAGATCGGA
53546 TATGAGTGTGGAGTCGTTGCACCATATGCATGGAAGAACTCGGNNNAGATCGGAAC
47046 TATGAGTGTGGAGTCGTTGCCAGTAATAAAACACAAGGGCGGCNNNAGATCGGAAC
39431 TATGAGTGTGGAGTCGTTGCAGAGAATCAAAGGAGTGGGTGGANNNAGATCGGAA
66247 TATGAGTGTGGAGTCGTTGCCCTGGATTGCCCTTCATTTCTGTNNNAGATCGGAAGA
71676 TATGAGTGTGGAGTCGTTGCACTGCTGCAACTACATCACAGAGNNNAGATCGGAAG
41680 TATGAGTGTGGAGTCGTTGCTTGTATGTCTCCGAGGGTGTGTANNNAGATCGGAAC
57869 TATGAGTGTGGAGTCGTTGCCCAGACATACCCGTGTTCACAATNNNAGATCGGAAGA
110069 TATGAGTGTGGAGTCGTTGCTTCAGAGAAAAAGAACCTGGGACCNNNAGATCGGAA
71623 TATGAGTGTGGAGTCGTTGCAGAGCCCCATTCAAAGGCTCCGNNNAGATCGGAAGA
51023 TATGAGTGTGGAGTCGTTGCAGGTTCAAGTACAAGAGCAGCAGNNNAGATCGGAA
50396 TATGAGTGTGGAGTCGTTGCCGGGAGGCTTTTCAGAAGAGAAGNNNAGATCGGAA
106683 TATGAGTGTGGAGTCGTTGCGTGTAGTGGTGCAATCCTGGTTTANNNAGATCGGAA
37769 TATGAGTGTGGAGTCGTTGCTGACAGAAGTCTGGTTCCACAGANNNAGATCGGAAC
103453 TATGAGTGTGGAGTCGTTGCTCTGACCACAAAAGGCTGAAGGNNNAGATCGGAAGA
116663 TATGAGTGTGGAGTCGTTGCCTCTCTTTCTTCTCCCTCAGTTGCNNNAGATCGGAAGA
71654 TATGAGTGTGGAGTCGTTGCCCTCTCCCCCTATGCCCTCTTNNNAGATCGGAAGAGCA
104272 TATGAGTGTGGAGTCGTTGCCACACCACTGTTGGCCTGTTTNNNAGATCGGAAGAGC
42936 TATGAGTGTGGAGTCGTTGCGATTGCATGCTTGTAGGTGCACNNNAGATCGGAAGA
44029 TATGAGTGTGGAGTCGTTGCAGTGAAGACTTAGACCCCAGACCNNNAGATCGGAAG
115961 TATGAGTGTGGAGTCGTTGCTTTTCTTAGAGATAGGTCTCGCTCTNNNAGATCGGAA
68849 TATGAGTGTGGAGTCGTTGCTGCCTGACTGTGTGAATGCATTTNNNAGATCGGAAG
51406 TATGAGTGTGGAGTCGTTGCTCTCTTTTCCTCCCTTCCCAGAGNNNAGATCGGAAGAC
41319 TATGAGTGTGGAGTCGTTGCTGTGTGGTCCCAGTGGTATTCTANNNAGATCGGAAG

60202 TATGAGTGTGGAGTCGTTGCTGTAGTCTGTCCAGATGCTCACTNNNAGATCGGAAG
42322 TATGAGTGTGGAGTCGTTGCGTGCAATCCTGGTTTAGTGCAACNNNAGATCGGAAG
51241 TATGAGTGTGGAGTCGTTGCGATACAAATGTCCCAGGGCTCAGNNNAGATCGGAA
96216 TATGAGTGTGGAGTCGTTGCCACAAAACTCAGCCCAGCCAGGNNNAGATCGGAAGA
100527 TATGAGTGTGGAGTCGTTGCTGTGTCTCCATCTGTGTTCACCNNNAGATCGGAAGA
58151 TATGAGTGTGGAGTCGTTGCGGTGCCAGGAATCGGGAAATAATNNNAGATCGGAA
54599 TATGAGTGTGGAGTCGTTGCTTATGTGTGTGCTTGATCCCTGGNNNAGATCGGAAG
45678 TATGAGTGTGGAGTCGTTGCAAGGGAAAAGGGCAAAAGTCTCCNNNAGATCGGAA
50317 TATGAGTGTGGAGTCGTTGCGCAAAACTCAAATGTGGTGCAAGNNNAGATCGGAA
108619 TATGAGTGTGGAGTCGTTGCGGATATTTTGACACTGCATTTTCCGNNNAGATCGGA
99820 TATGAGTGTGGAGTCGTTGCAACCCTTCTGGCTTTGTCCTTNNNAGATCGGAAGAG
54883 TATGAGTGTGGAGTCGTTGCGGTTCTGGTTTAGGGAAAGGTGGNNNAGATCGGA
107228 TATGAGTGTGGAGTCGTTGCCACAGGAAAAAGGGTGCTCACTTTNNNAGATCGGA
44589 TATGAGTGTGGAGTCGTTGCCCTTCACTAGAACACACTGCCCNNNAGATCGGAAGA
69323 TATGAGTGTGGAGTCGTTGCTGCTGGTAACTCATTGTGGGTTTNNNAGATCGGAAG
103474 TATGAGTGTGGAGTCGTTGCTTGGGGAGTGGTAGTAGAAAACACCNNNAGATCGG
45808 TATGAGTGTGGAGTCGTTGCGAGATCCAGACAACTCAGTGTCCNNNAGATCGGAAG
55766 TATGAGTGTGGAGTCGTTGCCATCATTGCTTTGCCAGGAACTGNNNAGATCGGAAG
95882 TATGAGTGTGGAGTCGTTGCCGCCCATTCCCCCATTCCACNNNAGATCGGAAGAGCA
99458 TATGAGTGTGGAGTCGTTGCTGAACCTGGTGGATCCCATTAGANNNAGATCGGAA
57210 TATGAGTGTGGAGTCGTTGCGTGCCAGCCAGTAAAATGGATTGNNNAGATCGGAA
117686 TATGAGTGTGGAGTCGTTGCACCCTAGCAATCTTCCCACCCCNNNAGATCGGAAGAG
40190 TATGAGTGTGGAGTCGTTGCCCAGAGTGAGAGTGATCAGGTGANNNAGATCGGA
99462 TATGAGTGTGGAGTCGTTGCCGGCTTGCTGAATCGTTGAGATGNNNAGATCGGAA
96948 TATGAGTGTGGAGTCGTTGCGGCACCTCTAGCCCGGGGATNNNAGATCGGAAGAG
97266 TATGAGTGTGGAGTCGTTGCGTTCAGGCCTCTCGTCGCCGNNNAGATCGGAAGAG
99815 TATGAGTGTGGAGTCGTTGCCGCACCCAATAAAGCGAATGGNNNAGATCGGAAGA
98580 TATGAGTGTGGAGTCGTTGCCGCCCCACTACACTAGGCTAATTNNNAGATCGGAAG
103445 TATGAGTGTGGAGTCGTTGCCTTTTCAACCCAACACATAGGTCNNNAGATCGGAAG
47248 TATGAGTGTGGAGTCGTTGCCCATCAGGAGAATAGCAACTGGCNNNAGATCGGAA
31026 TATGAGTGTGGAGTCGTTGCCCAGGATTTTAGACTTGCCCCAANNNAGATCGGAAG
101972 TATGAGTGTGGAGTCGTTGCAGGGTTTTGCTCTTGTGGCACNNNAGATCGGAAGA
37411 TATGAGTGTGGAGTCGTTGCGGCATTTGCACTGTCTGTTTTCANNNAGATCGGAAG
96219 TATGAGTGTGGAGTCGTTGCCACCCATTGTGACCTGGAGCTANNNAGATCGGAAGA
71637 TATGAGTGTGGAGTCGTTGCATTACAAGATGAGCAGTAAGGCCNNNAGATCGGAA
34669 TATGAGTGTGGAGTCGTTGCTCAGGAAAAGTTGGAAATGGCCANNNAGATCGGAA
47879 TATGAGTGTGGAGTCGTTGCAGGCCACCTTTCTATAACAGTGCNNNAGATCGGAAG
65088 TATGAGTGTGGAGTCGTTGCTAAATGCAGGTCACTCCTAGGGTNNNAGATCGGAA
51724 TATGAGTGTGGAGTCGTTGCCTTGTTGAGAATGGAGGCTGGAGNNNAGATCGGA
95885 TATGAGTGTGGAGTCGTTGCGTGCCAGCCATGGAGCCTCCNNNAGATCGGAAGAG
29842 TATGAGTGTGGAGTCGTTGCGACTGTACGGAGAAATGGAAGGCNNNAGATCGGA
71764 TATGAGTGTGGAGTCGTTGCAGGTGCAGTCTATCTGACTTTCANNNAGATCGGAAG
44873 TATGAGTGTGGAGTCGTTGCTTTTCTACTCCTTCTGGCAAGCCNNNAGATCGGAAGA

51918 TATGAGTGTGGAGTCGTTGCGACTGGAGCAGAGGGAAATTGAGNNNAGATCGGAA
101035 TATGAGTGTGGAGTCGTTGCGGTGGCTGATGTAGGAAGATTTNNNAGATCGGAAG
59760 TATGAGTGTGGAGTCGTTGCGAACAACAACCTCTCCCCAAACTNNNAGATCGGAAGA
48448 TATGAGTGTGGAGTCGTTGCCTTTGCAGACATTCCCCCAGATCNNNAGATCGGAAGA
67416 TATGAGTGTGGAGTCGTTGCCCCACTCCATAGGTCCCAGTATTNNNAGATCGGAAGA
62035 TATGAGTGTGGAGTCGTTGCTGGTATATGTTTCACAACGGGCTNNNAGATCGGAAG
63024 TATGAGTGTGGAGTCGTTGCACTGGCACTGAATCCATCTCTCTNNNAGATCGGAAGA
71672 TATGAGTGTGGAGTCGTTGCTTGCTGTGAAAAAGACGTTCACGNNNAGATCGGAAG
71737 TATGAGTGTGGAGTCGTTGCCCGTCCCCTGTACCTACTACATTNNNAGATCGGAAGA
45734 TATGAGTGTGGAGTCGTTGCCAGAGGAAACTTCAGCTCAGTCCNNNAGATCGGAAG
102834 TATGAGTGTGGAGTCGTTGCGCCTGGTGAAGCATCTGCTGAGNNNAGATCGGAAGA
41813 TATGAGTGTGGAGTCGTTGCCCAGGGTTCAACCAATTCCCTTANNNAGATCGGAAGA
71820 TATGAGTGTGGAGTCGTTGCGTCAAGGCCATGGGCAACTGNNNAGATCGGAAGAG
102013 TATGAGTGTGGAGTCGTTGCCAAAAGCAACGAGCCCTGTTAGTNNNAGATCGGAAG
104206 TATGAGTGTGGAGTCGTTGCTTCTCACCCATGAAATTGGCAAAAATTNNNAGATCGG
71812 TATGAGTGTGGAGTCGTTGCTGATGATGAGGGTGACAAGGACTNNNAGATCGGAA
101922 TATGAGTGTGGAGTCGTTGCCTGGTGACTGAATTTAGAGCAGTGNNNAGATCGGAA
100872 TATGAGTGTGGAGTCGTTGCTCCGAAAGCCTTTGCTGTGTTCNNNAGATCGGAAGAG
107227 TATGAGTGTGGAGTCGTTGCCCAGGATGATGTCATCGCCTCATNNNAGATCGGAAG
71815 TATGAGTGTGGAGTCGTTGCATGTGGCAGAGTGTTCAGTTAGCNNNAGATCGGAAG
100994 TATGAGTGTGGAGTCGTTGCCCTAAAAATCCTAGAGTAGGCCAGGNNNAGATCGGA
98581 TATGAGTGTGGAGTCGTTGCGAGACATCTTGGCGAGGCTCAANNNAGATCGGAAGA
101484 TATGAGTGTGGAGTCGTTGCATCACCCTCAGCTAATTTTTGCTTNNNAGATCGGAAG
102011 TATGAGTGTGGAGTCGTTGCCAGGCTCGATCTCCCAAGGTCNNNAGATCGGAAGAG
100576 TATGAGTGTGGAGTCGTTGCATGTTTGGACCCTGCCGATTTGANNNAGATCGGAAG
115769 TATGAGTGTGGAGTCGTTGCATCACAGTTTGACCAAGGCGAANNNAGATCGGAAGA
97260 TATGAGTGTGGAGTCGTTGCAACTGGAGAAATGTGGCAGAGGCNNNAGATCGGAA
109142 TATGAGTGTGGAGTCGTTGCCCCTGCCTCTGGAGTTTTCTTACNNNAGATCGGAAGA
103451 TATGAGTGTGGAGTCGTTGCAATGCCTGGACACTATACAGCAGNNNAGATCGGAAG
34578 TATGAGTGTGGAGTCGTTGCCGCCATCATCTATCAATCAGCCANNNAGATCGGAAGA
67190 TATGAGTGTGGAGTCGTTGCCGCCTACATGTTCTGGCATCATTNNNAGATCGGAAGA
45015 TATGAGTGTGGAGTCGTTGCTCACTCCAGAGCTAGTCTTAGCCNNNAGATCGGAAGA
56274 TATGAGTGTGGAGTCGTTGCGTAGAGGGAGGAGGATGTGCTGNNNAGATCGGAA
45205 TATGAGTGTGGAGTCGTTGCATGTAACCATGTCAGGTGATGCCNNNAGATCGGAAG
69162 TATGAGTGTGGAGTCGTTGCCCCCAACCCCATCCTCTATCTTTNNNAGATCGGAAGAG
58569 TATGAGTGTGGAGTCGTTGCGCTGGGCATTCCTTATTCTCCATNNNAGATCGGAAGA
71413 TATGAGTGTGGAGTCGTTGCGAGAGAGAGAAAACACGTGCTGTNNNAGATCGGAA
64224 TATGAGTGTGGAGTCGTTGCGCATCACATTGAACTCTTCATAGAGTNNNAGATCGGA
37129 TATGAGTGTGGAGTCGTTGCACCCTCACACCCATAGTCATTCANNNAGATCGGAAGA
65520 TATGAGTGTGGAGTCGTTGCTCTGCCTGTCTCTAGTTATTGGTNNNAGATCGGAAGA
51072 TATGAGTGTGGAGTCGTTGCATCTGCTGCACTGTTATCGCAGNNNAGATCGGAAGA
31131 TATGAGTGTGGAGTCGTTGCAGAGCACAAATTCACAGCTCCAANNNAGATCGGAAG
58742 TATGAGTGTGGAGTCGTTGCCTCATATGCTTTACGCTGTGCATNNNAGATCGGAAGA

56801 TATGAGTGTGGAGTCGTTGCATGTATGTGTGTGTCAGTGGGTGNNNAGATCGGAA
51520 TATGAGTGTGGAGTCGTTGCTTCCCGCTTTGTAACTGTGAGAGNNNAGATCGGAAG
47986 TATGAGTGTGGAGTCGTTGCTGTGTGTATGCTCATGGATGTGCNNNAGATCGGAAG
35413 TATGAGTGTGGAGTCGTTGCGAGATCCTTGTTGCTAATGAGCANNNAGATCGGAAG
56726 TATGAGTGTGGAGTCGTTGCCCCACAAAGGAACAATGTAAGGTGNNNAGATCGGAA
46010 TATGAGTGTGGAGTCGTTGCACATGCACCTCTGGTTTCTTTCCNNNAGATCGGAAGA
56700 TATGAGTGTGGAGTCGTTGCTGTATAATTGTTTATAGTGTGAGCGTGNNNAGATCG
30155 TATGAGTGTGGAGTCGTTGCAAAGCAAGAGAGGGCATGGAAAANNNAGATCGGAA
48573 TATGAGTGTGGAGTCGTTGCCCCTCTGTCCAAGACACAAACTCNNNAGATCGGAAGA
65528 TATGAGTGTGGAGTCGTTGCTTTTGAGATTTCTCACCTCCTTGGTNNNAGATCGGAA
58837 TATGAGTGTGGAGTCGTTGCAGCCTTGACTTCCTCACTCTCATNNNAGATCGGAAGA
52477 TATGAGTGTGGAGTCGTTGCATCGCAAATCTGATGAGGTTCCGNNNAGATCGGAAG
66358 TATGAGTGTGGAGTCGTTGCGCATGTATAGCATTTACACTATCGTGTNNNAGATCGG
62737 TATGAGTGTGGAGTCGTTGCCGCATTTTCCTGTACCTGTATCTNNNAGATCGGAAGA
47057 TATGAGTGTGGAGTCGTTGCCCAAAACTCGAGGATAGAAGGGCNNNAGATCGGAA
36002 TATGAGTGTGGAGTCGTTGCAGGAGTTGACATGTGTTTCTATATGCANNNAGATCG
62006 TATGAGTGTGGAGTCGTTGCCCCTTCCCATATTGTCCTAGGCTNNNAGATCGGAAGA
96631 TATGAGTGTGGAGTCGTTGCTTTCTTCCAGATAGGGTACACTTCTNNNAGATCGGAA
69583 TATGAGTGTGGAGTCGTTGCAGGGTTTCCACCACAGATGTTTTNNNAGATCGGAAG
39873 TATGAGTGTGGAGTCGTTGCTTTTCTGGGTCTCAGTCACCTGANNNAGATCGGAAGA
38203 TATGAGTGTGGAGTCGTTGCGGAGAGGAACTGGAAGAAGGAGANNNAGATCGGA
100235 TATGAGTGTGGAGTCGTTGCCTTCGTTACACAAATGTGAAGGTNNNAGATCGGAAG
32417 TATGAGTGTGGAGTCGTTGCGCTGCGTACACTTGGAACTCTAANNNAGATCGGAAG
68936 TATGAGTGTGGAGTCGTTGCCAGGTGCCAGGGTAGTTCTATTTNNNAGATCGGAAG
109968 TATGAGTGTGGAGTCGTTGCTGTTTATAGTGTGAGCGTGTATAATAGNNNAGATCG
116201 TATGAGTGTGGAGTCGTTGCTCTCTCCCCTTTCTCTCCAGTCNNNAGATCGGAAGAG
46979 TATGAGTGTGGAGTCGTTGCGAGAAACAGAGACAGAGTGAGGCNNNAGATCGGA
64928 TATGAGTGTGGAGTCGTTGCAAGGTCACTCTGCTTACTGAGGTNNNAGATCGGAAG
67338 TATGAGTGTGGAGTCGTTGCAGTTCATCCTGAGGCACCTGATTNNNAGATCGGAAG
57944 TATGAGTGTGGAGTCGTTGCCAGCTTGGATGCTAAGGAGCAATNNNAGATCGGAA
61430 TATGAGTGTGGAGTCGTTGCTACTTGCTGCCTACCTTTCTCCTNNNAGATCGGAAGA
49085 TATGAGTGTGGAGTCGTTGCGGATCTCTCCTAACAGGGGTCTCNNNAGATCGGAAG
35687 TATGAGTGTGGAGTCGTTGCTCTCCCTAGGCATTTCTTAGGCANNNAGATCGGAAGA
43804 TATGAGTGTGGAGTCGTTGCTGGCTCTCTTTCTTAGGAGAACCNNNAGATCGGAAG
37347 TATGAGTGTGGAGTCGTTGCTCACAGAGAAAGCACCACTTTCANNNAGATCGGAAG
40194 TATGAGTGTGGAGTCGTTGCGATGAAGAGAGTGGGAGAGGTGANNNAGATCGGA
52833 TATGAGTGTGGAGTCGTTGCTGGGATAGAGACAGAGCTGAAGGNNNAGATCGGA
101041 TATGAGTGTGGAGTCGTTGCAAAACAACAATAACCCACACACCAATNNNAGATCGGA
57130 TATGAGTGTGGAGTCGTTGCTGTACTGGGGAGTTGTGTTTGTGNNNAGATCGGAA
69404 TATGAGTGTGGAGTCGTTGCCCCCAGGAACTTCCCACAATTTTNNNAGATCGGAAGA
106216 TATGAGTGTGGAGTCGTTGCTTCGACTGAACACTTTGCTTTCCTNNNAGATCGGAAG
57309 TATGAGTGTGGAGTCGTTGCGCTACAGATGATAGATAGAATAGCTTGNNNAGATCG
40059 TATGAGTGTGGAGTCGTTGCTCTGCCCATTTTTGATCGTCTGANNNAGATCGGAAGA

52481 TATGAGTGTGGAGTCGTTGCTGGAGACAGAAGATGGAAAAGCGNNNAGATCGGA
52845 TATGAGTGTGGAGTCGTTGCTAAGGAGGCAAGCATGTTGAAGGNNNAGATCGGA
44402 TATGAGTGTGGAGTCGTTGCTACACACACAATCTCTCTCCCCCNNNAGATCGGAAGA
62889 TATGAGTGTGGAGTCGTTGCACTGACAATTCCTAACCCTCCTCTNNNAGATCGGAAG
44001 TATGAGTGTGGAGTCGTTGCAGACATCCTCATCCTCCTTCACCNNNAGATCGGAAGA
66700 TATGAGTGTGGAGTCGTTGCAGCCCATCAGAAAATCGAGTTGTNNNAGATCGGAA
49859 TATGAGTGTGGAGTCGTTGCGGCAGGGCAGATGACAAAATTTCNNNAGATCGGAA
68140 TATGAGTGTGGAGTCGTTGCCTGCATGAATTGGAAGATCATTCTTNNNAGATCGGA
37135 TATGAGTGTGGAGTCGTTGCTCTTGGCACCAACCTTTCATTCANNNAGATCGGAAGA
32004 TATGAGTGTGGAGTCGTTGCAACCCACACACCAATACTTGGAANNNAGATCGGAAG
38296 TATGAGTGTGGAGTCGTTGCTGTAGAGGATGGCAGTGATGAGANNNAGATCGGA
35574 TATGAGTGTGGAGTCGTTGCAGAGAGAGAGTAGAAAGAAAGGCANNNAGATCGG
47826 TATGAGTGTGGAGTCGTTGCGAAATCTGTGCTTTGGTGTCTGCNNNAGATCGGAAG
101460 TATGAGTGTGGAGTCGTTGCCCAAACTGCCCCTTTTTATAAGGAGANNNAGATCGG
50342 TATGAGTGTGGAGTCGTTGCTAGGTCAGGAGTGGTAGCTCAAGNNNAGATCGGAA
61207 TATGAGTGTGGAGTCGTTGCAGTAGTGCTTAGAGGAAAATGCCTNNNAGATCGGA
42105 TATGAGTGTGGAGTCGTTGCTCAACTAGGGCCCCTGTCTTTTANNNAGATCGGAAG
102810 TATGAGTGTGGAGTCGTTGCTTTAGATGGAGGCTTACGCTGTTNNNAGATCGGAA
62267 TATGAGTGTGGAGTCGTTGCCCAATGAGCTGTCCAGTTATGCTNNNAGATCGGAAG
67566 TATGAGTGTGGAGTCGTTGCTATCCCAAAGAGCTCCAGGACTTNNNAGATCGGAAG
57490 TATGAGTGTGGAGTCGTTGCTTAGATGGAGGCTTACGCTGTTGNNNAGATCGGAA
40856 TATGAGTGTGGAGTCGTTGCGGAGTGGATATTGGGGGTGGATANNNAGATCGGA
55821 TATGAGTGTGGAGTCGTTGCAAACCAGGAGAACTTCTGCACTGNNNAGATCGGAA
62414 TATGAGTGTGGAGTCGTTGCTGTGAGAGGACTGTATGTGTGCTNNNAGATCGGAA
44710 TATGAGTGTGGAGTCGTTGCATTTCTTTGCTTCTCCAGCTCCCNNNAGATCGGAAGA
98858 TATGAGTGTGGAGTCGTTGCGAAGTCCTAAGCTCCCGTACCTCNNNAGATCGGAAG
40572 TATGAGTGTGGAGTCGTTGCTAGCAAGCCTCTTTGCCTTTTGANNNAGATCGGAAG
65560 TATGAGTGTGGAGTCGTTGCATCAGACCTGTGAGTTGATTTGGTNNNAGATCGGA
69503 TATGAGTGTGGAGTCGTTGCTGTAGGGAAGCAAGTTGGCTTTTNNNAGATCGGAA
71699 TATGAGTGTGGAGTCGTTGCGGTAGTAGCTCTTGAAGCACCAANNNAGATCGGAA
39985 TATGAGTGTGGAGTCGTTGCGTGATCTTTGGGATCTTGGCTGANNNAGATCGGAA
60270 TATGAGTGTGGAGTCGTTGCCGCAGACTAGCAAAATCCAGACTNNNAGATCGGAAG
66987 TATGAGTGTGGAGTCGTTGCTGTTCCATTATCACTTCAAAGGAATTNNNAGATCGG
41509 TATGAGTGTGGAGTCGTTGCGCATTGTTGAGTTCTGGGAGGTANNNAGATCGGAA
54861 TATGAGTGTGGAGTCGTTGCAAGAAGCTGTGGTTCATGAGTGGNNNAGATCGGAA
49273 TATGAGTGTGGAGTCGTTGCCTGGGAAAATGCAGGAAAACGTCNNNAGATCGGAA
38823 TATGAGTGTGGAGTCGTTGCAAGGGGGAGATAGAAGAGGAGGANNNAGATCGG
58025 TATGAGTGTGGAGTCGTTGCACACTCTTTTGGGGGTCAAGAATNNNAGATCGGAA
34049 TATGAGTGTGGAGTCGTTGCACATCATGAGACTTACCCCACCANNNAGATCGGAAG
48768 TATGAGTGTGGAGTCGTTGCGGCAAGATCTCACTTACTGCCTCNNNAGATCGGAAG
101634 TATGAGTGTGGAGTCGTTGCATCTCACATCTCACTCTGTGGCCNNNAGATCGGAAGA
42791 TATGAGTGTGGAGTCGTTGCATGATGGAAGGCAACTTCTCCACNNNAGATCGGAAG
54710 TATGAGTGTGGAGTCGTTGCTGAGGAAAAGGACAGAATGCTGGNNNAGATCGGA

| | |
|---|---|
| 42439 | TATGAGTGTGGAGTCGTTGCAAACACCCCATCAGAGGTTGAACNNNAGATCGGAAG |
| 68449 | TATGAGTGTGGAGTCGTTGCATGTTCCAAATTGGTTCCGGGTTNNNAGATCGGAAG |
| 99854 | TATGAGTGTGGAGTCGTTGCCTGTCTTTGCTTATGTGTGCAGGNNNAGATCGGAAG |
| 97862 | TATGAGTGTGGAGTCGTTGCTTTCCAGCATGTGGGTGTATGTNNNAGATCGGAAG |
| 55082 | TATGAGTGTGGAGTCGTTGCAAGGAGAAAAACAGGAGGCTTGGNNNAGATCGGA |
| 56261 | TATGAGTGTGGAGTCGTTGCATGCTTCATGCTTCCTCTGCTGNNNAGATCGGAAGA |
| 58547 | TATGAGTGTGGAGTCGTTGCTCCCAGTGTATGTCACCATCCATNNNAGATCGGAAG |
| 35832 | TATGAGTGTGGAGTCGTTGCTCAAATCACATATGCAGGTGGCANNNAGATCGGAA |
| 33381 | TATGAGTGTGGAGTCGTTGCAGCGAGAAAGAGAGGATCTCACANNNAGATCGGAA |
| 62029 | TATGAGTGTGGAGTCGTTGCTCTCCTTTCGATCATACAGGGCTNNNAGATCGGAAG |
| 53557 | TATGAGTGTGGAGTCGTTGCTTGAGGATTGAATGAGCAGTCGGNNNAGATCGGAA |
| 56133 | TATGAGTGTGGAGTCGTTGCGAGTGAGGGTTGAAGAAGAGCTGNNNAGATCGGA |
| 66060 | TATGAGTGTGGAGTCGTTGCCTTTTGAATCCCATGGCAGCTGTNNNAGATCGGAAG |
| 50195 | TATGAGTGTGGAGTCGTTGCATGGTCACACCTGCCAAGTAAAGNNNAGATCGGAA |
| 39935 | TATGAGTGTGGAGTCGTTGCCCGTGAATTGTATGGCTTCCTGANNNAGATCGGAAG |
| 41041 | TATGAGTGTGGAGTCGTTGCCATTTCACTGCATTCCCACCCTANNNAGATCGGAAGA |
| 45864 | TATGAGTGTGGAGTCGTTGCGCAAGGACTACCAGACAACTTCCNNNAGATCGGAAG |
| 46283 | TATGAGTGTGGAGTCGTTGCTGGATGCTTTGCATTTCTACAGCNNNAGATCGGAAG |
| 66509 | TATGAGTGTGGAGTCGTTGCGCATAACTACTCCTGGTGTGTGTNNNAGATCGGAAG |
| 29841 | TATGAGTGTGGAGTCGTTGCGGGACCTCCAAATAATCTAAGCAGGNNNAGATCGG |
| 56883 | TATGAGTGTGGAGTCGTTGCAAGGACACTGCGGTACTAATGTGNNNAGATCGGAA |
| 45366 | TATGAGTGTGGAGTCGTTGCCTGCAACAGCTTCTAAGTTTGCCNNNAGATCGGAAG |
| 52920 | TATGAGTGTGGAGTCGTTGCAGAAAGGGTCTGTTGTTGACAGGNNNAGATCGGAA |
| 67136 | TATGAGTGTGGAGTCGTTGCCGAGTAGAAATCAGCCCAGCATTNNNAGATCGGAA |
| 60016 | TATGAGTGTGGAGTCGTTGCTCAGGACCTTCAGTAATGACACTNNNAGATCGGAAG |
| 35420 | TATGAGTGTGGAGTCGTTGCAACTCCACATGAAACACTGAGCANNNAGATCGGAAG |
| 108589 | TATGAGTGTGGAGTCGTTGCACCTATATGCTAGATGGCCCACTGNNNAGATCGGAA |
| 49616 | TATGAGTGTGGAGTCGTTGCGTATAGTGCAGGCCACTCACTTCNNNAGATCGGAAG |
| 48574 | TATGAGTGTGGAGTCGTTGCTATGGAGCAGAAGAGGCAAACTCNNNAGATCGGAA |
| 110078 | TATGAGTGTGGAGTCGTTGCTCTGTCTGTTGTATGTGTGCGANNNAGATCGGAAGA |
| 55662 | TATGAGTGTGGAGTCGTTGCGGATGGGAGACTGTGACTTGATGNNNAGATCGGAA |
| 33226 | TATGAGTGTGGAGTCGTTGCAGGCATGATTTCCCTTTATCCACANNNAGATCGGAA |
| 43071 | TATGAGTGTGGAGTCGTTGCGAGGAGGGGTGTGTACAATTCACNNNAGATCGGAA |
| 50670 | TATGAGTGTGGAGTCGTTGCCACAGAGAGACAGCTTTCCACAGNNNAGATCGGAA |
| 32814 | TATGAGTGTGGAGTCGTTGCCCAAACATTTTCTGAGCACCAACANNNAGATCGGAA |
| 33810 | TATGAGTGTGGAGTCGTTGCCCAGCCTAATCCCTGTACCTACANNNAGATCGGAAGA |
| 41051 | TATGAGTGTGGAGTCGTTGCGATACAAAATGGCCACAGCCCTANNNAGATCGGAAG |
| 38681 | TATGAGTGTGGAGTCGTTGCGAACAAGCACAGCTAAGGAAGGANNNAGATCGGAA |
| 33498 | TATGAGTGTGGAGTCGTTGCAGAAAGAACCACAGAGACAGACANNNAGATCGGAA |
| 64447 | TATGAGTGTGGAGTCGTTGCGCTACCAAACACAGCACTCTAGTNNNAGATCGGAAG |
| 51661 | TATGAGTGTGGAGTCGTTGCTGTGGTCAAAGTGGTTTAGGGAGNNNAGATCGGAA |
| 40213 | TATGAGTGTGGAGTCGTTGCACAGACCGAGAACTAATGGGTGANNNAGATCGGAA |
| 58751 | TATGAGTGTGGAGTCGTTGCTACAGTGTCTGTGCATGTTGCATNNNAGATCGGAAG |

46867 TATGAGTGTGGAGTCGTTGCAACTGTGAAGGATCAAGACAGGCNNNAGATCGGAA

68585 TATGAGTGTGGAGTCGTTGCCCTCACGGAGCATTTCGTATGTTNNNAGATCGGAAG

99595 TATGAGTGTGGAGTCGTTGCACTCCCTTCGTGAATTTCCCTTTCNNNAGATCGGAAG

66673 TATGAGTGTGGAGTCGTTGCCACTCATTCCTGCTTGGTCTTGTNNNAGATCGGAAG

33850 TATGAGTGTGGAGTCGTTGCCCAGGACCCGTAGGATTTCTACANNNAGATCGGAAG

58402 TATGAGTGTGGAGTCGTTGCTCCCGTGGTCATAGCTGATACATNNNAGATCGGAAG

39993 TATGAGTGTGGAGTCGTTGCCTGTAACACACATACACTGCTGANNNAGATCGGAAG

44725 TATGAGTGTGGAGTCGTTGCAGGAGGTTCTTCTTTCCTCTCCCNNNAGATCGGAAG

36825 TATGAGTGTGGAGTCGTTGCTTGAAATTGGCAGAGGCTTCTCANNNAGATCGGAA

60664 TATGAGTGTGGAGTCGTTGCTAGCTGCGTAGACTTGTACACCTNNNAGATCGGAAG

35047 TATGAGTGTGGAGTCGTTGCAGCAGAAAGATCCAATCCTTCCANNNAGATCGGAAG

35804 TATGAGTGTGGAGTCGTTGCTGACACTTCTAAACAACCTGGCANNNAGATCGGAAG

44999 TATGAGTGTGGAGTCGTTGCTAGAGGAGGCAGATAACGTAGCCNNNAGATCGGAA

99847 TATGAGTGTGGAGTCGTTGCAGAGTAGATTGGAGTAGGAAGAGGTNNNAGATCG

65505 TATGAGTGTGGAGTCGTTGCCTTGCTTTCCCTTGGGAATTGGTNNNAGATCGGAAG

47181 TATGAGTGTGGAGTCGTTGCACAAAAGTTTTCCTGAGTTGGGCNNNAGATCGGAA

64255 TATGAGTGTGGAGTCGTTGCGGAAATTGTTGAGGCTGAGGAGTNNNAGATCGGA

52008 TATGAGTGTGGAGTCGTTGCAATCCCCAAGAGCCCCTGTATAGNNNAGATCGGAAG

42674 TATGAGTGTGGAGTCGTTGCTGCCTTTCACCTTCTGTAACCACNNNAGATCGGAAGA

52987 TATGAGTGTGGAGTCGTTGCGTTTGTTTCCAGAAATGGCCAGGNNNAGATCGGAA

50319 TATGAGTGTGGAGTCGTTGCCAGAAACAGTGTGGATGTGCAAGNNNAGATCGGAA

57424 TATGAGTGTGGAGTCGTTGCCCCCTCCAGTGTTGTATCAGTTGNNNAGATCGGAAG

71774 TATGAGTGTGGAGTCGTTGCTTCCTTTGCATACGGTCTCTTCGNNNAGATCGGAAG

101455 TATGAGTGTGGAGTCGTTGCGTCCCTTTGGCCTTTGCCTTNNNAGATCGGAAGAGC

65209 TATGAGTGTGGAGTCGTTGCTATTTCATGCATGTGTGGTGGGTNNNAGATCGGAA

67173 TATGAGTGTGGAGTCGTTGCCCCTCCAAGCAGAATACTGCATTNNNAGATCGGAAG

46197 TATGAGTGTGGAGTCGTTGCATCACAGCAGAAGTGAAGGAAGCNNNAGATCGGAA

65338 TATGAGTGTGGAGTCGTTGCACAGTGAGAGTTACAAAACACTGGTNNNAGATCGG

40801 TATGAGTGTGGAGTCGTTGCCAATGTCAGCCACACACCAGATANNNAGATCGGAAG

42120 TATGAGTGTGGAGTCGTTGCAGAGGCCTGCCATCAGCTTTTTANNNAGATCGGAAG

59078 TATGAGTGTGGAGTCGTTGCGAGAGAGAGAGGCTGGTTGAGATNNNAGATCGGA

54738 TATGAGTGTGGAGTCGTTGCTTTCTCTCCCTCCAGTCATCTGGNNNAGATCGGAAGA

48812 TATGAGTGTGGAGTCGTTGCTCTCAGCATCTCTAGCTGTCCTCNNNAGATCGGAAGA

31542 TATGAGTGTGGAGTCGTTGCTCTGGGGACACGACTTACAAGAANNNAGATCGGAA

70675 TATGAGTGTGGAGTCGTTGCGAGAAGGGTCCATGGGTCTCTAGNNNAGATCGGAA

49370 TATGAGTGTGGAGTCGTTGCATCTGCAACTTGGAGGACATGTCNNNAGATCGGAA

64262 TATGAGTGTGGAGTCGTTGCCATTTGACGAAACGCTTAGGAGTNNNAGATCGGAA

51909 TATGAGTGTGGAGTCGTTGCTGGTGGATAGAGGTACGTGTGAGNNNAGATCGGA

99588 TATGAGTGTGGAGTCGTTGCAAAACAACCAACAACCCAGTGTTNNNAGATCGGAAG

36849 TATGAGTGTGGAGTCGTTGCGCTGCATATGGTCTCCAAAGTCANNNAGATCGGAAG

39597 TATGAGTGTGGAGTCGTTGCTGGATAGCTCTGAAAAACTGATAATGANNNAGATCG

57611 TATGAGTGTGGAGTCGTTGCTGAGGCACAGTGATAGTGGTTTGNNNAGATCGGAA

46513 TATGAGTGTGGAGTCGTTGCTAACCTTGGCTTTTGGGTAGAGCNNNAGATCGGAA

40146 TATGAGTGTGGAGTCGTTGCGAAAACGCAGATTCCATTCAGTGANNNAGATCGGAA

62508 TATGAGTGTGGAGTCGTTGCTACAAAGCCCATCACACTTTGCTNNNAGATCGGAAG

46574 TATGAGTGTGGAGTCGTTGCGGCAACTCCAGGTAAAAATGAGCNNNAGATCGGAA

66310 TATGAGTGTGGAGTCGTTGCGGAGCAGATTTGGAAGTCAGTGTNNNAGATCGGAA

64598 TATGAGTGTGGAGTCGTTGCCTATCCTGTGCTTTCGTTACCGTNNNAGATCGGAAG

68323 TATGAGTGTGGAGTCGTTGCGGTGCTTTGTACAGACTGGAGTTNNNAGATCGGAA

57206 TATGAGTGTGGAGTCGTTGCCAGAAGAGCTGAGTGAGGGATTGNNNAGATCGGA

51054 TATGAGTGTGGAGTCGTTGCCGTACTGCTAGAGAGTTGAGCAGNNNAGATCGGAA

54756 TATGAGTGTGGAGTCGTTGCTCATGCAAACTCATTTGCTCTGGNNNAGATCGGAAG

65366 TATGAGTGTGGAGTCGTTGCTGTCATAGACTGATGAGCCTGGTNNNAGATCGGAA

103327 TATGAGTGTGGAGTCGTTGCTACCTCGGTCTTCTCTTTCAGGNNNAGATCGGAAGA

53740 TATGAGTGTGGAGTCGTTGCTACCTCGGTCTTCTCTTTCAGGGNNNAGATCGGAAG

30897 TATGAGTGTGGAGTCGTTGCACGGAAGTTGTAAAGTCTCACAANNNAGATCGGAA

117312 TATGAGTGTGGAGTCGTTGCTCCACAGTTTTGAGCTTACTCCATNNNAGATCGGAAG

47114 TATGAGTGTGGAGTCGTTGCGTAGAGCTGACTTTATGAGGGGCNNNAGATCGGAA

44510 TATGAGTGTGGAGTCGTTGCTATGTCTGCTTTCTCAGAAGCCCNNNAGATCGGAAG

53611 TATGAGTGTGGAGTCGTTGCTCTGCTTTTGAGATGCTCAAGGGNNNAGATCGGAAG

30110 TATGAGTGTGGAGTCGTTGCGCCAGACTTTGGGGAGAAGAAAANNNAGATCGGA

46701 TATGAGTGTGGAGTCGTTGCCCCAGCTCAACCAGACTATTAGCNNNAGATCGGAAG

54777 TATGAGTGTGGAGTCGTTGCAAAGACACCCTGGGAAATTCTGGNNNAGATCGGAA

64388 TATGAGTGTGGAGTCGTTGCGGATCATCAGCTGGGCAAATAGTNNNAGATCGGAA

54743 TATGAGTGTGGAGTCGTTGCCCACACGTTCTGTAGCTATCTGGNNNAGATCGGAAG

101710 TATGAGTGTGGAGTCGTTGCTGTCTGTGTTGGTGGATTCAGGGNNNAGATCGGAA

33107 TATGAGTGTGGAGTCGTTGCCACAAACTCATCAGTAGATTTGACACANNNAGATCG

54171 TATGAGTGTGGAGTCGTTGCGAATAGAAGGCACAGACAGTGGGNNNAGATCGGA

99902 TATGAGTGTGGAGTCGTTGCCAAAAGAGTAAGTGCGGCCTGANNNAGATCGGAAG

96630 TATGAGTGTGGAGTCGTTGCCAGACATACACATCAGCGGATGCTCCNNNAGATCGG

67702 TATGAGTGTGGAGTCGTTGCTTCAGTGGTTTGTGTTGTCCCTTNNNAGATCGGAAG

37597 TATGAGTGTGGAGTCGTTGCTCCATCAGAGAGTGGCTTCAAGANNNAGATCGGAA

43080 TATGAGTGTGGAGTCGTTGCTTGCATGTGCGTGTTTACTTCACNNNAGATCGGAAG

56098 TATGAGTGTGGAGTCGTTGCCTGAGAAGCTACTGGACAAGCTGNNNAGATCGGAA

48564 TATGAGTGTGGAGTCGTTGCCCTGACAGTTGGCCCTTCTTATCNNNAGATCGGAAG

48783 TATGAGTGTGGAGTCGTTGCGCATGAACAGGTTCCAGATCCTCNNNAGATCGGAAG

42042 TATGAGTGTGGAGTCGTTGCCCCTGGCCATCATCTCCTCTTTANNNAGATCGGAAGA

37502 TATGAGTGTGGAGTCGTTGCTGCCTTTGTACATCCAGGAAAGANNNAGATCGGAAG

38991 TATGAGTGTGGAGTCGTTGCTGGACAAAATTTGGGGTAAGGGANNNAGATCGGA

30381 TATGAGTGTGGAGTCGTTGCATTTAGGTGAGGTTGCAGGCAAANNNAGATCGGAA

47999 TATGAGTGTGGAGTCGTTGCCTAAGCAAAGTATGCTGCTGTGCNNNAGATCGGAAG

61148 TATGAGTGTGGAGTCGTTGCTGCTTTCAACAAGTACAAGGCCTNNNAGATCGGAAG

29840 TATGAGTGTGGAGTCGTTGCGTAGGAAGCAATGACACCTTCCANNNAGATCGGAAG

68280 TATGAGTGTGGAGTCGTTGCCAGATGTTTGTCTCAGGGCAGTTNNNAGATCGGAAG

62827 TATGAGTGTGGAGTCGTTGCAAGTCTCCTGTAATCCCACCTCTNNNAGATCGGAAGA

62339 TATGAGTGTGGAGTCGTTGCATGCGTTCAGAAAACTCTCTGCTNNNAGATCGGAAG

62241 TATGAGTGTGGAGTCGTTGCAGTAGGAACATTACAACGATGCTNNNAGATCGGAA

65939 TATGAGTGTGGAGTCGTTGCGCAGTGGAAGATGGAATGACTGTNNNAGATCGGA

99898 TATGAGTGTGGAGTCGTTGCATGGTTTTAGCACTCAGGCTATGNNNAGATCGGAAG

50528 TATGAGTGTGGAGTCGTTGCGCGTGCGTGTATGTGAATTGAAGNNNAGATCGGAA

46824 TATGAGTGTGGAGTCGTTGCGATTTCAGGGCAAAGAACAAGGCNNNAGATCGGAA

44197 TATGAGTGTGGAGTCGTTGCCTCTCTTGGTTGCAGGACTTACCNNNAGATCGGAAG

104905 TATGAGTGTGGAGTCGTTGCAGTTGTAAAGTCTCACAATATGCCATCNNNAGATCG

105037 TATGAGTGTGGAGTCGTTGCCAGAGTAGTTCAGTGAGGCCATCANNNAGATCGGA

43494 TATGAGTGTGGAGTCGTTGCCTGCTCCTATTCCTGTGGCATACNNNAGATCGGAAG

44747 TATGAGTGTGGAGTCGTTGCATGCTGGAATGAATGGAAGTCCCNNNAGATCGGAA

34138 TATGAGTGTGGAGTCGTTGCACTCCCTTCCATGATGATGACCANNNAGATCGGAAG

52318 TATGAGTGTGGAGTCGTTGCACATATAGGCTCACCATTGAGACGNNNAGATCGGAA

31477 TATGAGTGTGGAGTCGTTGCACCCACATACATATAGGCATTCAANNNAGATCGGAA

42167 TATGAGTGTGGAGTCGTTGCCAAGGAATGGCAGACACACAAACNNNAGATCGGAA

108540 TATGAGTGTGGAGTCGTTGCGCCTGTAAGGAGACCCATACTTNNNAGATCGGAAG

40817 TATGAGTGTGGAGTCGTTGCGCACAAACCAGAGCAGTGAGATANNNAGATCGGAA

67087 TATGAGTGTGGAGTCGTTGCTAAGGGATCTGGGAGCGTACATTNNNAGATCGGAA

66437 TATGAGTGTGGAGTCGTTGCGTCCAGGTCTTTGGATTATGTGTNNNAGATCGGAAG

69508 TATGAGTGTGGAGTCGTTGCCACAATATGGGCAACCTGCTTTTNNNAGATCGGAAG

34836 TATGAGTGTGGAGTCGTTGCAGCAAATAATTCACCCGTGATCCANNNAGATCGGAA

58752 TATGAGTGTGGAGTCGTTGCCAGGGCCTTTGTTCTTGTTGCATNNNAGATCGGAAG

47490 TATGAGTGTGGAGTCGTTGCCAATTCAGTCCGTAGCAACATGCNNNAGATCGGAAG

38382 TATGAGTGTGGAGTCGTTGCTGTGTACCTCTCACTGGCATAGANNNAGATCGGAAG

55272 TATGAGTGTGGAGTCGTTGCAGAGTGGTGCAAGATAGCCAATGNNNAGATCGGAA

67277 TATGAGTGTGGAGTCGTTGCATCCAGACACATCCCCAGAGATTNNNAGATCGGAAG

32487 TATGAGTGTGGAGTCGTTGCACAGACAGACAGACAAGGGGTAANNNAGATCGGA

47920 TATGAGTGTGGAGTCGTTGCCCATTTCCTTGTCATTAGCGTGCNNNAGATCGGAAG

32947 TATGAGTGTGGAGTCGTTGCACAAGCCCCTAGGTTGAATAACANNNAGATCGGAAG

41362 TATGAGTGTGGAGTCGTTGCCACCCTGGTCTTCTCGTCAAGTANNNAGATCGGAAG

60123 TATGAGTGTGGAGTCGTTGCACAAATACTCAGAATCAGGAGCACTNNNAGATCGGA

37742 TATGAGTGTGGAGTCGTTGCTCCAAATAACTATGGCATCCAACAGANNNAGATCGG

45610 TATGAGTGTGGAGTCGTTGCCCCTACTTCCTTCTTCTTTCCTCCNNNAGATCGGAAGA

66810 TATGAGTGTGGAGTCGTTGCCTATGTGCCTCTGTCTGCTTTGTNNNAGATCGGAAG

101459 TATGAGTGTGGAGTCGTTGCGATGAAATCTTACTCTTGCCCAGNNNAGATCGGAAG

100506 TATGAGTGTGGAGTCGTTGCGTTATCTTCTCCAGTCTGCTACCANNNAGATCGGAAG

29951 TATGAGTGTGGAGTCGTTGCTGGACACAACAGGCATCTCTAGANNNAGATCGGAA

47284 TATGAGTGTGGAGTCGTTGCAAAATCTGTAGGGTATGGCTGGCNNNAGATCGGAA

56751 TATGAGTGTGGAGTCGTTGCCAAGAGACTGAAGCTGAGAGGTGNNNAGATCGGA

47424 TATGAGTGTGGAGTCGTTGCCCCACTTTCTTTTCGTCTTTGGCNNNAGATCGGAAGA

101709 TATGAGTGTGGAGTCGTTGCGAACACAGACTGGGCAAACTAGANNNAGATCGGAA

101056 TATGAGTGTGGAGTCGTTGCTGGCTTCTACTTCTTATGTCTTGGAAANNNAGATCGG

101471 TATGAGTGTGGAGTCGTTGCACCCATCCAGGCAGTTATGAAAGNNNAGATCGGAA

100750 TATGAGTGTGGAGTCGTTGCATTTGAGTGGAAGTCACGTGGATGNNNAGATCGGA

100948 TATGAGTGTGGAGTCGTTGCGAGTAGGGTCAGTTCTCGCTCTTTNNNAGATCGGAA
102879 TATGAGTGTGGAGTCGTTGCGCCCACCCAGAGAATGTTCTCGNNNAGATCGGAAGA
101050 TATGAGTGTGGAGTCGTTGCTTCCTGTTGAATCTGGACAAAGTTANNNAGATCGGA
117092 TATGAGTGTGGAGTCGTTGCGTGACATCGAGGACCTTAGGAAAGNNNAGATCGGA
102556 TATGAGTGTGGAGTCGTTGCCAGAAGGAAGCATTGCTAATATAGGANNNAGATCG
105036 TATGAGTGTGGAGTCGTTGCCCCTTTGAAAGTAGAGAGAGGTGTNNNAGATCGGA
106233 TATGAGTGTGGAGTCGTTGCTTCTTTGGGGTGTGACCATATGTTCNNNAGATCGGA
106251 TATGAGTGTGGAGTCGTTGCTATGGCAGTTCCCTCATGGTCCTNNNAGATCGGAAG
112456 TATGAGTGTGGAGTCGTTGCCATCCAGGGTCATTCAGTAATGCNNNAGATCGGAAG
101440 TATGAGTGTGGAGTCGTTGCTGTATGGGTGTGTGTTTATCCAGTTNNNAGATCGGA
102611 TATGAGTGTGGAGTCGTTGCACTGCCCTGATGTGCATAACTTTNNNAGATCGGAAG
104173 TATGAGTGTGGAGTCGTTGCCACACACACATACATCTCTCCATGTNNNAGATCGGAA
106278 TATGAGTGTGGAGTCGTTGCCAGTTCAAGTCGAAGTCCAAAGGNNNAGATCGGAA
103346 TATGAGTGTGGAGTCGTTGCGCTCTAACTTGTTTCACACCTGGCGNNNAGATCGGA
106640 TATGAGTGTGGAGTCGTTGCCAGGGGTGACAAAAACCTGGATANNNAGATCGGAA
117336 TATGAGTGTGGAGTCGTTGCAAGAAACCAACCTAAGAAGGAGGCNNNAGATCGGA
117280 TATGAGTGTGGAGTCGTTGCGACCATTTTTGTGGGCAGGGAAGNNNAGATCGGAA
104147 TATGAGTGTGGAGTCGTTGCAGGTAGTGGCATGGGTTATGTGTGNNNAGATCGG
97040 TATGAGTGTGGAGTCGTTGCGAGTAGAGGCTGAGGCCTGGANNNAGATCGGAAG
100078 TATGAGTGTGGAGTCGTTGCGCTATTTCCCAGTAACCACTGATCNNNAGATCGGAA
109999 TATGAGTGTGGAGTCGTTGCGTTGGGAGGAACCAGGGCTATNNNAGATCGGAAG
68829 TATGAGTGTGGAGTCGTTGCGTGTACTAGAGTGCCTGCCATTTNNNAGATCGGAAG
99604 TATGAGTGTGGAGTCGTTGCGAAACGCGAGTGACTGGGTCTGNNNAGATCGGAAG
98710 TATGAGTGTGGAGTCGTTGCGAGGGACTTCAGGCCACTGACNNNAGATCGGAAGA
69191 TATGAGTGTGGAGTCGTTGCGGATGACAGAAGCCACAGTCTTTNNNAGATCGGAA
48281 TATGAGTGTGGAGTCGTTGCCCTTTGCTTTTCCGCTGCAATCNNNAGATCGGAAGAG
102595 TATGAGTGTGGAGTCGTTGCGTGTGTTTGCTTTTAGCCTTTGAANNNAGATCGGAA
66273 TATGAGTGTGGAGTCGTTGCTCCAATTCCGTAACCTCAAGTGTNNNAGATCGGAAG
30182 TATGAGTGTGGAGTCGTTGCCCCCATGTGCTCCCATCATAAAANNNAGATCGGAAG
71661 TATGAGTGTGGAGTCGTTGCATCAGAGCCAAGCAGGACTTCTANNNAGATCGGAA
59191 TATGAGTGTGGAGTCGTTGCCTGAAGGATGGAAATTTGGGGATNNNAGATCGGA
64045 TATGAGTGTGGAGTCGTTGCTCTTTTTGATGGGTATTCTGGCAGTNNNAGATCGGA
102828 TATGAGTGTGGAGTCGTTGCATAGAGGGGTCTTTGAGTTTTCTAANNNAGATCGG
44515 TATGAGTGTGGAGTCGTTGCCCACACTAGAACACAGTAAGCCCNNNAGATCGGAAG
29839 TATGAGTGTGGAGTCGTTGCACCAGTTTTCATGTGGATGACCCNNNAGATCGGAAG
55994 TATGAGTGTGGAGTCGTTGCGGGGATTCACAAAACAAAGCCTGNNNAGATCGGAA
46864 TATGAGTGTGGAGTCGTTGCCACACACATATCCAAACACAGGCNNNAGATCGGAAG
61822 TATGAGTGTGGAGTCGTTGCGCTCTCAGCAAGTTCCAATAGCTNNNAGATCGGAAG
56425 TATGAGTGTGGAGTCGTTGCTATGGCAATCACTGTGTGGTCTGNNNAGATCGGAA
39369 TATGAGTGTGGAGTCGTTGCTGACAGAGTGAAAAGCATCTGGANNNAGATCGGAA
50563 TATGAGTGTGGAGTCGTTGCATGCTTCTGCTTGGGGTTCTAAGNNNAGATCGGAAG
66201 TATGAGTGTGGAGTCGTTGCTGGTCTCCCCTGTATTAGTCTGTNNNAGATCGGAAG
100566 TATGAGTGTGGAGTCGTTGCATGTGTGTTTGGGGTTTGTTTGTTNNNAGATCGGA

49838 TATGAGTGTGGAGTCGTTGCCCCTCTCACTCTTTCCAGTGTTCNNNAGATCGGAAGA
59842 TATGAGTGTGGAGTCGTTGCGCACCCATACATACATGGCAACTNNNAGATCGGAAG
41479 TATGAGTGTGGAGTCGTTGCGCCAGTAAGCCCAAGAAAAGGTANNNAGATCGGAA
48351 TATGAGTGTGGAGTCGTTGCCTGTGAGGAAGAAGCTCACCATCNNNAGATCGGAA
59097 TATGAGTGTGGAGTCGTTGCGGACCTACACTGCCCTGTTAGATNNNAGATCGGAAG
71799 TATGAGTGTGGAGTCGTTGCCAGCCAAAGAAGAAACCACTGGANNNAGATCGGAA
48830 TATGAGTGTGGAGTCGTTGCCCTACTGCTCACACACTTTCCTCNNNAGATCGGAAGA
64861 TATGAGTGTGGAGTCGTTGCTCTGGAAAACTGCATACTTCAGGTNNNAGATCGGAA
99915 TATGAGTGTGGAGTCGTTGCAAGACCGACAATAATTCTCTCCANNNAGATCGGAAG
48938 TATGAGTGTGGAGTCGTTGCCTCCAAAGACCTGCTATCTGCTCNNNAGATCGGAAG
38539 TATGAGTGTGGAGTCGTTGCTCCTAAAGCAACCAAAGATTTACGANNNAGATCGGA
29838 TATGAGTGTGGAGTCGTTGCCTCCGGGACACAGTTGAACAATTNNNAGATCGGAAG
60282 TATGAGTGTGGAGTCGTTGCTTCCCCTACAGTCCAGTCAGACTNNNAGATCGGAAG
41591 TATGAGTGTGGAGTCGTTGCACCACCATCCTGAATCCTTGGTANNNAGATCGGAAG
41830 TATGAGTGTGGAGTCGTTGCCCTGCACATTGGATCGTTCCTTANNNAGATCGGAAG
109901 TATGAGTGTGGAGTCGTTGCATGTATGTATGTATGTATGTTGAAGGCNNNAGATCG
56430 TATGAGTGTGGAGTCGTTGCCAAACCCCAGGTAAGACTGTCTGNNNAGATCGGAAG
63823 TATGAGTGTGGAGTCGTTGCCAGGGGAGGTGCTAAGAATATAAGTNNNAGATCGG
46957 TATGAGTGTGGAGTCGTTGCGTGAAAAAGAGAGAGGGAGAGGCNNNAGATCGGA
42728 TATGAGTGTGGAGTCGTTGCTTGCTTACCCCTAAAAAGCCCACNNNAGATCGGAAG
58545 TATGAGTGTGGAGTCGTTGCCATCTCCACACGTCCAAATCCATNNNAGATCGGAAGA
102865 TATGAGTGTGGAGTCGTTGCGGGTTGGGGATTGGGGTAATGNNNAGATCGGAAG
49139 TATGAGTGTGGAGTCGTTGCCTGCTCTCTGGACCCAATTTCTCNNNAGATCGGAAGA
55121 TATGAGTGTGGAGTCGTTGCACACACATTAGTACAGGGGTTGGNNNAGATCGGAA
44823 TATGAGTGTGGAGTCGTTGCCTGCTAACTCAAACCAGGTTCCCNNNAGATCGGAAG
48418 TATGAGTGTGGAGTCGTTGCGTTCACCTCCTTCAGAGCTCATCNNNAGATCGGAAGA
30235 TATGAGTGTGGAGTCGTTGCGAGGCAGATACCACACACACAAANNNAGATCGGAA
43152 TATGAGTGTGGAGTCGTTGCGCCTGGTCATGATGGATACAGACNNNAGATCGGAA
46609 TATGAGTGTGGAGTCGTTGCCACCACTTCACTTCATTGTGAGCNNNAGATCGGAAG
67603 TATGAGTGTGGAGTCGTTGCCGGGTCACAGGGAGTTACTACTTNNNAGATCGGAA
47065 TATGAGTGTGGAGTCGTTGCTATGCAGTTTCTGATGACAGGGCNNNAGATCGGAA
68185 TATGAGTGTGGAGTCGTTGCCAGTTTTGCCCCCTACCTTTCTTNNNAGATCGGAAGA
45773 TATGAGTGTGGAGTCGTTGCCTCCTCTCTAGGAAACCTGGTCCNNNAGATCGGAAG
58107 TATGAGTGTGGAGTCGTTGCGAAGGCAGGGACAATGATGGAATNNNAGATCGGA
49448 TATGAGTGTGGAGTCGTTGCGCAAAAGTAAAGCTGGACGTGTCNNNAGATCGGAA
97648 TATGAGTGTGGAGTCGTTGCGTCTAGTGATGCTTGGCTACCCNNNAGATCGGAAGA
34478 TATGAGTGTGGAGTCGTTGCACATAATATTCCTGGGGGTCCCANNNAGATCGGAAG
31323 TATGAGTGTGGAGTCGTTGCGGATCTGAATCTTGGCTCTGCAANNNAGATCGGAAG
42735 TATGAGTGTGGAGTCGTTGCACAAAAATGGCCCATAATCCCACNNNAGATCGGAAG
101896 TATGAGTGTGGAGTCGTTGCGGATTGGTAGATAGGTGAATGCAGGNNNAGATCG
104193 TATGAGTGTGGAGTCGTTGCGAAAACTGAGTACAAGCAGGACAGGNNNAGATCGG
59604 TATGAGTGTGGAGTCGTTGCTGTATGCGTGTGCATGTACGTATNNNAGATCGGAA
67296 TATGAGTGTGGAGTCGTTGCACTACTTTTCCCTCTCAGCGATTNNNAGATCGGAAGA

45009 TATGAGTGTGGAGTCGTTGCGAGAGAAGCTGTAGCCATTAGCCNNNAGATCGGAA
38959 TATGAGTGTGGAGTCGTTGCACATTGATGAGGGGAATATAAAGGGANNNAGATCG
117113 TATGAGTGTGGAGTCGTTGCTCTCGGTGGGAAAACCAAATGGNNNAGATCGGAAG
45642 TATGAGTGTGGAGTCGTTGCAGGTCAATTTCCACTCTTGCTCCNNNAGATCGGAAG
64707 TATGAGTGTGGAGTCGTTGCCCAGGTTTATCAGCCCAAAAGGTNNNAGATCGGAAG
67455 TATGAGTGTGGAGTCGTTGCGCACGCAATGAGAATGCTTTATTNNNAGATCGGAAG
98448 TATGAGTGTGGAGTCGTTGCAGCAGAGCTCCCAGTCACATCCNNNAGATCGGAAGA
52390 TATGAGTGTGGAGTCGTTGCTTGTGTATCTGAATCTGCACCCGNNNAGATCGGAAG
44281 TATGAGTGTGGAGTCGTTGCCTGTCTCTGCTCTTTTCTCACCCNNNAGATCGGAAGA
43269 TATGAGTGTGGAGTCGTTGCGGAAACCAGGAGAGAAACAGGACNNNAGATCGGA
55115 TATGAGTGTGGAGTCGTTGCTTATAGCTTCCCCAAGACGTTGGNNNAGATCGGAAG
36773 TATGAGTGTGGAGTCGTTGCGGTTGTAAGATGCAGCACTCTCANNNAGATCGGAA
56984 TATGAGTGTGGAGTCGTTGCACACCTGCTTTATTTGAACTCTGTGNNNAGATCGGA
30259 TATGAGTGTGGAGTCGTTGCCTGATTGACGTTGGGGAGACAAANNNAGATCGGAA
65030 TATGAGTGTGGAGTCGTTGCAAACAAGGTTACAAAATTGCGGTNNNAGATCGGAA
117789 TATGAGTGTGGAGTCGTTGCACATTTTATAGCCTACTTTTCGTCATNNNAGATCGGA
43244 TATGAGTGTGGAGTCGTTGCCAGTATGCCCCATGGTTGTAGACNNNAGATCGGAAG
59450 TATGAGTGTGGAGTCGTTGCGGGACAGAGGGACAGAGAGATATNNNAGATCGGA
58180 TATGAGTGTGGAGTCGTTGCCATACCATGGCCCACAAGCTAATNNNAGATCGGAAG
62548 TATGAGTGTGGAGTCGTTGCCAAGAACCAGTGGAGCGAAATCTNNNAGATCGGAA
48523 TATGAGTGTGGAGTCGTTGCCCTGACCCTTTGTCACCACTATCNNNAGATCGGAAGA
71342 TATGAGTGTGGAGTCGTTGCCACACACACACTTTGCAAAAGGTNNNAGATCGGAAG
34900 TATGAGTGTGGAGTCGTTGCGAAGGTCATTTAGGCAAGCTCCANNNAGATCGGAA
56632 TATGAGTGTGGAGTCGTTGCCTCACTCTTATCACCCTGGAGTGNNNAGATCGGAAG
71706 TATGAGTGTGGAGTCGTTGCCTGTCGGCCATCTACAAGTGGATNNNAGATCGGAAG
63425 TATGAGTGTGGAGTCGTTGCTACCTGCTTGCTACCTGTCTTCTNNNAGATCGGAAGA
116346 TATGAGTGTGGAGTCGTTGCTTCTGACAAAGTTTTGCTCCTGTGGNNNAGATCGGA
53522 TATGAGTGTGGAGTCGTTGCTTTAGTCAGGTCCTAGTAGGCGGNNNAGATCGGAA
109865 TATGAGTGTGGAGTCGTTGCAGAGGGTACTGAATAACTCACCTCANNNAGATCGGA
51206 TATGAGTGTGGAGTCGTTGCGAGCAGTAGGCACAGGTTATCAGNNNAGATCGGAA
41980 TATGAGTGTGGAGTCGTTGCGCCTCAGAGTTGCCCAGAATTTANNNAGATCGGAAG
59389 TATGAGTGTGGAGTCGTTGCCACGCACGCACATACACAAATATNNNAGATCGGAAG
59525 TATGAGTGTGGAGTCGTTGCGCACACGCAAACCCATATCCTATNNNAGATCGGAAG
43780 TATGAGTGTGGAGTCGTTGCAAAAGAAAGAACCCAACGCAACCNNNAGATCGGAA
97872 TATGAGTGTGGAGTCGTTGCCTCCTAACCTGTGCTCGTGTGNNNAGATCGGAAGAG
68772 TATGAGTGTGGAGTCGTTGCATGACCTCAGACAACGCCAATTTNNNAGATCGGAAG
33479 TATGAGTGTGGAGTCGTTGCTGGTGATAGGCAGAAGGAAGACANNNAGATCGGA
65243 TATGAGTGTGGAGTCGTTGCGGCTTTTGTAGGAGGGAAATGGTNNNAGATCGGA
41129 TATGAGTGTGGAGTCGTTGCAGAGCCCAGCAAGGTATTTCCTANNNAGATCGGAAG
108546 TATGAGTGTGGAGTCGTTGCCGTTGCCTCAGAGAAGATTCCCNNNAGATCGGAAGA
95946 TATGAGTGTGGAGTCGTTGCGCCTGTCATCCTGTGTAACAGTTNNNAGATCGGAAG
97043 TATGAGTGTGGAGTCGTTGCTGACATTCCGAGTTCCTGGAAGNNNAGATCGGAAG
95493 TATGAGTGTGGAGTCGTTGCTGGTCAAAATACCCCAATCCAGTAANNNAGATCGGA

| | |
|---|---|
| 57810 | TATGAGTGTGGAGTCGTTGCTGGGCAGCTGAATTTCCTGAAATNNNAGATCGGAA |
| 44945 | TATGAGTGTGGAGTCGTTGCCCCAGTCATACACATGAAGAGCCNNNAGATCGGAAG |
| 63336 | TATGAGTGTGGAGTCGTTGCATCCGCCTCTTTTCATGCATTCTNNNAGATCGGAAGA |
| 50800 | TATGAGTGTGGAGTCGTTGCCCTCCCTCTACCTACTTCACCAGNNNAGATCGGAAGA |
| 31933 | TATGAGTGTGGAGTCGTTGCACAAGGATGGGAACTAGTGGGAANNNAGATCGGA |
| 40230 | TATGAGTGTGGAGTCGTTGCCCTTGGGAAACAAGCTATGGTGANNNAGATCGGAA |
| 54487 | TATGAGTGTGGAGTCGTTGCAAACCTCCAAAGGGGAAAACTGGNNNAGATCGGAA |
| 69566 | TATGAGTGTGGAGTCGTTGCGTGTTTGAACTGCAGTGGGTTTTNNNAGATCGGAA |
| 104792 | TATGAGTGTGGAGTCGTTGCAGGAAAAGGTGGTGGACAAGGNNNAGATCGGAAG |
| 38650 | TATGAGTGTGGAGTCGTTGCTAGGAAAAGGTGGTGGACAAGGANNNAGATCGGA |
| 44186 | TATGAGTGTGGAGTCGTTGCCAAACACATGCACACTGTGTACCNNNAGATCGGAAG |
| 38483 | TATGAGTGTGGAGTCGTTGCAACTGGACCAGCAGCTTCTTAGANNNAGATCGGAAG |
| 39059 | TATGAGTGTGGAGTCGTTGCAGATGGAGCAGATATCAACGGGANNNAGATCGGA |
| 58141 | TATGAGTGTGGAGTCGTTGCTGTGTGCCCATGTGTATGTGAATNNNAGATCGGAA |
| 109885 | TATGAGTGTGGAGTCGTTGCATACAGGAAATGCAGCAACATAGANNNAGATCGGA |
| 95947 | TATGAGTGTGGAGTCGTTGCAAAACAACCCAACAGCTCAAAATNNNAGATCGGAAG |
| 101485 | TATGAGTGTGGAGTCGTTGCATGAAGGAATCCTCCCCTATATGGAGNNNAGATCGG |
| 33032 | TATGAGTGTGGAGTCGTTGCAAATGCCCACTGAGAAACACACANNNAGATCGGAAG |
| 35810 | TATGAGTGTGGAGTCGTTGCTGGACTGAACATTTTTCCTGGCANNNAGATCGGAAG |
| 37640 | TATGAGTGTGGAGTCGTTGCGGCTTTTGTGTCAAGAGGGAAGANNNAGATCGGAA |
| 107155 | TATGAGTGTGGAGTCGTTGCATGTGTGCGTTTTAGACATTAGTCCNNNAGATCGGA |
| 98330 | TATGAGTGTGGAGTCGTTGCAACACACCTGGCAGAAGACAGACNNNAGATCGGAA |
| 56866 | TATGAGTGTGGAGTCGTTGCCTGGGCTTGAGGTGTTGGTGNNNAGATCGGAAGA |
| 44618 | TATGAGTGTGGAGTCGTTGCTGTGTTTTGTCACTTTGTTGCCCNNNAGATCGGAAG |
| 106725 | TATGAGTGTGGAGTCGTTGCGTGGATTTTTCCATGAGGATGCCANNNAGATCGGA |
| 107089 | TATGAGTGTGGAGTCGTTGCGGACTGAACATTTTTCCTGGCAGNNNAGATCGGAAG |
| 53402 | TATGAGTGTGGAGTCGTTGCGGTAAGCTCTGCATGTGAGTAGGNNNAGATCGGAA |
| 117382 | TATGAGTGTGGAGTCGTTGCCTCCCATCTCTCAGAGTTCAGCTNNNAGATCGGAAG |
| 42711 | TATGAGTGTGGAGTCGTTGCCTTATTTCTGCTCCTCCACCCACNNNAGATCGGAAGA |
| 65831 | TATGAGTGTGGAGTCGTTGCGGGTGTCTCGCTCTGTTATATGTNNNAGATCGGAAG |
| 100245 | TATGAGTGTGGAGTCGTTGCAACAATAGTAGATAGAGGCCAGGTGCNNNAGATCG |
| 95100 | TATGAGTGTGGAGTCGTTGCGTCTACATTCGCTGAGAGGCAGNNNAGATCGGAAG |
| 67665 | TATGAGTGTGGAGTCGTTGCCTGTGGCTCCTACTGAGTACCTTNNNAGATCGGAAG |
| 42707 | TATGAGTGTGGAGTCGTTGCGCTCCCTCTTTTCACTAACCCACNNNAGATCGGAAGA |
| 46582 | TATGAGTGTGGAGTCGTTGCCACAAGGAGGGATAACACTGAGCNNNAGATCGGAA |
| 69259 | TATGAGTGTGGAGTCGTTGCAGTTTGAAGCACAGCCTCAGTTTNNNAGATCGGAAG |
| 106280 | TATGAGTGTGGAGTCGTTGCGGTGGAGCAGAAACTGAATTTGNNNAGATCGGAAG |
| 30560 | TATGAGTGTGGAGTCGTTGCTCCAGCAATGCTCTGTTACGAAANNNAGATCGGAAG |
| 44596 | TATGAGTGTGGAGTCGTTGCCTTGAACAGAGTACATCCTGCCCNNNAGATCGGAAG |
| 61625 | TATGAGTGTGGAGTCGTTGCTTTTTCCATGAGGATGCCAAGCTNNNAGATCGGAAG |
| 99912 | TATGAGTGTGGAGTCGTTGCCTACCCTCCGGGCTCACTTCGANNNAGATCGGAAGA |
| 31434 | TATGAGTGTGGAGTCGTTGCGTGCTGCAGATTCTTCCTCTCAANNNAGATCGGAAG |
| 69164 | TATGAGTGTGGAGTCGTTGCGGGGGTGGGGTTGTTTATCTTTNNNAGATCGGAA |

33979 TATGAGTGTGGAGTCGTTGCGATGAGAAGGCCTAAAGCAACCANNNAGATCGGAA
35199 TATGAGTGTGGAGTCGTTGCCAGAGAGACGGAGATAGGAAGCANNNAGATCGGA
41474 TATGAGTGTGGAGTCGTTGCCCTGTCGTTTGCAGTAAGTCGTANNNAGATCGGAAG
40390 TATGAGTGTGGAGTCGTTGCTGGGGGAAAATGATTGATACTATTGANNNAGATCG
31198 TATGAGTGTGGAGTCGTTGCCCACTCAGAAGATTTCCCAGCAANNNAGATCGGAAG
45575 TATGAGTGTGGAGTCGTTGCTCACATCAGTCCCATCCCTCCNNNAGATCGGAAGAGC
53694 TATGAGTGTGGAGTCGTTGCAAGAGGTTCAAAGAACAGCAGGGNNNAGATCGGA
116184 TATGAGTGTGGAGTCGTTGCACAGCTCACTACAGCCTGGATTCNNNAGATCGGAAG
100544 TATGAGTGTGGAGTCGTTGCGGCTGGAGTGCTGTAGTAGGATAANNNAGATCGGA
101493 TATGAGTGTGGAGTCGTTGCTCCTCTCTCAGGAAATAAAGGCTACANNNAGATCGG
58819 TATGAGTGTGGAGTCGTTGCACTGAAAGAGAGAGGTGCCTCATNNNAGATCGGAA
34686 TATGAGTGTGGAGTCGTTGCATGTCAGTTTGTCTTGTTGGCCANNNAGATCGGAAG
69431 TATGAGTGTGGAGTCGTTGCAGATTGTGGAAATACAAGGGATTTTNNNAGATCGG
102572 TATGAGTGTGGAGTCGTTGCACATGCCTGATTGAAAGTTTCCTGANNNAGATCGGA
96779 TATGAGTGTGGAGTCGTTGCCTGTGCCCGCACCCTCATCCNNNAGATCGGAAGAGC
55302 TATGAGTGTGGAGTCGTTGCCACCAACGTTGCCTTGTAGAATGNNNAGATCGGAAG
68662 TATGAGTGTGGAGTCGTTGCCATCCACTTTGGCTCACAGTGTTNNNAGATCGGAAG
51463 TATGAGTGTGGAGTCGTTGCCCAGGAGGTAGGACTGAAGAGAGNNNAGATCGGA
41664 TATGAGTGTGGAGTCGTTGCGCTGTCCCAAAAGGCTTAGTGTANNNAGATCGGAA
116569 TATGAGTGTGGAGTCGTTGCAAACCTCCCTGAGTACTCCCCNNNAGATCGGAAGAG
42625 TATGAGTGTGGAGTCGTTGCTATGAAGCGTTTCTCCAGGACACNNNAGATCGGAAG
50762 TATGAGTGTGGAGTCGTTGCCACAGCTAGGAGACAAGCTACAGNNNAGATCGGAA
32438 TATGAGTGTGGAGTCGTTGCAGAGGGGTCTTTGAGTTTTCTAANNNAGATCGGAA
109996 TATGAGTGTGGAGTCGTTGCGTCACTCAGCACCCAGGCTAGNNNAGATCGGAAGA
40814 TATGAGTGTGGAGTCGTTGCAACAACCACTGACCCAGGAGATANNNAGATCGGAA
100698 TATGAGTGTGGAGTCGTTGCATAAAATGTGTGTGTGTGTATACGTATNNNAGATCG
105047 TATGAGTGTGGAGTCGTTGCAAAAAGGCAATTCCGCAATAAGGTTATNNNAGATCG
52445 TATGAGTGTGGAGTCGTTGCACCACTTTCTAAAAGGCCATCCGNNNAGATCGGAAG
104838 TATGAGTGTGGAGTCGTTGCTCTATGGCTTCCAACTTACTGCANNNAGATCGGAAG
44887 TATGAGTGTGGAGTCGTTGCTGTCTATCTGAGGCTCCTAAGCCNNNAGATCGGAAG
41106 TATGAGTGTGGAGTCGTTGCGCTGCCTTGATTTGTCCATCCTANNNAGATCGGAAG
62772 TATGAGTGTGGAGTCGTTGCTTCAGCGGTTTCATTGACACTCTNNNAGATCGGAAG
48661 TATGAGTGTGGAGTCGTTGCTGGGTTGGCCTTGAAGTTTACTCNNNAGATCGGAAG
47256 TATGAGTGTGGAGTCGTTGCCTTCTTAGTGATCACCCACTGGCNNNAGATCGGAAG
41957 TATGAGTGTGGAGTCGTTGCTCCACACCTACCTTCAGCTGTTANNNAGATCGGAAGA
32215 TATGAGTGTGGAGTCGTTGCCCACTCACCGTTCTCACTTTGAANNNAGATCGGAAGA
46542 TATGAGTGTGGAGTCGTTGCGAGAGAGAGGAGAGCATAGGAGCNNNAGATCGGA
40838 TATGAGTGTGGAGTCGTTGCGATGGATTTTGACAGGCCGGATANNNAGATCGGAA
42281 TATGAGTGTGGAGTCGTTGCCCCTACCTACCTACCTACCTACAACNNNAGATCGGAA
45100 TATGAGTGTGGAGTCGTTGCTTAGTCTTAGAGACAGGTGGGCCNNNAGATCGGAA
64995 TATGAGTGTGGAGTCGTTGCTGAAAAGGGGCAATTTTGTAGGTNNNAGATCGGAA
97996 TATGAGTGTGGAGTCGTTGCAAAACCCAAAAACCGTTCGGCANNNAGATCGGAAGA
29950 TATGAGTGTGGAGTCGTTGCGAGAGCAGGGAATTCAGTTCAGGNNNAGATCGGA

68484 TATGAGTGTGGAGTCGTTGCCAGTTGAAGGGGTGGATATGGTTNNNAGATCGGAA
55217 TATGAGTGTGGAGTCGTTGCCTTGGGGTCATTGAGCTTTTTGGNNNAGATCGGAAG
116961 TATGAGTGTGGAGTCGTTGCTCAATAGTACCCCCTACCTCACCNNNAGATCGGAAGA
41449 TATGAGTGTGGAGTCGTTGCCAGGTACACACACGTGCTTAGTANNNAGATCGGAAG
117581 TATGAGTGTGGAGTCGTTGCTTGAGATGGAGTACAGTGTTGCAGNNNAGATCGGA
45601 TATGAGTGTGGAGTCGTTGCTGTGAATAAGGGGGTTCTCCTCCNNNAGATCGGAAG
117885 TATGAGTGTGGAGTCGTTGCCTCTGTTTTCAGTGAGGGAATTAGGTNNNAGATCGG
98783 TATGAGTGTGGAGTCGTTGCAGCCGTAAAGCATCATGGGCATANNNAGATCGGAA
98583 TATGAGTGTGGAGTCGTTGCAAGAGGAAGCCTTGACCCCCTTNNNAGATCGGAAGA
55549 TATGAGTGTGGAGTCGTTGCCGGGAGAAGGCTATGTGAAGATGNNNAGATCGGA
31595 TATGAGTGTGGAGTCGTTGCATGGTCAAACCAGATGCACAGAANNNAGATCGGAA
103330 TATGAGTGTGGAGTCGTTGCATCTTGGCTCACTGCTCATTTTNNNAGATCGGAAGAG
66020 TATGAGTGTGGAGTCGTTGCGCTTCCAACTTACTGCATCCTGTNNNAGATCGGAAGA
37694 TATGAGTGTGGAGTCGTTGCTTGGGAGAGAAGTGGCACTAAGANNNAGATCGGAA
116339 TATGAGTGTGGAGTCGTTGCCTTTGTAACCTGCTGTCCTACAGCNNNAGATCGGAAG
47194 TATGAGTGTGGAGTCGTTGCAAGAAGATGGGTGGTGAAATGGCNNNAGATCGGA
38676 TATGAGTGTGGAGTCGTTGCACACTGCTTCAGAGCTAGAAGGANNNAGATCGGAA
32486 TATGAGTGTGGAGTCGTTGCCTTATGTAACCGAGCCCGGGTAANNNAGATCGGAAG
41253 TATGAGTGTGGAGTCGTTGCTTCTTTCTCCCCAGTGGCATCTANNNAGATCGGAAGA
48563 TATGAGTGTGGAGTCGTTGCCACCTGCACATCCACCTCTTATCNNNAGATCGGAAGA
34763 TATGAGTGTGGAGTCGTTGCGAACCACCCCTCTTTTATTGCCANNNAGATCGGAAGA
57092 TATGAGTGTGGAGTCGTTGCTGGAGAACCCTAACACACATTGTGNNNAGATCGGAA
105038 TATGAGTGTGGAGTCGTTGCATTGGGGTCAGAGTGAAAAGAGCNNNAGATCGGAA
45117 TATGAGTGTGGAGTCGTTGCTGTTGCACTTGTGAATTTATGGCCNNNAGATCGGAA
62335 TATGAGTGTGGAGTCGTTGCGCTGAGAACAAATAGGCTCTGCTNNNAGATCGGAAG
53141 TATGAGTGTGGAGTCGTTGCAGAATGGTGGATTCATGCAGAGGNNNAGATCGGAA
34729 TATGAGTGTGGAGTCGTTGCCAGAAATAGAAGCTCTGCTGCCANNNAGATCGGAAG
67925 TATGAGTGTGGAGTCGTTGCGGTATGTTTTCTCTCCGTGGCTTNNNAGATCGGAAG
63920 TATGAGTGTGGAGTCGTTGCACAACTCAGAGGATGAAGATACAGTNNNAGATCGG
52773 TATGAGTGTGGAGTCGTTGCGCAGCGAGAGCTATAACTCAAGGNNNAGATCGGAA
69149 TATGAGTGTGGAGTCGTTGCTGTAATTCGGGACAGCTTGCTTTNNNAGATCGGAAG
109963 TATGAGTGTGGAGTCGTTGCAGCAGGCACCGTATATTTGTTCTGNNNAGATCGGAA
37584 TATGAGTGTGGAGTCGTTGCGGCTCAGCATCCTGAAATCAAGANNNAGATCGGAAG
32733 TATGAGTGTGGAGTCGTTGCACTTCTGCTATTCCAACAAAGAAACANNNAGATCGG
69166 TATGAGTGTGGAGTCGTTGCTGGGGGAGTGATACCAACTCTTTNNNAGATCGGAAG
59452 TATGAGTGTGGAGTCGTTGCGGTCCCTCCACATGGCTAGATATNNNAGATCGGAAG
50362 TATGAGTGTGGAGTCGTTGCCAGTGGTGCATCCTAGGTTCAAGNNNAGATCGGAA
49261 TATGAGTGTGGAGTCGTTGCGGGGCTGGGGAAGATATCTAGTCNNNAGATCGGAA
60844 TATGAGTGTGGAGTCGTTGCTAAGTTTCAGCACATTGCACCCTNNNAGATCGGAAG
49987 TATGAGTGTGGAGTCGTTGCCACTCTTGGCTCCTCTCCTTTTCNNNAGATCGGAAGA
33376 TATGAGTGTGGAGTCGTTGCAGTAGTAACCTGTCCAGCTCACANNNAGATCGGAAG
56209 TATGAGTGTGGAGTCGTTGCACAAAATAGGGTTGACTGGGCTGNNNAGATCGGAA
47360 TATGAGTGTGGAGTCGTTGCCCAGTAATAACACCCTCTGTGGCNNNAGATCGGAAG

41741 TATGAGTGTGGAGTCGTTGCCAGCCACTAAAATGGGGCCATTANNNAGATCGGAAG
41050 TATGAGTGTGGAGTCGTTGCACCTAGTCATGGAACAAGCCCTANNNAGATCGGAAG
58590 TATGAGTGTGGAGTCGTTGCACTGTGAATTGCCACCTTTCCATNNNAGATCGGAAGA
105048 TATGAGTGTGGAGTCGTTGCTTCTGGCCGTGTTCTGGTTAGNNNAGATCGGAAGAG
34973 TATGAGTGTGGAGTCGTTGCATTAGTCCATCCATAGCTGGTCCANNNAGATCGGAA
35323 TATGAGTGTGGAGTCGTTGCTCTTTCTTTGCTTCCCATCAGCANNNAGATCGGAAGA
107153 TATGAGTGTGGAGTCGTTGCTCAGATGGTGTCTCATTTTGTCGNNNAGATCGGAAG
52207 TATGAGTGTGGAGTCGTTGCGGGTCAGAAAAGAGGTGGCTTAGNNNAGATCGGAA
96422 TATGAGTGTGGAGTCGTTGCCTAACCAGGGCAGAAACCTAAAANNNAGATCGGAAG
71632 TATGAGTGTGGAGTCGTTGCAGGGGAATCAGGGCCTGCAAGNNNAGATCGGAAGA
44920 TATGAGTGTGGAGTCGTTGCTCATCTCTACCTGTAGTCCAGCCNNNAGATCGGAAGA
42319 TATGAGTGTGGAGTCGTTGCAAGGCTGTTAGAACACTGGCAACNNNAGATCGGAA
96419 TATGAGTGTGGAGTCGTTGCTCTTGGATTCCGGAAGGGGAAGGNNNAGATCGGAA
40104 TATGAGTGTGGAGTCGTTGCGCCTTATGGACCAACAGAAGTGANNNAGATCGGAA
38968 TATGAGTGTGGAGTCGTTGCTGAACAGGATGGAAAAGAAGGGANNNAGATCGGA
43671 TATGAGTGTGGAGTCGTTGCAGAGAGGGACAGAGAGCACTTACNNNAGATCGGAA
29484 TATGAGTGTGGAGTCGTTGCAGGCTCATGGATTTCCTGGCNNNAGATCGGAAGAG
49413 TATGAGTGTGGAGTCGTTGCAGAGTGAAAGAAAGCCAGCTGTCNNNAGATCGGAA
50729 TATGAGTGTGGAGTCGTTGCCGCTGGCATTTTCTTTGAGACAGNNNAGATCGGAAG
58992 TATGAGTGTGGAGTCGTTGCGGAAGGATCCGAGAGATCCAGATNNNAGATCGGAA
42220 TATGAGTGTGGAGTCGTTGCAAGGAAGTGGTGAGACACGAAACNNNAGATCGGAA
37092 TATGAGTGTGGAGTCGTTGCACCTCAATACCCCCAGGAATTCANNNAGATCGGAAGA
58695 TATGAGTGTGGAGTCGTTGCTTCTTGGAGACTGAGACTGGCATNNNAGATCGGAAG
44673 TATGAGTGTGGAGTCGTTGCCTTGTGGGAGCCTTTCTTATCCCNNNAGATCGGAAGA
32145 TATGAGTGTGGAGTCGTTGCAAAAGGCTGCAGTCGAGAGTGAANNNAGATCGGAA
56820 TATGAGTGTGGAGTCGTTGCTTGCTGCTTTTTATGCCATGGTGNNNAGATCGGAAGA
107191 TATGAGTGTGGAGTCGTTGCGAGGGACAGAGAGCACTTACTACANNNAGATCGGA
55033 TATGAGTGTGGAGTCGTTGCTCTGTCATCATCACCGTCATTGGNNNAGATCGGAAGA
55807 TATGAGTGTGGAGTCGTTGCATGCTATCTAGTAACTGGCCACTGNNNAGATCGGAA
37735 TATGAGTGTGGAGTCGTTGCCCCAACACACACAGAGAAACAGANNNAGATCGGAAG
49179 TATGAGTGTGGAGTCGTTGCCCTAGAGAACACGTGCAGAAGTCNNNAGATCGGAA
29485 TATGAGTGTGGAGTCGTTGCACCGAGTGACAATACACAGCANNNAGATCGGAAGA
35550 TATGAGTGTGGAGTCGTTGCAACCCAAACTTTTTCTCTCCGCANNNAGATCGGAAGA
52259 TATGAGTGTGGAGTCGTTGCTGTCTTTGAACCACCTGAGAACGNNNAGATCGGAAG
100765 TATGAGTGTGGAGTCGTTGCCCCTCTCTGTCCCAATCCCANNNAGATCGGAAGAGCA
100914 TATGAGTGTGGAGTCGTTGCGTGCTGCAGATTCTTCCTCTCAAAGNNNAGATCGGAA
71889 TATGAGTGTGGAGTCGTTGCCCCACCTCTTACCGATTTCTTCCNNNAGATCGGAAGA
71789 TATGAGTGTGGAGTCGTTGCGCAGCTCTTGGCCTGAGACTNNNAGATCGGAAGAG
101652 TATGAGTGTGGAGTCGTTGCTCAGGGATCAGCATACCATGGNNNAGATCGGAAGA
71694 TATGAGTGTGGAGTCGTTGCCGCATCTCTAAGGAGGAGTACCANNNAGATCGGAA
102589 TATGAGTGTGGAGTCGTTGCCTGAGTGGGGAATGAGTGGGTAANNNAGATCGGA
71728 TATGAGTGTGGAGTCGTTGCATTACCTGTTCCAGCACCATGTCNNNAGATCGGAAGA
71794 TATGAGTGTGGAGTCGTTGCCTTCCACACTCTGTCCCCATCTGNNNAGATCGGAAGA

| | |
|---|---|
| 101052 | TATGAGTGTGGAGTCGTTGCCCGTGCTCAGCCAATACATTATANNNAGATCGGAAG |
| 71890 | TATGAGTGTGGAGTCGTTGCGGTGCAGTTATGCCTCAGATTCANNNAGATCGGAAG |
| 107185 | TATGAGTGTGGAGTCGTTGCCTATCGGTGATGTAAGGGGGCNNNAGATCGGAAG |
| 100542 | TATGAGTGTGGAGTCGTTGCTATCGGTGATGTAAGGGGGCGNNNAGATCGGAAG |
| 101614 | TATGAGTGTGGAGTCGTTGCCAGCTAGAGCCCTCATGAAGGAGNNNAGATCGGAA |
| 102557 | TATGAGTGTGGAGTCGTTGCCACTGGCTGTCTCTTACTAATCCNNNAGATCGGAAG |
| 122003 | TATGAGTGTGGAGTCGTTGCGTTTCTCCTGGTGAGTTGCCATTTNNNAGATCGGAA |
| 71693 | TATGAGTGTGGAGTCGTTGCGCTGTACCTCATGAGTTGGTTGTNNNAGATCGGAAG |
| 100704 | TATGAGTGTGGAGTCGTTGCGTTTGGTGATGTTCCTCCAGATTTNNNAGATCGGAA |
| 103392 | TATGAGTGTGGAGTCGTTGCTGCCTGTATTTGCTGGAGAACTNNNAGATCGGAAG |
| 71888 | TATGAGTGTGGAGTCGTTGCCACTGATTGCTCTTAGGTCTGGCNNNAGATCGGAAG |
| 104982 | TATGAGTGTGGAGTCGTTGCTTCCTCCTCAACACTCTCCCACCNNNAGATCGGAAGA |
| 114796 | TATGAGTGTGGAGTCGTTGCGGGCAAGGTAAGAGCTGCAANNNAGATCGGAAGA |
| 105017 | TATGAGTGTGGAGTCGTTGCCTACCGCCAATGTTCACTCTAAGCNNNAGATCGGAA |
| 102854 | TATGAGTGTGGAGTCGTTGCCTCAGACCCCAAACTCTCCCAGNNNAGATCGGAAGA |
| 71892 | TATGAGTGTGGAGTCGTTGCGGTTGGAAAGCCATAGGATTCCANNNAGATCGGAA |
| 103394 | TATGAGTGTGGAGTCGTTGCTCAGATGGGAACTCACAGAAGTCTGNNNAGATCGG |
| 117584 | TATGAGTGTGGAGTCGTTGCCTGGAGAATCCTGACTGAAACACCTANNNAGATCGG |
| 98703 | TATGAGTGTGGAGTCGTTGCTAGTCCTGAACCTTGTGCTTTCTCNNNAGATCGGAAG |
| 101367 | TATGAGTGTGGAGTCGTTGCTACCTGCCCTCCTTGAATGACACNNNAGATCGGAAG |
| 106212 | TATGAGTGTGGAGTCGTTGCCTTCACATCACAGCTGGCGGANNNAGATCGGAAGAG |
| 101414 | TATGAGTGTGGAGTCGTTGCCAGTGAGCCATGTGAGTGTANNNAGATCGGAAGAG |
| 97405 | TATGAGTGTGGAGTCGTTGCCTCCCTTGACCTTCCTCCAGCTNNNAGATCGGAAGAG |
| 98640 | TATGAGTGTGGAGTCGTTGCCCCTGCTTTTTGACTTGTAGGCTNNNAGATCGGAAG |
| 58939 | TATGAGTGTGGAGTCGTTGCCGTGGTAAGAGCACCTCAAAGATNNNAGATCGGAA |
| 50723 | TATGAGTGTGGAGTCGTTGCAGAAGCATGAATGGCAGAGACAGNNNAGATCGGA |
| 69657 | TATGAGTGTGGAGTCGTTGCGTTCTTCCTTTGGTGGCGTTTTTNNNAGATCGGAAG |
| 68189 | TATGAGTGTGGAGTCGTTGCAAGTTTCCCTGTGCCTCTTTCTTNNNAGATCGGAAGA |
| 49006 | TATGAGTGTGGAGTCGTTGCTGGTGTGCTTTCTGCTGTATCTCNNNAGATCGGAAG |
| 54379 | TATGAGTGTGGAGTCGTTGCGCAACATGCTATACCTTTCCATGGNNNAGATCGGAA |
| 47759 | TATGAGTGTGGAGTCGTTGCAATGTGGAGAACTTACAGGCTGCNNNAGATCGGAA |
| 40718 | TATGAGTGTGGAGTCGTTGCGAGTTTCTGGGAGGGTAGCCATANNNAGATCGGAA |
| 46972 | TATGAGTGTGGAGTCGTTGCATTGTTTCCTCAGTTCTGGAGGCNNNAGATCGGAAG |
| 33188 | TATGAGTGTGGAGTCGTTGCCCACAACTTACTGAAACCCCACANNNAGATCGGAAG |
| 37409 | TATGAGTGTGGAGTCGTTGCTGGCTGCATCTTTCAATGTTTTCANNNAGATCGGAAG |
| 106695 | TATGAGTGTGGAGTCGTTGCTTCCTAGTTTACTTGTGGGGTGTNNNAGATCGGAAG |
| 46096 | TATGAGTGTGGAGTCGTTGCCCTTCGGCAACCATTTAGAAAGCNNNAGATCGGAAG |
| 39642 | TATGAGTGTGGAGTCGTTGCGGTGTGTAGGTGTAATGCCATGANNNAGATCGGAA |
| 68439 | TATGAGTGTGGAGTCGTTGCCCTGATCGTAAGTTGGTTCGGTTNNNAGATCGGAAG |
| 40703 | TATGAGTGTGGAGTCGTTGCACCATTCTCGACCAGAGACCATANNNAGATCGGAAG |
| 39611 | TATGAGTGTGGAGTCGTTGCCACCCATTGACCTGTTCACATGANNNAGATCGGAAG |
| 69048 | TATGAGTGTGGAGTCGTTGCGTGGGAGTTGTATGCCATCCTTTNNNAGATCGGAAG |
| 58778 | TATGAGTGTGGAGTCGTTGCTGAAGCAAACCCTGCAAGATCATNNNAGATCGGAAG |

53414 TATGAGTGTGGAGTCGTTGCAAAGAGTGGTGTGTGTGTAGGNNNAGATCGGA
60430 TATGAGTGTGGAGTCGTTGCTGGGAGAGTGATACGGTTTGACTNNNAGATCGGAA
31429 TATGAGTGTGGAGTCGTTGCGGGCCAGTCTAACTTGATCTCAANNNAGATCGGAAG
98970 TATGAGTGTGGAGTCGTTGCCACACAAATAGAAGCCCTCTGTGANNNAGATCGGAA
64178 TATGAGTGTGGAGTCGTTGCTGACACCTGAGTTGGAGAAGAGTNNNAGATCGGAA
52310 TATGAGTGTGGAGTCGTTGCTAGCAGGGAGATCTAGACTTCACGNNNAGATCGGA
48863 TATGAGTGTGGAGTCGTTGCACAAAGAGAGAGGGAGAGAGCTCNNNAGATCGGA
32912 TATGAGTGTGGAGTCGTTGCGTCAACAACTTGCTAGAAATGGAACANNNAGATCGG
29837 TATGAGTGTGGAGTCGTTGCAGGAAAATGACCCTCCCCTCCNNNAGATCGGAAGAG
32828 TATGAGTGTGGAGTCGTTGCAATAGAAATGGGCTTGGCCAACANNNAGATCGGAA
64081 TATGAGTGTGGAGTCGTTGCGTCACTCCTTGTATGTTTTGCAGTNNNAGATCGGAA
31764 TATGAGTGTGGAGTCGTTGCGCCCAAGCATCATCCCATTAGAANNNAGATCGGAAG
49730 TATGAGTGTGGAGTCGTTGCATGCTCACAGGAAAGCCTCTTCNNNAGATCGGAAGA
61075 TATGAGTGTGGAGTCGTTGCACTGGTGAAACCTGAATAAGCCTNNNAGATCGGAAG
104996 TATGAGTGTGGAGTCGTTGCAGCAAGATCCCCTAACCTGAGANNNAGATCGGAAGA
35399 TATGAGTGTGGAGTCGTTGCCAGTTGAGAGAGAGTAGGGAGCANNNAGATCGGA
39280 TATGAGTGTGGAGTCGTTGCTCTTTGGATGGTGGGATGATGGANNNAGATCGGAA
65812 TATGAGTGTGGAGTCGTTGCCACACCCTGCTGTAATATCTGATGTNNNAGATCGGA
57749 TATGAGTGTGGAGTCGTTGCCCCCCACAAACCCTAAGTCAAATNNNAGATCGGAAGA
58362 TATGAGTGTGGAGTCGTTGCATGCACAAGTGAGATGCAGACATNNNAGATCGGAA
38873 TATGAGTGTGGAGTCGTTGCTGTGGTATGGCTCTCAAATAGGANNNAGATCGGAA
66848 TATGAGTGTGGAGTCGTTGCTCCAAAGGGCAATCCTTGTTTGTNNNAGATCGGAAG
45787 TATGAGTGTGGAGTCGTTGCCCTGGGCATCTCTTTGTATGTCCNNNAGATCGGAAG
42554 TATGAGTGTGGAGTCGTTGCTGAGAGCTGGTTTACCAACACACNNNAGATCGGAAG
42144 TATGAGTGTGGAGTCGTTGCGAGAGAGAGAGACCCTGCAAAACNNNAGATCGGAA
67418 TATGAGTGTGGAGTCGTTGCGGGGCTTTTTACCCTGCAGTATTNNNAGATCGGAAG
57028 TATGAGTGTGGAGTCGTTGCTTGCTGGAGAACACTCTAGTGTGNNNAGATCGGAA
51517 TATGAGTGTGGAGTCGTTGCGCTGCTCAGACTTACAGTGAGAGNNNAGATCGGAA
46636 TATGAGTGTGGAGTCGTTGCATGGAATGCAGTGGTGAGATAGCNNNAGATCGGAA
59702 TATGAGTGTGGAGTCGTTGCGAGGGTGAGGGGAGGATTCTTATNNNAGATCGGA
96129 TATGAGTGTGGAGTCGTTGCGCTGTGACCCATCTACCAATTTGNNNAGATCGGAAG
58912 TATGAGTGTGGAGTCGTTGCGGATAACAACCCCTTCCGTTCATNNNAGATCGGAAG
71754 TATGAGTGTGGAGTCGTTGCCCGCTGGGAAGAAGAAAACATCTNNNAGATCGGAA
64354 TATGAGTGTGGAGTCGTTGCCCCTGAACTGTCTAAAAGTGAGTNNNAGATCGGAAG
42811 TATGAGTGTGGAGTCGTTGCGCACACATACATGCCATTTCCACNNNAGATCGGAAGA
55231 TATGAGTGTGGAGTCGTTGCACAAAAGCCAGAGTGCACAAATGNNNAGATCGGAA
36600 TATGAGTGTGGAGTCGTTGCCTGGGGGAGAAGGAACTTACTCANNNAGATCGGAA
56608 TATGAGTGTGGAGTCGTTGCGTCCTTGTACCTGTGTGTCAGTGNNNAGATCGGAAG
39498 TATGAGTGTGGAGTCGTTGCTGTGTCTAAGTAAGGGGCTTGGANNNAGATCGGAA
36946 TATGAGTGTGGAGTCGTTGCTATCTGTTGGCACTCTGAGGTCANNNAGATCGGAAG
99729 TATGAGTGTGGAGTCGTTGCTACCAGCCTCCTACTGTCCTGCNNNAGATCGGAAGAG
38733 TATGAGTGTGGAGTCGTTGCAAACCCAGTCTCTAACACCAGGANNNAGATCGGAAG
56581 TATGAGTGTGGAGTCGTTGCTCCAGCTCCAAATGTAAGCAGTGNNNAGATCGGAAG

40884 TATGAGTGTGGAGTCGTTGCGGACAGCCACACAGTTGCATATANNNAGATCGGAA
34555 TATGAGTGTGGAGTCGTTGCTTTGTTTTGATTCAGTGAAGCCANNNAGATCGGAAG
117755 TATGAGTGTGGAGTCGTTGCTTGTAATGCCAGGTCTTGCTCTGNNNAGATCGGAAG
55694 TATGAGTGTGGAGTCGTTGCGACCTCCAAACTGGGAAGCTATGNNNAGATCGGAA
57115 TATGAGTGTGGAGTCGTTGCTACTGATGGGCTTGGATGTTGTGNNNAGATCGGAA
67224 TATGAGTGTGGAGTCGTTGCAGGCTACTTTCAGATCATAGTCATTNNNAGATCGGA
52721 TATGAGTGTGGAGTCGTTGCACACTGGACACTGAGTTGAAAGGNNNAGATCGGAA
42329 TATGAGTGTGGAGTCGTTGCATTGTCACTGGCAGCTCATCAACNNNAGATCGGAAG
60447 TATGAGTGTGGAGTCGTTGCTTGGTGATACCCTGCTCCATACTNNNAGATCGGAAG
57530 TATGAGTGTGGAGTCGTTGCTTTCTACCCTGGCTCCACATTTGNNNAGATCGGAAGA
68802 TATGAGTGTGGAGTCGTTGCATGATGTGCACAGAGGCACATTTNNNAGATCGGAA
64867 TATGAGTGTGGAGTCGTTGCGGTTGCTATACAGGACAAGAGGTNNNAGATCGGAA
51629 TATGAGTGTGGAGTCGTTGCTTACTTGGGGCCTGAGATAGGAGNNNAGATCGGAA
38649 TATGAGTGTGGAGTCGTTGCTTTACTGGGCTCTGAGACAAGGANNNAGATCGGAA
42183 TATGAGTGTGGAGTCGTTGCGAGGGTCTAGCCATTTCCCAAACNNNAGATCGGAAG
103368 TATGAGTGTGGAGTCGTTGCAATGCCATGGTGAAAGAATTGGTNNNAGATCGGAA
68137 TATGAGTGTGGAGTCGTTGCCAGGGGACAAGACGGAAATTCTTNNNAGATCGGAA
56092 TATGAGTGTGGAGTCGTTGCAGAACTCACAGTGACCAAAGCTGNNNAGATCGGAA
71619 TATGAGTGTGGAGTCGTTGCTGAAGCAATTTAGGGCAAACTATCCNNNAGATCGGA
31069 TATGAGTGTGGAGTCGTTGCACAGTTCTGGCAAATACAGCCAANNNAGATCGGAAG
42759 TATGAGTGTGGAGTCGTTGCACATGAGTTGAGTAACTGGCCACNNNAGATCGGAA
97715 TATGAGTGTGGAGTCGTTGCTGTATATGAACCTGGTCTCCTTGGNNNAGATCGGAA
100396 TATGAGTGTGGAGTCGTTGCGTATTCTGTTAGACTTGGGCCATGNNNAGATCGGAA
98948 TATGAGTGTGGAGTCGTTGCATCAGAACTTGGTCTCTCCCACTNNNAGATCGGAAG
43151 TATGAGTGTGGAGTCGTTGCCCAAAGTGGGATCTCTGACAGACNNNAGATCGGAA
57616 TATGAGTGTGGAGTCGTTGCCAGTGGTGGTCCTAGTCTGTTTGNNNAGATCGGAA
33492 TATGAGTGTGGAGTCGTTGCAGTATCCAATACACACACAGACANNNAGATCGGAAG
57203 TATGAGTGTGGAGTCGTTGCACCATCTTCTGCCCATAGGATTGNNNAGATCGGAAG
39014 TATGAGTGTGGAGTCGTTGCTGTGGTCAACTAATAGGCAGGGANNNAGATCGGAA
103358 TATGAGTGTGGAGTCGTTGCAAATGCAAAAGACTGTAGGTCTCTGANNNAGATCG
34013 TATGAGTGTGGAGTCGTTGCTGTGTGAGTGAGTAGATATGTAACCANNNAGATCG
49334 TATGAGTGTGGAGTCGTTGCAAACCTTCAAACCTTCCTGGGTCNNNAGATCGGAAG
37313 TATGAGTGTGGAGTCGTTGCGGTACTTGGTCACAGATGTGTTCANNNAGATCGGA
101377 TATGAGTGTGGAGTCGTTGCTATGTGTTTTCAGAATGGGCGTGGNNNAGATCGGA
30148 TATGAGTGTGGAGTCGTTGCTGTGAAGTACCTGGGAGGGAAAANNNAGATCGGA
49823 TATGAGTGTGGAGTCGTTGCACATGCGCACACAATCATGTTCNNNAGATCGGAAGA
55340 TATGAGTGTGGAGTCGTTGCCCTTCTGGGGAAATGGGGTAATGNNNAGATCGGAA
61352 TATGAGTGTGGAGTCGTTGCAAGCAGCTAGAAGCCCTTATCCTNNNAGATCGGAAG
97082 TATGAGTGTGGAGTCGTTGCGGACGCTGATGTGCACAGTTNNNAGATCGGAAGAG
29835 TATGAGTGTGGAGTCGTTGCGTGGTGTGAAGGCCAAAAATCTGNNNAGATCGGAA
48431 TATGAGTGTGGAGTCGTTGCCATCTGTCTGCCTCTCTGTCATCNNNAGATCGGAAGA
58447 TATGAGTGTGGAGTCGTTGCACAGTTGGCAATAACCTGACCATNNNAGATCGGAAG
52433 TATGAGTGTGGAGTCGTTGCACATAAGAAACCACTCTTGGCCGNNNAGATCGGAAG

98729 TATGAGTGTGGAGTCGTTGCGACCATCCATGCCGTTGTCCNNNAGATCGGAAGAGC

98887 TATGAGTGTGGAGTCGTTGCTGATTTTTGTCCACAGACCCCANNNAGATCGGAAGA

49920 TATGAGTGTGGAGTCGTTGCGTAGTCCGTATCACTCGGCTTTCNNNAGATCGGAAG

64638 TATGAGTGTGGAGTCGTTGCTATGCCTTGAAATGTTGAGGCGTNNNAGATCGGAA

35743 TATGAGTGTGGAGTCGTTGCCCTTCATTCATCTGTAAGTGGGCANNNAGATCGGAA

115887 TATGAGTGTGGAGTCGTTGCTTCTTTCTTAGACAGGGTCTCCTGCNNNAGATCGGA

117385 TATGAGTGTGGAGTCGTTGCGGCAGTGGTACAAACACTGTTCNNNAGATCGGAAG

57396 TATGAGTGTGGAGTCGTTGCGCAATCTTCGCTCTGCTTTCTTGNNNAGATCGGAAGA

68448 TATGAGTGTGGAGTCGTTGCAAGACTGTAGGTCTCTGAGGGTTNNNAGATCGGAA

58998 TATGAGTGTGGAGTCGTTGCAGGGAGAGATTCACTTCGCAGATNNNAGATCGGAA

104941 TATGAGTGTGGAGTCGTTGCGTCAGTGTGCTGGCATATGTGTGNNNAGATCGGAA

37306 TATGAGTGTGGAGTCGTTGCGGCAGTGGTACAAACACTGTTCANNNAGATCGGAA

71675 TATGAGTGTGGAGTCGTTGCATCATCTGGTGCTACCTGAGGGNNNAGATCGGAAG

51355 TATGAGTGTGGAGTCGTTGCGCAAGGCACCCTATACACAAGAGNNNAGATCGGAA

31695 TATGAGTGTGGAGTCGTTGCGGCACATGAAGACACAAGGAGAANNNAGATCGGA

48932 TATGAGTGTGGAGTCGTTGCTTCTTAGACAGGGTCTCCTGCTCNNNAGATCGGAAG

50818 TATGAGTGTGGAGTCGTTGCAGAAGGGCAAGATGAAAACCCAGNNNAGATCGGA

100392 TATGAGTGTGGAGTCGTTGCGCCCCTTGAACTGAAAAGCACTATCNNNAGATCGG

33090 TATGAGTGTGGAGTCGTTGCCCACGAGAGAAGTTCAAGACACANNNAGATCGGAA

63763 TATGAGTGTGGAGTCGTTGCACTCTCATTCCACAAAGCAGAAGTNNNAGATCGGAA

58709 TATGAGTGTGGAGTCGTTGCAAATCCGAATAAGGCGGATGCATNNNAGATCGGAA

122016 TATGAGTGTGGAGTCGTTGCCCGAGATTTCAATCCAACTGAATGNNNAGATCGGAA

103362 TATGAGTGTGGAGTCGTTGCAACTAATAGGCAGGGATTTCCTCANNNAGATCGGAA

47330 TATGAGTGTGGAGTCGTTGCAAATGTCCAGAGCTAAGAGTGGCNNNAGATCGGAA

49734 TATGAGTGTGGAGTCGTTGCGGCTGAATACCCTGCTTCTCTTCNNNAGATCGGAAG

41607 TATGAGTGTGGAGTCGTTGCATCTGTCTTTCTGGGGGCATGTANNNAGATCGGAAG

48019 TATGAGTGTGGAGTCGTTGCGCTTCTCATGGACGAAATTGTGCNNNAGATCGGAAG

107178 TATGAGTGTGGAGTCGTTGCTCATGGTTTTGTGTTGTAGGTACACNNNAGATCGGA

45956 TATGAGTGTGGAGTCGTTGCTTGCTTGTAGCCATAGGTGTTCCNNNAGATCGGAAG

110003 TATGAGTGTGGAGTCGTTGCCAGAGACCTGGGAACCAGATATGNNNAGATCGGAA

29830 TATGAGTGTGGAGTCGTTGCCAGGGTTTGCATTAGCTTCAGGANNNAGATCGGAA

46273 TATGAGTGTGGAGTCGTTGCCGCATAACAATGTCTTGGACAGCNNNAGATCGGAAG

63371 TATGAGTGTGGAGTCGTTGCTCCTTGTTAGCCCAGTGACTTCTNNNAGATCGGAAG

67867 TATGAGTGTGGAGTCGTTGCTGTTCATTTGGCATCAGGAGCTTNNNAGATCGGAAG

31863 TATGAGTGTGGAGTCGTTGCGGACCCCACAATTCCTCTAGGAANNNAGATCGGAAG

52650 TATGAGTGTGGAGTCGTTGCGTTTGTATGCATGGCTGTCTTCGNNNAGATCGGAAG

40881 TATGAGTGTGGAGTCGTTGCACCTTGCACCTGCGTGAAATATANNNAGATCGGAAG

107129 TATGAGTGTGGAGTCGTTGCTGGAGAGAATACCTGAATCAGTGTGGNNNAGATCG

68844 TATGAGTGTGGAGTCGTTGCCCCAAGGCTAGAATCAGGCATTTNNNAGATCGGAAG

35739 TATGAGTGTGGAGTCGTTGCTATCAAGATCTACTGGCTGGGCANNNAGATCGGAA

109113 TATGAGTGTGGAGTCGTTGCCCAAGGCTAGAATCAGGCATTTCNNNAGATCGGAAG

109133 TATGAGTGTGGAGTCGTTGCAAGAGAAAGAAATCATGAGGAGACTNNNAGATCGG

117613 TATGAGTGTGGAGTCGTTGCATTAGCTTAGGGCTCCACCCTTANNNAGATCGGAAG

99724 TATGAGTGTGGAGTCGTTGCAAGGGATGAGGCTAGAATAAAGGGANNNAGATCG
69262 TATGAGTGTGGAGTCGTTGCGGGGAACATGTCTGCTTCAGTTTNNNAGATCGGAA
48791 TATGAGTGTGGAGTCGTTGCCTCATTTTTCCTCTGCCCTCCTCNNNAGATCGGAAGA
99322 TATGAGTGTGGAGTCGTTGCTGCTTTGATAACAGGGTGGTTTCCNNNAGATCGGAA
99399 TATGAGTGTGGAGTCGTTGCAGGCTTAGAGGGCAAGTGGAGNNNAGATCGGAAG
66086 TATGAGTGTGGAGTCGTTGCTCTCTTTCTCTCCTTTTGGCTGTNNNAGATCGGAAGA
51608 TATGAGTGTGGAGTCGTTGCTGTGAGTGTGTGTTTGAAGAGGAGNNNAGATCGG
97709 TATGAGTGTGGAGTCGTTGCAGAACGTACACCCCTTCTGTGATNNNAGATCGGAAG
44992 TATGAGTGTGGAGTCGTTGCCAGGCTAGTCATAACTCCTAGCCNNNAGATCGGAAG
49958 TATGAGTGTGGAGTCGTTGCCCTGACCTAACCCACTTGGTTTCNNNAGATCGGAAGA
45566 TATGAGTGTGGAGTCGTTGCATCTCCTATCAACCTCACCCTCCNNNAGATCGGAAGA
33664 TATGAGTGTGGAGTCGTTGCACACAGCTACCTGAAAGATGACANNNAGATCGGAA
115962 TATGAGTGTGGAGTCGTTGCACATCTCACACATGCACCTTCTGNNNAGATCGGAAGA
117793 TATGAGTGTGGAGTCGTTGCCTTTCGTCTCTCTCTCTTTCTGTGNNNAGATCGGAAG
57230 TATGAGTGTGGAGTCGTTGCCTTCCAGTCATCCTCCCAACTTGNNNAGATCGGAAGA
41336 TATGAGTGTGGAGTCGTTGCGTGAAACCCAAGCCACAGTTCTANNNAGATCGGAAG
62041 TATGAGTGTGGAGTCGTTGCCATTTATTGAGCATCCTGGGGCTNNNAGATCGGAAG
46753 TATGAGTGTGGAGTCGTTGCAATAGTTGAAGGCAGTCTACCGCNNNAGATCGGAA
41942 TATGAGTGTGGAGTCGTTGCCTTCTAGCCCCTGCTTTTGGTTANNNAGATCGGAAGA
44546 TATGAGTGTGGAGTCGTTGCAGGAACTTGATCATTGTGTAGCCCNNNAGATCGGAA
55748 TATGAGTGTGGAGTCGTTGCGGATAGAAGCTGCCCAGAAACTGNNNAGATCGGAA
67821 TATGAGTGTGGAGTCGTTGCATAGGTCTGGGACGTGTTTCCTTNNNAGATCGGAAG
67811 TATGAGTGTGGAGTCGTTGCTACGCACAGACTTTTGATTCCTTNNNAGATCGGAAG
97075 TATGAGTGTGGAGTCGTTGCATGGATTCCTCCTGTCTCTCTCTCNNNAGATCGGAAG
45882 TATGAGTGTGGAGTCGTTGCCTTTGTTGCCTTCATTGCCTTCCNNNAGATCGGAAGA
30129 TATGAGTGTGGAGTCGTTGCTCTGTCAAGAAACTGCCCGAAAANNNAGATCGGAAG
39183 TATGAGTGTGGAGTCGTTGCCCAGGAGGAGATTTATGGTGGGANNNAGATCGGA
59568 TATGAGTGTGGAGTCGTTGCAGCGTATGTCTGTTTCCGTCTATNNNAGATCGGAAG
96814 TATGAGTGTGGAGTCGTTGCTGGCTTTCTCTATCACTCCGTCANNNAGATCGGAAGA
105035 TATGAGTGTGGAGTCGTTGCACTGGGGAAGATCAAAAACTGGANNNAGATCGGAA
52866 TATGAGTGTGGAGTCGTTGCAATAAGGAGTGGGTGGGGTAAGGNNNAGATCGGA
58440 TATGAGTGTGGAGTCGTTGCAACAATTTATGACCGGGCACCATNNNAGATCGGAAG
36629 TATGAGTGTGGAGTCGTTGCCAAAGCACTAACCATTCCCCTCANNNAGATCGGAAGA
57042 TATGAGTGTGGAGTCGTTGCTTGTAGTTGTACGTGTGGGTGTGNNNAGATCGGAA
32494 TATGAGTGTGGAGTCGTTGCAGACTTTTCTGCCTCCCTGGTAANNNAGATCGGAAG
116231 TATGAGTGTGGAGTCGTTGCCAAAAACCTCAACTATTCCTCCTCNNNAGATCGGAAG
99677 TATGAGTGTGGAGTCGTTGCCCCCACCCTTGTCTCTACAAAATNNNAGATCGGAAGA
37193 TATGAGTGTGGAGTCGTTGCCCTTGAAGATGCCTCTCCCTTCANNNAGATCGGAAGA
50065 TATGAGTGTGGAGTCGTTGCCAGTTGGCCTGAAACACACAAAGNNNAGATCGGAA
40348 TATGAGTGTGGAGTCGTTGCCCTGCTCTTGAAGGAGTTTGTGANNNAGATCGGAA
30603 TATGAGTGTGGAGTCGTTGCTATGCCCATGAGAAGGAGGGAAANNNAGATCGGA
100395 TATGAGTGTGGAGTCGTTGCTCAACTGGGACATGGTTTGTTGGNNNAGATCGGAA
68774 TATGAGTGTGGAGTCGTTGCCCCCTCCCTTCGAAATCCAATTTNNNAGATCGGAAGA

| | |
|---|---|
| 51633 | TATGAGTGTGGAGTCGTTGCGCACCCACCTTTTTAGGTAGGAGNNNAGATCGGAAG |
| 43013 | TATGAGTGTGGAGTCGTTGCTTCTCTGTGTGTGAAGCTCTCACNNNAGATCGGAAG |
| 43014 | TATGAGTGTGGAGTCGTTGCCGCTCTCACTTTCACTCTCTCACNNNAGATCGGAAGA |
| 63003 | TATGAGTGTGGAGTCGTTGCACTCCTTTTCCCTTCCACTCTCTNNNAGATCGGAAGAG |
| 64646 | TATGAGTGTGGAGTCGTTGCACTGTGTGTACATGTGAATGCGTNNNAGATCGGAAG |
| 59868 | TATGAGTGTGGAGTCGTTGCCCAGCTACTAATGGCTTTCAACTNNNAGATCGGAAG |
| 99401 | TATGAGTGTGGAGTCGTTGCGCAGAGGGAAAGGAAGGTTGCANNNAGATCGGAA |
| 100990 | TATGAGTGTGGAGTCGTTGCCAACAGCTGCAATGGGCCAAATAANNNAGATCGGA |
| 107060 | TATGAGTGTGGAGTCGTTGCTTGAGCATCCTGGGGCTATGAATACNNNAGATCGG |
| 55044 | TATGAGTGTGGAGTCGTTGCTTGCACACCCTACACTTGATTGGNNNAGATCGGAAG |
| 109037 | TATGAGTGTGGAGTCGTTGCCCATCTCTTATCCCTTTGTCATGTNNNAGATCGGAAG |
| 99709 | TATGAGTGTGGAGTCGTTGCAGGGAAGCAGGATGGAGATTAGANNNAGATCGGA |
| 117529 | TATGAGTGTGGAGTCGTTGCGCTCTCCATATTCTGGGGTTCCANNNAGATCGGAAG |
| 104938 | TATGAGTGTGGAGTCGTTGCTGGGGGTTCTGGAAAATTCTACTGNNNAGATCGGA |
| 101407 | TATGAGTGTGGAGTCGTTGCTGCTGAAAAATACAACTCCAGGTGTNNNAGATCGGA |
| 30631 | TATGAGTGTGGAGTCGTTGCGCCATAGGGGAAAAACCTGGAAANNNAGATCGGAA |
| 109026 | TATGAGTGTGGAGTCGTTGCCACACCCTACACTTGATTGGCTNNNAGATCGGAAGA |
| 98943 | TATGAGTGTGGAGTCGTTGCGGGAAGCAGGATGGAGATTAGAATNNNAGATCGG |
| 96812 | TATGAGTGTGGAGTCGTTGCCACCTCCCAGGAATATTTTTGTCTNNNAGATCGGAAG |
| 63616 | TATGAGTGTGGAGTCGTTGCTTGTGGGCTGTGATCAGAAAAGTNNNAGATCGGAA |
| 41545 | TATGAGTGTGGAGTCGTTGCACTGTGACAACAACTTTGGGGTANNNAGATCGGAAG |
| 62818 | TATGAGTGTGGAGTCGTTGCGGGTGGGTAAGGGATTCTACTCTNNNAGATCGGAA |
| 41240 | TATGAGTGTGGAGTCGTTGCGCTGGATGTAGTTGGTGTTGCTANNNAGATCGGAA |
| 53328 | TATGAGTGTGGAGTCGTTGCGCTGCAATGGGCCAAATAATAGGNNNAGATCGGAA |
| 117168 | TATGAGTGTGGAGTCGTTGCCTGCAAGTGGCCAGATCTCTGTNNNAGATCGGAAGA |
| 98872 | TATGAGTGTGGAGTCGTTGCGAACCACCTTGTGAGCACAGATTCNNNAGATCGGAA |
| 100061 | TATGAGTGTGGAGTCGTTGCATTATCCACACACTCAGCTTTGAAGTNNNAGATCGGA |
| 61033 | TATGAGTGTGGAGTCGTTGCTGGTGTTCTGTATCAGTGTCCCTNNNAGATCGGAAG |
| 116352 | TATGAGTGTGGAGTCGTTGCAAATGAGGCCTCTGCCTCACAANNNAGATCGGAAGA |
| 67517 | TATGAGTGTGGAGTCGTTGCTCAACGTACTTACCTTTCCACTTNNNAGATCGGAAGA |
| 63698 | TATGAGTGTGGAGTCGTTGCGGACTGACTTTGGAAACCCAAGTNNNAGATCGGAAG |
| 43237 | TATGAGTGTGGAGTCGTTGCACCTCTTTTCTCCCCCACTAGACNNNAGATCGGAAGA |
| 57956 | TATGAGTGTGGAGTCGTTGCCCCACAAAGATCCAAGAGGCAATNNNAGATCGGAAG |
| 99052 | TATGAGTGTGGAGTCGTTGCACAGAATGTGCGCCAAAGATTCTNNNAGATCGGAAG |
| 53366 | TATGAGTGTGGAGTCGTTGCTCTGCAATGGGAAGGGTACTAGGNNNAGATCGGAA |
| 62640 | TATGAGTGTGGAGTCGTTGCCCCTGATAACTCAGCATGCATCTNNNAGATCGGAAG |
| 68522 | TATGAGTGTGGAGTCGTTGCCAGGCACATTCTCTCTTGTGGTTNNNAGATCGGAAG |
| 57563 | TATGAGTGTGGAGTCGTTGCTCCTCCATATACTGGCAGCTTTGNNNAGATCGGAAG |
| 63095 | TATGAGTGTGGAGTCGTTGCTGCCCTACTGGTTCTCTTTCTCTNNNAGATCGGAAGA |
| 51347 | TATGAGTGTGGAGTCGTTGCGAAGGGAGGGTAGAGGGAAAGAGNNNAGATCGG |
| 97401 | TATGAGTGTGGAGTCGTTGCTAACCAGGGCACAGAGACACANNNAGATCGGAAGA |
| 66347 | TATGAGTGTGGAGTCGTTGCAGTCTGTGTGTGTGTTCACGTGTNNNAGATCGGAAG |
| 60960 | TATGAGTGTGGAGTCGTTGCTGAGCCTCTAAACTATTGGCCCTNNNAGATCGGAAG |

| | |
|---|---|
| 56383 | TATGAGTGTGGAGTCGTTGCGGGGTAACATGCCAAACTCTCTGNNNAGATCGGAAG |
| 57961 | TATGAGTGTGGAGTCGTTGCAAATCAGTAACTGCAATGGCAATNNNAGATCGGAAG |
| 58249 | TATGAGTGTGGAGTCGTTGCCACTCCGTCACACAGAGAAACATNNNAGATCGGAAG |
| 35032 | TATGAGTGTGGAGTCGTTGCCTAACTCCCTGCACAAACTTCCANNNAGATCGGAAGA |
| 50777 | TATGAGTGTGGAGTCGTTGCTCAGAGCTGGGTGCCTTACAGNNNAGATCGGAAGA |
| 48067 | TATGAGTGTGGAGTCGTTGCTTGAATTCCTACTATTTGTCAGATTGCNNNAGATCGG |
| 29836 | TATGAGTGTGGAGTCGTTGCAGAAGCCTGGAGCTATTCAGGAANNNAGATCGGAA |
| 52556 | TATGAGTGTGGAGTCGTTGCTCATCAAGGCAGAACTGGAATCGNNNAGATCGGAA |
| 32725 | TATGAGTGTGGAGTCGTTGCAGAGAGAGAGACTGGCTCAAACANNNAGATCGGAA |
| 34377 | TATGAGTGTGGAGTCGTTGCCTCTACCTGGATGACTTAGCCCANNNAGATCGGAAG |
| 109144 | TATGAGTGTGGAGTCGTTGCCCTGTGTGCTTCCTGTGGTTTNNNAGATCGGAAGAG |
| 38792 | TATGAGTGTGGAGTCGTTGCATCCCTTTGTCATGTAACAGAGGANNNAGATCGGAA |
| 56175 | TATGAGTGTGGAGTCGTTGCTACATAGCTACGGGTACAGGCTGNNNAGATCGGAA |
| 117650 | TATGAGTGTGGAGTCGTTGCCAAGGAGATAGTTTGCAGACGAAAGNNNAGATCGG |
| 97145 | TATGAGTGTGGAGTCGTTGCCCATTTGTCTACCCACCTATCCACCNNNAGATCGGAA |
| 38258 | TATGAGTGTGGAGTCGTTGCTCAGACAAACTCTCCCATGGAGANNNAGATCGGAAG |
| 62502 | TATGAGTGTGGAGTCGTTGCAAATCATCAACTTCTCCATTTGCTNNNAGATCGGAAG |
| 30196 | TATGAGTGTGGAGTCGTTGCCCAACCCATCTTGGCCTCTAAAANNNAGATCGGAAG |
| 59642 | TATGAGTGTGGAGTCGTTGCTCCACTTTCCCACCTCTGTGTATNNNAGATCGGAAGA |
| 69663 | TATGAGTGTGGAGTCGTTGCCCTGTGTGCTTCCTGTGGTTTTTNNNAGATCGGAAG |
| 38391 | TATGAGTGTGGAGTCGTTGCTCGCCAGTCTCCAAGCATATAGANNNAGATCGGAAG |
| 35348 | TATGAGTGTGGAGTCGTTGCTCCCTAGACTAACAGCTTCAGCANNNAGATCGGAAG |
| 57389 | TATGAGTGTGGAGTCGTTGCAAGTTTGGTACTGGGGCTTCTTGNNNAGATCGGAA |
| 50356 | TATGAGTGTGGAGTCGTTGCACCATTCCCGAGTTTCTGTCAAGNNNAGATCGGAAG |
| 108568 | TATGAGTGTGGAGTCGTTGCCTGGGAGACAAAGCGGAAATCANNNAGATCGGAAG |
| 99725 | TATGAGTGTGGAGTCGTTGCGAATGAGGCACCCTGGAAGTGGNNNAGATCGGAAG |
| 65613 | TATGAGTGTGGAGTCGTTGCGTTGAGGGACAATGGGGAAATGTNNNAGATCGGA |
| 59033 | TATGAGTGTGGAGTCGTTGCAAGCTCTGAGAGAGAGAGAGAGATNNNAGATCGG |
| 41766 | TATGAGTGTGGAGTCGTTGCTGACCCCATGCCCACATAGATTANNNAGATCGGAAG |
| 65308 | TATGAGTGTGGAGTCGTTGCTTTGAGGATGCCACTTTGATGGTNNNAGATCGGAAG |
| 63653 | TATGAGTGTGGAGTCGTTGCAGGCTCCCATCACCATAGAAAGTNNNAGATCGGAAG |
| 48391 | TATGAGTGTGGAGTCGTTGCAGAGAGAGAGAGGAACTGGCATCNNNAGATCGGA |
| 51643 | TATGAGTGTGGAGTCGTTGCTTTGAGTGACTTTCTCTGCGGAGNNNAGATCGGAAG |
| 55478 | TATGAGTGTGGAGTCGTTGCCTGTCATACACACGTACTTGCATGNNNAGATCGGAA |
| 108159 | TATGAGTGTGGAGTCGTTGCGGGATTGTGGGGACCTATGTGTNNNAGATCGGAAG |
| 36189 | TATGAGTGTGGAGTCGTTGCTCAATCTTGTGAGGTTGTGTGCANNNAGATCGGAAG |
| 71522 | TATGAGTGTGGAGTCGTTGCCACAGTAATAGGAACGCTGCCTTNNNAGATCGGAAG |
| 56463 | TATGAGTGTGGAGTCGTTGCAAGAGGCAACCAAAGTCCTTCTGNNNAGATCGGAAG |
| 40577 | TATGAGTGTGGAGTCGTTGCCTCCTCCTCCACTCCTCTTTTGANNNAGATCGGAAGA |
| 34731 | TATGAGTGTGGAGTCGTTGCCCTCCTCATAGTTTATTCATCTGCCANNNAGATCGGA |
| 60853 | TATGAGTGTGGAGTCGTTGCAGAGCCCAATTACTTTAGACCCTNNNAGATCGGAAG |
| 107151 | TATGAGTGTGGAGTCGTTGCGAACCTGATGACCAATGGATTGATNNNAGATCGGA |
| 109105 | TATGAGTGTGGAGTCGTTGCGAGAGAGAGAGAAGAGAGCACTGTCNNNAGATCG |

55051 TATGAGTGTGGAGTCGTTGCAGAAGAAGGCAGGGCATTATTGGNNNAGATCGGA
99263 TATGAGTGTGGAGTCGTTGCCAGAACACACCCCCTACACACNNNAGATCGGAAGAG
31617 TATGAGTGTGGAGTCGTTGCATTCCTCTGGTCTTCAGCCAGAANNNAGATCGGAAG
60217 TATGAGTGTGGAGTCGTTGCTCGTCTACTCTCACACAGTCACTNNNAGATCGGAAGA
29833 TATGAGTGTGGAGTCGTTGCAGATTCAGATATCACGCTCCCCANNNAGATCGGAAG
56220 TATGAGTGTGGAGTCGTTGCACATCAGGTTTTCTAGCTGGCTGNNNAGATCGGAAG
49213 TATGAGTGTGGAGTCGTTGCGGACACCCATCATGCCAGTCNNNAGATCGGAAGAGC
37491 TATGAGTGTGGAGTCGTTGCAAGCAGTCTGCAAACCAGAAAGANNNAGATCGGAA
52352 TATGAGTGTGGAGTCGTTGCAGTAATTGGTCACAGCCAGTACGNNNAGATCGGAA
46487 TATGAGTGTGGAGTCGTTGCCCCTGAATTTCTCTGGTCAGAGCNNNAGATCGGAAG
101055 TATGAGTGTGGAGTCGTTGCATGCCTGATTTCTCAACTGCAAANNNAGATCGGAAG
101915 TATGAGTGTGGAGTCGTTGCTGCCATCCCATCCAAATACATTTNNNAGATCGGAAGA
46537 TATGAGTGTGGAGTCGTTGCACTAGAACCAGAACCAGAGGAGCNNNAGATCGGAA
30089 TATGAGTGTGGAGTCGTTGCACCCTCAGTTCTCACATGCAAAANNNAGATCGGAAG
100999 TATGAGTGTGGAGTCGTTGCCACAGTCAAGGACAGGGTTTACTTNNNAGATCGGA
71791 TATGAGTGTGGAGTCGTTGCACTGTCAGGCAGAAGACACTGGNNNAGATCGGAAG
101045 TATGAGTGTGGAGTCGTTGCCAGGGACTGAACCAGATGATCCTNNNAGATCGGAA
104175 TATGAGTGTGGAGTCGTTGCGTAGGCAAGCTGGTAACAAAAGAGNNNAGATCGG
100988 TATGAGTGTGGAGTCGTTGCGAGTTGGGAGAGAGGGGAAAGAANNNAGATCGG
101919 TATGAGTGTGGAGTCGTTGCCCTTGAAGATGCCTCTCCCTTCACNNNAGATCGGAAG
102594 TATGAGTGTGGAGTCGTTGCTTTCTGCCTGTGTGATACTGCATTNNNAGATCGGAA
105014 TATGAGTGTGGAGTCGTTGCCATACCCGACTTGCTTGCTTATTTANNNAGATCGGAA
101895 TATGAGTGTGGAGTCGTTGCCCACCTGAAATGGGAATCATTACCNNNAGATCGGAA
103391 TATGAGTGTGGAGTCGTTGCGAAGAGGAGGAGGAGACATTTGGNNNAGATCGGA
100813 TATGAGTGTGGAGTCGTTGCACATTCCTGCATAGCTCTGTGGNNNAGATCGGAAGA
109998 TATGAGTGTGGAGTCGTTGCCCTGGCAGTAAAAGAACGTATGANNNAGATCGGAA
114840 TATGAGTGTGGAGTCGTTGCTCCACCAAACTCCCCAGGAAANNNAGATCGGAAGAG
109063 TATGAGTGTGGAGTCGTTGCCATATAGAGTGGTGTGTGCACAGTANNNAGATCGG
122013 TATGAGTGTGGAGTCGTTGCGAAGGGCAAGAAAATTGAACCCTGNNNAGATCGGA
103350 TATGAGTGTGGAGTCGTTGCGGGAAACAGACCTCCCAGTTCNNNAGATCGGAAGA
100568 TATGAGTGTGGAGTCGTTGCGAATGGAGTGGCAGGAGTGCNNNAGATCGGAAGA
106161 TATGAGTGTGGAGTCGTTGCGGGCAGGGAGAGTAGTTACTAGGTNNNAGATCGG
104157 TATGAGTGTGGAGTCGTTGCGGAGCAGTGGGAAGTCTCATGATNNNAGATCGGAA
108587 TATGAGTGTGGAGTCGTTGCCGTGAGATTTCTAGGATGGTTCCCNNNAGATCGGAA
59141 TATGAGTGTGGAGTCGTTGCGAAACCTGGTCATGGGAGAGGATNNNAGATCGGA
99106 TATGAGTGTGGAGTCGTTGCCTATGAACAACGACATTCTCCGGNNNAGATCGGAAG
30912 TATGAGTGTGGAGTCGTTGCCCCTTTCAAATGTGTTCAGACAANNNAGATCGGAAG
29828 TATGAGTGTGGAGTCGTTGCACTCACACAAATCTGTAACAGCTNNNAGATCGGAAG
63019 TATGAGTGTGGAGTCGTTGCACGCTTACCCTCTTTTCCTCTCTNNNAGATCGGAAGA
34342 TATGAGTGTGGAGTCGTTGCAGTCATCTTTGTCATCCTCCCCANNNAGATCGGAAGA
55965 TATGAGTGTGGAGTCGTTGCAACATGAAAGGAGCATAGCCCTGNNNAGATCGGAA
33833 TATGAGTGTGGAGTCGTTGCTTCCTTCAAGCAGAGGTGCTACANNNAGATCGGAAG
34752 TATGAGTGTGGAGTCGTTGCAGTTGAATCACACAGAATTGCCANNNAGATCGGAAG

40337 TATGAGTGTGGAGTCGTTGCATGCAGACCAAAAATGGTTGTGANNNAGATCGGAA
110832 TATGAGTGTGGAGTCGTTGCTGTGTAGAAGCAGAGGGATATTTCTNNNAGATCGG
45760 TATGAGTGTGGAGTCGTTGCCTTCTTTCTCACCCCAAAGGTCCNNNAGATCGGAAGA
58426 TATGAGTGTGGAGTCGTTGCATGCACAGAAAAGAAACAACCATNNNAGATCGGAA
39678 TATGAGTGTGGAGTCGTTGCCCCGTAAGACGAAAGTTTCATGANNNAGATCGGAA
59307 TATGAGTGTGGAGTCGTTGCACATTGCAAAAGAGGAGGCTGATNNNAGATCGGAA
96421 TATGAGTGTGGAGTCGTTGCTGCTCATCTTAGGGACAAGTAAGAANNNAGATCGG
41302 TATGAGTGTGGAGTCGTTGCGTCTGAACCCTTACCCCTGTCTANNNAGATCGGAAGA
56009 TATGAGTGTGGAGTCGTTGCACAGTTTAGTTCATGGAGGCCTGNNNAGATCGGAA
51425 TATGAGTGTGGAGTCGTTGCTGCCATGTAAGATGAAGGCAGAGNNNAGATCGGAA
97885 TATGAGTGTGGAGTCGTTGCTGAAACCCCTTCTTCCAAAAAGANNNAGATCGGAAG
33080 TATGAGTGTGGAGTCGTTGCAGATGGAGGCTCACTATCACACANNNAGATCGGAA
97881 TATGAGTGTGGAGTCGTTGCTCAGATGTGGAGTTGCTGCTGNNNAGATCGGAAGA
69629 TATGAGTGTGGAGTCGTTGCGGGGGTGGGGAGTTTGTATTTTTNNNAGATCGGAA
65577 TATGAGTGTGGAGTCGTTGCTCAGATTTCGTGGCACTTTTGGTNNNAGATCGGAAG
48943 TATGAGTGTGGAGTCGTTGCTTGAAGTGTCTGGCATAGTGCTCNNNAGATCGGAA
65333 TATGAGTGTGGAGTCGTTGCAGACACAAAGAAGGGAAACTGGTNNNAGATCGGA
99292 TATGAGTGTGGAGTCGTTGCTCTACTGATTGGCCCATTTTACTGANNNAGATCGGA
58707 TATGAGTGTGGAGTCGTTGCCTCCTACTCAAGTCCCCATGCATNNNAGATCGGAAGA
101916 TATGAGTGTGGAGTCGTTGCGTTGTCTTTGTGCACATGAGGAGANNNAGATCGGA
33838 TATGAGTGTGGAGTCGTTGCATGCAGCAGAGGTTTACATCTACANNNAGATCGGAA
64889 TATGAGTGTGGAGTCGTTGCGTTGGAATGGAAATGAAACGAGGTNNNAGATCGG
103309 TATGAGTGTGGAGTCGTTGCAGGCTCACTATCACACAGGCTNNNAGATCGGAAGAG
31162 TATGAGTGTGGAGTCGTTGCGTGGTTTGAACGTGTGTTTCCAANNNAGATCGGAA
54120 TATGAGTGTGGAGTCGTTGCAAGACTAAAACACACAGGCTGGGNNNAGATCGGAA
64719 TATGAGTGTGGAGTCGTTGCATCTCACCATTTTGCCCAAAGGTNNNAGATCGGAAG
35914 TATGAGTGTGGAGTCGTTGCCCACAAAAGCTATGTCATGAATGCANNNAGATCGGA
51242 TATGAGTGTGGAGTCGTTGCCCTGCTTTCTTCTATGGGCTCAGNNNAGATCGGAAG
53638 TATGAGTGTGGAGTCGTTGCTGGTGTGCGTTAGTATTGAAGGGNNNAGATCGGAA
52231 TATGAGTGTGGAGTCGTTGCCCAGGGCACAACAAGCATTTTAGNNNAGATCGGAA
34917 TATGAGTGTGGAGTCGTTGCCTTGTTGCCAGCTATGATCTCCANNNAGATCGGAAG
52229 TATGAGTGTGGAGTCGTTGCATGCAGAGGCACACAGTCTTTAGNNNAGATCGGAA
66968 TATGAGTGTGGAGTCGTTGCGGCAGTGGGTAGGCTAATCAATTNNNAGATCGGAA
36685 TATGAGTGTGGAGTCGTTGCCTTTTTCCCTCTGTGTGAGCTCANNNAGATCGGAAGA
58777 TATGAGTGTGGAGTCGTTGCCGCACTTACTCACCCATCATCATNNNAGATCGGAAGA
34474 TATGAGTGTGGAGTCGTTGCTAGGTCAAGAAGCATCAGTCCCANNNAGATCGGAA
53302 TATGAGTGTGGAGTCGTTGCAATAGTCCACAGGTGAGGTGAGGNNNAGATCGGA
47691 TATGAGTGTGGAGTCGTTGCGGAAGGGTTGGCTCTATTACTGCNNNAGATCGGAA
55206 TATGAGTGTGGAGTCGTTGCAGCCTATGTCTGCCTCATTTTGGNNNAGATCGGAAG
52861 TATGAGTGTGGAGTCGTTGCGCCACACATGAAACATTCCTAAGGNNNAGATCGGAA
60194 TATGAGTGTGGAGTCGTTGCTGTCCCCGATATATGTCCTCACTNNNAGATCGGAAGA
33318 TATGAGTGTGGAGTCGTTGCCCTTTCCAGTCATCATCTGCACANNNAGATCGGAAGA
53613 TATGAGTGTGGAGTCGTTGCCTATAACAGGGCATGGTCAAGGGNNNAGATCGGAA

| | |
|---|---|
| 44133 | TATGAGTGTGGAGTCGTTGCTGCGTTTGTGTCACTTTATTCATACCNNNAGATCGGA |
| 32972 | TATGAGTGTGGAGTCGTTGCAGTCTGGTTCTGTCTGGGTAACANNNAGATCGGAA |
| 65288 | TATGAGTGTGGAGTCGTTGCCCAATCTTTTCCCTATGGGATGGTNNNAGATCGGAA |
| 104180 | TATGAGTGTGGAGTCGTTGCAAAGCTCATAACATTGGATTTGGCNNNAGATCGGAA |
| 34054 | TATGAGTGTGGAGTCGTTGCGCTCGGAAATAACTGATCCACCANNNAGATCGGAAG |
| 37897 | TATGAGTGTGGAGTCGTTGCTGCCTACAGTGAATCTCTCCAGANNNAGATCGGAAG |
| 41113 | TATGAGTGTGGAGTCGTTGCTCTGGGGGTTAGTTTTGCTCCTANNNAGATCGGAAG |
| 64277 | TATGAGTGTGGAGTCGTTGCGAGGCTATGAAAGGACTGGGAGTNNNAGATCGGA |
| 31141 | TATGAGTGTGGAGTCGTTGCCCACTTCCACCAGAAAAGTCCAANNNAGATCGGAAG |
| 109057 | TATGAGTGTGGAGTCGTTGCAAGCTCGGAAATAACTGATCCACCATNNNAGATCGG |
| 71647 | TATGAGTGTGGAGTCGTTGCCCATGCCTTTGCAGGTAAGTANNNAGATCGGAAGA |
| 63520 | TATGAGTGTGGAGTCGTTGCCGATGCCATGTGTCTTCCTTTCTNNNAGATCGGAAG |
| 40416 | TATGAGTGTGGAGTCGTTGCGGTGTTTGGGGAATTCACCTTGANNNAGATCGGAA |
| 104199 | TATGAGTGTGGAGTCGTTGCTCTTTATCTGTGTCCATAAAGCATTTNNNAGATCGGA |
| 39633 | TATGAGTGTGGAGTCGTTGCGGAAGGACATGTTTGGACCATGANNNAGATCGGAA |
| 39149 | TATGAGTGTGGAGTCGTTGCGGACCTATGTGAAAGACCTGGGANNNAGATCGGAA |
| 30305 | TATGAGTGTGGAGTCGTTGCTATTCAGGACATGCCAACCCAAANNNAGATCGGAAG |
| 60445 | TATGAGTGTGGAGTCGTTGCAGTCAAGACAGCCTTACCATACTNNNAGATCGGAAG |
| 61529 | TATGAGTGTGGAGTCGTTGCGGTGGGGATAGTAGAAGCTTCCTNNNAGATCGGAA |
| 69634 | TATGAGTGTGGAGTCGTTGCTTTGGCAACACCATGCACTTTTTNNNAGATCGGAAG |
| 30713 | TATGAGTGTGGAGTCGTTGCCCTTAGCTTGCCCAGAGCATAAANNNAGATCGGAAG |
| 108485 | TATGAGTGTGGAGTCGTTGCGTGGTGGATCTCCCTTGTTCTNNNAGATCGGAAGAG |
| 57217 | TATGAGTGTGGAGTCGTTGCCAGCCAGAGAGACAGAGTGATTGNNNAGATCGGAA |
| 43861 | TATGAGTGTGGAGTCGTTGCGAAAAGCGAGTAACAGCAACACCNNNAGATCGGAA |
| 100152 | TATGAGTGTGGAGTCGTTGCTGTGTGGTGGAAGCTTTGTCCTNNNAGATCGGAAG |
| 64341 | TATGAGTGTGGAGTCGTTGCATCCCCTTAGGCTTTTCCTGAGTNNNAGATCGGAAG |
| 68496 | TATGAGTGTGGAGTCGTTGCGCAGTTTCTCCAGTTTCCTGGTTNNNAGATCGGAAG |
| 50380 | TATGAGTGTGGAGTCGTTGCTGACTAACAAAAAGAATACCACAGAAGNNNAGATCG |
| 62823 | TATGAGTGTGGAGTCGTTGCAGTTCTCTGGTGGCTGTTACTCTNNNAGATCGGAAG |
| 52735 | TATGAGTGTGGAGTCGTTGCTTCATCCATCTTCCAGCACAAGGNNNAGATCGGAAG |
| 63862 | TATGAGTGTGGAGTCGTTGCTTGTTGCTGAACCATTCAACAGTNNNAGATCGGAAG |
| 39283 | TATGAGTGTGGAGTCGTTGCTTGTCCAGCTATCCATTGATGGANNNAGATCGGAAG |
| 45090 | TATGAGTGTGGAGTCGTTGCCTTTGGAAACAATTCAAGGGGCCNNNAGATCGGAA |
| 39083 | TATGAGTGTGGAGTCGTTGCAACTGAGGACTTTTCGTAGGGGANNNAGATCGGAA |
| 62333 | TATGAGTGTGGAGTCGTTGCCCTGTTTTCCTGTTTCCTCTGCTNNNAGATCGGAAGA |
| 66536 | TATGAGTGTGGAGTCGTTGCGGAGGGAAAAGAACCTGTTGTGTNNNAGATCGGA |
| 52387 | TATGAGTGTGGAGTCGTTGCTGAAATGCCCTTCCTCTTTACCGNNNAGATCGGAAGA |
| 38616 | TATGAGTGTGGAGTCGTTGCGCCTCAAAGTCACTTCCAAAGGANNNAGATCGGAAG |
| 71768 | TATGAGTGTGGAGTCGTTGCGGGATGTCAGGACACATGTCTTCNNNAGATCGGAA |
| 55602 | TATGAGTGTGGAGTCGTTGCGAGGGGCTAATATTGGCAGGATGNNNAGATCGGA |
| 65994 | TATGAGTGTGGAGTCGTTGCTGTCTGAACCTATCTCAGCCTGTNNNAGATCGGAAG |
| 42630 | TATGAGTGTGGAGTCGTTGCAGCCTTGCAGAGTTAGATGACACNNNAGATCGGAA |
| 47695 | TATGAGTGTGGAGTCGTTGCCTCTCTCTCTTTCACCAACCTGCNNNAGATCGGAAGA |

60591 TATGAGTGTGGAGTCGTTGCAGGGGCATCAGAGTTCTAAACCTNNNAGATCGGAA
62051 TATGAGTGTGGAGTCGTTGCTACCTTACAAAGATCCCTGGGCTNNNAGATCGGAAG
64703 TATGAGTGTGGAGTCGTTGCGGGGGCAGTGGATATATTTTCGTNNNAGATCGGAA
48356 TATGAGTGTGGAGTCGTTGCCCCTCCTGGATTATCTCCCCATCNNNAGATCGGAAGA
39178 TATGAGTGTGGAGTCGTTGCTTTGACCATACTCTGTCGTGGGANNNAGATCGGAAG
67088 TATGAGTGTGGAGTCGTTGCTCAAATCCACACTGGTGTACATTNNNAGATCGGAAG
57840 TATGAGTGTGGAGTCGTTGCAGTCGGGTCTCACAGAGCTAAATNNNAGATCGGAA
32244 TATGAGTGTGGAGTCGTTGCGCTAGAATGCGGTGGCTGAATAANNNAGATCGGAA
107895 TATGAGTGTGGAGTCGTTGCCCACTCCCAAGCTATGTGCTTATNNNAGATCGGAAG
45707 TATGAGTGTGGAGTCGTTGCTCCTTTTCCGGTTCTGTTTCTCCNNNAGATCGGAAGA
47306 TATGAGTGTGGAGTCGTTGCTTTGGGGTTTTCTCTCTTCTGGCNNNAGATCGGAAG
54071 TATGAGTGTGGAGTCGTTGCCTCCGGGTTCAAAGCTATGGGNNNAGATCGGAAGA
56983 TATGAGTGTGGAGTCGTTGCCAGCTTCATGGGTGTGATCTGTGNNNAGATCGGAA
104977 TATGAGTGTGGAGTCGTTGCTAGAATGCGGTGGCTGAATAACGNNNAGATCGGAA
106166 TATGAGTGTGGAGTCGTTGCTGTGTGTGTGTTCACCAAGAGATATNNNAGATCGG
59532 TATGAGTGTGGAGTCGTTGCTGTGGGGTCCATCATGTTCCTATNNNAGATCGGAAG
66781 TATGAGTGTGGAGTCGTTGCTTGTCCCTCCCTCCCTTATTTGTNNNAGATCGGAAGA
57708 TATGAGTGTGGAGTCGTTGCCAGGTGGTGCATGTAGACCAAATNNNAGATCGGAA
49568 TATGAGTGTGGAGTCGTTGCTGGCAAAGTTTGAAGGCTGATTCNNNAGATCGGAA
60243 TATGAGTGTGGAGTCGTTGCACCACTTTCTTGCATTCTTTCACTNNNAGATCGGAAG
38624 TATGAGTGTGGAGTCGTTGCAATAAGGCTGCATGCTCAAAGGANNNAGATCGGAA
47564 TATGAGTGTGGAGTCGTTGCAGATTGGTGTGGTAACGAGATGCNNNAGATCGGAA
51554 TATGAGTGTGGAGTCGTTGCGTTTCGGGACTGCAGCCGAGNNNAGATCGGAAGAG
31458 TATGAGTGTGGAGTCGTTGCTGCCAAGAAACAAGTCTGGTCAANNNAGATCGGAA
59359 TATGAGTGTGGAGTCGTTGCAAGAAAGGGCAATCAGCCTTGATNNNAGATCGGAA
48366 TATGAGTGTGGAGTCGTTGCTTATACCTAGCCCGAGCTCCATCNNNAGATCGGAAG
66535 TATGAGTGTGGAGTCGTTGCTGTGTTTGGATGTGATGTTGTGTNNNAGATCGGAA
117357 TATGAGTGTGGAGTCGTTGCCCCCTGATAATCCTCATACCCCTGANNNAGATCGGAA
117927 TATGAGTGTGGAGTCGTTGCCAATGAAGTCTCGCCTCGCTNNNAGATCGGAAGAGC
66988 TATGAGTGTGGAGTCGTTGCTGGGATGAGGAGGGAAAGGAATTNNNAGATCGGA
46712 TATGAGTGTGGAGTCGTTGCGGTATCTCCCAACTGCCTTTAGCNNNAGATCGGAAG
57986 TATGAGTGTGGAGTCGTTGCGGGGGACTGGTAACACAATCAATNNNAGATCGGAA
56124 TATGAGTGTGGAGTCGTTGCCCTGGCCATTACTTATCCAGCTGNNNAGATCGGAAG
59270 TATGAGTGTGGAGTCGTTGCCCCACCCATCCTTCTCTCATGATNNNAGATCGGAAGA
42017 TATGAGTGTGGAGTCGTTGCTGTCACCAATGTGATGCCCTTTANNNAGATCGGAAG
47547 TATGAGTGTGGAGTCGTTGCGGTGGCATTCATTCTGTTCATGCNNNAGATCGGAAG
67226 TATGAGTGTGGAGTCGTTGCGGGCAACTAAGCAGCTAGTCATTNNNAGATCGGAA
44703 TATGAGTGTGGAGTCGTTGCTGGCCTCTTCATGATATCCTCCCNNNAGATCGGAAGA
47171 TATGAGTGTGGAGTCGTTGCCATTCACAACTTCTTTGGTGGGCNNNAGATCGGAAG
33485 TATGAGTGTGGAGTCGTTGCCATCAAACCCCCACTTGAAGACANNNAGATCGGAAG
102304 TATGAGTGTGGAGTCGTTGCGCAGGGAACCAGAGGAAAATGAGNNNAGATCGGA
104741 TATGAGTGTGGAGTCGTTGCCCCAGTTCACTGAAAGGATCCTNNNAGATCGGAAGA
30651 TATGAGTGTGGAGTCGTTGCAGGTCATTGCTGGGATCATGAAANNNAGATCGGAA

43535 TATGAGTGTGGAGTCGTTGCCTTGGGGCAGGTCCTTTTACTACNNNAGATCGGAAG
36465 TATGAGTGTGGAGTCGTTGCACCTTTTGGTGCTCTGTTCATCANNNAGATCGGAAG
50565 TATGAGTGTGGAGTCGTTGCTGTCCCCTGATGGGGTAAGTAAGNNNAGATCGGAA
104908 TATGAGTGTGGAGTCGTTGCATCTCCCAACTGCCTTTAGCATCTNNNAGATCGGAAG
41201 TATGAGTGTGGAGTCGTTGCAAGCAATTCAGCATCTGTGGCTANNNAGATCGGAAG
109908 TATGAGTGTGGAGTCGTTGCTAGACCTTTTGGTGCTCTGTTCATCATNNNAGATCGG
116916 TATGAGTGTGGAGTCGTTGCGGTGCATGTAGACCAAATAGGAAAAGNNNAGATCG
96522 TATGAGTGTGGAGTCGTTGCTTTTAGATGATGGAGCCATGCATGCNNNAGATCGG
59538 TATGAGTGTGGAGTCGTTGCGCCAGGTGTCTCTAGTGAGCTATNNNAGATCGGAA
41901 TATGAGTGTGGAGTCGTTGCCAGGGTAGCAATGTGCTCAGTTANNNAGATCGGAA
33663 TATGAGTGTGGAGTCGTTGCACCTCTGCCTTTTGTTCATGACANNNAGATCGGAAG
66285 TATGAGTGTGGAGTCGTTGCACCGCTTCTAGGCTACTAAGTGTNNNAGATCGGAAG
30048 TATGAGTGTGGAGTCGTTGCATGCCTATAAAAGACCATCACAAAANNNAGATCGGA
51130 TATGAGTGTGGAGTCGTTGCTCTAGGGAAGATGCAAACTGCAGNNNAGATCGGAA
71662 TATGAGTGTGGAGTCGTTGCTCTTGCTCAGACTGACCTTCCATNNNAGATCGGAAG
61681 TATGAGTGTGGAGTCGTTGCATTTGGAAACAGGAAGGACAGCTNNNAGATCGGAA
57138 TATGAGTGTGGAGTCGTTGCGTGCTAAGGTCTCAGCGAAATTGNNNAGATCGGAA
47344 TATGAGTGTGGAGTCGTTGCAGGTCCAGATAGAAGTAGGTGGCNNNAGATCGGA
47951 TATGAGTGTGGAGTCGTTGCGAATGAAAACAAGAGCTGGGTGCNNNAGATCGGA
101877 TATGAGTGTGGAGTCGTTGCCGTCACTTATGGGTGTGAAATTCNNNAGATCGGAAG
46072 TATGAGTGTGGAGTCGTTGCATCATCCCCCTTTCATCCAAAGCNNNAGATCGGAAGA
56410 TATGAGTGTGGAGTCGTTGCCTGCAATGGGCGGTTATAGTCTGNNNAGATCGGAA
58428 TATGAGTGTGGAGTCGTTGCGCCCCTTACTGGAATTCAACCATNNNAGATCGGAAG
57162 TATGAGTGTGGAGTCGTTGCGCTGCATGGTATGCCTTACATTGNNNAGATCGGAAG
56301 TATGAGTGTGGAGTCGTTGCGTTTGTGTGTGTGCCAGAATCTGNNNAGATCGGAA
38062 TATGAGTGTGGAGTCGTTGCGTGTATGGGCGTATGTGTTCAGANNNAGATCGGAA
41343 TATGAGTGTGGAGTCGTTGCAAGGCAAACGCCTCAACTTTCTANNNAGATCGGAAG
67163 TATGAGTGTGGAGTCGTTGCGTGGGTCATAGGGTGAATGCATTNNNAGATCGGAA
30701 TATGAGTGTGGAGTCGTTGCCCATGCTAGGCCCTGCAAATAAANNNAGATCGGAAG
51725 TATGAGTGTGGAGTCGTTGCTACTTTTTGAGACAGGGCTGGAGNNNAGATCGGAA
32280 TATGAGTGTGGAGTCGTTGCGGCATGGGGAGAGATGGAGATAANNNAGATCGGA
41622 TATGAGTGTGGAGTCGTTGCCCCTAGAAAGTAGGCCCACTGTANNNAGATCGGAAG
30352 TATGAGTGTGGAGTCGTTGCCTGATACGCCAGCAAAAAGCAAANNNAGATCGGAA
45683 TATGAGTGTGGAGTCGTTGCAGACTACTTCAGCCTCTGTCTCCNNNAGATCGGAAG
57269 TATGAGTGTGGAGTCGTTGCCATTTCTCCCCAGGTTACCCTTGNNNAGATCGGAAGA
96874 TATGAGTGTGGAGTCGTTGCACTTTTCCAAGGACAGAGCTTCTNNNAGATCGGAAG
96181 TATGAGTGTGGAGTCGTTGCGCTGGCCTTCTCTCTAGCTCTNNNAGATCGGAAGAG
46688 TATGAGTGTGGAGTCGTTGCCCCAAATCAGACCTTCCTGTAGCNNNAGATCGGAAG
35500 TATGAGTGTGGAGTCGTTGCCTCCCTCTCAAATCCTGGTAGCANNNAGATCGGAAG
67020 TATGAGTGTGGAGTCGTTGCCTTGCCGAAGGTCAGAGCTAATTNNNAGATCGGAA
69256 TATGAGTGTGGAGTCGTTGCTGAGTTCCCTAAATTGGCAGTTTNNNAGATCGGAAG
38572 TATGAGTGTGGAGTCGTTGCTCCACAATATCTTCAGAAATGCGANNNAGATCGGAA
68881 TATGAGTGTGGAGTCGTTGCTCTCCATCCCACTCTCCAGATTTNNNAGATCGGAAGA

| | |
|---|---|
| 44739 | TATGAGTGTGGAGTCGTTGCTTGAACCAAACCAGTTGTCTCCCNNNAGATCGGAAG |
| 55086 | TATGAGTGTGGAGTCGTTGCCCTGCAAAAACAGACCATCTTGGNNNAGATCGGAAG |
| 49198 | TATGAGTGTGGAGTCGTTGCAGCAAAAGTAGCCACTGACAGTCNNNAGATCGGAA |
| 54116 | TATGAGTGTGGAGTCGTTGCAGAGAAAGAGAGAGTGAGCTGGGNNNAGATCGGA |
| 117520 | TATGAGTGTGGAGTCGTTGCATAAAGCCCCACTTTTCTGGGGNNNAGATCGGAAGA |
| 32300 | TATGAGTGTGGAGTCGTTGCAACAGTGGTCAGGGTCCTGATAANNNAGATCGGAA |
| 45711 | TATGAGTGTGGAGTCGTTGCGTGGGTACAGATCGTGAAAGTCCNNNAGATCGGAA |
| 97205 | TATGAGTGTGGAGTCGTTGCGTAAAAGCCATCAGGATGACACACNNNAGATCGGA |
| 57299 | TATGAGTGTGGAGTCGTTGCGACAGAGACATCCCAGAAGCTTGNNNAGATCGGAA |
| 45245 | TATGAGTGTGGAGTCGTTGCTACATTCTGATGCTTTCCCTGCCNNNAGATCGGAAGA |
| 51846 | TATGAGTGTGGAGTCGTTGCGTAAACAGAGGTTCTGGGCTGAGNNNAGATCGGAA |
| 51813 | TATGAGTGTGGAGTCGTTGCGTCAAGTGGCTACCAGAACTGAGNNNAGATCGGAA |
| 45125 | TATGAGTGTGGAGTCGTTGCCTTTCATCTGTCAGCATCTGGCCNNNAGATCGGAAG |
| 48819 | TATGAGTGTGGAGTCGTTGCGATGACTGCCTCCTCACTTCCTCNNNAGATCGGAAGA |
| 103266 | TATGAGTGTGGAGTCGTTGCGAAAGTCTGAAAGACTAGGCACTGCNNNAGATCGG |
| 58797 | TATGAGTGTGGAGTCGTTGCGCCCCACAAGATCAGAAACTCATNNNAGATCGGAAG |
| 36762 | TATGAGTGTGGAGTCGTTGCTCGTGTGATCTCCCTTCATCTCANNNAGATCGGAAGA |
| 38091 | TATGAGTGTGGAGTCGTTGCCAGTCGTTTGCTAGAGCAAGAGANNNAGATCGGAA |
| 47977 | TATGAGTGTGGAGTCGTTGCCCTCCCAGAATAGGAACATGTGCNNNAGATCGGAAG |
| 34045 | TATGAGTGTGGAGTCGTTGCTGGCACAAGGAAGTTTTACACCANNNAGATCGGAA |
| 98372 | TATGAGTGTGGAGTCGTTGCGACAGAACAGGTGAGGATGTGANNNAGATCGGAA |
| 53386 | TATGAGTGTGGAGTCGTTGCACCCTTCAGACCAATAATGCTAGGNNNAGATCGGAA |
| 35174 | TATGAGTGTGGAGTCGTTGCTAACTTCACAAAAGGACCAAGCANNNAGATCGGAAG |
| 32822 | TATGAGTGTGGAGTCGTTGCTCCAGTCAGTTTCTAAGCCAACANNNAGATCGGAAG |
| 67747 | TATGAGTGTGGAGTCGTTGCCAAGCACATACATACCCTGCCTTNNNAGATCGGAAG |
| 109862 | TATGAGTGTGGAGTCGTTGCCCTTGATGGACCAGAAATTATGGGGNNNAGATCGG |
| 53943 | TATGAGTGTGGAGTCGTTGCGAATAAAGAAGATGCCCTGGGGGNNNAGATCGGA |
| 38085 | TATGAGTGTGGAGTCGTTGCCACTGAAAGCCCACTGAAAGAGANNNAGATCGGAA |
| 43514 | TATGAGTGTGGAGTCGTTGCTGCCAAATCCCTGGCTGTTATACNNNAGATCGGAAG |
| 102830 | TATGAGTGTGGAGTCGTTGCTCTTTTCCCTATGGGATGGTACATACTNNNAGATCGG |
| 57640 | TATGAGTGTGGAGTCGTTGCCCTTTTGGCATTTTGGGCTTTTGNNNAGATCGGAAG |
| 52933 | TATGAGTGTGGAGTCGTTGCCTCTCACACACTAACACACCAGGNNNAGATCGGAAG |
| 49095 | TATGAGTGTGGAGTCGTTGCCGGTCATCTTCTGTCTGTGTCTCNNNAGATCGGAAG |
| 32263 | TATGAGTGTGGAGTCGTTGCCAGTGGTAGCTTTCGAGGCATAANNNAGATCGGAA |
| 67592 | TATGAGTGTGGAGTCGTTGCGAACATCAGAGGCATGGTGACTTNNNAGATCGGAA |
| 57068 | TATGAGTGTGGAGTCGTTGCACAAAGCACAAGACATGTGTGTGNNNAGATCGGAA |
| 69481 | TATGAGTGTGGAGTCGTTGCCAGTGTTCTATCGCTTGCCTTTTNNNAGATCGGAAGA |
| 68770 | TATGAGTGTGGAGTCGTTGCTTGATGGAGCACAGACCCAATTTNNNAGATCGGAAG |
| 102272 | TATGAGTGTGGAGTCGTTGCTGGTGTCTTTCCTTCACAGGGCTNNNAGATCGGAAG |
| 57173 | TATGAGTGTGGAGTCGTTGCCTTGGCTGGCTTAGAAAGCATTGNNNAGATCGGAA |
| 41787 | TATGAGTGTGGAGTCGTTGCGACAACCAGCAGCATACCACTTANNNAGATCGGAAG |
| 51639 | TATGAGTGTGGAGTCGTTGCAGGTAGAACTCTGATCACCGGAGNNNAGATCGGAA |
| 49648 | TATGAGTGTGGAGTCGTTGCATCTTGGTAAGACCCTCCCCTTCNNNAGATCGGAAGA |

106678 TATGAGTGTGGAGTCGTTGCTCAGCTCTGGTAAGGGCTACTAGTNNNAGATCGGAA

32731 TATGAGTGTGGAGTCGTTGCAGAAGGATCCTGAATAATTTCAAACANNNAGATCGG

51346 TATGAGTGTGGAGTCGTTGCTATGCAGGGAGAGGAGGAAAGAGNNNAGATCGGA

55685 TATGAGTGTGGAGTCGTTGCGCTCCCCTGTGTGTGTGTATATGNNNAGATCGGAAG

33037 TATGAGTGTGGAGTCGTTGCTTTCCAGCTTTGACACACACACANNNAGATCGGAAG

49866 TATGAGTGTGGAGTCGTTGCCAGCATGACAGCACTGACATTTCNNNAGATCGGAAG

51119 TATGAGTGTGGAGTCGTTGCAAAGCCATAAAGGCACAATGCAGNNNAGATCGGAA

50679 TATGAGTGTGGAGTCGTTGCAAGTAACCTGAAGGACTGCACAGNNNAGATCGGAA

50485 TATGAGTGTGGAGTCGTTGCCAGCCAAAACAGCTTCCATGAAGNNNAGATCGGAAG

104166 TATGAGTGTGGAGTCGTTGCTTCTATTTCACTTCTTGGAAGACTGCCNNNAGATCGG

64365 TATGAGTGTGGAGTCGTTGCACCAGAGTGTTGATGTTGTGAGTNNNAGATCGGAA

47197 TATGAGTGTGGAGTCGTTGCTACCCAGTGCCTAAAACAATGGCNNNAGATCGGAAG

67834 TATGAGTGTGGAGTCGTTGCGTGAGGGTGGAGAAAAGAAGCTTNNNAGATCGGA

110822 TATGAGTGTGGAGTCGTTGCTCACTTGGGACTTTCTCAGTTCAANNNAGATCGGAA

62860 TATGAGTGTGGAGTCGTTGCCAGACGCCATCTCTCCCTCTNNNAGATCGGAAGAGC

68642 TATGAGTGTGGAGTCGTTGCCCCAGGAGCATGAACTATCTGTTNNNAGATCGGAAG

51200 TATGAGTGTGGAGTCGTTGCCTCTCACTTCGCTTCCTGATCAGNNNAGATCGGAAGA

102586 TATGAGTGTGGAGTCGTTGCCTGTGTGTCTAAGCCTCACTCTANNNAGATCGGAAG

57359 TATGAGTGTGGAGTCGTTGCTCTTCATTTCACCCCTGCTCTTGNNNAGATCGGAAGA

61787 TATGAGTGTGGAGTCGTTGCATCAAAACCTGGGTCTAGGAGCTNNNAGATCGGAA

31670 TATGAGTGTGGAGTCGTTGCCTTGACCAGCAGAGCTTTCAGAANNNAGATCGGAAG

37705 TATGAGTGTGGAGTCGTTGCTCCCATGCCCTTCCAATCTAAGANNNAGATCGGAAGA

58810 TATGAGTGTGGAGTCGTTGCGAACACCATTGTCCCAACCTCATNNNAGATCGGAAG

39656 TATGAGTGTGGAGTCGTTGCAGCAAAGAATGGAACTGGCATGANNNAGATCGGAA

56650 TATGAGTGTGGAGTCGTTGCTCAAATGGCTACATCCGTGAGTGNNNAGATCGGAA

53306 TATGAGTGTGGAGTCGTTGCACAAAACAACCCTGTCTGTGAGGNNNAGATCGGAA

34526 TATGAGTGTGGAGTCGTTGCAGCACATGGATATTTGATAAAAAGCCANNNAGATCG

68380 TATGAGTGTGGAGTCGTTGCGGGAGAAACAGTAAGGAAGCGTTNNNAGATCGGA

38068 TATGAGTGTGGAGTCGTTGCCTGGGATTCTCTGAGCTTTCAGANNNAGATCGGAAG

36268 TATGAGTGTGGAGTCGTTGCGCACACCCTATTTTTGAGTTGCANNNAGATCGGAAG

39923 TATGAGTGTGGAGTCGTTGCGGGGGTAATTTTTGGTATCCTGANNNAGATCGGAA

45583 TATGAGTGTGGAGTCGTTGCTATGTAGGTTTTCCCTGGCCTCCNNNAGATCGGAAG

108518 TATGAGTGTGGAGTCGTTGCGGGTGTTCTCTCATTTTCTTGTTGNNNAGATCGGAA

108504 TATGAGTGTGGAGTCGTTGCTCCATCCACTCGTTCATCCAATNNNAGATCGGAAGAG

31423 TATGAGTGTGGAGTCGTTGCGGCAGAACACTAGAACTGCTCAANNNAGATCGGAA

37671 TATGAGTGTGGAGTCGTTGCCATTGCCCTCCGAATTCTGAAGANNNAGATCGGAAG

30160 TATGAGTGTGGAGTCGTTGCATGATGCCAGGTCTTGTGGAAAANNNAGATCGGAA

60150 TATGAGTGTGGAGTCGTTGCGGTCCCCTCTGGAATGCACTNNNAGATCGGAAGAGC

66962 TATGAGTGTGGAGTCGTTGCGGGCTTCTGAGGAATCTGCAATTNNNAGATCGGAA

35336 TATGAGTGTGGAGTCGTTGCTGAGCCTATTCTCCTTCTCAGCANNNAGATCGGAAGA

96179 TATGAGTGTGGAGTCGTTGCATAGCAGGCAGGAAATCTGGCNNNAGATCGGAAGA

48681 TATGAGTGTGGAGTCGTTGCGATGTAGCACTGACCATGACCTCNNNAGATCGGAAG

40258 TATGAGTGTGGAGTCGTTGCTGCTGAGACAAATTATCCATGTGANNNAGATCGGAA

| | |
|---|---|
| 39153 | TATGAGTGTGGAGTCGTTGCACTGATTAGCTGTAAGCCTGGGANNNAGATCGGAA |
| 32243 | TATGAGTGTGGAGTCGTTGCACCATGACCCTGACCTGGAATAANNNAGATCGGAAG |
| 48596 | TATGAGTGTGGAGTCGTTGCCCACCATAACCTACTGCACACTCNNNAGATCGGAAGA |
| 107107 | TATGAGTGTGGAGTCGTTGCCTGGCACTTCAGATCAAGGACANNNAGATCGGAAG |
| 58712 | TATGAGTGTGGAGTCGTTGCAACACACACTCACATGGATGCATNNNAGATCGGAAG |
| 61551 | TATGAGTGTGGAGTCGTTGCGAGGTTTTGGAACATGGGTTCCTNNNAGATCGGAA |
| 59233 | TATGAGTGTGGAGTCGTTGCTTTTTGTTGGAGTGGGTGTGGATNNNAGATCGGAA |
| 35595 | TATGAGTGTGGAGTCGTTGCAAGACCATTTGCTGATGAAGGCANNNAGATCGGAA |
| 52757 | TATGAGTGTGGAGTCGTTGCAGCAGGAATGACTTCATCCAAGGNNNAGATCGGAA |
| 52048 | TATGAGTGTGGAGTCGTTGCACAGATGAACAGGCACAAGCTAGNNNAGATCGGAA |
| 55353 | TATGAGTGTGGAGTCGTTGCGAGTTTCCCAACATGGCAACATGNNNAGATCGGAAG |
| 31860 | TATGAGTGTGGAGTCGTTGCGTGTGTATGTGCATGCCTAGGAANNNAGATCGGAA |
| 101061 | TATGAGTGTGGAGTCGTTGCTAAGAGGAGGAGAGGCCCGAACNNNAGATCGGAA |
| 102547 | TATGAGTGTGGAGTCGTTGCTAGCAAAGACACGGGGTTCACNNNAGATCGGAAGA |
| 41952 | TATGAGTGTGGAGTCGTTGCTTGTGATTGGGTTGCCACTGTTANNNAGATCGGAAG |
| 70540 | TATGAGTGTGGAGTCGTTGCCCCTTTCTCCACTTTCCCAAGTCNNNAGATCGGAAGA |
| 109033 | TATGAGTGTGGAGTCGTTGCTACCTACCCATGCGAATTCAGAGNNNAGATCGGAAG |
| 53807 | TATGAGTGTGGAGTCGTTGCCCAGACCACTCTGTCCTATAGGGNNNAGATCGGAAG |
| 32276 | TATGAGTGTGGAGTCGTTGCTGGAGGAAGGCTAAGGCAGATAANNNAGATCGGA |
| 62230 | TATGAGTGTGGAGTCGTTGCGGGTGTTACAGTGAGTTCATGCTNNNAGATCGGAA |
| 44539 | TATGAGTGTGGAGTCGTTGCAATCTGAAGCTCAATGCTAGCCCNNNAGATCGGAAG |
| 71804 | TATGAGTGTGGAGTCGTTGCGAGAAGATTGCTAAGACCCCTGCNNNAGATCGGAA |
| 42614 | TATGAGTGTGGAGTCGTTGCCACACACGATCCCAGATAGACACNNNAGATCGGAAG |
| 104723 | TATGAGTGTGGAGTCGTTGCAAGAATCCCTATTTGGCAGATCTGAAANNNAGATCG |
| 31582 | TATGAGTGTGGAGTCGTTGCGGTAGAGGGCTGTCAGGTAAGAANNNAGATCGGA |
| 65681 | TATGAGTGTGGAGTCGTTGCGGCACTTCAGATCAAGGACATGTNNNAGATCGGAA |
| 46970 | TATGAGTGTGGAGTCGTTGCTTGGATTTCAGATGTTGGGAGGCNNNAGATCGGAA |
| 117034 | TATGAGTGTGGAGTCGTTGCCATCAACCATGCAGGTAAAGACANNNAGATCGGAA |
| 99627 | TATGAGTGTGGAGTCGTTGCAATGGAACAAATAAGGCGCGGTNNNAGATCGGAAG |
| 34683 | TATGAGTGTGGAGTCGTTGCACACAGTTGATTCCATATATTGGCCANNNAGATCGG |
| 35141 | TATGAGTGTGGAGTCGTTGCAGAAAAGTGATCCTGCCAAAGCANNNAGATCGGAA |
| 65192 | TATGAGTGTGGAGTCGTTGCGGAAGGGAGAGAGAAATCTGGGTNNNAGATCGGA |
| 34635 | TATGAGTGTGGAGTCGTTGCTGCTATTCTACAGTCTACCGCCANNNAGATCGGAAG |
| 55579 | TATGAGTGTGGAGTCGTTGCCCTGACTTTGCCCTTAGGAGATGNNNAGATCGGAAG |
| 38238 | TATGAGTGTGGAGTCGTTGCAGATATTGATGAGTGCCGGGAGANNNAGATCGGA |
| 96871 | TATGAGTGTGGAGTCGTTGCCTAATAAACTCTGTTGCCAGGGGANNNAGATCGGA |
| 45087 | TATGAGTGTGGAGTCGTTGCATCACTCTTATTAGGGGAGGGCCNNNAGATCGGAA |
| 38236 | TATGAGTGTGGAGTCGTTGCTGGCAACAAGGAGAATAGGGAGANNNAGATCGGA |
| 52969 | TATGAGTGTGGAGTCGTTGCGTGATTTTGGAAGTTTAGTTCCCAGGNNNAGATCGG |
| 57088 | TATGAGTGTGGAGTCGTTGCTGTGTCATTGTGTGTGTTGTGTGNNNAGATCGGAA |
| 45979 | TATGAGTGTGGAGTCGTTGCAGATGGTTGCTGACGGATTTCCNNNAGATCGGAAG |
| 38718 | TATGAGTGTGGAGTCGTTGCGCAATAGTTGGAGGAAGACAGGANNNAGATCGGA |
| 70655 | TATGAGTGTGGAGTCGTTGCCTACCCATGCGAATTCAGAGGAGNNNAGATCGGAA |

| | |
|---|---|
| 101708 | TATGAGTGTGGAGTCGTTGCTGTATATGTGTGCTTGGTGCTTNNNAGATCGGAAG |
| 40512 | TATGAGTGTGGAGTCGTTGCACCTCCACATGTTTATCCAAAATTTGANNNAGATCGG |
| 43728 | TATGAGTGTGGAGTCGTTGCCCCTATTTGGCAGATCTGAAAACCNNNAGATCGGAA |
| 104185 | TATGAGTGTGGAGTCGTTGCATGCTTGAAAGTGGTCTCAGAGCNNNAGATCGGAA |
| 71755 | TATGAGTGTGGAGTCGTTGCCCCGTTAGGGTGTTTTTAGGCTTNNNAGATCGGAAG |
| 43836 | TATGAGTGTGGAGTCGTTGCCTGGCAATGGAGGTCGATAACCNNNAGATCGGAAG |
| 100726 | TATGAGTGTGGAGTCGTTGCAGTTGATTCCATATATTGGCCACTNNNAGATCGGAA |
| 68573 | TATGAGTGTGGAGTCGTTGCTTGGTTCTTGTCTGTCTCATGTTNNNAGATCGGAAG |
| 104152 | TATGAGTGTGGAGTCGTTGCGCTAGTTCAGGACTGGATTCACANNNAGATCGGAAG |
| 48117 | TATGAGTGTGGAGTCGTTGCGTATATGTGTGCTTGGTGCTTGCNNNAGATCGGAAG |
| 31196 | TATGAGTGTGGAGTCGTTGCGGTCCAGATGGTTTTACCAGCAANNNAGATCGGAAG |
| 59176 | TATGAGTGTGGAGTCGTTGCTGATTTTGGTGGAGGAGAGGGATNNNAGATCGGA |
| 29827 | TATGAGTGTGGAGTCGTTGCAGCCAAACAAAAATGTGTACTGTNNNAGATCGGAAG |
| 38295 | TATGAGTGTGGAGTCGTTGCAAGGACCCAGTGAATGGATGAGANNNAGATCGGAA |
| 29829 | TATGAGTGTGGAGTCGTTGCGCAATGACAAAAGCAACAGCAACNNNAGATCGGAA |
| 34958 | TATGAGTGTGGAGTCGTTGCAGGGGGAGATACCATCTAGTCCANNNAGATCGGAA |
| 52944 | TATGAGTGTGGAGTCGTTGCTTTAGAATCAGCCTTCACCCAGGNNNAGATCGGAAG |
| 43256 | TATGAGTGTGGAGTCGTTGCTAGAGAGCTTACCCTTAGGCGACNNNAGATCGGAAG |
| 41743 | TATGAGTGTGGAGTCGTTGCCAGTCACTCCCTCCTGTCCATTANNNAGATCGGAAGA |
| 62666 | TATGAGTGTGGAGTCGTTGCCCACACACACAGGAAGAAGATCTNNNAGATCGGAAG |
| 100530 | TATGAGTGTGGAGTCGTTGCAGGGTTCCAGTGTTCTTGTACCTCNNNAGATCGGAA |
| 29947 | TATGAGTGTGGAGTCGTTGCATTCCACAGCATACTTCTCCCCANNNAGATCGGAAGA |
| 29949 | TATGAGTGTGGAGTCGTTGCAGTGCTAAGTTTCCTACGCATGGNNNAGATCGGAAG |
| 31671 | TATGAGTGTGGAGTCGTTGCAGATCCTTACGGGCCAAAGAGAANNNAGATCGGAA |
| 101011 | TATGAGTGTGGAGTCGTTGCATTGGATCGTCTGAAGCCTTGGNNNAGATCGGAAG |
| 101650 | TATGAGTGTGGAGTCGTTGCGGCTATTATGAAGACTGTGGCAANNNAGATCGGAA |
| 102577 | TATGAGTGTGGAGTCGTTGCCCAGCCCTCCATTGATGAATCCTNNNAGATCGGAAG |
| 101639 | TATGAGTGTGGAGTCGTTGCGTGTTCTTTGGCTTGCATTCTCTTTCNNNAGATCGGA |
| 100779 | TATGAGTGTGGAGTCGTTGCCCAAGATGTTCCCTAGCAAATCAGANNNAGATCGGA |
| 106172 | TATGAGTGTGGAGTCGTTGCGTGTTTTATAAGTGCCTTTCTAAATGCNNNAGATCGG |
| 101374 | TATGAGTGTGGAGTCGTTGCTGAGCATATGTGTGTGTGTGGCNNNAGATCGGAAG |
| 107099 | TATGAGTGTGGAGTCGTTGCCCATTCAAATCTAGGAACGCTACACNNNAGATCGGA |
| 107908 | TATGAGTGTGGAGTCGTTGCTGCTAAAAAGTACATTGAGAGCCATNNNAGATCGGA |
| 102286 | TATGAGTGTGGAGTCGTTGCTGAAGGATGACCAGGTGGAAAGNNNAGATCGGAA |
| 101640 | TATGAGTGTGGAGTCGTTGCTTTGTTGGAGTGGGTGTGGATGTNNNAGATCGGAA |
| 71836 | TATGAGTGTGGAGTCGTTGCCCATCCTCTTGTCTCTGCAGTACNNNAGATCGGAAGA |
| 107890 | TATGAGTGTGGAGTCGTTGCAATGCCGGGTCACAGCAACCNNNAGATCGGAAGAG |
| 116115 | TATGAGTGTGGAGTCGTTGCTCTCCACTTTCCCAAGTCATTCATNNNAGATCGGAAG |
| 104197 | TATGAGTGTGGAGTCGTTGCGCACCCGTGATGGTGTACTTCNNNAGATCGGAAGAG |
| 100913 | TATGAGTGTGGAGTCGTTGCTTAGTCAATGTGCAGTGCTGTGATNNNAGATCGGAA |
| 102266 | TATGAGTGTGGAGTCGTTGCCGGACAGCTCAAGAACAAGGTNNNAGATCGGAAGA |
| 106686 | TATGAGTGTGGAGTCGTTGCTGTCATTCCTCTGTGTGTGGACATNNNAGATCGGAA |
| 108594 | TATGAGTGTGGAGTCGTTGCTATTGGGCAGTGTTGGAATCATTNNNAGATCGGAA |

| | |
|---|---|
| 116078 | TATGAGTGTGGAGTCGTTGCTCTGTCTCTGGCCAGTTCTCATNNNAGATCGGAAGA |
| 102244 | TATGAGTGTGGAGTCGTTGCCACCAACCTGGCCTGCTTTTTNNNAGATCGGAAGAG |
| 101019 | TATGAGTGTGGAGTCGTTGCTGAGAGGTGATCAGGAAAGCTAGANNNAGATCGG |
| 105559 | TATGAGTGTGGAGTCGTTGCTTGGTTTGCATTTCTCCCCTTCNNNAGATCGGAAGAG |
| 98369 | TATGAGTGTGGAGTCGTTGCAAACAGGGCTTCCAAGGGACAGTNNNAGATCGGAA |
| 107909 | TATGAGTGTGGAGTCGTTGCGTGTGTGTGTGTGATTCATAGGANNNAGATCGGAA |
| 108608 | TATGAGTGTGGAGTCGTTGCACACTCGCCATCTTACAAACATAGNNNAGATCGGAA |
| 51074 | TATGAGTGTGGAGTCGTTGCAAGGTCTGCAAAGGAAAAGGCAGNNNAGATCGGA |
| 50400 | TATGAGTGTGGAGTCGTTGCCATTTGGGGGATTGGGAGAGAAGNNNAGATCGGA |
| 98723 | TATGAGTGTGGAGTCGTTGCTTTGGCCATTCTTCGCCTGACNNNAGATCGGAAGAG |
| 99016 | TATGAGTGTGGAGTCGTTGCTGTGTCTCCTCTAAAATTCGGGTGNNNAGATCGGAA |
| 63690 | TATGAGTGTGGAGTCGTTGCTGTTTTTGACCCTGAGGACAAGTNNNAGATCGGAAG |
| 96954 | TATGAGTGTGGAGTCGTTGCGTATCCATCACCCTAGAAGTTCCTTNNNAGATCGGAA |
| 68961 | TATGAGTGTGGAGTCGTTGCATACACTGTCAGAGACACACTTTNNNAGATCGGAAG |
| 30962 | TATGAGTGTGGAGTCGTTGCATTGAGCTTCTTGAAATGGTACAANNNAGATCGGAA |
| 41568 | TATGAGTGTGGAGTCGTTGCATTGTGAGTTGGGGAGACTGGTANNNAGATCGGA |
| 51269 | TATGAGTGTGGAGTCGTTGCTTTTGGGGGTTCAGACAAGTCAGNNNAGATCGGAA |
| 33321 | TATGAGTGTGGAGTCGTTGCAAGCTGAGGAAATTGTCTGCACANNNAGATCGGAA |
| 30395 | TATGAGTGTGGAGTCGTTGCCACAGGAGTTTGGCAAATGCAAANNNAGATCGGAA |
| 71808 | TATGAGTGTGGAGTCGTTGCTGGTGACCCTCTTCCTCTTACAANNNAGATCGGAAGA |
| 36997 | TATGAGTGTGGAGTCGTTGCACACCTATCCCAAAGGGATGTCANNNAGATCGGAAG |
| 49976 | TATGAGTGTGGAGTCGTTGCGGGGGTGGATTTGGAGGATTTTCNNNAGATCGGA |
| 49814 | TATGAGTGTGGAGTCGTTGCTCCTTGTACCTTGCAAGTGGTTCNNNAGATCGGAAG |
| 60710 | TATGAGTGTGGAGTCGTTGCAGAGAGAGCCCCAATATTCACCTNNNAGATCGGAAG |
| 52160 | TATGAGTGTGGAGTCGTTGCTACACTCTCCCACCCTCCTGTAGNNNAGATCGGAAGA |
| 68128 | TATGAGTGTGGAGTCGTTGCGTCCCTGTAGTTGAGGGTGTCTTNNNAGATCGGAA |
| 31499 | TATGAGTGTGGAGTCGTTGCACTGCTTCAGCTTCTCTGTTCAANNNAGATCGGAAG |
| 57313 | TATGAGTGTGGAGTCGTTGCTTTTCCTGGATTCCTCAGGCTTGNNNAGATCGGAAG |
| 46729 | TATGAGTGTGGAGTCGTTGCAATAGTGTGGATGATTCCCACGCNNNAGATCGGAA |
| 62635 | TATGAGTGTGGAGTCGTTGCCCTCTTGGGAGATAAGGGCATCTNNNAGATCGGAA |
| 41149 | TATGAGTGTGGAGTCGTTGCTTCTTTTGCCATTCAGGCAGCTANNNAGATCGGAAG |
| 38688 | TATGAGTGTGGAGTCGTTGCACTGAGGATGTAGAGGATGAAGGANNNAGATCGG |
| 43847 | TATGAGTGTGGAGTCGTTGCATGGGTAGACCAGGGAAGTAACCNNNAGATCGGAA |
| 30320 | TATGAGTGTGGAGTCGTTGCATGTTCCCTCACTGTTGCCCAAANNNAGATCGGAAG |
| 38695 | TATGAGTGTGGAGTCGTTGCTGGTGGTGTTTGTGACAATAAGGANNNAGATCGGA |
| 30357 | TATGAGTGTGGAGTCGTTGCAAAAATGAAACGTTGGAAAGCAAANNNAGATCGGA |
| 37284 | TATGAGTGTGGAGTCGTTGCAAACAGATACTGGGGGAGGTTCANNNAGATCGGAA |
| 63255 | TATGAGTGTGGAGTCGTTGCAGCCAATGGTGATTTTGCTGTCTNNNAGATCGGAAG |
| 46696 | TATGAGTGTGGAGTCGTTGCCCAGCAACTGCAAGTCAATTAGCNNNAGATCGGAAG |
| 71634 | TATGAGTGTGGAGTCGTTGCGCTGACCAAAAATGTCTCAAAGTCAANNNAGATCGG |
| 43125 | TATGAGTGTGGAGTCGTTGCACAAACACCATATTTGGCCAAGACNNNAGATCGGAA |
| 50464 | TATGAGTGTGGAGTCGTTGCGTCTCCTTTGCTCTCCATGGAAGNNNAGATCGGAAG |
| 103901 | TATGAGTGTGGAGTCGTTGCCACATTGTATTAGTCCATTCCCACATNNNAGATCGGA |

31400 TATGAGTGTGGAGTCGTTGCTGCCCTCCTGATTAGAACCTCAANNNAGATCGGAAG
103871 TATGAGTGTGGAGTCGTTGCAAGCCTCTTCCCATGAAGAGAGAANNNAGATCGGAA
57241 TATGAGTGTGGAGTCGTTGCAGAAGGGGAAATTGGCTCACTTGNNNAGATCGGAA
63994 TATGAGTGTGGAGTCGTTGCCTGACCAAGTAAGGTTTATTCCAGTNNNAGATCGGA
97976 TATGAGTGTGGAGTCGTTGCACACACACACAAAAGAACTCTCCANNNAGATCGGAA
33659 TATGAGTGTGGAGTCGTTGCAGGGAGGGAGTGTTTACATGACANNNAGATCGGAA
39647 TATGAGTGTGGAGTCGTTGCGGGGGTCCATATGAGTTCCATGANNNAGATCGGAA
35998 TATGAGTGTGGAGTCGTTGCCGTACTCCCACAACTCATATGCANNNAGATCGGAAG
43462 TATGAGTGTGGAGTCGTTGCACTTTCCAGGCTATGTCCTTTGACNNNAGATCGGAAG
57454 TATGAGTGTGGAGTCGTTGCGGGGAGGGTTTTTCCTTAGGTTGNNNAGATCGGAA
69304 TATGAGTGTGGAGTCGTTGCGAGCCCCGTGATTTTGTAGGTTTNNNAGATCGGAAG
57582 TATGAGTGTGGAGTCGTTGCCTCAGACAACTCCTGGCTCTTTGNNNAGATCGGAAG
62976 TATGAGTGTGGAGTCGTTGCTGTTTTGCAGATGGTCTTGCTCTNNNAGATCGGAAG
42800 TATGAGTGTGGAGTCGTTGCGTTACACACAGACCTCTGTCCACNNNAGATCGGAAG
47295 TATGAGTGTGGAGTCGTTGCCTGGTGATCTTTGTCTCTCTGGCNNNAGATCGGAAG
106696 TATGAGTGTGGAGTCGTTGCGCCCCGTGATTTTGTAGGTTTAGANNNAGATCGGAA
104142 TATGAGTGTGGAGTCGTTGCCCTGAATGGATAAGGAGGGACANNNAGATCGGAAG
45830 TATGAGTGTGGAGTCGTTGCCAGGCTCTCCTCTTGGAAATTCCNNNAGATCGGAAG
45083 TATGAGTGTGGAGTCGTTGCCTTAGAGAAGACAACCAAGGGCCNNNAGATCGGAA
108488 TATGAGTGTGGAGTCGTTGCGCCGCATTTATCTAAGTAGCAGTNNNAGATCGGAAG
49793 TATGAGTGTGGAGTCGTTGCAAATCTCGCCTCCATGAAGGTTCNNNAGATCGGAAG
58363 TATGAGTGTGGAGTCGTTGCGAGAGGTGAGATGGGAGAGACATNNNAGATCGGA
30552 TATGAGTGTGGAGTCGTTGCGCCTGCTGGGAACTTCTTAGAAANNNAGATCGGAAG
65535 TATGAGTGTGGAGTCGTTGCTCCACTCATCATCCATATCTTGGTNNNAGATCGGAAG
61644 TATGAGTGTGGAGTCGTTGCGCTTGACATACCAGCATGAAGCTNNNAGATCGGAAG
39509 TATGAGTGTGGAGTCGTTGCTTTGGCTTATCTTCCCAGTTGGANNNAGATCGGAAG
36157 TATGAGTGTGGAGTCGTTGCGGTATTGGAAGCTGTATGGTGCANNNAGATCGGAA
49584 TATGAGTGTGGAGTCGTTGCATAGCCCTAGTGTGCCCCTATTCNNNAGATCGGAAG
31909 TATGAGTGTGGAGTCGTTGCTCCCTGACAAACATAGAGGGGAANNNAGATCGGAA
34941 TATGAGTGTGGAGTCGTTGCTCAGGGTCATAAAGTTGTTCTCCANNNAGATCGGAA
41863 TATGAGTGTGGAGTCGTTGCGACCTGGGTGCTAGTTTTGCTTANNNAGATCGGAAG
108479 TATGAGTGTGGAGTCGTTGCGTTATCCAAGCTGAACTCAAACTCCNNNAGATCGGAA
31048 TATGAGTGTGGAGTCGTTGCGACCAGCGATTTGTCTATCCCAANNNAGATCGGAAG
62284 TATGAGTGTGGAGTCGTTGCTGAAAGGAAAAGGTAAGACTGCTNNNAGATCGGAA
99283 TATGAGTGTGGAGTCGTTGCCAGTGATGCTGGGCAAATTTACACNNNAGATCGGAA
40082 TATGAGTGTGGAGTCGTTGCCGTTCTTGGAGCCTGATTTCTGANNNAGATCGGAAG
44622 TATGAGTGTGGAGTCGTTGCATTTCTGGAGACAAGCTTTGCCCNNNAGATCGGAAG
37460 TATGAGTGTGGAGTCGTTGCCACACCTGGAGTTGGAACAAAGANNNAGATCGGAA
39877 TATGAGTGTGGAGTCGTTGCACATTCCAACCTATGGGACCTGANNNAGATCGGAAG
38983 TATGAGTGTGGAGTCGTTGCGCTGCAAGATTCTTTTGAAGGGANNNAGATCGGAA
46561 TATGAGTGTGGAGTCGTTGCATAAGGTTGTCAGCAACTGGAGCNNNAGATCGGAA
65245 TATGAGTGTGGAGTCGTTGCGCAGGGAGGCTATTGTAAATGGTNNNAGATCGGAA
66045 TATGAGTGTGGAGTCGTTGCAACTGATGGGGAAAACAAGCTGTNNNAGATCGGAA

49599 TATGAGTGTGGAGTCGTTGCGAAGGCTCAATGTGGCAAACTTCNNNAGATCGGAA
39601 TATGAGTGTGGAGTCGTTGCAGGCTGGAGTACAGGAAACATGANNNAGATCGGA
116690 TATGAGTGTGGAGTCGTTGCATCCCCTGGACACGACCACNNNAGATCGGAAGAG
41306 TATGAGTGTGGAGTCGTTGCTCCAATCAGCCTTCTGGTGTCTANNNAGATCGGAAG
53562 TATGAGTGTGGAGTCGTTGCTCTTCCAAATCCCAGCAATTCGGNNNAGATCGGAAG
106180 TATGAGTGTGGAGTCGTTGCTTGGCACTACCCTCTCCTCTTTACNNNAGATCGGAAG
65497 TATGAGTGTGGAGTCGTTGCGTGTAGGTGTTGATGGTTGTGGTNNNAGATCGGAA
107861 TATGAGTGTGGAGTCGTTGCAAGAGAGCGAGAGAGAGAAAGCATANNNAGATCG
66483 TATGAGTGTGGAGTCGTTGCGAGATATGGCTGAATGGGTGTGTNNNAGATCGGA
69499 TATGAGTGTGGAGTCGTTGCTGCCCTTTGAGAATCTGGCTTTTNNNAGATCGGAAG
34180 TATGAGTGTGGAGTCGTTGCGGAGGAACTGAGACCTTCTACCANNNAGATCGGAA
38078 TATGAGTGTGGAGTCGTTGCATAGCATTGCTGGAGATAAAAGAGANNNAGATCGG
31958 TATGAGTGTGGAGTCGTTGCTCAGTACAGGTAGGGTGATGGAANNNAGATCGGA
45774 TATGAGTGTGGAGTCGTTGCTTTTCCCAGTTTCTCAGTGGTCCNNNAGATCGGAAGA
48531 TATGAGTGTGGAGTCGTTGCCATCCACACATGCCCTTCCTATCNNNAGATCGGAAGA
108531 TATGAGTGTGGAGTCGTTGCTAGTGGGTCTGCATGGAAAGAGNNNAGATCGGAAG
32353 TATGAGTGTGGAGTCGTTGCAAAGAGGAAGTGGGGAACCCTAANNNAGATCGGA
65064 TATGAGTGTGGAGTCGTTGCTAGAACCAGTGACTTTCCAGGGTNNNAGATCGGAAG
39144 TATGAGTGTGGAGTCGTTGCCCTCACTGTCACTTAGACTGGGANNNAGATCGGAAG
96332 TATGAGTGTGGAGTCGTTGCAAGTGCCTCCTGGTGCACCGNNNAGATCGGAAGAG
102479 TATGAGTGTGGAGTCGTTGCAGAGACAGAAAGAGCAAGAGCTNNNAGATCGGAA
49122 TATGAGTGTGGAGTCGTTGCCTCCCTTCACAGTCAAGCTTCTCNNNAGATCGGAAGA
30457 TATGAGTGTGGAGTCGTTGCATCCAATGACAGCATGTATTCAAANNNAGATCGGAA
64582 TATGAGTGTGGAGTCGTTGCAGCAAGCCTTAGGAATCTTACGTNNNAGATCGGAAG
108507 TATGAGTGTGGAGTCGTTGCATGTATGTGAGACAGAGCCTCACNNNAGATCGGAA
47395 TATGAGTGTGGAGTCGTTGCTGAGTGAGTACATCTTGCTTGGCNNNAGATCGGAAG
103861 TATGAGTGTGGAGTCGTTGCGCATCGTATATCCCTTTGGAAGTTTGNNNAGATCGG
62677 TATGAGTGTGGAGTCGTTGCCAAGCCAAGGAGAGAGGAGATCTNNNAGATCGGA
39025 TATGAGTGTGGAGTCGTTGCGTGGTGGCTAAAAGTTTCAGGGANNNAGATCGGAA
36563 TATGAGTGTGGAGTCGTTGCTGAGGATATTTAGGGCCCACTCANNNAGATCGGAAG
104895 TATGAGTGTGGAGTCGTTGCAGGGATGTTGTTGAGGAGGAGGNNNAGATCGGAA
53028 TATGAGTGTGGAGTCGTTGCCAGAAGCCAAATGTATGAGCAGGNNNAGATCGGAA
38844 TATGAGTGTGGAGTCGTTGCATAAAACCTGCAGAGGTGGAGGANNNAGATCGGA
49760 TATGAGTGTGGAGTCGTTGCTGTTTGTGTCTCTCCCGAAGTTCNNNAGATCGGAAG
64716 TATGAGTGTGGAGTCGTTGCTCCAGAAAAATCTCCCAACAAAGGTNNNAGATCGGA
60348 TATGAGTGTGGAGTCGTTGCGGAAGAGTACCAGACCATGGACTNNNAGATCGGAA
61581 TATGAGTGTGGAGTCGTTGCCACCCACACTAAGCCATTTTCCTNNNAGATCGGAAGA
108532 TATGAGTGTGGAGTCGTTGCGGTTTCAGAAAAATGCCGTGGAANNNAGATCGGAA
116439 TATGAGTGTGGAGTCGTTGCTGGCTTTTGTCAGTTCTCCATTCANNNAGATCGGAAG
101405 TATGAGTGTGGAGTCGTTGCCACGACATAATGGAAACAACTAATGTNNNAGATCGG
71478 TATGAGTGTGGAGTCGTTGCGTGTGTCTTCACCAGTCACCATTNNNAGATCGGAAG
58792 TATGAGTGTGGAGTCGTTGCAAAGAACGTGCCTGCCATATCATNNNAGATCGGAAG
41102 TATGAGTGTGGAGTCGTTGCCCTTGGTCTGCAAGATTTGCCTANNNAGATCGGAAG

| | |
|---|---|
| 49916 | TATGAGTGTGGAGTCGTTGCTCTGGGGCAAGTTCATAGCTTTCNNNAGATCGGAAG |
| 68008 | TATGAGTGTGGAGTCGTTGCAGGTAGATGGGGCCTCAAATCTTNNNAGATCGGAA |
| 30446 | TATGAGTGTGGAGTCGTTGCGAGCTTCCTGTACACACGTCAAANNNAGATCGGAAG |
| 46195 | TATGAGTGTGGAGTCGTTGCCAGAAAGGAGGTGAGAAGGAAGCNNNAGATCGGA |
| 101926 | TATGAGTGTGGAGTCGTTGCACCTTTTACCCCAGAAGACTTCAGTANNNAGATCGG |
| 32901 | TATGAGTGTGGAGTCGTTGCAACAACTACAACAGGCAGGAACANNNAGATCGGAA |
| 30652 | TATGAGTGTGGAGTCGTTGCACACCCACATGCATCTCATGAAANNNAGATCGGAAG |
| 66349 | TATGAGTGTGGAGTCGTTGCACAGCCAAGTTATGCATACGTGTNNNAGATCGGAAG |
| 43635 | TATGAGTGTGGAGTCGTTGCGCACACAGGGTACTAAGTGGTACNNNAGATCGGAA |
| 108533 | TATGAGTGTGGAGTCGTTGCGCAGATCCAGTGTGCTTCTCAAATATNNNAGATCGG |
| 38976 | TATGAGTGTGGAGTCGTTGCTGGCAGAATTCACTAGGAAGGGANNNAGATCGGAA |
| 65777 | TATGAGTGTGGAGTCGTTGCGCTAGAGGCAAAAGCAGAGATGTNNNAGATCGGA |
| 29824 | TATGAGTGTGGAGTCGTTGCTGACTGCCCAAAATACGACACATNNNAGATCGGAAG |
| 45789 | TATGAGTGTGGAGTCGTTGCTGGTTGATGGTGAGAAACTGTCCNNNAGATCGGAA |
| 34851 | TATGAGTGTGGAGTCGTTGCGCATCAATGAACACTAAGTTGTATCCANNNAGATCG |
| 49315 | TATGAGTGTGGAGTCGTTGCCACAGCTTCTGGTGTCTTAGGTCNNNAGATCGGAAG |
| 110752 | TATGAGTGTGGAGTCGTTGCTGTTTGCATGCTTTGATATATACTAGCNNNAGATCG |
| 46604 | TATGAGTGTGGAGTCGTTGCAGAAAAAGAGAGGAACGGTGAGCNNNAGATCGGA |
| 41697 | TATGAGTGTGGAGTCGTTGCACACACACACAACCTTCCTTGTANNNAGATCGGAAGA |
| 69920 | TATGAGTGTGGAGTCGTTGCGCCTCTTCTAATGACACATTGCANNNAGATCGGAAG |
| 51384 | TATGAGTGTGGAGTCGTTGCACACATGAACACAGGCTGAAGAGNNNAGATCGGAA |
| 32127 | TATGAGTGTGGAGTCGTTGCTAGCCAAGAGCTTAGCCTCTGAANNNAGATCGGAAG |
| 106709 | TATGAGTGTGGAGTCGTTGCAGTGCAGAAAAGGTGGCATTTATTNNNAGATCGGA |
| 116430 | TATGAGTGTGGAGTCGTTGCCTCTCCAGTATGGACCCATTTTTNNNAGATCGGAAGA |
| 34095 | TATGAGTGTGGAGTCGTTGCGCAACATGACATACATAAGACCANNNAGATCGGAAG |
| 48936 | TATGAGTGTGGAGTCGTTGCGGAATGACATTGTTGAGCTGCTCNNNAGATCGGAA |
| 38368 | TATGAGTGTGGAGTCGTTGCATTCATCCGGCTCCTCTTTGAGANNNAGATCGGAAG |
| 39664 | TATGAGTGTGGAGTCGTTGCACGCAAAGGAACACTTCTATCATGANNNAGATCGGA |
| 58138 | TATGAGTGTGGAGTCGTTGCGAGAGCGAGTGCACTAAGTGAATNNNAGATCGGAA |
| 45870 | TATGAGTGTGGAGTCGTTGCACTGATCCTCCATCCAGACTTCCNNNAGATCGGAAGA |
| 97694 | TATGAGTGTGGAGTCGTTGCCTCCCCATCACCCTAAGACCTGTNNNAGATCGGAAGA |
| 32177 | TATGAGTGTGGAGTCGTTGCATACTACCCAGCTCGGATGTGAANNNAGATCGGAAG |
| 39317 | TATGAGTGTGGAGTCGTTGCACTCACCATTTTCCCAGACTGGANNNAGATCGGAAG |
| 36957 | TATGAGTGTGGAGTCGTTGCAGCAATGACTATTTTGCAGGGTCANNNAGATCGGA |
| 103290 | TATGAGTGTGGAGTCGTTGCGAGGGATTTACTTTTCGGTGAGCNNNAGATCGGAA |
| 68675 | TATGAGTGTGGAGTCGTTGCAGAGGCTAAGATGAGTGGGTGTTNNNAGATCGGA |
| 47866 | TATGAGTGTGGAGTCGTTGCCTGTCTACCTGTGGGTAAAGTGCNNNAGATCGGAA |
| 35782 | TATGAGTGTGGAGTCGTTGCGTCACATTGAACTTGATAAGATGGCANNNAGATCGG |
| 37332 | TATGAGTGTGGAGTCGTTGCGGTGCATGTAGGTAGCCATTTCANNNAGATCGGAA |
| 41974 | TATGAGTGTGGAGTCGTTGCGGTGGTGGTGAGAAGTGTTGTTANNNAGATCGGA |
| 44327 | TATGAGTGTGGAGTCGTTGCCTGACTAGGGGAGTGGTACCCNNNAGATCGGAAGA |
| 44972 | TATGAGTGTGGAGTCGTTGCAGGGATTTACTTTTCGGTGAGCCNNNAGATCGGAA |
| 50945 | TATGAGTGTGGAGTCGTTGCACTCAAAAAGCTGCAACAATCCAGNNNAGATCGGAA |

29826 TATGAGTGTGGAGTCGTTGCGAGTGTGGTTCTTCCATGCTCTTNNNAGATCGGAAG
33721 TATGAGTGTGGAGTCGTTGCTGACAGAATCACCAGCATTGACANNNAGATCGGAAG
30956 TATGAGTGTGGAGTCGTTGCCCGAATGTGCATCCCTTCTACAANNNAGATCGGAAG
55004 TATGAGTGTGGAGTCGTTGCAGAGTGACATCAGCCAATTGTGGNNNAGATCGGAA
65631 TATGAGTGTGGAGTCGTTGCAACTGGAACAATCTGGGCAATGTNNNAGATCGGAA
69083 TATGAGTGTGGAGTCGTTGCGCTTCCTTTGCTTGTTCAGCTTTNNNAGATCGGAAGA
66684 TATGAGTGTGGAGTCGTTGCCCTCCTGGGAAAGCAAAAGTTGTNNNAGATCGGAA
116901 TATGAGTGTGGAGTCGTTGCTAAGGAATAGGGTGAGAGGGCACANNNAGATCGG
58144 TATGAGTGTGGAGTCGTTGCAATCTCACCCCTCCCACTTGAATNNNAGATCGGAAGA
39992 TATGAGTGTGGAGTCGTTGCGAGCATCCCTTGGTTTATGCTGANNNAGATCGGAAG
103842 TATGAGTGTGGAGTCGTTGCGCATAGTGATATTAGGGTAAAGCAAGTNNNAGATC
41679 TATGAGTGTGGAGTCGTTGCATGCCTCTATTTCAGCGTGTGTANNNAGATCGGAAG
52542 TATGAGTGTGGAGTCGTTGCTACTAAGAAAAACTGGTGGGTGCGNNNAGATCGGA
41095 TATGAGTGTGGAGTCGTTGCTCAAAATACTTCTGGAGTGCCTANNNAGATCGGAAG
34845 TATGAGTGTGGAGTCGTTGCATAACGTCTTCTGCTCCTATCCANNNAGATCGGAAGA
45065 TATGAGTGTGGAGTCGTTGCCATCAGTTCAGATTCTGGAGGCCNNNAGATCGGAAG
116724 TATGAGTGTGGAGTCGTTGCTACAGCAAGAAGGCAGTGGTCATNNNAGATCGGAA
71697 TATGAGTGTGGAGTCGTTGCGCATCTTCACATCACTGTCGGATNNNAGATCGGAAG
33313 TATGAGTGTGGAGTCGTTGCTCTAACGCTCTTTGTGATGCACANNNAGATCGGAAG
40757 TATGAGTGTGGAGTCGTTGCCACAGTTGGTGGCATACTGCATANNNAGATCGGAAG
63772 TATGAGTGTGGAGTCGTTGCACCAGATGGCCACTTTGAGAAGTNNNAGATCGGAA
33786 TATGAGTGTGGAGTCGTTGCCCAGACCCTAGCCATCTGATACANNNAGATCGGAAG
36003 TATGAGTGTGGAGTCGTTGCAAGCTACATGGCTATATGTATATGCANNNAGATCGG
67064 TATGAGTGTGGAGTCGTTGCCAGCAGTGTGATTTTGGCACATTNNNAGATCGGAAG
60009 TATGAGTGTGGAGTCGTTGCAAGTCGTGCATTTGTGAGACACTNNNAGATCGGAAG
117433 TATGAGTGTGGAGTCGTTGCCCTGGATCCTCTTGCTTAGATCCNNNAGATCGGAAG
96647 TATGAGTGTGGAGTCGTTGCGGTAGGTCCGTAGGTTTCAGCNNNAGATCGGAAGA
96462 TATGAGTGTGGAGTCGTTGCCCAAGGATCAGGAAGATCCAGAGCNNNAGATCGGA
104084 TATGAGTGTGGAGTCGTTGCCCTTTTCTAATCGATCATAGCTGTTCNNNAGATCGGA
57994 TATGAGTGTGGAGTCGTTGCAGCAATCGTCCTTTCACCTCAATNNNAGATCGGAAGA
57266 TATGAGTGTGGAGTCGTTGCCATCCCCACCATACTTGACCTTGNNNAGATCGGAAGA
62832 TATGAGTGTGGAGTCGTTGCTGTCATTCTTCAGGTGGACCTCTNNNAGATCGGAAG
44753 TATGAGTGTGGAGTCGTTGCAAACCACATGTATGGTAGTAGTCCCNNNAGATCGGA
102603 TATGAGTGTGGAGTCGTTGCAGTAGAGATGGTGTGTTGTGTGTGNNNAGATCGGA
53866 TATGAGTGTGGAGTCGTTGCTAGCTCAGATCTCTCCAAAGGGGNNNAGATCGGAAG
46041 TATGAGTGTGGAGTCGTTGCCTGCTGGGGTTTGGAATTTTTCCNNNAGATCGGAAG
108539 TATGAGTGTGGAGTCGTTGCACAACCTGCCCTCACCCAAATNNNAGATCGGAAGAG
67851 TATGAGTGTGGAGTCGTTGCTCTGTAAGGCAGAGGTACAGCTTNNNAGATCGGAA
31969 TATGAGTGTGGAGTCGTTGCTACCTTTTCCCTTGGACCTGGAANNNAGATCGGAAG
43048 TATGAGTGTGGAGTCGTTGCGGAATTTTGATGCAGCTGGTCACNNNAGATCGGAA
46097 TATGAGTGTGGAGTCGTTGCAGGCAAGGAAGATGAAGGAAAGCNNNAGATCGGA
64257 TATGAGTGTGGAGTCGTTGCACTCACAGGGTATTTTGAGGAGTNNNAGATCGGAA
66960 TATGAGTGTGGAGTCGTTGCTAAGCAACATGGGAACTGCAATTNNNAGATCGGAA

| | |
|---|---|
| 63654 | TATGAGTGTGGAGTCGTTGCTGAGCCATAGAAAGCCATAGAAAGTNNNAGATCGG |
| 39329 | TATGAGTGTGGAGTCGTTGCTATTACATTCGCAGCAACCTGGANNNAGATCGGAAG |
| 67344 | TATGAGTGTGGAGTCGTTGCTTTTCAGGGCATCCTTCCTGATTNNNAGATCGGAAG |
| 109070 | TATGAGTGTGGAGTCGTTGCAATCTGGATCATGTTGCATACACNNNAGATCGGAAG |
| 51414 | TATGAGTGTGGAGTCGTTGCAAATTGCTGTGGAGGATCCAGAGNNNAGATCGGAA |
| 44045 | TATGAGTGTGGAGTCGTTGCTTTTTCCTGGGTCAATGGAGACCNNNAGATCGGAAG |
| 44257 | TATGAGTGTGGAGTCGTTGCACCTCCCATGAGATACTAACACCCNNNAGATCGGAA |
| 34090 | TATGAGTGTGGAGTCGTTGCTGACAAGCACGGAATAAAGACCANNNAGATCGGAA |
| 33121 | TATGAGTGTGGAGTCGTTGCAATCTGGATCATGTTGCATACACANNNAGATCGGAA |
| 55204 | TATGAGTGTGGAGTCGTTGCCAGATGCAGATACGTGTGTTTGGNNNAGATCGGAA |
| 43877 | TATGAGTGTGGAGTCGTTGCGCCCTTTTGAGTTTCTTGACACCNNNAGATCGGAAG |
| 50052 | TATGAGTGTGGAGTCGTTGCGGGGAGCTGGGAGGTAATAAAAGNNNAGATCGGA |
| 34897 | TATGAGTGTGGAGTCGTTGCGGGAAAACCCACTTATTCCTCCANNNAGATCGGAAG |
| 107916 | TATGAGTGTGGAGTCGTTGCATCTTATCATCTGCAAGTGGAAACANNNAGATCGG |
| 66251 | TATGAGTGTGGAGTCGTTGCCCAGCATGCCTTACTCTTTCTGTNNNAGATCGGAAGA |
| 45476 | TATGAGTGTGGAGTCGTTGCCCTGTGTGTATCAACAATGTGATCCNNNAGATCGG |
| 53077 | TATGAGTGTGGAGTCGTTGCATTTGGGAACACAGTCACTCAGGNNNAGATCGGAA |
| 54083 | TATGAGTGTGGAGTCGTTGCTGATGTGCATTCTCAAGACTGGGNNNAGATCGGAA |
| 108464 | TATGAGTGTGGAGTCGTTGCTCTTCCCTGAAAACACTGATAGATCCNNNAGATCGGA |
| 39625 | TATGAGTGTGGAGTCGTTGCTGGGATGAGCTGAGTGTACATGANNNAGATCGGAA |
| 102231 | TATGAGTGTGGAGTCGTTGCCCATTCTCTCCTCAGCCTCACTNNNAGATCGGAAGAG |
| 53995 | TATGAGTGTGGAGTCGTTGCATGAAAAATCCACAGTGGTGGGGNNNAGATCGGAA |
| 63587 | TATGAGTGTGGAGTCGTTGCTTTGGCTCCCTGTTGAGTTTTCTNNNAGATCGGAAG |
| 107837 | TATGAGTGTGGAGTCGTTGCTGGCTTGGTAAACTGTTTCTTGGGNNNAGATCGGA |
| 45201 | TATGAGTGTGGAGTCGTTGCAGAACTGGTTGATCATGGATGCCNNNAGATCGGAA |
| 70926 | TATGAGTGTGGAGTCGTTGCGGACCTCTTTCTCTGTGTGTGTGNNNAGATCGGAAG |
| 98602 | TATGAGTGTGGAGTCGTTGCCACCACCACCACGAAAAATACAANNNAGATCGGAAG |
| 51677 | TATGAGTGTGGAGTCGTTGCGGCTTTGATGGTAAGGATGGGAGNNNAGATCGGA |
| 100771 | TATGAGTGTGGAGTCGTTGCACTGGAATATGAGCCCCTATATTTCNNNAGATCGGA |
| 64712 | TATGAGTGTGGAGTCGTTGCTCTCTCCCCTCACATGAAAAGGTNNNAGATCGGAAG |
| 56762 | TATGAGTGTGGAGTCGTTGCGGGGCTTCTCATGTGTATAGGTGNNNAGATCGGAA |
| 62996 | TATGAGTGTGGAGTCGTTGCATCATGGACTGCTGTGTATCTCTNNNAGATCGGAAG |
| 35260 | TATGAGTGTGGAGTCGTTGCTCCAGTGTCCATAATCATTACAGCANNNAGATCGGA |
| 55062 | TATGAGTGTGGAGTCGTTGCCGTTGGTTGTCACAAAGACTTGGNNNAGATCGGAA |
| 36379 | TATGAGTGTGGAGTCGTTGCAATGTCCCCTTCTCTGGGAATCANNNAGATCGGAAG |
| 48859 | TATGAGTGTGGAGTCGTTGCCCACAGGGTGTATTGAAGAGCTCNNNAGATCGGAA |
| 42651 | TATGAGTGTGGAGTCGTTGCAGGGAATGACCACTGAAAACCACNNNAGATCGGAA |
| 50836 | TATGAGTGTGGAGTCGTTGCGCTCTGTTCCAAATCCTACCCAGNNNAGATCGGAAG |
| 105574 | TATGAGTGTGGAGTCGTTGCAGTTTGGAGTGTTACATGCTAGAGTNNNAGATCGG |
| 36809 | TATGAGTGTGGAGTCGTTGCCAAGGAGAAGGTTTCTGTGTCTCANNNAGATCGGA |
| 54349 | TATGAGTGTGGAGTCGTTGCATCAGCGTCTAAGCATCACATGGNNNAGATCGGAAG |
| 57483 | TATGAGTGTGGAGTCGTTGCTGTATTCTGTCAAAGAAGGATATGTTGNNNAGATCG |
| 51431 | TATGAGTGTGGAGTCGTTGCAGGGCATAAATCACAGTGCAGAGNNNAGATCGGAA |

52923 TATGAGTGTGGAGTCGTTGCGTCAGCAGTGGAAAGACTACAGGNNNAGATCGGAA
55585 TATGAGTGTGGAGTCGTTGCTGTAGTAGGTGGAGGCTGAGATGNNNAGATCGGA
34761 TATGAGTGTGGAGTCGTTGCCTATGACCACCATGTGATTGCCANNNAGATCGGAAG
62884 TATGAGTGTGGAGTCGTTGCACATAAAGACCCCTCTTGCCTCTNNNAGATCGGAAG
66637 TATGAGTGTGGAGTCGTTGCGTGATGCCCAGTAACATCCTTGTNNNAGATCGGAAG
69645 TATGAGTGTGGAGTCGTTGCCAGTTTCCCCTCTCAGGCTTTTTNNNAGATCGGAAGA
35947 TATGAGTGTGGAGTCGTTGCTTTCCATGTGATTCCAGCATGCANNNAGATCGGAAG
64991 TATGAGTGTGGAGTCGTTGCATCTGCTTTTGGCATTGGTAGGTNNNAGATCGGAAG
58851 TATGAGTGTGGAGTCGTTGCAGCAACAGTGTCTTCCTAGTCATNNNAGATCGGAAG
69215 TATGAGTGTGGAGTCGTTGCCTCTCCTTCCCTCCACCTTCTTTNNNAGATCGGAAGAG
117451 TATGAGTGTGGAGTCGTTGCGTGTGTGTGTGTTGGACAGTTATNNNAGATCGGAA
52100 TATGAGTGTGGAGTCGTTGCGGAGGGGTGGTCGATTTTTCTAGNNNAGATCGGAA
42239 TATGAGTGTGGAGTCGTTGCTAGCATCCTTGCCTTCGCTAAACNNNAGATCGGAAG
57495 TATGAGTGTGGAGTCGTTGCTTTAGATGGAGCCTGCTCTGTTGNNNAGATCGGAAG
29946 TATGAGTGTGGAGTCGTTGCTTCCCCAGCTTTCCTCTCCTATCNNNAGATCGGAAGA
101867 TATGAGTGTGGAGTCGTTGCCTCAAAAATGGGACTAGAAGGGANNNAGATCGGAA
44179 TATGAGTGTGGAGTCGTTGCACAGATACAGAGGTGACTGGTACCNNNAGATCGGA
41985 TATGAGTGTGGAGTCGTTGCGGAGCCCATCACCTGTACATTTANNNAGATCGGAAG
37466 TATGAGTGTGGAGTCGTTGCTTGAAATCCACCAGCACCAAAGANNNAGATCGGAAG
98604 TATGAGTGTGGAGTCGTTGCAAGGGATTGTGTTTGTATGTGTGCANNNAGATCGG
46016 TATGAGTGTGGAGTCGTTGCAGCCTTTGGAGTATGTCGTTTCCNNNAGATCGGAAG
51035 TATGAGTGTGGAGTCGTTGCAACTTTGATTTGCGTGTCAGCAGNNNAGATCGGAAG
46379 TATGAGTGTGGAGTCGTTGCATTCTTTTTCCTCTCCCTGCAGCNNNAGATCGGAAGA
66383 TATGAGTGTGGAGTCGTTGCTATCAAGTGGGAGATGTGGGTGTNNNAGATCGGA
35427 TATGAGTGTGGAGTCGTTGCCCATTTTTCGCTTTCTCTGAGCANNNAGATCGGAAGA
60639 TATGAGTGTGGAGTCGTTGCACAATCTCAGCTGGCTCTAACCTNNNAGATCGGAAG
58956 TATGAGTGTGGAGTCGTTGCGCACCCTGTTGACAGCATAAGATNNNAGATCGGAAG
49974 TATGAGTGTGGAGTCGTTGCATCCTGGGTGCACTCTCATTTTCNNNAGATCGGAAG
37068 TATGAGTGTGGAGTCGTTGCCTTTGAGGGCAGGAAACTTGTCANNNAGATCGGAA
64143 TATGAGTGTGGAGTCGTTGCTTCCTCCTCATCTGTCCTTCAGTNNNAGATCGGAAGA
39806 TATGAGTGTGGAGTCGTTGCCACAGGTCAGATTCAAGCACTGANNNAGATCGGAA
56796 TATGAGTGTGGAGTCGTTGCAAAGAGAGGTGAGAAACTGGGTGNNNAGATCGGA
68697 TATGAGTGTGGAGTCGTTGCTGTGTGTGTAGTCTTGCTGTGTTNNNAGATCGGAAG
104743 TATGAGTGTGGAGTCGTTGCTGTACTCTTGCACAACTTTGCTNNNAGATCGGAAGA
51485 TATGAGTGTGGAGTCGTTGCAATAGAGCGCTGCAAAAGGAGAGNNNAGATCGGA
108474 TATGAGTGTGGAGTCGTTGCGAGGGGAAGTGAGGAGAGAGAGANNNAGATCGG
97041 TATGAGTGTGGAGTCGTTGCAAACTGCTGACAAGTGAAGTCTANNNAGATCGGAA
57712 TATGAGTGTGGAGTCGTTGCCCTTGGGATAAGATGGCCCAAATNNNAGATCGGAA
103866 TATGAGTGTGGAGTCGTTGCGCTAACCCAATCTTACAATCAGCAANNNAGATCGGA
34272 TATGAGTGTGGAGTCGTTGCATTATTGCCTGGGTCACTACCCANNNAGATCGGAAG
55414 TATGAGTGTGGAGTCGTTGCACTTATCTTGCAGCATCCCCATGNNNAGATCGGAAG
53519 TATGAGTGTGGAGTCGTTGCACATCATTCAATACATTGCGCGGNNNAGATCGGAAG
97384 TATGAGTGTGGAGTCGTTGCCCCTCCACCTTCTTTCACGTTTTNNNAGATCGGAAGA

29825 TATGAGTGTGGAGTCGTTGCCGTGTGTGTGTGTATGACACTNNNAGATCGGAA
48385 TATGAGTGTGGAGTCGTTGCTCAATGGCACAGTCACTAGCATCNNNAGATCGGAAG
66093 TATGAGTGTGGAGTCGTTGCATGTGGAAACACAGACATGCTGTNNNAGATCGGAA
55361 TATGAGTGTGGAGTCGTTGCCAGGGAAAAGACCACACCACATGNNNAGATCGGAA
48377 TATGAGTGTGGAGTCGTTGCCACCTAACCAGGAAGACAGCATCNNNAGATCGGAAG
57995 TATGAGTGTGGAGTCGTTGCCAGGTATGCAACAGGTGCTCAATNNNAGATCGGAA
40556 TATGAGTGTGGAGTCGTTGCTGGGACTAGAAGGGATGGTTTGANNNAGATCGGA
51486 TATGAGTGTGGAGTCGTTGCAGGTGCTGTGTTTTCAAGGAGAGNNNAGATCGGAA
104732 TATGAGTGTGGAGTCGTTGCCAGGTGCTGTGTTTTCAAGGAGAGNNNAGATCGGA
68740 TATGAGTGTGGAGTCGTTGCGGACAATGTGGTTAGGGGTTGTTNNNAGATCGGAA
34062 TATGAGTGTGGAGTCGTTGCGAGTGTAGAGCATTGATGCACCANNNAGATCGGAA
48032 TATGAGTGTGGAGTCGTTGCTTTGTGTGTCTGTCTGTTTGTGCNNNAGATCGGAAG
48976 TATGAGTGTGGAGTCGTTGCTGGAATGAGCAGCAGGAAATCTCNNNAGATCGGAA
69569 TATGAGTGTGGAGTCGTTGCTGTGTGTGTGCTTGTATGGTTTTNNNAGATCGGAAG
48393 TATGAGTGTGGAGTCGTTGCTCAAGGTTCACTCATGTGGCATCNNNAGATCGGAAG
34632 TATGAGTGTGGAGTCGTTGCGGACAGGGTCTTTGTTTTAGCCANNNAGATCGGAA
51906 TATGAGTGTGGAGTCGTTGCCTCTTGCACAACTTTGCTGTGAGNNNAGATCGGAAG
58082 TATGAGTGTGGAGTCGTTGCGAGACATACCCCTAGCCAGGAATNNNAGATCGGAA
58982 TATGAGTGTGGAGTCGTTGCCTGAAGGACGGCTACCCAGATNNNAGATCGGAAGA
51314 TATGAGTGTGGAGTCGTTGCTGGGTTCATGAGTTCTGCTTCAGNNNAGATCGGAAG
57793 TATGAGTGTGGAGTCGTTGCCTCCAAAGCCATCACCTGGAAATNNNAGATCGGAAG
32395 TATGAGTGTGGAGTCGTTGCCAAAATCTGACACGGGAGGCTAANNNAGATCGGAA
61008 TATGAGTGTGGAGTCGTTGCAACGTATTTTTCCTGACCTCCCTNNNAGATCGGAAGA
49474 TATGAGTGTGGAGTCGTTGCATTGTCTTAGCCAGCCACTTGTCNNNAGATCGGAAG
61526 TATGAGTGTGGAGTCGTTGCATCTTTCCTTCCACTGCCTTCCTNNNAGATCGGAAGA
32029 TATGAGTGTGGAGTCGTTGCGAGCAAGAGCAGACAGGAATGAANNNAGATCGGA
101267 TATGAGTGTGGAGTCGTTGCGGATGGTTTTGGTACCTGTGTAATGNNNAGATCGG
95479 TATGAGTGTGGAGTCGTTGCCCCACAAGCAGGAAACCTGAAANNNAGATCGGAAG
52010 TATGAGTGTGGAGTCGTTGCGGATGGGCTTGGAGACAAACTAGNNNAGATCGGA
65145 TATGAGTGTGGAGTCGTTGCACGCAGATGATACAGAAATGGGTNNNAGATCGGAA
71780 TATGAGTGTGGAGTCGTTGCTTCCCAAGTTCTTCACCTTTGGCNNNAGATCGGAAGA
102477 TATGAGTGTGGAGTCGTTGCCCTACTTGGGGAAGCCTAAAACTGNNNAGATCGGA
102545 TATGAGTGTGGAGTCGTTGCATACCAAGTCCACTCTGTTCAGTNNNAGATCGGAAG
104149 TATGAGTGTGGAGTCGTTGCTCTGGGCTTCAGGTTCTGAGAAATNNNAGATCGGA
71773 TATGAGTGTGGAGTCGTTGCAAATGCTCTTACAGCAGAGAGCCNNNAGATCGGAA
107074 TATGAGTGTGGAGTCGTTGCCAACACCATAGCACGTACTTACAAANNNAGATCGGA
105591 TATGAGTGTGGAGTCGTTGCGATGGCAACATAATAATGTGGGANNNAGATCGGAA
102220 TATGAGTGTGGAGTCGTTGCGACCTAGAGCTTGGCAAGAGCATANNNAGATCGGA
106669 TATGAGTGTGGAGTCGTTGCGAAAAGCGAATGTAGGGTGGGGANNNAGATCGGA
105555 TATGAGTGTGGAGTCGTTGCATAACTTACCCTGTCTCCTCAGTNNNAGATCGGAAGA
100958 TATGAGTGTGGAGTCGTTGCGAAATGCAACCATGTACTCCCTCNNNAGATCGGAAG
106199 TATGAGTGTGGAGTCGTTGCGACATTACAGGTGCATGCCATGGNNNAGATCGGAA
107829 TATGAGTGTGGAGTCGTTGCACCTGAATGGAATTGGAGACTATTGTCNNNAGATCG

| | |
|---|---|
| 109058 | TATGAGTGTGGAGTCGTTGCAACAGTCAGAATTGGATTACAGCCTNNNAGATCGGA |
| 100574 | TATGAGTGTGGAGTCGTTGCCACCACACCCAGACATACTTACTTTNNNAGATCGGAA |
| 103870 | TATGAGTGTGGAGTCGTTGCTGGTGTGCAAGTGTTGTTGGAGNNNAGATCGGAAG |
| 101864 | TATGAGTGTGGAGTCGTTGCGCCCTAGAGGAAGTCATTGTTTCCNNNAGATCGGAA |
| 101427 | TATGAGTGTGGAGTCGTTGCCCCAATTACACACACCAAGTCTANNNAGATCGGAAG |
| 102218 | TATGAGTGTGGAGTCGTTGCTCATGGTGATCGGCAAAGTACAGNNNAGATCGGAA |
| 97383 | TATGAGTGTGGAGTCGTTGCCTCAAGTTTTCAGATTTGGTGAAGTGNNNAGATCGG |
| 110745 | TATGAGTGTGGAGTCGTTGCTTTTTGTTTGTTTCCCAGTGCATTTNNNAGATCGGAA |
| 100125 | TATGAGTGTGGAGTCGTTGCGCAGGGAGTTTAGGAAAAGAGGCNNNAGATCGGA |
| 116318 | TATGAGTGTGGAGTCGTTGCCTCCTGCACTGATTTTACCTTCCANNNAGATCGGAAG |
| 112130 | TATGAGTGTGGAGTCGTTGCCTGGAGTAACTTGGAGTGATCATCTTCNNNAGATCG |
| 113336 | TATGAGTGTGGAGTCGTTGCTACATACCACCACGCCCATATTTTTANNNAGATCGGA |
| 101421 | TATGAGTGTGGAGTCGTTGCGCTAACTAGAGTTCTTACTGTTGTTTTNNNAGATCGG |
| 103296 | TATGAGTGTGGAGTCGTTGCAGGCCGTGTATCTGTCACTCTNNNAGATCGGAAGAG |
| 96440 | TATGAGTGTGGAGTCGTTGCTTTTCCATGATTGCACCAGTGCANNNAGATCGGAAG |
| 102197 | TATGAGTGTGGAGTCGTTGCCCAGATAATGAAGATGCTGCTGGNNNAGATCGGAA |
| 97385 | TATGAGTGTGGAGTCGTTGCTTTCCCCATTCGGTATGATGAGGNNNAGATCGGAAG |
| 110716 | TATGAGTGTGGAGTCGTTGCGTGTGTTGTTGGAAACTGGACCTTANNNAGATCGG |
| 103876 | TATGAGTGTGGAGTCGTTGCTACAGAGGAGTGGGATTACTGGANNNAGATCGGA |
| 30806 | TATGAGTGTGGAGTCGTTGCTTTCTGGCCATTGTATTGAAACAANNNAGATCGGAA |
| 71736 | TATGAGTGTGGAGTCGTTGCCAGCATCCCTTTTAGCCAAAACGNNNAGATCGGAAG |
| 60421 | TATGAGTGTGGAGTCGTTGCACAGTATTTTGACCTAGGGATTGACTNNNAGATCGG |
| 41549 | TATGAGTGTGGAGTCGTTGCGGGGTCAACCTTTTTGATGGGTANNNAGATCGGAA |
| 63002 | TATGAGTGTGGAGTCGTTGCTGAGAACACAAGACCCACTCTCTNNNAGATCGGAAG |
| 46963 | TATGAGTGTGGAGTCGTTGCGTTATTCCAGCAAGGAGGAGGCNNNAGATCGGAAG |
| 41841 | TATGAGTGTGGAGTCGTTGCCACTATGCATGCCCATCAGCTTANNNAGATCGGAAG |
| 51179 | TATGAGTGTGGAGTCGTTGCGCAGGCAGAAGGATAATTTGCAGNNNAGATCGGAA |
| 55075 | TATGAGTGTGGAGTCGTTGCTCCTCTTTCACATACACTCCTTGGNNNAGATCGGAAG |
| 39451 | TATGAGTGTGGAGTCGTTGCATGGGGTAGGAGAAAAGTGTGGANNNAGATCGGA |
| 49227 | TATGAGTGTGGAGTCGTTGCCTTCTCTTCTGCTCTCTGCAGTCNNNAGATCGGAAGA |
| 54014 | TATGAGTGTGGAGTCGTTGCTGCCCATTTTGAAATTGTTGGGGNNNAGATCGGAAG |
| 62540 | TATGAGTGTGGAGTCGTTGCCCCAAAAGACAATGGAGAAAAATCTNNNAGATCGG |
| 59968 | TATGAGTGTGGAGTCGTTGCACCAGTTTCAGTGAAGTATAGAACACTNNNAGATCG |
| 57423 | TATGAGTGTGGAGTCGTTGCGCATTTTATCCCTGGTGCAGTTGNNNAGATCGGAAG |
| 35961 | TATGAGTGTGGAGTCGTTGCTAGTTCTCTCCGTCTCTCATGCANNNAGATCGGAAGA |
| 40108 | TATGAGTGTGGAGTCGTTGCTTGAAGGAAGAAACGGGAAGTGANNNAGATCGGA |
| 69580 | TATGAGTGTGGAGTCGTTGCGTGGGGTGAGTGGATCATGTTTTNNNAGATCGGAA |
| 49045 | TATGAGTGTGGAGTCGTTGCGAAGGGTCCCTGGTCTATCTCTCNNNAGATCGGAAG |
| 41632 | TATGAGTGTGGAGTCGTTGCCAAGGCATTTTACCCAGCCTGTANNNAGATCGGAAG |
| 44407 | TATGAGTGTGGAGTCGTTGCGTTAGTGGGAGATACAATTCCCCCNNNAGATCGGAA |
| 63598 | TATGAGTGTGGAGTCGTTGCTGCCTTCCTCAGCAAGTTTTTCTNNNAGATCGGAAGA |
| 36727 | TATGAGTGTGGAGTCGTTGCAGCAGACCTTGTTTGTATGCTCANNNAGATCGGAAG |
| 71625 | TATGAGTGTGGAGTCGTTGCATTACCTCAAAGGAAAACTGGAAGAANNNAGATCG |

| | |
|---|---|
| 52590 | TATGAGTGTGGAGTCGTTGCTCCTTCTGGCATTTCTTTCCTCGNNNAGATCGGAAGA |
| 39409 | TATGAGTGTGGAGTCGTTGCGTGAAGACAGGCTTTTCAGTGGANNNAGATCGGAA |
| 32467 | TATGAGTGTGGAGTCGTTGCCCTGGTCCTCACTGCTCTAGTAANNNAGATCGGAAG |
| 54546 | TATGAGTGTGGAGTCGTTGCTGACTGAAACATAGGTGAGACTGGNNNAGATCGGA |
| 29822 | TATGAGTGTGGAGTCGTTGCATACACCTGGCTACTAAGCACCANNNAGATCGGAAG |
| 103894 | TATGAGTGTGGAGTCGTTGCAGCCACAGAAGACTGGAAGAAGTCNNNAGATCGGA |
| 68556 | TATGAGTGTGGAGTCGTTGCTGGGGGAGGGTTATGAGAATGTTNNNAGATCGGA |
| 31539 | TATGAGTGTGGAGTCGTTGCGGGTAACTACGTGGCAACAAGAANNNAGATCGGAA |
| 105564 | TATGAGTGTGGAGTCGTTGCCTGTGTGTGTGTATGTGAATCCTTANNNAGATCGGA |
| 33593 | TATGAGTGTGGAGTCGTTGCACCATAGAAATGACCAGAGGACANNNAGATCGGAA |
| 46063 | TATGAGTGTGGAGTCGTTGCCCCAAAAGACAGTGTGACAAAGCNNNAGATCGGAA |
| 121982 | TATGAGTGTGGAGTCGTTGCAAGTTGAAGCGCGGCACGAATCNNNAGATCGGAAG |
| 50426 | TATGAGTGTGGAGTCGTTGCACAGCAACTCACCAAGAAGGAAGNNNAGATCGGAA |
| 30258 | TATGAGTGTGGAGTCGTTGCCCCACCCACTTTTCTCAGACAAANNNAGATCGGAAGA |
| 66443 | TATGAGTGTGGAGTCGTTGCTAAGCTGCTCCTATGACCTGTGTNNNAGATCGGAAG |
| 40328 | TATGAGTGTGGAGTCGTTGCCATTGACCTAAACAGGCTTGTGANNNAGATCGGAAG |
| 68267 | TATGAGTGTGGAGTCGTTGCTCTCCCTTTCTTCCACTCCAGTTNNNAGATCGGAAGA |
| 61971 | TATGAGTGTGGAGTCGTTGCTCTATACCCAGACAAAGCAGGCTNNNAGATCGGAAG |
| 103304 | TATGAGTGTGGAGTCGTTGCGTTTGTATGCTCATCCTCTGAAGACNNNAGATCGGA |
| 50608 | TATGAGTGTGGAGTCGTTGCGCATCCCACCTTTACAGAAACAGNNNAGATCGGAAG |
| 64384 | TATGAGTGTGGAGTCGTTGCGTCAGCAACGTCCTACTTTGAGTNNNAGATCGGAAG |
| 67042 | TATGAGTGTGGAGTCGTTGCTAAGGCCACAGTTCTGCAACATTNNNAGATCGGAAG |
| 65708 | TATGAGTGTGGAGTCGTTGCGCACTCCATACTTTCCTCCATGTNNNAGATCGGAAGA |
| 69264 | TATGAGTGTGGAGTCGTTGCAGGGGAAGACAGGGAAAGAGTTTNNNAGATCGGA |
| 96666 | TATGAGTGTGGAGTCGTTGCCCCCCAAGAAAGTGACCCAAGNNNAGATCGGAAGA |
| 38635 | TATGAGTGTGGAGTCGTTGCGGAAACACGACACGTCTAAAGGANNNAGATCGGAA |
| 104089 | TATGAGTGTGGAGTCGTTGCGTGAGGGGCTATATCGTTTGTTANNNAGATCGGAA |
| 71309 | TATGAGTGTGGAGTCGTTGCAGAGAGAAGGGGTGTTGAAGAGTNNNAGATCGGA |
| 33846 | TATGAGTGTGGAGTCGTTGCACAGTAAGTCAGGGATAAAGTCTACANNNAGATCG |
| 41261 | TATGAGTGTGGAGTCGTTGCCAGCAGTGGAAGCAGCATATCTANNNAGATCGGAA |
| 67844 | TATGAGTGTGGAGTCGTTGCCATTTGGGACCTGAGCTAAGCTTNNNAGATCGGAAG |
| 49185 | TATGAGTGTGGAGTCGTTGCGCAGCTAGACGATCAGTGAAGTCNNNAGATCGGAA |
| 30233 | TATGAGTGTGGAGTCGTTGCGGGGGAATTCAGAACATAACAAANNNAGATCGGAA |
| 56697 | TATGAGTGTGGAGTCGTTGCTCATCTCTGTTCTTTGCTCCGTGNNNAGATCGGAAGA |
| 97928 | TATGAGTGTGGAGTCGTTGCGAGTACCATCTCGGCTCACCGNNNAGATCGGAAGAG |
| 44891 | TATGAGTGTGGAGTCGTTGCACATTTGAGGGGTGAAGTAAGCCNNNAGATCGGAA |
| 66298 | TATGAGTGTGGAGTCGTTGCGGGAATGAAAGTTGTCCCAGTGTNNNAGATCGGAA |
| 39167 | TATGAGTGTGGAGTCGTTGCAAGAGCTCAGACATGTTCTGGGANNNAGATCGGAA |
| 104701 | TATGAGTGTGGAGTCGTTGCCACTTATCTCCCTTTTTCCTTTACTATNNNAGATCGGA |
| 65034 | TATGAGTGTGGAGTCGTTGCCTCATCAATTGTGTCCCTTCGGTNNNAGATCGGAAG |
| 97960 | TATGAGTGTGGAGTCGTTGCCATTCTAATCACCCACCTGTGCANNNAGATCGGAAGA |
| 29821 | TATGAGTGTGGAGTCGTTGCATCTGCAGCCATTCTCACCTTTANNNAGATCGGAAGA |
| 100775 | TATGAGTGTGGAGTCGTTGCCTGAAAAACGGAAGGAAATCTTGTCNNNAGATCGG |

33507 TATGAGTGTGGAGTCGTTGCTGAATTTTGGGATAACCCAGACANNNAGATCGGAA
42877 TATGAGTGTGGAGTCGTTGCAGAACAGATCCGAAAAGAGGCACNNNAGATCGGAA
49619 TATGAGTGTGGAGTCGTTGCTAGGCTGGTCAGGATGAGACTTCNNNAGATCGGAA
39726 TATGAGTGTGGAGTCGTTGCTGTGTGTGTGTGTAAGCTGATGANNNAGATCGGAA
100784 TATGAGTGTGGAGTCGTTGCCTAATTTTTCACCGTTTGGCCAGNNNAGATCGGAAG
38759 TATGAGTGTGGAGTCGTTGCATGCAATAAAGTGAAGTGCAGGANNNAGATCGGAA
63872 TATGAGTGTGGAGTCGTTGCAGCAAACTAAAAGTATACAACACAGTNNNAGATCGG
37132 TATGAGTGTGGAGTCGTTGCTGAGTGTTCAGAGTTACTTCATTCANNNAGATCGGA
36153 TATGAGTGTGGAGTCGTTGCTGACATAGAGAGAGAGACGTGCANNNAGATCGGAA
32519 TATGAGTGTGGAGTCGTTGCTGTGTGCATCTATGTGCCTGTAANNNAGATCGGAAG
43785 TATGAGTGTGGAGTCGTTGCGTCTGTCTGTCTGTGTCTCAACCNNNAGATCGGAAG
37224 TATGAGTGTGGAGTCGTTGCAGTGCCTAAGGTTGCAAATCTTCANNNAGATCGGAA
67243 TATGAGTGTGGAGTCGTTGCCAGGCACTTACCCAGTGTTCATTNNNAGATCGGAAG
62703 TATGAGTGTGGAGTCGTTGCTTGATGTAGATCCTAGTGGTGATCTNNNAGATCGGA
66372 TATGAGTGTGGAGTCGTTGCCCCTGCTGTGTCAAAATAGGTGTNNNAGATCGGAAG
71614 TATGAGTGTGGAGTCGTTGCTGGTGGGACAAGCATTTAGTTTTNNNAGATCGGAA
58830 TATGAGTGTGGAGTCGTTGCAGGGAAAATACAGATGTGCTCATNNNAGATCGGAA
33873 TATGAGTGTGGAGTCGTTGCTGTTGTGCATAGATACTTGATAGTACANNNAGATCG
47343 TATGAGTGTGGAGTCGTTGCAAATCTGTTCCGTTCTAGGTGGCNNNAGATCGGAAG
59572 TATGAGTGTGGAGTCGTTGCATGGGTTTTTCAGCGCCTTCTATNNNAGATCGGAAG
98104 TATGAGTGTGGAGTCGTTGCACAGCCTTCCTCGAGACACACATNNNAGATCGGAAG
30024 TATGAGTGTGGAGTCGTTGCGGAGGAGTGTCATGGTGGAAAAANNNAGATCGGA
62185 TATGAGTGTGGAGTCGTTGCAACCTGGTATTCCCTTTTTGGCTNNNAGATCGGAAG
37312 TATGAGTGTGGAGTCGTTGCTGATTGGTGTTACAAATGGTGTTCANNNAGATCGGA
31618 TATGAGTGTGGAGTCGTTGCTTATGCATGTAAGTTCGCCAGAANNNAGATCGGAAG
55895 TATGAGTGTGGAGTCGTTGCTCACATCAAATCCTGATCAGTACTGNNNAGATCGGA
61184 TATGAGTGTGGAGTCGTTGCTATTCGTTGAGGAGAACTGGCCTNNNAGATCGGAA
52029 TATGAGTGTGGAGTCGTTGCGCAAACTCACACTGTTCCCCTAGNNNAGATCGGAAG
108491 TATGAGTGTGGAGTCGTTGCGCTTCTTGGATCTGTGAATTGAGTTNNNAGATCGGA
60460 TATGAGTGTGGAGTCGTTGCATGCCTTCACTTCTCCCACTACTNNNAGATCGGAAGA
41130 TATGAGTGTGGAGTCGTTGCCTCCACCTTCCCCTTCTTTCCTANNNAGATCGGAAGAG
105563 TATGAGTGTGGAGTCGTTGCGGCAGGTATATCTCATTGCTGACTTNNNAGATCGGA
107046 TATGAGTGTGGAGTCGTTGCCAAACTCACACTGTTCCCCTAGANNNAGATCGGAAG
59500 TATGAGTGTGGAGTCGTTGCTACCCACATTCACTGCCACCTATNNNAGATCGGAAGA
32898 TATGAGTGTGGAGTCGTTGCCATCACCCATTTCCCAAGGAACANNNAGATCGGAAG
38500 TATGAGTGTGGAGTCGTTGCGGAGCCATGCACATCTGTTTAGANNNAGATCGGAA
56309 TATGAGTGTGGAGTCGTTGCTTGCCTCCACTTCTGAACATCTGNNNAGATCGGAAGA
58560 TATGAGTGTGGAGTCGTTGCTTCGGTGTTTTGTTCACCTCCATNNNAGATCGGAAGA
64497 TATGAGTGTGGAGTCGTTGCAGTACATGGTGGGGCTCATTAGTNNNAGATCGGAA
56185 TATGAGTGTGGAGTCGTTGCGGTCCTAGACACTAGAGAGGCTGNNNAGATCGGAA
57800 TATGAGTGTGGAGTCGTTGCGTTGCCCCTAGGAACTTGGAAATNNNAGATCGGAA
59458 TATGAGTGTGGAGTCGTTGCGCCAGTGACAAACCCCATGATATNNNAGATCGGAAG
32237 TATGAGTGTGGAGTCGTTGCGCCTCTTGGGTAGGTTGCAATAANNNAGATCGGAA

106553 TATGAGTGTGGAGTCGTTGCGCATGTAAGTTCGCCAGAAATTTTCNNNAGATCGGA
56337 TATGAGTGTGGAGTCGTTGCTGCCTCCTTACCTCAAAAACTCTGNNNAGATCGGAAG
43470 TATGAGTGTGGAGTCGTTGCGTTACAGCATCGTGGCCAAATACNNNAGATCGGAAG
45492 TATGAGTGTGGAGTCGTTGCTGGAGATGATGCCCTAGCTATCCNNNAGATCGGAAG
46956 TATGAGTGTGGAGTCGTTGCGTGAGTGTGTGTATGAGAGAGGCNNNAGATCGGA
71703 TATGAGTGTGGAGTCGTTGCCTATGTTGTCAGGGGCAATGTGANNNAGATCGGAA
50302 TATGAGTGTGGAGTCGTTGCAGCATTTGCCTTATGTGGGCAAGNNNAGATCGGAA
40717 TATGAGTGTGGAGTCGTTGCGTTTTAGGCCAGTGTGAGCCATANNNAGATCGGAA
106178 TATGAGTGTGGAGTCGTTGCCACTTGTAGGTTGATGCGCATTTGNNNAGATCGGAA
57915 TATGAGTGTGGAGTCGTTGCCCTTCAGCTGATTACTGGCCAATNNNAGATCGGAAG
54174 TATGAGTGTGGAGTCGTTGCACCCAAGGAGTATCTATCAGTGGGNNNAGATCGGA
71769 TATGAGTGTGGAGTCGTTGCCACACATTGGAGAGGTGTTTTGCNNNAGATCGGAA
60926 TATGAGTGTGGAGTCGTTGCGCATCTGTGTGTGTATTTCCCCTNNNAGATCGGAAG
64161 TATGAGTGTGGAGTCGTTGCCCATCAGAGGAGAAATTTTTCAGTNNNAGATCGGAA
110737 TATGAGTGTGGAGTCGTTGCCCCCACAGCCATGTTGGTTGNNNAGATCGGAAGAGC
58536 TATGAGTGTGGAGTCGTTGCTCTGACAAGAAATGTGCTGCCATNNNAGATCGGAAG
107826 TATGAGTGTGGAGTCGTTGCATCAACTGAAATCAGGATCTTGAAGACNNNAGATCG
63284 TATGAGTGTGGAGTCGTTGCGGCTTTGCTGTCTGAGATTGTCTNNNAGATCGGAAG
35503 TATGAGTGTGGAGTCGTTGCCGTACACATACCAGCATGTAGCANNNAGATCGGAAG
45439 TATGAGTGTGGAGTCGTTGCTAAGGGAATGACACCATGCATCCNNNAGATCGGAA
42687 TATGAGTGTGGAGTCGTTGCGATTTCTGGTGGGAGAAGACCACNNNAGATCGGAA
56773 TATGAGTGTGGAGTCGTTGCTTTGAGTACTACCCATTCTCGGTGNNNAGATCGGAA
34120 TATGAGTGTGGAGTCGTTGCTCTTGTGTGAAATGCTTAGGACCANNNAGATCGGAA
45669 TATGAGTGTGGAGTCGTTGCAAAACAACTGGAAGGAGCTCTCCNNNAGATCGGAA
56429 TATGAGTGTGGAGTCGTTGCCTTCTCCACATTCCCACTGTCTGNNNAGATCGGAAGA
43911 TATGAGTGTGGAGTCGTTGCGTGATATGGGATAACCTCCCACCNNNAGATCGGAAG
54559 TATGAGTGTGGAGTCGTTGCCTTCCCTGAGGAGGTAGTACTGGNNNAGATCGGAA
61755 TATGAGTGTGGAGTCGTTGCTTACCTCCAACATGGAAAGAGCTNNNAGATCGGAAG
60024 TATGAGTGTGGAGTCGTTGCGTCATTTGGCCTATGCAATACACTNNNAGATCGGAA
111134 TATGAGTGTGGAGTCGTTGCTGCTTCTCTCTGGGCATTTCTCANNNAGATCGGAAGA
59533 TATGAGTGTGGAGTCGTTGCGGAGGTGACAGAGGCTAAGCTATNNNAGATCGGAA
106167 TATGAGTGTGGAGTCGTTGCGAGTTGGCTTATCCCTTAGATTTCATCNNNAGATCGG
33509 TATGAGTGTGGAGTCGTTGCATCACAATGGTCTCTCCCAGACANNNAGATCGGAAG
65349 TATGAGTGTGGAGTCGTTGCAGTACCCAACTCACTTCACTGGTNNNAGATCGGAAG
38612 TATGAGTGTGGAGTCGTTGCTTGGCTCACACGATTACAAAGGANNNAGATCGGAA
64364 TATGAGTGTGGAGTCGTTGCTCCTGCCACAAATTCTTGTGAGTNNNAGATCGGAAG
101682 TATGAGTGTGGAGTCGTTGCGTGCTCCTGCTTCAGAGGAACTANNNAGATCGGAAG
105560 TATGAGTGTGGAGTCGTTGCCTCCCAGAACACGTCAGTAACANNNAGATCGGAAGA
58796 TATGAGTGTGGAGTCGTTGCCCTGGGCAGCCTTGATTTATCATNNNAGATCGGAAG
55426 TATGAGTGTGGAGTCGTTGCGCTGCTACTTTTTGGAAGCCATGNNNAGATCGGAAG
32808 TATGAGTGTGGAGTCGTTGCATGACGCGGGAAAAGATACAACANNNAGATCGGAA
117738 TATGAGTGTGGAGTCGTTGCATGTCCTCATGAATAGCCATTCCANNNAGATCGGAA
68882 TATGAGTGTGGAGTCGTTGCTGGCCTTTGGACTTTGCAGATTTNNNAGATCGGAAG

55331 TATGAGTGTGGAGTCGTTGCGCAGTGGGTACATCCACCTAATGNNNAGATCGGAAG
52631 TATGAGTGTGGAGTCGTTGCCATTGAGCATCCAAGTTGTGTCGNNNAGATCGGAAG
69281 TATGAGTGTGGAGTCGTTGCGTGGCCCGTAGTCTGTTTAGTTTNNNAGATCGGAAG
50296 TATGAGTGTGGAGTCGTTGCGAAAGCAGAGAGAGACAGGCAAGNNNAGATCGGA
61229 TATGAGTGTGGAGTCGTTGCAGCAGCAACTAATTACTATGCCTNNNAGATCGGAAG
32924 TATGAGTGTGGAGTCGTTGCATGCAGACACACACAGATGAACANNNAGATCGGAAG
64763 TATGAGTGTGGAGTCGTTGCTCTGAAGGCAAACTCAAGAAGGTNNNAGATCGGAA
102840 TATGAGTGTGGAGTCGTTGCAGTAAAGTGAAACAATCGGCTTCACNNNAGATCGGA
64347 TATGAGTGTGGAGTCGTTGCGCAGCCTAGTAATCCATCTGAGTNNNAGATCGGAAG
31817 TATGAGTGTGGAGTCGTTGCATAAAAACCCAAGTGCCCAGGAANNNAGATCGGAAG
106580 TATGAGTGTGGAGTCGTTGCGTATTGCTCTAAGGAACTCCCTGANNNAGATCGGAA
43033 TATGAGTGTGGAGTCGTTGCCAACCCTACAGACACACAGTCACNNNAGATCGGAAG
30822 TATGAGTGTGGAGTCGTTGCGCGTGTCTTATTTGTGGCAACAANNNAGATCGGAAG
71787 TATGAGTGTGGAGTCGTTGCAAATAAGTGTGCCACCATCACGANNNAGATCGGAAG
67481 TATGAGTGTGGAGTCGTTGCACTGTCCTCCCAAGTCTCAACTTNNNAGATCGGAAGA
31601 TATGAGTGTGGAGTCGTTGCCACAGGGAAAGCGAACTACAGAANNNAGATCGGAA
117873 TATGAGTGTGGAGTCGTTGCCTCAAAGATGTAGGAACTGTGCTGNNNAGATCGGA
52236 TATGAGTGTGGAGTCGTTGCGACATTGCCCTTCCTCAAAAACGNNNAGATCGGAAG
53120 TATGAGTGTGGAGTCGTTGCGCAGGAGGTCAAAGGTAAAGAGGNNNAGATCGGA
61760 TATGAGTGTGGAGTCGTTGCCCCACTGTAGTACTGGAAGAGCTNNNAGATCGGAAG
54764 TATGAGTGTGGAGTCGTTGCGGTCTGTCCACTAAGAAAGTCTGGNNNAGATCGGA
103838 TATGAGTGTGGAGTCGTTGCCCCAACCCCGAAATAATCTCTTGAANNNAGATCGGA
66002 TATGAGTGTGGAGTCGTTGCTCCTTTTCATTTTGGATGCCTGTNNNAGATCGGAAGA
45473 TATGAGTGTGGAGTCGTTGCTGCTCTAAGGAACTCCCTGATCCNNNAGATCGGAAG
53129 TATGAGTGTGGAGTCGTTGCCCACCTCAACGAGAATGAAGAGGNNNAGATCGGAA
40912 TATGAGTGTGGAGTCGTTGCTTCCCACCCTCTTTGCCAGTATANNNAGATCGGAAGA
108481 TATGAGTGTGGAGTCGTTGCACATGCACAAAAGTGAAATAGCCTTNNNAGATCGGA
97281 TATGAGTGTGGAGTCGTTGCCCACCCTCCACCTGCAGATTTTANNNAGATCGGAAGA
97851 TATGAGTGTGGAGTCGTTGCCAAGGAGCAACAGTCTATGTGTCTNNNAGATCGGA
51140 TATGAGTGTGGAGTCGTTGCGAAACAATCGGCTTCACTGCAGNNNAGATCGGAAG
38912 TATGAGTGTGGAGTCGTTGCTGTTTTGCAATATCCGTGGTAGGANNNAGATCGGA
68999 TATGAGTGTGGAGTCGTTGCACTTATTTGGTCGGTGCACCTTTNNNAGATCGGAAG
69306 TATGAGTGTGGAGTCGTTGCTGAAGGGCTAGGTTTTTCGGTTTNNNAGATCGGAAG
110695 TATGAGTGTGGAGTCGTTGCTACACACCCACACAAATCCTATTGGTNNNAGATCGGA
31206 TATGAGTGTGGAGTCGTTGCCGAGGCTTTGAGAGAAAGAGCAANNNAGATCGGA
47105 TATGAGTGTGGAGTCGTTGCAAAATGCTTCCCCAATAAGGGGCNNNAGATCGGAAG
41624 TATGAGTGTGGAGTCGTTGCTGGTGTGATAGTCGCTCACTGTANNNAGATCGGAAG
116163 TATGAGTGTGGAGTCGTTGCGAAAAGGCCAGAGCTTCTGCCNNNAGATCGGAAGA
67336 TATGAGTGTGGAGTCGTTGCGGGTGGGCCAGAAAGATATGATTNNNAGATCGGA
61255 TATGAGTGTGGAGTCGTTGCATTATCTGTGGACCTGTCTGCCTNNNAGATCGGAAG
37551 TATGAGTGTGGAGTCGTTGCTCCAAAGACTAAAGAAGCCAAGANNNAGATCGGAA
59364 TATGAGTGTGGAGTCGTTGCGGCATATGGGAAGAGTGCTTGATNNNAGATCGGAA
104702 TATGAGTGTGGAGTCGTTGCTGCTGCCATTTGAATGATCTATCANNNAGATCGGAA

45515 TATGAGTGTGGAGTCGTTGCATCATTCTCTGCGGTTCAACTCCNNNAGATCGGAAGA
43921 TATGAGTGTGGAGTCGTTGCATCCTAGTTGATCCTCTGCCACCNNNAGATCGGAAGA
31742 TATGAGTGTGGAGTCGTTGCTGAGAATTGTGCGGCATATAGAANNNAGATCGGAA[illegible]
34505 TATGAGTGTGGAGTCGTTGCATGTACCAGGAGAATGGTTCCCANNNAGATCGGAAG
100020 TATGAGTGTGGAGTCGTTGCACTATCAACTTCTGCTGAGGCATAAANNNAGATCGGA
30230 TATGAGTGTGGAGTCGTTGCCCTGTCCTTGGCTCTACAACAAANNNAGATCGGAAGA
34667 TATGAGTGTGGAGTCGTTGCATAAGAGTGATGGAGTTGGGCCANNNAGATCGGAA
42252 TATGAGTGTGGAGTCGTTGCGCCAGAGACAAGAGCAAAACAACNNNAGATCGGAA[illegible]
57481 TATGAGTGTGGAGTCGTTGCCGTCAGGGATGTAGACGATGTTGNNNAGATCGGAA
58909 TATGAGTGTGGAGTCGTTGCCCCACTCTGCAGTTGTTCTTCATNNNAGATCGGAAGA
61302 TATGAGTGTGGAGTCGTTGCTTCTCCTCTTCCTGGGAAATCCTNNNAGATCGGAAGA
52384 TATGAGTGTGGAGTCGTTGCCTGATTGAAACCCCCAGATACCGNNNAGATCGGAAG
63618 TATGAGTGTGGAGTCGTTGCCCATCACCTTTCTGGGGAAAAGTNNNAGATCGGAAG
122020 TATGAGTGTGGAGTCGTTGCTGTGAAAACTCAGGGTACTGCCCNNNAGATCGGAAG
69171 TATGAGTGTGGAGTCGTTGCTCCCACCTCATCTTTCCCTCTTTNNNAGATCGGAAGAG
49921 TATGAGTGTGGAGTCGTTGCCTGCTTTTCTGCAGATGGCTTTCNNNAGATCGGAAGA
40321 TATGAGTGTGGAGTCGTTGCTCTTCACATAGAACATAGTGTGTGANNNAGATCGGA
108972 TATGAGTGTGGAGTCGTTGCTTTGGTGATGTGTTTAGCATGAAGCNNNAGATCGGA
29945 TATGAGTGTGGAGTCGTTGCTCAAGACTGCTTCTAACCTCTGTNNNAGATCGGAAGA
30408 TATGAGTGTGGAGTCGTTGCGGGTGGGGGAATAGAATTGCAAANNNAGATCGGAA
45423 TATGAGTGTGGAGTCGTTGCTCTCTGTCTGTACCCATCCATCCNNNAGATCGGAAGA[illegible]
64820 TATGAGTGTGGAGTCGTTGCAGAGGTTCAGATTTGACCCAGGTNNNAGATCGGAA[illegible]
62381 TATGAGTGTGGAGTCGTTGCGCTGTTCTTATGTGTTTGTAGTGCTNNNAGATCGGAA
53809 TATGAGTGTGGAGTCGTTGCTGAACTTTGCCTTTGTCCTAGGGNNNAGATCGGAAG
110734 TATGAGTGTGGAGTCGTTGCTCCCCTCAGCAAAGATGCCAATNNNAGATCGGAAGA[illegible]
53667 TATGAGTGTGGAGTCGTTGCGCAGGAAGATGAAAGGTACAGGGNNNAGATCGGAA
55436 TATGAGTGTGGAGTCGTTGCAGCCAGACTGTGATATTGCCATGNNNAGATCGGAAG
68949 TATGAGTGTGGAGTCGTTGCCCCCCAGAAGGGTGGATTTATTTNNNAGATCGGAAG
34812 TATGAGTGTGGAGTCGTTGCTGAACACACATTCACAGCATCCANNNAGATCGGAAGA
112147 TATGAGTGTGGAGTCGTTGCAGCAACTGTCTGCAAACCATGAANNNAGATCGGAAG
30501 TATGAGTGTGGAGTCGTTGCGTGATGGGTGGTGCTTACAGAAANNNAGATCGGAA
63934 TATGAGTGTGGAGTCGTTGCTCACAGATTGAAAAGGAACCAGTNNNAGATCGGAA[illegible]
48547 TATGAGTGTGGAGTCGTTGCGGTGAGAGACGATGCACAGTATCNNNAGATCGGAA
33778 TATGAGTGTGGAGTCGTTGCGCCTCATTCTCCCTGTAGATACANNNAGATCGGAAGA
51622 TATGAGTGTGGAGTCGTTGCTTACCTGTTCCTCACTGGAGGAGNNNAGATCGGAAG
106661 TATGAGTGTGGAGTCGTTGCCTTTGCCTTTGTCCTAGGGAGCNNNAGATCGGAAGA
32833 TATGAGTGTGGAGTCGTTGCAACCTGGTATGAAGCAAGCAACANNNAGATCGGAA[illegible]
45084 TATGAGTGTGGAGTCGTTGCATAAAAACTAACAGGCCAGGGCCNNNAGATCGGAA[illegible]
42627 TATGAGTGTGGAGTCGTTGCTTGCATAGTGTTTGGAGGGACACNNNAGATCGGAA[illegible]
38515 TATGAGTGTGGAGTCGTTGCCCATTATTCTTTCCCCAGCACGANNNAGATCGGAAGA
39565 TATGAGTGTGGAGTCGTTGCTTGCATTGCTGTCATCACAATGANNNAGATCGGAAGA
108519 TATGAGTGTGGAGTCGTTGCGCCACACCCTCTGATCTCTTTACNNNAGATCGGAAGA
101635 TATGAGTGTGGAGTCGTTGCCCTCATCTTTCCCTCTTTCTTTCAANNNAGATCGGAAG

| | |
|---|---|
| 39965 | TATGAGTGTGGAGTCGTTGCGTTCTCATAGTTCTGGACGCTGANNNAGATCGGAAG |
| 33989 | TATGAGTGTGGAGTCGTTGCCTGCAATATTCCCAACAGAACCANNNAGATCGGAAG |
| 40643 | TATGAGTGTGGAGTCGTTGCTGCGTTGGGGCAGAATTTCAATANNNAGATCGGAA |
| 58724 | TATGAGTGTGGAGTCGTTGCATTGTCTGTGTGATGAGCTGCATNNNAGATCGGAA |
| 42070 | TATGAGTGTGGAGTCGTTGCGCATTTGTGTGGGTGCATGTTTANNNAGATCGGAA |
| 59056 | TATGAGTGTGGAGTCGTTGCTGTATCCCAGTTGCTCTGGAGATNNNAGATCGGAAG |
| 32167 | TATGAGTGTGGAGTCGTTGCATAGCCTCTTCCCACATGGTGAANNNAGATCGGAAG |
| 99631 | TATGAGTGTGGAGTCGTTGCGGGTAGGAGAGTTGGGCATGANNNAGATCGGAAG |
| 52978 | TATGAGTGTGGAGTCGTTGCACAGATTATCAGAAAGCGCCAGGNNNAGATCGGAA |
| 62791 | TATGAGTGTGGAGTCGTTGCTTCTCTGAGCTGTTGGTCACTCTNNNAGATCGGAAG |
| 29823 | TATGAGTGTGGAGTCGTTGCTGGTCGGGGAGCAGTATATTCATNNNAGATCGGAA |
| 31326 | TATGAGTGTGGAGTCGTTGCCCAGTAGCAGTGTCTTTCTGCAANNNAGATCGGAAG |
| 53222 | TATGAGTGTGGAGTCGTTGCTATTTATTTCGGGGGAGGGGAGGNNNAGATCGGA |
| 34670 | TATGAGTGTGGAGTCGTTGCACAGTGCTAATTGCTAATGGCCANNNAGATCGGAAG |
| 37277 | TATGAGTGTGGAGTCGTTGCTCTGCAACATTTTCTTTATCCGTTCANNNAGATCGGA |
| 99515 | TATGAGTGTGGAGTCGTTGCCTGGCTTATTTCACCGACCTCATANNNAGATCGGAAG |
| 42718 | TATGAGTGTGGAGTCGTTGCCAATAGCTTTCCTCATCCCCCACNNNAGATCGGAAGA |
| 65437 | TATGAGTGTGGAGTCGTTGCAGCAGTTCCGAGTAGTTAGTGGTNNNAGATCGGAA |
| 52758 | TATGAGTGTGGAGTCGTTGCTACCCAGTCTCACATCTCCAAGGNNNAGATCGGAAG |
| 34714 | TATGAGTGTGGAGTCGTTGCTCCTGTTCCCAACTTTATATGCCANNNAGATCGGAAG |
| 51030 | TATGAGTGTGGAGTCGTTGCAAACAGATGAAGAGCCTCAGCAGNNNAGATCGGAA |
| 108492 | TATGAGTGTGGAGTCGTTGCTGGTATAAAAACCTTTTTGCGGCNNNAGATCGGAAG |
| 62176 | TATGAGTGTGGAGTCGTTGCGAAAACTAGTGCAGTCATTTGGCTNNNAGATCGGA |
| 50034 | TATGAGTGTGGAGTCGTTGCGGCAAGAGCTGACTTTGCAAAAGNNNAGATCGGAA |
| 56528 | TATGAGTGTGGAGTCGTTGCGCATTTGTTGTCCTCCATGAAGTGNNNAGATCGGAA |
| 67703 | TATGAGTGTGGAGTCGTTGCGTCATGCTCATTCCACTTCCCTTNNNAGATCGGAAGA |
| 53107 | TATGAGTGTGGAGTCGTTGCGAGGGAGCACAAGAGATTTCAGGNNNAGATCGGA |
| 32278 | TATGAGTGTGGAGTCGTTGCGCCCTTGCCATTGTCAGAGATAANNNAGATCGGAAG |
| 43773 | TATGAGTGTGGAGTCGTTGCAACAGACACACACACTACCAACCNNNAGATCGGAAG |
| 33685 | TATGAGTGTGGAGTCGTTGCGCTCCTTTGCTCTAATCCTGACANNNAGATCGGAAGA |
| 42244 | TATGAGTGTGGAGTCGTTGCTACCTCAAAGCAGGGCAGTAAACNNNAGATCGGAA |
| 100267 | TATGAGTGTGGAGTCGTTGCCTGGTGACAAAAAGGGTAAGGACTNNNAGATCGGA |
| 29820 | TATGAGTGTGGAGTCGTTGCGCCCTGTATAGACGTACACCAACNNNAGATCGGAAG |
| 29942 | TATGAGTGTGGAGTCGTTGCTCCCAGAAGCAGATGTAGAGGAGNNNAGATCGGAA |
| 68536 | TATGAGTGTGGAGTCGTTGCTTGCACGTTACTGAGGTTTGGTTNNNAGATCGGAAG |
| 102539 | TATGAGTGTGGAGTCGTTGCTTCATATTCTTTGCCCTTTCCCAAATTNNNAGATCGGA |
| 67251 | TATGAGTGTGGAGTCGTTGCCTGCTGTGGAAAGGGAGAAGATTNNNAGATCGGA |
| 109905 | TATGAGTGTGGAGTCGTTGCTTGTGTGTCTTGATGGATTCAGAGNNNAGATCGGA |
| 57643 | TATGAGTGTGGAGTCGTTGCGTGCTTTGCCATTGTTGCTTTTGNNNAGATCGGAAG |
| 101791 | TATGAGTGTGGAGTCGTTGCAGTGTCTCTAGCAGCTGAAAGTANNNAGATCGGAA |
| 52227 | TATGAGTGTGGAGTCGTTGCGAGTGAACCAGGAGGTGCTTTAGNNNAGATCGGAA |
| 101678 | TATGAGTGTGGAGTCGTTGCAGACAGACGGGAACAGACAAAGGNNNAGATCGGA |
| 40232 | TATGAGTGTGGAGTCGTTGCGGAAATGCTTGCTAGACTGGTGANNNAGATCGGAA |

62287 TATGAGTGTGGAGTCGTTGCATCTTATTCCCGGATGGACTGCTNNNAGATCGGAAG
30351 TATGAGTGTGGAGTCGTTGCGGCTGACTGGAGAGTTTTCCAAANNNAGATCGGAA
48589 TATGAGTGTGGAGTCGTTGCCATAAAGGGGCCAGTCTGAACTCNNNAGATCGGAA
36834 TATGAGTGTGGAGTCGTTGCCCACAACCTCCACAATGTTCTCANNNAGATCGGAAGA
95105 TATGAGTGTGGAGTCGTTGCTAGCTTCTTTCAGCTTCCTCCCTNNNAGATCGGAAGA
48290 TATGAGTGTGGAGTCGTTGCTTCCTGGATCTCCCCTGAGAATCNNNAGATCGGAAG
70469 TATGAGTGTGGAGTCGTTGCTTCTTATAAAGCTCCCCACGATGCNNNAGATCGGAA
64782 TATGAGTGTGGAGTCGTTGCACTGGAGATGTGGCTATTCTAAGGTNNNAGATCGG
48627 TATGAGTGTGGAGTCGTTGCTGGTAGGTGATTGTGCTTCACTCNNNAGATCGGAAG
51482 TATGAGTGTGGAGTCGTTGCACCCATCAATCTAAAGGCGAGAGNNNAGATCGGAA
62043 TATGAGTGTGGAGTCGTTGCGAGAAATCTGTAGGTTTGGGGCTNNNAGATCGGAA
104951 TATGAGTGTGGAGTCGTTGCCCGAGGGCCAACCATACTATGTNNNAGATCGGAAG
43459 TATGAGTGTGGAGTCGTTGCGGGTGGGTCAATTCTTTGTTGACNNNAGATCGGAA
42400 TATGAGTGTGGAGTCGTTGCTTGATTTTGCCTCTAGGGGGAACNNNAGATCGGAA
59623 TATGAGTGTGGAGTCGTTGCCATTTGGCTGGGGTCAATGGTATNNNAGATCGGAA
117536 TATGAGTGTGGAGTCGTTGCGCCTGTGGAACTGTTAGTCCTTANNNAGATCGGAAG
62971 TATGAGTGTGGAGTCGTTGCTCTAAACCTCATCCAATATTATGCTCTNNNAGATCGG
49258 TATGAGTGTGGAGTCGTTGCCAGGGAAGCACATGCCTATAGTCNNNAGATCGGAA
64888 TATGAGTGTGGAGTCGTTGCACTGAAGGAAGGGCATTAGAGGTNNNAGATCGGA
48518 TATGAGTGTGGAGTCGTTGCCGAATGGCTTTTGGTGCCATATCNNNAGATCGGAAG
43749 TATGAGTGTGGAGTCGTTGCAGACGTACAATGCTCCTCAAACCNNNAGATCGGAAG
63300 TATGAGTGTGGAGTCGTTGCCTTTCTGGCAACTCCCTTTGTCTNNNAGATCGGAAGA
67977 TATGAGTGTGGAGTCGTTGCATTCCACCTTGACTTTGGTGCTTNNNAGATCGGAAG
40173 TATGAGTGTGGAGTCGTTGCTGATCTACCCTGTGCTTACGTGANNNAGATCGGAAG
69537 TATGAGTGTGGAGTCGTTGCGGACAGACCCAACTCCAAGTTTTNNNAGATCGGAAG
67582 TATGAGTGTGGAGTCGTTGCGTCCTCACCACAAACTTGGACTTNNNAGATCGGAAG
117595 TATGAGTGTGGAGTCGTTGCAGTTTTAGGAGCCTGAGGGTTTTNNNAGATCGGAA
101051 TATGAGTGTGGAGTCGTTGCTGTGGGAGGTGGGAATACAAACNNNAGATCGGAA
48490 TATGAGTGTGGAGTCGTTGCTATGACACCAGCCGTATTGGATCNNNAGATCGGAAG
38344 TATGAGTGTGGAGTCGTTGCGGTATTGCTGTCTTGCTGTGAGANNNAGATCGGAA
63589 TATGAGTGTGGAGTCGTTGCACCCAAACTGATACACATGTTTTCTNNNAGATCGGAA
54590 TATGAGTGTGGAGTCGTTGCATAGGAGAGGAGAAACACCCTGGNNNAGATCGGA
47470 TATGAGTGTGGAGTCGTTGCTCGGAAGGCTAACAGAAGAATGCNNNAGATCGGAA
108202 TATGAGTGTGGAGTCGTTGCCAGACCATGAGTACCCTTCAAAGAANNNAGATCGGA
30632 TATGAGTGTGGAGTCGTTGCGCTTTATGGTAAGGGGCTGGAAANNNAGATCGGAA
52216 TATGAGTGTGGAGTCGTTGCGACCACCAATACAGGCAGGTTAGNNNAGATCGGAA
37745 TATGAGTGTGGAGTCGTTGCACCAACATACAGACAGTTCAACAGANNNAGATCGGA
44081 TATGAGTGTGGAGTCGTTGCTATTTCATTTGGGCAGCTGGACCNNNAGATCGGAAG
67746 TATGAGTGTGGAGTCGTTGCTTCACAAAGACAAGTACTGCCTTNNNAGATCGGAAG
34427 TATGAGTGTGGAGTCGTTGCAAGGAAAGCCAAAGTAGATCCCANNNAGATCGGAA
56856 TATGAGTGTGGAGTCGTTGCGGAGACAAATGTGTGTTGTGGTGNNNAGATCGGA
40478 TATGAGTGTGGAGTCGTTGCCCAATGTAGCACCAAGAGGTTGANNNAGATCGGAA
66473 TATGAGTGTGGAGTCGTTGCGGGTGAGAGAGAGAAGAGTGTGTNNNAGATCGGA

| | |
|---|---|
| 29944 | TATGAGTGTGGAGTCGTTGCCCATAGTAGCCTGGAACCACAGANNNAGATCGGAA |
| 47884 | TATGAGTGTGGAGTCGTTGCACACAGACATACTGAAGCAGTGCNNNAGATCGGAA |
| 37000 | TATGAGTGTGGAGTCGTTGCGTCACTTTCAATGTCATGATGTCANNNAGATCGGAA |
| 29943 | TATGAGTGTGGAGTCGTTGCGGCTGCACTTCAGAAATTTGGAGNNNAGATCGGAA |
| 29941 | TATGAGTGTGGAGTCGTTGCCTATTTGCCCCTAAAGCCTCTGGNNNAGATCGGAAG |
| 30279 | TATGAGTGTGGAGTCGTTGCACCCTAAAACTTAGTGGCTTACAAANNNAGATCGG |
| 71832 | TATGAGTGTGGAGTCGTTGCCCACCCATGCCTTCTTCTCTTTTNNNAGATCGGAAGA |
| 55544 | TATGAGTGTGGAGTCGTTGCTTAGCGATGCAGATGGGAAGATGNNNAGATCGGA |
| 71835 | TATGAGTGTGGAGTCGTTGCCGGCTCCTCAGATAATGAGTACTNNNAGATCGGAAG |
| 71834 | TATGAGTGTGGAGTCGTTGCTCACCATTTGTCTTTGCAGGGAANNNAGATCGGAAG |
| 52928 | TATGAGTGTGGAGTCGTTGCACCCTGCCTTTGGAAATTACAGGNNNAGATCGGAAG |
| 58355 | TATGAGTGTGGAGTCGTTGCAACACTTTGCAACTGCAAGACATNNNAGATCGGAAG |
| 71831 | TATGAGTGTGGAGTCGTTGCAAAATGGAGTGGATTGGGGTGTCNNNAGATCGGA |
| 30947 | TATGAGTGTGGAGTCGTTGCTCATGTGGGAGGTGGGAATACAANNNAGATCGGA |
| 71750 | TATGAGTGTGGAGTCGTTGCATAAATAAGCTGCAGGCAGGGTCNNNAGATCGGAA |
| 100687 | TATGAGTGTGGAGTCGTTGCCTGTAACCTCACACTGGGCTTANNNAGATCGGAAGA |
| 50798 | TATGAGTGTGGAGTCGTTGCTTAGCATGGAAGACACTCACCAGNNNAGATCGGAA |
| 102241 | TATGAGTGTGGAGTCGTTGCGAATGGCTAGAAATGTGACCAAANNNAGATCGGAA |
| 70456 | TATGAGTGTGGAGTCGTTGCGGTTTTGACAAACTGCATCTGGCNNNAGATCGGAAG |
| 55266 | TATGAGTGTGGAGTCGTTGCCTTTAGCAGGACACTGCACAATGNNNAGATCGGAAG |
| 100793 | TATGAGTGTGGAGTCGTTGCTCAGAGAAAGAGGCAGAGAAACCTNNNAGATCGGA |
| 103263 | TATGAGTGTGGAGTCGTTGCAACCTTTCCCCAGTGGTTTCNNNAGATCGGAAGAG |
| 106617 | TATGAGTGTGGAGTCGTTGCGTCTGCACTTTCATCCAGGAGTATTNNNAGATCGGA |
| 107828 | TATGAGTGTGGAGTCGTTGCTAAAATCACAAGCCATAGAGAACATGGNNNAGATCG |
| 114615 | TATGAGTGTGGAGTCGTTGCAGAAGATGAGACTAGACTGTGAGGANNNAGATCG |
| 103688 | TATGAGTGTGGAGTCGTTGCCCATTGAGGATCCATAGCTTGAATCNNNAGATCGGA |
| 102515 | TATGAGTGTGGAGTCGTTGCTGGATGGTAACAAATTGTCAGATGGANNNAGATCG |
| 101395 | TATGAGTGTGGAGTCGTTGCAGAGGAATGCAAAGCCAAAACATAANNNAGATCGG |
| 71905 | TATGAGTGTGGAGTCGTTGCCAAGCACCTTGAACTCAAAAGNNNAGATCGGAAGA |
| 103298 | TATGAGTGTGGAGTCGTTGCAACCTCCTATGCCTCAGCAATTNNNAGATCGGAAGA |
| 104096 | TATGAGTGTGGAGTCGTTGCGTCAACCGGTCATTAGGTATTTCTNNNAGATCGGAA |
| 106169 | TATGAGTGTGGAGTCGTTGCGAGTTCTTGGTCGTGCAGAATGTNNNAGATCGGAA |
| 103905 | TATGAGTGTGGAGTCGTTGCTAAGATATGTGTGTGCGCCTATGACNNNAGATCGGA |
| 103863 | TATGAGTGTGGAGTCGTTGCTTGGGGCTTAGCTCAGAATAGTTNNNAGATCGGAA |
| 107056 | TATGAGTGTGGAGTCGTTGCTGTCAAAGAATCAGCAGTGAGCAAGANNNAGATCG |
| 107834 | TATGAGTGTGGAGTCGTTGCAGTTGCATGGATAGTTGTTCAAATCGNNNAGATCGG |
| 113345 | TATGAGTGTGGAGTCGTTGCCTTTGTATTGTATTCAGGAGGCACANNNAGATCGGA |
| 107083 | TATGAGTGTGGAGTCGTTGCAAGATACATGAAACGGAGGCANNNAGATCGGAAGA |
| 108515 | TATGAGTGTGGAGTCGTTGCTCAGCGCATGTCTATTGGTTTAGNNNAGATCGGAAG |
| 115835 | TATGAGTGTGGAGTCGTTGCTTCTTTCTGCATGGTCTTGTTCTGTNNNAGATCGGAA |
| 96082 | TATGAGTGTGGAGTCGTTGCGATACGGGGTTCCAGGAAAGGNNNAGATCGGAAGA |
| 99345 | TATGAGTGTGGAGTCGTTGCAGAAAAGCCACCTCACCTCCCTNNNAGATCGGAAGA |
| 96359 | TATGAGTGTGGAGTCGTTGCCTTCCAATTTAGTTGGGCCAGCCNNNAGATCGGAAG |

| | |
|---|---|
| 108427 | TATGAGTGTGGAGTCGTTGCGTGTGTGTAAACAAGTCCTGTCTCNNNAGATCGGAA |
| 62779 | TATGAGTGTGGAGTCGTTGCACCAGTTACTTTGTATCCCACTCTNNNAGATCGGAAG |
| 60893 | TATGAGTGTGGAGTCGTTGCAGATCACCTAAAACCTACCCCCTNNNAGATCGGAAGA |
| 60719 | TATGAGTGTGGAGTCGTTGCTCACTTACACGGTGTTTTCACCTNNNAGATCGGAAGA |
| 45045 | TATGAGTGTGGAGTCGTTGCAGTTTCCCTATTTTTCCCAGGCCNNNAGATCGGAAGA |
| 68163 | TATGAGTGTGGAGTCGTTGCCCCTCTCTGTAACTCTGCTTCTTNNNAGATCGGAAGAC |
| 103269 | TATGAGTGTGGAGTCGTTGCCAGGGTATCTATCACCTAGGCTGNNNAGATCGGAAG |
| 48118 | TATGAGTGTGGAGTCGTTGCTGTAGGTCTGTGTATGTGCTTGCNNNAGATCGGAAG |
| 41293 | TATGAGTGTGGAGTCGTTGCGTGTTCAAGGCTAGGAGGGTCTANNNAGATCGGAA |
| 54125 | TATGAGTGTGGAGTCGTTGCGGGTATCTATCACCTAGGCTGGGNNNAGATCGGAAC |
| 39031 | TATGAGTGTGGAGTCGTTGCTGGAGTGGAGGTTAGTAGAGGGANNNAGATCGGA |
| 33429 | TATGAGTGTGGAGTCGTTGCCCCCAGAAATCTGACAGTTCACANNNAGATCGGAAGA |
| 96036 | TATGAGTGTGGAGTCGTTGCTCTATTTAGTCATGCCACCGCTCAANNNAGATCGGAA |
| 52111 | TATGAGTGTGGAGTCGTTGCAGTGTGGTAGTATGCGCCAGTAGNNNAGATCGGAA |
| 65726 | TATGAGTGTGGAGTCGTTGCGTAAGTGCATGTATAGGGGCATGTNNNAGATCGGA |
| 46208 | TATGAGTGTGGAGTCGTTGCAACTCCCTCTTCATGTTGGAAGCNNNAGATCGGAAGA |
| 40940 | TATGAGTGTGGAGTCGTTGCGGCACAGGCAGTAAGAACAACTANNNAGATCGGAAC |
| 107736 | TATGAGTGTGGAGTCGTTGCATGTGTGTAAACAAGGCCTGTCTNNNAGATCGGAAG |
| 29816 | TATGAGTGTGGAGTCGTTGCCACTCTACCTCCATGAAGAGCACNNNAGATCGGAAGA |
| 58159 | TATGAGTGTGGAGTCGTTGCAGGGAATGTGGGGGAGACATAATNNNAGATCGGAA |
| 32793 | TATGAGTGTGGAGTCGTTGCTACATACCACCTCCCACACAACANNNAGATCGGAAGA |
| 70852 | TATGAGTGTGGAGTCGTTGCTGGATTGGCCAGATGTTCTGATGNNNAGATCGGAAC |
| 40982 | TATGAGTGTGGAGTCGTTGCCAACCCAAAGTGGTGCAAGACTANNNAGATCGGAAG |
| 31921 | TATGAGTGTGGAGTCGTTGCGCAACCTTAGAGGCAAATGGGAANNNAGATCGGAA |
| 45200 | TATGAGTGTGGAGTCGTTGCAAGGATCATGTGAATGGGATGCCNNNAGATCGGAA |
| 37955 | TATGAGTGTGGAGTCGTTGCGCTGTGTAAAGATGAAGGGCAGANNNAGATCGGAA |
| 53030 | TATGAGTGTGGAGTCGTTGCCTGCCTTTGGTTTTGATAGCAGGNNNAGATCGGAAG |
| 37673 | TATGAGTGTGGAGTCGTTGCATGTGCTGGCAAGAAAGTGAAGANNNAGATCGGAA |
| 32379 | TATGAGTGTGGAGTCGTTGCGAGAGAGCGCGCTAAAAAGCTAANNNAGATCGGAA |
| 42242 | TATGAGTGTGGAGTCGTTGCAGGCAGGAGTTGCTGTTCTAAACNNNAGATCGGAAC |
| 41797 | TATGAGTGTGGAGTCGTTGCCCTGGCACAGTGTTGAGTACTTANNNAGATCGGAAG |
| 63558 | TATGAGTGTGGAGTCGTTGCGCACCTACCCTTTCTCTGTTTCTNNNAGATCGGAAGAC |
| 66658 | TATGAGTGTGGAGTCGTTGCAGACAGACTAGGAAAAAGTGCTTGTNNNAGATCGGA |
| 67398 | TATGAGTGTGGAGTCGTTGCGTGGGGGAGTAATTATTGGCTATTNNNAGATCGGAA |
| 54303 | TATGAGTGTGGAGTCGTTGCCCCTCCCTCCTAAAACCAAATGGNNNAGATCGGAAGA |
| 61248 | TATGAGTGTGGAGTCGTTGCTTGCATCTAAAACCATGCTGCCTNNNAGATCGGAAGA |
| 59365 | TATGAGTGTGGAGTCGTTGCGGTGGGTTCTGTGCTTTCTTGATNNNAGATCGGAAG |
| 37693 | TATGAGTGTGGAGTCGTTGCCAGGAAGGGTGGCATAACTAAGANNNAGATCGGAA |
| 50287 | TATGAGTGTGGAGTCGTTGCCACACACAAACACATCCAGCAAGNNNAGATCGGAAG |
| 42156 | TATGAGTGTGGAGTCGTTGCCTGAAGTTGGGAGAGGGGAAAACNNNAGATCGGAA |
| 30020 | TATGAGTGTGGAGTCGTTGCGTGGCCTGGTAGGGGGAAAAANNNAGATCGGAAGA |
| 34615 | TATGAGTGTGGAGTCGTTGCAGGACTAATTCCCTTGCTAGCCANNNAGATCGGAAGA |
| 37160 | TATGAGTGTGGAGTCGTTGCATGCACAATTTCCTTGCTTATTCANNNAGATCGGAAG |

68658 TATGAGTGTGGAGTCGTTGCTGTGTGTGTCCTGAAACAGTGTTNNNAGATCGGAAG
36421 TATGAGTGTGGAGTCGTTGCGCACACATCCATACATCCCATCANNNAGATCGGAAGA
63838 TATGAGTGTGGAGTCGTTGCGTAGTGCTTGAGCATCAGTAAGTNNNAGATCGGAA
60696 TATGAGTGTGGAGTCGTTGCACCATGACTGAAACATTGCACCTNNNAGATCGGAAG
56471 TATGAGTGTGGAGTCGTTGCCACAGACCCTTGGTGATCTTCTGNNNAGATCGGAAG
44523 TATGAGTGTGGAGTCGTTGCTCTTTTCACTGTAATCCCAGCCCNNNAGATCGGAAGA
38204 TATGAGTGTGGAGTCGTTGCAGAGAGGGATGAGGAGAAGGAGANNNAGATCGG
52940 TATGAGTGTGGAGTCGTTGCAGAGAGTTGTGAGAGGTACCAGGNNNAGATCGGA
37250 TATGAGTGTGGAGTCGTTGCGTGCATGCTGTAGTCATTTCTTCANNNAGATCGGAA
37738 TATGAGTGTGGAGTCGTTGCTGGAGTGGGAGATTAGAAACAGANNNAGATCGGA
30808 TATGAGTGTGGAGTCGTTGCGCTCTGGCATCAGGGATAAACAANNNAGATCGGAA
67487 TATGAGTGTGGAGTCGTTGCGCCAACAGGTCTTCTCAGAACTTNNNAGATCGGAAG
43116 TATGAGTGTGGAGTCGTTGCTCAGGAAGAGCAAGAGCAAAGACNNNAGATCGGAA
46832 TATGAGTGTGGAGTCGTTGCTCTGAGGAATTGTACTGCAAGGCNNNAGATCGGAA
38486 TATGAGTGTGGAGTCGTTGCTGGCTTTTGGTGGAACAGTTAGANNNAGATCGGAA
44175 TATGAGTGTGGAGTCGTTGCTCCCAAGAGCCTAGACTAGTACCNNNAGATCGGAAG
95149 TATGAGTGTGGAGTCGTTGCAAGGCCAATAGTAACTCCTGGTGNNNAGATCGGAAG
58910 TATGAGTGTGGAGTCGTTGCCTCTCTCTCTCCACCGAGTTCATNNNAGATCGGAAGA
107823 TATGAGTGTGGAGTCGTTGCATCACATCAGTTAGAAACCCCTGTTTCNNNAGATCGG
38179 TATGAGTGTGGAGTCGTTGCAGACACACATTCAGAGGCTAGAGANNNAGATCGGA
98036 TATGAGTGTGGAGTCGTTGCAGATGGAGCCTGTTGGTGTTTAGNNNAGATCGGAA
54271 TATGAGTGTGGAGTCGTTGCCCTCTTAGACCCCAACATTTGGGNNNAGATCGGAAG
49341 TATGAGTGTGGAGTCGTTGCTGAATTGCTGGAGGCTTATGGTCNNNAGATCGGAA
65931 TATGAGTGTGGAGTCGTTGCTTCCTTTCTCTGACCCAGACTGTNNNAGATCGGAAGA
29814 TATGAGTGTGGAGTCGTTGCTGATTGACAATGCTGTCAAGGCTNNNAGATCGGAAG
46833 TATGAGTGTGGAGTCGTTGCGGACATAAGCAAATCTGCAAGGCNNNAGATCGGAA
66847 TATGAGTGTGGAGTCGTTGCCAGCCGAAATATTCATTTGTGTTTGTNNNAGATCGG
34134 TATGAGTGTGGAGTCGTTGCCATAAGCAAGCAAGCAATGACCANNNAGATCGGAAG
62138 TATGAGTGTGGAGTCGTTGCAAGTCTGATTACCCTGTCTGGCTNNNAGATCGGAAG
66081 TATGAGTGTGGAGTCGTTGCTTCTAACTCTGGGCTATGGCTGTNNNAGATCGGAAG
41866 TATGAGTGTGGAGTCGTTGCCCTGCACAAGTGTGTCGATCTTANNNAGATCGGAAG
107079 TATGAGTGTGGAGTCGTTGCCATAAGCAAGCAAGCAATGACCAGNNNAGATCGGA
67644 TATGAGTGTGGAGTCGTTGCCCCTTCCTACCTCATAGCACCTTNNNAGATCGGAAGA
33427 TATGAGTGTGGAGTCGTTGCAGTGGGTTTTCATCTGTCTTCACANNNAGATCGGAA
104072 TATGAGTGTGGAGTCGTTGCAATCTTTTCTTCCTCCTTTCCTTCCNNNAGATCGGAAG
43162 TATGAGTGTGGAGTCGTTGCCCTACACCTTCTCCCTACCAGACNNNAGATCGGAAGA
116731 TATGAGTGTGGAGTCGTTGCCTCTCTCTCTCTCTCTGTTTGAAGAGNNNAGATCGGA
58684 TATGAGTGTGGAGTCGTTGCAACATGAAAAGAAGCCTGGGCATNNNAGATCGGAA
63619 TATGAGTGTGGAGTCGTTGCTTTAGGCATGTTTACTGGAAAAGTNNNAGATCGGAA
60334 TATGAGTGTGGAGTCGTTGCCTTACCCACCTACATACCGGACTNNNAGATCGGAAGA
105502 TATGAGTGTGGAGTCGTTGCTGGTCCTGTTGGAAGAGGAGTTANNNAGATCGGAA
39734 TATGAGTGTGGAGTCGTTGCACAAAAACAAAACCCTGTGATGANNNAGATCGGAAG
95127 TATGAGTGTGGAGTCGTTGCTAAGTACCCTCTTCTCCCTCCCTNNNAGATCGGAAGA

42111 TATGAGTGTGGAGTCGTTGCGCCTAGTGGGGAGAGAGGTTTTANNNAGATCGGAA
32664 TATGAGTGTGGAGTCGTTGCCCCTACCTCAAAAGGGCAAAACANNNAGATCGGAAG
68783 TATGAGTGTGGAGTCGTTGCCAGCAGCTAGGGAACAGGAATTTNNNAGATCGGAA
100942 TATGAGTGTGGAGTCGTTGCGGTTTTCATCTGTCTTCACATCANNNAGATCGGAAG
56648 TATGAGTGTGGAGTCGTTGCGGATGCACAGTCAGAGTGAGTGNNNAGATCGGAA
104070 TATGAGTGTGGAGTCGTTGCGTGTGTTCTTTGGCCTTGTTTTNNNAGATCGGAAGA
62892 TATGAGTGTGGAGTCGTTGCCTCACCTGGTCTATTGCTCCTCTNNNAGATCGGAAGA
47366 TATGAGTGTGGAGTCGTTGCTATGAAGAGGCAGTGTTTGTGGCNNNAGATCGGAA
36731 TATGAGTGTGGAGTCGTTGCCATACTGATCTCAGAGCTGCTCANNNAGATCGGAAG
43254 TATGAGTGTGGAGTCGTTGCTATGAAGACATGTGACCAGCGACNNNAGATCGGAA
31035 TATGAGTGTGGAGTCGTTGCTTTCAGGCAAATTACAAGCCCAANNNAGATCGGAAG
31993 TATGAGTGTGGAGTCGTTGCTGGGTGTGGGTTATATGGTGGAANNNAGATCGGA
50360 TATGAGTGTGGAGTCGTTGCTGGACCTCTACCCAGTCTTCAAGNNNAGATCGGAAG
44826 TATGAGTGTGGAGTCGTTGCTCGCTTCTTTTCTCACTGTTCCCNNNAGATCGGAAGA
35614 TATGAGTGTGGAGTCGTTGCTTTTCAAGATTCACAGCCAGGCANNNAGATCGGAAG
43584 TATGAGTGTGGAGTCGTTGCAGACCCCCAGCATACTGATCTACNNNAGATCGGAAG
33449 TATGAGTGTGGAGTCGTTGCTGGTGGTTGTGGTCAAAAAGACANNNAGATCGGAA
44730 TATGAGTGTGGAGTCGTTGCCTTACATGCTCACACAGTCTCCCNNNAGATCGGAAGA
45382 TATGAGTGTGGAGTCGTTGCTCCTCAGTTGCTCTTCTCAAATCCNNNAGATCGGAAG
46738 TATGAGTGTGGAGTCGTTGCGAGGATGATAAGCTGAAGGACGCNNNAGATCGGA
43201 TATGAGTGTGGAGTCGTTGCGACAAAAATACAGTTGTTGAGAGAGACNNNAGATC
104822 TATGAGTGTGGAGTCGTTGCTTGGAACCTTCAGAATAGTTACCTNNNAGATCGGAA
58085 TATGAGTGTGGAGTCGTTGCATCACTGAAATTTTGGCAGGAATNNNAGATCGGAA
66679 TATGAGTGTGGAGTCGTTGCCAGATGACGTGTGTCCTTCTTGTNNNAGATCGGAAG
56978 TATGAGTGTGGAGTCGTTGCCTGCACACCATTGTGAATCTGTGNNNAGATCGGAAG
44420 TATGAGTGTGGAGTCGTTGCTAGAATAGCAAAAAGCGAGCCCCNNNAGATCGGAA
33402 TATGAGTGTGGAGTCGTTGCAGTTCTGGAGAGATTTGGTCACANNNAGATCGGAA
66829 TATGAGTGTGGAGTCGTTGCTTTCTCAGCACCCAGTAGTTTGTNNNAGATCGGAAG
65979 TATGAGTGTGGAGTCGTTGCCATAAAAGACACTGCCACCCTGTNNNAGATCGGAAG
65391 TATGAGTGTGGAGTCGTTGCGAGCCCATCAAAAGACATCTGGTNNNAGATCGGAA
42290 TATGAGTGTGGAGTCGTTGCGAAGTCAGAAAAATGCCCCCAACNNNAGATCGGAA
31779 TATGAGTGTGGAGTCGTTGCAGTTAGCATGGGGAGATCTCGAANNNAGATCGGAA
33705 TATGAGTGTGGAGTCGTTGCCTGAGACTAGGAAGCAGGTGACANNNAGATCGGAA
33835 TATGAGTGTGGAGTCGTTGCTCCTATCACTTCCCTGTTGCTACANNNAGATCGGAAG
102896 TATGAGTGTGGAGTCGTTGCGCTGTGAAGTCAAGGTCACTATTCCNNNAGATCGGA
49282 TATGAGTGTGGAGTCGTTGCCAAACAGGAAAAGGAAAGCCGTCNNNAGATCGGAA
40536 TATGAGTGTGGAGTCGTTGCGGGTGCTGTATTACTGGCTTTGANNNAGATCGGAA
34778 TATGAGTGTGGAGTCGTTGCGCAGACATAAAGACCTTTTTGCCANNNAGATCGGAA
95132 TATGAGTGTGGAGTCGTTGCAAAGGATCGAATGGCCAAGTACCNNNAGATCGGAA
50569 TATGAGTGTGGAGTCGTTGCGGGGTGATGGTTTTCCAGGTAAGNNNAGATCGGAA
60873 TATGAGTGTGGAGTCGTTGCGTTTCCTTGAAGGTTTATTTTACCCTNNNAGATCGGA
38822 TATGAGTGTGGAGTCGTTGCTAACTACAGCAACGGAAGGAGGANNNAGATCGGAA
56741 TATGAGTGTGGAGTCGTTGCATTTTTGCCACATCATGCAGGTGNNNAGATCGGAAG

66941 TATGAGTGTGGAGTCGTTGCGAAGTACCAGGTTCCAGCCAATTNNNAGATCGGAAG
102483 TATGAGTGTGGAGTCGTTGCTGTTAGGTTCCGTGTGTGTGTGNNNAGATCGGAAGA
70912 TATGAGTGTGGAGTCGTTGCCTGTCTGTCTATTTGGGTGGGTGNNNAGATCGGAAG
42556 TATGAGTGTGGAGTCGTTGCCCTTTCGGTTATACACACACACACNNNAGATCGGAAG
58381 TATGAGTGTGGAGTCGTTGCTGCTACTTTTGCACTTGGGACATNNNAGATCGGAAGA
46136 TATGAGTGTGGAGTCGTTGCATGAAGTGCTACTATGGCCAAGCNNNAGATCGGAAG
64435 TATGAGTGTGGAGTCGTTGCGGTTGTTCCTCCCTTCTCCTAGTNNNAGATCGGAAGA
69082 TATGAGTGTGGAGTCGTTGCGCATACCTTGGGAGTTCAGCTTTNNNAGATCGGAAG
58425 TATGAGTGTGGAGTCGTTGCATGCCAACATACGGTGAAACCATNNNAGATCGGAAG
106131 TATGAGTGTGGAGTCGTTGCGTGCAGATCATGGTTTTGTTGTTGTNNNAGATCGGA
39182 TATGAGTGTGGAGTCGTTGCGAACAACGTGAATATGGGTGGGANNNAGATCGGAA
44329 TATGAGTGTGGAGTCGTTGCTTGCACACTCCAAAAATGTACCCNNNAGATCGGAAGA
57244 TATGAGTGTGGAGTCGTTGCGACGCAGGCCTATAAGAGACTTGNNNAGATCGGAAG
96409 TATGAGTGTGGAGTCGTTGCTTTCCCTAGAATGGTAACGGGCANNNAGATCGGAAG
53806 TATGAGTGTGGAGTCGTTGCTTACAGAGGCCAGATGGATAGGGNNNAGATCGGAA
56757 TATGAGTGTGGAGTCGTTGCGCACTTTGTGTCTTTGTGAGGTGNNNAGATCGGAAG
50077 TATGAGTGTGGAGTCGTTGCCTGAAAGCTGAACAGCACCAAAGNNNAGATCGGAAG
105479 TATGAGTGTGGAGTCGTTGCTTTGGTAGTGCTGTTTTCCAGGCNNNAGATCGGAAG
59148 TATGAGTGTGGAGTCGTTGCACCTTTCATAGTGGGCTGAGGATNNNAGATCGGAAG
30629 TATGAGTGTGGAGTCGTTGCATACCAGCAGCAAACACTGGAAANNNAGATCGGAAG
53135 TATGAGTGTGGAGTCGTTGCAGCATCTGCAACTAAACCAGAGGNNNAGATCGGAAG
101060 TATGAGTGTGGAGTCGTTGCCTTTATGCGATTGTGGGATCTGGNNNAGATCGGAAG
58824 TATGAGTGTGGAGTCGTTGCCTCTTAGCCATCATGCAGCTCATNNNAGATCGGAAGA
68457 TATGAGTGTGGAGTCGTTGCTGATGGGATGTTTTTCTGGGGTTNNNAGATCGGAAG
52605 TATGAGTGTGGAGTCGTTGCTGGACTGAGGTTACAAAGTCTCGNNNAGATCGGAAG
49885 TATGAGTGTGGAGTCGTTGCGTCTCGCTCTCCCCTGTTATTTCNNNAGATCGGAAGAG
37344 TATGAGTGTGGAGTCGTTGCCGTGTAGACCAGCAGTGATTTCANNNAGATCGGAAG
58728 TATGAGTGTGGAGTCGTTGCATCGTTTAGGTAGCTTGCTGCATNNNAGATCGGAAG
37006 TATGAGTGTGGAGTCGTTGCTCTGTTCCTCTGCAGGTATGTCANNNAGATCGGAAGA
52926 TATGAGTGTGGAGTCGTTGCAAAAGAGAGGGGAGGTCTACAGGNNNAGATCGGAA
67181 TATGAGTGTGGAGTCGTTGCCTGGAGAAAGGCCTACTTGCATTNNNAGATCGGAAG
51223 TATGAGTGTGGAGTCGTTGCGCTCTGGAACTTTCATCCCTCAGNNNAGATCGGAAGA
44848 TATGAGTGTGGAGTCGTTGCTTTCCAGTGCAATTCCAAAAGCCNNNAGATCGGAAGA
47472 TATGAGTGTGGAGTCGTTGCGAGATGACCCCAGATCAGAATGCNNNAGATCGGAAG
70748 TATGAGTGTGGAGTCGTTGCGGTATTCAGCACCTTCCCTTAGGNNNAGATCGGAAGA
63408 TATGAGTGTGGAGTCGTTGCTCTTGTGGATTCTTTGGGCTTCTNNNAGATCGGAAGA
36032 TATGAGTGTGGAGTCGTTGCAGTGAACACGTGAATGTTTATGCANNNAGATCGGAA
59823 TATGAGTGTGGAGTCGTTGCTCCCAGGCTATCAACTACCAACTNNNAGATCGGAAGA
59302 TATGAGTGTGGAGTCGTTGCCCCAGACCTTTCTCTCTCCTGATNNNAGATCGGAAGAG
64123 TATGAGTGTGGAGTCGTTGCACTTAGCAGCAAAAGCAGTCAGTNNNAGATCGGAAG
41949 TATGAGTGTGGAGTCGTTGCCGCATGGACCCAGAAAACTGTTANNNAGATCGGAAG
63067 TATGAGTGTGGAGTCGTTGCGCTTCTGTTGCCTGGAATTCTCTNNNAGATCGGAAGA
109019 TATGAGTGTGGAGTCGTTGCTGTATGACTGTGTGTGTGTCTCTATGNNNAGATCGG

69303 TATGAGTGTGGAGTCGTTGCTGTTTTTCCCAGCACGTAGGTTTNNNAGATCGGAAG
50327 TATGAGTGTGGAGTCGTTGCCAGTGAATCACCTCCCCATCAAGNNNAGATCGGAAG
35567 TATGAGTGTGGAGTCGTTGCGTGCTGGACAATTTCAAAAGGCANNNAGATCGGAA
68504 TATGAGTGTGGAGTCGTTGCCCAAGTGAGTAGAGGGTCTGGTTNNNAGATCGGAA
62572 TATGAGTGTGGAGTCGTTGCGTCACCTAGGCTGTAGGGAATCTNNNAGATCGGAA
44680 TATGAGTGTGGAGTCGTTGCATTATAGCTCCTCCCACACTCCCNNNAGATCGGAAGA
36591 TATGAGTGTGGAGTCGTTGCTCTTGGAGGAGATGCAGATACTCANNNAGATCGGA
59751 TATGAGTGTGGAGTCGTTGCTGGTTCTCTTGCTTTGGAAAACTNNNAGATCGGAAG
56834 TATGAGTGTGGAGTCGTTGCCTTCTTTTAGGCCAGCCTGGTGNNNAGATCGGAAGA
61512 TATGAGTGTGGAGTCGTTGCTTCACTTTCTGCTTGCACTTCCTNNNAGATCGGAAGA
61733 TATGAGTGTGGAGTCGTTGCCACTATTCCTAGGGAGCTCAGCTNNNAGATCGGAAG
105536 TATGAGTGTGGAGTCGTTGCAAATGCAGATGTGTCATGCATACTACNNNAGATCGG
49993 TATGAGTGTGGAGTCGTTGCCCCTAGCCAACCCTGATCTTTTCNNNAGATCGGAAGA
109031 TATGAGTGTGGAGTCGTTGCATAAAGACATATCAGTGTGGCCATNNNAGATCGGA
62281 TATGAGTGTGGAGTCGTTGCTCCAGCCTCAAAATATCACTGCTNNNAGATCGGAAG
39177 TATGAGTGTGGAGTCGTTGCGTGAGACTAAGGGATACGTGGGANNNAGATCGGA
57943 TATGAGTGTGGAGTCGTTGCGCATGTATCCCTTACCCAGCAATNNNAGATCGGAAG
95104 TATGAGTGTGGAGTCGTTGCCCCGTGATCATCTCTGACCTGATNNNAGATCGGAAG
58996 TATGAGTGTGGAGTCGTTGCGTTGATAGAGCAAGGCAGCAGATNNNAGATCGGAA
63859 TATGAGTGTGGAGTCGTTGCACCCTTATGATTTGACATCAACAGTNNNAGATCGGA
63975 TATGAGTGTGGAGTCGTTGCTCCTCAAAGATCTTGCAATCCAGTNNNAGATCGGAA
30115 TATGAGTGTGGAGTCGTTGCCCATTCCACCTGGTCACAGAAAANNNAGATCGGAAG
36643 TATGAGTGTGGAGTCGTTGCCATCTCTCACCACTTTTGCCTCANNNAGATCGGAAGA
97670 TATGAGTGTGGAGTCGTTGCAAGGGGGAAAGGATGAGAGGAAGNNNAGATCGG
29818 TATGAGTGTGGAGTCGTTGCGCTATCCCAGCCTCTTCTCTGAANNNAGATCGGAAG
104810 TATGAGTGTGGAGTCGTTGCATACGGTGAAACCATGAAGTATAAAGNNNAGATCG
33171 TATGAGTGTGGAGTCGTTGCTGACTTTGTTCTCTCTGACCACANNNAGATCGGAAG
39760 TATGAGTGTGGAGTCGTTGCACCTTGGGAGCCTCTACTTATGANNNAGATCGGAAG
31692 TATGAGTGTGGAGTCGTTGCACCCACATAAGCCAGGTAGAGAANNNAGATCGGAA
51301 TATGAGTGTGGAGTCGTTGCCGGCCTGAACAAAAGACATTCAGNNNAGATCGGAA
38212 TATGAGTGTGGAGTCGTTGCTATGTTAGGCGACCATCAGGAGANNNAGATCGGAA
44718 TATGAGTGTGGAGTCGTTGCCTTGTACACACATACCTGCTCCCNNNAGATCGGAAGA
43167 TATGAGTGTGGAGTCGTTGCGGCTCACTTGTAGTTTCCCAGACNNNAGATCGGAAG
70373 TATGAGTGTGGAGTCGTTGCGGGCATGCAAGATGAAACATTCCNNNAGATCGGAA
44990 TATGAGTGTGGAGTCGTTGCGTTACTGAATTGGCTCCCTAGCCNNNAGATCGGAAG
64845 TATGAGTGTGGAGTCGTTGCGATTTTTGAGCTTGCATGCAGGTNNNAGATCGGAA
106117 TATGAGTGTGGAGTCGTTGCGCTGCCATTTCTGTGTTTATACAGGNNNAGATCGGA
37290 TATGAGTGTGGAGTCGTTGCTTAGTGTGGAAACTTGGGGTTCANNNAGATCGGAA
53147 TATGAGTGTGGAGTCGTTGCGCTTCTCTGGGGAAATTCAGAGGNNNAGATCGGAA
41220 TATGAGTGTGGAGTCGTTGCTATGGCACAGACATTTGCTGCTANNNAGATCGGAAG
110633 TATGAGTGTGGAGTCGTTGCGAGGAACAGTTTGAAACACTTGGCNNNAGATCGGA
45047 TATGAGTGTGGAGTCGTTGCATTTACTGTGCATTCTCCAGGCCNNNAGATCGGAAG
71644 TATGAGTGTGGAGTCGTTGCGGGATATTACTCGGTCCAGGTNNNAGATCGGAAGA

| | |
|---|---|
| 55489 | TATGAGTGTGGAGTCGTTGCGGGAATGACAGGGTCTCATCATGNNNAGATCGGAA |
| 61117 | TATGAGTGTGGAGTCGTTGCAGACATTTCATCTCCCTGAGCCTNNNAGATCGGAAG |
| 45415 | TATGAGTGTGGAGTCGTTGCGGGAACTAATGTGACCACCATCCNNNAGATCGGAAG |
| 104782 | TATGAGTGTGGAGTCGTTGCTACAAAGTCTCGTGTGTCTGTCTNNNAGATCGGAAG |
| 115908 | TATGAGTGTGGAGTCGTTGCTGTGTGTGAGATAAAGTCTCACTGTGNNNAGATCG |
| 68926 | TATGAGTGTGGAGTCGTTGCCCCACCGCCTTTCTCTCATATTTNNNAGATCGGAAGA |
| 40951 | TATGAGTGTGGAGTCGTTGCGAGGATGGGCTCTAGTCGAACTANNNAGATCGGAA |
| 45391 | TATGAGTGTGGAGTCGTTGCGCTCTCTCTTTCTCTGGAGAATCCNNNAGATCGGAAG |
| 41675 | TATGAGTGTGGAGTCGTTGCGTGTGCTGGAGCCAATATGTGTANNNAGATCGGAA |
| 46565 | TATGAGTGTGGAGTCGTTGCTATTCTCTCTGGGTGAGTGGAGCNNNAGATCGGAA |
| 45082 | TATGAGTGTGGAGTCGTTGCCTAAGAACCTAAAACCTCCGGCCNNNAGATCGGAAG |
| 29938 | TATGAGTGTGGAGTCGTTGCATGATAGTCTTCAAGCCTGAGCCNNNAGATCGGAAG |
| 51150 | TATGAGTGTGGAGTCGTTGCTTAGAATAGCAAAGGGGCTGCAGNNNAGATCGGAA |
| 102520 | TATGAGTGTGGAGTCGTTGCTTGTTTTCATGTGGTTTTGGGTTTNNNAGATCGGAA |
| 103314 | TATGAGTGTGGAGTCGTTGCGAGACATAGAACAGACACTTTCAGGTNNNAGATCG |
| 65530 | TATGAGTGTGGAGTCGTTGCGGAAGCATCTAATCTGGCTTGGTNNNAGATCGGAA |
| 62201 | TATGAGTGTGGAGTCGTTGCAGAAGGTCGAGAGAGACAATGCTNNNAGATCGGAA |
| 30128 | TATGAGTGTGGAGTCGTTGCTGAGCAGTGGTTGTGGTAGAAAANNNAGATCGGAA |
| 59422 | TATGAGTGTGGAGTCGTTGCGCTGACGCTGTACACATCCATATNNNAGATCGGAAG |
| 35703 | TATGAGTGTGGAGTCGTTGCGCAACCTCTACAAACTAAGGGCANNNAGATCGGAAG |
| 56791 | TATGAGTGTGGAGTCGTTGCTAACATTGGTTGCCTATGGGGTGNNNAGATCGGAA |
| 43545 | TATGAGTGTGGAGTCGTTGCCACCTGCTGTTTTCTCTCCCTACNNNAGATCGGAAGA |
| 63240 | TATGAGTGTGGAGTCGTTGCCTTTTGCAGGACAGCTACTGTCTNNNAGATCGGAAG |
| 38455 | TATGAGTGTGGAGTCGTTGCGCTCAAATGCTCATGCCTGTAGANNNAGATCGGAAG |
| 39098 | TATGAGTGTGGAGTCGTTGCTCCAGCCTTTTATTTGAATGGGGANNNAGATCGGAA |
| 115921 | TATGAGTGTGGAGTCGTTGCTCTCTCTCTCCCTTGCCACCATTNNNAGATCGGAAGA |
| 34923 | TATGAGTGTGGAGTCGTTGCTGAAACAGACCTATGGCTCTCCANNNAGATCGGAAG |
| 54436 | TATGAGTGTGGAGTCGTTGCACTTGGGGACAGGTATAGGATGGNNNAGATCGGA |
| 41229 | TATGAGTGTGGAGTCGTTGCGAAGCGAAAAGTAGTGGGTGCTANNNAGATCGGA |
| 35095 | TATGAGTGTGGAGTCGTTGCTGTTGTTTCACACATAGTACTTTCCANNNAGATCGGA |
| 53568 | TATGAGTGTGGAGTCGTTGCAGGCTTTTGTTGTTGTTGTTCGGNNNAGATCGGAAG |
| 56988 | TATGAGTGTGGAGTCGTTGCTTTGTTTTCAGGCTTGCTCTGTGNNNAGATCGGAAG |
| 102213 | TATGAGTGTGGAGTCGTTGCTCCGTGTGTATTATGGGATGGCNNNAGATCGGAAG |
| 57176 | TATGAGTGTGGAGTCGTTGCAATCCAGTTTGTTGGCAGCATTGNNNAGATCGGAAG |
| 103267 | TATGAGTGTGGAGTCGTTGCCGGAAGACACTGCTACTAGCCTNNNAGATCGGAAGA |
| 51021 | TATGAGTGTGGAGTCGTTGCAAATATGAAGTCCCAGCCAGCAGNNNAGATCGGAA |
| 36087 | TATGAGTGTGGAGTCGTTGCACAATCACATATTCTCAGTAGCTGCANNNAGATCGG |
| 55052 | TATGAGTGTGGAGTCGTTGCTCACCATCACCACTCAAACTTGGNNNAGATCGGAAG |
| 49168 | TATGAGTGTGGAGTCGTTGCAGGCAGCCAGAGAAAAGAAAGTCNNNAGATCGGA |
| 49689 | TATGAGTGTGGAGTCGTTGCTCAGCTTTAGCCACAGTAGCTTCNNNAGATCGGAAG |
| 36028 | TATGAGTGTGGAGTCGTTGCTCAGTGTGGCCATAAAGTTATGCANNNAGATCGGA |
| 48055 | TATGAGTGTGGAGTCGTTGCATGCTTGAAAAAGGACCCATTGCNNNAGATCGGAA |
| 98025 | TATGAGTGTGGAGTCGTTGCTTTTGGTCCCCGAGTTTGTTTTTNNNAGATCGGAAG |

| | |
|---|---|
| 54299 | TATGAGTGTGGAGTCGTTGCAGATCCTGCCTTGGGAAAAATGGNNNAGATCGGAA |
| 58487 | TATGAGTGTGGAGTCGTTGCCACACAGAATCAGCACATCCCATNNNAGATCGGAAG |
| 56596 | TATGAGTGTGGAGTCGTTGCACCCAGAAGCTACAACATCAGTGNNNAGATCGGAAG |
| 46953 | TATGAGTGTGGAGTCGTTGCTGAGACTGGAAAGATAAAAGAGAGGCNNNAGATC |
| 62418 | TATGAGTGTGGAGTCGTTGCTTGGCAGGAGTAATTTGTGTGCTNNNAGATCGGAA |
| 60347 | TATGAGTGTGGAGTCGTTGCGTGCAGTGGTGCTATAATGGACTNNNAGATCGGAA |
| 51227 | TATGAGTGTGGAGTCGTTGCGGGTAACTGCTAAAGAGCCTCAGNNNAGATCGGAA |
| 56346 | TATGAGTGTGGAGTCGTTGCTGCTCAAGGCTTGTCTTACTCTGNNNAGATCGGAAG |
| 65305 | TATGAGTGTGGAGTCGTTGCTGAACAGCTTATACTTTGTGATGGTNNNAGATCGGA |
| 58097 | TATGAGTGTGGAGTCGTTGCTTGAAGGTGACAGAGGAGGGAATNNNAGATCGGA |
| 99127 | TATGAGTGTGGAGTCGTTGCCCAGGGAGGTGAGTTCAGACTNNNAGATCGGAAGA |
| 48781 | TATGAGTGTGGAGTCGTTGCGGTCTGGTTGTCCATGAATCCTCNNNAGATCGGAAG |
| 46792 | TATGAGTGTGGAGTCGTTGCGAAAGACAATCCCTCTCTGTCGCNNNAGATCGGAAG |
| 60118 | TATGAGTGTGGAGTCGTTGCTCTGCCAACTCTATCTCAGCACTNNNAGATCGGAAGA |
| 41433 | TATGAGTGTGGAGTCGTTGCTTGGCTCTCAAGGATGTGGAGTANNNAGATCGGAA |
| 39447 | TATGAGTGTGGAGTCGTTGCTCTTGCTCCTTGTATCCTGTGGANNNAGATCGGAAG |
| 56817 | TATGAGTGTGGAGTCGTTGCTTCATCTCTGCCCATGAATGGTGNNNAGATCGGAAG |
| 35968 | TATGAGTGTGGAGTCGTTGCCAGGAAGACCAAGTGAAGATGCANNNAGATCGGAA |
| 48951 | TATGAGTGTGGAGTCGTTGCTGGGACATGAACACTCATTGCTCNNNAGATCGGAAG |
| 67491 | TATGAGTGTGGAGTCGTTGCAAAGCAGGATGCAAAGGGAACTTNNNAGATCGGAA |
| 106120 | TATGAGTGTGGAGTCGTTGCCTGTGTGTAAGGTTTTGGACCCACNNNAGATCGGAA |
| 108477 | TATGAGTGTGGAGTCGTTGCACAGTGCCACAACAAACATGAGATNNNAGATCGGA |
| 42303 | TATGAGTGTGGAGTCGTTGCTTTGCTGTAGACAGGTGTCCAACNNNAGATCGGAAG |
| 117354 | TATGAGTGTGGAGTCGTTGCGACTGATAGGATCGGTGTGTACGGNNNAGATCGGA |
| 98043 | TATGAGTGTGGAGTCGTTGCAAGGCTCCGTCTGTTGATGCNNNAGATCGGAAGAG |
| 29817 | TATGAGTGTGGAGTCGTTGCAGGCTGGACCTGAAGATTAGTCANNNAGATCGGAA |
| 104724 | TATGAGTGTGGAGTCGTTGCGGGAACAGCAGATAGTGGGGAAAANNNAGATCGG |
| 30277 | TATGAGTGTGGAGTCGTTGCCTTCAGAGTGGAAGCGGTACAAANNNAGATCGGAA |
| 107762 | TATGAGTGTGGAGTCGTTGCTTGGCAGAGAGTAATAAGAGCCAGNNNAGATCGGA |
| 51786 | TATGAGTGTGGAGTCGTTGCCTGACATAGGTGGATCGCATGAGNNNAGATCGGAA |
| 44309 | TATGAGTGTGGAGTCGTTGCAATCCACTTCTCTACCCTGACCCNNNAGATCGGAAGA |
| 38746 | TATGAGTGTGGAGTCGTTGCATAGATCCTGTCCTCCTCCAGGANNNAGATCGGAAG |
| 56713 | TATGAGTGTGGAGTCGTTGCATTAGCCCATGCTTCACTTCGTGNNNAGATCGGAAG |
| 42004 | TATGAGTGTGGAGTCGTTGCCCTGAGCCTGGCCTTGTTATTTANNNAGATCGGAAG |
| 50842 | TATGAGTGTGGAGTCGTTGCTTATAGACATGGCAGGTCCCCAGNNNAGATCGGAA |
| 36159 | TATGAGTGTGGAGTCGTTGCAAAGTACAAATCGGTCTGGTGCANNNAGATCGGAA |
| 99960 | TATGAGTGTGGAGTCGTTGCTTGAGCCTTTATCCAACTTCAGCNNNAGATCGGAAG |
| 65533 | TATGAGTGTGGAGTCGTTGCGGACTAGGAGAGGGAATCTTGGTNNNAGATCGGA |
| 44038 | TATGAGTGTGGAGTCGTTGCTTCTCCCTGTGCTTACAGAGACCNNNAGATCGGAAG |
| 110715 | TATGAGTGTGGAGTCGTTGCTACGATAGCCACAAGGTACACGTNNNAGATCGGAA |
| 117212 | TATGAGTGTGGAGTCGTTGCCGGCTGCAGAGCTCAAACTACNNNAGATCGGAAGA |
| 101693 | TATGAGTGTGGAGTCGTTGCTGTGTGAAGAATTATGCTGACCTTNNNAGATCGGAA |
| 55660 | TATGAGTGTGGAGTCGTTGCCAGGAGAACAAGAGGGTGTGATGNNNAGATCGGA |

40834 TATGAGTGTGGAGTCGTTGCGATGAGCCCTGCTTAGGAGGATANNNAGATCGGAA
104090 TATGAGTGTGGAGTCGTTGCCCATGATTTCTCAGAGGGAGTGNNNAGATCGGAAG
51337 TATGAGTGTGGAGTCGTTGCACAGCAGATAGTGGGGAAAAGAGNNNAGATCGGA
48350 TATGAGTGTGGAGTCGTTGCATTACCGCTTTGAGAGCACCATCNNNAGATCGGAAG
62425 TATGAGTGTGGAGTCGTTGCTCTGTGCCATACTTTGTTGTGCTNNNAGATCGGAAG
42520 TATGAGTGTGGAGTCGTTGCCACCCATGCAAATACACCAACACNNNAGATCGGAAG
59499 TATGAGTGTGGAGTCGTTGCGCTGGGAAGGAGTAGGAACCTATNNNAGATCGGAA
106573 TATGAGTGTGGAGTCGTTGCATCCAGTAAGTGAGAAAAGATGTGANNNAGATCGG
107729 TATGAGTGTGGAGTCGTTGCGATTTTTCAGGTTTTTCTGGACCCTNNNAGATCGGAA
61223 TATGAGTGTGGAGTCGTTGCGGGTTTCATCCATCAAGATGCCTNNNAGATCGGAAG
45309 TATGAGTGTGGAGTCGTTGCTCAGGTTCTGCTAAAGATGTGCCNNNAGATCGGAAG
48707 TATGAGTGTGGAGTCGTTGCTTTGACTTAGAGGTAGCCCCCTCNNNAGATCGGAAG
104825 TATGAGTGTGGAGTCGTTGCTGACTTAGAGGTAGCCCCCTCNNNAGATCGGAAGAG
67804 TATGAGTGTGGAGTCGTTGCTGAGACCTGATTCCAGTGTCCTTNNNAGATCGGAAG
32108 TATGAGTGTGGAGTCGTTGCGAGGAGCATGTAAACCAGCTGAANNNAGATCGGAA
32279 TATGAGTGTGGAGTCGTTGCATTGTGGGAAGGGCAGGAGATAANNNAGATCGGA
116418 TATGAGTGTGGAGTCGTTGCTAAGGAACTGAAGCAGAGATCCTNNNAGATCGGAA
99340 TATGAGTGTGGAGTCGTTGCCCTCTTTTTCATCCTGCCTGATCAGNNNAGATCGGAA
37112 TATGAGTGTGGAGTCGTTGCACAGGAGGCCAGATTACCATTCANNNAGATCGGAAG
97668 TATGAGTGTGGAGTCGTTGCCGGATGTCGCCGGATTTTATGATCNNNAGATCGGAA
47627 TATGAGTGTGGAGTCGTTGCCCCAATGGTTGTGAGCTTTATGCNNNAGATCGGAAG
55681 TATGAGTGTGGAGTCGTTGCAGAGAGATTGGGGTGGGGATATGNNNAGATCGGA
48341 TATGAGTGTGGAGTCGTTGCCCCCCTGACAGCTGGAAATACATCNNNAGATCGGAAG
55047 TATGAGTGTGGAGTCGTTGCCCCCCCAAAACATCTGCTATTGGNNNAGATCGGAAGA
57655 TATGAGTGTGGAGTCGTTGCAACTGCATGTGGTCATGGTTTTGNNNAGATCGGAAG
43685 TATGAGTGTGGAGTCGTTGCTACAACAAATGCCCCCAGGTTACNNNAGATCGGAAG
48965 TATGAGTGTGGAGTCGTTGCGTCTGTTGCTTGTCAAGTTGCTCNNNAGATCGGAAG
58341 TATGAGTGTGGAGTCGTTGCCAAGGGCACTAATCCCATCACATNNNAGATCGGAAG
39066 TATGAGTGTGGAGTCGTTGCTGTTGGTAGCATAGGATTCGGGANNNAGATCGGAA
52269 TATGAGTGTGGAGTCGTTGCCACGTTTATGCTGTACACACACGNNNAGATCGGAAG
62395 TATGAGTGTGGAGTCGTTGCAGTCTGCCTCTAAATTTGGTGCTNNNAGATCGGAAG
110670 TATGAGTGTGGAGTCGTTGCATGCAATGATGGAAACTGGGAAGTNNNAGATCGGA
109932 TATGAGTGTGGAGTCGTTGCTGGTAGAAAATCAAAGGTGTATGTGNNNAGATCGG
68720 TATGAGTGTGGAGTCGTTGCCACTGCCTGGGTTGTAGATTGTTNNNAGATCGGAAG
102211 TATGAGTGTGGAGTCGTTGCGTTCCTCTATAGGCGAGATGTCCTNNNAGATCGGAA
64426 TATGAGTGTGGAGTCGTTGCACTTGCCAGCTTACAGGACTAGTNNNAGATCGGAAG
64984 TATGAGTGTGGAGTCGTTGCTCTGGACCATCCCTCTAGTAGGTNNNAGATCGGAAG
52647 TATGAGTGTGGAGTCGTTGCTCTGTTTTCTGAGGACTGCTTCGNNNAGATCGGAAG
99280 TATGAGTGTGGAGTCGTTGCAGTTCTCAGATCATGCGTCTGTNNNAGATCGGAAGA
43478 TATGAGTGTGGAGTCGTTGCTGGCAGCACACTGATGAGAATACNNNAGATCGGAA
40365 TATGAGTGTGGAGTCGTTGCCCGTTTAGGTGTCCAGACATTGANNNAGATCGGAAG
50123 TATGAGTGTGGAGTCGTTGCAAACCAGCAAGAGGGGAAGAAAGNNNAGATCGGA
105485 TATGAGTGTGGAGTCGTTGCACTGTCAATAGCCATCTTTGTTACCNNNAGATCGGAA

47692 TATGAGTGTGGAGTCGTTGCATACAACAGGCTGCTCTTACTGCNNNAGATCGGAAG
106576 TATGAGTGTGGAGTCGTTGCAGTTGTTTTTCATGAACTGCTCANNNAGATCGGAAG
58685 TATGAGTGTGGAGTCGTTGCTCACTCACTAACTAGCTGGGCATNNNAGATCGGAAG
58370 TATGAGTGTGGAGTCGTTGCACCTGTAGATGCAGTACCGACATNNNAGATCGGAAG
106115 TATGAGTGTGGAGTCGTTGCCTGACCCTCTATGTGTCGGGGNNNAGATCGGAAGA
69242 TATGAGTGTGGAGTCGTTGCGGACTAGGAGGTCAGGGAAGTTTNNNAGATCGGA
51503 TATGAGTGTGGAGTCGTTGCAGCAGAAATACCCAGCTGGAGAGNNNAGATCGGAA
99374 TATGAGTGTGGAGTCGTTGCGCCAGTCATAGAATTCTGCACACNNNAGATCGGAAG
42474 TATGAGTGTGGAGTCGTTGCTCATCCAGTCAAAATGTGAAGTAACNNNAGATCGGA
103857 TATGAGTGTGGAGTCGTTGCGTGTGTGTGTGTTGTGGTGTAGACNNNAGATCGGA
104813 TATGAGTGTGGAGTCGTTGCGAATGAATGCCTAATGTGTGCAAACANNNAGATCG
59378 TATGAGTGTGGAGTCGTTGCGGTCTTCTCACATCGCCTTTGATNNNAGATCGGAAG
58466 TATGAGTGTGGAGTCGTTGCCCAGCAAACACATACAGACCCATNNNAGATCGGAAG
60633 TATGAGTGTGGAGTCGTTGCAGCACTGCAAAGAGACATAACCTNNNAGATCGGAA
29939 TATGAGTGTGGAGTCGTTGCAGGCATTTCTCTGTTGCACTGAANNNAGATCGGAAG
103850 TATGAGTGTGGAGTCGTTGCTAGGCCGGAGTGCATGATCATAGNNNAGATCGGAA
32723 TATGAGTGTGGAGTCGTTGCTGAATGCCTAATGTGTGCAAACANNNAGATCGGAA
54729 TATGAGTGTGGAGTCGTTGCACGAGATTTCACTACCTTGCTGGNNNAGATCGGAAG
57300 TATGAGTGTGGAGTCGTTGCCATATTGCTGCTTGGGAAGCTTGNNNAGATCGGAA
58994 TATGAGTGTGGAGTCGTTGCCCTTTATGCCAAGTGCTCCAGATNNNAGATCGGAAG
54170 TATGAGTGTGGAGTCGTTGCTGGAAATGTGATGAAGACAGTGGGNNNAGATCGG
31402 TATGAGTGTGGAGTCGTTGCCCTAGAAGACAGGAGGACCTCAANNNAGATCGGAA
35945 TATGAGTGTGGAGTCGTTGCTATTCGCTAAGCCTTTCCATGCANNNAGATCGGAAG
43179 TATGAGTGTGGAGTCGTTGCAGTGTCACAGAATCACTGCAGACNNNAGATCGGAA
117338 TATGAGTGTGGAGTCGTTGCACCTATCTACCTATCTGTCTATTCATCNNNAGATCGG
66280 TATGAGTGTGGAGTCGTTGCGGTGTGATGGTCAGTATGAAGTGTNNNAGATCGGA
44076 TATGAGTGTGGAGTCGTTGCGGTAACACTGAAATGAGGGGACCNNNAGATCGGAA
51904 TATGAGTGTGGAGTCGTTGCTAAAACAGCACTCGGGATGTGAGNNNAGATCGGAA
38773 TATGAGTGTGGAGTCGTTGCTCATTACCTGGGAAAATGTCAGGANNNAGATCGGA
65449 TATGAGTGTGGAGTCGTTGCGGAAACAAAGAGAGAAGAGGTGGTNNNAGATCGG
39250 TATGAGTGTGGAGTCGTTGCTAGAAGATCATGGTGGCCATGGANNNAGATCGGAA
67210 TATGAGTGTGGAGTCGTTGCACACGAAGAGAAAGTTGCTCATTNNNAGATCGGAA
53790 TATGAGTGTGGAGTCGTTGCAGAGAACTTTGTGTTGCTGAGGGNNNAGATCGGAA
64038 TATGAGTGTGGAGTCGTTGCCCAAATAAGAGGCAAGAGGCAGTNNNAGATCGGAA
107749 TATGAGTGTGGAGTCGTTGCGTGTGTTGGGGGAGGGTTCAGNNNAGATCGGAAG
30297 TATGAGTGTGGAGTCGTTGCATCTGGCAATGTGATGGACCAAANNNAGATCGGAA
51569 TATGAGTGTGGAGTCGTTGCTCCAATCAGTTGCCTTCTTCGAGNNNAGATCGGAAG
110735 TATGAGTGTGGAGTCGTTGCTGGCTTTGACTAGCAGTCTATACTCNNNAGATCGGA
32812 TATGAGTGTGGAGTCGTTGCTCCCACGGACAACATTTACAACANNNAGATCGGAAG
37412 TATGAGTGTGGAGTCGTTGCTCTTGCATCTGTGGGTGTTTTCANNNAGATCGGAAG
51207 TATGAGTGTGGAGTCGTTGCGCCTGGTACACACTAACACTCAGNNNAGATCGGAAG
38137 TATGAGTGTGGAGTCGTTGCTCCTGGTAAGAAAGCAAAAAGAGAGANNNAGATCG
46648 TATGAGTGTGGAGTCGTTGCATTTCGGAAGACACTGCTACTAGCNNNAGATCGGAA

103302 TATGAGTGTGGAGTCGTTGCATGGACAACTCTGATGGGCTGGGNNNAGATCGGAA
117827 TATGAGTGTGGAGTCGTTGCAGCACCTAAACCCCTTTTGCTCNNNAGATCGGAAGAG
96744 TATGAGTGTGGAGTCGTTGCACTTCATCTTCCCCACCACTAAGANNNAGATCGGAAG
40014 TATGAGTGTGGAGTCGTTGCAAGGCTGTTGCTTTTCCATCTGANNNAGATCGGAAG
41770 TATGAGTGTGGAGTCGTTGCGGGGGAAAGAGGCAGATGGATTANNNAGATCGGA
65345 TATGAGTGTGGAGTCGTTGCGGGACTAAAGATGTATCACTGGTNNNAGATCGGAA
69479 TATGAGTGTGGAGTCGTTGCCCCTTCTGGTGTGTATGCCTTTTNNNAGATCGGAAG
53189 TATGAGTGTGGAGTCGTTGCAAACTCCCCTTAGAGCAAGGAGGNNNAGATCGGAA
60859 TATGAGTGTGGAGTCGTTGCTGTGTAAAAGTTTCCCTTGACCCTNNNAGATCGGAAG
37788 TATGAGTGTGGAGTCGTTGCAAAGAGAGCACTCCCAAGACAGANNNAGATCGGAA
47955 TATGAGTGTGGAGTCGTTGCTTATAGATCCTGTGCCATGGTGCNNNAGATCGGAAG
44632 TATGAGTGTGGAGTCGTTGCAGTCAGATATGGGTGGGAATCCCNNNAGATCGGAA
44438 TATGAGTGTGGAGTCGTTGCAACTCACAGGTTTTCTATGCCCCNNNAGATCGGAAG
56004 TATGAGTGTGGAGTCGTTGCGTCTTCACTTTTAGGGGAGCCTGNNNAGATCGGAAG
46201 TATGAGTGTGGAGTCGTTGCTGTAAGCTCCAAAAGAGGGAAGCNNNAGATCGGAA
34382 TATGAGTGTGGAGTCGTTGCGAGAAGTCTAAGATCAAAGGCCCANNNAGATCGGA
54911 TATGAGTGTGGAGTCGTTGCTCAACATCATGGAAACTGGGTGGNNNAGATCGGAA
32647 TATGAGTGTGGAGTCGTTGCTCCCCCTCACAAGATCAAAAACANNNAGATCGGAAG
60905 TATGAGTGTGGAGTCGTTGCATTCTCTTCACTTCAGAGCCCCTNNNAGATCGGAAGA
67497 TATGAGTGTGGAGTCGTTGCACAGGGCTCAAGGAGGATAACTTNNNAGATCGGAA
38518 TATGAGTGTGGAGTCGTTGCGCCATTGTCCTACAGAATCACGANNNAGATCGGAAG
30518 TATGAGTGTGGAGTCGTTGCAGAGCCTGCATGTAGAAGAGAAANNNAGATCGGAA
34068 TATGAGTGTGGAGTCGTTGCACATCTCTCTCTCTTGAACTCACCANNNAGATCGGAA
68891 TATGAGTGTGGAGTCGTTGCGAGGCTGCAGGACTTTAGGATTTNNNAGATCGGAA
71110 TATGAGTGTGGAGTCGTTGCGCCTGTCTGTCTCTTTCTGGACTNNNAGATCGGAAG
106085 TATGAGTGTGGAGTCGTTGCATGTGTTTGTTTCTTCCAGCCTTGNNNAGATCGGAAG
45479 TATGAGTGTGGAGTCGTTGCCTCCATGGTTTCCTCTGTGATCCNNNAGATCGGAAG
67158 TATGAGTGTGGAGTCGTTGCAAAAGCTACTTCAGCCTGGCATTNNNAGATCGGAAG
36254 TATGAGTGTGGAGTCGTTGCTTTCCCTGTGGAATCATCTTGCANNNAGATCGGAAG
61147 TATGAGTGTGGAGTCGTTGCTTCCTTGGTGTTTTGAAAGGCCTNNNAGATCGGAAG
66793 TATGAGTGTGGAGTCGTTGCGTGTGTGTGTCTACCACCTTTGTNNNAGATCGGAAG
117262 TATGAGTGTGGAGTCGTTGCAGGTTTCCTTCCGGTCAGTCANNNAGATCGGAAGAG
51312 TATGAGTGTGGAGTCGTTGCGTTTCACTGTTGTTGCCCTTCAGNNNAGATCGGAAG
34586 TATGAGTGTGGAGTCGTTGCAATGCGATCTATGAAAAGAGCCANNNAGATCGGAA
40745 TATGAGTGTGGAGTCGTTGCTCTAAGGATGGCTGAAGGGCATANNNAGATCGGAA
68810 TATGAGTGTGGAGTCGTTGCTCCATTCTCTGCAGTGGACATTTNNNAGATCGGAAG
35690 TATGAGTGTGGAGTCGTTGCTCTTGATCTGGTCATCTTTTAGGCANNNAGATCGGA
57279 TATGAGTGTGGAGTCGTTGCATTTGCTTCACCTCTGAGCCTTGNNNAGATCGGAAG
31282 TATGAGTGTGGAGTCGTTGCTGTGTACTTATGCTGCTTGGCAANNNAGATCGGAAG
32202 TATGAGTGTGGAGTCGTTGCAGATGTGACATGGTGCTCTTGAANNNAGATCGGAA
50316 TATGAGTGTGGAGTCGTTGCGTGATACGGAGGCTATCTGCAAGNNNAGATCGGAA
49049 TATGAGTGTGGAGTCGTTGCTGGGTATGCAACTCTCCTCTCTCNNNAGATCGGAAG
42345 TATGAGTGTGGAGTCGTTGCCCTTTGCCCCAAAACTCTTCAACNNNAGATCGGAAGA

| | |
|---|---|
| 32020 | TATGAGTGTGGAGTCGTTGCCGTGTTCTGAATGCCCAAATGAANNNAGATCGGAAG |
| 36147 | TATGAGTGTGGAGTCGTTGCTGGCTTTACCACTTACTAGTGCANNNAGATCGGAAG |
| 102882 | TATGAGTGTGGAGTCGTTGCTACGGTTGTGGGATAGAGGTCCTNNNAGATCGGAA |
| 64838 | TATGAGTGTGGAGTCGTTGCCAGGAAAAGAATGAGAGGCAGGTNNNAGATCGGA |
| 47812 | TATGAGTGTGGAGTCGTTGCGCCATTCATTTTGAAAAGTCTCTGCNNNAGATCGGA |
| 59629 | TATGAGTGTGGAGTCGTTGCGTGGTCGCATGTGTGTGATGTATNNNAGATCGGAA |
| 71496 | TATGAGTGTGGAGTCGTTGCGCTCCTGGAAACTGTTGCCTATTNNNAGATCGGAAG |
| 59088 | TATGAGTGTGGAGTCGTTGCTGCCACATGGATGGATGGTAGATNNNAGATCGGAA |
| 103258 | TATGAGTGTGGAGTCGTTGCTCAGGAAAAGAATGAGAGGCAGGTTNNNAGATCG |
| 67268 | TATGAGTGTGGAGTCGTTGCCGATCAGGGAGAGTAGGCAGATTNNNAGATCGGA |
| 107766 | TATGAGTGTGGAGTCGTTGCGTGTGTGGGTGTGAGTATGTGTACANNNAGATCGG |
| 51302 | TATGAGTGTGGAGTCGTTGCGCTTCTTACCCTCACCCATTCAGNNNAGATCGGAAGA |
| 42949 | TATGAGTGTGGAGTCGTTGCTGGTGAACAAAAGATGCTTGCACNNNAGATCGGAA |
| 59958 | TATGAGTGTGGAGTCGTTGCAGGATGGCAAGTGTCAAAACACTNNNAGATCGGAA |
| 100120 | TATGAGTGTGGAGTCGTTGCGTTTGTTTTCTTTGGTCTTCCCTCCNNNAGATCGGAA |
| 52005 | TATGAGTGTGGAGTCGTTGCGTCTCACAGCCAAGTGGCTATAGNNNAGATCGGAA |
| 105545 | TATGAGTGTGGAGTCGTTGCAAGGATATGGAGGAGAAACAGGAGGNNNAGATCG |
| 34393 | TATGAGTGTGGAGTCGTTGCCGTGGTAGGTTATGAAATGCCCANNNAGATCGGAA |
| 34073 | TATGAGTGTGGAGTCGTTGCAGCAAAATCTGCATAGCTCACCANNNAGATCGGAAG |
| 29940 | TATGAGTGTGGAGTCGTTGCTCCCCATCTAAGTTTCTGGAGGTNNNAGATCGGAAG |
| 33148 | TATGAGTGTGGAGTCGTTGCCTCTGGGCCTCTGTTTTTACACANNNAGATCGGAAG |
| 35870 | TATGAGTGTGGAGTCGTTGCATGAAACTGGGGAAAGTTTGGCANNNAGATCGGAA |
| 57467 | TATGAGTGTGGAGTCGTTGCCTAAGTATCCGCACCTCTGGTTGNNNAGATCGGAAG |
| 29815 | TATGAGTGTGGAGTCGTTGCTGCCCTCTACTGGCAACAATATCNNNAGATCGGAAG |
| 54287 | TATGAGTGTGGAGTCGTTGCCCTATGCTCGGTGTATCTTTGGGNNNAGATCGGAAG |
| 31900 | TATGAGTGTGGAGTCGTTGCTGACAAGAGTAGCCATGAGGGAANNNAGATCGGA |
| 56727 | TATGAGTGTGGAGTCGTTGCCAGAAAAGAGAGCCAAACAGGTGNNNAGATCGGA |
| 51108 | TATGAGTGTGGAGTCGTTGCACTGATAGGCAAAAGACTGGCAGNNNAGATCGGAA |
| 43988 | TATGAGTGTGGAGTCGTTGCGAACATGGGAAAGAGAGGTCACCNNNAGATCGGA |
| 36384 | TATGAGTGTGGAGTCGTTGCCTGTCCCTTTGGCCACATAATCANNNAGATCGGAAG |
| 71841 | TATGAGTGTGGAGTCGTTGCAGATATTCACCATTATAGGTGGGTTTNNNAGATCGG |
| 47077 | TATGAGTGTGGAGTCGTTGCTCAAGATAGTTGCAACAGAGGGCNNNAGATCGGAA |
| 71850 | TATGAGTGTGGAGTCGTTGCTCAACACCTACGAAGGAAACATCANNNAGATCGGAA |
| 100663 | TATGAGTGTGGAGTCGTTGCGAGAAAGATCCTGCCTTGGGAAAANNNAGATCGGA |
| 100724 | TATGAGTGTGGAGTCGTTGCTTCATCTCTCCAGTTCAAGTGATCNNNAGATCGGAAG |
| 71849 | TATGAGTGTGGAGTCGTTGCATAGTCCATTGGAAAGGGAGGCANNNAGATCGGAA |
| 100705 | TATGAGTGTGGAGTCGTTGCGATTTGCGAGTGAGGACAGAGTTNNNAGATCGGAA |
| 67347 | TATGAGTGTGGAGTCGTTGCGGACACTTTCTGGTCTGCTGATTNNNAGATCGGAAG |
| 52304 | TATGAGTGTGGAGTCGTTGCGTGTTTCCAAGTAATGGCTCACGNNNAGATCGGAAG |
| 71753 | TATGAGTGTGGAGTCGTTGCGTTCGGGAGCAAGAATTGGTTTGNNNAGATCGGAA |
| 100581 | TATGAGTGTGGAGTCGTTGCCAGTGGGTCTGCCCTTGTAGANNNAGATCGGAAGA |
| 47746 | TATGAGTGTGGAGTCGTTGCGTGTACTTGTGTCAAACAGCTGCNNNAGATCGGAAG |
| 104117 | TATGAGTGTGGAGTCGTTGCGGCTCATCATTGGAACTGCTTCCNNNAGATCGGAAG |

| | |
|---|---|
| 100936 | TATGAGTGTGGAGTCGTTGCGCATCGCAGGATTTTAGGCAAAGGNNNAGATCGGA |
| 106095 | TATGAGTGTGGAGTCGTTGCCCTAGACCTTTTCCTCCCTGTACANNNAGATCGGAAG |
| 100952 | TATGAGTGTGGAGTCGTTGCCTGTGGGCTCTGTTAGGACAGGANNNAGATCGGAA |
| 102262 | TATGAGTGTGGAGTCGTTGCTTAGTTGGGTGCTGTGGCAGANNNAGATCGGAAGA |
| 103912 | TATGAGTGTGGAGTCGTTGCATTCCTCCCTCCCCTTTTTGTCNNNAGATCGGAAGAG |
| 105517 | TATGAGTGTGGAGTCGTTGCACTCACTGCAATCTCAAATTCATGGNNNAGATCGGA |
| 107830 | TATGAGTGTGGAGTCGTTGCCACATACATATACACACGGAAGCANNNAGATCGGAA |
| 108439 | TATGAGTGTGGAGTCGTTGCACAGACCTATGGGGCCTGGAAGNNNAGATCGGAAG |
| 100003 | TATGAGTGTGGAGTCGTTGCGGAGGAGTTCACAGATCCGCTNNNAGATCGGAAGA |
| 102512 | TATGAGTGTGGAGTCGTTGCCAGGGGATAAGAGGAAAGCTGCANNNAGATCGGA |
| 104123 | TATGAGTGTGGAGTCGTTGCGGGGTGTGTATGTAGTAAGTAGGANNNAGATCGG |
| 106067 | TATGAGTGTGGAGTCGTTGCCTTCTGTGGAATAACGCTTGTGTNNNAGATCGGAAG |
| 100785 | TATGAGTGTGGAGTCGTTGCGCACCAGTCCACTGGAAAATCNNNAGATCGGAAGA |
| 104814 | TATGAGTGTGGAGTCGTTGCTGCCAACTCTATCTCAGCACTGNNNAGATCGGAAGA |
| 100549 | TATGAGTGTGGAGTCGTTGCGTCATAAACACAGGAGTTCAGCTATNNNAGATCGGA |
| 107831 | TATGAGTGTGGAGTCGTTGCGCATAGAACTGGGCTTCCCACACNNNAGATCGGAAG |
| 109871 | TATGAGTGTGGAGTCGTTGCCCTCTGGGAAAGAGACCTGGAAGNNNAGATCGGAA |
| 105478 | TATGAGTGTGGAGTCGTTGCCCCAGGATTGTAATGAGTGTCTCATANNNAGATCGG |
| 106996 | TATGAGTGTGGAGTCGTTGCCTCTACCTCCCAGGCTCAAAGGCNNNAGATCGGAAG |
| 104861 | TATGAGTGTGGAGTCGTTGCCCTGTTTTCATTCCCATTTTTCCCANNNAGATCGGAAG |
| 116228 | TATGAGTGTGGAGTCGTTGCACAGATGCTCAAGTTGCTCATACNNNAGATCGGAAG |
| 100760 | TATGAGTGTGGAGTCGTTGCACAATACACAGGACAGCCTTTCANNNAGATCGGAAG |
| 109857 | TATGAGTGTGGAGTCGTTGCGTGGGATGACAGGTATGTACCACNNNAGATCGGAA |
| 110653 | TATGAGTGTGGAGTCGTTGCTGCCCTAATGGAACTGGCCANNNAGATCGGAAGAG |
| 121967 | TATGAGTGTGGAGTCGTTGCAAAGGCATGAAATACAGCAACTCNNNAGATCGGAA |
| 108522 | TATGAGTGTGGAGTCGTTGCAGGCTTTGGAAACAAGCAGATGANNNAGATCGGAA |
| 71904 | TATGAGTGTGGAGTCGTTGCAAGGGGAAAGGAATTACATGATANNNAGATCGGA |
| 117443 | TATGAGTGTGGAGTCGTTGCAGTGTAGCCTTGACCTGTCAATCTNNNAGATCGGAA |
| 96762 | TATGAGTGTGGAGTCGTTGCGTGCTGAACTGCAGGTGTCTCTTNNNAGATCGGAAG |
| 105507 | TATGAGTGTGGAGTCGTTGCTCTTCTCTCTGTCACATTGCCTCTNNNAGATCGGAAG |
| 107809 | TATGAGTGTGGAGTCGTTGCCACACGCACCCACACAGTTATGNNNAGATCGGAAGA |
| 114529 | TATGAGTGTGGAGTCGTTGCATGCAAAGTACTGTTGTGGGAGCANNNAGATCGGA |
| 116592 | TATGAGTGTGGAGTCGTTGCAGTGTAGCCTTGACCTGTCAATCTCNNNAGATCGGA |
| 116116 | TATGAGTGTGGAGTCGTTGCCCTGACTAATGTGTGGTTTGCAGNNNAGATCGGAA |
| 108999 | TATGAGTGTGGAGTCGTTGCATTTTCCCACCCGTCAAATTCATTNNNAGATCGGAAG |
| 99341 | TATGAGTGTGGAGTCGTTGCTTGGTGACACAGCTCTTCTTGTTNNNAGATCGGAAG |
| 103841 | TATGAGTGTGGAGTCGTTGCATGCAATTGTTGAGGCTTGCAANNNAGATCGGAAG |
| 107008 | TATGAGTGTGGAGTCGTTGCTTTGGCAGGTCACTGGTTTGAANNNAGATCGGAAG |
| 99518 | TATGAGTGTGGAGTCGTTGCCTTGGTATACTGGAGAAGTCAGCANNNAGATCGGA |
| 65599 | TATGAGTGTGGAGTCGTTGCCCGTATCCATTCCAAGCAAATGTNNNAGATCGGAAG |
| 97634 | TATGAGTGTGGAGTCGTTGCCCAGGATACTTTCCCACTGAGCNNNAGATCGGAAGA |
| 95130 | TATGAGTGTGGAGTCGTTGCCAAGCCCTTTCTGGGAGTTTTCTNNNAGATCGGAAG |
| 45879 | TATGAGTGTGGAGTCGTTGCTCCATAGCTCTCTATGCCCTTCCNNNAGATCGGAAGA |

65869 TATGAGTGTGGAGTCGTTGCTTGCTATCGCTGCAGGTTTATGTNNNAGATCGGAAG
33642 TATGAGTGTGGAGTCGTTGCAAGAAAAGGCCATCTAGTGGACANNNAGATCGGAA
38485 TATGAGTGTGGAGTCGTTGCGGGGGATTTGGTGCTAAGTTAGANNNAGATCGGAA
96602 TATGAGTGTGGAGTCGTTGCAAGGCAGAAGGAGCTTGAGATCCNNNAGATCGGAA
46745 TATGAGTGTGGAGTCGTTGCTTTGGTGAACTGCCTATGTACGCNNNAGATCGGAAG
66943 TATGAGTGTGGAGTCGTTGCGTAAATGGGCATGGGAGCCAATTNNNAGATCGGAA
95122 TATGAGTGTGGAGTCGTTGCGGTAGTGCAAGGGGTCACAGATNNNAGATCGGAAG
49986 TATGAGTGTGGAGTCGTTGCCCCATCCAACTCACCTCCTTTTCNNNAGATCGGAAGAG
96172 TATGAGTGTGGAGTCGTTGCGAAAAATCACTGACTAAAGGCAGGTNNNAGATCGGA
48144 TATGAGTGTGGAGTCGTTGCAGACTGTGTTTCCCTCTTCTTGCNNNAGATCGGAAGA
56996 TATGAGTGTGGAGTCGTTGCGGGAGGGTCTAAAGATGTCTGTGNNNAGATCGGAA
34454 TATGAGTGTGGAGTCGTTGCAGAGAGAGAGAGAATCCCTCCCANNNAGATCGGAA
71734 TATGAGTGTGGAGTCGTTGCAATCCAGCAGAGAGCTATGAGGTNNNAGATCGGAA
46911 TATGAGTGTGGAGTCGTTGCCCATAATTGACTGAGGTCCAGGCNNNAGATCGGAAG
30245 TATGAGTGTGGAGTCGTTGCCACACTTATGCCATGTCCACAAANNNAGATCGGAAGA
96166 TATGAGTGTGGAGTCGTTGCTCCACGAGATCCCTCCCATGATANNNAGATCGGAAGA
60795 TATGAGTGTGGAGTCGTTGCTTGGTGTCAGAATGATGCTACCTNNNAGATCGGAAG
30339 TATGAGTGTGGAGTCGTTGCCGCCTTCTGAGACAATCTCCAAANNNAGATCGGAAGA
56226 TATGAGTGTGGAGTCGTTGCGCTATCTCCTTGTGACTTGGCTGNNNAGATCGGAAG
50368 TATGAGTGTGGAGTCGTTGCATGCTCCCGAAGTCAGAAAGAAGNNNAGATCGGAA
45831 TATGAGTGTGGAGTCGTTGCTGGCTGGTGTTTGTTGAAATTCCNNNAGATCGGAAG
55739 TATGAGTGTGGAGTCGTTGCAACCCAAACACCACAGAAAACTGNNNAGATCGGAAG
106546 TATGAGTGTGGAGTCGTTGCCTATCTCCTTGTGACTTGGCTGCNNNAGATCGGAAGA
41062 TATGAGTGTGGAGTCGTTGCACTCATGCCCAGTGTTATCCCTANNNAGATCGGAAGA
107010 TATGAGTGTGGAGTCGTTGCAATGGACAACACAGGTCAAGTCATCNNNAGATCGGA
30979 TATGAGTGTGGAGTCGTTGCGCAACCGAGGAATACACAACCAANNNAGATCGGAAG
71638 TATGAGTGTGGAGTCGTTGCCCAGAACAGACCACTATCCCAANNNAGATCGGAAGA
34577 TATGAGTGTGGAGTCGTTGCGAAACTGACTTGTTTTGCAGCCANNNAGATCGGAAG
38580 TATGAGTGTGGAGTCGTTGCGACAACACAGGTCAAGTCATCGANNNAGATCGGAAG
35883 TATGAGTGTGGAGTCGTTGCACACATATCCAATCCCAAATGCANNNAGATCGGAAGA
60494 TATGAGTGTGGAGTCGTTGCCCCACAGAACCTATGAGATTCTACTNNNAGATCGGAA
41372 TATGAGTGTGGAGTCGTTGCAGTGGGGATGTGTGAGAACAGTANNNAGATCGGAA
57120 TATGAGTGTGGAGTCGTTGCCCTCTCCATCCTGACGATTTGTGNNNAGATCGGAAGA
59625 TATGAGTGTGGAGTCGTTGCAACTCCTCAGGAAGGGATGGTATNNNAGATCGGAA
55569 TATGAGTGTGGAGTCGTTGCTTCATTTCCCCCACCAAGAGATGNNNAGATCGGAAGA
50394 TATGAGTGTGGAGTCGTTGCGCCCCTACTGTCCAAAAGAGAAGNNNAGATCGGAAG
48467 TATGAGTGTGGAGTCGTTGCACAACCACAAGTTCACCAGGATCNNNAGATCGGAAG
34891 TATGAGTGTGGAGTCGTTGCTGAGACTGTAAGTGAAGCCTCCANNNAGATCGGAA
41912 TATGAGTGTGGAGTCGTTGCATTCGCTGCTCATGAGTGAGTTANNNAGATCGGAAG
62738 TATGAGTGTGGAGTCGTTGCATTCCCTTCTTGGGCCTGTATCTNNNAGATCGGAAGA
47910 TATGAGTGTGGAGTCGTTGCTAGTAGGTGAGCCCAGATAGTGCNNNAGATCGGAA
58656 TATGAGTGTGGAGTCGTTGCAAAGTAAGGTGTTGGCAAGGCATNNNAGATCGGAA
96164 TATGAGTGTGGAGTCGTTGCCTCTCTCTAGGACAAGTGTGTGAGNNNAGATCGGAA

| | |
|---|---|
| 37161 | TATGAGTGTGGAGTCGTTGCGGACCGTGTGCTGTAGTTATTCANNNAGATCGGAAG |
| 42047 | TATGAGTGTGGAGTCGTTGCCACTACCACCACCACTGTCTTTANNNAGATCGGAAGA |
| 38302 | TATGAGTGTGGAGTCGTTGCTGCTACTTACAGGAAATGTATGAGANNNAGATCGGA |
| 105550 | TATGAGTGTGGAGTCGTTGCTTTGTGTGGATTGGGGCTGTTANNNAGATCGGAAGA |
| 30720 | TATGAGTGTGGAGTCGTTGCATGGGGAGGGGCAGTATGATAAANNNAGATCGGAA |
| 48313 | TATGAGTGTGGAGTCGTTGCCTTAGGGCTCTGCCACAAACATCNNNAGATCGGAAGA |
| 55512 | TATGAGTGTGGAGTCGTTGCGGCGAAAAGACCAACAAGTCATGNNNAGATCGGAA |
| 42903 | TATGAGTGTGGAGTCGTTGCCACTCACATTCACAAACTGGCACNNNAGATCGGAAGA |
| 62374 | TATGAGTGTGGAGTCGTTGCGAGAGAGAAAGGGAGAGAGTGCTNNNAGATCGGA |
| 56326 | TATGAGTGTGGAGTCGTTGCACTTTGTGCCAGGTACTGATCTGNNNAGATCGGAAG |
| 44018 | TATGAGTGTGGAGTCGTTGCCTGAAACTGCTTTTGCCAAGACCNNNAGATCGGAAGA |
| 108426 | TATGAGTGTGGAGTCGTTGCGTGTATGAGAGAGAGAAATGCTAACCNNNAGATCG |
| 52075 | TATGAGTGTGGAGTCGTTGCCACCACCAGCATGTTTTTGCTAGNNNAGATCGGAAGA |
| 101225 | TATGAGTGTGGAGTCGTTGCTCCCTATCCTTATGGGTTTTGGGNNNAGATCGGAAGA |
| 61520 | TATGAGTGTGGAGTCGTTGCTCCCTAACAAGAAATCCCTTCCTNNNAGATCGGAAGA |
| 62469 | TATGAGTGTGGAGTCGTTGCGGCTATGTTTGCTTTTGCTTGCTNNNAGATCGGAAGA |
| 37538 | TATGAGTGTGGAGTCGTTGCCCCTCAGCATCTTCTGTATACAAGANNNAGATCGGAA |
| 29812 | TATGAGTGTGGAGTCGTTGCTTCTCCCATCCTCCTAAAGGTGCNNNAGATCGGAAGA |
| 35019 | TATGAGTGTGGAGTCGTTGCACAAATCAAAAGGGCCCATTCCANNNAGATCGGAAG |
| 63398 | TATGAGTGTGGAGTCGTTGCAATGACAAACAGACGCAGCTTCTNNNAGATCGGAAG |
| 56037 | TATGAGTGTGGAGTCGTTGCGTGGGGAAACACTTGAGATCCTGNNNAGATCGGAA |
| 43775 | TATGAGTGTGGAGTCGTTGCTCCTTCAACTTTCTGTCCCAACCNNNAGATCGGAAGA |
| 44447 | TATGAGTGTGGAGTCGTTGCTTTTTGTGTATGTGCTTTGCCCCNNNAGATCGGAAGA |
| 35414 | TATGAGTGTGGAGTCGTTGCGGCATCTGAGAGTAAACATGAGCANNNAGATCGGAA |
| 57731 | TATGAGTGTGGAGTCGTTGCACGCAAGGATTGGTCTTCCAAATNNNAGATCGGAAG |
| 42642 | TATGAGTGTGGAGTCGTTGCGGCTTGAGAGAAGCAGAGTACACNNNAGATCGGAA |
| 54481 | TATGAGTGTGGAGTCGTTGCAAAGCCATGGTGACTTCTGTATGGNNNAGATCGGAA |
| 64592 | TATGAGTGTGGAGTCGTTGCGTGCAGTTACCAATATCCACCGTNNNAGATCGGAAGA |
| 36182 | TATGAGTGTGGAGTCGTTGCTGTAAATGTCTGTGAGAGTGTGCANNNAGATCGGAA |
| 52657 | TATGAGTGTGGAGTCGTTGCCCCCTCCTCCACTTATTTGTTCGNNNAGATCGGAAGA |
| 34167 | TATGAGTGTGGAGTCGTTGCAGTTGCTCCTTGAATTGGATACCANNNAGATCGGAA |
| 37737 | TATGAGTGTGGAGTCGTTGCAGAGAGCGAGAGCTAGAAACAGANNNAGATCGGAA |
| 58024 | TATGAGTGTGGAGTCGTTGCGTCACAGCCTCAGAGTAAAGAATNNNAGATCGGAAG |
| 31734 | TATGAGTGTGGAGTCGTTGCGGGGGAAATCTAGGGGTTGAGAANNNAGATCGGAA |
| 61139 | TATGAGTGTGGAGTCGTTGCAACATATACATACACCCCCGCCTNNNAGATCGGAAGA |
| 46940 | TATGAGTGTGGAGTCGTTGCATTGTGCTTATTGTGTGTCAGGCNNNAGATCGGAAG |
| 35552 | TATGAGTGTGGAGTCGTTGCGAGAGAGACAGAGATAGAGCGCANNNAGATCGGAA |
| 57271 | TATGAGTGTGGAGTCGTTGCGAAAGAAAAAGAGAGCGCCCTTGNNNAGATCGGAA |
| 98776 | TATGAGTGTGGAGTCGTTGCACAACCATTACCCAAAAAGGGAGANNNAGATCGGAA |
| 56717 | TATGAGTGTGGAGTCGTTGCCAGAAGCATTGTCTTATCCAAGGTGNNNAGATCGGA |
| 57227 | TATGAGTGTGGAGTCGTTGCTGAAGGAGAAGAGGCCAAACTTGNNNAGATCGGAA |
| 60419 | TATGAGTGTGGAGTCGTTGCCGCAAAAGCTGAAGTTTGTGACTNNNAGATCGGAAG |
| 41234 | TATGAGTGTGGAGTCGTTGCACCCTCCAGCAAGAAAATTGCTANNNAGATCGGAAG |

| | |
|---|---|
| 41927 | TATGAGTGTGGAGTCGTTGCGCAAGGGAGAGATCCTGAGGTTANNNAGATCGGAA |
| 71515 | TATGAGTGTGGAGTCGTTGCGAGAGAGACAGGCAGACTACCTTNNNAGATCGGAA |
| 50379 | TATGAGTGTGGAGTCGTTGCAGTGCTGGAAAGGAACACAGAAGNNNAGATCGGAA |
| 39061 | TATGAGTGTGGAGTCGTTGCAATCTCTTTCCTTCCTAGCGGGANNNAGATCGGAAG |
| 48396 | TATGAGTGTGGAGTCGTTGCAAGATTGAACCCAGGGATGCATCNNNAGATCGGAA |
| 43268 | TATGAGTGTGGAGTCGTTGCTTATTTCCTGCCACGTGAAGGACNNNAGATCGGAAG |
| 51390 | TATGAGTGTGGAGTCGTTGCGAACACAGCCCAGTCTAACAGAGNNNAGATCGGAA |
| 54924 | TATGAGTGTGGAGTCGTTGCGAACTCCTCATCTTCCATGGTGGNNNAGATCGGAAG |
| 117838 | TATGAGTGTGGAGTCGTTGCCCTCCCATCACATAACTCTGCATNNNAGATCGGAAGA |
| 36782 | TATGAGTGTGGAGTCGTTGCAGACACACAGATACCAGCTCTCANNNAGATCGGAAG |
| 63765 | TATGAGTGTGGAGTCGTTGCTCTTCTTTTCATAGCCAGCAGAAGTNNNAGATCGGAA |
| 44980 | TATGAGTGTGGAGTCGTTGCAGAGACAGTACCACCTCATAGCCNNNAGATCGGAAG |
| 105552 | TATGAGTGTGGAGTCGTTGCGTTTGAAGACTAAAGGATTTGGGGNNNAGATCGGA |
| 41966 | TATGAGTGTGGAGTCGTTGCTTTGAAGGGAGAGGAGGGTGTTANNNAGATCGGA |
| 57073 | TATGAGTGTGGAGTCGTTGCTGTATTGTAGCCCAGTGTGTGTGNNNAGATCGGAA |
| 67833 | TATGAGTGTGGAGTCGTTGCACACCAAATGGATGCTCAAGCTTNNNAGATCGGAAG |
| 59253 | TATGAGTGTGGAGTCGTTGCAGTTGCATTTTTCCAAAGAAATGATNNNAGATCGGA |
| 99393 | TATGAGTGTGGAGTCGTTGCGCATGTGTCAAGCAATTCTGAAANNNAGATCGGAA |
| 42558 | TATGAGTGTGGAGTCGTTGCGGGGAGAAAGAGTACACACACACNNNAGATCGGAA |
| 39180 | TATGAGTGTGGAGTCGTTGCGGTGTAGGTATGTGTAGGTGGGANNNAGATCGGA |
| 66042 | TATGAGTGTGGAGTCGTTGCCAGAAGGAACAGGACAAAGCTGTNNNAGATCGGAA |
| 40182 | TATGAGTGTGGAGTCGTTGCTCCACTTGCCATTTCAAAGGTGANNNAGATCGGAAG |
| 66708 | TATGAGTGTGGAGTCGTTGCACAAGCACCTAAACTTCCGTTGTNNNAGATCGGAAG |
| 32041 | TATGAGTGTGGAGTCGTTGCGCACATCTCTTCCAGAGCATGAANNNAGATCGGAAG |
| 106958 | TATGAGTGTGGAGTCGTTGCAGTTGCATTTTTCCAAAGAAATGATAANNNAGATCG |
| 34435 | TATGAGTGTGGAGTCGTTGCACAGACAGTGGTACACATATCCANNNAGATCGGAA |
| 61181 | TATGAGTGTGGAGTCGTTGCAGTAACAAAGATGGTTTGGGCCTNNNAGATCGGAA |
| 30274 | TATGAGTGTGGAGTCGTTGCACGCACACACAGAGCTATACAAANNNAGATCGGAAG |
| 69572 | TATGAGTGTGGAGTCGTTGCGGTAGGGCAACATCTCTGGTTTTNNNAGATCGGAA |
| 54088 | TATGAGTGTGGAGTCGTTGCCTTTTCTCTGTGGCTGTACTGGGNNNAGATCGGAAG |
| 57715 | TATGAGTGTGGAGTCGTTGCGAAAAAGCCCACATCAGCCAAATNNNAGATCGGAAG |
| 38097 | TATGAGTGTGGAGTCGTTGCACCTCTCTGTACCAAGATAGAAGAGANNNAGATCGG |
| 40155 | TATGAGTGTGGAGTCGTTGCAGAGAGAGAGGAGAGACTGAGTGANNNAGATCGG |
| 62842 | TATGAGTGTGGAGTCGTTGCTCAGAAAGAACTGAGCACCCTCTNNNAGATCGGAAG |
| 63609 | TATGAGTGTGGAGTCGTTGCAACGAGAATTGCTCCCCAAAAGTNNNAGATCGGAAG |
| 49492 | TATGAGTGTGGAGTCGTTGCGAAACAGCAGCAGGAGGAAATTCNNNAGATCGGAA |
| 47133 | TATGAGTGTGGAGTCGTTGCATCCCTAAACCATTCCAATGGGCNNNAGATCGGAAG |
| 109883 | TATGAGTGTGGAGTCGTTGCCAAAATCCAGAGGGGCAGTCCCNNNAGATCGGAAG |
| 30401 | TATGAGTGTGGAGTCGTTGCCCTGTCATTCCTCTTCCTGCAAANNNAGATCGGAAGA |
| 46845 | TATGAGTGTGGAGTCGTTGCCTAACAGGATGGTTCAGGAAGGCNNNAGATCGGAA |
| 102845 | TATGAGTGTGGAGTCGTTGCAAGCTCCAGGGCTATCCTTGAGAGNNNAGATCGGAA |
| 66015 | TATGAGTGTGGAGTCGTTGCGGTTGCTCTGTCGTAAATCCTGTNNNAGATCGGAAG |
| 45221 | TATGAGTGTGGAGTCGTTGCACCAAGATGTGACTGTAACTGCCNNNAGATCGGAAG |

| | |
|---|---|
| 55719 | TATGAGTGTGGAGTCGTTGCGAGTGTGGGCCAGTGTATGTATGNNNAGATCGGAA |
| 62536 | TATGAGTGTGGAGTCGTTGCTGTGTGTGTGTCTCAGTTTTGCTNNNAGATCGGAAG |
| 56578 | TATGAGTGTGGAGTCGTTGCGTTCAAAGTGACAGCTTCCAGTGNNNAGATCGGAA |
| 107012 | TATGAGTGTGGAGTCGTTGCACACAGTAGTATATGATTCGGCCATNNNAGATCGGA |
| 50402 | TATGAGTGTGGAGTCGTTGCCCATGCAGAGAGAACAGGAGAAGNNNAGATCGGA |
| 49609 | TATGAGTGTGGAGTCGTTGCCCAGAAGGACATTTCCCCACTTCNNNAGATCGGAAG |
| 62274 | TATGAGTGTGGAGTCGTTGCTGTTGTCTGATTGCTCAACTGCTNNNAGATCGGAAG |
| 34694 | TATGAGTGTGGAGTCGTTGCTGGGAAGTCCATAGGTAATGCCANNNAGATCGGAA |
| 40658 | TATGAGTGTGGAGTCGTTGCTTTGGCCACTCACTGGGTGAATANNNAGATCGGAAG |
| 31650 | TATGAGTGTGGAGTCGTTGCCCTGTGTGCATGAATCTGCAGAANNNAGATCGGAA |
| 32272 | TATGAGTGTGGAGTCGTTGCAGGCACTGGGGGTATGAAGATAANNNAGATCGGA |
| 105379 | TATGAGTGTGGAGTCGTTGCATAACTAAAAGGGGAGAAAAGCTTGNNNAGATCGG |
| 66232 | TATGAGTGTGGAGTCGTTGCGGAGGTGTTCTACCCTCTTCTGTNNNAGATCGGAAG |
| 110678 | TATGAGTGTGGAGTCGTTGCAGAGATTCATAGTTTGGATTGGCATGNNNAGATCG |
| 46124 | TATGAGTGTGGAGTCGTTGCATGATGACAAGAGGGTGACAAGCNNNAGATCGGA |
| 41294 | TATGAGTGTGGAGTCGTTGCCACCAATTGCCTTTCTGGGTCTANNNAGATCGGAAG |
| 104746 | TATGAGTGTGGAGTCGTTGCACATCTGGAAAGTGACCAGGGANNNAGATCGGAAG |
| 34561 | TATGAGTGTGGAGTCGTTGCTCTAGCTTTGGAAATGGTAAGCCANNNAGATCGGA |
| 44645 | TATGAGTGTGGAGTCGTTGCTCACTAACCTCTCTTCCCATCCCNNNAGATCGGAAGA |
| 55367 | TATGAGTGTGGAGTCGTTGCACACAGTTTCACCAATGCACATGNNNAGATCGGAAG |
| 71747 | TATGAGTGTGGAGTCGTTGCGCTGCCTCATCCTCATCGTTATTNNNAGATCGGAAGA |
| 51962 | TATGAGTGTGGAGTCGTTGCAGGGGGTCTGGGGAAAGAAATAGNNNAGATCGGA |
| 69057 | TATGAGTGTGGAGTCGTTGCACATCCACCATCTTCCCTCCTTTNNNAGATCGGAAGA |
| 48777 | TATGAGTGTGGAGTCGTTGCAACTTCCTTCTCCGAATTGCCTCNNNAGATCGGAAGA |
| 59660 | TATGAGTGTGGAGTCGTTGCGCAGGTGGACAGTGGTTCATTATNNNAGATCGGAA |
| 46202 | TATGAGTGTGGAGTCGTTGCTTTTCCAGGTACCAATGGGAAGCNNNAGATCGGAA |
| 57662 | TATGAGTGTGGAGTCGTTGCTCTGGCTGTGCTTTTGTGTTTTGNNNAGATCGGAAG |
| 50445 | TATGAGTGTGGAGTCGTTGCACAGATTTTCTGCCACAGGGAAGNNNAGATCGGAA |
| 116936 | TATGAGTGTGGAGTCGTTGCCCTGAATGGGGTGACCTCTTTGGNNNAGATCGGAA |
| 62207 | TATGAGTGTGGAGTCGTTGCACCTTTTCCTTCCTTTTCCTAATGCTNNNAGATCGGAA |
| 102253 | TATGAGTGTGGAGTCGTTGCCAGGGTGTTATTCTGTCATCCAAGNNNAGATCGGAA |
| 44312 | TATGAGTGTGGAGTCGTTGCTTCTGACAGTCCTCATCTGACCCNNNAGATCGGAAGA |
| 40799 | TATGAGTGTGGAGTCGTTGCGCCCTACTATGTACCCACAGATANNNAGATCGGAAG |
| 44285 | TATGAGTGTGGAGTCGTTGCTGCCAAAACTTGTTTACATTCACCCNNNAGATCGGAA |
| 108432 | TATGAGTGTGGAGTCGTTGCCCAATCTCCCTCTGCTTCCTTTTTATNNNAGATCGGAA |
| 57759 | TATGAGTGTGGAGTCGTTGCGGGGACTGGGAGGATGAAGAAATNNNAGATCGGA |
| 51020 | TATGAGTGTGGAGTCGTTGCCCACAGAATTTCAAAGCCAGCAGNNNAGATCGGAAG |
| 103881 | TATGAGTGTGGAGTCGTTGCGAGAGATGACTAGCAGGAATTGCAANNNAGATCGG |
| 51014 | TATGAGTGTGGAGTCGTTGCGCAAAGCAAAGAAAACACAGCAGNNNAGATCGGAA |
| 104501 | TATGAGTGTGGAGTCGTTGCTTCTTTGCATATGTTTTGCTAAGCCTNNNAGATCGGA |
| 32653 | TATGAGTGTGGAGTCGTTGCTGGTTGTGTATGTGTGCTAAAAACANNNAGATCGG |
| 117363 | TATGAGTGTGGAGTCGTTGCGACACAGTGAGAAGGCAAGCTAANNNAGATCGGAA |
| 98848 | TATGAGTGTGGAGTCGTTGCCCATACTGAGCTCCTCTGCTAAANNNAGATCGGAAG |

50694 TATGAGTGTGGAGTCGTTGCGCTCACTTGAAGGTGTCTCACAGNNNAGATCGGAAG
56320 TATGAGTGTGGAGTCGTTGCATGCCTGATCCCAAACTCATCTGNNNAGATCGGAAG
45429 TATGAGTGTGGAGTCGTTGCGGGTGTACCATCTGTTTCCATCCNNNAGATCGGAAG
97900 TATGAGTGTGGAGTCGTTGCGGTTATTTATCACTTCCGGCTGCNNNAGATCGGAAG
96635 TATGAGTGTGGAGTCGTTGCGTCAGAGCTAAGCAAAGCCCCCNNNAGATCGGAAG
39377 TATGAGTGTGGAGTCGTTGCTCTGTTGTCTCTGTCTCTCTGGANNNAGATCGGAAG
54907 TATGAGTGTGGAGTCGTTGCAGAGAGAGAAGGTAAAGGGGTGGNNNAGATCGG
50307 TATGAGTGTGGAGTCGTTGCCTGCACTTTCACTTCTTGGCAAGNNNAGATCGGAAG
36751 TATGAGTGTGGAGTCGTTGCATTCTCTAGGCAGGCCAATCTCANNNAGATCGGAAG
40841 TATGAGTGTGGAGTCGTTGCGGACACCCCTTAGTTGAGGGATANNNAGATCGGAA
63546 TATGAGTGTGGAGTCGTTGCGCACAAAAGTATTGCCAGGTTTCTNNNAGATCGGAA
107767 TATGAGTGTGGAGTCGTTGCGTGATGAGGAACTGACCTGACNNNAGATCGGAAGA
58145 TATGAGTGTGGAGTCGTTGCGGAGAAGAGGGAGGGACTTGAATNNNAGATCGGA
31681 TATGAGTGTGGAGTCGTTGCCAAGCACAGACACGATCAGAGAANNNAGATCGGAA
59292 TATGAGTGTGGAGTCGTTGCAAGTGAACCGTACTGGGACTGATNNNAGATCGGAA
67767 TATGAGTGTGGAGTCGTTGCTTTTTCCATGAGCCAGCATCCTTNNNAGATCGGAAG
35505 TATGAGTGTGGAGTCGTTGCTAATGTGGGGAAGGAGTGTAGCANNNAGATCGGA
103880 TATGAGTGTGGAGTCGTTGCGGGTTGGGGTGAGCAAAGTTTNNNAGATCGGAAG
59603 TATGAGTGTGGAGTCGTTGCGGGGAGAAAAGCTTGTCACGTATNNNAGATCGGAA
57618 TATGAGTGTGGAGTCGTTGCCTTGGTTTTCCTCGGTGTGTTTGNNNAGATCGGAAG
63971 TATGAGTGTGGAGTCGTTGCCAATAGAGCCTAGAGCTGCCAGTNNNAGATCGGAA
46443 TATGAGTGTGGAGTCGTTGCGTTCCTCTCCTCCCAATTTCAGCNNNAGATCGGAAGA
33678 TATGAGTGTGGAGTCGTTGCAGATCAGTGTGAGTAGGACTGACANNNAGATCGGA
44970 TATGAGTGTGGAGTCGTTGCGGATTTCTCGACTTCTCTGAGCCNNNAGATCGGAAG
61054 TATGAGTGTGGAGTCGTTGCGCTGACTTCCCAGTATTTTTCCCTNNNAGATCGGAAG
53293 TATGAGTGTGGAGTCGTTGCCAGTGTTTTCCCCTCAAGTGAGGNNNAGATCGGAAG
62612 TATGAGTGTGGAGTCGTTGCTCCCTTCAGATGTTGGACCATCTNNNAGATCGGAAG
103770 TATGAGTGTGGAGTCGTTGCCAGTATCCTCTCATGGAAGCAGCNNNAGATCGGAAG
105486 TATGAGTGTGGAGTCGTTGCGGAAACCAGGAAACCTATACTGTNNNAGATCGGAA
57760 TATGAGTGTGGAGTCGTTGCTGCTGGAACAAGTTTGAAGAAATNNNAGATCGGAA
51638 TATGAGTGTGGAGTCGTTGCAAAAAGAAGGGCAGAAGACGGAGNNNAGATCGGA
62141 TATGAGTGTGGAGTCGTTGCCGGATTTCTAAGGCTTTCTGGCTNNNAGATCGGAAG
69430 TATGAGTGTGGAGTCGTTGCGAAAGGGAGGCAGTCAGGATTTTNNNAGATCGGA
44585 TATGAGTGTGGAGTCGTTGCACGATACTTGCAGTTCTATGCCCNNNAGATCGGAAG
60583 TATGAGTGTGGAGTCGTTGCTTCAGGTTTCTGCTTCCAAACCTNNNAGATCGGAAG
55689 TATGAGTGTGGAGTCGTTGCCCACCCTACCCTACTCCACTATGNNNAGATCGGAAGA
63567 TATGAGTGTGGAGTCGTTGCGCCGTTGTCACCACTGATTTTCTNNNAGATCGGAAG
99775 TATGAGTGTGGAGTCGTTGCCTTTCCATGGCCTCTACTCCTCANNNAGATCGGAAGA
68158 TATGAGTGTGGAGTCGTTGCCCTAGCGGAATGCTTTCCTTCTTNNNAGATCGGAAG
65313 TATGAGTGTGGAGTCGTTGCGCTGCTTTGGCTAGTGATATGGTNNNAGATCGGAA
50329 TATGAGTGTGGAGTCGTTGCGACAGCAGGAAGCAGAGATCAAGNNNAGATCGGA
67632 TATGAGTGTGGAGTCGTTGCTGTGCATGTCTAAGTGCAACCTTNNNAGATCGGAAG
102353 TATGAGTGTGGAGTCGTTGCTCCCAGAAATCCAGCCCTGAGNNNAGATCGGAAGAG

54315 TATGAGTGTGGAGTCGTTGCTCCTTCCTACGCAGATACAATGGNNNAGATCGGAAG
55538 TATGAGTGTGGAGTCGTTGCCAGGCATAGAGACTGCCAAGATGNNNAGATCGGAA
35302 TATGAGTGTGGAGTCGTTGCTCAAATTGCAAAGAAAGGCAGCANNNAGATCGGAA
48576 TATGAGTGTGGAGTCGTTGCGCAGTCCCACAGACTAGAAACTCNNNAGATCGGAAG
60039 TATGAGTGTGGAGTCGTTGCATCTGGGGAAGCTGATAACCACTNNNAGATCGGAA
71085 TATGAGTGTGGAGTCGTTGCACAGGACAGGCTGGATAAGAACTNNNAGATCGGAA
55352 TATGAGTGTGGAGTCGTTGCGGGGTTTGGTAGAACCCAACATGNNNAGATCGGAA
31531 TATGAGTGTGGAGTCGTTGCGCTGGGTTTTGAGGGAGAAAGAANNNAGATCGGA
34402 TATGAGTGTGGAGTCGTTGCGGGAGTTTCTATTGTTCTGCCCANNNAGATCGGAAG
37542 TATGAGTGTGGAGTCGTTGCTGCTAAATTCCTTTGGAACCAAGANNNAGATCGGAA
66367 TATGAGTGTGGAGTCGTTGCAGATATGAGACAGTGCCAGGTGTNNNAGATCGGAA
102800 TATGAGTGTGGAGTCGTTGCGAGATATGAGACAGTGCCAGGTGTNNNAGATCGGA
65190 TATGAGTGTGGAGTCGTTGCGAGATAGCCATTTTTGGCTGGGTNNNAGATCGGAA
44717 TATGAGTGTGGAGTCGTTGCCCTCCCTATCCAGATATGCTCCCNNNAGATCGGAAGA
64530 TATGAGTGTGGAGTCGTTGCTTTCATTGTGGTGTGGAGAACGTNNNAGATCGGAA
104107 TATGAGTGTGGAGTCGTTGCGTAACTTCTAATTGGCCATCTTGCTGNNNAGATCGG
43397 TATGAGTGTGGAGTCGTTGCAAATACAAGTGGCCTCAGCTGACNNNAGATCGGAA
61932 TATGAGTGTGGAGTCGTTGCACTTTGGAAATGACCTGTTCGCTNNNAGATCGGAAG
42169 TATGAGTGTGGAGTCGTTGCTATGTATGCCTCTGGCCACAAACNNNAGATCGGAAG
40319 TATGAGTGTGGAGTCGTTGCGTGTTCAAGGGTCAACTGTGTGANNNAGATCGGAA
108403 TATGAGTGTGGAGTCGTTGCACAGTAGGTAACAAATGCTAGTTTGTNNNAGATCGG
67459 TATGAGTGTGGAGTCGTTGCCACACTCCTCCAGGCAAAAACTTNNNAGATCGGAAG
47316 TATGAGTGTGGAGTCGTTGCCTCTGTCTCACCATACAAGTGGCNNNAGATCGGAAG
59015 TATGAGTGTGGAGTCGTTGCAAAACAATTGCCCTGGGTCAGATNNNAGATCGGAA
55017 TATGAGTGTGGAGTCGTTGCGCATTGTGGGTTCTGACAATTGGNNNAGATCGGAA
108421 TATGAGTGTGGAGTCGTTGCTATTAGGATGACTTGATCCAGAGCTGANNNAGATCG
32056 TATGAGTGTGGAGTCGTTGCTGGAAAACGACACAGTGGATGAANNNAGATCGGAA
116701 TATGAGTGTGGAGTCGTTGCGGAAAACGACACAGTGGATGAANNNAGATCGGAA
43017 TATGAGTGTGGAGTCGTTGCCTGGCACTGTAATCCTATAGTCTCACNNNAGATCGG
59608 TATGAGTGTGGAGTCGTTGCACTTGCCCTGTATACTGTCGTATNNNAGATCGGAAG
45493 TATGAGTGTGGAGTCGTTGCACACCTGACAAAGGACTTCTATCCNNNAGATCGGAA
51772 TATGAGTGTGGAGTCGTTGCGATTGCCAGCTTTCGGAAATGAGNNNAGATCGGAA
59232 TATGAGTGTGGAGTCGTTGCCCACTTGAGGGTTTTCTGTGGATNNNAGATCGGAAG
49973 TATGAGTGTGGAGTCGTTGCTCATGTGTGCCTGTTGCATTTTCNNNAGATCGGAAG
46645 TATGAGTGTGGAGTCGTTGCAAAAACCTGCCAGCCTAACTAGCNNNAGATCGGAAG
62019 TATGAGTGTGGAGTCGTTGCTCTGAATAACTCTTATGCACGGCTNNNAGATCGGAA
61443 TATGAGTGTGGAGTCGTTGCTGCTAAGCCTCTTTCTCAGTCCTNNNAGATCGGAAGA
104113 TATGAGTGTGGAGTCGTTGCCATTTTGTCTAATGAGAGCCCCTCANNNAGATCGGA
47629 TATGAGTGTGGAGTCGTTGCACTGCAAGGATAAACTCAAAACTGCNNNAGATCGGA
40902 TATGAGTGTGGAGTCGTTGCGCTGAGGGATGGTACAGCCTATANNNAGATCGGAA
60474 TATGAGTGTGGAGTCGTTGCAGATCATCCTGGTTTTGCCTACTNNNAGATCGGAAG
35057 TATGAGTGTGGAGTCGTTGCCCTGAGAAGGAGGGAATCTTCCANNNAGATCGGAA
67620 TATGAGTGTGGAGTCGTTGCGGAAGGTGAGGGCATTTGTACTTNNNAGATCGGAA

45275 TATGAGTGTGGAGTCGTTGCTACAAAAGAAGCTGGGATCTGCCNNNAGATCGGAA
60313 TATGAGTGTGGAGTCGTTGCACATAGGGGACATGAAAAGGACTNNNAGATCGGAA
98212 TATGAGTGTGGAGTCGTTGCCAGAACTCTAAGATGCTTGGGCANNNAGATCGGAA
66806 TATGAGTGTGGAGTCGTTGCTTACTCTCTCCTGCTGGCTTTGTNNNAGATCGGAAGA
67525 TATGAGTGTGGAGTCGTTGCAAGCTCTATTACAGTGGGCACTTNNNAGATCGGAAG
47168 TATGAGTGTGGAGTCGTTGCTGCTAAGATAGCTCTGTCTGGGCNNNAGATCGGAA
102470 TATGAGTGTGGAGTCGTTGCTTTTGTGTCTGTGTGTCCAGCATNNNAGATCGGAAG
49665 TATGAGTGTGGAGTCGTTGCCCTCATGCTCTGACACATCCTTCNNNAGATCGGAAGA
51572 TATGAGTGTGGAGTCGTTGCAGGCAAAGGAAGACCAAAAGGAGNNNAGATCGGA
46464 TATGAGTGTGGAGTCGTTGCTAGGGGAAGCTGTTTTGAAGAGCNNNAGATCGGAA
108445 TATGAGTGTGGAGTCGTTGCATGTCACCAGTTTCTCCCCCTTNNNAGATCGGAAGAG
57613 TATGAGTGTGGAGTCGTTGCACATGCCAATACCACACTGTTTGNNNAGATCGGAAG
59210 TATGAGTGTGGAGTCGTTGCTGCCATATTTGTGATTTCATGGATNNNAGATCGGAA
67118 TATGAGTGTGGAGTCGTTGCACACAGTCATTCACAATGCCATTNNNAGATCGGAAG
52511 TATGAGTGTGGAGTCGTTGCTGATGTGATGCTTTTAGGAGGCGNNNAGATCGGAA
97631 TATGAGTGTGGAGTCGTTGCGCCGGGTTTTTGTTCCCCTAGATTTAGNNNAGATCG
47532 TATGAGTGTGGAGTCGTTGCAGAGAGAGAAAGGCTATGCATGCNNNAGATCGGAA
68482 TATGAGTGTGGAGTCGTTGCGACAAGTAGCGGTCTTCATGGTTNNNAGATCGGAA
38740 TATGAGTGTGGAGTCGTTGCGTTTTCATCATCCTCAGCCAGGANNNAGATCGGAAG
51582 TATGAGTGTGGAGTCGTTGCGGATTGGAAGGTTCGAGAAGGAGNNNAGATCGGA
43564 TATGAGTGTGGAGTCGTTGCCCATCCCCTGAATTGACAGCTACNNNAGATCGGAAG
69305 TATGAGTGTGGAGTCGTTGCCCCAGGTATTATGTGTGCGGTTTNNNAGATCGGAAG
39511 TATGAGTGTGGAGTCGTTGCACAGCTACTTGACGTTAGTTGGANNNAGATCGGAA
33874 TATGAGTGTGGAGTCGTTGCCCGTGAGTTTGCCCATTAGTACANNNAGATCGGAAG
42095 TATGAGTGTGGAGTCGTTGCGGGATGGGTGATGTTGCCTTTTANNNAGATCGGAA
31819 TATGAGTGTGGAGTCGTTGCAGTTTAGATAGGCTGGCCAGGAANNNAGATCGGAA
46571 TATGAGTGTGGAGTCGTTGCGAGGGAGAAAATCTGGTTGGAGCNNNAGATCGGA
51995 TATGAGTGTGGAGTCGTTGCGAGTAGTGCCAAGAAGCGGATAGNNNAGATCGGA
55220 TATGAGTGTGGAGTCGTTGCGCTCTCCTCTGGGTCTCAAAATGNNNAGATCGGAAG
106088 TATGAGTGTGGAGTCGTTGCCCTGTGCCAAATGTGTTATACAAATGANNNAGATCG
64256 TATGAGTGTGGAGTCGTTGCTGCTATACGTGTCAGTGAGGAGTNNNAGATCGGAA
32090 TATGAGTGTGGAGTCGTTGCTGCCCAGATAGACTTGACCTGAANNNAGATCGGAAG
67523 TATGAGTGTGGAGTCGTTGCTCCTTTTGGAAACTTCAGCACTTNNNAGATCGGAAG
38998 TATGAGTGTGGAGTCGTTGCCCAAGGTTTGATTTGGACAGGGANNNAGATCGGAA
104013 TATGAGTGTGGAGTCGTTGCGTGTGTGGTACATTTACTCCTTGNNNAGATCGGAAG
62047 TATGAGTGTGGAGTCGTTGCATTTATTGATGGGCACATGGGCTNNNAGATCGGAA
39140 TATGAGTGTGGAGTCGTTGCTGAGGGTAAAAAGAACAACTGGGANNNAGATCGG
44187 TATGAGTGTGGAGTCGTTGCCACACACGACTGAATTGTGTACCNNNAGATCGGAAG
121986 TATGAGTGTGGAGTCGTTGCGAGAGGGCTACGAAGTGAGAGTAGNNNAGATCGG
101379 TATGAGTGTGGAGTCGTTGCTTTGTGCTGTGGTATGTTGCAGTNNNAGATCGGAA
33319 TATGAGTGTGGAGTCGTTGCTGTTATGCATCTGTACTATCTGCACANNNAGATCGG
35949 TATGAGTGTGGAGTCGTTGCAGTTTTGGGTCTTTCTAGCATGCANNNAGATCGGAA
31049 TATGAGTGTGGAGTCGTTGCCCACTTGATGTTGGAACTCCCAANNNAGATCGGAAG

115968 TATGAGTGTGGAGTCGTTGCAGCCCTCCTAGCCACCTAAAGAATNNNAGATCGGAA
44223 TATGAGTGTGGAGTCGTTGCTAGCCAAACATTGCATCAAACCCNNNAGATCGGAAG
65400 TATGAGTGTGGAGTCGTTGCTCCAGTAGGTTCTGTTTCTCTGGTNNNAGATCGGAA
60707 TATGAGTGTGGAGTCGTTGCCTCAGCAACAGTGGAATTCACCTNNNAGATCGGAAG
52763 TATGAGTGTGGAGTCGTTGCTCAGGGAGTTCAAATCAGCAAGGNNNAGATCGGAA
110614 TATGAGTGTGGAGTCGTTGCAGGTAGGTAAGTGGAGAGAGTGAGANNNAGATCG
64625 TATGAGTGTGGAGTCGTTGCGAAGACAAGAGTATTGGCAGCGTNNNAGATCGGAA
49873 TATGAGTGTGGAGTCGTTGCCTGTGCCTTCCGGATTGCATTTCNNNAGATCGGAAG
36506 TATGAGTGTGGAGTCGTTGCATCTGGCTGGCTGAAGGATATCANNNAGATCGGAA
61546 TATGAGTGTGGAGTCGTTGCTGGTTTGTCAGTTCACAGTTCCTNNNAGATCGGAAG
57216 TATGAGTGTGGAGTCGTTGCCCTGTGGCTTTTTGAGCTGATTGNNNAGATCGGAAG
64936 TATGAGTGTGGAGTCGTTGCTTTACTAACCAGCAGGGTGAGGTNNNAGATCGGAA
50482 TATGAGTGTGGAGTCGTTGCACAAGAGGGCATGAGTTTGGAAGNNNAGATCGGA
99784 TATGAGTGTGGAGTCGTTGCGAACCTTGCCCTTTTCCTGTTGANNNAGATCGGAAG
29811 TATGAGTGTGGAGTCGTTGCTCCCATTCTGTTTTGATGCTCCANNNAGATCGGAAGA
63610 TATGAGTGTGGAGTCGTTGCCTAACTTGGGTGGAGCCAAAAGTNNNAGATCGGAA
38081 TATGAGTGTGGAGTCGTTGCAGAGACCAGAGGCAAGAAAGAGANNNAGATCGGA
66440 TATGAGTGTGGAGTCGTTGCTCTTGTGAAGGTTCTCACTGTGTNNNAGATCGGAAG
57069 TATGAGTGTGGAGTCGTTGCGTGTATGCTTGCAGATGTGTGTGNNNAGATCGGAA
105518 TATGAGTGTGGAGTCGTTGCAACAGTAGCAGGAACATGGCTCATNNNAGATCGGA
69311 TATGAGTGTGGAGTCGTTGCGATTACAGTCACTGGCAGGGTTTNNNAGATCGGAA
31874 TATGAGTGTGGAGTCGTTGCGGACATCACAGATGAAACCGGAANNNAGATCGGAA
63649 TATGAGTGTGGAGTCGTTGCTTGGTGGCAAAGGAACAGAAAGTNNNAGATCGGA
56626 TATGAGTGTGGAGTCGTTGCACACACAAAATTAGGCGGGAGTGNNNAGATCGGAA
109924 TATGAGTGTGGAGTCGTTGCATACACACAAAATTAGGCGGGAGTGNNNAGATCGG
46551 TATGAGTGTGGAGTCGTTGCGTAATGGGGTTTTAGGAGGGAGCNNNAGATCGGA
42032 TATGAGTGTGGAGTCGTTGCGAAATCAGGGTGAGCTGGCTTTANNNAGATCGGAA
59942 TATGAGTGTGGAGTCGTTGCGCGTGATTACATGCAGGGTAACTNNNAGATCGGAA
38278 TATGAGTGTGGAGTCGTTGCCAGTTGCTCATCCTTTGTGGAGANNNAGATCGGAAG
49206 TATGAGTGTGGAGTCGTTGCTGAGGACCAAACAGTTACCAGTCNNNAGATCGGAA
99780 TATGAGTGTGGAGTCGTTGCTGGTTGCCACTGTGAGTTCAGNNNAGATCGGAAGA
49786 TATGAGTGTGGAGTCGTTGCATACGCCTGCATGAAAAACGTTCNNNAGATCGGAAG
122012 TATGAGTGTGGAGTCGTTGCGAGCAACAGAAAGAACACAATTGCNNNAGATCGGA
70388 TATGAGTGTGGAGTCGTTGCAATGACCTCCTAGTTGCCAAAGCNNNAGATCGGAAG
63377 TATGAGTGTGGAGTCGTTGCCGGAGGACACATTTTCACCTTCTNNNAGATCGGAAG
30597 TATGAGTGTGGAGTCGTTGCTATGTAGCTGGGGTGAAGGGAAANNNAGATCGGA
68139 TATGAGTGTGGAGTCGTTGCGAGGTGCCCATTCTTCCATTCTTNNNAGATCGGAAG
52801 TATGAGTGTGGAGTCGTTGCTGGGTGTTTGGATTACAGGAAGGNNNAGATCGGA
41139 TATGAGTGTGGAGTCGTTGCATTGTGTGTGTTGCTTCAAGCTANNNAGATCGGAAG
56005 TATGAGTGTGGAGTCGTTGCAGCATAACTTTGAGTGGAGCCTGNNNAGATCGGAA
50443 TATGAGTGTGGAGTCGTTGCCTGGTTGGTATGGGAAAGGGAAGNNNAGATCGGA
61493 TATGAGTGTGGAGTCGTTGCGCACCCATTATTCACCCATTCCTNNNAGATCGGAAGA
36839 TATGAGTGTGGAGTCGTTGCTCAGGACAGAGAAGGATTTCTCANNNAGATCGGAA

| | |
|---|---|
| 48790 | TATGAGTGTGGAGTCGTTGCAGAAAGTCCCTGAAGACCTCCTCNNNAGATCGGAAG |
| 45288 | TATGAGTGTGGAGTCGTTGCGATGACCCACTGATACTTCTGCCNNNAGATCGGAAG |
| 106093 | TATGAGTGTGGAGTCGTTGCTGACCCACTGATACTTCTGCCNNNAGATCGGAAGAG |
| 50487 | TATGAGTGTGGAGTCGTTGCGACCCAAATGTTAGCGCATGAAGNNNAGATCGGAA |
| 29935 | TATGAGTGTGGAGTCGTTGCAGTTCATCTGAGCATTCCAAGCCNNNAGATCGGAAG |
| 121999 | TATGAGTGTGGAGTCGTTGCGTTCCCGTGGAGGAAGAGGTGGNNNAGATCGGAA |
| 62240 | TATGAGTGTGGAGTCGTTGCATTTCTGTGCTGCTGTAGATGCTNNNAGATCGGAAG |
| 55613 | TATGAGTGTGGAGTCGTTGCGGACAGAATGAATCCTGGGGATGNNNAGATCGGA |
| 35142 | TATGAGTGTGGAGTCGTTGCCCCCCAGGATAAAATCCAAAGCANNNAGATCGGAAG |
| 61190 | TATGAGTGTGGAGTCGTTGCTCCTGATATTCTGTGTCTGGCCTNNNAGATCGGAAG |
| 41669 | TATGAGTGTGGAGTCGTTGCCATCACATTGTACGCCTGGTGTANNNAGATCGGAAG |
| 63340 | TATGAGTGTGGAGTCGTTGCTCCTGAGGGGTTGATGTCATTCTNNNAGATCGGAAG |
| 59057 | TATGAGTGTGGAGTCGTTGCAACCATGACAATTCCGTGGAGATNNNAGATCGGAA |
| 35322 | TATGAGTGTGGAGTCGTTGCGGAAGCTTAGACACACATCAGCANNNAGATCGGAA |
| 40006 | TATGAGTGTGGAGTCGTTGCACACACAAGCAGGAGAAATCTGANNNAGATCGGAA |
| 48816 | TATGAGTGTGGAGTCGTTGCACCTCTCAGGAACAGGATTCCTCNNNAGATCGGAAG |
| 51871 | TATGAGTGTGGAGTCGTTGCTGGCTAAATCACAGCTGTCTGAGNNNAGATCGGAA |
| 68684 | TATGAGTGTGGAGTCGTTGCCTCAAAGGGGTGGGGAATGTGTTNNNAGATCGGA |
| 61455 | TATGAGTGTGGAGTCGTTGCCTTGTGGCTTAGAGGTAGGTCCTNNNAGATCGGAA |
| 38055 | TATGAGTGTGGAGTCGTTGCCAGCAGATTCTATACCCTTCAGANNNAGATCGGAAG |
| 36350 | TATGAGTGTGGAGTCGTTGCGCCAACCCTGGTACCATAAATCANNNAGATCGGAAG |
| 31967 | TATGAGTGTGGAGTCGTTGCAGGTCTTGTCTTGGCTACTGGAANNNAGATCGGAA |
| 49643 | TATGAGTGTGGAGTCGTTGCGCCTTCTCAGTTCAGCTACCTTCNNNAGATCGGAAGA |
| 59908 | TATGAGTGTGGAGTCGTTGCGTGGATGAAGCCATCTTGGAACTNNNAGATCGGAA |
| 40394 | TATGAGTGTGGAGTCGTTGCACTGGAGAAAGTAGGGTGTATTGANNNAGATCGGA |
| 51040 | TATGAGTGTGGAGTCGTTGCAAAGAAACCAGACTGCAGAGCAGNNNAGATCGGAA |
| 58153 | TATGAGTGTGGAGTCGTTGCACGCCACTTTCAGCCTCAATAATNNNAGATCGGAAG |
| 51186 | TATGAGTGTGGAGTCGTTGCTTCGGCCATCCTTTCTGAATCAGNNNAGATCGGAAG |
| 54711 | TATGAGTGTGGAGTCGTTGCTTTCTCCAAGGCTAGAATGCTGGNNNAGATCGGAAG |
| 57777 | TATGAGTGTGGAGTCGTTGCGGCACGTCTTTGTCCTGAGAAATNNNAGATCGGAAG |
| 117286 | TATGAGTGTGGAGTCGTTGCTTCTTGGCCTCGCTCTGTCATTCNNNAGATCGGAAGA |
| 97256 | TATGAGTGTGGAGTCGTTGCCAGACCTAGGGGCAGACAATTNNNAGATCGGAAGA |
| 29809 | TATGAGTGTGGAGTCGTTGCTAGACCAGTTGGGGGTGTGTGNNNAGATCGGAAGA |
| 33907 | TATGAGTGTGGAGTCGTTGCCTGTTCCACTGATTGCCCTTACANNNAGATCGGAAGA |
| 62886 | TATGAGTGTGGAGTCGTTGCGGTTGCATCTGTCCAAATCCTCTNNNAGATCGGAAG |
| 100318 | TATGAGTGTGGAGTCGTTGCCACACACTCACATCCACCTTCCNNNAGATCGGAAGAG |
| 102536 | TATGAGTGTGGAGTCGTTGCTGTCTTTCTCTGCCACCCAGGATANNNAGATCGGAA |
| 41136 | TATGAGTGTGGAGTCGTTGCAGTGGAAGTAGAGCCACAAGCTANNNAGATCGGAA |
| 48465 | TATGAGTGTGGAGTCGTTGCAGAGAGAACCCTGTCCAAGGATCNNNAGATCGGAA |
| 36672 | TATGAGTGTGGAGTCGTTGCCTGTGTGAACTCATCTTTTCCTCANNNAGATCGGAAG |
| 45197 | TATGAGTGTGGAGTCGTTGCTAGGGAAAGGGGATAGAGATGCCNNNAGATCGGA |
| 41630 | TATGAGTGTGGAGTCGTTGCGAAGGAAGGCAAGCAAACCTGTANNNAGATCGGAA |
| 69163 | TATGAGTGTGGAGTCGTTGCTGGCCCTGCAGATGTCTATCTTTNNNAGATCGGAAG |

106540 TATGAGTGTGGAGTCGTTGCAGGGGTTGATGTCATTCTCCCCNNNAGATCGGAAGA
31637 TATGAGTGTGGAGTCGTTGCCACACACAGAGACATACGCAGAANNNAGATCGGAA
57876 TATGAGTGTGGAGTCGTTGCAGTGATATGGGGCCAGGTACAATNNNAGATCGGAA
51557 TATGAGTGTGGAGTCGTTGCCTGATTCCAACTCAACCTCCGAGNNNAGATCGGAAG
61995 TATGAGTGTGGAGTCGTTGCTGGACCTAATTTTGTCTGAGGCTNNNAGATCGGAAG
43939 TATGAGTGTGGAGTCGTTGCAAAGGTCTCAAAAGGCAGCACCNNNAGATCGGAAG
40600 TATGAGTGTGGAGTCGTTGCGGCAGCATACCACACAGCAAATANNNAGATCGGAA
42485 TATGAGTGTGGAGTCGTTGCGCTTCTTTGGTGCCCTTTGTAACNNNAGATCGGAAG
96171 TATGAGTGTGGAGTCGTTGCCCTCCCCGCCGTCTCTAGTANNNAGATCGGAAGAGC
50805 TATGAGTGTGGAGTCGTTGCCATAACAAATGGAGGGGGACCAGNNNAGATCGGA
35931 TATGAGTGTGGAGTCGTTGCGTCTGTATAACAATATGCACATGCANNNAGATCGGA
66126 TATGAGTGTGGAGTCGTTGCTGTGGTTTTCTTCATGTTTGCTGTNNNAGATCGGAA
104517 TATGAGTGTGGAGTCGTTGCAAACACAAATGAACAATCTCAACGATNNNAGATCGG
38859 TATGAGTGTGGAGTCGTTGCTTCCCCTGTTTCATACCTGAGGANNNAGATCGGAAG
46412 TATGAGTGTGGAGTCGTTGCCTTTCTCACCAACCTCCTCAGCNNNAGATCGGAAGAG
47530 TATGAGTGTGGAGTCGTTGCATGCCACACATTTACATGCATGCNNNAGATCGGAAG
71748 TATGAGTGTGGAGTCGTTGCGGATACGTTCACCAGAAGGAAGGNNNAGATCGGAA
67189 TATGAGTGTGGAGTCGTTGCTGCCTCCTACCTTAGCCATCATTNNNAGATCGGAAGA
108440 TATGAGTGTGGAGTCGTTGCTTGTACCCATTTTTGAGCGTTCATNNNAGATCGGAA
52346 TATGAGTGTGGAGTCGTTGCCCTTTGTGAGTGGCTAGACTACGNNNAGATCGGAA
70210 TATGAGTGTGGAGTCGTTGCGTGTGTTTCTCCACTAAGCCAACNNNAGATCGGAAG
71517 TATGAGTGTGGAGTCGTTGCCCATGTTTTCTTCTGAGCCCCTTNNNAGATCGGAAGA
95931 TATGAGTGTGGAGTCGTTGCGATGGATGTGCGGTTGGATACGTANNNAGATCGGA
29810 TATGAGTGTGGAGTCGTTGCAATGGGAAAGAAGCCATGAGCAANNNAGATCGGA
32183 TATGAGTGTGGAGTCGTTGCAGATAGATCATGGGGGCTGTGAANNNAGATCGGA
102541 TATGAGTGTGGAGTCGTTGCTCTATTCCGCAGGGTAAGCACTTNNNAGATCGGAAG
49853 TATGAGTGTGGAGTCGTTGCGGAACTGACTGGAGGAGTTGTTCNNNAGATCGGAA
48092 TATGAGTGTGGAGTCGTTGCAATAGTTTTCTGCAGGTACTTGCNNNAGATCGGAAG
62607 TATGAGTGTGGAGTCGTTGCAAGAACGCCTGCCAAATACATCTNNNAGATCGGAAG
66964 TATGAGTGTGGAGTCGTTGCCTGAGGCGTGTGAATTTGCAATTNNNAGATCGGAA
57683 TATGAGTGTGGAGTCGTTGCTAGGAGTGCCGAAGCAAGAAAATNNNAGATCGGAA
110604 TATGAGTGTGGAGTCGTTGCGAGAACAGAGTAGGCCTCACCCNNNAGATCGGAAG
71258 TATGAGTGTGGAGTCGTTGCACATAGTCTCAATTATACCTCTGGTCTNNNAGATCGG
97905 TATGAGTGTGGAGTCGTTGCACACACACATACACCAAGCAACACNNNAGATCGGAA
51305 TATGAGTGTGGAGTCGTTGCAGCACCTTGTGATGTGGATTCAGNNNAGATCGGAA
48686 TATGAGTGTGGAGTCGTTGCAAAAAGCACAGACCTCTGACCTCNNNAGATCGGAAG
105898 TATGAGTGTGGAGTCGTTGCCTGGTCACTTGGATCACCTACACANNNAGATCGGAA
60143 TATGAGTGTGGAGTCGTTGCAAGAACTCATAGGGTCAGGCACTNNNAGATCGGAA
47979 TATGAGTGTGGAGTCGTTGCGGGAGCTATCTCTGTACATGTGCNNNAGATCGGAA
100325 TATGAGTGTGGAGTCGTTGCGGCTCAGGGCACTTTTCAAAAATNNNAGATCGGAAG
101623 TATGAGTGTGGAGTCGTTGCGATAGGAGATGTTCCTGGTTGTCCNNNAGATCGGA
41906 TATGAGTGTGGAGTCGTTGCGTGTGTGAGGGACAGGAGAGTTANNNAGATCGGA
102216 TATGAGTGTGGAGTCGTTGCAGCCAAATGTTTCTTCTGGCATNNNAGATCGGAAGA

41013 TATGAGTGTGGAGTCGTTGCGGGAAGGGTGAGGTGATGTACTANNNAGATCGGA
41131 TATGAGTGTGGAGTCGTTGCCCAGCATGAGGCAGTTTTTCCTANNNAGATCGGAAG
43102 TATGAGTGTGGAGTCGTTGCTTCCCTTCCAAGAGTGCTTTCACNNNAGATCGGAAGA
55611 TATGAGTGTGGAGTCGTTGCCGTCATGATTAGAGAGCGGGATGNNNAGATCGGAA
54319 TATGAGTGTGGAGTCGTTGCGCTTTCTGTTTCTTGGCCAATGGNNNAGATCGGAAG
57378 TATGAGTGTGGAGTCGTTGCAGGTAGGGGCTTTGTTTGTCTTGNNNAGATCGGAA
110401 TATGAGTGTGGAGTCGTTGCTTGGTGCCCTTTGTAACCCTTCNNNAGATCGGAAGA
31325 TATGAGTGTGGAGTCGTTGCATTCTTTGGGAAGCAGTCTGCAANNNAGATCGGAA
102248 TATGAGTGTGGAGTCGTTGCTGTACCTGGCAAGCTGTAGACNNNAGATCGGAAGA
64797 TATGAGTGTGGAGTCGTTGCACTATGAGACTGAAGCCACAGGTNNNAGATCGGAA
68572 TATGAGTGTGGAGTCGTTGCAGAAAGGGGATTGTGTGCATGTTNNNAGATCGGAA
52241 TATGAGTGTGGAGTCGTTGCAACGAATTGAGGCATGTCAAACGNNNAGATCGGAA
41700 TATGAGTGTGGAGTCGTTGCTACCTCCATGCCCAGTTCTTGTANNNAGATCGGAAGA
34213 TATGAGTGTGGAGTCGTTGCCGTTTCCAGATTCACAAAACCCANNNAGATCGGAAG
95928 TATGAGTGTGGAGTCGTTGCAAAGCTACACCCACTCCCCACNNNAGATCGGAAGAG
96746 TATGAGTGTGGAGTCGTTGCCAACAGGAGAAACAGAACAAAACGANNNAGATCGG
31167 TATGAGTGTGGAGTCGTTGCCAAATGCTGACACCCAAAAGCAANNNAGATCGGAAG
33207 TATGAGTGTGGAGTCGTTGCTTATGAATCCCCTCTGAGCCACANNNAGATCGGAAG
63294 TATGAGTGTGGAGTCGTTGCTTTCTGGTTGCATCGAGTTGTCTNNNAGATCGGAAG
109041 TATGAGTGTGGAGTCGTTGCTGGGAATAAAGACTGGTACAGAGACNNNAGATCGG
54403 TATGAGTGTGGAGTCGTTGCTCCTGATGAAAGCCATCTCATGGNNNAGATCGGAAG
56277 TATGAGTGTGGAGTCGTTGCTTCTTAGCATGCACTTTGTGCTGNNNAGATCGGAAG
96261 TATGAGTGTGGAGTCGTTGCGCCAGGGGCTCTAAGAGTGANNNAGATCGGAAGA
52442 TATGAGTGTGGAGTCGTTGCAGTTGGGGTGGAAAGTAAATCCGNNNAGATCGGA
31720 TATGAGTGTGGAGTCGTTGCTCCAGATTGAAAAACACCTGAGAANNNAGATCGGA
104765 TATGAGTGTGGAGTCGTTGCTCTACATATCAAGCCTGTGCCCNNNAGATCGGAAGA
104132 TATGAGTGTGGAGTCGTTGCTTTCTTGCTTTGGTCTTGCAATTCNNNAGATCGGAAG
44740 TATGAGTGTGGAGTCGTTGCATTGATGGTGCTGCTATTCTCCCNNNAGATCGGAAG
43917 TATGAGTGTGGAGTCGTTGCCCAAATCAGATTCAATGGCCACCNNNAGATCGGAAG
52834 TATGAGTGTGGAGTCGTTGCAGCAACATGAAGTGAGCTGAAGGNNNAGATCGGA
60858 TATGAGTGTGGAGTCGTTGCGTTACATAGGGCACATTGACCCTNNNAGATCGGAAG
55867 TATGAGTGTGGAGTCGTTGCTCACAGTGCAGATTTCCTGACTGNNNAGATCGGAAG
55408 TATGAGTGTGGAGTCGTTGCGCAGGAAGAGACAAGACCCATGNNNAGATCGGAAG
43233 TATGAGTGTGGAGTCGTTGCCCCAGACTTCAAAGGCCATAGACNNNAGATCGGAAG
68456 TATGAGTGTGGAGTCGTTGCGCAAGTCTATTTTCCAGGGGGTTNNNAGATCGGAA
44393 TATGAGTGTGGAGTCGTTGCTATGTTTGTCTCCTCACTCCCCCNNNAGATCGGAAGA
60971 TATGAGTGTGGAGTCGTTGCTCTACATATCAAGCCTGTGCCCTNNNAGATCGGAAG
44024 TATGAGTGTGGAGTCGTTGCTGAAAAAGGAGGCAGAACAGACCNNNAGATCGGA
54213 TATGAGTGTGGAGTCGTTGCGAGGTGAAATTCCTCAGTGTGGGNNNAGATCGGAA
46436 TATGAGTGTGGAGTCGTTGCAATCAATGAGAGCGCTATTCAGCNNNAGATCGGAA
66374 TATGAGTGTGGAGTCGTTGCAGCAGTACCAATGTAATTAGGTGTNNNAGATCGGA
116525 TATGAGTGTGGAGTCGTTGCCCAGATAAAGAGGAGGTGGAATGGGNNNAGATCG
48620 TATGAGTGTGGAGTCGTTGCTGGCTTCAGATGATTCTGCACTCNNNAGATCGGAAG

38840 TATGAGTGTGGAGTCGTTGCGAGAACATGAATGGATCTGGAGGANNNAGATCGGA
59790 TATGAGTGTGGAGTCGTTGCTGCCTGAAAAACAGAGGGAAACTNNNAGATCGGAA
60728 TATGAGTGTGGAGTCGTTGCTTGCTTTTCCATTCAGCAGACCTNNNAGATCGGAAGA
97084 TATGAGTGTGGAGTCGTTGCTCTTTGTCATGTCCTGTGGCTTTCNNNAGATCGGAAG
29808 TATGAGTGTGGAGTCGTTGCCACTCCCTCCTCCCCATTATCTTNNNAGATCGGAAGAG
34889 TATGAGTGTGGAGTCGTTGCTAGCAAAGAAGACTGTCCCTCCANNNAGATCGGAAG
37639 TATGAGTGTGGAGTCGTTGCCGTGCAAAATCCTGGTAGGAAGANNNAGATCGGAA
29813 TATGAGTGTGGAGTCGTTGCGAACCCATAAGCAGTGGGGTAAGNNNAGATCGGAA
55291 TATGAGTGTGGAGTCGTTGCTGCGTTGATACCAGGGAAGAATGNNNAGATCGGAA
71666 TATGAGTGTGGAGTCGTTGCGGCTGCTTCCTTCTCTAAAGCTTNNNAGATCGGAAGA
71894 TATGAGTGTGGAGTCGTTGCTGTGATGAGAGGGATTGGAAAAGACNNNAGATCGG
71749 TATGAGTGTGGAGTCGTTGCTTTTGGTTGATGAGCAGCTCCTTNNNAGATCGGAAG
71740 TATGAGTGTGGAGTCGTTGCATCCCAGAAACCTCATCAGGGAANNNAGATCGGAAG
47028 TATGAGTGTGGAGTCGTTGCTCACAATGTTGGGGTACTTTAGGCNNNAGATCGGAA
63419 TATGAGTGTGGAGTCGTTGCAGGAGGAGGATTTTTGCTCTTCTNNNAGATCGGAAG
121930 TATGAGTGTGGAGTCGTTGCTCAAGCTCCTTGGTAAACAGGCTNNNAGATCGGAAG
39323 TATGAGTGTGGAGTCGTTGCGCTCTCAGCTTTCTAGATACTGGANNNAGATCGGAA
48846 TATGAGTGTGGAGTCGTTGCTTACGATGCAGAACTGACAGCTCNNNAGATCGGAAG
102082 TATGAGTGTGGAGTCGTTGCGGAAGTAAAAGTCAGTTGACCCCANNNAGATCGGAA
102745 TATGAGTGTGGAGTCGTTGCCTGAAATAGTCCCAGACAAATGGGGNNNAGATCGGA
64206 TATGAGTGTGGAGTCGTTGCGGAGAAGGGATCACCAAGAGAGTNNNAGATCGGAA
100941 TATGAGTGTGGAGTCGTTGCTGGGTGGAAAAGAGAATAGATTGGAANNNAGATCG
59215 TATGAGTGTGGAGTCGTTGCCCTTCAACCTAGGACACCTGGATNNNAGATCGGAAGA
122017 TATGAGTGTGGAGTCGTTGCAGGAAATGTGAAGGAGGCAGGAANNNAGATCGGA
104068 TATGAGTGTGGAGTCGTTGCTAGGCTGGAGTGGCACAATCATAANNNAGATCGGAA
104511 TATGAGTGTGGAGTCGTTGCTACCAAAACAATGACCTCAAACTGTCNNNAGATCGGA
105480 TATGAGTGTGGAGTCGTTGCTCACCGTAAACGCCCCAAGATCNNNAGATCGGAAGA
101234 TATGAGTGTGGAGTCGTTGCATGAATTTGAGTTCTGAGAGTGGGTNNNAGATCGGA
100517 TATGAGTGTGGAGTCGTTGCGCCTCAGTTTGTGTCTGGGAATANNNAGATCGGAAG
103705 TATGAGTGTGGAGTCGTTGCCAAACCTCGACCACTGGTAAAAATNNNAGATCGGAA
103898 TATGAGTGTGGAGTCGTTGCTCAGGGTACATTCCCCACATTTTNNNAGATCGGAAGA
110656 TATGAGTGTGGAGTCGTTGCTGGTCTCTGGTTTCTGCTTGTGTNNNAGATCGGAAGA
107748 TATGAGTGTGGAGTCGTTGCAAGGCTGGAGTAAAAGCAATGTCCNNNAGATCGGAA
104110 TATGAGTGTGGAGTCGTTGCAGGAGACAAAACAGACCTCCTCTTTNNNAGATCGGAA
102206 TATGAGTGTGGAGTCGTTGCATCACTTTCTCAAATCATCCACCGNNNAGATCGGAAG
101422 TATGAGTGTGGAGTCGTTGCCATCAGTGGCATCATCTTGGCNNNAGATCGGAAGAG
115900 TATGAGTGTGGAGTCGTTGCTGTAAGCTTTGGGAGTGATGGACNNNAGATCGGAA
107740 TATGAGTGTGGAGTCGTTGCCACATCGGAGGTCAGCAAATTTGGNNNAGATCGGAA
107711 TATGAGTGTGGAGTCGTTGCCTCCTGACCTCACTCGATTCTCTCNNNAGATCGGAAGA
104061 TATGAGTGTGGAGTCGTTGCTTGCTCCTTTATCAATGATCTGTTGANNNAGATCGGA
107745 TATGAGTGTGGAGTCGTTGCCACCACTCACACACCTTCCTGNNNAGATCGGAAGAG
110643 TATGAGTGTGGAGTCGTTGCGGTGGCTATTACTGAAATCGCAATNNNAGATCGGAA
103250 TATGAGTGTGGAGTCGTTGCACAACCCAATAATTGAGAATAGGCAANNNAGATCGG

110663 TATGAGTGTGGAGTCGTTGCTGTGACGTGTATGTGAGTGGTGTGNNNAGATCGGA
105527 TATGAGTGTGGAGTCGTTGCGTGCGCCTCCATCGATGCATATNNNAGATCGGAAGA
104787 TATGAGTGTGGAGTCGTTGCCATTGACTAAAGGGGAGGCCAAANNNAGATCGGAA
110631 TATGAGTGTGGAGTCGTTGCGTGGGGTGGAGATGGACAGCAAANNNAGATCGGA
104105 TATGAGTGTGGAGTCGTTGCATGTTACCAAAAGACCCAGCAATTCNNNAGATCGGA
122021 TATGAGTGTGGAGTCGTTGCTTCCAGACCCTATCTCAGAGCCNNNAGATCGGAAGA
104843 TATGAGTGTGGAGTCGTTGCAGAAAGAAAATTAGTGGTGGGAGTGGNNNAGATC
114477 TATGAGTGTGGAGTCGTTGCCTGGTTAGTTTCTGCTTGTGCGNNNAGATCGGAAGA
103301 TATGAGTGTGGAGTCGTTGCTGGGTATCAAGGAAACCATCCTATCCNNNAGATCGG
101188 TATGAGTGTGGAGTCGTTGCGGTTTCCCTGCTTGCCTCATTCANNNAGATCGGAAGA
102490 TATGAGTGTGGAGTCGTTGCTTCCAGTTCTGAGTTGAGGGCAANNNAGATCGGAA
102258 TATGAGTGTGGAGTCGTTGCGCTCAAACAGTCCACCTACCCANNNAGATCGGAAGA
122018 TATGAGTGTGGAGTCGTTGCAGTAGCAGATGGCGCTAATTTCCTANNNAGATCGGA
102205 TATGAGTGTGGAGTCGTTGCTGCCCAGCCTTTCTTCTTCTTTTTNNNAGATCGGAAGA
103864 TATGAGTGTGGAGTCGTTGCTGGACTTTGAGGGACTTTCAGGNNNAGATCGGAAG
107759 TATGAGTGTGGAGTCGTTGCGTGGAGAGAAAACTGTGTGATTCAGNNNAGATCGG
103714 TATGAGTGTGGAGTCGTTGCGAGAATGATGCCATACACAGCTNNNAGATCGGAAG
117374 TATGAGTGTGGAGTCGTTGCTAGGGCTTCAGAGATGACTGATCTTNNNAGATCGGA
106113 TATGAGTGTGGAGTCGTTGCGGTCAGGTCTTTTGAGCCCTTGNNNAGATCGGAAGA
99785 TATGAGTGTGGAGTCGTTGCTTGCCCCATCTCAGTCTACTATCTGNNNAGATCGGAA
71903 TATGAGTGTGGAGTCGTTGCATATAAGCAATGAGCCACCAAATNNNAGATCGGAAG
109848 TATGAGTGTGGAGTCGTTGCGGATTGTGTAGAATTGGCGTTGANNNAGATCGGAA
110647 TATGAGTGTGGAGTCGTTGCCTTCTGTTCGTTCTTTCATTTGTTCNNNAGATCGGAA
100286 TATGAGTGTGGAGTCGTTGCAAATAGTCCGCAGGCTGGGTNNNAGATCGGAAGAG
96399 TATGAGTGTGGAGTCGTTGCCTTCCACATCCCCAGCATCCATNNNAGATCGGAAGAG
58378 TATGAGTGTGGAGTCGTTGCGCTTCATGGGACAATACGGACATNNNAGATCGGAA
55216 TATGAGTGTGGAGTCGTTGCCGTGTGCATGTATCTGGTTTTGGNNNAGATCGGAA
96947 TATGAGTGTGGAGTCGTTGCAGTCTGGCTAGAACAAATGCTAGAATTNNNAGATCG
50325 TATGAGTGTGGAGTCGTTGCACACATCCCCCTCACAAATCAAGNNNAGATCGGAAGA
30695 TATGAGTGTGGAGTCGTTGCCCCAGATGACCAGTTGGTTGAAANNNAGATCGGAA
51928 TATGAGTGTGGAGTCGTTGCTTCCTACCTGGGAGATCCTTGAGNNNAGATCGGAAG
62507 TATGAGTGTGGAGTCGTTGCGTCCCCGAGTTTCTGTATTTGCTNNNAGATCGGAAG
66209 TATGAGTGTGGAGTCGTTGCACATGCATACCTCTTCTGTCTGTNNNAGATCGGAAGA
98452 TATGAGTGTGGAGTCGTTGCCCACCCAATGCACATACCTGTTTCNNNAGATCGGAAG
71630 TATGAGTGTGGAGTCGTTGCTCTAAGCTTCTCGTCACTGGTNNNAGATCGGAAGAG
96728 TATGAGTGTGGAGTCGTTGCCCCCATTCATTCTTTCAAGCTGNNNAGATCGGAAGAG
105996 TATGAGTGTGGAGTCGTTGCAGAGAGAGGAAGATGGTGTGGGNNNAGATCGGAA
53022 TATGAGTGTGGAGTCGTTGCGAGAGAGAGGGAAACTAAGCAGGNNNAGATCGGA
32214 TATGAGTGTGGAGTCGTTGCACCACAACCACCACAACTTTGAANNNAGATCGGAAG
66070 TATGAGTGTGGAGTCGTTGCAAATGAAAATGAAGCTGCGCTGTNNNAGATCGGAA
117480 TATGAGTGTGGAGTCGTTGCCCAGCATGAGCACAGAGAAAAGNNNAGATCGGAAG
98617 TATGAGTGTGGAGTCGTTGCTTGGTCACTTTGTCACATTCTACTGTNNNAGATCGGA
62707 TATGAGTGTGGAGTCGTTGCAGCATGGCAGTAAATCCTTGATCTNNNAGATCGGAA

| | |
|---|---|
| 40587 | TATGAGTGTGGAGTCGTTGCGGGGACAGGGAACCTGAAAAATANNNAGATCGGAA |
| 95135 | TATGAGTGTGGAGTCGTTGCCTTAGACCTGGACGCCATGTTGNNNAGATCGGAAGA |
| 59310 | TATGAGTGTGGAGTCGTTGCATTTCCTAGCCCTGTCTGCTGATNNNAGATCGGAAGA |
| 65596 | TATGAGTGTGGAGTCGTTGCATGCCTGCTAGGAAGACAAATGTNNNAGATCGGAAC |
| 63898 | TATGAGTGTGGAGTCGTTGCGGACTTGTATCCAGATTGAGACAGTNNNAGATCGGA |
| 110558 | TATGAGTGTGGAGTCGTTGCCTGCATCTTGCTATAGTTGCCTAANNNAGATCGGAAG |
| 45926 | TATGAGTGTGGAGTCGTTGCTCATCTGCCTTTCACACAGTTCCNNNAGATCGGAAGA |
| 30177 | TATGAGTGTGGAGTCGTTGCTCGCCAATAGAAAACAGTTGAAAANNNAGATCGGAA |
| 44881 | TATGAGTGTGGAGTCGTTGCAACTTTGTGCTCAGAATGAAGCCNNNAGATCGGAAG |
| 61500 | TATGAGTGTGGAGTCGTTGCAGCCAGGTATGGTGCATATTCCTNNNAGATCGGAAG |
| 35508 | TATGAGTGTGGAGTCGTTGCGTCTGGAAGCATCCATTGTAGCANNNAGATCGGAAG |
| 33920 | TATGAGTGTGGAGTCGTTGCTTTCAAGAATTCATAGGACATGTTACANNNAGATCGG |
| 51616 | TATGAGTGTGGAGTCGTTGCGAGAATTTGGAGGCAAGGAGGAGNNNAGATCGGA |
| 58975 | TATGAGTGTGGAGTCGTTGCGCAGAACCCTGACTATCACAGATNNNAGATCGGAAG |
| 95134 | TATGAGTGTGGAGTCGTTGCGCTGTTGCCATCTGGTTGATCTANNNAGATCGGAAG |
| 57119 | TATGAGTGTGGAGTCGTTGCTGCCCTTCTCTCACTCATTTGTGNNNAGATCGGAAGA |
| 30427 | TATGAGTGTGGAGTCGTTGCTGTGCATGTAGTGCCAGATCAAANNNAGATCGGAAG |
| 40828 | TATGAGTGTGGAGTCGTTGCCTGGGTAAGGGAGCAGAAGGATANNNAGATCGGA |
| 106025 | TATGAGTGTGGAGTCGTTGCTGGGAAGAGACAACAAAAGCCTACNNNAGATCGGA |
| 97701 | TATGAGTGTGGAGTCGTTGCTTTCCAAGTCTCCTTTACCCTGTGANNNAGATCGGAA |
| 54871 | TATGAGTGTGGAGTCGTTGCAGGTTTTCACACCACATTCTAGTGGNNNAGATCGGA |
| 51863 | TATGAGTGTGGAGTCGTTGCTTGTCCGTGCATGTGTATCTGAGNNNAGATCGGAAG |
| 50008 | TATGAGTGTGGAGTCGTTGCCGCACGCACATATACCCTTTTTCNNNAGATCGGAAGA |
| 35282 | TATGAGTGTGGAGTCGTTGCTTGGAGGGCTATATGATCCAGCANNNAGATCGGAAC |
| 67379 | TATGAGTGTGGAGTCGTTGCGCGTGTGTGTGAGTTTGGATATTNNNAGATCGGAAC |
| 106548 | TATGAGTGTGGAGTCGTTGCCTTCTCTCACTCATTTGTGTAGCNNNAGATCGGAAGA |
| 31757 | TATGAGTGTGGAGTCGTTGCACCTGGCATCAGCAAAAGTAGAANNNAGATCGGAAC |
| 58530 | TATGAGTGTGGAGTCGTTGCCCGCTCTAAAAATCTGTGGCCATNNNAGATCGGAAGA |
| 44776 | TATGAGTGTGGAGTCGTTGCGAAGTCTCCTGTTAGTTCTGTCCCNNNAGATCGGAAG |
| 96697 | TATGAGTGTGGAGTCGTTGCTTACAGGGCCATCCATCTTCTTTGNNNAGATCGGAAG |
| 67082 | TATGAGTGTGGAGTCGTTGCGCAGTCAGCATACCACTGACATTNNNAGATCGGAAG |
| 29802 | TATGAGTGTGGAGTCGTTGCCTTGGGAAGCCTCTTTGCTTACTNNNAGATCGGAAGA |
| 40948 | TATGAGTGTGGAGTCGTTGCGCCACGTATCAAGCACAGAACTANNNAGATCGGAAG |
| 51403 | TATGAGTGTGGAGTCGTTGCTAATGCTGCCATGAACACCAGAGNNNAGATCGGAAG |
| 37305 | TATGAGTGTGGAGTCGTTGCCGTCCATAACAGTAAAGATTATGTTCANNNAGATCGG |
| 59726 | TATGAGTGTGGAGTCGTTGCTCTGGTGACCCTTGGACCTTTATNNNAGATCGGAAGA |
| 47990 | TATGAGTGTGGAGTCGTTGCTGCCAAGGCTTATAAAAACTGTGCNNNAGATCGGAA |
| 41430 | TATGAGTGTGGAGTCGTTGCCGTCTGTGTAGACTGTGGGAGTANNNAGATCGGAAC |
| 29803 | TATGAGTGTGGAGTCGTTGCAGTGTGTGAAGACAGAAGGCCTANNNAGATCGGAA |
| 29804 | TATGAGTGTGGAGTCGTTGCCAGACACGAGAAAGCATACCCAANNNAGATCGGAAG |
| 68909 | TATGAGTGTGGAGTCGTTGCCCCCCATCTTCCCATTCTGATTTNNNAGATCGGAAGAC |
| 61143 | TATGAGTGTGGAGTCGTTGCGAACTAGATATGCACTTGCGCCTNNNAGATCGGAAG |
| 63707 | TATGAGTGTGGAGTCGTTGCACTGAACATCACAATGGCCAAGTNNNAGATCGGAAG |

49803 TATGAGTGTGGAGTCGTTGCCCAATCTCTGCAGCTTAGGGTTCNNNAGATCGGAAG
34355 TATGAGTGTGGAGTCGTTGCGCTAGTCTACATCAGGTTCCCCANNNAGATCGGAAG
35975 TATGAGTGTGGAGTCGTTGCGAAACACTGTAGCAGGAGATGCANNNAGATCGGAA
30681 TATGAGTGTGGAGTCGTTGCCCCAAGCATGAAGAAACGTGAAANNNAGATCGGAA
37918 TATGAGTGTGGAGTCGTTGCAAAGCAACTAGGACAACAGCAGANNNAGATCGGAA
31372 TATGAGTGTGGAGTCGTTGCTTCCAAATTCCTGGCCACATCAANNNAGATCGGAAG
41992 TATGAGTGTGGAGTCGTTGCGGTGGACTGGATGCTTGCATTTANNNAGATCGGAA
97702 TATGAGTGTGGAGTCGTTGCCCGTCCCTAGTCAAATTTGCAANNNAGATCGGAAGA
47274 TATGAGTGTGGAGTCGTTGCAAATCACTTAGGACAAGCCTGGCNNNAGATCGGAA
50636 TATGAGTGTGGAGTCGTTGCTTAGAAGCGGATCGCCATAACAGNNNAGATCGGAA
51297 TATGAGTGTGGAGTCGTTGCTCCCCTTTCTATGAGGGTGTCAGNNNAGATCGGAAG
96731 TATGAGTGTGGAGTCGTTGCCCCACCGGTCTTTCATAAAGGANNNAGATCGGAAGA
97985 TATGAGTGTGGAGTCGTTGCGGAAAACAAATAGCAAGAGGTCCNNNAGATCGGAA
117255 TATGAGTGTGGAGTCGTTGCTGTGTGTATTTGGTTGGCTAGTTAGTNNNAGATCGG
65439 TATGAGTGTGGAGTCGTTGCGAGTATGTATGTGTGCACGTGGTNNNAGATCGGAA
47204 TATGAGTGTGGAGTCGTTGCAAGCCCTCAATGCTAGTAATGGCNNNAGATCGGAAG
117228 TATGAGTGTGGAGTCGTTGCGTCTCTCCCCCTTTCACATTTGTNNNAGATCGGAAGA
46147 TATGAGTGTGGAGTCGTTGCTAGAGAAGTCCTCAGAGGCAAGCNNNAGATCGGAA
32065 TATGAGTGTGGAGTCGTTGCGGAAGCAGGTTTGGTGGTATGAANNNAGATCGGA
48931 TATGAGTGTGGAGTCGTTGCAAAGGTGTGATATGAGCCTGCTCNNNAGATCGGAA
68331 TATGAGTGTGGAGTCGTTGCTTCCAGAGGACTGACAGTGAGTTNNNAGATCGGAA
42637 TATGAGTGTGGAGTCGTTGCTACCCAGTTTTCAGTGCCTACACNNNAGATCGGAAG
30523 TATGAGTGTGGAGTCGTTGCTAGGGTGAGCCTTGTGAGAGAAANNNAGATCGGA
107476 TATGAGTGTGGAGTCGTTGCTGTCCCCGAGTTTCTGTATTTGCNNNAGATCGGAAG
54220 TATGAGTGTGGAGTCGTTGCAACATCGGTTTCTACTGTGTGGGNNNAGATCGGAA
38431 TATGAGTGTGGAGTCGTTGCTGTGTGTAACCAGTGCTTCTAGANNNAGATCGGAAG
100248 TATGAGTGTGGAGTCGTTGCCCCACTTGGCTTCTTCTGTTTACNNNAGATCGGAAGA
50546 TATGAGTGTGGAGTCGTTGCGGTGGGGGCGGTGTTATTATAAGNNNAGATCGGA
54157 TATGAGTGTGGAGTCGTTGCACTGACTTCCTTTGCATTCTGGGNNNAGATCGGAAG
40522 TATGAGTGTGGAGTCGTTGCCCTGATGTTCTGCTGGGATTTGANNNAGATCGGAAG
64148 TATGAGTGTGGAGTCGTTGCCAGCATGGGAGGAAAAGTTCAGTNNNAGATCGGAA
56152 TATGAGTGTGGAGTCGTTGCTGACAAGGGAGAGAGACTAGCTGNNNAGATCGGA
46679 TATGAGTGTGGAGTCGTTGCAAGATCAGTGCCAAGAAGGTAGCNNNAGATCGGAA
53506 TATGAGTGTGGAGTCGTTGCTATGTGTTTGTCTTACCTGCCGGNNNAGATCGGAAG
35985 TATGAGTGTGGAGTCGTTGCCATCAGGAAAGGTGATGGATGCANNNAGATCGGAA
31164 TATGAGTGTGGAGTCGTTGCACACGCCACTGATTCTTTTCCAANNNAGATCGGAAGA
97021 TATGAGTGTGGAGTCGTTGCGTTATTATGCAAGCGGCATTGTGNNNAGATCGGAA
52753 TATGAGTGTGGAGTCGTTGCTAGGATTAGGCAACATGCCAAGGNNNAGATCGGAA
49754 TATGAGTGTGGAGTCGTTGCGAAGAGGGGAGTGCAGTTTCTTCNNNAGATCGGAA
67045 TATGAGTGTGGAGTCGTTGCACCACATTTTCTTTGTGGAACATTNNNAGATCGGAA
66930 TATGAGTGTGGAGTCGTTGCATGACGCCAATCTGCCTACAATTNNNAGATCGGAAG
66919 TATGAGTGTGGAGTCGTTGCCTGCTTTAGCCCCATGGGAAATTNNNAGATCGGAAG
51306 TATGAGTGTGGAGTCGTTGCGTCTCCTCTACCTCCCACTTCAGNNNAGATCGGAAGA

| | |
|---|---|
| 37964 | TATGAGTGTGGAGTCGTTGCAATGGTCTAACCAAACATGCAGANNNAGATCGGAAG |
| 101386 | TATGAGTGTGGAGTCGTTGCGCCACTGTCATCTGGTGGTTANNNAGATCGGAAGAG |
| 56589 | TATGAGTGTGGAGTCGTTGCTTTGAGATCTGTCCTTGGCAGTGNNNAGATCGGAAG |
| 38217 | TATGAGTGTGGAGTCGTTGCCAGATGGAGAGATGGAGAGGAGANNNAGATCGGA |
| 68939 | TATGAGTGTGGAGTCGTTGCGGGTGTGTGTGTCCTGAGTATTTNNNAGATCGGAA |
| 97323 | TATGAGTGTGGAGTCGTTGCCTATCGAGCACTCTTGGGGAGNNNAGATCGGAAGA |
| 62348 | TATGAGTGTGGAGTCGTTGCTTTCTGAGGTTTCAGCTTCTGCTNNNAGATCGGAAG |
| 55070 | TATGAGTGTGGAGTCGTTGCGGGGTAGCTGAAATAAGCCTTGGNNNAGATCGGAA |
| 33802 | TATGAGTGTGGAGTCGTTGCAGCCAGAAGAAAACACACACTACANNNAGATCGGAA |
| 57730 | TATGAGTGTGGAGTCGTTGCCGCTCTTCAGTGACCTTCCAAATNNNAGATCGGAAG |
| 41104 | TATGAGTGTGGAGTCGTTGCTCGCCCATGCAGATAGAATCCTANNNAGATCGGAAG |
| 59447 | TATGAGTGTGGAGTCGTTGCGGCAAAATCAGGCCCTCAGATATNNNAGATCGGAAG |
| 62308 | TATGAGTGTGGAGTCGTTGCAAAGTAGTGATCCTGGTCCTGCTNNNAGATCGGAAG |
| 42331 | TATGAGTGTGGAGTCGTTGCAGAACAAATGTTGACCACATATCAACNNNAGATCGG |
| 67983 | TATGAGTGTGGAGTCGTTGCCATCTCAGGCTCATCTTGTGCTTNNNAGATCGGAAG |
| 40441 | TATGAGTGTGGAGTCGTTGCTGTACCAGGCATGCAAATCTTGANNNAGATCGGAAG |
| 54176 | TATGAGTGTGGAGTCGTTGCTAAATTCCCTTCTGGGGAGTGGGNNNAGATCGGAA |
| 52182 | TATGAGTGTGGAGTCGTTGCATCCTTTCTGCCTTGCCTTGTAGNNNAGATCGGAAGA |
| 56986 | TATGAGTGTGGAGTCGTTGCCAAGCTACAAACCTGCCTCTGTGNNNAGATCGGAAG |
| 60916 | TATGAGTGTGGAGTCGTTGCATCCACTCTATATCCAGTCCCCTNNNAGATCGGAAGA |
| 40084 | TATGAGTGTGGAGTCGTTGCTGTCTGAAAGTTCCCGTTTCTGANNNAGATCGGAAG |
| 31254 | TATGAGTGTGGAGTCGTTGCAGATTTGAAAGAAGACAGGGCAANNNAGATCGGAA |
| 34720 | TATGAGTGTGGAGTCGTTGCTTGAAATGAAGCTGAGACTGCCANNNAGATCGGAA |
| 54285 | TATGAGTGTGGAGTCGTTGCGAAAAGTCCTGAATGGCTTTGGGNNNAGATCGGAA |
| 64775 | TATGAGTGTGGAGTCGTTGCCACCTACCTACCTCCTTGAAGGTNNNAGATCGGAAG |
| 46199 | TATGAGTGTGGAGTCGTTGCCTCTTCCTTTCAGCCTAGGAAGCNNNAGATCGGAAG |
| 34888 | TATGAGTGTGGAGTCGTTGCTCCATAAGTCTAAATGCCCTCCANNNAGATCGGAAG |
| 30205 | TATGAGTGTGGAGTCGTTGCCGTATTGCTGGGGATTGGTAAAANNNAGATCGGAA |
| 40381 | TATGAGTGTGGAGTCGTTGCTAGTGGAGGGGAGCTTTCATTGANNNAGATCGGAA |
| 38559 | TATGAGTGTGGAGTCGTTGCAGAGAGATGAGGAGTGAAAGCGANNNAGATCGGA |
| 44280 | TATGAGTGTGGAGTCGTTGCCTGTCTTGAGCATCTTCTCACCCNNNAGATCGGAAGA |
| 38594 | TATGAGTGTGGAGTCGTTGCGATACACATGCCTACGTCTTCGANNNAGATCGGAAG |
| 104011 | TATGAGTGTGGAGTCGTTGCATGTGAGAGAAAGAGAGAGGTGGGNNNAGATCGG |
| 50648 | TATGAGTGTGGAGTCGTTGCATGGGTAACACACAGACACACAGNNNAGATCGGAA |
| 106052 | TATGAGTGTGGAGTCGTTGCTGTGGTGCAATTCTGTGCTGTNNNAGATCGGAAGA |
| 59939 | TATGAGTGTGGAGTCGTTGCATGCCACAGAGGAAGAGGTAACTNNNAGATCGGAA |
| 62250 | TATGAGTGTGGAGTCGTTGCTTCATCTGACTTCAATGTTGATGCTNNNAGATCGGAA |
| 51180 | TATGAGTGTGGAGTCGTTGCTTAGTGGCATTTGGCATTTGCAGNNNAGATCGGAA |
| 96775 | TATGAGTGTGGAGTCGTTGCCATCCACTCTCCATGGTGTGGNNNAGATCGGAAGAG |
| 107721 | TATGAGTGTGGAGTCGTTGCCTTGGGTGTTTGTAGGACTTACTTCANNNAGATCGG |
| 34562 | TATGAGTGTGGAGTCGTTGCAAGCAGTGAACACTTGTAAGCCANNNAGATCGGAA |
| 51949 | TATGAGTGTGGAGTCGTTGCGGTTACAGGCTGCTCTCTTTGAGNNNAGATCGGAAG |
| 63087 | TATGAGTGTGGAGTCGTTGCGCATTCATGTGGGCAATTTCTCTNNNAGATCGGAAG |

48905 TATGAGTGTGGAGTCGTTGCATTCTCCTCAGGACTACTGGCTCNNNAGATCGGAAG
117102 TATGAGTGTGGAGTCGTTGCCTTGGAGTGCGGTGGCATGATTANNNAGATCGGAA
40125 TATGAGTGTGGAGTCGTTGCAGAGAGACCTTTTGTCCCAGTGANNNAGATCGGAAG
59714 TATGAGTGTGGAGTCGTTGCAGGCATTTGAAGCTGGCTGTTATNNNAGATCGGAAG
63875 TATGAGTGTGGAGTCGTTGCACTCTCACTCACTCACACACAGTNNNAGATCGGAAGA
35850 TATGAGTGTGGAGTCGTTGCGTCACAAAGGACTGTACTTGGCANNNAGATCGGAAG
33004 TATGAGTGTGGAGTCGTTGCTATCCCCTACCACATCCAACACANNNAGATCGGAAGA
54542 TATGAGTGTGGAGTCGTTGCACTAGAGGAAGCGAGAAGACTGGNNNAGATCGGA
65537 TATGAGTGTGGAGTCGTTGCACGATTCAGGTTCAAGTCTTGGTNNNAGATCGGAAG
105434 TATGAGTGTGGAGTCGTTGCTGTGTACTATGCCAAGTGTTGATANNNAGATCGGAA
60543 TATGAGTGTGGAGTCGTTGCAGAGAGAGAGAGGAGGGCTTACTNNNAGATCGGA
69249 TATGAGTGTGGAGTCGTTGCCCAAGAGCAAGAGAGCACAGTTTNNNAGATCGGAA
59007 TATGAGTGTGGAGTCGTTGCTTGTGCACTGTGTACTTGCAGATNNNAGATCGGAAG
35126 TATGAGTGTGGAGTCGTTGCACAGACAGACACTCACAAAAGCANNNAGATCGGAAG
117737 TATGAGTGTGGAGTCGTTGCCCCTTTGGAGAGGACAAAGTTGCNNNAGATCGGAA
48280 TATGAGTGTGGAGTCGTTGCTGAAGGCACAGATCAGAGCAATCNNNAGATCGGAA
59405 TATGAGTGTGGAGTCGTTGCTGAGAGGTTGTTCGGGGGAATATNNNAGATCGGAA
31368 TATGAGTGTGGAGTCGTTGCGGCAGGAAACTGGTCTGAATCAANNNAGATCGGAA
67973 TATGAGTGTGGAGTCGTTGCCCAATCAAGTGACTTTCGTGCTTNNNAGATCGGAAG
30508 TATGAGTGTGGAGTCGTTGCGTGGTGAGAAATCTGGGCAGAAANNNAGATCGGA
45656 TATGAGTGTGGAGTCGTTGCAGGCTCAGAGTTAGGTCATCTCCNNNAGATCGGAAG
107719 TATGAGTGTGGAGTCGTTGCTGTGTATGGTCTGGGTTGGCANNNAGATCGGAAGA
101418 TATGAGTGTGGAGTCGTTGCCTGGCTAAATTCTGGCGGCTTCTTNNNAGATCGGAA
101687 TATGAGTGTGGAGTCGTTGCATGGAATGGAACCCCAAAACTGCNNNAGATCGGAAG
54092 TATGAGTGTGGAGTCGTTGCCACCATCATTATCACCACCTGGGNNNAGATCGGAAG
46960 TATGAGTGTGGAGTCGTTGCTGAAACCCTCAGAGAGTAGAGGCNNNAGATCGGAA
103832 TATGAGTGTGGAGTCGTTGCGGAGGCCGAATCGCATGAGANNNAGATCGGAAGAG
60759 TATGAGTGTGGAGTCGTTGCGCCCATCATACACAAGTATGACCTNNNAGATCGGAA
50663 TATGAGTGTGGAGTCGTTGCATGATAAGCTGGTGAAGCCACAGNNNAGATCGGAA
44822 TATGAGTGTGGAGTCGTTGCCACCTCAGCATCACTTAGTTCCCNNNAGATCGGAAGA
30665 TATGAGTGTGGAGTCGTTGCGCAAAGTGGAGACACACCTGAAANNNAGATCGGAA
102108 TATGAGTGTGGAGTCGTTGCAGAGGTTGTTCGGGGGAATATAGANNNAGATCGGA
44294 TATGAGTGTGGAGTCGTTGCCCTATTCCAACCATGACAGACCCNNNAGATCGGAAG
60879 TATGAGTGTGGAGTCGTTGCTCTCTGTAGGGTGAGACCCCTNNNAGATCGGAAGAG
54567 TATGAGTGTGGAGTCGTTGCTCCCAAAGAAAACAAAGAAACCTGGNNNAGATCGGA
40564 TATGAGTGTGGAGTCGTTGCAGATCCTCAGTTGGTGTGTTTGANNNAGATCGGAAG
57440 TATGAGTGTGGAGTCGTTGCCTTCCAGTTTCTCCTACCCGTTGNNNAGATCGGAAGA
110599 TATGAGTGTGGAGTCGTTGCCGCTCAGAGGCTTTCACACAAANNNAGATCGGAAGA
59813 TATGAGTGTGGAGTCGTTGCTTGAGTGGTGAAATGCCACAACTNNNAGATCGGAAG
57459 TATGAGTGTGGAGTCGTTGCTGGAATAGAAGTCAGTGGGGTTGNNNAGATCGGA
54635 TATGAGTGTGGAGTCGTTGCCAGCAGTGATCTTGAACTCCTGGNNNAGATCGGAAG
57395 TATGAGTGTGGAGTCGTTGCCAATTTCCCTGGCTGCTTTCTTGNNNAGATCGGAAGA
44345 TATGAGTGTGGAGTCGTTGCACAAAGGCAGAAACTGTAACCCCNNNAGATCGGAAG

49557 TATGAGTGTGGAGTCGTTGCACCTGCTCAAACGTGGTAGATTCNNNAGATCGGAAG
55244 TATGAGTGTGGAGTCGTTGCGCAGGTCCGCACTAAATCAAATGNNNAGATCGGAA
52398 TATGAGTGTGGAGTCGTTGCTTTTTGAGACAACTCTGTCCCCGNNNAGATCGGAAG
44352 TATGAGTGTGGAGTCGTTGCAGCAAAATGAAAATGCAAGACCCCNNNAGATCGGA
41337 TATGAGTGTGGAGTCGTTGCTGGGTTATGGAGCCACAGTTCTANNNAGATCGGAA
103647 TATGAGTGTGGAGTCGTTGCAGTGAACAGGAATCACAGCATACANNNAGATCGGA
37828 TATGAGTGTGGAGTCGTTGCAACAGAAATGGAAGGACACCAGANNNAGATCGGAA
37029 TATGAGTGTGGAGTCGTTGCTGAAAACTGTAAAGCACTCTGTCANNNAGATCGGAA
47753 TATGAGTGTGGAGTCGTTGCTTTGTAGAGCTTCAGTGAGCTGCNNNAGATCGGAA
64336 TATGAGTGTGGAGTCGTTGCCGCTGAGAGAGAGAGTACTGAGTNNNAGATCGGA
41811 TATGAGTGTGGAGTCGTTGCCTTTCTCTTCCTTGCTGCCCTTANNNAGATCGGAAGA
51111 TATGAGTGTGGAGTCGTTGCATGTCTTGAGGTTATGCTGGCAGNNNAGATCGGAA
96734 TATGAGTGTGGAGTCGTTGCGGATGTCAGTATCACCCGTATCAANNNAGATCGGAA
42886 TATGAGTGTGGAGTCGTTGCGCATTAGCCTGAAAATACGGCACNNNAGATCGGAA
44096 TATGAGTGTGGAGTCGTTGCGAGGACACTTGGGAATGATGACCNNNAGATCGGAA
53190 TATGAGTGTGGAGTCGTTGCGGTTGGAACTGAAGAGAAGGAGGNNNAGATCGGA
102737 TATGAGTGTGGAGTCGTTGCTTGTCTGGGGGTAAATGAGAGGGNNNAGATCGGA
43076 TATGAGTGTGGAGTCGTTGCAGAGGAATGGATTGGCGATTCACNNNAGATCGGAA
46235 TATGAGTGTGGAGTCGTTGCCAACACTTGGGAGACCTCTAAGCNNNAGATCGGAAG
51123 TATGAGTGTGGAGTCGTTGCAGTGTCCTATGAACAGCATGCAGNNNAGATCGGAA
55944 TATGAGTGTGGAGTCGTTGCGAGAAAGACTTCAGACCACCCTGNNNAGATCGGAA
53843 TATGAGTGTGGAGTCGTTGCTAAGGATACCAGTCTGCTACGGGNNNAGATCGGAA
33671 TATGAGTGTGGAGTCGTTGCCCATAAACACCAGCTGGATGACANNNAGATCGGAAG
58376 TATGAGTGTGGAGTCGTTGCAGAGAAAGGAAGCAGGAGGACATNNNAGATCGGA
63338 TATGAGTGTGGAGTCGTTGCGCTTTCTCCTTTGTTACTCTCATTCTNNNAGATCGGAA
49459 TATGAGTGTGGAGTCGTTGCTCAGAGGCAATGCTGTATGTGTCNNNAGATCGGAA
33876 TATGAGTGTGGAGTCGTTGCTCTCAAAAGTCAATGAACCGTACANNNAGATCGGAA
50968 TATGAGTGTGGAGTCGTTGCTTCTCCTCCCCTTCTTTCTCCAGNNNAGATCGGAAGA
63495 TATGAGTGTGGAGTCGTTGCCTTGGTAAGGTGGTGGCATTTCTNNNAGATCGGAA
121977 TATGAGTGTGGAGTCGTTGCCAGAGACTTGACATGGAAACCTNNNAGATCGGAAG
64809 TATGAGTGTGGAGTCGTTGCAACAGGAATCACAGCATACAGGTNNNAGATCGGAA
53230 TATGAGTGTGGAGTCGTTGCTCTTACCTTCTGTGAGTGGGAGGNNNAGATCGGAA
62721 TATGAGTGTGGAGTCGTTGCGTGCAGGGAAAATGCACCTATCTNNNAGATCGGAA
41769 TATGAGTGTGGAGTCGTTGCACAAGGGGCTTTCTCTGGGATTANNNAGATCGGAA
44012 TATGAGTGTGGAGTCGTTGCGGCATATGGGATACAATGAAAAGACCNNNAGATCG
70282 TATGAGTGTGGAGTCGTTGCCTTGAGTTTGTTTGCCTGGAACCNNNAGATCGGAAG
66809 TATGAGTGTGGAGTCGTTGCCCCAAACAACCTTACTGCTTTGTNNNAGATCGGAAG
110660 TATGAGTGTGGAGTCGTTGCCTTGCGTCAAATCGAAAGCTGTNNNAGATCGGAAG
63829 TATGAGTGTGGAGTCGTTGCTCAGTAGAGGCAAGTCGCTAAGTNNNAGATCGGAA
29931 TATGAGTGTGGAGTCGTTGCTTTTGGCAGTTAGGCTTGTTCCTNNNAGATCGGAAG
63059 TATGAGTGTGGAGTCGTTGCTATCTGGGTGCATCTGTGTCTCTNNNAGATCGGAAG
68935 TATGAGTGTGGAGTCGTTGCGCACAGCGTCGTTTTGTCTATTTNNNAGATCGGAAG
30666 TATGAGTGTGGAGTCGTTGCTGAGGGTTTTCATTGCCCTGAAANNNAGATCGGAAG

| | |
|---|---|
| 45008 | TATGAGTGTGGAGTCGTTGCCTTGGCTCCTATGGACATTAGCCNNNAGATCGGAAG |
| 41490 | TATGAGTGTGGAGTCGTTGCAAACCTCTCGATTGGGGAAGGTANNNAGATCGGAA |
| 96369 | TATGAGTGTGGAGTCGTTGCTGCTGACCTCTGAATACAAAACAANNNAGATCGGAA |
| 100244 | TATGAGTGTGGAGTCGTTGCTCCTGGCATTCGAAGTTCCCNNNAGATCGGAAGAGC |
| 50230 | TATGAGTGTGGAGTCGTTGCCCCTGGCTGAGAATATCGACAAGNNNAGATCGGAA |
| 53243 | TATGAGTGTGGAGTCGTTGCAACCCCAAAACTGCTTATGGAGGNNNAGATCGGAA |
| 42705 | TATGAGTGTGGAGTCGTTGCAACCCACACATACAAGAACCCACNNNAGATCGGAAG |
| 34319 | TATGAGTGTGGAGTCGTTGCTCATTTCCCATGGATATCCCCCANNNAGATCGGAAGA |
| 117585 | TATGAGTGTGGAGTCGTTGCCGCTCACATAAATGACTGGTTTGNNNAGATCGGAAG |
| 97319 | TATGAGTGTGGAGTCGTTGCTAAAAACACCTTCTTCAGCCGGGNNNAGATCGGAAG |
| 55347 | TATGAGTGTGGAGTCGTTGCTCATGCAAGCCAACAGAAACATGNNNAGATCGGAA |
| 32408 | TATGAGTGTGGAGTCGTTGCATCGAGTGGGACAGATATGCTAANNNAGATCGGAA |
| 45647 | TATGAGTGTGGAGTCGTTGCCTTGAGCTGTGGGATCAATCTCCNNNAGATCGGAAG |
| 36965 | TATGAGTGTGGAGTCGTTGCGCACAGAACAAGGTACATGGTCANNNAGATCGGAA |
| 35204 | TATGAGTGTGGAGTCGTTGCTTTTTCTGCTGATGTGGGAAGCANNNAGATCGGAA |
| 35473 | TATGAGTGTGGAGTCGTTGCAGTGAACATCATTTGGCCTAGCANNNAGATCGGAAG |
| 67239 | TATGAGTGTGGAGTCGTTGCCTTCTGCCTGTGGCTAGTTCATTNNNAGATCGGAAG |
| 33315 | TATGAGTGTGGAGTCGTTGCATAAGCAGATTCACAGCTGCACANNNAGATCGGAAG |
| 41593 | TATGAGTGTGGAGTCGTTGCCAGGCAAGTTTTCGGCAAATGTANNNAGATCGGAA |
| 40032 | TATGAGTGTGGAGTCGTTGCGTCATATCATAGAAGCTGCCTCTGANNNAGATCGGA |
| 60575 | TATGAGTGTGGAGTCGTTGCCGAGGCACTGTGAATGAAAACCTNNNAGATCGGAA |
| 104021 | TATGAGTGTGGAGTCGTTGCTCCCCTCTCCTTACAAGGACACTNNNAGATCGGAAGA |
| 60811 | TATGAGTGTGGAGTCGTTGCCACCAAGGCTTCCCAAATTACCTNNNAGATCGGAAG |
| 49808 | TATGAGTGTGGAGTCGTTGCTTCTCACTCCAACATGTGGGTTCNNNAGATCGGAAG |
| 54076 | TATGAGTGTGGAGTCGTTGCGTCCTCTTCCTGTGCTTTATGGGNNNAGATCGGAAG |
| 117272 | TATGAGTGTGGAGTCGTTGCTCACTCAGCTGCCTCTTTAGACNNNAGATCGGAAGA |
| 117590 | TATGAGTGTGGAGTCGTTGCTCATGGCATTCACCAGAATCCTGNNNAGATCGGAAG |
| 100285 | TATGAGTGTGGAGTCGTTGCGACCCCCACACACAGACATGCNNNAGATCGGAAGAG |
| 64907 | TATGAGTGTGGAGTCGTTGCAAGGCATGAATAAGAGGGGAGGTNNNAGATCGGA |
| 34136 | TATGAGTGTGGAGTCGTTGCGGGGGAATGGAATATCATGACCANNNAGATCGGAA |
| 63390 | TATGAGTGTGGAGTCGTTGCGTTCTGGCGTTTCTTCTCCTTCTNNNAGATCGGAAGA |
| 57410 | TATGAGTGTGGAGTCGTTGCTGTGCCAGACAGACACTAAGTTGNNNAGATCGGAA |
| 32369 | TATGAGTGTGGAGTCGTTGCTCTCTACAGTCAGCCTTGCCTAANNNAGATCGGAAG |
| 104451 | TATGAGTGTGGAGTCGTTGCGGGGGCATCACTCAACATTCANNNAGATCGGAAGA |
| 56493 | TATGAGTGTGGAGTCGTTGCGTCCGTCAGTCCTATGCTTTCTGNNNAGATCGGAAG |
| 101081 | TATGAGTGTGGAGTCGTTGCGCTCAATGTCTTCTTGGTCTGTGNNNAGATCGGAAG |
| 60465 | TATGAGTGTGGAGTCGTTGCGGGCTCAGGAGAAGATGACTACTNNNAGATCGGAA |
| 50002 | TATGAGTGTGGAGTCGTTGCATTTTGCATGGGATGGCGTTTTCNNNAGATCGGAAG |
| 106007 | TATGAGTGTGGAGTCGTTGCGCTTGTTGTAGACTGGAGGTGCNNNAGATCGGAAG |
| 49464 | TATGAGTGTGGAGTCGTTGCGTGTCCCTGTGTATGTCTGTGTCNNNAGATCGGAAG |
| 52389 | TATGAGTGTGGAGTCGTTGCCAAAAGTTTCACCTGGAAACCCGNNNAGATCGGAAG |
| 52526 | TATGAGTGTGGAGTCGTTGCATTACTCATGGAAGCCATGGCGNNNAGATCGGAAG |
| 116942 | TATGAGTGTGGAGTCGTTGCCCTCTCTGTTTCTCTCCAACTCCCNNNAGATCGGAAGA |

40878 TATGAGTGTGGAGTCGTTGCGTGGAAAGTTGTGGGGGTTGATANNNAGATCGGA
51708 TATGAGTGTGGAGTCGTTGCCACCCTCTTTACACACACTGGAGNNNAGATCGGAAG
45012 TATGAGTGTGGAGTCGTTGCAAAAGAACACTCCCCACTTAGCCNNNAGATCGGAAG
60507 TATGAGTGTGGAGTCGTTGCGACGGACACAGGAGACTAGTACTNNNAGATCGGAA
45313 TATGAGTGTGGAGTCGTTGCTCAATGTCTTCTTGGTCTGTGCCNNNAGATCGGAAG
47557 TATGAGTGTGGAGTCGTTGCACTGGCCGCTTATATACAGATGCNNNAGATCGGAAG
38209 TATGAGTGTGGAGTCGTTGCGTGCTTCTGAGACCCAGGAGANNNAGATCGGAAGA
48606 TATGAGTGTGGAGTCGTTGCTGGTATAAATGTGCAGCCCACTCNNNAGATCGGAAG
36463 TATGAGTGTGGAGTCGTTGCGCCTCAGCCACAGAAATTCATCANNNAGATCGGAAG
115964 TATGAGTGTGGAGTCGTTGCAGATTCACAGCTGCACAGGAGAGNNNAGATCGGAA
61819 TATGAGTGTGGAGTCGTTGCGGGGGAAAATGGTAGTTTGAGCTNNNAGATCGGA
60433 TATGAGTGTGGAGTCGTTGCACTGACGCATCGAGAAAAATACTNNNAGATCGGAA
49960 TATGAGTGTGGAGTCGTTGCCCTTCTCTGGCTCTCCATGTTTCNNNAGATCGGAAGA
50550 TATGAGTGTGGAGTCGTTGCGGAGACATGCCAAGGAACCTAAGNNNAGATCGGAA
41230 TATGAGTGTGGAGTCGTTGCCCCTGATGGGTAGAAATGTGCTANNNAGATCGGAA
103830 TATGAGTGTGGAGTCGTTGCTAGTGTCTGTGCGGGTCTCTGNNNAGATCGGAAGA
47318 TATGAGTGTGGAGTCGTTGCTCCTCTCAAAGCAAGTAAGTGGCNNNAGATCGGAAG
65662 TATGAGTGTGGAGTCGTTGCCACAGAGTACCTTTCCAAACATGTNNNAGATCGGAA
65677 TATGAGTGTGGAGTCGTTGCGTGTATGTGGGTGTGTCACATGTNNNAGATCGGAA
57493 TATGAGTGTGGAGTCGTTGCTTGTCCACAGACACACTCTGTTGNNNAGATCGGAAG
68249 TATGAGTGTGGAGTCGTTGCTGAATGTAAGAGTTACAGGACAGTTNNNAGATCGG
65853 TATGAGTGTGGAGTCGTTGCGGGGGAAGGGGGATAATGTATGTNNNAGATCGGA
116079 TATGAGTGTGGAGTCGTTGCGGGGGAAAATGGTAGTTTGAGCTANNNAGATCGG
54024 TATGAGTGTGGAGTCGTTGCGAGAGAGAAAGCCAGACAATGGGNNNAGATCGGA
53797 TATGAGTGTGGAGTCGTTGCAATAAGAAGCCCTGGGTTGAGGGNNNAGATCGGA
43849 TATGAGTGTGGAGTCGTTGCATTCTTCACCTGTGAGCGTAACCNNNAGATCGGAAG
45693 TATGAGTGTGGAGTCGTTGCCTCCTTCTCCACTCTCCTTCTCCNNNAGATCGGAAGAG
39846 TATGAGTGTGGAGTCGTTGCGTGTGTGTGTGTGGCTATACTGANNNAGATCGGAA
117367 TATGAGTGTGGAGTCGTTGCAAGGGGCATAGATAAGGATGGTCNNNAGATCGGA
64966 TATGAGTGTGGAGTCGTTGCGGGATCTTATGCAGGACCTAGGTNNNAGATCGGAA
116019 TATGAGTGTGGAGTCGTTGCACCAAGGAAACCCTACTTTCAGANNNAGATCGGAAG
103792 TATGAGTGTGGAGTCGTTGCATTCTGAGTGCACAAAGGCAGCANNNAGATCGGAA
52673 TATGAGTGTGGAGTCGTTGCCCACCTGAAGAAGGAGGAAAAGGNNNAGATCGGA
60505 TATGAGTGTGGAGTCGTTGCCCCAAACAAAACCTTTGAGAGTACTNNNAGATCGGA
60297 TATGAGTGTGGAGTCGTTGCGTGCAGACCAAGATTCTGAGACTNNNAGATCGGAA
107491 TATGAGTGTGGAGTCGTTGCATAAATGTGCAGCCCACTCAGGNNNAGATCGGAAG
51289 TATGAGTGTGGAGTCGTTGCGAGAGCACCTCAGATTTGGTCAGNNNAGATCGGAA
43402 TATGAGTGTGGAGTCGTTGCGTTTGGGTACAGGGACATCTGACNNNAGATCGGAA
45137 TATGAGTGTGGAGTCGTTGCCCCAGATGATTCTAATGGTGGCCNNNAGATCGGAAG
55638 TATGAGTGTGGAGTCGTTGCAGAAAGGGCACCTCATGATGATGNNNAGATCGGAA
102109 TATGAGTGTGGAGTCGTTGCCCAGATGTAGGAAAAGTGGCAAANNNAGATCGGAA
33834 TATGAGTGTGGAGTCGTTGCTCTAGGGCAGTCTCATTGCTACANNNAGATCGGAAG
61051 TATGAGTGTGGAGTCGTTGCGGTGGGCACATGATATTTTCCCTNNNAGATCGGAAG

49534 TATGAGTGTGGAGTCGTTGCTGTCCTGCTAGTGTTGAGCATTCNNNAGATCGGAAG
49118 TATGAGTGTGGAGTCGTTGCGGCACTTTCCTTCTTCCCTTCTCNNNAGATCGGAAGA
62901 TATGAGTGTGGAGTCGTTGCTCATTCTCTGACAGCCTTCCTCTNNNAGATCGGAAGA
117446 TATGAGTGTGGAGTCGTTGCCCCCCAAAGTCAGCTACTCAANNNAGATCGGAAGAG
41579 TATGAGTGTGGAGTCGTTGCGACAACCAAGAATGCCAGTGGTANNNAGATCGGAA
56561 TATGAGTGTGGAGTCGTTGCTCCCATGTTTTTACCCACCAGTGNNNAGATCGGAAGA
63895 TATGAGTGTGGAGTCGTTGCTCACCCTCATTCTTCAAAGACAGTNNNAGATCGGAAG
44773 TATGAGTGTGGAGTCGTTGCCCAGAGTTAGGAAAAGCTGTCCCNNNAGATCGGAA
36264 TATGAGTGTGGAGTCGTTGCCCTCCACTTCAGTGATTCTTGCANNNAGATCGGAAGA
55103 TATGAGTGTGGAGTCGTTGCAAATGGCTGGGGATCATTCTTGGNNNAGATCGGAA
50635 TATGAGTGTGGAGTCGTTGCGCCTGTTCCTTCTCTCTGAACAGNNNAGATCGGAAGA
105541 TATGAGTGTGGAGTCGTTGCAGTGGCTCATGGGCTGTTTCNNNAGATCGGAAGAG
30072 TATGAGTGTGGAGTCGTTGCACCAGTGTCATAACCTGCCAAAANNNAGATCGGAAG
38804 TATGAGTGTGGAGTCGTTGCAGGAGAGAGACTGACAGAGAGGANNNAGATCGGA
116367 TATGAGTGTGGAGTCGTTGCTCTGTCTCTGTCTCTCTCTTATCTCTTNNNAGATCGGA
64099 TATGAGTGTGGAGTCGTTGCCACACACTTCATCTAGCTAACTCAGTNNNAGATCGGA
57461 TATGAGTGTGGAGTCGTTGCTGGTTCTGGTGATGAGATGGTTGNNNAGATCGGAA
32950 TATGAGTGTGGAGTCGTTGCTCCCCTCTGCCTTGTACATAACANNNAGATCGGAAGA
65431 TATGAGTGTGGAGTCGTTGCTCCCTGAACTTGTCTATCAGTGGTNNNAGATCGGAA
116156 TATGAGTGTGGAGTCGTTGCCCTTGCCTTTCTGTCTTGACCANNNAGATCGGAAGAG
68275 TATGAGTGTGGAGTCGTTGCGTTCACACCTAGCCATAGCAGTTNNNAGATCGGAAG
117597 TATGAGTGTGGAGTCGTTGCAGTTTTTCTGGCTCCAGCTGTCCNNNAGATCGGAAG
97704 TATGAGTGTGGAGTCGTTGCACTCCTACAAGAACGGCACACNNNAGATCGGAAGAG
50332 TATGAGTGTGGAGTCGTTGCGCCCATTCTCATCCACACTCAAGNNNAGATCGGAAGA
43192 TATGAGTGTGGAGTCGTTGCACATCTGGGATCCAGCAAGAGACNNNAGATCGGAA
49583 TATGAGTGTGGAGTCGTTGCTGATGAGTTTGCAGCCCCTATTCNNNAGATCGGAAG
59145 TATGAGTGTGGAGTCGTTGCGGGAATACTATGGGCTGGAGGATNNNAGATCGGA
45514 TATGAGTGTGGAGTCGTTGCTCTGTTGCCACATCTTCAACTCCNNNAGATCGGAAGA
49436 TATGAGTGTGGAGTCGTTGCTTTGTTCTGTGGTCCCTTCTGTCNNNAGATCGGAAGA
48610 TATGAGTGTGGAGTCGTTGCTCTAGGATAAGCTCTGGCCACTCNNNAGATCGGAAG
101106 TATGAGTGTGGAGTCGTTGCAACGGGCGTAGATTCAATCAGATNNNAGATCGGAA
56312 TATGAGTGTGGAGTCGTTGCAAGGGTGCTGAGTTTCACATCTGNNNAGATCGGAA
43115 TATGAGTGTGGAGTCGTTGCGGCTCCGAATAGATCCCAAAGACNNNAGATCGGAA
102759 TATGAGTGTGGAGTCGTTGCGGAGGGGGTGGTCTTACTAGGTCNNNAGATCGGA
49656 TATGAGTGTGGAGTCGTTGCTTGTTCTAGCACCTTCAGCCTTCNNNAGATCGGAAGA
57609 TATGAGTGTGGAGTCGTTGCAAGAGGTCACCTTGGATGGTTTGNNNAGATCGGAA
59037 TATGAGTGTGGAGTCGTTGCACATTTGGTGAGGGGTGAGAGATNNNAGATCGGA
62558 TATGAGTGTGGAGTCGTTGCTAGGAGAGAACCAAGGCCAATCTNNNAGATCGGAA
96691 TATGAGTGTGGAGTCGTTGCACATGCCAATGATATCAGGGGTCANNNAGATCGGA
121961 TATGAGTGTGGAGTCGTTGCTTCTTGTCTGTTGACCTGTTGATAGANNNAGATCGG
39102 TATGAGTGTGGAGTCGTTGCTCTAACAGACTGAAAGCTGGGGANNNAGATCGGAA
55039 TATGAGTGTGGAGTCGTTGCAGAAAAAGCCCCGTAAGGATTGGNNNAGATCGGAA
60102 TATGAGTGTGGAGTCGTTGCAGATCCCTAGCCTTTGTTCCACTNNNAGATCGGAAGA

36794 TATGAGTGTGGAGTCGTTGCACCCATATTCCTGGGAAGTCTCANNNAGATCGGAAG
32002 TATGAGTGTGGAGTCGTTGCCCCCCTACACATTCCACTTGGAANNNAGATCGGAAG
97325 TATGAGTGTGGAGTCGTTGCAAGAGACTTCACTGGGCACAGTNNNAGATCGGAAG
68114 TATGAGTGTGGAGTCGTTGCAGGGGGTGGTCTTACTAGGTCTTNNNAGATCGGAA
42393 TATGAGTGTGGAGTCGTTGCGGGCACGTCCTATTATCAGGAACNNNAGATCGGAA
105334 TATGAGTGTGGAGTCGTTGCTCTGTTATCATTGGGGATGGGAAGNNNAGATCGGA
43490 TATGAGTGTGGAGTCGTTGCCATTAGCCAGTCCTTGGCCATACNNNAGATCGGAAG
116012 TATGAGTGTGGAGTCGTTGCTACCACAATTGTAAGCCTCCTGAGNNNAGATCGGAA
38111 TATGAGTGTGGAGTCGTTGCAGAGGGAAAAGCACAGACAGAGANNNAGATCGGA
54055 TATGAGTGTGGAGTCGTTGCGGGCGTAGATTCAATCAGATGGGNNNAGATCGGAA
101114 TATGAGTGTGGAGTCGTTGCAAGCCCCGTAAGGATTGGGANNNAGATCGGAAGAG
37823 TATGAGTGTGGAGTCGTTGCACCAGCTAAATTCAGAAACCAGANNNAGATCGGAAG
54417 TATGAGTGTGGAGTCGTTGCAAATGCTCTAGGCAACGAGATGGNNNAGATCGGAA
46778 TATGAGTGTGGAGTCGTTGCTTCCATTTGAGCTTGATTTGCGCNNNAGATCGGAAG
39739 TATGAGTGTGGAGTCGTTGCAACAGGAGCTACAGGCCATATGANNNAGATCGGAA
40226 TATGAGTGTGGAGTCGTTGCAGTGGAGCATGTAGTAGATGGTGANNNAGATCGG
34056 TATGAGTGTGGAGTCGTTGCAGGATCAGAAATGAGAAGCACCANNNAGATCGGAA
63695 TATGAGTGTGGAGTCGTTGCCTGGGAGGCATTCAAAACCAAGTNNNAGATCGGAA
50582 TATGAGTGTGGAGTCGTTGCACATTTGGGAGACGGTGTGTAAGNNNAGATCGGAA
46040 TATGAGTGTGGAGTCGTTGCTGCTGTGTCTGGGTATGTTTTCCNNNAGATCGGAAG
64481 TATGAGTGTGGAGTCGTTGCGACAGAGAGAGAGAGACTGTAGTNNNAGATCGGA
34162 TATGAGTGTGGAGTCGTTGCTGAAAGACCCCATCCTCATACCANNNAGATCGGAAG
97310 TATGAGTGTGGAGTCGTTGCTCAGGACCTCCATTTCTGGACCCNNNAGATCGGAAG
68565 TATGAGTGTGGAGTCGTTGCGGTGGTGTCTTCTGAACCATGTTNNNAGATCGGAAG
61744 TATGAGTGTGGAGTCGTTGCGCCCTAAAATAGCCACTTCAGCTNNNAGATCGGAAG
65715 TATGAGTGTGGAGTCGTTGCAAAAACACTGGGTCTCAGCATGTNNNAGATCGGAA
44795 TATGAGTGTGGAGTCGTTGCACTGGAAGAGTAAGGAACTTCCCNNNAGATCGGAA
34957 TATGAGTGTGGAGTCGTTGCAGATGAAATGGCAATAGCTAGTCCANNNAGATCGG
30276 TATGAGTGTGGAGTCGTTGCACGCAGCTTTCTGGTTCTACAAANNNAGATCGGAAG
109000 TATGAGTGTGGAGTCGTTGCCTAAGCATCAGGAGAGGCAGTANNNAGATCGGAAG
52985 TATGAGTGTGGAGTCGTTGCTTGAAAACATATCTGGGGCCAGGNNNAGATCGGAA
65650 TATGAGTGTGGAGTCGTTGCACTCCCCCTAACTCCATGAATGTNNNAGATCGGAAGA
53785 TATGAGTGTGGAGTCGTTGCGTCACATTGGGAGAAACTGAGGGNNNAGATCGGAA
116140 TATGAGTGTGGAGTCGTTGCCTTCTTGCTTTCCTTCAACTAAGTANNNAGATCGGAA
31541 TATGAGTGTGGAGTCGTTGCCCCACCATGCAAGGATACAAGAANNNAGATCGGAA
66491 TATGAGTGTGGAGTCGTTGCTGAGGGAGAGTGAGTATGTGTGTNNNAGATCGGA
59118 TATGAGTGTGGAGTCGTTGCAGAGAGAGGACAGCAACAAGGATNNNAGATCGGA
46984 TATGAGTGTGGAGTCGTTGCGGGGTAGGACAATTTCTTGAGGCNNNAGATCGGAA
65206 TATGAGTGTGGAGTCGTTGCTGGAAAATAATTGCACCGTGGGTNNNAGATCGGAA
50568 TATGAGTGTGGAGTCGTTGCGGGAAGGTGTTACTCAGCGTAAGNNNAGATCGGAA
55419 TATGAGTGTGGAGTCGTTGCGTTAGTGGAGTAGCAGCCCATGNNNAGATCGGAAG
55945 TATGAGTGTGGAGTCGTTGCAGAGATAGAGACAGAGCACCCTGNNNAGATCGGAA
56917 TATGAGTGTGGAGTCGTTGCTGGTGTGACTACTTCAGGTATGTGNNNAGATCGGA

53986 TATGAGTGTGGAGTCGTTGCTCCCAAGTAAGAGAGAAGTGGGGNNNAGATCGGA
33605 TATGAGTGTGGAGTCGTTGCTGTGTGCCTACCATATTACTAGGACANNNAGATCGG
97022 TATGAGTGTGGAGTCGTTGCAGCCTAGTCAACGTAGAGAAAACCNNNAGATCGGA
97849 TATGAGTGTGGAGTCGTTGCAGGATGGATGTGAGATCAAGCAGNNNAGATCGGA
97033 TATGAGTGTGGAGTCGTTGCGTTAGAGTACAGTGGCAAGACCTCGNNNAGATCGG
96001 TATGAGTGTGGAGTCGTTGCTGTCCAGCAAGGAAGTCTTTGGGNNNAGATCGGAA
103791 TATGAGTGTGGAGTCGTTGCACCTGGCTCTGTCACCCATACTGNNNAGATCGGAAG
104054 TATGAGTGTGGAGTCGTTGCCTGAAAATGAGTGTTCCTCCTCCTNNNAGATCGGAA
40418 TATGAGTGTGGAGTCGTTGCTGGCTATGGTTTGAGACCCTTGANNNAGATCGGAA
42076 TATGAGTGTGGAGTCGTTGCCGAGGATCACGTGTGTGTGTTTANNNAGATCGGAA
110382 TATGAGTGTGGAGTCGTTGCAAACATATCTGGGGCCAGGCCNNNAGATCGGAAGA
61189 TATGAGTGTGGAGTCGTTGCTGTATGACTTCACTCTCTGGCCTNNNAGATCGGAAG
64295 TATGAGTGTGGAGTCGTTGCTCAGGTATCTCGTGTAAGTGGAGTNNNAGATCGGA
48714 TATGAGTGTGGAGTCGTTGCCTTTAGGCACCAAGAATCCCCTCNNNAGATCGGAAG
35472 TATGAGTGTGGAGTCGTTGCTAGAGTTTCCACAAGGCCTAGCANNNAGATCGGAAG
45271 TATGAGTGTGGAGTCGTTGCTATGGCATGAAAAATTATGCTGCCNNNAGATCGGAA
37787 TATGAGTGTGGAGTCGTTGCGCATGAAGGGATACCAAGACAGANNNAGATCGGAA
52434 TATGAGTGTGGAGTCGTTGCACAAAAGCATCATCAGAATGCCGNNNAGATCGGAA
40025 TATGAGTGTGGAGTCGTTGCTAGAACATGCCAGGACACTCTGANNNAGATCGGAA
54000 TATGAGTGTGGAGTCGTTGCTCTCTTTCTCCTTCTCTGTGGGGNNNAGATCGGAAGA
48843 TATGAGTGTGGAGTCGTTGCCAGAAACGTTCCCCTAGAAGCTCNNNAGATCGGAAG
34307 TATGAGTGTGGAGTCGTTGCTCCTCTCTCTCTGAAGTACCCCANNNAGATCGGAAGA
58276 TATGAGTGTGGAGTCGTTGCGAGCCCCCAAATCAGCATAACATNNNAGATCGGAAG
63102 TATGAGTGTGGAGTCGTTGCCCTCTGCTGCCATCTTTTTCTCTNNNAGATCGGAAGA
33068 TATGAGTGTGGAGTCGTTGCTAGTGACTGAGAAGTCCCACACANNNAGATCGGAA
46821 TATGAGTGTGGAGTCGTTGCGGAGAGCCTTGACTGATAAAGGCNNNAGATCGGAA
37631 TATGAGTGTGGAGTCGTTGCAACACCCTTACCATCCAGGAAGANNNAGATCGGAAG
43086 TATGAGTGTGGAGTCGTTGCACCCTCTATGGCTCTCTCTTCACNNNAGATCGGAAGA
39188 TATGAGTGTGGAGTCGTTGCTGACCTAGAGCTTGATTGTGGGANNNAGATCGGAA
41339 TATGAGTGTGGAGTCGTTGCGTGCAGATGCCAGTGATGTTCTANNNAGATCGGAA
96732 TATGAGTGTGGAGTCGTTGCTACTAGACACCATGCTTCCATCTTGNNNAGATCGGAA
57778 TATGAGTGTGGAGTCGTTGCTCTTTCCCCCACCCACTAGAAATNNNAGATCGGAAGA
62325 TATGAGTGTGGAGTCGTTGCCTGGGATTTGCCTTCAATCTGCTNNNAGATCGGAAG
104516 TATGAGTGTGGAGTCGTTGCTCAGAGGGACATGGGGTGATTGNNNAGATCGGAA
31556 TATGAGTGTGGAGTCGTTGCGACAGCCTGCAGTAACTCAAGAANNNAGATCGGAA
58903 TATGAGTGTGGAGTCGTTGCTTGAAAAAGAGGGGCTGCTTCATNNNAGATCGGAA
110538 TATGAGTGTGGAGTCGTTGCTTAGCATCAGGAGAAACAGAAACAANNNAGATCGG
47172 TATGAGTGTGGAGTCGTTGCCTCCTCTACATTAGCATGTGGGCNNNAGATCGGAAG
53559 TATGAGTGTGGAGTCGTTGCAACAATTGGCGAATCTAGGTCGGNNNAGATCGGAA
29932 TATGAGTGTGGAGTCGTTGCATCAGGGTCAGTGTGCAGACATNNNAGATCGGAAG
71727 TATGAGTGTGGAGTCGTTGCCTGACCGCCTGATGAGAATGAAANNNAGATCGGAA
102112 TATGAGTGTGGAGTCGTTGCAATAGCCACTTCAGCTTTCATCTNNNAGATCGGAAGA
62384 TATGAGTGTGGAGTCGTTGCGTTTCTACCTCTTGTGTCGTGCTNNNAGATCGGAAG

103622 TATGAGTGTGGAGTCGTTGCCTCATTCATTCTCTTGCTGGGTACNNNAGATCGGAAG
51822 TATGAGTGTGGAGTCGTTGCGTGCTTCCTAACCAGTCACTGAGNNNAGATCGGAAG
70451 TATGAGTGTGGAGTCGTTGCGAAGAAACGTAAAGCACACTGGCNNNAGATCGGAA
68018 TATGAGTGTGGAGTCGTTGCTCCTCAGTGACATTTCTAACATCTTNNNAGATCGGAA
47823 TATGAGTGTGGAGTCGTTGCTTAGAAAATCCCAACGGGTCTGCNNNAGATCGGAAG
59311 TATGAGTGTGGAGTCGTTGCGCCTGAAGTATGTGTGTGCTGATNNNAGATCGGAA
66518 TATGAGTGTGGAGTCGTTGCTGAGCCAGTTCCATATTGTGTGTNNNAGATCGGAAG
54281 TATGAGTGTGGAGTCGTTGCTTTGATGAGGCTTACCCTTTGGGNNNAGATCGGAAG
56811 TATGAGTGTGGAGTCGTTGCGGAAGGAATAGCTCTGTTGGGTGNNNAGATCGGAA
101172 TATGAGTGTGGAGTCGTTGCCTTAGTGAGGAGGTAGTTAGCAAGGNNNAGATCGG
55529 TATGAGTGTGGAGTCGTTGCCCCAGCTGACTTCTAGGTTCATGNNNAGATCGGAAG
63163 TATGAGTGTGGAGTCGTTGCAGAGATTGGAAGGGAAGACGTCTNNNAGATCGGAA
43549 TATGAGTGTGGAGTCGTTGCCCCCACACATATTTCAGGCCTACNNNAGATCGGAAGA
36490 TATGAGTGTGGAGTCGTTGCTTTTTCCCTTGGTTTCTGGATCANNNAGATCGGAAGA
116944 TATGAGTGTGGAGTCGTTGCATTCAGATGTTCCTAGGTGTGTGGNNNAGATCGGAA
96402 TATGAGTGTGGAGTCGTTGCTAGCAGGTGTCTCTGGCTTCTNNNAGATCGGAAGAG
29801 TATGAGTGTGGAGTCGTTGCGGGGAGACAGAGACAAAGACAACNNNAGATCGGAA
29806 TATGAGTGTGGAGTCGTTGCCTGACCATTACTGCACCAGTCAANNNAGATCGGAAG
49004 TATGAGTGTGGAGTCGTTGCCTTCCCGCTTCCCGTCTATCTCNNNAGATCGGAAGAG
31752 TATGAGTGTGGAGTCGTTGCGTCTGGTCTAGCTCCTGCTAGAANNNAGATCGGAAG
29805 TATGAGTGTGGAGTCGTTGCACCATGTGCTACAAGTGCATCATNNNAGATCGGAAG
29933 TATGAGTGTGGAGTCGTTGCTCCTACTGACTTCTCCCCTTTGGNNNAGATCGGAAGA
39168 TATGAGTGTGGAGTCGTTGCTGTTCGCTTTCTTGTTTCTGGGANNNAGATCGGAAG
52463 TATGAGTGTGGAGTCGTTGCAAATGGCAGTGAAGGTATCTCCGNNNAGATCGGAA
29934 TATGAGTGTGGAGTCGTTGCGCCACAGATTTCCAACATCCATGNNNAGATCGGAAG
71867 TATGAGTGTGGAGTCGTTGCAAAGAAATGGGGAGAGAGGTGGANNNAGATCGGA
71868 TATGAGTGTGGAGTCGTTGCTGATGTAAACTGGCTTCCTCGACNNNAGATCGGAAG
29929 TATGAGTGTGGAGTCGTTGCGTTGCCTCACTGAAAAGAAACCGNNNAGATCGGAAG
29930 TATGAGTGTGGAGTCGTTGCGCCATAACACGTTTACTGTTCCCNNNAGATCGGAAGA
63939 TATGAGTGTGGAGTCGTTGCAGGCCATCAACATTTTCACCAGTNNNAGATCGGAAG
71870 TATGAGTGTGGAGTCGTTGCTGCCCAAACTTATCAATCCTCCCNNNAGATCGGAAGA
62069 TATGAGTGTGGAGTCGTTGCTGACTTGAAACAAACATTTTGGGCTNNNAGATCGGA
100706 TATGAGTGTGGAGTCGTTGCTAGACATGTAACTGAAAGACCAGCANNNAGATCGGA
32449 TATGAGTGTGGAGTCGTTGCGTCCTGTCATGGCACTGAAGTAANNNAGATCGGAAG
71726 TATGAGTGTGGAGTCGTTGCCCCAGTGTAGGCAAATGGCTATCNNNAGATCGGAAG
122009 TATGAGTGTGGAGTCGTTGCAAGCAAAAATGCCATGTTCCACATCTNNNAGATCGGA
100696 TATGAGTGTGGAGTCGTTGCCCAGGACCACATCTGAAGAAGACNNNAGATCGGAAG
102501 TATGAGTGTGGAGTCGTTGCCCTGACCTCATTGCGCAAAGGNNNAGATCGGAAGAG
101193 TATGAGTGTGGAGTCGTTGCAGGAAGCAAGGAAGCAATCGGNNNAGATCGGAAG
49738 TATGAGTGTGGAGTCGTTGCCTTCGCTGCCTAGTATGGTCTTCNNNAGATCGGAAG
102356 TATGAGTGTGGAGTCGTTGCATAAGATTAGTCAAGGATTCAGGCCANNNAGATCGG
71865 TATGAGTGTGGAGTCGTTGCCTGCACACAGAAACATAAAGCCANNNAGATCGGAAG
102709 TATGAGTGTGGAGTCGTTGCGTGTCTCCCAATGTCATGGTGGNNNAGATCGGAAGA

104034 TATGAGTGTGGAGTCGTTGCTGGCAGCAAAACAAGTCACCTNNNAGATCGGAAGA
101177 TATGAGTGTGGAGTCGTTGCAGGATCAGAGAGGGCAGTCTCTNNNAGATCGGAAG
104014 TATGAGTGTGGAGTCGTTGCTCTCCATGAAACATGACCTTAGGAANNNAGATCGGA
101176 TATGAGTGTGGAGTCGTTGCGTTCTGGTGAGATTGCAGGACTNNNAGATCGGAAG
103662 TATGAGTGTGGAGTCGTTGCGACAGAGAACTGGACAGGAGGAGNNNAGATCGGA
102074 TATGAGTGTGGAGTCGTTGCTACTTTTAGATTGAGGTTTGCCCATCANNNAGATCG
100743 TATGAGTGTGGAGTCGTTGCAGTGGGGAAACTAAGGCCAAAATGNNNAGATCGG
102769 TATGAGTGTGGAGTCGTTGCGTGTGCTTTCAGTTCATCATGTCAATNNNAGATCGG
119535 TATGAGTGTGGAGTCGTTGCTCCCTGAGAAAGGCGATATATTTGNNNAGATCGGA
105514 TATGAGTGTGGAGTCGTTGCAGTCCACAGTTCCCAGGGTTTTNNNAGATCGGAAGA
105525 TATGAGTGTGGAGTCGTTGCACGGACAACCATATCCCTTACANNNAGATCGGAAGA
71866 TATGAGTGTGGAGTCGTTGCAGGTCTAGCATAAATGTTTTCTCAGGANNNAGATCG
117608 TATGAGTGTGGAGTCGTTGCGGAATAGAGAAGTCCAAGAAGCCTNNNAGATCGGA
101178 TATGAGTGTGGAGTCGTTGCAACTCACTGAAGCCTCTCCTGGANNNAGATCGGAAG
100720 TATGAGTGTGGAGTCGTTGCATTAGCTGGCATGAAGCACTGTNNNAGATCGGAAG
103995 TATGAGTGTGGAGTCGTTGCGAGGCCATTGCTGTAGAAGCAGGNNNAGATCGGAA
110550 TATGAGTGTGGAGTCGTTGCAGGAGAACAGAAGAAAGAGCAATCNNNAGATCGG
71690 TATGAGTGTGGAGTCGTTGCGCTGCCTCTTCTATCATCCAGACNNNAGATCGGAAG
100768 TATGAGTGTGGAGTCGTTGCCCTTCTCATAGCTGCCCCCGNNNAGATCGGAAGAGC
106119 TATGAGTGTGGAGTCGTTGCTGAAACGGCACAATCATAGGTCACNNNAGATCGGA
102264 TATGAGTGTGGAGTCGTTGCCCCCATGCTCTTGTACATACTTTTNNNAGATCGGAAG
108338 TATGAGTGTGGAGTCGTTGCATCCGTCTGCCTCAGGTTACCANNNAGATCGGAAGA
110547 TATGAGTGTGGAGTCGTTGCTAGACTTTATGTGGTATGTACACTTGANNNAGATCG
115953 TATGAGTGTGGAGTCGTTGCTCCTCCCTTTCTTGGATCTTGGTNNNAGATCGGAAGA
102199 TATGAGTGTGGAGTCGTTGCGCCACACCCAGCTAGTATTGTTNNNAGATCGGAAGA
104044 TATGAGTGTGGAGTCGTTGCGCTGCAATAAACATACAAGAGCAAGTANNNAGATCG
114360 TATGAGTGTGGAGTCGTTGCAGAGCAATCACATCTAAGGGACANNNAGATCGGAA
115990 TATGAGTGTGGAGTCGTTGCCTTTGGTATCAGGTTAGTGCTTGCTNNNAGATCGGA
104491 TATGAGTGTGGAGTCGTTGCGAAAGAGCATTAGAGCTATGCCTNNNAGATCGGAA
100675 TATGAGTGTGGAGTCGTTGCGCCTCCTGGCCTATTCTTTAGTNNNAGATCGGAAGA
95131 TATGAGTGTGGAGTCGTTGCAAAAGGGAGCAGGAGACAGGTAANNNAGATCGGA
71685 TATGAGTGTGGAGTCGTTGCGCTCAGGATTCCAAGTGCTTTCANNNAGATCGGAAG
97707 TATGAGTGTGGAGTCGTTGCCACACACACTCACAGCCATCTCNNNAGATCGGAAGA
57534 TATGAGTGTGGAGTCGTTGCGTGTGTGTAGGGAGGGTCATTTGNNNAGATCGGAA
96578 TATGAGTGTGGAGTCGTTGCCTTGACACAGCTGCCTGGGAATNNNAGATCGGAAGA
49115 TATGAGTGTGGAGTCGTTGCGTCCCCTTCTGACAACACTTCTCNNNAGATCGGAAGA
30078 TATGAGTGTGGAGTCGTTGCCCAAGCAGGCATGATAAGCAAAANNNAGATCGGAA
47214 TATGAGTGTGGAGTCGTTGCATGTTTAGTCTGATTGCCATGGCNNNAGATCGGAAG
110564 TATGAGTGTGGAGTCGTTGCGTGGTGTGTGGCATGAAAAATCTAGNNNAGATCGG
34898 TATGAGTGTGGAGTCGTTGCACACACAAACATTCCTTCCTCCANNNAGATCGGAAGA
39133 TATGAGTGTGGAGTCGTTGCTGAAGGTCAGGGAGTATATGGGANNNAGATCGGA
62102 TATGAGTGTGGAGTCGTTGCTATGGCTATGTGAGATGATGGCTNNNAGATCGGAA
40087 TATGAGTGTGGAGTCGTTGCATCTGGTGTCTCCTCCTTTTCTGANNNAGATCGGAAG

34713 TATGAGTGTGGAGTCGTTGCCGTACACACACACGTATATGCCANNNAGATCGGAAG
66270 TATGAGTGTGGAGTCGTTGCTGTGCATGTATGTGAGCAAGTGTNNNAGATCGGAA
30093 TATGAGTGTGGAGTCGTTGCCCTGACACAAAGGCACATCAAAANNNAGATCGGAAG
38243 TATGAGTGTGGAGTCGTTGCACTGTATTGCCAAATATGGGGAGANNNAGATCGGA
67663 TATGAGTGTGGAGTCGTTGCTTTAGCTCTTGCCTCCCTACCTTNNNAGATCGGAAGA
107655 TATGAGTGTGGAGTCGTTGCCACCATCATCCACTTAGCAGCTCNNNAGATCGGAAGA
43446 TATGAGTGTGGAGTCGTTGCTGGCAGACAGATTTCAGCTTGACNNNAGATCGGAA
97578 TATGAGTGTGGAGTCGTTGCCCCACCCATCCACATAGCCGNNNAGATCGGAAGAGC
61857 TATGAGTGTGGAGTCGTTGCGAGCTGTAGTCCTAGCAGTAGCTNNNAGATCGGAA
104420 TATGAGTGTGGAGTCGTTGCTGCCCATCATTATTATTGCGATCTTTNNNAGATCGGA
55576 TATGAGTGTGGAGTCGTTGCCACTGAGGCTGCTGTTAGAGATGNNNAGATCGGAA
40997 TATGAGTGTGGAGTCGTTGCTCCTTCTCTTCCTCTGGTGACTANNNAGATCGGAAGA
38699 TATGAGTGTGGAGTCGTTGCTCACTCCTAGCAAGAGCTAAGGANNNAGATCGGAA
41991 TATGAGTGTGGAGTCGTTGCGGGGGAAATGCTTTGTGCATTTANNNAGATCGGAA
96701 TATGAGTGTGGAGTCGTTGCAAGCCAGTTGAACAGATACAAAAANNNAGATCGGA
97126 TATGAGTGTGGAGTCGTTGCTCCTTTCATTTTTGGCCTTGCCTANNNAGATCGGAAG
32820 TATGAGTGTGGAGTCGTTGCTCCATGCTTATTGGTTACCCAACANNNAGATCGGAA
36089 TATGAGTGTGGAGTCGTTGCTTTCTTTAGATGAGCAGGCTGCANNNAGATCGGAAG
48836 TATGAGTGTGGAGTCGTTGCGTCAAGAGGGAGACACAAGCTCNNNAGATCGGAAG
30526 TATGAGTGTGGAGTCGTTGCGTGGAGCTCAGAGGAAGGAGAAANNNAGATCGGA
53426 TATGAGTGTGGAGTCGTTGCGAGGCACTTGTTCTGTCCTTAGGNNNAGATCGGAAG
51508 TATGAGTGTGGAGTCGTTGCTAATGGTTTCCGCAGGATGAGAGNNNAGATCGGAA
59694 TATGAGTGTGGAGTCGTTGCCCAAGCACTCTATGCAGGCTTATNNNAGATCGGAAG
35904 TATGAGTGTGGAGTCGTTGCACTGTATGGAATTGAACAGAATGCANNNAGATCGG
53295 TATGAGTGTGGAGTCGTTGCAGACGGGAGTAACTACAGTGAGGNNNAGATCGGA
51323 TATGAGTGTGGAGTCGTTGCTTCTTTAGGCAGAACTATTGTTCAGNNNAGATCGGA
35415 TATGAGTGTGGAGTCGTTGCTGAAACACCTGCTAACATGAGCANNNAGATCGGAAG
105282 TATGAGTGTGGAGTCGTTGCTCTTTTCCAACAGTCTCCTTCTGNNNAGATCGGAAGA
44608 TATGAGTGTGGAGTCGTTGCATTTCTGTTCCCCTAGAGTGCCCNNNAGATCGGAAG
101202 TATGAGTGTGGAGTCGTTGCTGTATGAACCTTCTTGTCTCTCGNNNAGATCGGAAG
47016 TATGAGTGTGGAGTCGTTGCTTCCAAAGGGCACAATTGTAGGCNNNAGATCGGAA
68136 TATGAGTGTGGAGTCGTTGCTTGGGCTTGTCTCTCTTTGTCTTNNNAGATCGGAAGA
62723 TATGAGTGTGGAGTCGTTGCGGAAGAGATCACCAGGCCTATCTNNNAGATCGGAA
66375 TATGAGTGTGGAGTCGTTGCTTTCTTTATCAAAAGCCCGGTGTNNNAGATCGGAAG
58016 TATGAGTGTGGAGTCGTTGCTGCTTCCCCTTCACCCTTTCAATNNNAGATCGGAAGA
59929 TATGAGTGTGGAGTCGTTGCTCTACAGCTTCTGGCCCATAACTNNNAGATCGGAAG
41694 TATGAGTGTGGAGTCGTTGCATGGGTGGTCTCAGGAACTTGTANNNAGATCGGAA
49190 TATGAGTGTGGAGTCGTTGCCAAGCACTGACAGGTCCTAAGTCNNNAGATCGGAAG
60671 TATGAGTGTGGAGTCGTTGCAAAAGGTAGAGGTAGCACCACCTNNNAGATCGGAA
38972 TATGAGTGTGGAGTCGTTGCCTGGTTAGGCACTAGAGAAGGGANNNAGATCGGA
48176 TATGAGTGTGGAGTCGTTGCTGTGGTGAAAAGAAATGTTGTTGCNNNAGATCGGA
41788 TATGAGTGTGGAGTCGTTGCCTTTCCCCTTTGTGAGCCACTTANNNAGATCGGAAGA
103784 TATGAGTGTGGAGTCGTTGCGCCACTCATTCATCTTTTTGTATGGGNNNAGATCGG

30907 TATGAGTGTGGAGTCGTTGCAGGATAGTGGTGGGAACAGACAANNNAGATCGGA
53900 TATGAGTGTGGAGTCGTTGCGACATTTGGGTAGGTTTGAGGGGNNNAGATCGGA
52061 TATGAGTGTGGAGTCGTTGCATGAAATGGGATTGGCTGGCTAGNNNAGATCGGAA
44514 TATGAGTGTGGAGTCGTTGCTGGAGTTACATTCAGATAAGCCCNNNAGATCGGAAG
97572 TATGAGTGTGGAGTCGTTGCACAGGTGGTACGATCATAGCTCATNNNAGATCGGA
68635 TATGAGTGTGGAGTCGTTGCAGAAGTGAGGTTAGACTGCTGTTNNNAGATCGGAA
51996 TATGAGTGTGGAGTCGTTGCTAGGGGGAGAGAGACAGGGATAGNNNAGATCGGA
40890 TATGAGTGTGGAGTCGTTGCACTGCCACCCAGGTCGATATATANNNAGATCGGAAG
69177 TATGAGTGTGGAGTCGTTGCTCATCAGCAAGATGCTGCTCTTTNNNAGATCGGAAG
39268 TATGAGTGTGGAGTCGTTGCGGAGGTTGTCAGAAAGTTCATGGANNNAGATCGGA
67411 TATGAGTGTGGAGTCGTTGCGATGGGGTGCCATGGTTTCTATTNNNAGATCGGAA
55863 TATGAGTGTGGAGTCGTTGCTTTTTCCCCAAGATGGTGGACTGNNNAGATCGGAAG
42404 TATGAGTGTGGAGTCGTTGCGGCGAATACCTGTATCTGGGAACNNNAGATCGGAA
34145 TATGAGTGTGGAGTCGTTGCAGTTGAAGCATTTGACATTGACCANNNAGATCGGAA
49426 TATGAGTGTGGAGTCGTTGCACACACACGAGTTTCACTCTGTCNNNAGATCGGAAG
37931 TATGAGTGTGGAGTCGTTGCTAAAAGCCCATCTGAGGAGCAGANNNAGATCGGAA
64401 TATGAGTGTGGAGTCGTTGCTCCCTTCCTTTCACCTGCATAGTNNNAGATCGGAAGA
67370 TATGAGTGTGGAGTCGTTGCGTCTAATGCCAAAGGGCAATATTNNNAGATCGGAAG
66573 TATGAGTGTGGAGTCGTTGCGCATGAAACTTGAAGGCCATTGTNNNAGATCGGAA
57048 TATGAGTGTGGAGTCGTTGCATTAGCTGAGACATGGTGGTGTGNNNAGATCGGAA
43149 TATGAGTGTGGAGTCGTTGCAGAGTGGTGAGGAGAAGACAGACNNNAGATCGGA
43036 TATGAGTGTGGAGTCGTTGCACCTCCTTACAAATCAGCAGTCACNNNAGATCGGAAG
36203 TATGAGTGTGGAGTCGTTGCAGTGGGAATTTTGAGGAAATTGCANNNAGATCGGA
42269 TATGAGTGTGGAGTCGTTGCAGCCTGTCCAGAATCAAGACAACNNNAGATCGGAAG
36252 TATGAGTGTGGAGTCGTTGCTGAGAAGCAATACACCATCTTGCANNNAGATCGGAA
107688 TATGAGTGTGGAGTCGTTGCCGGGCTATTTCTGGAAGGATTGAANNNAGATCGGA
52766 TATGAGTGTGGAGTCGTTGCAGAGGACAAAACTCAAGGCAAGGNNNAGATCGGA
43311 TATGAGTGTGGAGTCGTTGCCCTTTCTTTCTACCCTTGGGGACNNNAGATCGGAAGA
34746 TATGAGTGTGGAGTCGTTGCGAAACTACTTGAACCTGGTGCCANNNAGATCGGAAG
55857 TATGAGTGTGGAGTCGTTGCAGGCTTAGATAACCACAAGGACTGNNNAGATCGGA
57753 TATGAGTGTGGAGTCGTTGCCTTGTGTGTGCAGTGGTTCAAATNNNAGATCGGAA
106610 TATGAGTGTGGAGTCGTTGCATGTGTGTGCATTTTGGGAGGGTANNNAGATCGGA
51515 TATGAGTGTGGAGTCGTTGCAGCATTTGATGGATGGCTGAGAGNNNAGATCGGAA
41821 TATGAGTGTGGAGTCGTTGCAAATTCAGCCTGGGACATCCTTANNNAGATCGGAAG
71756 TATGAGTGTGGAGTCGTTGCCTGGGGTCTCATATACAGGGACCNNNAGATCGGAA
54099 TATGAGTGTGGAGTCGTTGCAATAGCCTTCCTAAAGGCCTGGGNNNAGATCGGAA
50732 TATGAGTGTGGAGTCGTTGCAGTTTTTGGGGGCAGTTAGACAGNNNAGATCGGAA
62135 TATGAGTGTGGAGTCGTTGCTTCCTCTTTTAGGGTCTCTGGCTNNNAGATCGGAAG
41018 TATGAGTGTGGAGTCGTTGCGATACTTGCTGAGCACCAACCTANNNAGATCGGAAG
33452 TATGAGTGTGGAGTCGTTGCTGAACCACCCACAACAAAAGACANNNAGATCGGAAG
46662 TATGAGTGTGGAGTCGTTGCATGAAAGAGTGAGTGCTGCTAGCNNNAGATCGGAA
34693 TATGAGTGTGGAGTCGTTGCACGAGATGACTTTCTGAATGCCANNNAGATCGGAAG
51700 TATGAGTGTGGAGTCGTTGCCCACACACTAACTCACCATGGAGNNNAGATCGGAAG

42481 TATGAGTGTGGAGTCGTTGCAGTTGCCCAGAATAGCCTGTAACNNNAGATCGGAAG
52875 TATGAGTGTGGAGTCGTTGCAGGTCATGTTGCCTGTAAACAGGNNNAGATCGGAA
29927 TATGAGTGTGGAGTCGTTGCGGAGGAGAGGACCAAACAGATGANNNAGATCGGA
56079 TATGAGTGTGGAGTCGTTGCTGGTAGGGAGTGAAGTCTTCCTGNNNAGATCGGAA
33547 TATGAGTGTGGAGTCGTTGCGGGGGTTGAAACTGAATGAGACANNNAGATCGGA
37936 TATGAGTGTGGAGTCGTTGCTCTGTAGCCTCAACTTCTAGCAGANNNAGATCGGAA
63047 TATGAGTGTGGAGTCGTTGCTAGAGATACCGAGATATGGTCTCTNNNAGATCGGA
97897 TATGAGTGTGGAGTCGTTGCACATATCACAGGTTGAAAAAGCGANNNAGATCGGA
61346 TATGAGTGTGGAGTCGTTGCAAGATAGGTGCTCGTGGTATCCTNNNAGATCGGAA
42566 TATGAGTGTGGAGTCGTTGCGAGAGAAACATGCATGCACACACNNNAGATCGGAA
69339 TATGAGTGTGGAGTCGTTGCTTTTACTTCATCTCCTTTGTTGGTTTNNNAGATCGGA
44035 TATGAGTGTGGAGTCGTTGCCTGGTCCTTTTACTCTGCAGACCNNNAGATCGGAAG
60216 TATGAGTGTGGAGTCGTTGCTTCACACACACACACAAGTCACTNNNAGATCGGAAG
58206 TATGAGTGTGGAGTCGTTGCCCAAAAAGGAGTGCCTGGGTAATNNNAGATCGGAA
32939 TATGAGTGTGGAGTCGTTGCACAAGGCAACATTTGATGTGAACANNNAGATCGGA
60746 TATGAGTGTGGAGTCGTTGCGTGTGTTTTCCACTCAAGGACCTNNNAGATCGGAAG
54309 TATGAGTGTGGAGTCGTTGCGGGCTCAGTAACTCCTCAAATGGNNNAGATCGGAA
42933 TATGAGTGTGGAGTCGTTGCACGAATAAGACCTGGTTCTGCACNNNAGATCGGAAG
63911 TATGAGTGTGGAGTCGTTGCTGTGTGTGCATGTATGTATGACAGTNNNAGATCGG
41933 TATGAGTGTGGAGTCGTTGCTCAGGGAATGAAGGACAGGGTTANNNAGATCGGA
55626 TATGAGTGTGGAGTCGTTGCGCTTGAGAGCTATGCAGTGGATGNNNAGATCGGAA
47643 TATGAGTGTGGAGTCGTTGCCCCCTCAGTGAATGAGTAACTGCNNNAGATCGGAAG
67610 TATGAGTGTGGAGTCGTTGCGGAACAAGTGGAAGGTGCTACTTNNNAGATCGGAA
30431 TATGAGTGTGGAGTCGTTGCCCTCCCAAACCCAAGAACTCAAANNNAGATCGGAAG
47621 TATGAGTGTGGAGTCGTTGCCACCTGAGCCTGTAGAGTTATGCNNNAGATCGGAAG
49182 TATGAGTGTGGAGTCGTTGCCCACAAGGATAGCTCAGGAAGTCNNNAGATCGGAA
35000 TATGAGTGTGGAGTCGTTGCCAGAGCTTCTTTCCACTTGTCCANNNAGATCGGAAG
30937 TATGAGTGTGGAGTCGTTGCTGCTCTTCCCTACTCTGTGACAANNNAGATCGGAAG
32152 TATGAGTGTGGAGTCGTTGCACAGCCAGTGTCTTCATAGTGAANNNAGATCGGAAG
58554 TATGAGTGTGGAGTCGTTGCAGACCTTGCCTTCCCTTATCCATNNNAGATCGGAAGA
63337 TATGAGTGTGGAGTCGTTGCGCCTGAATGGTCCAACTCATTCTNNNAGATCGGAAG
29926 TATGAGTGTGGAGTCGTTGCGCCTAACCTGATAGCAGGGTTTCNNNAGATCGGAAG
40120 TATGAGTGTGGAGTCGTTGCAAGGAAGGAAGGAAAACCAGTGANNNAGATCGGA
61299 TATGAGTGTGGAGTCGTTGCAGAAGGGAGAGAGAGTTTTGCCTNNNAGATCGGA
34942 TATGAGTGTGGAGTCGTTGCTCCAATTGCCTCCAATTTCTCCANNNAGATCGGAAGA
68310 TATGAGTGTGGAGTCGTTGCTGACATGAGTGGAAGGGAGAGTTNNNAGATCGGA
71713 TATGAGTGTGGAGTCGTTGCAGTACCTCAGCCCATGCCAAGNNNAGATCGGAAGAG
31959 TATGAGTGTGGAGTCGTTGCGCTGGCTTCCTTCCAATATGGAANNNAGATCGGAAG
31114 TATGAGTGTGGAGTCGTTGCGTGAACGAGTGAGAACCATCCAANNNAGATCGGAA
58057 TATGAGTGTGGAGTCGTTGCTCCCCTCTGTTTTCCTGGAGAATNNNAGATCGGAAG
71341 TATGAGTGTGGAGTCGTTGCTTTATGATCCAGACCAGGCTCGTNNNAGATCGGAAG
39770 TATGAGTGTGGAGTCGTTGCAGTTGACAGAAGCACCAAACTGANNNAGATCGGAA
43204 TATGAGTGTGGAGTCGTTGCTCTCACCCTCCCAATGTAGAGACNNNAGATCGGAAG

57258 TATGAGTGTGGAGTCGTTGCAAATGCGTGTGCTTGTGTACTTGNNNAGATCGGAA
30934 TATGAGTGTGGAGTCGTTGCTATGCTTTGGTTGACGGTGACAANNNAGATCGGAA
69175 TATGAGTGTGGAGTCGTTGCCAGTGTCACCTTGAAGGCTCTTTNNNAGATCGGAAG
34018 TATGAGTGTGGAGTCGTTGCAGCATTGCCTGAAGAAAACACCANNNAGATCGGAA
56685 TATGAGTGTGGAGTCGTTGCACAGGTTTTTCTGCATGTACGTGNNNAGATCGGAAG
104447 TATGAGTGTGGAGTCGTTGCTTTGATGCTGTCCCTTTGTCTCNNNAGATCGGAAGA
67520 TATGAGTGTGGAGTCGTTGCCCTCATACGTAACCACAGCACTTNNNAGATCGGAAG
39539 TATGAGTGTGGAGTCGTTGCTGGGAAAACAATTGGAAAAGTTTGGANNNAGATCG
102177 TATGAGTGTGGAGTCGTTGCATAGCTTGTTTCCTTCCACTCTATGNNNAGATCGGAA
41194 TATGAGTGTGGAGTCGTTGCATTTACCCCTCCACTGATGGCTANNNAGATCGGAAG
108375 TATGAGTGTGGAGTCGTTGCAAAGCTGGGAGAGAGGAGACTGNNNAGATCGGAA
116288 TATGAGTGTGGAGTCGTTGCCCCAGAATAGCCTGTAACCATCCTNNNAGATCGGAA
61575 TATGAGTGTGGAGTCGTTGCTGGAAAAACATAATAGAAGAGTTTCCTNNNAGATCG
67542 TATGAGTGTGGAGTCGTTGCAAGCCAACAGCAAGACTTCACTTNNNAGATCGGAAG
65995 TATGAGTGTGGAGTCGTTGCAAGGGAGAGAAATGAGAGCCTGTNNNAGATCGGA
105258 TATGAGTGTGGAGTCGTTGCTCCAGACCAGGCTCGTAATAAACANNNAGATCGGAA
106049 TATGAGTGTGGAGTCGTTGCTCAAGAGGACACATGGTCAGGGNNNAGATCGGAAG
39681 TATGAGTGTGGAGTCGTTGCAAAAAGCTGCCACTGGAAGATGANNNAGATCGGAA
70992 TATGAGTGTGGAGTCGTTGCTTTTTGTGCGTCCGTTTTGACATNNNAGATCGGAAG
65634 TATGAGTGTGGAGTCGTTGCTCCACTTTTGAGAGCTGCAATGTNNNAGATCGGAAG
65456 TATGAGTGTGGAGTCGTTGCTTTGTGATGAGATTGAGGGTGGTNNNAGATCGGAA
58128 TATGAGTGTGGAGTCGTTGCGGGTAGGCAGTGCATTACTGAATNNNAGATCGGAA
35529 TATGAGTGTGGAGTCGTTGCAGTGATGACAAAGAGAACACGCANNNAGATCGGAA
30833 TATGAGTGTGGAGTCGTTGCCAAGCCCTTTCAGGAGAGAACAANNNAGATCGGAA
29925 TATGAGTGTGGAGTCGTTGCCTTTGTCCTCACTCCTCGTGTCNNNAGATCGGAAGAG
37797 TATGAGTGTGGAGTCGTTGCGGGAGGACTGAGAGAGAGACAGANNNAGATCGGA
44041 TATGAGTGTGGAGTCGTTGCAGGGGCATATACATGGAGAGACCNNNAGATCGGAA
41185 TATGAGTGTGGAGTCGTTGCGAGACGAGCTTTTGCTTAGGCTANNNAGATCGGAA
34564 TATGAGTGTGGAGTCGTTGCCCTACTTAGCCTTCATAACAGCCANNNAGATCGGAAG
99241 TATGAGTGTGGAGTCGTTGCTCTTCCTTCATTGCACTGTCCACTCNNNAGATCGGAA
29982 TATGAGTGTGGAGTCGTTGCCACAAAAACAACTCCCTCTGGGANNNAGATCGGAAG
102173 TATGAGTGTGGAGTCGTTGCGAGACGAGCTTTTGCTTAGGCTATNNNAGATCGGA
66816 TATGAGTGTGGAGTCGTTGCTCCCAAACCCTGTTTTATTCTTTGTNNNAGATCGGAA
108371 TATGAGTGTGGAGTCGTTGCTCTGAACTGACAAAGAAATGTTTATTGNNNAGATCG
43699 TATGAGTGTGGAGTCGTTGCGCAGAGACAGCAACACCATTTACNNNAGATCGGAAG
31911 TATGAGTGTGGAGTCGTTGCCATTCTCTGGGGGAATAGGGGAANNNAGATCGGAA
55336 TATGAGTGTGGAGTCGTTGCGCAGCAGGAATAGCAAGGTAATGNNNAGATCGGA
53908 TATGAGTGTGGAGTCGTTGCCTGCCTAAAATCCAAGCTAGGGGNNNAGATCGGAA
102168 TATGAGTGTGGAGTCGTTGCTCCCTGTTGAGGTTGCTTTTCCATNNNAGATCGGAA
35544 TATGAGTGTGGAGTCGTTGCTCCTCTGATCTCTGCTATTACGCANNNAGATCGGAAG
63982 TATGAGTGTGGAGTCGTTGCTGTCTTGGTTCCCTTACTCCAGTNNNAGATCGGAAG
110578 TATGAGTGTGGAGTCGTTGCAATAGGAAGCCTACAGTGTTCTTTANNNAGATCGGA
110541 TATGAGTGTGGAGTCGTTGCGAAGGCACTGCTGGCACAATATNNNAGATCGGAAG

116468 TATGAGTGTGGAGTCGTTGCCTCACTCTGTTGTGCAGGCCTGNNNAGATCGGAAGA
59490 TATGAGTGTGGAGTCGTTGCGCAGGATACATGGGGGTGACTATNNNAGATCGGAA
63861 TATGAGTGTGGAGTCGTTGCATTCAGGCCAAACTGTCAACAGTNNNAGATCGGAAG
32883 TATGAGTGTGGAGTCGTTGCCCTTATCTGTGACTGGTCAGAACANNNAGATCGGAA
60652 TATGAGTGTGGAGTCGTTGCCACTATGAACACATGCCACACCTNNNAGATCGGAAG
37174 TATGAGTGTGGAGTCGTTGCGTAAATCTGCTGCCTCCACTTCANNNAGATCGGAAG
50881 TATGAGTGTGGAGTCGTTGCACAGTTTCAGCTTTGTTTCCCAGNNNAGATCGGAAG
30838 TATGAGTGTGGAGTCGTTGCTGGCATTTGTTATGGGGGAACAANNNAGATCGGAA
61541 TATGAGTGTGGAGTCGTTGCTCACCTTCCTGGTTTCTCTTCCTNNNAGATCGGAAGA
97954 TATGAGTGTGGAGTCGTTGCAGGCATCTTGACTTCACAACAGNNNAGATCGGAAGA
101308 TATGAGTGTGGAGTCGTTGCGTGTGTCTGTGTAGACATGGTCTANNNAGATCGGA
60994 TATGAGTGTGGAGTCGTTGCGGAACCTCTGTACCTTGATCCCTNNNAGATCGGAAG
50431 TATGAGTGTGGAGTCGTTGCTCCCTTTGTCTCTCTCCAGGAAGNNNAGATCGGAAG
30116 TATGAGTGTGGAGTCGTTGCGGGGTTCACAGGATTCCAGAAAANNNAGATCGGAA
116729 TATGAGTGTGGAGTCGTTGCTGGTTTCATTGATCGCTGTCCCTNNNAGATCGGAAG
53057 TATGAGTGTGGAGTCGTTGCCATCTCCTCAAACCCTATGCAGGNNNAGATCGGAAG
68552 TATGAGTGTGGAGTCGTTGCGAGGGAGGAGCAGAAAGAATGTTNNNAGATCGGA
52652 TATGAGTGTGGAGTCGTTGCTGTGTGTCTCATGTGTAGGTTCGNNNAGATCGGAA
99182 TATGAGTGTGGAGTCGTTGCAGGGGGCGACAGGCCAATCTNNNAGATCGGAAGA
58947 TATGAGTGTGGAGTCGTTGCACAGTCTTGGGAGAGTGCAAGATNNNAGATCGGAA
41990 TATGAGTGTGGAGTCGTTGCCTCTGGTGGAAGCCATGCATTTANNNAGATCGGAAG
48363 TATGAGTGTGGAGTCGTTGCCGCAGAAAGTAGGTGATGCCATCNNNAGATCGGAA
69342 TATGAGTGTGGAGTCGTTGCTGTATCCCTCGCCCAAAATGTTTNNNAGATCGGAAG
99569 TATGAGTGTGGAGTCGTTGCAGTTCCCCACTCATGCAATGTTGNNNAGATCGGAAG
56711 TATGAGTGTGGAGTCGTTGCCCTCGTTGTGGTTTCAATTCGTGNNNAGATCGGAAG
35012 TATGAGTGTGGAGTCGTTGCTCTTGGCCATATCCATAATTCCANNNAGATCGGAAGA
32184 TATGAGTGTGGAGTCGTTGCCCACTCCTTTTCCACTGTGTGAANNNAGATCGGAAGA
40285 TATGAGTGTGGAGTCGTTGCTGCTTACATGATTTAGCTACTGTGANNNAGATCGGA
99237 TATGAGTGTGGAGTCGTTGCGAGGTTTGCTCTCACACAGGCNNNAGATCGGAAGA
54806 TATGAGTGTGGAGTCGTTGCGTCCATGCTGCAGGTTTCTGGNNNAGATCGGAAGA
55666 TATGAGTGTGGAGTCGTTGCAGCCTGATTGCTGCATTTTGATGNNNAGATCGGAAG
102185 TATGAGTGTGGAGTCGTTGCTTATGCTGTCAGGAGTGAACTTGNNNAGATCGGAA
59874 TATGAGTGTGGAGTCGTTGCGGGAGTTCCAGGTAGACAGAACTNNNAGATCGGAA
41886 TATGAGTGTGGAGTCGTTGCTGGGGAGCTGGAGATTGTTCTTANNNAGATCGGAA
46333 TATGAGTGTGGAGTCGTTGCATTCATGTCAGCAGAGATCCAGCNNNAGATCGGAAG
99145 TATGAGTGTGGAGTCGTTGCTGCTCACTTTATAGGCCCTGCGANNNAGATCGGAAG
101571 TATGAGTGTGGAGTCGTTGCAAATCCAAGCTAGGGGAAACACAGNNNAGATCGGA
50168 TATGAGTGTGGAGTCGTTGCGTTAGACCACTGTGGGTGGAAAGNNNAGATCGGAA
110570 TATGAGTGTGGAGTCGTTGCAAAGTCATCTGTTGTTGGGCACNNNAGATCGGAAG
59695 TATGAGTGTGGAGTCGTTGCGCCAAGACCTCTTCCATGCTTATNNNAGATCGGAAG
43846 TATGAGTGTGGAGTCGTTGCACCCCAGAATGCAGAAAGTAACCNNNAGATCGGAA
65951 TATGAGTGTGGAGTCGTTGCATCGTTCTAGACAAAGACTACTGTNNNAGATCGGAA
41847 TATGAGTGTGGAGTCGTTGCAGGGTGCACAGATCTATGGCTTANNNAGATCGGAA

103988 TATGAGTGTGGAGTCGTTGCGGGCTGGTACACATTTTTGCAATTTNNNAGATCGGA
53748 TATGAGTGTGGAGTCGTTGCGATTTCAGCCTAACACCAGAGGGNNNAGATCGGAA
38131 TATGAGTGTGGAGTCGTTGCAAAAACTCTTGGAACCTTCAGAGANNNAGATCGGAA
56989 TATGAGTGTGGAGTCGTTGCAACCTCTGTGTAGTTGCTCTGTGNNNAGATCGGAAG
71362 TATGAGTGTGGAGTCGTTGCACAGATTCCTAACAACCAGGGGTNNNAGATCGGAA
105251 TATGAGTGTGGAGTCGTTGCTCTATCACCTCAAATGACACCACATNNNAGATCGGAA
49414 TATGAGTGTGGAGTCGTTGCAGCGGTAGGTATAGAGAGCTGTCNNNAGATCGGAA
33022 TATGAGTGTGGAGTCGTTGCCTCCTCCCAGTGCAAATAACACANNNAGATCGGAAG
65627 TATGAGTGTGGAGTCGTTGCAGATTGTGGTGGTCATTTTACAATGTNNNAGATCGG
111619 TATGAGTGTGGAGTCGTTGCCCTGGTTTCTCTTCCTTTACAGCANNNAGATCGGAAG
38484 TATGAGTGTGGAGTCGTTGCTCCCCCAAACCATTTGAAAGTTAGANNNAGATCGGA
39494 TATGAGTGTGGAGTCGTTGCGAGAGGGGTATGGTTACCTTGGANNNAGATCGGA
55939 TATGAGTGTGGAGTCGTTGCCATTGTCAGGGGTGTGTACCTGNNNAGATCGGAAG
117628 TATGAGTGTGGAGTCGTTGCTGGTGATTCTTTGTTTTCCTGCCTANNNAGATCGGAA
99961 TATGAGTGTGGAGTCGTTGCTTCAGTCACCATGATGTTTGTGANNNAGATCGGAAG
100671 TATGAGTGTGGAGTCGTTGCATCCCAAGAGACTGTGAATTCTCNNNAGATCGGAAG
62653 TATGAGTGTGGAGTCGTTGCCCTGCTCCAGTACGATCTCATCTNNNAGATCGGAAGA
43480 TATGAGTGTGGAGTCGTTGCGCGACACCAAACAGGATGAATACNNNAGATCGGAA
116289 TATGAGTGTGGAGTCGTTGCTGGGAGAGTGCAAGATAATTCACNNNAGATCGGAA
108941 TATGAGTGTGGAGTCGTTGCTGATTTTGGGGCAGAACTTTCTATNNNAGATCGGAA
50687 TATGAGTGTGGAGTCGTTGCCCCCTTCCCCCATTTTATCACAGNNNAGATCGGAAGA
36749 TATGAGTGTGGAGTCGTTGCCGATCATTCCACCAAGAAAAATCTCANNNAGATCGG
38425 TATGAGTGTGGAGTCGTTGCCACTCAGGAGAAGAGGTGCTAGANNNAGATCGGAA
55177 TATGAGTGTGGAGTCGTTGCTAGCTGAACAGAGTGAGCTTTGGNNNAGATCGGAA
59907 TATGAGTGTGGAGTCGTTGCGACAGGGACTTAGAGGTGGAACTNNNAGATCGGA
68148 TATGAGTGTGGAGTCGTTGCCCGTAACTTTGAGGGCACTTCTTNNNAGATCGGAAG
35418 TATGAGTGTGGAGTCGTTGCGAGCTACAGCATAACTGATGAGCANNNAGATCGGA
106631 TATGAGTGTGGAGTCGTTGCCCAGGCCTCTAATTTTGTCTTCCTNNNAGATCGGAAG
110331 TATGAGTGTGGAGTCGTTGCGCTGAAGGCACACAACAATTGGTNNNAGATCGGAA
57634 TATGAGTGTGGAGTCGTTGCTGGGTCTTTGCTTTTGCCTTTTGNNNAGATCGGAAG
60522 TATGAGTGTGGAGTCGTTGCGTATGCCCGAATGTGCATGTACTNNNAGATCGGAAG
60924 TATGAGTGTGGAGTCGTTGCTCAGCTCTTCTGTCATTTCCCCTNNNAGATCGGAAGA
53085 TATGAGTGTGGAGTCGTTGCGCTCATGAATGTTACAGCTCAGGNNNAGATCGGAA
42099 TATGAGTGTGGAGTCGTTGCGTCTGTCCTCACTTGGGCTTTTANNNAGATCGGAAG
34142 TATGAGTGTGGAGTCGTTGCACCAGATAAAAGAAAATTCCTGACCANNNAGATCGG
40602 TATGAGTGTGGAGTCGTTGCGAATAGGCTGTGGTGGGCAAATANNNAGATCGGA
62205 TATGAGTGTGGAGTCGTTGCTTGTTCAGAGGCCATAGAATGCTNNNAGATCGGAA
102105 TATGAGTGTGGAGTCGTTGCGCTCCAGTACGATCTCATCTTAGCNNNAGATCGGAA
49100 TATGAGTGTGGAGTCGTTGCCCTGTCTCACTTGGTGTTGTCTCNNNAGATCGGAAG
50121 TATGAGTGTGGAGTCGTTGCGGGGAGGGGAGAATCAAAGAAAGNNNAGATCGG
50055 TATGAGTGTGGAGTCGTTGCCCCAGCCACCATCTGAGTAAAAGNNNAGATCGGAAG
66424 TATGAGTGTGGAGTCGTTGCTGCTGTTGATGATGGTGATGTGTNNNAGATCGGAA
36522 TATGAGTGTGGAGTCGTTGCTTGCCTTACCCTGTCCTGTATCANNNAGATCGGAAGA

| | |
|---|---|
| 30283 | TATGAGTGTGGAGTCGTTGCTTGTGGTGGCTGGATAAACCAAANNNAGATCGGAA |
| 45574 | TATGAGTGTGGAGTCGTTGCGGATCAATTTTTGGTTGCCCTCCNNNAGATCGGAAG |
| 55620 | TATGAGTGTGGAGTCGTTGCAAGAAGAGCAGCAAGGATGGATGNNNAGATCGGA |
| 66516 | TATGAGTGTGGAGTCGTTGCCGTGGAGAATAGTGTGTGTGTGTNNNAGATCGGAA |
| 65862 | TATGAGTGTGGAGTCGTTGCGGGACAGAGAGACAATGCTTATGTNNNAGATCGGA |
| 55200 | TATGAGTGTGGAGTCGTTGCTGGGAAAACATGTTCCTGTTTGGNNNAGATCGGAA |
| 105259 | TATGAGTGTGGAGTCGTTGCGTTAGGGTTGCAATGAAACGTGTGNNNAGATCGGA |
| 30396 | TATGAGTGTGGAGTCGTTGCAGTTGTCCCTTGAACAATGCAAANNNAGATCGGAAG |
| 55298 | TATGAGTGTGGAGTCGTTGCGTGTTGTGTTGTGCAGGAGAATGNNNAGATCGGAA |
| 48580 | TATGAGTGTGGAGTCGTTGCTAAAGGCTGAGATGTCCCAACTCNNNAGATCGGAAG |
| 40609 | TATGAGTGTGGAGTCGTTGCCCTAATGTGGCCAGCTGGAAATANNNAGATCGGAA |
| 68324 | TATGAGTGTGGAGTCGTTGCAGATGCAACAGGTGCAATGAGTTNNNAGATCGGAA |
| 45795 | TATGAGTGTGGAGTCGTTGCCTCGATCATCTTACAGCCTGTCCNNNAGATCGGAAG |
| 51562 | TATGAGTGTGGAGTCGTTGCGGGAAGATAAAGAAGGGTGCGAGNNNAGATCGGA |
| 34539 | TATGAGTGTGGAGTCGTTGCTCAGCCTCAGAAAATGAAAGCCANNNAGATCGGAA |
| 30263 | TATGAGTGTGGAGTCGTTGCGGGCCAAACCTCATTAGGACAAANNNAGATCGGAA |
| 50704 | TATGAGTGTGGAGTCGTTGCTCTGTGCCTCTGTTTCTTCACAGNNNAGATCGGAAG |
| 68694 | TATGAGTGTGGAGTCGTTGCTGTGTTATGCCTTCCACTGTGTTNNNAGATCGGAAG |
| 36803 | TATGAGTGTGGAGTCGTTGCACCAAGGAGAATATCTGGTCTCANNNAGATCGGAA |
| 45598 | TATGAGTGTGGAGTCGTTGCTCACATTTTGCCTTTCCTCCTCCNNNAGATCGGAAGA |
| 59392 | TATGAGTGTGGAGTCGTTGCGGTGGAGTGGCTGTTCCAAATATNNNAGATCGGAA |
| 104439 | TATGAGTGTGGAGTCGTTGCATGGGATTTCTCTCTTGTCCAGGNNNAGATCGGAAG |
| 67357 | TATGAGTGTGGAGTCGTTGCGGGTTGCAATGAAACGTGTGATTNNNAGATCGGAA |
| 106551 | TATGAGTGTGGAGTCGTTGCATGCAGGAGGTATTGTTATCCCANNNAGATCGGAA |
| 50140 | TATGAGTGTGGAGTCGTTGCAGGACCAGGAGAAGACAGGAAAGNNNAGATCGGA |
| 108377 | TATGAGTGTGGAGTCGTTGCGTTTGTAGTGTGTGCGTGTTCNNNAGATCGGAAGA |
| 31781 | TATGAGTGTGGAGTCGTTGCAGGTAGTGCAGCATCAAAAGGAANNNAGATCGGAA |
| 44860 | TATGAGTGTGGAGTCGTTGCCGATTTTGCTCTGGTGTAAAGCCNNNAGATCGGAAG |
| 68916 | TATGAGTGTGGAGTCGTTGCCCGTGACCTGAGATGTGTGATTTNNNAGATCGGAA |
| 110576 | TATGAGTGTGGAGTCGTTGCGGGCTACTACCTCTGGTCTGTGGNNNAGATCGGAA |
| 34858 | TATGAGTGTGGAGTCGTTGCTTTTTCTGGGGCAATCAACTCCANNNAGATCGGAAG |
| 47724 | TATGAGTGTGGAGTCGTTGCCATAAAAAGCAGGCAACTCCTGCNNNAGATCGGAAG |
| 97892 | TATGAGTGTGGAGTCGTTGCCACACAAATACACTCCTTTATTCCACANNNAGATCGG |
| 43642 | TATGAGTGTGGAGTCGTTGCGACAGACAATCAGGAGGCTGTACNNNAGATCGGAA |
| 68222 | TATGAGTGTGGAGTCGTTGCCAGCACACACTCATGGAGAAGTTNNNAGATCGGAA |
| 68884 | TATGAGTGTGGAGTCGTTGCCCAAATGCCTGCTCCAGAGATTTNNNAGATCGGAAG |
| 41160 | TATGAGTGTGGAGTCGTTGCAAGGAGCCTCCAAACACTAGCTANNNAGATCGGAAG |
| 107546 | TATGAGTGTGGAGTCGTTGCGAGAGACTGAGAGTGTCAGGCANNNAGATCGGAA |
| 69156 | TATGAGTGTGGAGTCGTTGCCATACAGCCTCAGCCACATCTTTNNNAGATCGGAAG |
| 30402 | TATGAGTGTGGAGTCGTTGCGCTTTCCTTTCTGAAGCTGCAAANNNAGATCGGAAG |
| 49165 | TATGAGTGTGGAGTCGTTGCGGTGGCAAAAGGTGCTAAAAGTCNNNAGATCGGAA |
| 39152 | TATGAGTGTGGAGTCGTTGCTAGGTGTGGAAATAAGCCTGGGANNNAGATCGGAA |
| 104032 | TATGAGTGTGGAGTCGTTGCAGGTATCAGGAACAAAACAAAGGTTTTNNNAGATC |

57886 TATGAGTGTGGAGTCGTTGCTGCCCATTGCTCCAATAACCAATNNNAGATCGGAAGA
66282 TATGAGTGTGGAGTCGTTGCTGAAAAGCAGGATGGTGAAGTGTNNNAGATCGGA
97575 TATGAGTGTGGAGTCGTTGCGATTGGAGAAGGCTTCACGGGTNNNAGATCGGAAG
56031 TATGAGTGTGGAGTCGTTGCTATTCTAGGCCCTCCACATCCTGNNNAGATCGGAAGA
49020 TATGAGTGTGGAGTCGTTGCATCATTTGCAAGCGTACCCTCTCNNNAGATCGGAAGA
70713 TATGAGTGTGGAGTCGTTGCTCTACTCTGCTACTGGGAACAGGNNNAGATCGGAAG
46353 TATGAGTGTGGAGTCGTTGCGAAACTCAAAAGTCAGCAGCAGCNNNAGATCGGAA
54049 TATGAGTGTGGAGTCGTTGCAGATTGTGATGGCTAAGCATGGGNNNAGATCGGAA
52984 TATGAGTGTGGAGTCGTTGCATAGTTGTTGAAATGGGGCCAGGNNNAGATCGGAA
65435 TATGAGTGTGGAGTCGTTGCGATGTCTCTAGGAACCATAGTGGTNNNAGATCGGA
63628 TATGAGTGTGGAGTCGTTGCCCAAAGCAGTTAGGGGACAAAGTNNNAGATCGGAA
113107 TATGAGTGTGGAGTCGTTGCTGTGTGTTTCAGATGCAGTCTTGNNNAGATCGGAAG
41964 TATGAGTGTGGAGTCGTTGCGGTTGGTGTTGGTCTCAGTGTTANNNAGATCGGAA
49029 TATGAGTGTGGAGTCGTTGCCTACTCGCAGCTGATAAGCTCTCNNNAGATCGGAAG
51983 TATGAGTGTGGAGTCGTTGCATTGCTCCTGCATCTCTGCATAGNNNAGATCGGAAG
61496 TATGAGTGTGGAGTCGTTGCTTGAGAGGGACACCTTCATTCCTNNNAGATCGGAAG
58464 TATGAGTGTGGAGTCGTTGCCCATCTGTGTCACTTTCACCCATNNNAGATCGGAAGA
69575 TATGAGTGTGGAGTCGTTGCTGAATAGCCAGCACCTTGGTTTTNNNAGATCGGAAG
38314 TATGAGTGTGGAGTCGTTGCTGCCTGAACCCTAATTCCTGAGANNNAGATCGGAAG
105280 TATGAGTGTGGAGTCGTTGCTAATTCATGCTGCTTTCTCAACTGANNNAGATCGGAA
66884 TATGAGTGTGGAGTCGTTGCTTTGCACCTGGTCTGATTTTTGTNNNAGATCGGAAG
63208 TATGAGTGTGGAGTCGTTGCAACATAGAAACCGACCCTGGTCTNNNAGATCGGAAG
98343 TATGAGTGTGGAGTCGTTGCCAATGCAGCGATGTGATCAGCTCNNNAGATCGGAA
102175 TATGAGTGTGGAGTCGTTGCAAGGTTGAAAACGAATGGACGAAGATNNNAGATCG
34591 TATGAGTGTGGAGTCGTTGCGTTCCATTTTGTCAGGAGAGCCANNNAGATCGGAAG
65668 TATGAGTGTGGAGTCGTTGCACCCAATACATGGCTGAACATGTNNNAGATCGGAAG
116460 TATGAGTGTGGAGTCGTTGCATAAAATCACATGAGGCAATTCCTTNNNAGATCGGA
67611 TATGAGTGTGGAGTCGTTGCTGTCAAGGCTCTCAGCTCTACTTNNNAGATCGGAAG
41218 TATGAGTGTGGAGTCGTTGCAAGAGTTAGGGCAACAGCTGCTANNNAGATCGGAA
116700 TATGAGTGTGGAGTCGTTGCGTTGCCAGGCTGATCTCAAACCTNNNAGATCGGAAG
57627 TATGAGTGTGGAGTCGTTGCCAGCCACAGCACATCAGATTTTGNNNAGATCGGAAG
99006 TATGAGTGTGGAGTCGTTGCCCTCTTCTCTTTTTGGGAACGTGNNNAGATCGGAAG
44477 TATGAGTGTGGAGTCGTTGCTGAGAAGCATACCAGGTCCCCNNNAGATCGGAAGA
44131 TATGAGTGTGGAGTCGTTGCGCTGCCATTATACCCAGCATACCNNNAGATCGGAAG
30942 TATGAGTGTGGAGTCGTTGCAGTAGGAGGGTGTCAGTTGACAANNNAGATCGGAA
36022 TATGAGTGTGGAGTCGTTGCGCATGGGTGTGTGTATGTATGCANNNAGATCGGAA
71665 TATGAGTGTGGAGTCGTTGCAGATGTAAGTAGGTGCGGGTTTGNNNAGATCGGAA
47871 TATGAGTGTGGAGTCGTTGCCTGTTACCTCTGTGGAGAAGTGCNNNAGATCGGAA
66882 TATGAGTGTGGAGTCGTTGCAATGTCCTTTGGTGTAGATTTTTGTNNNAGATCGGA
59069 TATGAGTGTGGAGTCGTTGCTGTAGGGGGAAACCAAGTGAGATNNNAGATCGGA
32306 TATGAGTGTGGAGTCGTTGCGCACACTGACCTCACCCATATAANNNAGATCGGAAG
63344 TATGAGTGTGGAGTCGTTGCAAAGACCCATGAGCAGGGATTCTNNNAGATCGGAA
101372 TATGAGTGTGGAGTCGTTGCAAAGTACAAGACTTGGGGGTCTATCNNNAGATCGG

68723 TATGAGTGTGGAGTCGTTGCTGCTGCTTTCTCAACTGATTGTTNNNAGATCGGAAG
37974 TATGAGTGTGGAGTCGTTGCCATCGTGTTTCTGCTACTGCAGANNNAGATCGGAAG
42122 TATGAGTGTGGAGTCGTTGCACACACACCCTGCACAGTTTTANNNAGATCGGAAG
31973 TATGAGTGTGGAGTCGTTGCAAGCCAACATCATTAGCCTGGAANNNAGATCGGAAG
35451 TATGAGTGTGGAGTCGTTGCTTGCAGCATGTGTCATAATAGCANNNAGATCGGAAG
67583 TATGAGTGTGGAGTCGTTGCGGAGACAGCTCGATTTTGGACTTNNNAGATCGGAA
101338 TATGAGTGTGGAGTCGTTGCATGGCAGTGGGATCTGGATCTGNNNAGATCGGAAG
45483 TATGAGTGTGGAGTCGTTGCACCCCAAAAGCCACTCAATATCCNNNAGATCGGAAG
43678 TATGAGTGTGGAGTCGTTGCCTAACTGCCTCTCGAAGGCTTACNNNAGATCGGAAG
100615 TATGAGTGTGGAGTCGTTGCTCAGGAGGCTGTACATATAAGTCCGNNNAGATCGG
33167 TATGAGTGTGGAGTCGTTGCTGGAAATGTAATTGCTGGACCACANNNAGATCGGA
102466 TATGAGTGTGGAGTCGTTGCCCACTCCAAGGCAAAACAGCAGNNNAGATCGGAAG
70135 TATGAGTGTGGAGTCGTTGCCTCACCCAGTCTGCTGATGACTANNNAGATCGGAAG
57526 TATGAGTGTGGAGTCGTTGCAGAACTGTGGGATGGGGAATTTGNNNAGATCGGA
110542 TATGAGTGTGGAGTCGTTGCGGATAGTTCAGGGAAGGTGAGGTNNNAGATCGGA
97848 TATGAGTGTGGAGTCGTTGCCCCCCGCTACCACGGCTTTTNNNAGATCGGAAGAGC
42823 TATGAGTGTGGAGTCGTTGCAAATCAGTCTGGCTGAGAAGCACNNNAGATCGGAA
48579 TATGAGTGTGGAGTCGTTGCAAGGGAGACTGGAAAACCAACTCNNNAGATCGGAA
102201 TATGAGTGTGGAGTCGTTGCGGGATTACCTGGTCCAGCCTTCNNNAGATCGGAAGA
51415 TATGAGTGTGGAGTCGTTGCATGGGCTAGGCTCTTATCCAGAGNNNAGATCGGAA
46285 TATGAGTGTGGAGTCGTTGCTTCCCAAGAGCTTTCCTTACAGCNNNAGATCGGAAG
55505 TATGAGTGTGGAGTCGTTGCGGAAATTGGCTCATTGGCTCATGNNNAGATCGGAA
42900 TATGAGTGTGGAGTCGTTGCGGTCTAATAAGAGGGCATGGCACNNNAGATCGGAA
50376 TATGAGTGTGGAGTCGTTGCGTGGGCAGTAAAGAGGGAAGAAGNNNAGATCGGA
29923 TATGAGTGTGGAGTCGTTGCAGATTTCAGTGCAGCTGTGATGANNNAGATCGGAA
71729 TATGAGTGTGGAGTCGTTGCTTCATCACCACCAAGTCCAATGGNNNAGATCGGAAG
37505 TATGAGTGTGGAGTCGTTGCTGTAGATCTGTGCAGGGGAAAGANNNAGATCGGAA
56491 TATGAGTGTGGAGTCGTTGCCAGCTTTTCCACAGCACTTTCTGNNNAGATCGGAAG
29984 TATGAGTGTGGAGTCGTTGCAAGCTGGAAATCTCTCTCAGGGTNNNAGATCGGAA
38023 TATGAGTGTGGAGTCGTTGCACCATCTCAAGCTATAGGCTCAGANNNAGATCGGAA
29983 TATGAGTGTGGAGTCGTTGCTCAAAAATGGCAAGAAGGCAAGCNNNAGATCGGAA
46599 TATGAGTGTGGAGTCGTTGCTACGTTCATCAGACCAAGTGAGCNNNAGATCGGAAG
101221 TATGAGTGTGGAGTCGTTGCCCTGAAGCTCTGTTCATTTGTTTGGNNNAGATCGGA
101661 TATGAGTGTGGAGTCGTTGCCCTGGCAGTCATCTCTCCAACTCNNNAGATCGGAAG
70370 TATGAGTGTGGAGTCGTTGCGGGTAAATTCTGATAAGGGACAGTCCNNNAGATCG
71788 TATGAGTGTGGAGTCGTTGCCTTTCTCGGCATCTGGAAGCTTCNNNAGATCGGAAG
106009 TATGAGTGTGGAGTCGTTGCCGTGATGTCCAGATTCTCACAATNNNAGATCGGAAG
103985 TATGAGTGTGGAGTCGTTGCACAAGATCCACAAGGCAATATAGCANNNAGATCGG
102140 TATGAGTGTGGAGTCGTTGCTATTTGCACTGTCATGTTTATTGCAGTNNNAGATCGG
107701 TATGAGTGTGGAGTCGTTGCATGCCTGGCTGACAACTCAATTNNNAGATCGGAAGA
106005 TATGAGTGTGGAGTCGTTGCTCCACCTTTCTGCTTCTTCCTTTNNNAGATCGGAAGA
102071 TATGAGTGTGGAGTCGTTGCACTTCTGCAGCTTCATGTGCCNNNAGATCGGAAGAG
71710 TATGAGTGTGGAGTCGTTGCGGCTTCGGAAGTAGAAGCACTTTNNNAGATCGGAA

95487 TATGAGTGTGGAGTCGTTGCAAGACCAAAGTGCAATATGATAGGTTGNNNAGATC
121953 TATGAGTGTGGAGTCGTTGCATGGGCAAAAAGAGGAACTGAANNNAGATCGGAA
105238 TATGAGTGTGGAGTCGTTGCCTTTTCATCTCTGGTGTGCCACNNNAGATCGGAAGA
101164 TATGAGTGTGGAGTCGTTGCCCCAGCATTTCTCAGAGGAGAACNNNAGATCGGAAG
102458 TATGAGTGTGGAGTCGTTGCCATCTGTGCATTCAACCTTGGNNNAGATCGGAAGAG
103768 TATGAGTGTGGAGTCGTTGCGTAAGCATGCATGTGCAAGTATCANNNAGATCGGA
71801 TATGAGTGTGGAGTCGTTGCCATCGGTTACTACAAACTGGCCANNNAGATCGGAAG
104379 TATGAGTGTGGAGTCGTTGCGGGAGCTGGAGATTGTTCTTATGGNNNAGATCGGA
104026 TATGAGTGTGGAGTCGTTGCAAAAAGGAATGATTACAGCCAGGNNNAGATCGGAA
106040 TATGAGTGTGGAGTCGTTGCGCACTGGTGTTTCTCAAACTTCTCNNNAGATCGGAA
103599 TATGAGTGTGGAGTCGTTGCTAATTTCACATTTCAGTGAGGCTGNNNAGATCGGAA
107677 TATGAGTGTGGAGTCGTTGCTACACATGGTGGCTGTGTCTGNNNAGATCGGAAGA
103771 TATGAGTGTGGAGTCGTTGCTCTGGAATACAGTGATGCAGTCACNNNAGATCGGA
110925 TATGAGTGTGGAGTCGTTGCAGCCCTTGGAACTTCATAAAGGTNNNAGATCGGAAG
105998 TATGAGTGTGGAGTCGTTGCAGACAGCTAGAGGAGAGACAGAGNNNAGATCGGA
108373 TATGAGTGTGGAGTCGTTGCCTGACAACAACTAATCCGTTCTCCNNNAGATCGGAAG
106044 TATGAGTGTGGAGTCGTTGCTCCCTCTGCAAATTCTTAGGAATGANNNAGATCGGA
108957 TATGAGTGTGGAGTCGTTGCTAACTCCATACGCCCTCACTTTACNNNAGATCGGAAG
117785 TATGAGTGTGGAGTCGTTGCGCAGACTGGACATTTCAGGGGAAANNNAGATCGGA
104010 TATGAGTGTGGAGTCGTTGCACTAACAAGTTGATGACATGGATGTNNNAGATCGG
104428 TATGAGTGTGGAGTCGTTGCGTGTGTTTCCTTGTATGGCTATCTNNNAGATCGGAA
104000 TATGAGTGTGGAGTCGTTGCGTTTTCCTGTGATTGCCTCAGAGCNNNAGATCGGAA
101376 TATGAGTGTGGAGTCGTTGCAGTTTCTTACGGGGCTATTGACTNNNAGATCGGAAG
95099 TATGAGTGTGGAGTCGTTGCGTTGTTGTACGGAATGTGCAAGGNNNAGATCGGAA
61896 TATGAGTGTGGAGTCGTTGCACATCCTGCACATGATAAACGCTNNNAGATCGGAAG
61484 TATGAGTGTGGAGTCGTTGCGTCCGCAAAACAATTCTTGTCCTNNNAGATCGGAAG
53842 TATGAGTGTGGAGTCGTTGCTTTTTCTTTCCGTCGAGACGGGNNNAGATCGGAAGA
38399 TATGAGTGTGGAGTCGTTGCTGCTATTTGTCAAGAGTAAAGACTAGANNNAGATCG
69480 TATGAGTGTGGAGTCGTTGCGTTGTATTTGTGCCGTGCCTTTTNNNAGATCGGAAG
69209 TATGAGTGTGGAGTCGTTGCTGACTGGCTGAAAGGCATTCTTTNNNAGATCGGAAG
32530 TATGAGTGTGGAGTCGTTGCCAGGAATGGTGGTGCACTTGTAANNNAGATCGGAA
98398 TATGAGTGTGGAGTCGTTGCGCCCCGTTACACCGAGTTCTTNNNAGATCGGAAGAG
65820 TATGAGTGTGGAGTCGTTGCGGTTCTCATTGCAGGCAAATATGTNNNAGATCGGAA
29912 TATGAGTGTGGAGTCGTTGCGGAAAAGCATCAACCTTTGTGCANNNAGATCGGAA
42777 TATGAGTGTGGAGTCGTTGCGCACAAACAAACCAAACATCCACNNNAGATCGGAAG
66127 TATGAGTGTGGAGTCGTTGCTCCACCCACTGAAATTTTGCTGTNNNAGATCGGAAG
59018 TATGAGTGTGGAGTCGTTGCAAGGGTAGAGTGCCACTTCAGATNNNAGATCGGAA
42429 TATGAGTGTGGAGTCGTTGCAGTCAATTACATACTGATGAGTGAACNNNAGATCGG
110464 TATGAGTGTGGAGTCGTTGCTTGGCAGGGTTTCACAATGTCNNNAGATCGGAAGA
43653 TATGAGTGTGGAGTCGTTGCGGGGAGAGAGGGAGAGATTGTACNNNAGATCGGA
62968 TATGAGTGTGGAGTCGTTGCTGACTATCCCCTTGACATGCTCTNNNAGATCGGAAGA
59719 TATGAGTGTGGAGTCGTTGCCACGGACCTGGGTGAGAATTTATNNNAGATCGGAA
63490 TATGAGTGTGGAGTCGTTGCACCCCACTGAGTTTTAGGAATTTCTNNNAGATCGGA

64082 TATGAGTGTGGAGTCGTTGCATGGTGTTTGGCACTCAATCAGTNNNAGATCGGAAG
43889 TATGAGTGTGGAGTCGTTGCGTTGTTGTTGTTGTCCACCACCNNNAGATCGGAAGA
101542 TATGAGTGTGGAGTCGTTGCTGTTGTTGTTGTCCACCACCNNNAGATCGGAAGAGC
62200 TATGAGTGTGGAGTCGTTGCTTGGTGCTTAGGAAAACAATGCTNNNAGATCGGAA
59589 TATGAGTGTGGAGTCGTTGCCTGCCTTCCCTTTTCTGCAGTATNNNAGATCGGAAGA
49098 TATGAGTGTGGAGTCGTTGCACCACATGCAGATGTCTTGTCTCNNNAGATCGGAAG
44958 TATGAGTGTGGAGTCGTTGCAGATAAAATGAGGGGGAGGAGCCNNNAGATCGGA
60885 TATGAGTGTGGAGTCGTTGCTTATGAACCGAATGGGTACCCCTNNNAGATCGGAAG
66548 TATGAGTGTGGAGTCGTTGCTGATGAACATACGTAGCTTTTGTGTNNNAGATCGGA
49124 TATGAGTGTGGAGTCGTTGCCCTCCCATGACTTGATGCTTCTCNNNAGATCGGAAGA
37152 TATGAGTGTGGAGTCGTTGCAAGCAAATGTGGGACAATATTCANNNAGATCGGAA
64390 TATGAGTGTGGAGTCGTTGCTGTCAGAAGGCCTGGGAAATAGTNNNAGATCGGAA
110531 TATGAGTGTGGAGTCGTTGCCGGAAGGGAAGTAGGTTGTCATNNNAGATCGGAA
69091 TATGAGTGTGGAGTCGTTGCCTGTATGGGAGAGCAGTAGCTTTNNNAGATCGGAA
57337 TATGAGTGTGGAGTCGTTGCGTTTTGAGGCACAGTCTTGCTTGNNNAGATCGGAAG
68143 TATGAGTGTGGAGTCGTTGCCACTGACTGTTTCTGCTTATTCTTNNNAGATCGGAAG
58038 TATGAGTGTGGAGTCGTTGCGCTGTCACTGATTGGACCAGAATNNNAGATCGGAA
49387 TATGAGTGTGGAGTCGTTGCCTGCCTTTTGCCTTTGCTATGTCNNNAGATCGGAAGA
59139 TATGAGTGTGGAGTCGTTGCGAGGGAGGTGGTTGTAAGAGGATNNNAGATCGGA
40410 TATGAGTGTGGAGTCGTTGCCCACACTGATTTCGGTGACTTGANNNAGATCGGAAG
57818 TATGAGTGTGGAGTCGTTGCTGGCTGTGTGTCTTTGGTGAAATNNNAGATCGGAA
66713 TATGAGTGTGGAGTCGTTGCAGGGACTAACTAGAAGCGGTTGTNNNAGATCGGAA
58054 TATGAGTGTGGAGTCGTTGCGGGTGTGGGGTAAGGAGAGAATNNNAGATCGGAA
49872 TATGAGTGTGGAGTCGTTGCCTACACTAGGTCCCGTGCATTTCNNNAGATCGGAAG
55371 TATGAGTGTGGAGTCGTTGCATCAAGCACCAGAACCAGACATGNNNAGATCGGAA
37706 TATGAGTGTGGAGTCGTTGCCTGTGGAGCCAGCATTTCTAAGANNNAGATCGGAA
45322 TATGAGTGTGGAGTCGTTGCAGCATCATCCCTTCCTAATTGCCNNNAGATCGGAAGA
117517 TATGAGTGTGGAGTCGTTGCCCAACCCATTTACCCTTCTCCTTNNNAGATCGGAAGA
54246 TATGAGTGTGGAGTCGTTGCAGATGAGGCTAACACTCCTTGGGNNNAGATCGGAA
38622 TATGAGTGTGGAGTCGTTGCAAGCTGAGGAAGGGATCAAAGGANNNAGATCGGA
35430 TATGAGTGTGGAGTCGTTGCTGATTCTAGTGTCCCAGTGAGCANNNAGATCGGAAG
36817 TATGAGTGTGGAGTCGTTGCAGTCCTTTCAGTCCTGAGTATTCTCANNNAGATCGGA
29919 TATGAGTGTGGAGTCGTTGCCTAAGCTCTTCCCTGAAGGTCCTNNNAGATCGGAAG
51438 TATGAGTGTGGAGTCGTTGCTGAGGGGATACTGCAGATCAGAGNNNAGATCGGAA
38145 TATGAGTGTGGAGTCGTTGCATGAGAGAGAGCATGCAGAGAGANNNAGATCGGA
39688 TATGAGTGTGGAGTCGTTGCGCAGCCTGGACTAGAAGAGATGANNNAGATCGGAA
38221 TATGAGTGTGGAGTCGTTGCACACTAGTCCTTAGTTGAATAGGAGANNNAGATCGG
51979 TATGAGTGTGGAGTCGTTGCACCACTGCCTCCTCAGCATAGNNNAGATCGGAAGAG
57651 TATGAGTGTGGAGTCGTTGCTGTAAGGAGGGCTCTTCGTTTTGNNNAGATCGGAA
43933 TATGAGTGTGGAGTCGTTGCGCTACACTGCATTACAAAGCACCNNNAGATCGGAAG
56945 TATGAGTGTGGAGTCGTTGCGCTGTGAGGTTTCTCCTACTGTGNNNAGATCGGAAG
67555 TATGAGTGTGGAGTCGTTGCTTGAATGGGATTGCAGCAGACTTNNNAGATCGGAA
49781 TATGAGTGTGGAGTCGTTGCACCTGTGAGACGAACAGTAGTTCNNNAGATCGGAA

69625 TATGAGTGTGGAGTCGTTGCGTGCATGAGAGTGAGTGATTTTTNNNAGATCGGAA
36741 TATGAGTGTGGAGTCGTTGCAGCAGGAAAGAGAAACTTGCTCANNNAGATCGGAA
59899 TATGAGTGTGGAGTCGTTGCTTCATTGTGCAAATTGGGGAACTNNNAGATCGGAA
67624 TATGAGTGTGGAGTCGTTGCTGAAGCACCAACCAAGCTTACTTNNNAGATCGGAAG
41494 TATGAGTGTGGAGTCGTTGCTGAGGCTACATGAAACACAGGTANNNAGATCGGAA
53246 TATGAGTGTGGAGTCGTTGCCACTCATACACACTTCCTGGAGGNNNAGATCGGAAG
59313 TATGAGTGTGGAGTCGTTGCCCCCGCATTGTTGTTTCTCTGATNNNAGATCGGAAG
69601 TATGAGTGTGGAGTCGTTGCTTTTCCCAAGTGGAGCTTGTTTTNNNAGATCGGAAG
63405 TATGAGTGTGGAGTCGTTGCTGCTCACAGAGAAGTAGGCTTCTNNNAGATCGGAA
61363 TATGAGTGTGGAGTCGTTGCCCCCTACAAGAGTTCACACTCCTNNNAGATCGGAAG
34175 TATGAGTGTGGAGTCGTTGCAGGTTCAGAGTGAGATGCTACCANNNAGATCGGAA
32668 TATGAGTGTGGAGTCGTTGCAGCAGCCACTCAGAAAGAAAACANNNAGATCGGAA
35990 TATGAGTGTGGAGTCGTTGCTTCATTTGTTTGCTTGTGATGCANNNAGATCGGAAG
64848 TATGAGTGTGGAGTCGTTGCCAGGCCAACCAAATGAATCAGGTNNNAGATCGGAA
48335 TATGAGTGTGGAGTCGTTGCGCCTCCAGGACATATAGGACATCNNNAGATCGGAAG
52113 TATGAGTGTGGAGTCGTTGCAGCCCCTTCTTTTCTTCCAGTAGNNNAGATCGGAAG
52296 TATGAGTGTGGAGTCGTTGCCATACTCAACACTTCTTGGCACGNNNAGATCGGAAG
60483 TATGAGTGTGGAGTCGTTGCGAGGGAGAAGAGGACTTGCTACTNNNAGATCGGAA
101599 TATGAGTGTGGAGTCGTTGCCCAACAGCCTCACATTTGTTCTTNNNAGATCGGAAG
37601 TATGAGTGTGGAGTCGTTGCGGGCAACAAAAAGCGAAAGAAGANNNAGATCGGA
37979 TATGAGTGTGGAGTCGTTGCTTTGTAATTCAGCCAGCTGCAGANNNAGATCGGAAG
40351 TATGAGTGTGGAGTCGTTGCCAGGGCCTCCAATTCTTTTGTGANNNAGATCGGAAG
32344 TATGAGTGTGGAGTCGTTGCGCTTTCCTCCAAGGCCTTACTAANNNAGATCGGAAG
43140 TATGAGTGTGGAGTCGTTGCATAGGGTGGTAAAGGGGGAAGACNNNAGATCGGA
38761 TATGAGTGTGGAGTCGTTGCTACACAGGAGCTCCAAATCAGGANNNAGATCGGAA
67353 TATGAGTGTGGAGTCGTTGCGGAGCCACCTAATGAAGGTGATTNNNAGATCGGAA
60843 TATGAGTGTGGAGTCGTTGCGTAGGGTAGAAAGGTAGCACCCTNNNAGATCGGAA
33849 TATGAGTGTGGAGTCGTTGCACCTCTGGTATAACCGGTCTACANNNAGATCGGAAG
47626 TATGAGTGTGGAGTCGTTGCCCAGAACTGCCCAAACTTTATGCNNNAGATCGGAAG
57998 TATGAGTGTGGAGTCGTTGCGGTACATAAGCCCTGCAGTCAATNNNAGATCGGAAG
56586 TATGAGTGTGGAGTCGTTGCACACAGTGAGAGTAGAGGCAGTGNNNAGATCGGA
52503 TATGAGTGTGGAGTCGTTGCGTAAATGATTTGGGTTGGACGCGNNNAGATCGGAA
41514 TATGAGTGTGGAGTCGTTGCGGGGTGATGTTTCTGCCTAGGTANNNAGATCGGAA
52128 TATGAGTGTGGAGTCGTTGCTCATACACAGAGTGGGGACGTAGNNNAGATCGGAA
39479 TATGAGTGTGGAGTCGTTGCGGTGAGGTTTCTCAAAGGATTGGANNNAGATCGGA
108969 TATGAGTGTGGAGTCGTTGCGAGAGAGGAGTGTATGTTGTCAGCNNNAGATCGG
34106 TATGAGTGTGGAGTCGTTGCAGAGAGAGGAGAGATACTGAGACCANNNAGATCG
39335 TATGAGTGTGGAGTCGTTGCACACATACTGTAGCAACCCTGGANNNAGATCGGAAG
58816 TATGAGTGTGGAGTCGTTGCACCATGAGATACTCCAGCCTCATNNNAGATCGGAAG
43361 TATGAGTGTGGAGTCGTTGCCACACCAGCTTTGGGAAAATGACNNNAGATCGGAA
71397 TATGAGTGTGGAGTCGTTGCTCTTCTTCCATCCTTTCAGTCATGTNNNAGATCGGAA
49945 TATGAGTGTGGAGTCGTTGCGCACCACATCCACATAACGTTTCNNNAGATCGGAAG
52221 TATGAGTGTGGAGTCGTTGCGATGGGGTTTGGCTCTCTGTTAGNNNAGATCGGAA

| | |
|---|---|
| 39190 | TATGAGTGTGGAGTCGTTGCTCTCTCTCTCACTCTCAATTGGGANNNAGATCGGAAG |
| 95096 | TATGAGTGTGGAGTCGTTGCAACAATGGCCAGTCCGACTTCNNNAGATCGGAAGAG |
| 47625 | TATGAGTGTGGAGTCGTTGCCGTTTTCCCAGAGCAGATTTATGCNNNAGATCGGAA |
| 31564 | TATGAGTGTGGAGTCGTTGCGCAGTGGGTCAGAGAGAGAAGAANNNAGATCGGA |
| 55584 | TATGAGTGTGGAGTCGTTGCTCATGGGGCTTTTTCCTGAGATGNNNAGATCGGAAG |
| 63191 | TATGAGTGTGGAGTCGTTGCAGATGAGAGAGAGGTCTGGGTCTNNNAGATCGGA |
| 68742 | TATGAGTGTGGAGTCGTTGCCGTGGGAGGATGATTCTGTTGTTNNNAGATCGGAA |
| 67324 | TATGAGTGTGGAGTCGTTGCTCCCACAAATAACCCACTGGATTNNNAGATCGGAAG |
| 97477 | TATGAGTGTGGAGTCGTTGCGGACTAACCCAGAAGTTGCAGTCNNNAGATCGGAA |
| 40812 | TATGAGTGTGGAGTCGTTGCTGCGTTTGGATTTGCCAGAGATANNNAGATCGGAA |
| 39141 | TATGAGTGTGGAGTCGTTGCTCTAACAGATGAGGCAACTGGGANNNAGATCGGAA |
| 69109 | TATGAGTGTGGAGTCGTTGCTAGATGAGAAGGAGCAGGGCTTTNNNAGATCGGA |
| 69617 | TATGAGTGTGGAGTCGTTGCTAGCAGGGGTGGATTGCATTTTTNNNAGATCGGAA |
| 33913 | TATGAGTGTGGAGTCGTTGCGATGGACACCTAGCTTGCTTACANNNAGATCGGAAG |
| 39808 | TATGAGTGTGGAGTCGTTGCAGATGAGAGAGAGAGAGCACTGANNNAGATCGGA |
| 63770 | TATGAGTGTGGAGTCGTTGCGTAAACATGACCTTGGGAGAAGTNNNAGATCGGAA |
| 30016 | TATGAGTGTGGAGTCGTTGCCACCATGAGGTAGGCAGGAAAAANNNAGATCGGAA |
| 46158 | TATGAGTGTGGAGTCGTTGCTGCTTACCTTCTCACAGTCAAGCNNNAGATCGGAAG |
| 49952 | TATGAGTGTGGAGTCGTTGCGAAAACGTGAATGCCTGGGTTTCNNNAGATCGGAA |
| 107667 | TATGAGTGTGGAGTCGTTGCTAATGAGTTGGGGAGGAGTGTTNNNAGATCGGAA |
| 29921 | TATGAGTGTGGAGTCGTTGCTGGCTTTACGTAGCTCAATCCTGNNNAGATCGGAAG |
| 57446 | TATGAGTGTGGAGTCGTTGCGCGTGCACATTTATCAAAGGTTGNNNAGATCGGAA |
| 54111 | TATGAGTGTGGAGTCGTTGCCATTTGAATCTGCAACAGCTGGGNNNAGATCGGAA |
| 54568 | TATGAGTGTGGAGTCGTTGCGAGGGACAAGAATAGCAACCTGGNNNAGATCGGA |
| 58799 | TATGAGTGTGGAGTCGTTGCCATCCTGACCCCTCCTAACTCATNNNAGATCGGAAGA |
| 116363 | TATGAGTGTGGAGTCGTTGCGTTTTGTACAGAGGGGAAGGTGAGNNNAGATCGG |
| 34963 | TATGAGTGTGGAGTCGTTGCTTTCAGGATTTGAATGCCGTCCANNNAGATCGGAAG |
| 31357 | TATGAGTGTGGAGTCGTTGCGCCTGCCAAGTTTTTCTTTGCAANNNAGATCGGAAG |
| 103963 | TATGAGTGTGGAGTCGTTGCTTTGGCTTTTGCGCACATAATGGNNNAGATCGGAAG |
| 66908 | TATGAGTGTGGAGTCGTTGCTAGTGCTTGCAACAACGCAAATTNNNAGATCGGAAG |
| 53858 | TATGAGTGTGGAGTCGTTGCTAGAAAATTGCTTTGGGTCCGGGNNNAGATCGGAA |
| 35569 | TATGAGTGTGGAGTCGTTGCAGAATGACACATGGGAAAAGGCANNNAGATCGGA |
| 41586 | TATGAGTGTGGAGTCGTTGCGTTCCTGTAGTGGGAGTGTGGTANNNAGATCGGAA |
| 36211 | TATGAGTGTGGAGTCGTTGCTGACTTCTCCCATAGCTAATTGCANNNAGATCGGAA |
| 31456 | TATGAGTGTGGAGTCGTTGCTTGACAGGAGGAAAGAGGGTCAANNNAGATCGGA |
| 98685 | TATGAGTGTGGAGTCGTTGCTGTGAAATTGCTTTGTGGTGCANNNAGATCGGAAG |
| 29911 | TATGAGTGTGGAGTCGTTGCCGTACATTCTATTTTGGCTCCCANNNAGATCGGAAG |
| 56395 | TATGAGTGTGGAGTCGTTGCCGGTGACCACTCAACTTCTCTGNNNAGATCGGAAGA |
| 57742 | TATGAGTGTGGAGTCGTTGCGGAACCAGCCATTCCCATCAAATNNNAGATCGGAAG |
| 43200 | TATGAGTGTGGAGTCGTTGCCAGACAGACAGACAAGCAGAGACNNNAGATCGGAA |
| 37958 | TATGAGTGTGGAGTCGTTGCATGTTGTCAGCTTGAATGGCAGANNNAGATCGGAA |
| 30282 | TATGAGTGTGGAGTCGTTGCACTCTTAGATGCAGCAAACCAAANNNAGATCGGAAG |
| 110289 | TATGAGTGTGGAGTCGTTGCATAGAGACCTCAGGGGGCAACTNNNAGATCGGAAG |

99063 TATGAGTGTGGAGTCGTTGCGAACTTGCCTCCTCATGGACCTGNNNAGATCGGAAG
54205 TATGAGTGTGGAGTCGTTGCGAAGTAGAAATGGGGCATGTGGGNNNAGATCGGA
32277 TATGAGTGTGGAGTCGTTGCGCTGGGCTTGGGAATTCAGATAANNNAGATCGGAA
102193 TATGAGTGTGGAGTCGTTGCCAGAGGAGAGAAGAAGGGAGTGATNNNAGATCGG
58113 TATGAGTGTGGAGTCGTTGCGACAATGGGCAGAGATGTGGAATNNNAGATCGGA
48733 TATGAGTGTGGAGTCGTTGCGGGAATTTGGGTTACAGTCCCTCNNNAGATCGGAA
45546 TATGAGTGTGGAGTCGTTGCACAAACATTGCAGCTCTTTACTCCNNNAGATCGGAAG
70984 TATGAGTGTGGAGTCGTTGCGAAGTGGGGGCAGATGTTTGAATNNNAGATCGGA
117681 TATGAGTGTGGAGTCGTTGCGGCTTTTCAGTGACCACAGCTATNNNAGATCGGAAG
63499 TATGAGTGTGGAGTCGTTGCGGGGAATCCTGCCAAAGATTTCTNNNAGATCGGAA
67056 TATGAGTGTGGAGTCGTTGCTCATCATTCCCCTCTCCCACATTNNNAGATCGGAAGA
42557 TATGAGTGTGGAGTCGTTGCTTAGCTTCCTGTGACACACACACNNNAGATCGGAAG
49051 TATGAGTGTGGAGTCGTTGCCCCATGAAACAAGGCTCTCTCTCNNNAGATCGGAAG
29918 TATGAGTGTGGAGTCGTTGCAATGTGATTTTGCCACCCACAAANNNAGATCGGAAG
52307 TATGAGTGTGGAGTCGTTGCGGAAGAAAATCCCAGCATTCACGNNNAGATCGGAA
59485 TATGAGTGTGGAGTCGTTGCGCAAGCACTCTGACTTCCACTATNNNAGATCGGAAG
36144 TATGAGTGTGGAGTCGTTGCGAGAGAAAGACAGCATGAGTGCANNNAGATCGGA
58738 TATGAGTGTGGAGTCGTTGCGGTCTGAATGGTAAGTGGTGCATNNNAGATCGGAA
106705 TATGAGTGTGGAGTCGTTGCTTCTGCTGAGGAACCTTAGGCNNNAGATCGGAAGA
42935 TATGAGTGTGGAGTCGTTGCATGAAGTGACAAGGAAGGTGCACNNNAGATCGGA
55006 TATGAGTGTGGAGTCGTTGCTGGAAAAGCTTAGGGGATTGTGGNNNAGATCGGA
62760 TATGAGTGTGGAGTCGTTGCCCTGCCTAATTCCATGCAACTCTNNNAGATCGGAAGA
50377 TATGAGTGTGGAGTCGTTGCGTGGCTGAGGAAGAGTGAAGAAGNNNAGATCGGA
63661 TATGAGTGTGGAGTCGTTGCTCTCCAGTGAGGAAAGGGAAAGTNNNAGATCGGAA
29917 TATGAGTGTGGAGTCGTTGCGGGGAGCTTTGATCTGAGTTTGANNNAGATCGGAA
42359 TATGAGTGTGGAGTCGTTGCAGGGAGGATAAGGAACCCAGAACNNNAGATCGGA
67457 TATGAGTGTGGAGTCGTTGCGGCAGAAGGCAGACAGGTTTATTNNNAGATCGGAA
38377 TATGAGTGTGGAGTCGTTGCTAAGGTCATGCCATCTCTGAATAGANNNAGATCGGA
48446 TATGAGTGTGGAGTCGTTGCCAGGGGAGAGAGAGAGACAGATCNNNAGATCGGA
107672 TATGAGTGTGGAGTCGTTGCACCAAGAGACAGAAGGGTAATAGCANNNAGATCGG
68104 TATGAGTGTGGAGTCGTTGCGGCTGGGCTTTTAGGAAAGTCTTNNNAGATCGGAA
108258 TATGAGTGTGGAGTCGTTGCAGATAAAACGCAAGTCAGGGACTGNNNAGATCGGA
34052 TATGAGTGTGGAGTCGTTGCCCAAAGGTTTCTAAGAGCCACCANNNAGATCGGAAG
41862 TATGAGTGTGGAGTCGTTGCTCCCAACACACCACATTTGCTTANNNAGATCGGAAGA
69031 TATGAGTGTGGAGTCGTTGCCTCAGCCTTCCTTTTGTCCCTTTNNNAGATCGGAAGA
46282 TATGAGTGTGGAGTCGTTGCGCCCAGTCAGAGAAATCTACAGCNNNAGATCGGAA
58952 TATGAGTGTGGAGTCGTTGCAGCCCATTTGAAGAGAGGAAGATNNNAGATCGGAA
60086 TATGAGTGTGGAGTCGTTGCTCTCTCTCCCAGTCCTATCCACTNNNAGATCGGAAGA
66072 TATGAGTGTGGAGTCGTTGCATGTTCCCACCATTCTAGGCTGTNNNAGATCGGAAG
63030 TATGAGTGTGGAGTCGTTGCTCTGGTGTTTCCTTCTCTCTCTCTNNNAGATCGGAAG
46933 TATGAGTGTGGAGTCGTTGCGAGACAGAAAGTAGAGCTCAGGCNNNAGATCGGA
44785 TATGAGTGTGGAGTCGTTGCTACAACTTTCAGCCTGCATTCCCNNNAGATCGGAAGA
31491 TATGAGTGTGGAGTCGTTGCCAGGGCCATAATTCCGAGTTCAANNNAGATCGGAAG

32713 TATGAGTGTGGAGTCGTTGCAGTGTTTGTCGGTAGAGCAAACANNNAGATCGGAA
44792 TATGAGTGTGGAGTCGTTGCAGTGAGGAAGAGGTGACTATTCCCNNNAGATCGGA
39731 TATGAGTGTGGAGTCGTTGCTGTAGTTTGGTGGTATCAGTGATGANNNAGATCGG
46631 TATGAGTGTGGAGTCGTTGCATTGAAGGAGTATTCATTCTCCATAGCNNNAGATCG
50024 TATGAGTGTGGAGTCGTTGCGGGGTGCGATGATGCATAAAAAGNNNAGATCGGA
68638 TATGAGTGTGGAGTCGTTGCGTTGACCCCATACCTTTGCTGTTNNNAGATCGGAAG
66534 TATGAGTGTGGAGTCGTTGCACAGAATGTAGAGCATTGGTTGTGTNNNAGATCGG
46365 TATGAGTGTGGAGTCGTTGCATGTTTTCAGGGTAGAAGGCAGCNNNAGATCGGAA
58156 TATGAGTGTGGAGTCGTTGCACCTCAGGGGGCAACTGAATAATNNNAGATCGGAA
96804 TATGAGTGTGGAGTCGTTGCGGGAGGAGACACAACAACAAAAAGNNNAGATCGG
31168 TATGAGTGTGGAGTCGTTGCGAGAGTGAGCAGAGACAAAGCAANNNAGATCGGA
59722 TATGAGTGTGGAGTCGTTGCGTGTCCCTGTAGCAGCGATTTATNNNAGATCGGAAG
65358 TATGAGTGTGGAGTCGTTGCACTGGTACAGAGAAGAACCTGGTNNNAGATCGGAA
69637 TATGAGTGTGGAGTCGTTGCTGCCCTCAAGCAGTGTACTTTTTNNNAGATCGGAAG
102194 TATGAGTGTGGAGTCGTTGCAAGGTGAAAATGGAGAGCAGTGTCANNNAGATCG
57284 TATGAGTGTGGAGTCGTTGCTGTCCAAACTATCCTTGGCCTTGNNNAGATCGGAAG
51544 TATGAGTGTGGAGTCGTTGCCGTTGAGGTTCCCACTGTTAGAGNNNAGATCGGAA
39607 TATGAGTGTGGAGTCGTTGCTTCCACTTTCAAAGGCCACATGANNNAGATCGGAAG
67468 TATGAGTGTGGAGTCGTTGCTCACCACAGATCCTATTGAAACTTNNNAGATCGGAA
58701 TATGAGTGTGGAGTCGTTGCCACACACGCACAGATAAATGCATNNNAGATCGGAAG
58294 TATGAGTGTGGAGTCGTTGCCAACTGACAAGCAAAGCACACATNNNAGATCGGAAG
31339 TATGAGTGTGGAGTCGTTGCCATCTGATTGTGGAGGTGTGCAANNNAGATCGGAA
101516 TATGAGTGTGGAGTCGTTGCGGTCTTTCGTTCATGTACTCAGCTNNNAGATCGGAA
49904 TATGAGTGTGGAGTCGTTGCCCACAGAGGAAACGGCCTTTCNNNAGATCGGAAGA
30463 TATGAGTGTGGAGTCGTTGCTGCGAAGCATACGAGAGTTCAAANNNAGATCGGAA
105429 TATGAGTGTGGAGTCGTTGCGTTTAGTACAGGGTTCCACTCTGTTANNNAGATCGG
66758 TATGAGTGTGGAGTCGTTGCTGCCACAGAACAGAAAAACATTTGTNNNAGATCGGA
62359 TATGAGTGTGGAGTCGTTGCTCACTCCCATGAAATCAAGTGCTNNNAGATCGGAAG
59631 TATGAGTGTGGAGTCGTTGCTGGGACTGGTGTGTGTACTGTATNNNAGATCGGAA
51649 TATGAGTGTGGAGTCGTTGCAGTGTTGAGTAAGGGACAGGGAGNNNAGATCGGA
71667 TATGAGTGTGGAGTCGTTGCATCGAACTGTTCTCCCTCTTCCTNNNAGATCGGAAGA
48270 TATGAGTGTGGAGTCGTTGCCCTGGGCTGGAATTCTACCAATCNNNAGATCGGAAG
57929 TATGAGTGTGGAGTCGTTGCCCCATCTTCTGGGAGTTTCCAATNNNAGATCGGAAG
33162 TATGAGTGTGGAGTCGTTGCTCCCTTACAGCCTATTCACCACANNNAGATCGGAAGA
34070 TATGAGTGTGGAGTCGTTGCGTGATTTGTACAGGACCTCACCANNNAGATCGGAAG
35187 TATGAGTGTGGAGTCGTTGCCCTATGTGTGTGGTCAAGAAGCANNNAGATCGGAA
50604 TATGAGTGTGGAGTCGTTGCGCCAAGTGAAAAGAGCCAAACAGNNNAGATCGGAA
53627 TATGAGTGTGGAGTCGTTGCAGGACATCTGAGAAAGGGAAGGGNNNAGATCGGA
32436 TATGAGTGTGGAGTCGTTGCAGCCACCTTTGCTTTGGTTCTAANNNAGATCGGAAG
53289 TATGAGTGTGGAGTCGTTGCAAAATCTCTGCTCAGCTCTGAGGNNNAGATCGGAAG
100038 TATGAGTGTGGAGTCGTTGCGTGCTGATTCCCGGACATCTATGNNNAGATCGGAAG
61091 TATGAGTGTGGAGTCGTTGCAAGTCAACAATAAGCAGCAGCCTNNNAGATCGGAA
31032 TATGAGTGTGGAGTCGTTGCTCACTGTCTATTCAGGTCCCCAANNNAGATCGGAAG

59356 TATGAGTGTGGAGTCGTTGCTACACCCCTTTGGGAGGATTGATNNNAGATCGGAAG
101133 TATGAGTGTGGAGTCGTTGCTTCAGACTGGGAAGCCCCACTCTNNNAGATCGGAAG
41124 TATGAGTGTGGAGTCGTTGCCCGAGAGATCAGAAGCCTTCCTANNNAGATCGGAAG
58647 TATGAGTGTGGAGTCGTTGCACATTTTCTGCAAATCCACGCATNNNAGATCGGAAG
44397 TATGAGTGTGGAGTCGTTGCTTTGTCTTCAATATGCCTCCCCCNNNAGATCGGAAGA
48386 TATGAGTGTGGAGTCGTTGCCCAAGTGATCAAGAATGCGCATCNNNAGATCGGAA
31712 TATGAGTGTGGAGTCGTTGCCAGGAAGAAGGGTACCTGGAGAANNNAGATCGGA
62930 TATGAGTGTGGAGTCGTTGCTCTCTCCCAAATATCCTTCTAGCTCTNNNAGATCGGA
48796 TATGAGTGTGGAGTCGTTGCTATTGCCCCTGAAAATGCTCCTCNNNAGATCGGAAG
110458 TATGAGTGTGGAGTCGTTGCTGAATTGCTCTGGAATCACTGTCAAAANNNAGATCG
45899 TATGAGTGTGGAGTCGTTGCTCCATTCAGCCATCAATGCTTCCNNNAGATCGGAAGA
62531 TATGAGTGTGGAGTCGTTGCGTCCACCCTTATCTGTACTTTTGCTNNNAGATCGGAA
53367 TATGAGTGTGGAGTCGTTGCAACTACTGACAGTGGGACCTAGGNNNAGATCGGAA
58552 TATGAGTGTGGAGTCGTTGCCCGCCTTCTCCATCTCTATCCATNNNAGATCGGAAGA
45967 TATGAGTGTGGAGTCGTTGCATCCCCCAAGCTGTACAATTTCCNNNAGATCGGAAG
36347 TATGAGTGTGGAGTCGTTGCAGCCACAGATTCTGCTGAAATCANNNAGATCGGAAG
69547 TATGAGTGTGGAGTCGTTGCGGCATCATCTCTCAGCCAGTTTTNNNAGATCGGAAG
102772 TATGAGTGTGGAGTCGTTGCTCTTCTGGGAGTTTCCAATGGTATNNNAGATCGGAA
103813 TATGAGTGTGGAGTCGTTGCTAATGGCTCTCAGCTCCAAATCCANNNAGATCGGAA
68511 TATGAGTGTGGAGTCGTTGCAGAGTAGGAGAAAACAGGTGGTTNNNAGATCGGA
53571 TATGAGTGTGGAGTCGTTGCGCCTCAAATTATTCTGGCTTTCGGNNNAGATCGGAA
58167 TATGAGTGTGGAGTCGTTGCGCTCGCTTGCTCACACCTATAATNNNAGATCGGAAG
45163 TATGAGTGTGGAGTCGTTGCCTGAATCGTGTCCCTGAAATGCCNNNAGATCGGAAG
32373 TATGAGTGTGGAGTCGTTGCGCTTGTGGCTAACACAGTCCTAANNNAGATCGGAAG
106585 TATGAGTGTGGAGTCGTTGCTTGTTCGTAGTGTTTTGGTGGGANNNAGATCGGAA
42093 TATGAGTGTGGAGTCGTTGCGGAGGAAGTGGTGATGCCTTTTANNNAGATCGGAA
29981 TATGAGTGTGGAGTCGTTGCCCAAGACCCATTGAGAAACTTCCTNNNAGATCGGAA
54469 TATGAGTGTGGAGTCGTTGCAATCCTGCCATCTGTGACTATGGNNNAGATCGGAAG
102092 TATGAGTGTGGAGTCGTTGCAGTGAAAAGAGCCAAACAGGAAAAGNNNAGATCGG
63263 TATGAGTGTGGAGTCGTTGCGGTTGCCAAGAAGGATTCTGTCTNNNAGATCGGAA
47993 TATGAGTGTGGAGTCGTTGCTATTTCACTTCAGCCAGCTGTGCNNNAGATCGGAAG
65700 TATGAGTGTGGAGTCGTTGCAGGTAAAGACTATATGGCCCATGTNNNAGATCGGA
41181 TATGAGTGTGGAGTCGTTGCGAAGTGAATTGGCCTTGAGGCTANNNAGATCGGAA
66808 TATGAGTGTGGAGTCGTTGCCTGGGAAGCAAAAACTGCTTTGTNNNAGATCGGAA
68584 TATGAGTGTGGAGTCGTTGCAAGAGGTTGGGATCCCGTATGTTNNNAGATCGGAA
37752 TATGAGTGTGGAGTCGTTGCAGAGGGAAAAATGAAGCTAACAGANNNAGATCGG
65611 TATGAGTGTGGAGTCGTTGCTCCAGGTTTGGGTTAGGAAATGTNNNAGATCGGAA
59644 TATGAGTGTGGAGTCGTTGCGGTGGGTCCCCAGAAAATTGTATNNNAGATCGGAA
32959 TATGAGTGTGGAGTCGTTGCGCACATTATTTGTACAGGCACTAACANNNAGATCGG
66429 TATGAGTGTGGAGTCGTTGCTGGACCTGTGAAGACCTATGTGTNNNAGATCGGAA
68874 TATGAGTGTGGAGTCGTTGCACACTTTTGACTATGGCTTTCATTTNNNAGATCGGAA
37436 TATGAGTGTGGAGTCGTTGCGGGACACAGGAACCAACAAAAGANNNAGATCGGAA
68232 TATGAGTGTGGAGTCGTTGCGCCTGTTGTCTCTGGATGAAGTTNNNAGATCGGAAG

| | |
|---|---|
| 55595 | TATGAGTGTGGAGTCGTTGCAAATGGTTGTAGGGCCAAGGATGNNNAGATCGGAA |
| 60268 | TATGAGTGTGGAGTCGTTGCTAACTGATGGTATACCCCAGACTNNNAGATCGGAAG |
| 35519 | TATGAGTGTGGAGTCGTTGCGCACTCCTTAGGACTTGTTAGCANNNAGATCGGAAG |
| 55467 | TATGAGTGTGGAGTCGTTGCGGTAAGGTATCTTGGGTGGCATGNNNAGATCGGAA |
| 47761 | TATGAGTGTGGAGTCGTTGCGTCTTCACACAAAATCAGGCTGCNNNAGATCGGAAG |
| 43343 | TATGAGTGTGGAGTCGTTGCATCAGACCCTGCAATATCTGGACNNNAGATCGGAAG |
| 60502 | TATGAGTGTGGAGTCGTTGCTACAAAACAACCTGGCCAGTACTNNNAGATCGGAAG |
| 64877 | TATGAGTGTGGAGTCGTTGCCCTTCAGTAAAGCAACAGAGAGGTNNNAGATCGGAA |
| 43350 | TATGAGTGTGGAGTCGTTGCAGTGTGTGAAATGTCTGGTGGACNNNAGATCGGAA |
| 64741 | TATGAGTGTGGAGTCGTTGCTTCTGACTGACTGGCTACAAGGTNNNAGATCGGAAG |
| 35760 | TATGAGTGTGGAGTCGTTGCAAATGCTGTGGGTAAGAATGGCANNNAGATCGGAA |
| 35218 | TATGAGTGTGGAGTCGTTGCAGACAGTTTATTGCAGGATCTAAGCANNNAGATCGG |
| 45308 | TATGAGTGTGGAGTCGTTGCACCTCTCTTTTAGTTGTGGTGCCNNNAGATCGGAAGA |
| 51651 | TATGAGTGTGGAGTCGTTGCAGAATAAAATGGCTGGCAGGGAGNNNAGATCGGAA |
| 47023 | TATGAGTGTGGAGTCGTTGCGAGCACATAGCACTCTCTTAGGCNNNAGATCGGAAG |
| 116214 | TATGAGTGTGGAGTCGTTGCGCGGTGCCTCAATCTTGGACTTCNNNAGATCGGAAG |
| 64130 | TATGAGTGTGGAGTCGTTGCGCATAATGTGTGATTTCAAATGTCAGTNNNAGATCGG |
| 54391 | TATGAGTGTGGAGTCGTTGCAACAAGACTATAGCTGGGCATGGNNNAGATCGGAAG |
| 36241 | TATGAGTGTGGAGTCGTTGCTTCACTGAGCAGCAATACTTGCANNNAGATCGGAAG |
| 34695 | TATGAGTGTGGAGTCGTTGCACACTAAGGATAAGAACATGCCANNNAGATCGGAAG |
| 30660 | TATGAGTGTGGAGTCGTTGCCCTCCCAAATGACCCAACTGAAANNNAGATCGGAAGA |
| 51249 | TATGAGTGTGGAGTCGTTGCACCCAGTGATTCTCCCATCTCAGNNNAGATCGGAAGA |
| 117273 | TATGAGTGTGGAGTCGTTGCCAACATGATAAACTGCAGTGCAANNNAGATCGGAAG |
| 105305 | TATGAGTGTGGAGTCGTTGCTTTTCTTCCTCTGAGCCCTGAAAGNNNAGATCGGAAG |
| 41914 | TATGAGTGTGGAGTCGTTGCTTGAGTGAGGCTTGGGGTAGTTANNNAGATCGGAA |
| 30386 | TATGAGTGTGGAGTCGTTGCACACTCACATATTTCCGGGCAAANNNAGATCGGAAGA |
| 44651 | TATGAGTGTGGAGTCGTTGCTATTTGCAAGCTCCCATCATCCCNNNAGATCGGAAGA |
| 71802 | TATGAGTGTGGAGTCGTTGCACTTTTTGATCAGAGCACGGACANNNAGATCGGAAG |
| 37509 | TATGAGTGTGGAGTCGTTGCTTCTTCCTCTGAGCCCTGAAAGANNNAGATCGGAAGA |
| 114179 | TATGAGTGTGGAGTCGTTGCAAGCAGGCCCAACGTAAACATANNNAGATCGGAAGA |
| 100032 | TATGAGTGTGGAGTCGTTGCAGGGATGTCCTCATTTCCAAACTNNNAGATCGGAAGA |
| 38929 | TATGAGTGTGGAGTCGTTGCCAGTCAATTGAGAAGGACACGGANNNAGATCGGAA |
| 51659 | TATGAGTGTGGAGTCGTTGCGTGTGTAAGTTTGGGCTAGGGAGNNNAGATCGGAA |
| 71123 | TATGAGTGTGGAGTCGTTGCGACAAAAGAGGTTGCTGCCTACTNNNAGATCGGAAG |
| 68839 | TATGAGTGTGGAGTCGTTGCCTCAAGACGGGCTACAGCATTTNNNAGATCGGAAGA |
| 34666 | TATGAGTGTGGAGTCGTTGCAATGTAAATCAGGATGTGGGCCANNNAGATCGGAAG |
| 44765 | TATGAGTGTGGAGTCGTTGCAAACACTTAGATGCCTTGGTCCCNNNAGATCGGAAGA |
| 46686 | TATGAGTGTGGAGTCGTTGCCCAGGCTCTTGAGTTACTGTAGCNNNAGATCGGAAG |
| 65561 | TATGAGTGTGGAGTCGTTGCTTGGCCTCACAATCATATTTGGTNNNAGATCGGAAGA |
| 65746 | TATGAGTGTGGAGTCGTTGCTCTGCATCTCAATGTTTCTCATGTNNNAGATCGGAAG |
| 61952 | TATGAGTGTGGAGTCGTTGCAAACAGGAGACAGTAGGAAGGCTNNNAGATCGGAA |
| 60620 | TATGAGTGTGGAGTCGTTGCTGCTTCTGTTTCTTTTGAGAACCTNNNAGATCGGAAG |
| 69144 | TATGAGTGTGGAGTCGTTGCTCAACTCTGGTGGTTTGTGCTTTNNNAGATCGGAAGA |

| | |
|---|---|
| 43708 | TATGAGTGTGGAGTCGTTGCTATGTCCAGGCACTGTGCTTTACNNNAGATCGGAAG |
| 65751 | TATGAGTGTGGAGTCGTTGCTGGTCAGCCCCAGTAATTCATGTNNNAGATCGGAAG |
| 100819 | TATGAGTGTGGAGTCGTTGCCGAGATCACAGCTCAATGCAGACNNNAGATCGGAAG |
| 63457 | TATGAGTGTGGAGTCGTTGCTGTCTGCTTTCTTGCATCGTTCTNNNAGATCGGAAGA |
| 35068 | TATGAGTGTGGAGTCGTTGCCAGGGGAAATTTGGAAGGTTCCANNNAGATCGGAA |
| 64008 | TATGAGTGTGGAGTCGTTGCTGGCTAGTGTGAAAAACAGCAGTNNNAGATCGGAA |
| 55864 | TATGAGTGTGGAGTCGTTGCGGCATAGTCCTCCATTTGGACTGNNNAGATCGGAAG |
| 52328 | TATGAGTGTGGAGTCGTTGCTCAGAGAACAAACACAATTGGACGNNNAGATCGGA |
| 64561 | TATGAGTGTGGAGTCGTTGCCTGTCCTTTTCGTACTTGGACGTNNNAGATCGGAAG |
| 50141 | TATGAGTGTGGAGTCGTTGCACCCACAGCAGAATACAGGAAAGNNNAGATCGGAA |
| 45705 | TATGAGTGTGGAGTCGTTGCTCTGTCTTTTGGTGCCTTTCTCCNNNAGATCGGAAGA |
| 30120 | TATGAGTGTGGAGTCGTTGCCTATTGCTCTGGCTGGCAGAAAANNNAGATCGGAAG |
| 64240 | TATGAGTGTGGAGTCGTTGCGCCAGCTTCTAGGAAAAAGGAGTNNNAGATCGGAA |
| 58922 | TATGAGTGTGGAGTCGTTGCCACACAACGAGCAAAGCTTTCATNNNAGATCGGAAG |
| 97769 | TATGAGTGTGGAGTCGTTGCCAGCTTCACACTTCAGCCCATNNNAGATCGGAAGAG |
| 31633 | TATGAGTGTGGAGTCGTTGCCCATCCTGGTTTGAGAAGCAGAANNNAGATCGGAAG |
| 48728 | TATGAGTGTGGAGTCGTTGCGGTCAAATTCAGTCACCTCCCTCNNNAGATCGGAAG |
| 43095 | TATGAGTGTGGAGTCGTTGCGAGGGGTGGTTAGTCCATTTCACNNNAGATCGGAA |
| 108899 | TATGAGTGTGGAGTCGTTGCAAGGATAAGAACATGCCAAATTCANNNAGATCGGA |
| 31793 | TATGAGTGTGGAGTCGTTGCGGAAACAGAGCACACAGAAGGAANNNAGATCGGA |
| 61713 | TATGAGTGTGGAGTCGTTGCCTCTTGAAACCTTCTGAGCAGCTNNNAGATCGGAAG |
| 110503 | TATGAGTGTGGAGTCGTTGCGAAGAACCCTCACTAATAGCGGANNNAGATCGGAAG |
| 45281 | TATGAGTGTGGAGTCGTTGCACTAAGAGAACAAGTGTCTCTGCCNNNAGATCGGAA |
| 117208 | TATGAGTGTGGAGTCGTTGCCATTTGCCTAAAACAGTGGCTGTNNNAGATCGGAAG |
| 29915 | TATGAGTGTGGAGTCGTTGCTCTCTTTCTACTCCCATCTCCAGANNNAGATCGGAAG |
| 58512 | TATGAGTGTGGAGTCGTTGCCTGAATGAGCTGGAAAGAGCCATNNNAGATCGGAA |
| 50427 | TATGAGTGTGGAGTCGTTGCGGTTCTTGCCCTTGAGAAGGAAGNNNAGATCGGAA |
| 36949 | TATGAGTGTGGAGTCGTTGCGCTTCGGTATGAAGTAACGGTCANNNAGATCGGAA |
| 61072 | TATGAGTGTGGAGTCGTTGCTTAGCCTTTCCAATGTGAAGCCTNNNAGATCGGAAG |
| 61669 | TATGAGTGTGGAGTCGTTGCATTTCTCAAGGGCAGAAACAGCTNNNAGATCGGAAG |
| 36963 | TATGAGTGTGGAGTCGTTGCAAAGTGCGTCCTAATTTGGGTCANNNAGATCGGAAG |
| 32216 | TATGAGTGTGGAGTCGTTGCTAACTGTTGTTCCCCGCTTTGAANNNAGATCGGAAG |
| 64736 | TATGAGTGTGGAGTCGTTGCACATGGAGTTCAAGCAACAAGGTNNNAGATCGGAA |
| 63902 | TATGAGTGTGGAGTCGTTGCGTGAAAAAGGAAAGGGGGACAGTNNNAGATCGGA |
| 53164 | TATGAGTGTGGAGTCGTTGCTGGGATGACAGAGTAGTGAGAGGNNNAGATCGGA |
| 105953 | TATGAGTGTGGAGTCGTTGCCTGATTTGGATCAAGTGTCAAAGTGNNNAGATCGG |
| 107670 | TATGAGTGTGGAGTCGTTGCGTGTATGTGCGTTTATTAGTGAGGGTNNNAGATCG |
| 46214 | TATGAGTGTGGAGTCGTTGCAAGATGCAGAAGGTAGCTGAAGCNNNAGATCGGAA |
| 40391 | TATGAGTGTGGAGTCGTTGCACACACAGAAAACATCTCCCTATTGANNNAGATCGG |
| 55482 | TATGAGTGTGGAGTCGTTGCCCAGTTATTGCCCCATTTGCATGNNNAGATCGGAAG |
| 33862 | TATGAGTGTGGAGTCGTTGCTGCTCAATCCTTGTGTCAGTACANNNAGATCGGAAG |
| 67458 | TATGAGTGTGGAGTCGTTGCGGCTGGGTTTCACTGGGTTTATTNNNAGATCGGAAG |
| 59219 | TATGAGTGTGGAGTCGTTGCTGATGGTCTTTCCCATCCTGGATNNNAGATCGGAAG |

56698 TATGAGTGTGGAGTCGTTGCGCAGAGGTTACTAGAGAAGCGTGNNNAGATCGGA
46323 TATGAGTGTGGAGTCGTTGCGAAGTTTTGAAAAGTGGGCCAGCNNNAGATCGGAA
68932 TATGAGTGTGGAGTCGTTGCTGCCCCCTCTAATCAGGCTATTTNNNAGATCGGAAG
70359 TATGAGTGTGGAGTCGTTGCGTTTCCCATCTGTGATTCCCTCCNNNAGATCGGAAGA
101518 TATGAGTGTGGAGTCGTTGCTTTCTGCTTAGCATTTGACAGCTCNNNAGATCGGAA
108292 TATGAGTGTGGAGTCGTTGCGAGAAATTGGGGTATTGAGGAGGNNNAGATCGGA
50218 TATGAGTGTGGAGTCGTTGCCATAAACCCGCCCCCACAAGNNNAGATCGGAAGAGC
117214 TATGAGTGTGGAGTCGTTGCTTGTTCTTGCTCTCTCTTCCCTCTNNNAGATCGGAAG
48546 TATGAGTGTGGAGTCGTTGCCTGCTGTGTCCAGCTCAAGTATCNNNAGATCGGAAG
36995 TATGAGTGTGGAGTCGTTGCTTTGTCATGCATGTGTTCATGTCANNNAGATCGGAA
59265 TATGAGTGTGGAGTCGTTGCTTGGAAGGTGAGAGGTCCATGATNNNAGATCGGAA
104459 TATGAGTGTGGAGTCGTTGCGTGGGAGTAGGGGTATGAAATCNNNAGATCGGAA
54072 TATGAGTGTGGAGTCGTTGCTAAAGCGTTCCCAGTTCTATGGGNNNAGATCGGAAG
46952 TATGAGTGTGGAGTCGTTGCCCAAATCCCAGATTCTCAGAGGCNNNAGATCGGAAG
71006 TATGAGTGTGGAGTCGTTGCCATACACACTGAAGACAGGCCATNNNAGATCGGAAG
71648 TATGAGTGTGGAGTCGTTGCGTGCGCTTGGATCTGATAAACGAGGNNNAGATCGG
49790 TATGAGTGTGGAGTCGTTGCAGGTAGGAGAGGAACCAAGGTTCNNNAGATCGGA
45552 TATGAGTGTGGAGTCGTTGCGCTCAGTCAATAGCACACCTCCNNNAGATCGGAAGA
65963 TATGAGTGTGGAGTCGTTGCACACACAAACGTTTTCAAACCTGTNNNAGATCGGAA
51226 TATGAGTGTGGAGTCGTTGCAGAAAACATCAGAGCAGCCTCAGNNNAGATCGGAA
36426 TATGAGTGTGGAGTCGTTGCTTAGTCTGATACTGGGGCCATCANNNAGATCGGAAG
106604 TATGAGTGTGGAGTCGTTGCGTTTGATTGATGAGGTCGATGGTGNNNAGATCGGA
39053 TATGAGTGTGGAGTCGTTGCTTCATGCTGGGTGTAAGTAGGGANNNAGATCGGAA
43866 TATGAGTGTGGAGTCGTTGCAGACAGACACATGAACACACACCNNNAGATCGGAAG
58461 TATGAGTGTGGAGTCGTTGCCCTTTCAAACAGCTACCACCCATNNNAGATCGGAAGA
70975 TATGAGTGTGGAGTCGTTGCCAGGCAGGAAGAAGCCAAAGAATNNNAGATCGGA
67328 TATGAGTGTGGAGTCGTTGCAAATGCCACATTCTGGGTGGATTNNNAGATCGGAA
101186 TATGAGTGTGGAGTCGTTGCGATGGGAATGCCAGGAGGAGAAGNNNAGATCGGA
101305 TATGAGTGTGGAGTCGTTGCAGTCATCTTGAATTGGGGCCCCNNNAGATCGGAAG
102467 TATGAGTGTGGAGTCGTTGCATCAGAAACTTCTTGGGAGGCATAGNNNAGATCGG
54019 TATGAGTGTGGAGTCGTTGCCAGCGGCACAATAGAAAAATGGGNNNAGATCGGAA
108347 TATGAGTGTGGAGTCGTTGCTATGTGTGTGATATACTTGCGTGTCTCNNNAGATCG
65300 TATGAGTGTGGAGTCGTTGCAGAGAGAGACAAAAGCTGATGGTNNNAGATCGGA
116225 TATGAGTGTGGAGTCGTTGCCCTGCTTCCCCATCATATCCATCNNNAGATCGGAAGA
49163 TATGAGTGTGGAGTCGTTGCACCCCCTCCTCTCTCTTTTTCTCNNNAGATCGGAAGAG
116986 TATGAGTGTGGAGTCGTTGCTCTTTTTCATGGTGTCTGTCACCTAGNNNAGATCGGA
39118 TATGAGTGTGGAGTCGTTGCACCAGATGACAGAGTGAATGGGANNNAGATCGGA
55634 TATGAGTGTGGAGTCGTTGCGGAGTGGGTAGTAAGCCATGATGNNNAGATCGGA
35270 TATGAGTGTGGAGTCGTTGCATTTATGTTGCCAAGACCCAGCANNNAGATCGGAAG
67060 TATGAGTGTGGAGTCGTTGCCCCCTACAGAAAAGGAGCACATTNNNAGATCGGAAG
103670 TATGAGTGTGGAGTCGTTGCAGAGCCTAAATTCTCATCCTCATGCNNNAGATCGGA
49407 TATGAGTGTGGAGTCGTTGCTAAACTGCAGCCAAGTTCCTGTCNNNAGATCGGAAG
57045 TATGAGTGTGGAGTCGTTGCTGATTGATGAGGTCGATGGTGTGNNNAGATCGGAA

47773 TATGAGTGTGGAGTCGTTGCTTCAACAAGTAGCCGTATGCTGCNNNAGATCGGAAG
117321 TATGAGTGTGGAGTCGTTGCGATGTGGGTGGGGTGGTGACANNNAGATCGGAAG
54484 TATGAGTGTGGAGTCGTTGCTTTGCTCTCAGCTCTCCTTATGGNNNAGATCGGAAGA
46385 TATGAGTGTGGAGTCGTTGCACCTTTCTTCTTTCCTCTGCAGCNNNAGATCGGAAGA
57699 TATGAGTGTGGAGTCGTTGCGCCTGCAAATCCCAATCACAAATNNNAGATCGGAAG
64425 TATGAGTGTGGAGTCGTTGCATGCCTGATTCCCACAGACTAGTNNNAGATCGGAAG
65950 TATGAGTGTGGAGTCGTTGCATGATGCAAGACGTCACTACTGTNNNAGATCGGAAG
50152 TATGAGTGTGGAGTCGTTGCGCGATCCAAAAATGCAGGGAAAGNNNAGATCGGAA
50886 TATGAGTGTGGAGTCGTTGCGGGTTCTTACTCTGACAAGCCAGNNNAGATCGGAAG
59177 TATGAGTGTGGAGTCGTTGCGTGTGTATGGGTAGGTGAGGGATNNNAGATCGGA
62556 TATGAGTGTGGAGTCGTTGCCCTGCTTGTGGTGTTTACAATCTNNNAGATCGGAAG
65800 TATGAGTGTGGAGTCGTTGCAGGGGATGACTATAGCTTGGATGTNNNAGATCGGA
100561 TATGAGTGTGGAGTCGTTGCTTTCTTTAGGCAAATACCCAGTAGTNNNAGATCGGA
102075 TATGAGTGTGGAGTCGTTGCCACTTTTGATTTTGTCAATTGAACAGTNNNAGATCGG
63880 TATGAGTGTGGAGTCGTTGCATTTGTTCTCTTTGGCCCACAGTNNNAGATCGGAAGA
53686 TATGAGTGTGGAGTCGTTGCCTCACCCATACTAGGATCCAGGGNNNAGATCGGAAG
62986 TATGAGTGTGGAGTCGTTGCATGAGCACTAGCAGACCATCTCTNNNAGATCGGAAG
33026 TATGAGTGTGGAGTCGTTGCTTCCCCACCTCAGTTCTAACACANNNAGATCGGAAGA
69269 TATGAGTGTGGAGTCGTTGCACTGGCTGTGTAAAACTGGAGTTTNNNAGATCGGAA
59120 TATGAGTGTGGAGTCGTTGCTCTGTCCACATTTTCCCAAGGATNNNAGATCGGAAGA
101206 TATGAGTGTGGAGTCGTTGCTGTCCATTCCCTCATAAGACTTTCTNNNAGATCGGAA
70811 TATGAGTGTGGAGTCGTTGCAGTGCAAAAATTTGGAGAGCTGGNNNAGATCGGAA
41716 TATGAGTGTGGAGTCGTTGCGACACAAGAGTGCCACCCAATTANNNAGATCGGAAG
47379 TATGAGTGTGGAGTCGTTGCCTTCTGTATCCTTGGGGATTGGCNNNAGATCGGAAG
29920 TATGAGTGTGGAGTCGTTGCATAAAACCGCTGACACAACTGGTNNNAGATCGGAAG
51695 TATGAGTGTGGAGTCGTTGCAGTTGTGGTAGGCCAAAATGGAGNNNAGATCGGAA
103927 TATGAGTGTGGAGTCGTTGCCCTCTCAACCCACCTTTTGAAAATNNNAGATCGGAAG
42695 TATGAGTGTGGAGTCGTTGCTTAGGGAGTTCTCGATAGAAATACCACNNNAGATCG
46681 TATGAGTGTGGAGTCGTTGCGCAGAGGTGCTATTTAGGGTAGCNNNAGATCGGAA
52339 TATGAGTGTGGAGTCGTTGCATTCTGTCCATTCGGGACATACGNNNAGATCGGAAG
49300 TATGAGTGTGGAGTCGTTGCTGGCAGTGAAGAATTACCAGGTCNNNAGATCGGAA
67768 TATGAGTGTGGAGTCGTTGCGCATTTGACAGCTCCTCATCCTTNNNAGATCGGAAGA
43947 TATGAGTGTGGAGTCGTTGCTAGCAGGAAACATGCTTAGCACCNNNAGATCGGAAG
57962 TATGAGTGTGGAGTCGTTGCTTCTCTAGCTCCCTCATGGCAATNNNAGATCGGAAGA
57921 TATGAGTGTGGAGTCGTTGCACTCTTACTTTGCCAGCTCCAATNNNAGATCGGAAGA
107706 TATGAGTGTGGAGTCGTTGCGCGTCCCTTCAAAAAGATATGTTNNNAGATCGGAAG
107475 TATGAGTGTGGAGTCGTTGCCACATGAACACACACCCAGTATANNNAGATCGGAAG
45401 TATGAGTGTGGAGTCGTTGCGATGCCCTTCACACCTGTAATCCNNNAGATCGGAAGA
71803 TATGAGTGTGGAGTCGTTGCATGCCACCCAACTTTTCATACGTNNNAGATCGGAAGA
59755 TATGAGTGTGGAGTCGTTGCAGTTTCAGGCACCAGAACAAACTNNNAGATCGGAAG
102106 TATGAGTGTGGAGTCGTTGCTTACTGGGTCGGGGGATTGCGATNNNAGATCGGAA
107639 TATGAGTGTGGAGTCGTTGCTGTTGGTGTGTGTTGGGTATGTNNNAGATCGGAAG
50850 TATGAGTGTGGAGTCGTTGCAAAGCACACACTACTATGCCCAGNNNAGATCGGAAG

68652 TATGAGTGTGGAGTCGTTGCTTGCTTCTCTGTGCACTTCTGTTNNNAGATCGGAAGA
68202 TATGAGTGTGGAGTCGTTGCACAGGAGTTTGGCCAGAAAAGTTNNNAGATCGGAA
30658 TATGAGTGTGGAGTCGTTGCAGCAGTAATGGCAGTGGTATGAAANNNAGATCGGA
46076 TATGAGTGTGGAGTCGTTGCATCCTCTACTGGGAAAGCAAAGCNNNAGATCGGAA
60207 TATGAGTGTGGAGTCGTTGCTACCAACACTAGGAGATCTCACTNNNAGATCGGAAG
36161 TATGAGTGTGGAGTCGTTGCAACTGCCCTCTTATTTTGGTGCANNNAGATCGGAAG
39567 TATGAGTGTGGAGTCGTTGCCCCACCCAGAAAAGTGACAATGANNNAGATCGGAA
110521 TATGAGTGTGGAGTCGTTGCAGGAAGCATGGAAGCATAGCAGNNNAGATCGGAA
116672 TATGAGTGTGGAGTCGTTGCACAGCTACCACCCATTCTTGGATTNNNAGATCGGAA
29914 TATGAGTGTGGAGTCGTTGCATTTTGTTCCGTTTGCAATGCCANNNAGATCGGAAG
34477 TATGAGTGTGGAGTCGTTGCATACCCACACACACATAGTCCCANNNAGATCGGAAG
41231 TATGAGTGTGGAGTCGTTGCATGATCGCACTGGAAATGTGCTANNNAGATCGGAA
50486 TATGAGTGTGGAGTCGTTGCTCTCTCACACACACAGCATGAAGNNNAGATCGGAAG
106521 TATGAGTGTGGAGTCGTTGCAACGTCCAAGTGCAGGTATGTNNNAGATCGGAAGA
61024 TATGAGTGTGGAGTCGTTGCCAATCTGCCCAACTTTTCTCCCTNNNAGATCGGAAGA
111655 TATGAGTGTGGAGTCGTTGCTGCTTCTCTGTGCACTTCTGTTACNNNAGATCGGAAG
102434 TATGAGTGTGGAGTCGTTGCTACCAGATGTGGGTGCATCTTNNNAGATCGGAAGA
32686 TATGAGTGTGGAGTCGTTGCATGATTACCTGATGTTGCTAAAACANNNAGATCGGA
49898 TATGAGTGTGGAGTCGTTGCCCCACCCTCCCTTGTTTACTTTCNNNAGATCGGAAGA
30800 TATGAGTGTGGAGTCGTTGCAACACAGTTTCTGAGACAAACAANNNAGATCGGAAG
56299 TATGAGTGTGGAGTCGTTGCATGGACAAACCTGGGACAATCTGNNNAGATCGGAA
68716 TATGAGTGTGGAGTCGTTGCCCAGAGGACCACGATCAATTGTTNNNAGATCGGAAG
49368 TATGAGTGTGGAGTCGTTGCTCTGGGATGCAGTAGAACATGTCNNNAGATCGGAA
100039 TATGAGTGTGGAGTCGTTGCCCTACTTGGATTCTCCTGCTCGNNNAGATCGGAAGA
96517 TATGAGTGTGGAGTCGTTGCAAGCCATTTACATGCGAGGAAGTGANNNAGATCGG
58706 TATGAGTGTGGAGTCGTTGCCGTCTGCACCATAGACATGCATNNNAGATCGGAAGA
105326 TATGAGTGTGGAGTCGTTGCACTCTTTACTGGAATACATTGTGGTNNNAGATCGGA
30202 TATGAGTGTGGAGTCGTTGCCCTAGAGTAACTGTGTGAGGTAAAANNNAGATCGG
29922 TATGAGTGTGGAGTCGTTGCTGAATTTTCTGGAAGAGGGGCAGNNNAGATCGGAA
29978 TATGAGTGTGGAGTCGTTGCATAAACCCTGGAATGCAGAAGGCNNNAGATCGGAA
29916 TATGAGTGTGGAGTCGTTGCTGATGAGAAGGGTTTGAGGCTCTNNNAGATCGGAA
70464 TATGAGTGTGGAGTCGTTGCCACACATTCTCTGCTTGGAATGCNNNAGATCGGAAG
29977 TATGAGTGTGGAGTCGTTGCGATGTGCTGTGGATGGAGATCTGNNNAGATCGGAA
53525 TATGAGTGTGGAGTCGTTGCACATAACCCTCATTTTCTGGCGGNNNAGATCGGAAG
61433 TATGAGTGTGGAGTCGTTGCTGTCTTCCTGAGGCTTTTCTCCTNNNAGATCGGAAGA
29980 TATGAGTGTGGAGTCGTTGCCCTCCTATGCCTCTAGCTCAGTTNNNAGATCGGAAGA
34710 TATGAGTGTGGAGTCGTTGCAGACAAGAAGATCCCTGATGCCANNNAGATCGGAA
29979 TATGAGTGTGGAGTCGTTGCGTGATCCTCTAGTTCTGTGGGCTNNNAGATCGGAAG
46025 TATGAGTGTGGAGTCGTTGCTCTGTCATTGCTGAGGTGTTTCCNNNAGATCGGAAG
101309 TATGAGTGTGGAGTCGTTGCTGTGTGTGTTTGGGATGGTAGAGTANNNAGATCGG
103785 TATGAGTGTGGAGTCGTTGCACTACACACACTGTCTTGAATGCCNNNAGATCGGAA
105420 TATGAGTGTGGAGTCGTTGCGGTATTTTCACATTCTTCATGGTGTGTNNNAGATCG
102719 TATGAGTGTGGAGTCGTTGCCAGACCTTTGATCCTTTCCAGATNNNAGATCGGAAG

| | |
|---|---|
| 103680 | TATGAGTGTGGAGTCGTTGCCGCTTAGACTGCAGAATCACTTTANNNAGATCGGAA |
| 103946 | TATGAGTGTGGAGTCGTTGCCAGAAGCAAGTGCTGCCATTTGNNNAGATCGGAAG |
| 97214 | TATGAGTGTGGAGTCGTTGCCTGGCATGAGGATGTAGAAAGCTNNNAGATCGGAA |
| 101364 | TATGAGTGTGGAGTCGTTGCCATCCTGTCACCTTGTAGCGTTNNNAGATCGGAAGA |
| 104421 | TATGAGTGTGGAGTCGTTGCATAATACTGGTCACCTTCTTGCCTNNNAGATCGGAAG |
| 108260 | TATGAGTGTGGAGTCGTTGCTGATTGATGGCATTTGGAATGGTTCNNNAGATCGG |
| 105278 | TATGAGTGTGGAGTCGTTGCCAGGACCTCACCAGAAGATATAAATGANNNAGATCG |
| 105247 | TATGAGTGTGGAGTCGTTGCGATGAGAAGCCACTGTTTCACAANNNAGATCGGAA |
| 102135 | TATGAGTGTGGAGTCGTTGCGATGGTGTTGGTGGACATTTAGGTNNNAGATCGGA |
| 108328 | TATGAGTGTGGAGTCGTTGCTTTTATAGAGGAAAAGTCTCGCCANNNAGATCGGAA |
| 105235 | TATGAGTGTGGAGTCGTTGCCCATCCTGGTTTGAGAAGCAGAATNNNAGATCGGA |
| 105463 | TATGAGTGTGGAGTCGTTGCGAGAGGAGTATCACAGGATGGAATNNNAGATCGG |
| 104015 | TATGAGTGTGGAGTCGTTGCGGAGTTGTTCCCCAAAGGAAAACNNNAGATCGGAA |
| 107674 | TATGAGTGTGGAGTCGTTGCTGGTGTGGTGTGTGTGTGTAAAAANNNAGATCGGA |
| 116143 | TATGAGTGTGGAGTCGTTGCAGGAGAGACAGCATATCAATGGANNNAGATCGGAA |
| 103728 | TATGAGTGTGGAGTCGTTGCCTCTACCTCCCCAGGCTCACATCNNNAGATCGGAAGA |
| 103683 | TATGAGTGTGGAGTCGTTGCGAAGAGGCCTGGAATGGTCACANNNAGATCGGAAG |
| 107668 | TATGAGTGTGGAGTCGTTGCGGCTAATGGTTTCGGCATGGANNNAGATCGGAAGA |
| 116561 | TATGAGTGTGGAGTCGTTGCACCAATGCCTGGCTCATTTCGTANNNAGATCGGAAG |
| 105428 | TATGAGTGTGGAGTCGTTGCGTGGTGTGGAGCAATATACCGTTNNNAGATCGGAA |
| 110480 | TATGAGTGTGGAGTCGTTGCAACAATGGGCTGAATCAGGAGATANNNAGATCGGA |
| 103922 | TATGAGTGTGGAGTCGTTGCTTAGGCTCTGGACTGAAACCCANNNAGATCGGAAG |
| 105472 | TATGAGTGTGGAGTCGTTGCAGATGCATACCATCATGCCAGCTNNNAGATCGGAAG |
| 107669 | TATGAGTGTGGAGTCGTTGCAATACTGGTACACACCACCACACTNNNAGATCGGAA |
| 104056 | TATGAGTGTGGAGTCGTTGCTGAGGGGTTGGTAGAGGTGAGTGNNNAGATCGGA |
| 102150 | TATGAGTGTGGAGTCGTTGCTGCCTGGCCTCCAATTCTTAGTAANNNAGATCGGAA |
| 100827 | TATGAGTGTGGAGTCGTTGCTTCAAAAATGGCATTGACGGATACANNNAGATCGG |
| 117744 | TATGAGTGTGGAGTCGTTGCTCTTTCCTTGTCTCCCCTCTCTNNNAGATCGGAAGAG |
| 97080 | TATGAGTGTGGAGTCGTTGCTCCATGTCTTATTTCCAGTTTGGGNNNAGATCGGAA |
| 102191 | TATGAGTGTGGAGTCGTTGCCGCAAGTACCTTTTTGCTATAATGACTNNNAGATCGG |
| 114241 | TATGAGTGTGGAGTCGTTGCAGAGCAGCTGAGAAAGGCTAAANNNAGATCGGAA |
| 122001 | TATGAGTGTGGAGTCGTTGCGTGGCTATCTCCAACTTGCTGGGNNNAGATCGGAAG |
| 104055 | TATGAGTGTGGAGTCGTTGCAGACAAGACCTTCCTTTGGTCAANNNAGATCGGAAG |
| 110494 | TATGAGTGTGGAGTCGTTGCTGGCCTTACCTTGGGAGAATAGTNNNAGATCGGAA |
| 100805 | TATGAGTGTGGAGTCGTTGCCTTGATCTTGTGGCACCTTGATCTNNNAGATCGGAA |
| 104023 | TATGAGTGTGGAGTCGTTGCCTGGGACCATAATGGGTGCATGNNNAGATCGGAAG |
| 56906 | TATGAGTGTGGAGTCGTTGCGGTCGCTAAGTGAATTATGGATGTGNNNAGATCGG |
| 62897 | TATGAGTGTGGAGTCGTTGCTCAAGCTCTCTCATGTGTCCTCTNNNAGATCGGAAGA |
| 108279 | TATGAGTGTGGAGTCGTTGCGTATCCATGTGTTTGAATTACTGCCNNNAGATCGGA |
| 59196 | TATGAGTGTGGAGTCGTTGCCCTGTACTCGGGGATTATGGGATNNNAGATCGGAA |
| 40723 | TATGAGTGTGGAGTCGTTGCCTGTTCCTGGTTGTCATGCCATANNNAGATCGGAAG |
| 45599 | TATGAGTGTGGAGTCGTTGCATACTAGGGAAGTGTGCTCCTCCNNNAGATCGGAAG |
| 43767 | TATGAGTGTGGAGTCGTTGCGTGCATTTGTTGCAAGGTAAACCNNNAGATCGGAA |

69229 TATGAGTGTGGAGTCGTTGCTGGCTACAGAGCATGGTTTCTTTNNNAGATCGGAAG
39189 TATGAGTGTGGAGTCGTTGCTGTTAGCCTTCAGCAAATTGGGANNNAGATCGGAA
61936 TATGAGTGTGGAGTCGTTGCTGGCATCAGTTCTTGAAAAGGCTNNNAGATCGGAA
42209 TATGAGTGTGGAGTCGTTGCACTTTCAGCCCACTTGTTTCAAACNNNAGATCGGAAG
32213 TATGAGTGTGGAGTCGTTGCGCACAGAGCCATCCTGATTTGAANNNAGATCGGAAG
61217 TATGAGTGTGGAGTCGTTGCCTTGCCTCTCTTGTTGAATGCCTNNNAGATCGGAAG
36339 TATGAGTGTGGAGTCGTTGCTGTTGAAGCGAAATGGAAAATCANNNAGATCGGAA
39288 TATGAGTGTGGAGTCGTTGCTGAGGGACACACAGAACTATGGANNNAGATCGGAA
43859 TATGAGTGTGGAGTCGTTGCCGAAGAAACATAGCTGGAAAACACCNNNAGATCGG
58260 TATGAGTGTGGAGTCGTTGCGCTTTGTGGCTATTGCCAACATNNNAGATCGGAAGA
44368 TATGAGTGTGGAGTCGTTGCTTTTCTTGGCAGTTCCTTACCCCNNNAGATCGGAAGA
100405 TATGAGTGTGGAGTCGTTGCCCTTTGACTAGGTAGCTTTGAGCNNNAGATCGGAAG
56827 TATGAGTGTGGAGTCGTTGCTGCGTATGTGTATTATTGCTATGGTGNNNAGATCGG
31724 TATGAGTGTGGAGTCGTTGCGAATACGACCGTAACAGTGAGAANNNAGATCGGAA
46928 TATGAGTGTGGAGTCGTTGCTTTCCACAAAGGAGAGATCAGGCNNNAGATCGGAA
31474 TATGAGTGTGGAGTCGTTGCGACCTCCTGCCAGTCAAATTCAANNNAGATCGGAAG
67178 TATGAGTGTGGAGTCGTTGCGCTTGTGTGTGTTTGTCTGCATTNNNAGATCGGAAG
103753 TATGAGTGTGGAGTCGTTGCGAGAGAGAGAGACAGAGCATCTATNNNAGATCGG
35172 TATGAGTGTGGAGTCGTTGCTGGGAGAGCACTGTATACAAGCANNNAGATCGGAA
47734 TATGAGTGTGGAGTCGTTGCCTGGACCTGTTGACTATTCCTGCNNNAGATCGGAAG
32505 TATGAGTGTGGAGTCGTTGCTCCAAAGCCCAGATTTCATGTAANNNAGATCGGAAG
98670 TATGAGTGTGGAGTCGTTGCGTCTACTTAGTTGTGGAGTTCGTGTNNNAGATCGGA
41682 TATGAGTGTGGAGTCGTTGCCCAATCACCCCAAAGTGTGTGTANNNAGATCGGAAG
43239 TATGAGTGTGGAGTCGTTGCAGGGGTTGGGATGACTTCTAGACNNNAGATCGGAA
97730 TATGAGTGTGGAGTCGTTGCCCCAAGGAACACCCGCCCACNNNAGATCGGAAGAGC
36093 TATGAGTGTGGAGTCGTTGCTCCCTCTATTATTCCTGGCTGCANNNAGATCGGAAGA
32003 TATGAGTGTGGAGTCGTTGCGGAAGAGGCAACTTGACTTGGAANNNAGATCGGAA
60173 TATGAGTGTGGAGTCGTTGCAACCATGGAGGTGAAGGATCACTNNNAGATCGGAA
47392 TATGAGTGTGGAGTCGTTGCACATATCCTTGGAACAGCTTGGCNNNAGATCGGAAG
67667 TATGAGTGTGGAGTCGTTGCCACCCCCACAAATTCTGTACCTTNNNAGATCGGAAGA
33887 TATGAGTGTGGAGTCGTTGCTCAGGCATTCATGCACATGTACANNNAGATCGGAAG
97134 TATGAGTGTGGAGTCGTTGCTAACAAAAGCCCACGAAGAGTTTTCNNNAGATCGGA
55151 TATGAGTGTGGAGTCGTTGCAGCATTTCATTCTCTTTTCTTTGTTGGNNNAGATCGG
47087 TATGAGTGTGGAGTCGTTGCATTCCCTACTTCACTGATAGGGCNNNAGATCGGAAG
40722 TATGAGTGTGGAGTCGTTGCGCAAGGAATTCTCACTGGCCATANNNAGATCGGAAG
34305 TATGAGTGTGGAGTCGTTGCTTTACTCAACTTCCCTCACCCCANNNAGATCGGAAGA
57484 TATGAGTGTGGAGTCGTTGCTTCAGGGTGCATGCTCTATGTTGNNNAGATCGGAAG
41983 TATGAGTGTGGAGTCGTTGCCACACACACACGGCAAACATTTANNNAGATCGGAAG
60783 TATGAGTGTGGAGTCGTTGCGGGTCCTAGCTGATGAACTACCTNNNAGATCGGAAG
68293 TATGAGTGTGGAGTCGTTGCTAGCAGATTCCTCTGGCTCAGTTNNNAGATCGGAAG
39412 TATGAGTGTGGAGTCGTTGCGTGATTTGTGCACCTAGAGTGGANNNAGATCGGAA
57243 TATGAGTGTGGAGTCGTTGCCTCAGTTCAGGGTGTCAGACTTGNNNAGATCGGAA
64920 TATGAGTGTGGAGTCGTTGCAGTGCTCATGACAGAAATAATGAGGTNNNAGATCG

52266 TATGAGTGTGGAGTCGTTGCTACTTCAACCCAGTGACAACACGNNNAGATCGGAAG
32971 TATGAGTGTGGAGTCGTTGCTGGTGTATAATAGCAGACAGGTAACANNNAGATCG
59787 TATGAGTGTGGAGTCGTTGCGTCACATTCCCAGAACGGAAACTNNNAGATCGGAAG
43659 TATGAGTGTGGAGTCGTTGCGAAGTCCCTCTGCTGAGCATTACNNNAGATCGGAAG
65740 TATGAGTGTGGAGTCGTTGCATTTGTGGCCAGGTTCATCATGTNNNAGATCGGAAG
32370 TATGAGTGTGGAGTCGTTGCGACACATGCACACACACTCCTAANNNAGATCGGAAG
67099 TATGAGTGTGGAGTCGTTGCGACTTTCCACTCATCCACCCATTNNNAGATCGGAAGA
31980 TATGAGTGTGGAGTCGTTGCAAAAGGTGTCCGGTAATCTGGAANNNAGATCGGAA
41523 TATGAGTGTGGAGTCGTTGCCCACAAATGGTAAGCACAGGGTANNNAGATCGGAA
49142 TATGAGTGTGGAGTCGTTGCCGTAAGGCAAACAAGCCTTTCTCNNNAGATCGGAAG
57635 TATGAGTGTGGAGTCGTTGCCCCCTTCACAGACCTTCCTTTTGNNNAGATCGGAAGA
65753 TATGAGTGTGGAGTCGTTGCACCTCTCTACCTTGGAGTTCATGTNNNAGATCGGAA
63926 TATGAGTGTGGAGTCGTTGCCACACCATGAGGTTGTCTACAGTNNNAGATCGGAAG
29909 TATGAGTGTGGAGTCGTTGCTAATGTTGGAGGGAAGGACTGGTNNNAGATCGGA
47357 TATGAGTGTGGAGTCGTTGCTGCTCTGAATCATTACCTGTGGCNNNAGATCGGAAG
57485 TATGAGTGTGGAGTCGTTGCTTATGCGCCCGTTTGTTATGTTGNNNAGATCGGAAG
62379 TATGAGTGTGGAGTCGTTGCATGGAAATGTTGGGTCTAGTGCTNNNAGATCGGAA
38272 TATGAGTGTGGAGTCGTTGCAAAGAACCTCCTTCCAGTGGAGANNNAGATCGGAA
69272 TATGAGTGTGGAGTCGTTGCACCCCTTACCATGAGTTGAGTTTNNNAGATCGGAAG
39134 TATGAGTGTGGAGTCGTTGCAAATACCCACACGTCCTATGGGANNNAGATCGGAAG
35136 TATGAGTGTGGAGTCGTTGCACCAGAAGAAAAATTGCTAAAAGCANNNAGATCGG
32933 TATGAGTGTGGAGTCGTTGCTCCCCAAGGTATGTGTCTGAACANNNAGATCGGAAG
64340 TATGAGTGTGGAGTCGTTGCCCTGCTTATTTGTTTTGTTTCCTGAGTNNNAGATCGG
51907 TATGAGTGTGGAGTCGTTGCTGCCTTCGAGAGAAATCTGTGAGNNNAGATCGGAA
102196 TATGAGTGTGGAGTCGTTGCTTGCCTGTTATCAGTGTAGTCTTNNNAGATCGGAAG
29483 TATGAGTGTGGAGTCGTTGCGGCAAGTTTTACCTACAGCTCTNNNAGATCGGAAGA
67597 TATGAGTGTGGAGTCGTTGCAGTCCCCAGATGTGAGCATACTTNNNAGATCGGAAG
58924 TATGAGTGTGGAGTCGTTGCTGCCAAAGACCTCTTCGTTTCATNNNAGATCGGAAG
60560 TATGAGTGTGGAGTCGTTGCCTGTGCCTGAATCCTGCTTTACTNNNAGATCGGAAG
55110 TATGAGTGTGGAGTCGTTGCATGGCAGCTCTAAATGGAGTTGGNNNAGATCGGAA
62678 TATGAGTGTGGAGTCGTTGCAGGGTCAAACACAACTGAGATCTNNNAGATCGGAA
58703 TATGAGTGTGGAGTCGTTGCAAGTTGTGGCTTCCTGAATGCATNNNAGATCGGAAG
47419 TATGAGTGTGGAGTCGTTGCTGTGTGTGTATCTTCAACTTTGGCNNNAGATCGGAA
61737 TATGAGTGTGGAGTCGTTGCTGGTCCTAATAAGATCTCTCAGCTNNNAGATCGGAA
67285 TATGAGTGTGGAGTCGTTGCACACAATTCCAAAGTCCTGAGATTNNNAGATCGGAA
30563 TATGAGTGTGGAGTCGTTGCCCCTAACACTACCAGTAGCGAAANNNAGATCGGAAG
66900 TATGAGTGTGGAGTCGTTGCGGGGTGGTGGTGCTATGAAAATTNNNAGATCGGA
53847 TATGAGTGTGGAGTCGTTGCGAAAGTTATTTTGCTCTCCCGGGNNNAGATCGGAAG
61015 TATGAGTGTGGAGTCGTTGCCTTCATGCAACCATTTCCTCCCTNNNAGATCGGAAGA
40423 TATGAGTGTGGAGTCGTTGCCCTCTCTCTCACACACTCCTTGANNNAGATCGGAAGA
60323 TATGAGTGTGGAGTCGTTGCATGTTCTTTGAGTACAGCAGGACTNNNAGATCGGAA
66594 TATGAGTGTGGAGTCGTTGCTCTGAGTAAGGTCACTGGATTGTNNNAGATCGGAA
57629 TATGAGTGTGGAGTCGTTGCAGTTTCCCGTGGTCTGGATTTTGNNNAGATCGGAAG

55951 TATGAGTGTGGAGTCGTTGCCTGGGTTTTCTTCAGGTACCCTGNNNAGATCGGAAG
41073 TATGAGTGTGGAGTCGTTGCCTTTCCTGCCTTGCTTTTCCCTANNNAGATCGGAAGA
102740 TATGAGTGTGGAGTCGTTGCGAAGAAAAATTGCTAAAAGCATTGAAANNNAGATC
66624 TATGAGTGTGGAGTCGTTGCTGAGCAGCAGCATTTTACCTTGTNNNAGATCGGAAG
31603 TATGAGTGTGGAGTCGTTGCGGAAAGGCTGAAGGAAACCAGAANNNAGATCGGA
38675 TATGAGTGTGGAGTCGTTGCAGCCACATCCCTTAGAATAGAAGGANNNAGATCGG
37369 TATGAGTGTGGAGTCGTTGCTTCTCTGCATCTGGCTTCTTTCANNNAGATCGGAAGA
39382 TATGAGTGTGGAGTCGTTGCAGTGGACCTTAGCCAATTCTGGANNNAGATCGGAA
55848 TATGAGTGTGGAGTCGTTGCCACTCTGAAACCTCACCAGACTGNNNAGATCGGAAG
31135 TATGAGTGTGGAGTCGTTGCACATAGGCTTCCCTCATCTCCAANNNAGATCGGAAG
103724 TATGAGTGTGGAGTCGTTGCGGGGCAGTCCATTGATAAATACTTNNNAGATCGGA
64655 TATGAGTGTGGAGTCGTTGCAGTGGAATGCTGTTATATATGTGCGTNNNAGATCG
33462 TATGAGTGTGGAGTCGTTGCATGGGAGGTCATCTGGAAAGACANNNAGATCGGAA
46085 TATGAGTGTGGAGTCGTTGCACCTTGTCCATCACTGTCAAAGCNNNAGATCGGAAG
41805 TATGAGTGTGGAGTCGTTGCCCATTTCTCCAAAGCCACCCTTANNNAGATCGGAAGA
64887 TATGAGTGTGGAGTCGTTGCGCTCAGGAGTGTCTTCTAGAGGTNNNAGATCGGAA
41968 TATGAGTGTGGAGTCGTTGCTACTGGGGGACATGGATGTGTTANNNAGATCGGAA
43831 TATGAGTGTGGAGTCGTTGCTCAGCCAAAAGGTAAACTTGAACCNNNAGATCGGAA
58766 TATGAGTGTGGAGTCGTTGCGGGGAGCCTTTCCTTTCAATCATNNNAGATCGGAAG
50638 TATGAGTGTGGAGTCGTTGCTCTCCATTGCCCTTCACTAACAGNNNAGATCGGAAGA
65992 TATGAGTGTGGAGTCGTTGCAGGGACAGTTTTTCTTTTCCCTGTNNNAGATCGGAA
64399 TATGAGTGTGGAGTCGTTGCTAAGCAGTGTTGGGACCCATAGTNNNAGATCGGAA
29906 TATGAGTGTGGAGTCGTTGCGCAACGCAGGCTGAAGTATAATGNNNAGATCGGAA
33186 TATGAGTGTGGAGTCGTTGCCCTCCAAATTGTGAAATTACCCACANNNAGATCGGA
62341 TATGAGTGTGGAGTCGTTGCTAGGCAAAATCTGCTTCTCTGCTNNNAGATCGGAAG
33525 TATGAGTGTGGAGTCGTTGCGAGTGAGGAGACAGACAGAGACANNNAGATCGGA
52256 TATGAGTGTGGAGTCGTTGCCTGTGCTCCACTAAAATGCAACGNNNAGATCGGAAG
66655 TATGAGTGTGGAGTCGTTGCTTCAGTTGTTCCAGAACTGCTTGTNNNAGATCGGAA
46837 TATGAGTGTGGAGTCGTTGCAGGTGTTAGTGATGCTTCAAGGCNNNAGATCGGAA
58108 TATGAGTGTGGAGTCGTTGCCCACACACATGGGCTTATGGAATNNNAGATCGGAAG
66388 TATGAGTGTGGAGTCGTTGCACCCTTCAACATCACAATGGTGTNNNAGATCGGAAG
105958 TATGAGTGTGGAGTCGTTGCACCCTCCTTTCGGTCTGTTTGTGNNNAGATCGGAAG
66991 TATGAGTGTGGAGTCGTTGCGTATGGAGGGCCCTGAAGGAATTNNNAGATCGGAA
68895 TATGAGTGTGGAGTCGTTGCAGAGAAGCCAGCAGTTTGGATTTNNNAGATCGGAA
40651 TATGAGTGTGGAGTCGTTGCTTGCATGTTTGGTGGCAGGAATANNNAGATCGGAA
36781 TATGAGTGTGGAGTCGTTGCGCAATTCAGGAAGACAGCTCTCANNNAGATCGGAA
71622 TATGAGTGTGGAGTCGTTGCGCAATGACAAAGGTAACAATTCCAGNNNAGATCGG
71731 TATGAGTGTGGAGTCGTTGCATGTTGATGGGATTCCAGTAGCCNNNAGATCGGAA
69557 TATGAGTGTGGAGTCGTTGCAGTTCCAGAGGGGTGGTAGTTTTNNNAGATCGGAA
45953 TATGAGTGTGGAGTCGTTGCGTTGCTTCTGTTGCTTCTGTTCCNNNAGATCGGAAG
116242 TATGAGTGTGGAGTCGTTGCCTCTCTCCTCTCTCTCTCATCTCTTTNNNAGATCGGAA
29482 TATGAGTGTGGAGTCGTTGCTGTGGATGGTAAGAATTGAGGCTNNNAGATCGGAA
37281 TATGAGTGTGGAGTCGTTGCTGGAAGAGATTTTACTCAAGGTTCANNNAGATCGG

56498 TATGAGTGTGGAGTCGTTGCGTGGGTTTTTCTTTTCTGGTTTTCTGNNNAGATCGGA
67428 TATGAGTGTGGAGTCGTTGCTCTCTCCAGTGGTGTGTGGTATTNNNAGATCGGAAG
45403 TATGAGTGTGGAGTCGTTGCGCATTTGTGTTGCTTTCAACATCCNNNAGATCGGAAG
51287 TATGAGTGTGGAGTCGTTGCCCAGGTTCCAGTACAATGGTCAGNNNAGATCGGAAG
62345 TATGAGTGTGGAGTCGTTGCTGCCTTAGAACCTGAAATTCTGCTNNNAGATCGGAAG
55486 TATGAGTGTGGAGTCGTTGCAGCAGCAAAACCAAAGCATCATGNNNAGATCGGAAG
30125 TATGAGTGTGGAGTCGTTGCGCAGGTGGTTACCTTTGAGAAAANNNAGATCGGAAG
60028 TATGAGTGTGGAGTCGTTGCTCTGAATGGCTTGGGAGTACACTNNNAGATCGGAAG
106451 TATGAGTGTGGAGTCGTTGCATGTCTTTCACTCATGGAGGGAGNNNAGATCGGAAG
107527 TATGAGTGTGGAGTCGTTGCAAGGAAGTGCTGCTGTAGGTAAGNNNAGATCGGAA
51927 TATGAGTGTGGAGTCGTTGCGTCTACTGAAGATGGGCCTTGAGNNNAGATCGGAAG
64176 TATGAGTGTGGAGTCGTTGCATTGCTCCAGGAGAGAGAAGAGTNNNAGATCGGAA
52806 TATGAGTGTGGAGTCGTTGCAAAGCACATCACCATCAGGAAGGNNNAGATCGGAAG
64268 TATGAGTGTGGAGTCGTTGCTGTCTTTCACTCATGGAGGGAGTNNNAGATCGGAAG
66610 TATGAGTGTGGAGTCGTTGCGCACCAAATGCATTCCAACTTGTNNNAGATCGGAAGA
61311 TATGAGTGTGGAGTCGTTGCCTACAGGGAGCTGCTCTAATCCTNNNAGATCGGAAG
64131 TATGAGTGTGGAGTCGTTGCGAGAGAGAGAATCACCTTGTCAGTNNNAGATCGGA
68225 TATGAGTGTGGAGTCGTTGCAGCGCACATTTCCTTTAGAAGTTNNNAGATCGGAAGA
116784 TATGAGTGTGGAGTCGTTGCGTCCTGGGGAAATGGGACATCAANNNAGATCGGAA
99330 TATGAGTGTGGAGTCGTTGCGAGGGTGTATGACCTTTGCGATATTNNNAGATCGGA
57586 TATGAGTGTGGAGTCGTTGCGGAGAAGTAGGCAGGTGTCTTTGNNNAGATCGGAA
51435 TATGAGTGTGGAGTCGTTGCCCGAGGCAGATAGAACATCAGAGNNNAGATCGGAA
60477 TATGAGTGTGGAGTCGTTGCAGGTTTCTGAAGCAATGGCTACTNNNAGATCGGAAG
56838 TATGAGTGTGGAGTCGTTGCCCTAATATGGAAGCCTGCTGGTGNNNAGATCGGAAG
42054 TATGAGTGTGGAGTCGTTGCGCATGTGTGTTTCCCTAAGTTTANNNAGATCGGAAGA
51838 TATGAGTGTGGAGTCGTTGCTTTGGTCCATTGGTGTTCCTGAGNNNAGATCGGAAG
41537 TATGAGTGTGGAGTCGTTGCAGGAGGGTGACTACTAAGGGGTANNNAGATCGGAA
48994 TATGAGTGTGGAGTCGTTGCCGGGGTCTTAGCATTCTCATCTCNNNAGATCGGAAGA
47711 TATGAGTGTGGAGTCGTTGCGAAAAATATAACCGCTGCCCTGCNNNAGATCGGAAG
38122 TATGAGTGTGGAGTCGTTGCGTAATGTGCCCTTCAAGCAGAGANNNAGATCGGAAG
65307 TATGAGTGTGGAGTCGTTGCCTGCAAAGACAGACCTTGATGGTNNNAGATCGGAAG
95128 TATGAGTGTGGAGTCGTTGCTGGCCCTCTTTTGCCTACTATTGNNNAGATCGGAAGA
68690 TATGAGTGTGGAGTCGTTGCATTCCCCCATCTGTTCATGTGTTNNNAGATCGGAAGA
52368 TATGAGTGTGGAGTCGTTGCAAAGCGTAATGACACAAACACCGNNNAGATCGGAAG
67402 TATGAGTGTGGAGTCGTTGCCAGCTTCCCACCATCTTGCTATTNNNAGATCGGAAGA
40667 TATGAGTGTGGAGTCGTTGCCAGCACACCGCATGGATCTAATANNNAGATCGGAAG
42605 TATGAGTGTGGAGTCGTTGCGCACGCACTATGCTATTTCACACNNNAGATCGGAAGA
33760 TATGAGTGTGGAGTCGTTGCTGGTGACATAAGCACTCCATACANNNAGATCGGAAG
55105 TATGAGTGTGGAGTCGTTGCTAGTAGGGGTTGAGGGTTCTTGGNNNAGATCGGAA
48057 TATGAGTGTGGAGTCGTTGCAGAGGCAGAGAATACAGCATTGCNNNAGATCGGAA
47442 TATGAGTGTGGAGTCGTTGCTAGAAATGGCCAAGAGCAAATGCNNNAGATCGGAAG
37676 TATGAGTGTGGAGTCGTTGCCCAAAAAGTGAAGTCAATGTGAAGANNNAGATCGGA
31721 TATGAGTGTGGAGTCGTTGCAGGAGCTCAGAAAAGCCTGAGAANNNAGATCGGAA

| | |
|---|---|
| 117218 | TATGAGTGTGGAGTCGTTGCGTGCAAACTGCTTATAGGTGTGGTANNNAGATCGG |
| 47717 | TATGAGTGTGGAGTCGTTGCATTTACTGTGGAACTCAGCCTGCNNNAGATCGGAAG |
| 67707 | TATGAGTGTGGAGTCGTTGCGCTGTTTGCTCTCTCTTTCCCTTNNNAGATCGGAAGA |
| 42171 | TATGAGTGTGGAGTCGTTGCGCTCAGTCTGGAAAAGCACAAACNNNAGATCGGAA |
| 41815 | TATGAGTGTGGAGTCGTTGCATGAGAGCCCACAGATAGCCTTANNNAGATCGGAA |
| 68201 | TATGAGTGTGGAGTCGTTGCTTTTCCCAATATCCTACTCTTTTTCTTNNNAGATCGGA |
| 37668 | TATGAGTGTGGAGTCGTTGCACCCCTTTTACTTTACAGTCTGAAGANNNAGATCGGA |
| 33105 | TATGAGTGTGGAGTCGTTGCACTGGATCAGGCTAATTGACACANNNAGATCGGAA |
| 42147 | TATGAGTGTGGAGTCGTTGCGGGCAAGGATACATGCAGAAAACNNNAGATCGGAA |
| 44787 | TATGAGTGTGGAGTCGTTGCACAAACAGCTACTGTCTCATTCCCNNNAGATCGGAAG |
| 61602 | TATGAGTGTGGAGTCGTTGCACAGACATTCACGTGTCAAAGCTNNNAGATCGGAAG |
| 34388 | TATGAGTGTGGAGTCGTTGCATGATTCTTTTGTACTGGGCCCANNNAGATCGGAAG |
| 35351 | TATGAGTGTGGAGTCGTTGCGGCAGCTAATAAATGTGTTCAGCANNNAGATCGGA |
| 49088 | TATGAGTGTGGAGTCGTTGCGTCCTGGGAAATCCTTTGGTCTCNNNAGATCGGAAG |
| 62717 | TATGAGTGTGGAGTCGTTGCACACAGCCCTGCCATCTATATCTNNNAGATCGGAAGA |
| 116527 | TATGAGTGTGGAGTCGTTGCGTTGTGCTCTCTCTCTCTTACTCTNNNAGATCGGAAG |
| 48175 | TATGAGTGTGGAGTCGTTGCCCTAAACATTCAGCAGGTGTTGCNNNAGATCGGAAG |
| 35682 | TATGAGTGTGGAGTCGTTGCTCTTTTTCCATGGACTGTAGGCANNNAGATCGGAAG |
| 47590 | TATGAGTGTGGAGTCGTTGCATCCCTAACAACACTGGTGATGCNNNAGATCGGAAG |
| 57139 | TATGAGTGTGGAGTCGTTGCGGCAGGGGGTATACAGGAAATTGNNNAGATCGGA |
| 61315 | TATGAGTGTGGAGTCGTTGCGCATTTTTCTACCTTGCATACATCCTNNNAGATCGGA |
| 104445 | TATGAGTGTGGAGTCGTTGCTGTGTGTGTTTGTAGGAGAGAAGANNNAGATCGGA |
| 105944 | TATGAGTGTGGAGTCGTTGCGTGTGTGTGTCTTGAAAGCTCANNNAGATCGGAAG |
| 33555 | TATGAGTGTGGAGTCGTTGCGAAAGATGGAAGGCCTTGAGACANNNAGATCGGAA |
| 108244 | TATGAGTGTGGAGTCGTTGCCCTTTCCTTAGGGTTTGCTGTTCTNNNAGATCGGAAG |
| 65907 | TATGAGTGTGGAGTCGTTGCTAGCACTTTCTGCTTTCCACTGTNNNAGATCGGAAGA |
| 96460 | TATGAGTGTGGAGTCGTTGCCCCGCAAAAGTTTCTCCTACCTTGGNNNAGATCGGAA |
| 71770 | TATGAGTGTGGAGTCGTTGCTTGACTGCACTGGAAATCCTCTGNNNAGATCGGAAG |
| 62391 | TATGAGTGTGGAGTCGTTGCATTGAGTACTGGTAGTGGGTGCTNNNAGATCGGAA |
| 47555 | TATGAGTGTGGAGTCGTTGCTCAAGAGGAGGACAAACAGATGCNNNAGATCGGAA |
| 57826 | TATGAGTGTGGAGTCGTTGCGGAGTTCGACCTGGGTTTGAAATNNNAGATCGGAA |
| 68153 | TATGAGTGTGGAGTCGTTGCAAGCCACTCAGATCTCCCTTCTTNNNAGATCGGAAGA |
| 56169 | TATGAGTGTGGAGTCGTTGCGAGACAGATGGTTCTGAAGGCTGNNNAGATCGGAA |
| 105970 | TATGAGTGTGGAGTCGTTGCGGTATGTGTTGTATGTGCATGTGNNNAGATCGGAA |
| 71195 | TATGAGTGTGGAGTCGTTGCTGAGTTACTTCGATGTTTCAGCTNNNAGATCGGAAG |
| 108238 | TATGAGTGTGGAGTCGTTGCTCAATACCTCATAGTGCTAGTGCTACTNNNAGATCGG |
| 49956 | TATGAGTGTGGAGTCGTTGCTCTTGCTTGCTCCTTCTGGTTTCNNNAGATCGGAAGA |
| 52228 | TATGAGTGTGGAGTCGTTGCTGGGGGAGGTGGAGAATCTTTAGNNNAGATCGGA |
| 35147 | TATGAGTGTGGAGTCGTTGCTACAGGCATGCTCAAAGAAAGCANNNAGATCGGAA |
| 35306 | TATGAGTGTGGAGTCGTTGCAAGAGCATTTAGATGTGGCAGCANNNAGATCGGAA |
| 37499 | TATGAGTGTGGAGTCGTTGCGGGGCAGAAAAGGAAAGGAAAGANNNAGATCGGA |
| 54999 | TATGAGTGTGGAGTCGTTGCCTGTAGTGGAGGTTTCTGTGTGGNNNAGATCGGAA |
| 53746 | TATGAGTGTGGAGTCGTTGCAATAGCCAGGAGACTGAAGAGGGNNNAGATCGGA |

65188 TATGAGTGTGGAGTCGTTGCGTATAAAGTGAACTGGGCTGGGTNNNAGATCGGAA
57599 TATGAGTGTGGAGTCGTTGCTGGAAGAGGTGGTGTTGAGTTTGNNNAGATCGGA
48259 TATGAGTGTGGAGTCGTTGCCTGGAGCAGGGAAGAGTGAAATCNNNAGATCGGA
105954 TATGAGTGTGGAGTCGTTGCAGACGAGGGTTCCATGACAGAGNNNAGATCGGAAG
49907 TATGAGTGTGGAGTCGTTGCTCTGAGGGAAAAAGCCTCCTTTCNNNAGATCGGAAG
63231 TATGAGTGTGGAGTCGTTGCTCTGATTACTGGCAACGATGTCTNNNAGATCGGAAG
63894 TATGAGTGTGGAGTCGTTGCGGGTCAGATGTCACAAAGACAGTNNNAGATCGGAA
64290 TATGAGTGTGGAGTCGTTGCATCTGGTTCACAGTAGCTGGAGTNNNAGATCGGAA
52144 TATGAGTGTGGAGTCGTTGCTCTTATTGGCCAAAGCTGGGTAGNNNAGATCGGAA
33426 TATGAGTGTGGAGTCGTTGCCATAGCCTCCAAGCATCTTCACANNNAGATCGGAAG
65531 TATGAGTGTGGAGTCGTTGCTAGCCCAAATGGAAATGCTTGGTNNNAGATCGGAA
30945 TATGAGTGTGGAGTCGTTGCTCCCAAACCTCATGCTTTGACAANNNAGATCGGAAG
67389 TATGAGTGTGGAGTCGTTGCTGAAACAATGCTTAGGCATACTATTNNNAGATCGGA
46508 TATGAGTGTGGAGTCGTTGCCTGAACTCCAACCTCTTGAGAGCNNNAGATCGGAAG
71541 TATGAGTGTGGAGTCGTTGCTATGGCCGTGTGACTAAGGTCTTNNNAGATCGGAA
59788 TATGAGTGTGGAGTCGTTGCACGGCAAAGAAAGAAGGGAAACTNNNAGATCGGA
55209 TATGAGTGTGGAGTCGTTGCCCTGGGACCTGACGATATTTTGGNNNAGATCGGAA
116756 TATGAGTGTGGAGTCGTTGCGTTGCAGGGAAAAGACTGGGGTNNNAGATCGGAA
29905 TATGAGTGTGGAGTCGTTGCAGGCCGCATAAGATAATTAGGGTNNNAGATCGGAA
54699 TATGAGTGTGGAGTCGTTGCTTAGTGTACTGCAAACTGGCTGGNNNAGATCGGAA
54506 TATGAGTGTGGAGTCGTTGCTCTCGTTTGTCAGAACACACTGGNNNAGATCGGAAG
48878 TATGAGTGTGGAGTCGTTGCCTTGTCTGTCCCCACTCGCTCNNNAGATCGGAAGAG
45623 TATGAGTGTGGAGTCGTTGCGTCACCCGAAATACAGTAGCTCCNNNAGATCGGAAG
71651 TATGAGTGTGGAGTCGTTGCGGGACAGGTAAAGAGACGAGAAAGNNNAGATCGG
62924 TATGAGTGTGGAGTCGTTGCGAGAAGCTGAATGCAAGAGCTCTNNNAGATCGGAA
59768 TATGAGTGTGGAGTCGTTGCCCACATTCATGCACATGCAAACTNNNAGATCGGAAG
55137 TATGAGTGTGGAGTCGTTGCTCTAGGCACCTTGCATATGTTGGNNNAGATCGGAAG
49684 TATGAGTGTGGAGTCGTTGCGTGTGCTTTGTTTCCTCAGCTTCNNNAGATCGGAAG
34918 TATGAGTGTGGAGTCGTTGCATACAGCATGCACTTGATCTCCANNNAGATCGGAAG
39399 TATGAGTGTGGAGTCGTTGCACTCAAATCTCTTCCCCAGTGGANNNAGATCGGAAG
102777 TATGAGTGTGGAGTCGTTGCATGAGAAGCTGAATGCAAGAGCTCTANNNAGATCG
48795 TATGAGTGTGGAGTCGTTGCGCCAGTCTTCTTATTGGCTCCTCNNNAGATCGGAAGA
103620 TATGAGTGTGGAGTCGTTGCGTGTACTGCAAACTGGCTGGTTNNNAGATCGGAAG
39596 TATGAGTGTGGAGTCGTTGCCTGTAAGGGCACAGGCATAATGANNNAGATCGGAA
32512 TATGAGTGTGGAGTCGTTGCTCTCAGGTAGGGCAACACTGTAANNNAGATCGGAA
42306 TATGAGTGTGGAGTCGTTGCGTAAGTGCTGCTTCGTAAGCAACNNNAGATCGGAA
48045 TATGAGTGTGGAGTCGTTGCCTGTCACTTGCTAGGTCAATTGCNNNAGATCGGAAG
110467 TATGAGTGTGGAGTCGTTGCACAGAGGAAGTGACCAGAGCAANNNAGATCGGAA
39128 TATGAGTGTGGAGTCGTTGCACACCTTTCTGTGAAAAGATGGGANNNAGATCGGA
71830 TATGAGTGTGGAGTCGTTGCGACGTGAGAATTGGAACTCCCTTNNNAGATCGGAA
42037 TATGAGTGTGGAGTCGTTGCGGCAATCCACCAGGTATGCTTTANNNAGATCGGAAG
53115 TATGAGTGTGGAGTCGTTGCGGAGAGAACACAGACCAAAGAGGNNNAGATCGGA
54955 TATGAGTGTGGAGTCGTTGCCATTGGTGCAAAGCAGTATGTGGNNNAGATCGGAA

39426 TATGAGTGTGGAGTCGTTGCAAGTGGGGACATGATTAGGTGGANNNAGATCGGA

106483 TATGAGTGTGGAGTCGTTGCTACTTATTTTCTCTTGCCTGGCCTNNNAGATCGGAAG

62817 TATGAGTGTGGAGTCGTTGCTCTGGGAGATTTTCTGCTCTACTCTNNNAGATCGGA

55820 TATGAGTGTGGAGTCGTTGCGAGGACATTTTGAGTTGGCACTGNNNAGATCGGAA

47285 TATGAGTGTGGAGTCGTTGCAGTGAACAAGTTGAGATGCTGGCNNNAGATCGGAA

103715 TATGAGTGTGGAGTCGTTGCGATGCAGGGGGTTGGTTTCAGNNNAGATCGGAAG

33712 TATGAGTGTGGAGTCGTTGCCTGGGTAACTGCAATCTGTGACANNNAGATCGGAA

108235 TATGAGTGTGGAGTCGTTGCGGAACTCTCTCTGAAATGTGTGCTTNNNAGATCGGA

49662 TATGAGTGTGGAGTCGTTGCGGAAAGACCTCACTTCTGCCTTCNNNAGATCGGAAG

35972 TATGAGTGTGGAGTCGTTGCCTCAAACCCCAGAAAGAGATGCANNNAGATCGGAA

46465 TATGAGTGTGGAGTCGTTGCTGGGGAAATGAAAGACACAGAGCNNNAGATCGGA

40955 TATGAGTGTGGAGTCGTTGCACGGGACAGACAACCTGTAACTANNNAGATCGGAA

42305 TATGAGTGTGGAGTCGTTGCCCCAGGAAACTGACAAAAGCAACNNNAGATCGGAA

37244 TATGAGTGTGGAGTCGTTGCCCTGTCCACAAAACCTGTCTTCANNNAGATCGGAAGA

56702 TATGAGTGTGGAGTCGTTGCGATGATAGAGAAGCAGAGGCGTGNNNAGATCGGA

69366 TATGAGTGTGGAGTCGTTGCCTCCAAAACAGGCCTCTCTGTTTNNNAGATCGGAAG

56253 TATGAGTGTGGAGTCGTTGCTTTCCTCAATGAACCTCCTGCTGNNNAGATCGGAAGA

57044 TATGAGTGTGGAGTCGTTGCCCCAGATGAGAGTTCATGGTGTGNNNAGATCGGAA

31673 TATGAGTGTGGAGTCGTTGCTTGGCCTTGATTGCTAAAGAGAANNNAGATCGGAA

45498 TATGAGTGTGGAGTCGTTGCTATGGTGATCATGCGCTGTATCCNNNAGATCGGAAG

68440 TATGAGTGTGGAGTCGTTGCGGAAACACACCAGGTAAAGGGTTNNNAGATCGGAA

117485 TATGAGTGTGGAGTCGTTGCGCCCTCACCAAGAACCTAATTATACTGNNNAGATCGG

50420 TATGAGTGTGGAGTCGTTGCGGGATAGCAATGAATTGCCGAAGNNNAGATCGGAA

54279 TATGAGTGTGGAGTCGTTGCGCCCTGCTTTCAAATACTTTGGGNNNAGATCGGAAG

39034 TATGAGTGTGGAGTCGTTGCGCTGCTTGGAAAATAAGGAGGGANNNAGATCGGA

66650 TATGAGTGTGGAGTCGTTGCCGCTTTATGAAGTTTTGGCTTGTNNNAGATCGGAAG

69172 TATGAGTGTGGAGTCGTTGCATTTTCTCTTGCCTGGCCTCTTTNNNAGATCGGAAGA

65327 TATGAGTGTGGAGTCGTTGCGGAGGGAGGGAGGAAATTATGGTNNNAGATCGGA

63917 TATGAGTGTGGAGTCGTTGCTTACAAGTTCCCTGTGTGACAGTNNNAGATCGGAAG

48715 TATGAGTGTGGAGTCGTTGCCTGAGCAAATCTGGAATCCCCTCNNNAGATCGGAAG

50388 TATGAGTGTGGAGTCGTTGCAAATCCCTCCATGGGATCAGAAGNNNAGATCGGAA

37343 TATGAGTGTGGAGTCGTTGCTGTGAGACTGTGGTTCATGATTTCANNNAGATCGGA

38343 TATGAGTGTGGAGTCGTTGCTCAAAGGAAAAGGTAACTGTGAGANNNAGATCGGA

62429 TATGAGTGTGGAGTCGTTGCACAAGAGGGAGTATTATTTTGTGCTNNNAGATCGG

53233 TATGAGTGTGGAGTCGTTGCCTCTCTCTCTCTCAGTTGGGAGGNNNAGATCGGAAG

30564 TATGAGTGTGGAGTCGTTGCTATTCATTTTGGGGTGGGCGAAANNNAGATCGGAA

98341 TATGAGTGTGGAGTCGTTGCAACTACTGAACCGCCAGGCATNNNAGATCGGAAGA

100754 TATGAGTGTGGAGTCGTTGCACACACATACTGTGGAACATTTTCNNNAGATCGGAA

41589 TATGAGTGTGGAGTCGTTGCGCCTGGTTGTGTCTTGATTGGTANNNAGATCGGAA

54554 TATGAGTGTGGAGTCGTTGCCGGTTTCCTTGGATAGTGACTGGNNNAGATCGGAA

65434 TATGAGTGTGGAGTCGTTGCGAAAATGCCACAGAGTGAGTGGTNNNAGATCGGAA

112962 TATGAGTGTGGAGTCGTTGCGTGAGATGCTGCTCCTGTTTACCNNNAGATCGGAAG

32426 TATGAGTGTGGAGTCGTTGCAGAGAGAGAGGGAGACGCTCTAANNNAGATCGGA

| | |
|---|---|
| 108257 | TATGAGTGTGGAGTCGTTGCTGAGAGCCCCATAATATGCTGTCTNNNAGATCGGAA |
| 61028 | TATGAGTGTGGAGTCGTTGCTCGGAGAAATCAAGTCAGTCCCTNNNAGATCGGAAG |
| 36173 | TATGAGTGTGGAGTCGTTGCGGGAGGTGTACAATTCATGTGCANNNAGATCGGAA |
| 51740 | TATGAGTGTGGAGTCGTTGCGACACTCCTAAACAGCAGTGGAGNNNAGATCGGAA |
| 68987 | TATGAGTGTGGAGTCGTTGCTTGAAGGGTTACTATTTTGTGACTTTNNNAGATCGG |
| 31530 | TATGAGTGTGGAGTCGTTGCAAAATGAGCCACCAGAGAAAGAANNNAGATCGGAA |
| 57431 | TATGAGTGTGGAGTCGTTGCACAAAAGAAGGTGAGGGGAGTTGNNNAGATCGGA |
| 59609 | TATGAGTGTGGAGTCGTTGCGGGATATAGTCGGCCCTTCGTATNNNAGATCGGAAG |
| 108313 | TATGAGTGTGGAGTCGTTGCGCTTATCCAGGGTCCCTTTTGGNNNAGATCGGAAGA |
| 108306 | TATGAGTGTGGAGTCGTTGCGAGAAACCTGGAGGGCATTATGTTTNNNAGATCGG |
| 29908 | TATGAGTGTGGAGTCGTTGCGCTGGGGGTTTCAACATAAAACCNNNAGATCGGAAG |
| 35056 | TATGAGTGTGGAGTCGTTGCCCCGTAAGGATTCTTTGCTTCCANNNAGATCGGAAG |
| 63522 | TATGAGTGTGGAGTCGTTGCCCCCCACACTCTAAATGCTTTCTNNNAGATCGGAAGA |
| 38011 | TATGAGTGTGGAGTCGTTGCCACGTTGTTTGGGGGATATCAGANNNAGATCGGAA |
| 30842 | TATGAGTGTGGAGTCGTTGCTGTGAGCACTGACTTGTGAACAANNNAGATCGGAAG |
| 61457 | TATGAGTGTGGAGTCGTTGCATCCTGACAAGCTATTGGGTCCTNNNAGATCGGAAG |
| 40994 | TATGAGTGTGGAGTCGTTGCATTGTGGCTACCTCAGCTGACTANNNAGATCGGAAG |
| 30844 | TATGAGTGTGGAGTCGTTGCTGGAGGAAGTGGGGAAATAACAANNNAGATCGGA |
| 67329 | TATGAGTGTGGAGTCGTTGCTCAGTGGAGAATCCCTGTGGATTNNNAGATCGGAAG |
| 57430 | TATGAGTGTGGAGTCGTTGCAGGCACTTCTCATGTGAGAGTTGNNNAGATCGGAAG |
| 46336 | TATGAGTGTGGAGTCGTTGCTCCAAATGTATCAGGCTATCCAGCNNNAGATCGGAA |
| 34469 | TATGAGTGTGGAGTCGTTGCGCATATTAGAAGCGTAAAGTCCCANNNAGATCGGAA |
| 95133 | TATGAGTGTGGAGTCGTTGCATCAGGCCTTCCCTCTTCTGATTNNNAGATCGGAAGA |
| 58800 | TATGAGTGTGGAGTCGTTGCCCTCTCTGAACTTTGCCACTCATNNNAGATCGGAAGA |
| 48688 | TATGAGTGTGGAGTCGTTGCATTCTTGGTGAAGGCCATACCTCNNNAGATCGGAAG |
| 63407 | TATGAGTGTGGAGTCGTTGCGATTGATGGGGATTTGGGCTTCTNNNAGATCGGAA |
| 66458 | TATGAGTGTGGAGTCGTTGCTTTTCTTGAGGGTCTTGCTGTGTNNNAGATCGGAAG |
| 51409 | TATGAGTGTGGAGTCGTTGCAACTCCTCTTACCAGAGCCAGAGNNNAGATCGGAAG |
| 102028 | TATGAGTGTGGAGTCGTTGCACTCCTCTTACCAGAGCCAGAGNNNAGATCGGAAGA |
| 102091 | TATGAGTGTGGAGTCGTTGCATTGCTGTACCTTTTGCTGAATTCAANNNAGATCGGA |
| 45392 | TATGAGTGTGGAGTCGTTGCCCCCAGAAAAGTGGCTAGAATCCNNNAGATCGGAAG |
| 103738 | TATGAGTGTGGAGTCGTTGCTGTTTTCTCGTTTGAGACGGAATNNNAGATCGGAAG |
| 46316 | TATGAGTGTGGAGTCGTTGCTAAAAGACCTCTCCCTTCCCAGCNNNAGATCGGAAGA |
| 68741 | TATGAGTGTGGAGTCGTTGCAGGATTGTGGGTTGACTGTTGTTNNNAGATCGGAA |
| 67959 | TATGAGTGTGGAGTCGTTGCGAGGGCACTGTTTATAGCTGCTTNNNAGATCGGAAG |
| 32380 | TATGAGTGTGGAGTCGTTGCCACGGATTGCTTGCAAAAGCTAANNNAGATCGGAAG |
| 107526 | TATGAGTGTGGAGTCGTTGCTGATAGAAACATTTTGGCATTAGCTNNNAGATCGGA |
| 115965 | TATGAGTGTGGAGTCGTTGCCTATCTCTCTCTTTCTTGGAGGCTGNNNAGATCGGAA |
| 117500 | TATGAGTGTGGAGTCGTTGCCACCATCTTCCCCAAAGATCACATGNNNAGATCGGAA |
| 96870 | TATGAGTGTGGAGTCGTTGCCCCCTCGCCCAATCTCTACAAANNNAGATCGGAAGAG |
| 58375 | TATGAGTGTGGAGTCGTTGCGGAGCAGGAGACAAAGAGGACATNNNAGATCGGA |
| 101539 | TATGAGTGTGGAGTCGTTGCTTTCACTCACCCGTAGCTTGGNNNAGATCGGAAGAG |
| 52496 | TATGAGTGTGGAGTCGTTGCGCTATTCAACCCCTCTTAGAGCGNNNAGATCGGAAG |

53444 TATGAGTGTGGAGTCGTTGCTTAGGGGAGGATACAGGAAACGGNNNAGATCGGA
34123 TATGAGTGTGGAGTCGTTGCTAACTGCCTTCTGAAAGGGACCANNNAGATCGGAAG
54527 TATGAGTGTGGAGTCGTTGCTTCATTCTGTGAGGCATCACTGGNNNAGATCGGAAG
52815 TATGAGTGTGGAGTCGTTGCTTCTTGCATTTCCCTTGGGAAGGNNNAGATCGGAAG
53440 TATGAGTGTGGAGTCGTTGCATCCTTTTCACGGTGGCTTTAGGNNNAGATCGGAAG
99679 TATGAGTGTGGAGTCGTTGCATAAGGAAGTCTGGCCCTGTAAGNNNAGATCGGAA
51533 TATGAGTGTGGAGTCGTTGCGGATCAAGGGGCCTCTCTAGAGNNNAGATCGGAAG
58993 TATGAGTGTGGAGTCGTTGCTCCCAGAAGCATCGTATCCAGATNNNAGATCGGAAG
50758 TATGAGTGTGGAGTCGTTGCCCACCCCCTGTGAGATACTACAGNNNAGATCGGAAG
35726 TATGAGTGTGGAGTCGTTGCAGAGAGAGAATAATGTGGGGGCANNNAGATCGGA
57165 TATGAGTGTGGAGTCGTTGCGGAGGAGGAAACCTTAGCCATTGNNNAGATCGGAA
103600 TATGAGTGTGGAGTCGTTGCTAAGGATTCCAAACTACTCCAAACANNNAGATCGGA
34356 TATGAGTGTGGAGTCGTTGCTAAGTCCTCACACTTGTTCCCCANNNAGATCGGAAGA
49822 TATGAGTGTGGAGTCGTTGCAGTCAGCCACAACTACCATGTTCNNNAGATCGGAAG
71720 TATGAGTGTGGAGTCGTTGCATCAATCAAGACCTCGACGCTTTNNNAGATCGGAAG
45781 TATGAGTGTGGAGTCGTTGCTTATTACTCGCCTGGCAATGTCCNNNAGATCGGAAG
39867 TATGAGTGTGGAGTCGTTGCAGTCTCATTTCCTATTTCCAACCTGANNNAGATCGGA
68063 TATGAGTGTGGAGTCGTTGCACACAGGCTCTCCATCTACTCTTNNNAGATCGGAAGA
43833 TATGAGTGTGGAGTCGTTGCCTCACAGAGCTTGGGTTTGAACCNNNAGATCGGAAG
56653 TATGAGTGTGGAGTCGTTGCGTGAAAGTGGTGGCTGTGAGTGNNNAGATCGGAA
35705 TATGAGTGTGGAGTCGTTGCATTGGCTAACATTCTACAGGGCANNNAGATCGGAAG
114141 TATGAGTGTGGAGTCGTTGCTGTGTGTGTGAATGTGCTCTACANNNAGATCGGAA
44212 TATGAGTGTGGAGTCGTTGCTGCTGGCTCATTCTTCCTTTACCNNNAGATCGGAAGA
108310 TATGAGTGTGGAGTCGTTGCGGGTAGGGGCTGTAAGTAAACTNNNAGATCGGAA
50469 TATGAGTGTGGAGTCGTTGCTGCTTATGCCAGTTGACTGGAAGNNNAGATCGGAA
46315 TATGAGTGTGGAGTCGTTGCATTCCCTTGATGAGATTCCCAGCNNNAGATCGGAAG
98013 TATGAGTGTGGAGTCGTTGCAGAAACTTGAATTTACCTTGGGGGNNNAGATCGGA
59457 TATGAGTGTGGAGTCGTTGCTCACTCACCCGTAGCTTGGATATNNNAGATCGGAAG
42013 TATGAGTGTGGAGTCGTTGCGGACCCGTAGACCAATACCTTTANNNAGATCGGAAG
32990 TATGAGTGTGGAGTCGTTGCAGGATTCCAAACTACTCCAAACACANNNAGATCGGA
97425 TATGAGTGTGGAGTCGTTGCGTTACCACAAAAACTAGGCAACGNNNAGATCGGAA
68653 TATGAGTGTGGAGTCGTTGCTCCTGCATTCTACTGGTTCTGTTNNNAGATCGGAAGA
103928 TATGAGTGTGGAGTCGTTGCTGGGTATGTAGGTGGCTTAGGCNNNAGATCGGAAG
30884 TATGAGTGTGGAGTCGTTGCTCCCCTTTCTCCATTCTCCACAANNNAGATCGGAAGA
57046 TATGAGTGTGGAGTCGTTGCACAATAGGAAAGGCACTGGTGTGNNNAGATCGGAA
108873 TATGAGTGTGGAGTCGTTGCACGGTGGAAGCATGAAGATCTATNNNAGATCGGAA
57748 TATGAGTGTGGAGTCGTTGCGCAGAAGGGAGCAGAAGTCAAATNNNAGATCGGA
96528 TATGAGTGTGGAGTCGTTGCTTTTCCTGCTTCTTAGTGTGGTTTNNNAGATCGGAAG
102440 TATGAGTGTGGAGTCGTTGCGGATCTCTGGCTCTGGTACCNNNAGATCGGAAGAG
30798 TATGAGTGTGGAGTCGTTGCTGTATGCCTGGAGAAACAAACAANNNAGATCGGAA
33891 TATGAGTGTGGAGTCGTTGCACCTTCTCCAGGTTGTCTGTACANNNAGATCGGAAG
36459 TATGAGTGTGGAGTCGTTGCTGACCTCTCCTTCACAGTCATCANNNAGATCGGAAGA
64504 TATGAGTGTGGAGTCGTTGCCTCCTGGACTAGGGCCTCTTAGTNNNAGATCGGAAG

| | |
|---|---|
| 100341 | TATGAGTGTGGAGTCGTTGCGGTCCCCCTACCGGTGGAGGNNNAGATCGGAAGAG |
| 64621 | TATGAGTGTGGAGTCGTTGCTTCCTTACCTTCCTCCATTCCGTNNNAGATCGGAAGA |
| 50419 | TATGAGTGTGGAGTCGTTGCCAGAATTGAGTAGAGGGCCGAAGNNNAGATCGGA |
| 66889 | TATGAGTGTGGAGTCGTTGCTCCAACCAGTGTCTTATCTTTTTGTNNNAGATCGGAA |
| 107602 | TATGAGTGTGGAGTCGTTGCTTCTGCTAAACAACCTCAATGAAAATCNNNAGATCGG |
| 41409 | TATGAGTGTGGAGTCGTTGCTCATGTTGCTGGAGCTCAGAGTANNNAGATCGGAA |
| 56159 | TATGAGTGTGGAGTCGTTGCAGAGAAGTCAACAGCTGTAGCTGNNNAGATCGGAA |
| 37653 | TATGAGTGTGGAGTCGTTGCCCTAAGAGGCAGGAAGTGGAAGANNNAGATCGGA |
| 29910 | TATGAGTGTGGAGTCGTTGCCTCTCACTGGGGAAGGTGACTAGNNNAGATCGGAA |
| 37929 | TATGAGTGTGGAGTCGTTGCATGCAGGAAATTTGCAGAGCAGANNNAGATCGGAA |
| 71765 | TATGAGTGTGGAGTCGTTGCGCTTTTCTTCCCAGGAGCTACAGNNNAGATCGGAAG |
| 63196 | TATGAGTGTGGAGTCGTTGCTATCTCTGCTTCCTTGTGGGTCTNNNAGATCGGAAG |
| 47735 | TATGAGTGTGGAGTCGTTGCCACTCTACTCTGCAACTTCCTGCNNNAGATCGGAAGA |
| 68284 | TATGAGTGTGGAGTCGTTGCATTTTTGGAGGAGAGCTGCAGTTNNNAGATCGGAA |
| 55527 | TATGAGTGTGGAGTCGTTGCACTGTTCTTCCTACGAGCTTCATGNNNAGATCGGAA |
| 64247 | TATGAGTGTGGAGTCGTTGCATGATGCCACAACAAACAGGAGTNNNAGATCGGAA |
| 110462 | TATGAGTGTGGAGTCGTTGCGAGAGAAGGCTGATTTCTGCTGNNNAGATCGGAAG |
| 41627 | TATGAGTGTGGAGTCGTTGCATGGGCCAAAGATTGAGACTGTANNNAGATCGGAA |
| 39893 | TATGAGTGTGGAGTCGTTGCCACTAAGGCATCTTTTGCCCTGANNNAGATCGGAAG |
| 66453 | TATGAGTGTGGAGTCGTTGCGGGAAATGGAATGGTTCCTGTGTNNNAGATCGGAA |
| 39316 | TATGAGTGTGGAGTCGTTGCTATCAGCTATCACCCAGACTGGANNNAGATCGGAAG |
| 37486 | TATGAGTGTGGAGTCGTTGCGGTAGTGGGGAGGGAAAGAAAGANNNAGATCGG |
| 44471 | TATGAGTGTGGAGTCGTTGCACATGGCAAAATCATCTCTCCCCNNNAGATCGGAAG |
| 51105 | TATGAGTGTGGAGTCGTTGCTAGCAGGGTGTGATAAATGGCAGNNNAGATCGGA |
| 70408 | TATGAGTGTGGAGTCGTTGCTTCTTTGAAGTGGTGTCCTCAGCNNNAGATCGGAAG |
| 106536 | TATGAGTGTGGAGTCGTTGCTGAAGAAAAATTTTGGAGTAGGACCANNNAGATCG |
| 45129 | TATGAGTGTGGAGTCGTTGCTTCACAACTACTTTGGTCTGGCCNNNAGATCGGAAG |
| 110473 | TATGAGTGTGGAGTCGTTGCTTCAAAACCAACCTATTTGACAGCATNNNAGATCGG |
| 38887 | TATGAGTGTGGAGTCGTTGCAAGCCAAGAAAAGAACCACTAGGANNNAGATCGGA |
| 37259 | TATGAGTGTGGAGTCGTTGCCACACCCCACATCTTCAAGTTCANNNAGATCGGAAGA |
| 55118 | TATGAGTGTGGAGTCGTTGCAATGTCAACTGTACCGAGGTTGGNNNAGATCGGAA |
| 63350 | TATGAGTGTGGAGTCGTTGCCTCCTTCCGGGTTCAGCTATTCTNNNAGATCGGAAG |
| 45714 | TATGAGTGTGGAGTCGTTGCATGGTTGGCTCATTCTCAAGTCCNNNAGATCGGAAG |
| 50326 | TATGAGTGTGGAGTCGTTGCAGGGAATGGCCAGTTGAATCAAGNNNAGATCGGAA |
| 117583 | TATGAGTGTGGAGTCGTTGCCGTCGGCATCTACACCAATGTNNNAGATCGGAAGAG |
| 47751 | TATGAGTGTGGAGTCGTTGCTCCCAAGCTTTATTCTCAGCTGCNNNAGATCGGAAG |
| 102424 | TATGAGTGTGGAGTCGTTGCAGATGCTGTAGGCTCACCCTGNNNAGATCGGAAGA |
| 103719 | TATGAGTGTGGAGTCGTTGCCCCGACTTTCCTTTTATCCAAAGAGNNNAGATCGGAA |
| 32031 | TATGAGTGTGGAGTCGTTGCCTCACTCTGCGGAGAAACATGAANNNAGATCGGAA |
| 117886 | TATGAGTGTGGAGTCGTTGCCCCCAAATCAACAGTGGCTAATAANNNAGATCGGAA |
| 98419 | TATGAGTGTGGAGTCGTTGCATGGTGATTTGTGCCTGAAATCTCNNNAGATCGGAA |
| 49880 | TATGAGTGTGGAGTCGTTGCCACTTGCCATGTTCCTGGATTTCNNNAGATCGGAAG |
| 102449 | TATGAGTGTGGAGTCGTTGCCAAGTCCCAGGCTACAGTGGTTCNNNAGATCGGAAG |

65234 TATGAGTGTGGAGTCGTTGCTGGAAAACCTTGATTCTTTTGGGTNNNAGATCGGAA
107634 TATGAGTGTGGAGTCGTTGCAGTGATTTTCCTGCTTCTGCGNNNAGATCGGAAGAG
67985 TATGAGTGTGGAGTCGTTGCAAACCTGTGAGGTGAAATTGCTTNNNAGATCGGAA
52836 TATGAGTGTGGAGTCGTTGCCACAACTTAGACTGGGCTGAAGGNNNAGATCGGAA
56778 TATGAGTGTGGAGTCGTTGCTAGGATTGGGATTTTGCAGGGTGNNNAGATCGGAA
57741 TATGAGTGTGGAGTCGTTGCGCTCCACTGTGTGTTTTGCAAATNNNAGATCGGAAG
35124 TATGAGTGTGGAGTCGTTGCCTCTCTCTCCCCCTCTTTTTCCANNNAGATCGGAAGAG
57329 TATGAGTGTGGAGTCGTTGCAAGATGACTTCAGAGGCTGCTTGNNNAGATCGGAA
32236 TATGAGTGTGGAGTCGTTGCGCAAGCACACCTGAGAGCAATAANNNAGATCGGAA
104413 TATGAGTGTGGAGTCGTTGCTACTCTGCAACTTCCTGCTTTTTNNNAGATCGGAAGA
34443 TATGAGTGTGGAGTCGTTGCTGCCAAGTCCAGTTTTTATCCCANNNAGATCGGAAG
116013 TATGAGTGTGGAGTCGTTGCCTTTAGGGCTGTCCCTCCTGGNNNAGATCGGAAGAG
29907 TATGAGTGTGGAGTCGTTGCTCTAAAAATGAACTACCCCTGAACANNNAGATCGGA
52942 TATGAGTGTGGAGTCGTTGCTATTTGCACTCTGGAAACCCAGGNNNAGATCGGAAG
31639 TATGAGTGTGGAGTCGTTGCTACCAGGGCAGGAATAGGCAGAANNNAGATCGGA
54327 TATGAGTGTGGAGTCGTTGCTCTAGGGGAAGGGAACAGAATGGNNNAGATCGGA
103642 TATGAGTGTGGAGTCGTTGCCAGGGTGTGATAAATGGCAGACTACNNNAGATCGG
103694 TATGAGTGTGGAGTCGTTGCGTCCTTGAACTCCTCAGCTCAAACNNNAGATCGGAA
52846 TATGAGTGTGGAGTCGTTGCACAGTCCAGAGACATCTTTGAAGGNNNAGATCGGA
39367 TATGAGTGTGGAGTCGTTGCTGTATGTCCATTTATCTGTTGCTGGANNNAGATCGG
40192 TATGAGTGTGGAGTCGTTGCTCCATTGGTTGACACTCAGGTGANNNAGATCGGAAG
57975 TATGAGTGTGGAGTCGTTGCAGTTCTCCAAGTGGTTGTGCAATNNNAGATCGGAAG
36931 TATGAGTGTGGAGTCGTTGCAGGGATGGTATGAGATCAAGGTCANNNAGATCGGA
69362 TATGAGTGTGGAGTCGTTGCTGACAGTAGTCTCCATGCTGTTTNNNAGATCGGAAG
101331 TATGAGTGTGGAGTCGTTGCAATCAACCACCAATCTCTGGCCATNNNAGATCGGAA
66704 TATGAGTGTGGAGTCGTTGCATTTTGGAGTAGGACCACGTTGTNNNAGATCGGAA
105917 TATGAGTGTGGAGTCGTTGCAATCTCCATTGTTCACAGTAACCTNNNAGATCGGAAG
57515 TATGAGTGTGGAGTCGTTGCTCACCAGTAGGTTCCCTTTGTTGNNNAGATCGGAAG
46761 TATGAGTGTGGAGTCGTTGCGCTCTAATCTTCCTTAGCTCCGCNNNAGATCGGAAGA
39234 TATGAGTGTGGAGTCGTTGCTCTGAAAGGAAAGGTAGAGAATGGANNNAGATCG
37926 TATGAGTGTGGAGTCGTTGCTTGCTTTCTAACCATGCAGCAGANNNAGATCGGAAG
65821 TATGAGTGTGGAGTCGTTGCCAGCAGTCCCTCGAACAATATGTNNNAGATCGGAAG
29975 TATGAGTGTGGAGTCGTTGCATTCTCCCACACATGGAGTTCTGNNNAGATCGGAAG
29903 TATGAGTGTGGAGTCGTTGCATGTTAGTTGAGGAGGAAAGGGTNNNAGATCGGA
56750 TATGAGTGTGGAGTCGTTGCTTGAACCAAACAGTGGAGAGGTGNNNAGATCGGAA
48806 TATGAGTGTGGAGTCGTTGCATTGTCTGAAAAACGGGGTCCTCNNNAGATCGGAA
39044 TATGAGTGTGGAGTCGTTGCTAAAACCCTCTTCCAGTGAGGGANNNAGATCGGAAG
47503 TATGAGTGTGGAGTCGTTGCAGAGAGAGACTGCATTGACATGCNNNAGATCGGAA
59682 TATGAGTGTGGAGTCGTTGCCATCACCTCAGGGCTCCCTTATNNNAGATCGGAAGA
29976 TATGAGTGTGGAGTCGTTGCAGACTTTTCCTGCACCTTGTGAGNNNAGATCGGAAG
71829 TATGAGTGTGGAGTCGTTGCTTGCCATAAACTGCCTTTTCAGCNNNAGATCGGAAG
71761 TATGAGTGTGGAGTCGTTGCGCCCACAGCATAAACAAAGTTGGNNNAGATCGGAA
102138 TATGAGTGTGGAGTCGTTGCGAGAATCTTGCTTGTTACCCAGGNNNAGATCGGAAG

| | |
|---|---|
| 101590 | TATGAGTGTGGAGTCGTTGCGCTACCACCAATACATATTTGTAGGGNNNAGATCGG |
| 71681 | TATGAGTGTGGAGTCGTTGCAGTTCTCTAGCATTCTGTCTTTCGTNNNAGATCGGAA |
| 102457 | TATGAGTGTGGAGTCGTTGCGACCATAGACACATACCACTACACCNNNAGATCGGA |
| 106441 | TATGAGTGTGGAGTCGTTGCGCAAGGGCAATCCAGAAATCAGANNNAGATCGGAA |
| 102093 | TATGAGTGTGGAGTCGTTGCGCAGGCTTGTCACCATTATCAATNNNAGATCGGAAG |
| 95491 | TATGAGTGTGGAGTCGTTGCTTGGCCTTCACATTAGACTGTAGAGNNNAGATCGGA |
| 105962 | TATGAGTGTGGAGTCGTTGCCCTGGCCCACAGTAGATTGTTTCNNNAGATCGGAAG |
| 101224 | TATGAGTGTGGAGTCGTTGCCTATCAGCAGCTTTTACATACGGCTNNNAGATCGGA |
| 102442 | TATGAGTGTGGAGTCGTTGCGAGAGTAGCTGAGGTTCAGAGAGANNNAGATCGG |
| 102451 | TATGAGTGTGGAGTCGTTGCGAAAGCCAACATTTCTGTGAAATTGTNNNAGATCGG |
| 108284 | TATGAGTGTGGAGTCGTTGCTCTATCCCTCCCCTGACAAGACNNNAGATCGGAAGA |
| 71711 | TATGAGTGTGGAGTCGTTGCTGCTTCTTGGCAGGACTTTGAAANNNAGATCGGAAG |
| 103762 | TATGAGTGTGGAGTCGTTGCTCAACGTGTCCCCCTAAATTCATGNNNAGATCGGAA |
| 101112 | TATGAGTGTGGAGTCGTTGCGTTGCTTTCATTCAGTTGATCCAGACNNNAGATCGG |
| 102126 | TATGAGTGTGGAGTCGTTGCGTGCCTGGCCTATTGATCTAAATNNNAGATCGGAAG |
| 101137 | TATGAGTGTGGAGTCGTTGCGTTAGAAGAACTGGCGTGTCTGCNNNAGATCGGAA |
| 107579 | TATGAGTGTGGAGTCGTTGCCTTCAACACACAATTCCTAATATGCCNNNAGATCGGA |
| 116158 | TATGAGTGTGGAGTCGTTGCCGCACCTGTCTTAGCCTCTGAAANNNAGATCGGAAG |
| 103655 | TATGAGTGTGGAGTCGTTGCACATTGAGTAAGCTCTGGAAAGTNNNAGATCGGAA |
| 100741 | TATGAGTGTGGAGTCGTTGCCCAGGTAGTGTACATTTTACCCAGTNNNAGATCGGA |
| 107617 | TATGAGTGTGGAGTCGTTGCTCCCTAGGCCTTACCTAAACTACNNNAGATCGGAAGA |
| 102088 | TATGAGTGTGGAGTCGTTGCGGGCAATGCATAAGTGTTCACATTTNNNAGATCGGA |
| 101307 | TATGAGTGTGGAGTCGTTGCTTCGGTCTGTTTGTGGCATATCNNNAGATCGGAAGA |
| 110509 | TATGAGTGTGGAGTCGTTGCCGCCGGACCACTTGTTTGTTTTGTTTTNNNAGATCGG |
| 101523 | TATGAGTGTGGAGTCGTTGCTCAGAAACAGAAAGACAAAAGCCACANNNAGATCG |
| 102184 | TATGAGTGTGGAGTCGTTGCACCGTGCCTAGCTAATCCATTTNNNAGATCGGAAGA |
| 104391 | TATGAGTGTGGAGTCGTTGCGAAAGTTGATTTTGCCAAGGCTAATNNNAGATCGGA |
| 101354 | TATGAGTGTGGAGTCGTTGCTGGAAAGCTGGAGGTTCAGGANNNAGATCGGAAG |
| 103605 | TATGAGTGTGGAGTCGTTGCGGAGTTCAGTTCAAAGCCATCAGANNNAGATCGGA |
| 102409 | TATGAGTGTGGAGTCGTTGCCACTAAGCCCAGCTCAGCATGATNNNAGATCGGAAG |
| 112985 | TATGAGTGTGGAGTCGTTGCAAGAGGAAGCTAATCTGTGAGGATACNNNAGATCG |
| 108885 | TATGAGTGTGGAGTCGTTGCACAAGAGGCTTCTGCCTATCCCNNNAGATCGGAAGA |
| 100523 | TATGAGTGTGGAGTCGTTGCAGTTTCCTGAGGGTAGGCTTTTTNNNAGATCGGAAG |
| 105952 | TATGAGTGTGGAGTCGTTGCTTTGGGAGACGCATCTCGTGTATNNNAGATCGGAA |
| 110505 | TATGAGTGTGGAGTCGTTGCTAAGTGGAAAACACAGTTCTCCTGGNNNAGATCGGA |
| 100374 | TATGAGTGTGGAGTCGTTGCTTTCACATTGTCCCACCTTTCTGNNNAGATCGGAAGA |
| 96453 | TATGAGTGTGGAGTCGTTGCAAGCCTGGAACTAAGTGAAAACAGNNNAGATCGGA |
| 98128 | TATGAGTGTGGAGTCGTTGCGGAAGGACGGGTTACAGAGGAGANNNAGATCGGA |
| 39289 | TATGAGTGTGGAGTCGTTGCTCTGACACTGAGCACACTATGGANNNAGATCGGAAG |
| 62634 | TATGAGTGTGGAGTCGTTGCAGAAAGAGACCTGAAGGGCATCTNNNAGATCGGAA |
| 98619 | TATGAGTGTGGAGTCGTTGCTTACTCCTAACCCTCAGGAAGCCNNNAGATCGGAAG |
| 59577 | TATGAGTGTGGAGTCGTTGCGTGCGTGCATCTTTCTAAAGTATNNNAGATCGGAAG |
| 105846 | TATGAGTGTGGAGTCGTTGCTTCTGGGGAGTATGCAAATTTCTCNNNAGATCGGAA |

108906 TATGAGTGTGGAGTCGTTGCGAAGCTGGCATATATTCTCTCACAGNNNAGATCGGA
36767 TATGAGTGTGGAGTCGTTGCAGGGAGTGGCATAATGGATCTCANNNAGATCGGAA
30579 TATGAGTGTGGAGTCGTTGCTGTAGGTTCCGAATATTTCAGGAAANNNAGATCGGA
61388 TATGAGTGTGGAGTCGTTGCATGACTGGTTTAGGCTTCCTCCTNNNAGATCGGAAG
97759 TATGAGTGTGGAGTCGTTGCAATACAAGCATGTGAAAGACCTCNNNAGATCGGAA
50651 TATGAGTGTGGAGTCGTTGCGATGCACAGAGGTACAGACACAGNNNAGATCGGAA
48001 TATGAGTGTGGAGTCGTTGCTTCAAGTGTATCAGGCTCTGTGCNNNAGATCGGAAG
55031 TATGAGTGTGGAGTCGTTGCTCCAATCAGCCATTGATCCATTGGNNNAGATCGGAA
36481 TATGAGTGTGGAGTCGTTGCAAACGTCAAAATCCACAGGATCANNNAGATCGGAAG
63064 TATGAGTGTGGAGTCGTTGCATCAAGCCTGGAAGTTTGTCTCTNNNAGATCGGAAG
41870 TATGAGTGTGGAGTCGTTGCGAAAACCAGCCACACACCTCTTANNNAGATCGGAAG
116448 TATGAGTGTGGAGTCGTTGCGCCTCTCTCCTCCTTCCTTTCCANNNAGATCGGAAGA
96781 TATGAGTGTGGAGTCGTTGCATAAAAAGTAGAAGTTGGGGCTGGNNNAGATCGG
48142 TATGAGTGTGGAGTCGTTGCCAAGCCATTCCAATCCTTCTTGCNNNAGATCGGAAGA
116858 TATGAGTGTGGAGTCGTTGCCTCTCACTCTCGTTGGTAAGGTTTANNNAGATCGGA
97543 TATGAGTGTGGAGTCGTTGCGTCTCTCTAGTTTCCAGTTGACACCNNNAGATCGGAA
97542 TATGAGTGTGGAGTCGTTGCGGACATCATTTTCTTTCTTCAACGCNNNAGATCGGAA
53372 TATGAGTGTGGAGTCGTTGCCCCCTTTATGAAATCTGCCCTAGGNNNAGATCGGAA
32802 TATGAGTGTGGAGTCGTTGCTACAGCAAGAAGGCATGACAACANNNAGATCGGAA
41166 TATGAGTGTGGAGTCGTTGCCCCACCTCTCTGAACAGTAGCTANNNAGATCGGAAG
98741 TATGAGTGTGGAGTCGTTGCCTTGTGTTTGGGTCTTACTAGTTACAANNNAGATCG
35694 TATGAGTGTGGAGTCGTTGCGAGTAAGTGAATTTTGTCCGGCANNNAGATCGGAA
66255 TATGAGTGTGGAGTCGTTGCGGTAGCCACAGTTCTATTTTCTGTNNNAGATCGGAA
49812 TATGAGTGTGGAGTCGTTGCTTACATATTCCGTGGCCTGGTTCNNNAGATCGGAAG
117753 TATGAGTGTGGAGTCGTTGCCCCCAAAAGCCTTGTATAGACAGNNNAGATCGGAAG
62309 TATGAGTGTGGAGTCGTTGCTTGTTGACCTGTTCTTTCCTGCTNNNAGATCGGAAGA
107409 TATGAGTGTGGAGTCGTTGCTGAGGTCTTGCGCAAAATAAAGANNNAGATCGGAA
57175 TATGAGTGTGGAGTCGTTGCTACTCAATCTCAGGGCAGCATTGNNNAGATCGGAAG
31965 TATGAGTGTGGAGTCGTTGCGGGGATTTGAAGCTTGACTGGAANNNAGATCGGAA
64711 TATGAGTGTGGAGTCGTTGCGTGGGAAAGGTCATGGAAAAGGTNNNAGATCGGA
47437 TATGAGTGTGGAGTCGTTGCCCAGGCTGCTTATACGAAAATGCNNNAGATCGGAAG
116893 TATGAGTGTGGAGTCGTTGCCTCTCACATCCCTTGCTCTGCTNNNAGATCGGAAGAG
37195 TATGAGTGTGGAGTCGTTGCCCTGGTTGTTTCTGTTCCCTTCANNNAGATCGGAAGA
31701 TATGAGTGTGGAGTCGTTGCGAGGAGCTAGGTATGCAGGAGAANNNAGATCGGA
68885 TATGAGTGTGGAGTCGTTGCTTCATGGGCAGACACTGAGATTTNNNAGATCGGAAG
40998 TATGAGTGTGGAGTCGTTGCCGCTCAGTGGTTTCATGTGACTANNNAGATCGGAAG
47888 TATGAGTGTGGAGTCGTTGCTGTGTCTGTCTGTCTATCAGTGCNNNAGATCGGAAG
103692 TATGAGTGTGGAGTCGTTGCCTCTTGGAGACAGAGTGTCACTNNNAGATCGGAAG
59648 TATGAGTGTGGAGTCGTTGCCTGGATCCCTGTCCACTTTGTATNNNAGATCGGAAG
66339 TATGAGTGTGGAGTCGTTGCCCCTTCTTGAGTTGGTGTTAGTGTNNNAGATCGGAA
62998 TATGAGTGTGGAGTCGTTGCAAGGTCCCGTTGTCTTTATCTCTNNNAGATCGGAAG
59022 TATGAGTGTGGAGTCGTTGCCCCCTTCGAAGTCTGTTTCAGATNNNAGATCGGAAG
31065 TATGAGTGTGGAGTCGTTGCCGTGTCTATTTGGAATAAGCCAANNNAGATCGGAAG

36453 TATGAGTGTGGAGTCGTTGCTGAAAATGTCTTTGATGGCTCATCANNNAGATCGGA
61273 TATGAGTGTGGAGTCGTTGCACTATGTTCGTCTCCAATTGCCTNNNAGATCGGAAG
45840 TATGAGTGTGGAGTCGTTGCACCAGGGGTATGGGAGTAATTCCNNNAGATCGGAA
96823 TATGAGTGTGGAGTCGTTGCGGGTCACAAAAACATAGACACTGTCNNNAGATCGG
54110 TATGAGTGTGGAGTCGTTGCAACATTACCACTGTGAAGCTGGGNNNAGATCGGAA
103959 TATGAGTGTGGAGTCGTTGCATGTGTGTGTATCCTTTCAGCANNNAGATCGGAAGA
51118 TATGAGTGTGGAGTCGTTGCACCTTTCTTCTCCCATTTGGCAGNNNAGATCGGAAGA
50848 TATGAGTGTGGAGTCGTTGCCTTCCAAAAGAAACATGGCCCAGNNNAGATCGGAAG
40234 TATGAGTGTGGAGTCGTTGCAAAGGAACTGAAAATGCTGGTGANNNAGATCGGAA
38593 TATGAGTGTGGAGTCGTTGCTCTTGATTTGCCTCTCACTTCGANNNAGATCGGAAGA
34214 TATGAGTGTGGAGTCGTTGCAGGATTTGATCAAGGAAGAAAACCCANNNAGATCG
68575 TATGAGTGTGGAGTCGTTGCACACGAGAAACCTGTTAGATGTTNNNAGATCGGAA
42340 TATGAGTGTGGAGTCGTTGCACTTCCTGCTCTCATTCTCTCAACNNNAGATCGGAAG
30805 TATGAGTGTGGAGTCGTTGCCTGGGAAACATCCAGGGAAACAANNNAGATCGGAA
105887 TATGAGTGTGGAGTCGTTGCGGTGATGGCTTCTGTGTTATTACCTNNNAGATCGGA
56373 TATGAGTGTGGAGTCGTTGCCTGGGTAAGGTCTTTTGGCTCTGNNNAGATCGGAA
66913 TATGAGTGTGGAGTCGTTGCGCACGCAAGAGCTCATTCAAATTNNNAGATCGGAAG
51413 TATGAGTGTGGAGTCGTTGCCTTTATGGGACTGGATGCCAGAGNNNAGATCGGAA
56143 TATGAGTGTGGAGTCGTTGCAGACTTTGCAATACTGGGAGCTGNNNAGATCGGAA
63964 TATGAGTGTGGAGTCGTTGCACCAGAGCTATCACTTTCCCAGTNNNAGATCGGAAG
65555 TATGAGTGTGGAGTCGTTGCGTGTGATGTGCATGCAATTTGGTNNNAGATCGGAA
51916 TATGAGTGTGGAGTCGTTGCGAGCTTCCTACCACTGTTGTGAGNNNAGATCGGAAG
62080 TATGAGTGTGGAGTCGTTGCTACAAGCAACACTGTGAATGGCTNNNAGATCGGAA
110299 TATGAGTGTGGAGTCGTTGCTGATTTGCCTCTCACTTCGAATGTNNNAGATCGGAA
31089 TATGAGTGTGGAGTCGTTGCGGTGAGAAAGTTTATGGGGCCAANNNAGATCGGA
41677 TATGAGTGTGGAGTCGTTGCCTTCACCAGACATTGGCTGTGTANNNAGATCGGAAG
43104 TATGAGTGTGGAGTCGTTGCTATGCAGTGAGACAGGGTTTCACNNNAGATCGGAA
53929 TATGAGTGTGGAGTCGTTGCTTTTGGTTTGGCATTAGAGGGGGNNNAGATCGGAA
57779 TATGAGTGTGGAGTCGTTGCGGGAAACAGCCTGCACTAGAAATNNNAGATCGGAA
38428 TATGAGTGTGGAGTCGTTGCAGGTGTGTAGGACGTTTTCTCTAGANNNAGATCGG
37572 TATGAGTGTGGAGTCGTTGCGTCTGAGTTTTGCCAGAGCAAGANNNAGATCGGAA
33378 TATGAGTGTGGAGTCGTTGCAGCAGGTATTTCCTAGGCTCACANNNAGATCGGAAG
40016 TATGAGTGTGGAGTCGTTGCAGGGTCTCCTAAAATGTCATCTGANNNAGATCGGAA
49772 TATGAGTGTGGAGTCGTTGCGAGGAGGAGGTGTGTTTCAGTTCNNNAGATCGGAA
107594 TATGAGTGTGGAGTCGTTGCAAGGCCCCAAAGAGACCACTAANNNAGATCGGAAG
64314 TATGAGTGTGGAGTCGTTGCGGTGGTGATTCTAAAGGACATGAGTNNNAGATCGG
50305 TATGAGTGTGGAGTCGTTGCCAGAAAGCTGACAAAGTGGCAAGNNNAGATCGGAA
102366 TATGAGTGTGGAGTCGTTGCTCTGCTCTGGGTGTATATGTGTGATANNNAGATCGG
67578 TATGAGTGTGGAGTCGTTGCTTGAGTGAATCAGACATGGACTTNNNAGATCGGAA
64927 TATGAGTGTGGAGTCGTTGCTCACTCACATACACACTACTGAGGTNNNAGATCGGA
116894 TATGAGTGTGGAGTCGTTGCCCCCCTTCCAAAGATGGAGAGTANNNAGATCGGAAG
43967 TATGAGTGTGGAGTCGTTGCTCTCCCACTCCCATATCATCACCNNNAGATCGGAAGA
63236 TATGAGTGTGGAGTCGTTGCTGAAAGGAAAGCAGTAAAACTGTCTNNNAGATCGG

53134 TATGAGTGTGGAGTCGTTGCGATGGATGCCAAGGTTACAGAGGNNNAGATCGGAA
48605 TATGAGTGTGGAGTCGTTGCCTTGAGCTTCCCATTACCCACTCNNNAGATCGGAAGA
49711 TATGAGTGTGGAGTCGTTGCCTGAAGCCCCTGTATACTGCTTCNNNAGATCGGAAGA
110449 TATGAGTGTGGAGTCGTTGCATCTCTCTTTTCTGCTTCCTGGTNNNAGATCGGAAGA
51560 TATGAGTGTGGAGTCGTTGCATAAGTTTTAGCCCACAGGCGAGNNNAGATCGGAA
63559 TATGAGTGTGGAGTCGTTGCTTGGCTATGACTGCTGTGTTTCTNNNAGATCGGAAGA
38809 TATGAGTGTGGAGTCGTTGCTTTCTGCGTGTTTTGTGAGAGGANNNAGATCGGAA
38300 TATGAGTGTGGAGTCGTTGCCCAAGAGATGAGAGACCTCTATGAGANNNAGATCG
54865 TATGAGTGTGGAGTCGTTGCTGCTGCCTTTGGAATCATAGTGGNNNAGATCGGAA
49931 TATGAGTGTGGAGTCGTTGCCCTCATGCCCTCATGTCTCTTTCNNNAGATCGGAAGA
95125 TATGAGTGTGGAGTCGTTGCCTTCCCGCCCATGCTGTTATTATNNNAGATCGGAAGA
41546 TATGAGTGTGGAGTCGTTGCTTATGCTCCGGGATACATGGGTANNNAGATCGGAA
58404 TATGAGTGTGGAGTCGTTGCACTTACGGCACCATCATATACATNNNAGATCGGAAGA
42644 TATGAGTGTGGAGTCGTTGCAGCACCTTAGCTCTCTGGTACACNNNAGATCGGAAGA
54091 TATGAGTGTGGAGTCGTTGCGTCAGCTCTTATGTTACACCTGGGNNNAGATCGGAA
55271 TATGAGTGTGGAGTCGTTGCCTGATGGTGAGAGGTTCCCAATGNNNAGATCGGAA
107393 TATGAGTGTGGAGTCGTTGCTGGAGAAGGAGCAAAAGTCTACTNNNAGATCGGAA
38474 TATGAGTGTGGAGTCGTTGCAGTTTGTTTACCACATGTCCTTAGANNNAGATCGGAA
62032 TATGAGTGTGGAGTCGTTGCGCTAGAATATGAATGTCATGAGGGCTNNNAGATCG
45723 TATGAGTGTGGAGTCGTTGCAGCACCCCAGAGATATACAGTCCNNNAGATCGGAAG
100023 TATGAGTGTGGAGTCGTTGCTGGCACATGGATCTTGGACATCANNNAGATCGGAA
32015 TATGAGTGTGGAGTCGTTGCGGGAGACAATGGACAGTTTGGAANNNAGATCGGAA
66017 TATGAGTGTGGAGTCGTTGCCCCTCAAACAAAGGGAATCCTGTNNNAGATCGGAAG
64363 TATGAGTGTGGAGTCGTTGCTGGTTGGATGTAATGATTGTGAGTNNNAGATCGGAA
48173 TATGAGTGTGGAGTCGTTGCCTCTCTCTATGCCACAGTGTTGCNNNAGATCGGAAGA
43302 TATGAGTGTGGAGTCGTTGCACCTTGGGGGTTATCATAGGGACNNNAGATCGGAA
108209 TATGAGTGTGGAGTCGTTGCTGTGTGTGTGTCTTGAGGGAAAGNNNAGATCGGAA
67408 TATGAGTGTGGAGTCGTTGCGCCCATACACACAGCTGTCTATTNNNAGATCGGAAGA
71745 TATGAGTGTGGAGTCGTTGCGGCAGAAGAAGAGGATGTTCAGGNNNAGATCGGAA
49892 TATGAGTGTGGAGTCGTTGCCAGAACAGGGAGCTGAGACTTTCNNNAGATCGGAA
47263 TATGAGTGTGGAGTCGTTGCTAGAACCCTCCTTCCTTACTGGCNNNAGATCGGAAGA
32350 TATGAGTGTGGAGTCGTTGCCCCGCGTTCCTTTGAATACCTAANNNAGATCGGAAGA
32384 TATGAGTGTGGAGTCGTTGCCCCTGTGGAGTAGGAGAAGCTAANNNAGATCGGAA
32217 TATGAGTGTGGAGTCGTTGCCCCAAAGATCCTCCTGCTTTGAANNNAGATCGGAAGA
40435 TATGAGTGTGGAGTCGTTGCAGTGTGGTATCAAGGAGGCTTGANNNAGATCGGAA
68728 TATGAGTGTGGAGTCGTTGCTTGCTACAGTTCTGTGCCTTGTTNNNAGATCGGAAGA
96770 TATGAGTGTGGAGTCGTTGCAACTCCCTAACAGTTCTCGCTTCNNNAGATCGGAAGA
42447 TATGAGTGTGGAGTCGTTGCGCTCCCACAGCACCTAGAATAACNNNAGATCGGAAGA
65172 TATGAGTGTGGAGTCGTTGCTGCTATCCAGAGATCTTATGGGTNNNAGATCGGAAG
61693 TATGAGTGTGGAGTCGTTGCTGCTCTACTTCTGTAACCCAGCTNNNAGATCGGAAGA
51336 TATGAGTGTGGAGTCGTTGCGGGGGAGAAACCAAGGAAAAGAGNNNAGATCGGA
45937 TATGAGTGTGGAGTCGTTGCCATTGCTGTGAACAACTGGTTCCNNNAGATCGGAAG
53510 TATGAGTGTGGAGTCGTTGCATCTCTCTCTATGTGGTCTCCGGNNNAGATCGGAAGA

30638 TATGAGTGTGGAGTCGTTGCTGGCATCATAGAATGCGTGGAAANNNAGATCGGAA
39822 TATGAGTGTGGAGTCGTTGCTGGTAGTGAACTGTGTTTCACTGANNNAGATCGGA
105339 TATGAGTGTGGAGTCGTTGCTGATAAGGCCAGAAAAATCACCATNNNAGATCGGA
67360 TATGAGTGTGGAGTCGTTGCATGCAAACTGCCTAGGTGTGATTNNNAGATCGGAA
39254 TATGAGTGTGGAGTCGTTGCGGACTTGATGCTCTTATCCATGGANNNAGATCGGAA
64981 TATGAGTGTGGAGTCGTTGCTCAGTGGATTAGAGACTAAGTAGGTNNNAGATCGG
52159 TATGAGTGTGGAGTCGTTGCACCGGAAGGAAGTAGGACTGTAGNNNAGATCGGA
57504 TATGAGTGTGGAGTCGTTGCTTGGGGGAGTAACAATGGTGTTGNNNAGATCGGA
43975 TATGAGTGTGGAGTCGTTGCTCACTCTCCTGAACTCACTCACCNNNAGATCGGAAGA
50775 TATGAGTGTGGAGTCGTTGCGGGGATCACGAGGTGAATTACAGNNNAGATCGGA
102756 TATGAGTGTGGAGTCGTTGCGTTCATCATTTGGGGAAGAAAAAGTNNNAGATCGG
34603 TATGAGTGTGGAGTCGTTGCAGAGAGTATGTATCAGGTGAGCCANNNAGATCGGA
52063 TATGAGTGTGGAGTCGTTGCGCAGGATGAAGAACTGTGGCTAGNNNAGATCGGA
49676 TATGAGTGTGGAGTCGTTGCAGGCTGCACGAATCTTTTCCTTCNNNAGATCGGAAG
59259 TATGAGTGTGGAGTCGTTGCAGGAAAGGAGGAACGGGAATGATNNNAGATCGGA
66579 TATGAGTGTGGAGTCGTTGCGGATTTTGATCGGGTTGCATTGTNNNAGATCGGAA
69401 TATGAGTGTGGAGTCGTTGCAGATACAGCAAATTGTGGTAAAATTTTNNNAGATCG
47210 TATGAGTGTGGAGTCGTTGCCATATTACTCCCAAAGCCATGGCNNNAGATCGGAAG
49395 TATGAGTGTGGAGTCGTTGCGTGTGTAGGAACCTGTGACTGTCNNNAGATCGGAA
40588 TATGAGTGTGGAGTCGTTGCAGGAATGCCAAGCCCACAAAATANNNAGATCGGAA
54565 TATGAGTGTGGAGTCGTTGCATTCAAACCTGAGGCAAACCTGGNNNAGATCGGAA
33023 TATGAGTGTGGAGTCGTTGCGCCTGAGCAGTACCAATAACACANNNAGATCGGAAG
57633 TATGAGTGTGGAGTCGTTGCGGTGCAAGGAAATCACCCTTTTGNNNAGATCGGAA
60810 TATGAGTGTGGAGTCGTTGCTTGGGCTGATGGCAAAATTACCTNNNAGATCGGAA
51920 TATGAGTGTGGAGTCGTTGCGCCCTCTACTTGGACACATTGAGNNNAGATCGGAAG
47648 TATGAGTGTGGAGTCGTTGCCTGGTTTATGAAGGGGACACTGCNNNAGATCGGAA
62958 TATGAGTGTGGAGTCGTTGCCTTTTATCAGGGCAAGTGGCTCTNNNAGATCGGAAG
35043 TATGAGTGTGGAGTCGTTGCAGTTTTTACCTGTTCTTGACTTCCANNNAGATCGGAA
42088 TATGAGTGTGGAGTCGTTGCTCTTACTCCAGGGGGCACTTTTANNNAGATCGGAAG
65648 TATGAGTGTGGAGTCGTTGCTGCCAACTTTTGTGGTAGGAATGTNNNAGATCGGA
37791 TATGAGTGTGGAGTCGTTGCACAAGATTAGAAGCACAGACAGANNNAGATCGGAA
97826 TATGAGTGTGGAGTCGTTGCCCCTGCAGAACCTGAGTATACCNNNAGATCGGAAGA
67559 TATGAGTGTGGAGTCGTTGCGGACAGGGCATCATTCTAGACTTNNNAGATCGGAA
54862 TATGAGTGTGGAGTCGTTGCTAGAGCCAGGGAAATCTGAGTGGNNNAGATCGGA
59594 TATGAGTGTGGAGTCGTTGCAGGAAGTGAGTGCTTGGGAGTATNNNAGATCGGA
62402 TATGAGTGTGGAGTCGTTGCTGCCAGAAATTGTACACTGTGCTNNNAGATCGGAAG
47818 TATGAGTGTGGAGTCGTTGCGGCTTGTTCTAATAAATTTCAGTCTGCNNNAGATCG
116414 TATGAGTGTGGAGTCGTTGCTCTTTGTTTTTCTGTGTGCATTTGCTNNNAGATCGGA
54062 TATGAGTGTGGAGTCGTTGCGGATGAAGTTAGCCTTGGATGGGNNNAGATCGGA
35590 TATGAGTGTGGAGTCGTTGCATGTGCTGACAAATGAGAAGGCANNNAGATCGGAA
46345 TATGAGTGTGGAGTCGTTGCCTCCTTCCTTGAGTTCTTCCAGCNNNAGATCGGAAGA
53859 TATGAGTGTGGAGTCGTTGCAGTGCTATGAATTCTTCAGCGGGNNNAGATCGGAA
65593 TATGAGTGTGGAGTCGTTGCTCCCAAAGGACATACACTAAAATGTNNNAGATCGGA

| | |
|---|---|
| 34036 | TATGAGTGTGGAGTCGTTGCTCTCAAAGGGACAAAAAGACACCANNNAGATCGGA |
| 108186 | TATGAGTGTGGAGTCGTTGCATCCTAGAAAGAGGTGATTGTCAGGNNNAGATCGG |
| 42769 | TATGAGTGTGGAGTCGTTGCCTTCCATTTATGGGCATTGCCACNNNAGATCGGAAG |
| 48190 | TATGAGTGTGGAGTCGTTGCCTCTCTTGTGGTGACTGATTTGCNNNAGATCGGAAG |
| 42108 | TATGAGTGTGGAGTCGTTGCCAGTGCTCACCCTCTTAGTTTTANNNAGATCGGAAG |
| 117668 | TATGAGTGTGGAGTCGTTGCTTTTTGCTGGGGAACAAGATCTTNNNAGATCGGAAG |
| 29894 | TATGAGTGTGGAGTCGTTGCATTGTGTGCCTGTATTCCCAGTGNNNAGATCGGAAG |
| 29895 | TATGAGTGTGGAGTCGTTGCCAATTTCAGACAGGGTGGCTAGGNNNAGATCGGAA |
| 71687 | TATGAGTGTGGAGTCGTTGCTCTTCTCACTGAACCAGACCTGTNNNAGATCGGAAG |
| 48240 | TATGAGTGTGGAGTCGTTGCACGCTTTCTAAAGTACTATGTTTTTGCNNNAGATCGG |
| 36668 | TATGAGTGTGGAGTCGTTGCCAGATAGTTGCTGCAGTTCCTCANNNAGATCGGAAG |
| 49487 | TATGAGTGTGGAGTCGTTGCGGGGAGGGGTGTACTCAAAATTCNNNAGATCGGAA |
| 30992 | TATGAGTGTGGAGTCGTTGCGCAATCAAACAGCAGAACACCAANNNAGATCGGAA |
| 57212 | TATGAGTGTGGAGTCGTTGCAGTGTCTCTCCCACCTGGATTGNNNAGATCGGAAGA |
| 44936 | TATGAGTGTGGAGTCGTTGCACAATGACAGGAGACAATCAGCCNNNAGATCGGAA |
| 97541 | TATGAGTGTGGAGTCGTTGCACACATCAACCTCCACGAAAGGTNNNAGATCGGAAG |
| 99400 | TATGAGTGTGGAGTCGTTGCACTTGTCTGGTTCAGCATTTCACNNNAGATCGGAAG |
| 52785 | TATGAGTGTGGAGTCGTTGCGTATAAGCCTGAGCCCAAGAAGGNNNAGATCGGAA |
| 56423 | TATGAGTGTGGAGTCGTTGCTCCTCTCTGTAGCATCTGGTCTGNNNAGATCGGAAG |
| 54889 | TATGAGTGTGGAGTCGTTGCGCAAGAATACAGGGGTAAGGTGGNNNAGATCGGA |
| 65050 | TATGAGTGTGGAGTCGTTGCAGGTTTTGACTGTGAGGAAGGGTNNNAGATCGGA |
| 65233 | TATGAGTGTGGAGTCGTTGCTCTGCCCATTACATCTTTTGGGTNNNAGATCGGAAG |
| 106463 | TATGAGTGTGGAGTCGTTGCATTGATTTCTGCAACTGTGGAAACNNNAGATCGGAA |
| 38672 | TATGAGTGTGGAGTCGTTGCTAGAGAAGGACTGGGGAGAAGGANNNAGATCGGA |
| 97163 | TATGAGTGTGGAGTCGTTGCCTTAGTTCCTCCCTCCACCCCCNNNAGATCGGAAGAG |
| 29902 | TATGAGTGTGGAGTCGTTGCGAGATTCACCAGGGGGAAAAAGGNNNAGATCGGA |
| 46736 | TATGAGTGTGGAGTCGTTGCGTTTTGACTTGAGCTTGAGACGCNNNAGATCGGAA |
| 43022 | TATGAGTGTGGAGTCGTTGCGCCCTGAGCTTATCAGTTCTCACNNNAGATCGGAAG |
| 50331 | TATGAGTGTGGAGTCGTTGCGCTGGGGAAAGTGGTCTATCAAGNNNAGATCGGAA |
| 68283 | TATGAGTGTGGAGTCGTTGCTTCTTGTAACCACCACTGCAGTTNNNAGATCGGAAG |
| 56281 | TATGAGTGTGGAGTCGTTGCGCGTAAATGAATGCAACTTGCTGNNNAGATCGGAA |
| 108895 | TATGAGTGTGGAGTCGTTGCGTGTTTGAAATGTCTGTTCATATGCNNNAGATCGGA |
| 65196 | TATGAGTGTGGAGTCGTTGCATTAGTGATGGCTGTCTCTGGGTNNNAGATCGGAA |
| 57472 | TATGAGTGTGGAGTCGTTGCGGGCTCAGAGAAAGGATTGGTTGNNNAGATCGGA |
| 64831 | TATGAGTGTGGAGTCGTTGCTGGGGAAGAAAAAGTTTTATCCAGGTNNNAGATCG |
| 67859 | TATGAGTGTGGAGTCGTTGCTTTTCAGCCTGGTTGTTCAGCTTNNNAGATCGGAAG |
| 45387 | TATGAGTGTGGAGTCGTTGCTATCAGGAATGGTTGGCCAATCCNNNAGATCGGAA |
| 56129 | TATGAGTGTGGAGTCGTTGCGGGTGGGTGTTATAAGTCAGCTGNNNAGATCGGAA |
| 65960 | TATGAGTGTGGAGTCGTTGCTTTCCCATCAAGTGCTATTACTGTNNNAGATCGGAAG |
| 67367 | TATGAGTGTGGAGTCGTTGCCTTCAGGAGTTTGGGATACAAATATTNNNAGATCGG |
| 103613 | TATGAGTGTGGAGTCGTTGCAGGCGAAAAGGGAGAGAGAAAGNNNAGATCGGA |
| 42383 | TATGAGTGTGGAGTCGTTGCTTTTGCAGCCACCTGGATAGAACNNNAGATCGGAAG |
| 55219 | TATGAGTGTGGAGTCGTTGCTCCCATCATCTGCCCTCAAAATGNNNAGATCGGAAG |

| | |
|---|---|
| 42271 | TATGAGTGTGGAGTCGTTGCGTCCCACTCTCTGAACAGACAACNNNAGATCGGAAG |
| 108212 | TATGAGTGTGGAGTCGTTGCGGTGCTATGGAATGGAGACAGAATNNNAGATCGG |
| 58750 | TATGAGTGTGGAGTCGTTGCATCACTCTCTGCTTTGGTTGCATNNNAGATCGGAAG |
| 117770 | TATGAGTGTGGAGTCGTTGCTATTGGGGCTGGAATGCAATGGCNNNAGATCGGAA |
| 31774 | TATGAGTGTGGAGTCGTTGCTGAATACCCAAAGCAATCACGAANNNAGATCGGAAG |
| 102714 | TATGAGTGTGGAGTCGTTGCTGAGGCTAACAGGAGAGGGTAATNNNAGATCGGA |
| 61202 | TATGAGTGTGGAGTCGTTGCCCTTTTGTTGTTTGCTTTGGCCTNNNAGATCGGAAG |
| 43839 | TATGAGTGTGGAGTCGTTGCGCAGAGCTTTTTCGACACTAACCNNNAGATCGGAAG |
| 59149 | TATGAGTGTGGAGTCGTTGCGGGACAGTGGGAATAGTGAGGATNNNAGATCGGA |
| 64173 | TATGAGTGTGGAGTCGTTGCTGGGATGCTGAGATACAGAAGAGTNNNAGATCGG |
| 58987 | TATGAGTGTGGAGTCGTTGCCCAATTCCATTTGCCTCCCAGATNNNAGATCGGAAGA |
| 41119 | TATGAGTGTGGAGTCGTTGCCAAGGCTCTGCTGGAAATTCCTANNNAGATCGGAAG |
| 55650 | TATGAGTGTGGAGTCGTTGCTGCCCTTCACATTAGCTCTGATGNNNAGATCGGAAG |
| 32463 | TATGAGTGTGGAGTCGTTGCTTCTCTTGCCCAGACTGGAGTAANNNAGATCGGAAG |
| 32289 | TATGAGTGTGGAGTCGTTGCACACAGCCTAAGCAGAGGGATAANNNAGATCGGAA |
| 103644 | TATGAGTGTGGAGTCGTTGCTTCAGTGCAGTGGCATCATCCTTNNNAGATCGGAAG |
| 68702 | TATGAGTGTGGAGTCGTTGCGTTGTAAAGCCGGGAGTGTGTTNNNAGATCGGAAG |
| 31769 | TATGAGTGTGGAGTCGTTGCCAGCTTCTCTTCTTAGTCATTTAGAANNNAGATCGGA |
| 43377 | TATGAGTGTGGAGTCGTTGCTTGTTTCCAGGGGACATGATGACNNNAGATCGGAA |
| 69239 | TATGAGTGTGGAGTCGTTGCTCTGCAACTGTGGAAACAAGTTTNNNAGATCGGAAG |
| 55207 | TATGAGTGTGGAGTCGTTGCAGGCTGGGGAAGAAAGATTTTGGNNNAGATCGGA |
| 60720 | TATGAGTGTGGAGTCGTTGCTCATTGCCCTAAACAGAAAGACCTNNNAGATCGGAA |
| 97159 | TATGAGTGTGGAGTCGTTGCTGTCTCCCTCTCCACTTCTGGNNNAGATCGGAAGAG |
| 71508 | TATGAGTGTGGAGTCGTTGCTGACCTAACCTGTGTCCAGACTTNNNAGATCGGAAG |
| 52115 | TATGAGTGTGGAGTCGTTGCGATGGGTGAGGCAATGTCAGTAGNNNAGATCGGA |
| 69222 | TATGAGTGTGGAGTCGTTGCAATGTTTTGGCACCATGTTCTTTNNNAGATCGGAAG |
| 43873 | TATGAGTGTGGAGTCGTTGCACACAGTGAAGGTTTCAGACACCNNNAGATCGGAA |
| 31629 | TATGAGTGTGGAGTCGTTGCGGCAAAGGGTGGTAATTCCAGAANNNAGATCGGAA |
| 37214 | TATGAGTGTGGAGTCGTTGCTACTTTGAGCCTACCAGGCTTCANNNAGATCGGAAG |
| 70764 | TATGAGTGTGGAGTCGTTGCTTATGAGAGGAAATGGGGGAGGGNNNAGATCGGA |
| 98122 | TATGAGTGTGGAGTCGTTGCCTACCCCCTCAAGAGTATCACACNNNAGATCGGAAG |
| 49455 | TATGAGTGTGGAGTCGTTGCCCGAATACAATGTCCCTGGTGTCNNNAGATCGGAAG |
| 33252 | TATGAGTGTGGAGTCGTTGCCGAAAAGGGAGAGAGAAAGCACANNNAGATCGGA |
| 67197 | TATGAGTGTGGAGTCGTTGCAGCCTGTCACCCAAAAACTCATTNNNAGATCGGAAG |
| 102346 | TATGAGTGTGGAGTCGTTGCCTCACAAGTACCTAAGTGGCCTNNNAGATCGGAAGA |
| 63383 | TATGAGTGTGGAGTCGTTGCTCTAGCTCTTTTGGGAGCCTTCTNNNAGATCGGAAG |
| 49191 | TATGAGTGTGGAGTCGTTGCTTGTCCTGCCCAAACTGTAAGTCNNNAGATCGGAAG |
| 65396 | TATGAGTGTGGAGTCGTTGCGATGGTCAAATCAAATTACTCTGGTNNNAGATCGGA |
| 104403 | TATGAGTGTGGAGTCGTTGCGCTAACACAACCATGGCTTGNNNAGATCGGAAGAG |
| 36041 | TATGAGTGTGGAGTCGTTGCGGGAGGGAAGGAGAATAACTGCANNNAGATCGGA |
| 44699 | TATGAGTGTGGAGTCGTTGCCAAGAGAGCATGATTTCCCTCCCNNNAGATCGGAAG |
| 67519 | TATGAGTGTGGAGTCGTTGCTTGGCTATAAGGAAACAGCACTTNNNAGATCGGAA |
| 115011 | TATGAGTGTGGAGTCGTTGCATGCAGCCAGATTTCATGACAGTNNNAGATCGGAAG |

96987 TATGAGTGTGGAGTCGTTGCTTTTCTCTTTCAGGCCTCAACCTNNNAGATCGGAAGA
53254 TATGAGTGTGGAGTCGTTGCGCTGCTGTTTTGTTATGTGGAGGNNNAGATCGGAA
54759 TATGAGTGTGGAGTCGTTGCCCGTATTACACAAGCCTCTCTGGNNNAGATCGGAAG
40198 TATGAGTGTGGAGTCGTTGCGTAACCCTGACACTTGGAGGTGANNNAGATCGGAA
53262 TATGAGTGTGGAGTCGTTGCAAGTATAGGTGGCGTTTTGGAGGNNNAGATCGGAA
45183 TATGAGTGTGGAGTCGTTGCCCATTGTACTCTCCTAGCATGCCNNNAGATCGGAAGA
39207 TATGAGTGTGGAGTCGTTGCTGTGTGATGAATGCCATTTGGGANNNAGATCGGAA
47969 TATGAGTGTGGAGTCGTTGCCAGTGACTTACACAAAGTTGGTGCNNNAGATCGGAA
32940 TATGAGTGTGGAGTCGTTGCCCTTGATAGAAGTGCCTGTGAACANNNAGATCGGAA
35560 TATGAGTGTGGAGTCGTTGCGTTCATTGTTGTGCATTCTCGCANNNAGATCGGAAG
116396 TATGAGTGTGGAGTCGTTGCTCTCTCTCTTGAAAGGGGTGCCNNNAGATCGGAAGA
46523 TATGAGTGTGGAGTCGTTGCTGAAGAAGAGGAAGAATGCGAGCNNNAGATCGGA
112890 TATGAGTGTGGAGTCGTTGCATGGGAGGCTACATATAGGCTAGANNNAGATCGGA
117798 TATGAGTGTGGAGTCGTTGCCTCTTCCTCCCTTCTCCTGTTGATANNNAGATCGGAAG
101534 TATGAGTGTGGAGTCGTTGCAAAAATGGCACCGGAAAAGAATGAGAANNNAGATC
34407 TATGAGTGTGGAGTCGTTGCTAACAGGAGAGGGTAATTGCCCANNNAGATCGGAA
48365 TATGAGTGTGGAGTCGTTGCCTGCTCACAGCTCTTGATCCATCNNNAGATCGGAAGA
58178 TATGAGTGTGGAGTCGTTGCGGGGAGGGGTCTAAACTCCTAATNNNAGATCGGAA
103968 TATGAGTGTGGAGTCGTTGCCCTCTGTCCTCCCCTATGGGANNNAGATCGGAAGAG
64301 TATGAGTGTGGAGTCGTTGCGCTCTCATGACTTCCTTTGGAGTNNNAGATCGGAAG
107596 TATGAGTGTGGAGTCGTTGCTCTGTAGCCATTGGATGACCATTTNNNAGATCGGAA
53820 TATGAGTGTGGAGTCGTTGCCAGTCAAAGGGTGAAGTGTAGGGNNNAGATCGGA
30535 TATGAGTGTGGAGTCGTTGCTGGCACCGGAAAAGAATGAGAAANNNAGATCGGAA
57734 TATGAGTGTGGAGTCGTTGCATCCCAACAACACTGGAGCAAATNNNAGATCGGAAG
38356 TATGAGTGTGGAGTCGTTGCGCTAACACAACCATGGCTTGAGANNNAGATCGGAAG
35269 TATGAGTGTGGAGTCGTTGCTCTTCAATGCTTTCTAACCCAGCANNNAGATCGGAAG
67153 TATGAGTGTGGAGTCGTTGCGGAGAGGGGGTAATCTAGGCATTNNNAGATCGGAA
105975 TATGAGTGTGGAGTCGTTGCGTTGGTCAAATGATTATGGAGGCCNNNAGATCGGA
30803 TATGAGTGTGGAGTCGTTGCGAACCTGGCAACACTTCAAACAANNNAGATCGGAAG
31227 TATGAGTGTGGAGTCGTTGCAGTGAGCATTTGAAGGCTAGCAANNNAGATCGGAA
99004 TATGAGTGTGGAGTCGTTGCGGTAGAATCAGAGGAGGTAGTGGNNNAGATCGGA
42064 TATGAGTGTGGAGTCGTTGCGTGTGTGTTGTGAGGAGGGTTTANNNAGATCGGAA
67515 TATGAGTGTGGAGTCGTTGCTGGCTACGTGTATCTCTCCACTTNNNAGATCGGAAG
52157 TATGAGTGTGGAGTCGTTGCGGGCCTGTACTGGTAGCATGTAGNNNAGATCGGAA
37470 TATGAGTGTGGAGTCGTTGCATTCAGCACATACCTGCCAAAGANNNAGATCGGAAG
67218 TATGAGTGTGGAGTCGTTGCTTCTCCATGCATGCTGTCTCATTNNNAGATCGGAAGA
47475 TATGAGTGTGGAGTCGTTGCACCAAAGCTGTGACCTAGAATGCNNNAGATCGGAAG
45608 TATGAGTGTGGAGTCGTTGCCTCCTGCCATTCTACTTTCCTCCNNNAGATCGGAAGA
65347 TATGAGTGTGGAGTCGTTGCCCAGAAGCAAAACTCTCACTGGTNNNAGATCGGAAG
35481 TATGAGTGTGGAGTCGTTGCTGAACCAGGAATAATTTTGCTAGCANNNAGATCGGA
52564 TATGAGTGTGGAGTCGTTGCTGAGCAGAGAAAGGACAAGATCGNNNAGATCGGAA
53631 TATGAGTGTGGAGTCGTTGCCAATGGAAATCTGAGGGGAAGGGNNNAGATCGGA
58915 TATGAGTGTGGAGTCGTTGCGCTAGTTCCACTCTCATGTTCATNNNAGATCGGAAGA

107566 TATGAGTGTGGAGTCGTTGCAGTTGAGGAATTCAAAATTTATTCCAGNNNAGATCG
54069 TATGAGTGTGGAGTCGTTGCAACCCAGCCTTCTAGACTATGGGNNNAGATCGGAAG
68097 TATGAGTGTGGAGTCGTTGCCACTCCCTTCCCTATGCTCTCTTNNNAGATCGGAAGA
63461 TATGAGTGTGGAGTCGTTGCTGTTATCTTTTGGTGCCAGGTTCTNNNAGATCGGAA
110398 TATGAGTGTGGAGTCGTTGCGTGTGTGTTTCCACCTAAGTTGATAGNNNAGATCGG
31457 TATGAGTGTGGAGTCGTTGCTCATGTAGTCCCACCATGGTCAANNNAGATCGGAAG
61982 TATGAGTGTGGAGTCGTTGCTGTGAAAGGTAAGTAGATTCAGGCTNNNAGATCGG
40613 TATGAGTGTGGAGTCGTTGCTTCTTCGCTCCCTCCCTGAAATANNNAGATCGGAAGA
33693 TATGAGTGTGGAGTCGTTGCTCTACTGCACTTATCACTATCTGACANNNAGATCGGA
64207 TATGAGTGTGGAGTCGTTGCATGGGAAGAAGGCTGAAGAGAGTNNNAGATCGGA
56214 TATGAGTGTGGAGTCGTTGCCTCTCTCTCATTCGTGATGGCTGNNNAGATCGGAAG
68983 TATGAGTGTGGAGTCGTTGCAACTGTGTTCCCATGCTGACTTTNNNAGATCGGAAG
37765 TATGAGTGTGGAGTCGTTGCGGAACCCTTGCACTTTACACAGANNNAGATCGGAAG
105392 TATGAGTGTGGAGTCGTTGCAAGCTGTGTCAGGGCAGGAATACNNNAGATCGGAA
35005 TATGAGTGTGGAGTCGTTGCTACCTCTCCACTGGCAAATTCCANNNAGATCGGAAGA
41054 TATGAGTGTGGAGTCGTTGCCCACTGACATAGATGAGGCCCTANNNAGATCGGAAG
53826 TATGAGTGTGGAGTCGTTGCAGAGCAATGCACATGGATTAGGGNNNAGATCGGAA
40761 TATGAGTGTGGAGTCGTTGCCAAAGAGAACAGGCAGGTGCATANNNAGATCGGAA
55907 TATGAGTGTGGAGTCGTTGCACAGGTTCTCTGGTTGAAACCTGNNNAGATCGGAAG
62581 TATGAGTGTGGAGTCGTTGCAGGCACCCTAAGATGGTGAATCTNNNAGATCGGAA
56433 TATGAGTGTGGAGTCGTTGCAAAGTCCCATCAAGTGCTGTCTGNNNAGATCGGAAG
32450 TATGAGTGTGGAGTCGTTGCGGCATCATCCTTGCTCACAGTAANNNAGATCGGAAG
54172 TATGAGTGTGGAGTCGTTGCGCTTGTTTCTACTACCCAGTGGGNNNAGATCGGAAG
46570 TATGAGTGTGGAGTCGTTGCAAGTGAGAGCACCATATTGGAGCNNNAGATCGGAA
68549 TATGAGTGTGGAGTCGTTGCAGGGTGTGGATTCATGCAATGTTNNNAGATCGGAA
43595 TATGAGTGTGGAGTCGTTGCTTCTAGCTGGGTGTCTCGTCTACNNNAGATCGGAAG
46204 TATGAGTGTGGAGTCGTTGCGTAAGAGACCTGGGAATGGAAGCNNNAGATCGGA
41908 TATGAGTGTGGAGTCGTTGCTGGCAAGTCCTGAGTGAGAGTTANNNAGATCGGAA
52577 TATGAGTGTGGAGTCGTTGCACCTTCCAATGAGTGAGAACTCGNNNAGATCGGAAG
59492 TATGAGTGTGGAGTCGTTGCTGGAGGGGGCATATGTTGACTATNNNAGATCGGAA
42778 TATGAGTGTGGAGTCGTTGCAACAGGAGGTAAATGCCATCCACNNNAGATCGGAA
50401 TATGAGTGTGGAGTCGTTGCTGGTGTGTGCTTTCGTAGAGAAGNNNAGATCGGAA
48307 TATGAGTGTGGAGTCGTTGCACCCCATCCCACTCCATCTAATCNNNAGATCGGAAGA
31324 TATGAGTGTGGAGTCGTTGCACAGCCACAATACAAGTCTGCAANNNAGATCGGAAG
63764 TATGAGTGTGGAGTCGTTGCGCCACAGTTGACTAAGCAGAAGTNNNAGATCGGAA
49332 TATGAGTGTGGAGTCGTTGCGTTGGTCATTCCTCTGATGGGTCNNNAGATCGGAAG
43522 TATGAGTGTGGAGTCGTTGCCCACTGCTTCTGTAACCCACTACNNNAGATCGGAAGA
31036 TATGAGTGTGGAGTCGTTGCACTTTCAATCATGCAGAGCCCAANNNAGATCGGAAG
51343 TATGAGTGTGGAGTCGTTGCTGGGGGTACAGAGACAGAAAGAGNNNAGATCGGA
106534 TATGAGTGTGGAGTCGTTGCTGGGTCATTTGGCTTGATTCATTANNNAGATCGGAA
106527 TATGAGTGTGGAGTCGTTGCCACAAAAGTGAGGACCCCAGTATANNNAGATCGGA
46973 TATGAGTGTGGAGTCGTTGCAGGTACAGATATCAGGTGGAGGCNNNAGATCGGA
35707 TATGAGTGTGGAGTCGTTGCATGATAACTTCCTCGAGAGGGCANNNAGATCGGAA

99409 TATGAGTGTGGAGTCGTTGCCCTGGAGCTGCGCTGCTGATNNNAGATCGGAAGAG
56428 TATGAGTGTGGAGTCGTTGCCCCATTCCAATACCCACTGTCTGNNNAGATCGGAAGA
58679 TATGAGTGTGGAGTCGTTGCTTTAGGGATCTTCCAAGGGGCATNNNAGATCGGAA
43540 TATGAGTGTGGAGTCGTTGCTGCTGTATTCAGTTGCCACCTACNNNAGATCGGAAG
54043 TATGAGTGTGGAGTCGTTGCAGATCAACTCACCAATCCATGGGNNNAGATCGGAAG
56490 TATGAGTGTGGAGTCGTTGCGACAAAATTGCAAAGGCATTTCTGNNNAGATCGGA
34154 TATGAGTGTGGAGTCGTTGCAAAAGCACTGGCAGGAAATACCANNNAGATCGGAA
48183 TATGAGTGTGGAGTCGTTGCTCCCAAGAATGTGCAGAATTTGCNNNAGATCGGAAG
64458 TATGAGTGTGGAGTCGTTGCAGACAAGAAGCTGGGAAAGTAGTNNNAGATCGGA
97822 TATGAGTGTGGAGTCGTTGCTTAGCGCCCTCTCTTATGTGGGNNNAGATCGGAAGA
103943 TATGAGTGTGGAGTCGTTGCATAGGGATGATTTCAGCTTAGCATAANNNAGATCGG
63782 TATGAGTGTGGAGTCGTTGCAGGGCAAAATATGCAAAGGAAGTNNNAGATCGGA
64029 TATGAGTGTGGAGTCGTTGCAAGGATGTCTTGCATTACGCAGTNNNAGATCGGAA
42967 TATGAGTGTGGAGTCGTTGCTATGTACTGGCAAAGGCCATCACNNNAGATCGGAAG
100582 TATGAGTGTGGAGTCGTTGCGAGGCTAGTGTTGAACCCCTGGANNNAGATCGGAA
34706 TATGAGTGTGGAGTCGTTGCGTGAGATAGCCAAATGGATGCCANNNAGATCGGAA
59700 TATGAGTGTGGAGTCGTTGCGAAAGGAAGGCCAAAAGTCTTATNNNAGATCGGAA
61199 TATGAGTGTGGAGTCGTTGCTGGTTGTTGGAATTTCTTGGCCTNNNAGATCGGAAG
49718 TATGAGTGTGGAGTCGTTGCTGGTCCTTTCTCCTCATTGCTTCNNNAGATCGGAAGA
68246 TATGAGTGTGGAGTCGTTGCCCAAGATAGCAGCACAGACAGTTNNNAGATCGGAA
65334 TATGAGTGTGGAGTCGTTGCGCTGAGGATCACTTTCAACTGGTNNNAGATCGGAAG
40966 TATGAGTGTGGAGTCGTTGCAAGTAACCACTCTCCAGCCACTANNNAGATCGGAAG
44650 TATGAGTGTGGAGTCGTTGCATCATCCCCAAAGCAATCATCCCNNNAGATCGGAAGA
59260 TATGAGTGTGGAGTCGTTGCGGGGTGATGCAAGTGGTAATGATNNNAGATCGGA
34849 TATGAGTGTGGAGTCGTTGCTGTTGTCCTTCCCTTGGTATCCANNNAGATCGGAAGA
31897 TATGAGTGTGGAGTCGTTGCGTGAGAGAGAGACACAGAGGGAANNNAGATCGGA
40240 TATGAGTGTGGAGTCGTTGCTGACAGGGTGTGATAAGTGGTGANNNAGATCGGA
103630 TATGAGTGTGGAGTCGTTGCCAAGATAGCAGCACAGACAGTTCNNNAGATCGGAA
104450 TATGAGTGTGGAGTCGTTGCTTCTACTACCCAGTGGGAATTTGGNNNAGATCGGAA
60498 TATGAGTGTGGAGTCGTTGCTGGGTTGCAGGAGAGAAAGTACTNNNAGATCGGA
69055 TATGAGTGTGGAGTCGTTGCAGGCTCCCATAAGCTACTCCTTTNNNAGATCGGAAGA
116714 TATGAGTGTGGAGTCGTTGCCTGAGGAACAGTCATGTGAGTACATNNNAGATCGG
63411 TATGAGTGTGGAGTCGTTGCTGTGTTTCCCTTTATTGTGCTTCTNNNAGATCGGAAG
63018 TATGAGTGTGGAGTCGTTGCTCCTATGGGTCTCCTTCCTCTCTNNNAGATCGGAAGA
63543 TATGAGTGTGGAGTCGTTGCTCAAACTGCTACTACAAAATAGTTTCTNNNAGATCGG
105937 TATGAGTGTGGAGTCGTTGCTGGGTGCAGGTGATAAGGACANNNAGATCGGAAG
37993 TATGAGTGTGGAGTCGTTGCCTTTAGATGGGTCTGCTTGCAGANNNAGATCGGAA
30881 TATGAGTGTGGAGTCGTTGCGACAGACACACAGACATCCACAANNNAGATCGGAAG
29897 TATGAGTGTGGAGTCGTTGCAAAAGACCCCTCACCAGATGAGANNNAGATCGGAA
29973 TATGAGTGTGGAGTCGTTGCCACAAGAACCCTAGCTCAGTGTCNNNAGATCGGAAG
101528 TATGAGTGTGGAGTCGTTGCCTATTCCTATTCTATCTTCCTAGAGCTNNNAGATCGG
31352 TATGAGTGTGGAGTCGTTGCAACTTTCACCCTCATGGTTGCAANNNAGATCGGAAG
65815 TATGAGTGTGGAGTCGTTGCTGCCCCTGTATAAGTAGTGATGTNNNAGATCGGAAG

42817 TATGAGTGTGGAGTCGTTGCGCAATCCAATGGAGGAAAAGCACNNNAGATCGGAA
107368 TATGAGTGTGGAGTCGTTGCTTGTCCTTCCCTTGGTATCCATGGNNNAGATCGGAA
110434 TATGAGTGTGGAGTCGTTGCAGAAGTCACAGTGCTCCCATCTNNNAGATCGGAAGA
32269 TATGAGTGTGGAGTCGTTGCTGATGGATCCTGGGGCAAGATAANNNAGATCGGAA
57918 TATGAGTGTGGAGTCGTTGCGAAGTCCATCGCAGTGATCCAATNNNAGATCGGAAG
45418 TATGAGTGTGGAGTCGTTGCGAAGAAACTGGTGGTCACCATCCNNNAGATCGGAA
104435 TATGAGTGTGGAGTCGTTGCGATACTCCTCCACTTACGCTGGGNNNAGATCGGAAG
42654 TATGAGTGTGGAGTCGTTGCGCTATGGGGAAATGTCAAACCACNNNAGATCGGAA
65511 TATGAGTGTGGAGTCGTTGCAGATGAGCATGCCAAAGATTGGTNNNAGATCGGAA
48587 TATGAGTGTGGAGTCGTTGCCTGGTTCTGTCTCTGGAGAACTCNNNAGATCGGAAG
34446 TATGAGTGTGGAGTCGTTGCCACTCAATCAGCATGAACTCCCANNNAGATCGGAAG
64584 TATGAGTGTGGAGTCGTTGCGAGCATTTTTGCCGTTCTTACGTNNNAGATCGGAAG
35106 TATGAGTGTGGAGTCGTTGCCTGAGGTTTCCACAAGGTTTCCANNNAGATCGGAAG
35235 TATGAGTGTGGAGTCGTTGCAGATGAGGGGAGTACAAACAGCANNNAGATCGGA
57637 TATGAGTGTGGAGTCGTTGCGCAACCACCGTAAGTAGCTTTTGNNNAGATCGGAAG
47221 TATGAGTGTGGAGTCGTTGCCCATTCACTCACAGAATCATGGCNNNAGATCGGAAG
48653 TATGAGTGTGGAGTCGTTGCAGGATTTAGAGGCCACCCTACTCNNNAGATCGGAAG
68195 TATGAGTGTGGAGTCGTTGCGTGCATGCGTTTGTGTGTTTCTTNNNAGATCGGAAG
45234 TATGAGTGTGGAGTCGTTGCTTTTGATGGGTACAGAACCTGCCNNNAGATCGGAAG
63188 TATGAGTGTGGAGTCGTTGCTTCTGTGTCAGTAACTGGGGTCTNNNAGATCGGAAG
66870 TATGAGTGTGGAGTCGTTGCTGCTGTGGCTACTATACAGTTTTGTNNNAGATCGGA
107573 TATGAGTGTGGAGTCGTTGCTGTCCAAAAATGTTTCAGCTGGGANNNAGATCGGA
108232 TATGAGTGTGGAGTCGTTGCGTAAGCCAGACATTCTATGCAGCTNNNAGATCGGAA
117084 TATGAGTGTGGAGTCGTTGCCTGCTTCTGGTGTGGAGAGTTANNNAGATCGGAAG
29901 TATGAGTGTGGAGTCGTTGCGGGGAATCACAACTTCAGGTTGANNNAGATCGGAA
30471 TATGAGTGTGGAGTCGTTGCTTCCTGGTTACATTCATCAAAAGAAANNNAGATCGG
112916 TATGAGTGTGGAGTCGTTGCTCCAAAAATGTTTCAGCTGGGATNNNAGATCGGAAG
59074 TATGAGTGTGGAGTCGTTGCGCTTTCCAGATCCAGTGTGAGATNNNAGATCGGAAG
110417 TATGAGTGTGGAGTCGTTGCCTTTTTCAGCACCAACATGCTCAAAANNNAGATCGGA
30157 TATGAGTGTGGAGTCGTTGCGGAGAGTGAGGGCTACTGGAAAANNNAGATCGGA
38224 TATGAGTGTGGAGTCGTTGCAAGAAACTGGACCACTACGGAGANNNAGATCGGAA
103948 TATGAGTGTGGAGTCGTTGCGGTGGAGTTTCTTTTCCTCAATCTNNNAGATCGGAA
54597 TATGAGTGTGGAGTCGTTGCTTTTTCTGATGCTCTCCCCTGGNNNAGATCGGAAGA
57874 TATGAGTGTGGAGTCGTTGCATACGCGCAGACTTTCTGACAATNNNAGATCGGAAG
117183 TATGAGTGTGGAGTCGTTGCCAATGAAGAGATGAGCTGCGCCNNNAGATCGGAAG
51126 TATGAGTGTGGAGTCGTTGCGTTGGGAGAACAGACCTATGCAGNNNAGATCGGAA
50079 TATGAGTGTGGAGTCGTTGCACTGTAAACCAGCTGGACCAAAGNNNAGATCGGAA
42370 TATGAGTGTGGAGTCGTTGCGTGGTGGTGGGTAAGAAGAGAACNNNAGATCGGA
60481 TATGAGTGTGGAGTCGTTGCCCTTGTGTTGTAGAGCTGCTACTNNNAGATCGGAAG
53389 TATGAGTGTGGAGTCGTTGCCCTGCCCAAGATTTCTCTCTAGGNNNAGATCGGAAG
29974 TATGAGTGTGGAGTCGTTGCAGGGAAAACTCAGCCAAAACAGTNNNAGATCGGAA
66686 TATGAGTGTGGAGTCGTTGCTGATAGCCACAGTGAGAAAGTTGTNNNAGATCGGA
34330 TATGAGTGTGGAGTCGTTGCATTATGTGGTTTCTTTGGCCCCANNNAGATCGGAAG

44871 TATGAGTGTGGAGTCGTTGCCTGGAAATAAACCATGCCAAGCCNNNAGATCGGAAG
102743 TATGAGTGTGGAGTCGTTGCTTTACCAAGATTTTTGCATTGGCCATANNNAGATCGG
57317 TATGAGTGTGGAGTCGTTGCATGTATACCAAGTGCTGGGCTTGNNNAGATCGGAA
61359 TATGAGTGTGGAGTCGTTGCTCTTGAGTGTCAGTGGAACTCCTNNNAGATCGGAAG
36972 TATGAGTGTGGAGTCGTTGCGAGCTAACCAGTAAACCTGGTCANNNAGATCGGAA
37247 TATGAGTGTGGAGTCGTTGCCAGTGTGGCACAGATCTTCTTCANNNAGATCGGAAG
51159 TATGAGTGTGGAGTCGTTGCGTGGTTGTGGTGAAATTCTGCAGNNNAGATCGGAA
50436 TATGAGTGTGGAGTCGTTGCTGGGGAGAAAGAAGGAGAGGAAGNNNAGATCGGA
44478 TATGAGTGTGGAGTCGTTGCCAACCTGCTATACATTAGGTCCCCNNNAGATCGGAA
40986 TATGAGTGTGGAGTCGTTGCCCTGGTGTTGCACTTTGAGACTANNNAGATCGGAAG
47808 TATGAGTGTGGAGTCGTTGCAAAATGCTACCCCTCATCTCTGCNNNAGATCGGAAGA
42263 TATGAGTGTGGAGTCGTTGCGATAGGCTCTTGCTACCCACAACNNNAGATCGGAAG
56942 TATGAGTGTGGAGTCGTTGCCCCAGAAGAGAAGTATGTAAGACTGTGNNNAGATC
52359 TATGAGTGTGGAGTCGTTGCGTGGGTAGGTGGGAAAAGTTACGNNNAGATCGGA
47455 TATGAGTGTGGAGTCGTTGCTCTGCCTTGAGAAGAGACAATGCNNNAGATCGGAA
62383 TATGAGTGTGGAGTCGTTGCTGTAGAAACAACACCTACGTGCTNNNAGATCGGAAG
48309 TATGAGTGTGGAGTCGTTGCTCAGGGCAAGGCAAAGTCTAATCNNNAGATCGGAA
34674 TATGAGTGTGGAGTCGTTGCCTGAAGAAAGGATTTCCTGGCCANNNAGATCGGAA
46159 TATGAGTGTGGAGTCGTTGCCATTTACAATCTGCGGGTCAAGCNNNAGATCGGAAG
101189 TATGAGTGTGGAGTCGTTGCTGGTTTTGGACACCTAGGTTGATCNNNAGATCGGAA
54769 TATGAGTGTGGAGTCGTTGCATCTGGTTTTTGCAGAGGTCTGGNNNAGATCGGAA
50117 TATGAGTGTGGAGTCGTTGCGCAGCTCTTCTGTGACTTCAAAGNNNAGATCGGAAG
64177 TATGAGTGTGGAGTCGTTGCATGAGGAACAGCTGGAGAAGAGTNNNAGATCGGA
108224 TATGAGTGTGGAGTCGTTGCGCTATCAGACACAGACCTGGGAGNNNAGATCGGAA
67241 TATGAGTGTGGAGTCGTTGCAGAAAAACAGTCTGGAGGTTCATTNNNAGATCGGA
33265 TATGAGTGTGGAGTCGTTGCCATCCCTTTATAGTGCCAGCACANNNAGATCGGAAG
105906 TATGAGTGTGGAGTCGTTGCCAGTGTTGAGTGGAGCACATACNNNAGATCGGAAG
106497 TATGAGTGTGGAGTCGTTGCGAGATTACACAGCAAAGCCTGAGTNNNAGATCGGA
31841 TATGAGTGTGGAGTCGTTGCTGAGCTGGGATGTTGTAGAGGAANNNAGATCGGA
45053 TATGAGTGTGGAGTCGTTGCACCCCTCTCTTTTATCTCAGGCCNNNAGATCGGAAGA
48122 TATGAGTGTGGAGTCGTTGCTCCAGTGAAAAACATATTTTGCTTGCNNNAGATCGG
68032 TATGAGTGTGGAGTCGTTGCAGGCTCTCAGGTTCCAAGATCTTNNNAGATCGGAAG
32987 TATGAGTGTGGAGTCGTTGCATCTTCCTAGAGCTGTAAAACACANNNAGATCGGAA
39286 TATGAGTGTGGAGTCGTTGCTTCACACATCCACAGCATATGGANNNAGATCGGAAG
69457 TATGAGTGTGGAGTCGTTGCTGGTAGAATGTTGCACAACTTTTNNNAGATCGGAAG
104444 TATGAGTGTGGAGTCGTTGCCTTGGTTTGCAATATGGAGGCAGNNNAGATCGGAA
31399 TATGAGTGTGGAGTCGTTGCTACCTTCAGCAATGCAACCTCAANNNAGATCGGAAG
52007 TATGAGTGTGGAGTCGTTGCGGCTGGAGCAAATCCGGTATAGNNNAGATCGGAAG
47203 TATGAGTGTGGAGTCGTTGCGGGGAGAAACTGGAACTAATGGCNNNAGATCGGA
34188 TATGAGTGTGGAGTCGTTGCTTGTAGTGAAACTGCAGAGTACCANNNAGATCGGA
98738 TATGAGTGTGGAGTCGTTGCCACACATTTCCCGTCCAGCAAACNNNAGATCGGAAG
67755 TATGAGTGTGGAGTCGTTGCACAACCTCCTTTCAAAGTGCCTTNNNAGATCGGAAGA
53794 TATGAGTGTGGAGTCGTTGCTTTAGCTCACAGGACTAGTGAGGGNNNAGATCGGA

| | |
|---|---|
| 57854 | TATGAGTGTGGAGTCGTTGCGGGTCAACGAGGATTGGAACAATNNNAGATCGGAA |
| 43759 | TATGAGTGTGGAGTCGTTGCGAGAAGGTGGTCTGTCTGAAACCNNNAGATCGGAA |
| 63467 | TATGAGTGTGGAGTCGTTGCCACTTACGCTGGGGTTATGTTCTNNNAGATCGGAAG |
| 31072 | TATGAGTGTGGAGTCGTTGCAAAGGAAGATGGAAAGCAGCCAANNNAGATCGGA |
| 63907 | TATGAGTGTGGAGTCGTTGCATGCAGCCAGATTTCATGACAGTNNNAGATCGGAAG |
| 67528 | TATGAGTGTGGAGTCGTTGCGCCCCTGCCAATAACAATCACTTNNNAGATCGGAAG |
| 67521 | TATGAGTGTGGAGTCGTTGCAGCATTAGTACATGGCAGCACTTNNNAGATCGGAAG |
| 96047 | TATGAGTGTGGAGTCGTTGCACCACCACCACTGTCAGAGCNNNAGATCGGAAGAGC |
| 68616 | TATGAGTGTGGAGTCGTTGCAAGGTGGCTCTGAGTATCCTGTTNNNAGATCGGAA |
| 40012 | TATGAGTGTGGAGTCGTTGCTCTCACTCACTGAAGCCATCTGANNNAGATCGGAAG |
| 105936 | TATGAGTGTGGAGTCGTTGCGCTTGTTTTGGAGGTCTGCTGAANNNAGATCGGAA |
| 57849 | TATGAGTGTGGAGTCGTTGCACGGCCTGCAATTTTCAAACAATNNNAGATCGGAAG |
| 45522 | TATGAGTGTGGAGTCGTTGCAAAGAAGCAGACAGAGACACTCCNNNAGATCGGAA |
| 115952 | TATGAGTGTGGAGTCGTTGCCTCTTGTCTCCTTTTTGGAGGGGNNNAGATCGGAAG |
| 67963 | TATGAGTGTGGAGTCGTTGCCCCACAATGAGGACAATCTGCTTNNNAGATCGGAAG |
| 100371 | TATGAGTGTGGAGTCGTTGCCTTGGGTAAGCCTTGGTTTGTCNNNAGATCGGAAG |
| 29900 | TATGAGTGTGGAGTCGTTGCTTCAGCAAATGAGTATCCCGGTGNNNAGATCGGAA |
| 31647 | TATGAGTGTGGAGTCGTTGCTGAAGAAAGGTGTGGATGCAGAANNNAGATCGGA |
| 49108 | TATGAGTGTGGAGTCGTTGCTGCCTCTCTTGCACTTCATTCTCNNNAGATCGGAAGA |
| 41309 | TATGAGTGTGGAGTCGTTGCATGTGAGAGCCATTGTGTGTCTANNNAGATCGGAA |
| 56808 | TATGAGTGTGGAGTCGTTGCTTCTTCCCATTTCCACTTGGGTGNNNAGATCGGAAG |
| 44949 | TATGAGTGTGGAGTCGTTGCACTCCTTGTAAGTTTGCAGAGCCNNNAGATCGGAAG |
| 61771 | TATGAGTGTGGAGTCGTTGCCAGGAGGAGGAGAAAAGAGAGCTNNNAGATCGGA |
| 34496 | TATGAGTGTGGAGTCGTTGCTTTACACCAGGGATGACTTCCCANNNAGATCGGAAG |
| 52805 | TATGAGTGTGGAGTCGTTGCAGAGTAGCTGGGAATCAGGAAGGNNNAGATCGGA |
| 70173 | TATGAGTGTGGAGTCGTTGCCAGGGTCTCTTAGACCAGCTGTANNNAGATCGGAAG |
| 71758 | TATGAGTGTGGAGTCGTTGCGCCCTGGAAATAGAGGTAGATGGNNNAGATCGGA |
| 34661 | TATGAGTGTGGAGTCGTTGCTCCATTCATCTGTAGATGGGCCANNNAGATCGGAAG |
| 100912 | TATGAGTGTGGAGTCGTTGCTACCCATCATTTGCCTGAAAGACCNNNAGATCGGAA |
| 29993 | TATGAGTGTGGAGTCGTTGCGCCGGTCCAAGGGAATACAAAAANNNAGATCGGAA |
| 34297 | TATGAGTGTGGAGTCGTTGCGGCTCAATGTGCTTATAACCCCANNNAGATCGGAAG |
| 31902 | TATGAGTGTGGAGTCGTTGCTTTCTCAAGGCCTCACTAGGGAANNNAGATCGGAAG |
| 30744 | TATGAGTGTGGAGTCGTTGCCGTGACCCCAGAAGAAAGCTAAANNNAGATCGGAA |
| 34521 | TATGAGTGTGGAGTCGTTGCTCCTCCACTCTCAACTTTTCCCANNNAGATCGGAAGA |
| 69477 | TATGAGTGTGGAGTCGTTGCCCTAGTGGGAGGACTAGCCTTTTNNNAGATCGGAA |
| 96820 | TATGAGTGTGGAGTCGTTGCCCTGATTTCAAAGTCTCCCCTCCANNNAGATCGGAAG |
| 60721 | TATGAGTGTGGAGTCGTTGCCCCATCATTTGCCTGAAAGACCTNNNAGATCGGAAG |
| 43617 | TATGAGTGTGGAGTCGTTGCATCTTGTCTCAGGGCTGGAGTACNNNAGATCGGAA |
| 39754 | TATGAGTGTGGAGTCGTTGCAGCAGCAGACACAAACTGTATGANNNAGATCGGAA |
| 47429 | TATGAGTGTGGAGTCGTTGCTGCCTTGTTCTATCACAATTTTGGCNNNAGATCGGAA |
| 97447 | TATGAGTGTGGAGTCGTTGCATTCTCGGATGCTCAAGTTCCTTATTNNNAGATCGGA |
| 97780 | TATGAGTGTGGAGTCGTTGCTCTTCAAGTGTCGACCAAACTCCTNNNAGATCGGAA |
| 29896 | TATGAGTGTGGAGTCGTTGCAGTTGCCTCTGATCAAAGCTGTTNNNAGATCGGAAG |

61409 TATGAGTGTGGAGTCGTTGCTCTATGTCCTGGACCTCTCTCCTNNNAGATCGGAAGA
47251 TATGAGTGTGGAGTCGTTGCTACGCTGAATTGTCCTAACTGGCNNNAGATCGGAAG
32889 TATGAGTGTGGAGTCGTTGCTATGGAAAACAGCGTGGAGAACANNNAGATCGGAA
68512 TATGAGTGTGGAGTCGTTGCTGGTCTATGAAGACAGGGTGGTNNNAGATCGGAA
58966 TATGAGTGTGGAGTCGTTGCACTTTTAGAGCAGGGAACAGATNNNAGATCGGAA
37965 TATGAGTGTGGAGTCGTTGCTTTCTGTTGTGATAGCCATGCAGANNNAGATCGGAA
110424 TATGAGTGTGGAGTCGTTGCCCTGACCCCTGAAGCATGAAAGTNNNAGATCGGAAG
43443 TATGAGTGTGGAGTCGTTGCTCACTAGGTACTAGCCCCTTGACNNNAGATCGGAAG
35800 TATGAGTGTGGAGTCGTTGCCTATCGCCAATATGTGACTGGCANNNAGATCGGAAG
102747 TATGAGTGTGGAGTCGTTGCTAGCAGCAGACACAAACTGTATGAAANNNAGATCG
39116 TATGAGTGTGGAGTCGTTGCACAAACTGGGAAAAGGAATGGGANNNAGATCGGA
105859 TATGAGTGTGGAGTCGTTGCATGTGGCTTTTTCTGTCTACACTTGTNNNAGATCGGA
36815 TATGAGTGTGGAGTCGTTGCAGGGAAGATGGTGGTCATTCTCANNNAGATCGGAA
48215 TATGAGTGTGGAGTCGTTGCGGTCATCAGAATGCATGGTTTGCNNNAGATCGGAA
44573 TATGAGTGTGGAGTCGTTGCCTTTGCAGTGATGCTAAATGCCCNNNAGATCGGAAG
58678 TATGAGTGTGGAGTCGTTGCTCCAGTTTGAAGCTTTAGGGCATNNNAGATCGGAAG
57470 TATGAGTGTGGAGTCGTTGCGTTTTCCGGACATGATGTGGTTGNNNAGATCGGAA
46811 TATGAGTGTGGAGTCGTTGCAAAAGGTAGTGGTACCCAAAGGCNNNAGATCGGAA
29898 TATGAGTGTGGAGTCGTTGCGGTTTAGCCAAGTGATTTGCTGGNNNAGATCGGAA
70753 TATGAGTGTGGAGTCGTTGCTTTCCTTTTCCCTTTCTGGACGGNNNAGATCGGAAGA
65124 TATGAGTGTGGAGTCGTTGCTGTCTGTGGTGATAGATAGGGGTNNNAGATCGGAA
58356 TATGAGTGTGGAGTCGTTGCGGACCTGAGATGGGAGAAGACATNNNAGATCGGA
29972 TATGAGTGTGGAGTCGTTGCCTGGTCATCTGCCTGTAATTGCTNNNAGATCGGAAG
56203 TATGAGTGTGGAGTCGTTGCATCTGTCGAAATCACAAGGGCTGNNNAGATCGGAA
41086 TATGAGTGTGGAGTCGTTGCAAAGACGGAGGACCATATGCCTANNNAGATCGGAA
71717 TATGAGTGTGGAGTCGTTGCGGAAGAAGACAGTGAAGGGCATGNNNAGATCGGA
103746 TATGAGTGTGGAGTCGTTGCTGATTTTGGACTTTCTGGTCTCCAGNNNAGATCGGA
101077 TATGAGTGTGGAGTCGTTGCGGGAGGTTTTCGTTTTCTAAGACTGNNNAGATCGGA
116136 TATGAGTGTGGAGTCGTTGCCCTGTGTCCAGACTTTTCTCTGTTTNNNAGATCGGAA
101141 TATGAGTGTGGAGTCGTTGCTCACAAGTGGAGATATTTGCAACTTNNNAGATCGGA
58248 TATGAGTGTGGAGTCGTTGCCTAAGCTAACCGGGCAGAAACATNNNAGATCGGAA
71760 TATGAGTGTGGAGTCGTTGCTGACGCACTCACTGAGGATTTTTNNNAGATCGGAAG
104410 TATGAGTGTGGAGTCGTTGCTCCTTTCCATTTTGGGTGTTTTACTTTNNNAGATCGG
44226 TATGAGTGTGGAGTCGTTGCAGTATTGGCAGGATGCTAAACCCNNNAGATCGGAA
71715 TATGAGTGTGGAGTCGTTGCCACCTTTCTGATGTGGTTCTGCTNNNAGATCGGAAG
102181 TATGAGTGTGGAGTCGTTGCAAACCAAAAGTCCAACTCTAGTAACNNNAGATCGGA
103960 TATGAGTGTGGAGTCGTTGCGAACCCCAGGCAGACAGAATCNNNAGATCGGAAGA
103623 TATGAGTGTGGAGTCGTTGCATTCTGGAGCTAAGGAGGGCAANNNAGATCGGAAG
103745 TATGAGTGTGGAGTCGTTGCATAGTAATAAAGAGGTGCGGAGAGNNNAGATCGG
103980 TATGAGTGTGGAGTCGTTGCCTGAACCTATTGAGATGATCACATGATNNNAGATCG
102754 TATGAGTGTGGAGTCGTTGCGCTTCACCTCTAGGCATTGTATCCNNNAGATCGGAAG
100716 TATGAGTGTGGAGTCGTTGCACCAAGTCTGGAATACAGTGATGCANNNAGATCGGA
105267 TATGAGTGTGGAGTCGTTGCGCTAAACCCACCTATACAGAGGACNNNAGATCGGAA

| | |
|---|---|
| 105901 | TATGAGTGTGGAGTCGTTGCAACAATGAAAAAGTAGTCAGGCAGCCTNNNAGATCG |
| 105357 | TATGAGTGTGGAGTCGTTGCGCCAACGAGGGAGATAAGGTGNNNAGATCGGAAGA |
| 106539 | TATGAGTGTGGAGTCGTTGCGTGGCTGGCAAACAAGTGTTGGNNNAGATCGGAAG |
| 107578 | TATGAGTGTGGAGTCGTTGCATCTCTGTGAAGCAGTCAGATTGCNNNAGATCGGAA |
| 105389 | TATGAGTGTGGAGTCGTTGCCACAGGTACTTCACAAAAGAGGGNNNAGATCGGAA |
| 104437 | TATGAGTGTGGAGTCGTTGCCTCTCTGGTGGTAAAAGTTTGATGTGNNNAGATCGG |
| 108912 | TATGAGTGTGGAGTCGTTGCAGGGTTCCTTACTCTTTGATTTTCTGNNNAGATCGGA |
| 102190 | TATGAGTGTGGAGTCGTTGCCCACATCAAGGCTGGGGTTGATNNNAGATCGGAAGA |
| 105365 | TATGAGTGTGGAGTCGTTGCTAGCGGTTGCTTCCTGGAGAAANNNAGATCGGAAGA |
| 101134 | TATGAGTGTGGAGTCGTTGCCCTGTTTTGTGGTGGTTCGACGTTNNNAGATCGGAA |
| 116099 | TATGAGTGTGGAGTCGTTGCGGTCATGGACAAGGTGAGGGGTANNNAGATCGGAA |
| 107614 | TATGAGTGTGGAGTCGTTGCACAGGTGATCCTCCTCAGCCACNNNAGATCGGAAGA |
| 71744 | TATGAGTGTGGAGTCGTTGCGCAGCTCTCTACAGTCTCTTTCCNNNAGATCGGAAGA |
| 102788 | TATGAGTGTGGAGTCGTTGCACAACCCCGAAATGGAATAAGTAGGNNNAGATCGGA |
| 101210 | TATGAGTGTGGAGTCGTTGCTTGGGGAGATATCCAGTAATGCGNNNAGATCGGAA |
| 110394 | TATGAGTGTGGAGTCGTTGCGGACCATCTCTAGCCCATGTGANNNAGATCGGAAGA |
| 107589 | TATGAGTGTGGAGTCGTTGCGGCACACTGAAGCTGACAACNNNAGATCGGAAGAG |
| 107630 | TATGAGTGTGGAGTCGTTGCTACAAAGAGTGGTAAGTGAGCATTTNNNAGATCGGA |
| 108955 | TATGAGTGTGGAGTCGTTGCGGATTCTGAGTGTGTGCTTTGCNNNAGATCGGAAGA |
| 107574 | TATGAGTGTGGAGTCGTTGCAGGGTCACAATATTTATCAGGGCANNNAGATCGGAA |
| 101311 | TATGAGTGTGGAGTCGTTGCTCCTGCTGTCACTTCTGCATGTNNNAGATCGGAAGA |
| 105359 | TATGAGTGTGGAGTCGTTGCACTTTCCATCCTCAGGACAAGCANNNAGATCGGAAGA |
| 110431 | TATGAGTGTGGAGTCGTTGCTCAGCAAGAAGAGATTATACCAAGACANNNAGATCG |
| 100411 | TATGAGTGTGGAGTCGTTGCGGTGGTGGGCAGAAGATTGTATTANNNAGATCGGA |
| 105884 | TATGAGTGTGGAGTCGTTGCGCCTATTGTGCGGGACTACTCNNNAGATCGGAAGAG |
| 105340 | TATGAGTGTGGAGTCGTTGCCTTTCATACCAGGAAATCGAAGAAGNNNAGATCGGA |
| 110399 | TATGAGTGTGGAGTCGTTGCACTAGGCTCAAGCCATCTCCCTNNNAGATCGGAAGA |
| 112878 | TATGAGTGTGGAGTCGTTGCTCCTCCTCTCTCAGTATGCTGANNNAGATCGGAAGAG |
| 106454 | TATGAGTGTGGAGTCGTTGCTAGATTGTCCAGGTTCCAGGAAGNNNAGATCGGAA |
| 98708 | TATGAGTGTGGAGTCGTTGCGCCCACTGCACACTGTATTCAAANNNAGATCGGAAGA |
| 117621 | TATGAGTGTGGAGTCGTTGCCTGTGCTCCTCTTGCAGTTAGTTNNNAGATCGGAAGA |
| 117846 | TATGAGTGTGGAGTCGTTGCTCTTTCCCCTCTCTCTCTCTCCTGNNNAGATCGGAAGA |
| 63426 | TATGAGTGTGGAGTCGTTGCTCCCTCACACAAAACATTCTTCTNNNAGATCGGAAGA |
| 30497 | TATGAGTGTGGAGTCGTTGCCACAGGCACAGAAGAGACAGAAANNNAGATCGGAA |
| 52006 | TATGAGTGTGGAGTCGTTGCACTGTGCTGCGAGAATGCTATAGNNNAGATCGGAA |
| 100077 | TATGAGTGTGGAGTCGTTGCAGTCTCCCTCTCTTGTGGAGANNNAGATCGGAAGAG |
| 54242 | TATGAGTGTGGAGTCGTTGCTGTGTATACGAGTGACCCTTGGGNNNAGATCGGAA |
| 107462 | TATGAGTGTGGAGTCGTTGCAAACACAAGGGGGCTTCCTTTTNNNAGATCGGAAGA |
| 40205 | TATGAGTGTGGAGTCGTTGCGAGAGAAGAGACAGACAGGGTGANNNAGATCGGA |
| 109823 | TATGAGTGTGGAGTCGTTGCTAGAATGCAAGTCTGTGAGGGCANNNAGATCGGAA |
| 35402 | TATGAGTGTGGAGTCGTTGCACAATAGAAGGAAATTGGGAGCANNNAGATCGGAA |
| 41505 | TATGAGTGTGGAGTCGTTGCGCCTTTGGTGGAGAAGAGAGGTANNNAGATCGGAA |
| 39761 | TATGAGTGTGGAGTCGTTGCCCCCACCAACAATCACCTTATGANNNAGATCGGAAGA |

48585 TATGAGTGTGGAGTCGTTGCCGCACCAGAAAAACCTTCAACTCNNNAGATCGGAAG
106988 TATGAGTGTGGAGTCGTTGCAGTCAGGCCCTATGTAACAATGNNNAGATCGGAAG
48167 TATGAGTGTGGAGTCGTTGCGGTTGTATTGTTGTTGCTGTTGCNNNAGATCGGAAG
54893 TATGAGTGTGGAGTCGTTGCGATACCAAGTGCTTCTGAGGTGGNNNAGATCGGAA
97507 TATGAGTGTGGAGTCGTTGCTTTATTGGGAGAAGCTGGGTGCTNNNAGATCGGAA
40748 TATGAGTGTGGAGTCGTTGCGTTACACAGCTGAATGGGGCATANNNAGATCGGAA
33365 TATGAGTGTGGAGTCGTTGCCCTTGACAGGTTCTGGTATCACANNNAGATCGGAAG
32290 TATGAGTGTGGAGTCGTTGCGAACTCAGGGGATCCTGGGATAANNNAGATCGGAA
65977 TATGAGTGTGGAGTCGTTGCTGCTGACTTGAGGATATTACCTGTNNNAGATCGGAA
59897 TATGAGTGTGGAGTCGTTGCGGGGAGAGATTGAAGAGGGAACTNNNAGATCGGA
60623 TATGAGTGTGGAGTCGTTGCCCCAAAGCAAGAGAAAGGAACCTNNNAGATCGGAA
68785 TATGAGTGTGGAGTCGTTGCTGATTCCACGAGGACTGGAATTTNNNAGATCGGAA
69151 TATGAGTGTGGAGTCGTTGCTCACTCAGCATTTGCTTTGCTTTNNNAGATCGGAAGA
57934 TATGAGTGTGGAGTCGTTGCATGTCACAGATGCTCCAAGCAATNNNAGATCGGAAG
108149 TATGAGTGTGGAGTCGTTGCGGGGGAAGCTAGGAATGAACTGNNNAGATCGGAA
62545 TATGAGTGTGGAGTCGTTGCGCTGTCAATCCAGCATCAAATCTNNNAGATCGGAAG
62085 TATGAGTGTGGAGTCGTTGCAGAGAGACACAGAGAACATGGCTNNNAGATCGGAA
38499 TATGAGTGTGGAGTCGTTGCCTTAGTCTGGCAAGCTGTTTAGANNNAGATCGGAAG
103597 TATGAGTGTGGAGTCGTTGCTGGATCGCTGTAGAGTAGCATTGATNNNAGATCGG
52934 TATGAGTGTGGAGTCGTTGCAGACTATGAAGCCTCTCACCAGGNNNAGATCGGAAG
50632 TATGAGTGTGGAGTCGTTGCACACATCAGTCTGCAAGGAACAGNNNAGATCGGAA
66200 TATGAGTGTGGAGTCGTTGCGCAAAGATGAATACAGTACAGTCTGTNNNAGATCG
34807 TATGAGTGTGGAGTCGTTGCGAGCAAGCAAACTAGGACATCCANNNAGATCGGAA
41945 TATGAGTGTGGAGTCGTTGCGTGCCTGCCATACAGTCATGTTANNNAGATCGGAAG
55212 TATGAGTGTGGAGTCGTTGCTCAGCCAAGGAACAGACTTTTGGNNNAGATCGGAA
52690 TATGAGTGTGGAGTCGTTGCCGCCTGTTCTCTCTATCCAAAGGNNNAGATCGGAAG
42198 TATGAGTGTGGAGTCGTTGCTACAGATGTGGCTCTAGGCAAACNNNAGATCGGAA
36461 TATGAGTGTGGAGTCGTTGCACACATTTACACCTTGGTCATCANNNAGATCGGAAG
109791 TATGAGTGTGGAGTCGTTGCATGTGTGTGTTTGGGAGTAGAGTGTNNNAGATCGG
52396 TATGAGTGTGGAGTCGTTGCAAATAAACCTGTCACCATCCCCGNNNAGATCGGAAG
30461 TATGAGTGTGGAGTCGTTGCGAGCCCACCTTTTACTCTTCAAANNNAGATCGGAAG
48293 TATGAGTGTGGAGTCGTTGCCTTTTTGCTGGCCTTGAGGAATCNNNAGATCGGAAG
41504 TATGAGTGTGGAGTCGTTGCGCCAGGTTTGCCTCAGAGGTANNNAGATCGGAAGA
30119 TATGAGTGTGGAGTCGTTGCACAGGTATGCACATGGCAGAAAANNNAGATCGGAA
31947 TATGAGTGTGGAGTCGTTGCGAGAGAGAGAGGGGTGAATGGAANNNAGATCGG
59526 TATGAGTGTGGAGTCGTTGCACATTGGAGAACAGCCCTCCTATNNNAGATCGGAAG
65475 TATGAGTGTGGAGTCGTTGCGCAGTCTTTGAGATCACTGTGGTNNNAGATCGGAA
43488 TATGAGTGTGGAGTCGTTGCCAGAGCATAGGCAAGACCCATACNNNAGATCGGAA
63754 TATGAGTGTGGAGTCGTTGCGCAATTCAGGCACAAGAAAGAAGTNNNAGATCGGA
50771 TATGAGTGTGGAGTCGTTGCCACAAGAGCAAGAGCCTGTACAGNNNAGATCGGAA
36568 TATGAGTGTGGAGTCGTTGCAGAGAGAGAGAGATCATAGCACTCANNNAGATCGG
107036 TATGAGTGTGGAGTCGTTGCTATGTGATGATTGATGAGTGCACCAANNNAGATCG
40566 TATGAGTGTGGAGTCGTTGCTCTTCCATTTGGCTAAAGAGATTTTGANNNAGATCG

49490 TATGAGTGTGGAGTCGTTGCCGGCAAAGGATGGTTCCAAATTCNNNAGATCGGAA
69094 TATGAGTGTGGAGTCGTTGCGGCAGGGTGTCTTGATTAGCTTTNNNAGATCGGAA
36011 TATGAGTGTGGAGTCGTTGCACACGCACACTTAGTTCTATGCANNNAGATCGGAAG
31500 TATGAGTGTGGAGTCGTTGCACACATTAGAGGGAAATTTGTTCAANNNAGATCGG
98130 TATGAGTGTGGAGTCGTTGCCAAAGCAGCCTCTCCTTCCGNNNAGATCGGAAGAGC
46902 TATGAGTGTGGAGTCGTTGCAAAATAAAGGTCTGGGCCAGGCNNNAGATCGGAAG
39218 TATGAGTGTGGAGTCGTTGCATGAAAGAATGGCACTTTTGGGANNNAGATCGGAA
60378 TATGAGTGTGGAGTCGTTGCTGACTGGGTGGGGAAATATGACTNNNAGATCGGAA
47020 TATGAGTGTGGAGTCGTTGCGCAACCCTTATGTCTCCTTAGGCNNNAGATCGGAAG
34827 TATGAGTGTGGAGTCGTTGCTTGTCTCCTCAGCTTCAGATCCANNNAGATCGGAAGA
34445 TATGAGTGTGGAGTCGTTGCTTATCAGGTCATGGGAACTCCCANNNAGATCGGAAG
46185 TATGAGTGTGGAGTCGTTGCTGAGAATGAACAGGTGGAGAAGCNNNAGATCGGA
58500 TATGAGTGTGGAGTCGTTGCTCCTTCCTTCTTTACGGTCCCATNNNAGATCGGAAGA
42158 TATGAGTGTGGAGTCGTTGCGGTAGCAGGTAAGGGTGGAAAACNNNAGATCGGA
57166 TATGAGTGTGGAGTCGTTGCTTCCAGCTACCTTCTTGCCATTGNNNAGATCGGAAGA
47807 TATGAGTGTGGAGTCGTTGCTCAGAAATGAGGTCTCGCTCTGCNNNAGATCGGAAG
39387 TATGAGTGTGGAGTCGTTGCCCAGCACTTTCTTACTTTCTGGANNNAGATCGGAAGA
55417 TATGAGTGTGGAGTCGTTGCGAGGGCAGTATGATTTCCCCATGNNNAGATCGGAA
65063 TATGAGTGTGGAGTCGTTGCAGTGATACAAACTCTTCCAGGGTNNNAGATCGGAAG
48971 TATGAGTGTGGAGTCGTTGCGCGTGTATGTGTCTTGTTTGCTCNNNAGATCGGAAG
30372 TATGAGTGTGGAGTCGTTGCGTTCCATCCCCAACAGTAGCAAANNNAGATCGGAAG
64979 TATGAGTGTGGAGTCGTTGCGTTGATGCATTAGAAGTTTCTAGGTNNNAGATCGGA
101315 TATGAGTGTGGAGTCGTTGCGTGTAAGGATGATGGGGAGAGAANNNAGATCGGA
46482 TATGAGTGTGGAGTCGTTGCATTTTTGGCATTTCCTGCAGAGCNNNAGATCGGAAG
34514 TATGAGTGTGGAGTCGTTGCGGAATGAGTTGGGTGATTTCCCANNNAGATCGGAA
40582 TATGAGTGTGGAGTCGTTGCTGTCAGCAATAGTCTGCTTTTTGANNNAGATCGGAA
56905 TATGAGTGTGGAGTCGTTGCGAGATGAAAGGCTCTGGGATGTGNNNAGATCGGA
63967 TATGAGTGTGGAGTCGTTGCACCTATCTACATGAAGGGCCAGTNNNAGATCGGAAG
54881 TATGAGTGTGGAGTCGTTGCTTTCGCCACAATCCTTATCGTGGNNNAGATCGGAAG
58064 TATGAGTGTGGAGTCGTTGCAGGCAGGGGAGGTGAATTAGAATNNNAGATCGGA
68650 TATGAGTGTGGAGTCGTTGCCTTTCTTCCTGCTGCAGTCTGTTNNNAGATCGGAAGA
40974 TATGAGTGTGGAGTCGTTGCGGGCTGATGAGAAATTGGCACTANNNAGATCGGAA
39272 TATGAGTGTGGAGTCGTTGCTTGCCAGGATTCCCATAGATGGANNNAGATCGGAAG
59495 TATGAGTGTGGAGTCGTTGCCTGGCTCCTCTGCCACTTACTATNNNAGATCGGAAGA
42725 TATGAGTGTGGAGTCGTTGCTCCCTGTCTCTATTTCTCCCCACNNNAGATCGGAAGA
100082 TATGAGTGTGGAGTCGTTGCCCCCCGCCAATAGGTACATTTTNNNAGATCGGAAGA
48352 TATGAGTGTGGAGTCGTTGCAATGCCCTCTCCTTTTCACCATCNNNAGATCGGAAGA
56303 TATGAGTGTGGAGTCGTTGCTACAGTTGCAGGATGAGAATCTGNNNAGATCGGAA
104292 TATGAGTGTGGAGTCGTTGCCACCAGGCAGTTCTGCTTTCANNNAGATCGGAAGAG
61424 TATGAGTGTGGAGTCGTTGCAGACACCAGTGAATAGATTCTCCTNNNAGATCGGAA
56620 TATGAGTGTGGAGTCGTTGCGAAGAGATGTGGGGAAGAGAGTGNNNAGATCGGA
46536 TATGAGTGTGGAGTCGTTGCACACATACACACTTTCCAGGAGCNNNAGATCGGAAG
61711 TATGAGTGTGGAGTCGTTGCAAATGAACTTCCTAGCAGCAGCTNNNAGATCGGAAG

32083 TATGAGTGTGGAGTCGTTGCTATTTCAGGAAACGGCTACTGAANNNAGATCGGAAG

46437 TATGAGTGTGGAGTCGTTGCCACACAGACAACGATATCTTATTCAGCNNNAGATCGG

29886 TATGAGTGTGGAGTCGTTGCCAGAAGGATAAACTGGCTTGCCTNNNAGATCGGAAG

33735 TATGAGTGTGGAGTCGTTGCAAAGACTTGGGGCATCTTTGACANNNAGATCGGAAG

37204 TATGAGTGTGGAGTCGTTGCCCATCACACCTCATTTCTCCTTCANNNAGATCGGAAGA

51271 TATGAGTGTGGAGTCGTTGCTGTGATGTTATGCACCCAGTCAGNNNAGATCGGAAG

36357 TATGAGTGTGGAGTCGTTGCTTTCCTTTCACAGGACCACAATCANNNAGATCGGAAG

116142 TATGAGTGTGGAGTCGTTGCTAGCCATCTGCAACCCAGGAATANNNAGATCGGAAG

95108 TATGAGTGTGGAGTCGTTGCCGATTGGTCAGTGAGTTCCTGTTNNNAGATCGGAAG

59887 TATGAGTGTGGAGTCGTTGCAGGACTTAGCCCCATGTAGAACTNNNAGATCGGAAG

57150 TATGAGTGTGGAGTCGTTGCTTGGAGGCTGATGATGGGAATTGNNNAGATCGGAA

52490 TATGAGTGTGGAGTCGTTGCCCTGGCAATAGAAATTCACAGCGNNNAGATCGGAAG

50531 TATGAGTGTGGAGTCGTTGCTTAGGGACAGGGGGATCTTGAAGNNNAGATCGGAA

64761 TATGAGTGTGGAGTCGTTGCGAGACAGAGAAGCCTTTCAAGGTNNNAGATCGGAA

52947 TATGAGTGTGGAGTCGTTGCATTTTCTGTGCTGTGTACCCAGGNNNAGATCGGAAG

32704 TATGAGTGTGGAGTCGTTGCACCTCCATTTGCTTTCCCAAACANNNAGATCGGAAGA

34879 TATGAGTGTGGAGTCGTTGCAATCACTTCTTACACCACCTCCANNNAGATCGGAAGA

60780 TATGAGTGTGGAGTCGTTGCAGTGCCTGTTCAGGTAGATACCTNNNAGATCGGAAG

64259 TATGAGTGTGGAGTCGTTGCGCCAGAAGCATACATGTAGGAGTNNNAGATCGGAA

49461 TATGAGTGTGGAGTCGTTGCTAGGGGTCTCACTAAGCTGTGTCNNNAGATCGGAAG

54153 TATGAGTGTGGAGTCGTTGCCTTTGACATCAGATGGGTCTGGGNNNAGATCGGAA

40732 TATGAGTGTGGAGTCGTTGCCCGTCAGCCATCTTTGTTCCATANNNAGATCGGAAGA

66160 TATGAGTGTGGAGTCGTTGCCCTGCTTCATTCTCCAACTCTGTNNNAGATCGGAAGA

52676 TATGAGTGTGGAGTCGTTGCGCCAGGGGAGTATTACTGAAAAGGNNNAGATCGGA

47415 TATGAGTGTGGAGTCGTTGCTGGGGTTGAATGGAAAATTTGGCNNNAGATCGGAA

52849 TATGAGTGTGGAGTCGTTGCCCATCTGCAACCCAGGAATAAGGNNNAGATCGGAAG

52589 TATGAGTGTGGAGTCGTTGCTTCTGGGTTCTCTCTACTCCTCGNNNAGATCGGAAGA

64439 TATGAGTGTGGAGTCGTTGCCGAAAGACTGAGAATGCGCTAGTNNNAGATCGGAA

53480 TATGAGTGTGGAGTCGTTGCATTTTAGGCACACAACAGACCGGNNNAGATCGGAAG

50227 TATGAGTGTGGAGTCGTTGCGTATCCGAAAGGGCCAGACAAGNNNAGATCGGAAG

31276 TATGAGTGTGGAGTCGTTGCACAGAGAGAAGCAAATGTGGCAANNNAGATCGGAA

36411 TATGAGTGTGGAGTCGTTGCAGACAGAGAGGGAAAGAGTACATCANNNAGATCGG

36960 TATGAGTGTGGAGTCGTTGCAGTAGATTCCACATCGTGGGTCANNNAGATCGGAAG

67841 TATGAGTGTGGAGTCGTTGCGGGCCTTGGTCTAACATAAGCTTNNNAGATCGGAAG

65262 TATGAGTGTGGAGTCGTTGCTGGAGTAAAGGATGAAACATGGTNNNAGATCGGAA

52431 TATGAGTGTGGAGTCGTTGCACTGTAAGATTTATGCCTGGCCGNNNAGATCGGAAG

32526 TATGAGTGTGGAGTCGTTGCCCTGCACAATACCACAGGTGTAANNNAGATCGGAAG

60786 TATGAGTGTGGAGTCGTTGCCGCACGTGTCTATCTACCTACCTNNNAGATCGGAAGA

35953 TATGAGTGTGGAGTCGTTGCAAAAGTGTGTCTACCTGCATGCANNNAGATCGGAAG

51882 TATGAGTGTGGAGTCGTTGCTTTTGAATGAGGCCTCGAGTGAGNNNAGATCGGAA

67388 TATGAGTGTGGAGTCGTTGCGTGTTGGGAAGCCTGTGACTATTNNNAGATCGGAA

51801 TATGAGTGTGGAGTCGTTGCGAGCTACAAATGGTGGGGATGAGNNNAGATCGGAA

116388 TATGAGTGTGGAGTCGTTGCCACTCCTTCTCTCTCTCTTCCTANNNAGATCGGAAG

| | |
|---|---|
| 100404 | TATGAGTGTGGAGTCGTTGCTGCCTGGCCCACAAAGATACTTTANNNAGATCGGAA |
| 43509 | TATGAGTGTGGAGTCGTTGCCCTTTTCAGGTGGCCGGATATACNNNAGATCGGAAG |
| 52877 | TATGAGTGTGGAGTCGTTGCCAGTCTCATGGTAAAAAGTACAACAGGNNNAGATCG |
| 35398 | TATGAGTGTGGAGTCGTTGCGAAATGGAGAAAGGAAGGGAGCANNNAGATCGGA |
| 30757 | TATGAGTGTGGAGTCGTTGCACTGGTGCGTGTGTTTCTCTAAANNNAGATCGGAAG |
| 67403 | TATGAGTGTGGAGTCGTTGCCTGCCTTCTGACCCTTTGCTATTNNNAGATCGGAAGA |
| 62081 | TATGAGTGTGGAGTCGTTGCGATTTCCGGGTTCACTAATGGCTNNNAGATCGGAAG |
| 58818 | TATGAGTGTGGAGTCGTTGCTTCTGTCTGGTACTCTGCCTCATNNNAGATCGGAAGA |
| 101600 | TATGAGTGTGGAGTCGTTGCACCTTTTCCTTCTCCATGAAGCNNNAGATCGGAAGAG |
| 46448 | TATGAGTGTGGAGTCGTTGCAGTCGATCCCAAGTGAAAAGAGCNNNAGATCGGAA |
| 34436 | TATGAGTGTGGAGTCGTTGCAATGAGAGATGTTTTGGATATCCCANNNAGATCGGA |
| 35028 | TATGAGTGTGGAGTCGTTGCTTCTGGCTTCCTTCAGTATTCCANNNAGATCGGAAGA |
| 55303 | TATGAGTGTGGAGTCGTTGCCCAGGAAGACTATAGGCCGAATGNNNAGATCGGAA |
| 66006 | TATGAGTGTGGAGTCGTTGCTCCTCCCTTCTATATTTATGCCTGTNNNAGATCGGAA |
| 48463 | TATGAGTGTGGAGTCGTTGCCTCATCACACATTCCCCTCGATCNNNAGATCGGAAGA |
| 99279 | TATGAGTGTGGAGTCGTTGCCTAAAAAGGGGTCTTCCTGAGCTNNNAGATCGGAA |
| 68830 | TATGAGTGTGGAGTCGTTGCGCTCTCATGGTAAGCTGCCATTTNNNAGATCGGAAG |
| 54558 | TATGAGTGTGGAGTCGTTGCAGTTCTAGGCTGTCTGCTACTGGNNNAGATCGGAAG |
| 38928 | TATGAGTGTGGAGTCGTTGCAGGGGGAAAAATGACATATAACGGANNNAGATCG |
| 38893 | TATGAGTGTGGAGTCGTTGCTTCCTTTCCTACAGTCCCTAGGANNNAGATCGGAAGA |
| 30473 | TATGAGTGTGGAGTCGTTGCACGGGTGGGATAGTGGAAAGAAANNNAGATCGGA |
| 36726 | TATGAGTGTGGAGTCGTTGCCCAGAATGATTGGCCTATGCTCANNNAGATCGGAAG |
| 106491 | TATGAGTGTGGAGTCGTTGCCCCATTAGACTGTCCAGGCTCTTCNNNAGATCGGAA |
| 30006 | TATGAGTGTGGAGTCGTTGCACAAATACACTCTAGTGCTCAAAAANNNAGATCGGA |
| 61868 | TATGAGTGTGGAGTCGTTGCGCACTGATGTGATCATGGTAGCTNNNAGATCGGAA |
| 48904 | TATGAGTGTGGAGTCGTTGCGGCTTGGGTTTCAAATATGGCTCNNNAGATCGGAA |
| 40293 | TATGAGTGTGGAGTCGTTGCCAAGGCCAGTTAGTTAGCTGTGANNNAGATCGGAA |
| 44900 | TATGAGTGTGGAGTCGTTGCATTATGAAGTTGAGGCCACAGCCNNNAGATCGGAA |
| 53052 | TATGAGTGTGGAGTCGTTGCTGGAGAACTCTGATCAATGCAGGNNNAGATCGGAA |
| 96862 | TATGAGTGTGGAGTCGTTGCCCATTTTCTGTGTTGGGGTTTCCTNNNAGATCGGAA |
| 61899 | TATGAGTGTGGAGTCGTTGCCTTTTGTTCCTCTGGAGAACGCTNNNAGATCGGAAG |
| 32123 | TATGAGTGTGGAGTCGTTGCTAAAGGCCAAGAGCGAATCTGAANNNAGATCGGAA |
| 107006 | TATGAGTGTGGAGTCGTTGCGCTCACCATCCTCTGTTTGCANNNAGATCGGAAGAG |
| 41572 | TATGAGTGTGGAGTCGTTGCATCCATTTGCACAGATGCTGGTANNNAGATCGGAAG |
| 68665 | TATGAGTGTGGAGTCGTTGCGGGAAATGACCTTTGGGAGTGTTNNNAGATCGGAA |
| 115683 | TATGAGTGTGGAGTCGTTGCGAGGAAGGAGAAAGTCCCTTGTCANNNAGATCGGA |
| 100428 | TATGAGTGTGGAGTCGTTGCTTGACACATGAATATGGAGCACANNNAGATCGGAA |
| 98441 | TATGAGTGTGGAGTCGTTGCGAGTGTGTTGGTCGAGAGCAATCNNNAGATCGGAA |
| 99039 | TATGAGTGTGGAGTCGTTGCGACAGTTCACTTTATCCCCCAGGNNNAGATCGGAAG |
| 36883 | TATGAGTGTGGAGTCGTTGCTTTCTCCCAGGGATATGCAGTCANNNAGATCGGAAG |
| 58931 | TATGAGTGTGGAGTCGTTGCAGCATTTTCTTATCTTCGGTTTTCATNNNAGATCGGA |
| 37708 | TATGAGTGTGGAGTCGTTGCTGGGAAAGAAGAAAGGACAGTAAGANNNAGATCG |
| 41577 | TATGAGTGTGGAGTCGTTGCCCTGCTAGCTCTGTTCTCTGGTANNNAGATCGGAAG |

106980 TATGAGTGTGGAGTCGTTGCCAGACGTAAGCATATTCGTGGGGNNNAGATCGGAA
56649 TATGAGTGTGGAGTCGTTGCAGCCTGAGTGTTACGTGAGTGNNNAGATCGGAAGA
48115 TATGAGTGTGGAGTCGTTGCGGCTTCATCTTTCATTGGCTTGCNNNAGATCGGAAG
42516 TATGAGTGTGGAGTCGTTGCAACTTGTCCACCAGGCTAAACACNNNAGATCGGAAG
44483 TATGAGTGTGGAGTCGTTGCGGAGTAAGTCAAAGATGTGTCCCCNNNAGATCGGA
36372 TATGAGTGTGGAGTCGTTGCTGGGCTTCTAAACTGATGAAGAATCANNNAGATCGG
117408 TATGAGTGTGGAGTCGTTGCCTGTCTGCCCTTTTCTGTCTGGNNNAGATCGGAAGA
102030 TATGAGTGTGGAGTCGTTGCGCATTTTCTTATCTTCGGTTTTCATTTNNNAGATCGG
38149 TATGAGTGTGGAGTCGTTGCTCACGTAGTGGGAGAGAGAGAGANNNAGATCGGA
39155 TATGAGTGTGGAGTCGTTGCAGTCTCAGTACAAACTTCCTGGGANNNAGATCGGAA
56812 TATGAGTGTGGAGTCGTTGCCGAGATGTTGAGTTTACTTTGGGTGNNNAGATCGG
46968 TATGAGTGTGGAGTCGTTGCCTTTGCAACAAGATATGGGAGGCNNNAGATCGGAA
63824 TATGAGTGTGGAGTCGTTGCCACTTGCTTCTGTACCCCTAAGTNNNAGATCGGAAGA
67595 TATGAGTGTGGAGTCGTTGCCCTGACCTGGAGTGTTTTGACTTNNNAGATCGGAAG
29998 TATGAGTGTGGAGTCGTTGCCAGCTTGTGTCACTCCCCAAAAANNNAGATCGGAAG
39138 TATGAGTGTGGAGTCGTTGCGGAAGGGAAGGGAGATTATGGGANNNAGATCGGA
100085 TATGAGTGTGGAGTCGTTGCAAGTAGCAAGTACTGGCAAGGATNNNAGATCGGAA
29891 TATGAGTGTGGAGTCGTTGCGCCTGGTGCAAATAAGAGTGAGANNNAGATCGGAA
56156 TATGAGTGTGGAGTCGTTGCGAGAATGCTGCTGGTAGTAGCTGNNNAGATCGGAA
63600 TATGAGTGTGGAGTCGTTGCATTGTACCGGCCTTGGTTTTTCTNNNAGATCGGAAG
43897 TATGAGTGTGGAGTCGTTGCAGATTGGGTTACAACAACCCACCNNNAGATCGGAAG
55312 TATGAGTGTGGAGTCGTTGCATGCCAGTTTTGCAATGGGAATGNNNAGATCGGAA
69113 TATGAGTGTGGAGTCGTTGCTAGGAGCATAATCCCTGGGCTTTNNNAGATCGGAAG
50085 TATGAGTGTGGAGTCGTTGCGAATGAAGTGAAACGGCCCAAAGNNNAGATCGGAA
105869 TATGAGTGTGGAGTCGTTGCTGGTGGCAGGTCACATGGTTATTNNNAGATCGGAA
51022 TATGAGTGTGGAGTCGTTGCATGTACCAGCATTTCTCCAGCAGNNNAGATCGGAAG
106987 TATGAGTGTGGAGTCGTTGCGTGTGTGTGGAGATGGCTTATCTNNNAGATCGGAA
40682 TATGAGTGTGGAGTCGTTGCGCTCCCCTTCTCTCTCACACATANNNAGATCGGAAGA
39954 TATGAGTGTGGAGTCGTTGCTTTCTCCAACCACTTTCAGCTGANNNAGATCGGAAGA
49551 TATGAGTGTGGAGTCGTTGCAACTCCTCCCTAAGGGGTCATTCNNNAGATCGGAAG
30810 TATGAGTGTGGAGTCGTTGCTGGAAATCTATTTGGTTCCTAAACAANNNAGATCGG
48444 TATGAGTGTGGAGTCGTTGCTTCACACTGGGCTTCAACAGATCNNNAGATCGGAAG
45481 TATGAGTGTGGAGTCGTTGCTTGGCTGTTTCTCAGGTTGATCCNNNAGATCGGAAG
52995 TATGAGTGTGGAGTCGTTGCTATGCAGGAATTCATGTGCCAGGNNNAGATCGGAA
108839 TATGAGTGTGGAGTCGTTGCTTGGAGTTTCTCATGGTTAGGGCNNNAGATCGGAA
46617 TATGAGTGTGGAGTCGTTGCAGCCTCTTTTCAGTCAGTTGAGCNNNAGATCGGAAG
35878 TATGAGTGTGGAGTCGTTGCCTTGCTCTAGGAATTGCAAAATGCANNNAGATCGGA
100599 TATGAGTGTGGAGTCGTTGCTTGGGGAGCAAGCAAAGTTTCANNNAGATCGGAAG
38797 TATGAGTGTGGAGTCGTTGCTACATAAAGACAGCCCCAGAGGANNNAGATCGGAA
58867 TATGAGTGTGGAGTCGTTGCGTTCTGCACAGGTACTTGGTCATNNNAGATCGGAAG
109397 TATGAGTGTGGAGTCGTTGCTGGGGCAGGTGGTATTTCAAAANNNAGATCGGAAG
29885 TATGAGTGTGGAGTCGTTGCATTAGCTGTGGGAAAGGGAGACTNNNAGATCGGA
101108 TATGAGTGTGGAGTCGTTGCTGGGGACTTACTAGATTGTCCAAGCNNNAGATCGG

52791 TATGAGTGTGGAGTCGTTGCTTCACCCTGGTTTTGGAAGAAGGNNNAGATCGGAA
60604 TATGAGTGTGGAGTCGTTGCAGCATACATACATACAAGCAACCTNNNAGATCGGAA
42241 TATGAGTGTGGAGTCGTTGCTCTGAGGCCTCTTTCCTTCTAAACNNNAGATCGGAAG
107410 TATGAGTGTGGAGTCGTTGCGGCACCATGAGTGATATCTTCCTNNNAGATCGGAAG
53970 TATGAGTGTGGAGTCGTTGCGCACACAAGTAATTCAACTGGGGNNNAGATCGGAA
51100 TATGAGTGTGGAGTCGTTGCGGAGTTTCTCATGGTTAGGGCAGNNNAGATCGGAA
64229 TATGAGTGTGGAGTCGTTGCTTCCAAGGCCATGTCTCTAGAGTNNNAGATCGGAAG
42388 TATGAGTGTGGAGTCGTTGCTCATGCAAAGGAAAATGCCGAACNNNAGATCGGAA
64986 TATGAGTGTGGAGTCGTTGCACCAAAAAGTGCAGAAAGGTAGGTNNNAGATCGGA
71620 TATGAGTGTGGAGTCGTTGCTTGCTTTCAACTTTGGGTGGCTNNNAGATCGGAAGA
51894 TATGAGTGTGGAGTCGTTGCAAAGCAAGCTGAGATGAGGTGAGNNNAGATCGGA
60798 TATGAGTGTGGAGTCGTTGCTGTATGGGTTCAGTGCTCTACCTNNNAGATCGGAAG
106472 TATGAGTGTGGAGTCGTTGCTAGTTATTGCAGACGGGGCAAATNNNAGATCGGAA
54422 TATGAGTGTGGAGTCGTTGCAAAAGCAATGAGGGTGGAGATGGNNNAGATCGGA
47792 TATGAGTGTGGAGTCGTTGCCTTGTTGCGGACTTTGTATCTGCNNNAGATCGGAAG
57724 TATGAGTGTGGAGTCGTTGCAGGTCAAACATCCCATGCCAAATNNNAGATCGGAAG
65563 TATGAGTGTGGAGTCGTTGCTCTCTAGGCTTTTCCCCTTTGGTNNNAGATCGGAAGA
56474 TATGAGTGTGGAGTCGTTGCAGCACATACACAGAAACATTCTTCTGNNNAGATCGG
39830 TATGAGTGTGGAGTCGTTGCGCCTGTTTTCCATTTGGGACTGANNNAGATCGGAAG
53716 TATGAGTGTGGAGTCGTTGCGACAAGCCTTCAGATGATCAGGGNNNAGATCGGAA
43249 TATGAGTGTGGAGTCGTTGCTCTCCAGTTTCTTTGTGACCGACNNNAGATCGGAAG
34782 TATGAGTGTGGAGTCGTTGCTGAATTGTCTCCCTCTCAAATCCANNNAGATCGGAAG
109830 TATGAGTGTGGAGTCGTTGCGAATTGGAGGTAGCCAGTGTCAANNNAGATCGGAA
58286 TATGAGTGTGGAGTCGTTGCGGGGTGTGGTGATACAAACACATNNNAGATCGGAA
32387 TATGAGTGTGGAGTCGTTGCCTTAGTGCCATCTTGCCAGCTAANNNAGATCGGAAG
60618 TATGAGTGTGGAGTCGTTGCAGATGTAGCAATCCCCAGAACCTNNNAGATCGGAAG
67823 TATGAGTGTGGAGTCGTTGCTTCGTGGTAAGCAGCTTTTCCTTNNNAGATCGGAAG
48325 TATGAGTGTGGAGTCGTTGCGTTACAAGAACCCAACGCACATCNNNAGATCGGAAG
33816 TATGAGTGTGGAGTCGTTGCTACCTAACACACACAGGCCTACANNNAGATCGGAAG
64890 TATGAGTGTGGAGTCGTTGCCAACAAACCAATCGTAACGAGGTNNNAGATCGGAAG
48693 TATGAGTGTGGAGTCGTTGCACAGAAGGAGTGTGCTTTACCTCNNNAGATCGGAAG
107004 TATGAGTGTGGAGTCGTTGCAGAAAATGCTCAAGCAAGCAGGAGNNNAGATCGGA
46467 TATGAGTGTGGAGTCGTTGCTAGAAGCCCTGAGTAGACAGAGCNNNAGATCGGAA
56292 TATGAGTGTGGAGTCGTTGCTTGCTGCTGTTATTGTGTTGCTGNNNAGATCGGAAG
36322 TATGAGTGTGGAGTCGTTGCAAGTGAGAAGTCTGCTGTTTGCANNNAGATCGGAA
63716 TATGAGTGTGGAGTCGTTGCTTCTGAGACCCCTGATTCCAAGTNNNAGATCGGAAG
54029 TATGAGTGTGGAGTCGTTGCAACAATGAGGCTGAAGGAATGGGNNNAGATCGGA
66795 TATGAGTGTGGAGTCGTTGCTTGCTGGCCAATTATCCCTTTGTNNNAGATCGGAAG
103053 TATGAGTGTGGAGTCGTTGCCTTGGCTCAGTGACAGTCTCTAGTTNNNAGATCGGA
103289 TATGAGTGTGGAGTCGTTGCCTCTTACCATGCACTGTGGCTTTATNNNAGATCGGAA
37862 TATGAGTGTGGAGTCGTTGCTTTCTTCTGGGTAAAGTCCCAGANNNAGATCGGAAG
68858 TATGAGTGTGGAGTCGTTGCCAGCCAAGATGGGACCATCATTTNNNAGATCGGAAG
34439 TATGAGTGTGGAGTCGTTGCAAGCTCATCCTACACGTATCCCANNNAGATCGGAAG

| | |
|---|---|
| 38070 | TATGAGTGTGGAGTCGTTGCTGGTGCTATTGTGAGGTTTCAGANNNAGATCGGAA |
| 50029 | TATGAGTGTGGAGTCGTTGCGGGAGGAAGAGAGAAGCCAAAAGNNNAGATCGGA |
| 53226 | TATGAGTGTGGAGTCGTTGCTAATGGAGAAGAAGGTGGGGAGGNNNAGATCGG |
| 54763 | TATGAGTGTGGAGTCGTTGCAACCAGCAAACCTTCTTCTCTGGNNNAGATCGGAAG |
| 56449 | TATGAGTGTGGAGTCGTTGCGGGAGAGGGAGAGAGGAATTCTGNNNAGATCGGA |
| 54836 | TATGAGTGTGGAGTCGTTGCCTCTTCTCAATCCACTCCAGTGGNNNAGATCGGAAGA |
| 57828 | TATGAGTGTGGAGTCGTTGCCTGGAGCACCCCGACTAATAAATNNNAGATCGGAAG |
| 61691 | TATGAGTGTGGAGTCGTTGCGAATGCCAGACTTTACACCAGCTNNNAGATCGGAAG |
| 30124 | TATGAGTGTGGAGTCGTTGCTCAGTAACCAACGCTGGAGAAAANNNAGATCGGAA |
| 103560 | TATGAGTGTGGAGTCGTTGCGCCACTGAAGAGCAACCAAAATAATNNNAGATCGG |
| 62868 | TATGAGTGTGGAGTCGTTGCCATAACACCCCTCTTCAGCCTCTNNNAGATCGGAAGA |
| 106512 | TATGAGTGTGGAGTCGTTGCTTTGGCTGTTATTGCCTACGTTATCCNNNAGATCGGA |
| 49589 | TATGAGTGTGGAGTCGTTGCATTCGTGGGGTCACTTGCTATTCNNNAGATCGGAAG |
| 67474 | TATGAGTGTGGAGTCGTTGCGGAGCATGCATAAGGTCCAACTTNNNAGATCGGAA |
| 51636 | TATGAGTGTGGAGTCGTTGCGGTGCAGGTTTGCTTTTTAGGAGNNNAGATCGGAA |
| 38910 | TATGAGTGTGGAGTCGTTGCTGCTCATTATTTTTGAGGGTAGGANNNAGATCGGA |
| 32482 | TATGAGTGTGGAGTCGTTGCTGGCTGTGAGCTGTACTAGGTAANNNAGATCGGAA |
| 36308 | TATGAGTGTGGAGTCGTTGCTCTTCTGATGGAGTTGCTTTGCANNNAGATCGGAAG |
| 37882 | TATGAGTGTGGAGTCGTTGCATTCTGATCCCTGATCTGCCAGANNNAGATCGGAAG |
| 30057 | TATGAGTGTGGAGTCGTTGCCACACACGAACCCAAAACCAAAANNNAGATCGGAAG |
| 116752 | TATGAGTGTGGAGTCGTTGCACTAAATCAGCTTCAATCATCACAGCCNNNAGATCGG |
| 117511 | TATGAGTGTGGAGTCGTTGCTTACCAGCTCTTTGAAGGAATATTCANNNAGATCGG |
| 31506 | TATGAGTGTGGAGTCGTTGCGGGGTTGAGTACTAGCCTTTCAANNNAGATCGGAA |
| 39084 | TATGAGTGTGGAGTCGTTGCTAGCAATCACCATGTGTAGGGGANNNAGATCGGAA |
| 50616 | TATGAGTGTGGAGTCGTTGCCAGCCTAAGCCCTTCCTAAACAGNNNAGATCGGAAG |
| 32729 | TATGAGTGTGGAGTCGTTGCTGCTGTAACTTCTGGTTCAAACANNNAGATCGGAAG |
| 61193 | TATGAGTGTGGAGTCGTTGCGGATAGCTAGTGGTATGTGGCCTNNNAGATCGGAA |
| 39305 | TATGAGTGTGGAGTCGTTGCTGGCATGAGCTTATTTACAACTGGANNNAGATCGGA |
| 33113 | TATGAGTGTGGAGTCGTTGCACCCATGTACATACAACATACACANNNAGATCGGAA |
| 95115 | TATGAGTGTGGAGTCGTTGCGTTAGAGAGAAGCCCCATCACCTNNNAGATCGGAAG |
| 40184 | TATGAGTGTGGAGTCGTTGCTTTATGGAGAGGCCTCAAGGTGANNNAGATCGGAA |
| 101591 | TATGAGTGTGGAGTCGTTGCTCCTCATCCTTTTCAACTCTTGGAANNNAGATCGGAA |
| 104754 | TATGAGTGTGGAGTCGTTGCGAAGGAAGCTTGGACAATGAGTACANNNAGATCGG |
| 58510 | TATGAGTGTGGAGTCGTTGCGGTTGTTTGGAGTACTCAGCCATNNNAGATCGGAA |
| 105833 | TATGAGTGTGGAGTCGTTGCTTTAGTTGATGGACATTTAGGCTGGNNNAGATCGG |
| 56141 | TATGAGTGTGGAGTCGTTGCAATGCACACTGATGGTAGAGCTGNNNAGATCGGAA |
| 34184 | TATGAGTGTGGAGTCGTTGCTGAAAGAATGGTGTTGAAAGTACCANNNAGATCGG |
| 54779 | TATGAGTGTGGAGTCGTTGCGTAGAGATTGGGGCACATTCTGGNNNAGATCGGAA |
| 32309 | TATGAGTGTGGAGTCGTTGCGGATCTGCTGGGGTGGACTATAANNNAGATCGGAA |
| 47032 | TATGAGTGTGGAGTCGTTGCCAGGAGGGAAGGATGTTTTAGGCNNNAGATCGGA |
| 30040 | TATGAGTGTGGAGTCGTTGCCTACCTGCCAGCATGGGTAAAAANNNAGATCGGAA |
| 62865 | TATGAGTGTGGAGTCGTTGCCAGCTTCAATCATCACAGCCTCTNNNAGATCGGAAGA |
| 98799 | TATGAGTGTGGAGTCGTTGCAATGATGGTGGTGAAGACCAGTGNNNAGATCGGAA |

97789 TATGAGTGTGGAGTCGTTGCACCCAGAGTAGGAGGGAGGANNNAGATCGGAAGA
100052 TATGAGTGTGGAGTCGTTGCCAAAACACTCTTCAGAACCTCATCNNNAGATCGGAAG
98754 TATGAGTGTGGAGTCGTTGCGGCACTAAGCACTCTCTGTACCCNNNAGATCGGAAG
38429 TATGAGTGTGGAGTCGTTGCTACCTGCTACCTCCCCTTCTAGANNNAGATCGGAAGA
59859 TATGAGTGTGGAGTCGTTGCACTTCATGTGGGCAAAGTCAACTNNNAGATCGGAAG
54843 TATGAGTGTGGAGTCGTTGCATTACCCAGTTTGGATGCAGTGGNNNAGATCGGAA
62465 TATGAGTGTGGAGTCGTTGCTAACAGAGCCAATCATGCTTGCTNNNAGATCGGAAG
65854 TATGAGTGTGGAGTCGTTGCATGCAGGTGGTAGGAGTGTATGTNNNAGATCGGA
43766 TATGAGTGTGGAGTCGTTGCAGACGGGGCAAATTCTCTAAACCNNNAGATCGGAA
49022 TATGAGTGTGGAGTCGTTGCGAAGGGGCATTATTCAGCCTCTCNNNAGATCGGAAG
66633 TATGAGTGTGGAGTCGTTGCTCTCCTCCAGTGATTAGCCTTGTNNNAGATCGGAAG
49770 TATGAGTGTGGAGTCGTTGCTGCTTTGTTTCAAGCCTCAGTTCNNNAGATCGGAAG
60424 TATGAGTGTGGAGTCGTTGCCACTCCAAACTGTGCTCTTGACTNNNAGATCGGAAG
48751 TATGAGTGTGGAGTCGTTGCCACGATCTTCTGCTTCAGCCTCNNNAGATCGGAAGA
100642 TATGAGTGTGGAGTCGTTGCGCCAGGTTTTGGTATCAGGAATGCNNNAGATCGGA
37819 TATGAGTGTGGAGTCGTTGCTGGAAAAGTGGTAGAGGCTACAGANNNAGATCGG
66156 TATGAGTGTGGAGTCGTTGCTCCTGGCAGACCTTGATATCTGTNNNAGATCGGAAG
69556 TATGAGTGTGGAGTCGTTGCTGGGCTAGGATGCAAGTAGTTTTNNNAGATCGGAA
40385 TATGAGTGTGGAGTCGTTGCGCACTCGTGTTTTTAGTTTAGATTGANNNAGATCGG
44794 TATGAGTGTGGAGTCGTTGCATAACACCCCATGCAACTTCCCNNNAGATCGGAAGA
39354 TATGAGTGTGGAGTCGTTGCGAAAGATGCAGGAAAGAGCTGGANNNAGATCGGA
64523 TATGAGTGTGGAGTCGTTGCTGCCATCTATATTTGAAAGAAACGTNNNAGATCGGA
44426 TATGAGTGTGGAGTCGTTGCCTTCTTTTCAGGACACTAGCCCCNNNAGATCGGAAGA
64430 TATGAGTGTGGAGTCGTTGCGAGGTGACCATGAAGGACCTAGTNNNAGATCGGAA
31383 TATGAGTGTGGAGTCGTTGCGCCTCATTGGCCTTGGATATCAANNNAGATCGGAAG
60969 TATGAGTGTGGAGTCGTTGCGTCTCTCACAACAAATGTGCCCTNNNAGATCGGAAG
44080 TATGAGTGTGGAGTCGTTGCAAAACCTAAAAGTGCCCTGGACCNNNAGATCGGAAG
39070 TATGAGTGTGGAGTCGTTGCAGAGGATTTGAGCTAACAGGGGANNNAGATCGGA
46836 TATGAGTGTGGAGTCGTTGCCCATCCTCACATCATCTCAAGGCNNNAGATCGGAAGA
67773 TATGAGTGTGGAGTCGTTGCGGGTGTTGAAAGTGGGTATCCTTNNNAGATCGGAA
59104 TATGAGTGTGGAGTCGTTGCGCATGTATGTTACTCCTCACGATNNNAGATCGGAAG
43112 TATGAGTGTGGAGTCGTTGCACCTGCACTGTGGAAGAAAAGACNNNAGATCGGAA
63677 TATGAGTGTGGAGTCGTTGCCCCAGAGCTCAAAAAGGTAAAGTNNNAGATCGGAA
105838 TATGAGTGTGGAGTCGTTGCGTGCGGGTATGTTGACCAATAACCNNNAGATCGGA
50045 TATGAGTGTGGAGTCGTTGCGCATTGTTTGCCTCTGGGAAAAGNNNAGATCGGAA
42928 TATGAGTGTGGAGTCGTTGCGGATAAGTTTTACGTGGCTGCACNNNAGATCGGAA
64307 TATGAGTGTGGAGTCGTTGCAGCCAGGCAAATTGGAAATGAGTNNNAGATCGGAA
102056 TATGAGTGTGGAGTCGTTGCCAATTCTCCCCTTGTGTGGAGACNNNAGATCGGAAG
64957 TATGAGTGTGGAGTCGTTGCAGATTGTTGGGTGCAAGATAGGTNNNAGATCGGAA
34270 TATGAGTGTGGAGTCGTTGCATGGCCTCGTCATCATATACCCANNNAGATCGGAAG
68959 TATGAGTGTGGAGTCGTTGCACCTCTCCTGAGCTCTGAACTTTNNNAGATCGGAAGA
106984 TATGAGTGTGGAGTCGTTGCATCAGAAACCATTTCACTTCAGCCANNNAGATCGGA
42667 TATGAGTGTGGAGTCGTTGCTGGAAATAAAGGGCCAGAACCACNNNAGATCGGAA

| | |
|---|---|
| 41038 | TATGAGTGTGGAGTCGTTGCGGGGAACAACTGAGCCTTACCTANNNAGATCGGAA |
| 62621 | TATGAGTGTGGAGTCGTTGCTACCCCCAGACCTATCTCCATCTNNNAGATCGGAAGA |
| 51469 | TATGAGTGTGGAGTCGTTGCGGCAAGGGGAGAAAAAGAGAGAGNNNAGATCGG |
| 31040 | TATGAGTGTGGAGTCGTTGCACATTTATTGGCTTCAGGCCCAANNNAGATCGGAAG |
| 61865 | TATGAGTGTGGAGTCGTTGCATTACCAGTATGGACGGGTAGCTNNNAGATCGGAA |
| 58852 | TATGAGTGTGGAGTCGTTGCGGGGTAGTGGTCATGGTAGTCATNNNAGATCGGAA |
| 30162 | TATGAGTGTGGAGTCGTTGCGCCTGTCTGTGCACAAATGAAAANNNAGATCGGAA |
| 48254 | TATGAGTGTGGAGTCGTTGCGCAAGAGCACACTTCTGCAAATCNNNAGATCGGAAG |
| 68970 | TATGAGTGTGGAGTCGTTGCTCTCCCCATAGCTCTCTCACTTTNNNAGATCGGAAGA |
| 38293 | TATGAGTGTGGAGTCGTTGCTGTGAGAGCTCAGTGTCATGAGANNNAGATCGGAA |
| 116993 | TATGAGTGTGGAGTCGTTGCCAGCAATCAATGAGATAGGAGACATNNNAGATCGG |
| 100978 | TATGAGTGTGGAGTCGTTGCGGCGGAGCAAAATGTATATGAGANNNAGATCGGA |
| 53219 | TATGAGTGTGGAGTCGTTGCTGGGATGTGTTTTATGTAGGGAGGNNNAGATCGG |
| 52247 | TATGAGTGTGGAGTCGTTGCTAAGCTCTTCAAGGGTGGAAACGNNNAGATCGGAA |
| 33829 | TATGAGTGTGGAGTCGTTGCACTGCTTTTGGTTCTTGGCTACANNNAGATCGGAAG |
| 39161 | TATGAGTGTGGAGTCGTTGCTATGGCATGGAGAATAGCTGGGANNNAGATCGGA |
| 68668 | TATGAGTGTGGAGTCGTTGCATTTCTCTGCTAGTGTGCGTGTTNNNAGATCGGAAG |
| 33051 | TATGAGTGTGGAGTCGTTGCCCCACTTCCAATTCTGACACACANNNAGATCGGAAGA |
| 71757 | TATGAGTGTGGAGTCGTTGCCAGCCCAAAACCCTAATTTCAGCNNNAGATCGGAAG |
| 41582 | TATGAGTGTGGAGTCGTTGCAAGTCCCAGTCAATGAGGTGGTANNNAGATCGGAA |
| 58325 | TATGAGTGTGGAGTCGTTGCGCAAACAGAATTGACCAGCACATNNNAGATCGGAA |
| 105866 | TATGAGTGTGGAGTCGTTGCCCAGCACCCCAAAAAGATCCTGNNNAGATCGGAAGA |
| 64382 | TATGAGTGTGGAGTCGTTGCGACAGCATGTGCAATTGTTGAGTNNNAGATCGGAA |
| 62970 | TATGAGTGTGGAGTCGTTGCATGAAAGGCAAAGGTGATGCTCTNNNAGATCGGAA |
| 108882 | TATGAGTGTGGAGTCGTTGCGTCTCCATCCTCTTTCTCTTTGCNNNAGATCGGAAGA |
| 61785 | TATGAGTGTGGAGTCGTTGCCTCCATGGTCCTTGATAGGAGCTNNNAGATCGGAAG |
| 50734 | TATGAGTGTGGAGTCGTTGCGTGTAGGCAGTCTTGAAGGACAGNNNAGATCGGAA |
| 102096 | TATGAGTGTGGAGTCGTTGCTGTAGGCAGTCTTGAAGGACAGNNNAGATCGGAAG |
| 36238 | TATGAGTGTGGAGTCGTTGCAACAGGATACTGGAGGACTTGCANNNAGATCGGAA |
| 63189 | TATGAGTGTGGAGTCGTTGCTACAGGGAGGATCCTATGGGTCTNNNAGATCGGAA |
| 43931 | TATGAGTGTGGAGTCGTTGCACACAGTACTTAGCTCTTCCACCNNNAGATCGGAAG |
| 52004 | TATGAGTGTGGAGTCGTTGCCACAACCACACACCTGGCTATAGNNNAGATCGGAAG |
| 40728 | TATGAGTGTGGAGTCGTTGCGTGCCCATGGTTTAGTGTCCATANNNAGATCGGAAG |
| 39975 | TATGAGTGTGGAGTCGTTGCACCTTTCATACCTGTGAGGCTGANNNAGATCGGAAG |
| 71628 | TATGAGTGTGGAGTCGTTGCGCGGGGTCCTTCCCATGATACATNNNAGATCGGAAG |
| 51083 | TATGAGTGTGGAGTCGTTGCAGAGAAGGGATAAGTCCAGGCAGNNNAGATCGGA |
| 33201 | TATGAGTGTGGAGTCGTTGCAGGACTTCCAGTACATTCCCACANNNAGATCGGAAG |
| 47177 | TATGAGTGTGGAGTCGTTGCAAGCCTGAACATCGATATTGGGCNNNAGATCGGAA |
| 35884 | TATGAGTGTGGAGTCGTTGCTTTGTGTCTCCTTCCCAAATGCANNNAGATCGGAAG |
| 31246 | TATGAGTGTGGAGTCGTTGCTTTTGCGCAGTTTTATGAGGCAANNNAGATCGGAAG |
| 53062 | TATGAGTGTGGAGTCGTTGCTATTCTGTCTAGGGAGCTGCAGGNNNAGATCGGAA |
| 60544 | TATGAGTGTGGAGTCGTTGCCTTGGGAAATGGGTGTGCTTACTNNNAGATCGGAA |
| 35858 | TATGAGTGTGGAGTCGTTGCGCAGGTATAGAGACTAGTTGGCANNNAGATCGGAA |

109431 TATGAGTGTGGAGTCGTTGCCAGATAGCACTGGAACCAGGACNNNAGATCGGAAG
109790 TATGAGTGTGGAGTCGTTGCAGTTGTTTCCCACTTGAACTCCTGNNNAGATCGGAAG
51605 TATGAGTGTGGAGTCGTTGCCAGCTATGGCACTTTCTCAGGAGNNNAGATCGGAAG
117848 TATGAGTGTGGAGTCGTTGCGTCTGGAATGCAGTGTTGTGGCNNNAGATCGGAAG
100433 TATGAGTGTGGAGTCGTTGCTGTCGGAGAGCCAAGGAATGATNNNAGATCGGAAG
29882 TATGAGTGTGGAGTCGTTGCGGGGTACTGAAGAGCTTATTCTCCNNNAGATCGGAA
45757 TATGAGTGTGGAGTCGTTGCCCACTGTCTTCAATCAAAAGGTCCNNNAGATCGGAAG
50177 TATGAGTGTGGAGTCGTTGCTTATACCTGCCTGCTCCTGAAAGNNNAGATCGGAAG
68493 TATGAGTGTGGAGTCGTTGCTCTGACTAGGTTCTCACCTGGTTNNNAGATCGGAAG
54835 TATGAGTGTGGAGTCGTTGCACAGCAATAAGTTAGGCCAGTGGNNNAGATCGGAA
35527 TATGAGTGTGGAGTCGTTGCGAGAGAGAAACAAGGCTAACGCANNNAGATCGGAA
109381 TATGAGTGTGGAGTCGTTGCCAAGACTGGGTAATGTATGGAGANNNAGATCGGAA
49633 TATGAGTGTGGAGTCGTTGCGGGTACTTCCGTGTGTGTACTTCNNNAGATCGGAAG
52066 TATGAGTGTGGAGTCGTTGCAACTAGAGCCCTGAGGATGCTAGNNNAGATCGGAA
38331 TATGAGTGTGGAGTCGTTGCTTCTAACCTCAAGCCCAGTGAGANNNAGATCGGAAG
45028 TATGAGTGTGGAGTCGTTGCTCCCTCACATCAAAGAAAAGGCCNNNAGATCGGAAG
54592 TATGAGTGTGGAGTCGTTGCAGGAAGGATGTTTTTGACCCTGGNNNAGATCGGAA
47602 TATGAGTGTGGAGTCGTTGCGCATGAGAAGTCCCAGGATATGCNNNAGATCGGAA
38805 TATGAGTGTGGAGTCGTTGCAAGACATTCATCCTCAGAGAGGANNNAGATCGGAAG
67685 TATGAGTGTGGAGTCGTTGCTATTTGTCTCTCTGCCTCCCCTTNNNAGATCGGAAGA
29893 TATGAGTGTGGAGTCGTTGCGTTCAGAAAGGCGTGCACTTCNNNAGATCGGAAGA
30486 TATGAGTGTGGAGTCGTTGCCAGCCTTCTCATGTGGGAAGAAANNNAGATCGGAAG
42473 TATGAGTGTGGAGTCGTTGCATGGGGGTAGTGGGAGAAGTAACNNNAGATCGGA
41655 TATGAGTGTGGAGTCGTTGCGCAGTTTCTGGAGGCTTTCTGTANNNAGATCGGAAG
62154 TATGAGTGTGGAGTCGTTGCAATGTCATAATGGTGTGGTGGCTNNNAGATCGGAA
37968 TATGAGTGTGGAGTCGTTGCAGGAAAACGTCACATGATGCAGANNNAGATCGGAA
103685 TATGAGTGTGGAGTCGTTGCGCAATAAGTTAGGCCAGTGGTTNNNAGATCGGAAG
69464 TATGAGTGTGGAGTCGTTGCACAGCTTGACATTCCCGACTTTTNNNAGATCGGAAGA
52649 TATGAGTGTGGAGTCGTTGCCCGCTTGTAGTTCTCTCTCTTCGNNNAGATCGGAAGA
105171 TATGAGTGTGGAGTCGTTGCCCAGGACTGCATGGATTGGTACANNNAGATCGGAA
48821 TATGAGTGTGGAGTCGTTGCTTACCTCTGAGTTCCCCTTCCTCNNNAGATCGGAAGA
37480 TATGAGTGTGGAGTCGTTGCAGCCTGTGTACATTTGGCAAAGANNNAGATCGGAAG
52249 TATGAGTGTGGAGTCGTTGCCCAGATCTTACCCAAGACCAACGNNNAGATCGGAAG
69279 TATGAGTGTGGAGTCGTTGCCGCTCCCTTTCCAGTGTTAGTTTNNNAGATCGGAAGA
41602 TATGAGTGTGGAGTCGTTGCGGTCTTTCGTCTGTGCACATGTANNNAGATCGGAAG
116971 TATGAGTGTGGAGTCGTTGCAACCAACCCTTTTCCGTATCTATGAGNNNAGATCGGA
100926 TATGAGTGTGGAGTCGTTGCGGCTTACATCATGGCCAGAGTATNNNAGATCGGAAG
54586 TATGAGTGTGGAGTCGTTGCACTTCCTGTTCTCTCTCTACCTGGNNNAGATCGGAAG
40311 TATGAGTGTGGAGTCGTTGCACAGAAAGAGTGAGAAATCGTGTGANNNAGATCGG
55493 TATGAGTGTGGAGTCGTTGCGAGCTAAGCCAATGCTGATCATGNNNAGATCGGAAG
53460 TATGAGTGTGGAGTCGTTGCTGGTATTAGAAACCTGCAGACGGNNNAGATCGGAA
61201 TATGAGTGTGGAGTCGTTGCATCAGAACAAAGTGCTTTGGCCTNNNAGATCGGAAG
38831 TATGAGTGTGGAGTCGTTGCTGGAAAAATCCCATGAGGGAGGANNNAGATCGGAA

64631 TATGAGTGTGGAGTCGTTGCGGCACAGTATTTGAGATAGCGTNNNAGATCGGAA
98947 TATGAGTGTGGAGTCGTTGCCTGAAAATGCTATGGGCAGTCACNNNAGATCGGAA
29883 TATGAGTGTGGAGTCGTTGCCACAGCTGCTTCAGCATCATCTTNNNAGATCGGAAG
43007 TATGAGTGTGGAGTCGTTGCGCTATTACGAGGAACTGGCTCACNNNAGATCGGAA
101139 TATGAGTGTGGAGTCGTTGCTGATTTAGGTATGATGCCAGGAGCNNNAGATCGGA
31366 TATGAGTGTGGAGTCGTTGCAGCCAGTGGAGAAGGAGAATCAANNNAGATCGGA
58101 TATGAGTGTGGAGTCGTTGCTGTGTACTGATGGGAAGGGGAATNNNAGATCGGA
33990 TATGAGTGTGGAGTCGTTGCAGTCAAGAAAGCTTTGGAGAACCANNNAGATCGGA
54557 TATGAGTGTGGAGTCGTTGCTGCCAAAAGTTACCATTGACTGGNNNAGATCGGAAG
52268 TATGAGTGTGGAGTCGTTGCGCCGAAGTAACTAGGAGAACACGNNNAGATCGGAA
71741 TATGAGTGTGGAGTCGTTGCAGGCCGCATGATTGTATTTCCTNNNAGATCGGAAG
100694 TATGAGTGTGGAGTCGTTGCCCCCAGTACCCTATTCATTTGATGTGNNNAGATCGGA
63511 TATGAGTGTGGAGTCGTTGCCCATTCTGCCTGGAACACTTTCTNNNAGATCGGAAGA
61682 TATGAGTGTGGAGTCGTTGCTCTCTGAAGAACACTAACATACAGCTNNNAGATCGG
49750 TATGAGTGTGGAGTCGTTGCTGGACTGGGTTGAATGGTTCTTCNNNAGATCGGAA
50048 TATGAGTGTGGAGTCGTTGCTGACAGATTGCAGCACTGAAAAGNNNAGATCGGAA
46773 TATGAGTGTGGAGTCGTTGCCAAATAGATGACCCCTATGGCGCNNNAGATCGGAAG
41499 TATGAGTGTGGAGTCGTTGCACCATCCACAAAGACCTCAGGTANNNAGATCGGAAG
31628 TATGAGTGTGGAGTCGTTGCCCTTGGAACCAAACTGTCCAGAANNNAGATCGGAAG
50131 TATGAGTGTGGAGTCGTTGCACACACGCTTGCTTTTCAGAAAGNNNAGATCGGAAG
42973 TATGAGTGTGGAGTCGTTGCTGTGTCTGTAAAAGAAGGGAGATCACNNNAGATCG
48740 TATGAGTGTGGAGTCGTTGCATCCAGCACTCTTTGTTTCCCTCNNNAGATCGGAAGA
34023 TATGAGTGTGGAGTCGTTGCGTTCATTGTTGTGTCCAACACCANNNAGATCGGAAG
102066 TATGAGTGTGGAGTCGTTGCCATTGTTGTGTCCAACACCANNNAGATCGGAAGAGC
31796 TATGAGTGTGGAGTCGTTGCTTGGGATCCAGCAGTAGAAGGAANNNAGATCGGAA
52239 TATGAGTGTGGAGTCGTTGCTCTCTTTGGCTGGAAGACAAACGNNNAGATCGGAAG
30064 TATGAGTGTGGAGTCGTTGCAAAACAACCTAAGAACATTTACCAAAANNNAGATCG
42821 TATGAGTGTGGAGTCGTTGCAAGTTAGCTACCAACCCAAGCACNNNAGATCGGAAG
43883 TATGAGTGTGGAGTCGTTGCATTGTCATCTGGCCCATTACACCNNNAGATCGGAAG
36513 TATGAGTGTGGAGTCGTTGCGGCTGGTCTGCAATGTTCTATCANNNAGATCGGAAG
36145 TATGAGTGTGGAGTCGTTGCACAGTAAGGAACTGATGAGTGCANNNAGATCGGAA
54209 TATGAGTGTGGAGTCGTTGCGGAGGAGAAAAAGTCACTGTGGGNNNAGATCGGA
68841 TATGAGTGTGGAGTCGTTGCCTCAGCTTTGTCAGAGAGCATTTNNNAGATCGGAAG
102734 TATGAGTGTGGAGTCGTTGCTTTTTGACCCTGGGTACTCCNNNAGATCGGAAGAGC
40137 TATGAGTGTGGAGTCGTTGCTGGAGACTTTATCCTGGCAGTGANNNAGATCGGAA
103526 TATGAGTGTGGAGTCGTTGCGATAGAATCTCTGACCAAGACAATAATNNNAGATCG
39158 TATGAGTGTGGAGTCGTTGCCAATTCTTACGTGGAAGCTGGGANNNAGATCGGAA
44301 TATGAGTGTGGAGTCGTTGCTAAGCTCTTCAGCAAAAGGACCCNNNAGATCGGAAG
116676 TATGAGTGTGGAGTCGTTGCCCTCTCTTCCCTCCTTTTGAAGTNNNAGATCGGAAGA
97795 TATGAGTGTGGAGTCGTTGCCTCTGGCACGCTGCTTATCAGNNNAGATCGGAAGAG
63409 TATGAGTGTGGAGTCGTTGCGGACAATGACAACAGTGGCTTCTNNNAGATCGGAA
30862 TATGAGTGTGGAGTCGTTGCGCTCCTGATGCCAAGATACACAANNNAGATCGGAAG
34265 TATGAGTGTGGAGTCGTTGCTTTGTGGTTTCTTTGTGGACCCANNNAGATCGGAAG

| | |
|---|---|
| 47663 | TATGAGTGTGGAGTCGTTGCGACCTTTCCATGCATTTCACTGCNNNAGATCGGAAGA |
| 38155 | TATGAGTGTGGAGTCGTTGCAGCGTGTGTGTGACATAGAGAGANNNAGATCGGAA |
| 109447 | TATGAGTGTGGAGTCGTTGCGTAACTGAGGACAGGGAGCTATGCNNNAGATCGGA |
| 37448 | TATGAGTGTGGAGTCGTTGCTCTGGCATGGGGAGTAGAAAAGANNNAGATCGGAA |
| 40869 | TATGAGTGTGGAGTCGTTGCGACCACCTTCAGGGATTCTGATANNNAGATCGGAAG |
| 41695 | TATGAGTGTGGAGTCGTTGCGTTTCACCCCACAGTGACTTGTANNNAGATCGGAAGA |
| 53690 | TATGAGTGTGGAGTCGTTGCCCAAACTGCCCATTTCCAGGGNNNAGATCGGAAGAG |
| 50577 | TATGAGTGTGGAGTCGTTGCATGTATGCAAGTGGGCCTGTAAGNNNAGATCGGAA |
| 49953 | TATGAGTGTGGAGTCGTTGCTTTGGGTTTGCAGTTTGGGTTTCNNNAGATCGGAAG |
| 36881 | TATGAGTGTGGAGTCGTTGCTGGAGGTTGAGAAATCCCAGTCANNNAGATCGGAA |
| 64779 | TATGAGTGTGGAGTCGTTGCCCAAAGCCAAATGAAAAGATAAGGTNNNAGATCGGA |
| 64375 | TATGAGTGTGGAGTCGTTGCAAAACCTGCCAGATTGCTTGAGTNNNAGATCGGAAG |
| 39952 | TATGAGTGTGGAGTCGTTGCAGTAATCCTTACAACTCTCAGCTGANNNAGATCGGAA |
| 33836 | TATGAGTGTGGAGTCGTTGCGTGTGTGTAACTCCCAAATCTACANNNAGATCGGAA |
| 43128 | TATGAGTGTGGAGTCGTTGCATAAGACACTGGAACCGCAAGACNNNAGATCGGAA |
| 61126 | TATGAGTGTGGAGTCGTTGCTGCCTGTCTATCTATATCTAGCCTNNNAGATCGGAAG |
| 35315 | TATGAGTGTGGAGTCGTTGCACCCTATTCATTTGATGTGCAGCANNNAGATCGGAA |
| 57726 | TATGAGTGTGGAGTCGTTGCACACATGCCCTTTCCATCCAAATNNNAGATCGGAAGA |
| 42958 | TATGAGTGTGGAGTCGTTGCAGAGAGGGAGAGAGAGCAATCACNNNAGATCGGAA |
| 36395 | TATGAGTGTGGAGTCGTTGCGCGGCCAAAAATAGTCAACATCANNNAGATCGGAAG |
| 104646 | TATGAGTGTGGAGTCGTTGCCCATCAGTGGCAGGCTGGATATATNNNAGATCGGAA |
| 50111 | TATGAGTGTGGAGTCGTTGCTGACACCTCCCCTACTCTCAAAGNNNAGATCGGAAGA |
| 66151 | TATGAGTGTGGAGTCGTTGCATTTCCTGTGGCTGTTCATCTGTNNNAGATCGGAAGA |
| 96857 | TATGAGTGTGGAGTCGTTGCGGAGATGCACCACATGAGATGTGNNNAGATCGGAA |
| 38878 | TATGAGTGTGGAGTCGTTGCGCCTCCAGAAGGACTACATAGGANNNAGATCGGAA |
| 68977 | TATGAGTGTGGAGTCGTTGCTGAAGGTCAGAGAGTGGGACTTTNNNAGATCGGAA |
| 30604 | TATGAGTGTGGAGTCGTTGCAGGATCTGGGCCAATTAGGGAAANNNAGATCGGAA |
| 31810 | TATGAGTGTGGAGTCGTTGCAAGCTGACATGCAATCACAGGAANNNAGATCGGAA |
| 47599 | TATGAGTGTGGAGTCGTTGCACACACACTGCTTTTGCATATGCNNNAGATCGGAAGA |
| 47380 | TATGAGTGTGGAGTCGTTGCTCGGAATGAACAAACATGATTGGCNNNAGATCGGAA |
| 29888 | TATGAGTGTGGAGTCGTTGCGGCTCTGGATGTTTAGGTTCTCGNNNAGATCGGAAG |
| 46739 | TATGAGTGTGGAGTCGTTGCATTTGAAAGTACGGGTAGGACGCNNNAGATCGGAA |
| 33632 | TATGAGTGTGGAGTCGTTGCTATGATGCCAGGAGCTATGGACANNNAGATCGGAA |
| 58311 | TATGAGTGTGGAGTCGTTGCATGAACCTCTCCCTTCACCACATNNNAGATCGGAAGA |
| 103083 | TATGAGTGTGGAGTCGTTGCGTTCCCCTGCACCATCTCAGANNNAGATCGGAAGAG |
| 39215 | TATGAGTGTGGAGTCGTTGCAGAGAGACTACAGCTGTTTGGGANNNAGATCGGAA |
| 52696 | TATGAGTGTGGAGTCGTTGCTCCAGTACAAAAGTGGGCAAAGGNNNAGATCGGAA |
| 52039 | TATGAGTGTGGAGTCGTTGCCAGGCTCTCAGATCCCATCCTAGNNNAGATCGGAAGA |
| 34178 | TATGAGTGTGGAGTCGTTGCTTCACCAGACAGCAGATCTACCANNNAGATCGGAAGA |
| 69248 | TATGAGTGTGGAGTCGTTGCTCACATTTGGAACCCCACAGTTTNNNAGATCGGAAGA |
| 54587 | TATGAGTGTGGAGTCGTTGCACCAGATGATAAATCACAGTACCTGGNNNAGATCGG |
| 58322 | TATGAGTGTGGAGTCGTTGCTCAACCACCACACTTTTCCACATNNNAGATCGGAAGA |
| 116304 | TATGAGTGTGGAGTCGTTGCCAGAGAGACAGAGGTGTGTATGTATNNNAGATCGG |

59511 TATGAGTGTGGAGTCGTTGCATACCCTCACTGTCTCTCCCTATNNNAGATCGGAAGA
64461 TATGAGTGTGGAGTCGTTGCTGTATTTGAAAACGCCCAGTAGTNNNAGATCGGAAG
55531 TATGAGTGTGGAGTCGTTGCGGCTGTCATCATGGCATTTCATGNNNAGATCGGAAG
57644 TATGAGTGTGGAGTCGTTGCTGTACTCAGCGAGAGCTCTTTTGNNNAGATCGGAAG
32367 TATGAGTGTGGAGTCGTTGCGAATGGGGAGCTTTTCTGCCTAANNNAGATCGGAA
41836 TATGAGTGTGGAGTCGTTGCACGGGGTCATTACAGGAAGCTTANNNAGATCGGAA
64359 TATGAGTGTGGAGTCGTTGCGGATGAGGAGACAGACTGTGAGTNNNAGATCGGA
55164 TATGAGTGTGGAGTCGTTGCTCACATGCACACACACTTTATTTGGNNNAGATCGGA
52107 TATGAGTGTGGAGTCGTTGCGCCAACTGTGCTTCTACCAGTAGNNNAGATCGGAAG
109440 TATGAGTGTGGAGTCGTTGCAGTGAGGCAGAGATTTGCATACAANNNAGATCGGA
56089 TATGAGTGTGGAGTCGTTGCCGCAAGCCATAGAAGAAAAGCTGNNNAGATCGGAA
49386 TATGAGTGTGGAGTCGTTGCCTTTTTGCCTGTCGTGTGATGTCNNNAGATCGGAAG
53150 TATGAGTGTGGAGTCGTTGCCAGTTGGTGATGGACAAGAGAGGNNNAGATCGGA
38312 TATGAGTGTGGAGTCGTTGCTTTCTAACCAGTACGGCCTGAGANNNAGATCGGAAG
55230 TATGAGTGTGGAGTCGTTGCCCCCATCTTCTCTCCCACAAATGNNNAGATCGGAAGA
46013 TATGAGTGTGGAGTCGTTGCGTTTCCACTCCCCTGTAGTTTCCNNNAGATCGGAAGA
104698 TATGAGTGTGGAGTCGTTGCGCGATTCCCACATTTTCATTCAATNNNAGATCGGAAG
60755 TATGAGTGTGGAGTCGTTGCATCTTTAGACCCTACGTGGACCTNNNAGATCGGAAG
61347 TATGAGTGTGGAGTCGTTGCAATGCAGGGTCTCACTGTATCCTNNNAGATCGGAAG
51719 TATGAGTGTGGAGTCGTTGCGTGAAGTCTTGGAGAAGCTGGAGNNNAGATCGGA
96159 TATGAGTGTGGAGTCGTTGCCATACTCTGTCTACCCTACCAGGCNNNAGATCGGAAG
65178 TATGAGTGTGGAGTCGTTGCGTGGAAATATTAGGTCACTGGGTNNNAGATCGGAA
59039 TATGAGTGTGGAGTCGTTGCAAGTGTACTGATTCAGCTTAGAGATNNNAGATCGGA
55904 TATGAGTGTGGAGTCGTTGCCTCTGACCAAGACAATAATTTTTACTGNNNAGATCGG
64222 TATGAGTGTGGAGTCGTTGCAGACACAACCTCTGAATAATAGAGTNNNAGATCGGA
107450 TATGAGTGTGGAGTCGTTGCTGGGTGGGGAAATATGACTGAGAANNNAGATCGG
31756 TATGAGTGTGGAGTCGTTGCTGGGGCTACACACAGTTCTAGAANNNAGATCGGAA
59656 TATGAGTGTGGAGTCGTTGCCCTGGAAGGGTCCCTAGCATTATNNNAGATCGGAAG
34637 TATGAGTGTGGAGTCGTTGCGGAAAGGACTAACTTTCTCGCCANNNAGATCGGAAG
115832 TATGAGTGTGGAGTCGTTGCACTTCAGCAGGACATTTCTGTGCTNNNAGATCGGAA
59437 TATGAGTGTGGAGTCGTTGCCCAGTCCTGAGAACCTTGCATATNNNAGATCGGAAG
116458 TATGAGTGTGGAGTCGTTGCATCTATCCATCCTTCCACATATCTATTNNNAGATCGGA
44219 TATGAGTGTGGAGTCGTTGCTACTCCAAAGCCTAGCAAAACCCNNNAGATCGGAAG
30298 TATGAGTGTGGAGTCGTTGCCATAGATTCTGGGGTGGACCAAANNNAGATCGGAA
67922 TATGAGTGTGGAGTCGTTGCGGATCTGGGTACAAAAGTGGCTTNNNAGATCGGAA
32400 TATGAGTGTGGAGTCGTTGCCAGGAACCAAGAAGCATGGCTAANNNAGATCGGAA
64832 TATGAGTGTGGAGTCGTTGCAGCGATATGTGTTCTCTCCAGGTNNNAGATCGGAAG
71577 TATGAGTGTGGAGTCGTTGCAAACCCTTCAATGCCTCTGTGTTNNNAGATCGGAAG
109786 TATGAGTGTGGAGTCGTTGCCGTCGTAGAATCTGTATGATTGTTGTTNNNAGATCG
55073 TATGAGTGTGGAGTCGTTGCCCTCACATAATCTCCTGCCTTGGNNNAGATCGGAAGA
39692 TATGAGTGTGGAGTCGTTGCAGGTGACATTATGCTTGGAGATGANNNAGATCGGA
37487 TATGAGTGTGGAGTCGTTGCACCTAATATTTCTGGGGAAAGAAAGANNNAGATCG
53451 TATGAGTGTGGAGTCGTTGCTAGAATTTTGTGGGTGGACACGGNNNAGATCGGAA

| | |
|---|---|
| 47434 | TATGAGTGTGGAGTCGTTGCCTGTAGTGAAGTGCCCAAAATGCNNNAGATCGGAA |
| 108207 | TATGAGTGTGGAGTCGTTGCAGGAGAGGTGGCAAAGTGATAAGNNNAGATCGGA |
| 117851 | TATGAGTGTGGAGTCGTTGCGGACATCTCATTGTGGTTTGTGTNNNAGATCGGAAG |
| 61227 | TATGAGTGTGGAGTCGTTGCCTGGAACGTGCTTACATATGCCTNNNAGATCGGAAG |
| 52903 | TATGAGTGTGGAGTCGTTGCAAGGAAGCACCAAGTTTCACAGGNNNAGATCGGAA |
| 30952 | TATGAGTGTGGAGTCGTTGCTGTTAGAAGGTGTAAAAGGCTACAANNNAGATCGG |
| 47190 | TATGAGTGTGGAGTCGTTGCCTCCATTTCACCCATTTTTGGGCNNNAGATCGGAAGA |
| 63379 | TATGAGTGTGGAGTCGTTGCTTGAATCCCTTGTCTCCCCTTCTNNNAGATCGGAAGA |
| 48064 | TATGAGTGTGGAGTCGTTGCGGCTGTCATCTCAGAAAAGATTGCNNNAGATCGGA |
| 57574 | TATGAGTGTGGAGTCGTTGCACTGGGTCAGAATCTCTGCTTTGNNNAGATCGGAAG |
| 106530 | TATGAGTGTGGAGTCGTTGCATAGGCAAGACCCATACCAGTACTNNNAGATCGGAA |
| 62433 | TATGAGTGTGGAGTCGTTGCATGTCACAACACATGACAATTGCTNNNAGATCGGAA |
| 63183 | TATGAGTGTGGAGTCGTTGCTGGTTACAGCTCCAGTTAGGTCTNNNAGATCGGAAG |
| 41720 | TATGAGTGTGGAGTCGTTGCGGAGTGCAGCATCAAAGCAATTANNNAGATCGGAA |
| 42568 | TATGAGTGTGGAGTCGTTGCCGGCAAGACTGAAATTCACACACNNNAGATCGGAAG |
| 33743 | TATGAGTGTGGAGTCGTTGCGGAGCAGTGGTGTGGAAAATACANNNAGATCGGA |
| 34887 | TATGAGTGTGGAGTCGTTGCTCTTACATTCCCATTCCCCTCCANNNAGATCGGAAGA |
| 35952 | TATGAGTGTGGAGTCGTTGCTTCTTCTTGGAAACCTGCATGCANNNAGATCGGAAG |
| 98139 | TATGAGTGTGGAGTCGTTGCGCAAAGCACAACCTCAGCTCANNNAGATCGGAAGAG |
| 48004 | TATGAGTGTGGAGTCGTTGCCTTTCACACCAAGTGTTCTGTGCNNNAGATCGGAAG |
| 44063 | TATGAGTGTGGAGTCGTTGCAGTGTGTGGAACATTGAAGGACCNNNAGATCGGAA |
| 29892 | TATGAGTGTGGAGTCGTTGCTCTGAAATCACTCGGAATGTACAGTNNNAGATCGGA |
| 38482 | TATGAGTGTGGAGTCGTTGCGTAGGCTGAGCTCCTGTCTTAGANNNAGATCGGAA |
| 29970 | TATGAGTGTGGAGTCGTTGCACGGCAAATATTATGGCAGCCTTNNNAGATCGGAAG |
| 71752 | TATGAGTGTGGAGTCGTTGCGGTCGCTAAGGTCAGGTTGTATCNNNAGATCGGAA |
| 63438 | TATGAGTGTGGAGTCGTTGCGGGAGAGGGAAGAAGGAAGTTCTNNNAGATCGGA |
| 30131 | TATGAGTGTGGAGTCGTTGCAGAGACGAGGATGGGAAGGAAAANNNAGATCGGA |
| 29969 | TATGAGTGTGGAGTCGTTGCACTACAGAACCCCAGGATTCCTCNNNAGATCGGAAG |
| 44998 | TATGAGTGTGGAGTCGTTGCTTTTCCTCTTGCCATGAGTAGCCNNNAGATCGGAAG |
| 109401 | TATGAGTGTGGAGTCGTTGCATTTCTTTCTCTCTCTCCTGGAGCANNNAGATCGGAA |
| 102102 | TATGAGTGTGGAGTCGTTGCCACAGAGAAGGGAGAGCTTGCTNNNAGATCGGAAG |
| 38920 | TATGAGTGTGGAGTCGTTGCTACACAGCAACTCCAGGTTAGGANNNAGATCGGAA |
| 47170 | TATGAGTGTGGAGTCGTTGCGGAGTTTTTATCCTCAGGTGGGCNNNAGATCGGAA |
| 102803 | TATGAGTGTGGAGTCGTTGCACGGTTGGTTTCCTCTCTACACNNNAGATCGGAAGA |
| 100961 | TATGAGTGTGGAGTCGTTGCCACACGACTTGCCTGCCTCANNNAGATCGGAAGAGC |
| 103531 | TATGAGTGTGGAGTCGTTGCCTTAGCCCCATGTAGAACTTATAGTNNNAGATCGGA |
| 107020 | TATGAGTGTGGAGTCGTTGCTGTGTGTGTAGACAGTCTTGCTATGNNNAGATCGG |
| 104755 | TATGAGTGTGGAGTCGTTGCTATCTGAAAGTGGGGGCAGTTGTANNNAGATCGGA |
| 100991 | TATGAGTGTGGAGTCGTTGCTATTTACTGTGCTCAGCCAGCAGAGNNNAGATCGGA |
| 71704 | TATGAGTGTGGAGTCGTTGCCTGGAGGAAAAGTCTGGTAGGGANNNAGATCGGA |
| 103055 | TATGAGTGTGGAGTCGTTGCACATTTCCAGGTCATAGGGCATNNNAGATCGGAAGA |
| 101825 | TATGAGTGTGGAGTCGTTGCTTTCTTTTCCTCTGGGCACATACANNNAGATCGGAAG |
| 44316 | TATGAGTGTGGAGTCGTTGCTGAAAGTGGCGTATTGAAATACCCNNNAGATCGGA |

101840 TATGAGTGTGGAGTCGTTGCTTCTAGGTGTCAGAGGGCTATCTNNNAGATCGGAAG

33964 TATGAGTGTGGAGTCGTTGCTGTCAAGATGCTACAATGGAAACCANNNAGATCGGA

104665 TATGAGTGTGGAGTCGTTGCTCGCCAAGTTTAGTTCTACTTGACTTNNNAGATCGGA

102392 TATGAGTGTGGAGTCGTTGCAGAGAGAAGTAGCGAAGTCCTAGNNNAGATCGGAA

102350 TATGAGTGTGGAGTCGTTGCACCACACTCAGCTACCTTGTTGNNNAGATCGGAAGAG

105840 TATGAGTGTGGAGTCGTTGCAATATCCTTCCCATGGTGTCCATNNNAGATCGGAAGA

101147 TATGAGTGTGGAGTCGTTGCTTTCTCTCTAATGGCTAGGGGGNNNAGATCGGAAGA

121981 TATGAGTGTGGAGTCGTTGCATGTGCCTGGAGAGGAGCCTGAANNNAGATCGGAA

104298 TATGAGTGTGGAGTCGTTGCAGTCATTCAATCCTTCCACTGCAGCNNNAGATCGGAA

101841 TATGAGTGTGGAGTCGTTGCAAAGGGCAAGTGAATCAGAGGAANNNAGATCGGAA

103276 TATGAGTGTGGAGTCGTTGCGCCCAGCCCAGTTTATCCTTTTNNNAGATCGGAAGAG

103093 TATGAGTGTGGAGTCGTTGCTCCTTTTAGATACTCAGTAGTGGGANNNAGATCGGA

105256 TATGAGTGTGGAGTCGTTGCAACTCCCATTCATTCATCTGCCCNNNAGATCGGAAGAG

104283 TATGAGTGTGGAGTCGTTGCTTACAGTCTACCACAACCACACCNNNAGATCGGAAGA

109444 TATGAGTGTGGAGTCGTTGCGTTCCAGGCATAAGCTACCATGTNNNAGATCGGAAG

108132 TATGAGTGTGGAGTCGTTGCGAGAAAGACACTCAGAGGAAGGANNNAGATCGGAA

103579 TATGAGTGTGGAGTCGTTGCTGGGAACCAAATTGCTTGTATTCNNNAGATCGGAAG

102333 TATGAGTGTGGAGTCGTTGCCTGTGTCAGGAATGGGTAGGGANNNAGATCGGAAG

115940 TATGAGTGTGGAGTCGTTGCTTGGCACCTGTTTGTTTGTAACTNNNAGATCGGAAGA

102379 TATGAGTGTGGAGTCGTTGCATGTGGAATGTGCCTAGCAAAACTNNNAGATCGGAA

109471 TATGAGTGTGGAGTCGTTGCTCTCTGGATCATACGGTCATTCTNNNAGATCGGAAGA

102116 TATGAGTGTGGAGTCGTTGCGCTGCTGGCTAATCAGTTTACCCNNNAGATCGGAAGA

104622 TATGAGTGTGGAGTCGTTGCGTAGGTGGTAGGTGGTAGGTTTTCNNNAGATCGGA

108208 TATGAGTGTGGAGTCGTTGCCTTTACTATTCTGCCTAGTGGTCTNNNAGATCGGAAG

109513 TATGAGTGTGGAGTCGTTGCGGAGCTTTGCTGAATCTTTTGCAGNNNAGATCGGAA

109799 TATGAGTGTGGAGTCGTTGCAGTTCTTCCAAGCATCAACCCCNNNAGATCGGAAGAG

105879 TATGAGTGTGGAGTCGTTGCGGAGGTGATTATGGCTTGGATCTGNNNAGATCGGA

118244 TATGAGTGTGGAGTCGTTGCTCAGCCTCCCAAAGGTCTACAATNNNAGATCGGAAGA

110267 TATGAGTGTGGAGTCGTTGCACTCTCCCTTTTTCCCCTCTTCNNNAGATCGGAAGAGC

105839 TATGAGTGTGGAGTCGTTGCGGTTCAAGTCCAAAGGCAAGGAAANNNAGATCGGA

100643 TATGAGTGTGGAGTCGTTGCGCCCTTCTGGCTCTTTTTCTGATNNNAGATCGGAAGA

98443 TATGAGTGTGGAGTCGTTGCACCTCTCTCTTACAGGAACTCTTGTNNNAGATCGGAA

101781 TATGAGTGTGGAGTCGTTGCACAAGAAGTAACAAGTTTTGGCAAGGNNNAGATCGG

98383 TATGAGTGTGGAGTCGTTGCACAAGAGAAGAGCTGCTGTAAAANNNAGATCGGAA

101119 TATGAGTGTGGAGTCGTTGCACCAGGGCATCATTTTCCACCTNNNAGATCGGAAGAG

103064 TATGAGTGTGGAGTCGTTGCGGATTGCTGGACCGAAGGATAGAANNNAGATCGGA

104299 TATGAGTGTGGAGTCGTTGCGTGAGAAGGGGAATGTCCTTGAGNNNAGATCGGAA

104656 TATGAGTGTGGAGTCGTTGCCGTGCCTGGCCTGTGATACATANNNAGATCGGAAGA

105191 TATGAGTGTGGAGTCGTTGCCTCAGCCCTCCCTAGCCCTCNNNAGATCGGAAGAGCA

116809 TATGAGTGTGGAGTCGTTGCAACCTCGTTCTCTATACACACATACNNNAGATCGGAAG

104297 TATGAGTGTGGAGTCGTTGCCTCCTGTTTTCCACGTGGTTCTCNNNAGATCGGAAGA

101353 TATGAGTGTGGAGTCGTTGCCAGTTGGCCCAAAAAGTGTGTGTNNNAGATCGGAAG

108177 TATGAGTGTGGAGTCGTTGCTGAACCACCATGCTCAGCAATACTNNNAGATCGGAAG

98074 TATGAGTGTGGAGTCGTTGCGCACCTTGAGCCTTCCTCATTATGANNNAGATCGGAA
71635 TATGAGTGTGGAGTCGTTGCTCATCAGGGGTTGTCACAGTCACNNNAGATCGGAAG
64353 TATGAGTGTGGAGTCGTTGCCGTGCGTCTAATGGTTTCTGAGTNNNAGATCGGAAG
121951 TATGAGTGTGGAGTCGTTGCCTCCCCATAGCCAGGGATTGTATCNNNAGATCGGAA
65500 TATGAGTGTGGAGTCGTTGCATGCCTGTGTATGTGTTTGTGGTNNNAGATCGGAA
35944 TATGAGTGTGGAGTCGTTGCAAGGGAAAGAAGGATTTCCATGCANNNAGATCGGA
52712 TATGAGTGTGGAGTCGTTGCATGAGCACTCCCTCCTAGAAAGGNNNAGATCGGAAG
50698 TATGAGTGTGGAGTCGTTGCAGGGGTAGTTTTGCTTGTCACAGNNNAGATCGGAA
104687 TATGAGTGTGGAGTCGTTGCTGTCCATCCTTGTCTGGTTTTGATATNNNAGATCGGA
45996 TATGAGTGTGGAGTCGTTGCTGCCAGATACTTGCTTCCTTTCCNNNAGATCGGAAGA
30612 TATGAGTGTGGAGTCGTTGCCAGGCTTCTCTTTCTTGGGGAAANNNAGATCGGAAG
71698 TATGAGTGTGGAGTCGTTGCCGAAGCTTTTCCCTCATTTCTGCNNNAGATCGGAAGA
54313 TATGAGTGTGGAGTCGTTGCAACTTATGTGGCAGTCACAATGGNNNAGATCGGAA
50145 TATGAGTGTGGAGTCGTTGCAGACAGAGAGATGGGCAGGAAAGNNNAGATCGGA
52573 TATGAGTGTGGAGTCGTTGCAAATCCCATGAGAAGGCTGATCGNNNAGATCGGAA
61870 TATGAGTGTGGAGTCGTTGCACTTTTTGTGTGTCCTGGTAGCTNNNAGATCGGAAG
52380 TATGAGTGTGGAGTCGTTGCCTGTTTCTGAGGCATCTAGACCGNNNAGATCGGAAG
38322 TATGAGTGTGGAGTCGTTGCTGGCTTTGTTTATCATGCTGAGANNNAGATCGGAAG
57214 TATGAGTGTGGAGTCGTTGCAGGGTGTCTGTGTGAGATGATTGNNNAGATCGGAA
51395 TATGAGTGTGGAGTCGTTGCCCTACATCCAGCTTGTCACAGAGNNNAGATCGGAAG
50656 TATGAGTGTGGAGTCGTTGCACCATCAACATCAGTCACCACAGNNNAGATCGGAAG
30559 TATGAGTGTGGAGTCGTTGCGGGTATAGTGCACGCTTACGAAANNNAGATCGGAA
54063 TATGAGTGTGGAGTCGTTGCCCAAGATAACAATTTGAGGTGATGGGNNNAGATCG
60048 TATGAGTGTGGAGTCGTTGCACTTCTGTTCAATTTGCTACCACTNNNAGATCGGAAG
108097 TATGAGTGTGGAGTCGTTGCCACTACTCAGCCATGACGAAAATNNNAGATCGGAAG
44359 TATGAGTGTGGAGTCGTTGCCATACTGGCTTCATCTTGACCCCNNNAGATCGGAAGA
57106 TATGAGTGTGGAGTCGTTGCTATTCACATGCTGGCAGTTGTGNNNAGATCGGAAGA
56776 TATGAGTGTGGAGTCGTTGCGCAAGTGTGAAGTTTCAAGGGTGNNNAGATCGGAA
32719 TATGAGTGTGGAGTCGTTGCGGAGATATCATGTGCATGCAAACANNNAGATCGGAA
41710 TATGAGTGTGGAGTCGTTGCGCCCATGAATGCTGCTGAAATTANNNAGATCGGAAG
61545 TATGAGTGTGGAGTCGTTGCTCTGTGGATTCCTGGAAGTTCCTNNNAGATCGGAAG
42273 TATGAGTGTGGAGTCGTTGCAAGCAAAAACCAAGCCAGACAACNNNAGATCGGAA
38752 TATGAGTGTGGAGTCGTTGCTGCCAAACTTTCAATCAGCAGGANNNAGATCGGAAG
41403 TATGAGTGTGGAGTCGTTGCTTATTGTGACCTGGCCCAGAGTANNNAGATCGGAAG
57785 TATGAGTGTGGAGTCGTTGCACACACACACGATAGCAGGAAATNNNAGATCGGAA
44384 TATGAGTGTGGAGTCGTTGCAAGACTACCTAAAAACAGCCCCCNNNAGATCGGAAG
61715 TATGAGTGTGGAGTCGTTGCAATATGGAAAGTTTGGGGCAGCTNNNAGATCGGAA
116255 TATGAGTGTGGAGTCGTTGCTACTCCCCATTAGCCAAACCCAANNNAGATCGGAAGA
34848 TATGAGTGTGGAGTCGTTGCACACTATTCTGCATCCAGTATCCANNNAGATCGGAAG
41322 TATGAGTGTGGAGTCGTTGCGAGCATGTGAGTGCACACTTCTANNNAGATCGGAA
34199 TATGAGTGTGGAGTCGTTGCTCCCCTCAGGTTTTGTTATTACCANNNAGATCGGAAG
62158 TATGAGTGTGGAGTCGTTGCTGCCAGAAAAGTTTGAAATTGGCTNNNAGATCGGAA
65615 TATGAGTGTGGAGTCGTTGCTCTCTCTCCCCACTTGGAAATGTNNNAGATCGGAAGA

95113 TATGAGTGTGGAGTCGTTGCGACTTACGAGAGACGACCGAAAANNNAGATCGGAA
50794 TATGAGTGTGGAGTCGTTGCGCGATAGTTTTCTCAACCACCAGNNNAGATCGGAAG
30965 TATGAGTGTGGAGTCGTTGCGGCGATCTTGAGACTGTGTACAANNNAGATCGGAA
41486 TATGAGTGTGGAGTCGTTGCGGAGTTATTGGGTGTGCAAGGTANNNAGATCGGA
36329 TATGAGTGTGGAGTCGTTGCGGGCCTCTAAAGTAGCTTTTGCANNNAGATCGGAAG
32154 TATGAGTGTGGAGTCGTTGCTGGAGGGTGAAGAGAAAGGTGAANNNAGATCGGA
57311 TATGAGTGTGGAGTCGTTGCTTTGTAATGGTACCCCAGGCTTGNNNAGATCGGAAG
98772 TATGAGTGTGGAGTCGTTGCAGTCATGTGTGGCTTTTGATTGANNNAGATCGGAA
67587 TATGAGTGTGGAGTCGTTGCGGTCAAGAAGCAGAAGCTGACTTNNNAGATCGGAA
31488 TATGAGTGTGGAGTCGTTGCGGCTGGTGCTCAAATCTCTTCAANNNAGATCGGAAG
31566 TATGAGTGTGGAGTCGTTGCGGGGATGTTTGGGGTTAGAAGAANNNAGATCGGA
115877 TATGAGTGTGGAGTCGTTGCAGATTCCCATGCCATAGCTGCNNNAGATCGGAAGAG
56784 TATGAGTGTGGAGTCGTTGCTCTAAGTGGTCCAAGGTAGGGTGNNNAGATCGGAA
32932 TATGAGTGTGGAGTCGTTGCTCTTTCCCTAGAGGTGCTGAACANNNAGATCGGAAG
37144 TATGAGTGTGGAGTCGTTGCTTGACATTGTGCTTGGGGATTCANNNAGATCGGAA
109441 TATGAGTGTGGAGTCGTTGCAGGGTGACTGAAAAATGCTTGCTNNNAGATCGGAA
50701 TATGAGTGTGGAGTCGTTGCGCATTGTCTCTCCCCATTCACAGNNNAGATCGGAAGA
40200 TATGAGTGTGGAGTCGTTGCAATGTATGACAGGGGTGAGGTGANNNAGATCGGA
64696 TATGAGTGTGGAGTCGTTGCAGTTGTTCTAGGTCTGGGTTCGTNNNAGATCGGAA
66307 TATGAGTGTGGAGTCGTTGCTGGTTCTGCTTCTAAAATCTCAGTGTNNNAGATCGG
40397 TATGAGTGTGGAGTCGTTGCATGACAGCAGCACCATTTATTGANNNAGATCGGAAG
64000 TATGAGTGTGGAGTCGTTGCCATCAGAGACACACCAAAGCAGTNNNAGATCGGAA
65540 TATGAGTGTGGAGTCGTTGCACAGCCTCAGAATAAGTTTCTTGGTNNNAGATCGGA
68238 TATGAGTGTGGAGTCGTTGCACTCTGGCTACATCCTGTAAGTTNNNAGATCGGAAG
104306 TATGAGTGTGGAGTCGTTGCGGGTGAAGAGAAAGGTGAAAGACNNNAGATCGGA
64766 TATGAGTGTGGAGTCGTTGCAGAATCATCTGGATTTGTAGAAGGTNNNAGATCGG
40642 TATGAGTGTGGAGTCGTTGCAGTGTCTACCCACGGTGTCAATANNNAGATCGGAAG
53247 TATGAGTGTGGAGTCGTTGCAGAGAGAAATGGGTAGCTGGAGGNNNAGATCGGA
60293 TATGAGTGTGGAGTCGTTGCCCCATCCTGATGAAACTGAGACTNNNAGATCGGAAG
99492 TATGAGTGTGGAGTCGTTGCCTACTTGTCAGGGCTGGAACTATTNNNAGATCGGAA
55723 TATGAGTGTGGAGTCGTTGCGGCGGGGTCCGATAGTAATTATGNNNAGATCGGAA
68175 TATGAGTGTGGAGTCGTTGCAGAGGCTGTTGCCAATTGTTCTTNNNAGATCGGAAG
32533 TATGAGTGTGGAGTCGTTGCCTGCTGATGGTTCCCAGTTGTAANNNAGATCGGAAG
104695 TATGAGTGTGGAGTCGTTGCTCTTTGCCTGACTAGCCACCAANNNAGATCGGAAGA
38984 TATGAGTGTGGAGTCGTTGCAGAGACAGAGAGTGCATAAGGGANNNAGATCGGA
38246 TATGAGTGTGGAGTCGTTGCTTTTCTCCGAATGGAATGGGAGANNNAGATCGGAA
51296 TATGAGTGTGGAGTCGTTGCCTCTCTCTCTCTCGGAGTGTCAGNNNAGATCGGAAG
47303 TATGAGTGTGGAGTCGTTGCATGGTGTTCACTTGACTTCTGGCNNNAGATCGGAAG
64950 TATGAGTGTGGAGTCGTTGCAGTTATGACCTGGATACACATAGGTNNNAGATCGGA
69593 TATGAGTGTGGAGTCGTTGCCGTGTATGAGAGGCAGGTGTTTTNNNAGATCGGAA
36857 TATGAGTGTGGAGTCGTTGCGCATCCAGCCTGTAATCAAGTCANNNAGATCGGAAG
100139 TATGAGTGTGGAGTCGTTGCGTTCCCATATTATGTCCATAGCTGCNNNAGATCGGA
95117 TATGAGTGTGGAGTCGTTGCGAGTTCCTCCATCAGCCAAGATGNNNAGATCGGAAG

39530 TATGAGTGTGGAGTCGTTGCTGAGCTATGGTGAGACCTTTGGANNNAGATCGGAA
43532 TATGAGTGTGGAGTCGTTGCCTGGAAGACTTGCCCATGACTACNNNAGATCGGAAG
60288 TATGAGTGTGGAGTCGTTGCTTCCCAGCAATGGAGTAGAGACTNNNAGATCGGAA
56352 TATGAGTGTGGAGTCGTTGCGCTCCACACTCAATAACCCTCTGNNNAGATCGGAAG
34519 TATGAGTGTGGAGTCGTTGCACCTACACCAGATCAGTTTCCCANNNAGATCGGAAG
59784 TATGAGTGTGGAGTCGTTGCTCTATGTGGGTGGTCAGGAAACTNNNAGATCGGAA
63450 TATGAGTGTGGAGTCGTTGCTGTAAGAAACATTGGTTTGAGTTCTNNNAGATCGGA
50613 TATGAGTGTGGAGTCGTTGCATGCAGCTATCAGTGGGAAACAGNNNAGATCGGAA
31895 TATGAGTGTGGAGTCGTTGCCGCTAAAGACAACACTCAGGGAANNNAGATCGGAA
66921 TATGAGTGTGGAGTCGTTGCAAAGCCAGAGAGCAGTGGAAATTNNNAGATCGGAA
43063 TATGAGTGTGGAGTCGTTGCTCCATAAATGCCTTGGGTGTCACNNNAGATCGGAAG
69525 TATGAGTGTGGAGTCGTTGCTTCCCCAAAGAGCAAAGTCTTTTNNNAGATCGGAAG
53937 TATGAGTGTGGAGTCGTTGCATTGGCATGTGTGATTATGGGGGNNNAGATCGGAA
117910 TATGAGTGTGGAGTCGTTGCAGCCAACTTACAGAGAGGTCCACNNNAGATCGGAA
35030 TATGAGTGTGGAGTCGTTGCACTTGCCAGCTGTTTGTATTCCANNNAGATCGGAAG
33716 TATGAGTGTGGAGTCGTTGCGGTGGATCAGGAAATGTGTGACANNNAGATCGGAA
31932 TATGAGTGTGGAGTCGTTGCAAGCATTCAAGACACAGTGGGAANNNAGATCGGAA
69220 TATGAGTGTGGAGTCGTTGCACCCAGAGAGAAGGAGGTTCTTTNNNAGATCGGAA
50991 TATGAGTGTGGAGTCGTTGCCCTCCCTCTCATTGTGTTTCCAGNNNAGATCGGAAGA
52524 TATGAGTGTGGAGTCGTTGCGTTGCTGGAGTTGATAAATGGCGNNNAGATCGGAA
105822 TATGAGTGTGGAGTCGTTGCGGGAATAGCCTTGCATTCATTCATNNNAGATCGGAA
57958 TATGAGTGTGGAGTCGTTGCGGACCAAGAATAGCTGAGGCAATNNNAGATCGGAA
50336 TATGAGTGTGGAGTCGTTGCTGCTGCTTCAGGAACTACTCAAGNNNAGATCGGAAG
62039 TATGAGTGTGGAGTCGTTGCTGGCTGTCTAAAATAAATGGGGCTNNNAGATCGGA
58285 TATGAGTGTGGAGTCGTTGCGTTGGGGGAGAAGGAAAACACATNNNAGATCGGA
40550 TATGAGTGTGGAGTCGTTGCGGCTATGATTGAGAGACAGTTTGANNNAGATCGGA
31365 TATGAGTGTGGAGTCGTTGCTCTCCATCCTATGCTTCAATCAANNNAGATCGGAAGA
60166 TATGAGTGTGGAGTCGTTGCGTGCATATACTAGACTTCGAATCACTNNNAGATCGG
39973 TATGAGTGTGGAGTCGTTGCAGATATTGGTGTGTTTCAGGCTGANNNAGATCGGA
69433 TATGAGTGTGGAGTCGTTGCTGCTTGTTGAGGTTGGGGATTTTNNNAGATCGGAA
38495 TATGAGTGTGGAGTCGTTGCTGTCCCAAGAGTTGTTGCTTTAGANNNAGATCGGAA
103254 TATGAGTGTGGAGTCGTTGCCACTTTTGGAGGGAGGATGTTCATANNNAGATCGG
30723 TATGAGTGTGGAGTCGTTGCCCAAGGGCAGCAAGAGCTATAAANNNAGATCGGAA
67818 TATGAGTGTGGAGTCGTTGCACTAGGGGTCTTTGTGGTTCCTTNNNAGATCGGAAG
57064 TATGAGTGTGGAGTCGTTGCGGACACTTCTCTGTGAGTGTGTGNNNAGATCGGAA
63033 TATGAGTGTGGAGTCGTTGCTGCTTCCAGAGGTTTCTCTCTCTNNNAGATCGGAAG
60478 TATGAGTGTGGAGTCGTTGCGTTGTGTCCATAGCTTGGCTACTNNNAGATCGGAAG
51427 TATGAGTGTGGAGTCGTTGCCCCACATGATTACTGAGGCAGAGNNNAGATCGGAA
34852 TATGAGTGTGGAGTCGTTGCACAACTTCATCGGGTAACTTATCCANNNAGATCGGA
106917 TATGAGTGTGGAGTCGTTGCAAACAACTTCTCTTTTGAGTTCCCANNNAGATCGGAA
52444 TATGAGTGTGGAGTCGTTGCCTCAACCTCTGTGTTCACATCCGNNNAGATCGGAAG
55735 TATGAGTGTGGAGTCGTTGCGTGGGTGAGACTGGGAGTTTATGNNNAGATCGGA
41246 TATGAGTGTGGAGTCGTTGCGGAAGTTTGGCAGGCAGAATCTANNNAGATCGGAA

37455 TATGAGTGTGGAGTCGTTGCACGCATTCTGAGGACTTGAAAAGANNNAGATCGGA
34322 TATGAGTGTGGAGTCGTTGCTTTTCTTTCTCCTTCCTCCCCCANNNAGATCGGAAGAG
38726 TATGAGTGTGGAGTCGTTGCAGGGCAAGCTTTACTATACAGGANNNAGATCGGAA
31177 TATGAGTGTGGAGTCGTTGCCACAAGGCAAACATCACAAGCAANNNAGATCGGAA
59527 TATGAGTGTGGAGTCGTTGCTGCTTCCTTGAGTGCTCTCCTATNNNAGATCGGAAG
45157 TATGAGTGTGGAGTCGTTGCGTGTCAGCCTTTGTTTTGGCCNNNAGATCGGAAGAG
29880 TATGAGTGTGGAGTCGTTGCACAATGGAGCTGTAAGGTCCTCANNNAGATCGGAA
63670 TATGAGTGTGGAGTCGTTGCAGAGGTGGAAAGGCACATAAAGTNNNAGATCGGA
60200 TATGAGTGTGGAGTCGTTGCTGATGACTAGACAGTGGCTCACTNNNAGATCGGAA
53291 TATGAGTGTGGAGTCGTTGCCAAAGGGTTGGGATTCTCTGAGGNNNAGATCGGAA
55545 TATGAGTGTGGAGTCGTTGCTAGAGGGAGCAGGATGGAAGATGNNNAGATCGGA
46885 TATGAGTGTGGAGTCGTTGCGCTCATGCATATTTTACCCAGGCNNNAGATCGGAAG
43513 TATGAGTGTGGAGTCGTTGCGCCTCAGGTAGCCGTCATTATACNNNAGATCGGAAG
36276 TATGAGTGTGGAGTCGTTGCCATTATTTGCCTGCCATGTTGCANNNAGATCGGAAG
61317 TATGAGTGTGGAGTCGTTGCGTTGCTTTTCTCTCCACCATCCTNNNAGATCGGAAGA
52982 TATGAGTGTGGAGTCGTTGCAAAGAATAGAAGCAAGGGCCAGGNNNAGATCGGA
37235 TATGAGTGTGGAGTCGTTGCCATCTGGTTGACTTCGCTCTTCANNNAGATCGGAAG
37498 TATGAGTGTGGAGTCGTTGCGCAACCACAACTGAAACGAAAGANNNAGATCGGAA
33812 TATGAGTGTGGAGTCGTTGCAACCTAGAATGCTACACCCTACANNNAGATCGGAAG
45908 TATGAGTGTGGAGTCGTTGCCCACACAGAGCAGTAGATCTTCCNNNAGATCGGAAG
37287 TATGAGTGTGGAGTCGTTGCTTTGGCTTGGTTTCTAGGGTTCANNNAGATCGGAAG
59670 TATGAGTGTGGAGTCGTTGCGGTAGAGGGGCAGGGTAACTTATNNNAGATCGGA
58490 TATGAGTGTGGAGTCGTTGCCTTCCTCTCCTTCTGGATCCCATNNNAGATCGGAAGA
43145 TATGAGTGTGGAGTCGTTGCCCACCAAAGGGAAGAGGTAAGACNNNAGATCGGAA
66875 TATGAGTGTGGAGTCGTTGCTCTGTCTCAGTGCTTGGTTTTGTNNNAGATCGGAAG
58617 TATGAGTGTGGAGTCGTTGCTCTTCTACCGTATGAGCCAGCATNNNAGATCGGAAG
38875 TATGAGTGTGGAGTCGTTGCGCACATCTTTATGGGTCAATAGGANNNAGATCGGA
43858 TATGAGTGTGGAGTCGTTGCCAGGGCAAAAGGAAGAAAACACCNNNAGATCGGAA
46930 TATGAGTGTGGAGTCGTTGCAAGAAGGAAATCAGCACTCAGGCNNNAGATCGGAA
57510 TATGAGTGTGGAGTCGTTGCTACCACCAAATGCAGTCTTGTTGNNNAGATCGGAAG
61606 TATGAGTGTGGAGTCGTTGCTCCTTTGTTCTTTTCAAACTGAAAGCTNNNAGATCGG
69613 TATGAGTGTGGAGTCGTTGCCTTGCCACACGTGTCCATTTTTNNNAGATCGGAAGA
53829 TATGAGTGTGGAGTCGTTGCCAAGGGGATAAGGGGACTTAGGGNNNAGATCGGA
58252 TATGAGTGTGGAGTCGTTGCCGGCAAGGGGAAACAATAAACATNNNAGATCGGAA
70742 TATGAGTGTGGAGTCGTTGCTGTAGTCAAGCTGTTTGCTGAGGNNNAGATCGGAA
37804 TATGAGTGTGGAGTCGTTGCTGGTCTATGCTATTGGGGACAGANNNAGATCGGAA
46052 TATGAGTGTGGAGTCGTTGCTGTGTGTTTCCTAAGCAGAAAAGCNNNAGATCGGA
45142 TATGAGTGTGGAGTCGTTGCTCTATTTAGCATCTGCTGTGGCCNNNAGATCGGAAG
51873 TATGAGTGTGGAGTCGTTGCATCACACTTCCGTGGTTTCTGAGNNNAGATCGGAAG
53055 TATGAGTGTGGAGTCGTTGCTATTCCCTGATGCTTCATGCAGGNNNAGATCGGAAG
68786 TATGAGTGTGGAGTCGTTGCTGGGTAGGGGGAGAACTGAATTTNNNAGATCGGA
30067 TATGAGTGTGGAGTCGTTGCGCTTCATAGCTGTTCTCCCAAAANNNAGATCGGAAG
55382 TATGAGTGTGGAGTCGTTGCAAACATGCCTGGACCACTACATGNNNAGATCGGAAG

64332 TATGAGTGTGGAGTCGTTGCTTAGGTAGCCATCAGCACTGAGTNNNAGATCGGAAG
39741 TATGAGTGTGGAGTCGTTGCAAGAGACGTGTCAGGGAACTATGANNNAGATCGGA
43212 TATGAGTGTGGAGTCGTTGCTGAGAGGCATTGTTTGAGGAGACNNNAGATCGGAA
47664 TATGAGTGTGGAGTCGTTGCTCCCTCCCTTGATTTTTCACTGCNNNAGATCGGAAGA
32321 TATGAGTGTGGAGTCGTTGCCCCCTTCAGCTCCTTGAAACTAANNNAGATCGGAAGA
99491 TATGAGTGTGGAGTCGTTGCACAAACCAATCCTTCCCCTTTTTCANNNAGATCGGAAG
34788 TATGAGTGTGGAGTCGTTGCCTCTGCATCCCATTTCCAATCCANNNAGATCGGAAGA
63672 TATGAGTGTGGAGTCGTTGCCTGCAAGGGCCAGATCATAAAGTNNNAGATCGGAA
105817 TATGAGTGTGGAGTCGTTGCCTGCTTGGTCTTCCCAGAGTTCANNNAGATCGGAAG
30967 TATGAGTGTGGAGTCGTTGCTGTTCCTCCTGGTCACCTTACAANNNAGATCGGAAGA
68908 TATGAGTGTGGAGTCGTTGCCCTTGCCTGGACTTGTCTGATTTNNNAGATCGGAAG
117024 TATGAGTGTGGAGTCGTTGCGTCCTTCTCATCTTAGCACCTGNNNAGATCGGAAGAG
64531 TATGAGTGTGGAGTCGTTGCGGGAGGGCAATCAAATAGAACGTNNNAGATCGGAA
34996 TATGAGTGTGGAGTCGTTGCTATGTGTTGCAAAGGTGTGTCCANNNAGATCGGAAG
59703 TATGAGTGTGGAGTCGTTGCTGGGAAGGGGCTGAACTTCTTATNNNAGATCGGAA
68418 TATGAGTGTGGAGTCGTTGCAGTGCTCAGTGTAGTCTCAGGTTNNNAGATCGGAAG
43077 TATGAGTGTGGAGTCGTTGCAGTGGTTTCACGGGCATATTCACNNNAGATCGGAAG
42647 TATGAGTGTGGAGTCGTTGCAACTGCTGGAACTTTGCTTACACNNNAGATCGGAAG
56977 TATGAGTGTGGAGTCGTTGCCTTGCTTGAATGGAGTTGCTGTGNNNAGATCGGAAG
59073 TATGAGTGTGGAGTCGTTGCAGGGCAGAGAAAAGAGGTGAGATNNNAGATCGGA
67816 TATGAGTGTGGAGTCGTTGCTTCCTCCCATACTCCTCTTCCTTNNNAGATCGGAAGAG
99146 TATGAGTGTGGAGTCGTTGCGTTATCTGTGTGGGATACATTCCAAGNNNAGATCGG
100095 TATGAGTGTGGAGTCGTTGCCAAAAATGATCACCGAGGGGGTANNNAGATCGGAA
59478 TATGAGTGTGGAGTCGTTGCAGCCCCCACCCATGGAATTATATNNNAGATCGGAAG
53275 TATGAGTGTGGAGTCGTTGCTCAGGGAAGAGGGATAACTGAGGNNNAGATCGGA
104648 TATGAGTGTGGAGTCGTTGCATCCCAATACTATTTACCCAAAGGANNNAGATCGGAA
60877 TATGAGTGTGGAGTCGTTGCCACTATTTATAGCCCCCACCCCTNNNAGATCGGAAGA
61835 TATGAGTGTGGAGTCGTTGCACAAGTTTAGACCAAGGAAACTAGCTNNNAGATCGG
35857 TATGAGTGTGGAGTCGTTGCAGACAAAACAGAGATAGAGTTGGCANNNAGATCGG
67581 TATGAGTGTGGAGTCGTTGCCGAAGAACCCCATGATTGGACTTNNNAGATCGGAAG
110218 TATGAGTGTGGAGTCGTTGCGTAAGGATGGGGTAACTATTCTAGCNNNAGATCGGA
101832 TATGAGTGTGGAGTCGTTGCTGTTCAGGGCTTTGAGAAAAGCANNNAGATCGGAA
61152 TATGAGTGTGGAGTCGTTGCAGACAGAAAGGTAGGTAAGGCCTNNNAGATCGGAA
104691 TATGAGTGTGGAGTCGTTGCCCAGAGGTGGGATTTCTTTAGAATGNNNAGATCGGA
59040 TATGAGTGTGGAGTCGTTGCGCATTTCTCTCTCCCTACGAGATNNNAGATCGGAAGA
30721 TATGAGTGTGGAGTCGTTGCCCGTGTCTGGTACTGCTGATAAANNNAGATCGGAAG
62178 TATGAGTGTGGAGTCGTTGCTGGAAAAGAGACAAGCTTTGGCTNNNAGATCGGAA
39019 TATGAGTGTGGAGTCGTTGCTCTTCCAAGGAGTATTGCAGGGANNNAGATCGGAA
57158 TATGAGTGTGGAGTCGTTGCAGGTGGGTTGATGACAGACATTGNNNAGATCGGAA
109707 TATGAGTGTGGAGTCGTTGCGTGTTTGTGTGTTTGTGACTACAGNNNAGATCGGAA
116481 TATGAGTGTGGAGTCGTTGCTTCCTCTTTCTTGTGGGGATTTCTGNNNAGATCGGAA
67588 TATGAGTGTGGAGTCGTTGCTTCTCATCTCCAAAGGCTGACTTNNNAGATCGGAAGA
52341 TATGAGTGTGGAGTCGTTGCCTCTCTCTCTTTGTGGCCATACGNNNAGATCGGAAGA

| | |
|---|---|
| 68828 | TATGAGTGTGGAGTCGTTGCCAACGCTTTCATGATGGCCATTTNNNAGATCGGAAG |
| 34842 | TATGAGTGTGGAGTCGTTGCTCCAGGGTGTCTTTCCATATCCANNNAGATCGGAAG |
| 52795 | TATGAGTGTGGAGTCGTTGCGATGTGCTTGAAACTGCAGAAGGNNNAGATCGGAA |
| 106436 | TATGAGTGTGGAGTCGTTGCGTTCCTTTGAATTTTGGGGTTTGNNNAGATCGGAAG |
| 31526 | TATGAGTGTGGAGTCGTTGCGGATGGGATCAAGAGGCAAAGAANNNAGATCGGA |
| 58499 | TATGAGTGTGGAGTCGTTGCGGCAGTATCTCCTTAGGTCCCATNNNAGATCGGAAG |
| 37651 | TATGAGTGTGGAGTCGTTGCCTGATGTGTCCTTTGCTGGAAGANNNAGATCGGAA |
| 68660 | TATGAGTGTGGAGTCGTTGCTAGAGGCAAAAGGCAACAGTGTTNNNAGATCGGAA |
| 41989 | TATGAGTGTGGAGTCGTTGCTGCCATCTGTTTCCTGGCATTTANNNAGATCGGAAG |
| 29877 | TATGAGTGTGGAGTCGTTGCTCTGGCTCCATTTCTTGAAGAGACNNNAGATCGGAA |
| 37603 | TATGAGTGTGGAGTCGTTGCTCCAAAAAGAGTAGGAAACAAGAAGANNNAGATCG |
| 105766 | TATGAGTGTGGAGTCGTTGCTGTGCATGGTGAGTGACTTTGGANNNAGATCGGAA |
| 107357 | TATGAGTGTGGAGTCGTTGCGGATGGGATCAAGAGGCAAAGAATTNNNAGATCG |
| 108785 | TATGAGTGTGGAGTCGTTGCTGACTATTACCCAGTGTGTGTTTTTNNNAGATCGGA |
| 32324 | TATGAGTGTGGAGTCGTTGCGAGATGCTGGTGGCATCAACTAANNNAGATCGGAA |
| 35394 | TATGAGTGTGGAGTCGTTGCATTTCCTCATCTAGGCAGGAGCANNNAGATCGGAAG |
| 58448 | TATGAGTGTGGAGTCGTTGCGCCCCAAAAGAGAATGTGACCATNNNAGATCGGAA |
| 98153 | TATGAGTGTGGAGTCGTTGCGCTAAGGCTCTCAGAACTCCCGNNNAGATCGGAAGA |
| 37108 | TATGAGTGTGGAGTCGTTGCAGTTCTCCAAAGGGGAATACATTCANNNAGATCGGA |
| 31527 | TATGAGTGTGGAGTCGTTGCCAGCCAAAACAACCTGCAAAGAANNNAGATCGGAA |
| 44748 | TATGAGTGTGGAGTCGTTGCACTCACTCCTTCCTTGAAGTCCCNNNAGATCGGAAGA |
| 39142 | TATGAGTGTGGAGTCGTTGCACCACAGAAATAAAAGGAACTGGGANNNAGATCGG |
| 36393 | TATGAGTGTGGAGTCGTTGCGCAGCTAGAGGACTGAAAACATCANNNAGATCGGA |
| 53098 | TATGAGTGTGGAGTCGTTGCGGAGACCACTGCACTAATTCAGGNNNAGATCGGAA |
| 43396 | TATGAGTGTGGAGTCGTTGCGGATTGACAGAGGGTTTCCTGACNNNAGATCGGAA |
| 109429 | TATGAGTGTGGAGTCGTTGCTATCTCACTCCGGGCCTCCTGNNNAGATCGGAAGAG |
| 45109 | TATGAGTGTGGAGTCGTTGCTTTTCTCTCTCTCATGCATGGCCNNNAGATCGGAAGA |
| 56043 | TATGAGTGTGGAGTCGTTGCTATTTGCCAGCTCAGAACTCCTGNNNAGATCGGAAG |
| 116738 | TATGAGTGTGGAGTCGTTGCTCTCTCACTTCCCCATCACAGTAATNNNAGATCGGAA |
| 64834 | TATGAGTGTGGAGTCGTTGCTCAATTTCTCCAGGAAAGCAGGTNNNAGATCGGAAG |
| 55221 | TATGAGTGTGGAGTCGTTGCACTGCATCTGGGCATTCAAAATGNNNAGATCGGAAG |
| 47601 | TATGAGTGTGGAGTCGTTGCGCCAGCCCTTGATTTAGATATGCNNNAGATCGGAAG |
| 67541 | TATGAGTGTGGAGTCGTTGCGCAAATACCACAGCACTTCACTTNNNAGATCGGAAG |
| 62894 | TATGAGTGTGGAGTCGTTGCGTGTACCTAAAACCCCTCTCCTCTNNNAGATCGGAAG |
| 34394 | TATGAGTGTGGAGTCGTTGCAAGCAACAATGGTGATATGCCCANNNAGATCGGAA |
| 68497 | TATGAGTGTGGAGTCGTTGCTACCATGCCAGAAAAAGCTGGTTNNNAGATCGGAA |
| 118842 | TATGAGTGTGGAGTCGTTGCACATACACATCACACATACCATGANNNAGATCGGAA |
| 116370 | TATGAGTGTGGAGTCGTTGCGAATGTACGCTGATACGTTTCACTNNNAGATCGGAA |
| 102046 | TATGAGTGTGGAGTCGTTGCATGGGAGTAGACAGAAAGGAAAGAAGNNNAGATC |
| 62492 | TATGAGTGTGGAGTCGTTGCGCTATCCCCTTGAACATGTTGCTNNNAGATCGGAAG |
| 49480 | TATGAGTGTGGAGTCGTTGCACAGAGATCCACGATGAGTTGTCNNNAGATCGGAA |
| 39322 | TATGAGTGTGGAGTCGTTGCTCGTGGACAAAAAGAATACTGGANNNAGATCGGAA |
| 51348 | TATGAGTGTGGAGTCGTTGCGCGGGAAGGGTGATTCTAAAGAGNNNAGATCGGA |

| | |
|---|---|
| 60626 | TATGAGTGTGGAGTCGTTGCAAATGCGAACATTCAGGGAACCTNNNAGATCGGAA |
| 107017 | TATGAGTGTGGAGTCGTTGCCCCACATGGCAAGTTTACTTTTTCTNNNAGATCGGAA |
| 32117 | TATGAGTGTGGAGTCGTTGCGGTCTTGCTGTGTAGATGCTGAANNNAGATCGGAA |
| 109748 | TATGAGTGTGGAGTCGTTGCGTGTGTTTCTGTGCTATGAAAAACTNNNAGATCGGA |
| 46886 | TATGAGTGTGGAGTCGTTGCCTTTTCTGGTTAGTACCCCAGGCNNNAGATCGGAAG |
| 95114 | TATGAGTGTGGAGTCGTTGCATCAAAGGGGGAAATGGCATGAANNNAGATCGGA |
| 30521 | TATGAGTGTGGAGTCGTTGCGGATTGGAAAGGGCACAGAGAAANNNAGATCGGA |
| 34885 | TATGAGTGTGGAGTCGTTGCAAACTGATAAAGCAAACCCTCCANNNAGATCGGAAG |
| 48191 | TATGAGTGTGGAGTCGTTGCCTGGTAACAACGGTCTGATTTGCNNNAGATCGGAAG |
| 60654 | TATGAGTGTGGAGTCGTTGCCCAAATGCATAACCCTCACACCTNNNAGATCGGAAG |
| 42422 | TATGAGTGTGGAGTCGTTGCCCACAGACGTACTCTCACTGAACNNNAGATCGGAAG |
| 46573 | TATGAGTGTGGAGTCGTTGCTACCCAGTGTGTGTTTTTGGAGCNNNAGATCGGAAG |
| 65925 | TATGAGTGTGGAGTCGTTGCTGTACGCTGATACGTTTCACTGTNNNAGATCGGAAG |
| 99812 | TATGAGTGTGGAGTCGTTGCTTCTGGGTCTCACTTCCCTCATAANNNAGATCGGAAG |
| 36167 | TATGAGTGTGGAGTCGTTGCCAGACATTATGGGGGAATGTGCANNNAGATCGGAA |
| 56643 | TATGAGTGTGGAGTCGTTGCACAGAAGGATGGAGAATGATGAGTGNNNAGATCG |
| 52290 | TATGAGTGTGGAGTCGTTGCCAAAGTAGACTGACCCTTCCACGNNNAGATCGGAAG |
| 57456 | TATGAGTGTGGAGTCGTTGCAAACATCACTCACTCCAGGGTTGNNNAGATCGGAAG |
| 66500 | TATGAGTGTGGAGTCGTTGCGCTCTTTCTCCACATCTGTGTGTNNNAGATCGGAAG |
| 37105 | TATGAGTGTGGAGTCGTTGCTTCTGCCTAGGTTACGACATTCANNNAGATCGGAAG |
| 67368 | TATGAGTGTGGAGTCGTTGCGCCTCTTGAGCAGCTCCAATATTNNNAGATCGGAAG |
| 64874 | TATGAGTGTGGAGTCGTTGCAAATCTAGAGACCCTGCAGAGGTNNNAGATCGGAA |
| 44976 | TATGAGTGTGGAGTCGTTGCAGGTGGAAACGGATATTTGAGCCNNNAGATCGGAA |
| 101854 | TATGAGTGTGGAGTCGTTGCGAGTATTTGTTCCAGGACCCCTANNNAGATCGGAAG |
| 61817 | TATGAGTGTGGAGTCGTTGCTTGCAGGAAAACACATTTGAGCTNNNAGATCGGAA |
| 102694 | TATGAGTGTGGAGTCGTTGCTTCCTTACATTCCCACCAACACTNNNAGATCGGAAGA |
| 62106 | TATGAGTGTGGAGTCGTTGCCCTGGCTCATAACTCCTATGGCTNNNAGATCGGAAG |
| 30248 | TATGAGTGTGGAGTCGTTGCCAAACTGAATGGAGCAGCACAAANNNAGATCGGAA |
| 39024 | TATGAGTGTGGAGTCGTTGCTGTGCCTAGAATGAGTTCAGGGANNNAGATCGGAA |
| 109765 | TATGAGTGTGGAGTCGTTGCGATGGGTGCGAGGTGACATTTCANNNAGATCGGAA |
| 40985 | TATGAGTGTGGAGTCGTTGCGAGCCCTAGAAGCAGTGAGACTANNNAGATCGGAA |
| 57021 | TATGAGTGTGGAGTCGTTGCTTCAGCACATCCTCATCAGTGTGNNNAGATCGGAAG |
| 116354 | TATGAGTGTGGAGTCGTTGCAGGATACAGCAAGGTGTCTTTCCANNNAGATCGGAA |
| 54430 | TATGAGTGTGGAGTCGTTGCAATACATCTGGCTGGTACGATGGNNNAGATCGGAA |
| 32938 | TATGAGTGTGGAGTCGTTGCATCCAAAATCTGCAGGGTGAACANNNAGATCGGAA |
| 63421 | TATGAGTGTGGAGTCGTTGCACTTCCACATCATCCTCTCTTCTNNNAGATCGGAAGA |
| 31489 | TATGAGTGTGGAGTCGTTGCCCTTGGTTGAGCTCCTTCTTCAANNNAGATCGGAAG |
| 37628 | TATGAGTGTGGAGTCGTTGCCTCAACCAGATGCACAAGGAAGANNNAGATCGGAA |
| 55780 | TATGAGTGTGGAGTCGTTGCTCCTTACATTCCCACCAACACTGNNNAGATCGGAAGA |
| 42366 | TATGAGTGTGGAGTCGTTGCTTGGCTCTGTTTGTTTGCAGAACNNNAGATCGGAAG |
| 56875 | TATGAGTGTGGAGTCGTTGCGTCCACTAGGGCAAGGAAATGTGNNNAGATCGGAA |
| 31148 | TATGAGTGTGGAGTCGTTGCCACCTTCTCCTACCTCTGTCCAANNNAGATCGGAAGA |
| 108095 | TATGAGTGTGGAGTCGTTGCTACCAAGGGCAGAGACTGGATNNNAGATCGGAAGA |

44715 TATGAGTGTGGAGTCGTTGCATTATGGAGGTTGAGGGCTCCCNNNAGATCGGAAG
45884 TATGAGTGTGGAGTCGTTGCGTTGTGTGAAGTCCTCTCCTTCCNNNAGATCGGAAG
55156 TATGAGTGTGGAGTCGTTGCTGATGAGATTCCTGGGCATTTGGNNNAGATCGGAA
96208 TATGAGTGTGGAGTCGTTGCCCGTTTCAGCATTGTGTGTGTTCNNNAGATCGGAAG
49817 TATGAGTGTGGAGTCGTTGCTTGTTACCAGCCCAGATTGGTTCNNNAGATCGGAAG
35906 TATGAGTGTGGAGTCGTTGCTGTCTCTTTCTGTGCAGAATGCANNNAGATCGGAAG
101598 TATGAGTGTGGAGTCGTTGCAATCCAAAATCTGCAGGGTGAACAGNNNAGATCGG
38334 TATGAGTGTGGAGTCGTTGCACCAAATAGCCAAAGCAATAGTGAGANNNAGATCG
55916 TATGAGTGTGGAGTCGTTGCCTCAATGCAATGTCAACCACCTGNNNAGATCGGAAG
46959 TATGAGTGTGGAGTCGTTGCTCCCAATCTACGTTTTGAGAGGCNNNAGATCGGAAG
110185 TATGAGTGTGGAGTCGTTGCGTATGTATTTTCCTGAGGGATGTCCCNNNAGATCGG
49280 TATGAGTGTGGAGTCGTTGCAAAGACCATATTGGCAGACCGTCNNNAGATCGGAA
50237 TATGAGTGTGGAGTCGTTGCACAGGTTTTGCACCGTTTACAAGNNNAGATCGGAAG
34819 TATGAGTGTGGAGTCGTTGCACACCACCCTATGATGTCATCCANNNAGATCGGAAG
101155 TATGAGTGTGGAGTCGTTGCGTCCTACCTCGTAAACAAATGCTNNNAGATCGGAAG
37885 TATGAGTGTGGAGTCGTTGCTGTACTTTGGAATAAAGTTGCCAGANNNAGATCGG
46131 TATGAGTGTGGAGTCGTTGCGCAGAACAATCAGTTATCCCAAGCNNNAGATCGGAA
106907 TATGAGTGTGGAGTCGTTGCTTTCTTTGTACGATGCCATGTTCTGNNNAGATCGGA
53118 TATGAGTGTGGAGTCGTTGCTGAGGGTGAGAGAAGGAAAGAGGNNNAGATCGG
58279 TATGAGTGTGGAGTCGTTGCCCTTTTGAGGCTGCCACTAACATNNNAGATCGGAAG
30134 TATGAGTGTGGAGTCGTTGCGGCAATCTCTTCCTCCAGGAAAANNNAGATCGGAAG
67952 TATGAGTGTGGAGTCGTTGCGCCAGGGTTCATTTCTACTGCTTNNNAGATCGGAAG
58110 TATGAGTGTGGAGTCGTTGCAAGGAGCCTCACTCATCTGGAATNNNAGATCGGAAG
31163 TATGAGTGTGGAGTCGTTGCTGGTCAGTCATGAGAATTTTCCAANNNAGATCGGAA
59200 TATGAGTGTGGAGTCGTTGCCCTACAAAGAAGCATGCTGGGATNNNAGATCGGAA
68544 TATGAGTGTGGAGTCGTTGCACCAAAAGATGCCTGGAAATGTTNNNAGATCGGAA
35466 TATGAGTGTGGAGTCGTTGCAGGTTGGGAGGCATTCTATAGCANNNAGATCGGAA
66215 TATGAGTGTGGAGTCGTTGCAGTAAGAGAAATGCCCTTGTCTGTNNNAGATCGGA
51795 TATGAGTGTGGAGTCGTTGCGGGAAAGGCTTCACAGAGATGAGNNNAGATCGGA
47312 TATGAGTGTGGAGTCGTTGCCACACACACACTCTTTTTCTGGCNNNAGATCGGAAGA
109313 TATGAGTGTGGAGTCGTTGCGTGTGTGTATCATTTATTTTGACACAANNNAGATCG
49652 TATGAGTGTGGAGTCGTTGCGAAATGGATGCTTTGTGCCCTTCNNNAGATCGGAAG
30256 TATGAGTGTGGAGTCGTTGCAAGTTCCAATAGGCTGTCACAAANNNAGATCGGAAG
29878 TATGAGTGTGGAGTCGTTGCCTCTTACCACTCTTCTCCACCCANNNAGATCGGAAGA
95112 TATGAGTGTGGAGTCGTTGCCAGGTGTGGGGTAATTCTGTTCCNNNAGATCGGAA
49348 TATGAGTGTGGAGTCGTTGCTGATTTCCAAAACTGAGCTGGTCNNNAGATCGGAAG
67512 TATGAGTGTGGAGTCGTTGCAAAGTCTCACTCTGTCCCCACTTNNNAGATCGGAAGA
42106 TATGAGTGTGGAGTCGTTGCGAGAGCGAGCCAGGTTTCTTTTANNNAGATCGGAA
35326 TATGAGTGTGGAGTCGTTGCTGGCTTTCAGGTACATATCAGCANNNAGATCGGAAG
108103 TATGAGTGTGGAGTCGTTGCTCACTTCAGCCTCCCAAGATTTTNNNAGATCGGAAGA
46801 TATGAGTGTGGAGTCGTTGCCACTTCAGTATGCAAAGCTTCGCNNNAGATCGGAAG
116251 TATGAGTGTGGAGTCGTTGCTGCTCTTGTTTTCATTCTCTTGCTCTNNNAGATCGGA
42170 TATGAGTGTGGAGTCGTTGCCAGCAAGGTGTCTTTCCACAAACNNNAGATCGGAAG

| | |
|---|---|
| 29881 | TATGAGTGTGGAGTCGTTGCTCTGTTAGAAGGGGGAAGAGGTGNNNAGATCGGA |
| 63896 | TATGAGTGTGGAGTCGTTGCGGAAAAATATGAATTCTCAGAGACAGTNNNAGATCG |
| 62939 | TATGAGTGTGGAGTCGTTGCGCACATGAGATTCTTCAGGCTCTNNNAGATCGGAAG |
| 104659 | TATGAGTGTGGAGTCGTTGCCTCCCATTATTTACTACCTAGCTTGGTNNNAGATCGG |
| 56630 | TATGAGTGTGGAGTCGTTGCTGATGGACTAGGCCTATGGAGTGNNNAGATCGGAA |
| 35243 | TATGAGTGTGGAGTCGTTGCACAGAACACACACATACACAGCANNNAGATCGGAAG |
| 36651 | TATGAGTGTGGAGTCGTTGCAGGCTTACTTATGACAGATCCTCANNNAGATCGGAA |
| 37948 | TATGAGTGTGGAGTCGTTGCGGAGAGAGAGCTATTCAGGCAGANNNAGATCGGA |
| 53620 | TATGAGTGTGGAGTCGTTGCCACACACACATATGGGAGAAGGGNNNAGATCGGAA |
| 39663 | TATGAGTGTGGAGTCGTTGCACCCTGGGAAAATTTGATCATGANNNAGATCGGAA |
| 47346 | TATGAGTGTGGAGTCGTTGCTAATAGAGAGTAGCGAGGGTGGCNNNAGATCGGA |
| 58681 | TATGAGTGTGGAGTCGTTGCAGAGAGAAAGAGAGTTGGGGCATNNNAGATCGGA |
| 68748 | TATGAGTGTGGAGTCGTTGCTGGAGGTGGCTTTATGCTTTGTTNNNAGATCGGAA |
| 53738 | TATGAGTGTGGAGTCGTTGCACAGCCTCTTTTCACTTTCAGGGNNNAGATCGGAAG |
| 41665 | TATGAGTGTGGAGTCGTTGCGTTAGTGTCAGAAGCCACGTGTANNNAGATCGGAA |
| 52210 | TATGAGTGTGGAGTCGTTGCCACCACTGTGGCTTCCTTCTTAGNNNAGATCGGAAG |
| 37043 | TATGAGTGTGGAGTCGTTGCGCAGCAAGATACAGAAAAGTGTCANNNAGATCGGA |
| 43506 | TATGAGTGTGGAGTCGTTGCCCACCACCACAGAGCACATATACNNNAGATCGGAAG |
| 46632 | TATGAGTGTGGAGTCGTTGCAGGCAGTCTGTACTTTGCATAGCNNNAGATCGGAAG |
| 109346 | TATGAGTGTGGAGTCGTTGCGACAGAGAGAAAGAGGGGGATATNNNAGATCGGA |
| 43217 | TATGAGTGTGGAGTCGTTGCAAAGAGAGAAAGGGGAGGGAGACNNNAGATCGG |
| 65913 | TATGAGTGTGGAGTCGTTGCGAATGAAATGCTGCATGCACTGTNNNAGATCGGAA |
| 71837 | TATGAGTGTGGAGTCGTTGCGCTGCCAAGTCTCTGTGAATACANNNAGATCGGAAG |
| 30364 | TATGAGTGTGGAGTCGTTGCTTTACACTCCCACTTACAGCAAANNNAGATCGGAAGA |
| 54720 | TATGAGTGTGGAGTCGTTGCTTCAAGCACCTGCTATATGCTGGNNNAGATCGGAAG |
| 63931 | TATGAGTGTGGAGTCGTTGCACCATAATAGCAAAGACATCTTACAGTNNNAGATCG |
| 35417 | TATGAGTGTGGAGTCGTTGCTGAAAGAAGGCTCAAGATGAGCANNNAGATCGGAA |
| 41047 | TATGAGTGTGGAGTCGTTGCACTCTACTCCACACTCTCCCCTANNNAGATCGGAAGA |
| 52983 | TATGAGTGTGGAGTCGTTGCTAGAATATCTCACAGGGGCCAGGNNNAGATCGGAA |
| 52600 | TATGAGTGTGGAGTCGTTGCGCAGAAGGTAAGAAGACCTCTCGNNNAGATCGGAA |
| 54888 | TATGAGTGTGGAGTCGTTGCTCCCCAAGTTGTTTAGAAGGTGGNNNAGATCGGAA |
| 55149 | TATGAGTGTGGAGTCGTTGCGATACTACCACTGGGCTTGTTGGNNNAGATCGGAA |
| 54270 | TATGAGTGTGGAGTCGTTGCCTGCCAAAGAAAGGGAATTTGGGNNNAGATCGGAA |
| 58478 | TATGAGTGTGGAGTCGTTGCAGCAAAGCTGAAAAATAGCCCATNNNAGATCGGAA |
| 63484 | TATGAGTGTGGAGTCGTTGCTGTTGCTTTGTTCCTGTTTGTTCTNNNAGATCGGAAG |
| 34332 | TATGAGTGTGGAGTCGTTGCCCCTCCTACCTTATTATGCCCCANNNAGATCGGAAGA |
| 57221 | TATGAGTGTGGAGTCGTTGCCGCTCAGGGAGGGAAGAATATTGNNNAGATCGGAA |
| 116065 | TATGAGTGTGGAGTCGTTGCTAGAATCTCGCTCTGTCTCCCATTNNNAGATCGGAAG |
| 40074 | TATGAGTGTGGAGTCGTTGCTTCTGGTGAGCTCATCCTTCTGANNNAGATCGGAAG |
| 99087 | TATGAGTGTGGAGTCGTTGCCAGATTGAGCCAAATATCCTGCGNNNAGATCGGAAG |
| 54817 | TATGAGTGTGGAGTCGTTGCCCCTCATCTATTGGCTGAAGTGGNNNAGATCGGAAG |
| 71798 | TATGAGTGTGGAGTCGTTGCGGAGCACTTTACTATGGTGGTGGNNNAGATCGGAA |
| 103187 | TATGAGTGTGGAGTCGTTGCACTTTCCATCTTGTGTCTTCCCAAANNNAGATCGGAA |

36479 TATGAGTGTGGAGTCGTTGCTGGACAAACTGTTACCTATTAGATCANNNAGATCGG
67633 TATGAGTGTGGAGTCGTTGCTTCAAGTGGATTGGCTCAACCTTNNNAGATCGGAAG
66194 TATGAGTGTGGAGTCGTTGCTCCCACTCTGCTTCTTTCTCTGTNNNAGATCGGAAGA
46369 TATGAGTGTGGAGTCGTTGCAGAATGCATGTGGTATAGGCAGCNNNAGATCGGAA
61756 TATGAGTGTGGAGTCGTTGCGCACAGAACCAACTGAAAGAGCTNNNAGATCGGAA
66989 TATGAGTGTGGAGTCGTTGCGTTACAGAGCCGGGAAAGGAATTNNNAGATCGGAA
36414 TATGAGTGTGGAGTCGTTGCACTGTCCTCTTACTGGACCATCANNNAGATCGGAAG
48139 TATGAGTGTGGAGTCGTTGCTTTCAGTCACTCCATGGTCTTGCNNNAGATCGGAAG
41211 TATGAGTGTGGAGTCGTTGCGAAGCCAACTGAAGGAACTGCTANNNAGATCGGAA
66929 TATGAGTGTGGAGTCGTTGCCACAGTACAGCAGCCCTACAATTNNNAGATCGGAAG
31103 TATGAGTGTGGAGTCGTTGCTTGGGACTGTTTTTCTCTGCCAANNNAGATCGGAAG
71724 TATGAGTGTGGAGTCGTTGCAGAAAGGTTGAGGGGAAATCAGCNNNAGATCGGA
38358 TATGAGTGTGGAGTCGTTGCTTTCTTGGAACTCTAGTCTTGAGANNNAGATCGGAA
65556 TATGAGTGTGGAGTCGTTGCTCCAAGACTGTGACTGAATTTGGTNNNAGATCGGAA
68403 TATGAGTGTGGAGTCGTTGCTGGTCATCTCTGGTGGTAAGGTTNNNAGATCGGAA
30242 TATGAGTGTGGAGTCGTTGCGCACTATCACAAGGTCCCACAAANNNAGATCGGAAG
35211 TATGAGTGTGGAGTCGTTGCCCTCACAAGGTGATACTGAAGCANNNAGATCGGAAG
53399 TATGAGTGTGGAGTCGTTGCTCATGCTTTGTTCACCGAGTAGGNNNAGATCGGAAG
36543 TATGAGTGTGGAGTCGTTGCACAGTCATTATTCAGAGCAACTCANNNAGATCGGAA
107355 TATGAGTGTGGAGTCGTTGCAAACAGTCATTATTCAGAGCAACTCANNNAGATCGG
52213 TATGAGTGTGGAGTCGTTGCGAGAATGGGCAGGTGGAAGTTAGNNNAGATCGGA
50125 TATGAGTGTGGAGTCGTTGCGGAGCGGAAGACCACTAAGAAAGNNNAGATCGGA
32502 TATGAGTGTGGAGTCGTTGCGAGGGATGTCCCAGGTTTGGTAANNNAGATCGGAA
49871 TATGAGTGTGGAGTCGTTGCCCCATCTCTGATTGCAGCATTTCNNNAGATCGGAAG
65274 TATGAGTGTGGAGTCGTTGCTTGAAATGGAAATATGCTCATGGTNNNAGATCGGA
34174 TATGAGTGTGGAGTCGTTGCTAAGCAGTATGACAGCCCTACCANNNAGATCGGAAG
51849 TATGAGTGTGGAGTCGTTGCACTGCTCATGACTTCATGCTGAGNNNAGATCGGAAG
67038 TATGAGTGTGGAGTCGTTGCTCCTCATAGTAGAAGCACAACATTNNNAGATCGGAA
38369 TATGAGTGTGGAGTCGTTGCCTCTTCCTGCATGCCTTTTGAGANNNAGATCGGAAG
34089 TATGAGTGTGGAGTCGTTGCCAGTGAGGACATTTGGAAAGACCANNNAGATCGGA
33840 TATGAGTGTGGAGTCGTTGCGCTGGCAAGTCCCATTATCTACANNNAGATCGGAAG
109732 TATGAGTGTGGAGTCGTTGCAACTAGACATGTTGTGTAGCAGAGTAGNNNAGATC
109739 TATGAGTGTGGAGTCGTTGCAAGATACCATCTCACAGCATCAGAATGNNNAGATCG
58538 TATGAGTGTGGAGTCGTTGCCACTACATGATGGCTTCTGCCATNNNAGATCGGAAG
99451 TATGAGTGTGGAGTCGTTGCAATCCACTCACTACCCACCAGATNNNAGATCGGAAG
45668 TATGAGTGTGGAGTCGTTGCTTGAGACCCTTGATCTCCTCTCCNNNAGATCGGAAG
55608 TATGAGTGTGGAGTCGTTGCGCTCTCGCTCTCAAAAATAGGATGNNNAGATCGGAA
59243 TATGAGTGTGGAGTCGTTGCGGGGGAGCGATTTCTATTTGGATNNNAGATCGGAA
35168 TATGAGTGTGGAGTCGTTGCAAAATCTACAGGGTGCACAAGCANNNAGATCGGAA
52137 TATGAGTGTGGAGTCGTTGCTTGGAGAGGCATGACCTAGGTAGNNNAGATCGGAA
58358 TATGAGTGTGGAGTCGTTGCACACACACAGACATAACAGACATNNNAGATCGGAAG
38579 TATGAGTGTGGAGTCGTTGCCGGTCTGAAGACATCCTCATCGANNNAGATCGGAAG
52485 TATGAGTGTGGAGTCGTTGCACCTGACTACAATTGGAGAAGCGNNNAGATCGGAA

| | |
|---|---|
| 48286 | TATGAGTGTGGAGTCGTTGCGCCCATGCTGAGGTAGAAGAATCNNNAGATCGGAA |
| 44547 | TATGAGTGTGGAGTCGTTGCAGATTTTGAGACTGACAACGCCCNNNAGATCGGAAG |
| 67557 | TATGAGTGTGGAGTCGTTGCGGTGGGAATGGGTAGAGAGACTTNNNAGATCGGA |
| 44938 | TATGAGTGTGGAGTCGTTGCGAGGTTCTCACTGTTGATCAGCCNNNAGATCGGAAG |
| 36999 | TATGAGTGTGGAGTCGTTGCTCACTTAGCCTCTGTGGATGTCANNNAGATCGGAAG |
| 38680 | TATGAGTGTGGAGTCGTTGCTGAGGAGATACCTTGAAGGAAGGANNNAGATCGG |
| 48939 | TATGAGTGTGGAGTCGTTGCCAATTCTCCCACCTTCTCTGCTCNNNAGATCGGAAGA |
| 44939 | TATGAGTGTGGAGTCGTTGCATCACCACCATTTTCTCTCAGCCNNNAGATCGGAAGA |
| 64898 | TATGAGTGTGGAGTCGTTGCACATAGAGCAAAGGGAAGGAGGTNNNAGATCGGA |
| 55849 | TATGAGTGTGGAGTCGTTGCAGTATTCATCTGTTGCCCAGACTGNNNAGATCGGAA |
| 48666 | TATGAGTGTGGAGTCGTTGCCTCCCTTCCTCCATACCAACCTCNNNAGATCGGAAGA |
| 70796 | TATGAGTGTGGAGTCGTTGCAACCAAGAGGGTGAAAGGAATGGNNNAGATCGGA |
| 56806 | TATGAGTGTGGAGTCGTTGCAAAAAGAAACGTGTGTGTGGGTGNNNAGATCGGA |
| 66366 | TATGAGTGTGGAGTCGTTGCTTTCCCTGATCTAACCCAGGTGTNNNAGATCGGAAG |
| 37579 | TATGAGTGTGGAGTCGTTGCTCTAGCAGGGAAAAGTTGCAAGANNNAGATCGGAA |
| 64644 | TATGAGTGTGGAGTCGTTGCAGAAATGAGGCATATTGTGGCGTNNNAGATCGGAA |
| 68557 | TATGAGTGTGGAGTCGTTGCAGCGTTTTCCACATATGAATGTTNNNAGATCGGAAG |
| 65315 | TATGAGTGTGGAGTCGTTGCAGACTGTAGAATGCATTGTTATATGGTNNNAGATCG |
| 117472 | TATGAGTGTGGAGTCGTTGCTCTTTCAACGGACTTTTGCTCTGNNNAGATCGGAAG |
| 34079 | TATGAGTGTGGAGTCGTTGCTTTCTAGTCTCGCTCTGTCACCANNNAGATCGGAAGA |
| 117479 | TATGAGTGTGGAGTCGTTGCGTTTGGTTTTGTTGCTTGCAATNNNAGATCGGAAGA |
| 43312 | TATGAGTGTGGAGTCGTTGCAGACCCTTGAACAATTTGGGGACNNNAGATCGGAA |
| 49542 | TATGAGTGTGGAGTCGTTGCCCAGTGCAATCACTCATGCATTCNNNAGATCGGAAG |
| 38361 | TATGAGTGTGGAGTCGTTGCGGTTGGCTGAGTTCTTGTTGAGANNNAGATCGGAA |
| 37121 | TATGAGTGTGGAGTCGTTGCACAAGCATCAAAGAGGGCATTCANNNAGATCGGAA |
| 104685 | TATGAGTGTGGAGTCGTTGCATTTTCTCTTCCTCTCCCTCTCTCTGNNNAGATCGGAA |
| 48705 | TATGAGTGTGGAGTCGTTGCATACTCACACTCACACCCCCTCNNNAGATCGGAAGAG |
| 105785 | TATGAGTGTGGAGTCGTTGCCCTGAAATACACACATCCACTCTTCTGNNNAGATCGG |
| 45890 | TATGAGTGTGGAGTCGTTGCATAGGACACTCCCACTTCCTTCCNNNAGATCGGAAGA |
| 68542 | TATGAGTGTGGAGTCGTTGCGGGTAGGGGATCGAGGAAATGTTNNNAGATCGGA |
| 67830 | TATGAGTGTGGAGTCGTTGCATTTCAGATGGTGAGGCAAGCTTNNNAGATCGGAA |
| 34559 | TATGAGTGTGGAGTCGTTGCACACCAGACACCAATATAAGCCANNNAGATCGGAAG |
| 34865 | TATGAGTGTGGAGTCGTTGCTCCTAAAGAGGTGCTTCACTCCANNNAGATCGGAAG |
| 103194 | TATGAGTGTGGAGTCGTTGCTGGCTTGTGACTATAGAGACTTGGTNNNAGATCGG |
| 65169 | TATGAGTGTGGAGTCGTTGCGAAGGCTTGATAACTCCTATATGGGTNNNAGATCGG |
| 62844 | TATGAGTGTGGAGTCGTTGCCATTTTCCTTTGCCTGACCCTCTNNNAGATCGGAAGA |
| 97066 | TATGAGTGTGGAGTCGTTGCACAGGAGACTGCCAGAAGTTCTTNNNAGATCGGAA |
| 60761 | TATGAGTGTGGAGTCGTTGCGTGCTTGTTAGAAGTGCTGACCTNNNAGATCGGAAG |
| 55443 | TATGAGTGTGGAGTCGTTGCTTATTCCGTCCTGTTGCTCCATGNNNAGATCGGAAGA |
| 34623 | TATGAGTGTGGAGTCGTTGCAGGGTAGCAGTTCTGATTAGCCANNNAGATCGGAA |
| 46294 | TATGAGTGTGGAGTCGTTGCGGGAAATCTTTTCAATGGACCAGCNNNAGATCGGAA |
| 60435 | TATGAGTGTGGAGTCGTTGCCGAAGGCCCCACTTACAAATACTNNNAGATCGGAAG |
| 98462 | TATGAGTGTGGAGTCGTTGCCCAACCAGCCCTTCTTCCCATTNNNAGATCGGAAGAG |

53478 TATGAGTGTGGAGTCGTTGCAGGTGTCTCTGATATACTCACCGGNNNAGATCGGAA
46808 TATGAGTGTGGAGTCGTTGCAAAATCCCCCTTCCAGAAAAGGCNNNAGATCGGAAG
56172 TATGAGTGTGGAGTCGTTGCCTTTTCTGCTGTTGTGAAGGCTGNNNAGATCGGAAG
41174 TATGAGTGTGGAGTCGTTGCCACATTGCTCCAGGATAAGGCTANNNAGATCGGAAG
50310 TATGAGTGTGGAGTCGTTGCCCTCTTCTGCCATCTGATGCAAGNNNAGATCGGAAG
105770 TATGAGTGTGGAGTCGTTGCATGGAATATTGGGCTTGAGGACACNNNAGATCGGA
68825 TATGAGTGTGGAGTCGTTGCTCATCTCTGCACCAGTCCCATTTNNNAGATCGGAAGA
32103 TATGAGTGTGGAGTCGTTGCCAGTCCCAATAGCCAAAGCTGAANNNAGATCGGAAG
38935 TATGAGTGTGGAGTCGTTGCTTGTAACCATGGGTGATGACGGANNNAGATCGGAA
104632 TATGAGTGTGGAGTCGTTGCATGTGTGAGGCGTTTACAGGGNNNAGATCGGAAGA
105778 TATGAGTGTGGAGTCGTTGCGCTGGGCACTTTCCATTATCTAGNNNAGATCGGAAG
32000 TATGAGTGTGGAGTCGTTGCTCATAGATGCCGCAAAATTGGAANNNAGATCGGAA
108782 TATGAGTGTGGAGTCGTTGCTCATAGATAGATGGCAGCAGATATCNNNAGATCGG
109801 TATGAGTGTGGAGTCGTTGCTATTACCTTTCTAGTAGTTGCATGTTTNNNAGATCGG
99771 TATGAGTGTGGAGTCGTTGCTTGCCTATGTTTTCTTTCCGTAGTTNNNAGATCGGAA
55170 TATGAGTGTGGAGTCGTTGCCCACATTTTGTCTGTCTGACTTTGGNNNAGATCGGA
54524 TATGAGTGTGGAGTCGTTGCACTGGTCACTGTTTTAGCACTGGNNNAGATCGGAAG
46257 TATGAGTGTGGAGTCGTTGCATGGTATCGGGTTTCTCAACAGCNNNAGATCGGAAG
62236 TATGAGTGTGGAGTCGTTGCCTGAATTGGGACTGACAGATGCTNNNAGATCGGAA
104643 TATGAGTGTGGAGTCGTTGCAGTAGGTAAGGGCCAGATATCTTGTNNNAGATCGG
46150 TATGAGTGTGGAGTCGTTGCCAGTATGTATGCCCTTTGCAAGCNNNAGATCGGAAG
39603 TATGAGTGTGGAGTCGTTGCTCACACCTATACTTTGGCAACATGANNNAGATCGGA
63361 TATGAGTGTGGAGTCGTTGCCTGCACCATACAGCATCACTTCTNNNAGATCGGAAG
57142 TATGAGTGTGGAGTCGTTGCGCTCTAATGGAGGTTGGCAATTGNNNAGATCGGAA
53722 TATGAGTGTGGAGTCGTTGCTAGAATACCACGCCTACTCAGGGNNNAGATCGGAAG
69246 TATGAGTGTGGAGTCGTTGCTCCCCTCATCCTGTGGTAAGTTTNNNAGATCGGAAG
48894 TATGAGTGTGGAGTCGTTGCCCTATGGTTTGTGATGAGGGCTCNNNAGATCGGAA
108801 TATGAGTGTGGAGTCGTTGCCACCTATACTTTGGCAACATGATNNNAGATCGGAAG
37433 TATGAGTGTGGAGTCGTTGCAGAATGCAGCACAGGGAAAAAGANNNAGATCGGA
68373 TATGAGTGTGGAGTCGTTGCCTGAGAACACACTAAACACCGTTNNNAGATCGGAAG
103578 TATGAGTGTGGAGTCGTTGCTCCCTTTGGCAGTCAGTATAGCCNNNAGATCGGAAG
54274 TATGAGTGTGGAGTCGTTGCACAGCAAGACAGCTAGATTTGGGNNNAGATCGGAA
71578 TATGAGTGTGGAGTCGTTGCCTCCAAACACCAGACTGCTTGTTNNNAGATCGGAAG
52700 TATGAGTGTGGAGTCGTTGCAAGCCTCCCAAAACTCTCAAAGGNNNAGATCGGAAG
56704 TATGAGTGTGGAGTCGTTGCCTGAGCTTATTCTGTCTTGCGTGNNNAGATCGGAAG
55878 TATGAGTGTGGAGTCGTTGCAATCTGGGTGGGAGAGGATACTGNNNAGATCGGA
41825 TATGAGTGTGGAGTCGTTGCCTGTGTGTCTGTCGATGTCCTTANNNAGATCGGAAG
51821 TATGAGTGTGGAGTCGTTGCCACACCTTCTCCAAGTCACTGAGNNNAGATCGGAAG
64906 TATGAGTGTGGAGTCGTTGCGCTTTGTAGGGAGTTAGGGAGGTNNNAGATCGGA
58682 TATGAGTGTGGAGTCGTTGCTGATTTATGCATGGCATGGGCATNNNAGATCGGAA
61366 TATGAGTGTGGAGTCGTTGCTCTTAGGTCATACATCCACTCCTNNNAGATCGGAAGA
38721 TATGAGTGTGGAGTCGTTGCAGAAAGTATCAAGTATCTGGACAGGANNNAGATCG
63645 TATGAGTGTGGAGTCGTTGCCTCCAAGCCCCATTTCTCAAAGTNNNAGATCGGAAG

| | |
|---|---|
| 66244 | TATGAGTGTGGAGTCGTTGCTGGGACACTTAGGTTGGTTCTGTNNNAGATCGGAA |
| 106978 | TATGAGTGTGGAGTCGTTGCGTGTTCCTCAAATTTTGCTTTGAACNNNAGATCGGA |
| 57720 | TATGAGTGTGGAGTCGTTGCGCTAGAATCAGAGCAGGCCAAATNNNAGATCGGAA |
| 71057 | TATGAGTGTGGAGTCGTTGCCCCCAGGAGTAGTGCACAAATATNNNAGATCGGAA |
| 99769 | TATGAGTGTGGAGTCGTTGCAAGAGGGGCTGGCTGTCCCTNNNAGATCGGAAGAG |
| 45898 | TATGAGTGTGGAGTCGTTGCTTCTAGTTGCTCACTTGGCTTCCNNNAGATCGGAAG |
| 48848 | TATGAGTGTGGAGTCGTTGCGGCTGGAGTGAAGATTACAGCTCNNNAGATCGGAA |
| 52367 | TATGAGTGTGGAGTCGTTGCCCCTCTTCCCATTTCAGTAACCGNNNAGATCGGAAGA |
| 61087 | TATGAGTGTGGAGTCGTTGCCTGCATTCATGATATTCCCAGCCTNNNAGATCGGAAG |
| 50352 | TATGAGTGTGGAGTCGTTGCTTGTACCCACTACCACAGTCAAGNNNAGATCGGAAG |
| 44674 | TATGAGTGTGGAGTCGTTGCGCAGACATCAAGGCAATTTATCCCNNNAGATCGGAA |
| 96907 | TATGAGTGTGGAGTCGTTGCGCAAGTGAAGATGAGATTTGGGAGNNNAGATCGG |
| 58607 | TATGAGTGTGGAGTCGTTGCCTTGCCTCATGTGAAAACAGCATNNNAGATCGGAAG |
| 32129 | TATGAGTGTGGAGTCGTTGCCGCACAACCTCAGATTCTCTGAANNNAGATCGGAAG |
| 39752 | TATGAGTGTGGAGTCGTTGCGCTTTGCCTTTAGCCATGTATGANNNAGATCGGAAG |
| 59980 | TATGAGTGTGGAGTCGTTGCGATTTTCGCAAAGTGCACACACTNNNAGATCGGAAG |
| 57318 | TATGAGTGTGGAGTCGTTGCTGAACATCAGATCTCTGGGCTTGNNNAGATCGGAAG |
| 43625 | TATGAGTGTGGAGTCGTTGCAAGATTGCTAGGCTGCAAGGTACNNNAGATCGGAA |
| 62238 | TATGAGTGTGGAGTCGTTGCTCAAATCCAGCTGTAGAGATGCTNNNAGATCGGAAG |
| 58380 | TATGAGTGTGGAGTCGTTGCGGTGAACTCATTCTTGGGGACATNNNAGATCGGAA |
| 119251 | TATGAGTGTGGAGTCGTTGCGGTACTGTTCTGAGAGCTTTATGTGNNNAGATCGG |
| 64700 | TATGAGTGTGGAGTCGTTGCTCAGCCTCATTCACAAGTTTCGTNNNAGATCGGAAG |
| 55963 | TATGAGTGTGGAGTCGTTGCATTGTATTTCTGCTCCAGCCCTGNNNAGATCGGAAG |
| 36778 | TATGAGTGTGGAGTCGTTGCTAATTGCCCAGACATCCCTCTCANNNAGATCGGAAG |
| 70068 | TATGAGTGTGGAGTCGTTGCTGTTTCTACACCGCAAACCATGANNNAGATCGGAAG |
| 71879 | TATGAGTGTGGAGTCGTTGCACTGCTACCAAAAATTGGTGGGTNNNAGATCGGAA |
| 47728 | TATGAGTGTGGAGTCGTTGCTCTCTATCACCAACCTCTCCTGCNNNAGATCGGAAGA |
| 58626 | TATGAGTGTGGAGTCGTTGCAGCATTTGGTCTCCTAAGAGCATNNNAGATCGGAAG |
| 71838 | TATGAGTGTGGAGTCGTTGCGTGGCCCAGATGAGTTTAGTGTCNNNAGATCGGAA |
| 71881 | TATGAGTGTGGAGTCGTTGCCAAGGCTGCCCTCTAGTTGAATTNNNAGATCGGAAG |
| 38899 | TATGAGTGTGGAGTCGTTGCAAGATGGACGGTGTTAGCTAGGANNNAGATCGGA |
| 41900 | TATGAGTGTGGAGTCGTTGCACTAGCCAATCCCCCATCAGTTANNNAGATCGGAAG |
| 71882 | TATGAGTGTGGAGTCGTTGCTGAAGCTTGTTGTGACTTCTGCTNNNAGATCGGAAG |
| 101273 | TATGAGTGTGGAGTCGTTGCCATCAGAAACCATCCCTGCTGGCNNNAGATCGGAAG |
| 68436 | TATGAGTGTGGAGTCGTTGCAAGGTTGAACAGATTCTGCGGTTNNNAGATCGGAA |
| 71896 | TATGAGTGTGGAGTCGTTGCATTCAGATTTGTGTTGGTGGCCTNNNAGATCGGAAG |
| 71880 | TATGAGTGTGGAGTCGTTGCTGGCTGCTTTTCCTGACCATTAGNNNAGATCGGAAG |
| 55835 | TATGAGTGTGGAGTCGTTGCATTTGGTCACGATGAAGTCACTGNNNAGATCGGAA |
| 102389 | TATGAGTGTGGAGTCGTTGCGTTCAAAGGTGACTGGATCACAANNNAGATCGGAA |
| 103065 | TATGAGTGTGGAGTCGTTGCTTTTTCTGGGTGAGAGGATCCTANNNAGATCGGAA |
| 71887 | TATGAGTGTGGAGTCGTTGCCTTGCAGCAGATAAGGGCACTTANNNAGATCGGAA |
| 115573 | TATGAGTGTGGAGTCGTTGCCCAGGCTCAGCTTTAGTACCAGAANNNAGATCGGAA |
| 39569 | TATGAGTGTGGAGTCGTTGCTCTACTCAGTCCACAGCCAATGANNNAGATCGGAAG |

102644 TATGAGTGTGGAGTCGTTGCTATGTGCGTATCCATCCAGAGGGNNNAGATCGGAA
103233 TATGAGTGTGGAGTCGTTGCATGGGACAGAAACAATTTGCATTACTANNNAGATCG
104638 TATGAGTGTGGAGTCGTTGCTACATTGCGGTAAACTCAGGGCNNNAGATCGGAAG
103561 TATGAGTGTGGAGTCGTTGCGAGACAGGGAAATTCAGGCAACANNNAGATCGGAA
104635 TATGAGTGTGGAGTCGTTGCCTTGTGTGCATGTGTCTCTTTCANNNAGATCGGAAG
48683 TATGAGTGTGGAGTCGTTGCTAGAACCCTGCCCTTATGACCTCNNNAGATCGGAAG
106419 TATGAGTGTGGAGTCGTTGCAGGTTCAAGTGGATTGGCTCAACNNNAGATCGGAA
106884 TATGAGTGTGGAGTCGTTGCAAAGCAGGTAGCTTGAACAATGAGNNNAGATCGGA
71659 TATGAGTGTGGAGTCGTTGCCTAAAATGGATCCACCAGCCCACNNNAGATCGGAAG
104696 TATGAGTGTGGAGTCGTTGCGCACCTATAATCATGCCAGACCANNNAGATCGGAAG
105207 TATGAGTGTGGAGTCGTTGCAGAAAATCAAAGGGCTGCATGGANNNAGATCGGAA
106887 TATGAGTGTGGAGTCGTTGCCCAGCGCTCACGTTTTATTTGTACNNNAGATCGGAAG
109430 TATGAGTGTGGAGTCGTTGCTATGTGTATGGGTGTGTGTATATATCTNNNAGATCG
108115 TATGAGTGTGGAGTCGTTGCGAAAGGCATCCTCCCCTTGTGTCANNNAGATCGGAA
106912 TATGAGTGTGGAGTCGTTGCCAACAGCAACTCCGCTAAGGATNNNAGATCGGAAGA
107314 TATGAGTGTGGAGTCGTTGCTGTTTCCATGCCCCACTTCTTNNNAGATCGGAAGAGC
103182 TATGAGTGTGGAGTCGTTGCGAGATAGAATGCTGAATGTTGTCCCNNNAGATCGG
105814 TATGAGTGTGGAGTCGTTGCAGTTCAAAATCTCCTGGTCAGGTNNNAGATCGGAAG
117754 TATGAGTGTGGAGTCGTTGCAAAAACCTCCAGTAGGGCAGTGTNNNAGATCGGAA
108138 TATGAGTGTGGAGTCGTTGCAATACAGTGGGCCAGGCATACAGNNNAGATCGGAA
102369 TATGAGTGTGGAGTCGTTGCGCCTCTGAATTATCCTGACCTGCNNNAGATCGGAAG
99486 TATGAGTGTGGAGTCGTTGCACACACAAAAAGCAACCCTGCCNNNAGATCGGAAGA
103076 TATGAGTGTGGAGTCGTTGCCCACCATCCCCCACCCTTATCNNNAGATCGGAAGAGC
103073 TATGAGTGTGGAGTCGTTGCGAACTTGATTGGTTCTATTTCTCTGGNNNAGATCGG
97974 TATGAGTGTGGAGTCGTTGCTGTCTGCACTCCTGATTCCAATGNNNAGATCGGAAG
100515 TATGAGTGTGGAGTCGTTGCGGTCAATTTATGGCTGTGGTTTTGNNNAGATCGGAA
101545 TATGAGTGTGGAGTCGTTGCGTGGCTCAAAACAACACAAAAGANNNAGATCGGAA
106368 TATGAGTGTGGAGTCGTTGCCTTAGGAGGGTGAGGAATGGCCNNNAGATCGGAA
108810 TATGAGTGTGGAGTCGTTGCCGCACACATGCACAGAGTTATGTNNNAGATCGGAAG
116966 TATGAGTGTGGAGTCGTTGCAACACAATCTCAGCTTACGGCAGNNNAGATCGGAAG
105151 TATGAGTGTGGAGTCGTTGCCATCCTGCCAAATTCTCCCGAGNNNAGATCGGAAGA
46576 TATGAGTGTGGAGTCGTTGCAGTGGTAATGGAACCGAATGAGCNNNAGATCGGAA
35115 TATGAGTGTGGAGTCGTTGCCCCACAGACACATACGTATTTTCCANNNAGATCGGAA
30827 TATGAGTGTGGAGTCGTTGCGCATATCACTTGGGGGTCAACAANNNAGATCGGAA
31265 TATGAGTGTGGAGTCGTTGCATGCGTGTGTTTCATTATGGCAANNNAGATCGGAAG
31291 TATGAGTGTGGAGTCGTTGCGGTTTTGCATACTCTTACATGCAANNNAGATCGGAA
66922 TATGAGTGTGGAGTCGTTGCCCAGGTTTGAAGCTTCTGAAATTNNNAGATCGGAAG
109663 TATGAGTGTGGAGTCGTTGCACTGGAAGGCATGGGGTAATGANNNAGATCGGAA
34560 TATGAGTGTGGAGTCGTTGCAAACAGAGCAAGTGAGTAAGCCANNNAGATCGGAA
38865 TATGAGTGTGGAGTCGTTGCTTTGTCCTGTCCTTTTCTGAGGANNNAGATCGGAAG
95140 TATGAGTGTGGAGTCGTTGCGCTGGCAAAGGTAAACTACCAGANNNAGATCGGAA
58242 TATGAGTGTGGAGTCGTTGCCACGTATGCACAGAGACAAACATNNNAGATCGGAAG
58808 TATGAGTGTGGAGTCGTTGCCCCCACACCTGTTCTCTACTCATNNNAGATCGGAAGA

47534 TATGAGTGTGGAGTCGTTGCCCTCTGTGTATTTGTGTGCATGCNNNAGATCGGAAG
66095 TATGAGTGTGGAGTCGTTGCTCACATACACAGTAGCATGCTGTNNNAGATCGGAAG
59560 TATGAGTGTGGAGTCGTTGCCAGGCATCAGAGGTGTAATCTATNNNAGATCGGAA
41365 TATGAGTGTGGAGTCGTTGCGTGTGTGCGCGCATGTATAAGTANNNAGATCGGAA
42084 TATGAGTGTGGAGTCGTTGCTGCCCATGTTCCAAGGCATTTTANNNAGATCGGAAG
48078 TATGAGTGTGGAGTCGTTGCCCCACAATCCCTTAGGAGTTATTGCNNNAGATCGGA
106939 TATGAGTGTGGAGTCGTTGCTGTCTGTCTTTTCTCACTCAAGGNNNAGATCGGAAG
31525 TATGAGTGTGGAGTCGTTGCCGCCCAAAGACTAAGGCAAAGAANNNAGATCGGAA
100042 TATGAGTGTGGAGTCGTTGCGAAACTTTTGCTCTGTTGTCCAGGNNNAGATCGGAA
99561 TATGAGTGTGGAGTCGTTGCGTTCTTGAGGCTCTTGATGAGCTNNNAGATCGGAAG
32652 TATGAGTGTGGAGTCGTTGCGCCCTGGTCCAAACATAAAAACANNNAGATCGGAAG
44504 TATGAGTGTGGAGTCGTTGCACACGGACACAAATACAAAGCCCNNNAGATCGGAAG
59777 TATGAGTGTGGAGTCGTTGCTGATGGAAGAAGACGGAAGAAACTNNNAGATCGG
50870 TATGAGTGTGGAGTCGTTGCTTCTTTGTGGAGCATTGTCCCAGNNNAGATCGGAAG
51754 TATGAGTGTGGAGTCGTTGCCCCCTTTTAGCCTCTTTGTGGAGNNNAGATCGGAAG
52809 TATGAGTGTGGAGTCGTTGCATTGGCTCAAGCAGTTAGGAAGGNNNAGATCGGAA
54022 TATGAGTGTGGAGTCGTTGCTCAATCATCTGGGAGCAAATGGGNNNAGATCGGAA
71643 TATGAGTGTGGAGTCGTTGCTTTGCCTAGAGAAGTCACAGTGNNNAGATCGGAAG
71686 TATGAGTGTGGAGTCGTTGCAGACTTGCCTCCATATCGAGTGTNNNAGATCGGAAG
42543 TATGAGTGTGGAGTCGTTGCTGCTAAGTGCAGCAACAATAACACNNNAGATCGGAA
105760 TATGAGTGTGGAGTCGTTGCGGGAGAGACTGAAGAGGAAATGANNNAGATCGGA
44265 TATGAGTGTGGAGTCGTTGCGAGATATATGCATGCACCCACCCNNNAGATCGGAAG
32257 TATGAGTGTGGAGTCGTTGCCACACTATGTTTCCCTCCCATAANNNAGATCGGAAGA
48869 TATGAGTGTGGAGTCGTTGCCTCTTTGTTCTGGGACTGAGCTCNNNAGATCGGAAG
102370 TATGAGTGTGGAGTCGTTGCTGTAGAAGGGGTCTTTCTGTGTNNNAGATCGGAAG
48407 TATGAGTGTGGAGTCGTTGCTCTGCCAGGGTAAAAGCATCATCNNNAGATCGGAAG
61019 TATGAGTGTGGAGTCGTTGCACAAAGAAGTATCCCAATCTCCCTNNNAGATCGGAA
40428 TATGAGTGTGGAGTCGTTGCTCAGGCTTATACCCATTTCCTTGANNNAGATCGGAAG
105732 TATGAGTGTGGAGTCGTTGCCATAACAAACCTTACATTTTACCTGTTNNNAGATCGG
35603 TATGAGTGTGGAGTCGTTGCAGGGTAGAATACTTGGGTAAGGCANNNAGATCGGA
40975 TATGAGTGTGGAGTCGTTGCCATCACCCAAATGCCCATCACTANNNAGATCGGAAG
65774 TATGAGTGTGGAGTCGTTGCAGGGTTTCTAGATGTTCCAGATGTNNNAGATCGGA
52399 TATGAGTGTGGAGTCGTTGCACACACAACTTTCAAAAAGCCCGNNNAGATCGGAAG
50910 TATGAGTGTGGAGTCGTTGCTATCCAATTCCCAAACAGGCCAGNNNAGATCGGAAG
50628 TATGAGTGTGGAGTCGTTGCAATCTTTTCCAAGTGGGAGAACAGNNNAGATCGGA
43188 TATGAGTGTGGAGTCGTTGCCACTGCAAGGAGAGGAAAGAGACNNNAGATCGGA
51340 TATGAGTGTGGAGTCGTTGCAGAGGGTAGGGCAGATCAAAGAGNNNAGATCGGA
37872 TATGAGTGTGGAGTCGTTGCCATCTCCATTTGCACTAGCCAGANNNAGATCGGAAG
99564 TATGAGTGTGGAGTCGTTGCAATGTGTCCACTGATGCGCAGNNNAGATCGGAAGA
95118 TATGAGTGTGGAGTCGTTGCCCAGAAGTGACAACTGTCCAAAANNNAGATCGGAA
54341 TATGAGTGTGGAGTCGTTGCAGCAGCAGCAACAGTAGTAATGGNNNAGATCGGAA
33444 TATGAGTGTGGAGTCGTTGCGGAAGTATGTGCCCTGTTTCACANNNAGATCGGAAG
45679 TATGAGTGTGGAGTCGTTGCCCTTGATATTGTGCCCAGTCTCCNNNAGATCGGAAG

109680 TATGAGTGTGGAGTCGTTGCTGTGTCAGGTGGGTCTCACTCNNNAGATCGGAAGA
45345 TATGAGTGTGGAGTCGTTGCCCCAGGTACATACACTTCTTGCCNNNAGATCGGAAG
35311 TATGAGTGTGGAGTCGTTGCAGAGAGAGAGTGTGTATGCAGCANNNAGATCGGA
39658 TATGAGTGTGGAGTCGTTGCCCTTGTGCTTCTGGTATGCATGANNNAGATCGGAAG
101734 TATGAGTGTGGAGTCGTTGCGTGTGTGTGTGTCATTTTCCTTTTTCTNNNAGATCGG
32492 TATGAGTGTGGAGTCGTTGCAGAGAGTTGGTTTGTGTGGGTAANNNAGATCGGA
67320 TATGAGTGTGGAGTCGTTGCTTGCGTGAAGGTTTGTTGGGATTNNNAGATCGGAA
47410 TATGAGTGTGGAGTCGTTGCGCAGGGCATTATGAAATGTTGGCNNNAGATCGGAA
52500 TATGAGTGTGGAGTCGTTGCAGCATTCCAAAAGGAGAGTAGCGNNNAGATCGGAA
66021 TATGAGTGTGGAGTCGTTGCAAAACAAAAAGCCTCCCTCCTGTNNNAGATCGGAAG
43163 TATGAGTGTGGAGTCGTTGCTTCACCGAACCTCACCCAGACNNNAGATCGGAAGAG
39120 TATGAGTGTGGAGTCGTTGCAACACTGGTCCAGACTAATGGGANNNAGATCGGAA
54219 TATGAGTGTGGAGTCGTTGCGTTTTTCACCAGTGATGTGTGGGNNNAGATCGGAA
95136 TATGAGTGTGGAGTCGTTGCCACACGATCTACGTGGAAGGACNNNAGATCGGAAG
37372 TATGAGTGTGGAGTCGTTGCTGGAAAGTTGACTGGCAGTTTCANNNAGATCGGAA
51905 TATGAGTGTGGAGTCGTTGCCAGTAAAGTCAGGGCACTGTGAGNNNAGATCGGAA
59822 TATGAGTGTGGAGTCGTTGCAGAGAGAGATGCTGACACCAACTNNNAGATCGGAA
55703 TATGAGTGTGGAGTCGTTGCAGCATGTGGCAACTAGCAGTATGNNNAGATCGGAA
31204 TATGAGTGTGGAGTCGTTGCATGCAGTAGATTATGCTCAGCAANNNAGATCGGAA
32128 TATGAGTGTGGAGTCGTTGCACATGCTCACACAAAGCTCTGAANNNAGATCGGAAG
100655 TATGAGTGTGGAGTCGTTGCATTACCCCCTTCCTTCCTGCCTGNNNAGATCGGAAGA
55921 TATGAGTGTGGAGTCGTTGCGGCTCCGAAGAAAGAGACCTGNNNAGATCGGAAGA
43196 TATGAGTGTGGAGTCGTTGCCAACTGCCTCTTGGTACAGAGACNNNAGATCGGAAG
30734 TATGAGTGTGGAGTCGTTGCTCTCACACAAACAAATCACCTAAANNNAGATCGGAA
65619 TATGAGTGTGGAGTCGTTGCAGGTAACAAAACCCCTCAATAAATGTNNNAGATCGG
36328 TATGAGTGTGGAGTCGTTGCGCAGGAAATAAGCAGCTTTTGCANNNAGATCGGAA
46733 TATGAGTGTGGAGTCGTTGCATCTTCTCAGTGGTTCTTCACGCNNNAGATCGGAAG
95119 TATGAGTGTGGAGTCGTTGCTGATATTGGTTTCTGGGCACTGGNNNAGATCGGAA
71337 TATGAGTGTGGAGTCGTTGCAAAGCTGCTAAAACTCAGTCCGTNNNAGATCGGAAG
40690 TATGAGTGTGGAGTCGTTGCCAGGGAGAGGAAGAGCAGACATANNNAGATCGGA
41703 TATGAGTGTGGAGTCGTTGCGGGGGAGGAGTGGTGTATTTGTANNNAGATCGGA
38261 TATGAGTGTGGAGTCGTTGCCCCAACGCATGTATGTATGGAGANNNAGATCGGAA
69340 TATGAGTGTGGAGTCGTTGCTCCCAAGAATGTCAGTTTGGTTTNNNAGATCGGAAG
46187 TATGAGTGTGGAGTCGTTGCCGTCCCTAACAGCATCTAGAAGCNNNAGATCGGAAG
108869 TATGAGTGTGGAGTCGTTGCGCTGTCTATACCATTTCTGGACCTCNNNAGATCGGAA
68886 TATGAGTGTGGAGTCGTTGCTGGTCAACTTACTAGGCCGATTTNNNAGATCGGAAG
54634 TATGAGTGTGGAGTCGTTGCTGATTGGAAAGTGGTGATCCTGGNNNAGATCGGAA
99557 TATGAGTGTGGAGTCGTTGCACACACACATCTATCCAGTTCACNNNAGATCGGAAGA
50666 TATGAGTGTGGAGTCGTTGCGTCCTGGCACAGAATATCCACAGNNNAGATCGGAAG
67393 TATGAGTGTGGAGTCGTTGCGTGCTGATGAAGGCAGACCTATTNNNAGATCGGAA
32121 TATGAGTGTGGAGTCGTTGCGAGAGGTTCAGCTGATTGCTGAANNNAGATCGGAA
35866 TATGAGTGTGGAGTCGTTGCACTCCTCTACTTGTACTTTGGCANNNAGATCGGAAGA
44604 TATGAGTGTGGAGTCGTTGCTGTTACTTGGGGATTTTCTGCCCNNNAGATCGGAAG

| | |
|---|---|
| 44010 | TATGAGTGTGGAGTCGTTGCTACAAGCCTATCTGGGTTTCACCNNNAGATCGGAAG |
| 42573 | TATGAGTGTGGAGTCGTTGCTGCTATTGGTGCCTGATACACACNNNAGATCGGAAG |
| 61913 | TATGAGTGTGGAGTCGTTGCTGACATCCAAAGGTAATTCCGCTNNNAGATCGGAAG |
| 68423 | TATGAGTGTGGAGTCGTTGCAGAAACAGGATGGTCAGGAGGTTNNNAGATCGGA |
| 48862 | TATGAGTGTGGAGTCGTTGCCACTGTCTACCTTTGCAGAGCTCNNNAGATCGGAAG |
| 39018 | TATGAGTGTGGAGTCGTTGCAGAGAGAGAGATTTGTGCAGGGANNNAGATCGGA |
| 54928 | TATGAGTGTGGAGTCGTTGCGATTGGAGGGGATGTTATGGTGGNNNAGATCGGA |
| 40889 | TATGAGTGTGGAGTCGTTGCCAACCCTCCCACCCCCATATATANNNAGATCGGAAGA |
| 45692 | TATGAGTGTGGAGTCGTTGCAAGACATTCTCTGCTCCTTCTCCNNNAGATCGGAAGA |
| 31223 | TATGAGTGTGGAGTCGTTGCTGGAGGTGGAAAGTTCATAGCAANNNAGATCGGAA |
| 35157 | TATGAGTGTGGAGTCGTTGCATGCAACACACACTCCTAAAGCANNNAGATCGGAAG |
| 54458 | TATGAGTGTGGAGTCGTTGCTTGACTATGCTGGCTTGTGATGGNNNAGATCGGAA |
| 51061 | TATGAGTGTGGAGTCGTTGCGGAAGAAAGTTGGCCTTTAGCAGNNNAGATCGGAA |
| 39205 | TATGAGTGTGGAGTCGTTGCTTTGTCATCTCCAGATGTTGGGANNNAGATCGGAAG |
| 59535 | TATGAGTGTGGAGTCGTTGCGGAGGGGAGATGAAGAGAGCTATNNNAGATCGGA |
| 56615 | TATGAGTGTGGAGTCGTTGCAGGAACTGCAGCATGTAAGAGTGNNNAGATCGGAA |
| 43247 | TATGAGTGTGGAGTCGTTGCACTTAGAAAAAGAAACACACACGACNNNAGATCGG |
| 42425 | TATGAGTGTGGAGTCGTTGCTAAGGTGCACAGAAGCTCTGAACNNNAGATCGGAA |
| 99009 | TATGAGTGTGGAGTCGTTGCTGAAGTGGGTCATTTGATGGGGTNNNAGATCGGAA |
| 57789 | TATGAGTGTGGAGTCGTTGCAGCAATAGAGAGGTGGGGGAAATNNNAGATCGGA |
| 51167 | TATGAGTGTGGAGTCGTTGCGAGTTCTGACTGAGAAGGTGCAGNNNAGATCGGAA |
| 33165 | TATGAGTGTGGAGTCGTTGCACCACTTGATTTTGAAGACCACANNNAGATCGGAAG |
| 40755 | TATGAGTGTGGAGTCGTTGCATGCTCGTGTGGATATGCATANNNAGATCGGAAGA |
| 42911 | TATGAGTGTGGAGTCGTTGCGAGACAAGGTCACACAAATGCACNNNAGATCGGAA |
| 66820 | TATGAGTGTGGAGTCGTTGCAGTTGGAGGGAGCAAAAGTTTGTNNNAGATCGGA |
| 49502 | TATGAGTGTGGAGTCGTTGCAAGACAGAGTTGGGCCTGAATTCNNNAGATCGGAA |
| 56280 | TATGAGTGTGGAGTCGTTGCTTATTCCCACTGCCTGATTGCTGNNNAGATCGGAAG |
| 36334 | TATGAGTGTGGAGTCGTTGCGCTCATGTTGTATGTCTTTTTGCANNNAGATCGGAA |
| 62062 | TATGAGTGTGGAGTCGTTGCTTCTTGTTCTAGACCATTGGGCTNNNAGATCGGAAG |
| 53671 | TATGAGTGTGGAGTCGTTGCTTTTGTTTTAGGCCTCACCAGGGNNNAGATCGGAAG |
| 30353 | TATGAGTGTGGAGTCGTTGCGGCAAAAGCTGGGTAAAAGCAAANNNAGATCGGA |
| 56224 | TATGAGTGTGGAGTCGTTGCTCAGATGGAAAGGATTATGTGGCTGNNNAGATCGG |
| 66474 | TATGAGTGTGGAGTCGTTGCAAAGAAGGGGAAATAAGAGTGTGTNNNAGATCGG |
| 60403 | TATGAGTGTGGAGTCGTTGCAAGGTGGTGCTATGAGAGTGACTNNNAGATCGGAA |
| 68000 | TATGAGTGTGGAGTCGTTGCACACACTCTCACACACTTTGCTTNNNAGATCGGAAGA |
| 97300 | TATGAGTGTGGAGTCGTTGCAGAACCTAAATTGGATCGAGCCTTTNNNAGATCGGA |
| 62413 | TATGAGTGTGGAGTCGTTGCAATAAGGAATTGGCTCGTGTGCTNNNAGATCGGAA |
| 117283 | TATGAGTGTGGAGTCGTTGCAAAATAGCCTAGCATAGTGCTGCNNNAGATCGGAA |
| 117688 | TATGAGTGTGGAGTCGTTGCAAAGTGCAGAGGTGCAATGATAGCNNNAGATCGGA |
| 51658 | TATGAGTGTGGAGTCGTTGCATGTGGGAAGTGATCCTAGGGAGNNNAGATCGGA |
| 49802 | TATGAGTGTGGAGTCGTTGCCGATCTCCAAGCTGTAAGGGTTCNNNAGATCGGAAG |
| 35491 | TATGAGTGTGGAGTCGTTGCGTGACCATGGGTTGAAAGTAGCANNNAGATCGGAA |
| 38821 | TATGAGTGTGGAGTCGTTGCGTCCTGTGTCTGTAGAAGGAGGANNNAGATCGGAA |

34662 TATGAGTGTGGAGTCGTTGCTATTTGGTGTTTTCACTGGGCCANNNAGATCGGAAG
66304 TATGAGTGTGGAGTCGTTGCGTGCATTTTTATTGGAGGCAGTGTNNNAGATCGGA
60058 TATGAGTGTGGAGTCGTTGCAGGAAAGAGAAGGAAGACCCACTNNNAGATCGGA
45357 TATGAGTGTGGAGTCGTTGCATTACTTCAGTCCCAGTGTTGCCNNNAGATCGGAAG
36293 TATGAGTGTGGAGTCGTTGCTGCTGGGAGATGTTTCATTTGCANNNAGATCGGAA
40832 TATGAGTGTGGAGTCGTTGCACCCACTGATGGTGTTCAGGATANNNAGATCGGAA
59803 TATGAGTGTGGAGTCGTTGCTTGCCCAGAAAGCATGGTAAACTNNNAGATCGGAA
48669 TATGAGTGTGGAGTCGTTGCCCTCTGGAACTCCTCAGAACCTCNNNAGATCGGAAG
41007 TATGAGTGTGGAGTCGTTGCAGTTGGAGACAAGCAGGCTACTANNNAGATCGGAA
63671 TATGAGTGTGGAGTCGTTGCAACACACAGAACGGGCATAAAGTNNNAGATCGGAA
59237 TATGAGTGTGGAGTCGTTGCAGAGAACACAAAGATCCTTGGATNNNAGATCGGAA
98250 TATGAGTGTGGAGTCGTTGCACTGATTGGTGCCAGTTTACAATCNNNAGATCGGAA
32879 TATGAGTGTGGAGTCGTTGCCACTCTCTGTTTGTCCCAGAACANNNAGATCGGAAG
62493 TATGAGTGTGGAGTCGTTGCTTTCTGCCTTAGTCCATGTTGCTNNNAGATCGGAAG
37408 TATGAGTGTGGAGTCGTTGCGCACGATGTGGTAATTAGTTTTCANNNAGATCGGAA
63869 TATGAGTGTGGAGTCGTTGCAAGTAGAGCCCAGTGGTAACAGTNNNAGATCGGAA
41874 TATGAGTGTGGAGTCGTTGCCCAGGCACAAGATTCTGCTCTTANNNAGATCGGAAG
62126 TATGAGTGTGGAGTCGTTGCTTGCTTTCTTTTCTGTCCTGGCTNNNAGATCGGAAGA
66279 TATGAGTGTGGAGTCGTTGCTTCGGATCATGTTGTGGAAGTGTNNNAGATCGGAA
46514 TATGAGTGTGGAGTCGTTGCGCACAGGGAAAATATTGGTAGAGCNNNAGATCGGA
41371 TATGAGTGTGGAGTCGTTGCCTTTCCTGTGCCGATCAACAGTANNNAGATCGGAAG
67461 TATGAGTGTGGAGTCGTTGCCCACCGGAGCTATAGCAAAACTTNNNAGATCGGAAG
40616 TATGAGTGTGGAGTCGTTGCCTGCCTGCCTGCCTGTATAAATANNNAGATCGGAAG
37600 TATGAGTGTGGAGTCGTTGCCCCTTCACTGAAAAACATAAAAGAAGANNNAGATCG
95145 TATGAGTGTGGAGTCGTTGCGTTTCCTGGCAGAAGGCATTTACNNNAGATCGGAAG
37675 TATGAGTGTGGAGTCGTTGCGGGAAGGGAGAAAGGAGTGAAGANNNAGATCGG
33466 TATGAGTGTGGAGTCGTTGCTGAACTGCAGAGTAGCATAAAGACANNNAGATCGG
33257 TATGAGTGTGGAGTCGTTGCTGGTTCTCAACAAGATGAAGCACANNNAGATCGGA
61743 TATGAGTGTGGAGTCGTTGCTGGGACTGCATTATTGATTCAGCTNNNAGATCGGAA
68237 TATGAGTGTGGAGTCGTTGCAGGCAGAGACAGTCGAGTAAGTTNNNAGATCGGAA
62202 TATGAGTGTGGAGTCGTTGCTGTATGGATTTGAATCCCAATGCTNNNAGATCGGAA
46627 TATGAGTGTGGAGTCGTTGCAGGCTTTCAGGCCGAATAATAGCNNNAGATCGGAA
55604 TATGAGTGTGGAGTCGTTGCTAGTGATTAGGGCCAGGAGGATGNNNAGATCGGA
40839 TATGAGTGTGGAGTCGTTGCGATGTGGTAGGGGTGAAGGGATANNNAGATCGGA
43209 TATGAGTGTGGAGTCGTTGCAGGGGTAAGGAGACAAAGGAGACNNNAGATCGGA
60673 TATGAGTGTGGAGTCGTTGCGAAGGCTTGTGTGTATACCACCTNNNAGATCGGAAG
58716 TATGAGTGTGGAGTCGTTGCCACGCACTTACGTACATATGCATNNNAGATCGGAAG
55173 TATGAGTGTGGAGTCGTTGCAGCCAAACACTCTGGTACTTTGGNNNAGATCGGAAG
32297 TATGAGTGTGGAGTCGTTGCGGTCCAGAAAAGGCCACTGATAANNNAGATCGGAA
63014 TATGAGTGTGGAGTCGTTGCTGTTTTTCTCTCCCACCCTCTCTNNNAGATCGGAAGA
71684 TATGAGTGTGGAGTCGTTGCCCAATGAAGCCATTACCTCCCAANNNAGATCGGAAG
60255 TATGAGTGTGGAGTCGTTGCAGTACAGCCATAAGGTGAAAGACTNNNAGATCGGA
32776 TATGAGTGTGGAGTCGTTGCGCAACTCTTTAGACAGAAAGGTAAACANNNAGATCG

64155 TATGAGTGTGGAGTCGTTGCTGCTCTTTCTGTCTGTCTTTCAGTNNNAGATCGGAAG
54539 TATGAGTGTGGAGTCGTTGCTGCTACTTTTGCATTGATTTCACTGGNNNAGATCGGA
31303 TATGAGTGTGGAGTCGTTGCTATCTGTAGGGCAGGCTATGCAANNNAGATCGGAA
111450 TATGAGTGTGGAGTCGTTGCTACACAGCAAACATTCTAGCCGNNNAGATCGGAAGA
31310 TATGAGTGTGGAGTCGTTGCTGGCTACTCTGATAACACTGCAANNNAGATCGGAAG
59641 TATGAGTGTGGAGTCGTTGCCCCCACACACCATACAGGTGTATNNNAGATCGGAAG
43679 TATGAGTGTGGAGTCGTTGCTCCCTATGGCTAGAAGGGCTTACNNNAGATCGGAAG
71296 TATGAGTGTGGAGTCGTTGCACACACACACAAAGCCTAACAGTNNNAGATCGGAAG
53551 TATGAGTGTGGAGTCGTTGCATCCATGTTGTAGCCAATCTCGGNNNAGATCGGAAG
49468 TATGAGTGTGGAGTCGTTGCTATTTGGGGGATGTGTGTGTGTCNNNAGATCGGAA
53536 TATGAGTGTGGAGTCGTTGCGAGAGAGAGAGTAGTGTTTGCGGNNNAGATCGGA
53490 TATGAGTGTGGAGTCGTTGCTACACAGCAAACATTCTAGCCGGNNNAGATCGGAAG
98830 TATGAGTGTGGAGTCGTTGCTTTAGGGGTCCCAGACCTAGGNNNAGATCGGAAGA
58030 TATGAGTGTGGAGTCGTTGCCAGGACCCACAACTTGGAAGAATNNNAGATCGGAA
55991 TATGAGTGTGGAGTCGTTGCATGATGCTGTAAGTGCTTCCCTGNNNAGATCGGAAG
50975 TATGAGTGTGGAGTCGTTGCTTCCCTTGGACTAAGAGGTCCAGNNNAGATCGGAAG
47783 TATGAGTGTGGAGTCGTTGCCTCTGCTTATTGCTGATTGCTGCNNNAGATCGGAAG
54934 TATGAGTGTGGAGTCGTTGCAAATAACATCGATGGGGTGGTGGNNNAGATCGGA
62048 TATGAGTGTGGAGTCGTTGCACACAGGAAAAATGACATGGGCTNNNAGATCGGAA
109256 TATGAGTGTGGAGTCGTTGCGTCTCCCATACTTTCCCTATCTCTCTNNNAGATCGGAA
48085 TATGAGTGTGGAGTCGTTGCCCTTCTCTCTCATTCTCTCACTTGCNNNAGATCGGAAG
63259 TATGAGTGTGGAGTCGTTGCCACTGTATTTCTAGTTAGCTCTGTCTNNNAGATCGGA
98997 TATGAGTGTGGAGTCGTTGCGACCTCCAAACTAAAGGAACAGGAATNNNAGATCG
95121 TATGAGTGTGGAGTCGTTGCAACAAAGGTAGGTGCCAGAATGGNNNAGATCGGA
48127 TATGAGTGTGGAGTCGTTGCACTGCATGACAACAATGTATCTTGCNNNAGATCGGA
44837 TATGAGTGTGGAGTCGTTGCTACATGAGCAACACCAGTTTCCCNNNAGATCGGAAG
50181 TATGAGTGTGGAGTCGTTGCTTTGCTGAAGGCTTTGGTGAAAGNNNAGATCGGAA
44728 TATGAGTGTGGAGTCGTTGCCCTTGAATCCTTTGCTCTCTCCCNNNAGATCGGAAGA
45024 TATGAGTGTGGAGTCGTTGCCAAGTAAATACGGTCCTCTCGCCNNNAGATCGGAAG
52190 TATGAGTGTGGAGTCGTTGCTGCTCTACGCCATTCGTCATTAGNNNAGATCGGAAG
67863 TATGAGTGTGGAGTCGTTGCACAATCTCCAAATCGCAGAGCTTNNNAGATCGGAAG
41840 TATGAGTGTGGAGTCGTTGCGCCAGGAGGGAGTAATCAGCTTANNNAGATCGGAA
49972 TATGAGTGTGGAGTCGTTGCAGCCAAAACAGAGCAGCATTTTCNNNAGATCGGAAG
52917 TATGAGTGTGGAGTCGTTGCAGAAGTAGAGAGGCTCTGACAGGNNNAGATCGGA
64105 TATGAGTGTGGAGTCGTTGCACAGAGGGGAAAGAGATACTCAGTNNNAGATCGG
65331 TATGAGTGTGGAGTCGTTGCCTGAGGCCAAAAGGTGTTATGGTNNNAGATCGGAA
56560 TATGAGTGTGGAGTCGTTGCTCAGTCAACTTCTCTGCTTACAGTGNNNAGATCGGA
64012 TATGAGTGTGGAGTCGTTGCGCAAGGAAAGATCCATCAGCAGTNNNAGATCGGAA
62132 TATGAGTGTGGAGTCGTTGCTGATCTCCTGACCTAATCTGGCTNNNAGATCGGAAG
49515 TATGAGTGTGGAGTCGTTGCCTGACACACACAACAGCACATTCNNNAGATCGGAAG
102089 TATGAGTGTGGAGTCGTTGCAGCCATAAGGTGAAAGACTACTTTTNNNAGATCGGA
48410 TATGAGTGTGGAGTCGTTGCAGTAGCAGCAGCAGCATATCATCNNNAGATCGGAA
105218 TATGAGTGTGGAGTCGTTGCCTCACCTTTCCATGCAGGCACNNNAGATCGGAAGAG

108003 TATGAGTGTGGAGTCGTTGCATACACAAATATACACGCACATGCNNNAGATCGGAA
67626 TATGAGTGTGGAGTCGTTGCAAATGATTGTGCTACTGGTTTACTTNNNAGATCGGA
39961 TATGAGTGTGGAGTCGTTGCCCCTATACACACTCCTGAGCTGANNNAGATCGGAAG
58520 TATGAGTGTGGAGTCGTTGCCGTGTATGGGTGAAGTTAGCCATNNNAGATCGGAA
101152 TATGAGTGTGGAGTCGTTGCGACATTTTAGCTGTACCTGATTTTATCNNNAGATCGG
102065 TATGAGTGTGGAGTCGTTGCTTTTGTGTTGGGGGAGAATCAGAGNNNAGATCGGA
41668 TATGAGTGTGGAGTCGTTGCTGCTAGGTCATGCTAAGGGTGTANNNAGATCGGAA
59871 TATGAGTGTGGAGTCGTTGCACACACTATCCAACCTCAAGAACTNNNAGATCGGAAG
116815 TATGAGTGTGGAGTCGTTGCTTTCCTCTTTCTTTGTCTTGTTCAGNNNAGATCGGAA
96107 TATGAGTGTGGAGTCGTTGCTTGTAGGTTTCGTGGTCGGGNNNAGATCGGAAGAG
49593 TATGAGTGTGGAGTCGTTGCGGAATGTGCTTGGGAGGGTATTCNNNAGATCGGAA
40530 TATGAGTGTGGAGTCGTTGCTTGCTCTATTGTGCTGCCTTTGANNNAGATCGGAAG
51248 TATGAGTGTGGAGTCGTTGCCATCCCTTCATTTCGGTGCTCAGNNNAGATCGGAAG
69010 TATGAGTGTGGAGTCGTTGCTCAACTCAAGATGGATAAAATACCTTTNNNAGATCG
47323 TATGAGTGTGGAGTCGTTGCTGTCTAGGATATAGGGCAGTGGCNNNAGATCGGAA
66780 TATGAGTGTGGAGTCGTTGCATGTTTATGACAAGAGCCTATTTGTNNNAGATCGGA
44564 TATGAGTGTGGAGTCGTTGCAAAGTGGTGCCTAATAATGGCCCNNNAGATCGGAA
59475 TATGAGTGTGGAGTCGTTGCCACACACACCCCAACACGTATATNNNAGATCGGAAG
60224 TATGAGTGTGGAGTCGTTGCAGCCAGGGACTTATTCTGTCACTNNNAGATCGGAAG
33062 TATGAGTGTGGAGTCGTTGCACACACATACATAAACATCTACACACANNNAGATCGG
117195 TATGAGTGTGGAGTCGTTGCTCTAGGATATAGGGCAGTGGCTNNNAGATCGGAAG
98881 TATGAGTGTGGAGTCGTTGCATACCTTACAGACCGGTAGAGAGNNNAGATCGGAA
101284 TATGAGTGTGGAGTCGTTGCCCCACTGCAAGCTTTGTCTCTCNNNAGATCGGAAGA
38405 TATGAGTGTGGAGTCGTTGCTTCCCATGAAAGCTTCCCCTAGANNNAGATCGGAAG
108831 TATGAGTGTGGAGTCGTTGCGAAAGTAACAGATGCTGTGGCAAANNNAGATCGGA
65850 TATGAGTGTGGAGTCGTTGCTGTGTGTGTATGGTATGGTATGTNNNAGATCGGAA
116279 TATGAGTGTGGAGTCGTTGCCGTTATTTGTCTCCTCTGCAAGGNNNAGATCGGAAG
98236 TATGAGTGTGGAGTCGTTGCCCCACAACGGGAGTGGCATCNNNAGATCGGAAGAG
62928 TATGAGTGTGGAGTCGTTGCAGGCTGACTAGAATGTGAGCTCTNNNAGATCGGAA
62224 TATGAGTGTGGAGTCGTTGCTCCTTTACATGCTCATGCATGCTNNNAGATCGGAAGA
61407 TATGAGTGTGGAGTCGTTGCTTTTAGCTGTACCTGATTTTATCTCCTNNNAGATCGG
33383 TATGAGTGTGGAGTCGTTGCCCCCAAAGGTGTACATTCTCACANNNAGATCGGAAG
45834 TATGAGTGTGGAGTCGTTGCAATCCCATTGCCTGGAGAATTCCNNNAGATCGGAAG
68724 TATGAGTGTGGAGTCGTTGCAGGCATCACCCTGGACTATTGTTNNNAGATCGGAAG
109310 TATGAGTGTGGAGTCGTTGCTCTGGCTCACAGCTATGGACTAANNNAGATCGGAAG
41561 TATGAGTGTGGAGTCGTTGCCTCCCTCTCTCTCGGAAATGGTANNNAGATCGGAAG
104682 TATGAGTGTGGAGTCGTTGCTCAAAGATTCCAGAAGAGGAAGAGGNNNAGATCGG
61975 TATGAGTGTGGAGTCGTTGCCCAGTCTAAAACAGATGCAGGCTNNNAGATCGGAAG
60911 TATGAGTGTGGAGTCGTTGCAAAACAAGAGTAGCTCCTCCCCTNNNAGATCGGAAG
117843 TATGAGTGTGGAGTCGTTGCCGATCTCACCTTGTGGCAATCTCNNNAGATCGGAAG
62789 TATGAGTGTGGAGTCGTTGCAGAAAGAGATCCCTGTGCACTCTNNNAGATCGGAAG
34207 TATGAGTGTGGAGTCGTTGCTCCTTCTGGAATAAAACATGTTACCANNNAGATCGG
35360 TATGAGTGTGGAGTCGTTGCAAAGGAATGGACAACCAAGAGCANNNAGATCGGAA

53191 TATGAGTGTGGAGTCGTTGCTGGGGTTAGGGAACATAAGGAGGNNNAGATCGGA
106842 TATGAGTGTGGAGTCGTTGCAGTTTGAGACCCAAGGTACTGGGNNNAGATCGGAA
54089 TATGAGTGTGGAGTCGTTGCGGGGTAGAGGAAGAGTTACTGGGNNNAGATCGGA
36108 TATGAGTGTGGAGTCGTTGCACTGTAGACGTAGTCACTCTGCANNNAGATCGGAAG
96292 TATGAGTGTGGAGTCGTTGCGCCTAATCGTCCCCACAAAATCTTANNNAGATCGGAA
95913 TATGAGTGTGGAGTCGTTGCATGTTCTTCATCCCCTTCTCCCTGNNNAGATCGGAAG
52874 TATGAGTGTGGAGTCGTTGCAAACTTCCAGCCCTACAAACAGGNNNAGATCGGAAG
103546 TATGAGTGTGGAGTCGTTGCGTTTTCACTGGGCCAACATATTACNNNAGATCGGAA
104329 TATGAGTGTGGAGTCGTTGCCTTTGTCATTTTTAGGTGTTGCAANNNAGATCGGAA
30005 TATGAGTGTGGAGTCGTTGCATCTCCTCAGTGCCTTGCAAAAANNNAGATCGGAAG
58176 TATGAGTGTGGAGTCGTTGCCCAAAAACCAGGTGAGGCCTAATNNNAGATCGGAA
40317 TATGAGTGTGGAGTCGTTGCCTGTTACCCAGACAATGGTGTGANNNAGATCGGAA
97308 TATGAGTGTGGAGTCGTTGCGGTTGTTTACCCAGCAGGTGGNNNAGATCGGAAGA
95138 TATGAGTGTGGAGTCGTTGCAAATGCTACACATCCTCCTAGCTNNNAGATCGGAAG
37569 TATGAGTGTGGAGTCGTTGCGCACACCTGACTATGAAGCAAGANNNAGATCGGAA
39753 TATGAGTGTGGAGTCGTTGCACACGGCCTCTTTTCATGTATGANNNAGATCGGAAG
29873 TATGAGTGTGGAGTCGTTGCGCTTTGCCATCCTCTGACTAGACNNNAGATCGGAAG
42022 TATGAGTGTGGAGTCGTTGCTCCCAACAAAACCTTGGCCTTTANNNAGATCGGAAG
47162 TATGAGTGTGGAGTCGTTGCTTTTCTTAGAGAGGTTGCTGGGCNNNAGATCGGAA
49263 TATGAGTGTGGAGTCGTTGCCCCTGGTTCTTCTGACAGTAGTCNNNAGATCGGAAG
41924 TATGAGTGTGGAGTCGTTGCAGCGTTTAGTCACCTCCAGGTTANNNAGATCGGAAG
48548 TATGAGTGTGGAGTCGTTGCGCATGTTGCTGGTAAAGCGTATCNNNAGATCGGAA
98249 TATGAGTGTGGAGTCGTTGCTCACTCCTTTACCCACTGCATTCNNNAGATCGGAAGA
49306 TATGAGTGTGGAGTCGTTGCCTTGCTGGTTCTCTTCAGAGGTCNNNAGATCGGAAG
102624 TATGAGTGTGGAGTCGTTGCCATCAGTAGAAATGGCAACTGCCTNNNAGATCGGAA
31974 TATGAGTGTGGAGTCGTTGCTTCATTGAGACAACTGCCTGGAANNNAGATCGGAAG
53736 TATGAGTGTGGAGTCGTTGCATTGGATGAAAGCTCGTTCAGGGNNNAGATCGGAA
35903 TATGAGTGTGGAGTCGTTGCAGAAAGCCTGATGTGTCAATGCANNNAGATCGGAA
55588 TATGAGTGTGGAGTCGTTGCTGGGTGTCTTCAGCAACTAGATGNNNAGATCGGAA
62152 TATGAGTGTGGAGTCGTTGCTGTGTATAGCCTGTATGGTGGCTNNNAGATCGGAA
34267 TATGAGTGTGGAGTCGTTGCTTTTGGCTGATAGTACTGACCCANNNAGATCGGAAG
62385 TATGAGTGTGGAGTCGTTGCATCTTGACCCAAACTTTCGTGCTNNNAGATCGGAAG
67807 TATGAGTGTGGAGTCGTTGCTTTCCACTTGGCCAACATTCCTTNNNAGATCGGAAGA
46680 TATGAGTGTGGAGTCGTTGCGGAATATGCTCAGCAGAGGTAGCNNNAGATCGGAA
30772 TATGAGTGTGGAGTCGTTGCGCTGTGGAAGTCTTGAGGGTAAANNNAGATCGGAA
68454 TATGAGTGTGGAGTCGTTGCTTGAATACCATCCTGGAGGGGTTNNNAGATCGGAA
49369 TATGAGTGTGGAGTCGTTGCGGAGAATCAGAGAGGCACATGTCNNNAGATCGGAA
56804 TATGAGTGTGGAGTCGTTGCGGCATATACATACAGGGTGGGTGNNNAGATCGGAA
51029 TATGAGTGTGGAGTCGTTGCGTTGTGAAGTAGGACCTCAGCAGNNNAGATCGGAA
102077 TATGAGTGTGGAGTCGTTGCGCTTGGGAGGGTATTCAAATATTCAGTNNNAGATCG
102973 TATGAGTGTGGAGTCGTTGCTGGGAGTAATGACTGAGGAAAGGANNNAGATCGG
69200 TATGAGTGTGGAGTCGTTGCGAACCTCTACGCAACCTGTCTTTNNNAGATCGGAAG
51536 TATGAGTGTGGAGTCGTTGCCGTGTGTGTGAGGGTAAGTAGAGNNNAGATCGGA

| | |
|---|---|
| 70913 | TATGAGTGTGGAGTCGTTGCGCAAGTTCAAATGTAGCCTGGTGNNNAGATCGGAA |
| 41765 | TATGAGTGTGGAGTCGTTGCATGCAGGCACAGCTTTCAGATTANNNAGATCGGAAG |
| 109640 | TATGAGTGTGGAGTCGTTGCAGTAAAATGTCAAAGGCGACTGACNNNAGATCGGA |
| 98301 | TATGAGTGTGGAGTCGTTGCTGGTGGGTGAGAGTGGTTAGAATNNNAGATCGGA |
| 50790 | TATGAGTGTGGAGTCGTTGCCGATGAGGGATCTTTGGAACCAGNNNAGATCGGAA |
| 56039 | TATGAGTGTGGAGTCGTTGCAAGTGTTGGCTCACATGATCCTGNNNAGATCGGAAG |
| 34677 | TATGAGTGTGGAGTCGTTGCTGAAACTACCTACCATCTGGCCANNNAGATCGGAAG |
| 116811 | TATGAGTGTGGAGTCGTTGCTGAGAGTGTGTGTGTGTTTGTAGTNNNAGATCGGA |
| 31115 | TATGAGTGTGGAGTCGTTGCCCCCCTGGTTCAAATTCATCCAANNNAGATCGGAAGA |
| 41562 | TATGAGTGTGGAGTCGTTGCATGGTGAGATGGTGGGAATGGTANNNAGATCGGA |
| 69103 | TATGAGTGTGGAGTCGTTGCATTTTCCCTGTTGCTGAGGCTTTNNNAGATCGGAAG |
| 106879 | TATGAGTGTGGAGTCGTTGCAGATAGCCTCTTAGGATACAAAGGGACNNNAGATC |
| 55325 | TATGAGTGTGGAGTCGTTGCTCCCCTGAGTTCATAAGCTGAATGNNNAGATCGGAA |
| 45103 | TATGAGTGTGGAGTCGTTGCTCTTGATAACTGCCCAAATGGCCNNNAGATCGGAAG |
| 54477 | TATGAGTGTGGAGTCGTTGCGCCCTCTGGTTTTACTGCTATGGNNNAGATCGGAAG |
| 34121 | TATGAGTGTGGAGTCGTTGCAGAGAGAGAGTGAATAAAAGGGACCANNNAGATC |
| 65508 | TATGAGTGTGGAGTCGTTGCAATGACATCCCCTGAACATTGGTNNNAGATCGGAAG |
| 44735 | TATGAGTGTGGAGTCGTTGCTTCTTCCCATCTTCTGGTCTCCCNNNAGATCGGAAGA |
| 38790 | TATGAGTGTGGAGTCGTTGCGTCATGCTTTGGAGTGAAGAGGANNNAGATCGGAA |
| 40219 | TATGAGTGTGGAGTCGTTGCAGTGAGTGCATGTAACAATGGTGANNNAGATCGGA |
| 106942 | TATGAGTGTGGAGTCGTTGCGGCTTTCTAACACTTGGTAGGATATNNNAGATCGGA |
| 46477 | TATGAGTGTGGAGTCGTTGCAGGAATCTGTTTGTGAGCAGAGCNNNAGATCGGAA |
| 49905 | TATGAGTGTGGAGTCGTTGCACTGTCATGGGTATGTGCCTTTCNNNAGATCGGAAG |
| 61524 | TATGAGTGTGGAGTCGTTGCTAGTGAACAAAGCAAGCCTTCCTNNNAGATCGGAAG |
| 43904 | TATGAGTGTGGAGTCGTTGCTTTAGAGAAGTCTTTGCCCCACCNNNAGATCGGAAG |
| 104333 | TATGAGTGTGGAGTCGTTGCTCATCTTGTTTGCCTATAGAGAGGGNNNAGATCGGA |
| 59473 | TATGAGTGTGGAGTCGTTGCATGAGTGCCCAGGAAGGCTATATNNNAGATCGGAA |
| 31247 | TATGAGTGTGGAGTCGTTGCCCTGCTGCTTCCAATATAGGCAANNNAGATCGGAAG |
| 34038 | TATGAGTGTGGAGTCGTTGCCTCGACTACCTGTTAGGACACCANNNAGATCGGAAG |
| 54852 | TATGAGTGTGGAGTCGTTGCAGCACAGTTATGAAGCAGAGTGGNNNAGATCGGAA |
| 45489 | TATGAGTGTGGAGTCGTTGCTCAGAAGAGAGCATGCACTATCCNNNAGATCGGAA |
| 51097 | TATGAGTGTGGAGTCGTTGCCAGCTTCAAAATTCACAGGGCAGNNNAGATCGGAA |
| 30423 | TATGAGTGTGGAGTCGTTGCGAGGAGGTATCAGCAGCATCAAANNNAGATCGGAA |
| 44872 | TATGAGTGTGGAGTCGTTGCACTAGAGCTAAGGACAGCAAGCCNNNAGATCGGAA |
| 59600 | TATGAGTGTGGAGTCGTTGCTACTCCCCCAACAGAACACGTATNNNAGATCGGAAG |
| 56456 | TATGAGTGTGGAGTCGTTGCAACAGAGGGTCAAAGCACTTCTGNNNAGATCGGAA |
| 60151 | TATGAGTGTGGAGTCGTTGCACACTTGGGTATCTTTCATGCACTNNNAGATCGGAA |
| 60990 | TATGAGTGTGGAGTCGTTGCAAGATAACCCACTACAGATCCCTNNNAGATCGGAAG |
| 42576 | TATGAGTGTGGAGTCGTTGCATTCTGGTAGCCATTAGCCACACNNNAGATCGGAAG |
| 48342 | TATGAGTGTGGAGTCGTTGCAGAGGCCTGATCCAGGATACATCNNNAGATCGGAA |
| 30284 | TATGAGTGTGGAGTCGTTGCTGAAGAGATGGCCAACAACCAAANNNAGATCGGAA |
| 61825 | TATGAGTGTGGAGTCGTTGCTCACTCTCTCTCTTATCCATAGCTNNNAGATCGGAAG |
| 64551 | TATGAGTGTGGAGTCGTTGCTGGGACATAATTGAAGTGCACGTNNNAGATCGGAA |

54947 TATGAGTGTGGAGTCGTTGCAGCACAATCATAGCTCCATGTGGNNNAGATCGGAAG
39439 TATGAGTGTGGAGTCGTTGCATGGGAAATGAGAAGCATGTGGANNNAGATCGGA
44571 TATGAGTGTGGAGTCGTTGCGTTCTGTGTTGACAATTTGGCCCNNNAGATCGGAAG
39345 TATGAGTGTGGAGTCGTTGCAATGACACATGAGAGGTCCTGGANNNAGATCGGAA
59174 TATGAGTGTGGAGTCGTTGCTACATCTCCCTCCCATCAGGGATNNNAGATCGGAAG
30729 TATGAGTGTGGAGTCGTTGCGTTTCAAAGGGCAACGCACTAAANNNAGATCGGAA
44568 TATGAGTGTGGAGTCGTTGCTACTACGATTTCCTAGGTGGCCCNNNAGATCGGAAG
38432 TATGAGTGTGGAGTCGTTGCGCCTTCAGCCTGTGTTTTCTAGANNNAGATCGGAAG
66433 TATGAGTGTGGAGTCGTTGCTTAGTGTGCCTGTGTGTATGTGTNNNAGATCGGAA
33114 TATGAGTGTGGAGTCGTTGCTTCCCCAATTGCCAACATACACANNNAGATCGGAAGA
55295 TATGAGTGTGGAGTCGTTGCACACTAGCACTAACGGCAGAATGNNNAGATCGGAA
66526 TATGAGTGTGGAGTCGTTGCTCAGCAAACTTTAGTGCTTGTGTNNNAGATCGGAAG
115880 TATGAGTGTGGAGTCGTTGCACAACTTCTGGCTTCCTTTCCTCCNNNAGATCGGAAG
53732 TATGAGTGTGGAGTCGTTGCCACTTCTTCCTTCTGTGTCAGGGNNNAGATCGGAAG
67955 TATGAGTGTGGAGTCGTTGCCTACGTGAAGATTCTGCCTGCTTNNNAGATCGGAAG
29866 TATGAGTGTGGAGTCGTTGCTGAAAAAGCCTTCTTGGAGGAGGNNNAGATCGGAA
38378 TATGAGTGTGGAGTCGTTGCTCACGCAGAATGTTGCACATAGANNNAGATCGGAA
48301 TATGAGTGTGGAGTCGTTGCAGCAACAGGTGGTGAGTTGAATCNNNAGATCGGAA
57187 TATGAGTGTGGAGTCGTTGCAGAAGCGCTATGAAGGGTCATTGNNNAGATCGGAA
45516 TATGAGTGTGGAGTCGTTGCCTGCATTCACTCTGGAGAACTCCNNNAGATCGGAAG
102400 TATGAGTGTGGAGTCGTTGCAATGACAAGGCAAAATGGCAAGATNNNAGATCGGA
102383 TATGAGTGTGGAGTCGTTGCCCAGTTTTCCAGCTTTTCTGGCTNNNAGATCGGAAGA
57906 TATGAGTGTGGAGTCGTTGCAGGTGTTGCAAAGTTCTCCCAATNNNAGATCGGAAG
64445 TATGAGTGTGGAGTCGTTGCAAGACCACTGCTACACTGCTAGTNNNAGATCGGAAG
53815 TATGAGTGTGGAGTCGTTGCCCTTGTAACAGAGCAACGTAGGGNNNAGATCGGAA
71254 TATGAGTGTGGAGTCGTTGCACATGCTCTGCTAACTTTTTCTCTNNNAGATCGGAAG
35018 TATGAGTGTGGAGTCGTTGCGTTTCTGCCCTAACACCATTCCANNNAGATCGGAAGA
43439 TATGAGTGTGGAGTCGTTGCGAGGCGCAGAGAAATTTTGTGACNNNAGATCGGAA
109327 TATGAGTGTGGAGTCGTTGCAGACCCCTTCAATCCACAAACTTNNNAGATCGGAAGA
42981 TATGAGTGTGGAGTCGTTGCCCTTCAGTCTTCCGGGTTATCACNNNAGATCGGAAG
64668 TATGAGTGTGGAGTCGTTGCTGTTCCAAGACCCCTGATATCGTNNNAGATCGGAAG
55716 TATGAGTGTGGAGTCGTTGCTGCTCTGGACACTGTTTGGTATGNNNAGATCGGAAG
61400 TATGAGTGTGGAGTCGTTGCCATTGTGTATTCGCTTTGCTCCTNNNAGATCGGAAGA
33292 TATGAGTGTGGAGTCGTTGCAGATGTCAATATGCTAATTAGGCACANNNAGATCGG
55845 TATGAGTGTGGAGTCGTTGCGCCAAGACTGTTTTGTGAAGACTGNNNAGATCGGA
71297 TATGAGTGTGGAGTCGTTGCCGGGATGCTAATCTATGCACAGTNNNAGATCGGAA
43672 TATGAGTGTGGAGTCGTTGCATGTACGCAGACACACACCTTACNNNAGATCGGAAG
37650 TATGAGTGTGGAGTCGTTGCTGAAGAATAACATCAAAGCTGGAAGANNNAGATCG
32651 TATGAGTGTGGAGTCGTTGCAGGTAGCATATCCGTGGAAAAACANNNAGATCGGA
100901 TATGAGTGTGGAGTCGTTGCCCTTGTGTGTTCAGGTAGTACTCTACNNNAGATCGG
41539 TATGAGTGTGGAGTCGTTGCCATAGTTGAGTAGGGCAGGGGTANNNAGATCGGA
36416 TATGAGTGTGGAGTCGTTGCGCTGTATTGTACCAAACCCATCANNNAGATCGGAAG
61257 TATGAGTGTGGAGTCGTTGCTGGCTCTTTCAAAACTTCTGCCTNNNAGATCGGAAGA

| | |
|---|---|
| 62448 | TATGAGTGTGGAGTCGTTGCTGAAGCATTTGACCCTTATTGCTNNNAGATCGGAAG |
| 44968 | TATGAGTGTGGAGTCGTTGCATAAAGTATGCCTGCTCTGAGCCNNNAGATCGGAAG |
| 104558 | TATGAGTGTGGAGTCGTTGCAATATCTGTGAAGCCTAAAAAGCAATANNNAGATCG |
| 31287 | TATGAGTGTGGAGTCGTTGCGTGTGCTCTTGGGTCAATGCAANNNAGATCGGAAG |
| 108124 | TATGAGTGTGGAGTCGTTGCTGTGTTGGGATGAACAGGTCTGANNNAGATCGGAA |
| 46938 | TATGAGTGTGGAGTCGTTGCTATGTAATGCACCAAGGTCAGGCNNNAGATCGGAA |
| 29869 | TATGAGTGTGGAGTCGTTGCAAAACCCAAGAAACACAGACCCANNNAGATCGGAAG |
| 34980 | TATGAGTGTGGAGTCGTTGCGAATGCCACTTTTTCCATGTCCANNNAGATCGGAAG |
| 102655 | TATGAGTGTGGAGTCGTTGCAAAGGACTAGACTTGAGACCAGGCNNNAGATCGGA |
| 30938 | TATGAGTGTGGAGTCGTTGCGTGCCATACACACACATTGACAANNNAGATCGGAAG |
| 62557 | TATGAGTGTGGAGTCGTTGCAGGCTGAGTGAAAATAGCCAATCTNNNAGATCGGA |
| 53644 | TATGAGTGTGGAGTCGTTGCCAAAGAGAAGAGCAGGCTAAGGGNNNAGATCGGA |
| 35456 | TATGAGTGTGGAGTCGTTGCGCAAAGGAAAGAAAGACCATAGCANNNAGATCGGA |
| 116965 | TATGAGTGTGGAGTCGTTGCAGTCCTGTAAACATTGTCACTGAGNNNAGATCGGAA |
| 36271 | TATGAGTGTGGAGTCGTTGCTCAGACAGAAAGACTAGGTTGCANNNAGATCGGAA |
| 31697 | TATGAGTGTGGAGTCGTTGCTGGAAAGTGGGGAAGAAGGAGAANNNAGATCGGA |
| 104548 | TATGAGTGTGGAGTCGTTGCCTAATCGGGCAGTCTTCTGTGTTTTNNNAGATCGGA |
| 40875 | TATGAGTGTGGAGTCGTTGCACCTGACTGCTCCTCTTGTGATANNNAGATCGGAAG |
| 46847 | TATGAGTGTGGAGTCGTTGCAATACCTCTTCTAGGGAGGAAGGCNNNAGATCGGA |
| 33136 | TATGAGTGTGGAGTCGTTGCGACCCATTAGGGCTTTCTACACANNNAGATCGGAAG |
| 37519 | TATGAGTGTGGAGTCGTTGCGTGATGGGCATGGCTTCTAAAGANNNAGATCGGAA |
| 67139 | TATGAGTGTGGAGTCGTTGCTTTTACCAGAAGGTGGCAGCATTNNNAGATCGGAA |
| 44388 | TATGAGTGTGGAGTCGTTGCTTTGTTTTCTTCCCTCATCCCCCNNNAGATCGGAAGA |
| 49555 | TATGAGTGTGGAGTCGTTGCTTTGGAACCGTGTCCTCAGATTCNNNAGATCGGAAG |
| 99789 | TATGAGTGTGGAGTCGTTGCGTTCTCAAGTCCTGCCTGAAAGAGNNNAGATCGGAA |
| 55014 | TATGAGTGTGGAGTCGTTGCTCCCTCTGGTTCATTGTTTGTGGNNNAGATCGGAAG |
| 56639 | TATGAGTGTGGAGTCGTTGCTCTCTCAATAGTGGTACCATGAGTGNNNAGATCGGA |
| 58718 | TATGAGTGTGGAGTCGTTGCGCCTATAGAGAGGGGAACTGCATNNNAGATCGGAA |
| 36267 | TATGAGTGTGGAGTCGTTGCTGTGGAGGTTTTTCTGAGTTGCANNNAGATCGGAA |
| 32510 | TATGAGTGTGGAGTCGTTGCTCACCCCAATACCCCAACTGTAANNNAGATCGGAAGA |
| 63115 | TATGAGTGTGGAGTCGTTGCAGAGGCTGTGTATGGTCAAGTCTNNNAGATCGGAA |
| 62617 | TATGAGTGTGGAGTCGTTGCTGTCTTGACTAGCTTTCCCATCTNNNAGATCGGAAGA |
| 40736 | TATGAGTGTGGAGTCGTTGCCTGCATGGACTTGAGTGAGCATANNNAGATCGGAA |
| 110817 | TATGAGTGTGGAGTCGTTGCAATGAAGAACATCCATCCATCCATCCNNNAGATCGG |
| 56286 | TATGAGTGTGGAGTCGTTGCCTACATTGCCTTGCATCTTGCTGNNNAGATCGGAAG |
| 116375 | TATGAGTGTGGAGTCGTTGCCTCCCATAGTTGACAGAATGCCAGANNNAGATCGGA |
| 64463 | TATGAGTGTGGAGTCGTTGCATGTGGTCTTGGACTGGAGTAGTNNNAGATCGGAA |
| 68871 | TATGAGTGTGGAGTCGTTGCAGGGAAAGGAGGGGAGTTCATTTNNNAGATCGGA |
| 54850 | TATGAGTGTGGAGTCGTTGCCAGAAACTGGGTACACAGAGTGGNNNAGATCGGAA |
| 37715 | TATGAGTGTGGAGTCGTTGCTCCTCAAAGGGGAATTACTGTAAGANNNAGATCGG |
| 57501 | TATGAGTGTGGAGTCGTTGCCCCTCCAGATTGTAGCAGTGTTGNNNAGATCGGAAG |
| 41452 | TATGAGTGTGGAGTCGTTGCTCTGGCTGGCATATTTGAACGTANNNAGATCGGAAG |
| 109690 | TATGAGTGTGGAGTCGTTGCATGAACCAGCAGTTTCCAATTTNNNAGATCGGAAGA |

| | |
|---|---|
| 61305 | TATGAGTGTGGAGTCGTTGCTTTTTGTCCAAGTCTAGAGAATCCTNNNAGATCGGA |
| 46399 | TATGAGTGTGGAGTCGTTGCTGAACTTCTGTGTGTCCATCAGCNNNAGATCGGAAG |
| 37898 | TATGAGTGTGGAGTCGTTGCAGGAAGTACATGAAGGTCTCCAGANNNAGATCGGA |
| 30172 | TATGAGTGTGGAGTCGTTGCTTCCTGCCATACCAAAGTGAAAANNNAGATCGGAAG |
| 48944 | TATGAGTGTGGAGTCGTTGCATAGTCAGGGGTTCCTAGTGCTCNNNAGATCGGAA |
| 71280 | TATGAGTGTGGAGTCGTTGCTGCTCTCTCTTCTGTAGCAAAGTNNNAGATCGGAAG |
| 117161 | TATGAGTGTGGAGTCGTTGCCAAATGGTTGAATGAATGGTGAAAAAGNNNAGATC |
| 99940 | TATGAGTGTGGAGTCGTTGCAGAGTTCCTGCTGCCTCAGACNNNAGATCGGAAGAG |
| 30557 | TATGAGTGTGGAGTCGTTGCCCCAGGACCTGTAAGTAACGAAANNNAGATCGGAA |
| 56305 | TATGAGTGTGGAGTCGTTGCACTTGGAGTGCCTCTTGAATCTGNNNAGATCGGAAG |
| 45044 | TATGAGTGTGGAGTCGTTGCAGTAACATGTATGTCCCCAGGCCNNNAGATCGGAAG |
| 62124 | TATGAGTGTGGAGTCGTTGCCTAATGTGTGTATGTGCCTGGCTNNNAGATCGGAAG |
| 70755 | TATGAGTGTGGAGTCGTTGCCAGAGGACCAAAGAGGAAAAGGGNNNAGATCGGA |
| 60029 | TATGAGTGTGGAGTCGTTGCTAACACCATCAGCTGGGTACACTNNNAGATCGGAAG |
| 33248 | TATGAGTGTGGAGTCGTTGCAGGGTCTAGATGGTATTTTCCACANNNAGATCGGAA |
| 30538 | TATGAGTGTGGAGTCGTTGCCTTACCTGCTCTTGCCTGAGAAANNNAGATCGGAAG |
| 61174 | TATGAGTGTGGAGTCGTTGCGTAAACATTGTCACTGAGGGCCTNNNAGATCGGAA |
| 50046 | TATGAGTGTGGAGTCGTTGCGCAAGGAAACGTCAGTGGAAAAGNNNAGATCGGA |
| 41026 | TATGAGTGTGGAGTCGTTGCTTTCTGCTGCTACTCTCCACCTANNNAGATCGGAAGA |
| 45919 | TATGAGTGTGGAGTCGTTGCTCCTTTCATGTAGCAGCTTCTTCCNNNAGATCGGAAG |
| 46445 | TATGAGTGTGGAGTCGTTGCTGTTCCTCACCTTCTCTTTCAGCNNNAGATCGGAAGA |
| 40459 | TATGAGTGTGGAGTCGTTGCTCGTTTTCTGGGGGAAAAGTTGANNNAGATCGGAA |
| 63209 | TATGAGTGTGGAGTCGTTGCGTCCTTCCTAACTGTCCTGGTCTNNNAGATCGGAAGA |
| 37133 | TATGAGTGTGGAGTCGTTGCAGATGCCTTTGGCTGTTCATTCANNNAGATCGGAAG |
| 48457 | TATGAGTGTGGAGTCGTTGCGGGGACTGAAAAGTGCTGAGATCNNNAGATCGGA |
| 117057 | TATGAGTGTGGAGTCGTTGCCCCTTCTTTCCTGCCTTGGATATTNNNAGATCGGAAG |
| 68525 | TATGAGTGTGGAGTCGTTGCGTTTGCAGATGGCATTTGTGGTTNNNAGATCGGAA |
| 41982 | TATGAGTGTGGAGTCGTTGCGACCCCTGGTGCTTGTGAATTTANNNAGATCGGAAG |
| 103558 | TATGAGTGTGGAGTCGTTGCTACTTTGGGAGATCATAGCGTCTNNNAGATCGGAAG |
| 39404 | TATGAGTGTGGAGTCGTTGCAACATAGAGTAGGCAGCAGTGGANNNAGATCGGA |
| 31431 | TATGAGTGTGGAGTCGTTGCTGGCTATTTCTCCCCACTCTCAANNNAGATCGGAAGA |
| 49642 | TATGAGTGTGGAGTCGTTGCTGAAGAGCTTTGGGCATACCTTCNNNAGATCGGAAG |
| 65119 | TATGAGTGTGGAGTCGTTGCTTAGACGTTCAGTTACCAGGGGTNNNAGATCGGAA |
| 62775 | TATGAGTGTGGAGTCGTTGCCTTGAACCTCCAGTCACCACTCTNNNAGATCGGAAGA |
| 39683 | TATGAGTGTGGAGTCGTTGCATGCTATCAAAACTTATGCAGATGANNNAGATCGGA |
| 105221 | TATGAGTGTGGAGTCGTTGCTCCAGAAACCTTAGCTTCTCCAANNNAGATCGGAAG |
| 106951 | TATGAGTGTGGAGTCGTTGCTGTGAAATCTTCTGCCTGTCTCCANNNAGATCGGAA |
| 68389 | TATGAGTGTGGAGTCGTTGCACCCAGCCATCTTCAAATTCGTTNNNAGATCGGAAGA |
| 65808 | TATGAGTGTGGAGTCGTTGCGGACACCCCCAAATTCATGATGTNNNAGATCGGAAG |
| 107351 | TATGAGTGTGGAGTCGTTGCGGGCATGGCTTCTAAAGATTTCTCNNNAGATCGGAA |
| 107998 | TATGAGTGTGGAGTCGTTGCGGAGAGAGAGAGAAGCTTTGATGGNNNAGATCGG |
| 29867 | TATGAGTGTGGAGTCGTTGCGGGCCATGCATATGGTAAAAACCNNNAGATCGGAA |
| 35177 | TATGAGTGTGGAGTCGTTGCTCATTTCTGAAGGTTCCCAAGCANNNAGATCGGAAG |

50742 TATGAGTGTGGAGTCGTTGCAGTGCCATGAAGGCAAATGACAGNNNAGATCGGAA
44626 TATGAGTGTGGAGTCGTTGCAGCGGAAGTGATTGAGAAATCCCNNNAGATCGGAA
42341 TATGAGTGTGGAGTCGTTGCTCCTCTTCCATGTAGCTGTCAACNNNAGATCGGAAGA
106385 TATGAGTGTGGAGTCGTTGCAAAATCTCCAGGGCTCTTTACTAANNNAGATCGGAA
58439 TATGAGTGTGGAGTCGTTGCGGAGAGAAAGAAAGCAGCACCATNNNAGATCGGAA
53236 TATGAGTGTGGAGTCGTTGCCACCCACCTACAAATCATGGAGGNNNAGATCGGAAG
68822 TATGAGTGTGGAGTCGTTGCAGAACCAATCAGTTGGCCCATTTNNNAGATCGGAAG
64434 TATGAGTGTGGAGTCGTTGCGCAAGAGAGAAGGCAATCCTAGTNNNAGATCGGAA
99175 TATGAGTGTGGAGTCGTTGCATCAACAGCACCTGAGGGATTCTTNNNAGATCGGAA
29872 TATGAGTGTGGAGTCGTTGCAAAACCCTTCCTTGGGAGTAGCANNNAGATCGGAAG
100537 TATGAGTGTGGAGTCGTTGCAATCCAGACAGGGTAACAGCAACANNNAGATCGGAA
61609 TATGAGTGTGGAGTCGTTGCGGGGAAACAGACGGAATAAAGCTNNNAGATCGGAA
46799 TATGAGTGTGGAGTCGTTGCTACTTCTGGGTGGTGTAATTCGCNNNAGATCGGAAG
47760 TATGAGTGTGGAGTCGTTGCGAGTAAATTTTGGAGCAGGCTGCNNNAGATCGGAA
60195 TATGAGTGTGGAGTCGTTGCTGTGAACCTATTGTCAAGCTCACTNNNAGATCGGAA
102661 TATGAGTGTGGAGTCGTTGCCTGTGAACCTATTGTCAAGCTCACTNNNAGATCGGAA
43154 TATGAGTGTGGAGTCGTTGCACTGTCTCCCTCACCTTACAGACNNNAGATCGGAAGA
65180 TATGAGTGTGGAGTCGTTGCACAGAAAAGAAGGCCTACTGGGTNNNAGATCGGAA
108055 TATGAGTGTGGAGTCGTTGCGTGTAACCCAAGCAGTAAATTTCAGNNNAGATCGGA
65546 TATGAGTGTGGAGTCGTTGCTACGATTTCCTCTGGAGGTTGGTNNNAGATCGGAAG
71252 TATGAGTGTGGAGTCGTTGCGCTCTTCTCCCCTCCATTTCTCTNNNAGATCGGAAGAG
47667 TATGAGTGTGGAGTCGTTGCATCCCAACTCACAATCAGACTGCNNNAGATCGGAAGA
41444 TATGAGTGTGGAGTCGTTGCTGGCTACATGGTCACAGCTAGTANNNAGATCGGAAG
45456 TATGAGTGTGGAGTCGTTGCACGCACACTACATTTTGTTTCATCCNNNAGATCGGAA
42534 TATGAGTGTGGAGTCGTTGCAAAAACAGGGGACAGGAGAACACNNNAGATCGGAA
53130 TATGAGTGTGGAGTCGTTGCGTAGGAGGTGGGAGGATAAGAGGNNNAGATCGGA
117610 TATGAGTGTGGAGTCGTTGCTTTAGGAGTATAGCTGCAGTGGGNNNAGATCGGAA
69087 TATGAGTGTGGAGTCGTTGCGGACTGGGAAGAGTTTGAGCTTTNNNAGATCGGAA
100837 TATGAGTGTGGAGTCGTTGCGGCAGGGATGATTCTCAGGTTTATNNNAGATCGGAA
56380 TATGAGTGTGGAGTCGTTGCATGTCACGCGATCAAGATCTCTGNNNAGATCGGAAG
53421 TATGAGTGTGGAGTCGTTGCAGAGACAGCCACTCAGGATTAGGNNNAGATCGGAA
49798 TATGAGTGTGGAGTCGTTGCAAAACAATTGCCTCTGCAGGTTCNNNAGATCGGAAG
42459 TATGAGTGTGGAGTCGTTGCCCTACCTTCCATGGGCAACTAACNNNAGATCGGAAGA
46658 TATGAGTGTGGAGTCGTTGCAAAGACACACACATCCAGCTAGCNNNAGATCGGAAG
104863 TATGAGTGTGGAGTCGTTGCCTGTGATTTCCTATGGTGCTTTAGGGANNNAGATCG
55413 TATGAGTGTGGAGTCGTTGCTGCTTACTTTTAGAGGCCCCATGNNNAGATCGGAAGA
63621 TATGAGTGTGGAGTCGTTGCACTGGCATGTGCATCTGAAAAGTNNNAGATCGGAAG
49902 TATGAGTGTGGAGTCGTTGCCAAATACGGCAAAACCCCCTTTCNNNAGATCGGAAGA
60002 TATGAGTGTGGAGTCGTTGCCCAGGCTCAAGAACAAAGACACTNNNAGATCGGAAG
61406 TATGAGTGTGGAGTCGTTGCGTCAAATTCTTCTTCCGGTATCTCCTNNNAGATCGGAA
116863 TATGAGTGTGGAGTCGTTGCTGTGAGAAATTCTGTGAGGCTGCNNNAGATCGGAA
43609 TATGAGTGTGGAGTCGTTGCAGGTGCTAATTGGGCATCAGTACNNNAGATCGGAA
45356 TATGAGTGTGGAGTCGTTGCAGTTGCCTTACATCTCTGTTGCCNNNAGATCGGAAGA

36449 TATGAGTGTGGAGTCGTTGCTCCCAAAGGGCTTCTACTCATCANNNAGATCGGAAG
33962 TATGAGTGTGGAGTCGTTGCCACAGTCCCTTTGACAGAAACCANNNAGATCGGAAG
69384 TATGAGTGTGGAGTCGTTGCGACTTCCCAACTCCAGTGTGTTTNNNAGATCGGAAG
50242 TATGAGTGTGGAGTCGTTGCAGTAGTTCCATTCTGGCACCAAGNNNAGATCGGAAG
62148 TATGAGTGTGGAGTCGTTGCTAAAGGAAATGGGGAGAGTGGCTNNNAGATCGGA
51234 TATGAGTGTGGAGTCGTTGCTTGGTTGTGTCTACAGAGCTCAGNNNAGATCGGAA
64260 TATGAGTGTGGAGTCGTTGCATCAAAACCCTGGGTGTAGGAGTNNNAGATCGGAA
52937 TATGAGTGTGGAGTCGTTGCAAATGTAACCTGGGTTGACCAGGNNNAGATCGGAA
105201 TATGAGTGTGGAGTCGTTGCTACATCCAAACATATGTACAAAAACACNNNAGATCG
47873 TATGAGTGTGGAGTCGTTGCCTCCAGGGCTCTTTACTAAGTGCNNNAGATCGGAAG
50258 TATGAGTGTGGAGTCGTTGCCCCCCTCTATCCCTAATCCCAAGNNNAGATCGGAAGA
109620 TATGAGTGTGGAGTCGTTGCAGACTTATAAGGGCAAATGCCATGCNNNAGATCGG
68710 TATGAGTGTGGAGTCGTTGCCAGATCCAAAGGAGGCTTGTGTTNNNAGATCGGAA
35073 TATGAGTGTGGAGTCGTTGCCTATGGACGCATGGAATGTTCCANNNAGATCGGAA
66718 TATGAGTGTGGAGTCGTTGCTATGAGCCACTACACCTGGTTGTNNNAGATCGGAAG
37467 TATGAGTGTGGAGTCGTTGCACAAGAACGTTTACCCCCAAAGANNNAGATCGGAAG
63142 TATGAGTGTGGAGTCGTTGCCAGACTTTTATGTGCAAACTCAGTCTNNNAGATCGG
31740 TATGAGTGTGGAGTCGTTGCCACGCACGAAGGAGATCATAGAANNNAGATCGGAA
108838 TATGAGTGTGGAGTCGTTGCGAAACATCCTCTTGAAAACACTAGGTANNNAGATCG
51961 TATGAGTGTGGAGTCGTTGCTGCTGAACCAGCCTCTCAAATAGNNNAGATCGGAAG
55995 TATGAGTGTGGAGTCGTTGCAACACTGCCACTTAGAAAGCCTGNNNAGATCGGAAG
44053 TATGAGTGTGGAGTCGTTGCACCCGCACTGGAGTATATAGACCNNNAGATCGGAAG
97743 TATGAGTGTGGAGTCGTTGCAAATCTCTTCAGTTTCTGCAACACTGNNNAGATCGGA
48368 TATGAGTGTGGAGTCGTTGCATCTTCCACCAATGCTGTCCATCNNNAGATCGGAAGA
41384 TATGAGTGTGGAGTCGTTGCGAACCAGCATAAGGTTGCCAGTANNNAGATCGGAA
42809 TATGAGTGTGGAGTCGTTGCTGGAAATGCTCTGTTTTCTTCCACNNNAGATCGGAA
31665 TATGAGTGTGGAGTCGTTGCCTGGCACTGCTTTACTGTCAGAANNNAGATCGGAAG
43000 TATGAGTGTGGAGTCGTTGCTGGTAGTACAGTATAGGAAGTCCTCACNNNAGATCG
61629 TATGAGTGTGGAGTCGTTGCTGGAAATAGGTCTCCCTCAAGCTNNNAGATCGGAAG
59080 TATGAGTGTGGAGTCGTTGCGGTTTCCAAGCTGTTGAATAGATNNNAGATCGGAA
51077 TATGAGTGTGGAGTCGTTGCGAGAAGAGACAGTGTGAAGGCAGNNNAGATCGGA
68989 TATGAGTGTGGAGTCGTTGCGGCTGGTGGGTACCAAATACTTTNNNAGATCGGAA
51368 TATGAGTGTGGAGTCGTTGCTGGAAGGGAAGAGGTGAGAAGAGNNNAGATCGG
60482 TATGAGTGTGGAGTCGTTGCCCTGGAAGTCTTTCATTGCTACTNNNAGATCGGAAG
61069 TATGAGTGTGGAGTCGTTGCTGACAAACCAAGAGAAGAAGCCTNNNAGATCGGAA
34990 TATGAGTGTGGAGTCGTTGCCGAACTGCTCTCATTTCTGTCCANNNAGATCGGAAGA
31345 TATGAGTGTGGAGTCGTTGCCCCTTATGCTTGCCAAATTGCAANNNAGATCGGAAG
33028 TATGAGTGTGGAGTCGTTGCTCCAAACATATGTACAAAAACACACANNNAGATCGG
53503 TATGAGTGTGGAGTCGTTGCTAAAAATGTGCAACATTGGCCGGNNNAGATCGGAA
108088 TATGAGTGTGGAGTCGTTGCAATCTTTCCACCTAAGCCATCAAGTANNNAGATCGGA
34176 TATGAGTGTGGAGTCGTTGCGAATAACAGTGCCAGTGCTACCANNNAGATCGGAA
43801 TATGAGTGTGGAGTCGTTGCGCTACAACAAAAACCCCAGAACCNNNAGATCGGAAG
45957 TATGAGTGTGGAGTCGTTGCTGGGATTGTTGTATACGGTGTTCCNNNAGATCGGAA

45980 TATGAGTGTGGAGTCGTTGCATGCCTTTTGTCTGGGGATTTCCNNNAGATCGGAAG
61415 TATGAGTGTGGAGTCGTTGCGTCTTTCTATCCCCCTGTCTCCTNNNAGATCGGAAGA
30634 TATGAGTGTGGAGTCGTTGCAGCACCACTGTCAAATCTGGAAANNNAGATCGGAAG
54394 TATGAGTGTGGAGTCGTTGCCAATCTGCAAAAGGACTGCATGGNNNAGATCGGAA
47152 TATGAGTGTGGAGTCGTTGCAAATATACCCACAAAGGCTGGGCNNNAGATCGGAA
107347 TATGAGTGTGGAGTCGTTGCATCCTACTGAGCCCATCTGCAANNNAGATCGGAAGA
48084 TATGAGTGTGGAGTCGTTGCATGAATCTCCCAACCTCACTTGCNNNAGATCGGAAGA
48850 TATGAGTGTGGAGTCGTTGCCCTGCAAAGGTCAAAATCAGCTCNNNAGATCGGAAG
110265 TATGAGTGTGGAGTCGTTGCACAGGGTCTACTCAGGTTTCCCNNNAGATCGGAAGA
43182 TATGAGTGTGGAGTCGTTGCGTTTATTGGGTCAGGCCTCAGACNNNAGATCGGAA
101195 TATGAGTGTGGAGTCGTTGCACCAAGTATGACCAGCAGGCAATNNNAGATCGGAA
34937 TATGAGTGTGGAGTCGTTGCTGGCACATTTCTTCACTTCTCCANNNAGATCGGAAGA
58109 TATGAGTGTGGAGTCGTTGCCTTGTGATCTGAGCTGCTGGAATNNNAGATCGGAA
48434 TATGAGTGTGGAGTCGTTGCGTAAGACAGGCCTCGGATTCATCNNNAGATCGGAA
68247 TATGAGTGTGGAGTCGTTGCCATAGCACATACAATACAGACAGTTNNNAGATCGGA
59483 TATGAGTGTGGAGTCGTTGCCCACCACGACCAACAACAACTATNNNAGATCGGAAG
36179 TATGAGTGTGGAGTCGTTGCTGGGAATGTTACAAATCTGTGCANNNAGATCGGAA
109305 TATGAGTGTGGAGTCGTTGCGTTTTGTGAATGGCTCTAGCCTTNNNAGATCGGAAG
41310 TATGAGTGTGGAGTCGTTGCACTGCTTTGTTCCAGCTTGTCTANNNAGATCGGAAG
115977 TATGAGTGTGGAGTCGTTGCTCAGTTGGCCCATTTTTCCACAGCNNNAGATCGGAA
43923 TATGAGTGTGGAGTCGTTGCTGTCAAGGTCATGATTCTTATCCACCNNNAGATCGG
43791 TATGAGTGTGGAGTCGTTGCTCTAAAGCATGTAGCCTTCAACCNNNAGATCGGAAG
36766 TATGAGTGTGGAGTCGTTGCGAGCTATGGCAGAGAGGATCTCANNNAGATCGGAA
36828 TATGAGTGTGGAGTCGTTGCTTAGAATGGGGGCTGTCTTCTCANNNAGATCGGAA
37836 TATGAGTGTGGAGTCGTTGCTGCCATGAGATAACCTGACCAGANNNAGATCGGAA
42862 TATGAGTGTGGAGTCGTTGCCTAAAATACACACACCCACGCACNNNAGATCGGAAG
40952 TATGAGTGTGGAGTCGTTGCGGTCCCAGAGGAGAAAGGAACTANNNAGATCGGAA
45646 TATGAGTGTGGAGTCGTTGCTCTAGCATGAAGCTGTTTGCTCCNNNAGATCGGAAG
41807 TATGAGTGTGGAGTCGTTGCTGTCACTCCTTTGTGACCCCTTANNNAGATCGGAAGA
37280 TATGAGTGTGGAGTCGTTGCTGTGTGTGTGTGAAGAAAGGTTCANNNAGATCGGA
45155 TATGAGTGTGGAGTCGTTGCTGAGTTTACCTATCACTAAGTTGGCCNNNAGATCGG
48205 TATGAGTGTGGAGTCGTTGCAATATCATGCCCTCTCCCTTTGCNNNAGATCGGAAGA
109607 TATGAGTGTGGAGTCGTTGCTTTGTTCATCAGTGGACACTTAGCTTNNNAGATCGG
44719 TATGAGTGTGGAGTCGTTGCACTCATCAACACATCTTGCTCCCNNNAGATCGGAAGA
117346 TATGAGTGTGGAGTCGTTGCCTACCTACCTACCTATTCATCTCTATCNNNAGATCGGA
100200 TATGAGTGTGGAGTCGTTGCGTGCTCGCAGTAGATTCCAAGTNNNAGATCGGAAG
56298 TATGAGTGTGGAGTCGTTGCAGGCTGAGGAATCCAACAATCTGNNNAGATCGGAA
43160 TATGAGTGTGGAGTCGTTGCGGATAATGAGGCACAGACCAGACNNNAGATCGGAA
63123 TATGAGTGTGGAGTCGTTGCTCTTTGAAGGTTCAGGTCTAAGTCTNNNAGATCGGA
70865 TATGAGTGTGGAGTCGTTGCAGTGTAAAACACACACACCCCTGNNNAGATCGGAAG
109741 TATGAGTGTGGAGTCGTTGCATTTGCAGCAACCCAGTTGGAGTNNNAGATCGGAA
117432 TATGAGTGTGGAGTCGTTGCGAGCCGATCCTTCCACCATTTACNNNAGATCGGAAG
35662 TATGAGTGTGGAGTCGTTGCCAGCTCCATCTTCTGAATAGGCANNNAGATCGGAAG

59413 TATGAGTGTGGAGTCGTTGCCTTGACCTCCTGGGCCTACATATNNNAGATCGGAAGA
62863 TATGAGTGTGGAGTCGTTGCACAGACTACACATGAAAAAGCCTCTNNNAGATCGGAA
63277 TATGAGTGTGGAGTCGTTGCCGTACTCGTGTGTGTATGTGTCTNNNAGATCGGAAG
39203 TATGAGTGTGGAGTCGTTGCTCTTGCTACCTAGCTATCTTGGGANNNAGATCGGAAG
44901 TATGAGTGTGGAGTCGTTGCAGCAGAGAAAAACAGAGACAGCCNNNAGATCGGAA
52043 TATGAGTGTGGAGTCGTTGCTGGTGCTTTTTCCCAAGTCCTAGNNNAGATCGGAAGA
109654 TATGAGTGTGGAGTCGTTGCCAGTAACACAGCACTCAACCAAGNNNAGATCGGAAG
64349 TATGAGTGTGGAGTCGTTGCACATCCTGAGTTGAGCTCTGAGTNNNAGATCGGAAG
41108 TATGAGTGTGGAGTCGTTGCTCATTGTGGCTTCCTCACTCCTANNNAGATCGGAAGA
64670 TATGAGTGTGGAGTCGTTGCACCTACCTACCTATTCATCTCTATCGTNNNAGATCGGA
52703 TATGAGTGTGGAGTCGTTGCCCCAGGAATGGATTGTTTCAAAGGNNNAGATCGGAA
39129 TATGAGTGTGGAGTCGTTGCAGGCTTATCTGGTTTCTAGATGGGANNNAGATCGGA
102387 TATGAGTGTGGAGTCGTTGCTTTCCAACACCACAAGAAGCACTANNNAGATCGGAAG
59471 TATGAGTGTGGAGTCGTTGCGCTCTGGCATTTGCCACCTATATNNNAGATCGGAAGA
37182 TATGAGTGTGGAGTCGTTGCGCACAGTTCATCCTGCTACTTCANNNAGATCGGAAGA
35575 TATGAGTGTGGAGTCGTTGCTTGGTTGGACATTGAGAAAGGCANNNAGATCGGAA
63581 TATGAGTGTGGAGTCGTTGCCCCAAGTGGCATAGCTCTTTTCTNNNAGATCGGAAGA
60813 TATGAGTGTGGAGTCGTTGCACCACACCCTGTGTACATTACCTNNNAGATCGGAAGA
97065 TATGAGTGTGGAGTCGTTGCCACCGGCAAAACCATCACGANNNAGATCGGAAGAGC
98449 TATGAGTGTGGAGTCGTTGCCCCCAAGAACATATGCAGACCCTNNNAGATCGGAAGA
53018 TATGAGTGTGGAGTCGTTGCAAATGTGGTCCGAATCAAGCAGGNNNAGATCGGAAG
42696 TATGAGTGTGGAGTCGTTGCACCTTTTGGTCCATGACATACCACNNNAGATCGGAAG
48570 TATGAGTGTGGAGTCGTTGCAACATGCCAGTGGAGGAAAACTCNNNAGATCGGAAG
46993 TATGAGTGTGGAGTCGTTGCGTGATTGTGAGGGCTAACTAGGCNNNAGATCGGAA
49380 TATGAGTGTGGAGTCGTTGCAAAAGCCTTACCCTGCAGATGTCNNNAGATCGGAAG
59999 TATGAGTGTGGAGTCGTTGCGAGCATGATGATGTGGTATCACACTNNNAGATCGGA
37071 TATGAGTGTGGAGTCGTTGCGAAAAGCACCCCATCTCTTGTCANNNAGATCGGAAGA
117904 TATGAGTGTGGAGTCGTTGCTGGTGTTTCACTCTTGTGATCTTGNNNAGATCGGAAG
42470 TATGAGTGTGGAGTCGTTGCTATGTTGCACAGGGTGGTCTAACNNNAGATCGGAAG
61778 TATGAGTGTGGAGTCGTTGCCTCAAAGGAGGATTTGAAGGAGCTNNNAGATCGGA
105691 TATGAGTGTGGAGTCGTTGCCACACATACATCCCTACTCTTCACTTNNNAGATCGGAA
50032 TATGAGTGTGGAGTCGTTGCATGTCACAGCATTGGTGCAAAAGNNNAGATCGGAAG
40796 TATGAGTGTGGAGTCGTTGCCCTGCGGACCTGATCCTAAGATANNNAGATCGGAAG
46846 TATGAGTGTGGAGTCGTTGCTAGGAGCAGTATCAGAGGAAGGCNNNAGATCGGAA
51367 TATGAGTGTGGAGTCGTTGCTTCCTTTGCTCAAATCAGAGAAGAGNNNAGATCGGA
36865 TATGAGTGTGGAGTCGTTGCTTGCCATTTTTCCCCTGAAGTCANNNAGATCGGAAGA
52022 TATGAGTGTGGAGTCGTTGCGGGTGATCAGAGCAGAAACCTAGNNNAGATCGGAA
52363 TATGAGTGTGGAGTCGTTGCAGCGTCTTTCATTGCTCATTTACGNNNAGATCGGAAG
105703 TATGAGTGTGGAGTCGTTGCCACAACCGTGTCCATGTAAGTACNNNAGATCGGAAG
68923 TATGAGTGTGGAGTCGTTGCAGCCTGGGAGAGACCACATATTTNNNAGATCGGAAG
61011 TATGAGTGTGGAGTCGTTGCAGTTTCCTCTACTCTCCCTCCCTNNNAGATCGGAAGAG
62687 TATGAGTGTGGAGTCGTTGCAGTGGAACCATACCAGTGGATCTNNNAGATCGGAAG
50605 TATGAGTGTGGAGTCGTTGCTTTACTGGGGACGTTCCAAACAGNNNAGATCGGAAG

49944 TATGAGTGTGGAGTCGTTGCAGGGTGTAGCTGGAGAGAGTTTCNNNAGATCGGA
59372 TATGAGTGTGGAGTCGTTGCGGTTGCTGAAGAGCACTGTTGATNNNAGATCGGAA
29868 TATGAGTGTGGAGTCGTTGCACTTTATCTGCTCTTCCTGCTGCNNNAGATCGGAAGA
44434 TATGAGTGTGGAGTCGTTGCGAATCTTTGAAACCTCATGCCCCNNNAGATCGGAAG
29963 TATGAGTGTGGAGTCGTTGCCTACTATTCCAGTTGCTTGCCCANNNAGATCGGAAG
51858 TATGAGTGTGGAGTCGTTGCATTCTTACGCTGTGGCATCTGAGNNNAGATCGGAAG
46405 TATGAGTGTGGAGTCGTTGCCCATACCCAGCTCATGTATCAGCNNNAGATCGGAAG
52754 TATGAGTGTGGAGTCGTTGCCTGGATCATAAGCTCTGCCAAGGNNNAGATCGGAA
58753 TATGAGTGTGGAGTCGTTGCTTCCTGGCTTCTTGACTTTGCATNNNAGATCGGAAG
71701 TATGAGTGTGGAGTCGTTGCCCTCTGCCTCACAGTTCAATTCTNNNAGATCGGAAGA
29870 TATGAGTGTGGAGTCGTTGCGGAAAAGCCTAAGGCACTGGTTANNNAGATCGGAA
36103 TATGAGTGTGGAGTCGTTGCTTCTCTGGTCTTTTGCATCTGCANNNAGATCGGAAG
29965 TATGAGTGTGGAGTCGTTGCAGTTATTGCATCTGTCTGCCAGGNNNAGATCGGAAG
29871 TATGAGTGTGGAGTCGTTGCACAAGTAAAGACTTCCACCTGCCNNNAGATCGGAAG
29964 TATGAGTGTGGAGTCGTTGCGCTCTTTGCCCTGGACTGTAATTNNNAGATCGGAAG
56467 TATGAGTGTGGAGTCGTTGCTCTGGACCTACTACAGGCTTCTGNNNAGATCGGAAG
71722 TATGAGTGTGGAGTCGTTGCCTTGCGACGTTCTATGAAGCATGNNNAGATCGGAAG
40961 TATGAGTGTGGAGTCGTTGCTTACTGTCCTCTTGGCACCACTANNNAGATCGGAAG
50575 TATGAGTGTGGAGTCGTTGCTAGGCATGGGTGGACACTGTAAGNNNAGATCGGA
71671 TATGAGTGTGGAGTCGTTGCCAAGCAATGTGGATGATGACAGCNNNAGATCGGAA
62423 TATGAGTGTGGAGTCGTTGCGAGGAAGAAGACAGTGTTGTGCTNNNAGATCGGA
71806 TATGAGTGTGGAGTCGTTGCTCTTTGTGTACGTTATGCCTTGCNNNAGATCGGAAG
70679 TATGAGTGTGGAGTCGTTGCAGTTTGGACTGGTTGCTGTGTAGNNNAGATCGGAA
45590 TATGAGTGTGGAGTCGTTGCTTTCTTGTCTGTTGCCATCCTCCNNNAGATCGGAAGA
45797 TATGAGTGTGGAGTCGTTGCTTGAACTGCTTCTCATCCTGTCCNNNAGATCGGAAGA
101526 TATGAGTGTGGAGTCGTTGCTGATTGTATGAACACCTGGGATCTGNNNAGATCGG
33603 TATGAGTGTGGAGTCGTTGCGCTTGGTTTCCCTGAATAGGACANNNAGATCGGAAG
122002 TATGAGTGTGGAGTCGTTGCTGGAGTATGTCAAGAATGGCCAGAGNNNAGATCGG
102377 TATGAGTGTGGAGTCGTTGCCAAACAGATGAACACCACAAGGTGNNNAGATCGGA
105135 TATGAGTGTGGAGTCGTTGCATCTTGTCTGACACCTGCTACCANNNAGATCGGAAG
102640 TATGAGTGTGGAGTCGTTGCGTTGCTCTGACTAGGACTTCCAAAACNNNAGATCGG
49373 TATGAGTGTGGAGTCGTTGCTGTTCAACTCAGGACACCATGTCNNNAGATCGGAAG
103230 TATGAGTGTGGAGTCGTTGCGTTTTCTAATGAGCCTGATGCATGCNNNAGATCGGA
100627 TATGAGTGTGGAGTCGTTGCAAGAAATCCTGTGACCTGCTGTNNNAGATCGGAAG
102335 TATGAGTGTGGAGTCGTTGCCACACCCAGTACTCCAGCCTTATNNNAGATCGGAAG
100931 TATGAGTGTGGAGTCGTTGCTCTGTGTCTGGATAGCTGAGAGTGNNNAGATCGGA
103520 TATGAGTGTGGAGTCGTTGCTCAGCAGCATCAAAAATCCAAGGANNNAGATCGGA
106402 TATGAGTGTGGAGTCGTTGCGCTCCTTCTACATTCATTTCCAGTNNNAGATCGGAAG
101970 TATGAGTGTGGAGTCGTTGCCACCTCCCTCCAAGATCCTAGCNNNAGATCGGAAGA
104571 TATGAGTGTGGAGTCGTTGCTTTTGTATCAGCATATTCCTTGCCNNNAGATCGGAAG
105806 TATGAGTGTGGAGTCGTTGCTGTTTTCATCCATGGTTCGTGACTNNNAGATCGGAA
102360 TATGAGTGTGGAGTCGTTGCTATCCCAAAGCTGCTAAAACTCAGTNNNAGATCGGA
106952 TATGAGTGTGGAGTCGTTGCTTAGGTTCTTCCCACTCCACATCTNNNAGATCGGAAG

71657 TATGAGTGTGGAGTCGTTGCCACAGCAGAGAGCAACTTCCTAGNNNAGATCGGAAG

115529 TATGAGTGTGGAGTCGTTGCTCTTCCCACTCCACATCTCCAANNNAGATCGGAAGAG

104301 TATGAGTGTGGAGTCGTTGCATTATGTCTTGGGCTTCAAGGGGNNNAGATCGGAA

101800 TATGAGTGTGGAGTCGTTGCACATAATGAGCCCCAGTAATTGTNNNAGATCGGAA

103213 TATGAGTGTGGAGTCGTTGCAAAGTCAATATCTTGGTCTCGTTCCTANNNAGATCGG

101758 TATGAGTGTGGAGTCGTTGCTCCCTCATCCTCTTCACATCCCTTNNNAGATCGGAAGA

103003 TATGAGTGTGGAGTCGTTGCGTTTAGGCTCCAAAGGCAGAGGNNNAGATCGGAAG

106877 TATGAGTGTGGAGTCGTTGCATGAGCCCCAGTAATTGTTATCNNNAGATCGGAAG

105755 TATGAGTGTGGAGTCGTTGCTCCTGATTTTCTTCTGTGTTCCTTTNNNAGATCGGAAG

104567 TATGAGTGTGGAGTCGTTGCTGTGCTGCTGTGATTATGTGTGNNNAGATCGGAAG

109325 TATGAGTGTGGAGTCGTTGCGCCCCAAATGTAGCTGGAACTACNNNAGATCGGAAG

95481 TATGAGTGTGGAGTCGTTGCCATTACAGGCTTCTTCTGCTTGGAANNNAGATCGGA

113707 TATGAGTGTGGAGTCGTTGCGTGTGTGTGTAGTATCCTAGGCANNNAGATCGGAA

101473 TATGAGTGTGGAGTCGTTGCTTCATCAGTGCCTTTTTGACGTTANNNAGATCGGAAG

106913 TATGAGTGTGGAGTCGTTGCCTGTCTTTCACATAAAACATCATCTTTNNNAGATCGG

101789 TATGAGTGTGGAGTCGTTGCACAAGGAGGATTCTGGGGGATTNNNAGATCGGAAG

115980 TATGAGTGTGGAGTCGTTGCAACCCTGGCTAAAACAGTGAACTCNNNAGATCGGAA

103199 TATGAGTGTGGAGTCGTTGCTGAAAATGCCTACCACCTCCCTNNNAGATCGGAAGAG

106849 TATGAGTGTGGAGTCGTTGCTTTTATTTGACAGAGATGCCTCAGGNNNAGATCGGA

104279 TATGAGTGTGGAGTCGTTGCTACAGCTGCTTGACAACAGAANNNAGATCGGAAGAG

116324 TATGAGTGTGGAGTCGTTGCGAGGTTATGGTGAGATTACGCGCNNNAGATCGGAA

117440 TATGAGTGTGGAGTCGTTGCGCATAAACAACCACACATGGTTACNNNAGATCGGAA

116309 TATGAGTGTGGAGTCGTTGCTTGACCATTATTTTGGTGCTTTCCNNNAGATCGGAAG

108031 TATGAGTGTGGAGTCGTTGCGTCCAGTCATTCTCTCCAACACAGANNNAGATCGGAA

108017 TATGAGTGTGGAGTCGTTGCAAGAATGAGTTGATGGCATTTGCAACNNNAGATCGG

101255 TATGAGTGTGGAGTCGTTGCCATACTGATTCTGGCTCAAGGTGNNNAGATCGGAAG

122005 TATGAGTGTGGAGTCGTTGCACTTACCTGTTTGGATGCCTAGCNNNAGATCGGAAG

101283 TATGAGTGTGGAGTCGTTGCCCCAAAGAGCCGATGGTATAAATCCNNNAGATCGGA

103217 TATGAGTGTGGAGTCGTTGCGCAGCTATCTCTTCAACATACCAACCNNNAGATCGGA

100713 TATGAGTGTGGAGTCGTTGCGAAGTGTGTGTACTGCCAGCGTTNNNAGATCGGAA

107294 TATGAGTGTGGAGTCGTTGCAGAGGCTTGATCATTACAGGGTNNNAGATCGGAAG

101815 TATGAGTGTGGAGTCGTTGCTCTCTTTGGGTTCCTCTCTCTNNNAGATCGGAAGAGC

104354 TATGAGTGTGGAGTCGTTGCATGGGGGATAGATGTCTTGCCAANNNAGATCGGAA

101582 TATGAGTGTGGAGTCGTTGCCTTAGGCACACTGGGAGAATAGATGNNNAGATCGG

105211 TATGAGTGTGGAGTCGTTGCCATAATTGCTCCGACTAGGACATCCNNNAGATCGGA

110204 TATGAGTGTGGAGTCGTTGCTTTTGATGTTTAGTTTTGAACTGGCTNNNAGATCGG

109687 TATGAGTGTGGAGTCGTTGCACATACCAGGGAACAGAACAGAGANNNAGATCGGA

107336 TATGAGTGTGGAGTCGTTGCCACACACACAGTTTTCATAAGTGTTTNNNAGATCGGA

98260 TATGAGTGTGGAGTCGTTGCGCATTTCTCTTTAGATGGGGTTGGNNNAGATCGGAA

108002 TATGAGTGTGGAGTCGTTGCTGCCACAATCTAGTACCACAAACTAAANNNAGATCGG

115831 TATGAGTGTGGAGTCGTTGCATTGAAGGTGACTTCTGGCCAACNNNAGATCGGAAG

103175 TATGAGTGTGGAGTCGTTGCTCACTGCATTCCTCCTCTTTCAGGNNNAGATCGGAAG

101297 TATGAGTGTGGAGTCGTTGCGTCTACTCACTTCAGCCTCCCAGNNNAGATCGGAAGA

| | |
|---|---|
| 102968 | TATGAGTGTGGAGTCGTTGCGGAGAAGCAAGCAAGCAAGAAGNNNAGATCGGAA |
| 109072 | TATGAGTGTGGAGTCGTTGCACAGTCTGCTGGAGAAGTCCTTNNNAGATCGGAAG |
| 102976 | TATGAGTGTGGAGTCGTTGCCGGATCATGAGGTCGAGATAGCNNNAGATCGGAAG |
| 107310 | TATGAGTGTGGAGTCGTTGCAGATACATTTTGAGGCACAGAGCNNNAGATCGGAA |
| 104542 | TATGAGTGTGGAGTCGTTGCGTCCAGCCAAGTTTTGTTATTGTNNNAGATCGGAAG |
| 95148 | TATGAGTGTGGAGTCGTTGCGAGCACTTCAGAGCCAAGCTATTNNNAGATCGGAAG |
| 96200 | TATGAGTGTGGAGTCGTTGCACCCGCCAAAATTTATGTCTTTCTCNNNAGATCGGAA |
| 117307 | TATGAGTGTGGAGTCGTTGCTTTCCAGGGTCTTGCTCAATTGCNNNAGATCGGAAG |
| 95106 | TATGAGTGTGGAGTCGTTGCATCATGTAGAGCTGCTCGTTCACNNNAGATCGGAAG |
| 35656 | TATGAGTGTGGAGTCGTTGCAGGTGTGAGTTCAATATGAGGCANNNAGATCGGAA |
| 62951 | TATGAGTGTGGAGTCGTTGCGCCTCAAAGATAGCAATGGCTCTNNNAGATCGGAAG |
| 100101 | TATGAGTGTGGAGTCGTTGCAACACACACACTACACATAAATCATTTNNNAGATCGG |
| 50682 | TATGAGTGTGGAGTCGTTGCACGCTATCCACTACTTTGCACAGNNNAGATCGGAAG |
| 37435 | TATGAGTGTGGAGTCGTTGCACGCGCTGCCAAGTATAAAAAGANNNAGATCGGAA |
| 69032 | TATGAGTGTGGAGTCGTTGCACCCCCTTTTTCTCCTTCCCTTTNNNAGATCGGAAGAG |
| 30927 | TATGAGTGTGGAGTCGTTGCCCGCCTTGAAAGAGAGATGACAANNNAGATCGGAA |
| 31852 | TATGAGTGTGGAGTCGTTGCGGCACAAAAGTTCTCCTGAGGAANNNAGATCGGAA |
| 29856 | TATGAGTGTGGAGTCGTTGCACCTTCCTGTAGGATACACCCAGNNNAGATCGGAAG |
| 52677 | TATGAGTGTGGAGTCGTTGCGCACGCAAAAGAGAGTGAAAAGGNNNAGATCGGA |
| 55547 | TATGAGTGTGGAGTCGTTGCGCCAGTGTACTGAGTGGAAGATGNNNAGATCGGAA |
| 49511 | TATGAGTGTGGAGTCGTTGCTGCCCTCTGGATTGTGAACATTCNNNAGATCGGAAG |
| 46838 | TATGAGTGTGGAGTCGTTGCCCACACCATTTTCTCTTTCAAGGCNNNAGATCGGAAG |
| 36031 | TATGAGTGTGGAGTCGTTGCTGTGCCTGTGGAAGTATTTATGCANNNAGATCGGA |
| 42564 | TATGAGTGTGGAGTCGTTGCTTCTTTCAGCTCTGAGCACACACNNNAGATCGGAAG |
| 46891 | TATGAGTGTGGAGTCGTTGCATTTTCTGGCATGTATCCCAGGCNNNAGATCGGAAG |
| 54218 | TATGAGTGTGGAGTCGTTGCCTTCAGGATTCCACATGTGTGGGNNNAGATCGGAA |
| 60329 | TATGAGTGTGGAGTCGTTGCTGCCGAACTCATAACCTAGGACTNNNAGATCGGAAG |
| 58616 | TATGAGTGTGGAGTCGTTGCAGTTCTGGCATTTTTCCCAGCATNNNAGATCGGAAG |
| 48823 | TATGAGTGTGGAGTCGTTGCTTTTCCCTGTGTTGCTCTTCCTCNNNAGATCGGAAGA |
| 34129 | TATGAGTGTGGAGTCGTTGCCAGGTGTTTCTTCCATTGGACCANNNAGATCGGAAG |
| 31210 | TATGAGTGTGGAGTCGTTGCCAGGAGAGAGGGTAAGAGAGCAANNNAGATCGGA |
| 69100 | TATGAGTGTGGAGTCGTTGCGAGAGGTGAGGATGGAAGGCTTTNNNAGATCGGA |
| 66088 | TATGAGTGTGGAGTCGTTGCTCCAGAAAGCAAGACAATGCTGTNNNAGATCGGAA |
| 58759 | TATGAGTGTGGAGTCGTTGCACCCAGCTTCTACCCCAAATCATNNNAGATCGGAAGA |
| 55957 | TATGAGTGTGGAGTCGTTGCCCTATTCCTGTTTGTTCCCCCTGNNNAGATCGGAAGA |
| 35654 | TATGAGTGTGGAGTCGTTGCACTTCGTCTTTTCAGAATGAGGCANNNAGATCGGAA |
| 51650 | TATGAGTGTGGAGTCGTTGCCAGAAACAGAGCTAACCAGGGAGNNNAGATCGGAA |
| 41173 | TATGAGTGTGGAGTCGTTGCCCCTTTCTGTCTCTCTCTCGCTANNNAGATCGGAAGA |
| 117407 | TATGAGTGTGGAGTCGTTGCCAACAGCATCAAGGCTGAACATNNNAGATCGGAAG |
| 95876 | TATGAGTGTGGAGTCGTTGCGTGCTCAAGTTGTCTCTACAACCNNNAGATCGGAAG |
| 50175 | TATGAGTGTGGAGTCGTTGCCACCTTCCAGCTGTACCTGAAAGNNNAGATCGGAAG |
| 47516 | TATGAGTGTGGAGTCGTTGCGCCCACAATTATTTCTGCCATGCNNNAGATCGGAAG |
| 51326 | TATGAGTGTGGAGTCGTTGCCCTGTGTCTGCAGTACCTTTCAGNNNAGATCGGAAG |

| | |
|---|---|
| 56288 | TATGAGTGTGGAGTCGTTGCCTCAACACAGGCAATTCTTGCTGNNNAGATCGGAAG |
| 117678 | TATGAGTGTGGAGTCGTTGCTGTGTGTGTTTCTCTTTTCTTTTGGAANNNAGATCGG |
| 48893 | TATGAGTGTGGAGTCGTTGCGCATTGTGTTCTTGATACGGCTCNNNAGATCGGAAG |
| 45036 | TATGAGTGTGGAGTCGTTGCTTGTGGAACTGGTGAATAAGGCCNNNAGATCGGAA |
| 55056 | TATGAGTGTGGAGTCGTTGCGCTGTTTTCACCCAAGAACTTGGNNNAGATCGGAAG |
| 55193 | TATGAGTGTGGAGTCGTTGCCACACTGTCTTCTTACGGTTTGGNNNAGATCGGAAG |
| 47852 | TATGAGTGTGGAGTCGTTGCCCCTTAGTATTGGCAGTTTCTGCNNNAGATCGGAAG |
| 58160 | TATGAGTGTGGAGTCGTTGCCACTCCAAGGCAGATGCCATAATNNNAGATCGGAAG |
| 54497 | TATGAGTGTGGAGTCGTTGCTGGGGAGGAGAAAGAAGAACTGGNNNAGATCGGA |
| 106862 | TATGAGTGTGGAGTCGTTGCAAAAATATTAGCTGGGCGCCACCNNNAGATCGGAA |
| 99420 | TATGAGTGTGGAGTCGTTGCGCCTTTCTCTGCTTCACTCACTCNNNAGATCGGAAGA |
| 52129 | TATGAGTGTGGAGTCGTTGCACTCCAAATGCTGTAAGGCGTAGNNNAGATCGGAA |
| 45717 | TATGAGTGTGGAGTCGTTGCACACAGAGCAGAAAGCTAAGTCCNNNAGATCGGAA |
| 54535 | TATGAGTGTGGAGTCGTTGCTCCAGCCCAATAATTGTCACTGGNNNAGATCGGAAG |
| 97649 | TATGAGTGTGGAGTCGTTGCCAGCTACGTTTGCATGATAACGCNNNAGATCGGAAG |
| 44461 | TATGAGTGTGGAGTCGTTGCGAAGGCAGGACATTTTACTCCCCNNNAGATCGGAAG |
| 60568 | TATGAGTGTGGAGTCGTTGCTACCCACGTTGACAACAAAACCTNNNAGATCGGAAG |
| 32462 | TATGAGTGTGGAGTCGTTGCGCGTCTCTGTCAGTGGAGAGTAANNNAGATCGGAA |
| 48106 | TATGAGTGTGGAGTCGTTGCACCTTTCTGTAAGTGGTCCTTGCNNNAGATCGGAAG |
| 63758 | TATGAGTGTGGAGTCGTTGCGGAAGGGGGAGTAAGGAAGAAGTNNNAGATCGG |
| 41773 | TATGAGTGTGGAGTCGTTGCGCTTATGGGAACCCCACTGATTANNNAGATCGGAAG |
| 40731 | TATGAGTGTGGAGTCGTTGCGGGATTGTTCCTGCACTTCCATANNNAGATCGGAAG |
| 109537 | TATGAGTGTGGAGTCGTTGCGTATGATGCAAAGGTCCCATCTCANNNAGATCGGAA |
| 45952 | TATGAGTGTGGAGTCGTTGCTGCCTGAAATTTGACTTCTGTTCCNNNAGATCGGAA |
| 42041 | TATGAGTGTGGAGTCGTTGCTTCTTTGCGGTGGAGCCTCTTTANNNAGATCGGAAG |
| 53333 | TATGAGTGTGGAGTCGTTGCTTTGCGATCCCATCACTACATAGGNNNAGATCGGAA |
| 67456 | TATGAGTGTGGAGTCGTTGCAGGGCCAGTGGAGGATCTTTATTNNNAGATCGGAA |
| 44613 | TATGAGTGTGGAGTCGTTGCAGGGGTAAAGCACATTATTGCCCNNNAGATCGGAA |
| 106823 | TATGAGTGTGGAGTCGTTGCACCCACACAGACATTTATCTACATCTNNNAGATCGGA |
| 69335 | TATGAGTGTGGAGTCGTTGCAATGTAGCTTTCACGGGTGGTTTNNNAGATCGGAA |
| 49724 | TATGAGTGTGGAGTCGTTGCCCCCCTGTTCACTTGCTATCTTCNNNAGATCGGAAGA |
| 29854 | TATGAGTGTGGAGTCGTTGCAGGTGCAGGCTAATATACCTGATNNNAGATCGGAA |
| 55399 | TATGAGTGTGGAGTCGTTGCGGCACCAAACAGAATTCACCATGNNNAGATCGGAAG |
| 56414 | TATGAGTGTGGAGTCGTTGCCTCACTACCATTGCTTCCGTCTGNNNAGATCGGAAGA |
| 109233 | TATGAGTGTGGAGTCGTTGCTGTGTTCTTGCTCTCACTCTCTCACCCNNNAGATCGG |
| 65976 | TATGAGTGTGGAGTCGTTGCCCTCTGTCTCTCTGTGTACCTGTNNNAGATCGGAAGA |
| 105723 | TATGAGTGTGGAGTCGTTGCCCTTCCTTCTTTGTACCGGTTTCTNNNAGATCGGAAG |
| 66771 | TATGAGTGTGGAGTCGTTGCAGGAGGCACACAACTTCATTTGTNNNAGATCGGAAG |
| 33089 | TATGAGTGTGGAGTCGTTGCCACTTACCCACACACAAGACACANNNAGATCGGAAG |
| 30474 | TATGAGTGTGGAGTCGTTGCACCCATTGAGCTGCTGAAAGAAANNNAGATCGGAA |
| 40443 | TATGAGTGTGGAGTCGTTGCGAAACCCGGTCATCACATCTTGANNNAGATCGGAAG |
| 68931 | TATGAGTGTGGAGTCGTTGCCCACTGTAAGCCACAGGCTATTTNNNAGATCGGAAG |
| 69577 | TATGAGTGTGGAGTCGTTGCGAGAGAGAAGCTGGGTTGGTTTTNNNAGATCGGAA |

44061 TATGAGTGTGGAGTCGTTGCACAACCATCACCACTTGTAGACCNNNAGATCGGAAG
97512 TATGAGTGTGGAGTCGTTGCTGATTGGCACGCACCTCTTTTNNNAGATCGGAAGAG
49556 TATGAGTGTGGAGTCGTTGCGGATCAGTTCTGCCTGGAGATTCNNNAGATCGGAA
47447 TATGAGTGTGGAGTCGTTGCCTCTGTAGCCTAGCCTGAAATGCNNNAGATCGGAAG
100014 TATGAGTGTGGAGTCGTTGCGTGGAAGCATTCCTGTTTCATCTNNNAGATCGGAAG
64794 TATGAGTGTGGAGTCGTTGCCCAACCACCATTAGGAAACAGGTNNNAGATCGGAAG
49120 TATGAGTGTGGAGTCGTTGCCCTTTCTCCTCATTGGCCTTCTCNNNAGATCGGAAGA
62580 TATGAGTGTGGAGTCGTTGCACTGGAACCATCAGCCTGAATCTNNNAGATCGGAAG
36409 TATGAGTGTGGAGTCGTTGCATGCCTGATGCCTAGCTACATCANNNAGATCGGAAG
96554 TATGAGTGTGGAGTCGTTGCCGAGGGTACTGTGAACTTTACGTNNNAGATCGGAA
68796 TATGAGTGTGGAGTCGTTGCCGAGAACCAAGTTGGCAACATTTNNNAGATCGGAA
59970 TATGAGTGTGGAGTCGTTGCTACACAGGATACGTGGGAACACTNNNAGATCGGAA
67822 TATGAGTGTGGAGTCGTTGCAATGTGGGAAGCACCTTTTCCTTNNNAGATCGGAAG
103121 TATGAGTGTGGAGTCGTTGCTATTCTTCCAACCCAAATGTCCAACANNNAGATCGGA
32045 TATGAGTGTGGAGTCGTTGCCTGTGACCCACACACATCATGAANNNAGATCGGAAG
44205 TATGAGTGTGGAGTCGTTGCATGAACCAGTGGATGCTGTTACCNNNAGATCGGAA
110137 TATGAGTGTGGAGTCGTTGCTTCCAGATCCATGGTATTCAGTCNNNAGATCGGAAG
49164 TATGAGTGTGGAGTCGTTGCTGCCACCAAAAACCCAAAAAGTCNNNAGATCGGAAG
98804 TATGAGTGTGGAGTCGTTGCTCCACCTTCTAGAGAAGCCGCNNNAGATCGGAAGAG
48773 TATGAGTGTGGAGTCGTTGCTGCTTATGGACACATTCTGCCTCNNNAGATCGGAAG
33074 TATGAGTGTGGAGTCGTTGCCGTATCAGACAGTGATGCACACANNNAGATCGGAA
54490 TATGAGTGTGGAGTCGTTGCACAAGTGCAAGCATTCAAACTGGNNNAGATCGGAA
52716 TATGAGTGTGGAGTCGTTGCAAGTTAGGGAGGCTAGGGAAAGGNNNAGATCGGA
39461 TATGAGTGTGGAGTCGTTGCTGTGGATTATGAGCAACTTGTGGANNNAGATCGGA
108684 TATGAGTGTGGAGTCGTTGCCTGGAACCATCAGCCTGAATCTAANNNAGATCGGAA
54878 TATGAGTGTGGAGTCGTTGCGAGGTTACAGATCTTTCCCGTGGNNNAGATCGGAA
30678 TATGAGTGTGGAGTCGTTGCGTTGTGTCCAAGGCCTAGTGAAANNNAGATCGGAA
51376 TATGAGTGTGGAGTCGTTGCACAAAGGCCTGGAATTGGAAGAGNNNAGATCGGAA
60525 TATGAGTGTGGAGTCGTTGCGCATGATAGCACGTGACTGTACTNNNAGATCGGAA
34773 TATGAGTGTGGAGTCGTTGCCCTGAGAGAATGGAGTTTTGCCANNNAGATCGGAA
68147 TATGAGTGTGGAGTCGTTGCCAGGACACTCACAAGCACTTCTTNNNAGATCGGAAG
69666 TATGAGTGTGGAGTCGTTGCGCCATCAGCAGCAAGATGTTTTTNNNAGATCGGAAG
65647 TATGAGTGTGGAGTCGTTGCGGCTCATTCATCCTCAGGAATGTNNNAGATCGGAAG
39832 TATGAGTGTGGAGTCGTTGCTGTTTTGTTCAACAGTGGACTGANNNAGATCGGAAG
34989 TATGAGTGTGGAGTCGTTGCCTGTGAAACACCAGTTCTGTCCANNNAGATCGGAAG
58568 TATGAGTGTGGAGTCGTTGCTACACACATCCTGTGGTCTCCATNNNAGATCGGAAG
37727 TATGAGTGTGGAGTCGTTGCACACATGAGGACAAAGAAAACAGANNNAGATCGGA
68973 TATGAGTGTGGAGTCGTTGCCCCTTTAGCCTGGGTTTCACTTTNNNAGATCGGAAGA
35568 TATGAGTGTGGAGTCGTTGCTGAGCAGGAAGAAAGAAAAGGCANNNAGATCGGA
51473 TATGAGTGTGGAGTCGTTGCAGACATGCAAGGGAGAGAGAGAGNNNAGATCGGA
62537 TATGAGTGTGGAGTCGTTGCTAGGCGCTTCTCTTTGTTTTGCTNNNAGATCGGAAG
55429 TATGAGTGTGGAGTCGTTGCTATTTGGTTCCCAGATGGCCATGNNNAGATCGGAAG
58365 TATGAGTGTGGAGTCGTTGCCTCATATGCCTCCCCTGAGACATNNNAGATCGGAAG

| | |
|---|---|
| 116885 | TATGAGTGTGGAGTCGTTGCGTTTTTGCTAGGGTCTTGCTGTCNNNAGATCGGAAG |
| 30697 | TATGAGTGTGGAGTCGTTGCGAGCAACTGGGATCTGTTTGAAANNNAGATCGGAA |
| 30543 | TATGAGTGTGGAGTCGTTGCACCACTACCTGGGTCACTAGAAANNNAGATCGGAAG |
| 63515 | TATGAGTGTGGAGTCGTTGCTCCTGTTCCAGTTGCTTACTTTCTNNNAGATCGGAAG |
| 64593 | TATGAGTGTGGAGTCGTTGCCGTATCTTTGAAAACCAATCACCGTNNNAGATCGGA |
| 36232 | TATGAGTGTGGAGTCGTTGCTGGAGATGGTAGTTATGAACTTGCANNNAGATCGG |
| 30105 | TATGAGTGTGGAGTCGTTGCAAGCACTCTTCCTCCAAAGAAAANNNAGATCGGAAG |
| 56302 | TATGAGTGTGGAGTCGTTGCTTTATTAGCGTCACTCAGAATCTGNNNAGATCGGAA |
| 67944 | TATGAGTGTGGAGTCGTTGCCCTGTGGGATGAAAGAACTGCTTNNNAGATCGGAA |
| 59376 | TATGAGTGTGGAGTCGTTGCACACTCTTTGGGTGCTTGTTGATNNNAGATCGGAAG |
| 44046 | TATGAGTGTGGAGTCGTTGCTGCGAGGAAGACTGGAGACCNNNAGATCGGAAGA |
| 49962 | TATGAGTGTGGAGTCGTTGCGTTGCACATGGATGGACTGTTTCNNNAGATCGGAA |
| 51294 | TATGAGTGTGGAGTCGTTGCTTTTGCTAGGGTCTTGCTGTCAGNNNAGATCGGAAG |
| 58481 | TATGAGTGTGGAGTCGTTGCGTAGAAACAATCCAAGGGCCCATNNNAGATCGGAA |
| 45968 | TATGAGTGTGGAGTCGTTGCTGTGAGTTTCTTCCCAAGAATTTCCNNNAGATCGGA |
| 33809 | TATGAGTGTGGAGTCGTTGCTGGACTTACGAGACATACCTACANNNAGATCGGAAG |
| 69627 | TATGAGTGTGGAGTCGTTGCCTGGGCTCGGAGGTTTGATTTTTNNNAGATCGGAA |
| 40949 | TATGAGTGTGGAGTCGTTGCGAGTTTATGCTAGGGCAGAACTANNNAGATCGGAA |
| 34248 | TATGAGTGTGGAGTCGTTGCACCACAAATCTCCTAGTCACCCANNNAGATCGGAAG |
| 33567 | TATGAGTGTGGAGTCGTTGCGTTGTTGGGGGACACTTTAGACANNNAGATCGGAA |
| 100098 | TATGAGTGTGGAGTCGTTGCACCTCTGTTCCCTTCTGGAAGAGNNNAGATCGGAAG |
| 100342 | TATGAGTGTGGAGTCGTTGCTCTTGCTCAGTTTCCTGGCAGGNNNAGATCGGAAGA |
| 37712 | TATGAGTGTGGAGTCGTTGCGGCTGAGGGCTTAGAAGGTAAGANNNAGATCGGA |
| 46162 | TATGAGTGTGGAGTCGTTGCCTGGAAACAATGCAGCTTCAAGCNNNAGATCGGAA |
| 59384 | TATGAGTGTGGAGTCGTTGCCACCACCATGCTGAGATTTTGATNNNAGATCGGAAG |
| 35373 | TATGAGTGTGGAGTCGTTGCGCCTCTGATCAGTAAAGAGAGCANNNAGATCGGAA |
| 116121 | TATGAGTGTGGAGTCGTTGCTGTGTCTAGTTGTGCCTTCTCTGNNNAGATCGGAAG |
| 38734 | TATGAGTGTGGAGTCGTTGCATTCCAGTGACAGTACACCAGGANNNAGATCGGAA |
| 65269 | TATGAGTGTGGAGTCGTTGCTTTGGAGAAAGAAAGGCCATGGTNNNAGATCGGAA |
| 57977 | TATGAGTGTGGAGTCGTTGCCCCCCGAATACAGTCATTGCAATNNNAGATCGGAAG |
| 50259 | TATGAGTGTGGAGTCGTTGCGTTTTGGGGACTGAACTCCCAAGNNNAGATCGGAA |
| 46244 | TATGAGTGTGGAGTCGTTGCGAGAGAGAACTGGCCTTGTAAGCNNNAGATCGGAA |
| 66935 | TATGAGTGTGGAGTCGTTGCTGAGAATTCCTGCTTCCCCAATTNNNAGATCGGAAG |
| 34606 | TATGAGTGTGGAGTCGTTGCTCCTGGTCCTATACACATAGCCANNNAGATCGGAAG |
| 66328 | TATGAGTGTGGAGTCGTTGCAGCAGTCACCAAAATACTGAGTGTNNNAGATCGGA |
| 31299 | TATGAGTGTGGAGTCGTTGCTAATGATGGGAGGGAGGATGCAANNNAGATCGGA |
| 36086 | TATGAGTGTGGAGTCGTTGCACATTTTCAGTTGACCAGCTGCANNNAGATCGGAAG |
| 41244 | TATGAGTGTGGAGTCGTTGCTCCCTGCACCACCAGTAAATCTANNNAGATCGGAAG |
| 43386 | TATGAGTGTGGAGTCGTTGCCTGCTTGTCTAGAGGACCCTGACNNNAGATCGGAAG |
| 37776 | TATGAGTGTGGAGTCGTTGCAACAAATCTGCATTTCTGCACAGANNNAGATCGGAA |
| 66955 | TATGAGTGTGGAGTCGTTGCGGCTTGCAAAAGTGTGAGCAATTNNNAGATCGGAA |
| 42131 | TATGAGTGTGGAGTCGTTGCCACCACAGCATGGATGGAAAAACNNNAGATCGGAA |
| 30367 | TATGAGTGTGGAGTCGTTGCTGTAAGAGATGGGGAGGAGCAAANNNAGATCGGA |

| | |
|---|---|
| 49963 | TATGAGTGTGGAGTCGTTGCTGTGCTGGTATTGACCCTGTTTCNNNAGATCGGAAG |
| 57819 | TATGAGTGTGGAGTCGTTGCGCTGACTGTACAGGGTGTGAAATNNNAGATCGGAA |
| 34546 | TATGAGTGTGGAGTCGTTGCCCAGAAGAAATGTAATGCAAGCCANNNAGATCGGA |
| 65002 | TATGAGTGTGGAGTCGTTGCTGCTACCTTTACCTGCCTTAGGTNNNAGATCGGAAG |
| 71451 | TATGAGTGTGGAGTCGTTGCACACACACATACATATAGCCATTGTNNNAGATCGGA |
| 47367 | TATGAGTGTGGAGTCGTTGCGAGACAGAACGACGAAAATTGGCNNNAGATCGGAA |
| 61026 | TATGAGTGTGGAGTCGTTGCAGGCTGTACTCTTTCAAGTCCCTNNNAGATCGGAAG |
| 52168 | TATGAGTGTGGAGTCGTTGCCCCCTGCTCTCTACTCTCTGTAGNNNAGATCGGAAGA |
| 36050 | TATGAGTGTGGAGTCGTTGCGGAAGAGATCTATAAACTGCACTGCANNNAGATCG |
| 36440 | TATGAGTGTGGAGTCGTTGCGCTACTTTAGAGAGGGGGCATCANNNAGATCGGAA |
| 57223 | TATGAGTGTGGAGTCGTTGCTGCTTTGGGCTGGGTCTATATTGNNNAGATCGGAA |
| 52935 | TATGAGTGTGGAGTCGTTGCTTCAACTACCTATGGGGACCAGGNNNAGATCGGAA |
| 56658 | TATGAGTGTGGAGTCGTTGCACACCCCAAAGTTCACTCATAGTGNNNAGATCGGAA |
| 116913 | TATGAGTGTGGAGTCGTTGCTAGCCTTGGTAACCCCATTTTTACCNNNAGATCGGAA |
| 66755 | TATGAGTGTGGAGTCGTTGCGCAGGATGAATGTGGCAATTTGTNNNAGATCGGAA |
| 59027 | TATGAGTGTGGAGTCGTTGCACAGGTATCAGAGCCACAGAGATNNNAGATCGGAA |
| 68093 | TATGAGTGTGGAGTCGTTGCGGGTTTGGGGTGTTGAATCTCTTNNNAGATCGGAA |
| 30410 | TATGAGTGTGGAGTCGTTGCTGTAGGACCAAGCAACTTGCAAANNNAGATCGGAA |
| 65216 | TATGAGTGTGGAGTCGTTGCAAAGAGAGAGACAGTCTGTGGGTNNNAGATCGGA |
| 51583 | TATGAGTGTGGAGTCGTTGCTGTGTGTGAGTAGAGGGAAGGAGNNNAGATCGGA |
| 63578 | TATGAGTGTGGAGTCGTTGCCCAGGCTCATGTAGTGCTTTTCTNNNAGATCGGAAG |
| 30846 | TATGAGTGTGGAGTCGTTGCAGCAACAACAATAATGTCCTAACAANNNAGATCGGA |
| 117919 | TATGAGTGTGGAGTCGTTGCTCTCGCTCTCTCTTGTATGTGGGTNNNAGATCGGAA |
| 97516 | TATGAGTGTGGAGTCGTTGCTACAAGGGACCAGGCTTTTTCCNNNAGATCGGAAGA |
| 51019 | TATGAGTGTGGAGTCGTTGCCAGCAAAATCCTTTTCCCAGCAGNNNAGATCGGAAG |
| 39744 | TATGAGTGTGGAGTCGTTGCTGGGGTGGTATTTGATTCCTATGANNNAGATCGGA |
| 60244 | TATGAGTGTGGAGTCGTTGCTGTTTTGAATCCTGCTTCTTTCACTNNNAGATCGGAA |
| 33130 | TATGAGTGTGGAGTCGTTGCTCCCACTATTCCATGAAACTACACANNNAGATCGGAA |
| 100182 | TATGAGTGTGGAGTCGTTGCCCTTGGCTAAAAGGGACCAACATAANNNAGATCGG |
| 95110 | TATGAGTGTGGAGTCGTTGCCCTCTCCGAAACTGAGGAGTAGTNNNAGATCGGAA |
| 38516 | TATGAGTGTGGAGTCGTTGCATAGACAACTACAGAGGGCACGANNNAGATCGGAA |
| 59268 | TATGAGTGTGGAGTCGTTGCAGTTTCCCTACCCTATGCATGATNNNAGATCGGAAG |
| 46314 | TATGAGTGTGGAGTCGTTGCTGGTGTGTATTTCTTCTCCCAGCNNNAGATCGGAAG |
| 44290 | TATGAGTGTGGAGTCGTTGCAACAAGACATCAAGGCAAGACCCNNNAGATCGGAA |
| 54304 | TATGAGTGTGGAGTCGTTGCTACATAAGCCCACACCCAAATGGNNNAGATCGGAAG |
| 48885 | TATGAGTGTGGAGTCGTTGCCCCACTTTCACACTTAGAGGCTCNNNAGATCGGAAG |
| 55989 | TATGAGTGTGGAGTCGTTGCTGGGAATTGCATAAGTGTCCCTGNNNAGATCGGAA |
| 53438 | TATGAGTGTGGAGTCGTTGCGCCAGAAAGCAGGGAGATTTAGGNNNAGATCGGA |
| 45987 | TATGAGTGTGGAGTCGTTGCGCCCACAAACAAAACAACTTTCCNNNAGATCGGAAG |
| 64717 | TATGAGTGTGGAGTCGTTGCGAGAGAGAAGTGTGCACAAAGGTNNNAGATCGGA |
| 55508 | TATGAGTGTGGAGTCGTTGCGGAGAACCCTGATTGCTCTCATGNNNAGATCGGAAG |
| 42435 | TATGAGTGTGGAGTCGTTGCCACACAGGCACACAGAATTGAACNNNAGATCGGAA |
| 108732 | TATGAGTGTGGAGTCGTTGCGTGAGAGGCAAGAAGGAGTAAGGANNNAGATCGG |

59278 TATGAGTGTGGAGTCGTTGCGCCTGAATGTGTCCTTGGATGATNNNAGATCGGAA

100059 TATGAGTGTGGAGTCGTTGCAATGAAGAAGCTGAGGTATAGCCACNNNAGATCGG

68693 TATGAGTGTGGAGTCGTTGCCGTGCCTGGTGCATATATGTGTTNNNAGATCGGAAG

59303 TATGAGTGTGGAGTCGTTGCTGTTGCATTTGAGGCTTCCTGATNNNAGATCGGAAG

33730 TATGAGTGTGGAGTCGTTGCTGTTTGGCATTTCCCTCTTGACANNNAGATCGGAAG

33401 TATGAGTGTGGAGTCGTTGCATTCCATGTTTGGTGTGGTCACANNNAGATCGGAAG

32993 TATGAGTGTGGAGTCGTTGCACCCTGACCAATACAGAAACACANNNAGATCGGAAG

39986 TATGAGTGTGGAGTCGTTGCTGGGTTCCAATTCTATTTGGCTGANNNAGATCGGAA

70556 TATGAGTGTGGAGTCGTTGCACACACACCCCTTTTGAGACTTCNNNAGATCGGAAG

56587 TATGAGTGTGGAGTCGTTGCATTACCTGGTTTTGGAGGCAGTGNNNAGATCGGAA

37262 TATGAGTGTGGAGTCGTTGCCTTCCTCCCCAGATCTCAGTTCANNNAGATCGGAAGA

63106 TATGAGTGTGGAGTCGTTGCCTTTCCCCTCCCCATCAAAGTCTNNNAGATCGGAAGA

38323 TATGAGTGTGGAGTCGTTGCTTCTAGCTCATTTTCTCAATCTGAGANNNAGATCGGA

98105 TATGAGTGTGGAGTCGTTGCCATTCTGGGACTGCTTTCCAAAANNNAGATCGGAAG

43441 TATGAGTGTGGAGTCGTTGCACATGAACCAGAACCCACTTGACNNNAGATCGGAAG

64112 TATGAGTGTGGAGTCGTTGCTTGGTATGGGCTAAGAGCTCAGTNNNAGATCGGAA

66682 TATGAGTGTGGAGTCGTTGCTCTTGGTTCACGGAGTTTCTTGTNNNAGATCGGAAG

68009 TATGAGTGTGGAGTCGTTGCGCTGGCGTAGTAAGGGAAATCTTNNNAGATCGGAA

37587 TATGAGTGTGGAGTCGTTGCCAAGGGGCAAAGACAGATCAAGANNNAGATCGGAA

51853 TATGAGTGTGGAGTCGTTGCATAGGTAACCCCCAGTGCTGAGNNNAGATCGGAAG

62626 TATGAGTGTGGAGTCGTTGCTGTTCCATCTAAACCACAAGCATCTNNNAGATCGGAA

36038 TATGAGTGTGGAGTCGTTGCTTGGGCTGTAATTTCCAACTGCANNNAGATCGGAAG

30374 TATGAGTGTGGAGTCGTTGCTTCAGAGGGGAGAAAAAGGCAAANNNAGATCGGA

50719 TATGAGTGTGGAGTCGTTGCCCATGTTCTTCACCCTCAGACAGNNNAGATCGGAAG

41590 TATGAGTGTGGAGTCGTTGCAAGCAGGGAGGGTCATATTGGTANNNAGATCGGAA

95107 TATGAGTGTGGAGTCGTTGCGTGCAGGTAACTCTTCATCCGTTNNNAGATCGGAAG

101466 TATGAGTGTGGAGTCGTTGCGCTCACATGCTCTCAGTGAATAGTANNNAGATCGG

60548 TATGAGTGTGGAGTCGTTGCTGGGCACTGATGTATTGGTTACTNNNAGATCGGAA

54282 TATGAGTGTGGAGTCGTTGCTTCCAAAGACTTGTCCCTTTGGGNNNAGATCGGAAG

40777 TATGAGTGTGGAGTCGTTGCCTCCTGCAAATAGCTGGCTCATANNNAGATCGGAAG

102698 TATGAGTGTGGAGTCGTTGCTGGGACACACTGTGGTTGTCANNNAGATCGGAAGA

56508 TATGAGTGTGGAGTCGTTGCGAGGGGAGAGAGAGAGACAAGTGNNNAGATCGG

59031 TATGAGTGTGGAGTCGTTGCGGGCAAGGGAGAAAGAGAGAGATNNNAGATCGG

40481 TATGAGTGTGGAGTCGTTGCAGATGTGAAAGACTTATCGGTTGANNNAGATCGG

51690 TATGAGTGTGGAGTCGTTGCTGAATGAAGCCCTGAATTGGGAGNNNAGATCGGAA

64306 TATGAGTGTGGAGTCGTTGCAATACATGTCAGAAAAACGAAATGAGTNNNAGATC

54696 TATGAGTGTGGAGTCGTTGCTCTCTGTGTTGTAACATGGCTGGNNNAGATCGGAAG

50898 TATGAGTGTGGAGTCGTTGCTCCAAATCCACAGATTCAGCCAGNNNAGATCGGAAG

56396 TATGAGTGTGGAGTCGTTGCTCTCTCTCCAGCCATCTTCTCTGNNNAGATCGGAAGA

39713 TATGAGTGTGGAGTCGTTGCTTACACCTTCTACCGTGGGATGANNNAGATCGGAAG

100299 TATGAGTGTGGAGTCGTTGCACTCACCGGGGTAGCCAGAAAAGNNNAGATCGGAA

63633 TATGAGTGTGGAGTCGTTGCGTAAGCCCATGCTGATCCAAAGTNNNAGATCGGAAG

37230 TATGAGTGTGGAGTCGTTGCTCTGCCAAAAGAGACCTATCTTCANNNAGATCGGAA

| | |
|---|---|
| 52520 | TATGAGTGTGGAGTCGTTGCGTCTAAATCAAAATGCTGGGGCGNNNAGATCGGAA |
| 54819 | TATGAGTGTGGAGTCGTTGCCATCTGGGAGGAACTTGAAGTGGNNNAGATCGGAA |
| 47957 | TATGAGTGTGGAGTCGTTGCAATGAATGGAGGGATGATGGTGCNNNAGATCGGA |
| 54837 | TATGAGTGTGGAGTCGTTGCTTTTTCTTGCCTTGTTCCAGTGGNNNAGATCGGAAG |
| 108723 | TATGAGTGTGGAGTCGTTGCATGGGGACCAGGCAGACAATNNNAGATCGGAAGA |
| 63073 | TATGAGTGTGGAGTCGTTGCACCTACACAATTGCAAAACTTCTCTNNNAGATCGGAA |
| 56964 | TATGAGTGTGGAGTCGTTGCAGTCCCTTGAATCTTCAGCTGTGNNNAGATCGGAAG |
| 71646 | TATGAGTGTGGAGTCGTTGCGCAGAGCAAGAGTATGAGCATGTTTNNNAGATCGG |
| 67274 | TATGAGTGTGGAGTCGTTGCAGGGAGGAGAAGCACTTCAGATTNNNAGATCGGAA |
| 103183 | TATGAGTGTGGAGTCGTTGCCCCCAAGGCAACCATTTTCACATCNNNAGATCGGAAG |
| 30425 | TATGAGTGTGGAGTCGTTGCACACATAGACACACACTCATCAAANNNAGATCGGAA |
| 105681 | TATGAGTGTGGAGTCGTTGCTGCCATGTAGTCTCCAATTAGTGNNNAGATCGGAAG |
| 63477 | TATGAGTGTGGAGTCGTTGCTCTCCCAAAGTGTGTAGTGTTCTNNNAGATCGGAAG |
| 47495 | TATGAGTGTGGAGTCGTTGCAGGACGTAAATTGTGACACATGCNNNAGATCGGAA |
| 35971 | TATGAGTGTGGAGTCGTTGCGCAATCCCCATCTTACAGATGCANNNAGATCGGAAG |
| 61060 | TATGAGTGTGGAGTCGTTGCGAAACTGCAAAAACCCAAAGCCTNNNAGATCGGAAG |
| 70851 | TATGAGTGTGGAGTCGTTGCGGGAGAGCTGAGGGTAGATGATGNNNAGATCGGA |
| 29850 | TATGAGTGTGGAGTCGTTGCCAGTGTCAGGAGAGATTTGGTTTNNNAGATCGGAA |
| 67837 | TATGAGTGTGGAGTCGTTGCGTGTTGGGCTAAGAAGGAAGCTTNNNAGATCGGAA |
| 32974 | TATGAGTGTGGAGTCGTTGCCTTTCCTCTGTGTGGCTGTAACANNNAGATCGGAAG |
| 49494 | TATGAGTGTGGAGTCGTTGCTTCTAGGCAGCACAGACCAATTCNNNAGATCGGAAG |
| 36518 | TATGAGTGTGGAGTCGTTGCGTGCATACTAATGCCAGGGTATCANNNAGATCGGA |
| 104310 | TATGAGTGTGGAGTCGTTGCTACTGCAATGAAAATTTTTGGCTTGNNNAGATCGGA |
| 43906 | TATGAGTGTGGAGTCGTTGCCCTTACATGCTACTTTCCCCACCNNNAGATCGGAAGA |
| 47780 | TATGAGTGTGGAGTCGTTGCTTGGTGCCAGATTATTCTGCTGCNNNAGATCGGAAG |
| 47414 | TATGAGTGTGGAGTCGTTGCTTGATTGATTGCAAGGTGTTGGCNNNAGATCGGAA |
| 30836 | TATGAGTGTGGAGTCGTTGCTGAGCAATGATGGAGAGGAACAANNNAGATCGGA |
| 109541 | TATGAGTGTGGAGTCGTTGCTCTTCACTCTCATCCTGCAGTGGNNNAGATCGGAAG |
| 36010 | TATGAGTGTGGAGTCGTTGCAAACAAACCCAATGCTCTATGCANNNAGATCGGAAG |
| 41311 | TATGAGTGTGGAGTCGTTGCTGGGAGCAGAGTGGATTTGTCTANNNAGATCGGAA |
| 45250 | TATGAGTGTGGAGTCGTTGCAAGTTCCTTAGTCTTTGCCTGCCNNNAGATCGGAAG |
| 47988 | TATGAGTGTGGAGTCGTTGCAACGTCAGGCTTAGAGTATGTGCNNNAGATCGGAA |
| 102284 | TATGAGTGTGGAGTCGTTGCTCTTTCAGTATGTCTTCAAATTCACTANNNAGATCGG |
| 68607 | TATGAGTGTGGAGTCGTTGCGGAGCTTACTTTGTGAGCCTGTTNNNAGATCGGAAG |
| 44492 | TATGAGTGTGGAGTCGTTGCTCTCTTTGGGTTTCATCTTCCCCNNNAGATCGGAAGA |
| 30872 | TATGAGTGTGGAGTCGTTGCTGCTTGTATTCACACTCCCACAANNNAGATCGGAAGA |
| 34110 | TATGAGTGTGGAGTCGTTGCACATACACACACACATCATAGACCANNNAGATCGGA |
| 32095 | TATGAGTGTGGAGTCGTTGCATGTCTGAAGGCAAATGCCTGAANNNAGATCGGAA |
| 38010 | TATGAGTGTGGAGTCGTTGCGACCAGGGAGGGTTCATATCAGANNNAGATCGGAA |
| 32676 | TATGAGTGTGGAGTCGTTGCTGAGCACTGACTTCTGGAAAACANNNAGATCGGAAG |
| 65797 | TATGAGTGTGGAGTCGTTGCAAACATGCCCCATTTCTGGATGTNNNAGATCGGAAG |
| 52609 | TATGAGTGTGGAGTCGTTGCGGTCTCGAACTCTCTAGGTCTCGNNNAGATCGGAAG |
| 57132 | TATGAGTGTGGAGTCGTTGCTCAGAAAAATGCAGAGAAATTTTGTGNNNAGATCG |

106816 TATGAGTGTGGAGTCGTTGCATAAAACATATGCCTTGGACGGGNNNAGATCGGAA
63805 TATGAGTGTGGAGTCGTTGCACCTTTCCTACAGTCCCTGAAGTNNNAGATCGGAAG
57837 TATGAGTGTGGAGTCGTTGCAAGAGGGCACCGTTTCCCTAAATNNNAGATCGGAAG
59864 TATGAGTGTGGAGTCGTTGCTGCCCCTTACAACTTTGTCAACTNNNAGATCGGAAGA
62836 TATGAGTGTGGAGTCGTTGCCTCTTGCCTGTAGTCCTACCTCTNNNAGATCGGAAGA
116674 TATGAGTGTGGAGTCGTTGCGCAATGTTGTGTTCACAGTCTAATTNNNAGATCGGA
29849 TATGAGTGTGGAGTCGTTGCTTTCCTCTTTCAGCCTTACAGGGNNNAGATCGGAAG
51304 TATGAGTGTGGAGTCGTTGCCTCTCCCCGTCTTTGTCATTCAGNNNAGATCGGAAGA
51233 TATGAGTGTGGAGTCGTTGCCCACGTCTCTGAACAGCTCAGNNNAGATCGGAAGAG
40231 TATGAGTGTGGAGTCGTTGCTACTTGCAGGGGAGTTATGGTGANNNAGATCGGAA
60649 TATGAGTGTGGAGTCGTTGCTTCAATCCACTCACAGAACACCTNNNAGATCGGAAG
40308 TATGAGTGTGGAGTCGTTGCTCACTGCTGTTTGTTCAGTGTGANNNAGATCGGAAG
42995 TATGAGTGTGGAGTCGTTGCTAGTAACAGACTGGGACCCTCACNNNAGATCGGAAG
30306 TATGAGTGTGGAGTCGTTGCAGTGAAAGTCCTTGGAACCCAAANNNAGATCGGAA
46950 TATGAGTGTGGAGTCGTTGCTTGGGAAAATTCAGGGAAGAGGCNNNAGATCGGA
71109 TATGAGTGTGGAGTCGTTGCGGCATTTCCTGCTCTTATGGACTNNNAGATCGGAAG
50083 TATGAGTGTGGAGTCGTTGCTCTAAAGGCAGCTTTCCCCAAAGNNNAGATCGGAAG
63035 TATGAGTGTGGAGTCGTTGCGCTTTTTGAGATGGAGGTCTCTCTNNNAGATCGGAA
35534 TATGAGTGTGGAGTCGTTGCTCAAGCATATCAAATCCCACGCANNNAGATCGGAAG
31348 TATGAGTGTGGAGTCGTTGCGGTCTCCAAAGACATCCTTGCAANNNAGATCGGAAG
43829 TATGAGTGTGGAGTCGTTGCCTTCCTGATCTGGGTTGTGAACCNNNAGATCGGAAG
40507 TATGAGTGTGGAGTCGTTGCAAGCCAATCATCCAGCTTGTTGANNNAGATCGGAAG
34400 TATGAGTGTGGAGTCGTTGCCCTTCCTGTGTAAAATCTGCCCANNNAGATCGGAAG
67799 TATGAGTGTGGAGTCGTTGCCCCTAAGTTCTCTGACCGTCCTTNNNAGATCGGAAGA
43799 TATGAGTGTGGAGTCGTTGCACACACACCAGAGAAACAGAACCNNNAGATCGGAA
42475 TATGAGTGTGGAGTCGTTGCCTCCTCTCCCCTCCCTAAGTAACNNNAGATCGGAAGA
42831 TATGAGTGTGGAGTCGTTGCACACATATTCCACCCAACAGCACNNNAGATCGGAAG
51364 TATGAGTGTGGAGTCGTTGCAAGGGGTGGTTGAGGAAGAAGAGNNNAGATCGG
66951 TATGAGTGTGGAGTCGTTGCGAGTGAGAAGGCACTGTCCAATTNNNAGATCGGAA
48956 TATGAGTGTGGAGTCGTTGCATACACAAGGCAAGGACTTGCTCNNNAGATCGGAA
104368 TATGAGTGTGGAGTCGTTGCATAGGGTTGGGGAAAGATGGGANNNAGATCGGAA
34499 TATGAGTGTGGAGTCGTTGCAAAATAGAAGCTTTGGCTTCCCANNNAGATCGGAAG
66097 TATGAGTGTGGAGTCGTTGCTCTTCATTTTTGCAGGATGCTGTNNNAGATCGGAAG
55357 TATGAGTGTGGAGTCGTTGCTCTGTGGTGTACGTCGTAACATGNNNAGATCGGAA
45733 TATGAGTGTGGAGTCGTTGCTTGACTTGAAGTTGCCTCAGTCCNNNAGATCGGAAG
46668 TATGAGTGTGGAGTCGTTGCCTTTCCCCCTCTGTGTTTCTAGCNNNAGATCGGAAGA
109159 TATGAGTGTGGAGTCGTTGCTGTGTGTGTAGCAATCTCTCTCTCTANNNAGATCGG
109254 TATGAGTGTGGAGTCGTTGCGTGGAGAATGTTGGGGAGGTNNNAGATCGGAAGA
97233 TATGAGTGTGGAGTCGTTGCCCACCAAGGGAATTCAATGGGNNNAGATCGGAAGA
96081 TATGAGTGTGGAGTCGTTGCACCCTGCTCCTCTTTTCTCAGGNNNAGATCGGAAGAG
30531 TATGAGTGTGGAGTCGTTGCTTCTTATTCTGCATACTGGGAGAAANNNAGATCGGA
52435 TATGAGTGTGGAGTCGTTGCTGTGTGTTCAAATTGGAATGCCGNNNAGATCGGAA
38192 TATGAGTGTGGAGTCGTTGCGAGGAGAAATGGAAGCAAAGGAGANNNAGATCGG

60322 TATGAGTGTGGAGTCGTTGCACTGGAATGATGAAAGCAGGACTNNNAGATCGGAA
33331 TATGAGTGTGGAGTCGTTGCTCCTTTCACTTTCAGTGTGCACANNNAGATCGGAAG
44192 TATGAGTGTGGAGTCGTTGCTAACAAGACCAACAAGGTACTATTACCNNNAGATCG
64211 TATGAGTGTGGAGTCGTTGCAATGTATGGTCAGCCAGGAGAGTNNNAGATCGGAA
67531 TATGAGTGTGGAGTCGTTGCCTGTGAAACACCCTCCATCACTTNNNAGATCGGAAG
31580 TATGAGTGTGGAGTCGTTGCGTTCCTGACACTGTGGCTAAGAANNNAGATCGGAA
54840 TATGAGTGTGGAGTCGTTGCGGTGATGAATCTCTGAGCAGTGGNNNAGATCGGAA
49970 TATGAGTGTGGAGTCGTTGCCCCTCTAAAGTCTCTGGAAATTTTCNNNAGATCGGAA
40204 TATGAGTGTGGAGTCGTTGCTTGTGGAAGGTGTAGAAGGGTGANNNAGATCGGA
43518 TATGAGTGTGGAGTCGTTGCAGGACATTTGCCAGGGAAACTACNNNAGATCGGAA
50166 TATGAGTGTGGAGTCGTTGCGGGATAGGTGCTTCAGTGGAAAGNNNAGATCGGA
49204 TATGAGTGTGGAGTCGTTGCCACAGGCAAAAACTTCACCAGTCNNNAGATCGGAAG
37463 TATGAGTGTGGAGTCGTTGCAGCCAGAGATTAGTAAGCACAAAGANNNAGATCGG
37164 TATGAGTGTGGAGTCGTTGCTGGCACTCAGTTATCGAAACTTCANNNAGATCGGAA
57811 TATGAGTGTGGAGTCGTTGCGGCTGGAAGGAAGAAGCTGAAATNNNAGATCGGA
51877 TATGAGTGTGGAGTCGTTGCCCCGAACTGTGGAAGTAAGTGAGNNNAGATCGGAA
65021 TATGAGTGTGGAGTCGTTGCTGCTCTATGATGTCTCTTCCGGTNNNAGATCGGAAG
69422 TATGAGTGTGGAGTCGTTGCCAGTTACCAAACCTGCGCATTTTNNNAGATCGGAAG
55314 TATGAGTGTGGAGTCGTTGCTGTCTCTTCACCAACTGGGAATGNNNAGATCGGAAG
55871 TATGAGTGTGGAGTCGTTGCGAAGAGACACTGAGCTGTGACTGNNNAGATCGGAA
69125 TATGAGTGTGGAGTCGTTGCCATCCTTCTGCATTGCATGCTTTNNNAGATCGGAAGA
68945 TATGAGTGTGGAGTCGTTGCGTCACGCCCAGTCATCTGTATTTNNNAGATCGGAAG
57987 TATGAGTGTGGAGTCGTTGCTTGGCACCTGAAGCTGAATCAATNNNAGATCGGAAG
105178 TATGAGTGTGGAGTCGTTGCGGGGCACTGCTACCTTACTTTAGNNNAGATCGGAAG
59170 TATGAGTGTGGAGTCGTTGCTGGAGTTTCTTGGATCCAGGGATNNNAGATCGGAA
36953 TATGAGTGTGGAGTCGTTGCTGTATGCTTCATTGAAAGGGTCANNNAGATCGGAA
107288 TATGAGTGTGGAGTCGTTGCCCTGGACTGATGGAAAGTAAGAGGNNNAGATCGGA
108712 TATGAGTGTGGAGTCGTTGCAAGACCAACAAGGTACTATTACCATNNNAGATCGGA
45052 TATGAGTGTGGAGTCGTTGCTGATTATCCAAGAACTGCAGGCCNNNAGATCGGAAG
52667 TATGAGTGTGGAGTCGTTGCAAGGGGAGCATGAAGGAAAAAGGNNNAGATCGGA
116840 TATGAGTGTGGAGTCGTTGCACTGGGACATTGAGAAGAAGGATGNNNAGATCGG
57275 TATGAGTGTGGAGTCGTTGCATGGATTTGCTTCTCCTCCCTTGNNNAGATCGGAAGA
33095 TATGAGTGTGGAGTCGTTGCTGCGAAAGGATTTCAAGGACACANNNAGATCGGAA
31833 TATGAGTGTGGAGTCGTTGCGCTTCAAATGTGGTCCAGAGGAANNNAGATCGGAA
116395 TATGAGTGTGGAGTCGTTGCGTGTGGGGGTGCATTTTGTATNNNAGATCGGAAGA
46155 TATGAGTGTGGAGTCGTTGCGCAGCAACACATAATCCTCAAGCNNNAGATCGGAAG
39550 TATGAGTGTGGAGTCGTTGCTGTAGTGGCAGATGCAAAAATGANNNAGATCGGAA
63415 TATGAGTGTGGAGTCGTTGCCCCCACAGAGTGTAGGATCTTCTNNNAGATCGGAAG
49830 TATGAGTGTGGAGTCGTTGCAGGACATTGGATTGGAGCTGTTCNNNAGATCGGAA
106345 TATGAGTGTGGAGTCGTTGCGGGGGCAGGGAGCAATTGATTANNNAGATCGGAA
50126 TATGAGTGTGGAGTCGTTGCGGGGTGCAGAAAGGGTAAGAAAGNNNAGATCGGA
108768 TATGAGTGTGGAGTCGTTGCCACACCAGAGAAACAGAACCAATGTNNNAGATCGGA
45074 TATGAGTGTGGAGTCGTTGCGAAATAACCCCACATCCTAGGCCNNNAGATCGGAAG

29955 TATGAGTGTGGAGTCGTTGCTTGAATGGTGACAAAATTGGTGANNNAGATCGGAA
31268 TATGAGTGTGGAGTCGTTGCTCAGACCCTATTCTACCTGGCAANNNAGATCGGAAG
57508 TATGAGTGTGGAGTCGTTGCTATTGCAAGGGCTTTGCTTGTTGNNNAGATCGGAAG
100908 TATGAGTGTGGAGTCGTTGCCAGTCCCTCTACTCCTACCCTGGNNNAGATCGGAAGA
32191 TATGAGTGTGGAGTCGTTGCCACCATTGGCAATCAGCATTGAANNNAGATCGGAAG
31009 TATGAGTGTGGAGTCGTTGCTCTCCTGTGCTTCCTGATACCAANNNAGATCGGAAG
65362 TATGAGTGTGGAGTCGTTGCATCCTTCTGTTCTTTCCCCTGGTNNNAGATCGGAAGA
43687 TATGAGTGTGGAGTCGTTGCTCCTCGACTTCCAATGAGGTTACNNNAGATCGGAAG
71499 TATGAGTGTGGAGTCGTTGCTGTGTGGGGGTGCATTTTGTATTNNNAGATCGGAA
61764 TATGAGTGTGGAGTCGTTGCTTCAGTGAGTGGGATACAGAGCTNNNAGATCGGAA
58635 TATGAGTGTGGAGTCGTTGCGTCAGAGATTTGCTCCTGAGCATNNNAGATCGGAAG
35769 TATGAGTGTGGAGTCGTTGCGGGCTTGACTAGAAAACATGGCANNNAGATCGGAA
65876 TATGAGTGTGGAGTCGTTGCAGGACAAAAACTGCAGAAACTGTNNNAGATCGGAA
109579 TATGAGTGTGGAGTCGTTGCTCTATGTGTGTATTACTAGAGCCTTCTNNNAGATCGG
39228 TATGAGTGTGGAGTCGTTGCACTGTAGCATATAGGGTGAAATGGANNNAGATCGG
57291 TATGAGTGTGGAGTCGTTGCACCATTTCCACTTCTGCTCCTTGNNNAGATCGGAAGA
100969 TATGAGTGTGGAGTCGTTGCCAGAGAAGGCAGGGGGAGAGAAANNNAGATCGGA
61107 TATGAGTGTGGAGTCGTTGCTGTTCACTGCTGTACTAGAGCCTNNNAGATCGGAAG
34098 TATGAGTGTGGAGTCGTTGCATGAAAAACCAAGTCCCAGACCANNNAGATCGGAAG
64913 TATGAGTGTGGAGTCGTTGCGTTGGTTTTACAGGAGTGGAGGTNNNAGATCGGAA
106791 TATGAGTGTGGAGTCGTTGCGCCACTGTTTAGTAGACTGCTTNNNAGATCGGAAGA
53142 TATGAGTGTGGAGTCGTTGCTACCACCTCTAAGACTGCAGAGGNNNAGATCGGAAG
52385 TATGAGTGTGGAGTCGTTGCAATGACTTCAGCAGACTGTACCGNNNAGATCGGAAG
96908 TATGAGTGTGGAGTCGTTGCCACCCCTACTTCTGTCCAGTAGTNNNAGATCGGAAGA
46358 TATGAGTGTGGAGTCGTTGCAAGAATGAGTTGAGTGGAGCAGCNNNAGATCGGA
54929 TATGAGTGTGGAGTCGTTGCGTTTGCGAATCTTCAACTGGTGGNNNAGATCGGAAG
51748 TATGAGTGTGGAGTCGTTGCAAGAAAATGGACTTGGGGTGGAGNNNAGATCGGA
69607 TATGAGTGTGGAGTCGTTGCTTTTGCTCTCACTGAAACAATTTTTNNNAGATCGGA
30384 TATGAGTGTGGAGTCGTTGCTTGGTTCTGTTACTGTCGGCAAANNNAGATCGGAAG
34660 TATGAGTGTGGAGTCGTTGCATAGTGTCAGTTTGAATGGGCCANNNAGATCGGAA
42201 TATGAGTGTGGAGTCGTTGCAACTGGCCTTCTGGAGATCAAACNNNAGATCGGAAG
31230 TATGAGTGTGGAGTCGTTGCTCCAAACGTGCATACAGTAGCAANNNAGATCGGAAG
66690 TATGAGTGTGGAGTCGTTGCGCATTTTCTACTCTTTTGAACAGTTGTNNNAGATCGG
52081 TATGAGTGTGGAGTCGTTGCACATGCCCAAAAGAAGCCTCTAGNNNAGATCGGAAG
65432 TATGAGTGTGGAGTCGTTGCGGATCAAAGCGAGATTCAGTGGTNNNAGATCGGAA
43551 TATGAGTGTGGAGTCGTTGCCTCTGGGTCTACCAGTTGCCTACNNNAGATCGGAAG
66047 TATGAGTGTGGAGTCGTTGCCGCCTAAAAAGTCATCAAGCTGTNNNAGATCGGAAG
59900 TATGAGTGTGGAGTCGTTGCCACAGGAAGGAAACATGGGAACTNNNAGATCGGAA
40607 TATGAGTGTGGAGTCGTTGCTACCAGCCAAGCACAAGGAAATANNNAGATCGGAA
45132 TATGAGTGTGGAGTCGTTGCCTTGGGTTCTATGAGAAGTGGCCNNNAGATCGGAA
49954 TATGAGTGTGGAGTCGTTGCCCCTCTACTCCTACCCTGGTTTCNNNAGATCGGAAGA
48560 TATGAGTGTGGAGTCGTTGCGACCACTAGGCAACAGGCTTATCNNNAGATCGGAAG
101743 TATGAGTGTGGAGTCGTTGCAGAGGTTTTGTTGATTGCCTACCANNNAGATCGGAA

43338 TATGAGTGTGGAGTCGTTGCGAATAGCCCAAGCAATCCTGGACNNNAGATCGGAA
51236 TATGAGTGTGGAGTCGTTGCAGCGTATCACCTGTCTAGCTCAGNNNAGATCGGAAG
31868 TATGAGTGTGGAGTCGTTGCGGGGAAAGATGGGAGAGTAGGAANNNAGATCGG
38357 TATGAGTGTGGAGTCGTTGCGGATGTTGCATGAGCTCTTGAGANNNAGATCGGAA
53849 TATGAGTGTGGAGTCGTTGCTAAAGTTGTATACGAAGGCCGGGNNNAGATCGGAA
43410 TATGAGTGTGGAGTCGTTGCGTGTTTCAGAGAGGGGAAGTGACNNNAGATCGGA
40791 TATGAGTGTGGAGTCGTTGCCCGCAAGCTAAGAGGCAAAGATANNNAGATCGGAA
52952 TATGAGTGTGGAGTCGTTGCATAGAAAAGCAGATCAGCCCAGGNNNAGATCGGAA
51209 TATGAGTGTGGAGTCGTTGCCTGGCTTTCCAAAAAGCACTCAGNNNAGATCGGAAG
47042 TATGAGTGTGGAGTCGTTGCGTGCTTTTTGAGTACTTTCCGGCNNNAGATCGGAAG
38785 TATGAGTGTGGAGTCGTTGCAGTGCTTGAACATACCAAGAGGANNNAGATCGGAA
64028 TATGAGTGTGGAGTCGTTGCTGTCCATAATATGCTGTTGTTAGCAGTNNNAGATCG
35465 TATGAGTGTGGAGTCGTTGCATCCCCAAACCAGATCTATAGCANNNAGATCGGAAG
66802 TATGAGTGTGGAGTCGTTGCACATCCTCTTTGACCTTCCTTTGTNNNAGATCGGAAG
109217 TATGAGTGTGGAGTCGTTGCCAGAGAGAGGCTTAGGTCCGATTGNNNAGATCGGA
54605 TATGAGTGTGGAGTCGTTGCTTCTTGACAAACCACTTCCCTGGNNNAGATCGGAAG
35644 TATGAGTGTGGAGTCGTTGCATAGAGCAAGTAGCTGAGAGGCANNNAGATCGGA
50198 TATGAGTGTGGAGTCGTTGCTCCCCTGGACCTTGGTAGTAAAGNNNAGATCGGAAG
102314 TATGAGTGTGGAGTCGTTGCAGCACCTGCACCTTTTATGTTCCNNNAGATCGGAAG
45209 TATGAGTGTGGAGTCGTTGCTGGTGGAGGTTATAGCATATGCCNNNAGATCGGAA
64878 TATGAGTGTGGAGTCGTTGCAGAAGAAACAGGAGTTCAGAGAGGTNNNAGATCG
61489 TATGAGTGTGGAGTCGTTGCTTGGCTTGAGAAAGAGAAATTCCTNNNAGATCGGA
104584 TATGAGTGTGGAGTCGTTGCATGTGGTCAAAACTCCCAAAAGNNNAGATCGGAAG
60479 TATGAGTGTGGAGTCGTTGCAAGACCTCTAAGTGTCAATGCTACTNNNAGATCGGA
39687 TATGAGTGTGGAGTCGTTGCGGTGGGGTAAGGGAAAGAGATGANNNAGATCGG
30252 TATGAGTGTGGAGTCGTTGCGCACATACACGAAGATGCACAAANNNAGATCGGAA
39278 TATGAGTGTGGAGTCGTTGCAGTGAGGATGGGAACATGATGGANNNAGATCGGA
63301 TATGAGTGTGGAGTCGTTGCGGAAGGGAACCAGAAGTTTGTCTNNNAGATCGGAA
29846 TATGAGTGTGGAGTCGTTGCATAAAGCAGCTGGGTTCAGTAGGNNNAGATCGGAA
42342 TATGAGTGTGGAGTCGTTGCTGGTGGGATGTTTCTCTGTCAACNNNAGATCGGAAG
59576 TATGAGTGTGGAGTCGTTGCTTCGCTGTCACGTTGGTTTCTATNNNAGATCGGAAG
44967 TATGAGTGTGGAGTCGTTGCGCACGAAATCAGAAATCTGAGCCNNNAGATCGGAA
109622 TATGAGTGTGGAGTCGTTGCGTGGAGGGTAGACATGGAGAAAANNNAGATCGGA
63037 TATGAGTGTGGAGTCGTTGCGTGGAGGTCACATCCTTCTCTCTNNNAGATCGGAAG
121970 TATGAGTGTGGAGTCGTTGCTTCCAAGCACAGCTATCTCCTTTNNNAGATCGGAAGA
32281 TATGAGTGTGGAGTCGTTGCGCCCCATCCCCAGATTTAGATAANNNAGATCGGAAG
49756 TATGAGTGTGGAGTCGTTGCGCACTTGGGTTCTGTGTTTCTTCNNNAGATCGGAAG
45764 TATGAGTGTGGAGTCGTTGCCTTGTAAATGAGCCCAGAGGTCCNNNAGATCGGAA
61301 TATGAGTGTGGAGTCGTTGCATGGTTGTCAACTGCCAAATCCTNNNAGATCGGAAG
54774 TATGAGTGTGGAGTCGTTGCTTCCTAACAGTGTGCTTGTCTGGNNNAGATCGGAAG
65729 TATGAGTGTGGAGTCGTTGCGTGTGAGGAGTGTGTATGCATGTNNNAGATCGGAA
56725 TATGAGTGTGGAGTCGTTGCCCCCAGTGTTTCAGAGTAAGGTGNNNAGATCGGAA
42660 TATGAGTGTGGAGTCGTTGCTGTGGTAGTGGAGTAACAACCACNNNAGATCGGAA

68357 TATGAGTGTGGAGTCGTTGCGGGGGCATAGAGAGAGCTTAGTTNNNAGATCGGA
67423 TATGAGTGTGGAGTCGTTGCAGCGTTCCTTGGTCATTCGTATTNNNAGATCGGAAG
42206 TATGAGTGTGGAGTCGTTGCCAGCAGCAATGGTTCTCTCAAACNNNAGATCGGAAG
99690 TATGAGTGTGGAGTCGTTGCAGTTTTGAAGCAGACACCTGGAANNNAGATCGGAA
42231 TATGAGTGTGGAGTCGTTGCTGTTGCCCATACCAGACTGAAACNNNAGATCGGAAG
36813 TATGAGTGTGGAGTCGTTGCGTTTCTTGGGCTCACCATTCTCANNNAGATCGGAAG
101298 TATGAGTGTGGAGTCGTTGCTGGAATGTAATGGCAAAATCACAACNNNAGATCGG
43087 TATGAGTGTGGAGTCGTTGCACAGGAACTCAGCAGAAGTTCACNNNAGATCGGAA
64686 TATGAGTGTGGAGTCGTTGCTTTTCTCACATCTTGGCTGTCGTNNNAGATCGGAAG
103171 TATGAGTGTGGAGTCGTTGCCTAATGGGTTCACCTTTCTTGCTGNNNAGATCGGAA
47623 TATGAGTGTGGAGTCGTTGCTTGGGCACGAAGATTTGTTATGCNNNAGATCGGAA
106348 TATGAGTGTGGAGTCGTTGCTCCCCAAACCAGATCTATAGCATACANNNAGATCGG
50247 TATGAGTGTGGAGTCGTTGCAACAGCACCTTTCTGAGACCAAGNNNAGATCGGAAG
49175 TATGAGTGTGGAGTCGTTGCCACTCATCCCTTCTGCTCAAGTCNNNAGATCGGAAGA
47521 TATGAGTGTGGAGTCGTTGCAGAGAGAGAGCATACAAGCATGCNNNAGATCGGAA
29959 TATGAGTGTGGAGTCGTTGCAAGGCTCAGGTAATTTCTCTGCCNNNAGATCGGAAG
42097 TATGAGTGTGGAGTCGTTGCTCTCCTCCATGCCCTTCCTTTTANNNAGATCGGAAGA
71658 TATGAGTGTGGAGTCGTTGCACTGTGATGCTGAGGCATGAAAANNNAGATCGGAA
44188 TATGAGTGTGGAGTCGTTGCGCTTGCCTGTCCTTTATTGTACCNNNAGATCGGAAG
39418 TATGAGTGTGGAGTCGTTGCGGCTTCCTCATCTCTAAGGTGGANNNAGATCGGAAG
71428 TATGAGTGTGGAGTCGTTGCAGTGTGTATGTATGTAGCAGTGTNNNAGATCGGAA
30686 TATGAGTGTGGAGTCGTTGCACACGGCTTGAGAAGATGTGAAANNNAGATCGGAA
117462 TATGAGTGTGGAGTCGTTGCATGTGCTGAACTACAGAAGCANNNAGATCGGAAGA
117302 TATGAGTGTGGAGTCGTTGCCTCTGTCTCTCAGGACAACTAGANNNAGATCGGAAG
29851 TATGAGTGTGGAGTCGTTGCTGGAAAAGCAAAGGAAGGAAGCTNNNAGATCGGA
95109 TATGAGTGTGGAGTCGTTGCTGCCAAGGACACAGAAACATTCANNNAGATCGGAA
54098 TATGAGTGTGGAGTCGTTGCAAAAGAGATTTCAGCTCCCTGGGNNNAGATCGGAA
49192 TATGAGTGTGGAGTCGTTGCTGCTCTGCTTGTTTTGAACAGTCNNNAGATCGGAAG
54431 TATGAGTGTGGAGTCGTTGCTTATCCAGTCCACCATTCGATGGNNNAGATCGGAAG
65782 TATGAGTGTGGAGTCGTTGCGAGCTTTGCTAAAGGGGAGATGTNNNAGATCGGAA
40970 TATGAGTGTGGAGTCGTTGCCGTCCACCAAAGATTGAGCACTANNNAGATCGGAAG
52406 TATGAGTGTGGAGTCGTTGCCCAGCATAAGGTGTATGATCCCGNNNAGATCGGAA
35831 TATGAGTGTGGAGTCGTTGCTTGAAGCAATCTCTGTAGTGGCANNNAGATCGGAA
38034 TATGAGTGTGGAGTCGTTGCTACAGCAGTCTCACCAAGTCAGANNNAGATCGGAAG
44671 TATGAGTGTGGAGTCGTTGCACCACCTTACCTTACCCTTATCCCNNNAGATCGGAAG
32186 TATGAGTGTGGAGTCGTTGCTTCTGGTTGGGGTTTGTTGTGAANNNAGATCGGAA
71344 TATGAGTGTGGAGTCGTTGCACAAGGACAAGAAAAAGGAAGGTNNNAGATCGGA
68752 TATGAGTGTGGAGTCGTTGCGCTGCCTTCCTGTTTTGTTTGTTNNNAGATCGGAAG
117723 TATGAGTGTGGAGTCGTTGCCCTCTACAACCATTGCTTGGGATCTNNNAGATCGGA
100438 TATGAGTGTGGAGTCGTTGCAACCTTTGCCTTTCTACCGGACNNNAGATCGGAAGA
29857 TATGAGTGTGGAGTCGTTGCGCTCTCACTCATTCTCAGCTCAGNNNAGATCGGAAG
30512 TATGAGTGTGGAGTCGTTGCTCAAGCTCTCCTGTCCTCAGAAANNNAGATCGGAAG
69533 TATGAGTGTGGAGTCGTTGCGGGGCAGGGTATGTCTTTCTTTTNNNAGATCGGAA

| | |
|---|---|
| 32284 | TATGAGTGTGGAGTCGTTGCGTGGAAGCTGAAAGCCAGGATAANNNAGATCGGA |
| 57933 | TATGAGTGTGGAGTCGTTGCTCTGCTGGGAAAAGGAAAGCAATNNNAGATCGGAA |
| 64451 | TATGAGTGTGGAGTCGTTGCCTGCTCTTGCACATTGGTCTAGTNNNAGATCGGAAG |
| 65499 | TATGAGTGTGGAGTCGTTGCTCCAGTCTCTCTCAGTTTGTGGTNNNAGATCGGAAG |
| 52932 | TATGAGTGTGGAGTCGTTGCTGTAGGGCTTGCAAATAACCAGGNNNAGATCGGAA |
| 43999 | TATGAGTGTGGAGTCGTTGCCTGAGAATTACTGCCCCTTCACCNNNAGATCGGAAG |
| 61876 | TATGAGTGTGGAGTCGTTGCAAAACAGAAGTGCCAGTGTAGCTNNNAGATCGGAA |
| 106366 | TATGAGTGTGGAGTCGTTGCTTGAGGAAGTGGGTCTGGTGTTGNNNAGATCGGA |
| 56858 | TATGAGTGTGGAGTCGTTGCGGAAAACAAGATGCCCATTGGTGNNNAGATCGGAA |
| 67282 | TATGAGTGTGGAGTCGTTGCGCTGGTTCACATCCTTGGAGATTNNNAGATCGGAAG |
| 53307 | TATGAGTGTGGAGTCGTTGCGAGAAATGGAGGATCGTGTGAGGNNNAGATCGGA |
| 50476 | TATGAGTGTGGAGTCGTTGCATTTCTGCCTCAAAGGGTGGAAGNNNAGATCGGAA |
| 109277 | TATGAGTGTGGAGTCGTTGCATCCTTTCAAAGACCAAGCTTTTGNNNAGATCGGAA |
| 65302 | TATGAGTGTGGAGTCGTTGCTACACACTCCATGGTCTGATGGTNNNAGATCGGAAG |
| 47222 | TATGAGTGTGGAGTCGTTGCAAACTCCAAGTCCTGATCATGGCNNNAGATCGGAAG |
| 57968 | TATGAGTGTGGAGTCGTTGCGAGATGTGGGAGGAAACTGCAATNNNAGATCGGA |
| 41248 | TATGAGTGTGGAGTCGTTGCGGAGGCGGTAAGTCTGGAATCTANNNAGATCGGAA |
| 48268 | TATGAGTGTGGAGTCGTTGCGGAAGTGGGGATGTCTCACAATCNNNAGATCGGAA |
| 95875 | TATGAGTGTGGAGTCGTTGCCTTGTCGATAGCTTGCTGGTGTACNNNAGATCGGAA |
| 29853 | TATGAGTGTGGAGTCGTTGCTTCTCAGGGTGTGGGCTTCTAATNNNAGATCGGAAG |
| 29958 | TATGAGTGTGGAGTCGTTGCGTCATAGTGTGTGTCAAGTGCCTNNNAGATCGGAA |
| 50438 | TATGAGTGTGGAGTCGTTGCGAAATGGAGTGCTCAGGAGGAAGNNNAGATCGGA |
| 71824 | TATGAGTGTGGAGTCGTTGCCCTCCAGTGGTCTCCTTAGCTGNNNAGATCGGAAGA |
| 70043 | TATGAGTGTGGAGTCGTTGCAGTGTATCTTGAAGACCCTGGGANNNAGATCGGAA |
| 48995 | TATGAGTGTGGAGTCGTTGCCCCAGCGCATCTCTATTCATCTCNNNAGATCGGAAGA |
| 54833 | TATGAGTGTGGAGTCGTTGCAGAACACATATCTTGCCCAGTGGNNNAGATCGGAAG |
| 44800 | TATGAGTGTGGAGTCGTTGCCTGTCTTCTCTAGGAAACCTTCCCNNNAGATCGGAAG |
| 63436 | TATGAGTGTGGAGTCGTTGCCAATGGGCCTAAACCCAAGTTCTNNNAGATCGGAAG |
| 102699 | TATGAGTGTGGAGTCGTTGCAGCAGTGGAATCATGAGCAATTTTNNNAGATCGGA |
| 66044 | TATGAGTGTGGAGTCGTTGCAAACCACCCAGTTGTAAAGCTGTNNNAGATCGGAAG |
| 52405 | TATGAGTGTGGAGTCGTTGCAAATCTGCATAGATGCCATCCCGNNNAGATCGGAAG |
| 52692 | TATGAGTGTGGAGTCGTTGCAGCAGCAGGAGAATAAGCAAAGGNNNAGATCGGA |
| 31389 | TATGAGTGTGGAGTCGTTGCCTGAAGCCCATCCATGAACTCAANNNAGATCGGAAG |
| 61487 | TATGAGTGTGGAGTCGTTGCCCAGTCTCACAATGCTTTGTCCTNNNAGATCGGAAG |
| 56431 | TATGAGTGTGGAGTCGTTGCTAGTGAGGTTTTGTGCCTGTCTGNNNAGATCGGAA |
| 29957 | TATGAGTGTGGAGTCGTTGCAGTAAATCAGGCACTACCTGTCTNNNAGATCGGAAG |
| 71714 | TATGAGTGTGGAGTCGTTGCGGGGCTGTGTAGCGCTGATCNNNAGATCGGAAGAG |
| 67287 | TATGAGTGTGGAGTCGTTGCGGCATCAGACTGGCTTTGAGATTNNNAGATCGGAA |
| 68312 | TATGAGTGTGGAGTCGTTGCGGCAGTGGCATATGATGAGAGTTNNNAGATCGGAA |
| 37658 | TATGAGTGTGGAGTCGTTGCGACCAAAATGGTAAGAGTTGGAAGANNNAGATCGG |
| 57685 | TATGAGTGTGGAGTCGTTGCCCCACACCATCTCCAGTGAAAATNNNAGATCGGAAG |
| 62574 | TATGAGTGTGGAGTCGTTGCCATTCTGCTCTGCACTGGAATCTNNNAGATCGGAAG |
| 105689 | TATGAGTGTGGAGTCGTTGCTTGAAAGAAGAACCTCCCAACTAATANNNAGATCGG |

| | |
|---|---|
| 60336 | TATGAGTGTGGAGTCGTTGCTCAGTTTCAGATTTTTCAAGGGACTNNNAGATCGGA |
| 69660 | TATGAGTGTGGAGTCGTTGCTTCGGCAGCGTTAGGGTTTTTNNNAGATCGGAAGA |
| 56351 | TATGAGTGTGGAGTCGTTGCCATATCCATGGCTTCCACCTCTGNNNAGATCGGAAGA |
| 45078 | TATGAGTGTGGAGTCGTTGCTGGTCTTTTCCTGTGTTTAGGCCNNNAGATCGGAAG |
| 109282 | TATGAGTGTGGAGTCGTTGCGATTGTCTAGTGGGGAGGACTGNNNAGATCGGAAG |
| 35747 | TATGAGTGTGGAGTCGTTGCAGAGCTGAAAATTCTTGTGGGCANNNAGATCGGAA |
| 60388 | TATGAGTGTGGAGTCGTTGCATGTAGCTTTGTTTTGCCTGACTNNNAGATCGGAAG |
| 110115 | TATGAGTGTGGAGTCGTTGCCTGGTTGGGGTTTGTTGTGAATCANNNAGATCGGA |
| 71679 | TATGAGTGTGGAGTCGTTGCGGAGCAAGTCCTTCCCTTTCATTNNNAGATCGGAAG |
| 63743 | TATGAGTGTGGAGTCGTTGCACTTAGCCTTGTGGAACTCAAGTNNNAGATCGGAAG |
| 65813 | TATGAGTGTGGAGTCGTTGCGATTGCCCTGTTTCCTCTGATGTNNNAGATCGGAAG |
| 52011 | TATGAGTGTGGAGTCGTTGCATCGGCAGCTCAAACTCAACTAGNNNAGATCGGAAG |
| 52797 | TATGAGTGTGGAGTCGTTGCTACGCTGTGATTTGTGGAGAAGGNNNAGATCGGAA |
| 33933 | TATGAGTGTGGAGTCGTTGCACCAGCTTCTTCTCTGGTTTACANNNAGATCGGAAGA |
| 105700 | TATGAGTGTGGAGTCGTTGCTCCTTCCTTCCTGTCTTTCTTTGACTANNNAGATCGGA |
| 69157 | TATGAGTGTGGAGTCGTTGCCCCTCACATACACAGACATCTTTNNNAGATCGGAAGA |
| 38535 | TATGAGTGTGGAGTCGTTGCATCACCTTCCATCACCTCTACGANNNAGATCGGAAGA |
| 36617 | TATGAGTGTGGAGTCGTTGCCTGGGTCAAGCAATCCTACCTCANNNAGATCGGAAG |
| 40167 | TATGAGTGTGGAGTCGTTGCTGGAGGATATTTTTGTTTGGTTAGTGANNNAGATCG |
| 52464 | TATGAGTGTGGAGTCGTTGCAGGATGAAGAAGGATGGTCTCCGNNNAGATCGGAA |
| 54808 | TATGAGTGTGGAGTCGTTGCTGTGACTATGGCAGTGTTTCTGGNNNAGATCGGAA |
| 103406 | TATGAGTGTGGAGTCGTTGCTAGCACTGTCTGGTAACCACCATNNNAGATCGGAAG |
| 64875 | TATGAGTGTGGAGTCGTTGCACAGTCTGGGTTTCATCAGAGGTNNNAGATCGGAA |
| 40725 | TATGAGTGTGGAGTCGTTGCCCCCTAATGCATCCAGCTCCATANNNAGATCGGAAGA |
| 110116 | TATGAGTGTGGAGTCGTTGCGTCAGAACTTATGTGTTTGTGCGTNNNAGATCGGAA |
| 57936 | TATGAGTGTGGAGTCGTTGCCCACTTCCAGTCACAGAAGCAATNNNAGATCGGAAG |
| 67510 | TATGAGTGTGGAGTCGTTGCATTTCCCACACTTTGGACCACTTNNNAGATCGGAAGA |
| 58890 | TATGAGTGTGGAGTCGTTGCCGCACTAGGCATTGTACCTTCATNNNAGATCGGAAG |
| 57632 | TATGAGTGTGGAGTCGTTGCGTGAAATCAAGCCCCACCTTTTGNNNAGATCGGAAG |
| 38673 | TATGAGTGTGGAGTCGTTGCTAGGGAAGGCAACATGAGAAGGANNNAGATCGGA |
| 106354 | TATGAGTGTGGAGTCGTTGCGGCATTTGTCGGCTTCATGTTCNNNAGATCGGAAGA |
| 50877 | TATGAGTGTGGAGTCGTTGCAGGTGAGGTCTAAGAGTTCCCAGNNNAGATCGGAA |
| 30100 | TATGAGTGTGGAGTCGTTGCGCAAACAGCACCTGATGTCAAAANNNAGATCGGAA |
| 38408 | TATGAGTGTGGAGTCGTTGCACACTTTGGGGAGTTTCCCTAGANNNAGATCGGAAG |
| 110126 | TATGAGTGTGGAGTCGTTGCTGTAGCTTTGTTTTGCCTGACTAAANNNAGATCGGA |
| 45026 | TATGAGTGTGGAGTCGTTGCAGGGAATTTGTACTCCATTCGCCNNNAGATCGGAAG |
| 29845 | TATGAGTGTGGAGTCGTTGCAACTTTGCGCATCTCATTTCCTTNNNAGATCGGAAGA |
| 49422 | TATGAGTGTGGAGTCGTTGCCTTTTCAGCCTCTCCCATCTGTCNNNAGATCGGAAGA |
| 68121 | TATGAGTGTGGAGTCGTTGCGCCGACCTTGACTGTAATGTCTTNNNAGATCGGAAG |
| 67931 | TATGAGTGTGGAGTCGTTGCCCAAATTCAAGCACTGTTGGCTTNNNAGATCGGAAG |
| 35973 | TATGAGTGTGGAGTCGTTGCAATATGCTGTGTTACAGAGATGCANNNAGATCGGA |
| 37544 | TATGAGTGTGGAGTCGTTGCCAGAGCCACTATGTTGACCAAGANNNAGATCGGAA |
| 66387 | TATGAGTGTGGAGTCGTTGCTGGGAACTTTATCTTGTTGGGTGTNNNAGATCGGAA |

| | |
|---|---|
| 50501 | TATGAGTGTGGAGTCGTTGCGAAAGGACACCTGCTTCCTGAAGNNNAGATCGGAA |
| 105224 | TATGAGTGTGGAGTCGTTGCAATATGACAGGGCAGGGAAGGAGNNNAGATCGGA |
| 105198 | TATGAGTGTGGAGTCGTTGCCTTGCACATTGGTCTAGTGCTTTCNNNAGATCGGAA |
| 39247 | TATGAGTGTGGAGTCGTTGCCACAGCCTGAATTGGTACATGGANNNAGATCGGAA |
| 107289 | TATGAGTGTGGAGTCGTTGCCCTTTTCTGAGCAACTGATGACATTNNNAGATCGGA |
| 67825 | TATGAGTGTGGAGTCGTTGCGAGAGAGACAGCAACCAAAGCTTNNNAGATCGGAA |
| 44177 | TATGAGTGTGGAGTCGTTGCATGCGTTGAAATGAGGAGGTACCNNNAGATCGGAA |
| 70976 | TATGAGTGTGGAGTCGTTGCCCCAGGTGAAGAGAGGGAAGAATNNNAGATCGGA |
| 109625 | TATGAGTGTGGAGTCGTTGCGGTCACACTTGATTTTCACACTTGNNNAGATCGGAA |
| 50300 | TATGAGTGTGGAGTCGTTGCCGCTATGAAGTTTTTGGGGCAAGNNNAGATCGGAA |
| 36317 | TATGAGTGTGGAGTCGTTGCAGAGAGCACTGCATTAGTTTGCANNNAGATCGGAA |
| 43433 | TATGAGTGTGGAGTCGTTGCGTAACATCCAGGACACCTGTGACNNNAGATCGGAAG |
| 47299 | TATGAGTGTGGAGTCGTTGCCAAATGGCTAAAAGCTGTCTGGCNNNAGATCGGAA |
| 50621 | TATGAGTGTGGAGTCGTTGCTTTGTCACCTGGATTCCCAACAGNNNAGATCGGAAG |
| 56064 | TATGAGTGTGGAGTCGTTGCTCCTTTAGGTGAGTGAGGTCCTGNNNAGATCGGAA |
| 56991 | TATGAGTGTGGAGTCGTTGCTTAGAATGGTGGGCAAGTCTGTGNNNAGATCGGAA |
| 101496 | TATGAGTGTGGAGTCGTTGCATCAGACTGGCTTTGAGATTTAGCANNNAGATCGGA |
| 37289 | TATGAGTGTGGAGTCGTTGCCTCTAAGGGAAGAGTGGGGTTCANNNAGATCGGAA |
| 50832 | TATGAGTGTGGAGTCGTTGCGTTCTGCTTTAGGTCTGACCCAGNNNAGATCGGAAG |
| 105716 | TATGAGTGTGGAGTCGTTGCCTTGGTTGATGTTTGGCATCATCNNNAGATCGGAAG |
| 49137 | TATGAGTGTGGAGTCGTTGCGCATTTGTCGGCTTCATGTTCTCNNNAGATCGGAAG |
| 32453 | TATGAGTGTGGAGTCGTTGCTGCTGGGCAACTAAAAGCAGTAANNNAGATCGGAA |
| 65526 | TATGAGTGTGGAGTCGTTGCGCATGTTCTCTCTGAACCTTGGTNNNAGATCGGAAG |
| 61267 | TATGAGTGTGGAGTCGTTGCTTCAGCATTTCTAGGATGTGCCTNNNAGATCGGAAG |
| 44134 | TATGAGTGTGGAGTCGTTGCTCACTCCTGCCTACCTTCATACCNNNAGATCGGAAGA |
| 45019 | TATGAGTGTGGAGTCGTTGCCAGCATGTATTCCATTTCACGCCNNNAGATCGGAAG |
| 46814 | TATGAGTGTGGAGTCGTTGCGCAACAGAATAATGGGCAAAGGCNNNAGATCGGAA |
| 115957 | TATGAGTGTGGAGTCGTTGCGATTTACTTCTCAAGGCTACATTTTGGNNNAGATCG |
| 68349 | TATGAGTGTGGAGTCGTTGCCTGAGGACAAGCCAAAGCTAGTTNNNAGATCGGAA |
| 96544 | TATGAGTGTGGAGTCGTTGCCAGACAAAGAAGCAGGCACCTATNNNAGATCGGAA |
| 96510 | TATGAGTGTGGAGTCGTTGCCAGTGTGATGCATTTTCAAAGTTCCTNNNAGATCGG |
| 29855 | TATGAGTGTGGAGTCGTTGCCTCCAGCTGTTCCTTTCCTTCAANNNAGATCGGAAGA |
| 42702 | TATGAGTGTGGAGTCGTTGCAAGCAGTGTTCCATCTGTACCACNNNAGATCGGAAG |
| 65516 | TATGAGTGTGGAGTCGTTGCAACAGAGCAAAGGCAATATTGGTNNNAGATCGGAA |
| 41835 | TATGAGTGTGGAGTCGTTGCGGAGCATCTCATTCCCAAGCTTANNNAGATCGGAAG |
| 101765 | TATGAGTGTGGAGTCGTTGCCTGATTTTGAATGGCCAGTAGGTGTNNNAGATCGG |
| 101730 | TATGAGTGTGGAGTCGTTGCATTGTAACCTCCATCATCCCCACGNNNAGATCGGAAG |
| 37335 | TATGAGTGTGGAGTCGTTGCAAAACGGGTCCAGTTCCATTTCANNNAGATCGGAAG |
| 59190 | TATGAGTGTGGAGTCGTTGCCAGAAAAGAAACTGGCTGGGGATNNNAGATCGGA |
| 32397 | TATGAGTGTGGAGTCGTTGCGGTAATGAGGGGAGAAGGGCTAANNNAGATCGGA |
| 50108 | TATGAGTGTGGAGTCGTTGCTTGTTCTATTGGGGGCCTCAAAGNNNAGATCGGAA |
| 65680 | TATGAGTGTGGAGTCGTTGCTCTGTGTCTGTTATATGTAGACATGTNNNAGATCGG |
| 102617 | TATGAGTGTGGAGTCGTTGCGAGGTTAGGTGTAGCCACACTTNNNAGATCGGAAG |

62770 TATGAGTGTGGAGTCGTTGCTGGGGTCTATAGTAGGACACTCTNNNAGATCGGAA
64925 TATGAGTGTGGAGTCGTTGCGTTTGCTACTGGATTCACTGAGGTNNNAGATCGGAA
66823 TATGAGTGTGGAGTCGTTGCACAGCCACACATAGCAAGTTTGTNNNAGATCGGAAG
34510 TATGAGTGTGGAGTCGTTGCTGGCCTTGATATTCAAATTTCCCANNNAGATCGGAA
41849 TATGAGTGTGGAGTCGTTGCGGGGAAGAAACAGGTTTGGCTTANNNAGATCGGA
39094 TATGAGTGTGGAGTCGTTGCGGTTGAATCTGAAGTTTGGGGGANNNAGATCGGA
57085 TATGAGTGTGGAGTCGTTGCTAGGAGCCACATGAACTTGTGTGNNNAGATCGGAA
68049 TATGAGTGTGGAGTCGTTGCGTTGTGAGCCACAGGGTTATCTTNNNAGATCGGAA
55327 TATGAGTGTGGAGTCGTTGCCCTGGGTATGTGGATGGTGAATGNNNAGATCGGAA
109659 TATGAGTGTGGAGTCGTTGCAAGAATGAATTGATGGCACTTGCAGNNNAGATCGG
116853 TATGAGTGTGGAGTCGTTGCCTCCCTCCCTCCCATTTGTAGANNNAGATCGGAAGAG
69231 TATGAGTGTGGAGTCGTTGCACAGTGAAGGGCTGCTTTTCTTTNNNAGATCGGAAG
43407 TATGAGTGTGGAGTCGTTGCCTCTTCTGGGAAGCCTTTCTGACNNNAGATCGGAAG
59061 TATGAGTGTGGAGTCGTTGCTGGAGACCCAGGAAAGATGAGATNNNAGATCGGA
34826 TATGAGTGTGGAGTCGTTGCAGCAGTCTAAGATAGCCAGATCCANNNAGATCGGA
56171 TATGAGTGTGGAGTCGTTGCCGATTAGCCTAAGGTGAAGGCTGNNNAGATCGGAA
68220 TATGAGTGTGGAGTCGTTGCACATTTGGCAGTGAGCAGAAGTTNNNAGATCGGAA
59451 TATGAGTGTGGAGTCGTTGCTTTGACCCCTTTCGGGGAGATATNNNAGATCGGAAG
39641 TATGAGTGTGGAGTCGTTGCGGAGCATAATATGGTGGCCATGANNNAGATCGGAA
36738 TATGAGTGTGGAGTCGTTGCATCTAGCACAAAGGAGGTGCTCANNNAGATCGGAA
62408 TATGAGTGTGGAGTCGTTGCTGCTTCCTTATGCAATCTGTGCTNNNAGATCGGAAG
55891 TATGAGTGTGGAGTCGTTGCGGCAGAGTGAAGGCTTTCTACTGNNNAGATCGGAA
105102 TATGAGTGTGGAGTCGTTGCGGTAGCATAAAATACACACTGCTTTGNNNAGATCGG
50262 TATGAGTGTGGAGTCGTTGCTATTTGGCTCAAGGCTTCCCAAGNNNAGATCGGAAG
31160 TATGAGTGTGGAGTCGTTGCGGTCTTTTTCTCTGCCTTTCCAANNNAGATCGGAAGA
57963 TATGAGTGTGGAGTCGTTGCCGCCTCCATACTTACTTGGCAATNNNAGATCGGAAG
96072 TATGAGTGTGGAGTCGTTGCGCCTACACAGGTTCGTTTTCTGGTTGANNNAGATCG
71674 TATGAGTGTGGAGTCGTTGCTGGTACTTTACTGGGTCTGGAGGNNNAGATCGGAA
58170 TATGAGTGTGGAGTCGTTGCAAAGGCCAGGGAGGTGGTATAATNNNAGATCGGA
100701 TATGAGTGTGGAGTCGTTGCCAGTGAAGGGCTGCTTTTCTTTTNNNAGATCGGAAG
50634 TATGAGTGTGGAGTCGTTGCATAACCAGAGAGCAGCTGAACAGNNNAGATCGGAA
67726 TATGAGTGTGGAGTCGTTGCCAAAGACTGCAACTCCTAGCCTTNNNAGATCGGAAG
59339 TATGAGTGTGGAGTCGTTGCGCATCTTGCAAACTAGGGGTGATNNNAGATCGGAA
63899 TATGAGTGTGGAGTCGTTGCGAGCAAGCATATTCTCCGACAGTNNNAGATCGGAAG
107290 TATGAGTGTGGAGTCGTTGCGTTCTCTCTGAACCTTGGTTCCTNNNAGATCGGAAGA
47846 TATGAGTGTGGAGTCGTTGCGGGTGGTTGCTAAAATGTTCTGCNNNAGATCGGAA
53036 TATGAGTGTGGAGTCGTTGCCTCAATGTAGATAAGGGCGCAGGNNNAGATCGGAA
97805 TATGAGTGTGGAGTCGTTGCGTGTTCTTAGCCCCACTCGACNNNAGATCGGAAGAG
44067 TATGAGTGTGGAGTCGTTGCTTTTGACTAGGGTGACCAGGACCNNNAGATCGGAA
47446 TATGAGTGTGGAGTCGTTGCGGGAGGAGAGCATACTGAAATGCNNNAGATCGGA
55259 TATGAGTGTGGAGTCGTTGCCGGGTAGGGAGGTCTTTGAAATGNNNAGATCGGA
62427 TATGAGTGTGGAGTCGTTGCGTGTGCATATGCTTGTTTGTGCTNNNAGATCGGAAG
104589 TATGAGTGTGGAGTCGTTGCCTACCTTTCTGCTTCCTTCAGGANNNAGATCGGAAGA

| | |
|---|---|
| 53680 | TATGAGTGTGGAGTCGTTGCAATAACTTACGAGCTTCCCAGGGNNNAGATCGGAAG |
| 58668 | TATGAGTGTGGAGTCGTTGCCACAGGAGAAAGAAGTGAGGCATNNNAGATCGGAA |
| 47995 | TATGAGTGTGGAGTCGTTGCAGATAGGACAAGCTTAGCTGTGCNNNAGATCGGAA |
| 35775 | TATGAGTGTGGAGTCGTTGCTAAAGAAAGAAGCCACCATGGCANNNAGATCGGAA |
| 109239 | TATGAGTGTGGAGTCGTTGCATAATCCTCCCACCTCAGTGAAGNNNAGATCGGAAG |
| 39400 | TATGAGTGTGGAGTCGTTGCCTGAAGTGTTTTTCCCCAGTGGANNNAGATCGGAAG |
| 67616 | TATGAGTGTGGAGTCGTTGCTCCAGTTTCCTGAGCAGGTACTTNNNAGATCGGAAG |
| 45238 | TATGAGTGTGGAGTCGTTGCTTTGCTCATACTCTTCACCTGCCNNNAGATCGGAAGA |
| 59601 | TATGAGTGTGGAGTCGTTGCCTACCCAGCCCCCTCACACGTATNNNAGATCGGAAGA |
| 37263 | TATGAGTGTGGAGTCGTTGCTGTCACTTTGATTGATATGTCAGTTCANNNAGATCG |
| 64214 | TATGAGTGTGGAGTCGTTGCGAAACGAAACACATGGGGAGAGTNNNAGATCGGA |
| 41827 | TATGAGTGTGGAGTCGTTGCCAGAGTGTCAGGGACCTTCCTTANNNAGATCGGAAG |
| 56212 | TATGAGTGTGGAGTCGTTGCAAGTTTTGTCTGGAGGATGGCTGNNNAGATCGGAA |
| 108690 | TATGAGTGTGGAGTCGTTGCGTGTTTATTTTGGTAAAATGGGCAGGANNNAGATC |
| 44700 | TATGAGTGTGGAGTCGTTGCTGTATGTATGTGCTTTGCCTCCCNNNAGATCGGAAG |
| 116281 | TATGAGTGTGGAGTCGTTGCAGGCTTCTTCTGTTTCTCAGGCNNNAGATCGGAAGA |
| 59528 | TATGAGTGTGGAGTCGTTGCCTTCTCCAGCTTGGGTCTCCTATNNNAGATCGGAAGA |
| 41910 | TATGAGTGTGGAGTCGTTGCGTTGCATGGGAAACCCTGAGTTANNNAGATCGGAA |
| 36552 | TATGAGTGTGGAGTCGTTGCCCAATACCTCCTCCCCTAACTCANNNAGATCGGAAGA |
| 64258 | TATGAGTGTGGAGTCGTTGCGAAGGCTAGCTGGGTATAGGAGTNNNAGATCGGA |
| 55009 | TATGAGTGTGGAGTCGTTGCACTTCTGAAACCACACCTTGTGGNNNAGATCGGAAG |
| 67068 | TATGAGTGTGGAGTCGTTGCGGGAACACAGGGAAACTCACATTNNNAGATCGGAA |
| 50491 | TATGAGTGTGGAGTCGTTGCAGCACCCCTTGTCTTTGATGAAGNNNAGATCGGAAG |
| 33576 | TATGAGTGTGGAGTCGTTGCTGCTGATAGCTTCCTAAAGGACANNNAGATCGGAAG |
| 35072 | TATGAGTGTGGAGTCGTTGCTGTCTTCATTCCTGAAATGTTCCANNNAGATCGGAAG |
| 37253 | TATGAGTGTGGAGTCGTTGCGTCCCTGTAAGGCTGTTTTCTTCANNNAGATCGGAA |
| 62315 | TATGAGTGTGGAGTCGTTGCTCTCCTGGTAATTCTGAGCTGCTNNNAGATCGGAAG |
| 69155 | TATGAGTGTGGAGTCGTTGCAGGGGTTCATGCCTGTAATCTTTNNNAGATCGGAAG |
| 67614 | TATGAGTGTGGAGTCGTTGCGGGCCGCTAGAAATGAAGTACTTNNNAGATCGGAA |
| 36429 | TATGAGTGTGGAGTCGTTGCTCTTCTCCAAATCCATGCCATCANNNAGATCGGAAGA |
| 71777 | TATGAGTGTGGAGTCGTTGCACAGGCTATGGTTCATGTTTTGCNNNAGATCGGAAG |
| 100684 | TATGAGTGTGGAGTCGTTGCAGTTCCTGGTTCTGTAAGTCTGGNNNAGATCGGAAG |
| 44849 | TATGAGTGTGGAGTCGTTGCATCTCCTGTCTCATGGAAAAGCCNNNAGATCGGAAG |
| 33722 | TATGAGTGTGGAGTCGTTGCTTGTGCAGTCCATAGCATTGACANNNAGATCGGAAG |
| 52080 | TATGAGTGTGGAGTCGTTGCAAACAAAAACGGAGGCCCTCTAGNNNAGATCGGAA |
| 66582 | TATGAGTGTGGAGTCGTTGCCTCTTACGGATGCCAGTCATTGTNNNAGATCGGAAG |
| 103421 | TATGAGTGTGGAGTCGTTGCTTGTGGAGAAGGAAGAGGGAAGGANNNAGATCGG |
| 104555 | TATGAGTGTGGAGTCGTTGCTTGGGCTGCTTCCATATTATTACAANNNAGATCGGA |
| 70994 | TATGAGTGTGGAGTCGTTGCACCCAGTGAAAAGGTGAAACCATNNNAGATCGGAA |
| 34396 | TATGAGTGTGGAGTCGTTGCGTGGGAAAACAGAAGTATGCCCANNNAGATCGGAA |
| 69492 | TATGAGTGTGGAGTCGTTGCAGCATTGTGGTGAAGGAGCTTTTNNNAGATCGGAA |
| 107311 | TATGAGTGTGGAGTCGTTGCGGGTTCATGCCTGTAATCTTTTNNNAGATCGGAAGA |
| 45231 | TATGAGTGTGGAGTCGTTGCTACACACACACTCACATACTGCCNNNAGATCGGAAGA |

33788 TATGAGTGTGGAGTCGTTGCCAAGGCACACTGAAGGTGATACANNNAGATCGGAA
45302 TATGAGTGTGGAGTCGTTGCAACAGCAAGTCAAATTCAGTGCCNNNAGATCGGAAG
33117 TATGAGTGTGGAGTCGTTGCGGCTAGAAGTGCACACATACACANNNAGATCGGAA
65564 TATGAGTGTGGAGTCGTTGCTACTGTGGAAAGAGAGCTTTGGTNNNAGATCGGAA
56113 TATGAGTGTGGAGTCGTTGCCACAATGGCATGTTACACAGCTGNNNAGATCGGAAG
61064 TATGAGTGTGGAGTCGTTGCGAGAGAGAGAAGAGGGAAAGCCTNNNAGATCGGA
69341 TATGAGTGTGGAGTCGTTGCTGCTGGCTCTGGTATTTTGGTTTNNNAGATCGGAAG
62534 TATGAGTGTGGAGTCGTTGCAGCCCTGGTTACAATCTTTTGCTNNNAGATCGGAAG
64663 TATGAGTGTGGAGTCGTTGCGTGCTTTCCATGATCCAGATCGTNNNAGATCGGAAG
43004 TATGAGTGTGGAGTCGTTGCAAATAAGCAAGCCCCTAGCTCACNNNAGATCGGAAG
34884 TATGAGTGTGGAGTCGTTGCTCACCATGCTTTACATTACCTCCANNNAGATCGGAAG
102941 TATGAGTGTGGAGTCGTTGCATGTCTGTGGACCTGAATGTGCNNNAGATCGGAAG
59977 TATGAGTGTGGAGTCGTTGCGAAACTGAGGTGCTGCTAACACTNNNAGATCGGAA
70440 TATGAGTGTGGAGTCGTTGCTTGCCAGAGAATTGTGTTAGGGCNNNAGATCGGAA
53728 TATGAGTGTGGAGTCGTTGCTTGCTGGGTCTTTTTAGTCAGGGNNNAGATCGGAA
46490 TATGAGTGTGGAGTCGTTGCGTCATTGTTATCATTCCTTTTCAGAGCNNNAGATCGG
66530 TATGAGTGTGGAGTCGTTGCTTTCGCATTCCTACCAGTTGTGTNNNAGATCGGAAG
66967 TATGAGTGTGGAGTCGTTGCCGTGTGTGTGTACGTTTGCAATTNNNAGATCGGAAG
39970 TATGAGTGTGGAGTCGTTGCTCAAAGTAGTTTCTCAAGGCTGANNNAGATCGGAAG
117289 TATGAGTGTGGAGTCGTTGCCTTTCCAGTTGAGGTCTCACGTGNNNAGATCGGAAG
61826 TATGAGTGTGGAGTCGTTGCACTTTGTGCCAATGAGCATAGCTNNNAGATCGGAAG
42626 TATGAGTGTGGAGTCGTTGCGTATAAGGGAGCCCAGGGACACNNNAGATCGGAAG
100932 TATGAGTGTGGAGTCGTTGCTCAGCACCATGGAAAAGGTACCNNNAGATCGGAAG
48586 TATGAGTGTGGAGTCGTTGCTCAGCTAGCAACTCACAGAACTCNNNAGATCGGAAG
34875 TATGAGTGTGGAGTCGTTGCTGGTTTGGGCCTATTCTACTCCANNNAGATCGGAAG
58775 TATGAGTGTGGAGTCGTTGCACTGGCTTAGGAGTGTCCATCATNNNAGATCGGAAG
68566 TATGAGTGTGGAGTCGTTGCGAAGCCACTATGGACACCATGTTNNNAGATCGGAAG
67255 TATGAGTGTGGAGTCGTTGCTGTGATTGCCTGGGGGTAAGATTNNNAGATCGGAA
32404 TATGAGTGTGGAGTCGTTGCTCCTCACCAGTCCTCAATGCTAANNNAGATCGGAAGA
47917 TATGAGTGTGGAGTCGTTGCAGCCATTTGAGAATTACAAACGTGCNNNAGATCGGA
40333 TATGAGTGTGGAGTCGTTGCTGTCCAAAGTACAAGCGTTGTGANNNAGATCGGAA
60337 TATGAGTGTGGAGTCGTTGCACAAAAGAAGAATAATGACAGGGACTNNNAGATCG
42403 TATGAGTGTGGAGTCGTTGCAGTACGCAAATGTACCTGGGAACNNNAGATCGGAA
65277 TATGAGTGTGGAGTCGTTGCGTTTGGGATGCTGAGAAGATGGTNNNAGATCGGAA
121997 TATGAGTGTGGAGTCGTTGCCCCTGCCACTAGGAGACAGTANNNAGATCGGAAGA
67334 TATGAGTGTGGAGTCGTTGCGACTGGAGTACGTTGGCATGATTNNNAGATCGGAA
36732 TATGAGTGTGGAGTCGTTGCTGTCATCTGTCTTTTTCTTCTGCTCANNNAGATCGGA
45240 TATGAGTGTGGAGTCGTTGCGATTCTACTTTCCTGACCCTGCCNNNAGATCGGAAGA
62700 TATGAGTGTGGAGTCGTTGCACAGTCTCCAGAGCTTCTGATCTNNNAGATCGGAAG
35754 TATGAGTGTGGAGTCGTTGCTCTCTTTCTCTTTTCAGTTTGGGCANNNAGATCGGAA
30391 TATGAGTGTGGAGTCGTTGCGTTCTTCAACTGTGATTGGCAAANNNAGATCGGAAG
31654 TATGAGTGTGGAGTCGTTGCGTCTCCCCAAGTCCCTATCAGAANNNAGATCGGAAG
51198 TATGAGTGTGGAGTCGTTGCTGAATGAGGCACACTTGGATCAGNNNAGATCGGAA

| | |
|---|---|
| 67586 | TATGAGTGTGGAGTCGTTGCGACACAGGTTGGTTTCCTGACTTNNNAGATCGGAAG |
| 50163 | TATGAGTGTGGAGTCGTTGCTGTCTGGTACTTTGGCTGGAAAGNNNAGATCGGAA |
| 109276 | TATGAGTGTGGAGTCGTTGCTTTCACCCAGGATTGAGTGTACTNNNAGATCGGAAG |
| 109667 | TATGAGTGTGGAGTCGTTGCCTGTGGTGAATCAAGCAATTCTCNNNAGATCGGAAG |
| 61241 | TATGAGTGTGGAGTCGTTGCCATATAAAGCAGAGTGCCTGCCTNNNAGATCGGAAG |
| 36068 | TATGAGTGTGGAGTCGTTGCACTAGCAAATTGAATCCACCTGCANNNAGATCGGAA |
| 54838 | TATGAGTGTGGAGTCGTTGCGTGATTTTGGAACCAAGCAGTGGNNNAGATCGGAA |
| 37901 | TATGAGTGTGGAGTCGTTGCGAGAGGTGGAGGTATTGTCCAGANNNAGATCGGA |
| 31548 | TATGAGTGTGGAGTCGTTGCATCCAGAACAAAGGCAGCAAGAANNNAGATCGGAA |
| 104572 | TATGAGTGTGGAGTCGTTGCAAATTCAAGGCGTTCTCTCAACAANNNAGATCGGAA |
| 104566 | TATGAGTGTGGAGTCGTTGCTCTCTACAGGCACATATGCTTTAGTNNNAGATCGGA |
| 34117 | TATGAGTGTGGAGTCGTTGCATTGGAAGGAGATGTCAGGACCANNNAGATCGGAA |
| 37534 | TATGAGTGTGGAGTCGTTGCAACCTGAGGGAACAGAGACAAGANNNAGATCGGA |
| 43384 | TATGAGTGTGGAGTCGTTGCACTGGCTCTGCATGATTACTGACNNNAGATCGGAAG |
| 42226 | TATGAGTGTGGAGTCGTTGCAGAACGTGCATATCCCTGGAAACNNNAGATCGGAA |
| 50214 | TATGAGTGTGGAGTCGTTGCGTCACACCAAAGCATCACACAAGNNNAGATCGGAAG |
| 49564 | TATGAGTGTGGAGTCGTTGCGGCAAACACAGCTCATTGGATTCNNNAGATCGGAAG |
| 65625 | TATGAGTGTGGAGTCGTTGCTGGTAAAATGGGCAGGACAATGTNNNAGATCGGAA |
| 115289 | TATGAGTGTGGAGTCGTTGCCCAGTAAGTAGCCTAGAGCAGTTNNNAGATCGGAA |
| 50313 | TATGAGTGTGGAGTCGTTGCGTGAGAAGGTATCCACCTGCAAGNNNAGATCGGAA |
| 46191 | TATGAGTGTGGAGTCGTTGCGAGGGTGGTGACAAGTTAGAAGCNNNAGATCGGA |
| 43553 | TATGAGTGTGGAGTCGTTGCCTTCCAGGCAAATCACCTCCTACNNNAGATCGGAAG |
| 50961 | TATGAGTGTGGAGTCGTTGCTTCAAGTCCCAAATTCCCTCCAGNNNAGATCGGAAG |
| 30188 | TATGAGTGTGGAGTCGTTGCAGCCAGCCATCTTGAAGCTAAAANNNAGATCGGAAG |
| 102010 | TATGAGTGTGGAGTCGTTGCAACGACCTAAATTCTAGCACTGCNNNAGATCGGAAG |
| 101987 | TATGAGTGTGGAGTCGTTGCCGGGGTAGTACCTAAGACCATTCTCNNNAGATCGGA |
| 34365 | TATGAGTGTGGAGTCGTTGCTCTTCACTGACTGTATCAGCCCANNNAGATCGGAAG |
| 33274 | TATGAGTGTGGAGTCGTTGCTCCAGAGAAAAAGAGATAGCACANNNAGATCGGAA |
| 103104 | TATGAGTGTGGAGTCGTTGCCCGCAGCTTTACATTTGTGTTTGNNNAGATCGGAAG |
| 54787 | TATGAGTGTGGAGTCGTTGCTCTGCATGTCATTCACCTTCTGGNNNAGATCGGAAG |
| 56296 | TATGAGTGTGGAGTCGTTGCCTTCTGTTGGCTCTGTTTTGCTGNNNAGATCGGAAG |
| 64704 | TATGAGTGTGGAGTCGTTGCACACACTGCTTTGTACTTTTCGTNNNAGATCGGAAG |
| 37696 | TATGAGTGTGGAGTCGTTGCGTGCATAGCCCACAAACCTAAGANNNAGATCGGAAG |
| 54573 | TATGAGTGTGGAGTCGTTGCAACCTGGCTTTAGAAACACCTGGNNNAGATCGGAAG |
| 61236 | TATGAGTGTGGAGTCGTTGCTCTGGCATTTTACAAACCTGCCTNNNAGATCGGAAG |
| 35195 | TATGAGTGTGGAGTCGTTGCGTGTCTCCACAGCATTAGAAGCANNNAGATCGGAAG |
| 52252 | TATGAGTGTGGAGTCGTTGCGTTCTCAGGGAAGATCTCCAACGNNNAGATCGGAA |
| 38544 | TATGAGTGTGGAGTCGTTGCTGCCATACATCCACATATTACCGANNNAGATCGGAA |
| 33013 | TATGAGTGTGGAGTCGTTGCACTGAAACCACCAAAGGAACACANNNAGATCGGAA |
| 54155 | TATGAGTGTGGAGTCGTTGCTAGTGGAGGCTCTAATGTCTGGGNNNAGATCGGAA |
| 36208 | TATGAGTGTGGAGTCGTTGCTCACACATTTCCTGGGAATTGCANNNAGATCGGAAG |
| 115245 | TATGAGTGTGGAGTCGTTGCACGAGGGGTTGTGGATTACAGGANNNAGATCGGA |
| 99365 | TATGAGTGTGGAGTCGTTGCAAAAACAATTGTCCAGCTAGTGGANNNAGATCGGA |

| | |
|---|---|
| 33771 | TATGAGTGTGGAGTCGTTGCGGGGGTTGTATCTGCATCATACANNNAGATCGGAA |
| 63882 | TATGAGTGTGGAGTCGTTGCAAGTGAGGCACAAATTCCACAGTNNNAGATCGGAA |
| 67508 | TATGAGTGTGGAGTCGTTGCGTTTTTCCTCCTCCTGCACACTTNNNAGATCGGAAGA |
| 101990 | TATGAGTGTGGAGTCGTTGCCAACGGAGCAATCTCCCCTCANNNAGATCGGAAGAG |
| 36047 | TATGAGTGTGGAGTCGTTGCACGACCTAAATTCTAGCACTGCANNNAGATCGGAAG |
| 46135 | TATGAGTGTGGAGTCGTTGCTAAATGGAAAGACCAGGCCAAGCNNNAGATCGGAA |
| 53317 | TATGAGTGTGGAGTCGTTGCTGTAGGTGTGATAGGGGTTGAGGNNNAGATCGGA |
| 66409 | TATGAGTGTGGAGTCGTTGCTTATATTGGCCCATACTGGAATGTGTNNNAGATCGG |
| 109610 | TATGAGTGTGGAGTCGTTGCAATCTGACTGAGACAGGCACACAANNNAGATCGGA |
| 55632 | TATGAGTGTGGAGTCGTTGCAAACACAGCATCCGCAAATGATGNNNAGATCGGAA |
| 100937 | TATGAGTGTGGAGTCGTTGCGTGTATTTGTAGGAGTGCCATGAGANNNAGATCGG |
| 41859 | TATGAGTGTGGAGTCGTTGCCCCTTTCTGGGTGAATGTGCTTANNNAGATCGGAAG |
| 31161 | TATGAGTGTGGAGTCGTTGCCACTTCCTGGGGTTTCTTTCCAANNNAGATCGGAAG |
| 62317 | TATGAGTGTGGAGTCGTTGCTTGAAACAATTCCTCTGGCTGCTNNNAGATCGGAAG |
| 67350 | TATGAGTGTGGAGTCGTTGCGCCTTTGGGTACTGACAGTGATTNNNAGATCGGAA |
| 58761 | TATGAGTGTGGAGTCGTTGCCAGTGGCACAAAGCACAATCATNNNAGATCGGAAG |
| 31722 | TATGAGTGTGGAGTCGTTGCTAGGGGTTGCAGTTAGCTGAGAANNNAGATCGGAA |
| 57354 | TATGAGTGTGGAGTCGTTGCAACTCTGGCGCTTGATACTCTTGNNNAGATCGGAAG |
| 57098 | TATGAGTGTGGAGTCGTTGCGATGAAAGGTCCGTCTCCTTGTGNNNAGATCGGAA |
| 67275 | TATGAGTGTGGAGTCGTTGCCACACATGACCCTGCAAGAGATTNNNAGATCGGAAG |
| 109180 | TATGAGTGTGGAGTCGTTGCACATGCTTGGTGTTCCTTCATTTNNNAGATCGGAAG |
| 46164 | TATGAGTGTGGAGTCGTTGCTCTCTCTCCCAAAGAGTTCAAGCNNNAGATCGGAAG |
| 64159 | TATGAGTGTGGAGTCGTTGCACTGCCAGGGCTTTATTTTCAGTNNNAGATCGGAAG |
| 29848 | TATGAGTGTGGAGTCGTTGCAGTCCACAGTATGCTCTGATCCCNNNAGATCGGAAG |
| 59721 | TATGAGTGTGGAGTCGTTGCGGAGGGCAACGGATACGATTTATNNNAGATCGGAA |
| 38849 | TATGAGTGTGGAGTCGTTGCGGACCAAGATTGATTTTGGAGGANNNAGATCGGAA |
| 102686 | TATGAGTGTGGAGTCGTTGCTTCTGTGGAGGTCTGAGCCATCANNNAGATCGGAA |
| 60531 | TATGAGTGTGGAGTCGTTGCGAGGCAGCACAATTCCTTGTACTNNNAGATCGGAAG |
| 53146 | TATGAGTGTGGAGTCGTTGCACTACAGGAAGCTAGCTCAGAGGNNNAGATCGGAA |
| 67339 | TATGAGTGTGGAGTCGTTGCTAGCTGGGGTGGTATACCTGATTNNNAGATCGGAA |
| 107281 | TATGAGTGTGGAGTCGTTGCGGTGTCATCTAGGTAGTGGATTTGGNNNAGATCGG |
| 31873 | TATGAGTGTGGAGTCGTTGCTGGACTTCCTCTCAACTACGGAANNNAGATCGGAAG |
| 44534 | TATGAGTGTGGAGTCGTTGCTACCTTTGTAGCAATGTGAGCCCNNNAGATCGGAAG |
| 35735 | TATGAGTGTGGAGTCGTTGCAGTGTTTATGCAAAGGTATGGGCANNNAGATCGGA |
| 67238 | TATGAGTGTGGAGTCGTTGCCCATACCAAAGGGCCTCTTCATTNNNAGATCGGAAG |
| 116891 | TATGAGTGTGGAGTCGTTGCTTTGTGCTCCAGTTAGGCATATCTNNNAGATCGGAA |
| 55254 | TATGAGTGTGGAGTCGTTGCTGTCCCTCTAAGGGGAGGAAATGNNNAGATCGGAA |
| 65630 | TATGAGTGTGGAGTCGTTGCGAGCAGAACAGGATGAGCAATGTNNNAGATCGGAA |
| 103026 | TATGAGTGTGGAGTCGTTGCTAGGCAGGGAGAAGTAGATGAGGANNNAGATCGG |
| 58874 | TATGAGTGTGGAGTCGTTGCGCCCACTACAGTTCCTTTGTCATNNNAGATCGGAAGA |
| 54003 | TATGAGTGTGGAGTCGTTGCAATGGGCTGATGTAATTGTGGGGNNNAGATCGGAA |
| 68115 | TATGAGTGTGGAGTCGTTGCCAACACTTGGTCTAGTCGGTCTTNNNAGATCGGAAG |
| 44796 | TATGAGTGTGGAGTCGTTGCAATGTCCTGTTTTCCCACTTCCCNNNAGATCGGAAGA |

109272 TATGAGTGTGGAGTCGTTGCGGAGCAAGAACTGAAAGGATTAGCNNNAGATCGGA
65672 TATGAGTGTGGAGTCGTTGCCACAGGTTAGGATGCACACATGTNNNAGATCGGAA
116001 TATGAGTGTGGAGTCGTTGCGGTGACAAGTTAGAAGCGCTTGNNNAGATCGGAAG
116519 TATGAGTGTGGAGTCGTTGCCCCTATTCTCTGTGGATGTCACCANNNAGATCGGAA
117160 TATGAGTGTGGAGTCGTTGCTGCTTGATTCTACTTCTGCCCANNNAGATCGGAAGA
31217 TATGAGTGTGGAGTCGTTGCTACTAAACCCACAGAGGGAGCAANNNAGATCGGAA
29847 TATGAGTGTGGAGTCGTTGCAAAAACCCGTGTAATAAACCTGCNNNAGATCGGAAG
54610 TATGAGTGTGGAGTCGTTGCTAATGCTTTCCTTGTACAGCCTGGNNNAGATCGGAA
42965 TATGAGTGTGGAGTCGTTGCTGTGTGGATTAGAGTGACCATCACNNNAGATCGGA
29956 TATGAGTGTGGAGTCGTTGCTCCAAGATCCAGTGGTACATCTGTNNNAGATCGGAA
45778 TATGAGTGTGGAGTCGTTGCTGAGTTTCAGAAAGCCTTGGTCCNNNAGATCGGAAG
34317 TATGAGTGTGGAGTCGTTGCATGAAAAATTGTAGCAGCCCCCANNNAGATCGGAAG
56427 TATGAGTGTGGAGTCGTTGCTAGAGGGAGCCTTGAGATGTCTGNNNAGATCGGAA
37635 TATGAGTGTGGAGTCGTTGCTTCATGGTCTTCCAGGAGGAAGANNNAGATCGGAA
71825 TATGAGTGTGGAGTCGTTGCGCAACTCTGGCAAAACGAAATGANNNAGATCGGAA
71716 TATGAGTGTGGAGTCGTTGCTGGCTTTGTCACAGGTTGTTGATNNNAGATCGGAAG
71721 TATGAGTGTGGAGTCGTTGCATGGCCCAGAAATTTCCAACTTGNNNAGATCGGAAG
55881 TATGAGTGTGGAGTCGTTGCGATGCTTGAGACCACGACTACTGNNNAGATCGGAA
29961 TATGAGTGTGGAGTCGTTGCTGTGTTGGTTCAAGTTGTTTGGTNNNAGATCGGAA
34224 TATGAGTGTGGAGTCGTTGCCATAACCAACCAGTGTCAACCCANNNAGATCGGAAGA
71779 TATGAGTGTGGAGTCGTTGCTGGGACTAGAGGTACACACACAGNNNAGATCGGAA
51796 TATGAGTGTGGAGTCGTTGCCTGTGTTGGGAAAGGGAGATGAGNNNAGATCGGA
40422 TATGAGTGTGGAGTCGTTGCCACCAACCACAGTTCATCCTTGANNNAGATCGGAAGA
29954 TATGAGTGTGGAGTCGTTGCAAGCTCAAAAGGCAACAGGAAGANNNAGATCGGAA
49769 TATGAGTGTGGAGTCGTTGCCCAGAAAAGCCCCCATCAGTTCNNNAGATCGGAAGA
44931 TATGAGTGTGGAGTCGTTGCCTACACAGAAACAAACTGCAGCCNNNAGATCGGAAG
53215 TATGAGTGTGGAGTCGTTGCTACCTTTTTCAGTGCAAGGGAGGNNNAGATCGGAA
44233 TATGAGTGTGGAGTCGTTGCACCTGGCTTATTCTCTTCAACCCNNNAGATCGGAAGA
101261 TATGAGTGTGGAGTCGTTGCGGCTTATCCAGACTGAAAATCTCACNNNAGATCGGA
29953 TATGAGTGTGGAGTCGTTGCTTATTTCTCCAGCCAGTCACACCNNNAGATCGGAAGA
71778 TATGAGTGTGGAGTCGTTGCTTGGGCTACTGATTTGGGGAGATNNNAGATCGGAA
101244 TATGAGTGTGGAGTCGTTGCTAAATAGGATTATAGGTGCGTGCCNNNAGATCGGA
47898 TATGAGTGTGGAGTCGTTGCGTGTCTATCACAGGCTAGAGTGCNNNAGATCGGAA
43724 TATGAGTGTGGAGTCGTTGCCTCTCTCTCCTTCCAGCAAAACCNNNAGATCGGAAGA
29960 TATGAGTGTGGAGTCGTTGCTGTTGGTTTCTGGCCATATGAGGNNNAGATCGGAA
29962 TATGAGTGTGGAGTCGTTGCAGAGCTGAATCAGGTAGGCATGANNNAGATCGGAA
103030 TATGAGTGTGGAGTCGTTGCGCAGGAAAGCATGGTTTGATATGGNNNAGATCGGA
55305 TATGAGTGTGGAGTCGTTGCCGCTGCATGAATACACTCGAATGNNNAGATCGGAAG
109263 TATGAGTGTGGAGTCGTTGCTACTGGTGTGTGTGTAGATAGTGGTNNNAGATCGG
105727 TATGAGTGTGGAGTCGTTGCCTCCTCACTGTGTTGGCCAGGATNNNAGATCGGAAG
71823 TATGAGTGTGGAGTCGTTGCCTTGGATATTTCTGCCCTGGGACNNNAGATCGGAAG
71826 TATGAGTGTGGAGTCGTTGCGCGTTAGTGACAAGAGGGACATCNNNAGATCGGAA
117664 TATGAGTGTGGAGTCGTTGCGACACAGCTCTCTTCTTATGAGGACNNNAGATCGGA

71827 TATGAGTGTGGAGTCGTTGCGCCTCTGCTGAATGACTAGCTGNNNAGATCGGAAGA
102653 TATGAGTGTGGAGTCGTTGCCCACGGAATAGCCAAACCATAAGNNNAGATCGGAA
52581 TATGAGTGTGGAGTCGTTGCACTGCCCTATGCACTTTTACTCGNNNAGATCGGAAGA
104604 TATGAGTGTGGAGTCGTTGCGTGACTGTGAATTGTGCTGCTTNNNAGATCGGAAGA
104547 TATGAGTGTGGAGTCGTTGCGTCCCTGAAGCATATCCCAGCAGNNNAGATCGGAAG
106831 TATGAGTGTGGAGTCGTTGCAGTGGTTACTTTCCCAGGGTGANNNAGATCGGAAGA
104587 TATGAGTGTGGAGTCGTTGCTCCTAAAATCAGGCATAAGGCAGNNNAGATCGGAA
100859 TATGAGTGTGGAGTCGTTGCGGGGTTTTCATATAATTTGGGGTGTNNNAGATCGGA
104598 TATGAGTGTGGAGTCGTTGCGTGCATAGTCTACAACTCCATGTNNNAGATCGGAAGA
106317 TATGAGTGTGGAGTCGTTGCCCTTCATGGCTTTAGCCTTTCCAGNNNAGATCGGAAG
55824 TATGAGTGTGGAGTCGTTGCACTAAACACTTTGCCTTTGCACTGNNNAGATCGGAA
115265 TATGAGTGTGGAGTCGTTGCATGTCTACTTCAGGGTTCAGCTTTANNNAGATCGGAA
104308 TATGAGTGTGGAGTCGTTGCTAATTGGAATGGAGCCCAATGCNNNAGATCGGAAGA
102634 TATGAGTGTGGAGTCGTTGCCGTATTGTTTTGTGGCTGCTTTTGTNNNAGATCGGAA
103203 TATGAGTGTGGAGTCGTTGCGACTTTCCAAGGAGCTACTGCTNNNAGATCGGAAGA
107935 TATGAGTGTGGAGTCGTTGCTGCCTTCTTTGAGCTTGATGATNNNAGATCGGAAGA
110143 TATGAGTGTGGAGTCGTTGCCAGCCTTGATGTGTTCTAATGGCNNNAGATCGGAAG
100877 TATGAGTGTGGAGTCGTTGCGTCAGCAGCCCTATCTATAATGTNNNAGATCGGAAGA
106839 TATGAGTGTGGAGTCGTTGCACCCCAAAGTTACCAGATCACTTNNNAGATCGGAAGA
101965 TATGAGTGTGGAGTCGTTGCAATCTACATAAAACAAGGCAAAACGGGNNNAGATC
107363 TATGAGTGTGGAGTCGTTGCCTTGCCTACTTCATCTTTGCTGNNNAGATCGGAAGAG
109269 TATGAGTGTGGAGTCGTTGCAGGTATGGGAAGAACCCTCCTNNNAGATCGGAAGA
117757 TATGAGTGTGGAGTCGTTGCCTGAGGTGAAGAAATAGGGAGGGANNNAGATCGG
71776 TATGAGTGTGGAGTCGTTGCTTGTGGATGAGCAGCAGAAAGTTNNNAGATCGGAA
109558 TATGAGTGTGGAGTCGTTGCGTATTCTGGATCCCTTTCTGTTGTNNNAGATCGGAA
108049 TATGAGTGTGGAGTCGTTGCGAGTAGATGCCCAGTGGTGGAATNNNAGATCGGAA
108717 TATGAGTGTGGAGTCGTTGCTGATCTGTTTTCTACGGTATTCCATNNNAGATCGGAA
107951 TATGAGTGTGGAGTCGTTGCAGGAAAATGAATGGAGCTAGAGGTNNNAGATCGGA
104373 TATGAGTGTGGAGTCGTTGCGGAATGGAACCAAGTATCGTTATGTNNNAGATCGGA
101275 TATGAGTGTGGAGTCGTTGCTAGTTGTTTGCCATCTGAACAGGNNNAGATCGGAAG
101121 TATGAGTGTGGAGTCGTTGCCCTGCTGCCTTAGTATGAGCAANNNAGATCGGAAGA
102322 TATGAGTGTGGAGTCGTTGCCCCGGCTTTTAGCATCAAAANNNAGATCGGAAGAGC
100588 TATGAGTGTGGAGTCGTTGCTTGTTTCTTGGAAAACACCGGCTNNNAGATCGGAAG
105740 TATGAGTGTGGAGTCGTTGCTCCCACTCTCTTGGCTATTACAAATNNNAGATCGGAA
98150 TATGAGTGTGGAGTCGTTGCGCTCATACCTGCATTTGGTGGTNNNAGATCGGAAGA
100681 TATGAGTGTGGAGTCGTTGCTACATTGTCATACCAGCTCAGGCNNNAGATCGGAAGA
100639 TATGAGTGTGGAGTCGTTGCGGCTTTCTTTCCTTTTCTGCCATGNNNAGATCGGAAG
105686 TATGAGTGTGGAGTCGTTGCGTAGAGACGGGGGAACTCACCANNNAGATCGGAAG
101076 TATGAGTGTGGAGTCGTTGCCACTTCTTCACTAGATTTCTTCCAACANNNAGATCGGA
106401 TATGAGTGTGGAGTCGTTGCGGGCAGATGCTGGGAAATGGAANNNAGATCGGAA
108719 TATGAGTGTGGAGTCGTTGCCTGACCACACCACGTAATCATCTNNNAGATCGGAAGA
107928 TATGAGTGTGGAGTCGTTGCATTTACTCACCGATGCCCAGAANNNAGATCGGAAGA
101247 TATGAGTGTGGAGTCGTTGCGATGGCTATGAATCTGTGTGGTCNNNAGATCGGAA

| | |
|---|---|
| 109638 | TATGAGTGTGGAGTCGTTGCATTATGCCTGGGTTCCATCCCTNNNAGATCGGAAGA |
| 106825 | TATGAGTGTGGAGTCGTTGCGACCTTTAGCCTCTAGCTTCTCTCTNNNAGATCGGAA |
| 106398 | TATGAGTGTGGAGTCGTTGCAAAAGGTGTTTGATCCCTTCCCTNNNAGATCGGAAG |
| 71795 | TATGAGTGTGGAGTCGTTGCCTGCTCTTGGATACTGCACATGTNNNAGATCGGAAG |
| 117694 | TATGAGTGTGGAGTCGTTGCAGTGCCTCAAACTCCTCTACTAAACNNNAGATCGGAA |
| 102295 | TATGAGTGTGGAGTCGTTGCGAAGGAGCTTTGTGTGTGTTTTCTNNNAGATCGGAA |
| 113677 | TATGAGTGTGGAGTCGTTGCAAACACTAGCCCCATGCTCAGTTCNNNAGATCGGAA |
| 98263 | TATGAGTGTGGAGTCGTTGCCCACCCTTTTCTCAACCTTTGCNNNAGATCGGAAGAG |
| 102331 | TATGAGTGTGGAGTCGTTGCTCCTCTAGCCTCTGCAGTCTCNNNAGATCGGAAGAG |
| 109287 | TATGAGTGTGGAGTCGTTGCGTGAGTAGCTGGGATTACAAGCTTANNNAGATCGG |
| 103130 | TATGAGTGTGGAGTCGTTGCCTGGTAGGCCATAAAGCCAATGATNNNAGATCGGA |
| 95116 | TATGAGTGTGGAGTCGTTGCTACTTGGTGGCTAATTCAGCAGGNNNAGATCGGAA |
| 95147 | TATGAGTGTGGAGTCGTTGCTCCACCCTAAAGAGATAAGAGCANNNAGATCGGAAG |
| 100418 | TATGAGTGTGGAGTCGTTGCATTCTCCCTGGGGTAAGTCAACTTNNNAGATCGGAA |
| 96336 | TATGAGTGTGGAGTCGTTGCAGACTGGAGGAGTGTTACAGTGANNNAGATCGGAA |
| 105069 | TATGAGTGTGGAGTCGTTGCAAAAGGCAAATGTGGGGACCCNNNAGATCGGAAGA |
| 65836 | TATGAGTGTGGAGTCGTTGCGCACTCATATGCCTATCCCTATGTNNNAGATCGGAAG |
| 62116 | TATGAGTGTGGAGTCGTTGCTACGCCAAAGAAAAAGACTGGCTNNNAGATCGGAA |
| 32815 | TATGAGTGTGGAGTCGTTGCCCTTATCACCAACAGTGACCAACANNNAGATCGGAA |
| 59986 | TATGAGTGTGGAGTCGTTGCCAGACAAAGATGACAGCCACACTNNNAGATCGGAAG |
| 47745 | TATGAGTGTGGAGTCGTTGCAGAACCAGTTGACAGTAAGCTGCNNNAGATCGGAA |
| 106833 | TATGAGTGTGGAGTCGTTGCTACACCGCTGGGCATGGTAGNNNAGATCGGAAGAG |
| 32064 | TATGAGTGTGGAGTCGTTGCTGGTGGCCAGAGATGAGTATGAANNNAGATCGGAA |
| 58954 | TATGAGTGTGGAGTCGTTGCTGTAGCAGGGAGACATGGAAGATNNNAGATCGGAA |
| 65691 | TATGAGTGTGGAGTCGTTGCTGTATGTATGTTTTTGACAACCATGTNNNAGATCGG |
| 53527 | TATGAGTGTGGAGTCGTTGCTCATCAAGCCAAAATGTATGCGGNNNAGATCGGAAG |
| 37805 | TATGAGTGTGGAGTCGTTGCATCAATGACTGGAAATGGACAGANNNAGATCGGAA |
| 29800 | TATGAGTGTGGAGTCGTTGCTCTACACAGAGGGGAAACTAGGGNNNAGATCGGAA |
| 42579 | TATGAGTGTGGAGTCGTTGCTCTTTCCAGGACTTTGTCCACACNNNAGATCGGAAGA |
| 57547 | TATGAGTGTGGAGTCGTTGCTGTTCCTTTGTAGCAGCACTTTGNNNAGATCGGAAG |
| 50772 | TATGAGTGTGGAGTCGTTGCTGGCCGTTCTAGTAAGTGTACAGNNNAGATCGGAA |
| 99469 | TATGAGTGTGGAGTCGTTGCGAGCGGATGCTATTGCCTACCNNNAGATCGGAAGA |
| 29798 | TATGAGTGTGGAGTCGTTGCACCAAGCAGTGCCAAAAAGTACTNNNAGATCGGAA |
| 66788 | TATGAGTGTGGAGTCGTTGCTCTAGGAAAAAGGAGCACTTTGTNNNAGATCGGAA |
| 31480 | TATGAGTGTGGAGTCGTTGCCACAAGAGCTCCCGGTTATTCAANNNAGATCGGAAG |
| 36632 | TATGAGTGTGGAGTCGTTGCTCTCCATTTCCTTTTCTCCCTCANNNAGATCGGAAGAG |
| 70130 | TATGAGTGTGGAGTCGTTGCTGTCCACCTGTTCTGTGCCTATANNNAGATCGGAAGA |
| 57823 | TATGAGTGTGGAGTCGTTGCACACACATACATCATGGTTGAAATNNNAGATCGGAA |
| 35826 | TATGAGTGTGGAGTCGTTGCGCATGTGTTCTCATTTTCTGGCANNNAGATCGGAAG |
| 41279 | TATGAGTGTGGAGTCGTTGCGCTCCCACTGGTGTTGATCTCTANNNAGATCGGAAG |
| 59291 | TATGAGTGTGGAGTCGTTGCCAAAGCCATTTTCTGGCACTGATNNNAGATCGGAAG |
| 40608 | TATGAGTGTGGAGTCGTTGCGCACACACGTGAGGAAGGAAATANNNAGATCGGAA |
| 35432 | TATGAGTGTGGAGTCGTTGCTTGTATGTCTGGGGTAGTGAGCANNNAGATCGGAA |

64226 TATGAGTGTGGAGTCGTTGCCCCATACATTCACACACAACTAGAGTNNNAGATCGGA
45686 TATGAGTGTGGAGTCGTTGCCTCACTTCATCCCCGAATTCTCCNNNAGATCGGAAGA
36875 TATGAGTGTGGAGTCGTTGCGCATGTTCCTTCAGTCACAGTCANNNAGATCGGAAG
108681 TATGAGTGTGGAGTCGTTGCTGTGCATGTGTGCTTGTAGTGANNNAGATCGGAAG
60030 TATGAGTGTGGAGTCGTTGCCCCACACACATGCTCATTACACTNNNAGATCGGAAGA
54754 TATGAGTGTGGAGTCGTTGCTCCTTACCAGAACTCTGCTCTGGNNNAGATCGGAAG
59332 TATGAGTGTGGAGTCGTTGCTGTGCATGTGTGCTTGTAGTGATNNNAGATCGGAA
52271 TATGAGTGTGGAGTCGTTGCACACACACATTTTTCTGACACACGNNNAGATCGGAA
41771 TATGAGTGTGGAGTCGTTGCGGCAGGGGTAGGGGAAATGATTANNNAGATCGGA
30047 TATGAGTGTGGAGTCGTTGCCCCTCCCTACCAAAGACACAAAANNNAGATCGGAAG
67262 TATGAGTGTGGAGTCGTTGCCCCGCTTCGTGAGATTACAGATTNNNAGATCGGAAG
56072 TATGAGTGTGGAGTCGTTGCCAGCCATAGAATGCCTTGTCCTGNNNAGATCGGAAG
35986 TATGAGTGTGGAGTCGTTGCTTGGAACAAAAGTCCTGGATGCANNNAGATCGGAA
32765 TATGAGTGTGGAGTCGTTGCACTGTCTTTGGATCACAGATAAACANNNAGATCGGA
52199 TATGAGTGTGGAGTCGTTGCATTAGGGCTGGACAGGACCTTAGNNNAGATCGGAA
38509 TATGAGTGTGGAGTCGTTGCACCTGGCAAAACAATACTAACGANNNAGATCGGAAG
116865 TATGAGTGTGGAGTCGTTGCATGAGAGAGAGAGAGATACAGGGTNNNAGATCGG
95141 TATGAGTGTGGAGTCGTTGCTCCTGTCTATCCCCATGTTCCTGNNNAGATCGGAAGA
53838 TATGAGTGTGGAGTCGTTGCATAATTGCTTACCTGCCACGGGNNNAGATCGGAAGA
40631 TATGAGTGTGGAGTCGTTGCAAGAGCCAATTAGTGGGCCAATANNNAGATCGGAA
56146 TATGAGTGTGGAGTCGTTGCTAGGATTGCACAGAAGTGAGCTGNNNAGATCGGAA
36059 TATGAGTGTGGAGTCGTTGCAAAGGCAGGTAATACTGACTGCANNNAGATCGGAA
54927 TATGAGTGTGGAGTCGTTGCACAGTGAGGGAGTAGTATGGTGGNNNAGATCGGA
63230 TATGAGTGTGGAGTCGTTGCTTTCAACATGAGGGCAGATGTCTNNNAGATCGGAA
55451 TATGAGTGTGGAGTCGTTGCGGGGACTAAGAGAGAGAAGCATGNNNAGATCGGA
107941 TATGAGTGTGGAGTCGTTGCGGGTCTCAGTCTGGTGCTTTTNNNAGATCGGAAGA
40417 TATGAGTGTGGAGTCGTTGCTCTGATCATCTGGAGAACCCTTGANNNAGATCGGAA
34146 TATGAGTGTGGAGTCGTTGCCCCATACATTCCTGCATTGACCANNNAGATCGGAAGA
53812 TATGAGTGTGGAGTCGTTGCATAGGACTGAGCAAGGAGTAGGGNNNAGATCGGA
100060 TATGAGTGTGGAGTCGTTGCTAAGACCCCTCCTAAAAAGTGCCATNNNAGATCGGA
98973 TATGAGTGTGGAGTCGTTGCTCTTCCCTCCCACCTTGCACNNNAGATCGGAAGAGCA
32315 TATGAGTGTGGAGTCGTTGCAGGGTCTGCTGCTTCTCCTATAANNNAGATCGGAAG
41731 TATGAGTGTGGAGTCGTTGCGGTCCCAAAGTGTAGGCACATTANNNAGATCGGAA
38467 TATGAGTGTGGAGTCGTTGCGCAGGCACACACAGAAGATTAGANNNAGATCGGAA
48959 TATGAGTGTGGAGTCGTTGCATGTATTGAGACGGAGCTTGCTCNNNAGATCGGAA
62973 TATGAGTGTGGAGTCGTTGCACACTCAGGAGACTTTCTGCTCTNNNAGATCGGAAG
50726 TATGAGTGTGGAGTCGTTGCGGAAGACCATGTGAGGGAGACAGNNNAGATCGGA
68928 TATGAGTGTGGAGTCGTTGCCAGGCTCACTGCAGGTGATATTTNNNAGATCGGAAG
30144 TATGAGTGTGGAGTCGTTGCATGCAGAAGGCAATCAGGGAAAANNNAGATCGGAA
103426 TATGAGTGTGGAGTCGTTGCATGTATTGAGACGGAGCTTGCTCTNNNAGATCGGAA
46699 TATGAGTGTGGAGTCGTTGCCAGACATCAGGTCCCACATTAGCNNNAGATCGGAAG
42466 TATGAGTGTGGAGTCGTTGCCTCATGCCACACTGTCAGCTAACNNNAGATCGGAAG
67145 TATGAGTGTGGAGTCGTTGCCGTGGAAACAAGCATTCAGCATTNNNAGATCGGAA

49462 TATGAGTGTGGAGTCGTTGCCAGTGACAGAATCTTGCTGTGTCNNNAGATCGGAAG
68788 TATGAGTGTGGAGTCGTTGCCCTGGTGGAGGTTTCCTGAATTTNNNAGATCGGAAG
99869 TATGAGTGTGGAGTCGTTGCCTACAACACCTCCATCACTTTGCNNNAGATCGGAAGA
55814 TATGAGTGTGGAGTCGTTGCCAAACAATGGGGCAAGCACTGNNNAGATCGGAAGA
42365 TATGAGTGTGGAGTCGTTGCAACTACTGGCAAGGATGCAGAACNNNAGATCGGAA
55767 TATGAGTGTGGAGTCGTTGCAAATGGTCTGCAGCTAGGAACTGNNNAGATCGGAA
43835 TATGAGTGTGGAGTCGTTGCAACAGGCCTTGGCTTTCATAACCNNNAGATCGGAAG
116828 TATGAGTGTGGAGTCGTTGCATCTTCTGAGCACACAGTGAGACNNNAGATCGGAAG
43012 TATGAGTGTGGAGTCGTTGCCAGCCAGAGATAAGCACTCTCACNNNAGATCGGAAG
36981 TATGAGTGTGGAGTCGTTGCTGATTTTCACCTTGGGTTGGTCANNNAGATCGGAAG
69313 TATGAGTGTGGAGTCGTTGCTTCACTTTTTGGTCTCGGGGTTTNNNAGATCGGAAG
38951 TATGAGTGTGGAGTCGTTGCAAATCCTGTTGTCTGCAAAGGGANNNAGATCGGAA
49296 TATGAGTGTGGAGTCGTTGCGAAACGTCTTACTGGGGAAGGTCNNNAGATCGGAA
44009 TATGAGTGTGGAGTCGTTGCTGTGTGTCTCTCATTAGTTTCACCNNNAGATCGGAAG
98188 TATGAGTGTGGAGTCGTTGCGACCTTATTCCTTGGGGTCTGANNNAGATCGGAAGA
46491 TATGAGTGTGGAGTCGTTGCGTAGAGGCAGTTTTCAAAGAGAGCNNNAGATCGGA
30596 TATGAGTGTGGAGTCGTTGCCACTAAGGCAGCATCAAGGGAAANNNAGATCGGAA
69080 TATGAGTGTGGAGTCGTTGCTCAAATACATGCCATGCAGCTTTNNNAGATCGGAAG
30929 TATGAGTGTGGAGTCGTTGCGGCTTTTGATCTTCCCCTGACAANNNAGATCGGAAG
41712 TATGAGTGTGGAGTCGTTGCACGCTGTCACGCTCCATAAATTANNNAGATCGGAAG
54226 TATGAGTGTGGAGTCGTTGCTGCTAAGCTTCTGGATTTGTGGGNNNAGATCGGAA
105112 TATGAGTGTGGAGTCGTTGCTTCTAGCTCCGTCTCTTGTGAAGGNNNAGATCGGAA
42941 TATGAGTGTGGAGTCGTTGCGCAGGTTGAGAGACAAATTGCACNNNAGATCGGAA
117809 TATGAGTGTGGAGTCGTTGCCACACACACAGAGTTATGGTGATNNNAGATCGGAA
109165 TATGAGTGTGGAGTCGTTGCATGTGTGTTTGGCAGCATAAATTGNNNAGATCGGA
33119 TATGAGTGTGGAGTCGTTGCAACACAGAGTGTGGACATACACANNNAGATCGGAA
109171 TATGAGTGTGGAGTCGTTGCTTGCTGATGAACATTTCGGTTGCTNNNAGATCGGAA
31978 TATGAGTGTGGAGTCGTTGCTGCTATGTAATGTGGTGCTGGAANNNAGATCGGAA
29799 TATGAGTGTGGAGTCGTTGCAGAAAGGGGCAACTGACCAAATTNNNAGATCGGAA
38493 TATGAGTGTGGAGTCGTTGCCCTGCCTTTCCTCTCCCTTTAGANNNAGATCGGAAGA
48233 TATGAGTGTGGAGTCGTTGCCCCTCCCTAACCTCTTCTTTTGCNNNAGATCGGAAGA
64973 TATGAGTGTGGAGTCGTTGCGCAATGGTAAAGAAAGGGCTAGGTNNNAGATCGG
60888 TATGAGTGTGGAGTCGTTGCAAGAGGAGACAATTCAACCCCCTNNNAGATCGGAAG
71316 TATGAGTGTGGAGTCGTTGCGACAGTTGGAGAGAGACAGGAGTNNNAGATCGG
37072 TATGAGTGTGGAGTCGTTGCGAATCATGGCCGATCTCTTGTCANNNAGATCGGAAG
31403 TATGAGTGTGGAGTCGTTGCAACTCTCTGGAAGAACCCCTCAANNNAGATCGGAAG
62552 TATGAGTGTGGAGTCGTTGCAAGGCAAAGGACCCAAACAATCTNNNAGATCGGAA
61561 TATGAGTGTGGAGTCGTTGCACATCTCTCTTGTTTTACTTGTTCCTNNNAGATCGGA
29795 TATGAGTGTGGAGTCGTTGCCCCACACCAGAGAATTCCACAAANNNAGATCGGAAG
67162 TATGAGTGTGGAGTCGTTGCGCGTGCGTGTTTGAAATGCATTNNNAGATCGGAAG
60527 TATGAGTGTGGAGTCGTTGCTGGAGTTGGGGTTTCTCTGTACTNNNAGATCGGAA
48452 TATGAGTGTGGAGTCGTTGCGCTGGGCCATATTAGTGCAGATCNNNAGATCGGAA
30640 TATGAGTGTGGAGTCGTTGCATCATCAGGTCAGGCTGTGGAAANNNAGATCGGAA

30169 TATGAGTGTGGAGTCGTTGCGCCAAACCACATGATGCTGAAAANNNAGATCGGAAG
30858 TATGAGTGTGGAGTCGTTGCACCAGATACTGTGCTAGACACAANNNAGATCGGAAG
38656 TATGAGTGTGGAGTCGTTGCCAGCACTTGTCTAACAGCAAGGANNNAGATCGGAAG
41323 TATGAGTGTGGAGTCGTTGCTCTCCTCGTCCTCCTCACTTCTANNNAGATCGGAAGA
43930 TATGAGTGTGGAGTCGTTGCCTGGTCTACCTACCTCTTCCACCNNNAGATCGGAAGA
60341 TATGAGTGTGGAGTCGTTGCGGACAATCTACTCCATGGGGACTNNNAGATCGGAAG
107252 TATGAGTGTGGAGTCGTTGCCATTCAGCATTCTAGCCCGGGANNNAGATCGGAAGA
49140 TATGAGTGTGGAGTCGTTGCTTGGAGCCTCTGCAGAATTTCTCNNNAGATCGGAAG
48023 TATGAGTGTGGAGTCGTTGCTATTTGGGACCCTGAACTTGTGCNNNAGATCGGAAG
49199 TATGAGTGTGGAGTCGTTGCGGGGTAGGGTAGGGAATACAGTCNNNAGATCGGA
61129 TATGAGTGTGGAGTCGTTGCCTTGTTCAGGAAACCTGTAGCCTNNNAGATCGGAAG
32428 TATGAGTGTGGAGTCGTTGCACTGACAGAAGCTGCCTCTCTAANNNAGATCGGAAG
64446 TATGAGTGTGGAGTCGTTGCACTCTGGGTATGTAGCTCATCTAGTNNNAGATCGGA
40893 TATGAGTGTGGAGTCGTTGCGCGGAGAGGGAGGGATAACTATANNNAGATCGGA
63986 TATGAGTGTGGAGTCGTTGCGACCTGTTCCCCTTATCTCCAGTNNNAGATCGGAAGA
31471 TATGAGTGTGGAGTCGTTGCCCCAACACCTACATCCTTGTCAANNNAGATCGGAAGA
99481 TATGAGTGTGGAGTCGTTGCACTTGCCTCTTCTTTACTCAGGACNNNAGATCGGAAG
42731 TATGAGTGTGGAGTCGTTGCAATAAGGCAAAACACATGCCCACNNNAGATCGGAAG
30983 TATGAGTGTGGAGTCGTTGCCCAGCTACAACTTTTGCAACCAANNNAGATCGGAAG
44375 TATGAGTGTGGAGTCGTTGCAAATGAGATGCATCAGAACCCCCNNNAGATCGGAAG
51953 TATGAGTGTGGAGTCGTTGCCCTCCCTTGACCATCCTTTTGAGNNNAGATCGGAAGA
60982 TATGAGTGTGGAGTCGTTGCAGAAAGGGAGATTTGGAAATCCCTNNNAGATCGGA
51331 TATGAGTGTGGAGTCGTTGCCCCACCCCCAAATCATTTTTCAGNNNAGATCGGAAGA
38363 TATGAGTGTGGAGTCGTTGCTGTTTTTGTGCACCCATTTGAGANNNAGATCGGAAG
107995 TATGAGTGTGGAGTCGTTGCCATTGGGACAATTAGATGAAAAGATTTNNNAGATCG
64152 TATGAGTGTGGAGTCGTTGCCCCTCTCAATGAATAACTGTTCAGTNNNAGATCGGAA
56452 TATGAGTGTGGAGTCGTTGCCTTCTCCCACGTGTGGATTCTGNNNAGATCGGAAGA
42830 TATGAGTGTGGAGTCGTTGCATACACCTCCTTCACAACAGCACNNNAGATCGGAAGA
51447 TATGAGTGTGGAGTCGTTGCTCCTCCTCTCTTTCACGTCAGAGNNNAGATCGGAAGA
117724 TATGAGTGTGGAGTCGTTGCAGTTCAGTGGTGCAATCCTCCCNNNAGATCGGAAGA
34328 TATGAGTGTGGAGTCGTTGCCTCTTTTGGCAGAATAAGCCCCANNNAGATCGGAAG
44431 TATGAGTGTGGAGTCGTTGCTTATACACAATATGGCTGGCCCCNNNAGATCGGAAG
41219 TATGAGTGTGGAGTCGTTGCCTCTGCATGGTAAAAGGCTGCTANNNAGATCGGAAG
32474 TATGAGTGTGGAGTCGTTGCGGGGCACAGTTCCTAAATCGTAANNNAGATCGGAAG
34381 TATGAGTGTGGAGTCGTTGCTGTGTGTCTAGGACTTTAGCCCANNNAGATCGGAAG
34528 TATGAGTGTGGAGTCGTTGCACTCTAACAATGGGCAAAAGCCANNNAGATCGGAAG
96378 TATGAGTGTGGAGTCGTTGCTAGCACTTGGAAATGAGCCTCANNNAGATCGGAAGA
36790 TATGAGTGTGGAGTCGTTGCCTCCTCTCCTTCACTCTCTCTCANNNAGATCGGAAGAG
31925 TATGAGTGTGGAGTCGTTGCATGGAGCAGAGAGCATATGGGAANNNAGATCGGA
52959 TATGAGTGTGGAGTCGTTGCTTTCCTGATGCTCAAATCCCAGGNNNAGATCGGAAG
31145 TATGAGTGTGGAGTCGTTGCTGTATTAGGCCATAGGGGTCCAANNNAGATCGGAA
51675 TATGAGTGTGGAGTCGTTGCTAAAGCAGGTAAAGCCATGGGAGNNNAGATCGGAA
100167 TATGAGTGTGGAGTCGTTGCATTTCTTTGGCTCTTTGGGTACTTNNNAGATCGGAAG

40247 TATGAGTGTGGAGTCGTTGCAGGCTTGTTGCTAACTTTGGTGANNNAGATCGGAAG

69186 TATGAGTGTGGAGTCGTTGCGAATGGTCACCTGCTGTCTCTTTNNNAGATCGGAAG

40147 TATGAGTGTGGAGTCGTTGCTCCTTGGCTCATAACAAAGAGTGANNNAGATCGGAA

45746 TATGAGTGTGGAGTCGTTGCACACACCCCACAAACAATAGTCCNNNAGATCGGAAG

62630 TATGAGTGTGGAGTCGTTGCCCTGGGAGTATCTGTCTAGCATCTNNNAGATCGGAA

32347 TATGAGTGTGGAGTCGTTGCAAACCTTGAGGCAATGCACCTAANNNAGATCGGAAG

48080 TATGAGTGTGGAGTCGTTGCCCACACATCTTACTCTACAACTTGCNNNAGATCGGAA

56364 TATGAGTGTGGAGTCGTTGCCAATAAATGCTGGCACAGCTCTGNNNAGATCGGAAG

110128 TATGAGTGTGGAGTCGTTGCTTGGAGTGGTCTAATACAGGGTTNNNAGATCGGAA

109531 TATGAGTGTGGAGTCGTTGCTGACAGGGGACAGTGTTTCCAGNNNAGATCGGAAG

116063 TATGAGTGTGGAGTCGTTGCCTGGAGCTGAGACATTCTTCTCCANNNAGATCGGAA

55069 TATGAGTGTGGAGTCGTTGCGTGTGTGTGTCTGTATCCCTTGGNNNAGATCGGAAG

58181 TATGAGTGTGGAGTCGTTGCTCTCTTCTGCTCCCCAAGCTAATNNNAGATCGGAAGA

34607 TATGAGTGTGGAGTCGTTGCCAACTGTTTGGTCCAGATAGCCANNNAGATCGGAAG

65805 TATGAGTGTGGAGTCGTTGCACCAAATCCAAATAGGCATGATGTNNNAGATCGGAA

48590 TATGAGTGTGGAGTCGTTGCTGATGGAACAGGCTCTTGAACTCNNNAGATCGGAAG

52584 TATGAGTGTGGAGTCGTTGCCCAAAGACCAAATACAACCCTCGNNNAGATCGGAAG

109458 TATGAGTGTGGAGTCGTTGCATCTGTGGGAGTGCATTCATTTTTNNNAGATCGGAA

110100 TATGAGTGTGGAGTCGTTGCCACACTCATATCTGTCGGCAAACNNNAGATCGGAAG

53766 TATGAGTGTGGAGTCGTTGCTAGGTAAGGCTTCACTGAGAGGGNNNAGATCGGAA

43745 TATGAGTGTGGAGTCGTTGCACACTCATATCTGTCGGCAAACCNNNAGATCGGAAG

95142 TATGAGTGTGGAGTCGTTGCCAAGATGTATGTTACAGGATTAGTGGANNNAGATCG

45316 TATGAGTGTGGAGTCGTTGCCATTTATTGGATGCCTGTGTGCCNNNAGATCGGAAG

100960 TATGAGTGTGGAGTCGTTGCGAGAGAGAGATAGACCCAGTCTCACTNNNAGATCG

44017 TATGAGTGTGGAGTCGTTGCAGGTTCATGTACACACCAAGACCNNNAGATCGGAAG

48223 TATGAGTGTGGAGTCGTTGCGAGCCGTGGTATCTTTTGTTTGCNNNAGATCGGAAG

55956 TATGAGTGTGGAGTCGTTGCTCCTTCCACCTTATTACCCCCTGNNNAGATCGGAAGA

102962 TATGAGTGTGGAGTCGTTGCCAGGGAAGATTCCAGCAGTTACCNNNAGATCGGAAG

39599 TATGAGTGTGGAGTCGTTGCAGGGTGGAAAAACAGGCTAATGANNNAGATCGGAA

109532 TATGAGTGTGGAGTCGTTGCCTCAATAGTCTCTTCTGAATGTTGATANNNAGATCGG

109581 TATGAGTGTGGAGTCGTTGCCTACTGGTTCTGCTCCTTTAGAAANNNAGATCGGAAG

51830 TATGAGTGTGGAGTCGTTGCATGGCAGTTGTAAGCTCCCTGAGNNNAGATCGGAAG

69854 TATGAGTGTGGAGTCGTTGCAAAACTCCAAAGGTTCACTCCCANNNAGATCGGAAG

96964 TATGAGTGTGGAGTCGTTGCCCCCCGCAATAGAAATCTACACTNNNAGATCGGAAG

71719 TATGAGTGTGGAGTCGTTGCTGTAACGACCAGTGTCTTGAACCNNNAGATCGGAAG

36129 TATGAGTGTGGAGTCGTTGCAGAGAAAAGACACACCAAGTGCANNNAGATCGGAA

69161 TATGAGTGTGGAGTCGTTGCAGGCTGCGTTTATAGTTTTATATCTTTNNNAGATCGG

60461 TATGAGTGTGGAGTCGTTGCAGGAAGAAGGGATTGCCACTACTNNNAGATCGGAA

54337 TATGAGTGTGGAGTCGTTGCAGGTCACCAGGCTAAATGAATGGNNNAGATCGGAA

49180 TATGAGTGTGGAGTCGTTGCAGGTCTTCCAAGCTATAGAAGTCNNNAGATCGGAAG

37368 TATGAGTGTGGAGTCGTTGCCACCACCAGAACATGGTCTTTCANNNAGATCGGAAG

69529 TATGAGTGTGGAGTCGTTGCGCTCTCCTTCCACAGGTTCTTTTNNNAGATCGGAAGA

36444 TATGAGTGTGGAGTCGTTGCTGATTCTGGTGCTCTTTGCATCANNNAGATCGGAAG

64509 TATGAGTGTGGAGTCGTTGCTGGGTGAAAGTCATTGTGTTAGTNNNAGATCGGAA
63503 TATGAGTGTGGAGTCGTTGCGGGTGGGTGTGTTTGTGATTTCTNNNAGATCGGAA
44957 TATGAGTGTGGAGTCGTTGCCTACACAAGTAACACCAGGAGCCNNNAGATCGGAAG
33742 TATGAGTGTGGAGTCGTTGCAGTTCTGACCCATACCCAAAATACANNNAGATCGGA
49690 TATGAGTGTGGAGTCGTTGCCTTCTGGGGTCTTCGTAGCTTCNNNAGATCGGAAGA
67826 TATGAGTGTGGAGTCGTTGCACTTACTGCTGCCTTCAAAGCTTNNNAGATCGGAAG
48347 TATGAGTGTGGAGTCGTTGCTGGTTTGCCCTATTGGCTACATCNNNAGATCGGAAG
56635 TATGAGTGTGGAGTCGTTGCGTGAAGATCTTGGGCTTGGAGTGNNNAGATCGGAA
100553 TATGAGTGTGGAGTCGTTGCGTATGCTTCCAGAAGAGATCAGCATNNNAGATCGG
64024 TATGAGTGTGGAGTCGTTGCGAGCTTCTAGGTAATGTAGCAGTNNNAGATCGGAA
69611 TATGAGTGTGGAGTCGTTGCACACACACACTTGGTACATTTTTNNNAGATCGGAAG
37242 TATGAGTGTGGAGTCGTTGCCCAGTTTCACACAGTGGTCTTCANNNAGATCGGAAG
30355 TATGAGTGTGGAGTCGTTGCTCGGTCTTGACATCCAAAGCAAANNNAGATCGGAAG
47376 TATGAGTGTGGAGTCGTTGCGCTGGTTTTTCTGGTAAGATTGGCNNNAGATCGGA
54131 TATGAGTGTGGAGTCGTTGCGATATTTTTGAGCCTTGGCTGGGNNNAGATCGGAA
37555 TATGAGTGTGGAGTCGTTGCAGAAACAGCATCTATCATGCCAAGANNNAGATCGGA
115823 TATGAGTGTGGAGTCGTTGCTGTATGTGTGTGTTCTTCCTACCCTCNNNAGATCGGA
30330 TATGAGTGTGGAGTCGTTGCAGCCTGGAGAACATTATGCCAAANNNAGATCGGAA
67651 TATGAGTGTGGAGTCGTTGCACAGAAAACGAAGCCTTCACCTTNNNAGATCGGAAG
46597 TATGAGTGTGGAGTCGTTGCTGCTAAAAGTGGTGTCTCTGAGCNNNAGATCGGAA
49982 TATGAGTGTGGAGTCGTTGCCTTGTCAGCACAGCAGACTTTTCNNNAGATCGGAAG
40668 TATGAGTGTGGAGTCGTTGCTCCCCCATGCACCTTCAGTAATANNNAGATCGGAAGA
116945 TATGAGTGTGGAGTCGTTGCCTTGGTGAGTGGCTTGAGATTCCNNNAGATCGGAA
61382 TATGAGTGTGGAGTCGTTGCAGTAATGTGGGCTTCATCCTCCTNNNAGATCGGAAG
67929 TATGAGTGTGGAGTCGTTGCTGAGGGGACGATTATCTTGGCTTNNNAGATCGGAA
32681 TATGAGTGTGGAGTCGTTGCTGAATGAAACAGCTCTGAAAACANNNAGATCGGAA
104221 TATGAGTGTGGAGTCGTTGCACAGTCTTTTGTGATAGCTCCTGTTNNNAGATCGGA
67979 TATGAGTGTGGAGTCGTTGCAACAAAAATTGCCTGCTGTGCTTNNNAGATCGGAAG
52546 TATGAGTGTGGAGTCGTTGCGAAACTTGTCACACGGTATTGCGNNNAGATCGGAA
66009 TATGAGTGTGGAGTCGTTGCCCTACTCTTACCATTGTGCCTGTNNNAGATCGGAAGA
30963 TATGAGTGTGGAGTCGTTGCTGCCCATGTTGTGAACTGTACAANNNAGATCGGAAG
96963 TATGAGTGTGGAGTCGTTGCTGAGGTAGAATCTTGCTGTCACCNNNAGATCGGAAG
29793 TATGAGTGTGGAGTCGTTGCGAAAGGAAAGTTACCCTGGGACCNNNAGATCGGAA
46669 TATGAGTGTGGAGTCGTTGCCACGCTGGAGTACATACAGTAGCNNNAGATCGGAA
67910 TATGAGTGTGGAGTCGTTGCGGGAGGATAGCAGAAAATGGCTTNNNAGATCGGA
45453 TATGAGTGTGGAGTCGTTGCAAACAAGGCTTAGCCAGTCATCCNNNAGATCGGAAG
46243 TATGAGTGTGGAGTCGTTGCAGAGAGAGAGGGGTTCTGTAAGCNNNAGATCGGA
67105 TATGAGTGTGGAGTCGTTGCCCTGGAGTTTTAGGTTGCCCATTNNNAGATCGGAAG
70843 TATGAGTGTGGAGTCGTTGCACTTTTCACTCCCGTGTAGAATGNNNAGATCGGAAG
31091 TATGAGTGTGGAGTCGTTGCGCAGAGTATTTCGATTGGGCCAANNNAGATCGGAA
55299 TATGAGTGTGGAGTCGTTGCGCTGGGTGCCAAGAAGTAGAATGNNNAGATCGGA
107968 TATGAGTGTGGAGTCGTTGCATGACAACTGGCACAAAACAAACCTNNNAGATCGGA
43774 TATGAGTGTGGAGTCGTTGCATGAGAAAAATGGGACCCCAACCNNNAGATCGGAA

50808 TATGAGTGTGGAGTCGTTGCAGGAGCCACATTTTCTTGACCAGNNNAGATCGGAAG
65984 TATGAGTGTGGAGTCGTTGCTTTCCCCTAAGGATTACCCCTGTNNNAGATCGGAAGA
41965 TATGAGTGTGGAGTCGTTGCGCAGGCTGTGATCTTGAGTGTTANNNAGATCGGAA
109527 TATGAGTGTGGAGTCGTTGCAAAGACAGCACTTTTCCTGCCCNNNAGATCGGAAGA
101751 TATGAGTGTGGAGTCGTTGCCTGTGGTGTGAGGTATGTGTGGNNNAGATCGGAAG
51063 TATGAGTGTGGAGTCGTTGCATTATAGCCAGAGATGGACGCAGNNNAGATCGGAA
67734 TATGAGTGTGGAGTCGTTGCGTTTTCAATGAAGGGAGGGCCTTNNNAGATCGGAA
104266 TATGAGTGTGGAGTCGTTGCTACTCCCTACAGAGCCCAGAGTTNNNAGATCGGAAG
48664 TATGAGTGTGGAGTCGTTGCGTCAAAAGCGTCCTTGAAACCTCNNNAGATCGGAAG
106804 TATGAGTGTGGAGTCGTTGCCCTGACAGAAGACAGTGTTTCCAGAANNNAGATCGG
57383 TATGAGTGTGGAGTCGTTGCGCACCACACTGTCTTGATTCTTGNNNAGATCGGAAG
36846 TATGAGTGTGGAGTCGTTGCTCACCTCTAGCCTGGTTTTCTCANNNAGATCGGAAGA
57926 TATGAGTGTGGAGTCGTTGCGGTGAGTGGCTTGAGATTCCAATNNNAGATCGGAA
117249 TATGAGTGTGGAGTCGTTGCCCCACCCTCCAGAATGGTTCTATTNNNAGATCGGAAG
100886 TATGAGTGTGGAGTCGTTGCCCAAGCTTCTTTCTTCTGAAACATANNNAGATCGGAA
101084 TATGAGTGTGGAGTCGTTGCGAGGGGACGATTATCTTGGCTTTTANNNAGATCGG
66236 TATGAGTGTGGAGTCGTTGCTGCCTGACACCATAAAGTTCTGTNNNAGATCGGAAG
46143 TATGAGTGTGGAGTCGTTGCCAAGAAATCCCACAGAAGCAAGCNNNAGATCGGAA
61422 TATGAGTGTGGAGTCGTTGCGTCTACCTCTGCCAGATTCTCCTNNNAGATCGGAAGA
109585 TATGAGTGTGGAGTCGTTGCAAGGCAGATACCTCAGAAGCAAGNNNAGATCGGAA
39911 TATGAGTGTGGAGTCGTTGCACTTTCTCTGAAAACATGCCTGANNNAGATCGGAAG
33430 TATGAGTGTGGAGTCGTTGCACAAGCAAGCAAGAGAGTTCACANNNAGATCGGAA
96966 TATGAGTGTGGAGTCGTTGCTGTCTGGGATGGTGGAAGTCTNNNAGATCGGAAGA
65107 TATGAGTGTGGAGTCGTTGCTGAAAATTCTGTGAAGTCCGGGTNNNAGATCGGAA
66166 TATGAGTGTGGAGTCGTTGCGTTCTTCCTTTTGCCTCCTCTGTNNNAGATCGGAAGA
63562 TATGAGTGTGGAGTCGTTGCTTCAAGGCTGGTAAGAATAATTTTCTNNNAGATCGG
66463 TATGAGTGTGGAGTCGTTGCATAGCAGGAGTGTGTCTCTGTGTNNNAGATCGGAA
42236 TATGAGTGTGGAGTCGTTGCGACAACAAGCAGGGTGTTGAAACNNNAGATCGGAA
43503 TATGAGTGTGGAGTCGTTGCGCTGCCTCTCTGATTCTGGATACNNNAGATCGGAAG
55869 TATGAGTGTGGAGTCGTTGCTCTGCAACATGCTCTAGTGACTGNNNAGATCGGAAG
62778 TATGAGTGTGGAGTCGTTGCGTTTTGTTTAGGTGGCCCACTCTNNNAGATCGGAAG
107958 TATGAGTGTGGAGTCGTTGCCTGCAAATGGGTAGGAGTTGAACTANNNAGATCGG
56015 TATGAGTGTGGAGTCGTTGCAAGAGAATATGTGTGCTGGCCTGNNNAGATCGGAA
36160 TATGAGTGTGGAGTCGTTGCTCTTTGCAATATGGTGTGGTGCANNNAGATCGGAAG
54917 TATGAGTGTGGAGTCGTTGCAGTGAAATCTGAATAAGGTGGGTGGNNNAGATCGG
109562 TATGAGTGTGGAGTCGTTGCTAGGCATTTAGACTGGTTCCACACNNNAGATCGGAA
43811 TATGAGTGTGGAGTCGTTGCGGTACTGCATCTGGTTAGGAACCNNNAGATCGGAAG
30727 TATGAGTGTGGAGTCGTTGCGGGAAGGTCACAGCACAACTAAANNNAGATCGGAA
68567 TATGAGTGTGGAGTCGTTGCAAGGTGACATGTTCTGCCATGTTNNNAGATCGGAAG
62714 TATGAGTGTGGAGTCGTTGCTGGTTGATATGGGAAATGAGATATCTNNNAGATCG
55763 TATGAGTGTGGAGTCGTTGCTGAGACCCACTTCAGGAGAACTGNNNAGATCGGAAG
57428 TATGAGTGTGGAGTCGTTGCGAAGACCTGCAACAACAGAGTTGNNNAGATCGGAA
43156 TATGAGTGTGGAGTCGTTGCTTATGTGCTGACACCAACCAGACNNNAGATCGGAAG

102628 TATGAGTGTGGAGTCGTTGCTATGTGCTGACACCAACCAGACNNNAGATCGGAAGA

52027 TATGAGTGTGGAGTCGTTGCTCTTTGGGTAACAGTCCCCCTAGNNNAGATCGGAAG

47237 TATGAGTGTGGAGTCGTTGCGTGCTGGGTTATGCTTATATGGCNNNAGATCGGAA

51895 TATGAGTGTGGAGTCGTTGCGAAAATCCTGGGCACTAGGTGAGNNNAGATCGGAA

107933 TATGAGTGTGGAGTCGTTGCAACGGGGAGCAATGCTAAGATTTNNNAGATCGGAA

37020 TATGAGTGTGGAGTCGTTGCGCACAGCTTCATACTTCCTGTCANNNAGATCGGAAG

49704 TATGAGTGTGGAGTCGTTGCATAATCCCAGGCAATCTGGCTTCNNNAGATCGGAAG

37349 TATGAGTGTGGAGTCGTTGCAAATCAACTGAGCAGCCCTTTCANNNAGATCGGAAG

46334 TATGAGTGTGGAGTCGTTGCTTGCTGGTTTCTGTAGATCCAGCNNNAGATCGGAAG

69544 TATGAGTGTGGAGTCGTTGCAGGGAAAGAAATGCTAACAGTTTTNNNAGATCGGA

67193 TATGAGTGTGGAGTCGTTGCCCAACCCACCCATAACGATCATTNNNAGATCGGAAGA

50982 TATGAGTGTGGAGTCGTTGCTGACCATGCAGGTTTTATTCCAGNNNAGATCGGAAG

50020 TATGAGTGTGGAGTCGTTGCTGAGTGAGAGGGGTGAGAAAAAGNNNAGATCGGA

61984 TATGAGTGTGGAGTCGTTGCACCTCATCCTCTTTCTTCAGGCTNNNAGATCGGAAGA

99453 TATGAGTGTGGAGTCGTTGCACATGACAATTCTAAGCCCCTTTNNNAGATCGGAAG

55806 TATGAGTGTGGAGTCGTTGCGGGAAATTAGGAGACAGCCACTGNNNAGATCGGAA

40676 TATGAGTGTGGAGTCGTTGCCCCCTGACTTCCCAAAGAACATANNNAGATCGGAAG

101241 TATGAGTGTGGAGTCGTTGCTTTGATTTTTGACACATAAGAATTGCANNNAGATCG

56231 TATGAGTGTGGAGTCGTTGCAGGTTCCCTCTTCACAAATGCTGNNNAGATCGGAAG

45029 TATGAGTGTGGAGTCGTTGCCCTCAGTCTTCTTCAGAAAGGCCNNNAGATCGGAAG

39887 TATGAGTGTGGAGTCGTTGCCCTAGAACAATGAAACCCCCTGANNNAGATCGGAAG

106285 TATGAGTGTGGAGTCGTTGCTGTCACTACAGGATGATCTAGGCTCNNNAGATCGGA

45790 TATGAGTGTGGAGTCGTTGCATACACACAATCCCACACTGTCCNNNAGATCGGAAGA

47615 TATGAGTGTGGAGTCGTTGCTTGTCAGAAAGGCCTGAGTATGCNNNAGATCGGAA

61650 TATGAGTGTGGAGTCGTTGCAATCTCCCCTTCTCTGTGAAGCTNNNAGATCGGAAGA

44224 TATGAGTGTGGAGTCGTTGCCAGCCATGAAAATGAAGGAAACCCNNNAGATCGGA

45242 TATGAGTGTGGAGTCGTTGCTATTGTGTTCATATGCCCCTGCCNNNAGATCGGAAG

43628 TATGAGTGTGGAGTCGTTGCCGTTCTGGTCACAGCTTAGGTACNNNAGATCGGAAG

54941 TATGAGTGTGGAGTCGTTGCGTGAACAGACAGTTGCAATGTGGNNNAGATCGGAA

62386 TATGAGTGTGGAGTCGTTGCCCAACGGGAAAGATAAAGGTGCTNNNAGATCGGAA

38697 TATGAGTGTGGAGTCGTTGCCCGCTGCCTCTCTTCTATAAGGANNNAGATCGGAAG

101741 TATGAGTGTGGAGTCGTTGCTCTCTGTGTGTGGACATGAAGCTTNNNAGATCGGAA

56661 TATGAGTGTGGAGTCGTTGCCCTGACCACATTTGGGGATAGTGNNNAGATCGGAA

42172 TATGAGTGTGGAGTCGTTGCACCAGCTGCCTCTAAAGACAAACNNNAGATCGGAAG

30018 TATGAGTGTGGAGTCGTTGCACTTGCTGGGAACAAGGGAAAAANNNAGATCGGAA

55483 TATGAGTGTGGAGTCGTTGCCACCAGGCTCACATATTTGCATGNNNAGATCGGAAG

54526 TATGAGTGTGGAGTCGTTGCCAGATTTCAGAACTCATTGCACTGGNNNAGATCGGA

68424 TATGAGTGTGGAGTCGTTGCGCTTCTCCAGTTTTCTGGAGGTTNNNAGATCGGAAG

69451 TATGAGTGTGGAGTCGTTGCATGCACGCGAAAGGGATTATTTTNNNAGATCGGAA

62891 TATGAGTGTGGAGTCGTTGCTTTGCCCTAGATTCTCCTCCTCTNNNAGATCGGAAGA

56094 TATGAGTGTGGAGTCGTTGCTAGCTGAGTGCCTTGTAAAGCTGNNNAGATCGGAA

64814 TATGAGTGTGGAGTCGTTGCTGCTTTTTGATGGCATACCAGGTNNNAGATCGGAAG

40361 TATGAGTGTGGAGTCGTTGCATCACATTTGGGAGACTAGAATTGANNNAGATCGG

61659 TATGAGTGTGGAGTCGTTGCAACTTCCCAACAGCTACTAAGCTNNNAGATCGGAAG

43586 TATGAGTGTGGAGTCGTTGCCGTAGCAGGAGCATTCACTCTACNNNAGATCGGAAG

48712 TATGAGTGTGGAGTCGTTGCAAATGAAAGCAGAGAAGCCCCTCNNNAGATCGGAA

39780 TATGAGTGTGGAGTCGTTGCTAGCTCTGTGTGGGAAGAACTGANNNAGATCGGAA

104228 TATGAGTGTGGAGTCGTTGCTCTTAGCTCTGTGTGGGAAGAACNNNAGATCGGAA

56476 TATGAGTGTGGAGTCGTTGCATGCAGGTCTGTTTCCAGTTCTGNNNAGATCGGAAG

41238 TATGAGTGTGGAGTCGTTGCAGGAGTACATGAGGCAGTTGCTANNNAGATCGGAA

44444 TATGAGTGTGGAGTCGTTGCTGCTTCTAATTATGGTGTGCCCCNNNAGATCGGAAG

39959 TATGAGTGTGGAGTCGTTGCTCTGATGAGTGACATGGAGCTGANNNAGATCGGAA

46929 TATGAGTGTGGAGTCGTTGCCAGCTAAGGAAGTGGTATCAGGCNNNAGATCGGAA

62907 TATGAGTGTGGAGTCGTTGCTATCTTGCCCAGACCTTTCCTCTNNNAGATCGGAAGA

101713 TATGAGTGTGGAGTCGTTGCGTGACATACGAAAGAGCCATTCGNNNAGATCGGAA

56609 TATGAGTGTGGAGTCGTTGCTCAGTGCATTATGTACACTTCAGTGNNNAGATCGGA

29987 TATGAGTGTGGAGTCGTTGCCGCACACACATGAACAACAAAAANNNAGATCGGAAG

53983 TATGAGTGTGGAGTCGTTGCAGGCATGCAAAGATATTCTGGGGNNNAGATCGGAA

57116 TATGAGTGTGGAGTCGTTGCGGCAGCATAGTACACATTGTTGTGNNNAGATCGGA

35643 TATGAGTGTGGAGTCGTTGCATTCATCTAGAGAGAGAGAGAGGCANNNAGATCGG

65594 TATGAGTGTGGAGTCGTTGCGGAGGCCAGCAAGTCTAAAATGTNNNAGATCGGAA

63011 TATGAGTGTGGAGTCGTTGCATCCTGCTTGACTTGACCTCTCTNNNAGATCGGAAGA

32657 TATGAGTGTGGAGTCGTTGCGAATGCACTTGAGGAACAAAACANNNAGATCGGAA

104550 TATGAGTGTGGAGTCGTTGCAGAAATGGCGGAAGTTGGAGTATTNNNAGATCGG

59556 TATGAGTGTGGAGTCGTTGCAGATGAGGGGAAGGTTGTGCTATNNNAGATCGGA

59539 TATGAGTGTGGAGTCGTTGCCTATGCAGTTAGCCACACGCTATNNNAGATCGGAAG

30333 TATGAGTGTGGAGTCGTTGCTACTCATATTTGGTTTTAGTTGCCAAANNNAGATCGG

56848 TATGAGTGTGGAGTCGTTGCGAGAGAGAAGGTTCATGGTGGTGNNNAGATCGGA

109204 TATGAGTGTGGAGTCGTTGCTATGCACTTGGATGGAATTGTCCANNNAGATCGGAA

30915 TATGAGTGTGGAGTCGTTGCACCCAACTCTTCTCTGGAGACAANNNAGATCGGAAG

45933 TATGAGTGTGGAGTCGTTGCTAGTCCTGTTGGATCAGGGTTCCNNNAGATCGGAAG

96589 TATGAGTGTGGAGTCGTTGCTCTTCTTGAGGCGGAATCCATCTNNNAGATCGGAAG

65707 TATGAGTGTGGAGTCGTTGCGCAAAGGGAAAAACAACTCCATGTNNNAGATCGGA

54485 TATGAGTGTGGAGTCGTTGCTTCCATCAGCATGTCAGATTTATGGNNNAGATCGGA

53355 TATGAGTGTGGAGTCGTTGCTCTCAGGGTGCTTGAGGTATAGGNNNAGATCGGAA

70239 TATGAGTGTGGAGTCGTTGCGACATACACTCCAGGAGAAGCACNNNAGATCGGAA

67606 TATGAGTGTGGAGTCGTTGCCAAGCAGGGCTACAATCCTACTTNNNAGATCGGAAG

44813 TATGAGTGTGGAGTCGTTGCTGTGGCAGGAATTTCTTCTTCCCNNNAGATCGGAAG

53461 TATGAGTGTGGAGTCGTTGCAGGACAGAAGAGAAGAGAGACGGNNNAGATCGGA

31024 TATGAGTGTGGAGTCGTTGCGTGAGCTTGAAAGAAGACCCCAANNNAGATCGGAA

105647 TATGAGTGTGGAGTCGTTGCCAAACTCCTGGCTTGAGCAACCCNNNAGATCGGAAG

40916 TATGAGTGTGGAGTCGTTGCTGGGGCAGGAGTGATTACGTATANNNAGATCGGAA

35777 TATGAGTGTGGAGTCGTTGCGCACTTGAATCTTACCCATGGCANNNAGATCGGAAG

68023 TATGAGTGTGGAGTCGTTGCAACCCATTTCAACTGCCCATCTTNNNAGATCGGAAGA

99270 TATGAGTGTGGAGTCGTTGCTTCACAGAATGGCAGATGCCTGAANNNAGATCGGA

63181 TATGAGTGTGGAGTCGTTGCGTTTCAGGCCTAGAGTGAGGTCTNNNAGATCGGAA

48961 TATGAGTGTGGAGTCGTTGCGTGTAGTGGCAGGATCTTGCTCNNNAGATCGGAAG
54121 TATGAGTGTGGAGTCGTTGCTAGAAAGAGAAGGACAGGCTGGGNNNAGATCGGA
31203 TATGAGTGTGGAGTCGTTGCTCACATTGGAAACCCATCAGCAANNNAGATCGGAAG
60928 TATGAGTGTGGAGTCGTTGCTTGTTTTCCTCCTCTTTTCCCCTNNNAGATCGGAAGAG
61641 TATGAGTGTGGAGTCGTTGCACACCTACAGTGAGATGGAAGCTNNNAGATCGGAA
99973 TATGAGTGTGGAGTCGTTGCATTTCTAATGTCTGCAACTTTCATGANNNAGATCGGA
40053 TATGAGTGTGGAGTCGTTGCCTTTGGCCCTGGATCAGTCTGANNNAGATCGGAAGA
39856 TATGAGTGTGGAGTCGTTGCTCCATCTCCTGACCTTGTACTGANNNAGATCGGAAGA
105099 TATGAGTGTGGAGTCGTTGCATGGCGGTGCCTTAGGTTCCNNNAGATCGGAAGAG
34781 TATGAGTGTGGAGTCGTTGCGTTCAGGTTCAGGAGCAAATCCANNNAGATCGGAA
42960 TATGAGTGTGGAGTCGTTGCCCATCGCCCTTTTGCATAATCACNNNAGATCGGAAGA
30528 TATGAGTGTGGAGTCGTTGCGGGGCAAAGAACAAGAGGAGAAANNNAGATCGGA
43450 TATGAGTGTGGAGTCGTTGCATAAGCAGCCAAGTGCTCTTGACNNNAGATCGGAAG
38872 TATGAGTGTGGAGTCGTTGCAAAACAAAATCCGAGTTTTTGAGGANNNAGATCGG
40603 TATGAGTGTGGAGTCGTTGCATAGCGGCAGCAGCAAAGAAATANNNAGATCGGAA
95891 TATGAGTGTGGAGTCGTTGCCCCCATTTGTCTGAAGTGCTGNNNAGATCGGAAGAG
29486 TATGAGTGTGGAGTCGTTGCGCGAAGGCTTCCTCTGTGTANNNAGATCGGAAGAG
39788 TATGAGTGTGGAGTCGTTGCTGTTTCCTGGTTTGCTGAACTGANNNAGATCGGAAG
30870 TATGAGTGTGGAGTCGTTGCTGGTTTTCCAACTTAGCCCACAANNNAGATCGGAAG
63701 TATGAGTGTGGAGTCGTTGCATGTGAATGCATACAGCCCAAGTNNNAGATCGGAA
65237 TATGAGTGTGGAGTCGTTGCTGCCATTATTCTTTGGACAAATGGTNNNAGATCGGA
55693 TATGAGTGTGGAGTCGTTGCTTGCCCACACATCCATTCCTATGNNNAGATCGGAAGA
38424 TATGAGTGTGGAGTCGTTGCGGAGGCTCAGAATCTCTGCTAGANNNAGATCGGAA
40928 TATGAGTGTGGAGTCGTTGCGCCTGCATTTCTGTGCCCTTATANNNAGATCGGAAG
55198 TATGAGTGTGGAGTCGTTGCTCCTTGCTTGCTTGTTTATGTTTGGNNNAGATCGGA
106330 TATGAGTGTGGAGTCGTTGCAAACTACATGGAAATGGAAAAGTGCANNNAGATCG
48870 TATGAGTGTGGAGTCGTTGCCTATTTCCTGGGCTGTGAGCTCNNNAGATCGGAAGA
51344 TATGAGTGTGGAGTCGTTGCGGCGCATGTCTAGAGAGAAAGAGNNNAGATCGGA
37841 TATGAGTGTGGAGTCGTTGCAGTAAGCACATTGACAAACCCAGANNNAGATCGGA
42455 TATGAGTGTGGAGTCGTTGCCATACCTCCCTGCACACGATAACNNNAGATCGGAAG
48871 TATGAGTGTGGAGTCGTTGCGCTCATCTTTTTGCCTTGAGCTCNNNAGATCGGAAG
58585 TATGAGTGTGGAGTCGTTGCCAGGGCTTTGGCCTTTATTCCATNNNAGATCGGAAG
116250 TATGAGTGTGGAGTCGTTGCTCAACGGCTCACTATCTTGCCCANNNAGATCGGAAG
55659 TATGAGTGTGGAGTCGTTGCGGTGTTTGTGACAAGCTGTGATGNNNAGATCGGAA
59006 TATGAGTGTGGAGTCGTTGCCTCCCCAACTTGGAATTGCAGATNNNAGATCGGAAG
53775 TATGAGTGTGGAGTCGTTGCCTTGAGATAAGTCCCCAGGAGGGNNNAGATCGGAA
50921 TATGAGTGTGGAGTCGTTGCGAAATAGACCCAATCTGGGCCAGNNNAGATCGGAA
49941 TATGAGTGTGGAGTCGTTGCGTCCCCTTTTCAGCCTCAGTTTCNNNAGATCGGAAGA
31251 TATGAGTGTGGAGTCGTTGCGCTAACGGAGTAGAAAAGGGCAANNNAGATCGGA
69089 TATGAGTGTGGAGTCGTTGCTGTAAATCCCGCATGCTAGCTTTNNNAGATCGGAAG
39528 TATGAGTGTGGAGTCGTTGCTTCTTCCAGGCAAACACTTTGGANNNAGATCGGAAG
38879 TATGAGTGTGGAGTCGTTGCAGCCAAATGTGCAAACCATAGGANNNAGATCGGAA
30495 TATGAGTGTGGAGTCGTTGCCTTACCCTTGTCGAGCACAGAAANNNAGATCGGAAG

| | |
|---|---|
| 64397 | TATGAGTGTGGAGTCGTTGCACCCTAATGCCATTCCAATAATAGTNNNAGATCGGAA |
| 58327 | TATGAGTGTGGAGTCGTTGCAGAGGTATTCCATGTCAGCACATNNNAGATCGGAAG |
| 69345 | TATGAGTGTGGAGTCGTTGCCCACACTCACCCCTAGAATGTTTNNNAGATCGGAAGA |
| 102919 | TATGAGTGTGGAGTCGTTGCGCTCCTGGGCTTTTGAACTCCTNNNAGATCGGAAGA |
| 31019 | TATGAGTGTGGAGTCGTTGCTTGTTGGAAACTGAGTGACCCAANNNAGATCGGAA |
| 103488 | TATGAGTGTGGAGTCGTTGCCATTGTTGGAAACTGAGTGACCCAANNNAGATCGGA |
| 60961 | TATGAGTGTGGAGTCGTTGCTCTTACAATGTAGCTTTGGCCCTNNNAGATCGGAAG |
| 35966 | TATGAGTGTGGAGTCGTTGCATGGCACACCTAAACAAGATGCANNNAGATCGGAA |
| 64023 | TATGAGTGTGGAGTCGTTGCACCTTACTCTAGTGCCTAGCAGTNNNAGATCGGAAG |
| 116190 | TATGAGTGTGGAGTCGTTGCAGGTCATCCACACTGTCCACACNNNAGATCGGAAGA |
| 69527 | TATGAGTGTGGAGTCGTTGCCCACCTATGCCTCTGTGTCTTTTNNNAGATCGGAAGA |
| 41670 | TATGAGTGTGGAGTCGTTGCCAGGTGCATGGTGTATTGGTGTANNNAGATCGGAA |
| 69561 | TATGAGTGTGGAGTCGTTGCCCGATCCCACAAGATGAGGTTTTNNNAGATCGGAAG |
| 58562 | TATGAGTGTGGAGTCGTTGCAGCATTGGTTCTCTCTCCTCCATNNNAGATCGGAAGA |
| 65089 | TATGAGTGTGGAGTCGTTGCAGAAAGATGTGGCAGTCTAGGGTNNNAGATCGGAA |
| 61351 | TATGAGTGTGGAGTCGTTGCATGTGCTCAGACTTTTACATTATCCTNNNAGATCGGA |
| 35631 | TATGAGTGTGGAGTCGTTGCTGGTAATTGTGTATTTATTGTCAGGCANNNAGATCG |
| 53695 | TATGAGTGTGGAGTCGTTGCGAAAAGGGTACTACACAGCAGGGNNNAGATCGGAA |
| 62059 | TATGAGTGTGGAGTCGTTGCGCTGTAGAATACTTGTGTGGGCTNNNAGATCGGAA |
| 46002 | TATGAGTGTGGAGTCGTTGCAGGCAAAGTCCATCACTCTTTCCNNNAGATCGGAAG |
| 64035 | TATGAGTGTGGAGTCGTTGCTGGACTGAAGTAAAACAGGCAGTNNNAGATCGGAA |
| 30953 | TATGAGTGTGGAGTCGTTGCACAACTGCAATCAAGGGCTACAANNNAGATCGGAA |
| 35731 | TATGAGTGTGGAGTCGTTGCAAATTCTAGCAAATCCTGGGGCANNNAGATCGGAA |
| 41027 | TATGAGTGTGGAGTCGTTGCCTCCAGAAAGTCTTGTCCACCTANNNAGATCGGAAG |
| 116523 | TATGAGTGTGGAGTCGTTGCGATTGGAGTTCAGTGGCGGAGNNNAGATCGGAAG |
| 117173 | TATGAGTGTGGAGTCGTTGCTTTCCGCACCCTATTTGCTTGNNNAGATCGGAAGAG |
| 53186 | TATGAGTGTGGAGTCGTTGCAGAAAAAGGCATGGACAAGGAGGNNNAGATCGGA |
| 61468 | TATGAGTGTGGAGTCGTTGCAGAAAAGGAGCTGGAGATGTCCTNNNAGATCGGAA |
| 43467 | TATGAGTGTGGAGTCGTTGCGGGGAAGGTGGGGAGAAAAATACNNNAGATCGGA |
| 60745 | TATGAGTGTGGAGTCGTTGCTTGGCTGTAAATGGAAAGGACCTNNNAGATCGGAA |
| 64758 | TATGAGTGTGGAGTCGTTGCACAGGAGAGAGTATAACTTCAAGGTNNNAGATCGG |
| 46390 | TATGAGTGTGGAGTCGTTGCTCTTTGGTACTACCCATTGCAGCNNNAGATCGGAAG |
| 55617 | TATGAGTGTGGAGTCGTTGCGAGAGAGAAACAGTGCTGGGATGNNNAGATCGGA |
| 46407 | TATGAGTGTGGAGTCGTTGCTAGTGAGGGGTGGATAACTCAGCNNNAGATCGGAA |
| 34490 | TATGAGTGTGGAGTCGTTGCTTGGATCAGGGATCAGATTCCCANNNAGATCGGAA |
| 30747 | TATGAGTGTGGAGTCGTTGCAAGAGCGGGGAGACTTAGCTAAANNNAGATCGGAA |
| 48384 | TATGAGTGTGGAGTCGTTGCCTTTAGGTTGGTCCCTGAGCATCNNNAGATCGGAAG |
| 100290 | TATGAGTGTGGAGTCGTTGCACTTGGGTTCCACAGAATGTTAGTTTNNNAGATCGG |
| 53629 | TATGAGTGTGGAGTCGTTGCTTTCAAGGAGGGTAGGGAAGGGNNNAGATCGGAA |
| 30709 | TATGAGTGTGGAGTCGTTGCGATACGGTAGTGGGAGGAATAAANNNAGATCGGA |
| 40444 | TATGAGTGTGGAGTCGTTGCCTGTGGCTTGGATACCATCTTGANNNAGATCGGAAG |
| 65649 | TATGAGTGTGGAGTCGTTGCTGGCTCCTGAAGTTTTGGAATGTNNNAGATCGGAA |
| 104251 | TATGAGTGTGGAGTCGTTGCAGAGAAGAGAGACGGGAAAGGCANNNAGATCGGA |

| | |
|---|---|
| 40095 | TATGAGTGTGGAGTCGTTGCTGGGAAAGTGTCCACTAAAGTGANNNAGATCGGAA |
| 63374 | TATGAGTGTGGAGTCGTTGCACCCAGTAACTACTCTTCTTACTTCTNNNAGATCGGA |
| 117304 | TATGAGTGTGGAGTCGTTGCCGCAGTGAGTGGAAGTAGCCTGANNNAGATCGGAA |
| 67765 | TATGAGTGTGGAGTCGTTGCCACATCCCAACAGACACATCCTTNNNAGATCGGAAG |
| 98655 | TATGAGTGTGGAGTCGTTGCAATACACCCCTCCTCTTAGCCAAAGNNNAGATCGGAA |
| 62734 | TATGAGTGTGGAGTCGTTGCACTGAGCCTGGCAGTAAGTATCTNNNAGATCGGAA |
| 95144 | TATGAGTGTGGAGTCGTTGCTGACCAGGAGAGGAGAGTTAGTGNNNAGATCGGA |
| 33791 | TATGAGTGTGGAGTCGTTGCCGCACAATATATCTGGATACATATACANNNAGATCG |
| 55010 | TATGAGTGTGGAGTCGTTGCTATCTGTTCTTGGCTGCTTGTGGNNNAGATCGGAAG |
| 47660 | TATGAGTGTGGAGTCGTTGCACAGAGGGCTAAATTCTCACTGCNNNAGATCGGAAG |
| 48962 | TATGAGTGTGGAGTCGTTGCGGCTTTGGTTTGAACTCTTGCTCNNNAGATCGGAAG |
| 60703 | TATGAGTGTGGAGTCGTTGCTTTCCCACTCCATTCTCTCACCTNNNAGATCGGAAGA |
| 37778 | TATGAGTGTGGAGTCGTTGCATCTGGTCAGGGGAAATCACAGANNNAGATCGGAA |
| 40078 | TATGAGTGTGGAGTCGTTGCTAGTCCTGTGAAGTGGGTTCTGANNNAGATCGGAA |
| 105066 | TATGAGTGTGGAGTCGTTGCAGCGGGGAGACTTAGCTAAAAANNNAGATCGGAAG |
| 61423 | TATGAGTGTGGAGTCGTTGCGAGAGGCTGGAAGAGATTCTCCTNNNAGATCGGAA |
| 57114 | TATGAGTGTGGAGTCGTTGCATGTGTGCCCCTTCATGTTGTGNNNAGATCGGAAGA |
| 116040 | TATGAGTGTGGAGTCGTTGCTGAGGAGGAGACGGGTACGTACNNNAGATCGGAA |
| 71147 | TATGAGTGTGGAGTCGTTGCTCTCCCATCTCTAGATTTGCCCTNNNAGATCGGAAGA |
| 99047 | TATGAGTGTGGAGTCGTTGCCTCTATGCTCCCAGGCTCACCNNNAGATCGGAAGAG |
| 71695 | TATGAGTGTGGAGTCGTTGCGCTTGTTCTCGGCATTATGATGCNNNAGATCGGAAG |
| 50484 | TATGAGTGTGGAGTCGTTGCCCTGAAAAGCTGCCCCATGAAGNNNAGATCGGAAG |
| 45116 | TATGAGTGTGGAGTCGTTGCCTGTCCCTCTTCATGTTATGGCCNNNAGATCGGAAG |
| 38980 | TATGAGTGTGGAGTCGTTGCACCCAAGTACATGATGGAAGGGANNNAGATCGGAA |
| 109201 | TATGAGTGTGGAGTCGTTGCAGAGACGGAGACGGAGAAGGGAANNNAGATCGG |
| 45732 | TATGAGTGTGGAGTCGTTGCTTGTCTTCCTAGAGCCTCAGTCCNNNAGATCGGAAG |
| 59705 | TATGAGTGTGGAGTCGTTGCGGGGAGCAAAGCTGTACAGTTATNNNAGATCGGAA |
| 117003 | TATGAGTGTGGAGTCGTTGCCTGGATCTCTTCCTCTAAAACCCNNNAGATCGGAAGA |
| 53452 | TATGAGTGTGGAGTCGTTGCGATAATACAGGCAGGTGACACGGNNNAGATCGGAA |
| 59820 | TATGAGTGTGGAGTCGTTGCTCTCCCTTCTCTGAACACCAACTNNNAGATCGGAAGA |
| 61221 | TATGAGTGTGGAGTCGTTGCAGAGAGATACAATGTGCATGCCTNNNAGATCGGAA |
| 50779 | TATGAGTGTGGAGTCGTTGCAAGCCCCATAGCCATTGTTACAGNNNAGATCGGAAG |
| 51921 | TATGAGTGTGGAGTCGTTGCGGTTGGAACTGGAGGTCATTGAGNNNAGATCGGA |
| 64129 | TATGAGTGTGGAGTCGTTGCCCCTCAAAAGCTGAATGGTCAGTNNNAGATCGGAAG |
| 38645 | TATGAGTGTGGAGTCGTTGCCCAAGGTGGAGGAAAAACAAGGANNNAGATCGGA |
| 107983 | TATGAGTGTGGAGTCGTTGCTTGTTTTGAAGGAGAAAGAATAGTCTTNNNAGATCG |
| 108748 | TATGAGTGTGGAGTCGTTGCCAATGTGGCAAGGATTATGACAGGNNNAGATCGGA |
| 67585 | TATGAGTGTGGAGTCGTTGCCCAAGCCAGGTTCTGAATGACTTNNNAGATCGGAAG |
| 67425 | TATGAGTGTGGAGTCGTTGCAGGAGAAGCCCAAAGGAGGTATTNNNAGATCGGA |
| 56366 | TATGAGTGTGGAGTCGTTGCTTGCTGAATAGGATGCAGCTCTGNNNAGATCGGAA |
| 42678 | TATGAGTGTGGAGTCGTTGCGGGAGGAGAAATAACACCACCACNNNAGATCGGAA |
| 50210 | TATGAGTGTGGAGTCGTTGCCTGGTCACAGGGAGATGAACAAGNNNAGATCGGAA |
| 60075 | TATGAGTGTGGAGTCGTTGCCTCTCTCCCTTCTTACAGCCACTNNNAGATCGGAAGA |

56730 TATGAGTGTGGAGTCGTTGCAATGAAGCAGAGAACCACAGGTGNNNAGATCGGAA
41831 TATGAGTGTGGAGTCGTTGCCCCCCTTGGGATTCCTTTCCTTANNNAGATCGGAAGA
36533 TATGAGTGTGGAGTCGTTGCTGTGGCAAAACACATAACTTTATCANNNAGATCGGA
47363 TATGAGTGTGGAGTCGTTGCGGAATGCTATGATAGGGTGTGGCNNNAGATCGGAA
63529 TATGAGTGTGGAGTCGTTGCGCGTATACATTTTTCACCTCTTTCTNNNAGATCGGAA
100041 TATGAGTGTGGAGTCGTTGCCTCCAAGGTCAAGTCAGCCACNNNAGATCGGAAGAG
50276 TATGAGTGTGGAGTCGTTGCCTTACAGCCTCCTCCCTCCAAGNNNAGATCGGAAGAG
68817 TATGAGTGTGGAGTCGTTGCGGGTCAGAGGAGAGAGACCATTTNNNAGATCGGAA
67903 TATGAGTGTGGAGTCGTTGCCCTGGTGAAAAATCAGAGGGCTTNNNAGATCGGAA
40808 TATGAGTGTGGAGTCGTTGCTCCGCGTTGGAAAAACTCAGATANNNAGATCGGAAG
33557 TATGAGTGTGGAGTCGTTGCGGGCGTGTATGTATGTTGAGACANNNAGATCGGAA
31560 TATGAGTGTGGAGTCGTTGCACCGATACTCTGGCCAAGAAGAANNNAGATCGGAAG
46704 TATGAGTGTGGAGTCGTTGCTGGAGGTGCACAAATAGGTTAGCNNNAGATCGGAA
48544 TATGAGTGTGGAGTCGTTGCCACCACTTCACCCAGCTTCTATCNNNAGATCGGAAGA
59735 TATGAGTGTGGAGTCGTTGCGGGGTGGTGACAGAGCATTTTATNNNAGATCGGAA
68422 TATGAGTGTGGAGTCGTTGCGCAGACCACATCCTGTAGAGGTTNNNAGATCGGAAG
44167 TATGAGTGTGGAGTCGTTGCTGCTACCTTCCCCTTCTTCTACCNNNAGATCGGAAGAG
71016 TATGAGTGTGGAGTCGTTGCCACCATGCTAAGTGACAACGCATNNNAGATCGGAAG
61219 TATGAGTGTGGAGTCGTTGCTTCCCCTAAAGCAAAACATGCCTNNNAGATCGGAAGA
95862 TATGAGTGTGGAGTCGTTGCAAACAAGCCCTCTTCTCCGTAGTNNNAGATCGGAAGA
53059 TATGAGTGTGGAGTCGTTGCGGCATCCTTAGAGATACTGCAGGNNNAGATCGGAA
65323 TATGAGTGTGGAGTCGTTGCTTATCCCCAAGATACTTTTGGTATGGTNNNAGATCGG
65584 TATGAGTGTGGAGTCGTTGCATGGGTCTGTTTCTGGTTTTGGTNNNAGATCGGAAG
46251 TATGAGTGTGGAGTCGTTGCAGACGACCCATCATAACAACAGCNNNAGATCGGAAG
41939 TATGAGTGTGGAGTCGTTGCTCACCTCTGCCTTATCCTGGTTANNNAGATCGGAAGA
56852 TATGAGTGTGGAGTCGTTGCACTGCAAATAAGTCCCTGTGGTGNNNAGATCGGAAG
105618 TATGAGTGTGGAGTCGTTGCCAGCCTCAAACTCCTAGACAATCNNNAGATCGGAAGA
66909 TATGAGTGTGGAGTCGTTGCCCTGGCTGATTCCCACTCAAATTNNNAGATCGGAAGA
34403 TATGAGTGTGGAGTCGTTGCGCAATGAGTATGTGTAGTGCCCANNNAGATCGGAA
50521 TATGAGTGTGGAGTCGTTGCGAGGATCACTGGGGAGGTGAAGNNNAGATCGGAA
33524 TATGAGTGTGGAGTCGTTGCCACAGGAATGGAAAGGAAGAGACANNNAGATCGGA
41806 TATGAGTGTGGAGTCGTTGCCCCCTCCATCTGGTTTACCCTTANNNAGATCGGAAGA
116512 TATGAGTGTGGAGTCGTTGCCTCTCTTTCTTTCTCTGCTGGTCTTTNNNAGATCGGAA
100161 TATGAGTGTGGAGTCGTTGCTTATGAGAAAACAAAAGGCGCACTGANNNAGATCG
53989 TATGAGTGTGGAGTCGTTGCTCTATATTTGGGGCAGAGTGGGGNNNAGATCGGAA
70999 TATGAGTGTGGAGTCGTTGCTTAGAAAGACTGAGCCCACCCATNNNAGATCGGAAG
50540 TATGAGTGTGGAGTCGTTGCATGTGAAGGTGAGCAGGCATAAGNNNAGATCGGAA
52530 TATGAGTGTGGAGTCGTTGCTTGTTCCTTGGAATTGCAATGCGNNNAGATCGGAAG
105083 TATGAGTGTGGAGTCGTTGCATTGTATGTGTCCATGATGAGTGCNNNAGATCGGAA
50633 TATGAGTGTGGAGTCGTTGCGGGGTGTTACTGCTACTGAACAGNNNAGATCGGAA
106292 TATGAGTGTGGAGTCGTTGCAACTGATTCACTCCCACCTCTTTNNNAGATCGGAAGA
31020 TATGAGTGTGGAGTCGTTGCTTGGGGAATGCTTTTTGACCCAANNNAGATCGGAAG
68012 TATGAGTGTGGAGTCGTTGCAAACAATTGCACAAAGCAATCTTNNNAGATCGGAAG

55740 TATGAGTGTGGAGTCGTTGCATGACACGCTTGTTGACAAACTGNNNAGATCGGAAG
68434 TATGAGTGTGGAGTCGTTGCGGTGCATGGAGACACATTAGGTTNNNAGATCGGAA
99555 TATGAGTGTGGAGTCGTTGCGATTTTCTCCAAGGCCATGTGTCNNNAGATCGGAAG
98189 TATGAGTGTGGAGTCGTTGCTTGCTGCCATTACTGTGGCTCANNNAGATCGGAAGA
61559 TATGAGTGTGGAGTCGTTGCCCTGAAAATGCCTCTGTGTTCCTNNNAGATCGGAAG
53884 TATGAGTGTGGAGTCGTTGCTACTGGTGTATAGAGCTCAGGGGNNNAGATCGGAA
58468 TATGAGTGTGGAGTCGTTGCCATATTACCACCACCTCCCCCATNNNAGATCGGAAGA
62639 TATGAGTGTGGAGTCGTTGCGACATCATGGGTGACATGCATCTNNNAGATCGGAAG
50207 TATGAGTGTGGAGTCGTTGCGAGTTGGAGGCTGAGAGAACAAGNNNAGATCGGA
39154 TATGAGTGTGGAGTCGTTGCTATCTACTGAGAGAGGCCTGGGANNNAGATCGGAA
55111 TATGAGTGTGGAGTCGTTGCAAAAGCTGTGGGATTGGAGTTGGNNNAGATCGGA
30341 TATGAGTGTGGAGTCGTTGCCTTGGCATTTCCCTCTCTCCAAANNNAGATCGGAAGA
60449 TATGAGTGTGGAGTCGTTGCACAAAATGGGGTCATCTCATACTNNNAGATCGGAAG
108762 TATGAGTGTGGAGTCGTTGCAACAGGCAGTAAAGCAAACAATTNNNAGATCGGAA
31464 TATGAGTGTGGAGTCGTTGCCAGGTTGTCCTTTACCCTGTCAANNNAGATCGGAAG
64772 TATGAGTGTGGAGTCGTTGCTGCCAAACCAAAATCTGGAAGGTNNNAGATCGGAA
99449 TATGAGTGTGGAGTCGTTGCCCATGTAACCTCCCACTACCACCTNNNAGATCGGAAG
60803 TATGAGTGTGGAGTCGTTGCTCTTTCCATGCTGGCTAGTACCTNNNAGATCGGAAG
47576 TATGAGTGTGGAGTCGTTGCGAGAGAGAGATGCACATGGATGCNNNAGATCGGA
48820 TATGAGTGTGGAGTCGTTGCACAAAGTGACAAACCCCTTCCTCNNNAGATCGGAAG
71611 TATGAGTGTGGAGTCGTTGCGGCTGGCAAGATATGGAGCTTTTNNNAGATCGGAA
117197 TATGAGTGTGGAGTCGTTGCAAACTTCTGAGTGGGTAGAATCTTGNNNAGATCGG
37851 TATGAGTGTGGAGTCGTTGCTAGTCACCTTTTACGTCCCCAGANNNAGATCGGAAG
44355 TATGAGTGTGGAGTCGTTGCTAGGGGATTGATTTCTCGACCCCNNNAGATCGGAAG
34484 TATGAGTGTGGAGTCGTTGCCCAAATGTTCAGTCAATTGTCCCANNNAGATCGGAA
32529 TATGAGTGTGGAGTCGTTGCGTGCTCTCCGTATGTGTGTGTAANNNAGATCGGAAG
65044 TATGAGTGTGGAGTCGTTGCAGTAGAAGATGAAGGGCAAGGGTNNNAGATCGGA
33133 TATGAGTGTGGAGTCGTTGCCAGCCCAAACAGTTAGCTACACANNNAGATCGGAAG
71784 TATGAGTGTGGAGTCGTTGCCTGAGCCTCTTTCTCATCCCAAGNNNAGATCGGAAGA
54186 TATGAGTGTGGAGTCGTTGCTAGGTTTTATGGCTGTTCGTGGGNNNAGATCGGAA
57682 TATGAGTGTGGAGTCGTTGCGGGAGGGGAAGAGGTTTCAAAATNNNAGATCGGA
60462 TATGAGTGTGGAGTCGTTGCATACACACCTCACCAGCACTACTNNNAGATCGGAAGA
57283 TATGAGTGTGGAGTCGTTGCCATACAAACACACTGTGGCCTTGNNNAGATCGGAAG
34545 TATGAGTGTGGAGTCGTTGCTGAGAGAGAGAAAGAGCAAGCCANNNAGATCGGA
30296 TATGAGTGTGGAGTCGTTGCTCCCTCTTTTCTTCCAGACCAAANNNAGATCGGAAGA
67849 TATGAGTGTGGAGTCGTTGCCACCCACAGATTCCTAACAGCTTNNNAGATCGGAAG
53195 TATGAGTGTGGAGTCGTTGCCCATGAATGTGGTTCAGAGGAGGNNNAGATCGGAA
58854 TATGAGTGTGGAGTCGTTGCCTGCTATTCCACTGTGACGTCATNNNAGATCGGAAG
33195 TATGAGTGTGGAGTCGTTGCACCTACTGCCTAATAAGCCCACANNNAGATCGGAAG
40683 TATGAGTGTGGAGTCGTTGCTATCCCAAGTGGTGGCCCACATANNNAGATCGGAAG
66697 TATGAGTGTGGAGTCGTTGCCTCTTCCTCTGTCATGCAGTTGTNNNAGATCGGAAGA
41167 TATGAGTGTGGAGTCGTTGCCTGGTCCTTGTCTGGAGTAGCTANNNAGATCGGAAG
53170 TATGAGTGTGGAGTCGTTGCGACAGACTTGCTTGGTGTAGAGGNNNAGATCGGAA

55893 TATGAGTGTGGAGTCGTTGCGGGAATGAACTGAGGCAGTACTGNNNAGATCGGAA
99231 TATGAGTGTGGAGTCGTTGCTGCTGCCTCTCAAACACACATATCNNNAGATCGGAAG
29796 TATGAGTGTGGAGTCGTTGCACTAAGGATAAACCACTGGCCCANNNAGATCGGAAG
60018 TATGAGTGTGGAGTCGTTGCTGCCCATTTATCTGTTGATGACACTNNNAGATCGGAA
30505 TATGAGTGTGGAGTCGTTGCTTCCTAATGACCGTATAGCAGAAANNNAGATCGGAA
59020 TATGAGTGTGGAGTCGTTGCTGGCTACTCCTGGGATTTCAGATNNNAGATCGGAAG
52305 TATGAGTGTGGAGTCGTTGCCCTCATTCTGATAGGCAGTCACGNNNAGATCGGAAG
71170 TATGAGTGTGGAGTCGTTGCTCTGAGTGGGTAGAATCTTGCCTNNNAGATCGGAAG
51879 TATGAGTGTGGAGTCGTTGCGCAGGAAGACATACCACAGTGAGNNNAGATCGGAA
41090 TATGAGTGTGGAGTCGTTGCAGATTGCTTTTCTTGGCTGCCTANNNAGATCGGAAG
64041 TATGAGTGTGGAGTCGTTGCTGCCATCCTAGAAATAGGGCAGTNNNAGATCGGAAG
62992 TATGAGTGTGGAGTCGTTGCGGTGGAAGGTAGGTGTGATCTCTNNNAGATCGGAA
68129 TATGAGTGTGGAGTCGTTGCAACATTTCCAGCACTGGTGTCTTNNNAGATCGGAAG
62856 TATGAGTGTGGAGTCGTTGCACTAACATCAAGCACCTCCCTCTNNNAGATCGGAAGA
68576 TATGAGTGTGGAGTCGTTGCGCAATAACACAGATGAAGGATGTTNNNAGATCGGA
48321 TATGAGTGTGGAGTCGTTGCACCCTTCCTGTAGAGGACACATCNNNAGATCGGAAG
69562 TATGAGTGTGGAGTCGTTGCGGAGGGAAGTGTGTGTAGGTTTTNNNAGATCGGAA
47857 TATGAGTGTGGAGTCGTTGCGACTGGCTGTGTCATTTTTCTGCNNNAGATCGGAAG
52515 TATGAGTGTGGAGTCGTTGCAGAGAGAGAGAGATGAAAGGGCGNNNAGATCGGA
116273 TATGAGTGTGGAGTCGTTGCAAGGGCAAAAGGACATTAGTGGCANNNAGATCGGA
44191 TATGAGTGTGGAGTCGTTGCATACCTACCCTGAGGGCATTACCNNNAGATCGGAAG
66599 TATGAGTGTGGAGTCGTTGCCGGTGCCCTTCAAGGAATATTGTNNNAGATCGGAAG
51721 TATGAGTGTGGAGTCGTTGCGGAAAAGTCCATGACGCTGGAGNNNAGATCGGAAG
41621 TATGAGTGTGGAGTCGTTGCTTGAGTGACATAGCCCCACTGTANNNAGATCGGAAG
32124 TATGAGTGTGGAGTCGTTGCCTAAAAGGAGCGCCAACTCTGAANNNAGATCGGAAG
30055 TATGAGTGTGGAGTCGTTGCTACTGGCTTTGGCTGCTACAAAANNNAGATCGGAAG
55581 TATGAGTGTGGAGTCGTTGCCAAAAGGACCCAGACTGGAGATGNNNAGATCGGAA
103161 TATGAGTGTGGAGTCGTTGCGCTCCAAACTGTCTTCAAAGGGTNNNAGATCGGAAG
68551 TATGAGTGTGGAGTCGTTGCCCCAGTGGCTCTAGTTCAATGTTNNNAGATCGGAAG
107966 TATGAGTGTGGAGTCGTTGCCAGGAGCAACAAATATCTTCACCAAGANNNAGATCG
61957 TATGAGTGTGGAGTCGTTGCCAATTTCCAGTGAGGAACAGGCTNNNAGATCGGAAG
31693 TATGAGTGTGGAGTCGTTGCCTGCTAAAAGAGACAGCCGAGAANNNAGATCGGAA
70547 TATGAGTGTGGAGTCGTTGCACACACCCCAACACCTTTATGTCNNNAGATCGGAAGA
42804 TATGAGTGTGGAGTCGTTGCACCCAAGGAAACAGGAATTCCACNNNAGATCGGAAG
67995 TATGAGTGTGGAGTCGTTGCAATGGGGTCTTGCTTAGTTGCTTNNNAGATCGGAAG
66910 TATGAGTGTGGAGTCGTTGCGTGTTCCTTCCCACCCTCAAATTNNNAGATCGGAAGA
56321 TATGAGTGTGGAGTCGTTGCCGGGTTTGGTTTGATCTCATCTGNNNAGATCGGAAG
31563 TATGAGTGTGGAGTCGTTGCGACAAGCAGTACCTGCAGAAGAANNNAGATCGGAA
55804 TATGAGTGTGGAGTCGTTGCCTAACCAGGAACCAATCCCACTGNNNAGATCGGAAG
41289 TATGAGTGTGGAGTCGTTGCCCTGATTCCTTCCCCACAGTCTANNNAGATCGGAAGA
42736 TATGAGTGTGGAGTCGTTGCAGGGGGAACCACTATAATCCCACNNNAGATCGGAAG
47467 TATGAGTGTGGAGTCGTTGCGCAATCTTCCAGAGCTTCAATGCNNNAGATCGGAAG
117596 TATGAGTGTGGAGTCGTTGCTTCCGCCATCTGATCTTGGACTNNNAGATCGGAAGA

60346 TATGAGTGTGGAGTCGTTGCACAGAAGAAGGAAGAAAATTTGGGACTNNNAGATC
42694 TATGAGTGTGGAGTCGTTGCTGACTTACTGAAGCTGTGACCACNNNAGATCGGAAG
53471 TATGAGTGTGGAGTCGTTGCGGTGGCCCAATGTATATCTATACGGNNNAGATCGGA
54123 TATGAGTGTGGAGTCGTTGCGGTTTACAGTTCTTGAGGCTGGGNNNAGATCGGAAG
61068 TATGAGTGTGGAGTCGTTGCAAGTAGTGCATTCAAGCAAGCCTNNNAGATCGGAAG
33283 TATGAGTGTGGAGTCGTTGCCAGACATCATACACACAGGCACANNNAGATCGGAAG
33042 TATGAGTGTGGAGTCGTTGCCCCAAATATACCGGACACACACANNNAGATCGGAAGA
43893 TATGAGTGTGGAGTCGTTGCAAGAAATGAGCACACCTACCACCNNNAGATCGGAAG
54406 TATGAGTGTGGAGTCGTTGCACATGACAACCCTTCCTTCATGGNNNAGATCGGAAGA
66691 TATGAGTGTGGAGTCGTTGCAGAGAAATATACCTACGACAGTTGTNNNAGATCGGA
55631 TATGAGTGTGGAGTCGTTGCTGTGTAGGATTGGCCTTTGGATGNNNAGATCGGAAG
49319 TATGAGTGTGGAGTCGTTGCTCAGATTTGGTCTTGCAAGGGTCNNNAGATCGGAAG
98951 TATGAGTGTGGAGTCGTTGCATGTCACTGCCACTATACTGCAGNNNAGATCGGAAGA
68086 TATGAGTGTGGAGTCGTTGCGTAGGTGGAGTTAGGGAGCTCTTNNNAGATCGGAA
71670 TATGAGTGTGGAGTCGTTGCTCTGCATCCCTTGTTTCCTGATGNNNAGATCGGAAGA
41973 TATGAGTGTGGAGTCGTTGCAGTGAGGCCCAGAGAAGTTGTTANNNAGATCGGAA
50299 TATGAGTGTGGAGTCGTTGCACTTGGGAAAATATGGGGGCAAGNNNAGATCGGAA
38867 TATGAGTGTGGAGTCGTTGCTATGCCTGTGTCTATGGTGAGGANNNAGATCGGAAG
102280 TATGAGTGTGGAGTCGTTGCTTTCAGGGAAGAGGGAAACATTCTGTNNNAGATCGG
61102 TATGAGTGTGGAGTCGTTGCCCACACTGTCTTCAAAAGAGCCTNNNAGATCGGAAGA
102626 TATGAGTGTGGAGTCGTTGCATTACTCCAAGTCCCAAGTCCACNNNAGATCGGAAGA
56588 TATGAGTGTGGAGTCGTTGCAAGCATGTAAAGAGAGGGCAGTGNNNAGATCGGAA
30576 TATGAGTGTGGAGTCGTTGCGCCAGGACAGCTAAATCAGGAAANNNAGATCGGAAG
63879 TATGAGTGTGGAGTCGTTGCATTTTTGTGAGTGTCCCCACAGTNNNAGATCGGAAGA
70150 TATGAGTGTGGAGTCGTTGCTTGTGGAAGGGACCTGTATGCTANNNAGATCGGAAG
46453 TATGAGTGTGGAGTCGTTGCCCCACTCTTTCTCTTCCAAGAGCNNNAGATCGGAAGA
38313 TATGAGTGTGGAGTCGTTGCGTTAGCATTTGCTTTGCCTGAGANNNAGATCGGAAG
35333 TATGAGTGTGGAGTCGTTGCTAGATCCTTTCCTCTGCTCAGCANNNAGATCGGAAGA
43926 TATGAGTGTGGAGTCGTTGCTTACTCCAAGTCCCAAGTCCACCNNNAGATCGGAAGA
59638 TATGAGTGTGGAGTCGTTGCGGAATGGCTGCATCTGTCTGTATNNNAGATCGGAAG
104538 TATGAGTGTGGAGTCGTTGCAGAAGGGTGACTTGAGCTCTAGNNNAGATCGGAAG
56285 TATGAGTGTGGAGTCGTTGCATGTTTATTGCAGCATCTTGCTGNNNAGATCGGAAGA
41824 TATGAGTGTGGAGTCGTTGCAACTTCTCCCCAAGCACTCCTTANNNAGATCGGAAGA
46435 TATGAGTGTGGAGTCGTTGCCACACAGTGGAAAGCTATTCAGCNNNAGATCGGAAG
34929 TATGAGTGTGGAGTCGTTGCTGATTCCTGTTCCTCAGTCTCCANNNAGATCGGAAGA
30231 TATGAGTGTGGAGTCGTTGCGTTCTCTGTGGAACCCCAACAAANNNAGATCGGAAG
99559 TATGAGTGTGGAGTCGTTGCTGGGATATTTAGGACAACCAAGGGNNNAGATCGGA
97332 TATGAGTGTGGAGTCGTTGCCCCGAGACAGTATCTCACCATATNNNAGATCGGAAGA
34399 TATGAGTGTGGAGTCGTTGCAATAAGTGGCCAAAATCTGCCCANNNAGATCGGAAG
57199 TATGAGTGTGGAGTCGTTGCTCATCTCCTGGGTCCTGAGATTGNNNAGATCGGAAG
30163 TATGAGTGTGGAGTCGTTGCGGGCCTCTTTGAGGAGATGAAAANNNAGATCGGAA
37818 TATGAGTGTGGAGTCGTTGCAAAGCTAATGGGGAGAATACAGANNNAGATCGGAA
33163 TATGAGTGTGGAGTCGTTGCCAGTCGTACAGTTATTCACCACANNNAGATCGGAAGA

30795 TATGAGTGTGGAGTCGTTGCGGCACAGCTGTTGTGAAAAACAANNNAGATCGGAA
69410 TATGAGTGTGGAGTCGTTGCTGAAACCAATAAGGAAAAGTGAATTTTNNNAGATCG
44925 TATGAGTGTGGAGTCGTTGCGTAGGTAGTTCAGGAAAGCAGCCNNNAGATCGGAA
45417 TATGAGTGTGGAGTCGTTGCCTTATTGGATCAGGGCACCATCCNNNAGATCGGAAG
30419 TATGAGTGTGGAGTCGTTGCTCTCTGACACCAAACACATCAAANNNAGATCGGAAG
108767 TATGAGTGTGGAGTCGTTGCGGTGTGTTACAGTGTCAGTGGGNNNAGATCGGAAG
63414 TATGAGTGTGGAGTCGTTGCGCCTCCTCATCCCTTCATCTTCTNNNAGATCGGAAGA
60007 TATGAGTGTGGAGTCGTTGCACCTGCTTATCCACTCAGACACTNNNAGATCGGAAGA
39928 TATGAGTGTGGAGTCGTTGCTGGTAAGGAAATGAGGCTCCTGANNNAGATCGGAA
117673 TATGAGTGTGGAGTCGTTGCCACATGCACGCCTATATTAGTCTGNNNAGATCGGAA
99281 TATGAGTGTGGAGTCGTTGCATTGAGCCCTCAGAAAGCCTTAGNNNAGATCGGAAG
43263 TATGAGTGTGGAGTCGTTGCAGGAACTGTTTGGAGACAAGGACNNNAGATCGGAA
60691 TATGAGTGTGGAGTCGTTGCTGGAATGACCAAAGAAAGCACCTNNNAGATCGGAA
100909 TATGAGTGTGGAGTCGTTGCGGAAAGATGGATTTGTGTGGAGGTNNNAGATCGG
53277 TATGAGTGTGGAGTCGTTGCATTCGTAAGCTGTCACACTGAGGNNNAGATCGGAA
57335 TATGAGTGTGGAGTCGTTGCAAGGTTGATGGGAAGTGTGCTTGNNNAGATCGGA
32142 TATGAGTGTGGAGTCGTTGCCTGCATCCTCCAAAGTCAGTGAANNNAGATCGGAAG
43857 TATGAGTGTGGAGTCGTTGCGGATCAGAGTGTTGCAAAACACCNNNAGATCGGAA
34790 TATGAGTGTGGAGTCGTTGCCTGGCACTTCACGTATCAATCCANNNAGATCGGAAG
53655 TATGAGTGTGGAGTCGTTGCGGATCACTCCTTTCTTCACAGGGNNNAGATCGGAAG
61476 TATGAGTGTGGAGTCGTTGCTACTAAATGCCAGGCTCTGTCCTNNNAGATCGGAAG
107917 TATGAGTGTGGAGTCGTTGCCCATCTCACGATCTCAGCTCACTNNNAGATCGGAAGA
49052 TATGAGTGTGGAGTCGTTGCCCACTCCCATACCTCTCTCTCTCNNNAGATCGGAAGA
37905 TATGAGTGTGGAGTCGTTGCACTGTCTTGTCTTCCCTTCCAGANNNAGATCGGAAGA
43380 TATGAGTGTGGAGTCGTTGCTATTTGCAGTCACGCCCTATGACNNNAGATCGGAAG
39185 TATGAGTGTGGAGTCGTTGCTAGATGAAGGTGCTGATGTGGGANNNAGATCGGA
61367 TATGAGTGTGGAGTCGTTGCCCCTTATCCTCCTCTCCACTCCTNNNAGATCGGAAGAG
36922 TATGAGTGTGGAGTCGTTGCCTGCACCACATCAGTAATCGTCANNNAGATCGGAAG
60891 TATGAGTGTGGAGTCGTTGCTAAAATACAAGCCAGGACCCCCTNNNAGATCGGAAG
53786 TATGAGTGTGGAGTCGTTGCCTGAGGGTAGGGATTACTGAGGGNNNAGATCGGA
49890 TATGAGTGTGGAGTCGTTGCCTCTACCTTCCATGCCCACTTTCNNNAGATCGGAAGA
49825 TATGAGTGTGGAGTCGTTGCAGAAGGTCCCCTCACCTATGTTCNNNAGATCGGAAG
34876 TATGAGTGTGGAGTCGTTGCTTTCCCCAATTCTGGGTACTCCANNNAGATCGGAAGA
109549 TATGAGTGTGGAGTCGTTGCGTGTCAGTGTTGTAGGACTTTTTCCNNNAGATCGGA
68474 TATGAGTGTGGAGTCGTTGCTTGGAGAAGCACGAACAATGGTTNNNAGATCGGAA
31659 TATGAGTGTGGAGTCGTTGCCATTCACACAGCCACTCTCAGAANNNAGATCGGAAG
97737 TATGAGTGTGGAGTCGTTGCCCTATGGCATCTAAAGAAGGAGTGTANNNAGATCG
100898 TATGAGTGTGGAGTCGTTGCATTTGCATCTCCTTCCAAGTAACTCTNNNAGATCGGA
102691 TATGAGTGTGGAGTCGTTGCCAACAGCAACAATAAGAGCTAGCCNNNAGATCGGAA
56096 TATGAGTGTGGAGTCGTTGCACGAAAGTTGCATACACAAGCTGNNNAGATCGGAA
34811 TATGAGTGTGGAGTCGTTGCTTCCTTTCATGCTGAAGCATCCANNNAGATCGGAAG
41153 TATGAGTGTGGAGTCGTTGCTTGCCCTACTTGCATTTCAGCTANNNAGATCGGAAGA
67855 TATGAGTGTGGAGTCGTTGCAATGACTACAAACCTGGCAGCTTNNNAGATCGGAAG

30982 TATGAGTGTGGAGTCGTTGCATCTTTGATGGCTGTCCAACCAANNNAGATCGGAAG

52153 TATGAGTGTGGAGTCGTTGCCCCCCACCACCTTTTATTGGTAGNNNAGATCGGAAG

116958 TATGAGTGTGGAGTCGTTGCAGGAAACAGGTAGAAGGAGACTATATCNNNAGATC

69628 TATGAGTGTGGAGTCGTTGCCCCCTGGTTTCCACCCTATTTTTNNNAGATCGGAAGA

50221 TATGAGTGTGGAGTCGTTGCGACATAAACTCGCCAAGCACAAGNNNAGATCGGAA

53881 TATGAGTGTGGAGTCGTTGCAAATTCAGATTGAACCCCAGGGGNNNAGATCGGAA

36712 TATGAGTGTGGAGTCGTTGCAACAACTCTGTGTAACTGGCTCANNNAGATCGGAAG

61558 TATGAGTGTGGAGTCGTTGCAAATTCTTGCAGAGCCTGTTCCTNNNAGATCGGAAG

59872 TATGAGTGTGGAGTCGTTGCCTCTGGTGGCAAGAAGAAGAACTNNNAGATCGGAA

106767 TATGAGTGTGGAGTCGTTGCCTCCCAGGTTCAAGCGTGATCTCNNNAGATCGGAAG

42753 TATGAGTGTGGAGTCGTTGCGCTAAATTACAGTCAAGCGCCACNNNAGATCGGAAG

109499 TATGAGTGTGGAGTCGTTGCCGTGTCTTTCAGCATGGGGANNNAGATCGGAAGAG

41003 TATGAGTGTGGAGTCGTTGCACAATAGGCTGCCTGCAATACTANNNAGATCGGAAG

117835 TATGAGTGTGGAGTCGTTGCTAGCCTTGTCTCTCTGTGTCCATTNNNAGATCGGAAG

60094 TATGAGTGTGGAGTCGTTGCTGTCTCCAACTGACTTCTCCACTNNNAGATCGGAAGA

101492 TATGAGTGTGGAGTCGTTGCACGATGTACTGGCTGGAAGGAANNNAGATCGGAAG

66498 TATGAGTGTGGAGTCGTTGCTCTTCCCCATCCATACTGTGTGTNNNAGATCGGAAGA

56605 TATGAGTGTGGAGTCGTTGCTGAGCCAATCATGGAAGTCAGTGNNNAGATCGGAA

44190 TATGAGTGTGGAGTCGTTGCCAAGCCCAAGGCATCAAATTACCNNNAGATCGGAAG

36942 TATGAGTGTGGAGTCGTTGCAACAGATTCACACTTCGAGGTCANNNAGATCGGAAG

104474 TATGAGTGTGGAGTCGTTGCGTAGAAAAGTAGGTGAGAACACGTNNNAGATCGG

70953 TATGAGTGTGGAGTCGTTGCGGGGACAGCCTGTACTACATTTGNNNAGATCGGAA

30013 TATGAGTGTGGAGTCGTTGCAACCGTGTCCAATCCCAGAAAAANNNAGATCGGAAG

107932 TATGAGTGTGGAGTCGTTGCTTTATGGCATTTACAGCGACCNNNAGATCGGAAGAG

50328 TATGAGTGTGGAGTCGTTGCATGCCCGCACTCTATTCATCAAGNNNAGATCGGAAG

37166 TATGAGTGTGGAGTCGTTGCATAATGGCGCTACAGCAACTTCANNNAGATCGGAAG

32414 TATGAGTGTGGAGTCGTTGCCACAGCCAGTCAGAACTTGCTAANNNAGATCGGAAG

36376 TATGAGTGTGGAGTCGTTGCACTCAACTCCCAATAAACAGGAATCANNNAGATCGG

109551 TATGAGTGTGGAGTCGTTGCATGGGGATTTGGTTCTAGGATCCNNNAGATCGGAA

116923 TATGAGTGTGGAGTCGTTGCCTGTGGAACCCCAACAAATACAACTNNNAGATCGGA

67392 TATGAGTGTGGAGTCGTTGCCCCAACCATGGCCTAAACCTATTNNNAGATCGGAAG

68107 TATGAGTGTGGAGTCGTTGCGGCACCACTACTTTTCCAGTCTTNNNAGATCGGAAG

63574 TATGAGTGTGGAGTCGTTGCTTGGTCCTCATGATCCCTTTTCTNNNAGATCGGAAGA

34739 TATGAGTGTGGAGTCGTTGCCCATCCCCATATTGTTTCTGCCANNNAGATCGGAAGA

52104 TATGAGTGTGGAGTCGTTGCGCTTCTTTGGGAAGGGGAAGTAGNNNAGATCGGA

46181 TATGAGTGTGGAGTCGTTGCCAGAAAGAGAGAGAGCGAGAAGCNNNAGATCGGA

46258 TATGAGTGTGGAGTCGTTGCGAGAGAGCATTCCATTCAACAGCNNNAGATCGGAA

64857 TATGAGTGTGGAGTCGTTGCAATCACCCAGAATCCTCTCAGGTNNNAGATCGGAAG

109493 TATGAGTGTGGAGTCGTTGCTTACTAGCCATACCATCTGGCCTNNNAGATCGGAAG

66029 TATGAGTGTGGAGTCGTTGCGTCCAGAATGCTCACATTCCTGTNNNAGATCGGAAG

112575 TATGAGTGTGGAGTCGTTGCCCCCAACCATGGCCTAAACCTATTANNNAGATCGGAA

29794 TATGAGTGTGGAGTCGTTGCAACTTAGTCTGAGGGATTTGGCANNNAGATCGGAA

42669 TATGAGTGTGGAGTCGTTGCGCTTCTCTTTCCTATGGAACCACNNNAGATCGGAAG

71688 TATGAGTGTGGAGTCGTTGCGACAGGAAAGCCTCAAAGGGTTTNNNAGATCGGAA[illegible]
30717 TATGAGTGTGGAGTCGTTGCGATGGTGAGAGGGGGAGGATAAANNNAGATCGG[illegible]
29928 TATGAGTGTGGAGTCGTTGCGAGCTATTTCTTTGCTGACGTGCNNNAGATCGGAAG

AGCACACGTCTGAACTCTTTCCCTACACGACGCTCTTCCGATCTNNNGACCCGCATTGATAATCAGAG
GAGCACACGTCTGAACTCTTTCCCTACACGACGCTCTTCCGATCTNNNTATTTAGGCCCTGCTTTACCT
GAGCACACGTCTGAACTCTTTCCCTACACGACGCTCTTCCGATCTNNNTGGCTAAACTTTTCTAGGCAT
AGCACACGTCTGAACTCTTTCCCTACACGACGCTCTTCCGATCTNNNAGAGAATAAAGAGAGTATTT
AGCACACGTCTGAACTCTTTCCCTACACGACGCTCTTCCGATCTNNNCAAGGATTAGAGGAGGCATG
AGAGCACACGTCTGAACTCTTTCCCTACACGACGCTCTTCCGATCTNNNCAAATACCTGTGAGCGTTG
GAGCACACGTCTGAACTCTTTCCCTACACGACGCTCTTCCGATCTNNNTGCATAAAGAATAGGGAGG
AGAGCACACGTCTGAACTCTTTCCCTACACGACGCTCTTCCGATCTNNNTCCGAATACTGGGATTAGA
GAGCACACGTCTGAACTCTTTCCCTACACGACGCTCTTCCGATCTNNNTCAATGACAAATTTGATCTG
AGCACACGTCTGAACTCTTTCCCTACACGACGCTCTTCCGATCTNNNTTAGAATTCCAGGATGGTGTG
AGCACACGTCTGAACTCTTTCCCTACACGACGCTCTTCCGATCTNNNTCTGTCAAGGTCTGTCTAGGTT
GAGCACACGTCTGAACTCTTTCCCTACACGACGCTCTTCCGATCTNNNTTAGGAGCTCGTTCCCCTGAT
AGCACACGTCTGAACTCTTTCCCTACACGACGCTCTTCCGATCTNNNATCGCTGTTCATCAAGGATGC
AGCACACGTCTGAACTCTTTCCCTACACGACGCTCTTCCGATCTNNNAGTGAGCTGAGACCACTGACT
GCACACGTCTGAACTCTTTCCCTACACGACGCTCTTCCGATCTNNNGGCTTTTTGCACTAACAGCAACT
GCACACGTCTGAACTCTTTCCCTACACGACGCTCTTCCGATCTNNNTCAAAGATATGTGTGTCCGTGT
GCACACGTCTGAACTCTTTCCCTACACGACGCTCTTCCGATCTNNNAAGATGTGTAGCAGCTGTGGT
AGCACACGTCTGAACTCTTTCCCTACACGACGCTCTTCCGATCTNNNCTGGCCAGAGAAGTTTTTGAC
CACACGTCTGAACTCTTTCCCTACACGACGCTCTTCCGATCTNNNTGTCTCCCTTCCTCTCTTCTGTCCA
AGCACACGTCTGAACTCTTTCCCTACACGACGCTCTTCCGATCTNNNTTGTGTCAGTGTTCATCGTGG
GAGCACACGTCTGAACTCTTTCCCTACACGACGCTCTTCCGATCTNNNTGTCATCACTCTGACCACAGC
GCACACGTCTGAACTCTTTCCCTACACGACGCTCTTCCGATCTNNNTAGACACTGGGGCCAACTTGAT
AGCACACGTCTGAACTCTTTCCCTACACGACGCTCTTCCGATCTNNNGTTGGGAACTGTGGACAAAA
CACACGTCTGAACTCTTTCCCTACACGACGCTCTTCCGATCTNNNTGATGGCCAACTAGTCTAACATGG
AGCACACGTCTGAACTCTTTCCCTACACGACGCTCTTCCGATCTNNNCAAACAGCAGCCCTACAGAAG
GCACACGTCTGAACTCTTTCCCTACACGACGCTCTTCCGATCTNNNGACATCATGATCTACCTGCTTCC
GCACACGTCTGAACTCTTTCCCTACACGACGCTCTTCCGATCTNNNACATGGTGCATTATTCAAAACAA
CACACGTCTGAACTCTTTCCCTACACGACGCTCTTCCGATCTNNNAAAATGCCTTGGACACTACCCAG
CACACGTCTGAACTCTTTCCCTACACGACGCTCTTCCGATCTNNNAGCTGAGAACATAGGCATGAACT
AGCACACGTCTGAACTCTTTCCCTACACGACGCTCTTCCGATCTNNNTCTAGTACTGGACGATTTGTG
AGCACACGTCTGAACTCTTTCCCTACACGACGCTCTTCCGATCTNNNTCCTTTCTCTATCTCTCTTTCGCC
CACACGTCTGAACTCTTTCCCTACACGACGCTCTTCCGATCTNNNTAGGAGAAAACTTGAGCCCAGGA
GCACACGTCTGAACTCTTTCCCTACACGACGCTCTTCCGATCTNNNTGTCTATTCTCCACAGGCTTTGC
AGCACACGTCTGAACTCTTTCCCTACACGACGCTCTTCCGATCTNNNACCCCTGGCTTGAAAATTTGG
AGCACACGTCTGAACTCTTTCCCTACACGACGCTCTTCCGATCTNNNGCTTACCAGGTAGCTGGGAG
AGCACACGTCTGAACTCTTTCCCTACACGACGCTCTTCCGATCTNNNAGTGAGGCATGATGGTAGG
GAGCACACGTCTGAACTCTTTCCCTACACGACGCTCTTCCGATCTNNNACCTAGAGCAAGAGTTGATG
GCACACGTCTGAACTCTTTCCCTACACGACGCTCTTCCGATCTNNNTCTTTCTCAGCATTGCCAACCTT
GAGCACACGTCTGAACTCTTTCCCTACACGACGCTCTTCCGATCTNNNCTTTTTGCTTAGGATTGTCG
AGCACACGTCTGAACTCTTTCCCTACACGACGCTCTTCCGATCTNNNAGGGCCCAGCTAGAAAAACAA
AGCACACGTCTGAACTCTTTCCCTACACGACGCTCTTCCGATCTNNNGTGGAGATCCTGCCAATTCACT
GAGCACACGTCTGAACTCTTTCCCTACACGACGCTCTTCCGATCTNNNGCTGAGACAAGGGAGTGTT

AGAGCACACGTCTGAACTCTTTCCCTACACGACGCTCTTCCGATCTNNNAAGACGCATAAGTAAAGAAA
AAGAGCACACGTCTGAACTCTTTCCCTACACGACGCTCTTCCGATCTNNNTATTGCACCTTGCCAGGTA
GAAGAGCACACGTCTGAACTCTTTCCCTACACGACGCTCTTCCGATCTNNNATGGTGTTTTGTGATGTT
AGCACACGTCTGAACTCTTTCCCTACACGACGCTCTTCCGATCTNNNCTGAAGGAGGATGAAACAGTG
GAGCACACGTCTGAACTCTTTCCCTACACGACGCTCTTCCGATCTNNNTTCACCATGCCGGTGATAAAC
GCACACGTCTGAACTCTTTCCCTACACGACGCTCTTCCGATCTNNNTTGCCTTCTATCAGTAACCCCACA
AGCACACGTCTGAACTCTTTCCCTACACGACGCTCTTCCGATCTNNNCGCCAACATATCTGTGTATTTG
GAGCACACGTCTGAACTCTTTCCCTACACGACGCTCTTCCGATCTNNNTCATCTGGAATCTGTCGGTTC
AAGAGCACACGTCTGAACTCTTTCCCTACACGACGCTCTTCCGATCTNNNACAAATTCAAAGCCAGCAC
GCACACGTCTGAACTCTTTCCCTACACGACGCTCTTCCGATCTNNNCCGCAGAAATTTTGATGGGGAGA
AGAGCACACGTCTGAACTCTTTCCCTACACGACGCTCTTCCGATCTNNNAGAAGCTCACCTTACCACAG
AGCACACGTCTGAACTCTTTCCCTACACGACGCTCTTCCGATCTNNNTTGGACAAGGGACAGTAAAGCA
GAGCACACGTCTGAACTCTTTCCCTACACGACGCTCTTCCGATCTNNNTACAACTGTGGGGGTAAAACT
AGCACACGTCTGAACTCTTTCCCTACACGACGCTCTTCCGATCTNNNGCATGACAGTAGAAGAGCTGCA
GAGCACACGTCTGAACTCTTTCCCTACACGACGCTCTTCCGATCTNNNACAACAGAAGCAGAGGAACAC
AGCACACGTCTGAACTCTTTCCCTACACGACGCTCTTCCGATCTNNNTTGTTTTCAAGCACGAAACTGGC
AGAGCACACGTCTGAACTCTTTCCCTACACGACGCTCTTCCGATCTNNNCACATACCAGGAGATTGCCT
AGCACACGTCTGAACTCTTTCCCTACACGACGCTCTTCCGATCTNNNTATACACCCACATGCCCAGGTAA
AGCACACGTCTGAACTCTTTCCCTACACGACGCTCTTCCGATCTNNNCTTGCAGCTGCCGATATATCACT
AGCACACGTCTGAACTCTTTCCCTACACGACGCTCTTCCGATCTNNNATAGAACTGCACAAAACAGCTT
AGCACACGTCTGAACTCTTTCCCTACACGACGCTCTTCCGATCTNNNTGGGTAGAGAAGGGCATATGC
GAGCACACGTCTGAACTCTTTCCCTACACGACGCTCTTCCGATCTNNNCAGTTTGCTCAGATATGCCCT
CACACGTCTGAACTCTTTCCCTACACGACGCTCTTCCGATCTNNNTTTATTTTCCTCCTCCTCCTTCTTCTT
AGAGCACACGTCTGAACTCTTTCCCTACACGACGCTCTTCCGATCTNNNTTGGGAGTGCCATTAGTTCA
AGCACACGTCTGAACTCTTTCCCTACACGACGCTCTTCCGATCTNNNGGGGTTGGAAGCATCTTCTGTA
GAGCACACGTCTGAACTCTTTCCCTACACGACGCTCTTCCGATCTNNNGGGACTTCTGCAAAACCATACC
GAGCACACGTCTGAACTCTTTCCCTACACGACGCTCTTCCGATCTNNNCCATGCTGGTGTCAAACCTCTA
GCACACGTCTGAACTCTTTCCCTACACGACGCTCTTCCGATCTNNNTGAAGTCTTTTAGTTGAAATGCGT
GAGCACACGTCTGAACTCTTTCCCTACACGACGCTCTTCCGATCTNNNCTGGTCATTTGAGGCCAAGAG
GAGCACACGTCTGAACTCTTTCCCTACACGACGCTCTTCCGATCTNNNTGAAATCCAGTCCAGTCTGGG
AGCACACGTCTGAACTCTTTCCCTACACGACGCTCTTCCGATCTNNNTCTGATGGTGTTGACTTCTCTGA
AAGAGCACACGTCTGAACTCTTTCCCTACACGACGCTCTTCCGATCTNNNGCGTAAATCCACGATACCA
GAGCACACGTCTGAACTCTTTCCCTACACGACGCTCTTCCGATCTNNNTGGTTAGCTGCTTCCTCTCTTC
AGCACACGTCTGAACTCTTTCCCTACACGACGCTCTTCCGATCTNNNTGTTAGGATCTACACATCTGCC
AGAGCACACGTCTGAACTCTTTCCCTACACGACGCTCTTCCGATCTNNNTTTCCCAGTCAAGGGAAAGG
GAGCACACGTCTGAACTCTTTCCCTACACGACGCTCTTCCGATCTNNNTGCAACCTCTGCTTCCTTTGAAT
AGCACACGTCTGAACTCTTTCCCTACACGACGCTCTTCCGATCTNNNCCCCTCAGAACTTGAAGGAACTG
AGCACACGTCTGAACTCTTTCCCTACACGACGCTCTTCCGATCTNNNAGCTTTTGCATGTACTACTGGG
GCACACGTCTGAACTCTTTCCCTACACGACGCTCTTCCGATCTNNNTTGATGGAGATGGGTTGCTGAA
AAGAGCACACGTCTGAACTCTTTCCCTACACGACGCTCTTCCGATCTNNNTAAATTTGACCTGCCCTCCT
AGCACACGTCTGAACTCTTTCCCTACACGACGCTCTTCCGATCTNNNTGTGTCGGTCTAGGAGGTTATC
GAAGAGCACACGTCTGAACTCTTTCCCTACACGACGCTCTTCCGATCTNNNAAGCAGATCACTGGTTGA
AGCACACGTCTGAACTCTTTCCCTACACGACGCTCTTCCGATCTNNNAATGTGGATCAACTATGTGTGTC

GAGCACACGTCTGAACTCTTTCCCTACACGACGCTCTTCCGATCTNNNCTACAGTTCTGGGTTCTTGG
CACACGTCTGAACTCTTTCCCTACACGACGCTCTTCCGATCTNNNTGTAGTCCCAGTTACTTGTGGTTC
GCACACGTCTGAACTCTTTCCCTACACGACGCTCTTCCGATCTNNNGAAATTTCATGTTTCTGCGGGG
ACACGTCTGAACTCTTTCCCTACACGACGCTCTTCCGATCTNNNATTTCTCCCTAAGGCACCCAGATCA
ACACGTCTGAACTCTTTCCCTACACGACGCTCTTCCGATCTNNNAGCTGTGAAATTACTCAGATGTTG
GAGCACACGTCTGAACTCTTTCCCTACACGACGCTCTTCCGATCTNNNAATTAGTGCTGAATGAAAG
AAGAGCACACGTCTGAACTCTTTCCCTACACGACGCTCTTCCGATCTNNNACATTTGCACCGACCTAAG
AGCACACGTCTGAACTCTTTCCCTACACGACGCTCTTCCGATCTNNNGCAAAGATGAGGAGTGGAA
GCACACGTCTGAACTCTTTCCCTACACGACGCTCTTCCGATCTNNNAAAGCCCAAAAATTAGGGAGG
AAGAGCACACGTCTGAACTCTTTCCCTACACGACGCTCTTCCGATCTNNNTGCTGGGTCTTGATTTCC
CACACGTCTGAACTCTTTCCCTACACGACGCTCTTCCGATCTNNNGCAAATGCTCCTCATTTCCTCCTCA
GCACACGTCTGAACTCTTTCCCTACACGACGCTCTTCCGATCTNNNTACAGATGTAAGCCACGAGCCT
AGCACACGTCTGAACTCTTTCCCTACACGACGCTCTTCCGATCTNNNGAAGCTTAGCAAGCCCCAAGT
AGAGCACACGTCTGAACTCTTTCCCTACACGACGCTCTTCCGATCTNNNTGCACTCCATCACGACATG
AGCACACGTCTGAACTCTTTCCCTACACGACGCTCTTCCGATCTNNNAAATCCGAGGTCTGAAACCTG
AGAGCACACGTCTGAACTCTTTCCCTACACGACGCTCTTCCGATCTNNNTACACACATGCACACACCAT
AGCACACGTCTGAACTCTTTCCCTACACGACGCTCTTCCGATCTNNNGTGCTTCCCTCTTCTTTCACTCT
CACACGTCTGAACTCTTTCCCTACACGACGCTCTTCCGATCTNNNAAGGATGAGATTGCCCCTCTTCTC
CACACGTCTGAACTCTTTCCCTACACGACGCTCTTCCGATCTNNNAGACTTTCTCCACTTCAATTTTCCC
GAGCACACGTCTGAACTCTTTCCCTACACGACGCTCTTCCGATCTNNNTTCCTACTTTTGTTGGTTCCC
AGCACACGTCTGAACTCTTTCCCTACACGACGCTCTTCCGATCTNNNCAAGGGAGTTGCCAATCTAGC
CACACGTCTGAACTCTTTCCCTACACGACGCTCTTCCGATCTNNNAAAAACAGGCATGGAGTTTGCAG
GCACACGTCTGAACTCTTTCCCTACACGACGCTCTTCCGATCTNNNCTGTATTCCCCACTGTGACAAG
AGCACACGTCTGAACTCTTTCCCTACACGACGCTCTTCCGATCTNNNAGCTGAGGAAACTTACCTACCA
CACACGTCTGAACTCTTTCCCTACACGACGCTCTTCCGATCTNNNGCTGATCCAGAAAGAGGTAAGC
GAGCACACGTCTGAACTCTTTCCCTACACGACGCTCTTCCGATCTNNNTTTATCCTCTCTCCTTGTTGC
AAGAGCACACGTCTGAACTCTTTCCCTACACGACGCTCTTCCGATCTNNNACTGAAGCATTTAGGTC
GCACACGTCTGAACTCTTTCCCTACACGACGCTCTTCCGATCTNNNGGTGTGAGCTCTGAATCCAGTTT
GCACACGTCTGAACTCTTTCCCTACACGACGCTCTTCCGATCTNNNACGATGACCTCCAACATTGCTTT
GAGCACACGTCTGAACTCTTTCCCTACACGACGCTCTTCCGATCTNNNATTTGAAGGTACACAGTGT
AGAGCACACGTCTGAACTCTTTCCCTACACGACGCTCTTCCGATCTNNNATTGGAGATAGTGCTTTGT
GCACACGTCTGAACTCTTTCCCTACACGACGCTCTTCCGATCTNNNGGCTTCATCTTCAATATCATCCCT
AGCACACGTCTGAACTCTTTCCCTACACGACGCTCTTCCGATCTNNNCCTTTCCCTTGTGTTAGTGCC
AGCACACGTCTGAACTCTTTCCCTACACGACGCTCTTCCGATCTNNNGTGCAGTTCCAAACAAAATCCC
GAGCACACGTCTGAACTCTTTCCCTACACGACGCTCTTCCGATCTNNNTGAATGTTCTTTGAGCCTTG
AGAGCACACGTCTGAACTCTTTCCCTACACGACGCTCTTCCGATCTNNNTTCACTCACTCAACCAGCTA
GCACACGTCTGAACTCTTTCCCTACACGACGCTCTTCCGATCTNNNTAAAAGGCGTGAAATAGTAACG
GCACACGTCTGAACTCTTTCCCTACACGACGCTCTTCCGATCTNNNTTGAGGGTATATATGTGTGTGT
GAGCACACGTCTGAACTCTTTCCCTACACGACGCTCTTCCGATCTNNNATTTCACTGTCACACAGCAG
GAGCACACGTCTGAACTCTTTCCCTACACGACGCTCTTCCGATCTNNNAGAAACACACCCAAATCAGT
GAGCACACGTCTGAACTCTTTCCCTACACGACGCTCTTCCGATCTNNNTGCTTACTACATCCATCACTTC
AGCACACGTCTGAACTCTTTCCCTACACGACGCTCTTCCGATCTNNNTGCTTAGCTTTTGAATATGAAC
CACACGTCTGAACTCTTTCCCTACACGACGCTCTTCCGATCTNNNAAGATTGCTCATGGCCAGTGTTG

AGCACACGTCTGAACTCTTTCCCTACACGACGCTCTTCCGATCTNNNGTTTGATTTGGTGGCTGTTCTG
GCACACGTCTGAACTCTTTCCCTACACGACGCTCTTCCGATCTNNNCATCAATTCTCTTGGGGGTCCTG
GCACACGTCTGAACTCTTTCCCTACACGACGCTCTTCCGATCTNNNAATATAGTGAGACCGAGACCCC
GAGCACACGTCTGAACTCTTTCCCTACACGACGCTCTTCCGATCTNNNACTGTGTCTATATGCCTGGA
AGAGCACACGTCTGAACTCTTTCCCTACACGACGCTCTTCCGATCTNNNTCTATCTTGAAATGCGTGT
GAGCACACGTCTGAACTCTTTCCCTACACGACGCTCTTCCGATCTNNNCATGCTTCTCCTTTGTACGTG
GAGCACACGTCTGAACTCTTTCCCTACACGACGCTCTTCCGATCTNNNACAACTTGGTCCCAAATGGTT
GCACACGTCTGAACTCTTTCCCTACACGACGCTCTTCCGATCTNNNAACAATAACCGGCACACACAAA
GAGCACACGTCTGAACTCTTTCCCTACACGACGCTCTTCCGATCTNNNAGTGAGAGAAAAGAGAAC
AGCACACGTCTGAACTCTTTCCCTACACGACGCTCTTCCGATCTNNNACTAAAGCATGGAGAAGTCCC
GCACACGTCTGAACTCTTTCCCTACACGACGCTCTTCCGATCTNNNCCACAGGTGGCAAGAACTCTTA
AGCACACGTCTGAACTCTTTCCCTACACGACGCTCTTCCGATCTNNNTGTACCTGTTCACTCACACTTG
AGCACACGTCTGAACTCTTTCCCTACACGACGCTCTTCCGATCTNNNTAGGAATGTCATGTGCTCTCG
AGCACACGTCTGAACTCTTTCCCTACACGACGCTCTTCCGATCTNNNTGCACCAAACTAATATTTCCAC
GAGCACACGTCTGAACTCTTTCCCTACACGACGCTCTTCCGATCTNNNAAGATGAGAGAGAGGCAG
CACACGTCTGAACTCTTTCCCTACACGACGCTCTTCCGATCTNNNGTGTGGTCTTTTGTCCCTCATCAC
GAGCACACGTCTGAACTCTTTCCCTACACGACGCTCTTCCGATCTNNNTTCTACCCACTAGGCCATGT
AGCACACGTCTGAACTCTTTCCCTACACGACGCTCTTCCGATCTNNNACACCAAGGAGTGTCTCTCTCT
GAAGAGCACACGTCTGAACTCTTTCCCTACACGACGCTCTTCCGATCTNNNTCTGACAGAAGCGATG
GCACACGTCTGAACTCTTTCCCTACACGACGCTCTTCCGATCTNNNAACAATCCTTCTTGGCATATGGA
AGCACACGTCTGAACTCTTTCCCTACACGACGCTCTTCCGATCTNNNGTGTTTCACTATAGGCAGCTG
GCACACGTCTGAACTCTTTCCCTACACGACGCTCTTCCGATCTNNNTCTCAGACTTAGACTTCTTCCTG
GAGCACACGTCTGAACTCTTTCCCTACACGACGCTCTTCCGATCTNNNACAGATAAGAAGACTGAGT
GAGCACACGTCTGAACTCTTTCCCTACACGACGCTCTTCCGATCTNNNGACAGCGATAAAGACAAGA
GCACACGTCTGAACTCTTTCCCTACACGACGCTCTTCCGATCTNNNTGCACACATATATAGGCATCTA
AGAGCACACGTCTGAACTCTTTCCCTACACGACGCTCTTCCGATCTNNNCTGTGTTCAGGAACCTGGT
GAGCACACGTCTGAACTCTTTCCCTACACGACGCTCTTCCGATCTNNNACATGACCCCTTTCTTTGTGC
AGCACACGTCTGAACTCTTTCCCTACACGACGCTCTTCCGATCTNNNTGCTATGTGAATTTGGGGGA
GAGCACACGTCTGAACTCTTTCCCTACACGACGCTCTTCCGATCTNNNCTACTAGGCCCTACACACAC
AGCACACGTCTGAACTCTTTCCCTACACGACGCTCTTCCGATCTNNNACATCACCTACATCACACATAC
AGCACACGTCTGAACTCTTTCCCTACACGACGCTCTTCCGATCTNNNTCCCAATGTTTCAGGAACTGAC
GAGCACACGTCTGAACTCTTTCCCTACACGACGCTCTTCCGATCTNNNTGTCTGCTGTGCAGAAGTTT
GAGCACACGTCTGAACTCTTTCCCTACACGACGCTCTTCCGATCTNNNCAGTTTGTGCTTATGGATGG
GAGCACACGTCTGAACTCTTTCCCTACACGACGCTCTTCCGATCTNNNCTGTGCATGCGTGTTTTTCAT
AGAGCACACGTCTGAACTCTTTCCCTACACGACGCTCTTCCGATCTNNNCCAGGAATGTGTAAGGAA
GAGCACACGTCTGAACTCTTTCCCTACACGACGCTCTTCCGATCTNNNAAACTAAGGAGTTGTGCGT
AGAGCACACGTCTGAACTCTTTCCCTACACGACGCTCTTCCGATCTNNNGCTGATCCAGAAAGAGGT
AGCACACGTCTGAACTCTTTCCCTACACGACGCTCTTCCGATCTNNNACATATATGGGTATGGGAGT
GCACACGTCTGAACTCTTTCCCTACACGACGCTCTTCCGATCTNNNAGGGGGCAATTTTCAGTTTCAG
ACACGTCTGAACTCTTTCCCTACACGACGCTCTTCCGATCTNNNTCCGAAGGAAAGGCTTATTCACGC
AGCACACGTCTGAACTCTTTCCCTACACGACGCTCTTCCGATCTNNNGCAGAAGTGCCACCTTTACTTC
GAGCACACGTCTGAACTCTTTCCCTACACGACGCTCTTCCGATCTNNNCCAAAGTCAGCCCAGTGAGA
AGCACACGTCTGAACTCTTTCCCTACACGACGCTCTTCCGATCTNNNACCACGGAAAGAAAGAAAG

GAGCACACGTCTGAACTCTTTCCCTACACGACGCTCTTCCGATCTNNNAGACTGCATCTTTCCTCTTGG
AGCACACGTCTGAACTCTTTCCCTACACGACGCTCTTCCGATCTNNNTATGGGTTGGTAGAGGAACAG
CACACGTCTGAACTCTTTCCCTACACGACGCTCTTCCGATCTNNNTAAGAAACATCTGACCAGCCGGG
GCACACGTCTGAACTCTTTCCCTACACGACGCTCTTCCGATCTNNNCCATCCTTAGTCCCAGAAATGGC
AGAGCACACGTCTGAACTCTTTCCCTACACGACGCTCTTCCGATCTNNNCCTGTTGCATGGTTTTGTTT
AGAGCACACGTCTGAACTCTTTCCCTACACGACGCTCTTCCGATCTNNNGCCAGTGCTCCAACTTTTTC
GAGCACACGTCTGAACTCTTTCCCTACACGACGCTCTTCCGATCTNNNCAGAGACATTGACACGTAAGG
GCACACGTCTGAACTCTTTCCCTACACGACGCTCTTCCGATCTNNNCACCACACTGAGTGCCAATATCA
GAGCACACGTCTGAACTCTTTCCCTACACGACGCTCTTCCGATCTNNNTTTGTATTTCTCCCACCCAACCC
GAGCACACGTCTGAACTCTTTCCCTACACGACGCTCTTCCGATCTNNNTCCCTTACCTTCTCCCAAGCAAT
AAGAGCACACGTCTGAACTCTTTCCCTACACGACGCTCTTCCGATCTNNNGCCCACAAAATTGACAAAA
GAGCACACGTCTGAACTCTTTCCCTACACGACGCTCTTCCGATCTNNNATCTGCAGAAGAGGGGTGTG
GAGCACACGTCTGAACTCTTTCCCTACACGACGCTCTTCCGATCTNNNGGGGTGAAGAGATGGATATG
GAGCACACGTCTGAACTCTTTCCCTACACGACGCTCTTCCGATCTNNNGCCACATGCTGCTGAAAATG
AGAGCACACGTCTGAACTCTTTCCCTACACGACGCTCTTCCGATCTNNNTATATGCTGTTGCCATGGGT
AGCACACGTCTGAACTCTTTCCCTACACGACGCTCTTCCGATCTNNNTGTATGCATGTGACTTGTGTAT
GAAGAGCACACGTCTGAACTCTTTCCCTACACGACGCTCTTCCGATCTNNNCTTGAGCACTTTTTACTTG
GAGCACACGTCTGAACTCTTTCCCTACACGACGCTCTTCCGATCTNNNAAATGTTCTTACCACCCCTCTG
GAGCACACGTCTGAACTCTTTCCCTACACGACGCTCTTCCGATCTNNNCCATTTCGTGGCCTCTTCCATTA
AGCACACGTCTGAACTCTTTCCCTACACGACGCTCTTCCGATCTNNNAGCATAAGCTCCTTTGCCTTTGT
GAAGAGCACACGTCTGAACTCTTTCCCTACACGACGCTCTTCCGATCTNNNAATTTGCTAGCAGGGTA
GAGCACACGTCTGAACTCTTTCCCTACACGACGCTCTTCCGATCTNNNGGAGGGGAAAGAAGAGACA
AGCACACGTCTGAACTCTTTCCCTACACGACGCTCTTCCGATCTNNNACTCTTTGGTGAAACATTTCTCT
AGCACACGTCTGAACTCTTTCCCTACACGACGCTCTTCCGATCTNNNCGAAGTAGAGCATCCTGGAGT
GAGCACACGTCTGAACTCTTTCCCTACACGACGCTCTTCCGATCTNNNAAGCCTTCATTCTCATGTTGG
GAGCACACGTCTGAACTCTTTCCCTACACGACGCTCTTCCGATCTNNNACCCAGACTGGTTTTGTGGTT
GCACACGTCTGAACTCTTTCCCTACACGACGCTCTTCCGATCTNNNTCTCTTCCCAAACACAGTTCTGCC
AGCACACGTCTGAACTCTTTCCCTACACGACGCTCTTCCGATCTNNNGTGTGTCTGGCCAACATGAAT
GAGCACACGTCTGAACTCTTTCCCTACACGACGCTCTTCCGATCTNNNAGCAGGTGTATGTGTGCATG
AGCACACGTCTGAACTCTTTCCCTACACGACGCTCTTCCGATCTNNNTCCATTTCCTAAGTGAGACTTG
GAGCACACGTCTGAACTCTTTCCCTACACGACGCTCTTCCGATCTNNNCGAGGGGATGAGAGAGAGA
GAGCACACGTCTGAACTCTTTCCCTACACGACGCTCTTCCGATCTNNNTGTCATTCTAGCAGCCCTTTCA
AGAGCACACGTCTGAACTCTTTCCCTACACGACGCTCTTCCGATCTNNNTCTCCTTTCCCATCCTGCACAT
GAGCACACGTCTGAACTCTTTCCCTACACGACGCTCTTCCGATCTNNNGAGTAGAGTGACTGTGGGAA
GAGCACACGTCTGAACTCTTTCCCTACACGACGCTCTTCCGATCTNNNATGGCTTAGGGTGTAGGGTAT
AGAGCACACGTCTGAACTCTTTCCCTACACGACGCTCTTCCGATCTNNNGCATTGTTTTTGCTGCTGTTG
GAGCACACGTCTGAACTCTTTCCCTACACGACGCTCTTCCGATCTNNNGCCAATAGACACACAAAAGAT
GAGCACACGTCTGAACTCTTTCCCTACACGACGCTCTTCCGATCTNNNAACTGCTTGGTGTTTTCAGTC
AGAGCACACGTCTGAACTCTTTCCCTACACGACGCTCTTCCGATCTNNNCCCCAAGTAACCTCATAGGTT
GAGCACACGTCTGAACTCTTTCCCTACACGACGCTCTTCCGATCTNNNTCTAATAGGATTTGGAACACAC
GCACACGTCTGAACTCTTTCCCTACACGACGCTCTTCCGATCTNNNACTATATGCCTAGTTGTGTGCATG
GAGCACACGTCTGAACTCTTTCCCTACACGACGCTCTTCCGATCTNNNACGAGACAAATAGCCACACAC
AGCACACGTCTGAACTCTTTCCCTACACGACGCTCTTCCGATCTNNNTGCCAAAAATATTCTTTTCTTCTC

AGCACACGTCTGAACTCTTTCCCTACACGACGCTCTTCCGATCTNNNGAACTAGCTCCTGCATAGCACT
GAGCACACGTCTGAACTCTTTCCCTACACGACGCTCTTCCGATCTNNNTCCCTTGGTTTCCTTGAACTCA
GCACACGTCTGAACTCTTTCCCTACACGACGCTCTTCCGATCTNNNCGCTTGGCCTGCAATGTACTCCAT
GCACACGTCTGAACTCTTTCCCTACACGACGCTCTTCCGATCTNNNTTCCTCCAGTTGCTACTCTGTGACA
AGAGCACACGTCTGAACTCTTTCCCTACACGACGCTCTTCCGATCTNNNCCGAGTTCTTCCAAGAAACAG
AGCACACGTCTGAACTCTTTCCCTACACGACGCTCTTCCGATCTNNNGTATGGCTTCTGGGTCATTGCAT
GAGCACACGTCTGAACTCTTTCCCTACACGACGCTCTTCCGATCTNNNAGTTGTTCTGTGGTCTTCCTTT
GAGCACACGTCTGAACTCTTTCCCTACACGACGCTCTTCCGATCTNNNTACATGGCTTGGTAGGGATG
AGCACACGTCTGAACTCTTTCCCTACACGACGCTCTTCCGATCTNNNATCTGGATGGCTTCAAGCTCTTT
AGCACACGTCTGAACTCTTTCCCTACACGACGCTCTTCCGATCTNNNCCGGTAACTGAGCAAACACTAG
GAGCACACGTCTGAACTCTTTCCCTACACGACGCTCTTCCGATCTNNNTCCATCCTGATATGTGTGTGT
AGAGCACACGTCTGAACTCTTTCCCTACACGACGCTCTTCCGATCTNNNCGTTCTCCATCAAGGATATAG
GAGCACACGTCTGAACTCTTTCCCTACACGACGCTCTTCCGATCTNNNCTCTGCACCTTATTCAGAGCTG
AAGAGCACACGTCTGAACTCTTTCCCTACACGACGCTCTTCCGATCTNNNTAAAGTGGCTATTTGCACAG
GCACACGTCTGAACTCTTTCCCTACACGACGCTCTTCCGATCTNNNTCTTCACATCTTATTGCTCTTGCTG
GCACACGTCTGAACTCTTTCCCTACACGACGCTCTTCCGATCTNNNAGATCCAGTCTTCCACCACTCATCA
CACACGTCTGAACTCTTTCCCTACACGACGCTCTTCCGATCTNNNTGTCCATTCTCTTGGGAGCACTACA
GAGCACACGTCTGAACTCTTTCCCTACACGACGCTCTTCCGATCTNNNCCTTTTCTGAGCCTTTGTTGCT
GCACACGTCTGAACTCTTTCCCTACACGACGCTCTTCCGATCTNNNGGTTTGAAGAGGTGTCTTGCTGA
GAGCACACGTCTGAACTCTTTCCCTACACGACGCTCTTCCGATCTNNNCACTGGCCTACAGAGCATTTC
CACACGTCTGAACTCTTTCCCTACACGACGCTCTTCCGATCTNNNACTGTTAGTATTCCTGGATGATGGA
AGCACACGTCTGAACTCTTTCCCTACACGACGCTCTTCCGATCTNNNCTCTTGGGAATACTCCCCTGGTA
AGCACACGTCTGAACTCTTTCCCTACACGACGCTCTTCCGATCTNNNGTGGAACTCTCTCGCTCTTTCTG
AGCACACGTCTGAACTCTTTCCCTACACGACGCTCTTCCGATCTNNNTGTCCCCAGTATCTGTTTTGTGC
GAGCACACGTCTGAACTCTTTCCCTACACGACGCTCTTCCGATCTNNNATCTGAAACCAAACACCTCTCC
GAGCACACGTCTGAACTCTTTCCCTACACGACGCTCTTCCGATCTNNNCGCTTTGTGGAATTATCGTTG
AGCACACGTCTGAACTCTTTCCCTACACGACGCTCTTCCGATCTNNNTAGAGAGTTCTGTGCCCCAATCA
AGAGCACACGTCTGAACTCTTTCCCTACACGACGCTCTTCCGATCTNNNGATAGCTCTAGAGACACCGA
GAGCACACGTCTGAACTCTTTCCCTACACGACGCTCTTCCGATCTNNNCTATCTTTCTTCCCCACCCCCTC
AGCACACGTCTGAACTCTTTCCCTACACGACGCTCTTCCGATCTNNNCATGGCGAGACCCTATGTTTACC
GAGCACACGTCTGAACTCTTTCCCTACACGACGCTCTTCCGATCTNNNCTACAAGGAATGAGCCACCAT
AAGAGCACACGTCTGAACTCTTTCCCTACACGACGCTCTTCCGATCTNNNTTCCTGTCTACCTTTGCTGAT
GCACACGTCTGAACTCTTTCCCTACACGACGCTCTTCCGATCTNNNCTCTACCAGGCAACTTTCCCAAGC
GAAGAGCACACGTCTGAACTCTTTCCCTACACGACGCTCTTCCGATCTNNNTGCATCAGAAAAGCATTG
AGAGCACACGTCTGAACTCTTTCCCTACACGACGCTCTTCCGATCTNNNAGGTAGGCATTGGTATGTGG
AGAGCACACGTCTGAACTCTTTCCCTACACGACGCTCTTCCGATCTNNNTGCACTTGGGTTCATCAGGG
AGCACACGTCTGAACTCTTTCCCTACACGACGCTCTTCCGATCTNNNGCTAGTCTTACAGACCCATTGG
AGCACACGTCTGAACTCTTTCCCTACACGACGCTCTTCCGATCTNNNTGTAAACATATAACACCCCACAG
AGCACACGTCTGAACTCTTTCCCTACACGACGCTCTTCCGATCTNNNCCCATTTATTACAACGAACTTAT
AGCACACGTCTGAACTCTTTCCCTACACGACGCTCTTCCGATCTNNNGGACCATTTAGATAACAGCCCC
AGAGCACACGTCTGAACTCTTTCCCTACACGACGCTCTTCCGATCTNNNTATCCCCCAACTCAGTTTTGAG
AGCACACGTCTGAACTCTTTCCCTACACGACGCTCTTCCGATCTNNNACGGTTGAAGTTACACTGAGCT
GAGCACACGTCTGAACTCTTTCCCTACACGACGCTCTTCCGATCTNNNCCATCTGCAAAGACGGACAAT

AGCACACGTCTGAACTCTTTCCCTACACGACGCTCTTCCGATCTNNNACTGCCAAGAATAAAAGGAGC
GCACACGTCTGAACTCTTTCCCTACACGACGCTCTTCCGATCTNNNCCACCACCAAACAGTGACCATA
GCACACGTCTGAACTCTTTCCCTACACGACGCTCTTCCGATCTNNNGAAAGCACCAGGCTTCAAAATC
GAGCACACGTCTGAACTCTTTCCCTACACGACGCTCTTCCGATCTNNNAGCAACAGCGTGATCTAAG
AGCACACGTCTGAACTCTTTCCCTACACGACGCTCTTCCGATCTNNNGTTTTGCTGTTTGTTCCTGCC
AGCACACGTCTGAACTCTTTCCCTACACGACGCTCTTCCGATCTNNNTATTACATATCGGCCTGCAGT
GAGCACACGTCTGAACTCTTTCCCTACACGACGCTCTTCCGATCTNNNTTGCCTTGTATACGTGTGTG
AGCACACGTCTGAACTCTTTCCCTACACGACGCTCTTCCGATCTNNNCCTGGTTGCATTCATGGAGA
GCACACGTCTGAACTCTTTCCCTACACGACGCTCTTCCGATCTNNNATCACCCAACCTGTGCTTCTAC
GCACACGTCTGAACTCTTTCCCTACACGACGCTCTTCCGATCTNNNTCTCTTGCTGTTTTCAGTGTCCT
AGCACACGTCTGAACTCTTTCCCTACACGACGCTCTTCCGATCTNNNCTTTGCTTACCACCGTCTCTCT
AGAGCACACGTCTGAACTCTTTCCCTACACGACGCTCTTCCGATCTNNNATACGCAATTTGTGTGTGT
AGCACACGTCTGAACTCTTTCCCTACACGACGCTCTTCCGATCTNNNCCGAACTACGTCACTCTGTGTA
AGCACACGTCTGAACTCTTTCCCTACACGACGCTCTTCCGATCTNNNGGATGACCAGAATGAGAGA
GAGCACACGTCTGAACTCTTTCCCTACACGACGCTCTTCCGATCTNNNACACAAAGGGACACCTGTCA
GCACACGTCTGAACTCTTTCCCTACACGACGCTCTTCCGATCTNNNTTGGTGTAAGGCATGATGGTG
AAGAGCACACGTCTGAACTCTTTCCCTACACGACGCTCTTCCGATCTNNNCCTCAGGCCCAACATAGA
AGCACACGTCTGAACTCTTTCCCTACACGACGCTCTTCCGATCTNNNTTACTTGGTGCCATCGAAGTCT
AGCACACGTCTGAACTCTTTCCCTACACGACGCTCTTCCGATCTNNNCTAGGTGTGTTCCTAGGGGA
AGCACACGTCTGAACTCTTTCCCTACACGACGCTCTTCCGATCTNNNCCCCAGCTACCCTTCTAACATA
GCACACGTCTGAACTCTTTCCCTACACGACGCTCTTCCGATCTNNNCACGGATGGCAGAGTTTCTGTA
AGCACACGTCTGAACTCTTTCCCTACACGACGCTCTTCCGATCTNNNCACAGACATAAACCTCCTCCTG
GAGCACACGTCTGAACTCTTTCCCTACACGACGCTCTTCCGATCTNNNAAACCACATCCACACAGTGAC
GCACACGTCTGAACTCTTTCCCTACACGACGCTCTTCCGATCTNNNATCTTCAGGCCTTTGCTCAACTG
AGCACACGTCTGAACTCTTTCCCTACACGACGCTCTTCCGATCTNNNACGTACTTATCAGCTCACGTTC
GCACACGTCTGAACTCTTTCCCTACACGACGCTCTTCCGATCTNNNGGGAACAGTGTTGAGCTGTAT
GCACACGTCTGAACTCTTTCCCTACACGACGCTCTTCCGATCTNNNTGAAAAAGCAAGGCTAGGGGA
GAGCACACGTCTGAACTCTTTCCCTACACGACGCTCTTCCGATCTNNNGCTAACAATGCTGTTTGCGA
GAGCACACGTCTGAACTCTTTCCCTACACGACGCTCTTCCGATCTNNNGTAACAGAGCTAGATGCTGT
AAGAGCACACGTCTGAACTCTTTCCCTACACGACGCTCTTCCGATCTNNNCACGTGCTTTTCTTTGAG
AGCACACGTCTGAACTCTTTCCCTACACGACGCTCTTCCGATCTNNNCCGTTCCCTTCCTTTCTCTTCTC
AGAGCACACGTCTGAACTCTTTCCCTACACGACGCTCTTCCGATCTNNNGGCTTGACCTCTCTGAAGA
GCACACGTCTGAACTCTTTCCCTACACGACGCTCTTCCGATCTNNNTTATGTTGAGGGACACATGGCT
AGCACACGTCTGAACTCTTTCCCTACACGACGCTCTTCCGATCTNNNTGACATTCCCAACTAGGATCG
GAAGAGCACACGTCTGAACTCTTTCCCTACACGACGCTCTTCCGATCTNNNCTATTTTGGGAGGATGA
AGCACACGTCTGAACTCTTTCCCTACACGACGCTCTTCCGATCTNNNCCTGAGGATAAGCAGGGAGT
GCACACGTCTGAACTCTTTCCCTACACGACGCTCTTCCGATCTNNNGCTGAGTATTATATTTCCATATT
AGCACACGTCTGAACTCTTTCCCTACACGACGCTCTTCCGATCTNNNTGGGAATGTGTTATGATGCCT
GAGCACACGTCTGAACTCTTTCCCTACACGACGCTCTTCCGATCTNNNGAGAGAGATGTAGGCAGGA
AGCACACGTCTGAACTCTTTCCCTACACGACGCTCTTCCGATCTNNNACTTCCTAGGTGGATTCCTAG
AGCACACGTCTGAACTCTTTCCCTACACGACGCTCTTCCGATCTNNNGACAGAGCGAGACTGTAGGG
AGCACACGTCTGAACTCTTTCCCTACACGACGCTCTTCCGATCTNNNATGCCTGGTACCAATATGTCTG
AGCACACGTCTGAACTCTTTCCCTACACGACGCTCTTCCGATCTNNNGTGAAGGTACTTTGAGGTGG

AGCACACGTCTGAACTCTTTCCCTACACGACGCTCTTCCGATCTNNNAAAGGTGTGTATGTGTGGGG
AGCACACGTCTGAACTCTTTCCCTACACGACGCTCTTCCGATCTNNNTGTGCATTTTCACATGTATGA
CACACGTCTGAACTCTTTCCCTACACGACGCTCTTCCGATCTNNNAAGGCTTCATAATACATTTGTGG
GCACACGTCTGAACTCTTTCCCTACACGACGCTCTTCCGATCTNNNGGCTCATGATATGGCTACAGCT
GAGCACACGTCTGAACTCTTTCCCTACACGACGCTCTTCCGATCTNNNGACACCGTTGTTAGGTGGCT
CACACGTCTGAACTCTTTCCCTACACGACGCTCTTCCGATCTNNNTGCAAGTTTTCCACTAAAGGCAC
GAGCACACGTCTGAACTCTTTCCCTACACGACGCTCTTCCGATCTNNNCTAATTCAGGCAGGGCAAA
AGCACACGTCTGAACTCTTTCCCTACACGACGCTCTTCCGATCTNNNCTCTTTCTGGTTCTCACTGGCT
CACACGTCTGAACTCTTTCCCTACACGACGCTCTTCCGATCTNNNACCCAATCTAAAGTTGCAACCGAC
GAGCACACGTCTGAACTCTTTCCCTACACGACGCTCTTCCGATCTNNNGTGCCTCAGGCTGCTATCTT
AGCACACGTCTGAACTCTTTCCCTACACGACGCTCTTCCGATCTNNNTTGCAAGATCCAGGCAAAG
AGCACACGTCTGAACTCTTTCCCTACACGACGCTCTTCCGATCTNNNCTTCTTCGCACTAGACAATGAG
AAGAGCACACGTCTGAACTCTTTCCCTACACGACGCTCTTCCGATCTNNNAATGTGTCAGACGTTGTG
GCACACGTCTGAACTCTTTCCCTACACGACGCTCTTCCGATCTNNNCTAGTGCTGGGGAACTTTGAGA
AGCACACGTCTGAACTCTTTCCCTACACGACGCTCTTCCGATCTNNNCATGCCAATGGAGGTGCATTT
AGCACACGTCTGAACTCTTTCCCTACACGACGCTCTTCCGATCTNNNCATACGCACACAAACAACACA
AGCACACGTCTGAACTCTTTCCCTACACGACGCTCTTCCGATCTNNNACATGCAAAGTGCTTAGGGTA
GAAGAGCACACGTCTGAACTCTTTCCCTACACGACGCTCTTCCGATCTNNNAAGCTGTGGGATCTCTC
GAGCACACGTCTGAACTCTTTCCCTACACGACGCTCTTCCGATCTNNNAGAGTATGATCCCTGTTTG
AGCACACGTCTGAACTCTTTCCCTACACGACGCTCTTCCGATCTNNNAGCACTCCCTTATCTGCTTTGT
AGCACACGTCTGAACTCTTTCCCTACACGACGCTCTTCCGATCTNNNTTCTTGCATAACAGCACCTGCA
AGCACACGTCTGAACTCTTTCCCTACACGACGCTCTTCCGATCTNNNTTGGGCTGTCTAGGTAACAG
AGCACACGTCTGAACTCTTTCCCTACACGACGCTCTTCCGATCTNNNGCCTAAAAGGATGGGGAAG
AGCACACGTCTGAACTCTTTCCCTACACGACGCTCTTCCGATCTNNNGATTCTGGACAGACAGACAG
GAGCACACGTCTGAACTCTTTCCCTACACGACGCTCTTCCGATCTNNNGTTAGCTGCAGGAGTAGCTT
AGCACACGTCTGAACTCTTTCCCTACACGACGCTCTTCCGATCTNNNCCTGGGCAATTTAGCAAGAAA
GAGCACACGTCTGAACTCTTTCCCTACACGACGCTCTTCCGATCTNNNTCTTTACCCATCTGTCCTCCTG
AGCACACGTCTGAACTCTTTCCCTACACGACGCTCTTCCGATCTNNNCACAATTCCATAACTCCCCCAC
AGCACACGTCTGAACTCTTTCCCTACACGACGCTCTTCCGATCTNNNATTCAGTTTCTCTCGGTGTTCC
AGCACACGTCTGAACTCTTTCCCTACACGACGCTCTTCCGATCTNNNAATGTCTTGCTGCTAGGATTG
AGCACACGTCTGAACTCTTTCCCTACACGACGCTCTTCCGATCTNNNAGTCTCTGCACCTAATACTGCA
GCACACGTCTGAACTCTTTCCCTACACGACGCTCTTCCGATCTNNNACTCAGACTATGTCAGGGATTG
AGCACACGTCTGAACTCTTTCCCTACACGACGCTCTTCCGATCTNNNTCACTGAAGGGGAATCTGGG
GAGCACACGTCTGAACTCTTTCCCTACACGACGCTCTTCCGATCTNNNAGATGAGGATTGGCAGGAG
AGCACACGTCTGAACTCTTTCCCTACACGACGCTCTTCCGATCTNNNCCAGTCTTGCTGAGAGAGAG
AGAGCACACGTCTGAACTCTTTCCCTACACGACGCTCTTCCGATCTNNNTTATTCAATGCACATGGCCA
GAGCACACGTCTGAACTCTTTCCCTACACGACGCTCTTCCGATCTNNNACTTGAGAGAAAGAGAGGG
AGCACACGTCTGAACTCTTTCCCTACACGACGCTCTTCCGATCTNNNCCTCAGCATCTCAGACCAACAT
GAGCACACGTCTGAACTCTTTCCCTACACGACGCTCTTCCGATCTNNNCTGCCTTCTCATTTGACAGAC
GCACACGTCTGAACTCTTTCCCTACACGACGCTCTTCCGATCTNNNTAATTCACCCCTTCCCAACATGG
GAGCACACGTCTGAACTCTTTCCCTACACGACGCTCTTCCGATCTNNNACAGGGAGATAGGACTTGA
GAGCACACGTCTGAACTCTTTCCCTACACGACGCTCTTCCGATCTNNNCATCACTCCATAATTGCTGCC
AGCACACGTCTGAACTCTTTCCCTACACGACGCTCTTCCGATCTNNNTAGCAGATAGCACTGCAGTTT

GCACACGTCTGAACTCTTTCCCTACACGACGCTCTTCCGATCTNNNACTCACCTAGGAATATGGCTTCT
GAGCACACGTCTGAACTCTTTCCCTACACGACGCTCTTCCGATCTNNNACCACACCTTCTTGATTACTG
AGAGCACACGTCTGAACTCTTTCCCTACACGACGCTCTTCCGATCTNNNTTGCTAAAGGCAAATTCTAG
GAGCACACGTCTGAACTCTTTCCCTACACGACGCTCTTCCGATCTNNNGTCCTTACAAGACGGGGAGT
AGCACACGTCTGAACTCTTTCCCTACACGACGCTCTTCCGATCTNNNCGCTACCAGCCTGTCTCTTTATC
GCACACGTCTGAACTCTTTCCCTACACGACGCTCTTCCGATCTNNNATAGTTGGTGGAGGGGATAAG
AGCACACGTCTGAACTCTTTCCCTACACGACGCTCTTCCGATCTNNNTGCATCATCTGCATCACACACAT
AGCACACGTCTGAACTCTTTCCCTACACGACGCTCTTCCGATCTNNNTTCCGCTCATCAGATTTTCCAG
AGCACACGTCTGAACTCTTTCCCTACACGACGCTCTTCCGATCTNNNAGAGCCCTACTTTGGGCTACA
ACACGTCTGAACTCTTTCCCTACACGACGCTCTTCCGATCTNNNTGACAACTTCTCTCCCCTTCAGACAT
GCACACGTCTGAACTCTTTCCCTACACGACGCTCTTCCGATCTNNNGGAGGAGGAGGAACAGAACA
AGCACACGTCTGAACTCTTTCCCTACACGACGCTCTTCCGATCTNNNGTACACCGCGGCTTTTGCTGCA
GAGCACACGTCTGAACTCTTTCCCTACACGACGCTCTTCCGATCTNNNCATGCCTTATGGCGTTCAGA
AGCACACGTCTGAACTCTTTCCCTACACGACGCTCTTCCGATCTNNNTCCTTGTCCCTTCTTCTCCCTTTC
GCACACGTCTGAACTCTTTCCCTACACGACGCTCTTCCGATCTNNNCAGGCAAGTCAACAAATTACCCC
GAGCACACGTCTGAACTCTTTCCCTACACGACGCTCTTCCGATCTNNNAGCAGATAAAAGCAAGCTGG
AAGAGCACACGTCTGAACTCTTTCCCTACACGACGCTCTTCCGATCTNNNTATGATTACATCGGAACGG
GAGCACACGTCTGAACTCTTTCCCTACACGACGCTCTTCCGATCTNNNGTGTCCAAACATTTGAGGAAG
AGAGCACACGTCTGAACTCTTTCCCTACACGACGCTCTTCCGATCTNNNACTCAGCTCACAGGTAGAAT
GAGCACACGTCTGAACTCTTTCCCTACACGACGCTCTTCCGATCTNNNAGCTTTATTCGCCCTAACCAAG
AAGAGCACACGTCTGAACTCTTTCCCTACACGACGCTCTTCCGATCTNNNTCCCACTTCAGCCTCCTAAT
AGCACACGTCTGAACTCTTTCCCTACACGACGCTCTTCCGATCTNNNCGACAAAACGAGACACTCTGTC
AGCACACGTCTGAACTCTTTCCCTACACGACGCTCTTCCGATCTNNNAAGCCTATTCCTTCAACCTCTGC
AGCACACGTCTGAACTCTTTCCCTACACGACGCTCTTCCGATCTNNNTGTTCCACTGTTTGTTTTGGAG
AGAGCACACGTCTGAACTCTTTCCCTACACGACGCTCTTCCGATCTNNNAGCCAATCTTCATTCTGCTT
GAGCACACGTCTGAACTCTTTCCCTACACGACGCTCTTCCGATCTNNNGAGAGCAGGAAAGGAGAAG
AGCACACGTCTGAACTCTTTCCCTACACGACGCTCTTCCGATCTNNNATTCATTTTCCCCACAACCCTCC
AGCACACGTCTGAACTCTTTCCCTACACGACGCTCTTCCGATCTNNNAGGTCCCTCAAGGGAGAAATT
AGCACACGTCTGAACTCTTTCCCTACACGACGCTCTTCCGATCTNNNATGGGAAAATTCTCTCCCTCTT
GAGCACACGTCTGAACTCTTTCCCTACACGACGCTCTTCCGATCTNNNGCACTCCAATCTGAATGCCAG
AGAGCACACGTCTGAACTCTTTCCCTACACGACGCTCTTCCGATCTNNNCACTGTTGTAAGCCCTCTCA
GCACACGTCTGAACTCTTTCCCTACACGACGCTCTTCCGATCTNNNTGTTTTACGGTTACAAGGGCACA
AAGAGCACACGTCTGAACTCTTTCCCTACACGACGCTCTTCCGATCTNNNAAGTAATGAGCTTCAGCT
GCACACGTCTGAACTCTTTCCCTACACGACGCTCTTCCGATCTNNNATCTATGTGGAAGTGCAGCAAG
AGCACACGTCTGAACTCTTTCCCTACACGACGCTCTTCCGATCTNNNTGGGCACCTTAGAAGTTCCTCT
GAGCACACGTCTGAACTCTTTCCCTACACGACGCTCTTCCGATCTNNNATCATCTGGAGCAGTTTCTTG
AGCACACGTCTGAACTCTTTCCCTACACGACGCTCTTCCGATCTNNNAGATTTCTTGAAGATGACACA
GAGCACACGTCTGAACTCTTTCCCTACACGACGCTCTTCCGATCTNNNTCCATAAAGCCCTCTTGGTCT
AGCACACGTCTGAACTCTTTCCCTACACGACGCTCTTCCGATCTNNNCGTTCTGGATGACATTAGGAG
AGAGCACACGTCTGAACTCTTTCCCTACACGACGCTCTTCCGATCTNNNAAACGTTCAAACCATCGCA
AGAGCACACGTCTGAACTCTTTCCCTACACGACGCTCTTCCGATCTNNNATTGTACCTAGCAGGTGT
AGAGCACACGTCTGAACTCTTTCCCTACACGACGCTCTTCCGATCTNNNTGCATGTCATCACTTCCTTTC
AGCACACGTCTGAACTCTTTCCCTACACGACGCTCTTCCGATCTNNNCAGAAAGGTGGGAAGACTTG

GAGCACACGTCTGAACTCTTTCCCTACACGACGCTCTTCCGATCTNNNGTGGACTAAGGCAAAAGGT
AGCACACGTCTGAACTCTTTCCCTACACGACGCTCTTCCGATCTNNNGAATCCACAGTGTGATGGAT
GCACACGTCTGAACTCTTTCCCTACACGACGCTCTTCCGATCTNNNCCCACATAAATGCACCCAACTGA
AGCACACGTCTGAACTCTTTCCCTACACGACGCTCTTCCGATCTNNNCAGCAACAGAATGACACTCGT
GCACACGTCTGAACTCTTTCCCTACACGACGCTCTTCCGATCTNNNAGGGAAATGGGGAAAGGGTA
AGCACACGTCTGAACTCTTTCCCTACACGACGCTCTTCCGATCTNNNCTCCGAACAAAATACCCCTCT
AGCACACGTCTGAACTCTTTCCCTACACGACGCTCTTCCGATCTNNNATTGTTGCTCAAAGTGGTGG
AGCACACGTCTGAACTCTTTCCCTACACGACGCTCTTCCGATCTNNNTCTGAGAAAGACGGAAAACT
AGCACACGTCTGAACTCTTTCCCTACACGACGCTCTTCCGATCTNNNAAGCCACCTGATGCACAGATA
AGCACACGTCTGAACTCTTTCCCTACACGACGCTCTTCCGATCTNNNAGCTTGTCTGTGAGGAATGT
AGCACACGTCTGAACTCTTTCCCTACACGACGCTCTTCCGATCTNNNAGATGTTTTTCTCTGATGGTC
GAGCACACGTCTGAACTCTTTCCCTACACGACGCTCTTCCGATCTNNNGGAGCATTTGCTTTAGGGAC
GCACACGTCTGAACTCTTTCCCTACACGACGCTCTTCCGATCTNNNTGGCATGGCCATTATTTCATCCA
GAGCACACGTCTGAACTCTTTCCCTACACGACGCTCTTCCGATCTNNNTATTTCCCAACGCTGTTCTAG
AGCACACGTCTGAACTCTTTCCCTACACGACGCTCTTCCGATCTNNNACTGCAAGTGGTAAACGTGA
GAGCACACGTCTGAACTCTTTCCCTACACGACGCTCTTCCGATCTNNNAAAGAACCTCTTCACACACA
GCACACGTCTGAACTCTTTCCCTACACGACGCTCTTCCGATCTNNNGTTGAGCTGAGATCCTGCCATTA
GAAGAGCACACGTCTGAACTCTTTCCCTACACGACGCTCTTCCGATCTNNNCTATGTGGTGAGTTTCA
GCACACGTCTGAACTCTTTCCCTACACGACGCTCTTCCGATCTNNNGAGAGAGAGAGTGTGCAAGA
GAGCACACGTCTGAACTCTTTCCCTACACGACGCTCTTCCGATCTNNNGTGAGGGAACTGGGAATCA
GAGCACACGTCTGAACTCTTTCCCTACACGACGCTCTTCCGATCTNNNTTGCCAGGTGTAACTGCAAG
GAGCACACGTCTGAACTCTTTCCCTACACGACGCTCTTCCGATCTNNNACAGACTAAGACGAGGGTGT
AGAGCACACGTCTGAACTCTTTCCCTACACGACGCTCTTCCGATCTNNNCACCAACTGGACAAAGCCA
GAGCACACGTCTGAACTCTTTCCCTACACGACGCTCTTCCGATCTNNNTCCACTGCTCTGGCTTAGTGA
AGCACACGTCTGAACTCTTTCCCTACACGACGCTCTTCCGATCTNNNTCAGTGCCAAATCTTCTGTCTG
AGCACACGTCTGAACTCTTTCCCTACACGACGCTCTTCCGATCTNNNCTGAGTTCCTTAGAGGCAGCA
AAGAGCACACGTCTGAACTCTTTCCCTACACGACGCTCTTCCGATCTNNNTCTGAACACACTGAGAGC
AGAGCACACGTCTGAACTCTTTCCCTACACGACGCTCTTCCGATCTNNNACTGGCTAACAGTGAACTA
GCACACGTCTGAACTCTTTCCCTACACGACGCTCTTCCGATCTNNNTCTGAGACTTGGGCACAGATAC
AGCACACGTCTGAACTCTTTCCCTACACGACGCTCTTCCGATCTNNNCTTAGCAAAGCAGTTTGCCTCT
GCACACGTCTGAACTCTTTCCCTACACGACGCTCTTCCGATCTNNNAAAAATGAGCCTTTCTCCTCCCC
AGAGCACACGTCTGAACTCTTTCCCTACACGACGCTCTTCCGATCTNNNGGGTAGAGAGTTGGGGC
GCACACGTCTGAACTCTTTCCCTACACGACGCTCTTCCGATCTNNNCGGTCAACAAACATCAAGTCAG
GCACACGTCTGAACTCTTTCCCTACACGACGCTCTTCCGATCTNNNTAGATCTGGGTTCCCTGGAGTTG
AGCACACGTCTGAACTCTTTCCCTACACGACGCTCTTCCGATCTNNNGCCTGGCCCATAGCTTTCTTTT
GAGCACACGTCTGAACTCTTTCCCTACACGACGCTCTTCCGATCTNNNATGGGGATCACATGACAGGA
GAAGAGCACACGTCTGAACTCTTTCCCTACACGACGCTCTTCCGATCTNNNAGCTTGTAGTGAGCTGA
AGAGCACACGTCTGAACTCTTTCCCTACACGACGCTCTTCCGATCTNNNGCACACAAACACACAAGAC
GAGCACACGTCTGAACTCTTTCCCTACACGACGCTCTTCCGATCTNNNTCTTGGGACTTCTCCAGATTC
GCACACGTCTGAACTCTTTCCCTACACGACGCTCTTCCGATCTNNNTGATCCATGATCGGGCCATTGT
AGCACACGTCTGAACTCTTTCCCTACACGACGCTCTTCCGATCTNNNATTTATCCCTGGTGGCTCCAAA
AGCACACGTCTGAACTCTTTCCCTACACGACGCTCTTCCGATCTNNNTCGTCCATAGTCACACTACCCAA
GCACACGTCTGAACTCTTTCCCTACACGACGCTCTTCCGATCTNNNGGGAAATTTCTACGTGGTTGG

GCACACGTCTGAACTCTTTCCCTACACGACGCTCTTCCGATCTNNNGCTCCCTATATCTCCTCTCCCCTC
GCACACGTCTGAACTCTTTCCCTACACGACGCTCTTCCGATCTNNNACCTATAGGATCCTCTCCCTCCT
AGCACACGTCTGAACTCTTTCCCTACACGACGCTCTTCCGATCTNNNTCATCCTGTTCCTCTTTGCTGAA
GAGCACACGTCTGAACTCTTTCCCTACACGACGCTCTTCCGATCTNNNCCCAAAATTTCACTGACACAC
AGCACACGTCTGAACTCTTTCCCTACACGACGCTCTTCCGATCTNNNAGGGTTGAAGAGGATGTAGT
GCACACGTCTGAACTCTTTCCCTACACGACGCTCTTCCGATCTNNNACAAGTCCCACAGTGATTTCCTC
AGCACACGTCTGAACTCTTTCCCTACACGACGCTCTTCCGATCTNNNCTTCCCTCTCCCCCTTGATGTAG
AGCACACGTCTGAACTCTTTCCCTACACGACGCTCTTCCGATCTNNNTCCATCCCATCAGGACAGACTT
AGCACACGTCTGAACTCTTTCCCTACACGACGCTCTTCCGATCTNNNGCAACTGTGAGAACAAAAGG
AAGAGCACACGTCTGAACTCTTTCCCTACACGACGCTCTTCCGATCTNNNACTCAGGCATCATTCAGAT
AGCACACGTCTGAACTCTTTCCCTACACGACGCTCTTCCGATCTNNNAAAGGATGTGTCCGTCTGGT
GCACACGTCTGAACTCTTTCCCTACACGACGCTCTTCCGATCTNNNCACCTTTCAGTGCCTCCAAGAAT
CACACGTCTGAACTCTTTCCCTACACGACGCTCTTCCGATCTNNNTGTGTTCATGCCTTGATACCCTGC
AGCACACGTCTGAACTCTTTCCCTACACGACGCTCTTCCGATCTNNNGCTGAGTATTTCCTGTGGACC
GAGCACACGTCTGAACTCTTTCCCTACACGACGCTCTTCCGATCTNNNCTGTTCCATTGTATCACCCCAG
AGCACACGTCTGAACTCTTTCCCTACACGACGCTCTTCCGATCTNNNTCCTCAAAACTTAGGATACCC
GAGCACACGTCTGAACTCTTTCCCTACACGACGCTCTTCCGATCTNNNCCCCACCCCAGTTTAGGAGAT
AGCACACGTCTGAACTCTTTCCCTACACGACGCTCTTCCGATCTNNNAGCCCTTTCAGTGGTTCTTGAT
AGCACACGTCTGAACTCTTTCCCTACACGACGCTCTTCCGATCTNNNAGACCAAAACCACATGCAAAA
GAGCACACGTCTGAACTCTTTCCCTACACGACGCTCTTCCGATCTNNNAGTACTTCTGCCCCAGAGTTA
AGAGCACACGTCTGAACTCTTTCCCTACACGACGCTCTTCCGATCTNNNATCGTCCTTCATAACTGGG
AGCACACGTCTGAACTCTTTCCCTACACGACGCTCTTCCGATCTNNNTTATACACAAAGCAGCCCCTAC
AGCACACGTCTGAACTCTTTCCCTACACGACGCTCTTCCGATCTNNNACCTGGCTATTAGTGAGTTACT
AGAGCACACGTCTGAACTCTTTCCCTACACGACGCTCTTCCGATCTNNNAGCAAAAATGAACAAACTG
GCACACGTCTGAACTCTTTCCCTACACGACGCTCTTCCGATCTNNNGCACAGATCAATGCACAGCCAT
AGCACACGTCTGAACTCTTTCCCTACACGACGCTCTTCCGATCTNNNAGCAAAGATAGAGGGGTGTC
AGAGCACACGTCTGAACTCTTTCCCTACACGACGCTCTTCCGATCTNNNAGGATGAGTAAGAGTTTTC
AGAGCACACGTCTGAACTCTTTCCCTACACGACGCTCTTCCGATCTNNNAGGTTTCATGAAGCTGCCAA
GAAGAGCACACGTCTGAACTCTTTCCCTACACGACGCTCTTCCGATCTNNNCAGCACTGAGCCGAGAT
AGCACACGTCTGAACTCTTTCCCTACACGACGCTCTTCCGATCTNNNGTCCAGCCAGTATTGAGGCTTA
GAGCACACGTCTGAACTCTTTCCCTACACGACGCTCTTCCGATCTNNNACCCATAGCACTGGGAAAAA
GAGCACACGTCTGAACTCTTTCCCTACACGACGCTCTTCCGATCTNNNAGAAGTTATTTTGCCTCCCTG
AGCACACGTCTGAACTCTTTCCCTACACGACGCTCTTCCGATCTNNNTTTTGGAAAGGGAGCATCGGT
AGCACACGTCTGAACTCTTTCCCTACACGACGCTCTTCCGATCTNNNTCACACCTATGTTTGTGTGTGT
AGAGCACACGTCTGAACTCTTTCCCTACACGACGCTCTTCCGATCTNNNCTATGGCTCGCATCTTTCAC
GAGCACACGTCTGAACTCTTTCCCTACACGACGCTCTTCCGATCTNNNCTTAGGTCACGAACCCTACAG
AGCACACGTCTGAACTCTTTCCCTACACGACGCTCTTCCGATCTNNNGCAGGTTTGGGAAAGAGGAG
GCACACGTCTGAACTCTTTCCCTACACGACGCTCTTCCGATCTNNNCTTGAATGAGTTTACACGGCTG
GCACACGTCTGAACTCTTTCCCTACACGACGCTCTTCCGATCTNNNGCTGATATTCTGGGTATCTTTTC
GCACACGTCTGAACTCTTTCCCTACACGACGCTCTTCCGATCTNNNATGCTGGCAACTTCTCTCCTTTCC
GAGCACACGTCTGAACTCTTTCCCTACACGACGCTCTTCCGATCTNNNTCTGGCCAAACTTTTGTATG
AGCACACGTCTGAACTCTTTCCCTACACGACGCTCTTCCGATCTNNNTCTGAATTTATCCCACCCCTCA
AAGAGCACACGTCTGAACTCTTTCCCTACACGACGCTCTTCCGATCTNNNGAACTCTGTGGAACTTTG

GAGCACACGTCTGAACTCTTTCCCTACACGACGCTCTTCCGATCTNNNTGCCATATAACTGGAAATGCA

AGCACACGTCTGAACTCTTTCCCTACACGACGCTCTTCCGATCTNNNAAAACCCTGCTCCATTGTTTGC

AGCACACGTCTGAACTCTTTCCCTACACGACGCTCTTCCGATCTNNNTGTGACCAGGTTTCATCCAATCA

AGAGCACACGTCTGAACTCTTTCCCTACACGACGCTCTTCCGATCTNNNAATAAAGTGTGTGTGTGCAT

GAGCACACGTCTGAACTCTTTCCCTACACGACGCTCTTCCGATCTNNNTTGTCTCTTCCCCTCTTCCTGAC

GAGCACACGTCTGAACTCTTTCCCTACACGACGCTCTTCCGATCTNNNAAGCTTGTGGCACAGAGTAAA

AGCACACGTCTGAACTCTTTCCCTACACGACGCTCTTCCGATCTNNNAGTGGTTGCTCTAAGGTATCCAT

AGCACACGTCTGAACTCTTTCCCTACACGACGCTCTTCCGATCTNNNCCATCATAGTTCCATGGAGGGT

AGAGCACACGTCTGAACTCTTTCCCTACACGACGCTCTTCCGATCTNNNGGTGGAGGCTATGAGGGT

AGCACACGTCTGAACTCTTTCCCTACACGACGCTCTTCCGATCTNNNGCATTCCAATTTTCACCTCCAGCC

AGCACACGTCTGAACTCTTTCCCTACACGACGCTCTTCCGATCTNNNATCGCTGTATTCTTGACTGAGCC

AAGAGCACACGTCTGAACTCTTTCCCTACACGACGCTCTTCCGATCTNNNACTCACTTTGCTTTCCATTTG

GAGCACACGTCTGAACTCTTTCCCTACACGACGCTCTTCCGATCTNNNAAGCACTTGGAAAACATTGCA

AGAGCACACGTCTGAACTCTTTCCCTACACGACGCTCTTCCGATCTNNNCATAAAACTGCCTCCAATGCA

AGAGCACACGTCTGAACTCTTTCCCTACACGACGCTCTTCCGATCTNNNTGTATATCTCCTAATGCTGGG

GAGCACACGTCTGAACTCTTTCCCTACACGACGCTCTTCCGATCTNNNAGCCTTCTGCCTCTTCCTTCATA

AGCACACGTCTGAACTCTTTCCCTACACGACGCTCTTCCGATCTNNNGTCACATATCCCACTGCTTCAGC

AGCACACGTCTGAACTCTTTCCCTACACGACGCTCTTCCGATCTNNNGGCATAGCATCACCAAGTTGGA

AAGAGCACACGTCTGAACTCTTTCCCTACACGACGCTCTTCCGATCTNNNAGTGAGACACTTCAGGGAA

AGCACACGTCTGAACTCTTTCCCTACACGACGCTCTTCCGATCTNNNACTACATAGTCAGGCAAGGTCC

GAGCACACGTCTGAACTCTTTCCCTACACGACGCTCTTCCGATCTNNNTGCTACACTCAAAGTTTTGGCC

GAGCACACGTCTGAACTCTTTCCCTACACGACGCTCTTCCGATCTNNNCTGGCTTCAGACCTCACTCAAT

GAGCACACGTCTGAACTCTTTCCCTACACGACGCTCTTCCGATCTNNNTGAACTCTTGACTTGCAGTTTT

AGAGCACACGTCTGAACTCTTTCCCTACACGACGCTCTTCCGATCTNNNAGAAGTGGCCCAGATGACT

AAGAGCACACGTCTGAACTCTTTCCCTACACGACGCTCTTCCGATCTNNNTAGAGTTGATATGCACACA

GAGCACACGTCTGAACTCTTTCCCTACACGACGCTCTTCCGATCTNNNTTGCACCAAGCGAATAGAAGT

GAGCACACGTCTGAACTCTTTCCCTACACGACGCTCTTCCGATCTNNNACACATGGGAATGGGTAACTT

AGAGCACACGTCTGAACTCTTTCCCTACACGACGCTCTTCCGATCTNNNTAAGGTAAGGCTGTCCTTCTG

AGAGCACACGTCTGAACTCTTTCCCTACACGACGCTCTTCCGATCTNNNATACCTGGCATTTCTTAGCTG

AGCACACGTCTGAACTCTTTCCCTACACGACGCTCTTCCGATCTNNNGAATGTGGAAGAGACTGTGAC

AGAGCACACGTCTGAACTCTTTCCCTACACGACGCTCTTCCGATCTNNNGTTGGGTGTGTACATGTGTG

AGCACACGTCTGAACTCTTTCCCTACACGACGCTCTTCCGATCTNNNCTGTGCTGGCTTCTGTCTATTCA

AGCACACGTCTGAACTCTTTCCCTACACGACGCTCTTCCGATCTNNNGTGACAATTGGCGAGAGAGAGA

GAGCACACGTCTGAACTCTTTCCCTACACGACGCTCTTCCGATCTNNNTGGCCATAGAAACAGTAATGG

CACACGTCTGAACTCTTTCCCTACACGACGCTCTTCCGATCTNNNCCGCTTTGTCTCTTCACTTCTTGCAT

GAGCACACGTCTGAACTCTTTCCCTACACGACGCTCTTCCGATCTNNNATGGATTGAGCCAGATGCCAT

AGAGCACACGTCTGAACTCTTTCCCTACACGACGCTCTTCCGATCTNNNTTTCTATGTTCTTCCTCCTCC

AGAGCACACGTCTGAACTCTTTCCCTACACGACGCTCTTCCGATCTNNNACTGTCTCTTAGGGTCTGTG

AGCACACGTCTGAACTCTTTCCCTACACGACGCTCTTCCGATCTNNNTGCACACTGTGGATTCGTAAGA

GAGCACACGTCTGAACTCTTTCCCTACACGACGCTCTTCCGATCTNNNTGCCTGGTACACGTTTGTCATT

AGCACACGTCTGAACTCTTTCCCTACACGACGCTCTTCCGATCTNNNAATTACCGTGAGAGTGAGGGA

GAGCACACGTCTGAACTCTTTCCCTACACGACGCTCTTCCGATCTNNNGTCAAAGCCAAGAGGTCCTGA

GAGCACACGTCTGAACTCTTTCCCTACACGACGCTCTTCCGATCTNNNGCATGGATTGCAGGGAAACT

AGCACACGTCTGAACTCTTTCCCTACACGACGCTCTTCCGATCTNNNAACATGAGTTTGTCAGTGGAC
AGCACACGTCTGAACTCTTTCCCTACACGACGCTCTTCCGATCTNNNCAGGCTGTTCTGTTGACCTGT
GAGCACACGTCTGAACTCTTTCCCTACACGACGCTCTTCCGATCTNNNAGTTTCAATGAGACCCAAGC
AGCACACGTCTGAACTCTTTCCCTACACGACGCTCTTCCGATCTNNNAGCACTATTGCTGGATTGATG
AGCACACGTCTGAACTCTTTCCCTACACGACGCTCTTCCGATCTNNNAAGTAGCTGAGACTACATTGCG
AGAGCACACGTCTGAACTCTTTCCCTACACGACGCTCTTCCGATCTNNNACTAGAACCTGGGCGAGA
GAGCACACGTCTGAACTCTTTCCCTACACGACGCTCTTCCGATCTNNNTGTCGCATATTTCAGAATTCC
AGAGCACACGTCTGAACTCTTTCCCTACACGACGCTCTTCCGATCTNNNCGCAAATAGGCTCAAAACA
AGCACACGTCTGAACTCTTTCCCTACACGACGCTCTTCCGATCTNNNAATAAAGGTGTTGGCTGCTTC
GAGCACACGTCTGAACTCTTTCCCTACACGACGCTCTTCCGATCTNNNCAGCTAATGACCACAACCAG
GCACACGTCTGAACTCTTTCCCTACACGACGCTCTTCCGATCTNNNAGGAATGGCCATACTTCCTCACT
AGCACACGTCTGAACTCTTTCCCTACACGACGCTCTTCCGATCTNNNGTCTGCTCAGTTCCCGCTAAG
AGCACACGTCTGAACTCTTTCCCTACACGACGCTCTTCCGATCTNNNTGAACACCCACTCCATGAAGA
AGAGCACACGTCTGAACTCTTTCCCTACACGACGCTCTTCCGATCTNNNAGGAAGCAAACTCTTCTCA
GAGCACACGTCTGAACTCTTTCCCTACACGACGCTCTTCCGATCTNNNATAATAACCGTCCCCAACA
GAGCACACGTCTGAACTCTTTCCCTACACGACGCTCTTCCGATCTNNNTGTGTAGTGGGACCAATGA
GAGCACACGTCTGAACTCTTTCCCTACACGACGCTCTTCCGATCTNNNTTGGATTTGTGTCTCTGCATC
AGCACACGTCTGAACTCTTTCCCTACACGACGCTCTTCCGATCTNNNAATGCTCTACATGCAGGCTTAC
AGAGCACACGTCTGAACTCTTTCCCTACACGACGCTCTTCCGATCTNNNCCCAAACTGTGGAAATTGC
AGCACACGTCTGAACTCTTTCCCTACACGACGCTCTTCCGATCTNNNAACCTGCCAAACAAATCAATG
GCACACGTCTGAACTCTTTCCCTACACGACGCTCTTCCGATCTNNNATGGATTGGTGGAATGGTAGT
AGCACACGTCTGAACTCTTTCCCTACACGACGCTCTTCCGATCTNNNAAAGACATCCTGCCTAGTTCC
AGCACACGTCTGAACTCTTTCCCTACACGACGCTCTTCCGATCTNNNATTGGCAACAAACCATATGTG
AGCACACGTCTGAACTCTTTCCCTACACGACGCTCTTCCGATCTNNNGAACCTAGAATTGGGGCTGAG
CACACGTCTGAACTCTTTCCCTACACGACGCTCTTCCGATCTNNNATGATTATTTCATCTGAGCACTTG
CACACGTCTGAACTCTTTCCCTACACGACGCTCTTCCGATCTNNNCATGACACAGTGGCTATGAGGAG
GAGCACACGTCTGAACTCTTTCCCTACACGACGCTCTTCCGATCTNNNCGCGAGACGGAGGTTGCAA
AAGAGCACACGTCTGAACTCTTTCCCTACACGACGCTCTTCCGATCTNNNTGTGGTAGTTGCACACAA
GAGCACACGTCTGAACTCTTTCCCTACACGACGCTCTTCCGATCTNNNACCTTGGGCAAAACAAGCAA
GAGCACACGTCTGAACTCTTTCCCTACACGACGCTCTTCCGATCTNNNTGAAAGTAAGAGTGAGAGG
GAGCACACGTCTGAACTCTTTCCCTACACGACGCTCTTCCGATCTNNNCCTCTGGGGCTCAAATGTTTC
CACACGTCTGAACTCTTTCCCTACACGACGCTCTTCCGATCTNNNCCATGCCTTGCATCCATTTATCCCA
GCACACGTCTGAACTCTTTCCCTACACGACGCTCTTCCGATCTNNNACTTACGTAAATGTACGCAGCC
AGCACACGTCTGAACTCTTTCCCTACACGACGCTCTTCCGATCTNNNAGAACAGCGAAAGTCCATAGC
GAGCACACGTCTGAACTCTTTCCCTACACGACGCTCTTCCGATCTNNNTTCCCATGTTCTGGGTTGAC
AGCACACGTCTGAACTCTTTCCCTACACGACGCTCTTCCGATCTNNNACCTGGTAGTGGGTTTATGTC
AGCACACGTCTGAACTCTTTCCCTACACGACGCTCTTCCGATCTNNNCTGGGTTGCTGTTGAAGTTTT
GCACACGTCTGAACTCTTTCCCTACACGACGCTCTTCCGATCTNNNGAAGGCCTGGAGGTATGAAAC
AGCACACGTCTGAACTCTTTCCCTACACGACGCTCTTCCGATCTNNNTTAGTTGGCTCCCATGTCTCTC
GCACACGTCTGAACTCTTTCCCTACACGACGCTCTTCCGATCTNNNGCACATCCAGCTGTTAGTCTTG
AGCACACGTCTGAACTCTTTCCCTACACGACGCTCTTCCGATCTNNNGAGTTGGGGTGTCTACATTG
AGCACACGTCTGAACTCTTTCCCTACACGACGCTCTTCCGATCTNNNCAAGCCTTTGTAATGGTGAG
GAGCACACGTCTGAACTCTTTCCCTACACGACGCTCTTCCGATCTNNNGCCTGGCCAAGTATCACTTTT

GAGCACACGTCTGAACTCTTTCCCTACACGACGCTCTTCCGATCTNNNGGCTACAAGCATCGTATAGT
AGCACACGTCTGAACTCTTTCCCTACACGACGCTCTTCCGATCTNNNAAGAGAGGGAAGGGTAGTTG
GCACACGTCTGAACTCTTTCCCTACACGACGCTCTTCCGATCTNNNTACAGTTCATCTTGTGGAGGGG
AGCACACGTCTGAACTCTTTCCCTACACGACGCTCTTCCGATCTNNNTGTTCATCAGACCTTCCCTCCTT
GAGCACACGTCTGAACTCTTTCCCTACACGACGCTCTTCCGATCTNNNCTCACCAGGAAAGTATGTGG
AGCACACGTCTGAACTCTTTCCCTACACGACGCTCTTCCGATCTNNNCTCTGAACACGACTTGGGCTTT
GAGCACACGTCTGAACTCTTTCCCTACACGACGCTCTTCCGATCTNNNATTGAGTAGGAGTGAGCCA
AGCACACGTCTGAACTCTTTCCCTACACGACGCTCTTCCGATCTNNNTTTCACTTGCCTGTGTATCACCT
AAGAGCACACGTCTGAACTCTTTCCCTACACGACGCTCTTCCGATCTNNNTGGGCTGGTTACAACCTC
GAGCACACGTCTGAACTCTTTCCCTACACGACGCTCTTCCGATCTNNNAAGGCAAGTACCTCCTCTCTC
GCACACGTCTGAACTCTTTCCCTACACGACGCTCTTCCGATCTNNNACCACCAGTTCAGTACCATCTGT
AGAGCACACGTCTGAACTCTTTCCCTACACGACGCTCTTCCGATCTNNNGAGAAGAAGCAAGCATTC
CACACGTCTGAACTCTTTCCCTACACGACGCTCTTCCGATCTNNNCCCCTCAAACCATGCAATTTGTCC
GAGCACACGTCTGAACTCTTTCCCTACACGACGCTCTTCCGATCTNNNACTCATGGTGAGGACAAAGG
GAGCACACGTCTGAACTCTTTCCCTACACGACGCTCTTCCGATCTNNNTTGCTCTGATTTTACTGACTG
AGAGCACACGTCTGAACTCTTTCCCTACACGACGCTCTTCCGATCTNNNTTATCTCTGTCCACTGGTCC
AGCACACGTCTGAACTCTTTCCCTACACGACGCTCTTCCGATCTNNNCACCTGAGGAGATACACACAC
AGAGCACACGTCTGAACTCTTTCCCTACACGACGCTCTTCCGATCTNNNATCAAGCTGGAGGGAAAA
GAGCACACGTCTGAACTCTTTCCCTACACGACGCTCTTCCGATCTNNNGCAAAATCTGGCTCCTAGTG
GAGCACACGTCTGAACTCTTTCCCTACACGACGCTCTTCCGATCTNNNATTGGTTAGGGTTCCTCATTG
GAGCACACGTCTGAACTCTTTCCCTACACGACGCTCTTCCGATCTNNNGTCCCCACCAATACCACCAGAT
AAGAGCACACGTCTGAACTCTTTCCCTACACGACGCTCTTCCGATCTNNNGGCCAGAAGAGGCAACTT
AAGAGCACACGTCTGAACTCTTTCCCTACACGACGCTCTTCCGATCTNNNTTCAAATCCCACTCACTCG
AGAGCACACGTCTGAACTCTTTCCCTACACGACGCTCTTCCGATCTNNNACTTGCCTTTGGATAGCAAG
AGCACACGTCTGAACTCTTTCCCTACACGACGCTCTTCCGATCTNNNCTTTAGGTGAGAAGGGTTTCG
GAGCACACGTCTGAACTCTTTCCCTACACGACGCTCTTCCGATCTNNNTGGGCTTTCCAAGAATACCA
AGCACACGTCTGAACTCTTTCCCTACACGACGCTCTTCCGATCTNNNGCCCCAGGAGCCCTAAATAAAT
AGCACACGTCTGAACTCTTTCCCTACACGACGCTCTTCCGATCTNNNGACCCTGCATTACTGTACCACT
GAAGAGCACACGTCTGAACTCTTTCCCTACACGACGCTCTTCCGATCTNNNCCTGAAGCCTTGCTGAA
AGAGCACACGTCTGAACTCTTTCCCTACACGACGCTCTTCCGATCTNNNTTTGCCTAGTATTGCTTTGG
AGCACACGTCTGAACTCTTTCCCTACACGACGCTCTTCCGATCTNNNGCTCTGTCAAACTGTGTAGAGG
AGCACACGTCTGAACTCTTTCCCTACACGACGCTCTTCCGATCTNNNGGATTCTGATTCCCACTCCAAG
GAGCACACGTCTGAACTCTTTCCCTACACGACGCTCTTCCGATCTNNNTGAGGAGGGAATCCAAACTC
AGCACACGTCTGAACTCTTTCCCTACACGACGCTCTTCCGATCTNNNCCTAGGACATGCCTTCCAAAGA
GAGCACACGTCTGAACTCTTTCCCTACACGACGCTCTTCCGATCTNNNGGCTGAAAACTAGCCTTGGA
GAGCACACGTCTGAACTCTTTCCCTACACGACGCTCTTCCGATCTNNNTCAAAATGAGGGTGTCTGCT
AGCACACGTCTGAACTCTTTCCCTACACGACGCTCTTCCGATCTNNNCCCCTGTAAGAAGAAACACCG
AGCACACGTCTGAACTCTTTCCCTACACGACGCTCTTCCGATCTNNNCGACAGTTCAGCCTTATCCTCTG
GAGCACACGTCTGAACTCTTTCCCTACACGACGCTCTTCCGATCTNNNAATATCTGGGGTGCTGCCAT
GCACACGTCTGAACTCTTTCCCTACACGACGCTCTTCCGATCTNNNTTTACCATGTCCCACAGCAAAA
AAGAGCACACGTCTGAACTCTTTCCCTACACGACGCTCTTCCGATCTNNNTTGTGCAAAACTGGGTAA
AGAGCACACGTCTGAACTCTTTCCCTACACGACGCTCTTCCGATCTNNNCCATGCAGTAGCATAGATG
AAGAGCACACGTCTGAACTCTTTCCCTACACGACGCTCTTCCGATCTNNNTCATCTACAAGGAAGGCT

AGCACACGTCTGAACTCTTTCCCTACACGACGCTCTTCCGATCTNNNCAGGGAAGAGAGAGTGAGAG
GAGCACACGTCTGAACTCTTTCCCTACACGACGCTCTTCCGATCTNNNTGTCATTAGTTACACTGAATG
GAGCACACGTCTGAACTCTTTCCCTACACGACGCTCTTCCGATCTNNNGCGACAAAGCAAGATTGTGT
GAGCACACGTCTGAACTCTTTCCCTACACGACGCTCTTCCGATCTNNNAAATCTCTCCCTTCCTCCCTGTC
GAGCACACGTCTGAACTCTTTCCCTACACGACGCTCTTCCGATCTNNNATGAGCAGACATTTCAAAGCC
AGCACACGTCTGAACTCTTTCCCTACACGACGCTCTTCCGATCTNNNAGTGTCAGTTCATGTTGCCTCT
GCACACGTCTGAACTCTTTCCCTACACGACGCTCTTCCGATCTNNNGAAGTTGGAGAACTCATTGGGC
AGCACACGTCTGAACTCTTTCCCTACACGACGCTCTTCCGATCTNNNTTTAGCCACTCACCAACTGAAG
AGCACACGTCTGAACTCTTTCCCTACACGACGCTCTTCCGATCTNNNGTTGGTATGTCCATTCCCCTTG
AGCACACGTCTGAACTCTTTCCCTACACGACGCTCTTCCGATCTNNNGGGACTTTCAGGTTGGATGAG
AGCACACGTCTGAACTCTTTCCCTACACGACGCTCTTCCGATCTNNNGCATTGACCCAGCCAATACAAA
GAGCACACGTCTGAACTCTTTCCCTACACGACGCTCTTCCGATCTNNNTGAATACCTCTGCCCCACTTCAT
AGCACACGTCTGAACTCTTTCCCTACACGACGCTCTTCCGATCTNNNACATACCTACACACATTCATACA
AGAGCACACGTCTGAACTCTTTCCCTACACGACGCTCTTCCGATCTNNNACAACTGACATACCATCTCTG
GAGCACACGTCTGAACTCTTTCCCTACACGACGCTCTTCCGATCTNNNACTTCCCATGTCCAGACTAGGT
GCACACGTCTGAACTCTTTCCCTACACGACGCTCTTCCGATCTNNNCTCAGTATCTCCAGCTTGCCAACC
GAGCACACGTCTGAACTCTTTCCCTACACGACGCTCTTCCGATCTNNNGGTGAGATGAGTGAAACTCT
GAAGAGCACACGTCTGAACTCTTTCCCTACACGACGCTCTTCCGATCTNNNGCACAGACTAACCAGCTC
AGCACACGTCTGAACTCTTTCCCTACACGACGCTCTTCCGATCTNNNATACACACACGTATAATACACA
GAGCACACGTCTGAACTCTTTCCCTACACGACGCTCTTCCGATCTNNNGTGGGTTCAGCATTTTCTGTT
GCACACGTCTGAACTCTTTCCCTACACGACGCTCTTCCGATCTNNNGAGAGAGAGGAAGAGAGCTTG
GAGCACACGTCTGAACTCTTTCCCTACACGACGCTCTTCCGATCTNNNCACACATACCAACAGACAGGCA
AGCACACGTCTGAACTCTTTCCCTACACGACGCTCTTCCGATCTNNNTTCTAGGGAAGGCATGAGCAAT
AGCACACGTCTGAACTCTTTCCCTACACGACGCTCTTCCGATCTNNNAAGTCATGCAGAGTTCCACCATT
AGAGCACACGTCTGAACTCTTTCCCTACACGACGCTCTTCCGATCTNNNGAGGGTGGAATTCAAAGCC
AGCACACGTCTGAACTCTTTCCCTACACGACGCTCTTCCGATCTNNNTTACCTGGATTGCAAATGGACC
AGCACACGTCTGAACTCTTTCCCTACACGACGCTCTTCCGATCTNNNAGCACTCAGTGGTAGAGACTTC
GAGCACACGTCTGAACTCTTTCCCTACACGACGCTCTTCCGATCTNNNGTGATGGTGGTAGGAAGATC
AAGAGCACACGTCTGAACTCTTTCCCTACACGACGCTCTTCCGATCTNNNAGAAGACTTGTCCGGAGGA
AGAGCACACGTCTGAACTCTTTCCCTACACGACGCTCTTCCGATCTNNNTCCTGTGTCTTTACTTTTGC
AGAGCACACGTCTGAACTCTTTCCCTACACGACGCTCTTCCGATCTNNNACTTGAGCAGCCCGGAAGT
GAGCACACGTCTGAACTCTTTCCCTACACGACGCTCTTCCGATCTNNNCGGAGCTTGACACCTAGTAAA
AGCACACGTCTGAACTCTTTCCCTACACGACGCTCTTCCGATCTNNNTATTTTCCCAGACTGGAGTGGC
AGCACACGTCTGAACTCTTTCCCTACACGACGCTCTTCCGATCTNNNTCACTCGAATCTACACGGGATGA
GAGCACACGTCTGAACTCTTTCCCTACACGACGCTCTTCCGATCTNNNTGTCCCAAAAATGTATCATGCA
AGAGCACACGTCTGAACTCTTTCCCTACACGACGCTCTTCCGATCTNNNGAATCCCCAGCTCTCCTGCTA
AGAGCACACGTCTGAACTCTTTCCCTACACGACGCTCTTCCGATCTNNNGAAGCGAGAAAGGTCTGA
AGAGCACACGTCTGAACTCTTTCCCTACACGACGCTCTTCCGATCTNNNTTAGACTTCTGGGCAGCTAT
AGAGCACACGTCTGAACTCTTTCCCTACACGACGCTCTTCCGATCTNNNCCATCTGTTGTAGCAAACATC
AGCACACGTCTGAACTCTTTCCCTACACGACGCTCTTCCGATCTNNNTGGTCCCTTCTTTCAGCGTGTAT
AGAGCACACGTCTGAACTCTTTCCCTACACGACGCTCTTCCGATCTNNNGCTTGTTCTTTCTGCCTAAGG
GAGCACACGTCTGAACTCTTTCCCTACACGACGCTCTTCCGATCTNNNTGCAGTGATTCATTGGCTGTCT
GAGCACACGTCTGAACTCTTTCCCTACACGACGCTCTTCCGATCTNNNCACACTACACTTCTGTGGGAA

AGCACACGTCTGAACTCTTTCCCTACACGACGCTCTTCCGATCTNNNACAAGGTATAAGGTGCCTGG
GAGCACACGTCTGAACTCTTTCCCTACACGACGCTCTTCCGATCTNNNCACCAAGTGGAAAGATACCT
CACACGTCTGAACTCTTTCCCTACACGACGCTCTTCCGATCTNNNGCAGGTGCTAGTAAACCTGATCC
GCACACGTCTGAACTCTTTCCCTACACGACGCTCTTCCGATCTNNNATCTGCCAGAAAGGCTTAGAG
AGCACACGTCTGAACTCTTTCCCTACACGACGCTCTTCCGATCTNNNCACAGGAGAACCACTAAGGT
GAGCACACGTCTGAACTCTTTCCCTACACGACGCTCTTCCGATCTNNNCCCCCTCTACAATGCCTGTTT
AGCACACGTCTGAACTCTTTCCCTACACGACGCTCTTCCGATCTNNNTCCTGTTTCTCCCACTGTACCA
AGCACACGTCTGAACTCTTTCCCTACACGACGCTCTTCCGATCTNNNTTGCATTCATAGCTGGGCAGT
AGCACACGTCTGAACTCTTTCCCTACACGACGCTCTTCCGATCTNNNTTGCTCCTTGTCTTGAAGAGC
GCACACGTCTGAACTCTTTCCCTACACGACGCTCTTCCGATCTNNNGGACCCCCTCTAATCTGTTCCAT
GAGCACACGTCTGAACTCTTTCCCTACACGACGCTCTTCCGATCTNNNTCCCAAACTGATTTGGCCAT
AGCACACGTCTGAACTCTTTCCCTACACGACGCTCTTCCGATCTNNNGCCAGACCCCACAAATCAAGT
GCACACGTCTGAACTCTTTCCCTACACGACGCTCTTCCGATCTNNNACAGACTATGAGCGTGAGAAT
GAGCACACGTCTGAACTCTTTCCCTACACGACGCTCTTCCGATCTNNNCCATGTATTTGCTGTGTTTG
AGCACACGTCTGAACTCTTTCCCTACACGACGCTCTTCCGATCTNNNTGGGCTTTCAGCACAACAGAT
GCACACGTCTGAACTCTTTCCCTACACGACGCTCTTCCGATCTNNNTGAAATAAGGACCCCAAACGC
GAGCACACGTCTGAACTCTTTCCCTACACGACGCTCTTCCGATCTNNNTCTCTCTCTAGACAAATGA
AGCACACGTCTGAACTCTTTCCCTACACGACGCTCTTCCGATCTNNNTTAGCAAAGGACTTGGCACAG
AGCACACGTCTGAACTCTTTCCCTACACGACGCTCTTCCGATCTNNNTCCCTGGGGACTTTAGGAATT
GCACACGTCTGAACTCTTTCCCTACACGACGCTCTTCCGATCTNNNACCTACGTCTAGCTACCTACCCA
AGCACACGTCTGAACTCTTTCCCTACACGACGCTCTTCCGATCTNNNCTATCCCTCCACTAGCCTCCTCC
GAGCACACGTCTGAACTCTTTCCCTACACGACGCTCTTCCGATCTNNNCAATAGCTAAGCTCCTCACTG
GAGCACACGTCTGAACTCTTTCCCTACACGACGCTCTTCCGATCTNNNGCCTCAGTGAGCCATTATTTT
AGCACACGTCTGAACTCTTTCCCTACACGACGCTCTTCCGATCTNNNAGAGACGAGAGAATCACAGA
AGCACACGTCTGAACTCTTTCCCTACACGACGCTCTTCCGATCTNNNTCTGTACTAGACTGCACACCA
AGCACACGTCTGAACTCTTTCCCTACACGACGCTCTTCCGATCTNNNAGCCCCCAGAGAATCTGGTAA
AGCACACGTCTGAACTCTTTCCCTACACGACGCTCTTCCGATCTNNNAATCCATCTCACTGGGGGAAA
AGCACACGTCTGAACTCTTTCCCTACACGACGCTCTTCCGATCTNNNTCAAGGTTTTCGAGTTGCCCA
AGCACACGTCTGAACTCTTTCCCTACACGACGCTCTTCCGATCTNNNCTCTAGGGGATGCTGGGAATC
GAGCACACGTCTGAACTCTTTCCCTACACGACGCTCTTCCGATCTNNNCCTGTACCTATTGGCCTTTCC
GAGCACACGTCTGAACTCTTTCCCTACACGACGCTCTTCCGATCTNNNGGAGGCTTGACACTAGGAG
GAGCACACGTCTGAACTCTTTCCCTACACGACGCTCTTCCGATCTNNNTGTTCACAAGTGACATGGCA
GAGCACACGTCTGAACTCTTTCCCTACACGACGCTCTTCCGATCTNNNTTGGAGTGTGCTCTCTCTCT
AGCACACGTCTGAACTCTTTCCCTACACGACGCTCTTCCGATCTNNNGGAGAAAAGGGAAGGAACCC
AGCACACGTCTGAACTCTTTCCCTACACGACGCTCTTCCGATCTNNNATCCAATCCCGGATCCTACAGT
GAGCACACGTCTGAACTCTTTCCCTACACGACGCTCTTCCGATCTNNNAGAGAGGTTTAGGTGTCTG
AGCACACGTCTGAACTCTTTCCCTACACGACGCTCTTCCGATCTNNNTGAGAAGGAGGAAAGGGAA
AGCACACGTCTGAACTCTTTCCCTACACGACGCTCTTCCGATCTNNNCTAGGCAGACTACACTCCCTCTT
AGCACACGTCTGAACTCTTTCCCTACACGACGCTCTTCCGATCTNNNTGATACAAAGCTACCCTGCCTC
GAGCACACGTCTGAACTCTTTCCCTACACGACGCTCTTCCGATCTNNNTCACCTTGGAAGGTAGAGAG
AGCACACGTCTGAACTCTTTCCCTACACGACGCTCTTCCGATCTNNNTTCCCTTCCCATCTTTTTGGCAG
AGCACACGTCTGAACTCTTTCCCTACACGACGCTCTTCCGATCTNNNGCGTTATGCACAAAATTTCCC
GAGCACACGTCTGAACTCTTTCCCTACACGACGCTCTTCCGATCTNNNCAGAACTGCTGGAAAGACAA

GCACACGTCTGAACTCTTTCCCTACACGACGCTCTTCCGATCTNNNGGGAATTTCTAGGGTGTCTTTTT

AGCACACGTCTGAACTCTTTCCCTACACGACGCTCTTCCGATCTNNNTGGTCCAAAGGTTTTGTAAGTG

GAGCACACGTCTGAACTCTTTCCCTACACGACGCTCTTCCGATCTNNNTTATTTCTCCTCCCTCCCCATC

AGCACACGTCTGAACTCTTTCCCTACACGACGCTCTTCCGATCTNNNTTCCCTGAAGCGAGTACACATT

AAGAGCACACGTCTGAACTCTTTCCCTACACGACGCTCTTCCGATCTNNNGCGCCTGGCTGAAAATGA

GCACACGTCTGAACTCTTTCCCTACACGACGCTCTTCCGATCTNNNTAATCTGGCAGGTAGGTGAGG

AAGAGCACACGTCTGAACTCTTTCCCTACACGACGCTCTTCCGATCTNNNGTTTGGCCCATTATAGGC

AGAGCACACGTCTGAACTCTTTCCCTACACGACGCTCTTCCGATCTNNNGTGTGCAATTGGGTATGTT

AGAGCACACGTCTGAACTCTTTCCCTACACGACGCTCTTCCGATCTNNNGAGATAGGGACAGGGTGG

GAGCACACGTCTGAACTCTTTCCCTACACGACGCTCTTCCGATCTNNNAGTCAAATTACCTGGCTGAG

GAGCACACGTCTGAACTCTTTCCCTACACGACGCTCTTCCGATCTNNNGGACCCCATAGCCTGAAACA

AGCACACGTCTGAACTCTTTCCCTACACGACGCTCTTCCGATCTNNNGGAGGCCAGCATTCATACATT

AGCACACGTCTGAACTCTTTCCCTACACGACGCTCTTCCGATCTNNNTAAATGCTAGAGAACTGGAG

AGCACACGTCTGAACTCTTTCCCTACACGACGCTCTTCCGATCTNNNGCACCACCAAAACTGAGGAAG

GCACACGTCTGAACTCTTTCCCTACACGACGCTCTTCCGATCTNNNTTTAGCAAGGTCTCTGTGCAG

AGCACACGTCTGAACTCTTTCCCTACACGACGCTCTTCCGATCTNNNATCTGTGTGGAGGCAGCAAT

AGCACACGTCTGAACTCTTTCCCTACACGACGCTCTTCCGATCTNNNATCAGCAACTTCGCAGACTTAG

GCACACGTCTGAACTCTTTCCCTACACGACGCTCTTCCGATCTNNNCACCCCATTCTCACCAACACATAC

AGCACACGTCTGAACTCTTTCCCTACACGACGCTCTTCCGATCTNNNTGAAACGGAAATACCCTTTGC

AGCACACGTCTGAACTCTTTCCCTACACGACGCTCTTCCGATCTNNNTTTAGCCTCTGCCCTTCCCTTTA

AGCACACGTCTGAACTCTTTCCCTACACGACGCTCTTCCGATCTNNNTGGACACAGGATGGCAACAA

GCACACGTCTGAACTCTTTCCCTACACGACGCTCTTCCGATCTNNNTTGGGAAGGGGGTGTGTATAT

AGCACACGTCTGAACTCTTTCCCTACACGACGCTCTTCCGATCTNNNTTTTTACCTAAGATAATGGCCT

GAGCACACGTCTGAACTCTTTCCCTACACGACGCTCTTCCGATCTNNNAGACTGCATCTTTCCTCTTGG

AGAGCACACGTCTGAACTCTTTCCCTACACGACGCTCTTCCGATCTNNNGGCTATCTTCTTGGGCCTTC

GCACACGTCTGAACTCTTTCCCTACACGACGCTCTTCCGATCTNNNACATGTATCTCCTGTTATTGTGT

GAGCACACGTCTGAACTCTTTCCCTACACGACGCTCTTCCGATCTNNNGAAGTCTCTCCCACATCAAG

GAGCACACGTCTGAACTCTTTCCCTACACGACGCTCTTCCGATCTNNNTTTCACTGCTCAATGCTTCCT

GCACACGTCTGAACTCTTTCCCTACACGACGCTCTTCCGATCTNNNATTTCTGACAACCAAAGGGAAG

AGCACACGTCTGAACTCTTTCCCTACACGACGCTCTTCCGATCTNNNCAAAGAACACCCTTCCCCTTCT

AGCACACGTCTGAACTCTTTCCCTACACGACGCTCTTCCGATCTNNNCCTAGTTTTAGAAGTTTCAAT

GCACACGTCTGAACTCTTTCCCTACACGACGCTCTTCCGATCTNNNATGCACGTGGTAGACACTCAGT

AGCACACGTCTGAACTCTTTCCCTACACGACGCTCTTCCGATCTNNNGGACCAACAGTCAGCACTTAC

GCACACGTCTGAACTCTTTCCCTACACGACGCTCTTCCGATCTNNNTATTTGTCTTCGCCGCTTGTTG

AGCACACGTCTGAACTCTTTCCCTACACGACGCTCTTCCGATCTNNNCACCAAGAACCATAAACCGAG

GAGCACACGTCTGAACTCTTTCCCTACACGACGCTCTTCCGATCTNNNTTATTGCCTGCTCTCTCTCCTG

CACACGTCTGAACTCTTTCCCTACACGACGCTCTTCCGATCTNNNATACACAAGCCCCTTCACAATCCC

GAGCACACGTCTGAACTCTTTCCCTACACGACGCTCTTCCGATCTNNNAGCATACATAGTCATGGACA

GAGCACACGTCTGAACTCTTTCCCTACACGACGCTCTTCCGATCTNNNAGGAAAAGTCCAAACCCATC

AGCACACGTCTGAACTCTTTCCCTACACGACGCTCTTCCGATCTNNNAAATGATAGATAACGCCCCAC

AGCACACGTCTGAACTCTTTCCCTACACGACGCTCTTCCGATCTNNNTCTCTGGGTGTCTTTCTCCGAA

GAGCACACGTCTGAACTCTTTCCCTACACGACGCTCTTCCGATCTNNNAAAGAGAGGCTGTTAGACA

AAGAGCACACGTCTGAACTCTTTCCCTACACGACGCTCTTCCGATCTNNNCCTTTGTATCTGGTCTGG

GAGCACACGTCTGAACTCTTTCCCTACACGACGCTCTTCCGATCTNNNTCCACTTGGCTGTTAGAACAC
GCACACGTCTGAACTCTTTCCCTACACGACGCTCTTCCGATCTNNNGTCCAGTTGAAAACTTGATTCCCC
AGCACACGTCTGAACTCTTTCCCTACACGACGCTCTTCCGATCTNNNGAGAGAGAGACAGACCCAGTA
GCACACGTCTGAACTCTTTCCCTACACGACGCTCTTCCGATCTNNNCGGCAAGGTGATACTCCATCTCCA
AGCACACGTCTGAACTCTTTCCCTACACGACGCTCTTCCGATCTNNNTGAGAAGTGTGGAGGAAACTT
AGCACACGTCTGAACTCTTTCCCTACACGACGCTCTTCCGATCTNNNAGACATACTGTTGAGAAAACTG
AGCACACGTCTGAACTCTTTCCCTACACGACGCTCTTCCGATCTNNNTTTGCAAAGCCAAGAAGTGTG
AGCACACGTCTGAACTCTTTCCCTACACGACGCTCTTCCGATCTNNNACGGTTTCAAACAATGTGCTGA
AGCACACGTCTGAACTCTTTCCCTACACGACGCTCTTCCGATCTNNNTCTGGATCATCTCCAGCTCCATTC
AGCACACGTCTGAACTCTTTCCCTACACGACGCTCTTCCGATCTNNNTAGCACACACTTTTCCACCACAAC
AGCACACGTCTGAACTCTTTCCCTACACGACGCTCTTCCGATCTNNNGCATGGCTTCAGAGCTTGAGTA
GCACACGTCTGAACTCTTTCCCTACACGACGCTCTTCCGATCTNNNCATGCTCTTACCTGGGTGATGTCC
GAGCACACGTCTGAACTCTTTCCCTACACGACGCTCTTCCGATCTNNNACCCGAGGACGAAAACTTGTA
GAAGAGCACACGTCTGAACTCTTTCCCTACACGACGCTCTTCCGATCTNNNACAGTGCCTAGCCTAGAA
GCACACGTCTGAACTCTTTCCCTACACGACGCTCTTCCGATCTNNNCAGGCCCTAATCCAGGATGATCTC
GAGCACACGTCTGAACTCTTTCCCTACACGACGCTCTTCCGATCTNNNGAGTGCTGCTTCACATCCTACA
GAGCACACGTCTGAACTCTTTCCCTACACGACGCTCTTCCGATCTNNNCAAGGAACAGTTGGAGACCG
AGAGCACACGTCTGAACTCTTTCCCTACACGACGCTCTTCCGATCTNNNCCTTTGACCCATCCCATGTTTC
GAAGAGCACACGTCTGAACTCTTTCCCTACACGACGCTCTTCCGATCTNNNATGGTAGCACTAGAATC
GAGCACACGTCTGAACTCTTTCCCTACACGACGCTCTTCCGATCTNNNAGGGTGTTGCTTCCTCTACCAT
GAGCACACGTCTGAACTCTTTCCCTACACGACGCTCTTCCGATCTNNNCTTGCCATTCACAACAGTTGCA
GAGCACACGTCTGAACTCTTTCCCTACACGACGCTCTTCCGATCTNNNCGTTGTATTTGTTACAGCAGC
GAAGAGCACACGTCTGAACTCTTTCCCTACACGACGCTCTTCCGATCTNNNCTCCAGCAGCCAAAGTACT
GAGCACACGTCTGAACTCTTTCCCTACACGACGCTCTTCCGATCTNNNCTGTCCTTCAACTATTGGCCAC
GAGCACACGTCTGAACTCTTTCCCTACACGACGCTCTTCCGATCTNNNTCTACAACAGCCACCTTCCTAG
AGCACACGTCTGAACTCTTTCCCTACACGACGCTCTTCCGATCTNNNGATTGTGTCCAGGTCCTTGTTCA
AGCACACGTCTGAACTCTTTCCCTACACGACGCTCTTCCGATCTNNNTTACCTTTCATCTCCCACCACCTC
AGAGCACACGTCTGAACTCTTTCCCTACACGACGCTCTTCCGATCTNNNTGACACACAGTAATTGTAGC
AGAGCACACGTCTGAACTCTTTCCCTACACGACGCTCTTCCGATCTNNNCAGGAGTTTTTGCTGGGTGC
GAGCACACGTCTGAACTCTTTCCCTACACGACGCTCTTCCGATCTNNNTCACATGTAATCCCACCATCTTC
GCACACGTCTGAACTCTTTCCCTACACGACGCTCTTCCGATCTNNNAGGCATAAGCAGCAACTGACAAT
GCACACGTCTGAACTCTTTCCCTACACGACGCTCTTCCGATCTNNNTACAGCTATACTCCCACCTCAGCC
GAGCACACGTCTGAACTCTTTCCCTACACGACGCTCTTCCGATCTNNNCTCTGCCCCCAATTTAGAGTGA
GAGCACACGTCTGAACTCTTTCCCTACACGACGCTCTTCCGATCTNNNAGCAATTTCCGTAAGTGTGGT
GAGCACACGTCTGAACTCTTTCCCTACACGACGCTCTTCCGATCTNNNAAGTCCCTTTTGGGGAAGAAC
AGCACACGTCTGAACTCTTTCCCTACACGACGCTCTTCCGATCTNNNGTTTGCAGACAGCCACTTCTCTA
AGCACACGTCTGAACTCTTTCCCTACACGACGCTCTTCCGATCTNNNCTCAGCTTCAAGTGGCTCAGAAT
GCACACGTCTGAACTCTTTCCCTACACGACGCTCTTCCGATCTNNNAGCACCGTTTCTTCTTTCTCTGTCA
AAGAGCACACGTCTGAACTCTTTCCCTACACGACGCTCTTCCGATCTNNNCGGCTGTGAATCCATCATAT
AGCACACGTCTGAACTCTTTCCCTACACGACGCTCTTCCGATCTNNNCTGGACTAAAATCAAGGGGACC
AGCACACGTCTGAACTCTTTCCCTACACGACGCTCTTCCGATCTNNNCAGTAGCCTTGTTTGGGAAGG
AGAGCACACGTCTGAACTCTTTCCCTACACGACGCTCTTCCGATCTNNNCTTTGGCTTTTCCTGGCTTCT
AGCACACGTCTGAACTCTTTCCCTACACGACGCTCTTCCGATCTNNNATTATTGTTGACCAACGGGGCT

GAAGAGCACACGTCTGAACTCTTTCCCTACACGACGCTCTTCCGATCTNNNCACTGCATGTCTTTGTGCA
GAGCACACGTCTGAACTCTTTCCCTACACGACGCTCTTCCGATCTNNNGTGCCAGGAAAGATGAGACA
AGAGCACACGTCTGAACTCTTTCCCTACACGACGCTCTTCCGATCTNNNACAAGCTCTTCAGTTGCAAA
GCACACGTCTGAACTCTTTCCCTACACGACGCTCTTCCGATCTNNNGAAGTGTCCACCCTACTCCTCTT
AGCACACGTCTGAACTCTTTCCCTACACGACGCTCTTCCGATCTNNNTCCTACTATACTGGAGACACTG
AAGAGCACACGTCTGAACTCTTTCCCTACACGACGCTCTTCCGATCTNNNATGATCAGCCCACTACACT
GAAGAGCACACGTCTGAACTCTTTCCCTACACGACGCTCTTCCGATCTNNNAGATAACTTGAAGGGGA
CACACGTCTGAACTCTTTCCCTACACGACGCTCTTCCGATCTNNNCTTCCTCAGCAGGAGATAGACCACA
GAGCACACGTCTGAACTCTTTCCCTACACGACGCTCTTCCGATCTNNNTGCCCAGCTAGTGTGTATGTG
AGAGCACACGTCTGAACTCTTTCCCTACACGACGCTCTTCCGATCTNNNACATGCTAAGCTTTGTGATCA
AGCACACGTCTGAACTCTTTCCCTACACGACGCTCTTCCGATCTNNNAAAGTGGATGGAGCATACACA
GAGCACACGTCTGAACTCTTTCCCTACACGACGCTCTTCCGATCTNNNGGTAGTTTGTTCAGGGAACCT
GAGCACACGTCTGAACTCTTTCCCTACACGACGCTCTTCCGATCTNNNTTTGAAACCCCTCCCTTACCAGA
AGCACACGTCTGAACTCTTTCCCTACACGACGCTCTTCCGATCTNNNGTACTTGAGGTGGTGGAGCTA
AGCACACGTCTGAACTCTTTCCCTACACGACGCTCTTCCGATCTNNNAATCAATCTGTCTTTCTCCAGG
GAGCACACGTCTGAACTCTTTCCCTACACGACGCTCTTCCGATCTNNNCACAAACCACTCCCAGAGAACA
AGCACACGTCTGAACTCTTTCCCTACACGACGCTCTTCCGATCTNNNCAATGCATGAGTGTGCTGTGTT
AGCACACGTCTGAACTCTTTCCCTACACGACGCTCTTCCGATCTNNNAGCCATCTAGCAAAAAGCCAGA
AGCACACGTCTGAACTCTTTCCCTACACGACGCTCTTCCGATCTNNNCCAGCTTCCTCACTGTCAAACTT
GCACACGTCTGAACTCTTTCCCTACACGACGCTCTTCCGATCTNNNAGGATGGCTACTGATCCTAGCA
GCACACGTCTGAACTCTTTCCCTACACGACGCTCTTCCGATCTNNNCTGAACCCGACTGGCTATCATTT
GAGCACACGTCTGAACTCTTTCCCTACACGACGCTCTTCCGATCTNNNTCATGGACTTTACCTTGGGCAA
GAGCACACGTCTGAACTCTTTCCCTACACGACGCTCTTCCGATCTNNNCTGCCATGGTTGGTAACCCTAT
GAGCACACGTCTGAACTCTTTCCCTACACGACGCTCTTCCGATCTNNNTCAAGAATTAGGGACGCTGCA
AGCACACGTCTGAACTCTTTCCCTACACGACGCTCTTCCGATCTNNNAAAGGGGCAGATAACACCCAAA
AGCACACGTCTGAACTCTTTCCCTACACGACGCTCTTCCGATCTNNNCATTCCCAGGGAGCAGAAAAT
AGAGCACACGTCTGAACTCTTTCCCTACACGACGCTCTTCCGATCTNNNAGCACCATTGAGAGATCTG
GCACACGTCTGAACTCTTTCCCTACACGACGCTCTTCCGATCTNNNTGTACATGGATGGAAAGTGGTG
AGCACACGTCTGAACTCTTTCCCTACACGACGCTCTTCCGATCTNNNAGTAACAGTCTTACACTGCCAA
AGAGCACACGTCTGAACTCTTTCCCTACACGACGCTCTTCCGATCTNNNGGCATAGCATCACCAAGTTG
AGAGCACACGTCTGAACTCTTTCCCTACACGACGCTCTTCCGATCTNNNCCACTTGGTAAAAAGGATG
GAGCACACGTCTGAACTCTTTCCCTACACGACGCTCTTCCGATCTNNNACTGATTACCACTCTGTGCTTG
AGCACACGTCTGAACTCTTTCCCTACACGACGCTCTTCCGATCTNNNTCAACATCACACGGATAAGGGT
GAGCACACGTCTGAACTCTTTCCCTACACGACGCTCTTCCGATCTNNNAAGACTGACAGAAGAGCTGT
GAGCACACGTCTGAACTCTTTCCCTACACGACGCTCTTCCGATCTNNNAGTGTCCACCAGTGAATGGTA
AGCACACGTCTGAACTCTTTCCCTACACGACGCTCTTCCGATCTNNNGATCCCCACAGTTCTTCATTGCT
GAGCACACGTCTGAACTCTTTCCCTACACGACGCTCTTCCGATCTNNNTGTGAGGTTTTGTTCCCTCCAT
GAGCACACGTCTGAACTCTTTCCCTACACGACGCTCTTCCGATCTNNNTTGCAAATTTCAAGCCAAGGG
AGCACACGTCTGAACTCTTTCCCTACACGACGCTCTTCCGATCTNNNTAAAGCCCTTTCTTGGCTGCAAT
AGAGCACACGTCTGAACTCTTTCCCTACACGACGCTCTTCCGATCTNNNACATCAAGATAGCCACAGGA
GAGCACACGTCTGAACTCTTTCCCTACACGACGCTCTTCCGATCTNNNTCAAGTATGGCCCCATCTGAG
AGCACACGTCTGAACTCTTTCCCTACACGACGCTCTTCCGATCTNNNATCAGGATAACCCTCCTCCCTTG
AGCACACGTCTGAACTCTTTCCCTACACGACGCTCTTCCGATCTNNNACATCAGTCAATAGAAGAAAGA

AGCACACGTCTGAACTCTTTCCCTACACGACGCTCTTCCGATCTNNNCTCACTCTCACTCTCTCCTCTCCC
AGCACACGTCTGAACTCTTTCCCTACACGACGCTCTTCCGATCTNNNCAGAGGGTCCTACACCCTGAC
AAGAGCACACGTCTGAACTCTTTCCCTACACGACGCTCTTCCGATCTNNNTATTCATCCCTTGGTGGA
AGCACACGTCTGAACTCTTTCCCTACACGACGCTCTTCCGATCTNNNAGCTTTTGTTGGCTATAACAG
AGCACACGTCTGAACTCTTTCCCTACACGACGCTCTTCCGATCTNNNGCAAAGCAAGGTTTGTACTCA
GCACACGTCTGAACTCTTTCCCTACACGACGCTCTTCCGATCTNNNTGGACAGATATGTGTGTGCCTC
GCACACGTCTGAACTCTTTCCCTACACGACGCTCTTCCGATCTNNNGAGCCTACCTTTTCTTCTGAGGG
GCACACGTCTGAACTCTTTCCCTACACGACGCTCTTCCGATCTNNNCTCCCCACCCAAAGTGATGTTAG
AGCACACGTCTGAACTCTTTCCCTACACGACGCTCTTCCGATCTNNNGAGAGTTTCTTTCCCAAGGCC
AGCACACGTCTGAACTCTTTCCCTACACGACGCTCTTCCGATCTNNNAATCAAGGGAATGATCAACA
GAGCACACGTCTGAACTCTTTCCCTACACGACGCTCTTCCGATCTNNNTGTGGATCTTGTTGAGTGTG
AGCACACGTCTGAACTCTTTCCCTACACGACGCTCTTCCGATCTNNNATAGGCCCCAAAGCAGTAGT
GAGCACACGTCTGAACTCTTTCCCTACACGACGCTCTTCCGATCTNNNGTTGGGTCTTGTGGGCTAAT
GAGCACACGTCTGAACTCTTTCCCTACACGACGCTCTTCCGATCTNNNATGTCTTGTGTTCTCCATCCA
AGAGCACACGTCTGAACTCTTTCCCTACACGACGCTCTTCCGATCTNNNTGAGGACCCAAGGGCTAAA
AGCACACGTCTGAACTCTTTCCCTACACGACGCTCTTCCGATCTNNNCACTATTCACTTGGCCCACTCT
CACACGTCTGAACTCTTTCCCTACACGACGCTCTTCCGATCTNNNCACAGGCAAAAATGTACCCCTCTC
AGCACACGTCTGAACTCTTTCCCTACACGACGCTCTTCCGATCTNNNTGCATTTTGGAGTGGTCGTAT
AGAGCACACGTCTGAACTCTTTCCCTACACGACGCTCTTCCGATCTNNNCTGGAGTGTTAGGTTTTGG
GAGCACACGTCTGAACTCTTTCCCTACACGACGCTCTTCCGATCTNNNTTGCTGTTGTCATGTGTGTTT
AGCACACGTCTGAACTCTTTCCCTACACGACGCTCTTCCGATCTNNNTTTGGAACACAAAGGTGGTTT
GAGCACACGTCTGAACTCTTTCCCTACACGACGCTCTTCCGATCTNNNGTGGAAGCCTCCAAAAGCAG
AGCACACGTCTGAACTCTTTCCCTACACGACGCTCTTCCGATCTNNNCCCTCCTCCACAAAGCAAAAGAT
AGCACACGTCTGAACTCTTTCCCTACACGACGCTCTTCCGATCTNNNGACTTGTTTCCAAGGCTGACA
AGCACACGTCTGAACTCTTTCCCTACACGACGCTCTTCCGATCTNNNCCGTTTTGTGGTTACAGGAGG
GCACACGTCTGAACTCTTTCCCTACACGACGCTCTTCCGATCTNNNTCTTCTCACATGGTAACATTTTC
AGCACACGTCTGAACTCTTTCCCTACACGACGCTCTTCCGATCTNNNCCTCCTCAAGGGTTTAGGATG
AGCACACGTCTGAACTCTTTCCCTACACGACGCTCTTCCGATCTNNNGAGAGACCCTCAGATGCACAC
AGCACACGTCTGAACTCTTTCCCTACACGACGCTCTTCCGATCTNNNAGACATATGCAAACCTCCCAAA
GAAGAGCACACGTCTGAACTCTTTCCCTACACGACGCTCTTCCGATCTNNNTTCCTTATGTCCCATAG
AGAGCACACGTCTGAACTCTTTCCCTACACGACGCTCTTCCGATCTNNNGTGTCAGTGGCAATACTCC
AGCACACGTCTGAACTCTTTCCCTACACGACGCTCTTCCGATCTNNNTCAATGGAACACCCTCCTCCAA
AGCACACGTCTGAACTCTTTCCCTACACGACGCTCTTCCGATCTNNNTTCTTCTTGGGTCCTCCTGGTA
AAGAGCACACGTCTGAACTCTTTCCCTACACGACGCTCTTCCGATCTNNNACATGATGTTTGGCTGAA
AGCACACGTCTGAACTCTTTCCCTACACGACGCTCTTCCGATCTNNNGCATTTTCCCAGCTTCCACTTT
GAGCACACGTCTGAACTCTTTCCCTACACGACGCTCTTCCGATCTNNNTTGGCAGTACTCACTGTAGG
AGCACACGTCTGAACTCTTTCCCTACACGACGCTCTTCCGATCTNNNACCAAAGGAAAACCTTAGGC
AGCACACGTCTGAACTCTTTCCCTACACGACGCTCTTCCGATCTNNNCACATCATTGCTGGAGCCTTA
AGCACACGTCTGAACTCTTTCCCTACACGACGCTCTTCCGATCTNNNGGAACATCTCTCTCTCCAGTCA
GCACACGTCTGAACTCTTTCCCTACACGACGCTCTTCCGATCTNNNCCAATGACTGCCTTCTTGCTTTG
GCACACGTCTGAACTCTTTCCCTACACGACGCTCTTCCGATCTNNNACTTATGGGCACCATGTCAACTC
AGCACACGTCTGAACTCTTTCCCTACACGACGCTCTTCCGATCTNNNAAAGGAAAGGGCTTCTTGCTG
AGCACACGTCTGAACTCTTTCCCTACACGACGCTCTTCCGATCTNNNGGTTGATCCACTTTGTTGAGT

GAGCACACGTCTGAACTCTTTCCCTACACGACGCTCTTCCGATCTNNNAGTACTGTCTGTTTCGTTTCAC
GAGCACACGTCTGAACTCTTTCCCTACACGACGCTCTTCCGATCTNNNAGTTCTTCCAAAGAGAGTGCT
AGAGCACACGTCTGAACTCTTTCCCTACACGACGCTCTTCCGATCTNNNTGAAGTCAGGACTGTTAGG
GAGCACACGTCTGAACTCTTTCCCTACACGACGCTCTTCCGATCTNNNGCAGATTAGAGAGCAACGTC
GAGCACACGTCTGAACTCTTTCCCTACACGACGCTCTTCCGATCTNNNTTGTGTTTTCTCTGTCTCCCACC
GAGCACACGTCTGAACTCTTTCCCTACACGACGCTCTTCCGATCTNNNAACACAGGAACAACCTCTGAC
GAGCACACGTCTGAACTCTTTCCCTACACGACGCTCTTCCGATCTNNNCTCAGAACTAACAGGCAAGGG
GCACACGTCTGAACTCTTTCCCTACACGACGCTCTTCCGATCTNNNGAATGCAGAGTTTGCTTGACAGC
AGCACACGTCTGAACTCTTTCCCTACACGACGCTCTTCCGATCTNNNCAAATGCAGCATGCATGTAGGT
GAGCACACGTCTGAACTCTTTCCCTACACGACGCTCTTCCGATCTNNNGAAGAAAAGTAAGGATGCCA
AGAGCACACGTCTGAACTCTTTCCCTACACGACGCTCTTCCGATCTNNNCACCAGACAGGCCCATTTTTC
GAGCACACGTCTGAACTCTTTCCCTACACGACGCTCTTCCGATCTNNNGGAGGACCAACTTTCACATGC
AGCACACGTCTGAACTCTTTCCCTACACGACGCTCTTCCGATCTNNNCTGCAGGTAGTTGAGTTGGAAC
AGCACACGTCTGAACTCTTTCCCTACACGACGCTCTTCCGATCTNNNAGACTTGTTAGTGTTTGCAAAGT
AGCACACGTCTGAACTCTTTCCCTACACGACGCTCTTCCGATCTNNNTGTCCACGGACCAACATAGAGA
AGCACACGTCTGAACTCTTTCCCTACACGACGCTCTTCCGATCTNNNAGCCATAGTTGAAGGTTTAGGT
AGAGCACACGTCTGAACTCTTTCCCTACACGACGCTCTTCCGATCTNNNGGCCTTGGAAGTGTTAGATG
AGCACACGTCTGAACTCTTTCCCTACACGACGCTCTTCCGATCTNNNCTAGGGAAAAGCTGGTGTCACT
GAGCACACGTCTGAACTCTTTCCCTACACGACGCTCTTCCGATCTNNNTGAGGAGCAAAAGGAAGTGA
GAGCACACGTCTGAACTCTTTCCCTACACGACGCTCTTCCGATCTNNNAATTTGTGAGCACTAAGATTGA
GAGCACACGTCTGAACTCTTTCCCTACACGACGCTCTTCCGATCTNNNGTTTTCTGGCATCTTTCTGCCTC
AGCACACGTCTGAACTCTTTCCCTACACGACGCTCTTCCGATCTNNNCTTTCTGGGAGAACCAAAGGAC
AGCACACGTCTGAACTCTTTCCCTACACGACGCTCTTCCGATCTNNNATCATGTTTTAGTTGGAGGCGC
AGAGCACACGTCTGAACTCTTTCCCTACACGACGCTCTTCCGATCTNNNTCCTATTGTTATCCCTCCCCAA
AGCACACGTCTGAACTCTTTCCCTACACGACGCTCTTCCGATCTNNNCAGTAGCTCCCCATTTACACACCC
AGAGCACACGTCTGAACTCTTTCCCTACACGACGCTCTTCCGATCTNNNTTGCTTCCTTAGCATCTGGTT
AAGAGCACACGTCTGAACTCTTTCCCTACACGACGCTCTTCCGATCTNNNCAGAATGATGTCAGGACTT
GAGCACACGTCTGAACTCTTTCCCTACACGACGCTCTTCCGATCTNNNAGGCTGAAGCAGGTAGATAG
CACACGTCTGAACTCTTTCCCTACACGACGCTCTTCCGATCTNNNTATGTGTGAGAGCGTCTGTGAGAC
AGCACACGTCTGAACTCTTTCCCTACACGACGCTCTTCCGATCTNNNTTTCCTCCCTTCATACACACACACA
AGAGCACACGTCTGAACTCTTTCCCTACACGACGCTCTTCCGATCTNNNCACCAATCCTTGATCATGCCA
AGCACACGTCTGAACTCTTTCCCTACACGACGCTCTTCCGATCTNNNTGTTCCCTTCTAATAAGACAAAA
AGCACACGTCTGAACTCTTTCCCTACACGACGCTCTTCCGATCTNNNTTCAATCTTCCAGGGTGTGTGTC
AGCACACGTCTGAACTCTTTCCCTACACGACGCTCTTCCGATCTNNNAGGAGCAAGGTACAGGGAAGT
AGCACACGTCTGAACTCTTTCCCTACACGACGCTCTTCCGATCTNNNTGGAAGGCTGTGCATATGTGT
AGCACACGTCTGAACTCTTTCCCTACACGACGCTCTTCCGATCTNNNGATTTTCTCTCCCCCAAGAAGCA
AGCACACGTCTGAACTCTTTCCCTACACGACGCTCTTCCGATCTNNNGTGCTTTTACCTAGTAACTGCTCA
AGCACACGTCTGAACTCTTTCCCTACACGACGCTCTTCCGATCTNNNAAACATGGTGGGCACTTCAGAA
AGAGCACACGTCTGAACTCTTTCCCTACACGACGCTCTTCCGATCTNNNCAGCCACACAGTCCAAAACA
AGCACACGTCTGAACTCTTTCCCTACACGACGCTCTTCCGATCTNNNCCCTCTTGATAGGGACATCCTGT
AGCACACGTCTGAACTCTTTCCCTACACGACGCTCTTCCGATCTNNNACCCTTTAGTACGGAATTGGGG
AGCACACGTCTGAACTCTTTCCCTACACGACGCTCTTCCGATCTNNNACATTTAGGTTTGTCTGCTTTTG
AGAGCACACGTCTGAACTCTTTCCCTACACGACGCTCTTCCGATCTNNNTATCTTGGAAGGTTGAGACG

GAGCACACGTCTGAACTCTTTCCCTACACGACGCTCTTCCGATCTNNNTTTTTCAAGAAGCCATCTCCT

AGCACACGTCTGAACTCTTTCCCTACACGACGCTCTTCCGATCTNNNTCCCTTTATGCACTTCCTTGCTG

AGCACACGTCTGAACTCTTTCCCTACACGACGCTCTTCCGATCTNNNGGTTGGTACCCCAAACCACTA

AGCACACGTCTGAACTCTTTCCCTACACGACGCTCTTCCGATCTNNNACCATGAGCAAGAATCAGTG

AGAGCACACGTCTGAACTCTTTCCCTACACGACGCTCTTCCGATCTNNNGAAAATTGGTGCCCAGATA

AGCACACGTCTGAACTCTTTCCCTACACGACGCTCTTCCGATCTNNNGGACTTTAGACACACAGAACA

GCACACGTCTGAACTCTTTCCCTACACGACGCTCTTCCGATCTNNNCCAACCCAGCATCTTTCTAGTCC

AGCACACGTCTGAACTCTTTCCCTACACGACGCTCTTCCGATCTNNNCGAATGTAAGAGAACAGGGT

AGCACACGTCTGAACTCTTTCCCTACACGACGCTCTTCCGATCTNNNAAGAACTTTCAGGGTTTTGTGC

GAGCACACGTCTGAACTCTTTCCCTACACGACGCTCTTCCGATCTNNNGACCAAGGAAACCCAGAGAC

GCACACGTCTGAACTCTTTCCCTACACGACGCTCTTCCGATCTNNNTCCTTTGCATTTTCAATGTTGCC

CACACGTCTGAACTCTTTCCCTACACGACGCTCTTCCGATCTNNNCACTAACCAAGGTTATCTGCACA

GCACACGTCTGAACTCTTTCCCTACACGACGCTCTTCCGATCTNNNTGCAAACCCTAACTGATGTGGA

GAGCACACGTCTGAACTCTTTCCCTACACGACGCTCTTCCGATCTNNNAAAGCACCGCTTCTAAGAGG

AGAGCACACGTCTGAACTCTTTCCCTACACGACGCTCTTCCGATCTNNNTTGAATTTCTGCTCCACCAC

GAGCACACGTCTGAACTCTTTCCCTACACGACGCTCTTCCGATCTNNNTCTTGTACCCATCAGTCTTTG

AGCACACGTCTGAACTCTTTCCCTACACGACGCTCTTCCGATCTNNNGGCCACTTTGCTCTCATTTACCT

GCACACGTCTGAACTCTTTCCCTACACGACGCTCTTCCGATCTNNNGCTGGTAAATTGAAGAGCTGG

AGCACACGTCTGAACTCTTTCCCTACACGACGCTCTTCCGATCTNNNAGGCTCAATTGTTTTGAGCAG

GAGCACACGTCTGAACTCTTTCCCTACACGACGCTCTTCCGATCTNNNGTTCTGCTCTGGGGTACTCT

GCACACGTCTGAACTCTTTCCCTACACGACGCTCTTCCGATCTNNNATATTGTTGGGCCAGTCTCTCCT

GAGCACACGTCTGAACTCTTTCCCTACACGACGCTCTTCCGATCTNNNAAGACACTGCCCTGAAAATCA

AGCACACGTCTGAACTCTTTCCCTACACGACGCTCTTCCGATCTNNNTGTCTCTTTACTTGCCTGTGCTC

GCACACGTCTGAACTCTTTCCCTACACGACGCTCTTCCGATCTNNNGTGTGTGTCTGTCTCTCCATCAC

AGAGCACACGTCTGAACTCTTTCCCTACACGACGCTCTTCCGATCTNNNCTAGTCCCTGCCAGGTGAA

GCACACGTCTGAACTCTTTCCCTACACGACGCTCTTCCGATCTNNNACTACTTACTTCTGCATCAGACA

AGCACACGTCTGAACTCTTTCCCTACACGACGCTCTTCCGATCTNNNTGGCAGCTGCAATAGACAAAA

GAGCACACGTCTGAACTCTTTCCCTACACGACGCTCTTCCGATCTNNNAAATTCCCACAGAAGGGAAT

AGCACACGTCTGAACTCTTTCCCTACACGACGCTCTTCCGATCTNNNCCAGGATTTCTGTTATTTGAAT

GCACACGTCTGAACTCTTTCCCTACACGACGCTCTTCCGATCTNNNTTTCCTTCCTTGGGTCCTCTCTCC

AGCACACGTCTGAACTCTTTCCCTACACGACGCTCTTCCGATCTNNNTTAGACCTGAAATCGTGCCAC

AGAGCACACGTCTGAACTCTTTCCCTACACGACGCTCTTCCGATCTNNNGAAGAAGACCCTGTCCCTC

AGCACACGTCTGAACTCTTTCCCTACACGACGCTCTTCCGATCTNNNAAGTGCCTTCCAATTACGAGCC

AGAGCACACGTCTGAACTCTTTCCCTACACGACGCTCTTCCGATCTNNNTATGCTGCTGGACTTAGTT

AGAGCACACGTCTGAACTCTTTCCCTACACGACGCTCTTCCGATCTNNNAGAACATGCCTGGTGCATA

AGAGCACACGTCTGAACTCTTTCCCTACACGACGCTCTTCCGATCTNNNGAGGGATAATTTCACACAG

AGCACACGTCTGAACTCTTTCCCTACACGACGCTCTTCCGATCTNNNTGTGCTGTAAAGCAAGTTCCA

AGAGCACACGTCTGAACTCTTTCCCTACACGACGCTCTTCCGATCTNNNTAGCCCGGTCAACAAGAGC

GCACACGTCTGAACTCTTTCCCTACACGACGCTCTTCCGATCTNNNCCTCTTTTATCCAAGACTAGGAC

AGCACACGTCTGAACTCTTTCCCTACACGACGCTCTTCCGATCTNNNTCAGATGGCTATGTGTGTGTG

AGCACACGTCTGAACTCTTTCCCTACACGACGCTCTTCCGATCTNNNACTTCCGGCCTCTTTCTCTCTTT

AGCACACGTCTGAACTCTTTCCCTACACGACGCTCTTCCGATCTNNNGGCCCCTTCCTTCCAAGATTTT

GCACACGTCTGAACTCTTTCCCTACACGACGCTCTTCCGATCTNNNTGTGCTTTGTGACATGATGGT

AGCACACGTCTGAACTCTTTCCCTACACGACGCTCTTCCGATCTNNNTCCCTTTCCCTTGCTGATTTTGG
AGAGCACACGTCTGAACTCTTTCCCTACACGACGCTCTTCCGATCTNNNCAGTGTTCAATGTCCTCACA
AGCACACGTCTGAACTCTTTCCCTACACGACGCTCTTCCGATCTNNNCACTGCTTTCACTAACTGTTGG
AGAGCACACGTCTGAACTCTTTCCCTACACGACGCTCTTCCGATCTNNNTGATTGAGTAAGACCCTGCC
AGAGCACACGTCTGAACTCTTTCCCTACACGACGCTCTTCCGATCTNNNAGGGGATATATGTGATGG
AGCACACGTCTGAACTCTTTCCCTACACGACGCTCTTCCGATCTNNNAGAAAACGGTAGCATGACAGC
AGCACACGTCTGAACTCTTTCCCTACACGACGCTCTTCCGATCTNNNTTCGAACCATAGCAAGAAGGC
AGAGCACACGTCTGAACTCTTTCCCTACACGACGCTCTTCCGATCTNNNACAGGAGCCACACTATCTTC
GCACACGTCTGAACTCTTTCCCTACACGACGCTCTTCCGATCTNNNTAGAAGGGTTGGTGTATGTATG
GAGCACACGTCTGAACTCTTTCCCTACACGACGCTCTTCCGATCTNNNACACACAGAGACCCTTGAGT
AGCACACGTCTGAACTCTTTCCCTACACGACGCTCTTCCGATCTNNNGTGTCTGTTCACATCCTTTGGC
AGAGCACACGTCTGAACTCTTTCCCTACACGACGCTCTTCCGATCTNNNTGCAGTGAACCAAAATTGTG
ACACGTCTGAACTCTTTCCCTACACGACGCTCTTCCGATCTNNNCTCTCTGACGTTTCCCAAGTGAGCA
AGAGCACACGTCTGAACTCTTTCCCTACACGACGCTCTTCCGATCTNNNGAAGGCACCGGATAATTGA
GCACACGTCTGAACTCTTTCCCTACACGACGCTCTTCCGATCTNNNAGCCATATTTTTGGGAGGCAGA
AGCACACGTCTGAACTCTTTCCCTACACGACGCTCTTCCGATCTNNNGCTCCAGTTCAGTTAGCTAGA
AGCACACGTCTGAACTCTTTCCCTACACGACGCTCTTCCGATCTNNNTGTTTGATCTTCTCTGTGGGCT
GAGCACACGTCTGAACTCTTTCCCTACACGACGCTCTTCCGATCTNNNATGGAAAGCTTTTCCTGACCT
AGAGCACACGTCTGAACTCTTTCCCTACACGACGCTCTTCCGATCTNNNAATCGCTTGAACCCGAGATT
AGCACACGTCTGAACTCTTTCCCTACACGACGCTCTTCCGATCTNNNAATGCGGAAAGGTCCTCAAGA
AGAGCACACGTCTGAACTCTTTCCCTACACGACGCTCTTCCGATCTNNNAAAGGTAGAGCCATACCTCC
GAGCACACGTCTGAACTCTTTCCCTACACGACGCTCTTCCGATCTNNNCAAAGAGCAGACCTACCTTCC
AGCACACGTCTGAACTCTTTCCCTACACGACGCTCTTCCGATCTNNNGGTTTCCATATTGAGTTCCTGT
GCACACGTCTGAACTCTTTCCCTACACGACGCTCTTCCGATCTNNNATGAAATTTGGCTGCTCCCAGAC
GAGCACACGTCTGAACTCTTTCCCTACACGACGCTCTTCCGATCTNNNCCGCGATACCCAAAGTCAATA
AGCACACGTCTGAACTCTTTCCCTACACGACGCTCTTCCGATCTNNNGGGGTCGGATTTCATTCTTCT
GAGCACACGTCTGAACTCTTTCCCTACACGACGCTCTTCCGATCTNNNCCAAATGCCATCAATCAACAG
AGCACACGTCTGAACTCTTTCCCTACACGACGCTCTTCCGATCTNNNAGCCTTATAGGGATAACATTGA
AGCACACGTCTGAACTCTTTCCCTACACGACGCTCTTCCGATCTNNNCCCAGCTGGTGGAAAATATGT
GAAGAGCACACGTCTGAACTCTTTCCCTACACGACGCTCTTCCGATCTNNNTGGCCTTTGTATCTTGAC
AGAGCACACGTCTGAACTCTTTCCCTACACGACGCTCTTCCGATCTNNNTACGTATAGCTTGCCACCTT
AGCACACGTCTGAACTCTTTCCCTACACGACGCTCTTCCGATCTNNNCCCTCAGCCAAAGATGGAAAAG
AGAGCACACGTCTGAACTCTTTCCCTACACGACGCTCTTCCGATCTNNNTGAAAAGACAAGATGCATAC
AGCACACGTCTGAACTCTTTCCCTACACGACGCTCTTCCGATCTNNNGCAACAGCTACATGGCACAAA
GAGCACACGTCTGAACTCTTTCCCTACACGACGCTCTTCCGATCTNNNTAATGCAGCATCCTGTTTGGT
AGCACACGTCTGAACTCTTTCCCTACACGACGCTCTTCCGATCTNNNTTTACTGTCACACATGGGCACAA
AGCACACGTCTGAACTCTTTCCCTACACGACGCTCTTCCGATCTNNNTGCTAAAAATACAATAGCCTGG
GAGCACACGTCTGAACTCTTTCCCTACACGACGCTCTTCCGATCTNNNTCAAGGACGCCAGGAAAATCA
AGCACACGTCTGAACTCTTTCCCTACACGACGCTCTTCCGATCTNNNGAGGGATAATTTCACACAGCA
AGAGCACACGTCTGAACTCTTTCCCTACACGACGCTCTTCCGATCTNNNCCAGCAATATCCAGGACCAG
GAGCACACGTCTGAACTCTTTCCCTACACGACGCTCTTCCGATCTNNNACAATTGTATACCTAAAACTG
AGCACACGTCTGAACTCTTTCCCTACACGACGCTCTTCCGATCTNNNTCTGCAAAGATAAGCAGCTCT
AGCACACGTCTGAACTCTTTCCCTACACGACGCTCTTCCGATCTNNNGCTTTTATGTTCAGCCCCATCCT

AGCACACGTCTGAACTCTTTCCCTACACGACGCTCTTCCGATCTNNNTAAACCAGTTTCCATGTGGGTC
GAGCACACGTCTGAACTCTTTCCCTACACGACGCTCTTCCGATCTNNNTCATGTCCATGTGAACAGTA
GCACACGTCTGAACTCTTTCCCTACACGACGCTCTTCCGATCTNNNCTGTGACAGGTTGTAGAACCCA
AGCACACGTCTGAACTCTTTCCCTACACGACGCTCTTCCGATCTNNNTCCTGCCTCTGCATTCTCATTTC
AGCACACGTCTGAACTCTTTCCCTACACGACGCTCTTCCGATCTNNNCAAATGCTAACGAACCACCCCT
GCACACGTCTGAACTCTTTCCCTACACGACGCTCTTCCGATCTNNNTGCAAGCATAGTATAAGAAACT
AGCACACGTCTGAACTCTTTCCCTACACGACGCTCTTCCGATCTNNNCAACCTAGGTGGTCATGAATG
AAGAGCACACGTCTGAACTCTTTCCCTACACGACGCTCTTCCGATCTNNNTGAGATGGTCACAAGAA
GAAGAGCACACGTCTGAACTCTTTCCCTACACGACGCTCTTCCGATCTNNNCATATTCCTGGGTTTGG
GAGCACACGTCTGAACTCTTTCCCTACACGACGCTCTTCCGATCTNNNCCTGAGTGAAGAAGCCAAG
AGCACACGTCTGAACTCTTTCCCTACACGACGCTCTTCCGATCTNNNACTTTCAAAATGTTGCCTTGGG
AGCACACGTCTGAACTCTTTCCCTACACGACGCTCTTCCGATCTNNNATCAGGCATTTGAGCTCTTTGC
GCACACGTCTGAACTCTTTCCCTACACGACGCTCTTCCGATCTNNNCCCATGAGAAGAACTCAGTAG
GAGCACACGTCTGAACTCTTTCCCTACACGACGCTCTTCCGATCTNNNGTATAGAAGCGCGAAGATG
GCACACGTCTGAACTCTTTCCCTACACGACGCTCTTCCGATCTNNNGATGAACCCGATTCGTCTGTTGA
AGCACACGTCTGAACTCTTTCCCTACACGACGCTCTTCCGATCTNNNCCATGGATATGAAGGCCCAAC
AGCACACGTCTGAACTCTTTCCCTACACGACGCTCTTCCGATCTNNNGCACTCAGTAATCGTTTGGGAA
GCACACGTCTGAACTCTTTCCCTACACGACGCTCTTCCGATCTNNNTATTAGCATCTTGGTGGGAGGG
AGCACACGTCTGAACTCTTTCCCTACACGACGCTCTTCCGATCTNNNCCAAGTTGAGGTGGTGGGTTT
GCACACGTCTGAACTCTTTCCCTACACGACGCTCTTCCGATCTNNNTCACTTGCCATCAAGCTGATTCTC
CACACGTCTGAACTCTTTCCCTACACGACGCTCTTCCGATCTNNNTATTGCCATTCACACTCCACCAGC
GCACACGTCTGAACTCTTTCCCTACACGACGCTCTTCCGATCTNNNGCTCTTTCAGTTGTGCATTGTCT
GCACACGTCTGAACTCTTTCCCTACACGACGCTCTTCCGATCTNNNAGCAAGTCCTCTTGATACCAGAT
AGCACACGTCTGAACTCTTTCCCTACACGACGCTCTTCCGATCTNNNTGTGCCAATCAGTAATGCACAG
AGAGCACACGTCTGAACTCTTTCCCTACACGACGCTCTTCCGATCTNNNTGCTAACGTGGAGAGAGA
GAGCACACGTCTGAACTCTTTCCCTACACGACGCTCTTCCGATCTNNNTCTTGACCTGTGAAAGCAGT
GAAGAGCACACGTCTGAACTCTTTCCCTACACGACGCTCTTCCGATCTNNNGCTGTGTTGGTCGTTGT
GAGCACACGTCTGAACTCTTTCCCTACACGACGCTCTTCCGATCTNNNAGTGAGAGGTCCCCATGTGT
AGAGCACACGTCTGAACTCTTTCCCTACACGACGCTCTTCCGATCTNNNTTTGAGGCTTTGTTGGAGG
AGCACACGTCTGAACTCTTTCCCTACACGACGCTCTTCCGATCTNNNCCCTTTCTTTCTCCCGAACAGA
GAGCACACGTCTGAACTCTTTCCCTACACGACGCTCTTCCGATCTNNNTCGAGCGTGCTGTGTATTAC
AGCACACGTCTGAACTCTTTCCCTACACGACGCTCTTCCGATCTNNNCAGCAGGCTGTGTAGACTTC
AGAGCACACGTCTGAACTCTTTCCCTACACGACGCTCTTCCGATCTNNNTCAAACACCCATCACTTTCC
GAGCACACGTCTGAACTCTTTCCCTACACGACGCTCTTCCGATCTNNNTGACGTCCAAATTAGTGCCT
AGCACACGTCTGAACTCTTTCCCTACACGACGCTCTTCCGATCTNNNCAGGAATAATGGAGTTTACCC
AGCACACGTCTGAACTCTTTCCCTACACGACGCTCTTCCGATCTNNNAAGAAGTTTCCTCCACCAAATA
GCACACGTCTGAACTCTTTCCCTACACGACGCTCTTCCGATCTNNNGCTAATCACCAACCATGCCACTA
AGCACACGTCTGAACTCTTTCCCTACACGACGCTCTTCCGATCTNNNGCTCCAATCTCCCTCTCTCTCTCC
AGCACACGTCTGAACTCTTTCCCTACACGACGCTCTTCCGATCTNNNTGCCAGAATGATGCTTGTAAA
GAGCACACGTCTGAACTCTTTCCCTACACGACGCTCTTCCGATCTNNNTAGACATAAGGGGGCACTA
GAGCACACGTCTGAACTCTTTCCCTACACGACGCTCTTCCGATCTNNNATGAGCCTTAGGACAATGCT
AGCACACGTCTGAACTCTTTCCCTACACGACGCTCTTCCGATCTNNNAGTCAAGGTTTGGCCTGTGTA
GCACACGTCTGAACTCTTTCCCTACACGACGCTCTTCCGATCTNNNAATAACCGAGAGAGAGGAGG

AGCACACGTCTGAACTCTTTCCCTACACGACGCTCTTCCGATCTNNNGCTCCTCTGTTGTGTATGCTTG
AGCACACGTCTGAACTCTTTCCCTACACGACGCTCTTCCGATCTNNNAGCTTCACCCAGGAGAGCTAA
GAGCACACGTCTGAACTCTTTCCCTACACGACGCTCTTCCGATCTNNNCCACGGAATTGCCTTTGAAA
GCACACGTCTGAACTCTTTCCCTACACGACGCTCTTCCGATCTNNNGGCTTGCTCTCTGTGTCTCTTAG
AGAGCACACGTCTGAACTCTTTCCCTACACGACGCTCTTCCGATCTNNNACCAAACTCATCCAAAGAC
GAGCACACGTCTGAACTCTTTCCCTACACGACGCTCTTCCGATCTNNNTAGGTACTCATTGCTTCTGG
GAGCACACGTCTGAACTCTTTCCCTACACGACGCTCTTCCGATCTNNNACCAAAATCCTGCTGTGGTTT
GCACACGTCTGAACTCTTTCCCTACACGACGCTCTTCCGATCTNNNGCCACTGTTTACCAAGACACTG
AGCACACGTCTGAACTCTTTCCCTACACGACGCTCTTCCGATCTNNNCTTACCAAGTAGATGCCCCAAG
AGCACACGTCTGAACTCTTTCCCTACACGACGCTCTTCCGATCTNNNAGCCAGTTGACACAAAAGTCT
GAGCACACGTCTGAACTCTTTCCCTACACGACGCTCTTCCGATCTNNNTGGAGTGAGACTATGTCTCG
GAGCACACGTCTGAACTCTTTCCCTACACGACGCTCTTCCGATCTNNNCATTCAGTAATGGTTCTGGG
AGAGCACACGTCTGAACTCTTTCCCTACACGACGCTCTTCCGATCTNNNGCAGACTTGTTTGCTTTCAG
AGAGCACACGTCTGAACTCTTTCCCTACACGACGCTCTTCCGATCTNNNATGAGGAATGAGGACAAG
CACACGTCTGAACTCTTTCCCTACACGACGCTCTTCCGATCTNNNAGCGTAATCTAACGGAGGAGTG
AGCACACGTCTGAACTCTTTCCCTACACGACGCTCTTCCGATCTNNNGAAACCTTGCACATGTATCCCC
AGCACACGTCTGAACTCTTTCCCTACACGACGCTCTTCCGATCTNNNTCACAGATATCATCCCCCACTCC
GAGCACACGTCTGAACTCTTTCCCTACACGACGCTCTTCCGATCTNNNTCTGCAGATGTGGAATGTGA
GAGCACACGTCTGAACTCTTTCCCTACACGACGCTCTTCCGATCTNNNCCCCATGAGAAGAACTCAGTA
AGCACACGTCTGAACTCTTTCCCTACACGACGCTCTTCCGATCTNNNTACTCCAGACTGGGCAATATGA
AGCACACGTCTGAACTCTTTCCCTACACGACGCTCTTCCGATCTNNNCACTTTGGGAGGTTCTCTAAG
GAGCACACGTCTGAACTCTTTCCCTACACGACGCTCTTCCGATCTNNNGGAAGTAGTGTGTGCTTGG
AGCACACGTCTGAACTCTTTCCCTACACGACGCTCTTCCGATCTNNNTGATTGCCCTTCACCTGAAACA
AGAGCACACGTCTGAACTCTTTCCCTACACGACGCTCTTCCGATCTNNNACAAAAGGGCTTATTGGAA
AGCACACGTCTGAACTCTTTCCCTACACGACGCTCTTCCGATCTNNNGAGTTCACACAAGCCGTTCACT
GAGCACACGTCTGAACTCTTTCCCTACACGACGCTCTTCCGATCTNNNGGATGAGGAGGGAGAGAG
GAGCACACGTCTGAACTCTTTCCCTACACGACGCTCTTCCGATCTNNNGCTAATATGGAAGCCCTGCA
AGCACACGTCTGAACTCTTTCCCTACACGACGCTCTTCCGATCTNNNTCCCCTATTCCCTTGTGCTTGTA
AGCACACGTCTGAACTCTTTCCCTACACGACGCTCTTCCGATCTNNNCAGCAGTTTGACAATGATGTG
AGAGCACACGTCTGAACTCTTTCCCTACACGACGCTCTTCCGATCTNNNACCTGGAAACTCTCCCTAGT
GAGCACACGTCTGAACTCTTTCCCTACACGACGCTCTTCCGATCTNNNACCTGTAACTGTGAGATGGC
GAGCACACGTCTGAACTCTTTCCCTACACGACGCTCTTCCGATCTNNNTGGTGTACAGGCTAGAAAGC
AGCACACGTCTGAACTCTTTCCCTACACGACGCTCTTCCGATCTNNNACCCAATTAGGGTGTATTAGT
GAGCACACGTCTGAACTCTTTCCCTACACGACGCTCTTCCGATCTNNNTAGAAACATGCCAGTGTTGG
GAAGAGCACACGTCTGAACTCTTTCCCTACACGACGCTCTTCCGATCTNNNATCCCCTCCTGTACCTTGT
GAGCACACGTCTGAACTCTTTCCCTACACGACGCTCTTCCGATCTNNNCTTGATGCTCACCAGTACTCCA
GCACACGTCTGAACTCTTTCCCTACACGACGCTCTTCCGATCTNNNTCCTGAAGCCAAAAACCAAATTT
AGCACACGTCTGAACTCTTTCCCTACACGACGCTCTTCCGATCTNNNACTTAGGATTGGAGATAGCGC
GCACACGTCTGAACTCTTTCCCTACACGACGCTCTTCCGATCTNNNCACAAACAGCGCCTCAAAGAAT
AGCACACGTCTGAACTCTTTCCCTACACGACGCTCTTCCGATCTNNNATGGCAAACACCCACCTTTACA
AGCACACGTCTGAACTCTTTCCCTACACGACGCTCTTCCGATCTNNNGATGCTGCAAAAGGCAAGATT
GAGCACACGTCTGAACTCTTTCCCTACACGACGCTCTTCCGATCTNNNGGCAACAGCATCTGGACTAT
AGCACACGTCTGAACTCTTTCCCTACACGACGCTCTTCCGATCTNNNTACTCTCAGGTTCTCTGCTGTC

AGCACACGTCTGAACTCTTTCCCTACACGACGCTCTTCCGATCTNNNTGGGAACTAGTGACATTTTGCC
AGCACACGTCTGAACTCTTTCCCTACACGACGCTCTTCCGATCTNNNGCTGCAGTGAGCTGAGATAGAT
GCACACGTCTGAACTCTTTCCCTACACGACGCTCTTCCGATCTNNNAAATACCATAGTCTGGGGGTGGC
AGCACACGTCTGAACTCTTTCCCTACACGACGCTCTTCCGATCTNNNAAGCAGAAATAGATCCCGCCTT
AGCACACGTCTGAACTCTTTCCCTACACGACGCTCTTCCGATCTNNNTGCATGGTAGGGATACTCAGA
GAGCACACGTCTGAACTCTTTCCCTACACGACGCTCTTCCGATCTNNNGCTCACCTTGGAAAGTGACAT
GAGCACACGTCTGAACTCTTTCCCTACACGACGCTCTTCCGATCTNNNTGGATTTTGAGGTTGAACCCA
AGAGCACACGTCTGAACTCTTTCCCTACACGACGCTCTTCCGATCTNNNGATAATGTGAGGGCCCTTTG
AGCACACGTCTGAACTCTTTCCCTACACGACGCTCTTCCGATCTNNNATCAGAAATGACAGCCTGGTGA
AGAGCACACGTCTGAACTCTTTCCCTACACGACGCTCTTCCGATCTNNNGTATGCTCCACAATGCCAAG
AGCACACGTCTGAACTCTTTCCCTACACGACGCTCTTCCGATCTNNNCTAGTCAAGGTAGCGAGACACC
GAGCACACGTCTGAACTCTTTCCCTACACGACGCTCTTCCGATCTNNNTACCGTTGCTACAACAACAGCA
GAGCACACGTCTGAACTCTTTCCCTACACGACGCTCTTCCGATCTNNNCCTGGCCCATGTGAAGGTTTTA
GAAGAGCACACGTCTGAACTCTTTCCCTACACGACGCTCTTCCGATCTNNNAAGTGGCTTCTGTAGAG
AGCACACGTCTGAACTCTTTCCCTACACGACGCTCTTCCGATCTNNNTCCCAGACTGATGAGAAAGCCA
AGAGCACACGTCTGAACTCTTTCCCTACACGACGCTCTTCCGATCTNNNGCTGATCTGCTCCTTGATGTG
AGCACACGTCTGAACTCTTTCCCTACACGACGCTCTTCCGATCTNNNGTCCCTGGAAGATGCTGTTTTCA
GAGCACACGTCTGAACTCTTTCCCTACACGACGCTCTTCCGATCTNNNGTGCCTGAACTGGGATGTCAA
AGCACACGTCTGAACTCTTTCCCTACACGACGCTCTTCCGATCTNNNCACCCCCAACAAAGATTCCTTCA
AAGAGCACACGTCTGAACTCTTTCCCTACACGACGCTCTTCCGATCTNNNGAGGAAGGTCTCCCAACAA
AGCACACGTCTGAACTCTTTCCCTACACGACGCTCTTCCGATCTNNNTGGGAAATGCAGGTTTATGAAC
GAGCACACGTCTGAACTCTTTCCCTACACGACGCTCTTCCGATCTNNNTCTTTCCTGTAGATTGTGTTCT
GAGCACACGTCTGAACTCTTTCCCTACACGACGCTCTTCCGATCTNNNCGTTGTTCAGTGGGGGACTTT
GAGCACACGTCTGAACTCTTTCCCTACACGACGCTCTTCCGATCTNNNTAAGCCCTCCAAACTGTTCAACT
GAGCACACGTCTGAACTCTTTCCCTACACGACGCTCTTCCGATCTNNNGCTCCAAACACAGACACATACC
GAGCACACGTCTGAACTCTTTCCCTACACGACGCTCTTCCGATCTNNNGATGTGAGGTCCCCGTTCTTT
AGCACACGTCTGAACTCTTTCCCTACACGACGCTCTTCCGATCTNNNTGACTTGGAGGCACTGTTTTTC
GAGCACACGTCTGAACTCTTTCCCTACACGACGCTCTTCCGATCTNNNGTGAATTGAACCATCAGCATG
GAGCACACGTCTGAACTCTTTCCCTACACGACGCTCTTCCGATCTNNNGAGTTCTCACATATCTACGCCC
GAGCACACGTCTGAACTCTTTCCCTACACGACGCTCTTCCGATCTNNNGTCTGCAATCTGAAAGCAACA
AGAGCACACGTCTGAACTCTTTCCCTACACGACGCTCTTCCGATCTNNNTGCGATTGCCATCTCTACCAA
GCACACGTCTGAACTCTTTCCCTACACGACGCTCTTCCGATCTNNNTCGAATTCTACTCAGGGACGGAA
GAGCACACGTCTGAACTCTTTCCCTACACGACGCTCTTCCGATCTNNNTGCTCTGTTCTGTTCTCGTCTTC
GGAAGAGCACACGTCTGAACTCTTTCCCTACACGACGCTCTTCCGATCTNNNAATGGCTTGAAAGGCCT
CACACGTCTGAACTCTTTCCCTACACGACGCTCTTCCGATCTNNNCCGCCAATATTTAGGCTTTGCAGCA
GGAAGAGCACACGTCTGAACTCTTTCCCTACACGACGCTCTTCCGATCTNNNTCACTAGACAAGCTCAA
GAGCACACGTCTGAACTCTTTCCCTACACGACGCTCTTCCGATCTNNNGGCAGGAGAATTGAGCCAAC
GAGCACACGTCTGAACTCTTTCCCTACACGACGCTCTTCCGATCTNNNCATAACTGTGTGAGCTGGCTT
AGCACACGTCTGAACTCTTTCCCTACACGACGCTCTTCCGATCTNNNCTTCTTTGACATGTGGTCCTGCT
AGAGCACACGTCTGAACTCTTTCCCTACACGACGCTCTTCCGATCTNNNTCAGGTTGATTTTGTGCCTG
GAGCACACGTCTGAACTCTTTCCCTACACGACGCTCTTCCGATCTNNNACTGGGAAAGAAAACTCTCTA
GAGCACACGTCTGAACTCTTTCCCTACACGACGCTCTTCCGATCTNNNACAAGCACAATAGAAGCAGG
GAGCACACGTCTGAACTCTTTCCCTACACGACGCTCTTCCGATCTNNNAGAAGCTCAGAAGGCTTTGCT

AGCACACGTCTGAACTCTTTCCCTACACGACGCTCTTCCGATCTNNNCTCCTCTCTTAGCATTCGGTTCT
GAGCACACGTCTGAACTCTTTCCCTACACGACGCTCTTCCGATCTNNNTCCATTTAGTTGGTTCTGGG
GAGCACACGTCTGAACTCTTTCCCTACACGACGCTCTTCCGATCTNNNAGAAGCAGCAGGATAGTGA
GCACACGTCTGAACTCTTTCCCTACACGACGCTCTTCCGATCTNNNATTCTTTTTGTCCCTTCCCTCCCC
AGCACACGTCTGAACTCTTTCCCTACACGACGCTCTTCCGATCTNNNGTGCACATGGCATAAAATCCG
AGCACACGTCTGAACTCTTTCCCTACACGACGCTCTTCCGATCTNNNACAATTCATATGGGGTCTTAT
AGCACACGTCTGAACTCTTTCCCTACACGACGCTCTTCCGATCTNNNACCCATGATATACAATACACAT
AGAGCACACGTCTGAACTCTTTCCCTACACGACGCTCTTCCGATCTNNNGTTGGGTCAATCAGTCAG
GAGCACACGTCTGAACTCTTTCCCTACACGACGCTCTTCCGATCTNNNGATGGACTGGAGGTAGAGA
AGCACACGTCTGAACTCTTTCCCTACACGACGCTCTTCCGATCTNNNATGCATTGCTCAAGTACTCAG
GAGCACACGTCTGAACTCTTTCCCTACACGACGCTCTTCCGATCTNNNAGTCCAGGTTGAATTGGCTT
GAGCACACGTCTGAACTCTTTCCCTACACGACGCTCTTCCGATCTNNNGGGCTCGAGATAGAGGTTT
CACACGTCTGAACTCTTTCCCTACACGACGCTCTTCCGATCTNNNCTGGCCCCCTACTATGTTCTCTACA
AGCACACGTCTGAACTCTTTCCCTACACGACGCTCTTCCGATCTNNNACAGAAACCAGGATTTCAGG
AGCACACGTCTGAACTCTTTCCCTACACGACGCTCTTCCGATCTNNNGATTTCAGACTGACAGGGACA
AGAGCACACGTCTGAACTCTTTCCCTACACGACGCTCTTCCGATCTNNNAGTGGCTATTCAACTGGT
GAGCACACGTCTGAACTCTTTCCCTACACGACGCTCTTCCGATCTNNNAGTAAAGCACAGTGTGAGTA
GAGCACACGTCTGAACTCTTTCCCTACACGACGCTCTTCCGATCTNNNAAGAAGTTGAGAGACTGCTT
GAGCACACGTCTGAACTCTTTCCCTACACGACGCTCTTCCGATCTNNNGCACTATGCTGAGACACATG
AGCACACGTCTGAACTCTTTCCCTACACGACGCTCTTCCGATCTNNNTAGATATACCCACTCACCCAC
CACACGTCTGAACTCTTTCCCTACACGACGCTCTTCCGATCTNNNCCATCTGGTTCCTTCTCATCTGGC
AGCACACGTCTGAACTCTTTCCCTACACGACGCTCTTCCGATCTNNNTCTCCTTCTCCCCTAGGCAATT
AGCACACGTCTGAACTCTTTCCCTACACGACGCTCTTCCGATCTNNNGTCAGGGTTCTCTATGTGTGT
AGCACACGTCTGAACTCTTTCCCTACACGACGCTCTTCCGATCTNNNTCTGATCACTCTCACAGGCTAG
GCACACGTCTGAACTCTTTCCCTACACGACGCTCTTCCGATCTNNNCACCCAGCCTCTATCTTTCCTTCC
GAGCACACGTCTGAACTCTTTCCCTACACGACGCTCTTCCGATCTNNNTTCCACATTTGCATGTGCTCT
CACACGTCTGAACTCTTTCCCTACACGACGCTCTTCCGATCTNNNAAACAGACCCAAGCAGAGTTTCC
AGCACACGTCTGAACTCTTTCCCTACACGACGCTCTTCCGATCTNNNCCTCAGAGTGGTTCTCTCTCTC
GCACACGTCTGAACTCTTTCCCTACACGACGCTCTTCCGATCTNNNCGTGGACAAAATCACTTGAAG
AGCACACGTCTGAACTCTTTCCCTACACGACGCTCTTCCGATCTNNNGGTTCATCCATGTTGTATAGC
CACACGTCTGAACTCTTTCCCTACACGACGCTCTTCCGATCTNNNCCTCTCAGAGCCAAAAGTGGAAA
GCACACGTCTGAACTCTTTCCCTACACGACGCTCTTCCGATCTNNNAGGCTGGTTTCAAGTTTTGGGA
CACACGTCTGAACTCTTTCCCTACACGACGCTCTTCCGATCTNNNTTCACCTCACGATCTAGCCTTCTCA
AGAGCACACGTCTGAACTCTTTCCCTACACGACGCTCTTCCGATCTNNNCAAAACTTCGTCTCTCTCCC
GAGCACACGTCTGAACTCTTTCCCTACACGACGCTCTTCCGATCTNNNGTTGACAGGCTGTAGTTTG
CACACGTCTGAACTCTTTCCCTACACGACGCTCTTCCGATCTNNNAGCGGAATTTTCTCTCCAGACACC
GCACACGTCTGAACTCTTTCCCTACACGACGCTCTTCCGATCTNNNGATCAAACCAGCCTTCAGAGCT
GAGCACACGTCTGAACTCTTTCCCTACACGACGCTCTTCCGATCTNNNTTCTCATGAATTGGTCCTCGT
AGCACACGTCTGAACTCTTTCCCTACACGACGCTCTTCCGATCTNNNGAGGAAAGAGAAACCCACCC
GCACACGTCTGAACTCTTTCCCTACACGACGCTCTTCCGATCTNNNGCATAAGTTCATCCATGTTGTCA
GAGCACACGTCTGAACTCTTTCCCTACACGACGCTCTTCCGATCTNNNGGAAGGGCAGGCCTATTCTA
GAGCACACGTCTGAACTCTTTCCCTACACGACGCTCTTCCGATCTNNNATGCTTGTTTGCTGAGAAAG
GAGCACACGTCTGAACTCTTTCCCTACACGACGCTCTTCCGATCTNNNAGTAGCTGCCCTTAGAAAGT

AGCACACGTCTGAACTCTTTCCCTACACGACGCTCTTCCGATCTNNNTCAACTTATCTTTCAACACCCAG
GAGCACACGTCTGAACTCTTTCCCTACACGACGCTCTTCCGATCTNNNGAGGGGCTCAAGAGCAGATT
AGCACACGTCTGAACTCTTTCCCTACACGACGCTCTTCCGATCTNNNAGAATCAACATAAGGCCCACA
GAGCACACGTCTGAACTCTTTCCCTACACGACGCTCTTCCGATCTNNNAAAAAGGTGTGAATGAAGCC
AGCACACGTCTGAACTCTTTCCCTACACGACGCTCTTCCGATCTNNNTCAGTTGTCTTTCCTCACCCATC
GCACACGTCTGAACTCTTTCCCTACACGACGCTCTTCCGATCTNNNGCAAGATCTGACATCCAGAACC
GAGCACACGTCTGAACTCTTTCCCTACACGACGCTCTTCCGATCTNNNTGGTACCAGCATCTGGATCAT
GCACACGTCTGAACTCTTTCCCTACACGACGCTCTTCCGATCTNNNGCACATTCAGGAATCCCTAAGCA
GAGCACACGTCTGAACTCTTTCCCTACACGACGCTCTTCCGATCTNNNGTTTGAATTATTTCTAGTAC
AGCACACGTCTGAACTCTTTCCCTACACGACGCTCTTCCGATCTNNNCTGACCAGCTAGATCCCACTTC
GCACACGTCTGAACTCTTTCCCTACACGACGCTCTTCCGATCTNNNTCTTGCTCCTCAATGTGTGTGTG
AGAGCACACGTCTGAACTCTTTCCCTACACGACGCTCTTCCGATCTNNNCCTTCAGAAAGCTCCAATCA
GAGCACACGTCTGAACTCTTTCCCTACACGACGCTCTTCCGATCTNNNGGTCGCACTTTCCAATTCTCTC
AAGAGCACACGTCTGAACTCTTTCCCTACACGACGCTCTTCCGATCTNNNCTGAGGTGTGTACCATTC
AGCACACGTCTGAACTCTTTCCCTACACGACGCTCTTCCGATCTNNNTGGGAGTGATGCAACATTAG
GAGCACACGTCTGAACTCTTTCCCTACACGACGCTCTTCCGATCTNNNGAAAGATCATGGCTCTGGGA
AGCACACGTCTGAACTCTTTCCCTACACGACGCTCTTCCGATCTNNNAGAGGCAAGCAAACTTCAACC
GAGCACACGTCTGAACTCTTTCCCTACACGACGCTCTTCCGATCTNNNATAACTGGTAACCACAGGGA
GAGCACACGTCTGAACTCTTTCCCTACACGACGCTCTTCCGATCTNNNGAACAGAGGGTAGAAAATG
AGCACACGTCTGAACTCTTTCCCTACACGACGCTCTTCCGATCTNNNTCAATCTTTGCATGTTCCCACCA
AGCACACGTCTGAACTCTTTCCCTACACGACGCTCTTCCGATCTNNNAAGACTGAGAGTGAGGTGGA
GAGCACACGTCTGAACTCTTTCCCTACACGACGCTCTTCCGATCTNNNCTATACACTTTCCTGCGTGTG
AGCACACGTCTGAACTCTTTCCCTACACGACGCTCTTCCGATCTNNNTTGTGGAACAGGGAGCTGAG
GAGCACACGTCTGAACTCTTTCCCTACACGACGCTCTTCCGATCTNNNAAATACTAGTGAGGGTGGG
AGCACACGTCTGAACTCTTTCCCTACACGACGCTCTTCCGATCTNNNGAGAGCTGAACTAAACACAGG
AGCACACGTCTGAACTCTTTCCCTACACGACGCTCTTCCGATCTNNNGCTTTCTTGTTTGGTGGATGGT
CACACGTCTGAACTCTTTCCCTACACGACGCTCTTCCGATCTNNNGACTGAAAAGGCCAGAGCAAATG
AGCACACGTCTGAACTCTTTCCCTACACGACGCTCTTCCGATCTNNNGGATTTTGGGCTGCTCTTTTC
AGAGCACACGTCTGAACTCTTTCCCTACACGACGCTCTTCCGATCTNNNAGTGTTGGGAAATCCCCTG
GCACACGTCTGAACTCTTTCCCTACACGACGCTCTTCCGATCTNNNCTCCAGTGCCTAGCACATCTCTT
GAGCACACGTCTGAACTCTTTCCCTACACGACGCTCTTCCGATCTNNNAAAACAAATTCCTAACCAGT
GAGCACACGTCTGAACTCTTTCCCTACACGACGCTCTTCCGATCTNNNGTACAGCACCCTAGTTCAGA
GAGCACACGTCTGAACTCTTTCCCTACACGACGCTCTTCCGATCTNNNGATGGGTTATTTGCTTGGTT
AGCACACGTCTGAACTCTTTCCCTACACGACGCTCTTCCGATCTNNNCCTCAGCTAACAGCACTGAAC
AAGAGCACACGTCTGAACTCTTTCCCTACACGACGCTCTTCCGATCTNNNACCCGTAACAAGCAGCTA
GAGCACACGTCTGAACTCTTTCCCTACACGACGCTCTTCCGATCTNNNGCCCCTCTTCCTCTAGAAAT
AGCACACGTCTGAACTCTTTCCCTACACGACGCTCTTCCGATCTNNNGAAACAGTTGGGTGGCTAAC
AGCACACGTCTGAACTCTTTCCCTACACGACGCTCTTCCGATCTNNNAGGCCCTCACAGTAATCAAGG
GCACACGTCTGAACTCTTTCCCTACACGACGCTCTTCCGATCTNNNTAGTATACTCCTGTCCAGGCTC
GCACACGTCTGAACTCTTTCCCTACACGACGCTCTTCCGATCTNNNAGCTTTGAATGCATCAAGGTCT
CACACGTCTGAACTCTTTCCCTACACGACGCTCTTCCGATCTNNNATTTCCAGAGTGCCCATTGTGTG
GAGCACACGTCTGAACTCTTTCCCTACACGACGCTCTTCCGATCTNNNTCAGAGAGGATTTTGGTGT
GAAGAGCACACGTCTGAACTCTTTCCCTACACGACGCTCTTCCGATCTNNNTGGGGCAAAAACCCTAG

GCACACGTCTGAACTCTTTCCCTACACGACGCTCTTCCGATCTNNNCTGAGGGCTGTAAGTAGGAGT
GAGCACACGTCTGAACTCTTTCCCTACACGACGCTCTTCCGATCTNNNGAGGACAAGATGTGAGCTCC
AGCACACGTCTGAACTCTTTCCCTACACGACGCTCTTCCGATCTNNNGACGAATAGGGACAGGAAGA
GAGCACACGTCTGAACTCTTTCCCTACACGACGCTCTTCCGATCTNNNGTGGAATCTCAGAAACCCTG
AGCACACGTCTGAACTCTTTCCCTACACGACGCTCTTCCGATCTNNNTATGGGAAAGCACAACACATC
AGAGCACACGTCTGAACTCTTTCCCTACACGACGCTCTTCCGATCTNNNGAGCTGAGGTCACAGCAAC
CACACGTCTGAACTCTTTCCCTACACGACGCTCTTCCGATCTNNNGAGGAAGATCTGCAAGTGCTGT
GAGCACACGTCTGAACTCTTTCCCTACACGACGCTCTTCCGATCTNNNTCTTTCCTGCTCTCTTCTCCTC
AGCACACGTCTGAACTCTTTCCCTACACGACGCTCTTCCGATCTNNNTTCCTCAAGGTGTGCAAAGTT
AGAGCACACGTCTGAACTCTTTCCCTACACGACGCTCTTCCGATCTNNNGCAGACAATGGTTTCTTTTT
AGCACACGTCTGAACTCTTTCCCTACACGACGCTCTTCCGATCTNNNCACACACTCTAGACATTCACCC
AGCACACGTCTGAACTCTTTCCCTACACGACGCTCTTCCGATCTNNNACCATTCGCCAATGTCCTTGTT
GAGCACACGTCTGAACTCTTTCCCTACACGACGCTCTTCCGATCTNNNGGGGAGAAAGGCAAGTTAT
AGCACACGTCTGAACTCTTTCCCTACACGACGCTCTTCCGATCTNNNTACAATGCCTGTGCCACATTTC
GCACACGTCTGAACTCTTTCCCTACACGACGCTCTTCCGATCTNNNATAGTACTTTTCTCTCTCTCTCT
GAGCACACGTCTGAACTCTTTCCCTACACGACGCTCTTCCGATCTNNNTTACCTGGATCCACTGAAAG
AGCACACGTCTGAACTCTTTCCCTACACGACGCTCTTCCGATCTNNNCACACTCAAAGTTCCTGTCCTC
AGCACACGTCTGAACTCTTTCCCTACACGACGCTCTTCCGATCTNNNCCCTTTCATCAACCTGGGATCA
AGCACACGTCTGAACTCTTTCCCTACACGACGCTCTTCCGATCTNNNAACTGTTCTGTCTCATGACTG
GCACACGTCTGAACTCTTTCCCTACACGACGCTCTTCCGATCTNNNCCATTTGTTTGTGCTACCCACAG
GCACACGTCTGAACTCTTTCCCTACACGACGCTCTTCCGATCTNNNTCTCACTGCTGACTACTTGCTTTC
AGCACACGTCTGAACTCTTTCCCTACACGACGCTCTTCCGATCTNNNCTTCAAAATGCCCTGCTGCTTT
AGCACACGTCTGAACTCTTTCCCTACACGACGCTCTTCCGATCTNNNTGCCTAAATGTTCACAAGCCC
AGAGCACACGTCTGAACTCTTTCCCTACACGACGCTCTTCCGATCTNNNTTCCAGCAGGTGAAGGAAA
GCACACGTCTGAACTCTTTCCCTACACGACGCTCTTCCGATCTNNNCTAAATAGGCTGTGAATGGGGG
CACACGTCTGAACTCTTTCCCTACACGACGCTCTTCCGATCTNNNAGACAAGAGAGTAGCCAGCTGA
GCACACGTCTGAACTCTTTCCCTACACGACGCTCTTCCGATCTNNNTGCATGACACAAGTTCCAGTTC
GCACACGTCTGAACTCTTTCCCTACACGACGCTCTTCCGATCTNNNCGGTACATGGGTAAGAGTGGA
GCACACGTCTGAACTCTTTCCCTACACGACGCTCTTCCGATCTNNNGTTTGTATTCCTGCCTCCTCCAGC
AGAGCACACGTCTGAACTCTTTCCCTACACGACGCTCTTCCGATCTNNNCTCCATTCTGGTCCAACTC
AGCACACGTCTGAACTCTTTCCCTACACGACGCTCTTCCGATCTNNNAGGCCAGGAGAGAGGAAAT
AGCACACGTCTGAACTCTTTCCCTACACGACGCTCTTCCGATCTNNNTGTATTCTCCTTACCCCAACCC
AGCACACGTCTGAACTCTTTCCCTACACGACGCTCTTCCGATCTNNNAAGCTTTTCCATAGTCGACCA
GAGCACACGTCTGAACTCTTTCCCTACACGACGCTCTTCCGATCTNNNATCCAACTCCGTGTGTGGTT
CACACGTCTGAACTCTTTCCCTACACGACGCTCTTCCGATCTNNNTGCCCGTCTAAGAGGCTATTTTC
GAGCACACGTCTGAACTCTTTCCCTACACGACGCTCTTCCGATCTNNNCCCCTAAAATGTTGACAGGG
GAGCACACGTCTGAACTCTTTCCCTACACGACGCTCTTCCGATCTNNNCAAAATACTCACCTACCCACC
GAGCACACGTCTGAACTCTTTCCCTACACGACGCTCTTCCGATCTNNNTGGGGAAGAATCTGACAGC
AGCACACGTCTGAACTCTTTCCCTACACGACGCTCTTCCGATCTNNNACAGTCTTTCTGAAACATTCAA
GAGCACACGTCTGAACTCTTTCCCTACACGACGCTCTTCCGATCTNNNAGGCTCAGCACAACTTACAC
AGAGCACACGTCTGAACTCTTTCCCTACACGACGCTCTTCCGATCTNNNTTAGGGGTCACTACTCAAC
AGCACACGTCTGAACTCTTTCCCTACACGACGCTCTTCCGATCTNNNCACTTGAGTACGAAGATCGTC
CACACGTCTGAACTCTTTCCCTACACGACGCTCTTCCGATCTNNNAGGGAACACTAATTTTCTTTCCTG

GAGCACACGTCTGAACTCTTTCCCTACACGACGCTCTTCCGATCTNNNTTTTGGGGATGCTGCCAGAA
GCACACGTCTGAACTCTTTCCCTACACGACGCTCTTCCGATCTNNNTGTCCTACCCTGTACTGTCACTG
GCACACGTCTGAACTCTTTCCCTACACGACGCTCTTCCGATCTNNNTGACAAGGCCATGATACCATCA
AGCACACGTCTGAACTCTTTCCCTACACGACGCTCTTCCGATCTNNNTAAATCGCCCCTTCAGAAGTC
GAGCACACGTCTGAACTCTTTCCCTACACGACGCTCTTCCGATCTNNNGAGGCTCTGACCACTCTTGG
GAGCACACGTCTGAACTCTTTCCCTACACGACGCTCTTCCGATCTNNNGACCTGTGTGGAAACTGGC
GAGCACACGTCTGAACTCTTTCCCTACACGACGCTCTTCCGATCTNNNCACTCACAGTTTGGGTAGAG
GAGCACACGTCTGAACTCTTTCCCTACACGACGCTCTTCCGATCTNNNTCAGCATTGAAGTTGCTGGA
AGCACACGTCTGAACTCTTTCCCTACACGACGCTCTTCCGATCTNNNAAAAGTCTGCCCACTCAACAC
GCACACGTCTGAACTCTTTCCCTACACGACGCTCTTCCGATCTNNNATTATAAAGGGCTTGGGGCTG
GCACACGTCTGAACTCTTTCCCTACACGACGCTCTTCCGATCTNNNTCTTTGGATTACCAGGCATGGA
AGCACACGTCTGAACTCTTTCCCTACACGACGCTCTTCCGATCTNNNCAAGCTGAGCTAGCAAGGGAT
AGCACACGTCTGAACTCTTTCCCTACACGACGCTCTTCCGATCTNNNACTCTCATGAATGGAATCGGT
GAGCACACGTCTGAACTCTTTCCCTACACGACGCTCTTCCGATCTNNNATGAAAGCAGCAAAGGTCCA
AGCACACGTCTGAACTCTTTCCCTACACGACGCTCTTCCGATCTNNNACGGTTTTCAGCTACATTCCA
AAGAGCACACGTCTGAACTCTTTCCCTACACGACGCTCTTCCGATCTNNNATGATACCACTTTTCCCCTC
GAGCACACGTCTGAACTCTTTCCCTACACGACGCTCTTCCGATCTNNNGAGGACCCTATTGCCTCTTT
GAGCACACGTCTGAACTCTTTCCCTACACGACGCTCTTCCGATCTNNNGCACCAGGCATTATCAAAGT
CACACGTCTGAACTCTTTCCCTACACGACGCTCTTCCGATCTNNNCTTACCTTCCTCCCCCTCCTCTTCAT
GAGCACACGTCTGAACTCTTTCCCTACACGACGCTCTTCCGATCTNNNTTGCTTTCTGAGATGGGACA
GCACACGTCTGAACTCTTTCCCTACACGACGCTCTTCCGATCTNNNCCATCTAGCCTAGCCCCCATTTAC
AGCACACGTCTGAACTCTTTCCCTACACGACGCTCTTCCGATCTNNNTTACCAGCCAATGACAAACAC
GAGCACACGTCTGAACTCTTTCCCTACACGACGCTCTTCCGATCTNNNAGGGCTCAGAAGACCTACTC
AGCACACGTCTGAACTCTTTCCCTACACGACGCTCTTCCGATCTNNNTGTTTTGTGAAACACTGGCTTC
AGAGCACACGTCTGAACTCTTTCCCTACACGACGCTCTTCCGATCTNNNAAAGGCTTTGATAGGTTCTC
GAGCACACGTCTGAACTCTTTCCCTACACGACGCTCTTCCGATCTNNNTGGGGAAAGTCTCATCTTGC
GAGCACACGTCTGAACTCTTTCCCTACACGACGCTCTTCCGATCTNNNAGTAAGTTTTAGTGTGTTTC
GAGCACACGTCTGAACTCTTTCCCTACACGACGCTCTTCCGATCTNNNTCCAGTTATATGCCCAATGC
AGCACACGTCTGAACTCTTTCCCTACACGACGCTCTTCCGATCTNNNACCTCCACCTTCCTGAACACAT
AGCACACGTCTGAACTCTTTCCCTACACGACGCTCTTCCGATCTNNNGCCACAGATCTTATAGAGATG
AGAGCACACGTCTGAACTCTTTCCCTACACGACGCTCTTCCGATCTNNNTGGACTTGGCAGAAGAAT
GAGCACACGTCTGAACTCTTTCCCTACACGACGCTCTTCCGATCTNNNGAATCTTCAACCTCTGCCAGC
AGCACACGTCTGAACTCTTTCCCTACACGACGCTCTTCCGATCTNNNAAAGAAAACAAGGACATTTGC
AGCACACGTCTGAACTCTTTCCCTACACGACGCTCTTCCGATCTNNNAGGCAATACAGTAGGGTGTG
AGCACACGTCTGAACTCTTTCCCTACACGACGCTCTTCCGATCTNNNCACTCCTTCCCCATCAATACTGC
GAGCACACGTCTGAACTCTTTCCCTACACGACGCTCTTCCGATCTNNNCGTATATCTGGGGGAAGCA
CACACGTCTGAACTCTTTCCCTACACGACGCTCTTCCGATCTNNNGAGAGAGACCCCGTCTAAAGAG
AAGAGCACACGTCTGAACTCTTTCCCTACACGACGCTCTTCCGATCTNNNAGTGAGCCAATGTAGTGC
GAGCACACGTCTGAACTCTTTCCCTACACGACGCTCTTCCGATCTNNNACTATCCCCAGTCTCATCCA
AGAGCACACGTCTGAACTCTTTCCCTACACGACGCTCTTCCGATCTNNNCCACCACCATACCACTTTTG
GAGCACACGTCTGAACTCTTTCCCTACACGACGCTCTTCCGATCTNNNTCCTTTCTTCCCCTGTAGCAA
GAGCACACGTCTGAACTCTTTCCCTACACGACGCTCTTCCGATCTNNNCAATCACACTGAAAACCGT
GCACACGTCTGAACTCTTTCCCTACACGACGCTCTTCCGATCTNNNACACTTTTCACATGGTAGGTGC

GAGCACACGTCTGAACTCTTTCCCTACACGACGCTCTTCCGATCTNNNAAAACAGGATGGTGCTGGTG
AAGAGCACACGTCTGAACTCTTTCCCTACACGACGCTCTTCCGATCTNNNTGCAACTACACCTCAGAGA
GAAGAGCACACGTCTGAACTCTTTCCCTACACGACGCTCTTCCGATCTNNNTTGGACATAAGACAACCC
CACACGTCTGAACTCTTTCCCTACACGACGCTCTTCCGATCTNNNTTTTGTGGGTTGATCTTGTCCCCCAT
GAGCACACGTCTGAACTCTTTCCCTACACGACGCTCTTCCGATCTNNNAGTTAGCCAGGTGTGGTAAT
AGCACACGTCTGAACTCTTTCCCTACACGACGCTCTTCCGATCTNNNTTTGCTTGGGATTCTATTACTTG
AGAGCACACGTCTGAACTCTTTCCCTACACGACGCTCTTCCGATCTNNNAGCAGGGAAAAGAGGAAA
AGCACACGTCTGAACTCTTTCCCTACACGACGCTCTTCCGATCTNNNCAGCCTCAAGTAAGTCTCCACA
AGCACACGTCTGAACTCTTTCCCTACACGACGCTCTTCCGATCTNNNCTGTGCTTGACACACATGACGTA
GAGCACACGTCTGAACTCTTTCCCTACACGACGCTCTTCCGATCTNNNAAGCCAGAATCTTTGTTCAGC
GCACACGTCTGAACTCTTTCCCTACACGACGCTCTTCCGATCTNNNTCTGGCTGTGTAGAAATGAAAG
AGCACACGTCTGAACTCTTTCCCTACACGACGCTCTTCCGATCTNNNAGCATATCTGCGACTACCTAAG
AGCACACGTCTGAACTCTTTCCCTACACGACGCTCTTCCGATCTNNNATATCTGCCCAGTGCCATCTTCT
AGAGCACACGTCTGAACTCTTTCCCTACACGACGCTCTTCCGATCTNNNCAGTGTGTAAGTCTTTTACCT
GAGCACACGTCTGAACTCTTTCCCTACACGACGCTCTTCCGATCTNNNTTGAAGAGGATCCCACCTTCCT
AGAGCACACGTCTGAACTCTTTCCCTACACGACGCTCTTCCGATCTNNNAGATTGAGAGGCAATAACT
GAGCACACGTCTGAACTCTTTCCCTACACGACGCTCTTCCGATCTNNNTCACTCCATGTTTTCCCTTTTC
GAGCACACGTCTGAACTCTTTCCCTACACGACGCTCTTCCGATCTNNNTCCCCAAGGATTAGTTTTGAG
AGAGCACACGTCTGAACTCTTTCCCTACACGACGCTCTTCCGATCTNNNAGATCCTCTGCAGAAATGTA
GAGCACACGTCTGAACTCTTTCCCTACACGACGCTCTTCCGATCTNNNAAATCTCAGTGTGTGTCTGTC
GAGCACACGTCTGAACTCTTTCCCTACACGACGCTCTTCCGATCTNNNCATTCCCACTTTGCATTCTGCT
GAGCACACGTCTGAACTCTTTCCCTACACGACGCTCTTCCGATCTNNNAGGAAGGAGAAGGACCTAT
GAGCACACGTCTGAACTCTTTCCCTACACGACGCTCTTCCGATCTNNNAAATAGAGCAGGGCTGTTTG
AGCACACGTCTGAACTCTTTCCCTACACGACGCTCTTCCGATCTNNNAAGGTGAGAGAGAAGCCAGAT
GAGCACACGTCTGAACTCTTTCCCTACACGACGCTCTTCCGATCTNNNGTGTCTAATCGAAGTGTCTGC
GAAGAGCACACGTCTGAACTCTTTCCCTACACGACGCTCTTCCGATCTNNNAAGCTCAAGCTGTCAACC
AGCACACGTCTGAACTCTTTCCCTACACGACGCTCTTCCGATCTNNNTGGGAAGCTCCACCAAGAGATT
GAGCACACGTCTGAACTCTTTCCCTACACGACGCTCTTCCGATCTNNNTATTGTTCTGTGGGACGAGGG
AGCACACGTCTGAACTCTTTCCCTACACGACGCTCTTCCGATCTNNNCTGTCATGCTCCAGTCACAGTTT
AGAGCACACGTCTGAACTCTTTCCCTACACGACGCTCTTCCGATCTNNNTGCCTTGGTGGTCTTGAATT
AGCACACGTCTGAACTCTTTCCCTACACGACGCTCTTCCGATCTNNNAATACATGCTCAGGCATACCCA
AGAGCACACGTCTGAACTCTTTCCCTACACGACGCTCTTCCGATCTNNNTAATGAGGCATCAGTGGAG
AGCACACGTCTGAACTCTTTCCCTACACGACGCTCTTCCGATCTNNNAGAGCTGGTGATGTGTGATGT
AGCACACGTCTGAACTCTTTCCCTACACGACGCTCTTCCGATCTNNNAACACCTACATGTGCCAAACAGT
AGAGCACACGTCTGAACTCTTTCCCTACACGACGCTCTTCCGATCTNNNACAGAAGGGACATCACTCCA
GCACACGTCTGAACTCTTTCCCTACACGACGCTCTTCCGATCTNNNGCTTCCTTGTGCTTAGAGTTGGT
GAGCACACGTCTGAACTCTTTCCCTACACGACGCTCTTCCGATCTNNNAGTGCAGTCTTTGATTACTCCT
GAGCACACGTCTGAACTCTTTCCCTACACGACGCTCTTCCGATCTNNNAGATCAGAGGCATACATGTAA
AAGAGCACACGTCTGAACTCTTTCCCTACACGACGCTCTTCCGATCTNNNCTCCTGCAGTTCTCCTACAA
AGCACACGTCTGAACTCTTTCCCTACACGACGCTCTTCCGATCTNNNCTGAAGCAAAGGCTGGAGCAAT
AGCACACGTCTGAACTCTTTCCCTACACGACGCTCTTCCGATCTNNNCTCGCTTTGAGAGACAGGTATC
AGAGCACACGTCTGAACTCTTTCCCTACACGACGCTCTTCCGATCTNNNAGTGGTTTCCACAAGAAGGC
AGCACACGTCTGAACTCTTTCCCTACACGACGCTCTTCCGATCTNNNACATGCATGGACCTCCAGAGTTA

GCACACGTCTGAACTCTTTCCCTACACGACGCTCTTCCGATCTNNNTACTGACAGACTGGTGCTGGTAA
GCACACGTCTGAACTCTTTCCCTACACGACGCTCTTCCGATCTNNNGAGGCTACGGTGAACCAGGATAG
AGAGCACACGTCTGAACTCTTTCCCTACACGACGCTCTTCCGATCTNNNACTCAGGTGCTTTTCGAACTA
GAGCACACGTCTGAACTCTTTCCCTACACGACGCTCTTCCGATCTNNNATGCCAGTTTTGGCTTACTTC
AGCACACGTCTGAACTCTTTCCCTACACGACGCTCTTCCGATCTNNNGGATCACACAGCGTGTAGTCTTT
AGAGCACACGTCTGAACTCTTTCCCTACACGACGCTCTTCCGATCTNNNACCTACATCCTGTCTCCATCCA
AGCACACGTCTGAACTCTTTCCCTACACGACGCTCTTCCGATCTNNNTCATGGTTGAATCATTGTGGTTG
GAGCACACGTCTGAACTCTTTCCCTACACGACGCTCTTCCGATCTNNNGGCTCTATTCTTTCTCACAGG
AGCACACGTCTGAACTCTTTCCCTACACGACGCTCTTCCGATCTNNNCGGATTCCAGTCTGTCTCTCTCT
GAGCACACGTCTGAACTCTTTCCCTACACGACGCTCTTCCGATCTNNNACAAACAGCATACTTGAGTTG
AGCACACGTCTGAACTCTTTCCCTACACGACGCTCTTCCGATCTNNNTCAGATGGTTGTGAAGAAATTG
AGCACACGTCTGAACTCTTTCCCTACACGACGCTCTTCCGATCTNNNGGAAAGCCAACTCCTGGATTCA
AGCACACGTCTGAACTCTTTCCCTACACGACGCTCTTCCGATCTNNNTAGGTTACTTCAGCAGGGTGT
AGAGCACACGTCTGAACTCTTTCCCTACACGACGCTCTTCCGATCTNNNCCACAAAATGAGAAAGGCCT
AGCACACGTCTGAACTCTTTCCCTACACGACGCTCTTCCGATCTNNNAAGCCCAGTTGTACTCTCCAGAT
GAAGAGCACACGTCTGAACTCTTTCCCTACACGACGCTCTTCCGATCTNNNGGAATTGCCATGACTGG
AGCACACGTCTGAACTCTTTCCCTACACGACGCTCTTCCGATCTNNNGCAATTTTTGACTGGATGGCGA
AAGAGCACACGTCTGAACTCTTTCCCTACACGACGCTCTTCCGATCTNNNTAAGTGCTCATCATCCCACA
AGCACACGTCTGAACTCTTTCCCTACACGACGCTCTTCCGATCTNNNGCTCAACAATGATACACACACG
GAGCACACGTCTGAACTCTTTCCCTACACGACGCTCTTCCGATCTNNNTGCTTTGTATTTGGCCAAGACT
AGAGCACACGTCTGAACTCTTTCCCTACACGACGCTCTTCCGATCTNNNTGTATGAGTACTTGCACACA
AGCACACGTCTGAACTCTTTCCCTACACGACGCTCTTCCGATCTNNNAAAGAAGAAAGAACCAGAAAA
GAAGAGCACACGTCTGAACTCTTTCCCTACACGACGCTCTTCCGATCTNNNCACCTCACTTCTGTCTTCA
GAGCACACGTCTGAACTCTTTCCCTACACGACGCTCTTCCGATCTNNNCTATACAGGCATGGTTGCCT
AGAGCACACGTCTGAACTCTTTCCCTACACGACGCTCTTCCGATCTNNNACCGTTAGGATGCATAGGA
GCACACGTCTGAACTCTTTCCCTACACGACGCTCTTCCGATCTNNNGGAATTCCTTTGTGCAGGCTTTC
GAGCACACGTCTGAACTCTTTCCCTACACGACGCTCTTCCGATCTNNNGCCCACTTTCTGATGGAGATGA
AGCACACGTCTGAACTCTTTCCCTACACGACGCTCTTCCGATCTNNNTAAGGCAGACTTTCAGAGACC
AGAGCACACGTCTGAACTCTTTCCCTACACGACGCTCTTCCGATCTNNNTTGAGCTACTCAGGAGGCTG
AGAGCACACGTCTGAACTCTTTCCCTACACGACGCTCTTCCGATCTNNNTCATTCTTAGGGGATCCATG
AGCACACGTCTGAACTCTTTCCCTACACGACGCTCTTCCGATCTNNNATTCCACAGGTCCCTGTTCTCTT
AGCACACGTCTGAACTCTTTCCCTACACGACGCTCTTCCGATCTNNNTACTCGGCAAAGTCTCCTTTGT
GCACACGTCTGAACTCTTTCCCTACACGACGCTCTTCCGATCTNNNAGGGGAAACCAATGAAGAGTTG
AGAGCACACGTCTGAACTCTTTCCCTACACGACGCTCTTCCGATCTNNNTGCTGAAGAGAGAGAGGAT
GAGCACACGTCTGAACTCTTTCCCTACACGACGCTCTTCCGATCTNNNGGGATAAGTATGGAGAGGCA
GAGCACACGTCTGAACTCTTTCCCTACACGACGCTCTTCCGATCTNNNCTGCCACTCTGACATACATAG
AGAGCACACGTCTGAACTCTTTCCCTACACGACGCTCTTCCGATCTNNNAGCATGATTACTTCTATGGCT
AGCACACGTCTGAACTCTTTCCCTACACGACGCTCTTCCGATCTNNNACTAGATGATCTCTTCCTCTAGA
AGCACACGTCTGAACTCTTTCCCTACACGACGCTCTTCCGATCTNNNCGCCCTCACTCCTCTGTCCTTCAT
CACACGTCTGAACTCTTTCCCTACACGACGCTCTTCCGATCTNNNAATTTTTGTATTTGTATCGGCGGC
GAGCACACGTCTGAACTCTTTCCCTACACGACGCTCTTCCGATCTNNNGAGGAGTGGTCTCCTCACTCT
AAGAGCACACGTCTGAACTCTTTCCCTACACGACGCTCTTCCGATCTNNNCGAAGGCATAGAGGGCAA
GAAGAGCACACGTCTGAACTCTTTCCCTACACGACGCTCTTCCGATCTNNNCAGATACCCTGTTCCCCT

GAGCACACGTCTGAACTCTTTCCCTACACGACGCTCTTCCGATCTNNNTCCTTTCCCTGTCAGTCTCTCTC
GCACACGTCTGAACTCTTTCCCTACACGACGCTCTTCCGATCTNNNAGAAAAGGCAAAAGAATTGTTCC
AAGAGCACACGTCTGAACTCTTTCCCTACACGACGCTCTTCCGATCTNNNGCCAAATCCCATCTCTCTCG
GCACACGTCTGAACTCTTTCCCTACACGACGCTCTTCCGATCTNNNGAAAAGCACCACTTTCTTGGTGG
AGCACACGTCTGAACTCTTTCCCTACACGACGCTCTTCCGATCTNNNGCAGCATTACTAGGCTTTAGGC
GAGCACACGTCTGAACTCTTTCCCTACACGACGCTCTTCCGATCTNNNCCCTGTGTTGGCAAGAGTATG
AGCACACGTCTGAACTCTTTCCCTACACGACGCTCTTCCGATCTNNNTTCCTTGCCCTTTTCAGCTTTCTCA
GAGCACACGTCTGAACTCTTTCCCTACACGACGCTCTTCCGATCTNNNCCTTTCCAAGATGACGGGTTG
AGCACACGTCTGAACTCTTTCCCTACACGACGCTCTTCCGATCTNNNGACACTAAGCCCACCTACAACAG
AGAGCACACGTCTGAACTCTTTCCCTACACGACGCTCTTCCGATCTNNNTATCACCTCCTCCAAACAGACA
GAGCACACGTCTGAACTCTTTCCCTACACGACGCTCTTCCGATCTNNNTGCCATGTAGAAAAGGCAGTT
AGAGCACACGTCTGAACTCTTTCCCTACACGACGCTCTTCCGATCTNNNACATTGACCTCTCTCTTTCCCC
GAAGAGCACACGTCTGAACTCTTTCCCTACACGACGCTCTTCCGATCTNNNGTCGTTTCATATCTCCACCA
AGCACACGTCTGAACTCTTTCCCTACACGACGCTCTTCCGATCTNNNCAGGAAGGGAAAACCAATGGG
GAGCACACGTCTGAACTCTTTCCCTACACGACGCTCTTCCGATCTNNNGAGGGGCATTGACATAAGCA
AGCACACGTCTGAACTCTTTCCCTACACGACGCTCTTCCGATCTNNNCCTAGAACCATGTATGGAACAA
AGCACACGTCTGAACTCTTTCCCTACACGACGCTCTTCCGATCTNNNTCAAGCTTTTCTGCCTCATTTGCC
GAGCACACGTCTGAACTCTTTCCCTACACGACGCTCTTCCGATCTNNNTGGCCCATACATAAAAGGCAA
GAGCACACGTCTGAACTCTTTCCCTACACGACGCTCTTCCGATCTNNNATCCCTCACTTCTGCTTCCACTT
GAGCACACGTCTGAACTCTTTCCCTACACGACGCTCTTCCGATCTNNNATGTGGAACACAGAAGTTTCA
GAGCACACGTCTGAACTCTTTCCCTACACGACGCTCTTCCGATCTNNNTCAAGTGGAAGGGGCATACTA
GAGCACACGTCTGAACTCTTTCCCTACACGACGCTCTTCCGATCTNNNAGGAAAACGGGTATGAAGGA
AGAGCACACGTCTGAACTCTTTCCCTACACGACGCTCTTCCGATCTNNNAAAGTGGTTCACCACCTCTCT
AGCACACGTCTGAACTCTTTCCCTACACGACGCTCTTCCGATCTNNNATTTCCAAGTTTGGCCACTGAGA
GCACACGTCTGAACTCTTTCCCTACACGACGCTCTTCCGATCTNNNAGATTCATAGCATACCCCTCCCCC
AGCACACGTCTGAACTCTTTCCCTACACGACGCTCTTCCGATCTNNNGAATGCCGGATGCTTGACTTCT
AGCACACGTCTGAACTCTTTCCCTACACGACGCTCTTCCGATCTNNNGCCACCAGAACTCTTACTGCTAA
AGCACACGTCTGAACTCTTTCCCTACACGACGCTCTTCCGATCTNNNGGCATTACAAGCATTGACCTCAC
GCACACGTCTGAACTCTTTCCCTACACGACGCTCTTCCGATCTNNNGCCAGTATAGACAATAATTGGCC
AAGAGCACACGTCTGAACTCTTTCCCTACACGACGCTCTTCCGATCTNNNGAGATAGGAGGTACCCAG
GCACACGTCTGAACTCTTTCCCTACACGACGCTCTTCCGATCTNNNGAGGTGAGCGTCTATCCCTGGCA
AGCACACGTCTGAACTCTTTCCCTACACGACGCTCTTCCGATCTNNNGTTGTCCAAACCTGTTACCTCCA
GAGCACACGTCTGAACTCTTTCCCTACACGACGCTCTTCCGATCTNNNCCACATGAGGCAAGGAGTTCA
AGCACACGTCTGAACTCTTTCCCTACACGACGCTCTTCCGATCTNNNTGCCATCAGTAACACAAACCCA
AGAGCACACGTCTGAACTCTTTCCCTACACGACGCTCTTCCGATCTNNNAGTCAGCGTTAGTATGGAGC
GCACACGTCTGAACTCTTTCCCTACACGACGCTCTTCCGATCTNNNCATTTGCCTTCCACATTTGGGTCC
AGAGCACACGTCTGAACTCTTTCCCTACACGACGCTCTTCCGATCTNNNTGGGAAGTGCCATCATTTTC
GAGCACACGTCTGAACTCTTTCCCTACACGACGCTCTTCCGATCTNNNAAGCAAAGCAGGACCACAAA
AAGAGCACACGTCTGAACTCTTTCCCTACACGACGCTCTTCCGATCTNNNTTTTGGCAGAGCTTTAGCT
AGCACACGTCTGAACTCTTTCCCTACACGACGCTCTTCCGATCTNNNTTTTACTTACTTCCGCTCCCCACC
AGCACACGTCTGAACTCTTTCCCTACACGACGCTCTTCCGATCTNNNCCAGGTTCCTCCTACTTCTAGGG
AGAGCACACGTCTGAACTCTTTCCCTACACGACGCTCTTCCGATCTNNNGCCCCTTTGAACTCACCTCTTT
GAGCACACGTCTGAACTCTTTCCCTACACGACGCTCTTCCGATCTNNNGCTCACAATTGCTAAGTGAGG

AGCACACGTCTGAACTCTTTCCCTACACGACGCTCTTCCGATCTNNNTCCCCTCTCTCTCTTTACATGC
AGCACACGTCTGAACTCTTTCCCTACACGACGCTCTTCCGATCTNNNCAACCAGTTCAGCCTGGAGTA
GAGCACACGTCTGAACTCTTTCCCTACACGACGCTCTTCCGATCTNNNTCAGAGGCTAATCAGTCCTG
AGAGCACACGTCTGAACTCTTTCCCTACACGACGCTCTTCCGATCTNNNAAAAATTGGAGCAGCATTG
GAGCACACGTCTGAACTCTTTCCCTACACGACGCTCTTCCGATCTNNNTCAGACACCCAATATGGATC
GCACACGTCTGAACTCTTTCCCTACACGACGCTCTTCCGATCTNNNTTGTCATATTTGCGTACCCACCC
AGCACACGTCTGAACTCTTTCCCTACACGACGCTCTTCCGATCTNNNACCTCTGTGTATGCCTGTCTGA
AGCACACGTCTGAACTCTTTCCCTACACGACGCTCTTCCGATCTNNNTCATGCAAATGAATCCCCTGG
AAGAGCACACGTCTGAACTCTTTCCCTACACGACGCTCTTCCGATCTNNNTAGGAGCAAACTTCTCAG
AGCACACGTCTGAACTCTTTCCCTACACGACGCTCTTCCGATCTNNNTCATGCTGTCAAACTCTGGTG
GCACACGTCTGAACTCTTTCCCTACACGACGCTCTTCCGATCTNNNATGGCTTTGCATGTGTGTATGT
AGCACACGTCTGAACTCTTTCCCTACACGACGCTCTTCCGATCTNNNAGGGCAATATAAAGGAAAAT
GCACACGTCTGAACTCTTTCCCTACACGACGCTCTTCCGATCTNNNCAGATTCCCAAGCAAGGGAGA
AGCACACGTCTGAACTCTTTCCCTACACGACGCTCTTCCGATCTNNNAATTTTCTCCCATTCTGTTGTC
GCACACGTCTGAACTCTTTCCCTACACGACGCTCTTCCGATCTNNNTCAGGTTTTCTCATCTGTGTGG
AGCACACGTCTGAACTCTTTCCCTACACGACGCTCTTCCGATCTNNNGCCAACTAGACATAACAGTTT
AGCACACGTCTGAACTCTTTCCCTACACGACGCTCTTCCGATCTNNNCTGACATCCTAGCTCCTGACTG
GAGCACACGTCTGAACTCTTTCCCTACACGACGCTCTTCCGATCTNNNTGAGAAGTGCTCAATGCTC
AGCACACGTCTGAACTCTTTCCCTACACGACGCTCTTCCGATCTNNNAAGGGAAGTGTCTTAGTCCC
AGCACACGTCTGAACTCTTTCCCTACACGACGCTCTTCCGATCTNNNGCAATGCCTCCTCCTCATGTTA
AGCACACGTCTGAACTCTTTCCCTACACGACGCTCTTCCGATCTNNNAGTACCTGCTTTCATTCTAGGG
GAGCACACGTCTGAACTCTTTCCCTACACGACGCTCTTCCGATCTNNNAGGCATTATGTGGGTGCCAA
AGCACACGTCTGAACTCTTTCCCTACACGACGCTCTTCCGATCTNNNTTTCTTCTCTTTTCCACCCCACC
GAGCACACGTCTGAACTCTTTCCCTACACGACGCTCTTCCGATCTNNNCCCATTTTCTTTTGGTGACGC
CACACGTCTGAACTCTTTCCCTACACGACGCTCTTCCGATCTNNNGGTACAGAGCAAACAATGTGGG
AGCACACGTCTGAACTCTTTCCCTACACGACGCTCTTCCGATCTNNNGTCTCTCCAGCCAGAAAGAGT
GCACACGTCTGAACTCTTTCCCTACACGACGCTCTTCCGATCTNNNATGACAGAAGCGAGCAAGAAC
GCACACGTCTGAACTCTTTCCCTACACGACGCTCTTCCGATCTNNNTTCCTTCTCTCTGTTCCCTCACCC
GCACACGTCTGAACTCTTTCCCTACACGACGCTCTTCCGATCTNNNAGACTTACAGGAACTGCAGCCT
AGAGCACACGTCTGAACTCTTTCCCTACACGACGCTCTTCCGATCTNNNTGGGATTTCAGAAGAAGG
AGCACACGTCTGAACTCTTTCCCTACACGACGCTCTTCCGATCTNNNAAACAAGCAAAGCACAAGTCC
GCACACGTCTGAACTCTTTCCCTACACGACGCTCTTCCGATCTNNNGACCTGCCAGCCCACAATACAT
AGCACACGTCTGAACTCTTTCCCTACACGACGCTCTTCCGATCTNNNAACTCTGTGGGACAGATCTCA
AGCACACGTCTGAACTCTTTCCCTACACGACGCTCTTCCGATCTNNNATACATGAGAGTGCAGGTAC
GAGCACACGTCTGAACTCTTTCCCTACACGACGCTCTTCCGATCTNNNGTGCAGACTACACAGGTAG
AGCACACGTCTGAACTCTTTCCCTACACGACGCTCTTCCGATCTNNNTCTTTCTGTATTTCCAATTTTCT
AGCACACGTCTGAACTCTTTCCCTACACGACGCTCTTCCGATCTNNNGGCATCTGTTCTTGTTTGCAC
GCACACGTCTGAACTCTTTCCCTACACGACGCTCTTCCGATCTNNNCCTGGAGAACTTGTTCAAAGCA
AGCACACGTCTGAACTCTTTCCCTACACGACGCTCTTCCGATCTNNNGGAAGGGAATGGGGAGTAG
AGCACACGTCTGAACTCTTTCCCTACACGACGCTCTTCCGATCTNNNCACCCACTGCTCTAGATTTGG
GCACACGTCTGAACTCTTTCCCTACACGACGCTCTTCCGATCTNNNCTCTTCGTCGTTTACCAGCTCAA
GAGCACACGTCTGAACTCTTTCCCTACACGACGCTCTTCCGATCTNNNAAACTGCTCTAGCCTCTTTCA
GCACACGTCTGAACTCTTTCCCTACACGACGCTCTTCCGATCTNNNTGCCTTCTCCTGTCCTTCTCAATC

GAGCACACGTCTGAACTCTTTCCCTACACGACGCTCTTCCGATCTNNNACACAAGGCTCCTACAGAGTA
GAGCACACGTCTGAACTCTTTCCCTACACGACGCTCTTCCGATCTNNNTGTCACTCTAGGGAAAGGGTT
AGAGCACACGTCTGAACTCTTTCCCTACACGACGCTCTTCCGATCTNNNGCCCTAGAGGAAGTTGGAA
AGAGCACACGTCTGAACTCTTTCCCTACACGACGCTCTTCCGATCTNNNAGACATCCATTCCCAACAAAG
AGAGCACACGTCTGAACTCTTTCCCTACACGACGCTCTTCCGATCTNNNTCAAATTAGCTTTTGCTGGC
AGAGCACACGTCTGAACTCTTTCCCTACACGACGCTCTTCCGATCTNNNGTGTGAAAATGGAAGAGGG
GAGCACACGTCTGAACTCTTTCCCTACACGACGCTCTTCCGATCTNNNTGGACTGTTCTACCAGATCCTC
AGAGCACACGTCTGAACTCTTTCCCTACACGACGCTCTTCCGATCTNNNTGAGATGAATGAGTTCTGG
AGCACACGTCTGAACTCTTTCCCTACACGACGCTCTTCCGATCTNNNAAAAAGACACCTCCTGGCATAG
AGCACACGTCTGAACTCTTTCCCTACACGACGCTCTTCCGATCTNNNTTGACAGTTTGGGCTTCAACGT
GAGCACACGTCTGAACTCTTTCCCTACACGACGCTCTTCCGATCTNNNGAAGCTGATTAGAGGCTCTTC
ACACGTCTGAACTCTTTCCCTACACGACGCTCTTCCGATCTNNNGAGACCCGGAATGCGCAGGGCATC
GCACACGTCTGAACTCTTTCCCTACACGACGCTCTTCCGATCTNNNCAACAGAGCACATTTTAGGGGA
AGCACACGTCTGAACTCTTTCCCTACACGACGCTCTTCCGATCTNNNGGTGAGGTAGATTTCCGGAATG
AGAGCACACGTCTGAACTCTTTCCCTACACGACGCTCTTCCGATCTNNNTACCATGCGAGTAAGTGCCC
GAGCACACGTCTGAACTCTTTCCCTACACGACGCTCTTCCGATCTNNNTTGCACACGCAATTGCAAAAAT
AGAGCACACGTCTGAACTCTTTCCCTACACGACGCTCTTCCGATCTNNNTGAATTCTTCCCGGTTCTCTCC
AGAGCACACGTCTGAACTCTTTCCCTACACGACGCTCTTCCGATCTNNNGAACCAGTAGGATGTGTGTG
AGCACACGTCTGAACTCTTTCCCTACACGACGCTCTTCCGATCTNNNAGAGCTGAAAGACAGAAAGAT
AGAGCACACGTCTGAACTCTTTCCCTACACGACGCTCTTCCGATCTNNNGATCAGTGATGTACACGCAC
GAGCACACGTCTGAACTCTTTCCCTACACGACGCTCTTCCGATCTNNNTCTTCCTTTTGCTTACTGCACCA
AGAGCACACGTCTGAACTCTTTCCCTACACGACGCTCTTCCGATCTNNNTCTCCAGGTTACACAGCGTAG
GAGCACACGTCTGAACTCTTTCCCTACACGACGCTCTTCCGATCTNNNGAGGTTTCCAGACTGATTTGCC
AGCACACGTCTGAACTCTTTCCCTACACGACGCTCTTCCGATCTNNNTTGTTATGGGGCACAACACTGT
AGAGCACACGTCTGAACTCTTTCCCTACACGACGCTCTTCCGATCTNNNTCCTCTGCTTTGGTTACGTGA
AAGAGCACACGTCTGAACTCTTTCCCTACACGACGCTCTTCCGATCTNNNTCACTCCCATGAGTATTGAA
AGCACACGTCTGAACTCTTTCCCTACACGACGCTCTTCCGATCTNNNGTCTGCACACATAATTCACGCTC
GAGCACACGTCTGAACTCTTTCCCTACACGACGCTCTTCCGATCTNNNACACACAAACTTACATGGACAC
AGCACACGTCTGAACTCTTTCCCTACACGACGCTCTTCCGATCTNNNGCTGTGTTCAAGTGGGATGTG
AGCACACGTCTGAACTCTTTCCCTACACGACGCTCTTCCGATCTNNNTCTTTGGGATGAGTGCTCAACTC
AGAGCACACGTCTGAACTCTTTCCCTACACGACGCTCTTCCGATCTNNNGCAGACTTCAATGAGACTTG
GAGCACACGTCTGAACTCTTTCCCTACACGACGCTCTTCCGATCTNNNACTTATACCCTGATGAACTGG
GAGCACACGTCTGAACTCTTTCCCTACACGACGCTCTTCCGATCTNNNTTATGTGACTGTTGCCAGGA
AGCACACGTCTGAACTCTTTCCCTACACGACGCTCTTCCGATCTNNNCCTGACCCATTTCTACAACCCAGC
AGAGCACACGTCTGAACTCTTTCCCTACACGACGCTCTTCCGATCTNNNCTGCAAAGAGGCATTTTTCC
AGCACACGTCTGAACTCTTTCCCTACACGACGCTCTTCCGATCTNNNCCATCCCATGGTGGAAAGCATA
AGCACACGTCTGAACTCTTTCCCTACACGACGCTCTTCCGATCTNNNGTTGAGGCTGTAGCTGTAGTG
AAGAGCACACGTCTGAACTCTTTCCCTACACGACGCTCTTCCGATCTNNNTCAAGTCAAGAACTGAAAC
AGAGCACACGTCTGAACTCTTTCCCTACACGACGCTCTTCCGATCTNNNATCTAGTTAGTTCCCTCCAGC
GAGCACACGTCTGAACTCTTTCCCTACACGACGCTCTTCCGATCTNNNCTGCTGACATCCTTGACCTTGA
GAGCACACGTCTGAACTCTTTCCCTACACGACGCTCTTCCGATCTNNNTCCTGGGCATGACATGAGAAA
AGCACACGTCTGAACTCTTTCCCTACACGACGCTCTTCCGATCTNNNTCCAAGACAGTAGACTTTGGGT
AGCACACGTCTGAACTCTTTCCCTACACGACGCTCTTCCGATCTNNNTGTTGTCTGTGTACTCATAATTG

GAGCACACGTCTGAACTCTTTCCCTACACGACGCTCTTCCGATCTNNNCAGCTGAAGGGTATTTCTCT
AGCACACGTCTGAACTCTTTCCCTACACGACGCTCTTCCGATCTNNNCCCAAAATGTTGCCACAGCTTT
AGCACACGTCTGAACTCTTTCCCTACACGACGCTCTTCCGATCTNNNCACTCAGAATACGGCACACAT
AGCACACGTCTGAACTCTTTCCCTACACGACGCTCTTCCGATCTNNNCAAACCCTGAGTGTGGGTGT
AGCACACGTCTGAACTCTTTCCCTACACGACGCTCTTCCGATCTNNNTGGTCAGATTTCCTAGCCTCTG
GAGCACACGTCTGAACTCTTTCCCTACACGACGCTCTTCCGATCTNNNAGAACAGGTAAGTACGGG
AGAGCACACGTCTGAACTCTTTCCCTACACGACGCTCTTCCGATCTNNNGTAGAGGTGGAGATACAC
AGCACACGTCTGAACTCTTTCCCTACACGACGCTCTTCCGATCTNNNCCTTTGCAGCTTGGCATCATAT
AGCACACGTCTGAACTCTTTCCCTACACGACGCTCTTCCGATCTNNNCAGTCTGTTTTCTGTGTGTGTG
GCACACGTCTGAACTCTTTCCCTACACGACGCTCTTCCGATCTNNNTTGTATGTGGTGGAGCACAAC
AGCACACGTCTGAACTCTTTCCCTACACGACGCTCTTCCGATCTNNNGGACAGTGCTGAAGAGAAGT
AGCACACGTCTGAACTCTTTCCCTACACGACGCTCTTCCGATCTNNNGGTTCCAAGTCACTGCAGAAG
AGCACACGTCTGAACTCTTTCCCTACACGACGCTCTTCCGATCTNNNACTTTCTGGGAAATGCAACCT
CACACGTCTGAACTCTTTCCCTACACGACGCTCTTCCGATCTNNNTTTTTCTTACCCCTTGGGCTTCCCA
GCACACGTCTGAACTCTTTCCCTACACGACGCTCTTCCGATCTNNNGGAATCCATGTGTCTTGCTCCTT
GAGCACACGTCTGAACTCTTTCCCTACACGACGCTCTTCCGATCTNNNTTCCAAGTTTCCTCAGATCC
AGCACACGTCTGAACTCTTTCCCTACACGACGCTCTTCCGATCTNNNCAGTATTCTCAGGCAGTGTGG
GCACACGTCTGAACTCTTTCCCTACACGACGCTCTTCCGATCTNNNTGGTATCAGTCAGTCAATAAGC
GCACACGTCTGAACTCTTTCCCTACACGACGCTCTTCCGATCTNNNCCGGCTGATTTCTGGGTTGGCA
CACACGTCTGAACTCTTTCCCTACACGACGCTCTTCCGATCTNNNTCAACTCATTCTCGGTAAAGGTC
AGCACACGTCTGAACTCTTTCCCTACACGACGCTCTTCCGATCTNNNAAAGCTGGAATTCAGAGCGCA
AAGAGCACACGTCTGAACTCTTTCCCTACACGACGCTCTTCCGATCTNNNTGTGTGTATCCATCCATC
AGCACACGTCTGAACTCTTTCCCTACACGACGCTCTTCCGATCTNNNTTTTTCAGGCTCTCATGTTTCT
AGCACACGTCTGAACTCTTTCCCTACACGACGCTCTTCCGATCTNNNTAGCTTGCATCTTGGATCTTG
AGCACACGTCTGAACTCTTTCCCTACACGACGCTCTTCCGATCTNNNAGTCTCTGTAGGACAGTATTC
GAGCACACGTCTGAACTCTTTCCCTACACGACGCTCTTCCGATCTNNNGCATGGACGAACCTTGTAAG
GAGCACACGTCTGAACTCTTTCCCTACACGACGCTCTTCCGATCTNNNTACACTCTCACACTGTCCTGG
GCACACGTCTGAACTCTTTCCCTACACGACGCTCTTCCGATCTNNNTCTACCTGCCTAGTGGACTTTCC
AGCACACGTCTGAACTCTTTCCCTACACGACGCTCTTCCGATCTNNNGCACTGTTCGAAACACTATTA
AGCACACGTCTGAACTCTTTCCCTACACGACGCTCTTCCGATCTNNNCGAAACCATCCATCTTCCTCCTC
AGCACACGTCTGAACTCTTTCCCTACACGACGCTCTTCCGATCTNNNTATCCCTGGGAGAAAGCCAG
AGCACACGTCTGAACTCTTTCCCTACACGACGCTCTTCCGATCTNNNAGACCTAAGACAACAGGAACA
GAGCACACGTCTGAACTCTTTCCCTACACGACGCTCTTCCGATCTNNNATGGATGTTTTCTGGAGAGG
GCACACGTCTGAACTCTTTCCCTACACGACGCTCTTCCGATCTNNNAGTAAGCACATAGTGTCACGAG
AGCACACGTCTGAACTCTTTCCCTACACGACGCTCTTCCGATCTNNNGGAAGACAGCATCCTTTCACC
GCACACGTCTGAACTCTTTCCCTACACGACGCTCTTCCGATCTNNNCAACTGCAAATGGCTTTCGACAT
AGCACACGTCTGAACTCTTTCCCTACACGACGCTCTTCCGATCTNNNAGTGAAATTCCATCTCTCTCTC
AGCACACGTCTGAACTCTTTCCCTACACGACGCTCTTCCGATCTNNNTCCTCTAGCCAAAGGACCAGA
GAGCACACGTCTGAACTCTTTCCCTACACGACGCTCTTCCGATCTNNNCCTCTGAATTGGGAGCACAG
AGCACACGTCTGAACTCTTTCCCTACACGACGCTCTTCCGATCTNNNCAGAGGGAGAGAGAAGGAT
GCACACGTCTGAACTCTTTCCCTACACGACGCTCTTCCGATCTNNNAGCCCAATCACAGAAACCAGAA
AGCACACGTCTGAACTCTTTCCCTACACGACGCTCTTCCGATCTNNNAACATCTAACCAACAGGAAG
AGCACACGTCTGAACTCTTTCCCTACACGACGCTCTTCCGATCTNNNGATCATGGCCCTAATCCCAACC

AGCACACGTCTGAACTCTTTCCCTACACGACGCTCTTCCGATCTNNNCTGGCCAGGAGAGAAAAGAA
AGAGCACACGTCTGAACTCTTTCCCTACACGACGCTCTTCCGATCTNNNACCCAGTTTTATTTGCTTAG
GAGCACACGTCTGAACTCTTTCCCTACACGACGCTCTTCCGATCTNNNAAAACCACTTGTAGCCAGACA
AGCACACGTCTGAACTCTTTCCCTACACGACGCTCTTCCGATCTNNNACTACCATTTGCAAAAACATTC
AGAGCACACGTCTGAACTCTTTCCCTACACGACGCTCTTCCGATCTNNNACAGGTCCTCTTGGTACAAT
GCACACGTCTGAACTCTTTCCCTACACGACGCTCTTCCGATCTNNNCAGCACATGGGACTTTTTCCAGA
GAGCACACGTCTGAACTCTTTCCCTACACGACGCTCTTCCGATCTNNNCTGGTGCCTTACTCTGACATT
GAGCACACGTCTGAACTCTTTCCCTACACGACGCTCTTCCGATCTNNNTGTTTTGGGACTACCATGGA
GAGCACACGTCTGAACTCTTTCCCTACACGACGCTCTTCCGATCTNNNACTCTCATGATGGTACATTTT
GCACACGTCTGAACTCTTTCCCTACACGACGCTCTTCCGATCTNNNTACTCATTTGGCTCCGCATTTCAC
GCACACGTCTGAACTCTTTCCCTACACGACGCTCTTCCGATCTNNNTGAGTTCATTGTAATCAAACAAA
AGAGCACACGTCTGAACTCTTTCCCTACACGACGCTCTTCCGATCTNNNCTGCTGGTTTTCAGTGAAC
GAAGAGCACACGTCTGAACTCTTTCCCTACACGACGCTCTTCCGATCTNNNCAGTGGTTCCCAACCTTA
AGAGCACACGTCTGAACTCTTTCCCTACACGACGCTCTTCCGATCTNNNAAAGGTTATGAATGCCAGG
GAGCACACGTCTGAACTCTTTCCCTACACGACGCTCTTCCGATCTNNNAAACAAAAGCAGTTCCTGGA
AGAGCACACGTCTGAACTCTTTCCCTACACGACGCTCTTCCGATCTNNNAGCATCAAGCAAGTCCAAAA
AGCACACGTCTGAACTCTTTCCCTACACGACGCTCTTCCGATCTNNNCCCTTAGCCACTGAGTTTGTCT
GAGCACACGTCTGAACTCTTTCCCTACACGACGCTCTTCCGATCTNNNAGCAGCAAGAACAACTTTTG
AGAGCACACGTCTGAACTCTTTCCCTACACGACGCTCTTCCGATCTNNNAAGGAGAAGAGAGAGAC
GAAGAGCACACGTCTGAACTCTTTCCCTACACGACGCTCTTCCGATCTNNNTTCAAGCCACAGCATTTC
AGCACACGTCTGAACTCTTTCCCTACACGACGCTCTTCCGATCTNNNATGGACTACCATGCCTGGCTTC
AGCACACGTCTGAACTCTTTCCCTACACGACGCTCTTCCGATCTNNNAGAATTGAGGAGCTAATGCCT
AGCACACGTCTGAACTCTTTCCCTACACGACGCTCTTCCGATCTNNNGGGACATCTCCAAATTCCTGCA
GAGCACACGTCTGAACTCTTTCCCTACACGACGCTCTTCCGATCTNNNTTCTGAGAGACCTGGCCATTC
AGCACACGTCTGAACTCTTTCCCTACACGACGCTCTTCCGATCTNNNGCACTTTTCTAGGTCCTTTTCGT
AGCACACGTCTGAACTCTTTCCCTACACGACGCTCTTCCGATCTNNNGCCAAGAGACTGGGTGTTGA
AGCACACGTCTGAACTCTTTCCCTACACGACGCTCTTCCGATCTNNNAGGTCCATGTCACAGAATTGCT
AGCACACGTCTGAACTCTTTCCCTACACGACGCTCTTCCGATCTNNNCTGCTATCCACGCTCTGAAAGA
AGCACACGTCTGAACTCTTTCCCTACACGACGCTCTTCCGATCTNNNAGAGCCATCTGGGATGTTAG
GAGCACACGTCTGAACTCTTTCCCTACACGACGCTCTTCCGATCTNNNACACATGAGGGCTAAAGATG
GAGCACACGTCTGAACTCTTTCCCTACACGACGCTCTTCCGATCTNNNTGGAAGAAAAGGCTAAGGC
AGCACACGTCTGAACTCTTTCCCTACACGACGCTCTTCCGATCTNNNCTCTCAAGAAATAGGTGTGGC
AGCACACGTCTGAACTCTTTCCCTACACGACGCTCTTCCGATCTNNNAGTATCCATTTGCCCCTCATTG
GAGCACACGTCTGAACTCTTTCCCTACACGACGCTCTTCCGATCTNNNATGTAAGCACCCTATTGGTTC
AGAGCACACGTCTGAACTCTTTCCCTACACGACGCTCTTCCGATCTNNNAATTTCCAGAATGAATGAAT
AGCACACGTCTGAACTCTTTCCCTACACGACGCTCTTCCGATCTNNNGCTCCACATTCATGGCAGCTA
AGAGCACACGTCTGAACTCTTTCCCTACACGACGCTCTTCCGATCTNNNCACCTATCTGCTATAGTGAG
AGCACACGTCTGAACTCTTTCCCTACACGACGCTCTTCCGATCTNNNCTTCCTGAAGCTCAGGTCAATG
GCACACGTCTGAACTCTTTCCCTACACGACGCTCTTCCGATCTNNNGCTAAAAGCATGGACACTGGAG
AGCACACGTCTGAACTCTTTCCCTACACGACGCTCTTCCGATCTNNNTTAGGGCTGGGTTGGAGATTA
GAGCACACGTCTGAACTCTTTCCCTACACGACGCTCTTCCGATCTNNNGAAAAAGTGCTGACCCCAGA
AAGAGCACACGTCTGAACTCTTTCCCTACACGACGCTCTTCCGATCTNNNGAAATACTGTGCTCATGT
AGCACACGTCTGAACTCTTTCCCTACACGACGCTCTTCCGATCTNNNGACAATACAGCTGGGCAACAT

GAGCACACGTCTGAACTCTTTCCCTACACGACGCTCTTCCGATCTNNNTGTGCCCTCAGGTTATTCTAC
GAGCACACGTCTGAACTCTTTCCCTACACGACGCTCTTCCGATCTNNNTCTGCCAGAGAGAGATTGTG
GAGCACACGTCTGAACTCTTTCCCTACACGACGCTCTTCCGATCTNNNACACACACACATTCCCAGCTTA
GAGCACACGTCTGAACTCTTTCCCTACACGACGCTCTTCCGATCTNNNGAAAAGCAGAGCCAACGTTG
AGAGCACACGTCTGAACTCTTTCCCTACACGACGCTCTTCCGATCTNNNGGGTAGCAAAGAATTATGC
GAGCACACGTCTGAACTCTTTCCCTACACGACGCTCTTCCGATCTNNNAAATGGGTCAGCACCTCCAA
AGCACACGTCTGAACTCTTTCCCTACACGACGCTCTTCCGATCTNNNGGACGTCAGTCAGAAAGGATG
GAGCACACGTCTGAACTCTTTCCCTACACGACGCTCTTCCGATCTNNNAATGCTCCATGCACAGGATTA
GAGCACACGTCTGAACTCTTTCCCTACACGACGCTCTTCCGATCTNNNCAAGGAATGGCCAAGCTAGA
GAGCACACGTCTGAACTCTTTCCCTACACGACGCTCTTCCGATCTNNNAAATTCAGAGGCTACTGTCTG
GAGCACACGTCTGAACTCTTTCCCTACACGACGCTCTTCCGATCTNNNGAAATCGCCTAACAACGCATT
AGCACACGTCTGAACTCTTTCCCTACACGACGCTCTTCCGATCTNNNAATTTCCATGACCACCCAGACA
AGCACACGTCTGAACTCTTTCCCTACACGACGCTCTTCCGATCTNNNGTCTCCAGTGAGAATGCTTCCA
AGAGCACACGTCTGAACTCTTTCCCTACACGACGCTCTTCCGATCTNNNTTGGACCAAAGGAAACCTA
GAGCACACGTCTGAACTCTTTCCCTACACGACGCTCTTCCGATCTNNNGCTCTGCACATATCTGCCTAAC
GAGCACACGTCTGAACTCTTTCCCTACACGACGCTCTTCCGATCTNNNGACCTACAATTCTGAGGCAC
AAGAGCACACGTCTGAACTCTTTCCCTACACGACGCTCTTCCGATCTNNNGGGTTTCTCTCAGGTCTA
AGAGCACACGTCTGAACTCTTTCCCTACACGACGCTCTTCCGATCTNNNAGCTTTTGCAATTCTGATTCT
GAGCACACGTCTGAACTCTTTCCCTACACGACGCTCTTCCGATCTNNNAATCCCGGAAGCAAATATGC
AGAGCACACGTCTGAACTCTTTCCCTACACGACGCTCTTCCGATCTNNNACAGCAAACCAAATCCAAAG
AGCACACGTCTGAACTCTTTCCCTACACGACGCTCTTCCGATCTNNNACGTGTGAATTGGGGTAGGAA
AGCACACGTCTGAACTCTTTCCCTACACGACGCTCTTCCGATCTNNNGACTTCCCAAGCCTCCACAATAA
GAGCACACGTCTGAACTCTTTCCCTACACGACGCTCTTCCGATCTNNNAGTGCCTGCATTCCCTTAGGA
AGCACACGTCTGAACTCTTTCCCTACACGACGCTCTTCCGATCTNNNGCCAGGTGTAGAAGGCTTCAT
GAGCACACGTCTGAACTCTTTCCCTACACGACGCTCTTCCGATCTNNNAGGCAATCAGAGGTATGTTT
GAGCACACGTCTGAACTCTTTCCCTACACGACGCTCTTCCGATCTNNNCTTGAAAACAATGTGCGTCCT
GAGCACACGTCTGAACTCTTTCCCTACACGACGCTCTTCCGATCTNNNAGGTGTATGACCATCACCCAA
AGCACACGTCTGAACTCTTTCCCTACACGACGCTCTTCCGATCTNNNCAGCTGTGGTTATTTGGCTCCT
GAGCACACGTCTGAACTCTTTCCCTACACGACGCTCTTCCGATCTNNNTCGAGGGTGACTTTCTTCTCT
AGCACACGTCTGAACTCTTTCCCTACACGACGCTCTTCCGATCTNNNAGTTCTAATAAGCTTGTGACTG
AGCACACGTCTGAACTCTTTCCCTACACGACGCTCTTCCGATCTNNNATGTAGCTTGCACCATTCACTGT
AGCACACGTCTGAACTCTTTCCCTACACGACGCTCTTCCGATCTNNNCATCCTCTTTTGTGCTCTTCAG
GCACACGTCTGAACTCTTTCCCTACACGACGCTCTTCCGATCTNNNGCCCAGGGTCAAAATCAAACCAT
GAGCACACGTCTGAACTCTTTCCCTACACGACGCTCTTCCGATCTNNNCAGAACAGCTTGGAATGGCT
GAGCACACGTCTGAACTCTTTCCCTACACGACGCTCTTCCGATCTNNNTGCAGAGGGCTGAAAGAAA
AGCACACGTCTGAACTCTTTCCCTACACGACGCTCTTCCGATCTNNNATGGAACGCATTTTGGGTTGTT
GCACACGTCTGAACTCTTTCCCTACACGACGCTCTTCCGATCTNNNGTTCCAGCCACCTCCATTATGCCA
GAGCACACGTCTGAACTCTTTCCCTACACGACGCTCTTCCGATCTNNNTGCACTCTATGTGTTTTGTGG
AGAGCACACGTCTGAACTCTTTCCCTACACGACGCTCTTCCGATCTNNNCCACTCTGGGTAAGGGAAG
AGCACACGTCTGAACTCTTTCCCTACACGACGCTCTTCCGATCTNNNCCTCAAGATACCTTCCCACCTCA
AGCACACGTCTGAACTCTTTCCCTACACGACGCTCTTCCGATCTNNNCACATCAGGTGCGGTGAAGTC
GAGCACACGTCTGAACTCTTTCCCTACACGACGCTCTTCCGATCTNNNACCCCTAGGAAGACTGAAACA
GAGCACACGTCTGAACTCTTTCCCTACACGACGCTCTTCCGATCTNNNTCAAAGACATGCAAAACTGAG

AGCACACGTCTGAACTCTTTCCCTACACGACGCTCTTCCGATCTNNNGAGCTGGGGAGATGGATGA
AGCACACGTCTGAACTCTTTCCCTACACGACGCTCTTCCGATCTNNNAATACCTGAGGTGGAACAGG
GCACACGTCTGAACTCTTTCCCTACACGACGCTCTTCCGATCTNNNGGACAGCTCACCTGATTGTTCTA
GCACACGTCTGAACTCTTTCCCTACACGACGCTCTTCCGATCTNNNGGAGTTACAAACCAGTGCTCCAT
GCACACGTCTGAACTCTTTCCCTACACGACGCTCTTCCGATCTNNNACCTTTGAGCTATTGTACAACCT
GAGCACACGTCTGAACTCTTTCCCTACACGACGCTCTTCCGATCTNNNTGCCCGGCCCATATTATGTAT
GAGCACACGTCTGAACTCTTTCCCTACACGACGCTCTTCCGATCTNNNCCAGCAGATTCGGTGATAG
AGCACACGTCTGAACTCTTTCCCTACACGACGCTCTTCCGATCTNNNGCAGATCGAATATGGCAAAAG
GCACACGTCTGAACTCTTTCCCTACACGACGCTCTTCCGATCTNNNTAGCCAAGCAAAGTTCATAGG
AGCACACGTCTGAACTCTTTCCCTACACGACGCTCTTCCGATCTNNNGGCAAGTGGTCCCTTTATGTG
AGCACACGTCTGAACTCTTTCCCTACACGACGCTCTTCCGATCTNNNTAATATGAGGCACAGCGTGAT
GAGCACACGTCTGAACTCTTTCCCTACACGACGCTCTTCCGATCTNNNGATCCCAAACCGTTGAAGG
GAGCACACGTCTGAACTCTTTCCCTACACGACGCTCTTCCGATCTNNNTTGTCCCACATGGTCTAAGT
AGCACACGTCTGAACTCTTTCCCTACACGACGCTCTTCCGATCTNNNCTTGTGAAATTACCTGGCCCT
GCACACGTCTGAACTCTTTCCCTACACGACGCTCTTCCGATCTNNNTTAGGCACTGTGAGCTACATAT
GAGCACACGTCTGAACTCTTTCCCTACACGACGCTCTTCCGATCTNNNTCACTAACTGGCAGAAAATG
GAGCACACGTCTGAACTCTTTCCCTACACGACGCTCTTCCGATCTNNNTGTCAACAAAGCACAGTTTG
AGAGCACACGTCTGAACTCTTTCCCTACACGACGCTCTTCCGATCTNNNCTCGGGGCCAGTTTCCTTAT
AGCACACGTCTGAACTCTTTCCCTACACGACGCTCTTCCGATCTNNNTGCTCGCTAACACTTCTCATTG
AGCACACGTCTGAACTCTTTCCCTACACGACGCTCTTCCGATCTNNNGGATGCTGTCGACAAGTTTCT
AGCACACGTCTGAACTCTTTCCCTACACGACGCTCTTCCGATCTNNNTGCAAATGGACAAATGGGAA
GCACACGTCTGAACTCTTTCCCTACACGACGCTCTTCCGATCTNNNAGGGACTTGGGGTGATCATTT
AGCACACGTCTGAACTCTTTCCCTACACGACGCTCTTCCGATCTNNNGAATCCACCAAGGCAATTTAC
GAGCACACGTCTGAACTCTTTCCCTACACGACGCTCTTCCGATCTNNNAGGAAGTCCGACAAAGTCCT
AGCACACGTCTGAACTCTTTCCCTACACGACGCTCTTCCGATCTNNNTGCATGGGTATGGCTTTTATGT
AGCACACGTCTGAACTCTTTCCCTACACGACGCTCTTCCGATCTNNNGGGACACAGACAGGAATGGA
AGCACACGTCTGAACTCTTTCCCTACACGACGCTCTTCCGATCTNNNTGTAGACAGTCATGTACCCAC
ACACGTCTGAACTCTTTCCCTACACGACGCTCTTCCGATCTNNNGAGAAACACGTCCTCAAGCCCATC
AGCACACGTCTGAACTCTTTCCCTACACGACGCTCTTCCGATCTNNNCCAGCACATTCCCAAATGGTTT
GAGCACACGTCTGAACTCTTTCCCTACACGACGCTCTTCCGATCTNNNAGCTTCAATTGTGAATGCAG
AGCACACGTCTGAACTCTTTCCCTACACGACGCTCTTCCGATCTNNNGGTGGGTCCAGTCATAAGGAT
AGCACACGTCTGAACTCTTTCCCTACACGACGCTCTTCCGATCTNNNGTGGAGTGCTGTTTGTGGTG
AGCACACGTCTGAACTCTTTCCCTACACGACGCTCTTCCGATCTNNNCTGGTCTTTTCCTTTTCTGCCC
GCACACGTCTGAACTCTTTCCCTACACGACGCTCTTCCGATCTNNNAACGAATACTGCTCATGGTTGC
GAGCACACGTCTGAACTCTTTCCCTACACGACGCTCTTCCGATCTNNNATGAAATCAAGACGCTCCTG
GCACACGTCTGAACTCTTTCCCTACACGACGCTCTTCCGATCTNNNTGTGAGTGGCTCTCTACTTGTGA
AGCACACGTCTGAACTCTTTCCCTACACGACGCTCTTCCGATCTNNNAGGAATGAGGAGGATGTTTG
GAGCACACGTCTGAACTCTTTCCCTACACGACGCTCTTCCGATCTNNNAAATCTTCTCTTCCCCCACTG
AGCACACGTCTGAACTCTTTCCCTACACGACGCTCTTCCGATCTNNNAACTCCTGCTTCAACAAGCCTT
AGCACACGTCTGAACTCTTTCCCTACACGACGCTCTTCCGATCTNNNGACTTCTGCTCCAAAGGTTCC
GAGCACACGTCTGAACTCTTTCCCTACACGACGCTCTTCCGATCTNNNTAACAGGCCTGCTGTTAGCTT
AAGAGCACACGTCTGAACTCTTTCCCTACACGACGCTCTTCCGATCTNNNAAATTGAGCAGGGTGAG
AGCACACGTCTGAACTCTTTCCCTACACGACGCTCTTCCGATCTNNNGCACCACACTCTCAATACCA

GCACACGTCTGAACTCTTTCCCTACACGACGCTCTTCCGATCTNNNACGCCTCTTAGACCTCAACACTA
GAGCACACGTCTGAACTCTTTCCCTACACGACGCTCTTCCGATCTNNNAGTGGAGCTATATGGGTCCT
AGAGCACACGTCTGAACTCTTTCCCTACACGACGCTCTTCCGATCTNNNGCTCCTTTGGAAGAAAGTT
AGCACACGTCTGAACTCTTTCCCTACACGACGCTCTTCCGATCTNNNACTCTGTGCTAGGTTCCGAGT
GAGCACACGTCTGAACTCTTTCCCTACACGACGCTCTTCCGATCTNNNTGAGGGGCATGTATGAGTGG
AGCACACGTCTGAACTCTTTCCCTACACGACGCTCTTCCGATCTNNNCCAGCATTCAGTCAGAAACACC
GCACACGTCTGAACTCTTTCCCTACACGACGCTCTTCCGATCTNNNAGGCATCAATCCAAATGACTTGC
GCACACGTCTGAACTCTTTCCCTACACGACGCTCTTCCGATCTNNNCTGCCACCTTCCAGGATCATTTT
GAGCACACGTCTGAACTCTTTCCCTACACGACGCTCTTCCGATCTNNNTTTGTGCCTACGGTACAGTTC
GCACACGTCTGAACTCTTTCCCTACACGACGCTCTTCCGATCTNNNCTGCCTCCTCTTCAGCTTGATTTC
GAGCACACGTCTGAACTCTTTCCCTACACGACGCTCTTCCGATCTNNNTTGAGCTTTCATAACCTGCTG
AGCACACGTCTGAACTCTTTCCCTACACGACGCTCTTCCGATCTNNNCTAGCAATGGAGTGATGAGA
GCACACGTCTGAACTCTTTCCCTACACGACGCTCTTCCGATCTNNNTGTCAGAACTTTGTCACCTGTCT
AGCACACGTCTGAACTCTTTCCCTACACGACGCTCTTCCGATCTNNNAATCTTTAGCAGGGAAGTGCT
AGCACACGTCTGAACTCTTTCCCTACACGACGCTCTTCCGATCTNNNAAAAATCCTGCTTTTTCCTCTGC
GCACACGTCTGAACTCTTTCCCTACACGACGCTCTTCCGATCTNNNCTTGTTAGCTCGCTGGCTCTTTA
AGAGCACACGTCTGAACTCTTTCCCTACACGACGCTCTTCCGATCTNNNCCATTTGGGAGAGTGTCTC
AGCACACGTCTGAACTCTTTCCCTACACGACGCTCTTCCGATCTNNNCTGGCCTTGAACTTTTCTGGTT
GAGCACACGTCTGAACTCTTTCCCTACACGACGCTCTTCCGATCTNNNCCCTGAAGTTTGGGTGACA
GCACACGTCTGAACTCTTTCCCTACACGACGCTCTTCCGATCTNNNAGCTGGAATGATGATGCTGAG
GAGCACACGTCTGAACTCTTTCCCTACACGACGCTCTTCCGATCTNNNAGCTCCCTTCTGTGTAGGAAC
GAGCACACGTCTGAACTCTTTCCCTACACGACGCTCTTCCGATCTNNNATGGTGACAAGCACTACACA
AGCACACGTCTGAACTCTTTCCCTACACGACGCTCTTCCGATCTNNNAAGATCAACCTGGGAAGCGTT
GAGCACACGTCTGAACTCTTTCCCTACACGACGCTCTTCCGATCTNNNCCTGTCACCTCCCAGTATGGA
GAGCACACGTCTGAACTCTTTCCCTACACGACGCTCTTCCGATCTNNNTTGTGAGTGTGTTGAGACTT
GAGCACACGTCTGAACTCTTTCCCTACACGACGCTCTTCCGATCTNNNGCCAGATCTTCATTCCCTATG
AGCACACGTCTGAACTCTTTCCCTACACGACGCTCTTCCGATCTNNNAGGGAGTGGGGATGGTCAAT
AGCACACGTCTGAACTCTTTCCCTACACGACGCTCTTCCGATCTNNNCCCCTTCACATTGCCTCAACTTA
AGCACACGTCTGAACTCTTTCCCTACACGACGCTCTTCCGATCTNNNGACAAGTTCCATAGCTTGAGG
AGCACACGTCTGAACTCTTTCCCTACACGACGCTCTTCCGATCTNNNACAATTCTGGGAGAGCATCAA
AGCACACGTCTGAACTCTTTCCCTACACGACGCTCTTCCGATCTNNNGGAGCAATTGGCTGACTTCTA
GCACACGTCTGAACTCTTTCCCTACACGACGCTCTTCCGATCTNNNGCTCCCAAGTCAAGGATCTGTTT
AGCACACGTCTGAACTCTTTCCCTACACGACGCTCTTCCGATCTNNNTGAGGTGAATTTCCACCCTTA
AGCACACGTCTGAACTCTTTCCCTACACGACGCTCTTCCGATCTNNNAGGACAGAACATGGAAAGCCT
AGCACACGTCTGAACTCTTTCCCTACACGACGCTCTTCCGATCTNNNCTGGGCTAAGTTCAAGGTGTG
AGCACACGTCTGAACTCTTTCCCTACACGACGCTCTTCCGATCTNNNATGCTCCAGAACACGTTACACT
AGCACACGTCTGAACTCTTTCCCTACACGACGCTCTTCCGATCTNNNTGCCTTGGTGATCTTGGATAAT
AGAGCACACGTCTGAACTCTTTCCCTACACGACGCTCTTCCGATCTNNNTGATCATTAGATCTGGGGT
AGCACACGTCTGAACTCTTTCCCTACACGACGCTCTTCCGATCTNNNTGGGGAAAAATAAGTCCTAC
AGCACACGTCTGAACTCTTTCCCTACACGACGCTCTTCCGATCTNNNCCATAATCATGAGAGCCAGGT
AGCACACGTCTGAACTCTTTCCCTACACGACGCTCTTCCGATCTNNNAAACCATTACAGTAGCCAGTT
AGCACACGTCTGAACTCTTTCCCTACACGACGCTCTTCCGATCTNNNTAATCTGATAGGGACAGGGCT
GCACACGTCTGAACTCTTTCCCTACACGACGCTCTTCCGATCTNNNACTCCATACACAAGGCCCCTATT

AGCACACGTCTGAACTCTTTCCCTACACGACGCTCTTCCGATCTNNNAAAACAAAAGTGCATGCCACA
AGCACACGTCTGAACTCTTTCCCTACACGACGCTCTTCCGATCTNNNAATTCTAGAGCCAGAGACAGG
GAGCACACGTCTGAACTCTTTCCCTACACGACGCTCTTCCGATCTNNNAGTGATAAAAATCAGGCCCA
AGCACACGTCTGAACTCTTTCCCTACACGACGCTCTTCCGATCTNNNGCAGTGGTGTAATCACAGACC
GAGCACACGTCTGAACTCTTTCCCTACACGACGCTCTTCCGATCTNNNGAAGCAAAATGGGTCTCCCAT
GAGCACACGTCTGAACTCTTTCCCTACACGACGCTCTTCCGATCTNNNACCTGCTAAAAAGTTCTAGG
GAGCACACGTCTGAACTCTTTCCCTACACGACGCTCTTCCGATCTNNNCCTGCTCTTCATCAATCCTCCCT
AGCACACGTCTGAACTCTTTCCCTACACGACGCTCTTCCGATCTNNNACCCAATAACTGGTGATCTAA
GCACACGTCTGAACTCTTTCCCTACACGACGCTCTTCCGATCTNNNCAGCTGGATTTGAAGATAGTCC
AGAGCACACGTCTGAACTCTTTCCCTACACGACGCTCTTCCGATCTNNNTACTGTTCACACAGCATCTT
GAGCACACGTCTGAACTCTTTCCCTACACGACGCTCTTCCGATCTNNNACAATAGGGAACTTTGGAGG
GAGCACACGTCTGAACTCTTTCCCTACACGACGCTCTTCCGATCTNNNTGATGCCAGAAGCACAACTT
AGCACACGTCTGAACTCTTTCCCTACACGACGCTCTTCCGATCTNNNTGGCAAATGACAGGACTTCTT
GAGCACACGTCTGAACTCTTTCCCTACACGACGCTCTTCCGATCTNNNTGATACCCATTGAAGGATGC
AGCACACGTCTGAACTCTTTCCCTACACGACGCTCTTCCGATCTNNNAGCAAACATAAGATTGATGCA
AGCACACGTCTGAACTCTTTCCCTACACGACGCTCTTCCGATCTNNNAAGACATGCCGGAGTGTTGTA
AAGAGCACACGTCTGAACTCTTTCCCTACACGACGCTCTTCCGATCTNNNTCTCAAGAATTTCGGATA
AGCACACGTCTGAACTCTTTCCCTACACGACGCTCTTCCGATCTNNNTTTCCTCATTTTTGCCCAACTGC
GAGCACACGTCTGAACTCTTTCCCTACACGACGCTCTTCCGATCTNNNGGTAGTGACACACTCCCTTGA
AGCACACGTCTGAACTCTTTCCCTACACGACGCTCTTCCGATCTNNNCAAGGCTCATAGTGAGGAGG
AGCACACGTCTGAACTCTTTCCCTACACGACGCTCTTCCGATCTNNNAAAAGAAGGAGGCCCGTATG
AGCACACGTCTGAACTCTTTCCCTACACGACGCTCTTCCGATCTNNNTCAGCTTAGAGTTTTGAATGA
GAGCACACGTCTGAACTCTTTCCCTACACGACGCTCTTCCGATCTNNNCCTGGTCTGTTGGAACTCCAT
GAGCACACGTCTGAACTCTTTCCCTACACGACGCTCTTCCGATCTNNNAGAGGGACAGGAGGAGAG
AGCACACGTCTGAACTCTTTCCCTACACGACGCTCTTCCGATCTNNNGTCTGTGTGGGTGAATGCTTG
AGCACACGTCTGAACTCTTTCCCTACACGACGCTCTTCCGATCTNNNATTGCATCCTTTTGCTCCCATC
GAGCACACGTCTGAACTCTTTCCCTACACGACGCTCTTCCGATCTNNNATGCAATGTCCTTCTTAGTGC
AAGAGCACACGTCTGAACTCTTTCCCTACACGACGCTCTTCCGATCTNNNGAGAGTGGTCCAGAGGT
CACACGTCTGAACTCTTTCCCTACACGACGCTCTTCCGATCTNNNGTTTTGGGCTGAAAGATGGCGCA
GCACACGTCTGAACTCTTTCCCTACACGACGCTCTTCCGATCTNNNTGTTATGACGATTACACGAGCC
GAGCACACGTCTGAACTCTTTCCCTACACGACGCTCTTCCGATCTNNNTCAGTTCAAGTCCACTGAGGG
AGCACACGTCTGAACTCTTTCCCTACACGACGCTCTTCCGATCTNNNTCCTAACTTCAGTGTTGTCCCC
GCACACGTCTGAACTCTTTCCCTACACGACGCTCTTCCGATCTNNNGCTCTCCATTTTCGCTCACACATC
AGAGCACACGTCTGAACTCTTTCCCTACACGACGCTCTTCCGATCTNNNTAACGAGAGATTGGGGCA
GAGCACACGTCTGAACTCTTTCCCTACACGACGCTCTTCCGATCTNNNACAATACAGGGGACAAGAG
AGCACACGTCTGAACTCTTTCCCTACACGACGCTCTTCCGATCTNNNTCAACATGAGCTCTCTCTCCCA
GAGCACACGTCTGAACTCTTTCCCTACACGACGCTCTTCCGATCTNNNTTTCAAGGAAGAGCAACCTCT
AGCACACGTCTGAACTCTTTCCCTACACGACGCTCTTCCGATCTNNNTAAACCGCCAGTGGGAATGTA
GAGCACACGTCTGAACTCTTTCCCTACACGACGCTCTTCCGATCTNNNTGTGTCAGGAAAGAAAATAC
GAGCACACGTCTGAACTCTTTCCCTACACGACGCTCTTCCGATCTNNNAAGCTGGAGAGGGAAGGG
GAGCACACGTCTGAACTCTTTCCCTACACGACGCTCTTCCGATCTNNNGGTGTGATTGGGCTCAAAGG
GAGCACACGTCTGAACTCTTTCCCTACACGACGCTCTTCCGATCTNNNAAGCCCCCTAAAGATGGAGA
AGCACACGTCTGAACTCTTTCCCTACACGACGCTCTTCCGATCTNNNTGCATGTCTCCTTATGGGTCTC

GAGCACACGTCTGAACTCTTTCCCTACACGACGCTCTTCCGATCTNNNGACTTTACCGCCACCTTCCTG
AGCACACGTCTGAACTCTTTCCCTACACGACGCTCTTCCGATCTNNNGGCATACTTGTTTGTTCCGCAT
GAAGAGCACACGTCTGAACTCTTTCCCTACACGACGCTCTTCCGATCTNNNGCTACGAGTGAATACTT
AGCACACGTCTGAACTCTTTCCCTACACGACGCTCTTCCGATCTNNNGAACCTCAAAGGCCCTTATCCT
GAGCACACGTCTGAACTCTTTCCCTACACGACGCTCTTCCGATCTNNNGTTCAAGCCACAACAAAAGT
AGCACACGTCTGAACTCTTTCCCTACACGACGCTCTTCCGATCTNNNCGAGTGATGGCATGTGACTT
AGCACACGTCTGAACTCTTTCCCTACACGACGCTCTTCCGATCTNNNGACCGGTCTGTCCATTTCTACT
AGCACACGTCTGAACTCTTTCCCTACACGACGCTCTTCCGATCTNNNTGCTTAGGCCACTCTAGGAC
AGCACACGTCTGAACTCTTTCCCTACACGACGCTCTTCCGATCTNNNTTGCTATGGATTTGCCAGTCC
GAGCACACGTCTGAACTCTTTCCCTACACGACGCTCTTCCGATCTNNNTGGCACTACTCAAAAAGCCA
AGCACACGTCTGAACTCTTTCCCTACACGACGCTCTTCCGATCTNNNTCCAGGTAGTTCAGTTTCCTCG
CACACGTCTGAACTCTTTCCCTACACGACGCTCTTCCGATCTNNNTATACCTCTCGTGCATGAAGCCTC
AGCACACGTCTGAACTCTTTCCCTACACGACGCTCTTCCGATCTNNNTCTTTGGATTTCCAGGTGCAT
CACACGTCTGAACTCTTTCCCTACACGACGCTCTTCCGATCTNNNAGCTGCTGATCACTCATTTCAGGC
CACACGTCTGAACTCTTTCCCTACACGACGCTCTTCCGATCTNNNGCCCAAGCTACAGAAGCATTTTT
GCACACGTCTGAACTCTTTCCCTACACGACGCTCTTCCGATCTNNNCCCCCTTAGTTCCTCCCTACTTCC
AGCACACGTCTGAACTCTTTCCCTACACGACGCTCTTCCGATCTNNNAGCTGGCTACAGAATCCTGAA
AGCACACGTCTGAACTCTTTCCCTACACGACGCTCTTCCGATCTNNNAATTATTTGGGGGTAAAGGG
AGCACACGTCTGAACTCTTTCCCTACACGACGCTCTTCCGATCTNNNGGAGGTGGGGAGGCATATGT
GAGCACACGTCTGAACTCTTTCCCTACACGACGCTCTTCCGATCTNNNGAAATGGTAGCCAATTCTGG
AGCACACGTCTGAACTCTTTCCCTACACGACGCTCTTCCGATCTNNNGCAGGCAGTTTTACAGATGA
AGCACACGTCTGAACTCTTTCCCTACACGACGCTCTTCCGATCTNNNGGATCTGGTTTCCATTGCCTCT
AGAGCACACGTCTGAACTCTTTCCCTACACGACGCTCTTCCGATCTNNNAGCATAAATCAGATGCCAC
GAGCACACGTCTGAACTCTTTCCCTACACGACGCTCTTCCGATCTNNNACCAACAACCAGCAATCATTG
AGCACACGTCTGAACTCTTTCCCTACACGACGCTCTTCCGATCTNNNGAACCTGCCTTGCCCCATAAAT
GCACACGTCTGAACTCTTTCCCTACACGACGCTCTTCCGATCTNNNGGGTTGGCAATGGCTTTCTTTT
AGAGCACACGTCTGAACTCTTTCCCTACACGACGCTCTTCCGATCTNNNTTTGCCCACACCCTCACATA
AGCACACGTCTGAACTCTTTCCCTACACGACGCTCTTCCGATCTNNNACAACAGGAATGACAGCTAA
AGCACACGTCTGAACTCTTTCCCTACACGACGCTCTTCCGATCTNNNCCCATTTGAACCAGTTCAGGA
AGCACACGTCTGAACTCTTTCCCTACACGACGCTCTTCCGATCTNNNACTGACACACACATACACACA
AGCACACGTCTGAACTCTTTCCCTACACGACGCTCTTCCGATCTNNNATGGGGAATAGGGACACACT
AGCACACGTCTGAACTCTTTCCCTACACGACGCTCTTCCGATCTNNNGGATAATTCACCAGAGAGCTG
GCACACGTCTGAACTCTTTCCCTACACGACGCTCTTCCGATCTNNNTTACGGATACATTTTTGTGGGC
AGCACACGTCTGAACTCTTTCCCTACACGACGCTCTTCCGATCTNNNCCCAGTGCTAACAGCTATCCTG
AGCACACGTCTGAACTCTTTCCCTACACGACGCTCTTCCGATCTNNNTGTTTTGTATAGTGGCAGCAG
AGCACACGTCTGAACTCTTTCCCTACACGACGCTCTTCCGATCTNNNGATTACAGTGCCTGGCCATAG
CACACGTCTGAACTCTTTCCCTACACGACGCTCTTCCGATCTNNNAGTTTTCTTCCCAGGAAACCCAG
GAGCACACGTCTGAACTCTTTCCCTACACGACGCTCTTCCGATCTNNNACCAAGTGGGAGGACAAAT
GAAGAGCACACGTCTGAACTCTTTCCCTACACGACGCTCTTCCGATCTNNNAGCAGACATTTCGGTGA
GCACACGTCTGAACTCTTTCCCTACACGACGCTCTTCCGATCTNNNTAGATAGAACACTGGAACACAG
GAGCACACGTCTGAACTCTTTCCCTACACGACGCTCTTCCGATCTNNNATACCCAAGTTACCCTACCTG
GCACACGTCTGAACTCTTTCCCTACACGACGCTCTTCCGATCTNNNACTGATGAATGCTCAGACTGGG
AGCACACGTCTGAACTCTTTCCCTACACGACGCTCTTCCGATCTNNNTCTTGTACATGGGAAGAGAG

GCACACGTCTGAACTCTTTCCCTACACGACGCTCTTCCGATCTNNNTCTTTGGGATTTTGAGCCATGT
GAGCACACGTCTGAACTCTTTCCCTACACGACGCTCTTCCGATCTNNNGGTAGGTTCTGATGGCATCT
GCACACGTCTGAACTCTTTCCCTACACGACGCTCTTCCGATCTNNNATTGAAGCAGGGTCAGGGTTT
AGCACACGTCTGAACTCTTTCCCTACACGACGCTCTTCCGATCTNNNAGTAGTTCTCAACCTTGGCTT
GCACACGTCTGAACTCTTTCCCTACACGACGCTCTTCCGATCTNNNCTGAAAGTGCAGAGGAAAGCT
AGCACACGTCTGAACTCTTTCCCTACACGACGCTCTTCCGATCTNNNAGAGACATTTGGGGGTACTCA
AGCACACGTCTGAACTCTTTCCCTACACGACGCTCTTCCGATCTNNNCGCATCGCATGTTAGGTTTAT
GAGCACACGTCTGAACTCTTTCCCTACACGACGCTCTTCCGATCTNNNGCAGCAAAATAGCTGCATTC
GAGCACACGTCTGAACTCTTTCCCTACACGACGCTCTTCCGATCTNNNAGGCTCCTCTTTCTGATCCTC
AGCACACGTCTGAACTCTTTCCCTACACGACGCTCTTCCGATCTNNNCTCTCCATGTGAGCTCCTCCAT
AGCACACGTCTGAACTCTTTCCCTACACGACGCTCTTCCGATCTNNNGGAATATTTCTGTGCTTGCAG
GCACACGTCTGAACTCTTTCCCTACACGACGCTCTTCCGATCTNNNGCTTTTCACAAGTAAACAGGCC
AGCACACGTCTGAACTCTTTCCCTACACGACGCTCTTCCGATCTNNNTTGTAGAAAGGGTTGGCAGA
GCACACGTCTGAACTCTTTCCCTACACGACGCTCTTCCGATCTNNNACACATTGATTTAGAGCTGGGG
AGCACACGTCTGAACTCTTTCCCTACACGACGCTCTTCCGATCTNNNGGGCTGTAAGGTTTCTAGGC
GAGCACACGTCTGAACTCTTTCCCTACACGACGCTCTTCCGATCTNNNACATCCATTGAACACCAAGT
GCACACGTCTGAACTCTTTCCCTACACGACGCTCTTCCGATCTNNNAACAGCAGCACAGAAACAAAAC
GCACACGTCTGAACTCTTTCCCTACACGACGCTCTTCCGATCTNNNAGCCAACAGAACATTGCATTCT
AGCACACGTCTGAACTCTTTCCCTACACGACGCTCTTCCGATCTNNNGGAGAAGGGAAATCAGCCA
GCACACGTCTGAACTCTTTCCCTACACGACGCTCTTCCGATCTNNNACAACACTACCCTCCTCTCTTCCC
AGCACACGTCTGAACTCTTTCCCTACACGACGCTCTTCCGATCTNNNCAAGAAAGGGCAAATGCTTC
GAGCACACGTCTGAACTCTTTCCCTACACGACGCTCTTCCGATCTNNNCTTCCCAGCATGTGTGAGGT
AGAGCACACGTCTGAACTCTTTCCCTACACGACGCTCTTCCGATCTNNNGAGACTGGCTTTCTGCTTC
AGCACACGTCTGAACTCTTTCCCTACACGACGCTCTTCCGATCTNNNAAATTGTGTCTAGGGCTGAG
GAAGAGCACACGTCTGAACTCTTTCCCTACACGACGCTCTTCCGATCTNNNGAGACAACAGCCTTACA
AGAGCACACGTCTGAACTCTTTCCCTACACGACGCTCTTCCGATCTNNNCACCTTACTGATGGGCTAT
GCACACGTCTGAACTCTTTCCCTACACGACGCTCTTCCGATCTNNNTGCAGACATTGGAAGGTCATC
GAGCACACGTCTGAACTCTTTCCCTACACGACGCTCTTCCGATCTNNNCCCATAGAAGCCGTTATTGG
GAGCACACGTCTGAACTCTTTCCCTACACGACGCTCTTCCGATCTNNNTGGATTGTGGTCATGGTTAC
GAGCACACGTCTGAACTCTTTCCCTACACGACGCTCTTCCGATCTNNNAAATAGATGTGTGAGCTGG
AGCACACGTCTGAACTCTTTCCCTACACGACGCTCTTCCGATCTNNNAGAGAAGGGGCTGCTGATG
AGCACACGTCTGAACTCTTTCCCTACACGACGCTCTTCCGATCTNNNAATATCTGGCTTCGTGTGAGC
AGCACACGTCTGAACTCTTTCCCTACACGACGCTCTTCCGATCTNNNTGCCACTGTAGGCTTGAGATA
GAGCACACGTCTGAACTCTTTCCCTACACGACGCTCTTCCGATCTNNNTGGTATCTGATCCAAGTACCA
GCACACGTCTGAACTCTTTCCCTACACGACGCTCTTCCGATCTNNNTGTGTCAGAGAGAGAGCAAGG
GCACACGTCTGAACTCTTTCCCTACACGACGCTCTTCCGATCTNNNAGGGACTTGGCATCCATGAATT
AGCACACGTCTGAACTCTTTCCCTACACGACGCTCTTCCGATCTNNNCATATGGCAATTTGGGGGACT
AGCACACGTCTGAACTCTTTCCCTACACGACGCTCTTCCGATCTNNNTCCAATATGTCAAGTTGCCTTT
AGCACACGTCTGAACTCTTTCCCTACACGACGCTCTTCCGATCTNNNTATTTGTGACAAGGGTAGGT
AGCACACGTCTGAACTCTTTCCCTACACGACGCTCTTCCGATCTNNNTGTGTTCAAATTACTTTGTGGT
GCACACGTCTGAACTCTTTCCCTACACGACGCTCTTCCGATCTNNNCCTTAGCTCATTTCCGCCTATCCC
AGAGCACACGTCTGAACTCTTTCCCTACACGACGCTCTTCCGATCTNNNGGAGTTCGCAGAGTGGAA
AGCACACGTCTGAACTCTTTCCCTACACGACGCTCTTCCGATCTNNNCGTGCATGTTGAACCATTAT

GAGCACACGTCTGAACTCTTTCCCTACACGACGCTCTTCCGATCTNNNCAGAAGTTTGCTGGAGGAGT
AGCACACGTCTGAACTCTTTCCCTACACGACGCTCTTCCGATCTNNNACTTGACCTATTGGACTGTTGT
AGCACACGTCTGAACTCTTTCCCTACACGACGCTCTTCCGATCTNNNAGTGCGTTATGAACCCAGCTAT
GAGCACACGTCTGAACTCTTTCCCTACACGACGCTCTTCCGATCTNNNTGCATTCCAACATTTTTCTGA
AGAGCACACGTCTGAACTCTTTCCCTACACGACGCTCTTCCGATCTNNNGTGGAGAATCAACACCCTCT
AGCACACGTCTGAACTCTTTCCCTACACGACGCTCTTCCGATCTNNNCCAGGAAAGAAGCCAGTTCAT
GAGCACACGTCTGAACTCTTTCCCTACACGACGCTCTTCCGATCTNNNTTTCTACCATGGGCACCAGT
AGAGCACACGTCTGAACTCTTTCCCTACACGACGCTCTTCCGATCTNNNGAGTAGCTAGGACCGTGGG
AGAGCACACGTCTGAACTCTTTCCCTACACGACGCTCTTCCGATCTNNNTGCAGGAGAACGTTTTAGG
GAGCACACGTCTGAACTCTTTCCCTACACGACGCTCTTCCGATCTNNNCAGTGTGAAGAGGTCATGG
GAGCACACGTCTGAACTCTTTCCCTACACGACGCTCTTCCGATCTNNNAAGGTCAGCAGGGGAGAA
AGCACACGTCTGAACTCTTTCCCTACACGACGCTCTTCCGATCTNNNCAAGCTCCTATCAGGGTTCACA
AGCACACGTCTGAACTCTTTCCCTACACGACGCTCTTCCGATCTNNNGCCTGAGCAATGCAACAGATT
GAGCACACGTCTGAACTCTTTCCCTACACGACGCTCTTCCGATCTNNNGTAGGACCTGCATGGGATA
GAGCACACGTCTGAACTCTTTCCCTACACGACGCTCTTCCGATCTNNNCAGGATTTCTTTGTTTCTGTC
AGCACACGTCTGAACTCTTTCCCTACACGACGCTCTTCCGATCTNNNGGACCATTTTAGGGCTAGACA
AGCACACGTCTGAACTCTTTCCCTACACGACGCTCTTCCGATCTNNNCCTGAGTATTGGGTGTGTTTG
AAGAGCACACGTCTGAACTCTTTCCCTACACGACGCTCTTCCGATCTNNNTGTGACCTTGGGCTTTCA
GCACACGTCTGAACTCTTTCCCTACACGACGCTCTTCCGATCTNNNAGCGGGAGATTTTCCAGGTTTT
GCACACGTCTGAACTCTTTCCCTACACGACGCTCTTCCGATCTNNNAGAAGGTTCAATGACCTCCCAG
AGCACACGTCTGAACTCTTTCCCTACACGACGCTCTTCCGATCTNNNAAGGTGATGGCCTCCATTTTC
AGCACACGTCTGAACTCTTTCCCTACACGACGCTCTTCCGATCTNNNGGTTTACCCTGGGTGTCTTCTT
AGCACACGTCTGAACTCTTTCCCTACACGACGCTCTTCCGATCTNNNTTCTGTCCCTTCCAAAAGCACAA
GAGCACACGTCTGAACTCTTTCCCTACACGACGCTCTTCCGATCTNNNAGCTTGTACGAAGAACTGAG
GAGCACACGTCTGAACTCTTTCCCTACACGACGCTCTTCCGATCTNNNAAGAATGAGATCAGGGGGA
GCACACGTCTGAACTCTTTCCCTACACGACGCTCTTCCGATCTNNNAAGTCTGTCCCACATCCTCTCTCC
AGCACACGTCTGAACTCTTTCCCTACACGACGCTCTTCCGATCTNNNCATCAGCAAACTGAACAAACC
AGCACACGTCTGAACTCTTTCCCTACACGACGCTCTTCCGATCTNNNTTCGTGCCCTCTTATTGTGGAA
GAGCACACGTCTGAACTCTTTCCCTACACGACGCTCTTCCGATCTNNNCCTGGCTCATGGATGATGACA
AGCACACGTCTGAACTCTTTCCCTACACGACGCTCTTCCGATCTNNNTGAAACCAAATTATGCGACCAG
AGAGCACACGTCTGAACTCTTTCCCTACACGACGCTCTTCCGATCTNNNTCATCTAGGAGTGGGGAG
AGCACACGTCTGAACTCTTTCCCTACACGACGCTCTTCCGATCTNNNTGCCCTGTTGCATTAGTTCTTC
GAGCACACGTCTGAACTCTTTCCCTACACGACGCTCTTCCGATCTNNNGTACTACAGCCTGGGCGAGT
AGCACACGTCTGAACTCTTTCCCTACACGACGCTCTTCCGATCTNNNTCTTCTCTATGGGAATCACCGG
AGCACACGTCTGAACTCTTTCCCTACACGACGCTCTTCCGATCTNNNTACCCAGCCAATGAGGACACAT
GAGCACACGTCTGAACTCTTTCCCTACACGACGCTCTTCCGATCTNNNAGACACACACTGTCTAATAG
AGCACACGTCTGAACTCTTTCCCTACACGACGCTCTTCCGATCTNNNTTCCATGTGAACTTGACTTTT
AGCACACGTCTGAACTCTTTCCCTACACGACGCTCTTCCGATCTNNNAAGACCTGCCACATTAGGGAA
GAGCACACGTCTGAACTCTTTCCCTACACGACGCTCTTCCGATCTNNNCTTCTGGAGAGGCAAGAACT
AGAGCACACGTCTGAACTCTTTCCCTACACGACGCTCTTCCGATCTNNNAAAGAAGCGTAATAGCTGG
AGCACACGTCTGAACTCTTTCCCTACACGACGCTCTTCCGATCTNNNAACAACAAACTACTGATGAGG
AGAGCACACGTCTGAACTCTTTCCCTACACGACGCTCTTCCGATCTNNNTAGCTTACTTCCCCTGTAGC
GAGCACACGTCTGAACTCTTTCCCTACACGACGCTCTTCCGATCTNNNACGCAAATTCCACTAGGTACT

GAGCACACGTCTGAACTCTTTCCCTACACGACGCTCTTCCGATCTNNNTAAGGGAGAAAAGCAAGTGC
GCACACGTCTGAACTCTTTCCCTACACGACGCTCTTCCGATCTNNNCTGCAGTTTGGATAAGGCAGGAT
AGCACACGTCTGAACTCTTTCCCTACACGACGCTCTTCCGATCTNNNTCCAGAGGTGGCTAGAGTTGA
AGAGCACACGTCTGAACTCTTTCCCTACACGACGCTCTTCCGATCTNNNTTCAGGATCACACAGCAAGT
AGCACACGTCTGAACTCTTTCCCTACACGACGCTCTTCCGATCTNNNGAAGCAAAGCTGACATTGGAA
AGCACACGTCTGAACTCTTTCCCTACACGACGCTCTTCCGATCTNNNTCAAGGTGTCAAATCCCACGTTT
GAGCACACGTCTGAACTCTTTCCCTACACGACGCTCTTCCGATCTNNNCACTGGATTGCAAGTAGGCTT
GAGCACACGTCTGAACTCTTTCCCTACACGACGCTCTTCCGATCTNNNTATCAGACCCTTTCGACACACA
AGCACACGTCTGAACTCTTTCCCTACACGACGCTCTTCCGATCTNNNACATGTGTGTACATATACTTCTG
GAGCACACGTCTGAACTCTTTCCCTACACGACGCTCTTCCGATCTNNNCAAAAGGAGACACTCATGGCA
AGCACACGTCTGAACTCTTTCCCTACACGACGCTCTTCCGATCTNNNTGGATGTTTGATAGGATTGCA
AGCACACGTCTGAACTCTTTCCCTACACGACGCTCTTCCGATCTNNNAGGCCCACATCATTCATTTCCATC
GCACACGTCTGAACTCTTTCCCTACACGACGCTCTTCCGATCTNNNGGTTCGTTTTACGTTCCGTGAAG
AGCACACGTCTGAACTCTTTCCCTACACGACGCTCTTCCGATCTNNNACCCCTACTGTTTTAGTTTGGGC
AGCACACGTCTGAACTCTTTCCCTACACGACGCTCTTCCGATCTNNNACAAACTATTTGAGAGTGAGTTC
AGCACACGTCTGAACTCTTTCCCTACACGACGCTCTTCCGATCTNNNCTCCTACAGAGCAGTTCACTGGG
AGCACACGTCTGAACTCTTTCCCTACACGACGCTCTTCCGATCTNNNGGCATTTACAGATCATCCAGCC
GCACACGTCTGAACTCTTTCCCTACACGACGCTCTTCCGATCTNNNGGCCAAGCTAGTGATTGATGTGA
AGCACACGTCTGAACTCTTTCCCTACACGACGCTCTTCCGATCTNNNGACCAAGATGTGGAACAAGTGC
AGAGCACACGTCTGAACTCTTTCCCTACACGACGCTCTTCCGATCTNNNACTCATCGTCATGACAACCAC
AGAGCACACGTCTGAACTCTTTCCCTACACGACGCTCTTCCGATCTNNNAGCACATCATGGGACTTCTC
GAGCACACGTCTGAACTCTTTCCCTACACGACGCTCTTCCGATCTNNNCAGAGAAGTTAGGCTTGAAG
AGAGCACACGTCTGAACTCTTTCCCTACACGACGCTCTTCCGATCTNNNTGAGCAAAGGAATGAATGAC
AGAGCACACGTCTGAACTCTTTCCCTACACGACGCTCTTCCGATCTNNNCTGGAGGAAACCTGCAAAAC
AGCACACGTCTGAACTCTTTCCCTACACGACGCTCTTCCGATCTNNNGCTCTTGCCCTCTTCTCTGGTTAC
GAGCACACGTCTGAACTCTTTCCCTACACGACGCTCTTCCGATCTNNNGAGCCTGTGCCCATGAGTATA
AGCACACGTCTGAACTCTTTCCCTACACGACGCTCTTCCGATCTNNNCGCCCCTTCACATAAGGGATATG
AGCACACGTCTGAACTCTTTCCCTACACGACGCTCTTCCGATCTNNNTGACATAGGGCCTGTGAGAAAG
AGCACACGTCTGAACTCTTTCCCTACACGACGCTCTTCCGATCTNNNCCAAATGCTGTCGTTTTTGCCTCA
GAGCACACGTCTGAACTCTTTCCCTACACGACGCTCTTCCGATCTNNNGTGCCTGTGCAAAAATGTTCC
GCACACGTCTGAACTCTTTCCCTACACGACGCTCTTCCGATCTNNNCCCTGGAGCAAAGACTACAACAG
GAGCACACGTCTGAACTCTTTCCCTACACGACGCTCTTCCGATCTNNNTTTGGGCCTCCAGGAATACAAT
GAGCACACGTCTGAACTCTTTCCCTACACGACGCTCTTCCGATCTNNNGTGAAAAATGTCAAAGTGCA
GAGCACACGTCTGAACTCTTTCCCTACACGACGCTCTTCCGATCTNNNTTCCAACAGTGAACCCCTGTTTC
AGAGCACACGTCTGAACTCTTTCCCTACACGACGCTCTTCCGATCTNNNCCAAAGAATTCTGAACCAACA
AGAGCACACGTCTGAACTCTTTCCCTACACGACGCTCTTCCGATCTNNNCCACGGTTGTCTCAATCTCTC
AGCACACGTCTGAACTCTTTCCCTACACGACGCTCTTCCGATCTNNNTGGAGAAAGAGAAGCCATAGC
AGAGCACACGTCTGAACTCTTTCCCTACACGACGCTCTTCCGATCTNNNTCACTATGTTGGTGAGGGAT
GAGCACACGTCTGAACTCTTTCCCTACACGACGCTCTTCCGATCTNNNCTCAGAATGCTACCATAATTCA
AGCACACGTCTGAACTCTTTCCCTACACGACGCTCTTCCGATCTNNNGCCTGTGGTGTGTTAGAGAGAA
GCACACGTCTGAACTCTTTCCCTACACGACGCTCTTCCGATCTNNNCCCACAGGTTTTAGCTCCCTTTTCA
AGCACACGTCTGAACTCTTTCCCTACACGACGCTCTTCCGATCTNNNAAATGAACTCACAAGGTTCCAG
GAGCACACGTCTGAACTCTTTCCCTACACGACGCTCTTCCGATCTNNNAGACTGCTTTCGAACTGTAGCT

AAGAGCACACGTCTGAACTCTTTCCCTACACGACGCTCTTCCGATCTNNNATGGGAGAGAAAACACTA
GAGCACACGTCTGAACTCTTTCCCTACACGACGCTCTTCCGATCTNNNGCAAGAGGTAGCACAAAACA
GAGCACACGTCTGAACTCTTTCCCTACACGACGCTCTTCCGATCTNNNGCCAAATAAAATAAAAAGCAG
GCACACGTCTGAACTCTTTCCCTACACGACGCTCTTCCGATCTNNNCACAATAGGGCCAGGTAACAGA
GAGCACACGTCTGAACTCTTTCCCTACACGACGCTCTTCCGATCTNNNGGAGGCTGTTGGCTTCTGTAT
GAGCACACGTCTGAACTCTTTCCCTACACGACGCTCTTCCGATCTNNNTTCTCACTCCTGGACTACAGCA
GAAGAGCACACGTCTGAACTCTTTCCCTACACGACGCTCTTCCGATCTNNNATGGGGGAACCAGTTACA
AGCACACGTCTGAACTCTTTCCCTACACGACGCTCTTCCGATCTNNNTCCTCAGGGTCCCCTTTATTCAT
GAGCACACGTCTGAACTCTTTCCCTACACGACGCTCTTCCGATCTNNNAGGCCACTCACATCACACAAT
AGAGCACACGTCTGAACTCTTTCCCTACACGACGCTCTTCCGATCTNNNGGCTGTTTAGTGTTCAACCCA
AGCACACGTCTGAACTCTTTCCCTACACGACGCTCTTCCGATCTNNNTTTCGTTAGTCTAGCTAGTGGT
AGCACACGTCTGAACTCTTTCCCTACACGACGCTCTTCCGATCTNNNTCAATCTCCTGCTTCTCAGCTTCC
GAGCACACGTCTGAACTCTTTCCCTACACGACGCTCTTCCGATCTNNNATTGAAGAGATGGAGCTCAG
AGCACACGTCTGAACTCTTTCCCTACACGACGCTCTTCCGATCTNNNGCTGTGGTCTTCTGCTCCTAAA
AGCACACGTCTGAACTCTTTCCCTACACGACGCTCTTCCGATCTNNNAGAGGAGATAGTGAGGGTCA
AAGAGCACACGTCTGAACTCTTTCCCTACACGACGCTCTTCCGATCTNNNGAGAGGGTTGCTCAGTAG
AGCACACGTCTGAACTCTTTCCCTACACGACGCTCTTCCGATCTNNNGAGCTAATTCCACTCCTGACAGC
GCACACGTCTGAACTCTTTCCCTACACGACGCTCTTCCGATCTNNNTCCTCAAGCCATGTCCTCAATTCCA
GCACACGTCTGAACTCTTTCCCTACACGACGCTCTTCCGATCTNNNGCTGGGAGGTGGATAACTAGGA
GAGCACACGTCTGAACTCTTTCCCTACACGACGCTCTTCCGATCTNNNAAGTGCAGTGAACTAGGCCGT
AGCACACGTCTGAACTCTTTCCCTACACGACGCTCTTCCGATCTNNNAACTCACGTGTGCCAGAACTAT
AGCACACGTCTGAACTCTTTCCCTACACGACGCTCTTCCGATCTNNNTCCAGGATTTCAACATTCCAGGA
AGCACACGTCTGAACTCTTTCCCTACACGACGCTCTTCCGATCTNNNTTTGTCTGCCTCTGTCACACATC
AGCACACGTCTGAACTCTTTCCCTACACGACGCTCTTCCGATCTNNNACGATGACTGGATAAAAACAAT
GAGCACACGTCTGAACTCTTTCCCTACACGACGCTCTTCCGATCTNNNTTCCTCCTCACACTAGACAAG
AGCACACGTCTGAACTCTTTCCCTACACGACGCTCTTCCGATCTNNNGGGTCTCCATCTTGCAGACTAT
GAGCACACGTCTGAACTCTTTCCCTACACGACGCTCTTCCGATCTNNNGGAGGAACTACTAATTGCAC
GAGCACACGTCTGAACTCTTTCCCTACACGACGCTCTTCCGATCTNNNGTCTCCCGAGTTCTAAGCGAT
AGCACACGTCTGAACTCTTTCCCTACACGACGCTCTTCCGATCTNNNCATCCTGGTGGGTACAGAGTTT
AGCACACGTCTGAACTCTTTCCCTACACGACGCTCTTCCGATCTNNNTAGGATTGCTCACTGTTGGGAT
GAGCACACGTCTGAACTCTTTCCCTACACGACGCTCTTCCGATCTNNNATGCCCCAGAGAGAGGTAGG
AGCACACGTCTGAACTCTTTCCCTACACGACGCTCTTCCGATCTNNNAAGGTGTGTGCCTTTTACTGTG
GAGCACACGTCTGAACTCTTTCCCTACACGACGCTCTTCCGATCTNNNGCAGACTTGTAGATACCACCT
GAGCACACGTCTGAACTCTTTCCCTACACGACGCTCTTCCGATCTNNNTTGCCTTTGTTTTCCTCCTCC
GAGCACACGTCTGAACTCTTTCCCTACACGACGCTCTTCCGATCTNNNTTCTAGTGCCAAAATGCTGTG
AGCACACGTCTGAACTCTTTCCCTACACGACGCTCTTCCGATCTNNNGCCCCTGGTGTAGAGGAAAAT
AGCACACGTCTGAACTCTTTCCCTACACGACGCTCTTCCGATCTNNNAGTGGTCAGAAAGCACAAGTG
GCACACGTCTGAACTCTTTCCCTACACGACGCTCTTCCGATCTNNNTACCAGTCTTGACCCCAAACAAT
AGCACACGTCTGAACTCTTTCCCTACACGACGCTCTTCCGATCTNNNTGGAGAAAACATAGCCCTATTA
GAGCACACGTCTGAACTCTTTCCCTACACGACGCTCTTCCGATCTNNNAACGTACAAGGGCTCCAATGT
AGAGCACACGTCTGAACTCTTTCCCTACACGACGCTCTTCCGATCTNNNACGTGCCAAAGACATATTT
AGCACACGTCTGAACTCTTTCCCTACACGACGCTCTTCCGATCTNNNCAGAGCCTGACAAGGATTCAGA
AGCACACGTCTGAACTCTTTCCCTACACGACGCTCTTCCGATCTNNNTGGATTTTCAATGCTATGCCAG

AGCACACGTCTGAACTCTTTCCCTACACGACGCTCTTCCGATCTNNNGTGTCTCCTGTTACCAACCGA
AGCACACGTCTGAACTCTTTCCCTACACGACGCTCTTCCGATCTNNNAAATTCCTGTCAGTCCTCCAAG
AAGAGCACACGTCTGAACTCTTTCCCTACACGACGCTCTTCCGATCTNNNGCGACACCCTGTCCCTCTA
GAGCACACGTCTGAACTCTTTCCCTACACGACGCTCTTCCGATCTNNNCCTTACTGGTGTCATGAGCA
AGCACACGTCTGAACTCTTTCCCTACACGACGCTCTTCCGATCTNNNGGGTTATTGTGAGGACTGA
AGCACACGTCTGAACTCTTTCCCTACACGACGCTCTTCCGATCTNNNGCTTTACAAACCAGAAATGA
GAGCACACGTCTGAACTCTTTCCCTACACGACGCTCTTCCGATCTNNNACACACTGGTAGTCTCTGTA
AGCACACGTCTGAACTCTTTCCCTACACGACGCTCTTCCGATCTNNNTTGCTGAATCCAAACCCATTTG
AGCACACGTCTGAACTCTTTCCCTACACGACGCTCTTCCGATCTNNNACTTGGAAAGAAACCAAACAT
GCACACGTCTGAACTCTTTCCCTACACGACGCTCTTCCGATCTNNNCAAAGTGTGGAAGCACTTAGG
AGCACACGTCTGAACTCTTTCCCTACACGACGCTCTTCCGATCTNNNGTGTGGGGGAGTGTATTAG
ACACGTCTGAACTCTTTCCCTACACGACGCTCTTCCGATCTNNNAGAATTTTAGGTTGGTGGGCCTTC
AGCACACGTCTGAACTCTTTCCCTACACGACGCTCTTCCGATCTNNNACCATGCCCAGCCACTTATTA
AGAGCACACGTCTGAACTCTTTCCCTACACGACGCTCTTCCGATCTNNNGACCCTATTCTCCTGCTGCA
AGCACACGTCTGAACTCTTTCCCTACACGACGCTCTTCCGATCTNNNATGAACCATGGCAGGAGAAT
GAGCACACGTCTGAACTCTTTCCCTACACGACGCTCTTCCGATCTNNNTAGCTGTGCTTTCAGTTGAG
AGCACACGTCTGAACTCTTTCCCTACACGACGCTCTTCCGATCTNNNTAAGTGCTCCAGAAACCTACC
GCACACGTCTGAACTCTTTCCCTACACGACGCTCTTCCGATCTNNNAACCCAGCACCTCATGTCTAATG
AGCACACGTCTGAACTCTTTCCCTACACGACGCTCTTCCGATCTNNNAAAAGAAATAAGACACCTCC
GAGCACACGTCTGAACTCTTTCCCTACACGACGCTCTTCCGATCTNNNTGCCAACGTATACCCATTTG
GCACACGTCTGAACTCTTTCCCTACACGACGCTCTTCCGATCTNNNAGGAAAGCTGAGAAGCATGAG
GCACACGTCTGAACTCTTTCCCTACACGACGCTCTTCCGATCTNNNAATTCTAGACATTGGTCTCGCC
AGAGCACACGTCTGAACTCTTTCCCTACACGACGCTCTTCCGATCTNNNAGTCATGCTTCTGTTATAG
AGCACACGTCTGAACTCTTTCCCTACACGACGCTCTTCCGATCTNNNAAGGAACTGGCATTACTGGC
GAGCACACGTCTGAACTCTTTCCCTACACGACGCTCTTCCGATCTNNNACTCCAAGTTTTCTCCGAAGA
GAGCACACGTCTGAACTCTTTCCCTACACGACGCTCTTCCGATCTNNNGTGGTTGAAGTGAAAACGA
AGCACACGTCTGAACTCTTTCCCTACACGACGCTCTTCCGATCTNNNTGGGGAAAGTGATCCTGTCA
GAGCACACGTCTGAACTCTTTCCCTACACGACGCTCTTCCGATCTNNNCCACTAATACAGGCTTGCCC
AGCACACGTCTGAACTCTTTCCCTACACGACGCTCTTCCGATCTNNNTCCAGGACACTAAGCAAAACA
GCACACGTCTGAACTCTTTCCCTACACGACGCTCTTCCGATCTNNNGCCCTTCTTTCTTGTTCCCTTCAC
GCACACGTCTGAACTCTTTCCCTACACGACGCTCTTCCGATCTNNNTTGGATGAGTGGGCACATAAA
AGAGCACACGTCTGAACTCTTTCCCTACACGACGCTCTTCCGATCTNNNGTCTTGAGCCTGTCTCGAA
AGCACACGTCTGAACTCTTTCCCTACACGACGCTCTTCCGATCTNNNACTTGTTCATCCGTATTGTGG
GCACACGTCTGAACTCTTTCCCTACACGACGCTCTTCCGATCTNNNCCTGACTGGTAGAAAACTCCAG
AGCACACGTCTGAACTCTTTCCCTACACGACGCTCTTCCGATCTNNNCTGAGTGTAGAGCTTGGAAA
GAGCACACGTCTGAACTCTTTCCCTACACGACGCTCTTCCGATCTNNNCTGCCGGAGAAAAGGAACC
AGCACACGTCTGAACTCTTTCCCTACACGACGCTCTTCCGATCTNNNTACCCACAGCTAACAACATACC
AGCACACGTCTGAACTCTTTCCCTACACGACGCTCTTCCGATCTNNNCATGTCAGCTCATGCCTGTAA
AAGAGCACACGTCTGAACTCTTTCCCTACACGACGCTCTTCCGATCTNNNGCCATCCCCAGACTTCCC
GCACACGTCTGAACTCTTTCCCTACACGACGCTCTTCCGATCTNNNAAGTTCCTTTGTGCTTCCTGTGT
AGCACACGTCTGAACTCTTTCCCTACACGACGCTCTTCCGATCTNNNAATATCTATGAGTTGGGGCCA
AGCACACGTCTGAACTCTTTCCCTACACGACGCTCTTCCGATCTNNNGATAATCCAAAGGCTTCCCCAG
CACACGTCTGAACTCTTTCCCTACACGACGCTCTTCCGATCTNNNTTGAAGGCTTGATATGGTTTGG

AAGAGCACACGTCTGAACTCTTTCCCTACACGACGCTCTTCCGATCTNNNTGTTTTCCAAAGCTACTGT

GAGCACACGTCTGAACTCTTTCCCTACACGACGCTCTTCCGATCTNNNTTCCATAAGGTCTGAGTGTGT

GCACACGTCTGAACTCTTTCCCTACACGACGCTCTTCCGATCTNNNAGCATTTACTGGTGCTCATGTCA

AGCACACGTCTGAACTCTTTCCCTACACGACGCTCTTCCGATCTNNNTCCACAATTGCATGTCAGTTCCT

GCACACGTCTGAACTCTTTCCCTACACGACGCTCTTCCGATCTNNNCAGAAGAACCGCTTTGCTTGAAC

GAGCACACGTCTGAACTCTTTCCCTACACGACGCTCTTCCGATCTNNNCCCAGACACTGGACTATAGCG

AGAGCACACGTCTGAACTCTTTCCCTACACGACGCTCTTCCGATCTNNNATGGTTCCACATGAAAGCCA

AGAGCACACGTCTGAACTCTTTCCCTACACGACGCTCTTCCGATCTNNNCCATCAGCTTTGTGCTAGGT

GAGCACACGTCTGAACTCTTTCCCTACACGACGCTCTTCCGATCTNNNAAATGGCCACAGCAACTACAG

AAGAGCACACGTCTGAACTCTTTCCCTACACGACGCTCTTCCGATCTNNNGTAGAGGCAGAGAGAACA

AGAGCACACGTCTGAACTCTTTCCCTACACGACGCTCTTCCGATCTNNNTGAAGGTACATCCTCAGAA

GAGCACACGTCTGAACTCTTTCCCTACACGACGCTCTTCCGATCTNNNTCAACCTCGCCAAAAATCAAA

GAGCACACGTCTGAACTCTTTCCCTACACGACGCTCTTCCGATCTNNNCAGATGGTTATGTTCCTCCCT

AGCACACGTCTGAACTCTTTCCCTACACGACGCTCTTCCGATCTNNNTGGGACAAATCATGCACCTGTA

AGAGCACACGTCTGAACTCTTTCCCTACACGACGCTCTTCCGATCTNNNCCCTCCTTTCCTAGCTGTCTCT

GAGCACACGTCTGAACTCTTTCCCTACACGACGCTCTTCCGATCTNNNAAAAGGAGAGAGAGCGGCAA

GAGCACACGTCTGAACTCTTTCCCTACACGACGCTCTTCCGATCTNNNGCCCGTGGATATTATCTCCTT

GAGCACACGTCTGAACTCTTTCCCTACACGACGCTCTTCCGATCTNNNTGGGATTTGACTTAGGTGATC

AGCACACGTCTGAACTCTTTCCCTACACGACGCTCTTCCGATCTNNNGGCAAGTGACAGTGAGGAAGA

GAGCACACGTCTGAACTCTTTCCCTACACGACGCTCTTCCGATCTNNNAATGGAGGTGGAAGCTTGCC

GAGCACACGTCTGAACTCTTTCCCTACACGACGCTCTTCCGATCTNNNATGGACTGGTTTTTCTTTCAG

AAGAGCACACGTCTGAACTCTTTCCCTACACGACGCTCTTCCGATCTNNNCTAGCACAGAGCTTAGAAC

AGCACACGTCTGAACTCTTTCCCTACACGACGCTCTTCCGATCTNNNCCCAAACACTGGGAGACAGATT

GCACACGTCTGAACTCTTTCCCTACACGACGCTCTTCCGATCTNNNTCGACCATTCCAGTATCTCCTACT

AGAGCACACGTCTGAACTCTTTCCCTACACGACGCTCTTCCGATCTNNNTCCCTTCAGGTCCAAACAGA

GAGCACACGTCTGAACTCTTTCCCTACACGACGCTCTTCCGATCTNNNCCCCCATCAAAATTTATCCATC

AGCACACGTCTGAACTCTTTCCCTACACGACGCTCTTCCGATCTNNNCCATGACAGGATGTAGTGACCA

AGCACACGTCTGAACTCTTTCCCTACACGACGCTCTTCCGATCTNNNAGTTGAGCTACGGAGACAGA

AGCACACGTCTGAACTCTTTCCCTACACGACGCTCTTCCGATCTNNNCCCAAATCAAACACTCTAGCCCA

AGAGCACACGTCTGAACTCTTTCCCTACACGACGCTCTTCCGATCTNNNAAAGTCCAAATCTCTTGCAAT

GAGCACACGTCTGAACTCTTTCCCTACACGACGCTCTTCCGATCTNNNACAAAACTTCTAATCCCGCCCT

GAGCACACGTCTGAACTCTTTCCCTACACGACGCTCTTCCGATCTNNNGGATTTTCAGAACCAACTGG

GCACACGTCTGAACTCTTTCCCTACACGACGCTCTTCCGATCTNNNACAAAATGCCTCAGGTCCTTCAG

AGCACACGTCTGAACTCTTTCCCTACACGACGCTCTTCCGATCTNNNTCTGAGAGCTGCAGTTAGAGG

AGAGCACACGTCTGAACTCTTTCCCTACACGACGCTCTTCCGATCTNNNTCCTAGGTAGGAGTTTTGG

AGAGCACACGTCTGAACTCTTTCCCTACACGACGCTCTTCCGATCTNNNAAAGAAACTCAGGGTCCTCA

AGCACACGTCTGAACTCTTTCCCTACACGACGCTCTTCCGATCTNNNAAAATGGAGATGGTGTGTGGT

AGCACACGTCTGAACTCTTTCCCTACACGACGCTCTTCCGATCTNNNGGAGCTATGGCACAGATACTG

GCACACGTCTGAACTCTTTCCCTACACGACGCTCTTCCGATCTNNNAGTACTGATGAAGATCACCACG

AGCACACGTCTGAACTCTTTCCCTACACGACGCTCTTCCGATCTNNNGCTGTACCAAGTGATAGGGTG

GAGCACACGTCTGAACTCTTTCCCTACACGACGCTCTTCCGATCTNNNAAGCAACAGTGGTGTTTTCCA

AGCACACGTCTGAACTCTTTCCCTACACGACGCTCTTCCGATCTNNNACCTCCAAACCTCCTTTACCTCACA

AAGAGCACACGTCTGAACTCTTTCCCTACACGACGCTCTTCCGATCTNNNCAGGATTTCATTCCATTGTA

AGCACACGTCTGAACTCTTTCCCTACACGACGCTCTTCCGATCTNNNCACTGGCCTTCTGTTTCTCTCTC
AGAGCACACGTCTGAACTCTTTCCCTACACGACGCTCTTCCGATCTNNNCAAGAGGGAGGGACAGAG
AGCACACGTCTGAACTCTTTCCCTACACGACGCTCTTCCGATCTNNNCATCTTGCAAAAGCTGCAGTA
GAGCACACGTCTGAACTCTTTCCCTACACGACGCTCTTCCGATCTNNNCGTGCCTTTTGATAATGCAG
GAGCACACGTCTGAACTCTTTCCCTACACGACGCTCTTCCGATCTNNNTGGAGGTACCAACCTATCCA
AGAGCACACGTCTGAACTCTTTCCCTACACGACGCTCTTCCGATCTNNNTCTATACCAGGGTTCATTCC
CACACGTCTGAACTCTTTCCCTACACGACGCTCTTCCGATCTNNNGAGACTGAGGTGAGAGTTTTGC
GAGCACACGTCTGAACTCTTTCCCTACACGACGCTCTTCCGATCTNNNAGTATTGGCAAGAGAAGTG
AGCACACGTCTGAACTCTTTCCCTACACGACGCTCTTCCGATCTNNNGCTGCTATTTGAGAAACTGTG
GCACACGTCTGAACTCTTTCCCTACACGACGCTCTTCCGATCTNNNACAGTGACACAACACACACATC
AGCACACGTCTGAACTCTTTCCCTACACGACGCTCTTCCGATCTNNNACATTGACCTGTGCCACATAC
AGCACACGTCTGAACTCTTTCCCTACACGACGCTCTTCCGATCTNNNCATGTAACTCCAGGCAGCAAT
GCACACGTCTGAACTCTTTCCCTACACGACGCTCTTCCGATCTNNNACACTGCCTTTGACATACTGCAT
GAGCACACGTCTGAACTCTTTCCCTACACGACGCTCTTCCGATCTNNNCAGTCTGAAGGGCACATCTT
AGAGCACACGTCTGAACTCTTTCCCTACACGACGCTCTTCCGATCTNNNTGGCCATTGACTAGGTCAA
AAGAGCACACGTCTGAACTCTTTCCCTACACGACGCTCTTCCGATCTNNNTCTTGAGACAGCTCAAAAT
GCACACGTCTGAACTCTTTCCCTACACGACGCTCTTCCGATCTNNNGAGCAACACTCATTCTCCCATGA
CACACGTCTGAACTCTTTCCCTACACGACGCTCTTCCGATCTNNNTGCATAGGCTTTGTATGGTAGG
GAGCACACGTCTGAACTCTTTCCCTACACGACGCTCTTCCGATCTNNNCCAGAAGAAGCAGTTTGCA
GAGCACACGTCTGAACTCTTTCCCTACACGACGCTCTTCCGATCTNNNTACCTGAGCACTCCTTCATTT
CACACGTCTGAACTCTTTCCCTACACGACGCTCTTCCGATCTNNNCTTCTCCCTTAGAACCGCGTGCAT
AGCACACGTCTGAACTCTTTCCCTACACGACGCTCTTCCGATCTNNNAGTCTGTTGCCAAACTTTTCAT
AGCACACGTCTGAACTCTTTCCCTACACGACGCTCTTCCGATCTNNNGAATCACTAGAGCCCACAAGG
GCACACGTCTGAACTCTTTCCCTACACGACGCTCTTCCGATCTNNNACCTATAAGACGGTGTGCTAAT
AGAGCACACGTCTGAACTCTTTCCCTACACGACGCTCTTCCGATCTNNNTGATGACATCCCTCTCAACG
AGCACACGTCTGAACTCTTTCCCTACACGACGCTCTTCCGATCTNNNGGCATCTTTTGAGTGGGAACT
GCACACGTCTGAACTCTTTCCCTACACGACGCTCTTCCGATCTNNNGCAGGCTTACATGCCTGTTTGA
AGCACACGTCTGAACTCTTTCCCTACACGACGCTCTTCCGATCTNNNACATGTTGCCGAACAGACAT
AAGAGCACACGTCTGAACTCTTTCCCTACACGACGCTCTTCCGATCTNNNAGAGTAGCACCCAGGAAT
AGAGCACACGTCTGAACTCTTTCCCTACACGACGCTCTTCCGATCTNNNATCAGAGAGTGGAGAAAG
AGCACACGTCTGAACTCTTTCCCTACACGACGCTCTTCCGATCTNNNCGTGCCCAGCCTCATAGATATT
GCACACGTCTGAACTCTTTCCCTACACGACGCTCTTCCGATCTNNNCCATGTCAATTCCTCTCACTGGG
GCACACGTCTGAACTCTTTCCCTACACGACGCTCTTCCGATCTNNNAAGACGGATCTTCTGTCCTCTGG
CACACGTCTGAACTCTTTCCCTACACGACGCTCTTCCGATCTNNNACTGTGCAGCAAAGAAGACTGAA
AGAGCACACGTCTGAACTCTTTCCCTACACGACGCTCTTCCGATCTNNNGTTGTTAGAGCATAAGCTG
AGCACACGTCTGAACTCTTTCCCTACACGACGCTCTTCCGATCTNNNTTTGCCCTTTTGGAGCCATAGT
CACACGTCTGAACTCTTTCCCTACACGACGCTCTTCCGATCTNNNCCTTCAAACCTCCACTCCAAGTTC
AGCACACGTCTGAACTCTTTCCCTACACGACGCTCTTCCGATCTNNNGAGGTACTGCCTTGGTGATAA
AGCACACGTCTGAACTCTTTCCCTACACGACGCTCTTCCGATCTNNNAGATTCATATTTCCCAATAAG
CACACGTCTGAACTCTTTCCCTACACGACGCTCTTCCGATCTNNNATCGTTTTCCGATGGAAGAGGTC
AGAGCACACGTCTGAACTCTTTCCCTACACGACGCTCTTCCGATCTNNNCACTGCAATCATGGGATTCT
AAGAGCACACGTCTGAACTCTTTCCCTACACGACGCTCTTCCGATCTNNNTTGATCGTGAGTTTGCTCA
AGCACACGTCTGAACTCTTTCCCTACACGACGCTCTTCCGATCTNNNTGTGGTGGCTAGAAACTAGGA

AGAGCACACGTCTGAACTCTTTCCCTACACGACGCTCTTCCGATCTNNNCATTCCCTATGCTGCCTAAT
GAGCACACGTCTGAACTCTTTCCCTACACGACGCTCTTCCGATCTNNNGAACTCACTGAGCCAGAACA
AGCACACGTCTGAACTCTTTCCCTACACGACGCTCTTCCGATCTNNNGGGAAAGCGGGAAAAACAAA
AGCACACGTCTGAACTCTTTCCCTACACGACGCTCTTCCGATCTNNNCCATGAGGACAGTAGCAATCC
AGCACACGTCTGAACTCTTTCCCTACACGACGCTCTTCCGATCTNNNTGTAGGACACCGTCATGGTAC
GAGCACACGTCTGAACTCTTTCCCTACACGACGCTCTTCCGATCTNNNTCAGGCATGCTCAAACAGTA
GCACACGTCTGAACTCTTTCCCTACACGACGCTCTTCCGATCTNNNCTCAGAAGCACTACAAGAAGCG
GAGCACACGTCTGAACTCTTTCCCTACACGACGCTCTTCCGATCTNNNTGCTGGAGGCAAATCACATA
AGAGCACACGTCTGAACTCTTTCCCTACACGACGCTCTTCCGATCTNNNAAGTTGGGATGCAGGGTAC
GAGCACACGTCTGAACTCTTTCCCTACACGACGCTCTTCCGATCTNNNCAGAGCCGAGATCACGTTCA
AGCACACGTCTGAACTCTTTCCCTACACGACGCTCTTCCGATCTNNNTGAGAGCACCTACTAGAAGCC
AGCACACGTCTGAACTCTTTCCCTACACGACGCTCTTCCGATCTNNNTGTCTGAGAGTAGGCCGTAAG
GAGCACACGTCTGAACTCTTTCCCTACACGACGCTCTTCCGATCTNNNTCTAGCCCCTCCATGATTTATG
AGCACACGTCTGAACTCTTTCCCTACACGACGCTCTTCCGATCTNNNCCAACTTTTTGTCTTGTTGGTC
GCACACGTCTGAACTCTTTCCCTACACGACGCTCTTCCGATCTNNNCTCTGCAATTTCCCTGGAACTGA
GAGCACACGTCTGAACTCTTTCCCTACACGACGCTCTTCCGATCTNNNAGGTAAAAGCTCAAGAAAG
GAGCACACGTCTGAACTCTTTCCCTACACGACGCTCTTCCGATCTNNNAACAGCACTGCAAAAGCATGA
AGCACACGTCTGAACTCTTTCCCTACACGACGCTCTTCCGATCTNNNAGTCTGTCTCTCACTGCCATTCT
GAGCACACGTCTGAACTCTTTCCCTACACGACGCTCTTCCGATCTNNNCAGGACCCTTTTGAGAGTGAA
GAAGAGCACACGTCTGAACTCTTTCCCTACACGACGCTCTTCCGATCTNNNACAATCAGCCCTCTATAA
GAGCACACGTCTGAACTCTTTCCCTACACGACGCTCTTCCGATCTNNNCCCTGGAATGTTAGTTGCAGC
AGCACACGTCTGAACTCTTTCCCTACACGACGCTCTTCCGATCTNNNCAGAGAAAACAACCGGAGAG
AAGAGCACACGTCTGAACTCTTTCCCTACACGACGCTCTTCCGATCTNNNTGAGATCTCCGCAGGAAC
AAGAGCACACGTCTGAACTCTTTCCCTACACGACGCTCTTCCGATCTNNNTGGGGGAAAATTCCCAAC
AGCACACGTCTGAACTCTTTCCCTACACGACGCTCTTCCGATCTNNNGGTGTTTAGGGGTGCTTGTTG
GAAGAGCACACGTCTGAACTCTTTCCCTACACGACGCTCTTCCGATCTNNNGGCCCATCATTTCTTGTG
AGCACACGTCTGAACTCTTTCCCTACACGACGCTCTTCCGATCTNNNAACTTTCTGCCCAGTTTTCCTGT
AAGAGCACACGTCTGAACTCTTTCCCTACACGACGCTCTTCCGATCTNNNGCATCTTCTGGTTGATGCC
AGCACACGTCTGAACTCTTTCCCTACACGACGCTCTTCCGATCTNNNGGATTGAGGTGAATCAGTCCA
GCACACGTCTGAACTCTTTCCCTACACGACGCTCTTCCGATCTNNNAGAAAAAGCAAAGCAGTGTAG
AGCACACGTCTGAACTCTTTCCCTACACGACGCTCTTCCGATCTNNNCTGGCTTTGTGCTAGGGGAGA
AAGAGCACACGTCTGAACTCTTTCCCTACACGACGCTCTTCCGATCTNNNCCATTCCTCTGTATTGGTG
GCACACGTCTGAACTCTTTCCCTACACGACGCTCTTCCGATCTNNNCGTGTGGTAATCTGAGCTGCAT
GAGCACACGTCTGAACTCTTTCCCTACACGACGCTCTTCCGATCTNNNGCATGGACCTTATTGGCTCTT
GAGCACACGTCTGAACTCTTTCCCTACACGACGCTCTTCCGATCTNNNCGGGAAACAGCAAGGACTAA
AGCACACGTCTGAACTCTTTCCCTACACGACGCTCTTCCGATCTNNNCCTCTCGCTGAGTTCACAAAGA
GCACACGTCTGAACTCTTTCCCTACACGACGCTCTTCCGATCTNNNGAAAACTGCCATGTTGATCTCCG
GCACACGTCTGAACTCTTTCCCTACACGACGCTCTTCCGATCTNNNACGTGGATTGCCTTTACTTTGGT
GCACACGTCTGAACTCTTTCCCTACACGACGCTCTTCCGATCTNNNTGTTGAAGAGCCTTCGTTGAGA
GAGCACACGTCTGAACTCTTTCCCTACACGACGCTCTTCCGATCTNNNGACCCCGTCTCTGCATTTTACA
GAGCACACGTCTGAACTCTTTCCCTACACGACGCTCTTCCGATCTNNNGACTGGCTTATTTCACATCAC
GAGCACACGTCTGAACTCTTTCCCTACACGACGCTCTTCCGATCTNNNCTCCGGTAGGACAGGAGGG
AGAGCACACGTCTGAACTCTTTCCCTACACGACGCTCTTCCGATCTNNNAGTATTTTGGCATGCTGAG

GAGCACACGTCTGAACTCTTTCCCTACACGACGCTCTTCCGATCTNNNGATCTCTGATGATCCGCTCTCC
GAAGAGCACACGTCTGAACTCTTTCCCTACACGACGCTCTTCCGATCTNNNTTCCAGGGTTGGATATGT
AGCACACGTCTGAACTCTTTCCCTACACGACGCTCTTCCGATCTNNNAAGAGCAGAAGAAGCTAGAGG
GAGCACACGTCTGAACTCTTTCCCTACACGACGCTCTTCCGATCTNNNCATCCATGTGCACGTTTATGCC
AGCACACGTCTGAACTCTTTCCCTACACGACGCTCTTCCGATCTNNNCATACATATAGGGCCCCTGCTCT
GAGCACACGTCTGAACTCTTTCCCTACACGACGCTCTTCCGATCTNNNTGATGGCTTCCTTCCATTCTGTC
AAGAGCACACGTCTGAACTCTTTCCCTACACGACGCTCTTCCGATCTNNNATCTTGGGGAGGAGTACAC
GCACACGTCTGAACTCTTTCCCTACACGACGCTCTTCCGATCTNNNACACATACGCGTACACACTTCTCC
GAGCACACGTCTGAACTCTTTCCCTACACGACGCTCTTCCGATCTNNNTCTGATAGGAACTTGCATTGC
GAGCACACGTCTGAACTCTTTCCCTACACGACGCTCTTCCGATCTNNNGCCTAGGTCGAGTAGGAATG
GAGCACACGTCTGAACTCTTTCCCTACACGACGCTCTTCCGATCTNNNCTCTTCTCGCTCCGTCTATCTCT
AGCACACGTCTGAACTCTTTCCCTACACGACGCTCTTCCGATCTNNNCCATGTGAGGATTGCATACTTG
GAGCACACGTCTGAACTCTTTCCCTACACGACGCTCTTCCGATCTNNNGGAAGCAGGACAGACTAAAA
AGAGCACACGTCTGAACTCTTTCCCTACACGACGCTCTTCCGATCTNNNCCTAACACTAACTGAATGGG
AGAGCACACGTCTGAACTCTTTCCCTACACGACGCTCTTCCGATCTNNNCACCATGTTTGTGAAGTAGG
AAGAGCACACGTCTGAACTCTTTCCCTACACGACGCTCTTCCGATCTNNNGAATACATTGCTCTGGGG
GAGCACACGTCTGAACTCTTTCCCTACACGACGCTCTTCCGATCTNNNCCACGGTAGAGTTGCAGATAC
GAGCACACGTCTGAACTCTTTCCCTACACGACGCTCTTCCGATCTNNNCCCTCATAGAGGACACATCCAC
GAGCACACGTCTGAACTCTTTCCCTACACGACGCTCTTCCGATCTNNNTCCCTTACTCCCTTTGGGTCTTT
AGCACACGTCTGAACTCTTTCCCTACACGACGCTCTTCCGATCTNNNTATCCTGTGCATGGCTCAGTCAG
AGAGCACACGTCTGAACTCTTTCCCTACACGACGCTCTTCCGATCTNNNGAAAAGGGAGAGAAACCAA
AGAGCACACGTCTGAACTCTTTCCCTACACGACGCTCTTCCGATCTNNNATAGCAGGAACACAAAAGG
GAGCACACGTCTGAACTCTTTCCCTACACGACGCTCTTCCGATCTNNNTACTCGGCTGGATTCAGTTTGT
GAGCACACGTCTGAACTCTTTCCCTACACGACGCTCTTCCGATCTNNNTTACCATGGGACCTGAACTGC
GAGCACACGTCTGAACTCTTTCCCTACACGACGCTCTTCCGATCTNNNCAGTGGAGCTGAATTGCTCAA
AGCACACGTCTGAACTCTTTCCCTACACGACGCTCTTCCGATCTNNNTTTGCATTCCTTTGTGTCTCCCA
AGCACACGTCTGAACTCTTTCCCTACACGACGCTCTTCCGATCTNNNTGACTTTCTCCTCACCTCCATCAC
AGCACACGTCTGAACTCTTTCCCTACACGACGCTCTTCCGATCTNNNGTACTTGGGTTTCAGCTCCAGGC
GAGCACACGTCTGAACTCTTTCCCTACACGACGCTCTTCCGATCTNNNATCAAGGCATTAGGCAAGTAC
AGCACACGTCTGAACTCTTTCCCTACACGACGCTCTTCCGATCTNNNTCAACAACCTGAATTTCCTCCAG
AAGAGCACACGTCTGAACTCTTTCCCTACACGACGCTCTTCCGATCTNNNTCCAAGTTATCTTCCCAAAC
GAGCACACGTCTGAACTCTTTCCCTACACGACGCTCTTCCGATCTNNNAACAGAGGAACCAAACTGCCT
AGAGCACACGTCTGAACTCTTTCCCTACACGACGCTCTTCCGATCTNNNGTCAAAATCAATTGCTGAAG
GAGCACACGTCTGAACTCTTTCCCTACACGACGCTCTTCCGATCTNNNTCAGATGTCGCCAAGTTTCAA
AGCACACGTCTGAACTCTTTCCCTACACGACGCTCTTCCGATCTNNNAGTAGAGGGGCTCAACTGACA
CACACGTCTGAACTCTTTCCCTACACGACGCTCTTCCGATCTNNNGGCCTCTGCTGGATTTTGAGATTCA
AGAGCACACGTCTGAACTCTTTCCCTACACGACGCTCTTCCGATCTNNNTGAGGAACCATGGACTTTTC
GAGCACACGTCTGAACTCTTTCCCTACACGACGCTCTTCCGATCTNNNTGGCCTACACAAATGTACCCA
GAGCACACGTCTGAACTCTTTCCCTACACGACGCTCTTCCGATCTNNNCATACTGGAAGGATGTGGGA
GAGCACACGTCTGAACTCTTTCCCTACACGACGCTCTTCCGATCTNNNGGCCTAGTTTCAAAGAAGATA
AGAGCACACGTCTGAACTCTTTCCCTACACGACGCTCTTCCGATCTNNNTTACAGGTGGATGACACAAT
AGCACACGTCTGAACTCTTTCCCTACACGACGCTCTTCCGATCTNNNAACTTACCAAGACCTCCTGGGGC
AGCACACGTCTGAACTCTTTCCCTACACGACGCTCTTCCGATCTNNNAAAAATTGGGAACCAGCTTGG

GAGCACACGTCTGAACTCTTTCCCTACACGACGCTCTTCCGATCTNNNTATGCCCACAGTCCAAACTTGA
GAAGAGCACACGTCTGAACTCTTTCCCTACACGACGCTCTTCCGATCTNNNAGCTTTTGCCTATGCTTC
AGCACACGTCTGAACTCTTTCCCTACACGACGCTCTTCCGATCTNNNCTAGGCTAAGGGTGTCCTTCACT
GCACACGTCTGAACTCTTTCCCTACACGACGCTCTTCCGATCTNNNATAGGAGGTTTGGAGATCAGGC
AGAGCACACGTCTGAACTCTTTCCCTACACGACGCTCTTCCGATCTNNNAGGAGGGAAATTTGTCCACA
AGAGCACACGTCTGAACTCTTTCCCTACACGACGCTCTTCCGATCTNNNAAAAATAGCCGTGTGTGGCT
AGCACACGTCTGAACTCTTTCCCTACACGACGCTCTTCCGATCTNNNCGGACATTTGGAATCCCAGCTA
GAGCACACGTCTGAACTCTTTCCCTACACGACGCTCTTCCGATCTNNNAGATAGAACTCTGGGACTGA
AGAGCACACGTCTGAACTCTTTCCCTACACGACGCTCTTCCGATCTNNNGGGATCTTCCTCTTGGGATC
AGCACACGTCTGAACTCTTTCCCTACACGACGCTCTTCCGATCTNNNAACAAAGCAATGAGGTTGGCA
GAAGAGCACACGTCTGAACTCTTTCCCTACACGACGCTCTTCCGATCTNNNGCCCCTGCAATCATCAATC
GAGCACACGTCTGAACTCTTTCCCTACACGACGCTCTTCCGATCTNNNAGTAGCACAATCATGGCTGAA
GAGCACACGTCTGAACTCTTTCCCTACACGACGCTCTTCCGATCTNNNTGTGCTATGACCTTCTTTACCT
AGAGCACACGTCTGAACTCTTTCCCTACACGACGCTCTTCCGATCTNNNAAAACACCACAAATCAGCTG
GAGCACACGTCTGAACTCTTTCCCTACACGACGCTCTTCCGATCTNNNTGATTATTTGAAAAGCTTCGC
AGAGCACACGTCTGAACTCTTTCCCTACACGACGCTCTTCCGATCTNNNCTCTCCTCGATTTGCACGGTC
AGCACACGTCTGAACTCTTTCCCTACACGACGCTCTTCCGATCTNNNTTCACCTTTGGAGGATACTAGC
AGCACACGTCTGAACTCTTTCCCTACACGACGCTCTTCCGATCTNNNTTCCTCTCTCGAGCCAGTCTATG
GCACACGTCTGAACTCTTTCCCTACACGACGCTCTTCCGATCTNNNCAATCAATCTCCTGCCTCAGCTCC
AGCACACGTCTGAACTCTTTCCCTACACGACGCTCTTCCGATCTNNNCCCCTGACTTTACGGCCACGCATC
AGAGCACACGTCTGAACTCTTTCCCTACACGACGCTCTTCCGATCTNNNACTCTGGGATTTGAGTTGGC
AAGAGCACACGTCTGAACTCTTTCCCTACACGACGCTCTTCCGATCTNNNGGAACAATATAGGGGAAA
CACACGTCTGAACTCTTTCCCTACACGACGCTCTTCCGATCTNNNACCGGTGTTGCAAATAACGTTTCC
AGCACACGTCTGAACTCTTTCCCTACACGACGCTCTTCCGATCTNNNTGTGTGAATCTTGTGATTGTGT
GCACACGTCTGAACTCTTTCCCTACACGACGCTCTTCCGATCTNNNCAGGAGGGGCACTGTAGCTGCA
AAGAGCACACGTCTGAACTCTTTCCCTACACGACGCTCTTCCGATCTNNNATACATTGAAAACTCCCCCT
GAGCACACGTCTGAACTCTTTCCCTACACGACGCTCTTCCGATCTNNNTGAAATGAAGAAGTGTAGGA
AGAGCACACGTCTGAACTCTTTCCCTACACGACGCTCTTCCGATCTNNNTGGTCCTCCCCTTTTGTCTTTT
AGCACACGTCTGAACTCTTTCCCTACACGACGCTCTTCCGATCTNNNCCTTGGTCTCTTCGGTCTCCTCAT
AGCACACGTCTGAACTCTTTCCCTACACGACGCTCTTCCGATCTNNNAGGCAAGTCAGTCACTTCAGTT
AGAGCACACGTCTGAACTCTTTCCCTACACGACGCTCTTCCGATCTNNNTCCCTACACCACGAAATATGC
AGAGCACACGTCTGAACTCTTTCCCTACACGACGCTCTTCCGATCTNNNGGAGGTACAAACAGCAGAG
AGCACACGTCTGAACTCTTTCCCTACACGACGCTCTTCCGATCTNNNCCAGGGCAGGTAAGTCAAACA
AGCACACGTCTGAACTCTTTCCCTACACGACGCTCTTCCGATCTNNNGCCCCTAACTTAGACAGCAAGT
GAGCACACGTCTGAACTCTTTCCCTACACGACGCTCTTCCGATCTNNNTATTAGGGGGTGGGGACTTG
GAGCACACGTCTGAACTCTTTCCCTACACGACGCTCTTCCGATCTNNNTCTCAGCTGGATAATGTCCCCA
GAGCACACGTCTGAACTCTTTCCCTACACGACGCTCTTCCGATCTNNNTCTCCTTTCAGGGTGGCAAGA
CACACGTCTGAACTCTTTCCCTACACGACGCTCTTCCGATCTNNNTTTCGGTTCCATCTGTGCCTTAGCA
AAGAGCACACGTCTGAACTCTTTCCCTACACGACGCTCTTCCGATCTNNNCAGTGCATGCTCCATCAAGT
AGCACACGTCTGAACTCTTTCCCTACACGACGCTCTTCCGATCTNNNAGGAGAATTGCTGCTTGAACTC
CACACGTCTGAACTCTTTCCCTACACGACGCTCTTCCGATCTNNNGTCATCTCATGCCCTTTCACTCCCAT
AGCACACGTCTGAACTCTTTCCCTACACGACGCTCTTCCGATCTNNNTTTTGTGCATTGAGGTAGGCTG
AGAGCACACGTCTGAACTCTTTCCCTACACGACGCTCTTCCGATCTNNNCTACCCCTGTTTTTCACTTGGC

CACACGTCTGAACTCTTTCCCTACACGACGCTCTTCCGATCTNNNTGGTGTGATGATTATATCTGCCAC
AGAGCACACGTCTGAACTCTTTCCCTACACGACGCTCTTCCGATCTNNNTGAGAACGGACTAATCCAC
GAGCACACGTCTGAACTCTTTCCCTACACGACGCTCTTCCGATCTNNNTCTACATGCTATCAAGGGCCA
GCACACGTCTGAACTCTTTCCCTACACGACGCTCTTCCGATCTNNNAACTGTGGATGCCCTTGTTACAG
ACACGTCTGAACTCTTTCCCTACACGACGCTCTTCCGATCTNNNCTTCTCGCCACTCTTACTAGGGTCAT
AAGAGCACACGTCTGAACTCTTTCCCTACACGACGCTCTTCCGATCTNNNTTAGAGCCAGGAGATAGA
GAGCACACGTCTGAACTCTTTCCCTACACGACGCTCTTCCGATCTNNNCACACCAGTTTTCTTCCTGCTC
AGAGCACACGTCTGAACTCTTTCCCTACACGACGCTCTTCCGATCTNNNTGTCAGCTGCTGTTGACTTT
GAAGAGCACACGTCTGAACTCTTTCCCTACACGACGCTCTTCCGATCTNNNTTTGTCACTGGTTGCTGT
GAGCACACGTCTGAACTCTTTCCCTACACGACGCTCTTCCGATCTNNNTGGATGTTTCATGGCATCCTC
AGAGCACACGTCTGAACTCTTTCCCTACACGACGCTCTTCCGATCTNNNATAAAAGCATTGGTATGGC
GAGCACACGTCTGAACTCTTTCCCTACACGACGCTCTTCCGATCTNNNTGTGGCAATGCTTTCTTGGAT
GCACACGTCTGAACTCTTTCCCTACACGACGCTCTTCCGATCTNNNTCCAACATCTTGTTGCCATTGGAC
AAGAGCACACGTCTGAACTCTTTCCCTACACGACGCTCTTCCGATCTNNNCCGGTTGCTGCATCCATTAT
GCACACGTCTGAACTCTTTCCCTACACGACGCTCTTCCGATCTNNNTTTTGGGGTGGGGACACAGTCT
GAGCACACGTCTGAACTCTTTCCCTACACGACGCTCTTCCGATCTNNNCTCTCTTGAAGCTGCCCTGAA
AGCACACGTCTGAACTCTTTCCCTACACGACGCTCTTCCGATCTNNNGAGTAGGTGGGATTTCAGGCA
GCACACGTCTGAACTCTTTCCCTACACGACGCTCTTCCGATCTNNNTCAAGGCCCCGTCTCTACAAAAG
AGAGCACACGTCTGAACTCTTTCCCTACACGACGCTCTTCCGATCTNNNCCCTTTTGCCTAACAGTGGG
AGCACACGTCTGAACTCTTTCCCTACACGACGCTCTTCCGATCTNNNATACCTCAAGCTGCCTTTGACA
GAGCACACGTCTGAACTCTTTCCCTACACGACGCTCTTCCGATCTNNNCAGTGTGTGCCATTCTGTAAT
AGCACACGTCTGAACTCTTTCCCTACACGACGCTCTTCCGATCTNNNTTCCTCAAGAAGGCTTGATTGA
AAGAGCACACGTCTGAACTCTTTCCCTACACGACGCTCTTCCGATCTNNNGCTCGCCCTCTTTCTATTCT
GAGCACACGTCTGAACTCTTTCCCTACACGACGCTCTTCCGATCTNNNTGGATAGCACTCATAATGTGG
AAGAGCACACGTCTGAACTCTTTCCCTACACGACGCTCTTCCGATCTNNNTGGAGGGTGGTTGCCTATA
GAGCACACGTCTGAACTCTTTCCCTACACGACGCTCTTCCGATCTNNNAGAAATGAAGAGGCAGAGGT
CACACGTCTGAACTCTTTCCCTACACGACGCTCTTCCGATCTNNNAAATCTCGAGACCCCAGATCTCCCAT
GCACACGTCTGAACTCTTTCCCTACACGACGCTCTTCCGATCTNNNTGCTTTGCACACATTATCTCCCACA
AGAGCACACGTCTGAACTCTTTCCCTACACGACGCTCTTCCGATCTNNNTATCCAGGAGCCCACCAAAA
GAGCACACGTCTGAACTCTTTCCCTACACGACGCTCTTCCGATCTNNNACCTGGTAGGTGCTCCATAAA
GAGCACACGTCTGAACTCTTTCCCTACACGACGCTCTTCCGATCTNNNGTCCCCAGTTACTTTGGAGAC
GAAGAGCACACGTCTGAACTCTTTCCCTACACGACGCTCTTCCGATCTNNNACGCATCCTTATTGCTTTT
AGCACACGTCTGAACTCTTTCCCTACACGACGCTCTTCCGATCTNNNCTTTCCAAACTCTCTCCGCTCTCC
CACACGTCTGAACTCTTTCCCTACACGACGCTCTTCCGATCTNNNGTAAAATGCTTGGTTCAGTGCCCC
GAGCACACGTCTGAACTCTTTCCCTACACGACGCTCTTCCGATCTNNNAAGAGATGTGGTCTTGCCGT
AAGAGCACACGTCTGAACTCTTTCCCTACACGACGCTCTTCCGATCTNNNCCCCCAATGGTAACATCCT
GAGCACACGTCTGAACTCTTTCCCTACACGACGCTCTTCCGATCTNNNCCTTGTCCTTATTTCTCCAGG
GCACACGTCTGAACTCTTTCCCTACACGACGCTCTTCCGATCTNNNAAACTCATGGAGCGTATTCCCAC
GGAAGAGCACACGTCTGAACTCTTTCCCTACACGACGCTCTTCCGATCTNNNGGTGAGCCTGACTGCA
AGCACACGTCTGAACTCTTTCCCTACACGACGCTCTTCCGATCTNNNGAAAGTCACTACTCCTGCTTGC
AAGAGCACACGTCTGAACTCTTTCCCTACACGACGCTCTTCCGATCTNNNCTGAAAAATGAACAGAGCT
GAGCACACGTCTGAACTCTTTCCCTACACGACGCTCTTCCGATCTNNNCCAAACACTGTACTGACGTCCA
GCACACGTCTGAACTCTTTCCCTACACGACGCTCTTCCGATCTNNNCCTCAACCCTGCCAGAAAAGTAAC

AGCACACGTCTGAACTCTTTCCCTACACGACGCTCTTCCGATCTNNNTAGTTTGGGTCCTGTTGGTTTG
AGCACACGTCTGAACTCTTTCCCTACACGACGCTCTTCCGATCTNNNCCAGTGTTTTAGCCCAGAACCT
AAGAGCACACGTCTGAACTCTTTCCCTACACGACGCTCTTCCGATCTNNNAATCACTCTTTGCCTAACCC
GAGCACACGTCTGAACTCTTTCCCTACACGACGCTCTTCCGATCTNNNCAACAGAAGAAAGGTTGGTG
AGCACACGTCTGAACTCTTTCCCTACACGACGCTCTTCCGATCTNNNCTATGTATTTTCCGGGGTTGGC
GAGCACACGTCTGAACTCTTTCCCTACACGACGCTCTTCCGATCTNNNTCTTACCCACAGCAGCCATAAA
GAGCACACGTCTGAACTCTTTCCCTACACGACGCTCTTCCGATCTNNNGGGGGTGAGGGGATAAAGA
AGCACACGTCTGAACTCTTTCCCTACACGACGCTCTTCCGATCTNNNTTGCACCTGAACACTAAAGGAC
GCACACGTCTGAACTCTTTCCCTACACGACGCTCTTCCGATCTNNNGCCACAGTTGAAAAACCCTGTTCC
AGAGCACACGTCTGAACTCTTTCCCTACACGACGCTCTTCCGATCTNNNAGGGTTTCAGGGAGACACTG
CACACGTCTGAACTCTTTCCCTACACGACGCTCTTCCGATCTNNNTCATATCAAGAGCCGTGTGTGTGC
AGCACACGTCTGAACTCTTTCCCTACACGACGCTCTTCCGATCTNNNGCCTGGCCAATATTCTGTGTGTA
AGAGCACACGTCTGAACTCTTTCCCTACACGACGCTCTTCCGATCTNNNAGTACAAGTGGAGGCCATC
GAAGAGCACACGTCTGAACTCTTTCCCTACACGACGCTCTTCCGATCTNNNAAACAATCATGGCAAGG
AGCACACGTCTGAACTCTTTCCCTACACGACGCTCTTCCGATCTNNNCGCTGTCCTTTGATAACTCCCTCC
GAGCACACGTCTGAACTCTTTCCCTACACGACGCTCTTCCGATCTNNNTGCAAATTTGGAGGGAGGAA
AGCACACGTCTGAACTCTTTCCCTACACGACGCTCTTCCGATCTNNNCCTGAGCCATAGCACCTTTTCAT
GCACACGTCTGAACTCTTTCCCTACACGACGCTCTTCCGATCTNNNTGTTGACACATCATACTCACTTGA
AGAGCACACGTCTGAACTCTTTCCCTACACGACGCTCTTCCGATCTNNNTTCCAGTCCTTGCCAGATCAA
GGAAGAGCACACGTCTGAACTCTTTCCCTACACGACGCTCTTCCGATCTNNNCTACAGCCTGGGAGTTC
AGAGCACACGTCTGAACTCTTTCCCTACACGACGCTCTTCCGATCTNNNGAGAAGGCTGATGTCTTTGC
AGCACACGTCTGAACTCTTTCCCTACACGACGCTCTTCCGATCTNNNATGGTCTTCAGTTGCCTATGCTC
AAGAGCACACGTCTGAACTCTTTCCCTACACGACGCTCTTCCGATCTNNNGAAGATGCAGTGAGCCAC
AGCACACGTCTGAACTCTTTCCCTACACGACGCTCTTCCGATCTNNNCTCTAGATCAATCACTGCCTGCC
AGCACACGTCTGAACTCTTTCCCTACACGACGCTCTTCCGATCTNNNGAAATCCAGTCCGGCTTTTCCAG
GCACACGTCTGAACTCTTTCCCTACACGACGCTCTTCCGATCTNNNAATCCCACATACTCGCTACTTGGC
GAGCACACGTCTGAACTCTTTCCCTACACGACGCTCTTCCGATCTNNNCAGGCTCATGATTCACAACAG
AGAGCACACGTCTGAACTCTTTCCCTACACGACGCTCTTCCGATCTNNNGTGTTACAGAGGCAATGAGC
AGAGCACACGTCTGAACTCTTTCCCTACACGACGCTCTTCCGATCTNNNATCTCCTAGTGTGCAGGTCT
AGAGCACACGTCTGAACTCTTTCCCTACACGACGCTCTTCCGATCTNNNTCTGATAGGAACTTGCATTC
GCACACGTCTGAACTCTTTCCCTACACGACGCTCTTCCGATCTNNNCTGAGGAGGAGGGGAGAGCGCA
GAGCACACGTCTGAACTCTTTCCCTACACGACGCTCTTCCGATCTNNNGCAATTATTCACAACTGGGGC
AGCACACGTCTGAACTCTTTCCCTACACGACGCTCTTCCGATCTNNNTTTTCCTGGGCACATACAACCTCC
GCACACGTCTGAACTCTTTCCCTACACGACGCTCTTCCGATCTNNNATTTGCTCATCGTGGCGGAGCATC
AAGAGCACACGTCTGAACTCTTTCCCTACACGACGCTCTTCCGATCTNNNTCTGTTGTCAAGGCTGAAG
AGAGCACACGTCTGAACTCTTTCCCTACACGACGCTCTTCCGATCTNNNAGTGGGGGAAGGAGATGTT
GAGCACACGTCTGAACTCTTTCCCTACACGACGCTCTTCCGATCTNNNTAGTGTCTACTGCTTGTTGGGC
AGCACACGTCTGAACTCTTTCCCTACACGACGCTCTTCCGATCTNNNCATGTGTCTTTGAACCCATGCAC
GAGCACACGTCTGAACTCTTTCCCTACACGACGCTCTTCCGATCTNNNCGTTACTAACTAGAGGGGAGC
AGCACACGTCTGAACTCTTTCCCTACACGACGCTCTTCCGATCTNNNCAACCCTCCCGACTTCTTCTAACC
AAGAGCACACGTCTGAACTCTTTCCCTACACGACGCTCTTCCGATCTNNNGGGGGTGTCTGTGAATACA
AGCACACGTCTGAACTCTTTCCCTACACGACGCTCTTCCGATCTNNNCCATCCTACTGGTGTAAGGTGC
CACACGTCTGAACTCTTTCCCTACACGACGCTCTTCCGATCTNNNGTTTACCTGTCTGCCTCCTCATTCAT

GCACACGTCTGAACTCTTTCCCTACACGACGCTCTTCCGATCTNNNTCCAGATCCTCACCAATCTCTCCCA
GCACACGTCTGAACTCTTTCCCTACACGACGCTCTTCCGATCTNNNTTGCTGATAATCTCATGGGAGCG
AGAGCACACGTCTGAACTCTTTCCCTACACGACGCTCTTCCGATCTNNNTTAGGGCACCTGCAGTTTGA
AGCACACGTCTGAACTCTTTCCCTACACGACGCTCTTCCGATCTNNNTGGCTGACGTGATATTCAAGCT
AGAGCACACGTCTGAACTCTTTCCCTACACGACGCTCTTCCGATCTNNNGAATCTGGCTCAGGGCAACT
GCACACGTCTGAACTCTTTCCCTACACGACGCTCTTCCGATCTNNNCAAGACCCATACACGATTCAGCAC
GCACACGTCTGAACTCTTTCCCTACACGACGCTCTTCCGATCTNNNCCTTCATCTCCATACCCTCGACACA
AGCACACGTCTGAACTCTTTCCCTACACGACGCTCTTCCGATCTNNNCCAGGTGTTTCTGTGCCTGTAAA
AGCACACGTCTGAACTCTTTCCCTACACGACGCTCTTCCGATCTNNNTCTGTTCATGTAAGCAGGAAGG
AGCACACGTCTGAACTCTTTCCCTACACGACGCTCTTCCGATCTNNNCTAGGAAATGAAGCGAGGCTGT
AGCACACGTCTGAACTCTTTCCCTACACGACGCTCTTCCGATCTNNNGCAGTAGCAAGACCTTGTCTCA
AGCACACGTCTGAACTCTTTCCCTACACGACGCTCTTCCGATCTNNNAGTCCAATTTCTTCAGAGGCTGA
AGAGCACACGTCTGAACTCTTTCCCTACACGACGCTCTTCCGATCTNNNATCTGACTTCCCACTGAGAGA
GCACACGTCTGAACTCTTTCCCTACACGACGCTCTTCCGATCTNNNAAAAAGTGAGGGAAAAGGCTGG
GAGCACACGTCTGAACTCTTTCCCTACACGACGCTCTTCCGATCTNNNAGATGAACATTCCCATTCCTGG
AGCACACGTCTGAACTCTTTCCCTACACGACGCTCTTCCGATCTNNNAGCTCCTGGGGGTAATCTTGAA
GCACACGTCTGAACTCTTTCCCTACACGACGCTCTTCCGATCTNNNAACCTGTTCTTCCATGAGGCATTC
AGCACACGTCTGAACTCTTTCCCTACACGACGCTCTTCCGATCTNNNGAACATGCTCATACACACCCACA
GAGCACACGTCTGAACTCTTTCCCTACACGACGCTCTTCCGATCTNNNTAGGTGGTAACCAGGAAGTG
GCACACGTCTGAACTCTTTCCCTACACGACGCTCTTCCGATCTNNNGTCTCTTCCTAACAAGGGCACTCC
AGCACACGTCTGAACTCTTTCCCTACACGACGCTCTTCCGATCTNNNGAGCCAGTAGGGAACAACTCAA
GGAAGAGCACACGTCTGAACTCTTTCCCTACACGACGCTCTTCCGATCTNNNTCTTGCTTGGACCTCTCA
AAGAGCACACGTCTGAACTCTTTCCCTACACGACGCTCTTCCGATCTNNNGGAGTGTAGTGAACTCCTG
GCACACGTCTGAACTCTTTCCCTACACGACGCTCTTCCGATCTNNNTGAGCAGGAAGGGAGAATGAAC
GCACACGTCTGAACTCTTTCCCTACACGACGCTCTTCCGATCTNNNGTCAGAAGGGGTTCTAAAGCCAC
GCACACGTCTGAACTCTTTCCCTACACGACGCTCTTCCGATCTNNNCAGCGATCACAACAGCTTCAGATG
AAGAGCACACGTCTGAACTCTTTCCCTACACGACGCTCTTCCGATCTNNNAGAAAGAGACTGAGGGA
GAGCACACGTCTGAACTCTTTCCCTACACGACGCTCTTCCGATCTNNNCAGCAAGGAGAAAACTGTTC
ACACGTCTGAACTCTTTCCCTACACGACGCTCTTCCGATCTNNNACCCATCCTCCCGTAGGCCCCATCGG
GCACACGTCTGAACTCTTTCCCTACACGACGCTCTTCCGATCTNNNGAAGCAGCTTTCAGATCTGGTGA
AGAGCACACGTCTGAACTCTTTCCCTACACGACGCTCTTCCGATCTNNNCCAAACTGTGCCCTAACCTTT
GAGCACACGTCTGAACTCTTTCCCTACACGACGCTCTTCCGATCTNNNTTGGAGTTGCAGTTGATGTGT
GAGCACACGTCTGAACTCTTTCCCTACACGACGCTCTTCCGATCTNNNGGTTTGATGTGGGTTGGGTT
GAGCACACGTCTGAACTCTTTCCCTACACGACGCTCTTCCGATCTNNNTGGGAGTGGGAGGGGAAAA
GAGCACACGTCTGAACTCTTTCCCTACACGACGCTCTTCCGATCTNNNTACATAGCGGTTGTTGATGGC
AGAGCACACGTCTGAACTCTTTCCCTACACGACGCTCTTCCGATCTNNNGGGGTTTGGCAGAGATGTG
AGCACACGTCTGAACTCTTTCCCTACACGACGCTCTTCCGATCTNNNTATGATGTCTGTGTGGAAAGC
GAGCACACGTCTGAACTCTTTCCCTACACGACGCTCTTCCGATCTNNNTCACTTTAGCTACCCCCAAGTG
GAGCACACGTCTGAACTCTTTCCCTACACGACGCTCTTCCGATCTNNNATGGGGACGAGAAGGGATTT
AGAGCACACGTCTGAACTCTTTCCCTACACGACGCTCTTCCGATCTNNNACCGTAGTATACCCCCGGTCC
GAGCACACGTCTGAACTCTTTCCCTACACGACGCTCTTCCGATCTNNNCTCAGGTAGGTGTCCTTATCCC
AGAGCACACGTCTGAACTCTTTCCCTACACGACGCTCTTCCGATCTNNNCAAAAGCGAGACCCTGTCCTC
CACACGTCTGAACTCTTTCCCTACACGACGCTCTTCCGATCTNNNCAATCTCTTGGGCTTGGGTGATTCA

AGCACACGTCTGAACTCTTTCCCTACACGACGCTCTTCCGATCTNNNATGAAATTGCAGGCAGTGTTT
AGCACACGTCTGAACTCTTTCCCTACACGACGCTCTTCCGATCTNNNGCCATCTCTAGTGGACACACCT
GCACACGTCTGAACTCTTTCCCTACACGACGCTCTTCCGATCTNNNTGCAGGGAATTTAGGAAAGAG
AGCACACGTCTGAACTCTTTCCCTACACGACGCTCTTCCGATCTNNNCTGGAGAGAGCTACATACCCA
AGAGCACACGTCTGAACTCTTTCCCTACACGACGCTCTTCCGATCTNNNTACCTCTCTCCATGTCCCCTC
GAGCACACGTCTGAACTCTTTCCCTACACGACGCTCTTCCGATCTNNNACTGGATCCATCATCCCTCAT
GCACACGTCTGAACTCTTTCCCTACACGACGCTCTTCCGATCTNNNACACATTCATATGCAGTTGCCCA
AGCACACGTCTGAACTCTTTCCCTACACGACGCTCTTCCGATCTNNNCTTGAGGGTGCTCTTGTCTTG
CACACGTCTGAACTCTTTCCCTACACGACGCTCTTCCGATCTNNNGGAGGCTCAAGTCTGTGTATGTG
GCACACGTCTGAACTCTTTCCCTACACGACGCTCTTCCGATCTNNNAGCTGCTGTGTGGTGTATACTTG
GAGCACACGTCTGAACTCTTTCCCTACACGACGCTCTTCCGATCTNNNGGTAGGAGGAGGAATGCAG
GCACACGTCTGAACTCTTTCCCTACACGACGCTCTTCCGATCTNNNGTTTCTTCCTCCAGGAGTGCTTC
AGAGCACACGTCTGAACTCTTTCCCTACACGACGCTCTTCCGATCTNNNGTTGGGCGCTCTTTCTTACA
GAGCACACGTCTGAACTCTTTCCCTACACGACGCTCTTCCGATCTNNNCTGCACCTGGAAAGTTCTAG
AGCACACGTCTGAACTCTTTCCCTACACGACGCTCTTCCGATCTNNNGGTACACATTCACATGTAGTTG
GCACACGTCTGAACTCTTTCCCTACACGACGCTCTTCCGATCTNNNGTGGTTTCTGTAGCGTGTGTAC
GCACACGTCTGAACTCTTTCCCTACACGACGCTCTTCCGATCTNNNTCTTGTATCCTACAACCTTATTGA
GAGCACACGTCTGAACTCTTTCCCTACACGACGCTCTTCCGATCTNNNAGGCAGAAGGAAGAGAAGA
CACACGTCTGAACTCTTTCCCTACACGACGCTCTTCCGATCTNNNGTGGTTTCTGTAGCGTGTGTACT
AGCACACGTCTGAACTCTTTCCCTACACGACGCTCTTCCGATCTNNNATTTAGGGTGTTCTTCCACCCC
GAGCACACGTCTGAACTCTTTCCCTACACGACGCTCTTCCGATCTNNNTTGTGTTGATGGTAACCCAA
GAGCACACGTCTGAACTCTTTCCCTACACGACGCTCTTCCGATCTNNNTCTCTCTTACGCACACACACA
AAGAGCACACGTCTGAACTCTTTCCCTACACGACGCTCTTCCGATCTNNNTCAAAATCAGGGCTCAAC
AGCACACGTCTGAACTCTTTCCCTACACGACGCTCTTCCGATCTNNNCCGGGTATTTACTGTGTAGAG
AGCACACGTCTGAACTCTTTCCCTACACGACGCTCTTCCGATCTNNNGGGGGTCATGTCATGTGTCAA
GAGCACACGTCTGAACTCTTTCCCTACACGACGCTCTTCCGATCTNNNGATCAGGGAGGAAGCAACA
CACACGTCTGAACTCTTTCCCTACACGACGCTCTTCCGATCTNNNCTGGGTACCCTGTCTTCCTCCATCA
AGCACACGTCTGAACTCTTTCCCTACACGACGCTCTTCCGATCTNNNTGAAGTGTGTGTGTGTGTGT
AGCACACGTCTGAACTCTTTCCCTACACGACGCTCTTCCGATCTNNNACCATGCTAACGTGTCTACTT
GCACACGTCTGAACTCTTTCCCTACACGACGCTCTTCCGATCTNNNTTGCACCTGTGAGTATGTCTGT
AAGAGCACACGTCTGAACTCTTTCCCTACACGACGCTCTTCCGATCTNNNGTGAGTTTCCAAGCTCCAA
GAGCACACGTCTGAACTCTTTCCCTACACGACGCTCTTCCGATCTNNNCATTCTGGTCATCCAGTCCTG
AGCACACGTCTGAACTCTTTCCCTACACGACGCTCTTCCGATCTNNNCAGCCCAGATTCAAGGAGGT
GAGCACACGTCTGAACTCTTTCCCTACACGACGCTCTTCCGATCTNNNGTGAGTTTCCAAGCTCCAAAG
AGCACACGTCTGAACTCTTTCCCTACACGACGCTCTTCCGATCTNNNTAACAGCAGCTGGATGGATGA
GAGCACACGTCTGAACTCTTTCCCTACACGACGCTCTTCCGATCTNNNTAATACAAATGGGCTGACTG
AGAGCACACGTCTGAACTCTTTCCCTACACGACGCTCTTCCGATCTNNNGTGAGTGGAGATTGCATT
AAGAGCACACGTCTGAACTCTTTCCCTACACGACGCTCTTCCGATCTNNNTCCCAACTTTATCTAGAG
AGCACACGTCTGAACTCTTTCCCTACACGACGCTCTTCCGATCTNNNTGCAAAATAAAATCCCAGCACC
AGAGCACACGTCTGAACTCTTTCCCTACACGACGCTCTTCCGATCTNNNGTGTGTCAGAATGAGTAGC
AGAGCACACGTCTGAACTCTTTCCCTACACGACGCTCTTCCGATCTNNNAGCAAGAACAGCAGAGGAA
AGCACACGTCTGAACTCTTTCCCTACACGACGCTCTTCCGATCTNNNGCATGGCTCAGGAGAAATGA
GCACACGTCTGAACTCTTTCCCTACACGACGCTCTTCCGATCTNNNGAAAGGCACATGTGACAAAGA

GAGCACACGTCTGAACTCTTTCCCTACACGACGCTCTTCCGATCTNNNACTAGATGATTCCCAAGGGGC
AAGAGCACACGTCTGAACTCTTTCCCTACACGACGCTCTTCCGATCTNNNTGGTGGTCCCAACCACTAC
AGCACACGTCTGAACTCTTTCCCTACACGACGCTCTTCCGATCTNNNTTAGTCTCAAGGCTTCGGTCTCT
GCACACGTCTGAACTCTTTCCCTACACGACGCTCTTCCGATCTNNNACCTAACCTGCTACTAGCTTGACA
AGAGCACACGTCTGAACTCTTTCCCTACACGACGCTCTTCCGATCTNNNCTATTGAGGTCCCAGAGAGG
AGCACACGTCTGAACTCTTTCCCTACACGACGCTCTTCCGATCTNNNCCTGTCACTGTGTGTGTCTCTGT
GAGCACACGTCTGAACTCTTTCCCTACACGACGCTCTTCCGATCTNNNTGCACTGTGTTGAATTTTGGTC
AGAGCACACGTCTGAACTCTTTCCCTACACGACGCTCTTCCGATCTNNNTTCTTGTGCATCCTCTATGTG
AGCACACGTCTGAACTCTTTCCCTACACGACGCTCTTCCGATCTNNNACTTTCCTTGTCTCTAGCCCTGTC
AGAGCACACGTCTGAACTCTTTCCCTACACGACGCTCTTCCGATCTNNNACAGCACCACACACACACATA
AGCACACGTCTGAACTCTTTCCCTACACGACGCTCTTCCGATCTNNNGAATGCTACATATGACCCTGCCT
AGAGCACACGTCTGAACTCTTTCCCTACACGACGCTCTTCCGATCTNNNCCTCTCTTCTGAGCTGTTGTC
AGCACACGTCTGAACTCTTTCCCTACACGACGCTCTTCCGATCTNNNGTTTCAGCAACACCAACTTGCTTC
CACACGTCTGAACTCTTTCCCTACACGACGCTCTTCCGATCTNNNCCGTGCCTAGCCAGAAATTGTATCA
AGCACACGTCTGAACTCTTTCCCTACACGACGCTCTTCCGATCTNNNATAGGAGGGGGTTTCCCTTTCT
AGAGCACACGTCTGAACTCTTTCCCTACACGACGCTCTTCCGATCTNNNGGTTAGACTTACCCTCCTCCC
GAGCACACGTCTGAACTCTTTCCCTACACGACGCTCTTCCGATCTNNNTTTGTAGAATTGCCTGACCCAG
AGCACACGTCTGAACTCTTTCCCTACACGACGCTCTTCCGATCTNNNAAAAGATGTGTGTGGGGAGTC
GAGCACACGTCTGAACTCTTTCCCTACACGACGCTCTTCCGATCTNNNACAACAGTTGAGGGTTTTTGC
AGCACACGTCTGAACTCTTTCCCTACACGACGCTCTTCCGATCTNNNAGCGGAAACATTGCAAACACAA
GAGCACACGTCTGAACTCTTTCCCTACACGACGCTCTTCCGATCTNNNGTGTGGTAGTCTGGGTAGTCT
GAGCACACGTCTGAACTCTTTCCCTACACGACGCTCTTCCGATCTNNNTGTGTACCCAGAGAGAAGAGA
GAGCACACGTCTGAACTCTTTCCCTACACGACGCTCTTCCGATCTNNNCTTTGTGACTGTGAGAACCCT
ACACGTCTGAACTCTTTCCCTACACGACGCTCTTCCGATCTNNNTAGAAGGTGTTTGTGAAGTCGGCCA
AGAGCACACGTCTGAACTCTTTCCCTACACGACGCTCTTCCGATCTNNNTTGGGCATTTTTCCCCAGTAG
GAGCACACGTCTGAACTCTTTCCCTACACGACGCTCTTCCGATCTNNNCTTACCCTCAGTGTTCTCCATGC
AGCACACGTCTGAACTCTTTCCCTACACGACGCTCTTCCGATCTNNNAAACTCAGCCTCAAGGGAAAGA
GCACACGTCTGAACTCTTTCCCTACACGACGCTCTTCCGATCTNNNCCCATGGAAATGACTGAACAGGT
AGAGCACACGTCTGAACTCTTTCCCTACACGACGCTCTTCCGATCTNNNTACAGCTCTGACTGTGCTGA
AGCACACGTCTGAACTCTTTCCCTACACGACGCTCTTCCGATCTNNNAGACAGTTTGGAACAACCCAAG
AGAGCACACGTCTGAACTCTTTCCCTACACGACGCTCTTCCGATCTNNNACCACCCTTATTTTCTTCCTCT
GCACACGTCTGAACTCTTTCCCTACACGACGCTCTTCCGATCTNNNAAGCCCACAGAAGAACAGGTTTC
AGCACACGTCTGAACTCTTTCCCTACACGACGCTCTTCCGATCTNNNTGCACTAGGATGTCACTGCTTTC
GAGCACACGTCTGAACTCTTTCCCTACACGACGCTCTTCCGATCTNNNCCCATTCAGCTTGTCACTTGTC
AGCACACGTCTGAACTCTTTCCCTACACGACGCTCTTCCGATCTNNNCCCAGCAGCTTATGTGAATGTCT
AGCACACGTCTGAACTCTTTCCCTACACGACGCTCTTCCGATCTNNNCCTCTGTGACTGTAGGAACAAC
AGCACACGTCTGAACTCTTTCCCTACACGACGCTCTTCCGATCTNNNGCCAGTTCACGGTCTAGTTCAAG
AGCACACGTCTGAACTCTTTCCCTACACGACGCTCTTCCGATCTNNNAGGAGAGGGAGAAAGGGAAGA
AGAGCACACGTCTGAACTCTTTCCCTACACGACGCTCTTCCGATCTNNNAAGGAAACCTCCAGCTTTGTC
AAGAGCACACGTCTGAACTCTTTCCCTACACGACGCTCTTCCGATCTNNNCAAGGTCACAACAGATACTC
GAGCACACGTCTGAACTCTTTCCCTACACGACGCTCTTCCGATCTNNNCCCAGCAGCTTATGTGAATGTC
GAGCACACGTCTGAACTCTTTCCCTACACGACGCTCTTCCGATCTNNNGCCAGGTTGCTAGAAAACTAC
AGCACACGTCTGAACTCTTTCCCTACACGACGCTCTTCCGATCTNNNATAGCCTTTACTTACGTGCCAGC

GAGCACACGTCTGAACTCTTTCCCTACACGACGCTCTTCCGATCTNNNGGTTGGTCAGGCTTTGTCTCTT
GAGCACACGTCTGAACTCTTTCCCTACACGACGCTCTTCCGATCTNNNAGTGGCTGTCTTCTTCATGTGT
GAGCACACGTCTGAACTCTTTCCCTACACGACGCTCTTCCGATCTNNNCAGGTGCACACGTTATGACTA
AGCACACGTCTGAACTCTTTCCCTACACGACGCTCTTCCGATCTNNNGAGAACTTCTACATTCCCCAGC
GAGCACACGTCTGAACTCTTTCCCTACACGACGCTCTTCCGATCTNNNACTTCAAATCCAACTGGTGGGT
AGCACACGTCTGAACTCTTTCCCTACACGACGCTCTTCCGATCTNNNCCTTCTGGGGCTGATTTCAGTT
GAGCACACGTCTGAACTCTTTCCCTACACGACGCTCTTCCGATCTNNNTAACAGGCGACAGACGACGA
AGCACACGTCTGAACTCTTTCCCTACACGACGCTCTTCCGATCTNNNCAAAGTCTTGGCTGCTCTGTCT
AGCACACGTCTGAACTCTTTCCCTACACGACGCTCTTCCGATCTNNNATTCAACTCCAGGTAATGGTGG
AGCACACGTCTGAACTCTTTCCCTACACGACGCTCTTCCGATCTNNNAGCATATCTAGTCATAGAAAAG
AAGAGCACACGTCTGAACTCTTTCCCTACACGACGCTCTTCCGATCTNNNGCAGCATGTCTCAGGTAT
AGAGCACACGTCTGAACTCTTTCCCTACACGACGCTCTTCCGATCTNNNAATGATGACCTACGGTTCCAT
GAGCACACGTCTGAACTCTTTCCCTACACGACGCTCTTCCGATCTNNNCAATAAGCCACAAACACACAC
AGCACACGTCTGAACTCTTTCCCTACACGACGCTCTTCCGATCTNNNAGACCCATCAGAGATTTGTCCC
GAGCACACGTCTGAACTCTTTCCCTACACGACGCTCTTCCGATCTNNNTGCAAAATAAAATCCCAGCAC
AGCACACGTCTGAACTCTTTCCCTACACGACGCTCTTCCGATCTNNNGGGAGTGTATACGTGGGAGAG
AGCACACGTCTGAACTCTTTCCCTACACGACGCTCTTCCGATCTNNNGGGAGTGTATACGTGGGAGA
GAGCACACGTCTGAACTCTTTCCCTACACGACGCTCTTCCGATCTNNNCCCGTGAGATGAAAGACACA
AGCACACGTCTGAACTCTTTCCCTACACGACGCTCTTCCGATCTNNNTCATTTTGAAACCAAACAACAC
AGCACACGTCTGAACTCTTTCCCTACACGACGCTCTTCCGATCTNNNGGTCAGACGTAGAATCCAGAC
GCACACGTCTGAACTCTTTCCCTACACGACGCTCTTCCGATCTNNNACAAACCCCAGCTCCACTCTCATC
AGAGCACACGTCTGAACTCTTTCCCTACACGACGCTCTTCCGATCTNNNTTCAAGGTCTTTCCACTGTGC
GAGCACACGTCTGAACTCTTTCCCTACACGACGCTCTTCCGATCTNNNAGGCAAAATCAGTGAAGCTG
AGCACACGTCTGAACTCTTTCCCTACACGACGCTCTTCCGATCTNNNATTCTTGGCAAAGAGGTGCTCT
AGCACACGTCTGAACTCTTTCCCTACACGACGCTCTTCCGATCTNNNGTCGAAGTTGCAGAGAGCAAG
GAGCACACGTCTGAACTCTTTCCCTACACGACGCTCTTCCGATCTNNNGTGGATTTGTAACAGTGTCCC
AGAGCACACGTCTGAACTCTTTCCCTACACGACGCTCTTCCGATCTNNNGTGAAAATACCTCACCCCTG
AGCACACGTCTGAACTCTTTCCCTACACGACGCTCTTCCGATCTNNNGTCAGTACCCCTAATCTCTGCCT
AGCACACGTCTGAACTCTTTCCCTACACGACGCTCTTCCGATCTNNNAGCATTGCTGTGAACTTGTGA
AGCACACGTCTGAACTCTTTCCCTACACGACGCTCTTCCGATCTNNNAGCAAATCTTCCATGTTCCTCG
GAGCACACGTCTGAACTCTTTCCCTACACGACGCTCTTCCGATCTNNNCTGATTACAGAAACGACGGT
GAGCACACGTCTGAACTCTTTCCCTACACGACGCTCTTCCGATCTNNNGACATTTTGGGGAAGGGCTT
AGCACACGTCTGAACTCTTTCCCTACACGACGCTCTTCCGATCTNNNTTGCCTATAGGTGTGTGAGGT
AGAGCACACGTCTGAACTCTTTCCCTACACGACGCTCTTCCGATCTNNNCCACTCTCTAACACTTCTGTG
AGAGCACACGTCTGAACTCTTTCCCTACACGACGCTCTTCCGATCTNNNTTTCCCAGTCATGTAGACCC
AGCACACGTCTGAACTCTTTCCCTACACGACGCTCTTCCGATCTNNNGCAGGATGGGAATCTCTTCTGA
AGAGCACACGTCTGAACTCTTTCCCTACACGACGCTCTTCCGATCTNNNTACTGCGTTGTAAAGCACTGA
AGCACACGTCTGAACTCTTTCCCTACACGACGCTCTTCCGATCTNNNACCTGTAAAACATGAGACTGCA
GAGCACACGTCTGAACTCTTTCCCTACACGACGCTCTTCCGATCTNNNATGATGCTTTGGTTATTGCGG
AGCACACGTCTGAACTCTTTCCCTACACGACGCTCTTCCGATCTNNNTGCCCACAATCATAGTTCACTGC
AGCACACGTCTGAACTCTTTCCCTACACGACGCTCTTCCGATCTNNNTAACAGATTTCACCTCACTGGG
AGCACACGTCTGAACTCTTTCCCTACACGACGCTCTTCCGATCTNNNTCTACGGAGGACTGGATGAAG
AGCACACGTCTGAACTCTTTCCCTACACGACGCTCTTCCGATCTNNNATAAGCTTGCGAAAGGGAAGA

GAGCACACGTCTGAACTCTTTCCCTACACGACGCTCTTCCGATCTNNNCAAGTGTAGCAATGGCACTT
CACACGTCTGAACTCTTTCCCTACACGACGCTCTTCCGATCTNNNTTAGCAAAAGCCCCAGAGATTCC
AGCACACGTCTGAACTCTTTCCCTACACGACGCTCTTCCGATCTNNNTCATTGAGAGACAAGGTCCTG
AAGAGCACACGTCTGAACTCTTTCCCTACACGACGCTCTTCCGATCTNNNGGAATGGAAGGGAAATG
CACACGTCTGAACTCTTTCCCTACACGACGCTCTTCCGATCTNNNAGACCCATCAGAGATTTGTCCCA
GAGCACACGTCTGAACTCTTTCCCTACACGACGCTCTTCCGATCTNNNCAGGTTTGAGCTGTCACTTC
GAGCACACGTCTGAACTCTTTCCCTACACGACGCTCTTCCGATCTNNNATGAGAGGATCGCTTGTGTC
AGCACACGTCTGAACTCTTTCCCTACACGACGCTCTTCCGATCTNNNGCAGTGAGACCTCTCTCTCTCT
CACACGTCTGAACTCTTTCCCTACACGACGCTCTTCCGATCTNNNCTGTGTCCAGCTCTCCTTGCCATCG
AGCACACGTCTGAACTCTTTCCCTACACGACGCTCTTCCGATCTNNNAGTTTACAGCCTAAGCCCCTGT
GAGCACACGTCTGAACTCTTTCCCTACACGACGCTCTTCCGATCTNNNGAGAGCTCTGATTGGCCAA
GAGCACACGTCTGAACTCTTTCCCTACACGACGCTCTTCCGATCTNNNACAGATTCATACACACCTGC
AGCACACGTCTGAACTCTTTCCCTACACGACGCTCTTCCGATCTNNNTCTTTGGATAGATTTGCCTCTC
GAGCACACGTCTGAACTCTTTCCCTACACGACGCTCTTCCGATCTNNNCAATCCCCTTGCATTGCTCAA
GAGCACACGTCTGAACTCTTTCCCTACACGACGCTCTTCCGATCTNNNAGCACAGTACAGCAAACTTG
AGAGCACACGTCTGAACTCTTTCCCTACACGACGCTCTTCCGATCTNNNAAACTCTTTCCTGCCCCTTCT
AGCACACGTCTGAACTCTTTCCCTACACGACGCTCTTCCGATCTNNNATGTCCAGGCTACTCTCTCATC
GAAGAGCACACGTCTGAACTCTTTCCCTACACGACGCTCTTCCGATCTNNNTGGGTCCCAGAAGAAA
AGCACACGTCTGAACTCTTTCCCTACACGACGCTCTTCCGATCTNNNTTTCCCTCTACAAATCAGCGTG
AGAGCACACGTCTGAACTCTTTCCCTACACGACGCTCTTCCGATCTNNNATCCCCTTTGGTGACAGGA
AGCACACGTCTGAACTCTTTCCCTACACGACGCTCTTCCGATCTNNNATTCTTGTGCTTGAGGTGTGC
AGCACACGTCTGAACTCTTTCCCTACACGACGCTCTTCCGATCTNNNTTGGTATTCTTCATCACATTGC
AGAGCACACGTCTGAACTCTTTCCCTACACGACGCTCTTCCGATCTNNNTTTTATTGGGGCTACTCTCT
AGCACACGTCTGAACTCTTTCCCTACACGACGCTCTTCCGATCTNNNATAGAGGAGAGGTTGAGGA
AGCACACGTCTGAACTCTTTCCCTACACGACGCTCTTCCGATCTNNNTCAATCTCCTCTTCCCCATCCTC
GCACACGTCTGAACTCTTTCCCTACACGACGCTCTTCCGATCTNNNTGTGGGTACTAGGAGTTGGTT
AGCACACGTCTGAACTCTTTCCCTACACGACGCTCTTCCGATCTNNNATTTTTCCAGAACCCTCTCACCC
AGCACACGTCTGAACTCTTTCCCTACACGACGCTCTTCCGATCTNNNGCTACAGGGGAAAAGGTGTG
AAGAGCACACGTCTGAACTCTTTCCCTACACGACGCTCTTCCGATCTNNNATCCCCTTTGGTGACAGG
GAGCACACGTCTGAACTCTTTCCCTACACGACGCTCTTCCGATCTNNNGCACATGATCCAACGTTAGC
GCACACGTCTGAACTCTTTCCCTACACGACGCTCTTCCGATCTNNNCCCCTTATGGCATCTTTGAGCAT
AGAGCACACGTCTGAACTCTTTCCCTACACGACGCTCTTCCGATCTNNNTCCAGGCCTCAGAGACATC
AGCACACGTCTGAACTCTTTCCCTACACGACGCTCTTCCGATCTNNNTCCGATACAGCACAGTGATTAC
GCACACGTCTGAACTCTTTCCCTACACGACGCTCTTCCGATCTNNNACTTTGGAAGTCAATATTCCCA
AGCACACGTCTGAACTCTTTCCCTACACGACGCTCTTCCGATCTNNNTTGGTCTGGGTACATTGTGTG
GCACACGTCTGAACTCTTTCCCTACACGACGCTCTTCCGATCTNNNAAACCTACTACGATCAGCAGCA
GCACACGTCTGAACTCTTTCCCTACACGACGCTCTTCCGATCTNNNGAAAAGGCCAACACCTCACAAT
GCACACGTCTGAACTCTTTCCCTACACGACGCTCTTCCGATCTNNNGGAACACTCAGTCTCTTCAGCAG
AGCACACGTCTGAACTCTTTCCCTACACGACGCTCTTCCGATCTNNNGTATGATTTTGGGGTGTGTG
AGCACACGTCTGAACTCTTTCCCTACACGACGCTCTTCCGATCTNNNTTTCGTATTTCACCCAGCCTTGT
GCACACGTCTGAACTCTTTCCCTACACGACGCTCTTCCGATCTNNNTGTGCCATGCATTCATGTTCTCT
GCACACGTCTGAACTCTTTCCCTACACGACGCTCTTCCGATCTNNNTGAGGGGAAATGGATGGGGT
GCACACGTCTGAACTCTTTCCCTACACGACGCTCTTCCGATCTNNNACAACAAAAGCCCAGTGAGACA

GAGCACACGTCTGAACTCTTTCCCTACACGACGCTCTTCCGATCTNNNTAGCCCCAGCTCTTTATTTTG

AGCACACGTCTGAACTCTTTCCCTACACGACGCTCTTCCGATCTNNNAATAAAGAGCACACACCAGAG

CACACGTCTGAACTCTTTCCCTACACGACGCTCTTCCGATCTNNNTGCTGGGTCCCAAAATTATCGTT

AGCACACGTCTGAACTCTTTCCCTACACGACGCTCTTCCGATCTNNNAGAATTTCAAAGTCGGGAGC

AGCACACGTCTGAACTCTTTCCCTACACGACGCTCTTCCGATCTNNNGTTTCCCCTGCAGATAAGGAA

AGCACACGTCTGAACTCTTTCCCTACACGACGCTCTTCCGATCTNNNCCAAGCAATACTGTCTCCCTT

AGCACACGTCTGAACTCTTTCCCTACACGACGCTCTTCCGATCTNNNCTTGCATGATACAGGGAGGC

GAGCACACGTCTGAACTCTTTCCCTACACGACGCTCTTCCGATCTNNNGTTTCCCCTGCAGATAAGGA

GAGCACACGTCTGAACTCTTTCCCTACACGACGCTCTTCCGATCTNNNGCTGTTGCCACTGAGTAAAA

GAGCACACGTCTGAACTCTTTCCCTACACGACGCTCTTCCGATCTNNNTTTCCCTGTTGTCTTCGAGCA

AGCACACGTCTGAACTCTTTCCCTACACGACGCTCTTCCGATCTNNNTTTCTGTCTTTTTCCTGCTGCCT

GAGCACACGTCTGAACTCTTTCCCTACACGACGCTCTTCCGATCTNNNGGAAAAAGCAGCTATCTGAT

GAGCACACGTCTGAACTCTTTCCCTACACGACGCTCTTCCGATCTNNNTTTGTATCCTGCCCTTTACTG

AGAGCACACGTCTGAACTCTTTCCCTACACGACGCTCTTCCGATCTNNNTGCTCAGCATATCAAAGCA

AGCACACGTCTGAACTCTTTCCCTACACGACGCTCTTCCGATCTNNNCACTGTAAATCCCTTGGCCTCT

AGCACACGTCTGAACTCTTTCCCTACACGACGCTCTTCCGATCTNNNTCTAGAATCACAGTACATGTT

GCACACGTCTGAACTCTTTCCCTACACGACGCTCTTCCGATCTNNNGGTGAAACACCTGTGATGGGA

AGAGCACACGTCTGAACTCTTTCCCTACACGACGCTCTTCCGATCTNNNACAATGGTCACTACGAAGT

AGCACACGTCTGAACTCTTTCCCTACACGACGCTCTTCCGATCTNNNTTCTGGGTGTGTTGTTTGTCT

AGCACACGTCTGAACTCTTTCCCTACACGACGCTCTTCCGATCTNNNTATGGCTCTACAGAGAGAAGG

AGCACACGTCTGAACTCTTTCCCTACACGACGCTCTTCCGATCTNNNCTGTCTGCAGCTTCTATCTTCC

AGAGCACACGTCTGAACTCTTTCCCTACACGACGCTCTTCCGATCTNNNTTATGCATGCACACAAAGC

AGCACACGTCTGAACTCTTTCCCTACACGACGCTCTTCCGATCTNNNAGACCCAAGCAGACATAGAC

AAGAGCACACGTCTGAACTCTTTCCCTACACGACGCTCTTCCGATCTNNNAGAATTTCAAAGTCGGGA

AGCACACGTCTGAACTCTTTCCCTACACGACGCTCTTCCGATCTNNNAGCTCCTACACTCCCTTCTTACA

CACACGTCTGAACTCTTTCCCTACACGACGCTCTTCCGATCTNNNTCAAAACCAGAAACGGAACTACA

AGCACACGTCTGAACTCTTTCCCTACACGACGCTCTTCCGATCTNNNGATTCTACCTGCCCATCTTAGC

GCACACGTCTGAACTCTTTCCCTACACGACGCTCTTCCGATCTNNNACTTCTGTCTTAGAATTTTTGCT

GAGCACACGTCTGAACTCTTTCCCTACACGACGCTCTTCCGATCTNNNCTTGATTCTTGGGTCTGGG

AGAGCACACGTCTGAACTCTTTCCCTACACGACGCTCTTCCGATCTNNNAGTTCAGTGTTTCATGGAC

CACGTCTGAACTCTTTCCCTACACGACGCTCTTCCGATCTNNNCAGGACAAGCACGAGTACTACAGC

GCACACGTCTGAACTCTTTCCCTACACGACGCTCTTCCGATCTNNNACCTCCTTTATCCTGTCAGCCTTC

AGCACACGTCTGAACTCTTTCCCTACACGACGCTCTTCCGATCTNNNGGCAGCACTAACCTCATTCTAG

GAGCACACGTCTGAACTCTTTCCCTACACGACGCTCTTCCGATCTNNNCCAGAGTGAGACCCTGAAA

AGCACACGTCTGAACTCTTTCCCTACACGACGCTCTTCCGATCTNNNATAGCACACATGCCTCCAAACT

GCACACGTCTGAACTCTTTCCCTACACGACGCTCTTCCGATCTNNNGGTTGAGATGATTACCTCGGAC

AAGAGCACACGTCTGAACTCTTTCCCTACACGACGCTCTTCCGATCTNNNCTTATTAGGGCTGGCAAA

GAGCACACGTCTGAACTCTTTCCCTACACGACGCTCTTCCGATCTNNNGAAAGGAAGTTTGAGCTCCC

AGCACACGTCTGAACTCTTTCCCTACACGACGCTCTTCCGATCTNNNACAGGATCTGGTCCTACAGGA

AGCACACGTCTGAACTCTTTCCCTACACGACGCTCTTCCGATCTNNNAGGTCTTTAGCTGAAATGTAC

AGCACACGTCTGAACTCTTTCCCTACACGACGCTCTTCCGATCTNNNTTCCCATGTCCTCCTCTTCTCTC

GAGCACACGTCTGAACTCTTTCCCTACACGACGCTCTTCCGATCTNNNGTCAGGATTGAAACTGGCT

AGCACACGTCTGAACTCTTTCCCTACACGACGCTCTTCCGATCTNNNGCCAAAAAGGCAAGTCATGG

AGCACACGTCTGAACTCTTTCCCTACACGACGCTCTTCCGATCTNNNTGCTCTGCTTACTTACCTACCCC
GAGCACACGTCTGAACTCTTTCCCTACACGACGCTCTTCCGATCTNNNAATGTCAGGAAAAGGACATG
AGAGCACACGTCTGAACTCTTTCCCTACACGACGCTCTTCCGATCTNNNCGGCCTGCTTGAATCACAAA
AGCACACGTCTGAACTCTTTCCCTACACGACGCTCTTCCGATCTNNNAATTTATTCTGAGGGGGCTGT
GAGCACACGTCTGAACTCTTTCCCTACACGACGCTCTTCCGATCTNNNACTGACACTGTACATCTGGTG
GAGCACACGTCTGAACTCTTTCCCTACACGACGCTCTTCCGATCTNNNGCTTACCATGCATGTGGACCTA
AGCACACGTCTGAACTCTTTCCCTACACGACGCTCTTCCGATCTNNNTCTCATAAGAACCTACCTGGCCC
AGCACACGTCTGAACTCTTTCCCTACACGACGCTCTTCCGATCTNNNGGGAGGGGGAGAGTTATCAT
AGCACACGTCTGAACTCTTTCCCTACACGACGCTCTTCCGATCTNNNATGAATCTCTGGCTGGACATAC
AGCACACGTCTGAACTCTTTCCCTACACGACGCTCTTCCGATCTNNNTTAGGCCAGTTTCACTCCACCTT
GCACACGTCTGAACTCTTTCCCTACACGACGCTCTTCCGATCTNNNCTCACCTGAGCTCTGCCTTTATCC
GAGCACACGTCTGAACTCTTTCCCTACACGACGCTCTTCCGATCTNNNTTCTTGCCTAGAAAGCTTAGC
GAGCACACGTCTGAACTCTTTCCCTACACGACGCTCTTCCGATCTNNNGTCCTTCCTGTGGTCTCTACCT
GAGCACACGTCTGAACTCTTTCCCTACACGACGCTCTTCCGATCTNNNGGGGACAATTACAGGCAGGA
GCACACGTCTGAACTCTTTCCCTACACGACGCTCTTCCGATCTNNNAAGTACCCATCCCAGTGTCTTGTC
GAGCACACGTCTGAACTCTTTCCCTACACGACGCTCTTCCGATCTNNNAGATCAATAGGGCCAGAAGTG
AGCACACGTCTGAACTCTTTCCCTACACGACGCTCTTCCGATCTNNNTAAAGCACCCAAGAAAAACCACC
GAGCACACGTCTGAACTCTTTCCCTACACGACGCTCTTCCGATCTNNNTCGTTGGGTGACACTAGCATA
AAGAGCACACGTCTGAACTCTTTCCCTACACGACGCTCTTCCGATCTNNNTGCAGTATGATTCCAGGAC
GAGCACACGTCTGAACTCTTTCCCTACACGACGCTCTTCCGATCTNNNCATTCCTCTCTGCCTCTACCAG
AGAGCACACGTCTGAACTCTTTCCCTACACGACGCTCTTCCGATCTNNNTGCTGCTCTCTTAGACATAGC
GCACACGTCTGAACTCTTTCCCTACACGACGCTCTTCCGATCTNNNCACATTAGAAATCCAAGGAGGG
GAGCACACGTCTGAACTCTTTCCCTACACGACGCTCTTCCGATCTNNNTGTGTGATTTAGAGATAACCA
GAGCACACGTCTGAACTCTTTCCCTACACGACGCTCTTCCGATCTNNNTGAGTGTGTGTGTGTTACATA
GAGCACACGTCTGAACTCTTTCCCTACACGACGCTCTTCCGATCTNNNAGTCACTTAGCCTGTGTGCTTA
GAGCACACGTCTGAACTCTTTCCCTACACGACGCTCTTCCGATCTNNNGCTATTTTGGCCACACAACCAA
AAGAGCACACGTCTGAACTCTTTCCCTACACGACGCTCTTCCGATCTNNNAATCACAACAACGTCTTCCT
AGCACACGTCTGAACTCTTTCCCTACACGACGCTCTTCCGATCTNNNCTGTGAAATGCCCAAAAGCACAT
AGCACACGTCTGAACTCTTTCCCTACACGACGCTCTTCCGATCTNNNTCATCTAATTCCACAGTGCCACG
GAGCACACGTCTGAACTCTTTCCCTACACGACGCTCTTCCGATCTNNNAATCACAACAACGTCTTCCTG
GAGCACACGTCTGAACTCTTTCCCTACACGACGCTCTTCCGATCTNNNATGACAGTGGAGATGGCATT
GAGCACACGTCTGAACTCTTTCCCTACACGACGCTCTTCCGATCTNNNAGCCAAAAACCAGAATTAGCCT
GAGCACACGTCTGAACTCTTTCCCTACACGACGCTCTTCCGATCTNNNAGAAATTACTCCAGGAGCAC
GAGCACACGTCTGAACTCTTTCCCTACACGACGCTCTTCCGATCTNNNCAAGGCAGGACCTTCTGACTG
AGCACACGTCTGAACTCTTTCCCTACACGACGCTCTTCCGATCTNNNGGATATTGGAAAGGCATCAGG
GCACACGTCTGAACTCTTTCCCTACACGACGCTCTTCCGATCTNNNACTGACCTGTGAAGTAGATGATC
AGCACACGTCTGAACTCTTTCCCTACACGACGCTCTTCCGATCTNNNTTCAGTCCTGTCTCCCAAGGTAA
AGCACACGTCTGAACTCTTTCCCTACACGACGCTCTTCCGATCTNNNCACTGGTAGATGTGCCATAGCT
GAGCACACGTCTGAACTCTTTCCCTACACGACGCTCTTCCGATCTNNNTTTCTACTTGGCTTTCCTAGGT
GAGCACACGTCTGAACTCTTTCCCTACACGACGCTCTTCCGATCTNNNGAAGGCAGCTTCCTGTAAGTC
GAGCACACGTCTGAACTCTTTCCCTACACGACGCTCTTCCGATCTNNNTGGGCCAGAGTGATACTCTAA
AGCACACGTCTGAACTCTTTCCCTACACGACGCTCTTCCGATCTNNNATCATATTCCAGTGTGCCTACCT
GAGCACACGTCTGAACTCTTTCCCTACACGACGCTCTTCCGATCTNNNGCAAAATCCCCAATACATGTCA

AAGAGCACACGTCTGAACTCTTTCCCTACACGACGCTCTTCCGATCTNNNAGAAGTGAGGAAGCTGT
AGCACACGTCTGAACTCTTTCCCTACACGACGCTCTTCCGATCTNNNCTCACAGGGACACACACACAG
AGCACACGTCTGAACTCTTTCCCTACACGACGCTCTTCCGATCTNNNATGGGATGCCTCTGCTGAGA
GCACACGTCTGAACTCTTTCCCTACACGACGCTCTTCCGATCTNNNCCATTCTGCTCATTCAGTCCTGT
GAGCACACGTCTGAACTCTTTCCCTACACGACGCTCTTCCGATCTNNNATTGATACCTAGGGCCTCTG
GCACACGTCTGAACTCTTTCCCTACACGACGCTCTTCCGATCTNNNAAATTGCTGGTGGAAACACTTG
GCACACGTCTGAACTCTTTCCCTACACGACGCTCTTCCGATCTNNNAGTTTACTGCGTAGTGCTTTGG
AGAGCACACGTCTGAACTCTTTCCCTACACGACGCTCTTCCGATCTNNNAAACTGTCCAGATGGTTCT
CACACGTCTGAACTCTTTCCCTACACGACGCTCTTCCGATCTNNNAGTCAAAAGTTGAGGTCGGTGTC
CACACGTCTGAACTCTTTCCCTACACGACGCTCTTCCGATCTNNNTATTCTGCAGTTCAAAGGCCTGCC
GCACACGTCTGAACTCTTTCCCTACACGACGCTCTTCCGATCTNNNACTAAACTGATTTTTGCCATGG
GCACACGTCTGAACTCTTTCCCTACACGACGCTCTTCCGATCTNNNAGCTAGAATGGTTCTCAGAAA
AGCACACGTCTGAACTCTTTCCCTACACGACGCTCTTCCGATCTNNNAGAAGCTGGGTTGGCAATAA
GAGCACACGTCTGAACTCTTTCCCTACACGACGCTCTTCCGATCTNNNACAGAGCAGCAGCAATTACA
AGCACACGTCTGAACTCTTTCCCTACACGACGCTCTTCCGATCTNNNAGTCACTTAGCCTGTGTGCTTA
AGCACACGTCTGAACTCTTTCCCTACACGACGCTCTTCCGATCTNNNACACATAGATCACATGGTACTC
AGCACACGTCTGAACTCTTTCCCTACACGACGCTCTTCCGATCTNNNCTAGCTTGGTGTCTCCTTCAAC
AGCACACGTCTGAACTCTTTCCCTACACGACGCTCTTCCGATCTNNNGGCAATGATGAAAACTACTCG
GCACACGTCTGAACTCTTTCCCTACACGACGCTCTTCCGATCTNNNGACAAGGATTCAACCATTGCCGC
GAGCACACGTCTGAACTCTTTCCCTACACGACGCTCTTCCGATCTNNNCTTCTTTGGGGCTCTGAAAAT
CACACGTCTGAACTCTTTCCCTACACGACGCTCTTCCGATCTNNNCGCAGCATGGTGGAGGTGCTCAT
GCACACGTCTGAACTCTTTCCCTACACGACGCTCTTCCGATCTNNNGCCACCAACCTCATTCTGTTTGC
GAGCACACGTCTGAACTCTTTCCCTACACGACGCTCTTCCGATCTNNNGGCCCAAAATGGAAAACATA
CACACGTCTGAACTCTTTCCCTACACGACGCTCTTCCGATCTNNNTTTGCAGGGTCCTAACTCAATCGC
AGAGCACACGTCTGAACTCTTTCCCTACACGACGCTCTTCCGATCTNNNCCCCGCATTCACCTCTTCTA
AAGAGCACACGTCTGAACTCTTTCCCTACACGACGCTCTTCCGATCTNNNCCATGAATTAGATGCAG
AGCACACGTCTGAACTCTTTCCCTACACGACGCTCTTCCGATCTNNNGGGAAGGTTGTTGCCATAGA
CACACGTCTGAACTCTTTCCCTACACGACGCTCTTCCGATCTNNNCAAGAGGATTGGTGGATGGATG
CACACGTCTGAACTCTTTCCCTACACGACGCTCTTCCGATCTNNNCATCAGTGTGAACCAGATGGACC
GCACACGTCTGAACTCTTTCCCTACACGACGCTCTTCCGATCTNNNCTCACTGTCCTTCCCTGGCTACC
GCACACGTCTGAACTCTTTCCCTACACGACGCTCTTCCGATCTNNNTCGTTGGGTGACACTAGCATAT
GAGCACACGTCTGAACTCTTTCCCTACACGACGCTCTTCCGATCTNNNGGGACTGTGTATGTGTGTC
GAGCACACGTCTGAACTCTTTCCCTACACGACGCTCTTCCGATCTNNNTCAGTGGAAACATTCCCTTCA
AAGAGCACACGTCTGAACTCTTTCCCTACACGACGCTCTTCCGATCTNNNTATTTTGTCTCGCCCATG
GCACACGTCTGAACTCTTTCCCTACACGACGCTCTTCCGATCTNNNCTTGTGTGGTTTGGAGTGAGT
GAGCACACGTCTGAACTCTTTCCCTACACGACGCTCTTCCGATCTNNNGTAATTTCCTCTTGCTCCCCC
GCACACGTCTGAACTCTTTCCCTACACGACGCTCTTCCGATCTNNNCGTCTACTCAGTGTCACGGTTT
GCACACGTCTGAACTCTTTCCCTACACGACGCTCTTCCGATCTNNNAATGAAAAGCATCACCTCCTCCA
AGCACACGTCTGAACTCTTTCCCTACACGACGCTCTTCCGATCTNNNTATGGTGGAATATGCTGGCCC
GCACACGTCTGAACTCTTTCCCTACACGACGCTCTTCCGATCTNNNCCATCCTGCCCTACTACATCACG
GCACACGTCTGAACTCTTTCCCTACACGACGCTCTTCCGATCTNNNACCTAAACCTGATGTCTCCACCA
GAGCACACGTCTGAACTCTTTCCCTACACGACGCTCTTCCGATCTNNNAGCAGTTTCTACTGCACCCTC
AGCACACGTCTGAACTCTTTCCCTACACGACGCTCTTCCGATCTNNNAGCTAATGAAGCCTGACCATG

AGCACACGTCTGAACTCTTTCCCTACACGACGCTCTTCCGATCTNNNTACCTGTAATGGCATGTGTGC
AAGAGCACACGTCTGAACTCTTTCCCTACACGACGCTCTTCCGATCTNNNGGCAACAGCTAATTCACA
AGCACACGTCTGAACTCTTTCCCTACACGACGCTCTTCCGATCTNNNGAATACTTACAGGGCTCCAGT
GCACACGTCTGAACTCTTTCCCTACACGACGCTCTTCCGATCTNNNGTGAATGAAATGCTTGGAGCAG
AGCACACGTCTGAACTCTTTCCCTACACGACGCTCTTCCGATCTNNNTTGTGACAGTTTGGAGTAGA
AGCACACGTCTGAACTCTTTCCCTACACGACGCTCTTCCGATCTNNNCTTTACTGGTGGTGATGGTGG
AGCACACGTCTGAACTCTTTCCCTACACGACGCTCTTCCGATCTNNNTACATGCACACATAAATGCAGG
AGCACACGTCTGAACTCTTTCCCTACACGACGCTCTTCCGATCTNNNAAAAACAAGCCTGCTTGAGCA
AGCACACGTCTGAACTCTTTCCCTACACGACGCTCTTCCGATCTNNNATCAGAAAGACACTCCTACCC
AGCACACGTCTGAACTCTTTCCCTACACGACGCTCTTCCGATCTNNNACCCGTTCTTCCAGAATAGGTC
GCACACGTCTGAACTCTTTCCCTACACGACGCTCTTCCGATCTNNNACCTCTGGTATAGTTCTTTTCTG
AGCACACGTCTGAACTCTTTCCCTACACGACGCTCTTCCGATCTNNNCTCTGCTCTCACTTTCATCACC
GCACACGTCTGAACTCTTTCCCTACACGACGCTCTTCCGATCTNNNGAACAAATGGGAGGAACTCCT
AGCACACGTCTGAACTCTTTCCCTACACGACGCTCTTCCGATCTNNNCCCAACTCTTGTTTGGGGGTA
AAGAGCACACGTCTGAACTCTTTCCCTACACGACGCTCTTCCGATCTNNNTCAAAAGGCTCTTCAGTG
GCACACGTCTGAACTCTTTCCCTACACGACGCTCTTCCGATCTNNNTGTAAATCCAGATTCCAAAACC
AGCACACGTCTGAACTCTTTCCCTACACGACGCTCTTCCGATCTNNNGTTACATGTGAAACAGAGCAC
AGCACACGTCTGAACTCTTTCCCTACACGACGCTCTTCCGATCTNNNTCTCTCTTTTGCTTTCTCTTGCA
AGCACACGTCTGAACTCTTTCCCTACACGACGCTCTTCCGATCTNNNTGTGCCCTTAGGCAGTTTCTTT
GAGCACACGTCTGAACTCTTTCCCTACACGACGCTCTTCCGATCTNNNCATCTCTCTTCCATCCCCTAGC
GAGCACACGTCTGAACTCTTTCCCTACACGACGCTCTTCCGATCTNNNGCAAATGCCATTATTTCACGC
GAGCACACGTCTGAACTCTTTCCCTACACGACGCTCTTCCGATCTNNNATTCTGAGCTCCTTCTTTGCA
GAGCACACGTCTGAACTCTTTCCCTACACGACGCTCTTCCGATCTNNNAAGAAAGGGTGGTGAGACA
GAGCACACGTCTGAACTCTTTCCCTACACGACGCTCTTCCGATCTNNNCATGAGGGAGACTTTGAGG
GAGCACACGTCTGAACTCTTTCCCTACACGACGCTCTTCCGATCTNNNGTGCCCTCAATAAGGCACAT
AGCACACGTCTGAACTCTTTCCCTACACGACGCTCTTCCGATCTNNNTTGTTGTTCAACCCACGATGTC
AGAGCACACGTCTGAACTCTTTCCCTACACGACGCTCTTCCGATCTNNNATTCTGCTGCTGTCCCCTAG
AGCACACGTCTGAACTCTTTCCCTACACGACGCTCTTCCGATCTNNNCCCAATATTTCTGCAGTGCAAC
GCACACGTCTGAACTCTTTCCCTACACGACGCTCTTCCGATCTNNNGTGGTGATGGAGCAAGACCCT
GCACACGTCTGAACTCTTTCCCTACACGACGCTCTTCCGATCTNNNTCCTTTCCTTCTGTCTTCCATGCC
AGCACACGTCTGAACTCTTTCCCTACACGACGCTCTTCCGATCTNNNATCAATATATACATGCACACAC
GCACACGTCTGAACTCTTTCCCTACACGACGCTCTTCCGATCTNNNTAAAAATGTCCCTTTTGGCACG
AGCACACGTCTGAACTCTTTCCCTACACGACGCTCTTCCGATCTNNNTTTCTCCACTACCAAAACTGG
AGCACACGTCTGAACTCTTTCCCTACACGACGCTCTTCCGATCTNNNGTCTTTCAGTTCCATCCTCTCC
AGCACACGTCTGAACTCTTTCCCTACACGACGCTCTTCCGATCTNNNTAGAAACCCAGGTCAGACACA
AGCACACGTCTGAACTCTTTCCCTACACGACGCTCTTCCGATCTNNNTCATTCTGTTATGATTCTCTACA
AGAGCACACGTCTGAACTCTTTCCCTACACGACGCTCTTCCGATCTNNNAGAGACAAGGCTCACTGAG
AGCACACGTCTGAACTCTTTCCCTACACGACGCTCTTCCGATCTNNNGTTTCTGTGGCTCCATCTATCC
AGCACACGTCTGAACTCTTTCCCTACACGACGCTCTTCCGATCTNNNCACCATAGCATGTATAGTAGG
AGCACACGTCTGAACTCTTTCCCTACACGACGCTCTTCCGATCTNNNTGCACGTCTGTGTATTTGTGT
AGAGCACACGTCTGAACTCTTTCCCTACACGACGCTCTTCCGATCTNNNGTTTGGCCTGAGTTTCTGG
GAGCACACGTCTGAACTCTTTCCCTACACGACGCTCTTCCGATCTNNNAAATTTCAGATGCTGGAAAC
AGAGCACACGTCTGAACTCTTTCCCTACACGACGCTCTTCCGATCTNNNCCCCCAAGAGTGTCAATTTC

GCACACGTCTGAACTCTTTCCCTACACGACGCTCTTCCGATCTNNNTCTCCTTCCTCATCTCCTACCCTCAT
AAGAGCACACGTCTGAACTCTTTCCCTACACGACGCTCTTCCGATCTNNNATTCTGTCACTTCCCTTCTG
GAGCACACGTCTGAACTCTTTCCCTACACGACGCTCTTCCGATCTNNNCCGCCTCACTTTCTTTCTCTCTC
AGAGCACACGTCTGAACTCTTTCCCTACACGACGCTCTTCCGATCTNNNGGCCTGTTTCCTCTAGCTTTT
AGCACACGTCTGAACTCTTTCCCTACACGACGCTCTTCCGATCTNNNAGATGTGGCAAAAACCAAACTG
GAGCACACGTCTGAACTCTTTCCCTACACGACGCTCTTCCGATCTNNNTCAAAAGCCATATACGCCCACA
GAGCACACGTCTGAACTCTTTCCCTACACGACGCTCTTCCGATCTNNNAGTCACTTAGCTGGTCTTCCTG
AGCACACGTCTGAACTCTTTCCCTACACGACGCTCTTCCGATCTNNNTGGGGCCAAAGGAAAATAACC
GAGCACACGTCTGAACTCTTTCCCTACACGACGCTCTTCCGATCTNNNCTAGCTGGGTGGGTTACACTT
GAGCACACGTCTGAACTCTTTCCCTACACGACGCTCTTCCGATCTNNNACATGAAGCTCATGCAGTACC
AGCACACGTCTGAACTCTTTCCCTACACGACGCTCTTCCGATCTNNNCCTGCTGGTCTCCCAATTCTTTGC
AGCACACGTCTGAACTCTTTCCCTACACGACGCTCTTCCGATCTNNNTACGTCTTCTAGAGAGCTTGTG
AGCACACGTCTGAACTCTTTCCCTACACGACGCTCTTCCGATCTNNNACATGGGGCTTTGGTAAAGAAC
AGCACACGTCTGAACTCTTTCCCTACACGACGCTCTTCCGATCTNNNGACCAAAAGCTAGTCAGCACTC
AGCACACGTCTGAACTCTTTCCCTACACGACGCTCTTCCGATCTNNNCAGTGCCAAGGTTCTGAGACTA
GAGCACACGTCTGAACTCTTTCCCTACACGACGCTCTTCCGATCTNNNATCTTCCCCTGTGCTCTTTCTCT
AGCACACGTCTGAACTCTTTCCCTACACGACGCTCTTCCGATCTNNNAGTTGGATCCCAAAATGGATCG
AGCACACGTCTGAACTCTTTCCCTACACGACGCTCTTCCGATCTNNNTTACACTACATGCACACACATGC
AGCACACGTCTGAACTCTTTCCCTACACGACGCTCTTCCGATCTNNNATGATTTGCTTCGCATAGACAGC
GAGCACACGTCTGAACTCTTTCCCTACACGACGCTCTTCCGATCTNNNGAATGGTGTCCTGAGCTTCTG
GAGCACACGTCTGAACTCTTTCCCTACACGACGCTCTTCCGATCTNNNACAGAATGAGCACAAAAGCCA
AGCACACGTCTGAACTCTTTCCCTACACGACGCTCTTCCGATCTNNNTAATAATGCATGGTGCCCACAG
GAGCACACGTCTGAACTCTTTCCCTACACGACGCTCTTCCGATCTNNNTCGAGATAGAGCTCGTCTAGT
AGCACACGTCTGAACTCTTTCCCTACACGACGCTCTTCCGATCTNNNATCCAGGATGTGATGTATGGGC
GAGCACACGTCTGAACTCTTTCCCTACACGACGCTCTTCCGATCTNNNGCATATGCTTCCTGGTTGAAA
GAGCACACGTCTGAACTCTTTCCCTACACGACGCTCTTCCGATCTNNNAGAGGGAATGTGCTATGTGT
GAGCACACGTCTGAACTCTTTCCCTACACGACGCTCTTCCGATCTNNNGGTAATTGGCCCAGTGATTTT
AGAGCACACGTCTGAACTCTTTCCCTACACGACGCTCTTCCGATCTNNNTGAAGAGCTGACCCATAGTG
GCACACGTCTGAACTCTTTCCCTACACGACGCTCTTCCGATCTNNNTTCCCATCACTGCTTCCCTCTAACA
AGCACACGTCTGAACTCTTTCCCTACACGACGCTCTTCCGATCTNNNCTCCTTCTCCCACTTCCTCTCTTCA
GAGCACACGTCTGAACTCTTTCCCTACACGACGCTCTTCCGATCTNNNGGAGGATTTGTTTTGTGAAG
AGCACACGTCTGAACTCTTTCCCTACACGACGCTCTTCCGATCTNNNACATGGGGCTTTGGTAAAGAAG
GAGCACACGTCTGAACTCTTTCCCTACACGACGCTCTTCCGATCTNNNCCTTCTGTGCACTTCCTACTCTG
AGAGCACACGTCTGAACTCTTTCCCTACACGACGCTCTTCCGATCTNNNTAGCAATACCCCCTCGGTACA
GAGCACACGTCTGAACTCTTTCCCTACACGACGCTCTTCCGATCTNNNTGTACACTGTGCAGAGGTAA
GAGCACACGTCTGAACTCTTTCCCTACACGACGCTCTTCCGATCTNNNTGTACACTGTGCAGAGGTAAG
AGCACACGTCTGAACTCTTTCCCTACACGACGCTCTTCCGATCTNNNAGGTGATTCTCCACCTCTCAGTC
AGCACACGTCTGAACTCTTTCCCTACACGACGCTCTTCCGATCTNNNATAACAAGACAGGGGATGGCT
GAGCACACGTCTGAACTCTTTCCCTACACGACGCTCTTCCGATCTNNNACCTCTTCCAAATCACACACAC
AGAGCACACGTCTGAACTCTTTCCCTACACGACGCTCTTCCGATCTNNNTAGCAATACCCCCTCGGTACA
ACACGTCTGAACTCTTTCCCTACACGACGCTCTTCCGATCTNNNAGGTGATTCTCCACCTCTCAGTCCAT
AGCACACGTCTGAACTCTTTCCCTACACGACGCTCTTCCGATCTNNNGCTCCTGTCAATTTCCACAGTCA
GAGCACACGTCTGAACTCTTTCCCTACACGACGCTCTTCCGATCTNNNACTGGCTCAGACAAGACTACAG

GCACACGTCTGAACTCTTTCCCTACACGACGCTCTTCCGATCTNNNATAAATGCCTCTTCTCATCCGCCC
GAGCACACGTCTGAACTCTTTCCCTACACGACGCTCTTCCGATCTNNNATGTGAGCCTTCCTCTGATTC
AGAGCACACGTCTGAACTCTTTCCCTACACGACGCTCTTCCGATCTNNNACATATTTGCTTAGCAGGG
AGAGCACACGTCTGAACTCTTTCCCTACACGACGCTCTTCCGATCTNNNTCATCATTGGCTTTGTTTTG
GAGCACACGTCTGAACTCTTTCCCTACACGACGCTCTTCCGATCTNNNGTGTTGTGTGCACATGCT
AGCACACGTCTGAACTCTTTCCCTACACGACGCTCTTCCGATCTNNNGGGCATGCTATATGCAAGTGA
AGCACACGTCTGAACTCTTTCCCTACACGACGCTCTTCCGATCTNNNCTCCTACCATATTCCCCTCCCTG
AGCACACGTCTGAACTCTTTCCCTACACGACGCTCTTCCGATCTNNNATTTGGTTTGCAATGGGAGCT
GAGCACACGTCTGAACTCTTTCCCTACACGACGCTCTTCCGATCTNNNACATAGAACAGGGAGGAGCT
GAGCACACGTCTGAACTCTTTCCCTACACGACGCTCTTCCGATCTNNNAAAACTCCACAAGCCCCAGAA
AGCACACGTCTGAACTCTTTCCCTACACGACGCTCTTCCGATCTNNNCAGACACAGACCCACTCACAAA
AGAGCACACGTCTGAACTCTTTCCCTACACGACGCTCTTCCGATCTNNNCCCCAAGAGTGTCAATTTC
GCACACGTCTGAACTCTTTCCCTACACGACGCTCTTCCGATCTNNNCTAGTGGTCCTACGTCCTCATCT
GAGCACACGTCTGAACTCTTTCCCTACACGACGCTCTTCCGATCTNNNCCCAGCCAAGAACTTTCACTT
AGCACACGTCTGAACTCTTTCCCTACACGACGCTCTTCCGATCTNNNTCACTCTATGATCCTGGGCAAC
GAGCACACGTCTGAACTCTTTCCCTACACGACGCTCTTCCGATCTNNNGGAGAATGGGGGAAGAAGC
AGCACACGTCTGAACTCTTTCCCTACACGACGCTCTTCCGATCTNNNAAAGTCAGAGACAGGAAGCGG
AAGAGCACACGTCTGAACTCTTTCCCTACACGACGCTCTTCCGATCTNNNCCAAGACCGAAAGGCCTA
GAGCACACGTCTGAACTCTTTCCCTACACGACGCTCTTCCGATCTNNNAGCAGAAACCACAATCTCTCT
AGAGCACACGTCTGAACTCTTTCCCTACACGACGCTCTTCCGATCTNNNACTGTTGCAGTCCGGAAAC
GAGCACACGTCTGAACTCTTTCCCTACACGACGCTCTTCCGATCTNNNAGCTTTTGACTTTTGGCCACA
AGCACACGTCTGAACTCTTTCCCTACACGACGCTCTTCCGATCTNNNGACTTGCTCAAGGTCTCTGCAT
AGAGCACACGTCTGAACTCTTTCCCTACACGACGCTCTTCCGATCTNNNCTGCACCTGGACAACTTTCT
AGCACACGTCTGAACTCTTTCCCTACACGACGCTCTTCCGATCTNNNTACTGTGTCTCACAGAGAAGG
GCACACGTCTGAACTCTTTCCCTACACGACGCTCTTCCGATCTNNNACTCACTTGAACAGATTTGTAAG
GAGCACACGTCTGAACTCTTTCCCTACACGACGCTCTTCCGATCTNNNGTTTACGTTAGTCCTGTGGG
AGAGCACACGTCTGAACTCTTTCCCTACACGACGCTCTTCCGATCTNNNATTTGGTTGGCTCTGGAAT
GAGCACACGTCTGAACTCTTTCCCTACACGACGCTCTTCCGATCTNNNTTTTCTCCTGCAGCTTCCCTC
GCACACGTCTGAACTCTTTCCCTACACGACGCTCTTCCGATCTNNNTTATCCCAGAATCAGCTGGCAAC
GAGCACACGTCTGAACTCTTTCCCTACACGACGCTCTTCCGATCTNNNTGTACATGGACTTCCCTCTCC
CACACGTCTGAACTCTTTCCCTACACGACGCTCTTCCGATCTNNNTCCCCTGGCTTAGAAACAGATGAC
AGCACACGTCTGAACTCTTTCCCTACACGACGCTCTTCCGATCTNNNCTCTGCTCACAAGTCCTTCCTTC
AGCACACGTCTGAACTCTTTCCCTACACGACGCTCTTCCGATCTNNNCATATTTGCACACAGGCACATG
GAGCACACGTCTGAACTCTTTCCCTACACGACGCTCTTCCGATCTNNNATAGTAACCCACAAACCCCTG
GCACACGTCTGAACTCTTTCCCTACACGACGCTCTTCCGATCTNNNCAAAGAAAGGAAGATGGCGAG
GCACACGTCTGAACTCTTTCCCTACACGACGCTCTTCCGATCTNNNGGTTAGCTAGTGGTTTGGCAGA
AGAGCACACGTCTGAACTCTTTCCCTACACGACGCTCTTCCGATCTNNNCCTGAGGGTCGATCTCATC
GAGCACACGTCTGAACTCTTTCCCTACACGACGCTCTTCCGATCTNNNGCCACAACGCTAGTCATAGT
GAGCACACGTCTGAACTCTTTCCCTACACGACGCTCTTCCGATCTNNNTGACCTACTCAGGTTTCACCAG
GAGCACACGTCTGAACTCTTTCCCTACACGACGCTCTTCCGATCTNNNACAAGACACAGACATCTTCCTG
GCACACGTCTGAACTCTTTCCCTACACGACGCTCTTCCGATCTNNNTGACATCTAGCAAGAAGTTGGG
AGCACACGTCTGAACTCTTTCCCTACACGACGCTCTTCCGATCTNNNAATCCAGGCTTTCCCAAATGTG
GAGCACACGTCTGAACTCTTTCCCTACACGACGCTCTTCCGATCTNNNGACAGCGATCAACGTGTAGT

AAGAGCACACGTCTGAACTCTTTCCCTACACGACGCTCTTCCGATCTNNNCCAAGACCGAAAGGCCT
AGAGCACACGTCTGAACTCTTTCCCTACACGACGCTCTTCCGATCTNNNAAGGACCCTCCTTTGTTTCT
AGCACACGTCTGAACTCTTTCCCTACACGACGCTCTTCCGATCTNNNCCTGGCCCACTAATGTACTTTC
GAGCACACGTCTGAACTCTTTCCCTACACGACGCTCTTCCGATCTNNNTTCAATGGCTTCAGCTCCCATT
AGCACACGTCTGAACTCTTTCCCTACACGACGCTCTTCCGATCTNNNTGTTGGTATTTCAGGTGGTGC
GAGCACACGTCTGAACTCTTTCCCTACACGACGCTCTTCCGATCTNNNGCTTCACTGGAATGAGAAGG
AGAGCACACGTCTGAACTCTTTCCCTACACGACGCTCTTCCGATCTNNNAAAAGTCCCTCTTTGGAAT
AGCACACGTCTGAACTCTTTCCCTACACGACGCTCTTCCGATCTNNNACTGTTGACTGCCTACCCAAAA
AGCACACGTCTGAACTCTTTCCCTACACGACGCTCTTCCGATCTNNNCGTTTCAATGCCTCTCCTGAAT
AGCACACGTCTGAACTCTTTCCCTACACGACGCTCTTCCGATCTNNNACCTGAACATTATAGCCCCACC
AGAGCACACGTCTGAACTCTTTCCCTACACGACGCTCTTCCGATCTNNNTTATCTGTTGGATGCTCCCT
GCACACGTCTGAACTCTTTCCCTACACGACGCTCTTCCGATCTNNNTAGACAAGCAAGACAGCTTCCA
AAGAGCACACGTCTGAACTCTTTCCCTACACGACGCTCTTCCGATCTNNNCTTGCGTACTTTGAGTTT
GAGCACACGTCTGAACTCTTTCCCTACACGACGCTCTTCCGATCTNNNGTGACACAGCCAAACAATCA
AGCACACGTCTGAACTCTTTCCCTACACGACGCTCTTCCGATCTNNNACCTCGTCTGAAAAACAATGA
GCACACGTCTGAACTCTTTCCCTACACGACGCTCTTCCGATCTNNNGGACTTTTTGACCTCCAGCCTAC
GAGCACACGTCTGAACTCTTTCCCTACACGACGCTCTTCCGATCTNNNGGGGAGAGTTTTGTTGTCCC
AAGAGCACACGTCTGAACTCTTTCCCTACACGACGCTCTTCCGATCTNNNCCTGGCCCACTAATGTACT
GAGCACACGTCTGAACTCTTTCCCTACACGACGCTCTTCCGATCTNNNACCTGAACATTATAGCCCCA
AGCACACGTCTGAACTCTTTCCCTACACGACGCTCTTCCGATCTNNNGTCTTGGTGGATCTGGCTTGG
AGCACACGTCTGAACTCTTTCCCTACACGACGCTCTTCCGATCTNNNAAGGGTATCACATGAAACCTG
GAGCACACGTCTGAACTCTTTCCCTACACGACGCTCTTCCGATCTNNNACAAAATGGTCCATGCCTCTG
GAGCACACGTCTGAACTCTTTCCCTACACGACGCTCTTCCGATCTNNNTTTGTACAATGAGGAGGTG
GCACACGTCTGAACTCTTTCCCTACACGACGCTCTTCCGATCTNNNACTCAGACAGGAACACACACAA
GAGCACACGTCTGAACTCTTTCCCTACACGACGCTCTTCCGATCTNNNCTTCTCCAGCCTAGCAGTGTT
GAGCACACGTCTGAACTCTTTCCCTACACGACGCTCTTCCGATCTNNNTAAAGCTAATGCAGACACAC
AGCACACGTCTGAACTCTTTCCCTACACGACGCTCTTCCGATCTNNNGCCTGGCCTTGTATGTCAATTC
GAGCACACGTCTGAACTCTTTCCCTACACGACGCTCTTCCGATCTNNNAGTTCCAACTCCTAGTCTCCT
GAGCACACGTCTGAACTCTTTCCCTACACGACGCTCTTCCGATCTNNNCCTTCCACCTTTGCTGTTGAG
AGCACACGTCTGAACTCTTTCCCTACACGACGCTCTTCCGATCTNNNTGTTGGCTTGTAACCAATGAC
GAGCACACGTCTGAACTCTTTCCCTACACGACGCTCTTCCGATCTNNNTAACTTGGCAGGAAACCTGT
AGCACACGTCTGAACTCTTTCCCTACACGACGCTCTTCCGATCTNNNAAACCAAAACCGAGACACACA
GAGCACACGTCTGAACTCTTTCCCTACACGACGCTCTTCCGATCTNNNAGAGCAGAGATGGCAAATT
GCACACGTCTGAACTCTTTCCCTACACGACGCTCTTCCGATCTNNNATGAGGGGCCCTGAAGAAGTA
AGCACACGTCTGAACTCTTTCCCTACACGACGCTCTTCCGATCTNNNCGGGATTTCAAACACAGACAG
CACACGTCTGAACTCTTTCCCTACACGACGCTCTTCCGATCTNNNACCTGATATCCAGGGCCATTTCTC
AGCACACGTCTGAACTCTTTCCCTACACGACGCTCTTCCGATCTNNNTAAGGCCTTGAATGGGAGAGT
AGCACACGTCTGAACTCTTTCCCTACACGACGCTCTTCCGATCTNNNTTCGACCTTTGATGACTGGAC
AGCACACGTCTGAACTCTTTCCCTACACGACGCTCTTCCGATCTNNNAAAGGGGCTCTCTTTCTCTGTC
AGCACACGTCTGAACTCTTTCCCTACACGACGCTCTTCCGATCTNNNTACAGGGGCTATGGCTATGAA
AGCACACGTCTGAACTCTTTCCCTACACGACGCTCTTCCGATCTNNNAATCTCATTGTTTCCCCAATGC
AGCACACGTCTGAACTCTTTCCCTACACGACGCTCTTCCGATCTNNNGACAGCGATTGACGTGTAGTT
AGCACACGTCTGAACTCTTTCCCTACACGACGCTCTTCCGATCTNNNACTGAATGCTTGCTCTCAAAG

AGCACACGTCTGAACTCTTTCCCTACACGACGCTCTTCCGATCTNNNAGTGTCTTGTACCCAGCCTATG
AGCACACGTCTGAACTCTTTCCCTACACGACGCTCTTCCGATCTNNNAGAGAGAAAGAAGACACCAG
GCACACGTCTGAACTCTTTCCCTACACGACGCTCTTCCGATCTNNNGGCTCGAGCATCTGTATCATCTG
GCACACGTCTGAACTCTTTCCCTACACGACGCTCTTCCGATCTNNNCCACCCAATCTGCCTCTATCACT
GAGCACACGTCTGAACTCTTTCCCTACACGACGCTCTTCCGATCTNNNCGAACTCATACTGCAGATCCT
AGAGCACACGTCTGAACTCTTTCCCTACACGACGCTCTTCCGATCTNNNTTTCTGCCTGCTCAGTGATT
GAGCACACGTCTGAACTCTTTCCCTACACGACGCTCTTCCGATCTNNNATTCATGCTAGCAAGCTCTCC
GAGCACACGTCTGAACTCTTTCCCTACACGACGCTCTTCCGATCTNNNAATACATGACTTCAGGGTGG
GAGCACACGTCTGAACTCTTTCCCTACACGACGCTCTTCCGATCTNNNAGAGTGTATAAGAGGGGG
AGCACACGTCTGAACTCTTTCCCTACACGACGCTCTTCCGATCTNNNTAAAGGTTCCCATGTGTCTGG
GAGCACACGTCTGAACTCTTTCCCTACACGACGCTCTTCCGATCTNNNACTTCAGCCCTGAACTATGG
AGCACACGTCTGAACTCTTTCCCTACACGACGCTCTTCCGATCTNNNCCTGAGGCTGGGAGAACATT
AGCACACGTCTGAACTCTTTCCCTACACGACGCTCTTCCGATCTNNNCTCCTTCCCCAGGTTATCATTG
AAGAGCACACGTCTGAACTCTTTCCCTACACGACGCTCTTCCGATCTNNNGCTGTGAGCTCCAAGAA
CACACGTCTGAACTCTTTCCCTACACGACGCTCTTCCGATCTNNNGCACCTGGTTTTTCCTATGCAACC
AGAGCACACGTCTGAACTCTTTCCCTACACGACGCTCTTCCGATCTNNNTTAGGGAGGCAGAAGCAA
AGAGCACACGTCTGAACTCTTTCCCTACACGACGCTCTTCCGATCTNNNTTCCCCAAACTTTTGAAGA
GCACACGTCTGAACTCTTTCCCTACACGACGCTCTTCCGATCTNNNGGATTCCGAAGGCAAAGAGCT
AGCACACGTCTGAACTCTTTCCCTACACGACGCTCTTCCGATCTNNNCTGGGCTGAACAACCACAGA
AAGAGCACACGTCTGAACTCTTTCCCTACACGACGCTCTTCCGATCTNNNATAGCTGAGGTAGAGGT
GAGCACACGTCTGAACTCTTTCCCTACACGACGCTCTTCCGATCTNNNTGTGTAAGAGCATGGGGGA
GAGCACACGTCTGAACTCTTTCCCTACACGACGCTCTTCCGATCTNNNAAGGGGTAACTTAGCAGG
AGCACACGTCTGAACTCTTTCCCTACACGACGCTCTTCCGATCTNNNGAGTTTGGAGTACAGCTGTG
GCACACGTCTGAACTCTTTCCCTACACGACGCTCTTCCGATCTNNNTAAGTCTCAGCCAGACTTGGACA
AGCACACGTCTGAACTCTTTCCCTACACGACGCTCTTCCGATCTNNNTTTCTGCCATTTCTGACTGAG
GCACACGTCTGAACTCTTTCCCTACACGACGCTCTTCCGATCTNNNCTGACAATTCCAGATCCTCAGG
AGCACACGTCTGAACTCTTTCCCTACACGACGCTCTTCCGATCTNNNGGACCTACTCGGTACTTGGAT
AGCACACGTCTGAACTCTTTCCCTACACGACGCTCTTCCGATCTNNNGATCTAGGAGGCACGCATGAT
GCACACGTCTGAACTCTTTCCCTACACGACGCTCTTCCGATCTNNNATTGGGTACACTGAATGCTTGC
GAGCACACGTCTGAACTCTTTCCCTACACGACGCTCTTCCGATCTNNNAGGTTCAGCAGCTATTGTGA
GCACACGTCTGAACTCTTTCCCTACACGACGCTCTTCCGATCTNNNCCAGTGGTGAGAAACATGGTG
GAGCACACGTCTGAACTCTTTCCCTACACGACGCTCTTCCGATCTNNNGCCATGGAAAACAGGTCATC
GAGCACACGTCTGAACTCTTTCCCTACACGACGCTCTTCCGATCTNNNAGAGAATTTCAGACTTGAGC
AAGAGCACACGTCTGAACTCTTTCCCTACACGACGCTCTTCCGATCTNNNCAACAAAGTGAGACCTCTG
GAGCACACGTCTGAACTCTTTCCCTACACGACGCTCTTCCGATCTNNNGGGATGGGTAGGATCTAACT
GAGCACACGTCTGAACTCTTTCCCTACACGACGCTCTTCCGATCTNNNTGTGCCTTCAGAGCTTCTCTT
AGCACACGTCTGAACTCTTTCCCTACACGACGCTCTTCCGATCTNNNGATTGAGCAATGCCAGGCTAA
AGCACACGTCTGAACTCTTTCCCTACACGACGCTCTTCCGATCTNNNAGAGGGAAAAGCAGGAGTA
AGCACACGTCTGAACTCTTTCCCTACACGACGCTCTTCCGATCTNNNTTTGAAACCTTGCCTAGGGTC
CACACGTCTGAACTCTTTCCCTACACGACGCTCTTCCGATCTNNNCTGTGATGTGGAATGAAAGCAG
AGCACACGTCTGAACTCTTTCCCTACACGACGCTCTTCCGATCTNNNTGTAAGGGACAGAGAGGGAG
AGCACACGTCTGAACTCTTTCCCTACACGACGCTCTTCCGATCTNNNAATGACCTTTCAGGTGGTGCA
AAGAGCACACGTCTGAACTCTTTCCCTACACGACGCTCTTCCGATCTNNNGGTCCTCATAAGCTTTCAC

AGCACACGTCTGAACTCTTTCCCTACACGACGCTCTTCCGATCTNNNTGTCCTTGTTGTCCTTGTTCTG
GCACACGTCTGAACTCTTTCCCTACACGACGCTCTTCCGATCTNNNTGCATCTCCAGATTCACACCCTA
AGCACACGTCTGAACTCTTTCCCTACACGACGCTCTTCCGATCTNNNACCAAACACCTTCATGCCTAGA
GAGCACACGTCTGAACTCTTTCCCTACACGACGCTCTTCCGATCTNNNCACTGGGCTATTATCTACCGC
GAGCACACGTCTGAACTCTTTCCCTACACGACGCTCTTCCGATCTNNNGAAGGTAGCTGACATTGGC
AGCACACGTCTGAACTCTTTCCCTACACGACGCTCTTCCGATCTNNNCCAGAAACCTGCTCTCTCATTC
AGCACACGTCTGAACTCTTTCCCTACACGACGCTCTTCCGATCTNNNATCAAGGGAATCCTGGTTTCA
GAGCACACGTCTGAACTCTTTCCCTACACGACGCTCTTCCGATCTNNNAAATAGGTTTGGGAGGTTG
AGCACACGTCTGAACTCTTTCCCTACACGACGCTCTTCCGATCTNNNACATCGTCCTCAACATTGACCT
AGCACACGTCTGAACTCTTTCCCTACACGACGCTCTTCCGATCTNNNCCCTACTAGATCAGCACCAAGG
AGCACACGTCTGAACTCTTTCCCTACACGACGCTCTTCCGATCTNNNCAGGGGCGATATAGCATTCA
GAGCACACGTCTGAACTCTTTCCCTACACGACGCTCTTCCGATCTNNNGGCTATGCTCTGGAGAGAG
AGCACACGTCTGAACTCTTTCCCTACACGACGCTCTTCCGATCTNNNGTGGTCCTCAAATCCCTGCATA
GCACACGTCTGAACTCTTTCCCTACACGACGCTCTTCCGATCTNNNAAAGTCAGAGACAGGAAGCGG
GCACACGTCTGAACTCTTTCCCTACACGACGCTCTTCCGATCTNNNGGAAGACTGGCTTTGCATTCTC
AGCACACGTCTGAACTCTTTCCCTACACGACGCTCTTCCGATCTNNNCTACCACCACACCCAGCTCTATC
AAGAGCACACGTCTGAACTCTTTCCCTACACGACGCTCTTCCGATCTNNNCTCTGGTTCCCAGGAGGC
AGCACACGTCTGAACTCTTTCCCTACACGACGCTCTTCCGATCTNNNCATTTCAGAGCTGAGTTCCTG
AGCACACGTCTGAACTCTTTCCCTACACGACGCTCTTCCGATCTNNNAGCAGGATGCAGAAGCAAAT
CACACGTCTGAACTCTTTCCCTACACGACGCTCTTCCGATCTNNNCGTGTGTCTATCGACTGGGTTTT
ACACGTCTGAACTCTTTCCCTACACGACGCTCTTCCGATCTNNNCCTGCTGGTCTCCCAATTCTTTGCA
AGCACACGTCTGAACTCTTTCCCTACACGACGCTCTTCCGATCTNNNAGCACCTTCACCTCCTCAAGTA
GCACACGTCTGAACTCTTTCCCTACACGACGCTCTTCCGATCTNNNTCGTCCCTCTCCTCCATACCTAAC
AGCACACGTCTGAACTCTTTCCCTACACGACGCTCTTCCGATCTNNNGAATTGTGGTGGAGGTTGTC
AGCACACGTCTGAACTCTTTCCCTACACGACGCTCTTCCGATCTNNNGCCAGCACAAAAATGAGTTAC
AGCACACGTCTGAACTCTTTCCCTACACGACGCTCTTCCGATCTNNNAGTGATGGTATGCACCTGTAT
GAGCACACGTCTGAACTCTTTCCCTACACGACGCTCTTCCGATCTNNNTAGTTACAGGGGTGGAGCA
AGCACACGTCTGAACTCTTTCCCTACACGACGCTCTTCCGATCTNNNAGAAGGGCCTTATCCACATCC
AGAGCACACGTCTGAACTCTTTCCCTACACGACGCTCTTCCGATCTNNNGCTCGGTTCCTTCAACCATT
GAGCACACGTCTGAACTCTTTCCCTACACGACGCTCTTCCGATCTNNNTCGTGCTGCGTGTATTATTT
AGCACACGTCTGAACTCTTTCCCTACACGACGCTCTTCCGATCTNNNCAGCAGAGGTGATGTTGTACC
GCACACGTCTGAACTCTTTCCCTACACGACGCTCTTCCGATCTNNNCAGCACTGTCACCTACATGATG
GCACACGTCTGAACTCTTTCCCTACACGACGCTCTTCCGATCTNNNATCAACCCCCAGCATTCTGATTCC
GCACACGTCTGAACTCTTTCCCTACACGACGCTCTTCCGATCTNNNACGAAAGCAAAATGGTCAAAG
GCACACGTCTGAACTCTTTCCCTACACGACGCTCTTCCGATCTNNNCAGGGGAATGCCCAATACTTCT
GAGCACACGTCTGAACTCTTTCCCTACACGACGCTCTTCCGATCTNNNACATGAGTTTTGGAAGGGAC
GAGCACACGTCTGAACTCTTTCCCTACACGACGCTCTTCCGATCTNNNTGAAGTGTTCCAGGCAGAAA
AGCACACGTCTGAACTCTTTCCCTACACGACGCTCTTCCGATCTNNNTTCCACCATCACCGAAAGTTCT
CACACGTCTGAACTCTTTCCCTACACGACGCTCTTCCGATCTNNNCCAGGTCTGTTTCTTCTTCTCCCC
AGAGCACACGTCTGAACTCTTTCCCTACACGACGCTCTTCCGATCTNNNTCTAGTTCCTGTAGGTGTG
AGCACACGTCTGAACTCTTTCCCTACACGACGCTCTTCCGATCTNNNAGGGAGAACGGAAAGGTGTG
GCACACGTCTGAACTCTTTCCCTACACGACGCTCTTCCGATCTNNNATCATCCAGCAAAGCACTAGAC
GCACACGTCTGAACTCTTTCCCTACACGACGCTCTTCCGATCTNNNCAGTTTCCCTAAAGGGGAGAGG

GAGCACACGTCTGAACTCTTTCCCTACACGACGCTCTTCCGATCTNNNTGGCAGATACTCTTAGGACG
AGCACACGTCTGAACTCTTTCCCTACACGACGCTCTTCCGATCTNNNCAGAACACAGGAAGGGTTTTG
CACACGTCTGAACTCTTTCCCTACACGACGCTCTTCCGATCTNNNCACAGACAGACCTGGACTTGAACC
GCACACGTCTGAACTCTTTCCCTACACGACGCTCTTCCGATCTNNNCTCCACCAGGTCTGTGATGTTTT
AGCACACGTCTGAACTCTTTCCCTACACGACGCTCTTCCGATCTNNNCATGATCAATTCCAGCTTCCTGC
GCACACGTCTGAACTCTTTCCCTACACGACGCTCTTCCGATCTNNNCTACTGCACTTCAGCAAGACACT
AGCACACGTCTGAACTCTTTCCCTACACGACGCTCTTCCGATCTNNNTCATCTTACAGCTTCAAGTTAC
AGCACACGTCTGAACTCTTTCCCTACACGACGCTCTTCCGATCTNNNGGTTGCCGTGGGTTTGTTTG
CACACGTCTGAACTCTTTCCCTACACGACGCTCTTCCGATCTNNNTCCAATTAGATCTGACCAGGCCCC
GAGCACACGTCTGAACTCTTTCCCTACACGACGCTCTTCCGATCTNNNGCCCAGCCTAAGATCCTATGT
GCACACGTCTGAACTCTTTCCCTACACGACGCTCTTCCGATCTNNNCACTCATCAACTCCAGGCAACTT
GAGCACACGTCTGAACTCTTTCCCTACACGACGCTCTTCCGATCTNNNCTCTGAGTACAGTCGTGTGT
GCACACGTCTGAACTCTTTCCCTACACGACGCTCTTCCGATCTNNNAGAAGATATGGCAGTTCCAGTG
CACACGTCTGAACTCTTTCCCTACACGACGCTCTTCCGATCTNNNCGAGATCCCATCCGAACATACACC
GCACACGTCTGAACTCTTTCCCTACACGACGCTCTTCCGATCTNNNCTGAAACTCTGCCCCGTGAAAT
GAGCACACGTCTGAACTCTTTCCCTACACGACGCTCTTCCGATCTNNNGCAGGCCCCTCATCTTGAATA
AGAGCACACGTCTGAACTCTTTCCCTACACGACGCTCTTCCGATCTNNNGACAAATGAAACTGGCAG
AGCACACGTCTGAACTCTTTCCCTACACGACGCTCTTCCGATCTNNNCTGGCCAACAGAGACAGTTAC
AGCACACGTCTGAACTCTTTCCCTACACGACGCTCTTCCGATCTNNNGTTTGCCTTTTTATCCCTGTGG
GAGCACACGTCTGAACTCTTTCCCTACACGACGCTCTTCCGATCTNNNTTCAAAGAGATGCCTTCTTCG
GCACACGTCTGAACTCTTTCCCTACACGACGCTCTTCCGATCTNNNTTCATTCTTGCTGAGCGAGTAC
AGAGCACACGTCTGAACTCTTTCCCTACACGACGCTCTTCCGATCTNNNAGGCCACATGGAGGAGTT
AGCACACGTCTGAACTCTTTCCCTACACGACGCTCTTCCGATCTNNNCCATTCTAAGCTCACATTCGCC
AGCACACGTCTGAACTCTTTCCCTACACGACGCTCTTCCGATCTNNNGCTGAGACCCTACACGTTTTTC
AGAGCACACGTCTGAACTCTTTCCCTACACGACGCTCTTCCGATCTNNNGTGTGGGTTAGAATCCTGA
GAGCACACGTCTGAACTCTTTCCCTACACGACGCTCTTCCGATCTNNNGTGTGGGTTAGAATCCTGA
AGCACACGTCTGAACTCTTTCCCTACACGACGCTCTTCCGATCTNNNCACAGAGCAACAGCATCATAG
GCACACGTCTGAACTCTTTCCCTACACGACGCTCTTCCGATCTNNNTTTCATCCTTCCCTCCTCCTTCTCA
AGCACACGTCTGAACTCTTTCCCTACACGACGCTCTTCCGATCTNNNTATGACCTTCACTCTGGGACAG
GAGCACACGTCTGAACTCTTTCCCTACACGACGCTCTTCCGATCTNNNGTATTGGCTGCTGTGGAAG
AGCACACGTCTGAACTCTTTCCCTACACGACGCTCTTCCGATCTNNNATGAGGGGCTTTCCTATCCTGT
CACACGTCTGAACTCTTTCCCTACACGACGCTCTTCCGATCTNNNCGCCTTCTTCTCTGCTCTGTAAGC
GCACACGTCTGAACTCTTTCCCTACACGACGCTCTTCCGATCTNNNAGTCTAGGAATAAGGGCCTTGG
GAGCACACGTCTGAACTCTTTCCCTACACGACGCTCTTCCGATCTNNNTTCTATACAGTTGCACAATGC
GAGCACACGTCTGAACTCTTTCCCTACACGACGCTCTTCCGATCTNNNACCAAGCAAGGGTGAAAGC
CACACGTCTGAACTCTTTCCCTACACGACGCTCTTCCGATCTNNNCTCGCATTAGAGGAGAAGAAGG
AGCACACGTCTGAACTCTTTCCCTACACGACGCTCTTCCGATCTNNNGCAGAGGTGGGGTAGAATT
CACACGTCTGAACTCTTTCCCTACACGACGCTCTTCCGATCTNNNTGTATGGGAAAGTGTCCACAACG
AGCACACGTCTGAACTCTTTCCCTACACGACGCTCTTCCGATCTNNNCTAGACATGCGTAGAGGCACA
AGCACACGTCTGAACTCTTTCCCTACACGACGCTCTTCCGATCTNNNCTATGGTTCAGTTACAGCAGG
GAGCACACGTCTGAACTCTTTCCCTACACGACGCTCTTCCGATCTNNNCCCACCTACCTCACCCTTCTTA
GAGCACACGTCTGAACTCTTTCCCTACACGACGCTCTTCCGATCTNNNTTATCATGGGTGACTTTACC
AGCACACGTCTGAACTCTTTCCCTACACGACGCTCTTCCGATCTNNNTAGATGGGCGAGTTTGTGGA

AGCACACGTCTGAACTCTTTCCCTACACGACGCTCTTCCGATCTNNNCTTTTGGACACACACTCTCCAG

CACACGTCTGAACTCTTTCCCTACACGACGCTCTTCCGATCTNNNCCTTCTCAAGGAAGGAGGTGAG

AGCACACGTCTGAACTCTTTCCCTACACGACGCTCTTCCGATCTNNNGCGCCTTATCTCTGTCTCTCTCT

GAGCACACGTCTGAACTCTTTCCCTACACGACGCTCTTCCGATCTNNNTCATTCACAGGGAATCTATG

CACACGTCTGAACTCTTTCCCTACACGACGCTCTTCCGATCTNNNCACTGGCACCAGACCATCAAAGCA

AGAGCACACGTCTGAACTCTTTCCCTACACGACGCTCTTCCGATCTNNNGAATAAATGCCTGCTCCCA

AGCACACGTCTGAACTCTTTCCCTACACGACGCTCTTCCGATCTNNNGCCAGCAGGAAAGTGCAATAA

AGCACACGTCTGAACTCTTTCCCTACACGACGCTCTTCCGATCTNNNCGTACTCATGTGGGGAAACAA

AGCACACGTCTGAACTCTTTCCCTACACGACGCTCTTCCGATCTNNNACTCTCTACATACCAGGCACAG

AGCACACGTCTGAACTCTTTCCCTACACGACGCTCTTCCGATCTNNNCAGTTGAGTGAGGAACAGA

GCACACGTCTGAACTCTTTCCCTACACGACGCTCTTCCGATCTNNNTCATTGCTTTGTTCTTGCTGCTGC

GCACACGTCTGAACTCTTTCCCTACACGACGCTCTTCCGATCTNNNTGGGGAATTCTAATGCACCCAAC

AGCACACGTCTGAACTCTTTCCCTACACGACGCTCTTCCGATCTNNNAATTACTACCGTGACTGGGAT

AGCACACGTCTGAACTCTTTCCCTACACGACGCTCTTCCGATCTNNNTGAGAGAAAAAGATGTCAAA

AGCACACGTCTGAACTCTTTCCCTACACGACGCTCTTCCGATCTNNNCCAGGTTTATGTGCCTTTCTG

AGCACACGTCTGAACTCTTTCCCTACACGACGCTCTTCCGATCTNNNTGGAATGCATGTTTGTTTCAT

GCACACGTCTGAACTCTTTCCCTACACGACGCTCTTCCGATCTNNNTTCAGAGGTGTTGAAGGAATG

CACACGTCTGAACTCTTTCCCTACACGACGCTCTTCCGATCTNNNCTTTCCTCCATGGCATTCACAGAC

GAGCACACGTCTGAACTCTTTCCCTACACGACGCTCTTCCGATCTNNNGGGGACGCAGATGACATTC

AGCACACGTCTGAACTCTTTCCCTACACGACGCTCTTCCGATCTNNNCTCATGTGGTCCTCCTTAGGTG

GAGCACACGTCTGAACTCTTTCCCTACACGACGCTCTTCCGATCTNNNCCTGGGGGACACTATTACAG

AGCACACGTCTGAACTCTTTCCCTACACGACGCTCTTCCGATCTNNNGCATTAGGGTGGAAAAATGG

AGCACACGTCTGAACTCTTTCCCTACACGACGCTCTTCCGATCTNNNCTCTGCCTAAAAGATGGGAGC

AGAGCACACGTCTGAACTCTTTCCCTACACGACGCTCTTCCGATCTNNNCACTCCCTTCACCCATCACA

AGCACACGTCTGAACTCTTTCCCTACACGACGCTCTTCCGATCTNNNGAGACAAGTGAAAGCCGAA

AGCACACGTCTGAACTCTTTCCCTACACGACGCTCTTCCGATCTNNNGGCGCACAGACAAAGAAAGT

AGCACACGTCTGAACTCTTTCCCTACACGACGCTCTTCCGATCTNNNAAATATCCACTGCCCTTGGACA

AGCACACGTCTGAACTCTTTCCCTACACGACGCTCTTCCGATCTNNNATAATCTCCCCCACCACACTTCT

AGCACACGTCTGAACTCTTTCCCTACACGACGCTCTTCCGATCTNNNCTGAAGAGGCTGCAAGAAG

AGCACACGTCTGAACTCTTTCCCTACACGACGCTCTTCCGATCTNNNAGTAGAATTTCAGGCCTGTCT

GCACACGTCTGAACTCTTTCCCTACACGACGCTCTTCCGATCTNNNTTTGGTTGGCTTTCACTGTCTGA

AGAGCACACGTCTGAACTCTTTCCCTACACGACGCTCTTCCGATCTNNNAGGGATAGTGGGAAGTT

GAGCACACGTCTGAACTCTTTCCCTACACGACGCTCTTCCGATCTNNNCAGGCCACTCTCAACTAAGT

AGCACACGTCTGAACTCTTTCCCTACACGACGCTCTTCCGATCTNNNGTCAAAATAACAACTCCGCCC

AGAGCACACGTCTGAACTCTTTCCCTACACGACGCTCTTCCGATCTNNNGTGATTGTACCACCACACAT

AGCACACGTCTGAACTCTTTCCCTACACGACGCTCTTCCGATCTNNNATCCATAGTTGTGTACTGGGG

AGCACACGTCTGAACTCTTTCCCTACACGACGCTCTTCCGATCTNNNGCTGATCTTGAAGCCCTTGAG

AGCACACGTCTGAACTCTTTCCCTACACGACGCTCTTCCGATCTNNNACCCACAGAAACTGAAAGACA

AGCACACGTCTGAACTCTTTCCCTACACGACGCTCTTCCGATCTNNNTCTCTCGCTGTCTTCCTATGTGA

GCACACGTCTGAACTCTTTCCCTACACGACGCTCTTCCGATCTNNNAGAAAACAGGTCTGTGTGTGTG

AGCACACGTCTGAACTCTTTCCCTACACGACGCTCTTCCGATCTNNNGTGCAGCACAGAACATCCTCA

AGCACACGTCTGAACTCTTTCCCTACACGACGCTCTTCCGATCTNNNGCTGCAGTAACCCATGATTCA

AGCACACGTCTGAACTCTTTCCCTACACGACGCTCTTCCGATCTNNNTGAAGAAGTCCTCTCTGCTCCA

AGCACACGTCTGAACTCTTTCCCTACACGACGCTCTTCCGATCTNNNAAGGATGAATCCCAGTTGCTT
AGCACACGTCTGAACTCTTTCCCTACACGACGCTCTTCCGATCTNNNCTGTCCTTGAGTAACCATCCTG
AGCACACGTCTGAACTCTTTCCCTACACGACGCTCTTCCGATCTNNNAACTTCCCTAGCCCCTGAAAT
AGCACACGTCTGAACTCTTTCCCTACACGACGCTCTTCCGATCTNNNGGAAGAACTCACACTGGGGA
AGCACACGTCTGAACTCTTTCCCTACACGACGCTCTTCCGATCTNNNGACCCCAAGTCAGCAAGTCAG
GAGCACACGTCTGAACTCTTTCCCTACACGACGCTCTTCCGATCTNNNCCTGGGACAAAGCAGAACCT
GCACACGTCTGAACTCTTTCCCTACACGACGCTCTTCCGATCTNNNTTCGGCGAAATGATTTTCCACAC
AGCACACGTCTGAACTCTTTCCCTACACGACGCTCTTCCGATCTNNNCCTGCCATCTAAACCCATAACCC
AGCACACGTCTGAACTCTTTCCCTACACGACGCTCTTCCGATCTNNNGGAGGTACTGAGGTGAATTC
AGCACACGTCTGAACTCTTTCCCTACACGACGCTCTTCCGATCTNNNAATGCGTTCACCATGTAACAGG
CACACGTCTGAACTCTTTCCCTACACGACGCTCTTCCGATCTNNNCTGTGGTTTAGCCCAGGAATCTC
AGCACACGTCTGAACTCTTTCCCTACACGACGCTCTTCCGATCTNNNATTCCATCCCCTCTGTCACTCCC
GAGCACACGTCTGAACTCTTTCCCTACACGACGCTCTTCCGATCTNNNAAGCAGGTTGCAGAATACC
GAGCACACGTCTGAACTCTTTCCCTACACGACGCTCTTCCGATCTNNNTTCTACTGGATTAGGGTGG
AGCACACGTCTGAACTCTTTCCCTACACGACGCTCTTCCGATCTNNNTGAAGAAGTCCTCTCTGCTCC
GCACACGTCTGAACTCTTTCCCTACACGACGCTCTTCCGATCTNNNGCTGCTTCAGCACATCATGGAG
GAGCACACGTCTGAACTCTTTCCCTACACGACGCTCTTCCGATCTNNNTGTCACTTTTGTCCTCTTCCAC
GAGCACACGTCTGAACTCTTTCCCTACACGACGCTCTTCCGATCTNNNGTGAGTGCACACGAGTATG
GCACACGTCTGAACTCTTTCCCTACACGACGCTCTTCCGATCTNNNAGTGTTCATGTTACTGGGAAG
AGAGCACACGTCTGAACTCTTTCCCTACACGACGCTCTTCCGATCTNNNCTGGATGGGAATCCTGCTT
AGCACACGTCTGAACTCTTTCCCTACACGACGCTCTTCCGATCTNNNTTGCTCAGGGTCATGTGTCTG
GAGCACACGTCTGAACTCTTTCCCTACACGACGCTCTTCCGATCTNNNTCACAAGTCTCTGACACAAC
AGAGCACACGTCTGAACTCTTTCCCTACACGACGCTCTTCCGATCTNNNAGTTGGCAAGATTTTGGGG
GCACACGTCTGAACTCTTTCCCTACACGACGCTCTTCCGATCTNNNTAACTCCAGCCTCATCTCTGTCCC
AGCACACGTCTGAACTCTTTCCCTACACGACGCTCTTCCGATCTNNNTCGGCAAGTACCCCAAGGAGC
AGCACACGTCTGAACTCTTTCCCTACACGACGCTCTTCCGATCTNNNGACTAGCTGCTCTGGGTGTTT
GCACACGTCTGAACTCTTTCCCTACACGACGCTCTTCCGATCTNNNACACATCTTACATGCAGAGGCA
GAGCACACGTCTGAACTCTTTCCCTACACGACGCTCTTCCGATCTNNNGAAGAGCAGAGGAGTAAC
GCACACGTCTGAACTCTTTCCCTACACGACGCTCTTCCGATCTNNNCCTCTCTGGTCTGGCCTCGCCAT
GAGCACACGTCTGAACTCTTTCCCTACACGACGCTCTTCCGATCTNNNGGTGATGCCTTGAATGAGG
GAGCACACGTCTGAACTCTTTCCCTACACGACGCTCTTCCGATCTNNNTTGAGTGACACAAAAGAGT
GAGCACACGTCTGAACTCTTTCCCTACACGACGCTCTTCCGATCTNNNAATCATCACTGCACTTTGAG
AGCACACGTCTGAACTCTTTCCCTACACGACGCTCTTCCGATCTNNNAGAGCCAACCCCTCACTTCTTA
AGCACACGTCTGAACTCTTTCCCTACACGACGCTCTTCCGATCTNNNTGTTTGTGTGCATGCATTGTGT
GCACACGTCTGAACTCTTTCCCTACACGACGCTCTTCCGATCTNNNACCAGTGTTCTCCTTCCTTCCTTC
CACGTCTGAACTCTTTCCCTACACGACGCTCTTCCGATCTNNNGACAGCCGGACCAGGCATACTGCAT
CACACGTCTGAACTCTTTCCCTACACGACGCTCTTCCGATCTNNNACCATATGAGCTTGTGTGTGCATC
GCACACGTCTGAACTCTTTCCCTACACGACGCTCTTCCGATCTNNNAGGGAAGAGAAGGTCCAAGG
AGCACACGTCTGAACTCTTTCCCTACACGACGCTCTTCCGATCTNNNGAATCCAGCAACCAACCAGTA
GAGCACACGTCTGAACTCTTTCCCTACACGACGCTCTTCCGATCTNNNATATTCTCCCTATCCTGCTGC
AGCACACGTCTGAACTCTTTCCCTACACGACGCTCTTCCGATCTNNNGCTTGGTGTATGTCTGTGGTC
CACACGTCTGAACTCTTTCCCTACACGACGCTCTTCCGATCTNNNAAGAGCCTTGTGGGAAGAGAG
AGCACACGTCTGAACTCTTTCCCTACACGACGCTCTTCCGATCTNNNCATTTCAGGCTTCCAGCAGAA

AGCACACGTCTGAACTCTTTCCCTACACGACGCTCTTCCGATCTNNNGTTCAGAAGTCTCAGAAACCCC
AGCACACGTCTGAACTCTTTCCCTACACGACGCTCTTCCGATCTNNNAATCATCACTGCACTTTGAGG
AGCACACGTCTGAACTCTTTCCCTACACGACGCTCTTCCGATCTNNNCCAAAGCCTGGGGTTTTCATC
GCACACGTCTGAACTCTTTCCCTACACGACGCTCTTCCGATCTNNNGAGCAATAAGGGCTGGGTTTG
CACACGTCTGAACTCTTTCCCTACACGACGCTCTTCCGATCTNNNTGGAATCCACCTACTTCCAGGAG
GAGCACACGTCTGAACTCTTTCCCTACACGACGCTCTTCCGATCTNNNACATGAGGCAATTCAGACAT
AGCACACGTCTGAACTCTTTCCCTACACGACGCTCTTCCGATCTNNNGAACGAGAGGCGTGTCAGAA
GAGCACACGTCTGAACTCTTTCCCTACACGACGCTCTTCCGATCTNNNTCAGGTCTGACCCAGTCATA
GAGCACACGTCTGAACTCTTTCCCTACACGACGCTCTTCCGATCTNNNCCAATGGAGGGTCAGAGAA
AGAGCACACGTCTGAACTCTTTCCCTACACGACGCTCTTCCGATCTNNNTCTGTATACACGTCTCTCTCT
ACACGTCTGAACTCTTTCCCTACACGACGCTCTTCCGATCTNNNGTTTTCCTCCTGGCAGGTAGATGC
AGAGCACACGTCTGAACTCTTTCCCTACACGACGCTCTTCCGATCTNNNAACCCCCTAACGTGGATGAA
AGAGCACACGTCTGAACTCTTTCCCTACACGACGCTCTTCCGATCTNNNGCGATTGAGTGACTGCCTCA
CACACGTCTGAACTCTTTCCCTACACGACGCTCTTCCGATCTNNNTGCCACCTATGAACCAAGGACTA
AGCACACGTCTGAACTCTTTCCCTACACGACGCTCTTCCGATCTNNNTTGACTGGATATCTGGATTGC
AAGAGCACACGTCTGAACTCTTTCCCTACACGACGCTCTTCCGATCTNNNAGGTCACTGACAGGCAG
AGCACACGTCTGAACTCTTTCCCTACACGACGCTCTTCCGATCTNNNAAGTGATGTGGGGCAAGTTA
AGCACACGTCTGAACTCTTTCCCTACACGACGCTCTTCCGATCTNNNCGACTTTCTGGAAGAGGAAGC
CACGTCTGAACTCTTTCCCTACACGACGCTCTTCCGATCTNNNCCCTTGGAGCATTTTGTCACTGGCAT
AGCACACGTCTGAACTCTTTCCCTACACGACGCTCTTCCGATCTNNNACTCTGTCTGTCCCCTAAAGGT
GAGCACACGTCTGAACTCTTTCCCTACACGACGCTCTTCCGATCTNNNTAATGGATCTACACTCCGGCG
CACACGTCTGAACTCTTTCCCTACACGACGCTCTTCCGATCTNNNAGAAGGGCTGGACACCTTACTTA
GAGCACACGTCTGAACTCTTTCCCTACACGACGCTCTTCCGATCTNNNGGTGATTTCATACACGCACA
AGCACACGTCTGAACTCTTTCCCTACACGACGCTCTTCCGATCTNNNACAATTTCTTGGCATCTCTCCC
CACACGTCTGAACTCTTTCCCTACACGACGCTCTTCCGATCTNNNCTCCAGTTCCCGATCCACCCCATCG
ACACGTCTGAACTCTTTCCCTACACGACGCTCTTCCGATCTNNNGAGTTTCTGGACCCCCTCATGGCAT
CACACGTCTGAACTCTTTCCCTACACGACGCTCTTCCGATCTNNNTCTCCCCGTTTTCTCTTTTGGTGC
GCACACGTCTGAACTCTTTCCCTACACGACGCTCTTCCGATCTNNNGTTTGATTGTGTCTTGCTCTGC
GCACACGTCTGAACTCTTTCCCTACACGACGCTCTTCCGATCTNNNAGGACAGTAGGACACACACAG
AGCACACGTCTGAACTCTTTCCCTACACGACGCTCTTCCGATCTNNNATAAGGAGTGCTGAGGAAG
AGCACACGTCTGAACTCTTTCCCTACACGACGCTCTTCCGATCTNNNATATAACTAGAGGCCGAGCCA
CACACGTCTGAACTCTTTCCCTACACGACGCTCTTCCGATCTNNNATGTCATAGCTCAAGTGGCAAGG
GCACACGTCTGAACTCTTTCCCTACACGACGCTCTTCCGATCTNNNTAGAGTAGTGGTCTCCAACTCC
GCACACGTCTGAACTCTTTCCCTACACGACGCTCTTCCGATCTNNNCATCACTGGGAAGAGAGAGGG
AGCACACGTCTGAACTCTTTCCCTACACGACGCTCTTCCGATCTNNNAGACAAAACTTGTGCCTGAA
GAGCACACGTCTGAACTCTTTCCCTACACGACGCTCTTCCGATCTNNNTAGCACATGCCCATAGTCCCA
AGCACACGTCTGAACTCTTTCCCTACACGACGCTCTTCCGATCTNNNGGATTCGGATCAAAGGAGCTG
AGCACACGTCTGAACTCTTTCCCTACACGACGCTCTTCCGATCTNNNATGAGGGTCTCTGCTGTTCTT
GAGCACACGTCTGAACTCTTTCCCTACACGACGCTCTTCCGATCTNNNCCTTGGGACACTTTTGTTTG
ACACGTCTGAACTCTTTCCCTACACGACGCTCTTCCGATCTNNNAGTCCCCCAACTCCCTGGTCCATCG
GAGCACACGTCTGAACTCTTTCCCTACACGACGCTCTTCCGATCTNNNAATGGAATTTGCCACAGAG
AGCACACGTCTGAACTCTTTCCCTACACGACGCTCTTCCGATCTNNNACAAAAGCTTTCTATCCCCCAG
GCACACGTCTGAACTCTTTCCCTACACGACGCTCTTCCGATCTNNNTACACAAACACAAAAACAGGCC

AGAGCACACGTCTGAACTCTTTCCCTACACGACGCTCTTCCGATCTNNNCCAGGAGGTAGACTTAGCA
AGCACACGTCTGAACTCTTTCCCTACACGACGCTCTTCCGATCTNNNGCCCAGATTCATTTGTTTCTGC
AGCACACGTCTGAACTCTTTCCCTACACGACGCTCTTCCGATCTNNNGGTGACATCTTTCACAATCGG
AGCACACGTCTGAACTCTTTCCCTACACGACGCTCTTCCGATCTNNNCTTCCAGCATCTCGGACAACAG
AGCACACGTCTGAACTCTTTCCCTACACGACGCTCTTCCGATCTNNNAGGTCCAAGCTGTTGAAGGAA
AGCACACGTCTGAACTCTTTCCCTACACGACGCTCTTCCGATCTNNNCAGCCCGGCTCTAACAATAAG
AGCACACGTCTGAACTCTTTCCCTACACGACGCTCTTCCGATCTNNNCCTTATCCCTCTGTTTCTCTGCG
GAGCACACGTCTGAACTCTTTCCCTACACGACGCTCTTCCGATCTNNNACCCCTCCTCCTTTATTACCCAG
GCACACGTCTGAACTCTTTCCCTACACGACGCTCTTCCGATCTNNNACCTGGGACACATCAATCACAAC
AGCACACGTCTGAACTCTTTCCCTACACGACGCTCTTCCGATCTNNNATACTGCCATGAAAGCCAGTC
AGCACACGTCTGAACTCTTTCCCTACACGACGCTCTTCCGATCTNNNGTGAGTGCACACGAGTATGTG
AGCACACGTCTGAACTCTTTCCCTACACGACGCTCTTCCGATCTNNNAAGCACTGTCTGTGAGTAATGG
CACACGTCTGAACTCTTTCCCTACACGACGCTCTTCCGATCTNNNAAGAAGTCGGTGGACAAGAACA
AGCACACGTCTGAACTCTTTCCCTACACGACGCTCTTCCGATCTNNNTATTAGTAGAGACTGGGCCTC
AAGAGCACACGTCTGAACTCTTTCCCTACACGACGCTCTTCCGATCTNNNAATGCCAAAGTCTGGATG
GAGCACACGTCTGAACTCTTTCCCTACACGACGCTCTTCCGATCTNNNACCGATGTGATGTGTGTGG
AGAGCACACGTCTGAACTCTTTCCCTACACGACGCTCTTCCGATCTNNNGCCCTCAGGTCATTTTAGGA
GCACACGTCTGAACTCTTTCCCTACACGACGCTCTTCCGATCTNNNCAGAGACATTCCAGGGGTCCAG
AGCACACGTCTGAACTCTTTCCCTACACGACGCTCTTCCGATCTNNNATAGTCACACAAACACCCAACC
GAGCACACGTCTGAACTCTTTCCCTACACGACGCTCTTCCGATCTNNNTGGGAAGGTCTTCGGTCTAA
AGAGCACACGTCTGAACTCTTTCCCTACACGACGCTCTTCCGATCTNNNTGTGGATAACACCCTGGAC
AGCACACGTCTGAACTCTTTCCCTACACGACGCTCTTCCGATCTNNNCGACTTTCTGGAAGAGGAAGC
AGCACACGTCTGAACTCTTTCCCTACACGACGCTCTTCCGATCTNNNCCTATGGACTTGGGAAAGCAC
CACACGTCTGAACTCTTTCCCTACACGACGCTCTTCCGATCTNNNGGAGGAAGGGAGAGAAGATCTG
AGCACACGTCTGAACTCTTTCCCTACACGACGCTCTTCCGATCTNNNCCAGAAGGAGAGAAGTGTGT
AGCACACGTCTGAACTCTTTCCCTACACGACGCTCTTCCGATCTNNNGCCCTATCTGAATCCCTTTGCC
AGAGCACACGTCTGAACTCTTTCCCTACACGACGCTCTTCCGATCTNNNGAATCCATTCAGGTCCGACT
AGCACACGTCTGAACTCTTTCCCTACACGACGCTCTTCCGATCTNNNTTCTTATCTCGGGTCACCACCTT
AGCACACGTCTGAACTCTTTCCCTACACGACGCTCTTCCGATCTNNNCCGCTTTCTTTTCTCATGGTTG
AGCACACGTCTGAACTCTTTCCCTACACGACGCTCTTCCGATCTNNNCAAGGCATTACTTGCCAACTGT
AGCACACGTCTGAACTCTTTCCCTACACGACGCTCTTCCGATCTNNNACTAATGTCGGTGTCTAGAACC
AGCACACGTCTGAACTCTTTCCCTACACGACGCTCTTCCGATCTNNNCCCACCTGACTTGAAAACAATG
GAGCACACGTCTGAACTCTTTCCCTACACGACGCTCTTCCGATCTNNNCTGTACTGACTGCAGACACG
AGCACACGTCTGAACTCTTTCCCTACACGACGCTCTTCCGATCTNNNTTGCTGAGAGAGTAGGTGTC
GCACACGTCTGAACTCTTTCCCTACACGACGCTCTTCCGATCTNNNAGATGTACCTTTGGATGTCTGTC
AGCACACGTCTGAACTCTTTCCCTACACGACGCTCTTCCGATCTNNNTTGACACTGTATTCCCTCTGCCT
GAGCACACGTCTGAACTCTTTCCCTACACGACGCTCTTCCGATCTNNNTTTAGGTACTGAGGGGCAA
AGAGCACACGTCTGAACTCTTTCCCTACACGACGCTCTTCCGATCTNNNTAGCTTTTGAAATCAGCCC
GCACACGTCTGAACTCTTTCCCTACACGACGCTCTTCCGATCTNNNAGGATGGTAAGGATGAAAAAG
AGCACACGTCTGAACTCTTTCCCTACACGACGCTCTTCCGATCTNNNACATCCAAATCAGGAAATCGGC
GCACACGTCTGAACTCTTTCCCTACACGACGCTCTTCCGATCTNNNCTTATGCTGTGTTTAGCATGCAC
AGCACACGTCTGAACTCTTTCCCTACACGACGCTCTTCCGATCTNNNGCCAGACTACGTCTCTCTCTCT
AGCACACGTCTGAACTCTTTCCCTACACGACGCTCTTCCGATCTNNNTCTGGATCTGGGTGGTTGATA

GAGCACACGTCTGAACTCTTTCCCTACACGACGCTCTTCCGATCTNNNCTGGCTGGTAGAACTGTGAA
AGCACACGTCTGAACTCTTTCCCTACACGACGCTCTTCCGATCTNNNTTCTTCTAAAGCCCAAGGTAGG
AGCACACGTCTGAACTCTTTCCCTACACGACGCTCTTCCGATCTNNNGCATGGTGGTAGGTATGATT
AGCACACGTCTGAACTCTTTCCCTACACGACGCTCTTCCGATCTNNNATGCAGTGCCAAATAACATGG
AGAGCACACGTCTGAACTCTTTCCCTACACGACGCTCTTCCGATCTNNNACCTTGCTAAGAGGAGGTC
GCACACGTCTGAACTCTTTCCCTACACGACGCTCTTCCGATCTNNNCGACAGACACCCTGTCATACAAA
GAAGAGCACACGTCTGAACTCTTTCCCTACACGACGCTCTTCCGATCTNNNACACTCCCAGATTTGGG
AGAGCACACGTCTGAACTCTTTCCCTACACGACGCTCTTCCGATCTNNNTTTCTCCTGCCTCTCAGCCTA
AGCACACGTCTGAACTCTTTCCCTACACGACGCTCTTCCGATCTNNNTACCTACAGTCAACCCTGGACAC
GAGCACACGTCTGAACTCTTTCCCTACACGACGCTCTTCCGATCTNNNTGCAATACATTACGTGGCAC
GCACACGTCTGAACTCTTTCCCTACACGACGCTCTTCCGATCTNNNATTGTCCCTTTCCGCTTTCCTTTC
AGCACACGTCTGAACTCTTTCCCTACACGACGCTCTTCCGATCTNNNACAGAGCAAGACACCAGAAA
AAGAGCACACGTCTGAACTCTTTCCCTACACGACGCTCTTCCGATCTNNNCCTTGTTCTAACATTGGG
GCACACGTCTGAACTCTTTCCCTACACGACGCTCTTCCGATCTNNNCCTAGTTTGGGAAGCAGAGACA
GAGCACACGTCTGAACTCTTTCCCTACACGACGCTCTTCCGATCTNNNACTGGATTCAGTCACGCCTTT
GAAGAGCACACGTCTGAACTCTTTCCCTACACGACGCTCTTCCGATCTNNNAGAGGTAAAGGTGAAC
GCACACGTCTGAACTCTTTCCCTACACGACGCTCTTCCGATCTNNNTTTTCCTCAGGTGAGGCTGGA
GAGCACACGTCTGAACTCTTTCCCTACACGACGCTCTTCCGATCTNNNAGCGTTTCTTACATCAGCTCA
AGCACACGTCTGAACTCTTTCCCTACACGACGCTCTTCCGATCTNNNACGGGTTTTCTTTTTGCATTGG
AGCACACGTCTGAACTCTTTCCCTACACGACGCTCTTCCGATCTNNNAAATAAGACAACATGGGCAGC
AGAGCACACGTCTGAACTCTTTCCCTACACGACGCTCTTCCGATCTNNNCCTATACCTAGAGCCGGAG
AGCACACGTCTGAACTCTTTCCCTACACGACGCTCTTCCGATCTNNNAGTCCGTGCATATCCAAGCATA
AGCACACGTCTGAACTCTTTCCCTACACGACGCTCTTCCGATCTNNNTCCCTAGCCGATATCCACAGAA
AGCACACGTCTGAACTCTTTCCCTACACGACGCTCTTCCGATCTNNNCACTGAGATCAAAGCATAGCG
GAAGAGCACACGTCTGAACTCTTTCCCTACACGACGCTCTTCCGATCTNNNACACTCCCAGATTTGGG
CACACGTCTGAACTCTTTCCCTACACGACGCTCTTCCGATCTNNNACCCAACTGAACTTGAGAACTGGC
AGAGCACACGTCTGAACTCTTTCCCTACACGACGCTCTTCCGATCTNNNCAAATCAAGCAGAGGACAT
AGCACACGTCTGAACTCTTTCCCTACACGACGCTCTTCCGATCTNNNCAGCTGGTGGGATTCTAACCT
AGCACACGTCTGAACTCTTTCCCTACACGACGCTCTTCCGATCTNNNCCACATATGCGCACACATCTCT
GAGCACACGTCTGAACTCTTTCCCTACACGACGCTCTTCCGATCTNNNTGGGTTCCCACAGGTAAACC
GCACACGTCTGAACTCTTTCCCTACACGACGCTCTTCCGATCTNNNTGGGTTCTTACCTCTCCTCTGTGC
AGCACACGTCTGAACTCTTTCCCTACACGACGCTCTTCCGATCTNNNTGCTGCAAAATGATTTGGTCT
AGCACACGTCTGAACTCTTTCCCTACACGACGCTCTTCCGATCTNNNTCTGTGCTTTGACGAAGTCATG
AGCACACGTCTGAACTCTTTCCCTACACGACGCTCTTCCGATCTNNNGGTCTGTTGTTGACTGGTACT
AGCACACGTCTGAACTCTTTCCCTACACGACGCTCTTCCGATCTNNNGGCTAACCTGTGTATGCCACTG
AGAGCACACGTCTGAACTCTTTCCCTACACGACGCTCTTCCGATCTNNNAAGATTGCTTGTAGCCAAA
AGAGCACACGTCTGAACTCTTTCCCTACACGACGCTCTTCCGATCTNNNAGTCTAGCCAAGAGAGCTC
AAGAGCACACGTCTGAACTCTTTCCCTACACGACGCTCTTCCGATCTNNNCACAAGATCTGCCCTCAAG
GAGCACACGTCTGAACTCTTTCCCTACACGACGCTCTTCCGATCTNNNCAGACACTGTGACAGATCAT
GCACACGTCTGAACTCTTTCCCTACACGACGCTCTTCCGATCTNNNCCCTGGTCTCAGATATGTCTCCC
AGCACACGTCTGAACTCTTTCCCTACACGACGCTCTTCCGATCTNNNAACAGTCTGATGTGTGTGTGT
GAAGAGCACACGTCTGAACTCTTTCCCTACACGACGCTCTTCCGATCTNNNCCCCTCTAGCTTGCAA
GCACACGTCTGAACTCTTTCCCTACACGACGCTCTTCCGATCTNNNAATGTGCAAACTCCAGCATTTG

AGAGCACACGTCTGAACTCTTTCCCTACACGACGCTCTTCCGATCTNNNAGCATTGAAAAATTGGTGC
AGAGCACACGTCTGAACTCTTTCCCTACACGACGCTCTTCCGATCTNNNATGCCAGTCCTACATACATG
GCACACGTCTGAACTCTTTCCCTACACGACGCTCTTCCGATCTNNNCCACCAACATTTTGATTCATTCTC
AGCACACGTCTGAACTCTTTCCCTACACGACGCTCTTCCGATCTNNNTTAGGTTAGGTGCTCTTGGCTC
GCACACGTCTGAACTCTTTCCCTACACGACGCTCTTCCGATCTNNNACTTAGTGGTGGTAATGGTGGG
GAGCACACGTCTGAACTCTTTCCCTACACGACGCTCTTCCGATCTNNNGGTGGTGCTCTCCTGTAGTTT
GAGCACACGTCTGAACTCTTTCCCTACACGACGCTCTTCCGATCTNNNCCTTGTAGTCAAGTGGAACCC
AGAGCACACGTCTGAACTCTTTCCCTACACGACGCTCTTCCGATCTNNNTCCTCTCCCCTGATGTCCTAT
GCACACGTCTGAACTCTTTCCCTACACGACGCTCTTCCGATCTNNNCCAGCCCTTTTCTTGTGCAATTTC
AGCACACGTCTGAACTCTTTCCCTACACGACGCTCTTCCGATCTNNNCACAAGATCTGCCCTCAAGATC
AGAGCACACGTCTGAACTCTTTCCCTACACGACGCTCTTCCGATCTNNNTACATGTAGCTGCATTGAAC
AAGAGCACACGTCTGAACTCTTTCCCTACACGACGCTCTTCCGATCTNNNACACAGCTCAGTACCTAG
GAGCACACGTCTGAACTCTTTCCCTACACGACGCTCTTCCGATCTNNNGTCTCTGGTTTTGTGCAGTTTG
AAGAGCACACGTCTGAACTCTTTCCCTACACGACGCTCTTCCGATCTNNNCACTTCAACCTCTGCCTTCA
GAGCACACGTCTGAACTCTTTCCCTACACGACGCTCTTCCGATCTNNNTGCACATATTTAGTGTGCTAT
AGAGCACACGTCTGAACTCTTTCCCTACACGACGCTCTTCCGATCTNNNAGTGCTGAAAGACAAGCAA
AGCACACGTCTGAACTCTTTCCCTACACGACGCTCTTCCGATCTNNNCAGGGGAGCTCTCTGTAAAGA
GAGCACACGTCTGAACTCTTTCCCTACACGACGCTCTTCCGATCTNNNAGCAAGGTCAATCCTGTCTCTC
AGCACACGTCTGAACTCTTTCCCTACACGACGCTCTTCCGATCTNNNACACATGCAGACAAATAAATG
AGCACACGTCTGAACTCTTTCCCTACACGACGCTCTTCCGATCTNNNATTTCTCCCATGGTATGTGCGT
GAGCACACGTCTGAACTCTTTCCCTACACGACGCTCTTCCGATCTNNNAGCAAGGTCAATCCTGTCTCT
AGAGCACACGTCTGAACTCTTTCCCTACACGACGCTCTTCCGATCTNNNTCTGGTTTGTAATGTCACAG
GAGCACACGTCTGAACTCTTTCCCTACACGACGCTCTTCCGATCTNNNATGTGGGGTGGTATAAAGGG
GAGCACACGTCTGAACTCTTTCCCTACACGACGCTCTTCCGATCTNNNTTGCTGTCACAACTACCACCAT
GCACACGTCTGAACTCTTTCCCTACACGACGCTCTTCCGATCTNNNTCGCCCTCCTCTACCCATATAGAC
AGCACACGTCTGAACTCTTTCCCTACACGACGCTCTTCCGATCTNNNGAGACCTCTCAGCATTCCTGAA
AGCACACGTCTGAACTCTTTCCCTACACGACGCTCTTCCGATCTNNNAACCTAAAATATGCCTGCCCCTG
AGAGCACACGTCTGAACTCTTTCCCTACACGACGCTCTTCCGATCTNNNCACCAATCGCTCACCTAGTTT
GAGCACACGTCTGAACTCTTTCCCTACACGACGCTCTTCCGATCTNNNATATGTATCACACATGCCCCC
GAGCACACGTCTGAACTCTTTCCCTACACGACGCTCTTCCGATCTNNNCCGAGATATAAAGGAGCCAG
GAGCACACGTCTGAACTCTTTCCCTACACGACGCTCTTCCGATCTNNNTCACTTCACTTCACCTCAACCCT
GAGCACACGTCTGAACTCTTTCCCTACACGACGCTCTTCCGATCTNNNTGGCAACAAAATCACTCCTGA
AGAGCACACGTCTGAACTCTTTCCCTACACGACGCTCTTCCGATCTNNNAAGAGGAGAGACGAAGA
GAGCACACGTCTGAACTCTTTCCCTACACGACGCTCTTCCGATCTNNNTCCCAGCCACTAATGCACTTA
GAGCACACGTCTGAACTCTTTCCCTACACGACGCTCTTCCGATCTNNNTTTGGGCCATGTCTGTTTCAG
GAGCACACGTCTGAACTCTTTCCCTACACGACGCTCTTCCGATCTNNNCCCTCACTCATACTTCTCTCCG
AAGAGCACACGTCTGAACTCTTTCCCTACACGACGCTCTTCCGATCTNNNTTCTGAAAGCTCCCAATCT
GAGCACACGTCTGAACTCTTTCCCTACACGACGCTCTTCCGATCTNNNACAACCCTGATTCCATTGCTCT
AGCACACGTCTGAACTCTTTCCCTACACGACGCTCTTCCGATCTNNNTTTCAGCGGTAATACTTGCAGT
AGCACACGTCTGAACTCTTTCCCTACACGACGCTCTTCCGATCTNNNATCTTTGGGCTGTGATTGTTCC
GCACACGTCTGAACTCTTTCCCTACACGACGCTCTTCCGATCTNNNCCCCATGGAAGCCATGAAACAT
GAGCACACGTCTGAACTCTTTCCCTACACGACGCTCTTCCGATCTNNNAGAACAGACATGTCCATAGCC
AGAGCACACGTCTGAACTCTTTCCCTACACGACGCTCTTCCGATCTNNNAAGAATAGAGATGGGCTGT

AGCACACGTCTGAACTCTTTCCCTACACGACGCTCTTCCGATCTNNNGGGTGTGTTTTGATGGACCTG
AGCACACGTCTGAACTCTTTCCCTACACGACGCTCTTCCGATCTNNNCCTTTGTGTGGCTCCTGATTTT
AGCACACGTCTGAACTCTTTCCCTACACGACGCTCTTCCGATCTNNNTAAGTGAAGACGGGGTGCTG
GCACACGTCTGAACTCTTTCCCTACACGACGCTCTTCCGATCTNNNCTCATTGCAGACTACCCTATGCC
AGAGCACACGTCTGAACTCTTTCCCTACACGACGCTCTTCCGATCTNNNGAGGGACAGTTCATATTGG
CACACGTCTGAACTCTTTCCCTACACGACGCTCTTCCGATCTNNNCTCTCTCTCGTGGACACCTCACCA
GCACACGTCTGAACTCTTTCCCTACACGACGCTCTTCCGATCTNNNGTCCCAGAATGCTTTGCAATGT
AGCACACGTCTGAACTCTTTCCCTACACGACGCTCTTCCGATCTNNNTTCACCCGGATCTCAATTTTGC
GAGCACACGTCTGAACTCTTTCCCTACACGACGCTCTTCCGATCTNNNTCCATTTCTAGTGCTAGTCCC
GCACACGTCTGAACTCTTTCCCTACACGACGCTCTTCCGATCTNNNAAGCAAACCACAGAAAACCAAG
GAGCACACGTCTGAACTCTTTCCCTACACGACGCTCTTCCGATCTNNNAGCCTCGTCTCAGTAACTACC
AGAGCACACGTCTGAACTCTTTCCCTACACGACGCTCTTCCGATCTNNNGACTTCAGAAAATTCTCTCT
AGCACACGTCTGAACTCTTTCCCTACACGACGCTCTTCCGATCTNNNTGCTAAGAATTGTGAACAGGC
AGCACACGTCTGAACTCTTTCCCTACACGACGCTCTTCCGATCTNNNCAATATGCTTGCCTCAAGGGT
AGCACACGTCTGAACTCTTTCCCTACACGACGCTCTTCCGATCTNNNCCCTATGCATCTCTTTGGCCAT
GCACACGTCTGAACTCTTTCCCTACACGACGCTCTTCCGATCTNNNAATCCTTACTGGATCATGTGCCCC
AGCACACGTCTGAACTCTTTCCCTACACGACGCTCTTCCGATCTNNNTGCTTCCCATGTAGTGTGTTTC
AGCACACGTCTGAACTCTTTCCCTACACGACGCTCTTCCGATCTNNNTGGGTATGTTAGAACAAAACT
AGCACACGTCTGAACTCTTTCCCTACACGACGCTCTTCCGATCTNNNTCATGTTGAACGATTGAACTG
AGAGCACACGTCTGAACTCTTTCCCTACACGACGCTCTTCCGATCTNNNTAATGTCACAGAGCTCCAC
GAGCACACGTCTGAACTCTTTCCCTACACGACGCTCTTCCGATCTNNNTGGTTTTCCAGAGGCTATGA
GCACACGTCTGAACTCTTTCCCTACACGACGCTCTTCCGATCTNNNTTGCTGGAAGAACTCATGGCTA
GAGCACACGTCTGAACTCTTTCCCTACACGACGCTCTTCCGATCTNNNACCCCTAAGAAGTCCATAGA
AGCACACGTCTGAACTCTTTCCCTACACGACGCTCTTCCGATCTNNNCCCTTCTATAGCAGCTACACAC
AGCACACGTCTGAACTCTTTCCCTACACGACGCTCTTCCGATCTNNNATTGGGGACTAACTTGGATTG
AGCACACGTCTGAACTCTTTCCCTACACGACGCTCTTCCGATCTNNNACAAAGGCATAAGAAATTGTG
GAGCACACGTCTGAACTCTTTCCCTACACGACGCTCTTCCGATCTNNNAATTAGTGTGTGTGTGTGCA
AGCACACGTCTGAACTCTTTCCCTACACGACGCTCTTCCGATCTNNNATTCCTCCCACAGCAAAGAACA
GAGCACACGTCTGAACTCTTTCCCTACACGACGCTCTTCCGATCTNNNCACTGCTGTCAGTAAGCATA
GCACACGTCTGAACTCTTTCCCTACACGACGCTCTTCCGATCTNNNATGCTCTTGGTATGAGTGATGG
GAGCACACGTCTGAACTCTTTCCCTACACGACGCTCTTCCGATCTNNNTGGCCAAAAACAGGGTCAA
AGCACACGTCTGAACTCTTTCCCTACACGACGCTCTTCCGATCTNNNAGTTCTGTCCCTTTCTCTCTCCT
GAGCACACGTCTGAACTCTTTCCCTACACGACGCTCTTCCGATCTNNNGAAGCACTTCTCCTTGCTTTC
AGCACACGTCTGAACTCTTTCCCTACACGACGCTCTTCCGATCTNNNGTGTGGCATGTGTAAGAGAG
AGCACACGTCTGAACTCTTTCCCTACACGACGCTCTTCCGATCTNNNTCTAATCTCTGTTTGGGTTGC
GCACACGTCTGAACTCTTTCCCTACACGACGCTCTTCCGATCTNNNTGGTAGGTGTATTGCAAGTGG
AGCACACGTCTGAACTCTTTCCCTACACGACGCTCTTCCGATCTNNNAGGAACAGAAAACTGGGGAG
GAGCACACGTCTGAACTCTTTCCCTACACGACGCTCTTCCGATCTNNNGGCAAGAGAGCACACCAAG
GAGCACACGTCTGAACTCTTTCCCTACACGACGCTCTTCCGATCTNNNTGCATCTATTCTTCAAACTCA
AGCACACGTCTGAACTCTTTCCCTACACGACGCTCTTCCGATCTNNNTACAGTGGCCAGTCAGATTTTG
GAGCACACGTCTGAACTCTTTCCCTACACGACGCTCTTCCGATCTNNNCGCACATTCCCTCTCTCTCTTT
GAGCACACGTCTGAACTCTTTCCCTACACGACGCTCTTCCGATCTNNNTTACAGGATGGTGGTTGCGA
AGCACACGTCTGAACTCTTTCCCTACACGACGCTCTTCCGATCTNNNAGTGTTTTGCTTTCGTTCTTCC

GAGCACACGTCTGAACTCTTTCCCTACACGACGCTCTTCCGATCTNNNTGGGTATTGTTGAGTGAGA
AGCACACGTCTGAACTCTTTCCCTACACGACGCTCTTCCGATCTNNNGATACTTGTTCCAAAGCACACG
AGCACACGTCTGAACTCTTTCCCTACACGACGCTCTTCCGATCTNNNGAAGGTCAAAGTCCCTTTTCGG
GCACACGTCTGAACTCTTTCCCTACACGACGCTCTTCCGATCTNNNAGCCAACTAAGCCTTGTAGGTA
AGCACACGTCTGAACTCTTTCCCTACACGACGCTCTTCCGATCTNNNGAGGATCCCTTAGGCTGAGAA
AGCACACGTCTGAACTCTTTCCCTACACGACGCTCTTCCGATCTNNNAAACAAAGATTGGCATGCAGG
AGCACACGTCTGAACTCTTTCCCTACACGACGCTCTTCCGATCTNNNGATGCCACTATACAACCACTGG
GCACACGTCTGAACTCTTTCCCTACACGACGCTCTTCCGATCTNNNGTAAACTCAGTTGCACCCATTGC
AGCACACGTCTGAACTCTTTCCCTACACGACGCTCTTCCGATCTNNNGAATCATCCCACTCAACACAGC
AGCACACGTCTGAACTCTTTCCCTACACGACGCTCTTCCGATCTNNNTTTGGGTGGTGTCACTTTTGG
AGCACACGTCTGAACTCTTTCCCTACACGACGCTCTTCCGATCTNNNGCCTTCATTGATGCGTTCAGAT
AGCACACGTCTGAACTCTTTCCCTACACGACGCTCTTCCGATCTNNNAACAACACAGAATTACCCACCC
GAGCACACGTCTGAACTCTTTCCCTACACGACGCTCTTCCGATCTNNNGAAACTGGCCTTACCGGAGT
GAAGAGCACACGTCTGAACTCTTTCCCTACACGACGCTCTTCCGATCTNNNCCCGGTCTCATTGAAGG
AGCACACGTCTGAACTCTTTCCCTACACGACGCTCTTCCGATCTNNNATTTCCTGTACCTCATGGAGCC
AGCACACGTCTGAACTCTTTCCCTACACGACGCTCTTCCGATCTNNNTAAAGCTCACACTTGGAAGCA
GAGCACACGTCTGAACTCTTTCCCTACACGACGCTCTTCCGATCTNNNGCATCCTGATCCAGGTACAA
AGCACACGTCTGAACTCTTTCCCTACACGACGCTCTTCCGATCTNNNCAGCAAGCTGAGCTATGTCCTT
GCACACGTCTGAACTCTTTCCCTACACGACGCTCTTCCGATCTNNNGGAAGATTGAGAAGCTACTGG
AGCACACGTCTGAACTCTTTCCCTACACGACGCTCTTCCGATCTNNNTTGGCTTAGTGGGGAAGCAT
GAGCACACGTCTGAACTCTTTCCCTACACGACGCTCTTCCGATCTNNNTTGTTGCCATGTGGTAATGC
AGCACACGTCTGAACTCTTTCCCTACACGACGCTCTTCCGATCTNNNGCTTCTCCTGCCTTTCTCAAAGT
GCACACGTCTGAACTCTTTCCCTACACGACGCTCTTCCGATCTNNNAGGCCCTGTTTACTTGGTGATT
CACGTCTGAACTCTTTCCCTACACGACGCTCTTCCGATCTNNNTTTAGCTGCCCCCAACTTCATTTCATC
AGCACACGTCTGAACTCTTTCCCTACACGACGCTCTTCCGATCTNNNGGACCATGATTATCCCATGCC
AGCACACGTCTGAACTCTTTCCCTACACGACGCTCTTCCGATCTNNNAATTCCCTTCACCCACTGCATTT
GAGCACACGTCTGAACTCTTTCCCTACACGACGCTCTTCCGATCTNNNGCTCTCTTTACGGTGCTTACC
AAGAGCACACGTCTGAACTCTTTCCCTACACGACGCTCTTCCGATCTNNNTAAGAGCCAAGCCAAGAA
AGCACACGTCTGAACTCTTTCCCTACACGACGCTCTTCCGATCTNNNTATGAAAAAGAAACCGGGGG
AGCACACGTCTGAACTCTTTCCCTACACGACGCTCTTCCGATCTNNNAGGAAGGAATTTGGGCTGAG
AGAGCACACGTCTGAACTCTTTCCCTACACGACGCTCTTCCGATCTNNNTATGCCTTTTACCTCACCATG
GCACACGTCTGAACTCTTTCCCTACACGACGCTCTTCCGATCTNNNACACCAACAAGACCACAGGAGT
GCACACGTCTGAACTCTTTCCCTACACGACGCTCTTCCGATCTNNNCCTTTATTCTCATAGGTTGCTAG
GAGCACACGTCTGAACTCTTTCCCTACACGACGCTCTTCCGATCTNNNCACAGAATCACTCAGGATGCC
GAGCACACGTCTGAACTCTTTCCCTACACGACGCTCTTCCGATCTNNNGGCTGCTAACTTGTCAGTTCC
AGCACACGTCTGAACTCTTTCCCTACACGACGCTCTTCCGATCTNNNAGGCATCCAAATGTTTCCTCCT
AGCACACGTCTGAACTCTTTCCCTACACGACGCTCTTCCGATCTNNNTCTCAACACCATGGAAAAGTT
GAGCACACGTCTGAACTCTTTCCCTACACGACGCTCTTCCGATCTNNNGGGAGATTCTGTATAGCCAT
AGCACACGTCTGAACTCTTTCCCTACACGACGCTCTTCCGATCTNNNGGTGCTTTCCTGTAATCCCACT
AGCACACGTCTGAACTCTTTCCCTACACGACGCTCTTCCGATCTNNNAGGGTTTGGTTTTGTTGTTTG
GAAGAGCACACGTCTGAACTCTTTCCCTACACGACGCTCTTCCGATCTNNNTCCAGGACTACTCTTGG
GAGCACACGTCTGAACTCTTTCCCTACACGACGCTCTTCCGATCTNNNCCAAATGGGTACCCCTCAATC
AGCACACGTCTGAACTCTTTCCCTACACGACGCTCTTCCGATCTNNNCCTTCCTCCTCTGAAACTGCAT

GAGCACACGTCTGAACTCTTTCCCTACACGACGCTCTTCCGATCTNNNCAAAGGACCCCATGTGCTGA
AGCACACGTCTGAACTCTTTCCCTACACGACGCTCTTCCGATCTNNNCCTTGAACTTCTGAGGAGGTGA
GAGCACACGTCTGAACTCTTTCCCTACACGACGCTCTTCCGATCTNNNTTCTTTTGGTTTAGGGGCTTG
GAGCACACGTCTGAACTCTTTCCCTACACGACGCTCTTCCGATCTNNNAGAAAGAAGAAGTCAAGGG
GCACACGTCTGAACTCTTTCCCTACACGACGCTCTTCCGATCTNNNAGTTTCCTTTCTTTTTCACTTGCC
GAGCACACGTCTGAACTCTTTCCCTACACGACGCTCTTCCGATCTNNNTTATAGTCCTGGCTTTGGCAC
AGAGCACACGTCTGAACTCTTTCCCTACACGACGCTCTTCCGATCTNNNCTCAGCACAATGCTACTGGG
GAGCACACGTCTGAACTCTTTCCCTACACGACGCTCTTCCGATCTNNNTCAACCCTAACACTTTTTCCCC
AGCACACGTCTGAACTCTTTCCCTACACGACGCTCTTCCGATCTNNNTCCAGACCATCCTTCACCTTTCA
GAGCACACGTCTGAACTCTTTCCCTACACGACGCTCTTCCGATCTNNNCCAAGCCAGAGCATTGATACC
GCACACGTCTGAACTCTTTCCCTACACGACGCTCTTCCGATCTNNNCCTCCTTTGGTTTCTAGGAGCTGC
AGCACACGTCTGAACTCTTTCCCTACACGACGCTCTTCCGATCTNNNCCTCCTTTGGTTTCTAGGAGCTG
GAGCACACGTCTGAACTCTTTCCCTACACGACGCTCTTCCGATCTNNNATGTATATCTCTGTGCGTGTG
AGCACACGTCTGAACTCTTTCCCTACACGACGCTCTTCCGATCTNNNTGGGTTCCCACAGGTAAACCA
GAGCACACGTCTGAACTCTTTCCCTACACGACGCTCTTCCGATCTNNNAGTTGTGGTGCTTAGGGAA
AGCACACGTCTGAACTCTTTCCCTACACGACGCTCTTCCGATCTNNNTGGCTTTCCATGGTAATTTGGT
GAGCACACGTCTGAACTCTTTCCCTACACGACGCTCTTCCGATCTNNNCATGTCCTGGATAGCTGCATG
GAGCACACGTCTGAACTCTTTCCCTACACGACGCTCTTCCGATCTNNNCCCCAACTCTGTGCCAATAAA
AGCACACGTCTGAACTCTTTCCCTACACGACGCTCTTCCGATCTNNNCCTCTCTTTTTCTTTCACGCCCTC
GAGCACACGTCTGAACTCTTTCCCTACACGACGCTCTTCCGATCTNNNATTGTACTGTGTATGCTCCA
GAGCACACGTCTGAACTCTTTCCCTACACGACGCTCTTCCGATCTNNNAGCCTGCAATAAACTTGGTGT
AGCACACGTCTGAACTCTTTCCCTACACGACGCTCTTCCGATCTNNNCAAGAATGCCAAGGTATTTCTT
GAGCACACGTCTGAACTCTTTCCCTACACGACGCTCTTCCGATCTNNNTCCCTGCAGTACTGTTTACCA
GAAGAGCACACGTCTGAACTCTTTCCCTACACGACGCTCTTCCGATCTNNNACACTCAAGTCCAGAGGA
AGAGCACACGTCTGAACTCTTTCCCTACACGACGCTCTTCCGATCTNNNCACACACAGACCTTCTCTTCA
AGCACACGTCTGAACTCTTTCCCTACACGACGCTCTTCCGATCTNNNGAACGATGTTATGCCCTCCAGA
AAGAGCACACGTCTGAACTCTTTCCCTACACGACGCTCTTCCGATCTNNNCTGTGGATGCACCCCACAC
AGCACACGTCTGAACTCTTTCCCTACACGACGCTCTTCCGATCTNNNAAACCATGATACAGGTGCCAGA
GAGCACACGTCTGAACTCTTTCCCTACACGACGCTCTTCCGATCTNNNTGGACAATAGGCAACTGGGT
AGCACACGTCTGAACTCTTTCCCTACACGACGCTCTTCCGATCTNNNCTCTTTCCTTGTCGGAGCTTGTA
GAGCACACGTCTGAACTCTTTCCCTACACGACGCTCTTCCGATCTNNNAGGTTCTCCAGAAAAACCAAA
AGCACACGTCTGAACTCTTTCCCTACACGACGCTCTTCCGATCTNNNTCTCTCTCAGAAATCAACCCCCG
AGCACACGTCTGAACTCTTTCCCTACACGACGCTCTTCCGATCTNNNCTGCTGGTCAACCACTATGTGT
GCACACGTCTGAACTCTTTCCCTACACGACGCTCTTCCGATCTNNNCCTCAGGTTACTCAGGTTTCTGC
GAGCACACGTCTGAACTCTTTCCCTACACGACGCTCTTCCGATCTNNNCTATTAGTCCAAGGTTGGGCA
AGAGCACACGTCTGAACTCTTTCCCTACACGACGCTCTTCCGATCTNNNTTTTGAACCCACTAAGGAGC
GAGCACACGTCTGAACTCTTTCCCTACACGACGCTCTTCCGATCTNNNCAGTCCTACCAAGACACAACC
GAGCACACGTCTGAACTCTTTCCCTACACGACGCTCTTCCGATCTNNNAGTGCACTAATAGGGGACACA
AGCACACGTCTGAACTCTTTCCCTACACGACGCTCTTCCGATCTNNNACGCTTCAAAGGGACAAAGAA
GAGCACACGTCTGAACTCTTTCCCTACACGACGCTCTTCCGATCTNNNAGGTTGATGCTACCTGGGGTA
GAGCACACGTCTGAACTCTTTCCCTACACGACGCTCTTCCGATCTNNNCCTATCTCTGGGTGGATGCTG
GCACACGTCTGAACTCTTTCCCTACACGACGCTCTTCCGATCTNNNACAAAAGCAATTCATTCGCACTG
AGCACACGTCTGAACTCTTTCCCTACACGACGCTCTTCCGATCTNNNTCATTCATGTTCCCGAGCAATC

AGCACACGTCTGAACTCTTTCCCTACACGACGCTCTTCCGATCTNNNCTGAATGGAAGCTATCCTGG
GAGCACACGTCTGAACTCTTTCCCTACACGACGCTCTTCCGATCTNNNAGGTGTCAGAGGTTTTGAT
AGCACACGTCTGAACTCTTTCCCTACACGACGCTCTTCCGATCTNNNAGTTCCCTAGGTCCCTTTCACT
GAGCACACGTCTGAACTCTTTCCCTACACGACGCTCTTCCGATCTNNNAGGGTTTAGGATGACAGGG
GAGCACACGTCTGAACTCTTTCCCTACACGACGCTCTTCCGATCTNNNAGAAGACTGTAAGAAAAGC
AGCACACGTCTGAACTCTTTCCCTACACGACGCTCTTCCGATCTNNNTGCAACCAAGTTCAAGGTATGC
AAGAGCACACGTCTGAACTCTTTCCCTACACGACGCTCTTCCGATCTNNNATGTATATCTCTGTGCGT
GAGCACACGTCTGAACTCTTTCCCTACACGACGCTCTTCCGATCTNNNACCTCATGCATTTCCCTCTCT
GCACACGTCTGAACTCTTTCCCTACACGACGCTCTTCCGATCTNNNTTCTGGGCAGGGACTGTATTGT
AGCACACGTCTGAACTCTTTCCCTACACGACGCTCTTCCGATCTNNNAATGAGGCAAACAGATCAAT
AGCACACGTCTGAACTCTTTCCCTACACGACGCTCTTCCGATCTNNNTTGTGCTAGGCTGTACTTCACA
GAGCACACGTCTGAACTCTTTCCCTACACGACGCTCTTCCGATCTNNNTCAGAGCCTCTCTCTCTCAAC
AGCACACGTCTGAACTCTTTCCCTACACGACGCTCTTCCGATCTNNNTAAGCACTGTGTACAGCAAG
AGCACACGTCTGAACTCTTTCCCTACACGACGCTCTTCCGATCTNNNAGTAGGCCTCATTGGTCATTC
GCACACGTCTGAACTCTTTCCCTACACGACGCTCTTCCGATCTNNNAGGCACACGAAGGGACATTTAT
GAGCACACGTCTGAACTCTTTCCCTACACGACGCTCTTCCGATCTNNNATGACAGAACGGAACTCTGA
GAGCACACGTCTGAACTCTTTCCCTACACGACGCTCTTCCGATCTNNNGCTCTAGACTTGCTTCTGCTT
AGCACACGTCTGAACTCTTTCCCTACACGACGCTCTTCCGATCTNNNCAGGCACACACAAAGAAAGAT
AGCACACGTCTGAACTCTTTCCCTACACGACGCTCTTCCGATCTNNNTTCCCAGGTCAATCTCTCTCTC
AGCACACGTCTGAACTCTTTCCCTACACGACGCTCTTCCGATCTNNNGCAGGATCTTCTTGAGGTGT
AGCACACGTCTGAACTCTTTCCCTACACGACGCTCTTCCGATCTNNNACCTCCCACATCAAAGATTTAC
AGCACACGTCTGAACTCTTTCCCTACACGACGCTCTTCCGATCTNNNATAGTCACATAGGGGCTTAGG
AGCACACGTCTGAACTCTTTCCCTACACGACGCTCTTCCGATCTNNNCCGAGATTCCCTTAGGTGTGTT
GAGCACACGTCTGAACTCTTTCCCTACACGACGCTCTTCCGATCTNNNTCAAGACCCTACTTTGTATGA
AGCACACGTCTGAACTCTTTCCCTACACGACGCTCTTCCGATCTNNNGCCAGGTTCTTCCAAAGATGTC
GAGCACACGTCTGAACTCTTTCCCTACACGACGCTCTTCCGATCTNNNCCATTCCAGGCAATCTGAAG
GCACACGTCTGAACTCTTTCCCTACACGACGCTCTTCCGATCTNNNTGACTATTTGCACATCTGGCTCC
AGCACACGTCTGAACTCTTTCCCTACACGACGCTCTTCCGATCTNNNGGTAGAGCTAGGACTGGTTC
GCACACGTCTGAACTCTTTCCCTACACGACGCTCTTCCGATCTNNNAGATGCATGACAGCTGGTGAA
AGAGCACACGTCTGAACTCTTTCCCTACACGACGCTCTTCCGATCTNNNAGGCTCTGGCTGTACTCGA
AGAGCACACGTCTGAACTCTTTCCCTACACGACGCTCTTCCGATCTNNNTGTGTTCTTAGTTTTTGCC
GCACACGTCTGAACTCTTTCCCTACACGACGCTCTTCCGATCTNNNTCAGTCTGGGTGGCAGAATAAG
GCACACGTCTGAACTCTTTCCCTACACGACGCTCTTCCGATCTNNNTCTAACAAGCTCCTGGAAGGTG
AGCACACGTCTGAACTCTTTCCCTACACGACGCTCTTCCGATCTNNNTAGCTTCTTCCTCCTTCCCCTTCC
AGCACACGTCTGAACTCTTTCCCTACACGACGCTCTTCCGATCTNNNCTGCCTAACCATCAGACCATCA
AGCACACGTCTGAACTCTTTCCCTACACGACGCTCTTCCGATCTNNNCAGAGAGTGGCACCCTTATAG
GAGCACACGTCTGAACTCTTTCCCTACACGACGCTCTTCCGATCTNNNCAGGTATCAGATAAGTGTGG
GAGCACACGTCTGAACTCTTTCCCTACACGACGCTCTTCCGATCTNNNGTACACTTCTGGGGTCAGTC
GCACACGTCTGAACTCTTTCCCTACACGACGCTCTTCCGATCTNNNCTATGGTGCCAGATTTAGGCAG
GAGCACACGTCTGAACTCTTTCCCTACACGACGCTCTTCCGATCTNNNCAGTCCTACCAAGACACAACC
AAGAGCACACGTCTGAACTCTTTCCCTACACGACGCTCTTCCGATCTNNNCGCAATTGTTCCTTAGTGT
AGCACACGTCTGAACTCTTTCCCTACACGACGCTCTTCCGATCTNNNACTTATCCCTTGACTTCCGGTG
AGAGCACACGTCTGAACTCTTTCCCTACACGACGCTCTTCCGATCTNNNGAGGATCCCTTAGGCTGAG

GAGCACACGTCTGAACTCTTTCCCTACACGACGCTCTTCCGATCTNNNGTAAGAGGCAGGAAAGGTG
AGCACACGTCTGAACTCTTTCCCTACACGACGCTCTTCCGATCTNNNGTTGTGGTGCTTAGGGAAGAGA
AGAGCACACGTCTGAACTCTTTCCCTACACGACGCTCTTCCGATCTNNNACAAATGTGTGTGTGAGAG
AGAGCACACGTCTGAACTCTTTCCCTACACGACGCTCTTCCGATCTNNNTCCTCTTTCTGTCTCTGCTGT
GAAGAGCACACGTCTGAACTCTTTCCCTACACGACGCTCTTCCGATCTNNNCGTTTGTGGAGAGCTGG
AGAGCACACGTCTGAACTCTTTCCCTACACGACGCTCTTCCGATCTNNNATTGTGGTGGCATGGAAAC
AGAGCACACGTCTGAACTCTTTCCCTACACGACGCTCTTCCGATCTNNNACTGGGATTGGCTGATTTCA
AGCACACGTCTGAACTCTTTCCCTACACGACGCTCTTCCGATCTNNNTTGAAGTGCAACAGAACACACC
AGCACACGTCTGAACTCTTTCCCTACACGACGCTCTTCCGATCTNNNTTCTAGCAGTGCAATGTTCCAG
AAGAGCACACGTCTGAACTCTTTCCCTACACGACGCTCTTCCGATCTNNNATCCAGGTCCATTCGTTCCA
AGCACACGTCTGAACTCTTTCCCTACACGACGCTCTTCCGATCTNNNGAGGGTAAAGTCAAACCAGGC
GAGCACACGTCTGAACTCTTTCCCTACACGACGCTCTTCCGATCTNNNAGAATCTGGGCATGAAATGG
GAGCACACGTCTGAACTCTTTCCCTACACGACGCTCTTCCGATCTNNNGGGCAATTCTTCTGTTCCACAT
AGAGCACACGTCTGAACTCTTTCCCTACACGACGCTCTTCCGATCTNNNAACTAACTTAGCTGCTCACCG
GAGCACACGTCTGAACTCTTTCCCTACACGACGCTCTTCCGATCTNNNACCAACTCCAAAGCATGTGTT
AGAGCACACGTCTGAACTCTTTCCCTACACGACGCTCTTCCGATCTNNNCGGGGAGACAAGAGAAAA
GAGCACACGTCTGAACTCTTTCCCTACACGACGCTCTTCCGATCTNNNGAAGGTCCAGGAACAGAGGT
AAGAGCACACGTCTGAACTCTTTCCCTACACGACGCTCTTCCGATCTNNNTCAGGGAAAGAGCACTGGA
AGCACACGTCTGAACTCTTTCCCTACACGACGCTCTTCCGATCTNNNAGTGCTGTGACTGGGATAGTT
GAGCACACGTCTGAACTCTTTCCCTACACGACGCTCTTCCGATCTNNNTTCTTACGCCCCTCTATAGCCAA
AGCACACGTCTGAACTCTTTCCCTACACGACGCTCTTCCGATCTNNNTGAATGTGAGGAGGTCTACAG
AGCACACGTCTGAACTCTTTCCCTACACGACGCTCTTCCGATCTNNNACCTGCATATTTCACGAAGAG
GAGCACACGTCTGAACTCTTTCCCTACACGACGCTCTTCCGATCTNNNCGCAGGCTCCAATTCAATCCAC
GCACACGTCTGAACTCTTTCCCTACACGACGCTCTTCCGATCTNNNCATAGAGGGAACCCCTACCACTG
GAGCACACGTCTGAACTCTTTCCCTACACGACGCTCTTCCGATCTNNNCATGGCAACTGGTTTCTCCCAT
GCACACGTCTGAACTCTTTCCCTACACGACGCTCTTCCGATCTNNNCTCGCTCCCTCCATCTCTCTTATCAT
GAGCACACGTCTGAACTCTTTCCCTACACGACGCTCTTCCGATCTNNNACAAAAACAAAAACAGGTGA
AGCACACGTCTGAACTCTTTCCCTACACGACGCTCTTCCGATCTNNNATACCTGAATGGCTGCTTATGC
AGCACACGTCTGAACTCTTTCCCTACACGACGCTCTTCCGATCTNNNGTAAGCCAGGATTCACCAGGAA
GAGCACACGTCTGAACTCTTTCCCTACACGACGCTCTTCCGATCTNNNTCCTTGGTCTCCCTACAGGAAA
AGAGCACACGTCTGAACTCTTTCCCTACACGACGCTCTTCCGATCTNNNAATCTTCTGTCCATCGCCCTTT
AGAGCACACGTCTGAACTCTTTCCCTACACGACGCTCTTCCGATCTNNNCCTTCCTCCCTCCAATCTCTCT
AAGAGCACACGTCTGAACTCTTTCCCTACACGACGCTCTTCCGATCTNNNATATCAGGGTTTCAGAAGG
AGCACACGTCTGAACTCTTTCCCTACACGACGCTCTTCCGATCTNNNTATCCTAGCACTCTGCCTTCCAA
AGCACACGTCTGAACTCTTTCCCTACACGACGCTCTTCCGATCTNNNTGTAAAGCATTTGGAGACCAG
GAGCACACGTCTGAACTCTTTCCCTACACGACGCTCTTCCGATCTNNNCCCACTACATACCCACCGTCTTA
AGCACACGTCTGAACTCTTTCCCTACACGACGCTCTTCCGATCTNNNCACCCCAAAAACTCACACCAGTA
AGCACACGTCTGAACTCTTTCCCTACACGACGCTCTTCCGATCTNNNCCAACCCCTGTATCACACAAACA
GAGCACACGTCTGAACTCTTTCCCTACACGACGCTCTTCCGATCTNNNGACTGGAGCACGTAGGATGTT
GAGCACACGTCTGAACTCTTTCCCTACACGACGCTCTTCCGATCTNNNTTCGACCAAGACGTCATTACG
GAGCACACGTCTGAACTCTTTCCCTACACGACGCTCTTCCGATCTNNNTGATGTACCAGGGAGACTGTG
AGCACACGTCTGAACTCTTTCCCTACACGACGCTCTTCCGATCTNNNTTTGGAAGCACTGCAATGACA
AGAGCACACGTCTGAACTCTTTCCCTACACGACGCTCTTCCGATCTNNNAAAGGCAATTTCTGTGTATG

GCACACGTCTGAACTCTTTCCCTACACGACGCTCTTCCGATCTNNNCTGTGATCCTCCTTTTCCAACCACA
AGCACACGTCTGAACTCTTTCCCTACACGACGCTCTTCCGATCTNNNCCTTCTGCCTCAGGTGGAATTTT
GAGCACACGTCTGAACTCTTTCCCTACACGACGCTCTTCCGATCTNNNAGACCCTGCAGCTATCATAAG
GAGCACACGTCTGAACTCTTTCCCTACACGACGCTCTTCCGATCTNNNACACCCTGCTGCATCATTTGTA
AGCACACGTCTGAACTCTTTCCCTACACGACGCTCTTCCGATCTNNNATGGAATCTCCCTGTCCCATCTC
GAGCACACGTCTGAACTCTTTCCCTACACGACGCTCTTCCGATCTNNNGGTGCAGTTATGCCTCAGATT
GCACACGTCTGAACTCTTTCCCTACACGACGCTCTTCCGATCTNNNTTTGTTCTTTCCCAGCCCTACACCA
GAGCACACGTCTGAACTCTTTCCCTACACGACGCTCTTCCGATCTNNNTTTGGACTTCAAGGATGCCAA
AGCACACGTCTGAACTCTTTCCCTACACGACGCTCTTCCGATCTNNNCATCTCAGGTTCTGACAGTTGG
AGCACACGTCTGAACTCTTTCCCTACACGACGCTCTTCCGATCTNNNTTTGTGGAACTGACAAAGGCTG
AGAGCACACGTCTGAACTCTTTCCCTACACGACGCTCTTCCGATCTNNNGCTACCTTCTCCTTTTGTGAG
GAGCACACGTCTGAACTCTTTCCCTACACGACGCTCTTCCGATCTNNNTCATGAGAAGAGTGTTGCAGA
AGCACACGTCTGAACTCTTTCCCTACACGACGCTCTTCCGATCTNNNTTGTGTGCCCTTCCATGAATGTT
AGCACACGTCTGAACTCTTTCCCTACACGACGCTCTTCCGATCTNNNCTCATGATAGGATTGGGGGTCC
AGCACACGTCTGAACTCTTTCCCTACACGACGCTCTTCCGATCTNNNGCCCTCATCTCTTTCTCTCCTCTCA
GGAAGAGCACACGTCTGAACTCTTTCCCTACACGACGCTCTTCCGATCTNNNAGACCTAGCCCTCAGATA
GAGCACACGTCTGAACTCTTTCCCTACACGACGCTCTTCCGATCTNNNCAACAAGGAATCCAGTCTGACC
GAAGAGCACACGTCTGAACTCTTTCCCTACACGACGCTCTTCCGATCTNNNTCAGCCTGTTGACTACAGT
AAGAGCACACGTCTGAACTCTTTCCCTACACGACGCTCTTCCGATCTNNNACCTACTTTTGGATGGTTTG
AGCACACGTCTGAACTCTTTCCCTACACGACGCTCTTCCGATCTNNNTGAAAGGACCAAGTGAATAAAA
AGCACACGTCTGAACTCTTTCCCTACACGACGCTCTTCCGATCTNNNTCAATCTTTTCTCCTTGCCCCACCA
AGCACACGTCTGAACTCTTTCCCTACACGACGCTCTTCCGATCTNNNAGGATTTAGGGATGGAGGTTC
AGCACACGTCTGAACTCTTTCCCTACACGACGCTCTTCCGATCTNNNAACTCTAGGACTGGGGCAAATG
GAGCACACGTCTGAACTCTTTCCCTACACGACGCTCTTCCGATCTNNNTTAGGGAGTTTATTCATTCCCC
AGCACACGTCTGAACTCTTTCCCTACACGACGCTCTTCCGATCTNNNCTTCCTGGGAGAGGACTGAAGT
AGCACACGTCTGAACTCTTTCCCTACACGACGCTCTTCCGATCTNNNCACAGCCACTCTAGCTGATTCTCC
GAGCACACGTCTGAACTCTTTCCCTACACGACGCTCTTCCGATCTNNNCACACAGAGACCCATTCAGACA
GAGCACACGTCTGAACTCTTTCCCTACACGACGCTCTTCCGATCTNNNTGGGGCTGGACTTCTACACTT
AGCACACGTCTGAACTCTTTCCCTACACGACGCTCTTCCGATCTNNNTATTGCTAACAGGTCAGCATGG
GAGCACACGTCTGAACTCTTTCCCTACACGACGCTCTTCCGATCTNNNTAAGGATGGTAGCTAAAGGG
GAGCACACGTCTGAACTCTTTCCCTACACGACGCTCTTCCGATCTNNNTGGAGTGATTCTTTTCAGCAG
GCACACGTCTGAACTCTTTCCCTACACGACGCTCTTCCGATCTNNNATGGCTGTCTCATTCCAACCATCC
AGCACACGTCTGAACTCTTTCCCTACACGACGCTCTTCCGATCTNNNAAGCTTGCACTTCTGAGCCTTTT
AGAGCACACGTCTGAACTCTTTCCCTACACGACGCTCTTCCGATCTNNNTGCAAGTTGATCAGCCTAGT
GAGCACACGTCTGAACTCTTTCCCTACACGACGCTCTTCCGATCTNNNAAGCCACATACATTCCAAAGC
AGCACACGTCTGAACTCTTTCCCTACACGACGCTCTTCCGATCTNNNTTTCAGGGGGAAGGTTGGTTTT
AGCACACGTCTGAACTCTTTCCCTACACGACGCTCTTCCGATCTNNNTCAATTACCAGATGCACTCCTGG
GAGCACACGTCTGAACTCTTTCCCTACACGACGCTCTTCCGATCTNNNACATGAGGATACAGAGGTGGT
AGCACACGTCTGAACTCTTTCCCTACACGACGCTCTTCCGATCTNNNATGGATTAGGTTTTGTTGTACA
GAAGAGCACACGTCTGAACTCTTTCCCTACACGACGCTCTTCCGATCTNNNATCTCTACAACCCTAGGGG
GAAGAGCACACGTCTGAACTCTTTCCCTACACGACGCTCTTCCGATCTNNNACTGTACCTGCCCCCTCACT
GAGCACACGTCTGAACTCTTTCCCTACACGACGCTCTTCCGATCTNNNATTCCCTTCCATGCCTCCTTAGT
GCACACGTCTGAACTCTTTCCCTACACGACGCTCTTCCGATCTNNNGGGGTCTCACTCCGAATCATTAC

GAGCACACGTCTGAACTCTTTCCCTACACGACGCTCTTCCGATCTNNNTACCAGAGTGACTTAGAGCTG
GGAAGAGCACACGTCTGAACTCTTTCCCTACACGACGCTCTTCCGATCTNNNGCCAACTGGACAAATAG
AGCACACGTCTGAACTCTTTCCCTACACGACGCTCTTCCGATCTNNNGAGAAGGGGAGGAAGAAAGC
AGCACACGTCTGAACTCTTTCCCTACACGACGCTCTTCCGATCTNNNGACACTGAGGCTTTGTGTGATG
GAGCACACGTCTGAACTCTTTCCCTACACGACGCTCTTCCGATCTNNNCAACATTCACAGTCACGTTCAC
GAGCACACGTCTGAACTCTTTCCCTACACGACGCTCTTCCGATCTNNNGAACCCCAACAAGATACAGAG
GCACACGTCTGAACTCTTTCCCTACACGACGCTCTTCCGATCTNNNCCCCAAGGTACAATCTACAGGGA
AGCACACGTCTGAACTCTTTCCCTACACGACGCTCTTCCGATCTNNNTTAGGCTACCTGAGGCTGAAGA
GCACACGTCTGAACTCTTTCCCTACACGACGCTCTTCCGATCTNNNTCTTTGGGTACCTGACCTCTTCCCA
AGAGCACACGTCTGAACTCTTTCCCTACACGACGCTCTTCCGATCTNNNCAAAACACTCTGCAACACCAC
AGCACACGTCTGAACTCTTTCCCTACACGACGCTCTTCCGATCTNNNTGGGCACACATTTGTGTTACAG
GAGCACACGTCTGAACTCTTTCCCTACACGACGCTCTTCCGATCTNNNCACCTGTAGAACTGGTAAACCC
AGAGCACACGTCTGAACTCTTTCCCTACACGACGCTCTTCCGATCTNNNGAGCGAGCAGCAGATAAGA
GAGCACACGTCTGAACTCTTTCCCTACACGACGCTCTTCCGATCTNNNATCCAGACCATGACCCACTTTC
GAGCACACGTCTGAACTCTTTCCCTACACGACGCTCTTCCGATCTNNNTCTGCTTTATTCCTCTCTGGCCT
AGAGCACACGTCTGAACTCTTTCCCTACACGACGCTCTTCCGATCTNNNACACAGATAACCAGTCCTCCC
GAGCACACGTCTGAACTCTTTCCCTACACGACGCTCTTCCGATCTNNNGAACAGTTGTAGAGGGCTTGT
AGAGCACACGTCTGAACTCTTTCCCTACACGACGCTCTTCCGATCTNNNAGATAAAGAGCATTATGTCC
AGCACACGTCTGAACTCTTTCCCTACACGACGCTCTTCCGATCTNNNTTTCTCTTTGCCTCTGCTTTCGTC
GAGCACACGTCTGAACTCTTTCCCTACACGACGCTCTTCCGATCTNNNCACCTCTCTGCCCTCATGTTTTT
GCACACGTCTGAACTCTTTCCCTACACGACGCTCTTCCGATCTNNNATTGAAGGGGATAGGGTGCTTC
AGCACACGTCTGAACTCTTTCCCTACACGACGCTCTTCCGATCTNNNCTCTCTTCCATTGGTGCTCAGTC
GAGCACACGTCTGAACTCTTTCCCTACACGACGCTCTTCCGATCTNNNGCATCCTTTTGTGTGACTGTGT
AGCACACGTCTGAACTCTTTCCCTACACGACGCTCTTCCGATCTNNNTTGAACACTTTGTAAGTACCTGT
GAGCACACGTCTGAACTCTTTCCCTACACGACGCTCTTCCGATCTNNNCCCCCGACGACGTGGACTACCA
GCACACGTCTGAACTCTTTCCCTACACGACGCTCTTCCGATCTNNNAAAAAGTCTGAGGGCCTGGTTA
AGAGCACACGTCTGAACTCTTTCCCTACACGACGCTCTTCCGATCTNNNACTTCCGAGTTTTCTCTCTCTC
GAGCACACGTCTGAACTCTTTCCCTACACGACGCTCTTCCGATCTNNNTGCTGACATACCTGGCAACAA
AAGAGCACACGTCTGAACTCTTTCCCTACACGACGCTCTTCCGATCTNNNGGGACGGGAAAGGAAGAA
GAGCACACGTCTGAACTCTTTCCCTACACGACGCTCTTCCGATCTNNNATTCCTTCTGGGCCCTTTCAAT
AGCACACGTCTGAACTCTTTCCCTACACGACGCTCTTCCGATCTNNNACCTGATCTGCAAACAATCCTG
AGCACACGTCTGAACTCTTTCCCTACACGACGCTCTTCCGATCTNNNGTGTTCACTCAGAACCCTTGTCC
AGCACACGTCTGAACTCTTTCCCTACACGACGCTCTTCCGATCTNNNCCACTAATCTAAGGGCAACCCAC
GAGCACACGTCTGAACTCTTTCCCTACACGACGCTCTTCCGATCTNNNCCATCGTGAGGGTTTTGAGGT
CACACGTCTGAACTCTTTCCCTACACGACGCTCTTCCGATCTNNNGCACAAGGAGATAGGGTTCCTCTC
AGCACACGTCTGAACTCTTTCCCTACACGACGCTCTTCCGATCTNNNTCTCCATCACCCCTTTGTTTTGGC
AGAGCACACGTCTGAACTCTTTCCCTACACGACGCTCTTCCGATCTNNNCAGTTGGCTACTCTTGCTTCC
AGAGCACACGTCTGAACTCTTTCCCTACACGACGCTCTTCCGATCTNNNATCTGTTCAGCAGGGAGAGA
GAGCACACGTCTGAACTCTTTCCCTACACGACGCTCTTCCGATCTNNNCTGAAACCGAGAGTTCTCCCAA
AGCACACGTCTGAACTCTTTCCCTACACGACGCTCTTCCGATCTNNNTCTGCCTGGGGGAAAAATAGTT
AGCACACGTCTGAACTCTTTCCCTACACGACGCTCTTCCGATCTNNNCAGTCTGACACCTGCTCTTTTCA
AGCACACGTCTGAACTCTTTCCCTACACGACGCTCTTCCGATCTNNNCGTGACTATTTGTGCTTGCATCC
AGAGCACACGTCTGAACTCTTTCCCTACACGACGCTCTTCCGATCTNNNAACACCCAATATGTGCAGGT

AGCACACGTCTGAACTCTTTCCCTACACGACGCTCTTCCGATCTNNNTCTTGAACTACTGGACTGTGG
AGCACACGTCTGAACTCTTTCCCTACACGACGCTCTTCCGATCTNNNAACTTGTACACGGAAACACCTG
GCACACGTCTGAACTCTTTCCCTACACGACGCTCTTCCGATCTNNNGAGAAAACTATGGCTGGAGACC
CACACGTCTGAACTCTTTCCCTACACGACGCTCTTCCGATCTNNNCAATCATTGACCAGCCATGGAGA
GAGCACACGTCTGAACTCTTTCCCTACACGACGCTCTTCCGATCTNNNAGCTCATAAGTGCTTGGGC
GAGCACACGTCTGAACTCTTTCCCTACACGACGCTCTTCCGATCTNNNCTGTTCTGTGAGGTTGTTCAG
GAGCACACGTCTGAACTCTTTCCCTACACGACGCTCTTCCGATCTNNNGCACGTTCCCTTATAGACTGA
AGCACACGTCTGAACTCTTTCCCTACACGACGCTCTTCCGATCTNNNTGCATTCAGAAGCTTGGCATT
AGCACACGTCTGAACTCTTTCCCTACACGACGCTCTTCCGATCTNNNATGTTGGGTGAAAGCTCCTTG
AGAGCACACGTCTGAACTCTTTCCCTACACGACGCTCTTCCGATCTNNNATGTTGGGTGAAAGCTCCT
AGCACACGTCTGAACTCTTTCCCTACACGACGCTCTTCCGATCTNNNTTCCTTCCCTCATGGAGGTCTTT
AGCACACGTCTGAACTCTTTCCCTACACGACGCTCTTCCGATCTNNNTGATTACCCTCAACACTCCCCA
GAGCACACGTCTGAACTCTTTCCCTACACGACGCTCTTCCGATCTNNNCTGGACTACAGGAGGCCCTA
AGCACACGTCTGAACTCTTTCCCTACACGACGCTCTTCCGATCTNNNAGTACAACCGTAGGAGTAGT
AGAGCACACGTCTGAACTCTTTCCCTACACGACGCTCTTCCGATCTNNNTCAGAAGCTGCAGTTGTAA
AGCACACGTCTGAACTCTTTCCCTACACGACGCTCTTCCGATCTNNNTGTGGGGAGAGTACCTCTTAG
AAGAGCACACGTCTGAACTCTTTCCCTACACGACGCTCTTCCGATCTNNNGCCCGGCCTTGAGTTTTA
AGCACACGTCTGAACTCTTTCCCTACACGACGCTCTTCCGATCTNNNCGGGTGTTACAAACACTGCAA
AGCACACGTCTGAACTCTTTCCCTACACGACGCTCTTCCGATCTNNNGGCAGAGCAGAAGAAAGGAA
GAGCACACGTCTGAACTCTTTCCCTACACGACGCTCTTCCGATCTNNNAATTGACTAGCTGGACTGTG
AGAGCACACGTCTGAACTCTTTCCCTACACGACGCTCTTCCGATCTNNNCCGTGAATACATTCGCAAAG
GAGCACACGTCTGAACTCTTTCCCTACACGACGCTCTTCCGATCTNNNGGGAATAACTGGCTTCCAGA
CACACGTCTGAACTCTTTCCCTACACGACGCTCTTCCGATCTNNNGTACCTCTGCTTCCACACATACCC
GCACACGTCTGAACTCTTTCCCTACACGACGCTCTTCCGATCTNNNCTGGACTAGTTGAGGCCATTCA
GAGCACACGTCTGAACTCTTTCCCTACACGACGCTCTTCCGATCTNNNATAGACCTACTGAGCCTCACC
AGCACACGTCTGAACTCTTTCCCTACACGACGCTCTTCCGATCTNNNGGCAGAGCAGAAGAAAGGAA
GAGCACACGTCTGAACTCTTTCCCTACACGACGCTCTTCCGATCTNNNATAGCCACCACGTCTCTCTGT
GCACACGTCTGAACTCTTTCCCTACACGACGCTCTTCCGATCTNNNTTCTGGAAAGTTCCTCCTCCC
CACACGTCTGAACTCTTTCCCTACACGACGCTCTTCCGATCTNNNTTTAGCTCTGGGGCTGGTGTTAT
AGCACACGTCTGAACTCTTTCCCTACACGACGCTCTTCCGATCTNNNAGTCTCTTCTACTTCAAATAAA
GAAGAGCACACGTCTGAACTCTTTCCCTACACGACGCTCTTCCGATCTNNNAAAGTCACCTGGGATGA
AGCACACGTCTGAACTCTTTCCCTACACGACGCTCTTCCGATCTNNNTATGAGGAAGAGCTGACCGAC
AGCACACGTCTGAACTCTTTCCCTACACGACGCTCTTCCGATCTNNNACTTCTGATGACCACTCCCTCAA
GCACACGTCTGAACTCTTTCCCTACACGACGCTCTTCCGATCTNNNTCTGAGTTCCTAACCACTGAGCA
GAGCACACGTCTGAACTCTTTCCCTACACGACGCTCTTCCGATCTNNNCAGACTGGGTGTGTAATCAA
AGCACACGTCTGAACTCTTTCCCTACACGACGCTCTTCCGATCTNNNATGGTCACTGGAACATGGGTA
AGCACACGTCTGAACTCTTTCCCTACACGACGCTCTTCCGATCTNNNAGAGTGGAAAGGGGTATGTG
AGCACACGTCTGAACTCTTTCCCTACACGACGCTCTTCCGATCTNNNCATGCATGGACACACACAATA
AGCACACGTCTGAACTCTTTCCCTACACGACGCTCTTCCGATCTNNNGGGAAACCAGTGTTCTGGGTT
AGCACACGTCTGAACTCTTTCCCTACACGACGCTCTTCCGATCTNNNATAGACCTACTGAGCCTCACCT
CACACGTCTGAACTCTTTCCCTACACGACGCTCTTCCGATCTNNNCCAGGAAGACTGAGGTGTGTGT
GCACACGTCTGAACTCTTTCCCTACACGACGCTCTTCCGATCTNNNCTTGCCTTCAAGGAATTATGGC
GAGCACACGTCTGAACTCTTTCCCTACACGACGCTCTTCCGATCTNNNATTCAGCTGCTCAGTGTCATC

GAGCACACGTCTGAACTCTTTCCCTACACGACGCTCTTCCGATCTNNNTCATCTATACATGGGTTAGCA
AGAGCACACGTCTGAACTCTTTCCCTACACGACGCTCTTCCGATCTNNNTGTGGGGGAAAGGTAGAA
AGCACACGTCTGAACTCTTTCCCTACACGACGCTCTTCCGATCTNNNCCTCCCTGGGACTATGTCTCAA
GAAGAGCACACGTCTGAACTCTTTCCCTACACGACGCTCTTCCGATCTNNNGAGATGCCCCAGATAAA
AGCACACGTCTGAACTCTTTCCCTACACGACGCTCTTCCGATCTNNNCCACATGCTCAAAGGGTACATC
ACACGTCTGAACTCTTTCCCTACACGACGCTCTTCCGATCTNNNGATCTCTTTCTGGAAAGGGGGTGC
AGAGCACACGTCTGAACTCTTTCCCTACACGACGCTCTTCCGATCTNNNGAGATGTCTGGGGTGAATG
AGCACACGTCTGAACTCTTTCCCTACACGACGCTCTTCCGATCTNNNCCAGGGTGTTGATGTCTCCCTT
AGAGCACACGTCTGAACTCTTTCCCTACACGACGCTCTTCCGATCTNNNCCAGTACACAGCCAACATA
AAGAGCACACGTCTGAACTCTTTCCCTACACGACGCTCTTCCGATCTNNNGGCTTTGATCAACCACCAG
GAGCACACGTCTGAACTCTTTCCCTACACGACGCTCTTCCGATCTNNNGGATACCGGGAAGGTTTGT
AGCACACGTCTGAACTCTTTCCCTACACGACGCTCTTCCGATCTNNNGCTGGTTAGACAGCATGAGTG
AAGAGCACACGTCTGAACTCTTTCCCTACACGACGCTCTTCCGATCTNNNAGCAGTATTCATAGTAGG
AGAGCACACGTCTGAACTCTTTCCCTACACGACGCTCTTCCGATCTNNNTTCTCAGGAGATCTGGGGT
ACACGTCTGAACTCTTTCCCTACACGACGCTCTTCCGATCTNNNCCAACCCCGTTGTCCCCCTGCATCGG
AGCACACGTCTGAACTCTTTCCCTACACGACGCTCTTCCGATCTNNNCTTGAAGGGTTGAGTGAAGG
AGCACACGTCTGAACTCTTTCCCTACACGACGCTCTTCCGATCTNNNCCTTATGAAACTCCAGCGAGGA
AGAGCACACGTCTGAACTCTTTCCCTACACGACGCTCTTCCGATCTNNNAGATTGAGATCTCTGCGGA
GAGCACACGTCTGAACTCTTTCCCTACACGACGCTCTTCCGATCTNNNATTTGGCATTTGGGATGTGTG
CACACGTCTGAACTCTTTCCCTACACGACGCTCTTCCGATCTNNNGGTTGTTTGAAAGTGTGTGACTGA
AGCACACGTCTGAACTCTTTCCCTACACGACGCTCTTCCGATCTNNNAGGATTTAGGGATGGAGGTT
GAGCACACGTCTGAACTCTTTCCCTACACGACGCTCTTCCGATCTNNNGGGACGGGAAAGGAAGAAA
GAGCACACGTCTGAACTCTTTCCCTACACGACGCTCTTCCGATCTNNNCCGGAGAAGGTCCTCAAAGTA
GCACACGTCTGAACTCTTTCCCTACACGACGCTCTTCCGATCTNNNCTCAAGCTACCAAAGACCACTGTC
GAGCACACGTCTGAACTCTTTCCCTACACGACGCTCTTCCGATCTNNNTCCAACTGTCAGAGTGTCAGG
GAAGAGCACACGTCTGAACTCTTTCCCTACACGACGCTCTTCCGATCTNNNGATCATCCTGTCTCCGC
GAAGAGCACACGTCTGAACTCTTTCCCTACACGACGCTCTTCCGATCTNNNGCAGAGCTCTCACGGTT
AGCACACGTCTGAACTCTTTCCCTACACGACGCTCTTCCGATCTNNNAACACACAGACTTGTCCTACTG
AGCACACGTCTGAACTCTTTCCCTACACGACGCTCTTCCGATCTNNNTTGGATCCATCTGCAACTGAAG
AGCACACGTCTGAACTCTTTCCCTACACGACGCTCTTCCGATCTNNNAGTTGGGCACCCATATTTCACT
AGCACACGTCTGAACTCTTTCCCTACACGACGCTCTTCCGATCTNNNCCTCACCCTCCTCCAAGTACATA
AGCACACGTCTGAACTCTTTCCCTACACGACGCTCTTCCGATCTNNNGGTAGGTTGCAGAGCATAGGA
GAGCACACGTCTGAACTCTTTCCCTACACGACGCTCTTCCGATCTNNNTTAGTAGGGTCAGAGGCAGT
AGCACACGTCTGAACTCTTTCCCTACACGACGCTCTTCCGATCTNNNAAGGAAGGTCCAAGTCTTTGCT
GCACACGTCTGAACTCTTTCCCTACACGACGCTCTTCCGATCTNNNAATAGTCAGCCTGGTCCACAGA
GCACACGTCTGAACTCTTTCCCTACACGACGCTCTTCCGATCTNNNACGTTCAGCCATGGAGATGATT
AGAGCACACGTCTGAACTCTTTCCCTACACGACGCTCTTCCGATCTNNNTTATCTGGGCTGGAGCTAC
GAGCACACGTCTGAACTCTTTCCCTACACGACGCTCTTCCGATCTNNNATCCTCTGTTTCATCTGGCAA
GAGCACACGTCTGAACTCTTTCCCTACACGACGCTCTTCCGATCTNNNGATCACGAGAGGCCATGAAG
GAGCACACGTCTGAACTCTTTCCCTACACGACGCTCTTCCGATCTNNNGGTAGTTAGATGTCCCATCCT
GAGCACACGTCTGAACTCTTTCCCTACACGACGCTCTTCCGATCTNNNGTGGGGAGATGGTTTCTCTC
AGCACACGTCTGAACTCTTTCCCTACACGACGCTCTTCCGATCTNNNTGACGGCAGAAGGGAAAAGAA
AGAGCACACGTCTGAACTCTTTCCCTACACGACGCTCTTCCGATCTNNNATGCCCAAAGTACCATACCA

GAGCACACGTCTGAACTCTTTCCCTACACGACGCTCTTCCGATCTNNNTCAAATGTAGTGTATCCCTTT
AGCACACGTCTGAACTCTTTCCCTACACGACGCTCTTCCGATCTNNNTGAGATCACTTTGGGCAACAA
GCACACGTCTGAACTCTTTCCCTACACGACGCTCTTCCGATCTNNNAGGTCTCTTTTATACACACACAC
AGCACACGTCTGAACTCTTTCCCTACACGACGCTCTTCCGATCTNNNTGCTGGATGTGGATTGAGCTT
AGAGCACACGTCTGAACTCTTTCCCTACACGACGCTCTTCCGATCTNNNGGAACCTGGCAAGACCTCA
AGCACACGTCTGAACTCTTTCCCTACACGACGCTCTTCCGATCTNNNCTACAGTGAAGGAGTGAAGG
AAGAGCACACGTCTGAACTCTTTCCCTACACGACGCTCTTCCGATCTNNNGAGGTCCAGGCAAAGCA
AGCACACGTCTGAACTCTTTCCCTACACGACGCTCTTCCGATCTNNNCCCAGGATTGAGTTCTCTGCT
GCACACGTCTGAACTCTTTCCCTACACGACGCTCTTCCGATCTNNNTGCCCCAAGAAGACATGGTAGT
GAGCACACGTCTGAACTCTTTCCCTACACGACGCTCTTCCGATCTNNNTTCGAGATCATCCTGGGTCT
AGCACACGTCTGAACTCTTTCCCTACACGACGCTCTTCCGATCTNNNAAACAGAAGTCTACCACCCC
CACACGTCTGAACTCTTTCCCTACACGACGCTCTTCCGATCTNNNTTTTCTTTGGCAGCTGATCCTGC
GCACACGTCTGAACTCTTTCCCTACACGACGCTCTTCCGATCTNNNTTGGATCCATCTGCAACTGAAG
GAGCACACGTCTGAACTCTTTCCCTACACGACGCTCTTCCGATCTNNNCAGGTGCTGCCTATGAGTA
AGCACACGTCTGAACTCTTTCCCTACACGACGCTCTTCCGATCTNNNAATAGACAACGAGTTGCCTGA
GAGCACACGTCTGAACTCTTTCCCTACACGACGCTCTTCCGATCTNNNTGAACTTTGGCCATCTACAG
AGCACACGTCTGAACTCTTTCCCTACACGACGCTCTTCCGATCTNNNTGTGGGATAATGTGTGGAAA
AGCACACGTCTGAACTCTTTCCCTACACGACGCTCTTCCGATCTNNNGAGGTGCTGGGATTCAGGGT
GCACACGTCTGAACTCTTTCCCTACACGACGCTCTTCCGATCTNNNACGTAAAAACTAATTGCGAAAC
GCACACGTCTGAACTCTTTCCCTACACGACGCTCTTCCGATCTNNNGGTTTTCCTAATCTCCATGCCCC
GCACACGTCTGAACTCTTTCCCTACACGACGCTCTTCCGATCTNNNCTAAAAATCTCCAGGCACCCTG
GAGCACACGTCTGAACTCTTTCCCTACACGACGCTCTTCCGATCTNNNCTAGGACTTCTCGGCCCTCAT
GAGCACACGTCTGAACTCTTTCCCTACACGACGCTCTTCCGATCTNNNGGTTCTGTAGCCAAATGGG
GAGCACACGTCTGAACTCTTTCCCTACACGACGCTCTTCCGATCTNNNGGCGACAATAGCAGAAACT
AGCACACGTCTGAACTCTTTCCCTACACGACGCTCTTCCGATCTNNNAGAGTGATGGAATGTAGCCC
GAGCACACGTCTGAACTCTTTCCCTACACGACGCTCTTCCGATCTNNNGTGCCTTGTAGAAGGAAGG
AGCACACGTCTGAACTCTTTCCCTACACGACGCTCTTCCGATCTNNNAGGCAGTTGAGAGGAAAACC
AGCACACGTCTGAACTCTTTCCCTACACGACGCTCTTCCGATCTNNNTCTGCTTCTGTGCAACTTCTGTT
AGAGCACACGTCTGAACTCTTTCCCTACACGACGCTCTTCCGATCTNNNGAAAGGGTAAAGAGCTG
GAGCACACGTCTGAACTCTTTCCCTACACGACGCTCTTCCGATCTNNNCCCCCTCTTCTGTGATCAGTCT
AGCACACGTCTGAACTCTTTCCCTACACGACGCTCTTCCGATCTNNNTTCCCATCTTGCTGCTTCTTCTC
GAGCACACGTCTGAACTCTTTCCCTACACGACGCTCTTCCGATCTNNNCTCTTTCCCTCTCCCCACAATC
AGCACACGTCTGAACTCTTTCCCTACACGACGCTCTTCCGATCTNNNATGCAGGTATATGCAGGGAC
AGAGCACACGTCTGAACTCTTTCCCTACACGACGCTCTTCCGATCTNNNATTATGCCGTCCTGGGTAT
AGCACACGTCTGAACTCTTTCCCTACACGACGCTCTTCCGATCTNNNTCTGCTTGGATGGCAGATGTA
AGCACACGTCTGAACTCTTTCCCTACACGACGCTCTTCCGATCTNNNCTTCATTCAGTGTCTGGGGTCT
AGCACACGTCTGAACTCTTTCCCTACACGACGCTCTTCCGATCTNNNAGCTCTTGATTTTGGCACAGT
GAGCACACGTCTGAACTCTTTCCCTACACGACGCTCTTCCGATCTNNNCAAACATGAGGCATTTGTTC
GCACACGTCTGAACTCTTTCCCTACACGACGCTCTTCCGATCTNNNCCGAGTGAAAACTTCCTGCCAT
CACACGTCTGAACTCTTTCCCTACACGACGCTCTTCCGATCTNNNTTGTGCTTGTAGAACCTGCTGTT
AGCACACGTCTGAACTCTTTCCCTACACGACGCTCTTCCGATCTNNNAAACTGAAATGCTCATCAGGG
GAGCACACGTCTGAACTCTTTCCCTACACGACGCTCTTCCGATCTNNNAATGTGCTTGACAGTCACAC
AGCACACGTCTGAACTCTTTCCCTACACGACGCTCTTCCGATCTNNNTTGATTTTCTCTCTCCCCCACC

GAGCACACGTCTGAACTCTTTCCCTACACGACGCTCTTCCGATCTNNNAACTCTCCATTTTGGAAGCAGC

AGCACACGTCTGAACTCTTTCCCTACACGACGCTCTTCCGATCTNNNAGCAACGACAAAAGCCAAAACA

AGCACACGTCTGAACTCTTTCCCTACACGACGCTCTTCCGATCTNNNTGATGCCTTTTGCATCTCTCTCTC

GCACACGTCTGAACTCTTTCCCTACACGACGCTCTTCCGATCTNNNGGATGACGAACTTGGCATTCTCTC

GAGCACACGTCTGAACTCTTTCCCTACACGACGCTCTTCCGATCTNNNCCGTGAATACATTCGCAAAGT

GCACACGTCTGAACTCTTTCCCTACACGACGCTCTTCCGATCTNNNTCAGAAGCTGCAGTTGTAAGAAT

AGCACACGTCTGAACTCTTTCCCTACACGACGCTCTTCCGATCTNNNGGTCCTTCTTTCCTTCTCCTTTCC

AGAGCACACGTCTGAACTCTTTCCCTACACGACGCTCTTCCGATCTNNNGGGGTTCTCTGGGTTTTCTT

GAGCACACGTCTGAACTCTTTCCCTACACGACGCTCTTCCGATCTNNNACACCCATGCACACACTTTGTA

AGCACACGTCTGAACTCTTTCCCTACACGACGCTCTTCCGATCTNNNGCAAAGGAGACCCCTCCCATTCA

AGCACACGTCTGAACTCTTTCCCTACACGACGCTCTTCCGATCTNNNGTCAAACAGCTGAAGCCAAAGA

GAGCACACGTCTGAACTCTTTCCCTACACGACGCTCTTCCGATCTNNNTGATCCTCTACCGAGAGAGCA

AGAGCACACGTCTGAACTCTTTCCCTACACGACGCTCTTCCGATCTNNNTAAGTGTGAGGTAAGGTGG

GAGCACACGTCTGAACTCTTTCCCTACACGACGCTCTTCCGATCTNNNGCTTGTCAAGAGATTCCTTCGC

AGAGCACACGTCTGAACTCTTTCCCTACACGACGCTCTTCCGATCTNNNATCTCTACAACCCTAGGGCTC

AGAGCACACGTCTGAACTCTTTCCCTACACGACGCTCTTCCGATCTNNNCCCGTGTTATTCAAGGGTTAC

CACACGTCTGAACTCTTTCCCTACACGACGCTCTTCCGATCTNNNTGACATGGAAAATTTTGATAATCCT

GAGCACACGTCTGAACTCTTTCCCTACACGACGCTCTTCCGATCTNNNAAGAAAGATGCCATGACGTG

AGCACACGTCTGAACTCTTTCCCTACACGACGCTCTTCCGATCTNNNTCTTGGCACCTCAATACCAATCC

AGAGCACACGTCTGAACTCTTTCCCTACACGACGCTCTTCCGATCTNNNAAGTGATTTGGAAAACATGC

AGAGCACACGTCTGAACTCTTTCCCTACACGACGCTCTTCCGATCTNNNTCCCAGGACATTGGACCTAG

AGCACACGTCTGAACTCTTTCCCTACACGACGCTCTTCCGATCTNNNTTCTGCCCACCCAATATCCTTCTC

GAGCACACGTCTGAACTCTTTCCCTACACGACGCTCTTCCGATCTNNNTCATGAATACCTAGCATGTGCC

AGCACACGTCTGAACTCTTTCCCTACACGACGCTCTTCCGATCTNNNGCCTTCAGAACCTCCTTCATGTTC

AGAGCACACGTCTGAACTCTTTCCCTACACGACGCTCTTCCGATCTNNNCTAGAACCCGATCCACGTCTT

AGCACACGTCTGAACTCTTTCCCTACACGACGCTCTTCCGATCTNNNGATTTCTGCCCAGAAGAGGAGT

AGAGCACACGTCTGAACTCTTTCCCTACACGACGCTCTTCCGATCTNNNCCCGTGTTATTCAAGGGTTA

AGCACACGTCTGAACTCTTTCCCTACACGACGCTCTTCCGATCTNNNGTAGCCAAGGCTGATAGAGAG

GAGCACACGTCTGAACTCTTTCCCTACACGACGCTCTTCCGATCTNNNAGAAGACAACGCAAAGAGGT

GAGCACACGTCTGAACTCTTTCCCTACACGACGCTCTTCCGATCTNNNGGAAAAAGCCCAGTTCTCAATC

GAGCACACGTCTGAACTCTTTCCCTACACGACGCTCTTCCGATCTNNNCCTTTGCACTTGGACGGGAAT

GCACACGTCTGAACTCTTTCCCTACACGACGCTCTTCCGATCTNNNACCAAAGCAGCAGGAATAGGACA

GCACACGTCTGAACTCTTTCCCTACACGACGCTCTTCCGATCTNNNCCAACAAGATATTTTGCAAGCCA

AGCACACGTCTGAACTCTTTCCCTACACGACGCTCTTCCGATCTNNNTTCCTCCCTGCATGGATTCTGTA

CACGTCTGAACTCTTTCCCTACACGACGCTCTTCCGATCTNNNCTAGGCAATTCTGCCTCCATAGCCATC

AGAGCACACGTCTGAACTCTTTCCCTACACGACGCTCTTCCGATCTNNNCTTGGGCTATAAACACTCCCC

GCACACGTCTGAACTCTTTCCCTACACGACGCTCTTCCGATCTNNNATCTTGGGCCTGTGTTATCTCCTC

CACACGTCTGAACTCTTTCCCTACACGACGCTCTTCCGATCTNNNCTCACCTGCAATGATGAACCAGTCA

GCACACGTCTGAACTCTTTCCCTACACGACGCTCTTCCGATCTNNNCTGGATCTCTCTGTTAGAGGGGC

GAGCACACGTCTGAACTCTTTCCCTACACGACGCTCTTCCGATCTNNNCACCAAGATCTCCATCCTGGAA

AGAGCACACGTCTGAACTCTTTCCCTACACGACGCTCTTCCGATCTNNNTCCCCTATCTGTGTGGGTTTT

AGCACACGTCTGAACTCTTTCCCTACACGACGCTCTTCCGATCTNNNGCCTGGTGTGACATAAGCTGAA

GCACACGTCTGAACTCTTTCCCTACACGACGCTCTTCCGATCTNNNCTGAGGTGAGCCTGTGTTCTCTC

AGCACACGTCTGAACTCTTTCCCTACACGACGCTCTTCCGATCTNNNAGAGATTCTCATACTCGGCCTCC
AGCACACGTCTGAACTCTTTCCCTACACGACGCTCTTCCGATCTNNNAGGACCTGATTTCCTTACTGCC
GCACACGTCTGAACTCTTTCCCTACACGACGCTCTTCCGATCTNNNGATAGCTAGTCCCGGCGTGCTA
AGCACACGTCTGAACTCTTTCCCTACACGACGCTCTTCCGATCTNNNGATAGCTAGTCCCGGCGTGCTA
GAGCACACGTCTGAACTCTTTCCCTACACGACGCTCTTCCGATCTNNNCTGTTTCCTGAGGCCCAACAT
GCACACGTCTGAACTCTTTCCCTACACGACGCTCTTCCGATCTNNNATGTGGATGGGAGAGTGTGAA
GAGCACACGTCTGAACTCTTTCCCTACACGACGCTCTTCCGATCTNNNGGTAGGAATTCCGGCTTGTC
AGCACACGTCTGAACTCTTTCCCTACACGACGCTCTTCCGATCTNNNTCTTGGTTGGCACCAGTGTTT
GAGCACACGTCTGAACTCTTTCCCTACACGACGCTCTTCCGATCTNNNACTACTAAAAGGAGGCTGAG
GCACACGTCTGAACTCTTTCCCTACACGACGCTCTTCCGATCTNNNAAATAGCTTAGGGCAGAAGCA
AGCACACGTCTGAACTCTTTCCCTACACGACGCTCTTCCGATCTNNNGCCCTCAACAAGATGTTTTGCC
GCACACGTCTGAACTCTTTCCCTACACGACGCTCTTCCGATCTNNNACGCTCTAATCCTAGCTCCACTTC
GCACACGTCTGAACTCTTTCCCTACACGACGCTCTTCCGATCTNNNCCGCCTGCATCTCCTAATAAACAC
AGCACACGTCTGAACTCTTTCCCTACACGACGCTCTTCCGATCTNNNCCACTTTGGTGCTCTTGGAGA
GCACACGTCTGAACTCTTTCCCTACACGACGCTCTTCCGATCTNNNGCTGTACCATACACCCATCTACCC
GAGCACACGTCTGAACTCTTTCCCTACACGACGCTCTTCCGATCTNNNGCATGTTGCTTTTGTACCGTC
AAGAGCACACGTCTGAACTCTTTCCCTACACGACGCTCTTCCGATCTNNNGGGCAAGCTTGATCCCTAT
AAGAGCACACGTCTGAACTCTTTCCCTACACGACGCTCTTCCGATCTNNNCAGAACTCCAGGCTCTCTG
AGCACACGTCTGAACTCTTTCCCTACACGACGCTCTTCCGATCTNNNGTGGTGTTCAGGGATTTAGG
AGCACACGTCTGAACTCTTTCCCTACACGACGCTCTTCCGATCTNNNGTGCTTATGGCAGCAGAGAGG
GCACACGTCTGAACTCTTTCCCTACACGACGCTCTTCCGATCTNNNTTGCTGGACATGGATGGATTCTG
CACACGTCTGAACTCTTTCCCTACACGACGCTCTTCCGATCTNNNGTGTATATATGTATGTGTGAGAC
CACACGTCTGAACTCTTTCCCTACACGACGCTCTTCCGATCTNNNGATGACAGGATGTCCTCTTCAGG
AGCACACGTCTGAACTCTTTCCCTACACGACGCTCTTCCGATCTNNNTGACAGGTTTTATTCCGCTCCTG
GAGCACACGTCTGAACTCTTTCCCTACACGACGCTCTTCCGATCTNNNATTCATCATCTTCCACACGCAC
AGAGCACACGTCTGAACTCTTTCCCTACACGACGCTCTTCCGATCTNNNGCTAATCTAGCATGCCTGGC
AGCACACGTCTGAACTCTTTCCCTACACGACGCTCTTCCGATCTNNNGTTTTGTGTGCGTAAGTGCATG
GCACACGTCTGAACTCTTTCCCTACACGACGCTCTTCCGATCTNNNCTCTCTCTGTTCCATGGTCAGGT
AGCACACGTCTGAACTCTTTCCCTACACGACGCTCTTCCGATCTNNNATCTTATGCACACACACACGGA
GAGCACACGTCTGAACTCTTTCCCTACACGACGCTCTTCCGATCTNNNTTCCATTTGAAACCCTCTGGG
GAGCACACGTCTGAACTCTTTCCCTACACGACGCTCTTCCGATCTNNNGTAATACAGGCATGTGCCAC
GAGCACACGTCTGAACTCTTTCCCTACACGACGCTCTTCCGATCTNNNTTTTTACACCTGCCACCATTG
AGCACACGTCTGAACTCTTTCCCTACACGACGCTCTTCCGATCTNNNCGGAGAGGGACCATGTCTCTA
GCACACGTCTGAACTCTTTCCCTACACGACGCTCTTCCGATCTNNNCATAAGCTGTTTCGGCCTTTGG
AGCACACGTCTGAACTCTTTCCCTACACGACGCTCTTCCGATCTNNNACCAATGTCAGCTCAATAGCA
AGCACACGTCTGAACTCTTTCCCTACACGACGCTCTTCCGATCTNNNTGCAAACTGGCTTGTGATTTCT
AGCACACGTCTGAACTCTTTCCCTACACGACGCTCTTCCGATCTNNNTGATAACACCAGTAAGCTGTG
GAGCACACGTCTGAACTCTTTCCCTACACGACGCTCTTCCGATCTNNNTGTAAATGCGTGAGAAGAG
AGCACACGTCTGAACTCTTTCCCTACACGACGCTCTTCCGATCTNNNCCATGAAGATTCAGCACTGTG
AGCACACGTCTGAACTCTTTCCCTACACGACGCTCTTCCGATCTNNNACAGACATAGATACAGGCAGG
AGCACACGTCTGAACTCTTTCCCTACACGACGCTCTTCCGATCTNNNACCAACACTCCTCCCTCTATGTT
AGCACACGTCTGAACTCTTTCCCTACACGACGCTCTTCCGATCTNNNAAAGCTCACCTATTGCCCTCAA
AGCACACGTCTGAACTCTTTCCCTACACGACGCTCTTCCGATCTNNNTCTATACTAACCCAACCTCTACC

GAGCACACGTCTGAACTCTTTCCCTACACGACGCTCTTCCGATCTNNNAGCCATGAACATTCTAAAGA
GAGCACACGTCTGAACTCTTTCCCTACACGACGCTCTTCCGATCTNNNACAGGGATGTAGGACGTAG
AGCACACGTCTGAACTCTTTCCCTACACGACGCTCTTCCGATCTNNNAACATTACCCCTCTCATACCTG
GAGCACACGTCTGAACTCTTTCCCTACACGACGCTCTTCCGATCTNNNAGTGGAATCTAGGCAGGGA
GAGCACACGTCTGAACTCTTTCCCTACACGACGCTCTTCCGATCTNNNACCTTGATCATGAAGAGTAA
GAGCACACGTCTGAACTCTTTCCCTACACGACGCTCTTCCGATCTNNNCTAGAGTTGACAGTGAGGA
GAGCACACGTCTGAACTCTTTCCCTACACGACGCTCTTCCGATCTNNNAAGCAAGGACAATATGTGG
AAGAGCACACGTCTGAACTCTTTCCCTACACGACGCTCTTCCGATCTNNNAGAGCAAGAATCAAGTG
CACACGTCTGAACTCTTTCCCTACACGACGCTCTTCCGATCTNNNGAAGCAGAGCAAAAATGGCAGA
AGCACACGTCTGAACTCTTTCCCTACACGACGCTCTTCCGATCTNNNCCATGGTCGGCAGTCTTGAT
AGCACACGTCTGAACTCTTTCCCTACACGACGCTCTTCCGATCTNNNGGAGGCAAACATGATTCACA
AGCACACGTCTGAACTCTTTCCCTACACGACGCTCTTCCGATCTNNNAGAGCTTGCAGTGAGTAGACA
GCACACGTCTGAACTCTTTCCCTACACGACGCTCTTCCGATCTNNNAGTAAATAACGAAAGCCCCAGC
AGCACACGTCTGAACTCTTTCCCTACACGACGCTCTTCCGATCTNNNACCCAATCCACGCATGTGAT
GCACACGTCTGAACTCTTTCCCTACACGACGCTCTTCCGATCTNNNTCCATCACCTCTAAAAGCTTCAT
GAGCACACGTCTGAACTCTTTCCCTACACGACGCTCTTCCGATCTNNNAGAAATGGGACACAAAGA
GAGCACACGTCTGAACTCTTTCCCTACACGACGCTCTTCCGATCTNNNAGCTGAGTAGTCACACAGCA
GAGCACACGTCTGAACTCTTTCCCTACACGACGCTCTTCCGATCTNNNACATGATCCAGGGACAGTG
GCACACGTCTGAACTCTTTCCCTACACGACGCTCTTCCGATCTNNNCCAAAGTCTCACAGCCAGTTAG
AGCACACGTCTGAACTCTTTCCCTACACGACGCTCTTCCGATCTNNNAATCGGGAGAAAGAGACAG
AGCACACGTCTGAACTCTTTCCCTACACGACGCTCTTCCGATCTNNNCGGCACATATTTACCCAGAGG
AGCACACGTCTGAACTCTTTCCCTACACGACGCTCTTCCGATCTNNNATATCTGAAGAACTCGCTGTG
GCACACGTCTGAACTCTTTCCCTACACGACGCTCTTCCGATCTNNNCACTCTGTACTCACATGGGCTTT
AGCACACGTCTGAACTCTTTCCCTACACGACGCTCTTCCGATCTNNNTTGGTAAACACTACCAACCAG
GAGCACACGTCTGAACTCTTTCCCTACACGACGCTCTTCCGATCTNNNATACACCAATGTGATGGTG
AGCACACGTCTGAACTCTTTCCCTACACGACGCTCTTCCGATCTNNNCACATTTGAGTTTTGCAGGGT
AGCACACGTCTGAACTCTTTCCCTACACGACGCTCTTCCGATCTNNNCATATTGAGCCAAGGGAAGG
AGCACACGTCTGAACTCTTTCCCTACACGACGCTCTTCCGATCTNNNTCAAGGATGTGTAGGTGCAT
GAGCACACGTCTGAACTCTTTCCCTACACGACGCTCTTCCGATCTNNNTTCTCACCACTCTCTCTCTTCC
GAGCACACGTCTGAACTCTTTCCCTACACGACGCTCTTCCGATCTNNNCCGCTTCTGGTACTCTGATG
GCACACGTCTGAACTCTTTCCCTACACGACGCTCTTCCGATCTNNNGCCTAAAATTCACCCCCTTTGG
GCACACGTCTGAACTCTTTCCCTACACGACGCTCTTCCGATCTNNNTCTCTGCTCTCTCTCCTCTCTCAC
AGCACACGTCTGAACTCTTTCCCTACACGACGCTCTTCCGATCTNNNTGAGGAAAAGGAGGTGCCT
AGCACACGTCTGAACTCTTTCCCTACACGACGCTCTTCCGATCTNNNGTAAGCGTTTCTGTCTCTCCCT
GCACACGTCTGAACTCTTTCCCTACACGACGCTCTTCCGATCTNNNTATCAGCACAGGTCCTGGAACA
AGCACACGTCTGAACTCTTTCCCTACACGACGCTCTTCCGATCTNNNAGCCCACATAGACATGAACA
GAGCACACGTCTGAACTCTTTCCCTACACGACGCTCTTCCGATCTNNNTTCAATTATGAACAGGAGGG
AGCACACGTCTGAACTCTTTCCCTACACGACGCTCTTCCGATCTNNNAAAAGGAGGAGTGTGTGTGT
GAGCACACGTCTGAACTCTTTCCCTACACGACGCTCTTCCGATCTNNNGCTTCAGTTTTGGGGAAGG
AGCACACGTCTGAACTCTTTCCCTACACGACGCTCTTCCGATCTNNNAGAGTGGATAGAAAAGGGCA
CACACGTCTGAACTCTTTCCCTACACGACGCTCTTCCGATCTNNNGTTGAGTGCAACTCTGGCTTCTA
AGCACACGTCTGAACTCTTTCCCTACACGACGCTCTTCCGATCTNNNAAGAGGATCTTGTAGCCCAGG
AGCACACGTCTGAACTCTTTCCCTACACGACGCTCTTCCGATCTNNNCCCTACAATACACAGGACACCC

GAGCACACGTCTGAACTCTTTCCCTACACGACGCTCTTCCGATCTNNNGCTTACCCACACATGTACACTCA
AGCACACGTCTGAACTCTTTCCCTACACGACGCTCTTCCGATCTNNNAGAGTTCTGTAGGCCAAGGTA
AGCACACGTCTGAACTCTTTCCCTACACGACGCTCTTCCGATCTNNNCTGTTACAACACCCCAGCTTCAT
GAGCACACGTCTGAACTCTTTCCCTACACGACGCTCTTCCGATCTNNNGCAAAGCCATCTATTTCCATGC
GAGCACACGTCTGAACTCTTTCCCTACACGACGCTCTTCCGATCTNNNTTTCAGTCTTCATGTGTGTGG
AGAGCACACGTCTGAACTCTTTCCCTACACGACGCTCTTCCGATCTNNNAAGTGAAAAATCCATCCCCCT
GAGCACACGTCTGAACTCTTTCCCTACACGACGCTCTTCCGATCTNNNGAGGGAGCACCAAGAATGAT
AGCACACGTCTGAACTCTTTCCCTACACGACGCTCTTCCGATCTNNNAACCCCATTTCCTACACACATGC
AGCACACGTCTGAACTCTTTCCCTACACGACGCTCTTCCGATCTNNNTTTGATAGTCTGCTGTTGAGCCC
AGCACACGTCTGAACTCTTTCCCTACACGACGCTCTTCCGATCTNNNAGAGAAGTTGCTTTGGCCATTCA
GAGCACACGTCTGAACTCTTTCCCTACACGACGCTCTTCCGATCTNNNCCCTCAAGGGGGAAATCAAAA
GAGCACACGTCTGAACTCTTTCCCTACACGACGCTCTTCCGATCTNNNATTTACCAACTTGCATTTGTTA
GAGCACACGTCTGAACTCTTTCCCTACACGACGCTCTTCCGATCTNNNCTGGACAACATCGTAAAGCTC
GAGCACACGTCTGAACTCTTTCCCTACACGACGCTCTTCCGATCTNNNTGCTTGCATAGATGGACATTC
AGCACACGTCTGAACTCTTTCCCTACACGACGCTCTTCCGATCTNNNTCCCATACTCACTGATGCTCCAT
GAGCACACGTCTGAACTCTTTCCCTACACGACGCTCTTCCGATCTNNNAGTCTTGGCAGGTATGGACA
GAGCACACGTCTGAACTCTTTCCCTACACGACGCTCTTCCGATCTNNNGAAATGGACCCTTGGAGATG
GAGCACACGTCTGAACTCTTTCCCTACACGACGCTCTTCCGATCTNNNAGATCCAGAAGGTAAGGAAG
AGAGCACACGTCTGAACTCTTTCCCTACACGACGCTCTTCCGATCTNNNCTTTTCTTTCAGGCTTCCTAT
GAGCACACGTCTGAACTCTTTCCCTACACGACGCTCTTCCGATCTNNNTCACATCCACAGTACACATGCA
GAGCACACGTCTGAACTCTTTCCCTACACGACGCTCTTCCGATCTNNNCACATCCTACGCTCACATACCTC
GAGCACACGTCTGAACTCTTTCCCTACACGACGCTCTTCCGATCTNNNTGGAGGAGATAGAGAGGGG
AGAGCACACGTCTGAACTCTTTCCCTACACGACGCTCTTCCGATCTNNNCCAAGCATTCTGGCAAGTTTC
AGCACACGTCTGAACTCTTTCCCTACACGACGCTCTTCCGATCTNNNCCTGTCGACGGAGTGAGACTTA
GAGCACACGTCTGAACTCTTTCCCTACACGACGCTCTTCCGATCTNNNGGTCACAATCACCATGATTCGC
GAGCACACGTCTGAACTCTTTCCCTACACGACGCTCTTCCGATCTNNNCCTGGCCCTCTACCTTCATTTTT
AGCACACGTCTGAACTCTTTCCCTACACGACGCTCTTCCGATCTNNNTGCATTGTGGATCAATATTCTTC
AGCACACGTCTGAACTCTTTCCCTACACGACGCTCTTCCGATCTNNNCCAGGAATGGGGATTGCCAATA
GAGCACACGTCTGAACTCTTTCCCTACACGACGCTCTTCCGATCTNNNTGCTTGTTGTGTTATGACTTG
GAAGAGCACACGTCTGAACTCTTTCCCTACACGACGCTCTTCCGATCTNNNTGGAATATGTGGAGTGTC
GAAGAGCACACGTCTGAACTCTTTCCCTACACGACGCTCTTCCGATCTNNNATGCCTGCTAAGGAATTT
AGCACACGTCTGAACTCTTTCCCTACACGACGCTCTTCCGATCTNNNGCCTGCTTAGAGTAGCAGTAAC
AGAGCACACGTCTGAACTCTTTCCCTACACGACGCTCTTCCGATCTNNNACAGTTTCTCATGTTCTAGTC
AGAGCACACGTCTGAACTCTTTCCCTACACGACGCTCTTCCGATCTNNNGAGAACAACTGCTCTGGGG
AGAGCACACGTCTGAACTCTTTCCCTACACGACGCTCTTCCGATCTNNNTCCTTGCCTTTCTTACCCCTGT
GCACACGTCTGAACTCTTTCCCTACACGACGCTCTTCCGATCTNNNGGACTGGGAGGCAAATAACTCTC
GAGCACACGTCTGAACTCTTTCCCTACACGACGCTCTTCCGATCTNNNTATAAACGTGGCCAGTTCTCT
AGCACACGTCTGAACTCTTTCCCTACACGACGCTCTTCCGATCTNNNCCCCAGAATCCAGGACTTCCTAT
CACACGTCTGAACTCTTTCCCTACACGACGCTCTTCCGATCTNNNCACAGAGCGCCCTTTGAGGAGCAT
GAGCACACGTCTGAACTCTTTCCCTACACGACGCTCTTCCGATCTNNNATATGCTAGTGTGTCCTCCCCA
GCACACGTCTGAACTCTTTCCCTACACGACGCTCTTCCGATCTNNNAGCACTATTTGGATTCTGCTTGTC
AGCACACGTCTGAACTCTTTCCCTACACGACGCTCTTCCGATCTNNNTGACTGTCATCTGGAATTTCTCC
AGCACACGTCTGAACTCTTTCCCTACACGACGCTCTTCCGATCTNNNAGCATGCTATCATTTGTGTGGA

ACACGTCTGAACTCTTTCCCTACACGACGCTCTTCCGATCTNNNATCTGGGAGGACTTCAACTCCAACA
GCACACGTCTGAACTCTTTCCCTACACGACGCTCTTCCGATCTNNNAGGTCACATTCATACCCGAAGGA
AGCACACGTCTGAACTCTTTCCCTACACGACGCTCTTCCGATCTNNNGCCATGCTCTCTTCATGCAATTC
GAGCACACGTCTGAACTCTTTCCCTACACGACGCTCTTCCGATCTNNNATGAGGCTCACCAAATGTGTG
GAGCACACGTCTGAACTCTTTCCCTACACGACGCTCTTCCGATCTNNNGTAATGTCCTCTGGGTTCAT
AGAGCACACGTCTGAACTCTTTCCCTACACGACGCTCTTCCGATCTNNNCATCCCTTCATTACCTGTGGT
AGCACACGTCTGAACTCTTTCCCTACACGACGCTCTTCCGATCTNNNATGTTTTCTTCCAACCTCTGGCT
AGCACACGTCTGAACTCTTTCCCTACACGACGCTCTTCCGATCTNNNAGGGGTCTATACATACCATTTG
GAGCACACGTCTGAACTCTTTCCCTACACGACGCTCTTCCGATCTNNNTGGAATATGTGGAGTGTGTG
GAGCACACGTCTGAACTCTTTCCCTACACGACGCTCTTCCGATCTNNNCGAAGAGCCTGACATAAACA
GAGCACACGTCTGAACTCTTTCCCTACACGACGCTCTTCCGATCTNNNAAAAGGAGGAGTGTGTGTGT
GAGCACACGTCTGAACTCTTTCCCTACACGACGCTCTTCCGATCTNNNATGTTTTCTTCCAACCTCTGGCT
AGCACACGTCTGAACTCTTTCCCTACACGACGCTCTTCCGATCTNNNGAGTTTTCCTCCAGCAAAGCCAA
GAGCACACGTCTGAACTCTTTCCCTACACGACGCTCTTCCGATCTNNNTCTGATGCAAAACTCGACCTT
AGAGCACACGTCTGAACTCTTTCCCTACACGACGCTCTTCCGATCTNNNGAGTTGAATGAGGTCATTG
AGCACACGTCTGAACTCTTTCCCTACACGACGCTCTTCCGATCTNNNCATCCCTTCATTACCTGTGGTGT
AGAGCACACGTCTGAACTCTTTCCCTACACGACGCTCTTCCGATCTNNNTTGTCTTGTTTCCCAGAGAA
AGAGCACACGTCTGAACTCTTTCCCTACACGACGCTCTTCCGATCTNNNTCACCCTTGGCTTATTTTGCA
GAGCACACGTCTGAACTCTTTCCCTACACGACGCTCTTCCGATCTNNNATCTGGAGGACAATTGGGACA
GAGCACACGTCTGAACTCTTTCCCTACACGACGCTCTTCCGATCTNNNGCCTTTCTCTTTCTTGCTGTCTT
GAGCACACGTCTGAACTCTTTCCCTACACGACGCTCTTCCGATCTNNNTGATGTATGTTGGCTATGGT
GAGCACACGTCTGAACTCTTTCCCTACACGACGCTCTTCCGATCTNNNCCCAAGACGCTGACAACATGA
AGAGCACACGTCTGAACTCTTTCCCTACACGACGCTCTTCCGATCTNNNTGCTTGTTGTGTTATGACTTG
GAGCACACGTCTGAACTCTTTCCCTACACGACGCTCTTCCGATCTNNNGAAATGCTGCTCTGATAGGG
AGCACACGTCTGAACTCTTTCCCTACACGACGCTCTTCCGATCTNNNTCTTCCAAGCTGTGTTTCAAATG
GAGCACACGTCTGAACTCTTTCCCTACACGACGCTCTTCCGATCTNNNTTGTGACAAATGCCTCAGTGT
GAGCACACGTCTGAACTCTTTCCCTACACGACGCTCTTCCGATCTNNNGAGCTCATGGGTGTTTGAGGA
AGAGCACACGTCTGAACTCTTTCCCTACACGACGCTCTTCCGATCTNNNCTGCAAAGACCTAAGTCACA
AGCACACGTCTGAACTCTTTCCCTACACGACGCTCTTCCGATCTNNNTGAACTAACACAAGCAGCCAA
GAGCACACGTCTGAACTCTTTCCCTACACGACGCTCTTCCGATCTNNNCTCACTCCAGAGGCTTCATTT
GAGCACACGTCTGAACTCTTTCCCTACACGACGCTCTTCCGATCTNNNGATCACCAGTGGCCAATAACG
GAGCACACGTCTGAACTCTTTCCCTACACGACGCTCTTCCGATCTNNNATCCTAATTTCTCCCTCTTGTT
AGCACACGTCTGAACTCTTTCCCTACACGACGCTCTTCCGATCTNNNCATGTAGGAGTGGAAGCTTTC
AGCACACGTCTGAACTCTTTCCCTACACGACGCTCTTCCGATCTNNNCCAACGTACCACACACATACACA
AGCACACGTCTGAACTCTTTCCCTACACGACGCTCTTCCGATCTNNNAACCTCAGAGGAAGAATGGGT
AGCACACGTCTGAACTCTTTCCCTACACGACGCTCTTCCGATCTNNNGAGGACAGACAGAAAGCAAG
AGCACACGTCTGAACTCTTTCCCTACACGACGCTCTTCCGATCTNNNGGCCATTACAGGAAGTGCTTTT
GAAGAGCACACGTCTGAACTCTTTCCCTACACGACGCTCTTCCGATCTNNNTGCTGCATAGAGTGGCT
GAGCACACGTCTGAACTCTTTCCCTACACGACGCTCTTCCGATCTNNNTGACATACCCCCAACTCTCTCT
GAGCACACGTCTGAACTCTTTCCCTACACGACGCTCTTCCGATCTNNNAGGCAAATTCAGAAATACAGT
GAGCACACGTCTGAACTCTTTCCCTACACGACGCTCTTCCGATCTNNNTGACATACCCCCAACTCTCTCT
GAAGAGCACACGTCTGAACTCTTTCCCTACACGACGCTCTTCCGATCTNNNTGTCGAGCTTTGGATGCT
AGCACACGTCTGAACTCTTTCCCTACACGACGCTCTTCCGATCTNNNAACAAAGGCCATGTGTCCTCAC

GAAGAGCACACGTCTGAACTCTTTCCCTACACGACGCTCTTCCGATCTNNNAGCCTCCTCTGGTAACTG
GAGCACACGTCTGAACTCTTTCCCTACACGACGCTCTTCCGATCTNNNCACAAAGACTGCACCATGTTT
GCACACGTCTGAACTCTTTCCCTACACGACGCTCTTCCGATCTNNNAACCAGGCTGCTTCTTTGAGAC
GAGCACACGTCTGAACTCTTTCCCTACACGACGCTCTTCCGATCTNNNCCACCATCTTTCCCCAGTAAG
AGCACACGTCTGAACTCTTTCCCTACACGACGCTCTTCCGATCTNNNCCAGGTGGGCCAAATAAACAG
GCACACGTCTGAACTCTTTCCCTACACGACGCTCTTCCGATCTNNNACAGGATTTACCGGAAGAAGCA
AAGAGCACACGTCTGAACTCTTTCCCTACACGACGCTCTTCCGATCTNNNCCTCGAGAGAAACAGACC
AGCACACGTCTGAACTCTTTCCCTACACGACGCTCTTCCGATCTNNNTCCAGAGGGGAAGAGGATATT
AGCACACGTCTGAACTCTTTCCCTACACGACGCTCTTCCGATCTNNNTGCTTTTACATGTCCCAAGCTC
GCACACGTCTGAACTCTTTCCCTACACGACGCTCTTCCGATCTNNNGGTGTTTAGGTGAGACAACCTC
GCACACGTCTGAACTCTTTCCCTACACGACGCTCTTCCGATCTNNNCCATCTCTCCCAACCTCCATACACA
GAGCACACGTCTGAACTCTTTCCCTACACGACGCTCTTCCGATCTNNNTCCTCAATGACAAGGCAATTT
GCACACGTCTGAACTCTTTCCCTACACGACGCTCTTCCGATCTNNNGTGGCTTCATGCTGTTATCATG
AGCACACGTCTGAACTCTTTCCCTACACGACGCTCTTCCGATCTNNNCCAGAAACTACTGGCAGGAAC
GCACACGTCTGAACTCTTTCCCTACACGACGCTCTTCCGATCTNNNAGCTGGGACTACATGTGAATGT
AGCACACGTCTGAACTCTTTCCCTACACGACGCTCTTCCGATCTNNNTGAGGAAGTGTAGCTTCAAG
AGCACACGTCTGAACTCTTTCCCTACACGACGCTCTTCCGATCTNNNTGCTTTGTGTCAGATATGGCT
GAGCACACGTCTGAACTCTTTCCCTACACGACGCTCTTCCGATCTNNNAGTAGAAAGATGAAGTGGGT
GCACACGTCTGAACTCTTTCCCTACACGACGCTCTTCCGATCTNNNAGTTGTCATGCTGGTGGATGTT
GAGCACACGTCTGAACTCTTTCCCTACACGACGCTCTTCCGATCTNNNTGTTTGCTAGGTAGTTGTCT
GAGCACACGTCTGAACTCTTTCCCTACACGACGCTCTTCCGATCTNNNGGACAAGTGTCAGCAGGATT
AGCACACGTCTGAACTCTTTCCCTACACGACGCTCTTCCGATCTNNNTTTCAGTTTCTGCTCTGGAGCT
AGCACACGTCTGAACTCTTTCCCTACACGACGCTCTTCCGATCTNNNCTCTTCGAGGCCCATCTACAATG
AGCACACGTCTGAACTCTTTCCCTACACGACGCTCTTCCGATCTNNNAGATTTCACCATCTTGAGCAGG
GCACACGTCTGAACTCTTTCCCTACACGACGCTCTTCCGATCTNNNCCACTTGTTCAACTCCCGAAGAA
AGCACACGTCTGAACTCTTTCCCTACACGACGCTCTTCCGATCTNNNATGTGCCAGGAACTACCTTAT
AGAGCACACGTCTGAACTCTTTCCCTACACGACGCTCTTCCGATCTNNNGAGGCTGGAAAACTGGCAA
AGCACACGTCTGAACTCTTTCCCTACACGACGCTCTTCCGATCTNNNAATCACACACAAAGGGAAAGC
GCACACGTCTGAACTCTTTCCCTACACGACGCTCTTCCGATCTNNNAGAACCCAGAAGCCTCTTCTCTT
GAGCACACGTCTGAACTCTTTCCCTACACGACGCTCTTCCGATCTNNNGGGCCTTGGATGACATTATG
AGAGCACACGTCTGAACTCTTTCCCTACACGACGCTCTTCCGATCTNNNTAACCACCTGGTCTTGCTCT
AGCACACGTCTGAACTCTTTCCCTACACGACGCTCTTCCGATCTNNNATGAACATTTCGATCTGAGGG
GCACACGTCTGAACTCTTTCCCTACACGACGCTCTTCCGATCTNNNGGTTCAGATTCTCGTAGCTCTG
GAGCACACGTCTGAACTCTTTCCCTACACGACGCTCTTCCGATCTNNNTCGTTCAGGGTGTGTTTTGT
AGCACACGTCTGAACTCTTTCCCTACACGACGCTCTTCCGATCTNNNCATCTGCCAGGAATCCCTTTCT
AGCACACGTCTGAACTCTTTCCCTACACGACGCTCTTCCGATCTNNNCTCCCCAACAGTGGTTGAAAAG
GCACACGTCTGAACTCTTTCCCTACACGACGCTCTTCCGATCTNNNTTTCTTCGTCTTGACGGTACAGG
GCACACGTCTGAACTCTTTCCCTACACGACGCTCTTCCGATCTNNNAAACCTAGAGTTGGATGTGCGT
AGCACACGTCTGAACTCTTTCCCTACACGACGCTCTTCCGATCTNNNGGTCTCCCTCTCTCTTCACACA
AGCACACGTCTGAACTCTTTCCCTACACGACGCTCTTCCGATCTNNNCAGAGTATCCACTCCCAACAA
GAGCACACGTCTGAACTCTTTCCCTACACGACGCTCTTCCGATCTNNNCACACAATCTGTCCTCACCTC
AGCACACGTCTGAACTCTTTCCCTACACGACGCTCTTCCGATCTNNNCTGTAGCTTCAGACCCATTGA
GCACACGTCTGAACTCTTTCCCTACACGACGCTCTTCCGATCTNNNAATAGCTCAGAATGTGCCAGAC

GAGCACACGTCTGAACTCTTTCCCTACACGACGCTCTTCCGATCTNNNTGATGTAACAGTGAATAAAGA
AGCACACGTCTGAACTCTTTCCCTACACGACGCTCTTCCGATCTNNNAATCCTTTCAGGTGGCCCTTTTT
GCACACGTCTGAACTCTTTCCCTACACGACGCTCTTCCGATCTNNNGGCTAGGCACAGATGTGAGAAA
GCACACGTCTGAACTCTTTCCCTACACGACGCTCTTCCGATCTNNNTTCCCAGGAAGGTAGGAAAGAGC
GAGCACACGTCTGAACTCTTTCCCTACACGACGCTCTTCCGATCTNNNGGAGGAGGAGGAGTATGAG
AGCACACGTCTGAACTCTTTCCCTACACGACGCTCTTCCGATCTNNNTCAGTTGAGGCATCTTCTGCTTC
GAGCACACGTCTGAACTCTTTCCCTACACGACGCTCTTCCGATCTNNNTTGTAAAGTGGCAATGCAAG
AGAGCACACGTCTGAACTCTTTCCCTACACGACGCTCTTCCGATCTNNNGGTTGATTTTGGTGTTACCA
AAGAGCACACGTCTGAACTCTTTCCCTACACGACGCTCTTCCGATCTNNNTGCTTTGTGTCAGATATGG
AGCACACGTCTGAACTCTTTCCCTACACGACGCTCTTCCGATCTNNNGCCGAGGCATCTGTCTCAAAAT
AGCACACGTCTGAACTCTTTCCCTACACGACGCTCTTCCGATCTNNNCCATGTGACCAGTTACAGGTAA
AGAGCACACGTCTGAACTCTTTCCCTACACGACGCTCTTCCGATCTNNNCTTCCTTTCCCCTCCCGCTTAC
AGCACACGTCTGAACTCTTTCCCTACACGACGCTCTTCCGATCTNNNTGAGGAAGAGGTAGCTGGAAA
AGAGCACACGTCTGAACTCTTTCCCTACACGACGCTCTTCCGATCTNNNAGAGAAGTTGCTTTGGCCAT
AAGAGCACACGTCTGAACTCTTTCCCTACACGACGCTCTTCCGATCTNNNCCATGTGCTTCAGGGAAGA
AGAGCACACGTCTGAACTCTTTCCCTACACGACGCTCTTCCGATCTNNNGAAGGAAGATTCCAGGCAGA
GCACACGTCTGAACTCTTTCCCTACACGACGCTCTTCCGATCTNNNGCCGAGGCATCTGTCTCAAAATAC
AAGAGCACACGTCTGAACTCTTTCCCTACACGACGCTCTTCCGATCTNNNCTTCCTTTCCCCTCCCGCTTA
GAGCACACGTCTGAACTCTTTCCCTACACGACGCTCTTCCGATCTNNNCAGGAAACACTCCTTTGGGCTA
GAGCACACGTCTGAACTCTTTCCCTACACGACGCTCTTCCGATCTNNNTGATCAGAGTACATCCCAGAT
GAGCACACGTCTGAACTCTTTCCCTACACGACGCTCTTCCGATCTNNNAGTGAATAATGCATTGCTTCTC
GAGCACACGTCTGAACTCTTTCCCTACACGACGCTCTTCCGATCTNNNCTGCACTGAACCTGCATAGAT
GAGCACACGTCTGAACTCTTTCCCTACACGACGCTCTTCCGATCTNNNAGAGAGCCCCCAAGTTGATAC
GAGCACACGTCTGAACTCTTTCCCTACACGACGCTCTTCCGATCTNNNGGTTGATTTTGGTGTTACCAG
AGCACACGTCTGAACTCTTTCCCTACACGACGCTCTTCCGATCTNNNTCCTTGCCTCTGACTTCCCATTAC
GAGCACACGTCTGAACTCTTTCCCTACACGACGCTCTTCCGATCTNNNGTGTCCCCTGGAGAAGATCAA
AAGAGCACACGTCTGAACTCTTTCCCTACACGACGCTCTTCCGATCTNNNTTCTGTCCCGTTTTTGCCATC
AGCACACGTCTGAACTCTTTCCCTACACGACGCTCTTCCGATCTNNNGCAAAGCTCCTCTCTACTCTTCC
AGCACACGTCTGAACTCTTTCCCTACACGACGCTCTTCCGATCTNNNAGTTCAGAAAAATGCACGCACTC
GCACACGTCTGAACTCTTTCCCTACACGACGCTCTTCCGATCTNNNAGTTGGCAGAAGATAACCACTGT
GAGCACACGTCTGAACTCTTTCCCTACACGACGCTCTTCCGATCTNNNCACCAACACACAATTCCCCTGTA
GCACACGTCTGAACTCTTTCCCTACACGACGCTCTTCCGATCTNNNGTACCTCAGAGGACACATCTCCAC
GAGCACACGTCTGAACTCTTTCCCTACACGACGCTCTTCCGATCTNNNCTTTGCTCCCTCTCTCCAAGAC
GAGCACACGTCTGAACTCTTTCCCTACACGACGCTCTTCCGATCTNNNTCTGAGCACCAACTCATTCAAG
AGAGCACACGTCTGAACTCTTTCCCTACACGACGCTCTTCCGATCTNNNCAATCCTGAGATTCCCAACCCT
AGCACACGTCTGAACTCTTTCCCTACACGACGCTCTTCCGATCTNNNACAGCAATTTCTGCTCTCTCAGG
AGCACACGTCTGAACTCTTTCCCTACACGACGCTCTTCCGATCTNNNACAGGCTTGAAAGTCTCTGTCTC
AGCACACGTCTGAACTCTTTCCCTACACGACGCTCTTCCGATCTNNNTTCAGGTGAAAGTGGCTAAGTC
GCACACGTCTGAACTCTTTCCCTACACGACGCTCTTCCGATCTNNNGGTCTCCTTCTTGCTGACTGTAGC
AAGAGCACACGTCTGAACTCTTTCCCTACACGACGCTCTTCCGATCTNNNCCAAACAGATAGAAGGGC
GCACACGTCTGAACTCTTTCCCTACACGACGCTCTTCCGATCTNNNCTCATCCATCTTGTTCCCTGCTCCA
GAGCACACGTCTGAACTCTTTCCCTACACGACGCTCTTCCGATCTNNNTCCTAGGAAGCCACCTTCTCTG
AGCACACGTCTGAACTCTTTCCCTACACGACGCTCTTCCGATCTNNNCTCCTCCTCGTTCACAAAGTCAG

AGCACACGTCTGAACTCTTTCCCTACACGACGCTCTTCCGATCTNNNCAGCTCACACTTCAGTCACTCT
AGCACACGTCTGAACTCTTTCCCTACACGACGCTCTTCCGATCTNNNCACATGCCAATATGCTAATATA
AGCACACGTCTGAACTCTTTCCCTACACGACGCTCTTCCGATCTNNNAGAACCCAGAAGCCTCTTCTCT
GCACACGTCTGAACTCTTTCCCTACACGACGCTCTTCCGATCTNNNGTTTCTGAGTTCATCCTGGCCTT
CACACGTCTGAACTCTTTCCCTACACGACGCTCTTCCGATCTNNNCGCACATATTGCAACAGAACCCC
AAGAGCACACGTCTGAACTCTTTCCCTACACGACGCTCTTCCGATCTNNNTGATCCCCAATTCATGTCT
GAGCACACGTCTGAACTCTTTCCCTACACGACGCTCTTCCGATCTNNNCCCAAAAGTATGACCATGTG
AGCACACGTCTGAACTCTTTCCCTACACGACGCTCTTCCGATCTNNNAAGGGACAAGGACTGGGAA
GAGCACACGTCTGAACTCTTTCCCTACACGACGCTCTTCCGATCTNNNACTCCACAACATAGGATGGG
GCACACGTCTGAACTCTTTCCCTACACGACGCTCTTCCGATCTNNNCTGGATTTAGCCTCAGACCAAC
CACACGTCTGAACTCTTTCCCTACACGACGCTCTTCCGATCTNNNCTGGGAAAGAGAAATGGTAGGG
GAGCACACGTCTGAACTCTTTCCCTACACGACGCTCTTCCGATCTNNNCCATGTGACCAGTTACAGGT
AGCACACGTCTGAACTCTTTCCCTACACGACGCTCTTCCGATCTNNNCCTGGCCCCCTAATTTCACATA
AAGAGCACACGTCTGAACTCTTTCCCTACACGACGCTCTTCCGATCTNNNTGTTTCTTTGCTGAGCCCA
AGCACACGTCTGAACTCTTTCCCTACACGACGCTCTTCCGATCTNNNCCTGCCCACTCATAAATCCATC
AGCACACGTCTGAACTCTTTCCCTACACGACGCTCTTCCGATCTNNNAGTGCAGGGTGATTTGCAGTT
AGCACACGTCTGAACTCTTTCCCTACACGACGCTCTTCCGATCTNNNTTATTGATGCCTAGCGTGTCTC
GCACACGTCTGAACTCTTTCCCTACACGACGCTCTTCCGATCTNNNCTGAGACACAGGCATATGCAAC
GCACACGTCTGAACTCTTTCCCTACACGACGCTCTTCCGATCTNNNTGCAAGCCCTTTATTCTGGCATT
GCACACGTCTGAACTCTTTCCCTACACGACGCTCTTCCGATCTNNNCTGGGAAAGAGAAATGGTAGG
AGCACACGTCTGAACTCTTTCCCTACACGACGCTCTTCCGATCTNNNTGCAACCTCAAGCCCTGTATAA
GCACACGTCTGAACTCTTTCCCTACACGACGCTCTTCCGATCTNNNGCCACCAATATATACACCTGGT
AGCACACGTCTGAACTCTTTCCCTACACGACGCTCTTCCGATCTNNNGTGACAACAAACCACACACAC
GCACACGTCTGAACTCTTTCCCTACACGACGCTCTTCCGATCTNNNTCAGCCTCTGAAAGCCCTGATT
AGCACACGTCTGAACTCTTTCCCTACACGACGCTCTTCCGATCTNNNGTCAGATCCACAAGACCCCTTT
AGCACACGTCTGAACTCTTTCCCTACACGACGCTCTTCCGATCTNNNTCTTAGATCACTTGCTTGGGG
GAGCACACGTCTGAACTCTTTCCCTACACGACGCTCTTCCGATCTNNNCTCAGATGCTTCCCTGTTCTT
AGAGCACACGTCTGAACTCTTTCCCTACACGACGCTCTTCCGATCTNNNTGCAGAAATTAGAGCAAC
GCACACGTCTGAACTCTTTCCCTACACGACGCTCTTCCGATCTNNNCGAAAAGCACAAGATTCAACCC
AGCACACGTCTGAACTCTTTCCCTACACGACGCTCTTCCGATCTNNNTTATCAGAAGGAGGGATGGG
AGCACACGTCTGAACTCTTTCCCTACACGACGCTCTTCCGATCTNNNCCTGAAGTGTGTGAAGAAACC
GAGCACACGTCTGAACTCTTTCCCTACACGACGCTCTTCCGATCTNNNGTCAACACTGCTGAAGTTG
AGCACACGTCTGAACTCTTTCCCTACACGACGCTCTTCCGATCTNNNACCTACCATAAGCTGTACTCAG
AGCACACGTCTGAACTCTTTCCCTACACGACGCTCTTCCGATCTNNNCAGCTTCACTCTGTCTCTGTCT
AGCACACGTCTGAACTCTTTCCCTACACGACGCTCTTCCGATCTNNNGGTATCCATGAGTGTGTACCT
AGCACACGTCTGAACTCTTTCCCTACACGACGCTCTTCCGATCTNNNGGTGTGATTGTGCTACTGTAC
AGCACACGTCTGAACTCTTTCCCTACACGACGCTCTTCCGATCTNNNGATTGTGGGGACCTATGTGT
AGCACACGTCTGAACTCTTTCCCTACACGACGCTCTTCCGATCTNNNCACCTCCCAGAAAATGTTTGTT
CACACGTCTGAACTCTTTCCCTACACGACGCTCTTCCGATCTNNNCCCAAATGTCATTATGCAGCACCC
GAGCACACGTCTGAACTCTTTCCCTACACGACGCTCTTCCGATCTNNNCTCCCCAACAGTGGTTGAAA
GCACACGTCTGAACTCTTTCCCTACACGACGCTCTTCCGATCTNNNCTGGAACCAGATCAATACCAGG
GAGCACACGTCTGAACTCTTTCCCTACACGACGCTCTTCCGATCTNNNGAGTTCAAGCACGTCAAAAC
AAGAGCACACGTCTGAACTCTTTCCCTACACGACGCTCTTCCGATCTNNNGTTCTTTCAGAGAATGC

AGAGCACACGTCTGAACTCTTTCCCTACACGACGCTCTTCCGATCTNNNGCCTGTATCCCTTTCCATAAG
CACACGTCTGAACTCTTTCCCTACACGACGCTCTTCCGATCTNNNGACGAGGAGTATGAGCCCCACCAT
AGCACACGTCTGAACTCTTTCCCTACACGACGCTCTTCCGATCTNNNCAACCTTATTTGTCCTGACAGCC
AGCACACGTCTGAACTCTTTCCCTACACGACGCTCTTCCGATCTNNNGAGGGGGTACTCATTCAGTCCAT
AGCACACGTCTGAACTCTTTCCCTACACGACGCTCTTCCGATCTNNNATTTTTAGGAAGAGCTGGCTGC
AGCACACGTCTGAACTCTTTCCCTACACGACGCTCTTCCGATCTNNNGAACATGTACCTGCACTGTCCT
CACACGTCTGAACTCTTTCCCTACACGACGCTCTTCCGATCTNNNCTACCACTGCTCAGAGGAACAGACA
GAGCACACGTCTGAACTCTTTCCCTACACGACGCTCTTCCGATCTNNNTCATGAGGGATGGTTACCCAT
GAGCACACGTCTGAACTCTTTCCCTACACGACGCTCTTCCGATCTNNNTGTCTTGGTTCCTGAATGGTAC
AGCACACGTCTGAACTCTTTCCCTACACGACGCTCTTCCGATCTNNNAGAGATGGCACATTCTGCAAAC
AGCACACGTCTGAACTCTTTCCCTACACGACGCTCTTCCGATCTNNNTGGGAGCAAAACACCATTACAC
AGCACACGTCTGAACTCTTTCCCTACACGACGCTCTTCCGATCTNNNTGGCATTGTCCTCAAAACATCCAC
GAGCACACGTCTGAACTCTTTCCCTACACGACGCTCTTCCGATCTNNNGACAAAGTCCACTGGAAGGA
AGCACACGTCTGAACTCTTTCCCTACACGACGCTCTTCCGATCTNNNGACTAAAACTCAGCCAGCAGGA
AGAGCACACGTCTGAACTCTTTCCCTACACGACGCTCTTCCGATCTNNNGAGTGAGGTTTGCTGCTTTC
AGCACACGTCTGAACTCTTTCCCTACACGACGCTCTTCCGATCTNNNACGGCCGTGTAGATGATGAAG
GAGCACACGTCTGAACTCTTTCCCTACACGACGCTCTTCCGATCTNNNATTATGTTTGCAATGTGTGGG
AAGAGCACACGTCTGAACTCTTTCCCTACACGACGCTCTTCCGATCTNNNGCCATATTAGTCGGTGTTCT
AAGAGCACACGTCTGAACTCTTTCCCTACACGACGCTCTTCCGATCTNNNCCACTTGTTCAACTCCCGAA
AGCACACGTCTGAACTCTTTCCCTACACGACGCTCTTCCGATCTNNNTGGAATGTCTTCTGACCCCAGA
GAGCACACGTCTGAACTCTTTCCCTACACGACGCTCTTCCGATCTNNNAGCCATGATTGCTTCCTCATTG
GAGCACACGTCTGAACTCTTTCCCTACACGACGCTCTTCCGATCTNNNACCTAACAGTTCCCTTTCTTCA
GAGCACACGTCTGAACTCTTTCCCTACACGACGCTCTTCCGATCTNNNTCAGGACATTTCCTTCCTGTCC
AGAGCACACGTCTGAACTCTTTCCCTACACGACGCTCTTCCGATCTNNNTGCTGAACTAGTTGCTTGCA
GCACACGTCTGAACTCTTTCCCTACACGACGCTCTTCCGATCTNNNGAGTACTAGGATTACAGGCAAG
GAGCACACGTCTGAACTCTTTCCCTACACGACGCTCTTCCGATCTNNNAGCTCAGCTCACTCCAATCTCA
CACACGTCTGAACTCTTTCCCTACACGACGCTCTTCCGATCTNNNGCGTGTGGGGTGTGTGTGTAGAC
AAGAGCACACGTCTGAACTCTTTCCCTACACGACGCTCTTCCGATCTNNNGTCCATGAGTGTGCATGGC
AGAGCACACGTCTGAACTCTTTCCCTACACGACGCTCTTCCGATCTNNNGGAATGGGTTTGAGGTGG
GCACACGTCTGAACTCTTTCCCTACACGACGCTCTTCCGATCTNNNCAAAGCAGCTCTGTTTCCCTACTCA
GCACACGTCTGAACTCTTTCCCTACACGACGCTCTTCCGATCTNNNCACTATGCAGTTATCTGAATGGC
AAGAGCACACGTCTGAACTCTTTCCCTACACGACGCTCTTCCGATCTNNNGATCAAGCCAGGATTTGTT
AGAGCACACGTCTGAACTCTTTCCCTACACGACGCTCTTCCGATCTNNNCCTGAGGTGGGAAGGAATTT
AGAGCACACGTCTGAACTCTTTCCCTACACGACGCTCTTCCGATCTNNNTGTCTCCTGTCACTGAGGCTG
AGAGCACACGTCTGAACTCTTTCCCTACACGACGCTCTTCCGATCTNNNGCATAAGTAGGTCACAGGGA
AGCACACGTCTGAACTCTTTCCCTACACGACGCTCTTCCGATCTNNNGTTGCCGATATACCCTATGACC
AGCACACGTCTGAACTCTTTCCCTACACGACGCTCTTCCGATCTNNNTTAGTAATACCACCACACCACCA
AGCACACGTCTGAACTCTTTCCCTACACGACGCTCTTCCGATCTNNNAGGAAAGCTTTTAGCCCTCATG
GCACACGTCTGAACTCTTTCCCTACACGACGCTCTTCCGATCTNNNATATGTGCTCTCCCCCTCTCCTACA
GCACACGTCTGAACTCTTTCCCTACACGACGCTCTTCCGATCTNNNCAAACTCACACAACCAGGAAGGA
GAGCACACGTCTGAACTCTTTCCCTACACGACGCTCTTCCGATCTNNNTTTACTATGCTGGGCGACAAG
AGCACACGTCTGAACTCTTTCCCTACACGACGCTCTTCCGATCTNNNGTTTGAGCTCTGGATGCTTCTG
GAGCACACGTCTGAACTCTTTCCCTACACGACGCTCTTCCGATCTNNNATCTCTTTCTCTCTCGCTCTGGC

GAGCACACGTCTGAACTCTTTCCCTACACGACGCTCTTCCGATCTNNNAGGAACAAGTTTTCTGGGTC
AAGAGCACACGTCTGAACTCTTTCCCTACACGACGCTCTTCCGATCTNNNTCATCAATCAGGTGTGTGT
GCACACGTCTGAACTCTTTCCCTACACGACGCTCTTCCGATCTNNNGAACATGTCTTCCCTTGTAGGGC
GAGCACACGTCTGAACTCTTTCCCTACACGACGCTCTTCCGATCTNNNAAATGGCATGGTGTTTGTGT
GAGCACACGTCTGAACTCTTTCCCTACACGACGCTCTTCCGATCTNNNTACACCTCAACCCAGAACATAC
GAGCACACGTCTGAACTCTTTCCCTACACGACGCTCTTCCGATCTNNNAGTGGTGGAATAGATGCCTT
AAGAGCACACGTCTGAACTCTTTCCCTACACGACGCTCTTCCGATCTNNNCTCCACCATCAGATTTGGG
GCACACGTCTGAACTCTTTCCCTACACGACGCTCTTCCGATCTNNNAGCAGTCATGGACCACAATTCT
GAGCACACGTCTGAACTCTTTCCCTACACGACGCTCTTCCGATCTNNNAGAGGAAACGTGTACACACA
GAGCACACGTCTGAACTCTTTCCCTACACGACGCTCTTCCGATCTNNNAAAAGGGAAATAGGGGGA
AGCACACGTCTGAACTCTTTCCCTACACGACGCTCTTCCGATCTNNNCACATACACACTCTCGCTCTCAA
GAGCACACGTCTGAACTCTTTCCCTACACGACGCTCTTCCGATCTNNNGTTCATGCGATTCTCCTGGTTC
GCACACGTCTGAACTCTTTCCCTACACGACGCTCTTCCGATCTNNNCTTAGCTGGCTACCAGGCTCTCA
GAGCACACGTCTGAACTCTTTCCCTACACGACGCTCTTCCGATCTNNNCGGCCGTTCTCATGTATCTTG
AGCACACGTCTGAACTCTTTCCCTACACGACGCTCTTCCGATCTNNNATAGACCTCCTTTCCTCTCCTGC
GAGCACACGTCTGAACTCTTTCCCTACACGACGCTCTTCCGATCTNNNAGCTCAAGAAGTGCAGAAGT
AGAGCACACGTCTGAACTCTTTCCCTACACGACGCTCTTCCGATCTNNNACAAATATCCTTGCACATCTC
AGAGCACACGTCTGAACTCTTTCCCTACACGACGCTCTTCCGATCTNNNGGTTGCTACTGCTGCTTCAG
GCACACGTCTGAACTCTTTCCCTACACGACGCTCTTCCGATCTNNNCTGCCTTTTTCCTCCCTCTCTTCCA
AGAGCACACGTCTGAACTCTTTCCCTACACGACGCTCTTCCGATCTNNNGACATTCTCTCTGAGGCCAG
GAGCACACGTCTGAACTCTTTCCCTACACGACGCTCTTCCGATCTNNNTTGTTCCCAGGCCCTCTTAGA
AAGAGCACACGTCTGAACTCTTTCCCTACACGACGCTCTTCCGATCTNNNTCTGCCTTGTCTTTTTCAAG
CACACGTCTGAACTCTTTCCCTACACGACGCTCTTCCGATCTNNNGTTCATGCGATTCTCCTGGTTCTC
GAGCACACGTCTGAACTCTTTCCCTACACGACGCTCTTCCGATCTNNNTTTTCTAACTGGCATGTGGAG
GAGCACACGTCTGAACTCTTTCCCTACACGACGCTCTTCCGATCTNNNGCAGAGTGAGGCTTTGTGTC
AGCACACGTCTGAACTCTTTCCCTACACGACGCTCTTCCGATCTNNNCCAGAGAAATTGGCCTCTTTGG
AGAGCACACGTCTGAACTCTTTCCCTACACGACGCTCTTCCGATCTNNNGCACATGGGAGAAACAAA
AGCACACGTCTGAACTCTTTCCCTACACGACGCTCTTCCGATCTNNNTCTTCTCTCTCTCTCCCTGTCCTC
GAGCACACGTCTGAACTCTTTCCCTACACGACGCTCTTCCGATCTNNNTCTCCTTAGTCCACTGATGCC
GAGCACACGTCTGAACTCTTTCCCTACACGACGCTCTTCCGATCTNNNTCTGCTTCCTTCCCACTCTTCTT
AGCACACGTCTGAACTCTTTCCCTACACGACGCTCTTCCGATCTNNNAAAAGAGGAGGTGATAGCTCA
GAGCACACGTCTGAACTCTTTCCCTACACGACGCTCTTCCGATCTNNNCTGGAATTCCCAGCTATGCAT
GAGCACACGTCTGAACTCTTTCCCTACACGACGCTCTTCCGATCTNNNACTCCTACATGTTTGTGTGTG
GCACACGTCTGAACTCTTTCCCTACACGACGCTCTTCCGATCTNNNAGCATGCTAACTTCTGACTTGGT
GCACACGTCTGAACTCTTTCCCTACACGACGCTCTTCCGATCTNNNTTTCCACCCAAAAGGAATCCTCC
GAGCACACGTCTGAACTCTTTCCCTACACGACGCTCTTCCGATCTNNNGTTGCTTATTGGAAGGACATG
AGAGCACACGTCTGAACTCTTTCCCTACACGACGCTCTTCCGATCTNNNACGGCAATGAAACCAGGTA
GAGCACACGTCTGAACTCTTTCCCTACACGACGCTCTTCCGATCTNNNGTCCTCTGCTTCTTCGTGTTCA
AGCACACGTCTGAACTCTTTCCCTACACGACGCTCTTCCGATCTNNNACCCTCACAACTTCCTCACACTC
GAGCACACGTCTGAACTCTTTCCCTACACGACGCTCTTCCGATCTNNNAGTAGGTGACCCAGTGTTTT
GCACACGTCTGAACTCTTTCCCTACACGACGCTCTTCCGATCTNNNTGTTGTGGTAAAACCTGTGTCT
GCACACGTCTGAACTCTTTCCCTACACGACGCTCTTCCGATCTNNNTTTATTCTGGGCTCCTAGCACCT
GAGCACACGTCTGAACTCTTTCCCTACACGACGCTCTTCCGATCTNNNGTATGTGGACAGCATGAGA

AGAGCACACGTCTGAACTCTTTCCCTACACGACGCTCTTCCGATCTNNNACAGCAGGAAGCCATTACT
AGCACACGTCTGAACTCTTTCCCTACACGACGCTCTTCCGATCTNNNGTGTGTGTGTTCACAAGCA
AGCACACGTCTGAACTCTTTCCCTACACGACGCTCTTCCGATCTNNNTAAAAGAATCACCCACAGCAC
GAGCACACGTCTGAACTCTTTCCCTACACGACGCTCTTCCGATCTNNNTGACAGAGTAAGGTCCCTTC
AGCACACGTCTGAACTCTTTCCCTACACGACGCTCTTCCGATCTNNNTCTCAGAGAGCTTCTTCGGTCA
AGCACACGTCTGAACTCTTTCCCTACACGACGCTCTTCCGATCTNNNGTAAGGTTACTGTGTTAGGCC
AGCACACGTCTGAACTCTTTCCCTACACGACGCTCTTCCGATCTNNNTTTCAGTGTCTCTCCCTCTCT
GAGCACACGTCTGAACTCTTTCCCTACACGACGCTCTTCCGATCTNNNCACTCGTGTGTCCTTCATCAT
GCACACGTCTGAACTCTTTCCCTACACGACGCTCTTCCGATCTNNNGCTTCCACCAAACACCACATTT
AAGAGCACACGTCTGAACTCTTTCCCTACACGACGCTCTTCCGATCTNNNTCTCAGAGAGCTTCTTCGG
CACACGTCTGAACTCTTTCCCTACACGACGCTCTTCCGATCTNNNTGCTAAAGCGAATCCTGGGCTCA
GCACACGTCTGAACTCTTTCCCTACACGACGCTCTTCCGATCTNNNAGTTGCATCATTTAGGGTACCA
AGCACACGTCTGAACTCTTTCCCTACACGACGCTCTTCCGATCTNNNTAAGGTAAGGGTTGTGTGTG
AGAGCACACGTCTGAACTCTTTCCCTACACGACGCTCTTCCGATCTNNNACACACACACACAGATCTAT
GAGCACACGTCTGAACTCTTTCCCTACACGACGCTCTTCCGATCTNNNTGCTCTGACAGTGTCTAACCC
GAGCACACGTCTGAACTCTTTCCCTACACGACGCTCTTCCGATCTNNNGCATGTGAAACTGCAAAAGC
AGCACACGTCTGAACTCTTTCCCTACACGACGCTCTTCCGATCTNNNTAGTCATAGGGAATGTGTTGT
AGCACACGTCTGAACTCTTTCCCTACACGACGCTCTTCCGATCTNNNAGTTTCAAGTAGCCAACTGAG
GAGCACACGTCTGAACTCTTTCCCTACACGACGCTCTTCCGATCTNNNCATCTGGGAAACCAAGCTGT
GCACACGTCTGAACTCTTTCCCTACACGACGCTCTTCCGATCTNNNTGTGAAGTTCCACAGTATCTGA
AGCACACGTCTGAACTCTTTCCCTACACGACGCTCTTCCGATCTNNNTGCCTCTCACCCTTTACTAAGGT
CACACGTCTGAACTCTTTCCCTACACGACGCTCTTCCGATCTNNNATGCCCTGTACTGAGGAAAGAGA
GAGCACACGTCTGAACTCTTTCCCTACACGACGCTCTTCCGATCTNNNAATCAGTGAACAGACACACA
GAGCACACGTCTGAACTCTTTCCCTACACGACGCTCTTCCGATCTNNNGCTAACTGGCTGCAAGAAAC
GCACACGTCTGAACTCTTTCCCTACACGACGCTCTTCCGATCTNNNGAAACAGAATATGGGTGGGG
GCACACGTCTGAACTCTTTCCCTACACGACGCTCTTCCGATCTNNNGAAGAGGGGGAGAAAGGAGA
GCACACGTCTGAACTCTTTCCCTACACGACGCTCTTCCGATCTNNNCCTAGATTGCACCACACTACTGC
GAAGAGCACACGTCTGAACTCTTTCCCTACACGACGCTCTTCCGATCTNNNAGGTTACGTGAGCTAG
AGCACACGTCTGAACTCTTTCCCTACACGACGCTCTTCCGATCTNNNGACATTCTCTCTGAGGCCAGTT
GCACACGTCTGAACTCTTTCCCTACACGACGCTCTTCCGATCTNNNTGTTCATTATCTCTGGTGTGTG
AGCACACGTCTGAACTCTTTCCCTACACGACGCTCTTCCGATCTNNNTCCACATTTACATTCTCTCCCTCC
AGCACACGTCTGAACTCTTTCCCTACACGACGCTCTTCCGATCTNNNTCCATCTATGTTGCCGCATGAA
AGCACACGTCTGAACTCTTTCCCTACACGACGCTCTTCCGATCTNNNAAATTCTTTGCACTGTCCACCC
GAGCACACGTCTGAACTCTTTCCCTACACGACGCTCTTCCGATCTNNNGGATAACTCATGCTGCAGGG
GCACACGTCTGAACTCTTTCCCTACACGACGCTCTTCCGATCTNNNAGAGCCTGCAATGTTGATGTTG
GAGCACACGTCTGAACTCTTTCCCTACACGACGCTCTTCCGATCTNNNTGGGCTCTTTTAGTGTGTGT
GCACACGTCTGAACTCTTTCCCTACACGACGCTCTTCCGATCTNNNCACACAATGAGCAGCATGGAAA
AGCACACGTCTGAACTCTTTCCCTACACGACGCTCTTCCGATCTNNNAATAGGAAATGTCAAAGCGG
AGCACACGTCTGAACTCTTTCCCTACACGACGCTCTTCCGATCTNNNTTGCAGAGCATGGGTAAGAT
GAGCACACGTCTGAACTCTTTCCCTACACGACGCTCTTCCGATCTNNNAAAAAGTCCATTGTTGAGTA
GCACACGTCTGAACTCTTTCCCTACACGACGCTCTTCCGATCTNNNCTTGGGTTCAGACTGTCTCCTTG
AGCACACGTCTGAACTCTTTCCCTACACGACGCTCTTCCGATCTNNNCTGACAAAGAGCTCATACCCA
CACACGTCTGAACTCTTTCCCTACACGACGCTCTTCCGATCTNNNAGGCAGGCCTGTAAATTATCTGC

AGCACACGTCTGAACTCTTTCCCTACACGACGCTCTTCCGATCTNNNCGTCCAGCTGAAACGTGTGTA
AGCACACGTCTGAACTCTTTCCCTACACGACGCTCTTCCGATCTNNNTATAGACGCTTGAGGCCAGGA
GAGCACACGTCTGAACTCTTTCCCTACACGACGCTCTTCCGATCTNNNCTACACACCCAATCACATGGG
GCACACGTCTGAACTCTTTCCCTACACGACGCTCTTCCGATCTNNNTGCTTCCTTTACGTGGCTTTCTTC
AGCACACGTCTGAACTCTTTCCCTACACGACGCTCTTCCGATCTNNNAATACATGCACAGCACACACA
GCACACGTCTGAACTCTTTCCCTACACGACGCTCTTCCGATCTNNNAAAAGCCTGTAAACCACCAACC
AGCACACGTCTGAACTCTTTCCCTACACGACGCTCTTCCGATCTNNNCTACATGAAGGAAACAAACG
GAGCACACGTCTGAACTCTTTCCCTACACGACGCTCTTCCGATCTNNNGGGCTCTCCCATTCTGTTTCA
GCACACGTCTGAACTCTTTCCCTACACGACGCTCTTCCGATCTNNNAGCAACTGTATCAATCAGCTGT
CACACGTCTGAACTCTTTCCCTACACGACGCTCTTCCGATCTNNNGACAATGTATTCCAGAGGCAGG
GCACACGTCTGAACTCTTTCCCTACACGACGCTCTTCCGATCTNNNTGTGAAGGAAAAACCTGGTGA
CACACGTCTGAACTCTTTCCCTACACGACGCTCTTCCGATCTNNNCCCAGCAGTCATGAAAGAAGGA
AGCACACGTCTGAACTCTTTCCCTACACGACGCTCTTCCGATCTNNNATGGGGACTTCTCTCTCTCTCC
GAGCACACGTCTGAACTCTTTCCCTACACGACGCTCTTCCGATCTNNNGGGCTCTCCCATTCTGTTTCA
AGAGCACACGTCTGAACTCTTTCCCTACACGACGCTCTTCCGATCTNNNTGAACAGTCGACACATGTA
AGCACACGTCTGAACTCTTTCCCTACACGACGCTCTTCCGATCTNNNTCCACATACAGACAAACGTGCA
CACACGTCTGAACTCTTTCCCTACACGACGCTCTTCCGATCTNNNCTGGCCCTTTGGAGAAACTGAAA
AGCACACGTCTGAACTCTTTCCCTACACGACGCTCTTCCGATCTNNNTATACTCCTCAAAACACACAC
AGCACACGTCTGAACTCTTTCCCTACACGACGCTCTTCCGATCTNNNCAAGAGCAGAGTTGTAAACT
GCACACGTCTGAACTCTTTCCCTACACGACGCTCTTCCGATCTNNNTGCCTAGTTTGTTTGGCTTCCAT
AGCACACGTCTGAACTCTTTCCCTACACGACGCTCTTCCGATCTNNNTTAGGGAAGCAATGCAACTC
GAGCACACGTCTGAACTCTTTCCCTACACGACGCTCTTCCGATCTNNNATGGGCTAATCCTTATCTGG
CACACGTCTGAACTCTTTCCCTACACGACGCTCTTCCGATCTNNNGTAACGCATCATCTACCGTCCTCCA
AGCACACGTCTGAACTCTTTCCCTACACGACGCTCTTCCGATCTNNNACATTACAAGGTTTGAAAGG
AGCACACGTCTGAACTCTTTCCCTACACGACGCTCTTCCGATCTNNNTGTGCGTCTGGAAAGTGAAT
GCACACGTCTGAACTCTTTCCCTACACGACGCTCTTCCGATCTNNNATCTCTCCCTCACACACAGAAGT
AGCACACGTCTGAACTCTTTCCCTACACGACGCTCTTCCGATCTNNNATGGGTAAGTGTTGAAGAG
GAGCACACGTCTGAACTCTTTCCCTACACGACGCTCTTCCGATCTNNNATACCATCCCACCACTCACTCA
ACACGTCTGAACTCTTTCCCTACACGACGCTCTTCCGATCTNNNCGTGGTGGCGAAGAGAGTAAAAT
AGAGCACACGTCTGAACTCTTTCCCTACACGACGCTCTTCCGATCTNNNGCTTCCAGGAATCTCAAAT
GCACACGTCTGAACTCTTTCCCTACACGACGCTCTTCCGATCTNNNTTCACTGCAAATCAACTGGGTC
GAGCACACGTCTGAACTCTTTCCCTACACGACGCTCTTCCGATCTNNNGGGAAAGAATGAACAAAGC
GCACACGTCTGAACTCTTTCCCTACACGACGCTCTTCCGATCTNNNCTTGTGTATGTGTGTGTCCAGG
GCACACGTCTGAACTCTTTCCCTACACGACGCTCTTCCGATCTNNNGTGGGCAGATATTTGAAGGCTG
GCACACGTCTGAACTCTTTCCCTACACGACGCTCTTCCGATCTNNNATTTGATGCAGTGACATGGGT
AGCACACGTCTGAACTCTTTCCCTACACGACGCTCTTCCGATCTNNNGGGGATGGAGGCAGATGAAT
AGCACACGTCTGAACTCTTTCCCTACACGACGCTCTTCCGATCTNNNTAGTGTCCTTGAAGGCTCTGG
GCACACGTCTGAACTCTTTCCCTACACGACGCTCTTCCGATCTNNNCGACTACAGGAGCTCGTCACTAT
AGCACACGTCTGAACTCTTTCCCTACACGACGCTCTTCCGATCTNNNGAGAACTTCTTCCACTCCCCCAC
GCACACGTCTGAACTCTTTCCCTACACGACGCTCTTCCGATCTNNNTGCTGTCTACCAAGGTGACTTT
GAGCACACGTCTGAACTCTTTCCCTACACGACGCTCTTCCGATCTNNNTGGGAGGACTTAGGATTGT
GCACACGTCTGAACTCTTTCCCTACACGACGCTCTTCCGATCTNNNTGTGGGGATTAGGAGTTCCAGT
AGCACACGTCTGAACTCTTTCCCTACACGACGCTCTTCCGATCTNNNACAGCAGCTTCTCTCTGTCTCT

AGCACACGTCTGAACTCTTTCCCTACACGACGCTCTTCCGATCTNNNGAGGCTCTCAACAGTGTAATG
GCACACGTCTGAACTCTTTCCCTACACGACGCTCTTCCGATCTNNNAGGCAAGGATGAAAACAGCCA
AGCACACGTCTGAACTCTTTCCCTACACGACGCTCTTCCGATCTNNNACCAACATCAAGGTGAGGAA
AGCACACGTCTGAACTCTTTCCCTACACGACGCTCTTCCGATCTNNNTTCACTGCAAATCAACTGGGTC
AGCACACGTCTGAACTCTTTCCCTACACGACGCTCTTCCGATCTNNNTCCCATTGTGATTTCTCTCCTA
AAGAGCACACGTCTGAACTCTTTCCCTACACGACGCTCTTCCGATCTNNNAGGCAAGGATGAAAACA
GAAGAGCACACGTCTGAACTCTTTCCCTACACGACGCTCTTCCGATCTNNNTATACTCCTCAAAACACA
AGAGCACACGTCTGAACTCTTTCCCTACACGACGCTCTTCCGATCTNNNCCTGTTGGCCATATACCTTT
AGCACACGTCTGAACTCTTTCCCTACACGACGCTCTTCCGATCTNNNCTGGGCAACACTCCAGTGAAA
AGCACACGTCTGAACTCTTTCCCTACACGACGCTCTTCCGATCTNNNTTTTCTTTCCCTTGCCTCTCACC
GCACACGTCTGAACTCTTTCCCTACACGACGCTCTTCCGATCTNNNCCTAAGCAACGGAATGAAACCC
AGCACACGTCTGAACTCTTTCCCTACACGACGCTCTTCCGATCTNNNCAGTTATCTGTGAGCGTATGG
AGAGCACACGTCTGAACTCTTTCCCTACACGACGCTCTTCCGATCTNNNGCACTCCCATCAGGAGAGT
GAGCACACGTCTGAACTCTTTCCCTACACGACGCTCTTCCGATCTNNNCCCCTTGGACCCAAGATTACA
GCACACGTCTGAACTCTTTCCCTACACGACGCTCTTCCGATCTNNNTTTGGGAAACAAGACCCTTCTG
GAGCACACGTCTGAACTCTTTCCCTACACGACGCTCTTCCGATCTNNNAACCGATGGTTGTTTCCTGA
AGCACACGTCTGAACTCTTTCCCTACACGACGCTCTTCCGATCTNNNGCACTCTGTTTCAGTCTGGAG
GAGCACACGTCTGAACTCTTTCCCTACACGACGCTCTTCCGATCTNNNTTCCTTCCAAGACAAAGCCCA
GAGCACACGTCTGAACTCTTTCCCTACACGACGCTCTTCCGATCTNNNTTCCAGCCACTGACAGAGTA
AGCACACGTCTGAACTCTTTCCCTACACGACGCTCTTCCGATCTNNNGTGATGATCAGGGATGCAGT
GCACACGTCTGAACTCTTTCCCTACACGACGCTCTTCCGATCTNNNGCTGTAGTGGGGTGGTTTTTAC
AGCACACGTCTGAACTCTTTCCCTACACGACGCTCTTCCGATCTNNNTGGACACATAGGAAGTGTGT
GCACACGTCTGAACTCTTTCCCTACACGACGCTCTTCCGATCTNNNTGAGTTCCACCCATGAAAAAGC
AGCACACGTCTGAACTCTTTCCCTACACGACGCTCTTCCGATCTNNNAGAGAAGTCTCCAATAAGGG
AGCACACGTCTGAACTCTTTCCCTACACGACGCTCTTCCGATCTNNNCATCTGGCCCTTTATTCTTCCC
GAGCACACGTCTGAACTCTTTCCCTACACGACGCTCTTCCGATCTNNNTACAGCACTCCAAGTCTCTGA
AGCACACGTCTGAACTCTTTCCCTACACGACGCTCTTCCGATCTNNNACCAGAGGGATGAAAAGGAA
GAGCACACGTCTGAACTCTTTCCCTACACGACGCTCTTCCGATCTNNNAGGCTGCTTCTAGTCTTCTGC
AGCACACGTCTGAACTCTTTCCCTACACGACGCTCTTCCGATCTNNNGCCTTGTGAGATCTGTTCATTG
AGCACACGTCTGAACTCTTTCCCTACACGACGCTCTTCCGATCTNNNGTTTACAAGGGTGGAGACAG
AGAGCACACGTCTGAACTCTTTCCCTACACGACGCTCTTCCGATCTNNNTGGCTTCTTATGCCTAACAC
AGCACACGTCTGAACTCTTTCCCTACACGACGCTCTTCCGATCTNNNAATCCCTCCCCACATTCTCTCTC
AGCACACGTCTGAACTCTTTCCCTACACGACGCTCTTCCGATCTNNNAGTTTTGGGGGTTGTTTGAC
GCACACGTCTGAACTCTTTCCCTACACGACGCTCTTCCGATCTNNNCTTCTGTTCTCCTTCAAGCCACAC
GCACACGTCTGAACTCTTTCCCTACACGACGCTCTTCCGATCTNNNAGCTAATCACAGGCAGGACCTA
AGCACACGTCTGAACTCTTTCCCTACACGACGCTCTTCCGATCTNNNTTGCCACCTGTGGCTTTTTCTAT
ACACGTCTGAACTCTTTCCCTACACGACGCTCTTCCGATCTNNNGTGGATGGCTAGAGAAGGGAAG
GCACACGTCTGAACTCTTTCCCTACACGACGCTCTTCCGATCTNNNGAACAGAAGCAGACCACACAG
GCACACGTCTGAACTCTTTCCCTACACGACGCTCTTCCGATCTNNNTTTCCTTCCCTCTTGTCCCTTCCC
AGCACACGTCTGAACTCTTTCCCTACACGACGCTCTTCCGATCTNNNAATGTCAGGGCACAGAATGT
AGCACACGTCTGAACTCTTTCCCTACACGACGCTCTTCCGATCTNNNCCAAAGAGTTTGAGAACTGAT
AGCACACGTCTGAACTCTTTCCCTACACGACGCTCTTCCGATCTNNNGGTAGCCTGATGAAAGATGT
CACACGTCTGAACTCTTTCCCTACACGACGCTCTTCCGATCTNNNTGGCTTGCAATCATGTGCAGAAT

AGCACACGTCTGAACTCTTTCCCTACACGACGCTCTTCCGATCTNNNTCTGGGAGCTATTTTGTGGCTT
AGCACACGTCTGAACTCTTTCCCTACACGACGCTCTTCCGATCTNNNAGTGTGCCCATTTCTCTCTCTCT
GAGCACACGTCTGAACTCTTTCCCTACACGACGCTCTTCCGATCTNNNGGGGGATATAAAGGCTTGGT
AGAGCACACGTCTGAACTCTTTCCCTACACGACGCTCTTCCGATCTNNNAAAGAAGAGCTGGGGCAA
GCACACGTCTGAACTCTTTCCCTACACGACGCTCTTCCGATCTNNNAGCTACTTGTACCCCAAAAGCTG
GAGCACACGTCTGAACTCTTTCCCTACACGACGCTCTTCCGATCTNNNTTAGGGGATTCTGGGGAGA
GAGCACACGTCTGAACTCTTTCCCTACACGACGCTCTTCCGATCTNNNTGTGATACAGCGTCTACTGG
AGAGCACACGTCTGAACTCTTTCCCTACACGACGCTCTTCCGATCTNNNCCTCTCGATGTTGTAGTTGG
GAGCACACGTCTGAACTCTTTCCCTACACGACGCTCTTCCGATCTNNNTGGAGGAAGCTCTGATCCAA
GCACACGTCTGAACTCTTTCCCTACACGACGCTCTTCCGATCTNNNTTTGGCTCCTGCTTCATCTTCTCC
GAGCACACGTCTGAACTCTTTCCCTACACGACGCTCTTCCGATCTNNNATTGTTACTGTTTTCGCAGG
GAGCACACGTCTGAACTCTTTCCCTACACGACGCTCTTCCGATCTNNNCTCCTCCAGGAATCTTTGGCT
AGCACACGTCTGAACTCTTTCCCTACACGACGCTCTTCCGATCTNNNTCTGTTAGTTTTGAATTGGCCTG
GCACACGTCTGAACTCTTTCCCTACACGACGCTCTTCCGATCTNNNGAGTGAGATCCACTTTGGAGAC
AAGAGCACACGTCTGAACTCTTTCCCTACACGACGCTCTTCCGATCTNNNAAAGCATGGCTCTACCCAC
AGCACACGTCTGAACTCTTTCCCTACACGACGCTCTTCCGATCTNNNTGGATTCTAATCTCTGGTGCAT
GCACACGTCTGAACTCTTTCCCTACACGACGCTCTTCCGATCTNNNCTTGTTCATCAAAAGCTGCACCA
GAGCACACGTCTGAACTCTTTCCCTACACGACGCTCTTCCGATCTNNNTCTTCACATTGAGCATATGTA
AGCACACGTCTGAACTCTTTCCCTACACGACGCTCTTCCGATCTNNNCACATCCACGTTCTAGGCAGA
AGCACACGTCTGAACTCTTTCCCTACACGACGCTCTTCCGATCTNNNACGAGGTTCAGAAAGTGGCT
GAGCACACGTCTGAACTCTTTCCCTACACGACGCTCTTCCGATCTNNNCTGAGCCCAAGAGTTCAAGTA
GAGCACACGTCTGAACTCTTTCCCTACACGACGCTCTTCCGATCTNNNTGCGTTATCTCTCTCTCTCTC
AGCACACGTCTGAACTCTTTCCCTACACGACGCTCTTCCGATCTNNNTGTCAAGAGAAAGACCACTTC
AGCACACGTCTGAACTCTTTCCCTACACGACGCTCTTCCGATCTNNNTTGCTGCATTCCAGCCTAAAAG
AGCACACGTCTGAACTCTTTCCCTACACGACGCTCTTCCGATCTNNNCCTTGACATGTTAGTGTGGAGG
AAGAGCACACGTCTGAACTCTTTCCCTACACGACGCTCTTCCGATCTNNNTGTGCGTCTGGAAAGTGA
AGAGCACACGTCTGAACTCTTTCCCTACACGACGCTCTTCCGATCTNNNGGAAACCAGAGACACAAAA
GAGCACACGTCTGAACTCTTTCCCTACACGACGCTCTTCCGATCTNNNTGGATTTGATTGTGTTGGAT
AGCACACGTCTGAACTCTTTCCCTACACGACGCTCTTCCGATCTNNNCTGGAAGCATCTGCATCCTTCTT
AAGAGCACACGTCTGAACTCTTTCCCTACACGACGCTCTTCCGATCTNNNGGAGCTGCAGGGTAAAA
AGCACACGTCTGAACTCTTTCCCTACACGACGCTCTTCCGATCTNNNAAAACTTTGAATAAAGTGCTCA
AGCACACGTCTGAACTCTTTCCCTACACGACGCTCTTCCGATCTNNNACTGCCTTTGCCTAGTCACAAA
AGCACACGTCTGAACTCTTTCCCTACACGACGCTCTTCCGATCTNNNCTATGCTTCCTCTGAAGGCACTG
GAGCACACGTCTGAACTCTTTCCCTACACGACGCTCTTCCGATCTNNNACCACACCCTACCACTTCACTA
GAGCACACGTCTGAACTCTTTCCCTACACGACGCTCTTCCGATCTNNNCTTGCAGCTACACATGGACTT
GAGCACACGTCTGAACTCTTTCCCTACACGACGCTCTTCCGATCTNNNAGCGCCAGGAAAAGTAAATC
GCACACGTCTGAACTCTTTCCCTACACGACGCTCTTCCGATCTNNNCCTCCAACCCAACCAATATTGTC
AGCACACGTCTGAACTCTTTCCCTACACGACGCTCTTCCGATCTNNNTCCAGTTCTATCCACGATGCAA
AGCACACGTCTGAACTCTTTCCCTACACGACGCTCTTCCGATCTNNNTGGTGTGTGGTGTGTATCTGT
GAGCACACGTCTGAACTCTTTCCCTACACGACGCTCTTCCGATCTNNNCCCTTTGAGGATGGCCTGAAA
AGCACACGTCTGAACTCTTTCCCTACACGACGCTCTTCCGATCTNNNGACAGAGTGGCTTGAACACTA
GAGCACACGTCTGAACTCTTTCCCTACACGACGCTCTTCCGATCTNNNCTTTTCTCTCCCTCCATCCCCTT
GCACACGTCTGAACTCTTTCCCTACACGACGCTCTTCCGATCTNNNTCTTCCTCTTCTCTGCTGAAGTTC

GAGCACACGTCTGAACTCTTTCCCTACACGACGCTCTTCCGATCTNNNTTAGGCCTCTCTGATGAGGT
AAGAGCACACGTCTGAACTCTTTCCCTACACGACGCTCTTCCGATCTNNNTGTACATTCTGTTTGAGG
AGAGCACACGTCTGAACTCTTTCCCTACACGACGCTCTTCCGATCTNNNGAATTCTCTTTTCCTGGCTT
AGCACACGTCTGAACTCTTTCCCTACACGACGCTCTTCCGATCTNNNTTCCTTCTCAACAGGGAGAGG
GCACACGTCTGAACTCTTTCCCTACACGACGCTCTTCCGATCTNNNTTTCCAGAGCTGAGTACAGTG
AGCACACGTCTGAACTCTTTCCCTACACGACGCTCTTCCGATCTNNNAACTGAAATGTACACCGGTGA
GAGCACACGTCTGAACTCTTTCCCTACACGACGCTCTTCCGATCTNNNTCAGCATCTGCTTTAGTGAAA
GAGCACACGTCTGAACTCTTTCCCTACACGACGCTCTTCCGATCTNNNCCCTTTTGCTGGGCATAAAGT
AGCACACGTCTGAACTCTTTCCCTACACGACGCTCTTCCGATCTNNNGTCTGATGGATTTTTGCTCTG
AAGAGCACACGTCTGAACTCTTTCCCTACACGACGCTCTTCCGATCTNNNGGCAGCTTTACTCTCCTTT
GAGCACACGTCTGAACTCTTTCCCTACACGACGCTCTTCCGATCTNNNGTAGACCCCAAAGCACTATGT
AGCACACGTCTGAACTCTTTCCCTACACGACGCTCTTCCGATCTNNNGAGGCAGGTACAGCACAGAA
AGAGCACACGTCTGAACTCTTTCCCTACACGACGCTCTTCCGATCTNNNTAAAATCTGCCCCGATTGGG
GAGCACACGTCTGAACTCTTTCCCTACACGACGCTCTTCCGATCTNNNATTACATGATTGGCGTTAGGC
CACGTCTGAACTCTTTCCCTACACGACGCTCTTCCGATCTNNNCGACGCTTTCCTTTCATTTTTGCCAT
AGCACACGTCTGAACTCTTTCCCTACACGACGCTCTTCCGATCTNNNTATCCTGAGGTTGCCTACAAG
GCACACGTCTGAACTCTTTCCCTACACGACGCTCTTCCGATCTNNNCTGGAGGGCAACTCCTCCTTTCA
GCACACGTCTGAACTCTTTCCCTACACGACGCTCTTCCGATCTNNNCTCTGCCTTCGAATACCAGCTTT
GCACACGTCTGAACTCTTTCCCTACACGACGCTCTTCCGATCTNNNCTTTGACCCTGGGTGGTTCTTTC
GAGCACACGTCTGAACTCTTTCCCTACACGACGCTCTTCCGATCTNNNTACATTCTGCTTTCCTGGTGGT
AGCACACGTCTGAACTCTTTCCCTACACGACGCTCTTCCGATCTNNNATTCCTTCCTAGGAGCCCATCA
GCACACGTCTGAACTCTTTCCCTACACGACGCTCTTCCGATCTNNNTGAGCTAAGTACAGGACCCACA
GCACACGTCTGAACTCTTTCCCTACACGACGCTCTTCCGATCTNNNCTTTCTGGTGTGCTACCTTCTCC
GAGCACACGTCTGAACTCTTTCCCTACACGACGCTCTTCCGATCTNNNCTTGGAATATCCGGGTTGCT
AGCACACGTCTGAACTCTTTCCCTACACGACGCTCTTCCGATCTNNNGGGGGAGCTTAGAGTACATG
AGCACACGTCTGAACTCTTTCCCTACACGACGCTCTTCCGATCTNNNCCTGCTGATCTACAGCAGTGT
GAAGAGCACACGTCTGAACTCTTTCCCTACACGACGCTCTTCCGATCTNNNGGAGGGCCACCTTGAT
GAGCACACGTCTGAACTCTTTCCCTACACGACGCTCTTCCGATCTNNNCAGGAAGGTGGCTTTGGTA
AGCACACGTCTGAACTCTTTCCCTACACGACGCTCTTCCGATCTNNNTGACTTACAGAGTTTCAGACT
GCACACGTCTGAACTCTTTCCCTACACGACGCTCTTCCGATCTNNNCAGAGTTGTGTAAGAGCCTTG
GAGCACACGTCTGAACTCTTTCCCTACACGACGCTCTTCCGATCTNNNTATTGCAATGAGTGGGGGAG
GCACACGTCTGAACTCTTTCCCTACACGACGCTCTTCCGATCTNNNCCAAAAGTCCCCCAAAGCATAG
GAGCACACGTCTGAACTCTTTCCCTACACGACGCTCTTCCGATCTNNNGGTGTGGAATGGTGTCCCAT
CACACGTCTGAACTCTTTCCCTACACGACGCTCTTCCGATCTNNNGACAGGAAGGAAGAAAGAGGGT
GAGCACACGTCTGAACTCTTTCCCTACACGACGCTCTTCCGATCTNNNCTAGACGGAAGCTTGGAGCA
AGCACACGTCTGAACTCTTTCCCTACACGACGCTCTTCCGATCTNNNTTGAAGAAGGGACTGAGCGA
GAGCACACGTCTGAACTCTTTCCCTACACGACGCTCTTCCGATCTNNNACAATGGAGTGGGTCATGAA
ACACGTCTGAACTCTTTCCCTACACGACGCTCTTCCGATCTNNNCCTCCCTGTATCACCCCCTCCATCGG
GAGCACACGTCTGAACTCTTTCCCTACACGACGCTCTTCCGATCTNNNAAACCTCTGTACAGGTATGTG
GCACACGTCTGAACTCTTTCCCTACACGACGCTCTTCCGATCTNNNAGCTTGGGCTGAATAATGAGC
GCACACGTCTGAACTCTTTCCCTACACGACGCTCTTCCGATCTNNNGGCAAACTGTAGGTATTCAGC
AGCACACGTCTGAACTCTTTCCCTACACGACGCTCTTCCGATCTNNNAACATCCTGGGTACATGCACTT
GAGCACACGTCTGAACTCTTTCCCTACACGACGCTCTTCCGATCTNNNAGGCCAATGGAAAATAACA

AGCACACGTCTGAACTCTTTCCCTACACGACGCTCTTCCGATCTNNNGCCCAGTATGGGTCATAGGA
AGCACACGTCTGAACTCTTTCCCTACACGACGCTCTTCCGATCTNNNGTGTGTGTAAGTGATGCCAGT
GCACACGTCTGAACTCTTTCCCTACACGACGCTCTTCCGATCTNNNCCATGAAGATGATGTCCCACAG
GCACACGTCTGAACTCTTTCCCTACACGACGCTCTTCCGATCTNNNAGGATGCACAGGGCTAAGTAG
AGCACACGTCTGAACTCTTTCCCTACACGACGCTCTTCCGATCTNNNAGCCTGGATCAAAAGTCAGGA
AGCACACGTCTGAACTCTTTCCCTACACGACGCTCTTCCGATCTNNNACCACCAAACGACAGTTGAGA
GAGCACACGTCTGAACTCTTTCCCTACACGACGCTCTTCCGATCTNNNACACAAAGTCCAAACAATCTG
AGCACACGTCTGAACTCTTTCCCTACACGACGCTCTTCCGATCTNNNACACAGAACATATCCACACAC
AGCACACGTCTGAACTCTTTCCCTACACGACGCTCTTCCGATCTNNNGGTCCCAGGGAGAATGCAAT
GAGCACACGTCTGAACTCTTTCCCTACACGACGCTCTTCCGATCTNNNCCAATCAAGGACTTCAGAGT
AGCACACGTCTGAACTCTTTCCCTACACGACGCTCTTCCGATCTNNNATGTACCTTACCAGTGGCACT
AGCACACGTCTGAACTCTTTCCCTACACGACGCTCTTCCGATCTNNNTGTGTACAATGAAATACTCAG
AGCACACGTCTGAACTCTTTCCCTACACGACGCTCTTCCGATCTNNNTTAGGGGATTCTGGGGAGAA
GCACACGTCTGAACTCTTTCCCTACACGACGCTCTTCCGATCTNNNCTCCCGCTCTAAGTGATTCTCCC
AGCACACGTCTGAACTCTTTCCCTACACGACGCTCTTCCGATCTNNNGCTGGGGGTAGAAGAAAAAC
CACACGTCTGAACTCTTTCCCTACACGACGCTCTTCCGATCTNNNGCCCATAGCTACTCTGTTCTTGGC
AGCACACGTCTGAACTCTTTCCCTACACGACGCTCTTCCGATCTNNNTAAAATCGCTCAAGACACAGCC
AGCACACGTCTGAACTCTTTCCCTACACGACGCTCTTCCGATCTNNNCCATACAAACATCCAAAGCCTG
GAGCACACGTCTGAACTCTTTCCCTACACGACGCTCTTCCGATCTNNNAGTGATTGCTGACTGTTCAG
AGCACACGTCTGAACTCTTTCCCTACACGACGCTCTTCCGATCTNNNGGCTGCAGTTGAGCTATGGT
AGCACACGTCTGAACTCTTTCCCTACACGACGCTCTTCCGATCTNNNGAGCCACTGAATAGAGGAGCT
AGCACACGTCTGAACTCTTTCCCTACACGACGCTCTTCCGATCTNNNTGTCTTGAACCCAGGACAGTT
AGCACACGTCTGAACTCTTTCCCTACACGACGCTCTTCCGATCTNNNGTGGTTTATCCATGGAAGGGC
GAAGAGCACACGTCTGAACTCTTTCCCTACACGACGCTCTTCCGATCTNNNTGAACAACAACTGATCCC
GAGCACACGTCTGAACTCTTTCCCTACACGACGCTCTTCCGATCTNNNAATGGAGACTACCATGACGT
AGCACACGTCTGAACTCTTTCCCTACACGACGCTCTTCCGATCTNNNAGGATGCACAGGGCTAAGTA
GAGCACACGTCTGAACTCTTTCCCTACACGACGCTCTTCCGATCTNNNTTTTCTTGCGTGGGAGTGTT
AGCACACGTCTGAACTCTTTCCCTACACGACGCTCTTCCGATCTNNNAGCCTGAGTGATGGAGTAAG
AGCACACGTCTGAACTCTTTCCCTACACGACGCTCTTCCGATCTNNNCAGAAACCTGAGTGCTGACAT
AGAGCACACGTCTGAACTCTTTCCCTACACGACGCTCTTCCGATCTNNNCCATGTTCCTGTGAATGAC
AGCACACGTCTGAACTCTTTCCCTACACGACGCTCTTCCGATCTNNNAATGATATTCTCCCCACCTCCC
AGAGCACACGTCTGAACTCTTTCCCTACACGACGCTCTTCCGATCTNNNTCTCATTTCCCAGAGCTTTC
GCACACGTCTGAACTCTTTCCCTACACGACGCTCTTCCGATCTNNNGCAGTGACCTTGTGTATTGTGT
AGCACACGTCTGAACTCTTTCCCTACACGACGCTCTTCCGATCTNNNGAAGTAGTGCAGGTGATCAAG
GAGCACACGTCTGAACTCTTTCCCTACACGACGCTCTTCCGATCTNNNGATGTCCCTATTGCCATCACC
GAGCACACGTCTGAACTCTTTCCCTACACGACGCTCTTCCGATCTNNNACCCTTTTTGCCGTGTAACAG
AGCACACGTCTGAACTCTTTCCCTACACGACGCTCTTCCGATCTNNNCACTGTTCCAATGCCCATTTCA
GAGCACACGTCTGAACTCTTTCCCTACACGACGCTCTTCCGATCTNNNGGGCTGCAAAACTGACCTAT
AAGAGCACACGTCTGAACTCTTTCCCTACACGACGCTCTTCCGATCTNNNATGTGTGTGTGCACTCTC
AGCACACGTCTGAACTCTTTCCCTACACGACGCTCTTCCGATCTNNNGCCCTCCCTTGATGGTATGTAT
GCACACGTCTGAACTCTTTCCCTACACGACGCTCTTCCGATCTNNNTCTGCAAGAAGTAAGTTGGCTC
GAGCACACGTCTGAACTCTTTCCCTACACGACGCTCTTCCGATCTNNNAGAGACCCAGAGAGACAGT
AGCACACGTCTGAACTCTTTCCCTACACGACGCTCTTCCGATCTNNNTAAAATCGCTCAAGACACAGC

AGCACACGTCTGAACTCTTTCCCTACACGACGCTCTTCCGATCTNNNTGTGAGGAAGACATTACACATT
AAGAGCACACGTCTGAACTCTTTCCCTACACGACGCTCTTCCGATCTNNNCTTGGGTCCCATGGAGGA
GAGCACACGTCTGAACTCTTTCCCTACACGACGCTCTTCCGATCTNNNTGAACAACAACTGATCCCTCCA
GAGCACACGTCTGAACTCTTTCCCTACACGACGCTCTTCCGATCTNNNGGTGTGGGATATGGTACTTG
AGCACACGTCTGAACTCTTTCCCTACACGACGCTCTTCCGATCTNNNCCTGGACACAGTGAAGAACCTT
AGCACACGTCTGAACTCTTTCCCTACACGACGCTCTTCCGATCTNNNATCTGCCCACAGAGCCTACAATA
GAGCACACGTCTGAACTCTTTCCCTACACGACGCTCTTCCGATCTNNNTCCATGTTCCTGTGAATGACGA
AGCACACGTCTGAACTCTTTCCCTACACGACGCTCTTCCGATCTNNNATGCTCTTTTGGCCATGCTTTTC
GAGCACACGTCTGAACTCTTTCCCTACACGACGCTCTTCCGATCTNNNTATAGGGCTGTGTGGGAACAA
GAGCACACGTCTGAACTCTTTCCCTACACGACGCTCTTCCGATCTNNNTGTGAGGAAGACATTACACAT
GAGCACACGTCTGAACTCTTTCCCTACACGACGCTCTTCCGATCTNNNGGCGGGTCACCTAATCTCTCTA
AGAGCACACGTCTGAACTCTTTCCCTACACGACGCTCTTCCGATCTNNNCAAGTGCCTGTGTCAAGTAA
GAGCACACGTCTGAACTCTTTCCCTACACGACGCTCTTCCGATCTNNNCGTGTGCATGTTTCAAATGTT
AGAGCACACGTCTGAACTCTTTCCCTACACGACGCTCTTCCGATCTNNNTTATCTTGCATAGGCTATCCC
GAGCACACGTCTGAACTCTTTCCCTACACGACGCTCTTCCGATCTNNNATATGCTCATCTGGCCTTTCCC
GAGCACACGTCTGAACTCTTTCCCTACACGACGCTCTTCCGATCTNNNGGATGCCTCAGTTCTCTCTC
AGCACACGTCTGAACTCTTTCCCTACACGACGCTCTTCCGATCTNNNTTCAAGTGGCTACCTGAGAGTT
GAGCACACGTCTGAACTCTTTCCCTACACGACGCTCTTCCGATCTNNNGAAACTCAGGCTGGGTAACA
GCACACGTCTGAACTCTTTCCCTACACGACGCTCTTCCGATCTNNNAATCTCACAGCTCTAAGTGGCAG
GAGCACACGTCTGAACTCTTTCCCTACACGACGCTCTTCCGATCTNNNGAAGCCCCAGACCCATCTGTCA
GAGCACACGTCTGAACTCTTTCCCTACACGACGCTCTTCCGATCTNNNCGCCTGGTCAGAATCGTCATT
GCACACGTCTGAACTCTTTCCCTACACGACGCTCTTCCGATCTNNNGGCAGGGTGATCGTGTCATATT
GAGCACACGTCTGAACTCTTTCCCTACACGACGCTCTTCCGATCTNNNGGAAAATCTGGCAACATTGAG
GAGCACACGTCTGAACTCTTTCCCTACACGACGCTCTTCCGATCTNNNGCTGCTGTCCTGATGTCTTGTA
AGCACACGTCTGAACTCTTTCCCTACACGACGCTCTTCCGATCTNNNTTTGGCTCCTGCTTCATCTTCTC
GAGCACACGTCTGAACTCTTTCCCTACACGACGCTCTTCCGATCTNNNATGTTTGCGTGTCTGGCTTAT
AGCACACGTCTGAACTCTTTCCCTACACGACGCTCTTCCGATCTNNNGTTTACTATGTGCCCCTTCCCAG
AGAGCACACGTCTGAACTCTTTCCCTACACGACGCTCTTCCGATCTNNNGAACAGCCTAAGCACATCCT
AGAGCACACGTCTGAACTCTTTCCCTACACGACGCTCTTCCGATCTNNNGGTGGTTTTCCAGGTGGTA
GAAGAGCACACGTCTGAACTCTTTCCCTACACGACGCTCTTCCGATCTNNNATCATTCTGCGCTTACAGA
AGCACACGTCTGAACTCTTTCCCTACACGACGCTCTTCCGATCTNNNTTTCCATAAGGCTAGTGTGCAG
AGAGCACACGTCTGAACTCTTTCCCTACACGACGCTCTTCCGATCTNNNCTATGGCACTCCCTTTCAAGT
AGAGCACACGTCTGAACTCTTTCCCTACACGACGCTCTTCCGATCTNNNTAAAAATGTCACTGCTCCCC
GAGCACACGTCTGAACTCTTTCCCTACACGACGCTCTTCCGATCTNNNAAAAATTTAGCCAGGTGGGT
AGAGCACACGTCTGAACTCTTTCCCTACACGACGCTCTTCCGATCTNNNGCAAGAGAGCTCCTTCAGGT
AGCACACGTCTGAACTCTTTCCCTACACGACGCTCTTCCGATCTNNNCTCTGTCCTCACAGAGTTCAAGC
CACACGTCTGAACTCTTTCCCTACACGACGCTCTTCCGATCTNNNCTTTTTGCCAACATGATGCCTCGCA
AGCACACGTCTGAACTCTTTCCCTACACGACGCTCTTCCGATCTNNNCTCCCTAAGCAAGACAGGAAAC
GCACACGTCTGAACTCTTTCCCTACACGACGCTCTTCCGATCTNNNTCCATTGCTACATGTGCCTTTTTCA
AGAGCACACGTCTGAACTCTTTCCCTACACGACGCTCTTCCGATCTNNNCTGCTCTCTTTTGGGACCATC
GCACACGTCTGAACTCTTTCCCTACACGACGCTCTTCCGATCTNNNGAACATGTGTGGCCAGAACTAC
GAGCACACGTCTGAACTCTTTCCCTACACGACGCTCTTCCGATCTNNNAATGGTCACGTTCACTTCCTCA
GAGCACACGTCTGAACTCTTTCCCTACACGACGCTCTTCCGATCTNNNAGATTTCTGCCTTGGGGGAAA

GCACACGTCTGAACTCTTTCCCTACACGACGCTCTTCCGATCTNNNGAAGTAGTGCAGGTGATCAAGCA
CACACGTCTGAACTCTTTCCCTACACGACGCTCTTCCGATCTNNNAAAGATCCAAAACACTTCTGCTTCCC
AAGAGCACACGTCTGAACTCTTTCCCTACACGACGCTCTTCCGATCTNNNAGAGAAGAGGTGGTTGGT
CACACGTCTGAACTCTTTCCCTACACGACGCTCTTCCGATCTNNNCCCAGGATTCCAGATTCTGAGTTCA
GAGCACACGTCTGAACTCTTTCCCTACACGACGCTCTTCCGATCTNNNTCCTCCCTCTTTTTCCACACTCA
GAGCACACGTCTGAACTCTTTCCCTACACGACGCTCTTCCGATCTNNNATTCGGGACCCCAGTGAATTT
GAGCACACGTCTGAACTCTTTCCCTACACGACGCTCTTCCGATCTNNNGCCTCTTAGCAGAGCCTTCAAA
AGAGCACACGTCTGAACTCTTTCCCTACACGACGCTCTTCCGATCTNNNGACAGTCTTTCTCTCTCCTTGC
AGAGCACACGTCTGAACTCTTTCCCTACACGACGCTCTTCCGATCTNNNTCCCTATTTGTAAGTGCAGCA
CACACGTCTGAACTCTTTCCCTACACGACGCTCTTCCGATCTNNNACAGTTTACCAGCAGTCTAAGCGCA
GAGCACACGTCTGAACTCTTTCCCTACACGACGCTCTTCCGATCTNNNTTCAGTACCCTTCAGTACCCCT
GAGCACACGTCTGAACTCTTTCCCTACACGACGCTCTTCCGATCTNNNACACAATGAGGGGATTGGAGA
AGAGCACACGTCTGAACTCTTTCCCTACACGACGCTCTTCCGATCTNNNATTCTGAAAATGGCTGCACCT
AGCACACGTCTGAACTCTTTCCCTACACGACGCTCTTCCGATCTNNNAAAAGTAGAGATTATCAGAGTC
AGAGCACACGTCTGAACTCTTTCCCTACACGACGCTCTTCCGATCTNNNTGTCCTCAGTGTCATTTTCCTC
AGAGCACACGTCTGAACTCTTTCCCTACACGACGCTCTTCCGATCTNNNTTTCCTTCTTGCCTTCTTCCCAC
GAGCACACGTCTGAACTCTTTCCCTACACGACGCTCTTCCGATCTNNNAGGGTGGCTAAAAATGTGGA
GAGCACACGTCTGAACTCTTTCCCTACACGACGCTCTTCCGATCTNNNAGGTGGTGATAGGCTACCATT
GAGCACACGTCTGAACTCTTTCCCTACACGACGCTCTTCCGATCTNNNATCCTCACCAAAAATGGTGCTC
AGCACACGTCTGAACTCTTTCCCTACACGACGCTCTTCCGATCTNNNGATTTGATGGGGATCAGTGCCT
AGCACACGTCTGAACTCTTTCCCTACACGACGCTCTTCCGATCTNNNTATGGGAGAGACCCAGTATGC
AGAGCACACGTCTGAACTCTTTCCCTACACGACGCTCTTCCGATCTNNNTGTATTACCTCGAGTAGGGC
AGCACACGTCTGAACTCTTTCCCTACACGACGCTCTTCCGATCTNNNCTAGGGTTGTGTGGGTAAAGC
AGCACACGTCTGAACTCTTTCCCTACACGACGCTCTTCCGATCTNNNGTGTGAGGCCCAACAAATGAA
GCACACGTCTGAACTCTTTCCCTACACGACGCTCTTCCGATCTNNNTTTTGTGACCTGTGAAGGTGGT
GAGCACACGTCTGAACTCTTTCCCTACACGACGCTCTTCCGATCTNNNTCACATTTATACAGAGCCAAAC
AGCACACGTCTGAACTCTTTCCCTACACGACGCTCTTCCGATCTNNNTGTCCAGTTCCCAACAATTGCATC
AGCACACGTCTGAACTCTTTCCCTACACGACGCTCTTCCGATCTNNNATTCACTACACACACATGTGGAC
GAGCACACGTCTGAACTCTTTCCCTACACGACGCTCTTCCGATCTNNNTCTCTGACCACACAGAAAGTTG
GAGCACACGTCTGAACTCTTTCCCTACACGACGCTCTTCCGATCTNNNTTCTCAGATGTGTGGGAAAAG
AGCACACGTCTGAACTCTTTCCCTACACGACGCTCTTCCGATCTNNNTTTCTGTCTTTGCTTGGCTCCATC
AAGAGCACACGTCTGAACTCTTTCCCTACACGACGCTCTTCCGATCTNNNTCCAGTCTGGGAAAGACTT
AGAGCACACGTCTGAACTCTTTCCCTACACGACGCTCTTCCGATCTNNNGTGTCTGGTACAAAGGGTAA
AGCACACGTCTGAACTCTTTCCCTACACGACGCTCTTCCGATCTNNNCGCACCTGGCCATCTTTTATCTTC
AGAGCACACGTCTGAACTCTTTCCCTACACGACGCTCTTCCGATCTNNNACAAGGTTGCAATTAGAGG
AGAGCACACGTCTGAACTCTTTCCCTACACGACGCTCTTCCGATCTNNNTTCATAATGAGGTGCAAGG
AGAGCACACGTCTGAACTCTTTCCCTACACGACGCTCTTCCGATCTNNNTTGCCTGGGGTTTTTAGAGTT
AGCACACGTCTGAACTCTTTCCCTACACGACGCTCTTCCGATCTNNNTCCCATGAGAAGACACTTACCT
AGCACACGTCTGAACTCTTTCCCTACACGACGCTCTTCCGATCTNNNGCTATAAGGCTGATGTGTGGG
AAGAGCACACGTCTGAACTCTTTCCCTACACGACGCTCTTCCGATCTNNNCGTCTCTGGCCTATGTGAA
GAGCACACGTCTGAACTCTTTCCCTACACGACGCTCTTCCGATCTNNNGCTTAGTGATTCCCAACCTCCT
AGCACACGTCTGAACTCTTTCCCTACACGACGCTCTTCCGATCTNNNGACAAACCAGCAAATGAGCAAC
AGAGCACACGTCTGAACTCTTTCCCTACACGACGCTCTTCCGATCTNNNCCATCAGGTGTTTAGCCTCA

GCACACGTCTGAACTCTTTCCCTACACGACGCTCTTCCGATCTNNNAGCTTCTCTCATTACTTTTCACT
GAGCACACGTCTGAACTCTTTCCCTACACGACGCTCTTCCGATCTNNNCCTGGAACCAATCCTCCTACA
GAGCACACGTCTGAACTCTTTCCCTACACGACGCTCTTCCGATCTNNNTCTAATCTATCTGCCTGAGGG
AGAGCACACGTCTGAACTCTTTCCCTACACGACGCTCTTCCGATCTNNNAGCAAATGAAATCTACCTG
AGCACACGTCTGAACTCTTTCCCTACACGACGCTCTTCCGATCTNNNACAACCCTATGGAAGAGTGA
GAGCACACGTCTGAACTCTTTCCCTACACGACGCTCTTCCGATCTNNNGTGTGAATGGGCATGTGTT
AGCACACGTCTGAACTCTTTCCCTACACGACGCTCTTCCGATCTNNNTGCTCATGATTCCCTTGGCTA
GCACACGTCTGAACTCTTTCCCTACACGACGCTCTTCCGATCTNNNTACTGGGTCGGTCATGGTTTCT
AGCACACGTCTGAACTCTTTCCCTACACGACGCTCTTCCGATCTNNNAGCAACTGACCAAATTGCTAA
GAGCACACGTCTGAACTCTTTCCCTACACGACGCTCTTCCGATCTNNNTTCTGAGAATACACCACTGG
AGCACACGTCTGAACTCTTTCCCTACACGACGCTCTTCCGATCTNNNCCCTCCCATTACTGTGGGGAA
GCACACGTCTGAACTCTTTCCCTACACGACGCTCTTCCGATCTNNNAGATGGAGGAAGAGCCAATCC
AGCACACGTCTGAACTCTTTCCCTACACGACGCTCTTCCGATCTNNNTGAGGAGGGAGGTAACTTCCA
GCACACGTCTGAACTCTTTCCCTACACGACGCTCTTCCGATCTNNNGCCTATATCTCCAGGATGGCTC
AGCACACGTCTGAACTCTTTCCCTACACGACGCTCTTCCGATCTNNNTGGTTTGAAATTGCAAGTGTC
GAGCACACGTCTGAACTCTTTCCCTACACGACGCTCTTCCGATCTNNNCCCTCCCATTACTGTGGGGA
AGCACACGTCTGAACTCTTTCCCTACACGACGCTCTTCCGATCTNNNTGCATCCTGAATGGATAAGGT
GCACACGTCTGAACTCTTTCCCTACACGACGCTCTTCCGATCTNNNAATGTGAGGGTGCTTGATTCT
AGCACACGTCTGAACTCTTTCCCTACACGACGCTCTTCCGATCTNNNAGAAACTGGGGAGGTGAGG
AGCACACGTCTGAACTCTTTCCCTACACGACGCTCTTCCGATCTNNNGCACTTGGCAGGATTTGTATG
AGCACACGTCTGAACTCTTTCCCTACACGACGCTCTTCCGATCTNNNATGTGGTGTTTTCTGCCTGGA
AGAGCACACGTCTGAACTCTTTCCCTACACGACGCTCTTCCGATCTNNNGAAAAGATAAGCTCCCTGC
AGCACACGTCTGAACTCTTTCCCTACACGACGCTCTTCCGATCTNNNCTGTGCCTGTTCCTCTCTTTCT
AGCACACGTCTGAACTCTTTCCCTACACGACGCTCTTCCGATCTNNNTAGGCTGAACATCATTCCAGG
AGCACACGTCTGAACTCTTTCCCTACACGACGCTCTTCCGATCTNNNAAACAGATCAGACGAGAAGC
AGCACACGTCTGAACTCTTTCCCTACACGACGCTCTTCCGATCTNNNCCTACAGTTTCCTACAAAGCGT
GAGCACACGTCTGAACTCTTTCCCTACACGACGCTCTTCCGATCTNNNAGTGGTCATTCTCACCTTTTC
AGCACACGTCTGAACTCTTTCCCTACACGACGCTCTTCCGATCTNNNTCTTCAACCTGCTGCTGTCATA
GAGCACACGTCTGAACTCTTTCCCTACACGACGCTCTTCCGATCTNNNTTCTGCACAATCAGCATTACC
GAGCACACGTCTGAACTCTTTCCCTACACGACGCTCTTCCGATCTNNNAGCTGAATTTATCTATTGCCT
AGCACACGTCTGAACTCTTTCCCTACACGACGCTCTTCCGATCTNNNGCCTAAAAGGTGAAACCCCCA
AGAGCACACGTCTGAACTCTTTCCCTACACGACGCTCTTCCGATCTNNNCGCAGTCTTTTGAGTAGCTG
AGCACACGTCTGAACTCTTTCCCTACACGACGCTCTTCCGATCTNNNTGCACCACACAAATACACTCAC
GAGCACACGTCTGAACTCTTTCCCTACACGACGCTCTTCCGATCTNNNTCAAATTCCCCATTTGTTTCT
GAGCACACGTCTGAACTCTTTCCCTACACGACGCTCTTCCGATCTNNNGTTGTGCTGCCAATATCTTCT
AGCACACGTCTGAACTCTTTCCCTACACGACGCTCTTCCGATCTNNNGTTGAGTGGCTCTGATCCCAA
AGCACACGTCTGAACTCTTTCCCTACACGACGCTCTTCCGATCTNNNGCACATGCAATGCAGAGATCC
GAGCACACGTCTGAACTCTTTCCCTACACGACGCTCTTCCGATCTNNNTCCCCAGGTGAGAAAGAATG
AGCACACGTCTGAACTCTTTCCCTACACGACGCTCTTCCGATCTNNNCTTGGGCAAAGGAGCAAGACT
GAGCACACGTCTGAACTCTTTCCCTACACGACGCTCTTCCGATCTNNNGTTATGCTCTTTGTCCACCCGA
GAGCACACGTCTGAACTCTTTCCCTACACGACGCTCTTCCGATCTNNNCAGGGACAGTGTAATCTTGG
GAGCACACGTCTGAACTCTTTCCCTACACGACGCTCTTCCGATCTNNNCCGTGTAGAGATATTTTCACC
GAGCACACGTCTGAACTCTTTCCCTACACGACGCTCTTCCGATCTNNNTGTTCCAAAAGCCATAAGCCT

GAGCACACGTCTGAACTCTTTCCCTACACGACGCTCTTCCGATCTNNNCAGAGGTTGCAGGAGAGATT
AGAGCACACGTCTGAACTCTTTCCCTACACGACGCTCTTCCGATCTNNNGGCAATTTACAAACAAAACA
ACACGTCTGAACTCTTTCCCTACACGACGCTCTTCCGATCTNNNAGCCTTAGGGCCCACACATCCATCGC
AGCACACGTCTGAACTCTTTCCCTACACGACGCTCTTCCGATCTNNNCAAATAGTCTCACAGGGAAGCC
AGCACACGTCTGAACTCTTTCCCTACACGACGCTCTTCCGATCTNNNAAGGTGAGCCAGCTTCTCATTC
AGCACACGTCTGAACTCTTTCCCTACACGACGCTCTTCCGATCTNNNATCTCAGCCAAGTTGTGCCTATC
AGAGCACACGTCTGAACTCTTTCCCTACACGACGCTCTTCCGATCTNNNAAGATGCCATGAGAGCAAAG
GAAGAGCACACGTCTGAACTCTTTCCCTACACGACGCTCTTCCGATCTNNNTCTGTACTGAAGCTACAT
AGAGCACACGTCTGAACTCTTTCCCTACACGACGCTCTTCCGATCTNNNACTCAGAGTTTCCCTGATTCC
AGCACACGTCTGAACTCTTTCCCTACACGACGCTCTTCCGATCTNNNCCCCAACTGACATTCTAAGGAC
GAGCACACGTCTGAACTCTTTCCCTACACGACGCTCTTCCGATCTNNNCGGTCCTGGACACTAAACAAA
AAGAGCACACGTCTGAACTCTTTCCCTACACGACGCTCTTCCGATCTNNNTGCCAATGTAGCCATGATA
AGAGCACACGTCTGAACTCTTTCCCTACACGACGCTCTTCCGATCTNNNGAATTGCCTGTGAATGCCCAC
AGCACACGTCTGAACTCTTTCCCTACACGACGCTCTTCCGATCTNNNGCAAAGGACAGTTCTGCCTAAG
GCACACGTCTGAACTCTTTCCCTACACGACGCTCTTCCGATCTNNNCAGCTACCACCCTTCATCAATCCCA
GAGCACACGTCTGAACTCTTTCCCTACACGACGCTCTTCCGATCTNNNATGATGTGGTATGTATGTGTG
AGAGCACACGTCTGAACTCTTTCCCTACACGACGCTCTTCCGATCTNNNCAAAGAAGGTTCTGTGCTTA
GAGCACACGTCTGAACTCTTTCCCTACACGACGCTCTTCCGATCTNNNCCTATTTCCCAACTCCACTGCTC
AGCACACGTCTGAACTCTTTCCCTACACGACGCTCTTCCGATCTNNNCCATTCTATTGCAGGGCTCTCTC
CACACGTCTGAACTCTTTCCCTACACGACGCTCTTCCGATCTNNNGGAGCAGAAACTTTGCGTCTCTACA
GAGCACACGTCTGAACTCTTTCCCTACACGACGCTCTTCCGATCTNNNTATTTGATGCCAAACAGAGGG
GCACACGTCTGAACTCTTTCCCTACACGACGCTCTTCCGATCTNNNCCTTTCCCCTCAACGTGTACAATC
GAGCACACGTCTGAACTCTTTCCCTACACGACGCTCTTCCGATCTNNNTTGAAGGTGAACCCAGAAGA
GAGCACACGTCTGAACTCTTTCCCTACACGACGCTCTTCCGATCTNNNGGTGACAGAGCTGGACACTGT
GAGCACACGTCTGAACTCTTTCCCTACACGACGCTCTTCCGATCTNNNTATTGCCCTGAGCAGAGATGA
GAGCACACGTCTGAACTCTTTCCCTACACGACGCTCTTCCGATCTNNNTGCCAGAACAAATGCACTAGG
AAGAGCACACGTCTGAACTCTTTCCCTACACGACGCTCTTCCGATCTNNNTCAATTATGGTGCTCTCGA
AGAGCACACGTCTGAACTCTTTCCCTACACGACGCTCTTCCGATCTNNNGCAAAGAAACACTCGGAGTT
AGAGCACACGTCTGAACTCTTTCCCTACACGACGCTCTTCCGATCTNNNTCTTCGTCTTATGCTTTCACCT
GAGCACACGTCTGAACTCTTTCCCTACACGACGCTCTTCCGATCTNNNCAGTATCTGCCTCCATCACCTTC
GAGCACACGTCTGAACTCTTTCCCTACACGACGCTCTTCCGATCTNNNGCTTGGAAGAAAGTTCCGTG
GAGCACACGTCTGAACTCTTTCCCTACACGACGCTCTTCCGATCTNNNTGACCTGTCCATTTTGATGCA
AGAGCACACGTCTGAACTCTTTCCCTACACGACGCTCTTCCGATCTNNNAGGGGAAAGTGAGAGTTGA
AGCACACGTCTGAACTCTTTCCCTACACGACGCTCTTCCGATCTNNNAAGTCCTCTTTGGCAGATACAG
AGAGCACACGTCTGAACTCTTTCCCTACACGACGCTCTTCCGATCTNNNTCCTGTTTCTTCCTCTTCTCTG
GAGCACACGTCTGAACTCTTTCCCTACACGACGCTCTTCCGATCTNNNCCAAATTACCTTAGGTGAGGT
GCACACGTCTGAACTCTTTCCCTACACGACGCTCTTCCGATCTNNNAGAAGGATTGGGTCAGGTGATC
GAGCACACGTCTGAACTCTTTCCCTACACGACGCTCTTCCGATCTNNNTCAAGTGTGTCCCAAAGAACA
GAGCACACGTCTGAACTCTTTCCCTACACGACGCTCTTCCGATCTNNNAGACAGACACCTCCTTATTGGC
AAGAGCACACGTCTGAACTCTTTCCCTACACGACGCTCTTCCGATCTNNNACCTCCAAAAGCTTCCTCCT
AGCACACGTCTGAACTCTTTCCCTACACGACGCTCTTCCGATCTNNNCCACCTCTTTGGTCCACTTCTCTC
AGCACACGTCTGAACTCTTTCCCTACACGACGCTCTTCCGATCTNNNAAACGTTTTCTGGAGGTCCTTG
AGCACACGTCTGAACTCTTTCCCTACACGACGCTCTTCCGATCTNNNAGCTCATCTCCAGTCTATACCCA

AGCACACGTCTGAACTCTTTCCCTACACGACGCTCTTCCGATCTNNNGTCTATGAAGGTCTGCGGATT
AGCACACGTCTGAACTCTTTCCCTACACGACGCTCTTCCGATCTNNNACAAATAATGCACCCACATCC
AGCACACGTCTGAACTCTTTCCCTACACGACGCTCTTCCGATCTNNNTACAAAGAGCAGCACGAGATA
AGAGCACACGTCTGAACTCTTTCCCTACACGACGCTCTTCCGATCTNNNTGCTGAGTGAAACCAAATC
AGAGCACACGTCTGAACTCTTTCCCTACACGACGCTCTTCCGATCTNNNAATTTTGCCGCATTTGCTTT
AGCACACGTCTGAACTCTTTCCCTACACGACGCTCTTCCGATCTNNNGCTAATGGAAGAGACTTCTG
GCACACGTCTGAACTCTTTCCCTACACGACGCTCTTCCGATCTNNNGCCTAAGTCCAGAAGCAGGTT
AGCACACGTCTGAACTCTTTCCCTACACGACGCTCTTCCGATCTNNNGGTTGTAAACTCAACTCGGAG
AGCACACGTCTGAACTCTTTCCCTACACGACGCTCTTCCGATCTNNNACCAATAACCCAAACTGTATC
AAGAGCACACGTCTGAACTCTTTCCCTACACGACGCTCTTCCGATCTNNNTGTCTTGGGTGTTTGGTT
GAGCACACGTCTGAACTCTTTCCCTACACGACGCTCTTCCGATCTNNNAAGCTCTGAATTGGACTGTC
GAGCACACGTCTGAACTCTTTCCCTACACGACGCTCTTCCGATCTNNNATTGACCATGTTGAGTTCGG
AGCACACGTCTGAACTCTTTCCCTACACGACGCTCTTCCGATCTNNNTATATTCCATACCCCTCCCACC
AGCACACGTCTGAACTCTTTCCCTACACGACGCTCTTCCGATCTNNNCCTAAATGTGCTTTTTGCATG
AAGAGCACACGTCTGAACTCTTTCCCTACACGACGCTCTTCCGATCTNNNTCAATTTCATCTACCTGC
AGCACACGTCTGAACTCTTTCCCTACACGACGCTCTTCCGATCTNNNCCTCCTCCTTCCATTGCTTTCTC
AAGAGCACACGTCTGAACTCTTTCCCTACACGACGCTCTTCCGATCTNNNTTTCACTTTGCTTGGGAG
AGAGCACACGTCTGAACTCTTTCCCTACACGACGCTCTTCCGATCTNNNGCAGAAACATCCTCCTTCAG
GCACACGTCTGAACTCTTTCCCTACACGACGCTCTTCCGATCTNNNTGGTTTCCAGTACTCCTGGTTCT
GCACACGTCTGAACTCTTTCCCTACACGACGCTCTTCCGATCTNNNGAGCCATGTTTGCAACTGAGGT
GAGCACACGTCTGAACTCTTTCCCTACACGACGCTCTTCCGATCTNNNACCCACTACAGGATATCTGGC
AGCACACGTCTGAACTCTTTCCCTACACGACGCTCTTCCGATCTNNNAGCCTAGTCTAACATCACCACA
GAGCACACGTCTGAACTCTTTCCCTACACGACGCTCTTCCGATCTNNNCTGGAATGCACTGTGAAGC
GCACACGTCTGAACTCTTTCCCTACACGACGCTCTTCCGATCTNNNCTTGGATCTCTCCCTCTCTCCC
AGCACACGTCTGAACTCTTTCCCTACACGACGCTCTTCCGATCTNNNCTGGTGGTCACATAGGGAAT
AGCACACGTCTGAACTCTTTCCCTACACGACGCTCTTCCGATCTNNNCCATGACTTGACCAGCCTAAC
GCACACGTCTGAACTCTTTCCCTACACGACGCTCTTCCGATCTNNNTGGTAGGTGCACAGTGAATGT
AGAGCACACGTCTGAACTCTTTCCCTACACGACGCTCTTCCGATCTNNNTCTCCATTGATTCTTGATTC
GAGCACACGTCTGAACTCTTTCCCTACACGACGCTCTTCCGATCTNNNTGGGGGATTGTTTTTGTGCT
GCACACGTCTGAACTCTTTCCCTACACGACGCTCTTCCGATCTNNNAGCACAGGCAGAATTTCTTCAA
GCACACGTCTGAACTCTTTCCCTACACGACGCTCTTCCGATCTNNNCATTCCAGTGCATCATCTTGTGC
AGCACACGTCTGAACTCTTTCCCTACACGACGCTCTTCCGATCTNNNCGTCAGCACAAGAAATTGCTC
GCACACGTCTGAACTCTTTCCCTACACGACGCTCTTCCGATCTNNNATCCCCCTGTATTCTGTAGGCAT
GAGCACACGTCTGAACTCTTTCCCTACACGACGCTCTTCCGATCTNNNAGAAAAGTACTCCACGACAA
AGCACACGTCTGAACTCTTTCCCTACACGACGCTCTTCCGATCTNNNTGGCTTCCTCCTGGTTATGTA
GAGCACACGTCTGAACTCTTTCCCTACACGACGCTCTTCCGATCTNNNTGCATCTAATTCTGAGGGAC
AGCACACGTCTGAACTCTTTCCCTACACGACGCTCTTCCGATCTNNNGTCATTTTGTACACGGATGTG
AAGAGCACACGTCTGAACTCTTTCCCTACACGACGCTCTTCCGATCTNNNGGCCTCTGATTATCTTCTC
AGCACACGTCTGAACTCTTTCCCTACACGACGCTCTTCCGATCTNNNTTTCTGGTGGGATATATCTGG
GAGCACACGTCTGAACTCTTTCCCTACACGACGCTCTTCCGATCTNNNCAAGGAAGAAGGGACAGG
GCACACGTCTGAACTCTTTCCCTACACGACGCTCTTCCGATCTNNNGGCGACAGAGCTACTCTCAAAA
AGCACACGTCTGAACTCTTTCCCTACACGACGCTCTTCCGATCTNNNTGGCTTCCTCCTGGTTATGTAG
GAGCACACGTCTGAACTCTTTCCCTACACGACGCTCTTCCGATCTNNNACTGGAATGGTGGAAAAAG

AGCACACGTCTGAACTCTTTCCCTACACGACGCTCTTCCGATCTNNNTCAATCACTGAAACGAGTTACT
AGCACACGTCTGAACTCTTTCCCTACACGACGCTCTTCCGATCTNNNTTAGGGGATGCTCCTATCACT
AGCACACGTCTGAACTCTTTCCCTACACGACGCTCTTCCGATCTNNNTGGTAAGTGGTCTCTGGGTTA
GAGCACACGTCTGAACTCTTTCCCTACACGACGCTCTTCCGATCTNNNGCTACCCTAGTTTGGCTTTCT
GAGCACACGTCTGAACTCTTTCCCTACACGACGCTCTTCCGATCTNNNGCTACAGAGTTGTGATCATG
GCACACGTCTGAACTCTTTCCCTACACGACGCTCTTCCGATCTNNNCATCTGCAGCACCTCTATTTCCA
GAGCACACGTCTGAACTCTTTCCCTACACGACGCTCTTCCGATCTNNNAGGGAGGGATGATCAAGTTG
AAGAGCACACGTCTGAACTCTTTCCCTACACGACGCTCTTCCGATCTNNNGTGAACACTCCCAGGCTAT
GCACACGTCTGAACTCTTTCCCTACACGACGCTCTTCCGATCTNNNTCAAAGTTTGGTGGGGAAAACC
AGCACACGTCTGAACTCTTTCCCTACACGACGCTCTTCCGATCTNNNCTGCCTCTTGAATCCAGAACAG
GGAAGAGCACACGTCTGAACTCTTTCCCTACACGACGCTCTTCCGATCTNNNAAGTTTCCTAGGAGGG
AGCACACGTCTGAACTCTTTCCCTACACGACGCTCTTCCGATCTNNNCCATATCACTCCCCAAATGCGA
AGAGCACACGTCTGAACTCTTTCCCTACACGACGCTCTTCCGATCTNNNGCTAGAAGACCTGGGGGC
AGCACACGTCTGAACTCTTTCCCTACACGACGCTCTTCCGATCTNNNTGGCTTCTCTTCTCTCAATCTCT
GCACACGTCTGAACTCTTTCCCTACACGACGCTCTTCCGATCTNNNTGGTTTCAATGGTCTAAGGCTG
AGCACACGTCTGAACTCTTTCCCTACACGACGCTCTTCCGATCTNNNTTTGGAATCCAGATCACCCTCC
GAGCACACGTCTGAACTCTTTCCCTACACGACGCTCTTCCGATCTNNNAAAGAAACTCTCTCCTCCCTCC
AGCACACGTCTGAACTCTTTCCCTACACGACGCTCTTCCGATCTNNNTACTTCTGGGAGAAGCGCTTG
AGCACACGTCTGAACTCTTTCCCTACACGACGCTCTTCCGATCTNNNAGCTTCAGAGAAGATCTAGGC
AGCACACGTCTGAACTCTTTCCCTACACGACGCTCTTCCGATCTNNNTCCCGAGAGGCAACCAGTATT
GAGCACACGTCTGAACTCTTTCCCTACACGACGCTCTTCCGATCTNNNTCAAAGGAACATTGACCTGAG
AGCACACGTCTGAACTCTTTCCCTACACGACGCTCTTCCGATCTNNNAGAAACATTGAGAGATAGACA
AAGAGCACACGTCTGAACTCTTTCCCTACACGACGCTCTTCCGATCTNNNGAGCCTGGACAAATTCCC
AGCACACGTCTGAACTCTTTCCCTACACGACGCTCTTCCGATCTNNNTGGCGACACCAATTTCCAACTT
AGCACACGTCTGAACTCTTTCCCTACACGACGCTCTTCCGATCTNNNGACAAATACCCCAGTTTCCCTC
AGCACACGTCTGAACTCTTTCCCTACACGACGCTCTTCCGATCTNNNGTCTGCCTCCCCATCATACATTC
GCACACGTCTGAACTCTTTCCCTACACGACGCTCTTCCGATCTNNNCCTCCTGGAATGACACGTTTCTC
AGAGCACACGTCTGAACTCTTTCCCTACACGACGCTCTTCCGATCTNNNTGAAGCAGGCTCTTGTTTC
AGAGCACACGTCTGAACTCTTTCCCTACACGACGCTCTTCCGATCTNNNGGTGTTTTTCCAGCAGCAA
GCACACGTCTGAACTCTTTCCCTACACGACGCTCTTCCGATCTNNNAGGAAATGTACAAGTTGCTTTG
GCACACGTCTGAACTCTTTCCCTACACGACGCTCTTCCGATCTNNNACATTGGAAACACGATTGGATT
AGCACACGTCTGAACTCTTTCCCTACACGACGCTCTTCCGATCTNNNATGCTAGCTCAGCCTCTAGACA
AGAGCACACGTCTGAACTCTTTCCCTACACGACGCTCTTCCGATCTNNNCAGGGCTAACCAGAAAAG
AGAGCACACGTCTGAACTCTTTCCCTACACGACGCTCTTCCGATCTNNNACTTCCGCCTTTCCATTTCA
AGCACACGTCTGAACTCTTTCCCTACACGACGCTCTTCCGATCTNNNATATCCTGACAACACACACCCT
AGCACACGTCTGAACTCTTTCCCTACACGACGCTCTTCCGATCTNNNGCACTTATGTCAAGCAGGGAT
CACACGTCTGAACTCTTTCCCTACACGACGCTCTTCCGATCTNNNCCATTTGCGTTATCAGGCTCTCTCA
GAGCACACGTCTGAACTCTTTCCCTACACGACGCTCTTCCGATCTNNNGGACATCTTGTGTTGGCATTT
AGCACACGTCTGAACTCTTTCCCTACACGACGCTCTTCCGATCTNNNTGGAAGAGATCTTCGAGGGAA
GAGCACACGTCTGAACTCTTTCCCTACACGACGCTCTTCCGATCTNNNGCCACTCAAGCAGAATCTCTA
AGAGCACACGTCTGAACTCTTTCCCTACACGACGCTCTTCCGATCTNNNGCTTGGAAGAAAGTTCCGT
GAGCACACGTCTGAACTCTTTCCCTACACGACGCTCTTCCGATCTNNNCAGCTCTTTCCTGTTACCCTTC
AGCACACGTCTGAACTCTTTCCCTACACGACGCTCTTCCGATCTNNNACAGCTATAGGTACGGAAGT

AAGAGCACACGTCTGAACTCTTTCCCTACACGACGCTCTTCCGATCTNNNTTTGCACAGCACTCCTCTAA

AGCACACGTCTGAACTCTTTCCCTACACGACGCTCTTCCGATCTNNNTATGAAGCCAGCCTATATGCCC

AGCACACGTCTGAACTCTTTCCCTACACGACGCTCTTCCGATCTNNNGCTCTCTATCTGGGCAGCAAAA

AGCACACGTCTGAACTCTTTCCCTACACGACGCTCTTCCGATCTNNNATATTTGGGACACAGAACCTGC

GAGCACACGTCTGAACTCTTTCCCTACACGACGCTCTTCCGATCTNNNATGTTGGTCATGTTCTGCTTC

AGCACACGTCTGAACTCTTTCCCTACACGACGCTCTTCCGATCTNNNGCCAAGTTCTCATGTGCTCTGA

GAGCACACGTCTGAACTCTTTCCCTACACGACGCTCTTCCGATCTNNNAAAGCATAAAGGAGCCTTGCA

GAGCACACGTCTGAACTCTTTCCCTACACGACGCTCTTCCGATCTNNNCAGGGCTCTGCTACAAATGTG

GAGCACACGTCTGAACTCTTTCCCTACACGACGCTCTTCCGATCTNNNATATTTGGGACACAGAACCTG

GAGCACACGTCTGAACTCTTTCCCTACACGACGCTCTTCCGATCTNNNTGACTTTCTTTTCATCTTGCAT

AGCACACGTCTGAACTCTTTCCCTACACGACGCTCTTCCGATCTNNNACATTCTTCCCTGACTGTAAAGC

AGAGCACACGTCTGAACTCTTTCCCTACACGACGCTCTTCCGATCTNNNTTGATTGATGCACCTTCTCA

AGCACACGTCTGAACTCTTTCCCTACACGACGCTCTTCCGATCTNNNTCAGCAAACCCCAAAAAGAATG

AGAGCACACGTCTGAACTCTTTCCCTACACGACGCTCTTCCGATCTNNNACACAACTGATTTTGTGGGT

AGCACACGTCTGAACTCTTTCCCTACACGACGCTCTTCCGATCTNNNCCTCAACTCACCTCACTCCCTATCA

AGAGCACACGTCTGAACTCTTTCCCTACACGACGCTCTTCCGATCTNNNCTGCGCTGATAAGATGCCAT

GAGCACACGTCTGAACTCTTTCCCTACACGACGCTCTTCCGATCTNNNGGTCCATGAAGGCCAAGTAAC

GAGCACACGTCTGAACTCTTTCCCTACACGACGCTCTTCCGATCTNNNTTCCTCTCTCTTCTCCTCCTGGT

AAGAGCACACGTCTGAACTCTTTCCCTACACGACGCTCTTCCGATCTNNNTGGGCTAAGACTTCCTAGG

AGAGCACACGTCTGAACTCTTTCCCTACACGACGCTCTTCCGATCTNNNGTTGCCTACTTAGGGTTCTGA

CACACGTCTGAACTCTTTCCCTACACGACGCTCTTCCGATCTNNNTCAAACTCCACTCTGCTTCTGACCAT

AGAGCACACGTCTGAACTCTTTCCCTACACGACGCTCTTCCGATCTNNNATTCAGCTGACAGCCAATTCC

AGCACACGTCTGAACTCTTTCCCTACACGACGCTCTTCCGATCTNNNTATCTGGTCTCTGGGGTCATTGC

AGAGCACACGTCTGAACTCTTTCCCTACACGACGCTCTTCCGATCTNNNGTTTGGAGCCACAGAGGATT

GAGCACACGTCTGAACTCTTTCCCTACACGACGCTCTTCCGATCTNNNACTGCTGATTACAGGTTGTGT

AGCACACGTCTGAACTCTTTCCCTACACGACGCTCTTCCGATCTNNNGAGGTCCAAACGGAAGAAAGC

AGCACACGTCTGAACTCTTTCCCTACACGACGCTCTTCCGATCTNNNATAAACTTGAGGTAGGCCAGG

AGAGCACACGTCTGAACTCTTTCCCTACACGACGCTCTTCCGATCTNNNTCTGTAAGTACTTTGCCTGTC

AGAGCACACGTCTGAACTCTTTCCCTACACGACGCTCTTCCGATCTNNNGTGGTTTTCCAGGGACAAAA

AGCACACGTCTGAACTCTTTCCCTACACGACGCTCTTCCGATCTNNNCACCCAGGCCTTGTGCATATAAT

GAGCACACGTCTGAACTCTTTCCCTACACGACGCTCTTCCGATCTNNNTTTCAGCCTCTCTACTCTGCTG

AGCACACGTCTGAACTCTTTCCCTACACGACGCTCTTCCGATCTNNNAGTTTTGCTATTTTCTTCTGTGT

AGAGCACACGTCTGAACTCTTTCCCTACACGACGCTCTTCCGATCTNNNTGCAAGTTCGTCCCTGTAACT

GAGCACACGTCTGAACTCTTTCCCTACACGACGCTCTTCCGATCTNNNTTCAGCATAAGGCAGTTTGGG

AGCACACGTCTGAACTCTTTCCCTACACGACGCTCTTCCGATCTNNNAGGCATGACAGTTCTAGCCACT

GAGCACACGTCTGAACTCTTTCCCTACACGACGCTCTTCCGATCTNNNTAAGCTCCACCCCCTAAAGATC

GAGCACACGTCTGAACTCTTTCCCTACACGACGCTCTTCCGATCTNNNTGGATGGTTCAGGGATAGTTC

AGCACACGTCTGAACTCTTTCCCTACACGACGCTCTTCCGATCTNNNTCAGGCCCCTTCTGTCAATGTAC

AAGAGCACACGTCTGAACTCTTTCCCTACACGACGCTCTTCCGATCTNNNATAGTTTGTGAGCCACTGC

AGAGCACACGTCTGAACTCTTTCCCTACACGACGCTCTTCCGATCTNNNATGGATAAGTCTGCAAGTAC

GAGCACACGTCTGAACTCTTTCCCTACACGACGCTCTTCCGATCTNNNATGATGGCACTGGAGTTCTTTC

GAAGAGCACACGTCTGAACTCTTTCCCTACACGACGCTCTTCCGATCTNNNGTGGGGACACAGTAACA

GAGCACACGTCTGAACTCTTTCCCTACACGACGCTCTTCCGATCTNNNAGAATAGGGAAGGGGAAGC

GAGCACACGTCTGAACTCTTTCCCTACACGACGCTCTTCCGATCTNNNTATGGCATGATTTGAGTGCT
AGAGCACACGTCTGAACTCTTTCCCTACACGACGCTCTTCCGATCTNNNCCTAACTTGCAGAGGGACA
AGCACACGTCTGAACTCTTTCCCTACACGACGCTCTTCCGATCTNNNCAATTTGAATCTGGGCTTGGCT
GCACACGTCTGAACTCTTTCCCTACACGACGCTCTTCCGATCTNNNAATCAAGCTGGAATTAGGCCTG
AGCACACGTCTGAACTCTTTCCCTACACGACGCTCTTCCGATCTNNNTTCAACTTGGCCCACTTGGAAA
GCACACGTCTGAACTCTTTCCCTACACGACGCTCTTCCGATCTNNNTGCTTAGAAGTCCTTGGTGTTCT
AGCACACGTCTGAACTCTTTCCCTACACGACGCTCTTCCGATCTNNNCTGGAATGCACTGTGAAGCTA
AGCACACGTCTGAACTCTTTCCCTACACGACGCTCTTCCGATCTNNNTTTCTTTTGGTGGCAGGGAGT
AGCACACGTCTGAACTCTTTCCCTACACGACGCTCTTCCGATCTNNNACAGTACCATTTCACAGCCCAA
CACACGTCTGAACTCTTTCCCTACACGACGCTCTTCCGATCTNNNCCATCAGCTTGCTTTGGTCTGTAC
GAGCACACGTCTGAACTCTTTCCCTACACGACGCTCTTCCGATCTNNNAGTATTCACGGACATTGCATC
GAGCACACGTCTGAACTCTTTCCCTACACGACGCTCTTCCGATCTNNNATGCCCCAGAGATCTGGTTTT
GCACACGTCTGAACTCTTTCCCTACACGACGCTCTTCCGATCTNNNACCTACTTTTCTGCTTTCCCCAC
AGCACACGTCTGAACTCTTTCCCTACACGACGCTCTTCCGATCTNNNTGGGTAAACCTTTCTTTAGAA
GCACACGTCTGAACTCTTTCCCTACACGACGCTCTTCCGATCTNNNCAAGTCTTGACCTACAGCTGACC
GAGCACACGTCTGAACTCTTTCCCTACACGACGCTCTTCCGATCTNNNAGTGGAGATTTTGAAGGCA
AGAGCACACGTCTGAACTCTTTCCCTACACGACGCTCTTCCGATCTNNNTGTGTTCTACAAATTTTCCA
AGCACACGTCTGAACTCTTTCCCTACACGACGCTCTTCCGATCTNNNGTGCTTAGGCTTACTCAGTGCT
AGCACACGTCTGAACTCTTTCCCTACACGACGCTCTTCCGATCTNNNGGGGTAGCGATCCTGTCATTT
AAGAGCACACGTCTGAACTCTTTCCCTACACGACGCTCTTCCGATCTNNNAGTCCTGTCTCCGATACTT
AGCACACGTCTGAACTCTTTCCCTACACGACGCTCTTCCGATCTNNNTATAATGCTTACGGACCCAGTG
AGCACACGTCTGAACTCTTTCCCTACACGACGCTCTTCCGATCTNNNTAGGTACTCACACCTCCCTTTC
GCACACGTCTGAACTCTTTCCCTACACGACGCTCTTCCGATCTNNNGGCAAGTAGAATGCTGATGGGA
GAGCACACGTCTGAACTCTTTCCCTACACGACGCTCTTCCGATCTNNNGGGGAAGATATCTGCATCTG
GCACACGTCTGAACTCTTTCCCTACACGACGCTCTTCCGATCTNNNACTTTACTATTTGGAAGAAACGT
AGCACACGTCTGAACTCTTTCCCTACACGACGCTCTTCCGATCTNNNGTGGTTGGCTATCTGCCTTTCTT
AGCACACGTCTGAACTCTTTCCCTACACGACGCTCTTCCGATCTNNNGTTGGGCCTTAGGAGTAAATG
AGCACACGTCTGAACTCTTTCCCTACACGACGCTCTTCCGATCTNNNCAGCTCAGAGTGTAGGTTAGG
GAGCACACGTCTGAACTCTTTCCCTACACGACGCTCTTCCGATCTNNNTATGTTTCCCCTTCACCCTAGC
GCACACGTCTGAACTCTTTCCCTACACGACGCTCTTCCGATCTNNNGGACCTCAGAGAGACCTGAAAG
GAGCACACGTCTGAACTCTTTCCCTACACGACGCTCTTCCGATCTNNNGGGTTTGGCTTGGAGCTAATG
AGAGCACACGTCTGAACTCTTTCCCTACACGACGCTCTTCCGATCTNNNTGAGGAATTTCTGGATGTG
AGCACACGTCTGAACTCTTTCCCTACACGACGCTCTTCCGATCTNNNTGAAGCATGCCTACAAGATTT
GCACACGTCTGAACTCTTTCCCTACACGACGCTCTTCCGATCTNNNGGGTTGTAAGGGTGTGTTGATG
AGAGCACACGTCTGAACTCTTTCCCTACACGACGCTCTTCCGATCTNNNCCACATGGCTTAGTACTCAAT
AAGAGCACACGTCTGAACTCTTTCCCTACACGACGCTCTTCCGATCTNNNCCAGCCCCTGAAAAGATTT
GAGCACACGTCTGAACTCTTTCCCTACACGACGCTCTTCCGATCTNNNTAGCCATGGTTTCCTTCATCTG
GAGCACACGTCTGAACTCTTTCCCTACACGACGCTCTTCCGATCTNNNGCTCGGGTTTCTAAGGGAGA
AGAGCACACGTCTGAACTCTTTCCCTACACGACGCTCTTCCGATCTNNNTGAAGTTGGCTCCTGTATCC
AGCACACGTCTGAACTCTTTCCCTACACGACGCTCTTCCGATCTNNNAGGCTTTTAGGCAGTCTAGTCC
AGCACACGTCTGAACTCTTTCCCTACACGACGCTCTTCCGATCTNNNCTGTTCCCACCAGATCACAAGAG
AGCACACGTCTGAACTCTTTCCCTACACGACGCTCTTCCGATCTNNNTATTTGCCTGGCACGAACAAAA
GCACACGTCTGAACTCTTTCCCTACACGACGCTCTTCCGATCTNNNCCATCAGCTTGCTTTGGTCTGTAC

GAGCACACGTCTGAACTCTTTCCCTACACGACGCTCTTCCGATCTNNNTCGGTTTTATAGGCCAACAAG
AGCACACGTCTGAACTCTTTCCCTACACGACGCTCTTCCGATCTNNNTATTCCTGTGTGCATGCCTACCC
GAGCACACGTCTGAACTCTTTCCCTACACGACGCTCTTCCGATCTNNNTAAAGTTGCACTGTGACTGGG
GAGCACACGTCTGAACTCTTTCCCTACACGACGCTCTTCCGATCTNNNGAACAGGATTCAGGCTCTGTC
GAGCACACGTCTGAACTCTTTCCCTACACGACGCTCTTCCGATCTNNNTTTGTTTATGTTATGACGCATG
GAGCACACGTCTGAACTCTTTCCCTACACGACGCTCTTCCGATCTNNNAGAAACATGTAGGAGACGAA
AGAGCACACGTCTGAACTCTTTCCCTACACGACGCTCTTCCGATCTNNNAGTGGAGATTTTGAAGGCAG
GAGCACACGTCTGAACTCTTTCCCTACACGACGCTCTTCCGATCTNNNTTTTGGGTTGATGGGTTCCAG
AGAGCACACGTCTGAACTCTTTCCCTACACGACGCTCTTCCGATCTNNNTTTTGGGTTGATGGGTTCCA
AGAGCACACGTCTGAACTCTTTCCCTACACGACGCTCTTCCGATCTNNNTGAGGTGATTGGTTATGAAT
GAGCACACGTCTGAACTCTTTCCCTACACGACGCTCTTCCGATCTNNNGTGAACCTCATACAAAACCCCC
AGCACACGTCTGAACTCTTTCCCTACACGACGCTCTTCCGATCTNNNCTATGCCAAGCACTCATTCAGGT
GAGCACACGTCTGAACTCTTTCCCTACACGACGCTCTTCCGATCTNNNCTGCTGCCTCATAACCCTACAT
GAGCACACGTCTGAACTCTTTCCCTACACGACGCTCTTCCGATCTNNNTCCTGCTTCTTGTTTGACTTCCC
AGCACACGTCTGAACTCTTTCCCTACACGACGCTCTTCCGATCTNNNGTCCCTGGCAAGCATGAATTTA
GAGCACACGTCTGAACTCTTTCCCTACACGACGCTCTTCCGATCTNNNACAAGACTATCCCTGATGTCTC
AGCACACGTCTGAACTCTTTCCCTACACGACGCTCTTCCGATCTNNNGGGTTGTAAGGGTGTGTTGAT
GAGCACACGTCTGAACTCTTTCCCTACACGACGCTCTTCCGATCTNNNTGAACAGAACCTCTCCTCTTCG
GCACACGTCTGAACTCTTTCCCTACACGACGCTCTTCCGATCTNNNAGGCTCAAAGTCTAACGCCAATCG
GAGCACACGTCTGAACTCTTTCCCTACACGACGCTCTTCCGATCTNNNTAGGGGTGAAGGACATCTGAA
AGCACACGTCTGAACTCTTTCCCTACACGACGCTCTTCCGATCTNNNTGCTAACACTTGGCATCAAGTCA
GAGCACACGTCTGAACTCTTTCCCTACACGACGCTCTTCCGATCTNNNAATAGGATGGATGTGTGCGT
GCACACGTCTGAACTCTTTCCCTACACGACGCTCTTCCGATCTNNNAGCTGAGGAGCTCACAGACTAAA
AGCACACGTCTGAACTCTTTCCCTACACGACGCTCTTCCGATCTNNNCCGACTTAGGAGCAAACAATCCA
GCACACGTCTGAACTCTTTCCCTACACGACGCTCTTCCGATCTNNNTGGAACCATGGTCTTTTCCCTTGC
AGAGCACACGTCTGAACTCTTTCCCTACACGACGCTCTTCCGATCTNNNACCTTGTTGGAGTACAGAAG
AAGAGCACACGTCTGAACTCTTTCCCTACACGACGCTCTTCCGATCTNNNTGGATGTTGAAATGGCCTC
AGCACACGTCTGAACTCTTTCCCTACACGACGCTCTTCCGATCTNNNGCAACTCCCTTTCCTTACCTAGCC
AGAGCACACGTCTGAACTCTTTCCCTACACGACGCTCTTCCGATCTNNNCCAAGAGCCTTTTGTTCAGGT
GAGCACACGTCTGAACTCTTTCCCTACACGACGCTCTTCCGATCTNNNACAATCAAGAGTGCCATTAGC
AGCACACGTCTGAACTCTTTCCCTACACGACGCTCTTCCGATCTNNNAAACTGGCTGAATGAGACAGAG
AGAGCACACGTCTGAACTCTTTCCCTACACGACGCTCTTCCGATCTNNNACATACACAGAGTGAGCGAG
AGCACACGTCTGAACTCTTTCCCTACACGACGCTCTTCCGATCTNNNAGATTCCAGAGCCAGAGGTGAA
AGAGCACACGTCTGAACTCTTTCCCTACACGACGCTCTTCCGATCTNNNGCTTTGGGGAAGTTTGTTGC
GAGCACACGTCTGAACTCTTTCCCTACACGACGCTCTTCCGATCTNNNTTTGGACAGCATCAAGGAGCT
AGAGCACACGTCTGAACTCTTTCCCTACACGACGCTCTTCCGATCTNNNTGGTGTTTTAGCTTTCCTCTT
GAGCACACGTCTGAACTCTTTCCCTACACGACGCTCTTCCGATCTNNNAGTCCAACAGGAAAATATCAG
AGAGCACACGTCTGAACTCTTTCCCTACACGACGCTCTTCCGATCTNNNCTACTTCCACGTCCAAACTTC
AGAGCACACGTCTGAACTCTTTCCCTACACGACGCTCTTCCGATCTNNNTGGCAATGGTGAGGAAACA
AGCACACGTCTGAACTCTTTCCCTACACGACGCTCTTCCGATCTNNNAAGAGTTACTGGGTCACCACTGA
AGCACACGTCTGAACTCTTTCCCTACACGACGCTCTTCCGATCTNNNCCATTTTCCTGTGGGTAGAGGT
GAGCACACGTCTGAACTCTTTCCCTACACGACGCTCTTCCGATCTNNNCTTTCCTCAAGTTCCACCAACCA
GAAGAGCACACGTCTGAACTCTTTCCCTACACGACGCTCTTCCGATCTNNNTGAGGCTGTTCTTCCTGG

AGAGCACACGTCTGAACTCTTTCCCTACACGACGCTCTTCCGATCTNNNTAACAGAGTACCTTAGCCCC
GAGCACACGTCTGAACTCTTTCCCTACACGACGCTCTTCCGATCTNNNTAACTTCTGGCCTCCCAGGTT
AGCACACGTCTGAACTCTTTCCCTACACGACGCTCTTCCGATCTNNNGCCAGCAGTGTACTGTATCAC
GAGCACACGTCTGAACTCTTTCCCTACACGACGCTCTTCCGATCTNNNATCTAATCTCCTGGCTGGAA
GCACACGTCTGAACTCTTTCCCTACACGACGCTCTTCCGATCTNNNTGGAAGTAGAGCTGTCTCTCTG
AGCACACGTCTGAACTCTTTCCCTACACGACGCTCTTCCGATCTNNNTCCCACTATGGAATGATGGG
AAGAGCACACGTCTGAACTCTTTCCCTACACGACGCTCTTCCGATCTNNNTGTAGTTGACAAGGAGA
GAGCACACGTCTGAACTCTTTCCCTACACGACGCTCTTCCGATCTNNNCTATAGCTGCTGGAGCGACT
GAGCACACGTCTGAACTCTTTCCCTACACGACGCTCTTCCGATCTNNNCTTGAGTGACCTGTACCCAA
AGCACACGTCTGAACTCTTTCCCTACACGACGCTCTTCCGATCTNNNTGGAAGAGTGGGTTGGAACA
GAAGAGCACACGTCTGAACTCTTTCCCTACACGACGCTCTTCCGATCTNNNGGATTTGTGGGAGTTG
GAGCACACGTCTGAACTCTTTCCCTACACGACGCTCTTCCGATCTNNNCCTAAAGGTGTCCTGTTTGA
AAGAGCACACGTCTGAACTCTTTCCCTACACGACGCTCTTCCGATCTNNNGTTATTACTGGCTGTTGC
CACACGTCTGAACTCTTTCCCTACACGACGCTCTTCCGATCTNNNAGACCATGGATAAATGGCAACCG
GCACACGTCTGAACTCTTTCCCTACACGACGCTCTTCCGATCTNNNACACAGAATAAACAGCCAGGC
AGCACACGTCTGAACTCTTTCCCTACACGACGCTCTTCCGATCTNNNGGTATCCAATTACATCCACGT
AGCACACGTCTGAACTCTTTCCCTACACGACGCTCTTCCGATCTNNNACTCATTTTGGCATGTTGATTT
AGAGCACACGTCTGAACTCTTTCCCTACACGACGCTCTTCCGATCTNNNTGTAGCACAATTCACAATT
GAGCACACGTCTGAACTCTTTCCCTACACGACGCTCTTCCGATCTNNNTTACCACACATCTGGTTCCTC
GAGCACACGTCTGAACTCTTTCCCTACACGACGCTCTTCCGATCTNNNTAGATACACATGCCCACATGC
GCACACGTCTGAACTCTTTCCCTACACGACGCTCTTCCGATCTNNNCCTCAGTTCCTCATTGTCAAGG
AAGAGCACACGTCTGAACTCTTTCCCTACACGACGCTCTTCCGATCTNNNTGGCAATTTACATACACAC
AGCACACGTCTGAACTCTTTCCCTACACGACGCTCTTCCGATCTNNNATCTGTTTCAGCAATGACACC
AGCACACGTCTGAACTCTTTCCCTACACGACGCTCTTCCGATCTNNNTATCCATCACCTCACACAGGGA
AGCACACGTCTGAACTCTTTCCCTACACGACGCTCTTCCGATCTNNNGGGAAGGGTATGGTGACCAG
GCACACGTCTGAACTCTTTCCCTACACGACGCTCTTCCGATCTNNNTAAACAGTACTGGTCAGTGGC
GAGCACACGTCTGAACTCTTTCCCTACACGACGCTCTTCCGATCTNNNTCCACATGACAAAACACTGG
AGCACACGTCTGAACTCTTTCCCTACACGACGCTCTTCCGATCTNNNTTATGTTCCTCCAAACAGCACC
AGAGCACACGTCTGAACTCTTTCCCTACACGACGCTCTTCCGATCTNNNAGATAGCCCCAGGAAAAG
GCACACGTCTGAACTCTTTCCCTACACGACGCTCTTCCGATCTNNNAAGAAGGGAAATTCCTACGCAC
AGCACACGTCTGAACTCTTTCCCTACACGACGCTCTTCCGATCTNNNTCTCAGTTTACTCACAGAAACG
AGAGCACACGTCTGAACTCTTTCCCTACACGACGCTCTTCCGATCTNNNACAACGGACACCTTATTGA
GAAGAGCACACGTCTGAACTCTTTCCCTACACGACGCTCTTCCGATCTNNNTTGTATACAGGTTGCGC
AGCACACGTCTGAACTCTTTCCCTACACGACGCTCTTCCGATCTNNNACAAGAGCTGCCCTACTTACAG
GCACACGTCTGAACTCTTTCCCTACACGACGCTCTTCCGATCTNNNTAATGGTGATGCCATCTGTGTG
GAGCACACGTCTGAACTCTTTCCCTACACGACGCTCTTCCGATCTNNNTGATGGCATTGAGCAGAAG
GAGCACACGTCTGAACTCTTTCCCTACACGACGCTCTTCCGATCTNNNCTTGAATTCACTGGGTTTGT
AGCACACGTCTGAACTCTTTCCCTACACGACGCTCTTCCGATCTNNNAGTTTATAGTCAGGGTACCTC
AGCACACGTCTGAACTCTTTCCCTACACGACGCTCTTCCGATCTNNNAGAAGACATACACGCACACACT
GAGCACACGTCTGAACTCTTTCCCTACACGACGCTCTTCCGATCTNNNAAGCCTGGTCCATAAGGTTT
GCACACGTCTGAACTCTTTCCCTACACGACGCTCTTCCGATCTNNNTCTATTCATGACCTACCCCCGACC
AGCACACGTCTGAACTCTTTCCCTACACGACGCTCTTCCGATCTNNNCTGCGATCTCCTTGTCATAGAG
AAGAGCACACGTCTGAACTCTTTCCCTACACGACGCTCTTCCGATCTNNNGCAATGGATGTCACAAC

GCACACGTCTGAACTCTTTCCCTACACGACGCTCTTCCGATCTNNNGCTTCAAGGATTGTTGTGCCTAC
GAGCACACGTCTGAACTCTTTCCCTACACGACGCTCTTCCGATCTNNNTCCCATAGAGCCACAGAAGTT
AGCACACGTCTGAACTCTTTCCCTACACGACGCTCTTCCGATCTNNNTAGGAGTTGAAAACACACGCAC
AGAGCACACGTCTGAACTCTTTCCCTACACGACGCTCTTCCGATCTNNNAAGAACAGTGATGGTGAAC
AGCACACGTCTGAACTCTTTCCCTACACGACGCTCTTCCGATCTNNNATTGCTCTTTAGCCACACTGCTT
AGAGCACACGTCTGAACTCTTTCCCTACACGACGCTCTTCCGATCTNNNAGAATTATGGGTGTGTGTG
AGAGCACACGTCTGAACTCTTTCCCTACACGACGCTCTTCCGATCTNNNAGCAGAATACCAACAGTGCT
GAGCACACGTCTGAACTCTTTCCCTACACGACGCTCTTCCGATCTNNNTGCTAAGAGATTAGATTTAC
AGAGCACACGTCTGAACTCTTTCCCTACACGACGCTCTTCCGATCTNNNTAAAAAGCACTGCACAAACC
GAGCACACGTCTGAACTCTTTCCCTACACGACGCTCTTCCGATCTNNNCAAAAACGATATATCAGAGAC
GAGCACACGTCTGAACTCTTTCCCTACACGACGCTCTTCCGATCTNNNTGCCAGTTTATCTTACAGGCAC
AGCACACGTCTGAACTCTTTCCCTACACGACGCTCTTCCGATCTNNNCGTCTTCTGGATGTGGAGGAA
GAGCACACGTCTGAACTCTTTCCCTACACGACGCTCTTCCGATCTNNNCTAAGCTCCACTGATGTGCTCT
GCACACGTCTGAACTCTTTCCCTACACGACGCTCTTCCGATCTNNNAACCTGGACTGCAAATCACACAT
AGCACACGTCTGAACTCTTTCCCTACACGACGCTCTTCCGATCTNNNGTCACCTTCCAGAGAGAGAGAC
AGCACACGTCTGAACTCTTTCCCTACACGACGCTCTTCCGATCTNNNACATGAAAGAACCAATTCAAGT
GCACACGTCTGAACTCTTTCCCTACACGACGCTCTTCCGATCTNNNTGTGGTAATTGTGTACAGCAGGC
AGCACACGTCTGAACTCTTTCCCTACACGACGCTCTTCCGATCTNNNTGTGTGTATAGTTCACAGTGCC
AGAGCACACGTCTGAACTCTTTCCCTACACGACGCTCTTCCGATCTNNNCTGCGATCTCCTTGTCATAGA
AGCACACGTCTGAACTCTTTCCCTACACGACGCTCTTCCGATCTNNNAGGAACTGTGAACACTCCAAAC
AGCACACGTCTGAACTCTTTCCCTACACGACGCTCTTCCGATCTNNNGATGTGGGAGGAAAACATGAC
AGCACACGTCTGAACTCTTTCCCTACACGACGCTCTTCCGATCTNNNAAAAAGCCAGACAGAGGAAAG
GCACACGTCTGAACTCTTTCCCTACACGACGCTCTTCCGATCTNNNCTCCTCCCTCATATCAGCCTCAACA
AGAGCACACGTCTGAACTCTTTCCCTACACGACGCTCTTCCGATCTNNNAATGCATTCACTCAGCAAGT
GCACACGTCTGAACTCTTTCCCTACACGACGCTCTTCCGATCTNNNCAAACCTAGTGCAGACCCAAAGTC
GAGCACACGTCTGAACTCTTTCCCTACACGACGCTCTTCCGATCTNNNATGACTGTTCCACTGGTATGC
GAGCACACGTCTGAACTCTTTCCCTACACGACGCTCTTCCGATCTNNNTGATTACAGCACTGTGTGTGT
AGAGCACACGTCTGAACTCTTTCCCTACACGACGCTCTTCCGATCTNNNCTGATGCCCTTGATGGTTAC
GAAGAGCACACGTCTGAACTCTTTCCCTACACGACGCTCTTCCGATCTNNNTGATGAAGATTCCACTCA
GAGCACACGTCTGAACTCTTTCCCTACACGACGCTCTTCCGATCTNNNAACCAGCTTAGAAGACAAAAG
GAGCACACGTCTGAACTCTTTCCCTACACGACGCTCTTCCGATCTNNNGGAGACAAGATTGAGAGCCAC
GAGCACACGTCTGAACTCTTTCCCTACACGACGCTCTTCCGATCTNNNGTAGGGAAAGGAGACAGAAA
GAGCACACGTCTGAACTCTTTCCCTACACGACGCTCTTCCGATCTNNNAGAGGTCTACTCATTGCTCAC
GCACACGTCTGAACTCTTTCCCTACACGACGCTCTTCCGATCTNNNCCCAGTCTTAGTCCCCACACTCAT
CACACGTCTGAACTCTTTCCCTACACGACGCTCTTCCGATCTNNNTGTCTGTAGGCTTTGTACTGAAAC
GAGCACACGTCTGAACTCTTTCCCTACACGACGCTCTTCCGATCTNNNTCCTACACATTTTCATGTTTTT
GAGCACACGTCTGAACTCTTTCCCTACACGACGCTCTTCCGATCTNNNAAGAAATGTGCTCTACAGCTG
GAGCACACGTCTGAACTCTTTCCCTACACGACGCTCTTCCGATCTNNNAGAGCCACTCCCTGCTTCTAAT
AGAGCACACGTCTGAACTCTTTCCCTACACGACGCTCTTCCGATCTNNNTTGTATACAGGTTGCGCTCT
AGCACACGTCTGAACTCTTTCCCTACACGACGCTCTTCCGATCTNNNGCCTACAACATAGACCATGTCCC
GCACACGTCTGAACTCTTTCCCTACACGACGCTCTTCCGATCTNNNACTTTTTCGGTTCTGGCTGTCTTC
GCACACGTCTGAACTCTTTCCCTACACGACGCTCTTCCGATCTNNNTGAGTTTCTGGTAGGAATTGGA
AAGAGCACACGTCTGAACTCTTTCCCTACACGACGCTCTTCCGATCTNNNTGAATGGAAAAAGAAAAT

AGCACACGTCTGAACTCTTTCCCTACACGACGCTCTTCCGATCTNNNATTTGACCCATGGTCCCTTTAG
GAGCACACGTCTGAACTCTTTCCCTACACGACGCTCTTCCGATCTNNNGACACACCAGTACACACACAC
AGCACACGTCTGAACTCTTTCCCTACACGACGCTCTTCCGATCTNNNCAACAATCTGACTTGCCTCAG
GAGCACACGTCTGAACTCTTTCCCTACACGACGCTCTTCCGATCTNNNTCTGAAATACTAGGGTAGGG
GCACACGTCTGAACTCTTTCCCTACACGACGCTCTTCCGATCTNNNGAACACAGGTATGTGCCATCAC
GAGCACACGTCTGAACTCTTTCCCTACACGACGCTCTTCCGATCTNNNAACTTGCCTGACATAGGATT
AAGAGCACACGTCTGAACTCTTTCCCTACACGACGCTCTTCCGATCTNNNTCAGTTAGCTCTACATGG
GAGCACACGTCTGAACTCTTTCCCTACACGACGCTCTTCCGATCTNNNTGCCCACAATTACACTTCTA
GAGCACACGTCTGAACTCTTTCCCTACACGACGCTCTTCCGATCTNNNCCATGGACAGTCTACCTCCTT
AGCACACGTCTGAACTCTTTCCCTACACGACGCTCTTCCGATCTNNNGGAATGGATGGATACCTCCTT
GCACACGTCTGAACTCTTTCCCTACACGACGCTCTTCCGATCTNNNAGGAGACACGAGAAAGCTTAG
GAGCACACGTCTGAACTCTTTCCCTACACGACGCTCTTCCGATCTNNNACCATCTCTGTTTCTACTGAG
GCACACGTCTGAACTCTTTCCCTACACGACGCTCTTCCGATCTNNNGTTCTTTTGGTTGTTGGTGGTG
AGAGCACACGTCTGAACTCTTTCCCTACACGACGCTCTTCCGATCTNNNTCCCAGTGATCAAAAATAC
AGCACACGTCTGAACTCTTTCCCTACACGACGCTCTTCCGATCTNNNCAAGGAAGGAGGTGATGATG
AGCACACGTCTGAACTCTTTCCCTACACGACGCTCTTCCGATCTNNNTCATGTTGCCATGTGAATGTG
AGCACACGTCTGAACTCTTTCCCTACACGACGCTCTTCCGATCTNNNACATAGTTGCCAAGGCTTTGT
GAAGAGCACACGTCTGAACTCTTTCCCTACACGACGCTCTTCCGATCTNNNTATCTACCATTTCAGCAG
AGCACACGTCTGAACTCTTTCCCTACACGACGCTCTTCCGATCTNNNCTGAGCACAGTCTCTCTCTCTC
GCACACGTCTGAACTCTTTCCCTACACGACGCTCTTCCGATCTNNNGTGCAATAGTATGGCTGGGTC
GCACACGTCTGAACTCTTTCCCTACACGACGCTCTTCCGATCTNNNCTTCCCCGAACAGCAGTTCATA
GAGCACACGTCTGAACTCTTTCCCTACACGACGCTCTTCCGATCTNNNAGGTTTGTCAGCAACTTGT
GCACACGTCTGAACTCTTTCCCTACACGACGCTCTTCCGATCTNNNCTCAGATTGTTCAGCGGTTCCA
AGAGCACACGTCTGAACTCTTTCCCTACACGACGCTCTTCCGATCTNNNAGGAGGAAACTACCAACA
AGCACACGTCTGAACTCTTTCCCTACACGACGCTCTTCCGATCTNNNACCAGGAAACTTTTGAGATGG
GAGCACACGTCTGAACTCTTTCCCTACACGACGCTCTTCCGATCTNNNTGCACTCTAGGTAGTTGCTC
AGCACACGTCTGAACTCTTTCCCTACACGACGCTCTTCCGATCTNNNGACTTGGCTGTTAGGATGGG
GCACACGTCTGAACTCTTTCCCTACACGACGCTCTTCCGATCTNNNAGTGAAGCAACCCTTCTCTCTC
AGAGCACACGTCTGAACTCTTTCCCTACACGACGCTCTTCCGATCTNNNTGACCATGAAGTGCTTAGG
GCACACGTCTGAACTCTTTCCCTACACGACGCTCTTCCGATCTNNNGGCCAGGATTCATAGGCTCTCTA
CACACGTCTGAACTCTTTCCCTACACGACGCTCTTCCGATCTNNNTCCTCCACATTGTAAACCCTCCTCA
AGAGCACACGTCTGAACTCTTTCCCTACACGACGCTCTTCCGATCTNNNACCTGGAATTTTCTCTCTCT
GCACACGTCTGAACTCTTTCCCTACACGACGCTCTTCCGATCTNNNAGTGAAGCAACCCTTCTCTCTCT
GCACACGTCTGAACTCTTTCCCTACACGACGCTCTTCCGATCTNNNTCATCAACTCATGGCCAATCCTG
GCACACGTCTGAACTCTTTCCCTACACGACGCTCTTCCGATCTNNNATATGTGACACACAGACACGTA
AGCACACGTCTGAACTCTTTCCCTACACGACGCTCTTCCGATCTNNNAGATCATGAGGGTAGGAGAG
GCACACGTCTGAACTCTTTCCCTACACGACGCTCTTCCGATCTNNNTGTCTTACTGGTGTGGTTGTAG
GCACACGTCTGAACTCTTTCCCTACACGACGCTCTTCCGATCTNNNTTGGTCATCCTTCCTCACCATCTC
AGCACACGTCTGAACTCTTTCCCTACACGACGCTCTTCCGATCTNNNGTGACTTGGATAACTCATCAG
GAGCACACGTCTGAACTCTTTCCCTACACGACGCTCTTCCGATCTNNNCGAAACCCTGTCTCTGCAATG
AGCACACGTCTGAACTCTTTCCCTACACGACGCTCTTCCGATCTNNNGATGGTGACGTGGGATGTCA
AGCACACGTCTGAACTCTTTCCCTACACGACGCTCTTCCGATCTNNNTATAAACCGTGCACTCTCCCTC
AGCACACGTCTGAACTCTTTCCCTACACGACGCTCTTCCGATCTNNNGAGCTCTGTGATTAGCAGGA

AGAGCACACGTCTGAACTCTTTCCCTACACGACGCTCTTCCGATCTNNNACCAGGAAACTTTTGAGAT
AGCACACGTCTGAACTCTTTCCCTACACGACGCTCTTCCGATCTNNNCCTCCCTCCCCCTTCTACATATC
GAGCACACGTCTGAACTCTTTCCCTACACGACGCTCTTCCGATCTNNNATCAGTCACTTCTCTGGCCTCTT
GAGCACACGTCTGAACTCTTTCCCTACACGACGCTCTTCCGATCTNNNTAAGTTCATTCAGTGTACGTG
AGAGCACACGTCTGAACTCTTTCCCTACACGACGCTCTTCCGATCTNNNATTGATGTGGACAAGACTGT
GAGCACACGTCTGAACTCTTTCCCTACACGACGCTCTTCCGATCTNNNTTGACAGTGTGTTCACAGAGA
GAGCACACGTCTGAACTCTTTCCCTACACGACGCTCTTCCGATCTNNNTTTCACATTGACCACTCCACCAT
GAGCACACGTCTGAACTCTTTCCCTACACGACGCTCTTCCGATCTNNNTGCAGTGTATTTTTCCCCTCAC
AGAGCACACGTCTGAACTCTTTCCCTACACGACGCTCTTCCGATCTNNNTGCAAATGCCATTACTCTGTG
AGCACACGTCTGAACTCTTTCCCTACACGACGCTCTTCCGATCTNNNAGGGGTTTGGAGAAGTCACAG
AGAGCACACGTCTGAACTCTTTCCCTACACGACGCTCTTCCGATCTNNNTGTCAAAGCAGACAGATGGA
GAGCACACGTCTGAACTCTTTCCCTACACGACGCTCTTCCGATCTNNNGAGGCTCTGAAGTTTTCCCTGA
AGCACACGTCTGAACTCTTTCCCTACACGACGCTCTTCCGATCTNNNAAAAATTACTGCCTGGGGTGAG
AGAGCACACGTCTGAACTCTTTCCCTACACGACGCTCTTCCGATCTNNNTCCCACTTCTTGGAATCACTCA
CACACGTCTGAACTCTTTCCCTACACGACGCTCTTCCGATCTNNNACTGTTTTCATCAGTTTTCATGCGA
GAGCACACGTCTGAACTCTTTCCCTACACGACGCTCTTCCGATCTNNNTCCGTGTCATGTCATTCCTGAA
GAAGAGCACACGTCTGAACTCTTTCCCTACACGACGCTCTTCCGATCTNNNACAGCAGTCCAACTTCTG
AGCACACGTCTGAACTCTTTCCCTACACGACGCTCTTCCGATCTNNNCTTTCAGAAGGAGGATGGTTC
AGCACACGTCTGAACTCTTTCCCTACACGACGCTCTTCCGATCTNNNTAAGAAGAGAGAGAGTGGGA
GAGCACACGTCTGAACTCTTTCCCTACACGACGCTCTTCCGATCTNNNTCAGCTCTCTCTTACCTAGTGC
GAGCACACGTCTGAACTCTTTCCCTACACGACGCTCTTCCGATCTNNNTACCAGGAATACGGCAGCAA
GAGCACACGTCTGAACTCTTTCCCTACACGACGCTCTTCCGATCTNNNAGCACAATAACATATTACCTTG
AGAGCACACGTCTGAACTCTTTCCCTACACGACGCTCTTCCGATCTNNNTGTGACGGAGATGGGTATTT
GAGCACACGTCTGAACTCTTTCCCTACACGACGCTCTTCCGATCTNNNTTGATCTTGGGTCATCTGTGT
GCACACGTCTGAACTCTTTCCCTACACGACGCTCTTCCGATCTNNNCCTCTCTCCCCTCTGTCTCTCTACA
AGCACACGTCTGAACTCTTTCCCTACACGACGCTCTTCCGATCTNNNGGACTGTTCAGTTTATGTCCCC
GAGCACACGTCTGAACTCTTTCCCTACACGACGCTCTTCCGATCTNNNAAAAGTGCCCTGAAGGAAACT
AGCACACGTCTGAACTCTTTCCCTACACGACGCTCTTCCGATCTNNNTCGATTCTGTGCTTTTTATATCT
GAGCACACGTCTGAACTCTTTCCCTACACGACGCTCTTCCGATCTNNNTCCATCTGTCTCAGAGTTTAGA
AGCACACGTCTGAACTCTTTCCCTACACGACGCTCTTCCGATCTNNNGGCCTGGGTAATCTCTCTCTCTC
AGAGCACACGTCTGAACTCTTTCCCTACACGACGCTCTTCCGATCTNNNCATCCAACCTCAATCCTCAGCT
GAAGAGCACACGTCTGAACTCTTTCCCTACACGACGCTCTTCCGATCTNNNGGTGATGAAGAGGTGCT
AGCACACGTCTGAACTCTTTCCCTACACGACGCTCTTCCGATCTNNNAGACTTCTGCAATGTCAAGTGG
AGCACACGTCTGAACTCTTTCCCTACACGACGCTCTTCCGATCTNNNGATTTCTGCAGGAGGAACCAGA
GAGCACACGTCTGAACTCTTTCCCTACACGACGCTCTTCCGATCTNNNATTCTCCAGAGTGACAGAGCCA
AGCACACGTCTGAACTCTTTCCCTACACGACGCTCTTCCGATCTNNNGTTTGCCAGTTGGCAGTCAAAA
GAGCACACGTCTGAACTCTTTCCCTACACGACGCTCTTCCGATCTNNNAGCTAGAATTCAAGGGAGAG
GCACACGTCTGAACTCTTTCCCTACACGACGCTCTTCCGATCTNNNTTCCAGACTTTCTCCTGTGCCTAC
AGCACACGTCTGAACTCTTTCCCTACACGACGCTCTTCCGATCTNNNGCTGAGCAGAAAATTCAGGCA
AGCACACGTCTGAACTCTTTCCCTACACGACGCTCTTCCGATCTNNNTTGTTCTTTGAGTTAGCCAGG
GAGCACACGTCTGAACTCTTTCCCTACACGACGCTCTTCCGATCTNNNCAATATGCAGCTTGCTTCACCA
GAGCACACGTCTGAACTCTTTCCCTACACGACGCTCTTCCGATCTNNNCTAGTCATGAGCTGCCATCTGT
AGCACACGTCTGAACTCTTTCCCTACACGACGCTCTTCCGATCTNNNGCTTTCCCCTGCAAATACCCATA

GAGCACACGTCTGAACTCTTTCCCTACACGACGCTCTTCCGATCTNNNGGATGGGGGCAGTGATAATG
GAGCACACGTCTGAACTCTTTCCCTACACGACGCTCTTCCGATCTNNNACATCCTCAAACCAGGCCTTCTA
AGCACACGTCTGAACTCTTTCCCTACACGACGCTCTTCCGATCTNNNCAATTTGATGCAGTCTGATGCC
AGAGCACACGTCTGAACTCTTTCCCTACACGACGCTCTTCCGATCTNNNATGGGAAGTGCACTGTACTA
AGCACACGTCTGAACTCTTTCCCTACACGACGCTCTTCCGATCTNNNATCAAATCCAGTAGAGCCAAGC
GAGCACACGTCTGAACTCTTTCCCTACACGACGCTCTTCCGATCTNNNACGTTTTACCCCAGTTGCTTCA
GAGCACACGTCTGAACTCTTTCCCTACACGACGCTCTTCCGATCTNNNAATGTGAGTGGGCTGTCAATG
AAGAGCACACGTCTGAACTCTTTCCCTACACGACGCTCTTCCGATCTNNNTCCGGTTCCCGAAAGTTTCT
AGCACACGTCTGAACTCTTTCCCTACACGACGCTCTTCCGATCTNNNACTGCCCCTGGATCATCTGTATT
GAGCACACGTCTGAACTCTTTCCCTACACGACGCTCTTCCGATCTNNNGCAAGGTTGCTGGATATTGAA
GAGCACACGTCTGAACTCTTTCCCTACACGACGCTCTTCCGATCTNNNGCAGAAGCACAACTTCCACAA
AGCACACGTCTGAACTCTTTCCCTACACGACGCTCTTCCGATCTNNNGAATGCACCAGAACAGGTCTCA
GAGCACACGTCTGAACTCTTTCCCTACACGACGCTCTTCCGATCTNNNTACCTGTTCACTTCTGTCACCTG
GAGCACACGTCTGAACTCTTTCCCTACACGACGCTCTTCCGATCTNNNTTTACACTCCCAAACTCCAGTGC
AGCACACGTCTGAACTCTTTCCCTACACGACGCTCTTCCGATCTNNNGGTGAGAAGTCTACAGGGCTTT
GAGCACACGTCTGAACTCTTTCCCTACACGACGCTCTTCCGATCTNNNACACATAGGCAACATAGGCAC
AGAGCACACGTCTGAACTCTTTCCCTACACGACGCTCTTCCGATCTNNNGGACCTATGGTGGGACTTCT
AGCACACGTCTGAACTCTTTCCCTACACGACGCTCTTCCGATCTNNNAACCAAGAACCAACAGGAGACA
AGAGCACACGTCTGAACTCTTTCCCTACACGACGCTCTTCCGATCTNNNAAAAAGGCCAAACTGCTAAC
GAGCACACGTCTGAACTCTTTCCCTACACGACGCTCTTCCGATCTNNNACAAGACTGTGTGACTTCTTGC
AGAGCACACGTCTGAACTCTTTCCCTACACGACGCTCTTCCGATCTNNNGGAGGTATTGGTGTGAAGT
AGAGCACACGTCTGAACTCTTTCCCTACACGACGCTCTTCCGATCTNNNTAGTGTAGCGTGGGTGAATT
AGCACACGTCTGAACTCTTTCCCTACACGACGCTCTTCCGATCTNNNTTGTTTATTTGTGCAGACAGCTT
AGCACACGTCTGAACTCTTTCCCTACACGACGCTCTTCCGATCTNNNGCAGAAGCACAACTTCCACAATG
GAGCACACGTCTGAACTCTTTCCCTACACGACGCTCTTCCGATCTNNNCTGGGCCAGTAAAGAGATGC
GCACACGTCTGAACTCTTTCCCTACACGACGCTCTTCCGATCTNNNTAATCAAACCCTGGCTGAGGAGA
AGAGCACACGTCTGAACTCTTTCCCTACACGACGCTCTTCCGATCTNNNCAGATGCCAAGCTTTCTCTC
AGCACACGTCTGAACTCTTTCCCTACACGACGCTCTTCCGATCTNNNGTTTCAGCCATGTCAGCCAGGCA
AGAGCACACGTCTGAACTCTTTCCCTACACGACGCTCTTCCGATCTNNNCAGTGAGAGGTGCTTATTTC
AGCACACGTCTGAACTCTTTCCCTACACGACGCTCTTCCGATCTNNNTCCGGTTCCCGAAAGTTTCTTTTG
AGAGCACACGTCTGAACTCTTTCCCTACACGACGCTCTTCCGATCTNNNGGTGTTGTCAACCTTTACTCC
AGCACACGTCTGAACTCTTTCCCTACACGACGCTCTTCCGATCTNNNATTGCACCTTTAGGTAACCCAGC
GAGCACACGTCTGAACTCTTTCCCTACACGACGCTCTTCCGATCTNNNGCTCTTCTCTGCTAGTTCTGCTG
AGAGCACACGTCTGAACTCTTTCCCTACACGACGCTCTTCCGATCTNNNCCCTCTCTCTCTGGAGATATG
AGAGCACACGTCTGAACTCTTTCCCTACACGACGCTCTTCCGATCTNNNAATCTAAGGAGAAGCCAGGC
GAGCACACGTCTGAACTCTTTCCCTACACGACGCTCTTCCGATCTNNNGATCACCTACCTTCTGGTGACC
GAGCACACGTCTGAACTCTTTCCCTACACGACGCTCTTCCGATCTNNNGTCACCATCATGTGGCAATTTG
GCACACGTCTGAACTCTTTCCCTACACGACGCTCTTCCGATCTNNNGGACTGCATGTCTTCCATCCATTC
AGAGCACACGTCTGAACTCTTTCCCTACACGACGCTCTTCCGATCTNNNACACATACACAATTTTGCAAA
AGCACACGTCTGAACTCTTTCCCTACACGACGCTCTTCCGATCTNNNAGGCTGGGATGGATACTACTGA
GAGCACACGTCTGAACTCTTTCCCTACACGACGCTCTTCCGATCTNNNACTACCATGAACTAGCAGACA
AGAGCACACGTCTGAACTCTTTCCCTACACGACGCTCTTCCGATCTNNNAAAGTCACTGATCATGCCCCT
GAGCACACGTCTGAACTCTTTCCCTACACGACGCTCTTCCGATCTNNNTCCGTGTCATGTCATTCCTGAA

GCACACGTCTGAACTCTTTCCCTACACGACGCTCTTCCGATCTNNNTACATGGCCCAGTGTTGATAAG

GCACACGTCTGAACTCTTTCCCTACACGACGCTCTTCCGATCTNNNCCGTACACTTACAAGTGAGGCA

GAGCACACGTCTGAACTCTTTCCCTACACGACGCTCTTCCGATCTNNNTGCAAATGTTTTCCTCAGTTG

AGCACACGTCTGAACTCTTTCCCTACACGACGCTCTTCCGATCTNNNACTCCACACATGCACGATTCTT

AAGAGCACACGTCTGAACTCTTTCCCTACACGACGCTCTTCCGATCTNNNGCATCTAACATGGTACCG

GCACACGTCTGAACTCTTTCCCTACACGACGCTCTTCCGATCTNNNAGAAAAGAATCCAGCCCTCATG

GAGCACACGTCTGAACTCTTTCCCTACACGACGCTCTTCCGATCTNNNGGGAGGGATGTAGGAGAG

GAGCACACGTCTGAACTCTTTCCCTACACGACGCTCTTCCGATCTNNNGCATGAATGAAGTGGTAGC

GAGCACACGTCTGAACTCTTTCCCTACACGACGCTCTTCCGATCTNNNACTCCCTTTCTCATCTTCTCTT

GCACACGTCTGAACTCTTTCCCTACACGACGCTCTTCCGATCTNNNGAACTTGGTACCTTTCCTGCCTC

GCACACGTCTGAACTCTTTCCCTACACGACGCTCTTCCGATCTNNNCACTGCCACCACAATTTCATCTG

AGCACACGTCTGAACTCTTTCCCTACACGACGCTCTTCCGATCTNNNTGCAAGCTGGTCAGAAACTCA

AGCACACGTCTGAACTCTTTCCCTACACGACGCTCTTCCGATCTNNNAACAGTCCCAGAAAGGAAAAG

AGCACACGTCTGAACTCTTTCCCTACACGACGCTCTTCCGATCTNNNTTCAGACAGCCACAAATGTCT

CACACGTCTGAACTCTTTCCCTACACGACGCTCTTCCGATCTNNNTGAACTTTTTGAGGACCCCACACC

GCACACGTCTGAACTCTTTCCCTACACGACGCTCTTCCGATCTNNNGTCCTTACTGTCAGCATGAACC

AGAGCACACGTCTGAACTCTTTCCCTACACGACGCTCTTCCGATCTNNNACAAACAGAAGAGCACAG

AGAGCACACGTCTGAACTCTTTCCCTACACGACGCTCTTCCGATCTNNNAGTTGCTCCTGAAGAAGAT

GCACACGTCTGAACTCTTTCCCTACACGACGCTCTTCCGATCTNNNTCAGTGTATACTATCAGAAGG

GAGCACACGTCTGAACTCTTTCCCTACACGACGCTCTTCCGATCTNNNACTCCAAGACTTTCCCCAGA

GCACACGTCTGAACTCTTTCCCTACACGACGCTCTTCCGATCTNNNTGCCAAGTGTGTTTCTTGCTACT

AGCACACGTCTGAACTCTTTCCCTACACGACGCTCTTCCGATCTNNNAGCATGCACATAACCCAGAG

GAGCACACGTCTGAACTCTTTCCCTACACGACGCTCTTCCGATCTNNNCCGAAGTAGTTGCAGCATTT

AGCACACGTCTGAACTCTTTCCCTACACGACGCTCTTCCGATCTNNNACGTATTTGTGTGTGGCTGTT

GCACACGTCTGAACTCTTTCCCTACACGACGCTCTTCCGATCTNNNGATACGGCTTTGCACAATCTTGG

AGAGCACACGTCTGAACTCTTTCCCTACACGACGCTCTTCCGATCTNNNCCTAAGTCAGTCCCATTGAA

AGCACACGTCTGAACTCTTTCCCTACACGACGCTCTTCCGATCTNNNGCTAGTAGGAGTGTTTGCCTG

AGCACACGTCTGAACTCTTTCCCTACACGACGCTCTTCCGATCTNNNTAAGCTTGCCAAGTCACTGTC

GCACACGTCTGAACTCTTTCCCTACACGACGCTCTTCCGATCTNNNAAGAGAAAAGAAGGGGCATG

AGCACACGTCTGAACTCTTTCCCTACACGACGCTCTTCCGATCTNNNAGGAGACACGAGAAAGCTTA

GAGCACACGTCTGAACTCTTTCCCTACACGACGCTCTTCCGATCTNNNAGTGGGGAGAACTAAGACA

AGCACACGTCTGAACTCTTTCCCTACACGACGCTCTTCCGATCTNNNAGAAGTTTTACTTGGGTCAAT

GAGCACACGTCTGAACTCTTTCCCTACACGACGCTCTTCCGATCTNNNCACAGAGAATCACCTGCACA

GCACACGTCTGAACTCTTTCCCTACACGACGCTCTTCCGATCTNNNAGTTTTTGTTGGGGCTTGAAG

AGCACACGTCTGAACTCTTTCCCTACACGACGCTCTTCCGATCTNNNAGAAAACACTATGCCCACTCTC

GCACACGTCTGAACTCTTTCCCTACACGACGCTCTTCCGATCTNNNACGTATTTGTGTGTGGCTGTT

AGCACACGTCTGAACTCTTTCCCTACACGACGCTCTTCCGATCTNNNACGGCGTCATTAGTACACAAC

AGCACACGTCTGAACTCTTTCCCTACACGACGCTCTTCCGATCTNNNAGTCTCCCAGTCTTGGGTATG

AGCACACGTCTGAACTCTTTCCCTACACGACGCTCTTCCGATCTNNNAAGCCCAGAGAGTATTGTTC

GCACACGTCTGAACTCTTTCCCTACACGACGCTCTTCCGATCTNNNGTCAGCAATGGCAAAATCAGCA

AGCACACGTCTGAACTCTTTCCCTACACGACGCTCTTCCGATCTNNNAAACCAGAACACGTAACTGCCT

GCACACGTCTGAACTCTTTCCCTACACGACGCTCTTCCGATCTNNNGAGGAGACAAGGTGAGAGGA

AGCACACGTCTGAACTCTTTCCCTACACGACGCTCTTCCGATCTNNNTGAACTTTTTGAGGACCCCACA

AGCACACGTCTGAACTCTTTCCCTACACGACGCTCTTCCGATCTNNNCGACAGACAGAGCAAAGCTCA
GCACACGTCTGAACTCTTTCCCTACACGACGCTCTTCCGATCTNNNTAACTGTTCAGGATAGGGCGT
AGCACACGTCTGAACTCTTTCCCTACACGACGCTCTTCCGATCTNNNTGTGCATGAGCTGGACATCCT
AGCACACGTCTGAACTCTTTCCCTACACGACGCTCTTCCGATCTNNNCCTGCCCTCCTTATGATATGCA
GAGCACACGTCTGAACTCTTTCCCTACACGACGCTCTTCCGATCTNNNGTGCAGCAAGCTTGTGTAATA
AGCACACGTCTGAACTCTTTCCCTACACGACGCTCTTCCGATCTNNNACATCTTGCATGGATAAGGTG
AGCACACGTCTGAACTCTTTCCCTACACGACGCTCTTCCGATCTNNNATAGACAGCCTACAGCCAGACA
AGCACACGTCTGAACTCTTTCCCTACACGACGCTCTTCCGATCTNNNGAATGAGCAAAAACAGCCAGA
GAGCACACGTCTGAACTCTTTCCCTACACGACGCTCTTCCGATCTNNNTGTGACCAAGAAGCCAACGA
GCACACGTCTGAACTCTTTCCCTACACGACGCTCTTCCGATCTNNNAGTCCTATGTCAACATTCAGCG
GAGCACACGTCTGAACTCTTTCCCTACACGACGCTCTTCCGATCTNNNATCTAGAATCCCAGCCTAGTG
AGCACACGTCTGAACTCTTTCCCTACACGACGCTCTTCCGATCTNNNTATGGACAGTAAGCAAGCCCA
GAGCACACGTCTGAACTCTTTCCCTACACGACGCTCTTCCGATCTNNNCCAGAGTTGGAAAGTAAAC
AGCACACGTCTGAACTCTTTCCCTACACGACGCTCTTCCGATCTNNNCAGAATTTCCTCGTGGGTGGA
GAGCACACGTCTGAACTCTTTCCCTACACGACGCTCTTCCGATCTNNNTCATGTTGTTGCAAATGACG
AGCACACGTCTGAACTCTTTCCCTACACGACGCTCTTCCGATCTNNNAATAACACTCCACTTTTCCCCG
GCACACGTCTGAACTCTTTCCCTACACGACGCTCTTCCGATCTNNNGCAGGGTCTTGTCACATCTTCAA
GAGCACACGTCTGAACTCTTTCCCTACACGACGCTCTTCCGATCTNNNCCACCGCACCAGCTGTAAAAT
GCACACGTCTGAACTCTTTCCCTACACGACGCTCTTCCGATCTNNNGAAGGGAAAACGGAGGAAGA
AGCACACGTCTGAACTCTTTCCCTACACGACGCTCTTCCGATCTNNNTGCCAATTTGCCAATAACAACA
GAGCACACGTCTGAACTCTTTCCCTACACGACGCTCTTCCGATCTNNNCTACGTTCTGGGAAGGGAAG
AGCACACGTCTGAACTCTTTCCCTACACGACGCTCTTCCGATCTNNNACCTGCATGGCACGAATAAGA
GAGCACACGTCTGAACTCTTTCCCTACACGACGCTCTTCCGATCTNNNCTTCAGCCTGTTTGACAGAG
GAGCACACGTCTGAACTCTTTCCCTACACGACGCTCTTCCGATCTNNNGCCTGCCTGTCTCCATATCATT
GAGCACACGTCTGAACTCTTTCCCTACACGACGCTCTTCCGATCTNNNTCAAACAGTGCCACCAAGAA
GCACACGTCTGAACTCTTTCCCTACACGACGCTCTTCCGATCTNNNGTTCAGGTTGATGGGAGGGAA
GAGCACACGTCTGAACTCTTTCCCTACACGACGCTCTTCCGATCTNNNTTAGCACTCAACTGCTGTCT
AGCACACGTCTGAACTCTTTCCCTACACGACGCTCTTCCGATCTNNNCCATGTGCACTGTAGGTAGCA
GCACACGTCTGAACTCTTTCCCTACACGACGCTCTTCCGATCTNNNTGGTAGTTGAGTGGCAAAATG
GCACACGTCTGAACTCTTTCCCTACACGACGCTCTTCCGATCTNNNTTCCCTCACTCTTTGGATGCTCT
AGCACACGTCTGAACTCTTTCCCTACACGACGCTCTTCCGATCTNNNGGGTCAAAGGAGAAAAGGCA
AGAGCACACGTCTGAACTCTTTCCCTACACGACGCTCTTCCGATCTNNNAACTCTGCATGAGGGGGA
AGCACACGTCTGAACTCTTTCCCTACACGACGCTCTTCCGATCTNNNTTGTGGAAAGGAGCGTATCTG
GAGCACACGTCTGAACTCTTTCCCTACACGACGCTCTTCCGATCTNNNCTGTGCTCCTGGAAGTTGTT
AGCACACGTCTGAACTCTTTCCCTACACGACGCTCTTCCGATCTNNNCCCCCTTCCCCTCTATTTGTTTG
AGAGCACACGTCTGAACTCTTTCCCTACACGACGCTCTTCCGATCTNNNTGCAAAACCTGTACTGTGT
GAGCACACGTCTGAACTCTTTCCCTACACGACGCTCTTCCGATCTNNNACAAACTGAGGGAGAGGAA
GAGCACACGTCTGAACTCTTTCCCTACACGACGCTCTTCCGATCTNNNGTTTGCCAGTTGGCAGTCAA
GCACACGTCTGAACTCTTTCCCTACACGACGCTCTTCCGATCTNNNTATCCTGGAACTGGAAGATGG
AGCACACGTCTGAACTCTTTCCCTACACGACGCTCTTCCGATCTNNNAGGGTTCCCTCACAGATGAAT
GAGCACACGTCTGAACTCTTTCCCTACACGACGCTCTTCCGATCTNNNCCCTCTGGGTCAGACAATCA
GAGCACACGTCTGAACTCTTTCCCTACACGACGCTCTTCCGATCTNNNAATAGCCCAAGGGGTCATCT
GAGCACACGTCTGAACTCTTTCCCTACACGACGCTCTTCCGATCTNNNCAATCCCACTCCTTAGCAGGA

AGCACACGTCTGAACTCTTTCCCTACACGACGCTCTTCCGATCTNNNTGGGTATGTCTTTATTTTGGGC
GAGCACACGTCTGAACTCTTTCCCTACACGACGCTCTTCCGATCTNNNACCCTCAATAACCAGGACCAT
GAGCACACGTCTGAACTCTTTCCCTACACGACGCTCTTCCGATCTNNNGCAATTTTTGTTGTCCCATGG
AGCACACGTCTGAACTCTTTCCCTACACGACGCTCTTCCGATCTNNNGCACAGTCTCCGATTTCTGGATT
GCACACGTCTGAACTCTTTCCCTACACGACGCTCTTCCGATCTNNNACTCGCCGATGATAACTTTGTCCC
AGCACACGTCTGAACTCTTTCCCTACACGACGCTCTTCCGATCTNNNTGAGACACACAAAGCACTGTGA
GAGCACACGTCTGAACTCTTTCCCTACACGACGCTCTTCCGATCTNNNTGTAGTGGTCTCATCCCAGAC
AAGAGCACACGTCTGAACTCTTTCCCTACACGACGCTCTTCCGATCTNNNATTACCCATTTCAGAAGCG
GAGCACACGTCTGAACTCTTTCCCTACACGACGCTCTTCCGATCTNNNATCAATCCACTGCCTATGGGGT
GAGCACACGTCTGAACTCTTTCCCTACACGACGCTCTTCCGATCTNNNTCCCAGAACTCAAGGGTAAT
GAGCACACGTCTGAACTCTTTCCCTACACGACGCTCTTCCGATCTNNNCCAGCTTCAGCAGCTATAAAC
AGCACACGTCTGAACTCTTTCCCTACACGACGCTCTTCCGATCTNNNTGCAAGCTGGTCAGAAACTCAAT
GAGCACACGTCTGAACTCTTTCCCTACACGACGCTCTTCCGATCTNNNATGCCACCTTTGAAATCACTTG
GAGCACACGTCTGAACTCTTTCCCTACACGACGCTCTTCCGATCTNNNTTCTGATTGATGGCCTGCCTTA
GAGCACACGTCTGAACTCTTTCCCTACACGACGCTCTTCCGATCTNNNTGTAGATGAAGACAGCACTG
AGCACACGTCTGAACTCTTTCCCTACACGACGCTCTTCCGATCTNNNACAGAACATGTGTAGCACTTCC
AAGAGCACACGTCTGAACTCTTTCCCTACACGACGCTCTTCCGATCTNNNTCACCTAATGCTGTGGCTT
GAGCACACGTCTGAACTCTTTCCCTACACGACGCTCTTCCGATCTNNNAAATCATGCGTCTACACTCCAG
AGAGCACACGTCTGAACTCTTTCCCTACACGACGCTCTTCCGATCTNNNACTACTGAGCATACCTAGGT
AGCACACGTCTGAACTCTTTCCCTACACGACGCTCTTCCGATCTNNNTCGTACAAACCAGCCACTAAAG
AGCACACGTCTGAACTCTTTCCCTACACGACGCTCTTCCGATCTNNNGTTAGACAGCATTAGCTCCCAG
GCACACGTCTGAACTCTTTCCCTACACGACGCTCTTCCGATCTNNNACTCTCCTGTCCTCATCTCACTGC
AGCACACGTCTGAACTCTTTCCCTACACGACGCTCTTCCGATCTNNNTGATCATGGCAATCATATGGGA
AGCACACGTCTGAACTCTTTCCCTACACGACGCTCTTCCGATCTNNNAACCCAATTCAGCAAGGCCATT
AGCACACGTCTGAACTCTTTCCCTACACGACGCTCTTCCGATCTNNNACAGCTACACTCACTTCCTCCTG
AGCACACGTCTGAACTCTTTCCCTACACGACGCTCTTCCGATCTNNNGCTGTTATGGCAGGAATGGGT
GAGCACACGTCTGAACTCTTTCCCTACACGACGCTCTTCCGATCTNNNCAAGATCCACTCCAGGCTTTCT
GAGCACACGTCTGAACTCTTTCCCTACACGACGCTCTTCCGATCTNNNTTGCTCTACCCTTCTGTCTCTCC
AGCACACGTCTGAACTCTTTCCCTACACGACGCTCTTCCGATCTNNNCTGGAAAGATGTTGGTAGGG
GAGCACACGTCTGAACTCTTTCCCTACACGACGCTCTTCCGATCTNNNAGTGGTAGAGAGGGATCAAA
AGAGCACACGTCTGAACTCTTTCCCTACACGACGCTCTTCCGATCTNNNACCCCACAGATTTGCCTAAAT
AGAGCACACGTCTGAACTCTTTCCCTACACGACGCTCTTCCGATCTNNNCTGGCCTTTGGAAGTAGGTT
GAGCACACGTCTGAACTCTTTCCCTACACGACGCTCTTCCGATCTNNNATATAGGAGGGACTGAGTG
AGAGCACACGTCTGAACTCTTTCCCTACACGACGCTCTTCCGATCTNNNTCACCAGGCACTCTGGAGA
AGAGCACACGTCTGAACTCTTTCCCTACACGACGCTCTTCCGATCTNNNAGTTGCTCCTGAAGAAGATG
GAGCACACGTCTGAACTCTTTCCCTACACGACGCTCTTCCGATCTNNNGAGCAAATTCTGCAAGCTGA
AGAGCACACGTCTGAACTCTTTCCCTACACGACGCTCTTCCGATCTNNNATGGATTTCAGAGGCCCCA
AGCACACGTCTGAACTCTTTCCCTACACGACGCTCTTCCGATCTNNNATCTGGGCTCAAGAAATCCAC
AGCACACGTCTGAACTCTTTCCCTACACGACGCTCTTCCGATCTNNNGTCAGATCCCTCCCTCATTGTCT
GAGCACACGTCTGAACTCTTTCCCTACACGACGCTCTTCCGATCTNNNGGAATCCGATCTGAAAGAC
AGAGCACACGTCTGAACTCTTTCCCTACACGACGCTCTTCCGATCTNNNGTCATCTCATTGGTGGTCAC
GAGCACACGTCTGAACTCTTTCCCTACACGACGCTCTTCCGATCTNNNCAGCAATTCTGTGAGAGGTA
AAGAGCACACGTCTGAACTCTTTCCCTACACGACGCTCTTCCGATCTNNNTTGAAGTAATCCTCCTGCCT

AGCACACGTCTGAACTCTTTCCCTACACGACGCTCTTCCGATCTNNNGATTACAGACAACCTTGGTGC

AGCACACGTCTGAACTCTTTCCCTACACGACGCTCTTCCGATCTNNNCAGAAAAAGGACACTCTGAG

AGCACACGTCTGAACTCTTTCCCTACACGACGCTCTTCCGATCTNNNAGGCCCCACTATCTAACCTTG

AGCACACGTCTGAACTCTTTCCCTACACGACGCTCTTCCGATCTNNNTTGAGACACATATCAGATCCC

GCACACGTCTGAACTCTTTCCCTACACGACGCTCTTCCGATCTNNNGCTCTTCTACGTTTCACTGTCGT

GAGCACACGTCTGAACTCTTTCCCTACACGACGCTCTTCCGATCTNNNTGTTAGAAGGGACAAGTTT

CACACGTCTGAACTCTTTCCCTACACGACGCTCTTCCGATCTNNNTCACCATTTGTCTTTGCAGGGAA

GAGCACACGTCTGAACTCTTTCCCTACACGACGCTCTTCCGATCTNNNAAACAGGAGGCTCTCTTTGT

AGCACACGTCTGAACTCTTTCCCTACACGACGCTCTTCCGATCTNNNTGTTTGTCTCCTCTTCATTGTCG

AGCACACGTCTGAACTCTTTCCCTACACGACGCTCTTCCGATCTNNNCAGCTACAGATGGTACAGGTG

AGCACACGTCTGAACTCTTTCCCTACACGACGCTCTTCCGATCTNNNCTAACACAACCCTTTCCCAGGA

AGCACACGTCTGAACTCTTTCCCTACACGACGCTCTTCCGATCTNNNTTCCTCTCCTCACTCTCCTTGAC

GAGCACACGTCTGAACTCTTTCCCTACACGACGCTCTTCCGATCTNNNGTGATGAACGCAACAGCTTA

GAGCACACGTCTGAACTCTTTCCCTACACGACGCTCTTCCGATCTNNNGACCTACCAGTGGGACTTTC

GAGCACACGTCTGAACTCTTTCCCTACACGACGCTCTTCCGATCTNNNATTGTCACCAGTTCCATGGC

GCACACGTCTGAACTCTTTCCCTACACGACGCTCTTCCGATCTNNNGGTGCACCTGGCCTTTATTCTTA

AGCACACGTCTGAACTCTTTCCCTACACGACGCTCTTCCGATCTNNNTTGAGTGACTGAATTGCTAGG

GAGCACACGTCTGAACTCTTTCCCTACACGACGCTCTTCCGATCTNNNTTAGCACTCAACTGCTGTCTC

AGCACACGTCTGAACTCTTTCCCTACACGACGCTCTTCCGATCTNNNAAAGGTAGAACCCAACCATCC

AGCACACGTCTGAACTCTTTCCCTACACGACGCTCTTCCGATCTNNNCAAGCCAAGGCTGCAATAAGC

AGAGCACACGTCTGAACTCTTTCCCTACACGACGCTCTTCCGATCTNNNATGAAAGGCCCTGGAAATG

ACACGTCTGAACTCTTTCCCTACACGACGCTCTTCCGATCTNNNCCTTTCCCCCTATATGTGTGTGTGC

GAGCACACGTCTGAACTCTTTCCCTACACGACGCTCTTCCGATCTNNNGCTGTGCAGCAGGATTTTC

GAAGAGCACACGTCTGAACTCTTTCCCTACACGACGCTCTTCCGATCTNNNATTTCCAAAAAGTATGG

AAGAGCACACGTCTGAACTCTTTCCCTACACGACGCTCTTCCGATCTNNNTTTGACTGCAAGGCAGA

AGAGCACACGTCTGAACTCTTTCCCTACACGACGCTCTTCCGATCTNNNCCTTTCCCCCTATATGTGTG

AAGAGCACACGTCTGAACTCTTTCCCTACACGACGCTCTTCCGATCTNNNCCAGCAGTGTACAAAAG

AAGAGCACACGTCTGAACTCTTTCCCTACACGACGCTCTTCCGATCTNNNACACAAAGATTCCCAGCA

CACACGTCTGAACTCTTTCCCTACACGACGCTCTTCCGATCTNNNGGGTCGATCTCAAACAGGTGCAT

CACACGTCTGAACTCTTTCCCTACACGACGCTCTTCCGATCTNNNTCACTCTCTATTGCTCCCCCAAACA

AGCACACGTCTGAACTCTTTCCCTACACGACGCTCTTCCGATCTNNNAAAGAAGAGAGGGGGAAG

AGCACACGTCTGAACTCTTTCCCTACACGACGCTCTTCCGATCTNNNCCAGCAAATGTCCTATCCTGA

GAGCACACGTCTGAACTCTTTCCCTACACGACGCTCTTCCGATCTNNNTGGCAAGACTGGATTCATA

AGCACACGTCTGAACTCTTTCCCTACACGACGCTCTTCCGATCTNNNTGTCTCAAAAAGAGCAAAGG

AAGAGCACACGTCTGAACTCTTTCCCTACACGACGCTCTTCCGATCTNNNCCAAGCACAGAAGAGAG

AAGAGCACACGTCTGAACTCTTTCCCTACACGACGCTCTTCCGATCTNNNTCCTTCCCCCACATGAAAG

AGAGCACACGTCTGAACTCTTTCCCTACACGACGCTCTTCCGATCTNNNAAGATCATCGTCTGGTGAT

GCACACGTCTGAACTCTTTCCCTACACGACGCTCTTCCGATCTNNNTAGAGTGCGATTTGGAAATGG

AGCACACGTCTGAACTCTTTCCCTACACGACGCTCTTCCGATCTNNNTTATGCAACCTAAGAAATGCC

GAGCACACGTCTGAACTCTTTCCCTACACGACGCTCTTCCGATCTNNNGCCCGGCCTATGTATCACAAT

GCACACGTCTGAACTCTTTCCCTACACGACGCTCTTCCGATCTNNNCGGCACTCCCTAAATGATCTCTC

CACACGTCTGAACTCTTTCCCTACACGACGCTCTTCCGATCTNNNTGAATGAAGCCATCTTGGACACC

AGCACACGTCTGAACTCTTTCCCTACACGACGCTCTTCCGATCTNNNAAAAATCTCCCCAGACCTGCCC

GAGCACACGTCTGAACTCTTTCCCTACACGACGCTCTTCCGATCTNNNCGCATGTGTCCTCTGGAATTG
AGCACACGTCTGAACTCTTTCCCTACACGACGCTCTTCCGATCTNNNAGCCTGTTTCTCCTAGCATGTT
GCACACGTCTGAACTCTTTCCCTACACGACGCTCTTCCGATCTNNNATATCACCTTCCAGAGTTACGTT
GCACACGTCTGAACTCTTTCCCTACACGACGCTCTTCCGATCTNNNTCCTTAGAAATAAAGTGGAAAG
GCACACGTCTGAACTCTTTCCCTACACGACGCTCTTCCGATCTNNNCATCCTCCCAAGTTCATGGACAC
GCACACGTCTGAACTCTTTCCCTACACGACGCTCTTCCGATCTNNNCTTTGGTCTCTGTTCCTCCTCCCC
AGCACACGTCTGAACTCTTTCCCTACACGACGCTCTTCCGATCTNNNGGATGTTCCTCCCTCTGTATG
AGCACACGTCTGAACTCTTTCCCTACACGACGCTCTTCCGATCTNNNTCATCTTGTGTTTGGACAGTG
GAGCACACGTCTGAACTCTTTCCCTACACGACGCTCTTCCGATCTNNNACAGAGACACAACACAGGG
GAGCACACGTCTGAACTCTTTCCCTACACGACGCTCTTCCGATCTNNNGGATGTTCCTCCCTCTGTAT
AGAGCACACGTCTGAACTCTTTCCCTACACGACGCTCTTCCGATCTNNNTCATATTGTCACATTGGAA
AGCACACGTCTGAACTCTTTCCCTACACGACGCTCTTCCGATCTNNNCCAGTTATCTCTGTCTCTCTCG
GAGCACACGTCTGAACTCTTTCCCTACACGACGCTCTTCCGATCTNNNTGCTTGTGAAATGGAGAGTT
GAGCACACGTCTGAACTCTTTCCCTACACGACGCTCTTCCGATCTNNNGCCCAGGCAACAATGGTAAA
AGAGCACACGTCTGAACTCTTTCCCTACACGACGCTCTTCCGATCTNNNAGGGTGGCTAGCTGTAGC
AGCACACGTCTGAACTCTTTCCCTACACGACGCTCTTCCGATCTNNNATCTTTTGCTCCCTGCATTGTTT
GAGCACACGTCTGAACTCTTTCCCTACACGACGCTCTTCCGATCTNNNAGCTGTCAGATTGTTAGCTCT
AGCACACGTCTGAACTCTTTCCCTACACGACGCTCTTCCGATCTNNNCGTTTCCAACAGTTCCATGTGT
AGCACACGTCTGAACTCTTTCCCTACACGACGCTCTTCCGATCTNNNATGGGAAAGACTGGTGTCTGG
AGAGCACACGTCTGAACTCTTTCCCTACACGACGCTCTTCCGATCTNNNTGTGGTACTATTCTGGGAG
GCACACGTCTGAACTCTTTCCCTACACGACGCTCTTCCGATCTNNNAGTCTCCCAAGTCTGATAAAGT
GAGCACACGTCTGAACTCTTTCCCTACACGACGCTCTTCCGATCTNNNCACATCCCACCACTATCACCAA
AGCACACGTCTGAACTCTTTCCCTACACGACGCTCTTCCGATCTNNNAAAAACATGAAGGGCCACAC
GAGCACACGTCTGAACTCTTTCCCTACACGACGCTCTTCCGATCTNNNCATATTCATACACACGCACGC
GAGCACACGTCTGAACTCTTTCCCTACACGACGCTCTTCCGATCTNNNGGTGCTGAGGTGTTTCTTCT
AGAGCACACGTCTGAACTCTTTCCCTACACGACGCTCTTCCGATCTNNNAAAAGAGGGGGTCAGGGA
AGCACACGTCTGAACTCTTTCCCTACACGACGCTCTTCCGATCTNNNTGATCTGTAGTGGTGTGTGTG
AGAGCACACGTCTGAACTCTTTCCCTACACGACGCTCTTCCGATCTNNNGGAAACACAGCTACTACTTC
GAGCACACGTCTGAACTCTTTCCCTACACGACGCTCTTCCGATCTNNNACTGTAAACTCTAGAACAAC
GAGCACACGTCTGAACTCTTTCCCTACACGACGCTCTTCCGATCTNNNGGAATTTGTGTGTGTGTACC
AGCACACGTCTGAACTCTTTCCCTACACGACGCTCTTCCGATCTNNNTGGAGTAGGGATAATCACAC
GCACACGTCTGAACTCTTTCCCTACACGACGCTCTTCCGATCTNNNTCTACTGTTGTGTCAGGGTCACT
AAGAGCACACGTCTGAACTCTTTCCCTACACGACGCTCTTCCGATCTNNNCCTTGGATAATTGAGACC
AGAGCACACGTCTGAACTCTTTCCCTACACGACGCTCTTCCGATCTNNNCTTTACCTGATGCCCACCTA
AGCACACGTCTGAACTCTTTCCCTACACGACGCTCTTCCGATCTNNNGCACACTCTCATGCATTTTCTCT
AGCACACGTCTGAACTCTTTCCCTACACGACGCTCTTCCGATCTNNNTTCAAAAGACCCCAGACCCAAT
AGCACACGTCTGAACTCTTTCCCTACACGACGCTCTTCCGATCTNNNAATCAAAATGCCCCTGGAAGT
AGAGCACACGTCTGAACTCTTTCCCTACACGACGCTCTTCCGATCTNNNACTCTATTAGGAGGAGTGA
AGCACACGTCTGAACTCTTTCCCTACACGACGCTCTTCCGATCTNNNCCAGTTGAATTTTGGAAATCTG
AGAGCACACGTCTGAACTCTTTCCCTACACGACGCTCTTCCGATCTNNNTCATTCTTTGAAAAATTGGG
AGCACACGTCTGAACTCTTTCCCTACACGACGCTCTTCCGATCTNNNAAATTGCTGGGAACAAGGACT
AGCACACGTCTGAACTCTTTCCCTACACGACGCTCTTCCGATCTNNNTTCCCTGCCAACTACATCAACTC
AGCACACGTCTGAACTCTTTCCCTACACGACGCTCTTCCGATCTNNNAGGAGGAGATGTTAGTGACT

AGCACACGTCTGAACTCTTTCCCTACACGACGCTCTTCCGATCTNNNATATTCTTTCCCAGTCCCTGGC
GCACACGTCTGAACTCTTTCCCTACACGACGCTCTTCCGATCTNNNGGATCACCAGCACCTTATTGTT
GAGCACACGTCTGAACTCTTTCCCTACACGACGCTCTTCCGATCTNNNGCTCTGCTTGTTTCATTTGGT
AGCACACGTCTGAACTCTTTCCCTACACGACGCTCTTCCGATCTNNNTCTGGAAAACTGGAATTTGGG
AGCACACGTCTGAACTCTTTCCCTACACGACGCTCTTCCGATCTNNNCTCCTGGTTCCCATTACTAGCCT
GCACACGTCTGAACTCTTTCCCTACACGACGCTCTTCCGATCTNNNTCCAAGTGTTTTCAGAAGTCCCC
AGAGCACACGTCTGAACTCTTTCCCTACACGACGCTCTTCCGATCTNNNATGCCACTGCATGAAGCA
GAGCACACGTCTGAACTCTTTCCCTACACGACGCTCTTCCGATCTNNNAAGAAAGGAGTGTTCACAG
AGCACACGTCTGAACTCTTTCCCTACACGACGCTCTTCCGATCTNNNAAATGCATAGCTCCATGACCA
GAGCACACGTCTGAACTCTTTCCCTACACGACGCTCTTCCGATCTNNNTCCTATTGTTCGATAGAAGT
AGCACACGTCTGAACTCTTTCCCTACACGACGCTCTTCCGATCTNNNGCAGAAAGAGCAAGGAAGA
AGCACACGTCTGAACTCTTTCCCTACACGACGCTCTTCCGATCTNNNGCTTACCGCAGAGATAAGACA
GAGCACACGTCTGAACTCTTTCCCTACACGACGCTCTTCCGATCTNNNCAGACTAGAGTGGAGGAAG
AGCACACGTCTGAACTCTTTCCCTACACGACGCTCTTCCGATCTNNNACTTTCCATTGCATATTGTTGC
AGCACACGTCTGAACTCTTTCCCTACACGACGCTCTTCCGATCTNNNTGTGTCTCAGAGAATTGGGA
AGCACACGTCTGAACTCTTTCCCTACACGACGCTCTTCCGATCTNNNCTCTGGAAAGGCAGGGTTT
AGCACACGTCTGAACTCTTTCCCTACACGACGCTCTTCCGATCTNNNGCCAGCATCACCAGCAAAAGA
GCACACGTCTGAACTCTTTCCCTACACGACGCTCTTCCGATCTNNNCCTGTGGTTCTGTGATGTCTGTT
AAGAGCACACGTCTGAACTCTTTCCCTACACGACGCTCTTCCGATCTNNNCGTATTTGCCACTTTCCAG
GAGCACACGTCTGAACTCTTTCCCTACACGACGCTCTTCCGATCTNNNGTTGGTAAAACAGTACACG
GAGCACACGTCTGAACTCTTTCCCTACACGACGCTCTTCCGATCTNNNCAAGTGCTGTGGTTTGGATG
GCACACGTCTGAACTCTTTCCCTACACGACGCTCTTCCGATCTNNNGGCGTGTAGTTCTGGAATTCT
AGCACACGTCTGAACTCTTTCCCTACACGACGCTCTTCCGATCTNNNTTTTTCCCTACCTTTTCCCCAGC
GCACACGTCTGAACTCTTTCCCTACACGACGCTCTTCCGATCTNNNCACTGCTAAAGACAGAGCCAGA
AGCACACGTCTGAACTCTTTCCCTACACGACGCTCTTCCGATCTNNNTTCCAGGGAACAGACACTAGA
AGCACACGTCTGAACTCTTTCCCTACACGACGCTCTTCCGATCTNNNGTCCCCTAGTCTCTAGTTCCCT
AGAGCACACGTCTGAACTCTTTCCCTACACGACGCTCTTCCGATCTNNNGCCTTCTCTGTCTTTCTGTC
AGCACACGTCTGAACTCTTTCCCTACACGACGCTCTTCCGATCTNNNGCCTGGGCAATAGAAGTGAG
AGCACACGTCTGAACTCTTTCCCTACACGACGCTCTTCCGATCTNNNATTCTCTCCCTGTTGAATGCTCC
AGCACACGTCTGAACTCTTTCCCTACACGACGCTCTTCCGATCTNNNTGCTGTATCTGCTTATAATCTC
AGCACACGTCTGAACTCTTTCCCTACACGACGCTCTTCCGATCTNNNTGATGACATTTTGGTTGTGCG
GAGCACACGTCTGAACTCTTTCCCTACACGACGCTCTTCCGATCTNNNGCCTGGGCAATAGAAGTGA
CACACGTCTGAACTCTTTCCCTACACGACGCTCTTCCGATCTNNNGAGGTGCAGGAATGTAGGACA
AGCACACGTCTGAACTCTTTCCCTACACGACGCTCTTCCGATCTNNNGGAGGAGGCTGAGGAAAAT
AGCACACGTCTGAACTCTTTCCCTACACGACGCTCTTCCGATCTNNNAGTAAGGACCTACTCTTGTCAT
GCACACGTCTGAACTCTTTCCCTACACGACGCTCTTCCGATCTNNNCCCTCTACCCTCCTTCTGGAAAC
AGAGCACACGTCTGAACTCTTTCCCTACACGACGCTCTTCCGATCTNNNAATTCAGGTTCTGTAGGTC
GAGCACACGTCTGAACTCTTTCCCTACACGACGCTCTTCCGATCTNNNACAAAACCACAAAACGCAGA
GAGCACACGTCTGAACTCTTTCCCTACACGACGCTCTTCCGATCTNNNAACATGCTACCAGCCAATAA
GCACACGTCTGAACTCTTTCCCTACACGACGCTCTTCCGATCTNNNAAAACCTGTCTCCAAACAGACCA
GAGCACACGTCTGAACTCTTTCCCTACACGACGCTCTTCCGATCTNNNCTGACCTGGCCAACAGAGAA
AGCACACGTCTGAACTCTTTCCCTACACGACGCTCTTCCGATCTNNNGTTGTGTCAACTGCTGATATG
CACACGTCTGAACTCTTTCCCTACACGACGCTCTTCCGATCTNNNCCGCTCTCATCACTTTCCAACTTCA

GAGCACACGTCTGAACTCTTTCCCTACACGACGCTCTTCCGATCTNNNTGTAAATGGGATTGGAGGC
AGCACACGTCTGAACTCTTTCCCTACACGACGCTCTTCCGATCTNNNCCAGTGGCATAATGGCAAATG
GAGCACACGTCTGAACTCTTTCCCTACACGACGCTCTTCCGATCTNNNAGAATAGAGAGCCAGAAGG
GCACACGTCTGAACTCTTTCCCTACACGACGCTCTTCCGATCTNNNGGAGGAGGCTGAGGAAAATTC
AGCACACGTCTGAACTCTTTCCCTACACGACGCTCTTCCGATCTNNNGTACTACCAGCTTGGAAACCA
CACACGTCTGAACTCTTTCCCTACACGACGCTCTTCCGATCTNNNTGCCTTTGACAACAGCTTTCATTC
GCACACGTCTGAACTCTTTCCCTACACGACGCTCTTCCGATCTNNNTCCCCTTGACTGTCTCTCAAAGTC
GAGCACACGTCTGAACTCTTTCCCTACACGACGCTCTTCCGATCTNNNCGTTTCCCACCCAGAGAAGAT
AGCACACGTCTGAACTCTTTCCCTACACGACGCTCTTCCGATCTNNNTCATTGTCCTGGGCTACAGTTT
AGCACACGTCTGAACTCTTTCCCTACACGACGCTCTTCCGATCTNNNCCTCTGCAAAAATCACAGCCT
AGCACACGTCTGAACTCTTTCCCTACACGACGCTCTTCCGATCTNNNAAGAATGGTTGGAACGGTTG
GAGCACACGTCTGAACTCTTTCCCTACACGACGCTCTTCCGATCTNNNAGTTTGTTGTCTGGCTCTTC
AGCACACGTCTGAACTCTTTCCCTACACGACGCTCTTCCGATCTNNNTCATGCCAGTCCCTCTATTGGA
CACACGTCTGAACTCTTTCCCTACACGACGCTCTTCCGATCTNNNTTCTGGCACCTCTCACTCCATTAC
AGCACACGTCTGAACTCTTTCCCTACACGACGCTCTTCCGATCTNNNGCACATACACATGAGTTTTGG
AGCACACGTCTGAACTCTTTCCCTACACGACGCTCTTCCGATCTNNNGCCTGAGTGATAGAGAGCATC
GAGCACACGTCTGAACTCTTTCCCTACACGACGCTCTTCCGATCTNNNTGGTGCAACTTTGTGTCAGG
GCACACGTCTGAACTCTTTCCCTACACGACGCTCTTCCGATCTNNNAAGGCCTTCCCTCACTACATCTG
AGCACACGTCTGAACTCTTTCCCTACACGACGCTCTTCCGATCTNNNTTATCGCTTCTCAGCATTTGGC
GAGCACACGTCTGAACTCTTTCCCTACACGACGCTCTTCCGATCTNNNCCCATAAATGTGCATGGGG
GAAGAGCACACGTCTGAACTCTTTCCCTACACGACGCTCTTCCGATCTNNNACAGACTGCATGGTAT
AGCACACGTCTGAACTCTTTCCCTACACGACGCTCTTCCGATCTNNNGTACACAAGCTGGTATGTGT
AGCACACGTCTGAACTCTTTCCCTACACGACGCTCTTCCGATCTNNNTGCACTAGTTTTTGTAATTTTC
AGCACACGTCTGAACTCTTTCCCTACACGACGCTCTTCCGATCTNNNATCTGGTCTTCCCAAACTCATG
AGCACACGTCTGAACTCTTTCCCTACACGACGCTCTTCCGATCTNNNACGTTCTTCTCACTCCCTCATCA
AGCACACGTCTGAACTCTTTCCCTACACGACGCTCTTCCGATCTNNNTTTCTGTCCCCTAGAATCCTGC
AGCACACGTCTGAACTCTTTCCCTACACGACGCTCTTCCGATCTNNNCTCCTCTGATCTGGAACAGTG
AGCACACGTCTGAACTCTTTCCCTACACGACGCTCTTCCGATCTNNNATTTTCCACCCTCATGATGCAGT
AGCACACGTCTGAACTCTTTCCCTACACGACGCTCTTCCGATCTNNNAGCAACAGGCAGAATGGTTTG
AGCACACGTCTGAACTCTTTCCCTACACGACGCTCTTCCGATCTNNNGGATGGAGTTGGGAGGATG
AGCACACGTCTGAACTCTTTCCCTACACGACGCTCTTCCGATCTNNNCTGGACCACATCTGGAATGAA
GAGCACACGTCTGAACTCTTTCCCTACACGACGCTCTTCCGATCTNNNGCAAGAACTTTATCCATCCTT
GAGCACACGTCTGAACTCTTTCCCTACACGACGCTCTTCCGATCTNNNAGGATGCTCTAAGTACAGCC
AGCACACGTCTGAACTCTTTCCCTACACGACGCTCTTCCGATCTNNNTACTTTCTTGCAGCCTGTGGTT
AGAGCACACGTCTGAACTCTTTCCCTACACGACGCTCTTCCGATCTNNNCAGCTCTCAGGCATTCGAA
GAGCACACGTCTGAACTCTTTCCCTACACGACGCTCTTCCGATCTNNNAGGGTTGGTAAATGCTAGG
AGCACACGTCTGAACTCTTTCCCTACACGACGCTCTTCCGATCTNNNCCTGACATGAAGGAGAAGTG
GAGCACACGTCTGAACTCTTTCCCTACACGACGCTCTTCCGATCTNNNAGGGGCCGACTGTGTATGTA
AGCACACGTCTGAACTCTTTCCCTACACGACGCTCTTCCGATCTNNNTTTGGATGCTTCCAGGTCAGT
GAGCACACGTCTGAACTCTTTCCCTACACGACGCTCTTCCGATCTNNNTGGTTAGTCAATGTGTGCTC
AGAGCACACGTCTGAACTCTTTCCCTACACGACGCTCTTCCGATCTNNNCTCCTTCAACAGAAGATGCT
GAGCACACGTCTGAACTCTTTCCCTACACGACGCTCTTCCGATCTNNNAAGGGCTTAGGTGTGATTTG
AGCACACGTCTGAACTCTTTCCCTACACGACGCTCTTCCGATCTNNNCTAGTGGCTCTAAAGCATGAC

AGCACACGTCTGAACTCTTTCCCTACACGACGCTCTTCCGATCTNNNCCATTTTGTGCAAGTCACAGA
AGCACACGTCTGAACTCTTTCCCTACACGACGCTCTTCCGATCTNNNAAACACTGTGGATGCTTACCAG
GAGCACACGTCTGAACTCTTTCCCTACACGACGCTCTTCCGATCTNNNTCAGTTTGATGTGTTTCTGGG
AGCACACGTCTGAACTCTTTCCCTACACGACGCTCTTCCGATCTNNNGCAGTGTTTTGTGCAAAGTCT
AGCACACGTCTGAACTCTTTCCCTACACGACGCTCTTCCGATCTNNNTTGTTCACACAGAAAAACGAA
GAGCACACGTCTGAACTCTTTCCCTACACGACGCTCTTCCGATCTNNNTAACAGGGGTGACTTAGAGG
GCACACGTCTGAACTCTTTCCCTACACGACGCTCTTCCGATCTNNNCTTATAAGCATCCAGCCTCTGCC
AGCACACGTCTGAACTCTTTCCCTACACGACGCTCTTCCGATCTNNNCCTGACAGGTTTTACTACCCCT
AGCACACGTCTGAACTCTTTCCCTACACGACGCTCTTCCGATCTNNNCAACGTTGACTAAAACATTGC
AGAGCACACGTCTGAACTCTTTCCCTACACGACGCTCTTCCGATCTNNNTAAACCAGGAGCACTTTGT
AGAGCACACGTCTGAACTCTTTCCCTACACGACGCTCTTCCGATCTNNNCGGTGGAGAGAAACCTAAA
AGCACACGTCTGAACTCTTTCCCTACACGACGCTCTTCCGATCTNNNTGATTTGTCACTGGGGTATTGC
GAGCACACGTCTGAACTCTTTCCCTACACGACGCTCTTCCGATCTNNNAGGACGGAGAGAATGTGAA
AGCACACGTCTGAACTCTTTCCCTACACGACGCTCTTCCGATCTNNNTCTACAACGGCAACCACTTTCA
AGAGCACACGTCTGAACTCTTTCCCTACACGACGCTCTTCCGATCTNNNAGGATATGAACAGGAATTG
AGCACACGTCTGAACTCTTTCCCTACACGACGCTCTTCCGATCTNNNAATCACTTAGCACAACACCTG
GAGCACACGTCTGAACTCTTTCCCTACACGACGCTCTTCCGATCTNNNTGTCCACTGCAGCATCATATT
AGCACACGTCTGAACTCTTTCCCTACACGACGCTCTTCCGATCTNNNCCTGCATAATTGTTCCGTGAAG
GAGCACACGTCTGAACTCTTTCCCTACACGACGCTCTTCCGATCTNNNATGCTCTTGAGTTGCTGATGG
GAGCACACGTCTGAACTCTTTCCCTACACGACGCTCTTCCGATCTNNNAGGGTATCCACAAAGCAATG
GAGCACACGTCTGAACTCTTTCCCTACACGACGCTCTTCCGATCTNNNGTGTATTTCCTTGACACCCTC
GAGCACACGTCTGAACTCTTTCCCTACACGACGCTCTTCCGATCTNNNCAGCATTCCATCAGAGAAGCA
AGCACACGTCTGAACTCTTTCCCTACACGACGCTCTTCCGATCTNNNAGGCACTATCCTAGTTGTTGGG
GAGCACACGTCTGAACTCTTTCCCTACACGACGCTCTTCCGATCTNNNGGAACTAACATTGCTGGAGA
GAGCACACGTCTGAACTCTTTCCCTACACGACGCTCTTCCGATCTNNNACCTTGGAATTTGGACCAGTG
GCACACGTCTGAACTCTTTCCCTACACGACGCTCTTCCGATCTNNNTACAGAGTATAGCCACTTGCGTG
AGCACACGTCTGAACTCTTTCCCTACACGACGCTCTTCCGATCTNNNAGAAAACCAAAGGATGCCTTC
AGCACACGTCTGAACTCTTTCCCTACACGACGCTCTTCCGATCTNNNCCTCATATCGGGTTTCAGCAG
AGCACACGTCTGAACTCTTTCCCTACACGACGCTCTTCCGATCTNNNACCCTCTTCTTTCCTTAGCACA
AGAGCACACGTCTGAACTCTTTCCCTACACGACGCTCTTCCGATCTNNNTGATAAGAAATCGGTGAGC
AGCACACGTCTGAACTCTTTCCCTACACGACGCTCTTCCGATCTNNNTTTAGCCTTCCTGTACACTCTG
AGCACACGTCTGAACTCTTTCCCTACACGACGCTCTTCCGATCTNNNCCCTCTCAAACCATTCCCTTCTGC
AGCACACGTCTGAACTCTTTCCCTACACGACGCTCTTCCGATCTNNNAACCTGGGGTCCAAGGATAGA
GAGCACACGTCTGAACTCTTTCCCTACACGACGCTCTTCCGATCTNNNGCTGACTACCAAGCCATCATT
AGCACACGTCTGAACTCTTTCCCTACACGACGCTCTTCCGATCTNNNATCTGAATCTGAACAGTGGGT
AGCACACGTCTGAACTCTTTCCCTACACGACGCTCTTCCGATCTNNNTTCCCGGTGTACAGAAACACAA
AGAGCACACGTCTGAACTCTTTCCCTACACGACGCTCTTCCGATCTNNNTACCCTGTGGAACTGTCCTT
AGCACACGTCTGAACTCTTTCCCTACACGACGCTCTTCCGATCTNNNTGTGGATTTGCAGGCATTCCTT
GCACACGTCTGAACTCTTTCCCTACACGACGCTCTTCCGATCTNNNTGTTGGCTTCTCTCTATGCTCCTC
AGCACACGTCTGAACTCTTTCCCTACACGACGCTCTTCCGATCTNNNAGATAACATTGACCATGTGCA
AGCACACGTCTGAACTCTTTCCCTACACGACGCTCTTCCGATCTNNNAGTGCAGACATTGCATTTGAG
AGCACACGTCTGAACTCTTTCCCTACACGACGCTCTTCCGATCTNNNACACTAAAAGGAAATGGTGGG
AAGAGCACACGTCTGAACTCTTTCCCTACACGACGCTCTTCCGATCTNNNGGCCCATAGTCCAAGTGA

AGCACACGTCTGAACTCTTTCCCTACACGACGCTCTTCCGATCTNNNCTATTGCCTTTGCAATCCCGAAC
AGCACACGTCTGAACTCTTTCCCTACACGACGCTCTTCCGATCTNNNGAAGCTCAAGTGGCAAGGAAC
GAGCACACGTCTGAACTCTTTCCCTACACGACGCTCTTCCGATCTNNNTGACTTACAAGGCAAAGAAA
GAGCACACGTCTGAACTCTTTCCCTACACGACGCTCTTCCGATCTNNNTGGAATTGCTGGAGGCATGT
GAGCACACGTCTGAACTCTTTCCCTACACGACGCTCTTCCGATCTNNNGAGGGCTTTGCTCTATACATG
GCACACGTCTGAACTCTTTCCCTACACGACGCTCTTCCGATCTNNNGAGGGCGCTAGTTCTAAAGATG
AGAGCACACGTCTGAACTCTTTCCCTACACGACGCTCTTCCGATCTNNNTGAATGAAAAGGCAGGAAG
AGCACACGTCTGAACTCTTTCCCTACACGACGCTCTTCCGATCTNNNCACTCAGTTGAGGGCCAATTCT
GCACACGTCTGAACTCTTTCCCTACACGACGCTCTTCCGATCTNNNGCCTCTATCTCTTTCTGCACTGGC
GCACACGTCTGAACTCTTTCCCTACACGACGCTCTTCCGATCTNNNTCCAAATCTCTACTAAGCCGGGG
AGCACACGTCTGAACTCTTTCCCTACACGACGCTCTTCCGATCTNNNCAGCTCTCAGGCATTCGAAAAA
AAGAGCACACGTCTGAACTCTTTCCCTACACGACGCTCTTCCGATCTNNNTCTTGTTTGCCTCCTGTGT
GCACACGTCTGAACTCTTTCCCTACACGACGCTCTTCCGATCTNNNAAACTCTTGCATCCTGGGAAACC
AGAGCACACGTCTGAACTCTTTCCCTACACGACGCTCTTCCGATCTNNNTTTTCTCACACACCCATGTCAC
AGCACACGTCTGAACTCTTTCCCTACACGACGCTCTTCCGATCTNNNAGGCAATAGACCAGAGCCCATT
AGAGCACACGTCTGAACTCTTTCCCTACACGACGCTCTTCCGATCTNNNAGGTTTTCATGTTTCCAAGC
AGCACACGTCTGAACTCTTTCCCTACACGACGCTCTTCCGATCTNNNTGTTTTATTTGGAGTAGAAAGT
AGCACACGTCTGAACTCTTTCCCTACACGACGCTCTTCCGATCTNNNAGGATCACATGTTCGGTGTTCA
AGAGCACACGTCTGAACTCTTTCCCTACACGACGCTCTTCCGATCTNNNTTCCTCTCCTCTTCTCTGACA
AGAGCACACGTCTGAACTCTTTCCCTACACGACGCTCTTCCGATCTNNNCCAGTGCACGAAAGGTCTAG
GAGCACACGTCTGAACTCTTTCCCTACACGACGCTCTTCCGATCTNNNATACATATCTCTGTGAGCTGCC
AGCACACGTCTGAACTCTTTCCCTACACGACGCTCTTCCGATCTNNNTCAGCAGCTCTGGAGTATTCCT
GCACACGTCTGAACTCTTTCCCTACACGACGCTCTTCCGATCTNNNTCCTGTTCACTTGATCCCTTCCACA
AAGAGCACACGTCTGAACTCTTTCCCTACACGACGCTCTTCCGATCTNNNCTCCATAACCCCTCCCCCATT
AGCACACGTCTGAACTCTTTCCCTACACGACGCTCTTCCGATCTNNNCAGCCTGCAAAACCTTTCAGTGC
GAAGAGCACACGTCTGAACTCTTTCCCTACACGACGCTCTTCCGATCTNNNCAACGTTGACTAAAACAT
AGCACACGTCTGAACTCTTTCCCTACACGACGCTCTTCCGATCTNNNCAGCATGATTCCTAAGGCTTCCA
AGCACACGTCTGAACTCTTTCCCTACACGACGCTCTTCCGATCTNNNGTACACAAGCTGGTATGTGTG
GAGCACACGTCTGAACTCTTTCCCTACACGACGCTCTTCCGATCTNNNGGTGAGGATTTGGGGACAAC
GAGCACACGTCTGAACTCTTTCCCTACACGACGCTCTTCCGATCTNNNCGACAATGGGTGATAGTGGG
GAGCACACGTCTGAACTCTTTCCCTACACGACGCTCTTCCGATCTNNNCCAAGTGCCTTGTTACAGTAG
AGCACACGTCTGAACTCTTTCCCTACACGACGCTCTTCCGATCTNNNGCAACAGTGACCAAGACACAAA
AGCACACGTCTGAACTCTTTCCCTACACGACGCTCTTCCGATCTNNNCTCCATCTACCTTTCTGGTCCCTC
GAGCACACGTCTGAACTCTTTCCCTACACGACGCTCTTCCGATCTNNNGTGTGATTGCCTAGAGATTGC
AGCACACGTCTGAACTCTTTCCCTACACGACGCTCTTCCGATCTNNNCTCCGCAAGTATAAGGAGGAA
GAGCACACGTCTGAACTCTTTCCCTACACGACGCTCTTCCGATCTNNNGTGAACACCCCAGTGGTCAATA
AGAGCACACGTCTGAACTCTTTCCCTACACGACGCTCTTCCGATCTNNNGCCTTTCCACTGAGTTTTCCT
GAGCACACGTCTGAACTCTTTCCCTACACGACGCTCTTCCGATCTNNNTCAGTGTATGATCTAAAGACA
GAGCACACGTCTGAACTCTTTCCCTACACGACGCTCTTCCGATCTNNNTTTCCTTGGCCCATACAGTGTT
AGCACACGTCTGAACTCTTTCCCTACACGACGCTCTTCCGATCTNNNTAGCTACCTGTCCGTCCCTTATC
AGAGCACACGTCTGAACTCTTTCCCTACACGACGCTCTTCCGATCTNNNCGCATGTAAGGTTAGTCACC
AGCACACGTCTGAACTCTTTCCCTACACGACGCTCTTCCGATCTNNNCCTGGTAGCAAGCAAGAAGACA
GCACACGTCTGAACTCTTTCCCTACACGACGCTCTTCCGATCTNNNTTTGGCCCAACAGGCTATTTCATC

GAGCACACGTCTGAACTCTTTCCCTACACGACGCTCTTCCGATCTNNNGGCAAAGTCTGACCTCATAG
AGCACACGTCTGAACTCTTTCCCTACACGACGCTCTTCCGATCTNNNCAGTCTCCCTTCCTTAGTCCCTTC
AGCACACGTCTGAACTCTTTCCCTACACGACGCTCTTCCGATCTNNNAGCCTGAAATTGGCTGAACTT
AGCACACGTCTGAACTCTTTCCCTACACGACGCTCTTCCGATCTNNNACCTTGGAATTTGGACCAGTG
AAGAGCACACGTCTGAACTCTTTCCCTACACGACGCTCTTCCGATCTNNNGCCTTTGGGCATAGTCAT
GCACACGTCTGAACTCTTTCCCTACACGACGCTCTTCCGATCTNNNAGCCTTACTTCCTCATCAGGGTTC
GAGCACACGTCTGAACTCTTTCCCTACACGACGCTCTTCCGATCTNNNGGTAACTCACACTCATCCACC
AGCACACGTCTGAACTCTTTCCCTACACGACGCTCTTCCGATCTNNNCCAGCGAGCCAACAGTTCATA
GAGCACACGTCTGAACTCTTTCCCTACACGACGCTCTTCCGATCTNNNGCCAATGAGCTGTGAATGAA
GAGCACACGTCTGAACTCTTTCCCTACACGACGCTCTTCCGATCTNNNAGTGGACAAAACACGGGAAC
AGCACACGTCTGAACTCTTTCCCTACACGACGCTCTTCCGATCTNNNGAGGATTTGCGCTTGTGTATG
AGCACACGTCTGAACTCTTTCCCTACACGACGCTCTTCCGATCTNNNCTTGTGCCTTCTGGGAAAGAA
GAGCACACGTCTGAACTCTTTCCCTACACGACGCTCTTCCGATCTNNNCACCCAATACTGAGGAGGC
GAGCACACGTCTGAACTCTTTCCCTACACGACGCTCTTCCGATCTNNNACTGTCACTGTGGGGTTTTCT
AAGAGCACACGTCTGAACTCTTTCCCTACACGACGCTCTTCCGATCTNNNATGAGTGATTCTGGGTG
GAGCACACGTCTGAACTCTTTCCCTACACGACGCTCTTCCGATCTNNNCCCCACAAAGATCCCAATAGA
GAGCACACGTCTGAACTCTTTCCCTACACGACGCTCTTCCGATCTNNNAGCAGAAAACCTATCCCCAA
GAGCACACGTCTGAACTCTTTCCCTACACGACGCTCTTCCGATCTNNNTGTTCATCTACCATTCCCTTC
AGCACACGTCTGAACTCTTTCCCTACACGACGCTCTTCCGATCTNNNTAACTAGGGGAGGAAGCCTTA
AGCACACGTCTGAACTCTTTCCCTACACGACGCTCTTCCGATCTNNNTGTAGTGTCTGTCCTTTTGGC
GAGCACACGTCTGAACTCTTTCCCTACACGACGCTCTTCCGATCTNNNTGACACCACCTATGAAAGAG
GCACACGTCTGAACTCTTTCCCTACACGACGCTCTTCCGATCTNNNCTCCTCTTACCCCCTCCTTCCTCAT
AGCACACGTCTGAACTCTTTCCCTACACGACGCTCTTCCGATCTNNNCTGAGTGTGTGTGTCTGGA
AGCACACGTCTGAACTCTTTCCCTACACGACGCTCTTCCGATCTNNNCTCTAATCCAGGCAAAGAGGC
GAGCACACGTCTGAACTCTTTCCCTACACGACGCTCTTCCGATCTNNNTGCATTGCATGCTGATACCA
GCACACGTCTGAACTCTTTCCCTACACGACGCTCTTCCGATCTNNNCCCCTAGTGCTGTCCTTTGATAGA
AGCACACGTCTGAACTCTTTCCCTACACGACGCTCTTCCGATCTNNNTACCCCATTGGCTTTGTTGTTG
GAGCACACGTCTGAACTCTTTCCCTACACGACGCTCTTCCGATCTNNNAAGATATCCTGAGAGAGGT
GAGCACACGTCTGAACTCTTTCCCTACACGACGCTCTTCCGATCTNNNCGTGGTCACTCAGGGTCATA
AGAGCACACGTCTGAACTCTTTCCCTACACGACGCTCTTCCGATCTNNNCCAAAGTCATCCTTATCTGT
AGCACACGTCTGAACTCTTTCCCTACACGACGCTCTTCCGATCTNNNCTGCCTCCATGATCCTCTTCAA
AGCACACGTCTGAACTCTTTCCCTACACGACGCTCTTCCGATCTNNNGTTTTCCCCTTGTCTCTCCTTGG
AGCACACGTCTGAACTCTTTCCCTACACGACGCTCTTCCGATCTNNNTTTGAGATGTCAGCAGTCTCTG
AGCACACGTCTGAACTCTTTCCCTACACGACGCTCTTCCGATCTNNNATGAGAATAGGTCTCATGGG
GCACACGTCTGAACTCTTTCCCTACACGACGCTCTTCCGATCTNNNCCTAGTGCCTGGCATATGTGTA
GAGCACACGTCTGAACTCTTTCCCTACACGACGCTCTTCCGATCTNNNGGGTCCATGAAGGCAGAGAT
AAGAGCACACGTCTGAACTCTTTCCCTACACGACGCTCTTCCGATCTNNNTCAGTCCTTTTCCAAGTCCA
GCACACGTCTGAACTCTTTCCCTACACGACGCTCTTCCGATCTNNNAGCATCAGGCCTATAGCAGACT
GAGCACACGTCTGAACTCTTTCCCTACACGACGCTCTTCCGATCTNNNTCCTTGAAAAGTTCAGCAAC
AGCACACGTCTGAACTCTTTCCCTACACGACGCTCTTCCGATCTNNNTCCACAATGCCTTACATTTTGCA
GAGCACACGTCTGAACTCTTTCCCTACACGACGCTCTTCCGATCTNNNTTTTCTCACACACCCATGTCA
AGCACACGTCTGAACTCTTTCCCTACACGACGCTCTTCCGATCTNNNACTCAGTGATTGCCAAGGAAG
GCACACGTCTGAACTCTTTCCCTACACGACGCTCTTCCGATCTNNNACTCCACAGATGTACTGTAGCG

GAGCACACGTCTGAACTCTTTCCCTACACGACGCTCTTCCGATCTNNNTGAGTGGTTTTGCTCCCTTTT
AGCACACGTCTGAACTCTTTCCCTACACGACGCTCTTCCGATCTNNNTGAGCCTAAGGGAAAAATCTG
AGCACACGTCTGAACTCTTTCCCTACACGACGCTCTTCCGATCTNNNTGTTGGGAGCTACACCAATAG
GAAGAGCACACGTCTGAACTCTTTCCCTACACGACGCTCTTCCGATCTNNNAAAGAAAGGCAGCAG
GAGCACACGTCTGAACTCTTTCCCTACACGACGCTCTTCCGATCTNNNATGGCAAACCCAGAAATCAG
GAGCACACGTCTGAACTCTTTCCCTACACGACGCTCTTCCGATCTNNNTTGCCTGATTGAGAACCAAC
GAGCACACGTCTGAACTCTTTCCCTACACGACGCTCTTCCGATCTNNNTAAGCTTGGTTTTCTGACGCC
AGCACACGTCTGAACTCTTTCCCTACACGACGCTCTTCCGATCTNNNCACATTGAGAACACGTTCGGTT
AGAGCACACGTCTGAACTCTTTCCCTACACGACGCTCTTCCGATCTNNNAGAGGTATGGGGACAAAC
AGAGCACACGTCTGAACTCTTTCCCTACACGACGCTCTTCCGATCTNNNCAGTCACCTATTCCTGCTGC
GCACACGTCTGAACTCTTTCCCTACACGACGCTCTTCCGATCTNNNAATCAGCAAACACAGATGACCC
AGCACACGTCTGAACTCTTTCCCTACACGACGCTCTTCCGATCTNNNAGGGAAGGCATCAGTTCCTTA
AGCACACGTCTGAACTCTTTCCCTACACGACGCTCTTCCGATCTNNNCATTTGCTGCAACACAGACACT
GCACACGTCTGAACTCTTTCCCTACACGACGCTCTTCCGATCTNNNAAGGGATTTGGTCACTGGGATT
GAGCACACGTCTGAACTCTTTCCCTACACGACGCTCTTCCGATCTNNNTTTCTTCATCCCGCCATCCTTA
AGCACACGTCTGAACTCTTTCCCTACACGACGCTCTTCCGATCTNNNTCAAGGAGAAGAAAGCAGTTC
AGCACACGTCTGAACTCTTTCCCTACACGACGCTCTTCCGATCTNNNTGACTTCAGTGATCCTCCAAGG
GAGCACACGTCTGAACTCTTTCCCTACACGACGCTCTTCCGATCTNNNAGCTGGGGAGAGGGATATG
AGCACACGTCTGAACTCTTTCCCTACACGACGCTCTTCCGATCTNNNGCCAAAGTTGGAACAAAGTGT
GAGCACACGTCTGAACTCTTTCCCTACACGACGCTCTTCCGATCTNNNCACCGTATTGATAGGATAAG
GAGCACACGTCTGAACTCTTTCCCTACACGACGCTCTTCCGATCTNNNATTTCTGCTGGGCAGGATAC
GAGCACACGTCTGAACTCTTTCCCTACACGACGCTCTTCCGATCTNNNTGTGGCTGTATAGTACTCCA
AGCACACGTCTGAACTCTTTCCCTACACGACGCTCTTCCGATCTNNNTGAGGGATTGGAACTGAGGT
AGAGCACACGTCTGAACTCTTTCCCTACACGACGCTCTTCCGATCTNNNATCTTTTGCTCCCTGCATTG
CACACGTCTGAACTCTTTCCCTACACGACGCTCTTCCGATCTNNNCTGGGAGTTCACAGCTGTATGTGC
GAGCACACGTCTGAACTCTTTCCCTACACGACGCTCTTCCGATCTNNNAAGGGCAAGGACTGATATAC
AAGAGCACACGTCTGAACTCTTTCCCTACACGACGCTCTTCCGATCTNNNTGCAAGCATCTGTTTATGT
GAGCACACGTCTGAACTCTTTCCCTACACGACGCTCTTCCGATCTNNNGTCTAGTCTCCCTGTTCCCAA
AGAGCACACGTCTGAACTCTTTCCCTACACGACGCTCTTCCGATCTNNNACCTACTAGCTCTGTGTGT
GAGCACACGTCTGAACTCTTTCCCTACACGACGCTCTTCCGATCTNNNACACAGTGGCTAAAGAGAG
GCACACGTCTGAACTCTTTCCCTACACGACGCTCTTCCGATCTNNNTCTAGCTTTGAGGACAGAGGGT
AGCACACGTCTGAACTCTTTCCCTACACGACGCTCTTCCGATCTNNNTACACAAGCTAAAGCTGCACCA
AGCACACGTCTGAACTCTTTCCCTACACGACGCTCTTCCGATCTNNNTGTATTCCTAACCCTACCCCAGC
AGCACACGTCTGAACTCTTTCCCTACACGACGCTCTTCCGATCTNNNTGAAAAATTAGTTATTGCTTGG
AGCACACGTCTGAACTCTTTCCCTACACGACGCTCTTCCGATCTNNNCTTCAGTCTTTGCCTCCCCATA
GAGCACACGTCTGAACTCTTTCCCTACACGACGCTCTTCCGATCTNNNACTTGGAAGAGGCCTTCAAA
AGCACACGTCTGAACTCTTTCCCTACACGACGCTCTTCCGATCTNNNTAGCAAAGCATTCCTTTTGCCAC
AGAGCACACGTCTGAACTCTTTCCCTACACGACGCTCTTCCGATCTNNNTGTGCAACCCAGTTTGTGGT
AGCACACGTCTGAACTCTTTCCCTACACGACGCTCTTCCGATCTNNNTATCCAGGATCCCCAAGGACAA
AGCACACGTCTGAACTCTTTCCCTACACGACGCTCTTCCGATCTNNNGAGTTCTCTCTGTGTGTGCAT
GCACACGTCTGAACTCTTTCCCTACACGACGCTCTTCCGATCTNNNAGGCTTCATGTTGCTCCATTTCTC
GAGCACACGTCTGAACTCTTTCCCTACACGACGCTCTTCCGATCTNNNGGCCTGTAAGTGGGAATGA
AGAGCACACGTCTGAACTCTTTCCCTACACGACGCTCTTCCGATCTNNNGGTTATGGGTCTTTGCTTTG

GAGCACACGTCTGAACTCTTTCCCTACACGACGCTCTTCCGATCTNNNAGCCAGAAAATGAGATGAG
AGCACACGTCTGAACTCTTTCCCTACACGACGCTCTTCCGATCTNNNAAACTTGACAGTGGCTTTCCTG
AGAGCACACGTCTGAACTCTTTCCCTACACGACGCTCTTCCGATCTNNNTGCAAGCATCTGTTTATGTG
AGCACACGTCTGAACTCTTTCCCTACACGACGCTCTTCCGATCTNNNATGTGAAAGAGATGCTGTGAC
AGCACACGTCTGAACTCTTTCCCTACACGACGCTCTTCCGATCTNNNGGGGTCCACATGCCAATAATTT
AGCACACGTCTGAACTCTTTCCCTACACGACGCTCTTCCGATCTNNNAAACTGCTATCTCACCTCCCACT
GAGCACACGTCTGAACTCTTTCCCTACACGACGCTCTTCCGATCTNNNTATTTCAGCATCTGCCTCTTC
AAGAGCACACGTCTGAACTCTTTCCCTACACGACGCTCTTCCGATCTNNNATACATATCTCTGTGAGCT
GAGCACACGTCTGAACTCTTTCCCTACACGACGCTCTTCCGATCTNNNGATTTCTGGTTCAGTGACCC
AGCACACGTCTGAACTCTTTCCCTACACGACGCTCTTCCGATCTNNNGCTGAGTGGATCAGCTCAAAG
AGCACACGTCTGAACTCTTTCCCTACACGACGCTCTTCCGATCTNNNGCCCATTGCAGTATTTTGTTGC
AGCACACGTCTGAACTCTTTCCCTACACGACGCTCTTCCGATCTNNNTGCAATGTTCATTCCTGGTTGA
CACACGTCTGAACTCTTTCCCTACACGACGCTCTTCCGATCTNNNTGCAATGTTCATTCCTGGTTGAGA
AGCACACGTCTGAACTCTTTCCCTACACGACGCTCTTCCGATCTNNNTCTTCTCTTTGCTCCATTGGCAT
GAGCACACGTCTGAACTCTTTCCCTACACGACGCTCTTCCGATCTNNNGTCCAGGTGACTTGTGTATG
AGAGCACACGTCTGAACTCTTTCCCTACACGACGCTCTTCCGATCTNNNATTTTCCCCAATGCCCACAA
GAGCACACGTCTGAACTCTTTCCCTACACGACGCTCTTCCGATCTNNNACTCTTTCTCTCTCCTGTGGCT
GAGCACACGTCTGAACTCTTTCCCTACACGACGCTCTTCCGATCTNNNACACGCCAGGACCTCTAAAAA
AGCACACGTCTGAACTCTTTCCCTACACGACGCTCTTCCGATCTNNNTGCAGTATTGGACACGACAGA
GAGCACACGTCTGAACTCTTTCCCTACACGACGCTCTTCCGATCTNNNAGGCTGAAAATGCCTTGAGT
AGCACACGTCTGAACTCTTTCCCTACACGACGCTCTTCCGATCTNNNTCAAGATCCAGCCTGTGGTTTA
AGAGCACACGTCTGAACTCTTTCCCTACACGACGCTCTTCCGATCTNNNCCTGAGAATGGAGGGTAA
AGCACACGTCTGAACTCTTTCCCTACACGACGCTCTTCCGATCTNNNATTCCACTTCTCTGCCTCCAGTA
GCACACGTCTGAACTCTTTCCCTACACGACGCTCTTCCGATCTNNNTTCCAGACTGTTGCTGATTTCTTC
AGCACACGTCTGAACTCTTTCCCTACACGACGCTCTTCCGATCTNNNTGCTTCCTAGGTATGTAAGGC
AGCACACGTCTGAACTCTTTCCCTACACGACGCTCTTCCGATCTNNNCAAGAACTTCCAGGAATCAGTG
AGCACACGTCTGAACTCTTTCCCTACACGACGCTCTTCCGATCTNNNTCTTGGTCCCAACACAGAGGTT
AGCACACGTCTGAACTCTTTCCCTACACGACGCTCTTCCGATCTNNNTAGCCACTTCCTGGCTTGTAGA
GAGCACACGTCTGAACTCTTTCCCTACACGACGCTCTTCCGATCTNNNGGATACAAGGCATCTCAAGC
AGCACACGTCTGAACTCTTTCCCTACACGACGCTCTTCCGATCTNNNAAGGGTCTACGGACTTCAACA
AGCACACGTCTGAACTCTTTCCCTACACGACGCTCTTCCGATCTNNNTGCCCTGCTGTCACTCAATAAA
AGAGCACACGTCTGAACTCTTTCCCTACACGACGCTCTTCCGATCTNNNTAGTGTATTGGTCAGGATT
AAGAGCACACGTCTGAACTCTTTCCCTACACGACGCTCTTCCGATCTNNNGAGGTTCTAAGTGGTGG
AGCACACGTCTGAACTCTTTCCCTACACGACGCTCTTCCGATCTNNNTGCATCACAGGGTAGCATTTC
GAGCACACGTCTGAACTCTTTCCCTACACGACGCTCTTCCGATCTNNNAGAGTGAGAGAGAGATGTG
AGCACACGTCTGAACTCTTTCCCTACACGACGCTCTTCCGATCTNNNTATCTATAAGTGCGTGTGTGC
AGCACACGTCTGAACTCTTTCCCTACACGACGCTCTTCCGATCTNNNTGATCTGAGCTGTAAGGGAGG
AGCACACGTCTGAACTCTTTCCCTACACGACGCTCTTCCGATCTNNNGAAGACAGAGAGGCAGGAGA
GCACACGTCTGAACTCTTTCCCTACACGACGCTCTTCCGATCTNNNTAAAACCTGCCAGGGAACTCAA
GAGCACACGTCTGAACTCTTTCCCTACACGACGCTCTTCCGATCTNNNTCAGGCATGTGCTATGTGAAT
AGCACACGTCTGAACTCTTTCCCTACACGACGCTCTTCCGATCTNNNCCCAATATAGCAACAGGGGAA
AGAGCACACGTCTGAACTCTTTCCCTACACGACGCTCTTCCGATCTNNNGCATCACGGTAACCATAAA
GAGCACACGTCTGAACTCTTTCCCTACACGACGCTCTTCCGATCTNNNAAAGCCTTCTCACTACTCCCA

AGCACACGTCTGAACTCTTTCCCTACACGACGCTCTTCCGATCTNNNTGTGTATGACATGATCCTTTTG
AGCACACGTCTGAACTCTTTCCCTACACGACGCTCTTCCGATCTNNNAGGGTGGCTAGCTGTAGCTAT
AGCACACGTCTGAACTCTTTCCCTACACGACGCTCTTCCGATCTNNNTTGTGCACAGCCTCTGTAAATC
AGCACACGTCTGAACTCTTTCCCTACACGACGCTCTTCCGATCTNNNAGAAGATGCAAACAGCTGGG
CACACGTCTGAACTCTTTCCCTACACGACGCTCTTCCGATCTNNNCACTTGTCAGTTCCTGCACCATAC
GAGCACACGTCTGAACTCTTTCCCTACACGACGCTCTTCCGATCTNNNCCCTCAAGGAGTTTACACTC
GAGCACACGTCTGAACTCTTTCCCTACACGACGCTCTTCCGATCTNNNGTGAGGAGTGAGAAAGAG
AGCACACGTCTGAACTCTTTCCCTACACGACGCTCTTCCGATCTNNNCATGTGGGACCAAGTCTAGA
AGAGCACACGTCTGAACTCTTTCCCTACACGACGCTCTTCCGATCTNNNGGCCAACATGTCACCTAAC
AGAGCACACGTCTGAACTCTTTCCCTACACGACGCTCTTCCGATCTNNNACAGAACAGGCCTTCCTTC
AAGAGCACACGTCTGAACTCTTTCCCTACACGACGCTCTTCCGATCTNNNAGATGTTTCAGCACCCTC
GCACACGTCTGAACTCTTTCCCTACACGACGCTCTTCCGATCTNNNGGAAAGAGATGCAGAAAAGCC
AGCACACGTCTGAACTCTTTCCCTACACGACGCTCTTCCGATCTNNNTGGGACCTACAACTCAAGATG
AGCACACGTCTGAACTCTTTCCCTACACGACGCTCTTCCGATCTNNNTACCTTGAGGTATGACTGTG
AGCACACGTCTGAACTCTTTCCCTACACGACGCTCTTCCGATCTNNNTCATCTTGTTGTCTAACGCCTG
GAGCACACGTCTGAACTCTTTCCCTACACGACGCTCTTCCGATCTNNNCTCCCTAAAGTGTCTGCCACA
AGCACACGTCTGAACTCTTTCCCTACACGACGCTCTTCCGATCTNNNAAGATGTTTCAGCACCCTCCA
AGCACACGTCTGAACTCTTTCCCTACACGACGCTCTTCCGATCTNNNTTTGACAAGGCCTCTTACTGTC
AGCACACGTCTGAACTCTTTCCCTACACGACGCTCTTCCGATCTNNNCTGCATATCATCCTCCCACTCC
GCACACGTCTGAACTCTTTCCCTACACGACGCTCTTCCGATCTNNNGCACTCTGTGTGTGTGAGAGA
AGAGCACACGTCTGAACTCTTTCCCTACACGACGCTCTTCCGATCTNNNTGGTTTTATTCCTCTGGCA
GAGCACACGTCTGAACTCTTTCCCTACACGACGCTCTTCCGATCTNNNAACATCTCCTTCCACTCCTCC
GCACACGTCTGAACTCTTTCCCTACACGACGCTCTTCCGATCTNNNATTTGAAGCCCTTGCTGGGTAA
AGCACACGTCTGAACTCTTTCCCTACACGACGCTCTTCCGATCTNNNCATTTTGGTCATGCACATCCCT
AGAGCACACGTCTGAACTCTTTCCCTACACGACGCTCTTCCGATCTNNNAGATCTTTTATCCTGCTGC
AGAGCACACGTCTGAACTCTTTCCCTACACGACGCTCTTCCGATCTNNNAAGCAAAGTGTTGTGGTA
GAGCACACGTCTGAACTCTTTCCCTACACGACGCTCTTCCGATCTNNNACACAGATCCAGGCACAAAG
GAGCACACGTCTGAACTCTTTCCCTACACGACGCTCTTCCGATCTNNNGCAGTCACGGGATTTGCTAC
GAGCACACGTCTGAACTCTTTCCCTACACGACGCTCTTCCGATCTNNNACCAGTCAGGTTCATACCAG
AAGAGCACACGTCTGAACTCTTTCCCTACACGACGCTCTTCCGATCTNNNCACATGTCCTCTCCGAATC
GAGCACACGTCTGAACTCTTTCCCTACACGACGCTCTTCCGATCTNNNCCACTAATAGGATGAGGCAG
AGCACACGTCTGAACTCTTTCCCTACACGACGCTCTTCCGATCTNNNTTGCGACGAGTTCTGGAGTA
GAGCACACGTCTGAACTCTTTCCCTACACGACGCTCTTCCGATCTNNNTGCTGGGTAATCTTGTAGAA
GAGCACACGTCTGAACTCTTTCCCTACACGACGCTCTTCCGATCTNNNGTGGGCAGTAAAACAGGAC
AGCACACGTCTGAACTCTTTCCCTACACGACGCTCTTCCGATCTNNNTTCATGAACCAATCTCAGGAGC
AGCACACGTCTGAACTCTTTCCCTACACGACGCTCTTCCGATCTNNNGCAGTCCATGGTTCACCAAAA
GCACACGTCTGAACTCTTTCCCTACACGACGCTCTTCCGATCTNNNAGAGGCATGTGTACTTACGGC
GAGCACACGTCTGAACTCTTTCCCTACACGACGCTCTTCCGATCTNNNTCCTTTCTCTTCCTTGTTGGG
GCACACGTCTGAACTCTTTCCCTACACGACGCTCTTCCGATCTNNNCAGCATGTGAGAGTGTGAGAG
AGCACACGTCTGAACTCTTTCCCTACACGACGCTCTTCCGATCTNNNAGAGACAGAGCACCATGACTT
AGCACACGTCTGAACTCTTTCCCTACACGACGCTCTTCCGATCTNNNATTCCACATCTGTTTTCCCCACC
GAAGAGCACACGTCTGAACTCTTTCCCTACACGACGCTCTTCCGATCTNNNAAGGAACAGCTGGGAT
GAGCACACGTCTGAACTCTTTCCCTACACGACGCTCTTCCGATCTNNNTCCTTGTATCCCTCAGTGTCT

GAGCACACGTCTGAACTCTTTCCCTACACGACGCTCTTCCGATCTNNNTTGTTCCAGGATGTGTCTGTC
GCACACGTCTGAACTCTTTCCCTACACGACGCTCTTCCGATCTNNNAGGATTTGGCTAGCAGACTGAGA
AGCACACGTCTGAACTCTTTCCCTACACGACGCTCTTCCGATCTNNNTTGTGACTTGGGGAGTTTTCCT
AGCACACGTCTGAACTCTTTCCCTACACGACGCTCTTCCGATCTNNNCAGCATTCCATCAGAGAAGCAG
AGAGCACACGTCTGAACTCTTTCCCTACACGACGCTCTTCCGATCTNNNTAGATTGTGGCTTTCCAGAC
GAGCACACGTCTGAACTCTTTCCCTACACGACGCTCTTCCGATCTNNNAGCCAACTTGTCTTCTAGTACC
AGCACACGTCTGAACTCTTTCCCTACACGACGCTCTTCCGATCTNNNTGTATCCTGAGACTGGGTTGAA
GAGCACACGTCTGAACTCTTTCCCTACACGACGCTCTTCCGATCTNNNATGTACAAGGCAAGCAGCATC
GAGCACACGTCTGAACTCTTTCCCTACACGACGCTCTTCCGATCTNNNGGAGGGGTCCATAGGCTTATC
GAGCACACGTCTGAACTCTTTCCCTACACGACGCTCTTCCGATCTNNNTCCTTCTGTGTTCCTCTTTGTG
AGCACACGTCTGAACTCTTTCCCTACACGACGCTCTTCCGATCTNNNAGTTTTCACCGTCTGCTTCAGTT
GAGCACACGTCTGAACTCTTTCCCTACACGACGCTCTTCCGATCTNNNGACTTAGCATGTCCTCAGCCA
AGCACACGTCTGAACTCTTTCCCTACACGACGCTCTTCCGATCTNNNTTGTATATATCCCACCCCTGCCCC
GAGCACACGTCTGAACTCTTTCCCTACACGACGCTCTTCCGATCTNNNCAGACCCACAGGGAAATTCTGT
GAGCACACGTCTGAACTCTTTCCCTACACGACGCTCTTCCGATCTNNNTGCATCCTGCACTTTTTCTCAG
AGAGCACACGTCTGAACTCTTTCCCTACACGACGCTCTTCCGATCTNNNGCAAATATTGTCAACTGTGT
GAGCACACGTCTGAACTCTTTCCCTACACGACGCTCTTCCGATCTNNNTAGGTAAGGTTCCCCCAGACA
AGCACACGTCTGAACTCTTTCCCTACACGACGCTCTTCCGATCTNNNTGGTGAAAGGCACCTAGAACAA
GCACACGTCTGAACTCTTTCCCTACACGACGCTCTTCCGATCTNNNCCAAGCACCAAGGATGGGTAATA
GAGCACACGTCTGAACTCTTTCCCTACACGACGCTCTTCCGATCTNNNTAATGGAAGCTGGTCATCAGC
AGCACACGTCTGAACTCTTTCCCTACACGACGCTCTTCCGATCTNNNTCTACTTGCCTTTTTCTCTCTCTC
AGAGCACACGTCTGAACTCTTTCCCTACACGACGCTCTTCCGATCTNNNTGCAGAAGAAATTCTACCCTC
GAGCACACGTCTGAACTCTTTCCCTACACGACGCTCTTCCGATCTNNNAACCAGGCATATTGTTGGAAC
GAGCACACGTCTGAACTCTTTCCCTACACGACGCTCTTCCGATCTNNNCGCGTGCTGTAAGCTACCTAA
AGCACACGTCTGAACTCTTTCCCTACACGACGCTCTTCCGATCTNNNCTGGTAAGCAGCAGCATAGTTG
GAGCACACGTCTGAACTCTTTCCCTACACGACGCTCTTCCGATCTNNNTCTGAGAGGGCTGTTTTGTCCT
AGCACACGTCTGAACTCTTTCCCTACACGACGCTCTTCCGATCTNNNAACAGTAAGTTACCTTGCACCCG
AGCACACGTCTGAACTCTTTCCCTACACGACGCTCTTCCGATCTNNNGTTCAGGCACAGCTAGAATCCA
AGCACACGTCTGAACTCTTTCCCTACACGACGCTCTTCCGATCTNNNAATTCTCTCAAGGGTAGTGGGG
AGCACACGTCTGAACTCTTTCCCTACACGACGCTCTTCCGATCTNNNTATGAACAGGGCCTACCAGAAC
AGCACACGTCTGAACTCTTTCCCTACACGACGCTCTTCCGATCTNNNTCCAGGTAATGAGGCTTTAGGC
CACACGTCTGAACTCTTTCCCTACACGACGCTCTTCCGATCTNNNCAGGTCTCAGGGAGAGGGAACCA
AGCACACGTCTGAACTCTTTCCCTACACGACGCTCTTCCGATCTNNNTCCCCCACTCAAGCATTTGAATC
GAGCACACGTCTGAACTCTTTCCCTACACGACGCTCTTCCGATCTNNNTGTGAAGCTACAGTTTTCTGG
GAGCACACGTCTGAACTCTTTCCCTACACGACGCTCTTCCGATCTNNNAGGTCCTTCAAACAGACACAC
AGCACACGTCTGAACTCTTTCCCTACACGACGCTCTTCCGATCTNNNAATGTCCTCCAGCATTGTCTGTG
AGAGCACACGTCTGAACTCTTTCCCTACACGACGCTCTTCCGATCTNNNATGTGTTCACCAACTTCCTCTC
AGCACACGTCTGAACTCTTTCCCTACACGACGCTCTTCCGATCTNNNGCTTCAGGTCTCACATGTACCAA
AGCACACGTCTGAACTCTTTCCCTACACGACGCTCTTCCGATCTNNNAGGCTCCTCATCAGATTCCCATT
GAGCACACGTCTGAACTCTTTCCCTACACGACGCTCTTCCGATCTNNNTGTTGGTGGTCTCAAGTCTGT
GAGCACACGTCTGAACTCTTTCCCTACACGACGCTCTTCCGATCTNNNATCCGTATTAGTTGGGGTCCT
AGCACACGTCTGAACTCTTTCCCTACACGACGCTCTTCCGATCTNNNTTGTACCCAATTACAGAGCTTCA
GCACACGTCTGAACTCTTTCCCTACACGACGCTCTTCCGATCTNNNTACCATTTGGGTTTGTTGGGTGT

AGCACACGTCTGAACTCTTTCCCTACACGACGCTCTTCCGATCTNNNTCTCTAACAGCTTGCAGGTCAA
AGCACACGTCTGAACTCTTTCCCTACACGACGCTCTTCCGATCTNNNAGAAATGTTCTCTGCAGTTTTG
GAGCACACGTCTGAACTCTTTCCCTACACGACGCTCTTCCGATCTNNNATGAGAATAGGTCTCATGGG
AGAGCACACGTCTGAACTCTTTCCCTACACGACGCTCTTCCGATCTNNNTGCGTTTGTCTCAGTAAGCA
GAGCACACGTCTGAACTCTTTCCCTACACGACGCTCTTCCGATCTNNNACATTGGAGGGGGAAAAG
GAGCACACGTCTGAACTCTTTCCCTACACGACGCTCTTCCGATCTNNNGGCTAGATCAATGGGCTCTA
AGCACACGTCTGAACTCTTTCCCTACACGACGCTCTTCCGATCTNNNTCTCTACATTCCACCAGCACAGT
GAGCACACGTCTGAACTCTTTCCCTACACGACGCTCTTCCGATCTNNNCGTCTTGCCTATCCTGAAACC
AAGAGCACACGTCTGAACTCTTTCCCTACACGACGCTCTTCCGATCTNNNTGCGTTTGTCTCAGTAAG
GAGCACACGTCTGAACTCTTTCCCTACACGACGCTCTTCCGATCTNNNCCGCCCACCACTTTACCAAAT
AAGAGCACACGTCTGAACTCTTTCCCTACACGACGCTCTTCCGATCTNNNACCATCAAATTCCTTGGG
GCACACGTCTGAACTCTTTCCCTACACGACGCTCTTCCGATCTNNNGCATAGTACATGGGTGTCCTCA
GAGCACACGTCTGAACTCTTTCCCTACACGACGCTCTTCCGATCTNNNTCCAGCAAGGCTGCTAAAAGT
GAGCACACGTCTGAACTCTTTCCCTACACGACGCTCTTCCGATCTNNNTCCCTGCCCTCAAGAAACAAA
GAGCACACGTCTGAACTCTTTCCCTACACGACGCTCTTCCGATCTNNNTTTACGTTCCGGGGATTATGA
AGCACACGTCTGAACTCTTTCCCTACACGACGCTCTTCCGATCTNNNTCCCAACAATGTTTTGAGATA
GAAGAGCACACGTCTGAACTCTTTCCCTACACGACGCTCTTCCGATCTNNNGCCAAGGATGGCAAAG
GAGCACACGTCTGAACTCTTTCCCTACACGACGCTCTTCCGATCTNNNAGGAATAGTGCCCATTGTGT
AGCACACGTCTGAACTCTTTCCCTACACGACGCTCTTCCGATCTNNNTACTTGCCCAATGAGGAGCTTT
AGCACACGTCTGAACTCTTTCCCTACACGACGCTCTTCCGATCTNNNGGATCTTCTCTGAGCATGCCTA
GCACACGTCTGAACTCTTTCCCTACACGACGCTCTTCCGATCTNNNATGTACCAGCTACCATCCATCCA
GAGCACACGTCTGAACTCTTTCCCTACACGACGCTCTTCCGATCTNNNATGGGGTGAGAGTGACTG
AGCACACGTCTGAACTCTTTCCCTACACGACGCTCTTCCGATCTNNNAGAGACCCATTTCCTCCAATCC
GCACACGTCTGAACTCTTTCCCTACACGACGCTCTTCCGATCTNNNGAGTGTGTGCGCATGTTCTTT
AGCACACGTCTGAACTCTTTCCCTACACGACGCTCTTCCGATCTNNNATTTATGGCTGACTGGCATAC
GAGCACACGTCTGAACTCTTTCCCTACACGACGCTCTTCCGATCTNNNGAAGTCAGGTGGGAAAGG
GAGCACACGTCTGAACTCTTTCCCTACACGACGCTCTTCCGATCTNNNTCAGCTCCCTAGACAGAGTA
GAGCACACGTCTGAACTCTTTCCCTACACGACGCTCTTCCGATCTNNNTGTGCCCTGTGTGAAAACAG
GAGCACACGTCTGAACTCTTTCCCTACACGACGCTCTTCCGATCTNNNGTCTCCAGACCTGCCCAGAT
GCACACGTCTGAACTCTTTCCCTACACGACGCTCTTCCGATCTNNNGTTATGGTGGGATGCGACTTC
AAGAGCACACGTCTGAACTCTTTCCCTACACGACGCTCTTCCGATCTNNNGACTGTGTTTCTCCCTTCT
GAGCACACGTCTGAACTCTTTCCCTACACGACGCTCTTCCGATCTNNNAAAAATGAAGCTTTGCAGCA
GAGCACACGTCTGAACTCTTTCCCTACACGACGCTCTTCCGATCTNNNCCAAGAGGCCTAGCAAGAG
AGAGCACACGTCTGAACTCTTTCCCTACACGACGCTCTTCCGATCTNNNCCTTTGATTGTCCCCACACA
AGCACACGTCTGAACTCTTTCCCTACACGACGCTCTTCCGATCTNNNAAAGTGCTGGGACTACACAAG
GAGCACACGTCTGAACTCTTTCCCTACACGACGCTCTTCCGATCTNNNGACCTTTCAGAGGTAGGATC
GAGCACACGTCTGAACTCTTTCCCTACACGACGCTCTTCCGATCTNNNCCTTCAGCTCTTTGGTCTTGT
AGCACACGTCTGAACTCTTTCCCTACACGACGCTCTTCCGATCTNNNATTGATTCTGAGGAGGCAGG
AGCACACGTCTGAACTCTTTCCCTACACGACGCTCTTCCGATCTNNNAGAAGTCCTCTTCCCTATCAGC
GAGCACACGTCTGAACTCTTTCCCTACACGACGCTCTTCCGATCTNNNACACCTATCCACAAGTGTCAA
CACACGTCTGAACTCTTTCCCTACACGACGCTCTTCCGATCTNNNTTCTTCTCCCTAGTGGATTTGCGC
AGCACACGTCTGAACTCTTTCCCTACACGACGCTCTTCCGATCTNNNCTGGCACATGTCTACTAGAGG
AGCACACGTCTGAACTCTTTCCCTACACGACGCTCTTCCGATCTNNNGTGATATCAGACTGGCCCTTG

AGAGCACACGTCTGAACTCTTTCCCTACACGACGCTCTTCCGATCTNNNCAGGAAACTGGCATTACGT
AGCACACGTCTGAACTCTTTCCCTACACGACGCTCTTCCGATCTNNNAATCTGGAATTCGCTGTGTCT
GAGCACACGTCTGAACTCTTTCCCTACACGACGCTCTTCCGATCTNNNACATGAACTTCTCAGAGCCAC
GCACACGTCTGAACTCTTTCCCTACACGACGCTCTTCCGATCTNNNCTTCCATCCTCCCTCCTCTTCTCCA
ACACGTCTGAACTCTTTCCCTACACGACGCTCTTCCGATCTNNNAGGTCTCTGTATGCTTTGTGCATC
GAGCACACGTCTGAACTCTTTCCCTACACGACGCTCTTCCGATCTNNNAGAAGCGCAGGGTGGAATT
GAGCACACGTCTGAACTCTTTCCCTACACGACGCTCTTCCGATCTNNNGTTTCAGGCTGAGATGGGA
AGCACACGTCTGAACTCTTTCCCTACACGACGCTCTTCCGATCTNNNCCCACTTTCTCTTCTCTGTGTCG
GCACACGTCTGAACTCTTTCCCTACACGACGCTCTTCCGATCTNNNAATAACAAACGGGAGGAGGAA
GAGCACACGTCTGAACTCTTTCCCTACACGACGCTCTTCCGATCTNNNTCTGAGAGGGCTGTTTTGT
AGAGCACACGTCTGAACTCTTTCCCTACACGACGCTCTTCCGATCTNNNTGCTTCTGTGTCAGGTATA
AGCACACGTCTGAACTCTTTCCCTACACGACGCTCTTCCGATCTNNNGTATTTCTTCCCTTCCCTCCCCT
CACACGTCTGAACTCTTTCCCTACACGACGCTCTTCCGATCTNNNCCAGATTGAGGGAAGCTTCTTGG
CACACGTCTGAACTCTTTCCCTACACGACGCTCTTCCGATCTNNNCTAGAAGGCCTTGTTGCCTGTAG
AGAGCACACGTCTGAACTCTTTCCCTACACGACGCTCTTCCGATCTNNNACATGCTATGTTTGGCTAT
AGCACACGTCTGAACTCTTTCCCTACACGACGCTCTTCCGATCTNNNGTTGTGACCACAAGGTCTGGA
GAGCACACGTCTGAACTCTTTCCCTACACGACGCTCTTCCGATCTNNNTCCAGCAAGGCTGCTAAAAG
AAGAGCACACGTCTGAACTCTTTCCCTACACGACGCTCTTCCGATCTNNNACTGTTGTGTTGGGGAAT
GCACACGTCTGAACTCTTTCCCTACACGACGCTCTTCCGATCTNNNCTGCTTCACCTCAGTTCTCAACA
AGCACACGTCTGAACTCTTTCCCTACACGACGCTCTTCCGATCTNNNGGAACTCCTCAGGGACAACTA
AGCACACGTCTGAACTCTTTCCCTACACGACGCTCTTCCGATCTNNNTACTCAGCTCTTCTGGGTTTGG
AGCACACGTCTGAACTCTTTCCCTACACGACGCTCTTCCGATCTNNNTGAGCAGACAGAGAAGTCTG
GAGCACACGTCTGAACTCTTTCCCTACACGACGCTCTTCCGATCTNNNCTGCCCTCTGTCACTTCCAATT
ACACGTCTGAACTCTTTCCCTACACGACGCTCTTCCGATCTNNNCTCCCACGTTCTGCTTGAATAACA
AGCACACGTCTGAACTCTTTCCCTACACGACGCTCTTCCGATCTNNNATAATGACGGCTGTGATCTGC
GAGCACACGTCTGAACTCTTTCCCTACACGACGCTCTTCCGATCTNNNGGATTGATGTTCACTGTCAG
GAGCACACGTCTGAACTCTTTCCCTACACGACGCTCTTCCGATCTNNNCCGTACAATCTATTCTGTGA
AGCACACGTCTGAACTCTTTCCCTACACGACGCTCTTCCGATCTNNNTTTGAGACAAGGTCGGTCTC
AGCACACGTCTGAACTCTTTCCCTACACGACGCTCTTCCGATCTNNNCCTGCATTCTTACCTTTGCAAG
GCACACGTCTGAACTCTTTCCCTACACGACGCTCTTCCGATCTNNNCTCATTACAACTCTGCCTCTGGG
GCACACGTCTGAACTCTTTCCCTACACGACGCTCTTCCGATCTNNNCTGGACTTCCAACCTGCTTTGTA
AGAGCACACGTCTGAACTCTTTCCCTACACGACGCTCTTCCGATCTNNNTGGTTGATGGGAAAAACA
GAGCACACGTCTGAACTCTTTCCCTACACGACGCTCTTCCGATCTNNNTTTGAGACAAGGTCGGTCT
AGCACACGTCTGAACTCTTTCCCTACACGACGCTCTTCCGATCTNNNCAGGAGAGGCCAGAAGTTTG
AGCACACGTCTGAACTCTTTCCCTACACGACGCTCTTCCGATCTNNNTTGGAGCTCAGAAGTTTTCTG
GCACACGTCTGAACTCTTTCCCTACACGACGCTCTTCCGATCTNNNGTTCCCAAACAAGTATGCTCCC
GCACACGTCTGAACTCTTTCCCTACACGACGCTCTTCCGATCTNNNTTGGCATGGCATTCTCTCAGTT
GCACACGTCTGAACTCTTTCCCTACACGACGCTCTTCCGATCTNNNGTTTTGAAGGTGGTGAGGACG
GAGCACACGTCTGAACTCTTTCCCTACACGACGCTCTTCCGATCTNNNCTTTCCTTCTTTCCTTCCACCC
GCACACGTCTGAACTCTTTCCCTACACGACGCTCTTCCGATCTNNNCCACATGGGGCATGTCCATACA
GCACACGTCTGAACTCTTTCCCTACACGACGCTCTTCCGATCTNNNACACATTAGCTGTCACCAAAGCA
GCACACGTCTGAACTCTTTCCCTACACGACGCTCTTCCGATCTNNNTGTTCCTATGCTGAAAGCACCT
GCACACGTCTGAACTCTTTCCCTACACGACGCTCTTCCGATCTNNNACATTCAGGTTTTGCTGGTTTAG

AGCACACGTCTGAACTCTTTCCCTACACGACGCTCTTCCGATCTNNNACTTCAGGGTGACCAAAACTTC
GAGCACACGTCTGAACTCTTTCCCTACACGACGCTCTTCCGATCTNNNCCTAATGGTAGAAAGGCTTCC
AGAGCACACGTCTGAACTCTTTCCCTACACGACGCTCTTCCGATCTNNNCTGGAACTTTTCTCCACAGGC
AGAGCACACGTCTGAACTCTTTCCCTACACGACGCTCTTCCGATCTNNNCGGTGAGGAATTCAAATGTG
AGCACACGTCTGAACTCTTTCCCTACACGACGCTCTTCCGATCTNNNTCTTCTCAAGTTTAGCGGCCATG
GAGCACACGTCTGAACTCTTTCCCTACACGACGCTCTTCCGATCTNNNTGCAGATGCTAGAAAGTGCT
GAGCACACGTCTGAACTCTTTCCCTACACGACGCTCTTCCGATCTNNNCCACTGAAAACCCTGAGAACCA
GCACACGTCTGAACTCTTTCCCTACACGACGCTCTTCCGATCTNNNCATTCTCCAAGAGGGAGTGCAAA
AAGAGCACACGTCTGAACTCTTTCCCTACACGACGCTCTTCCGATCTNNNATTGCAGATGGCTCTTGT
AGCACACGTCTGAACTCTTTCCCTACACGACGCTCTTCCGATCTNNNGAGGAAGATCAGGAAATCCCAG
AGAGCACACGTCTGAACTCTTTCCCTACACGACGCTCTTCCGATCTNNNAAAAAGCATGGGGTGTGTG
GAGCACACGTCTGAACTCTTTCCCTACACGACGCTCTTCCGATCTNNNCACACGCTGACATACCGACATG
GAGCACACGTCTGAACTCTTTCCCTACACGACGCTCTTCCGATCTNNNTTTTCTCCCCTTCTGGAAGACC
GAGCACACGTCTGAACTCTTTCCCTACACGACGCTCTTCCGATCTNNNATCCTAATGCACACACTTGCAC
AGCACACGTCTGAACTCTTTCCCTACACGACGCTCTTCCGATCTNNNCAGTAACTGTTCATTTGGCAATT
AGCACACGTCTGAACTCTTTCCCTACACGACGCTCTTCCGATCTNNNGAGAATGAGAGTGGGGCAAAC
GAGCACACGTCTGAACTCTTTCCCTACACGACGCTCTTCCGATCTNNNTCAGGGACATTGATGGGCTTG
AGCACACGTCTGAACTCTTTCCCTACACGACGCTCTTCCGATCTNNNATCCACCTGGGTTTTGATTTGCT
AGCACACGTCTGAACTCTTTCCCTACACGACGCTCTTCCGATCTNNNGAAACAATTGCAGCCTGATTCC
GAGCACACGTCTGAACTCTTTCCCTACACGACGCTCTTCCGATCTNNNATTTGGAAGGCCACCATGTGA
GAGCACACGTCTGAACTCTTTCCCTACACGACGCTCTTCCGATCTNNNATGTTGTACCTTGGTGGCTTCA
AGCACACGTCTGAACTCTTTCCCTACACGACGCTCTTCCGATCTNNNATAGCACACCTTGGCTGGATTT
AGCACACGTCTGAACTCTTTCCCTACACGACGCTCTTCCGATCTNNNGAAACAATTGCAGCCTGATTCCC
AGCACACGTCTGAACTCTTTCCCTACACGACGCTCTTCCGATCTNNNAGTGAACACCACTGTTGACTAGC
AGAGCACACGTCTGAACTCTTTCCCTACACGACGCTCTTCCGATCTNNNTAATTATTGCCACCCCTTGG
GAGCACACGTCTGAACTCTTTCCCTACACGACGCTCTTCCGATCTNNNTACATTGGAGGAGCAGACTGC
GCACACGTCTGAACTCTTTCCCTACACGACGCTCTTCCGATCTNNNATTTCAGGACTTGCTACATTTTCC
AGCACACGTCTGAACTCTTTCCCTACACGACGCTCTTCCGATCTNNNGTCATTGTTCTTGGTACTTTGTT
GAGCACACGTCTGAACTCTTTCCCTACACGACGCTCTTCCGATCTNNNTAATTATTGCCACCCCTTGGCT
AGCACACGTCTGAACTCTTTCCCTACACGACGCTCTTCCGATCTNNNGCTCACTACATATAGACACCCTG
AGAGCACACGTCTGAACTCTTTCCCTACACGACGCTCTTCCGATCTNNNCACTAAAGCCAAACTTCTGCA
AGAGCACACGTCTGAACTCTTTCCCTACACGACGCTCTTCCGATCTNNNGGTGAAGGTGTTCAGGTAG
AGCACACGTCTGAACTCTTTCCCTACACGACGCTCTTCCGATCTNNNATGAAGATTTGGCCATCTTCCCC
GAGCACACGTCTGAACTCTTTCCCTACACGACGCTCTTCCGATCTNNNACCAGGATGCTGGAATAGAG
AGCACACGTCTGAACTCTTTCCCTACACGACGCTCTTCCGATCTNNNCCATGGATACCAAGGAATGAGC
GAGCACACGTCTGAACTCTTTCCCTACACGACGCTCTTCCGATCTNNNTCACCTTGTCTACACACACATGC
AGCACACGTCTGAACTCTTTCCCTACACGACGCTCTTCCGATCTNNNCCATCTCTCCTGGAGCCTAGTTT
GAGCACACGTCTGAACTCTTTCCCTACACGACGCTCTTCCGATCTNNNTGGATTGGAACTGCAATTTTG
GAGCACACGTCTGAACTCTTTCCCTACACGACGCTCTTCCGATCTNNNCAGAGCCCACAAACAATCTTCC
AGCACACGTCTGAACTCTTTCCCTACACGACGCTCTTCCGATCTNNNCCATTCTGGAAAGCAATTGGCAC
GAGCACACGTCTGAACTCTTTCCCTACACGACGCTCTTCCGATCTNNNTATGTCTCCGTGCTTACCAGTT
AGAGCACACGTCTGAACTCTTTCCCTACACGACGCTCTTCCGATCTNNNAACCTCTAATTTGGTTTGCCA
AGAGCACACGTCTGAACTCTTTCCCTACACGACGCTCTTCCGATCTNNNCTTTGCACAATCCTGTGACCA

AGCACACGTCTGAACTCTTTCCCTACACGACGCTCTTCCGATCTNNNAGGGAAGATAGTCGAGGAG
GCACACGTCTGAACTCTTTCCCTACACGACGCTCTTCCGATCTNNNATGTCAAGAATTTTTGCTGGTGT
AGAGCACACGTCTGAACTCTTTCCCTACACGACGCTCTTCCGATCTNNNAGAGCCCTTTGTATTTTGG
AGCACACGTCTGAACTCTTTCCCTACACGACGCTCTTCCGATCTNNNCCAGCTGGAGTTCAGAATAAC
AGAGCACACGTCTGAACTCTTTCCCTACACGACGCTCTTCCGATCTNNNTGGAGTTAGATCCAAGATA
AGCACACGTCTGAACTCTTTCCCTACACGACGCTCTTCCGATCTNNNCCTCCCAGTACCTACCTTTCTCC
GAGCACACGTCTGAACTCTTTCCCTACACGACGCTCTTCCGATCTNNNGACAATGAGAACACCTCTGG
AGCACACGTCTGAACTCTTTCCCTACACGACGCTCTTCCGATCTNNNTTTCAGATGCAGGCACATGTTT
AGAGCACACGTCTGAACTCTTTCCCTACACGACGCTCTTCCGATCTNNNGCTAAACCAGGGTTCTTTG
AGCACACGTCTGAACTCTTTCCCTACACGACGCTCTTCCGATCTNNNTAATAGCACAACAGCAGCAGC
AGCACACGTCTGAACTCTTTCCCTACACGACGCTCTTCCGATCTNNNACCCACATCATTGTGTCTTCCAC
GAAGAGCACACGTCTGAACTCTTTCCCTACACGACGCTCTTCCGATCTNNNTGTTGTATGGCTGAGCA
AGCACACGTCTGAACTCTTTCCCTACACGACGCTCTTCCGATCTNNNGCTCAGTGTGGACATGGAAAT
AGCACACGTCTGAACTCTTTCCCTACACGACGCTCTTCCGATCTNNNACTTTTGTGATGTGGAAAGTG
GCACACGTCTGAACTCTTTCCCTACACGACGCTCTTCCGATCTNNNAGCTGTAAATGTTTCTCTCTCTC
AGCACACGTCTGAACTCTTTCCCTACACGACGCTCTTCCGATCTNNNCATTCACATCCTTGTCCCCAGAG
GAGCACACGTCTGAACTCTTTCCCTACACGACGCTCTTCCGATCTNNNTAGTCCTTGTGTTCTGCACTC
GCACACGTCTGAACTCTTTCCCTACACGACGCTCTTCCGATCTNNNTGTGCCTCCTCCTTCTCATTTTCC
AGCACACGTCTGAACTCTTTCCCTACACGACGCTCTTCCGATCTNNNTATCCCAGACAGGGAGAACAG
AGCACACGTCTGAACTCTTTCCCTACACGACGCTCTTCCGATCTNNNGGTGCTGTCTTGTGTGGTAG
GAGCACACGTCTGAACTCTTTCCCTACACGACGCTCTTCCGATCTNNNAAAACGATGTCTGTCCACTCA
CACACGTCTGAACTCTTTCCCTACACGACGCTCTTCCGATCTNNNAACTCCCTTACAGGGGAGAAAA
GCACACGTCTGAACTCTTTCCCTACACGACGCTCTTCCGATCTNNNGAAAGCTCATGAAGGGTGGGA
AGAGCACACGTCTGAACTCTTTCCCTACACGACGCTCTTCCGATCTNNNTCATCCCACAGATTCAAGAG
AGCACACGTCTGAACTCTTTCCCTACACGACGCTCTTCCGATCTNNNACTTCTACACAAGACCACATGA
GAGCACACGTCTGAACTCTTTCCCTACACGACGCTCTTCCGATCTNNNATGCTTCCATGTTATAGCCCA
GAGCACACGTCTGAACTCTTTCCCTACACGACGCTCTTCCGATCTNNNTACCATCCTGGTGGGTGTGT
AGCACACGTCTGAACTCTTTCCCTACACGACGCTCTTCCGATCTNNNTCGATATTATTCAGATTTTCCG
GAGCACACGTCTGAACTCTTTCCCTACACGACGCTCTTCCGATCTNNNTCAGCAGGAAATGAATAAG
CACACGTCTGAACTCTTTCCCTACACGACGCTCTTCCGATCTNNNCCCAAGAACCGGGAACTCTAATG
GAGCACACGTCTGAACTCTTTCCCTACACGACGCTCTTCCGATCTNNNATGAATTGCTCTTCACCGACC
GAGCACACGTCTGAACTCTTTCCCTACACGACGCTCTTCCGATCTNNNAGGAGAAATTCAACAAGCC
AGCACACGTCTGAACTCTTTCCCTACACGACGCTCTTCCGATCTNNNAACCCATCAAAAGACAGTTCG
AGAGCACACGTCTGAACTCTTTCCCTACACGACGCTCTTCCGATCTNNNCATCTTTGAGCTTTCTGCAG
GCACACGTCTGAACTCTTTCCCTACACGACGCTCTTCCGATCTNNNTGACAACACTCAAGCTGAGAAC
AGCACACGTCTGAACTCTTTCCCTACACGACGCTCTTCCGATCTNNNCTTCCCAGGGTTGTTGTGAAG
GAGCACACGTCTGAACTCTTTCCCTACACGACGCTCTTCCGATCTNNNAGCCATTGCGCCATCTATAT
GAGCACACGTCTGAACTCTTTCCCTACACGACGCTCTTCCGATCTNNNACCGAGCTGATCAAAAGACA
GAGCACACGTCTGAACTCTTTCCCTACACGACGCTCTTCCGATCTNNNATTCACGGACCACTTTGATA
AGAGCACACGTCTGAACTCTTTCCCTACACGACGCTCTTCCGATCTNNNCCCTGTCCTGAACAGAGAA
GAGCACACGTCTGAACTCTTTCCCTACACGACGCTCTTCCGATCTNNNTTATCCCATGGTGGAAGGAG
AGCACACGTCTGAACTCTTTCCCTACACGACGCTCTTCCGATCTNNNTTGGCCCCAAAAAGTACACTT
AGCACACGTCTGAACTCTTTCCCTACACGACGCTCTTCCGATCTNNNCACCTGTGTCTCCTTGCTAGAT

AGAGCACACGTCTGAACTCTTTCCCTACACGACGCTCTTCCGATCTNNNTCCCACAAACCATCCATAATG
GAGCACACGTCTGAACTCTTTCCCTACACGACGCTCTTCCGATCTNNNATGTGTACCCAAGGGCAGAG
AGAGCACACGTCTGAACTCTTTCCCTACACGACGCTCTTCCGATCTNNNTGAAGCTGTAAAATTACAAG
AGCACACGTCTGAACTCTTTCCCTACACGACGCTCTTCCGATCTNNNGCACAATGAAAGCATTGCTCAG
GAGCACACGTCTGAACTCTTTCCCTACACGACGCTCTTCCGATCTNNNACGTGCACATATGGGAAAATC
AGCACACGTCTGAACTCTTTCCCTACACGACGCTCTTCCGATCTNNNACATTTTCTGACCCCCACAACTC
GAGCACACGTCTGAACTCTTTCCCTACACGACGCTCTTCCGATCTNNNTGGAGCAGAGGTGCACTATCT
AGCACACGTCTGAACTCTTTCCCTACACGACGCTCTTCCGATCTNNNAGAAATGGGAAGTGTTTGGGT
GCACACGTCTGAACTCTTTCCCTACACGACGCTCTTCCGATCTNNNTGGTGCTTTGGTTATCCTTGGTC
AGCACACGTCTGAACTCTTTCCCTACACGACGCTCTTCCGATCTNNNCCTCCCCATCCCTTACTTATCCCC
AGAGCACACGTCTGAACTCTTTCCCTACACGACGCTCTTCCGATCTNNNAAGGAAATTACAGACCAGCA
AGCACACGTCTGAACTCTTTCCCTACACGACGCTCTTCCGATCTNNNAATTCCCAGGTTCTTTCCAGTG
AAGAGCACACGTCTGAACTCTTTCCCTACACGACGCTCTTCCGATCTNNNCTAAACAAAGGGCATGGTG
AGAGCACACGTCTGAACTCTTTCCCTACACGACGCTCTTCCGATCTNNNTGTGCTGATAGAAACACCTG
GAGCACACGTCTGAACTCTTTCCCTACACGACGCTCTTCCGATCTNNNCCAGTGTGTTCATGGGCATCA
AGCACACGTCTGAACTCTTTCCCTACACGACGCTCTTCCGATCTNNNCTGAAGCAGTACCCTCACTTAG
AGAGCACACGTCTGAACTCTTTCCCTACACGACGCTCTTCCGATCTNNNCAGGGGAAATTTTTCAAGA
GAGCACACGTCTGAACTCTTTCCCTACACGACGCTCTTCCGATCTNNNCCAAAATGAGATACGGCTGCT
AGAGCACACGTCTGAACTCTTTCCCTACACGACGCTCTTCCGATCTNNNCTCCCAGGGGAACTTATTCAA
AGAGCACACGTCTGAACTCTTTCCCTACACGACGCTCTTCCGATCTNNNCACTAGGCAGAGGAAAAAG
AGAGCACACGTCTGAACTCTTTCCCTACACGACGCTCTTCCGATCTNNNACACTCTTACAACCACACACA
AGCACACGTCTGAACTCTTTCCCTACACGACGCTCTTCCGATCTNNNATCCCTTTTCCTCCTGTTCCTCAC
AGCACACGTCTGAACTCTTTCCCTACACGACGCTCTTCCGATCTNNNTCCCCATCCGTCTAGAAGTGAA
AGCACACGTCTGAACTCTTTCCCTACACGACGCTCTTCCGATCTNNNAGTGTCTGAGCATAGAGTTGG
GAAGAGCACACGTCTGAACTCTTTCCCTACACGACGCTCTTCCGATCTNNNCAGGGGAAATTTTTCAA
GAGCACACGTCTGAACTCTTTCCCTACACGACGCTCTTCCGATCTNNNGGAGTGTTTAGCTCAGTTAG
GAGCACACGTCTGAACTCTTTCCCTACACGACGCTCTTCCGATCTNNNGGACAACAGAAAGCACCCACT
GAGCACACGTCTGAACTCTTTCCCTACACGACGCTCTTCCGATCTNNNGTCCTTCAAGCTCCGTTTTCTCT
GAGCACACGTCTGAACTCTTTCCCTACACGACGCTCTTCCGATCTNNNTGGAATCCAGGAGCTGGTCAA
AGCACACGTCTGAACTCTTTCCCTACACGACGCTCTTCCGATCTNNNCAAAGATTTCCGTCCTTCTGCCT
GAGCACACGTCTGAACTCTTTCCCTACACGACGCTCTTCCGATCTNNNCAGAAAGCAAGGAAGTGCTTG
AAGAGCACACGTCTGAACTCTTTCCCTACACGACGCTCTTCCGATCTNNNAGTCATGTCACCTTTGGCT
AAGAGCACACGTCTGAACTCTTTCCCTACACGACGCTCTTCCGATCTNNNTGTCTCCCTTTTCTTAGCAT
AGCACACGTCTGAACTCTTTCCCTACACGACGCTCTTCCGATCTNNNCTTACAGACACCACACGCACATA
GAGCACACGTCTGAACTCTTTCCCTACACGACGCTCTTCCGATCTNNNACAAGGACAGACTGGAGTTTG
AGAGCACACGTCTGAACTCTTTCCCTACACGACGCTCTTCCGATCTNNNCACTGCACACAGACATATCCA
AGCACACGTCTGAACTCTTTCCCTACACGACGCTCTTCCGATCTNNNACAGATTGCTGTATAGTAGGAC
AGCACACGTCTGAACTCTTTCCCTACACGACGCTCTTCCGATCTNNNAGAAAAGTAGGCCAGTGTGTG
GCACACGTCTGAACTCTTTCCCTACACGACGCTCTTCCGATCTNNNTATGCAGTCTGAGGTTCAAAGCC
GAGCACACGTCTGAACTCTTTCCCTACACGACGCTCTTCCGATCTNNNATGGAGAAAATAACTGGGCT
AGAGCACACGTCTGAACTCTTTCCCTACACGACGCTCTTCCGATCTNNNGCAGTTCCAAGACCAATCAAT
AGCACACGTCTGAACTCTTTCCCTACACGACGCTCTTCCGATCTNNNGAAATCCTTTACCTTCCCCCGAC
GAGCACACGTCTGAACTCTTTCCCTACACGACGCTCTTCCGATCTNNNGTGACACCCCATCACTTTAGTC

GAGCACACGTCTGAACTCTTTCCCTACACGACGCTCTTCCGATCTNNNAGGCTTTCTTCTGTTCTCTGT
AGCACACGTCTGAACTCTTTCCCTACACGACGCTCTTCCGATCTNNNATAGGGCCCTGCTTTGTCATCT
AGCACACGTCTGAACTCTTTCCCTACACGACGCTCTTCCGATCTNNNTCTATTCTGGAAAGCTGAGGC
AGAGCACACGTCTGAACTCTTTCCCTACACGACGCTCTTCCGATCTNNNTACGAATACCCTACTGGTTG
GAGCACACGTCTGAACTCTTTCCCTACACGACGCTCTTCCGATCTNNNGGGAAGAAGGACAAGGAG
GCACACGTCTGAACTCTTTCCCTACACGACGCTCTTCCGATCTNNNTTCTCCTATCAGGTAGTGTGCG
GCACACGTCTGAACTCTTTCCCTACACGACGCTCTTCCGATCTNNNGACCTGTTTGATTTTCCCACCCT
GAGCACACGTCTGAACTCTTTCCCTACACGACGCTCTTCCGATCTNNNATGATAAGGGACCATGAAGG
AGCACACGTCTGAACTCTTTCCCTACACGACGCTCTTCCGATCTNNNTTGTGAGAAAGAGCCAACTG
AGCACACGTCTGAACTCTTTCCCTACACGACGCTCTTCCGATCTNNNTGTTTGAGGGGAAGCAAGTA
GAGCACACGTCTGAACTCTTTCCCTACACGACGCTCTTCCGATCTNNNAGATGGCCTAGTAAAGCAT
AAGAGCACACGTCTGAACTCTTTCCCTACACGACGCTCTTCCGATCTNNNAAGAACCCATCCCCCTACA
AGCACACGTCTGAACTCTTTCCCTACACGACGCTCTTCCGATCTNNNTGAGGGACAACAGTCTATAGG
AAGAGCACACGTCTGAACTCTTTCCCTACACGACGCTCTTCCGATCTNNNCTCATTCTAGCCCAGGTG
GAGCACACGTCTGAACTCTTTCCCTACACGACGCTCTTCCGATCTNNNCCACGGATCATACTGCCTCA
GCACACGTCTGAACTCTTTCCCTACACGACGCTCTTCCGATCTNNNACAATTACGAGAGAGGGGGA
AGCACACGTCTGAACTCTTTCCCTACACGACGCTCTTCCGATCTNNNAGCAAATTGCAGAGCCGTATG
GAGCACACGTCTGAACTCTTTCCCTACACGACGCTCTTCCGATCTNNNTTTTTGTGTGAATACCCTGG
CACACGTCTGAACTCTTTCCCTACACGACGCTCTTCCGATCTNNNTTCCCCTCCAATATCTGTCTGCTCA
AGCACACGTCTGAACTCTTTCCCTACACGACGCTCTTCCGATCTNNNAACCACTGCCCTGGTAAAGAA
GCACACGTCTGAACTCTTTCCCTACACGACGCTCTTCCGATCTNNNACATTGCCTACTTCTTCTGCGAC
AGAGCACACGTCTGAACTCTTTCCCTACACGACGCTCTTCCGATCTNNNGGTTGCAAAGTTTCTCGAG
CACACGTCTGAACTCTTTCCCTACACGACGCTCTTCCGATCTNNNTGGTTTACTTTGTGGTTCTCGGG
GCACACGTCTGAACTCTTTCCCTACACGACGCTCTTCCGATCTNNNGCAGTAGAGGTTTCCTGGTTTC
GAGCACACGTCTGAACTCTTTCCCTACACGACGCTCTTCCGATCTNNNCTGTGTCGCAGCACATGAAG
AGCACACGTCTGAACTCTTTCCCTACACGACGCTCTTCCGATCTNNNACTTCCAGTTGTTGGATTTTG
AGCACACGTCTGAACTCTTTCCCTACACGACGCTCTTCCGATCTNNNTTGTCTTGAGCTCACACAGTGG
AGCACACGTCTGAACTCTTTCCCTACACGACGCTCTTCCGATCTNNNACATGCAGGATGGGGTAGAA
AGCACACGTCTGAACTCTTTCCCTACACGACGCTCTTCCGATCTNNNAGCTGGTTTGTACATCAAAGG
GAGCACACGTCTGAACTCTTTCCCTACACGACGCTCTTCCGATCTNNNTTTTGGTTCTTTCCGAGTTGT
GAGCACACGTCTGAACTCTTTCCCTACACGACGCTCTTCCGATCTNNNAAGAATCAACTTTTCGCTTTG
GCACACGTCTGAACTCTTTCCCTACACGACGCTCTTCCGATCTNNNTTGACTGTCAGAGTGTTGATGG
GCACACGTCTGAACTCTTTCCCTACACGACGCTCTTCCGATCTNNNCCTGGCCCTGATGTGACTTTTA
GAGCACACGTCTGAACTCTTTCCCTACACGACGCTCTTCCGATCTNNNTATTTGGAGGAGGATGTGG
GAGCACACGTCTGAACTCTTTCCCTACACGACGCTCTTCCGATCTNNNGCCTCACCACTCCAATTTTCA
AGAGCACACGTCTGAACTCTTTCCCTACACGACGCTCTTCCGATCTNNNGAGTTGATCTCCTGGAGGT
AGCACACGTCTGAACTCTTTCCCTACACGACGCTCTTCCGATCTNNNTTTGTAGCCTTGCCTTTATGCC
AAGAGCACACGTCTGAACTCTTTCCCTACACGACGCTCTTCCGATCTNNNTGGGTGAAAGAATGAGA
GAGCACACGTCTGAACTCTTTCCCTACACGACGCTCTTCCGATCTNNNAGTGGCAAAACCACAATTAC
AGAGCACACGTCTGAACTCTTTCCCTACACGACGCTCTTCCGATCTNNNCAAAATCCAGAATCCCAGG
AGAGCACACGTCTGAACTCTTTCCCTACACGACGCTCTTCCGATCTNNNACATGTCCTCCTGTAGTCC
GAGCACACGTCTGAACTCTTTCCCTACACGACGCTCTTCCGATCTNNNGTGCATGAAGATAGGGCCT
GCACACGTCTGAACTCTTTCCCTACACGACGCTCTTCCGATCTNNNGCGAAGATCACATCACTGCATT

AGCACACGTCTGAACTCTTTCCCTACACGACGCTCTTCCGATCTNNNTTTTCCCCCAAATTCCTTCCCTG
AGCACACGTCTGAACTCTTTCCCTACACGACGCTCTTCCGATCTNNNGGTTCCTAAACACGCTCTCTCTC
AGCACACGTCTGAACTCTTTCCCTACACGACGCTCTTCCGATCTNNNGCATGAGTGCTTAGCTGTCTCT
AGCACACGTCTGAACTCTTTCCCTACACGACGCTCTTCCGATCTNNNCTCCCATTTTTGGCGAAGACAA
AGCACACGTCTGAACTCTTTCCCTACACGACGCTCTTCCGATCTNNNCAACCCCCTCCCTGACTGCATCA
AGCACACGTCTGAACTCTTTCCCTACACGACGCTCTTCCGATCTNNNCAAGGACTTGTCAGCCTCCATA
AGAGCACACGTCTGAACTCTTTCCCTACACGACGCTCTTCCGATCTNNNCCCTTTCTTGTTTTAGGGTG
AGCACACGTCTGAACTCTTTCCCTACACGACGCTCTTCCGATCTNNNGAGGGCCTGAGAGAGAAAAAG
AGCACACGTCTGAACTCTTTCCCTACACGACGCTCTTCCGATCTNNNGAGCACTGGAGGCTGTCTATA
GAGCACACGTCTGAACTCTTTCCCTACACGACGCTCTTCCGATCTNNNCAATCGAGATTGCCCTTGATG
GAGCACACGTCTGAACTCTTTCCCTACACGACGCTCTTCCGATCTNNNACCAAATGTCACTGAATTGTA
GCACACGTCTGAACTCTTTCCCTACACGACGCTCTTCCGATCTNNNAAAACAGTGCTGGAACACCTAG
AGAGCACACGTCTGAACTCTTTCCCTACACGACGCTCTTCCGATCTNNNAGAATGCAGGGGATATGC
GAGCACACGTCTGAACTCTTTCCCTACACGACGCTCTTCCGATCTNNNTCCCTTGATAGCCACTTTGAG
GAGCACACGTCTGAACTCTTTCCCTACACGACGCTCTTCCGATCTNNNATTGTGACAACTTACCCCACC
GCACACGTCTGAACTCTTTCCCTACACGACGCTCTTCCGATCTNNNACCTTGGCAGTGATTTCTTCTCC
AGAGCACACGTCTGAACTCTTTCCCTACACGACGCTCTTCCGATCTNNNACAACAATTTTGTTTCCTCCT
AGCACACGTCTGAACTCTTTCCCTACACGACGCTCTTCCGATCTNNNTTTGTTACCCCAACTCAAGAGG
GAGCACACGTCTGAACTCTTTCCCTACACGACGCTCTTCCGATCTNNNAAGAACCCATCCCCCTACAAC
AGCACACGTCTGAACTCTTTCCCTACACGACGCTCTTCCGATCTNNNCCTGCAAACACACAAACACAGA
GAGCACACGTCTGAACTCTTTCCCTACACGACGCTCTTCCGATCTNNNACTTTGCCTAAAGGAGCATTC
GCACACGTCTGAACTCTTTCCCTACACGACGCTCTTCCGATCTNNNTCTAGAGATGGAGGCACAACCA
AGAGCACACGTCTGAACTCTTTCCCTACACGACGCTCTTCCGATCTNNNCCAGGAAACAGGGGTTGA
AGCACACGTCTGAACTCTTTCCCTACACGACGCTCTTCCGATCTNNNTAAATGGACTTGAAGGGGCAT
AGCACACGTCTGAACTCTTTCCCTACACGACGCTCTTCCGATCTNNNTTAGGGTCCTCCACAGTGTCAT
AGCACACGTCTGAACTCTTTCCCTACACGACGCTCTTCCGATCTNNNCTGGCACGAGCATTTTGAGTA
AGCACACGTCTGAACTCTTTCCCTACACGACGCTCTTCCGATCTNNNTTAGCCGTCTTTCCCCTATTTGC
AGCACACGTCTGAACTCTTTCCCTACACGACGCTCTTCCGATCTNNNTACCTGCAAAAACTCTCCACAC
GAGCACACGTCTGAACTCTTTCCCTACACGACGCTCTTCCGATCTNNNGAGGCAAGTGGAGAAGGAA
GAGCACACGTCTGAACTCTTTCCCTACACGACGCTCTTCCGATCTNNNCCCAGACCACCTCAGTATGAA
AGAGCACACGTCTGAACTCTTTCCCTACACGACGCTCTTCCGATCTNNNCATGCCTGACATGCAGCAA
AGCACACGTCTGAACTCTTTCCCTACACGACGCTCTTCCGATCTNNNAAATGGGCTTCAAGTGATCAG
GAGCACACGTCTGAACTCTTTCCCTACACGACGCTCTTCCGATCTNNNTGCTCTGTGTAGACCCTGAT
AGCACACGTCTGAACTCTTTCCCTACACGACGCTCTTCCGATCTNNNATGTCTGCAAACACCTTCCATCA
GCACACGTCTGAACTCTTTCCCTACACGACGCTCTTCCGATCTNNNTTTACCATCTTCTGAGCAGCCAC
GCACACGTCTGAACTCTTTCCCTACACGACGCTCTTCCGATCTNNNGCCTTCATCTTCAAATGGAGTGT
AGCACACGTCTGAACTCTTTCCCTACACGACGCTCTTCCGATCTNNNTGTCCAACTCTCTCTCTCACACAC
GCACACGTCTGAACTCTTTCCCTACACGACGCTCTTCCGATCTNNNAAAAAGCCCTCCATCCATGTCTCC
GCACACGTCTGAACTCTTTCCCTACACGACGCTCTTCCGATCTNNNAGCTAGTACAAAGCACGCAGAA
GAGCACACGTCTGAACTCTTTCCCTACACGACGCTCTTCCGATCTNNNTCCTGGCTTCTACATGGTCAA
GAGCACACGTCTGAACTCTTTCCCTACACGACGCTCTTCCGATCTNNNGCGTCATACTTGCATTTTATT
AGCACACGTCTGAACTCTTTCCCTACACGACGCTCTTCCGATCTNNNTTGTTCCAAAAAGCTGTGAAG
CACACGTCTGAACTCTTTCCCTACACGACGCTCTTCCGATCTNNNCATAAGCTGTGCATAGCAGGCAT

AGCACACGTCTGAACTCTTTCCCTACACGACGCTCTTCCGATCTNNNGAGAATGGGGAGGTGGT
GAGCACACGTCTGAACTCTTTCCCTACACGACGCTCTTCCGATCTNNNCTAGGAGTTCTGGGCTTAGT
GAGCACACGTCTGAACTCTTTCCCTACACGACGCTCTTCCGATCTNNNGCCCTATGCTAATCACAAAA
AGCACACGTCTGAACTCTTTCCCTACACGACGCTCTTCCGATCTNNNAGAAAAGTAGGCCAGTGTGT
GAGCACACGTCTGAACTCTTTCCCTACACGACGCTCTTCCGATCTNNNGGAAGACTGAAGGGGAGAG
GAGCACACGTCTGAACTCTTTCCCTACACGACGCTCTTCCGATCTNNNCCCCTCTACTTGTCTGGAGA
GAGCACACGTCTGAACTCTTTCCCTACACGACGCTCTTCCGATCTNNNGGACAATGGATTTTGGTCT
AGAGCACACGTCTGAACTCTTTCCCTACACGACGCTCTTCCGATCTNNNGGGATGGAGTACACAGA
AGCACACGTCTGAACTCTTTCCCTACACGACGCTCTTCCGATCTNNNAGCAAAGTGAGAAAGAGCTG
GAGCACACGTCTGAACTCTTTCCCTACACGACGCTCTTCCGATCTNNNACGATGGAGTGCCATGATCA
GAGCACACGTCTGAACTCTTTCCCTACACGACGCTCTTCCGATCTNNNAGAGCTGGAAGAAGTGGAG
AGAGCACACGTCTGAACTCTTTCCCTACACGACGCTCTTCCGATCTNNNAGAGCTGGAAGAAGTGGA
GAGCACACGTCTGAACTCTTTCCCTACACGACGCTCTTCCGATCTNNNGAGAATTTGCTCAGTGGGAG
GCACACGTCTGAACTCTTTCCCTACACGACGCTCTTCCGATCTNNNAAAAACTCTGCAGGAGGTAGG
GAGCACACGTCTGAACTCTTTCCCTACACGACGCTCTTCCGATCTNNNACTTTTTCTTGGAGGCTGACA
AAGAGCACACGTCTGAACTCTTTCCCTACACGACGCTCTTCCGATCTNNNGCTGTTTTCTGGAGCTCTT
GAGCACACGTCTGAACTCTTTCCCTACACGACGCTCTTCCGATCTNNNGGCAAAGCTTGCCTGCTAAAG
AGCACACGTCTGAACTCTTTCCCTACACGACGCTCTTCCGATCTNNNATGAGCTAGGGGTTCATGAGA
AGCACACGTCTGAACTCTTTCCCTACACGACGCTCTTCCGATCTNNNAGAAAGTAAAGCTGCCTTGCC
GAGCACACGTCTGAACTCTTTCCCTACACGACGCTCTTCCGATCTNNNAACTTTTCCGTGAGCAAACAA
GAAGAGCACACGTCTGAACTCTTTCCCTACACGACGCTCTTCCGATCTNNNAGAAAATGACCCATCTGC
AGCACACGTCTGAACTCTTTCCCTACACGACGCTCTTCCGATCTNNNGATTCCCTGAGCCACTTTCAGA
AGCACACGTCTGAACTCTTTCCCTACACGACGCTCTTCCGATCTNNNATGGGACTTTCAGTGGTAAAC
GAGCACACGTCTGAACTCTTTCCCTACACGACGCTCTTCCGATCTNNNAGCATTTCCCTTTGTGTTTAGC
GAGCACACGTCTGAACTCTTTCCCTACACGACGCTCTTCCGATCTNNNCCCTCAGTTCAGAAGATGTA
GAAGAGCACACGTCTGAACTCTTTCCCTACACGACGCTCTTCCGATCTNNNAGAGTAAGTGTCATGG
AGAGCACACGTCTGAACTCTTTCCCTACACGACGCTCTTCCGATCTNNNTGAAGCTGTGTTGTTCAGAG
GAGCACACGTCTGAACTCTTTCCCTACACGACGCTCTTCCGATCTNNNTGAAGCTGTGTTGTTCAGAGA
AAGAGCACACGTCTGAACTCTTTCCCTACACGACGCTCTTCCGATCTNNNAGAAAGAGTGAAACGCAG
AGCACACGTCTGAACTCTTTCCCTACACGACGCTCTTCCGATCTNNNGACAATTACATACACGCACGCA
GAGCACACGTCTGAACTCTTTCCCTACACGACGCTCTTCCGATCTNNNAGGCAAAAATTAGTCCTAAG
GAGCACACGTCTGAACTCTTTCCCTACACGACGCTCTTCCGATCTNNNGCCTCCAGGTTTCCTTGTAGA
GAGCACACGTCTGAACTCTTTCCCTACACGACGCTCTTCCGATCTNNNCACTTTTATTGTACCAGCCACG
AGCACACGTCTGAACTCTTTCCCTACACGACGCTCTTCCGATCTNNNTTGGAATCAATTGCTCTGGCAC
AGCACACGTCTGAACTCTTTCCCTACACGACGCTCTTCCGATCTNNNTTCCATGCATTTCTCCAAGTGC
GAGCACACGTCTGAACTCTTTCCCTACACGACGCTCTTCCGATCTNNNTAAACCTCCCGAATCATGTGA
GCACACGTCTGAACTCTTTCCCTACACGACGCTCTTCCGATCTNNNCAAAATCCAGAATCCCAGGGTG
AGAGCACACGTCTGAACTCTTTCCCTACACGACGCTCTTCCGATCTNNNGCTTGTGGGAGCACAGTTTT
AGAGCACACGTCTGAACTCTTTCCCTACACGACGCTCTTCCGATCTNNNGAGACTATGCCCAAAGCCA
GAGCACACGTCTGAACTCTTTCCCTACACGACGCTCTTCCGATCTNNNACACGAAACAAATAGACATG
AGCACACGTCTGAACTCTTTCCCTACACGACGCTCTTCCGATCTNNNCCTGAGCCAGTGTACCATTTGA
GAGCACACGTCTGAACTCTTTCCCTACACGACGCTCTTCCGATCTNNNTCTTGGATGTGTATGTGCTTG
GAGCACACGTCTGAACTCTTTCCCTACACGACGCTCTTCCGATCTNNNAGCAGTGTTTCCCAGGAATG

GAGCACACGTCTGAACTCTTTCCCTACACGACGCTCTTCCGATCTNNNGTGGATCTACACATACATGGA
AGAGCACACGTCTGAACTCTTTCCCTACACGACGCTCTTCCGATCTNNNTCCATAGATGTGGATGGATA
GAGCACACGTCTGAACTCTTTCCCTACACGACGCTCTTCCGATCTNNNGGAGCCTGATGAATTGTCTGT
AGCACACGTCTGAACTCTTTCCCTACACGACGCTCTTCCGATCTNNNTGCAGCACATGACTGAATTGGTA
AGCACACGTCTGAACTCTTTCCCTACACGACGCTCTTCCGATCTNNNCCAACATCCTTCCCTACTTCAGC
GAGCACACGTCTGAACTCTTTCCCTACACGACGCTCTTCCGATCTNNNTTTGTCTTTTGCAGGCACTCTCT
AGCACACGTCTGAACTCTTTCCCTACACGACGCTCTTCCGATCTNNNAAGGACACTCGGAGGAGAAGA
GCACACGTCTGAACTCTTTCCCTACACGACGCTCTTCCGATCTNNNCTCCTCTTTACTGGGGCTTTCACC
AGAGCACACGTCTGAACTCTTTCCCTACACGACGCTCTTCCGATCTNNNAAACCACTAACTCTCACGCAT
AGAGCACACGTCTGAACTCTTTCCCTACACGACGCTCTTCCGATCTNNNACATGTCACATTTGAGACTAG
CACACGTCTGAACTCTTTCCCTACACGACGCTCTTCCGATCTNNNCCCTTCCCTTGGATTGGATTGAGCA
AGCACACGTCTGAACTCTTTCCCTACACGACGCTCTTCCGATCTNNNTCTTCAGAGAATTGTCTTTGTAC
GAGCACACGTCTGAACTCTTTCCCTACACGACGCTCTTCCGATCTNNNAATCTTAGGTGGTGCAGAGG
AGCACACGTCTGAACTCTTTCCCTACACGACGCTCTTCCGATCTNNNGGCAATAGGAAAGTTTGAAGG
GAGCACACGTCTGAACTCTTTCCCTACACGACGCTCTTCCGATCTNNNGGGTGGGTGGAAATGTGTG
GAAGAGCACACGTCTGAACTCTTTCCCTACACGACGCTCTTCCGATCTNNNGGAGAAGGAAGGTGGG
AGAGCACACGTCTGAACTCTTTCCCTACACGACGCTCTTCCGATCTNNNAGCATGCTGGCATATATCCTT
GAGCACACGTCTGAACTCTTTCCCTACACGACGCTCTTCCGATCTNNNCTCTCGTCACTACCCTCCAGTTC
AGCACACGTCTGAACTCTTTCCCTACACGACGCTCTTCCGATCTNNNAATGGCTACAGTCTTTGTGTTG
AGAGCACACGTCTGAACTCTTTCCCTACACGACGCTCTTCCGATCTNNNCTTGTGTTTTCAGATGGGTG
AGCACACGTCTGAACTCTTTCCCTACACGACGCTCTTCCGATCTNNNAAGGCAGTTATCAGCAGGCAAA
GAGCACACGTCTGAACTCTTTCCCTACACGACGCTCTTCCGATCTNNNCACTTCCTGGTTGGAGAGGAG
GAGCACACGTCTGAACTCTTTCCCTACACGACGCTCTTCCGATCTNNNAGGAATCATCTGGAAAGAGAT
AGCACACGTCTGAACTCTTTCCCTACACGACGCTCTTCCGATCTNNNTTGTTCATTGCCCTCTCCTTTCCC
GAGCACACGTCTGAACTCTTTCCCTACACGACGCTCTTCCGATCTNNNATGCAACCATGCTATACTTGG
GAGCACACGTCTGAACTCTTTCCCTACACGACGCTCTTCCGATCTNNNATGGAAGGGGAAGACGGAC
AGAGCACACGTCTGAACTCTTTCCCTACACGACGCTCTTCCGATCTNNNCACTTAGCTGAAGGGATGGA
AGAGCACACGTCTGAACTCTTTCCCTACACGACGCTCTTCCGATCTNNNACAAAGATCCATTCCCTCCCA
AGCACACGTCTGAACTCTTTCCCTACACGACGCTCTTCCGATCTNNNATGTGTGAGCCTGATGGCTTAT
GAAGAGCACACGTCTGAACTCTTTCCCTACACGACGCTCTTCCGATCTNNNTTGTTGTTGAAACCATGA
GAGCACACGTCTGAACTCTTTCCCTACACGACGCTCTTCCGATCTNNNACCATGATTTTTGCTGTTTTCC
GAGCACACGTCTGAACTCTTTCCCTACACGACGCTCTTCCGATCTNNNCAGTAGTGTTGAGCAGAGCTG
AGCACACGTCTGAACTCTTTCCCTACACGACGCTCTTCCGATCTNNNCCACTCCCTTCTCTCCCAAAATGC
GAGCACACGTCTGAACTCTTTCCCTACACGACGCTCTTCCGATCTNNNTTTGTGTTGGAACTTGGGTCA
AGCACACGTCTGAACTCTTTCCCTACACGACGCTCTTCCGATCTNNNATCTGTGTGTAGTGTGGGGGT
GAGCACACGTCTGAACTCTTTCCCTACACGACGCTCTTCCGATCTNNNACAGTAGTTCCCAGTTCCATCC
AAGAGCACACGTCTGAACTCTTTCCCTACACGACGCTCTTCCGATCTNNNTGAAAGTAGTGGTTTCGGC
AGCACACGTCTGAACTCTTTCCCTACACGACGCTCTTCCGATCTNNNTGCTTCTGGGGACAGGAATTAC
AAGAGCACACGTCTGAACTCTTTCCCTACACGACGCTCTTCCGATCTNNNAACGTCTTCTCTTCCCTCCTC
GAGCACACGTCTGAACTCTTTCCCTACACGACGCTCTTCCGATCTNNNATAATTTCCCTAGCCCCAGATG
AAGAGCACACGTCTGAACTCTTTCCCTACACGACGCTCTTCCGATCTNNNGGCATGGCTGTTGCTTCATA
GAGCACACGTCTGAACTCTTTCCCTACACGACGCTCTTCCGATCTNNNAGCCTAGGAACTGGAGAAGT
AGCACACGTCTGAACTCTTTCCCTACACGACGCTCTTCCGATCTNNNAAGAGTGGGGCTGTGATCTTA

GAGCACACGTCTGAACTCTTTCCCTACACGACGCTCTTCCGATCTNNNCCCTTCACAAACTAGGCCAAA
AGCACACGTCTGAACTCTTTCCCTACACGACGCTCTTCCGATCTNNNAACATGCTCATGCTTCCATTACT
GAGCACACGTCTGAACTCTTTCCCTACACGACGCTCTTCCGATCTNNNTTCAGATTTGCTAGCCCCTTCA
AGCACACGTCTGAACTCTTTCCCTACACGACGCTCTTCCGATCTNNNTGTTGTTTTTCATCCTTGGCAG
GAGCACACGTCTGAACTCTTTCCCTACACGACGCTCTTCCGATCTNNNGGGATTAGGAGCCAGTAGG
GAAGAGCACACGTCTGAACTCTTTCCCTACACGACGCTCTTCCGATCTNNNTGGGAAGATGCCTTGCT
AGAGCACACGTCTGAACTCTTTCCCTACACGACGCTCTTCCGATCTNNNAGATGTGTCCCGTCTCCTCTTA
AGCACACGTCTGAACTCTTTCCCTACACGACGCTCTTCCGATCTNNNCCCAGAAGCCACTTCATACAGA
GAGCACACGTCTGAACTCTTTCCCTACACGACGCTCTTCCGATCTNNNTGAATTTGGAGAAGGCACCA
AGCACACGTCTGAACTCTTTCCCTACACGACGCTCTTCCGATCTNNNCTTTTCCTCCAGGGCTTTGTGAT
AGCACACGTCTGAACTCTTTCCCTACACGACGCTCTTCCGATCTNNNACAGCTGTAGGAAAACCAAAC
GAGCACACGTCTGAACTCTTTCCCTACACGACGCTCTTCCGATCTNNNCCTGGACCAGACAGCAAAGAA
AGAGCACACGTCTGAACTCTTTCCCTACACGACGCTCTTCCGATCTNNNATGGCAGCCTCATTAGAGGA
AGCACACGTCTGAACTCTTTCCCTACACGACGCTCTTCCGATCTNNNTTATACCCCGTTACATCCCTCCCC
AGCACACGTCTGAACTCTTTCCCTACACGACGCTCTTCCGATCTNNNGCATGGGGTCTGTGAAACTGTA
GAGCACACGTCTGAACTCTTTCCCTACACGACGCTCTTCCGATCTNNNTTTGTATGCCCCTCTTTGCTTC
AGAGCACACGTCTGAACTCTTTCCCTACACGACGCTCTTCCGATCTNNNTGCCATTGATACACATTGCCT
AGCACACGTCTGAACTCTTTCCCTACACGACGCTCTTCCGATCTNNNAGGGCAATAGCCAGTTTTGTA
GAGCACACGTCTGAACTCTTTCCCTACACGACGCTCTTCCGATCTNNNTTCTCAGGCCCACTTCCTGTAT
AGAGCACACGTCTGAACTCTTTCCCTACACGACGCTCTTCCGATCTNNNTGTCCCCCTTATGGAGATGA
GAGCACACGTCTGAACTCTTTCCCTACACGACGCTCTTCCGATCTNNNCAAGGCTACCACTTTCTCTCCA
GAGCACACGTCTGAACTCTTTCCCTACACGACGCTCTTCCGATCTNNNACAACCTGTGTTCGTCAATGG
AGAGCACACGTCTGAACTCTTTCCCTACACGACGCTCTTCCGATCTNNNTAAACATGTACACAACCCTGC
GAGCACACGTCTGAACTCTTTCCCTACACGACGCTCTTCCGATCTNNNTGGGAATCCCTGTCTCTGCAA
AAGAGCACACGTCTGAACTCTTTCCCTACACGACGCTCTTCCGATCTNNNTGGGAATCCCTGTCTCTGC
AGAGCACACGTCTGAACTCTTTCCCTACACGACGCTCTTCCGATCTNNNGACTGGGATTGGTTCCACTC
GAGCACACGTCTGAACTCTTTCCCTACACGACGCTCTTCCGATCTNNNGGACCATGGAATGTGCTTTGA
GAGCACACGTCTGAACTCTTTCCCTACACGACGCTCTTCCGATCTNNNGAAGACACTTCCCACCTTCATG
AGCACACGTCTGAACTCTTTCCCTACACGACGCTCTTCCGATCTNNNTGCAAACCCTCTGAAAAACATG
GAGCACACGTCTGAACTCTTTCCCTACACGACGCTCTTCCGATCTNNNGGAATCACATCACACACATGCA
GCACACGTCTGAACTCTTTCCCTACACGACGCTCTTCCGATCTNNNTGGTCTCGTTGGCTTAGTAGTGA
AGCACACGTCTGAACTCTTTCCCTACACGACGCTCTTCCGATCTNNNAGACAATGAGGGCTTTGGATT
AGAGCACACGTCTGAACTCTTTCCCTACACGACGCTCTTCCGATCTNNNTGAGGCATTGGAGATGGTG
AGCACACGTCTGAACTCTTTCCCTACACGACGCTCTTCCGATCTNNNGTGATCCAAGTGACCAGAAGG
AGCACACGTCTGAACTCTTTCCCTACACGACGCTCTTCCGATCTNNNCTGTTGTGTTCCTGACATTGTG
AGAGCACACGTCTGAACTCTTTCCCTACACGACGCTCTTCCGATCTNNNGCATCATGCTGTGTCTTGAA
AGCACACGTCTGAACTCTTTCCCTACACGACGCTCTTCCGATCTNNNCTTCCCACCTTGGATATGCGTAG
AGAGCACACGTCTGAACTCTTTCCCTACACGACGCTCTTCCGATCTNNNTACATCGTTCCAAAACAGGT
AGCACACGTCTGAACTCTTTCCCTACACGACGCTCTTCCGATCTNNNTCTGTAACTACACCACCAGCCT
GAGCACACGTCTGAACTCTTTCCCTACACGACGCTCTTCCGATCTNNNCTGAGTGGGTGAGAGTTGAG
AGAGCACACGTCTGAACTCTTTCCCTACACGACGCTCTTCCGATCTNNNCATGAAAAGGTGGAGCAGA
GCACACGTCTGAACTCTTTCCCTACACGACGCTCTTCCGATCTNNNCCTGGGAATAACTCCCTCCTCTTC
GAGCACACGTCTGAACTCTTTCCCTACACGACGCTCTTCCGATCTNNNAACATCTTAGGCTCTGCTTGA

AGCACACGTCTGAACTCTTTCCCTACACGACGCTCTTCCGATCTNNNCTGTCTGTTTTCCGTGACTGTCT
AGCACACGTCTGAACTCTTTCCCTACACGACGCTCTTCCGATCTNNNTGGATGGGGGCAAATAATCACA
CACACGTCTGAACTCTTTCCCTACACGACGCTCTTCCGATCTNNNTGGATGGGGGCAAATAATCACATG
GAGCACACGTCTGAACTCTTTCCCTACACGACGCTCTTCCGATCTNNNAGCTGGTTTGTACATCAAAGG
AGCACACGTCTGAACTCTTTCCCTACACGACGCTCTTCCGATCTNNNTCTCATGCCTGGCCAGATAAAA
GAGCACACGTCTGAACTCTTTCCCTACACGACGCTCTTCCGATCTNNNTGAACAGGTAGAGGATGGTG
AGCACACGTCTGAACTCTTTCCCTACACGACGCTCTTCCGATCTNNNTGGAAGGTACTGGAGTTGGGA
AGAGCACACGTCTGAACTCTTTCCCTACACGACGCTCTTCCGATCTNNNTTTGTTCTGTGAAGCAAGGT
AGCACACGTCTGAACTCTTTCCCTACACGACGCTCTTCCGATCTNNNGCAAACAACATTGACATGGCTG
AGCACACGTCTGAACTCTTTCCCTACACGACGCTCTTCCGATCTNNNTTAGTATCCACGGAAGGACCTG
AGCACACGTCTGAACTCTTTCCCTACACGACGCTCTTCCGATCTNNNTGCTAACAGAGTAGATCTTGTG
GAGCACACGTCTGAACTCTTTCCCTACACGACGCTCTTCCGATCTNNNCGTGCACAGACACGCAGAAAA
GAGCACACGTCTGAACTCTTTCCCTACACGACGCTCTTCCGATCTNNNACAAAAGGGAAACTAAGGTCA
GAGCACACGTCTGAACTCTTTCCCTACACGACGCTCTTCCGATCTNNNGTGTCTGCCTGGGTAGTAACT
GAGCACACGTCTGAACTCTTTCCCTACACGACGCTCTTCCGATCTNNNTATAAAGGGATAGCACCTGG
AGCACACGTCTGAACTCTTTCCCTACACGACGCTCTTCCGATCTNNNTGCAGAACTGTGAAGTCCTCCT
GAGCACACGTCTGAACTCTTTCCCTACACGACGCTCTTCCGATCTNNNAGAATCAGTTCTAGGGTGCCT
AGAGCACACGTCTGAACTCTTTCCCTACACGACGCTCTTCCGATCTNNNCCCCCATTTCTCCACAACTTCT
GAGCACACGTCTGAACTCTTTCCCTACACGACGCTCTTCCGATCTNNNGATCCTTGATTTGGTCCCTGTG
AGCACACGTCTGAACTCTTTCCCTACACGACGCTCTTCCGATCTNNNACCTGATAGAACTAGTATGGCT
AGCACACGTCTGAACTCTTTCCCTACACGACGCTCTTCCGATCTNNNGGAACGTGAAGTACCACTCTCA
GAGCACACGTCTGAACTCTTTCCCTACACGACGCTCTTCCGATCTNNNACCTCCCTCTCTCTTCTCCCTTAC
GCACACGTCTGAACTCTTTCCCTACACGACGCTCTTCCGATCTNNNAAAGAGAGACCACATGGAAAGC
GAGCACACGTCTGAACTCTTTCCCTACACGACGCTCTTCCGATCTNNNGTAGCAAAAGACCATGCAGGA
AGAGCACACGTCTGAACTCTTTCCCTACACGACGCTCTTCCGATCTNNNTGGGTTCAGTCTGATGATG
GAGCACACGTCTGAACTCTTTCCCTACACGACGCTCTTCCGATCTNNNAGAGAAGCAAACGCAAATCC
AGCACACGTCTGAACTCTTTCCCTACACGACGCTCTTCCGATCTNNNTTTCTCTGCCTCCTCACCATACAC
AGCACACGTCTGAACTCTTTCCCTACACGACGCTCTTCCGATCTNNNTTATTGTCAAGAAGCAGAGCAG
GAGCACACGTCTGAACTCTTTCCCTACACGACGCTCTTCCGATCTNNNCTCTCCTCTTGTAGATGGCAGG
GAGCACACGTCTGAACTCTTTCCCTACACGACGCTCTTCCGATCTNNNTACCTGGAACATGGGTTAGGA
GCACACGTCTGAACTCTTTCCCTACACGACGCTCTTCCGATCTNNNAAGCTAAGTCAGTAAGCCACAG
GCACACGTCTGAACTCTTTCCCTACACGACGCTCTTCCGATCTNNNTACCTGCAGTTCCCAGGAAAAAC
AGCACACGTCTGAACTCTTTCCCTACACGACGCTCTTCCGATCTNNNGTGACAATCCTCAACCCTTTCCAG
GCACACGTCTGAACTCTTTCCCTACACGACGCTCTTCCGATCTNNNGTTTATAGCCTCTCCTCCCCTCCCA
AGCACACGTCTGAACTCTTTCCCTACACGACGCTCTTCCGATCTNNNCGTGGTCATAACAGAGCACTGA
CACACGTCTGAACTCTTTCCCTACACGACGCTCTTCCGATCTNNNCCAAACTCTGTGTCTGGGTTTCACA
GAGCACACGTCTGAACTCTTTCCCTACACGACGCTCTTCCGATCTNNNACGGGGACAAAGCTTCAAAGA
GAGCACACGTCTGAACTCTTTCCCTACACGACGCTCTTCCGATCTNNNCGGGTTGGATCAAAAATACG
GAGCACACGTCTGAACTCTTTCCCTACACGACGCTCTTCCGATCTNNNAGGCTCCAGGTACATAGAGTT
AGCACACGTCTGAACTCTTTCCCTACACGACGCTCTTCCGATCTNNNTTGCATTCTTGTCTTCACAGGCT
AGAGCACACGTCTGAACTCTTTCCCTACACGACGCTCTTCCGATCTNNNCTTGGGGCTTGCATGTTGTT
GAGCACACGTCTGAACTCTTTCCCTACACGACGCTCTTCCGATCTNNNGGAAGCAGAGATTGCAGAG
AGCACACGTCTGAACTCTTTCCCTACACGACGCTCTTCCGATCTNNNTGGTTTTGGGAGAGGAGGTAG

GCACACGTCTGAACTCTTTCCCTACACGACGCTCTTCCGATCTNNNCGTGCACAGACACGCAGAAAAT
AGCACACGTCTGAACTCTTTCCCTACACGACGCTCTTCCGATCTNNNTCTGGAGTTTTCATGGACTTT
GAGCACACGTCTGAACTCTTTCCCTACACGACGCTCTTCCGATCTNNNTTTTCACCAGAGCTAAACCCC
GCACACGTCTGAACTCTTTCCCTACACGACGCTCTTCCGATCTNNNTCAGCCCATACAAAACTGACCT
AGCACACGTCTGAACTCTTTCCCTACACGACGCTCTTCCGATCTNNNCCATGTGGTATGTAGAGATGT
GCACACGTCTGAACTCTTTCCCTACACGACGCTCTTCCGATCTNNNGACACACCTCATCAAGATGGG
AGCACACGTCTGAACTCTTTCCCTACACGACGCTCTTCCGATCTNNNTAATGTTGTGCAGAGTTGCTG
GCACACGTCTGAACTCTTTCCCTACACGACGCTCTTCCGATCTNNNTGTGTTTTGGCCAAGAGGTCA
CACGTCTGAACTCTTTCCCTACACGACGCTCTTCCGATCTNNNGCCCATCAGAAATAACCAACAGCCAT
GAGCACACGTCTGAACTCTTTCCCTACACGACGCTCTTCCGATCTNNNAGCAATGGAGGCTGTCAAA
AGAGCACACGTCTGAACTCTTTCCCTACACGACGCTCTTCCGATCTNNNACAGACTGGCCGTTTCTCAA
AGCACACGTCTGAACTCTTTCCCTACACGACGCTCTTCCGATCTNNNATCTCCCTGGGTGTAGAGAG
AAGAGCACACGTCTGAACTCTTTCCCTACACGACGCTCTTCCGATCTNNNTAGATTGGCATTCGTCAG
GCACACGTCTGAACTCTTTCCCTACACGACGCTCTTCCGATCTNNNCCAACTTGACCAAACTCACAGG
CACACGTCTGAACTCTTTCCCTACACGACGCTCTTCCGATCTNNNCATACACATACCTGCCACAACCACA
GCACACGTCTGAACTCTTTCCCTACACGACGCTCTTCCGATCTNNNTAATGATTGGGCTCTGGCTGA
GAGCACACGTCTGAACTCTTTCCCTACACGACGCTCTTCCGATCTNNNTGTGATATTGCACCTCTGTTA
GCACACGTCTGAACTCTTTCCCTACACGACGCTCTTCCGATCTNNNATAAGCAACCACAGATCCACACC
AGCACACGTCTGAACTCTTTCCCTACACGACGCTCTTCCGATCTNNNAAGAAAAACCATCTCCCTTCCC
AGCACACGTCTGAACTCTTTCCCTACACGACGCTCTTCCGATCTNNNCACGAACTCACACACACAACTT
GCACACGTCTGAACTCTTTCCCTACACGACGCTCTTCCGATCTNNNTGGATTCCAAATGGAGCCAAC
GCACACGTCTGAACTCTTTCCCTACACGACGCTCTTCCGATCTNNNGAGCCTAAGAGCAAGCAGTCAA
AGAGCACACGTCTGAACTCTTTCCCTACACGACGCTCTTCCGATCTNNNGATCGAGACCAGGGTTTTG
GAGCACACGTCTGAACTCTTTCCCTACACGACGCTCTTCCGATCTNNNAAGACTGAATGTGGTGTGT
GAGCACACGTCTGAACTCTTTCCCTACACGACGCTCTTCCGATCTNNNAGCTCGTGAGAATACTGCTT
AGCACACGTCTGAACTCTTTCCCTACACGACGCTCTTCCGATCTNNNCCCTCCTCTGTCTCCCTTTTCAT
GAGCACACGTCTGAACTCTTTCCCTACACGACGCTCTTCCGATCTNNNACACAGAGGTTGTGTTTCTC
AGCACACGTCTGAACTCTTTCCCTACACGACGCTCTTCCGATCTNNNAGCATGCTTTTTGTATGAGGC
AGCACACGTCTGAACTCTTTCCCTACACGACGCTCTTCCGATCTNNNAGTTCAACTGATCATGGCTGG
AGCACACGTCTGAACTCTTTCCCTACACGACGCTCTTCCGATCTNNNCATGCCTGATGTGCAACTACT
GAGCACACGTCTGAACTCTTTCCCTACACGACGCTCTTCCGATCTNNNATCTCAAGTGGCTTGTCTCTC
GCACACGTCTGAACTCTTTCCCTACACGACGCTCTTCCGATCTNNNAACTCCTACCTCCTCTGGGAAA
AAGAGCACACGTCTGAACTCTTTCCCTACACGACGCTCTTCCGATCTNNNTCTCAAAAGCCCAACTGA
AGCACACGTCTGAACTCTTTCCCTACACGACGCTCTTCCGATCTNNNCGCTGTACCCAAAGTGCTATT
AGCACACGTCTGAACTCTTTCCCTACACGACGCTCTTCCGATCTNNNAGCCCCAACAAAAGTTTGTCT
AGCACACGTCTGAACTCTTTCCCTACACGACGCTCTTCCGATCTNNNGCAGTTCCAAGACCAATCAATG
AGCACACGTCTGAACTCTTTCCCTACACGACGCTCTTCCGATCTNNNAGGTGACTTGCTCCCTGTAGT
AGCACACGTCTGAACTCTTTCCCTACACGACGCTCTTCCGATCTNNNTCATAGGAAGTGTTGCAGGG
AGCACACGTCTGAACTCTTTCCCTACACGACGCTCTTCCGATCTNNNTGGGTTTCAGCCTAAAGCTAC
AGCACACGTCTGAACTCTTTCCCTACACGACGCTCTTCCGATCTNNNGTGGATGATGTGTCCTTTCAC
GAGCACACGTCTGAACTCTTTCCCTACACGACGCTCTTCCGATCTNNNACCTTGGCAGTGATTTCTTC
GAGCACACGTCTGAACTCTTTCCCTACACGACGCTCTTCCGATCTNNNCAACCATACAGGGGCCAAA
GCACACGTCTGAACTCTTTCCCTACACGACGCTCTTCCGATCTNNNGAGGTTGAGGGTACAGCGAGT

AGAGCACACGTCTGAACTCTTTCCCTACACGACGCTCTTCCGATCTNNNTGATCTTCACACCAGGAGGA

AGCACACGTCTGAACTCTTTCCCTACACGACGCTCTTCCGATCTNNNGCAAGGAACAGGAACTAGATC

AGCACACGTCTGAACTCTTTCCCTACACGACGCTCTTCCGATCTNNNGGCCCATGAATGCGTATTTTCAG

AGAGCACACGTCTGAACTCTTTCCCTACACGACGCTCTTCCGATCTNNNCTGCTAATCCAGAAGACTTGC

AGCACACGTCTGAACTCTTTCCCTACACGACGCTCTTCCGATCTNNNATTTCCCTGTGCCCTTTGTTAGC

GAGCACACGTCTGAACTCTTTCCCTACACGACGCTCTTCCGATCTNNNAGGGCAAGTGACTGCATTCTA

GCACACGTCTGAACTCTTTCCCTACACGACGCTCTTCCGATCTNNNTGTGTTTTGGCCAAGAGGTCAAT

GAGCACACGTCTGAACTCTTTCCCTACACGACGCTCTTCCGATCTNNNGCTCCTACTGAATCCAGAAACC

GCACACGTCTGAACTCTTTCCCTACACGACGCTCTTCCGATCTNNNCAGGAAACAGAACCAACAAGATC

GAGCACACGTCTGAACTCTTTCCCTACACGACGCTCTTCCGATCTNNNAGCAAATTGCAGAGCCGTATG

AGAGCACACGTCTGAACTCTTTCCCTACACGACGCTCTTCCGATCTNNNCTTTGTGTGAAAAGGACAGG

GAGCACACGTCTGAACTCTTTCCCTACACGACGCTCTTCCGATCTNNNCCGTGATTGTACACAACGCTC

AGCACACGTCTGAACTCTTTCCCTACACGACGCTCTTCCGATCTNNNTGGTCCCACAGTTCTAGCTCATA

AGCACACGTCTGAACTCTTTCCCTACACGACGCTCTTCCGATCTNNNAGCCTGCTTTGTTTTGCATGTTT

CACACGTCTGAACTCTTTCCCTACACGACGCTCTTCCGATCTNNNTAACTGTACCGGGAGCACCTAAACA

AAGAGCACACGTCTGAACTCTTTCCCTACACGACGCTCTTCCGATCTNNNTCTTCACCCTTGTAGTAGG

GAGCACACGTCTGAACTCTTTCCCTACACGACGCTCTTCCGATCTNNNTGTTTACTATCTGTCCTTTTCAC

AGCACACGTCTGAACTCTTTCCCTACACGACGCTCTTCCGATCTNNNCAACTGGACCTCCTTGTGGAAA

AGCACACGTCTGAACTCTTTCCCTACACGACGCTCTTCCGATCTNNNAGTCTGATATTGAAGTTGGCCC

AAGAGCACACGTCTGAACTCTTTCCCTACACGACGCTCTTCCGATCTNNNTGAAAGTAGTGGTTTCGG

AGCACACGTCTGAACTCTTTCCCTACACGACGCTCTTCCGATCTNNNGGTCTGAGTGCAATTCCCAATC

AGCACACGTCTGAACTCTTTCCCTACACGACGCTCTTCCGATCTNNNTTTTCATTGCCTCGATCGTCCTC

GCACACGTCTGAACTCTTTCCCTACACGACGCTCTTCCGATCTNNNAAGCAGACATAGAGGAGGGGAA

AGAGCACACGTCTGAACTCTTTCCCTACACGACGCTCTTCCGATCTNNNAACTTCTCCAAGAGCAAGCAC

AGAGCACACGTCTGAACTCTTTCCCTACACGACGCTCTTCCGATCTNNNAGTTGAGAAAAGGGTGGCG

GCACACGTCTGAACTCTTTCCCTACACGACGCTCTTCCGATCTNNNATGTGAGTTTCATACCCAGGTGC

AGCACACGTCTGAACTCTTTCCCTACACGACGCTCTTCCGATCTNNNAAAGTCGGCTCCACAAGAACTT

AGCACACGTCTGAACTCTTTCCCTACACGACGCTCTTCCGATCTNNNCTCCCTAATGTAGGTCAGCCTCA

AGCACACGTCTGAACTCTTTCCCTACACGACGCTCTTCCGATCTNNNAAATTGAAAACCAGCATCTCTG

AGAGCACACGTCTGAACTCTTTCCCTACACGACGCTCTTCCGATCTNNNTACGAATACCCTACTGGTTGC

AGCACACGTCTGAACTCTTTCCCTACACGACGCTCTTCCGATCTNNNCACTAATATTTGGTTTACATGCT

AGCACACGTCTGAACTCTTTCCCTACACGACGCTCTTCCGATCTNNNGATTTTGGTGCAACCTTGGGAA

GAGCACACGTCTGAACTCTTTCCCTACACGACGCTCTTCCGATCTNNNGCCTCACCTTCAGGGGATAAAA

AGCACACGTCTGAACTCTTTCCCTACACGACGCTCTTCCGATCTNNNACTATACACACACTCCCCTTCCC

GAGCACACGTCTGAACTCTTTCCCTACACGACGCTCTTCCGATCTNNNAAGCCACATTTGTCCCCATGAA

GCACACGTCTGAACTCTTTCCCTACACGACGCTCTTCCGATCTNNNGGCTCCCCTTCTGGATCCTTCATC

AGCACACGTCTGAACTCTTTCCCTACACGACGCTCTTCCGATCTNNNATGTGAGTTTCATACCCAGGTGC

AGAGCACACGTCTGAACTCTTTCCCTACACGACGCTCTTCCGATCTNNNCTAAATGGGTTGTGGGGAG

GAGCACACGTCTGAACTCTTTCCCTACACGACGCTCTTCCGATCTNNNGAAAGCGGCTGACCATTTGTT

GAGCACACGTCTGAACTCTTTCCCTACACGACGCTCTTCCGATCTNNNTGTTTGTCATGGACTTGGTACC

AGAGCACACGTCTGAACTCTTTCCCTACACGACGCTCTTCCGATCTNNNAGGAAGTAAAGGCCACAGTA

GAAGAGCACACGTCTGAACTCTTTCCCTACACGACGCTCTTCCGATCTNNNTTGTTTGTCTCTTCCACC

AGCACACGTCTGAACTCTTTCCCTACACGACGCTCTTCCGATCTNNNGGGTACTTGGGTATCTTCACCC

AGAGCACACGTCTGAACTCTTTCCCTACACGACGCTCTTCCGATCTNNNTGGCACTTCCATTTTCATTG
GAGCACACGTCTGAACTCTTTCCCTACACGACGCTCTTCCGATCTNNNAGTGCTCTTATTCAGGATCAG
GCACACGTCTGAACTCTTTCCCTACACGACGCTCTTCCGATCTNNNGAAAGAAAGCCGATCTTTGTGC
AGCACACGTCTGAACTCTTTCCCTACACGACGCTCTTCCGATCTNNNCATTCACACAAAGGTGACTGG
CACACGTCTGAACTCTTTCCCTACACGACGCTCTTCCGATCTNNNACCAACTTGTCTTTGTCATCGGTC
AGCACACGTCTGAACTCTTTCCCTACACGACGCTCTTCCGATCTNNNTCAAGTACCAACCTAGAGGCTT
GAGCACACGTCTGAACTCTTTCCCTACACGACGCTCTTCCGATCTNNNAGGCTGGGTAAGTATGAAG
GAGCACACGTCTGAACTCTTTCCCTACACGACGCTCTTCCGATCTNNNTGAGCTCTGCAAAGTGAGTA
GAGCACACGTCTGAACTCTTTCCCTACACGACGCTCTTCCGATCTNNNGGCAGGGACCAGATCATACA
GCACACGTCTGAACTCTTTCCCTACACGACGCTCTTCCGATCTNNNTGAGTGTCATACATTGGTCCCCT
AAGAGCACACGTCTGAACTCTTTCCCTACACGACGCTCTTCCGATCTNNNGAGGCTAGACCTTCTCTG
AGCACACGTCTGAACTCTTTCCCTACACGACGCTCTTCCGATCTNNNAGGATGCTTTCTGTCTCCTCCAT
AGCACACGTCTGAACTCTTTCCCTACACGACGCTCTTCCGATCTNNNTACTTACGTCGGCAGACCTTAC
GAGCACACGTCTGAACTCTTTCCCTACACGACGCTCTTCCGATCTNNNGCTGTTGTAACTCCACTCAG
AGCACACGTCTGAACTCTTTCCCTACACGACGCTCTTCCGATCTNNNCCAGAAGAGCTCTGTGCTGTT
AGCACACGTCTGAACTCTTTCCCTACACGACGCTCTTCCGATCTNNNACACTCTCCTTTTCTGAAGCCT
GAGCACACGTCTGAACTCTTTCCCTACACGACGCTCTTCCGATCTNNNCCATCCACAGCCTTTCCTGAA
AGCACACGTCTGAACTCTTTCCCTACACGACGCTCTTCCGATCTNNNTTCTCCCAGAGGCAGTGACTTT
GAGCACACGTCTGAACTCTTTCCCTACACGACGCTCTTCCGATCTNNNTCCCTTTTCCTCCTGTTCCTCA
AGAGCACACGTCTGAACTCTTTCCCTACACGACGCTCTTCCGATCTNNNAGTTGAGAAAAGGGTGG
GAGCACACGTCTGAACTCTTTCCCTACACGACGCTCTTCCGATCTNNNTAGTGACTTCTTCAGGGGTA
GAAGAGCACACGTCTGAACTCTTTCCCTACACGACGCTCTTCCGATCTNNNTGCCATTGATACACATT
AGCACACGTCTGAACTCTTTCCCTACACGACGCTCTTCCGATCTNNNTGGTTTCTGGGAGAGGATGTT
AGAGCACACGTCTGAACTCTTTCCCTACACGACGCTCTTCCGATCTNNNAAACTCCTGAGACTGGCTTT
GAGCACACGTCTGAACTCTTTCCCTACACGACGCTCTTCCGATCTNNNGGAGACGGGGTTTTGTGGT
AGAGCACACGTCTGAACTCTTTCCCTACACGACGCTCTTCCGATCTNNNGGGATTAGGAGCCAGTAG
GCACACGTCTGAACTCTTTCCCTACACGACGCTCTTCCGATCTNNNTGGGCACTTCCAGTATCATCTCA
AAGAGCACACGTCTGAACTCTTTCCCTACACGACGCTCTTCCGATCTNNNGCTAATGGTGAAATGGA
AGCACACGTCTGAACTCTTTCCCTACACGACGCTCTTCCGATCTNNNCTCCCTCTGGGAGATAGTGTG
GAGCACACGTCTGAACTCTTTCCCTACACGACGCTCTTCCGATCTNNNGCCTCCTCTGACCTACACACAT
GCACACGTCTGAACTCTTTCCCTACACGACGCTCTTCCGATCTNNNTTGATGTTTCAGGCACACTGTC
AGCACACGTCTGAACTCTTTCCCTACACGACGCTCTTCCGATCTNNNCAGCGTCTCACTCTGGTTTCTA
AGAGCACACGTCTGAACTCTTTCCCTACACGACGCTCTTCCGATCTNNNGCAGCAGAACTAATATGGG
AGAGCACACGTCTGAACTCTTTCCCTACACGACGCTCTTCCGATCTNNNTGTGGGGAAGAGGAATCT
AGCACACGTCTGAACTCTTTCCCTACACGACGCTCTTCCGATCTNNNATGTGATTTTTGTTGGGCACT
CACACGTCTGAACTCTTTCCCTACACGACGCTCTTCCGATCTNNNTCAAGAGGTGGTTTTGTGTTGCT
GAGCACACGTCTGAACTCTTTCCCTACACGACGCTCTTCCGATCTNNNGCAAGAGTAAAGCACTGTTC
GAGCACACGTCTGAACTCTTTCCCTACACGACGCTCTTCCGATCTNNNACGCTTCGTTTTGGCATGAA
GCACACGTCTGAACTCTTTCCCTACACGACGCTCTTCCGATCTNNNTTTATTGGAGACAGGCTCACG
AGAGCACACGTCTGAACTCTTTCCCTACACGACGCTCTTCCGATCTNNNAGTCATCACCAAAGCCAAA
ACACGTCTGAACTCTTTCCCTACACGACGCTCTTCCGATCTNNNTCTTCCGTGCTTGGCATCTAAAACA
GAGCACACGTCTGAACTCTTTCCCTACACGACGCTCTTCCGATCTNNNAAGAAGAGCAGCTGGTGTT
AAGAGCACACGTCTGAACTCTTTCCCTACACGACGCTCTTCCGATCTNNNAGCTAAGACTACAGTGCC

AGAGCACACGTCTGAACTCTTTCCCTACACGACGCTCTTCCGATCTNNNCAAGTGTGTATGTGGTGT
GCACACGTCTGAACTCTTTCCCTACACGACGCTCTTCCGATCTNNNTGTCCATGGTCTGACTTGTCTCT
GAGCACACGTCTGAACTCTTTCCCTACACGACGCTCTTCCGATCTNNNGCTAGAAGAGCCCAGACTAC
AGAGCACACGTCTGAACTCTTTCCCTACACGACGCTCTTCCGATCTNNNTAGAGCCCTTGAACAAAGC
AGAGCACACGTCTGAACTCTTTCCCTACACGACGCTCTTCCGATCTNNNGCTTGGGAGGCTGAGTTAT
CACACGTCTGAACTCTTTCCCTACACGACGCTCTTCCGATCTNNNACATCAAACACCTACTTCTGGCTC
GAAGAGCACACGTCTGAACTCTTTCCCTACACGACGCTCTTCCGATCTNNNTCCCAGTGAGTAAGT
GCACACGTCTGAACTCTTTCCCTACACGACGCTCTTCCGATCTNNNGACAGGGTCTCTCTCTGGTTTCT
AAGAGCACACGTCTGAACTCTTTCCCTACACGACGCTCTTCCGATCTNNNAGGAAAACACATCAGGCT
GCACACGTCTGAACTCTTTCCCTACACGACGCTCTTCCGATCTNNNGAGATTCCTTCCAGGCCAATGA
AGCACACGTCTGAACTCTTTCCCTACACGACGCTCTTCCGATCTNNNGGTTGCTAATGTCTTCGGGA
CACACGTCTGAACTCTTTCCCTACACGACGCTCTTCCGATCTNNNTCATCAGGTTTGGAAGTTGTGG
AGAGCACACGTCTGAACTCTTTCCCTACACGACGCTCTTCCGATCTNNNCAGCATCTTGGTTCTGCAG
AGCACACGTCTGAACTCTTTCCCTACACGACGCTCTTCCGATCTNNNCCGATCTCACCTCACGGCAAGC
AGCACACGTCTGAACTCTTTCCCTACACGACGCTCTTCCGATCTNNNGCATTGCAAAAATATGGAACT
AAGAGCACACGTCTGAACTCTTTCCCTACACGACGCTCTTCCGATCTNNNCACCCCGGAGAATTTCAA
AGCACACGTCTGAACTCTTTCCCTACACGACGCTCTTCCGATCTNNNGCTGAAGACACTTTGGTAGTG
AGAGCACACGTCTGAACTCTTTCCCTACACGACGCTCTTCCGATCTNNNGAGGTCAAAGCTCCCATCT
GCACACGTCTGAACTCTTTCCCTACACGACGCTCTTCCGATCTNNNTGCTTGCATGCTGTGATGTCTAT
GAGCACACGTCTGAACTCTTTCCCTACACGACGCTCTTCCGATCTNNNGCTTACCTTGTGGGGTTGTT
AGCACACGTCTGAACTCTTTCCCTACACGACGCTCTTCCGATCTNNNGTTGTTGGAAAACATGAGGG
GAGCACACGTCTGAACTCTTTCCCTACACGACGCTCTTCCGATCTNNNACCATGTGCAAGAATAAGCT
AGCACACGTCTGAACTCTTTCCCTACACGACGCTCTTCCGATCTNNNTGAATACTCACCTCCAAAACAT
CACACGTCTGAACTCTTTCCCTACACGACGCTCTTCCGATCTNNNACACAAAGGGCAAACTCCTGATTC
CACACGTCTGAACTCTTTCCCTACACGACGCTCTTCCGATCTNNNAAGAAAGACAGCTTCCACACTGG
AGCACACGTCTGAACTCTTTCCCTACACGACGCTCTTCCGATCTNNNTACAGGTGTGTTGTGGAGTT
AGCACACGTCTGAACTCTTTCCCTACACGACGCTCTTCCGATCTNNNGAGACTAAGCTGTCTGGCTTG
GAAGAGCACACGTCTGAACTCTTTCCCTACACGACGCTCTTCCGATCTNNNATGAGATGGCTTTTTAC
AGCACACGTCTGAACTCTTTCCCTACACGACGCTCTTCCGATCTNNNTCCTCCCCTCTTTTTCTGCTACAC
AGCACACGTCTGAACTCTTTCCCTACACGACGCTCTTCCGATCTNNNTCCAGGCAAATGTGTAGGTG
AGCACACGTCTGAACTCTTTCCCTACACGACGCTCTTCCGATCTNNNAATCTGAATCCCTTGGCTGTGA
AGCACACGTCTGAACTCTTTCCCTACACGACGCTCTTCCGATCTNNNGAATGGTTCTGCTCTGTTTGCT
GCACACGTCTGAACTCTTTCCCTACACGACGCTCTTCCGATCTNNNATCGGCTAATGAAGCCTGTGAT
AGCACACGTCTGAACTCTTTCCCTACACGACGCTCTTCCGATCTNNNGGAGGATCCTTAGGTCAGGA
CACACGTCTGAACTCTTTCCCTACACGACGCTCTTCCGATCTNNNCTGTGGATTACATGGACCTGCCTT
CACACGTCTGAACTCTTTCCCTACACGACGCTCTTCCGATCTNNNCAAAGTTTAGCTTTCCCATGCCCC
GAGCACACGTCTGAACTCTTTCCCTACACGACGCTCTTCCGATCTNNNAACTTTGTGACTGTGTGCTT
AGAGCACACGTCTGAACTCTTTCCCTACACGACGCTCTTCCGATCTNNNTGAAACACATTACAGAGGG
AGCACACGTCTGAACTCTTTCCCTACACGACGCTCTTCCGATCTNNNGGAGGAAGGGTTGCATTTAG
GAGCACACGTCTGAACTCTTTCCCTACACGACGCTCTTCCGATCTNNNTCCTGATTGTTTACACACACA
AGCACACGTCTGAACTCTTTCCCTACACGACGCTCTTCCGATCTNNNAAAAAGAGGGAGAGGTAGCA
AGAGCACACGTCTGAACTCTTTCCCTACACGACGCTCTTCCGATCTNNNGCTCACATGGCACCTCTCAT
GAGCACACGTCTGAACTCTTTCCCTACACGACGCTCTTCCGATCTNNNTTTTCTGTTGAAGCTGTTGTT

AGAGCACACGTCTGAACTCTTTCCCTACACGACGCTCTTCCGATCTNNNTGTTTGACAGAGACAGAGAA
AGCACACGTCTGAACTCTTTCCCTACACGACGCTCTTCCGATCTNNNTTCTTTTCCTCTTCCTCGTCGCCA
AGCACACGTCTGAACTCTTTCCCTACACGACGCTCTTCCGATCTNNNCAGCATGGATGGAAAGACAGA
GAGCACACGTCTGAACTCTTTCCCTACACGACGCTCTTCCGATCTNNNAGTTAGATGAGGTGGAGGAA
AAGAGCACACGTCTGAACTCTTTCCCTACACGACGCTCTTCCGATCTNNNTGGCAAGATGCAGACTAGA
GAGCACACGTCTGAACTCTTTCCCTACACGACGCTCTTCCGATCTNNNTGCCTTGATTCTGCATCTACTG
GCACACGTCTGAACTCTTTCCCTACACGACGCTCTTCCGATCTNNNACATGAAAAACCTGACAGATACA
AGAGCACACGTCTGAACTCTTTCCCTACACGACGCTCTTCCGATCTNNNATGCCCAGTCCATCCAAGAGA
GAGCACACGTCTGAACTCTTTCCCTACACGACGCTCTTCCGATCTNNNATACACAACCCAAGGAAGTCT
GAGCACACGTCTGAACTCTTTCCCTACACGACGCTCTTCCGATCTNNNAGTGGGACCTCATGTGTACCAA
GAGCACACGTCTGAACTCTTTCCCTACACGACGCTCTTCCGATCTNNNGCTGTTGCAGGTCCATCAACTA
GAAGAGCACACGTCTGAACTCTTTCCCTACACGACGCTCTTCCGATCTNNNAGATACCGTCTGTTTGTT
AGAGCACACGTCTGAACTCTTTCCCTACACGACGCTCTTCCGATCTNNNATTGCGGTAACTCTGAGTGT
AGCACACGTCTGAACTCTTTCCCTACACGACGCTCTTCCGATCTNNNCTGCCTGCCTCTCTTCCTACATA
AGCACACGTCTGAACTCTTTCCCTACACGACGCTCTTCCGATCTNNNTGCTTTCTTCCTTTCGTTCGTCA
GCACACGTCTGAACTCTTTCCCTACACGACGCTCTTCCGATCTNNNTGTTTTGTGTACACCCTGGTCA
GAGCACACGTCTGAACTCTTTCCCTACACGACGCTCTTCCGATCTNNNCGCCTATTTGCACTGGGAATA
AGAGCACACGTCTGAACTCTTTCCCTACACGACGCTCTTCCGATCTNNNCCACAAGATTGCCGTGATGA
AGAGCACACGTCTGAACTCTTTCCCTACACGACGCTCTTCCGATCTNNNTCACTACAGGGTGTGTGTGT
GAGCACACGTCTGAACTCTTTCCCTACACGACGCTCTTCCGATCTNNNCGTAACAGAGCGAGACTATG
AGAGCACACGTCTGAACTCTTTCCCTACACGACGCTCTTCCGATCTNNNCAAAACAAAACGAAATGCCC
GAGCACACGTCTGAACTCTTTCCCTACACGACGCTCTTCCGATCTNNNGTAAGAGCAAGCAGAGACCCA
AGCACACGTCTGAACTCTTTCCCTACACGACGCTCTTCCGATCTNNNTAGAGGCTATCACATCCGAAGCA
GAGCACACGTCTGAACTCTTTCCCTACACGACGCTCTTCCGATCTNNNGCCAATAAAGCGTGGTTCTCT
GAGCACACGTCTGAACTCTTTCCCTACACGACGCTCTTCCGATCTNNNCCTTTGTTCTCTCCTACTCCCCA
AGCACACGTCTGAACTCTTTCCCTACACGACGCTCTTCCGATCTNNNTGTTTTGTGTACACCCTGGTCA
GAGCACACGTCTGAACTCTTTCCCTACACGACGCTCTTCCGATCTNNNTGGTACATACCATGACCTATA
AGCACACGTCTGAACTCTTTCCCTACACGACGCTCTTCCGATCTNNNACCCTGGTTCTCTCAACTGGTTA
GAGCACACGTCTGAACTCTTTCCCTACACGACGCTCTTCCGATCTNNNACACATACATACGTGTAGACAT
GAGCACACGTCTGAACTCTTTCCCTACACGACGCTCTTCCGATCTNNNGCGTGGTGCCTTATTTGTTGAA
AGCACACGTCTGAACTCTTTCCCTACACGACGCTCTTCCGATCTNNNGAGCTTTACCCTCATCTCCCTCT
AGCACACGTCTGAACTCTTTCCCTACACGACGCTCTTCCGATCTNNNAAATGCCTCCTAAGGGTGTTCT
GAGCACACGTCTGAACTCTTTCCCTACACGACGCTCTTCCGATCTNNNTTTCCAGTGTCTACCACCTCTGT
GAGCACACGTCTGAACTCTTTCCCTACACGACGCTCTTCCGATCTNNNCCCTGGACTGGAACACAAATT
GAAGAGCACACGTCTGAACTCTTTCCCTACACGACGCTCTTCCGATCTNNNAGCTACTTCTTTGTGTAT
AGCACACGTCTGAACTCTTTCCCTACACGACGCTCTTCCGATCTNNNTCAAGCCTCACACCTATTTTGGG
AGAGCACACGTCTGAACTCTTTCCCTACACGACGCTCTTCCGATCTNNNCTTGGCTACTGTGATAGCCCT
GAGCACACGTCTGAACTCTTTCCCTACACGACGCTCTTCCGATCTNNNACACAACCAATCATGAGACAG
AGAGCACACGTCTGAACTCTTTCCCTACACGACGCTCTTCCGATCTNNNTGGCAGCCTCTTAGGAAAAT
GAGCACACGTCTGAACTCTTTCCCTACACGACGCTCTTCCGATCTNNNCTTCACTATGTCCAATGCAGG
GCACACGTCTGAACTCTTTCCCTACACGACGCTCTTCCGATCTNNNACCTCTCCTATCCTCTTTGCCACCA
AGCACACGTCTGAACTCTTTCCCTACACGACGCTCTTCCGATCTNNNTCTGCACTTATTCTTTGTCAGC
GAGCACACGTCTGAACTCTTTCCCTACACGACGCTCTTCCGATCTNNNGAGTAGTCTGCACTCCCATGTT

AGCACACGTCTGAACTCTTTCCCTACACGACGCTCTTCCGATCTNNNTAGCCTGGAGGTGTGTCATTTT
AGCACACGTCTGAACTCTTTCCCTACACGACGCTCTTCCGATCTNNNCACACATGCCTTGATAAAACGG
GAGCACACGTCTGAACTCTTTCCCTACACGACGCTCTTCCGATCTNNNTAATGCTAGGAAGAGTGGG
GAGCACACGTCTGAACTCTTTCCCTACACGACGCTCTTCCGATCTNNNGAGGGAAGACATTCTCTCTCC
GAGCACACGTCTGAACTCTTTCCCTACACGACGCTCTTCCGATCTNNNTCTATACATAGACACACACAC
AGCACACGTCTGAACTCTTTCCCTACACGACGCTCTTCCGATCTNNNAGCCTTTTTCATCACAGGGAAG
GAGCACACGTCTGAACTCTTTCCCTACACGACGCTCTTCCGATCTNNNTGCACCCACATACACACACATT
GCACACGTCTGAACTCTTTCCCTACACGACGCTCTTCCGATCTNNNACTCAACCTCAAGAGAGAGGAG
GAGCACACGTCTGAACTCTTTCCCTACACGACGCTCTTCCGATCTNNNCCGGCCTATCAGTGGGTTTTT
GAGCACACGTCTGAACTCTTTCCCTACACGACGCTCTTCCGATCTNNNTGCTCGGACCCTGATTACTAT
AGCACACGTCTGAACTCTTTCCCTACACGACGCTCTTCCGATCTNNNAATCCGTTGGGTTCACTCCTTT
GCACACGTCTGAACTCTTTCCCTACACGACGCTCTTCCGATCTNNNTGATGCAGGAGCACAACTAAGA
GAGCACACGTCTGAACTCTTTCCCTACACGACGCTCTTCCGATCTNNNATTGTGTTCGAGTATCACCCA
AAGAGCACACGTCTGAACTCTTTCCCTACACGACGCTCTTCCGATCTNNNTATATGTGAAATCGCCCT
GAGCACACGTCTGAACTCTTTCCCTACACGACGCTCTTCCGATCTNNNTGGGGACAAGAAGAAACCAT
AGCACACGTCTGAACTCTTTCCCTACACGACGCTCTTCCGATCTNNNTACACAAGCATGTATGCACAG
GCACACGTCTGAACTCTTTCCCTACACGACGCTCTTCCGATCTNNNCTAGCACCTCCATCCTACCTCTCCA
GAGCACACGTCTGAACTCTTTCCCTACACGACGCTCTTCCGATCTNNNGACACATACATGCGGGTCTG
AGAGCACACGTCTGAACTCTTTCCCTACACGACGCTCTTCCGATCTNNNTGGCTGTCATGTAGCAATCC
GAGCACACGTCTGAACTCTTTCCCTACACGACGCTCTTCCGATCTNNNTACATTTCAGGAGACCAAGCA
GAGCACACGTCTGAACTCTTTCCCTACACGACGCTCTTCCGATCTNNNTGCATAAAACATGGCACTGCT
AGCACACGTCTGAACTCTTTCCCTACACGACGCTCTTCCGATCTNNNCAGAATAGCTCTCATGGGTTTG
AGAGCACACGTCTGAACTCTTTCCCTACACGACGCTCTTCCGATCTNNNAGTGGTAATGCAAGAGGAG
AGCACACGTCTGAACTCTTTCCCTACACGACGCTCTTCCGATCTNNNTCTGCTGCATCTCTGTGTTTCTC
GAGCACACGTCTGAACTCTTTCCCTACACGACGCTCTTCCGATCTNNNAAAGGGCCTACTCAGGAAAA
GAGCACACGTCTGAACTCTTTCCCTACACGACGCTCTTCCGATCTNNNAACAGTACCACAGTACCATGT
GCACACGTCTGAACTCTTTCCCTACACGACGCTCTTCCGATCTNNNTTGTTTTGGTTTGCCATTTCCCCC
AGAGCACACGTCTGAACTCTTTCCCTACACGACGCTCTTCCGATCTNNNAGCCAGCAGCTCATCTTTTTC
AGCACACGTCTGAACTCTTTCCCTACACGACGCTCTTCCGATCTNNNCCCCCTCCCATATTGTGGTAAGA
GAGCACACGTCTGAACTCTTTCCCTACACGACGCTCTTCCGATCTNNNATCCCTGACAGCCATTAGTAG
GAGCACACGTCTGAACTCTTTCCCTACACGACGCTCTTCCGATCTNNNACAGGTATTGCATACCAAGC
AGAGCACACGTCTGAACTCTTTCCCTACACGACGCTCTTCCGATCTNNNGCAAGAAACCAGATTGGAT
GAGCACACGTCTGAACTCTTTCCCTACACGACGCTCTTCCGATCTNNNTTGATGCAGTTGATGTAAAT
AGCACACGTCTGAACTCTTTCCCTACACGACGCTCTTCCGATCTNNNACGTGTGTGTAAGTAGCAATC
GAGCACACGTCTGAACTCTTTCCCTACACGACGCTCTTCCGATCTNNNCTCCCTGTCAAAGAGGTTCCC
AGCACACGTCTGAACTCTTTCCCTACACGACGCTCTTCCGATCTNNNACATGAGGGTTTTAGTGTCCTG
GAGCACACGTCTGAACTCTTTCCCTACACGACGCTCTTCCGATCTNNNCGTCTACAACAACACCCTCAAC
GAGCACACGTCTGAACTCTTTCCCTACACGACGCTCTTCCGATCTNNNTCCAGGCATGTCCATGTAGGT
GAGCACACGTCTGAACTCTTTCCCTACACGACGCTCTTCCGATCTNNNAAACCAGAAGGAGTTGAGCA
GAGCACACGTCTGAACTCTTTCCCTACACGACGCTCTTCCGATCTNNNCCTCCAGCTCTATTCAAGTGGT
AGCACACGTCTGAACTCTTTCCCTACACGACGCTCTTCCGATCTNNNCCTCTCAAGCTTCATCTCCAACC
AGCACACGTCTGAACTCTTTCCCTACACGACGCTCTTCCGATCTNNNTCAACTACCTCCATCCTAGGTCA
GCACACGTCTGAACTCTTTCCCTACACGACGCTCTTCCGATCTNNNATGAGCATGTACACACACACAGT

AGCACACGTCTGAACTCTTTCCCTACACGACGCTCTTCCGATCTNNNAGAGTTTATCTGGAGGAACA
CACACGTCTGAACTCTTTCCCTACACGACGCTCTTCCGATCTNNNCATCAGTGTCTGGAGGCTTTTCA
AGCACACGTCTGAACTCTTTCCCTACACGACGCTCTTCCGATCTNNNCATTTATATTGGCTGCCTGGT
AGAGCACACGTCTGAACTCTTTCCCTACACGACGCTCTTCCGATCTNNNAGCAGTGAGCTCTAACAGA
GAGCACACGTCTGAACTCTTTCCCTACACGACGCTCTTCCGATCTNNNAAGTGTTCCCAGTTTAGACCC
GCACACGTCTGAACTCTTTCCCTACACGACGCTCTTCCGATCTNNNAGAATTCCAGGGTGCAAGATGT
AGCACACGTCTGAACTCTTTCCCTACACGACGCTCTTCCGATCTNNNGTCTCTTGGAAGGGGTGTAGG
GAGCACACGTCTGAACTCTTTCCCTACACGACGCTCTTCCGATCTNNNCAGTTTATGGACAGAAGCTG
GAGCACACGTCTGAACTCTTTCCCTACACGACGCTCTTCCGATCTNNNTTCCCCACTCCTAACTTACA
AGCACACGTCTGAACTCTTTCCCTACACGACGCTCTTCCGATCTNNNTTCACTCCATTCCCCACACACATC
AGCACACGTCTGAACTCTTTCCCTACACGACGCTCTTCCGATCTNNNTCCAAGCACTCCAAAATCTCAC
AGCACACGTCTGAACTCTTTCCCTACACGACGCTCTTCCGATCTNNNTACATTTCAAGTGCTGTCCCTC
AGCACACGTCTGAACTCTTTCCCTACACGACGCTCTTCCGATCTNNNAAAAGGGACAGCAGTTTGGC
AAGAGCACACGTCTGAACTCTTTCCCTACACGACGCTCTTCCGATCTNNNTATGCTGGAGGCTTTCGT
AGCACACGTCTGAACTCTTTCCCTACACGACGCTCTTCCGATCTNNNGGAGTTTCCTTTCTTCTTCCACC
AGCACACGTCTGAACTCTTTCCCTACACGACGCTCTTCCGATCTNNNCTTTGGTGCTCCTGCTGTTTAG
GAGCACACGTCTGAACTCTTTCCCTACACGACGCTCTTCCGATCTNNNTCAGGTAGATCACACACACAC
GCACACGTCTGAACTCTTTCCCTACACGACGCTCTTCCGATCTNNNTGGTGTTGAAAGTGCTTCTTGA
AGCACACGTCTGAACTCTTTCCCTACACGACGCTCTTCCGATCTNNNCCTTACTTTCTCAGCTCAGTGG
GCACACGTCTGAACTCTTTCCCTACACGACGCTCTTCCGATCTNNNATGCTTTCAGGTCTTCAAGCCAA
AGCACACGTCTGAACTCTTTCCCTACACGACGCTCTTCCGATCTNNNAGATATAACACTGCTGGCTGG
GAGCACACGTCTGAACTCTTTCCCTACACGACGCTCTTCCGATCTNNNACCACTTCCCCAATCTCTCTCT
GAGCACACGTCTGAACTCTTTCCCTACACGACGCTCTTCCGATCTNNNACATATCCATAGTCAGGAGA
GAGCACACGTCTGAACTCTTTCCCTACACGACGCTCTTCCGATCTNNNGGCTGACCTCAACTCAGAAC
AGCACACGTCTGAACTCTTTCCCTACACGACGCTCTTCCGATCTNNNGGGGCATTATCCTTCACCACAA
AGCACACGTCTGAACTCTTTCCCTACACGACGCTCTTCCGATCTNNNATGTGAAGGAGACCCTGACAT
AGCACACGTCTGAACTCTTTCCCTACACGACGCTCTTCCGATCTNNNCTTCCAGGGGATTCTCATGTTG
GAGCACACGTCTGAACTCTTTCCCTACACGACGCTCTTCCGATCTNNNAAAATACATGCGTTGGAGGT
GAGCACACGTCTGAACTCTTTCCCTACACGACGCTCTTCCGATCTNNNAGCTCACATGAAAACATCCCC
AGAGCACACGTCTGAACTCTTTCCCTACACGACGCTCTTCCGATCTNNNGTCAGTCCCCCATGGGATA
GCACACGTCTGAACTCTTTCCCTACACGACGCTCTTCCGATCTNNNTCTGGGGAGTGTGTGTCTGTAA
AGCACACGTCTGAACTCTTTCCCTACACGACGCTCTTCCGATCTNNNACTTTGTCCTTGATATCCTTAG
AAGAGCACACGTCTGAACTCTTTCCCTACACGACGCTCTTCCGATCTNNNTCTGGTAAGGCCATGGG
GAGCACACGTCTGAACTCTTTCCCTACACGACGCTCTTCCGATCTNNNACAGATCACAACCCTGCCTCT
CACACGTCTGAACTCTTTCCCTACACGACGCTCTTCCGATCTNNNCACTGTATCCCAACTGCACATCAC
GAGCACACGTCTGAACTCTTTCCCTACACGACGCTCTTCCGATCTNNNATAGCACAGCTAGCATCAAG
GAGCACACGTCTGAACTCTTTCCCTACACGACGCTCTTCCGATCTNNNACAACTCTACCATCCCCTAAC
AGCACACGTCTGAACTCTTTCCCTACACGACGCTCTTCCGATCTNNNTAACCAAATGCCCACCAGTCA
CACACGTCTGAACTCTTTCCCTACACGACGCTCTTCCGATCTNNNTAGGCTGAGGTTGGTAAAGGTC
AAGAGCACACGTCTGAACTCTTTCCCTACACGACGCTCTTCCGATCTNNNTGAATTTCCCTTTGCCCCT
AGCACACGTCTGAACTCTTTCCCTACACGACGCTCTTCCGATCTNNNTTGGTGTTTCCCCTCACTCCTT
AGCACACGTCTGAACTCTTTCCCTACACGACGCTCTTCCGATCTNNNAGACAGGGTCTTCCAATAGGC
AGCACACGTCTGAACTCTTTCCCTACACGACGCTCTTCCGATCTNNNCCTCAGGTTCTGGAGGAATAC

AGCACACGTCTGAACTCTTTCCCTACACGACGCTCTTCCGATCTNNNCCAGGTAGGAAGTGATTTCTGG
GAGCACACGTCTGAACTCTTTCCCTACACGACGCTCTTCCGATCTNNNGGTCTGCGAAAGATGAGTTA
GAGCACACGTCTGAACTCTTTCCCTACACGACGCTCTTCCGATCTNNNCAGGTACCTAACCTGTCTCTGG
GAGCACACGTCTGAACTCTTTCCCTACACGACGCTCTTCCGATCTNNNCCTGTTGCTGCAGTTGATTTCT
AGCACACGTCTGAACTCTTTCCCTACACGACGCTCTTCCGATCTNNNAAAGATAAAGCCCCAAAGGCCA
GAGCACACGTCTGAACTCTTTCCCTACACGACGCTCTTCCGATCTNNNAGGACAGACTGAACGTCCTTA
GCACACGTCTGAACTCTTTCCCTACACGACGCTCTTCCGATCTNNNTCAGATCCACATTTTCCCAGTTTC
AGAGCACACGTCTGAACTCTTTCCCTACACGACGCTCTTCCGATCTNNNTGTTTACATAGAACCTTCCAC
GAGCACACGTCTGAACTCTTTCCCTACACGACGCTCTTCCGATCTNNNAGTCCTAATCCCCATCCTCAGAA
AGAGCACACGTCTGAACTCTTTCCCTACACGACGCTCTTCCGATCTNNNGGTGCTCAGTTTGCCATTTTG
AGAGCACACGTCTGAACTCTTTCCCTACACGACGCTCTTCCGATCTNNNGAAGTTGTGGATCTTCTCCC
GAGCACACGTCTGAACTCTTTCCCTACACGACGCTCTTCCGATCTNNNGTCCCTAAAGGAATGCTGTCA
AGCACACGTCTGAACTCTTTCCCTACACGACGCTCTTCCGATCTNNNGTCACTGAATCACATGGGTAG
GAGCACACGTCTGAACTCTTTCCCTACACGACGCTCTTCCGATCTNNNGACGGGTGACATTTTCATCTCC
GAGCACACGTCTGAACTCTTTCCCTACACGACGCTCTTCCGATCTNNNACTTCTATCTCAGGCTCGTCCT
GAGCACACGTCTGAACTCTTTCCCTACACGACGCTCTTCCGATCTNNNGTCCAATGCCACTGGTGTTCTT
AGAGCACACGTCTGAACTCTTTCCCTACACGACGCTCTTCCGATCTNNNCTTACACAGAGCCCAGACTTT
GAGCACACGTCTGAACTCTTTCCCTACACGACGCTCTTCCGATCTNNNAGGAAACAAGGCTTACCTAGA
AGCACACGTCTGAACTCTTTCCCTACACGACGCTCTTCCGATCTNNNATGGAAGGGAAGGGCAACTTT
AGAGCACACGTCTGAACTCTTTCCCTACACGACGCTCTTCCGATCTNNNTGCAGTGGTAGCAAAAGCAA
GAGCACACGTCTGAACTCTTTCCCTACACGACGCTCTTCCGATCTNNNCCCGGCCAGTTCTGATTCTTATT
GCACACGTCTGAACTCTTTCCCTACACGACGCTCTTCCGATCTNNNCAGCTCTAGAATCCTGGCTTTCCC
GAGCACACGTCTGAACTCTTTCCCTACACGACGCTCTTCCGATCTNNNACGACCAGAGAAAAGAGTGA
GAGCACACGTCTGAACTCTTTCCCTACACGACGCTCTTCCGATCTNNNGTCACGTTTTCTGTTCCCTCCTT
AGCACACGTCTGAACTCTTTCCCTACACGACGCTCTTCCGATCTNNNTGGCCTGCCAGGATTTAGAGAA
GAGCACACGTCTGAACTCTTTCCCTACACGACGCTCTTCCGATCTNNNCCCATATGACTAGTTTCCCAGC
GCACACGTCTGAACTCTTTCCCTACACGACGCTCTTCCGATCTNNNACATGTTTACTTGCTGCCTGAGAC
GAGCACACGTCTGAACTCTTTCCCTACACGACGCTCTTCCGATCTNNNTGCTTGAAGAATCATGGTCCC
GAGCACACGTCTGAACTCTTTCCCTACACGACGCTCTTCCGATCTNNNTCACTGCATGGTGACCACAAT
AGCACACGTCTGAACTCTTTCCCTACACGACGCTCTTCCGATCTNNNTACTGTTGCTACCTTTTGCATCTC
GAGCACACGTCTGAACTCTTTCCCTACACGACGCTCTTCCGATCTNNNCCCTGTAACTGCACCAAACTTC
AAGAGCACACGTCTGAACTCTTTCCCTACACGACGCTCTTCCGATCTNNNCTTACACAGAGCCCAGACTT
AGCACACGTCTGAACTCTTTCCCTACACGACGCTCTTCCGATCTNNNCTTTATGCAAATTGGCCTCGCTG
GCACACGTCTGAACTCTTTCCCTACACGACGCTCTTCCGATCTNNNCTGAGAGTTGCGGTGGCTATGCA
AAGAGCACACGTCTGAACTCTTTCCCTACACGACGCTCTTCCGATCTNNNTGCATTCTGAGATCAGATTA
GAGCACACGTCTGAACTCTTTCCCTACACGACGCTCTTCCGATCTNNNTCGCAGGGAGTAAAATGTGTC
GCACACGTCTGAACTCTTTCCCTACACGACGCTCTTCCGATCTNNNACTGTTCCAGGGCATAGTCTCAT
AGCACACGTCTGAACTCTTTCCCTACACGACGCTCTTCCGATCTNNNCAGCTAATGCTGCCTCCTTAGAG
GAGCACACGTCTGAACTCTTTCCCTACACGACGCTCTTCCGATCTNNNAAAGGTTCTTGGTGGTACTGC
AGAGCACACGTCTGAACTCTTTCCCTACACGACGCTCTTCCGATCTNNNTCAATCCTTAGATCAGTTCTT
GAGCACACGTCTGAACTCTTTCCCTACACGACGCTCTTCCGATCTNNNTGAATATGCCAGTTTGGATGCA
AGCACACGTCTGAACTCTTTCCCTACACGACGCTCTTCCGATCTNNNTTATGGCCTTCCTCAGCCTAACTC
GAGCACACGTCTGAACTCTTTCCCTACACGACGCTCTTCCGATCTNNNTGTCTTTGGAGGGATTTGAGT

AGCACACGTCTGAACTCTTTCCCTACACGACGCTCTTCCGATCTNNNACCTCAGAGCTCCTGTCATTAG
GAGCACACGTCTGAACTCTTTCCCTACACGACGCTCTTCCGATCTNNNACAGCAGGACATGTGTGCTAA
AGCACACGTCTGAACTCTTTCCCTACACGACGCTCTTCCGATCTNNNACGTCTACCAGAGTTCATTTGGA
AGAGCACACGTCTGAACTCTTTCCCTACACGACGCTCTTCCGATCTNNNATCACTCATTAGCAATCCACC
GAGCACACGTCTGAACTCTTTCCCTACACGACGCTCTTCCGATCTNNNTTTCACTAGCACCTTCCAAAGC
AGAGCACACGTCTGAACTCTTTCCCTACACGACGCTCTTCCGATCTNNNTACTAGGTCTGAGGTTCAGG
AGAGCACACGTCTGAACTCTTTCCCTACACGACGCTCTTCCGATCTNNNTGTGTGTGTGTAGATTCCAG
GAGCACACGTCTGAACTCTTTCCCTACACGACGCTCTTCCGATCTNNNACTAGAATCAGCCAATCTGGC
GAGCACACGTCTGAACTCTTTCCCTACACGACGCTCTTCCGATCTNNNCACAGTCATCCCTCCCTAAGACA
AGAGCACACGTCTGAACTCTTTCCCTACACGACGCTCTTCCGATCTNNNCCCCAGCCTTTCATGCTCTATT
GCACACGTCTGAACTCTTTCCCTACACGACGCTCTTCCGATCTNNNTTCCTGTGAAAAACTGTTTGGGTG
GAGCACACGTCTGAACTCTTTCCCTACACGACGCTCTTCCGATCTNNNATGGAACGTGGCAGAAGTGA
GAGCACACGTCTGAACTCTTTCCCTACACGACGCTCTTCCGATCTNNNTCAATACCTCATTGCCCACCTT
GAGCACACGTCTGAACTCTTTCCCTACACGACGCTCTTCCGATCTNNNCTGTCTCTCTGGCGTCATTTGT
GAGCACACGTCTGAACTCTTTCCCTACACGACGCTCTTCCGATCTNNNGGTAGATTCTGCAGTTTGGG
AGAGCACACGTCTGAACTCTTTCCCTACACGACGCTCTTCCGATCTNNNCATGGCATCCTGAGATGGA
AGAGCACACGTCTGAACTCTTTCCCTACACGACGCTCTTCCGATCTNNNCACTTGTCTACAGCATCAGCA
GAGCACACGTCTGAACTCTTTCCCTACACGACGCTCTTCCGATCTNNNAGGGAATCTCCTCATGGAACA
GAGCACACGTCTGAACTCTTTCCCTACACGACGCTCTTCCGATCTNNNCTGGAAAGCTTGGAAGATTGG
GAGCACACGTCTGAACTCTTTCCCTACACGACGCTCTTCCGATCTNNNCCTGCTTACACACTATCACCACC
GAGCACACGTCTGAACTCTTTCCCTACACGACGCTCTTCCGATCTNNNCCTAGAACATGGGTCTGGGTT
GAGCACACGTCTGAACTCTTTCCCTACACGACGCTCTTCCGATCTNNNTGGCACAGAGACAGAGAGAA
GAGCACACGTCTGAACTCTTTCCCTACACGACGCTCTTCCGATCTNNNGAGGGCAAAGAAGAGATCACA
AGAGCACACGTCTGAACTCTTTCCCTACACGACGCTCTTCCGATCTNNNACTTGAACTCACTTCTGTCCA
AGAGCACACGTCTGAACTCTTTCCCTACACGACGCTCTTCCGATCTNNNACAGGGCACTGTACTTGATTG
GAGCACACGTCTGAACTCTTTCCCTACACGACGCTCTTCCGATCTNNNACACAGGAAGGCATTTCTTAG
GAGCACACGTCTGAACTCTTTCCCTACACGACGCTCTTCCGATCTNNNACAAATGGGCATGAGGGAAA
GCACACGTCTGAACTCTTTCCCTACACGACGCTCTTCCGATCTNNNCTTTGTTCCTTTCCTGGTCTGGGCA
GAGCACACGTCTGAACTCTTTCCCTACACGACGCTCTTCCGATCTNNNCTCCCCAAGGTTTTGGAAAAG
AGCACACGTCTGAACTCTTTCCCTACACGACGCTCTTCCGATCTNNNAGGCCACACTAACTTTGAAAGCA
GAGCACACGTCTGAACTCTTTCCCTACACGACGCTCTTCCGATCTNNNTACTAGGTCTGAGGTTCAGGG
GAGCACACGTCTGAACTCTTTCCCTACACGACGCTCTTCCGATCTNNNTCCTCCTGGAAGCTCATTGAGA
GAGCACACGTCTGAACTCTTTCCCTACACGACGCTCTTCCGATCTNNNAGGCAAATGGATTTCTTGAGG
GAGCACACGTCTGAACTCTTTCCCTACACGACGCTCTTCCGATCTNNNCACAAATGTGCTTGCCTCTCTT
GAAGAGCACACGTCTGAACTCTTTCCCTACACGACGCTCTTCCGATCTNNNACTAAATGCCAGGTCAGT
AGCACACGTCTGAACTCTTTCCCTACACGACGCTCTTCCGATCTNNNTTATGACAGTGGTGCATCCATG
AGCACACGTCTGAACTCTTTCCCTACACGACGCTCTTCCGATCTNNNCCCCTCATTGAGAATCACTACGCC
AGCACACGTCTGAACTCTTTCCCTACACGACGCTCTTCCGATCTNNNACGTTCCTCTGGTATATTCGGGT
GAGCACACGTCTGAACTCTTTCCCTACACGACGCTCTTCCGATCTNNNCTGGCTTCCTCCAAAGTTAGCA
AGCACACGTCTGAACTCTTTCCCTACACGACGCTCTTCCGATCTNNNATATATGCCAGTCACACTCCCCC
AGCACACGTCTGAACTCTTTCCCTACACGACGCTCTTCCGATCTNNNAGCCACTGTGTACTTAGCATGT
AGCACACGTCTGAACTCTTTCCCTACACGACGCTCTTCCGATCTNNNCCCAAGGGAGGGAAGGTGTAA
GAGCACACGTCTGAACTCTTTCCCTACACGACGCTCTTCCGATCTNNNTGCTAAATCTCATGGGGCCATT

AGCACACGTCTGAACTCTTTCCCTACACGACGCTCTTCCGATCTNNNTTTGTGTTCTTGTCTGGCCTTGT
GAGCACACGTCTGAACTCTTTCCCTACACGACGCTCTTCCGATCTNNNTGCTTGTGGCTGGTACAAGTT
GAGCACACGTCTGAACTCTTTCCCTACACGACGCTCTTCCGATCTNNNAGAAAATACACGTCTGCTGCC
ACACGTCTGAACTCTTTCCCTACACGACGCTCTTCCGATCTNNNGCATAAAATCTGGAAAGCCACCGCA
GAGCACACGTCTGAACTCTTTCCCTACACGACGCTCTTCCGATCTNNNGAAGGATCCCTTCCCTCATCCT
GAGCACACGTCTGAACTCTTTCCCTACACGACGCTCTTCCGATCTNNNGTCACGTTTTCTGTTCCCTCCTT
AGCACACGTCTGAACTCTTTCCCTACACGACGCTCTTCCGATCTNNNCCCAGCCTCCTTCTTCTTTTTCTCA
AGCACACGTCTGAACTCTTTCCCTACACGACGCTCTTCCGATCTNNNATCTGTGTCATCCTTTGGAAGCA
AGCACACGTCTGAACTCTTTCCCTACACGACGCTCTTCCGATCTNNNTCCTGTTCTAAGGCAAGGCAAA
AGCACACGTCTGAACTCTTTCCCTACACGACGCTCTTCCGATCTNNNACGAACAAACAACCAACCAACA
GAGCACACGTCTGAACTCTTTCCCTACACGACGCTCTTCCGATCTNNNATTGCAATGTATGAGGCATCG
GAGCACACGTCTGAACTCTTTCCCTACACGACGCTCTTCCGATCTNNNTCACATCTCATTGTATCCAGGC
AGCACACGTCTGAACTCTTTCCCTACACGACGCTCTTCCGATCTNNNAGGCATCATCCAAGGGATTTAC
GAGCACACGTCTGAACTCTTTCCCTACACGACGCTCTTCCGATCTNNNAAGCATTCCTTGGTCTGGTTCT
GAGCACACGTCTGAACTCTTTCCCTACACGACGCTCTTCCGATCTNNNACCTAGGACCTCTCTAACTCCCT
GAGCACACGTCTGAACTCTTTCCCTACACGACGCTCTTCCGATCTNNNTCAAAGTGTGTGTGTGTATGC
AGCACACGTCTGAACTCTTTCCCTACACGACGCTCTTCCGATCTNNNCAACCTCTACTGGTCATCTGCAC
GAGCACACGTCTGAACTCTTTCCCTACACGACGCTCTTCCGATCTNNNCCTCCCTCCCTTTCCTAACTTCAC
GAGCACACGTCTGAACTCTTTCCCTACACGACGCTCTTCCGATCTNNNGAAAGTGGAAACAGCCTCCTG
AGAGCACACGTCTGAACTCTTTCCCTACACGACGCTCTTCCGATCTNNNTCCCTGTCTCACCACTGTTAC
GAGCACACGTCTGAACTCTTTCCCTACACGACGCTCTTCCGATCTNNNCCTCTGGAGGAACTCTGACCTT
AGCACACGTCTGAACTCTTTCCCTACACGACGCTCTTCCGATCTNNNTTGTCTTGTGTCTTGTTCCCTCCC
AGCACACGTCTGAACTCTTTCCCTACACGACGCTCTTCCGATCTNNNTCACTCTCCCAAATGCAGTAAAA
AGCACACGTCTGAACTCTTTCCCTACACGACGCTCTTCCGATCTNNNTCACAGCTTTTGGAGCTTCCATT
AGCACACGTCTGAACTCTTTCCCTACACGACGCTCTTCCGATCTNNNCTATGCCAGACATCATGCTAGG
GCACACGTCTGAACTCTTTCCCTACACGACGCTCTTCCGATCTNNNCCGGTTTGGTGTTTGATTGACAG
AGCACACGTCTGAACTCTTTCCCTACACGACGCTCTTCCGATCTNNNTCCAGCATGTTTACACACACACA
CACACGTCTGAACTCTTTCCCTACACGACGCTCTTCCGATCTNNNTGCAGAGACACAGACGTAGAGAG
AGAGCACACGTCTGAACTCTTTCCCTACACGACGCTCTTCCGATCTNNNAACTGACCCCTGTTTGCACTA
AGAGCACACGTCTGAACTCTTTCCCTACACGACGCTCTTCCGATCTNNNTTGCCATTCTCTTCCACAACAC
GCACACGTCTGAACTCTTTCCCTACACGACGCTCTTCCGATCTNNNTCAGGGTACTAGCACTGTTGGTT
GAGCACACGTCTGAACTCTTTCCCTACACGACGCTCTTCCGATCTNNNAAGTAGCAGAGCTGGAATTTC
AGCACACGTCTGAACTCTTTCCCTACACGACGCTCTTCCGATCTNNNCCATGGTGAGTGGAATGTTGGA
GCACACGTCTGAACTCTTTCCCTACACGACGCTCTTCCGATCTNNNGTCTTCCCAACTGCTCTTCTCTGCA
AGCACACGTCTGAACTCTTTCCCTACACGACGCTCTTCCGATCTNNNCTTCTGGGGAGAAAGAAGGCTC
AGCACACGTCTGAACTCTTTCCCTACACGACGCTCTTCCGATCTNNNCAATCCCACACCGATACCAAGAC
GAGCACACGTCTGAACTCTTTCCCTACACGACGCTCTTCCGATCTNNNATCTTTGGTCTGTTCAGCGCT
GAGCACACGTCTGAACTCTTTCCCTACACGACGCTCTTCCGATCTNNNACCACATAACTCAGAGTAGGG
AGCACACGTCTGAACTCTTTCCCTACACGACGCTCTTCCGATCTNNNTATTGACATGTGTGTGTGTGGC
AGCACACGTCTGAACTCTTTCCCTACACGACGCTCTTCCGATCTNNNGCTGGCCTTGGTATTAGTAGTC
AGCACACGTCTGAACTCTTTCCCTACACGACGCTCTTCCGATCTNNNTAGGGTGGCCTAAAAGACACT
AGCACACGTCTGAACTCTTTCCCTACACGACGCTCTTCCGATCTNNNCTGCAGGCAGTGATGCTATCTA
AGCACACGTCTGAACTCTTTCCCTACACGACGCTCTTCCGATCTNNNTCATGAGCCTGACATAATCAGGT

AGAGCACACGTCTGAACTCTTTCCCTACACGACGCTCTTCCGATCTNNNACAATGTCTATTCATGCGTG
AGCACACGTCTGAACTCTTTCCCTACACGACGCTCTTCCGATCTNNNCCTTGCCCATTTTGCCCTACTATC
AGCACACGTCTGAACTCTTTCCCTACACGACGCTCTTCCGATCTNNNTTCTCTCTTGCCTATCCCTACCGC
GAGCACACGTCTGAACTCTTTCCCTACACGACGCTCTTCCGATCTNNNGGGAGAGACAGGGAGAAAC
AGCACACGTCTGAACTCTTTCCCTACACGACGCTCTTCCGATCTNNNCAATCCCACACCGATACCAAGAC
AGCACACGTCTGAACTCTTTCCCTACACGACGCTCTTCCGATCTNNNAAAATGATAGCCGTCCTGTTGC
GCACACGTCTGAACTCTTTCCCTACACGACGCTCTTCCGATCTNNNAGCCACACGCAAAATGAAACAAA
AGCACACGTCTGAACTCTTTCCCTACACGACGCTCTTCCGATCTNNNAAGTGGGTCATTCCCACAGACT
AGCACACGTCTGAACTCTTTCCCTACACGACGCTCTTCCGATCTNNNGCCTGTTGTTCAAAGACACTTCC
GAGCACACGTCTGAACTCTTTCCCTACACGACGCTCTTCCGATCTNNNCCTCCCTCCCTTTCCTAACTTCA
AGAGCACACGTCTGAACTCTTTCCCTACACGACGCTCTTCCGATCTNNNACCTTCCAAGGACATTCAACT
GAGCACACGTCTGAACTCTTTCCCTACACGACGCTCTTCCGATCTNNNGGAGGGCACTAGTGAAGTCCT
GCACACGTCTGAACTCTTTCCCTACACGACGCTCTTCCGATCTNNNTATCTCTCATGCTCACCTCTGGGC
GAGCACACGTCTGAACTCTTTCCCTACACGACGCTCTTCCGATCTNNNTCCTAGCATCTCAGGGCATAA
GAGCACACGTCTGAACTCTTTCCCTACACGACGCTCTTCCGATCTNNNCCTGTGCCCTCTCATCATTCTCT
GCACACGTCTGAACTCTTTCCCTACACGACGCTCTTCCGATCTNNNAAATCACGTGTCAGTCTCCTTCCCA
GAGCACACGTCTGAACTCTTTCCCTACACGACGCTCTTCCGATCTNNNCATGCACCATCACCCTGGATAA
GAGCACACGTCTGAACTCTTTCCCTACACGACGCTCTTCCGATCTNNNTGGAGGTGTGAGTAAGATAG
GAGCACACGTCTGAACTCTTTCCCTACACGACGCTCTTCCGATCTNNNGTGTTAGCATGTCCACCTGAG
AGCACACGTCTGAACTCTTTCCCTACACGACGCTCTTCCGATCTNNNCCCACAGATTGAACTGAAATCTT
AAGAGCACACGTCTGAACTCTTTCCCTACACGACGCTCTTCCGATCTNNNAAACATACACACACAGCCTC
AGAGCACACGTCTGAACTCTTTCCCTACACGACGCTCTTCCGATCTNNNTTGTCCTCTGCAAAGAACCT
AAGAGCACACGTCTGAACTCTTTCCCTACACGACGCTCTTCCGATCTNNNACCTTCCAAGGACATTCAAC
AGAGCACACGTCTGAACTCTTTCCCTACACGACGCTCTTCCGATCTNNNATAACTGAGAAACAGTGGCT
AGAGCACACGTCTGAACTCTTTCCCTACACGACGCTCTTCCGATCTNNNTTTTCCTCCTCCCCAAACATGA
AGCACACGTCTGAACTCTTTCCCTACACGACGCTCTTCCGATCTNNNCCTGATACCCTCCTCTTATGGCA
CACACGTCTGAACTCTTTCCCTACACGACGCTCTTCCGATCTNNNGAAAGCACCACATCAAAATGGCAC
GAGCACACGTCTGAACTCTTTCCCTACACGACGCTCTTCCGATCTNNNAACGATGGAAACTGTACCCTA
AGAGCACACGTCTGAACTCTTTCCCTACACGACGCTCTTCCGATCTNNNCTGGAGAGTAGGAAGTGAG
GAGCACACGTCTGAACTCTTTCCCTACACGACGCTCTTCCGATCTNNNTGTCCCATCAATCAGTGTAGTC
AGCACACGTCTGAACTCTTTCCCTACACGACGCTCTTCCGATCTNNNTGCACCACTGTCATAAACACACA
GAGCACACGTCTGAACTCTTTCCCTACACGACGCTCTTCCGATCTNNNGTCTGTGAAGTGTGTGCCTGT
AGAGCACACGTCTGAACTCTTTCCCTACACGACGCTCTTCCGATCTNNNGTCATTTTGGCTTCTTTCCCTC
AGAGCACACGTCTGAACTCTTTCCCTACACGACGCTCTTCCGATCTNNNCAATGCCCAGGGAAATTGAC
GAGCACACGTCTGAACTCTTTCCCTACACGACGCTCTTCCGATCTNNNCCTCCCTATCGCTCTACTCCATT
AGCACACGTCTGAACTCTTTCCCTACACGACGCTCTTCCGATCTNNNAAGTGGGTCATTCCCACAGACTA
GAGCACACGTCTGAACTCTTTCCCTACACGACGCTCTTCCGATCTNNNAATTTTAGGCTTTGGGGCTGA
GAGCACACGTCTGAACTCTTTCCCTACACGACGCTCTTCCGATCTNNNTCTCCTCTACTGTCCCAGAAAG
GAGCACACGTCTGAACTCTTTCCCTACACGACGCTCTTCCGATCTNNNTTGCCAGCCATTCTGTTTGTAG
GAGCACACGTCTGAACTCTTTCCCTACACGACGCTCTTCCGATCTNNNGAGGTGATTATCCTCACTGCT
AGAGCACACGTCTGAACTCTTTCCCTACACGACGCTCTTCCGATCTNNNCCAGAGCAGTAGTTCCAAGA
AGCACACGTCTGAACTCTTTCCCTACACGACGCTCTTCCGATCTNNNACTCTCAGGTACTAAGAAGAAA
AGCACACGTCTGAACTCTTTCCCTACACGACGCTCTTCCGATCTNNNTACACTCAAACCACTCGGCAAA

AGCACACGTCTGAACTCTTTCCCTACACGACGCTCTTCCGATCTNNNAAAACACTAGTAACCCCACTGC
GCACACGTCTGAACTCTTTCCCTACACGACGCTCTTCCGATCTNNNTTCCATGGTAAACACTGTCAGCCC
GCACACGTCTGAACTCTTTCCCTACACGACGCTCTTCCGATCTNNNACAGATGGATGTAGGGGCTCAA
CACACGTCTGAACTCTTTCCCTACACGACGCTCTTCCGATCTNNNCCACACTTGGTTATGTCCTTCGTCA
GAGCACACGTCTGAACTCTTTCCCTACACGACGCTCTTCCGATCTNNNTAAGGGGAGGAGAGAGAGA
GCACACGTCTGAACTCTTTCCCTACACGACGCTCTTCCGATCTNNNCAAACTAGCGTCTTCTCTGGGTC
AGCACACGTCTGAACTCTTTCCCTACACGACGCTCTTCCGATCTNNNCAGGGACTTTGGAACACTTTG
GAGCACACGTCTGAACTCTTTCCCTACACGACGCTCTTCCGATCTNNNGGCTGTCTGCTGTCCATATTTC
GCACACGTCTGAACTCTTTCCCTACACGACGCTCTTCCGATCTNNNGCATGCCTACTAAATGCCAGCTA
GAGCACACGTCTGAACTCTTTCCCTACACGACGCTCTTCCGATCTNNNAGACTGAGATTTGCCTGTGTC
AGCACACGTCTGAACTCTTTCCCTACACGACGCTCTTCCGATCTNNNCTGAGAGAGGGAGGGATTGAG
CACACGTCTGAACTCTTTCCCTACACGACGCTCTTCCGATCTNNNTTAGGATCTGGCTCACTTGGGTTC
AGCACACGTCTGAACTCTTTCCCTACACGACGCTCTTCCGATCTNNNTCAAAGTACACGGTAGGTAGG
AGAGCACACGTCTGAACTCTTTCCCTACACGACGCTCTTCCGATCTNNNTGAACAAGCCCAAAATACTG
AGAGCACACGTCTGAACTCTTTCCCTACACGACGCTCTTCCGATCTNNNTAAGCTTACAATGCCAGGG
AGAGCACACGTCTGAACTCTTTCCCTACACGACGCTCTTCCGATCTNNNTTAGCTCCTCAAATGCACCA
GAGCACACGTCTGAACTCTTTCCCTACACGACGCTCTTCCGATCTNNNAATGTGACACAGTTCCAGTG
GCACACGTCTGAACTCTTTCCCTACACGACGCTCTTCCGATCTNNNATACTGGTGGCCAAAGGTTTCTC
GAGCACACGTCTGAACTCTTTCCCTACACGACGCTCTTCCGATCTNNNTGCTTTCAGCTATCTCAGTCTG
CACACGTCTGAACTCTTTCCCTACACGACGCTCTTCCGATCTNNNTTACAGTGCTTCAGGCCTCAGATC
AGCACACGTCTGAACTCTTTCCCTACACGACGCTCTTCCGATCTNNNTGTACTCTGTTACCAGCTTTTCA
AGCACACGTCTGAACTCTTTCCCTACACGACGCTCTTCCGATCTNNNGCTTTGCTTGGTGCTTCATTAGC
CACACGTCTGAACTCTTTCCCTACACGACGCTCTTCCGATCTNNNTAGCGGAGAGCTAAGGTACTCAC
GCACACGTCTGAACTCTTTCCCTACACGACGCTCTTCCGATCTNNNTCCAAGATGGATTCGTTAGCAG
GAGCACACGTCTGAACTCTTTCCCTACACGACGCTCTTCCGATCTNNNGACTTTCCACTTGTTCTCCTCC
AGCACACGTCTGAACTCTTTCCCTACACGACGCTCTTCCGATCTNNNGTAGAGAGTGGTGGAAATGG
GAGCACACGTCTGAACTCTTTCCCTACACGACGCTCTTCCGATCTNNNCACTTGTCTACAGCATCAGCA
GCACACGTCTGAACTCTTTCCCTACACGACGCTCTTCCGATCTNNNAGCTGTTAGCTCAGTGATCCTCT
GCACACGTCTGAACTCTTTCCCTACACGACGCTCTTCCGATCTNNNGCTGGGGCAAATGGCTACTAAA
AGCACACGTCTGAACTCTTTCCCTACACGACGCTCTTCCGATCTNNNATGTGTGAGAGACAGCGAGTA
GAGCACACGTCTGAACTCTTTCCCTACACGACGCTCTTCCGATCTNNNATCTAACCCCCTCTTCCTGTCAT
GAGCACACGTCTGAACTCTTTCCCTACACGACGCTCTTCCGATCTNNNTCCATCTACTCTCTTGCTCCCAA
GAGCACACGTCTGAACTCTTTCCCTACACGACGCTCTTCCGATCTNNNGAACCCTCCAGGAGACCATAA
GAGCACACGTCTGAACTCTTTCCCTACACGACGCTCTTCCGATCTNNNTTTGCTGTTGCTGAAAATGG
GCACACGTCTGAACTCTTTCCCTACACGACGCTCTTCCGATCTNNNCCTTTGCCCTCAGGACTATCTCTC
GAGCACACGTCTGAACTCTTTCCCTACACGACGCTCTTCCGATCTNNNTGGTAGCAGCATGAGTATCTC
GAGCACACGTCTGAACTCTTTCCCTACACGACGCTCTTCCGATCTNNNCGGCCTCTTCTTGCACATTTC
GAGCACACGTCTGAACTCTTTCCCTACACGACGCTCTTCCGATCTNNNTGGGTGTCCATGGTAGGGAA
GAGCACACGTCTGAACTCTTTCCCTACACGACGCTCTTCCGATCTNNNCTTCAAGGCAGAAACCAGCT
AAGAGCACACGTCTGAACTCTTTCCCTACACGACGCTCTTCCGATCTNNNTTGGTATGGCTTCTTGCTC
GAGCACACGTCTGAACTCTTTCCCTACACGACGCTCTTCCGATCTNNNTTGGTCCCTGGAAGTGAACA
GAGCACACGTCTGAACTCTTTCCCTACACGACGCTCTTCCGATCTNNNACATTCTCACACAGCTTCTGTC
GCACACGTCTGAACTCTTTCCCTACACGACGCTCTTCCGATCTNNNAGGCAGATGGAACAGAAAGGT

AGCACACGTCTGAACTCTTTCCCTACACGACGCTCTTCCGATCTNNNAACAGGGAAAAGGAGAGATG

GCACACGTCTGAACTCTTTCCCTACACGACGCTCTTCCGATCTNNNGATGAGTTCTGGGAACACTTGG

AGCACACGTCTGAACTCTTTCCCTACACGACGCTCTTCCGATCTNNNCTCTTCCTCTAGTTTTCCCCGCC

GAGCACACGTCTGAACTCTTTCCCTACACGACGCTCTTCCGATCTNNNTTTGCTGTTGCTGAAAATGGT

AGCACACGTCTGAACTCTTTCCCTACACGACGCTCTTCCGATCTNNNATAACCACATCACACGCCATAC

AGAGCACACGTCTGAACTCTTTCCCTACACGACGCTCTTCCGATCTNNNGCAGGCCCTTTCTTAGTACA

AGCACACGTCTGAACTCTTTCCCTACACGACGCTCTTCCGATCTNNNAAAGGGAGATAGTTGCATCCA

GAGCACACGTCTGAACTCTTTCCCTACACGACGCTCTTCCGATCTNNNCTATTTGAAGGCATGGCACCT

AGAGCACACGTCTGAACTCTTTCCCTACACGACGCTCTTCCGATCTNNNGGGAACTCAGACAAATGGG

GAGCACACGTCTGAACTCTTTCCCTACACGACGCTCTTCCGATCTNNNATCTAACCCCCTCTTCCTGTCA

CACACGTCTGAACTCTTTCCCTACACGACGCTCTTCCGATCTNNNACATTCTCACACAGCTTCTGTCCCA

AGCACACGTCTGAACTCTTTCCCTACACGACGCTCTTCCGATCTNNNGCACCCCCTTTACTCTCTATCCT

GAGCACACGTCTGAACTCTTTCCCTACACGACGCTCTTCCGATCTNNNGCCAATGCCTGTTTGGATTAC

AGCACACGTCTGAACTCTTTCCCTACACGACGCTCTTCCGATCTNNNAAGGGAAGGAGGAATCTAAG

GAGCACACGTCTGAACTCTTTCCCTACACGACGCTCTTCCGATCTNNNCCTCCTCAACTTAGACAGTGG

AGAGCACACGTCTGAACTCTTTCCCTACACGACGCTCTTCCGATCTNNNACCCCACATCTACAAAGAA

GAGCACACGTCTGAACTCTTTCCCTACACGACGCTCTTCCGATCTNNNCCCGTCACTCTACATATTCAG

GAGCACACGTCTGAACTCTTTCCCTACACGACGCTCTTCCGATCTNNNCTTCCTGAACACACACAGGAT

GAGCACACGTCTGAACTCTTTCCCTACACGACGCTCTTCCGATCTNNNACAGGTGCATTAGGATCTCT

AGCACACGTCTGAACTCTTTCCCTACACGACGCTCTTCCGATCTNNNACATGCAACTCACACACACAGA

AGAGCACACGTCTGAACTCTTTCCCTACACGACGCTCTTCCGATCTNNNTCCCGCCCCCAAATCAAATAT

AGCACACGTCTGAACTCTTTCCCTACACGACGCTCTTCCGATCTNNNGTACACTGTGATCCCACAGTCA

AGCACACGTCTGAACTCTTTCCCTACACGACGCTCTTCCGATCTNNNTAATCCCTGAGCTGCATCCAAA

AGCACACGTCTGAACTCTTTCCCTACACGACGCTCTTCCGATCTNNNTGGGTCATCACCAACATCATA

AGCACACGTCTGAACTCTTTCCCTACACGACGCTCTTCCGATCTNNNTATTTCTGGTGACAGCCTTTTG

AGCACACGTCTGAACTCTTTCCCTACACGACGCTCTTCCGATCTNNNCTTGCCTTCATTTGCATAGGG

GAGCACACGTCTGAACTCTTTCCCTACACGACGCTCTTCCGATCTNNNTGCAATTCAAACTCCTACAAA

AGAGCACACGTCTGAACTCTTTCCCTACACGACGCTCTTCCGATCTNNNCATTGTAATACCCTTCTCCC

AGCACACGTCTGAACTCTTTCCCTACACGACGCTCTTCCGATCTNNNACACACACAGATAAGTGGAAG

AGCACACGTCTGAACTCTTTCCCTACACGACGCTCTTCCGATCTNNNACTGTTCCAGGGCATAGTCTCA

GAGCACACGTCTGAACTCTTTCCCTACACGACGCTCTTCCGATCTNNNATTTTGTCTGCCATACATGCT

GCACACGTCTGAACTCTTTCCCTACACGACGCTCTTCCGATCTNNNCCACCTTCCTGTCAACTTGTCTCC

GAGCACACGTCTGAACTCTTTCCCTACACGACGCTCTTCCGATCTNNNCCACACATGTGATAAACCTG

GAGCACACGTCTGAACTCTTTCCCTACACGACGCTCTTCCGATCTNNNAGACTTTCTCACAGGGTAGA

AGCACACGTCTGAACTCTTTCCCTACACGACGCTCTTCCGATCTNNNGACGTTATTTGGAGGTAACG

AGAGCACACGTCTGAACTCTTTCCCTACACGACGCTCTTCCGATCTNNNTGACTTCCCAACAATCATGT

AGAGCACACGTCTGAACTCTTTCCCTACACGACGCTCTTCCGATCTNNNAGCCAAGCATGGTTTCTCCT

GAGCACACGTCTGAACTCTTTCCCTACACGACGCTCTTCCGATCTNNNATGTCTTGGAGAAAGCAAG

GAGCACACGTCTGAACTCTTTCCCTACACGACGCTCTTCCGATCTNNNCTGTGACACTTGGCATGGTTT

GAGCACACGTCTGAACTCTTTCCCTACACGACGCTCTTCCGATCTNNNAGATTGCTTCTGACAGCACA

AGCACACGTCTGAACTCTTTCCCTACACGACGCTCTTCCGATCTNNNTCACGCGTGGTCCTAAATGTCC

GAGCACACGTCTGAACTCTTTCCCTACACGACGCTCTTCCGATCTNNNAGCAGTGTTTCGTTTCTGGT

GAGCACACGTCTGAACTCTTTCCCTACACGACGCTCTTCCGATCTNNNCCTCCAGGTACTGAGAGATG

GAGCACACGTCTGAACTCTTTCCCTACACGACGCTCTTCCGATCTNNNCAAAGCAGCATGAGATCTCC
AAGAGCACACGTCTGAACTCTTTCCCTACACGACGCTCTTCCGATCTNNNGTTACTTGAGCTTTGGAG
GAGCACACGTCTGAACTCTTTCCCTACACGACGCTCTTCCGATCTNNNGGTGACCTGGGGATTCATTT
GAGCACACGTCTGAACTCTTTCCCTACACGACGCTCTTCCGATCTNNNGCAGGACAACTTTGTGCTTT
AAGAGCACACGTCTGAACTCTTTCCCTACACGACGCTCTTCCGATCTNNNCTTTCTTGTCCTCTCCTCC
AGCACACGTCTGAACTCTTTCCCTACACGACGCTCTTCCGATCTNNNGACTACTGGCTTTGTATGTG
GCACACGTCTGAACTCTTTCCCTACACGACGCTCTTCCGATCTNNNAATGCCTGCTTGTGTCATTTTCC
AGCACACGTCTGAACTCTTTCCCTACACGACGCTCTTCCGATCTNNNGATCGTGAAAGCCCCAAAAAT
AGCACACGTCTGAACTCTTTCCCTACACGACGCTCTTCCGATCTNNNTGGATGGATAAACTCATACTA
AGCACACGTCTGAACTCTTTCCCTACACGACGCTCTTCCGATCTNNNCCCAGATAACCCCTATTGCCA
CACACGTCTGAACTCTTTCCCTACACGACGCTCTTCCGATCTNNNATTTGTCTGCCATACATGCTGC
GCACACGTCTGAACTCTTTCCCTACACGACGCTCTTCCGATCTNNNATCCTGCGCTCACATTTCTTCTT
GAGCACACGTCTGAACTCTTTCCCTACACGACGCTCTTCCGATCTNNNAGTGAAAAACAGGTGAGA
GCACACGTCTGAACTCTTTCCCTACACGACGCTCTTCCGATCTNNNCACCAGGCATCAGCAATAAGCT
AGCACACGTCTGAACTCTTTCCCTACACGACGCTCTTCCGATCTNNNGAGGCTTCAAAGAATTCAGAG
GAGCACACGTCTGAACTCTTTCCCTACACGACGCTCTTCCGATCTNNNGGCCTCCAAATACTTGGCTTT
GAGCACACGTCTGAACTCTTTCCCTACACGACGCTCTTCCGATCTNNNAAACTGAGCACAGCTGATGA
AGCACACGTCTGAACTCTTTCCCTACACGACGCTCTTCCGATCTNNNTTGTTTAGCCAGTCACAAGAG
AGCACACGTCTGAACTCTTTCCCTACACGACGCTCTTCCGATCTNNNCACCAGAACACACTCACACATG
GCACACGTCTGAACTCTTTCCCTACACGACGCTCTTCCGATCTNNNAGCTCTAAGAGGTGGACATGA
AGCACACGTCTGAACTCTTTCCCTACACGACGCTCTTCCGATCTNNNATACTGTGTGTGGATGTAGGG
GCACACGTCTGAACTCTTTCCCTACACGACGCTCTTCCGATCTNNNTCAATTTTGAAAGCCCTGGTGGT
AGCACACGTCTGAACTCTTTCCCTACACGACGCTCTTCCGATCTNNNGCACACAGCACTCTCTGTTTCT
CACACGTCTGAACTCTTTCCCTACACGACGCTCTTCCGATCTNNNCAAGGGATTTCTCCCCTTGTAGC
AGCACACGTCTGAACTCTTTCCCTACACGACGCTCTTCCGATCTNNNACCTTCTCCAGAGAGACAGAA
GAGCACACGTCTGAACTCTTTCCCTACACGACGCTCTTCCGATCTNNNGTGGTCGTGTCCAGTTATTC
AGCACACGTCTGAACTCTTTCCCTACACGACGCTCTTCCGATCTNNNCATGAGGTGGATGAAAGGAC
GCACACGTCTGAACTCTTTCCCTACACGACGCTCTTCCGATCTNNNTCAGTCCAGGGATTGAGAGAAC
GAGCACACGTCTGAACTCTTTCCCTACACGACGCTCTTCCGATCTNNNAGACCAAGTGTCTTGTGTTCC
AGCACACGTCTGAACTCTTTCCCTACACGACGCTCTTCCGATCTNNNAATCACTGAATTCATGCTGGG
GAGCACACGTCTGAACTCTTTCCCTACACGACGCTCTTCCGATCTNNNGACTGCTTACACCTTAGCTGC
GCACACGTCTGAACTCTTTCCCTACACGACGCTCTTCCGATCTNNNAAGAGTGGGGACAAAACACTGT
AGCACACGTCTGAACTCTTTCCCTACACGACGCTCTTCCGATCTNNNTCTCTCTGTGTTGTCTTATGCCT
AGCACACGTCTGAACTCTTTCCCTACACGACGCTCTTCCGATCTNNNACACAACTTCTTCCACCACCTT
AGCACACGTCTGAACTCTTTCCCTACACGACGCTCTTCCGATCTNNNATCCCTACTGCCTTCTGGACTT
GAGCACACGTCTGAACTCTTTCCCTACACGACGCTCTTCCGATCTNNNGTCTCCACAGTCATTCTTGGC
AAGAGCACACGTCTGAACTCTTTCCCTACACGACGCTCTTCCGATCTNNNACCTGTCAACAAACATGTG
AGCACACGTCTGAACTCTTTCCCTACACGACGCTCTTCCGATCTNNNAAACTTTCTCTCTCACCCCGGTA
GAGCACACGTCTGAACTCTTTCCCTACACGACGCTCTTCCGATCTNNNAATATACACTGAGGGCCATG
AGCACACGTCTGAACTCTTTCCCTACACGACGCTCTTCCGATCTNNNTCAAACCCGGGTGTATTTCAAG
AGCACACGTCTGAACTCTTTCCCTACACGACGCTCTTCCGATCTNNNTGTGGAAACAAAACCACCCTT
GCACACGTCTGAACTCTTTCCCTACACGACGCTCTTCCGATCTNNNTATTTCTGGTGACAGCCTTTTGC
GCACACGTCTGAACTCTTTCCCTACACGACGCTCTTCCGATCTNNNTGCATGTTCACCTCCTTTACACA

AGCACACGTCTGAACTCTTTCCCTACACGACGCTCTTCCGATCTNNNAATGCACATAAACTGCATGGC
AGCACACGTCTGAACTCTTTCCCTACACGACGCTCTTCCGATCTNNNAGTGGAAGAAAAGAAAACCTC
GAGCACACGTCTGAACTCTTTCCCTACACGACGCTCTTCCGATCTNNNCCAGAATGATGCCAGGATT
GAGCACACGTCTGAACTCTTTCCCTACACGACGCTCTTCCGATCTNNNCCCTACAGCGTTTCCAGAAAG
AGCACACGTCTGAACTCTTTCCCTACACGACGCTCTTCCGATCTNNNCACCCACCGTACTCCATATACCT
AGCACACGTCTGAACTCTTTCCCTACACGACGCTCTTCCGATCTNNNTGACAGAGCTGTATCAGTTG
AGCACACGTCTGAACTCTTTCCCTACACGACGCTCTTCCGATCTNNNTTCTGAAGCTCTTGGCTGAAGT
AGCACACGTCTGAACTCTTTCCCTACACGACGCTCTTCCGATCTNNNATTCTGCCAAAGAGAAAGTGG
GAGCACACGTCTGAACTCTTTCCCTACACGACGCTCTTCCGATCTNNNTATGGCCAGGTGGTTCAATT
AAGAGCACACGTCTGAACTCTTTCCCTACACGACGCTCTTCCGATCTNNNAAACCGGAAAGTGTGTT
AGCACACGTCTGAACTCTTTCCCTACACGACGCTCTTCCGATCTNNNTGAATGTTTTTCAGGACCTGTG
GAGCACACGTCTGAACTCTTTCCCTACACGACGCTCTTCCGATCTNNNAGGCACCATCAAAAATGACC
GCACACGTCTGAACTCTTTCCCTACACGACGCTCTTCCGATCTNNNCGTGGGGACTATGGTGACTAC
GCACACGTCTGAACTCTTTCCCTACACGACGCTCTTCCGATCTNNNGGTGGGGTAGTGGCTTATCCT
GAGCACACGTCTGAACTCTTTCCCTACACGACGCTCTTCCGATCTNNNTTGCATTCCACATGCTCCTC
CACACGTCTGAACTCTTTCCCTACACGACGCTCTTCCGATCTNNNCTGCTCCCAACTTCTTGAAACCTCA
GAGCACACGTCTGAACTCTTTCCCTACACGACGCTCTTCCGATCTNNNAGCTTCCTCAGATTGAGGCT
AGCACACGTCTGAACTCTTTCCCTACACGACGCTCTTCCGATCTNNNTGGTCTCTCTCTCTCTTCTCTGTT
CACACGTCTGAACTCTTTCCCTACACGACGCTCTTCCGATCTNNNGATGGATCCAAGGGCTCAAACAA
AGCACACGTCTGAACTCTTTCCCTACACGACGCTCTTCCGATCTNNNCTGAGGCACATTCAGATCTTCC
AGCACACGTCTGAACTCTTTCCCTACACGACGCTCTTCCGATCTNNNCAACTTCTCCACTCAACCCTCAG
GCACACGTCTGAACTCTTTCCCTACACGACGCTCTTCCGATCTNNNATTTCTAGCCCAGTCAACCTGGA
AGCACACGTCTGAACTCTTTCCCTACACGACGCTCTTCCGATCTNNNTGGAGTTCCAGGTAATAGTTT
GAGCACACGTCTGAACTCTTTCCCTACACGACGCTCTTCCGATCTNNNCCCACCCCCAATCAGAAAGAA
AGCACACGTCTGAACTCTTTCCCTACACGACGCTCTTCCGATCTNNNAGAAGTGGCTTCCAAACTGTGT
AGAGCACACGTCTGAACTCTTTCCCTACACGACGCTCTTCCGATCTNNNTACCTCATCAAGTTTGGCAC
AGCACACGTCTGAACTCTTTCCCTACACGACGCTCTTCCGATCTNNNTCTTCTGAAAGATTTTTACAAA
AGCACACGTCTGAACTCTTTCCCTACACGACGCTCTTCCGATCTNNNTTGAGACAGCCTTTAGCACAG
GCACACGTCTGAACTCTTTCCCTACACGACGCTCTTCCGATCTNNNAGCTCTTGCTGGAATGGATAG
AGCACACGTCTGAACTCTTTCCCTACACGACGCTCTTCCGATCTNNNCGTCTGCTTCCAATTATCCTGG
GCACACGTCTGAACTCTTTCCCTACACGACGCTCTTCCGATCTNNNCAGGATAGGCTGTATCCCTGAC
AGAGCACACGTCTGAACTCTTTCCCTACACGACGCTCTTCCGATCTNNNGTGATCAAACCAAAGCAGC
AAGAGCACACGTCTGAACTCTTTCCCTACACGACGCTCTTCCGATCTNNNTTTGTACCAGGCATTGGGC
AGCACACGTCTGAACTCTTTCCCTACACGACGCTCTTCCGATCTNNNTAGGTGTTTGGTGGGAAAAC
AGCACACGTCTGAACTCTTTCCCTACACGACGCTCTTCCGATCTNNNCCGAGGTCTACAAGGCTGGAT
AAGAGCACACGTCTGAACTCTTTCCCTACACGACGCTCTTCCGATCTNNNGAACCTGCCTAACAGTGA
AGCACACGTCTGAACTCTTTCCCTACACGACGCTCTTCCGATCTNNNGCATGGGGTTTGCTTTCCTAA
CACACGTCTGAACTCTTTCCCTACACGACGCTCTTCCGATCTNNNCCATTCTTTATTTGGGGAGCGGA
AGCACACGTCTGAACTCTTTCCCTACACGACGCTCTTCCGATCTNNNTGTCACACCACTATAGACAGAC
AGCACACGTCTGAACTCTTTCCCTACACGACGCTCTTCCGATCTNNNCGCTATTGGCTCTTGACCTGAA
AAGAGCACACGTCTGAACTCTTTCCCTACACGACGCTCTTCCGATCTNNNCTGAGTTTGCTTGGGGTT
AGCACACGTCTGAACTCTTTCCCTACACGACGCTCTTCCGATCTNNNGCAGCAATCTGTCTTTTCTTGT
GCACACGTCTGAACTCTTTCCCTACACGACGCTCTTCCGATCTNNNTGCTCGGACCCTGATTACTATGT

GCACACGTCTGAACTCTTTCCCTACACGACGCTCTTCCGATCTNNNGAAATCTGGCATAGGAGCCTGA
AGCACACGTCTGAACTCTTTCCCTACACGACGCTCTTCCGATCTNNNCCTGAACAACTGAATAAAACC
AGAGCACACGTCTGAACTCTTTCCCTACACGACGCTCTTCCGATCTNNNGTCAACATGCAACCAAAAA
AGCACACGTCTGAACTCTTTCCCTACACGACGCTCTTCCGATCTNNNCAACCTCTTGTCCAGTGTTGG
AGAGCACACGTCTGAACTCTTTCCCTACACGACGCTCTTCCGATCTNNNACCCTTGCGATGAAAGGTG
GAAGAGCACACGTCTGAACTCTTTCCCTACACGACGCTCTTCCGATCTNNNGAAGGGCTGAGAAACCA
AGAGCACACGTCTGAACTCTTTCCCTACACGACGCTCTTCCGATCTNNNTAGGTGTTTGGTGGGAAA
AAGAGCACACGTCTGAACTCTTTCCCTACACGACGCTCTTCCGATCTNNNACCCTAGACAGACACCTTC
AGAGCACACGTCTGAACTCTTTCCCTACACGACGCTCTTCCGATCTNNNTCTTCTGGCCTAATGTACTG
GCACACGTCTGAACTCTTTCCCTACACGACGCTCTTCCGATCTNNNTCCCAGCCTAACCTAGTTCAACA
GCACACGTCTGAACTCTTTCCCTACACGACGCTCTTCCGATCTNNNATTGTGTGCATGTATTTGGTGTG
GAAGAGCACACGTCTGAACTCTTTCCCTACACGACGCTCTTCCGATCTNNNAATTGAGCTCCTCGGAC
AGAGCACACGTCTGAACTCTTTCCCTACACGACGCTCTTCCGATCTNNNACTTCCCTGATGCACACTTT
AGCACACGTCTGAACTCTTTCCCTACACGACGCTCTTCCGATCTNNNGTGTCTTGACAAGTGAAGTGT
AGCACACGTCTGAACTCTTTCCCTACACGACGCTCTTCCGATCTNNNGTGGTGTGATCTTAGGTCATTG
AGAGCACACGTCTGAACTCTTTCCCTACACGACGCTCTTCCGATCTNNNCACTGCTGGTAGCCATCTAC
AAGAGCACACGTCTGAACTCTTTCCCTACACGACGCTCTTCCGATCTNNNCCCTCAAGCGATTCACCCAC
AGCACACGTCTGAACTCTTTCCCTACACGACGCTCTTCCGATCTNNNCTCGGAGGGTCTCAGTTTCAAA
ACACGTCTGAACTCTTTCCCTACACGACGCTCTTCCGATCTNNNGTACATGCAGAGTGAGTAGCTGGC
AGAGCACACGTCTGAACTCTTTCCCTACACGACGCTCTTCCGATCTNNNTTACTGACAGTGGAGTTTGC
AGCACACGTCTGAACTCTTTCCCTACACGACGCTCTTCCGATCTNNNGAAACGAGAGGATCGCTTGAC
GCACACGTCTGAACTCTTTCCCTACACGACGCTCTTCCGATCTNNNTTTCTGAGGCGGGTTCTCACTTT
GAAGAGCACACGTCTGAACTCTTTCCCTACACGACGCTCTTCCGATCTNNNGTGCTCAGCAAAAGTCA
AGCACACGTCTGAACTCTTTCCCTACACGACGCTCTTCCGATCTNNNATGCCCAAAATCTCTGCTTTGGA
GCACACGTCTGAACTCTTTCCCTACACGACGCTCTTCCGATCTNNNTGCAATTGTCAGATGAATGAGA
GGAAGAGCACACGTCTGAACTCTTTCCCTACACGACGCTCTTCCGATCTNNNTGAGTGAGATTTTTCAC
GAGCACACGTCTGAACTCTTTCCCTACACGACGCTCTTCCGATCTNNNCCCTCAAGCGATTCACCCACTT
AGAGCACACGTCTGAACTCTTTCCCTACACGACGCTCTTCCGATCTNNNAGAGGGGCTATAATCCCAG
GAGCACACGTCTGAACTCTTTCCCTACACGACGCTCTTCCGATCTNNNCACATTCTTAGGGGCTTCCAC
GCACACGTCTGAACTCTTTCCCTACACGACGCTCTTCCGATCTNNNCTCTGGCTTCTGTCACTGCAACCA
AGAGCACACGTCTGAACTCTTTCCCTACACGACGCTCTTCCGATCTNNNTCAATGGTTCCTGTTGGCTC
AGCACACGTCTGAACTCTTTCCCTACACGACGCTCTTCCGATCTNNNCAAGGAGTTCTTGGCTTTCCTC
GCACACGTCTGAACTCTTTCCCTACACGACGCTCTTCCGATCTNNNTTTTGCCTTGTTTTCAAGCCTCAC
AGAGCACACGTCTGAACTCTTTCCCTACACGACGCTCTTCCGATCTNNNTCATTTCACTACCGTATGCCG
AGCACACGTCTGAACTCTTTCCCTACACGACGCTCTTCCGATCTNNNCTCCCCACTCCATAAAGCACATC
GCACACGTCTGAACTCTTTCCCTACACGACGCTCTTCCGATCTNNNAGTAAGTACTCAAACGCCTGCTG
GAGCACACGTCTGAACTCTTTCCCTACACGACGCTCTTCCGATCTNNNTAGGAGCCAGACCTGTCTCAA
AGCACACGTCTGAACTCTTTCCCTACACGACGCTCTTCCGATCTNNNTCAATTCTTGCATTGAATCTAT
AAGAGCACACGTCTGAACTCTTTCCCTACACGACGCTCTTCCGATCTNNNTAAGGTAATTGTGTGTGG
GCACACGTCTGAACTCTTTCCCTACACGACGCTCTTCCGATCTNNNGCAACCCTCTCTCTTTCTACAGGC
AGAGCACACGTCTGAACTCTTTCCCTACACGACGCTCTTCCGATCTNNNAGTGCCTAGTGCAGGATGT
GAGCACACGTCTGAACTCTTTCCCTACACGACGCTCTTCCGATCTNNNAGAAGATTTTAGGTGCTGTT
AGCACACGTCTGAACTCTTTCCCTACACGACGCTCTTCCGATCTNNNCCTCTCAGTGTATGCTCACAAG

AGCACACGTCTGAACTCTTTCCCTACACGACGCTCTTCCGATCTNNNTCTCAACTTATTCTTTCTCCTCGC
GAGCACACGTCTGAACTCTTTCCCTACACGACGCTCTTCCGATCTNNNAGATTCGATGTGTACTCTTGC
GAGCACACGTCTGAACTCTTTCCCTACACGACGCTCTTCCGATCTNNNAGAGCTCAATAAACCCAAGGCA
AGAGCACACGTCTGAACTCTTTCCCTACACGACGCTCTTCCGATCTNNNACCAGCTGAGATCCGTCTAT
GCACACGTCTGAACTCTTTCCCTACACGACGCTCTTCCGATCTNNNAGTAGTGCTGATTTCCAAATGCCA
AGCACACGTCTGAACTCTTTCCCTACACGACGCTCTTCCGATCTNNNTCTCTAAGTCAGCAGACCCCAAA
GAGCACACGTCTGAACTCTTTCCCTACACGACGCTCTTCCGATCTNNNAGTGTTGGCTTATTGCTTTGCA
ACACGTCTGAACTCTTTCCCTACACGACGCTCTTCCGATCTNNNCGCACAAGAACAAATGACAGTCCCAT
GAGCACACGTCTGAACTCTTTCCCTACACGACGCTCTTCCGATCTNNNACCCCAGAACCTAAAGGACTG
AGAGCACACGTCTGAACTCTTTCCCTACACGACGCTCTTCCGATCTNNNTGCCCTCTTTCCTGTACTCTG
GAGCACACGTCTGAACTCTTTCCCTACACGACGCTCTTCCGATCTNNNACTCCTTCCTTACCAAAGTGGC
GCACACGTCTGAACTCTTTCCCTACACGACGCTCTTCCGATCTNNNTACCTATTAGCCAGTGTCTCCCCC
GAGCACACGTCTGAACTCTTTCCCTACACGACGCTCTTCCGATCTNNNCAGCTGTAGGAAAAGCCAGTC
GAGCACACGTCTGAACTCTTTCCCTACACGACGCTCTTCCGATCTNNNACTGCATCACAAACCTATGACA
AGAGCACACGTCTGAACTCTTTCCCTACACGACGCTCTTCCGATCTNNNTGGTTGTAATGTGTTTCCGG
AGCACACGTCTGAACTCTTTCCCTACACGACGCTCTTCCGATCTNNNACCACCTAATAGTCAAAGGCAG
GAGCACACGTCTGAACTCTTTCCCTACACGACGCTCTTCCGATCTNNNAGAGGTGTGTGTGTGTCTGTT
GAGCACACGTCTGAACTCTTTCCCTACACGACGCTCTTCCGATCTNNNTTTTCAAATATTCCCCTCGCCCC
AGCACACGTCTGAACTCTTTCCCTACACGACGCTCTTCCGATCTNNNTGTTCTGGAGTTCCTCGACTTCT
AGAGCACACGTCTGAACTCTTTCCCTACACGACGCTCTTCCGATCTNNNTCCTGAAGACAAAGCAGTGC
GAGCACACGTCTGAACTCTTTCCCTACACGACGCTCTTCCGATCTNNNCTGCATCATCCCCAGCACATAAA
GAGCACACGTCTGAACTCTTTCCCTACACGACGCTCTTCCGATCTNNNAGCAAGATTCCAAACAGTAAA
AGCACACGTCTGAACTCTTTCCCTACACGACGCTCTTCCGATCTNNNCAAAGTGTGGCCACTCCTCAATA
AAGAGCACACGTCTGAACTCTTTCCCTACACGACGCTCTTCCGATCTNNNTCAGTATGCAGTCAGTGTG
AGAGCACACGTCTGAACTCTTTCCCTACACGACGCTCTTCCGATCTNNNTTCTGGTCCAAAATGCAACCA
AGAGCACACGTCTGAACTCTTTCCCTACACGACGCTCTTCCGATCTNNNAGATTCTTACTTGAAATCCCC
GAGCACACGTCTGAACTCTTTCCCTACACGACGCTCTTCCGATCTNNNAAAGGTCAGCTGAGTGATTTG
GCACACGTCTGAACTCTTTCCCTACACGACGCTCTTCCGATCTNNNGATTTTTGGCAGATGCCTTCCACC
AGCACACGTCTGAACTCTTTCCCTACACGACGCTCTTCCGATCTNNNTGGCCTGCAGATTTTGAGACTTT
AGCACACGTCTGAACTCTTTCCCTACACGACGCTCTTCCGATCTNNNGCACAGAACCAATCAGAGGGTA
GAGCACACGTCTGAACTCTTTCCCTACACGACGCTCTTCCGATCTNNNATGGGCTCCCAACAGATTGAA
AGCACACGTCTGAACTCTTTCCCTACACGACGCTCTTCCGATCTNNNTCCCATAGGAATGAAACCTCAAA
GAGCACACGTCTGAACTCTTTCCCTACACGACGCTCTTCCGATCTNNNAGCCAGTTATCTTGGGTCTTGC
AGCACACGTCTGAACTCTTTCCCTACACGACGCTCTTCCGATCTNNNATGTGTTTGCATGCCAGGTCTA
GCACACGTCTGAACTCTTTCCCTACACGACGCTCTTCCGATCTNNNCATCTGACTGTGTAGCACTTCCCC
AGCACACGTCTGAACTCTTTCCCTACACGACGCTCTTCCGATCTNNNTACATACATATACGGGGACGGG
GAGCACACGTCTGAACTCTTTCCCTACACGACGCTCTTCCGATCTNNNGCAGTTCTGTTCTCAACATCTG
GAGCACACGTCTGAACTCTTTCCCTACACGACGCTCTTCCGATCTNNNACAAGATCACTGGAAAGAGCC
GAGCACACGTCTGAACTCTTTCCCTACACGACGCTCTTCCGATCTNNNGGCAACTGTCATCATCAACTG
AGAGCACACGTCTGAACTCTTTCCCTACACGACGCTCTTCCGATCTNNNCTATTGTCATCCCTTGGGCAC
AGAGCACACGTCTGAACTCTTTCCCTACACGACGCTCTTCCGATCTNNNAGTCTCAAATATAGACAAGC
AGCACACGTCTGAACTCTTTCCCTACACGACGCTCTTCCGATCTNNNTCATCTTTTGGAAATGGAATAAG
AAGAGCACACGTCTGAACTCTTTCCCTACACGACGCTCTTCCGATCTNNNCCATGTGAAATTTGCTTCTG

GAGCACACGTCTGAACTCTTTCCCTACACGACGCTCTTCCGATCTNNNAGGCTCTACCACATAGACAA
GAGCACACGTCTGAACTCTTTCCCTACACGACGCTCTTCCGATCTNNNACTGCTGCTATAGCTATCCTT
GAGCACACGTCTGAACTCTTTCCCTACACGACGCTCTTCCGATCTNNNGGCATCTGAGTGTTCCATGC
AGCACACGTCTGAACTCTTTCCCTACACGACGCTCTTCCGATCTNNNTGGTATACATGGCTCTATGCG
GAGCACACGTCTGAACTCTTTCCCTACACGACGCTCTTCCGATCTNNNATGCCCTTTATTAGGAGTTC
GAGCACACGTCTGAACTCTTTCCCTACACGACGCTCTTCCGATCTNNNAAAAGTACATAACCCCTCTGC
AGAGCACACGTCTGAACTCTTTCCCTACACGACGCTCTTCCGATCTNNNTCACTCTGGAATGACTTTTT
AGCACACGTCTGAACTCTTTCCCTACACGACGCTCTTCCGATCTNNNTAATTGCTTGATGGCCTTGAG
AGCACACGTCTGAACTCTTTCCCTACACGACGCTCTTCCGATCTNNNTAGGGAAGTAGGGCGTTGAAT
AGAGCACACGTCTGAACTCTTTCCCTACACGACGCTCTTCCGATCTNNNACAGTAAACACAAACGAAC
AGCACACGTCTGAACTCTTTCCCTACACGACGCTCTTCCGATCTNNNAGGAGATGCAGGCAAGGTA
AGCACACGTCTGAACTCTTTCCCTACACGACGCTCTTCCGATCTNNNTGAGTGAAAGATCCTAGACAC
AGCACACGTCTGAACTCTTTCCCTACACGACGCTCTTCCGATCTNNNAGCAGTAATTTTCACAAATTC
GAGCACACGTCTGAACTCTTTCCCTACACGACGCTCTTCCGATCTNNNCAACAGTAAACTGTGGTAAG
GCACACGTCTGAACTCTTTCCCTACACGACGCTCTTCCGATCTNNNGCAAAATTTGTTTCAGCAGAGC
AGCACACGTCTGAACTCTTTCCCTACACGACGCTCTTCCGATCTNNNCTGGACAATAGAGCGAGTAA
GCACACGTCTGAACTCTTTCCCTACACGACGCTCTTCCGATCTNNNCAGAGCATGTCTGTCTGTCTGTC
AGCACACGTCTGAACTCTTTCCCTACACGACGCTCTTCCGATCTNNNCACACTAGGCCCAGGTCTTTTT
AGCACACGTCTGAACTCTTTCCCTACACGACGCTCTTCCGATCTNNNAGATCTGTACTCAATGGAAG
GAGCACACGTCTGAACTCTTTCCCTACACGACGCTCTTCCGATCTNNNCTGAGAAGTGAGGCAGGAG
AGAGCACACGTCTGAACTCTTTCCCTACACGACGCTCTTCCGATCTNNNTGAATTCATAGATGGCTCC
AGCACACGTCTGAACTCTTTCCCTACACGACGCTCTTCCGATCTNNNACTGCACAGCATTGATACAGA
AGAGCACACGTCTGAACTCTTTCCCTACACGACGCTCTTCCGATCTNNNGCACTCTGGCCTGGGTAAT
AGCACACGTCTGAACTCTTTCCCTACACGACGCTCTTCCGATCTNNNACCCAGCAAAATGTACAAACA
GAGCACACGTCTGAACTCTTTCCCTACACGACGCTCTTCCGATCTNNNGTTGGCTGTGCTGCTTATTTT
GAGCACACGTCTGAACTCTTTCCCTACACGACGCTCTTCCGATCTNNNGATTTTGTGTGCAGGGAG
GAGCACACGTCTGAACTCTTTCCCTACACGACGCTCTTCCGATCTNNNGAACTTGTAGCTGTCCACAC
GCACACGTCTGAACTCTTTCCCTACACGACGCTCTTCCGATCTNNNTGCATGCTCATTCACTTTAGGCA
AGCACACGTCTGAACTCTTTCCCTACACGACGCTCTTCCGATCTNNNAACTGACCATCTCAACCTACCC
GAGCACACGTCTGAACTCTTTCCCTACACGACGCTCTTCCGATCTNNNCCAGCCTGGTCATAGAGAG
AGCACACGTCTGAACTCTTTCCCTACACGACGCTCTTCCGATCTNNNTGCCGGCATTCTTCTTTCCTAT
AGAGCACACGTCTGAACTCTTTCCCTACACGACGCTCTTCCGATCTNNNCACCAGTCCCACGTTAGAAC
GAGCACACGTCTGAACTCTTTCCCTACACGACGCTCTTCCGATCTNNNTTACTGTTGTGAGCCAGTCT
AGCACACGTCTGAACTCTTTCCCTACACGACGCTCTTCCGATCTNNNTTATACTTACATACTATTCCCTG
AGCACACGTCTGAACTCTTTCCCTACACGACGCTCTTCCGATCTNNNATCAACCCAACAAAGCCTACC
AGCACACGTCTGAACTCTTTCCCTACACGACGCTCTTCCGATCTNNNCCTTCTAGCTCTTCCTTTCTTCT
GAGCACACGTCTGAACTCTTTCCCTACACGACGCTCTTCCGATCTNNNTTTCACTGGAGCAAAGAGTT
GCACACGTCTGAACTCTTTCCCTACACGACGCTCTTCCGATCTNNNAGTGTATTTATTGTTCCTGGG
AGCACACGTCTGAACTCTTTCCCTACACGACGCTCTTCCGATCTNNNTTGGTAACACTCATAGCACCTG
AGCACACGTCTGAACTCTTTCCCTACACGACGCTCTTCCGATCTNNNTGTCTTTCCAACTCCAAAGCCA
GAGCACACGTCTGAACTCTTTCCCTACACGACGCTCTTCCGATCTNNNAGGCAGAGCAGAAACGTAG
AGCACACGTCTGAACTCTTTCCCTACACGACGCTCTTCCGATCTNNNCTAACTGGGGGTTCTTTCTGC
AGCACACGTCTGAACTCTTTCCCTACACGACGCTCTTCCGATCTNNNAACGGTATGTTATATGGGTGC

AGCACACGTCTGAACTCTTTCCCTACACGACGCTCTTCCGATCTNNNTTTGGTAGGTTGGCATGTTTC
AGCACACGTCTGAACTCTTTCCCTACACGACGCTCTTCCGATCTNNNTCCTGCAACTTTACTAAACCAA
GAGCACACGTCTGAACTCTTTCCCTACACGACGCTCTTCCGATCTNNNTGGGAGAGGATCAATTTCT
GAGCACACGTCTGAACTCTTTCCCTACACGACGCTCTTCCGATCTNNNAGTCCATAAATGCAGGGAGA
GAGCACACGTCTGAACTCTTTCCCTACACGACGCTCTTCCGATCTNNNAGCAATCAGAGGGTTTTGG
AGCACACGTCTGAACTCTTTCCCTACACGACGCTCTTCCGATCTNNNTACAAACACAGAATGTCACCC
GAGCACACGTCTGAACTCTTTCCCTACACGACGCTCTTCCGATCTNNNTGTTGAATGAGGCAAATCA
GAGCACACGTCTGAACTCTTTCCCTACACGACGCTCTTCCGATCTNNNAGTTGATGGAGTTTGGCAT
AGCACACGTCTGAACTCTTTCCCTACACGACGCTCTTCCGATCTNNNTTACCAGGTACGGTGATAAG
AGCACACGTCTGAACTCTTTCCCTACACGACGCTCTTCCGATCTNNNTCACTCCTCCCTTTTCTTCGTCT
GAGCACACGTCTGAACTCTTTCCCTACACGACGCTCTTCCGATCTNNNTAGAACCTCCTATTCCACCAC
GCACACGTCTGAACTCTTTCCCTACACGACGCTCTTCCGATCTNNNAATGAGAAGGTTGCTTCTGCCT
GCACACGTCTGAACTCTTTCCCTACACGACGCTCTTCCGATCTNNNATGCATCCAGAAAAGTGCCTTG
GAGCACACGTCTGAACTCTTTCCCTACACGACGCTCTTCCGATCTNNNCAGAAAGCCTGGAAGCTGAA
GAGCACACGTCTGAACTCTTTCCCTACACGACGCTCTTCCGATCTNNNAGTGTATGCTCTCACACTAG
AGCACACGTCTGAACTCTTTCCCTACACGACGCTCTTCCGATCTNNNTGTGTTGACTGTTGAATGAGG
AGCACACGTCTGAACTCTTTCCCTACACGACGCTCTTCCGATCTNNNATCTCCACTGCTCCCATACACTT
AGCACACGTCTGAACTCTTTCCCTACACGACGCTCTTCCGATCTNNNCGCAGTGGTTACACACATCTA
AGAGCACACGTCTGAACTCTTTCCCTACACGACGCTCTTCCGATCTNNNCCCAGATGCACCATTTCTCT
GAGCACACGTCTGAACTCTTTCCCTACACGACGCTCTTCCGATCTNNNCATGAGCAGTGAATCTCACC
GAGCACACGTCTGAACTCTTTCCCTACACGACGCTCTTCCGATCTNNNGTCTCTCTTGTGCGTTCTCTCT
AGCACACGTCTGAACTCTTTCCCTACACGACGCTCTTCCGATCTNNNTCCTGTCACTTACAGAAGCTCC
GAGCACACGTCTGAACTCTTTCCCTACACGACGCTCTTCCGATCTNNNTAACCCTAAGGAGGGCTTAC
GCACACGTCTGAACTCTTTCCCTACACGACGCTCTTCCGATCTNNNACATATCCCCATAACAGCTCCCT
AGCACACGTCTGAACTCTTTCCCTACACGACGCTCTTCCGATCTNNNTGCCCTCACATCCATCCTATCTT
AGCACACGTCTGAACTCTTTCCCTACACGACGCTCTTCCGATCTNNNGCCAAGCTCTTTAGAGCCTAC
GCACACGTCTGAACTCTTTCCCTACACGACGCTCTTCCGATCTNNNAGGATGTCCTAACAGGACCTAG
GCACACGTCTGAACTCTTTCCCTACACGACGCTCTTCCGATCTNNNAAGGCCCAGTTCTCCTCTCATTA
AGCACACGTCTGAACTCTTTCCCTACACGACGCTCTTCCGATCTNNNTGCCAGGGAGTGAAAATTAG
GCACACGTCTGAACTCTTTCCCTACACGACGCTCTTCCGATCTNNNCTGACAATAGCCTCCTCTGTGCT
GCACACGTCTGAACTCTTTCCCTACACGACGCTCTTCCGATCTNNNTCTCACCCAGATGTTCTGCTGTA
AGCACACGTCTGAACTCTTTCCCTACACGACGCTCTTCCGATCTNNNGAGCTCTGCCCATGAGAGACT
GAGCACACGTCTGAACTCTTTCCCTACACGACGCTCTTCCGATCTNNNTCATTGAGAAGAAACCTTA
GAGCACACGTCTGAACTCTTTCCCTACACGACGCTCTTCCGATCTNNNTGCTGATACGCTGATGGAAT
GCACACGTCTGAACTCTTTCCCTACACGACGCTCTTCCGATCTNNNTGCAAGTCACAAGCTGAATCCTT
GAGCACACGTCTGAACTCTTTCCCTACACGACGCTCTTCCGATCTNNNTTGGGTGAAAAATAAAGGG
CACACGTCTGAACTCTTTCCCTACACGACGCTCTTCCGATCTNNNCTGTACCCTTGTTGGGTTGTCTGC
AGCACACGTCTGAACTCTTTCCCTACACGACGCTCTTCCGATCTNNNTCATCTCATAACCCTCTGACCCC
AGCACACGTCTGAACTCTTTCCCTACACGACGCTCTTCCGATCTNNNACACAGTGAGACCCCCTTTAC
GCACACGTCTGAACTCTTTCCCTACACGACGCTCTTCCGATCTNNNCACAATCTCCTGAGCCAATTCCA
AGCACACGTCTGAACTCTTTCCCTACACGACGCTCTTCCGATCTNNNCCATTCATCCTGCACGAACAGA
AGCACACGTCTGAACTCTTTCCCTACACGACGCTCTTCCGATCTNNNGTTCCAAGGAACACATCCTCC
GCACACGTCTGAACTCTTTCCCTACACGACGCTCTTCCGATCTNNNAGTGGGTGTTGCTTTTCACTCA

AGCACACGTCTGAACTCTTTCCCTACACGACGCTCTTCCGATCTNNNTTTCCTGAATGATGGCAACCCT
AGCACACGTCTGAACTCTTTCCCTACACGACGCTCTTCCGATCTNNNCACACTATGACAGCCAAGAAG
AGCACACGTCTGAACTCTTTCCCTACACGACGCTCTTCCGATCTNNNTGTCCACATCATTTCCTTGCACA
AGCACACGTCTGAACTCTTTCCCTACACGACGCTCTTCCGATCTNNNGATGATGTGGTGGGCATAAG
AGCACACGTCTGAACTCTTTCCCTACACGACGCTCTTCCGATCTNNNTTGGTCAGTAGCTACACAAAT
CACACGTCTGAACTCTTTCCCTACACGACGCTCTTCCGATCTNNNGCTGTTCCCCTTACCTAGAATGCC
GCACACGTCTGAACTCTTTCCCTACACGACGCTCTTCCGATCTNNNTGGTATGACAGAGGGAAACAG
GAAGAGCACACGTCTGAACTCTTTCCCTACACGACGCTCTTCCGATCTNNNATATGCCTTGGACTTTTT
GAGCACACGTCTGAACTCTTTCCCTACACGACGCTCTTCCGATCTNNNTCTGTTTTTGAGTTACAGCAG
GCACACGTCTGAACTCTTTCCCTACACGACGCTCTTCCGATCTNNNGCTTCCAGTTCTGTTCACATTGC
GAGCACACGTCTGAACTCTTTCCCTACACGACGCTCTTCCGATCTNNNTTACAGTTGCAGTTATCCTGC
AGCACACGTCTGAACTCTTTCCCTACACGACGCTCTTCCGATCTNNNGCTTCAGTTTTGGTAGGTTGG
GAAGAGCACACGTCTGAACTCTTTCCCTACACGACGCTCTTCCGATCTNNNTAGAGCTGAACCAAAG
AGCACACGTCTGAACTCTTTCCCTACACGACGCTCTTCCGATCTNNNTGATTGACAGTGGAGTAGCCA
AGAGCACACGTCTGAACTCTTTCCCTACACGACGCTCTTCCGATCTNNNGGAACTTTAGATCATGAAG
GAGCACACGTCTGAACTCTTTCCCTACACGACGCTCTTCCGATCTNNNGTTCGTGCAGGATGAATGG
AGCACACGTCTGAACTCTTTCCCTACACGACGCTCTTCCGATCTNNNGAGTATGCCTGGGAACCTTTG
GAGCACACGTCTGAACTCTTTCCCTACACGACGCTCTTCCGATCTNNNTCATGGGCAACACATACACA
GCACACGTCTGAACTCTTTCCCTACACGACGCTCTTCCGATCTNNNGGGCTTTGAGGAGCAATTCAA
AGCACACGTCTGAACTCTTTCCCTACACGACGCTCTTCCGATCTNNNGCTAATTCTGGGACTGTAGGC
GAGCACACGTCTGAACTCTTTCCCTACACGACGCTCTTCCGATCTNNNTGGTGTATCTATTGTCTGGC
GAGCACACGTCTGAACTCTTTCCCTACACGACGCTCTTCCGATCTNNNCACCAGTCCCACGTTAGAACA
GAGCACACGTCTGAACTCTTTCCCTACACGACGCTCTTCCGATCTNNNAACTCACTCAAAGTAACACCG
AGCACACGTCTGAACTCTTTCCCTACACGACGCTCTTCCGATCTNNNTTGAGCTAGTAGGAGAGCCA
CACACGTCTGAACTCTTTCCCTACACGACGCTCTTCCGATCTNNNCTTCCCCGCTGTCCATTCATTCATC
AGAGCACACGTCTGAACTCTTTCCCTACACGACGCTCTTCCGATCTNNNGGCCCTGAAGAGAAAAGA
GAGCACACGTCTGAACTCTTTCCCTACACGACGCTCTTCCGATCTNNNTGATCCACTGATACAGAACCT
AGCACACGTCTGAACTCTTTCCCTACACGACGCTCTTCCGATCTNNNTATCCAGCTAGGGACTTGGA
AGCACACGTCTGAACTCTTTCCCTACACGACGCTCTTCCGATCTNNNGCCAGGCCCTATTCTAAGTTCT
AGCACACGTCTGAACTCTTTCCCTACACGACGCTCTTCCGATCTNNNCTCCACACCTCACCTCCTCCAG
AGCACACGTCTGAACTCTTTCCCTACACGACGCTCTTCCGATCTNNNCCTGTTAGTTTCAGAGGCTGT
GAGCACACGTCTGAACTCTTTCCCTACACGACGCTCTTCCGATCTNNNTATCCAGCTAGGGACTTGGA
AGCACACGTCTGAACTCTTTCCCTACACGACGCTCTTCCGATCTNNNCAGTAGGTCACTGCCTATCAA
GAAGAGCACACGTCTGAACTCTTTCCCTACACGACGCTCTTCCGATCTNNNGAAGTCTTCCCTGGGTG
AGCACACGTCTGAACTCTTTCCCTACACGACGCTCTTCCGATCTNNNTGGGTTCACTGCCTTGTTTACT
GAGCACACGTCTGAACTCTTTCCCTACACGACGCTCTTCCGATCTNNNGGGTAAACAGGCCTGAAAA
GAGCACACGTCTGAACTCTTTCCCTACACGACGCTCTTCCGATCTNNNTCTGGTATTCTTCATACCACT
AGCACACGTCTGAACTCTTTCCCTACACGACGCTCTTCCGATCTNNNTCTGTAATTCTTGCCCAGACAG
AGCACACGTCTGAACTCTTTCCCTACACGACGCTCTTCCGATCTNNNCCTCAAATCATCCAGAAAGGC
AGCACACGTCTGAACTCTTTCCCTACACGACGCTCTTCCGATCTNNNTGGTGATTTTAGGGTTGTTGG
GCACACGTCTGAACTCTTTCCCTACACGACGCTCTTCCGATCTNNNTTTCAGGCTCTGATATTTGGCCC
AGAGCACACGTCTGAACTCTTTCCCTACACGACGCTCTTCCGATCTNNNTTTTATAACATCGGCGTGTG
GCACACGTCTGAACTCTTTCCCTACACGACGCTCTTCCGATCTNNNACCAGTTATGGGGTACTGGAC

GCACACGTCTGAACTCTTTCCCTACACGACGCTCTTCCGATCTNNNCCAGCTTAGCACAGAATGGGAA
GAGCACACGTCTGAACTCTTTCCCTACACGACGCTCTTCCGATCTNNNGCGCAGAACCATCTATTTTG
AGCACACGTCTGAACTCTTTCCCTACACGACGCTCTTCCGATCTNNNAGGTCTTAGCTTTGTTTATGTG
GAGCACACGTCTGAACTCTTTCCCTACACGACGCTCTTCCGATCTNNNTTCTCAGTACATCCAGTCCCCT
GCACACGTCTGAACTCTTTCCCTACACGACGCTCTTCCGATCTNNNTCCATTTATCCAGGGAGCAAAG
GCACACGTCTGAACTCTTTCCCTACACGACGCTCTTCCGATCTNNNCCCTTCTTTTGCCCTCTTCAACTC
GCACACGTCTGAACTCTTTCCCTACACGACGCTCTTCCGATCTNNNATTCCAAGCTGGGGAAGGAAG
GAGCACACGTCTGAACTCTTTCCCTACACGACGCTCTTCCGATCTNNNTGGATGCTACACTGCATCTTC
CACACGTCTGAACTCTTTCCCTACACGACGCTCTTCCGATCTNNNCTGACCCAGGAGATTTACCAAGC
GCACACGTCTGAACTCTTTCCCTACACGACGCTCTTCCGATCTNNNTCTTTCTTTCCAAGCAGGTTGAC
GAGCACACGTCTGAACTCTTTCCCTACACGACGCTCTTCCGATCTNNNTCAAACACCCAGTCATTGCA
GCACACGTCTGAACTCTTTCCCTACACGACGCTCTTCCGATCTNNNGCACTCCCTTTTCAGATTCTCCA
GCACACGTCTGAACTCTTTCCCTACACGACGCTCTTCCGATCTNNNGCTGTTCCCCTTACCTAGAATGC
GAGCACACGTCTGAACTCTTTCCCTACACGACGCTCTTCCGATCTNNNGTCGAGAGAGTGGTGACTA
GCACACGTCTGAACTCTTTCCCTACACGACGCTCTTCCGATCTNNNCTAACTCTCTGAGCTCTCCCTCC
GCACACGTCTGAACTCTTTCCCTACACGACGCTCTTCCGATCTNNNTTCAGGTGCCATCACTTTTTCCA
AGAGCACACGTCTGAACTCTTTCCCTACACGACGCTCTTCCGATCTNNNAGCCTCAGAAACTCAATGG
AGCACACGTCTGAACTCTTTCCCTACACGACGCTCTTCCGATCTNNNGACTTGATACAAGCCACAGGG
CACACGTCTGAACTCTTTCCCTACACGACGCTCTTCCGATCTNNNAGGGAGAATCCAGCCTGTAAGT
GAGCACACGTCTGAACTCTTTCCCTACACGACGCTCTTCCGATCTNNNTCGCTTTATAACAGCCAGAGT
AGCACACGTCTGAACTCTTTCCCTACACGACGCTCTTCCGATCTNNNTCATAGCTCTACAGCTTACTGG
GAGCACACGTCTGAACTCTTTCCCTACACGACGCTCTTCCGATCTNNNTTTCCTAGTCCTGGTACTTGC
GCACACGTCTGAACTCTTTCCCTACACGACGCTCTTCCGATCTNNNGGCTAAATCATACATACCAGGC
AGCACACGTCTGAACTCTTTCCCTACACGACGCTCTTCCGATCTNNNTAGGCACAGAAGCCTCCTTTTT
AGCACACGTCTGAACTCTTTCCCTACACGACGCTCTTCCGATCTNNNACAGGCTTCCACTTTTTCCACAT
GCACACGTCTGAACTCTTTCCCTACACGACGCTCTTCCGATCTNNNAGAATGATTATTCCCACACAGG
GCACACGTCTGAACTCTTTCCCTACACGACGCTCTTCCGATCTNNNGCTTTGCGAAACTTTTGTTTCAG
AGAGCACACGTCTGAACTCTTTCCCTACACGACGCTCTTCCGATCTNNNCTGAGTTCACCCGTGTTGTC
CACACGTCTGAACTCTTTCCCTACACGACGCTCTTCCGATCTNNNCAGCACTTCATACCCATGGTCTCC
CACACGTCTGAACTCTTTCCCTACACGACGCTCTTCCGATCTNNNGGGTGGATTTGGGACACTTATG
AGCACACGTCTGAACTCTTTCCCTACACGACGCTCTTCCGATCTNNNAATTTTGCTATGGGTTGCTGG
AGCACACGTCTGAACTCTTTCCCTACACGACGCTCTTCCGATCTNNNACCGTGTGAGAAATTGTTTGG
GAGCACACGTCTGAACTCTTTCCCTACACGACGCTCTTCCGATCTNNNGAATGAAGGATGCCCTGGA
GAGCACACGTCTGAACTCTTTCCCTACACGACGCTCTTCCGATCTNNNAGAAATGGGCACCTTAGTAC
AGCACACGTCTGAACTCTTTCCCTACACGACGCTCTTCCGATCTNNNTCAGTGATTCCATCCTCCCACTT
AGCACACGTCTGAACTCTTTCCCTACACGACGCTCTTCCGATCTNNNCTTACGCTTCCTTTTGGTGGAG
AGAGCACACGTCTGAACTCTTTCCCTACACGACGCTCTTCCGATCTNNNTCTGTAATTCTTGCCCAGAC
GAGCACACGTCTGAACTCTTTCCCTACACGACGCTCTTCCGATCTNNNGAGAGAACCTGAAAGGAAG
GCACACGTCTGAACTCTTTCCCTACACGACGCTCTTCCGATCTNNNACAGATTCATTCATGATGCCAC
GCACACGTCTGAACTCTTTCCCTACACGACGCTCTTCCGATCTNNNGGTCACATCCTGCACTATACC
AGCACACGTCTGAACTCTTTCCCTACACGACGCTCTTCCGATCTNNNGTTGGGATATGTGGGAAAGT
GAGCACACGTCTGAACTCTTTCCCTACACGACGCTCTTCCGATCTNNNTGGAGCTAGTATCGATGCAG
AGCACACGTCTGAACTCTTTCCCTACACGACGCTCTTCCGATCTNNNAGCCTGGGGATCACTGTATTT

AGAGCACACGTCTGAACTCTTTCCCTACACGACGCTCTTCCGATCTNNNCCTTTTGGGGCTGAGTACT
GAGCACACGTCTGAACTCTTTCCCTACACGACGCTCTTCCGATCTNNNTATTCGCCCATCCTTCAACACA
AGAGCACACGTCTGAACTCTTTCCCTACACGACGCTCTTCCGATCTNNNGTCAAAACAATGAGCACTCCT
AGCACACGTCTGAACTCTTTCCCTACACGACGCTCTTCCGATCTNNNCAACCTGGCTATGTCAGAGCAT
GAGCACACGTCTGAACTCTTTCCCTACACGACGCTCTTCCGATCTNNNTGCCTGGCCCACAGATAGATA
GAGCACACGTCTGAACTCTTTCCCTACACGACGCTCTTCCGATCTNNNAAGGCACTTTCCTGTTTTCCCT
AGAGCACACGTCTGAACTCTTTCCCTACACGACGCTCTTCCGATCTNNNAGTGAAGCCTTCTTACTCTTC
AGCACACGTCTGAACTCTTTCCCTACACGACGCTCTTCCGATCTNNNAGTGATTACCATAGGTAGCTTT
AAGAGCACACGTCTGAACTCTTTCCCTACACGACGCTCTTCCGATCTNNNTAGCATGGTGACCACGGT
AGCACACGTCTGAACTCTTTCCCTACACGACGCTCTTCCGATCTNNNCTGACCCAGGAGATTTACCAAG
AGAGCACACGTCTGAACTCTTTCCCTACACGACGCTCTTCCGATCTNNNGCCTTTACCCAGGTTCAGTCA
AGAGCACACGTCTGAACTCTTTCCCTACACGACGCTCTTCCGATCTNNNGTAATTTCCCAGGCAGTGAC
AGCACACGTCTGAACTCTTTCCCTACACGACGCTCTTCCGATCTNNNGGCTTGTGTGGAGGTAAGTCA
AGAGCACACGTCTGAACTCTTTCCCTACACGACGCTCTTCCGATCTNNNTACCTGGCACTCATATTCACC
AGCACACGTCTGAACTCTTTCCCTACACGACGCTCTTCCGATCTNNNTTACTACATTGCAGAAGCCCAG
AGCACACGTCTGAACTCTTTCCCTACACGACGCTCTTCCGATCTNNNTAACCCTAAGGAGGGCTTACGA
AGCACACGTCTGAACTCTTTCCCTACACGACGCTCTTCCGATCTNNNCCCTCTCTTCATCTTCACATGGCC
GAGCACACGTCTGAACTCTTTCCCTACACGACGCTCTTCCGATCTNNNATAAACCAAACCCCAGTGACG
GAGCACACGTCTGAACTCTTTCCCTACACGACGCTCTTCCGATCTNNNTCGCTTTATAACAGCCAGAGT
GAGCACACGTCTGAACTCTTTCCCTACACGACGCTCTTCCGATCTNNNTGCCATTATCTCTTCACAGCA
GCACACGTCTGAACTCTTTCCCTACACGACGCTCTTCCGATCTNNNTACAGTGCCTTGATTCACAACCCC
AAGAGCACACGTCTGAACTCTTTCCCTACACGACGCTCTTCCGATCTNNNGGCCTAACCTGTGATAGCC
AGAGCACACGTCTGAACTCTTTCCCTACACGACGCTCTTCCGATCTNNNTAAAACCTGGATCCCGTTCAG
GAGCACACGTCTGAACTCTTTCCCTACACGACGCTCTTCCGATCTNNNACAAAACACCACTATGCCCACA
AGAGCACACGTCTGAACTCTTTCCCTACACGACGCTCTTCCGATCTNNNCAAGAAACAGAAATGCGCTC
AGCACACGTCTGAACTCTTTCCCTACACGACGCTCTTCCGATCTNNNTAGGAAGAACAACCCTCCACTC
AGAGCACACGTCTGAACTCTTTCCCTACACGACGCTCTTCCGATCTNNNGGAGATGGTCAACAAATGCA
AGCACACGTCTGAACTCTTTCCCTACACGACGCTCTTCCGATCTNNNAGATTGCCCTCACATCCATCCTA
GAGCACACGTCTGAACTCTTTCCCTACACGACGCTCTTCCGATCTNNNTGGTCTTCCCTGCTTTTCCTTT
AGCACACGTCTGAACTCTTTCCCTACACGACGCTCTTCCGATCTNNNTAGCTGCTTCTCTATCCACTCCCC
GAGCACACGTCTGAACTCTTTCCCTACACGACGCTCTTCCGATCTNNNAACCACAAGAGGACTGCAGTA
GCACACGTCTGAACTCTTTCCCTACACGACGCTCTTCCGATCTNNNGGGAAAAAGCAAACTGTTTCCA
GAGCACACGTCTGAACTCTTTCCCTACACGACGCTCTTCCGATCTNNNACTATACTATGGTAGCATTTCC
AGCACACGTCTGAACTCTTTCCCTACACGACGCTCTTCCGATCTNNNATCTGATTGCCAGGTTGATCTG
AAGAGCACACGTCTGAACTCTTTCCCTACACGACGCTCTTCCGATCTNNNTGAGGTTGTATAATGCAT
AGAGCACACGTCTGAACTCTTTCCCTACACGACGCTCTTCCGATCTNNNGAGTGTGCAGTGAATCGAG
GAGCACACGTCTGAACTCTTTCCCTACACGACGCTCTTCCGATCTNNNTAGACTTACCCCAGGAAAGGC
GAGCACACGTCTGAACTCTTTCCCTACACGACGCTCTTCCGATCTNNNCCCTCTCTTCATCTTCACATGG
AGCACACGTCTGAACTCTTTCCCTACACGACGCTCTTCCGATCTNNNGCTTCCTTTCAGGGTTTGTGGA
AAGAGCACACGTCTGAACTCTTTCCCTACACGACGCTCTTCCGATCTNNNGATTTTTGGCAGATGCCTT
AGCACACGTCTGAACTCTTTCCCTACACGACGCTCTTCCGATCTNNNCCACCTACCTAAACCAGTGTGT
GAGCACACGTCTGAACTCTTTCCCTACACGACGCTCTTCCGATCTNNNCTCATTCATCTCCCTCTCCCAGT
AGCACACGTCTGAACTCTTTCCCTACACGACGCTCTTCCGATCTNNNAGAGGAGAGAATCAGCAAACA

GAGCACACGTCTGAACTCTTTCCCTACACGACGCTCTTCCGATCTNNNGGTGTCACAGCTCTCCAGAAA
AGCACACGTCTGAACTCTTTCCCTACACGACGCTCTTCCGATCTNNNTGACCGAGTTCTGAGCTCTATG
AGAGCACACGTCTGAACTCTTTCCCTACACGACGCTCTTCCGATCTNNNTGCAAGGAGAACCTTGGGA
GAGCACACGTCTGAACTCTTTCCCTACACGACGCTCTTCCGATCTNNNGATTTCTGCCTCAGCTCTTCTC
AGAGCACACGTCTGAACTCTTTCCCTACACGACGCTCTTCCGATCTNNNGAAACAGTGGGCTTTGTTTC
GAGCACACGTCTGAACTCTTTCCCTACACGACGCTCTTCCGATCTNNNAGGAACCAGAGGCTGTGTAG
AGAGCACACGTCTGAACTCTTTCCCTACACGACGCTCTTCCGATCTNNNGTATCAGCCTCTTGCTTCTAC
AGCACACGTCTGAACTCTTTCCCTACACGACGCTCTTCCGATCTNNNGAGGTTGTGAGCCGAAATCGT
GAGCACACGTCTGAACTCTTTCCCTACACGACGCTCTTCCGATCTNNNAGCTTTCTCCAAGGCTGTGTT
GAGCACACGTCTGAACTCTTTCCCTACACGACGCTCTTCCGATCTNNNTGTATATATTTGCCAGGGTTCT
GAGCACACGTCTGAACTCTTTCCCTACACGACGCTCTTCCGATCTNNNCCACAGCATTAGCTCTTGGCTT
GAGCACACGTCTGAACTCTTTCCCTACACGACGCTCTTCCGATCTNNNACCATCATCCTGGAGTTTTTCC
AGCACACGTCTGAACTCTTTCCCTACACGACGCTCTTCCGATCTNNNGGGGGTGGTTACACAGATGTAC
AGAGCACACGTCTGAACTCTTTCCCTACACGACGCTCTTCCGATCTNNNAGATGTTTCTGTGCATGTGC
GAGCACACGTCTGAACTCTTTCCCTACACGACGCTCTTCCGATCTNNNTCATTCTTTTCCCCCATCCTGA
GAGCACACGTCTGAACTCTTTCCCTACACGACGCTCTTCCGATCTNNNTAACTAATAGCTCCCTCGTGGC
AGCACACGTCTGAACTCTTTCCCTACACGACGCTCTTCCGATCTNNNATGCGGCTATTTCCAAGAGTGTT
AGCACACGTCTGAACTCTTTCCCTACACGACGCTCTTCCGATCTNNNCTCTGGAATTACACACAGGAGC
GAGCACACGTCTGAACTCTTTCCCTACACGACGCTCTTCCGATCTNNNATTTCTTCCTCCATGTGTGGGT
GCACACGTCTGAACTCTTTCCCTACACGACGCTCTTCCGATCTNNNAAACTGTTTTTATGGACCGTGCCC
GAGCACACGTCTGAACTCTTTCCCTACACGACGCTCTTCCGATCTNNNCAACTTGCAACAAGAGGAATC
AGCACACGTCTGAACTCTTTCCCTACACGACGCTCTTCCGATCTNNNAGTCCGAACAGATAAGCAGGG
GAGCACACGTCTGAACTCTTTCCCTACACGACGCTCTTCCGATCTNNNGTATCAGCCTCTTGCTTCTAGC
GAGCACACGTCTGAACTCTTTCCCTACACGACGCTCTTCCGATCTNNNTTCAGGTGCCATCACTTTTTCCA
AGAGCACACGTCTGAACTCTTTCCCTACACGACGCTCTTCCGATCTNNNGCTCTTTTGGAAAGCCCTCTC
GCACACGTCTGAACTCTTTCCCTACACGACGCTCTTCCGATCTNNNAGCTTATGGAGGATCATTTTGTTC
GAGCACACGTCTGAACTCTTTCCCTACACGACGCTCTTCCGATCTNNNCTGGGCTTTGGTTTGGAGTA
AGCACACGTCTGAACTCTTTCCCTACACGACGCTCTTCCGATCTNNNTTAGCTTTCCAACATCCCCCTTG
GAGCACACGTCTGAACTCTTTCCCTACACGACGCTCTTCCGATCTNNNAGCCACTCAAAGGAACTTTCTG
GAGCACACGTCTGAACTCTTTCCCTACACGACGCTCTTCCGATCTNNNGAACTGTGGCCTTCTTCCCTAA
AGCACACGTCTGAACTCTTTCCCTACACGACGCTCTTCCGATCTNNNATATGAATCACTTGGATGGCTG
GAGCACACGTCTGAACTCTTTCCCTACACGACGCTCTTCCGATCTNNNTACAACCCACTGACATCCCTGA
AAGAGCACACGTCTGAACTCTTTCCCTACACGACGCTCTTCCGATCTNNNTCCCTCCTCTTTGTCCTCATC
GAGCACACGTCTGAACTCTTTCCCTACACGACGCTCTTCCGATCTNNNGTACACGTATGAACAGTGCTC
GAGCACACGTCTGAACTCTTTCCCTACACGACGCTCTTCCGATCTNNNACCGTAATACCGTCTCCCAATG
AGCACACGTCTGAACTCTTTCCCTACACGACGCTCTTCCGATCTNNNCCACACCTGTTACTGAGGGTAG
GAGCACACGTCTGAACTCTTTCCCTACACGACGCTCTTCCGATCTNNNCCTCCATTACATTGTGACCAGG
GAGCACACGTCTGAACTCTTTCCCTACACGACGCTCTTCCGATCTNNNGCAGTTTTCATGTGCCTTTCAG
AGCACACGTCTGAACTCTTTCCCTACACGACGCTCTTCCGATCTNNNGTGGTGCATTCCTCTCTTCCAAA
AGCACACGTCTGAACTCTTTCCCTACACGACGCTCTTCCGATCTNNNTGGGAGAGACTAGCATCTCTGT
GAGCACACGTCTGAACTCTTTCCCTACACGACGCTCTTCCGATCTNNNTGCTTGGTTTCTCCTTGATGC
AGAGCACACGTCTGAACTCTTTCCCTACACGACGCTCTTCCGATCTNNNACATCAGGCCACTAAGGAAA
GGAAGAGCACACGTCTGAACTCTTTCCCTACACGACGCTCTTCCGATCTNNNATACATGATGTGGCCCT

GCACACGTCTGAACTCTTTCCCTACACGACGCTCTTCCGATCTNNNACTGACCCTTCCTTCTCTACCTGC
GAGCACACGTCTGAACTCTTTCCCTACACGACGCTCTTCCGATCTNNNTGTCCTCACAAGGTAGGAG
AGCACACGTCTGAACTCTTTCCCTACACGACGCTCTTCCGATCTNNNCAGCCCAGCACTGTATCTGTTT
GCACACGTCTGAACTCTTTCCCTACACGACGCTCTTCCGATCTNNNAAAGCCCTGAGGATCTGTAGG
GCACACGTCTGAACTCTTTCCCTACACGACGCTCTTCCGATCTNNNCCATCCACGACCTCTCTCTCTAAC
AGCACACGTCTGAACTCTTTCCCTACACGACGCTCTTCCGATCTNNNGTGTGTGCACGTGTGAAGAA
GAGCACACGTCTGAACTCTTTCCCTACACGACGCTCTTCCGATCTNNNTCAGATTCTCCTGAATCACCCC
GAGCACACGTCTGAACTCTTTCCCTACACGACGCTCTTCCGATCTNNNCAGGGCTGAAACCTTCTCTC
GAGCACACGTCTGAACTCTTTCCCTACACGACGCTCTTCCGATCTNNNTGTTGATGATCTAGTCAGCA
GAGCACACGTCTGAACTCTTTCCCTACACGACGCTCTTCCGATCTNNNCAGTACAGCATTCTGTGCAT
GAGCACACGTCTGAACTCTTTCCCTACACGACGCTCTTCCGATCTNNNAGAGAAAGTCAGGAGAATG
AGCACACGTCTGAACTCTTTCCCTACACGACGCTCTTCCGATCTNNNGAGGGAGGTGGACAAATGG
GAGCACACGTCTGAACTCTTTCCCTACACGACGCTCTTCCGATCTNNNAAAGGGAGAAGAAGAGGTG
AGCACACGTCTGAACTCTTTCCCTACACGACGCTCTTCCGATCTNNNAGCTTTGAGTTTGTGTATTGCT
AGCACACGTCTGAACTCTTTCCCTACACGACGCTCTTCCGATCTNNNGGTGGTCTCACAGGTCTTTGTT
AGCACACGTCTGAACTCTTTCCCTACACGACGCTCTTCCGATCTNNNCTGAGATTTCACCAGGCTGCAC
GCACACGTCTGAACTCTTTCCCTACACGACGCTCTTCCGATCTNNNTGTTTTCCACAAGCTCAAGTTCCC
AGCACACGTCTGAACTCTTTCCCTACACGACGCTCTTCCGATCTNNNTTCCTCCTTGACCCCTAAAGCAT
AGCACACGTCTGAACTCTTTCCCTACACGACGCTCTTCCGATCTNNNAGAATCAGAAATGTTCCTCGG
GAGCACACGTCTGAACTCTTTCCCTACACGACGCTCTTCCGATCTNNNCGTCTAATCCCATGGCTCAG
AGCACACGTCTGAACTCTTTCCCTACACGACGCTCTTCCGATCTNNNATGGAGGGGTATCACAACCAA
AGCACACGTCTGAACTCTTTCCCTACACGACGCTCTTCCGATCTNNNAGCCTGGGCAATAGCATAAGA
AGCACACGTCTGAACTCTTTCCCTACACGACGCTCTTCCGATCTNNNCCAAAAGATAATCCCAGCCCT
GAAGAGCACACGTCTGAACTCTTTCCCTACACGACGCTCTTCCGATCTNNNTTAGTGGGATCTCTCTC
AGCACACGTCTGAACTCTTTCCCTACACGACGCTCTTCCGATCTNNNAGGAATCAGCTTTGAGGAGA
AGCACACGTCTGAACTCTTTCCCTACACGACGCTCTTCCGATCTNNNACGGATATTGGACCTTCAAAC
AGAGCACACGTCTGAACTCTTTCCCTACACGACGCTCTTCCGATCTNNNTCAGCTTCTGAGCAAGAGTT
AGCACACGTCTGAACTCTTTCCCTACACGACGCTCTTCCGATCTNNNATTCTTTGGATTCTGGGGTCCC
GAGCACACGTCTGAACTCTTTCCCTACACGACGCTCTTCCGATCTNNNGATTTTTCCCGAGACCCTACC
AGCACACGTCTGAACTCTTTCCCTACACGACGCTCTTCCGATCTNNNCTTCCATGAGCTCAGCTTGAC
AGCACACGTCTGAACTCTTTCCCTACACGACGCTCTTCCGATCTNNNTGTTCGGTCTACTCCATGAACA
AGCACACGTCTGAACTCTTTCCCTACACGACGCTCTTCCGATCTNNNAAGTCCTAGGGTTCCCAAACCT
GCACACGTCTGAACTCTTTCCCTACACGACGCTCTTCCGATCTNNNGTCAACTCAAAGGGCTCCTCATTC
AGCACACGTCTGAACTCTTTCCCTACACGACGCTCTTCCGATCTNNNTGTTCGCAACAGAAAAACACGA
GAGCACACGTCTGAACTCTTTCCCTACACGACGCTCTTCCGATCTNNNACCGAAGTCAGACACCAAGG
GAGCACACGTCTGAACTCTTTCCCTACACGACGCTCTTCCGATCTNNNAGTGCTGTGTCTTGACTTTTG
GAGCACACGTCTGAACTCTTTCCCTACACGACGCTCTTCCGATCTNNNATCAAGGACATCAAGGAGG
AGCACACGTCTGAACTCTTTCCCTACACGACGCTCTTCCGATCTNNNGCACCATGCATACTCTCAAGTC
AGAGCACACGTCTGAACTCTTTCCCTACACGACGCTCTTCCGATCTNNNGGAAATGATGTATGACAGC
AGAGCACACGTCTGAACTCTTTCCCTACACGACGCTCTTCCGATCTNNNGACATGGGGTGAAGATGTC
AGCACACGTCTGAACTCTTTCCCTACACGACGCTCTTCCGATCTNNNACATCAAATCTAGGGGATTCTT
GAGCACACGTCTGAACTCTTTCCCTACACGACGCTCTTCCGATCTNNNTCAGCAACACAGCCAAATCTT
AAGAGCACACGTCTGAACTCTTTCCCTACACGACGCTCTTCCGATCTNNNAGCAATGCTTTGGATGAC

GCACACGTCTGAACTCTTTCCCTACACGACGCTCTTCCGATCTNNNCGTCTAATCCCATGGCTCAGAG
AGCACACGTCTGAACTCTTTCCCTACACGACGCTCTTCCGATCTNNNCTTCCAGCCTTTGGACATGAAC
GCACACGTCTGAACTCTTTCCCTACACGACGCTCTTCCGATCTNNNCCCAAACTGACACCTTTGTGTGT
AGCACACGTCTGAACTCTTTCCCTACACGACGCTCTTCCGATCTNNNAGGATCCAAGGGTGACAGGT
AGCACACGTCTGAACTCTTTCCCTACACGACGCTCTTCCGATCTNNNTTGAATCCAGCAAACCCATCAC
AGCACACGTCTGAACTCTTTCCCTACACGACGCTCTTCCGATCTNNNGAGGAAAGACACAGATGCAC
AGCACACGTCTGAACTCTTTCCCTACACGACGCTCTTCCGATCTNNNTTAGGGGGTTAGTTCTTTCCT
GCACACGTCTGAACTCTTTCCCTACACGACGCTCTTCCGATCTNNNAACTTCAACTGCTACCCAGACAC
GCACACGTCTGAACTCTTTCCCTACACGACGCTCTTCCGATCTNNNCTTGCAGATGGTTCACATGACC
AGAGCACACGTCTGAACTCTTTCCCTACACGACGCTCTTCCGATCTNNNAGTACACGTATGAACAGT
GAGCACACGTCTGAACTCTTTCCCTACACGACGCTCTTCCGATCTNNNTGCAATGGAAGTAGCATAA
GCACACGTCTGAACTCTTTCCCTACACGACGCTCTTCCGATCTNNNGACTAGACTTGTGTCAGGGCTT
GCACACGTCTGAACTCTTTCCCTACACGACGCTCTTCCGATCTNNNCAAAGAGCAGTGAACATGAG
GAGCACACGTCTGAACTCTTTCCCTACACGACGCTCTTCCGATCTNNNAGCAATTGTACATGGGTGT
GAGCACACGTCTGAACTCTTTCCCTACACGACGCTCTTCCGATCTNNNTCAGCTTTACCTTACAGGCA
CACGTCTGAACTCTTTCCCTACACGACGCTCTTCCGATCTNNNCCGTGGCTTTTACCCTCCACATCGG
AGCACACGTCTGAACTCTTTCCCTACACGACGCTCTTCCGATCTNNNCTATTCTCTGGGTTTGTGCTG
GAGCACACGTCTGAACTCTTTCCCTACACGACGCTCTTCCGATCTNNNGTGTTCCCTATCTCTTTCCTCG
GCACACGTCTGAACTCTTTCCCTACACGACGCTCTTCCGATCTNNNGCATTTCAGGACCTGTCCCTATG
AGCACACGTCTGAACTCTTTCCCTACACGACGCTCTTCCGATCTNNNATCCAGGAACTCAGATCTACC
GCACACGTCTGAACTCTTTCCCTACACGACGCTCTTCCGATCTNNNTCCTGCAAATCTGAACTCTTCCG
AGCACACGTCTGAACTCTTTCCCTACACGACGCTCTTCCGATCTNNNACACACTCACACCAAACACAAA
GAGCACACGTCTGAACTCTTTCCCTACACGACGCTCTTCCGATCTNNNACAGATGACGGACTGTAGG
AGAGCACACGTCTGAACTCTTTCCCTACACGACGCTCTTCCGATCTNNNTGCCACTGGTGGAAATTGT
AGCACACGTCTGAACTCTTTCCCTACACGACGCTCTTCCGATCTNNNAAGTGTAATGCACAGACACCT
GCACACGTCTGAACTCTTTCCCTACACGACGCTCTTCCGATCTNNNTCAAGGATTTCCATAGCACTGC
GAGCACACGTCTGAACTCTTTCCCTACACGACGCTCTTCCGATCTNNNACCAGACCTGAACCTACCAAT
GCACACGTCTGAACTCTTTCCCTACACGACGCTCTTCCGATCTNNNTTTGTCCACTGTTATGCCTTGCT
AGCACACGTCTGAACTCTTTCCCTACACGACGCTCTTCCGATCTNNNCCCAAAAACAGTCATGCAAGT
GAGCACACGTCTGAACTCTTTCCCTACACGACGCTCTTCCGATCTNNNCTGACTACAGGTCAGGCAGA
GAGCACACGTCTGAACTCTTTCCCTACACGACGCTCTTCCGATCTNNNACACAGCCAGATTTGGAACC
AGCACACGTCTGAACTCTTTCCCTACACGACGCTCTTCCGATCTNNNATATTTACTGGCTTGCCCCAAG
GAGCACACGTCTGAACTCTTTCCCTACACGACGCTCTTCCGATCTNNNTTCTTTGTGTACTTTCCGCAC
GCACACGTCTGAACTCTTTCCCTACACGACGCTCTTCCGATCTNNNATGTCATACAAGTGCCAGAGCA
AAGAGCACACGTCTGAACTCTTTCCCTACACGACGCTCTTCCGATCTNNNTCTCTAACAGCCATTCTG
GCACACGTCTGAACTCTTTCCCTACACGACGCTCTTCCGATCTNNNCAGACCTTGCCCGTGATGAGCA
AGCACACGTCTGAACTCTTTCCCTACACGACGCTCTTCCGATCTNNNAATTCTCCCAGCCTCGCCTCTTC
AGAGCACACGTCTGAACTCTTTCCCTACACGACGCTCTTCCGATCTNNNCAAGTCAGGTCATGAGGC
AAGAGCACACGTCTGAACTCTTTCCCTACACGACGCTCTTCCGATCTNNNAAAACCCAAATGCTGTCG
GCACACGTCTGAACTCTTTCCCTACACGACGCTCTTCCGATCTNNNTCCAATAAAGAACATGCATCCAG
CACACGTCTGAACTCTTTCCCTACACGACGCTCTTCCGATCTNNNGTGTGTGTGTGTCAATAAAAGGC
CACACGTCTGAACTCTTTCCCTACACGACGCTCTTCCGATCTNNNGATCCATCTGCAGAGCTGTTCTTC
AGCACACGTCTGAACTCTTTCCCTACACGACGCTCTTCCGATCTNNNGATCACTGACAGGCTAAAAG

GGAAGAGCACACGTCTGAACTCTTTCCCTACACGACGCTCTTCCGATCTNNNATCCACTTGAGGAGCCA
GAGCACACGTCTGAACTCTTTCCCTACACGACGCTCTTCCGATCTNNNACCTAAGCTTGATGATGGCAA
GCACACGTCTGAACTCTTTCCCTACACGACGCTCTTCCGATCTNNNATTCTTTGGATTCTGGGGTCCCTC
GAGCACACGTCTGAACTCTTTCCCTACACGACGCTCTTCCGATCTNNNCAGCCATTACGCATCCTGTTTT
CACACGTCTGAACTCTTTCCCTACACGACGCTCTTCCGATCTNNNAAGCTAGAACTTGAACTTGCCCCA
AGAGCACACGTCTGAACTCTTTCCCTACACGACGCTCTTCCGATCTNNNTCATCCAGGTTGTTGTGAAG
AGCACACGTCTGAACTCTTTCCCTACACGACGCTCTTCCGATCTNNNCCGAGAAGAAGTGGCTAATCT
AGAGCACACGTCTGAACTCTTTCCCTACACGACGCTCTTCCGATCTNNNCAAATCTGCCTTCAGACCTCA
GAGCACACGTCTGAACTCTTTCCCTACACGACGCTCTTCCGATCTNNNTAAAGGGGCATGCAGTTTAC
GAGCACACGTCTGAACTCTTTCCCTACACGACGCTCTTCCGATCTNNNTGCATTCAAAAGGACGGGAAA
GAGCACACGTCTGAACTCTTTCCCTACACGACGCTCTTCCGATCTNNNGGCCAGTTTTAGAAAGTTTG
GCACACGTCTGAACTCTTTCCCTACACGACGCTCTTCCGATCTNNNGTAAGCAGATGATGCCAGAATG
AGAGCACACGTCTGAACTCTTTCCCTACACGACGCTCTTCCGATCTNNNGCCAGTGAGGATAGAGTGA
GAGCACACGTCTGAACTCTTTCCCTACACGACGCTCTTCCGATCTNNNTGGACAACACTGCTCTACATTC
AGAGCACACGTCTGAACTCTTTCCCTACACGACGCTCTTCCGATCTNNNTTTCTGTTGACAGCTCACAT
GAGCACACGTCTGAACTCTTTCCCTACACGACGCTCTTCCGATCTNNNAGGTACAACCCAAGATCATCA
AGAGCACACGTCTGAACTCTTTCCCTACACGACGCTCTTCCGATCTNNNGTGCTATCAGGCCCAAAATC
AGCACACGTCTGAACTCTTTCCCTACACGACGCTCTTCCGATCTNNNTGTCCTCGATATAGCTGGGTGG
AGAGCACACGTCTGAACTCTTTCCCTACACGACGCTCTTCCGATCTNNNTCAGTCACTCTCTCCCACTTG
AAGAGCACACGTCTGAACTCTTTCCCTACACGACGCTCTTCCGATCTNNNAAACATTTAGTGGTCCGTT
GAGCACACGTCTGAACTCTTTCCCTACACGACGCTCTTCCGATCTNNNGTCACTAAGCTGCCAAGTTCT
GAGCACACGTCTGAACTCTTTCCCTACACGACGCTCTTCCGATCTNNNCATTGCTGCATTGCTTTCCAA
AGCACACGTCTGAACTCTTTCCCTACACGACGCTCTTCCGATCTNNNACTCTCGTTGCTTCAGGATTCA
GAGCACACGTCTGAACTCTTTCCCTACACGACGCTCTTCCGATCTNNNTTTGAGTGTCGAACTTGTGG
AGCACACGTCTGAACTCTTTCCCTACACGACGCTCTTCCGATCTNNNAGTCAGATGTGTTTTTGTGCAG
AGCACACGTCTGAACTCTTTCCCTACACGACGCTCTTCCGATCTNNNAAAATGAGTATACGCGCACCTG
GCACACGTCTGAACTCTTTCCCTACACGACGCTCTTCCGATCTNNNTCCCAGTTTGTCATGCAATTAGA
GAAGAGCACACGTCTGAACTCTTTCCCTACACGACGCTCTTCCGATCTNNNGGAGATCCTAGTCTTTA
AGCACACGTCTGAACTCTTTCCCTACACGACGCTCTTCCGATCTNNNGGCAGAGCTCCAAAGTATGTCA
GAGCACACGTCTGAACTCTTTCCCTACACGACGCTCTTCCGATCTNNNTGTGTTGTTGCACCTACCCAAA
GAGCACACGTCTGAACTCTTTCCCTACACGACGCTCTTCCGATCTNNNCTGGGCCACATAGGAAGATT
CACACGTCTGAACTCTTTCCCTACACGACGCTCTTCCGATCTNNNGCTCCAGTAGTGCCCTTTATCTGCA
AGAGCACACGTCTGAACTCTTTCCCTACACGACGCTCTTCCGATCTNNNACCTTACTATGTTCCCAAAAT
GAGCACACGTCTGAACTCTTTCCCTACACGACGCTCTTCCGATCTNNNTACCTGAAGTGGCATTCGATG
AGCACACGTCTGAACTCTTTCCCTACACGACGCTCTTCCGATCTNNNTGGGAAAATAATGGAATGGAA
AGCACACGTCTGAACTCTTTCCCTACACGACGCTCTTCCGATCTNNNTTAGTGCAGAAGACTTGCTTGC
GAGCACACGTCTGAACTCTTTCCCTACACGACGCTCTTCCGATCTNNNAGAGTGTCCAGAAATGGCTT
AAGAGCACACGTCTGAACTCTTTCCCTACACGACGCTCTTCCGATCTNNNGCTTATGCTACCCCTGTGT
GCACACGTCTGAACTCTTTCCCTACACGACGCTCTTCCGATCTNNNCTGTGTTCTGCCTCCTCAAATCTCA
AGAGCACACGTCTGAACTCTTTCCCTACACGACGCTCTTCCGATCTNNNTGACTATCGGCCATTGTAA
AGCACACGTCTGAACTCTTTCCCTACACGACGCTCTTCCGATCTNNNGGTCCTCACTTTCCTTGAACTC
GAGCACACGTCTGAACTCTTTCCCTACACGACGCTCTTCCGATCTNNNCAGAGTGAGCTAGAAATACG
AGAGCACACGTCTGAACTCTTTCCCTACACGACGCTCTTCCGATCTNNNTGTACAGACATGCACACAGA

AGCACACGTCTGAACTCTTTCCCTACACGACGCTCTTCCGATCTNNNGTGTGTGTGTATGTGAACCCT
GCACACGTCTGAACTCTTTCCCTACACGACGCTCTTCCGATCTNNNTTTGGTGCTAATGCTGATGGTCT
ACACGTCTGAACTCTTTCCCTACACGACGCTCTTCCGATCTNNNTTTGGTGCTAATGCTGATGGTCTCA
AGCACACGTCTGAACTCTTTCCCTACACGACGCTCTTCCGATCTNNNGCCTGGGGTCCTAGAGATTT
GCACACGTCTGAACTCTTTCCCTACACGACGCTCTTCCGATCTNNNGGGGCGGAAGCTAGTTTTACTT
AGCACACGTCTGAACTCTTTCCCTACACGACGCTCTTCCGATCTNNNTCCTATTCCTACAGTGCGTCCAT
AGAGCACACGTCTGAACTCTTTCCCTACACGACGCTCTTCCGATCTNNNAGGGTTCTTTCTACACTCCC
AGCACACGTCTGAACTCTTTCCCTACACGACGCTCTTCCGATCTNNNTCTTGCTTCCTTGCTACACAAC
AGAGCACACGTCTGAACTCTTTCCCTACACGACGCTCTTCCGATCTNNNATTGAATGCACTGTGTTCTT
GCACACGTCTGAACTCTTTCCCTACACGACGCTCTTCCGATCTNNNCCGTGTTTCACATTTGGCTTTGT
AGCACACGTCTGAACTCTTTCCCTACACGACGCTCTTCCGATCTNNNTCTCAAATGGTTCAGCAAAAA
GAGCACACGTCTGAACTCTTTCCCTACACGACGCTCTTCCGATCTNNNCTTCAAAATGTTACTGTCAGG
AGCACACGTCTGAACTCTTTCCCTACACGACGCTCTTCCGATCTNNNTGTGTGAGTGTACGAGAGAG
AGCACACGTCTGAACTCTTTCCCTACACGACGCTCTTCCGATCTNNNTGGTCTCCTTGTGTATTGTCA
AGCACACGTCTGAACTCTTTCCCTACACGACGCTCTTCCGATCTNNNACCATTCTTCACTTTCCTCAGG
AGCACACGTCTGAACTCTTTCCCTACACGACGCTCTTCCGATCTNNNCTTTCTCTCTCTCTGTCTGTGGG
AGCACACGTCTGAACTCTTTCCCTACACGACGCTCTTCCGATCTNNNACTGGTCTGACTACACCCCTTA
GCACACGTCTGAACTCTTTCCCTACACGACGCTCTTCCGATCTNNNCCAGCTTTGTCAGCAACATTCAG
AGAGCACACGTCTGAACTCTTTCCCTACACGACGCTCTTCCGATCTNNNACATGATAACATCTGGGTG
AGCACACGTCTGAACTCTTTCCCTACACGACGCTCTTCCGATCTNNNGCAAAGAGCCACTCATACATCC
AGCACACGTCTGAACTCTTTCCCTACACGACGCTCTTCCGATCTNNNTGACATTTCCATAATGGTTAG
GAGCACACGTCTGAACTCTTTCCCTACACGACGCTCTTCCGATCTNNNCACCATCATCCCAAGGTATCC
GAGCACACGTCTGAACTCTTTCCCTACACGACGCTCTTCCGATCTNNNCAGAGAGAATATATATGTGT
AGCACACGTCTGAACTCTTTCCCTACACGACGCTCTTCCGATCTNNNGCTTGAAATCAGGTAGTGTTAG
AGCACACGTCTGAACTCTTTCCCTACACGACGCTCTTCCGATCTNNNATGCGTGCTCTCTCTCCATGTA
AGCACACGTCTGAACTCTTTCCCTACACGACGCTCTTCCGATCTNNNACAAGGGCCATGCTTTTGATT
GCACACGTCTGAACTCTTTCCCTACACGACGCTCTTCCGATCTNNNTGCCTGCCAGACATTTTTCTTTG
GCACACGTCTGAACTCTTTCCCTACACGACGCTCTTCCGATCTNNNTATCTCATCTCCTTTGCCCTCCCCA
AGCACACGTCTGAACTCTTTCCCTACACGACGCTCTTCCGATCTNNNTCCATTACATCTATCCCCCACC
AGAGCACACGTCTGAACTCTTTCCCTACACGACGCTCTTCCGATCTNNNAGATTCTACTGCTGTTGGCA
AGCACACGTCTGAACTCTTTCCCTACACGACGCTCTTCCGATCTNNNAGCAATATTTCTCCACGTGTGT
AGAGCACACGTCTGAACTCTTTCCCTACACGACGCTCTTCCGATCTNNNAACCTGTGTGACCATGGAA
AGCACACGTCTGAACTCTTTCCCTACACGACGCTCTTCCGATCTNNNGGCAATGCCAACCCTAAAGACT
GAGCACACGTCTGAACTCTTTCCCTACACGACGCTCTTCCGATCTNNNGCTACACTCAGCACCTGTTAC
AGAGCACACGTCTGAACTCTTTCCCTACACGACGCTCTTCCGATCTNNNAATGCAAGGTGTATTAGCC
GAGCACACGTCTGAACTCTTTCCCTACACGACGCTCTTCCGATCTNNNAGGAAAACTCATATGTCCCA
AAGAGCACACGTCTGAACTCTTTCCCTACACGACGCTCTTCCGATCTNNNGGATTTATACACATGCAC
CACACGTCTGAACTCTTTCCCTACACGACGCTCTTCCGATCTNNNCAGGCTTTGTAAAAGCAAGGACC
AGCACACGTCTGAACTCTTTCCCTACACGACGCTCTTCCGATCTNNNGAACACGCTAGGATCTGTACG
AGCACACGTCTGAACTCTTTCCCTACACGACGCTCTTCCGATCTNNNGGACTGTTAGTATTGCTTGGG
AGCACACGTCTGAACTCTTTCCCTACACGACGCTCTTCCGATCTNNNGGGTCATGAAGGAGAAATGT
AGCACACGTCTGAACTCTTTCCCTACACGACGCTCTTCCGATCTNNNTCCACTTTCTCCTGAAACAGCT
AGCACACGTCTGAACTCTTTCCCTACACGACGCTCTTCCGATCTNNNGCTAGCTCCTGTGTTTCCATCA

GAGCACACGTCTGAACTCTTTCCCTACACGACGCTCTTCCGATCTNNNCTACTGCCTCGCAATACCAAA
GAGCACACGTCTGAACTCTTTCCCTACACGACGCTCTTCCGATCTNNNAGTTCCCATTTCCTCCTTCCTCA
GAGCACACGTCTGAACTCTTTCCCTACACGACGCTCTTCCGATCTNNNATCCACTAAGCCCAGCAGAAAT
AGCACACGTCTGAACTCTTTCCCTACACGACGCTCTTCCGATCTNNNTTGCTCAACTACAGCAATCCCAT
GAGCACACGTCTGAACTCTTTCCCTACACGACGCTCTTCCGATCTNNNTCAAGGTATTAGATGCACCCA
AGCACACGTCTGAACTCTTTCCCTACACGACGCTCTTCCGATCTNNNAAGTCAGGTGGTGGTTTTGTT
AGCACACGTCTGAACTCTTTCCCTACACGACGCTCTTCCGATCTNNNGACTTGCAGAACAGTAGGTGTC
AGCACACGTCTGAACTCTTTCCCTACACGACGCTCTTCCGATCTNNNACAGGCTTGTGGTGTATATGT
GAGCACACGTCTGAACTCTTTCCCTACACGACGCTCTTCCGATCTNNNCTCACTCTTTTCCCACTGCTGAA
AGCACACGTCTGAACTCTTTCCCTACACGACGCTCTTCCGATCTNNNTGGAGGAAGATGCTAGTAGAG
GAGCACACGTCTGAACTCTTTCCCTACACGACGCTCTTCCGATCTNNNATCTAACCAACCCTTCTCCCTA
GAGCACACGTCTGAACTCTTTCCCTACACGACGCTCTTCCGATCTNNNCCCAACAGACAGAGGTTGGA
AGCACACGTCTGAACTCTTTCCCTACACGACGCTCTTCCGATCTNNNATTTATCACTGCAAGCACCTCCA
GAGCACACGTCTGAACTCTTTCCCTACACGACGCTCTTCCGATCTNNNGGTCTATCTTTCACCCCTCCCTT
GAGCACACGTCTGAACTCTTTCCCTACACGACGCTCTTCCGATCTNNNCTGAACCTGAAAACAGGTCTG
AGCACACGTCTGAACTCTTTCCCTACACGACGCTCTTCCGATCTNNNACCAAGTGTTTGAATAGAAATA
AGCACACGTCTGAACTCTTTCCCTACACGACGCTCTTCCGATCTNNNATCTAAGTACCCCTTATACACGC
AGAGCACACGTCTGAACTCTTTCCCTACACGACGCTCTTCCGATCTNNNACTATGCTTTGCCTTCAACTC
AGCACACGTCTGAACTCTTTCCCTACACGACGCTCTTCCGATCTNNNGCTGAAGGCTGTTCTATGGACA
AGAGCACACGTCTGAACTCTTTCCCTACACGACGCTCTTCCGATCTNNNATGGCATTGGTGAGTTGTT
AGCACACGTCTGAACTCTTTCCCTACACGACGCTCTTCCGATCTNNNCCTCTGCATTAGTTTGGTCAGG
AGCACACGTCTGAACTCTTTCCCTACACGACGCTCTTCCGATCTNNNCCTTGGAAATATGTCCCCCTGT
AGCACACGTCTGAACTCTTTCCCTACACGACGCTCTTCCGATCTNNNTGTCCCAGAATGTAGGTCAATG
AGAGCACACGTCTGAACTCTTTCCCTACACGACGCTCTTCCGATCTNNNTCACCCGTCTTCTTCTAAGGT
GAGCACACGTCTGAACTCTTTCCCTACACGACGCTCTTCCGATCTNNNTGCTTGTATAATTCTGGGTGT
GAGCACACGTCTGAACTCTTTCCCTACACGACGCTCTTCCGATCTNNNATTGTTGGTAGGAATCACAG
GAGCACACGTCTGAACTCTTTCCCTACACGACGCTCTTCCGATCTNNNTAGGAGAGTGAGCTGGAAG
AGCACACGTCTGAACTCTTTCCCTACACGACGCTCTTCCGATCTNNNCAATGTGATGATCCAGGCACA
AGCACACGTCTGAACTCTTTCCCTACACGACGCTCTTCCGATCTNNNTGATGCCATAATCTCCAAAGCC
GAGCACACGTCTGAACTCTTTCCCTACACGACGCTCTTCCGATCTNNNCCACTTCCACACTCCTAACTCAC
AGAGCACACGTCTGAACTCTTTCCCTACACGACGCTCTTCCGATCTNNNGTGCTCCAGGACTTATCTTT
GAGCACACGTCTGAACTCTTTCCCTACACGACGCTCTTCCGATCTNNNTCCTCCTCTACCTTCCTGTAAG
GAGCACACGTCTGAACTCTTTCCCTACACGACGCTCTTCCGATCTNNNAAGCTGCTGTTTGGAGAGCTA
GAGCACACGTCTGAACTCTTTCCCTACACGACGCTCTTCCGATCTNNNTTCCTCATGATCTCTATTCCCC
AGAGCACACGTCTGAACTCTTTCCCTACACGACGCTCTTCCGATCTNNNGAGAGGAAAGGCCTGAAA
AAGAGCACACGTCTGAACTCTTTCCCTACACGACGCTCTTCCGATCTNNNAGCTTGATGGGGGATGTA
GAAGAGCACACGTCTGAACTCTTTCCCTACACGACGCTCTTCCGATCTNNNACTAAGTGCTAAGTCCCA
AGCACACGTCTGAACTCTTTCCCTACACGACGCTCTTCCGATCTNNNGCAGTATTTCTCACCAACTTGC
AGCACACGTCTGAACTCTTTCCCTACACGACGCTCTTCCGATCTNNNCTCCTGTGTCTCTTTGACAACCC
GAGCACACGTCTGAACTCTTTCCCTACACGACGCTCTTCCGATCTNNNGGTAAAAGTAGGGCTCTTGGA
GAGCACACGTCTGAACTCTTTCCCTACACGACGCTCTTCCGATCTNNNTTGAGAATTGAGGAGGAGG
AGCACACGTCTGAACTCTTTCCCTACACGACGCTCTTCCGATCTNNNAGAAGACCAATTATTGAAAAAC
GAGCACACGTCTGAACTCTTTCCCTACACGACGCTCTTCCGATCTNNNGGCCAATAACTACCTGGGGA

AGCACACGTCTGAACTCTTTCCCTACACGACGCTCTTCCGATCTNNNAACTTGTGGAGAAAAGCCAGA
CACACGTCTGAACTCTTTCCCTACACGACGCTCTTCCGATCTNNNAGTGTGCCATGGAGGAGAACTCA
GAGCACACGTCTGAACTCTTTCCCTACACGACGCTCTTCCGATCTNNNAGACCTTACCCTGTCATGCAG
AGAGCACACGTCTGAACTCTTTCCCTACACGACGCTCTTCCGATCTNNNGATTCTGACTGTAGTGCTTC
AGCACACGTCTGAACTCTTTCCCTACACGACGCTCTTCCGATCTNNNGCGGGTATATGCAACTTTGGG
GAGCACACGTCTGAACTCTTTCCCTACACGACGCTCTTCCGATCTNNNATGGCCAGAAGCAATGACTA
GAGCACACGTCTGAACTCTTTCCCTACACGACGCTCTTCCGATCTNNNTACCCTCTCAGCTTACAGCCAT
GCACACGTCTGAACTCTTTCCCTACACGACGCTCTTCCGATCTNNNAGAGCACACTATCCAGGTGATG
GAGCACACGTCTGAACTCTTTCCCTACACGACGCTCTTCCGATCTNNNCTTAGCCAGCCTGATGATGTC
GAGCACACGTCTGAACTCTTTCCCTACACGACGCTCTTCCGATCTNNNAATCAATGAAGGGAGGGAAA
GAGCACACGTCTGAACTCTTTCCCTACACGACGCTCTTCCGATCTNNNCTGAATGGTTCTCTGATCCTG
GAGCACACGTCTGAACTCTTTCCCTACACGACGCTCTTCCGATCTNNNGCTAGACTGGGCCTCCTACA
GAGCACACGTCTGAACTCTTTCCCTACACGACGCTCTTCCGATCTNNNCAATTGCCATTCGTTCCAGTG
AGCACACGTCTGAACTCTTTCCCTACACGACGCTCTTCCGATCTNNNTCTTTTTCTTCTTTCTCTCAAATT
AGAGCACACGTCTGAACTCTTTCCCTACACGACGCTCTTCCGATCTNNNGGCAAGTGTTTCAGATAGC
GAGCACACGTCTGAACTCTTTCCCTACACGACGCTCTTCCGATCTNNNGGGGATCAAACTGCCAACT
GAGCACACGTCTGAACTCTTTCCCTACACGACGCTCTTCCGATCTNNNCTTTGACAAATGTAGGGGCC
GCACACGTCTGAACTCTTTCCCTACACGACGCTCTTCCGATCTNNNCTTGATGCATGGGCTGAAATTG
GAGCACACGTCTGAACTCTTTCCCTACACGACGCTCTTCCGATCTNNNGCACAAAAGCACCTTCTCTCTC
GAGCACACGTCTGAACTCTTTCCCTACACGACGCTCTTCCGATCTNNNTTGGCCTTGAAGTGTGTATGG
AGCACACGTCTGAACTCTTTCCCTACACGACGCTCTTCCGATCTNNNGCTACAGCCTTCTAAGGAAACA
AGCACACGTCTGAACTCTTTCCCTACACGACGCTCTTCCGATCTNNNTTGTGCTTTTTCCCCAGTTTGT
AGCACACGTCTGAACTCTTTCCCTACACGACGCTCTTCCGATCTNNNACAAGGTGGAGGTGGATTAG
GAGCACACGTCTGAACTCTTTCCCTACACGACGCTCTTCCGATCTNNNGAAGGCTTTGCTGTCTGGAA
GCACACGTCTGAACTCTTTCCCTACACGACGCTCTTCCGATCTNNNCCCATTTCCTGCCTTCTACAACAC
AAGAGCACACGTCTGAACTCTTTCCCTACACGACGCTCTTCCGATCTNNNATGTGCAGATGTTCTGGG
AGCACACGTCTGAACTCTTTCCCTACACGACGCTCTTCCGATCTNNNGTAGAAAGAACTGACCAGCCA
AGCACACGTCTGAACTCTTTCCCTACACGACGCTCTTCCGATCTNNNCCAGAATTTTCTTCACTGTGGG
AGCACACGTCTGAACTCTTTCCCTACACGACGCTCTTCCGATCTNNNAAACGTAGTTCAGACCCAGAT
AGCACACGTCTGAACTCTTTCCCTACACGACGCTCTTCCGATCTNNNTCAACATGGTAAGTTGGGGAC
GAGCACACGTCTGAACTCTTTCCCTACACGACGCTCTTCCGATCTNNNATGGTACCCGTTGAGCCATTT
GAGCACACGTCTGAACTCTTTCCCTACACGACGCTCTTCCGATCTNNNGGGGCCTAGTGATGGTGAT
GAGCACACGTCTGAACTCTTTCCCTACACGACGCTCTTCCGATCTNNNAGAGATGAGACTTTGGAGT
AGCACACGTCTGAACTCTTTCCCTACACGACGCTCTTCCGATCTNNNTGGGTCCCATAAAGGGGAAT
GAGCACACGTCTGAACTCTTTCCCTACACGACGCTCTTCCGATCTNNNTAGCCATATTTGGAGGATG
GCACACGTCTGAACTCTTTCCCTACACGACGCTCTTCCGATCTNNNCCTGTGAAGGGACATTACTAAG
GCACACGTCTGAACTCTTTCCCTACACGACGCTCTTCCGATCTNNNACTAAGGTAACCAAATATCCCTC
GCACACGTCTGAACTCTTTCCCTACACGACGCTCTTCCGATCTNNNAGCACTGGCTGACATTTAGTAGC
AGCACACGTCTGAACTCTTTCCCTACACGACGCTCTTCCGATCTNNNAAATTCCCCTGAACTCACTGTG
GAGCACACGTCTGAACTCTTTCCCTACACGACGCTCTTCCGATCTNNNGCCAGGATAGGATGCACATT
GAGCACACGTCTGAACTCTTTCCCTACACGACGCTCTTCCGATCTNNNGCTTGATGGGGGATGTAGAG
AGCACACGTCTGAACTCTTTCCCTACACGACGCTCTTCCGATCTNNNGGGTCATGCATCTTTCAACAAC
AGCACACGTCTGAACTCTTTCCCTACACGACGCTCTTCCGATCTNNNGCACTGGCTCACTGAATGGAA

AGCACACGTCTGAACTCTTTCCCTACACGACGCTCTTCCGATCTNNNCCTGAGTTGGCCTTTGACTGAA
AGAGCACACGTCTGAACTCTTTCCCTACACGACGCTCTTCCGATCTNNNATGGATGGATAGTTGGGA
GAGCACACGTCTGAACTCTTTCCCTACACGACGCTCTTCCGATCTNNNACTTTTCCTCTTCCCTTTAGG
AAGAGCACACGTCTGAACTCTTTCCCTACACGACGCTCTTCCGATCTNNNTTGACCTGGCTTCATAGG
GAGCACACGTCTGAACTCTTTCCCTACACGACGCTCTTCCGATCTNNNTCCATGGGATGTATCAGTGA
AGCACACGTCTGAACTCTTTCCCTACACGACGCTCTTCCGATCTNNNACCACCTTACCCCAGTAACGTA
AGCACACGTCTGAACTCTTTCCCTACACGACGCTCTTCCGATCTNNNGCTTTATTGTCCTGGCCTCCT
GAGCACACGTCTGAACTCTTTCCCTACACGACGCTCTTCCGATCTNNNCCAATTCCCTGCTTCCCTTCA
AGCACACGTCTGAACTCTTTCCCTACACGACGCTCTTCCGATCTNNNCCCCACATGGTTTGAAAGAG
AGCACACGTCTGAACTCTTTCCCTACACGACGCTCTTCCGATCTNNNGATTAGGCATACGCCATCCTG
GCACACGTCTGAACTCTTTCCCTACACGACGCTCTTCCGATCTNNNCGTTGGTGGACATTTAGCTTCCT
AGCACACGTCTGAACTCTTTCCCTACACGACGCTCTTCCGATCTNNNTAACAGGTTGAATGCAGAGGC
AGCACACGTCTGAACTCTTTCCCTACACGACGCTCTTCCGATCTNNNGGCTCTTGCTTCCATACCTTCTC
AGCACACGTCTGAACTCTTTCCCTACACGACGCTCTTCCGATCTNNNTGATTGGGCCAGATTTTGTGT
GAGCACACGTCTGAACTCTTTCCCTACACGACGCTCTTCCGATCTNNNTCCTATCTGAATGGCTACGCT
AGCACACGTCTGAACTCTTTCCCTACACGACGCTCTTCCGATCTNNNTGCCAAAGCTCAAGTCTTCAAC
GAGCACACGTCTGAACTCTTTCCCTACACGACGCTCTTCCGATCTNNNGGACTTGTGGTTTGCTCTTC
GAGCACACGTCTGAACTCTTTCCCTACACGACGCTCTTCCGATCTNNNACCCCTCACACTGGTATGTTT
GCACACGTCTGAACTCTTTCCCTACACGACGCTCTTCCGATCTNNNTACACACACACATACAGAGCCAG
GAGCACACGTCTGAACTCTTTCCCTACACGACGCTCTTCCGATCTNNNCAGAACATAGCATCAGCACG
GAGCACACGTCTGAACTCTTTCCCTACACGACGCTCTTCCGATCTNNNCACCCACACACATACTCATGC
GCACACGTCTGAACTCTTTCCCTACACGACGCTCTTCCGATCTNNNCAAAACAGGAAATGGACAGTGG
AGAGCACACGTCTGAACTCTTTCCCTACACGACGCTCTTCCGATCTNNNGATTCAATAGCACACCAGC
GAGCACACGTCTGAACTCTTTCCCTACACGACGCTCTTCCGATCTNNNGAGCTCAAGTAGAGTGGTT
GAGCACACGTCTGAACTCTTTCCCTACACGACGCTCTTCCGATCTNNNGCAACACAATCCTCTTCATCCC
GAGCACACGTCTGAACTCTTTCCCTACACGACGCTCTTCCGATCTNNNGCCTGAATAGCCCATTTTTCT
GAGCACACGTCTGAACTCTTTCCCTACACGACGCTCTTCCGATCTNNNCTTGGCTCCATCTTCAGTCTA
AGAGCACACGTCTGAACTCTTTCCCTACACGACGCTCTTCCGATCTNNNATCTGCATTGTGCAACTTTC
AGAGCACACGTCTGAACTCTTTCCCTACACGACGCTCTTCCGATCTNNNATGTCTGCAGGCTACATTCA
AAGAGCACACGTCTGAACTCTTTCCCTACACGACGCTCTTCCGATCTNNNGGAACACAGAACCAAACC
GAGCACACGTCTGAACTCTTTCCCTACACGACGCTCTTCCGATCTNNNAACCTTTCCCCAGCAAAGAGA
AGAGCACACGTCTGAACTCTTTCCCTACACGACGCTCTTCCGATCTNNNGTGCTGAGTAAAACTGTGT
AGCACACGTCTGAACTCTTTCCCTACACGACGCTCTTCCGATCTNNNGAGGATCCCTTGGAATTCACCT
GCACACGTCTGAACTCTTTCCCTACACGACGCTCTTCCGATCTNNNTAAAACACCAACCTTTCACCCCAC
GCACACGTCTGAACTCTTTCCCTACACGACGCTCTTCCGATCTNNNTCACTGGCTCATTTCAACTGCATC
AGCACACGTCTGAACTCTTTCCCTACACGACGCTCTTCCGATCTNNNTTCTGCTCATATCCTGTGCTGT
GAGCACACGTCTGAACTCTTTCCCTACACGACGCTCTTCCGATCTNNNACTTTTTGAGTTTTTGCTTAG
GCACACGTCTGAACTCTTTCCCTACACGACGCTCTTCCGATCTNNNTTTGCTGTTGTGTTGGAGAGGT
AGCACACGTCTGAACTCTTTCCCTACACGACGCTCTTCCGATCTNNNGTAGAAAGCTGGGTGTTACAG
AGCACACGTCTGAACTCTTTCCCTACACGACGCTCTTCCGATCTNNNTCTCCTCTTTCTGGCTTGTGGAT
GAGCACACGTCTGAACTCTTTCCCTACACGACGCTCTTCCGATCTNNNTTATGGACTGGGAGGAAGA
GCACACGTCTGAACTCTTTCCCTACACGACGCTCTTCCGATCTNNNACCTCATGCAAGGCCAAATGTA
AGCACACGTCTGAACTCTTTCCCTACACGACGCTCTTCCGATCTNNNCCTTGGCAGTGACTTTGAAAC

GAGCACACGTCTGAACTCTTTCCCTACACGACGCTCTTCCGATCTNNNTGGCCATAAGCCAAATCCAAC
AGAGCACACGTCTGAACTCTTTCCCTACACGACGCTCTTCCGATCTNNNAGGCTACCACACACTTTTTA
AAGAGCACACGTCTGAACTCTTTCCCTACACGACGCTCTTCCGATCTNNNGTGCCATCTATGAACTTCA
GAAGAGCACACGTCTGAACTCTTTCCCTACACGACGCTCTTCCGATCTNNNCAAGAGGTGTCTCCTCCT
AGAGCACACGTCTGAACTCTTTCCCTACACGACGCTCTTCCGATCTNNNTCTCAATGCATGGTCACTGG
AGCACACGTCTGAACTCTTTCCCTACACGACGCTCTTCCGATCTNNNTCTGTGGCTAGATAGTATTCCA
AAGAGCACACGTCTGAACTCTTTCCCTACACGACGCTCTTCCGATCTNNNCTCCAGTCCTATTCTCCTTC
GAGCACACGTCTGAACTCTTTCCCTACACGACGCTCTTCCGATCTNNNCCACCGGATGAACTCCAAAAT
GAGCACACGTCTGAACTCTTTCCCTACACGACGCTCTTCCGATCTNNNGCACTGGCTCACTGAATGGA
AAGAGCACACGTCTGAACTCTTTCCCTACACGACGCTCTTCCGATCTNNNACAAAGAGGAGCTGGTAT
AGAGCACACGTCTGAACTCTTTCCCTACACGACGCTCTTCCGATCTNNNCTTGTGTCTTCCTGATCCCAC
AGCACACGTCTGAACTCTTTCCCTACACGACGCTCTTCCGATCTNNNTGTTGTTAGTGCATAGTTTGA
GAGCACACGTCTGAACTCTTTCCCTACACGACGCTCTTCCGATCTNNNCACTGAGTTTAGTTGGGGAT
AGCACACGTCTGAACTCTTTCCCTACACGACGCTCTTCCGATCTNNNGTACTGAAGGCCTGCTATGAAC
GAAGAGCACACGTCTGAACTCTTTCCCTACACGACGCTCTTCCGATCTNNNAGAAGTAAAAGCCTAAG
AGCACACGTCTGAACTCTTTCCCTACACGACGCTCTTCCGATCTNNNTCTTTGAAGGAAGCCTCAGCAT
GAGCACACGTCTGAACTCTTTCCCTACACGACGCTCTTCCGATCTNNNGATAGCGTCAGATCCAGACAC
AGCACACGTCTGAACTCTTTCCCTACACGACGCTCTTCCGATCTNNNTGGCTTTGTGATTTCTGCTTGA
GAGCACACGTCTGAACTCTTTCCCTACACGACGCTCTTCCGATCTNNNAACAGAGAGAAGGGAAGAG
AGCACACGTCTGAACTCTTTCCCTACACGACGCTCTTCCGATCTNNNGGACACTTCTGCTTCCTGATTG
GAGCACACGTCTGAACTCTTTCCCTACACGACGCTCTTCCGATCTNNNTCTGGGATGCTATCAAGAGTG
GAGCACACGTCTGAACTCTTTCCCTACACGACGCTCTTCCGATCTNNNAGGTTCTCTTGAGAGACAGA
GAGCACACGTCTGAACTCTTTCCCTACACGACGCTCTTCCGATCTNNNCTCTCCCCTTTTTCTGGCTACAC
GCACACGTCTGAACTCTTTCCCTACACGACGCTCTTCCGATCTNNNACTCATTTCACATGGTTTCCCGTC
GAGCACACGTCTGAACTCTTTCCCTACACGACGCTCTTCCGATCTNNNATGTTGGGAAACAGCTGGGT
AAGAGCACACGTCTGAACTCTTTCCCTACACGACGCTCTTCCGATCTNNNAGTCTCGACAGATAGGAT
AGAGCACACGTCTGAACTCTTTCCCTACACGACGCTCTTCCGATCTNNNCCACCCTACCTGTTCTCACCT
AGCACACGTCTGAACTCTTTCCCTACACGACGCTCTTCCGATCTNNNTGGCTGCTCAATTTACCACCTTA
GAGCACACGTCTGAACTCTTTCCCTACACGACGCTCTTCCGATCTNNNGCTCTCCCGGTTTTTGTTTTTG
AGAGCACACGTCTGAACTCTTTCCCTACACGACGCTCTTCCGATCTNNNGACGGCAGAATGAAACCTT
GCACACGTCTGAACTCTTTCCCTACACGACGCTCTTCCGATCTNNNGACCCTCCTGGAAACTCACCTCAT
AGCACACGTCTGAACTCTTTCCCTACACGACGCTCTTCCGATCTNNNGAGAAGTTTCCCCAAGGTTGG
AGCACACGTCTGAACTCTTTCCCTACACGACGCTCTTCCGATCTNNNTTCTTCTCCCATAGGGCTCTGTC
GCACACGTCTGAACTCTTTCCCTACACGACGCTCTTCCGATCTNNNTAATGTAGCGCATGACTGTGTG
AGCACACGTCTGAACTCTTTCCCTACACGACGCTCTTCCGATCTNNNCATGTGAGACAGCACCTTGTC
GAGCACACGTCTGAACTCTTTCCCTACACGACGCTCTTCCGATCTNNNAATTTCATGATGCGGAAGGC
GAGCACACGTCTGAACTCTTTCCCTACACGACGCTCTTCCGATCTNNNTCCATCTTCTCACATAGCTAG
AGAGCACACGTCTGAACTCTTTCCCTACACGACGCTCTTCCGATCTNNNCAAGAGTTTCCTGAGGCTA
AGCACACGTCTGAACTCTTTCCCTACACGACGCTCTTCCGATCTNNNTAGCTAGCAGCAGGATCAACTC
AGCACACGTCTGAACTCTTTCCCTACACGACGCTCTTCCGATCTNNNGCATTCCTGCAGCTCTAGTTTC
AGCACACGTCTGAACTCTTTCCCTACACGACGCTCTTCCGATCTNNNAACCTACTCCAAGCATCCTTCCT
GAGCACACGTCTGAACTCTTTCCCTACACGACGCTCTTCCGATCTNNNAGGCCTGAAGTATTGCTGACC
AGCACACGTCTGAACTCTTTCCCTACACGACGCTCTTCCGATCTNNNTGTTCCCCTCCTTCTTTTTCTGCC

AGCACACGTCTGAACTCTTTCCCTACACGACGCTCTTCCGATCTNNNCCTTCGCTTCTGTTCAATGCTT
AGCACACGTCTGAACTCTTTCCCTACACGACGCTCTTCCGATCTNNNACTATGCTTTGCCTTCAACTCCG
AGCACACGTCTGAACTCTTTCCCTACACGACGCTCTTCCGATCTNNNGCACACTTACCTCTCACTACAG
GCACACGTCTGAACTCTTTCCCTACACGACGCTCTTCCGATCTNNNTGGGGTCATGAGAAGACAAAT
GCACACGTCTGAACTCTTTCCCTACACGACGCTCTTCCGATCTNNNGCTGAAGGCTGTTCTATGGACA
AGCACACGTCTGAACTCTTTCCCTACACGACGCTCTTCCGATCTNNNAGCAGGAAATCCTGTGCATTC
GAGCACACGTCTGAACTCTTTCCCTACACGACGCTCTTCCGATCTNNNTGGGAAGCTATAATTGGAG
GAGCACACGTCTGAACTCTTTCCCTACACGACGCTCTTCCGATCTNNNCAGGGCTCAAAGAGAGTAA
GCACACGTCTGAACTCTTTCCCTACACGACGCTCTTCCGATCTNNNCACAGATGATCCACGTGTTGCT
GAAGAGCACACGTCTGAACTCTTTCCCTACACGACGCTCTTCCGATCTNNNTATGTGTGGTGCAAGAA
GCACACGTCTGAACTCTTTCCCTACACGACGCTCTTCCGATCTNNNGTCCTGTAGGATCATGCACACT
GAGCACACGTCTGAACTCTTTCCCTACACGACGCTCTTCCGATCTNNNCAGAAAATGCTGCCTTTCCC
GAGCACACGTCTGAACTCTTTCCCTACACGACGCTCTTCCGATCTNNNTTCCAGCCTCTCAGCAAAATA
GCACACGTCTGAACTCTTTCCCTACACGACGCTCTTCCGATCTNNNGCCCAGAGACAACTTGATTGTGA
GCACACGTCTGAACTCTTTCCCTACACGACGCTCTTCCGATCTNNNGCAAGGACAAAAGTAACACAG
AGCACACGTCTGAACTCTTTCCCTACACGACGCTCTTCCGATCTNNNAAACAACCAACCAACCACCCAA
GCACACGTCTGAACTCTTTCCCTACACGACGCTCTTCCGATCTNNNAACATGGTGTTCTCAGAGGCA
GAGCACACGTCTGAACTCTTTCCCTACACGACGCTCTTCCGATCTNNNTTCTTCTCCCATAGGGCTCTG
AGCACACGTCTGAACTCTTTCCCTACACGACGCTCTTCCGATCTNNNTTTGACATATGAAGGGGTGT
GAGCACACGTCTGAACTCTTTCCCTACACGACGCTCTTCCGATCTNNNAACTGTAGGATACCCCTTGC
GAGCACACGTCTGAACTCTTTCCCTACACGACGCTCTTCCGATCTNNNCATCTCAGAGGGTGGGTGGA
GCACACGTCTGAACTCTTTCCCTACACGACGCTCTTCCGATCTNNNTGCGGATACAAAACCTTCTAGC
AGCACACGTCTGAACTCTTTCCCTACACGACGCTCTTCCGATCTNNNAGTGTTGGGGTAGTTTAGCAA
AGCACACGTCTGAACTCTTTCCCTACACGACGCTCTTCCGATCTNNNGTCACCTGCACACATACAGACA
GAGCACACGTCTGAACTCTTTCCCTACACGACGCTCTTCCGATCTNNNCCTTATTCTTCCTCCAAGGTGG
GAGCACACGTCTGAACTCTTTCCCTACACGACGCTCTTCCGATCTNNNTGAATCTCACCATTGATGGG
GAGCACACGTCTGAACTCTTTCCCTACACGACGCTCTTCCGATCTNNNTCAGTTGGATGTAGTGGTCC
AGCACACGTCTGAACTCTTTCCCTACACGACGCTCTTCCGATCTNNNAATACGTGCCCATTGATGGAT
AAGAGCACACGTCTGAACTCTTTCCCTACACGACGCTCTTCCGATCTNNNTCCATCTTCTCACATAGCT
GAGCACACGTCTGAACTCTTTCCCTACACGACGCTCTTCCGATCTNNNGAGGGAAAGGAGGTGAGAG
GCACACGTCTGAACTCTTTCCCTACACGACGCTCTTCCGATCTNNNCTAGTAAGCAGTGGACTGGCA
GAGCACACGTCTGAACTCTTTCCCTACACGACGCTCTTCCGATCTNNNCTACATCCCAACTGAGGGGT
GAGCACACGTCTGAACTCTTTCCCTACACGACGCTCTTCCGATCTNNNCAGGAAGGCTTCATCCTGCA
GAGCACACGTCTGAACTCTTTCCCTACACGACGCTCTTCCGATCTNNNGCTTACCCTGCTGGAATATGC
GAGCACACGTCTGAACTCTTTCCCTACACGACGCTCTTCCGATCTNNNACAGGAATTCACATGAGAGA
AGAGCACACGTCTGAACTCTTTCCCTACACGACGCTCTTCCGATCTNNNACATTTAGTCAGTGAGGG
GAGCACACGTCTGAACTCTTTCCCTACACGACGCTCTTCCGATCTNNNTGTATACACAAAACTGTATA
AGCACACGTCTGAACTCTTTCCCTACACGACGCTCTTCCGATCTNNNTGGTGTCTCCATGACCTACCA
AAGAGCACACGTCTGAACTCTTTCCCTACACGACGCTCTTCCGATCTNNNCAAGACACACAGTAGGCA
AGCACACGTCTGAACTCTTTCCCTACACGACGCTCTTCCGATCTNNNAGGCTGTTGCAGAGTAGAAT
GAGCACACGTCTGAACTCTTTCCCTACACGACGCTCTTCCGATCTNNNTATTCATAAGCCAGCCCTGTG
GAGCACACGTCTGAACTCTTTCCCTACACGACGCTCTTCCGATCTNNNATTAGCCTTGGCGATCTTGT
AGCACACGTCTGAACTCTTTCCCTACACGACGCTCTTCCGATCTNNNATACCTAGTGGCTCTCTCCCTC

GAGCACACGTCTGAACTCTTTCCCTACACGACGCTCTTCCGATCTNNNAATATCACAGGGCACCCACA
AGCACACGTCTGAACTCTTTCCCTACACGACGCTCTTCCGATCTNNNTCCTTTGAGGAAAAAGACCC
AGCACACGTCTGAACTCTTTCCCTACACGACGCTCTTCCGATCTNNNCCTGCTGTACTTCCCCTGTTTT
GAGCACACGTCTGAACTCTTTCCCTACACGACGCTCTTCCGATCTNNNGGAATCAGTTCAGCAAGAG
AGCACACGTCTGAACTCTTTCCCTACACGACGCTCTTCCGATCTNNNTCGTCTCTTGTATCCTACACGC
AGCACACGTCTGAACTCTTTCCCTACACGACGCTCTTCCGATCTNNNAGGACAGCTTCCTCATTCTTT
AGCACACGTCTGAACTCTTTCCCTACACGACGCTCTTCCGATCTNNNCAATTTCACTCCGTGGTCTTCC
GAGCACACGTCTGAACTCTTTCCCTACACGACGCTCTTCCGATCTNNNCTGTGGGGAACCAAAGAGT
GAGCACACGTCTGAACTCTTTCCCTACACGACGCTCTTCCGATCTNNNTAAACTTTGCCAGAACTAGC
AGCACACGTCTGAACTCTTTCCCTACACGACGCTCTTCCGATCTNNNTACCTGAAGATAGTGCCAGTC
GAGCACACGTCTGAACTCTTTCCCTACACGACGCTCTTCCGATCTNNNGATGGGAAATGCTTGCCTGT
AAGAGCACACGTCTGAACTCTTTCCCTACACGACGCTCTTCCGATCTNNNGAGGTTGCAGTGAGACG
GCACACGTCTGAACTCTTTCCCTACACGACGCTCTTCCGATCTNNNTAACCATGGGAATTTCCTGGA
GAGCACACGTCTGAACTCTTTCCCTACACGACGCTCTTCCGATCTNNNTGCTTATAATTTTGTCAGCG
AGCACACGTCTGAACTCTTTCCCTACACGACGCTCTTCCGATCTNNNGCAGGAGAATGCTTGATCCC
AGCACACGTCTGAACTCTTTCCCTACACGACGCTCTTCCGATCTNNNGGATTATTTGGTCCCTCTGCC
GAAGAGCACACGTCTGAACTCTTTCCCTACACGACGCTCTTCCGATCTNNNGTACTCTGTGAGTGTGT
GAGCACACGTCTGAACTCTTTCCCTACACGACGCTCTTCCGATCTNNNCAACTCCAGCAGAGAAAGGG
AGCACACGTCTGAACTCTTTCCCTACACGACGCTCTTCCGATCTNNNGGCATCAGGTTTTCTTTCATG
AGCACACGTCTGAACTCTTTCCCTACACGACGCTCTTCCGATCTNNNTAAACACTAAAACCTCCGTGG
GCACACGTCTGAACTCTTTCCCTACACGACGCTCTTCCGATCTNNNGCCTGGAGCATCTTATAGTGCA
GCACACGTCTGAACTCTTTCCCTACACGACGCTCTTCCGATCTNNNACACAAAGAATGCACAAGCATG
AGCACACGTCTGAACTCTTTCCCTACACGACGCTCTTCCGATCTNNNGGGCAAAGCTTATCTACTGGC
AGCACACGTCTGAACTCTTTCCCTACACGACGCTCTTCCGATCTNNNATCAGGGTGATGTGTGTGTG
GAGCACACGTCTGAACTCTTTCCCTACACGACGCTCTTCCGATCTNNNCATTCACTGCTCCACATCATC
AGCACACGTCTGAACTCTTTCCCTACACGACGCTCTTCCGATCTNNNACCTCCAAGTCTCTCAGCTTCAT
GCACACGTCTGAACTCTTTCCCTACACGACGCTCTTCCGATCTNNNAAAAGGTGGGGGACCACTACAT
AGCACACGTCTGAACTCTTTCCCTACACGACGCTCTTCCGATCTNNNCACCCACCTTTTGGGTTGTTAC
GCACACGTCTGAACTCTTTCCCTACACGACGCTCTTCCGATCTNNNATCAGGGTGATGTGTGTGTGT
GCACACGTCTGAACTCTTTCCCTACACGACGCTCTTCCGATCTNNNTTTGGGGTAAAAATGCGTGTG
AGCACACGTCTGAACTCTTTCCCTACACGACGCTCTTCCGATCTNNNCTGCCTCACACTTACTGTGGAA
GAGCACACGTCTGAACTCTTTCCCTACACGACGCTCTTCCGATCTNNNAGAATGTCTCCAAGGCTGTT
GAGCACACGTCTGAACTCTTTCCCTACACGACGCTCTTCCGATCTNNNCCTTACATAAATTTTGCAGC
AGCACACGTCTGAACTCTTTCCCTACACGACGCTCTTCCGATCTNNNTGATTTTGACTGGGGACATTG
GCACACGTCTGAACTCTTTCCCTACACGACGCTCTTCCGATCTNNNACAGAGCTATATGCCTCCCAGA
GAGCACACGTCTGAACTCTTTCCCTACACGACGCTCTTCCGATCTNNNACCAAAATCCAGATGAGAGG
AGCACACGTCTGAACTCTTTCCCTACACGACGCTCTTCCGATCTNNNTTCTTTCCTGCTTCAGTTGTCCC
AGCACACGTCTGAACTCTTTCCCTACACGACGCTCTTCCGATCTNNNTCTTTCAATGATTTCTGGTGTG
GCACACGTCTGAACTCTTTCCCTACACGACGCTCTTCCGATCTNNNATCAAGGAAATAAAAGAGGAC
AGCACACGTCTGAACTCTTTCCCTACACGACGCTCTTCCGATCTNNNACACTCTGCTCAACTTTGCCTTT
GAGCACACGTCTGAACTCTTTCCCTACACGACGCTCTTCCGATCTNNNTTTTTGGCATGGTCTGCCTT
AGCACACGTCTGAACTCTTTCCCTACACGACGCTCTTCCGATCTNNNTTCTCCAGCTCTCAGTCCTTCA
AGCACACGTCTGAACTCTTTCCCTACACGACGCTCTTCCGATCTNNNATGTCTGGCACTTTTGGGTATG

AGCACACGTCTGAACTCTTTCCCTACACGACGCTCTTCCGATCTNNNTCAAAAGGGAGGAAGGGAGA

AGCACACGTCTGAACTCTTTCCCTACACGACGCTCTTCCGATCTNNNTTGTTGCACTGTCCTGAATGTG

AGCACACGTCTGAACTCTTTCCCTACACGACGCTCTTCCGATCTNNNAGTATACTACGCTGCTTCCCAG

GCACACGTCTGAACTCTTTCCCTACACGACGCTCTTCCGATCTNNNTATTTGCATGGGCTCCTTAGCTG

GAGCACACGTCTGAACTCTTTCCCTACACGACGCTCTTCCGATCTNNNTAGGACCTTGCTCTTACCCAC

GAGCACACGTCTGAACTCTTTCCCTACACGACGCTCTTCCGATCTNNNCCCAATAACCTGATGCACATG

AGCACACGTCTGAACTCTTTCCCTACACGACGCTCTTCCGATCTNNNAGATACCTAGCTTTGTGCGTG

GAGCACACGTCTGAACTCTTTCCCTACACGACGCTCTTCCGATCTNNNCACAGGGTGGGAACAAGAT

GCACACGTCTGAACTCTTTCCCTACACGACGCTCTTCCGATCTNNNGTTGTCTTTGCAAGTTCTTGGC

AGCACACGTCTGAACTCTTTCCCTACACGACGCTCTTCCGATCTNNNCAAACTCCATCCCATCCTTCCT

GCACACGTCTGAACTCTTTCCCTACACGACGCTCTTCCGATCTNNNGGGCAAGAATATGGCTGTGCTA

AGCACACGTCTGAACTCTTTCCCTACACGACGCTCTTCCGATCTNNNAACTTTGGGGAATACCAACCC

GAGCACACGTCTGAACTCTTTCCCTACACGACGCTCTTCCGATCTNNNTAATTTGCACCTAGTCCTGTG

AGCACACGTCTGAACTCTTTCCCTACACGACGCTCTTCCGATCTNNNACTTCTGAACTCAATCAAATAC

ACACGTCTGAACTCTTTCCCTACACGACGCTCTTCCGATCTNNNTCTCTAGCCTGAAAGAGTGCTGTC

AGCACACGTCTGAACTCTTTCCCTACACGACGCTCTTCCGATCTNNNTGGCTCCCCTGATAAACCATCT

GCACACGTCTGAACTCTTTCCCTACACGACGCTCTTCCGATCTNNNACGCACATACATATATACCTGCT

AGCACACGTCTGAACTCTTTCCCTACACGACGCTCTTCCGATCTNNNAACATCTTCCAGATTAGTGCG

GAGCACACGTCTGAACTCTTTCCCTACACGACGCTCTTCCGATCTNNNTAAACACTAAAACCTCCGTG

GAGCACACGTCTGAACTCTTTCCCTACACGACGCTCTTCCGATCTNNNTGTTCAAGAGGAGTTTTGG

GCACACGTCTGAACTCTTTCCCTACACGACGCTCTTCCGATCTNNNTGGCTGGGGCAATTAGAAAGA

AGCACACGTCTGAACTCTTTCCCTACACGACGCTCTTCCGATCTNNNGGGCCTCATTCAATCAGAGAA

GAGCACACGTCTGAACTCTTTCCCTACACGACGCTCTTCCGATCTNNNCAGTGGTGGTAGCAACAGA

AGCACACGTCTGAACTCTTTCCCTACACGACGCTCTTCCGATCTNNNCTGTGCGTACAGCTACTCTCAT

GCACACGTCTGAACTCTTTCCCTACACGACGCTCTTCCGATCTNNNCCAATGCTACTAGATAGGGGG

GAGCACACGTCTGAACTCTTTCCCTACACGACGCTCTTCCGATCTNNNCAGAGCCGTAATGAAGACAC

AGCACACGTCTGAACTCTTTCCCTACACGACGCTCTTCCGATCTNNNTACTCAGCACACACAAACACA

AGCACACGTCTGAACTCTTTCCCTACACGACGCTCTTCCGATCTNNNTCAAAGACATGAAGATAGTC

GCACACGTCTGAACTCTTTCCCTACACGACGCTCTTCCGATCTNNNTGATCTCGTATCTTGTGATTTTT

AGCACACGTCTGAACTCTTTCCCTACACGACGCTCTTCCGATCTNNNTGTCTCAAAGAGCCTTTCTCTG

AGCACACGTCTGAACTCTTTCCCTACACGACGCTCTTCCGATCTNNNGGCACTTTCAAAAGTCATGCT

GCACACGTCTGAACTCTTTCCCTACACGACGCTCTTCCGATCTNNNCTCTCCCCTTTTTCTGGCTACAC

AGCACACGTCTGAACTCTTTCCCTACACGACGCTCTTCCGATCTNNNGGCAACTTCCTCATGCTTTTT

AGAGCACACGTCTGAACTCTTTCCCTACACGACGCTCTTCCGATCTNNNCCGGCCTCGTGTATTGATTT

AGAGCACACGTCTGAACTCTTTCCCTACACGACGCTCTTCCGATCTNNNAGTCATGCTGCAAAGTTAT

AGAGCACACGTCTGAACTCTTTCCCTACACGACGCTCTTCCGATCTNNNCCATCTTTGACGGCAAACT

AAGAGCACACGTCTGAACTCTTTCCCTACACGACGCTCTTCCGATCTNNNGTTGTGTGGTGTGCAAT

GAGCACACGTCTGAACTCTTTCCCTACACGACGCTCTTCCGATCTNNNAGGAAGGGGCTCCGTTAGA

AGAGCACACGTCTGAACTCTTTCCCTACACGACGCTCTTCCGATCTNNNATAGCATGTGGCTACTGCA

GCACACGTCTGAACTCTTTCCCTACACGACGCTCTTCCGATCTNNNCCCTCCTGCATCACTCATTTTCA

GCACACGTCTGAACTCTTTCCCTACACGACGCTCTTCCGATCTNNNATGACAAATGAGCTTCTGGGC

AGCACACGTCTGAACTCTTTCCCTACACGACGCTCTTCCGATCTNNNGATTGGGACCTCAAAGTGAG

GCACACGTCTGAACTCTTTCCCTACACGACGCTCTTCCGATCTNNNTAGTTAGAGCTGCCTGGCTAA

GAGCACACGTCTGAACTCTTTCCCTACACGACGCTCTTCCGATCTNNNGGCTGCTGTTCTTACACAAT

GAGCACACGTCTGAACTCTTTCCCTACACGACGCTCTTCCGATCTNNNGTTGGCATGCAAGGATGTAT

GCACACGTCTGAACTCTTTCCCTACACGACGCTCTTCCGATCTNNNAAGGAGAAGGAAGGACTGGG

GCACACGTCTGAACTCTTTCCCTACACGACGCTCTTCCGATCTNNNAGGAGCAATGGAGCAGATACT

AGCACACGTCTGAACTCTTTCCCTACACGACGCTCTTCCGATCTNNNCGGATTCAGAATAGAGGGAG

AGAGCACACGTCTGAACTCTTTCCCTACACGACGCTCTTCCGATCTNNNAGTTGGATTGGAAGGTGG

ACACGTCTGAACTCTTTCCCTACACGACGCTCTTCCGATCTNNNGTGGCTATGCCTTCGGAGTTTCCAT

GCACACGTCTGAACTCTTTCCCTACACGACGCTCTTCCGATCTNNNGAAATCATCTACTCCATCCGCCC

AGCACACGTCTGAACTCTTTCCCTACACGACGCTCTTCCGATCTNNNAGGATTTCAATGCCAAAGTCCC

AGCACACGTCTGAACTCTTTCCCTACACGACGCTCTTCCGATCTNNNGCCAAAGATCAGGGAACTCAT

GAGCACACGTCTGAACTCTTTCCCTACACGACGCTCTTCCGATCTNNNTGGAAAACTGAGGGATTC

AGCACACGTCTGAACTCTTTCCCTACACGACGCTCTTCCGATCTNNNGGCATCTATTCCAAGAGGTAG

AGCACACGTCTGAACTCTTTCCCTACACGACGCTCTTCCGATCTNNNTAGATCTCTTCCTCCAACCCCTG

AGCACACGTCTGAACTCTTTCCCTACACGACGCTCTTCCGATCTNNNTGTTGAGCTGGTATGCAGAAA

AGCACACGTCTGAACTCTTTCCCTACACGACGCTCTTCCGATCTNNNTCGATGGCACCAACTATCTGA

AAGAGCACACGTCTGAACTCTTTCCCTACACGACGCTCTTCCGATCTNNNAGGCGGGAGGGTCACAT

GAGCACACGTCTGAACTCTTTCCCTACACGACGCTCTTCCGATCTNNNTGTGAGGAAGCTACTCCAAG

GCACACGTCTGAACTCTTTCCCTACACGACGCTCTTCCGATCTNNNTTCTTGCTGAATTCAAGCTGGGT

AGCACACGTCTGAACTCTTTCCCTACACGACGCTCTTCCGATCTNNNTGGTTTCCTGCCTTCAGAACAT

AGCACACGTCTGAACTCTTTCCCTACACGACGCTCTTCCGATCTNNNGAACTTAGAGTGGTAAGCTG

AGCACACGTCTGAACTCTTTCCCTACACGACGCTCTTCCGATCTNNNAGCTAACCTAAGAGAGCAGG

AGAGCACACGTCTGAACTCTTTCCCTACACGACGCTCTTCCGATCTNNNTCAGCGATGCATAACTCTGT

GAGCACACGTCTGAACTCTTTCCCTACACGACGCTCTTCCGATCTNNNGCAGTCTTTTGTTTGCTGAG

AGCACACGTCTGAACTCTTTCCCTACACGACGCTCTTCCGATCTNNNCCCTTTCTCTAGCCTCTTCACCT

GCACACGTCTGAACTCTTTCCCTACACGACGCTCTTCCGATCTNNNCTCCAGCAATTCTTCCCCATCTCC

GAGCACACGTCTGAACTCTTTCCCTACACGACGCTCTTCCGATCTNNNCCAACTGACTTCCTGCATCAA

AGCACACGTCTGAACTCTTTCCCTACACGACGCTCTTCCGATCTNNNCCTCTGAGGTTCCTTTTGGTA

AGAGCACACGTCTGAACTCTTTCCCTACACGACGCTCTTCCGATCTNNNAAAAAGCTCCACTTTTTGAG

GCACACGTCTGAACTCTTTCCCTACACGACGCTCTTCCGATCTNNNAGGGCCTCACTCAGCTTATAGAT

AGAGCACACGTCTGAACTCTTTCCCTACACGACGCTCTTCCGATCTNNNTTCTATCTGCAGTGCCAAT

GAGCACACGTCTGAACTCTTTCCCTACACGACGCTCTTCCGATCTNNNAGGCCAGACTGGAAGAAGA

AAGAGCACACGTCTGAACTCTTTCCCTACACGACGCTCTTCCGATCTNNNACTGGAAAATCTCCAAG

GAGCACACGTCTGAACTCTTTCCCTACACGACGCTCTTCCGATCTNNNGCCAGGATTTTTATCTGTGT

GCACACGTCTGAACTCTTTCCCTACACGACGCTCTTCCGATCTNNNCATGCTGTTCTCATGGTGGAGTT

CACACGTCTGAACTCTTTCCCTACACGACGCTCTTCCGATCTNNNGTCTTTGATCTGCCATTCTGGGAC

AGAGCACACGTCTGAACTCTTTCCCTACACGACGCTCTTCCGATCTNNNTGAGCCCACCACTAAGAGA

GAGCACACGTCTGAACTCTTTCCCTACACGACGCTCTTCCGATCTNNNATTCTCTGCAGTGTCCTTTGG

GAGCACACGTCTGAACTCTTTCCCTACACGACGCTCTTCCGATCTNNNCCAATGCTGTTCCCTTCTCTT

AGCACACGTCTGAACTCTTTCCCTACACGACGCTCTTCCGATCTNNNGCATTTCCAAAGCATGTTGTGT

AGCACACGTCTGAACTCTTTCCCTACACGACGCTCTTCCGATCTNNNAATCAATGGCTCAGCAGGAAG

GAGCACACGTCTGAACTCTTTCCCTACACGACGCTCTTCCGATCTNNNATAATGGCAGGGAAAAAGG

AGCACACGTCTGAACTCTTTCCCTACACGACGCTCTTCCGATCTNNNAACAATATTCTCAGGGCATCGG

GAGCACACGTCTGAACTCTTTCCCTACACGACGCTCTTCCGATCTNNNTCAGCGATGCATAACTCTGT

AGCACACGTCTGAACTCTTTCCCTACACGACGCTCTTCCGATCTNNNCTATGCTCATTGGCGTAGCACA

AGCACACGTCTGAACTCTTTCCCTACACGACGCTCTTCCGATCTNNNACAACCAGGGCACAGGGAATC

GCACACGTCTGAACTCTTTCCCTACACGACGCTCTTCCGATCTNNNTTTCTCTCCTCAACCCATCGTCCA

GCACACGTCTGAACTCTTTCCCTACACGACGCTCTTCCGATCTNNNGTTCAGCTGTCATTGCTCTCTGTC

AGCACACGTCTGAACTCTTTCCCTACACGACGCTCTTCCGATCTNNNTTTAGGGTCTCTCATGCTGCCAT

AGCACACGTCTGAACTCTTTCCCTACACGACGCTCTTCCGATCTNNNGCATGCAATTTTCTGCAAAGCT

AGCACACGTCTGAACTCTTTCCCTACACGACGCTCTTCCGATCTNNNTGGGACTAAATTCCTCCCCTCA

GAGCACACGTCTGAACTCTTTCCCTACACGACGCTCTTCCGATCTNNNCTGTTCATGTACACACACCCA

AGCACACGTCTGAACTCTTTCCCTACACGACGCTCTTCCGATCTNNNATCTGGAGCTAAAATGAGCCT

AGAGCACACGTCTGAACTCTTTCCCTACACGACGCTCTTCCGATCTNNNTCTGTTGGCTTGATTTTCAT

AGCACACGTCTGAACTCTTTCCCTACACGACGCTCTTCCGATCTNNNAGACTGCTGGATCAGGTGAAAT

AGAGCACACGTCTGAACTCTTTCCCTACACGACGCTCTTCCGATCTNNNAACACAGAGAGCTGGTTTCT

AGAGCACACGTCTGAACTCTTTCCCTACACGACGCTCTTCCGATCTNNNTGGGTTGATAGACCCTTAGA

AAGAGCACACGTCTGAACTCTTTCCCTACACGACGCTCTTCCGATCTNNNACTAAGTGCTAAGTCCCA

AGCACACGTCTGAACTCTTTCCCTACACGACGCTCTTCCGATCTNNNACACCTGGGAATTAGAAGGGGA

AGCACACGTCTGAACTCTTTCCCTACACGACGCTCTTCCGATCTNNNTATTCCAGTTTTGGGTTGTGG

AGCACACGTCTGAACTCTTTCCCTACACGACGCTCTTCCGATCTNNNCACAACAGGATGCAAGAAAAC

GCACACGTCTGAACTCTTTCCCTACACGACGCTCTTCCGATCTNNNAAAAGAGAACCCAGAAGCAGCA

AGAGCACACGTCTGAACTCTTTCCCTACACGACGCTCTTCCGATCTNNNAAACTCAGAGGAAGGATGT

AGCACACGTCTGAACTCTTTCCCTACACGACGCTCTTCCGATCTNNNCTGCCACAGAAGCAAATTCACTG

GAGCACACGTCTGAACTCTTTCCCTACACGACGCTCTTCCGATCTNNNTGCTTTGAACACAGGTGGAT

GAGCACACGTCTGAACTCTTTCCCTACACGACGCTCTTCCGATCTNNNTGTGTGTGTAAGCAGGAA

AGCACACGTCTGAACTCTTTCCCTACACGACGCTCTTCCGATCTNNNTATGGAGCTCAGAGAGGTGAA

AGCACACGTCTGAACTCTTTCCCTACACGACGCTCTTCCGATCTNNNCTGCCTTCAAAGGCCTACAATCT

AGCACACGTCTGAACTCTTTCCCTACACGACGCTCTTCCGATCTNNNAGGCCTTTTCCATACTCACACA

GAGCACACGTCTGAACTCTTTCCCTACACGACGCTCTTCCGATCTNNNGGCAATGTTTGCCATGAAAG

AGCACACGTCTGAACTCTTTCCCTACACGACGCTCTTCCGATCTNNNAACAGGAAATTTCTCCCAAGG

GAAGAGCACACGTCTGAACTCTTTCCCTACACGACGCTCTTCCGATCTNNNCACTTTTTCCTCCCGCTTT

GAGCACACGTCTGAACTCTTTCCCTACACGACGCTCTTCCGATCTNNNTGGATCATGGAATCACTTTCT

AGCACACGTCTGAACTCTTTCCCTACACGACGCTCTTCCGATCTNNNGGAAGTGAACACGGCTGATTA

GAGCACACGTCTGAACTCTTTCCCTACACGACGCTCTTCCGATCTNNNTTCTTCCCTCCCACACTCTGTA

GCACACGTCTGAACTCTTTCCCTACACGACGCTCTTCCGATCTNNNCGGATTCAGAATAGAGGGAGG

GAGCACACGTCTGAACTCTTTCCCTACACGACGCTCTTCCGATCTNNNTGTGCCCGGTATGGTTTTCAAT

GCACACGTCTGAACTCTTTCCCTACACGACGCTCTTCCGATCTNNNTGCAATCATTTTCATGACAGGG

GCACACGTCTGAACTCTTTCCCTACACGACGCTCTTCCGATCTNNNTGTCAGGCAAAAATAGCAAAAA

AGCACACGTCTGAACTCTTTCCCTACACGACGCTCTTCCGATCTNNNAAATCAGGTTCCTTCTCCCCAGA

AGCACACGTCTGAACTCTTTCCCTACACGACGCTCTTCCGATCTNNNCCCTTTCTCTAGCCTCTTCACCTC

AGCACACGTCTGAACTCTTTCCCTACACGACGCTCTTCCGATCTNNNCACCATGTAATCTCCCAAGAGCA

GCACACGTCTGAACTCTTTCCCTACACGACGCTCTTCCGATCTNNNGCCAAAGCAATGGTACCAGAGT

AGCACACGTCTGAACTCTTTCCCTACACGACGCTCTTCCGATCTNNNATGAGGAAGAAAAGGGGGA

GAGCACACGTCTGAACTCTTTCCCTACACGACGCTCTTCCGATCTNNNCTGTGGGGAACCAAAGAGTG

AGCACACGTCTGAACTCTTTCCCTACACGACGCTCTTCCGATCTNNNACATTCATTTCAGGCACACAGG

AGCACACGTCTGAACTCTTTCCCTACACGACGCTCTTCCGATCTNNNATGCTGTGTTCTCCATCTTTGCT

GCACACGTCTGAACTCTTTCCCTACACGACGCTCTTCCGATCTNNNAGAAGCTAGGAAGCCATCTGG
GAGCACACGTCTGAACTCTTTCCCTACACGACGCTCTTCCGATCTNNNCCAGATCCAAGCCAGACAGA
AGAGCACACGTCTGAACTCTTTCCCTACACGACGCTCTTCCGATCTNNNCCTCTCAACTACACAACCTA
AGCACACGTCTGAACTCTTTCCCTACACGACGCTCTTCCGATCTNNNTGCCTGACCATTCAAATCCATG
AGCACACGTCTGAACTCTTTCCCTACACGACGCTCTTCCGATCTNNNCTTCCCTCCCTTCTCCTTTTCTGC
GCACACGTCTGAACTCTTTCCCTACACGACGCTCTTCCGATCTNNNGGGTGGAATGTTCACACACACT
AGCACACGTCTGAACTCTTTCCCTACACGACGCTCTTCCGATCTNNNATCACACCAAACACCAGCAGA
AGCACACGTCTGAACTCTTTCCCTACACGACGCTCTTCCGATCTNNNCCATGTGAATTGAAGAGGGG
AGCACACGTCTGAACTCTTTCCCTACACGACGCTCTTCCGATCTNNNCTCCAGCAATTCTTCCCCATCT
GCACACGTCTGAACTCTTTCCCTACACGACGCTCTTCCGATCTNNNCCTGTAGTAAGAACCTAGCGC
GCACACGTCTGAACTCTTTCCCTACACGACGCTCTTCCGATCTNNNCACACCTTGCTGGTTCCTTCAGC
AGCACACGTCTGAACTCTTTCCCTACACGACGCTCTTCCGATCTNNNTGTGGAATTCTTGTCCTAGCA
AGCACACGTCTGAACTCTTTCCCTACACGACGCTCTTCCGATCTNNNGGAGGTGGGTATTGGTCAAG
CACACGTCTGAACTCTTTCCCTACACGACGCTCTTCCGATCTNNNTCTGTGGCTAGATAGTATTCCAC
CACACGTCTGAACTCTTTCCCTACACGACGCTCTTCCGATCTNNNTGCTACTGCCATTACTGCATTGT
AGCACACGTCTGAACTCTTTCCCTACACGACGCTCTTCCGATCTNNNAGAAGCTAGGAAGCCATCTGG
CACACGTCTGAACTCTTTCCCTACACGACGCTCTTCCGATCTNNNTTCCTTTTCTCTTTCCAGCCACCCA
AGAGCACACGTCTGAACTCTTTCCCTACACGACGCTCTTCCGATCTNNNGCTCTATGTGCTTGTTATTG
CACACGTCTGAACTCTTTCCCTACACGACGCTCTTCCGATCTNNNTTATCCCCTTGATCTTGCCTCCACA
AGCACACGTCTGAACTCTTTCCCTACACGACGCTCTTCCGATCTNNNCCAACTTCTCAAAAACCACACA
AGCACACGTCTGAACTCTTTCCCTACACGACGCTCTTCCGATCTNNNAAGCATCCCACAGTGACAGA
AGCACACGTCTGAACTCTTTCCCTACACGACGCTCTTCCGATCTNNNGACAGGTTTGGAATCAGGCTT
AGCACACGTCTGAACTCTTTCCCTACACGACGCTCTTCCGATCTNNNATATAGGCATATGCTCACCTA
CACACGTCTGAACTCTTTCCCTACACGACGCTCTTCCGATCTNNNAATTCCTCATTCTCTGGGCCCTTC
AAGAGCACACGTCTGAACTCTTTCCCTACACGACGCTCTTCCGATCTNNNGCCCAACTTTTCCTCCTTCC
GCACACGTCTGAACTCTTTCCCTACACGACGCTCTTCCGATCTNNNGTGCTTCCTGTAGACAGAGTTG
GAGCACACGTCTGAACTCTTTCCCTACACGACGCTCTTCCGATCTNNNACCCTTCTAACAACCTGGCA
AAGAGCACACGTCTGAACTCTTTCCCTACACGACGCTCTTCCGATCTNNNGCAAACACACTCACTCTCT
GAGCACACGTCTGAACTCTTTCCCTACACGACGCTCTTCCGATCTNNNGAAACAGAGTAGTAGCAGC
GAGCACACGTCTGAACTCTTTCCCTACACGACGCTCTTCCGATCTNNNAGAGAAGGTTTCCTTGCATTT
GAGCACACGTCTGAACTCTTTCCCTACACGACGCTCTTCCGATCTNNNCAGCTGGAGGAATTGTGCTC
GCACACGTCTGAACTCTTTCCCTACACGACGCTCTTCCGATCTNNNAAAGTGCTCAACTTGGTACCAG
GAGCACACGTCTGAACTCTTTCCCTACACGACGCTCTTCCGATCTNNNCTAGATCCTGTCTTGGAGCC
GCACACGTCTGAACTCTTTCCCTACACGACGCTCTTCCGATCTNNNCGGAAGCTGTGTTTCACTGTAG
GCACACGTCTGAACTCTTTCCCTACACGACGCTCTTCCGATCTNNNGCTGTTTCTAGCAGGCACCTATT
GCACACGTCTGAACTCTTTCCCTACACGACGCTCTTCCGATCTNNNTGAGGGACCACAGGGGAATTT
AGCACACGTCTGAACTCTTTCCCTACACGACGCTCTTCCGATCTNNNCCTGGTGATGAAATCTGTCCC
AGCACACGTCTGAACTCTTTCCCTACACGACGCTCTTCCGATCTNNNATTGCCAGTCAGTGTCTGTCTT
AGCACACGTCTGAACTCTTTCCCTACACGACGCTCTTCCGATCTNNNTGGTGCATGTCCTATATCAAAG
AAGAGCACACGTCTGAACTCTTTCCCTACACGACGCTCTTCCGATCTNNNCTCAGGGATGTTTCCCTC
AAGAGCACACGTCTGAACTCTTTCCCTACACGACGCTCTTCCGATCTNNNATTGGCTGAAAAGACTACCC
AAGAGCACACGTCTGAACTCTTTCCCTACACGACGCTCTTCCGATCTNNNATGCTGGTTTGCCTTCTG
GCACACGTCTGAACTCTTTCCCTACACGACGCTCTTCCGATCTNNNCTGTACTCCTTGTTCTGCACCTC

GAGCACACGTCTGAACTCTTTCCCTACACGACGCTCTTCCGATCTNNNATCAGGGCAAGTAGAAAGCA
AGCACACGTCTGAACTCTTTCCCTACACGACGCTCTTCCGATCTNNNTGAAATCCAGCAACCCCATTTCT
GAGCACACGTCTGAACTCTTTCCCTACACGACGCTCTTCCGATCTNNNGAGAAGTTTCCCCAAGGTTG
GCACACGTCTGAACTCTTTCCCTACACGACGCTCTTCCGATCTNNNTCCTTTCCCACATGAGATTCCCTCA
GAGCACACGTCTGAACTCTTTCCCTACACGACGCTCTTCCGATCTNNNTCAACATACCTGTACACACACA
AAGAGCACACGTCTGAACTCTTTCCCTACACGACGCTCTTCCGATCTNNNTGTGTCCTTATCCTTAGCTC
GAAGAGCACACGTCTGAACTCTTTCCCTACACGACGCTCTTCCGATCTNNNATGTCATTGGCTTTCTG
GAGCACACGTCTGAACTCTTTCCCTACACGACGCTCTTCCGATCTNNNTAAACTTTGCCAGAACTAGCG
AGAGCACACGTCTGAACTCTTTCCCTACACGACGCTCTTCCGATCTNNNAGGTCACATTTGTGAGAACA
AGAGCACACGTCTGAACTCTTTCCCTACACGACGCTCTTCCGATCTNNNTGTCACAAATTACCACAAAGT
AGAGCACACGTCTGAACTCTTTCCCTACACGACGCTCTTCCGATCTNNNCGTAGGCCGGTAAACGAAGT
AAGAGCACACGTCTGAACTCTTTCCCTACACGACGCTCTTCCGATCTNNNCTTACCAGCCAAGTTTCTG
GAGCACACGTCTGAACTCTTTCCCTACACGACGCTCTTCCGATCTNNNACGGACAGCTTCTAAGAGTG
AAGAGCACACGTCTGAACTCTTTCCCTACACGACGCTCTTCCGATCTNNNTGAGGTGGAGGATCTAAG
AGAGCACACGTCTGAACTCTTTCCCTACACGACGCTCTTCCGATCTNNNCATTGAGGAAGTGGGGGAA
GCACACGTCTGAACTCTTTCCCTACACGACGCTCTTCCGATCTNNNTCTCCTCATCTGTGAAATGCACC
GAGCACACGTCTGAACTCTTTCCCTACACGACGCTCTTCCGATCTNNNTAGCTTGTTCCTGTGTCCTCTG
GCACACGTCTGAACTCTTTCCCTACACGACGCTCTTCCGATCTNNNTCAGGCAATTACAGGAGGATTG
AGCACACGTCTGAACTCTTTCCCTACACGACGCTCTTCCGATCTNNNTGGTGTCACTCAAATTTTGCCCT
GAGCACACGTCTGAACTCTTTCCCTACACGACGCTCTTCCGATCTNNNAGGTCCCTCTTCCATTGTTCTC
AGAGCACACGTCTGAACTCTTTCCCTACACGACGCTCTTCCGATCTNNNGTGAGAATCTAACTGGTGG
AGCACACGTCTGAACTCTTTCCCTACACGACGCTCTTCCGATCTNNNAAGTGAAACTCACTGGGGCAAA
AGCACACGTCTGAACTCTTTCCCTACACGACGCTCTTCCGATCTNNNTCGTTGCCAAATCATAGACACT
GAGCACACGTCTGAACTCTTTCCCTACACGACGCTCTTCCGATCTNNNGGCTAGCATGTTTAGCATCAC
AGAGCACACGTCTGAACTCTTTCCCTACACGACGCTCTTCCGATCTNNNAAGACAGGGTCCCATTCAT
GAGCACACGTCTGAACTCTTTCCCTACACGACGCTCTTCCGATCTNNNCCCATCAGAGCTCCCTGTAG
AAGAGCACACGTCTGAACTCTTTCCCTACACGACGCTCTTCCGATCTNNNGTCTCCATGCAATACAGTG
CACACGTCTGAACTCTTTCCCTACACGACGCTCTTCCGATCTNNNGTTGTTTGCTAGGAGATGGCAGA
GAGCACACGTCTGAACTCTTTCCCTACACGACGCTCTTCCGATCTNNNTGTTTTACTATCCAGCAGTCTT
GAAGAGCACACGTCTGAACTCTTTCCCTACACGACGCTCTTCCGATCTNNNACACTCTCCAATTCCATCCA
GAGCACACGTCTGAACTCTTTCCCTACACGACGCTCTTCCGATCTNNNGTGTAGAAAAACAAGCCACAG
GAGCACACGTCTGAACTCTTTCCCTACACGACGCTCTTCCGATCTNNNCTTGCCTAGATTAGCTGCTGCA
AGCACACGTCTGAACTCTTTCCCTACACGACGCTCTTCCGATCTNNNACATTGTTCACAGTAGCCAAGA
GAGCACACGTCTGAACTCTTTCCCTACACGACGCTCTTCCGATCTNNNAGGAAGCTTCTGGTTCCTACCA
GAGCACACGTCTGAACTCTTTCCCTACACGACGCTCTTCCGATCTNNNGTCCCAACTACTACTCCCAAAG
AGCACACGTCTGAACTCTTTCCCTACACGACGCTCTTCCGATCTNNNGCTGTAGTCCTAAAACTCAACC
AAGAGCACACGTCTGAACTCTTTCCCTACACGACGCTCTTCCGATCTNNNTGCATTCCAAGTGTTCTCA
AGCACACGTCTGAACTCTTTCCCTACACGACGCTCTTCCGATCTNNNGATGCCTTCTTGTGGCTTTTTCC
AGAGCACACGTCTGAACTCTTTCCCTACACGACGCTCTTCCGATCTNNNCTTTTACCATTGGGGGTGTG
GAGCACACGTCTGAACTCTTTCCCTACACGACGCTCTTCCGATCTNNNGTTTTGCCTGGTTTGGCTCTT
AGCACACGTCTGAACTCTTTCCCTACACGACGCTCTTCCGATCTNNNAGATGAATCACTTTCATAACGG
GAGCACACGTCTGAACTCTTTCCCTACACGACGCTCTTCCGATCTNNNCTGAAGAATATTTCCCTGCCCC
GAGCACACGTCTGAACTCTTTCCCTACACGACGCTCTTCCGATCTNNNCAAGTACATGGGTCTCACGAA

AGCACACGTCTGAACTCTTTCCCTACACGACGCTCTTCCGATCTNNNAACAACTGCTCAGTTCCTTCCT
AGCACACGTCTGAACTCTTTCCCTACACGACGCTCTTCCGATCTNNNTCCTTTCTCCCTTACCTGTCCTG
AGCACACGTCTGAACTCTTTCCCTACACGACGCTCTTCCGATCTNNNGTGCGAGGTATGGATAGAAG
AGCACACGTCTGAACTCTTTCCCTACACGACGCTCTTCCGATCTNNNACAGTTGTTATGAAAATACTC
AGCACACGTCTGAACTCTTTCCCTACACGACGCTCTTCCGATCTNNNATCATGTTCTGGACTTCCTACG
GCACACGTCTGAACTCTTTCCCTACACGACGCTCTTCCGATCTNNNGTGACAGGGCCTATATGTGCAT
GAGCACACGTCTGAACTCTTTCCCTACACGACGCTCTTCCGATCTNNNACTTGAATCCTGCTTTTGCCT
GAGCACACGTCTGAACTCTTTCCCTACACGACGCTCTTCCGATCTNNNAAATTTGGAGAAGCCAACAT
AGAGCACACGTCTGAACTCTTTCCCTACACGACGCTCTTCCGATCTNNNTCTTCTTATCTGTGGGGAG
GCACACGTCTGAACTCTTTCCCTACACGACGCTCTTCCGATCTNNNTTTACAGATCAGGACCCCCTACG
GCACACGTCTGAACTCTTTCCCTACACGACGCTCTTCCGATCTNNNCAAAACCAAATTCCCGACCTTCG
AGCACACGTCTGAACTCTTTCCCTACACGACGCTCTTCCGATCTNNNGGCTCTTCTCAAGCATTTTCCC
AAGAGCACACGTCTGAACTCTTTCCCTACACGACGCTCTTCCGATCTNNNCCCAGCCGATATCCTGTTC
GAGCACACGTCTGAACTCTTTCCCTACACGACGCTCTTCCGATCTNNNTTCCAAAAGTTCAAACAGAAT
AGCACACGTCTGAACTCTTTCCCTACACGACGCTCTTCCGATCTNNNTGCAACAAGCAAAGAGATTG
GCACACGTCTGAACTCTTTCCCTACACGACGCTCTTCCGATCTNNNTGCTCTCTTTCACTCTTCCTCTCT
AGCACACGTCTGAACTCTTTCCCTACACGACGCTCTTCCGATCTNNNACCCTGTGTTGAATGATCATG
AGAGCACACGTCTGAACTCTTTCCCTACACGACGCTCTTCCGATCTNNNTCTTTGTGGCTGAATGGTA
GAGCACACGTCTGAACTCTTTCCCTACACGACGCTCTTCCGATCTNNNAAACCTCCAACAAGGAGGGA
GCACACGTCTGAACTCTTTCCCTACACGACGCTCTTCCGATCTNNNTTATCCATACACACCATGCCAGC
AGCACACGTCTGAACTCTTTCCCTACACGACGCTCTTCCGATCTNNNTAACATATTCCCCAAGTTGCCCC
AGAGCACACGTCTGAACTCTTTCCCTACACGACGCTCTTCCGATCTNNNCTGTACCTGTAATTGGACC
AGCACACGTCTGAACTCTTTCCCTACACGACGCTCTTCCGATCTNNNTGGCCTACATGGAGGTAACAG
GAGCACACGTCTGAACTCTTTCCCTACACGACGCTCTTCCGATCTNNNTGAGGCAGAATTGCCAGAAC
ACACGTCTGAACTCTTTCCCTACACGACGCTCTTCCGATCTNNNCGAAAGCCCACCTGACCATTGCATC
GCACACGTCTGAACTCTTTCCCTACACGACGCTCTTCCGATCTNNNATTCTTGTCTGAGCAACCCACTC
GAGCACACGTCTGAACTCTTTCCCTACACGACGCTCTTCCGATCTNNNAAAAACGCCTGGCTAGAGTT
GAGCACACGTCTGAACTCTTTCCCTACACGACGCTCTTCCGATCTNNNTGCAGACCTATTCACAATTG
AGCACACGTCTGAACTCTTTCCCTACACGACGCTCTTCCGATCTNNNCCTTGGGGGTTTTGCTAGGAT
GCACACGTCTGAACTCTTTCCCTACACGACGCTCTTCCGATCTNNNTGGGTTGGAAGAGGTGTTTTTG
AGCACACGTCTGAACTCTTTCCCTACACGACGCTCTTCCGATCTNNNGAAACCGGAATTCACCTTCCTG
AGAGCACACGTCTGAACTCTTTCCCTACACGACGCTCTTCCGATCTNNNCAACAAGAGTGAACCTCAG
AAGAGCACACGTCTGAACTCTTTCCCTACACGACGCTCTTCCGATCTNNNTTATTGGTCCTCTTTGTTC
GCACACGTCTGAACTCTTTCCCTACACGACGCTCTTCCGATCTNNNAATATATGTGTTTGTATGTGTG
AGCACACGTCTGAACTCTTTCCCTACACGACGCTCTTCCGATCTNNNCCACACTACAACTGCCCATATC
CACACGTCTGAACTCTTTCCCTACACGACGCTCTTCCGATCTNNNAGCTGACAACAACTAACCTTTGG
AGCACACGTCTGAACTCTTTCCCTACACGACGCTCTTCCGATCTNNNTCAATGAGTTCACCAGACAAC
AGCACACGTCTGAACTCTTTCCCTACACGACGCTCTTCCGATCTNNNGGCAATCGGACAATGTAGGAC
AGCACACGTCTGAACTCTTTCCCTACACGACGCTCTTCCGATCTNNNACCAGCACTTCCAAGAACACA
AGCACACGTCTGAACTCTTTCCCTACACGACGCTCTTCCGATCTNNNTGTATGGGGTTGCAGTCAGAG
AGCACACGTCTGAACTCTTTCCCTACACGACGCTCTTCCGATCTNNNCCTTTCCATAGCCGAGAAAGC
AGCACACGTCTGAACTCTTTCCCTACACGACGCTCTTCCGATCTNNNTAATCCAAGGCGATCCACAAA
AAGAGCACACGTCTGAACTCTTTCCCTACACGACGCTCTTCCGATCTNNNATTGGTGCTTGCAAAGT

AGCACACGTCTGAACTCTTTCCCTACACGACGCTCTTCCGATCTNNNAGAAGAAAATCCTCCAAATGAA
GAAGAGCACACGTCTGAACTCTTTCCCTACACGACGCTCTTCCGATCTNNNCTTGAGGTAGGGGTGAG
AGCACACGTCTGAACTCTTTCCCTACACGACGCTCTTCCGATCTNNNTTTTTCTCGCCGGGTAGTAGTT
AGCACACGTCTGAACTCTTTCCCTACACGACGCTCTTCCGATCTNNNTTTCTAGAGTGTTTGCCTCCTGC
AGCACACGTCTGAACTCTTTCCCTACACGACGCTCTTCCGATCTNNNACGGAAACCCACACACATTGTT
AGCACACGTCTGAACTCTTTCCCTACACGACGCTCTTCCGATCTNNNAGAATGTTCAATGGTTGGCTGA
GCACACGTCTGAACTCTTTCCCTACACGACGCTCTTCCGATCTNNNTTTCCTTCTCTTCCCCATGTGAGCA
GAGCACACGTCTGAACTCTTTCCCTACACGACGCTCTTCCGATCTNNNAGTTCTAGTATCTCTCTCTCTC
GAGCACACGTCTGAACTCTTTCCCTACACGACGCTCTTCCGATCTNNNGAATCCATCACCACAGCAGAC
AGCACACGTCTGAACTCTTTCCCTACACGACGCTCTTCCGATCTNNNCATAAGGCTTTGGGATCCTGTG
GCACACGTCTGAACTCTTTCCCTACACGACGCTCTTCCGATCTNNNTCTGCTGCATAAATTGCACCTCTC
GAGCACACGTCTGAACTCTTTCCCTACACGACGCTCTTCCGATCTNNNTACACCAAGCAACTCAATCCCA
AGCACACGTCTGAACTCTTTCCCTACACGACGCTCTTCCGATCTNNNGTTTCTACACTTAGGCCTGGAC
AGAGCACACGTCTGAACTCTTTCCCTACACGACGCTCTTCCGATCTNNNGCTTTTGAAAGGCAAGCAAA
AGCACACGTCTGAACTCTTTCCCTACACGACGCTCTTCCGATCTNNNTCCTAGAAGCAGCTTTCCCAATC
AGCACACGTCTGAACTCTTTCCCTACACGACGCTCTTCCGATCTNNNTCATGAGTGTTCCTGCTGTTTCC
GAGCACACGTCTGAACTCTTTCCCTACACGACGCTCTTCCGATCTNNNAGTGCTCTGGCTCTGAAGACT
GAGCACACGTCTGAACTCTTTCCCTACACGACGCTCTTCCGATCTNNNCTCCTGCCATTCTCTTGTAAGG
AGCACACGTCTGAACTCTTTCCCTACACGACGCTCTTCCGATCTNNNAGCAAATACTGAAGCTGTCTCCA
AGCACACGTCTGAACTCTTTCCCTACACGACGCTCTTCCGATCTNNNTGACAACCTGGAAGTCATTGGT
AAGAGCACACGTCTGAACTCTTTCCCTACACGACGCTCTTCCGATCTNNNAAAGACATGTGGAAGCGA
AGCACACGTCTGAACTCTTTCCCTACACGACGCTCTTCCGATCTNNNTTGAGTAGCCAAGTCAAGCAAG
AAGAGCACACGTCTGAACTCTTTCCCTACACGACGCTCTTCCGATCTNNNTTTCACCGTAGTGTGTGTC
GAGCACACGTCTGAACTCTTTCCCTACACGACGCTCTTCCGATCTNNNACCCTGTAATTATTCGGTAAGC
AGCACACGTCTGAACTCTTTCCCTACACGACGCTCTTCCGATCTNNNTTGGCGTCAGTTACTGGAACTT
GCACACGTCTGAACTCTTTCCCTACACGACGCTCTTCCGATCTNNNACCCGTAACACAAACAGGTTCATC
AGCACACGTCTGAACTCTTTCCCTACACGACGCTCTTCCGATCTNNNCCTGGTTGGCTCAAAACTACCT
AGCACACGTCTGAACTCTTTCCCTACACGACGCTCTTCCGATCTNNNCTCTTAGAGACCATCCACTGCAC
AGCACACGTCTGAACTCTTTCCCTACACGACGCTCTTCCGATCTNNNCCAAGCAGCCTTCCCTAATGAAT
GAGCACACGTCTGAACTCTTTCCCTACACGACGCTCTTCCGATCTNNNCTGTTGGCCTGTCTATTGGTC
GAGCACACGTCTGAACTCTTTCCCTACACGACGCTCTTCCGATCTNNNCTTGCCTCATCCCCCATTACTTC
GAGCACACGTCTGAACTCTTTCCCTACACGACGCTCTTCCGATCTNNNACTATCTTTCTAGAGGCATCCA
GAGCACACGTCTGAACTCTTTCCCTACACGACGCTCTTCCGATCTNNNTTTTTCCGTAGGATAACGATTC
GAGCACACGTCTGAACTCTTTCCCTACACGACGCTCTTCCGATCTNNNAAAGACACAAAATGTGGCTGC
GAGCACACGTCTGAACTCTTTCCCTACACGACGCTCTTCCGATCTNNNCCAGAGCTGGCAGTAGGATA
AGCACACGTCTGAACTCTTTCCCTACACGACGCTCTTCCGATCTNNNTCCCATTCAGTCCATTGTAGGG
AGAGCACACGTCTGAACTCTTTCCCTACACGACGCTCTTCCGATCTNNNTTATTCAACACACCTCTGGTC
AGCACACGTCTGAACTCTTTCCCTACACGACGCTCTTCCGATCTNNNGCCTCAGCCAACCAAGTAAATG
GCACACGTCTGAACTCTTTCCCTACACGACGCTCTTCCGATCTNNNTATCCCTACACTCCTACACCCACCA
GCACACGTCTGAACTCTTTCCCTACACGACGCTCTTCCGATCTNNNGTGCACCAGGTAATTCAGACCAA
AGAGCACACGTCTGAACTCTTTCCCTACACGACGCTCTTCCGATCTNNNAACGAAGAACTTGTTTGTGG
GAGCACACGTCTGAACTCTTTCCCTACACGACGCTCTTCCGATCTNNNGTGGCCACACGAATCTAACATC
AGCACACGTCTGAACTCTTTCCCTACACGACGCTCTTCCGATCTNNNTTAGCCTACAGGCTCTGTCTCA

AGCACACGTCTGAACTCTTTCCCTACACGACGCTCTTCCGATCTNNNAACTCAATTCTTCAAGACGCCAC
GCACACGTCTGAACTCTTTCCCTACACGACGCTCTTCCGATCTNNNTCCCTCCTCACCAACACATACATCA
GGAAGAGCACACGTCTGAACTCTTTCCCTACACGACGCTCTTCCGATCTNNNAAAGACATGTGGAAGC
AGCACACGTCTGAACTCTTTCCCTACACGACGCTCTTCCGATCTNNNAAAGTCATTCCCAGTCAAGGCA
AGAGCACACGTCTGAACTCTTTCCCTACACGACGCTCTTCCGATCTNNNGGAGAAAAATAATGCGAAG
AAGAGCACACGTCTGAACTCTTTCCCTACACGACGCTCTTCCGATCTNNNTCTCAGAGCCCGCCTAATTC
GCACACGTCTGAACTCTTTCCCTACACGACGCTCTTCCGATCTNNNCCGAGAAGCGACTGGTCATTATT
GAGCACACGTCTGAACTCTTTCCCTACACGACGCTCTTCCGATCTNNNAGGCCTTATCTGGAAAAAGCA
AGCACACGTCTGAACTCTTTCCCTACACGACGCTCTTCCGATCTNNNTGGATAAGGAATACTTGCCCTG
AGCACACGTCTGAACTCTTTCCCTACACGACGCTCTTCCGATCTNNNTTACTGAAAACAGCCCAGAGGA
AGAGCACACGTCTGAACTCTTTCCCTACACGACGCTCTTCCGATCTNNNAGGGACACTCTGTGTCTTTTT
GAAGAGCACACGTCTGAACTCTTTCCCTACACGACGCTCTTCCGATCTNNNTTTCCCCCTCCACATGCATC
AGAGCACACGTCTGAACTCTTTCCCTACACGACGCTCTTCCGATCTNNNAAATAAGCCAGCAAGGAGGA
AGCACACGTCTGAACTCTTTCCCTACACGACGCTCTTCCGATCTNNNTGGGAATGGTGCTTCAGAGAAA
GCACACGTCTGAACTCTTTCCCTACACGACGCTCTTCCGATCTNNNTCCTGTGTATCTGGCTCTGGTTTC
GAGCACACGTCTGAACTCTTTCCCTACACGACGCTCTTCCGATCTNNNGGGAGCCGTACACAAATCTCA
GAGCACACGTCTGAACTCTTTCCCTACACGACGCTCTTCCGATCTNNNGAGGAAGGAATGCTGGAGA
AGAGCACACGTCTGAACTCTTTCCCTACACGACGCTCTTCCGATCTNNNTGCTGAGTTACATTTACTCCA
AGCACACGTCTGAACTCTTTCCCTACACGACGCTCTTCCGATCTNNNTTCTGTTGCATCTGACAGGCTTT
AGCACACGTCTGAACTCTTTCCCTACACGACGCTCTTCCGATCTNNNGCCTGGCTTCATATCCAAAAGTG
GAGCACACGTCTGAACTCTTTCCCTACACGACGCTCTTCCGATCTNNNGTCACTTATTAGCCACAGGGCT
GAGCACACGTCTGAACTCTTTCCCTACACGACGCTCTTCCGATCTNNNGACAGTGACAGCTGTAGCAGA
GAGCACACGTCTGAACTCTTTCCCTACACGACGCTCTTCCGATCTNNNAGAAGGGTATTCTGATCTGG
AGAGCACACGTCTGAACTCTTTCCCTACACGACGCTCTTCCGATCTNNNACTGAAGGAGAGCATCATGA
AGCACACGTCTGAACTCTTTCCCTACACGACGCTCTTCCGATCTNNNATAAATCACAGCCAGCACTGTG
AGAGCACACGTCTGAACTCTTTCCCTACACGACGCTCTTCCGATCTNNNCTATGTCTCACATGGCCTTTC
AGCACACGTCTGAACTCTTTCCCTACACGACGCTCTTCCGATCTNNNTATTTCCAGTCTGCCAGGCATTC
GAGCACACGTCTGAACTCTTTCCCTACACGACGCTCTTCCGATCTNNNAGGGTCTAAGTACTCCCAAGT
GAGCACACGTCTGAACTCTTTCCCTACACGACGCTCTTCCGATCTNNNATTCCATTTAGCATCTCCCACCC
GAGCACACGTCTGAACTCTTTCCCTACACGACGCTCTTCCGATCTNNNACCTTCCCAAGGTGTCAAAAAG
AGCACACGTCTGAACTCTTTCCCTACACGACGCTCTTCCGATCTNNNTATCTGCTCAAATTGCGTGTGTC
AGCACACGTCTGAACTCTTTCCCTACACGACGCTCTTCCGATCTNNNCTAGACAAGCCCAAAGTGTGTG
AGAGCACACGTCTGAACTCTTTCCCTACACGACGCTCTTCCGATCTNNNCTGGGTCTGGAATCTGTGTCT
GAGCACACGTCTGAACTCTTTCCCTACACGACGCTCTTCCGATCTNNNAGCCAACTCCTGACCTGTTTTT
GAGCACACGTCTGAACTCTTTCCCTACACGACGCTCTTCCGATCTNNNTTTCTAGTGGATGGGGCTTCA
GAGCACACGTCTGAACTCTTTCCCTACACGACGCTCTTCCGATCTNNNGAAGAGGCATCTGACAGCTTG
AAGAGCACACGTCTGAACTCTTTCCCTACACGACGCTCTTCCGATCTNNNCCTGAGGTATATCTGTGCTC
GAGCACACGTCTGAACTCTTTCCCTACACGACGCTCTTCCGATCTNNNTGTCCTTTCCTTCTCACTGCTTC
GAGCACACGTCTGAACTCTTTCCCTACACGACGCTCTTCCGATCTNNNGAATTGAGACCACAGCCCTCA
AGCACACGTCTGAACTCTTTCCCTACACGACGCTCTTCCGATCTNNNTACACATGCCTGAGTGTAAGCAC
GAGCACACGTCTGAACTCTTTCCCTACACGACGCTCTTCCGATCTNNNCAGTAAGCCATGATAGCACCTC
AGAGCACACGTCTGAACTCTTTCCCTACACGACGCTCTTCCGATCTNNNTTCTTCATGAAGACCTCACAG
AAGAGCACACGTCTGAACTCTTTCCCTACACGACGCTCTTCCGATCTNNNGAAACGCAAATACAGGTGC

AGAGCACACGTCTGAACTCTTTCCCTACACGACGCTCTTCCGATCTNNNTATCCACCACTACCTCAGCT
AGCACACGTCTGAACTCTTTCCCTACACGACGCTCTTCCGATCTNNNTGTAGCAACTCTGGATGCCAAT
AGCACACGTCTGAACTCTTTCCCTACACGACGCTCTTCCGATCTNNNTGGCACATATGAATACACACA
AGCACACGTCTGAACTCTTTCCCTACACGACGCTCTTCCGATCTNNNGACATTCCGAGTATGCGTGTG
GAGCACACGTCTGAACTCTTTCCCTACACGACGCTCTTCCGATCTNNNGGAAATGCTTACTGTGCCAAC
AGCACACGTCTGAACTCTTTCCCTACACGACGCTCTTCCGATCTNNNGCCCAAAGAGCAGGTGAGAT
GAGCACACGTCTGAACTCTTTCCCTACACGACGCTCTTCCGATCTNNNAGGGAACTTCTTTTGCTCCCT
AGCACACGTCTGAACTCTTTCCCTACACGACGCTCTTCCGATCTNNNCTTCAGCTTCCCTCACTTTACC
AGCACACGTCTGAACTCTTTCCCTACACGACGCTCTTCCGATCTNNNCCATGACTGCTCATGCCATTTT
GAGCACACGTCTGAACTCTTTCCCTACACGACGCTCTTCCGATCTNNNTTTTGTTCCACTGGGAAAGC
GAGCACACGTCTGAACTCTTTCCCTACACGACGCTCTTCCGATCTNNNAGAGTATTTTCCTTCTCACTCC
GAGCACACGTCTGAACTCTTTCCCTACACGACGCTCTTCCGATCTNNNTTGTTCCCCATCCACCACATA
GAGCACACGTCTGAACTCTTTCCCTACACGACGCTCTTCCGATCTNNNAGCTTCACTCAGGAAGTGGC
AGCACACGTCTGAACTCTTTCCCTACACGACGCTCTTCCGATCTNNNCCATGACTGCTCATGCCATTTT
AGCACACGTCTGAACTCTTTCCCTACACGACGCTCTTCCGATCTNNNGGGCTCAGAAAGGTAACATAG
AGCACACGTCTGAACTCTTTCCCTACACGACGCTCTTCCGATCTNNNACTCCCCACTCTCATTTACTCCC
AGAGCACACGTCTGAACTCTTTCCCTACACGACGCTCTTCCGATCTNNNACTATTTACATTCTTTGTACA
AGCACACGTCTGAACTCTTTCCCTACACGACGCTCTTCCGATCTNNNTCTAAGAGGATGGAAGCAGTA
GAGCACACGTCTGAACTCTTTCCCTACACGACGCTCTTCCGATCTNNNTTTTCTTCTTTCTTTCCCCCTTC
AGAGCACACGTCTGAACTCTTTCCCTACACGACGCTCTTCCGATCTNNNACCTCCTCTGTGATATGGT
GAGCACACGTCTGAACTCTTTCCCTACACGACGCTCTTCCGATCTNNNTTCAGTAAACTTCATGGGGC
GAGCACACGTCTGAACTCTTTCCCTACACGACGCTCTTCCGATCTNNNCAAGAGCGACACTCTGTCGA
AGCACACGTCTGAACTCTTTCCCTACACGACGCTCTTCCGATCTNNNCTCCACAGCAGTGAGACCTTT
GAGCACACGTCTGAACTCTTTCCCTACACGACGCTCTTCCGATCTNNNTGCCTGGCTGAAATTTCTCCTT
AGCACACGTCTGAACTCTTTCCCTACACGACGCTCTTCCGATCTNNNCCTCTACAAGCTTCTGATGCGT
AGCACACGTCTGAACTCTTTCCCTACACGACGCTCTTCCGATCTNNNCCCAACATCATCAGAAGAGCC
AGAGCACACGTCTGAACTCTTTCCCTACACGACGCTCTTCCGATCTNNNATAGTGGTTTCCTGGAACTG
AGCACACGTCTGAACTCTTTCCCTACACGACGCTCTTCCGATCTNNNTCCATACTGAGACCCTTCACGTT
AGCACACGTCTGAACTCTTTCCCTACACGACGCTCTTCCGATCTNNNTTCTGCCCACCATACAGCTACT
GAGCACACGTCTGAACTCTTTCCCTACACGACGCTCTTCCGATCTNNNAGTGGAGACAGACCCACAGA
AGCACACGTCTGAACTCTTTCCCTACACGACGCTCTTCCGATCTNNNCTCCAGGAGGAAAACCAGAG
GCACACGTCTGAACTCTTTCCCTACACGACGCTCTTCCGATCTNNNCCTGGTACATCCCCTACATCAGT
GCACACGTCTGAACTCTTTCCCTACACGACGCTCTTCCGATCTNNNCTGTGGCCTTGCCTATACTTTGA
GAGCACACGTCTGAACTCTTTCCCTACACGACGCTCTTCCGATCTNNNTCTCAAATGGTTCAGGAGAAA
GCACACGTCTGAACTCTTTCCCTACACGACGCTCTTCCGATCTNNNTCTCCAGCACTGTATCCACTTGA
AGCACACGTCTGAACTCTTTCCCTACACGACGCTCTTCCGATCTNNNTGCAATAAATTCCAATCTATCG
AGCACACGTCTGAACTCTTTCCCTACACGACGCTCTTCCGATCTNNNCAGCTGCATCATCTGTTATTCCC
AGCACACGTCTGAACTCTTTCCCTACACGACGCTCTTCCGATCTNNNCATCGGGTTGAGCTGAGAAG
GAGCACACGTCTGAACTCTTTCCCTACACGACGCTCTTCCGATCTNNNATGCAAGTATGCTGTCTTGA
GAGCACACGTCTGAACTCTTTCCCTACACGACGCTCTTCCGATCTNNNTCCAGTGACACATATGCACTC
GAGCACACGTCTGAACTCTTTCCCTACACGACGCTCTTCCGATCTNNNGTAGACCTGAGGGACCATTC
AAGAGCACACGTCTGAACTCTTTCCCTACACGACGCTCTTCCGATCTNNNACCAGTCCAGGAATAGAG
GAGCACACGTCTGAACTCTTTCCCTACACGACGCTCTTCCGATCTNNNGCCCATCTTCCATTGGTGTAT

AAGAGCACACGTCTGAACTCTTTCCCTACACGACGCTCTTCCGATCTNNNTTCTCTAGCTTCCCCTAAGC
AGCACACGTCTGAACTCTTTCCCTACACGACGCTCTTCCGATCTNNNTTTGCCAGTTCCCTAGTTTCCAC
GCACACGTCTGAACTCTTTCCCTACACGACGCTCTTCCGATCTNNNGCAGACCATGACTTCCTTTGCTAC
AGCACACGTCTGAACTCTTTCCCTACACGACGCTCTTCCGATCTNNNTTCAAGCAGCACACCAAGTAA
AGCACACGTCTGAACTCTTTCCCTACACGACGCTCTTCCGATCTNNNATGGTACTGCACACAGTAGTT
GAGCACACGTCTGAACTCTTTCCCTACACGACGCTCTTCCGATCTNNNCTTCTCCATAACCAGAGAGG
AGAGCACACGTCTGAACTCTTTCCCTACACGACGCTCTTCCGATCTNNNTATACACATGTGCACTTGG
AGCACACGTCTGAACTCTTTCCCTACACGACGCTCTTCCGATCTNNNCATCATGTAGAACACAATGTA
GAGCACACGTCTGAACTCTTTCCCTACACGACGCTCTTCCGATCTNNNACAGTGGGTACAGTTACTCC
AGCACACGTCTGAACTCTTTCCCTACACGACGCTCTTCCGATCTNNNACAGAGGTAGGGCAGCTAAC
AGCACACGTCTGAACTCTTTCCCTACACGACGCTCTTCCGATCTNNNTGATACAATATGGCTTGTGCC
AGCACACGTCTGAACTCTTTCCCTACACGACGCTCTTCCGATCTNNNTGTCCGAAGTAATTTGAAAGG
GAGCACACGTCTGAACTCTTTCCCTACACGACGCTCTTCCGATCTNNNCTGTCACAGCAACTGTCCCA
AGCACACGTCTGAACTCTTTCCCTACACGACGCTCTTCCGATCTNNNTGAAGGAGGCCACATTTTACT
GCACACGTCTGAACTCTTTCCCTACACGACGCTCTTCCGATCTNNNTATACTCAAGAGAGACCCCAGG
GCACACGTCTGAACTCTTTCCCTACACGACGCTCTTCCGATCTNNNAGAAAATGCAGCATCCCTTGCT
AGCACACGTCTGAACTCTTTCCCTACACGACGCTCTTCCGATCTNNNAGACAACCACAATTCTCTGAGC
AGCACACGTCTGAACTCTTTCCCTACACGACGCTCTTCCGATCTNNNGGCTGTCCCTATAAGGGGGTA
AGCACACGTCTGAACTCTTTCCCTACACGACGCTCTTCCGATCTNNNAATCCTTCTTTCTGTGCTCTGCC
GAGCACACGTCTGAACTCTTTCCCTACACGACGCTCTTCCGATCTNNNAACATAGGTTCATCGGCTGT
GAGCACACGTCTGAACTCTTTCCCTACACGACGCTCTTCCGATCTNNNGCTCCGTTCCACTTATTTCTC
AGAGCACACGTCTGAACTCTTTCCCTACACGACGCTCTTCCGATCTNNNGCTAGTTTGAACCAGGCAT
GCACACGTCTGAACTCTTTCCCTACACGACGCTCTTCCGATCTNNNTAATACACATTCCTGGGCCATG
GAGCACACGTCTGAACTCTTTCCCTACACGACGCTCTTCCGATCTNNNTTGCCAAAACTAAATGCTGG
AGCACACGTCTGAACTCTTTCCCTACACGACGCTCTTCCGATCTNNNTGTACTTTGTATGCCTTGGGG
GCACACGTCTGAACTCTTTCCCTACACGACGCTCTTCCGATCTNNNATGTGTCTTGGGGTTTCATTGC
GAGCACACGTCTGAACTCTTTCCCTACACGACGCTCTTCCGATCTNNNCAGCTCCTGTAACACCTTGA
AGCACACGTCTGAACTCTTTCCCTACACGACGCTCTTCCGATCTNNNAGAACTGGTAAACAGCACTAT
GAGCACACGTCTGAACTCTTTCCCTACACGACGCTCTTCCGATCTNNNAGAGCAGACAGGTTGCTTCT
GAGCACACGTCTGAACTCTTTCCCTACACGACGCTCTTCCGATCTNNNAAGTTCAGTTCTTTTTCAATG
GAGCACACGTCTGAACTCTTTCCCTACACGACGCTCTTCCGATCTNNNCAATGGGTACAGTCAGACAG
GCACACGTCTGAACTCTTTCCCTACACGACGCTCTTCCGATCTNNNTCTTGGAAACAATGCCCATGTG
GAGCACACGTCTGAACTCTTTCCCTACACGACGCTCTTCCGATCTNNNGACCTATGTTGCCAGCTTCTC
GAGCACACGTCTGAACTCTTTCCCTACACGACGCTCTTCCGATCTNNNCACCACAAAAGAACACAAGG
AGCACACGTCTGAACTCTTTCCCTACACGACGCTCTTCCGATCTNNNTTGCTTATCCATTCAGCCTGTCC
AAGAGCACACGTCTGAACTCTTTCCCTACACGACGCTCTTCCGATCTNNNAAATGACTGTTCCAGTTC
GCACACGTCTGAACTCTTTCCCTACACGACGCTCTTCCGATCTNNNAACCCTGCTTCTTGGCATTTTG
GAGCACACGTCTGAACTCTTTCCCTACACGACGCTCTTCCGATCTNNNTCGAAGGTGTACTTTTCCCT
GAGCACACGTCTGAACTCTTTCCCTACACGACGCTCTTCCGATCTNNNTGCAAAATCAAGAGGCCATCC
GAGCACACGTCTGAACTCTTTCCCTACACGACGCTCTTCCGATCTNNNCCCTAGTCTTCTCAGTAGCCA
AGAGCACACGTCTGAACTCTTTCCCTACACGACGCTCTTCCGATCTNNNGAAGGCCATGCAGACAAA
GAGCACACGTCTGAACTCTTTCCCTACACGACGCTCTTCCGATCTNNNGATGCCTGGATACCCAGAGT
GAGCACACGTCTGAACTCTTTCCCTACACGACGCTCTTCCGATCTNNNTTCTGTGGTGCTCTCTTGCA

GAGCACACGTCTGAACTCTTTCCCTACACGACGCTCTTCCGATCTNNNCTGCAGAGAAACCAGTAGG
GAGCACACGTCTGAACTCTTTCCCTACACGACGCTCTTCCGATCTNNNAATGTGAGAGGAATCACGC
GAGCACACGTCTGAACTCTTTCCCTACACGACGCTCTTCCGATCTNNNCCCCTTCTTCCAAAGTTCCTA
AGCACACGTCTGAACTCTTTCCCTACACGACGCTCTTCCGATCTNNNAGCAGTGTGAATCATGTGTGT
AGCACACGTCTGAACTCTTTCCCTACACGACGCTCTTCCGATCTNNNATGGTTGTGGGTCTTTTCTGC
AGCACACGTCTGAACTCTTTCCCTACACGACGCTCTTCCGATCTNNNACACACACAGATATGTACTCAC
GAGCACACGTCTGAACTCTTTCCCTACACGACGCTCTTCCGATCTNNNAGCAAAGTAATGGCTCCCTC
AGCACACGTCTGAACTCTTTCCCTACACGACGCTCTTCCGATCTNNNTTAGCCTAGCACAACAGTACA
AGCACACGTCTGAACTCTTTCCCTACACGACGCTCTTCCGATCTNNNAAGTGCCAAACTCTCTCTCTCT
GCACACGTCTGAACTCTTTCCCTACACGACGCTCTTCCGATCTNNNTCATGCAAGAGTAATCAATGTA
AGCACACGTCTGAACTCTTTCCCTACACGACGCTCTTCCGATCTNNNGTAGGTGTTTTGCCATCGCAT
GCACACGTCTGAACTCTTTCCCTACACGACGCTCTTCCGATCTNNNTGTACAGATGTATACACACACA
AGAGCACACGTCTGAACTCTTTCCCTACACGACGCTCTTCCGATCTNNNACTTTTTGGACTACTGCTCA
AGCACACGTCTGAACTCTTTCCCTACACGACGCTCTTCCGATCTNNNTGGAGAGAGCTCACCTATTGC
AGCACACGTCTGAACTCTTTCCCTACACGACGCTCTTCCGATCTNNNTGAGAGCAAAACTAATGTGG
GAGCACACGTCTGAACTCTTTCCCTACACGACGCTCTTCCGATCTNNNTTGTTAGAAACCGACAGGT
AGCACACGTCTGAACTCTTTCCCTACACGACGCTCTTCCGATCTNNNAGAGCCAGCATTATTTGTGCC
AGCACACGTCTGAACTCTTTCCCTACACGACGCTCTTCCGATCTNNNGTCTGCAAATTTTGGACAGCT
GAGCACACGTCTGAACTCTTTCCCTACACGACGCTCTTCCGATCTNNNTAGGCTCAGAAAGACTTCCC
AGCACACGTCTGAACTCTTTCCCTACACGACGCTCTTCCGATCTNNNTTTGTCTGCCTTGTGCTCTCA
AGCACACGTCTGAACTCTTTCCCTACACGACGCTCTTCCGATCTNNNTCCCCTTTATGTGTACGTCCCTT
ACACGTCTGAACTCTTTCCCTACACGACGCTCTTCCGATCTNNNAAGACACAGTTTTCCGAAGTCCCCA
AGCACACGTCTGAACTCTTTCCCTACACGACGCTCTTCCGATCTNNNAAGCCTGTATCAGTCTGGTGG
AAGAGCACACGTCTGAACTCTTTCCCTACACGACGCTCTTCCGATCTNNNAGGCCATAAAACCTGCCC
GAGCACACGTCTGAACTCTTTCCCTACACGACGCTCTTCCGATCTNNNAGCTTTCTTCTGGTCCGACTT
GCACACGTCTGAACTCTTTCCCTACACGACGCTCTTCCGATCTNNNCATCTAGGGACTGCAGAGGAG
AGCACACGTCTGAACTCTTTCCCTACACGACGCTCTTCCGATCTNNNGCTTTTACATGGGTGTTCCCAC
GCACACGTCTGAACTCTTTCCCTACACGACGCTCTTCCGATCTNNNCCCACATACTTATCCATGGCAGG
AGCACACGTCTGAACTCTTTCCCTACACGACGCTCTTCCGATCTNNNTTTCCTCCCTATTATCGCCAGGT
GCACACGTCTGAACTCTTTCCCTACACGACGCTCTTCCGATCTNNNGATGCAGTCTACTCCCTGCTACT
AAGAGCACACGTCTGAACTCTTTCCCTACACGACGCTCTTCCGATCTNNNAGCTTTCTTCTGGTCCGA
GCACACGTCTGAACTCTTTCCCTACACGACGCTCTTCCGATCTNNNTCTCTGATCACACCCCACTTCC
GCACACGTCTGAACTCTTTCCCTACACGACGCTCTTCCGATCTNNNTTTGTCTGCCTTGTGCTCTCAT
GAGCACACGTCTGAACTCTTTCCCTACACGACGCTCTTCCGATCTNNNACAGTAGTGAATGTTCTCTCT
AGCACACGTCTGAACTCTTTCCCTACACGACGCTCTTCCGATCTNNNGATGCTCTCCTTTAGGTCTGT
AGCACACGTCTGAACTCTTTCCCTACACGACGCTCTTCCGATCTNNNTTTGTTCCACTGGGAAAGCT
AGCACACGTCTGAACTCTTTCCCTACACGACGCTCTTCCGATCTNNNTTGCTTCCAAGATTTCCAGCAG
AGCACACGTCTGAACTCTTTCCCTACACGACGCTCTTCCGATCTNNNGTGCTGTTTTGTGTGTTGTGT
GAGCACACGTCTGAACTCTTTCCCTACACGACGCTCTTCCGATCTNNNACTCTGCCCATTGACTGATCT
AGCACACGTCTGAACTCTTTCCCTACACGACGCTCTTCCGATCTNNNTCACTGGGCTGTGCTTACTTTT
AGCACACGTCTGAACTCTTTCCCTACACGACGCTCTTCCGATCTNNNATTGAGTGTGGAGAGGCAAA
GAGCACACGTCTGAACTCTTTCCCTACACGACGCTCTTCCGATCTNNNTACTGATTTGCCCTGTGTGT
GAGCACACGTCTGAACTCTTTCCCTACACGACGCTCTTCCGATCTNNNTGGATAAAAGTGTCTCTCTCT

GAGCACACGTCTGAACTCTTTCCCTACACGACGCTCTTCCGATCTNNNAGAGACCACACACACATGCA
AGCACACGTCTGAACTCTTTCCCTACACGACGCTCTTCCGATCTNNNACGTCGCTCCAGTGTAAACAAA
GAGCACACGTCTGAACTCTTTCCCTACACGACGCTCTTCCGATCTNNNACCAAAGGCAGAAAGAAGA
AGCACACGTCTGAACTCTTTCCCTACACGACGCTCTTCCGATCTNNNATCTTGGGTTCCTTGGTTTCA
GAGCACACGTCTGAACTCTTTCCCTACACGACGCTCTTCCGATCTNNNTTATTGATGGAGGGGTGGA
GCACACGTCTGAACTCTTTCCCTACACGACGCTCTTCCGATCTNNNAATGGTGATACCATGTGTGCCT
AGCACACGTCTGAACTCTTTCCCTACACGACGCTCTTCCGATCTNNNAGTCTCCTAAAAAGAGCGTTG
AGAGCACACGTCTGAACTCTTTCCCTACACGACGCTCTTCCGATCTNNNTAAGTCCCAAGCTGTCTGAG
GCACACGTCTGAACTCTTTCCCTACACGACGCTCTTCCGATCTNNNTCAGATGGTTGGGGTACTGTC
AGCACACGTCTGAACTCTTTCCCTACACGACGCTCTTCCGATCTNNNCCTACTGCTTACATCCCAAGCT
GAGCACACGTCTGAACTCTTTCCCTACACGACGCTCTTCCGATCTNNNTGTGATGGGGAGGTTATTC
AGCACACGTCTGAACTCTTTCCCTACACGACGCTCTTCCGATCTNNNCAGAACGAGACCCTATCTCCC
AGCACACGTCTGAACTCTTTCCCTACACGACGCTCTTCCGATCTNNNTCCATAGCTTCCCCACTGTAGT
GCACACGTCTGAACTCTTTCCCTACACGACGCTCTTCCGATCTNNNACTCCTACTCTACTGAGACACGC
GAGCACACGTCTGAACTCTTTCCCTACACGACGCTCTTCCGATCTNNNTACCCTCACAACATGGCAGT
GCACACGTCTGAACTCTTTCCCTACACGACGCTCTTCCGATCTNNNGCCAGATGCTCCTTTATCTGGT
GCACACGTCTGAACTCTTTCCCTACACGACGCTCTTCCGATCTNNNGGAGATGTACTGTGGAAAGCA
GAGCACACGTCTGAACTCTTTCCCTACACGACGCTCTTCCGATCTNNNTAGATCATGTGGTAGGGTTT
AGCACACGTCTGAACTCTTTCCCTACACGACGCTCTTCCGATCTNNNGACCTTCTGAGTTTGAGGCAG
AGCACACGTCTGAACTCTTTCCCTACACGACGCTCTTCCGATCTNNNATATCCAGCTTGTGTGTGTGT
GAGCACACGTCTGAACTCTTTCCCTACACGACGCTCTTCCGATCTNNNGGTGGCACCTACATTGCAAA
AAGAGCACACGTCTGAACTCTTTCCCTACACGACGCTCTTCCGATCTNNNGTGCCCTCATGATGGATT
GAGCACACGTCTGAACTCTTTCCCTACACGACGCTCTTCCGATCTNNNGGAGATAGGTCATCTACTGT
AGCACACGTCTGAACTCTTTCCCTACACGACGCTCTTCCGATCTNNNTCTGGGATCCTCTTTTAGGGCA
GAGCACACGTCTGAACTCTTTCCCTACACGACGCTCTTCCGATCTNNNGCAGCCCTATATGTGTAGCT
AGCACACGTCTGAACTCTTTCCCTACACGACGCTCTTCCGATCTNNNATGGACCCCAGGAATGGTTAC
GCACACGTCTGAACTCTTTCCCTACACGACGCTCTTCCGATCTNNNACGTCGCTCCAGTGTAAACAAAA
AGAGCACACGTCTGAACTCTTTCCCTACACGACGCTCTTCCGATCTNNNTAAGAAAACGACGACGAC
AGCACACGTCTGAACTCTTTCCCTACACGACGCTCTTCCGATCTNNNATCTGTGACGAAGCTGTGATG
AGCACACGTCTGAACTCTTTCCCTACACGACGCTCTTCCGATCTNNNTCACTTCCTTCTGACCCTCCTTT
AGCACACGTCTGAACTCTTTCCCTACACGACGCTCTTCCGATCTNNNATTCTGACAGGGCCAATTCAT
AGAGCACACGTCTGAACTCTTTCCCTACACGACGCTCTTCCGATCTNNNTGTGTCCACTGATCACATTT
AGAGCACACGTCTGAACTCTTTCCCTACACGACGCTCTTCCGATCTNNNGACCTAAGGTATGGCATGG
AAGAGCACACGTCTGAACTCTTTCCCTACACGACGCTCTTCCGATCTNNNACCAAGTCATCCTATTCTTC
AGCACACGTCTGAACTCTTTCCCTACACGACGCTCTTCCGATCTNNNCAGACACTGAGAATGGCCATG
GAGCACACGTCTGAACTCTTTCCCTACACGACGCTCTTCCGATCTNNNTGTTTTCTGTGTGGAAAGGG
CACACGTCTGAACTCTTTCCCTACACGACGCTCTTCCGATCTNNNGAAGATGTGCAGTCCAAGGCTAA
GAGCACACGTCTGAACTCTTTCCCTACACGACGCTCTTCCGATCTNNNCCAGGGAAAGGACATGATCT
AGCACACGTCTGAACTCTTTCCCTACACGACGCTCTTCCGATCTNNNTATTCAGTGCCAGCAGCTATCT
AGCACACGTCTGAACTCTTTCCCTACACGACGCTCTTCCGATCTNNNAGCCATGGCATAATGGGAAA
GAGCACACGTCTGAACTCTTTCCCTACACGACGCTCTTCCGATCTNNNAGAGATGATCCTTAGGCAGA
AGCACACGTCTGAACTCTTTCCCTACACGACGCTCTTCCGATCTNNNGCAACGATGCCATTGGAAAGA
GAGCACACGTCTGAACTCTTTCCCTACACGACGCTCTTCCGATCTNNNGGTTTTCACCACGGTGTTTC

GAGCACACGTCTGAACTCTTTCCCTACACGACGCTCTTCCGATCTNNNTCCTGACCTTCTTCCTAATGG
AGCACACGTCTGAACTCTTTCCCTACACGACGCTCTTCCGATCTNNNCATCCCTTTTCCTTCCTCCT
GAGCACACGTCTGAACTCTTTCCCTACACGACGCTCTTCCGATCTNNNAGTGAAAGAGGAGGTGATG
AGCACACGTCTGAACTCTTTCCCTACACGACGCTCTTCCGATCTNNNGCCAAGTAGAAGAGTACTAT
AAGAGCACACGTCTGAACTCTTTCCCTACACGACGCTCTTCCGATCTNNNCCTTTGAAGTAGGTCCCAT
GAGCACACGTCTGAACTCTTTCCCTACACGACGCTCTTCCGATCTNNNAAGTTCAAAGCCTGGTGAGA
AGAGCACACGTCTGAACTCTTTCCCTACACGACGCTCTTCCGATCTNNNAGACCCATAGCAAACTTGG
AGCACACGTCTGAACTCTTTCCCTACACGACGCTCTTCCGATCTNNNTGCCTTTAGTGTCCAAGGCAT
GCACACGTCTGAACTCTTTCCCTACACGACGCTCTTCCGATCTNNNTCTGTCCACCCAAACATCTACTC
AAGAGCACACGTCTGAACTCTTTCCCTACACGACGCTCTTCCGATCTNNNCATGTGCCTGTCAGTGAT
AGCACACGTCTGAACTCTTTCCCTACACGACGCTCTTCCGATCTNNNAAATCCAAAAGCCCCATCCTCT
GCACACGTCTGAACTCTTTCCCTACACGACGCTCTTCCGATCTNNNATTTTGTGTCTTTCCCAAACGCC
GCACACGTCTGAACTCTTTCCCTACACGACGCTCTTCCGATCTNNNTACACCTAATGGGCAAACAAGC
AGCACACGTCTGAACTCTTTCCCTACACGACGCTCTTCCGATCTNNNTCTGACTTTCCAAGTTGGCTG
AGCACACGTCTGAACTCTTTCCCTACACGACGCTCTTCCGATCTNNNCACACGTGACAAACTCCACCT
AGCACACGTCTGAACTCTTTCCCTACACGACGCTCTTCCGATCTNNNGGCAGCACAGACACTTAGGAT
AGCACACGTCTGAACTCTTTCCCTACACGACGCTCTTCCGATCTNNNCCACAGGGATGAAGTCAGTGA
GAGCACACGTCTGAACTCTTTCCCTACACGACGCTCTTCCGATCTNNNGCAGAAAGTAAGGCAAGA
AGCACACGTCTGAACTCTTTCCCTACACGACGCTCTTCCGATCTNNNGCCTTATCCCAAATGTTGTCAC
AGCACACGTCTGAACTCTTTCCCTACACGACGCTCTTCCGATCTNNNACAACTGTTAGAAGTGCCCTG
AGCACACGTCTGAACTCTTTCCCTACACGACGCTCTTCCGATCTNNNTTCCAACATCTAGACCCTGCT
GAGCACACGTCTGAACTCTTTCCCTACACGACGCTCTTCCGATCTNNNACCAAAGGACACATGGTAA
GCACACGTCTGAACTCTTTCCCTACACGACGCTCTTCCGATCTNNNTCAACCACAATGCAGGTCTTGA
GCACACGTCTGAACTCTTTCCCTACACGACGCTCTTCCGATCTNNNACATGTGTTTCACTAGTCTGCCA
AGCACACGTCTGAACTCTTTCCCTACACGACGCTCTTCCGATCTNNNATCTCATTGCATTACGGCCAGT
AGCACACGTCTGAACTCTTTCCCTACACGACGCTCTTCCGATCTNNNCCAGTACCTTTGCTGTGAATG
AGCACACGTCTGAACTCTTTCCCTACACGACGCTCTTCCGATCTNNNATTCAAGGGATCACTGTTCACC
AGCACACGTCTGAACTCTTTCCCTACACGACGCTCTTCCGATCTNNNATAGATGGGGGCCAAAATGTT
GCACACGTCTGAACTCTTTCCCTACACGACGCTCTTCCGATCTNNNATAGATGGGGGCCAAAATGTTC
AGAGCACACGTCTGAACTCTTTCCCTACACGACGCTCTTCCGATCTNNNAACATAGGTTCATCGGCTG
AGCACACGTCTGAACTCTTTCCCTACACGACGCTCTTCCGATCTNNNAACCTCACAAACCAGCTTCACT
AGCACACGTCTGAACTCTTTCCCTACACGACGCTCTTCCGATCTNNNACCATAATTGAGCTTGAGCTG
GCACACGTCTGAACTCTTTCCCTACACGACGCTCTTCCGATCTNNNTTTCTAAGTGCTTACGTGCTCAA
AGCACACGTCTGAACTCTTTCCCTACACGACGCTCTTCCGATCTNNNTGAGAAAATGACTTCTGATT
AGCACACGTCTGAACTCTTTCCCTACACGACGCTCTTCCGATCTNNNTGGATTGTGGAATGCTTAGCT
AGCACACGTCTGAACTCTTTCCCTACACGACGCTCTTCCGATCTNNNGCTAATGTCCTGACAAGAAGG
AGAGCACACGTCTGAACTCTTTCCCTACACGACGCTCTTCCGATCTNNNTGGTTGAAAACAAGCCTGT
GAGCACACGTCTGAACTCTTTCCCTACACGACGCTCTTCCGATCTNNNGCCCCTGAACCTGTAGTCTCT
GAGCACACGTCTGAACTCTTTCCCTACACGACGCTCTTCCGATCTNNNCTGATCTCCTCAACTTTCCTG
CACACGTCTGAACTCTTTCCCTACACGACGCTCTTCCGATCTNNNAATTGTAAAGCCTCGGCTAGGTG
GAGCACACGTCTGAACTCTTTCCCTACACGACGCTCTTCCGATCTNNNGGGGACAGAGTGAGATCC
CACACGTCTGAACTCTTTCCCTACACGACGCTCTTCCGATCTNNNTCCTCTATCCCCACAACCAACTTCAT
GCACACGTCTGAACTCTTTCCCTACACGACGCTCTTCCGATCTNNNTTGCTATTCCACTCTGCAGTCAC

AGAGCACACGTCTGAACTCTTTCCCTACACGACGCTCTTCCGATCTNNNACGGGAGGAGAGTGTTTCT
AGCACACGTCTGAACTCTTTCCCTACACGACGCTCTTCCGATCTNNNACTGTCAGCTCAACTGGGTTTT
AGCACACGTCTGAACTCTTTCCCTACACGACGCTCTTCCGATCTNNNACAATGATGGTAGGGCTTCAA
AGCACACGTCTGAACTCTTTCCCTACACGACGCTCTTCCGATCTNNNCGGACACACACACATACATCCA
AGCACACGTCTGAACTCTTTCCCTACACGACGCTCTTCCGATCTNNNATCAACAGTGAGAAATGCCCT
GAGCACACGTCTGAACTCTTTCCCTACACGACGCTCTTCCGATCTNNNTCTGACCTTAGGTAATTCGCC
AGCACACGTCTGAACTCTTTCCCTACACGACGCTCTTCCGATCTNNNAACTGACAACAGAACAGTGGA
AGCACACGTCTGAACTCTTTCCCTACACGACGCTCTTCCGATCTNNNAGTCAGACCCTAAAGACTATGT
AGCACACGTCTGAACTCTTTCCCTACACGACGCTCTTCCGATCTNNNGGTAATGTCATTGAACCATCCT
AGAGCACACGTCTGAACTCTTTCCCTACACGACGCTCTTCCGATCTNNNTCAGTGTAGTCAAGCCTTCC
GAGCACACGTCTGAACTCTTTCCCTACACGACGCTCTTCCGATCTNNNTCTTACTGGAGCTACCATAGG
AGAGCACACGTCTGAACTCTTTCCCTACACGACGCTCTTCCGATCTNNNAAGTCTTCAGGGTCTAGAG
GCACACGTCTGAACTCTTTCCCTACACGACGCTCTTCCGATCTNNNGGTCATTTCACACTTCTGCTTGG
AGCACACGTCTGAACTCTTTCCCTACACGACGCTCTTCCGATCTNNNAAGGCCTCTGGAGAGATCAGT
AGCACACGTCTGAACTCTTTCCCTACACGACGCTCTTCCGATCTNNNGAACTTTTCTCTGAGGGGGCT
AGCACACGTCTGAACTCTTTCCCTACACGACGCTCTTCCGATCTNNNAGTCTAATGGTTTCTAAACCCC
AGAGCACACGTCTGAACTCTTTCCCTACACGACGCTCTTCCGATCTNNNACGGCAATAGTCCACTGAG
GCACACGTCTGAACTCTTTCCCTACACGACGCTCTTCCGATCTNNNGAACAGCAGTATTTTCCTTGCGT
AGCACACGTCTGAACTCTTTCCCTACACGACGCTCTTCCGATCTNNNGTGGCATGTCTGTGTGTGTGA
GAGCACACGTCTGAACTCTTTCCCTACACGACGCTCTTCCGATCTNNNGCATGTATGATTGACAGCAG
AGCACACGTCTGAACTCTTTCCCTACACGACGCTCTTCCGATCTNNNTAAGCCACTAGTGAGGGAACT
GAGCACACGTCTGAACTCTTTCCCTACACGACGCTCTTCCGATCTNNNTCCCTATGTCTGCATTGCTCCT
GCACACGTCTGAACTCTTTCCCTACACGACGCTCTTCCGATCTNNNAGCAACAAATCCAATCTGTCCGT
GAGCACACGTCTGAACTCTTTCCCTACACGACGCTCTTCCGATCTNNNCGGACTTTTATCCAGATGCCTC
GAGCACACGTCTGAACTCTTTCCCTACACGACGCTCTTCCGATCTNNNACGCTATAGAGCCTGGAGTTT
AGCACACGTCTGAACTCTTTCCCTACACGACGCTCTTCCGATCTNNNTGGTTACTGTAGTTTGCTGAGC
AGAGCACACGTCTGAACTCTTTCCCTACACGACGCTCTTCCGATCTNNNGCTCATTTGTAATTAGGCCC
AGCACACGTCTGAACTCTTTCCCTACACGACGCTCTTCCGATCTNNNTCCTCTATCCCCACAACCAACTTC
AGCACACGTCTGAACTCTTTCCCTACACGACGCTCTTCCGATCTNNNTACTTACAGATCCTGGAGGGA
AGAGCACACGTCTGAACTCTTTCCCTACACGACGCTCTTCCGATCTNNNAAGTCTTCAGGGTCTAGAG
AGCACACGTCTGAACTCTTTCCCTACACGACGCTCTTCCGATCTNNNCCTTTCCCTATTTTGGGCACAC
GCACACGTCTGAACTCTTTCCCTACACGACGCTCTTCCGATCTNNNGTCACCCCTACAACACATAAACC
AGCACACGTCTGAACTCTTTCCCTACACGACGCTCTTCCGATCTNNNGTTTTGTGTGTGGCTGGAGA
GCACACGTCTGAACTCTTTCCCTACACGACGCTCTTCCGATCTNNNTGTGGCAAGGCAGTATGATTGTT
GAGCACACGTCTGAACTCTTTCCCTACACGACGCTCTTCCGATCTNNNCTTACCTCATTGCAGCTGTCT
GAGCACACGTCTGAACTCTTTCCCTACACGACGCTCTTCCGATCTNNNCAGTTCCTGGCACAAACTAAT
AGAGCACACGTCTGAACTCTTTCCCTACACGACGCTCTTCCGATCTNNNACACAAATGGGATCAGGAC
AGCACACGTCTGAACTCTTTCCCTACACGACGCTCTTCCGATCTNNNAGATTCTGCAGGCATCTTCACA
CACACGTCTGAACTCTTTCCCTACACGACGCTCTTCCGATCTNNNATAGTGAAAGTGTGTGTGTGTGCC
GAGCACACGTCTGAACTCTTTCCCTACACGACGCTCTTCCGATCTNNNTCTCAGACTTTTGGTTTTTACC
AGCACACGTCTGAACTCTTTCCCTACACGACGCTCTTCCGATCTNNNTTTCAGTGATGTATGTGGAGAA
GCACACGTCTGAACTCTTTCCCTACACGACGCTCTTCCGATCTNNNCACAGGAATGAGTGACTGTGGA
AGCACACGTCTGAACTCTTTCCCTACACGACGCTCTTCCGATCTNNNAAAGTGTCCCACATCCTTTTCCC

CACACGTCTGAACTCTTTCCCTACACGACGCTCTTCCGATCTNNNCTGCTCTCTAGACCTCCCGTGCATC
CACACGTCTGAACTCTTTCCCTACACGACGCTCTTCCGATCTNNNCAGAGCCAGAGTTAGTAGGGAG
GAGCACACGTCTGAACTCTTTCCCTACACGACGCTCTTCCGATCTNNNTCCACCCTTTTGCTTTTGCAT
GAGCACACGTCTGAACTCTTTCCCTACACGACGCTCTTCCGATCTNNNTCTGATGCTACACATCCTTCCC
AAGAGCACACGTCTGAACTCTTTCCCTACACGACGCTCTTCCGATCTNNNTTTCTGTCTTTCTGCGTAT
AGCACACGTCTGAACTCTTTCCCTACACGACGCTCTTCCGATCTNNNTCCATAGGAGACCAGTGTATA
GAGCACACGTCTGAACTCTTTCCCTACACGACGCTCTTCCGATCTNNNGTCCGTTGCTCATTCTCTCTCT
GAGCACACGTCTGAACTCTTTCCCTACACGACGCTCTTCCGATCTNNNACTTACACGTCTGGTACCTC
GAGCACACGTCTGAACTCTTTCCCTACACGACGCTCTTCCGATCTNNNTCTAAGCCGTACGCTCCAAAA
GAGCACACGTCTGAACTCTTTCCCTACACGACGCTCTTCCGATCTNNNACAAGAAGACAGGGATACG
AGCACACGTCTGAACTCTTTCCCTACACGACGCTCTTCCGATCTNNNGTCCAATTTCAGTGGAAGCAG
GCACACGTCTGAACTCTTTCCCTACACGACGCTCTTCCGATCTNNNATACCCATTCACTGCACATCAG
GCACACGTCTGAACTCTTTCCCTACACGACGCTCTTCCGATCTNNNTCTGGGATTATATCCTTCCATGC
GAGCACACGTCTGAACTCTTTCCCTACACGACGCTCTTCCGATCTNNNATCTCTGTCCCTCTTCCCCATT
AGCACACGTCTGAACTCTTTCCCTACACGACGCTCTTCCGATCTNNNTTTGAATGTTGACCTGGGTG
AGCACACGTCTGAACTCTTTCCCTACACGACGCTCTTCCGATCTNNNACAGCACCTGGTCCAATTTTC
AGCACACGTCTGAACTCTTTCCCTACACGACGCTCTTCCGATCTNNNGCTCTTTCTTAGGGGTTTCTGG
GAGCACACGTCTGAACTCTTTCCCTACACGACGCTCTTCCGATCTNNNGGATGTGACTAGGAGGGAG
GCACACGTCTGAACTCTTTCCCTACACGACGCTCTTCCGATCTNNNGTTTCCTAATGCCTGGGAGATG
GAGCACACGTCTGAACTCTTTCCCTACACGACGCTCTTCCGATCTNNNTCAGGAATGGGACCTCCTTC
AGCACACGTCTGAACTCTTTCCCTACACGACGCTCTTCCGATCTNNNAAGTGGACTACTAAAGCAAGG
AGAGCACACGTCTGAACTCTTTCCCTACACGACGCTCTTCCGATCTNNNGTGTGTAAACCAAGTCCCA
GCACACGTCTGAACTCTTTCCCTACACGACGCTCTTCCGATCTNNNCCCTGAATATTTTGCCCACACG
GAGCACACGTCTGAACTCTTTCCCTACACGACGCTCTTCCGATCTNNNTGATGATTTGGGAGTAGGG
AGCACACGTCTGAACTCTTTCCCTACACGACGCTCTTCCGATCTNNNACACACACACAGCTTCATCTGT
AAGAGCACACGTCTGAACTCTTTCCCTACACGACGCTCTTCCGATCTNNNGCAAGGTACTGCCTGAT
GCACACGTCTGAACTCTTTCCCTACACGACGCTCTTCCGATCTNNNGCCTAGAAACCTGTCACATGAG
AGAGCACACGTCTGAACTCTTTCCCTACACGACGCTCTTCCGATCTNNNCAGTCCCAATGCTCACGAA
AGAGCACACGTCTGAACTCTTTCCCTACACGACGCTCTTCCGATCTNNNTGTGGTGTAACACTCAAGA
GCACACGTCTGAACTCTTTCCCTACACGACGCTCTTCCGATCTNNNTGCATGAAAACCTTCAGGTGGA
AGCACACGTCTGAACTCTTTCCCTACACGACGCTCTTCCGATCTNNNAGCAATGGCTGGAAACACTT
GCACACGTCTGAACTCTTTCCCTACACGACGCTCTTCCGATCTNNNTTTCCAGATGCAGAAGCCAAAG
AGCACACGTCTGAACTCTTTCCCTACACGACGCTCTTCCGATCTNNNCTTTTCTTGCTCTCGGCATCCTT
GAGCACACGTCTGAACTCTTTCCCTACACGACGCTCTTCCGATCTNNNGAGTTGGAGCAGGACTAAG
CACACGTCTGAACTCTTTCCCTACACGACGCTCTTCCGATCTNNNGGACACTAGGGTTCCTGTGAGAA
GCACACGTCTGAACTCTTTCCCTACACGACGCTCTTCCGATCTNNNCTGCCCTTTTCTCACTTCCTCTCC
CACACGTCTGAACTCTTTCCCTACACGACGCTCTTCCGATCTNNNCAAATTTTTCCAGCTTTGGCCTCT
GAGCACACGTCTGAACTCTTTCCCTACACGACGCTCTTCCGATCTNNNAAAAGAAGGTCTGCTCAGCT
AGCACACGTCTGAACTCTTTCCCTACACGACGCTCTTCCGATCTNNNTTTTAGCCTGATTCCACAGCCA
GAGCACACGTCTGAACTCTTTCCCTACACGACGCTCTTCCGATCTNNNAGTCCCACATTTCATTGGTAG
GAGCACACGTCTGAACTCTTTCCCTACACGACGCTCTTCCGATCTNNNCTCATGGTGGCTCACACTTAT
GCACACGTCTGAACTCTTTCCCTACACGACGCTCTTCCGATCTNNNCTGAAATCCCACCACCCAAAGAT
AGCACACGTCTGAACTCTTTCCCTACACGACGCTCTTCCGATCTNNNTCTGTATTGCTGCAGTCTTTGT

GAGCACACGTCTGAACTCTTTCCCTACACGACGCTCTTCCGATCTNNNTGCATTTTATCATGTTGGATG
CACACGTCTGAACTCTTTCCCTACACGACGCTCTTCCGATCTNNNCGGCCAGGTCTCATGTGTTTTCAT
AGCACACGTCTGAACTCTTTCCCTACACGACGCTCTTCCGATCTNNNTCCAAGAACGGCTCAAAGATT
AGCACACGTCTGAACTCTTTCCCTACACGACGCTCTTCCGATCTNNNGCCTGATTGGTGAATCACTCT
GAGCACACGTCTGAACTCTTTCCCTACACGACGCTCTTCCGATCTNNNTACCCAGCCATCCTTGCCTAT
AGCACACGTCTGAACTCTTTCCCTACACGACGCTCTTCCGATCTNNNGGCTTGAAATGAACAGGGAAT
CACACGTCTGAACTCTTTCCCTACACGACGCTCTTCCGATCTNNNTCTCTGGGTCTCTGACGGTTTCCA
AGCACACGTCTGAACTCTTTCCCTACACGACGCTCTTCCGATCTNNNTGGGAAGGCAAATGTGTTTT
AGCACACGTCTGAACTCTTTCCCTACACGACGCTCTTCCGATCTNNNAGGACAGTTCAGAGATGCTG
GCACACGTCTGAACTCTTTCCCTACACGACGCTCTTCCGATCTNNNTCTGGGATTATATCCTTCCATGC
GCACACGTCTGAACTCTTTCCCTACACGACGCTCTTCCGATCTNNNACTATCATGGTTCCCAGCAAAA
CACACGTCTGAACTCTTTCCCTACACGACGCTCTTCCGATCTNNNATCAACAGTGAGAAATGCCCTCCC
GAGCACACGTCTGAACTCTTTCCCTACACGACGCTCTTCCGATCTNNNCCCTGACTAGGTTGAAGTG
AGCACACGTCTGAACTCTTTCCCTACACGACGCTCTTCCGATCTNNNGCTCTGTATCAGTCACGGTTCT
GAGCACACGTCTGAACTCTTTCCCTACACGACGCTCTTCCGATCTNNNTCATGAGGCTGCTTTCTCAG
GAGCACACGTCTGAACTCTTTCCCTACACGACGCTCTTCCGATCTNNNAGAACCCACGAACACAATTC
AAGAGCACACGTCTGAACTCTTTCCCTACACGACGCTCTTCCGATCTNNNTGATGATTTGGGAGTAG
AGCACACGTCTGAACTCTTTCCCTACACGACGCTCTTCCGATCTNNNCACACGTTACCATGTCTGCCTA
GAGCACACGTCTGAACTCTTTCCCTACACGACGCTCTTCCGATCTNNNTCCCTGAAATTAGTATTGGG
AGAGCACACGTCTGAACTCTTTCCCTACACGACGCTCTTCCGATCTNNNACATTTTCCATTTCCCTCAC
AGCACACGTCTGAACTCTTTCCCTACACGACGCTCTTCCGATCTNNNTGCCTGACTTAGCATAGTGTC
AGCACACGTCTGAACTCTTTCCCTACACGACGCTCTTCCGATCTNNNTGAATAAGGCTCCTGTGTTCTT
AGAGCACACGTCTGAACTCTTTCCCTACACGACGCTCTTCCGATCTNNNGCTCACTGAAATCGGAAG
AGCACACGTCTGAACTCTTTCCCTACACGACGCTCTTCCGATCTNNNTTTCTACAGTCAAGCCAGTTTG
AGCACACGTCTGAACTCTTTCCCTACACGACGCTCTTCCGATCTNNNGAAGAGCAACAAGAGCAACA
AGCACACGTCTGAACTCTTTCCCTACACGACGCTCTTCCGATCTNNNGCAAGGTACTGCCTGATGTA
AGCACACGTCTGAACTCTTTCCCTACACGACGCTCTTCCGATCTNNNCTGTGCCCAGCCTACATAAGA
GCACACGTCTGAACTCTTTCCCTACACGACGCTCTTCCGATCTNNNCCAATCTTATTCAGAGCCCTCCA
GCACACGTCTGAACTCTTTCCCTACACGACGCTCTTCCGATCTNNNACATTGTACGTGCTTTGGGACA
AAGAGCACACGTCTGAACTCTTTCCCTACACGACGCTCTTCCGATCTNNNGTCAATACTACAGCGAG
AGCACACGTCTGAACTCTTTCCCTACACGACGCTCTTCCGATCTNNNAGCGTTTTGACTTCTTGTTCCA
AGCACACGTCTGAACTCTTTCCCTACACGACGCTCTTCCGATCTNNNGGAAAATGGTTCTCTCCCTGA
GCACACGTCTGAACTCTTTCCCTACACGACGCTCTTCCGATCTNNNGGCTTTTCCTCTCCATACCTCCT
GAGCACACGTCTGAACTCTTTCCCTACACGACGCTCTTCCGATCTNNNCCCAGAATACCCCAAGATGG
GAGCACACGTCTGAACTCTTTCCCTACACGACGCTCTTCCGATCTNNNAATGCTAAATGCTAAAGGCG
AGCACACGTCTGAACTCTTTCCCTACACGACGCTCTTCCGATCTNNNAATGTACTGATGCCACTGAAC
AGCACACGTCTGAACTCTTTCCCTACACGACGCTCTTCCGATCTNNNTGGGTATGAGCACTGTGACTT
GAGCACACGTCTGAACTCTTTCCCTACACGACGCTCTTCCGATCTNNNGGTCCTATTTTGGCTCCTTCT
CACACGTCTGAACTCTTTCCCTACACGACGCTCTTCCGATCTNNNTGTATTACAGATTGTATTTTGCCA
AGCACACGTCTGAACTCTTTCCCTACACGACGCTCTTCCGATCTNNNGTATACCTGGTGAAAGCAGCA
GCACACGTCTGAACTCTTTCCCTACACGACGCTCTTCCGATCTNNNCTAGCATCTATTGCTGGCACCAT
AGCACACGTCTGAACTCTTTCCCTACACGACGCTCTTCCGATCTNNNCAGCAGATGTGTCTCATTCAC
AGCACACGTCTGAACTCTTTCCCTACACGACGCTCTTCCGATCTNNNGCCAGTGTACCCAGAGTGATT

AAGAGCACACGTCTGAACTCTTTCCCTACACGACGCTCTTCCGATCTNNNAGCCCTACCTTGTGCATGT
GAGCACACGTCTGAACTCTTTCCCTACACGACGCTCTTCCGATCTNNNTGCACTGGTTTTTGATCTTGC
GAGCACACGTCTGAACTCTTTCCCTACACGACGCTCTTCCGATCTNNNCAGGAAGACTGGTAATGGG
GAGCACACGTCTGAACTCTTTCCCTACACGACGCTCTTCCGATCTNNNGTGGTGTCTCAACTCTCAAGT
AGCACACGTCTGAACTCTTTCCCTACACGACGCTCTTCCGATCTNNNACAGCATTCTGGTCCCATCTTTC
AGAGCACACGTCTGAACTCTTTCCCTACACGACGCTCTTCCGATCTNNNCCACCATGTCTTGGTTCCTA
AGCACACGTCTGAACTCTTTCCCTACACGACGCTCTTCCGATCTNNNTCTTCCCAGGTTCAACAGATTTT
GAGCACACGTCTGAACTCTTTCCCTACACGACGCTCTTCCGATCTNNNTCATCTGGAGATGTTGGCAC
AGAGCACACGTCTGAACTCTTTCCCTACACGACGCTCTTCCGATCTNNNAGCTGACCCATACTGCAGA
AAGAGCACACGTCTGAACTCTTTCCCTACACGACGCTCTTCCGATCTNNNAACTTTTCCTCAAGAGCAC
CACACGTCTGAACTCTTTCCCTACACGACGCTCTTCCGATCTNNNAATCAAAGGTACTCGGAGGCTGA
AGCACACGTCTGAACTCTTTCCCTACACGACGCTCTTCCGATCTNNNGCCTCCAGTATATGTTAGCAG
AGCACACGTCTGAACTCTTTCCCTACACGACGCTCTTCCGATCTNNNAAGGGGAGGAACGGCTATAA
AGAGCACACGTCTGAACTCTTTCCCTACACGACGCTCTTCCGATCTNNNCTGTGGCCTTGCCTATACTT
AGCACACGTCTGAACTCTTTCCCTACACGACGCTCTTCCGATCTNNNACAACACCGAATTCAAGCGATT
GAGCACACGTCTGAACTCTTTCCCTACACGACGCTCTTCCGATCTNNNCCATCTAGAAACCACCAAGAG
AGAGCACACGTCTGAACTCTTTCCCTACACGACGCTCTTCCGATCTNNNACTGTCTTACTTTGCCTTGA
AGCACACGTCTGAACTCTTTCCCTACACGACGCTCTTCCGATCTNNNACAGTCCTTTGTTGCCTGATGT
GAGCACACGTCTGAACTCTTTCCCTACACGACGCTCTTCCGATCTNNNAACTTGCCTAGTCACCCCTACA
AGAGCACACGTCTGAACTCTTTCCCTACACGACGCTCTTCCGATCTNNNTGTGTAGACTTCAGGGACA
GCACACGTCTGAACTCTTTCCCTACACGACGCTCTTCCGATCTNNNAGATGAAGGAGGGCAGAGAA
AGAGCACACGTCTGAACTCTTTCCCTACACGACGCTCTTCCGATCTNNNATACCCATTCACTGCACATC
GCACACGTCTGAACTCTTTCCCTACACGACGCTCTTCCGATCTNNNTGTTACGCAAATTGAGCGAGAG
AAGAGCACACGTCTGAACTCTTTCCCTACACGACGCTCTTCCGATCTNNNGTGTCTCGCGCTTTCTCGG
GAAGAGCACACGTCTGAACTCTTTCCCTACACGACGCTCTTCCGATCTNNNTTCCTGATTCAGAACAGC
GCACACGTCTGAACTCTTTCCCTACACGACGCTCTTCCGATCTNNNTTTGGCAAGTTGACCAGGTTGAT
AGAGCACACGTCTGAACTCTTTCCCTACACGACGCTCTTCCGATCTNNNGCTATGACTGTGCACTTCTC
AGCACACGTCTGAACTCTTTCCCTACACGACGCTCTTCCGATCTNNNATAAGCAAGGGCCCAATTTCT
AGAGCACACGTCTGAACTCTTTCCCTACACGACGCTCTTCCGATCTNNNAACGTTTCCGCCACTTTATCT
AGCACACGTCTGAACTCTTTCCCTACACGACGCTCTTCCGATCTNNNTCTGGAGTGTGGTGGTGCAAA
GAAGAGCACACGTCTGAACTCTTTCCCTACACGACGCTCTTCCGATCTNNNAGCGAGTGGAATAAAAT
GCACACGTCTGAACTCTTTCCCTACACGACGCTCTTCCGATCTNNNTGACGTTGTATGTTTCGCTGTCT
AGCACACGTCTGAACTCTTTCCCTACACGACGCTCTTCCGATCTNNNCTTCCAGTAACTCCAGACCTTG
AGCACACGTCTGAACTCTTTCCCTACACGACGCTCTTCCGATCTNNNACAACTCTGTTGTGTGTGTGT
AGAGCACACGTCTGAACTCTTTCCCTACACGACGCTCTTCCGATCTNNNCACTCCCAAATTGTGGTTCT
GCACACGTCTGAACTCTTTCCCTACACGACGCTCTTCCGATCTNNNAAATCTACCTGTGAGCTGGAACC
AGAGCACACGTCTGAACTCTTTCCCTACACGACGCTCTTCCGATCTNNNGATAAAGAACTGCTGTGCT
AGAGCACACGTCTGAACTCTTTCCCTACACGACGCTCTTCCGATCTNNNCAGGTCCCCGAAAGAATGA
AGCACACGTCTGAACTCTTTCCCTACACGACGCTCTTCCGATCTNNNTTGTAGCCCCTACCAACAGAAG
GAGCACACGTCTGAACTCTTTCCCTACACGACGCTCTTCCGATCTNNNGATAGGATGTCCAGGATGG
AGCACACGTCTGAACTCTTTCCCTACACGACGCTCTTCCGATCTNNNAGAAATGTTACTGCTAAGCACC
AGCACACGTCTGAACTCTTTCCCTACACGACGCTCTTCCGATCTNNNGGGTAAGCTAGTGGTAGTTCT
GAGCACACGTCTGAACTCTTTCCCTACACGACGCTCTTCCGATCTNNNAATTCTGCATCTGTGTGTGTG

AGAGCACACGTCTGAACTCTTTCCCTACACGACGCTCTTCCGATCTNNNCGAAGGATATATGTGTGTG
GAGCACACGTCTGAACTCTTTCCCTACACGACGCTCTTCCGATCTNNNTGTGCTGTTTTCAGCTCTCTCT
AGAGCACACGTCTGAACTCTTTCCCTACACGACGCTCTTCCGATCTNNNAAATAATTGTGTGCTTGGG
AGCACACGTCTGAACTCTTTCCCTACACGACGCTCTTCCGATCTNNNGAAGAGCAGATGGTTGGCTAC
AGCACACGTCTGAACTCTTTCCCTACACGACGCTCTTCCGATCTNNNAGATCAAGCTGGGAAGCAAAT
GAGCACACGTCTGAACTCTTTCCCTACACGACGCTCTTCCGATCTNNNAGGAAAGTGTGCTTACAGAA
AGCACACGTCTGAACTCTTTCCCTACACGACGCTCTTCCGATCTNNNGGCAGCCAAAGGGTGAATCTA
GAGCACACGTCTGAACTCTTTCCCTACACGACGCTCTTCCGATCTNNNGGTGAGTAGCATTCCATAG
AGCACACGTCTGAACTCTTTCCCTACACGACGCTCTTCCGATCTNNNGCCTACCTGATCCAAGTAAAG
AGCACACGTCTGAACTCTTTCCCTACACGACGCTCTTCCGATCTNNNAATGTGGAAATTAGTTCCTCTG
AGCACACGTCTGAACTCTTTCCCTACACGACGCTCTTCCGATCTNNNAGCTGAACAAACCTGAGAGGG
CACACGTCTGAACTCTTTCCCTACACGACGCTCTTCCGATCTNNNTCCTTTCTGCTCAATTACAGGGCCA
AGAGCACACGTCTGAACTCTTTCCCTACACGACGCTCTTCCGATCTNNNGTGGTTTTCAAATCTTCCCC
GCACACGTCTGAACTCTTTCCCTACACGACGCTCTTCCGATCTNNNTTCTTTTCTGCCTTCCCCAACTCCA
GAGCACACGTCTGAACTCTTTCCCTACACGACGCTCTTCCGATCTNNNTGCATGGTTTTCTCTCTCACA
GAGCACACGTCTGAACTCTTTCCCTACACGACGCTCTTCCGATCTNNNTCCTAGGGACAAGAGGAAGT
GAGCACACGTCTGAACTCTTTCCCTACACGACGCTCTTCCGATCTNNNCAGGACCTTACTGCCACTTTCT
GAGCACACGTCTGAACTCTTTCCCTACACGACGCTCTTCCGATCTNNNGTCCATGCATCAGAAACAACA
GAGCACACGTCTGAACTCTTTCCCTACACGACGCTCTTCCGATCTNNNTCCTGCTAAAGTCTCCATCTTC
AGCACACGTCTGAACTCTTTCCCTACACGACGCTCTTCCGATCTNNNATGCACCATGCAGGATTGCATA
GAAGAGCACACGTCTGAACTCTTTCCCTACACGACGCTCTTCCGATCTNNNATAAAGCCCTGTGCAGGA
GAGCACACGTCTGAACTCTTTCCCTACACGACGCTCTTCCGATCTNNNAGGAGTTCAAGGTCATGGGT
GAGCACACGTCTGAACTCTTTCCCTACACGACGCTCTTCCGATCTNNNGCACCCTTGATCTATGTCTCTC
AGCACACGTCTGAACTCTTTCCCTACACGACGCTCTTCCGATCTNNNATGCCACACACATATCTATGCAC
AGCACACGTCTGAACTCTTTCCCTACACGACGCTCTTCCGATCTNNNCCCAGCTGCCTATTCTGAGTTTA
GCACACGTCTGAACTCTTTCCCTACACGACGCTCTTCCGATCTNNNCTATCTGGTTGGTGCTACTGCAA
GAGCACACGTCTGAACTCTTTCCCTACACGACGCTCTTCCGATCTNNNTTCTCCCTCTGGAATTTGCAAC
AGCACACGTCTGAACTCTTTCCCTACACGACGCTCTTCCGATCTNNNTCCAAAGTCATTCCAAAAGTAT
GAGCACACGTCTGAACTCTTTCCCTACACGACGCTCTTCCGATCTNNNCCCCATCTCCCAAAACAAGAA
AGAGCACACGTCTGAACTCTTTCCCTACACGACGCTCTTCCGATCTNNNGCATGCAGCAATTCATCTAG
AGCACACGTCTGAACTCTTTCCCTACACGACGCTCTTCCGATCTNNNATTCTTCACGGTGCTGTATCCT
CACACGTCTGAACTCTTTCCCTACACGACGCTCTTCCGATCTNNNAAGTTGGCTTGTGGTTACAACTTC
GCACACGTCTGAACTCTTTCCCTACACGACGCTCTTCCGATCTNNNAGGCAAATGACCCTCATACCTCT
AGAGCACACGTCTGAACTCTTTCCCTACACGACGCTCTTCCGATCTNNNCGAGCCAAATAATGGAAACT
AGCACACGTCTGAACTCTTTCCCTACACGACGCTCTTCCGATCTNNNTGGAACACTTTCCTTGACCGAT
AGCACACGTCTGAACTCTTTCCCTACACGACGCTCTTCCGATCTNNNGACAAACTCTCACTCTTGGGCT
AGCACACGTCTGAACTCTTTCCCTACACGACGCTCTTCCGATCTNNNGTCCTTGCCTAATGACCACAGA
AGCACACGTCTGAACTCTTTCCCTACACGACGCTCTTCCGATCTNNNTGAAACCCACTCCTCTCTCTCTCC
AGCACACGTCTGAACTCTTTCCCTACACGACGCTCTTCCGATCTNNNTCTTATTTCTGCCCACCCACTTCC
GAGCACACGTCTGAACTCTTTCCCTACACGACGCTCTTCCGATCTNNNCATGGCGTATTTTACAGCCAC
AGCACACGTCTGAACTCTTTCCCTACACGACGCTCTTCCGATCTNNNTCTGTGGTTGATATTGCTGCCTC
AGCACACGTCTGAACTCTTTCCCTACACGACGCTCTTCCGATCTNNNTTAGGAGGCCCCAAACTTCAAA
AGCACACGTCTGAACTCTTTCCCTACACGACGCTCTTCCGATCTNNNATACTGTGCTGTTTCTGGAGC

AGAGCACACGTCTGAACTCTTTCCCTACACGACGCTCTTCCGATCTNNNAAAATGCTAACACCACCACC
GCACACGTCTGAACTCTTTCCCTACACGACGCTCTTCCGATCTNNNAAAATGGAAAACAAATGTGCAT
GAGCACACGTCTGAACTCTTTCCCTACACGACGCTCTTCCGATCTNNNGACCTCCATAAAGTTCACCGC
AGAGCACACGTCTGAACTCTTTCCCTACACGACGCTCTTCCGATCTNNNGACACCCATTCACACACCAA
AGCACACGTCTGAACTCTTTCCCTACACGACGCTCTTCCGATCTNNNATACACAGCAAGGCAATGGG
GCACACGTCTGAACTCTTTCCCTACACGACGCTCTTCCGATCTNNNCCTACTTGTTTCTTTCCGTCCCCC
GCACACGTCTGAACTCTTTCCCTACACGACGCTCTTCCGATCTNNNTCAAAGCAGGCGACTAGTAGAG
AGCACACGTCTGAACTCTTTCCCTACACGACGCTCTTCCGATCTNNNTCTGGAAAAGGCAAACAATAC
GAGCACACGTCTGAACTCTTTCCCTACACGACGCTCTTCCGATCTNNNATTAGAAGGTAGCTGGAGG
GCACACGTCTGAACTCTTTCCCTACACGACGCTCTTCCGATCTNNNACCTCCTTGGTAGGATGTATTG
AAGAGCACACGTCTGAACTCTTTCCCTACACGACGCTCTTCCGATCTNNNTAGGTACTCAGTGAACA
AGCACACGTCTGAACTCTTTCCCTACACGACGCTCTTCCGATCTNNNTTTGTAAAAAGCACACACACA
AGCACACGTCTGAACTCTTTCCCTACACGACGCTCTTCCGATCTNNNAGGTCTCTTCTATTTTCCACCTC
GAGCACACGTCTGAACTCTTTCCCTACACGACGCTCTTCCGATCTNNNTCTAATGATGTGTGTGTGTG
AGAGCACACGTCTGAACTCTTTCCCTACACGACGCTCTTCCGATCTNNNAAGTACTGCATTGGCTTAG
AGCACACGTCTGAACTCTTTCCCTACACGACGCTCTTCCGATCTNNNTTCTTTTATCTCCCACTGCACCC
AGCACACGTCTGAACTCTTTCCCTACACGACGCTCTTCCGATCTNNNGCAGAAGGTAGGCTTTTCCAT
GAGCACACGTCTGAACTCTTTCCCTACACGACGCTCTTCCGATCTNNNTGAGCTGTGCATGGAGAAAA
GAGCACACGTCTGAACTCTTTCCCTACACGACGCTCTTCCGATCTNNNGGGAGACCTAGCTTCCTTTTC
AGCACACGTCTGAACTCTTTCCCTACACGACGCTCTTCCGATCTNNNTAGGGTCTTGTGTGTGCTCTCA
GAGCACACGTCTGAACTCTTTCCCTACACGACGCTCTTCCGATCTNNNGGGTTAGCGCTGTCTCTTTTG
AGCACACGTCTGAACTCTTTCCCTACACGACGCTCTTCCGATCTNNNCTCACTTGTTCTGTTCTCCCCTG
AGCACACGTCTGAACTCTTTCCCTACACGACGCTCTTCCGATCTNNNCCACAGGACGCACCTGTAATA
AGCACACGTCTGAACTCTTTCCCTACACGACGCTCTTCCGATCTNNNACCTCCTTGGTAGGATGTATT
GAGCACACGTCTGAACTCTTTCCCTACACGACGCTCTTCCGATCTNNNATCTATGCGAGTGTGTTTGT
AGCACACGTCTGAACTCTTTCCCTACACGACGCTCTTCCGATCTNNNATGTTTGGTCCCAGAGATGGG
GAGCACACGTCTGAACTCTTTCCCTACACGACGCTCTTCCGATCTNNNGCTGGATTTATAGGACCATG
GAGCACACGTCTGAACTCTTTCCCTACACGACGCTCTTCCGATCTNNNTGAGGCACCTTCCAGAGAAG
GAGCACACGTCTGAACTCTTTCCCTACACGACGCTCTTCCGATCTNNNACTAAGAATACTCCACCAAGG
AGAGCACACGTCTGAACTCTTTCCCTACACGACGCTCTTCCGATCTNNNTTCTCTGGGGACTGTGATG
GAGCACACGTCTGAACTCTTTCCCTACACGACGCTCTTCCGATCTNNNTACATGTGTGTTTCTGGGTG
AGCACACGTCTGAACTCTTTCCCTACACGACGCTCTTCCGATCTNNNACACCTCTATGTGAGCAAACA
GAGCACACGTCTGAACTCTTTCCCTACACGACGCTCTTCCGATCTNNNCTCACTTAGCAATGACACCCT
GAGCACACGTCTGAACTCTTTCCCTACACGACGCTCTTCCGATCTNNNCCCCAATCCAATTCCCCTTGA
AGCACACGTCTGAACTCTTTCCCTACACGACGCTCTTCCGATCTNNNTCTCACAAGTGCCTTGGCTATT
AAGAGCACACGTCTGAACTCTTTCCCTACACGACGCTCTTCCGATCTNNNTGGAAATCCTTAGATTGC
GAGCACACGTCTGAACTCTTTCCCTACACGACGCTCTTCCGATCTNNNCATGGCAGGCTTGTATGCAA
AAGAGCACACGTCTGAACTCTTTCCCTACACGACGCTCTTCCGATCTNNNAATCAGACTGGGCGCAG
AGCACACGTCTGAACTCTTTCCCTACACGACGCTCTTCCGATCTNNNCAAAGAAAAGACCAGGCACA
GAGCACACGTCTGAACTCTTTCCCTACACGACGCTCTTCCGATCTNNNTTGCATGTAGTCCCTTTCTCC
AGCACACGTCTGAACTCTTTCCCTACACGACGCTCTTCCGATCTNNNTACAGTTGAAAATATGGGCCC
CACACGTCTGAACTCTTTCCCTACACGACGCTCTTCCGATCTNNNACACACAGCTTTTCCCAAAGTTAC
AAGAGCACACGTCTGAACTCTTTCCCTACACGACGCTCTTCCGATCTNNNGGAACATAGGAAGTTCCA

GAGCACACGTCTGAACTCTTTCCCTACACGACGCTCTTCCGATCTNNNGCTGTAGTGAGGAAAGGAG
GCACACGTCTGAACTCTTTCCCTACACGACGCTCTTCCGATCTNNNGAGGCTCTCCTAGACGAAGAGT
AGCACACGTCTGAACTCTTTCCCTACACGACGCTCTTCCGATCTNNNCTTCAAACCAATCAGGGTGGCT
GCACACGTCTGAACTCTTTCCCTACACGACGCTCTTCCGATCTNNNAAATATGGATGCAGGGTGTGTG
AGCACACGTCTGAACTCTTTCCCTACACGACGCTCTTCCGATCTNNNCCTTCCTTTCACAACCCCTTCTC
AGCACACGTCTGAACTCTTTCCCTACACGACGCTCTTCCGATCTNNNCTGCATCTAGAATGGGAGTGT
AGCACACGTCTGAACTCTTTCCCTACACGACGCTCTTCCGATCTNNNCCCTTTCTACTGCCTCTTCCTCT
AAGAGCACACGTCTGAACTCTTTCCCTACACGACGCTCTTCCGATCTNNNCCAAAGCAAGAATGCAA
AGCACACGTCTGAACTCTTTCCCTACACGACGCTCTTCCGATCTNNNAACCTGGCAACATTTGGTAAA
GCACACGTCTGAACTCTTTCCCTACACGACGCTCTTCCGATCTNNNTTTGCAGCTGTCTTCTCCTTACGC
GCACACGTCTGAACTCTTTCCCTACACGACGCTCTTCCGATCTNNNGCTTCGGCAGGACAGAAATTA
AGCACACGTCTGAACTCTTTCCCTACACGACGCTCTTCCGATCTNNNAGCTCTAGTACACCTTACACAT
AGCACACGTCTGAACTCTTTCCCTACACGACGCTCTTCCGATCTNNNCTGGGAAGCCATTATTCTGTCC
AGCACACGTCTGAACTCTTTCCCTACACGACGCTCTTCCGATCTNNNCATAGGAAAACGCTGGAAGCA
GAGCACACGTCTGAACTCTTTCCCTACACGACGCTCTTCCGATCTNNNAATGAAACTTGCCCTTTGGCA
AGCACACGTCTGAACTCTTTCCCTACACGACGCTCTTCCGATCTNNNGGGTACAATTTGGGTACTGG
GAGCACACGTCTGAACTCTTTCCCTACACGACGCTCTTCCGATCTNNNAAGGAAAAACTGTGTGTGT
GAGCACACGTCTGAACTCTTTCCCTACACGACGCTCTTCCGATCTNNNCCCGGCCTCAAATCTTTATCT
AAGAGCACACGTCTGAACTCTTTCCCTACACGACGCTCTTCCGATCTNNNCGCATAGCTCCTATCACC
AGCACACGTCTGAACTCTTTCCCTACACGACGCTCTTCCGATCTNNNCCCGGCCCTGGGTATACATATT
AGCACACGTCTGAACTCTTTCCCTACACGACGCTCTTCCGATCTNNNCTCTGCTCTCCAACCTATGAGG
GAGCACACGTCTGAACTCTTTCCCTACACGACGCTCTTCCGATCTNNNTCTGAAGAGCTCAATACTGT
AGCACACGTCTGAACTCTTTCCCTACACGACGCTCTTCCGATCTNNNGCGTTCATTGGTGAATGAATG
AGAGCACACGTCTGAACTCTTTCCCTACACGACGCTCTTCCGATCTNNNTCAGAGCTGTGGTGCTCTT
GCACACGTCTGAACTCTTTCCCTACACGACGCTCTTCCGATCTNNNACCACTAATCCACTTCCTTGACC
AGCACACGTCTGAACTCTTTCCCTACACGACGCTCTTCCGATCTNNNGTATAGCTGGAGTGGAGGC
GAGCACACGTCTGAACTCTTTCCCTACACGACGCTCTTCCGATCTNNNCTTGTGTGTATCTTGGTCGT
GCACACGTCTGAACTCTTTCCCTACACGACGCTCTTCCGATCTNNNGTTAGTGGCCAGAATGTGGGA
GAGCACACGTCTGAACTCTTTCCCTACACGACGCTCTTCCGATCTNNNATCCGACTGTCGAAATGAG
AGCACACGTCTGAACTCTTTCCCTACACGACGCTCTTCCGATCTNNNCAAGCTGTAACAAGGTGCAT
GCACACGTCTGAACTCTTTCCCTACACGACGCTCTTCCGATCTNNNTTGTGTTTTGTTTTGAGGGCCA
GCACACGTCTGAACTCTTTCCCTACACGACGCTCTTCCGATCTNNNCACAGGTCCACATGTACACACA
GAGCACACGTCTGAACTCTTTCCCTACACGACGCTCTTCCGATCTNNNCTCTGCCTCCTCCACTATGAA
GCACACGTCTGAACTCTTTCCCTACACGACGCTCTTCCGATCTNNNAGAGACAACATGGAAAGGAAG
GAGCACACGTCTGAACTCTTTCCCTACACGACGCTCTTCCGATCTNNNAACCTAAGCCTTCACCTCTCA
GCACACGTCTGAACTCTTTCCCTACACGACGCTCTTCCGATCTNNNACCAAACCTTAGTGCCATATGA
GCACACGTCTGAACTCTTTCCCTACACGACGCTCTTCCGATCTNNNCTCCGCCCAAGTTCTTCTAACTCC
AGCACACGTCTGAACTCTTTCCCTACACGACGCTCTTCCGATCTNNNCTGTACAATGGGGAAAGTGAG
AGCACACGTCTGAACTCTTTCCCTACACGACGCTCTTCCGATCTNNNGACAAATCCCATGAGATGGCA
GCACACGTCTGAACTCTTTCCCTACACGACGCTCTTCCGATCTNNNGCACACACACACACCTTGAGATA
AGAGCACACGTCTGAACTCTTTCCCTACACGACGCTCTTCCGATCTNNNCTTCCCCTTCACTTCTTCAGT
AGCACACGTCTGAACTCTTTCCCTACACGACGCTCTTCCGATCTNNNGATGAATAAGGACAGTGACG
GCACACGTCTGAACTCTTTCCCTACACGACGCTCTTCCGATCTNNNAGCTGTACCAGTGGAGATGGA

GAGCACACGTCTGAACTCTTTCCCTACACGACGCTCTTCCGATCTNNNATCGGAAATGTTGGCCTGTA
GAGCACACGTCTGAACTCTTTCCCTACACGACGCTCTTCCGATCTNNNTCACTTGCACCGAGGCTAAG
GAGCACACGTCTGAACTCTTTCCCTACACGACGCTCTTCCGATCTNNNGAGAGAGAAGAACAGGCA
AGCACACGTCTGAACTCTTTCCCTACACGACGCTCTTCCGATCTNNNAGTTCTTTTGGGGGACCCTTT
GAGCACACGTCTGAACTCTTTCCCTACACGACGCTCTTCCGATCTNNNACCATGAATAGCATGACCAC
AAGAGCACACGTCTGAACTCTTTCCCTACACGACGCTCTTCCGATCTNNNCACAGTGGGAAAGAAAA
GAGCACACGTCTGAACTCTTTCCCTACACGACGCTCTTCCGATCTNNNGCAAGATCCATGTTGCTGAG
GAGCACACGTCTGAACTCTTTCCCTACACGACGCTCTTCCGATCTNNNTCAAAGTCAACGCAAAGAG
GCACACGTCTGAACTCTTTCCCTACACGACGCTCTTCCGATCTNNNACAGTCAACAGAGCACGTAACAT
GAGCACACGTCTGAACTCTTTCCCTACACGACGCTCTTCCGATCTNNNTCACTTTCTGTATCCCACAGG
AAGAGCACACGTCTGAACTCTTTCCCTACACGACGCTCTTCCGATCTNNNTCTAATTCTTGGGGGCTTA
AGAGCACACGTCTGAACTCTTTCCCTACACGACGCTCTTCCGATCTNNNTGCTCTTCACTGTGTAATGG
GAGCACACGTCTGAACTCTTTCCCTACACGACGCTCTTCCGATCTNNNTTTTCCAGCTCCCATTTTGC
GCACACGTCTGAACTCTTTCCCTACACGACGCTCTTCCGATCTNNNTCAAATGCCTCCTCCTCCACTAA
AGAGCACACGTCTGAACTCTTTCCCTACACGACGCTCTTCCGATCTNNNTGCATACAACTTGTTCTCAG
AGCACACGTCTGAACTCTTTCCCTACACGACGCTCTTCCGATCTNNNGCTATTATAGGTCCTTTTTACT
GAAGAGCACACGTCTGAACTCTTTCCCTACACGACGCTCTTCCGATCTNNNACTACCCAATTGTTCCAA
GCACACGTCTGAACTCTTTCCCTACACGACGCTCTTCCGATCTNNNTGTAACTAATACTGTACATCAC
AGCACACGTCTGAACTCTTTCCCTACACGACGCTCTTCCGATCTNNNGTGGTAGTAAAGGAAGCCCC
AGCACACGTCTGAACTCTTTCCCTACACGACGCTCTTCCGATCTNNNAATCTGTGTGGTCTGTCATGC
AGCACACGTCTGAACTCTTTCCCTACACGACGCTCTTCCGATCTNNNATAAAAGTCCAAGGGAGGTG
AGCACACGTCTGAACTCTTTCCCTACACGACGCTCTTCCGATCTNNNCCCCTCACCCGATACATAGACA
AGCACACGTCTGAACTCTTTCCCTACACGACGCTCTTCCGATCTNNNGTGGCCAGGAATTGAAAGAG
AGCACACGTCTGAACTCTTTCCCTACACGACGCTCTTCCGATCTNNNTTCTAGCTGGCAAACACACCAT
AGCACACGTCTGAACTCTTTCCCTACACGACGCTCTTCCGATCTNNNAAATGTATGCGTTCTACCCCA
GAGCACACGTCTGAACTCTTTCCCTACACGACGCTCTTCCGATCTNNNTCCCACAGATCCTCCATGAAG
AGCACACGTCTGAACTCTTTCCCTACACGACGCTCTTCCGATCTNNNTCTTCCCCTACACAGACACACAT
GAGCACACGTCTGAACTCTTTCCCTACACGACGCTCTTCCGATCTNNNTGTGCCCCTTCCCAATCAATA
AGCACACGTCTGAACTCTTTCCCTACACGACGCTCTTCCGATCTNNNTCGCAGGTACTTGGTGGTAGT
GAGCACACGTCTGAACTCTTTCCCTACACGACGCTCTTCCGATCTNNNACGGGCCTCTAAAAGAAATC
GAGCACACGTCTGAACTCTTTCCCTACACGACGCTCTTCCGATCTNNNTCAAAGTCTTTTCCTGGGAAA
GCACACGTCTGAACTCTTTCCCTACACGACGCTCTTCCGATCTNNNTACAGTTGTCTCTTGGTGTGTGC
AGCACACGTCTGAACTCTTTCCCTACACGACGCTCTTCCGATCTNNNAAAAAGTCCTGTTAGATGACA
GAGCACACGTCTGAACTCTTTCCCTACACGACGCTCTTCCGATCTNNNCTCTCTGCTTTGCATCTGGAC
GAGCACACGTCTGAACTCTTTCCCTACACGACGCTCTTCCGATCTNNNCCTGTGCGAAGACTTTAGGT
AGCACACGTCTGAACTCTTTCCCTACACGACGCTCTTCCGATCTNNNACTCCTGTTCCTTCCTCAGTTAC
AAGAGCACACGTCTGAACTCTTTCCCTACACGACGCTCTTCCGATCTNNNGAATCCACGGATGCAGA
AGAGCACACGTCTGAACTCTTTCCCTACACGACGCTCTTCCGATCTNNNAGGAACCCACCCAAGAAAA
GAGCACACGTCTGAACTCTTTCCCTACACGACGCTCTTCCGATCTNNNATCCACCCATCATCCTCAGCA
GAGCACACGTCTGAACTCTTTCCCTACACGACGCTCTTCCGATCTNNNATACTCTAAGTCGTGCAGCC
GCACACGTCTGAACTCTTTCCCTACACGACGCTCTTCCGATCTNNNCATGACGCCTGTCATAACTGCTA
AGCACACGTCTGAACTCTTTCCCTACACGACGCTCTTCCGATCTNNNGAAGTTCCTCTCTGGGCTAAT
AGAGCACACGTCTGAACTCTTTCCCTACACGACGCTCTTCCGATCTNNNAGAAATCATGTGAGGGGG

GAGCACACGTCTGAACTCTTTCCCTACACGACGCTCTTCCGATCTNNNTCCAAAGCAGGGCAAAACA
AAGAGCACACGTCTGAACTCTTTCCCTACACGACGCTCTTCCGATCTNNNTCCCCATGGGATATCTTC
GCACACGTCTGAACTCTTTCCCTACACGACGCTCTTCCGATCTNNNTGCCCACAAATATAGAGGGGA
AGCACACGTCTGAACTCTTTCCCTACACGACGCTCTTCCGATCTNNNAACTCAGTCTTTGGAGGGGCA
GCACACGTCTGAACTCTTTCCCTACACGACGCTCTTCCGATCTNNNCCAGGTGGAGAAGGGAGAATG
AGCACACGTCTGAACTCTTTCCCTACACGACGCTCTTCCGATCTNNNGTTTCTGGTTCTTGCTGAAGT
AGCACACGTCTGAACTCTTTCCCTACACGACGCTCTTCCGATCTNNNCTGGGCAACATAGAGACAAGG
GAGCACACGTCTGAACTCTTTCCCTACACGACGCTCTTCCGATCTNNNTGTGACGGAGGAAAATAAG
GCACACGTCTGAACTCTTTCCCTACACGACGCTCTTCCGATCTNNNTCAGGGTTATCTAGGGATGGC
AAGAGCACACGTCTGAACTCTTTCCCTACACGACGCTCTTCCGATCTNNNAAAGCCATTTGCATTCCAT
AGCACACGTCTGAACTCTTTCCCTACACGACGCTCTTCCGATCTNNNTGGGGGCTCTATATTTAGTCG
GAGCACACGTCTGAACTCTTTCCCTACACGACGCTCTTCCGATCTNNNAGCTCCTAAAGAGTTGTGAG
AGCACACGTCTGAACTCTTTCCCTACACGACGCTCTTCCGATCTNNNTGGACTGCAATCTTTAGTTTC
GCACACGTCTGAACTCTTTCCCTACACGACGCTCTTCCGATCTNNNCTGCCCATTTACCTGTCTGATGG
GAGCACACGTCTGAACTCTTTCCCTACACGACGCTCTTCCGATCTNNNTCCCAATAATGAGCCTTTCTG
AGCACACGTCTGAACTCTTTCCCTACACGACGCTCTTCCGATCTNNNGGGGAGAACTTGCATTCTGGA
GCACACGTCTGAACTCTTTCCCTACACGACGCTCTTCCGATCTNNNAAATGAGCTCGCAGAACCTACA
GAGCACACGTCTGAACTCTTTCCCTACACGACGCTCTTCCGATCTNNNATCCCTCTTCCCATCAAGAGC
GCACACGTCTGAACTCTTTCCCTACACGACGCTCTTCCGATCTNNNAAGCCACTCTCCGAAAACAAGA
GAGCACACGTCTGAACTCTTTCCCTACACGACGCTCTTCCGATCTNNNGCTGGAGCAAAAAGCAATG
GAGCACACGTCTGAACTCTTTCCCTACACGACGCTCTTCCGATCTNNNTTGTCTAGGCAGAAGTGGT
GCACACGTCTGAACTCTTTCCCTACACGACGCTCTTCCGATCTNNNAAGTGACATAAGAGTGAGGGC
GAGCACACGTCTGAACTCTTTCCCTACACGACGCTCTTCCGATCTNNNGTGTGTTAGTGAAGGTTCTC
AGAGCACACGTCTGAACTCTTTCCCTACACGACGCTCTTCCGATCTNNNGTGTTACCTCAGGCTATGT
ACACGTCTGAACTCTTTCCCTACACGACGCTCTTCCGATCTNNNATGACAGGTTATGGTAGCCCCTCC
GAGCACACGTCTGAACTCTTTCCCTACACGACGCTCTTCCGATCTNNNTCAAACATGGAAGGTAAGC
AGCACACGTCTGAACTCTTTCCCTACACGACGCTCTTCCGATCTNNNCCGTCAAACCCCTTCCTTACTT
GCACACGTCTGAACTCTTTCCCTACACGACGCTCTTCCGATCTNNNCTCATGAGAATGCTCTTGCATG
GAGCACACGTCTGAACTCTTTCCCTACACGACGCTCTTCCGATCTNNNCCTACTGCCACTACTACTACT
GCACACGTCTGAACTCTTTCCCTACACGACGCTCTTCCGATCTNNNCTACAGCACAGTAGCAAGACCA
AGCACACGTCTGAACTCTTTCCCTACACGACGCTCTTCCGATCTNNNCAAGGGGGTGGGCCAATAAA
AGAGCACACGTCTGAACTCTTTCCCTACACGACGCTCTTCCGATCTNNNTGGAAATCCTTAGATTGCCT
AGCACACGTCTGAACTCTTTCCCTACACGACGCTCTTCCGATCTNNNTTTACGAGTGCATGTGTCTGT
GAGCACACGTCTGAACTCTTTCCCTACACGACGCTCTTCCGATCTNNNTAGCTTCCATAAAAGCACCA
GAAGAGCACACGTCTGAACTCTTTCCCTACACGACGCTCTTCCGATCTNNNTCTAATTCTTGGGGGCTT
GCACACGTCTGAACTCTTTCCCTACACGACGCTCTTCCGATCTNNNGAAGCTCTTTCTGACTGGCCTA
AGCACACGTCTGAACTCTTTCCCTACACGACGCTCTTCCGATCTNNNCAGGCCAGGGACAAATAACT
GAGCACACGTCTGAACTCTTTCCCTACACGACGCTCTTCCGATCTNNNAGTGACTGTTTTCACCATTT
AGCACACGTCTGAACTCTTTCCCTACACGACGCTCTTCCGATCTNNNTCCCCCTTCTCCAAGAAATAGC
AAGAGCACACGTCTGAACTCTTTCCCTACACGACGCTCTTCCGATCTNNNCAAAATGAAGAGCTCTGG
GAGCACACGTCTGAACTCTTTCCCTACACGACGCTCTTCCGATCTNNNCAGATCTCTGGGATTCCTGT
AGCACACGTCTGAACTCTTTCCCTACACGACGCTCTTCCGATCTNNNAAGCCCACCAATTGATGAGTG
GCACACGTCTGAACTCTTTCCCTACACGACGCTCTTCCGATCTNNNAATGTTCTTGTGGGGACGGTAT

AGCACACGTCTGAACTCTTTCCCTACACGACGCTCTTCCGATCTNNNTCTGACATTCCTTGCCACAAACA
AGAGCACACGTCTGAACTCTTTCCCTACACGACGCTCTTCCGATCTNNNAATCACATGGTTGCTGTGG
GCACACGTCTGAACTCTTTCCCTACACGACGCTCTTCCGATCTNNNCATGCCAATCCCAGAAAGGTGT
GAGCACACGTCTGAACTCTTTCCCTACACGACGCTCTTCCGATCTNNNAAGGTTTATCAAAAGCACTG
AGCACACGTCTGAACTCTTTCCCTACACGACGCTCTTCCGATCTNNNCAACACAGTTCAAAATCTCTGC
GAGCACACGTCTGAACTCTTTCCCTACACGACGCTCTTCCGATCTNNNCAGTGGTGTATTTGGCTACA
GCACACGTCTGAACTCTTTCCCTACACGACGCTCTTCCGATCTNNNACGTTTTAGCAGATGATGTGGT
AGCACACGTCTGAACTCTTTCCCTACACGACGCTCTTCCGATCTNNNATGCTCAGTCTAGGCCAAATGT
AGAGCACACGTCTGAACTCTTTCCCTACACGACGCTCTTCCGATCTNNNTCCTTCAGGACATGTTTGTG
AGAGCACACGTCTGAACTCTTTCCCTACACGACGCTCTTCCGATCTNNNAAGATAGACAGAAGGAGC
GCACACGTCTGAACTCTTTCCCTACACGACGCTCTTCCGATCTNNNTTCTGGCTGATGAAGGTGTTCT
AGCACACGTCTGAACTCTTTCCCTACACGACGCTCTTCCGATCTNNNCTCCTGGTGAGATAGATGCTT
AGCACACGTCTGAACTCTTTCCCTACACGACGCTCTTCCGATCTNNNACTCCTTTGGACAGCCATAGTC
AGCACACGTCTGAACTCTTTCCCTACACGACGCTCTTCCGATCTNNNTTCTCCTGTAGGTACAGAATTG
GAGCACACGTCTGAACTCTTTCCCTACACGACGCTCTTCCGATCTNNNATATTCTTGAGAGCTTGGCC
AGCACACGTCTGAACTCTTTCCCTACACGACGCTCTTCCGATCTNNNTGTATGTGTGTGCGGTAGTGT
AGCACACGTCTGAACTCTTTCCCTACACGACGCTCTTCCGATCTNNNATCCAAATTCACTGGGTGCAT
AGAGCACACGTCTGAACTCTTTCCCTACACGACGCTCTTCCGATCTNNNTCTTTATACTCACCTGTGCC
AGCACACGTCTGAACTCTTTCCCTACACGACGCTCTTCCGATCTNNNTTAGCCGCCTTATTTGGCAG
AGCACACGTCTGAACTCTTTCCCTACACGACGCTCTTCCGATCTNNNGTGTGTTAGTGAAGGTTCTC
GAGCACACGTCTGAACTCTTTCCCTACACGACGCTCTTCCGATCTNNNCCCACTATCCCCAATCAGCTT
AGCACACGTCTGAACTCTTTCCCTACACGACGCTCTTCCGATCTNNNATCACATTACTCTCCACTCCCC
ACACGTCTGAACTCTTTCCCTACACGACGCTCTTCCGATCTNNNGGGTGTATGCACTCCACAATACACA
AGCACACGTCTGAACTCTTTCCCTACACGACGCTCTTCCGATCTNNNAGAGAGAGAGAGAGATGAGT
GAGCACACGTCTGAACTCTTTCCCTACACGACGCTCTTCCGATCTNNNCTCCACAGAGTAGCATTTGA
GCACACGTCTGAACTCTTTCCCTACACGACGCTCTTCCGATCTNNNTCAAATCATCTTCCAGTTTCCCT
AGCACACGTCTGAACTCTTTCCCTACACGACGCTCTTCCGATCTNNNTGTTTCCTCTGCCTTCATTGCA
GAGCACACGTCTGAACTCTTTCCCTACACGACGCTCTTCCGATCTNNNCTTCAGTTCAGGCACCACTTC
AGCACACGTCTGAACTCTTTCCCTACACGACGCTCTTCCGATCTNNNATCCAGGTTGGTAGTTCTGTG
AGAGCACACGTCTGAACTCTTTCCCTACACGACGCTCTTCCGATCTNNNACCCTTTACATCCAGTGACC
GCACACGTCTGAACTCTTTCCCTACACGACGCTCTTCCGATCTNNNCAATCCCTGCCTCAGTAAACTCC
AGCACACGTCTGAACTCTTTCCCTACACGACGCTCTTCCGATCTNNNTAGGGTCTTGTCAGCCTTCATC
GAGCACACGTCTGAACTCTTTCCCTACACGACGCTCTTCCGATCTNNNCAGATCTCTGGGATTCCTGT
AGCACACGTCTGAACTCTTTCCCTACACGACGCTCTTCCGATCTNNNCAAGACTGAAACTCCGTGTCAA
GCACACGTCTGAACTCTTTCCCTACACGACGCTCTTCCGATCTNNNAGCCAGGAACTGTGATGAAAAC
AGCACACGTCTGAACTCTTTCCCTACACGACGCTCTTCCGATCTNNNCATATCTGTAGGCATGAGCCAC
AGCACACGTCTGAACTCTTTCCCTACACGACGCTCTTCCGATCTNNNAGACAGAGCCACTGGAGAAGA
AGAGCACACGTCTGAACTCTTTCCCTACACGACGCTCTTCCGATCTNNNTTTTTGCTTTGGCTGAGTTT
ACACGTCTGAACTCTTTCCCTACACGACGCTCTTCCGATCTNNNAGGGAGTAGAATGAGTTCTTGGC
AGAGCACACGTCTGAACTCTTTCCCTACACGACGCTCTTCCGATCTNNNCGTTGGCTTCTCAGATGAA
AGCACACGTCTGAACTCTTTCCCTACACGACGCTCTTCCGATCTNNNGGAAGGCAACAGATATTGCAG
AGCACACGTCTGAACTCTTTCCCTACACGACGCTCTTCCGATCTNNNTCACACAACAGCAGGATCATC
AGCACACGTCTGAACTCTTTCCCTACACGACGCTCTTCCGATCTNNNATTGCGGGCTCTTATCTCTCTC

GCACACGTCTGAACTCTTTCCCTACACGACGCTCTTCCGATCTNNNAGAGAAGAAAACCCTGAACCAG
GAGCACACGTCTGAACTCTTTCCCTACACGACGCTCTTCCGATCTNNNAAAGACAGGGTGAGAGGAG
AGCACACGTCTGAACTCTTTCCCTACACGACGCTCTTCCGATCTNNNGGCAGCAGGATGATAGGTAG
GAGCACACGTCTGAACTCTTTCCCTACACGACGCTCTTCCGATCTNNNTGCCTTTCCTCAACAATTAGTG
AGAGCACACGTCTGAACTCTTTCCCTACACGACGCTCTTCCGATCTNNNGCCACGTTTGTTCAGAGGAC
AGCACACGTCTGAACTCTTTCCCTACACGACGCTCTTCCGATCTNNNTTTCAAATGCTTGCTGCTCTTGC
GAGCACACGTCTGAACTCTTTCCCTACACGACGCTCTTCCGATCTNNNCACAAGATGTAACAGCCATTG
AGAGCACACGTCTGAACTCTTTCCCTACACGACGCTCTTCCGATCTNNNTTTGCTGGTTCTATCTTGG
AGAGCACACGTCTGAACTCTTTCCCTACACGACGCTCTTCCGATCTNNNAGTCCCTTTGGTAAGCAATC
AGCACACGTCTGAACTCTTTCCCTACACGACGCTCTTCCGATCTNNNTTGATTCCTCACTGGCTTTGCTT
AGCACACGTCTGAACTCTTTCCCTACACGACGCTCTTCCGATCTNNNTTACAGACAGCATATCGTGGTG
AGAGCACACGTCTGAACTCTTTCCCTACACGACGCTCTTCCGATCTNNNACCCCCAAATTAGGAGACTT
AGAGCACACGTCTGAACTCTTTCCCTACACGACGCTCTTCCGATCTNNNTCAGGCAGTGAAGTTTTTC
GAGCACACGTCTGAACTCTTTCCCTACACGACGCTCTTCCGATCTNNNCCTAACCAGAACTTCTCCCCTG
GGAAGAGCACACGTCTGAACTCTTTCCCTACACGACGCTCTTCCGATCTNNNTTGATTGGAGCCTCCT
GAGCACACGTCTGAACTCTTTCCCTACACGACGCTCTTCCGATCTNNNCAGTGGTGTATTTGGCTACA
AGCACACGTCTGAACTCTTTCCCTACACGACGCTCTTCCGATCTNNNTACCTAAGGTCCATCGTCAGGT
GAGCACACGTCTGAACTCTTTCCCTACACGACGCTCTTCCGATCTNNNAGGCACGCAGAAATATACCT
CACACGTCTGAACTCTTTCCCTACACGACGCTCTTCCGATCTNNNTGGCTTTGGAAGTCAGAAGGAAG
AGCACACGTCTGAACTCTTTCCCTACACGACGCTCTTCCGATCTNNNAACTCCTCAGCAAAGTCGATAG
GAGCACACGTCTGAACTCTTTCCCTACACGACGCTCTTCCGATCTNNNATGGATTCATGGGTGTGTGT
AGAGCACACGTCTGAACTCTTTCCCTACACGACGCTCTTCCGATCTNNNTGAGGCTGGATTCAATATCG
AGAGCACACGTCTGAACTCTTTCCCTACACGACGCTCTTCCGATCTNNNTTGATTGGAGCCTCCTGATT
AGCACACGTCTGAACTCTTTCCCTACACGACGCTCTTCCGATCTNNNTTTCTCCTCCACAACTCACCTCT
GAGCACACGTCTGAACTCTTTCCCTACACGACGCTCTTCCGATCTNNNAGGGAGTAGAATGAGTTCTT
AGCACACGTCTGAACTCTTTCCCTACACGACGCTCTTCCGATCTNNNTCCTTTTCTTGCTGGTAGACAC
AGAGCACACGTCTGAACTCTTTCCCTACACGACGCTCTTCCGATCTNNNGATGTGGCTCAGGAAGCTT
AGAGCACACGTCTGAACTCTTTCCCTACACGACGCTCTTCCGATCTNNNTGCAGGAGGTGTACTAGTT
AGCACACGTCTGAACTCTTTCCCTACACGACGCTCTTCCGATCTNNNAGCAATAAGCACTAGAAGGAG
GAGCACACGTCTGAACTCTTTCCCTACACGACGCTCTTCCGATCTNNNTCAAGTTTGGTTCAGCACTAG
AGAGCACACGTCTGAACTCTTTCCCTACACGACGCTCTTCCGATCTNNNTAGTAGTGTCATTGCACTCC
AGAGCACACGTCTGAACTCTTTCCCTACACGACGCTCTTCCGATCTNNNGGTACACACCTTGGCTAAGA
AGCACACGTCTGAACTCTTTCCCTACACGACGCTCTTCCGATCTNNNTCAGTAGGAACAGTAATTTCAT
GAGCACACGTCTGAACTCTTTCCCTACACGACGCTCTTCCGATCTNNNGTAAAATGAACCTGTTGGCA
AGCACACGTCTGAACTCTTTCCCTACACGACGCTCTTCCGATCTNNNCACACACAATGGAATGGCTTCT
AGCACACGTCTGAACTCTTTCCCTACACGACGCTCTTCCGATCTNNNTAAATGATGGCTGACAAGCCTC
GAGCACACGTCTGAACTCTTTCCCTACACGACGCTCTTCCGATCTNNNATCCCAAAGGCCTCTCTTGACT
GCACACGTCTGAACTCTTTCCCTACACGACGCTCTTCCGATCTNNNTCCGGATCCAGTATCCTTACAGTC
AGCACACGTCTGAACTCTTTCCCTACACGACGCTCTTCCGATCTNNNCCTTAGTAACTTTCTGGTGGCC
AAGAGCACACGTCTGAACTCTTTCCCTACACGACGCTCTTCCGATCTNNNTCTGGTAATGTAGGGAAG
AGAGCACACGTCTGAACTCTTTCCCTACACGACGCTCTTCCGATCTNNNGAAGTTGAGGCAAACAGCA
AGAGCACACGTCTGAACTCTTTCCCTACACGACGCTCTTCCGATCTNNNTGTAAATTTCCAGGCCCTTC
AGCACACGTCTGAACTCTTTCCCTACACGACGCTCTTCCGATCTNNNTGGATTGTGTGCGTATGCAGTT

GAAGAGCACACGTCTGAACTCTTTCCCTACACGACGCTCTTCCGATCTNNNAGCAAGGAATAGGCAG
AGCACACGTCTGAACTCTTTCCCTACACGACGCTCTTCCGATCTNNNCAAGCTCAGGTTGGACACTACT
GCACACGTCTGAACTCTTTCCCTACACGACGCTCTTCCGATCTNNNTCCTCTTTGTCCCTTGCAAAACACA
GAGCACACGTCTGAACTCTTTCCCTACACGACGCTCTTCCGATCTNNNGGGGAAATATGACAGGACTG
AAGAGCACACGTCTGAACTCTTTCCCTACACGACGCTCTTCCGATCTNNNTGTTCATTCCACAGATCAG
AGCACACGTCTGAACTCTTTCCCTACACGACGCTCTTCCGATCTNNNTCTAAGGTTGAGCAAATGCATA
AAGAGCACACGTCTGAACTCTTTCCCTACACGACGCTCTTCCGATCTNNNTGTGCTCTGAACCACCAGA
GCACACGTCTGAACTCTTTCCCTACACGACGCTCTTCCGATCTNNNAGGGCAAAAATCACAATCCCTGT
AGAGCACACGTCTGAACTCTTTCCCTACACGACGCTCTTCCGATCTNNNACTAACCCACTACCTTCCCAG
AGAGCACACGTCTGAACTCTTTCCCTACACGACGCTCTTCCGATCTNNNTGGGAAGGTGCATCTCAGT
AGCACACGTCTGAACTCTTTCCCTACACGACGCTCTTCCGATCTNNNTGTAAGTGGAAGACAGGCAGA
AGCACACGTCTGAACTCTTTCCCTACACGACGCTCTTCCGATCTNNNTACATACCAAATCAGGGTGGG
AGCACACGTCTGAACTCTTTCCCTACACGACGCTCTTCCGATCTNNNTCACTCTGCTTGAAAGATCAGC
GAGCACACGTCTGAACTCTTTCCCTACACGACGCTCTTCCGATCTNNNTTTTCCTCGGAGATGTGTGT
GCACACGTCTGAACTCTTTCCCTACACGACGCTCTTCCGATCTNNNTGCCCTCATTTGTATCCAGTCTC
AGCACACGTCTGAACTCTTTCCCTACACGACGCTCTTCCGATCTNNNGAAGGGGGTTGTAAAGTGTG
GAGCACACGTCTGAACTCTTTCCCTACACGACGCTCTTCCGATCTNNNAGCAAACCATGAAAAGAAGA
GAGCACACGTCTGAACTCTTTCCCTACACGACGCTCTTCCGATCTNNNAAGAGGGGCAGACAGAAAA
AGCACACGTCTGAACTCTTTCCCTACACGACGCTCTTCCGATCTNNNTCCTTTCCTGACTGCTTGACCTA
GAGCACACGTCTGAACTCTTTCCCTACACGACGCTCTTCCGATCTNNNAACTTGCCACCTCTCACAAACA
AGCACACGTCTGAACTCTTTCCCTACACGACGCTCTTCCGATCTNNNGCCAGGGTTCTATGGAGAAAC
AGCACACGTCTGAACTCTTTCCCTACACGACGCTCTTCCGATCTNNNTGTATGTGTGTGCGGTAGTGT
AGCACACGTCTGAACTCTTTCCCTACACGACGCTCTTCCGATCTNNNATGTGCAGTGTGGGATGAGAT
GAGCACACGTCTGAACTCTTTCCCTACACGACGCTCTTCCGATCTNNNTCATGTTCAATTGGTGCTTCC
GAGCACACGTCTGAACTCTTTCCCTACACGACGCTCTTCCGATCTNNNAAGATTTCCTCCTGCCCTGTA
AGCACACGTCTGAACTCTTTCCCTACACGACGCTCTTCCGATCTNNNATTGGGGAAAAATGCCCATCT
AGAGCACACGTCTGAACTCTTTCCCTACACGACGCTCTTCCGATCTNNNAGACTGTCAACTAAAAGAGT
GAGCACACGTCTGAACTCTTTCCCTACACGACGCTCTTCCGATCTNNNCCGTACTCCTAACACACATACA
AGCACACGTCTGAACTCTTTCCCTACACGACGCTCTTCCGATCTNNNTCTCTCCCACTTTGGAGTTAGC
GAGCACACGTCTGAACTCTTTCCCTACACGACGCTCTTCCGATCTNNNCATTAGGAGCTGTAGACTGT
GAGCACACGTCTGAACTCTTTCCCTACACGACGCTCTTCCGATCTNNNCTGAGCAGCTGGAAAGTACA
GAGCACACGTCTGAACTCTTTCCCTACACGACGCTCTTCCGATCTNNNAGCCACTAAGTTTGCAGTAAT
GCACACGTCTGAACTCTTTCCCTACACGACGCTCTTCCGATCTNNNAGGACAAATATGTGCCTTGAGCA
AGCACACGTCTGAACTCTTTCCCTACACGACGCTCTTCCGATCTNNNAGCAAGTAGGGGAGAAGGAT
GAGCACACGTCTGAACTCTTTCCCTACACGACGCTCTTCCGATCTNNNACTACGCTAAGCACTTTACCT
AGCACACGTCTGAACTCTTTCCCTACACGACGCTCTTCCGATCTNNNGCAAAACTGAGGTTGACAAAG
GCACACGTCTGAACTCTTTCCCTACACGACGCTCTTCCGATCTNNNTGGGAAGTAACAGTAGTTGGT
AGCACACGTCTGAACTCTTTCCCTACACGACGCTCTTCCGATCTNNNATGAAGCTGTACTGGGATTTG
AGAGCACACGTCTGAACTCTTTCCCTACACGACGCTCTTCCGATCTNNNAGTACCCTATTCAGCAGAG
GAGCACACGTCTGAACTCTTTCCCTACACGACGCTCTTCCGATCTNNNAAGTGACATAAGAGTGAGG
AGAGCACACGTCTGAACTCTTTCCCTACACGACGCTCTTCCGATCTNNNGTTTGTCTTGGCTTCCCAGA
AGAGCACACGTCTGAACTCTTTCCCTACACGACGCTCTTCCGATCTNNNGGGTTAGTGGTGGTGGAGA
GAGCACACGTCTGAACTCTTTCCCTACACGACGCTCTTCCGATCTNNNTTGGGCAATTCTGTTCTGCAT

CACACGTCTGAACTCTTTCCCTACACGACGCTCTTCCGATCTNNNGCCGAAGGGATGTGCTCCTGCATC
GCACACGTCTGAACTCTTTCCCTACACGACGCTCTTCCGATCTNNNTTGCAAAACAAACCCCTGGACTA
GAGCACACGTCTGAACTCTTTCCCTACACGACGCTCTTCCGATCTNNNTCGACCTGGTCATTTGTACATG
AGCACACGTCTGAACTCTTTCCCTACACGACGCTCTTCCGATCTNNNGCAAATGATAATTTTGACCAGA
AGCACACGTCTGAACTCTTTCCCTACACGACGCTCTTCCGATCTNNNCCAGGAATGATAGACTTTGGG
GAGCACACGTCTGAACTCTTTCCCTACACGACGCTCTTCCGATCTNNNTGAATCTGCATGTGTGGTAT
GAGCACACGTCTGAACTCTTTCCCTACACGACGCTCTTCCGATCTNNNGATTGCTAGTGAGGCTTGTG
AGCACACGTCTGAACTCTTTCCCTACACGACGCTCTTCCGATCTNNNAGAGGTAATGCATACATGGAA
AGAGCACACGTCTGAACTCTTTCCCTACACGACGCTCTTCCGATCTNNNTAGAAAGGCTGGTGACTTAC
GCACACGTCTGAACTCTTTCCCTACACGACGCTCTTCCGATCTNNNCGACTGTCTGTAAAGTGGACCAA
AAGAGCACACGTCTGAACTCTTTCCCTACACGACGCTCTTCCGATCTNNNAGGGAGGGGTCCAAAAT
GAGCACACGTCTGAACTCTTTCCCTACACGACGCTCTTCCGATCTNNNCCTTCAATAGGCTTCTTGCCCA
GAGCACACGTCTGAACTCTTTCCCTACACGACGCTCTTCCGATCTNNNCATGAGTGAAGACCACTCACG
AGCACACGTCTGAACTCTTTCCCTACACGACGCTCTTCCGATCTNNNATGCAACCAACACCTCTTGTCTT
GAGCACACGTCTGAACTCTTTCCCTACACGACGCTCTTCCGATCTNNNGAGTAGCTTGGACTTTAGGG
GAGCACACGTCTGAACTCTTTCCCTACACGACGCTCTTCCGATCTNNNCTGCATTGCCTTCTGTGGTAT
GAGCACACGTCTGAACTCTTTCCCTACACGACGCTCTTCCGATCTNNNAAGGTTTGTGTGAGAATCAA
AGCACACGTCTGAACTCTTTCCCTACACGACGCTCTTCCGATCTNNNAACCTGCCTCTGGGTTTTTGTT
GCACACGTCTGAACTCTTTCCCTACACGACGCTCTTCCGATCTNNNGGCCATCCAAGACACATACACAT
GAGCACACGTCTGAACTCTTTCCCTACACGACGCTCTTCCGATCTNNNGCTTTGAATCCAGCTGCAACAT
AGCACACGTCTGAACTCTTTCCCTACACGACGCTCTTCCGATCTNNNAACTAATGGTGGCCTCTCTGTC
AGCACACGTCTGAACTCTTTCCCTACACGACGCTCTTCCGATCTNNNGTTCTCTCCCCACCTCCAAAAAG
GCACACGTCTGAACTCTTTCCCTACACGACGCTCTTCCGATCTNNNTCACCTATTTGGTAGTTCCCGGT
AGAGCACACGTCTGAACTCTTTCCCTACACGACGCTCTTCCGATCTNNNCCTGTCTCAGAAAACATGGC
GCACACGTCTGAACTCTTTCCCTACACGACGCTCTTCCGATCTNNNACATAACCTAACTGGCCCCTTGA
AGAGCACACGTCTGAACTCTTTCCCTACACGACGCTCTTCCGATCTNNNTTTGAAAGGTTCTCAGGAC
AGAGCACACGTCTGAACTCTTTCCCTACACGACGCTCTTCCGATCTNNNCTGCAGAAGAGTTGGGAGA
GAGCACACGTCTGAACTCTTTCCCTACACGACGCTCTTCCGATCTNNNGCTTTGAATCCAGCTGCAACAT
GAGCACACGTCTGAACTCTTTCCCTACACGACGCTCTTCCGATCTNNNATGTGCAGTGTGGGATGAGA
AGAGCACACGTCTGAACTCTTTCCCTACACGACGCTCTTCCGATCTNNNTGGATAAAATCACTGCCAACA
AGCACACGTCTGAACTCTTTCCCTACACGACGCTCTTCCGATCTNNNAGCCAGGTTTCACCAGTTTTATG
AAGAGCACACGTCTGAACTCTTTCCCTACACGACGCTCTTCCGATCTNNNGGTAGGGCCTTTGGAAGC
AGCACACGTCTGAACTCTTTCCCTACACGACGCTCTTCCGATCTNNNTCTTGGTCAGTTGGATTTTGGC
GCACACGTCTGAACTCTTTCCCTACACGACGCTCTTCCGATCTNNNTGATAAATAGTAAAACCTCTCTG
AAGAGCACACGTCTGAACTCTTTCCCTACACGACGCTCTTCCGATCTNNNGCTCTCAGAGTGGTGATT
AGCACACGTCTGAACTCTTTCCCTACACGACGCTCTTCCGATCTNNNAAGAGAGAAGGGGATGTGTGT
GAGCACACGTCTGAACTCTTTCCCTACACGACGCTCTTCCGATCTNNNTCTCACTTCTTCTTGATATCCCA
GAGCACACGTCTGAACTCTTTCCCTACACGACGCTCTTCCGATCTNNNCAGGAGGCATCGAGGTAGAG
GAGCACACGTCTGAACTCTTTCCCTACACGACGCTCTTCCGATCTNNNATGGTCAGATTGATGGTCTGC
AGCACACGTCTGAACTCTTTCCCTACACGACGCTCTTCCGATCTNNNTTGCATAGCACATGTCCTCTCTG
AAGAGCACACGTCTGAACTCTTTCCCTACACGACGCTCTTCCGATCTNNNCTGGCTGACCTAGCCAAAG
AGCACACGTCTGAACTCTTTCCCTACACGACGCTCTTCCGATCTNNNTGGATGTGGACAGACTAGAGCT
GAGCACACGTCTGAACTCTTTCCCTACACGACGCTCTTCCGATCTNNNACACACAATTTCTGATGGACAA

GAGCACACGTCTGAACTCTTTCCCTACACGACGCTCTTCCGATCTNNNAAGGAAAAACTGTGTGTGTG
AGCACACGTCTGAACTCTTTCCCTACACGACGCTCTTCCGATCTNNNACATAACCTAACTGGCCCCTTG
GCACACGTCTGAACTCTTTCCCTACACGACGCTCTTCCGATCTNNNGGAGTGGCAACTTTCTCTTTCCA
GAGCACACGTCTGAACTCTTTCCCTACACGACGCTCTTCCGATCTNNNAAACCACCTAGTGTTACAGG
AGCACACGTCTGAACTCTTTCCCTACACGACGCTCTTCCGATCTNNNGCAAGCATGTCCTCTTATGGT
AGCACACGTCTGAACTCTTTCCCTACACGACGCTCTTCCGATCTNNNAGTCCACAGAGCATCAGAACA
AGCACACGTCTGAACTCTTTCCCTACACGACGCTCTTCCGATCTNNNCATGTTCAGTACGCACGCAATT
AGCACACGTCTGAACTCTTTCCCTACACGACGCTCTTCCGATCTNNNCCCTGCAGAGACAGACTAGT
GAGCACACGTCTGAACTCTTTCCCTACACGACGCTCTTCCGATCTNNNGGCAGACCCCTACTCCAATAT
AGCACACGTCTGAACTCTTTCCCTACACGACGCTCTTCCGATCTNNNAAAATAAGATGTGTATACGTT
AGCACACGTCTGAACTCTTTCCCTACACGACGCTCTTCCGATCTNNNCTCCCACTCCCTAAAAAGCAAA
AGCACACGTCTGAACTCTTTCCCTACACGACGCTCTTCCGATCTNNNTTGCCAAAGCCTTAGGACTTCT
AGCACACGTCTGAACTCTTTCCCTACACGACGCTCTTCCGATCTNNNCGGAAACTTCGGAATAGACCA
GAGCACACGTCTGAACTCTTTCCCTACACGACGCTCTTCCGATCTNNNTTGTGGTGCACGTACTATTC
AGCACACGTCTGAACTCTTTCCCTACACGACGCTCTTCCGATCTNNNTCCCTAGCTGCCACATTCTACTT
AGCACACGTCTGAACTCTTTCCCTACACGACGCTCTTCCGATCTNNNTTGATATTTGTTTGGCAGGCAG
AGCACACGTCTGAACTCTTTCCCTACACGACGCTCTTCCGATCTNNNCATTCCTTGGCTTGCAAAGCAT
AGCACACGTCTGAACTCTTTCCCTACACGACGCTCTTCCGATCTNNNTCCACCTTCTCTGTTTCTGACCT
AGCACACGTCTGAACTCTTTCCCTACACGACGCTCTTCCGATCTNNNGGACAAACCTTACATCCCCTTC
AGCACACGTCTGAACTCTTTCCCTACACGACGCTCTTCCGATCTNNNTATTCAGGAGGCTTAGGCGG
GAGCACACGTCTGAACTCTTTCCCTACACGACGCTCTTCCGATCTNNNGCTCTCAGAACTCAGCACAT
GAGCACACGTCTGAACTCTTTCCCTACACGACGCTCTTCCGATCTNNNTACTCCAATGAAAGGCAAGA
GAGCACACGTCTGAACTCTTTCCCTACACGACGCTCTTCCGATCTNNNTTTTGCTAACCACAGTCCCA
AGCACACGTCTGAACTCTTTCCCTACACGACGCTCTTCCGATCTNNNTTGCAATTGTGTGTGAATGTG
AGCACACGTCTGAACTCTTTCCCTACACGACGCTCTTCCGATCTNNNTAATAATGGGAGAGAGCCTG
AGAGCACACGTCTGAACTCTTTCCCTACACGACGCTCTTCCGATCTNNNAGCGAATGTTGAGTTGAC
AGCACACGTCTGAACTCTTTCCCTACACGACGCTCTTCCGATCTNNNTACTCCAATGAAAGGCAAGAG
AGCACACGTCTGAACTCTTTCCCTACACGACGCTCTTCCGATCTNNNTCCCTCTCTGCCCTTTCAACTAT
AGAGCACACGTCTGAACTCTTTCCCTACACGACGCTCTTCCGATCTNNNGGGCAAGGAAAAGTGTGT
GAGCACACGTCTGAACTCTTTCCCTACACGACGCTCTTCCGATCTNNNAACTGTCTCATGCCAAGCTG
GAGCACACGTCTGAACTCTTTCCCTACACGACGCTCTTCCGATCTNNNTAGGTCTACCTCTTTCCCAGC
AGCACACGTCTGAACTCTTTCCCTACACGACGCTCTTCCGATCTNNNTGAGGAGTTTTGCACTGACG
CACACGTCTGAACTCTTTCCCTACACGACGCTCTTCCGATCTNNNGGTCCATGTGCAGGTTGTATGTA
AGCACACGTCTGAACTCTTTCCCTACACGACGCTCTTCCGATCTNNNTACAGTGAGCTGATACTGCGC
AGCACACGTCTGAACTCTTTCCCTACACGACGCTCTTCCGATCTNNNAAGAGGAAGCAGTGCATAGA
GAGCACACGTCTGAACTCTTTCCCTACACGACGCTCTTCCGATCTNNNTCCACCCCAAAGTAGATAGA
AGCACACGTCTGAACTCTTTCCCTACACGACGCTCTTCCGATCTNNNCCACGAGCTTCCTGAAGTCTA
AGAGCACACGTCTGAACTCTTTCCCTACACGACGCTCTTCCGATCTNNNCCAGAATTCAGCCTTGGAG
AGCACACGTCTGAACTCTTTCCCTACACGACGCTCTTCCGATCTNNNTGGGTGTATATTCATCCCCAAA
AGCACACGTCTGAACTCTTTCCCTACACGACGCTCTTCCGATCTNNNTGCACCAATTTCTTGCTAACCCT
AGCACACGTCTGAACTCTTTCCCTACACGACGCTCTTCCGATCTNNNAACCTTTGAGTGCCATAGCAA
GAGCACACGTCTGAACTCTTTCCCTACACGACGCTCTTCCGATCTNNNCAGCAATGAGCTTGCTACCT
AGCACACGTCTGAACTCTTTCCCTACACGACGCTCTTCCGATCTNNNACGCAATTCCTTTTCTGGGTAT

AGCACACGTCTGAACTCTTTCCCTACACGACGCTCTTCCGATCTNNNTGATGGTCTGTTTTCTTTTGCC
AAGAGCACACGTCTGAACTCTTTCCCTACACGACGCTCTTCCGATCTNNNACACACAGCTTTTCCCAAA
AGCACACGTCTGAACTCTTTCCCTACACGACGCTCTTCCGATCTNNNCCAAGCAAAAGTTGAAGAGC
AGCACACGTCTGAACTCTTTCCCTACACGACGCTCTTCCGATCTNNNGCATTTATGTGACCAGCCAAA
AGCACACGTCTGAACTCTTTCCCTACACGACGCTCTTCCGATCTNNNTTTCTACCAAAGGGCTAAGTG
AGCACACGTCTGAACTCTTTCCCTACACGACGCTCTTCCGATCTNNNACCTACAGAGGAATCATACTG
GAGCACACGTCTGAACTCTTTCCCTACACGACGCTCTTCCGATCTNNNGTGTTAGTCAAGTCTGACCC
AGAGCACACGTCTGAACTCTTTCCCTACACGACGCTCTTCCGATCTNNNGCAGCATCCTTTGTCCTCT
AGCACACGTCTGAACTCTTTCCCTACACGACGCTCTTCCGATCTNNNAACACATCCTCTGTCACCCATA
AGCACACGTCTGAACTCTTTCCCTACACGACGCTCTTCCGATCTNNNTGGCTTTCACTCGTTTTACAT
GCACACGTCTGAACTCTTTCCCTACACGACGCTCTTCCGATCTNNNAGGAAATCACCTCACCCCAAGTA
AGCACACGTCTGAACTCTTTCCCTACACGACGCTCTTCCGATCTNNNAGGGTGTCTAAATGGTCCCAG
GAAGAGCACACGTCTGAACTCTTTCCCTACACGACGCTCTTCCGATCTNNNGGCAGCTTTGTGGTGA
GAGCACACGTCTGAACTCTTTCCCTACACGACGCTCTTCCGATCTNNNCCCCTAAGGCCTTCAGTCTA
AGCACACGTCTGAACTCTTTCCCTACACGACGCTCTTCCGATCTNNNTGGCCCTGAAGATCCAGCTAT
AGCACACGTCTGAACTCTTTCCCTACACGACGCTCTTCCGATCTNNNAGAGTGTGGGAGAGGAAGG
AGCACACGTCTGAACTCTTTCCCTACACGACGCTCTTCCGATCTNNNATGGGTTGGAATTTGTCTGC
AGCACACGTCTGAACTCTTTCCCTACACGACGCTCTTCCGATCTNNNGGAAGTAGAGGGCTAAGCG
AGCACACGTCTGAACTCTTTCCCTACACGACGCTCTTCCGATCTNNNTGGCCAAAGCATTGAAAAGAT
GAGCACACGTCTGAACTCTTTCCCTACACGACGCTCTTCCGATCTNNNGTGTCCTCCAGTTTCATCCAC
AGCACACGTCTGAACTCTTTCCCTACACGACGCTCTTCCGATCTNNNCATACCAAGTGTCTCGATGCT
AGCACACGTCTGAACTCTTTCCCTACACGACGCTCTTCCGATCTNNNCAATATGTCATGCTTCACGCAC
GAGCACACGTCTGAACTCTTTCCCTACACGACGCTCTTCCGATCTNNNAAATGATTTGGGGCAGGT
AGCACACGTCTGAACTCTTTCCCTACACGACGCTCTTCCGATCTNNNGTTGGGCATTTGTGTGTGTT
GAGCACACGTCTGAACTCTTTCCCTACACGACGCTCTTCCGATCTNNNTATCTCGACCCAATTAGCAT
GCACACGTCTGAACTCTTTCCCTACACGACGCTCTTCCGATCTNNNTGAAGGAAGTTTTGCTTTGGG
AGCACACGTCTGAACTCTTTCCCTACACGACGCTCTTCCGATCTNNNGGTCAGTAACTACGTCCCTTTT
GAGCACACGTCTGAACTCTTTCCCTACACGACGCTCTTCCGATCTNNNATTGAAGGCTTCCCCTTCAC
GAGCACACGTCTGAACTCTTTCCCTACACGACGCTCTTCCGATCTNNNTACTGACATGGCAACATTTA
GCACACGTCTGAACTCTTTCCCTACACGACGCTCTTCCGATCTNNNTCTGCTCCTGGAGTAAGAAAGG
AGAGCACACGTCTGAACTCTTTCCCTACACGACGCTCTTCCGATCTNNNTTTCTGGGTAACTTTGAGC
AGCACACGTCTGAACTCTTTCCCTACACGACGCTCTTCCGATCTNNNACAAGAGAAGACCTTCATGAC
AGCACACGTCTGAACTCTTTCCCTACACGACGCTCTTCCGATCTNNNCTGGCTGACCTAGCCAAAGAT
AGCACACGTCTGAACTCTTTCCCTACACGACGCTCTTCCGATCTNNNGTCCAGAAAAGAAATCATAAA
AGCACACGTCTGAACTCTTTCCCTACACGACGCTCTTCCGATCTNNNTGTGTTAGGAGGAAAAAGAT
AGCACACGTCTGAACTCTTTCCCTACACGACGCTCTTCCGATCTNNNCCTGTGCGAAGACTTTAGGT
GCACACGTCTGAACTCTTTCCCTACACGACGCTCTTCCGATCTNNNGCCAGACAGGAGGTTACTTAG
AGCACACGTCTGAACTCTTTCCCTACACGACGCTCTTCCGATCTNNNAGCACAAAATTGACAAAAAGA
GAGCACACGTCTGAACTCTTTCCCTACACGACGCTCTTCCGATCTNNNTACGCCAACTTCTTATGCTGT
GAGCACACGTCTGAACTCTTTCCCTACACGACGCTCTTCCGATCTNNNAGGGAAAGGCAAAATTCAA
GCACACGTCTGAACTCTTTCCCTACACGACGCTCTTCCGATCTNNNCCAAGAGCACTTTGAGCTCAGT
GCACACGTCTGAACTCTTTCCCTACACGACGCTCTTCCGATCTNNNGTCCTCATGATGCATTGCCCTAA
GAGCACACGTCTGAACTCTTTCCCTACACGACGCTCTTCCGATCTNNNCTTTCCCTTTCCCCAACACTC

GAGCACACGTCTGAACTCTTTCCCTACACGACGCTCTTCCGATCTNNNTTCACTGAAGTGATGTAAAG
GAGCACACGTCTGAACTCTTTCCCTACACGACGCTCTTCCGATCTNNNATGAAACCAGAGAGCTTGTG
AGCACACGTCTGAACTCTTTCCCTACACGACGCTCTTCCGATCTNNNCATGTTCAGTACGCACGCAATT
GAGCACACGTCTGAACTCTTTCCCTACACGACGCTCTTCCGATCTNNNTGGAGACTACCAGAGAAAC
AGCACACGTCTGAACTCTTTCCCTACACGACGCTCTTCCGATCTNNNGGTCAGTAAGAAGGGAGACT
GCACACGTCTGAACTCTTTCCCTACACGACGCTCTTCCGATCTNNNGTTCAGGTCCTGTCGCCTAAAA
AGCACACGTCTGAACTCTTTCCCTACACGACGCTCTTCCGATCTNNNCCATGGCATGTAGGTAGGGT
ACACGTCTGAACTCTTTCCCTACACGACGCTCTTCCGATCTNNNAAGAGAAATTACAGGCCACAGCCC
AGCACACGTCTGAACTCTTTCCCTACACGACGCTCTTCCGATCTNNNGGGAAAATGTCCATGATCCCA
GCACACGTCTGAACTCTTTCCCTACACGACGCTCTTCCGATCTNNNCACGCAGCCCTATGAGGTAAAA
AGCACACGTCTGAACTCTTTCCCTACACGACGCTCTTCCGATCTNNNAATTTGGGCAGGACATGGAC
AGCACACGTCTGAACTCTTTCCCTACACGACGCTCTTCCGATCTNNNTGACGTCTGTAGCACACACAAA
GAGCACACGTCTGAACTCTTTCCCTACACGACGCTCTTCCGATCTNNNGCTGCAAAGTCTTAGAACCCA
AGCACACGTCTGAACTCTTTCCCTACACGACGCTCTTCCGATCTNNNTTCGGGTTTGGGTTTCACAAG
AGCACACGTCTGAACTCTTTCCCTACACGACGCTCTTCCGATCTNNNGTAAGCTCCAGTCCATGTCCAA
GCACACGTCTGAACTCTTTCCCTACACGACGCTCTTCCGATCTNNNACAAAACAGAAGCTCTTTGGGG
AGCACACGTCTGAACTCTTTCCCTACACGACGCTCTTCCGATCTNNNACTCACAGGTCTTACTGGTCC
GAGCACACGTCTGAACTCTTTCCCTACACGACGCTCTTCCGATCTNNNTGATCTTTCTGCTGTCTAGC
GCACACGTCTGAACTCTTTCCCTACACGACGCTCTTCCGATCTNNNGGCCTCCTCCTTCAAGACTTCTAC
AGCACACGTCTGAACTCTTTCCCTACACGACGCTCTTCCGATCTNNNCTGGACAGGAGGAACAGTCT
GCACACGTCTGAACTCTTTCCCTACACGACGCTCTTCCGATCTNNNGGCCATCTCTCCCTGGATAATCT
AGCACACGTCTGAACTCTTTCCCTACACGACGCTCTTCCGATCTNNNTCCCTTCTCAATTCCCATACCAC
AGAGCACACGTCTGAACTCTTTCCCTACACGACGCTCTTCCGATCTNNNCCACAGAAGCCAACAATCA
AGCACACGTCTGAACTCTTTCCCTACACGACGCTCTTCCGATCTNNNACTTTTTCTCAGGTGGCAAGTG
GAGCACACGTCTGAACTCTTTCCCTACACGACGCTCTTCCGATCTNNNCAATGGAGGAAGGTGACA
AGCACACGTCTGAACTCTTTCCCTACACGACGCTCTTCCGATCTNNNAGAGTAATGCTAGCTGTTGG
GAGCACACGTCTGAACTCTTTCCCTACACGACGCTCTTCCGATCTNNNTTACATGGTGGAAGCCTCTT
AGCACACGTCTGAACTCTTTCCCTACACGACGCTCTTCCGATCTNNNTATAATGTCCTCCGAGTTTGCC
AGCACACGTCTGAACTCTTTCCCTACACGACGCTCTTCCGATCTNNNTAGAGGAACTGTCAGGCTTCC
GAGCACACGTCTGAACTCTTTCCCTACACGACGCTCTTCCGATCTNNNGATCTGCCGGGAGAGGAGA
AGCACACGTCTGAACTCTTTCCCTACACGACGCTCTTCCGATCTNNNACACCAATCCTCCCAAAGTAAC
AGCACACGTCTGAACTCTTTCCCTACACGACGCTCTTCCGATCTNNNCTTTTTCCTTTGCTTTGAGGGC
AGCACACGTCTGAACTCTTTCCCTACACGACGCTCTTCCGATCTNNNGCGTGTCTATGTCGATTTGCA
GCACACGTCTGAACTCTTTCCCTACACGACGCTCTTCCGATCTNNNTCTTCCAAAATCAAAGCCCACTGC
AGCACACGTCTGAACTCTTTCCCTACACGACGCTCTTCCGATCTNNNGGCAAGAAAATCACATTGCCC
AGCACACGTCTGAACTCTTTCCCTACACGACGCTCTTCCGATCTNNNAGGTCACTCAGATGTGAACCA
GCACACGTCTGAACTCTTTCCCTACACGACGCTCTTCCGATCTNNNTAGAGGAACTGTCAGGCTTCCA
AGCACACGTCTGAACTCTTTCCCTACACGACGCTCTTCCGATCTNNNGACTTGAAATCCTGCCTGGAA
AGCACACGTCTGAACTCTTTCCCTACACGACGCTCTTCCGATCTNNNTCTCCCCAAGTAACTCACGTCA
GAGCACACGTCTGAACTCTTTCCCTACACGACGCTCTTCCGATCTNNNATGAAATGACCATCAAAGGC
AGAGCACACGTCTGAACTCTTTCCCTACACGACGCTCTTCCGATCTNNNGCCAGAAATGAGGGACAA
AGCACACGTCTGAACTCTTTCCCTACACGACGCTCTTCCGATCTNNNGCGATTTTCAGAACTTACTGCC
AGCACACGTCTGAACTCTTTCCCTACACGACGCTCTTCCGATCTNNNTCAACATGATGCACTAGAGGG

GCACACGTCTGAACTCTTTCCCTACACGACGCTCTTCCGATCTNNNGTTTCCCAGGTAACCTTCAGCTA
AGCACACGTCTGAACTCTTTCCCTACACGACGCTCTTCCGATCTNNNGTATTGCTAATCTCTGGGGTG
AGAGCACACGTCTGAACTCTTTCCCTACACGACGCTCTTCCGATCTNNNACCAAATACACAGACAGGAA
GAGCACACGTCTGAACTCTTTCCCTACACGACGCTCTTCCGATCTNNNTTCTTCAGAGGATGACCACCA
GAGCACACGTCTGAACTCTTTCCCTACACGACGCTCTTCCGATCTNNNTGAGGGTCCCTGAAAGAAAT
GAGCACACGTCTGAACTCTTTCCCTACACGACGCTCTTCCGATCTNNNTGCCTGTTGTACGTATATTCT
GAGCACACGTCTGAACTCTTTCCCTACACGACGCTCTTCCGATCTNNNAGAAAAGGTGGCTTTCAGACT
AGCACACGTCTGAACTCTTTCCCTACACGACGCTCTTCCGATCTNNNACATTCTCAGGATAGTGGATTT
AGCACACGTCTGAACTCTTTCCCTACACGACGCTCTTCCGATCTNNNTCATCTCCCGTTAGGTTAGCCA
GAAGAGCACACGTCTGAACTCTTTCCCTACACGACGCTCTTCCGATCTNNNTCTCCCAAGTAACTCACG
AGAGCACACGTCTGAACTCTTTCCCTACACGACGCTCTTCCGATCTNNNCCAGGAATTTGAGACCACCT
AAGAGCACACGTCTGAACTCTTTCCCTACACGACGCTCTTCCGATCTNNNGGGGTGAAACCACAGTTTT
GAGCACACGTCTGAACTCTTTCCCTACACGACGCTCTTCCGATCTNNNGTCCAGAGCAAGAATGGTAC
GAGCACACGTCTGAACTCTTTCCCTACACGACGCTCTTCCGATCTNNNAGTCAAGCATCCAAAGACAGC
AGCACACGTCTGAACTCTTTCCCTACACGACGCTCTTCCGATCTNNNAGTTGGGAAGGAGAGGTAAG
GAGCACACGTCTGAACTCTTTCCCTACACGACGCTCTTCCGATCTNNNTCCATATTGGGGCTGTGAACT
GAGCACACGTCTGAACTCTTTCCCTACACGACGCTCTTCCGATCTNNNGTCGACAAAAAGCTACACTGC
GAGCACACGTCTGAACTCTTTCCCTACACGACGCTCTTCCGATCTNNNTAACCCCATTTGAGCATGACC
GAGCACACGTCTGAACTCTTTCCCTACACGACGCTCTTCCGATCTNNNCACAGGCTCAGAGAGCAATGA
AGCACACGTCTGAACTCTTTCCCTACACGACGCTCTTCCGATCTNNNACATGCTCCTCAGTCAACAACTC
AGAGCACACGTCTGAACTCTTTCCCTACACGACGCTCTTCCGATCTNNNTGACCCTAAGATTTTCGCCAG
AGAGCACACGTCTGAACTCTTTCCCTACACGACGCTCTTCCGATCTNNNCCCAGAGAAAGAGCCATTAG
AGCACACGTCTGAACTCTTTCCCTACACGACGCTCTTCCGATCTNNNTCAGTTCTTTGTGAAGCCACGT
GAGCACACGTCTGAACTCTTTCCCTACACGACGCTCTTCCGATCTNNNTGCCCTAATCTTGGATTTCTG
GAGCACACGTCTGAACTCTTTCCCTACACGACGCTCTTCCGATCTNNNGGACCAAAAGGAGCAATTCT
AGAGCACACGTCTGAACTCTTTCCCTACACGACGCTCTTCCGATCTNNNAGCTGGACTTTTAGCCAACT
AGAGCACACGTCTGAACTCTTTCCCTACACGACGCTCTTCCGATCTNNNTGAGGGTGGATCCCCTATA
AAGAGCACACGTCTGAACTCTTTCCCTACACGACGCTCTTCCGATCTNNNTGTCAGGTGATGTGTGAG
GAGCACACGTCTGAACTCTTTCCCTACACGACGCTCTTCCGATCTNNNGGCATTGCTTTTGTCCTGACT
AGAGCACACGTCTGAACTCTTTCCCTACACGACGCTCTTCCGATCTNNNTGTGCTCTGAACCACCAGAT
GAGCACACGTCTGAACTCTTTCCCTACACGACGCTCTTCCGATCTNNNGGAAGAAGGTGACCTGGACC
AGCACACGTCTGAACTCTTTCCCTACACGACGCTCTTCCGATCTNNNAAGCTCTGGCTTATGGGTTTA
AAGAGCACACGTCTGAACTCTTTCCCTACACGACGCTCTTCCGATCTNNNGAAGAAACACCATGTTGCA
GAGCACACGTCTGAACTCTTTCCCTACACGACGCTCTTCCGATCTNNNGTGGGCAGTGCTATGATTTCT
AGCACACGTCTGAACTCTTTCCCTACACGACGCTCTTCCGATCTNNNCCACATTGAAGGAGGTGATCCA
AGAGCACACGTCTGAACTCTTTCCCTACACGACGCTCTTCCGATCTNNNGGAACTGGGGGAACAGGA
GCACACGTCTGAACTCTTTCCCTACACGACGCTCTTCCGATCTNNNGTGAGGGACTGTGCTATCTTTCC
GAGCACACGTCTGAACTCTTTCCCTACACGACGCTCTTCCGATCTNNNATGTTTCAGGCAAAGGCCATA
AAGAGCACACGTCTGAACTCTTTCCCTACACGACGCTCTTCCGATCTNNNAGCAGACAACTGATCACCC
GAAGAGCACACGTCTGAACTCTTTCCCTACACGACGCTCTTCCGATCTNNNCCATCCAGGTAATTGTGA
GAGCACACGTCTGAACTCTTTCCCTACACGACGCTCTTCCGATCTNNNGCTGTAACAGGAATCCCTACG
AAGAGCACACGTCTGAACTCTTTCCCTACACGACGCTCTTCCGATCTNNNTTACTGTTGCCTTGTGTCCA
GAGCACACGTCTGAACTCTTTCCCTACACGACGCTCTTCCGATCTNNNGTGGGCAGTGCTATGATTTC

GGAAGAGCACACGTCTGAACTCTTTCCCTACACGACGCTCTTCCGATCTNNNACCCTGAAGAAGCATA
AGCACACGTCTGAACTCTTTCCCTACACGACGCTCTTCCGATCTNNNACTCCTTCATTGCACACATGGTT
GAGCACACGTCTGAACTCTTTCCCTACACGACGCTCTTCCGATCTNNNTTTGCTGCCAGAATTTGAGAC
AGAGCACACGTCTGAACTCTTTCCCTACACGACGCTCTTCCGATCTNNNTCATGTCCTCCTGATTCCATC
GAGCACACGTCTGAACTCTTTCCCTACACGACGCTCTTCCGATCTNNNTCTCACTTTGGGCTTCAGCTTT
GAAGAGCACACGTCTGAACTCTTTCCCTACACGACGCTCTTCCGATCTNNNAGAGTTGGAAGCTTTGAA
AAGAGCACACGTCTGAACTCTTTCCCTACACGACGCTCTTCCGATCTNNNGGCCTCAAGAAATCCTCATT
AGCACACGTCTGAACTCTTTCCCTACACGACGCTCTTCCGATCTNNNGCATTTCAACCTCAACTTGCTG
AGCACACGTCTGAACTCTTTCCCTACACGACGCTCTTCCGATCTNNNCTTGTGTTGTGTATGTGTGCGT
GAGCACACGTCTGAACTCTTTCCCTACACGACGCTCTTCCGATCTNNNCATCCATCTCCCCATCCCTCTA
AGAGCACACGTCTGAACTCTTTCCCTACACGACGCTCTTCCGATCTNNNTGGTGGGAAAGTCAGAAAA
GCACACGTCTGAACTCTTTCCCTACACGACGCTCTTCCGATCTNNNGTTTCACTCACCCCATTTCCTGTCA
AGCACACGTCTGAACTCTTTCCCTACACGACGCTCTTCCGATCTNNNCAGTATGGTCGATGATGCTGGT
AAGAGCACACGTCTGAACTCTTTCCCTACACGACGCTCTTCCGATCTNNNGAAGTCTGCATGCTCTCCC
GAGCACACGTCTGAACTCTTTCCCTACACGACGCTCTTCCGATCTNNNGCTAATGTCTTCCAGCTCCTT
AGCACACGTCTGAACTCTTTCCCTACACGACGCTCTTCCGATCTNNNGCTTAGTGCATAGGACAGGCTTA
CACACGTCTGAACTCTTTCCCTACACGACGCTCTTCCGATCTNNNAGGAGATAAAGACACAGGCCATT
AAGAGCACACGTCTGAACTCTTTCCCTACACGACGCTCTTCCGATCTNNNCCAATCCAAGAGGAGCAT
AGCACACGTCTGAACTCTTTCCCTACACGACGCTCTTCCGATCTNNNCGCTTTCCAACACTGAAACCAAT
AGAGCACACGTCTGAACTCTTTCCCTACACGACGCTCTTCCGATCTNNNATACTGTGGGTTGGGCAGAA
GCACACGTCTGAACTCTTTCCCTACACGACGCTCTTCCGATCTNNNTTGTGGTCACCTTGCTCTCTTTACA
AGCACACGTCTGAACTCTTTCCCTACACGACGCTCTTCCGATCTNNNTTCTGTGAGTGGGTCAAACTTG
GGAAGAGCACACGTCTGAACTCTTTCCCTACACGACGCTCTTCCGATCTNNNCAGTTCTTAGAGCAAGT
AAGAGCACACGTCTGAACTCTTTCCCTACACGACGCTCTTCCGATCTNNNACACTGTCCTGTGATATTGT
GCACACGTCTGAACTCTTTCCCTACACGACGCTCTTCCGATCTNNNTTTAGGGCCACTTACTGGCTTCCA
AAGAGCACACGTCTGAACTCTTTCCCTACACGACGCTCTTCCGATCTNNNACTATAGGTCTGTCTGGCCA
GCACACGTCTGAACTCTTTCCCTACACGACGCTCTTCCGATCTNNNGGATAGAGTAGCTAGGACTACA
GCACACGTCTGAACTCTTTCCCTACACGACGCTCTTCCGATCTNNNCTGGCCCTGGTTTTCTTATCCATCA
GAGCACACGTCTGAACTCTTTCCCTACACGACGCTCTTCCGATCTNNNCCCCACTGCTTCATGTCTTACA
AGCACACGTCTGAACTCTTTCCCTACACGACGCTCTTCCGATCTNNNTGCAATTGGCAATCCATCTGGTA
AGCACACGTCTGAACTCTTTCCCTACACGACGCTCTTCCGATCTNNNAACTTTGGTGGCTGTCCTAAGT
GCACACGTCTGAACTCTTTCCCTACACGACGCTCTTCCGATCTNNNTAAAAGTGGCAGTTTCCCTTGAA
GCACACGTCTGAACTCTTTCCCTACACGACGCTCTTCCGATCTNNNTGCATCCTGACTAATAAAGCCCA
AGAGCACACGTCTGAACTCTTTCCCTACACGACGCTCTTCCGATCTNNNAGAAAGAGACAACTCTGTTC
GAGCACACGTCTGAACTCTTTCCCTACACGACGCTCTTCCGATCTNNNAGTATACCCATGATTCCAGCCT
CACACGTCTGAACTCTTTCCCTACACGACGCTCTTCCGATCTNNNTTATTATAGCTCCCCCTCCCCCTCAT
GAGCACACGTCTGAACTCTTTCCCTACACGACGCTCTTCCGATCTNNNTTTCATGCAAGGTTTGTGTCA
AGCACACGTCTGAACTCTTTCCCTACACGACGCTCTTCCGATCTNNNTTAGGTTACCAAGGGATGGGA
AGAGCACACGTCTGAACTCTTTCCCTACACGACGCTCTTCCGATCTNNNAAACACACATCTCTGTGCCT
GAGCACACGTCTGAACTCTTTCCCTACACGACGCTCTTCCGATCTNNNATGTACCCGTTTTGTGTGTGT
AGAGCACACGTCTGAACTCTTTCCCTACACGACGCTCTTCCGATCTNNNCCGGCCTGTAGTCTCTAGTTT
AGAGCACACGTCTGAACTCTTTCCCTACACGACGCTCTTCCGATCTNNNTCTGAACTGTTTGAGTAGAT
AGCACACGTCTGAACTCTTTCCCTACACGACGCTCTTCCGATCTNNNAACTATCCCAAGACACACTCTGC

AGCACACGTCTGAACTCTTTCCCTACACGACGCTCTTCCGATCTNNNATTCTTTCCTACACCAGGGGTGA
AGCACACGTCTGAACTCTTTCCCTACACGACGCTCTTCCGATCTNNNAGGACAGGGCATATGAGAGAG
GAGCACACGTCTGAACTCTTTCCCTACACGACGCTCTTCCGATCTNNNCACACATTTTCCCAGCACATTCA
GAGCACACGTCTGAACTCTTTCCCTACACGACGCTCTTCCGATCTNNNTGTGAACATAACATCAGGGAC
GAGCACACGTCTGAACTCTTTCCCTACACGACGCTCTTCCGATCTNNNATGTATGAAGGGGGAAAAAC
GAGCACACGTCTGAACTCTTTCCCTACACGACGCTCTTCCGATCTNNNTTACACCCTGGACCTCATCTCT
AGCACACGTCTGAACTCTTTCCCTACACGACGCTCTTCCGATCTNNNGAGGTTGTTCTGTTGGTTGTTC
GAGCACACGTCTGAACTCTTTCCCTACACGACGCTCTTCCGATCTNNNAAGGTCACTTTTCTCTCACAGC
GAGCACACGTCTGAACTCTTTCCCTACACGACGCTCTTCCGATCTNNNGAATCCAGGTGGGTAAACTG
AGAGCACACGTCTGAACTCTTTCCCTACACGACGCTCTTCCGATCTNNNCAGAAAGCAAAGGACCTCA
GAGCACACGTCTGAACTCTTTCCCTACACGACGCTCTTCCGATCTNNNGTCATTCTGAAGGGGAGTAG
GAGCACACGTCTGAACTCTTTCCCTACACGACGCTCTTCCGATCTNNNTACTGCTCCTTTCCAACACACAC
AGCACACGTCTGAACTCTTTCCCTACACGACGCTCTTCCGATCTNNNGCTGATTCTATCTCTCCCCACCTCA
AGCACACGTCTGAACTCTTTCCCTACACGACGCTCTTCCGATCTNNNTCCAGGACATAGGTCAAAGACC
GAGCACACGTCTGAACTCTTTCCCTACACGACGCTCTTCCGATCTNNNCATCGCTGTTTCTGAGTGAGT
AGCACACGTCTGAACTCTTTCCCTACACGACGCTCTTCCGATCTNNNAAAAACAGTCAGTGGCTCATTG
AGAGCACACGTCTGAACTCTTTCCCTACACGACGCTCTTCCGATCTNNNGGGGTAAAGGGCCATGATTT
GAGCACACGTCTGAACTCTTTCCCTACACGACGCTCTTCCGATCTNNNTGATTTTATCTGGGTCTCCTGG
AAGAGCACACGTCTGAACTCTTTCCCTACACGACGCTCTTCCGATCTNNNAAATTTCTTGAGGCCAGAG
GAGCACACGTCTGAACTCTTTCCCTACACGACGCTCTTCCGATCTNNNGCCACCGCACATAAATGTTCAA
GAGCACACGTCTGAACTCTTTCCCTACACGACGCTCTTCCGATCTNNNCAACTCTGTCCCTTTGAGGAAC
GAAGAGCACACGTCTGAACTCTTTCCCTACACGACGCTCTTCCGATCTNNNGCAGATGTAACACCCTGC
GAGCACACGTCTGAACTCTTTCCCTACACGACGCTCTTCCGATCTNNNAAGATCCAACTGGCATGATTG
AGCACACGTCTGAACTCTTTCCCTACACGACGCTCTTCCGATCTNNNTGAGATTTACAAGAGGGCCTG
GAGCACACGTCTGAACTCTTTCCCTACACGACGCTCTTCCGATCTNNNGCTGGTGTATAAGTCAGTAAG
AGCACACGTCTGAACTCTTTCCCTACACGACGCTCTTCCGATCTNNNTGATGATGGCAACAACCTGGAA
GAGCACACGTCTGAACTCTTTCCCTACACGACGCTCTTCCGATCTNNNTGTCTGCAAAACAAGGTAAAT
AGCACACGTCTGAACTCTTTCCCTACACGACGCTCTTCCGATCTNNNACTGTGTATTTATGTGTTTGTGC
AAGAGCACACGTCTGAACTCTTTCCCTACACGACGCTCTTCCGATCTNNNTACTTCCACGCAAAGGATC
AGCACACGTCTGAACTCTTTCCCTACACGACGCTCTTCCGATCTNNNGAAGGTAAGCGATCCTTCTGAC
AGCACACGTCTGAACTCTTTCCCTACACGACGCTCTTCCGATCTNNNTTTCTGCTCCAACCAGCAAGAAT
AGCACACGTCTGAACTCTTTCCCTACACGACGCTCTTCCGATCTNNNGGAAAGTACGTGGCATGGAAC
GCACACGTCTGAACTCTTTCCCTACACGACGCTCTTCCGATCTNNNCCTGGGGCCAAGGAAATTGTGCA
GAGCACACGTCTGAACTCTTTCCCTACACGACGCTCTTCCGATCTNNNTGTTCAGAAGTGGAACAGCA
AAGAGCACACGTCTGAACTCTTTCCCTACACGACGCTCTTCCGATCTNNNTGGAGTTGGAAAATGAAA
AGCACACGTCTGAACTCTTTCCCTACACGACGCTCTTCCGATCTNNNGGGATGCTCCCATCCTACTATG
GAGCACACGTCTGAACTCTTTCCCTACACGACGCTCTTCCGATCTNNNTCAAGTACTAACCATGGGCTTC
GAGCACACGTCTGAACTCTTTCCCTACACGACGCTCTTCCGATCTNNNGTCTGCAGAGACATAAAGTGC
AGAGCACACGTCTGAACTCTTTCCCTACACGACGCTCTTCCGATCTNNNACCTAGTAAAGCAGTTGAAT
GAGCACACGTCTGAACTCTTTCCCTACACGACGCTCTTCCGATCTNNNCCTCTATCTGCAAGAAGGACA
AAGAGCACACGTCTGAACTCTTTCCCTACACGACGCTCTTCCGATCTNNNTGCTTTATTGAGGTGGAG
GAAGAGCACACGTCTGAACTCTTTCCCTACACGACGCTCTTCCGATCTNNNACAAGTGGGTAAAAGAA
GAAGAGCACACGTCTGAACTCTTTCCCTACACGACGCTCTTCCGATCTNNNTCATGTAGTCACACACATC

GAGCACACGTCTGAACTCTTTCCCTACACGACGCTCTTCCGATCTNNNCCCTCTCCCCTGATTTCACATCT
GAGCACACGTCTGAACTCTTTCCCTACACGACGCTCTTCCGATCTNNNTGCCTTGTTTGGAAGACCTTG
AGCACACGTCTGAACTCTTTCCCTACACGACGCTCTTCCGATCTNNNGCCTTGCTACTCAGATTTCACCA
AAGAGCACACGTCTGAACTCTTTCCCTACACGACGCTCTTCCGATCTNNNGGCATGCCAATACCTAGGA
ACACGTCTGAACTCTTTCCCTACACGACGCTCTTCCGATCTNNNCTAAAAAGGGAGTTGGACCCTCGCA
AGCACACGTCTGAACTCTTTCCCTACACGACGCTCTTCCGATCTNNNACCAAGTCCCAATGTCTTCAGT
GAGCACACGTCTGAACTCTTTCCCTACACGACGCTCTTCCGATCTNNNGAGACTGTACACTACCAAGG
GAGCACACGTCTGAACTCTTTCCCTACACGACGCTCTTCCGATCTNNNCCAATGCAAGATACACACACA
AGCACACGTCTGAACTCTTTCCCTACACGACGCTCTTCCGATCTNNNCAAAGGCTGCAAAATGAGTGT
AGCACACGTCTGAACTCTTTCCCTACACGACGCTCTTCCGATCTNNNATGGTCAGGATGTGCTCTGGTT
AGCACACGTCTGAACTCTTTCCCTACACGACGCTCTTCCGATCTNNNCTCTTGCCTGAACTATCCTACG
AGAGCACACGTCTGAACTCTTTCCCTACACGACGCTCTTCCGATCTNNNGCAGACTGCACATGTTCTGA
GCACACGTCTGAACTCTTTCCCTACACGACGCTCTTCCGATCTNNNACCCAGTGCCCCTTTGATATACTC
AGAGCACACGTCTGAACTCTTTCCCTACACGACGCTCTTCCGATCTNNNAATTTCCAGTTGGGAAAGG
AGCACACGTCTGAACTCTTTCCCTACACGACGCTCTTCCGATCTNNNAGGAGTAAATGGGAGTTTTCTG
GAGCACACGTCTGAACTCTTTCCCTACACGACGCTCTTCCGATCTNNNTGGGCAATAGCAAGTCCCTAT
AGCACACGTCTGAACTCTTTCCCTACACGACGCTCTTCCGATCTNNNTTGCTGGTTCCTATGTCCCTTTG
GAGCACACGTCTGAACTCTTTCCCTACACGACGCTCTTCCGATCTNNNCTGGTGAGGACACGTTACACA
GAGCACACGTCTGAACTCTTTCCCTACACGACGCTCTTCCGATCTNNNTAATCATGTTGTGCTCACCCCA
AGCACACGTCTGAACTCTTTCCCTACACGACGCTCTTCCGATCTNNNCATGGTGATGTCCAGATTCCCA
AGCACACGTCTGAACTCTTTCCCTACACGACGCTCTTCCGATCTNNNTGTCCCAATCCCACACAGATACT
AAGAGCACACGTCTGAACTCTTTCCCTACACGACGCTCTTCCGATCTNNNTTGATGAGGAATCCTGAAT
AGAGCACACGTCTGAACTCTTTCCCTACACGACGCTCTTCCGATCTNNNCGTCTGGAAACCAGGATGT
AGCACACGTCTGAACTCTTTCCCTACACGACGCTCTTCCGATCTNNNTCTGCTATAGTTTCCCTTCAGTC
GAGCACACGTCTGAACTCTTTCCCTACACGACGCTCTTCCGATCTNNNACTAGATGTCTAATAAGAGG
GAGCACACGTCTGAACTCTTTCCCTACACGACGCTCTTCCGATCTNNNCTGCCACTGTCACAAGAAAGG
AGAGCACACGTCTGAACTCTTTCCCTACACGACGCTCTTCCGATCTNNNATGACACACACAAGCAGACA
GAGCACACGTCTGAACTCTTTCCCTACACGACGCTCTTCCGATCTNNNACAACTGACCCTTCACGTAGGT
AGCACACGTCTGAACTCTTTCCCTACACGACGCTCTTCCGATCTNNNTCAGATAGGCTTCCCCTACCTTC
AGAGCACACGTCTGAACTCTTTCCCTACACGACGCTCTTCCGATCTNNNGCATAGGACAAGGACCCAA
AGCACACGTCTGAACTCTTTCCCTACACGACGCTCTTCCGATCTNNNAAGTGATTGGATCTCTGAGGCT
GAGCACACGTCTGAACTCTTTCCCTACACGACGCTCTTCCGATCTNNNTTAGGTTACCAAGGGATGG
GAGCACACGTCTGAACTCTTTCCCTACACGACGCTCTTCCGATCTNNNTCAACATACTCATCTTCCTCCTG
AGCACACGTCTGAACTCTTTCCCTACACGACGCTCTTCCGATCTNNNTGCTGAGGTGCTTTGTAAACTCA
GCACACGTCTGAACTCTTTCCCTACACGACGCTCTTCCGATCTNNNTTTCTTCCTCCTGCTGCTTTGAACA
GCACACGTCTGAACTCTTTCCCTACACGACGCTCTTCCGATCTNNNAGCAAGTAAAGACATCTGAGCC
GCACACGTCTGAACTCTTTCCCTACACGACGCTCTTCCGATCTNNNCATGCCACCATGTTTTTGAACTGT
GAGCACACGTCTGAACTCTTTCCCTACACGACGCTCTTCCGATCTNNNTCCACTGGAAGAGACATAGT
CACACGTCTGAACTCTTTCCCTACACGACGCTCTTCCGATCTNNNGCCACCGCACATAAATGTTCAAGC
GAGCACACGTCTGAACTCTTTCCCTACACGACGCTCTTCCGATCTNNNACCAACTCACAGCTTCTCTCTC
AAGAGCACACGTCTGAACTCTTTCCCTACACGACGCTCTTCCGATCTNNNTTTGACCTCAGATAATAGA
AGCACACGTCTGAACTCTTTCCCTACACGACGCTCTTCCGATCTNNNATTCATTCCTTCCCCTCCACCTCC
AGCACACGTCTGAACTCTTTCCCTACACGACGCTCTTCCGATCTNNNTGTAAGGAAGATCCAGGGCTT

AGCACACGTCTGAACTCTTTCCCTACACGACGCTCTTCCGATCTNNNTGACATTATCGTTTGCCAGTC
AAGAGCACACGTCTGAACTCTTTCCCTACACGACGCTCTTCCGATCTNNNAGTACCCATGCTGTTCAA
AGCACACGTCTGAACTCTTTCCCTACACGACGCTCTTCCGATCTNNNTTGGTTGTGTAGCAATCCCTC
AGCACACGTCTGAACTCTTTCCCTACACGACGCTCTTCCGATCTNNNGTCCCTCTGTCCCTCTATTCCT
GCACACGTCTGAACTCTTTCCCTACACGACGCTCTTCCGATCTNNNTTGTTGGCAGCACATTCTAACCT
AGCACACGTCTGAACTCTTTCCCTACACGACGCTCTTCCGATCTNNNAAGTGCTTCTCTGAGGAGGG
AGCACACGTCTGAACTCTTTCCCTACACGACGCTCTTCCGATCTNNNAGTCCAACCATCACTTGATTTA
AGCACACGTCTGAACTCTTTCCCTACACGACGCTCTTCCGATCTNNNAGTACCCTTACAGGAAGAGCC
AGCACACGTCTGAACTCTTTCCCTACACGACGCTCTTCCGATCTNNNTTTAGCACCCAGCTGTCCTGAA
AGCACACGTCTGAACTCTTTCCCTACACGACGCTCTTCCGATCTNNNCCTATCACAATGCAAGCTCCAG
GAGCACACGTCTGAACTCTTTCCCTACACGACGCTCTTCCGATCTNNNGGTGTCCGTGGCTTATCTTT
AGCACACGTCTGAACTCTTTCCCTACACGACGCTCTTCCGATCTNNNCTGTTCTGATCCTCCAAGCTGT
GAGCACACGTCTGAACTCTTTCCCTACACGACGCTCTTCCGATCTNNNACGATCAGGAATGCAGAGG
GAGCACACGTCTGAACTCTTTCCCTACACGACGCTCTTCCGATCTNNNATCAGAATCCCAACAGAGGG
AGCACACGTCTGAACTCTTTCCCTACACGACGCTCTTCCGATCTNNNATGAACAGTCTGGTGCATGTC
GCACACGTCTGAACTCTTTCCCTACACGACGCTCTTCCGATCTNNNATTGCATGTCTGTGTGTATGGG
GCACACGTCTGAACTCTTTCCCTACACGACGCTCTTCCGATCTNNNTTTCAGCTTTATTTGCGTGGCAG
AGCACACGTCTGAACTCTTTCCCTACACGACGCTCTTCCGATCTNNNTGCTCATCACTCAAACACAACC
GAGCACACGTCTGAACTCTTTCCCTACACGACGCTCTTCCGATCTNNNGAAATTGGCATGGCAAAGG
AGCACACGTCTGAACTCTTTCCCTACACGACGCTCTTCCGATCTNNNCTGCAACAAATTGCCATCGAC
AGCACACGTCTGAACTCTTTCCCTACACGACGCTCTTCCGATCTNNNAGAGACATCCTGAGGCTAAG
GCACACGTCTGAACTCTTTCCCTACACGACGCTCTTCCGATCTNNNGCGGATGGAAAACACCACTTCT
GCACACGTCTGAACTCTTTCCCTACACGACGCTCTTCCGATCTNNNTGGCTAACCCCTCACTTCCAATAC
AGAGCACACGTCTGAACTCTTTCCCTACACGACGCTCTTCCGATCTNNNGTGGGGGTGAGGAAATT
GAGCACACGTCTGAACTCTTTCCCTACACGACGCTCTTCCGATCTNNNGCTGAATTTCAACCATCCCC
AGCACACGTCTGAACTCTTTCCCTACACGACGCTCTTCCGATCTNNNAGTACCCTTACAGGAAGAGCC
GCACACGTCTGAACTCTTTCCCTACACGACGCTCTTCCGATCTNNNAAGCTTGGAACACAAGAGTGG
GAGCACACGTCTGAACTCTTTCCCTACACGACGCTCTTCCGATCTNNNGAGGGTTAGCTGTCCTGAAA
AGCACACGTCTGAACTCTTTCCCTACACGACGCTCTTCCGATCTNNNTGCTAAGGAGGTTGGGGGA
AGCACACGTCTGAACTCTTTCCCTACACGACGCTCTTCCGATCTNNNAGTGACCTTTCACTTTTTCCCC
GAGCACACGTCTGAACTCTTTCCCTACACGACGCTCTTCCGATCTNNNAGGGCTTGATATGTGCAAA
AAGAGCACACGTCTGAACTCTTTCCCTACACGACGCTCTTCCGATCTNNNTGACTTCAAGCCCATGTT
AGCACACGTCTGAACTCTTTCCCTACACGACGCTCTTCCGATCTNNNGAGTTACACAGTTGGCAACAG
AGCACACGTCTGAACTCTTTCCCTACACGACGCTCTTCCGATCTNNNGGGCATGGAAAGGAATAG
GAGCACACGTCTGAACTCTTTCCCTACACGACGCTCTTCCGATCTNNNTGGGATCTGTAAAGTTAGA
AGCACACGTCTGAACTCTTTCCCTACACGACGCTCTTCCGATCTNNNTGTAAGGTTGCTGGGGAGAG
AGCACACGTCTGAACTCTTTCCCTACACGACGCTCTTCCGATCTNNNGCAACTTCTGCACTTGAACTGA
GCACACGTCTGAACTCTTTCCCTACACGACGCTCTTCCGATCTNNNCCCTTCCAATGCAGAAACTCAGA
AGCACACGTCTGAACTCTTTCCCTACACGACGCTCTTCCGATCTNNNATCCAAGAGGTAGAGGGGAA
GAGCACACGTCTGAACTCTTTCCCTACACGACGCTCTTCCGATCTNNNGCCACAGAGTGGAGATTTTC
AGCACACGTCTGAACTCTTTCCCTACACGACGCTCTTCCGATCTNNNTGGCCACATACAGAACTGCTT
GAGCACACGTCTGAACTCTTTCCCTACACGACGCTCTTCCGATCTNNNCCACTGCTAGCTCTCTCCTCA
AGCACACGTCTGAACTCTTTCCCTACACGACGCTCTTCCGATCTNNNAGTGTCCATAACTGTGGGAGC

GAGCACACGTCTGAACTCTTTCCCTACACGACGCTCTTCCGATCTNNNTTCCTTGGAATTAGCCATTCCC
AGCACACGTCTGAACTCTTTCCCTACACGACGCTCTTCCGATCTNNNTTTGAGGAGTTCCAGAGGTAG
GGAAGAGCACACGTCTGAACTCTTTCCCTACACGACGCTCTTCCGATCTNNNTGCCACTTTCTAAAAAGA
AAGAGCACACGTCTGAACTCTTTCCCTACACGACGCTCTTCCGATCTNNNACATGTTGCCTGTACCAGTT
AGCACACGTCTGAACTCTTTCCCTACACGACGCTCTTCCGATCTNNNCATTGCCACTATACCCTCAACTC
GCACACGTCTGAACTCTTTCCCTACACGACGCTCTTCCGATCTNNNTCCCTTTACAAGTGGTGGAGCTA
AGCACACGTCTGAACTCTTTCCCTACACGACGCTCTTCCGATCTNNNTTCTGCTGTTCAAGTTTGGCTC
AGCACACGTCTGAACTCTTTCCCTACACGACGCTCTTCCGATCTNNNCCTGCTCTAGTAAAGGAACCCCT
GCACACGTCTGAACTCTTTCCCTACACGACGCTCTTCCGATCTNNNAGACCATGTTCTCAACAGGACACC
GAGCACACGTCTGAACTCTTTCCCTACACGACGCTCTTCCGATCTNNNGAAGACCGCAGTAAACACTGA
AAGAGCACACGTCTGAACTCTTTCCCTACACGACGCTCTTCCGATCTNNNTTGAGGACTGCTGCCTTATA
AGCACACGTCTGAACTCTTTCCCTACACGACGCTCTTCCGATCTNNNCAATCCTCATATGTGCATGAGTG
GAGCACACGTCTGAACTCTTTCCCTACACGACGCTCTTCCGATCTNNNGGGTACGTTGATCCTTGTTTT
GAGCACACGTCTGAACTCTTTCCCTACACGACGCTCTTCCGATCTNNNTGGAGGAGCAAAAAGGCAAA
GCACACGTCTGAACTCTTTCCCTACACGACGCTCTTCCGATCTNNNGCAGTTTCGTCAGAGGTAAAGG
GAGCACACGTCTGAACTCTTTCCCTACACGACGCTCTTCCGATCTNNNCGTGGATGACCTTGGACATCA
AGCACACGTCTGAACTCTTTCCCTACACGACGCTCTTCCGATCTNNNACACTTTCAGTTTAGTGCACACA
GAGCACACGTCTGAACTCTTTCCCTACACGACGCTCTTCCGATCTNNNTGCTTACCTGTCCCACTTCACTA
GAAGAGCACACGTCTGAACTCTTTCCCTACACGACGCTCTTCCGATCTNNNGCCAGGACGGAACTTTTC
AGCACACGTCTGAACTCTTTCCCTACACGACGCTCTTCCGATCTNNNTCTTCTTATGTAACTGGGCACAA
AGAGCACACGTCTGAACTCTTTCCCTACACGACGCTCTTCCGATCTNNNTCCCAGATCAGCAAACATTTT
AGCACACGTCTGAACTCTTTCCCTACACGACGCTCTTCCGATCTNNNACAGCAGATAATGTTCAAATGC
GAGCACACGTCTGAACTCTTTCCCTACACGACGCTCTTCCGATCTNNNACCCAGTGCCCCTTTGATATAC
AGAGCACACGTCTGAACTCTTTCCCTACACGACGCTCTTCCGATCTNNNCAATGAAGAATTGCCCTCCA
GAGCACACGTCTGAACTCTTTCCCTACACGACGCTCTTCCGATCTNNNACACATGGCCTTCTGTTAGTGT
GAGCACACGTCTGAACTCTTTCCCTACACGACGCTCTTCCGATCTNNNGGGAAAAGGGGACGAAATGC
GAGCACACGTCTGAACTCTTTCCCTACACGACGCTCTTCCGATCTNNNTTTTCCAAACTTTCAACTGGTT
GAGCACACGTCTGAACTCTTTCCCTACACGACGCTCTTCCGATCTNNNAAGAGTGTATCAGTCAGGGTC
GAGCACACGTCTGAACTCTTTCCCTACACGACGCTCTTCCGATCTNNNTTTTGGGACTTTACCCAACTCC
GAGCACACGTCTGAACTCTTTCCCTACACGACGCTCTTCCGATCTNNNAAATGAGTGAGACCCTCCCTAC
AGCACACGTCTGAACTCTTTCCCTACACGACGCTCTTCCGATCTNNNCCTCCCCATTGTGGTTTAGGATC
GAGCACACGTCTGAACTCTTTCCCTACACGACGCTCTTCCGATCTNNNTGCTAGACAAAGGCAATTAGA
CACACGTCTGAACTCTTTCCCTACACGACGCTCTTCCGATCTNNNTACCATACAGGCCTTCTGCACATCAT
AGCACACGTCTGAACTCTTTCCCTACACGACGCTCTTCCGATCTNNNCTTAGACAGGAGACCTACACCC
GAGCACACGTCTGAACTCTTTCCCTACACGACGCTCTTCCGATCTNNNCGTACACACACACATCCCTTAC
AGAGCACACGTCTGAACTCTTTCCCTACACGACGCTCTTCCGATCTNNNACCACACACATTGGAGAGAC
GAGCACACGTCTGAACTCTTTCCCTACACGACGCTCTTCCGATCTNNNTCCTCTAAATTGCAATACGCCC
GAGCACACGTCTGAACTCTTTCCCTACACGACGCTCTTCCGATCTNNNGCCCTACTTCTTGGCTCCTCAC
AGCACACGTCTGAACTCTTTCCCTACACGACGCTCTTCCGATCTNNNCAAGCTCAGCGAATCCTTCTCTT
AGCACACGTCTGAACTCTTTCCCTACACGACGCTCTTCCGATCTNNNTGAAAGTTGAACTGAACTCCCC
AGAGCACACGTCTGAACTCTTTCCCTACACGACGCTCTTCCGATCTNNNATGTCATTCTCTCTCTGTCCCC
GAAGAGCACACGTCTGAACTCTTTCCCTACACGACGCTCTTCCGATCTNNNAGGAAAGATGGGGATAA
AGCACACGTCTGAACTCTTTCCCTACACGACGCTCTTCCGATCTNNNAGATCCTTTTGTAGTTGGTGCAC

GAGCACACGTCTGAACTCTTTCCCTACACGACGCTCTTCCGATCTNNNGAGCATTGTAGGTATTGCCC
GCACACGTCTGAACTCTTTCCCTACACGACGCTCTTCCGATCTNNNAACTGTTCCTAATATGTTGTTTG
AGCACACGTCTGAACTCTTTCCCTACACGACGCTCTTCCGATCTNNNTCTGTTACTCTAAGCAGGGTTA
AGCACACGTCTGAACTCTTTCCCTACACGACGCTCTTCCGATCTNNNACATTCAAACATGCTCCCTCTCT
AGAGCACACGTCTGAACTCTTTCCCTACACGACGCTCTTCCGATCTNNNCTCTCTCCAGCAACATCTG
AAGAGCACACGTCTGAACTCTTTCCCTACACGACGCTCTTCCGATCTNNNGCAGTGATGAACAGGAA
CACACGTCTGAACTCTTTCCCTACACGACGCTCTTCCGATCTNNNTTGGACCAGTGTCCCATGAGTAT
AAGAGCACACGTCTGAACTCTTTCCCTACACGACGCTCTTCCGATCTNNNATGTCATTCTCTCTCTGTCC
AGAGCACACGTCTGAACTCTTTCCCTACACGACGCTCTTCCGATCTNNNCGGTGTCCTTACCACAGAAT
GAGCACACGTCTGAACTCTTTCCCTACACGACGCTCTTCCGATCTNNNAGAAGGGCTCACCATTTACT
AAGAGCACACGTCTGAACTCTTTCCCTACACGACGCTCTTCCGATCTNNNTCTATTGAGATGGGGTAG
GAGCACACGTCTGAACTCTTTCCCTACACGACGCTCTTCCGATCTNNNAGCATTTCTCACTGACAGCAA
GCACACGTCTGAACTCTTTCCCTACACGACGCTCTTCCGATCTNNNACCTTTTCTGAGCTGTAGTCCTC
AGCACACGTCTGAACTCTTTCCCTACACGACGCTCTTCCGATCTNNNTGAGGTCTGTCGTTGACCAAA
AGCACACGTCTGAACTCTTTCCCTACACGACGCTCTTCCGATCTNNNCTACCACATACAAACGCCACTGT
AGAGCACACGTCTGAACTCTTTCCCTACACGACGCTCTTCCGATCTNNNAACCAGCCAGCTTGCTCTAT
GAGCACACGTCTGAACTCTTTCCCTACACGACGCTCTTCCGATCTNNNACATGCAAAAGAGTGCTAGG
GAGCACACGTCTGAACTCTTTCCCTACACGACGCTCTTCCGATCTNNNTCCGTCTTGACAACATTTCAA
GAGCACACGTCTGAACTCTTTCCCTACACGACGCTCTTCCGATCTNNNATCTGCAGTGTGAGGCAATG
AGCACACGTCTGAACTCTTTCCCTACACGACGCTCTTCCGATCTNNNGTAGCTGGGCAACGTCTAGTTT
AGCACACGTCTGAACTCTTTCCCTACACGACGCTCTTCCGATCTNNNCAGAGTGTGATGGTTCGTCTG
GAGCACACGTCTGAACTCTTTCCCTACACGACGCTCTTCCGATCTNNNGCATTTCTGGAGTAGGATGG
AGCACACGTCTGAACTCTTTCCCTACACGACGCTCTTCCGATCTNNNTACCATGGAGTTAGGCACAAG
GAGCACACGTCTGAACTCTTTCCCTACACGACGCTCTTCCGATCTNNNCCCTTTTGGTCTCTCTTGTGT
GAGCACACGTCTGAACTCTTTCCCTACACGACGCTCTTCCGATCTNNNAAGGCCAGTAACACTGAGAG
AGCACACGTCTGAACTCTTTCCCTACACGACGCTCTTCCGATCTNNNGTCCCACAGAGGTTTCTCTCTCT
GAGCACACGTCTGAACTCTTTCCCTACACGACGCTCTTCCGATCTNNNAGTCCTCCCTTCTCAGCCTCAT
GCACACGTCTGAACTCTTTCCCTACACGACGCTCTTCCGATCTNNNTGGCTGAGGACTTGGATAGAAC
GCACACGTCTGAACTCTTTCCCTACACGACGCTCTTCCGATCTNNNAAAAGAAGTGAGCCCTCTTGTG
AGCACACGTCTGAACTCTTTCCCTACACGACGCTCTTCCGATCTNNNACCATAAGTGTTGGTTTGGGT
AAGAGCACACGTCTGAACTCTTTCCCTACACGACGCTCTTCCGATCTNNNAAAGTGCATTGCTGGTAA
AGCACACGTCTGAACTCTTTCCCTACACGACGCTCTTCCGATCTNNNTGTGTGTAAGTCATTGCCAACA
AGCACACGTCTGAACTCTTTCCCTACACGACGCTCTTCCGATCTNNNCAGCGCCTGTAACTACATAAGC
GAGCACACGTCTGAACTCTTTCCCTACACGACGCTCTTCCGATCTNNNACCTTGACAATCTGCCTCCAA
GAGCACACGTCTGAACTCTTTCCCTACACGACGCTCTTCCGATCTNNNTTTGTAGGGAAAACCAGCAT
AGCACACGTCTGAACTCTTTCCCTACACGACGCTCTTCCGATCTNNNAACATGTTCGAAGCAGAAACC
AGAGCACACGTCTGAACTCTTTCCCTACACGACGCTCTTCCGATCTNNNCCCGTCTCCTTCCCTCTTATC
AGCACACGTCTGAACTCTTTCCCTACACGACGCTCTTCCGATCTNNNAGGTTTGCCAGTTTTCTTCCCA
GAGCACACGTCTGAACTCTTTCCCTACACGACGCTCTTCCGATCTNNNTGTCTGTGCAAAACACAAAAC
AGCACACGTCTGAACTCTTTCCCTACACGACGCTCTTCCGATCTNNNTTTGGGGTAAAAATGCCACAT
AGCACACGTCTGAACTCTTTCCCTACACGACGCTCTTCCGATCTNNNGGTCATCAGTTAGGCTTGCAG
GAGCACACGTCTGAACTCTTTCCCTACACGACGCTCTTCCGATCTNNNCAAACATGTCAGGAATTTGG
AGAGCACACGTCTGAACTCTTTCCCTACACGACGCTCTTCCGATCTNNNTCCGGAATAGACTGGACTA

GAGCACACGTCTGAACTCTTTCCCTACACGACGCTCTTCCGATCTNNNAAATTTCTAGGATGCTTGCGG
GAGCACACGTCTGAACTCTTTCCCTACACGACGCTCTTCCGATCTNNNGCACGAACAAGCAAAACCAGA
AGCACACGTCTGAACTCTTTCCCTACACGACGCTCTTCCGATCTNNNAGCCGTTGTATTCTGGAAACCT
AGCACACGTCTGAACTCTTTCCCTACACGACGCTCTTCCGATCTNNNTGGGAAAACACAGTGCAGTTCA
GAGCACACGTCTGAACTCTTTCCCTACACGACGCTCTTCCGATCTNNNTTACTTCCCATCACCTGTCCCAT
GAGCACACGTCTGAACTCTTTCCCTACACGACGCTCTTCCGATCTNNNTTTGTAGGGAAAACCAGCAT
AGCACACGTCTGAACTCTTTCCCTACACGACGCTCTTCCGATCTNNNCCACTAGGCCTCTTTTGAATGCT
GAGCACACGTCTGAACTCTTTCCCTACACGACGCTCTTCCGATCTNNNCACTGGCTTGCACTCACTATGA
AAGAGCACACGTCTGAACTCTTTCCCTACACGACGCTCTTCCGATCTNNNAGAGAGGGCGACAGAGTA
GCACACGTCTGAACTCTTTCCCTACACGACGCTCTTCCGATCTNNNCACCTCGGCTTTCACTGGAGACA
AGAGCACACGTCTGAACTCTTTCCCTACACGACGCTCTTCCGATCTNNNGGAGACCGAAGCTACTTTTT
AGCACACGTCTGAACTCTTTCCCTACACGACGCTCTTCCGATCTNNNTGTCCATCCTCTCTCTGTCTCTG
GAGCACACGTCTGAACTCTTTCCCTACACGACGCTCTTCCGATCTNNNTACTCAGGGCAGCTTACTTAG
AGAGCACACGTCTGAACTCTTTCCCTACACGACGCTCTTCCGATCTNNNGCTGCTTCTCTCTAAGCTTGA
AGCACACGTCTGAACTCTTTCCCTACACGACGCTCTTCCGATCTNNNCAGAATCCAGAGAGGAAGGGT
AGCACACGTCTGAACTCTTTCCCTACACGACGCTCTTCCGATCTNNNCACGGCCCAAGAGTGATGAATA
AGCACACGTCTGAACTCTTTCCCTACACGACGCTCTTCCGATCTNNNTCTGCAGTTTGAATATGGTTTCC
AAGAGCACACGTCTGAACTCTTTCCCTACACGACGCTCTTCCGATCTNNNAAAATCCCAATGTCTCTGCA
AGCACACGTCTGAACTCTTTCCCTACACGACGCTCTTCCGATCTNNNGGATGGGGTGAAAGGAAGA
GAGCACACGTCTGAACTCTTTCCCTACACGACGCTCTTCCGATCTNNNGAGAGTGATTCGTGCCAAGA
AGCACACGTCTGAACTCTTTCCCTACACGACGCTCTTCCGATCTNNNTCAAACGATACCTTGAACACTCA
AGCACACGTCTGAACTCTTTCCCTACACGACGCTCTTCCGATCTNNNGGCATACTCTGGGATAGTAGG
GAGCACACGTCTGAACTCTTTCCCTACACGACGCTCTTCCGATCTNNNAAACCCCAAAACAAACCAACC
GAGCACACGTCTGAACTCTTTCCCTACACGACGCTCTTCCGATCTNNNTCTGGGTTACCAGGTAGAGA
AGCACACGTCTGAACTCTTTCCCTACACGACGCTCTTCCGATCTNNNACCATTATTGTTTTGTGTTCCCCT
AGCACACGTCTGAACTCTTTCCCTACACGACGCTCTTCCGATCTNNNTCATTGGTTCATCAGTGGGAAA
AGCACACGTCTGAACTCTTTCCCTACACGACGCTCTTCCGATCTNNNCAGACTTTGAGCCCTGCTAACAT
AGCACACGTCTGAACTCTTTCCCTACACGACGCTCTTCCGATCTNNNACTGTTTGTTGGTCTGAATGT
AGCACACGTCTGAACTCTTTCCCTACACGACGCTCTTCCGATCTNNNCACATTGCCACAAAGCCTGAAAA
GAGCACACGTCTGAACTCTTTCCCTACACGACGCTCTTCCGATCTNNNACCAAAGCACTGTACACTTCAA
GAGCACACGTCTGAACTCTTTCCCTACACGACGCTCTTCCGATCTNNNCCTATCCCACTTCAGGGAAAG
AAGAGCACACGTCTGAACTCTTTCCCTACACGACGCTCTTCCGATCTNNNTTGCATGGTCTCTATGAGT
GAGCACACGTCTGAACTCTTTCCCTACACGACGCTCTTCCGATCTNNNAAATTGCCCTCTCTCTCCCTCT
AGCACACGTCTGAACTCTTTCCCTACACGACGCTCTTCCGATCTNNNCCTCTTTGTCTTTGTTTTTGGCA
GAGCACACGTCTGAACTCTTTCCCTACACGACGCTCTTCCGATCTNNNGAATGGTTGTCATCAGCCCTA
AGCACACGTCTGAACTCTTTCCCTACACGACGCTCTTCCGATCTNNNTCAGAGTTGCTGGGCTATTTCA
AGAGCACACGTCTGAACTCTTTCCCTACACGACGCTCTTCCGATCTNNNTGGCATAGTTTTGGCTCTAG
AGCACACGTCTGAACTCTTTCCCTACACGACGCTCTTCCGATCTNNNAGTCCCTTCCAAGGAATTAGCCA
AGAGCACACGTCTGAACTCTTTCCCTACACGACGCTCTTCCGATCTNNNAAACCAAATGCCCAGTTATG
AGAGCACACGTCTGAACTCTTTCCCTACACGACGCTCTTCCGATCTNNNGTGGCTCCAATGACTTTGCT
AGCACACGTCTGAACTCTTTCCCTACACGACGCTCTTCCGATCTNNNACTTCCCAGTATGAATGCACCAA
GAGCACACGTCTGAACTCTTTCCCTACACGACGCTCTTCCGATCTNNNACCCCATAAACCACACTCTCTCA
AGCACACGTCTGAACTCTTTCCCTACACGACGCTCTTCCGATCTNNNGTTTGCAAAGAGGTGGAGCAAA

GAGCACACGTCTGAACTCTTTCCCTACACGACGCTCTTCCGATCTNNNTTAGACAAGCTGCAGGAAGA
AGAGCACACGTCTGAACTCTTTCCCTACACGACGCTCTTCCGATCTNNNTCTCAGGTTCTCTTTCCCTCT
AGAGCACACGTCTGAACTCTTTCCCTACACGACGCTCTTCCGATCTNNNCCATGCCTCCCAAGGGATAT
AGCACACGTCTGAACTCTTTCCCTACACGACGCTCTTCCGATCTNNNCTTCTCTTGCTCATCCCATCCTGC
AGAGCACACGTCTGAACTCTTTCCCTACACGACGCTCTTCCGATCTNNNAAGGGGTGAAAACTGCTTA
GCACACGTCTGAACTCTTTCCCTACACGACGCTCTTCCGATCTNNNAGCCACAACCATGTGATTTCGAT
AGCACACGTCTGAACTCTTTCCCTACACGACGCTCTTCCGATCTNNNACAGACAGCAGATGGTAGGAC
AGCACACGTCTGAACTCTTTCCCTACACGACGCTCTTCCGATCTNNNAAACCAACAACCCATAATCCCCC
GAGCACACGTCTGAACTCTTTCCCTACACGACGCTCTTCCGATCTNNNTGTTTCCAAAGTCACAAAATGA
AGAGCACACGTCTGAACTCTTTCCCTACACGACGCTCTTCCGATCTNNNTGTAAGGAAGATCCAGGG
GCACACGTCTGAACTCTTTCCCTACACGACGCTCTTCCGATCTNNNTTTGAATGTACACGGTGTCAGGC
AGAGCACACGTCTGAACTCTTTCCCTACACGACGCTCTTCCGATCTNNNGCTGATAGAGGATGAGCTC
AGCACACGTCTGAACTCTTTCCCTACACGACGCTCTTCCGATCTNNNGAGCATTGTAGGTATTGCCCCA
GAGCACACGTCTGAACTCTTTCCCTACACGACGCTCTTCCGATCTNNNTCAGGGCTTTATGTCTCAAATC
GAGCACACGTCTGAACTCTTTCCCTACACGACGCTCTTCCGATCTNNNAAAATGCAGAGTAATGGTGG
AGAGCACACGTCTGAACTCTTTCCCTACACGACGCTCTTCCGATCTNNNTAGCAGGGTGAGACCTTGTC
GAGCACACGTCTGAACTCTTTCCCTACACGACGCTCTTCCGATCTNNNCTTTTCTAGAACCTGCCCCCA
AGCACACGTCTGAACTCTTTCCCTACACGACGCTCTTCCGATCTNNNGTACTGAGCGCAAATACCAAAC
AGCACACGTCTGAACTCTTTCCCTACACGACGCTCTTCCGATCTNNNTGATTTGGCCCTTGTTTTTCTTCA
AGCACACGTCTGAACTCTTTCCCTACACGACGCTCTTCCGATCTNNNAACAAAACCCCGTCTCAAAATG
AAGAGCACACGTCTGAACTCTTTCCCTACACGACGCTCTTCCGATCTNNNATATCTCAGTTTCTGCGGC
AAGAGCACACGTCTGAACTCTTTCCCTACACGACGCTCTTCCGATCTNNNATGCATCCACTAAATGACCC
GAGCACACGTCTGAACTCTTTCCCTACACGACGCTCTTCCGATCTNNNACCTAGACTGACTTCAGGGAA
GAGCACACGTCTGAACTCTTTCCCTACACGACGCTCTTCCGATCTNNNGCAACAATGGAGAGGGTTTC
AGCACACGTCTGAACTCTTTCCCTACACGACGCTCTTCCGATCTNNNGTACATGTCTCTAGGCAGAGGC
AGCACACGTCTGAACTCTTTCCCTACACGACGCTCTTCCGATCTNNNAGGGTCAGCAAAGTCAGGAAA
GAGCACACGTCTGAACTCTTTCCCTACACGACGCTCTTCCGATCTNNNTGTACCAGTCTCTCTCTGCAAC
AGAGCACACGTCTGAACTCTTTCCCTACACGACGCTCTTCCGATCTNNNGACATAATGGCTGTAGGGC
GAGCACACGTCTGAACTCTTTCCCTACACGACGCTCTTCCGATCTNNNCCAAATCTGGGCTTTACTTGTG
GAGCACACGTCTGAACTCTTTCCCTACACGACGCTCTTCCGATCTNNNTGCACAACAGTTCATTGTTTCTT
GAGCACACGTCTGAACTCTTTCCCTACACGACGCTCTTCCGATCTNNNTAGGGTAGAAGAGGTGGATT
GAGCACACGTCTGAACTCTTTCCCTACACGACGCTCTTCCGATCTNNNGTCATGTTGGTGGGTCAGAA
AAGAGCACACGTCTGAACTCTTTCCCTACACGACGCTCTTCCGATCTNNNTGAACTGTACAGGAAGGC
GAGCACACGTCTGAACTCTTTCCCTACACGACGCTCTTCCGATCTNNNCCCGCCTGTATTCCATAAACAG
AAGAGCACACGTCTGAACTCTTTCCCTACACGACGCTCTTCCGATCTNNNCCAGGTTGGCGTGTATGAC
GAGCACACGTCTGAACTCTTTCCCTACACGACGCTCTTCCGATCTNNNGGTCAATTTCTTGGGGGATTC
AAGAGCACACGTCTGAACTCTTTCCCTACACGACGCTCTTCCGATCTNNNCCTTGGGTTAGGCATCTCA
GAGCACACGTCTGAACTCTTTCCCTACACGACGCTCTTCCGATCTNNNCATTACTGCGATGAGATCCTG
GAGCACACGTCTGAACTCTTTCCCTACACGACGCTCTTCCGATCTNNNCCATCATTAGCCACAATGCCA
AGAGCACACGTCTGAACTCTTTCCCTACACGACGCTCTTCCGATCTNNNAGGAATTCAATCCCACTCCCA
GAGCACACGTCTGAACTCTTTCCCTACACGACGCTCTTCCGATCTNNNAGGGCATAAGGATGGATGGA
GCACACGTCTGAACTCTTTCCCTACACGACGCTCTTCCGATCTNNNATATCTCAGTTTCTGCGGGTCCA
AGAGCACACGTCTGAACTCTTTCCCTACACGACGCTCTTCCGATCTNNNAAACTGCTCAGGGTCTCATTT

GCACACGTCTGAACTCTTTCCCTACACGACGCTCTTCCGATCTNNNGGATGGTTGGGTTTTTCCGTACT

AGCACACGTCTGAACTCTTTCCCTACACGACGCTCTTCCGATCTNNNTCTGTCTGGTTTTATCCTTCAC

AGCACACGTCTGAACTCTTTCCCTACACGACGCTCTTCCGATCTNNNCTCTTGTTCACCACCCGTTTTTC

GCACACGTCTGAACTCTTTCCCTACACGACGCTCTTCCGATCTNNNCGTGAGGCCTGAATGCAATATCT

AGCACACGTCTGAACTCTTTCCCTACACGACGCTCTTCCGATCTNNNGGTAGTTGTTGTGCTGTGGGT

AGCACACGTCTGAACTCTTTCCCTACACGACGCTCTTCCGATCTNNNTGACACAGTACTTTCTCTCACT

AGCACACGTCTGAACTCTTTCCCTACACGACGCTCTTCCGATCTNNNTCCTTTCAATAGTTTTTGACCCT

GAGCACACGTCTGAACTCTTTCCCTACACGACGCTCTTCCGATCTNNNTGTACATTGACATTGGGGTT

AGCACACGTCTGAACTCTTTCCCTACACGACGCTCTTCCGATCTNNNTACTCAGGGCAGCTTACTTAG

AGCACACGTCTGAACTCTTTCCCTACACGACGCTCTTCCGATCTNNNCACCTGAAAAATCCCTTTTGCC

GCACACGTCTGAACTCTTTCCCTACACGACGCTCTTCCGATCTNNNCCTATTCTCCAGAGTGGTGGGA

GCACACGTCTGAACTCTTTCCCTACACGACGCTCTTCCGATCTNNNCTCAGTATGGGTGTATGAGCA

GCACACGTCTGAACTCTTTCCCTACACGACGCTCTTCCGATCTNNNATTTACACACATGCACGCTCTCT

CACACGTCTGAACTCTTTCCCTACACGACGCTCTTCCGATCTNNNATATAGCCCTCTGTGTCTGGTCCC

GAGCACACGTCTGAACTCTTTCCCTACACGACGCTCTTCCGATCTNNNCCGGCCTTGCCAGTATTTAT

AGAGCACACGTCTGAACTCTTTCCCTACACGACGCTCTTCCGATCTNNNAGTCCAGTGAGGCTGTGAT

AGCACACGTCTGAACTCTTTCCCTACACGACGCTCTTCCGATCTNNNAGTTTCCTCTCTCTTTCCCTCCG

GAGCACACGTCTGAACTCTTTCCCTACACGACGCTCTTCCGATCTNNNACAACACCTAATCCTCACATG

AAGAGCACACGTCTGAACTCTTTCCCTACACGACGCTCTTCCGATCTNNNAAGATTGAGGCTGCAGTG

CACACGTCTGAACTCTTTCCCTACACGACGCTCTTCCGATCTNNNTTCCACCTAGCAGATTGGGTACA

GAGCACACGTCTGAACTCTTTCCCTACACGACGCTCTTCCGATCTNNNCTATGCATGGAGCAGAACA

AGAGCACACGTCTGAACTCTTTCCCTACACGACGCTCTTCCGATCTNNNTTTTGATCTGCAGTTGCGT

GCACACGTCTGAACTCTTTCCCTACACGACGCTCTTCCGATCTNNNCTTCCTGGTCACTTTCTTGCTGT

GAGCACACGTCTGAACTCTTTCCCTACACGACGCTCTTCCGATCTNNNTGACATGGGCCCCTTTGATAT

AGCACACGTCTGAACTCTTTCCCTACACGACGCTCTTCCGATCTNNNTTGCTTTTCCCTTTCTTTTGCCCC

GCACACGTCTGAACTCTTTCCCTACACGACGCTCTTCCGATCTNNNGAAAGGCCACTATATCACCAGG

AGCACACGTCTGAACTCTTTCCCTACACGACGCTCTTCCGATCTNNNAAATTTCTTGAGGCCAGAGTT

GAGCACACGTCTGAACTCTTTCCCTACACGACGCTCTTCCGATCTNNNAGATGATAGCCCAAGGACA

GCACACGTCTGAACTCTTTCCCTACACGACGCTCTTCCGATCTNNNATCAAAGAAAGCTTTGGCCCTCT

GAGCACACGTCTGAACTCTTTCCCTACACGACGCTCTTCCGATCTNNNACAGTTTGGGATGGCATTTA

AGCACACGTCTGAACTCTTTCCCTACACGACGCTCTTCCGATCTNNNCAACAGCATCCTCACATAGGCTT

GAGCACACGTCTGAACTCTTTCCCTACACGACGCTCTTCCGATCTNNNTAAGTGGAAAAACCCCAGCT

AGCACACGTCTGAACTCTTTCCCTACACGACGCTCTTCCGATCTNNNTAACCATCATGCAAGGGCAAA

GAGCACACGTCTGAACTCTTTCCCTACACGACGCTCTTCCGATCTNNNCTTGGGTCTTCTTGTACCTGT

AGCACACGTCTGAACTCTTTCCCTACACGACGCTCTTCCGATCTNNNTTTGTGTTCCACACAGCTTGGT

AGCACACGTCTGAACTCTTTCCCTACACGACGCTCTTCCGATCTNNNGACAGTGCCTTTTCCCCTTCAG

GAGCACACGTCTGAACTCTTTCCCTACACGACGCTCTTCCGATCTNNNTGTTCAGTCTCTGTCTGTCTCT

AGCACACGTCTGAACTCTTTCCCTACACGACGCTCTTCCGATCTNNNATACAGCAGCATCTCTCTACGG

GAGCACACGTCTGAACTCTTTCCCTACACGACGCTCTTCCGATCTNNNATGTGGGTTTATGGATTGCT

GCACACGTCTGAACTCTTTCCCTACACGACGCTCTTCCGATCTNNNTAAGTTGGCCACGTTTAGACAG

GCACACGTCTGAACTCTTTCCCTACACGACGCTCTTCCGATCTNNNTCCCTGGGCAATTGTACTCATT

GAGCACACGTCTGAACTCTTTCCCTACACGACGCTCTTCCGATCTNNNCATTATTTGGCAGGGGGA

GAGCACACGTCTGAACTCTTTCCCTACACGACGCTCTTCCGATCTNNNTGGTTTCAGGTGGTTTTGGC

AGCACACGTCTGAACTCTTTCCCTACACGACGCTCTTCCGATCTNNNAATGGGTGGCTAGGAGAATA
GCACACGTCTGAACTCTTTCCCTACACGACGCTCTTCCGATCTNNNGCCCGAGAGCTGAAGTTTCATTA
AGAGCACACGTCTGAACTCTTTCCCTACACGACGCTCTTCCGATCTNNNAAGCATTTTATTGAACGGG
AGCACACGTCTGAACTCTTTCCCTACACGACGCTCTTCCGATCTNNNTTGCATTCTGAAGACAAGCCAG
AGCACACGTCTGAACTCTTTCCCTACACGACGCTCTTCCGATCTNNNACTTTGCTAAGCTAATAACA
GAGCACACGTCTGAACTCTTTCCCTACACGACGCTCTTCCGATCTNNNTAAAGGAAAGGTTCCAGCCC
AGCACACGTCTGAACTCTTTCCCTACACGACGCTCTTCCGATCTNNNTCTGTGTACATGGTGATGAAC
AGCACACGTCTGAACTCTTTCCCTACACGACGCTCTTCCGATCTNNNGCAGGCCTTCTTACTGAATCCT
CACACGTCTGAACTCTTTCCCTACACGACGCTCTTCCGATCTNNNTTGCCTTCACCAACACACATACAC
GAGCACACGTCTGAACTCTTTCCCTACACGACGCTCTTCCGATCTNNNTAATGTGGCAGAAAGTGAAG
AGAGCACACGTCTGAACTCTTTCCCTACACGACGCTCTTCCGATCTNNNTGTATATATGGAACACCAT
GAGCACACGTCTGAACTCTTTCCCTACACGACGCTCTTCCGATCTNNNTGACAAGCCACACTAAGCTC
AGAGCACACGTCTGAACTCTTTCCCTACACGACGCTCTTCCGATCTNNNCTCCATGTGCCAGATTCTGT
AGCACACGTCTGAACTCTTTCCCTACACGACGCTCTTCCGATCTNNNGCTGCTGAATGCCTTCCAACT
AGCACACGTCTGAACTCTTTCCCTACACGACGCTCTTCCGATCTNNNTCCTCTTCACATCAATAGGCTGA
GAGCACACGTCTGAACTCTTTCCCTACACGACGCTCTTCCGATCTNNNGTGGAGGGAAAGCAAATGC
AGCACACGTCTGAACTCTTTCCCTACACGACGCTCTTCCGATCTNNNTGTGTCCTGCATCTTACATGTG
GCACACGTCTGAACTCTTTCCCTACACGACGCTCTTCCGATCTNNNTACAGTTGACCTGGGACCTTTGT
AGCACACGTCTGAACTCTTTCCCTACACGACGCTCTTCCGATCTNNNGGGTCAGAGAAGCCATATTTCC
GAGCACACGTCTGAACTCTTTCCCTACACGACGCTCTTCCGATCTNNNCAGAATTTGCACTCCAGGCTA
AGCACACGTCTGAACTCTTTCCCTACACGACGCTCTTCCGATCTNNNCCAAAGCCTGACAGATAGACA
GCACACGTCTGAACTCTTTCCCTACACGACGCTCTTCCGATCTNNNTCTTGTGGTTCTGGATTACTGCC
AGCACACGTCTGAACTCTTTCCCTACACGACGCTCTTCCGATCTNNNAACAAGGCCTGCAAATAGAA
GAGCACACGTCTGAACTCTTTCCCTACACGACGCTCTTCCGATCTNNNAGTAGCTGGGATTCTGGCAT
CACACGTCTGAACTCTTTCCCTACACGACGCTCTTCCGATCTNNNCTTTGCAAGAACCTCTGTCTGCC
AGCACACGTCTGAACTCTTTCCCTACACGACGCTCTTCCGATCTNNNTTTGCGCATGTGTCCACTTTTA
GAGCACACGTCTGAACTCTTTCCCTACACGACGCTCTTCCGATCTNNNGGGAGAAATCATGGGAACC
AGCACACGTCTGAACTCTTTCCCTACACGACGCTCTTCCGATCTNNNGGGAGAAATCATGGGAACCC
GAGCACACGTCTGAACTCTTTCCCTACACGACGCTCTTCCGATCTNNNTCTTGAGCATGCACACATTTT
AGCACACGTCTGAACTCTTTCCCTACACGACGCTCTTCCGATCTNNNTTCTGTATCTGTCATCGGCAC
GCACACGTCTGAACTCTTTCCCTACACGACGCTCTTCCGATCTNNNACTCTGCTGTATGTATCTGTGGT
AGCACACGTCTGAACTCTTTCCCTACACGACGCTCTTCCGATCTNNNCTCTTTAGGTCAGCGGTGATTC
AGCACACGTCTGAACTCTTTCCCTACACGACGCTCTTCCGATCTNNNTGGAGGAGAATTTTCCAAAAA
AGCACACGTCTGAACTCTTTCCCTACACGACGCTCTTCCGATCTNNNCAGCTGTGAAGTCTCTCTCACT
AGCACACGTCTGAACTCTTTCCCTACACGACGCTCTTCCGATCTNNNTTGTTGGGAAATGGTAGTGAA
AGAGCACACGTCTGAACTCTTTCCCTACACGACGCTCTTCCGATCTNNNATGGGGAGAGTTGGTTATG
AGCACACGTCTGAACTCTTTCCCTACACGACGCTCTTCCGATCTNNNTAGGGCATGTTCTTTTGCTGAG
AGCACACGTCTGAACTCTTTCCCTACACGACGCTCTTCCGATCTNNNCAGCTCTTAGAAAAACACAGC
GCACACGTCTGAACTCTTTCCCTACACGACGCTCTTCCGATCTNNNCAGCCCTTGAAGCTATCTGGTTT
AGCACACGTCTGAACTCTTTCCCTACACGACGCTCTTCCGATCTNNNAATCTAACTGTCCATCAAGGG
AGCACACGTCTGAACTCTTTCCCTACACGACGCTCTTCCGATCTNNNAAAAATGCACTTGTGCCCTTA
AGCACACGTCTGAACTCTTTCCCTACACGACGCTCTTCCGATCTNNNTGGGAAAACACAGTGCAGTT
GAGCACACGTCTGAACTCTTTCCCTACACGACGCTCTTCCGATCTNNNTCTAAAATGAGCTGGCCAGG

AGCACACGTCTGAACTCTTTCCCTACACGACGCTCTTCCGATCTNNNGGCATCTGATCTGGGAAACAG
AGCACACGTCTGAACTCTTTCCCTACACGACGCTCTTCCGATCTNNNGAGGATGCCTAGATACGTCCA
AGCACACGTCTGAACTCTTTCCCTACACGACGCTCTTCCGATCTNNNCATGGAGGGCATCTCTTTCTCT
AGCACACGTCTGAACTCTTTCCCTACACGACGCTCTTCCGATCTNNNGTTGTAAGCTGGTCCCCCTAAT
AGCACACGTCTGAACTCTTTCCCTACACGACGCTCTTCCGATCTNNNCACTGGAGATGCATGAAGTTG
GAGCACACGTCTGAACTCTTTCCCTACACGACGCTCTTCCGATCTNNNGGGGAATTCCAGCTGATAG
GAGCACACGTCTGAACTCTTTCCCTACACGACGCTCTTCCGATCTNNNGGCAACATTGGCAAGATGAA
AGCACACGTCTGAACTCTTTCCCTACACGACGCTCTTCCGATCTNNNTTGCACACAGCCAGGATTATGT
AGCACACGTCTGAACTCTTTCCCTACACGACGCTCTTCCGATCTNNNTACTGAACTGTGTCCTCCAACC
GAGCACACGTCTGAACTCTTTCCCTACACGACGCTCTTCCGATCTNNNAGCTCTGTGCTGGAAATTCT
GAGCACACGTCTGAACTCTTTCCCTACACGACGCTCTTCCGATCTNNNCTATTCCTGGGGAAAACTCA
GAGCACACGTCTGAACTCTTTCCCTACACGACGCTCTTCCGATCTNNNCCACAGTGAATTTTGGTGGT
AGCACACGTCTGAACTCTTTCCCTACACGACGCTCTTCCGATCTNNNTTGTTCCAATGGTTCCAGCAAC
GAGCACACGTCTGAACTCTTTCCCTACACGACGCTCTTCCGATCTNNNTGTGCTTGTTTTTGCAGATGT
AGCACACGTCTGAACTCTTTCCCTACACGACGCTCTTCCGATCTNNNGAACACCCTCGCTAGTTTCAAG
AGCACACGTCTGAACTCTTTCCCTACACGACGCTCTTCCGATCTNNNTCGTTTATGGGACTGGACAGC
GAGCACACGTCTGAACTCTTTCCCTACACGACGCTCTTCCGATCTNNNATACACACTCTCTTGAGCCAA
GAGCACACGTCTGAACTCTTTCCCTACACGACGCTCTTCCGATCTNNNAGTCAGGGTTTCTCAGACAG
AGCACACGTCTGAACTCTTTCCCTACACGACGCTCTTCCGATCTNNNGTGAGGCTTTGGGAGGAAAA
GCACACGTCTGAACTCTTTCCCTACACGACGCTCTTCCGATCTNNNAGTCTTCACAAGCGCTGATCATT
GCACACGTCTGAACTCTTTCCCTACACGACGCTCTTCCGATCTNNNCTCGCAACGCTAAACAGTGAAAA
AGCACACGTCTGAACTCTTTCCCTACACGACGCTCTTCCGATCTNNNGCTGACATTGAGACAGACAG
AGAGCACACGTCTGAACTCTTTCCCTACACGACGCTCTTCCGATCTNNNTGCATGTTAGCACCCAAAA
AGCACACGTCTGAACTCTTTCCCTACACGACGCTCTTCCGATCTNNNCTGACGTGTCTCATGGAAAGG
GAGCACACGTCTGAACTCTTTCCCTACACGACGCTCTTCCGATCTNNNAGGGGAAGAAGGCATATGG
GAGCACACGTCTGAACTCTTTCCCTACACGACGCTCTTCCGATCTNNNTTCTTGCATTGTCTCCTTGCTG
AGCACACGTCTGAACTCTTTCCCTACACGACGCTCTTCCGATCTNNNAGCTCTGTGCTGGAAATTCTG
GCACACGTCTGAACTCTTTCCCTACACGACGCTCTTCCGATCTNNNCATGCCGCTGGATAAGATTCCA
GCACACGTCTGAACTCTTTCCCTACACGACGCTCTTCCGATCTNNNGCTCTATGAGGGGGAGTGTTTT
GAGCACACGTCTGAACTCTTTCCCTACACGACGCTCTTCCGATCTNNNATGGCATATGAGGTGTGTGT
GCACACGTCTGAACTCTTTCCCTACACGACGCTCTTCCGATCTNNNAGTGAAATGTGAAGCAGCAAGG
AGCACACGTCTGAACTCTTTCCCTACACGACGCTCTTCCGATCTNNNTTGCATGGTCTCTATGAGTGT
AGCACACGTCTGAACTCTTTCCCTACACGACGCTCTTCCGATCTNNNGCTTGTGTCACCCTACTCTCAA
GCACACGTCTGAACTCTTTCCCTACACGACGCTCTTCCGATCTNNNGGCTCCAGCATCTTTCCTTCTTT
AGCACACGTCTGAACTCTTTCCCTACACGACGCTCTTCCGATCTNNNTTTGAGGCTGGAGTAAGCAG
AGAGCACACGTCTGAACTCTTTCCCTACACGACGCTCTTCCGATCTNNNATGCATGCTAAGATCATGC
AGCACACGTCTGAACTCTTTCCCTACACGACGCTCTTCCGATCTNNNTTGCAAGTTTATTGCACGCAT
AGCACACGTCTGAACTCTTTCCCTACACGACGCTCTTCCGATCTNNNATCATCCTTACCCTTGCTGAGA
AAGAGCACACGTCTGAACTCTTTCCCTACACGACGCTCTTCCGATCTNNNGAGGCGTTTTGCCCATTA
GAGCACACGTCTGAACTCTTTCCCTACACGACGCTCTTCCGATCTNNNATTCCCAGGCTTTTTCTCTTGG
GAGCACACGTCTGAACTCTTTCCCTACACGACGCTCTTCCGATCTNNNCCCAAGGACATTCATCACTGG
AGCACACGTCTGAACTCTTTCCCTACACGACGCTCTTCCGATCTNNNGCCCTTTAGAGGCAAACACAT
GAGCACACGTCTGAACTCTTTCCCTACACGACGCTCTTCCGATCTNNNTATCTGGGGAGGCTGATTA

GAGCACACGTCTGAACTCTTTCCCTACACGACGCTCTTCCGATCTNNNCACCACTGTACAATGAGACCCT
GAGCACACGTCTGAACTCTTTCCCTACACGACGCTCTTCCGATCTNNNTAGTTTACCCCTAACTCCCCTG
AGCACACGTCTGAACTCTTTCCCTACACGACGCTCTTCCGATCTNNNTCCAGCATTGGTCATCAAGAACT
GAGCACACGTCTGAACTCTTTCCCTACACGACGCTCTTCCGATCTNNNCATGTGGCTCACCGTTTTCTTA
AGAGCACACGTCTGAACTCTTTCCCTACACGACGCTCTTCCGATCTNNNAGACACCTGCCTATCTCACCA
AGAGCACACGTCTGAACTCTTTCCCTACACGACGCTCTTCCGATCTNNNCTAAGTCAAGGGAGTGATG
AGAGCACACGTCTGAACTCTTTCCCTACACGACGCTCTTCCGATCTNNNGTCCAGGGGTCCATATACAC
AGAGCACACGTCTGAACTCTTTCCCTACACGACGCTCTTCCGATCTNNNTCTTTGCATGTTCACCACTGA
GAGCACACGTCTGAACTCTTTCCCTACACGACGCTCTTCCGATCTNNNTGAAGACTGTGTGTGTGTGTC
GAGCACACGTCTGAACTCTTTCCCTACACGACGCTCTTCCGATCTNNNCTAGGTTGGCCTCTCTCTCTCT
AGCACACGTCTGAACTCTTTCCCTACACGACGCTCTTCCGATCTNNNGAAAGTAAGTCCTCACCTCGTC
AGAGCACACGTCTGAACTCTTTCCCTACACGACGCTCTTCCGATCTNNNCACAACACATTTCTTCTTGCC
AGCACACGTCTGAACTCTTTCCCTACACGACGCTCTTCCGATCTNNNATGAGAAAGGGGTGAGGATGT
AAGAGCACACGTCTGAACTCTTTCCCTACACGACGCTCTTCCGATCTNNNGCAGATCATGAGACTGCTA
GAGCACACGTCTGAACTCTTTCCCTACACGACGCTCTTCCGATCTNNNATGTACTTACAAGCTCCATGT
GAGCACACGTCTGAACTCTTTCCCTACACGACGCTCTTCCGATCTNNNATGGAGAATGGGAAATCGT
GAGCACACGTCTGAACTCTTTCCCTACACGACGCTCTTCCGATCTNNNCCCTTCTCTGTCTGCATACTTCC
AGCACACGTCTGAACTCTTTCCCTACACGACGCTCTTCCGATCTNNNTACCTGTGTGCTACTCATTGTC
AGCACACGTCTGAACTCTTTCCCTACACGACGCTCTTCCGATCTNNNTCCTCACTGCCATTCCTCTATAT
AGCACACGTCTGAACTCTTTCCCTACACGACGCTCTTCCGATCTNNNAGGGGAAAGAGGGCAAGATA
GAAGAGCACACGTCTGAACTCTTTCCCTACACGACGCTCTTCCGATCTNNNACCATATTGTGCAGTACA
GCACACGTCTGAACTCTTTCCCTACACGACGCTCTTCCGATCTNNNCTTGGGCTGAACTATCCTCAGTG
AGCACACGTCTGAACTCTTTCCCTACACGACGCTCTTCCGATCTNNNAGAAAACAGTGAAGCCATGGT
ACACGTCTGAACTCTTTCCCTACACGACGCTCTTCCGATCTNNNAGAAAACAGTGAAGCCATGGTAGA
AGAGCACACGTCTGAACTCTTTCCCTACACGACGCTCTTCCGATCTNNNGTACAGCTTTACTGAAGCACC
GAGCACACGTCTGAACTCTTTCCCTACACGACGCTCTTCCGATCTNNNCAAACAAGCCAACAAACCCAA
GAAGAGCACACGTCTGAACTCTTTCCCTACACGACGCTCTTCCGATCTNNNAATAAGAGGTAGCCCCT
AGCACACGTCTGAACTCTTTCCCTACACGACGCTCTTCCGATCTNNNTTGTGCCCATTGTTTCTTGTCCT
AGCACACGTCTGAACTCTTTCCCTACACGACGCTCTTCCGATCTNNNTGATGTAATATTTTACCCATCCC
AGCACACGTCTGAACTCTTTCCCTACACGACGCTCTTCCGATCTNNNATATATACAGGGTGCGTGCGT
GAGCACACGTCTGAACTCTTTCCCTACACGACGCTCTTCCGATCTNNNAGGAACTCTGGACGAAGACT
AGAGCACACGTCTGAACTCTTTCCCTACACGACGCTCTTCCGATCTNNNGGTCAGCAAAGGAGAACCT
AGCACACGTCTGAACTCTTTCCCTACACGACGCTCTTCCGATCTNNNTGTGAAGAGTAGCACTTCTGT
ACACGTCTGAACTCTTTCCCTACACGACGCTCTTCCGATCTNNNATACACACTCTCTTGAGCCAAATCAT
GAGCACACGTCTGAACTCTTTCCCTACACGACGCTCTTCCGATCTNNNCCAGACTGCTTGTAACAGTGC
GAAGAGCACACGTCTGAACTCTTTCCCTACACGACGCTCTTCCGATCTNNNTAGTCAAGCCTCATTTTT
GAGCACACGTCTGAACTCTTTCCCTACACGACGCTCTTCCGATCTNNNTCCTCACTCCTTCCTCTCTCTCT
AGCACACGTCTGAACTCTTTCCCTACACGACGCTCTTCCGATCTNNNCCCCAAGTTTTCAACAGACATG
GCACACGTCTGAACTCTTTCCCTACACGACGCTCTTCCGATCTNNNCTCTCCATCCCATACAGCCAAACCA
CACACGTCTGAACTCTTTCCCTACACGACGCTCTTCCGATCTNNNAAGAAAACTCACCCTGTTGTGCTCA
GAGCACACGTCTGAACTCTTTCCCTACACGACGCTCTTCCGATCTNNNATAGCAACAACCAAAGTCCCCT
AGCACACGTCTGAACTCTTTCCCTACACGACGCTCTTCCGATCTNNNTGCAGCAATAACTGGAAATAG
AGCACACGTCTGAACTCTTTCCCTACACGACGCTCTTCCGATCTNNNAGCAAAAATGCAAGCCCTGAAT

GCACACGTCTGAACTCTTTCCCTACACGACGCTCTTCCGATCTNNNCAAACTTTAGCCTGTTTGGGCA
GAGCACACGTCTGAACTCTTTCCCTACACGACGCTCTTCCGATCTNNNCATGGAGGCCATACTGAAGT
AGAGCACACGTCTGAACTCTTTCCCTACACGACGCTCTTCCGATCTNNNTCCAAACACACAAAGTGGA
GAGCACACGTCTGAACTCTTTCCCTACACGACGCTCTTCCGATCTNNNAAACACTAGGGGAATTCAG
AGCACACGTCTGAACTCTTTCCCTACACGACGCTCTTCCGATCTNNNTCAGCGGATTAGAGTGGGAT
AGCACACGTCTGAACTCTTTCCCTACACGACGCTCTTCCGATCTNNNAGACCATGTTCTCAACAGGACA
CACACGTCTGAACTCTTTCCCTACACGACGCTCTTCCGATCTNNNAGCTCGAACTGCCTGGATAAAAG
GAGCACACGTCTGAACTCTTTCCCTACACGACGCTCTTCCGATCTNNNATTATCATACCACCCAGAGGC
AGCACACGTCTGAACTCTTTCCCTACACGACGCTCTTCCGATCTNNNTGAACCCAGATTGTAGCACAG
GAGCACACGTCTGAACTCTTTCCCTACACGACGCTCTTCCGATCTNNNCACTGCATTAGGGTTCTCCAG
AAGAGCACACGTCTGAACTCTTTCCCTACACGACGCTCTTCCGATCTNNNGTGGAACACACTTCAAGC
AGCACACGTCTGAACTCTTTCCCTACACGACGCTCTTCCGATCTNNNAAACTCACACACACTCTCTCCCT
AGAGCACACGTCTGAACTCTTTCCCTACACGACGCTCTTCCGATCTNNNCAAACTAGCTCAGCTTGACT
GAGCACACGTCTGAACTCTTTCCCTACACGACGCTCTTCCGATCTNNNATGGGGTGGGTTGATCTTA
GAGCACACGTCTGAACTCTTTCCCTACACGACGCTCTTCCGATCTNNNGTTCTGTAATTCGTGCGTGTG
AGCACACGTCTGAACTCTTTCCCTACACGACGCTCTTCCGATCTNNNGCACTCTTTACCTAGATTCACCT
GAGCACACGTCTGAACTCTTTCCCTACACGACGCTCTTCCGATCTNNNCACAACACATTTCTTCTTGCCC
GCACACGTCTGAACTCTTTCCCTACACGACGCTCTTCCGATCTNNNGTGCCTGCCTATGGGTGAAAAT
AGAGCACACGTCTGAACTCTTTCCCTACACGACGCTCTTCCGATCTNNNGCAAATTGCCCAAGGTAAC
AGCACACGTCTGAACTCTTTCCCTACACGACGCTCTTCCGATCTNNNAAAGACATTTACAGGGCCAG
AGAGCACACGTCTGAACTCTTTCCCTACACGACGCTCTTCCGATCTNNNTTGTTTGCTTCAGGTAATGG
GCACACGTCTGAACTCTTTCCCTACACGACGCTCTTCCGATCTNNNGCTGATAGAGGATGAGCTGCA
AGCACACGTCTGAACTCTTTCCCTACACGACGCTCTTCCGATCTNNNAACTTCTGTGCTTGATACCCTG
GAGCACACGTCTGAACTCTTTCCCTACACGACGCTCTTCCGATCTNNNGCACAGAGGCAGACACTGA
GAGCACACGTCTGAACTCTTTCCCTACACGACGCTCTTCCGATCTNNNCCTCAGCTTTGCAATCTTAGG
GAGCACACGTCTGAACTCTTTCCCTACACGACGCTCTTCCGATCTNNNTCATAAATACACACAGCGGC
GAGCACACGTCTGAACTCTTTCCCTACACGACGCTCTTCCGATCTNNNCCTGGATGAAAAGGCAGAG
GAGCACACGTCTGAACTCTTTCCCTACACGACGCTCTTCCGATCTNNNTGCTTTGGAAATACATCAGG
AGCACACGTCTGAACTCTTTCCCTACACGACGCTCTTCCGATCTNNNCCAGCTGGTCCTAAACAGAAAC
AGAGCACACGTCTGAACTCTTTCCCTACACGACGCTCTTCCGATCTNNNTCATCTCCAATCTACCTGCCC
AGCACACGTCTGAACTCTTTCCCTACACGACGCTCTTCCGATCTNNNCTACCATGCCTAAGCTGCTTTC
GAGCACACGTCTGAACTCTTTCCCTACACGACGCTCTTCCGATCTNNNACTCTTTCTGATCTCAAGGGA
GAGCACACGTCTGAACTCTTTCCCTACACGACGCTCTTCCGATCTNNNAGACACCTGCCTATCTCACCAT
GCACACGTCTGAACTCTTTCCCTACACGACGCTCTTCCGATCTNNNCTTTCCCATCAGCCCTTCAGTTTC
CACACGTCTGAACTCTTTCCCTACACGACGCTCTTCCGATCTNNNACAAAGAGAGGCCCTGTATCAAA
GAGCACACGTCTGAACTCTTTCCCTACACGACGCTCTTCCGATCTNNNGTCTGATCCAGCACAGACAC
GAGCACACGTCTGAACTCTTTCCCTACACGACGCTCTTCCGATCTNNNAGCAAGCTATGAACCAGGAG
AGCACACGTCTGAACTCTTTCCCTACACGACGCTCTTCCGATCTNNNTAAGGTAATACCAGTGTGCAG
AGCACACGTCTGAACTCTTTCCCTACACGACGCTCTTCCGATCTNNNTCTTTATACACAGCCCAGGCTC
AGCACACGTCTGAACTCTTTCCCTACACGACGCTCTTCCGATCTNNNGGATTCAACGTACCCTTCATGCT
AAGAGCACACGTCTGAACTCTTTCCCTACACGACGCTCTTCCGATCTNNNGAAGACAGAGCAAGCTG
GCACACGTCTGAACTCTTTCCCTACACGACGCTCTTCCGATCTNNNTAGGCTTCCTGTCCCACCTAAAA
AAGAGCACACGTCTGAACTCTTTCCCTACACGACGCTCTTCCGATCTNNNCAAAAAGCTTAGTGGCTT

GCACACGTCTGAACTCTTTCCCTACACGACGCTCTTCCGATCTNNNCAGGTGCACATATTAGCCAGAGC
GAGCACACGTCTGAACTCTTTCCCTACACGACGCTCTTCCGATCTNNNAGCATGAGAGCTGCTAAAACC
AGCACACGTCTGAACTCTTTCCCTACACGACGCTCTTCCGATCTNNNAACAGCCAGATACTCCTCACGT
AGCACACGTCTGAACTCTTTCCCTACACGACGCTCTTCCGATCTNNNCCCTGAAGGAAGCCAAACTACT
GAGCACACGTCTGAACTCTTTCCCTACACGACGCTCTTCCGATCTNNNAACAAGCCTCCATCCTGAAAAC
GAGCACACGTCTGAACTCTTTCCCTACACGACGCTCTTCCGATCTNNNGAGAGGACACACTCTAGTCCA
AGAGCACACGTCTGAACTCTTTCCCTACACGACGCTCTTCCGATCTNNNTGAAGCCCTACGTATCCAAAG
AGAGCACACGTCTGAACTCTTTCCCTACACGACGCTCTTCCGATCTNNNGATAACTGGCAAAACACTGT
AGCACACGTCTGAACTCTTTCCCTACACGACGCTCTTCCGATCTNNNACATGAGACAGAGGGGGTAGA
AGAGCACACGTCTGAACTCTTTCCCTACACGACGCTCTTCCGATCTNNNAATAGCAATTGCACATCCCCC
GAGCACACGTCTGAACTCTTTCCCTACACGACGCTCTTCCGATCTNNNGGATAACAGGCCTTGATATTC
AGCACACGTCTGAACTCTTTCCCTACACGACGCTCTTCCGATCTNNNACCAAACACCAAGTCCCTCTTTC
AGAGCACACGTCTGAACTCTTTCCCTACACGACGCTCTTCCGATCTNNNTCTGTATCAATATTCATGCCT
AGCACACGTCTGAACTCTTTCCCTACACGACGCTCTTCCGATCTNNNGTGTGTGTGTTCTGGTCTCT
GCACACGTCTGAACTCTTTCCCTACACGACGCTCTTCCGATCTNNNGGAGGATGCAATATTACCAGGG
GCACACGTCTGAACTCTTTCCCTACACGACGCTCTTCCGATCTNNNTTTTCCTTCTTCGCTCCTCTAGCCA
AGCACACGTCTGAACTCTTTCCCTACACGACGCTCTTCCGATCTNNNCAAATGGAAACCGTCAGAGTTC
AGCACACGTCTGAACTCTTTCCCTACACGACGCTCTTCCGATCTNNNTGAGAGAGATCTTCCACCGAAC
AGCACACGTCTGAACTCTTTCCCTACACGACGCTCTTCCGATCTNNNGCATGTATACAGGGATGTATTT
AGAGCACACGTCTGAACTCTTTCCCTACACGACGCTCTTCCGATCTNNNTTCCAAAAGGAATGATGGGA
GAGCACACGTCTGAACTCTTTCCCTACACGACGCTCTTCCGATCTNNNCAGGAGAATCTTGCAAAGCAG
GAGCACACGTCTGAACTCTTTCCCTACACGACGCTCTTCCGATCTNNNGGCATCTGGGTGTTTCCTAAG
AAGAGCACACGTCTGAACTCTTTCCCTACACGACGCTCTTCCGATCTNNNTCCCAGAGAGGACTGCTAC
GAAGAGCACACGTCTGAACTCTTTCCCTACACGACGCTCTTCCGATCTNNNTAGTCAAGCCTCATTTTG
AAGAGCACACGTCTGAACTCTTTCCCTACACGACGCTCTTCCGATCTNNNGTTGTGCGTTCCTAAACTCA
AAGAGCACACGTCTGAACTCTTTCCCTACACGACGCTCTTCCGATCTNNNGCTCTCTGCATCTCAATTCC
GAGCACACGTCTGAACTCTTTCCCTACACGACGCTCTTCCGATCTNNNGCTGCTGACTTGGAAAAACTC
GAGCACACGTCTGAACTCTTTCCCTACACGACGCTCTTCCGATCTNNNCATACATCAAACCCCACAACACC
GAGCACACGTCTGAACTCTTTCCCTACACGACGCTCTTCCGATCTNNNCCTCAAAAGTGCAGTGAGTTCC
GAGCACACGTCTGAACTCTTTCCCTACACGACGCTCTTCCGATCTNNNAACAAAAACTGGACGGTAGTC
AGCACACGTCTGAACTCTTTCCCTACACGACGCTCTTCCGATCTNNNCATAAACTGCCCAGATGGACAC
AAGAGCACACGTCTGAACTCTTTCCCTACACGACGCTCTTCCGATCTNNNTCTCATGGGAACAAGACCAT
AGCACACGTCTGAACTCTTTCCCTACACGACGCTCTTCCGATCTNNNTTGTGCCATTTCATGTCCTTCCAG
GAGCACACGTCTGAACTCTTTCCCTACACGACGCTCTTCCGATCTNNNCTGACTTGGCCTCTTGTTCTCA
GAGCACACGTCTGAACTCTTTCCCTACACGACGCTCTTCCGATCTNNNCCTGACAAGTTTGACACACAC
GAGCACACGTCTGAACTCTTTCCCTACACGACGCTCTTCCGATCTNNNTGTTGGATGTCTCCCTGCTCTA
AGCACACGTCTGAACTCTTTCCCTACACGACGCTCTTCCGATCTNNNTTTGGATGCCTTTTCTGAGGTC
AGCACACGTCTGAACTCTTTCCCTACACGACGCTCTTCCGATCTNNNCTTTCCCCACTCCCCTCTTTCTACA
GAAGAGCACACGTCTGAACTCTTTCCCTACACGACGCTCTTCCGATCTNNNTGATTTCTCTGTCTCATCA
AGCACACGTCTGAACTCTTTCCCTACACGACGCTCTTCCGATCTNNNTCCGGAATAGACTGGACTACAG
AGAGCACACGTCTGAACTCTTTCCCTACACGACGCTCTTCCGATCTNNNTGCCTTCACTCCTGAAGGAC
GAAGAGCACACGTCTGAACTCTTTCCCTACACGACGCTCTTCCGATCTNNNATCAGAAAGGACATGGA
GAGCACACGTCTGAACTCTTTCCCTACACGACGCTCTTCCGATCTNNNAATGGCAGATGGGGACTTGT

AGCACACGTCTGAACTCTTTCCCTACACGACGCTCTTCCGATCTNNNACCTCCAGTCCTTCTTTACCCT
AGAGCACACGTCTGAACTCTTTCCCTACACGACGCTCTTCCGATCTNNNTTCAAACCATGTCTGGACAA
AGCACACGTCTGAACTCTTTCCCTACACGACGCTCTTCCGATCTNNNGTGCTTGCACCATTTACAATCC
AGCACACGTCTGAACTCTTTCCCTACACGACGCTCTTCCGATCTNNNCTGAGACTGACCCATGTCATGT
AGCACACGTCTGAACTCTTTCCCTACACGACGCTCTTCCGATCTNNNAGTCAAGACATTTGAGGAAA
AAGAGCACACGTCTGAACTCTTTCCCTACACGACGCTCTTCCGATCTNNNAGCATTGTTACAGGCATTT
GCACACGTCTGAACTCTTTCCCTACACGACGCTCTTCCGATCTNNNAATGATGGGGGACAGACATTG
CACACGTCTGAACTCTTTCCCTACACGACGCTCTTCCGATCTNNNAATTTCATTGGAAGGAGGGGT
GAGCACACGTCTGAACTCTTTCCCTACACGACGCTCTTCCGATCTNNNGGAAATGTTCTCATGGGCA
AGCACACGTCTGAACTCTTTCCCTACACGACGCTCTTCCGATCTNNNAGGCTGTAGAGTTCTTTGCAT
GAGCACACGTCTGAACTCTTTCCCTACACGACGCTCTTCCGATCTNNNAGAGGAAAGTCTGTGAATG
AGCACACGTCTGAACTCTTTCCCTACACGACGCTCTTCCGATCTNNNCTGAGTACCCTGAGGAAGAG
AGCACACGTCTGAACTCTTTCCCTACACGACGCTCTTCCGATCTNNNGTACTTTTGAAACAGGGCTTG
AGCACACGTCTGAACTCTTTCCCTACACGACGCTCTTCCGATCTNNNACCTTCCAAAGTCGTACAGCT
AGCACACGTCTGAACTCTTTCCCTACACGACGCTCTTCCGATCTNNNCTGTGGCTGGAGTTGAAATA
GAGCACACGTCTGAACTCTTTCCCTACACGACGCTCTTCCGATCTNNNTGAGAACCAGTCTGAACAA
CACACGTCTGAACTCTTTCCCTACACGACGCTCTTCCGATCTNNNGATGCAGATGGCCAATAGTGG
GCACACGTCTGAACTCTTTCCCTACACGACGCTCTTCCGATCTNNNGAGGCATCCAAACCCACTTCAA
CACACGTCTGAACTCTTTCCCTACACGACGCTCTTCCGATCTNNNCATTGGAAAACGTTAGGTGGTGC
GCACACGTCTGAACTCTTTCCCTACACGACGCTCTTCCGATCTNNNAGCAAAACAAAACCCTGCAAGA
GAGCACACGTCTGAACTCTTTCCCTACACGACGCTCTTCCGATCTNNNACAATCAAGCAAATCTGGAG
AGAGCACACGTCTGAACTCTTTCCCTACACGACGCTCTTCCGATCTNNNTCTGCGCCTGAATTTCCTTA
AGCACACGTCTGAACTCTTTCCCTACACGACGCTCTTCCGATCTNNNTACTGTCTCTCACCTTGATGGC
AGCACACGTCTGAACTCTTTCCCTACACGACGCTCTTCCGATCTNNNGCCTTGCTGTACTTTAGAAGG
AGCACACGTCTGAACTCTTTCCCTACACGACGCTCTTCCGATCTNNNCCAGGTACAGTCTTGAAGCA
AGAGCACACGTCTGAACTCTTTCCCTACACGACGCTCTTCCGATCTNNNGTGGGTTTTACGACAGCAC
AGAGCACACGTCTGAACTCTTTCCCTACACGACGCTCTTCCGATCTNNNTGCCATATCTCAATGAAAG
GCACACGTCTGAACTCTTTCCCTACACGACGCTCTTCCGATCTNNNCTTAGCAAGTGGCTTCCAACTC
GCACACGTCTGAACTCTTTCCCTACACGACGCTCTTCCGATCTNNNTGTATTTCAACTTGCACCCACCT
AGCACACGTCTGAACTCTTTCCCTACACGACGCTCTTCCGATCTNNNATCCAAGGGGAAGGAGTAG
AGCACACGTCTGAACTCTTTCCCTACACGACGCTCTTCCGATCTNNNTTAGTGGAGGTCAGTAAGTG
AGCACACGTCTGAACTCTTTCCCTACACGACGCTCTTCCGATCTNNNAGCAGCTGCACATGAACATTT
AGCACACGTCTGAACTCTTTCCCTACACGACGCTCTTCCGATCTNNNACACAGGCTCGTTCATTCACA
CACACGTCTGAACTCTTTCCCTACACGACGCTCTTCCGATCTNNNTTCTTACCTGGCCTTTGCTTCAGC
CACGTCTGAACTCTTTCCCTACACGACGCTCTTCCGATCTNNNACAAAGACACCTAACTTCTCAGGCA
GAGCACACGTCTGAACTCTTTCCCTACACGACGCTCTTCCGATCTNNNTCCTGCAAATCCTATGTCTCC
AGAGCACACGTCTGAACTCTTTCCCTACACGACGCTCTTCCGATCTNNNACTTGAGCCTACCAAGCAG
AGAGCACACGTCTGAACTCTTTCCCTACACGACGCTCTTCCGATCTNNNGGGGAGACTGACTATCAAT
AGAGCACACGTCTGAACTCTTTCCCTACACGACGCTCTTCCGATCTNNNAGTGTTTCTCTCTCTCTCTCT
GAGCACACGTCTGAACTCTTTCCCTACACGACGCTCTTCCGATCTNNNATCCTTTTCTACCTCTGCTCCC
GCACACGTCTGAACTCTTTCCCTACACGACGCTCTTCCGATCTNNNCAGCACGAGATTTCTAAGTCCCC
AGCACACGTCTGAACTCTTTCCCTACACGACGCTCTTCCGATCTNNNGCTGTGAATGCCTTTGCTTCAT
GAGCACACGTCTGAACTCTTTCCCTACACGACGCTCTTCCGATCTNNNGCTGAAGCTTGTTTTCCCC

AGCACACGTCTGAACTCTTTCCCTACACGACGCTCTTCCGATCTNNNCCCTTGCCATACAACAACCTAG

AGAGCACACGTCTGAACTCTTTCCCTACACGACGCTCTTCCGATCTNNNTGGCCTGGTTGTTTCCGTA

AGAGCACACGTCTGAACTCTTTCCCTACACGACGCTCTTCCGATCTNNNCCTGCAAGGAAAGACTATG

GAGCACACGTCTGAACTCTTTCCCTACACGACGCTCTTCCGATCTNNNTGCAATTCTTTTAGCCTGG

CACACGTCTGAACTCTTTCCCTACACGACGCTCTTCCGATCTNNNACATTTGACTGAAAAGCTCGGGT

GCACACGTCTGAACTCTTTCCCTACACGACGCTCTTCCGATCTNNNAGGAAGTCATCTCTCTCTCGCTT

GCACACGTCTGAACTCTTTCCCTACACGACGCTCTTCCGATCTNNNCATTGACGCTTGGAAGGGAAA

GAGCACACGTCTGAACTCTTTCCCTACACGACGCTCTTCCGATCTNNNGCATAGAGCTCGAGGTTGAT

GAGCACACGTCTGAACTCTTTCCCTACACGACGCTCTTCCGATCTNNNAAGCAAAACTACCAGGTTCG

GAGCACACGTCTGAACTCTTTCCCTACACGACGCTCTTCCGATCTNNNATCCAAGGGGAAGGAGTA

CACACGTCTGAACTCTTTCCCTACACGACGCTCTTCCGATCTNNNCAACCTTTGGTGCTCAAGTGATT

AGAGCACACGTCTGAACTCTTTCCCTACACGACGCTCTTCCGATCTNNNTGATTACCAGGGTCTGAGG

GCACACGTCTGAACTCTTTCCCTACACGACGCTCTTCCGATCTNNNGATGCAGATGGCCAATAGTGGA

GCACACGTCTGAACTCTTTCCCTACACGACGCTCTTCCGATCTNNNAGAATTGGCAGGCATCACTTCT

AGCACACGTCTGAACTCTTTCCCTACACGACGCTCTTCCGATCTNNNCTTGAGTGCAGTGTCCCTATCA

GAGCACACGTCTGAACTCTTTCCCTACACGACGCTCTTCCGATCTNNNGTCATAAGCCTCTGTCCACA

AGCACACGTCTGAACTCTTTCCCTACACGACGCTCTTCCGATCTNNNGCTCTGTGGTCAAGTCTAGGA

AGCACACGTCTGAACTCTTTCCCTACACGACGCTCTTCCGATCTNNNAAGGGACTTCTGTTCCTCCTA

GCACACGTCTGAACTCTTTCCCTACACGACGCTCTTCCGATCTNNNGGGTATGATAGCGGGTTCCTG

GAGCACACGTCTGAACTCTTTCCCTACACGACGCTCTTCCGATCTNNNGTATACAGAGTTTGGGGCA

GCACACGTCTGAACTCTTTCCCTACACGACGCTCTTCCGATCTNNNTGTTGAACGTTCCTCCATACTGT

AGCACACGTCTGAACTCTTTCCCTACACGACGCTCTTCCGATCTNNNCCCTAAGCCCCCAACTAGAAAA

AGAGCACACGTCTGAACTCTTTCCCTACACGACGCTCTTCCGATCTNNNCTATGAATTGCTCCCCTACC

AGCACACGTCTGAACTCTTTCCCTACACGACGCTCTTCCGATCTNNNGGTGTTGGTCTACTCAGGGG

GAGCACACGTCTGAACTCTTTCCCTACACGACGCTCTTCCGATCTNNNAATCCTGGATGACCTGGCAA

CACACGTCTGAACTCTTTCCCTACACGACGCTCTTCCGATCTNNNGGGTTCCTATTTATCAGGCAGGC

AGAGCACACGTCTGAACTCTTTCCCTACACGACGCTCTTCCGATCTNNNGACGCTAGTGAGCAAGGA

GCACACGTCTGAACTCTTTCCCTACACGACGCTCTTCCGATCTNNNAGGAATGACAAGAGTGGGAA

ACACGTCTGAACTCTTTCCCTACACGACGCTCTTCCGATCTNNNTCCAAAAGCTAGTGTGTGTGTGTC

AGAGCACACGTCTGAACTCTTTCCCTACACGACGCTCTTCCGATCTNNNGGAACACCTCCATTCCTCTT

GCACACGTCTGAACTCTTTCCCTACACGACGCTCTTCCGATCTNNNGACTTGGCCTCTGGTTCATTTCA

GAGCACACGTCTGAACTCTTTCCCTACACGACGCTCTTCCGATCTNNNATCTCCCAGGTCCAAGCAATT

GAAGAGCACACGTCTGAACTCTTTCCCTACACGACGCTCTTCCGATCTNNNATACGCCTGTAGTCCCTA

GAGCACACGTCTGAACTCTTTCCCTACACGACGCTCTTCCGATCTNNNAAACATGTACTTACTCTAAAG

CACACGTCTGAACTCTTTCCCTACACGACGCTCTTCCGATCTNNNGATTGGTGCCCTCTGGAAACATC

AGAGCACACGTCTGAACTCTTTCCCTACACGACGCTCTTCCGATCTNNNAGTGAGAACACTTATTAGT

GAGCACACGTCTGAACTCTTTCCCTACACGACGCTCTTCCGATCTNNNCTATGTAGAACAGAGCTGC

GCACACGTCTGAACTCTTTCCCTACACGACGCTCTTCCGATCTNNNGACTGGAGTGTAGTGGTACCG

CACGTCTGAACTCTTTCCCTACACGACGCTCTTCCGATCTNNNTAATGGGTTGTTCCTGCTTGGAGCA

GAGCACACGTCTGAACTCTTTCCCTACACGACGCTCTTCCGATCTNNNATGAAAGTCTCCTTGCCTCTC

GCACACGTCTGAACTCTTTCCCTACACGACGCTCTTCCGATCTNNNCAGCCCTTGAAGCTATCTGGTTT

AGCACACGTCTGAACTCTTTCCCTACACGACGCTCTTCCGATCTNNNCATACTGCGAGGATGACTTCT

AGCACACGTCTGAACTCTTTCCCTACACGACGCTCTTCCGATCTNNNAGTAGTAGGGCAAGCATACC

GAGCACACGTCTGAACTCTTTCCCTACACGACGCTCTTCCGATCTNNNTAGTGAAGACAGGACCCAG
AGCACACGTCTGAACTCTTTCCCTACACGACGCTCTTCCGATCTNNNATTTCTCCCACCTTGCTATAGG
AGCACACGTCTGAACTCTTTCCCTACACGACGCTCTTCCGATCTNNNTGATGTTGTGGTGACGATGA
GAGCACACGTCTGAACTCTTTCCCTACACGACGCTCTTCCGATCTNNNGGTCAGACATGGACTATTGG
AGCACACGTCTGAACTCTTTCCCTACACGACGCTCTTCCGATCTNNNTGCATATACCATCGCCTCCAAA
AGAGCACACGTCTGAACTCTTTCCCTACACGACGCTCTTCCGATCTNNNGTTTGTTCTAAGGTTGCGT
AGCACACGTCTGAACTCTTTCCCTACACGACGCTCTTCCGATCTNNNGAACAGAAACTTGGGTGACA
GAGCACACGTCTGAACTCTTTCCCTACACGACGCTCTTCCGATCTNNNTTAGGAGCTGGTTGAAAAGG
AGAGCACACGTCTGAACTCTTTCCCTACACGACGCTCTTCCGATCTNNNTCAGGAATATTGGCCTATA
GCACACGTCTGAACTCTTTCCCTACACGACGCTCTTCCGATCTNNNGGGAGAAGGGATGTTCAGCTT
AGCACACGTCTGAACTCTTTCCCTACACGACGCTCTTCCGATCTNNNTATATCCTAATGGGAGGCGAA
GCACACGTCTGAACTCTTTCCCTACACGACGCTCTTCCGATCTNNNCGCTTTCCCTTTCTGAGATCTGG
AGCACACGTCTGAACTCTTTCCCTACACGACGCTCTTCCGATCTNNNTGAATCTCCTGCTTTTTCCTGG
AGAGCACACGTCTGAACTCTTTCCCTACACGACGCTCTTCCGATCTNNNAGGCTGTTGTGTGAGGAA
GAGCACACGTCTGAACTCTTTCCCTACACGACGCTCTTCCGATCTNNNATGTTCCTCTCCTCTCAGCATG
AGCACACGTCTGAACTCTTTCCCTACACGACGCTCTTCCGATCTNNNAAACAAGAGCAAGAAGAACTT
AGCACACGTCTGAACTCTTTCCCTACACGACGCTCTTCCGATCTNNNCTGTGCTTCACTCTCTCATCACA
AGCACACGTCTGAACTCTTTCCCTACACGACGCTCTTCCGATCTNNNTGCAGAGAGGGAAAGAAACA
GAGCACACGTCTGAACTCTTTCCCTACACGACGCTCTTCCGATCTNNNTGGCATTTTTGTGGAGAGAG
AGCACACGTCTGAACTCTTTCCCTACACGACGCTCTTCCGATCTNNNTTTAGACACCACCTCCCACAGAT
AGCACACGTCTGAACTCTTTCCCTACACGACGCTCTTCCGATCTNNNCCCCATATACCCCATATCCCAG
GAGCACACGTCTGAACTCTTTCCCTACACGACGCTCTTCCGATCTNNNCCTGAGGATTCATCCACCTGA
GAAGAGCACACGTCTGAACTCTTTCCCTACACGACGCTCTTCCGATCTNNNAGTTCTCAGTTGTTGTCA
AGCACACGTCTGAACTCTTTCCCTACACGACGCTCTTCCGATCTNNNTTATGTTGCCTGTCACAGTGTC
AGCACACGTCTGAACTCTTTCCCTACACGACGCTCTTCCGATCTNNNGCCCATCAATCAACAAGAGTGG
GCACACGTCTGAACTCTTTCCCTACACGACGCTCTTCCGATCTNNNACCTGGACCAATTGGAAAGGAT
GCACACGTCTGAACTCTTTCCCTACACGACGCTCTTCCGATCTNNNTCTACAACTGCCAGCATGCAAAT
GAGCACACGTCTGAACTCTTTCCCTACACGACGCTCTTCCGATCTNNNGATTTCTGTCAGACTGGTGA
AGAGCACACGTCTGAACTCTTTCCCTACACGACGCTCTTCCGATCTNNNAAGAAAGCAGGCACTCTTTT
AGCACACGTCTGAACTCTTTCCCTACACGACGCTCTTCCGATCTNNNACCAAAACATGGACATTTTCCC
AGCACACGTCTGAACTCTTTCCCTACACGACGCTCTTCCGATCTNNNTCCTAGAAGGCAATGCACCATT
AGCACACGTCTGAACTCTTTCCCTACACGACGCTCTTCCGATCTNNNGGGTAGATGCTTTGAAATGG
AGCACACGTCTGAACTCTTTCCCTACACGACGCTCTTCCGATCTNNNACATTGCACAATGCACATAAG
AGCACACGTCTGAACTCTTTCCCTACACGACGCTCTTCCGATCTNNNGAAGCACATGGGCAAGGTAT
AGCACACGTCTGAACTCTTTCCCTACACGACGCTCTTCCGATCTNNNTCTGATGCCACTTCCCAAAACA
AGCACACGTCTGAACTCTTTCCCTACACGACGCTCTTCCGATCTNNNTGGCATCAAAGACTACTGGAA
GAGCACACGTCTGAACTCTTTCCCTACACGACGCTCTTCCGATCTNNNCCCAGGAAGTATGTTTGGTA
GCACACGTCTGAACTCTTTCCCTACACGACGCTCTTCCGATCTNNNGGCAGACATACTCAGACCTCTCT
AGCACACGTCTGAACTCTTTCCCTACACGACGCTCTTCCGATCTNNNGTCTCCAACTTTACTGCAGAAG
AGCACACGTCTGAACTCTTTCCCTACACGACGCTCTTCCGATCTNNNTGTTGAAAATCAGGCCACCTA
AGCACACGTCTGAACTCTTTCCCTACACGACGCTCTTCCGATCTNNNAATGTTTTTGTTTGTGGGCCTC
AGAGCACACGTCTGAACTCTTTCCCTACACGACGCTCTTCCGATCTNNNTGAGCCTGTTATTACCTTTCC
GCACACGTCTGAACTCTTTCCCTACACGACGCTCTTCCGATCTNNNCTCTAGCACAGTGGGCTTTCTTC

GAGCACACGTCTGAACTCTTTCCCTACACGACGCTCTTCCGATCTNNNAGGAGTACTATCATGGCTCCT
AGCACACGTCTGAACTCTTTCCCTACACGACGCTCTTCCGATCTNNNATCATGTGTGTGGGTAGGAAG
GAGCACACGTCTGAACTCTTTCCCTACACGACGCTCTTCCGATCTNNNTAGCTTCAAACAGAAGTCGGC
GAGCACACGTCTGAACTCTTTCCCTACACGACGCTCTTCCGATCTNNNTGACATGGTCAATACTTCTGC
AGCACACGTCTGAACTCTTTCCCTACACGACGCTCTTCCGATCTNNNAGGCACTGAGAAATCTTATAG
AGAGCACACGTCTGAACTCTTTCCCTACACGACGCTCTTCCGATCTNNNATGATTAGTTGTGCCAGAT
AGCACACGTCTGAACTCTTTCCCTACACGACGCTCTTCCGATCTNNNTCTCTTCCACCCCCAAAAGAA
AGCACACGTCTGAACTCTTTCCCTACACGACGCTCTTCCGATCTNNNTGCAGATTATGCCACTGCACTC
GAGCACACGTCTGAACTCTTTCCCTACACGACGCTCTTCCGATCTNNNAGAGCTTAGATCTGTCTTCCC
AGCACACGTCTGAACTCTTTCCCTACACGACGCTCTTCCGATCTNNNAGATAACAAGTATGTGTATGT
AGAGCACACGTCTGAACTCTTTCCCTACACGACGCTCTTCCGATCTNNNTGAAGGGAAAACACAAAAA
CACACGTCTGAACTCTTTCCCTACACGACGCTCTTCCGATCTNNNAGTGCAGCTGGATACAAGTTCAA
GAGCACACGTCTGAACTCTTTCCCTACACGACGCTCTTCCGATCTNNNGTGTCGAGTTCTCCAGCTGTT
AGCACACGTCTGAACTCTTTCCCTACACGACGCTCTTCCGATCTNNNGAAGGATAAAATAAAAGTTGG
AGCACACGTCTGAACTCTTTCCCTACACGACGCTCTTCCGATCTNNNTTTGGCTGTTTTTGGATGCTAC
GAGCACACGTCTGAACTCTTTCCCTACACGACGCTCTTCCGATCTNNNGAAACTCAGCCTCCTCTGTAA
GCACACGTCTGAACTCTTTCCCTACACGACGCTCTTCCGATCTNNNCAGTGGGTGGGTACTCAAATAG
AGAGCACACGTCTGAACTCTTTCCCTACACGACGCTCTTCCGATCTNNNCCACCATTTGTACAATGCTAC
AGCACACGTCTGAACTCTTTCCCTACACGACGCTCTTCCGATCTNNNCACCACCTTCTGTCATGCTGAT
AAGAGCACACGTCTGAACTCTTTCCCTACACGACGCTCTTCCGATCTNNNTACTATGCCCAGCCGAAAA
AGCACACGTCTGAACTCTTTCCCTACACGACGCTCTTCCGATCTNNNCCTTATAGTCTTTAGGATTGCG
AGCACACGTCTGAACTCTTTCCCTACACGACGCTCTTCCGATCTNNNACCAACATTTCTGAAGAAGCA
AGAGCACACGTCTGAACTCTTTCCCTACACGACGCTCTTCCGATCTNNNTGTTGATGGTAATGCTGGT
AGCACACGTCTGAACTCTTTCCCTACACGACGCTCTTCCGATCTNNNTTCCATCCCGGGTGACAAACAT
AGAGCACACGTCTGAACTCTTTCCCTACACGACGCTCTTCCGATCTNNNATGATTAGTTGTGCCAGAT
AAGAGCACACGTCTGAACTCTTTCCCTACACGACGCTCTTCCGATCTNNNAATATGACTTACAGCTGTG
AGCACACGTCTGAACTCTTTCCCTACACGACGCTCTTCCGATCTNNNTCTATGGCCACAGCTCTAAGAC
AGAGCACACGTCTGAACTCTTTCCCTACACGACGCTCTTCCGATCTNNNAGCCTGATTGGGGTAAGTT
AGCACACGTCTGAACTCTTTCCCTACACGACGCTCTTCCGATCTNNNACAAAGTTGCTCCTCTCGTTCT
GAGCACACGTCTGAACTCTTTCCCTACACGACGCTCTTCCGATCTNNNGCTTGACAGGTTTCTAGAGT
GAGCACACGTCTGAACTCTTTCCCTACACGACGCTCTTCCGATCTNNNTCTTAGGACTGGGCCAAGGA
AGCACACGTCTGAACTCTTTCCCTACACGACGCTCTTCCGATCTNNNTGACACTCCCTTTTGAACTTGT
AGCACACGTCTGAACTCTTTCCCTACACGACGCTCTTCCGATCTNNNCCAAATTCTCCTCTCCTGCATCC
GCACACGTCTGAACTCTTTCCCTACACGACGCTCTTCCGATCTNNNCCGGCAGCTATACACTGGTCCAT
AGAGCACACGTCTGAACTCTTTCCCTACACGACGCTCTTCCGATCTNNNGTGTCCATTATCAGTGGTCT
GAGCACACGTCTGAACTCTTTCCCTACACGACGCTCTTCCGATCTNNNATGAGAGCATGTGAGTGTGA
AGCACACGTCTGAACTCTTTCCCTACACGACGCTCTTCCGATCTNNNGCAAGGGAAATTCACACACACA
AGCACACGTCTGAACTCTTTCCCTACACGACGCTCTTCCGATCTNNNTCAATGAGTGACCTTTTTGCAG
AGAGCACACGTCTGAACTCTTTCCCTACACGACGCTCTTCCGATCTNNNTTCCCAACATTCCAGATCCC
GAGCACACGTCTGAACTCTTTCCCTACACGACGCTCTTCCGATCTNNNGCACTGTGAATCTCCAAGAA
AGCACACGTCTGAACTCTTTCCCTACACGACGCTCTTCCGATCTNNNTGAAGTCGAAAGAGGACCACA
GAGCACACGTCTGAACTCTTTCCCTACACGACGCTCTTCCGATCTNNNCCAGATAGTGAACACGTACC
AGCACACGTCTGAACTCTTTCCCTACACGACGCTCTTCCGATCTNNNAGACACAGGAAGCAAACGATC

GAGCACACGTCTGAACTCTTTCCCTACACGACGCTCTTCCGATCTNNNCCCGGATCTCTCTGTTTCTCTCT
AGCACACGTCTGAACTCTTTCCCTACACGACGCTCTTCCGATCTNNNTGAACCTTACTAGGAAACACAC
GAGCACACGTCTGAACTCTTTCCCTACACGACGCTCTTCCGATCTNNNATGCCATCTCCCCAAATTCTTC
AGCACACGTCTGAACTCTTTCCCTACACGACGCTCTTCCGATCTNNNGTTGCCAAGCTTTTGGTGGAAA
AGCACACGTCTGAACTCTTTCCCTACACGACGCTCTTCCGATCTNNNATTCTAACCACTCCCTCTCCACCA
GAGCACACGTCTGAACTCTTTCCCTACACGACGCTCTTCCGATCTNNNGTCATCACCCTTCCAATCTCCTC
AAGAGCACACGTCTGAACTCTTTCCCTACACGACGCTCTTCCGATCTNNNACAATGTAAGCTTCCAGGTA
GAGCACACGTCTGAACTCTTTCCCTACACGACGCTCTTCCGATCTNNNACACAGCAAAACCTTGTCTGG
GAGCACACGTCTGAACTCTTTCCCTACACGACGCTCTTCCGATCTNNNTGACGGAACGAAGAGGGATT
AGAGCACACGTCTGAACTCTTTCCCTACACGACGCTCTTCCGATCTNNNTAAATGGAGCAGTGAGTGAC
AGCACACGTCTGAACTCTTTCCCTACACGACGCTCTTCCGATCTNNNATTATACCTTTCCCCGTCCCAGT
AGCACACGTCTGAACTCTTTCCCTACACGACGCTCTTCCGATCTNNNGCAACACAAAATCACCACACACA
AGAGCACACGTCTGAACTCTTTCCCTACACGACGCTCTTCCGATCTNNNTTGAGGGATGCCAGGATAAC
GAGCACACGTCTGAACTCTTTCCCTACACGACGCTCTTCCGATCTNNNGAATGGACAGCTTCTGTATGG
AGCACACGTCTGAACTCTTTCCCTACACGACGCTCTTCCGATCTNNNTCTCTGGGTGGTCTTGCTCAAAT
AGAGCACACGTCTGAACTCTTTCCCTACACGACGCTCTTCCGATCTNNNTGTGGTGTATTTGACACTATT
GAGCACACGTCTGAACTCTTTCCCTACACGACGCTCTTCCGATCTNNNATGGGAGGATGTGACAGAAT
GAGCACACGTCTGAACTCTTTCCCTACACGACGCTCTTCCGATCTNNNTTCTGAAAGTGTTTTGATTTCT
GCACACGTCTGAACTCTTTCCCTACACGACGCTCTTCCGATCTNNNAGCCCTTACTACCCACATACTCCCA
GAGCACACGTCTGAACTCTTTCCCTACACGACGCTCTTCCGATCTNNNTCAGGCCAGATTTGACCAACA
GAGCACACGTCTGAACTCTTTCCCTACACGACGCTCTTCCGATCTNNNTCCAAGGTTCTGTGAGGTAGA
GAGCACACGTCTGAACTCTTTCCCTACACGACGCTCTTCCGATCTNNNACAATCTTCACCAAACTCTAGC
AGCACACGTCTGAACTCTTTCCCTACACGACGCTCTTCCGATCTNNNTGCAAAGTCCTCCATCTTGAGT
AGAGCACACGTCTGAACTCTTTCCCTACACGACGCTCTTCCGATCTNNNGAACAACCCAAGACAATGTG
AGAGCACACGTCTGAACTCTTTCCCTACACGACGCTCTTCCGATCTNNNCTCAAAACTTTGGGCCAGTCA
AGAGCACACGTCTGAACTCTTTCCCTACACGACGCTCTTCCGATCTNNNCAAAATCAACCAGCTTGGCT
GCACACGTCTGAACTCTTTCCCTACACGACGCTCTTCCGATCTNNNCCTTAGCTGTTGTTGGCATCTTCC
AAGAGCACACGTCTGAACTCTTTCCCTACACGACGCTCTTCCGATCTNNNTGCAACATTCTTTCATCTCA
AGAGCACACGTCTGAACTCTTTCCCTACACGACGCTCTTCCGATCTNNNGAGGAACATGGATTTTCCGC
GAGCACACGTCTGAACTCTTTCCCTACACGACGCTCTTCCGATCTNNNAGATCCAACATCCAACAACAAC
GAGCACACGTCTGAACTCTTTCCCTACACGACGCTCTTCCGATCTNNNTGAAAGGTAATGGCTCTCAG
AAGAGCACACGTCTGAACTCTTTCCCTACACGACGCTCTTCCGATCTNNNGAACATCAGAAGTGCTGAC
GAGCACACGTCTGAACTCTTTCCCTACACGACGCTCTTCCGATCTNNNGCCTTCTTGCTCATGAAACTGA
AGCACACGTCTGAACTCTTTCCCTACACGACGCTCTTCCGATCTNNNTACATTTGTAGGCCCCTGTCAG
GAGCACACGTCTGAACTCTTTCCCTACACGACGCTCTTCCGATCTNNNTAAATCTGGGCCACAGAGCAA
AGCACACGTCTGAACTCTTTCCCTACACGACGCTCTTCCGATCTNNNAGGTTTTGCCATAACACTCAGTC
AGCACACGTCTGAACTCTTTCCCTACACGACGCTCTTCCGATCTNNNTATATGTACACTGCTGACCCCCA
GAGCACACGTCTGAACTCTTTCCCTACACGACGCTCTTCCGATCTNNNTGAACCATCATAAGTCAGGTTC
AGAGCACACGTCTGAACTCTTTCCCTACACGACGCTCTTCCGATCTNNNAGTCCCCGTGTCAAGATCTAC
GAGCACACGTCTGAACTCTTTCCCTACACGACGCTCTTCCGATCTNNNGGAAGAAACACCATAGTAAG
AGCACACGTCTGAACTCTTTCCCTACACGACGCTCTTCCGATCTNNNTTTATTGAGTGTTGCTGTTGGCT
AGAGCACACGTCTGAACTCTTTCCCTACACGACGCTCTTCCGATCTNNNTGGAGTACTGCAAAACCCAC
GAGCACACGTCTGAACTCTTTCCCTACACGACGCTCTTCCGATCTNNNGAAGACAGCCATAAGAAGGC

AGAGCACACGTCTGAACTCTTTCCCTACACGACGCTCTTCCGATCTNNNTAGAATCTGGCAGGGGAGG

CACACGTCTGAACTCTTTCCCTACACGACGCTCTTCCGATCTNNNCTCTAAGGATGTCCCCACCTTCCCA

GAGCACACGTCTGAACTCTTTCCCTACACGACGCTCTTCCGATCTNNNTGACATTGGTTGGAGAAACCC

GAGCACACGTCTGAACTCTTTCCCTACACGACGCTCTTCCGATCTNNNCATTTATAGGCCCCATACCCCCT

AGCACACGTCTGAACTCTTTCCCTACACGACGCTCTTCCGATCTNNNACATCCCACACACACATCTCTCTC

CACACGTCTGAACTCTTTCCCTACACGACGCTCTTCCGATCTNNNTAGTTGAAAGGACACTTGGGGGA

GAGCACACGTCTGAACTCTTTCCCTACACGACGCTCTTCCGATCTNNNGCTTTGCCTTCTGTGCTCTATG

AGAGCACACGTCTGAACTCTTTCCCTACACGACGCTCTTCCGATCTNNNAAATATATATACATAGGTGT

GAGCACACGTCTGAACTCTTTCCCTACACGACGCTCTTCCGATCTNNNAGATGGAATGTAAACCATGCC

GAGCACACGTCTGAACTCTTTCCCTACACGACGCTCTTCCGATCTNNNATCTGGACAGGTTAGACCAG

AGAGCACACGTCTGAACTCTTTCCCTACACGACGCTCTTCCGATCTNNNTATTGGTACTTACCTGCCCCC

AGCACACGTCTGAACTCTTTCCCTACACGACGCTCTTCCGATCTNNNTCTTTTCATCCTTTGGGTGGTTT

AGCACACGTCTGAACTCTTTCCCTACACGACGCTCTTCCGATCTNNNTTAGCCAACCCCTCTTTATCCCAG

AGCACACGTCTGAACTCTTTCCCTACACGACGCTCTTCCGATCTNNNAGGCCTAATACACATACACACAC

GCACACGTCTGAACTCTTTCCCTACACGACGCTCTTCCGATCTNNNCCACTACCAGTACCTGTCTCCTCCA

AGAGCACACGTCTGAACTCTTTCCCTACACGACGCTCTTCCGATCTNNNAACCAACCCCTAAATGATGG

AGCACACGTCTGAACTCTTTCCCTACACGACGCTCTTCCGATCTNNNAAGGTAAAGCCAAAGAAAGGC

GAGCACACGTCTGAACTCTTTCCCTACACGACGCTCTTCCGATCTNNNCCAGTGGCCTTCACACTTTTCT

AGCACACGTCTGAACTCTTTCCCTACACGACGCTCTTCCGATCTNNNGCAAGCAAGAAGTTAGTGCAG

AGCACACGTCTGAACTCTTTCCCTACACGACGCTCTTCCGATCTNNNTCCTAGTCGTGTTATTTTTCCAG

AGCACACGTCTGAACTCTTTCCCTACACGACGCTCTTCCGATCTNNNTGTCTTTGCATCCATCATCCAAT

AGAGCACACGTCTGAACTCTTTCCCTACACGACGCTCTTCCGATCTNNNTAGATCCCTCGTTTGTTTTGG

GCACACGTCTGAACTCTTTCCCTACACGACGCTCTTCCGATCTNNNGACACACACTCTTGGTTCCAGTTC

AGAGCACACGTCTGAACTCTTTCCCTACACGACGCTCTTCCGATCTNNNAGGGAGGAATGTGTCAGCTA

AGCACACGTCTGAACTCTTTCCCTACACGACGCTCTTCCGATCTNNNACTCTTGCTCCTCTCACTACCAGC

AGCACACGTCTGAACTCTTTCCCTACACGACGCTCTTCCGATCTNNNTCACCTCAAACTTGCTATACGAG

AGAGCACACGTCTGAACTCTTTCCCTACACGACGCTCTTCCGATCTNNNCTCTGAGACCTATGGGCAGG

AGAGCACACGTCTGAACTCTTTCCCTACACGACGCTCTTCCGATCTNNNCTCTTGGCTGCTGCAGTCTCC

GAGCACACGTCTGAACTCTTTCCCTACACGACGCTCTTCCGATCTNNNAGTGTCCATCAACCAATGAAG

AGCACACGTCTGAACTCTTTCCCTACACGACGCTCTTCCGATCTNNNATGGAAGGGAATTCAACTGTG

AAGAGCACACGTCTGAACTCTTTCCCTACACGACGCTCTTCCGATCTNNNCGTTTGGTTAGTGAAGTTC

GCACACGTCTGAACTCTTTCCCTACACGACGCTCTTCCGATCTNNNAAGAAAGAACAGCTAAGATGGCA

AGAGCACACGTCTGAACTCTTTCCCTACACGACGCTCTTCCGATCTNNNACAGACAATCCAGTCTTCTTC

GAGCACACGTCTGAACTCTTTCCCTACACGACGCTCTTCCGATCTNNNAAGAGACCTGCTAACACACAC

GAGCACACGTCTGAACTCTTTCCCTACACGACGCTCTTCCGATCTNNNTATGACCCAGAGCCACCTAGCT

GAGCACACGTCTGAACTCTTTCCCTACACGACGCTCTTCCGATCTNNNTATGCCCATGCTCCAAAAACCT

AGAGCACACGTCTGAACTCTTTCCCTACACGACGCTCTTCCGATCTNNNAGGCACTGATTCCAAGAAGA

AGCACACGTCTGAACTCTTTCCCTACACGACGCTCTTCCGATCTNNNATTTGGTTCCTTGCAAAGTGCTC

AGCACACGTCTGAACTCTTTCCCTACACGACGCTCTTCCGATCTNNNTCCTGGGTTTGTGTTGTTGCAT

AGCACACGTCTGAACTCTTTCCCTACACGACGCTCTTCCGATCTNNNGCAGGAGCTGGGTCTTTTACAT

AGAGCACACGTCTGAACTCTTTCCCTACACGACGCTCTTCCGATCTNNNCACAACTGCCCAGATACCTAA

AGCACACGTCTGAACTCTTTCCCTACACGACGCTCTTCCGATCTNNNTCTAGTAGATTTTCCTTTACATCT

AGCACACGTCTGAACTCTTTCCCTACACGACGCTCTTCCGATCTNNNACCAAGGAGAGAAGTGTCCTT

AGCACACGTCTGAACTCTTTCCCTACACGACGCTCTTCCGATCTNNNACAGTTACAAAGCACTAACGT
AGAGCACACGTCTGAACTCTTTCCCTACACGACGCTCTTCCGATCTNNNGGGAATGCCTTCATCAGAA
GAGCACACGTCTGAACTCTTTCCCTACACGACGCTCTTCCGATCTNNNAACATGACATAGGAGACACA
GCACACGTCTGAACTCTTTCCCTACACGACGCTCTTCCGATCTNNNTGGAGAACTGCAAGACAGTAGG
GCACACGTCTGAACTCTTTCCCTACACGACGCTCTTCCGATCTNNNCAGTGTCCTGAGAGCTACTACC
AGCACACGTCTGAACTCTTTCCCTACACGACGCTCTTCCGATCTNNNGTGTTCCTAGACACCTTCTGCT
GCACACGTCTGAACTCTTTCCCTACACGACGCTCTTCCGATCTNNNCCTGTCCCAAATACTTACCCAGCC
AGCACACGTCTGAACTCTTTCCCTACACGACGCTCTTCCGATCTNNNATCCATGGTGGATGATGCTA
GCACACGTCTGAACTCTTTCCCTACACGACGCTCTTCCGATCTNNNGGGTGTAGGGTATGTTCAGGG
GCACACGTCTGAACTCTTTCCCTACACGACGCTCTTCCGATCTNNNTATTTGCTTCCCATGTCCCCAG
GCACACGTCTGAACTCTTTCCCTACACGACGCTCTTCCGATCTNNNTGACCTCTCCTTCCATCCTCAAG
CACACGTCTGAACTCTTTCCCTACACGACGCTCTTCCGATCTNNNTTCAATGTCTCTAACACACGGCTC
GAGCACACGTCTGAACTCTTTCCCTACACGACGCTCTTCCGATCTNNNTGTATCCCAAAGAACAGGTT
AGCACACGTCTGAACTCTTTCCCTACACGACGCTCTTCCGATCTNNNATGGCAGTTGTTTCGTCTCTT
GAGCACACGTCTGAACTCTTTCCCTACACGACGCTCTTCCGATCTNNNAAAGCCTGATACACACCAAG
AGCACACGTCTGAACTCTTTCCCTACACGACGCTCTTCCGATCTNNNGCTACTTCATGCTACAGCCAGT
AGCACACGTCTGAACTCTTTCCCTACACGACGCTCTTCCGATCTNNNCCACCTCTGAAATGCCAGTGAT
AGCACACGTCTGAACTCTTTCCCTACACGACGCTCTTCCGATCTNNNTCCTCAGTGGAAACTGTGTTAG
AGCACACGTCTGAACTCTTTCCCTACACGACGCTCTTCCGATCTNNNTCTCAGGATGAGTTCAAGGTC
AGAGCACACGTCTGAACTCTTTCCCTACACGACGCTCTTCCGATCTNNNGCCCTTATGAAAGAGGAGA
CACACGTCTGAACTCTTTCCCTACACGACGCTCTTCCGATCTNNNCTACCTTACTCACCTTTGCCTGCCA
AAGAGCACACGTCTGAACTCTTTCCCTACACGACGCTCTTCCGATCTNNNTGGGATTCATTCATATTG
GAGCACACGTCTGAACTCTTTCCCTACACGACGCTCTTCCGATCTNNNGCTCTGGCTAATGGAAGTGA
GCACACGTCTGAACTCTTTCCCTACACGACGCTCTTCCGATCTNNNAAGGTGCTGGACATCTTTTCAC
GAGCACACGTCTGAACTCTTTCCCTACACGACGCTCTTCCGATCTNNNGGCATTTTGGGGGTGAAG
GAGCACACGTCTGAACTCTTTCCCTACACGACGCTCTTCCGATCTNNNAGTTTGAAGGACAAAGGGC
GCACACGTCTGAACTCTTTCCCTACACGACGCTCTTCCGATCTNNNACAGGTAAAAAGAACATAACGA
AAGAGCACACGTCTGAACTCTTTCCCTACACGACGCTCTTCCGATCTNNNCAGGTCTTTCTCAATAGC
AAGAGCACACGTCTGAACTCTTTCCCTACACGACGCTCTTCCGATCTNNNTTTGTGGGTAAAGAGCC
AGCACACGTCTGAACTCTTTCCCTACACGACGCTCTTCCGATCTNNNTGATCCCAGAATTTCAAGGCC
AAGAGCACACGTCTGAACTCTTTCCCTACACGACGCTCTTCCGATCTNNNCTGAGGCTTCTGAGCTGA
GAGCACACGTCTGAACTCTTTCCCTACACGACGCTCTTCCGATCTNNNAGCAAAGACTTGGAACCAAA
GAGCACACGTCTGAACTCTTTCCCTACACGACGCTCTTCCGATCTNNNTTTTACGTGGCAAAATGTAC
AAGAGCACACGTCTGAACTCTTTCCCTACACGACGCTCTTCCGATCTNNNGCTCTCCTAGGACACTGGG
GCACACGTCTGAACTCTTTCCCTACACGACGCTCTTCCGATCTNNNTGTACACTTCATGTCCCAGACCT
AGCACACGTCTGAACTCTTTCCCTACACGACGCTCTTCCGATCTNNNTAGCTCTGAGAACATAGTGGC
GAGCACACGTCTGAACTCTTTCCCTACACGACGCTCTTCCGATCTNNNACTGGTGCTCTTGAAAGACA
AGCACACGTCTGAACTCTTTCCCTACACGACGCTCTTCCGATCTNNNGCATCTCAGAACAAAAATCCC
GAGCACACGTCTGAACTCTTTCCCTACACGACGCTCTTCCGATCTNNNGGGCACTGGTTTGTGATACT
GAGCACACGTCTGAACTCTTTCCCTACACGACGCTCTTCCGATCTNNNAGTCTACTGTTCTTTACCCAC
GAGCACACGTCTGAACTCTTTCCCTACACGACGCTCTTCCGATCTNNNGCATTTGGAATCAGGGACTA
GCACACGTCTGAACTCTTTCCCTACACGACGCTCTTCCGATCTNNNGGAACCTTCTTGAGGCCATACA
GCACACGTCTGAACTCTTTCCCTACACGACGCTCTTCCGATCTNNNTGTTCAAGAGACCAGGCTACTC

AGCACACGTCTGAACTCTTTCCCTACACGACGCTCTTCCGATCTNNNACATTTTCTCTCAACTCATTGTG
AGCACACGTCTGAACTCTTTCCCTACACGACGCTCTTCCGATCTNNNTGCACTCTATTCTCTCTCTCTG
GAGCACACGTCTGAACTCTTTCCCTACACGACGCTCTTCCGATCTNNNATGTAAAGTGCTTACCCTGTG
AGCACACGTCTGAACTCTTTCCCTACACGACGCTCTTCCGATCTNNNTACATTTGTAGGCCCCTGTCAG
AGAGCACACGTCTGAACTCTTTCCCTACACGACGCTCTTCCGATCTNNNAGAGAAAAGAAACAGCAGA
AGCACACGTCTGAACTCTTTCCCTACACGACGCTCTTCCGATCTNNNTGGCATGTGCCAATCACCTATT
GAGCACACGTCTGAACTCTTTCCCTACACGACGCTCTTCCGATCTNNNAGTTTGGAAAAGAGATCTGCT
GAGCACACGTCTGAACTCTTTCCCTACACGACGCTCTTCCGATCTNNNACTCTGTGAGCCAATTCCATG
AGCACACGTCTGAACTCTTTCCCTACACGACGCTCTTCCGATCTNNNGACTTTGACAGCAAGTGTGTCTC
GAGCACACGTCTGAACTCTTTCCCTACACGACGCTCTTCCGATCTNNNTTGTAGATGGTGAGAGATGG
GAAGAGCACACGTCTGAACTCTTTCCCTACACGACGCTCTTCCGATCTNNNCCTTCACAGATTCCAGGA
GAGCACACGTCTGAACTCTTTCCCTACACGACGCTCTTCCGATCTNNNCTCTCCTCTATCCCTGCCACTTT
GAAGAGCACACGTCTGAACTCTTTCCCTACACGACGCTCTTCCGATCTNNNGAGAAAAGAAACAGCAG
AGAGCACACGTCTGAACTCTTTCCCTACACGACGCTCTTCCGATCTNNNCTACCAGTTCATCCATGTTGC
GAGCACACGTCTGAACTCTTTCCCTACACGACGCTCTTCCGATCTNNNGCTTCATAAGCATTGGATGAG
AGCACACGTCTGAACTCTTTCCCTACACGACGCTCTTCCGATCTNNNTCAGTGACCGAAATGAACCATC
GAGCACACGTCTGAACTCTTTCCCTACACGACGCTCTTCCGATCTNNNTGCCCAATCCCAACACAAATAC
AGCACACGTCTGAACTCTTTCCCTACACGACGCTCTTCCGATCTNNNGGAAAAATGGCCAATGGAACGA
AAGAGCACACGTCTGAACTCTTTCCCTACACGACGCTCTTCCGATCTNNNCTTCTGCCCCTACCCATAAAC
GAGCACACGTCTGAACTCTTTCCCTACACGACGCTCTTCCGATCTNNNATGTTTGGGTATGGAGCTTGT
GCACACGTCTGAACTCTTTCCCTACACGACGCTCTTCCGATCTNNNTGGCTCACATTAGTACCTTGCATC
AAGAGCACACGTCTGAACTCTTTCCCTACACGACGCTCTTCCGATCTNNNAGCTGTGACACATTCCAAA
AGAGCACACGTCTGAACTCTTTCCCTACACGACGCTCTTCCGATCTNNNTTTTCTGAAAGTCTTTGCACT
GAGCACACGTCTGAACTCTTTCCCTACACGACGCTCTTCCGATCTNNNTGTGTCTGTAATTCCCACCCCTA
GAGCACACGTCTGAACTCTTTCCCTACACGACGCTCTTCCGATCTNNNCCCTCTGTTCTCTCATTCCCCAT
CACACGTCTGAACTCTTTCCCTACACGACGCTCTTCCGATCTNNNATATTTCTCCTAGTGCCACAGCCCAT
GCACACGTCTGAACTCTTTCCCTACACGACGCTCTTCCGATCTNNNAATCCAGTGTTACCTGCCACTTCC
AGCACACGTCTGAACTCTTTCCCTACACGACGCTCTTCCGATCTNNNTATTCTCTCTTGTGCACTCGTGG
GAGCACACGTCTGAACTCTTTCCCTACACGACGCTCTTCCGATCTNNNAGAACTGAAGCCTCAAATGGC
GAGCACACGTCTGAACTCTTTCCCTACACGACGCTCTTCCGATCTNNNGGCCATCTGCTCAGTAGGTCTA
GAGCACACGTCTGAACTCTTTCCCTACACGACGCTCTTCCGATCTNNNTGCCAAAACTTCTCACTGAAGA
AGCACACGTCTGAACTCTTTCCCTACACGACGCTCTTCCGATCTNNNCCTAGCTTGGGGTGCAGAAATA
AGCACACGTCTGAACTCTTTCCCTACACGACGCTCTTCCGATCTNNNCCACCCAAACTATCAATTGTTGC
AGCACACGTCTGAACTCTTTCCCTACACGACGCTCTTCCGATCTNNNTTTACCTTCTGCCAAATGGGGT
GAGCACACGTCTGAACTCTTTCCCTACACGACGCTCTTCCGATCTNNNCATGACCCATTCTCTGTTCCCTC
GAGCACACGTCTGAACTCTTTCCCTACACGACGCTCTTCCGATCTNNNAGAGCCAGGCAATGTGTTTGT
GAGCACACGTCTGAACTCTTTCCCTACACGACGCTCTTCCGATCTNNNTCAATGGATGGTTGGATGAAC
GAGCACACGTCTGAACTCTTTCCCTACACGACGCTCTTCCGATCTNNNGAGAACATAAAGCAGATGCG
GGAAGAGCACACGTCTGAACTCTTTCCCTACACGACGCTCTTCCGATCTNNNACCCTTTGCTTTCCCCAT
AGCACACGTCTGAACTCTTTCCCTACACGACGCTCTTCCGATCTNNNGTTTGGGGAGGCTAAAGCAGT
GAGCACACGTCTGAACTCTTTCCCTACACGACGCTCTTCCGATCTNNNACTGGAAGACGAACTTGGAGA
GAGCACACGTCTGAACTCTTTCCCTACACGACGCTCTTCCGATCTNNNCTACCTCCAGGGAAGAGACAC
AGAGCACACGTCTGAACTCTTTCCCTACACGACGCTCTTCCGATCTNNNCTCTGGGAAAGAATCTGGCA

GAGCACACGTCTGAACTCTTTCCCTACACGACGCTCTTCCGATCTNNNTAGGATGAAGGGGCATCATG

GAGCACACGTCTGAACTCTTTCCCTACACGACGCTCTTCCGATCTNNNAACCTGCTAGCCTCATCATTCC

GAGCACACGTCTGAACTCTTTCCCTACACGACGCTCTTCCGATCTNNNCCAGCCAGAAGATGACCACAT

AGAGCACACGTCTGAACTCTTTCCCTACACGACGCTCTTCCGATCTNNNTTCTTATGAGCATGGGGCTT

AGCACACGTCTGAACTCTTTCCCTACACGACGCTCTTCCGATCTNNNTGAGGAGAAACAAGTGCAAGG

AGAGCACACGTCTGAACTCTTTCCCTACACGACGCTCTTCCGATCTNNNGGTACCATTTTCTTCAAATCT

AGAGCACACGTCTGAACTCTTTCCCTACACGACGCTCTTCCGATCTNNNCTGCCAGAACTACCAGTACTA

AGCACACGTCTGAACTCTTTCCCTACACGACGCTCTTCCGATCTNNNTCAAGTGGACAAAATACCCAGA

GAGCACACGTCTGAACTCTTTCCCTACACGACGCTCTTCCGATCTNNNGAAAAACAACGGGGCCTGTAT

AGCACACGTCTGAACTCTTTCCCTACACGACGCTCTTCCGATCTNNNGTTCCCTTTGCAATGGATGTGAC

AGCACACGTCTGAACTCTTTCCCTACACGACGCTCTTCCGATCTNNNCCACAGAACAGAGCGTCAAAAA

GAGCACACGTCTGAACTCTTTCCCTACACGACGCTCTTCCGATCTNNNCTACCAGTTCATCCATGTTGCCA

AGCACACGTCTGAACTCTTTCCCTACACGACGCTCTTCCGATCTNNNGAGAACATAAAGCAGATGCGT

AGCACACGTCTGAACTCTTTCCCTACACGACGCTCTTCCGATCTNNNGTACTGCCAGGCAAAGATGAG

GAGCACACGTCTGAACTCTTTCCCTACACGACGCTCTTCCGATCTNNNTTCCAGGAGCTGAATTTTCCA

GAAGAGCACACGTCTGAACTCTTTCCCTACACGACGCTCTTCCGATCTNNNTAGTTTCTTGCTGAGGGC

AGCACACGTCTGAACTCTTTCCCTACACGACGCTCTTCCGATCTNNNCACATGGCTTTCAGGAGGTCAA

AGCACACGTCTGAACTCTTTCCCTACACGACGCTCTTCCGATCTNNNGAACACACCTCAGAGTTACCAC

AGCACACGTCTGAACTCTTTCCCTACACGACGCTCTTCCGATCTNNNGTAGCTCTGTCTCTCTCCCTGTCC

AGCACACGTCTGAACTCTTTCCCTACACGACGCTCTTCCGATCTNNNAGGCTAAACTGTGTAGTGTTTG

AGCACACGTCTGAACTCTTTCCCTACACGACGCTCTTCCGATCTNNNCACTGCTTCCCCTTGCCTTATAG

GAGCACACGTCTGAACTCTTTCCCTACACGACGCTCTTCCGATCTNNNGCCTCTGCAGTCCAGATTTGAA

GAGCACACGTCTGAACTCTTTCCCTACACGACGCTCTTCCGATCTNNNGATTGAGCAACATTGCCTTAG

AGCACACGTCTGAACTCTTTCCCTACACGACGCTCTTCCGATCTNNNCGCCACCAATGGATCTGATTAC

GAGCACACGTCTGAACTCTTTCCCTACACGACGCTCTTCCGATCTNNNCAGGTCCTTTCTCAATATCATT

GCACACGTCTGAACTCTTTCCCTACACGACGCTCTTCCGATCTNNNATTGCACATTCTTGTGCTATGCCC

AGCACACGTCTGAACTCTTTCCCTACACGACGCTCTTCCGATCTNNNTGTGGATGAAGGTTGTCTGGG

GAGCACACGTCTGAACTCTTTCCCTACACGACGCTCTTCCGATCTNNNATCCTGACACCAAAACTGTGCA

GAGCACACGTCTGAACTCTTTCCCTACACGACGCTCTTCCGATCTNNNGGAATTGTTCTTCTCTCAGAG

GAGCACACGTCTGAACTCTTTCCCTACACGACGCTCTTCCGATCTNNNTATTCCTAACAGCGTACAAGG

GAGCACACGTCTGAACTCTTTCCCTACACGACGCTCTTCCGATCTNNNGTACACATGCACACAAGGAGA

AGCACACGTCTGAACTCTTTCCCTACACGACGCTCTTCCGATCTNNNACTTGGTTCAAATGACAGCTGA

AGAGCACACGTCTGAACTCTTTCCCTACACGACGCTCTTCCGATCTNNNTAATGTCAGGTGAGGGAAG

GAGCACACGTCTGAACTCTTTCCCTACACGACGCTCTTCCGATCTNNNAAGCTGAGAATGAAGTGCAG

GAGCACACGTCTGAACTCTTTCCCTACACGACGCTCTTCCGATCTNNNGGGTTTCCCCAGAGAAAGAGA

GCACACGTCTGAACTCTTTCCCTACACGACGCTCTTCCGATCTNNNATCTTGCTTCTCATTCTGTGGGCCA

AGCACACGTCTGAACTCTTTCCCTACACGACGCTCTTCCGATCTNNNCACCTCTACCTGTCTGAACTTGCC

GAGCACACGTCTGAACTCTTTCCCTACACGACGCTCTTCCGATCTNNNATTCTCATGGGTGCATTGCTTC

GCACACGTCTGAACTCTTTCCCTACACGACGCTCTTCCGATCTNNNATTGCACATTCTTGTGCTATGCCC

AGCACACGTCTGAACTCTTTCCCTACACGACGCTCTTCCGATCTNNNTGGGACTTAGCTTCCACAAACA

GAGCACACGTCTGAACTCTTTCCCTACACGACGCTCTTCCGATCTNNNGGCACAAATTGCAGTGTACAT

GCACACGTCTGAACTCTTTCCCTACACGACGCTCTTCCGATCTNNNAGGTCACAGTCATAAGCCTCCAAC

AGCACACGTCTGAACTCTTTCCCTACACGACGCTCTTCCGATCTNNNTCTGGAAGTGATGGAAGTGAAT

AGCACACGTCTGAACTCTTTCCCTACACGACGCTCTTCCGATCTNNNGGCAAAACCACCTAATGATGCA
AGCACACGTCTGAACTCTTTCCCTACACGACGCTCTTCCGATCTNNNAGTAGTATCAACCCCCACTTCAC
GAGCACACGTCTGAACTCTTTCCCTACACGACGCTCTTCCGATCTNNNGACACAGAGAGCTGAAAGAC
AGCACACGTCTGAACTCTTTCCCTACACGACGCTCTTCCGATCTNNNTCCTCCTATTCCATTACCACCGCC
AGCACACGTCTGAACTCTTTCCCTACACGACGCTCTTCCGATCTNNNTGAAAGGTCTGTTGGACAAGA
AGCACACGTCTGAACTCTTTCCCTACACGACGCTCTTCCGATCTNNNTACCTAATGTTTGCCCCTAAGGC
AAGAGCACACGTCTGAACTCTTTCCCTACACGACGCTCTTCCGATCTNNNGGGTTTCCCCAGAGAAAGA
GAAGAGCACACGTCTGAACTCTTTCCCTACACGACGCTCTTCCGATCTNNNGTTCCTCTCAAAAGTCTTA
AGCACACGTCTGAACTCTTTCCCTACACGACGCTCTTCCGATCTNNNGTGTGTGTACCTGGCTAAGAAC
AGCACACGTCTGAACTCTTTCCCTACACGACGCTCTTCCGATCTNNNTGTTGTGAGATGGCCACAAGT
AAGAGCACACGTCTGAACTCTTTCCCTACACGACGCTCTTCCGATCTNNNAAGCCCTGTCAAGATTTCT
GAGCACACGTCTGAACTCTTTCCCTACACGACGCTCTTCCGATCTNNNCCTGAGCCTCACTACCTTCCTAA
AGCACACGTCTGAACTCTTTCCCTACACGACGCTCTTCCGATCTNNNTGGATTTCTCCTTTTAGTTCTGT
AGCACACGTCTGAACTCTTTCCCTACACGACGCTCTTCCGATCTNNNGTCATTTTCCTGGCTTGCAGATG
AGCACACGTCTGAACTCTTTCCCTACACGACGCTCTTCCGATCTNNNTCTTAGGACTGGGCCAAGGAA
AAGAGCACACGTCTGAACTCTTTCCCTACACGACGCTCTTCCGATCTNNNTTCTACCCTCATCACCCCCAT
GAGCACACGTCTGAACTCTTTCCCTACACGACGCTCTTCCGATCTNNNGGGAATCCCAGCTGCCTAAA
AGAGCACACGTCTGAACTCTTTCCCTACACGACGCTCTTCCGATCTNNNGTCATGCTCACCAAACATGCA
AAGAGCACACGTCTGAACTCTTTCCCTACACGACGCTCTTCCGATCTNNNTCCTGTAATTTGTATTACTC
AGCACACGTCTGAACTCTTTCCCTACACGACGCTCTTCCGATCTNNNTCTTTGGCCATAAGGGGAGACA
AGCACACGTCTGAACTCTTTCCCTACACGACGCTCTTCCGATCTNNNTTGCAGAGATAGAGCCATCGG
AGCACACGTCTGAACTCTTTCCCTACACGACGCTCTTCCGATCTNNNGAACAGTTCTTGCCTGACAATG
GAGCACACGTCTGAACTCTTTCCCTACACGACGCTCTTCCGATCTNNNTGATGATCCTCTCCTGTTGCTG
GAGCACACGTCTGAACTCTTTCCCTACACGACGCTCTTCCGATCTNNNTACCCAACAGCTATGTGACTA
GAGCACACGTCTGAACTCTTTCCCTACACGACGCTCTTCCGATCTNNNTGGCTATTTTAGGATCTGTGC
GAGCACACGTCTGAACTCTTTCCCTACACGACGCTCTTCCGATCTNNNTGGAATGCCGCATATGTAAAA
GAGCACACGTCTGAACTCTTTCCCTACACGACGCTCTTCCGATCTNNNTCGTTTCTGTCTCCCTCTCTACC
AGAGCACACGTCTGAACTCTTTCCCTACACGACGCTCTTCCGATCTNNNCTTCCTTCAGGAACTGGTGCA
AGAGCACACGTCTGAACTCTTTCCCTACACGACGCTCTTCCGATCTNNNTAAGAGGGCCCAGTACTCAC
AGCACACGTCTGAACTCTTTCCCTACACGACGCTCTTCCGATCTNNNTGTATGACAGGATGCAGCTAGG
GAAGAGCACACGTCTGAACTCTTTCCCTACACGACGCTCTTCCGATCTNNNTGCAGTCTTTGAAGAACA
AGCACACGTCTGAACTCTTTCCCTACACGACGCTCTTCCGATCTNNNGACATAGTCAGGGAAGCCTGT
GAGCACACGTCTGAACTCTTTCCCTACACGACGCTCTTCCGATCTNNNTGAAACTTGAGACGGTGGTTC
GCACACGTCTGAACTCTTTCCCTACACGACGCTCTTCCGATCTNNNGCAGGACCCTACATTATTTGCCAC
GAGCACACGTCTGAACTCTTTCCCTACACGACGCTCTTCCGATCTNNNGCAGGCATCTCCGCTATAAAAC
AGCACACGTCTGAACTCTTTCCCTACACGACGCTCTTCCGATCTNNNTTGAAACACGATGCTAAATGAA
AGCACACGTCTGAACTCTTTCCCTACACGACGCTCTTCCGATCTNNNTGGGTAGTAAAACTGTGTCAACT
GAGCACACGTCTGAACTCTTTCCCTACACGACGCTCTTCCGATCTNNNCCGGAGGATCAGAATCAAGTC
AGCACACGTCTGAACTCTTTCCCTACACGACGCTCTTCCGATCTNNNCGTTGCTGATTCTGTCAAATGG
AGCACACGTCTGAACTCTTTCCCTACACGACGCTCTTCCGATCTNNNACAGTTACATTCCACCTTGCCTGC
AGCACACGTCTGAACTCTTTCCCTACACGACGCTCTTCCGATCTNNNTGCATTCTGATGCTCTGCTGAAA
AGAGCACACGTCTGAACTCTTTCCCTACACGACGCTCTTCCGATCTNNNCTGTGACTTTAGGATGCTGTC
AGCACACGTCTGAACTCTTTCCCTACACGACGCTCTTCCGATCTNNNTAATGTCAGGTGAGGGAAGAC

AGAGCACACGTCTGAACTCTTTCCCTACACGACGCTCTTCCGATCTNNNTGTCCAGTATGAACAGAACT
GGAAGAGCACACGTCTGAACTCTTTCCCTACACGACGCTCTTCCGATCTNNNCCCTCTAGTGTGAGTTG
AGCACACGTCTGAACTCTTTCCCTACACGACGCTCTTCCGATCTNNNAGCCATTACAAGAAGGGAAGC
AAGAGCACACGTCTGAACTCTTTCCCTACACGACGCTCTTCCGATCTNNNATTGGGCCATGGTTTAGTC
GAGCACACGTCTGAACTCTTTCCCTACACGACGCTCTTCCGATCTNNNTGCACATAGACCAGTAATCCC
AGCACACGTCTGAACTCTTTCCCTACACGACGCTCTTCCGATCTNNNAAAGGTGATTGTAAGCAAGCC
GAGCACACGTCTGAACTCTTTCCCTACACGACGCTCTTCCGATCTNNNAAGGGCCTAACACTTCAGATT
AGAGCACACGTCTGAACTCTTTCCCTACACGACGCTCTTCCGATCTNNNTCCCTTCAACCACAGCTCTTAC
GAGCACACGTCTGAACTCTTTCCCTACACGACGCTCTTCCGATCTNNNCCCTCCCTGTCATGTTTCTCTC
GAGCACACGTCTGAACTCTTTCCCTACACGACGCTCTTCCGATCTNNNTTGTGCTGAAGTCTGCCTGTAT
AGAGCACACGTCTGAACTCTTTCCCTACACGACGCTCTTCCGATCTNNNGGAGACAGAAGGAGAAAGT
AGAGCACACGTCTGAACTCTTTCCCTACACGACGCTCTTCCGATCTNNNTTGATAGCACAGTCTCCTGT
GAGCACACGTCTGAACTCTTTCCCTACACGACGCTCTTCCGATCTNNNACTTTGTTTTGTACTGGTTGTT
AGCACACGTCTGAACTCTTTCCCTACACGACGCTCTTCCGATCTNNNTGCTTCTCTGGTACAGGTTTCCT
GAGCACACGTCTGAACTCTTTCCCTACACGACGCTCTTCCGATCTNNNACATAGATCATCCACAAGCTGT
AGCACACGTCTGAACTCTTTCCCTACACGACGCTCTTCCGATCTNNNATGTTTTCTCCTTGCAAACGCTG
AGAGCACACGTCTGAACTCTTTCCCTACACGACGCTCTTCCGATCTNNNTACGTGAACTGATACACGTT
AGCACACGTCTGAACTCTTTCCCTACACGACGCTCTTCCGATCTNNNTCAAAGTTGGTGTCAAGGTCAG
GAGCACACGTCTGAACTCTTTCCCTACACGACGCTCTTCCGATCTNNNACATTCTGAGAGAGAGGGACA
AGAGCACACGTCTGAACTCTTTCCCTACACGACGCTCTTCCGATCTNNNAGCTAGCACCATGGGAAAA
AAGAGCACACGTCTGAACTCTTTCCCTACACGACGCTCTTCCGATCTNNNGAATTGGGACTTTGGGTGA
GAGCACACGTCTGAACTCTTTCCCTACACGACGCTCTTCCGATCTNNNAAACTCCTAAGAGGGAGGCTC
AGCACACGTCTGAACTCTTTCCCTACACGACGCTCTTCCGATCTNNNCAGGACTGTCAAGCAGAGAAT
AGCACACGTCTGAACTCTTTCCCTACACGACGCTCTTCCGATCTNNNATCTTTCTTGCACAGGTCCACTCC
AGCACACGTCTGAACTCTTTCCCTACACGACGCTCTTCCGATCTNNNCCTGCATCTTCTGCTACCATCTC
GAAGAGCACACGTCTGAACTCTTTCCCTACACGACGCTCTTCCGATCTNNNACATCTTCATTGCCCAATA
AGAGCACACGTCTGAACTCTTTCCCTACACGACGCTCTTCCGATCTNNNCCTGACACAAAGACCTCACAC
GCACACGTCTGAACTCTTTCCCTACACGACGCTCTTCCGATCTNNNTGCAGTTCTTGTCCACCTAAAAGT
GAGCACACGTCTGAACTCTTTCCCTACACGACGCTCTTCCGATCTNNNTATTACTGGCCTCCCGTCCTAA
GAGCACACGTCTGAACTCTTTCCCTACACGACGCTCTTCCGATCTNNNAATGGCATAAAACACACACGC
GAGCACACGTCTGAACTCTTTCCCTACACGACGCTCTTCCGATCTNNNTGGTGAAGGCATTAGAACAAC
GAGCACACGTCTGAACTCTTTCCCTACACGACGCTCTTCCGATCTNNNCTGAATCAGCCATCAGGGAAG
AGAGCACACGTCTGAACTCTTTCCCTACACGACGCTCTTCCGATCTNNNTCAAATTTAGGTCAGATTCAC
GAGCACACGTCTGAACTCTTTCCCTACACGACGCTCTTCCGATCTNNNGCAATTGGAAAATGCTAAGGC
AGCACACGTCTGAACTCTTTCCCTACACGACGCTCTTCCGATCTNNNCCAAGGGGAAGCCAAGAGATT
GCACACGTCTGAACTCTTTCCCTACACGACGCTCTTCCGATCTNNNGTCTACGAAAATCACCGAGCACTC
AGAGCACACGTCTGAACTCTTTCCCTACACGACGCTCTTCCGATCTNNNGGAAGGCTTTCCTGGAGAAC
AGCACACGTCTGAACTCTTTCCCTACACGACGCTCTTCCGATCTNNNGTGGGTAAATTCTCAGCCTTGC
AGCACACGTCTGAACTCTTTCCCTACACGACGCTCTTCCGATCTNNNATTTCTCTACCTTCCCAGGCCATC
GAGCACACGTCTGAACTCTTTCCCTACACGACGCTCTTCCGATCTNNNTGTCATGGGGGTCTGTCTTGTA
AGCACACGTCTGAACTCTTTCCCTACACGACGCTCTTCCGATCTNNNGAGGGCCAATTCAGAAGCCAT
GAGCACACGTCTGAACTCTTTCCCTACACGACGCTCTTCCGATCTNNNAGCAAGAGAGTACGACATTG
GAGCACACGTCTGAACTCTTTCCCTACACGACGCTCTTCCGATCTNNNCCTAAAACAACGTGTGTGTGT

AAGAGCACACGTCTGAACTCTTTCCCTACACGACGCTCTTCCGATCTNNNAGTCTTCTCTCCTGTCCCTTG
AGCACACGTCTGAACTCTTTCCCTACACGACGCTCTTCCGATCTNNNTAGTGAGCTTCCTACGTTGGCT
GCACACGTCTGAACTCTTTCCCTACACGACGCTCTTCCGATCTNNNTTTTCTGTTTCCTCAGTGAGCCCCA
GAGCACACGTCTGAACTCTTTCCCTACACGACGCTCTTCCGATCTNNNAAAGTCTGAAGCCAAAGAGG
GAGCACACGTCTGAACTCTTTCCCTACACGACGCTCTTCCGATCTNNNATGGAACAGAAAATGCACAAG
AGAGCACACGTCTGAACTCTTTCCCTACACGACGCTCTTCCGATCTNNNTGCAGGCTAAGTATGTTCCA
AGCACACGTCTGAACTCTTTCCCTACACGACGCTCTTCCGATCTNNNCTGGGTAAAGAGTGAGGCTGTG
AGCACACGTCTGAACTCTTTCCCTACACGACGCTCTTCCGATCTNNNTGTAGCTGAGGGAGAATGCTA
GAGCACACGTCTGAACTCTTTCCCTACACGACGCTCTTCCGATCTNNNGAAAGAGTTCCAATGGCCAA
AGCACACGTCTGAACTCTTTCCCTACACGACGCTCTTCCGATCTNNNACTTTGCATAAGCACACCACAA
AGCACACGTCTGAACTCTTTCCCTACACGACGCTCTTCCGATCTNNNTAGATCAATGCGCAGTCTGGTT
AGAGCACACGTCTGAACTCTTTCCCTACACGACGCTCTTCCGATCTNNNATACTGATCTCCATCCTCCTG
GAGCACACGTCTGAACTCTTTCCCTACACGACGCTCTTCCGATCTNNNCCAATTTGGCACTCTGTAGATA
AGAGCACACGTCTGAACTCTTTCCCTACACGACGCTCTTCCGATCTNNNTATCAGAAGCCTAAGGAGAG
GAGCACACGTCTGAACTCTTTCCCTACACGACGCTCTTCCGATCTNNNGGGGAGAACCTGATATAGT
AGCACACGTCTGAACTCTTTCCCTACACGACGCTCTTCCGATCTNNNTCCTCTAGTCACCAAAGAAACCA
GAGCACACGTCTGAACTCTTTCCCTACACGACGCTCTTCCGATCTNNNTCAGGACCCATGGTTTTCAGA
AGCACACGTCTGAACTCTTTCCCTACACGACGCTCTTCCGATCTNNNGTTTGCTGCTAAGTGTACTCCC
GAGCACACGTCTGAACTCTTTCCCTACACGACGCTCTTCCGATCTNNNGGCAGCAGCTAATGCTGAATA
AAGAGCACACGTCTGAACTCTTTCCCTACACGACGCTCTTCCGATCTNNNGGCAGCAGCTAATGCTGA
AGAGCACACGTCTGAACTCTTTCCCTACACGACGCTCTTCCGATCTNNNAGCCCATGCTAGTAGACCTC
AGAGCACACGTCTGAACTCTTTCCCTACACGACGCTCTTCCGATCTNNNAACTACTCTATGCGTGCCTCT
GAGCACACGTCTGAACTCTTTCCCTACACGACGCTCTTCCGATCTNNNAAGCCCTGTCAAGATTTCTGA
AGCACACGTCTGAACTCTTTCCCTACACGACGCTCTTCCGATCTNNNATTATTCTGCACAAGCAGAGGCA
AGAGCACACGTCTGAACTCTTTCCCTACACGACGCTCTTCCGATCTNNNAATGCATCATGGCACATCTAG
AGCACACGTCTGAACTCTTTCCCTACACGACGCTCTTCCGATCTNNNATCCTCAAAGCCAAATGTGGAG
AGCACACGTCTGAACTCTTTCCCTACACGACGCTCTTCCGATCTNNNGGGACCATGAGAAGCATGAAT
GAGCACACGTCTGAACTCTTTCCCTACACGACGCTCTTCCGATCTNNNTCCTTCTGCCTTTCAGGTTCAA
AGCACACGTCTGAACTCTTTCCCTACACGACGCTCTTCCGATCTNNNTGCAAAATTGGTATCCTGCTCTC
AGAGCACACGTCTGAACTCTTTCCCTACACGACGCTCTTCCGATCTNNNATGAAGCCAGGGTCCTAACT
AGCACACGTCTGAACTCTTTCCCTACACGACGCTCTTCCGATCTNNNATGTTAGGGCCCTCCAGAGAAA
GGAAGAGCACACGTCTGAACTCTTTCCCTACACGACGCTCTTCCGATCTNNNCATGACTGGCTTGCTTT
GAAGAGCACACGTCTGAACTCTTTCCCTACACGACGCTCTTCCGATCTNNNGCATTAGATTCTCACTGG
AGCACACGTCTGAACTCTTTCCCTACACGACGCTCTTCCGATCTNNNTGGTTTCCACATGTTCCCCAAAA
AGCACACGTCTGAACTCTTTCCCTACACGACGCTCTTCCGATCTNNNTAGGCACATCTTACATGGCAACT
AGCACACGTCTGAACTCTTTCCCTACACGACGCTCTTCCGATCTNNNCCTGGGTGTCCAGATTATGGATG
GAGCACACGTCTGAACTCTTTCCCTACACGACGCTCTTCCGATCTNNNTCCTGAAATTCTTGTTTACTGT
GAGCACACGTCTGAACTCTTTCCCTACACGACGCTCTTCCGATCTNNNTCTGTCCAGCCTCAAAGACCAT
GAGCACACGTCTGAACTCTTTCCCTACACGACGCTCTTCCGATCTNNNTTCACCATCTGCTAAGTGATGG
AGAGCACACGTCTGAACTCTTTCCCTACACGACGCTCTTCCGATCTNNNGCTCAAACCCAGATCTTCCAA
AGCACACGTCTGAACTCTTTCCCTACACGACGCTCTTCCGATCTNNNAGTGCCAATCTTCTCTTATGCTG
AGCACACGTCTGAACTCTTTCCCTACACGACGCTCTTCCGATCTNNNACAGTGGTGTGGACATTCGAA
GAGCACACGTCTGAACTCTTTCCCTACACGACGCTCTTCCGATCTNNNACTTCCTCCTCCCTCCAGAAAA

GAGCACACGTCTGAACTCTTTCCCTACACGACGCTCTTCCGATCTNNNTGAGGCATGTGAATGTCTTGA
GAGCACACGTCTGAACTCTTTCCCTACACGACGCTCTTCCGATCTNNNCAATCTGTCTATCTGGCCCCCTA
AGAGCACACGTCTGAACTCTTTCCCTACACGACGCTCTTCCGATCTNNNCTGCAGCTTCCCAGAAGCAA
AGCACACGTCTGAACTCTTTCCCTACACGACGCTCTTCCGATCTNNNCAAGGGTCAGTACAGAAAAGG
AGCACACGTCTGAACTCTTTCCCTACACGACGCTCTTCCGATCTNNNTCCTGTGACATAAGAAATGCGT
AAGAGCACACGTCTGAACTCTTTCCCTACACGACGCTCTTCCGATCTNNNGGCAGAGGCTAAAGAGGA
GCACACGTCTGAACTCTTTCCCTACACGACGCTCTTCCGATCTNNNTATGTTGACGCTCAAACCACACTC
GCACACGTCTGAACTCTTTCCCTACACGACGCTCTTCCGATCTNNNCCACAGAAGACAGTGTCCACAAA
AGAGCACACGTCTGAACTCTTTCCCTACACGACGCTCTTCCGATCTNNNGATGCAGTTTGATCCCTGGC
GAGCACACGTCTGAACTCTTTCCCTACACGACGCTCTTCCGATCTNNNAGTTTTGAATTCCTGGTTCTTC
GCACACGTCTGAACTCTTTCCCTACACGACGCTCTTCCGATCTNNNAAGCAGTGGTGCCTAAAGCTATC
AGAGCACACGTCTGAACTCTTTCCCTACACGACGCTCTTCCGATCTNNNGGCTGAGAACCCATGTGTGT
AGAGCACACGTCTGAACTCTTTCCCTACACGACGCTCTTCCGATCTNNNCTCTATTCTGCCACCCTCCTAC
AGCACACGTCTGAACTCTTTCCCTACACGACGCTCTTCCGATCTNNNACCATGTGGACTTGATGTTCATC
GAGCACACGTCTGAACTCTTTCCCTACACGACGCTCTTCCGATCTNNNTTCTTGACCTTGAGATCCGAC
GAGCACACGTCTGAACTCTTTCCCTACACGACGCTCTTCCGATCTNNNGTGAGCGTTCCTCCTCTTTTCT
AGCACACGTCTGAACTCTTTCCCTACACGACGCTCTTCCGATCTNNNTCATTCCCTTTTGAATGGTTGGT
GAAGAGCACACGTCTGAACTCTTTCCCTACACGACGCTCTTCCGATCTNNNACAAGGTACATCATGCCA
AGCACACGTCTGAACTCTTTCCCTACACGACGCTCTTCCGATCTNNNACAAAAGTCTCAGAGGAAAGCA
AGCACACGTCTGAACTCTTTCCCTACACGACGCTCTTCCGATCTNNNAAAGCTCTGCTTGGTCAGTTTGA
GCACACGTCTGAACTCTTTCCCTACACGACGCTCTTCCGATCTNNNAAAGTCAGTCTCTCTCTCTCCCTTC
GAGCACACGTCTGAACTCTTTCCCTACACGACGCTCTTCCGATCTNNNTGCAAGTACACTCTAATGTTCG
AGCACACGTCTGAACTCTTTCCCTACACGACGCTCTTCCGATCTNNNTCCCCATCAATCTCAATTGCTGTT
GAGCACACGTCTGAACTCTTTCCCTACACGACGCTCTTCCGATCTNNNCCTGGCTACATTTTGACACACC
GAGCACACGTCTGAACTCTTTCCCTACACGACGCTCTTCCGATCTNNNATTGTTTTCTGATGGGTGAGG
GAGCACACGTCTGAACTCTTTCCCTACACGACGCTCTTCCGATCTNNNGGGTGGGATAACAAGCAATA
CACACGTCTGAACTCTTTCCCTACACGACGCTCTTCCGATCTNNNTGCTTCTTCCCTCTTCTTCTTCACATC
AAGAGCACACGTCTGAACTCTTTCCCTACACGACGCTCTTCCGATCTNNNCAGAGATAGTGGGGCAGT
AGCACACGTCTGAACTCTTTCCCTACACGACGCTCTTCCGATCTNNNACCTTCTTCTGACCGCATTTGTTC
AGAGCACACGTCTGAACTCTTTCCCTACACGACGCTCTTCCGATCTNNNAGGTTTTAGGGAACTGGGT
GAGCACACGTCTGAACTCTTTCCCTACACGACGCTCTTCCGATCTNNNGCTTGATTGTTGTAAGGCCTG
AGCACACGTCTGAACTCTTTCCCTACACGACGCTCTTCCGATCTNNNCTGTAGACCTTGGGATTTGGCA
AGCACACGTCTGAACTCTTTCCCTACACGACGCTCTTCCGATCTNNNGAAATTTCTAGCCTCCCTGTGGC
AAGAGCACACGTCTGAACTCTTTCCCTACACGACGCTCTTCCGATCTNNNACAGAACCAACAGAGACAG
GAAGAGCACACGTCTGAACTCTTTCCCTACACGACGCTCTTCCGATCTNNNTTCCACAATCCCAGCTTTTA
GCACACGTCTGAACTCTTTCCCTACACGACGCTCTTCCGATCTNNNCACAGTGCCTGGCTTATAGACAA
GAGCACACGTCTGAACTCTTTCCCTACACGACGCTCTTCCGATCTNNNAAATGAGGCTGCCTTCTAGGG
GAGCACACGTCTGAACTCTTTCCCTACACGACGCTCTTCCGATCTNNNAGGGGTTTTTAGACCAATGCA
AGCACACGTCTGAACTCTTTCCCTACACGACGCTCTTCCGATCTNNNCTGCCTTGAATTCCATCAAGCCTC
GAGCACACGTCTGAACTCTTTCCCTACACGACGCTCTTCCGATCTNNNTCTTTGTCTTTCTGGCTGGCTT
AGAGCACACGTCTGAACTCTTTCCCTACACGACGCTCTTCCGATCTNNNTTTTTGATGCCTGCTGGATCT
AGCACACGTCTGAACTCTTTCCCTACACGACGCTCTTCCGATCTNNNCTGGTTTACAGATGGCAGTCTC
CACACGTCTGAACTCTTTCCCTACACGACGCTCTTCCGATCTNNNCAACTACCATTACACCCCAGTCCCAT

GAGCACACGTCTGAACTCTTTCCCTACACGACGCTCTTCCGATCTNNNTTTCTCCGTTTTTCAAACTGC
AGCACACGTCTGAACTCTTTCCCTACACGACGCTCTTCCGATCTNNNACACAGCAAGGAAAAGAACTG
AGCACACGTCTGAACTCTTTCCCTACACGACGCTCTTCCGATCTNNNTATGGCCTTTAGGCAAATGGC
AGCACACGTCTGAACTCTTTCCCTACACGACGCTCTTCCGATCTNNNTATCCTACAGGGAACAGGATG
GCACACGTCTGAACTCTTTCCCTACACGACGCTCTTCCGATCTNNNCCCATGCAACATAGGGGATGA
AGAGCACACGTCTGAACTCTTTCCCTACACGACGCTCTTCCGATCTNNNCCCACCAAAGAGTGGGAAA
GCACACGTCTGAACTCTTTCCCTACACGACGCTCTTCCGATCTNNNGTGGATACACTGTTGTTTGGTG
AGCACACGTCTGAACTCTTTCCCTACACGACGCTCTTCCGATCTNNNGGGGCCAGAAAATGACAAGA
GAGCACACGTCTGAACTCTTTCCCTACACGACGCTCTTCCGATCTNNNGGGGCTCTAACATATAACAT
GCACACGTCTGAACTCTTTCCCTACACGACGCTCTTCCGATCTNNNCCTGGTATCCATCTGGGTGACA
AGCACACGTCTGAACTCTTTCCCTACACGACGCTCTTCCGATCTNNNTGTAGCTCTCTGTGTATGTGTG
AGCACACGTCTGAACTCTTTCCCTACACGACGCTCTTCCGATCTNNNGCCAAATCAGGAATGCAATA
AGAGCACACGTCTGAACTCTTTCCCTACACGACGCTCTTCCGATCTNNNAACCTCTGGCTTTCAATCTC
AAGAGCACACGTCTGAACTCTTTCCCTACACGACGCTCTTCCGATCTNNNCTATACTCATGGCTCTGCA
GAGCACACGTCTGAACTCTTTCCCTACACGACGCTCTTCCGATCTNNNTGATTGATTCATTTGCTGTGC
GAGCACACGTCTGAACTCTTTCCCTACACGACGCTCTTCCGATCTNNNATGCCTGACAGGTTACCAAA
AGCACACGTCTGAACTCTTTCCCTACACGACGCTCTTCCGATCTNNNCCCTCTTTCCTCTCTTATGCTGC
AGCACACGTCTGAACTCTTTCCCTACACGACGCTCTTCCGATCTNNNGTCTCTGCCATACTTTTCCTGG
AGCACACGTCTGAACTCTTTCCCTACACGACGCTCTTCCGATCTNNNTTCATTTTGCTAGAGGCTCAG
AAGAGCACACGTCTGAACTCTTTCCCTACACGACGCTCTTCCGATCTNNNTGCTGCTGGGTATGTACC
AGCACACGTCTGAACTCTTTCCCTACACGACGCTCTTCCGATCTNNNAGTTAGCATTCAGCAGTCTCCA
GAGCACACGTCTGAACTCTTTCCCTACACGACGCTCTTCCGATCTNNNTGCCCTTGCCATTTATGTTTC
GCACACGTCTGAACTCTTTCCCTACACGACGCTCTTCCGATCTNNNAGGGTAGCAGGTAGCTGTATG
GAGCACACGTCTGAACTCTTTCCCTACACGACGCTCTTCCGATCTNNNGGCTCATAGTACATTTCCCCA
AGCACACGTCTGAACTCTTTCCCTACACGACGCTCTTCCGATCTNNNAGCATGAGAAAGTGAGAACC
GCACACGTCTGAACTCTTTCCCTACACGACGCTCTTCCGATCTNNNGAGTTCCATGGACACATGCACA
AGCACACGTCTGAACTCTTTCCCTACACGACGCTCTTCCGATCTNNNTGTGGTAGGAGGATCCTTGA
GCACACGTCTGAACTCTTTCCCTACACGACGCTCTTCCGATCTNNNTGCCCTTGCCATTTATGTTTCCT
GAGCACACGTCTGAACTCTTTCCCTACACGACGCTCTTCCGATCTNNNAGAGAGGAACTATGCCTCA
AGCACACGTCTGAACTCTTTCCCTACACGACGCTCTTCCGATCTNNNTGAAGGCAGCTAAATGCTGAA
AGCACACGTCTGAACTCTTTCCCTACACGACGCTCTTCCGATCTNNNGAAAGCAGCATCTGGGAAGG
GAGCACACGTCTGAACTCTTTCCCTACACGACGCTCTTCCGATCTNNNCACATTGGTTGCCCAGGAGA
GCACACGTCTGAACTCTTTCCCTACACGACGCTCTTCCGATCTNNNAACAGTCCACAAAACTGTACAC
AGCACACGTCTGAACTCTTTCCCTACACGACGCTCTTCCGATCTNNNAGGGATTCCCCAAAGGAATCA
AGCACACGTCTGAACTCTTTCCCTACACGACGCTCTTCCGATCTNNNTCACAAGAAAGATTCTGGTCTC
GAGCACACGTCTGAACTCTTTCCCTACACGACGCTCTTCCGATCTNNNTGCAGAATCAACTTGAAGCA
AGCACACGTCTGAACTCTTTCCCTACACGACGCTCTTCCGATCTNNNTAAAATCCAGTGCCCTCCCACT
GCACACGTCTGAACTCTTTCCCTACACGACGCTCTTCCGATCTNNNCTGATGCTGAAGGGTAGAGGA
GAGCACACGTCTGAACTCTTTCCCTACACGACGCTCTTCCGATCTNNNGGTGTTGTAGGATTCGGTT
AGCACACGTCTGAACTCTTTCCCTACACGACGCTCTTCCGATCTNNNGCTTAGGCCAAAGAAGTAAG
GCACACGTCTGAACTCTTTCCCTACACGACGCTCTTCCGATCTNNNGGTGGGCTAAGTTTCTGTCTCT
AAGAGCACACGTCTGAACTCTTTCCCTACACGACGCTCTTCCGATCTNNNGTCAAGATTTCAGGGAT
GCACACGTCTGAACTCTTTCCCTACACGACGCTCTTCCGATCTNNNACGCATCTCCCAAAATACACACA

GAGCACACGTCTGAACTCTTTCCCTACACGACGCTCTTCCGATCTNNNTCAGTCAGTTCTCAGAAGGGC
AGAGCACACGTCTGAACTCTTTCCCTACACGACGCTCTTCCGATCTNNNTCATGCAGAAGAGATGCCAA
GAGCACACGTCTGAACTCTTTCCCTACACGACGCTCTTCCGATCTNNNAAGCTCATCACCAGTGAGTAG
GAGCACACGTCTGAACTCTTTCCCTACACGACGCTCTTCCGATCTNNNTGTGTTTGGGAATAACCCAGT
CACACGTCTGAACTCTTTCCCTACACGACGCTCTTCCGATCTNNNCTCTTTACTCCCCGCAGCAAGCATC
AGCACACGTCTGAACTCTTTCCCTACACGACGCTCTTCCGATCTNNNAGTGAAGGAAAAACAGCAGGG
GAGCACACGTCTGAACTCTTTCCCTACACGACGCTCTTCCGATCTNNNCACCCAGGCCATAAACCTCAAA
GCACACGTCTGAACTCTTTCCCTACACGACGCTCTTCCGATCTNNNCCCACTCCTGTAATCTCAGCTTCCA
GAGCACACGTCTGAACTCTTTCCCTACACGACGCTCTTCCGATCTNNNCGGCATACCCTCCCTAAAGAAG
AGCACACGTCTGAACTCTTTCCCTACACGACGCTCTTCCGATCTNNNCAGCTTCACAACAAGGATGTTG
GAGCACACGTCTGAACTCTTTCCCTACACGACGCTCTTCCGATCTNNNTGTTCATATTTTGCCCCACTTG
AAGAGCACACGTCTGAACTCTTTCCCTACACGACGCTCTTCCGATCTNNNGTGAAGTGAGGGAAACCC
AGCACACGTCTGAACTCTTTCCCTACACGACGCTCTTCCGATCTNNNCGCTTCATGTCTCAGATTTGTCC
AGCACACGTCTGAACTCTTTCCCTACACGACGCTCTTCCGATCTNNNCTGGATCTTGGACACTGTCTGG
AGCACACGTCTGAACTCTTTCCCTACACGACGCTCTTCCGATCTNNNAGGAGGTATTTGTACTGGGTG
AGCACACGTCTGAACTCTTTCCCTACACGACGCTCTTCCGATCTNNNGGTAAACCATCATCACCATTGCC
GAGCACACGTCTGAACTCTTTCCCTACACGACGCTCTTCCGATCTNNNGCAAGTTACCATATTTATCATCT
GAGCACACGTCTGAACTCTTTCCCTACACGACGCTCTTCCGATCTNNNCCCCGACCCTACATAGTACACA
GAGCACACGTCTGAACTCTTTCCCTACACGACGCTCTTCCGATCTNNNGCAGTGCTTGGCAATTGAACTA
GAGCACACGTCTGAACTCTTTCCCTACACGACGCTCTTCCGATCTNNNGCAAACTCAGGGTGATCAAGC
AGAGCACACGTCTGAACTCTTTCCCTACACGACGCTCTTCCGATCTNNNTATAGCTCACACTCTCTCTCT
AGCACACGTCTGAACTCTTTCCCTACACGACGCTCTTCCGATCTNNNCATGCTTCCATTGCACCAAACATG
AGCACACGTCTGAACTCTTTCCCTACACGACGCTCTTCCGATCTNNNGGGCTTTGAACAGTGTCTACTG
AGCACACGTCTGAACTCTTTCCCTACACGACGCTCTTCCGATCTNNNATCTGAGAATCCCAAATGGCACC
AGCACACGTCTGAACTCTTTCCCTACACGACGCTCTTCCGATCTNNNAGCAAAGAGCATAACTAGGGCA
GAGCACACGTCTGAACTCTTTCCCTACACGACGCTCTTCCGATCTNNNTTTAGATGGGATTCCGCTTGG
GCACACGTCTGAACTCTTTCCCTACACGACGCTCTTCCGATCTNNNCAAGGGCATTCTTTTGCCTTCAGC
AGCACACGTCTGAACTCTTTCCCTACACGACGCTCTTCCGATCTNNNTGCTTAGATTCAGTGGACAATG
GAGCACACGTCTGAACTCTTTCCCTACACGACGCTCTTCCGATCTNNNAGACAGAGCCTAAACATCCCCT
AGCACACGTCTGAACTCTTTCCCTACACGACGCTCTTCCGATCTNNNTGCCACTTGGTGTCTCCATTTAC
AGAGCACACGTCTGAACTCTTTCCCTACACGACGCTCTTCCGATCTNNNTGTTTCCAGGGGGTAAGGAA
AGCACACGTCTGAACTCTTTCCCTACACGACGCTCTTCCGATCTNNNTGGAACTGCTCTTTGTCTTGTTC
AGCACACGTCTGAACTCTTTCCCTACACGACGCTCTTCCGATCTNNNTTGCTTCTGTAAGAGATGCCAC
AGCACACGTCTGAACTCTTTCCCTACACGACGCTCTTCCGATCTNNNGGTGGGGCTCTATACTACAGGA
GAGCACACGTCTGAACTCTTTCCCTACACGACGCTCTTCCGATCTNNNCTCACCCTCCTCAGTTTCAGAG
GAGCACACGTCTGAACTCTTTCCCTACACGACGCTCTTCCGATCTNNNCTTCCACACTCCCAAACTCATCC
AGCACACGTCTGAACTCTTTCCCTACACGACGCTCTTCCGATCTNNNGATCCTGGTGGGGTGATTCCTA
GAGCACACGTCTGAACTCTTTCCCTACACGACGCTCTTCCGATCTNNNACCAGTCAAATTAGAAGCGCTG
GAGCACACGTCTGAACTCTTTCCCTACACGACGCTCTTCCGATCTNNNAAGCTCTCCTAGCGTCAGAGT
GAGCACACGTCTGAACTCTTTCCCTACACGACGCTCTTCCGATCTNNNTCCACTGGGATCTCAGAAATGG
GAGCACACGTCTGAACTCTTTCCCTACACGACGCTCTTCCGATCTNNNCGGTAAGCTCCGAGTAGAGTT
GAGCACACGTCTGAACTCTTTCCCTACACGACGCTCTTCCGATCTNNNCAGCTAGGACCACAGGTAAG
AGCACACGTCTGAACTCTTTCCCTACACGACGCTCTTCCGATCTNNNTAAAGCTCCTCTCACCAGATGGC

AGCACACGTCTGAACTCTTTCCCTACACGACGCTCTTCCGATCTNNNCCTCCGTGAGGAAGATTATGC
GAAGAGCACACGTCTGAACTCTTTCCCTACACGACGCTCTTCCGATCTNNNCTACCTCCAGGGAAGAG
GCACACGTCTGAACTCTTTCCCTACACGACGCTCTTCCGATCTNNNGCTGAGAATCAGAGGCAAAAT
GAGCACACGTCTGAACTCTTTCCCTACACGACGCTCTTCCGATCTNNNGTTCCCTTTGCAATGGATGT
GCACACGTCTGAACTCTTTCCCTACACGACGCTCTTCCGATCTNNNTGAAACAATGGCCTTTGGAGC
GCACACGTCTGAACTCTTTCCCTACACGACGCTCTTCCGATCTNNNAAAGCTACAAGTCCCAGTGCAA
AGCACACGTCTGAACTCTTTCCCTACACGACGCTCTTCCGATCTNNNCATCTGCATTATCTCAGGGTC
AGAGCACACGTCTGAACTCTTTCCCTACACGACGCTCTTCCGATCTNNNACGAAACTGGACTGACGAA
AGCACACGTCTGAACTCTTTCCCTACACGACGCTCTTCCGATCTNNNTCCGACTTCCACTTGCAATACA
AGCACACGTCTGAACTCTTTCCCTACACGACGCTCTTCCGATCTNNNGTATGCCCAAGTCAGCAAAAT
GAGCACACGTCTGAACTCTTTCCCTACACGACGCTCTTCCGATCTNNNCAGAATCCTGGTCTCTGAAG
AGCACACGTCTGAACTCTTTCCCTACACGACGCTCTTCCGATCTNNNGTTAGTTCCAGCTTGGAGGTC
GAAGAGCACACGTCTGAACTCTTTCCCTACACGACGCTCTTCCGATCTNNNTTGCAAACACATGACTT
AGCACACGTCTGAACTCTTTCCCTACACGACGCTCTTCCGATCTNNNCTCCCTCCTGGGGGAAAAGA
GCACACGTCTGAACTCTTTCCCTACACGACGCTCTTCCGATCTNNNGATACCCTATAGGCGGTTTCCA
ACACGTCTGAACTCTTTCCCTACACGACGCTCTTCCGATCTNNNTGGCCAATCATAGATCCTTCCTGCA
AGAGCACACGTCTGAACTCTTTCCCTACACGACGCTCTTCCGATCTNNNTGTCACTATCACTGGTTTT
AGCACACGTCTGAACTCTTTCCCTACACGACGCTCTTCCGATCTNNNCCTTCTCCTCTCTCTCTGTCTGG
AGCACACGTCTGAACTCTTTCCCTACACGACGCTCTTCCGATCTNNNCTCCAGCTCCCACTGTAGACT
GAGCACACGTCTGAACTCTTTCCCTACACGACGCTCTTCCGATCTNNNCCTGGGTCTCCAGCTTACTGA
GCACACGTCTGAACTCTTTCCCTACACGACGCTCTTCCGATCTNNNTGCAGACAGAATGAAGCCAAG
GCACACGTCTGAACTCTTTCCCTACACGACGCTCTTCCGATCTNNNCTGTGAACTGCACACGTGAGTG
CACGTCTGAACTCTTTCCCTACACGACGCTCTTCCGATCTNNNAATACCTCATCACAGCCTCAACCCAT
AGAGCACACGTCTGAACTCTTTCCCTACACGACGCTCTTCCGATCTNNNCCTGACAGTTGGACTGGG
GCACACGTCTGAACTCTTTCCCTACACGACGCTCTTCCGATCTNNNCTCATTCAGGTCCCCAAGGGAG
GAGCACACGTCTGAACTCTTTCCCTACACGACGCTCTTCCGATCTNNNAAGGGATCTAGGGTCCTGT
AGCACACGTCTGAACTCTTTCCCTACACGACGCTCTTCCGATCTNNNACGTGGATTTTATGCATTTGT
AGCACACGTCTGAACTCTTTCCCTACACGACGCTCTTCCGATCTNNNGGGGTAAGGCAAATCAAAA
AGCACACGTCTGAACTCTTTCCCTACACGACGCTCTTCCGATCTNNNGGCTGGTGTCACCTCTTACATT
AGCACACGTCTGAACTCTTTCCCTACACGACGCTCTTCCGATCTNNNAGCATGGTCCTAGAATTTCCTC
CACACGTCTGAACTCTTTCCCTACACGACGCTCTTCCGATCTNNNTCATTAGACTTCTCGTGGTTGCG
GAGCACACGTCTGAACTCTTTCCCTACACGACGCTCTTCCGATCTNNNTGTGTGCATGTGTGAGCATA
GAGCACACGTCTGAACTCTTTCCCTACACGACGCTCTTCCGATCTNNNAAGTCCCTAAGGAGGGCTA
AGCACACGTCTGAACTCTTTCCCTACACGACGCTCTTCCGATCTNNNAGAGCCTTGTGTGTTATGGC
AGCACACGTCTGAACTCTTTCCCTACACGACGCTCTTCCGATCTNNNTAATCAGAACATCCCACAGTG
AGCACACGTCTGAACTCTTTCCCTACACGACGCTCTTCCGATCTNNNTGCCATGCCTTACACACACTA
AGCACACGTCTGAACTCTTTCCCTACACGACGCTCTTCCGATCTNNNAGTTGAAAGAGATAGATATA
AGCACACGTCTGAACTCTTTCCCTACACGACGCTCTTCCGATCTNNNATTCCACATTCATGCCCACAGT
GAGCACACGTCTGAACTCTTTCCCTACACGACGCTCTTCCGATCTNNNTGGATGTGAGCAACTAGAG
GCACACGTCTGAACTCTTTCCCTACACGACGCTCTTCCGATCTNNNCAACATTGCAGCAAACAGCCAT
AGCACACGTCTGAACTCTTTCCCTACACGACGCTCTTCCGATCTNNNCTGAACCTGAATGTCTGGCAT
AGCACACGTCTGAACTCTTTCCCTACACGACGCTCTTCCGATCTNNNCCTCCTTTGAGCAACCTTTGAG
CACACGTCTGAACTCTTTCCCTACACGACGCTCTTCCGATCTNNNTGTGTAGTACTCCATTGGCTGGA

AGCACACGTCTGAACTCTTTCCCTACACGACGCTCTTCCGATCTNNNAAATAGCCCATAGTGTGTGTGG
AGCACACGTCTGAACTCTTTCCCTACACGACGCTCTTCCGATCTNNNAGAGAACCACTGCTGCCAGTAT
AGCACACGTCTGAACTCTTTCCCTACACGACGCTCTTCCGATCTNNNGGCAGGAAAAGGAGAGAAA
GAGCACACGTCTGAACTCTTTCCCTACACGACGCTCTTCCGATCTNNNTTACAAGCCACAGTACCACGT
AGCACACGTCTGAACTCTTTCCCTACACGACGCTCTTCCGATCTNNNAACCTCAACAAATATGCCTTCAG
AGAGCACACGTCTGAACTCTTTCCCTACACGACGCTCTTCCGATCTNNNTTTTGATACCCCTACCCCTCT
AGCACACGTCTGAACTCTTTCCCTACACGACGCTCTTCCGATCTNNNTCATAAGTGTGTGCCTATGTGT
GAGCACACGTCTGAACTCTTTCCCTACACGACGCTCTTCCGATCTNNNCTATCAAACAAATGCCGAGGT
GAGCACACGTCTGAACTCTTTCCCTACACGACGCTCTTCCGATCTNNNATCCTCTTGCCTCGGCCTATAA
AGCACACGTCTGAACTCTTTCCCTACACGACGCTCTTCCGATCTNNNCCATCCTACACACAGCAACACA
AGCACACGTCTGAACTCTTTCCCTACACGACGCTCTTCCGATCTNNNTGTAAGTGTAGGCCTCTCTCTG
AGCACACGTCTGAACTCTTTCCCTACACGACGCTCTTCCGATCTNNNGCACACGCTGTCCTGTTGTTTA
AAGAGCACACGTCTGAACTCTTTCCCTACACGACGCTCTTCCGATCTNNNGGCTTTGTTTCTTACTCTTC
AGCACACGTCTGAACTCTTTCCCTACACGACGCTCTTCCGATCTNNNGTGTCAAAGAGAGCCAAGGCT
AGCACACGTCTGAACTCTTTCCCTACACGACGCTCTTCCGATCTNNNAGAAAATATGAGTGCACATGCC
GAGCACACGTCTGAACTCTTTCCCTACACGACGCTCTTCCGATCTNNNGCCAAAGAATCAAAACCGAC
GAGCACACGTCTGAACTCTTTCCCTACACGACGCTCTTCCGATCTNNNCCTCTTTTGCGCTCTCTCTGTA
AGCACACGTCTGAACTCTTTCCCTACACGACGCTCTTCCGATCTNNNTCAATGCTACCTTTGATGTGGA
AGCACACGTCTGAACTCTTTCCCTACACGACGCTCTTCCGATCTNNNACATACCTCTGTTCTGGCTCTGT
GAGCACACGTCTGAACTCTTTCCCTACACGACGCTCTTCCGATCTNNNACCAGGTACCCCATTCAACAT
AGAGCACACGTCTGAACTCTTTCCCTACACGACGCTCTTCCGATCTNNNGGACTATAAAAGCCCTGGG
AGCACACGTCTGAACTCTTTCCCTACACGACGCTCTTCCGATCTNNNAGAACCAGATTGAGAATGTAA
GCACACGTCTGAACTCTTTCCCTACACGACGCTCTTCCGATCTNNNATCCCTAGTGGACGACTGAAACT
AGCACACGTCTGAACTCTTTCCCTACACGACGCTCTTCCGATCTNNNCTGAGCCTTGCCAGAAGAGTAA
AGCACACGTCTGAACTCTTTCCCTACACGACGCTCTTCCGATCTNNNTGAGTAGCTAGTACCATGCCCT
AGCACACGTCTGAACTCTTTCCCTACACGACGCTCTTCCGATCTNNNCGTTGGAATTCACACATCTCAG
GAGCACACGTCTGAACTCTTTCCCTACACGACGCTCTTCCGATCTNNNCCACACCAGCCACTAGTAAAG
AGCACACGTCTGAACTCTTTCCCTACACGACGCTCTTCCGATCTNNNACAGTCTCTCTCTTTCTCACCCC
AAGAGCACACGTCTGAACTCTTTCCCTACACGACGCTCTTCCGATCTNNNGACCATCCTCTTTAGCTCAG
AGAGCACACGTCTGAACTCTTTCCCTACACGACGCTCTTCCGATCTNNNCACATAATTCCCAGAAGGT
GCACACGTCTGAACTCTTTCCCTACACGACGCTCTTCCGATCTNNNTCTCAAAGAGGCACACAAAGAC
GAGCACACGTCTGAACTCTTTCCCTACACGACGCTCTTCCGATCTNNNACCTTGGAGTGGTTTTGGTA
AGCACACGTCTGAACTCTTTCCCTACACGACGCTCTTCCGATCTNNNTTCCCAGAGGTAGCTGATTTGG
AGCACACGTCTGAACTCTTTCCCTACACGACGCTCTTCCGATCTNNNGGAGAGAAAAAGGGAGTAGG
AGAGCACACGTCTGAACTCTTTCCCTACACGACGCTCTTCCGATCTNNNCCTTGGCCCAATCAAGCCATT
AGAGCACACGTCTGAACTCTTTCCCTACACGACGCTCTTCCGATCTNNNAGTTTCTTGTGAGAATCTAG
AGAGCACACGTCTGAACTCTTTCCCTACACGACGCTCTTCCGATCTNNNAAAGGAACCAGCACATGAG
AGCACACGTCTGAACTCTTTCCCTACACGACGCTCTTCCGATCTNNNACATTCGAAATCAAGCTGTTGG
GCACACGTCTGAACTCTTTCCCTACACGACGCTCTTCCGATCTNNNGTCTGTGTCTCTCAGTTGTGCATC
AGCACACGTCTGAACTCTTTCCCTACACGACGCTCTTCCGATCTNNNACATGCATACACTTCTGTCTTCA
GAGCACACGTCTGAACTCTTTCCCTACACGACGCTCTTCCGATCTNNNAGGGACCAGTATTGTCACTCC
AGCACACGTCTGAACTCTTTCCCTACACGACGCTCTTCCGATCTNNNTTTGAACAGAACAGGGCTTGT
GCACACGTCTGAACTCTTTCCCTACACGACGCTCTTCCGATCTNNNACTCTCCCTCTTTTTCCTTCCCCCA

CACACGTCTGAACTCTTTCCCTACACGACGCTCTTCCGATCTNNNTCCTCATTTCTACCTGCAACGTGC
GCACACGTCTGAACTCTTTCCCTACACGACGCTCTTCCGATCTNNNTGCAAACTAACCTCACACCCTTC
GAGCACACGTCTGAACTCTTTCCCTACACGACGCTCTTCCGATCTNNNAGTGCCCAGGAAAATACTTC
AGCACACGTCTGAACTCTTTCCCTACACGACGCTCTTCCGATCTNNNTCATAAAGAGGGAGAAGCCTT
AAGAGCACACGTCTGAACTCTTTCCCTACACGACGCTCTTCCGATCTNNNCTTCGCACAGAGAGAAAG
GAGCACACGTCTGAACTCTTTCCCTACACGACGCTCTTCCGATCTNNNTACTCTGAAATGGTTCCCAA
AGCACACGTCTGAACTCTTTCCCTACACGACGCTCTTCCGATCTNNNCCCAGCTGTTTGATTGTTAT
GAGCACACGTCTGAACTCTTTCCCTACACGACGCTCTTCCGATCTNNNAATTTGGGTTGACTACCACCA
GAGCACACGTCTGAACTCTTTCCCTACACGACGCTCTTCCGATCTNNNTTGACAGAAAAGGGACAAA
GCACACGTCTGAACTCTTTCCCTACACGACGCTCTTCCGATCTNNNTGGCATTGTTTGTTTGCCTAAT
ACACGTCTGAACTCTTTCCCTACACGACGCTCTTCCGATCTNNNCACACATTTTACTCCGCTCTCCACAT
AGCACACGTCTGAACTCTTTCCCTACACGACGCTCTTCCGATCTNNNACTACTGGGTCCTGGGATGT
AGCACACGTCTGAACTCTTTCCCTACACGACGCTCTTCCGATCTNNNGTAGTCCCACATGTACACCAA
GCACACGTCTGAACTCTTTCCCTACACGACGCTCTTCCGATCTNNNCTCCACCGTCTGTGTATCGGTA
AGCACACGTCTGAACTCTTTCCCTACACGACGCTCTTCCGATCTNNNAGCCTAATCAAAGCCATTTGT
GAGCACACGTCTGAACTCTTTCCCTACACGACGCTCTTCCGATCTNNNAAAGCCATCTGTTACAGCACT
AGCACACGTCTGAACTCTTTCCCTACACGACGCTCTTCCGATCTNNNAGGCCACTCAACCATTTCATT
AGCACACGTCTGAACTCTTTCCCTACACGACGCTCTTCCGATCTNNNGGCTATTGCCACACGTTTTTC
AGCACACGTCTGAACTCTTTCCCTACACGACGCTCTTCCGATCTNNNAGAACATAGATCTCTCCCCCTG
AGCACACGTCTGAACTCTTTCCCTACACGACGCTCTTCCGATCTNNNAGAGGTGTATTCCCCTTTGAG
GCACACGTCTGAACTCTTTCCCTACACGACGCTCTTCCGATCTNNNATTAGGCTTTGTTTGTCCCAGCA
GCACACGTCTGAACTCTTTCCCTACACGACGCTCTTCCGATCTNNNTCAAGGGTATTTGAAGTTTGGG
AGCACACGTCTGAACTCTTTCCCTACACGACGCTCTTCCGATCTNNNAGGGATCTGCTTCTGAACTGG
AGCACACGTCTGAACTCTTTCCCTACACGACGCTCTTCCGATCTNNNCTCAATAAGCTCTCACTGCCTC
AAGAGCACACGTCTGAACTCTTTCCCTACACGACGCTCTTCCGATCTNNNATCCAAAATGCCCAAGCA
AGCACACGTCTGAACTCTTTCCCTACACGACGCTCTTCCGATCTNNNAGCTGATTTTTACTGCGTCTC
GCACACGTCTGAACTCTTTCCCTACACGACGCTCTTCCGATCTNNNTCCTTACAGCAAAGCAAAACGG
AGCACACGTCTGAACTCTTTCCCTACACGACGCTCTTCCGATCTNNNTGATGTCCTCGTAAGCAAACT
AGCACACGTCTGAACTCTTTCCCTACACGACGCTCTTCCGATCTNNNAGCAGTTGGATTTGTCTTCCCA
GAGCACACGTCTGAACTCTTTCCCTACACGACGCTCTTCCGATCTNNNAAGTCATGGCACATAGATG
GAGCACACGTCTGAACTCTTTCCCTACACGACGCTCTTCCGATCTNNNGCTCAACATTGGCCTGCTTT
AGCACACGTCTGAACTCTTTCCCTACACGACGCTCTTCCGATCTNNNTGCTTCCTTCCCAATCACTTTC
AGCACACGTCTGAACTCTTTCCCTACACGACGCTCTTCCGATCTNNNAAACCATGACTGATCTATCCAG
AGCACACGTCTGAACTCTTTCCCTACACGACGCTCTTCCGATCTNNNACTGCCCTCATACCAACTCTCAA
GAGCACACGTCTGAACTCTTTCCCTACACGACGCTCTTCCGATCTNNNAGAGAACCACTGCTGCCAGT
AGCACACGTCTGAACTCTTTCCCTACACGACGCTCTTCCGATCTNNNATCTGGTAACTCTGGTGTGAG
GAGCACACGTCTGAACTCTTTCCCTACACGACGCTCTTCCGATCTNNNAGCTTCAAGTCCTTCAAGACA
GCACACGTCTGAACTCTTTCCCTACACGACGCTCTTCCGATCTNNNAAGTTGGAGAGGAGGGAAACT
AGCACACGTCTGAACTCTTTCCCTACACGACGCTCTTCCGATCTNNNATGGAAAATGGGATACCCCAG
AGCACACGTCTGAACTCTTTCCCTACACGACGCTCTTCCGATCTNNNCCTTATCTCCTACCCATCCCCAC
GAGCACACGTCTGAACTCTTTCCCTACACGACGCTCTTCCGATCTNNNGGATGAGGACACACATGCTA
GCACACGTCTGAACTCTTTCCCTACACGACGCTCTTCCGATCTNNNGGTAAGCAGCCTGAAAGCATTT
AGCACACGTCTGAACTCTTTCCCTACACGACGCTCTTCCGATCTNNNTCAGATGCCCCAGCTGAATGTA

AGCACACGTCTGAACTCTTTCCCTACACGACGCTCTTCCGATCTNNNGGTTTCTCTTGTCCCCCAACATA
AGCACACGTCTGAACTCTTTCCCTACACGACGCTCTTCCGATCTNNNTACCCTCAACCTACAGAAGCACA
AGCACACGTCTGAACTCTTTCCCTACACGACGCTCTTCCGATCTNNNTTTACTCCTTAGACACCACCCCC
AGAGCACACGTCTGAACTCTTTCCCTACACGACGCTCTTCCGATCTNNNCCAAATGTTGCCCTTTGGTTC
AGCACACGTCTGAACTCTTTCCCTACACGACGCTCTTCCGATCTNNNATTACCAACATTGCCTTTTGCCCC
AGAGCACACGTCTGAACTCTTTCCCTACACGACGCTCTTCCGATCTNNNTTTGTCATAGAGACAGGGCA
AGAGCACACGTCTGAACTCTTTCCCTACACGACGCTCTTCCGATCTNNNGACTGGTACCACAGGAGTTC
GCACACGTCTGAACTCTTTCCCTACACGACGCTCTTCCGATCTNNNGTACAAGGGGCTCAAGTCGTTTA
GCACACGTCTGAACTCTTTCCCTACACGACGCTCTTCCGATCTNNNTGCCTATTTGGGCTTCTCTCTCTC
AGAGCACACGTCTGAACTCTTTCCCTACACGACGCTCTTCCGATCTNNNTGAGATGTACCAAATCCCTGT
AGCACACGTCTGAACTCTTTCCCTACACGACGCTCTTCCGATCTNNNGCGGGCATACATATTCAGTCCA
GAGCACACGTCTGAACTCTTTCCCTACACGACGCTCTTCCGATCTNNNGCCTATATTCACAGACGCCTGA
GAGCACACGTCTGAACTCTTTCCCTACACGACGCTCTTCCGATCTNNNTTGGGTGATGTAGCCAGAAC
AGCACACGTCTGAACTCTTTCCCTACACGACGCTCTTCCGATCTNNNTTTCCTGTAAGGATGTGAGTGA
AGAGCACACGTCTGAACTCTTTCCCTACACGACGCTCTTCCGATCTNNNTCTCTGGCTCATCTTCTCCTA
AGAGCACACGTCTGAACTCTTTCCCTACACGACGCTCTTCCGATCTNNNAAAGCAGACAAACCATTTGCC
AAGAGCACACGTCTGAACTCTTTCCCTACACGACGCTCTTCCGATCTNNNGCAACACAGAAGACAGTCT
GAGCACACGTCTGAACTCTTTCCCTACACGACGCTCTTCCGATCTNNNCGACAAAGCAAAACACTGTCT
AGAGCACACGTCTGAACTCTTTCCCTACACGACGCTCTTCCGATCTNNNATGTGTGTAAAGGGGGCTTT
AGAGCACACGTCTGAACTCTTTCCCTACACGACGCTCTTCCGATCTNNNACGAGGCAAGAATGAAAGT
AGAGCACACGTCTGAACTCTTTCCCTACACGACGCTCTTCCGATCTNNNACAGGTCAGTAGGAATCGTT
AGCACACGTCTGAACTCTTTCCCTACACGACGCTCTTCCGATCTNNNGAACCCCCTGACATTCCACAATA
GCACACGTCTGAACTCTTTCCCTACACGACGCTCTTCCGATCTNNNCCCAGCTGAAAGATAACTGGTGA
GAGCACACGTCTGAACTCTTTCCCTACACGACGCTCTTCCGATCTNNNGTCTGTAAGGGGTCACCAATG
AGAGCACACGTCTGAACTCTTTCCCTACACGACGCTCTTCCGATCTNNNGGTTTCTTCCAAGGCTAGTGG
GAGCACACGTCTGAACTCTTTCCCTACACGACGCTCTTCCGATCTNNNGCACGAGTTTTGCTTTGTAGG
AGCACACGTCTGAACTCTTTCCCTACACGACGCTCTTCCGATCTNNNGATTGATGAGATCCTGCAAGGC
GAGCACACGTCTGAACTCTTTCCCTACACGACGCTCTTCCGATCTNNNTGGAGTGAAATCCTGGTACTC
AGCACACGTCTGAACTCTTTCCCTACACGACGCTCTTCCGATCTNNNCAGCCTTCTGCTTTCAGTCTACA
AGCACACGTCTGAACTCTTTCCCTACACGACGCTCTTCCGATCTNNNAGTGGGTGACAGAACCTAGAC
AGAGCACACGTCTGAACTCTTTCCCTACACGACGCTCTTCCGATCTNNNAACTTCCCCGTGTAACTACCTC
AAGAGCACACGTCTGAACTCTTTCCCTACACGACGCTCTTCCGATCTNNNTTTTCATGTGAGGACAGTC
AAGAGCACACGTCTGAACTCTTTCCCTACACGACGCTCTTCCGATCTNNNAGAGCAAAAGCCAAAATCA
AGAGCACACGTCTGAACTCTTTCCCTACACGACGCTCTTCCGATCTNNNTGATTGAATCCAACAATGCGT
GCACACGTCTGAACTCTTTCCCTACACGACGCTCTTCCGATCTNNNCCAGGAACATTACTGCCTTGGAA
AGAGCACACGTCTGAACTCTTTCCCTACACGACGCTCTTCCGATCTNNNATGAAGGACCCTTGTTCATG
GAGCACACGTCTGAACTCTTTCCCTACACGACGCTCTTCCGATCTNNNAAGGTGGTGTGTTAGGGTTC
GAGCACACGTCTGAACTCTTTCCCTACACGACGCTCTTCCGATCTNNNGGAGCACTCTGGAATTCTCTCT
AAGAGCACACGTCTGAACTCTTTCCCTACACGACGCTCTTCCGATCTNNNTGGTGGCAACAGAAAGTAC
AGAGCACACGTCTGAACTCTTTCCCTACACGACGCTCTTCCGATCTNNNTCTGTCAAGCATCATCTGCTT
GAGCACACGTCTGAACTCTTTCCCTACACGACGCTCTTCCGATCTNNNGTAGGGATGGGGGATTCTGA
GAGCACACGTCTGAACTCTTTCCCTACACGACGCTCTTCCGATCTNNNGTGGACTGGAAAATTCAGCC
GAGCACACGTCTGAACTCTTTCCCTACACGACGCTCTTCCGATCTNNNGAGCTCAGGACAATAACATG

AGCACACGTCTGAACTCTTTCCCTACACGACGCTCTTCCGATCTNNNTAGGGACTGAGACCAAAGGA
GAGCACACGTCTGAACTCTTTCCCTACACGACGCTCTTCCGATCTNNNTGAATGGCTTCAGAAGGTC
GAGCACACGTCTGAACTCTTTCCCTACACGACGCTCTTCCGATCTNNNTCATCTCCTCCTGAGAGATCC
AGCACACGTCTGAACTCTTTCCCTACACGACGCTCTTCCGATCTNNNTCCTTGGCCAATTCCACAACTTT
AGCACACGTCTGAACTCTTTCCCTACACGACGCTCTTCCGATCTNNNGCAGGGTCCTAGAGAAAACC
AGCACACGTCTGAACTCTTTCCCTACACGACGCTCTTCCGATCTNNNTGCAACAGTAAGGATCCATTT
AGCACACGTCTGAACTCTTTCCCTACACGACGCTCTTCCGATCTNNNACTCATTCAGTACATATCACTG
AGCACACGTCTGAACTCTTTCCCTACACGACGCTCTTCCGATCTNNNGATGAGTGTGAGTAGTGCTGC
GAGCACACGTCTGAACTCTTTCCCTACACGACGCTCTTCCGATCTNNNGAAGCAAAGGGTCATCAGG
GAGCACACGTCTGAACTCTTTCCCTACACGACGCTCTTCCGATCTNNNTGTCCAATGTTCCATAAGGG
GAGCACACGTCTGAACTCTTTCCCTACACGACGCTCTTCCGATCTNNNTGTGGCATTATGAAGAGTGT
GAGCACACGTCTGAACTCTTTCCCTACACGACGCTCTTCCGATCTNNNCCTCCAGGCAGCTTGCTTTTA
AGCACACGTCTGAACTCTTTCCCTACACGACGCTCTTCCGATCTNNNTATAAAAGAGGCTCCCCAAGG
GCACACGTCTGAACTCTTTCCCTACACGACGCTCTTCCGATCTNNNCTTCCCTCATTTACAGACCCTGC
GAGCACACGTCTGAACTCTTTCCCTACACGACGCTCTTCCGATCTNNNAGGCACACCTAAAATCCTAC
GAGCACACGTCTGAACTCTTTCCCTACACGACGCTCTTCCGATCTNNNTGTGTTTTTGTTTCTTGCCAA
AGCACACGTCTGAACTCTTTCCCTACACGACGCTCTTCCGATCTNNNCCCATCCTATCCCATCTTGTCCT
GCACACGTCTGAACTCTTTCCCTACACGACGCTCTTCCGATCTNNNATGGCGATGAAGTGAAGACAG
GAGCACACGTCTGAACTCTTTCCCTACACGACGCTCTTCCGATCTNNNGCAAGGTTCAAAAACTATGT
AGAGCACACGTCTGAACTCTTTCCCTACACGACGCTCTTCCGATCTNNNTGCTCAGATCTACAGGCCT
AGCACACGTCTGAACTCTTTCCCTACACGACGCTCTTCCGATCTNNNTACATACACACAAAAGGGTGC
AGCACACGTCTGAACTCTTTCCCTACACGACGCTCTTCCGATCTNNNGGGAGCCAAACTCGAAAAATA
AGCACACGTCTGAACTCTTTCCCTACACGACGCTCTTCCGATCTNNNTGCTTTGCTCATTATTTGTCTC
GAAGAGCACACGTCTGAACTCTTTCCCTACACGACGCTCTTCCGATCTNNNGACAAGACCTATAGCGC
AGCACACGTCTGAACTCTTTCCCTACACGACGCTCTTCCGATCTNNNGGACACTTCTCTTACCTGCCAA
AAGAGCACACGTCTGAACTCTTTCCCTACACGACGCTCTTCCGATCTNNNATGAGGAACCTGCAGATA
AAGAGCACACGTCTGAACTCTTTCCCTACACGACGCTCTTCCGATCTNNNTGCGCAATTGATCTGTTTG
GAGCACACGTCTGAACTCTTTCCCTACACGACGCTCTTCCGATCTNNNGGGCTCTGGAACTTTGAATT
GAGCACACGTCTGAACTCTTTCCCTACACGACGCTCTTCCGATCTNNNTGTAGGTCTTTTACCTCCCTG
GAGCACACGTCTGAACTCTTTCCCTACACGACGCTCTTCCGATCTNNNATACAGGCCTCTTGTGTGTA
GAGCACACGTCTGAACTCTTTCCCTACACGACGCTCTTCCGATCTNNNTTCTCCCATAGCCTTCTACCAC
GAGCACACGTCTGAACTCTTTCCCTACACGACGCTCTTCCGATCTNNNATTCCCCATACACACACACAAG
AGAGCACACGTCTGAACTCTTTCCCTACACGACGCTCTTCCGATCTNNNGCTGGTGGAGGTAGCATAG
AGAGCACACGTCTGAACTCTTTCCCTACACGACGCTCTTCCGATCTNNNCTCCACTCAAACCACTGGAG
AGAGCACACGTCTGAACTCTTTCCCTACACGACGCTCTTCCGATCTNNNTCATGCACTGCAAGGCATA
AGCACACGTCTGAACTCTTTCCCTACACGACGCTCTTCCGATCTNNNACTTATCTTGTTCAAATTTCCC
AGCACACGTCTGAACTCTTTCCCTACACGACGCTCTTCCGATCTNNNGCAGATGCAAAAAGGGTATGG
AGCACACGTCTGAACTCTTTCCCTACACGACGCTCTTCCGATCTNNNCTAGTGTGCAGATCCAAGAGA
GAGCACACGTCTGAACTCTTTCCCTACACGACGCTCTTCCGATCTNNNGCCTATATACATGCGCACACA
CACACGTCTGAACTCTTTCCCTACACGACGCTCTTCCGATCTNNNTGAATGAGGCTGTACCTTGCTTC
AGCACACGTCTGAACTCTTTCCCTACACGACGCTCTTCCGATCTNNNTTGTGGCTCAAACATTTGGCTT
AGAGCACACGTCTGAACTCTTTCCCTACACGACGCTCTTCCGATCTNNNGCACCCCCTTTTAGCATGTA
GAAGAGCACACGTCTGAACTCTTTCCCTACACGACGCTCTTCCGATCTNNNCTGTCATACAGGAAGA

AGCACACGTCTGAACTCTTTCCCTACACGACGCTCTTCCGATCTNNNCATGGAGCCACCTATTGCTTACA
AAGAGCACACGTCTGAACTCTTTCCCTACACGACGCTCTTCCGATCTNNNAGAACAGTCTCTGCATAACA
GAGCACACGTCTGAACTCTTTCCCTACACGACGCTCTTCCGATCTNNNGGTGCAGTTCTGGTCTTCTCAT
GCACACGTCTGAACTCTTTCCCTACACGACGCTCTTCCGATCTNNNAGGCTTTACATTCCCTGTGCTTC
AGCACACGTCTGAACTCTTTCCCTACACGACGCTCTTCCGATCTNNNTCGATGACATTCTCACCTCTGTC
AGCACACGTCTGAACTCTTTCCCTACACGACGCTCTTCCGATCTNNNTCACGTACATCCTCCTTCATCCAC
GAGCACACGTCTGAACTCTTTCCCTACACGACGCTCTTCCGATCTNNNTACTCATCCTGCAGACTTTGGG
AGCACACGTCTGAACTCTTTCCCTACACGACGCTCTTCCGATCTNNNACCAAGGAACTAGGGATTGGG
AGCACACGTCTGAACTCTTTCCCTACACGACGCTCTTCCGATCTNNNATTCCCTGGCAACCACATTCATA
GAGCACACGTCTGAACTCTTTCCCTACACGACGCTCTTCCGATCTNNNCGCAGCATTGTGTCATATGGA
AGAGCACACGTCTGAACTCTTTCCCTACACGACGCTCTTCCGATCTNNNTGAAGTAAAACATTCAAGGC
AGCACACGTCTGAACTCTTTCCCTACACGACGCTCTTCCGATCTNNNAGGCTTTACATTCCCTGTGCTT
GCACACGTCTGAACTCTTTCCCTACACGACGCTCTTCCGATCTNNNGTAACACAAAGCAGGGCTGAAG
GAGCACACGTCTGAACTCTTTCCCTACACGACGCTCTTCCGATCTNNNCCAAGTCGAAGTTGCCAAAGA
AGCACACGTCTGAACTCTTTCCCTACACGACGCTCTTCCGATCTNNNTAATGCCTTGGAGGGATTTGG
GAGCACACGTCTGAACTCTTTCCCTACACGACGCTCTTCCGATCTNNNCTTGCTCGGTCTCACTGTATTG
AGCACACGTCTGAACTCTTTCCCTACACGACGCTCTTCCGATCTNNNCCACACCTTGGAAATGGATTTGG
AGAGCACACGTCTGAACTCTTTCCCTACACGACGCTCTTCCGATCTNNNAGTACTAGAAGAAACACACT
GAGCACACGTCTGAACTCTTTCCCTACACGACGCTCTTCCGATCTNNNTCTATTGGAGAAGCCTGCCTA
GAGCACACGTCTGAACTCTTTCCCTACACGACGCTCTTCCGATCTNNNTGGATGCATCTTTGCTTTTCCC
AGCACACGTCTGAACTCTTTCCCTACACGACGCTCTTCCGATCTNNNTGTGGGATAGACACCAAAGTT
AGAGCACACGTCTGAACTCTTTCCCTACACGACGCTCTTCCGATCTNNNAGGACATTTTGAGTACTGCT
GAAGAGCACACGTCTGAACTCTTTCCCTACACGACGCTCTTCCGATCTNNNGAACCTGTGAGGTGCTAT
AGAGCACACGTCTGAACTCTTTCCCTACACGACGCTCTTCCGATCTNNNTAGGAATAGCCAGGCCAGAA
AGAGCACACGTCTGAACTCTTTCCCTACACGACGCTCTTCCGATCTNNNCTTCTCACATATGTCCAGCCT
AGCACACGTCTGAACTCTTTCCCTACACGACGCTCTTCCGATCTNNNAGAGAATCCTGTGTCATGATTG
GAGCACACGTCTGAACTCTTTCCCTACACGACGCTCTTCCGATCTNNNAGAAGCTTCAGATCCTGGAAA
GCACACGTCTGAACTCTTTCCCTACACGACGCTCTTCCGATCTNNNAAATATTTGCTGACGGCCTCCCCA
AGCACACGTCTGAACTCTTTCCCTACACGACGCTCTTCCGATCTNNNAATATCCTCTGTGTCCCTCTCCCC
AGCACACGTCTGAACTCTTTCCCTACACGACGCTCTTCCGATCTNNNACATGAACAAGGACTTCAATACA
AGCACACGTCTGAACTCTTTCCCTACACGACGCTCTTCCGATCTNNNGGGAGATAGGATGGAGACACC
GAGCACACGTCTGAACTCTTTCCCTACACGACGCTCTTCCGATCTNNNACCTCCGGAGACTTGATTCAC
GAGCACACGTCTGAACTCTTTCCCTACACGACGCTCTTCCGATCTNNNTACCTGTTGGATTATGAAGGG
AGAGCACACGTCTGAACTCTTTCCCTACACGACGCTCTTCCGATCTNNNATGCACATGTGTGTATGTGT
AAGAGCACACGTCTGAACTCTTTCCCTACACGACGCTCTTCCGATCTNNNTGGGGACCATTGTTCAACCA
GAGCACACGTCTGAACTCTTTCCCTACACGACGCTCTTCCGATCTNNNGGTCATCTCTCAGTCTGATT
AGAGCACACGTCTGAACTCTTTCCCTACACGACGCTCTTCCGATCTNNNACAATCTGTTACAATGGTGG
GAGCACACGTCTGAACTCTTTCCCTACACGACGCTCTTCCGATCTNNNTTTTCACCTTCCTGAGACTTG
AGCACACGTCTGAACTCTTTCCCTACACGACGCTCTTCCGATCTNNNTGTAGGCAGTCTCTCTCTCTCAG
AGCACACGTCTGAACTCTTTCCCTACACGACGCTCTTCCGATCTNNNAAACTAAAGTCAGTCACTGGCC
AAGAGCACACGTCTGAACTCTTTCCCTACACGACGCTCTTCCGATCTNNNGCACCCCCTTTTAGCATGTA
GAGCACACGTCTGAACTCTTTCCCTACACGACGCTCTTCCGATCTNNNCCGCAACCTCCTTTATTACTGGT
CACACGTCTGAACTCTTTCCCTACACGACGCTCTTCCGATCTNNNGTCAAGCACCAAATGGAAACCAAC

GAGCACACGTCTGAACTCTTTCCCTACACGACGCTCTTCCGATCTNNNAGTTGTGTGCGGATTTTGAC
AGAGCACACGTCTGAACTCTTTCCCTACACGACGCTCTTCCGATCTNNNACCAGAGGACTACAACCCAC
AGCACACGTCTGAACTCTTTCCCTACACGACGCTCTTCCGATCTNNNTTCTCATTTTCCCACTCCCCTGAC
GAGCACACGTCTGAACTCTTTCCCTACACGACGCTCTTCCGATCTNNNGGAATATGGAGACCCAAGTC
AAGAGCACACGTCTGAACTCTTTCCCTACACGACGCTCTTCCGATCTNNNTTTTCCCTTCCTTCCTTTG
AGAGCACACGTCTGAACTCTTTCCCTACACGACGCTCTTCCGATCTNNNTTGAGTTCCCCTGCCAATCA
GAGCACACGTCTGAACTCTTTCCCTACACGACGCTCTTCCGATCTNNNGGAAAGTCCCTGTCCCAACAA
GAGCACACGTCTGAACTCTTTCCCTACACGACGCTCTTCCGATCTNNNGTCCTCTTGGCATTTTGGGGTT
GAGCACACGTCTGAACTCTTTCCCTACACGACGCTCTTCCGATCTNNNTCACCAGTGCTCATGAATTCTT
CACACGTCTGAACTCTTTCCCTACACGACGCTCTTCCGATCTNNNCTCTGCCCTCTTCCTTCCCCCCATCG
AGAGCACACGTCTGAACTCTTTCCCTACACGACGCTCTTCCGATCTNNNAAACGCCTTTCAGGAACTAGC
AGCACACGTCTGAACTCTTTCCCTACACGACGCTCTTCCGATCTNNNTGTGCAGGGTATAAGGTCTTTG
AGCACACGTCTGAACTCTTTCCCTACACGACGCTCTTCCGATCTNNNGCTGACTGCCAGGTTAGAATGA
GAAGAGCACACGTCTGAACTCTTTCCCTACACGACGCTCTTCCGATCTNNNACAGGCTCAACCTAATTG
GAGCACACGTCTGAACTCTTTCCCTACACGACGCTCTTCCGATCTNNNCTAAGGATGCTTGTGTTGCCT
AGAGCACACGTCTGAACTCTTTCCCTACACGACGCTCTTCCGATCTNNNTTGCAAGTCTCTGCATATTC
GAGCACACGTCTGAACTCTTTCCCTACACGACGCTCTTCCGATCTNNNAATCTTCACACACAGGATGAG
AGCACACGTCTGAACTCTTTCCCTACACGACGCTCTTCCGATCTNNNAACTTACTGTGTTTCAGGGCCAC
AGCACACGTCTGAACTCTTTCCCTACACGACGCTCTTCCGATCTNNNGTATGTGGGAGGGAAGTAGG
AAGAGCACACGTCTGAACTCTTTCCCTACACGACGCTCTTCCGATCTNNNTTCTCTCTTCTCCTGCTCTC
AGCACACGTCTGAACTCTTTCCCTACACGACGCTCTTCCGATCTNNNCTAAGGATGCTTGTGTTGCCTC
GAGCACACGTCTGAACTCTTTCCCTACACGACGCTCTTCCGATCTNNNACTTCCGAAGGCCAAGGATAA
GCACACGTCTGAACTCTTTCCCTACACGACGCTCTTCCGATCTNNNTGCTTGAGCTTTCCTTCTTCCTCCA
AGCACACGTCTGAACTCTTTCCCTACACGACGCTCTTCCGATCTNNNAATTTTCCCGCCTTGTATGGTGT
AGAGCACACGTCTGAACTCTTTCCCTACACGACGCTCTTCCGATCTNNNAAAATCTCTGCCCTTAGGGA
GAGCACACGTCTGAACTCTTTCCCTACACGACGCTCTTCCGATCTNNNTAGTTACCAGGTATCAACCATC
AGCACACGTCTGAACTCTTTCCCTACACGACGCTCTTCCGATCTNNNAAGTTGGTATGCCAGCAAAAAC
CACACGTCTGAACTCTTTCCCTACACGACGCTCTTCCGATCTNNNGAGAGAGGGAACTGGGGAAGTG
GAGCACACGTCTGAACTCTTTCCCTACACGACGCTCTTCCGATCTNNNCCTACCACATGACGTTCCTTTG
AGCACACGTCTGAACTCTTTCCCTACACGACGCTCTTCCGATCTNNNCTGTCTTTCTCTGGGGCTGATTC
AAGAGCACACGTCTGAACTCTTTCCCTACACGACGCTCTTCCGATCTNNNTTTTTCCCTTCCTTCCTTTGG
AGCACACGTCTGAACTCTTTCCCTACACGACGCTCTTCCGATCTNNNGGAAGGTCAGTCTCTGACAATG
AGCACACGTCTGAACTCTTTCCCTACACGACGCTCTTCCGATCTNNNAAAAGGGTGCAGTCTGTAGGA
AGCACACGTCTGAACTCTTTCCCTACACGACGCTCTTCCGATCTNNNAGGATGACTGTAATACCCAGCA
AGCACACGTCTGAACTCTTTCCCTACACGACGCTCTTCCGATCTNNNGGAAACTGCTAGTGCGTGATA
AGCACACGTCTGAACTCTTTCCCTACACGACGCTCTTCCGATCTNNNACTCTCACATGAAGAGAACCCCA
GAAGAGCACACGTCTGAACTCTTTCCCTACACGACGCTCTTCCGATCTNNNTTTTGGCTTCAAAAGGCTC
GAGCACACGTCTGAACTCTTTCCCTACACGACGCTCTTCCGATCTNNNACATTGCAGTTCTCTCACTGGT
AGCACACGTCTGAACTCTTTCCCTACACGACGCTCTTCCGATCTNNNACAGGTATGCACAACATCTCTCC
AGCACACGTCTGAACTCTTTCCCTACACGACGCTCTTCCGATCTNNNAGCTTGCAGGGAATATCTGTTC
GAGCACACGTCTGAACTCTTTCCCTACACGACGCTCTTCCGATCTNNNAATACAGGTCTGAAAGTCGTG
AAGAGCACACGTCTGAACTCTTTCCCTACACGACGCTCTTCCGATCTNNNTTTGGAGTGGGGGTTGTTT
AGAGCACACGTCTGAACTCTTTCCCTACACGACGCTCTTCCGATCTNNNCAAGTGAGCATGCATTGACCA

AGAGCACACGTCTGAACTCTTTCCCTACACGACGCTCTTCCGATCTNNNTCAGAGCTGACATTTGCTCC
GAGCACACGTCTGAACTCTTTCCCTACACGACGCTCTTCCGATCTNNNGCTGAATCCTGAGTAACAGAC
AGAGCACACGTCTGAACTCTTTCCCTACACGACGCTCTTCCGATCTNNNTTTGTGTAAAGCCTCAGAGC
AGCACACGTCTGAACTCTTTCCCTACACGACGCTCTTCCGATCTNNNAGCATACTAGCGTGTACATGTA
AGAGCACACGTCTGAACTCTTTCCCTACACGACGCTCTTCCGATCTNNNCAACATGAAATTCGGTTGTGC
AGCACACGTCTGAACTCTTTCCCTACACGACGCTCTTCCGATCTNNNCACTCAGACCCCATCTTACCCATC
AGCACACGTCTGAACTCTTTCCCTACACGACGCTCTTCCGATCTNNNAATTCCCAACCACCACTTTGGAA
GAGCACACGTCTGAACTCTTTCCCTACACGACGCTCTTCCGATCTNNNAAGTTTGCCTCTAGGGACTGA
GAGCACACGTCTGAACTCTTTCCCTACACGACGCTCTTCCGATCTNNNACTGTTCATGTCTTCACCCACT
AGCACACGTCTGAACTCTTTCCCTACACGACGCTCTTCCGATCTNNNACGGAAGCAAGGAAGGAATGT
GAGCACACGTCTGAACTCTTTCCCTACACGACGCTCTTCCGATCTNNNATACTCCACCAATAGTCCCCAC
GAGCACACGTCTGAACTCTTTCCCTACACGACGCTCTTCCGATCTNNNGGCCAAGTTTAGAAACATTCCT
GCACACGTCTGAACTCTTTCCCTACACGACGCTCTTCCGATCTNNNTCCCTCTCCTCTTTCCAATCCTGCA
AGCACACGTCTGAACTCTTTCCCTACACGACGCTCTTCCGATCTNNNGCTGGACATTGAGGAGGCTTTA
GAGCACACGTCTGAACTCTTTCCCTACACGACGCTCTTCCGATCTNNNCCTCCTCACAGGTCTCCCTAAA
AGCACACGTCTGAACTCTTTCCCTACACGACGCTCTTCCGATCTNNNCAAGAACTACTTGTCAACCTCCA
AGCACACGTCTGAACTCTTTCCCTACACGACGCTCTTCCGATCTNNNCTGCCCTGAGGCTTTTCACTAGC
AGCACACGTCTGAACTCTTTCCCTACACGACGCTCTTCCGATCTNNNGCCACTGCAAACATAGTAATCTT
GAGCACACGTCTGAACTCTTTCCCTACACGACGCTCTTCCGATCTNNNTTACAAGCAGTCCAAGAGCAT
AGCACACGTCTGAACTCTTTCCCTACACGACGCTCTTCCGATCTNNNTAACTCCCACTCCCATTCCTGAGC
AGCACACGTCTGAACTCTTTCCCTACACGACGCTCTTCCGATCTNNNACACTGTTTGTGGTAGGAAAG
GAGCACACGTCTGAACTCTTTCCCTACACGACGCTCTTCCGATCTNNNTCTCCTTCTTGAGCACCTGATG
GCACACGTCTGAACTCTTTCCCTACACGACGCTCTTCCGATCTNNNGAAAGTCACCCTCTCCCCACTATCA
AGCACACGTCTGAACTCTTTCCCTACACGACGCTCTTCCGATCTNNNATCTTGGGGATCAGCAAGGTTT
AGAGCACACGTCTGAACTCTTTCCCTACACGACGCTCTTCCGATCTNNNGTGTCCCTTCCTACTCACTTCC
GAGCACACGTCTGAACTCTTTCCCTACACGACGCTCTTCCGATCTNNNCCAGCTTGCATTCTAAGTGCTT
GAGCACACGTCTGAACTCTTTCCCTACACGACGCTCTTCCGATCTNNNGGCTATGTCTCAAACACCCCTT
GAGCACACGTCTGAACTCTTTCCCTACACGACGCTCTTCCGATCTNNNTTCATGGCAAAAGGCAGCAAA
GAGCACACGTCTGAACTCTTTCCCTACACGACGCTCTTCCGATCTNNNGTGACAGGTATGGCAGACAG
GAGCACACGTCTGAACTCTTTCCCTACACGACGCTCTTCCGATCTNNNGTGACATTGACAGAAGCCACT
AGCACACGTCTGAACTCTTTCCCTACACGACGCTCTTCCGATCTNNNCCAGTATGAAGTGGTTGAACTC
AGCACACGTCTGAACTCTTTCCCTACACGACGCTCTTCCGATCTNNNGCTATGTAATTTTAGGCAACACA
GCACACGTCTGAACTCTTTCCCTACACGACGCTCTTCCGATCTNNNCAGCCCCACATCAGTGACATCCAT
GAGCACACGTCTGAACTCTTTCCCTACACGACGCTCTTCCGATCTNNNGTGGGGGTGATACTCTCCACA
AGAGCACACGTCTGAACTCTTTCCCTACACGACGCTCTTCCGATCTNNNCACAGATACTCTGGGCACGTA
GAGCACACGTCTGAACTCTTTCCCTACACGACGCTCTTCCGATCTNNNCACACTTCTATTCTCCTTGCCCC
AGAGCACACGTCTGAACTCTTTCCCTACACGACGCTCTTCCGATCTNNNTTGAGTTCCCCTGCCAATCAA
AGAGCACACGTCTGAACTCTTTCCCTACACGACGCTCTTCCGATCTNNNCATGTGGTGTTTGAACATGG
GAGCACACGTCTGAACTCTTTCCCTACACGACGCTCTTCCGATCTNNNATCACTTCCTCCAGGTTGAAAC
GGAAGAGCACACGTCTGAACTCTTTCCCTACACGACGCTCTTCCGATCTNNNAAACGCCTTTCAGGAAC
AAGAGCACACGTCTGAACTCTTTCCCTACACGACGCTCTTCCGATCTNNNTGTAAAGTTACGTGCGTGA
AGCACACGTCTGAACTCTTTCCCTACACGACGCTCTTCCGATCTNNNTTTATCCAGCTGAGAAGCAGG
AGAGCACACGTCTGAACTCTTTCCCTACACGACGCTCTTCCGATCTNNNTGCTGCTTCATGAATGACCCT

GAGCACACGTCTGAACTCTTTCCCTACACGACGCTCTTCCGATCTNNNTCCTATTGGTTCTGGTTCAACC
GAGCACACGTCTGAACTCTTTCCCTACACGACGCTCTTCCGATCTNNNGTCAAGCACCAAATGGAAACCA
AGAGCACACGTCTGAACTCTTTCCCTACACGACGCTCTTCCGATCTNNNTGGAAAACTGCCTTATGACA
AGAGCACACGTCTGAACTCTTTCCCTACACGACGCTCTTCCGATCTNNNCCTTCCGTCCTTTTCTCTCCCC
GAGCACACGTCTGAACTCTTTCCCTACACGACGCTCTTCCGATCTNNNAAGGTTGGCATTCCATTTACC
GAGCACACGTCTGAACTCTTTCCCTACACGACGCTCTTCCGATCTNNNAAGCACTCAGTGACCATTAGCA
AGCACACGTCTGAACTCTTTCCCTACACGACGCTCTTCCGATCTNNNTAAAATACCTAGTACGGGCCGG
AGAGCACACGTCTGAACTCTTTCCCTACACGACGCTCTTCCGATCTNNNTAGAGGGTCTTCTCACCTTG
AGCACACGTCTGAACTCTTTCCCTACACGACGCTCTTCCGATCTNNNAAGGACTTTGTCTGCAAGCATCC
AGAGCACACGTCTGAACTCTTTCCCTACACGACGCTCTTCCGATCTNNNTGTTTACTGCATCTGTATGG
AGCACACGTCTGAACTCTTTCCCTACACGACGCTCTTCCGATCTNNNCCACTCTGCCTGCTTATGTCATTG
GGAAGAGCACACGTCTGAACTCTTTCCCTACACGACGCTCTTCCGATCTNNNCTTCCTTTGGGAAGGGA
GAGCACACGTCTGAACTCTTTCCCTACACGACGCTCTTCCGATCTNNNATACCGTGTTCCCATCTCTGAC
AGCACACGTCTGAACTCTTTCCCTACACGACGCTCTTCCGATCTNNNAGTGAAAGGTGCTATCTTGCTG
AGCACACGTCTGAACTCTTTCCCTACACGACGCTCTTCCGATCTNNNTCTGTAAAACATACCCACCCAGA
GGAAGAGCACACGTCTGAACTCTTTCCCTACACGACGCTCTTCCGATCTNNNGTTACCTCCCAATAGAC
AGCACACGTCTGAACTCTTTCCCTACACGACGCTCTTCCGATCTNNNGATGTGGACATCCCAAACCAAG
GCACACGTCTGAACTCTTTCCCTACACGACGCTCTTCCGATCTNNNTAAAACACAATCCCAACCTCCCCA
AGAGCACACGTCTGAACTCTTTCCCTACACGACGCTCTTCCGATCTNNNGAGCTCTAACTCTGAGCCAGC
AGAGCACACGTCTGAACTCTTTCCCTACACGACGCTCTTCCGATCTNNNACCTTGTGATAGAGCTGTCC
AGAGCACACGTCTGAACTCTTTCCCTACACGACGCTCTTCCGATCTNNNTTATCTCTCAGACCCTGTGTC
GAGCACACGTCTGAACTCTTTCCCTACACGACGCTCTTCCGATCTNNNGAGGCAGACAAAATCGCACTA
AGCACACGTCTGAACTCTTTCCCTACACGACGCTCTTCCGATCTNNNTAGTGGGAAGGAAGGAGGAA
AGCACACGTCTGAACTCTTTCCCTACACGACGCTCTTCCGATCTNNNAGTTTCTTTGGGAATGCAGGA
AGCACACGTCTGAACTCTTTCCCTACACGACGCTCTTCCGATCTNNNCTGTTCTCTCTCTCTTCCCCCTTCA
AGAGCACACGTCTGAACTCTTTCCCTACACGACGCTCTTCCGATCTNNNAGCTGTGCCTATGGTATGCT
GAGCACACGTCTGAACTCTTTCCCTACACGACGCTCTTCCGATCTNNNCTAACTCCTCACCTCCCTCCATC
AGCACACGTCTGAACTCTTTCCCTACACGACGCTCTTCCGATCTNNNGCTTGCATTTTGATGGAGCATG
AGCACACGTCTGAACTCTTTCCCTACACGACGCTCTTCCGATCTNNNGAAACAGGGCTCTTTATCCAGG
AGAGCACACGTCTGAACTCTTTCCCTACACGACGCTCTTCCGATCTNNNATCTCTCTCTTTCTCCCATCCGC
GAGCACACGTCTGAACTCTTTCCCTACACGACGCTCTTCCGATCTNNNACATTGTGATTTTCCCTAAAAC
GAGCACACGTCTGAACTCTTTCCCTACACGACGCTCTTCCGATCTNNNGGTTCAACCAAACCAGAGACA
GAGCACACGTCTGAACTCTTTCCCTACACGACGCTCTTCCGATCTNNNATGTGCTCGGCTAACAGTTTT
GAGCACACGTCTGAACTCTTTCCCTACACGACGCTCTTCCGATCTNNNATGGTCCATGAGAGACAGAA
AGCACACGTCTGAACTCTTTCCCTACACGACGCTCTTCCGATCTNNNAGAAACTGTCCTTGAGAGGGTC
GAGCACACGTCTGAACTCTTTCCCTACACGACGCTCTTCCGATCTNNNAAGGAGGTGATGTGAGAGTC
GAGCACACGTCTGAACTCTTTCCCTACACGACGCTCTTCCGATCTNNNGGGCCTTGTGATAGTGTGAGT
AGCACACGTCTGAACTCTTTCCCTACACGACGCTCTTCCGATCTNNNACACACAACTTAGGCTTCCTTCC
AGCACACGTCTGAACTCTTTCCCTACACGACGCTCTTCCGATCTNNNTACACACATAACACCCAGACACC
GAGCACACGTCTGAACTCTTTCCCTACACGACGCTCTTCCGATCTNNNAGTGAGAGCCACAAATTCCTCA
GAGCACACGTCTGAACTCTTTCCCTACACGACGCTCTTCCGATCTNNNGCTTGAAACTTGAAACAATGT
AGCACACGTCTGAACTCTTTCCCTACACGACGCTCTTCCGATCTNNNCAGGATCAGGTCAGAAATGGAC
GAGCACACGTCTGAACTCTTTCCCTACACGACGCTCTTCCGATCTNNNAGCTTCAAAGACACAGAAGGC

GAGCACACGTCTGAACTCTTTCCCTACACGACGCTCTTCCGATCTNNNGGTGAGAATGCTGTCTGTCTG
AGAGCACACGTCTGAACTCTTTCCCTACACGACGCTCTTCCGATCTNNNAAGGTGATGATCGACAAGTT
AGCACACGTCTGAACTCTTTCCCTACACGACGCTCTTCCGATCTNNNGGAAGCTTGCCACTGAAAGACT
GAGCACACGTCTGAACTCTTTCCCTACACGACGCTCTTCCGATCTNNNACAGTCCCACACACTCTCTCTT
AGCACACGTCTGAACTCTTTCCCTACACGACGCTCTTCCGATCTNNNAACTCTTAGGAGGATGCCCAACA
GAGCACACGTCTGAACTCTTTCCCTACACGACGCTCTTCCGATCTNNNGAGATCTCGTGCTCTCTCTCTC
AGCACACGTCTGAACTCTTTCCCTACACGACGCTCTTCCGATCTNNNGAGGCTGAGGTGAACCGTAAT
AGCACACGTCTGAACTCTTTCCCTACACGACGCTCTTCCGATCTNNNGTGAAGAAAGCTTGAGGTTTG
GAGCACACGTCTGAACTCTTTCCCTACACGACGCTCTTCCGATCTNNNCTCCAGGGAGAGATGATGTCC
AGCACACGTCTGAACTCTTTCCCTACACGACGCTCTTCCGATCTNNNAGGTGGGTTACTGAAAGGAGA
GAGCACACGTCTGAACTCTTTCCCTACACGACGCTCTTCCGATCTNNNACGAACTACAGAAAGGGTAAA
AGCACACGTCTGAACTCTTTCCCTACACGACGCTCTTCCGATCTNNNCTGCTCAGTGGATGGTTGTATC
AGCACACGTCTGAACTCTTTCCCTACACGACGCTCTTCCGATCTNNNCTTTAGCTTGCACAGCAGTCTCT
AGCACACGTCTGAACTCTTTCCCTACACGACGCTCTTCCGATCTNNNTTCTGAAGCAAGTGATTGTCCCC
AGCACACGTCTGAACTCTTTCCCTACACGACGCTCTTCCGATCTNNNTTGGGTCGTAAGGGTGGATTT
GAGCACACGTCTGAACTCTTTCCCTACACGACGCTCTTCCGATCTNNNAGCAGATAGTGTCCTGGTAG
GAGCACACGTCTGAACTCTTTCCCTACACGACGCTCTTCCGATCTNNNAAATCCCCATGTGGCCAAAAAC
GAGCACACGTCTGAACTCTTTCCCTACACGACGCTCTTCCGATCTNNNTCAGCTCCACTTCATATGTTTT
GAGCACACGTCTGAACTCTTTCCCTACACGACGCTCTTCCGATCTNNNTGAGGGACATACAGATTGCT
AGCACACGTCTGAACTCTTTCCCTACACGACGCTCTTCCGATCTNNNTGTCCCAGCTTTCAATGTCATCA
AGAGCACACGTCTGAACTCTTTCCCTACACGACGCTCTTCCGATCTNNNTTTTCTAACTTCAGCTGCCCC
AGCACACGTCTGAACTCTTTCCCTACACGACGCTCTTCCGATCTNNNACTGGAGATGGTCATACTGGCT
AGCACACGTCTGAACTCTTTCCCTACACGACGCTCTTCCGATCTNNNACTGTTCATGTCTTCACCCACTT
AGCACACGTCTGAACTCTTTCCCTACACGACGCTCTTCCGATCTNNNCCATGAATCCCAGCAATTAGCCT
GAGCACACGTCTGAACTCTTTCCCTACACGACGCTCTTCCGATCTNNNGAGCCCCACCACAAAAACAAA
AGCACACGTCTGAACTCTTTCCCTACACGACGCTCTTCCGATCTNNNTCACTGTGTCCTTATCTGGCCTT
AGCACACGTCTGAACTCTTTCCCTACACGACGCTCTTCCGATCTNNNCCATGATAACAAGGAGAAGCGC
GAGCACACGTCTGAACTCTTTCCCTACACGACGCTCTTCCGATCTNNNTGTCTGGCCTTGTTCTCATCTT
AGCACACGTCTGAACTCTTTCCCTACACGACGCTCTTCCGATCTNNNAGCTTTTCATTAGCTCAGCCTCGG
AGCACACGTCTGAACTCTTTCCCTACACGACGCTCTTCCGATCTNNNAGATCCTTCCTCATCTGCCACAAC
AGCACACGTCTGAACTCTTTCCCTACACGACGCTCTTCCGATCTNNNAGGGAAGTAAGGAAGTAGCCA
GAGCACACGTCTGAACTCTTTCCCTACACGACGCTCTTCCGATCTNNNCGGGTCTATCTGTGCTTGTTGA
AGCACACGTCTGAACTCTTTCCCTACACGACGCTCTTCCGATCTNNNCTTGGTCATACCTGCAATCTGCTC
AAGAGCACACGTCTGAACTCTTTCCCTACACGACGCTCTTCCGATCTNNNCCCTCTGTCTCTCTCTGGCT
AGAGCACACGTCTGAACTCTTTCCCTACACGACGCTCTTCCGATCTNNNCTCAAATCTGCCTTTCCTCCTC
GAGCACACGTCTGAACTCTTTCCCTACACGACGCTCTTCCGATCTNNNTGGCAAAATTCAAAATGCCTGA
AGCACACGTCTGAACTCTTTCCCTACACGACGCTCTTCCGATCTNNNTCGTACCTGTCACCTCCAAGATT
GAAGAGCACACGTCTGAACTCTTTCCCTACACGACGCTCTTCCGATCTNNNTGCACAATCCTTTCAAATT
AGAGCACACGTCTGAACTCTTTCCCTACACGACGCTCTTCCGATCTNNNAACATATTGCAGACATCACCC
AAGAGCACACGTCTGAACTCTTTCCCTACACGACGCTCTTCCGATCTNNNCTCTTCAAGTGGTGGCCAG
AGAGCACACGTCTGAACTCTTTCCCTACACGACGCTCTTCCGATCTNNNGATGCCAGAAAAATCCCCTG
AGCACACGTCTGAACTCTTTCCCTACACGACGCTCTTCCGATCTNNNTTGCTTGTTCTTTTTCTGTGCCCC
AGCACACGTCTGAACTCTTTCCCTACACGACGCTCTTCCGATCTNNNATTGCTACTCTTGTGGGAGAGG

AGCACACGTCTGAACTCTTTCCCTACACGACGCTCTTCCGATCTNNNCCACTAAGAGAATGCCCAATGG
AGCACACGTCTGAACTCTTTCCCTACACGACGCTCTTCCGATCTNNNAGAACAATGAACAGAAAAGAA
GAAGAGCACACGTCTGAACTCTTTCCCTACACGACGCTCTTCCGATCTNNNCCCAGCCAATGTAACTGC
AGCACACGTCTGAACTCTTTCCCTACACGACGCTCTTCCGATCTNNNACATGAAGATCACTTCAAACTTA
GAGCACACGTCTGAACTCTTTCCCTACACGACGCTCTTCCGATCTNNNTATGGGCTTTCAGGGGATAG
GAGCACACGTCTGAACTCTTTCCCTACACGACGCTCTTCCGATCTNNNTGGTAGATAGGATCTGCTGGT
GAGCACACGTCTGAACTCTTTCCCTACACGACGCTCTTCCGATCTNNNCTGGATCTACATACAGCCTTCC
AGCACACGTCTGAACTCTTTCCCTACACGACGCTCTTCCGATCTNNNTGTTCTTGTCACCTAAGTTTCTGA
AGAGCACACGTCTGAACTCTTTCCCTACACGACGCTCTTCCGATCTNNNTCCACCACTATTTGGGGCTT
AGCACACGTCTGAACTCTTTCCCTACACGACGCTCTTCCGATCTNNNTGCTTTGTTTGTATGTTAGATT
AGAGCACACGTCTGAACTCTTTCCCTACACGACGCTCTTCCGATCTNNNTCATGGTATATGCACAGGAC
AGAGCACACGTCTGAACTCTTTCCCTACACGACGCTCTTCCGATCTNNNAAAATCTCTGCCCTTAGGGA
AGAGCACACGTCTGAACTCTTTCCCTACACGACGCTCTTCCGATCTNNNAATTCACTCTCCTGCCATTGT
GAGCACACGTCTGAACTCTTTCCCTACACGACGCTCTTCCGATCTNNNCCAGAGCCCCTGGAATTTATG
GAGCACACGTCTGAACTCTTTCCCTACACGACGCTCTTCCGATCTNNNGTTCCTTCCTCCATGGCATTCT
AAGAGCACACGTCTGAACTCTTTCCCTACACGACGCTCTTCCGATCTNNNTTACTGCCCTCTCCTCACTTG
AGAGCACACGTCTGAACTCTTTCCCTACACGACGCTCTTCCGATCTNNNATGATGGAGGTCTGTCCAGT
AGCACACGTCTGAACTCTTTCCCTACACGACGCTCTTCCGATCTNNNTATGGCCTATCAACGTCTACTCG
AGAGCACACGTCTGAACTCTTTCCCTACACGACGCTCTTCCGATCTNNNCCAGAAGAGGGGGAAGGG
AGCACACGTCTGAACTCTTTCCCTACACGACGCTCTTCCGATCTNNNTCAGTACACAGACATGCAACTG
GAGCACACGTCTGAACTCTTTCCCTACACGACGCTCTTCCGATCTNNNCAGTTCTGCCATATCAGTGGC
GAGCACACGTCTGAACTCTTTCCCTACACGACGCTCTTCCGATCTNNNTCATGCACATGTAAGTGTTTGC
GCACACGTCTGAACTCTTTCCCTACACGACGCTCTTCCGATCTNNNAGCCAAATGGCCAGTTACAGAAA
AGCACACGTCTGAACTCTTTCCCTACACGACGCTCTTCCGATCTNNNCCTGTGAGATAGATGGCCAGAT
AGAGCACACGTCTGAACTCTTTCCCTACACGACGCTCTTCCGATCTNNNCAGTGATGCTTCCTTCAACCA
AGCACACGTCTGAACTCTTTCCCTACACGACGCTCTTCCGATCTNNNTCTGGAGCTCTGAAGTAAGTGT
AGAGCACACGTCTGAACTCTTTCCCTACACGACGCTCTTCCGATCTNNNAGGTTTGCTATGTCCCCATT
GAGCACACGTCTGAACTCTTTCCCTACACGACGCTCTTCCGATCTNNNAGCTGTGCCTATGGTATGCTA
GAGCACACGTCTGAACTCTTTCCCTACACGACGCTCTTCCGATCTNNNAGTGAAATAACCCATTTAGAA
AGCACACGTCTGAACTCTTTCCCTACACGACGCTCTTCCGATCTNNNTGTGTCACTTCTCTCAGGTCCTCC
GAGCACACGTCTGAACTCTTTCCCTACACGACGCTCTTCCGATCTNNNTGCTGGAGCAACTTGATATCT
AGCACACGTCTGAACTCTTTCCCTACACGACGCTCTTCCGATCTNNNAGCTGAACCCTTTGGTTCTCAAG
GAGCACACGTCTGAACTCTTTCCCTACACGACGCTCTTCCGATCTNNNGAAGTAGCTTCTGCTAAGGGC
AAGAGCACACGTCTGAACTCTTTCCCTACACGACGCTCTTCCGATCTNNNCTGGTCCACATGTGTGAGC
AGCACACGTCTGAACTCTTTCCCTACACGACGCTCTTCCGATCTNNNTAGTGGTAAGAGGGAAGGAGC
AGAGCACACGTCTGAACTCTTTCCCTACACGACGCTCTTCCGATCTNNNGAACTTTGGTATGGTAGCCT
GAGCACACGTCTGAACTCTTTCCCTACACGACGCTCTTCCGATCTNNNGAAAGTCATGTAGTCCCCACA
AGAGCACACGTCTGAACTCTTTCCCTACACGACGCTCTTCCGATCTNNNAGATGTCTAGAACCTGCTCTC
AGAGCACACGTCTGAACTCTTTCCCTACACGACGCTCTTCCGATCTNNNCTAGCCATGAGAAAGTCAAG
AAGAGCACACGTCTGAACTCTTTCCCTACACGACGCTCTTCCGATCTNNNTCCATGATTATGTTACCCCC
AGCACACGTCTGAACTCTTTCCCTACACGACGCTCTTCCGATCTNNNTGGACTTCCACAAGTTTCAGCT
GAGCACACGTCTGAACTCTTTCCCTACACGACGCTCTTCCGATCTNNNACATTGGGACTAAGTGTGGCA
GCACACGTCTGAACTCTTTCCCTACACGACGCTCTTCCGATCTNNNTCACTCACACTAGCAAATGAAAG

GAGCACACGTCTGAACTCTTTCCCTACACGACGCTCTTCCGATCTNNNCAATGTCTGGAAGAGAGAA
AGCACACGTCTGAACTCTTTCCCTACACGACGCTCTTCCGATCTNNNAGCAATAGATGCTGTGGCTTG
AGAGCACACGTCTGAACTCTTTCCCTACACGACGCTCTTCCGATCTNNNATCCTATCATTTTCTTCTGG
AGCACACGTCTGAACTCTTTCCCTACACGACGCTCTTCCGATCTNNNAACAACTTTACCTGAAGCACCG
AGCACACGTCTGAACTCTTTCCCTACACGACGCTCTTCCGATCTNNNTGTTTCCCATAGGTTGGTCTC
GCACACGTCTGAACTCTTTCCCTACACGACGCTCTTCCGATCTNNNAGGCCCCTTTACTTTGCTTGTTT
GGAAGAGCACACGTCTGAACTCTTTCCCTACACGACGCTCTTCCGATCTNNNTGCTGCAGGCAGGAT
CACACGTCTGAACTCTTTCCCTACACGACGCTCTTCCGATCTNNNAATGTCCATTTCCTCTTCCCCACCA
AGCACACGTCTGAACTCTTTCCCTACACGACGCTCTTCCGATCTNNNGGTTTCTCTGTCTCTCCACACA
AGCACACGTCTGAACTCTTTCCCTACACGACGCTCTTCCGATCTNNNGGCTTCATCACCAGACCAGAA
AGCACACGTCTGAACTCTTTCCCTACACGACGCTCTTCCGATCTNNNGCTGAGCTTATTGTGAGGAAG
AGCACACGTCTGAACTCTTTCCCTACACGACGCTCTTCCGATCTNNNAAATGGTCGAATTATGGGCGT
AGAGCACACGTCTGAACTCTTTCCCTACACGACGCTCTTCCGATCTNNNTGACTGCTGCACTCTCTTTT
AGCACACGTCTGAACTCTTTCCCTACACGACGCTCTTCCGATCTNNNTTCCACCACAGTACACTGTTTC
GAGCACACGTCTGAACTCTTTCCCTACACGACGCTCTTCCGATCTNNNGAAACAACACGGCCACTAGG
GCACACGTCTGAACTCTTTCCCTACACGACGCTCTTCCGATCTNNNTTCTGCAATACAACTTCCTGCCA
AGCACACGTCTGAACTCTTTCCCTACACGACGCTCTTCCGATCTNNNCCAGAGCCCCTGGAATTTATG
GAGCACACGTCTGAACTCTTTCCCTACACGACGCTCTTCCGATCTNNNACCAAGAGAGTAGATGAGG
GCACACGTCTGAACTCTTTCCCTACACGACGCTCTTCCGATCTNNNCACCTGGGGTTTTGGAAATCAG
AGCACACGTCTGAACTCTTTCCCTACACGACGCTCTTCCGATCTNNNAACCAGCGAGTACTGGGTATG
GCACACGTCTGAACTCTTTCCCTACACGACGCTCTTCCGATCTNNNGTTTCCCTCTCACCAGCCAATAGC
GAGCACACGTCTGAACTCTTTCCCTACACGACGCTCTTCCGATCTNNNATCTCTCATAGGCTCTACAGG
GCACACGTCTGAACTCTTTCCCTACACGACGCTCTTCCGATCTNNNATGTGTTTAGTCTCACCCACTGC
AGCACACGTCTGAACTCTTTCCCTACACGACGCTCTTCCGATCTNNNGCAACTCCAGGAAATTGATGTT
GAGCACACGTCTGAACTCTTTCCCTACACGACGCTCTTCCGATCTNNNTCCTTGGTTGCTAACCTCCTG
AGCACACGTCTGAACTCTTTCCCTACACGACGCTCTTCCGATCTNNNCCAATTGGCTGAAGACCTGGA
AGCACACGTCTGAACTCTTTCCCTACACGACGCTCTTCCGATCTNNNACAGGTACCATCTGTAACCAC
AGCACACGTCTGAACTCTTTCCCTACACGACGCTCTTCCGATCTNNNTGATCTTCTTGGCTCATTGTCT
AGCACACGTCTGAACTCTTTCCCTACACGACGCTCTTCCGATCTNNNATTTTCAGAGGAGATGGGGG
GAGCACACGTCTGAACTCTTTCCCTACACGACGCTCTTCCGATCTNNNGTGTGGCCTGGTTCTCTGTA
GCACACGTCTGAACTCTTTCCCTACACGACGCTCTTCCGATCTNNNGCATTGCATTCAGGAGGTTAGG
AGCACACGTCTGAACTCTTTCCCTACACGACGCTCTTCCGATCTNNNTTCCACCGATGGTAAACCAGA
AGCACACGTCTGAACTCTTTCCCTACACGACGCTCTTCCGATCTNNNTTTCTCCTTCTGTCTCCTCCCTG
GCACACGTCTGAACTCTTTCCCTACACGACGCTCTTCCGATCTNNNCCTACTCAACCAGAGGGGAAGA
AGAGCACACGTCTGAACTCTTTCCCTACACGACGCTCTTCCGATCTNNNACTTTTCAACCCATGAGGAA
GCACACGTCTGAACTCTTTCCCTACACGACGCTCTTCCGATCTNNNCTTGCTCGGTCTCACTGTATTGT
AGCACACGTCTGAACTCTTTCCCTACACGACGCTCTTCCGATCTNNNTTTTGGTATTTTGGTGGCTGC
GCACACGTCTGAACTCTTTCCCTACACGACGCTCTTCCGATCTNNNACGTTTACCCCTGCACATTTTCT
AGCACACGTCTGAACTCTTTCCCTACACGACGCTCTTCCGATCTNNNTTTGATAGCACAGAAGGGTGA
GAGCACACGTCTGAACTCTTTCCCTACACGACGCTCTTCCGATCTNNNCAGTTCTGCCATATCAGTGG
AAGAGCACACGTCTGAACTCTTTCCCTACACGACGCTCTTCCGATCTNNNTATGAGATTGAGCCAATG
GAGCACACGTCTGAACTCTTTCCCTACACGACGCTCTTCCGATCTNNNTCTCAGTGAAGTGCCCGATTT
AGCACACGTCTGAACTCTTTCCCTACACGACGCTCTTCCGATCTNNNCCTTTGTTCCATCGTGTGACTG

GAGCACACGTCTGAACTCTTTCCCTACACGACGCTCTTCCGATCTNNNTCTAGTTGCTAGTTACACAGC
GAGCACACGTCTGAACTCTTTCCCTACACGACGCTCTTCCGATCTNNNCCACTGAACCACCCAAGATTG
AGCACACGTCTGAACTCTTTCCCTACACGACGCTCTTCCGATCTNNNCCCATTTGCTTCCTCATTTGCTCCA
GAGCACACGTCTGAACTCTTTCCCTACACGACGCTCTTCCGATCTNNNTTGAGAGCAGGGCCTCATATA
AGAGCACACGTCTGAACTCTTTCCCTACACGACGCTCTTCCGATCTNNNGATGGGGTAGTTTGAGCAT
AGCACACGTCTGAACTCTTTCCCTACACGACGCTCTTCCGATCTNNNTGCAGAGGGCTCATCTCTAAT
AGCACACGTCTGAACTCTTTCCCTACACGACGCTCTTCCGATCTNNNTTCTAAGTCAGAATCCCCAGGC
GAGCACACGTCTGAACTCTTTCCCTACACGACGCTCTTCCGATCTNNNAGAGAATCCTAATGAGAGAC
GAGCACACGTCTGAACTCTTTCCCTACACGACGCTCTTCCGATCTNNNCACCTGTCTTTCTTGCTGAAG
GAGCACACGTCTGAACTCTTTCCCTACACGACGCTCTTCCGATCTNNNTGCCTCACTACTCCTGGAATTG
AGAGCACACGTCTGAACTCTTTCCCTACACGACGCTCTTCCGATCTNNNCATGGATGTCTGCTTTAGGC
AGAGCACACGTCTGAACTCTTTCCCTACACGACGCTCTTCCGATCTNNNTTTCTCTGATCAACAGCCACA
GAGCACACGTCTGAACTCTTTCCCTACACGACGCTCTTCCGATCTNNNCTTCTTTGTCTCTGTCCCTCCCT
GAGCACACGTCTGAACTCTTTCCCTACACGACGCTCTTCCGATCTNNNCACTGAACACAGGGAAGTCAT
GAGCACACGTCTGAACTCTTTCCCTACACGACGCTCTTCCGATCTNNNTTGTGGTTTTGGGTGTTGGTT
AGAGCACACGTCTGAACTCTTTCCCTACACGACGCTCTTCCGATCTNNNTTGTGGTTTTGGGTGTTGGT
GCACACGTCTGAACTCTTTCCCTACACGACGCTCTTCCGATCTNNNCCCATTGTCTGAATGGGATAAG
GAGCACACGTCTGAACTCTTTCCCTACACGACGCTCTTCCGATCTNNNGAACAAAGGCCATGGGGAAA
AGAGCACACGTCTGAACTCTTTCCCTACACGACGCTCTTCCGATCTNNNTCATGCTGTTTTCCCAGCTA
GAGCACACGTCTGAACTCTTTCCCTACACGACGCTCTTCCGATCTNNNTGGAAAGTCTCAACTGCTACA
GCACACGTCTGAACTCTTTCCCTACACGACGCTCTTCCGATCTNNNGCTCAGAGCTGGAGTTGACTTAG
AGCACACGTCTGAACTCTTTCCCTACACGACGCTCTTCCGATCTNNNAGACCTTTGTACACAGCCTACCT
GAGCACACGTCTGAACTCTTTCCCTACACGACGCTCTTCCGATCTNNNTGGTAAGAAAAGAGAGAAAA
GAGCACACGTCTGAACTCTTTCCCTACACGACGCTCTTCCGATCTNNNAAAAGTTTGGGGAAGGGAAA
AGAGCACACGTCTGAACTCTTTCCCTACACGACGCTCTTCCGATCTNNNATGAAGGACCCTTGTTCATG
AAGAGCACACGTCTGAACTCTTTCCCTACACGACGCTCTTCCGATCTNNNCCCATTGGCATTTCCCAAAT
GAGCACACGTCTGAACTCTTTCCCTACACGACGCTCTTCCGATCTNNNACAAGACCCTAGCAACTCTCT
GAGCACACGTCTGAACTCTTTCCCTACACGACGCTCTTCCGATCTNNNATGATTGCACTATGTGCTCCA
AGAGCACACGTCTGAACTCTTTCCCTACACGACGCTCTTCCGATCTNNNCCTACCACATGACGTTCCTTT
AGCACACGTCTGAACTCTTTCCCTACACGACGCTCTTCCGATCTNNNGCATCTAACGTATTTGCTACTT
GAGCACACGTCTGAACTCTTTCCCTACACGACGCTCTTCCGATCTNNNGGAGTGTTTGCCTAACTGTCC
AGCACACGTCTGAACTCTTTCCCTACACGACGCTCTTCCGATCTNNNTATGTGACCTTCACCTGACATC
AGCACACGTCTGAACTCTTTCCCTACACGACGCTCTTCCGATCTNNNTGTTGGAGACAACCACAAGTCA
AGCACACGTCTGAACTCTTTCCCTACACGACGCTCTTCCGATCTNNNATCGTGTCACCCACAACTTCTCT
GAAGAGCACACGTCTGAACTCTTTCCCTACACGACGCTCTTCCGATCTNNNTTTTAGTGGATCTGTAC
AGCACACGTCTGAACTCTTTCCCTACACGACGCTCTTCCGATCTNNNTGGACAGCGGTAAGATGAAG
GAGCACACGTCTGAACTCTTTCCCTACACGACGCTCTTCCGATCTNNNGAGCAAGCTTACCTGATGTCC
AGCACACGTCTGAACTCTTTCCCTACACGACGCTCTTCCGATCTNNNACAGTGGCAGAGCAAAACTAT
GAGCACACGTCTGAACTCTTTCCCTACACGACGCTCTTCCGATCTNNNGAAGCCAAAACTGTGAACTGC
AGAGCACACGTCTGAACTCTTTCCCTACACGACGCTCTTCCGATCTNNNTGTTAGCCCTCTATGATGTC
GAGCACACGTCTGAACTCTTTCCCTACACGACGCTCTTCCGATCTNNNTTCCTGTAGGAACTCTCCAGG
GAGCACACGTCTGAACTCTTTCCCTACACGACGCTCTTCCGATCTNNNTTCCAAACACTTGAGGTCAGA
AGCACACGTCTGAACTCTTTCCCTACACGACGCTCTTCCGATCTNNNAGGGTATTGCTTTCCTAGCCTGA

AGCACACGTCTGAACTCTTTCCCTACACGACGCTCTTCCGATCTNNNCTGTTCCTAGAGCCTCGTTTTCC
AGCACACGTCTGAACTCTTTCCCTACACGACGCTCTTCCGATCTNNNACTGATTTCCCCAGCTTTCACTG
AGCACACGTCTGAACTCTTTCCCTACACGACGCTCTTCCGATCTNNNCAAAGTCATCATGCCTCCCTCAT
AGCACACGTCTGAACTCTTTCCCTACACGACGCTCTTCCGATCTNNNTGATATGTCACTGCTCCCCAGA
AGAGCACACGTCTGAACTCTTTCCCTACACGACGCTCTTCCGATCTNNNCATGGGTTATCTGTGACGA
GAGCACACGTCTGAACTCTTTCCCTACACGACGCTCTTCCGATCTNNNTGCAAACCCTTCATTCACAAGG
GAGCACACGTCTGAACTCTTTCCCTACACGACGCTCTTCCGATCTNNNAGTTCACAAGTGTGCTCTGGT
GAGCACACGTCTGAACTCTTTCCCTACACGACGCTCTTCCGATCTNNNTTCTTACTGCTCTCCTCTACCC
GAAGAGCACACGTCTGAACTCTTTCCCTACACGACGCTCTTCCGATCTNNNGTGGAAGAGGTACCCCA
AGCACACGTCTGAACTCTTTCCCTACACGACGCTCTTCCGATCTNNNTTGAGAGCAGGGCCTCATATA
GCACACGTCTGAACTCTTTCCCTACACGACGCTCTTCCGATCTNNNCGTATGCACAAGCTCACACAATC
AAGAGCACACGTCTGAACTCTTTCCCTACACGACGCTCTTCCGATCTNNNTGAGAGAGAAGCTAGAGA
GAGCACACGTCTGAACTCTTTCCCTACACGACGCTCTTCCGATCTNNNGATGCTTGTGGGCTTAGAAA
GAGCACACGTCTGAACTCTTTCCCTACACGACGCTCTTCCGATCTNNNCTTGGCTAATCAGTACCACAC
AGCACACGTCTGAACTCTTTCCCTACACGACGCTCTTCCGATCTNNNAGCTTTCATTGTGTTCTACTGG
AGCACACGTCTGAACTCTTTCCCTACACGACGCTCTTCCGATCTNNNCCTTGCTACTGGAACTCAGCTT
AAGAGCACACGTCTGAACTCTTTCCCTACACGACGCTCTTCCGATCTNNNCACTGGAAAATAACCCAGC
GAGCACACGTCTGAACTCTTTCCCTACACGACGCTCTTCCGATCTNNNTAATGGAGGAGGGTGGTTT
GAAGAGCACACGTCTGAACTCTTTCCCTACACGACGCTCTTCCGATCTNNNAGGATGACTGTAATACC
AGCACACGTCTGAACTCTTTCCCTACACGACGCTCTTCCGATCTNNNCAGAAATCCTACCTTCCTTGCCA
AGCACACGTCTGAACTCTTTCCCTACACGACGCTCTTCCGATCTNNNGTGGCTTACCAGGAAGAACTC
GAGCACACGTCTGAACTCTTTCCCTACACGACGCTCTTCCGATCTNNNTATACACACTGCACTCCAAGCA
AGAGCACACGTCTGAACTCTTTCCCTACACGACGCTCTTCCGATCTNNNTACTGTGCCTAAGCCATTTG
GCACACGTCTGAACTCTTTCCCTACACGACGCTCTTCCGATCTNNNGCTGAAGGCTGTCTCTCTCTTTC
GAGCACACGTCTGAACTCTTTCCCTACACGACGCTCTTCCGATCTNNNTGGAGGTCTTAGAGTACAGA
GAGCACACGTCTGAACTCTTTCCCTACACGACGCTCTTCCGATCTNNNGGCTTTCTTCCCTTTGCTCTCAT
AGCACACGTCTGAACTCTTTCCCTACACGACGCTCTTCCGATCTNNNACTTACTGATGTGTTTCCCCTG
GAAGAGCACACGTCTGAACTCTTTCCCTACACGACGCTCTTCCGATCTNNNGTGTGGGTCTGTGCAT
AGCACACGTCTGAACTCTTTCCCTACACGACGCTCTTCCGATCTNNNTGCTGTAAGTGCCTCTCTAAGT
GAGCACACGTCTGAACTCTTTCCCTACACGACGCTCTTCCGATCTNNNTTCTTATATCTGCGTGTGTG
AGAGCACACGTCTGAACTCTTTCCCTACACGACGCTCTTCCGATCTNNNATTAGGTCATAAGGGTGG
GAGCACACGTCTGAACTCTTTCCCTACACGACGCTCTTCCGATCTNNNTTGATCCCTCAAGTCCTGAAT
AAGAGCACACGTCTGAACTCTTTCCCTACACGACGCTCTTCCGATCTNNNTTTTGAGTAAACCAGGCCC
AGCACACGTCTGAACTCTTTCCCTACACGACGCTCTTCCGATCTNNNTAGCCTGTGGGCCTTTTCTCTAT
GAGCACACGTCTGAACTCTTTCCCTACACGACGCTCTTCCGATCTNNNTGGGTTTGACCACTTTCTCCT
GCACACGTCTGAACTCTTTCCCTACACGACGCTCTTCCGATCTNNNTATCTGCTCGGTCAATCAGGCTA
AGCACACGTCTGAACTCTTTCCCTACACGACGCTCTTCCGATCTNNNTGGCTTGCGAATAAACAAATTT
AAGAGCACACGTCTGAACTCTTTCCCTACACGACGCTCTTCCGATCTNNNGTGGAAGAGGTACCCCAA
GAGCACACGTCTGAACTCTTTCCCTACACGACGCTCTTCCGATCTNNNGCTCCTTTGAACACACTCCAGT
AGAGCACACGTCTGAACTCTTTCCCTACACGACGCTCTTCCGATCTNNNTTTTGCCTTCCACTCTTCTTC
GAGCACACGTCTGAACTCTTTCCCTACACGACGCTCTTCCGATCTNNNTGTCACTAAACGTCATGCATCT
AGCACACGTCTGAACTCTTTCCCTACACGACGCTCTTCCGATCTNNNAGAGAGAGAGAAAGAGGAA
GAGCACACGTCTGAACTCTTTCCCTACACGACGCTCTTCCGATCTNNNCTGCTGCAACCTGGACTAGA

AGCACACGTCTGAACTCTTTCCCTACACGACGCTCTTCCGATCTNNNGCAAAGATAGAAGCAGAGAG
CACACGTCTGAACTCTTTCCCTACACGACGCTCTTCCGATCTNNNCTTTGGCTCTGTTTCTTTCTGCGC
AGCACACGTCTGAACTCTTTCCCTACACGACGCTCTTCCGATCTNNNTTCCGTCTTTCCTCTAATGTGG
AGCACACGTCTGAACTCTTTCCCTACACGACGCTCTTCCGATCTNNNATCTCACAGGTAGGGAGCAA
AGCACACGTCTGAACTCTTTCCCTACACGACGCTCTTCCGATCTNNNTGGTATGAAAAGCTCATTGAG
CACACGTCTGAACTCTTTCCCTACACGACGCTCTTCCGATCTNNNTACCCTCAACCTACAGAAGCACA
GCACACGTCTGAACTCTTTCCCTACACGACGCTCTTCCGATCTNNNCCTGAGGATTCTTTCTCTCTACC
AGCACACGTCTGAACTCTTTCCCTACACGACGCTCTTCCGATCTNNNTCAAGAAGACATTTTCCCCTTG
GCACACGTCTGAACTCTTTCCCTACACGACGCTCTTCCGATCTNNNGGGGTGGGGATACAAACATCAT
AGCACACGTCTGAACTCTTTCCCTACACGACGCTCTTCCGATCTNNNTCTCAAGCCTATCCTCTCCATCC
AGCACACGTCTGAACTCTTTCCCTACACGACGCTCTTCCGATCTNNNTGGAGATGAACAGGCAGGAC
GCACACGTCTGAACTCTTTCCCTACACGACGCTCTTCCGATCTNNNTACTTCTGTGAGCAAGAAGGAA
AGCACACGTCTGAACTCTTTCCCTACACGACGCTCTTCCGATCTNNNATAGCCTGTCCAAATTGTCCCA
AGCACACGTCTGAACTCTTTCCCTACACGACGCTCTTCCGATCTNNNTCAACGCCTTTCTCTCTCTCTCT
AGAGCACACGTCTGAACTCTTTCCCTACACGACGCTCTTCCGATCTNNNACCTCCCATACAACAGATGC
GCACACGTCTGAACTCTTTCCCTACACGACGCTCTTCCGATCTNNNGCTCTGGCTTCACTCTAAACCTC
AGCACACGTCTGAACTCTTTCCCTACACGACGCTCTTCCGATCTNNNCGTGTCTGTTTGTGAGCATGA
AGCACACGTCTGAACTCTTTCCCTACACGACGCTCTTCCGATCTNNNTAAATGAAGCGAGACAACACC
GCACACGTCTGAACTCTTTCCCTACACGACGCTCTTCCGATCTNNNTATCATCATCTCCTCTGCCCACCC
AGCACACGTCTGAACTCTTTCCCTACACGACGCTCTTCCGATCTNNNTTCTAAGTCAGAATCCCCAGG
AGAGCACACGTCTGAACTCTTTCCCTACACGACGCTCTTCCGATCTNNNGAGAAGCAGAGCCCAGCA
GCACACGTCTGAACTCTTTCCCTACACGACGCTCTTCCGATCTNNNTACCTACAAGATTTCCCCCAGCA
GAGCACACGTCTGAACTCTTTCCCTACACGACGCTCTTCCGATCTNNNAGGGAGACTGGGAATCAGG
AGCACACGTCTGAACTCTTTCCCTACACGACGCTCTTCCGATCTNNNTTGCATTGAGGCCTGGGAAAT
AGCACACGTCTGAACTCTTTCCCTACACGACGCTCTTCCGATCTNNNTAGCTCTTTGCTCCATTCAGTG
GCACACGTCTGAACTCTTTCCCTACACGACGCTCTTCCGATCTNNNCGAAGAAACCTTTTTCCCTGCAC
GAGCACACGTCTGAACTCTTTCCCTACACGACGCTCTTCCGATCTNNNTCCTTGACAAGTGTTGATGG
AGCACACGTCTGAACTCTTTCCCTACACGACGCTCTTCCGATCTNNNCATGCAGACATTGGATTCTTCT
GCACACGTCTGAACTCTTTCCCTACACGACGCTCTTCCGATCTNNNTCCTCTGGAACCTAACGTTTTCA
AGAGCACACGTCTGAACTCTTTCCCTACACGACGCTCTTCCGATCTNNNATTGAAACGAGACGGCACA
AAGAGCACACGTCTGAACTCTTTCCCTACACGACGCTCTTCCGATCTNNNGGGACCATCTTTTGCCTCA
GCACACGTCTGAACTCTTTCCCTACACGACGCTCTTCCGATCTNNNCCTTGTTCACTCAAAGAGCACCT
AGAGCACACGTCTGAACTCTTTCCCTACACGACGCTCTTCCGATCTNNNTGGGTCCATTCATTTTGTG
GCACACGTCTGAACTCTTTCCCTACACGACGCTCTTCCGATCTNNNTCTGAACCCATCCCACTCCTAAGC
AGAGCACACGTCTGAACTCTTTCCCTACACGACGCTCTTCCGATCTNNNAAGATCCTAGAGGGTCAGA
GCACACGTCTGAACTCTTTCCCTACACGACGCTCTTCCGATCTNNNCTCCATGTTACTTCTGCTTCGCT
AGCACACGTCTGAACTCTTTCCCTACACGACGCTCTTCCGATCTNNNCTAACAGTAGCTTGGGCAAG
GAGCACACGTCTGAACTCTTTCCCTACACGACGCTCTTCCGATCTNNNGCTGCCCTAGCAAAGGTCTA
AGAGCACACGTCTGAACTCTTTCCCTACACGACGCTCTTCCGATCTNNNACCCTCCTTGCATTCTCTTTA
AGCACACGTCTGAACTCTTTCCCTACACGACGCTCTTCCGATCTNNNGCTATCTAGGCACTCTGTGTGT
AGCACACGTCTGAACTCTTTCCCTACACGACGCTCTTCCGATCTNNNTCCTGATGGAACCAGCCTAAA
GCACACGTCTGAACTCTTTCCCTACACGACGCTCTTCCGATCTNNNTATTTGAATGAACGGGCCCTCA
AGCACACGTCTGAACTCTTTCCCTACACGACGCTCTTCCGATCTNNNGCCATCAGGGTGATGTTCTTAC

AGCACACGTCTGAACTCTTTCCCTACACGACGCTCTTCCGATCTNNNAAACAAAGTTCAAAACAGGCAG
AGAGCACACGTCTGAACTCTTTCCCTACACGACGCTCTTCCGATCTNNNATGGCTGACCATTTCAGGAC
AGCACACGTCTGAACTCTTTCCCTACACGACGCTCTTCCGATCTNNNAGGCTGGGGGTGTTCTAATAT
GAGCACACGTCTGAACTCTTTCCCTACACGACGCTCTTCCGATCTNNNCTGGTGGTCTTTGACTTGTGG
AGAGCACACGTCTGAACTCTTTCCCTACACGACGCTCTTCCGATCTNNNTTGTTTTTCCTCTGAGCTGCCT
AGCACACGTCTGAACTCTTTCCCTACACGACGCTCTTCCGATCTNNNTTCACACACCTGCATGTAAGCAT
AGCACACGTCTGAACTCTTTCCCTACACGACGCTCTTCCGATCTNNNATGCCTCTGTCTCATCCTCTCTC
GAGCACACGTCTGAACTCTTTCCCTACACGACGCTCTTCCGATCTNNNCCTGGCATCCTTCCTGTATCCTA
AGCACACGTCTGAACTCTTTCCCTACACGACGCTCTTCCGATCTNNNTGCCAAAGGTGTTCAATGAGTG
AAGAGCACACGTCTGAACTCTTTCCCTACACGACGCTCTTCCGATCTNNNAAGATCCTAGAGGGTCAGA
AGAGCACACGTCTGAACTCTTTCCCTACACGACGCTCTTCCGATCTNNNACACAAAGTACTGAAAGAGT
AAGAGCACACGTCTGAACTCTTTCCCTACACGACGCTCTTCCGATCTNNNGGTGGAGAGAAATCCCAG
GAGCACACGTCTGAACTCTTTCCCTACACGACGCTCTTCCGATCTNNNTTATTTCAGGTTTGGCCACTGC
AGCACACGTCTGAACTCTTTCCCTACACGACGCTCTTCCGATCTNNNTCCACCACTATTTGGGGCTTTTT
AGCACACGTCTGAACTCTTTCCCTACACGACGCTCTTCCGATCTNNNTTTGGGTTTGTATGTCTTGCCCT
GAGCACACGTCTGAACTCTTTCCCTACACGACGCTCTTCCGATCTNNNTCACTCAACATTGTAGTCAGT
AGCACACGTCTGAACTCTTTCCCTACACGACGCTCTTCCGATCTNNNTTAGCGCATCGATTTCTTCCTCAC
AGCACACGTCTGAACTCTTTCCCTACACGACGCTCTTCCGATCTNNNCTTTGGCCAACTCAGGGACTTAG
CACACGTCTGAACTCTTTCCCTACACGACGCTCTTCCGATCTNNNCTACTAGACTGATACCCAGGGGCCA
AGCACACGTCTGAACTCTTTCCCTACACGACGCTCTTCCGATCTNNNTCACACGCACACAGAAATACACT
GAGCACACGTCTGAACTCTTTCCCTACACGACGCTCTTCCGATCTNNNGGAGCTGTTTCTTTGGAGAG
GAGCACACGTCTGAACTCTTTCCCTACACGACGCTCTTCCGATCTNNNAGAAGCCACAAAAGCAAATGA
GAGCACACGTCTGAACTCTTTCCCTACACGACGCTCTTCCGATCTNNNGTGTCCCTTCCTACTCACTTCCT
GAGCACACGTCTGAACTCTTTCCCTACACGACGCTCTTCCGATCTNNNTAACAGTCAGGGTACCACCAC
AGCACACGTCTGAACTCTTTCCCTACACGACGCTCTTCCGATCTNNNGAAAGTCACAAGTGCCTCCTCA
AAGAGCACACGTCTGAACTCTTTCCCTACACGACGCTCTTCCGATCTNNNAGGCCTTTACCTCTACAACT
AGCACACGTCTGAACTCTTTCCCTACACGACGCTCTTCCGATCTNNNAGAAGCTGAATGCTTCTGAATG
GAGCACACGTCTGAACTCTTTCCCTACACGACGCTCTTCCGATCTNNNATTTCAAACTGCCATTGGATGC
GAGCACACGTCTGAACTCTTTCCCTACACGACGCTCTTCCGATCTNNNAAATCATGGCTTCAGTTAGTG
AGAGCACACGTCTGAACTCTTTCCCTACACGACGCTCTTCCGATCTNNNAACCATGTCTCAGCCTTCACT
AGAGCACACGTCTGAACTCTTTCCCTACACGACGCTCTTCCGATCTNNNCCGTGTAGGCTCACTGTAAG
GAGCACACGTCTGAACTCTTTCCCTACACGACGCTCTTCCGATCTNNNTTGCCTCCTCTCCTTCCCATAAA
AGCACACGTCTGAACTCTTTCCCTACACGACGCTCTTCCGATCTNNNAGTCTTGTTTCCCAAATGCTGG
AGAGCACACGTCTGAACTCTTTCCCTACACGACGCTCTTCCGATCTNNNAGAAGTCGAGCCAAAACCAA
AGAGCACACGTCTGAACTCTTTCCCTACACGACGCTCTTCCGATCTNNNTAAAGAGTTGCTTTGGAGC
AGCACACGTCTGAACTCTTTCCCTACACGACGCTCTTCCGATCTNNNGTACAATCACCTTCCAGTGGCAT
AGAGCACACGTCTGAACTCTTTCCCTACACGACGCTCTTCCGATCTNNNCTGTGGCTCTCTGTTTGCCTTA
GAGCACACGTCTGAACTCTTTCCCTACACGACGCTCTTCCGATCTNNNATTTTCAGAGGAGATGGGGG
AGCACACGTCTGAACTCTTTCCCTACACGACGCTCTTCCGATCTNNNCTTGTGTGTATCTGTATGTGCG
GAGCACACGTCTGAACTCTTTCCCTACACGACGCTCTTCCGATCTNNNTCTTTTGATCCCCCTCCTTGGAT
GCACACGTCTGAACTCTTTCCCTACACGACGCTCTTCCGATCTNNNCACCATTGATTCTACCCCCGATGCA
GAGCACACGTCTGAACTCTTTCCCTACACGACGCTCTTCCGATCTNNNGCCTGTACATCAAAACAAAGCC
GAGCACACGTCTGAACTCTTTCCCTACACGACGCTCTTCCGATCTNNNTCACATGAGGAAATTAGGCAA

GAGCACACGTCTGAACTCTTTCCCTACACGACGCTCTTCCGATCTNNNGGTGGCAGGTGGTGATAAT
AGCACACGTCTGAACTCTTTCCCTACACGACGCTCTTCCGATCTNNNGGTTGAAGATACTGGGTGGT
GCACACGTCTGAACTCTTTCCCTACACGACGCTCTTCCGATCTNNNCTATCCACAGGACACAAACCAGG
GCACACGTCTGAACTCTTTCCCTACACGACGCTCTTCCGATCTNNNGAACTCAAGTGATCCTGCCACT
GCACACGTCTGAACTCTTTCCCTACACGACGCTCTTCCGATCTNNNGCTGTGCTTCTCTCAGTCTTCAA
AGCACACGTCTGAACTCTTTCCCTACACGACGCTCTTCCGATCTNNNCCTTACACGCTCTGTGAACATC
GCACACGTCTGAACTCTTTCCCTACACGACGCTCTTCCGATCTNNNATTCCATTGTCACCAAACTGCCT
AGCACACGTCTGAACTCTTTCCCTACACGACGCTCTTCCGATCTNNNTGAGAGGGGACTTACAGGGA
GCACACGTCTGAACTCTTTCCCTACACGACGCTCTTCCGATCTNNNGGAAGATGGCATTTCATAGCCC
GCACACGTCTGAACTCTTTCCCTACACGACGCTCTTCCGATCTNNNCCAATGACAAAAGCGTCTTTCCC
GAGCACACGTCTGAACTCTTTCCCTACACGACGCTCTTCCGATCTNNNAGGGAGAGGAGATTGCTCA
GCACACGTCTGAACTCTTTCCCTACACGACGCTCTTCCGATCTNNNAACCTATACCCACCCTACAACCCC
AGCACACGTCTGAACTCTTTCCCTACACGACGCTCTTCCGATCTNNNAACATCGTGCCATTTTCCAATG
AGCACACGTCTGAACTCTTTCCCTACACGACGCTCTTCCGATCTNNNAGGCCCAGGTCGTCATACTTAT
GCACACGTCTGAACTCTTTCCCTACACGACGCTCTTCCGATCTNNNGTCTTAGAGGGGCATGATGGT
AGCACACGTCTGAACTCTTTCCCTACACGACGCTCTTCCGATCTNNNGCAGGGGCAGAGGAAAATAG
AGCACACGTCTGAACTCTTTCCCTACACGACGCTCTTCCGATCTNNNTCAATTCAGAGAACTGCACGG
GCACACGTCTGAACTCTTTCCCTACACGACGCTCTTCCGATCTNNNAAACAGGCAGAAATCCCCTGAA
GAGCACACGTCTGAACTCTTTCCCTACACGACGCTCTTCCGATCTNNNAGTAGGGGATGAGGGAATA
GAGCACACGTCTGAACTCTTTCCCTACACGACGCTCTTCCGATCTNNNGCTCCGATCGTGTACAAATGA
GAGCACACGTCTGAACTCTTTCCCTACACGACGCTCTTCCGATCTNNNTTTGCTGCAAGACAAGGTTT
GCACACGTCTGAACTCTTTCCCTACACGACGCTCTTCCGATCTNNNAAACACTATGGAGTGTTCTGGG
AGCACACGTCTGAACTCTTTCCCTACACGACGCTCTTCCGATCTNNNCCCAAACTGTCTGCCAGTAAT
GCACACGTCTGAACTCTTTCCCTACACGACGCTCTTCCGATCTNNNGACCCAGAGGATCTGGTTAGAG
GCACACGTCTGAACTCTTTCCCTACACGACGCTCTTCCGATCTNNNGGAACTATCCCCTCTGACTGGAG
AGAGCACACGTCTGAACTCTTTCCCTACACGACGCTCTTCCGATCTNNNTGGCTCATTAGAAGGCCAA
AGCACACGTCTGAACTCTTTCCCTACACGACGCTCTTCCGATCTNNNGGAAGGCAAAGGAATATGAA
AAGAGCACACGTCTGAACTCTTTCCCTACACGACGCTCTTCCGATCTNNNCCACTCTTTCTTTCTCCAGC
GAGCACACGTCTGAACTCTTTCCCTACACGACGCTCTTCCGATCTNNNGCAATTTGGTTCCTTCAAGC
GCACACGTCTGAACTCTTTCCCTACACGACGCTCTTCCGATCTNNNCACACTCTTTCTACTGGCTGACC
AGAGCACACGTCTGAACTCTTTCCCTACACGACGCTCTTCCGATCTNNNAGCAACTGATTTTGGTATG
AGCACACGTCTGAACTCTTTCCCTACACGACGCTCTTCCGATCTNNNCCAGCAAGGGTTTTAGAGCCT
AGAGCACACGTCTGAACTCTTTCCCTACACGACGCTCTTCCGATCTNNNTGGCTAGGTCAGCATTTTTC
GCACACGTCTGAACTCTTTCCCTACACGACGCTCTTCCGATCTNNNGTAGGAAGAGGAGGGGTGGA
GCACACGTCTGAACTCTTTCCCTACACGACGCTCTTCCGATCTNNNCCAGCCAGGTAAGGGGTGTAT
GAGCACACGTCTGAACTCTTTCCCTACACGACGCTCTTCCGATCTNNNAGACCCTGGTACTTCCAGAA
GAGCACACGTCTGAACTCTTTCCCTACACGACGCTCTTCCGATCTNNNGCCCACATAAACCACTTACAC
AGCACACGTCTGAACTCTTTCCCTACACGACGCTCTTCCGATCTNNNTAACCACAGAAAATGGTGCCCT
CACACGTCTGAACTCTTTCCCTACACGACGCTCTTCCGATCTNNNAGGTACTAGCCTCTCTCTCTCTCTC
AGCACACGTCTGAACTCTTTCCCTACACGACGCTCTTCCGATCTNNNGCACTTTTTGGCCTCATAAGCA
AGCACACGTCTGAACTCTTTCCCTACACGACGCTCTTCCGATCTNNNAACACCTCCATTTTGCTTTCCC
GAGCACACGTCTGAACTCTTTCCCTACACGACGCTCTTCCGATCTNNNTCCAAAACACCAGATGCAAA
AGCACACGTCTGAACTCTTTCCCTACACGACGCTCTTCCGATCTNNNCGCGTGGCCCACTATTTCTTAT

AGCACACGTCTGAACTCTTTCCCTACACGACGCTCTTCCGATCTNNNTTGGAGTATACCCAGGCAACA
CACACGTCTGAACTCTTTCCCTACACGACGCTCTTCCGATCTNNNCGCGTGGCCCACTATTTTCTTATCA
GAGCACACGTCTGAACTCTTTCCCTACACGACGCTCTTCCGATCTNNNAAAACCACAGTTAGCGGAAC
GAGCACACGTCTGAACTCTTTCCCTACACGACGCTCTTCCGATCTNNNAACATGCTATTTGGGGGTTT
AAGAGCACACGTCTGAACTCTTTCCCTACACGACGCTCTTCCGATCTNNNTCCTGATACTCTGCCAACT
GCACACGTCTGAACTCTTTCCCTACACGACGCTCTTCCGATCTNNNAACCCAGGAATGCTTTTCCCAAG
AGCACACGTCTGAACTCTTTCCCTACACGACGCTCTTCCGATCTNNNGTGTATTGCCTGTGTGTATGT
AGCACACGTCTGAACTCTTTCCCTACACGACGCTCTTCCGATCTNNNAACATGCTATTTGGGGGTTTC
GAGCACACGTCTGAACTCTTTCCCTACACGACGCTCTTCCGATCTNNNGCACATGGCAGGTACTTGAT
AGCACACGTCTGAACTCTTTCCCTACACGACGCTCTTCCGATCTNNNAACGCTGCTAATTCATTCCTTT
AGCACACGTCTGAACTCTTTCCCTACACGACGCTCTTCCGATCTNNNGGCTGTTGAGGCTGAAACATC
AGAGCACACGTCTGAACTCTTTCCCTACACGACGCTCTTCCGATCTNNNTCCCCGTGGGAAATACAGA
GAGCACACGTCTGAACTCTTTCCCTACACGACGCTCTTCCGATCTNNNAAGCTTACTAGGGAATGGAG
AGCACACGTCTGAACTCTTTCCCTACACGACGCTCTTCCGATCTNNNTTGCTTGTTCTTTTTCTGTGCC
AAGAGCACACGTCTGAACTCTTTCCCTACACGACGCTCTTCCGATCTNNNGGCTACTTGCCAGTCTTCA
AGCACACGTCTGAACTCTTTCCCTACACGACGCTCTTCCGATCTNNNGATCAAGCTTCTGCTCTGCACT
GAGCACACGTCTGAACTCTTTCCCTACACGACGCTCTTCCGATCTNNNCTACTGCCTTCCCTAGTTCTCC
GCACACGTCTGAACTCTTTCCCTACACGACGCTCTTCCGATCTNNNAGCCAAAGTGTCCTGATATGCTG
AGAGCACACGTCTGAACTCTTTCCCTACACGACGCTCTTCCGATCTNNNCACATCTGAGGATCTGGAG
GCACACGTCTGAACTCTTTCCCTACACGACGCTCTTCCGATCTNNNGTAGCTTCAGGTTTGACCGCTAA
GAGCACACGTCTGAACTCTTTCCCTACACGACGCTCTTCCGATCTNNNACTGCAGTCTTCATTGCTGC
GAGCACACGTCTGAACTCTTTCCCTACACGACGCTCTTCCGATCTNNNCTCCCTGGGTTGCTTTTTCAG
AGCACACGTCTGAACTCTTTCCCTACACGACGCTCTTCCGATCTNNNCCCAAACTGTCTGCCAGTAATG
GAGCACACGTCTGAACTCTTTCCCTACACGACGCTCTTCCGATCTNNNAAAATACTTCAGGCTGGGTT
GAAGAGCACACGTCTGAACTCTTTCCCTACACGACGCTCTTCCGATCTNNNACAAAAACGATACTTGCA
AGCACACGTCTGAACTCTTTCCCTACACGACGCTCTTCCGATCTNNNTCTGGACAAAGTTGATCCTCT
AGCACACGTCTGAACTCTTTCCCTACACGACGCTCTTCCGATCTNNNCGCTTTCACTCACTATCCACCAT
AGAGCACACGTCTGAACTCTTTCCCTACACGACGCTCTTCCGATCTNNNTAGCCTTGCCAAGTTGACA
AGAGCACACGTCTGAACTCTTTCCCTACACGACGCTCTTCCGATCTNNNAGGAAGGCCTTTGTCTTTC
GAGCACACGTCTGAACTCTTTCCCTACACGACGCTCTTCCGATCTNNNCTTTAGGAAGAGGACTGTGG
AGCACACGTCTGAACTCTTTCCCTACACGACGCTCTTCCGATCTNNNAATGCCTCCCATCAAAGCATCA
ACACGTCTGAACTCTTTCCCTACACGACGCTCTTCCGATCTNNNAAGTCATTATTCATACTCCCCTCTCC
GAGCACACGTCTGAACTCTTTCCCTACACGACGCTCTTCCGATCTNNNGGCTTTCTTCCCTTTGCTCTCA
AAGAGCACACGTCTGAACTCTTTCCCTACACGACGCTCTTCCGATCTNNNTCTTTGTTCCTCATGTGTG
AGAGCACACGTCTGAACTCTTTCCCTACACGACGCTCTTCCGATCTNNNTTGACATGCACCTGTGGTTG
AAGAGCACACGTCTGAACTCTTTCCCTACACGACGCTCTTCCGATCTNNNACAAGAAAGACCAACAAC
AGAGCACACGTCTGAACTCTTTCCCTACACGACGCTCTTCCGATCTNNNTGTGTAGCAAAGACAATTC
AGAGCACACGTCTGAACTCTTTCCCTACACGACGCTCTTCCGATCTNNNCCTCCTGAACCTTGCTTTCT
GAGCACACGTCTGAACTCTTTCCCTACACGACGCTCTTCCGATCTNNNCCACAGTGAGAACTGAATG
AAGAGCACACGTCTGAACTCTTTCCCTACACGACGCTCTTCCGATCTNNNCCATAGTGCACAGTGATT
AGCACACGTCTGAACTCTTTCCCTACACGACGCTCTTCCGATCTNNNGAGCCTCTCCCCTTTACAATGT
AGCACACGTCTGAACTCTTTCCCTACACGACGCTCTTCCGATCTNNNATCGGTGTTGATGGGGAAAG
GCACACGTCTGAACTCTTTCCCTACACGACGCTCTTCCGATCTNNNCTCACCCATTACCAGGTTCAAGG

GAGCACACGTCTGAACTCTTTCCCTACACGACGCTCTTCCGATCTNNNGCACATTTCAGTTCCTTTCAGG
AGCACACGTCTGAACTCTTTCCCTACACGACGCTCTTCCGATCTNNNCAATAGGATGAGGGGTGTGTG
AGCACACGTCTGAACTCTTTCCCTACACGACGCTCTTCCGATCTNNNAATTAGCCAGATGTTGTGACCC
GAGCACACGTCTGAACTCTTTCCCTACACGACGCTCTTCCGATCTNNNCAACCCATCTAAGGGGATTCA
AGCACACGTCTGAACTCTTTCCCTACACGACGCTCTTCCGATCTNNNTTAGGATGGAGTGAAGGCGA
GAGCACACGTCTGAACTCTTTCCCTACACGACGCTCTTCCGATCTNNNCAGATCTGTGGAAGAAAAAG
GAGCACACGTCTGAACTCTTTCCCTACACGACGCTCTTCCGATCTNNNTATTCAGACCACAGCATTCAG
AGCACACGTCTGAACTCTTTCCCTACACGACGCTCTTCCGATCTNNNTTTATCCGCACACATTCACCCA
AGCACACGTCTGAACTCTTTCCCTACACGACGCTCTTCCGATCTNNNGCACATGACATCTCACCACAGT
GAGCACACGTCTGAACTCTTTCCCTACACGACGCTCTTCCGATCTNNNTGTTTAGCCACAGTCACAGCT
GAGCACACGTCTGAACTCTTTCCCTACACGACGCTCTTCCGATCTNNNATCAAAGGGCAAAGCATTTCA
AAGAGCACACGTCTGAACTCTTTCCCTACACGACGCTCTTCCGATCTNNNGGTAACAAGAAACACTCC
AGCACACGTCTGAACTCTTTCCCTACACGACGCTCTTCCGATCTNNNTCTCCCTCACATTTCAGTACAAC
GAGCACACGTCTGAACTCTTTCCCTACACGACGCTCTTCCGATCTNNNACACCTACTAGAATTGCAACG
CACACGTCTGAACTCTTTCCCTACACGACGCTCTTCCGATCTNNNTGCAGGTTTTCTTTCAATGGGTGC
GAGCACACGTCTGAACTCTTTCCCTACACGACGCTCTTCCGATCTNNNACCTGAGATGTCCTCCTTTCA
GAGCACACGTCTGAACTCTTTCCCTACACGACGCTCTTCCGATCTNNNATCTTTCTCCCTCTCTCTCATCC
AGCACACGTCTGAACTCTTTCCCTACACGACGCTCTTCCGATCTNNNGCCAGCTGAGCCTACTCTTTTT
AGAGCACACGTCTGAACTCTTTCCCTACACGACGCTCTTCCGATCTNNNACCCAGGTTTAGGATTTATG
AGAGCACACGTCTGAACTCTTTCCCTACACGACGCTCTTCCGATCTNNNGTAGTGGCTTGGTCCTAAGG
AGCACACGTCTGAACTCTTTCCCTACACGACGCTCTTCCGATCTNNNACCAACAAACAAGTGGCTCGTA
AGCACACGTCTGAACTCTTTCCCTACACGACGCTCTTCCGATCTNNNCAGAACCAACAAGTTTGATCAC
AGAGCACACGTCTGAACTCTTTCCCTACACGACGCTCTTCCGATCTNNNTACTTCAGCTTATCTTTGGCG
AGCACACGTCTGAACTCTTTCCCTACACGACGCTCTTCCGATCTNNNCTGGAGAAGTGGGTACCTTTGT
AGCACACGTCTGAACTCTTTCCCTACACGACGCTCTTCCGATCTNNNTGCCGTCAATATGGGTTTTCCT
GAGCACACGTCTGAACTCTTTCCCTACACGACGCTCTTCCGATCTNNNCTTTGGTCTAAGGGTCACCAT
AGCACACGTCTGAACTCTTTCCCTACACGACGCTCTTCCGATCTNNNATATCATCAAATGACTGTGGG
AGCACACGTCTGAACTCTTTCCCTACACGACGCTCTTCCGATCTNNNAAATGCCAGGGTTGCAATATCA
GCACACGTCTGAACTCTTTCCCTACACGACGCTCTTCCGATCTNNNACTGTCATCTCCTTGGTATGGGG
AGCACACGTCTGAACTCTTTCCCTACACGACGCTCTTCCGATCTNNNCCTTTTTGTGACATCCTTGGTG
AGAGCACACGTCTGAACTCTTTCCCTACACGACGCTCTTCCGATCTNNNAAGAAAGCACAACACTAAG
AGAGCACACGTCTGAACTCTTTCCCTACACGACGCTCTTCCGATCTNNNGAAAGATGAATTGTGTGCG
GAGCACACGTCTGAACTCTTTCCCTACACGACGCTCTTCCGATCTNNNAGACCGATATAGCACTCACAT
AGCACACGTCTGAACTCTTTCCCTACACGACGCTCTTCCGATCTNNNCAGTTCTCCCACCTCAACCTATGC
GCACACGTCTGAACTCTTTCCCTACACGACGCTCTTCCGATCTNNNTGGATTCAAATCCCCTTGGATCA
GAGCACACGTCTGAACTCTTTCCCTACACGACGCTCTTCCGATCTNNNTCTCACACGTCTCCTTCTCTCT
AGAGCACACGTCTGAACTCTTTCCCTACACGACGCTCTTCCGATCTNNNTATCCAAGCTAAGAAGCACG
GCACACGTCTGAACTCTTTCCCTACACGACGCTCTTCCGATCTNNNTCCCTAAAATTGCACACCAGACCC
AGCACACGTCTGAACTCTTTCCCTACACGACGCTCTTCCGATCTNNNCCCATGCATTTGAACTAACTGG
AGCACACGTCTGAACTCTTTCCCTACACGACGCTCTTCCGATCTNNNTGGAGCCCTGATTTCACTCAAT
AGCACACGTCTGAACTCTTTCCCTACACGACGCTCTTCCGATCTNNNTTGCACATACACACACACAAAG
AGCACACGTCTGAACTCTTTCCCTACACGACGCTCTTCCGATCTNNNCCCCTCCCTTGTTTCATTAGCTCC
AGCACACGTCTGAACTCTTTCCCTACACGACGCTCTTCCGATCTNNNTAACACAGCTCTCACCACCATTC

AGCACACGTCTGAACTCTTTCCCTACACGACGCTCTTCCGATCTNNNGAGGCAGGTTTCTCTCTCTCT
AAGAGCACACGTCTGAACTCTTTCCCTACACGACGCTCTTCCGATCTNNNAAATGGGCTCTGTCTCTGT
AGCACACGTCTGAACTCTTTCCCTACACGACGCTCTTCCGATCTNNNCCTAGGTTTTGCGTGAGAGTC
GAGCACACGTCTGAACTCTTTCCCTACACGACGCTCTTCCGATCTNNNGGTGGGGGTATGAATCTAC
AGCACACGTCTGAACTCTTTCCCTACACGACGCTCTTCCGATCTNNNTGTGTCCGTGCATATGTGTGTA
AGCACACGTCTGAACTCTTTCCCTACACGACGCTCTTCCGATCTNNNTTCAGTAGGACACACCACCTGAT
AGCACACGTCTGAACTCTTTCCCTACACGACGCTCTTCCGATCTNNNATAAAAATATGTGAAGCTTGAT
AGCACACGTCTGAACTCTTTCCCTACACGACGCTCTTCCGATCTNNNATCACCAGAAGTCTGTCTTCCCC
AGAGCACACGTCTGAACTCTTTCCCTACACGACGCTCTTCCGATCTNNNTAACTTTCAGGACAGGTAG
AGCACACGTCTGAACTCTTTCCCTACACGACGCTCTTCCGATCTNNNTTTCTCAATAGGGGTGTGTGT
GCACACGTCTGAACTCTTTCCCTACACGACGCTCTTCCGATCTNNNGCTGGAGAGGGAAAGAATGACA
AGCACACGTCTGAACTCTTTCCCTACACGACGCTCTTCCGATCTNNNTAGGATTGCAAGTGAGTGGTC
AGCACACGTCTGAACTCTTTCCCTACACGACGCTCTTCCGATCTNNNGGGACTTAGAGGGTCTACTGC
AGCACACGTCTGAACTCTTTCCCTACACGACGCTCTTCCGATCTNNNTTTCATGACGCATCACTCTACCT
AGCACACGTCTGAACTCTTTCCCTACACGACGCTCTTCCGATCTNNNTATTGATCCGCCCCAATTGTCTC
AGCACACGTCTGAACTCTTTCCCTACACGACGCTCTTCCGATCTNNNTAAAGCCTGGTGAGAACTGGG
AGCACACGTCTGAACTCTTTCCCTACACGACGCTCTTCCGATCTNNNTTACAAATTGGGGGAGAAGGG
AGCACACGTCTGAACTCTTTCCCTACACGACGCTCTTCCGATCTNNNACCTGGGACAGTAACCACATT
AGCACACGTCTGAACTCTTTCCCTACACGACGCTCTTCCGATCTNNNGGAAAAGGGAGGTGAAGGAA
AGAGCACACGTCTGAACTCTTTCCCTACACGACGCTCTTCCGATCTNNNATTGGTGCAGCATGACATG
AGCACACGTCTGAACTCTTTCCCTACACGACGCTCTTCCGATCTNNNGGCTCAGGTATGACTGGTGAA
AGCACACGTCTGAACTCTTTCCCTACACGACGCTCTTCCGATCTNNNGATCACCTGTGTGTGATCTGCT
GAGCACACGTCTGAACTCTTTCCCTACACGACGCTCTTCCGATCTNNNGAGGGTATGATGGAAAAGCA
AGCACACGTCTGAACTCTTTCCCTACACGACGCTCTTCCGATCTNNNGTTCAGCCAGTAGCTAGAAGA
AGCACACGTCTGAACTCTTTCCCTACACGACGCTCTTCCGATCTNNNGTCCGAAGACCTCTTTTGTCTCC
AGCACACGTCTGAACTCTTTCCCTACACGACGCTCTTCCGATCTNNNTCCATCATGTAACCCTGTTGCA
AGCACACGTCTGAACTCTTTCCCTACACGACGCTCTTCCGATCTNNNAATGCAACTCTCTTTTCTCCGG
AGCACACGTCTGAACTCTTTCCCTACACGACGCTCTTCCGATCTNNNTCATTTCTCCCCTCTACTTCCCC
AGAGCACACGTCTGAACTCTTTCCCTACACGACGCTCTTCCGATCTNNNCTTTCAGTGGTCACATCCAG
AGCACACGTCTGAACTCTTTCCCTACACGACGCTCTTCCGATCTNNNTGTTTGTGCTGAGTTGTTTGCA
GCACACGTCTGAACTCTTTCCCTACACGACGCTCTTCCGATCTNNNATTAGCCTCAGGCCGTTAGTTTG
AGCACACGTCTGAACTCTTTCCCTACACGACGCTCTTCCGATCTNNNAGAGATTGCATGGGTAGAGC
AGCACACGTCTGAACTCTTTCCCTACACGACGCTCTTCCGATCTNNNTGCGTATGCATCCACATGAAAA
GCACACGTCTGAACTCTTTCCCTACACGACGCTCTTCCGATCTNNNATTGACATTCCAGGAAGATGGG
AGAGCACACGTCTGAACTCTTTCCCTACACGACGCTCTTCCGATCTNNNCCTCTGCCTCTCTCAGGTCTC
GCACACGTCTGAACTCTTTCCCTACACGACGCTCTTCCGATCTNNNACAGTTTACACTGGCATCTGGAA
AGCACACGTCTGAACTCTTTCCCTACACGACGCTCTTCCGATCTNNNTCCTTCCTTGTGTGTGTGTGTTT
AGCACACGTCTGAACTCTTTCCCTACACGACGCTCTTCCGATCTNNNGCAAAGGTTTTGAACATGGGC
AGCACACGTCTGAACTCTTTCCCTACACGACGCTCTTCCGATCTNNNTGAGCTTCATGCAAGAGAGTGA
AGCACACGTCTGAACTCTTTCCCTACACGACGCTCTTCCGATCTNNNTCCTGGTGTCCAAGGAAAGAA
AGCACACGTCTGAACTCTTTCCCTACACGACGCTCTTCCGATCTNNNTGACTTAGCCTGAGTTGGTCTC
AGCACACGTCTGAACTCTTTCCCTACACGACGCTCTTCCGATCTNNNCAGTTATTGAGCTAATGGCCAG
GAGCACACGTCTGAACTCTTTCCCTACACGACGCTCTTCCGATCTNNNTACAGCCTAGTGAGGATCCA

AGCACACGTCTGAACTCTTTCCCTACACGACGCTCTTCCGATCTNNNTAAAGGCAACTATTGCCCACTC
CACACGTCTGAACTCTTTCCCTACACGACGCTCTTCCGATCTNNNGAACTAAGCTACATGTGTGGGCA
AGCACACGTCTGAACTCTTTCCCTACACGACGCTCTTCCGATCTNNNTAGATAGAGTTGTGTGTGTG
AGCACACGTCTGAACTCTTTCCCTACACGACGCTCTTCCGATCTNNNAGGCCTTAGCCAAACAAATCCA
AGCACACGTCTGAACTCTTTCCCTACACGACGCTCTTCCGATCTNNNTAATGTGCATGCTCTCTCTCTCT
AGCACACGTCTGAACTCTTTCCCTACACGACGCTCTTCCGATCTNNNGCAAAATAGGTCTTATGACTT
GCACACGTCTGAACTCTTTCCCTACACGACGCTCTTCCGATCTNNNAGCTATTCATGCACACTCACACCC
AGCACACGTCTGAACTCTTTCCCTACACGACGCTCTTCCGATCTNNNTGGATTAGCATGTCCAATGCCT
AGCACACGTCTGAACTCTTTCCCTACACGACGCTCTTCCGATCTNNNAGTTGCACTGCTGTGAGAAAA
GAGCACACGTCTGAACTCTTTCCCTACACGACGCTCTTCCGATCTNNNGGGGGTCCTGTCTTCTTCTGT
AGCACACGTCTGAACTCTTTCCCTACACGACGCTCTTCCGATCTNNNGTTCCCATTCTGCTTTTTGAGC
GAGCACACGTCTGAACTCTTTCCCTACACGACGCTCTTCCGATCTNNNTGGATTTCAGGTGATTGCGA
AGCACACGTCTGAACTCTTTCCCTACACGACGCTCTTCCGATCTNNNTCTGCTGGTTCTCACTGATGGA
AGAGCACACGTCTGAACTCTTTCCCTACACGACGCTCTTCCGATCTNNNAACCAGTGTCACTTTCTGAA
AGCACACGTCTGAACTCTTTCCCTACACGACGCTCTTCCGATCTNNNTAGAATCCCAAGCCAAGTCAG
AGCACACGTCTGAACTCTTTCCCTACACGACGCTCTTCCGATCTNNNAGTGTTCCTCAGACCTGGTAA
AGCACACGTCTGAACTCTTTCCCTACACGACGCTCTTCCGATCTNNNATGGCTTGCTATTGCAGGGTTT
AGCACACGTCTGAACTCTTTCCCTACACGACGCTCTTCCGATCTNNNCTTCTTGGCCATGACATCTTCC
CACACGTCTGAACTCTTTCCCTACACGACGCTCTTCCGATCTNNNACTTATACTTCCACCTGGGGCTTC
AGCACACGTCTGAACTCTTTCCCTACACGACGCTCTTCCGATCTNNNAACACGCAACCTTCTTCTTGTCT
AGCACACGTCTGAACTCTTTCCCTACACGACGCTCTTCCGATCTNNNTGGCATCTGGGAGAGAACAT
GAGCACACGTCTGAACTCTTTCCCTACACGACGCTCTTCCGATCTNNNTGCTTCCAAACAAGAATGTGA
AGAGCACACGTCTGAACTCTTTCCCTACACGACGCTCTTCCGATCTNNNGGCTGCTGTTTTGTGTACG
AGAGCACACGTCTGAACTCTTTCCCTACACGACGCTCTTCCGATCTNNNGAAGAAGAGCATTAGGGC
GAGCACACGTCTGAACTCTTTCCCTACACGACGCTCTTCCGATCTNNNTGGATTAGCATGTCCAATGC
AGCACACGTCTGAACTCTTTCCCTACACGACGCTCTTCCGATCTNNNAAGTCATCCATTTGCAGGGAA
GAGCACACGTCTGAACTCTTTCCCTACACGACGCTCTTCCGATCTNNNTAGAACTGGCCCCTTGGGATA
GAGCACACGTCTGAACTCTTTCCCTACACGACGCTCTTCCGATCTNNNACCTATCTGTCACCTTCCATA
AGCACACGTCTGAACTCTTTCCCTACACGACGCTCTTCCGATCTNNNCCCAGAGATGGAACCCTCTAC
GAGCACACGTCTGAACTCTTTCCCTACACGACGCTCTTCCGATCTNNNCCCTCAGTACGCAGTAGCTTT
AGCACACGTCTGAACTCTTTCCCTACACGACGCTCTTCCGATCTNNNTAAAGTGGCCATTCTTCCTCAG
AGCACACGTCTGAACTCTTTCCCTACACGACGCTCTTCCGATCTNNNGTGGAACCCAATGCACTCTTGT
AGCACACGTCTGAACTCTTTCCCTACACGACGCTCTTCCGATCTNNNATGTTTATGGTCTCTCGTCCCCT
AGCACACGTCTGAACTCTTTCCCTACACGACGCTCTTCCGATCTNNNTTGCTGATGCCAGTGATTCAA
AGCACACGTCTGAACTCTTTCCCTACACGACGCTCTTCCGATCTNNNCCAGCAGCCAAAGAGAAAACA
AGCACACGTCTGAACTCTTTCCCTACACGACGCTCTTCCGATCTNNNTCCCTCTCTCTCTGAATCTCCCTC
AGAGCACACGTCTGAACTCTTTCCCTACACGACGCTCTTCCGATCTNNNCTTCCTAAAGCCCCCACCAA
GCACACGTCTGAACTCTTTCCCTACACGACGCTCTTCCGATCTNNNGTTGGCTCTGTTCGTTCACTAGA
AGAGCACACGTCTGAACTCTTTCCCTACACGACGCTCTTCCGATCTNNNCTGGACCCTAAACGAGTGT
AGAGCACACGTCTGAACTCTTTCCCTACACGACGCTCTTCCGATCTNNNACATACATGTTGAGATCCT
GCACACGTCTGAACTCTTTCCCTACACGACGCTCTTCCGATCTNNNCGTAATTTCCCCATGCCCATGTA
AGCACACGTCTGAACTCTTTCCCTACACGACGCTCTTCCGATCTNNNACACACTCAATTTGAAACCGCA
GCACACGTCTGAACTCTTTCCCTACACGACGCTCTTCCGATCTNNNACAGACCACAAGCTGAATTACG

AGCACACGTCTGAACTCTTTCCCTACACGACGCTCTTCCGATCTNNNTAGCTGTATCAGGTACTGCAG
GAGCACACGTCTGAACTCTTTCCCTACACGACGCTCTTCCGATCTNNNATCCTTCTTTCCCAACCATCAG
AGCACACGTCTGAACTCTTTCCCTACACGACGCTCTTCCGATCTNNNCTAATTACTACCCCCACTCCCCA
AGCACACGTCTGAACTCTTTCCCTACACGACGCTCTTCCGATCTNNNTGAGACTTCAAAAGGGATTGA
AGAGCACACGTCTGAACTCTTTCCCTACACGACGCTCTTCCGATCTNNNTTGGATTGTTGTTGCTCCTGT
AAGAGCACACGTCTGAACTCTTTCCCTACACGACGCTCTTCCGATCTNNNTGGGTTTGGGTAAACATA
AGCACACGTCTGAACTCTTTCCCTACACGACGCTCTTCCGATCTNNNTACCAGCGCTAGGATCACTATG
GAGCACACGTCTGAACTCTTTCCCTACACGACGCTCTTCCGATCTNNNATTCTCAGTTCTTTGCCTTCCAT
GAGCACACGTCTGAACTCTTTCCCTACACGACGCTCTTCCGATCTNNNCAGCTGGTTACTCATGGTTCT
AGCACACGTCTGAACTCTTTCCCTACACGACGCTCTTCCGATCTNNNTGCTTGTCAGTTTTCTCTCTTCCT
GCACACGTCTGAACTCTTTCCCTACACGACGCTCTTCCGATCTNNNAGGTATCTTGGCGTGACAACTTC
GAGCACACGTCTGAACTCTTTCCCTACACGACGCTCTTCCGATCTNNNAGGGGCACCTCCCTGTCTATAA
AGCACACGTCTGAACTCTTTCCCTACACGACGCTCTTCCGATCTNNNTAGCTGTATCAGGTACTGCAG
GAGCACACGTCTGAACTCTTTCCCTACACGACGCTCTTCCGATCTNNNACACTCAGGAGAATAACAAGT
AGAGCACACGTCTGAACTCTTTCCCTACACGACGCTCTTCCGATCTNNNACAGCACATGCAAATGGATT
AGCACACGTCTGAACTCTTTCCCTACACGACGCTCTTCCGATCTNNNGCATTACTAAATAACTTGCAAA
GAGCACACGTCTGAACTCTTTCCCTACACGACGCTCTTCCGATCTNNNAGGGATAGGGCCAGTGAGAA
GAGCACACGTCTGAACTCTTTCCCTACACGACGCTCTTCCGATCTNNNCCACCTATTGAATTGTGCCAG
AGCACACGTCTGAACTCTTTCCCTACACGACGCTCTTCCGATCTNNNAGTGGCATTTATGTGTTTCTCTT
GAGCACACGTCTGAACTCTTTCCCTACACGACGCTCTTCCGATCTNNNACATCCTGGTCCTCTCTCTCTTT
AGCACACGTCTGAACTCTTTCCCTACACGACGCTCTTCCGATCTNNNCAGGGCAGATGCAAGGAATAC
GCACACGTCTGAACTCTTTCCCTACACGACGCTCTTCCGATCTNNNGCAGGCAATCAGAGATTGTGGA
AGAGCACACGTCTGAACTCTTTCCCTACACGACGCTCTTCCGATCTNNNGACATCCTTCATTCCTAAGG
GAGCACACGTCTGAACTCTTTCCCTACACGACGCTCTTCCGATCTNNNAACTCTCCAGGTAAGTTGGCT
AGCACACGTCTGAACTCTTTCCCTACACGACGCTCTTCCGATCTNNNCTTGGCTATTCTTGGAGCTGTG
GAGCACACGTCTGAACTCTTTCCCTACACGACGCTCTTCCGATCTNNNAGAGGTGGAAAATAATGTG
AGCACACGTCTGAACTCTTTCCCTACACGACGCTCTTCCGATCTNNNACTTCTAGGCAGTTTTGTGGAG
GAGCACACGTCTGAACTCTTTCCCTACACGACGCTCTTCCGATCTNNNTCAGGAGAGAGGGAGGAATA
AGAGCACACGTCTGAACTCTTTCCCTACACGACGCTCTTCCGATCTNNNAATACCCTGTGTGTGTGGAT
AGCACACGTCTGAACTCTTTCCCTACACGACGCTCTTCCGATCTNNNTCCTCATAGTAGTTTTAGGTTAT
GAGCACACGTCTGAACTCTTTCCCTACACGACGCTCTTCCGATCTNNNCTCAAGTAAGAAGTTCCGGAG
GAGCACACGTCTGAACTCTTTCCCTACACGACGCTCTTCCGATCTNNNAGACACCAACCCTGTATGAA
AGCACACGTCTGAACTCTTTCCCTACACGACGCTCTTCCGATCTNNNTTGCCGGGTAAAGATCTAAGCA
AGCACACGTCTGAACTCTTTCCCTACACGACGCTCTTCCGATCTNNNGTGGGAGAAGATGGTACAAGA
AGAGCACACGTCTGAACTCTTTCCCTACACGACGCTCTTCCGATCTNNNACCCCAGTTTACATAAGTGTT
AGAGCACACGTCTGAACTCTTTCCCTACACGACGCTCTTCCGATCTNNNGGAGGAGTTGAGTGGAAAT
AGCACACGTCTGAACTCTTTCCCTACACGACGCTCTTCCGATCTNNNTCTTTTGCTGTTCTGGTTCCTCC
AGCACACGTCTGAACTCTTTCCCTACACGACGCTCTTCCGATCTNNNAGAAACCAAGTCCAGTAGAGG
AGCACACGTCTGAACTCTTTCCCTACACGACGCTCTTCCGATCTNNNTGTTGTTATGCATATGTGGCT
GAGCACACGTCTGAACTCTTTCCCTACACGACGCTCTTCCGATCTNNNAGTAGAAAGGCAGGAGAGCA
GAGCACACGTCTGAACTCTTTCCCTACACGACGCTCTTCCGATCTNNNACACAGGAGAAATCAGGAAC
GAGCACACGTCTGAACTCTTTCCCTACACGACGCTCTTCCGATCTNNNTATTATGGCTGCTGCACTGGT
AGAGCACACGTCTGAACTCTTTCCCTACACGACGCTCTTCCGATCTNNNCCTATTCCAGGGAGGGAAA

AGCACACGTCTGAACTCTTTCCCTACACGACGCTCTTCCGATCTNNNTTCTTTCGTTTTCAGCAGTTTGG

GAGCACACGTCTGAACTCTTTCCCTACACGACGCTCTTCCGATCTNNNTGGATTAGAGTAAGGGGAG

AGAGCACACGTCTGAACTCTTTCCCTACACGACGCTCTTCCGATCTNNNGGGGCCTTTTACAGGACACT

AGCACACGTCTGAACTCTTTCCCTACACGACGCTCTTCCGATCTNNNACACATATCTACTGGATCGTGGT

AGAGCACACGTCTGAACTCTTTCCCTACACGACGCTCTTCCGATCTNNNAAAGTCCTGCAACAGTAAGA

AGAGCACACGTCTGAACTCTTTCCCTACACGACGCTCTTCCGATCTNNNTGAACCTGAGCTCTGATCCTG

AGCACACGTCTGAACTCTTTCCCTACACGACGCTCTTCCGATCTNNNCTAAGCTCTGGCAGGGAGAAAT

GCACACGTCTGAACTCTTTCCCTACACGACGCTCTTCCGATCTNNNCTGAGACCCTGTCTCTAAACTGC

GAAGAGCACACGTCTGAACTCTTTCCCTACACGACGCTCTTCCGATCTNNNGCATAGGAGATTGACAG

GAGCACACGTCTGAACTCTTTCCCTACACGACGCTCTTCCGATCTNNNTCCTCTAACTCACTCGCTCTCT

GAGCACACGTCTGAACTCTTTCCCTACACGACGCTCTTCCGATCTNNNCAAAGAAAGGCTGGAGCACA

GAGCACACGTCTGAACTCTTTCCCTACACGACGCTCTTCCGATCTNNNTTGAGGAGGAAAGGCATGTT

GAGCACACGTCTGAACTCTTTCCCTACACGACGCTCTTCCGATCTNNNTGAGTGGTGATCATGCAGTAT

GAGCACACGTCTGAACTCTTTCCCTACACGACGCTCTTCCGATCTNNNTTTCTCTTAGGTGACCTCTCC

GAGCACACGTCTGAACTCTTTCCCTACACGACGCTCTTCCGATCTNNNTGGGGCATAGTAGAAACTGC

GAGCACACGTCTGAACTCTTTCCCTACACGACGCTCTTCCGATCTNNNCTAATGCCAGCATGATCCCTG

GAGCACACGTCTGAACTCTTTCCCTACACGACGCTCTTCCGATCTNNNCTCATCTCTTTTCAGCCCAGTC

GAGCACACGTCTGAACTCTTTCCCTACACGACGCTCTTCCGATCTNNNAACTCAGAAGTGAAGATGCCT

AGAGCACACGTCTGAACTCTTTCCCTACACGACGCTCTTCCGATCTNNNAGTAACCGCTATTTGTTGAG

AGAGCACACGTCTGAACTCTTTCCCTACACGACGCTCTTCCGATCTNNNTTGCATTTGCAACAAAGGA

AGCACACGTCTGAACTCTTTCCCTACACGACGCTCTTCCGATCTNNNCCTGCTATCTTGTTGCAGGACTT

AGAGCACACGTCTGAACTCTTTCCCTACACGACGCTCTTCCGATCTNNNCAAAAGGTAGGGGGACAA

GAGCACACGTCTGAACTCTTTCCCTACACGACGCTCTTCCGATCTNNNTCTCTTTTACCCCCATTCCCCAA

AGCACACGTCTGAACTCTTTCCCTACACGACGCTCTTCCGATCTNNNTACACTTCAAGTCCAGCCCTACAC

AGCACACGTCTGAACTCTTTCCCTACACGACGCTCTTCCGATCTNNNGAGTGTACCTGAAACCAGTCTG

AGAGCACACGTCTGAACTCTTTCCCTACACGACGCTCTTCCGATCTNNNCAATAGTTTTGGGATGGCTG

GAGCACACGTCTGAACTCTTTCCCTACACGACGCTCTTCCGATCTNNNTTTGGTGTCTCTCCTTCCAGTT

GAGCACACGTCTGAACTCTTTCCCTACACGACGCTCTTCCGATCTNNNATAGAAGTAAGGGAGGAGG

AAGAGCACACGTCTGAACTCTTTCCCTACACGACGCTCTTCCGATCTNNNATCACAAGCCTGGGGGCCT

GAGCACACGTCTGAACTCTTTCCCTACACGACGCTCTTCCGATCTNNNAGAGGACGTCAGGATTCACTC

GAGCACACGTCTGAACTCTTTCCCTACACGACGCTCTTCCGATCTNNNACAGGCATACTTGTCCATTTTT

AGCACACGTCTGAACTCTTTCCCTACACGACGCTCTTCCGATCTNNNACCTGTACTTCTGCCTGGAATCT

AGCACACGTCTGAACTCTTTCCCTACACGACGCTCTTCCGATCTNNNAGGATGAGAACAATAGAATGA

GAGCACACGTCTGAACTCTTTCCCTACACGACGCTCTTCCGATCTNNNAGTAAGCAGCCACTGACATCA

AGCACACGTCTGAACTCTTTCCCTACACGACGCTCTTCCGATCTNNNACCTAGAAAGAGGGACCATCAG

AGCACACGTCTGAACTCTTTCCCTACACGACGCTCTTCCGATCTNNNTTGAGAATCACTGCGTTAGTGG

GAGCACACGTCTGAACTCTTTCCCTACACGACGCTCTTCCGATCTNNNTTACAATGTGCTGCTAGGCAG

AGAGCACACGTCTGAACTCTTTCCCTACACGACGCTCTTCCGATCTNNNAGCCTTTGTGTTTGGATGGA

AGCACACGTCTGAACTCTTTCCCTACACGACGCTCTTCCGATCTNNNACACATAAGGCTGAAGGAAGTC

AGAGCACACGTCTGAACTCTTTCCCTACACGACGCTCTTCCGATCTNNNTTCAACAGCTGGAGCATGAA

GAGCACACGTCTGAACTCTTTCCCTACACGACGCTCTTCCGATCTNNNCTACACACCACACACACTTACCC

GAGCACACGTCTGAACTCTTTCCCTACACGACGCTCTTCCGATCTNNNAGATGTCTCTGTCTTCTGCCAG

AAGAGCACACGTCTGAACTCTTTCCCTACACGACGCTCTTCCGATCTNNNATTGGTGCAGCATGACAT

AGCACACGTCTGAACTCTTTCCCTACACGACGCTCTTCCGATCTNNNGACCCTCACAATGTCGTTGTT
AAGAGCACACGTCTGAACTCTTTCCCTACACGACGCTCTTCCGATCTNNNGAGCTGTTAGGCCTTGTG
AGCACACGTCTGAACTCTTTCCCTACACGACGCTCTTCCGATCTNNNCAACTCTGCTGGTTCTCTCCTT
AGCACACGTCTGAACTCTTTCCCTACACGACGCTCTTCCGATCTNNNTCTTTGAGCACTATGGTCTTGC
GCACACGTCTGAACTCTTTCCCTACACGACGCTCTTCCGATCTNNNAATTAGAGGCTAGATGCCCCAC
AGCACACGTCTGAACTCTTTCCCTACACGACGCTCTTCCGATCTNNNCTGCTGTTTGCTCTCTCTCTCT
AGCACACGTCTGAACTCTTTCCCTACACGACGCTCTTCCGATCTNNNTCAGCCAATCAATCAATCTGTC
GAGCACACGTCTGAACTCTTTCCCTACACGACGCTCTTCCGATCTNNNCACATTCAGCTTCAGAAATGC
GCACACGTCTGAACTCTTTCCCTACACGACGCTCTTCCGATCTNNNGGCAAAGACAGATCAATCTCCC
AGCACACGTCTGAACTCTTTCCCTACACGACGCTCTTCCGATCTNNNAGCAGTCAGGGCAAGATAGA
GCACACGTCTGAACTCTTTCCCTACACGACGCTCTTCCGATCTNNNGTCTGGAATGTTCCGCAATCTCT
CACACGTCTGAACTCTTTCCCTACACGACGCTCTTCCGATCTNNNACACAGCACTTATGTGTTTGCAC
AGAGCACACGTCTGAACTCTTTCCCTACACGACGCTCTTCCGATCTNNNACATTCCATTCTGTGTGTGT
AGCACACGTCTGAACTCTTTCCCTACACGACGCTCTTCCGATCTNNNCGACTTGAACTTCTCAAATGCC
AGCACACGTCTGAACTCTTTCCCTACACGACGCTCTTCCGATCTNNNTTGCTAACTTACCTGCAATGCC
AGCACACGTCTGAACTCTTTCCCTACACGACGCTCTTCCGATCTNNNAGTCATAATGTGGTAAGGAA
AGCACACGTCTGAACTCTTTCCCTACACGACGCTCTTCCGATCTNNNACAAGAATGTTTCAAAATGACT
GAGCACACGTCTGAACTCTTTCCCTACACGACGCTCTTCCGATCTNNNGTTTCCATCTTTCAGGCAACA
GAGCACACGTCTGAACTCTTTCCCTACACGACGCTCTTCCGATCTNNNGCCACCTTTTAGCATCCACCT
GCACACGTCTGAACTCTTTCCCTACACGACGCTCTTCCGATCTNNNTCCATTTCCCCTCCCTCATCAATC
GAGCACACGTCTGAACTCTTTCCCTACACGACGCTCTTCCGATCTNNNTCAATTTGGCCCGAATAGTA
AGCACACGTCTGAACTCTTTCCCTACACGACGCTCTTCCGATCTNNNACCTCAGAAAATTTGGCTTACA
GAGCACACGTCTGAACTCTTTCCCTACACGACGCTCTTCCGATCTNNNAAAAAGCACATCAACCACAT
GCACACGTCTGAACTCTTTCCCTACACGACGCTCTTCCGATCTNNNAGTTCCAGCCAACAGAACATGA
GAGCACACGTCTGAACTCTTTCCCTACACGACGCTCTTCCGATCTNNNTGATTACGTTTTTCCAAGGCT
AGCACACGTCTGAACTCTTTCCCTACACGACGCTCTTCCGATCTNNNTTGGCCAAAAGTACTGCATGG
AGAGCACACGTCTGAACTCTTTCCCTACACGACGCTCTTCCGATCTNNNGATACCAGGATAACAGTGC
AGCACACGTCTGAACTCTTTCCCTACACGACGCTCTTCCGATCTNNNTCAGAGCTTACTTTGGTGGTT
AGCACACGTCTGAACTCTTTCCCTACACGACGCTCTTCCGATCTNNNAATGTTTTGCAGAGAGGGACT
AGCACACGTCTGAACTCTTTCCCTACACGACGCTCTTCCGATCTNNNTGAAGCTTAGGAAACCAACCT
CACACGTCTGAACTCTTTCCCTACACGACGCTCTTCCGATCTNNNAAACCATGAAACCTCCATGTCCC
AGAGCACACGTCTGAACTCTTTCCCTACACGACGCTCTTCCGATCTNNNCCCTCTGGACAAAAAGCTG
AGAGCACACGTCTGAACTCTTTCCCTACACGACGCTCTTCCGATCTNNNACACAAAAAGCAAAGCTTG
GAGCACACGTCTGAACTCTTTCCCTACACGACGCTCTTCCGATCTNNNTGCATGCAAGGCCATCATTT
GAGCACACGTCTGAACTCTTTCCCTACACGACGCTCTTCCGATCTNNNGTGCTGCCCAAAGACAGAAT
GAAGAGCACACGTCTGAACTCTTTCCCTACACGACGCTCTTCCGATCTNNNGGATGTATTCCCTCAGG
AGCACACGTCTGAACTCTTTCCCTACACGACGCTCTTCCGATCTNNNAGTAAATGAGGAGGGCATCC
AGCACACGTCTGAACTCTTTCCCTACACGACGCTCTTCCGATCTNNNCTTCTAGTCTGGTGGTTGACAC
CACACGTCTGAACTCTTTCCCTACACGACGCTCTTCCGATCTNNNCTGAAGCCTCAGACCTTGGAAAG
AGCACACGTCTGAACTCTTTCCCTACACGACGCTCTTCCGATCTNNNAGGGTATGGCTTTGTTGTCAT
GAGCACACGTCTGAACTCTTTCCCTACACGACGCTCTTCCGATCTNNNGAGAAAGTTGTCTGGGCTTG
AGCACACGTCTGAACTCTTTCCCTACACGACGCTCTTCCGATCTNNNCATGGACTTTCCCCTGACTTAG
AGCACACGTCTGAACTCTTTCCCTACACGACGCTCTTCCGATCTNNNTCTATTTGGCCCTTTAGAATG

GAGCACACGTCTGAACTCTTTCCCTACACGACGCTCTTCCGATCTNNNCACACTCCAGCAAAACTCTGT
GAGCACACGTCTGAACTCTTTCCCTACACGACGCTCTTCCGATCTNNNCTTTGCTAGTGGGCAGTGT
AGAGCACACGTCTGAACTCTTTCCCTACACGACGCTCTTCCGATCTNNNTCCACTTTCACTGCTAATCAC
AGCACACGTCTGAACTCTTTCCCTACACGACGCTCTTCCGATCTNNNCAGATTTGCCACTTACACCTG
GCACACGTCTGAACTCTTTCCCTACACGACGCTCTTCCGATCTNNNTTGAGGAGGAAAGGCATGTTAG
GAGCACACGTCTGAACTCTTTCCCTACACGACGCTCTTCCGATCTNNNCACAGTGGGAACAGTTCTTG
AGCACACGTCTGAACTCTTTCCCTACACGACGCTCTTCCGATCTNNNGGTGTGCAACGAGGTGAGAA
AAGAGCACACGTCTGAACTCTTTCCCTACACGACGCTCTTCCGATCTNNNCCAGAAACCTCTGATTAT
GAGCACACGTCTGAACTCTTTCCCTACACGACGCTCTTCCGATCTNNNGCTTTTCAAAGAGGTGGTTT
AGCACACGTCTGAACTCTTTCCCTACACGACGCTCTTCCGATCTNNNTCTTGTTTTGGTAGGGGTGTG
GAGCACACGTCTGAACTCTTTCCCTACACGACGCTCTTCCGATCTNNNACTGTTCTCATTTCACAGGCCT
AGCACACGTCTGAACTCTTTCCCTACACGACGCTCTTCCGATCTNNNTGACACTGAACAGGTCATTTG
AGCACACGTCTGAACTCTTTCCCTACACGACGCTCTTCCGATCTNNNAGAGAACCACACTGACTATTTC
GAGCACACGTCTGAACTCTTTCCCTACACGACGCTCTTCCGATCTNNNCCAAATGTCCACCGACAGATA
AGCACACGTCTGAACTCTTTCCCTACACGACGCTCTTCCGATCTNNNGGCTCTATAGGTGCTCTCTCT
AGCACACGTCTGAACTCTTTCCCTACACGACGCTCTTCCGATCTNNNGAGAGGGAAGTGAATTGCAG
AGAGCACACGTCTGAACTCTTTCCCTACACGACGCTCTTCCGATCTNNNGCAGACTTGTTCAGAGAGC
GAGCACACGTCTGAACTCTTTCCCTACACGACGCTCTTCCGATCTNNNCAAATGGTCCACTTTTTCCCGT
GAGCACACGTCTGAACTCTTTCCCTACACGACGCTCTTCCGATCTNNNCCACCCCATTCAAAACAAATA
AGCACACGTCTGAACTCTTTCCCTACACGACGCTCTTCCGATCTNNNAAAACAGGCCAGTGGACATTA
AGCACACGTCTGAACTCTTTCCCTACACGACGCTCTTCCGATCTNNNTCCCAATGTCTCTGGGGTTTAG
GAGCACACGTCTGAACTCTTTCCCTACACGACGCTCTTCCGATCTNNNTGCAGCCATGCCTAAAGATA
AGAGCACACGTCTGAACTCTTTCCCTACACGACGCTCTTCCGATCTNNNACCATGACTAGCCAGTTCAG
GCACACGTCTGAACTCTTTCCCTACACGACGCTCTTCCGATCTNNNCGCTTTCCCTCTAGCTTTCTCTCCA
AGCACACGTCTGAACTCTTTCCCTACACGACGCTCTTCCGATCTNNNTTTCCCAATGCCTACACACAGA
GAGCACACGTCTGAACTCTTTCCCTACACGACGCTCTTCCGATCTNNNGGGCTCCAACAGAGAAACAG
AGAGCACACGTCTGAACTCTTTCCCTACACGACGCTCTTCCGATCTNNNTGTAGCTTTGAGTTTGGTGA
AGCACACGTCTGAACTCTTTCCCTACACGACGCTCTTCCGATCTNNNCTTCATCTGGGGAATAAGGTGC
AGCACACGTCTGAACTCTTTCCCTACACGACGCTCTTCCGATCTNNNATGGCTTTGGTTCTTAGGTGG
GAGCACACGTCTGAACTCTTTCCCTACACGACGCTCTTCCGATCTNNNTAGGTGTGAAACAGCTTCTG
AGCACACGTCTGAACTCTTTCCCTACACGACGCTCTTCCGATCTNNNGAAAAGAGTGTGTGTGTGTCC
GAGCACACGTCTGAACTCTTTCCCTACACGACGCTCTTCCGATCTNNNTGGAGACGAGTTCATGAGTT
AAGAGCACACGTCTGAACTCTTTCCCTACACGACGCTCTTCCGATCTNNNCATGGTGTCATGGTGTTTT
AGCACACGTCTGAACTCTTTCCCTACACGACGCTCTTCCGATCTNNNAAACCACGTACACAAGAACCT
GCACACGTCTGAACTCTTTCCCTACACGACGCTCTTCCGATCTNNNCTGGGAGAGGCATGGCAAAAAT
GCACACGTCTGAACTCTTTCCCTACACGACGCTCTTCCGATCTNNNTCTGGCTAACAGCTTTCTGAACT
AGAGCACACGTCTGAACTCTTTCCCTACACGACGCTCTTCCGATCTNNNCCACTTTTGGGCTTTAGCTA
GAGCACACGTCTGAACTCTTTCCCTACACGACGCTCTTCCGATCTNNNGTTTACTTCTCCCTCTGCCTGT
GAGCACACGTCTGAACTCTTTCCCTACACGACGCTCTTCCGATCTNNNAAAGCAGTTTGGTTGATGAG
AGCACACGTCTGAACTCTTTCCCTACACGACGCTCTTCCGATCTNNNTGAGCTTGGAGGTTACATGTT
AGCACACGTCTGAACTCTTTCCCTACACGACGCTCTTCCGATCTNNNACTTCTCTCTGCCTGTGTCTTCT
AGCACACGTCTGAACTCTTTCCCTACACGACGCTCTTCCGATCTNNNTAACTACCATGGTCCATTGCCT
GAAGAGCACACGTCTGAACTCTTTCCCTACACGACGCTCTTCCGATCTNNNAAAGTAGCAAAGGAAAG

GAGCACACGTCTGAACTCTTTCCCTACACGACGCTCTTCCGATCTNNNCTGTGTACTCTGGTACTGTCCC
AGCACACGTCTGAACTCTTTCCCTACACGACGCTCTTCCGATCTNNNTCCTCCCTCTCTATTCCTCTCTGTC
GAGCACACGTCTGAACTCTTTCCCTACACGACGCTCTTCCGATCTNNNTGGAGTTAGAGAGCTCCTTGT
AGCACACGTCTGAACTCTTTCCCTACACGACGCTCTTCCGATCTNNNACCTTGTTATTTATATTTGCGTCT
GCACACGTCTGAACTCTTTCCCTACACGACGCTCTTCCGATCTNNNTCACCTCTTCTCAGACAACCACTCA
AGAGCACACGTCTGAACTCTTTCCCTACACGACGCTCTTCCGATCTNNNTGTGGGTTTCATCACTAACCC
AGCACACGTCTGAACTCTTTCCCTACACGACGCTCTTCCGATCTNNNTGAATGACCTCAATAGAGCCCCA
GCACACGTCTGAACTCTTTCCCTACACGACGCTCTTCCGATCTNNNTGACCATACAACTAATCTTGTTCC
CACACGTCTGAACTCTTTCCCTACACGACGCTCTTCCGATCTNNNAGCCATCTTCATGTTTGAGGGTCC
GAGCACACGTCTGAACTCTTTCCCTACACGACGCTCTTCCGATCTNNNATACCCATGGGGAAATAAGG
AGCACACGTCTGAACTCTTTCCCTACACGACGCTCTTCCGATCTNNNAGTCTTTCCCTGTCTCTGTCTCTC
AGCACACGTCTGAACTCTTTCCCTACACGACGCTCTTCCGATCTNNNTTCCCCAAACAACAACAGCAATC
GAGCACACGTCTGAACTCTTTCCCTACACGACGCTCTTCCGATCTNNNCAATATGAACAAACGCACATGC
GAGCACACGTCTGAACTCTTTCCCTACACGACGCTCTTCCGATCTNNNCAACTATATGTGTCACCTCCCG
AGAGCACACGTCTGAACTCTTTCCCTACACGACGCTCTTCCGATCTNNNTGCCAAACCCTGATAGGTAG
AGCACACGTCTGAACTCTTTCCCTACACGACGCTCTTCCGATCTNNNGCTGACATCCCAATTGGATCACA
AGCACACGTCTGAACTCTTTCCCTACACGACGCTCTTCCGATCTNNNAAGGTAGAGAAGTGTTCCTCCC
GAGCACACGTCTGAACTCTTTCCCTACACGACGCTCTTCCGATCTNNNTGCAAAGTATCATGGCTGCAC
AGCACACGTCTGAACTCTTTCCCTACACGACGCTCTTCCGATCTNNNTACAGGATGGCTGAGAAACTTC
AGCACACGTCTGAACTCTTTCCCTACACGACGCTCTTCCGATCTNNNGACAATGCCCCCTCTAGACAATC
AGCACACGTCTGAACTCTTTCCCTACACGACGCTCTTCCGATCTNNNAAAACACCATGAGTGCTTCTGC
AGCACACGTCTGAACTCTTTCCCTACACGACGCTCTTCCGATCTNNNAGGGAAGTTCCAACATTTTCCC
AGCACACGTCTGAACTCTTTCCCTACACGACGCTCTTCCGATCTNNNCATGCAAATGAGCATTTTGTGG
AGCACACGTCTGAACTCTTTCCCTACACGACGCTCTTCCGATCTNNNTCTTGCTATAAATCACCAAACCAC
GAGCACACGTCTGAACTCTTTCCCTACACGACGCTCTTCCGATCTNNNTCTATGGTCGTGTGAGCCAAA
GCACACGTCTGAACTCTTTCCCTACACGACGCTCTTCCGATCTNNNTGGGAACACATAAAAGTCTGTCA
AGCACACGTCTGAACTCTTTCCCTACACGACGCTCTTCCGATCTNNNCGACATAGATGTAAAAATCCTCA
AAGAGCACACGTCTGAACTCTTTCCCTACACGACGCTCTTCCGATCTNNNTGATGTGAGGAATGTCCCA
GAGCACACGTCTGAACTCTTTCCCTACACGACGCTCTTCCGATCTNNNGATGATCTGCCAGACACCAAA
GAGCACACGTCTGAACTCTTTCCCTACACGACGCTCTTCCGATCTNNNGGCCATGTGAGCCAATTCCTTA
GAGCACACGTCTGAACTCTTTCCCTACACGACGCTCTTCCGATCTNNNATGAAATCAGTGTTGCCTTGC
AGCACACGTCTGAACTCTTTCCCTACACGACGCTCTTCCGATCTNNNGGTGGGGTATTATGGCCACTT
AGCACACGTCTGAACTCTTTCCCTACACGACGCTCTTCCGATCTNNNAAAGACATCCGAGACAGAACAC
GAGCACACGTCTGAACTCTTTCCCTACACGACGCTCTTCCGATCTNNNCCACACATACACATTCCCAACCA
AGCACACGTCTGAACTCTTTCCCTACACGACGCTCTTCCGATCTNNNATTGGTGGCTAGAGCTTCAGAA
GCACACGTCTGAACTCTTTCCCTACACGACGCTCTTCCGATCTNNNCTCCTCTGCCTGTTTTCGAGTTTCA
AAGAGCACACGTCTGAACTCTTTCCCTACACGACGCTCTTCCGATCTNNNGCAAATTTGTATGCAGAAG
GCACACGTCTGAACTCTTTCCCTACACGACGCTCTTCCGATCTNNNCGCTTCATGCTGACGAATGTTTTC
GCACACGTCTGAACTCTTTCCCTACACGACGCTCTTCCGATCTNNNCCTGTGAAAGGAATGTTTGTGGC
GCACACGTCTGAACTCTTTCCCTACACGACGCTCTTCCGATCTNNNCTGGGGAACTCCTAACCTCTTCACA
AGAGCACACGTCTGAACTCTTTCCCTACACGACGCTCTTCCGATCTNNNCACCCTGTCAATGTCTTCACT
GAGCACACGTCTGAACTCTTTCCCTACACGACGCTCTTCCGATCTNNNAATGAGAGAGAGAGAGCAGC
AAGAGCACACGTCTGAACTCTTTCCCTACACGACGCTCTTCCGATCTNNNCATTAGGGAGGAAGTCAC

GAGCACACGTCTGAACTCTTTCCCTACACGACGCTCTTCCGATCTNNNAGCATGGAATGAGCGGTTTT

GCACACGTCTGAACTCTTTCCCTACACGACGCTCTTCCGATCTNNNAGGCAAGAACTAAAATCCACAG

AAGAGCACACGTCTGAACTCTTTCCCTACACGACGCTCTTCCGATCTNNNAGCCACTAGGACCAGTA

GAGCACACGTCTGAACTCTTTCCCTACACGACGCTCTTCCGATCTNNNCCCTACCACAGGCTCATCATT

GCACACGTCTGAACTCTTTCCCTACACGACGCTCTTCCGATCTNNNAGGGCTTACCAAGCTGAATGAT

AGCACACGTCTGAACTCTTTCCCTACACGACGCTCTTCCGATCTNNNACCAAGTTAGTGAGCACAGCA

GAGCACACGTCTGAACTCTTTCCCTACACGACGCTCTTCCGATCTNNNGATGGCAGGACAGATTTGG

GAGCACACGTCTGAACTCTTTCCCTACACGACGCTCTTCCGATCTNNNTCTTCTCCCTGGAAAAGCAA

AGAGCACACGTCTGAACTCTTTCCCTACACGACGCTCTTCCGATCTNNNCATTCCTTTCCTGTCCCTTTG

AGCACACGTCTGAACTCTTTCCCTACACGACGCTCTTCCGATCTNNNGTAAATTGCCCAGCAGAATG

GAGCACACGTCTGAACTCTTTCCCTACACGACGCTCTTCCGATCTNNNGACCTAAAGGCCAAGCAAG

AGCACACGTCTGAACTCTTTCCCTACACGACGCTCTTCCGATCTNNNATGTCCATCCATCCCGTCCTAAT

AAGAGCACACGTCTGAACTCTTTCCCTACACGACGCTCTTCCGATCTNNNACCCACACACATACATTTA

AGCACACGTCTGAACTCTTTCCCTACACGACGCTCTTCCGATCTNNNGCAGCACTGAAGACTTTAGA

GAGCACACGTCTGAACTCTTTCCCTACACGACGCTCTTCCGATCTNNNTGGTCAACCATGTCATCATC

GAGCACACGTCTGAACTCTTTCCCTACACGACGCTCTTCCGATCTNNNCTGTTGCAGCCAATTCTTTCA

GCACACGTCTGAACTCTTTCCCTACACGACGCTCTTCCGATCTNNNTTCCCATTCCAGCCTCACAAAAA

GCACACGTCTGAACTCTTTCCCTACACGACGCTCTTCCGATCTNNNCAAAGCTGGAAACAGCCCTAG

AGCACACGTCTGAACTCTTTCCCTACACGACGCTCTTCCGATCTNNNTGGGGATGACAGAATGATGA

GAGCACACGTCTGAACTCTTTCCCTACACGACGCTCTTCCGATCTNNNCTCTTGCTGTCTCCTTACACA

GCACACGTCTGAACTCTTTCCCTACACGACGCTCTTCCGATCTNNNGTCAATGACACCAACAGAAACA

GCACACGTCTGAACTCTTTCCCTACACGACGCTCTTCCGATCTNNNTGCGCAGCTTCATCACATACAT

AGCACACGTCTGAACTCTTTCCCTACACGACGCTCTTCCGATCTNNNGCATGTTAGGTGGTTTTGTGT

AGCACACGTCTGAACTCTTTCCCTACACGACGCTCTTCCGATCTNNNGCTTCTCTGCCAAGTTCACAAA

AGCACACGTCTGAACTCTTTCCCTACACGACGCTCTTCCGATCTNNNTATGAGAACTAGGGCACACT

AGCACACGTCTGAACTCTTTCCCTACACGACGCTCTTCCGATCTNNNATTCAGTAGGGTTGGGACAG

AGAGCACACGTCTGAACTCTTTCCCTACACGACGCTCTTCCGATCTNNNCTGTGGGTGTGTTGTATTT

AGAGCACACGTCTGAACTCTTTCCCTACACGACGCTCTTCCGATCTNNNAGCCACTAGGACCAGTAAT

AGCACACGTCTGAACTCTTTCCCTACACGACGCTCTTCCGATCTNNNCAAGCAAACAAGTTTCGCCTC

AGAGCACACGTCTGAACTCTTTCCCTACACGACGCTCTTCCGATCTNNNGGAAAAGGCTGAAGG

AGAGCACACGTCTGAACTCTTTCCCTACACGACGCTCTTCCGATCTNNNCCAGGAAAGCAGCTCTTCA

GCACACGTCTGAACTCTTTCCCTACACGACGCTCTTCCGATCTNNNAGATAGATTAGCTAGACAAACA

AGCACACGTCTGAACTCTTTCCCTACACGACGCTCTTCCGATCTNNNACCAGAACAGAAACACTTTGT

AGCACACGTCTGAACTCTTTCCCTACACGACGCTCTTCCGATCTNNNTGAGTTGGAGAGACAGCCAA

GCACACGTCTGAACTCTTTCCCTACACGACGCTCTTCCGATCTNNNAGTATTGCAGTTGGAAAAACCT

AGCACACGTCTGAACTCTTTCCCTACACGACGCTCTTCCGATCTNNNGAACCCCCATTCCCATAGGTTT

GAGCACACGTCTGAACTCTTTCCCTACACGACGCTCTTCCGATCTNNNAATGCTTCCTTTCTCTTTTCCC

AGCACACGTCTGAACTCTTTCCCTACACGACGCTCTTCCGATCTNNNCTCTCCCTCCCACTTCTTTCAC

AGCACACGTCTGAACTCTTTCCCTACACGACGCTCTTCCGATCTNNNTGGCTGAGATGCATATGAGA

AGCACACGTCTGAACTCTTTCCCTACACGACGCTCTTCCGATCTNNNGGGCCATATTCACATGCACAA

AGAGCACACGTCTGAACTCTTTCCCTACACGACGCTCTTCCGATCTNNNAGGACAAAAGCTATAGGT

AGAGCACACGTCTGAACTCTTTCCCTACACGACGCTCTTCCGATCTNNNTCTATGGTCGTGTGAGCCA

AGCACACGTCTGAACTCTTTCCCTACACGACGCTCTTCCGATCTNNNTGAAATACGGTAAGCTCCCTC

GAGCACACGTCTGAACTCTTTCCCTACACGACGCTCTTCCGATCTNNNTGACTGACTACAACTATGGC
GCACACGTCTGAACTCTTTCCCTACACGACGCTCTTCCGATCTNNNTCATTGAGTGGTGGGATTTCAC
AGCACACGTCTGAACTCTTTCCCTACACGACGCTCTTCCGATCTNNNCATGCCCCTTGGAATGTAGAC
GCACACGTCTGAACTCTTTCCCTACACGACGCTCTTCCGATCTNNNATTTGGACAAATGTCTCCCCCAC
AGCACACGTCTGAACTCTTTCCCTACACGACGCTCTTCCGATCTNNNTCTCTTTGTAGGGGCAGCATTT
GCACACGTCTGAACTCTTTCCCTACACGACGCTCTTCCGATCTNNNTTCCAACCTTGCAGAGAGGGTA
GCACACGTCTGAACTCTTTCCCTACACGACGCTCTTCCGATCTNNNCTGTCCCCCAAAAATGCATACCTC
GCACACGTCTGAACTCTTTCCCTACACGACGCTCTTCCGATCTNNNAGGTAAACCATTCTTATCAGAC
AGCACACGTCTGAACTCTTTCCCTACACGACGCTCTTCCGATCTNNNAGTATTGCAGTTGGAAAAAC
GAGCACACGTCTGAACTCTTTCCCTACACGACGCTCTTCCGATCTNNNAATTTTTGCTCCCTCCCCATCT
AGCACACGTCTGAACTCTTTCCCTACACGACGCTCTTCCGATCTNNNTTTCCAGTGCTAAGGCATTCCA
GAGCACACGTCTGAACTCTTTCCCTACACGACGCTCTTCCGATCTNNNTGTGGAAGGAAGGGAGAAT
AGCACACGTCTGAACTCTTTCCCTACACGACGCTCTTCCGATCTNNNCGTTTTACTTGCTAGTTTTCAA
AAGAGCACACGTCTGAACTCTTTCCCTACACGACGCTCTTCCGATCTNNNTTTACCTGCCTTGGCCTAC
AAGAGCACACGTCTGAACTCTTTCCCTACACGACGCTCTTCCGATCTNNNTACAATCTACCGAGAGGG
GCACACGTCTGAACTCTTTCCCTACACGACGCTCTTCCGATCTNNNAAGGATTGGGACCTCCAAAGCT
AGAGCACACGTCTGAACTCTTTCCCTACACGACGCTCTTCCGATCTNNNAGTCAAAGTCCACACCTATT
AGCACACGTCTGAACTCTTTCCCTACACGACGCTCTTCCGATCTNNNTTTGTTTTGGTGTCTGTCTCCCC
AGCACACGTCTGAACTCTTTCCCTACACGACGCTCTTCCGATCTNNNCAACAGCAAGAGATGGTCTCC
GAGCACACGTCTGAACTCTTTCCCTACACGACGCTCTTCCGATCTNNNATGTTTGCAGTTGACAGACT
GCACACGTCTGAACTCTTTCCCTACACGACGCTCTTCCGATCTNNNATGGCTGTTTGAACCAGTGACA
AGCACACGTCTGAACTCTTTCCCTACACGACGCTCTTCCGATCTNNNACTGTTTCTCAAACAAGAGGG
AGCACACGTCTGAACTCTTTCCCTACACGACGCTCTTCCGATCTNNNCAACCCAGGTTGCCAAAGTTC
GCACACGTCTGAACTCTTTCCCTACACGACGCTCTTCCGATCTNNNATGTACAGAAAGATGACGCTG
GAGCACACGTCTGAACTCTTTCCCTACACGACGCTCTTCCGATCTNNNCCAAACCAAACAGTACTTCCC
AGCACACGTCTGAACTCTTTCCCTACACGACGCTCTTCCGATCTNNNTCTCTCCCCTTTCTCATTCAAC
GAGCACACGTCTGAACTCTTTCCCTACACGACGCTCTTCCGATCTNNNTCAATCGTGACGGTTTTATG
GAGCACACGTCTGAACTCTTTCCCTACACGACGCTCTTCCGATCTNNNCCCCTCAAGATCCTTCCTCGA
AGCACACGTCTGAACTCTTTCCCTACACGACGCTCTTCCGATCTNNNTCATCTTGGTGTGAACCAATCC
AGCACACGTCTGAACTCTTTCCCTACACGACGCTCTTCCGATCTNNNAAGGGGAGAGATTTATTTCC
AGCACACGTCTGAACTCTTTCCCTACACGACGCTCTTCCGATCTNNNAAGATCAGGCTTCCCTCAGAA
AGCACACGTCTGAACTCTTTCCCTACACGACGCTCTTCCGATCTNNNCCCTGAAATACTCACATTCCCA
AAGAGCACACGTCTGAACTCTTTCCCTACACGACGCTCTTCCGATCTNNNTCCTCCTACCTCTCTCTCCT
AGCACACGTCTGAACTCTTTCCCTACACGACGCTCTTCCGATCTNNNTCCTAGGCAAAATATTAGGTG
GAGCACACGTCTGAACTCTTTCCCTACACGACGCTCTTCCGATCTNNNGGGGATATGTTCTCAGGCTT
GCACACGTCTGAACTCTTTCCCTACACGACGCTCTTCCGATCTNNNGGCACTGTCTGTGTTTCTCTCTC
AGCACACGTCTGAACTCTTTCCCTACACGACGCTCTTCCGATCTNNNGTATTGGCTGGAAGAAACGC
GAGCACACGTCTGAACTCTTTCCCTACACGACGCTCTTCCGATCTNNNCCATGATAGCTGTGACATAG
AGCACACGTCTGAACTCTTTCCCTACACGACGCTCTTCCGATCTNNNACACTGCTTTTAGAGAATGCA
AGCACACGTCTGAACTCTTTCCCTACACGACGCTCTTCCGATCTNNNTTCCATTGTGAGGAAACTGG
CACACGTCTGAACTCTTTCCCTACACGACGCTCTTCCGATCTNNNATTTGGACAAATGTCTCCCCCACC
AGCACACGTCTGAACTCTTTCCCTACACGACGCTCTTCCGATCTNNNGGCCTATGGGAAAGAGTTGT
GAGCACACGTCTGAACTCTTTCCCTACACGACGCTCTTCCGATCTNNNAGGCTGAGTGGCTTACTTAC

AGCACACGTCTGAACTCTTTCCCTACACGACGCTCTTCCGATCTNNNAACACAACTGCTATCAAAATAC
AGCACACGTCTGAACTCTTTCCCTACACGACGCTCTTCCGATCTNNNCCAAATGTCCCTACCTCTAACAC
AGAGCACACGTCTGAACTCTTTCCCTACACGACGCTCTTCCGATCTNNNATGAAATCAGTGTTGCCTT
GAGCACACGTCTGAACTCTTTCCCTACACGACGCTCTTCCGATCTNNNTATAAATCAGATCCCCGACC
GAGCACACGTCTGAACTCTTTCCCTACACGACGCTCTTCCGATCTNNNGGTGAAGTAGAGGAAGTGA
GAGCACACGTCTGAACTCTTTCCCTACACGACGCTCTTCCGATCTNNNACTAAGCTGCATTGTAAGCG
GAGCACACGTCTGAACTCTTTCCCTACACGACGCTCTTCCGATCTNNNACTGGCACTTAGGAAATTCAG
GAGCACACGTCTGAACTCTTTCCCTACACGACGCTCTTCCGATCTNNNGAGCTCCAAGTGCATTTTCAG
AGCACACGTCTGAACTCTTTCCCTACACGACGCTCTTCCGATCTNNNTGCCTCTGCTGTCCTGTATCTAT
AGCACACGTCTGAACTCTTTCCCTACACGACGCTCTTCCGATCTNNNTTTCAGGTTGGTTGCTATGTCC
GAGCACACGTCTGAACTCTTTCCCTACACGACGCTCTTCCGATCTNNNAAAATTATACCCCTGCCCCCAC
GAAGAGCACACGTCTGAACTCTTTCCCTACACGACGCTCTTCCGATCTNNNTGACTTCCATTCAGGGAA
AGCACACGTCTGAACTCTTTCCCTACACGACGCTCTTCCGATCTNNNTCCCAGCCTTAGAGAGAACTAG
AGCACACGTCTGAACTCTTTCCCTACACGACGCTCTTCCGATCTNNNCAGTTCCATGCTTCCATGTGAG
AGAGCACACGTCTGAACTCTTTCCCTACACGACGCTCTTCCGATCTNNNGCAGAAGGAAGGCACGTAT
AGCACACGTCTGAACTCTTTCCCTACACGACGCTCTTCCGATCTNNNCAACTGTTCCTAGGCTGAATCCC
GAGCACACGTCTGAACTCTTTCCCTACACGACGCTCTTCCGATCTNNNTTCTCTCTCTTCTGCCTGTTGG
AGCACACGTCTGAACTCTTTCCCTACACGACGCTCTTCCGATCTNNNTGTCTTCCTTCCAGTGCTGTCT
GCACACGTCTGAACTCTTTCCCTACACGACGCTCTTCCGATCTNNNTGCTGAGAGGGAAGAAAACAA
GAGCACACGTCTGAACTCTTTCCCTACACGACGCTCTTCCGATCTNNNGCCCTTATGTGACATCATTCG
GAAGAGCACACGTCTGAACTCTTTCCCTACACGACGCTCTTCCGATCTNNNTATGTGCTGATCAGTGTA
GAGCACACGTCTGAACTCTTTCCCTACACGACGCTCTTCCGATCTNNNACCATGACTAGCCAGTTCAG
AGCACACGTCTGAACTCTTTCCCTACACGACGCTCTTCCGATCTNNNATAGACGTGTGTCTGTGTGTGT
AGAGCACACGTCTGAACTCTTTCCCTACACGACGCTCTTCCGATCTNNNAGACATTCTGATACCCTAAA
AGAGCACACGTCTGAACTCTTTCCCTACACGACGCTCTTCCGATCTNNNTCTAAGCATGTTACGTGAA
GAGCACACGTCTGAACTCTTTCCCTACACGACGCTCTTCCGATCTNNNTTCAACTCTGGAGGCAAACCA
AGAGCACACGTCTGAACTCTTTCCCTACACGACGCTCTTCCGATCTNNNTTTGACATGACACTCCCAC
GAGCACACGTCTGAACTCTTTCCCTACACGACGCTCTTCCGATCTNNNTACACCACACCTGAAGGAGA
GAGCACACGTCTGAACTCTTTCCCTACACGACGCTCTTCCGATCTNNNGTACAGAGGACATCACATGC
AGCACACGTCTGAACTCTTTCCCTACACGACGCTCTTCCGATCTNNNAAGAGGGCAAACCAAACAGG
AGCACACGTCTGAACTCTTTCCCTACACGACGCTCTTCCGATCTNNNTACATACCTCAAAACCCAGAGG
GAGCACACGTCTGAACTCTTTCCCTACACGACGCTCTTCCGATCTNNNCCTCTTTTGGCGCTTCTCTTTC
AGAGCACACGTCTGAACTCTTTCCCTACACGACGCTCTTCCGATCTNNNGTTGGGCAGAGGGATTAG
AGCACACGTCTGAACTCTTTCCCTACACGACGCTCTTCCGATCTNNNTTTGCCATGTGGGAGTTGGTA
GCACACGTCTGAACTCTTTCCCTACACGACGCTCTTCCGATCTNNNAAAATGCTCCTCTCTCCTCCCTTC
AGCACACGTCTGAACTCTTTCCCTACACGACGCTCTTCCGATCTNNNCACACCACCACACCCTGATTTAT
AGCACACGTCTGAACTCTTTCCCTACACGACGCTCTTCCGATCTNNNACATATGTGATATGACCCCAAT
AGCACACGTCTGAACTCTTTCCCTACACGACGCTCTTCCGATCTNNNACTCTTTCCCACAGTCAAGCTC
AGCACACGTCTGAACTCTTTCCCTACACGACGCTCTTCCGATCTNNNTCCTAGTGGAAGATGATAAGC
AGCACACGTCTGAACTCTTTCCCTACACGACGCTCTTCCGATCTNNNGTAGGAACCTATAGACCGCACC
GAGCACACGTCTGAACTCTTTCCCTACACGACGCTCTTCCGATCTNNNGCAAGAAGCCATGAGAAAG
AGCACACGTCTGAACTCTTTCCCTACACGACGCTCTTCCGATCTNNNAACTATTGTGTGTGTGCGTGT
GAGCACACGTCTGAACTCTTTCCCTACACGACGCTCTTCCGATCTNNNTGAAATTTCCTGATATTTTGG

AGAGCACACGTCTGAACTCTTTCCCTACACGACGCTCTTCCGATCTNNNCGAGTTCATAATCCAGGGGA
AGCACACGTCTGAACTCTTTCCCTACACGACGCTCTTCCGATCTNNNCCTGTCTTTTTCCACCAGTGTCA
AGCACACGTCTGAACTCTTTCCCTACACGACGCTCTTCCGATCTNNNCCACTGAGAGTAAATAAGGGC
GAGCACACGTCTGAACTCTTTCCCTACACGACGCTCTTCCGATCTNNNTAAATGCACCATGGAGGTGA
AGCACACGTCTGAACTCTTTCCCTACACGACGCTCTTCCGATCTNNNATCTTTGCAGATTGGCTGTCTG
AGCACACGTCTGAACTCTTTCCCTACACGACGCTCTTCCGATCTNNNAAGACCTCTCCTTCCCTTCTCTGC
AAGAGCACACGTCTGAACTCTTTCCCTACACGACGCTCTTCCGATCTNNNAAAAAGGCTTATTGGGGT
GAGCACACGTCTGAACTCTTTCCCTACACGACGCTCTTCCGATCTNNNACACACGCGGTTTAGCAATAT
AGCACACGTCTGAACTCTTTCCCTACACGACGCTCTTCCGATCTNNNTCGCGAATGGATCTCCTTCTTTC
GAGCACACGTCTGAACTCTTTCCCTACACGACGCTCTTCCGATCTNNNTGTTTCTCTCCCTCCTCTTTCTT
GAGCACACGTCTGAACTCTTTCCCTACACGACGCTCTTCCGATCTNNNCTTGAGGTTTGGGGATGAGT
AAGAGCACACGTCTGAACTCTTTCCCTACACGACGCTCTTCCGATCTNNNTCCCAGCCTTAGAGAGAAC
AGCACACGTCTGAACTCTTTCCCTACACGACGCTCTTCCGATCTNNNAGCCTCGAATGATTTCTCTCCTT
GCACACGTCTGAACTCTTTCCCTACACGACGCTCTTCCGATCTNNNCCAGACCCTTGTAAGTGCTCAAA
AGAGCACACGTCTGAACTCTTTCCCTACACGACGCTCTTCCGATCTNNNTTGTAGGAGACCACCTTCTCG
AGCACACGTCTGAACTCTTTCCCTACACGACGCTCTTCCGATCTNNNACATGTTTTATCATCTAGAGCC
GCACACGTCTGAACTCTTTCCCTACACGACGCTCTTCCGATCTNNNAAGAATCCGGAAGACCAGGAGA
GAGCACACGTCTGAACTCTTTCCCTACACGACGCTCTTCCGATCTNNNGCTTTTCATAGGGAAAGGCA
AGCACACGTCTGAACTCTTTCCCTACACGACGCTCTTCCGATCTNNNAAGGGCCAAGACAGCTATTTTC
AGCACACGTCTGAACTCTTTCCCTACACGACGCTCTTCCGATCTNNNTTTCAGGTTGCAGGTTTGGAA
GAGCACACGTCTGAACTCTTTCCCTACACGACGCTCTTCCGATCTNNNGGGTTGCATAGAAAAGCAAG
GAGCACACGTCTGAACTCTTTCCCTACACGACGCTCTTCCGATCTNNNCCGGGCTTCCTGTTTAGTCTC
GCACACGTCTGAACTCTTTCCCTACACGACGCTCTTCCGATCTNNNGAACACTGTTACCCATCAACTGCC
AGCACACGTCTGAACTCTTTCCCTACACGACGCTCTTCCGATCTNNNAGCTGTTGCCACAAATTCTGAA
AGAGCACACGTCTGAACTCTTTCCCTACACGACGCTCTTCCGATCTNNNGGATTGCAGTCAGCTTGTTT
GAGCACACGTCTGAACTCTTTCCCTACACGACGCTCTTCCGATCTNNNTTACCTGGCCTTTTCCTTTTCGT
GAGCACACGTCTGAACTCTTTCCCTACACGACGCTCTTCCGATCTNNNCGTCTTCATTTTTGGTGGTGG
AGCACACGTCTGAACTCTTTCCCTACACGACGCTCTTCCGATCTNNNAAGAAGTCTCCCACCCCTAGGT
AGCACACGTCTGAACTCTTTCCCTACACGACGCTCTTCCGATCTNNNCCAGCTGTATCCCTGTTGCTAG
AGAGCACACGTCTGAACTCTTTCCCTACACGACGCTCTTCCGATCTNNNTTTATTGCCTCACCTAGTTAC
AGCACACGTCTGAACTCTTTCCCTACACGACGCTCTTCCGATCTNNNGGATCACCAAAAGTCCTGAGA
GAGCACACGTCTGAACTCTTTCCCTACACGACGCTCTTCCGATCTNNNTAGCTAGAAATAGGCCTGGA
GAGCACACGTCTGAACTCTTTCCCTACACGACGCTCTTCCGATCTNNNTGTCCTAACTGACTGGCAGAA
AGCACACGTCTGAACTCTTTCCCTACACGACGCTCTTCCGATCTNNNTTCCCATGTTAGAAGAGAGGG
AGCACACGTCTGAACTCTTTCCCTACACGACGCTCTTCCGATCTNNNTCTCTGTAATGCCTATGGAACTT
GAGCACACGTCTGAACTCTTTCCCTACACGACGCTCTTCCGATCTNNNTTCTCTCCACCCCTAGAGTGTT
AGAGCACACGTCTGAACTCTTTCCCTACACGACGCTCTTCCGATCTNNNTCATAACCTGTGACATAGGA
AGCACACGTCTGAACTCTTTCCCTACACGACGCTCTTCCGATCTNNNTTCCCCATCAAATGACAAGGTG
AGAGCACACGTCTGAACTCTTTCCCTACACGACGCTCTTCCGATCTNNNCCAGAGACAGAGCAAAACA
GCACACGTCTGAACTCTTTCCCTACACGACGCTCTTCCGATCTNNNTTCAAGAAACTTCCCCTGCCTTCC
AGCACACGTCTGAACTCTTTCCCTACACGACGCTCTTCCGATCTNNNTGCAGGAAGAAGGAACAAAGG
AGCACACGTCTGAACTCTTTCCCTACACGACGCTCTTCCGATCTNNNTTTTCGGAGCAAGTTGAGGTT
GAGCACACGTCTGAACTCTTTCCCTACACGACGCTCTTCCGATCTNNNTTTCTTCCCTCAATTTCTGCCCA

GAGCACACGTCTGAACTCTTTCCCTACACGACGCTCTTCCGATCTNNNAAGCTAAGCCAAATAAGAG
GAGCACACGTCTGAACTCTTTCCCTACACGACGCTCTTCCGATCTNNNGCTCATACATCGTGTGTTCTT
AGCACACGTCTGAACTCTTTCCCTACACGACGCTCTTCCGATCTNNNGCAGGATTGACTGAAGGAGGT
AGCACACGTCTGAACTCTTTCCCTACACGACGCTCTTCCGATCTNNNACCACCTGCTATCTCATACCTCC
GAGCACACGTCTGAACTCTTTCCCTACACGACGCTCTTCCGATCTNNNAATCATCATGAAACCCTCTGG
AGCACACGTCTGAACTCTTTCCCTACACGACGCTCTTCCGATCTNNNGAGATCTCTGCTTGGAATGACC
GAGCACACGTCTGAACTCTTTCCCTACACGACGCTCTTCCGATCTNNNAAGTCTCTTCTCCCAGGCTGAT
GAGCACACGTCTGAACTCTTTCCCTACACGACGCTCTTCCGATCTNNNCTTTGTTCATAGCTTTGGGC
GAGCACACGTCTGAACTCTTTCCCTACACGACGCTCTTCCGATCTNNNGTCACCAGGTCACCATTCAAA
AGCACACGTCTGAACTCTTTCCCTACACGACGCTCTTCCGATCTNNNACCAGTGTATAATCAACGGAA
AGAGCACACGTCTGAACTCTTTCCCTACACGACGCTCTTCCGATCTNNNGAAGGGGTTTTCGGGAGCT
GAGCACACGTCTGAACTCTTTCCCTACACGACGCTCTTCCGATCTNNNCAGGTCAAGTCCACAACCTTG
GAGCACACGTCTGAACTCTTTCCCTACACGACGCTCTTCCGATCTNNNGGGGTCATTGTAATGCCACCT
AGCACACGTCTGAACTCTTTCCCTACACGACGCTCTTCCGATCTNNNAGTGCATTTTTATTTCCTCTCTC
AGCACACGTCTGAACTCTTTCCCTACACGACGCTCTTCCGATCTNNNATGGCTTGCTATTGCAGGGTTT
GAGCACACGTCTGAACTCTTTCCCTACACGACGCTCTTCCGATCTNNNCAGCCCAGCCCTATATCTTGG
GAGCACACGTCTGAACTCTTTCCCTACACGACGCTCTTCCGATCTNNNTCTGTGTTAGTCACGTTCTCC
GAGCACACGTCTGAACTCTTTCCCTACACGACGCTCTTCCGATCTNNNTGGGGAGGGACATAAACTTC
GAGCACACGTCTGAACTCTTTCCCTACACGACGCTCTTCCGATCTNNNCAGAACCTCCTATGACAAGG
AGCACACGTCTGAACTCTTTCCCTACACGACGCTCTTCCGATCTNNNTACGCTGCTTCTACCAAGTGAT
AGCACACGTCTGAACTCTTTCCCTACACGACGCTCTTCCGATCTNNNAACAGCAGCTCATTCCTTGTCT
AGAGCACACGTCTGAACTCTTTCCCTACACGACGCTCTTCCGATCTNNNGCAAGGCAGAATGAGCAA
GCACACGTCTGAACTCTTTCCCTACACGACGCTCTTCCGATCTNNNTAGAACTGCTTTCTGGAGTGTGC
GAGCACACGTCTGAACTCTTTCCCTACACGACGCTCTTCCGATCTNNNGGGCCAAAGAAGGTGGTAA
GCACACGTCTGAACTCTTTCCCTACACGACGCTCTTCCGATCTNNNGATGTGACTTCACAGGCACAGA
AGCACACGTCTGAACTCTTTCCCTACACGACGCTCTTCCGATCTNNNTGCAACCGTTGGGTTGAGATA
AGCACACGTCTGAACTCTTTCCCTACACGACGCTCTTCCGATCTNNNCCCAGCAATTCATCCCATCTCTC
AGCACACGTCTGAACTCTTTCCCTACACGACGCTCTTCCGATCTNNNGGGCCCTGGCTTTTCATTAGTT
GAGCACACGTCTGAACTCTTTCCCTACACGACGCTCTTCCGATCTNNNGATGGCAGGACAGATTTGG
AGCACACGTCTGAACTCTTTCCCTACACGACGCTCTTCCGATCTNNNCTGTGGTAAATGGTACTGGGT
AGCACACGTCTGAACTCTTTCCCTACACGACGCTCTTCCGATCTNNNGCACAGCACAGGAATGAATAG
AGAGCACACGTCTGAACTCTTTCCCTACACGACGCTCTTCCGATCTNNNATGCATCTGTCTTTCGTGGA
AGCACACGTCTGAACTCTTTCCCTACACGACGCTCTTCCGATCTNNNATGGACCCTCTTGAGCAACTGA
GCACACGTCTGAACTCTTTCCCTACACGACGCTCTTCCGATCTNNNAGAGCCAATCTCACATGCTTTGT
AGCACACGTCTGAACTCTTTCCCTACACGACGCTCTTCCGATCTNNNTCCCTTCAGAGGGAGAGAAT
AGCACACGTCTGAACTCTTTCCCTACACGACGCTCTTCCGATCTNNNAGACATGATTGCTAAGGTCCA
CACACGTCTGAACTCTTTCCCTACACGACGCTCTTCCGATCTNNNGGCCCTTCCTTTAGTTCAAGTCCC
AGCACACGTCTGAACTCTTTCCCTACACGACGCTCTTCCGATCTNNNCAATTCAGTGCTCCTTTCCCTC
GAGCACACGTCTGAACTCTTTCCCTACACGACGCTCTTCCGATCTNNNTGACTAGTGGGTGGATAGTT
AAGAGCACACGTCTGAACTCTTTCCCTACACGACGCTCTTCCGATCTNNNATTGAGTGAAAAAGGCA
AGCACACGTCTGAACTCTTTCCCTACACGACGCTCTTCCGATCTNNNCCTGAGAATGCCCTTGACTGTT
AGCACACGTCTGAACTCTTTCCCTACACGACGCTCTTCCGATCTNNNATTTATCTTTCCCAGAGCCCAGC
AAGAGCACACGTCTGAACTCTTTCCCTACACGACGCTCTTCCGATCTNNNAAAACAGCTCTGGCGTATT

AGAGCACACGTCTGAACTCTTTCCCTACACGACGCTCTTCCGATCTNNNCAAGGCTTCTGTACACCTTCT
GCACACGTCTGAACTCTTTCCCTACACGACGCTCTTCCGATCTNNNCGCTGTTGCTCTCTCTTTCTGTTCA
GCACACGTCTGAACTCTTTCCCTACACGACGCTCTTCCGATCTNNNGGCCACAGAAGATTAGACTAAC
AGCACACGTCTGAACTCTTTCCCTACACGACGCTCTTCCGATCTNNNGCCAAAGATGCTGTATCCTGCT
GAGCACACGTCTGAACTCTTTCCCTACACGACGCTCTTCCGATCTNNNCTGTGGACCAAAACTGACAGG
GAGCACACGTCTGAACTCTTTCCCTACACGACGCTCTTCCGATCTNNNGCTTTGCTTTTGTGAGGCTTT
AGCACACGTCTGAACTCTTTCCCTACACGACGCTCTTCCGATCTNNNCTGGTTACACCCCCTTCCCTAAC
AGAGCACACGTCTGAACTCTTTCCCTACACGACGCTCTTCCGATCTNNNTTCTCTCCACCCCTAGAGTGT
AGCACACGTCTGAACTCTTTCCCTACACGACGCTCTTCCGATCTNNNACAATGATGGTCGCTTTCCTCTC
AGCACACGTCTGAACTCTTTCCCTACACGACGCTCTTCCGATCTNNNTCATATTGCAGTTGTCCCTCGA
AGCACACGTCTGAACTCTTTCCCTACACGACGCTCTTCCGATCTNNNTAAAACACTCTCCCAGCAAACCC
AGCACACGTCTGAACTCTTTCCCTACACGACGCTCTTCCGATCTNNNATTGGGCATGAGTGGTTACAC
GAGCACACGTCTGAACTCTTTCCCTACACGACGCTCTTCCGATCTNNNGCTCTGGGTAACACACTTCTT
GCACACGTCTGAACTCTTTCCCTACACGACGCTCTTCCGATCTNNNACACTGGTCCAAACCATTCTCAA
AGAGCACACGTCTGAACTCTTTCCCTACACGACGCTCTTCCGATCTNNNCTGGTCCATTTGCAATCAAT
GCACACGTCTGAACTCTTTCCCTACACGACGCTCTTCCGATCTNNNAAGGTGAGAAGCCGTTATTGAC
GCACACGTCTGAACTCTTTCCCTACACGACGCTCTTCCGATCTNNNGGGGGTAAGAGGTGGTTAGAT
AGCACACGTCTGAACTCTTTCCCTACACGACGCTCTTCCGATCTNNNCTCTAGGGGAACCACAAAAAG
GAAGAGCACACGTCTGAACTCTTTCCCTACACGACGCTCTTCCGATCTNNNACTGAAAGACTGTCCTT
GAGCACACGTCTGAACTCTTTCCCTACACGACGCTCTTCCGATCTNNNCCAGGAAAGCAGCTCTTCAT
GAGCACACGTCTGAACTCTTTCCCTACACGACGCTCTTCCGATCTNNNGCAAGGTTGTAGAGATCCCA
AGCACACGTCTGAACTCTTTCCCTACACGACGCTCTTCCGATCTNNNTAGAACTGGCCCCTTGGGATAT
GAGCACACGTCTGAACTCTTTCCCTACACGACGCTCTTCCGATCTNNNAGGGGATGAATGTGTGCTTT
GCACACGTCTGAACTCTTTCCCTACACGACGCTCTTCCGATCTNNNGTGGGTGACCTTGATTCTCTCTG
GAGCACACGTCTGAACTCTTTCCCTACACGACGCTCTTCCGATCTNNNGGTTGACATGGGCAGTAGG
AGCACACGTCTGAACTCTTTCCCTACACGACGCTCTTCCGATCTNNNGGACAATCTGTGTCGATTGAG
GCACACGTCTGAACTCTTTCCCTACACGACGCTCTTCCGATCTNNNATTTACCTACCACCTTGTAGTGG
AGCACACGTCTGAACTCTTTCCCTACACGACGCTCTTCCGATCTNNNAGGAAGTGCTAACAAATGGTG
AGCACACGTCTGAACTCTTTCCCTACACGACGCTCTTCCGATCTNNNCCAGGGTGAGCAATCCAAAAG
AGAGCACACGTCTGAACTCTTTCCCTACACGACGCTCTTCCGATCTNNNTTTTCTCTTTCTCACGCACA
GCACACGTCTGAACTCTTTCCCTACACGACGCTCTTCCGATCTNNNGGGAAGAGTTTTTCTCCTCTGG
GAGCACACGTCTGAACTCTTTCCCTACACGACGCTCTTCCGATCTNNNGATCCATGCAGATCCAGTCTC
GAGCACACGTCTGAACTCTTTCCCTACACGACGCTCTTCCGATCTNNNTGTCCAACAGAGGAAACACAT
GAGCACACGTCTGAACTCTTTCCCTACACGACGCTCTTCCGATCTNNNCCCTGAGGATACAGCCCTAAAT
AGAGCACACGTCTGAACTCTTTCCCTACACGACGCTCTTCCGATCTNNNCTGGTTACACCCCCTTCCCTA
AGCACACGTCTGAACTCTTTCCCTACACGACGCTCTTCCGATCTNNNAGCCCCTGCTTTTTCCTTTCTAC
GCACACGTCTGAACTCTTTCCCTACACGACGCTCTTCCGATCTNNNTCTGCCTACTTTCTCCAGCTTCTC
GCACACGTCTGAACTCTTTCCCTACACGACGCTCTTCCGATCTNNNTGTCTGTGTGTGTGTTGAGTGA
AGCACACGTCTGAACTCTTTCCCTACACGACGCTCTTCCGATCTNNNTAACGAGGGCTCTGGGAGATA
AGCACACGTCTGAACTCTTTCCCTACACGACGCTCTTCCGATCTNNNACAGGCTTTGCATATTTCACCG
AGAGCACACGTCTGAACTCTTTCCCTACACGACGCTCTTCCGATCTNNNGCCACAAGTAGCTAGCAAAG
GAGCACACGTCTGAACTCTTTCCCTACACGACGCTCTTCCGATCTNNNGGCCTGACGGGTAAAGCTATT
GAGCACACGTCTGAACTCTTTCCCTACACGACGCTCTTCCGATCTNNNTCACTGTAGCAGTCTTTCTTC

GAGCACACGTCTGAACTCTTTCCCTACACGACGCTCTTCCGATCTNNNTCTCTGTCCAGCAGGAAGAAG
AGAGCACACGTCTGAACTCTTTCCCTACACGACGCTCTTCCGATCTNNNGCATGTTAGGTGGTTTTGTC
GAGCACACGTCTGAACTCTTTCCCTACACGACGCTCTTCCGATCTNNNGCAGGATTGACTGAAGGAGG
GAGCACACGTCTGAACTCTTTCCCTACACGACGCTCTTCCGATCTNNNCCTCATGAGCAACATTCTTCCCA
AGAGCACACGTCTGAACTCTTTCCCTACACGACGCTCTTCCGATCTNNNTGAATGAATGACGAACAGCA
GAGCACACGTCTGAACTCTTTCCCTACACGACGCTCTTCCGATCTNNNACATGCCCCTAGCTTACTAGCT
GAGCACACGTCTGAACTCTTTCCCTACACGACGCTCTTCCGATCTNNNACCCGGCACGATTTATTTGGT
AGAGCACACGTCTGAACTCTTTCCCTACACGACGCTCTTCCGATCTNNNGGCTGCAGGGTCATGTAAA
GAGCACACGTCTGAACTCTTTCCCTACACGACGCTCTTCCGATCTNNNAGCTACCACCCAAAACATAGCA
GAGCACACGTCTGAACTCTTTCCCTACACGACGCTCTTCCGATCTNNNTCTTCCCAGACCTAGACTGTGT
GAGCACACGTCTGAACTCTTTCCCTACACGACGCTCTTCCGATCTNNNAACAGCAACAGAGCAATATCC
GAGCACACGTCTGAACTCTTTCCCTACACGACGCTCTTCCGATCTNNNGCGACTTTTCTCTGCTTTCCATT
GAGCACACGTCTGAACTCTTTCCCTACACGACGCTCTTCCGATCTNNNCCTTTCGGGATAGTCGCAAGA
AGCACACGTCTGAACTCTTTCCCTACACGACGCTCTTCCGATCTNNNCCTTCATGCTGATGACTTCCAGT
GAGCACACGTCTGAACTCTTTCCCTACACGACGCTCTTCCGATCTNNNGCGAATACACACGCCCTGACCA
GAGCACACGTCTGAACTCTTTCCCTACACGACGCTCTTCCGATCTNNNAGTGTTCAGGGAAAACAGTC
AGAGCACACGTCTGAACTCTTTCCCTACACGACGCTCTTCCGATCTNNNCAATTCAGTGCTCCTTTCCCT
GAGCACACGTCTGAACTCTTTCCCTACACGACGCTCTTCCGATCTNNNTTTACCAAGGAACAGAGCCAT
AGCACACGTCTGAACTCTTTCCCTACACGACGCTCTTCCGATCTNNNATGGATAACATGTCCTCACTCCT
AGCACACGTCTGAACTCTTTCCCTACACGACGCTCTTCCGATCTNNNTCCCCAACCTGAGAAATTGTGT
AGCACACGTCTGAACTCTTTCCCTACACGACGCTCTTCCGATCTNNNGGGAAGAGTTTTTCTCCTCTGG
GAGCACACGTCTGAACTCTTTCCCTACACGACGCTCTTCCGATCTNNNTGGATGGCAAAGAACACATCC
AGCACACGTCTGAACTCTTTCCCTACACGACGCTCTTCCGATCTNNNAGCCCTTCTACAGTGGCATAAG
AGCACACGTCTGAACTCTTTCCCTACACGACGCTCTTCCGATCTNNNTGCCTGGCAGTAATGATTGGAT
GCACACGTCTGAACTCTTTCCCTACACGACGCTCTTCCGATCTNNNTGGTGTCTTTTATTCCACAGAGCC
AGCACACGTCTGAACTCTTTCCCTACACGACGCTCTTCCGATCTNNNGTGGGTGTCATAGCAACTAGAG
AGAGCACACGTCTGAACTCTTTCCCTACACGACGCTCTTCCGATCTNNNTGTGACTTACTGGCCCAACTT
GAAGAGCACACGTCTGAACTCTTTCCCTACACGACGCTCTTCCGATCTNNNAATGCATTCTCTCCCTCTCT
GAGCACACGTCTGAACTCTTTCCCTACACGACGCTCTTCCGATCTNNNTGTTCACAAGGTACTGCATACC
GAGCACACGTCTGAACTCTTTCCCTACACGACGCTCTTCCGATCTNNNCAAAAAGGGACAGCCAAAGTC
AAGAGCACACGTCTGAACTCTTTCCCTACACGACGCTCTTCCGATCTNNNCTCAAGGAGAAAGGGCAT
GCACACGTCTGAACTCTTTCCCTACACGACGCTCTTCCGATCTNNNCCTAGTTTGATATGGCTGGAGGC
AGCACACGTCTGAACTCTTTCCCTACACGACGCTCTTCCGATCTNNNAGCACTTCTGACTGATCCTCCTT
GAGCACACGTCTGAACTCTTTCCCTACACGACGCTCTTCCGATCTNNNAACCCAATATGTTCTGCATCCA
AGCACACGTCTGAACTCTTTCCCTACACGACGCTCTTCCGATCTNNNGAGCCTTCTCAGTTCTTTCCCAG
AAGAGCACACGTCTGAACTCTTTCCCTACACGACGCTCTTCCGATCTNNNCTTTGCTTTTATGCGTGTGT
AGCACACGTCTGAACTCTTTCCCTACACGACGCTCTTCCGATCTNNNACCTTGTGAAAGATCTGTGGCT
AGCACACGTCTGAACTCTTTCCCTACACGACGCTCTTCCGATCTNNNGATGCTTCCAGTCAGATGCTAC
AGAGCACACGTCTGAACTCTTTCCCTACACGACGCTCTTCCGATCTNNNCATCTACCCCTTTGTGAAGGA
AGAGCACACGTCTGAACTCTTTCCCTACACGACGCTCTTCCGATCTNNNTACCCCAATGAATGTGCCAT
GAGCACACGTCTGAACTCTTTCCCTACACGACGCTCTTCCGATCTNNNGAAGGCCTGAGAACCAAGAAC
AAGAGCACACGTCTGAACTCTTTCCCTACACGACGCTCTTCCGATCTNNNATAAAACAGGCAAAGGTC
AGCACACGTCTGAACTCTTTCCCTACACGACGCTCTTCCGATCTNNNTCAATCGTGACGGTTTTATGCCT

GAGCACACGTCTGAACTCTTTCCCTACACGACGCTCTTCCGATCTNNNCACACACAGACATATAAACGT
AGAGCACACGTCTGAACTCTTTCCCTACACGACGCTCTTCCGATCTNNNTGCAACTCTGTGATGTTGTTA
AGCACACGTCTGAACTCTTTCCCTACACGACGCTCTTCCGATCTNNNGGACGTAGGGTCCAACTTGAA
GAGCACACGTCTGAACTCTTTCCCTACACGACGCTCTTCCGATCTNNNACTCGAATGAAGCCTGTTTGG
AGAGCACACGTCTGAACTCTTTCCCTACACGACGCTCTTCCGATCTNNNATGGTATATCTGAGCCCTCCA
AGAGCACACGTCTGAACTCTTTCCCTACACGACGCTCTTCCGATCTNNNACAGCTGGAGGAGACTGAA
GAGCACACGTCTGAACTCTTTCCCTACACGACGCTCTTCCGATCTNNNAAAAAGTCCCCATGCTACACA
GAGCACACGTCTGAACTCTTTCCCTACACGACGCTCTTCCGATCTNNNATTTGGGGTGGACACAAACAC
AGAGCACACGTCTGAACTCTTTCCCTACACGACGCTCTTCCGATCTNNNTATCTTCTCGGCATGGCTTTT
AAGAGCACACGTCTGAACTCTTTCCCTACACGACGCTCTTCCGATCTNNNGCCAAAATGCCCATCAAAC
CACACGTCTGAACTCTTTCCCTACACGACGCTCTTCCGATCTNNNTTCTTGGGGTCAAGTTATGCACTC
AGCACACGTCTGAACTCTTTCCCTACACGACGCTCTTCCGATCTNNNATATGCACAGGTTTTGCATCCC
AGCACACGTCTGAACTCTTTCCCTACACGACGCTCTTCCGATCTNNNGTGCTTCATGTATCCTCTGGCTT
AGAGCACACGTCTGAACTCTTTCCCTACACGACGCTCTTCCGATCTNNNCGGTTCTCCAGAGAAACAAC
AGAGCACACGTCTGAACTCTTTCCCTACACGACGCTCTTCCGATCTNNNGGCTTTGGGCTAAGAAGGA
GAGCACACGTCTGAACTCTTTCCCTACACGACGCTCTTCCGATCTNNNAGATTTCCTTGGATGCCAGAA
GAGCACACGTCTGAACTCTTTCCCTACACGACGCTCTTCCGATCTNNNCCAGTTTTGCAGTCTGACACC
AGCACACGTCTGAACTCTTTCCCTACACGACGCTCTTCCGATCTNNNGAACCATCATAAGTTGGGACC
GAGCACACGTCTGAACTCTTTCCCTACACGACGCTCTTCCGATCTNNNAGAGGAGAGAGTGGATTGG
GAGCACACGTCTGAACTCTTTCCCTACACGACGCTCTTCCGATCTNNNACACTTTGCTTACTTACACCCA
AGCACACGTCTGAACTCTTTCCCTACACGACGCTCTTCCGATCTNNNTATGTCAGGAAACACACACACG
GAGCACACGTCTGAACTCTTTCCCTACACGACGCTCTTCCGATCTNNNAGCATTGCCATGGAAGTTGG
AAGAGCACACGTCTGAACTCTTTCCCTACACGACGCTCTTCCGATCTNNNCTTTCACCTTATCCCCATC
AGCACACGTCTGAACTCTTTCCCTACACGACGCTCTTCCGATCTNNNCTTTTTCTCCTACGTGCCCTTCA
GCACACGTCTGAACTCTTTCCCTACACGACGCTCTTCCGATCTNNNCTCCACAAAATTAGGCCATGCTG
AGCACACGTCTGAACTCTTTCCCTACACGACGCTCTTCCGATCTNNNACGTGAATATTGACAGGGGCTT
GAAGAGCACACGTCTGAACTCTTTCCCTACACGACGCTCTTCCGATCTNNNCGGAGGACAGCAGAGG
GAGCACACGTCTGAACTCTTTCCCTACACGACGCTCTTCCGATCTNNNCTCTTGCCTAAGTCTCAGGAT
AGAGCACACGTCTGAACTCTTTCCCTACACGACGCTCTTCCGATCTNNNGCCCACAACGACAGTCTTTTT
AGCACACGTCTGAACTCTTTCCCTACACGACGCTCTTCCGATCTNNNATATGCACAGGTTTTGCATCCC
GAGCACACGTCTGAACTCTTTCCCTACACGACGCTCTTCCGATCTNNNTCTTGCAGGGAGAGTAAGGT
AGCACACGTCTGAACTCTTTCCCTACACGACGCTCTTCCGATCTNNNTTGGATTAGTCAGGGTTCAGG
GAGCACACGTCTGAACTCTTTCCCTACACGACGCTCTTCCGATCTNNNCTGAGAATTTGGCAAAGGGC
GAGCACACGTCTGAACTCTTTCCCTACACGACGCTCTTCCGATCTNNNCAGAGAACAGGACCCTAGGA
AGCACACGTCTGAACTCTTTCCCTACACGACGCTCTTCCGATCTNNNAGCCCTTCTACAGTGGCATAAG
GAGCACACGTCTGAACTCTTTCCCTACACGACGCTCTTCCGATCTNNNGGAGACACCCTAAATGTGTCC
GAGCACACGTCTGAACTCTTTCCCTACACGACGCTCTTCCGATCTNNNAACAGTCTACTTTGGAAGGC
CACACGTCTGAACTCTTTCCCTACACGACGCTCTTCCGATCTNNNCCCTTCCTTATGAGGTTGGTGTGC
GAGCACACGTCTGAACTCTTTCCCTACACGACGCTCTTCCGATCTNNNCACCTAGCAAATAACAGGGGT
AGAGCACACGTCTGAACTCTTTCCCTACACGACGCTCTTCCGATCTNNNTCCCAACTTTTCCTTTGTCCCT
AGAGCACACGTCTGAACTCTTTCCCTACACGACGCTCTTCCGATCTNNNCAATCTCAGACCCTTCACAGC
AGCACACGTCTGAACTCTTTCCCTACACGACGCTCTTCCGATCTNNNAGTCTCTCATGTGTGTGAGGG
AGCACACGTCTGAACTCTTTCCCTACACGACGCTCTTCCGATCTNNNCCAGTAAAATGGCAGCAACAAG

AGCACACGTCTGAACTCTTTCCCTACACGACGCTCTTCCGATCTNNNGAAAATGACTTTCAAACAGAA
GAGCACACGTCTGAACTCTTTCCCTACACGACGCTCTTCCGATCTNNNCACAACCTCTGACATGGCAA
GAGCACACGTCTGAACTCTTTCCCTACACGACGCTCTTCCGATCTNNNAGCTGTGAGAGGGTGGTAA
GAGCACACGTCTGAACTCTTTCCCTACACGACGCTCTTCCGATCTNNNTGCAGACATGAAGGCAAGTC
AGCACACGTCTGAACTCTTTCCCTACACGACGCTCTTCCGATCTNNNTGAAGCTGCCCCTGTTTTGTAT
AGCACACGTCTGAACTCTTTCCCTACACGACGCTCTTCCGATCTNNNAGTCTTTTGGGGTTGGCATAG
AGCACACGTCTGAACTCTTTCCCTACACGACGCTCTTCCGATCTNNNTGTAAACATGCGGGAATTTGG
GCACACGTCTGAACTCTTTCCCTACACGACGCTCTTCCGATCTNNNCGCACTATAACTGCCCTACTTCCC
GCACACGTCTGAACTCTTTCCCTACACGACGCTCTTCCGATCTNNNGTGACCACGCCCCTTAGTCTCCA
AAGAGCACACGTCTGAACTCTTTCCCTACACGACGCTCTTCCGATCTNNNATGATCGTACACTACACT
GAGCACACGTCTGAACTCTTTCCCTACACGACGCTCTTCCGATCTNNNAAAACAACTCAGCATTCTGG
AGCACACGTCTGAACTCTTTCCCTACACGACGCTCTTCCGATCTNNNCAGGAACCAGTGCATACATTC
GAGCACACGTCTGAACTCTTTCCCTACACGACGCTCTTCCGATCTNNNGTCCAGCTGCACAAGAAGTA
GAAGAGCACACGTCTGAACTCTTTCCCTACACGACGCTCTTCCGATCTNNNTTGAGCTTCCAAATCAG
GCACACGTCTGAACTCTTTCCCTACACGACGCTCTTCCGATCTNNNGCAGCATTAGAAAGAGGCAGA
GCACACGTCTGAACTCTTTCCCTACACGACGCTCTTCCGATCTNNNCTGAAGCCTCAGACCTTGGAAAG
GCACACGTCTGAACTCTTTCCCTACACGACGCTCTTCCGATCTNNNCCTCTGGGGTCTTTGAGATTGT
AGCACACGTCTGAACTCTTTCCCTACACGACGCTCTTCCGATCTNNNGGGGATTGGAACAATGCCTG
CACACGTCTGAACTCTTTCCCTACACGACGCTCTTCCGATCTNNNTTTCCCCAAGATGTCCTGGTTTG
GAGCACACGTCTGAACTCTTTCCCTACACGACGCTCTTCCGATCTNNNGCATCAGATAAGCCAGAGG
AGCACACGTCTGAACTCTTTCCCTACACGACGCTCTTCCGATCTNNNATGCTAAGCTGAGGTTCCCAA
GAGCACACGTCTGAACTCTTTCCCTACACGACGCTCTTCCGATCTNNNTATCTGGCCCACCTATTCCTAG
AGCACACGTCTGAACTCTTTCCCTACACGACGCTCTTCCGATCTNNNATGCATCTGGAAAGCAAAAAG
AGCACACGTCTGAACTCTTTCCCTACACGACGCTCTTCCGATCTNNNATGCTGTGCAAGAGAAGTGA
AGCACACGTCTGAACTCTTTCCCTACACGACGCTCTTCCGATCTNNNTTGCAATTCCCTGTCCTTTGTA
AGCACACGTCTGAACTCTTTCCCTACACGACGCTCTTCCGATCTNNNTTTACATGTTTGGGTGGGTG
AGCACACGTCTGAACTCTTTCCCTACACGACGCTCTTCCGATCTNNNTGAAAATCCAAATGTGTATCCC
AGCACACGTCTGAACTCTTTCCCTACACGACGCTCTTCCGATCTNNNTTATTGCATCCCCATCAGTGTC
AGCACACGTCTGAACTCTTTCCCTACACGACGCTCTTCCGATCTNNNTAACGCTGGAGATGGAGGTT
GCACACGTCTGAACTCTTTCCCTACACGACGCTCTTCCGATCTNNNTGGTCTGAGATCTGTGTTCAAC
AGCACACGTCTGAACTCTTTCCCTACACGACGCTCTTCCGATCTNNNTGGGACTAGAGGTACACACAC
AGCACACGTCTGAACTCTTTCCCTACACGACGCTCTTCCGATCTNNNTCTGCTGTAGTGGTTCCCAGA
AGCACACGTCTGAACTCTTTCCCTACACGACGCTCTTCCGATCTNNNACAAGAACACTAAATGCTGTG
AGCACACGTCTGAACTCTTTCCCTACACGACGCTCTTCCGATCTNNNTCTCATGCACACGAGTGTTTCA
AGCACACGTCTGAACTCTTTCCCTACACGACGCTCTTCCGATCTNNNGAAGGAGCTTGCAGAGCATA
AGCACACGTCTGAACTCTTTCCCTACACGACGCTCTTCCGATCTNNNGCTGTGAAAGCATGACACCAA
AAGAGCACACGTCTGAACTCTTTCCCTACACGACGCTCTTCCGATCTNNNGCTCTGGGTAACACACTT
GAGCACACGTCTGAACTCTTTCCCTACACGACGCTCTTCCGATCTNNNAATTCATTCCAGGTTGCTGT
AGCACACGTCTGAACTCTTTCCCTACACGACGCTCTTCCGATCTNNNACACAACACAGATCACAGACC
GAGCACACGTCTGAACTCTTTCCCTACACGACGCTCTTCCGATCTNNNAGAATAAGCAGTATTGTGG
AGCACACGTCTGAACTCTTTCCCTACACGACGCTCTTCCGATCTNNNCCCAACTAAGGGTACAAATG
GCACACGTCTGAACTCTTTCCCTACACGACGCTCTTCCGATCTNNNTTATTGCATCCCCATCAGTGTCC
GCACACGTCTGAACTCTTTCCCTACACGACGCTCTTCCGATCTNNNCAGGGCAAGGGGTGTCTTTTA

GAGCACACGTCTGAACTCTTTCCCTACACGACGCTCTTCCGATCTNNNATCCCCACAGCAACCCTATAAG
GAGCACACGTCTGAACTCTTTCCCTACACGACGCTCTTCCGATCTNNNAAGTCAAAACAGGTCCAAGAG
GAGCACACGTCTGAACTCTTTCCCTACACGACGCTCTTCCGATCTNNNTGGAAACGCTATTCTCAACCTG
GAGCACACGTCTGAACTCTTTCCCTACACGACGCTCTTCCGATCTNNNAGAGAGACCTCATCTCTGTAG
GAGCACACGTCTGAACTCTTTCCCTACACGACGCTCTTCCGATCTNNNATACAGATGCAACCAAAGCAC
AGAGCACACGTCTGAACTCTTTCCCTACACGACGCTCTTCCGATCTNNNGGACTTCCACACTCGCCTATT
AGCACACGTCTGAACTCTTTCCCTACACGACGCTCTTCCGATCTNNNACCGAAGAACATGAACACACA
AGCACACGTCTGAACTCTTTCCCTACACGACGCTCTTCCGATCTNNNTGGACTTGTGGGTTATTTATG
AGCACACGTCTGAACTCTTTCCCTACACGACGCTCTTCCGATCTNNNGCCCATAAAGAGAATTTGCTGG
AGCACACGTCTGAACTCTTTCCCTACACGACGCTCTTCCGATCTNNNTACACAGAGACAGAAGGTCAG
AGCACACGTCTGAACTCTTTCCCTACACGACGCTCTTCCGATCTNNNACTGACACATCTATCACCACACA
AGCACACGTCTGAACTCTTTCCCTACACGACGCTCTTCCGATCTNNNTGGGTAAAGAGAACCCTCACCA
GAGCACACGTCTGAACTCTTTCCCTACACGACGCTCTTCCGATCTNNNCCTAGCTCTTCTGTCTAGCCCT
GAGCACACGTCTGAACTCTTTCCCTACACGACGCTCTTCCGATCTNNNGGTTCTCCAAGAGGTATCCAG
GAGCACACGTCTGAACTCTTTCCCTACACGACGCTCTTCCGATCTNNNTTCTGATGTTCCACCCTTCTGGA
AAGAGCACACGTCTGAACTCTTTCCCTACACGACGCTCTTCCGATCTNNNTGTCATAGACACCACCACT
AGCACACGTCTGAACTCTTTCCCTACACGACGCTCTTCCGATCTNNNGGGCGTCTAGGTTGATTCTATCT
GAGCACACGTCTGAACTCTTTCCCTACACGACGCTCTTCCGATCTNNNCTTGACGTTTACCACATGTCCCT
GAGCACACGTCTGAACTCTTTCCCTACACGACGCTCTTCCGATCTNNNTATCAGTTTGGGGAGGGAAGT
AGCACACGTCTGAACTCTTTCCCTACACGACGCTCTTCCGATCTNNNTTATTCTGTCACCAAGGTGTGC
AGCACACGTCTGAACTCTTTCCCTACACGACGCTCTTCCGATCTNNNAGGCAGGTAACAGTAGTCCTCA
GAGCACACGTCTGAACTCTTTCCCTACACGACGCTCTTCCGATCTNNNAAGGCAAGTAGCTGGGTTAGA
AGCACACGTCTGAACTCTTTCCCTACACGACGCTCTTCCGATCTNNNCTGGACCTGCAATTGCTCTACTT
AGCACACGTCTGAACTCTTTCCCTACACGACGCTCTTCCGATCTNNNTGACCCACAGATCATAGCCTTCA
AGCACACGTCTGAACTCTTTCCCTACACGACGCTCTTCCGATCTNNNCCCTTCTACTGCAGCCAAATGAT
GAGCACACGTCTGAACTCTTTCCCTACACGACGCTCTTCCGATCTNNNTCCATTGTTTGTCTGGTGCTGT
GAGCACACGTCTGAACTCTTTCCCTACACGACGCTCTTCCGATCTNNNCCCTGCTCATTAGTTTGTGGGA
GAGCACACGTCTGAACTCTTTCCCTACACGACGCTCTTCCGATCTNNNTGGGAGAAAATAATTGCAGT
AGCACACGTCTGAACTCTTTCCCTACACGACGCTCTTCCGATCTNNNATCTTGTGGACAGCTTGGGAAT
AAGAGCACACGTCTGAACTCTTTCCCTACACGACGCTCTTCCGATCTNNNTGCTTCATCATTGGCACTCT
GAGCACACGTCTGAACTCTTTCCCTACACGACGCTCTTCCGATCTNNNTGCAGCCAGTCCAAGATTTCAT
GAAGAGCACACGTCTGAACTCTTTCCCTACACGACGCTCTTCCGATCTNNNTCTAGAGAGCCCAAGTCA
GAGCACACGTCTGAACTCTTTCCCTACACGACGCTCTTCCGATCTNNNTGATTTGGATAATTGTTCCGG
AGAGCACACGTCTGAACTCTTTCCCTACACGACGCTCTTCCGATCTNNNATCCTGTGTAAAGACAGGCA
GCACACGTCTGAACTCTTTCCCTACACGACGCTCTTCCGATCTNNNTCCTCTTGGTGATGGGGTTGTAT
GAGCACACGTCTGAACTCTTTCCCTACACGACGCTCTTCCGATCTNNNAGATGTGCTTTGAAGGCTTTT
AGAGCACACGTCTGAACTCTTTCCCTACACGACGCTCTTCCGATCTNNNAATCTGGCCCACATGCACAAA
AGCACACGTCTGAACTCTTTCCCTACACGACGCTCTTCCGATCTNNNTTCATCTCCTGGATTATGTGCTTT
AGCACACGTCTGAACTCTTTCCCTACACGACGCTCTTCCGATCTNNNAAACCATGAAACCTCCATGTCC
GAGCACACGTCTGAACTCTTTCCCTACACGACGCTCTTCCGATCTNNNTGTTTTTGTCTGGCTGCCTTCA
AGCACACGTCTGAACTCTTTCCCTACACGACGCTCTTCCGATCTNNNAGCTTTATTTAGCACACACACA
AGCACACGTCTGAACTCTTTCCCTACACGACGCTCTTCCGATCTNNNATCAATTGCCAAATGTACCCACG
GAGCACACGTCTGAACTCTTTCCCTACACGACGCTCTTCCGATCTNNNGTGAAAGGCACTGGAGGTTG

AGCACACGTCTGAACTCTTTCCCTACACGACGCTCTTCCGATCTNNNGCTGCAAAGAGGCTGATTTCA
GAGCACACGTCTGAACTCTTTCCCTACACGACGCTCTTCCGATCTNNNATAATCCCAAAATGCAGTGGC
AGCACACGTCTGAACTCTTTCCCTACACGACGCTCTTCCGATCTNNNTACTCTGGAACATTCTGGCATC
AGCACACGTCTGAACTCTTTCCCTACACGACGCTCTTCCGATCTNNNTGGACTTTTGCACTACAGGTG
AGCACACGTCTGAACTCTTTCCCTACACGACGCTCTTCCGATCTNNNGAAATCAGAAAGGGCTGGTA
GAGCACACGTCTGAACTCTTTCCCTACACGACGCTCTTCCGATCTNNNGAGGTGCTTCACCCTAACATC
GAGCACACGTCTGAACTCTTTCCCTACACGACGCTCTTCCGATCTNNNTAAGGACAATCGAAACCCTCC
GAGCACACGTCTGAACTCTTTCCCTACACGACGCTCTTCCGATCTNNNGCACTCAATTGCTTACTTCCA
GAGCACACGTCTGAACTCTTTCCCTACACGACGCTCTTCCGATCTNNNGTTCAGTAGGAATTCGCTCA
GAGCACACGTCTGAACTCTTTCCCTACACGACGCTCTTCCGATCTNNNGAGGCTCTGTCTCAAAAGAA
GAGCACACGTCTGAACTCTTTCCCTACACGACGCTCTTCCGATCTNNNATCACTACCCCACAAAAGCA
GAGCACACGTCTGAACTCTTTCCCTACACGACGCTCTTCCGATCTNNNACTTCTAGGAACAAGGAGTA
GAGCACACGTCTGAACTCTTTCCCTACACGACGCTCTTCCGATCTNNNGATGATATATGGGCGGGCA
AGCACACGTCTGAACTCTTTCCCTACACGACGCTCTTCCGATCTNNNACAGGCAATCATTTAGCCTCCT
AGCACACGTCTGAACTCTTTCCCTACACGACGCTCTTCCGATCTNNNTGAGGTAGGAACCTGCAAGT
AGCACACGTCTGAACTCTTTCCCTACACGACGCTCTTCCGATCTNNNACCCATAGCTCAGAACACATGA
AGCACACGTCTGAACTCTTTCCCTACACGACGCTCTTCCGATCTNNNAAAAGGGAGACTGGGGAGA
GAGCACACGTCTGAACTCTTTCCCTACACGACGCTCTTCCGATCTNNNTTGAGCTTCCAAATCAGGTG
GAGCACACGTCTGAACTCTTTCCCTACACGACGCTCTTCCGATCTNNNGATCCTGTCTCTCTCTCTCCT
GAGCACACGTCTGAACTCTTTCCCTACACGACGCTCTTCCGATCTNNNAGTCCCTGTGATAGCTCCTGA
GAGCACACGTCTGAACTCTTTCCCTACACGACGCTCTTCCGATCTNNNCTCCATACAGCTTTCCATGG
GCACACGTCTGAACTCTTTCCCTACACGACGCTCTTCCGATCTNNNGCAGGAAGCACATCTACTAGCT
GCACACGTCTGAACTCTTTCCCTACACGACGCTCTTCCGATCTNNNCTCTATGTGAGCATCTTGGGGT
AGCACACGTCTGAACTCTTTCCCTACACGACGCTCTTCCGATCTNNNGGGTTTTCTGTCACTCTGCTGT
AGCACACGTCTGAACTCTTTCCCTACACGACGCTCTTCCGATCTNNNGGGCACCAGAGGCAATCTTAT
AGAGCACACGTCTGAACTCTTTCCCTACACGACGCTCTTCCGATCTNNNAGGGAAATGGAAGTAGTC
GCACACGTCTGAACTCTTTCCCTACACGACGCTCTTCCGATCTNNNACCCAATTTCCTTCATGCTCACA
GAGCACACGTCTGAACTCTTTCCCTACACGACGCTCTTCCGATCTNNNACAACACAGTACAAACACACT
AGCACACGTCTGAACTCTTTCCCTACACGACGCTCTTCCGATCTNNNTTGGTTTTCCATTCCATCCAGGT
GCACACGTCTGAACTCTTTCCCTACACGACGCTCTTCCGATCTNNNAGTTGGCACAGAAGGTGAAGA
AGCACACGTCTGAACTCTTTCCCTACACGACGCTCTTCCGATCTNNNTGGGATTTGTCATAATGAAGC
GCACACGTCTGAACTCTTTCCCTACACGACGCTCTTCCGATCTNNNCTTTCACCTTATCCCCCATCTGCC
AGCACACGTCTGAACTCTTTCCCTACACGACGCTCTTCCGATCTNNNTCACCCACATTTGAAACAGGA
AGCACACGTCTGAACTCTTTCCCTACACGACGCTCTTCCGATCTNNNGACCTTTCTGTCATGGCTCACA
GCACACGTCTGAACTCTTTCCCTACACGACGCTCTTCCGATCTNNNCTGCTCAGTTAGCTTAGTCGTC
AGCACACGTCTGAACTCTTTCCCTACACGACGCTCTTCCGATCTNNNGGTTTGGTTTGGTTGGTTTTG
AGCACACGTCTGAACTCTTTCCCTACACGACGCTCTTCCGATCTNNNCTCACCACCTAGGACATGGAG
AGCACACGTCTGAACTCTTTCCCTACACGACGCTCTTCCGATCTNNNAGTGACTGCAAATCAAATTTT
AGCACACGTCTGAACTCTTTCCCTACACGACGCTCTTCCGATCTNNNTCGATGACCTTACATCCACCCA
GAGCACACGTCTGAACTCTTTCCCTACACGACGCTCTTCCGATCTNNNTTCAAACCTCCTGAACACACCT
AGCACACGTCTGAACTCTTTCCCTACACGACGCTCTTCCGATCTNNNAAAGGGATGGAGAAACAGAG
GAGCACACGTCTGAACTCTTTCCCTACACGACGCTCTTCCGATCTNNNGAGACATGGAGGAGTTTGG
GAGCACACGTCTGAACTCTTTCCCTACACGACGCTCTTCCGATCTNNNCATGGAGACAGAGGCCAAG

GAGCACACGTCTGAACTCTTTCCCTACACGACGCTCTTCCGATCTNNNAGGCAGAAGTCTTATAGCTGG
GAGCACACGTCTGAACTCTTTCCCTACACGACGCTCTTCCGATCTNNNTCCTTTCTCATTGTGCCAGTTCC
GCACACGTCTGAACTCTTTCCCTACACGACGCTCTTCCGATCTNNNTCTGACACATCATGGCCACAAACC
CACACGTCTGAACTCTTTCCCTACACGACGCTCTTCCGATCTNNNTATTCCTCCTAGGCTGTGTGTGTCA
AGCACACGTCTGAACTCTTTCCCTACACGACGCTCTTCCGATCTNNNGGGCACCAGAGGCAATCTTATA
GAGCACACGTCTGAACTCTTTCCCTACACGACGCTCTTCCGATCTNNNAACTTTACCTGCCTCTGCTTTC
AGAGCACACGTCTGAACTCTTTCCCTACACGACGCTCTTCCGATCTNNNACAAAATAGTGGGGGTTCTG
AAGAGCACACGTCTGAACTCTTTCCCTACACGACGCTCTTCCGATCTNNNGGCAGTTTGAAGACCAATG
AGAGCACACGTCTGAACTCTTTCCCTACACGACGCTCTTCCGATCTNNNTTGGTCAGTGCAACGTAACT
GAGCACACGTCTGAACTCTTTCCCTACACGACGCTCTTCCGATCTNNNACTCTTGTTACAGGTACCCTCC
AAGAGCACACGTCTGAACTCTTTCCCTACACGACGCTCTTCCGATCTNNNTCTTCTCTTGTCAGTCAGTG
AGCACACGTCTGAACTCTTTCCCTACACGACGCTCTTCCGATCTNNNGGGAGGCTGAAGGAACAAAG
AGCACACGTCTGAACTCTTTCCCTACACGACGCTCTTCCGATCTNNNTCCTGGAATAGACTGCATGTCAG
AGCACACGTCTGAACTCTTTCCCTACACGACGCTCTTCCGATCTNNNCTTACAGAGGGACTGGTGTGTT
GAGCACACGTCTGAACTCTTTCCCTACACGACGCTCTTCCGATCTNNNTTCATTACCATCACTGCCACAGG
AGCACACGTCTGAACTCTTTCCCTACACGACGCTCTTCCGATCTNNNAGTCTCATATCACACACACACATG
AGAGCACACGTCTGAACTCTTTCCCTACACGACGCTCTTCCGATCTNNNGGCAGCATTGTGAAAACCCA
AGCACACGTCTGAACTCTTTCCCTACACGACGCTCTTCCGATCTNNNTGCTCAAGCATGAAAGAGAGCA
GAGCACACGTCTGAACTCTTTCCCTACACGACGCTCTTCCGATCTNNNCTGTGGGTGTGTTGTATTTGT
AGCACACGTCTGAACTCTTTCCCTACACGACGCTCTTCCGATCTNNNATTCACAGCCTCTTCCTGAATGG
AGCACACGTCTGAACTCTTTCCCTACACGACGCTCTTCCGATCTNNNGGCTAGACACACTGAAGGAAAG
AGCACACGTCTGAACTCTTTCCCTACACGACGCTCTTCCGATCTNNNCAGGTGCAATGATGTTACCTGG
AGCACACGTCTGAACTCTTTCCCTACACGACGCTCTTCCGATCTNNNAGTAATTCAAAAAGGCCAATGT
AGCACACGTCTGAACTCTTTCCCTACACGACGCTCTTCCGATCTNNNCTTGTCTCATGGGGCAGACATT
GAGCACACGTCTGAACTCTTTCCCTACACGACGCTCTTCCGATCTNNNGACTTCACCCATGATGTACCCA
GAGCACACGTCTGAACTCTTTCCCTACACGACGCTCTTCCGATCTNNNAGAGCTTGGTGACATGCATTG
GAGCACACGTCTGAACTCTTTCCCTACACGACGCTCTTCCGATCTNNNTGTTCCTTCTCAACACCTTGTGG
AGCACACGTCTGAACTCTTTCCCTACACGACGCTCTTCCGATCTNNNGCTGCATTGAGAACCAAGATG
GAGCACACGTCTGAACTCTTTCCCTACACGACGCTCTTCCGATCTNNNTTCGGTTTCTGATTTGGAGGT
GAGCACACGTCTGAACTCTTTCCCTACACGACGCTCTTCCGATCTNNNTCCAACTATTCTCTCCCCCAACC
AAGAGCACACGTCTGAACTCTTTCCCTACACGACGCTCTTCCGATCTNNNCTCCTCTGCCTGTTTTCGAG
AGCACACGTCTGAACTCTTTCCCTACACGACGCTCTTCCGATCTNNNGGGAATGGGGAGCTGAAAAG
AGCACACGTCTGAACTCTTTCCCTACACGACGCTCTTCCGATCTNNNCCCAGGGAAATGAGGCAAGTT
AGAGCACACGTCTGAACTCTTTCCCTACACGACGCTCTTCCGATCTNNNGAGAACTGCTCAGACATGG
AGCACACGTCTGAACTCTTTCCCTACACGACGCTCTTCCGATCTNNNTTGCCTTAGTTTTTGGTTTGGAA
GAGCACACGTCTGAACTCTTTCCCTACACGACGCTCTTCCGATCTNNNCCTGTTATCCTTTTCAGGGCCT
GAGCACACGTCTGAACTCTTTCCCTACACGACGCTCTTCCGATCTNNNCTCCACTGCAGAAAGGAGACT
AGAGCACACGTCTGAACTCTTTCCCTACACGACGCTCTTCCGATCTNNNCTAAAGCCTTCATCCCACTGG
AAGAGCACACGTCTGAACTCTTTCCCTACACGACGCTCTTCCGATCTNNNACTGGACTTGAGGAAACA
AGCACACGTCTGAACTCTTTCCCTACACGACGCTCTTCCGATCTNNNCAGCTCTCCTCAGCAAATTCAGT
AGAGCACACGTCTGAACTCTTTCCCTACACGACGCTCTTCCGATCTNNNCCAGAGAAACCAAACCTGTG
AGCACACGTCTGAACTCTTTCCCTACACGACGCTCTTCCGATCTNNNTCTTGGTAAATGTGAGTAGAAG
GCACACGTCTGAACTCTTTCCCTACACGACGCTCTTCCGATCTNNNCACATGACCATGCCAGCTAAGAT

AGAGCACACGTCTGAACTCTTTCCCTACACGACGCTCTTCCGATCTNNNTAGCTTCAGGCCAAAAGAC
AGCACACGTCTGAACTCTTTCCCTACACGACGCTCTTCCGATCTNNNGGCAAGGAGAAAGGACTAG
AGCACACGTCTGAACTCTTTCCCTACACGACGCTCTTCCGATCTNNNCTCCATACAGCTTTCCATGGGT
AGCACACGTCTGAACTCTTTCCCTACACGACGCTCTTCCGATCTNNNACAAAACTGTGTGTGCCTTCT
CACACGTCTGAACTCTTTCCCTACACGACGCTCTTCCGATCTNNNTTAGCCTAGCATACCTGAAGGGG
AGCACACGTCTGAACTCTTTCCCTACACGACGCTCTTCCGATCTNNNCATTCATCATTGGTGGAACGC
AGCACACGTCTGAACTCTTTCCCTACACGACGCTCTTCCGATCTNNNTAACAGCAAGATATAGGCCCT
AGCACACGTCTGAACTCTTTCCCTACACGACGCTCTTCCGATCTNNNAGTGCTTGACATCCTAGTTGG
GAGCACACGTCTGAACTCTTTCCCTACACGACGCTCTTCCGATCTNNNACACATGGTCAAACCAGGA
GAGCACACGTCTGAACTCTTTCCCTACACGACGCTCTTCCGATCTNNNGAATCTCCACCAACCATTCTG
AGCACACGTCTGAACTCTTTCCCTACACGACGCTCTTCCGATCTNNNGGAAATTCCCTCTCCTGCCACC
AGCACACGTCTGAACTCTTTCCCTACACGACGCTCTTCCGATCTNNNACTTGTGGATAAAAGTTACTA
GAGCACACGTCTGAACTCTTTCCCTACACGACGCTCTTCCGATCTNNNACAGCATGTGGTCATGTTGG
AGCACACGTCTGAACTCTTTCCCTACACGACGCTCTTCCGATCTNNNGAGGATGTGAGGTTAGGAG
AGCACACGTCTGAACTCTTTCCCTACACGACGCTCTTCCGATCTNNNCATCTGACCTGTTGTGCTCACT
AGCACACGTCTGAACTCTTTCCCTACACGACGCTCTTCCGATCTNNNAAGGAAAAGCCAGTACAGCAT
AGCACACGTCTGAACTCTTTCCCTACACGACGCTCTTCCGATCTNNNATCACCAAATGGCACCATACGA
AGCACACGTCTGAACTCTTTCCCTACACGACGCTCTTCCGATCTNNNAACCTCACTTCCCAAAAATGCC
AGCACACGTCTGAACTCTTTCCCTACACGACGCTCTTCCGATCTNNNCTGAGTTTCTGGATTGCAGGG
AGCACACGTCTGAACTCTTTCCCTACACGACGCTCTTCCGATCTNNNCCTGTATTCAGGGAGCTTGCTT
GAGCACACGTCTGAACTCTTTCCCTACACGACGCTCTTCCGATCTNNNGCTATTCTTATGGGCTGTGCC
AGAGCACACGTCTGAACTCTTTCCCTACACGACGCTCTTCCGATCTNNNGGTTGGCTATCACAAAACTG
AGCACACGTCTGAACTCTTTCCCTACACGACGCTCTTCCGATCTNNNTGCATTAGGGAACAAATGGGA
GAGCACACGTCTGAACTCTTTCCCTACACGACGCTCTTCCGATCTNNNTGCAGTTTCAGTGGGACATA
GCACACGTCTGAACTCTTTCCCTACACGACGCTCTTCCGATCTNNNCAGCACACAAGAACCAGGTCATC
AGCACACGTCTGAACTCTTTCCCTACACGACGCTCTTCCGATCTNNNGAAAATGAAATTTGCCAGCGG
AGCACACGTCTGAACTCTTTCCCTACACGACGCTCTTCCGATCTNNNCTTCTACCCAATGTGCCAAGGT
GCACACGTCTGAACTCTTTCCCTACACGACGCTCTTCCGATCTNNNCTGCAGGTGTATGTTACTGGCAT
AGAGCACACGTCTGAACTCTTTCCCTACACGACGCTCTTCCGATCTNNNGCATGCTCCTACTGTGCTGT
GCACACGTCTGAACTCTTTCCCTACACGACGCTCTTCCGATCTNNNAAATCCTCTTGGTGCAAGCTGA
GAGCACACGTCTGAACTCTTTCCCTACACGACGCTCTTCCGATCTNNNTGAAATCATTGAACATGGTA
AGAGCACACGTCTGAACTCTTTCCCTACACGACGCTCTTCCGATCTNNNTGAGTAGTTTTGTGTGTCTG
AGCACACGTCTGAACTCTTTCCCTACACGACGCTCTTCCGATCTNNNACACACAGCTTCTAACAACCA
GCACACGTCTGAACTCTTTCCCTACACGACGCTCTTCCGATCTNNNGTTTTGGAGATTTCTTGGCTGG
AGCACACGTCTGAACTCTTTCCCTACACGACGCTCTTCCGATCTNNNGGGATTCCAACAGGAGCAAT
GAGCACACGTCTGAACTCTTTCCCTACACGACGCTCTTCCGATCTNNNTTACAGTTCCGGCCTCTCAGT
AGAGCACACGTCTGAACTCTTTCCCTACACGACGCTCTTCCGATCTNNNTGTGAGATTGTGTTGGTGT
AGCACACGTCTGAACTCTTTCCCTACACGACGCTCTTCCGATCTNNNCTCGGCTGTGTTTATGATGGC
AAGAGCACACGTCTGAACTCTTTCCCTACACGACGCTCTTCCGATCTNNNTCCTCCCTAGGGTCAGATA
AGCACACGTCTGAACTCTTTCCCTACACGACGCTCTTCCGATCTNNNCATCATGCTCGGCTAAGAGTGT
AGCACACGTCTGAACTCTTTCCCTACACGACGCTCTTCCGATCTNNNCTTTCTTCCTGTCTGCCTCTGTC
GAGCACACGTCTGAACTCTTTCCCTACACGACGCTCTTCCGATCTNNNTATTCCCGATGACCCTGTCTT
AGAGCACACGTCTGAACTCTTTCCCTACACGACGCTCTTCCGATCTNNNATCCAGTAAGGGCTCTGGA

AGCACACGTCTGAACTCTTTCCCTACACGACGCTCTTCCGATCTNNNCTTGGCTCATCTTCAAACCGTCT
GAGCACACGTCTGAACTCTTTCCCTACACGACGCTCTTCCGATCTNNNAGTCTTTCCCTGTCTCTGTCTCT
AGCACACGTCTGAACTCTTTCCCTACACGACGCTCTTCCGATCTNNNCTTGTGGTAAGGAGTGGTTTG
AGCACACGTCTGAACTCTTTCCCTACACGACGCTCTTCCGATCTNNNCCTGAGTTTGAGTTCCATCCTG
AGCACACGTCTGAACTCTTTCCCTACACGACGCTCTTCCGATCTNNNTTGGAAGGAGGACCCTGAGAT
AGCACACGTCTGAACTCTTTCCCTACACGACGCTCTTCCGATCTNNNTCCTCCCTAGGGTCAGATAAAG
GAGCACACGTCTGAACTCTTTCCCTACACGACGCTCTTCCGATCTNNNCTCAGGAGGCTGAACAAGGA
AAGAGCACACGTCTGAACTCTTTCCCTACACGACGCTCTTCCGATCTNNNTTGATCTTCCCAGCTTCAGT
AGCACACGTCTGAACTCTTTCCCTACACGACGCTCTTCCGATCTNNNCTAGACCCCAATCAAAAGCCTC
GAGCACACGTCTGAACTCTTTCCCTACACGACGCTCTTCCGATCTNNNCACAGGCTGCTTCATCTCTTTCT
GAGCACACGTCTGAACTCTTTCCCTACACGACGCTCTTCCGATCTNNNTTGCTCCCGTGAACATCACATA
AGAGCACACGTCTGAACTCTTTCCCTACACGACGCTCTTCCGATCTNNNGGCTGAATTTGGGTCAGGT
AGCACACGTCTGAACTCTTTCCCTACACGACGCTCTTCCGATCTNNNAACTCTCCAGGTAAGTTGGCTG
AGAGCACACGTCTGAACTCTTTCCCTACACGACGCTCTTCCGATCTNNNCCAAATGTCCCTACCTCTAAC
AGCACACGTCTGAACTCTTTCCCTACACGACGCTCTTCCGATCTNNNGAAGGGTCACTTGAGCCTGAAA
GCACACGTCTGAACTCTTTCCCTACACGACGCTCTTCCGATCTNNNGAGATCAATCAAAGCGGTCCTCA
AGCACACGTCTGAACTCTTTCCCTACACGACGCTCTTCCGATCTNNNTTCATTTCCTAATGTCCTGGGG
AGCACACGTCTGAACTCTTTCCCTACACGACGCTCTTCCGATCTNNNAGGAGGCATATTGAAAGACTT
AGCACACGTCTGAACTCTTTCCCTACACGACGCTCTTCCGATCTNNNATGAAAAGCCAAAGAGGTGCT
GGAAGAGCACACGTCTGAACTCTTTCCCTACACGACGCTCTTCCGATCTNNNTGCTCCCTACTTCTACCT
GCACACGTCTGAACTCTTTCCCTACACGACGCTCTTCCGATCTNNNTATAAATCAGATCCCCCGACCCCCA
GCACACGTCTGAACTCTTTCCCTACACGACGCTCTTCCGATCTNNNGCCACCTCTTTGGCATTTGTATG
AAGAGCACACGTCTGAACTCTTTCCCTACACGACGCTCTTCCGATCTNNNCACTCTGCAGACGGTGTTA
GAGCACACGTCTGAACTCTTTCCCTACACGACGCTCTTCCGATCTNNNTTGGGCTACTGATTTGGGGA
GAGCACACGTCTGAACTCTTTCCCTACACGACGCTCTTCCGATCTNNNTTTTGCCCAGTGAACAGATGT
AGAGCACACGTCTGAACTCTTTCCCTACACGACGCTCTTCCGATCTNNNATCTCCAACCCCATTCAGATT
AGAGCACACGTCTGAACTCTTTCCCTACACGACGCTCTTCCGATCTNNNTTCTGATGTGAGGCTACCTTT
AAGAGCACACGTCTGAACTCTTTCCCTACACGACGCTCTTCCGATCTNNNTAAGACGCATGCATGTGAA
AAGAGCACACGTCTGAACTCTTTCCCTACACGACGCTCTTCCGATCTNNNGGAACAAGCCTAAATGCCC
GAGCACACGTCTGAACTCTTTCCCTACACGACGCTCTTCCGATCTNNNGGTATGTGTGTGAAGTGGGA
AGCACACGTCTGAACTCTTTCCCTACACGACGCTCTTCCGATCTNNNCTTCTACCCAATGTGCCAAGGTT
ACACGTCTGAACTCTTTCCCTACACGACGCTCTTCCGATCTNNNTCCTCACCTGTCACGAGGATAAACA
AGCACACGTCTGAACTCTTTCCCTACACGACGCTCTTCCGATCTNNNCTTTTCTTCAGTGTGGAGCCTG
GAGCACACGTCTGAACTCTTTCCCTACACGACGCTCTTCCGATCTNNNTGTCTGGAATCTTTCACTTGG
AGCACACGTCTGAACTCTTTCCCTACACGACGCTCTTCCGATCTNNNGTCTGAGGATGAGAAACCTTGG
AGCACACGTCTGAACTCTTTCCCTACACGACGCTCTTCCGATCTNNNAGTTTTCCTAAATGCCTGGAGC
AGCACACGTCTGAACTCTTTCCCTACACGACGCTCTTCCGATCTNNNCATTCTCTCACTTCAGCCTCCAA
GAGCACACGTCTGAACTCTTTCCCTACACGACGCTCTTCCGATCTNNNCATTGCTGCCATAGCAAGTGTT
AAGAGCACACGTCTGAACTCTTTCCCTACACGACGCTCTTCCGATCTNNNTGGTGTACGAAAAACTTGG
GAGCACACGTCTGAACTCTTTCCCTACACGACGCTCTTCCGATCTNNNATGTCAATGAGGGGGTTGAA
AGCACACGTCTGAACTCTTTCCCTACACGACGCTCTTCCGATCTNNNAGATAATAGTCTCTGCAGCTGT
GCACACGTCTGAACTCTTTCCCTACACGACGCTCTTCCGATCTNNNTTCTATCTATCCCTGGAGGCTGC
GAGCACACGTCTGAACTCTTTCCCTACACGACGCTCTTCCGATCTNNNCGAGGTGCTGCTGGATATGC

GCACACGTCTGAACTCTTTCCCTACACGACGCTCTTCCGATCTNNNTCAGAATCCTTTCAGGGGTGTGT
GAGCACACGTCTGAACTCTTTCCCTACACGACGCTCTTCCGATCTNNNCCTCACCTCCACTTCCCAAGTAC
AGCACACGTCTGAACTCTTTCCCTACACGACGCTCTTCCGATCTNNNTCGTTTTCTTTCTCCCACCACAAC
AGCACACGTCTGAACTCTTTCCCTACACGACGCTCTTCCGATCTNNNCAGGGGAAAGAGATGTTTGGC
AGAGCACACGTCTGAACTCTTTCCCTACACGACGCTCTTCCGATCTNNNCTGCCTATGTTTCCTTCATGG
AGAGCACACGTCTGAACTCTTTCCCTACACGACGCTCTTCCGATCTNNNGCTAGTCTCAGCTCAGCAGA
GAGCACACGTCTGAACTCTTTCCCTACACGACGCTCTTCCGATCTNNNGTTTCTTGTACTCTCCTGGAGG
CACACGTCTGAACTCTTTCCCTACACGACGCTCTTCCGATCTNNNATTCCTCAACCATTGACACACCCCAT
CACACGTCTGAACTCTTTCCCTACACGACGCTCTTCCGATCTNNNAGCTTTGAACTCCTAGGCTCTCTCAT
AAGAGCACACGTCTGAACTCTTTCCCTACACGACGCTCTTCCGATCTNNNATAGTGTTTCCTCTGATGC
AGAGCACACGTCTGAACTCTTTCCCTACACGACGCTCTTCCGATCTNNNCCCATCTCACAGCAGTTACTC
GAGCACACGTCTGAACTCTTTCCCTACACGACGCTCTTCCGATCTNNNTTGCAAGCTCCCTCTGAAAAGT
GAGCACACGTCTGAACTCTTTCCCTACACGACGCTCTTCCGATCTNNNGTGAGCTTCAGCAGGACCAAT
GAGCACACGTCTGAACTCTTTCCCTACACGACGCTCTTCCGATCTNNNCTGGGCTAGTGCTGATCATCT
AGAGCACACGTCTGAACTCTTTCCCTACACGACGCTCTTCCGATCTNNNCAAAACTCCCATGGCAGCATC
AGCACACGTCTGAACTCTTTCCCTACACGACGCTCTTCCGATCTNNNTAGTCAAGTGGTGTTGAGAGAC
GAGCACACGTCTGAACTCTTTCCCTACACGACGCTCTTCCGATCTNNNATGTAGAGGCTGCTTTATGTG
GAGCACACGTCTGAACTCTTTCCCTACACGACGCTCTTCCGATCTNNNTGAGTAAATGCATGCACAAGT
GAGCACACGTCTGAACTCTTTCCCTACACGACGCTCTTCCGATCTNNNTTGCCCTCCTCTCTACACACAAA
GAGCACACGTCTGAACTCTTTCCCTACACGACGCTCTTCCGATCTNNNCAAAGTGCCACCAAAAGCTCAA
GAGCACACGTCTGAACTCTTTCCCTACACGACGCTCTTCCGATCTNNNTTGATTACGCTCTCCTGCCTTTG
CACACGTCTGAACTCTTTCCCTACACGACGCTCTTCCGATCTNNNTAGGCCTCTCTTCTCTAGACCATCA
AGAGCACACGTCTGAACTCTTTCCCTACACGACGCTCTTCCGATCTNNNGGGTGAAAGTTCTGTGTACG
AGAGCACACGTCTGAACTCTTTCCCTACACGACGCTCTTCCGATCTNNNAAATAGCTCAGCCCCTTTTTG
AAGAGCACACGTCTGAACTCTTTCCCTACACGACGCTCTTCCGATCTNNNGTTTAGACCAATCTGAGTG
GAGCACACGTCTGAACTCTTTCCCTACACGACGCTCTTCCGATCTNNNACTCTAAACCATAGCAGACTGC
GAGCACACGTCTGAACTCTTTCCCTACACGACGCTCTTCCGATCTNNNTCTCTCCTCATTGTGTACAACC
AGAGCACACGTCTGAACTCTTTCCCTACACGACGCTCTTCCGATCTNNNCTCAGGAAGTCTCCCTCCTTCT
AGCACACGTCTGAACTCTTTCCCTACACGACGCTCTTCCGATCTNNNTGGCATGAAGATGGCACATTCTA
AGCACACGTCTGAACTCTTTCCCTACACGACGCTCTTCCGATCTNNNATGCACCCATAGTGTGCTAGAG
GAGCACACGTCTGAACTCTTTCCCTACACGACGCTCTTCCGATCTNNNTGCTCTGCATCTTTGTCACCAA
GCACACGTCTGAACTCTTTCCCTACACGACGCTCTTCCGATCTNNNCAACACAGGTTGCCTGTTCCCCAT
GAGCACACGTCTGAACTCTTTCCCTACACGACGCTCTTCCGATCTNNNTGTGACTAGTGCATGAGTTTCT
GAGCACACGTCTGAACTCTTTCCCTACACGACGCTCTTCCGATCTNNNAGACAGAAGAAAACTCTCCAA
AGCACACGTCTGAACTCTTTCCCTACACGACGCTCTTCCGATCTNNNCTGGTCACATATGCAGAAGCCA
GCACACGTCTGAACTCTTTCCCTACACGACGCTCTTCCGATCTNNNGATGGAGATGGGCAGAATCAAGC
AGCACACGTCTGAACTCTTTCCCTACACGACGCTCTTCCGATCTNNNGGGTGGGAGACAGTAAAAGGA
GAGCACACGTCTGAACTCTTTCCCTACACGACGCTCTTCCGATCTNNNTGTCCCATTCCTCTACCTACCAT
AGCACACGTCTGAACTCTTTCCCTACACGACGCTCTTCCGATCTNNNTGCCCTTTTGGCATAACCTCATT
AGCACACGTCTGAACTCTTTCCCTACACGACGCTCTTCCGATCTNNNTCAACCCTACCCAGAAATCTCCA
AGCACACGTCTGAACTCTTTCCCTACACGACGCTCTTCCGATCTNNNTCTAAAGGGTGTTGGAGCCCAT
AGAGCACACGTCTGAACTCTTTCCCTACACGACGCTCTTCCGATCTNNNTACAGTCAAGTGGAGGCATC
GAGCACACGTCTGAACTCTTTCCCTACACGACGCTCTTCCGATCTNNNAAGCAGCTCAGTTTCCCATTCT

AGAGCACACGTCTGAACTCTTTCCCTACACGACGCTCTTCCGATCTNNNGGCACACAATTGTCCCTAC
CACACGTCTGAACTCTTTCCCTACACGACGCTCTTCCGATCTNNNAGCTGTCTCGTACACACCCATCAT
GCACACGTCTGAACTCTTTCCCTACACGACGCTCTTCCGATCTNNNGCTACAAAGTGCTTTACTGACA
GCACACGTCTGAACTCTTTCCCTACACGACGCTCTTCCGATCTNNNTGTAGATCCCCCTACACTTCCTC
GCACACGTCTGAACTCTTTCCCTACACGACGCTCTTCCGATCTNNNGCTTGTGTTGAGAGAGAGTGA
GCACACGTCTGAACTCTTTCCCTACACGACGCTCTTCCGATCTNNNACATACATGCACATACACCCTG
AGCACACGTCTGAACTCTTTCCCTACACGACGCTCTTCCGATCTNNNTGTAGATCCCCCTACACTTCCT
AGCACACGTCTGAACTCTTTCCCTACACGACGCTCTTCCGATCTNNNCGCTCTGCCTCCTGCTCTATCAT
AGAGCACACGTCTGAACTCTTTCCCTACACGACGCTCTTCCGATCTNNNGAGGCCTGGTACTGTGACT
GCACACGTCTGAACTCTTTCCCTACACGACGCTCTTCCGATCTNNNAAGTTTGGCATTTCCTTGCCTCT
AGCACACGTCTGAACTCTTTCCCTACACGACGCTCTTCCGATCTNNNAAATAAAGGAAGGGTGGTGG
AGCACACGTCTGAACTCTTTCCCTACACGACGCTCTTCCGATCTNNNGGAGGACCCTTGTTGTTCAACT
AGCACACGTCTGAACTCTTTCCCTACACGACGCTCTTCCGATCTNNNAATAGCCTGCTACATGGCAAG
AGAGCACACGTCTGAACTCTTTCCCTACACGACGCTCTTCCGATCTNNNCATCTGCCACTGAACCATAC
GAGCACACGTCTGAACTCTTTCCCTACACGACGCTCTTCCGATCTNNNCTAACCTCCCCTCAGGCTATGT
AGCACACGTCTGAACTCTTTCCCTACACGACGCTCTTCCGATCTNNNCCACACATAACTCAGTCCGTGA
AAGAGCACACGTCTGAACTCTTTCCCTACACGACGCTCTTCCGATCTNNNTGACAGAGCGAAACCCTAT
GCACACGTCTGAACTCTTTCCCTACACGACGCTCTTCCGATCTNNNCGAGTACAAGTTCTTCTGGACGG
GCACACGTCTGAACTCTTTCCCTACACGACGCTCTTCCGATCTNNNGCTTACACAGAAACAGCTGCAA
AGCACACGTCTGAACTCTTTCCCTACACGACGCTCTTCCGATCTNNNATTCAATAGATATATATGCGT
GAGCACACGTCTGAACTCTTTCCCTACACGACGCTCTTCCGATCTNNNAGGGAGCAGCAGATGTGTAT
AGCACACGTCTGAACTCTTTCCCTACACGACGCTCTTCCGATCTNNNACTACACACCCCTTCTATCCCA
GAGCACACGTCTGAACTCTTTCCCTACACGACGCTCTTCCGATCTNNNCTTGGCCATGTGTAGCCTTT
AGCACACGTCTGAACTCTTTCCCTACACGACGCTCTTCCGATCTNNNGGGCTACCCCAAACAATATCA
AGAGCACACGTCTGAACTCTTTCCCTACACGACGCTCTTCCGATCTNNNACCAGCCCAATTCTACTCTCC
CACACGTCTGAACTCTTTCCCTACACGACGCTCTTCCGATCTNNNCAGCTGGTGCTTTTCTCGAACTTC
GAGCACACGTCTGAACTCTTTCCCTACACGACGCTCTTCCGATCTNNNAAAGAGAGAGAGTTCCTGAG
GCACACGTCTGAACTCTTTCCCTACACGACGCTCTTCCGATCTNNNTCATCACTCAACCTCACTGTTGCC
GAGCACACGTCTGAACTCTTTCCCTACACGACGCTCTTCCGATCTNNNCATAAGCACAGGGTTACCG
GAGCACACGTCTGAACTCTTTCCCTACACGACGCTCTTCCGATCTNNNGAACTCAAAGAAGTACTCCC
CACGTCTGAACTCTTTCCCTACACGACGCTCTTCCGATCTNNNGTTGTGTTATGTCTCCCTCCACCCATC
AGCACACGTCTGAACTCTTTCCCTACACGACGCTCTTCCGATCTNNNACAGACACCAAGCAATATGCTC
AGCACACGTCTGAACTCTTTCCCTACACGACGCTCTTCCGATCTNNNTCTCCATGAAGACAGGGAAG
GAGCACACGTCTGAACTCTTTCCCTACACGACGCTCTTCCGATCTNNNACAGTGCCCATCCTTAGTTCT
AGCACACGTCTGAACTCTTTCCCTACACGACGCTCTTCCGATCTNNNCCACCGTGACCAGCAGTTATT
GCACACGTCTGAACTCTTTCCCTACACGACGCTCTTCCGATCTNNNTGTGAGTTGTGCATGTAGTCT
GAGCACACGTCTGAACTCTTTCCCTACACGACGCTCTTCCGATCTNNNGCCTAATCCAATCCGACTGT
AGCACACGTCTGAACTCTTTCCCTACACGACGCTCTTCCGATCTNNNAGCATCAAGACACAGAGATGC
GAGCACACGTCTGAACTCTTTCCCTACACGACGCTCTTCCGATCTNNNTGACAAAGGTAAGCTCAGG
GAGCACACGTCTGAACTCTTTCCCTACACGACGCTCTTCCGATCTNNNCCACCGTGACCAGCAGTTAT
AGCACACGTCTGAACTCTTTCCCTACACGACGCTCTTCCGATCTNNNGGTAGGAAACAGCAACAGCA
GCACACGTCTGAACTCTTTCCCTACACGACGCTCTTCCGATCTNNNCTACCTGAGTTGTCACCGGAAA
AGCACACGTCTGAACTCTTTCCCTACACGACGCTCTTCCGATCTNNNCTCTACAGTGTGATGCTGGCT

AGCACACGTCTGAACTCTTTCCCTACACGACGCTCTTCCGATCTNNNGATTGCAAGCATTTGCCAGAA
AGCACACGTCTGAACTCTTTCCCTACACGACGCTCTTCCGATCTNNNATAAGAATCCATGTGGCCCAA
GCACACGTCTGAACTCTTTCCCTACACGACGCTCTTCCGATCTNNNGCTATTGCTGTGATGATGGTTG
GCACACGTCTGAACTCTTTCCCTACACGACGCTCTTCCGATCTNNNATTTCCGCAAGTACAGACTGAG
GAGCACACGTCTGAACTCTTTCCCTACACGACGCTCTTCCGATCTNNNACAACTACACATGTAAAAGA
GAGCACACGTCTGAACTCTTTCCCTACACGACGCTCTTCCGATCTNNNTGGAGGAGATCCTAATGGG
AGCACACGTCTGAACTCTTTCCCTACACGACGCTCTTCCGATCTNNNCAAGCACATATCCCACAATGCA
AGCACACGTCTGAACTCTTTCCCTACACGACGCTCTTCCGATCTNNNTGTAAGAAGAGGAAAGGAG
AGCACACGTCTGAACTCTTTCCCTACACGACGCTCTTCCGATCTNNNTCCAGGCCAAAAAGAAGTTGA
AGCACACGTCTGAACTCTTTCCCTACACGACGCTCTTCCGATCTNNNAAAGGTAACCAAGTGCTGCTA
AGCACACGTCTGAACTCTTTCCCTACACGACGCTCTTCCGATCTNNNCAGCCAAGTTTGGAGATCCCAT
AGCACACGTCTGAACTCTTTCCCTACACGACGCTCTTCCGATCTNNNTTTGTCTATCAGTTCCAAGAG
GAGCACACGTCTGAACTCTTTCCCTACACGACGCTCTTCCGATCTNNNACTGAGCTCTAATCTGGGCT
AGCACACGTCTGAACTCTTTCCCTACACGACGCTCTTCCGATCTNNNCCCTTTTTCACTCCTGATAGCG
GCACACGTCTGAACTCTTTCCCTACACGACGCTCTTCCGATCTNNNGGACAGTGACTTGGGTCCATTT
AGAGCACACGTCTGAACTCTTTCCCTACACGACGCTCTTCCGATCTNNNAGGCATATTAGCAGATGAG
GAGCACACGTCTGAACTCTTTCCCTACACGACGCTCTTCCGATCTNNNCAATCCTGAGAAACTTGGAG
AGCACACGTCTGAACTCTTTCCCTACACGACGCTCTTCCGATCTNNNACAAAGTCTCTACCCTACTGGA
AGCACACGTCTGAACTCTTTCCCTACACGACGCTCTTCCGATCTNNNTGAAGGAAAAAGAAGTGGGC
AGCACACGTCTGAACTCTTTCCCTACACGACGCTCTTCCGATCTNNNGTCAATATCTTGTGTTGGCGCT
AGCACACGTCTGAACTCTTTCCCTACACGACGCTCTTCCGATCTNNNTTACGTGTTTTGGTGATGGTG
GAGCACACGTCTGAACTCTTTCCCTACACGACGCTCTTCCGATCTNNNAGTATGGCTTTTACAGACTG
GAGCACACGTCTGAACTCTTTCCCTACACGACGCTCTTCCGATCTNNNAACCATTCATGGCAACAGGT
AGCACACGTCTGAACTCTTTCCCTACACGACGCTCTTCCGATCTNNNGTCTTGGGCTAAACCGATGAA
GAGCACACGTCTGAACTCTTTCCCTACACGACGCTCTTCCGATCTNNNAGTGAACTGTCTTTCTGGTT
GAGCACACGTCTGAACTCTTTCCCTACACGACGCTCTTCCGATCTNNNGCACTGACAAACACACGCATA
GAGCACACGTCTGAACTCTTTCCCTACACGACGCTCTTCCGATCTNNNATAATATTCCAGGTGTGTGT
GAGCACACGTCTGAACTCTTTCCCTACACGACGCTCTTCCGATCTNNNTGATAGCCTGCCTCTCTTCTCT
GAGCACACGTCTGAACTCTTTCCCTACACGACGCTCTTCCGATCTNNNACTGAAAAACTACAAGCCCAC
GCACACGTCTGAACTCTTTCCCTACACGACGCTCTTCCGATCTNNNACCGCCTTTGTCTTTGTTTTTCCC
GCACACGTCTGAACTCTTTCCCTACACGACGCTCTTCCGATCTNNNTCCTGTCTTATCTCCTTGGCTCCCA
AGAGCACACGTCTGAACTCTTTCCCTACACGACGCTCTTCCGATCTNNNGTTATCAGCCAGCCTCATT
AGCACACGTCTGAACTCTTTCCCTACACGACGCTCTTCCGATCTNNNAAATGCAGTCTGTGCAGAGTT
GAGCACACGTCTGAACTCTTTCCCTACACGACGCTCTTCCGATCTNNNACACACACACTTCTTACCAAC
AGCACACGTCTGAACTCTTTCCCTACACGACGCTCTTCCGATCTNNNTTGCTGACTTTGCCAACAATTC
AGCACACGTCTGAACTCTTTCCCTACACGACGCTCTTCCGATCTNNNAGGTGGCTGGAATTCTCTCTCT
AGCACACGTCTGAACTCTTTCCCTACACGACGCTCTTCCGATCTNNNTCGCTATCAAAATCCCAGGAG
GAGCACACGTCTGAACTCTTTCCCTACACGACGCTCTTCCGATCTNNNTGCAAGATGAAAGTGTGTG
AGCACACGTCTGAACTCTTTCCCTACACGACGCTCTTCCGATCTNNNTCAATGAACCTGAGGTAGGGT
AGCACACGTCTGAACTCTTTCCCTACACGACGCTCTTCCGATCTNNNGCATATGTCCTCTGGGGTGTAT
AGCACACGTCTGAACTCTTTCCCTACACGACGCTCTTCCGATCTNNNGTTGATGTCCTCCTGTGGCTA
AGCACACGTCTGAACTCTTTCCCTACACGACGCTCTTCCGATCTNNNGCACACAGGTCCACTTCTTAG
GAGCACACGTCTGAACTCTTTCCCTACACGACGCTCTTCCGATCTNNNGATCTTCCTTGCGTTTCCTTCC

AGCACACGTCTGAACTCTTTCCCTACACGACGCTCTTCCGATCTNNNCTGGCTGAATGTAATGTGTCC
AGCACACGTCTGAACTCTTTCCCTACACGACGCTCTTCCGATCTNNNAAAACACCCAAACCCCACAAAA
AGCACACGTCTGAACTCTTTCCCTACACGACGCTCTTCCGATCTNNNTCACGTTGCCACCTTACTTTCA
CACACGTCTGAACTCTTTCCCTACACGACGCTCTTCCGATCTNNNCACCTTGTCTGACATCTCCCTTGC
GCACACGTCTGAACTCTTTCCCTACACGACGCTCTTCCGATCTNNNCATAGCAGAGGGTAAACCACCA
AGCACACGTCTGAACTCTTTCCCTACACGACGCTCTTCCGATCTNNNTGGTTCTGTCTGTTACATCCCC
GCACACGTCTGAACTCTTTCCCTACACGACGCTCTTCCGATCTNNNCTCTACAGTGTGATGCTGGCTTT
AGCACACGTCTGAACTCTTTCCCTACACGACGCTCTTCCGATCTNNNATCTCCATGCCTATCCTCAGCTC
AGCACACGTCTGAACTCTTTCCCTACACGACGCTCTTCCGATCTNNNGGATATGCACTCAAAACACTG
GAGCACACGTCTGAACTCTTTCCCTACACGACGCTCTTCCGATCTNNNTAATTCTTGGCACCCAGAAC
AGCACACGTCTGAACTCTTTCCCTACACGACGCTCTTCCGATCTNNNTATTCCCCTGCACAGAGATGTC
AGCACACGTCTGAACTCTTTCCCTACACGACGCTCTTCCGATCTNNNGCTTTGCTCCCAATCTCTCTCTC
AGAGCACACGTCTGAACTCTTTCCCTACACGACGCTCTTCCGATCTNNNCAGATCTGGTGAAAGCTAG
GAGCACACGTCTGAACTCTTTCCCTACACGACGCTCTTCCGATCTNNNAGAAAGATGGCAAGACAA
GCACACGTCTGAACTCTTTCCCTACACGACGCTCTTCCGATCTNNNTTTATTTGAGGGAAAGCCGCCA
GCACACGTCTGAACTCTTTCCCTACACGACGCTCTTCCGATCTNNNTCATGCAGGGTAGGAAAGACA
AGCACACGTCTGAACTCTTTCCCTACACGACGCTCTTCCGATCTNNNCTTCAACTCCAGACCACCAACA
AGCACACGTCTGAACTCTTTCCCTACACGACGCTCTTCCGATCTNNNTGTTATCTTCTCCCCCTCATTGC
GCACACGTCTGAACTCTTTCCCTACACGACGCTCTTCCGATCTNNNTTGGGAGAAAGGGGTCAGTGT
AGCACACGTCTGAACTCTTTCCCTACACGACGCTCTTCCGATCTNNNGATGAACCACGGATCCTGCTA
GCACACGTCTGAACTCTTTCCCTACACGACGCTCTTCCGATCTNNNAGCATTGAAAAGCTGTGTGGGT
AGAGCACACGTCTGAACTCTTTCCCTACACGACGCTCTTCCGATCTNNNTTTGATCAAGTATTGAAAT
GCACACGTCTGAACTCTTTCCCTACACGACGCTCTTCCGATCTNNNGGCATCTTGGTCTCCCATAGGTA
AGCACACGTCTGAACTCTTTCCCTACACGACGCTCTTCCGATCTNNNCCCTCTCCCCTACTCTTCCTTTCC
GAAGAGCACACGTCTGAACTCTTTCCCTACACGACGCTCTTCCGATCTNNNAGCTCCAGAAACTGATG
GAGCACACGTCTGAACTCTTTCCCTACACGACGCTCTTCCGATCTNNNTTGCAAGACTGAGTTGTAGC
CACACGTCTGAACTCTTTCCCTACACGACGCTCTTCCGATCTNNNGAGATTAGGGCTCCCACCTATGA
GCACACGTCTGAACTCTTTCCCTACACGACGCTCTTCCGATCTNNNTGAGAGAGGCGAAGAATGAAG
GCACACGTCTGAACTCTTTCCCTACACGACGCTCTTCCGATCTNNNTCAGCAAGAGGGGAAAAGAGA
GCACACGTCTGAACTCTTTCCCTACACGACGCTCTTCCGATCTNNNAACTCTCCTGTCTTCCTTTCCCAC
AGCACACGTCTGAACTCTTTCCCTACACGACGCTCTTCCGATCTNNNCTGTACTGGCTGACCCCTTAGT
AGCACACGTCTGAACTCTTTCCCTACACGACGCTCTTCCGATCTNNNGTCCTTGTTTCCAGCTTTTCTGG
AGCACACGTCTGAACTCTTTCCCTACACGACGCTCTTCCGATCTNNNCACATTGTGTGGATCAAAGCC
AGCACACGTCTGAACTCTTTCCCTACACGACGCTCTTCCGATCTNNNATATGGCATTGTTGGACCCTG
AGCACACGTCTGAACTCTTTCCCTACACGACGCTCTTCCGATCTNNNACTGGACATGGAAGAGTTGTC
AGCACACGTCTGAACTCTTTCCCTACACGACGCTCTTCCGATCTNNNTTCTGAGGGGTCTACCTCCTA
GCACACGTCTGAACTCTTTCCCTACACGACGCTCTTCCGATCTNNNTAAAGTCACCCCAGTCCTCAGTT
CACACGTCTGAACTCTTTCCCTACACGACGCTCTTCCGATCTNNNCTCTTTGAATATTGGCAGGCTGG
GAGCACACGTCTGAACTCTTTCCCTACACGACGCTCTTCCGATCTNNNTCTTGTTTGCTTCCCGGCTTT
GCACACGTCTGAACTCTTTCCCTACACGACGCTCTTCCGATCTNNNACGTTGTTATAGGAGTATGGA
AGCACACGTCTGAACTCTTTCCCTACACGACGCTCTTCCGATCTNNNGCACTAATGTTTAGTTCTGTG
GAGCACACGTCTGAACTCTTTCCCTACACGACGCTCTTCCGATCTNNNGTCCCCATGATAAGATTAGC
AGCACACGTCTGAACTCTTTCCCTACACGACGCTCTTCCGATCTNNNTGTACCTGTACCCTCCTGTTTT

AGCACACGTCTGAACTCTTTCCCTACACGACGCTCTTCCGATCTNNNCCCACAGGACAAACACAAAAG
AGCACACGTCTGAACTCTTTCCCTACACGACGCTCTTCCGATCTNNNTGAATGATCTGCATTTGGGTG
GAGCACACGTCTGAACTCTTTCCCTACACGACGCTCTTCCGATCTNNNTTCTGCCAATTTGGAGTGTG
AGCACACGTCTGAACTCTTTCCCTACACGACGCTCTTCCGATCTNNNCATGGTTCATCAGGGAAGCTGT
GAGCACACGTCTGAACTCTTTCCCTACACGACGCTCTTCCGATCTNNNTTGATATTTTGCACTGCGGTG
AGCACACGTCTGAACTCTTTCCCTACACGACGCTCTTCCGATCTNNNCAGGCATGGTTTTTGAATGGT
GAGCACACGTCTGAACTCTTTCCCTACACGACGCTCTTCCGATCTNNNTATACCAGCTCACAGAGGTCC
AGCACACGTCTGAACTCTTTCCCTACACGACGCTCTTCCGATCTNNNTTTGACAGAACAGGTCAGCAC
GAGCACACGTCTGAACTCTTTCCCTACACGACGCTCTTCCGATCTNNNAGCAAGTGTCATAATCACGG
AGCACACGTCTGAACTCTTTCCCTACACGACGCTCTTCCGATCTNNNATATATGTGCGTTCATGGATG
GAGCACACGTCTGAACTCTTTCCCTACACGACGCTCTTCCGATCTNNNAATCCAATCACCTGGGGCAA
AGCACACGTCTGAACTCTTTCCCTACACGACGCTCTTCCGATCTNNNAAAGAGCAGATGGAGCAGAA
GCACACGTCTGAACTCTTTCCCTACACGACGCTCTTCCGATCTNNNTCCCCTACAGAATAGAAAGCCCC
AGCACACGTCTGAACTCTTTCCCTACACGACGCTCTTCCGATCTNNNGGTCAGCAGGTTTCTGAGAAC
GAGCACACGTCTGAACTCTTTCCCTACACGACGCTCTTCCGATCTNNNTGTTCAGTCTTCCATCTTGAA
AGCACACGTCTGAACTCTTTCCCTACACGACGCTCTTCCGATCTNNNTTTCATGGATGGAAAAAGCCA
AGCACACGTCTGAACTCTTTCCCTACACGACGCTCTTCCGATCTNNNCAAGGTCTTTGCCAGAGTGAG
GAGCACACGTCTGAACTCTTTCCCTACACGACGCTCTTCCGATCTNNNCCCATCTGAGGGTATTTCTGT
AGCACACGTCTGAACTCTTTCCCTACACGACGCTCTTCCGATCTNNNGGAGAAGAGAGCTGTTGTGTG
GAGCACACGTCTGAACTCTTTCCCTACACGACGCTCTTCCGATCTNNNAGTGACTGTGCTAAGGAGG
AGCACACGTCTGAACTCTTTCCCTACACGACGCTCTTCCGATCTNNNGGAGAAGAGAGCTGTTGTGTG
GGAAGAGCACACGTCTGAACTCTTTCCCTACACGACGCTCTTCCGATCTNNNCCCCAGTCTTATTCAGG
AGCACACGTCTGAACTCTTTCCCTACACGACGCTCTTCCGATCTNNNAACCCTCTGGTGCTGTCAAAAA
AAGAGCACACGTCTGAACTCTTTCCCTACACGACGCTCTTCCGATCTNNNCTCTATGTGACCAGGCAG
AGCACACGTCTGAACTCTTTCCCTACACGACGCTCTTCCGATCTNNNTATACCTGTCAGGGGTGGTGT
AGCACACGTCTGAACTCTTTCCCTACACGACGCTCTTCCGATCTNNNCCAGGTATCATTTTCTGAGCCC
GCACACGTCTGAACTCTTTCCCTACACGACGCTCTTCCGATCTNNNGAAGCCACAGGTAGAAGGAAGT
GAGCACACGTCTGAACTCTTTCCCTACACGACGCTCTTCCGATCTNNNAGAAGTCTTGTGATGGTTGG
GAGCACACGTCTGAACTCTTTCCCTACACGACGCTCTTCCGATCTNNNTCCCTAGTTCCTGGGTTGTG
AAGAGCACACGTCTGAACTCTTTCCCTACACGACGCTCTTCCGATCTNNNTATATGGTGCCCAACGAA
GAGCACACGTCTGAACTCTTTCCCTACACGACGCTCTTCCGATCTNNNGTGGGAATTCTCAGCCTCTGT
GAGCACACGTCTGAACTCTTTCCCTACACGACGCTCTTCCGATCTNNNTAAAAGAGTGTGGCTCCTCCT
AGCACACGTCTGAACTCTTTCCCTACACGACGCTCTTCCGATCTNNNAGCCTTTGAGCTCCTAGAAAAA
AGCACACGTCTGAACTCTTTCCCTACACGACGCTCTTCCGATCTNNNTGAGCAACCACTCAAGAGCAAA
AGCACACGTCTGAACTCTTTCCCTACACGACGCTCTTCCGATCTNNNACCACAGCTGCATTATTTGCTT
GAGCACACGTCTGAACTCTTTCCCTACACGACGCTCTTCCGATCTNNNAGTTCAAAAGCCCTCAATCAC
AAGAGCACACGTCTGAACTCTTTCCCTACACGACGCTCTTCCGATCTNNNAGCAAGACCTCATCTCATC
GAGCACACGTCTGAACTCTTTCCCTACACGACGCTCTTCCGATCTNNNAACCTTCCCTAGCTGCTCACAT
GAGCACACGTCTGAACTCTTTCCCTACACGACGCTCTTCCGATCTNNNCGTGAACCAAAGATTGGAAA
AGCACACGTCTGAACTCTTTCCCTACACGACGCTCTTCCGATCTNNNCCTTGGACCAGTATTGAAATG
AGCACACGTCTGAACTCTTTCCCTACACGACGCTCTTCCGATCTNNNTCCTTTTAGCCAAATCCTCAAAG
GCACACGTCTGAACTCTTTCCCTACACGACGCTCTTCCGATCTNNNATCCATCACACCATTTCAGAGCCC
AGCACACGTCTGAACTCTTTCCCTACACGACGCTCTTCCGATCTNNNCCCAAGCTTGATAGGAGCTTTC

AGCACACGTCTGAACTCTTTCCCTACACGACGCTCTTCCGATCTNNNACTGACATGCCCTTGACAAGT
AGCACACGTCTGAACTCTTTCCCTACACGACGCTCTTCCGATCTNNNTTGTCTGACAGTTTCCTCAGC
AGCACACGTCTGAACTCTTTCCCTACACGACGCTCTTCCGATCTNNNGTCTACAGAAGCCCCCTGCTA
GAGCACACGTCTGAACTCTTTCCCTACACGACGCTCTTCCGATCTNNNCATTCCTCAGCCCTCTATCCT
GCACACGTCTGAACTCTTTCCCTACACGACGCTCTTCCGATCTNNNCACCCTAGTTGGCTGATGTTGAA
GCACACGTCTGAACTCTTTCCCTACACGACGCTCTTCCGATCTNNNGCTTTAGAGTTCACAGTGAGAC
GCACACGTCTGAACTCTTTCCCTACACGACGCTCTTCCGATCTNNNAACAAGCCTTTTAGAGCAGGT
GAGCACACGTCTGAACTCTTTCCCTACACGACGCTCTTCCGATCTNNNCACTCTCTCTAGCCTGAGTGT
AGAGCACACGTCTGAACTCTTTCCCTACACGACGCTCTTCCGATCTNNNAACTTGACTGGGCTAAGG
AGCACACGTCTGAACTCTTTCCCTACACGACGCTCTTCCGATCTNNNGACCCTACCTCTCAGTCTCTCA
GCACACGTCTGAACTCTTTCCCTACACGACGCTCTTCCGATCTNNNTCTCTGAATTTAGCTCAATGGT
GCACACGTCTGAACTCTTTCCCTACACGACGCTCTTCCGATCTNNNCTTCATCGGGCTTCAGTGGTTT
AGCACACGTCTGAACTCTTTCCCTACACGACGCTCTTCCGATCTNNNGGAAGTCTACCCTGCTCTCTCA
GAGCACACGTCTGAACTCTTTCCCTACACGACGCTCTTCCGATCTNNNTTTCCAGACATGAGGTTGCT
AGCACACGTCTGAACTCTTTCCCTACACGACGCTCTTCCGATCTNNNAAGTGCTATGCAAATCAACTG
AGAGCACACGTCTGAACTCTTTCCCTACACGACGCTCTTCCGATCTNNNAGTCCTTCCTCCTCCATTCTT
AGAGCACACGTCTGAACTCTTTCCCTACACGACGCTCTTCCGATCTNNNACTTGAGCCACTGTGTCTG
AGCACACGTCTGAACTCTTTCCCTACACGACGCTCTTCCGATCTNNNTCCCCGTTCTTATGAGCCTTAC
AGCACACGTCTGAACTCTTTCCCTACACGACGCTCTTCCGATCTNNNTTCAGTGTGGAGCTTCTGGAC
AGCACACGTCTGAACTCTTTCCCTACACGACGCTCTTCCGATCTNNNAAACAAAGGATGCTTCCCTCC
GCACACGTCTGAACTCTTTCCCTACACGACGCTCTTCCGATCTNNNCAAGCTCCCATTTTCCCTCTCTC
GCACACGTCTGAACTCTTTCCCTACACGACGCTCTTCCGATCTNNNATTCTATGTGCACAGTGTCAGG
AGCACACGTCTGAACTCTTTCCCTACACGACGCTCTTCCGATCTNNNTTATCCGGGCTGAGAGATCCT
AGCACACGTCTGAACTCTTTCCCTACACGACGCTCTTCCGATCTNNNTTCCCATCTCTGTGTTCCCATCT
AGCACACGTCTGAACTCTTTCCCTACACGACGCTCTTCCGATCTNNNAGCAATTCAACGCTTCTCCTCA
AGCACACGTCTGAACTCTTTCCCTACACGACGCTCTTCCGATCTNNNTGCCTTCAGTCTTAGGTAATC
GAGCACACGTCTGAACTCTTTCCCTACACGACGCTCTTCCGATCTNNNGCAGTTACTGGGCCTCATAC
AGCACACGTCTGAACTCTTTCCCTACACGACGCTCTTCCGATCTNNNTGTCTGTGTACAACACTGATAG
AGCACACGTCTGAACTCTTTCCCTACACGACGCTCTTCCGATCTNNNATCCATGAAACCTTCCTCAGTG
GCACACGTCTGAACTCTTTCCCTACACGACGCTCTTCCGATCTNNNGGGTCCCTCATATGTGCATAGT
AGCACACGTCTGAACTCTTTCCCTACACGACGCTCTTCCGATCTNNNTAAGCACAGAGCCAAACACCTT
AGCACACGTCTGAACTCTTTCCCTACACGACGCTCTTCCGATCTNNNATTCTGAGGCATGCTACCTTTG
GAGCACACGTCTGAACTCTTTCCCTACACGACGCTCTTCCGATCTNNNATCAGTCATTGCTTGGCTAGT
AGCACACGTCTGAACTCTTTCCCTACACGACGCTCTTCCGATCTNNNAAATCCTGTGATCCTGTGAAT
GAGCACACGTCTGAACTCTTTCCCTACACGACGCTCTTCCGATCTNNNACAGACAGTCGTTTCTCTCCT
AGCACACGTCTGAACTCTTTCCCTACACGACGCTCTTCCGATCTNNNTACCCTTGACTACCTGGGAAG
GAGCACACGTCTGAACTCTTTCCCTACACGACGCTCTTCCGATCTNNNATGGCCTCCTAAAAGCTTCC
AGCACACGTCTGAACTCTTTCCCTACACGACGCTCTTCCGATCTNNNACAGATGTTCAGGGAGGAAT
GCACACGTCTGAACTCTTTCCCTACACGACGCTCTTCCGATCTNNNAATGGGCAAAGTGGTGTTTTT
AGCACACGTCTGAACTCTTTCCCTACACGACGCTCTTCCGATCTNNNAAAGCTAACCGGAACATCACA
GAGCACACGTCTGAACTCTTTCCCTACACGACGCTCTTCCGATCTNNNTCACATAGCGGCAAGGTGTA
AGAGCACACGTCTGAACTCTTTCCCTACACGACGCTCTTCCGATCTNNNAGTGTCTCCACTGTTGATTG
AGCACACGTCTGAACTCTTTCCCTACACGACGCTCTTCCGATCTNNNTTCTATGCACACCTCACATCAC

AGCACACGTCTGAACTCTTTCCCTACACGACGCTCTTCCGATCTNNNTCTTCAGAAAACAGGAATACA
GCACACGTCTGAACTCTTTCCCTACACGACGCTCTTCCGATCTNNNACCCACATATATACAGCATTGAA
GAGCACACGTCTGAACTCTTTCCCTACACGACGCTCTTCCGATCTNNNCTGTGTTCCAGTGTTACAACC
GCACACGTCTGAACTCTTTCCCTACACGACGCTCTTCCGATCTNNNAGCCCTCTTCACACCCATCTAATC
AGCACACGTCTGAACTCTTTCCCTACACGACGCTCTTCCGATCTNNNTGCCCATGCAATTTGACAGAT
GAGCACACGTCTGAACTCTTTCCCTACACGACGCTCTTCCGATCTNNNTAGAATACATGCACCCACTGC
GAGCACACGTCTGAACTCTTTCCCTACACGACGCTCTTCCGATCTNNNGGATTGGATGGGATAAATG
AGCACACGTCTGAACTCTTTCCCTACACGACGCTCTTCCGATCTNNNGTTGATGTCCTCCTGTGGCTAT
AGCACACGTCTGAACTCTTTCCCTACACGACGCTCTTCCGATCTNNNTGTAACATTAGGCCTGTTCCAG
AAGAGCACACGTCTGAACTCTTTCCCTACACGACGCTCTTCCGATCTNNNTCGTATATGTGTGTTCAT
GCACACGTCTGAACTCTTTCCCTACACGACGCTCTTCCGATCTNNNTGAGTGGTCTTGATAACCTTGT
GCACACGTCTGAACTCTTTCCCTACACGACGCTCTTCCGATCTNNNACTAAGAGCAGTCAGTCCCACTT
GAGCACACGTCTGAACTCTTTCCCTACACGACGCTCTTCCGATCTNNNTTATCCGGGCTGAGAGATCCT
AGCACACGTCTGAACTCTTTCCCTACACGACGCTCTTCCGATCTNNNGGAGAAAGGGAGGGAGAAG
GAGCACACGTCTGAACTCTTTCCCTACACGACGCTCTTCCGATCTNNNAGTTACCTCTGAATGAAACA
AAGAGCACACGTCTGAACTCTTTCCCTACACGACGCTCTTCCGATCTNNNAGCAATTCAACGCTTCTCCT
AGCACACGTCTGAACTCTTTCCCTACACGACGCTCTTCCGATCTNNNGCCAATATTGCGCCACTAAAGT
GAGCACACGTCTGAACTCTTTCCCTACACGACGCTCTTCCGATCTNNNGGAAGCTCCAATGAGGACTG
GCACACGTCTGAACTCTTTCCCTACACGACGCTCTTCCGATCTNNNGCCTAGCACATTGGAGAATGAC
GAGCACACGTCTGAACTCTTTCCCTACACGACGCTCTTCCGATCTNNNCCAAGAGGCTCTAAGACGAA
AGCACACGTCTGAACTCTTTCCCTACACGACGCTCTTCCGATCTNNNGAAGAAAGGAGAAGGAGAGC
GAGCACACGTCTGAACTCTTTCCCTACACGACGCTCTTCCGATCTNNNTAGAAGCCTAGAGCTGTTGG
GCACACGTCTGAACTCTTTCCCTACACGACGCTCTTCCGATCTNNNTCCCCATTCAAAGAGTACTACCC
GAGCACACGTCTGAACTCTTTCCCTACACGACGCTCTTCCGATCTNNNGATCTGGCAGCAGGTTTTGTT
GCACACGTCTGAACTCTTTCCCTACACGACGCTCTTCCGATCTNNNTCTAAAGGCATCACTCTCCCCATC
AAGAGCACACGTCTGAACTCTTTCCCTACACGACGCTCTTCCGATCTNNNCAGTATGCTCTACAAGCA
GAGCACACGTCTGAACTCTTTCCCTACACGACGCTCTTCCGATCTNNNCACTTCCTCACCATCCTTTCCTG
GAGCACACGTCTGAACTCTTTCCCTACACGACGCTCTTCCGATCTNNNAAGCCTCCCGGAAATGATTCT
AGCACACGTCTGAACTCTTTCCCTACACGACGCTCTTCCGATCTNNNCACTGGCTGACACACCCTATATT
AGCACACGTCTGAACTCTTTCCCTACACGACGCTCTTCCGATCTNNNTCATGGCTCTGCAGATTTTACC
AGCACACGTCTGAACTCTTTCCCTACACGACGCTCTTCCGATCTNNNTGCCTCATTTTGTCAGTAGTTT
AAGAGCACACGTCTGAACTCTTTCCCTACACGACGCTCTTCCGATCTNNNCGCACATTTTATATGCGCA
GAGCACACGTCTGAACTCTTTCCCTACACGACGCTCTTCCGATCTNNNATTCCTTGGTAGAAGTGCAA
AGCACACGTCTGAACTCTTTCCCTACACGACGCTCTTCCGATCTNNNTTCTACAACAGGCACTGGGGA
AAGAGCACACGTCTGAACTCTTTCCCTACACGACGCTCTTCCGATCTNNNACTATGGTAGGGTGGTT
GAGCACACGTCTGAACTCTTTCCCTACACGACGCTCTTCCGATCTNNNTCGTTCCTTTTTGTTTGTGACT
GAGCACACGTCTGAACTCTTTCCCTACACGACGCTCTTCCGATCTNNNGCCATTGTGCAACATACACAA
GAGCACACGTCTGAACTCTTTCCCTACACGACGCTCTTCCGATCTNNNTAGAATCATCCATTCCCATCG
AGCACACGTCTGAACTCTTTCCCTACACGACGCTCTTCCGATCTNNNATATCTGAGGTCTCCCAATCCC
GAAGAGCACACGTCTGAACTCTTTCCCTACACGACGCTCTTCCGATCTNNNACCACGTATTTCTCACTCT
GAGCACACGTCTGAACTCTTTCCCTACACGACGCTCTTCCGATCTNNNTCCCTGCAGCTAGGGTTTATCT
GAGCACACGTCTGAACTCTTTCCCTACACGACGCTCTTCCGATCTNNNGGAGAGATCAGCCACCTTTTC
AGCACACGTCTGAACTCTTTCCCTACACGACGCTCTTCCGATCTNNNAGTTCAGTTGGCACTGTTGTT

GCACACGTCTGAACTCTTTCCCTACACGACGCTCTTCCGATCTNNNAGTTCAGTTGGCACTGTTGTTG
AGCACACGTCTGAACTCTTTCCCTACACGACGCTCTTCCGATCTNNNCATGGTACGCAACCAGTGTGT
AGCACACGTCTGAACTCTTTCCCTACACGACGCTCTTCCGATCTNNNAAGCTATGGAGGTTGGAGTG
GAGCACACGTCTGAACTCTTTCCCTACACGACGCTCTTCCGATCTNNNCCTTAGCCATCTTAGGGAAG
GAGCACACGTCTGAACTCTTTCCCTACACGACGCTCTTCCGATCTNNNGCCGGAGATCAGTAAGTTGG
GCACACGTCTGAACTCTTTCCCTACACGACGCTCTTCCGATCTNNNGTGGGAGTAAGAAGAGGGAG
AGCACACGTCTGAACTCTTTCCCTACACGACGCTCTTCCGATCTNNNACTGCTCTAATGGACAGAAAC
AGCACACGTCTGAACTCTTTCCCTACACGACGCTCTTCCGATCTNNNAGGCAAGTAGCCTGTTTTTGCC
AGCACACGTCTGAACTCTTTCCCTACACGACGCTCTTCCGATCTNNNATCACCAGCGAGGAGACAGAA
AGAGCACACGTCTGAACTCTTTCCCTACACGACGCTCTTCCGATCTNNNACAATGCACTAGGAGATCA
GCACACGTCTGAACTCTTTCCCTACACGACGCTCTTCCGATCTNNNAAATCCCCTCTTCTCACTCTCCCC
AGCACACGTCTGAACTCTTTCCCTACACGACGCTCTTCCGATCTNNNAACATAAAGGAAAACACAGGT
AGAGCACACGTCTGAACTCTTTCCCTACACGACGCTCTTCCGATCTNNNAGGGACTGCTTGGATACTG
GCACACGTCTGAACTCTTTCCCTACACGACGCTCTTCCGATCTNNNTGACAAGTCATCCAAATGCACCA
AGCACACGTCTGAACTCTTTCCCTACACGACGCTCTTCCGATCTNNNGAGTGTCACTTGAGCTGGGAA
GAGCACACGTCTGAACTCTTTCCCTACACGACGCTCTTCCGATCTNNNTCACAAGAATGATTCCAGCC
AGCACACGTCTGAACTCTTTCCCTACACGACGCTCTTCCGATCTNNNGTTCCACCAATCCTGAGGGAAA
GAAGAGCACACGTCTGAACTCTTTCCCTACACGACGCTCTTCCGATCTNNNATGGCCTGATTCAATTCT
AGCACACGTCTGAACTCTTTCCCTACACGACGCTCTTCCGATCTNNNAGATGAGGGTTTGGAATCTTG
AGCACACGTCTGAACTCTTTCCCTACACGACGCTCTTCCGATCTNNNGGGTTGCAGGTAAAAACACCT
AGCACACGTCTGAACTCTTTCCCTACACGACGCTCTTCCGATCTNNNATACCTGAAGTCCTGAAAGGT
AGAGCACACGTCTGAACTCTTTCCCTACACGACGCTCTTCCGATCTNNNTCCAGTTGGCAGCTTCTGTG
GCACACGTCTGAACTCTTTCCCTACACGACGCTCTTCCGATCTNNNCCTACATAATCCATCACTGGGCG
AGCACACGTCTGAACTCTTTCCCTACACGACGCTCTTCCGATCTNNNTGACTTCTGTGTGAGTGTGTT
GCACACGTCTGAACTCTTTCCCTACACGACGCTCTTCCGATCTNNNCCTCAGTACCTTTACCCATTGCCC
AGAGCACACGTCTGAACTCTTTCCCTACACGACGCTCTTCCGATCTNNNACCTATTAGGAGTGCACAG
AGCACACGTCTGAACTCTTTCCCTACACGACGCTCTTCCGATCTNNNGGTATTCACCCTGGAGAAGCAA
AGCACACGTCTGAACTCTTTCCCTACACGACGCTCTTCCGATCTNNNTCCTAGAATCAGTCCTCCCAAC
GAGCACACGTCTGAACTCTTTCCCTACACGACGCTCTTCCGATCTNNNTAGCTAACGAGGATGCAGTG
GAGCACACGTCTGAACTCTTTCCCTACACGACGCTCTTCCGATCTNNNATGTTGCTGGCATGTAAACT
AGCACACGTCTGAACTCTTTCCCTACACGACGCTCTTCCGATCTNNNCACCTTTTCCTCTGTTCCAGTCA
GAGCACACGTCTGAACTCTTTCCCTACACGACGCTCTTCCGATCTNNNGAATTCCCCAAAGCAGCCTTA
AGCACACGTCTGAACTCTTTCCCTACACGACGCTCTTCCGATCTNNNCTTCAGTTTCAGGCACAGTGC
AGCACACGTCTGAACTCTTTCCCTACACGACGCTCTTCCGATCTNNNTGCCCTATCCTGGTCTCTACCTA
GAGCACACGTCTGAACTCTTTCCCTACACGACGCTCTTCCGATCTNNNCAGACAGCGTCTGGTCAGCT
AGCACACGTCTGAACTCTTTCCCTACACGACGCTCTTCCGATCTNNNTTCTCATATGCCTTTCCCCTCCA
AGAGCACACGTCTGAACTCTTTCCCTACACGACGCTCTTCCGATCTNNNGAACTAAGGCACCCCAAAG
AGCACACGTCTGAACTCTTTCCCTACACGACGCTCTTCCGATCTNNNACTTTTACTCAGTGCAGTTTGG
AGCACACGTCTGAACTCTTTCCCTACACGACGCTCTTCCGATCTNNNTAACTTGGTATGCGTTGCAAA
GCACACGTCTGAACTCTTTCCCTACACGACGCTCTTCCGATCTNNNCAACCTTCCTACCTCCTGCAAAAC
AGCACACGTCTGAACTCTTTCCCTACACGACGCTCTTCCGATCTNNNTACCATGGAGTTCTATCAGGG
AGCACACGTCTGAACTCTTTCCCTACACGACGCTCTTCCGATCTNNNCAGAGGAGGCAAACTGTGAT
AGAGCACACGTCTGAACTCTTTCCCTACACGACGCTCTTCCGATCTNNNTTTATGTTGGAGCGACAG

AGCACACGTCTGAACTCTTTCCCTACACGACGCTCTTCCGATCTNNNGCAAAGAAAAGAGAAGGGCAA
AGCACACGTCTGAACTCTTTCCCTACACGACGCTCTTCCGATCTNNNATATACTCTTCATGCCCCCACCA
GAGCACACGTCTGAACTCTTTCCCTACACGACGCTCTTCCGATCTNNNTTTCTCTCTCTCTCCCCCAGATG
GAGCACACGTCTGAACTCTTTCCCTACACGACGCTCTTCCGATCTNNNAGAGAGAAGAAGAGCTCAGT
GAGCACACGTCTGAACTCTTTCCCTACACGACGCTCTTCCGATCTNNNTGGAGAGAGAAGAAGAGCTC
AGCACACGTCTGAACTCTTTCCCTACACGACGCTCTTCCGATCTNNNTGGGTACAACAGGAGGAGAGA
GAGCACACGTCTGAACTCTTTCCCTACACGACGCTCTTCCGATCTNNNTCTGGGGATGGAGGTGGAA
AGCACACGTCTGAACTCTTTCCCTACACGACGCTCTTCCGATCTNNNTCTTTTCACCTGGCTCCTGAAAC
GAGCACACGTCTGAACTCTTTCCCTACACGACGCTCTTCCGATCTNNNGATCTGACAGTGGAAGGCAA
GAGCACACGTCTGAACTCTTTCCCTACACGACGCTCTTCCGATCTNNNGGGACTGATCATAAATGCAG
GCACACGTCTGAACTCTTTCCCTACACGACGCTCTTCCGATCTNNNCCTGGCTACCACCTCTCTTTCTACA
GAGCACACGTCTGAACTCTTTCCCTACACGACGCTCTTCCGATCTNNNGCCATACCAAAGAGCCATTCG
AGAGCACACGTCTGAACTCTTTCCCTACACGACGCTCTTCCGATCTNNNACACAGGTAGGTTAGAAGG
GAGCACACGTCTGAACTCTTTCCCTACACGACGCTCTTCCGATCTNNNGGCATCTCTGGAAGCCATATGA
GAGCACACGTCTGAACTCTTTCCCTACACGACGCTCTTCCGATCTNNNGAAGAGGCTAAATGAGGCAC
AGAGCACACGTCTGAACTCTTTCCCTACACGACGCTCTTCCGATCTNNNCAAAATGAATGTCTCTCGCTC
AAGAGCACACGTCTGAACTCTTTCCCTACACGACGCTCTTCCGATCTNNNAACAAGACAAGGATGTCA
GAGCACACGTCTGAACTCTTTCCCTACACGACGCTCTTCCGATCTNNNTCTCCAGGGTTTTCCAAGAAA
AGCACACGTCTGAACTCTTTCCCTACACGACGCTCTTCCGATCTNNNGTCTCCATGGTTTGCTCACAGAT
GAGCACACGTCTGAACTCTTTCCCTACACGACGCTCTTCCGATCTNNNACAATTAGGACCTTGCACCCAT
AAGAGCACACGTCTGAACTCTTTCCCTACACGACGCTCTTCCGATCTNNNTCTTTGTGGTAGTTGTTGG
AGAGCACACGTCTGAACTCTTTCCCTACACGACGCTCTTCCGATCTNNNAAGCTAACCCCCAGAACTTCC
AGCACACGTCTGAACTCTTTCCCTACACGACGCTCTTCCGATCTNNNCACACAACACTTACATGCCACAC
GAAGAGCACACGTCTGAACTCTTTCCCTACACGACGCTCTTCCGATCTNNNGCATGACTTACTGTGAGG
AGAGCACACGTCTGAACTCTTTCCCTACACGACGCTCTTCCGATCTNNNGCTTGGCTCAAACTCTCTCTC
AGAGCACACGTCTGAACTCTTTCCCTACACGACGCTCTTCCGATCTNNNTGCTCCATCCTACTTGCTACTG
AGCACACGTCTGAACTCTTTCCCTACACGACGCTCTTCCGATCTNNNTAGGAATGCTACCTCCCACTCAC
GAGCACACGTCTGAACTCTTTCCCTACACGACGCTCTTCCGATCTNNNTTGCAGATAGACCCCTTCTCACT
AGCACACGTCTGAACTCTTTCCCTACACGACGCTCTTCCGATCTNNNTCCTCTGAGGGCCTGAATACAA
AGAGCACACGTCTGAACTCTTTCCCTACACGACGCTCTTCCGATCTNNNCACTGGAGCTTCCCAAATCAT
AGAGCACACGTCTGAACTCTTTCCCTACACGACGCTCTTCCGATCTNNNCCTGACAGGTCTAAGCACAT
GAGCACACGTCTGAACTCTTTCCCTACACGACGCTCTTCCGATCTNNNAGTGAGAAGTGAAGTGTTAG
GAGCACACGTCTGAACTCTTTCCCTACACGACGCTCTTCCGATCTNNNAGTGAAGCTCACCTCACCCATA
AGCACACGTCTGAACTCTTTCCCTACACGACGCTCTTCCGATCTNNNCCCACTTTCAGAACCTGCTCTTT
AGCACACGTCTGAACTCTTTCCCTACACGACGCTCTTCCGATCTNNNTGGAATTCAACCAGCCCTAACTC
AGAGCACACGTCTGAACTCTTTCCCTACACGACGCTCTTCCGATCTNNNCTGCACAAGGGAGATGAAC
GAGCACACGTCTGAACTCTTTCCCTACACGACGCTCTTCCGATCTNNNCTTCCACTTCCAATAGCTGGCA
AGCACACGTCTGAACTCTTTCCCTACACGACGCTCTTCCGATCTNNNCCAAGTATCTGGCATGTACCAC
AGAGCACACGTCTGAACTCTTTCCCTACACGACGCTCTTCCGATCTNNNTTGGCTAGGAGACCAGGATT
AGCACACGTCTGAACTCTTTCCCTACACGACGCTCTTCCGATCTNNNGTTTGGATCTTCCTCCAACCTCC
GCACACGTCTGAACTCTTTCCCTACACGACGCTCTTCCGATCTNNNTTACTTACGATGCACCCCTCTCTCA
AGAGCACACGTCTGAACTCTTTCCCTACACGACGCTCTTCCGATCTNNNGCAGCCAGTCTGTGTATTAG
GAGCACACGTCTGAACTCTTTCCCTACACGACGCTCTTCCGATCTNNNTCAAGAGAAGACCCCTCTCACA

AGCACACGTCTGAACTCTTTCCCTACACGACGCTCTTCCGATCTNNNAATATGCCCTGTTGTCCCTCTGA
AGAGCACACGTCTGAACTCTTTCCCTACACGACGCTCTTCCGATCTNNNGACCTGAAGAACCAGGTAG
AGCACACGTCTGAACTCTTTCCCTACACGACGCTCTTCCGATCTNNNTGGCCTCAGTTACTGCAAACAA
GCACACGTCTGAACTCTTTCCCTACACGACGCTCTTCCGATCTNNNGGTGCTTGCTGAAGATAAGGGA
GAGCACACGTCTGAACTCTTTCCCTACACGACGCTCTTCCGATCTNNNCTCATGGAGCAGATTCCAGGA
AGAGCACACGTCTGAACTCTTTCCCTACACGACGCTCTTCCGATCTNNNTGAGGAGAAGAAAGCTAT
AGCACACGTCTGAACTCTTTCCCTACACGACGCTCTTCCGATCTNNNGCTGGTTGTTCAAAGCAAACTCT
AGCACACGTCTGAACTCTTTCCCTACACGACGCTCTTCCGATCTNNNACAAATGGTACACAGGCCAAAC
CACACGTCTGAACTCTTTCCCTACACGACGCTCTTCCGATCTNNNCAATGTGTGTAAGCGTGTGTTGTC
GAGCACACGTCTGAACTCTTTCCCTACACGACGCTCTTCCGATCTNNNTGTTTGACTATGTGCACTAGG
AGCACACGTCTGAACTCTTTCCCTACACGACGCTCTTCCGATCTNNNATAGGTGGGTTTTGAACATGGG
AGAGCACACGTCTGAACTCTTTCCCTACACGACGCTCTTCCGATCTNNNTTACCACCTTACATGCTGCCT
GAGCACACGTCTGAACTCTTTCCCTACACGACGCTCTTCCGATCTNNNACAAGAAGGCAGAAAGAGGA
AAGAGCACACGTCTGAACTCTTTCCCTACACGACGCTCTTCCGATCTNNNAGTAACTCATTGTGAAGGT
GAGCACACGTCTGAACTCTTTCCCTACACGACGCTCTTCCGATCTNNNTGTTATTTCCAGGCCAACCAC
CACACGTCTGAACTCTTTCCCTACACGACGCTCTTCCGATCTNNNTTCAGTGTCGAATAAGGGCACAG
CACACGTCTGAACTCTTTCCCTACACGACGCTCTTCCGATCTNNNTCCACACCCCCAGGATTCTTCATCG
AGCACACGTCTGAACTCTTTCCCTACACGACGCTCTTCCGATCTNNNACAAATAGGACCAAATGCAAC
AGCACACGTCTGAACTCTTTCCCTACACGACGCTCTTCCGATCTNNNGGGCATAGAAAGACCCCATCTC
GAGCACACGTCTGAACTCTTTCCCTACACGACGCTCTTCCGATCTNNNAGGAGATCAGAGCTGTCACTG
AGAGCACACGTCTGAACTCTTTCCCTACACGACGCTCTTCCGATCTNNNTCTTCCTGTTCCTTGTATTGT
AGCACACGTCTGAACTCTTTCCCTACACGACGCTCTTCCGATCTNNNGTGTATGCTAAGAGAGGGCATG
GAGCACACGTCTGAACTCTTTCCCTACACGACGCTCTTCCGATCTNNNCCAGATTCTGCAATAAAGCGG
AGCACACGTCTGAACTCTTTCCCTACACGACGCTCTTCCGATCTNNNACAAAACCTATGTGCTGTGCCTA
AGAGCACACGTCTGAACTCTTTCCCTACACGACGCTCTTCCGATCTNNNCATGTCTGTCCCTTTAGCAGC
GAAGAGCACACGTCTGAACTCTTTCCCTACACGACGCTCTTCCGATCTNNNTCCCTAAACACCTTCCCA
GCACACGTCTGAACTCTTTCCCTACACGACGCTCTTCCGATCTNNNCTGTGATGCAGTTCATCTCCTCCC
AGAGCACACGTCTGAACTCTTTCCCTACACGACGCTCTTCCGATCTNNNCATGCTCCTCAACTTGTGACA
AGAGCACACGTCTGAACTCTTTCCCTACACGACGCTCTTCCGATCTNNNTTCAATCTGAAGAGAAACCG
AGCACACGTCTGAACTCTTTCCCTACACGACGCTCTTCCGATCTNNNGCTTCAACCTCCTTCTAGCTGTCC
AGCACACGTCTGAACTCTTTCCCTACACGACGCTCTTCCGATCTNNNACACTGTGTGCTTTCTTGCTTTCC
AGCACACGTCTGAACTCTTTCCCTACACGACGCTCTTCCGATCTNNNAAGAGCTTCAGCAAAGGGTCA
AGCACACGTCTGAACTCTTTCCCTACACGACGCTCTTCCGATCTNNNGCAGATTTTAGGCAGCTCTCCTT
GAGCACACGTCTGAACTCTTTCCCTACACGACGCTCTTCCGATCTNNNTAAATGCTTTTGGAGCATCAG
AGCACACGTCTGAACTCTTTCCCTACACGACGCTCTTCCGATCTNNNACAAAGGATCATACTGCAGCCA
GAGCACACGTCTGAACTCTTTCCCTACACGACGCTCTTCCGATCTNNNAATTTGCAAGCCCTTTACTGT
GAGCACACGTCTGAACTCTTTCCCTACACGACGCTCTTCCGATCTNNNTATGATCCCAAAGCATTCCAG
GCACACGTCTGAACTCTTTCCCTACACGACGCTCTTCCGATCTNNNTAAGACAGGGAAAGGGGACTG
AGAGCACACGTCTGAACTCTTTCCCTACACGACGCTCTTCCGATCTNNNAAGTCTTCCTGGAGCAGAG
AGCACACGTCTGAACTCTTTCCCTACACGACGCTCTTCCGATCTNNNTTTCTCATCACCACTCACCACAC
AGCACACGTCTGAACTCTTTCCCTACACGACGCTCTTCCGATCTNNNTTCCCAATAATGCTGACACTGG
GAGCACACGTCTGAACTCTTTCCCTACACGACGCTCTTCCGATCTNNNCCTTTGGTAAACTGTAAAGGC
AGCACACGTCTGAACTCTTTCCCTACACGACGCTCTTCCGATCTNNNCCTTCCAGGACTGTGAGAAGA

GAGCACACGTCTGAACTCTTTCCCTACACGACGCTCTTCCGATCTNNNTTTGATATAAGTCCATGGT
AGCACACGTCTGAACTCTTTCCCTACACGACGCTCTTCCGATCTNNNATGCCAGATTCACAGTCCTCCT
GCACACGTCTGAACTCTTTCCCTACACGACGCTCTTCCGATCTNNNACCCAGTCTCATTTCTGTCTCCAC
GCACACGTCTGAACTCTTTCCCTACACGACGCTCTTCCGATCTNNNGGACCACAGCCCATAGCTAATTT
AGCACACGTCTGAACTCTTTCCCTACACGACGCTCTTCCGATCTNNNCAGTGGGGAGAGACAACAAT
AGAGCACACGTCTGAACTCTTTCCCTACACGACGCTCTTCCGATCTNNNCAGTGGGGAGAGACAACA
GCACACGTCTGAACTCTTTCCCTACACGACGCTCTTCCGATCTNNNAAAATAGACACACATATGCACG
AGCACACGTCTGAACTCTTTCCCTACACGACGCTCTTCCGATCTNNNTGGCTGGCTTGTGTTTATGTT
GCACACGTCTGAACTCTTTCCCTACACGACGCTCTTCCGATCTNNNTGCCAATAAAGACGCTTGTCAA
GCACACGTCTGAACTCTTTCCCTACACGACGCTCTTCCGATCTNNNGAGGAGAGCAGCAGTCTAGTTG
GCACACGTCTGAACTCTTTCCCTACACGACGCTCTTCCGATCTNNNCACTACAGTCATATCTGGCCCCT
GAGCACACGTCTGAACTCTTTCCCTACACGACGCTCTTCCGATCTNNNTCTACTGGGAACAGAATCCT
AGCACACGTCTGAACTCTTTCCCTACACGACGCTCTTCCGATCTNNNGGTCACTGGGTCTCTCTCTTTC
GCACACGTCTGAACTCTTTCCCTACACGACGCTCTTCCGATCTNNNTTTCAAGTTGGCTGTTCTCCACA
GAGCACACGTCTGAACTCTTTCCCTACACGACGCTCTTCCGATCTNNNGGAATTTTAGCCTGCTTCCC
AGAGCACACGTCTGAACTCTTTCCCTACACGACGCTCTTCCGATCTNNNTGAGTTTGGGGCATTTTTC
GAAGAGCACACGTCTGAACTCTTTCCCTACACGACGCTCTTCCGATCTNNNGCCTGTACCTGTCAACTT
GAGCACACGTCTGAACTCTTTCCCTACACGACGCTCTTCCGATCTNNNTAGTCAAGTGGTGTTGAGA
AGCACACGTCTGAACTCTTTCCCTACACGACGCTCTTCCGATCTNNNGGCCTCAGAATAGATCTCCCT
GCACACGTCTGAACTCTTTCCCTACACGACGCTCTTCCGATCTNNNATCTGCTCATGTGTTCCTTCTGC
GAGCACACGTCTGAACTCTTTCCCTACACGACGCTCTTCCGATCTNNNGCTAGCTGAACCAGGAAACA
AGCACACGTCTGAACTCTTTCCCTACACGACGCTCTTCCGATCTNNNCCTGGCCTGATTTGGATTTTC
AGCACACGTCTGAACTCTTTCCCTACACGACGCTCTTCCGATCTNNNGGAGTGCAGAATAAGGGTG
GCACACGTCTGAACTCTTTCCCTACACGACGCTCTTCCGATCTNNNTGTATAAACATATAGTGTCTGT
GCACACGTCTGAACTCTTTCCCTACACGACGCTCTTCCGATCTNNNTAACTCTTCCTTCCCTCCCTTCCC
ACACGTCTGAACTCTTTCCCTACACGACGCTCTTCCGATCTNNNAAACATACAGATTTTGGGGGCGAC
AGAGCACACGTCTGAACTCTTTCCCTACACGACGCTCTTCCGATCTNNNATACCAAGTGGACCGAAGG
GAGCACACGTCTGAACTCTTTCCCTACACGACGCTCTTCCGATCTNNNAAGGAAGAACTCCATCTTCC
AGAGCACACGTCTGAACTCTTTCCCTACACGACGCTCTTCCGATCTNNNAAATTTGGGGGTGTAGCT
GAGCACACGTCTGAACTCTTTCCCTACACGACGCTCTTCCGATCTNNNCCTGTGGTGATGTCAGGCTT
AAGAGCACACGTCTGAACTCTTTCCCTACACGACGCTCTTCCGATCTNNNTTGCTCTGCCCTGTAAGAA
GCACACGTCTGAACTCTTTCCCTACACGACGCTCTTCCGATCTNNNTTATGGAGCCGTTCAGACACAA
GAGCACACGTCTGAACTCTTTCCCTACACGACGCTCTTCCGATCTNNNTGAGACACAAACTTGGAAA
GAGCACACGTCTGAACTCTTTCCCTACACGACGCTCTTCCGATCTNNNATGTTCTGGATACGGGAAG
AGCACACGTCTGAACTCTTTCCCTACACGACGCTCTTCCGATCTNNNACACTTACATAAGTTGTGTGG
GAGCACACGTCTGAACTCTTTCCCTACACGACGCTCTTCCGATCTNNNTGGATGTGGTAGGATGAAG
AGCACACGTCTGAACTCTTTCCCTACACGACGCTCTTCCGATCTNNNACCCTGAGTGAATTGCCTAAA
AAGAGCACACGTCTGAACTCTTTCCCTACACGACGCTCTTCCGATCTNNNAACAAAAATCCCCAGGGA
GAGCACACGTCTGAACTCTTTCCCTACACGACGCTCTTCCGATCTNNNTGGATCTAAGCTACCTCCAGT
GAGCACACGTCTGAACTCTTTCCCTACACGACGCTCTTCCGATCTNNNAACCTCATTGATATGCATCA
AGCACACGTCTGAACTCTTTCCCTACACGACGCTCTTCCGATCTNNNAATCTCTTTTGTGCGCACTCTC
AGCACACGTCTGAACTCTTTCCCTACACGACGCTCTTCCGATCTNNNCCAATAGGATGTGTGTGAGT
AGAGCACACGTCTGAACTCTTTCCCTACACGACGCTCTTCCGATCTNNNCTGCACAAGGGAGATGAA

GAGCACACGTCTGAACTCTTTCCCTACACGACGCTCTTCCGATCTNNNGCAAGCACTCACTCTCCATTCT
AGAGCACACGTCTGAACTCTTTCCCTACACGACGCTCTTCCGATCTNNNTCAAAGCACTCTCCAAAGCCA
AGAGCACACGTCTGAACTCTTTCCCTACACGACGCTCTTCCGATCTNNNTAAAAAGAGCCTGTGGTTGC
AGCACACGTCTGAACTCTTTCCCTACACGACGCTCTTCCGATCTNNNCCATGGCCTAGTTGTTGATTCCA
AGAGCACACGTCTGAACTCTTTCCCTACACGACGCTCTTCCGATCTNNNGCTGCAGCTTTCTTATCTGGA
GAGCACACGTCTGAACTCTTTCCCTACACGACGCTCTTCCGATCTNNNGCAGTGATACTTTGGACGTCA
AGAGCACACGTCTGAACTCTTTCCCTACACGACGCTCTTCCGATCTNNNCTCACCTCGCTCATCTCCGCCA
GAAGAGCACACGTCTGAACTCTTTCCCTACACGACGCTCTTCCGATCTNNNAACCCAGATCCAAATGCCA
AGCACACGTCTGAACTCTTTCCCTACACGACGCTCTTCCGATCTNNNCCTTGGAGTTTGTAACTCACTTG
GAGCACACGTCTGAACTCTTTCCCTACACGACGCTCTTCCGATCTNNNAAGCAAGGTTGTGATTTGACT
AGCACACGTCTGAACTCTTTCCCTACACGACGCTCTTCCGATCTNNNTGAGGGTCAACTCTCTACAGAC
GCACACGTCTGAACTCTTTCCCTACACGACGCTCTTCCGATCTNNNCCTTCCTTACCATTGCTCCCAAGCA
GAGCACACGTCTGAACTCTTTCCCTACACGACGCTCTTCCGATCTNNNATAGGGTTGACAGGTCTTCCC
GAGCACACGTCTGAACTCTTTCCCTACACGACGCTCTTCCGATCTNNNTATGCCATCCACAGAAGTCTCT
GAGCACACGTCTGAACTCTTTCCCTACACGACGCTCTTCCGATCTNNNTGGATGTGGTAGGATGAAGG
GAGCACACGTCTGAACTCTTTCCCTACACGACGCTCTTCCGATCTNNNGGATATGCACATGCACACAAC
GCACACGTCTGAACTCTTTCCCTACACGACGCTCTTCCGATCTNNNGTGCCAGGGTATATGTGACTGTG
GAGCACACGTCTGAACTCTTTCCCTACACGACGCTCTTCCGATCTNNNATGTAACACCAACACACCAGCA
GCACACGTCTGAACTCTTTCCCTACACGACGCTCTTCCGATCTNNNAGTGCTGAATCCCCTGTAATCCTC
CACACGTCTGAACTCTTTCCCTACACGACGCTCTTCCGATCTNNNCTTGCCCAAGGTCATAAAAGGGTCA
AGCACACGTCTGAACTCTTTCCCTACACGACGCTCTTCCGATCTNNNAGAGTTGGGAGAAAGGGATG
AGCACACGTCTGAACTCTTTCCCTACACGACGCTCTTCCGATCTNNNAGAGAATGAGCACTGGCTCTGA
AGCACACGTCTGAACTCTTTCCCTACACGACGCTCTTCCGATCTNNNAGATAGATGATAAAGACCCATG
GAGCACACGTCTGAACTCTTTCCCTACACGACGCTCTTCCGATCTNNNCACAGGGAAAACCTAAACTGG
AAGAGCACACGTCTGAACTCTTTCCCTACACGACGCTCTTCCGATCTNNNTAGGAAGGGTTGAGTGAC
AGCACACGTCTGAACTCTTTCCCTACACGACGCTCTTCCGATCTNNNTGTTTTGACAAATGCTTGCTTTG
GAGCACACGTCTGAACTCTTTCCCTACACGACGCTCTTCCGATCTNNNTCTTCTTTCCTGACTTCCCAGG
AGCACACGTCTGAACTCTTTCCCTACACGACGCTCTTCCGATCTNNNAAAGAGCAGATGGAGCAGAATG
AGAGCACACGTCTGAACTCTTTCCCTACACGACGCTCTTCCGATCTNNNCAAATTTGAGGCTGCAGTGA
GCACACGTCTGAACTCTTTCCCTACACGACGCTCTTCCGATCTNNNCCTATCTTTATGCCCATGCCTCCCA
GAGCACACGTCTGAACTCTTTCCCTACACGACGCTCTTCCGATCTNNNGGATTCAGCATGCTCCTAGCA
AGCACACGTCTGAACTCTTTCCCTACACGACGCTCTTCCGATCTNNNTGGTAGGCTATTTGGCTCTACA
AGAGCACACGTCTGAACTCTTTCCCTACACGACGCTCTTCCGATCTNNNTCAGGTGCCCATCAGTGGAT
GAGCACACGTCTGAACTCTTTCCCTACACGACGCTCTTCCGATCTNNNGATGGCTTGGATGGAAGGAA
AGAGCACACGTCTGAACTCTTTCCCTACACGACGCTCTTCCGATCTNNNAAGTCCCCACCTTCATACAGC
GGAAGAGCACACGTCTGAACTCTTTCCCTACACGACGCTCTTCCGATCTNNNTACCGGCCATTTCAAGCA
AGAGCACACGTCTGAACTCTTTCCCTACACGACGCTCTTCCGATCTNNNTAGCTAACGAGGATGCAGT
AGCACACGTCTGAACTCTTTCCCTACACGACGCTCTTCCGATCTNNNATTGACAGGAGCCTGGGAAGT
AGAGCACACGTCTGAACTCTTTCCCTACACGACGCTCTTCCGATCTNNNCAGGTGCCAATTCCCCTTTAG
GAGCACACGTCTGAACTCTTTCCCTACACGACGCTCTTCCGATCTNNNTGAGATAGAGTGCCATGGAAG
GAGCACACGTCTGAACTCTTTCCCTACACGACGCTCTTCCGATCTNNNCCAAATAGTTCCTGCATGTGT
AGAGCACACGTCTGAACTCTTTCCCTACACGACGCTCTTCCGATCTNNNGAAAGGCTTATCAGTGAGGC
GCACACGTCTGAACTCTTTCCCTACACGACGCTCTTCCGATCTNNNATTAGGTAGTCCCAGCACTTTGG

GAGCACACGTCTGAACTCTTTCCCTACACGACGCTCTTCCGATCTNNNTGATCTCATGGGGTTACATT
GCACACGTCTGAACTCTTTCCCTACACGACGCTCTTCCGATCTNNNCCCCTGTTATAACCACCTACTCCA
AGAGCACACGTCTGAACTCTTTCCCTACACGACGCTCTTCCGATCTNNNGAATTGGGATTCTCTCACTT
GAGCACACGTCTGAACTCTTTCCCTACACGACGCTCTTCCGATCTNNNTACGGGCTAAGAAGAGATG
AGCACACGTCTGAACTCTTTCCCTACACGACGCTCTTCCGATCTNNNTTTTTGTGTATTCTTGCCCATG
CACACGTCTGAACTCTTTCCCTACACGACGCTCTTCCGATCTNNNATCCACCACACAGGTGTGAAAAA
CACACGTCTGAACTCTTTCCCTACACGACGCTCTTCCGATCTNNNCCTCGACAGTTCAGTCCCCATAGC
GAGCACACGTCTGAACTCTTTCCCTACACGACGCTCTTCCGATCTNNNTTTCTTAGGTGACCCTGGGG
GAGCACACGTCTGAACTCTTTCCCTACACGACGCTCTTCCGATCTNNNGACACAAACTCCTCCCACTCTC
AGCACACGTCTGAACTCTTTCCCTACACGACGCTCTTCCGATCTNNNGTTTTAGTGGCACCTCCTCCAT
GAGCACACGTCTGAACTCTTTCCCTACACGACGCTCTTCCGATCTNNNTGAAAAGAACGTGAGAGGT
AGCACACGTCTGAACTCTTTCCCTACACGACGCTCTTCCGATCTNNNGAATCCTTTTCACAACAGGCA
GAGCACACGTCTGAACTCTTTCCCTACACGACGCTCTTCCGATCTNNNCCATGCCATGTGAAGGACAT
GCACACGTCTGAACTCTTTCCCTACACGACGCTCTTCCGATCTNNNAGACTGACCCCTAAGTGCAAGA
GAGCACACGTCTGAACTCTTTCCCTACACGACGCTCTTCCGATCTNNNTTAGTGAGGCTTGAGACCCC
AGCACACGTCTGAACTCTTTCCCTACACGACGCTCTTCCGATCTNNNAAGTGGGAGGTAGAGGGAA
GCACACGTCTGAACTCTTTCCCTACACGACGCTCTTCCGATCTNNNTGTGTGACTGCAAGTAAGCATGT
AGCACACGTCTGAACTCTTTCCCTACACGACGCTCTTCCGATCTNNNTAATTCCACAACAAAGGCACCC
GCACACGTCTGAACTCTTTCCCTACACGACGCTCTTCCGATCTNNNCGTTAGCATGGTGGAGTATGT
GCACACGTCTGAACTCTTTCCCTACACGACGCTCTTCCGATCTNNNCTGAAAACTTGGTGGGAACTT
AGCACACGTCTGAACTCTTTCCCTACACGACGCTCTTCCGATCTNNNGTCAAACAGAGAGCAATTGA
AAGAGCACACGTCTGAACTCTTTCCCTACACGACGCTCTTCCGATCTNNNTTATGTGGGTCCAAGCA
AGCACACGTCTGAACTCTTTCCCTACACGACGCTCTTCCGATCTNNNTGTGTTATTGAGTTTGGATT
AGCACACGTCTGAACTCTTTCCCTACACGACGCTCTTCCGATCTNNNAGTTCTCTAGGATGGAGCTGT
GCACACGTCTGAACTCTTTCCCTACACGACGCTCTTCCGATCTNNNGTATACCAAGCCATCCACCTTCCC
AGCACACGTCTGAACTCTTTCCCTACACGACGCTCTTCCGATCTNNNAGGAAAGGAAATGCTGTCTC
GCACACGTCTGAACTCTTTCCCTACACGACGCTCTTCCGATCTNNNTGATCTCTGAGGGATGTGTGT
GCACACGTCTGAACTCTTTCCCTACACGACGCTCTTCCGATCTNNNCTCCAGCCATTGCTAACTGCTTA
AGCACACGTCTGAACTCTTTCCCTACACGACGCTCTTCCGATCTNNNAGTATGGCTTTTACAGACTGG
GAGCACACGTCTGAACTCTTTCCCTACACGACGCTCTTCCGATCTNNNCAGCTTGGGTAACTTGCTGA
AAGAGCACACGTCTGAACTCTTTCCCTACACGACGCTCTTCCGATCTNNNGCCCAGGGGTTCAAGTTA
GCACACGTCTGAACTCTTTCCCTACACGACGCTCTTCCGATCTNNNTCCAGCAGATCAGAACTCTTTGC
GAGCACACGTCTGAACTCTTTCCCTACACGACGCTCTTCCGATCTNNNACTTCCCTCTCTCTGTCTTTCT
AAGAGCACACGTCTGAACTCTTTCCCTACACGACGCTCTTCCGATCTNNNCCAGGAAGGAAGGCTTT
GAGCACACGTCTGAACTCTTTCCCTACACGACGCTCTTCCGATCTNNNTGGCTATAAGAAGTGTGTG
AGCACACGTCTGAACTCTTTCCCTACACGACGCTCTTCCGATCTNNNATGCTAATGGCACTCAAGTGT
GAGCACACGTCTGAACTCTTTCCCTACACGACGCTCTTCCGATCTNNNACACTTTCCTCCATCTCTCCTC
AGCACACGTCTGAACTCTTTCCCTACACGACGCTCTTCCGATCTNNNAAGCACAGGTTACACAGACAG
GAGCACACGTCTGAACTCTTTCCCTACACGACGCTCTTCCGATCTNNNCTGTGCCACTATGGCAACAA
AGCACACGTCTGAACTCTTTCCCTACACGACGCTCTTCCGATCTNNNTGCTGCCTACTCAGTCTCTTTC
GCACACGTCTGAACTCTTTCCCTACACGACGCTCTTCCGATCTNNNCTCTGCTCCAGTTCCTCAACATCC
AGCACACGTCTGAACTCTTTCCCTACACGACGCTCTTCCGATCTNNNGGCTATAGTGAGCCTCAAAGG
AGCACACGTCTGAACTCTTTCCCTACACGACGCTCTTCCGATCTNNNCCTCTAAATCTGAATGTAAGCC

AGCACACGTCTGAACTCTTTCCCTACACGACGCTCTTCCGATCTNNNTGCTCTGCTTTTCTGCTTTACCT
GAGCACACGTCTGAACTCTTTCCCTACACGACGCTCTTCCGATCTNNNCATCACCTGAGCAGTTTCTAC
AGCACACGTCTGAACTCTTTCCCTACACGACGCTCTTCCGATCTNNNGATCTCCTTCAGCTGTCAACTCC
GCACACGTCTGAACTCTTTCCCTACACGACGCTCTTCCGATCTNNNCTTTCCTATGGGGAGAGGGAGA
AGCACACGTCTGAACTCTTTCCCTACACGACGCTCTTCCGATCTNNNCAATGTCCCGGAAACAAAGTCC
GAGCACACGTCTGAACTCTTTCCCTACACGACGCTCTTCCGATCTNNNATGAAATCCAGGCCTGTGGA
GCACACGTCTGAACTCTTTCCCTACACGACGCTCTTCCGATCTNNNTCTCCCTTCCTTTTCTCTTCCCACAT
AGCACACGTCTGAACTCTTTCCCTACACGACGCTCTTCCGATCTNNNGTGCCCTAACCTTTTCTCTCCTG
AGAGCACACGTCTGAACTCTTTCCCTACACGACGCTCTTCCGATCTNNNAGAAGCCCTCCCAGAGATG
GAGCACACGTCTGAACTCTTTCCCTACACGACGCTCTTCCGATCTNNNGGATATTTTTGGCCCGTTGTT
AGAGCACACGTCTGAACTCTTTCCCTACACGACGCTCTTCCGATCTNNNGTGTGAATATTGGTCTGCG
GCACACGTCTGAACTCTTTCCCTACACGACGCTCTTCCGATCTNNNGGTAAGTAGGAGTTGCATGGG
AGCACACGTCTGAACTCTTTCCCTACACGACGCTCTTCCGATCTNNNGTCTAGCATTCTTTCACTGCTG
GAGCACACGTCTGAACTCTTTCCCTACACGACGCTCTTCCGATCTNNNAGCTGAACCAGGAAACAACTT
AGCACACGTCTGAACTCTTTCCCTACACGACGCTCTTCCGATCTNNNATCATTTGCCCATTTTGAATTGC
GAGCACACGTCTGAACTCTTTCCCTACACGACGCTCTTCCGATCTNNNCAGCAACAGCTTCCTTCCTGTA
AGCACACGTCTGAACTCTTTCCCTACACGACGCTCTTCCGATCTNNNGGAGAGCAAAACCCTGTCTAC
AGCACACGTCTGAACTCTTTCCCTACACGACGCTCTTCCGATCTNNNTTTCCTGTTGACTGAATGGTGG
AGAGCACACGTCTGAACTCTTTCCCTACACGACGCTCTTCCGATCTNNNCTCCCTGAAGTTGCTTGGGA
AGCACACGTCTGAACTCTTTCCCTACACGACGCTCTTCCGATCTNNNATTTACCACACTACAGGGGTG
GAGCACACGTCTGAACTCTTTCCCTACACGACGCTCTTCCGATCTNNNATTTTCTGTCTGCACGTGTGT
AAGAGCACACGTCTGAACTCTTTCCCTACACGACGCTCTTCCGATCTNNNGCTGCTGGTTTTGAAAGTG
AGCACACGTCTGAACTCTTTCCCTACACGACGCTCTTCCGATCTNNNAAACATCTGGAGGACCTCTCTG
AGCACACGTCTGAACTCTTTCCCTACACGACGCTCTTCCGATCTNNNGAGCTGTGTTAGTCTGGGTTC
GAGCACACGTCTGAACTCTTTCCCTACACGACGCTCTTCCGATCTNNNGCCTGCGATCTTCCAGATTGT
AGCACACGTCTGAACTCTTTCCCTACACGACGCTCTTCCGATCTNNNACTTCTGTGCCATCATGTGTAC
AGAGCACACGTCTGAACTCTTTCCCTACACGACGCTCTTCCGATCTNNNGCTGTCAACATGTATCCCTT
AGCACACGTCTGAACTCTTTCCCTACACGACGCTCTTCCGATCTNNNGTGTCTCTCTCCTGCTTCCAGTA
GCACACGTCTGAACTCTTTCCCTACACGACGCTCTTCCGATCTNNNCTACTTCCCTTCTTGCAGGCATT
GAGCACACGTCTGAACTCTTTCCCTACACGACGCTCTTCCGATCTNNNGCTTGGCTATTTCTAGGCAGA
AGAGCACACGTCTGAACTCTTTCCCTACACGACGCTCTTCCGATCTNNNGTTGGTTTCCGAAGCTCCAA
GCACACGTCTGAACTCTTTCCCTACACGACGCTCTTCCGATCTNNNTCCACTCATTCATGCTTCCTGTG
AGCACACGTCTGAACTCTTTCCCTACACGACGCTCTTCCGATCTNNNACATGGGAATTTTCTGGGGAC
AGAGCACACGTCTGAACTCTTTCCCTACACGACGCTCTTCCGATCTNNNCACATATGGGAGGAACTCT
GCACACGTCTGAACTCTTTCCCTACACGACGCTCTTCCGATCTNNNGGTGGCAATTGGTGGTCTTCTAT
AGCACACGTCTGAACTCTTTCCCTACACGACGCTCTTCCGATCTNNNGAGCCATTCACTTATTCCTGCCA
GAGCACACGTCTGAACTCTTTCCCTACACGACGCTCTTCCGATCTNNNTGTTTAGAGAACAGTGTCCC
AGCACACGTCTGAACTCTTTCCCTACACGACGCTCTTCCGATCTNNNAGGATTGAAGCCTCTCTCTCTGT
AGCACACGTCTGAACTCTTTCCCTACACGACGCTCTTCCGATCTNNNCGCAGCCTACACAATCAATAGC
AGCACACGTCTGAACTCTTTCCCTACACGACGCTCTTCCGATCTNNNATGGGGCTCTATGATGGTGTC
GCACACGTCTGAACTCTTTCCCTACACGACGCTCTTCCGATCTNNNACCTCCTGTATCAAACCCAGAGA
AGCACACGTCTGAACTCTTTCCCTACACGACGCTCTTCCGATCTNNNAACCTACTAGAAACCGTGGGT
GAGCACACGTCTGAACTCTTTCCCTACACGACGCTCTTCCGATCTNNNGAACCAAGAATGCATCATCG

GAGCACACGTCTGAACTCTTTCCCTACACGACGCTCTTCCGATCTNNNACTCAATGTTCAAGTTCTCCC
AGCACACGTCTGAACTCTTTCCCTACACGACGCTCTTCCGATCTNNNACACTCACACTCAAACACTTGC
AGCACACGTCTGAACTCTTTCCCTACACGACGCTCTTCCGATCTNNNTGGGGTGTGTGTGTGTAGAG
GAGCACACGTCTGAACTCTTTCCCTACACGACGCTCTTCCGATCTNNNACATGATGTGTTCCAGGCTC
GAGCACACGTCTGAACTCTTTCCCTACACGACGCTCTTCCGATCTNNNTGGAATGGAACAGAGCAAA
AGCACACGTCTGAACTCTTTCCCTACACGACGCTCTTCCGATCTNNNCTGCTCAAAATGACTGATGGG
AGCACACGTCTGAACTCTTTCCCTACACGACGCTCTTCCGATCTNNNTGCTGAGTGTATCCAAATCCTG
AGCACACGTCTGAACTCTTTCCCTACACGACGCTCTTCCGATCTNNNGCTGCTGGTTTTGAAAGTGG
GAGCACACGTCTGAACTCTTTCCCTACACGACGCTCTTCCGATCTNNNATCCCTGGCTCCTAGTTCAAT
AGCACACGTCTGAACTCTTTCCCTACACGACGCTCTTCCGATCTNNNGAGATGCTGGCCTGTGTAAGT
AGCACACGTCTGAACTCTTTCCCTACACGACGCTCTTCCGATCTNNNTGGAGCAGGAATTTGAACTG
GAGCACACGTCTGAACTCTTTCCCTACACGACGCTCTTCCGATCTNNNACACACATTGTCCTCTCTCTCT
AGCACACGTCTGAACTCTTTCCCTACACGACGCTCTTCCGATCTNNNCCCTCTGCACTTGGCTGAATTTT
GCACACGTCTGAACTCTTTCCCTACACGACGCTCTTCCGATCTNNNTCCTGGCAGTCCTAAAACAAACC
AGAGCACACGTCTGAACTCTTTCCCTACACGACGCTCTTCCGATCTNNNAACCTAAATGCCGGTCAACA
AGCACACGTCTGAACTCTTTCCCTACACGACGCTCTTCCGATCTNNNTGAGACTGACAGAGAAGTCCC
GAGCACACGTCTGAACTCTTTCCCTACACGACGCTCTTCCGATCTNNNCCTCATCTTCGTCACTTTCAG
AGCACACGTCTGAACTCTTTCCCTACACGACGCTCTTCCGATCTNNNGTAAGAGGTAGCTCTGCCCTA
AGAGCACACGTCTGAACTCTTTCCCTACACGACGCTCTTCCGATCTNNNAGGTAGGTGGACAATCCA
AGAGCACACGTCTGAACTCTTTCCCTACACGACGCTCTTCCGATCTNNNATACATGTTATGCAGCCCC
AGCACACGTCTGAACTCTTTCCCTACACGACGCTCTTCCGATCTNNNGTTTGGGGATCACTGAAGAGG
AGCACACGTCTGAACTCTTTCCCTACACGACGCTCTTCCGATCTNNNAACCACAGCTGCTGATTTCTCA
AGCACACGTCTGAACTCTTTCCCTACACGACGCTCTTCCGATCTNNNCACACACAGGCTCACATGCCTC
AGCACACGTCTGAACTCTTTCCCTACACGACGCTCTTCCGATCTNNNCCATTTGCACCACTTTCCAAGT
AGCACACGTCTGAACTCTTTCCCTACACGACGCTCTTCCGATCTNNNTTGGAAGTTTGGTGAGGTGT
AGCACACGTCTGAACTCTTTCCCTACACGACGCTCTTCCGATCTNNNACCAGGAGATCCAAAGACTGA
GAGCACACGTCTGAACTCTTTCCCTACACGACGCTCTTCCGATCTNNNGTCAGGCGGGATATTTGAGT
AGCACACGTCTGAACTCTTTCCCTACACGACGCTCTTCCGATCTNNNATCCACTAGCCTCAGGGAACAT
AGCACACGTCTGAACTCTTTCCCTACACGACGCTCTTCCGATCTNNNAACAAACGAATGTGACTTCTGC
GAAGAGCACACGTCTGAACTCTTTCCCTACACGACGCTCTTCCGATCTNNNTCCTTGATAGCCTTGCCA
AGCACACGTCTGAACTCTTTCCCTACACGACGCTCTTCCGATCTNNNTACCAGAAGTTTAGGCCTAGG
GAGCACACGTCTGAACTCTTTCCCTACACGACGCTCTTCCGATCTNNNTCACTCTGGTGTCTGCTATCT
GCACACGTCTGAACTCTTTCCCTACACGACGCTCTTCCGATCTNNNCGGGTTCTTGAGTTCTTGTCTCT
AGCACACGTCTGAACTCTTTCCCTACACGACGCTCTTCCGATCTNNNAAGAATTGGCTCCTTCACCTTC
AGCACACGTCTGAACTCTTTCCCTACACGACGCTCTTCCGATCTNNNTCATGGAAGACTTCTTCAAGC
GCACACGTCTGAACTCTTTCCCTACACGACGCTCTTCCGATCTNNNGCCTTTGTGGTCTGTTTGCTTTT
AGCACACGTCTGAACTCTTTCCCTACACGACGCTCTTCCGATCTNNNAGGAGGTTTACAAACTCAGAT
GAGCACACGTCTGAACTCTTTCCCTACACGACGCTCTTCCGATCTNNNTGTTTGAGTTGTAGCTTACTG
AGCACACGTCTGAACTCTTTCCCTACACGACGCTCTTCCGATCTNNNGGGATGGAGGTGAGTAGGTC
GCACACGTCTGAACTCTTTCCCTACACGACGCTCTTCCGATCTNNNACTGTGTTCTATGAGTTCTCTGC
AGCACACGTCTGAACTCTTTCCCTACACGACGCTCTTCCGATCTNNNGGCTAAAACCCAGCTGATTTC
AGCACACGTCTGAACTCTTTCCCTACACGACGCTCTTCCGATCTNNNCCCACTAGATGCTCTTGGAGAG
GCACACGTCTGAACTCTTTCCCTACACGACGCTCTTCCGATCTNNNAGAGAGAGAGATCAGAGTGTG

GGAAGAGCACACGTCTGAACTCTTTCCCTACACGACGCTCTTCCGATCTNNNATGCAACTGAGAATAT
AGCACACGTCTGAACTCTTTCCCTACACGACGCTCTTCCGATCTNNNTTCATGCTAGTCTGACCACAGG
AGAGCACACGTCTGAACTCTTTCCCTACACGACGCTCTTCCGATCTNNNAAAATCACTGCTCTAGCCA
GAGCACACGTCTGAACTCTTTCCCTACACGACGCTCTTCCGATCTNNNGCAACAATCCCATTCCCAATA
AGCACACGTCTGAACTCTTTCCCTACACGACGCTCTTCCGATCTNNNGGAAACCAAAGGGGGTCAAA
AGCACACGTCTGAACTCTTTCCCTACACGACGCTCTTCCGATCTNNNGGAGAGATCCTTGTGTGTCTT
AGCACACGTCTGAACTCTTTCCCTACACGACGCTCTTCCGATCTNNNAATCAGTCAGGGCTTTCCTTCA
AGCACACGTCTGAACTCTTTCCCTACACGACGCTCTTCCGATCTNNNTGAGAGACCTAGCTACAAACA
GCACACGTCTGAACTCTTTCCCTACACGACGCTCTTCCGATCTNNNGCAGTTACTGGGCCTCATACAA
AGAGCACACGTCTGAACTCTTTCCCTACACGACGCTCTTCCGATCTNNNCAGAGACATAGGAAGACC
GAGCACACGTCTGAACTCTTTCCCTACACGACGCTCTTCCGATCTNNNACATATGTGTCCTCTTTCCCC
AGCACACGTCTGAACTCTTTCCCTACACGACGCTCTTCCGATCTNNNCCTGGCCTTGTGTCCTTATCTT
AGCACACGTCTGAACTCTTTCCCTACACGACGCTCTTCCGATCTNNNTCACTTCCACAAATCAGGCACT
GAGCACACGTCTGAACTCTTTCCCTACACGACGCTCTTCCGATCTNNNACTCCTCTTCGGCATGAGACT
GCACACGTCTGAACTCTTTCCCTACACGACGCTCTTCCGATCTNNNCCAATGTCAGCCTTCTTCAGCGC
GAGCACACGTCTGAACTCTTTCCCTACACGACGCTCTTCCGATCTNNNTATAAATGGGAAGTGGCTG
GAGCACACGTCTGAACTCTTTCCCTACACGACGCTCTTCCGATCTNNNAGAGAAGAAAATGTGGAAA
GAGCACACGTCTGAACTCTTTCCCTACACGACGCTCTTCCGATCTNNNCAGGCTTCATCCTCAAACCTTG
GAAGAGCACACGTCTGAACTCTTTCCCTACACGACGCTCTTCCGATCTNNNGAAATCAGAACCCTAGTT
AGCACACGTCTGAACTCTTTCCCTACACGACGCTCTTCCGATCTNNNCCTCCCTCCCTCAATCTTACCTTC
GCACACGTCTGAACTCTTTCCCTACACGACGCTCTTCCGATCTNNNCTAGGCTTTTCCAATGGGACCT
GAGCACACGTCTGAACTCTTTCCCTACACGACGCTCTTCCGATCTNNNTTGATTCAAGGCAGGATTCA
GAGCACACGTCTGAACTCTTTCCCTACACGACGCTCTTCCGATCTNNNTAGAAATGAGCCCCCAAGTG
AGCACACGTCTGAACTCTTTCCCTACACGACGCTCTTCCGATCTNNNGGACTGGTGTTGACTTTGGAT
GAGCACACGTCTGAACTCTTTCCCTACACGACGCTCTTCCGATCTNNNTGTGTGTGTAGGCTGTGAAT
GCACACGTCTGAACTCTTTCCCTACACGACGCTCTTCCGATCTNNNTCTGAGTTTCCATCACTGCTTGG
AGCACACGTCTGAACTCTTTCCCTACACGACGCTCTTCCGATCTNNNCAGTGTCCGAAATGAGGAACT
GCACACGTCTGAACTCTTTCCCTACACGACGCTCTTCCGATCTNNNTGTGGTTTATTTAGAATGACCA
GCACACGTCTGAACTCTTTCCCTACACGACGCTCTTCCGATCTNNNCTAGGCTTTTCCAATGGGACCTC
AGCACACGTCTGAACTCTTTCCCTACACGACGCTCTTCCGATCTNNNAGAGCTCTATGTGGGTCCATG
AGCACACGTCTGAACTCTTTCCCTACACGACGCTCTTCCGATCTNNNCTATATGTGGTTTGTGTGTGT
AGCACACGTCTGAACTCTTTCCCTACACGACGCTCTTCCGATCTNNNTAGTTGTCCAAGGATGTTTTCC
GCACACGTCTGAACTCTTTCCCTACACGACGCTCTTCCGATCTNNNACTTTTTCTCCCTCTCTCCTGCTCA
AGCACACGTCTGAACTCTTTCCCTACACGACGCTCTTCCGATCTNNNATTTGCCCTCTCATGTTCAATG
GCACACGTCTGAACTCTTTCCCTACACGACGCTCTTCCGATCTNNNTTTTCTCCAGCCTTTGCCTTCTTC
AGCACACGTCTGAACTCTTTCCCTACACGACGCTCTTCCGATCTNNNATCAGAGGTGAGATTTGGGT
AGAGCACACGTCTGAACTCTTTCCCTACACGACGCTCTTCCGATCTNNNATCTCCTAGGTTCATGCCTC
AGCACACGTCTGAACTCTTTCCCTACACGACGCTCTTCCGATCTNNNTCCATTGTCTCCTTTCTTCCAGT
AGCACACGTCTGAACTCTTTCCCTACACGACGCTCTTCCGATCTNNNATCCATAATCAACAGAGCTGG
AGCACACGTCTGAACTCTTTCCCTACACGACGCTCTTCCGATCTNNNTCGTAATGGGGATACAGAAAG
GAGCACACGTCTGAACTCTTTCCCTACACGACGCTCTTCCGATCTNNNCAGTCAACATGACAGTCACCA
GAGCACACGTCTGAACTCTTTCCCTACACGACGCTCTTCCGATCTNNNAGCATGTCTTTCTTTCCTCAT
AGCACACGTCTGAACTCTTTCCCTACACGACGCTCTTCCGATCTNNNGCCTCCTTTACACAACCACCAAT

AGCACACGTCTGAACTCTTTCCCTACACGACGCTCTTCCGATCTNNNACCATCACACCCTTTAGGACA
GAAGAGCACACGTCTGAACTCTTTCCCTACACGACGCTCTTCCGATCTNNNCAACAAAAATGGACCCA
GAGCACACGTCTGAACTCTTTCCCTACACGACGCTCTTCCGATCTNNNTACACAAACATGCATGCCTTT
AGCACACGTCTGAACTCTTTCCCTACACGACGCTCTTCCGATCTNNNTGCATAGGGCCACCTTCTTACT
GCACACGTCTGAACTCTTTCCCTACACGACGCTCTTCCGATCTNNNACAGATCAATGGAAGAGTATG
AGCACACGTCTGAACTCTTTCCCTACACGACGCTCTTCCGATCTNNNGGATGCTGAAGTGTGAGTGG
CACACGTCTGAACTCTTTCCCTACACGACGCTCTTCCGATCTNNNAGAGGAAAACGTAAGCAGAACC
GCACACGTCTGAACTCTTTCCCTACACGACGCTCTTCCGATCTNNNGAGGAGAGCAGCAGTCTAGTT
GAGCACACGTCTGAACTCTTTCCCTACACGACGCTCTTCCGATCTNNNCATTGTGGCTTCCATCTTGGT
AGCACACGTCTGAACTCTTTCCCTACACGACGCTCTTCCGATCTNNNTCTTTCCATTGTGCCATCAACC
AGCACACGTCTGAACTCTTTCCCTACACGACGCTCTTCCGATCTNNNGTCTCAATGAAGACGATGGG
GAGCACACGTCTGAACTCTTTCCCTACACGACGCTCTTCCGATCTNNNAGACCTTCCAAAACAGGATT
AGCACACGTCTGAACTCTTTCCCTACACGACGCTCTTCCGATCTNNNCCCATTTCCAACTAACATACTG
AGAGCACACGTCTGAACTCTTTCCCTACACGACGCTCTTCCGATCTNNNTAGTAGGGATGCAGTAGC
AGCACACGTCTGAACTCTTTCCCTACACGACGCTCTTCCGATCTNNNTTAGAACGCCCTACTCTCTTCC
GAGCACACGTCTGAACTCTTTCCCTACACGACGCTCTTCCGATCTNNNACACAGTCCTTACTTCCTGA
AGCACACGTCTGAACTCTTTCCCTACACGACGCTCTTCCGATCTNNNTTAGGGTCAGACCAGTGCCTTT
AGCACACGTCTGAACTCTTTCCCTACACGACGCTCTTCCGATCTNNNGCATAGCAATGGGGGTGGA
GCACACGTCTGAACTCTTTCCCTACACGACGCTCTTCCGATCTNNNAGACTGAGGCCCTACTTTGCTT
AGCACACGTCTGAACTCTTTCCCTACACGACGCTCTTCCGATCTNNNACGCTCTGAGTTTATCATCGCT
GCACACGTCTGAACTCTTTCCCTACACGACGCTCTTCCGATCTNNNGCATGAGAGTTTGCTCTCTGTG
GCACACGTCTGAACTCTTTCCCTACACGACGCTCTTCCGATCTNNNGAAAGGCAGTTGGAACTTGTGT
CACACGTCTGAACTCTTTCCCTACACGACGCTCTTCCGATCTNNNTGGTGGGAGTAGTTATGTGGGA
GCACACGTCTGAACTCTTTCCCTACACGACGCTCTTCCGATCTNNNAGTCCGGTAACTCATCTGTTTGC
GCACACGTCTGAACTCTTTCCCTACACGACGCTCTTCCGATCTNNNAAAGAGGGTTAGGAGGTCTTG
GAGCACACGTCTGAACTCTTTCCCTACACGACGCTCTTCCGATCTNNNGCCATCTCCAAGGGCCTTTT
CACACGTCTGAACTCTTTCCCTACACGACGCTCTTCCGATCTNNNAAAGTCTTCCCCTGAATGCCTTGC
GCACACGTCTGAACTCTTTCCCTACACGACGCTCTTCCGATCTNNNGGTCTATGGAGAGAAGGAGC
AGCACACGTCTGAACTCTTTCCCTACACGACGCTCTTCCGATCTNNNTTGTCATAAATGCCTGGCAATG
GAGCACACGTCTGAACTCTTTCCCTACACGACGCTCTTCCGATCTNNNAATTCAGGCTCAGATGCTCT
CACACGTCTGAACTCTTTCCCTACACGACGCTCTTCCGATCTNNNTAGAAACCCTGCCTTCAGCTAGT
GCACACGTCTGAACTCTTTCCCTACACGACGCTCTTCCGATCTNNNGCCCAGGACAGTTGTGTATTA
GCACACGTCTGAACTCTTTCCCTACACGACGCTCTTCCGATCTNNNGACCAGAAGGGAGGAAAATGG
AGAGCACACGTCTGAACTCTTTCCCTACACGACGCTCTTCCGATCTNNNGGGTGTGTATGAGTGTGT
GAGCACACGTCTGAACTCTTTCCCTACACGACGCTCTTCCGATCTNNNGGAGTTGATCTGGTTTCAAT
GCACACGTCTGAACTCTTTCCCTACACGACGCTCTTCCGATCTNNNCAGAGGCAACTTTAGCCTCCAA
AAGAGCACACGTCTGAACTCTTTCCCTACACGACGCTCTTCCGATCTNNNTGTTCTGCTTCCACTTTCA
GAGCACACGTCTGAACTCTTTCCCTACACGACGCTCTTCCGATCTNNNGATCTGGCAGCAGGTTTTG
GAGCACACGTCTGAACTCTTTCCCTACACGACGCTCTTCCGATCTNNNAGTTCTCTAGGATGGAGCTG
AGCACACGTCTGAACTCTTTCCCTACACGACGCTCTTCCGATCTNNNTTGGGATGAAGCTCTGTGTT
GCACACGTCTGAACTCTTTCCCTACACGACGCTCTTCCGATCTNNNTCTCTACTTCTGCTTCTTGGGCT
GCACACGTCTGAACTCTTTCCCTACACGACGCTCTTCCGATCTNNNTGCAAAGTCATTCATATACTGCC
AGCACACGTCTGAACTCTTTCCCTACACGACGCTCTTCCGATCTNNNAAGGCATTTTGGCAATTCACT

AGCACACGTCTGAACTCTTTCCCTACACGACGCTCTTCCGATCTNNNTCCAACCATTCCTTCTTCTCTGCC

GCACACGTCTGAACTCTTTCCCTACACGACGCTCTTCCGATCTNNNGCTAAGGGACAGACTTGTTACG

GGAAGAGCACACGTCTGAACTCTTTCCCTACACGACGCTCTTCCGATCTNNNAACAAGACAAGGATGT

CACACGTCTGAACTCTTTCCCTACACGACGCTCTTCCGATCTNNNTATCTCCTGCATCAGTTCCTCCCCA

AGCACACGTCTGAACTCTTTCCCTACACGACGCTCTTCCGATCTNNNCCATCAGGCCTACGAAGAGTA

AGCACACGTCTGAACTCTTTCCCTACACGACGCTCTTCCGATCTNNNTGCTTCCAGACCTTTCAGTTGA

AGCACACGTCTGAACTCTTTCCCTACACGACGCTCTTCCGATCTNNNAAACCCAAGGTGAAACTGTAG

AGCACACGTCTGAACTCTTTCCCTACACGACGCTCTTCCGATCTNNNCCCAACCAGGAAACTAGAGTGA

GAGCACACGTCTGAACTCTTTCCCTACACGACGCTCTTCCGATCTNNNAGTCAGTCACAAACAAGTCAC

AGCACACGTCTGAACTCTTTCCCTACACGACGCTCTTCCGATCTNNNTCCACTTCTCAAGCTGGAACAA

AGCACACGTCTGAACTCTTTCCCTACACGACGCTCTTCCGATCTNNNACCCAACTCAGACAACCACTGA

CACACGTCTGAACTCTTTCCCTACACGACGCTCTTCCGATCTNNNAGGTTAGATCAGAAGAGCCCCAG

AGCACACGTCTGAACTCTTTCCCTACACGACGCTCTTCCGATCTNNNAGTTGAAATTTAGTTTTTCTTC

AGCACACGTCTGAACTCTTTCCCTACACGACGCTCTTCCGATCTNNNACTTCTTTCTCCTGGAACACAG

GCACACGTCTGAACTCTTTCCCTACACGACGCTCTTCCGATCTNNNATCGCCAGTGTTTCTCTCTTTGCC

AGCACACGTCTGAACTCTTTCCCTACACGACGCTCTTCCGATCTNNNTATTTCCACCGTCCCTGTCATCT

GCACACGTCTGAACTCTTTCCCTACACGACGCTCTTCCGATCTNNNCTTGCTCACCCTTCTCTTAGCAGC

AGCACACGTCTGAACTCTTTCCCTACACGACGCTCTTCCGATCTNNNTGAAGAGGCTGTTCTGAGGT

AGCACACGTCTGAACTCTTTCCCTACACGACGCTCTTCCGATCTNNNAGGTGAACAGAGAAGAGCTG

AGCACACGTCTGAACTCTTTCCCTACACGACGCTCTTCCGATCTNNNAGGTGATAAAGTCACTGAAGG

AGAGCACACGTCTGAACTCTTTCCCTACACGACGCTCTTCCGATCTNNNCCCAGGTTCTGATGATTGC

AGCACACGTCTGAACTCTTTCCCTACACGACGCTCTTCCGATCTNNNGAAGAAAGCACTCCATAAGG

AGCACACGTCTGAACTCTTTCCCTACACGACGCTCTTCCGATCTNNNATCGAAGAGTTCAACGTAGCC

CACACGTCTGAACTCTTTCCCTACACGACGCTCTTCCGATCTNNNACCAAAGAAACTGTGGTGTGTGT

AGCACACGTCTGAACTCTTTCCCTACACGACGCTCTTCCGATCTNNNATGAATGGTCTTGGGGAACAT

AGCACACGTCTGAACTCTTTCCCTACACGACGCTCTTCCGATCTNNNGGCCTAAACTGCCATCACCATT

AGCACACGTCTGAACTCTTTCCCTACACGACGCTCTTCCGATCTNNNATGTTAGACACACATGCACTG

AAGAGCACACGTCTGAACTCTTTCCCTACACGACGCTCTTCCGATCTNNNTTGACGGCTCCAGAAGACA

AGCACACGTCTGAACTCTTTCCCTACACGACGCTCTTCCGATCTNNNCATATCCGTGCCTGTGTATGT

AGAGCACACGTCTGAACTCTTTCCCTACACGACGCTCTTCCGATCTNNNATCAGAGGTGAGATTTGGG

GCACACGTCTGAACTCTTTCCCTACACGACGCTCTTCCGATCTNNNGCACAAGAATTCCAAAAGGCAC

GCACACGTCTGAACTCTTTCCCTACACGACGCTCTTCCGATCTNNNTAGCTCTTTCTCTCTCTCGGTCCC

GCACACGTCTGAACTCTTTCCCTACACGACGCTCTTCCGATCTNNNACAATAGGTGAGGGCCCCATTCA

GCACACGTCTGAACTCTTTCCCTACACGACGCTCTTCCGATCTNNNCCATCTGGAAACAGGTTGAGGT

GAGCACACGTCTGAACTCTTTCCCTACACGACGCTCTTCCGATCTNNNGAAGAAGTCCTGATGTCACT

AGAGCACACGTCTGAACTCTTTCCCTACACGACGCTCTTCCGATCTNNNCCTTTGTATACGGGGTGCTA

AGCACACGTCTGAACTCTTTCCCTACACGACGCTCTTCCGATCTNNNTGAAGCAAGCATTGACAGACT

AGCACACGTCTGAACTCTTTCCCTACACGACGCTCTTCCGATCTNNNCTGTTTCTCCTAACATCCTGGGC

GCACACGTCTGAACTCTTTCCCTACACGACGCTCTTCCGATCTNNNACACAGCTTGGGCTAAAGTGAA

AGCACACGTCTGAACTCTTTCCCTACACGACGCTCTTCCGATCTNNNGGCCAATGAACAAACTCTACAT

GAGCACACGTCTGAACTCTTTCCCTACACGACGCTCTTCCGATCTNNNGCACAAGAATTCCAAAAGGC

AGCACACGTCTGAACTCTTTCCCTACACGACGCTCTTCCGATCTNNNATTTGTTATATATGGTGTGTT

GCACACGTCTGAACTCTTTCCCTACACGACGCTCTTCCGATCTNNNTGAGATATTCTCTGTGAGGACT

GAGCACACGTCTGAACTCTTTCCCTACACGACGCTCTTCCGATCTNNNTATGAAGAGCCTGGGATGA[illegible]

[illegible]AGAGCACACGTCTGAACTCTTTCCCTACACGACGCTCTTCCGATCTNNNCATACATCTGTCCCCAACAA

AGCACACGTCTGAACTCTTTCCCTACACGACGCTCTTCCGATCTNNNGCCAGAGCCCCATCTAAAATC

GACATCGGACTCATCAGGAAGC
GCCATCGGACTCATCAGGAAGC
TGGGCATCGGACTCATCAGGAAGC
GCGTGACATCGGACTCATCAGGAAGC
CAACATCGGACTCATCAGGAAGC
GGTACATCGGACTCATCAGGAAGC
CAGCCATCGGACTCATCAGGAAGC
GGCACATCGGACTCATCAGGAAGC
GCGCATCGGACTCATCAGGAAGC
GCGCATCGGACTCATCAGGAAGC
TCATCGGACTCATCAGGAAGC
ACCATCGGACTCATCAGGAAGC
TGCATCGGACTCATCAGGAAGC
TGCATCGGACTCATCAGGAAGC
TCATCGGACTCATCAGGAAGC
GTCATCGGACTCATCAGGAAGC
TCCATCGGACTCATCAGGAAGC
CCTCATCGGACTCATCAGGAAGC
TCGGACTCATCAGGAAGC
ACATCGGACTCATCAGGAAGC
TACATCGGACTCATCAGGAAGC
ACATCGGACTCATCAGGAAGC
ACCCATCGGACTCATCAGGAAGC
GCATCGGACTCATCAGGAAGC
AGCATCGGACTCATCAGGAAGC
CCCATCGGACTCATCAGGAAGC
GTCATCGGACTCATCAGGAAGC
CATCGGACTCATCAGGAAGC
TCATCGGACTCATCAGGAAGC
GTCATCGGACTCATCAGGAAGC
CATCGGACTCATCAGGAAGC
ACATCGGACTCATCAGGAAGC
CATCGGACTCATCAGGAAGC
ATCATCGGACTCATCAGGAAGC
TAACATCGGACTCATCAGGAAGC
CTATCATCGGACTCATCAGGAAGC
GGTCATCGGACTCATCAGGAAGC
CATCGGACTCATCAGGAAGC
CTTCATCGGACTCATCAGGAAGC
AACATCGGACTCATCAGGAAGC
CCATCGGACTCATCAGGAAGC
CTAACATCGGACTCATCAGGAAGC

AGAACGCATCGGACTCATCAGGAAGC
AGTTCCATCGGACTCATCAGGAAGC
TCCGTCATCGGACTCATCAGGAAGC
GGCATCGGACTCATCAGGAAGC
ATCATCGGACTCATCAGGAAGC
ATCGGACTCATCAGGAAGC
TGTCATCGGACTCATCAGGAAGC
CTCATCGGACTCATCAGGAAGC
ATGGCATCGGACTCATCAGGAAGC
ACATCGGACTCATCAGGAAGC
GAGACATCGGACTCATCAGGAAGC
ACCATCGGACTCATCAGGAAGC
TGCCATCGGACTCATCAGGAAGC
ATCATCGGACTCATCAGGAAGC
CACCATCGGACTCATCAGGAAGC
GCATCGGACTCATCAGGAAGC
TCAGCATCGGACTCATCAGGAAGC
ACATCGGACTCATCAGGAAGC
TCATCGGACTCATCAGGAAGC
TGACATCGGACTCATCAGGAAGC
GAGCATCGGACTCATCAGGAAGC
CTCATCGGACTCATCAGGAAGC
TCATCGGACTCATCAGGAAGC
ACCCATCGGACTCATCAGGAAGC
AACATCGGACTCATCAGGAAGC
CACATCGGACTCATCAGGAAGC
AACATCGGACTCATCAGGAAGC
TGTCATCGGACTCATCAGGAAGC
GTTCATCGGACTCATCAGGAAGC
AACATCGGACTCATCAGGAAGC
ACATCGGACTCATCAGGAAGC
ACCATCATCGGACTCATCAGGAAGC
CTCATCGGACTCATCAGGAAGC
ACATCGGACTCATCAGGAAGC
GGTACATCGGACTCATCAGGAAGC
TCATCGGACTCATCAGGAAGC
GCATCGGACTCATCAGGAAGC
GTCATCGGACTCATCAGGAAGC
GCATCGGACTCATCAGGAAGC
GTCCATCGGACTCATCAGGAAGC
GGCATCGGACTCATCAGGAAGC
AAAGGGCATCGGACTCATCAGGAAGC
GCATCGGACTCATCAGGAAGC

GGACATCGGACTCATCAGGAAGC
CATCGGACTCATCAGGAAGC
GCATCGGACTCATCAGGAAGC
TCGGACTCATCAGGAAGC
ACATCGGACTCATCAGGAAGC
AGCGCATCGGACTCATCAGGAAGC
ATGTCATCGGACTCATCAGGAAGC
AGGTCATCGGACTCATCAGGAAGC
CACATCGGACTCATCAGGAAGC
TTCACCATCGGACTCATCAGGAAGC
TCGGACTCATCAGGAAGC
TGCATCGGACTCATCAGGAAGC
AACATCGGACTCATCAGGAAGC
ATACCATCGGACTCATCAGGAAGC
ACCATCGGACTCATCAGGAAGC
TGACATCGGACTCATCAGGAAGC
TCATCGGACTCATCAGGAAGC
ATCGGACTCATCAGGAAGC
TCATCGGACTCATCAGGAAGC
CCATCGGACTCATCAGGAAGC
CAGCATCGGACTCATCAGGAAGC
GCATCGGACTCATCAGGAAGC
GCATCGGACTCATCAGGAAGC
TCATCGGACTCATCAGGAAGC
ACATCGGACTCATCAGGAAGC
AGCATCGGACTCATCAGGAAGC
GTGTCCCATCGGACTCATCAGGAAGC
TCATCGGACTCATCAGGAAGC
CATCGGACTCATCAGGAAGC
GGGCATCGGACTCATCAGGAAGC
GGCTCATCGGACTCATCAGGAAGC
CATCGGACTCATCAGGAAGC
TCATCGGACTCATCAGGAAGC
CACATCGGACTCATCAGGAAGC
CCTCATCGGACTCATCAGGAAGC
ACCCATCGGACTCATCAGGAAGC
CTCATCGGACTCATCAGGAAGC
TGTGTCATCGGACTCATCAGGAAGC
GAGCATCGGACTCATCAGGAAGC
ATCACATCGGACTCATCAGGAAGC
ACACATCGGACTCATCAGGAAGC
CACACACATCGGACTCATCAGGAAGC
CATCGGACTCATCAGGAAGC

GGCATCGGACTCATCAGGAAGC
CATCGGACTCATCAGGAAGC
CCATCGGACTCATCAGGAAGC
AATGCATCGGACTCATCAGGAAGC
GGGACATCGGACTCATCAGGAAGC
TGCATCGGACTCATCAGGAAGC
CACATCGGACTCATCAGGAAGC
CCATCGGACTCATCAGGAAGC
AACTGTGCATCGGACTCATCAGGAAGC
TCCATCGGACTCATCAGGAAGC
TCATCGGACTCATCAGGAAGC
CCATCGGACTCATCAGGAAGC
TGCATCGGACTCATCAGGAAGC
CACATCGGACTCATCAGGAAGC
GAAGGCATCGGACTCATCAGGAAGC
ATCGGACTCATCAGGAAGC
CAACATCGGACTCATCAGGAAGC
CCATCGGACTCATCAGGAAGC
GCCCCATCGGACTCATCAGGAAGC
CATCGGACTCATCAGGAAGC
TGCATCGGACTCATCAGGAAGC
CCATCGGACTCATCAGGAAGC
TCCATGACATCGGACTCATCAGGAAGC
GGCACATCGGACTCATCAGGAAGC
TCCACATCGGACTCATCAGGAAGC
AAGGCATCGGACTCATCAGGAAGC
ATCATCGGACTCATCAGGAAGC
AGGCATCGGACTCATCAGGAAGC
TCACATCGGACTCATCAGGAAGC
CTGCATCGGACTCATCAGGAAGC
CCATCGGACTCATCAGGAAGC
TATCATCGGACTCATCAGGAAGC
GACGCATCGGACTCATCAGGAAGC
CTCATCGGACTCATCAGGAAGC
AGAGCACATCGGACTCATCAGGAAGC
GTGTCATCGGACTCATCAGGAAGC
AAGCACATCGGACTCATCAGGAAGC
GCATCATCGGACTCATCAGGAAGC
TCATCGGACTCATCAGGAAGC
ATCGGACTCATCAGGAAGC
CACATCGGACTCATCAGGAAGC
AAACATCGGACTCATCAGGAAGC
GAGGCATCGGACTCATCAGGAAGC

GACATCGGACTCATCAGGAAGC
GGGCATCGGACTCATCAGGAAGC
CATCGGACTCATCAGGAAGC
CATCGGACTCATCAGGAAGC
GGTCATCGGACTCATCAGGAAGC
TTTCATCGGACTCATCAGGAAGC
GGGCATCGGACTCATCAGGAAGC
ACATCGGACTCATCAGGAAGC
CCATCGGACTCATCAGGAAGC
TCATCGGACTCATCAGGAAGC
ACGCTCATCGGACTCATCAGGAAGC
TATCATCGGACTCATCAGGAAGC
GGGACATCGGACTCATCAGGAAGC
TAACATCGGACTCATCAGGAAGC
TGAGCATCGGACTCATCAGGAAGC
TGCCATCGGACTCATCAGGAAGC
GCCCACATCGGACTCATCAGGAAGC
GCCATCGGACTCATCAGGAAGC
ACATCGGACTCATCAGGAAGC
TCATCGGACTCATCAGGAAGC
GATCCGCATCGGACTCATCAGGAAGC
AGAGACATCGGACTCATCAGGAAGC
TCTCATCGGACTCATCAGGAAGC
TGCATCGGACTCATCAGGAAGC
CACATCGGACTCATCAGGAAGC
TTTCATCGGACTCATCAGGAAGC
CATCGGACTCATCAGGAAGC
CTCATCGGACTCATCAGGAAGC
TTTCATCGGACTCATCAGGAAGC
GTCATCGGACTCATCAGGAAGC
AGAAGCATCGGACTCATCAGGAAGC
CCATCGGACTCATCAGGAAGC
TACATCGGACTCATCAGGAAGC
AAGCCATCGGACTCATCAGGAAGC
TGGCATCGGACTCATCAGGAAGC
GTGCATCGGACTCATCAGGAAGC
GCTCATCGGACTCATCAGGAAGC
CTCATCGGACTCATCAGGAAGC
TTGTAACATCGGACTCATCAGGAAGC
CACACATCGGACTCATCAGGAAGC
CATCGGACTCATCAGGAAGC
CAACATCGGACTCATCAGGAAGC
GTCCATCGGACTCATCAGGAAGC

GCATCGGACTCATCAGGAAGC
ATCATCGGACTCATCAGGAAGC
TCGGACTCATCAGGAAGC
ATCGGACTCATCAGGAAGC
GAACCATCGGACTCATCAGGAAGC
TCATCGGACTCATCAGGAAGC
TTCATCGGACTCATCAGGAAGC
CTACATCGGACTCATCAGGAAGC
TCATCGGACTCATCAGGAAGC
GACATCGGACTCATCAGGAAGC
GTCATCGGACTCATCAGGAAGC
CGACATCGGACTCATCAGGAAGC
GGCATCGGACTCATCAGGAAGC
CCCGCATCGGACTCATCAGGAAGC
CATCGGACTCATCAGGAAGC
ATCGGACTCATCAGGAAGC
ATCGGACTCATCAGGAAGC
GCATCGGACTCATCAGGAAGC
ACATCGGACTCATCAGGAAGC
AACATCGGACTCATCAGGAAGC
ACATCGGACTCATCAGGAAGC
ACATCGGACTCATCAGGAAGC
CCATCGGACTCATCAGGAAGC
CATCGGACTCATCAGGAAGC
CCATCGGACTCATCAGGAAGC
GGCATCGGACTCATCAGGAAGC
ACATCGGACTCATCAGGAAGC
CCACATCGGACTCATCAGGAAGC
CATCGGACTCATCAGGAAGC
CATCGGACTCATCAGGAAGC
TGTCATCGGACTCATCAGGAAGC
TGCCATCGGACTCATCAGGAAGC
CATCGGACTCATCAGGAAGC
GTGTGTCATCGGACTCATCAGGAAGC
GTCTCATCGGACTCATCAGGAAGC
GATACATCGGACTCATCAGGAAGC
CCATCGGACTCATCAGGAAGC
GTCATCGGACTCATCAGGAAGC
GAGCCATCGGACTCATCAGGAAGC
CCATCGGACTCATCAGGAAGC
GCCATCGGACTCATCAGGAAGC
TACATCGGACTCATCAGGAAGC
TTCCATCGGACTCATCAGGAAGC

CACATCGGACTCATCAGGAAGC
ACATCGGACTCATCAGGAAGC
CCCATCGGACTCATCAGGAAGC
GAAACATCGGACTCATCAGGAAGC
TCATCGGACTCATCAGGAAGC
GGCATCGGACTCATCAGGAAGC
TGTCATCGGACTCATCAGGAAGC
TGCATCGGACTCATCAGGAAGC
CATCGGACTCATCAGGAAGC
CATCGGACTCATCAGGAAGC
CCATCGGACTCATCAGGAAGC
GTACATCGGACTCATCAGGAAGC
TCATCGGACTCATCAGGAAGC
GGGACATCGGACTCATCAGGAAGC
GTACATCGGACTCATCAGGAAGC
AGCATCGGACTCATCAGGAAGC
ACATTCCATCGGACTCATCAGGAAGC
TGCATCGGACTCATCAGGAAGC
AGGCATCGGACTCATCAGGAAGC
CCATCGGACTCATCAGGAAGC
ACATCGGACTCATCAGGAAGC
CCATCGGACTCATCAGGAAGC
AACATCGGACTCATCAGGAAGC
CATCGGACTCATCAGGAAGC
GCATCGGACTCATCAGGAAGC
CACATCGGACTCATCAGGAAGC
AGCATCGGACTCATCAGGAAGC
CATCATCGGACTCATCAGGAAGC
TCCGCATCGGACTCATCAGGAAGC
CTGTTCATCGGACTCATCAGGAAGC
CATCGGACTCATCAGGAAGC
ACAGCATCGGACTCATCAGGAAGC
GCATCGGACTCATCAGGAAGC
GTCATCGGACTCATCAGGAAGC
AGGGCAGCATCGGACTCATCAGGAAGC
ACACATCGGACTCATCAGGAAGC
GCCACATCGGACTCATCAGGAAGC
TGTCATCGGACTCATCAGGAAGC
AGGAGCATCGGACTCATCAGGAAGC
CCCATCGGACTCATCAGGAAGC
GCATCGGACTCATCAGGAAGC
GCCATCGGACTCATCAGGAAGC
CAACATCGGACTCATCAGGAAGC

AAACATCGGACTCATCAGGAAGC
CCATCGGACTCATCAGGAAGC
AACATCGGACTCATCAGGAAGC
CCATCGGACTCATCAGGAAGC
AATCATCGGACTCATCAGGAAGC
ATCGGACTCATCAGGAAGC
CAACCATCGGACTCATCAGGAAGC
CATCGGACTCATCAGGAAGC
ATCGGACTCATCAGGAAGC
TATCATCGGACTCATCAGGAAGC
TCTCATCGGACTCATCAGGAAGC
CCATCGGACTCATCAGGAAGC
CTAGGCATCGGACTCATCAGGAAGC
CCATCGGACTCATCAGGAAGC
ACATCGGACTCATCAGGAAGC
CACATCGGACTCATCAGGAAGC
CTCATCGGACTCATCAGGAAGC
TCTCTCCATCGGACTCATCAGGAAGC
GGACATCGGACTCATCAGGAAGC
GCATCGGACTCATCAGGAAGC
CCATCGGACTCATCAGGAAGC
AACATCGGACTCATCAGGAAGC
AAACATCGGACTCATCAGGAAGC
ACACCATCGGACTCATCAGGAAGC
CATCATCGGACTCATCAGGAAGC
CTGCATCGGACTCATCAGGAAGC
GCATCGGACTCATCAGGAAGC
ACATCGGACTCATCAGGAAGC
TCATCGGACTCATCAGGAAGC
CCCATCGGACTCATCAGGAAGC
CCATCGGACTCATCAGGAAGC
TCATCGGACTCATCAGGAAGC
TAGCATCGGACTCATCAGGAAGC
AGTTCATCGGACTCATCAGGAAGC
GACATCGGACTCATCAGGAAGC
CACCATCGGACTCATCAGGAAGC
ACGGCATCGGACTCATCAGGAAGC
CCATCGGACTCATCAGGAAGC
CACATCGGACTCATCAGGAAGC
CATCGGACTCATCAGGAAGC
ACAGTCATCGGACTCATCAGGAAGC
CACATCGGACTCATCAGGAAGC
CCCATCGGACTCATCAGGAAGC

GACATCGGACTCATCAGGAAGC
GCCATCGGACTCATCAGGAAGC
TCAACATCGGACTCATCAGGAAGC
TTGCATCGGACTCATCAGGAAGC
CATCGGACTCATCAGGAAGC
GGCATCGGACTCATCAGGAAGC
CATCGGACTCATCAGGAAGC
GCATCGGACTCATCAGGAAGC
AGCATCGGACTCATCAGGAAGC
CGGACTCATCAGGAAGC
AACCATCGGACTCATCAGGAAGC
TCGGACTCATCAGGAAGC
AACCATCGGACTCATCAGGAAGC
ATCGGACTCATCAGGAAGC
CATCGGACTCATCAGGAAGC
AGGCATCGGACTCATCAGGAAGC
GCACCCATCGGACTCATCAGGAAGC
GCCCATCGGACTCATCAGGAAGC
TGGTCATCGGACTCATCAGGAAGC
GGCATCGGACTCATCAGGAAGC
TAAACCATCGGACTCATCAGGAAGC
ACATCGGACTCATCAGGAAGC
CATCGGACTCATCAGGAAGC
GCATCGGACTCATCAGGAAGC
GGACATCGGACTCATCAGGAAGC
TGACCATCGGACTCATCAGGAAGC
CATCGGACTCATCAGGAAGC
GACATCGGACTCATCAGGAAGC
TCATCGGACTCATCAGGAAGC
AACATCGGACTCATCAGGAAGC
TCCCATCGGACTCATCAGGAAGC
CATCGGACTCATCAGGAAGC
GTGGCATCGGACTCATCAGGAAGC
GCATCGGACTCATCAGGAAGC
TCATCGGACTCATCAGGAAGC
GCCATCGGACTCATCAGGAAGC
CGCCATCGGACTCATCAGGAAGC
GTCATCGGACTCATCAGGAAGC
GTCATCGGACTCATCAGGAAGC
CATCCATCGGACTCATCAGGAAGC
GTGCCATCGGACTCATCAGGAAGC
CTCATCGGACTCATCAGGAAGC
AGGCATCGGACTCATCAGGAAGC

CACTCATCGGACTCATCAGGAAGC
GGCCATCGGACTCATCAGGAAGC
CATCGGACTCATCAGGAAGC
CTCATCGGACTCATCAGGAAGC
TGACATCGGACTCATCAGGAAGC
CCATCGGACTCATCAGGAAGC
GTACATCGGACTCATCAGGAAGC
GGTCATCGGACTCATCAGGAAGC
AGCATCGGACTCATCAGGAAGC
AGTCATCGGACTCATCAGGAAGC
TGCTCATCGGACTCATCAGGAAGC
ACTCATCGGACTCATCAGGAAGC
ACATCGGACTCATCAGGAAGC
GCCATCGGACTCATCAGGAAGC
GTTCATCGGACTCATCAGGAAGC
CCCATCGGACTCATCAGGAAGC
ACATCGGACTCATCAGGAAGC
GGCTGTCATCGGACTCATCAGGAAGC
ACCCATCGGACTCATCAGGAAGC
ACTCCATCGGACTCATCAGGAAGC
GAATCATCGGACTCATCAGGAAGC
GTACATCGGACTCATCAGGAAGC
GAATCATCGGACTCATCAGGAAGC
TACATCGGACTCATCAGGAAGC
GTCATCGGACTCATCAGGAAGC
ATCATCGGACTCATCAGGAAGC
CACTACATCGGACTCATCAGGAAGC
AGGCACATCGGACTCATCAGGAAGC
ACATCGGACTCATCAGGAAGC
GCATCGGACTCATCAGGAAGC
CATCGGACTCATCAGGAAGC
ATTTACATCGGACTCATCAGGAAGC
CCATCGGACTCATCAGGAAGC
GCATCGGACTCATCAGGAAGC
TCATCGGACTCATCAGGAAGC
GAGCATCGGACTCATCAGGAAGC
ATATCTCGCATCGGACTCATCAGGAAGC
AGACCATCGGACTCATCAGGAAGC
CGTCATCGGACTCATCAGGAAGC
ACATCGGACTCATCAGGAAGC
GCATCGGACTCATCAGGAAGC
ACATCGGACTCATCAGGAAGC
GACATCGGACTCATCAGGAAGC

ATCGGACTCATCAGGAAGC
CATCGGACTCATCAGGAAGC
ACATCGGACTCATCAGGAAGC
ACACATCGGACTCATCAGGAAGC
TGACACATCGGACTCATCAGGAAGC
CCATCGGACTCATCAGGAAGC
GCATCGGACTCATCAGGAAGC
TCATCGGACTCATCAGGAAGC
GTGCATCGGACTCATCAGGAAGC
TGGACATCGGACTCATCAGGAAGC
GTCATCGGACTCATCAGGAAGC
TCATCGGACTCATCAGGAAGC
ATCGGACTCATCAGGAAGC
AGCATCGGACTCATCAGGAAGC
GGCATCGGACTCATCAGGAAGC
CATCATCGGACTCATCAGGAAGC
TTACATCGGACTCATCAGGAAGC
TGCATCGGACTCATCAGGAAGC
CCCATCGGACTCATCAGGAAGC
AGGCATCGGACTCATCAGGAAGC
CAACCATCGGACTCATCAGGAAGC
ACATCGGACTCATCAGGAAGC
CTGCATCGGACTCATCAGGAAGC
GGGGCCATCGGACTCATCAGGAAGC
ACATCGGACTCATCAGGAAGC
AGACATCGGACTCATCAGGAAGC
TAGGCACATCGGACTCATCAGGAAGC
ATCACATCGGACTCATCAGGAAGC
TGAACATCGGACTCATCAGGAAGC
GCATCGGACTCATCAGGAAGC
CAACATCGGACTCATCAGGAAGC
GCCATCGGACTCATCAGGAAGC
TAGCATCGGACTCATCAGGAAGC
ACATCGGACTCATCAGGAAGC
ACTCATCGGACTCATCAGGAAGC
GACATCGGACTCATCAGGAAGC
ATTCATCGGACTCATCAGGAAGC
CCATCGGACTCATCAGGAAGC
CAGGTCATCGGACTCATCAGGAAGC
ATCGGACTCATCAGGAAGC
CCTCATCGGACTCATCAGGAAGC
GTCATCGGACTCATCAGGAAGC
ACCCTCATCGGACTCATCAGGAAGC

ACACATCGGACTCATCAGGAAGC
CCATCGGACTCATCAGGAAGC
ACATCGGACTCATCAGGAAGC
TGGGCATCGGACTCATCAGGAAGC
CCATCGGACTCATCAGGAAGC
ACACATCGGACTCATCAGGAAGC
TCATCGGACTCATCAGGAAGC
TGCATCGGACTCATCAGGAAGC
ACTTACATCGGACTCATCAGGAAGC
CATCGGACTCATCAGGAAGC
CCATCGGACTCATCAGGAAGC
GCACCATCGGACTCATCAGGAAGC
GGCATCGGACTCATCAGGAAGC
AGCCATCGGACTCATCAGGAAGC
CCCTCATCGGACTCATCAGGAAGC
ACATCGGACTCATCAGGAAGC
GCATCGGACTCATCAGGAAGC
AACATCGGACTCATCAGGAAGC
AGTCACATCGGACTCATCAGGAAGC
CCCATCGGACTCATCAGGAAGC
CTCATCGGACTCATCAGGAAGC
TGCATCGGACTCATCAGGAAGC
TTCATCGGACTCATCAGGAAGC
TTTGCATCGGACTCATCAGGAAGC
ACGCACATCGGACTCATCAGGAAGC
GTCATCGGACTCATCAGGAAGC
TCCCATCGGACTCATCAGGAAGC
CCCCATCGGACTCATCAGGAAGC
GCCCATCGGACTCATCAGGAAGC
CCCATCGGACTCATCAGGAAGC
GTCTCATCGGACTCATCAGGAAGC
ACATCGGACTCATCAGGAAGC
AGCATCGGACTCATCAGGAAGC
GTCCATCGGACTCATCAGGAAGC
TCGGACTCATCAGGAAGC
TACCATCGGACTCATCAGGAAGC
CGCATCGGACTCATCAGGAAGC
GTGTCATCGGACTCATCAGGAAGC
AGCATCGGACTCATCAGGAAGC
TTCATCGGACTCATCAGGAAGC
AGGCATCGGACTCATCAGGAAGC
AATCATCGGACTCATCAGGAAGC
TCTACATCGGACTCATCAGGAAGC

CCCATCGGACTCATCAGGAAGC
AGCATCGGACTCATCAGGAAGC
AGGCATCGGACTCATCAGGAAGC
GGCCATCGGACTCATCAGGAAGC
GGCATCGGACTCATCAGGAAGC
GCCATCGGACTCATCAGGAAGC
CTCATCGGACTCATCAGGAAGC
AATGGCATCGGACTCATCAGGAAGC
CCCATCGGACTCATCAGGAAGC
TGACATCGGACTCATCAGGAAGC
CATCGGACTCATCAGGAAGC
CATCGGACTCATCAGGAAGC
ACCATCGGACTCATCAGGAAGC
CAGCCATCGGACTCATCAGGAAGC
CACCATCGGACTCATCAGGAAGC
GGATCATCGGACTCATCAGGAAGC
CCCATCGGACTCATCAGGAAGC
CTCATCGGACTCATCAGGAAGC
TCCCATCGGACTCATCAGGAAGC
CCCATCGGACTCATCAGGAAGC
GCCATCGGACTCATCAGGAAGC
CACATCGGACTCATCAGGAAGC
CGCATCGGACTCATCAGGAAGC
TGCATCGGACTCATCAGGAAGC
TCATCGGACTCATCAGGAAGC
CATCGGACTCATCAGGAAGC
TACATCGGACTCATCAGGAAGC
TGGAACATCGGACTCATCAGGAAGC
AAAACATCGGACTCATCAGGAAGC
TGCTCATCGGACTCATCAGGAAGC
CTTCATCGGACTCATCAGGAAGC
TCGGACTCATCAGGAAGC
ACATCGGACTCATCAGGAAGC
AACATCGGACTCATCAGGAAGC
ATTCATCGGACTCATCAGGAAGC
CTGACATCGGACTCATCAGGAAGC
CCCATCGGACTCATCAGGAAGC
AGCATCGGACTCATCAGGAAGC
TCATCGGACTCATCAGGAAGC
ACATCGGACTCATCAGGAAGC
TCCCATCGGACTCATCAGGAAGC
ACCCATCGGACTCATCAGGAAGC
TCCATCGGACTCATCAGGAAGC

GTGACATCGGACTCATCAGGAAGC
GTTCATCGGACTCATCAGGAAGC
ACATCGGACTCATCAGGAAGC
CATCGGACTCATCAGGAAGC
TGGCATCGGACTCATCAGGAAGC
TCATCGGACTCATCAGGAAGC
GGTTCATCGGACTCATCAGGAAGC
CATCGGACTCATCAGGAAGC
TATGACATCGGACTCATCAGGAAGC
TCCATCGGACTCATCAGGAAGC
CATCGGACTCATCAGGAAGC
AGCACCATCGGACTCATCAGGAAGC
ATCGGACTCATCAGGAAGC
GAGACATCGGACTCATCAGGAAGC
GCCATCGGACTCATCAGGAAGC
CACCATCGGACTCATCAGGAAGC
ACCATCGGACTCATCAGGAAGC
AGCAACATCGGACTCATCAGGAAGC
TCACATCGGACTCATCAGGAAGC
CGCATCGGACTCATCAGGAAGC
TACATCGGACTCATCAGGAAGC
TATGACATCGGACTCATCAGGAAGC
TGTCCATCGGACTCATCAGGAAGC
ACCCATCGGACTCATCAGGAAGC
GGCATCGGACTCATCAGGAAGC
GTGCATCGGACTCATCAGGAAGC
TCCATCGGACTCATCAGGAAGC
GCATCGGACTCATCAGGAAGC
ATTTGCCATCGGACTCATCAGGAAGC
GGGCATCGGACTCATCAGGAAGC
GGCATCGGACTCATCAGGAAGC
GCATCGGACTCATCAGGAAGC
ATTCATCGGACTCATCAGGAAGC
CCATCGGACTCATCAGGAAGC
TGTCATCGGACTCATCAGGAAGC
GAGCATCGGACTCATCAGGAAGC
AGCATCGGACTCATCAGGAAGC
GCATCGGACTCATCAGGAAGC
AACCATCGGACTCATCAGGAAGC
CATCGGACTCATCAGGAAGC
AGTCCACATCGGACTCATCAGGAAGC
ACAGGCATCGGACTCATCAGGAAGC
GGGAACATCGGACTCATCAGGAAGC

AGGCATCGGACTCATCAGGAAGC
GCTCTCATCGGACTCATCAGGAAGC
CAACATCGGACTCATCAGGAAGC
CATCGGACTCATCAGGAAGC
CACATCGGACTCATCAGGAAGC
CCATCGGACTCATCAGGAAGC
ACATCGGACTCATCAGGAAGC
GCATCGGACTCATCAGGAAGC
GCATCGGACTCATCAGGAAGC
TGCATCGGACTCATCAGGAAGC
GCATCGGACTCATCAGGAAGC
TCATCGGACTCATCAGGAAGC
CACATCGGACTCATCAGGAAGC
GGCCATCGGACTCATCAGGAAGC
TTCATCGGACTCATCAGGAAGC
ATCGGACTCATCAGGAAGC
GCCCATCGGACTCATCAGGAAGC
TGAAACATCGGACTCATCAGGAAGC
CGCATCGGACTCATCAGGAAGC
TGTCATCGGACTCATCAGGAAGC
GACATCGGACTCATCAGGAAGC
ATCATCGGACTCATCAGGAAGC
TGCATCGGACTCATCAGGAAGC
TCATCGGACTCATCAGGAAGC
TCATCATCGGACTCATCAGGAAGC
CCATCGGACTCATCAGGAAGC
CCCATCGGACTCATCAGGAAGC
GTCGCATCGGACTCATCAGGAAGC
AAGAGCATCGGACTCATCAGGAAGC
GAGCCATCGGACTCATCAGGAAGC
TTCATCGGACTCATCAGGAAGC
AGGCATCGGACTCATCAGGAAGC
CTCATCGGACTCATCAGGAAGC
ACATCGGACTCATCAGGAAGC
AACATCGGACTCATCAGGAAGC
ATCATCGGACTCATCAGGAAGC
GGAAGCATCGGACTCATCAGGAAGC
GGACATCGGACTCATCAGGAAGC
GCCCATCGGACTCATCAGGAAGC
TCATCGGACTCATCAGGAAGC
GGTCATCGGACTCATCAGGAAGC
TTCATCGGACTCATCAGGAAGC
AGGCATCGGACTCATCAGGAAGC

GATCATCGGACTCATCAGGAAGC
TGGGCATCGGACTCATCAGGAAGC
CATCGGACTCATCAGGAAGC
GACATCGGACTCATCAGGAAGC
GCCATCGGACTCATCAGGAAGC
TTCATCGGACTCATCAGGAAGC
TCATCGGACTCATCAGGAAGC
AGCATCGGACTCATCAGGAAGC
TGCATCGGACTCATCAGGAAGC
CATCGGACTCATCAGGAAGC
AGACATCGGACTCATCAGGAAGC
AACATCGGACTCATCAGGAAGC
GCCATCGGACTCATCAGGAAGC
CCACATCGGACTCATCAGGAAGC
AACATCGGACTCATCAGGAAGC
ACATCGGACTCATCAGGAAGC
TTAGGCATCGGACTCATCAGGAAGC
AACATCGGACTCATCAGGAAGC
GTCATCGGACTCATCAGGAAGC
CATCGGACTCATCAGGAAGC
CATCGGACTCATCAGGAAGC
GCCATCGGACTCATCAGGAAGC
TGCCATCGGACTCATCAGGAAGC
GGGCATCGGACTCATCAGGAAGC
GCCATCGGACTCATCAGGAAGC
AACATCGGACTCATCAGGAAGC
AACATCGGACTCATCAGGAAGC
TTCATCGGACTCATCAGGAAGC
ATCATCGGACTCATCAGGAAGC
ACCATCGGACTCATCAGGAAGC
TTTGCATCGGACTCATCAGGAAGC
GAACATCGGACTCATCAGGAAGC
GTTCATCGGACTCATCAGGAAGC
CTTGCATCGGACTCATCAGGAAGC
TTCATCGGACTCATCAGGAAGC
AGGACATCGGACTCATCAGGAAGC
GGAGCATCGGACTCATCAGGAAGC
TCATCGGACTCATCAGGAAGC
TCATCGGACTCATCAGGAAGC
GTGGCATCGGACTCATCAGGAAGC
GCATCGGACTCATCAGGAAGC
CACATCGGACTCATCAGGAAGC
ACCTCATCGGACTCATCAGGAAGC

TGCCATCGGACTCATCAGGAAGC
GACATCGGACTCATCAGGAAGC
CCATCGGACTCATCAGGAAGC
GCATCGGACTCATCAGGAAGC
ATGACATCGGACTCATCAGGAAGC
AGCATCGGACTCATCAGGAAGC
ATCCACATCGGACTCATCAGGAAGC
CGTGCATCGGACTCATCAGGAAGC
GAAAGACATCGGACTCATCAGGAAGC
GGACATCGGACTCATCAGGAAGC
TCTCATCGGACTCATCAGGAAGC
CACATCGGACTCATCAGGAAGC
CCGCATCGGACTCATCAGGAAGC
TACATCGGACTCATCAGGAAGC
GCATCGGACTCATCAGGAAGC
AAGCATCGGACTCATCAGGAAGC
GCATCGGACTCATCAGGAAGC
ATCGGACTCATCAGGAAGC
CTCATCGGACTCATCAGGAAGC
CATCGGACTCATCAGGAAGC
TAGCATCGGACTCATCAGGAAGC
TGTCATCGGACTCATCAGGAAGC
TCCAGTCATCGGACTCATCAGGAAGC
GGACATCGGACTCATCAGGAAGC
CTTGCATCGGACTCATCAGGAAGC
GCCATCGGACTCATCAGGAAGC
AGCCATCGGACTCATCAGGAAGC
GACATCGGACTCATCAGGAAGC
TCATCGGACTCATCAGGAAGC
CCATCGGACTCATCAGGAAGC
AGCAGTCATCGGACTCATCAGGAAGC
ACATCGGACTCATCAGGAAGC
AACATCGGACTCATCAGGAAGC
CATCGGACTCATCAGGAAGC
GACATCGGACTCATCAGGAAGC
GTCATCGGACTCATCAGGAAGC
ATCGGACTCATCAGGAAGC
CAACCATCGGACTCATCAGGAAGC
CACCATCGGACTCATCAGGAAGC
TGCATCGGACTCATCAGGAAGC
TCATCGGACTCATCAGGAAGC
GTGACATCGGACTCATCAGGAAGC
GGTCTCATCGGACTCATCAGGAAGC

CTGCATCGGACTCATCAGGAAGC
CCATCGGACTCATCAGGAAGC
AGGCATCGGACTCATCAGGAAGC
ATCGGACTCATCAGGAAGC
TGCCATCGGACTCATCAGGAAGC
GCAACATCGGACTCATCAGGAAGC
TGCATCGGACTCATCAGGAAGC
ACCATCGGACTCATCAGGAAGC
CATCGGACTCATCAGGAAGC
CATCGGACTCATCAGGAAGC
ATCATCGGACTCATCAGGAAGC
CATCGGACTCATCAGGAAGC
ATGCATCGGACTCATCAGGAAGC
ATGCTTCATCGGACTCATCAGGAAGC
CATCGGACTCATCAGGAAGC
AACATCGGACTCATCAGGAAGC
GATTCATCGGACTCATCAGGAAGC
GACATCGGACTCATCAGGAAGC
AGTGAACATCGGACTCATCAGGAAGC
TTCATCGGACTCATCAGGAAGC
TCATCGGACTCATCAGGAAGC
CCCCATCGGACTCATCAGGAAGC
TGAATCATCGGACTCATCAGGAAGC
CCCATCGGACTCATCAGGAAGC
GTCATCGGACTCATCAGGAAGC
ACATCGGACTCATCAGGAAGC
CATCGGACTCATCAGGAAGC
GTCTCCCATCGGACTCATCAGGAAGC
GATACATCGGACTCATCAGGAAGC
GACATCGGACTCATCAGGAAGC
TCATCGGACTCATCAGGAAGC
CATCGGACTCATCAGGAAGC
ACCATCGGACTCATCAGGAAGC
TGTCATCGGACTCATCAGGAAGC
CTGCATCGGACTCATCAGGAAGC
ACATCGGACTCATCAGGAAGC
TCATCGGACTCATCAGGAAGC
ATCGGACTCATCAGGAAGC
TCCTCATCGGACTCATCAGGAAGC
GGCATCGGACTCATCAGGAAGC
AGCATCGGACTCATCAGGAAGC
TTGCATCGGACTCATCAGGAAGC
TTCATCGGACTCATCAGGAAGC

ATGAGCATCGGACTCATCAGGAAGC
AGACCATCGGACTCATCAGGAAGC
ACACATCGGACTCATCAGGAAGC
CATCGGACTCATCAGGAAGC
CTCATCGGACTCATCAGGAAGC
CTAGCCATCGGACTCATCAGGAAGC
AGGTGGGCATCGGACTCATCAGGAAGC
ATCGGACTCATCAGGAAGC
TACATCGGACTCATCAGGAAGC
ATTCATCGGACTCATCAGGAAGC
CACATCGGACTCATCAGGAAGC
TCCATCGGACTCATCAGGAAGC
ACATCGGACTCATCAGGAAGC
GACATCGGACTCATCAGGAAGC
GCATCGGACTCATCAGGAAGC
AACATCGGACTCATCAGGAAGC
TTCATCGGACTCATCAGGAAGC
ACATCGGACTCATCAGGAAGC
CATCGGACTCATCAGGAAGC
GCATCGGACTCATCAGGAAGC
CATCGGACTCATCAGGAAGC
ACCATCGGACTCATCAGGAAGC
TCCATCGGACTCATCAGGAAGC
AAACATCGGACTCATCAGGAAGC
AGCATCGGACTCATCAGGAAGC
CACATCGGACTCATCAGGAAGC
GGATCATCGGACTCATCAGGAAGC
GCATCGGACTCATCAGGAAGC
AGCCATCGGACTCATCAGGAAGC
GAACATCGGACTCATCAGGAAGC
GCCTCATCGGACTCATCAGGAAGC
GCATCGGACTCATCAGGAAGC
TCCATCGGACTCATCAGGAAGC
GTGCATCGGACTCATCAGGAAGC
TGCATCGGACTCATCAGGAAGC
CATCGGACTCATCAGGAAGC
TCCATCGGACTCATCAGGAAGC
TGCATCGGACTCATCAGGAAGC
TCATCGGACTCATCAGGAAGC
GGACATCGGACTCATCAGGAAGC
AACATCGGACTCATCAGGAAGC
CATCGGACTCATCAGGAAGC
AGAAACATCGGACTCATCAGGAAGC

ATCGGACTCATCAGGAAGC
CATCGGACTCATCAGGAAGC
CACTTGCATCGGACTCATCAGGAAGC
GGTCATCGGACTCATCAGGAAGC
ACTCATCGGACTCATCAGGAAGC
CATCGGACTCATCAGGAAGC
CATCGGACTCATCAGGAAGC
CATCGGACTCATCAGGAAGC
AACATCGGACTCATCAGGAAGC
CCACATCGGACTCATCAGGAAGC
GTGTCATCGGACTCATCAGGAAGC
ACACATCGGACTCATCAGGAAGC
TCACATCGGACTCATCAGGAAGC
CGCATCGGACTCATCAGGAAGC
GTAACATCGGACTCATCAGGAAGC
GCATCGGACTCATCAGGAAGC
ATCGGACTCATCAGGAAGC
GTCATCGGACTCATCAGGAAGC
TGCTCATCGGACTCATCAGGAAGC
TGGCATCGGACTCATCAGGAAGC
TGCCATCGGACTCATCAGGAAGC
TAGCATCGGACTCATCAGGAAGC
TCATCGGACTCATCAGGAAGC
ACCATCGGACTCATCAGGAAGC
AACATCGGACTCATCAGGAAGC
CTCATCGGACTCATCAGGAAGC
GACATCGGACTCATCAGGAAGC
TACATCGGACTCATCAGGAAGC
GTCATCGGACTCATCAGGAAGC
GCAACCCATCGGACTCATCAGGAAGC
CTTTCATCGGACTCATCAGGAAGC
ACATCGGACTCATCAGGAAGC
GCATCGGACTCATCAGGAAGC
CTGGTCATCGGACTCATCAGGAAGC
CCATCGGACTCATCAGGAAGC
TCACATCGGACTCATCAGGAAGC
CAGCATCGGACTCATCAGGAAGC
GTCATCGGACTCATCAGGAAGC
CCATCGGACTCATCAGGAAGC
CATCGGACTCATCAGGAAGC
CATCGGACTCATCAGGAAGC
AACATCGGACTCATCAGGAAGC
GCCATCGGACTCATCAGGAAGC

TGTCATCGGACTCATCAGGAAGC
GCCATCGGACTCATCAGGAAGC
GTGTCATCGGACTCATCAGGAAGC
CAACATCGGACTCATCAGGAAGC
CATCGGACTCATCAGGAAGC
ACCATCGGACTCATCAGGAAGC
TCCATCGGACTCATCAGGAAGC
GCATCGGACTCATCAGGAAGC
ACATCGGACTCATCAGGAAGC
TTCCACATCGGACTCATCAGGAAGC
ATCATCGGACTCATCAGGAAGC
CTTCATCGGACTCATCAGGAAGC
CACATCGGACTCATCAGGAAGC
TGACATCGGACTCATCAGGAAGC
TCATCGGACTCATCAGGAAGC
CACATCGGACTCATCAGGAAGC
AGGCATCGGACTCATCAGGAAGC
CCATCGGACTCATCAGGAAGC
AAAGACATCGGACTCATCAGGAAGC
AGACCATCGGACTCATCAGGAAGC
GCATCGGACTCATCAGGAAGC
CCATCGGACTCATCAGGAAGC
GGCATCGGACTCATCAGGAAGC
GTCATCGGACTCATCAGGAAGC
ATCGGACTCATCAGGAAGC
CTCATCGGACTCATCAGGAAGC
GGGGCATCGGACTCATCAGGAAGC
AGGCATCGGACTCATCAGGAAGC
ATCGGACTCATCAGGAAGC
ATCGGACTCATCAGGAAGC
CTCATCGGACTCATCAGGAAGC
GAAACATCGGACTCATCAGGAAGC
GCATCGGACTCATCAGGAAGC
AACATCGGACTCATCAGGAAGC
TTCATCGGACTCATCAGGAAGC
CATCGGACTCATCAGGAAGC
ACATCGGACTCATCAGGAAGC
TCATCGGACTCATCAGGAAGC
AATCATCGGACTCATCAGGAAGC
CATCGGACTCATCAGGAAGC
TCATCGGACTCATCAGGAAGC
ACCATCGGACTCATCAGGAAGC
GGACATCGGACTCATCAGGAAGC

GTCATCGGACTCATCAGGAAGC
CATCGGACTCATCAGGAAGC
TGCATCGGACTCATCAGGAAGC
TCTCATCGGACTCATCAGGAAGC
GTGGCATCGGACTCATCAGGAAGC
GGGCATCGGACTCATCAGGAAGC
CATCGGACTCATCAGGAAGC
CCGCATCGGACTCATCAGGAAGC
CGCATCGGACTCATCAGGAAGC
AATCATCGGACTCATCAGGAAGC
ACATCGGACTCATCAGGAAGC
CCATCGGACTCATCAGGAAGC
CCATCGGACTCATCAGGAAGC
AAGCATCGGACTCATCAGGAAGC
CAACATCGGACTCATCAGGAAGC
GCCATCGGACTCATCAGGAAGC
TCATCGGACTCATCAGGAAGC
GACATCGGACTCATCAGGAAGC
CACATCGGACTCATCAGGAAGC
GTACATCGGACTCATCAGGAAGC
CATCGGACTCATCAGGAAGC
AGCCATCGGACTCATCAGGAAGC
CATCGGACTCATCAGGAAGC
CATCGGACTCATCAGGAAGC
ACTTCATCGGACTCATCAGGAAGC
ACATCGGACTCATCAGGAAGC
ATCATCGGACTCATCAGGAAGC
GGGCATCGGACTCATCAGGAAGC
TTGGCATCGGACTCATCAGGAAGC
ATCGGACTCATCAGGAAGC
TGCATCGGACTCATCAGGAAGC
CTGCATCGGACTCATCAGGAAGC
CATCGGACTCATCAGGAAGC
TGCGCATCGGACTCATCAGGAAGC
AGAGCATCGGACTCATCAGGAAGC
CAGCCATCGGACTCATCAGGAAGC
CTCATCGGACTCATCAGGAAGC
AAAAACATCGGACTCATCAGGAAGC
TGCCATCGGACTCATCAGGAAGC
TGCATCGGACTCATCAGGAAGC
CATCGGACTCATCAGGAAGC
ACATCGGACTCATCAGGAAGC
GACATCGGACTCATCAGGAAGC

GCATCGGACTCATCAGGAAGC
GGTCATCGGACTCATCAGGAAGC
CCATCGGACTCATCAGGAAGC
CAGACATCGGACTCATCAGGAAGC
AGACACATCGGACTCATCAGGAAGC
CAACATCGGACTCATCAGGAAGC
TCCATCGGACTCATCAGGAAGC
CTTGCATCGGACTCATCAGGAAGC
CCATCGGACTCATCAGGAAGC
ACACATCGGACTCATCAGGAAGC
TATCATCGGACTCATCAGGAAGC
GAGACATCGGACTCATCAGGAAGC
TCGGACTCATCAGGAAGC
GTGTCATCGGACTCATCAGGAAGC
GCATCGGACTCATCAGGAAGC
GTTCATCGGACTCATCAGGAAGC
GCATCGGACTCATCAGGAAGC
CCCATCGGACTCATCAGGAAGC
GTCCATCGGACTCATCAGGAAGC
AGCATCGGACTCATCAGGAAGC
CGAACATCGGACTCATCAGGAAGC
ACCATCGGACTCATCAGGAAGC
GCCATCGGACTCATCAGGAAGC
CATCGGACTCATCAGGAAGC
GTCATCGGACTCATCAGGAAGC
GACATCGGACTCATCAGGAAGC
GTCATCGGACTCATCAGGAAGC
AACCTCATCGGACTCATCAGGAAGC
GACATCGGACTCATCAGGAAGC
CTGGTGCATCGGACTCATCAGGAAGC
TGGCATCGGACTCATCAGGAAGC
ACATCGGACTCATCAGGAAGC
CTGGGACATCGGACTCATCAGGAAGC
ATCATCGGACTCATCAGGAAGC
GTCATCGGACTCATCAGGAAGC
ACATCGGACTCATCAGGAAGC
GCGCATCGGACTCATCAGGAAGC
AAACATCGGACTCATCAGGAAGC
GCCATCGGACTCATCAGGAAGC
ATGCATCGGACTCATCAGGAAGC
GTGTAACATCGGACTCATCAGGAAGC
GGCATCGGACTCATCAGGAAGC
CATCGGACTCATCAGGAAGC

CCATCGGACTCATCAGGAAGC
GTGACATCGGACTCATCAGGAAGC
ACATCGGACTCATCAGGAAGC
CATCGGACTCATCAGGAAGC
TACATCGGACTCATCAGGAAGC
CTCACATCGGACTCATCAGGAAGC
CCATCGGACTCATCAGGAAGC
ATGCAGACATCGGACTCATCAGGAAGC
CCACTTCATCGGACTCATCAGGAAGC
AACACATCGGACTCATCAGGAAGC
CCATCGGACTCATCAGGAAGC
TCATCGGACTCATCAGGAAGC
GACATCGGACTCATCAGGAAGC
GGAGCATCGGACTCATCAGGAAGC
ACATCGGACTCATCAGGAAGC
TCCATCGGACTCATCAGGAAGC
ACCATCGGACTCATCAGGAAGC
GCATCGGACTCATCAGGAAGC
TACATCGGACTCATCAGGAAGC
CATCGGACTCATCAGGAAGC
ATCGGACTCATCAGGAAGC
CATCGGACTCATCAGGAAGC
TCATCGGACTCATCAGGAAGC
GACATCGGACTCATCAGGAAGC
GACATCATCGGACTCATCAGGAAGC
CTCCATCGGACTCATCAGGAAGC
TTGTAGCATCGGACTCATCAGGAAGC
TATGCATCGGACTCATCAGGAAGC
CTTACATCGGACTCATCAGGAAGC
GCATCGGACTCATCAGGAAGC
TTTCATCGGACTCATCAGGAAGC
AATCATCGGACTCATCAGGAAGC
AGGCATCGGACTCATCAGGAAGC
GAGCATCGGACTCATCAGGAAGC
CAGGCATCGGACTCATCAGGAAGC
ACCCATCGGACTCATCAGGAAGC
CATCGGACTCATCAGGAAGC
CATCGGACTCATCAGGAAGC
CGCATCGGACTCATCAGGAAGC
CAGGCATCGGACTCATCAGGAAGC
GTGCATCGGACTCATCAGGAAGC
AGCATCGGACTCATCAGGAAGC
AGGCATCGGACTCATCAGGAAGC

ACATCGGACTCATCAGGAAGC
ATCATCGGACTCATCAGGAAGC
CATCATCGGACTCATCAGGAAGC
CATCGGACTCATCAGGAAGC
ACCACATCGGACTCATCAGGAAGC
GGTCATCGGACTCATCAGGAAGC
CTCATCGGACTCATCAGGAAGC
ACATCGGACTCATCAGGAAGC
CCATCGGACTCATCAGGAAGC
TCCATCGGACTCATCAGGAAGC
TGAACATCGGACTCATCAGGAAGC
GCACATCGGACTCATCAGGAAGC
TCCCATCGGACTCATCAGGAAGC
CACACCATCGGACTCATCAGGAAGC
ACATCGGACTCATCAGGAAGC
TCATCGGACTCATCAGGAAGC
CATCGGACTCATCAGGAAGC
CTGCATCGGACTCATCAGGAAGC
AGGACATCGGACTCATCAGGAAGC
GCCATCGGACTCATCAGGAAGC
CCCATCGGACTCATCAGGAAGC
ACTTCATCGGACTCATCAGGAAGC
AACATCGGACTCATCAGGAAGC
GCCACATCGGACTCATCAGGAAGC
ACATCGGACTCATCAGGAAGC
AAGACCATCGGACTCATCAGGAAGC
CAACATCGGACTCATCAGGAAGC
CATCGGACTCATCAGGAAGC
GTCATCGGACTCATCAGGAAGC
TCCCATCGGACTCATCAGGAAGC
AAACATCGGACTCATCAGGAAGC
AAGCATCGGACTCATCAGGAAGC
CAGACATCGGACTCATCAGGAAGC
GTCCATCGGACTCATCAGGAAGC
TTTCACATCGGACTCATCAGGAAGC
AGCATCGGACTCATCAGGAAGC
CCCATCGGACTCATCAGGAAGC
CTCATCGGACTCATCAGGAAGC
ACATCGGACTCATCAGGAAGC
ACATCGGACTCATCAGGAAGC
TGTCATCGGACTCATCAGGAAGC
CTTCATCGGACTCATCAGGAAGC
CCATCGGACTCATCAGGAAGC

TCATCGGACTCATCAGGAAGC
TTCATCGGACTCATCAGGAAGC
GCATCGGACTCATCAGGAAGC
CCATCGGACTCATCAGGAAGC
CACATCGGACTCATCAGGAAGC
TCTCATCGGACTCATCAGGAAGC
GGCATCGGACTCATCAGGAAGC
GAGCATCGGACTCATCAGGAAGC
ACCATCGGACTCATCAGGAAGC
CTACATCGGACTCATCAGGAAGC
CCCATCGGACTCATCAGGAAGC
GCATCGGACTCATCAGGAAGC
TCATCGGACTCATCAGGAAGC
GCTTTCCATCGGACTCATCAGGAAGC
TCATCGGACTCATCAGGAAGC
GTACATCGGACTCATCAGGAAGC
CATCGGACTCATCAGGAAGC
TACATCGGACTCATCAGGAAGC
CATCGGACTCATCAGGAAGC
ACTCCATCGGACTCATCAGGAAGC
CAACATCGGACTCATCAGGAAGC
TGTCATCGGACTCATCAGGAAGC
TACATCGGACTCATCAGGAAGC
TCATCGGACTCATCAGGAAGC
CTCATCGGACTCATCAGGAAGC
TTCATCGGACTCATCAGGAAGC
TCATCGGACTCATCAGGAAGC
GACACATCGGACTCATCAGGAAGC
CCATCGGACTCATCAGGAAGC
GGCATCGGACTCATCAGGAAGC
ATCATCGGACTCATCAGGAAGC
CATCGGACTCATCAGGAAGC
GCATCGGACTCATCAGGAAGC
TAGAAGGCATCGGACTCATCAGGAAGC
TCGGACTCATCAGGAAGC
CAGCATCATCGGACTCATCAGGAAGC
GATTCATCGGACTCATCAGGAAGC
CTCATCGGACTCATCAGGAAGC
TCATCGGACTCATCAGGAAGC
GGTTCATCGGACTCATCAGGAAGC
TGTGGCATCGGACTCATCAGGAAGC
TGACATCGGACTCATCAGGAAGC
TTTCATCGGACTCATCAGGAAGC

CCATCGGACTCATCAGGAAGC
TGCATCGGACTCATCAGGAAGC
GGACATCGGACTCATCAGGAAGC
TCGGACTCATCAGGAAGC
TTCATCGGACTCATCAGGAAGC
CTCACATCGGACTCATCAGGAAGC
AACCCCATCGGACTCATCAGGAAGC
GTTTCATCGGACTCATCAGGAAGC
GAGCCATCGGACTCATCAGGAAGC
GCATCGGACTCATCAGGAAGC
GTCATCGGACTCATCAGGAAGC
AGGACATCGGACTCATCAGGAAGC
TCGGACTCATCAGGAAGC
ACCATCGGACTCATCAGGAAGC
CCCATCGGACTCATCAGGAAGC
TTCGCATCGGACTCATCAGGAAGC
GCACATCGGACTCATCAGGAAGC
GGGCATCGGACTCATCAGGAAGC
GAACATCGGACTCATCAGGAAGC
CCATCGGACTCATCAGGAAGC
TCGGACTCATCAGGAAGC
CATCGGACTCATCAGGAAGC
GTCATCGGACTCATCAGGAAGC
GCATCGGACTCATCAGGAAGC
TCGGACTCATCAGGAAGC
TCATCGGACTCATCAGGAAGC
CATCGGACTCATCAGGAAGC
TCATCGGACTCATCAGGAAGC
ACATCGGACTCATCAGGAAGC
CACATCGGACTCATCAGGAAGC
ATCGGACTCATCAGGAAGC
TCATCGGACTCATCAGGAAGC
TCGGACTCATCAGGAAGC
ACCATCGGACTCATCAGGAAGC
TGACATCGGACTCATCAGGAAGC
TCGGACTCATCAGGAAGC
TCATCGGACTCATCAGGAAGC
GCCATCGGACTCATCAGGAAGC
TACATCGGACTCATCAGGAAGC
CCATCGGACTCATCAGGAAGC
AACCATCGGACTCATCAGGAAGC
GCTCATCGGACTCATCAGGAAGC
GCCATCGGACTCATCAGGAAGC

ACATCGGACTCATCAGGAAGC
TACCATCGGACTCATCAGGAAGC
TCATCGGACTCATCAGGAAGC
TGCCATCGGACTCATCAGGAAGC
CATCGGACTCATCAGGAAGC
CATCGGACTCATCAGGAAGC
CTCATCGGACTCATCAGGAAGC
CATCGGACTCATCAGGAAGC
TGCTGGCATCGGACTCATCAGGAAGC
TCATCGGACTCATCAGGAAGC
ATCGGACTCATCAGGAAGC
GGACATCGGACTCATCAGGAAGC
TCATCGGACTCATCAGGAAGC
CCTGCATCGGACTCATCAGGAAGC
TCCATCGGACTCATCAGGAAGC
GTGCATCGGACTCATCAGGAAGC
TCATCGGACTCATCAGGAAGC
CCCATCGGACTCATCAGGAAGC
GGGGCATCGGACTCATCAGGAAGC
CATCGGACTCATCAGGAAGC
GACCATCGGACTCATCAGGAAGC
TCATCGGACTCATCAGGAAGC
TACATCGGACTCATCAGGAAGC
TGGCATCGGACTCATCAGGAAGC
TCATCGGACTCATCAGGAAGC
GCATCGGACTCATCAGGAAGC
CATCGGACTCATCAGGAAGC
CCATCGGACTCATCAGGAAGC
CTAACATCGGACTCATCAGGAAGC
ATCGGACTCATCAGGAAGC
GGCATCGGACTCATCAGGAAGC
TGCCATCGGACTCATCAGGAAGC
CCCATCGGACTCATCAGGAAGC
ACATCGGACTCATCAGGAAGC
AATGACATCGGACTCATCAGGAAGC
ACATCGGACTCATCAGGAAGC
TCACATCGGACTCATCAGGAAGC
ACATCGGACTCATCAGGAAGC
CATCGGACTCATCAGGAAGC
CTCATCGGACTCATCAGGAAGC
CATCGGACTCATCAGGAAGC
TCCCATCGGACTCATCAGGAAGC
TAACATCATCGGACTCATCAGGAAGC

GACATCGGACTCATCAGGAAGC
CTGCATCGGACTCATCAGGAAGC
GGACATCGGACTCATCAGGAAGC
ACCCATCGGACTCATCAGGAAGC
CCCATCGGACTCATCAGGAAGC
CATTCATCGGACTCATCAGGAAGC
ACATCGGACTCATCAGGAAGC
CATCGGACTCATCAGGAAGC
GTCATCGGACTCATCAGGAAGC
CTGGACATCGGACTCATCAGGAAGC
GCATCGGACTCATCAGGAAGC
TCATCGGACTCATCAGGAAGC
TTGGCATCGGACTCATCAGGAAGC
TCATCGGACTCATCAGGAAGC
CTCCATCGGACTCATCAGGAAGC
GCTCATCGGACTCATCAGGAAGC
ACATCGGACTCATCAGGAAGC
ACATCGGACTCATCAGGAAGC
CGCATCGGACTCATCAGGAAGC
CATCGGACTCATCAGGAAGC
ATCGGACTCATCAGGAAGC
TCATCGGACTCATCAGGAAGC
ATCATCGGACTCATCAGGAAGC
GTATCATCGGACTCATCAGGAAGC
CCATCGGACTCATCAGGAAGC
ACATCGGACTCATCAGGAAGC
ACATCGGACTCATCAGGAAGC
TGCATCGGACTCATCAGGAAGC
ATCGGACTCATCAGGAAGC
ACCCATCGGACTCATCAGGAAGC
GTAGCATCGGACTCATCAGGAAGC
CATCGGACTCATCAGGAAGC
CCATCGGACTCATCAGGAAGC
TTTCATCGGACTCATCAGGAAGC
CATCGGACTCATCAGGAAGC
TGACATCGGACTCATCAGGAAGC
CCATCGGACTCATCAGGAAGC
GTACATCGGACTCATCAGGAAGC
GGTCCATCGGACTCATCAGGAAGC
AATCATCGGACTCATCAGGAAGC
CAGCATCGGACTCATCAGGAAGC
CCCATCGGACTCATCAGGAAGC
ACATCGGACTCATCAGGAAGC

AATCATCGGACTCATCAGGAAGC
CATCGGACTCATCAGGAAGC
ACATCGGACTCATCAGGAAGC
ACCATCGGACTCATCAGGAAGC
ATACATCGGACTCATCAGGAAGC
AATACATCGGACTCATCAGGAAGC
GCAACATCGGACTCATCAGGAAGC
AGTCATCGGACTCATCAGGAAGC
ATCATCGGACTCATCAGGAAGC
TGCATCGGACTCATCAGGAAGC
CCATCGGACTCATCAGGAAGC
AGCATCGGACTCATCAGGAAGC
GCCATCGGACTCATCAGGAAGC
AGACATCGGACTCATCAGGAAGC
CACATCGGACTCATCAGGAAGC
ACCCATCGGACTCATCAGGAAGC
GTGCATCGGACTCATCAGGAAGC
GGACATCGGACTCATCAGGAAGC
CGGACTCATCAGGAAGC
CAACATCGGACTCATCAGGAAGC
CATCGGACTCATCAGGAAGC
CTCATCGGACTCATCAGGAAGC
TTGCATCGGACTCATCAGGAAGC
AGCATCGGACTCATCAGGAAGC
GCAACATCGGACTCATCAGGAAGC
CTACATCGGACTCATCAGGAAGC
TCCTTCACATCGGACTCATCAGGAAGC
AACATCGGACTCATCAGGAAGC
ACATCGGACTCATCAGGAAGC
TCCCACATCGGACTCATCAGGAAGC
CAGTGCATCGGACTCATCAGGAAGC
TTCATCGGACTCATCAGGAAGC
GACATCGGACTCATCAGGAAGC
TGTCATCGGACTCATCAGGAAGC
CATCGGACTCATCAGGAAGC
GGTACATCGGACTCATCAGGAAGC
ACATCGGACTCATCAGGAAGC
AACTGCATCGGACTCATCAGGAAGC
CTGCATCGGACTCATCAGGAAGC
TGTCATCGGACTCATCAGGAAGC
GACATCGGACTCATCAGGAAGC
CAGCCATCGGACTCATCAGGAAGC
CCATCGGACTCATCAGGAAGC

GCATCGGACTCATCAGGAAGC
ATGCCATCGGACTCATCAGGAAGC
TCAGCCATCGGACTCATCAGGAAGC
TCGGACTCATCAGGAAGC
TGGTCATCGGACTCATCAGGAAGC
GCCATCGGACTCATCAGGAAGC
GCAAACATCGGACTCATCAGGAAGC
CCATCGGACTCATCAGGAAGC
ACATCGGACTCATCAGGAAGC
CACATCGGACTCATCAGGAAGC
CACATCGGACTCATCAGGAAGC
GCCATCGGACTCATCAGGAAGC
TCATCGGACTCATCAGGAAGC
TGTGCCATCGGACTCATCAGGAAGC
TCCATCGGACTCATCAGGAAGC
GGCACATCGGACTCATCAGGAAGC
CACATCGGACTCATCAGGAAGC
GCCATCGGACTCATCAGGAAGC
AAAAGTCATCGGACTCATCAGGAAGC
GGCATCGGACTCATCAGGAAGC
GCATCGGACTCATCAGGAAGC
CCAACCATCGGACTCATCAGGAAGC
CTTCATCGGACTCATCAGGAAGC
TGTCATCGGACTCATCAGGAAGC
CACATCGGACTCATCAGGAAGC
TGAAACATCGGACTCATCAGGAAGC
TACATCGGACTCATCAGGAAGC
GATCATCGGACTCATCAGGAAGC
TCATCGGACTCATCAGGAAGC
TTGACATCGGACTCATCAGGAAGC
ACCATCGGACTCATCAGGAAGC
GTCACATCGGACTCATCAGGAAGC
TAACATCGGACTCATCAGGAAGC
TCATCGGACTCATCAGGAAGC
AGAACATCGGACTCATCAGGAAGC
CATCGGACTCATCAGGAAGC
TTGACATCGGACTCATCAGGAAGC
AAAGCCATCGGACTCATCAGGAAGC
AACCCATCGGACTCATCAGGAAGC
TACATCGGACTCATCAGGAAGC
CGCATCGGACTCATCAGGAAGC
CAAACATCGGACTCATCAGGAAGC
ACATCGGACTCATCAGGAAGC

CATCGGACTCATCAGGAAGC
GCATCGGACTCATCAGGAAGC
ACCCATCGGACTCATCAGGAAGC
CACATCGGACTCATCAGGAAGC
TCATCGGACTCATCAGGAAGC
AGCATCGGACTCATCAGGAAGC
GTCATCGGACTCATCAGGAAGC
GCCATCGGACTCATCAGGAAGC
CATCGGACTCATCAGGAAGC
GGACATCGGACTCATCAGGAAGC
CGCATCGGACTCATCAGGAAGC
ATCATCGGACTCATCAGGAAGC
GTCATCGGACTCATCAGGAAGC
TTCAACATCGGACTCATCAGGAAGC
TCATCGGACTCATCAGGAAGC
CTTAGACATCGGACTCATCAGGAAGC
AACATCGGACTCATCAGGAAGC
AAGCCATCGGACTCATCAGGAAGC
CCATCGGACTCATCAGGAAGC
TTTCATCGGACTCATCAGGAAGC
AAGCATCGGACTCATCAGGAAGC
ATCCTCCATCGGACTCATCAGGAAGC
CACCCATCGGACTCATCAGGAAGC
CAGCATCGGACTCATCAGGAAGC
GGATCATCGGACTCATCAGGAAGC
CATCGGACTCATCAGGAAGC
ATCATCGGACTCATCAGGAAGC
CACATCGGACTCATCAGGAAGC
GACCCATCGGACTCATCAGGAAGC
CTGCATCGGACTCATCAGGAAGC
CATCGGACTCATCAGGAAGC
GCATCGGACTCATCAGGAAGC
CCATCGGACTCATCAGGAAGC
GAGGCATCGGACTCATCAGGAAGC
AGGTCATCGGACTCATCAGGAAGC
CCCATCGGACTCATCAGGAAGC
TCCTCATCGGACTCATCAGGAAGC
ACTGTCATCGGACTCATCAGGAAGC
TCGGACTCATCAGGAAGC
CATCGGACTCATCAGGAAGC
CATCGGACTCATCAGGAAGC
ATGTAACATCGGACTCATCAGGAAGC
TCTCACATCGGACTCATCAGGAAGC

CCATCGGACTCATCAGGAAGC
CACATCGGACTCATCAGGAAGC
TTTCATCGGACTCATCAGGAAGC
CATCGGACTCATCAGGAAGC
GGCATCGGACTCATCAGGAAGC
GAACATCGGACTCATCAGGAAGC
ATCGGACTCATCAGGAAGC
ATCATCGGACTCATCAGGAAGC
GCATCGGACTCATCAGGAAGC
ACCATCGGACTCATCAGGAAGC
TGTCATCGGACTCATCAGGAAGC
CACATCGGACTCATCAGGAAGC
AACCTCATCGGACTCATCAGGAAGC
GAACATCGGACTCATCAGGAAGC
GAACATCGGACTCATCAGGAAGC
CCACATCGGACTCATCAGGAAGC
CATCGGACTCATCAGGAAGC
ACACATCGGACTCATCAGGAAGC
CATCGGACTCATCAGGAAGC
ACTCATCGGACTCATCAGGAAGC
ACACATCGGACTCATCAGGAAGC
AGGACATCGGACTCATCAGGAAGC
TCCCATCGGACTCATCAGGAAGC
ACATCGGACTCATCAGGAAGC
CATCGGACTCATCAGGAAGC
TACATCGGACTCATCAGGAAGC
ACATCGGACTCATCAGGAAGC
CCATCGGACTCATCAGGAAGC
CTGACATCGGACTCATCAGGAAGC
CTTGACATCGGACTCATCAGGAAGC
ATCGGACTCATCAGGAAGC
ACATCGGACTCATCAGGAAGC
AATCATCGGACTCATCAGGAAGC
TCATCGGACTCATCAGGAAGC
GGATCATCGGACTCATCAGGAAGC
CATCGGACTCATCAGGAAGC
TATCATCGGACTCATCAGGAAGC
AACCATCGGACTCATCAGGAAGC
AGGACATCGGACTCATCAGGAAGC
CATCGGACTCATCAGGAAGC
CATCGGACTCATCAGGAAGC
TTCATCGGACTCATCAGGAAGC
GGACATCGGACTCATCAGGAAGC

CATCGGACTCATCAGGAAGC
AGCATCGGACTCATCAGGAAGC
TCCCATCGGACTCATCAGGAAGC
GACACATCGGACTCATCAGGAAGC
GGACATCGGACTCATCAGGAAGC
CATCGGACTCATCAGGAAGC
ACATCGGACTCATCAGGAAGC
AGCATCGGACTCATCAGGAAGC
CAGGACATCGGACTCATCAGGAAGC
AGCATCGGACTCATCAGGAAGC
GCATCGGACTCATCAGGAAGC
TGGTCATCGGACTCATCAGGAAGC
AGCATCGGACTCATCAGGAAGC
TTTTCATCGGACTCATCAGGAAGC
GCATCGGACTCATCAGGAAGC
ATACATCATCGGACTCATCAGGAAGC
ACATCGGACTCATCAGGAAGC
TGACCATCGGACTCATCAGGAAGC
TGGCATCGGACTCATCAGGAAGC
TCATCGGACTCATCAGGAAGC
TCATCGGACTCATCAGGAAGC
TAGCATCGGACTCATCAGGAAGC
CATCGGACTCATCAGGAAGC
TTCATCGGACTCATCAGGAAGC
ACATCGGACTCATCAGGAAGC
TTCATCGGACTCATCAGGAAGC
ATCATCGGACTCATCAGGAAGC
ATCGGACTCATCAGGAAGC
ACATCGGACTCATCAGGAAGC
ACTGTCATCGGACTCATCAGGAAGC
ACCATCGGACTCATCAGGAAGC
ACATCGGACTCATCAGGAAGC
CTCATCGGACTCATCAGGAAGC
CCCCATCGGACTCATCAGGAAGC
GAGACATCGGACTCATCAGGAAGC
CCCTCATCGGACTCATCAGGAAGC
ATCATCGGACTCATCAGGAAGC
GCATCGGACTCATCAGGAAGC
AGGCATCGGACTCATCAGGAAGC
ACCATCGGACTCATCAGGAAGC
CATCGGACTCATCAGGAAGC
GCCATCGGACTCATCAGGAAGC
ATCGGACTCATCAGGAAGC

ACACATCGGACTCATCAGGAAGC
TCTCATCGGACTCATCAGGAAGC
AACACATCGGACTCATCAGGAAGC
GCCCATCGGACTCATCAGGAAGC
GTTGACATCGGACTCATCAGGAAGC
GAGGGCATCGGACTCATCAGGAAGC
CACATCGGACTCATCAGGAAGC
GGGGTCATCGGACTCATCAGGAAGC
GCCATCGGACTCATCAGGAAGC
TACATCGGACTCATCAGGAAGC
CGGCATCGGACTCATCAGGAAGC
GGACTCATCAGGAAGC
ACATCGGACTCATCAGGAAGC
GGCATCGGACTCATCAGGAAGC
CTACATCGGACTCATCAGGAAGC
TGCATCGGACTCATCAGGAAGC
CTCATCGGACTCATCAGGAAGC
GACCCATCGGACTCATCAGGAAGC
TTGTCATCGGACTCATCAGGAAGC
CGCCATCGGACTCATCAGGAAGC
ACATCGGACTCATCAGGAAGC
GATCATCGGACTCATCAGGAAGC
CTCATCGGACTCATCAGGAAGC
ACATCGGACTCATCAGGAAGC
AAACATCGGACTCATCAGGAAGC
AGCACCATCGGACTCATCAGGAAGC
CCATCGGACTCATCAGGAAGC
GGCATCGGACTCATCAGGAAGC
GTACATCGGACTCATCAGGAAGC
GCATCGGACTCATCAGGAAGC
GCCTCATCGGACTCATCAGGAAGC
TAGTCATCGGACTCATCAGGAAGC
AGGCATCGGACTCATCAGGAAGC
CATCGGACTCATCAGGAAGC
CCAACATCGGACTCATCAGGAAGC
ACATCGGACTCATCAGGAAGC
AGCATCGGACTCATCAGGAAGC
CCCGTCATCGGACTCATCAGGAAGC
CCCCATCGGACTCATCAGGAAGC
AACATCGGACTCATCAGGAAGC
AACCATCGGACTCATCAGGAAGC
TGCATCGGACTCATCAGGAAGC
GCCACATCGGACTCATCAGGAAGC

GGCATCGGACTCATCAGGAAGC
ATCATCGGACTCATCAGGAAGC
ACATCGGACTCATCAGGAAGC
TACATCGGACTCATCAGGAAGC
TCATCGGACTCATCAGGAAGC
CCAACATCGGACTCATCAGGAAGC
TGGGCATCGGACTCATCAGGAAGC
TCATCGGACTCATCAGGAAGC
CCATCGGACTCATCAGGAAGC
GCATCGGACTCATCAGGAAGC
AGCATCGGACTCATCAGGAAGC
ACATCGGACTCATCAGGAAGC
TCATCGGACTCATCAGGAAGC
TCGGACTCATCAGGAAGC
CATCGGACTCATCAGGAAGC
CTCATCGGACTCATCAGGAAGC
ACCATCGGACTCATCAGGAAGC
ACATCGGACTCATCAGGAAGC
TCGGACTCATCAGGAAGC
CATCGGACTCATCAGGAAGC
TACATCGGACTCATCAGGAAGC
TCCTCATCGGACTCATCAGGAAGC
TCATCGGACTCATCAGGAAGC
GCATCGGACTCATCAGGAAGC
AACATCGGACTCATCAGGAAGC
CATCATCGGACTCATCAGGAAGC
TTCATCGGACTCATCAGGAAGC
ATCGGACTCATCAGGAAGC
GGCCATCGGACTCATCAGGAAGC
CATCGGACTCATCAGGAAGC
AAACATCGGACTCATCAGGAAGC
AAACATCGGACTCATCAGGAAGC
TGCCATCGGACTCATCAGGAAGC
CCATCGGACTCATCAGGAAGC
AACATCGGACTCATCAGGAAGC
CATCGGACTCATCAGGAAGC
TCCCATCGGACTCATCAGGAAGC
AACATCGGACTCATCAGGAAGC
TTTCATCGGACTCATCAGGAAGC
GGAGCATCGGACTCATCAGGAAGC
GCATCGGACTCATCAGGAAGC
CCCATCGGACTCATCAGGAAGC
ATCGGACTCATCAGGAAGC

CAGCATCGGACTCATCAGGAAGC
GTAGCCATCGGACTCATCAGGAAGC
AGGCATCGGACTCATCAGGAAGC
CGTCATCGGACTCATCAGGAAGC
GTAATCATCGGACTCATCAGGAAGC
CATCGGACTCATCAGGAAGC
CCATCGGACTCATCAGGAAGC
TAAACATCGGACTCATCAGGAAGC
CATGCCATCGGACTCATCAGGAAGC
ATCGGACTCATCAGGAAGC
ACAACATCGGACTCATCAGGAAGC
CATTCATCGGACTCATCAGGAAGC
AGTGACCATCGGACTCATCAGGAAGC
GGACCATCGGACTCATCAGGAAGC
CTTCATCGGACTCATCAGGAAGC
AGGACATCGGACTCATCAGGAAGC
TCATCGGACTCATCAGGAAGC
TGACATCGGACTCATCAGGAAGC
CAGAGCATCGGACTCATCAGGAAGC
AGTCTCATCGGACTCATCAGGAAGC
CATCGGACTCATCAGGAAGC
GTCATCGGACTCATCAGGAAGC
ACATCGGACTCATCAGGAAGC
ATCATCGGACTCATCAGGAAGC
CATCGGACTCATCAGGAAGC
ATACATCGGACTCATCAGGAAGC
TCCATCGGACTCATCAGGAAGC
ACATCGGACTCATCAGGAAGC
GTTCATCGGACTCATCAGGAAGC
AACACATCGGACTCATCAGGAAGC
AGAGCATCGGACTCATCAGGAAGC
CCCATCGGACTCATCAGGAAGC
GCATCGGACTCATCAGGAAGC
GGCATCGGACTCATCAGGAAGC
TGGTCACATCGGACTCATCAGGAAGC
ATCATCGGACTCATCAGGAAGC
CTGTCATCGGACTCATCAGGAAGC
TCATCGGACTCATCAGGAAGC
ACATCGGACTCATCAGGAAGC
ACACATCGGACTCATCAGGAAGC
GATCATCGGACTCATCAGGAAGC
GTTGGCATCGGACTCATCAGGAAGC
TCTCATCGGACTCATCAGGAAGC

CTCATCGGACTCATCAGGAAGC
TGTCATCGGACTCATCAGGAAGC
GCATCGGACTCATCAGGAAGC
AAACATCGGACTCATCAGGAAGC
CTGCCATCGGACTCATCAGGAAGC
AAACATCGGACTCATCAGGAAGC
AGCATCGGACTCATCAGGAAGC
GGCATCGGACTCATCAGGAAGC
CAGCATCGGACTCATCAGGAAGC
GCCATCGGACTCATCAGGAAGC
CCCATCGGACTCATCAGGAAGC
GCATCGGACTCATCAGGAAGC
AGCATCGGACTCATCAGGAAGC
AGCCCATCGGACTCATCAGGAAGC
CTCATCGGACTCATCAGGAAGC
CATCATCGGACTCATCAGGAAGC
AAAGCCATCGGACTCATCAGGAAGC
TTGGTCATCGGACTCATCAGGAAGC
CAACATCGGACTCATCAGGAAGC
GTAAGCCATCGGACTCATCAGGAAGC
AGACATCGGACTCATCAGGAAGC
ACATCGGACTCATCAGGAAGC
CATCGGACTCATCAGGAAGC
TTCATCGGACTCATCAGGAAGC
GTTCATCGGACTCATCAGGAAGC
TCCCATCGGACTCATCAGGAAGC
GACATCGGACTCATCAGGAAGC
ACATCGGACTCATCAGGAAGC
GTCATCGGACTCATCAGGAAGC
GGACATCGGACTCATCAGGAAGC
TCATCGGACTCATCAGGAAGC
GCATCGGACTCATCAGGAAGC
TCATCGGACTCATCAGGAAGC
TTGCATCGGACTCATCAGGAAGC
AGAGCATCGGACTCATCAGGAAGC
TCATCGGACTCATCAGGAAGC
TCGGACTCATCAGGAAGC
TCACATCGGACTCATCAGGAAGC
GTAACATCGGACTCATCAGGAAGC
CATCGGACTCATCAGGAAGC
CATCGGACTCATCAGGAAGC
CACATCGGACTCATCAGGAAGC
GCACATCGGACTCATCAGGAAGC

TTTGCATCGGACTCATCAGGAAGC
TGACATCGGACTCATCAGGAAGC
CATCGGACTCATCAGGAAGC
CATCGGACTCATCAGGAAGC
CATCGGACTCATCAGGAAGC
TCCATCGGACTCATCAGGAAGC
TGGCATCGGACTCATCAGGAAGC
AGACCATCGGACTCATCAGGAAGC
CACATCGGACTCATCAGGAAGC
TTCATCGGACTCATCAGGAAGC
GACATCGGACTCATCAGGAAGC
AACTCATCGGACTCATCAGGAAGC
CTCATCGGACTCATCAGGAAGC
CACATCGGACTCATCAGGAAGC
CCATCGGACTCATCAGGAAGC
AGACACATCGGACTCATCAGGAAGC
GAACATCGGACTCATCAGGAAGC
CTACATCGGACTCATCAGGAAGC
TCATCGGACTCATCAGGAAGC
CTCATCGGACTCATCAGGAAGC
GGACATCGGACTCATCAGGAAGC
CTCATCGGACTCATCAGGAAGC
CACATCGGACTCATCAGGAAGC
GATCATCGGACTCATCAGGAAGC
GACATCGGACTCATCAGGAAGC
AAACATCGGACTCATCAGGAAGC
CTCATCGGACTCATCAGGAAGC
GGACTCATCAGGAAGC
CCATCGGACTCATCAGGAAGC
CTACATCGGACTCATCAGGAAGC
CTCATCGGACTCATCAGGAAGC
ATACATCGGACTCATCAGGAAGC
TCATCGGACTCATCAGGAAGC
CCATCGGACTCATCAGGAAGC
CAACATCGGACTCATCAGGAAGC
CATCGGACTCATCAGGAAGC
TGCCATCGGACTCATCAGGAAGC
GACATCGGACTCATCAGGAAGC
CCATCGGACTCATCAGGAAGC
ACATCGGACTCATCAGGAAGC
GCATCGGACTCATCAGGAAGC
AAGACCCATCGGACTCATCAGGAAGC
ACATCGGACTCATCAGGAAGC

CATCGGACTCATCAGGAAGC
GTACATCGGACTCATCAGGAAGC
TGGCCATCGGACTCATCAGGAAGC
TACATCGGACTCATCAGGAAGC
GATACATCGGACTCATCAGGAAGC
TCATCGGACTCATCAGGAAGC
CATCGGACTCATCAGGAAGC
CATCGGACTCATCAGGAAGC
AGCATCGGACTCATCAGGAAGC
CATCGGACTCATCAGGAAGC
CTCATCGGACTCATCAGGAAGC
CGGCATCGGACTCATCAGGAAGC
CATCGGACTCATCAGGAAGC
GGCATCGGACTCATCAGGAAGC
CATCGGACTCATCAGGAAGC
CATCGGACTCATCAGGAAGC
ATGGCATCGGACTCATCAGGAAGC
GCATCGGACTCATCAGGAAGC
GATACATCGGACTCATCAGGAAGC
TGCATCGGACTCATCAGGAAGC
CTTCATCGGACTCATCAGGAAGC
CAACATCGGACTCATCAGGAAGC
TTCATCGGACTCATCAGGAAGC
TCATCGGACTCATCAGGAAGC
CCACATCGGACTCATCAGGAAGC
GTGCATCGGACTCATCAGGAAGC
AAACATCGGACTCATCAGGAAGC
CATCGGACTCATCAGGAAGC
CACATCGGACTCATCAGGAAGC
CTCATCGGACTCATCAGGAAGC
ACACATCGGACTCATCAGGAAGC
CATCGGACTCATCAGGAAGC
GCCATCGGACTCATCAGGAAGC
TTGCATCGGACTCATCAGGAAGC
TTCATCGGACTCATCAGGAAGC
CCATCGGACTCATCAGGAAGC
TGTCATCGGACTCATCAGGAAGC
CTTGACATCGGACTCATCAGGAAGC
AAGACATCGGACTCATCAGGAAGC
CCCATCGGACTCATCAGGAAGC
CACATCGGACTCATCAGGAAGC
GGCATCGGACTCATCAGGAAGC
CATCGGACTCATCAGGAAGC

CACATCGGACTCATCAGGAAGC
CACATCGGACTCATCAGGAAGC
GCACATCGGACTCATCAGGAAGC
TTCATCGGACTCATCAGGAAGC
TCACATCGGACTCATCAGGAAGC
AAGAAACATCGGACTCATCAGGAAGC
CATCGGACTCATCAGGAAGC
CCCATCGGACTCATCAGGAAGC
CACATCGGACTCATCAGGAAGC
GGCCATCGGACTCATCAGGAAGC
GTGCATCGGACTCATCAGGAAGC
GATCATCGGACTCATCAGGAAGC
ATGTCATCGGACTCATCAGGAAGC
ACACATCGGACTCATCAGGAAGC
AATAGCCATCGGACTCATCAGGAAGC
AACATCGGACTCATCAGGAAGC
TGGGGCATCGGACTCATCAGGAAGC
CATCGGACTCATCAGGAAGC
GACATCGGACTCATCAGGAAGC
GTACATCGGACTCATCAGGAAGC
AAGCATCGGACTCATCAGGAAGC
TGGTCATCGGACTCATCAGGAAGC
TTCATCGGACTCATCAGGAAGC
TGATACATCGGACTCATCAGGAAGC
GAGCATCGGACTCATCAGGAAGC
TCATCGGACTCATCAGGAAGC
CGTCATCGGACTCATCAGGAAGC
AAATGCCATCGGACTCATCAGGAAGC
TCGGACTCATCAGGAAGC
ACATCGGACTCATCAGGAAGC
ATCATCGGACTCATCAGGAAGC
TCATCGGACTCATCAGGAAGC
ATCGGACTCATCAGGAAGC
AAACCCATCGGACTCATCAGGAAGC
CCTACATCGGACTCATCAGGAAGC
ACATCGGACTCATCAGGAAGC
TCGCATCGGACTCATCAGGAAGC
AACATCGGACTCATCAGGAAGC
AGGACACATCGGACTCATCAGGAAGC
TATTTCATCGGACTCATCAGGAAGC
TTACATCGGACTCATCAGGAAGC
CTGCATCGGACTCATCAGGAAGC
TCATCGGACTCATCAGGAAGC

CATCGGACTCATCAGGAAGC
CCATCGGACTCATCAGGAAGC
GGGTGACATCGGACTCATCAGGAAGC
GCATCGGACTCATCAGGAAGC
GCTCATCGGACTCATCAGGAAGC
GAACATCGGACTCATCAGGAAGC
GCATCGGACTCATCAGGAAGC
TTCATCGGACTCATCAGGAAGC
TCCATCGGACTCATCAGGAAGC
CTTCATCGGACTCATCAGGAAGC
GTCATCGGACTCATCAGGAAGC
ATCGGACTCATCAGGAAGC
TCCATCGGACTCATCAGGAAGC
ATCGGACTCATCAGGAAGC
CATCGGACTCATCAGGAAGC
ATCGGACTCATCAGGAAGC
AACATCGGACTCATCAGGAAGC
GCCACATCGGACTCATCAGGAAGC
TATTCATCGGACTCATCAGGAAGC
CCTCATCGGACTCATCAGGAAGC
GGACATCGGACTCATCAGGAAGC
TCATCGGACTCATCAGGAAGC
TGGGCATCGGACTCATCAGGAAGC
GACATCGGACTCATCAGGAAGC
ACATCGGACTCATCAGGAAGC
TCATCGGACTCATCAGGAAGC
TACCATCGGACTCATCAGGAAGC
CTTCATCGGACTCATCAGGAAGC
GACATCGGACTCATCAGGAAGC
GTCCATCGGACTCATCAGGAAGC
GAACATCGGACTCATCAGGAAGC
GGGCATCGGACTCATCAGGAAGC
ACATCGGACTCATCAGGAAGC
GACATCGGACTCATCAGGAAGC
CACATCGGACTCATCAGGAAGC
TCCATCGGACTCATCAGGAAGC
CATCGGACTCATCAGGAAGC
TATGCCATCGGACTCATCAGGAAGC
AAACCTCATCGGACTCATCAGGAAGC
GCCATCGGACTCATCAGGAAGC
CCCATCGGACTCATCAGGAAGC
ACATCGGACTCATCAGGAAGC
TTCACATCGGACTCATCAGGAAGC

GACATCGGACTCATCAGGAAGC
TCCCATCGGACTCATCAGGAAGC
TTCATCGGACTCATCAGGAAGC
GCCATCGGACTCATCAGGAAGC
GACATCGGACTCATCAGGAAGC
GGCATCGGACTCATCAGGAAGC
CGCATCGGACTCATCAGGAAGC
TCTACATCGGACTCATCAGGAAGC
TGCATCGGACTCATCAGGAAGC
ACATCGGACTCATCAGGAAGC
CCCATCGGACTCATCAGGAAGC
CCATCGGACTCATCAGGAAGC
AGGCATCGGACTCATCAGGAAGC
GCATCGGACTCATCAGGAAGC
TTCCATCGGACTCATCAGGAAGC
TCCACATCGGACTCATCAGGAAGC
CCATCGGACTCATCAGGAAGC
GCATCGGACTCATCAGGAAGC
AGAGCATCGGACTCATCAGGAAGC
ATCGGACTCATCAGGAAGC
AGGCATCGGACTCATCAGGAAGC
ATGCATCGGACTCATCAGGAAGC
TCTCCATCGGACTCATCAGGAAGC
AGCCATCGGACTCATCAGGAAGC
GGCACCATCGGACTCATCAGGAAGC
CACCCATCGGACTCATCAGGAAGC
ATCATCGGACTCATCAGGAAGC
TGGCATCGGACTCATCAGGAAGC
CACAGCATCGGACTCATCAGGAAGC
TGCCCATCGGACTCATCAGGAAGC
ATTACATCGGACTCATCAGGAAGC
TGCATCGGACTCATCAGGAAGC
GACATCGGACTCATCAGGAAGC
CAACATCGGACTCATCAGGAAGC
AACATCGGACTCATCAGGAAGC
TCATCGGACTCATCAGGAAGC
GTCATCGGACTCATCAGGAAGC
CTCATCGGACTCATCAGGAAGC
GGCCATCGGACTCATCAGGAAGC
TCCTCATCGGACTCATCAGGAAGC
ATCGGACTCATCAGGAAGC
GTTTCATCGGACTCATCAGGAAGC
GCCATCGGACTCATCAGGAAGC

TGAACATCGGACTCATCAGGAAGC
GTCATCGGACTCATCAGGAAGC
GCATCGGACTCATCAGGAAGC
CCTCATCGGACTCATCAGGAAGC
GAACATCGGACTCATCAGGAAGC
GTCATCGGACTCATCAGGAAGC
TTCATCGGACTCATCAGGAAGC
GTGAGCATCGGACTCATCAGGAAGC
GAAATGCATCGGACTCATCAGGAAGC
ACAGCATCGGACTCATCAGGAAGC
TAGACATCGGACTCATCAGGAAGC
ACCATCGGACTCATCAGGAAGC
TTCATCGGACTCATCAGGAAGC
AAGCCATCGGACTCATCAGGAAGC
CCACATCGGACTCATCAGGAAGC
GCCATCGGACTCATCAGGAAGC
CTCATCGGACTCATCAGGAAGC
CTTACCATCGGACTCATCAGGAAGC
TCATCGGACTCATCAGGAAGC
ACATCGGACTCATCAGGAAGC
ATCATCGGACTCATCAGGAAGC
GCATCGGACTCATCAGGAAGC
ACATCGGACTCATCAGGAAGC
ACACATCGGACTCATCAGGAAGC
CAGGCATCGGACTCATCAGGAAGC
ATCGGACTCATCAGGAAGC
CCCATCGGACTCATCAGGAAGC
GCATCGGACTCATCAGGAAGC
ATTCATCGGACTCATCAGGAAGC
ACATCGGACTCATCAGGAAGC
AATGCCATCGGACTCATCAGGAAGC
ACATCGGACTCATCAGGAAGC
AAGCATCGGACTCATCAGGAAGC
GCATCGGACTCATCAGGAAGC
TACATCGGACTCATCAGGAAGC
GGTTCATCGGACTCATCAGGAAGC
CACATCGGACTCATCAGGAAGC
GACATCGGACTCATCAGGAAGC
GAGCATCGGACTCATCAGGAAGC
AGCCCATCGGACTCATCAGGAAGC
CTCATCGGACTCATCAGGAAGC
ACCCATCGGACTCATCAGGAAGC
GACATCGGACTCATCAGGAAGC

CTGGCATCGGACTCATCAGGAAGC
TCATCGGACTCATCAGGAAGC
AAACATCGGACTCATCAGGAAGC
TCAGCATCGGACTCATCAGGAAGC
ACCCATCGGACTCATCAGGAAGC
TCATCGGACTCATCAGGAAGC
TCACATCGGACTCATCAGGAAGC
ACCATCGGACTCATCAGGAAGC
GGACATCGGACTCATCAGGAAGC
AGACATCGGACTCATCAGGAAGC
TGACCATCGGACTCATCAGGAAGC
CATCGGACTCATCAGGAAGC
CATCGGACTCATCAGGAAGC
CATCGGACTCATCAGGAAGC
GCAGCATCGGACTCATCAGGAAGC
GCATCGGACTCATCAGGAAGC
ACATCGGACTCATCAGGAAGC
ACATCGGACTCATCAGGAAGC
GTCATCGGACTCATCAGGAAGC
CTGCATCGGACTCATCAGGAAGC
CAATCATCGGACTCATCAGGAAGC
GCACATCGGACTCATCAGGAAGC
GGGGCATCGGACTCATCAGGAAGC
CAGACATCGGACTCATCAGGAAGC
CATCGGACTCATCAGGAAGC
AAACATCGGACTCATCAGGAAGC
GCATCGGACTCATCAGGAAGC
GTCATCGGACTCATCAGGAAGC
ATCGGACTCATCAGGAAGC
CTTCATCGGACTCATCAGGAAGC
GCATCGGACTCATCAGGAAGC
TGCATCGGACTCATCAGGAAGC
AAAGGTCATCGGACTCATCAGGAAGC
CCATCGGACTCATCAGGAAGC
AGCCCATCGGACTCATCAGGAAGC
CTCCATCGGACTCATCAGGAAGC
GGTCATCGGACTCATCAGGAAGC
TGGTCATCGGACTCATCAGGAAGC
ATTTCACATCGGACTCATCAGGAAGC
AGCATCGGACTCATCAGGAAGC
ATCGGACTCATCAGGAAGC
ACATCGGACTCATCAGGAAGC
TGCATCGGACTCATCAGGAAGC

TAGCACATCGGACTCATCAGGAAGC
CACCATCGGACTCATCAGGAAGC
TGACCCCATCGGACTCATCAGGAAGC
CCATCGGACTCATCAGGAAGC
TTCATCGGACTCATCAGGAAGC
ACATCGGACTCATCAGGAAGC
AAACCCATCGGACTCATCAGGAAGC
CATCGGACTCATCAGGAAGC
ACCATCGGACTCATCAGGAAGC
AAATGCATCGGACTCATCAGGAAGC
GTCATCGGACTCATCAGGAAGC
ATCGGACTCATCAGGAAGC
GGTCATCGGACTCATCAGGAAGC
ACATCGGACTCATCAGGAAGC
GGACATCGGACTCATCAGGAAGC
CTAGACATCGGACTCATCAGGAAGC
CATCGGACTCATCAGGAAGC
ATCGGACTCATCAGGAAGC
ATCATCGGACTCATCAGGAAGC
ATCATCGGACTCATCAGGAAGC
CCATCGGACTCATCAGGAAGC
TCATCGGACTCATCAGGAAGC
ATCGGACTCATCAGGAAGC
TGTGCCATCGGACTCATCAGGAAGC
GGCATCGGACTCATCAGGAAGC
GCATCGGACTCATCAGGAAGC
CGCATCGGACTCATCAGGAAGC
TTCATCGGACTCATCAGGAAGC
GCATCGGACTCATCAGGAAGC
GCATCGGACTCATCAGGAAGC
TAGCATCGGACTCATCAGGAAGC
TCATCGGACTCATCAGGAAGC
TGGCATCGGACTCATCAGGAAGC
CATCGGACTCATCAGGAAGC
GTCATCGGACTCATCAGGAAGC
CACATCGGACTCATCAGGAAGC
AACATCGGACTCATCAGGAAGC
CATCGGACTCATCAGGAAGC
CTCACATCGGACTCATCAGGAAGC
TTCATCGGACTCATCAGGAAGC
GCTTACCATCGGACTCATCAGGAAGC
TCATCGGACTCATCAGGAAGC
CCATCGGACTCATCAGGAAGC

ACATCGGACTCATCAGGAAGC
GCATCGGACTCATCAGGAAGC
ATACATCGGACTCATCAGGAAGC
GTTCATCGGACTCATCAGGAAGC
GTGCATCGGACTCATCAGGAAGC
GCGACATCGGACTCATCAGGAAGC
GGGCATCGGACTCATCAGGAAGC
GCCATCGGACTCATCAGGAAGC
CCCTCATCGGACTCATCAGGAAGC
GACATCGGACTCATCAGGAAGC
GGATCATCGGACTCATCAGGAAGC
ATCGGACTCATCAGGAAGC
TTATCATCGGACTCATCAGGAAGC
CTCCATCGGACTCATCAGGAAGC
CACCATCGGACTCATCAGGAAGC
GTCCATCGGACTCATCAGGAAGC
CCATCGGACTCATCAGGAAGC
CATCGGACTCATCAGGAAGC
TGACATCGGACTCATCAGGAAGC
CTCATCGGACTCATCAGGAAGC
TACATCGGACTCATCAGGAAGC
CCATCGGACTCATCAGGAAGC
GTCCCCATCGGACTCATCAGGAAGC
ACACATCGGACTCATCAGGAAGC
AGCCATCGGACTCATCAGGAAGC
CCATCATCGGACTCATCAGGAAGC
TCTCATCGGACTCATCAGGAAGC
AATCATCGGACTCATCAGGAAGC
TCACATCGGACTCATCAGGAAGC
ATCGGACTCATCAGGAAGC
GGCATCGGACTCATCAGGAAGC
CTTGCATCGGACTCATCAGGAAGC
CACATCGGACTCATCAGGAAGC
CCATCGGACTCATCAGGAAGC
CCACATCGGACTCATCAGGAAGC
TAAACATCGGACTCATCAGGAAGC
GCATCGGACTCATCAGGAAGC
GTCATCGGACTCATCAGGAAGC
CGCATCGGACTCATCAGGAAGC
CATCGGACTCATCAGGAAGC
GGCATCGGACTCATCAGGAAGC
GCATCGGACTCATCAGGAAGC
CTCATCGGACTCATCAGGAAGC

CCCACATCGGACTCATCAGGAAGC
TGTCATCGGACTCATCAGGAAGC
CATCGGACTCATCAGGAAGC
TCATCGGACTCATCAGGAAGC
CCATCGGACTCATCAGGAAGC
AACATCGGACTCATCAGGAAGC
CTTCATCGGACTCATCAGGAAGC
GAACATCGGACTCATCAGGAAGC
GAACATCGGACTCATCAGGAAGC
ATCCCTCATCGGACTCATCAGGAAGC
GGCACATCGGACTCATCAGGAAGC
GACATCGGACTCATCAGGAAGC
GGCATCGGACTCATCAGGAAGC
AGCATCGGACTCATCAGGAAGC
TTCATCGGACTCATCAGGAAGC
AAAACATCGGACTCATCAGGAAGC
GTCATCGGACTCATCAGGAAGC
CTGTCATCGGACTCATCAGGAAGC
AAACATCGGACTCATCAGGAAGC
CTATCATCGGACTCATCAGGAAGC
GACATCGGACTCATCAGGAAGC
CCAGGCATCGGACTCATCAGGAAGC
TTCATCGGACTCATCAGGAAGC
CATCGGACTCATCAGGAAGC
AGAACATCGGACTCATCAGGAAGC
CCTCATCGGACTCATCAGGAAGC
AACATCGGACTCATCAGGAAGC
CTTCATCGGACTCATCAGGAAGC
ACATCGGACTCATCAGGAAGC
TCACATCGGACTCATCAGGAAGC
CCATCGGACTCATCAGGAAGC
GGTCATCGGACTCATCAGGAAGC
CATCGGACTCATCAGGAAGC
GAGCATCGGACTCATCAGGAAGC
GTGGCATCGGACTCATCAGGAAGC
ACCTCATCGGACTCATCAGGAAGC
TCCCATCGGACTCATCAGGAAGC
CCATCGGACTCATCAGGAAGC
GCATCGGACTCATCAGGAAGC
TGCATCGGACTCATCAGGAAGC
AGACATCGGACTCATCAGGAAGC
ATCGGACTCATCAGGAAGC
ATGGGGCATCGGACTCATCAGGAAGC

CATCGGACTCATCAGGAAGC
GAGAATCATCGGACTCATCAGGAAGC
CACATCGGACTCATCAGGAAGC
TGACATCGGACTCATCAGGAAGC
GTGCATCGGACTCATCAGGAAGC
GGGCATCGGACTCATCAGGAAGC
CCATCGGACTCATCAGGAAGC
GCTGCATCGGACTCATCAGGAAGC
GCCATCGGACTCATCAGGAAGC
ACATCGGACTCATCAGGAAGC
TCCATCGGACTCATCAGGAAGC
CTCATCGGACTCATCAGGAAGC
TCATCGGACTCATCAGGAAGC
ACCCATCGGACTCATCAGGAAGC
AAAACATCGGACTCATCAGGAAGC
TCCCTCATCGGACTCATCAGGAAGC
ACATCGGACTCATCAGGAAGC
CCATCGGACTCATCAGGAAGC
CTTGCATCGGACTCATCAGGAAGC
GCCATCGGACTCATCAGGAAGC
CGGACTCATCAGGAAGC
TGTCATCGGACTCATCAGGAAGC
TCCATCGGACTCATCAGGAAGC
TTGTCATCGGACTCATCAGGAAGC
TCACATCGGACTCATCAGGAAGC
CTCATCGGACTCATCAGGAAGC
TCATCGGACTCATCAGGAAGC
GAACATCGGACTCATCAGGAAGC
TCATGTCATCGGACTCATCAGGAAGC
GGTGCTCATCGGACTCATCAGGAAGC
TTTCATCGGACTCATCAGGAAGC
CATCGGACTCATCAGGAAGC
GCATCGGACTCATCAGGAAGC
ACATCGGACTCATCAGGAAGC
GCCTCATCGGACTCATCAGGAAGC
TTCATCGGACTCATCAGGAAGC
ATCGGACTCATCAGGAAGC
ATGGCATCGGACTCATCAGGAAGC
ACAGCACATCGGACTCATCAGGAAGC
CATCGGACTCATCAGGAAGC
TGGACATCGGACTCATCAGGAAGC
AGAGGCATCGGACTCATCAGGAAGC
AGGCATCGGACTCATCAGGAAGC

GGGCATCGGACTCATCAGGAAGC
CTTCATCGGACTCATCAGGAAGC
ACACATCGGACTCATCAGGAAGC
CTCATCGGACTCATCAGGAAGC
TTCATCGGACTCATCAGGAAGC
AGTCATCGGACTCATCAGGAAGC
TCATCGGACTCATCAGGAAGC
ACTGCATCGGACTCATCAGGAAGC
CAAAGCATCGGACTCATCAGGAAGC
CCATCGGACTCATCAGGAAGC
AACATCGGACTCATCAGGAAGC
TTCATCGGACTCATCAGGAAGC
TCATCGGACTCATCAGGAAGC
CCCATCGGACTCATCAGGAAGC
CATCGGACTCATCAGGAAGC
CGACACATCGGACTCATCAGGAAGC
AACATCGGACTCATCAGGAAGC
CATCGGACTCATCAGGAAGC
GGCATCGGACTCATCAGGAAGC
CCATGTCATCGGACTCATCAGGAAGC
ATCATCGGACTCATCAGGAAGC
AGGCATCGGACTCATCAGGAAGC
TGTAACATCGGACTCATCAGGAAGC
CATTTCATCGGACTCATCAGGAAGC
AACATCGGACTCATCAGGAAGC
TAACTCATCGGACTCATCAGGAAGC
TCATCGGACTCATCAGGAAGC
CTAAGCATCGGACTCATCAGGAAGC
GCCATCGGACTCATCAGGAAGC
GACATCGGACTCATCAGGAAGC
ATACATCGGACTCATCAGGAAGC
CCTCCATCGGACTCATCAGGAAGC
CATCGGACTCATCAGGAAGC
GACATCGGACTCATCAGGAAGC
ATAGACATCGGACTCATCAGGAAGC
AACATCGGACTCATCAGGAAGC
CATCGGACTCATCAGGAAGC
CATCGGACTCATCAGGAAGC
GCATCGGACTCATCAGGAAGC
ATCATCGGACTCATCAGGAAGC
ATTCATCGGACTCATCAGGAAGC
GCATCGGACTCATCAGGAAGC
GGAGCATCGGACTCATCAGGAAGC

CTCATCGGACTCATCAGGAAGC
TGTGTGCATCGGACTCATCAGGAAGC
GGGCATCGGACTCATCAGGAAGC
CCATCGGACTCATCAGGAAGC
TCATCGGACTCATCAGGAAGC
GCATCGGACTCATCAGGAAGC
GTGTCCATCGGACTCATCAGGAAGC
CATCGGACTCATCAGGAAGC
CCCATCGGACTCATCAGGAAGC
GGAACATCGGACTCATCAGGAAGC
TCATCGGACTCATCAGGAAGC
CCATCGGACTCATCAGGAAGC
AGGGCATCGGACTCATCAGGAAGC
GGGCCATCGGACTCATCAGGAAGC
GGGTCATCGGACTCATCAGGAAGC
CTCAACATCGGACTCATCAGGAAGC
CACCATCGGACTCATCAGGAAGC
CACATCGGACTCATCAGGAAGC
TCATCGGACTCATCAGGAAGC
GCATCGGACTCATCAGGAAGC
GGGGCATCGGACTCATCAGGAAGC
GGAGGCATCGGACTCATCAGGAAGC
TGCATCGGACTCATCAGGAAGC
TACATCGGACTCATCAGGAAGC
AAGCATCGGACTCATCAGGAAGC
CATCGGACTCATCAGGAAGC
CATCGGACTCATCAGGAAGC
CATCGGACTCATCAGGAAGC
GGCCATCGGACTCATCAGGAAGC
GACATCGGACTCATCAGGAAGC
GCCACATCGGACTCATCAGGAAGC
TTCATCGGACTCATCAGGAAGC
GGCTCATCGGACTCATCAGGAAGC
AGACATCGGACTCATCAGGAAGC
AAACATCGGACTCATCAGGAAGC
ATCGGACTCATCAGGAAGC
GAGCCATCGGACTCATCAGGAAGC
AACCATCGGACTCATCAGGAAGC
ATGCCATCGGACTCATCAGGAAGC
AAATCGCATCGGACTCATCAGGAAGC
TGCCCATCGGACTCATCAGGAAGC
CATCGGACTCATCAGGAAGC
CCATCGGACTCATCAGGAAGC

AGCATCGGACTCATCAGGAAGC
CCATGACATCGGACTCATCAGGAAGC
TCATCGGACTCATCAGGAAGC
TCATCGGACTCATCAGGAAGC
AACCCATCGGACTCATCAGGAAGC
TGTCATCGGACTCATCAGGAAGC
CCATCGGACTCATCAGGAAGC
GGGCATCGGACTCATCAGGAAGC
CCATCATCGGACTCATCAGGAAGC
AAGCATCGGACTCATCAGGAAGC
CATGTCATCGGACTCATCAGGAAGC
AGGCATCGGACTCATCAGGAAGC
GGCATCGGACTCATCAGGAAGC
GGCCATCGGACTCATCAGGAAGC
AGGCATCGGACTCATCAGGAAGC
CCATCGGACTCATCAGGAAGC
GGACATCGGACTCATCAGGAAGC
CATCGGACTCATCAGGAAGC
ATCGGACTCATCAGGAAGC
CGGACTCATCAGGAAGC
GAGACATCGGACTCATCAGGAAGC
ACAGCCCCATCGGACTCATCAGGAAGC
ATCGGACTCATCAGGAAGC
TGCATCGGACTCATCAGGAAGC
ATCGGACTCATCAGGAAGC
TCCCCATCGGACTCATCAGGAAGC
AGTGACACATCGGACTCATCAGGAAGC
TCCATCGGACTCATCAGGAAGC
TCGGACTCATCAGGAAGC
TTCATCGGACTCATCAGGAAGC
AACATCGGACTCATCAGGAAGC
TGAGCATCGGACTCATCAGGAAGC
ATTCATCGGACTCATCAGGAAGC
CCATCGGACTCATCAGGAAGC
GGAACATCGGACTCATCAGGAAGC
AACATCGGACTCATCAGGAAGC
ATACATCGGACTCATCAGGAAGC
ATCGGACTCATCAGGAAGC
TACACATCGGACTCATCAGGAAGC
CGCATCGGACTCATCAGGAAGC
TCGGACTCATCAGGAAGC
GTCATCGGACTCATCAGGAAGC
CACATCGGACTCATCAGGAAGC

CATCGGACTCATCAGGAAGC
CAGACATCGGACTCATCAGGAAGC
GACATCGGACTCATCAGGAAGC
GCATCGGACTCATCAGGAAGC
CGGACTCATCAGGAAGC
AGGCTGCATCGGACTCATCAGGAAGC
TCATCGGACTCATCAGGAAGC
TGACATCGGACTCATCAGGAAGC
GAACTCATCGGACTCATCAGGAAGC
TGCATCGGACTCATCAGGAAGC
CAGGCATCGGACTCATCAGGAAGC
TACCATCGGACTCATCAGGAAGC
CATCGGACTCATCAGGAAGC
TTTTCATCGGACTCATCAGGAAGC
TCATCGGACTCATCAGGAAGC
ATCATCGGACTCATCAGGAAGC
ATTTGCATCGGACTCATCAGGAAGC
CATCGGACTCATCAGGAAGC
AATCATCGGACTCATCAGGAAGC
GCATCGGACTCATCAGGAAGC
TCGCATCGGACTCATCAGGAAGC
ACATCGGACTCATCAGGAAGC
CTGCATCGGACTCATCAGGAAGC
TCCCATCGGACTCATCAGGAAGC
AATGACATCGGACTCATCAGGAAGC
TGCCATCGGACTCATCAGGAAGC
TCGGACTCATCAGGAAGC
ATCGGACTCATCAGGAAGC
ACTACATCGGACTCATCAGGAAGC
AGACATCGGACTCATCAGGAAGC
TGAGCATCGGACTCATCAGGAAGC
TGGGACATCGGACTCATCAGGAAGC
CATCGGACTCATCAGGAAGC
ATCGGACTCATCAGGAAGC
CTCCATCGGACTCATCAGGAAGC
GGTACATCGGACTCATCAGGAAGC
CACATCGGACTCATCAGGAAGC
CCATCGGACTCATCAGGAAGC
ACCACCATCGGACTCATCAGGAAGC
TCATCGGACTCATCAGGAAGC
TGCCCCATCGGACTCATCAGGAAGC
AACATCGGACTCATCAGGAAGC
ATCGGACTCATCAGGAAGC

GGCATCGGACTCATCAGGAAGC
TCATCGGACTCATCAGGAAGC
CAGCATCGGACTCATCAGGAAGC
GGGACATCGGACTCATCAGGAAGC
GGCATCGGACTCATCAGGAAGC
ACCATCGGACTCATCAGGAAGC
AAGATCATCGGACTCATCAGGAAGC
CCATCGGACTCATCAGGAAGC
CATCGGACTCATCAGGAAGC
GAGCATCGGACTCATCAGGAAGC
CATCGGACTCATCAGGAAGC
ACATCGGACTCATCAGGAAGC
TGAGCATCGGACTCATCAGGAAGC
GTTGCAGCATCGGACTCATCAGGAAGC
CATCGGACTCATCAGGAAGC
ACACATCGGACTCATCAGGAAGC
CATCGGACTCATCAGGAAGC
ACATCGGACTCATCAGGAAGC
AGCATCGGACTCATCAGGAAGC
CCGTTCCATCGGACTCATCAGGAAGC
CTCACATCGGACTCATCAGGAAGC
GCATCGGACTCATCAGGAAGC
CAGTTCATCGGACTCATCAGGAAGC
CCATCGGACTCATCAGGAAGC
GCATCGGACTCATCAGGAAGC
CATCGGACTCATCAGGAAGC
GACATCGGACTCATCAGGAAGC
GAGGCATCGGACTCATCAGGAAGC
CTCCATCGGACTCATCAGGAAGC
GCCCCATCGGACTCATCAGGAAGC
ATCGGACTCATCAGGAAGC
GTGCATCGGACTCATCAGGAAGC
CATCGGACTCATCAGGAAGC
CGGACTCATCAGGAAGC
GTACGCATCGGACTCATCAGGAAGC
TCAATCATCGGACTCATCAGGAAGC
GACATCGGACTCATCAGGAAGC
CCATCGGACTCATCAGGAAGC
CCACATCGGACTCATCAGGAAGC
CATCGGACTCATCAGGAAGC
ACTTTCATCGGACTCATCAGGAAGC
GTCATCGGACTCATCAGGAAGC
TCGGACTCATCAGGAAGC

TCGGACTCATCAGGAAGC
CATCGGACTCATCAGGAAGC
TACCATCGGACTCATCAGGAAGC
GCATCGGACTCATCAGGAAGC
TTTCATCGGACTCATCAGGAAGC
ATCGGACTCATCAGGAAGC
TCGGACTCATCAGGAAGC
CATCGGACTCATCAGGAAGC
ACATCGGACTCATCAGGAAGC
TTCATCGGACTCATCAGGAAGC
ACATCGGACTCATCAGGAAGC
AGCATCGGACTCATCAGGAAGC
ACCCATCGGACTCATCAGGAAGC
GCATCGGACTCATCAGGAAGC
CCATCGGACTCATCAGGAAGC
TCATCGGACTCATCAGGAAGC
ATCGGACTCATCAGGAAGC
CATCGGACTCATCAGGAAGC
GTCCATCGGACTCATCAGGAAGC
CATCGGACTCATCAGGAAGC
AGCATCGGACTCATCAGGAAGC
AAAGTTCATCGGACTCATCAGGAAGC
ACCTCATCGGACTCATCAGGAAGC
GACATCGGACTCATCAGGAAGC
GCATCGGACTCATCAGGAAGC
CATCGGACTCATCAGGAAGC
CCAGAGCATCGGACTCATCAGGAAGC
CCACATCGGACTCATCAGGAAGC
ACTCATCAGGAAGC
CATCGGACTCATCAGGAAGC
GACATCGGACTCATCAGGAAGC
GACATCGGACTCATCAGGAAGC
TTTCATCGGACTCATCAGGAAGC
ATAATCATCGGACTCATCAGGAAGC
TGCATCGGACTCATCAGGAAGC
TTTACATCGGACTCATCAGGAAGC
GGCATCGGACTCATCAGGAAGC
TCATCGGACTCATCAGGAAGC
TGACATCGGACTCATCAGGAAGC
ATCGGACTCATCAGGAAGC
ACATCGGACTCATCAGGAAGC
GCATCGGACTCATCAGGAAGC
TCGGACTCATCAGGAAGC

GGCATCGGACTCATCAGGAAGC
TACATCGGACTCATCAGGAAGC
CCCATCGGACTCATCAGGAAGC
GACATCGGACTCATCAGGAAGC
TACATCGGACTCATCAGGAAGC
CCATCGGACTCATCAGGAAGC
CATCGGACTCATCAGGAAGC
TTCATCGGACTCATCAGGAAGC
CATCGGACTCATCAGGAAGC
CATCGGACTCATCAGGAAGC
GTATCATCGGACTCATCAGGAAGC
CATCGGACTCATCAGGAAGC
AACCATCGGACTCATCAGGAAGC
CATCATCGGACTCATCAGGAAGC
CCCATCGGACTCATCAGGAAGC
TCATCGGACTCATCAGGAAGC
AACTCATCGGACTCATCAGGAAGC
AGGTCATCGGACTCATCAGGAAGC
CATCGGACTCATCAGGAAGC
TCATCGGACTCATCAGGAAGC
AGCCATCGGACTCATCAGGAAGC
AGCATCGGACTCATCAGGAAGC
ACCAACATCGGACTCATCAGGAAGC
CGCATCGGACTCATCAGGAAGC
TCATCGGACTCATCAGGAAGC
GTTTCATCGGACTCATCAGGAAGC
TCGGACTCATCAGGAAGC
CATCATCGGACTCATCAGGAAGC
CCATCGGACTCATCAGGAAGC
GCATCGGACTCATCAGGAAGC
AGGACATCGGACTCATCAGGAAGC
TGCATCGGACTCATCAGGAAGC
AAGCATCGGACTCATCAGGAAGC
GACATCGGACTCATCAGGAAGC
ACCATCGGACTCATCAGGAAGC
GGCCATCGGACTCATCAGGAAGC
GTTGCCATCGGACTCATCAGGAAGC
ATTCCACATCGGACTCATCAGGAAGC
ACATCGGACTCATCAGGAAGC
CTGGCATCGGACTCATCAGGAAGC
CAAACATCGGACTCATCAGGAAGC
AGACATCGGACTCATCAGGAAGC
GCCATCGGACTCATCAGGAAGC

CTTCATCGGACTCATCAGGAAGC
CAATACATCGGACTCATCAGGAAGC
TCATCGGACTCATCAGGAAGC
ACATCGGACTCATCAGGAAGC
GAGGCATCGGACTCATCAGGAAGC
TCATCGGACTCATCAGGAAGC
GTCATCGGACTCATCAGGAAGC
CCCATCGGACTCATCAGGAAGC
CATCGGACTCATCAGGAAGC
ATTCATCGGACTCATCAGGAAGC
TCATCGGACTCATCAGGAAGC
CAGCATCGGACTCATCAGGAAGC
CATCGGACTCATCAGGAAGC
ATCGGACTCATCAGGAAGC
CCATCGGACTCATCAGGAAGC
CTCCATCGGACTCATCAGGAAGC
GCCATCGGACTCATCAGGAAGC
GACCATCGGACTCATCAGGAAGC
CTACATCGGACTCATCAGGAAGC
AACATCGGACTCATCAGGAAGC
TGTCATCGGACTCATCAGGAAGC
AGGCATCGGACTCATCAGGAAGC
TGTCATCGGACTCATCAGGAAGC
ATCGGACTCATCAGGAAGC
GACCATCGGACTCATCAGGAAGC
CCATCGGACTCATCAGGAAGC
ACCATCGGACTCATCAGGAAGC
TCATCGGACTCATCAGGAAGC
ACCATCGGACTCATCAGGAAGC
GACATCGGACTCATCAGGAAGC
TGCCATCGGACTCATCAGGAAGC
CCATCGGACTCATCAGGAAGC
CCATCGGACTCATCAGGAAGC
CTCATCGGACTCATCAGGAAGC
TCATCGGACTCATCAGGAAGC
GCCATCGGACTCATCAGGAAGC
GCATCGGACTCATCAGGAAGC
AATCATCGGACTCATCAGGAAGC
CTGCATCGGACTCATCAGGAAGC
GGCACATCGGACTCATCAGGAAGC
CTCATCGGACTCATCAGGAAGC
CGACATCGGACTCATCAGGAAGC
CATCGGACTCATCAGGAAGC

TACATCGGACTCATCAGGAAGC
TCCATCGGACTCATCAGGAAGC
GACATCGGACTCATCAGGAAGC
CCATCGGACTCATCAGGAAGC
TCCATCGGACTCATCAGGAAGC
TCATCGGACTCATCAGGAAGC
AAACATCGGACTCATCAGGAAGC
ACATCGGACTCATCAGGAAGC
CCATCGGACTCATCAGGAAGC
GGACACATCGGACTCATCAGGAAGC
GTCTTCATCGGACTCATCAGGAAGC
TCCCATCGGACTCATCAGGAAGC
CCCATCGGACTCATCAGGAAGC
CACATCGGACTCATCAGGAAGC
CACATCGGACTCATCAGGAAGC
AACATCGGACTCATCAGGAAGC
GAACATCGGACTCATCAGGAAGC
CATCATCGGACTCATCAGGAAGC
CTCATCGGACTCATCAGGAAGC
CTCATCGGACTCATCAGGAAGC
GGACTCATCAGGAAGC
TCCATCGGACTCATCAGGAAGC
GAACATCGGACTCATCAGGAAGC
TCCATCGGACTCATCAGGAAGC
ATCATCGGACTCATCAGGAAGC
CACATCGGACTCATCAGGAAGC
CATCATCGGACTCATCAGGAAGC
TCATCGGACTCATCAGGAAGC
GACATCGGACTCATCAGGAAGC
GCATCGGACTCATCAGGAAGC
CCACATCGGACTCATCAGGAAGC
TCTCATCGGACTCATCAGGAAGC
GACATCGGACTCATCAGGAAGC
GCCATCGGACTCATCAGGAAGC
CTTCATCGGACTCATCAGGAAGC
ACCATCGGACTCATCAGGAAGC
ACACATCGGACTCATCAGGAAGC
ACCATCGGACTCATCAGGAAGC
GTTCATCGGACTCATCAGGAAGC
CATCGGACTCATCAGGAAGC
CCATCGGACTCATCAGGAAGC
ACCATCGGACTCATCAGGAAGC
AGGCATCGGACTCATCAGGAAGC

ACGCATCGGACTCATCAGGAAGC
ATCGGACTCATCAGGAAGC
CTCATCGGACTCATCAGGAAGC
TCAACCTCATCGGACTCATCAGGAAGC
ATCGGACTCATCAGGAAGC
TTCATCGGACTCATCAGGAAGC
TGACATCGGACTCATCAGGAAGC
CATCGGACTCATCAGGAAGC
GACTCATCAGGAAGC
ACATCGGACTCATCAGGAAGC
ACTCATCGGACTCATCAGGAAGC
ACACATCGGACTCATCAGGAAGC
CCATCGGACTCATCAGGAAGC
ACATCGGACTCATCAGGAAGC
TCACATCGGACTCATCAGGAAGC
TCCATCGGACTCATCAGGAAGC
CCATCGGACTCATCAGGAAGC
ATATGCGCATCGGACTCATCAGGAAGC
CCATCGGACTCATCAGGAAGC
GATACATCGGACTCATCAGGAAGC
ATCATCGGACTCATCAGGAAGC
CCATCGGACTCATCAGGAAGC
GCCCATCGGACTCATCAGGAAGC
TGGCATCGGACTCATCAGGAAGC
ATCGGACTCATCAGGAAGC
GTCATCGGACTCATCAGGAAGC
CATCGGACTCATCAGGAAGC
TAGCATCGGACTCATCAGGAAGC
AGATACATCGGACTCATCAGGAAGC
ACTCATCGGACTCATCAGGAAGC
CATCGGACTCATCAGGAAGC
AAACATCGGACTCATCAGGAAGC
ACATCGGACTCATCAGGAAGC
ATAACATCGGACTCATCAGGAAGC
AACATCGGACTCATCAGGAAGC
CCATCGGACTCATCAGGAAGC
GCATCGGACTCATCAGGAAGC
CATCGGACTCATCAGGAAGC
TGCCATCGGACTCATCAGGAAGC
CATCGGACTCATCAGGAAGC
CATCGGACTCATCAGGAAGC
TATCATCGGACTCATCAGGAAGC
TCATCGGACTCATCAGGAAGC

CCCATCGGACTCATCAGGAAGC
GGCATCGGACTCATCAGGAAGC
CATCGGACTCATCAGGAAGC
AGACATCGGACTCATCAGGAAGC
GGCATCGGACTCATCAGGAAGC
CCATCGGACTCATCAGGAAGC
AATCATCGGACTCATCAGGAAGC
AGGCATCGGACTCATCAGGAAGC
ACCCATCGGACTCATCAGGAAGC
AACATCGGACTCATCAGGAAGC
CATCGGACTCATCAGGAAGC
GCCCATCGGACTCATCAGGAAGC
GACATCGGACTCATCAGGAAGC
CGATCATCGGACTCATCAGGAAGC
CCATCGGACTCATCAGGAAGC
CCATCCATCGGACTCATCAGGAAGC
TTCATCGGACTCATCAGGAAGC
CAGACATCGGACTCATCAGGAAGC
CTCATCGGACTCATCAGGAAGC
GCCATCGGACTCATCAGGAAGC
CCATCGGACTCATCAGGAAGC
TACGCATCGGACTCATCAGGAAGC
ACTCATCGGACTCATCAGGAAGC
AGCAGACATCGGACTCATCAGGAAGC
CATCGGACTCATCAGGAAGC
CATCGGACTCATCAGGAAGC
GCATCGGACTCATCAGGAAGC
GCCATCGGACTCATCAGGAAGC
ACAGCATCGGACTCATCAGGAAGC
ATCCTCATCGGACTCATCAGGAAGC
ATCGGACTCATCAGGAAGC
ATCGGACTCATCAGGAAGC
GCATCGGACTCATCAGGAAGC
CAAGCATCGGACTCATCAGGAAGC
CCATCGGACTCATCAGGAAGC
GCATCGGACTCATCAGGAAGC
GCTGCATCGGACTCATCAGGAAGC
CCCATCGGACTCATCAGGAAGC
TGCATCGGACTCATCAGGAAGC
AAAACACATCGGACTCATCAGGAAGC
CATCGGACTCATCAGGAAGC
CCATCATCGGACTCATCAGGAAGC
AAGCATCGGACTCATCAGGAAGC

CATCGGACTCATCAGGAAGC
CCACATCGGACTCATCAGGAAGC
AGCATCGGACTCATCAGGAAGC
ACATCGGACTCATCAGGAAGC
GTCATCGGACTCATCAGGAAGC
ATCATCGGACTCATCAGGAAGC
CCATCGGACTCATCAGGAAGC
AAGGCATCGGACTCATCAGGAAGC
CTCATCGGACTCATCAGGAAGC
TCATCGGACTCATCAGGAAGC
ATCGGACTCATCAGGAAGC
CGCATCGGACTCATCAGGAAGC
TCATCGGACTCATCAGGAAGC
ATTTCATCGGACTCATCAGGAAGC
CATCGGACTCATCAGGAAGC
GGACATCGGACTCATCAGGAAGC
CCATCGGACTCATCAGGAAGC
TCCATCGGACTCATCAGGAAGC
CCTATCATCGGACTCATCAGGAAGC
ACATCGGACTCATCAGGAAGC
CCTCATCGGACTCATCAGGAAGC
GGCATCGGACTCATCAGGAAGC
AGTGGCCATCGGACTCATCAGGAAGC
ATGAGCATCGGACTCATCAGGAAGC
ATCATCGGACTCATCAGGAAGC
AACATCGGACTCATCAGGAAGC
GGGCATCGGACTCATCAGGAAGC
TCATCGGACTCATCAGGAAGC
GCATCGGACTCATCAGGAAGC
GGCATCGGACTCATCAGGAAGC
GACCATCGGACTCATCAGGAAGC
TGACATCGGACTCATCAGGAAGC
CCTCATCGGACTCATCAGGAAGC
GTACATCGGACTCATCAGGAAGC
GCACATCGGACTCATCAGGAAGC
GCCATCGGACTCATCAGGAAGC
ACATCGGACTCATCAGGAAGC
TCATCGGACTCATCAGGAAGC
TGCATCGGACTCATCAGGAAGC
CACCATCGGACTCATCAGGAAGC
ACAACATCGGACTCATCAGGAAGC
TCATCGGACTCATCAGGAAGC
AAGACATCGGACTCATCAGGAAGC

GAGAACCATCGGACTCATCAGGAAGC
TCCATCGGACTCATCAGGAAGC
TAACATCGGACTCATCAGGAAGC
CATCGGACTCATCAGGAAGC
AGCCATCGGACTCATCAGGAAGC
ACATCGGACTCATCAGGAAGC
TCATCGGACTCATCAGGAAGC
GACGCATCGGACTCATCAGGAAGC
CCATCGGACTCATCAGGAAGC
ATCGGACTCATCAGGAAGC
GGCATCGGACTCATCAGGAAGC
AGTCATCGGACTCATCAGGAAGC
ACTCATCGGACTCATCAGGAAGC
ACAACATCGGACTCATCAGGAAGC
ATCATCGGACTCATCAGGAAGC
CCACATCGGACTCATCAGGAAGC
CCCATCGGACTCATCAGGAAGC
GCCATCGGACTCATCAGGAAGC
CATCGGACTCATCAGGAAGC
GCCATCGGACTCATCAGGAAGC
TCGGACTCATCAGGAAGC
CATCGGACTCATCAGGAAGC
AGGCCATCGGACTCATCAGGAAGC
CATCGGACTCATCAGGAAGC
ATCCATCGGACTCATCAGGAAGC
GCTGGTCATCGGACTCATCAGGAAGC
AGAGCATCGGACTCATCAGGAAGC
GGCATCGGACTCATCAGGAAGC
CCATCGGACTCATCAGGAAGC
ATCGGACTCATCAGGAAGC
TCCATCGGACTCATCAGGAAGC
TCTGCATCGGACTCATCAGGAAGC
ACACATCGGACTCATCAGGAAGC
TGTGTCATCGGACTCATCAGGAAGC
GACATCGGACTCATCAGGAAGC
TGCATCGGACTCATCAGGAAGC
TCATCGGACTCATCAGGAAGC
ACATCGGACTCATCAGGAAGC
TTCATCGGACTCATCAGGAAGC
CATCGGACTCATCAGGAAGC
CATCGGACTCATCAGGAAGC
TACATCGGACTCATCAGGAAGC
TGCATCGGACTCATCAGGAAGC

ATCATCGGACTCATCAGGAAGC
ACACCATCGGACTCATCAGGAAGC
GCCATCGGACTCATCAGGAAGC
GCATCGGACTCATCAGGAAGC
GCACATCGGACTCATCAGGAAGC
TGCATCGGACTCATCAGGAAGC
CCATCGGACTCATCAGGAAGC
AACATCGGACTCATCAGGAAGC
CCATCGGACTCATCAGGAAGC
ACATCGGACTCATCAGGAAGC
TTCATCGGACTCATCAGGAAGC
CATCGGACTCATCAGGAAGC
GGCATCGGACTCATCAGGAAGC
ACATCGGACTCATCAGGAAGC
ACAGGCATCGGACTCATCAGGAAGC
ACATCGGACTCATCAGGAAGC
GCCATCGGACTCATCAGGAAGC
GCATCGGACTCATCAGGAAGC
TCATCGGACTCATCAGGAAGC
ACATCGGACTCATCAGGAAGC
CACATCGGACTCATCAGGAAGC
GGCATCGGACTCATCAGGAAGC
GAGACATCGGACTCATCAGGAAGC
TTGGCATCGGACTCATCAGGAAGC
AACATCGGACTCATCAGGAAGC
TCATCGGACTCATCAGGAAGC
TATCATCGGACTCATCAGGAAGC
CCATCGGACTCATCAGGAAGC
TCATCGGACTCATCAGGAAGC
ATCGGACTCATCAGGAAGC
ATACACATCGGACTCATCAGGAAGC
CATCGGACTCATCAGGAAGC
ACATCGGACTCATCAGGAAGC
CATCGGACTCATCAGGAAGC
TCATCGGACTCATCAGGAAGC
AGCCATCGGACTCATCAGGAAGC
ACAACATCGGACTCATCAGGAAGC
CCATCGGACTCATCAGGAAGC
AGGGCATCGGACTCATCAGGAAGC
GACATCGGACTCATCAGGAAGC
AAGACATCGGACTCATCAGGAAGC
GCCATCGGACTCATCAGGAAGC
TGACATCGGACTCATCAGGAAGC

TCGGACTCATCAGGAAGC
GGTCATCGGACTCATCAGGAAGC
CATCGGACTCATCAGGAAGC
TCACATCGGACTCATCAGGAAGC
ACATCGGACTCATCAGGAAGC
ATCATCGGACTCATCAGGAAGC
GCATCGGACTCATCAGGAAGC
AGCATCGGACTCATCAGGAAGC
TCCATCGGACTCATCAGGAAGC
AGCATCGGACTCATCAGGAAGC
CATCGGACTCATCAGGAAGC
CCATCGGACTCATCAGGAAGC
GGCATCGGACTCATCAGGAAGC
ACATCGGACTCATCAGGAAGC
CCATCGGACTCATCAGGAAGC
TCATCGGACTCATCAGGAAGC
TCATCGGACTCATCAGGAAGC
CATCGGACTCATCAGGAAGC
CCATCGGACTCATCAGGAAGC
TCCATCGGACTCATCAGGAAGC
GACATCGGACTCATCAGGAAGC
GCATCGGACTCATCAGGAAGC
TGCCATCGGACTCATCAGGAAGC
GTCATCGGACTCATCAGGAAGC
ATCATCGGACTCATCAGGAAGC
TGTGCATCGGACTCATCAGGAAGC
TGCCATCGGACTCATCAGGAAGC
GAACATCGGACTCATCAGGAAGC
ATCGGACTCATCAGGAAGC
ATCGGACTCATCAGGAAGC
CCCCATCGGACTCATCAGGAAGC
GGCATCGGACTCATCAGGAAGC
CATCGGACTCATCAGGAAGC
AACATCGGACTCATCAGGAAGC
GGACATCGGACTCATCAGGAAGC
GACATCGGACTCATCAGGAAGC
CCATCGGACTCATCAGGAAGC
CTCATCGGACTCATCAGGAAGC
CCATCGGACTCATCAGGAAGC
AACATCGGACTCATCAGGAAGC
CGGACTCATCAGGAAGC
ACATCGGACTCATCAGGAAGC
CTCATCGGACTCATCAGGAAGC

CATCGGACTCATCAGGAAGC
CTCATCGGACTCATCAGGAAGC
GAGCCATCGGACTCATCAGGAAGC
GGCCATCGGACTCATCAGGAAGC
ATCCATCGGACTCATCAGGAAGC
GACATCGGACTCATCAGGAAGC
CATCGGACTCATCAGGAAGC
CTCATCGGACTCATCAGGAAGC
TGTCATCGGACTCATCAGGAAGC
AACATCGGACTCATCAGGAAGC
GCATCGGACTCATCAGGAAGC
TGACATCGGACTCATCAGGAAGC
CATCGGACTCATCAGGAAGC
GTCATCGGACTCATCAGGAAGC
CACATCGGACTCATCAGGAAGC
TAGACATCGGACTCATCAGGAAGC
GAACATCGGACTCATCAGGAAGC
TGAAACATCGGACTCATCAGGAAGC
TCTCATCGGACTCATCAGGAAGC
TAAGCATCGGACTCATCAGGAAGC
TCCATCGGACTCATCAGGAAGC
CATCGGACTCATCAGGAAGC
TCTCATCGGACTCATCAGGAAGC
CACATCGGACTCATCAGGAAGC
GACACATCGGACTCATCAGGAAGC
GAGCATCGGACTCATCAGGAAGC
GGAGCATCGGACTCATCAGGAAGC
CATCGGACTCATCAGGAAGC
CATCGGACTCATCAGGAAGC
TCCATCGGACTCATCAGGAAGC
ATCGGACTCATCAGGAAGC
CATCGGACTCATCAGGAAGC
GCATCGGACTCATCAGGAAGC
GTCATCGGACTCATCAGGAAGC
GACATCGGACTCATCAGGAAGC
TCATCGGACTCATCAGGAAGC
AAGACATCGGACTCATCAGGAAGC
GAACATCGGACTCATCAGGAAGC
GTCATCGGACTCATCAGGAAGC
GCCATCGGACTCATCAGGAAGC
GACATCGGACTCATCAGGAAGC
TCATCGGACTCATCAGGAAGC
TCTCATCGGACTCATCAGGAAGC

ATGAAACATCGGACTCATCAGGAAGC
GGACATCGGACTCATCAGGAAGC
TCATCGGACTCATCAGGAAGC
TTCATCGGACTCATCAGGAAGC
ATCATCGGACTCATCAGGAAGC
CTCCATCGGACTCATCAGGAAGC
GCCCCATCGGACTCATCAGGAAGC
TCATCGGACTCATCAGGAAGC
GCATCGGACTCATCAGGAAGC
CTCATCGGACTCATCAGGAAGC
TGCCATCGGACTCATCAGGAAGC
CCATCGGACTCATCAGGAAGC
CAGGCCATCGGACTCATCAGGAAGC
AGGCATCGGACTCATCAGGAAGC
CGCATCGGACTCATCAGGAAGC
CATCGGACTCATCAGGAAGC
TTTCATCGGACTCATCAGGAAGC
TTCTCATCGGACTCATCAGGAAGC
CCTCATCGGACTCATCAGGAAGC
CATCGGACTCATCAGGAAGC
GGCATCGGACTCATCAGGAAGC
TTCATCGGACTCATCAGGAAGC
CAGGCATCGGACTCATCAGGAAGC
GCATCGGACTCATCAGGAAGC
TTCATCGGACTCATCAGGAAGC
GACATCGGACTCATCAGGAAGC
TCATCGGACTCATCAGGAAGC
GTCATCGGACTCATCAGGAAGC
ATCATCGGACTCATCAGGAAGC
CACATCGGACTCATCAGGAAGC
GACCATCGGACTCATCAGGAAGC
CACATCGGACTCATCAGGAAGC
CCAGCATCGGACTCATCAGGAAGC
ATCATCGGACTCATCAGGAAGC
GACATCGGACTCATCAGGAAGC
ATCGGACTCATCAGGAAGC
TGACATCGGACTCATCAGGAAGC
AACATCGGACTCATCAGGAAGC
ACATCGGACTCATCAGGAAGC
CACATCGGACTCATCAGGAAGC
CCATCGGACTCATCAGGAAGC
TCTCATCGGACTCATCAGGAAGC
GTCATCGGACTCATCAGGAAGC

TCATCGGACTCATCAGGAAGC
GGCACATCGGACTCATCAGGAAGC
GCATCGGACTCATCAGGAAGC
CATCGGACTCATCAGGAAGC
TGGCATCGGACTCATCAGGAAGC
CTCCATCGGACTCATCAGGAAGC
ACCATCGGACTCATCAGGAAGC
GGACATCGGACTCATCAGGAAGC
CTTCCCATCGGACTCATCAGGAAGC
GGTCATCGGACTCATCAGGAAGC
CTACATCGGACTCATCAGGAAGC
TTTCATCGGACTCATCAGGAAGC
CCATCGGACTCATCAGGAAGC
ATCTGACATCGGACTCATCAGGAAGC
ATCGGACTCATCAGGAAGC
GAGTACATCGGACTCATCAGGAAGC
AGGCATCGGACTCATCAGGAAGC
AACATCGGACTCATCAGGAAGC
AATCATCGGACTCATCAGGAAGC
TGCAGTCATCGGACTCATCAGGAAGC
AAAGCATCGGACTCATCAGGAAGC
GATGTCATCGGACTCATCAGGAAGC
TGTCATCGGACTCATCAGGAAGC
CATCGGACTCATCAGGAAGC
GGCATCGGACTCATCAGGAAGC
CCATCGGACTCATCAGGAAGC
GACATCGGACTCATCAGGAAGC
TACCATCGGACTCATCAGGAAGC
TCATCGGACTCATCAGGAAGC
GTGCATCGGACTCATCAGGAAGC
AACATCGGACTCATCAGGAAGC
TCTCATCGGACTCATCAGGAAGC
TGGCATCGGACTCATCAGGAAGC
GCCTGCATCGGACTCATCAGGAAGC
TGGGCATCGGACTCATCAGGAAGC
TGCATCGGACTCATCAGGAAGC
ACCATCGGACTCATCAGGAAGC
GAGGCATCGGACTCATCAGGAAGC
TGCATCGGACTCATCAGGAAGC
CCATCGGACTCATCAGGAAGC
ATGCATCGGACTCATCAGGAAGC
AACATCGGACTCATCAGGAAGC
AGGGCATCGGACTCATCAGGAAGC

GCATCGGACTCATCAGGAAGC
CATCGGACTCATCAGGAAGC
ATCATCGGACTCATCAGGAAGC
CACCATCGGACTCATCAGGAAGC
AGAACATCGGACTCATCAGGAAGC
CCATCGGACTCATCAGGAAGC
GCCATCGGACTCATCAGGAAGC
AGGCCATCGGACTCATCAGGAAGC
TGCATCGGACTCATCAGGAAGC
ACATCGGACTCATCAGGAAGC
GTCCATCGGACTCATCAGGAAGC
ATGACATCGGACTCATCAGGAAGC
GCATCGGACTCATCAGGAAGC
GAACATCGGACTCATCAGGAAGC
TCATCGGACTCATCAGGAAGC
ATCGGACTCATCAGGAAGC
TAATACATCGGACTCATCAGGAAGC
CTCATCGGACTCATCAGGAAGC
GTACATCGGACTCATCAGGAAGC
CATCGGACTCATCAGGAAGC
TCGGACTCATCAGGAAGC
TCATCGGACTCATCAGGAAGC
ATCGGACTCATCAGGAAGC
AGACATCGGACTCATCAGGAAGC
CCCATCGGACTCATCAGGAAGC
TCTCATCGGACTCATCAGGAAGC
TTGTCATCGGACTCATCAGGAAGC
ATCATCGGACTCATCAGGAAGC
CTTCATCGGACTCATCAGGAAGC
GGACATCGGACTCATCAGGAAGC
AGCATCGGACTCATCAGGAAGC
ACATCGGACTCATCAGGAAGC
ATCGGACTCATCAGGAAGC
GGCATCGGACTCATCAGGAAGC
GCATCGGACTCATCAGGAAGC
CGGACATCGGACTCATCAGGAAGC
ACACATCGGACTCATCAGGAAGC
CCATCGGACTCATCAGGAAGC
ATCGGACTCATCAGGAAGC
ACGTACATCGGACTCATCAGGAAGC
ATCATCGGACTCATCAGGAAGC
CCATCGGACTCATCAGGAAGC
GACATCGGACTCATCAGGAAGC

GAGCATCGGACTCATCAGGAAGC
GACATCGGACTCATCAGGAAGC
ATCGGACTCATCAGGAAGC
CATCGGACTCATCAGGAAGC
CATCGGACTCATCAGGAAGC
CATCGGACTCATCAGGAAGC
TGTCATCGGACTCATCAGGAAGC
CATCGGACTCATCAGGAAGC
CATCGGACTCATCAGGAAGC
CACATCGGACTCATCAGGAAGC
CATCGGACTCATCAGGAAGC
GTGCATCGGACTCATCAGGAAGC
CCATCGGACTCATCAGGAAGC
CATCGGACTCATCAGGAAGC
ACATCGGACTCATCAGGAAGC
CCATCGGACTCATCAGGAAGC
CTCACCATCGGACTCATCAGGAAGC
AACATCGGACTCATCAGGAAGC
CCATCGGACTCATCAGGAAGC
GGCATCGGACTCATCAGGAAGC
ACATCGGACTCATCAGGAAGC
AGATTCATCGGACTCATCAGGAAGC
CTCATCGGACTCATCAGGAAGC
TCATCGGACTCATCAGGAAGC
AGCTGCATCGGACTCATCAGGAAGC
GCTGCATCGGACTCATCAGGAAGC
GTCATCGGACTCATCAGGAAGC
TCGGACTCATCAGGAAGC
GCATCGGACTCATCAGGAAGC
ATCACATCGGACTCATCAGGAAGC
TCATCGGACTCATCAGGAAGC
ATCGGACTCATCAGGAAGC
GCATCGGACTCATCAGGAAGC
ACCATCGGACTCATCAGGAAGC
ATTACATCGGACTCATCAGGAAGC
CCATCGGACTCATCAGGAAGC
GAAGCATCGGACTCATCAGGAAGC
GCATCGGACTCATCAGGAAGC
CCATCGGACTCATCAGGAAGC
GGCATCGGACTCATCAGGAAGC
TCATCGGACTCATCAGGAAGC
GCCCATCGGACTCATCAGGAAGC
AAGCATCGGACTCATCAGGAAGC

CCATCGGACTCATCAGGAAGC
ACATCGGACTCATCAGGAAGC
CATCGGACTCATCAGGAAGC
ACACATCGGACTCATCAGGAAGC
TCGGACTCATCAGGAAGC
CAGACATCGGACTCATCAGGAAGC
AACATCGGACTCATCAGGAAGC
CTCATCGGACTCATCAGGAAGC
GCATCGGACTCATCAGGAAGC
GACGCATCGGACTCATCAGGAAGC
ATCGGACTCATCAGGAAGC
CATCGGACTCATCAGGAAGC
GCCATCGGACTCATCAGGAAGC
GGGCCATCGGACTCATCAGGAAGC
GTCATCGGACTCATCAGGAAGC
GTCATCGGACTCATCAGGAAGC
GGCATCGGACTCATCAGGAAGC
ATCGGACTCATCAGGAAGC
ATAACATCGGACTCATCAGGAAGC
TCATCGGACTCATCAGGAAGC
AGACATCGGACTCATCAGGAAGC
GGTCATCGGACTCATCAGGAAGC
AGCATCGGACTCATCAGGAAGC
TTTCATCGGACTCATCAGGAAGC
GTCTCATCGGACTCATCAGGAAGC
TAAGCATCGGACTCATCAGGAAGC
CCATCGGACTCATCAGGAAGC
CATCGGACTCATCAGGAAGC
AGAGCATCGGACTCATCAGGAAGC
GCCATCGGACTCATCAGGAAGC
CATCGGACTCATCAGGAAGC
GCTTCTCATCGGACTCATCAGGAAGC
GACCATCGGACTCATCAGGAAGC
ACCATCGGACTCATCAGGAAGC
TTGCCATCGGACTCATCAGGAAGC
CTCATCGGACTCATCAGGAAGC
AACATCGGACTCATCAGGAAGC
CGCATCGGACTCATCAGGAAGC
CATCGGACTCATCAGGAAGC
TACATCGGACTCATCAGGAAGC
TACATCGGACTCATCAGGAAGC
GTCATCGGACTCATCAGGAAGC
ACATCGGACTCATCAGGAAGC

GGCATCGGACTCATCAGGAAGC
CCATCGGACTCATCAGGAAGC
GCATCGGACTCATCAGGAAGC
AAACATCGGACTCATCAGGAAGC
CATCGGACTCATCAGGAAGC
ATCCATCGGACTCATCAGGAAGC
CATCGGACTCATCAGGAAGC
CATCGGACTCATCAGGAAGC
ATGGCATCGGACTCATCAGGAAGC
GACATCGGACTCATCAGGAAGC
CATCGGACTCATCAGGAAGC
CATCGGACTCATCAGGAAGC
TTGGCATCGGACTCATCAGGAAGC
GTCCCATCGGACTCATCAGGAAGC
ACATCGGACTCATCAGGAAGC
CATCGGACTCATCAGGAAGC
CGCATCGGACTCATCAGGAAGC
TGAACATCGGACTCATCAGGAAGC
TCACATCGGACTCATCAGGAAGC
TCAGCATCGGACTCATCAGGAAGC
TACATCGGACTCATCAGGAAGC
GCCATCGGACTCATCAGGAAGC
GCATACATCGGACTCATCAGGAAGC
CATCGGACTCATCAGGAAGC
ATCGGACTCATCAGGAAGC
GCATCGGACTCATCAGGAAGC
GCATCGGACTCATCAGGAAGC
ACAGCCATCGGACTCATCAGGAAGC
CGGACTCATCAGGAAGC
GTAGCATCGGACTCATCAGGAAGC
GGGGCATCGGACTCATCAGGAAGC
GGGCATCGGACTCATCAGGAAGC
TCATCGGACTCATCAGGAAGC
TACATCGGACTCATCAGGAAGC
ATCGGACTCATCAGGAAGC
TCGGACTCATCAGGAAGC
ATCGGACTCATCAGGAAGC
AAGCATCGGACTCATCAGGAAGC
CACATCGGACTCATCAGGAAGC
GTCATCGGACTCATCAGGAAGC
TGCATCGGACTCATCAGGAAGC
ATCATCGGACTCATCAGGAAGC
GACATCGGACTCATCAGGAAGC

CCATCGGACTCATCAGGAAGC
GGCATCGGACTCATCAGGAAGC
TACATCGGACTCATCAGGAAGC
TCCATCGGACTCATCAGGAAGC
CATCGGACTCATCAGGAAGC
AGAGACATCGGACTCATCAGGAAGC
AAACATCGGACTCATCAGGAAGC
GTGCATCGGACTCATCAGGAAGC
ACAGCATCGGACTCATCAGGAAGC
CTCTCATCGGACTCATCAGGAAGC
ATCGGACTCATCAGGAAGC
TCTCATCGGACTCATCAGGAAGC
AAAACATCGGACTCATCAGGAAGC
CATCGGACTCATCAGGAAGC
CTCATCGGACTCATCAGGAAGC
CTATAACATCGGACTCATCAGGAAGC
CAGCATCGGACTCATCAGGAAGC
AACATCGGACTCATCAGGAAGC
CGGACTCATCAGGAAGC
TCCATCGGACTCATCAGGAAGC
ATCATCGGACTCATCAGGAAGC
CATCGGACTCATCAGGAAGC
CACCATCGGACTCATCAGGAAGC
CCATCGGACTCATCAGGAAGC
GACTCATCAGGAAGC
CGGACTCATCAGGAAGC
ATCGGACTCATCAGGAAGC
CATCGGACTCATCAGGAAGC
AACATCGGACTCATCAGGAAGC
GCTGCATCGGACTCATCAGGAAGC
CCATCGGACTCATCAGGAAGC
GCATCGGACTCATCAGGAAGC
GCATCGGACTCATCAGGAAGC
AACATCGGACTCATCAGGAAGC
GACATCGGACTCATCAGGAAGC
TACATCGGACTCATCAGGAAGC
TTCATCGGACTCATCAGGAAGC
CTCATCGGACTCATCAGGAAGC
ACGCATCGGACTCATCAGGAAGC
GACTCATCAGGAAGC
GAGGCATCGGACTCATCAGGAAGC
ACATCGGACTCATCAGGAAGC
CATCGGACTCATCAGGAAGC

GTGACATCGGACTCATCAGGAAGC
TCATCGGACTCATCAGGAAGC
ACATCGGACTCATCAGGAAGC
CATCGGACTCATCAGGAAGC
TGCATCGGACTCATCAGGAAGC
TCCATCGGACTCATCAGGAAGC
CATCGGACTCATCAGGAAGC
GCATCGGACTCATCAGGAAGC
CATCGGACTCATCAGGAAGC
TCCATCGGACTCATCAGGAAGC
AACATCGGACTCATCAGGAAGC
GCCATCGGACTCATCAGGAAGC
GCATCGGACTCATCAGGAAGC
ACCATCGGACTCATCAGGAAGC
GCTAGGCATCGGACTCATCAGGAAGC
TAAGCATCGGACTCATCAGGAAGC
ACAGCATCGGACTCATCAGGAAGC
CATCGGACTCATCAGGAAGC
CCATCGGACTCATCAGGAAGC
ACTCATCGGACTCATCAGGAAGC
CTAACATCGGACTCATCAGGAAGC
AACATCGGACTCATCAGGAAGC
TCCATCGGACTCATCAGGAAGC
GGCATCGGACTCATCAGGAAGC
GTGCATCGGACTCATCAGGAAGC
CATCGGACTCATCAGGAAGC
GTGCATCGGACTCATCAGGAAGC
CATCGGACTCATCAGGAAGC
GCATCGGACTCATCAGGAAGC
GCATCGGACTCATCAGGAAGC
TTTCATCGGACTCATCAGGAAGC
CCATCGGACTCATCAGGAAGC
ATGCATCGGACTCATCAGGAAGC
AAACATCGGACTCATCAGGAAGC
TGCATCGGACTCATCAGGAAGC
CATCGGACTCATCAGGAAGC
CTGTCATCGGACTCATCAGGAAGC
AGCACATCGGACTCATCAGGAAGC
CACATCGGACTCATCAGGAAGC
ACATCGGACTCATCAGGAAGC
CATCGGACTCATCAGGAAGC
CCATCGGACTCATCAGGAAGC
CACATCGGACTCATCAGGAAGC

ACACATCGGACTCATCAGGAAGC
TCATCGGACTCATCAGGAAGC
GGGCATCGGACTCATCAGGAAGC
GACATCGGACTCATCAGGAAGC
TCAACATCGGACTCATCAGGAAGC
CATCGGACTCATCAGGAAGC
GATAGACATCGGACTCATCAGGAAGC
TTCATCGGACTCATCAGGAAGC
CATCGGACTCATCAGGAAGC
TTCATCGGACTCATCAGGAAGC
ATCGGACTCATCAGGAAGC
AGACATCGGACTCATCAGGAAGC
GACCACATCGGACTCATCAGGAAGC
GCATCGGACTCATCAGGAAGC
TTCATCGGACTCATCAGGAAGC
TGAATCCCATCGGACTCATCAGGAAGC
GCATCGGACTCATCAGGAAGC
GACATCGGACTCATCAGGAAGC
TCATCGGACTCATCAGGAAGC
CTCATCGGACTCATCAGGAAGC
TGAGCATCGGACTCATCAGGAAGC
CCATCGGACTCATCAGGAAGC
GCATCGGACTCATCAGGAAGC
GACATCGGACTCATCAGGAAGC
GATAGACATCGGACTCATCAGGAAGC
ATCGGACTCATCAGGAAGC
TGCCCATCGGACTCATCAGGAAGC
TTCATCGGACTCATCAGGAAGC
ACATCGGACTCATCAGGAAGC
TACATCGGACTCATCAGGAAGC
ATCGGACTCATCAGGAAGC
AGCATCGGACTCATCAGGAAGC
TCATCGGACTCATCAGGAAGC
CCATCGGACTCATCAGGAAGC
CATCGGACTCATCAGGAAGC
GGGACATCGGACTCATCAGGAAGC
ACAACATCGGACTCATCAGGAAGC
ATCCCATCGGACTCATCAGGAAGC
GGCATCGGACTCATCAGGAAGC
CATCGGACTCATCAGGAAGC
GTCATCGGACTCATCAGGAAGC
AATGCATCGGACTCATCAGGAAGC
GCATCGGACTCATCAGGAAGC

AGGGCATCGGACTCATCAGGAAGC
GGTCATCGGACTCATCAGGAAGC
TGTCATCGGACTCATCAGGAAGC
CTCATCGGACTCATCAGGAAGC
GTCATCGGACTCATCAGGAAGC
TTCATCGGACTCATCAGGAAGC
CTGCATCGGACTCATCAGGAAGC
CTCATCGGACTCATCAGGAAGC
CATCGGACTCATCAGGAAGC
CCATCGGACTCATCAGGAAGC
CCCCATCGGACTCATCAGGAAGC
GACATTCATCGGACTCATCAGGAAGC
GGCATCGGACTCATCAGGAAGC
TGTCATCGGACTCATCAGGAAGC
TTTGGTCATCGGACTCATCAGGAAGC
ACAACATCGGACTCATCAGGAAGC
ATCATCGGACTCATCAGGAAGC
CTCATCGGACTCATCAGGAAGC
CCCCATCGGACTCATCAGGAAGC
GCATCGGACTCATCAGGAAGC
CTCATCGGACTCATCAGGAAGC
GGCCATCGGACTCATCAGGAAGC
GTGACATCGGACTCATCAGGAAGC
TCCATCGGACTCATCAGGAAGC
ATCGGACTCATCAGGAAGC
CCATCGGACTCATCAGGAAGC
GCATCGGACTCATCAGGAAGC
CTCATCGGACTCATCAGGAAGC
ACCATCGGACTCATCAGGAAGC
TGCCATCGGACTCATCAGGAAGC
TCATCGGACTCATCAGGAAGC
CACATCGGACTCATCAGGAAGC
GAGTGTCATCGGACTCATCAGGAAGC
CACATCGGACTCATCAGGAAGC
GTTCATCGGACTCATCAGGAAGC
TCATCGGACTCATCAGGAAGC
CTGCCATCGGACTCATCAGGAAGC
CCATCGGACTCATCAGGAAGC
TCATCGGACTCATCAGGAAGC
ACATCGGACTCATCAGGAAGC
ACATCGGACTCATCAGGAAGC
CTCATCGGACTCATCAGGAAGC
TGGGTCATCGGACTCATCAGGAAGC

ATCATCGGACTCATCAGGAAGC
GCATCGGACTCATCAGGAAGC
ATACATCGGACTCATCAGGAAGC
CATCGGACTCATCAGGAAGC
AGGCCATCGGACTCATCAGGAAGC
TCGGACTCATCAGGAAGC
TCATCGGACTCATCAGGAAGC
TCATCGGACTCATCAGGAAGC
CCCATCGGACTCATCAGGAAGC
CCATCGGACTCATCAGGAAGC
AACATCGGACTCATCAGGAAGC
GGCTCATCGGACTCATCAGGAAGC
AACATCGGACTCATCAGGAAGC
CTCATCGGACTCATCAGGAAGC
TCATCGGACTCATCAGGAAGC
CATCGGACTCATCAGGAAGC
CCATCGGACTCATCAGGAAGC
GGGTCATCGGACTCATCAGGAAGC
AGCCATCGGACTCATCAGGAAGC
ATGCCATCGGACTCATCAGGAAGC
GATCATCGGACTCATCAGGAAGC
CCATCGGACTCATCAGGAAGC
TCACATCGGACTCATCAGGAAGC
ACCATCGGACTCATCAGGAAGC
GGCATCGGACTCATCAGGAAGC
TGGCATCGGACTCATCAGGAAGC
GGTCATCGGACTCATCAGGAAGC
GCATCGGACTCATCAGGAAGC
CCACATCGGACTCATCAGGAAGC
CCATCGGACTCATCAGGAAGC
AATGCATCGGACTCATCAGGAAGC
CATCGGACTCATCAGGAAGC
CTCATCGGACTCATCAGGAAGC
AGGCATCGGACTCATCAGGAAGC
ATCATCGGACTCATCAGGAAGC
TGCATCGGACTCATCAGGAAGC
GAGCATCGGACTCATCAGGAAGC
ATTACATCGGACTCATCAGGAAGC
AGGGACATCGGACTCATCAGGAAGC
GCATCGGACTCATCAGGAAGC
CCATCGGACTCATCAGGAAGC
AAACATCGGACTCATCAGGAAGC
TCATCGGACTCATCAGGAAGC

GAGCCATCGGACTCATCAGGAAGC
CCATCGGACTCATCAGGAAGC
TCATCGGACTCATCAGGAAGC
GACATCGGACTCATCAGGAAGC
GTCATCGGACTCATCAGGAAGC
TGCATCGGACTCATCAGGAAGC
ACATCGGACTCATCAGGAAGC
CATCGGACTCATCAGGAAGC
ACATCGGACTCATCAGGAAGC
TGCATCGGACTCATCAGGAAGC
TCATCGGACTCATCAGGAAGC
CCATCGGACTCATCAGGAAGC
ATCCATCGGACTCATCAGGAAGC
GTTTATCATCGGACTCATCAGGAAGC
TCATCGGACTCATCAGGAAGC
GGCATCGGACTCATCAGGAAGC
CAGCATCGGACTCATCAGGAAGC
GCATCGGACTCATCAGGAAGC
GCCATCGGACTCATCAGGAAGC
AAGCATCGGACTCATCAGGAAGC
AAGCATCGGACTCATCAGGAAGC
CATCGGACTCATCAGGAAGC
GCATCGGACTCATCAGGAAGC
GGACTCATCAGGAAGC
TCCATCGGACTCATCAGGAAGC
CATCGGACTCATCAGGAAGC
TCCATCGGACTCATCAGGAAGC
CACAACATCGGACTCATCAGGAAGC
TGGCATCGGACTCATCAGGAAGC
TTGCATCGGACTCATCAGGAAGC
CCCATCGGACTCATCAGGAAGC
ACATCGGACTCATCAGGAAGC
ATGCCATCGGACTCATCAGGAAGC
AACATCGGACTCATCAGGAAGC
ATCATCGGACTCATCAGGAAGC
CCATCGGACTCATCAGGAAGC
GACATCGGACTCATCAGGAAGC
TGCGCATCGGACTCATCAGGAAGC
TCATCGGACTCATCAGGAAGC
CCCATCGGACTCATCAGGAAGC
TGAGTGCATCGGACTCATCAGGAAGC
TGCATCGGACTCATCAGGAAGC
CCATCGGACTCATCAGGAAGC

TTACATCGGACTCATCAGGAAGC
CCATCGGACTCATCAGGAAGC
GGCATCGGACTCATCAGGAAGC
CTGTCATCGGACTCATCAGGAAGC
ACATCGGACTCATCAGGAAGC
TGCATCGGACTCATCAGGAAGC
TAACATCGGACTCATCAGGAAGC
ACATCGGACTCATCAGGAAGC
CATCGGACTCATCAGGAAGC
ATCATCGGACTCATCAGGAAGC
CATCGGACTCATCAGGAAGC
GCATCGGACTCATCAGGAAGC
CCCATCGGACTCATCAGGAAGC
TACATCGGACTCATCAGGAAGC
GAGACATCGGACTCATCAGGAAGC
CACATCGGACTCATCAGGAAGC
TTCATCGGACTCATCAGGAAGC
GACATCGGACTCATCAGGAAGC
ATCGGACTCATCAGGAAGC
CTCATCGGACTCATCAGGAAGC
TCACATCGGACTCATCAGGAAGC
CCTCATCGGACTCATCAGGAAGC
GACATCGGACTCATCAGGAAGC
ATGACACATCGGACTCATCAGGAAGC
GGCATCGGACTCATCAGGAAGC
ATCATCGGACTCATCAGGAAGC
CACCATCGGACTCATCAGGAAGC
ACCATCGGACTCATCAGGAAGC
TTCCATCGGACTCATCAGGAAGC
ACATCGGACTCATCAGGAAGC
ACCACATCGGACTCATCAGGAAGC
CATCGGACTCATCAGGAAGC
TACATCGGACTCATCAGGAAGC
CATCGGACTCATCAGGAAGC
GGCATCGGACTCATCAGGAAGC
CACCATCGGACTCATCAGGAAGC
ACCATCGGACTCATCAGGAAGC
AGGCATCGGACTCATCAGGAAGC
AACATCGGACTCATCAGGAAGC
AATCATCGGACTCATCAGGAAGC
TACATCGGACTCATCAGGAAGC
TCATCGGACTCATCAGGAAGC
GCATCGGACTCATCAGGAAGC

TCCATCGGACTCATCAGGAAGC
AGCCATCGGACTCATCAGGAAGC
CATCGGACTCATCAGGAAGC
TAACACATCGGACTCATCAGGAAGC
GAGGGCATCGGACTCATCAGGAAGC
TCATCGGACTCATCAGGAAGC
GTGCCCATCGGACTCATCAGGAAGC
TCATCGGACTCATCAGGAAGC
ACATCGGACTCATCAGGAAGC
CACATCGGACTCATCAGGAAGC
CATCGGACTCATCAGGAAGC
GCATCGGACTCATCAGGAAGC
ACATCGGACTCATCAGGAAGC
CTCATCGGACTCATCAGGAAGC
CCATCGGACTCATCAGGAAGC
AGCCATCGGACTCATCAGGAAGC
TCATCGGACTCATCAGGAAGC
GCCATCGGACTCATCAGGAAGC
ATCGGACTCATCAGGAAGC
GACATCGGACTCATCAGGAAGC
CCCATCGGACTCATCAGGAAGC
GCCATCGGACTCATCAGGAAGC
GCATCGGACTCATCAGGAAGC
AGTGTCATCGGACTCATCAGGAAGC
ACATCGGACTCATCAGGAAGC
AACATCGGACTCATCAGGAAGC
CATCGGACTCATCAGGAAGC
ATCATCGGACTCATCAGGAAGC
ACATCGGACTCATCAGGAAGC
TAATCATCGGACTCATCAGGAAGC
GTCACATCGGACTCATCAGGAAGC
TCATCGGACTCATCAGGAAGC
ACATCGGACTCATCAGGAAGC
ATCGGACTCATCAGGAAGC
CATCGGACTCATCAGGAAGC
GACATCGGACTCATCAGGAAGC
GGGCATCGGACTCATCAGGAAGC
TCTCATCGGACTCATCAGGAAGC
CATCGGACTCATCAGGAAGC
ACCATCGGACTCATCAGGAAGC
GTGTCATCGGACTCATCAGGAAGC
ACATCGGACTCATCAGGAAGC
AAGTCATCGGACTCATCAGGAAGC

ACAGCATCGGACTCATCAGGAAGC<br>
CCATCGGACTCATCAGGAAGC<br>
AGACCATCGGACTCATCAGGAAGC<br>
CTCATCGGACTCATCAGGAAGC<br>
TGTAAACATCGGACTCATCAGGAAGC<br>
AAACCATCGGACTCATCAGGAAGC<br>
AAGTCATCGGACTCATCAGGAAGC<br>
TCATCGGACTCATCAGGAAGC<br>
ACATCGGACTCATCAGGAAGC<br>
TTGCATCGGACTCATCAGGAAGC<br>
AACATCGGACTCATCAGGAAGC<br>
CAACATCGGACTCATCAGGAAGC<br>
TTCATCGGACTCATCAGGAAGC<br>
CCATCGGACTCATCAGGAAGC<br>
CTCATCGGACTCATCAGGAAGC<br>
CTCCTCATCGGACTCATCAGGAAGC<br>
TACTCATCGGACTCATCAGGAAGC<br>
ACTAACATCGGACTCATCAGGAAGC<br>
CTCATCGGACTCATCAGGAAGC<br>
CATCGGACTCATCAGGAAGC<br>
GGCATCGGACTCATCAGGAAGC<br>
GGCATCGGACTCATCAGGAAGC<br>
ATCGGACTCATCAGGAAGC<br>
CATCGGACTCATCAGGAAGC<br>
ACATCGGACTCATCAGGAAGC<br>
CGGACTCATCAGGAAGC<br>
GGGACATCGGACTCATCAGGAAGC<br>
CCATCGGACTCATCAGGAAGC<br>
CCATCGGACTCATCAGGAAGC<br>
GCATCGGACTCATCAGGAAGC<br>
TCCATCGGACTCATCAGGAAGC<br>
TCCATCGGACTCATCAGGAAGC<br>
GGCTCATCGGACTCATCAGGAAGC<br>
CATCGGACTCATCAGGAAGC<br>
GGCATCGGACTCATCAGGAAGC<br>
ACATCGGACTCATCAGGAAGC<br>
CATCGGACTCATCAGGAAGC<br>
CATCGGACTCATCAGGAAGC<br>
TTTCATCGGACTCATCAGGAAGC<br>
CACATCGGACTCATCAGGAAGC<br>
AACATCGGACTCATCAGGAAGC<br>
CCATCGGACTCATCAGGAAGC<br>
CGGCATCGGACTCATCAGGAAGC

ATCGGACTCATCAGGAAGC
TCATCGGACTCATCAGGAAGC
GACATCGGACTCATCAGGAAGC
ACATCGGACTCATCAGGAAGC
CATCGGACTCATCAGGAAGC
TCACATCGGACTCATCAGGAAGC
ATCGGACTCATCAGGAAGC
AGTCATCGGACTCATCAGGAAGC
GCATCGGACTCATCAGGAAGC
ACATCGGACTCATCAGGAAGC
GGACATCGGACTCATCAGGAAGC
AGCCATCGGACTCATCAGGAAGC
TCATCGGACTCATCAGGAAGC
CTCATCGGACTCATCAGGAAGC
ATCGGACTCATCAGGAAGC
ATGCTCCATCGGACTCATCAGGAAGC
CCATCGGACTCATCAGGAAGC
TGTTTCATCGGACTCATCAGGAAGC
GGCACATCGGACTCATCAGGAAGC
AGGTCATCGGACTCATCAGGAAGC
ATCGGACTCATCAGGAAGC
CCCATCGGACTCATCAGGAAGC
TCATCGGACTCATCAGGAAGC
CCATCGGACTCATCAGGAAGC
TGCATCGGACTCATCAGGAAGC
CATCGGACTCATCAGGAAGC
AGCATCGGACTCATCAGGAAGC
TACATCGGACTCATCAGGAAGC
GCATCGGACTCATCAGGAAGC
CGACATCGGACTCATCAGGAAGC
CACATCGGACTCATCAGGAAGC
ATCGGACTCATCAGGAAGC
CATCGGACTCATCAGGAAGC
TCCCATCGGACTCATCAGGAAGC
CACATCGGACTCATCAGGAAGC
TCATCGGACTCATCAGGAAGC
CCATCGGACTCATCAGGAAGC
TGACATCGGACTCATCAGGAAGC
AGTCCATCGGACTCATCAGGAAGC
CTCACCATCGGACTCATCAGGAAGC
TGCATCGGACTCATCAGGAAGC
TCATCGGACTCATCAGGAAGC
CATCGGACTCATCAGGAAGC

GGGCATCGGACTCATCAGGAAGC
GAGGCATCATCGGACTCATCAGGAAGC
GGATCATCGGACTCATCAGGAAGC
ACATCGGACTCATCAGGAAGC
CCATCGGACTCATCAGGAAGC
GCCATCGGACTCATCAGGAAGC
ACATCGGACTCATCAGGAAGC
AGCATCGGACTCATCAGGAAGC
ATCGGACTCATCAGGAAGC
GTCCATCGGACTCATCAGGAAGC
GTCATCGGACTCATCAGGAAGC
CCCATCGGACTCATCAGGAAGC
TGAACATCGGACTCATCAGGAAGC
CTCATCGGACTCATCAGGAAGC
TCATCGGACTCATCAGGAAGC
CTCCATCGGACTCATCAGGAAGC
TGTCATCGGACTCATCAGGAAGC
GGCGCATCGGACTCATCAGGAAGC
CATCGGACTCATCAGGAAGC
TCATCGGACTCATCAGGAAGC
CTCATCGGACTCATCAGGAAGC
CCATCGGACTCATCAGGAAGC
TGCATCGGACTCATCAGGAAGC
TGCATCGGACTCATCAGGAAGC
ATCGGACTCATCAGGAAGC
CCATCGGACTCATCAGGAAGC
CTCTCATCGGACTCATCAGGAAGC
ATCATCGGACTCATCAGGAAGC
AAAAGTCATCGGACTCATCAGGAAGC
TGCATCGGACTCATCAGGAAGC
GCATCGGACTCATCAGGAAGC
CCATCGGACTCATCAGGAAGC
CCATCGGACTCATCAGGAAGC
TATCATCGGACTCATCAGGAAGC
CATCGGACTCATCAGGAAGC
CATCGGACTCATCAGGAAGC
CTTCATCGGACTCATCAGGAAGC
AGAGCATCGGACTCATCAGGAAGC
AGCATCGGACTCATCAGGAAGC
GCATCGGACTCATCAGGAAGC
ACATCGGACTCATCAGGAAGC
CCATCGGACTCATCAGGAAGC
TCTCCATCGGACTCATCAGGAAGC

GGCATCGGACTCATCAGGAAGC
GCATCGGACTCATCAGGAAGC
CCATCGGACTCATCAGGAAGC
CATCGGACTCATCAGGAAGC
AAACATCGGACTCATCAGGAAGC
GTCATCGGACTCATCAGGAAGC
GCCATCGGACTCATCAGGAAGC
TGCATCGGACTCATCAGGAAGC
AGCATCGGACTCATCAGGAAGC
TGAGCATCGGACTCATCAGGAAGC
CATCGGACTCATCAGGAAGC
CATCGGACTCATCAGGAAGC
AATCATCGGACTCATCAGGAAGC
TACATCGGACTCATCAGGAAGC
AATCATCGGACTCATCAGGAAGC
AATCATCGGACTCATCAGGAAGC
ATTTCATCGGACTCATCAGGAAGC
AGCATCGGACTCATCAGGAAGC
GAACATCGGACTCATCAGGAAGC
AGGCATCGGACTCATCAGGAAGC
TCGCATCGGACTCATCAGGAAGC
TGCATCGGACTCATCAGGAAGC
TCGGACTCATCAGGAAGC
CCATCGGACTCATCAGGAAGC
TGCATCGGACTCATCAGGAAGC
GAACATCGGACTCATCAGGAAGC
TTCATCGGACTCATCAGGAAGC
ATCGGACTCATCAGGAAGC
CATCGGACTCATCAGGAAGC
TGCCACATCGGACTCATCAGGAAGC
GAGTGACATCGGACTCATCAGGAAGC
AACATCGGACTCATCAGGAAGC
CATCGGACTCATCAGGAAGC
ATCGGACTCATCAGGAAGC
GGGCATCGGACTCATCAGGAAGC
GACATCGGACTCATCAGGAAGC
TGTCATCGGACTCATCAGGAAGC
GCATCGGACTCATCAGGAAGC
ATCATCGGACTCATCAGGAAGC
GCATCGGACTCATCAGGAAGC
ACATCGGACTCATCAGGAAGC
GCATCGGACTCATCAGGAAGC
GCATCGGACTCATCAGGAAGC

CACACATCGGACTCATCAGGAAGC
CTTCACATCGGACTCATCAGGAAGC
ACATCGGACTCATCAGGAAGC
CTGCAACATCGGACTCATCAGGAAGC
ACATCGGACTCATCAGGAAGC
ATCGGACTCATCAGGAAGC
TTCCCATCGGACTCATCAGGAAGC
ACATCGGACTCATCAGGAAGC
CACACATCGGACTCATCAGGAAGC
AAGACATCGGACTCATCAGGAAGC
GAAACATCGGACTCATCAGGAAGC
TTCATCGGACTCATCAGGAAGC
ACCCCTCATCGGACTCATCAGGAAGC
GTGCATCGGACTCATCAGGAAGC
ACTCATCAGGAAGC
AGGCATCGGACTCATCAGGAAGC
CCATCGGACTCATCAGGAAGC
GGAACATCGGACTCATCAGGAAGC
GTGCATCGGACTCATCAGGAAGC
ACATCGGACTCATCAGGAAGC
CCCCATCGGACTCATCAGGAAGC
AAGTCATCGGACTCATCAGGAAGC
AAACATCGGACTCATCAGGAAGC
ATCGGACTCATCAGGAAGC
ATCATCGGACTCATCAGGAAGC
CTCATCATCGGACTCATCAGGAAGC
CAAATACATCGGACTCATCAGGAAGC
CCATCGGACTCATCAGGAAGC
GCATCGGACTCATCAGGAAGC
GCATCGGACTCATCAGGAAGC
CATCGGACTCATCAGGAAGC
TGCATCGGACTCATCAGGAAGC
CAGCATCGGACTCATCAGGAAGC
TGCATCGGACTCATCAGGAAGC
ACATCGGACTCATCAGGAAGC
CCATCGGACTCATCAGGAAGC
AAGACATCGGACTCATCAGGAAGC
CCCATCGGACTCATCAGGAAGC
TTGCATCGGACTCATCAGGAAGC
GCCATCGGACTCATCAGGAAGC
TCTCATCGGACTCATCAGGAAGC
AACATCGGACTCATCAGGAAGC
CACCATCGGACTCATCAGGAAGC

TCACTCATCGGACTCATCAGGAAGC
CTCATCGGACTCATCAGGAAGC
ACACCATCGGACTCATCAGGAAGC
TTCATCGGACTCATCAGGAAGC
TTTTCATCGGACTCATCAGGAAGC
CTGCATCGGACTCATCAGGAAGC
ATATGACATCGGACTCATCAGGAAGC
TCCATCGGACTCATCAGGAAGC
ACATCGGACTCATCAGGAAGC
AGCCATCGGACTCATCAGGAAGC
TTCATCGGACTCATCAGGAAGC
ATCGGACTCATCAGGAAGC
GCATCGGACTCATCAGGAAGC
GTTGCATCGGACTCATCAGGAAGC
ACCCATCGGACTCATCAGGAAGC
CTCCATCGGACTCATCAGGAAGC
AGTCCCATCGGACTCATCAGGAAGC
TCATCGGACTCATCAGGAAGC
CTCATCGGACTCATCAGGAAGC
ATCGGACTCATCAGGAAGC
CATCGGACTCATCAGGAAGC
AACATCGGACTCATCAGGAAGC
TCTGCATCGGACTCATCAGGAAGC
CATCATCGGACTCATCAGGAAGC
AGACATCGGACTCATCAGGAAGC
TCAGCATCGGACTCATCAGGAAGC
TTGCATCGGACTCATCAGGAAGC
CATCGGACTCATCAGGAAGC
ATGCCACATCGGACTCATCAGGAAGC
TCATCGGACTCATCAGGAAGC
CATCGGACTCATCAGGAAGC
TCATCGGACTCATCAGGAAGC
TGACATCGGACTCATCAGGAAGC
GTCTCATCGGACTCATCAGGAAGC
ATGCATCGGACTCATCAGGAAGC
GCATCGGACTCATCAGGAAGC
GACATCGGACTCATCAGGAAGC
CCCCATCGGACTCATCAGGAAGC
ACATCGGACTCATCAGGAAGC
CATCGGACTCATCAGGAAGC
GGCATCGGACTCATCAGGAAGC
TGGCATCGGACTCATCAGGAAGC
CATCGGACTCATCAGGAAGC

GGCATCGGACTCATCAGGAAGC
AACATCGGACTCATCAGGAAGC
CATCGGACTCATCAGGAAGC
CATCGGACTCATCAGGAAGC
CGCATCGGACTCATCAGGAAGC
TCATCGGACTCATCAGGAAGC
CCATCGGACTCATCAGGAAGC
TGTCATCGGACTCATCAGGAAGC
ATCATCGGACTCATCAGGAAGC
ATCGGACTCATCAGGAAGC
ACCATCGGACTCATCAGGAAGC
AAGCATCGGACTCATCAGGAAGC
CTGACATCGGACTCATCAGGAAGC
GTCATCGGACTCATCAGGAAGC
CACCATCGGACTCATCAGGAAGC
CCTCATCGGACTCATCAGGAAGC
TGACATCGGACTCATCAGGAAGC
GCTACATCGGACTCATCAGGAAGC
GCATCGGACTCATCAGGAAGC
CACACATCGGACTCATCAGGAAGC
AAGCATCGGACTCATCAGGAAGC
CATCGGACTCATCAGGAAGC
CCCATCGGACTCATCAGGAAGC
CATCGGACTCATCAGGAAGC
TCCATCGGACTCATCAGGAAGC
TGCATCGGACTCATCAGGAAGC
ACCTCATCGGACTCATCAGGAAGC
GTCATCGGACTCATCAGGAAGC
TTGTCATCGGACTCATCAGGAAGC
GCCATCGGACTCATCAGGAAGC
TACATCGGACTCATCAGGAAGC
ATCGGACTCATCAGGAAGC
AGCATCGGACTCATCAGGAAGC
ACATCGGACTCATCAGGAAGC
CGGACTCATCAGGAAGC
GTCCATCGGACTCATCAGGAAGC
CATCGGACTCATCAGGAAGC
ATCGGACTCATCAGGAAGC
GCATCGGACTCATCAGGAAGC
ACCATCGGACTCATCAGGAAGC
TCTCATCGGACTCATCAGGAAGC
ATCATCGGACTCATCAGGAAGC
CCATCGGACTCATCAGGAAGC

CCATCGGACTCATCAGGAAGC
TCATCGGACTCATCAGGAAGC
CCATCGGACTCATCAGGAAGC
ACCATCGGACTCATCAGGAAGC
TACATCGGACTCATCAGGAAGC
AGACATCGGACTCATCAGGAAGC
ATTCATCGGACTCATCAGGAAGC
ATCATCGGACTCATCAGGAAGC
GCACATCGGACTCATCAGGAAGC
CCCATCGGACTCATCAGGAAGC
CATCGGACTCATCAGGAAGC
ATCGGACTCATCAGGAAGC
ATCGGACTCATCAGGAAGC
ATCATCGGACTCATCAGGAAGC
ATCGGACTCATCAGGAAGC
ATCATCGGACTCATCAGGAAGC
TGATTCATCGGACTCATCAGGAAGC
GTGTACATCGGACTCATCAGGAAGC
GACCATCGGACTCATCAGGAAGC
CATCGGACTCATCAGGAAGC
CATCGGACTCATCAGGAAGC
CCTCATCGGACTCATCAGGAAGC
CATCGGACTCATCAGGAAGC
GCATCGGACTCATCAGGAAGC
ACATCGGACTCATCAGGAAGC
TATTCATCGGACTCATCAGGAAGC
GTCATCGGACTCATCAGGAAGC
CATCGGACTCATCAGGAAGC
CCATCGGACTCATCAGGAAGC
TGCATCGGACTCATCAGGAAGC
AGACATCGGACTCATCAGGAAGC
AACATCGGACTCATCAGGAAGC
AACATCGGACTCATCAGGAAGC
GCATCGGACTCATCAGGAAGC
CACATCGGACTCATCAGGAAGC
TCCATCGGACTCATCAGGAAGC
GTCATCGGACTCATCAGGAAGC
CAAGTCATCGGACTCATCAGGAAGC
GACATCGGACTCATCAGGAAGC
CACATCGGACTCATCAGGAAGC
CATCGGACTCATCAGGAAGC
GCATCGGACTCATCAGGAAGC
CCACATCGGACTCATCAGGAAGC

AGAAGACATCGGACTCATCAGGAAGC
AAGCATCGGACTCATCAGGAAGC
CATCGGACTCATCAGGAAGC
ATTCATCGGACTCATCAGGAAGC
CAGTCATCGGACTCATCAGGAAGC
CAGCCATCGGACTCATCAGGAAGC
GCAGCATCGGACTCATCAGGAAGC
TTGGCACATCGGACTCATCAGGAAGC
CATCGGACTCATCAGGAAGC
TTCATCGGACTCATCAGGAAGC
GTCATCGGACTCATCAGGAAGC
GACATCGGACTCATCAGGAAGC
CATCGGACTCATCAGGAAGC
ACATCGGACTCATCAGGAAGC
GTCATCGGACTCATCAGGAAGC
GCAGCATCGGACTCATCAGGAAGC
AAGCATCGGACTCATCAGGAAGC
TAACATCGGACTCATCAGGAAGC
CATCGGACTCATCAGGAAGC
GAACATCGGACTCATCAGGAAGC
AGCATCGGACTCATCAGGAAGC
CCATCGGACTCATCAGGAAGC
CATCGGACTCATCAGGAAGC
GCATCGGACTCATCAGGAAGC
CTCCATCGGACTCATCAGGAAGC
GCATCGGACTCATCAGGAAGC
CATCATCGGACTCATCAGGAAGC
GTGCATCGGACTCATCAGGAAGC
TCATCGGACTCATCAGGAAGC
AAACATCGGACTCATCAGGAAGC
CATCGGACTCATCAGGAAGC
ATCGGACTCATCAGGAAGC
TGACATCGGACTCATCAGGAAGC
CCATCGGACTCATCAGGAAGC
TCATCGGACTCATCAGGAAGC
ACCATCGGACTCATCAGGAAGC
CGGCATCGGACTCATCAGGAAGC
CTGCATCGGACTCATCAGGAAGC
AGCATCGGACTCATCAGGAAGC
AGTCATCGGACTCATCAGGAAGC
ATCGGACTCATCAGGAAGC
AGCATCGGACTCATCAGGAAGC
TCATCGGACTCATCAGGAAGC

ACATCGGACTCATCAGGAAGC
AGACATCGGACTCATCAGGAAGC
TCATCGGACTCATCAGGAAGC
CTCATCGGACTCATCAGGAAGC
CACATCGGACTCATCAGGAAGC
TCCCATCGGACTCATCAGGAAGC
TCTGCATCGGACTCATCAGGAAGC
CATCGGACTCATCAGGAAGC
CCATCGGACTCATCAGGAAGC
ACATCGGACTCATCAGGAAGC
ACACATCGGACTCATCAGGAAGC
AGTGCATCGGACTCATCAGGAAGC
CCACATCGGACTCATCAGGAAGC
CTCATCGGACTCATCAGGAAGC
TCATCGGACTCATCAGGAAGC
ATAGCATCGGACTCATCAGGAAGC
TGGCATCGGACTCATCAGGAAGC
TGACATCGGACTCATCAGGAAGC
TCCACATCGGACTCATCAGGAAGC
GCATCGGACTCATCAGGAAGC
GCATCGGACTCATCAGGAAGC
GATGTCATCGGACTCATCAGGAAGC
CTGCATCGGACTCATCAGGAAGC
AGCATCGGACTCATCAGGAAGC
GTCATCGGACTCATCAGGAAGC
TCATCGGACTCATCAGGAAGC
GTCATCGGACTCATCAGGAAGC
AACATCGGACTCATCAGGAAGC
GCCCATCGGACTCATCAGGAAGC
GTGTGCCATCGGACTCATCAGGAAGC
TCAGGGCATCGGACTCATCAGGAAGC
CCCATCGGACTCATCAGGAAGC
AGGCCATCGGACTCATCAGGAAGC
AAAACATCGGACTCATCAGGAAGC
TACATCGGACTCATCAGGAAGC
GCATCGGACTCATCAGGAAGC
TGCCATCGGACTCATCAGGAAGC
TCATCGGACTCATCAGGAAGC
TCGGACTCATCAGGAAGC
ACCATCGGACTCATCAGGAAGC
CATCGGACTCATCAGGAAGC
GTCATCGGACTCATCAGGAAGC
AACATCGGACTCATCAGGAAGC

TCGGACTCATCAGGAAGC
CATCGGACTCATCAGGAAGC
CATCGGACTCATCAGGAAGC
TTCATCGGACTCATCAGGAAGC
CACCATCGGACTCATCAGGAAGC
GTCATCGGACTCATCAGGAAGC
CATCGGACTCATCAGGAAGC
GGCATCGGACTCATCAGGAAGC
TGCCATCGGACTCATCAGGAAGC
CCACATCGGACTCATCAGGAAGC
TCTGCATCGGACTCATCAGGAAGC
TCATCGGACTCATCAGGAAGC
ACATCGGACTCATCAGGAAGC
CACATCGGACTCATCAGGAAGC
GGGTCATCGGACTCATCAGGAAGC
TCATCGGACTCATCAGGAAGC
GCCCATCGGACTCATCAGGAAGC
AGTCATCGGACTCATCAGGAAGC
AGACATCGGACTCATCAGGAAGC
TCCATCGGACTCATCAGGAAGC
GGCCATCGGACTCATCAGGAAGC
ATACATCGGACTCATCAGGAAGC
GCCCATCGGACTCATCAGGAAGC
AGGCATCGGACTCATCAGGAAGC
TCATCGGACTCATCAGGAAGC
GACATCGGACTCATCAGGAAGC
AACATCGGACTCATCAGGAAGC
AGGACATCGGACTCATCAGGAAGC
CTCATCGGACTCATCAGGAAGC
CCCATCGGACTCATCAGGAAGC
TACATCGGACTCATCAGGAAGC
CTCATCGGACTCATCAGGAAGC
GTCATCGGACTCATCAGGAAGC
ACATCGGACTCATCAGGAAGC
TCACATCGGACTCATCAGGAAGC
GAGCATCGGACTCATCAGGAAGC
TGCATCGGACTCATCAGGAAGC
TATGTGCATCGGACTCATCAGGAAGC
CCATCGGACTCATCAGGAAGC
TCATGTCATCGGACTCATCAGGAAGC
CCATCGGACTCATCAGGAAGC
TAGTCCATCGGACTCATCAGGAAGC
CATCGGACTCATCAGGAAGC

TTCATCATCGGACTCATCAGGAAGC
CCCATCGGACTCATCAGGAAGC
CATCGGACTCATCAGGAAGC
TGCATCGGACTCATCAGGAAGC
TACATCGGACTCATCAGGAAGC
CCATCGGACTCATCAGGAAGC
AACAGCATCGGACTCATCAGGAAGC
TGTCATCGGACTCATCAGGAAGC
ACATCGGACTCATCAGGAAGC
CCCATCGGACTCATCAGGAAGC
ATCGGACTCATCAGGAAGC
CAGACATCGGACTCATCAGGAAGC
GCATCGGACTCATCAGGAAGC
AGCATCGGACTCATCAGGAAGC
GCATCGGACTCATCAGGAAGC
GGACATCGGACTCATCAGGAAGC
CCCATCGGACTCATCAGGAAGC
TGCGCATCGGACTCATCAGGAAGC
TCATCGGACTCATCAGGAAGC
CCCCATCGGACTCATCAGGAAGC
TGGCATCGGACTCATCAGGAAGC
TCATCGGACTCATCAGGAAGC
CATCGGACTCATCAGGAAGC
CCATCGGACTCATCAGGAAGC
CATCGGACTCATCAGGAAGC
GCCATCGGACTCATCAGGAAGC
AAAGCATCGGACTCATCAGGAAGC
CTCATCGGACTCATCAGGAAGC
CATCGGACTCATCAGGAAGC
AGGCATCGGACTCATCAGGAAGC
TCTCATCGGACTCATCAGGAAGC
CTCATCGGACTCATCAGGAAGC
CCATCGGACTCATCAGGAAGC
GAACATCGGACTCATCAGGAAGC
CATCGGACTCATCAGGAAGC
GACATCGGACTCATCAGGAAGC
CATCGGACTCATCAGGAAGC
TGCATCGGACTCATCAGGAAGC
TCATCGGACTCATCAGGAAGC
GCCATCGGACTCATCAGGAAGC
AGCATCGGACTCATCAGGAAGC
CCCATCGGACTCATCAGGAAGC
CCATCGGACTCATCAGGAAGC

TTGCCTCATCGGACTCATCAGGAAGC
CATCGGACTCATCAGGAAGC
TCATCGGACTCATCAGGAAGC
CATCGGACTCATCAGGAAGC
TGTCCATCGGACTCATCAGGAAGC
CATCGGACTCATCAGGAAGC
CTGCATCGGACTCATCAGGAAGC
GCGCATCGGACTCATCAGGAAGC
CTCCCATCGGACTCATCAGGAAGC
ACATCGGACTCATCAGGAAGC
GGCATCGGACTCATCAGGAAGC
ATCGGACTCATCAGGAAGC
CTCATCGGACTCATCAGGAAGC
TCACATCGGACTCATCAGGAAGC
TGAGCATCGGACTCATCAGGAAGC
TGGCATCGGACTCATCAGGAAGC
CATCGGACTCATCAGGAAGC
CATCGGACTCATCAGGAAGC
ACATCGGACTCATCAGGAAGC
GGTCATCGGACTCATCAGGAAGC
TAACATCGGACTCATCAGGAAGC
GTCATCGGACTCATCAGGAAGC
AACATCGGACTCATCAGGAAGC
CGCATCGGACTCATCAGGAAGC
ATCGGACTCATCAGGAAGC
ATGCATCGGACTCATCAGGAAGC
GTACATCGGACTCATCAGGAAGC
CATCGGACTCATCAGGAAGC
CATCGGACTCATCAGGAAGC
TCATCGGACTCATCAGGAAGC
ACATCGGACTCATCAGGAAGC
CATCGGACTCATCAGGAAGC
CATCGGACTCATCAGGAAGC
GCATCGGACTCATCAGGAAGC
TGCATCGGACTCATCAGGAAGC
CATCGGACTCATCAGGAAGC
CCATCGGACTCATCAGGAAGC
GCATCGGACTCATCAGGAAGC
CATCGGACTCATCAGGAAGC
AGGTCCATCGGACTCATCAGGAAGC
TCGGACTCATCAGGAAGC
CATCGGACTCATCAGGAAGC
CATCGGACTCATCAGGAAGC

GCATCGGACTCATCAGGAAGC
ACATGCCATCGGACTCATCAGGAAGC
TCATCGGACTCATCAGGAAGC
CATCGGACTCATCAGGAAGC
ATCGGACTCATCAGGAAGC
GCTGCATCGGACTCATCAGGAAGC
GGGCATCGGACTCATCAGGAAGC
GAATCATCGGACTCATCAGGAAGC
GTTCATCGGACTCATCAGGAAGC
CCATCGGACTCATCAGGAAGC
CCATCGGACTCATCAGGAAGC
AAGGCATCGGACTCATCAGGAAGC
ACATCGGACTCATCAGGAAGC
ATGTACATCGGACTCATCAGGAAGC
CCATCGGACTCATCAGGAAGC
TACATCGGACTCATCAGGAAGC
ACATCGGACTCATCAGGAAGC
CCATCGGACTCATCAGGAAGC
CATCGGACTCATCAGGAAGC
GGCCATCGGACTCATCAGGAAGC
CCATCGGACTCATCAGGAAGC
TGTTCATCGGACTCATCAGGAAGC
ACCATCGGACTCATCAGGAAGC
ACATCGGACTCATCAGGAAGC
CCATCGGACTCATCAGGAAGC
GTCATCGGACTCATCAGGAAGC
CCCATCGGACTCATCAGGAAGC
CAGGGCATCGGACTCATCAGGAAGC
CCATCGGACTCATCAGGAAGC
GAGCATCGGACTCATCAGGAAGC
CACATCGGACTCATCAGGAAGC
ACACCATCGGACTCATCAGGAAGC
ACATCGGACTCATCAGGAAGC
CCATCGGACTCATCAGGAAGC
GCCATCGGACTCATCAGGAAGC
CTCATCGGACTCATCAGGAAGC
GTGCATCGGACTCATCAGGAAGC
TGTCATCGGACTCATCAGGAAGC
ATCGGACTCATCAGGAAGC
AGACATCGGACTCATCAGGAAGC
GCCATCGGACTCATCAGGAAGC
CAGGCATCGGACTCATCAGGAAGC
CCCTGTCATCGGACTCATCAGGAAGC

CCCATCGGACTCATCAGGAAGC
CGGACTCATCAGGAAGC
CATCGGACTCATCAGGAAGC
TCATCGGACTCATCAGGAAGC
GCATCGGACTCATCAGGAAGC
CCATCGGACTCATCAGGAAGC
TCGGACTCATCAGGAAGC
GATCATCGGACTCATCAGGAAGC
GCCATCGGACTCATCAGGAAGC
ACATCGGACTCATCAGGAAGC
ACATCGGACTCATCAGGAAGC
CATCGGACTCATCAGGAAGC
CTGCATCGGACTCATCAGGAAGC
ACATCGGACTCATCAGGAAGC
TCTCATCGGACTCATCAGGAAGC
ACATCGGACTCATCAGGAAGC
GCCATCGGACTCATCAGGAAGC
TCCACATCGGACTCATCAGGAAGC
GAACATCGGACTCATCAGGAAGC
TCATCGGACTCATCAGGAAGC
TCATCGGACTCATCAGGAAGC
GCATCGGACTCATCAGGAAGC
TCATCGGACTCATCAGGAAGC
TCACATCGGACTCATCAGGAAGC
CGCATCGGACTCATCAGGAAGC
GCATCGGACTCATCAGGAAGC
ATCGGACTCATCAGGAAGC
GATTCATCGGACTCATCAGGAAGC
TATCTCATCGGACTCATCAGGAAGC
ATCGGACTCATCAGGAAGC
TAACATCGGACTCATCAGGAAGC
TGCCCATCGGACTCATCAGGAAGC
TAGCCATCGGACTCATCAGGAAGC
CATCGGACTCATCAGGAAGC
ACACCATCGGACTCATCAGGAAGC
GCATCGGACTCATCAGGAAGC
CATCGGACTCATCAGGAAGC
TCATCGGACTCATCAGGAAGC
TCGGACTCATCAGGAAGC
ACATCGGACTCATCAGGAAGC
AGCATCGGACTCATCAGGAAGC
TCATCGGACTCATCAGGAAGC
CATCGGACTCATCAGGAAGC

ATCCCATCGGACTCATCAGGAAGC
TGTGTCATCGGACTCATCAGGAAGC
CCATCGGACTCATCAGGAAGC
CAGCATCGGACTCATCAGGAAGC
GCATCGGACTCATCAGGAAGC
TCTCATCGGACTCATCAGGAAGC
ACACCATCGGACTCATCAGGAAGC
CCATCGGACTCATCAGGAAGC
CACCATCGGACTCATCAGGAAGC
CATGGCATCGGACTCATCAGGAAGC
CATCGGACTCATCAGGAAGC
CTCATCGGACTCATCAGGAAGC
TCGGACTCATCAGGAAGC
TTCATCGGACTCATCAGGAAGC
CATCGGACTCATCAGGAAGC
AAACATCGGACTCATCAGGAAGC
GTGTCATCGGACTCATCAGGAAGC
GCATCGGACTCATCAGGAAGC
TCGGACTCATCAGGAAGC
TTGCATCGGACTCATCAGGAAGC
ATCATCGGACTCATCAGGAAGC
CCTCATCGGACTCATCAGGAAGC
ATCGGACTCATCAGGAAGC
GGCATCGGACTCATCAGGAAGC
TAACATCGGACTCATCAGGAAGC
TCATCGGACTCATCAGGAAGC
CACAGCATCGGACTCATCAGGAAGC
ATCGGACTCATCAGGAAGC
AACATCGGACTCATCAGGAAGC
CATCGGACTCATCAGGAAGC
CGCATCGGACTCATCAGGAAGC
GTCATCGGACTCATCAGGAAGC
CTCATCGGACTCATCAGGAAGC
TCATCGGACTCATCAGGAAGC
CATCGGACTCATCAGGAAGC
GCGCATCGGACTCATCAGGAAGC
GAAGCATCGGACTCATCAGGAAGC
TCATCGGACTCATCAGGAAGC
CATCGGACTCATCAGGAAGC
CACCATCGGACTCATCAGGAAGC
TCTCATCGGACTCATCAGGAAGC
CATCGGACTCATCAGGAAGC
GTGACATCGGACTCATCAGGAAGC

GTTTCATCGGACTCATCAGGAAGC
ATCATCGGACTCATCAGGAAGC
CCCATCGGACTCATCAGGAAGC
TCAAACATCGGACTCATCAGGAAGC
TCATCGGACTCATCAGGAAGC
CCCATCGGACTCATCAGGAAGC
TCATCGGACTCATCAGGAAGC
TCACATCGGACTCATCAGGAAGC
CATCGGACTCATCAGGAAGC
GTCATCATCGGACTCATCAGGAAGC
TCGGACTCATCAGGAAGC
GTCATCGGACTCATCAGGAAGC
TGCCATCGGACTCATCAGGAAGC
GCAACATCGGACTCATCAGGAAGC
TTCATCGGACTCATCAGGAAGC
TGACATCGGACTCATCAGGAAGC
CCCATCGGACTCATCAGGAAGC
ATCATCGGACTCATCAGGAAGC
CTTCATCGGACTCATCAGGAAGC
CACATCGGACTCATCAGGAAGC
TCATCGGACTCATCAGGAAGC
CATCGGACTCATCAGGAAGC
CGCCATCGGACTCATCAGGAAGC
TCTCATCGGACTCATCAGGAAGC
ACAATCATCGGACTCATCAGGAAGC
GAGCATCGGACTCATCAGGAAGC
CATCGGACTCATCAGGAAGC
TGAGGTTCATCGGACTCATCAGGAAGC
GCATCGGACTCATCAGGAAGC
CCATCGGACTCATCAGGAAGC
CTCATCGGACTCATCAGGAAGC
ACATCGGACTCATCAGGAAGC
TCATCGGACTCATCAGGAAGC
AAACATCGGACTCATCAGGAAGC
ACATCGGACTCATCAGGAAGC
ACTCATCGGACTCATCAGGAAGC
ACATCGGACTCATCAGGAAGC
GCACATCGGACTCATCAGGAAGC
CACCATCGGACTCATCAGGAAGC
GTACGCATCGGACTCATCAGGAAGC
GCATCGGACTCATCAGGAAGC
CACATCGGACTCATCAGGAAGC
CATCGGACTCATCAGGAAGC

TTCATCGGACTCATCAGGAAGC
TTCATCGGACTCATCAGGAAGC
TCCATCGGACTCATCAGGAAGC
ATCGGACTCATCAGGAAGC
GACATCGGACTCATCAGGAAGC
ACATCGGACTCATCAGGAAGC
CGGCATCGGACTCATCAGGAAGC
TTCATCGGACTCATCAGGAAGC
GCATCGGACTCATCAGGAAGC
GCATCGGACTCATCAGGAAGC
GGCATCGGACTCATCAGGAAGC
GCATCGGACTCATCAGGAAGC
TCATCGGACTCATCAGGAAGC
TTCATCGGACTCATCAGGAAGC
CACCCATCGGACTCATCAGGAAGC
ACATCGGACTCATCAGGAAGC
CATCGGACTCATCAGGAAGC
GCCATCGGACTCATCAGGAAGC
GCCCATCGGACTCATCAGGAAGC
TCATCGGACTCATCAGGAAGC
CTCCATCGGACTCATCAGGAAGC
GCACATCGGACTCATCAGGAAGC
TCGGACTCATCAGGAAGC
GAGACATCGGACTCATCAGGAAGC
GAACATCGGACTCATCAGGAAGC
CATCGGACTCATCAGGAAGC
GGACATCGGACTCATCAGGAAGC
ACATCGGACTCATCAGGAAGC
CATCGGACTCATCAGGAAGC
GGCTCATCGGACTCATCAGGAAGC
ACATCGGACTCATCAGGAAGC
AGCCCATCGGACTCATCAGGAAGC
TCATCGGACTCATCAGGAAGC
ACATCGGACTCATCAGGAAGC
TGCATCGGACTCATCAGGAAGC
AACCATCGGACTCATCAGGAAGC
TTCATCGGACTCATCAGGAAGC
TCATCGGACTCATCAGGAAGC
ATCGGACTCATCAGGAAGC
CCATCGGACTCATCAGGAAGC
TGGAGCATCGGACTCATCAGGAAGC
TCATCGGACTCATCAGGAAGC
TCCATCGGACTCATCAGGAAGC

GGCATCGGACTCATCAGGAAGC
TACATCGGACTCATCAGGAAGC
ACCCATCGGACTCATCAGGAAGC
ACATCGGACTCATCAGGAAGC
GTCATCGGACTCATCAGGAAGC
GCCATACATCGGACTCATCAGGAAGC
CACGCCATCGGACTCATCAGGAAGC
GCTCATCGGACTCATCAGGAAGC
TCCATCGGACTCATCAGGAAGC
CATCGGACTCATCAGGAAGC
ACATCGGACTCATCAGGAAGC
GGCATCGGACTCATCAGGAAGC
ATTTCATCGGACTCATCAGGAAGC
GACATCGGACTCATCAGGAAGC
GCATCGGACTCATCAGGAAGC
CAACATCGGACTCATCAGGAAGC
AGCATCGGACTCATCAGGAAGC
AACATCGGACTCATCAGGAAGC
AGACATCGGACTCATCAGGAAGC
GTTCATCGGACTCATCAGGAAGC
ACATCGGACTCATCAGGAAGC
GTGCATCGGACTCATCAGGAAGC
ACATCGGACTCATCAGGAAGC
GCACATCGGACTCATCAGGAAGC
CCATCGGACTCATCAGGAAGC
GGCATCGGACTCATCAGGAAGC
GGACATCGGACTCATCAGGAAGC
TACATCGGACTCATCAGGAAGC
GGCATCGGACTCATCAGGAAGC
GCAACATCGGACTCATCAGGAAGC
CACCATCGGACTCATCAGGAAGC
CATCGGACTCATCAGGAAGC
CATCATCGGACTCATCAGGAAGC
CATCGGACTCATCAGGAAGC
ACATCGGACTCATCAGGAAGC
TCATCGGACTCATCAGGAAGC
ACATCGGACTCATCAGGAAGC
ATCATCGGACTCATCAGGAAGC
ATCGGACTCATCAGGAAGC
CAGCATCGGACTCATCAGGAAGC
GGGTCATCGGACTCATCAGGAAGC
GTGTCATCGGACTCATCAGGAAGC
TCATCGGACTCATCAGGAAGC

CCATCGGACTCATCAGGAAGC
CATCGGACTCATCAGGAAGC
GTGCATCGGACTCATCAGGAAGC
CATAGCATCGGACTCATCAGGAAGC
CATCGGACTCATCAGGAAGC
AGTGCATCGGACTCATCAGGAAGC
GAACATCGGACTCATCAGGAAGC
AGGCATCGGACTCATCAGGAAGC
AGACATCGGACTCATCAGGAAGC
ATCGGACTCATCAGGAAGC
GGACATCGGACTCATCAGGAAGC
TCCATCGGACTCATCAGGAAGC
GACATCGGACTCATCAGGAAGC
GCATCGGACTCATCAGGAAGC
AAAACATCGGACTCATCAGGAAGC
AGTGCATCGGACTCATCAGGAAGC
CATCGGACTCATCAGGAAGC
TTGTTTCATCGGACTCATCAGGAAGC
AACATCGGACTCATCAGGAAGC
TTTCATCGGACTCATCAGGAAGC
ACAACATCGGACTCATCAGGAAGC
CCATCGGACTCATCAGGAAGC
AAGACATCGGACTCATCAGGAAGC
ACATCGGACTCATCAGGAAGC
GGCATCGGACTCATCAGGAAGC
ATGAACATCGGACTCATCAGGAAGC
GCACCATCGGACTCATCAGGAAGC
CTGACATCGGACTCATCAGGAAGC
CATCGGACTCATCAGGAAGC
CACTAACATCGGACTCATCAGGAAGC
TGCCATCGGACTCATCAGGAAGC
ACATCGGACTCATCAGGAAGC
GCATCGGACTCATCAGGAAGC
ACATCGGACTCATCAGGAAGC
CTCATCGGACTCATCAGGAAGC
TAGGCATCGGACTCATCAGGAAGC
ACATCGGACTCATCAGGAAGC
ACATCGGACTCATCAGGAAGC
GCCATCGGACTCATCAGGAAGC
ATCATCGGACTCATCAGGAAGC
GGCATCGGACTCATCAGGAAGC
CATCGGACTCATCAGGAAGC
CATCGGACTCATCAGGAAGC

CAGCATCGGACTCATCAGGAAGC
TAGGCACATCGGACTCATCAGGAAGC
GCGCATCGGACTCATCAGGAAGC
TTCATCGGACTCATCAGGAAGC
GCATCGGACTCATCAGGAAGC
GGCATCGGACTCATCAGGAAGC
ACTCATCGGACTCATCAGGAAGC
TGCATCGGACTCATCAGGAAGC
CCATCGGACTCATCAGGAAGC
CGTTCATCGGACTCATCAGGAAGC
GTCATCGGACTCATCAGGAAGC
TTTCATCGGACTCATCAGGAAGC
ATGCATCGGACTCATCAGGAAGC
GACATCGGACTCATCAGGAAGC
GGACTCATCAGGAAGC
GGCATCGGACTCATCAGGAAGC
TCGGACTCATCAGGAAGC
CATCGGACTCATCAGGAAGC
ATCGGACTCATCAGGAAGC
GCATCGGACTCATCAGGAAGC
GCATCGGACTCATCAGGAAGC
ACATCGGACTCATCAGGAAGC
ATCGGACTCATCAGGAAGC
TGACATCGGACTCATCAGGAAGC
GATCATCGGACTCATCAGGAAGC
TTCATCGGACTCATCAGGAAGC
AGTATGTCATCGGACTCATCAGGAAGC
AGAACATCGGACTCATCAGGAAGC
CCATCGGACTCATCAGGAAGC
CTCATCGGACTCATCAGGAAGC
GACATCGGACTCATCAGGAAGC
TCATCGGACTCATCAGGAAGC
TTACATCGGACTCATCAGGAAGC
GCATCGGACTCATCAGGAAGC
TTGCATCGGACTCATCAGGAAGC
ATGCATCGGACTCATCAGGAAGC
TAACATCGGACTCATCAGGAAGC
ACTCATCAGGAAGC
CGCATCGGACTCATCAGGAAGC
GCATCGGACTCATCAGGAAGC
ACATCGGACTCATCAGGAAGC
GCATCGGACTCATCAGGAAGC
CCTCATCGGACTCATCAGGAAGC

GATCATCGGACTCATCAGGAAGC
CTCATCGGACTCATCAGGAAGC
GCATCGGACTCATCAGGAAGC
AACATCGGACTCATCAGGAAGC
GACATCGGACTCATCAGGAAGC
AACATCGGACTCATCAGGAAGC
ACCCATCGGACTCATCAGGAAGC
ACACATCGGACTCATCAGGAAGC
TAGCATCGGACTCATCAGGAAGC
GGCATCGGACTCATCAGGAAGC
TCATCGGACTCATCAGGAAGC
CAGCCATCGGACTCATCAGGAAGC
GTGCATCGGACTCATCAGGAAGC
ATCGGACTCATCAGGAAGC
CTCCATCGGACTCATCAGGAAGC
ATCGGACTCATCAGGAAGC
ACATCGGACTCATCAGGAAGC
CCATCGGACTCATCAGGAAGC
GTGCATCGGACTCATCAGGAAGC
ATCCATCGGACTCATCAGGAAGC
TGCATCGGACTCATCAGGAAGC
TGCATCGGACTCATCAGGAAGC
ATCATCGGACTCATCAGGAAGC
TCCACCATCGGACTCATCAGGAAGC
TGGACATCGGACTCATCAGGAAGC
GAACATCGGACTCATCAGGAAGC
GTACATCGGACTCATCAGGAAGC
ATCACATCGGACTCATCAGGAAGC
GACATCGGACTCATCAGGAAGC
GAGACATCGGACTCATCAGGAAGC
CCATCGGACTCATCAGGAAGC
AGCCATCGGACTCATCAGGAAGC
GCATCGGACTCATCAGGAAGC
GCACATCGGACTCATCAGGAAGC
ACCATCGGACTCATCAGGAAGC
ATCATCGGACTCATCAGGAAGC
CCATCGGACTCATCAGGAAGC
TACACATCGGACTCATCAGGAAGC
TCTCTCATCGGACTCATCAGGAAGC
TGCATCGGACTCATCAGGAAGC
CCATCGGACTCATCAGGAAGC
ATGCCATCGGACTCATCAGGAAGC
CACATCGGACTCATCAGGAAGC

GGACATCGGACTCATCAGGAAGC
AAAAGCATCGGACTCATCAGGAAGC
ACCATCGGACTCATCAGGAAGC
CACCATCGGACTCATCAGGAAGC
TTCATCGGACTCATCAGGAAGC
ACATCGGACTCATCAGGAAGC
AGCATCGGACTCATCAGGAAGC
CATCGGACTCATCAGGAAGC
AGACATCGGACTCATCAGGAAGC
TGGACATCGGACTCATCAGGAAGC
AACATCGGACTCATCAGGAAGC
ACCTCATCGGACTCATCAGGAAGC
CCCATCGGACTCATCAGGAAGC
CACATCGGACTCATCAGGAAGC
ACATCGGACTCATCAGGAAGC
CTCATCGGACTCATCAGGAAGC
TGCATCGGACTCATCAGGAAGC
GAGCATCGGACTCATCAGGAAGC
GCATCGGACTCATCAGGAAGC
ATCGGACTCATCAGGAAGC
TCCATCGGACTCATCAGGAAGC
TCATCGGACTCATCAGGAAGC
GCCATCGGACTCATCAGGAAGC
AACATCGGACTCATCAGGAAGC
CATCGGACTCATCAGGAAGC
TTCATCGGACTCATCAGGAAGC
CATCGGACTCATCAGGAAGC
TCACATCGGACTCATCAGGAAGC
TGAGCATCGGACTCATCAGGAAGC
ACACACATCGGACTCATCAGGAAGC
GCCATCGGACTCATCAGGAAGC
TGTCATCGGACTCATCAGGAAGC
CGAGCATCGGACTCATCAGGAAGC
GTGCATCGGACTCATCAGGAAGC
AACCATCGGACTCATCAGGAAGC
CATCGGACTCATCAGGAAGC
TCGGACTCATCAGGAAGC
CCCATCGGACTCATCAGGAAGC
ATCGGACTCATCAGGAAGC
TCCATCGGACTCATCAGGAAGC
TCATCGGACTCATCAGGAAGC
ACCATCGGACTCATCAGGAAGC
AAGCATCGGACTCATCAGGAAGC

ACATCGGACTCATCAGGAAGC
CATCGGACTCATCAGGAAGC
TGATGACATCGGACTCATCAGGAAGC
ATCGGACTCATCAGGAAGC
ACATCGGACTCATCAGGAAGC
TGCATCGGACTCATCAGGAAGC
ATCATCGGACTCATCAGGAAGC
CCCATCGGACTCATCAGGAAGC
AGGCATCGGACTCATCAGGAAGC
ATCGGACTCATCAGGAAGC
TCCATCGGACTCATCAGGAAGC
AAGCATCGGACTCATCAGGAAGC
TGTCATCGGACTCATCAGGAAGC
GGCTCATCGGACTCATCAGGAAGC
CACATCGGACTCATCAGGAAGC
GCATCGGACTCATCAGGAAGC
AGTGCATCGGACTCATCAGGAAGC
TGACATCGGACTCATCAGGAAGC
GTCATCGGACTCATCAGGAAGC
TCATCGGACTCATCAGGAAGC
CAGCATCGGACTCATCAGGAAGC
CAGGCATCGGACTCATCAGGAAGC
TGCATCGGACTCATCAGGAAGC
AGCATCGGACTCATCAGGAAGC
TCATCGGACTCATCAGGAAGC
CGCCATCGGACTCATCAGGAAGC
CATCGGACTCATCAGGAAGC
CCATCGGACTCATCAGGAAGC
GCCATCGGACTCATCAGGAAGC
GGTCATCGGACTCATCAGGAAGC
CATCGGACTCATCAGGAAGC
TTGTGCATCGGACTCATCAGGAAGC
AGGGCATCGGACTCATCAGGAAGC
CATCGGACTCATCAGGAAGC
GTGGGCATCGGACTCATCAGGAAGC
GAGGCATCGGACTCATCAGGAAGC
TCCCATCGGACTCATCAGGAAGC
TCATCGGACTCATCAGGAAGC
GAGCATCGGACTCATCAGGAAGC
AAACCACATCGGACTCATCAGGAAGC
TGCATCGGACTCATCAGGAAGC
CACATCGGACTCATCAGGAAGC
CAGCATCGGACTCATCAGGAAGC

GGGCATCGGACTCATCAGGAAGC
AACATCGGACTCATCAGGAAGC
GGCCATCGGACTCATCAGGAAGC
ACTATCCATCGGACTCATCAGGAAGC
CCACATCGGACTCATCAGGAAGC
TCTGCATCGGACTCATCAGGAAGC
TGCATCGGACTCATCAGGAAGC
ACATCGGACTCATCAGGAAGC
AACACATCGGACTCATCAGGAAGC
GTCCATCGGACTCATCAGGAAGC
TACATCGGACTCATCAGGAAGC
TTCATCGGACTCATCAGGAAGC
AAACATCGGACTCATCAGGAAGC
ACATCGGACTCATCAGGAAGC
GGCATCGGACTCATCAGGAAGC
ATACATCGGACTCATCAGGAAGC
TGTGCATCGGACTCATCAGGAAGC
GGCATCGGACTCATCAGGAAGC
TTAGCATCGGACTCATCAGGAAGC
TGCATCGGACTCATCAGGAAGC
ATCATCGGACTCATCAGGAAGC
CCTCCATCGGACTCATCAGGAAGC
CCATCGGACTCATCAGGAAGC
GTCATCGGACTCATCAGGAAGC
CAGCATCGGACTCATCAGGAAGC
CACATCGGACTCATCAGGAAGC
GTCATCGGACTCATCAGGAAGC
CATCGGACTCATCAGGAAGC
CTCATCGGACTCATCAGGAAGC
CCACATCGGACTCATCAGGAAGC
TCCATCGGACTCATCAGGAAGC
AGACATCGGACTCATCAGGAAGC
ACATCGGACTCATCAGGAAGC
CAACATCGGACTCATCAGGAAGC
TGCCATCGGACTCATCAGGAAGC
ATCATCGGACTCATCAGGAAGC
TACATCGGACTCATCAGGAAGC
GATCATCGGACTCATCAGGAAGC
TTCATCGGACTCATCAGGAAGC
ATCATCGGACTCATCAGGAAGC
TGGCATCGGACTCATCAGGAAGC
CCCCCATCGGACTCATCAGGAAGC
TGACATCGGACTCATCAGGAAGC

GTCCATCGGACTCATCAGGAAGC
AGTGACCATCGGACTCATCAGGAAGC
GACTCATCAGGAAGC
CCCATCGGACTCATCAGGAAGC
CCATCGGACTCATCAGGAAGC
GCATCGGACTCATCAGGAAGC
GGAGCATCGGACTCATCAGGAAGC
TGGGGTCATCGGACTCATCAGGAAGC
CCACATCGGACTCATCAGGAAGC
CCATCGGACTCATCAGGAAGC
CACATCGGACTCATCAGGAAGC
AATGTTCATCGGACTCATCAGGAAGC
GATCATCGGACTCATCAGGAAGC
ACATCGGACTCATCAGGAAGC
ATCGGACTCATCAGGAAGC
GTGCCATCGGACTCATCAGGAAGC
AGTTTCATCGGACTCATCAGGAAGC
CCATCGGACTCATCAGGAAGC
CCATCGGACTCATCAGGAAGC
ATCGGACTCATCAGGAAGC
GCACATCGGACTCATCAGGAAGC
ATCGGACTCATCAGGAAGC
ATCACATCGGACTCATCAGGAAGC
TATCATCGGACTCATCAGGAAGC
ATGCATCGGACTCATCAGGAAGC
AACATCGGACTCATCAGGAAGC
TCCCCATCGGACTCATCAGGAAGC
TTGGCATCGGACTCATCAGGAAGC
TGTCATCGGACTCATCAGGAAGC
CCATCGGACTCATCAGGAAGC
AGACATCGGACTCATCAGGAAGC
TTGCATCGGACTCATCAGGAAGC
AGGAACATCGGACTCATCAGGAAGC
CCATCGGACTCATCAGGAAGC
GTCTCATCGGACTCATCAGGAAGC
TCCACATCGGACTCATCAGGAAGC
GACATCGGACTCATCAGGAAGC
AGGCATCGGACTCATCAGGAAGC
CTCATCGGACTCATCAGGAAGC
TAGACATCGGACTCATCAGGAAGC
CATCGGACTCATCAGGAAGC
GGCATCGGACTCATCAGGAAGC
ACATCGGACTCATCAGGAAGC

GGCATCGGACTCATCAGGAAGC
CACATCGGACTCATCAGGAAGC
GGCATCGGACTCATCAGGAAGC
CACACATCGGACTCATCAGGAAGC
CCCCATCGGACTCATCAGGAAGC
TGGCCATCGGACTCATCAGGAAGC
AGCATCGGACTCATCAGGAAGC
GGCATCGGACTCATCAGGAAGC
CCTCATCGGACTCATCAGGAAGC
TTGGTCATCGGACTCATCAGGAAGC
CCTCATCGGACTCATCAGGAAGC
TGTCATCGGACTCATCAGGAAGC
CATCGGACTCATCAGGAAGC
AGCCATCGGACTCATCAGGAAGC
CGCAACATCGGACTCATCAGGAAGC
CATCGGACTCATCAGGAAGC
AACCACATCGGACTCATCAGGAAGC
ACCCATCGGACTCATCAGGAAGC
CATCGGACTCATCAGGAAGC
TACATCGGACTCATCAGGAAGC
CATCATCGGACTCATCAGGAAGC
ACATCGGACTCATCAGGAAGC
TAGACATCGGACTCATCAGGAAGC
ATCGGACTCATCAGGAAGC
GAACATCGGACTCATCAGGAAGC
AACATCGGACTCATCAGGAAGC
TGCATCGGACTCATCAGGAAGC
CTGCTCATCGGACTCATCAGGAAGC
TCTCATCGGACTCATCAGGAAGC
ACATCGGACTCATCAGGAAGC
CATCGGACTCATCAGGAAGC
TCCATCGGACTCATCAGGAAGC
TCATCGGACTCATCAGGAAGC
GGACATCGGACTCATCAGGAAGC
GTCATCGGACTCATCAGGAAGC
CCACATCGGACTCATCAGGAAGC
CCCATCGGACTCATCAGGAAGC
TTGCTCATCGGACTCATCAGGAAGC
CGCATCGGACTCATCAGGAAGC
GAAAGCATCGGACTCATCAGGAAGC
ACATCGGACTCATCAGGAAGC
GTCATCGGACTCATCAGGAAGC
GACCCATCGGACTCATCAGGAAGC

GCCATCGGACTCATCAGGAAGC
CCCATCGGACTCATCAGGAAGC
GGCATCGGACTCATCAGGAAGC
CCCATCGGACTCATCAGGAAGC
CCACATCGGACTCATCAGGAAGC
CATCGGACTCATCAGGAAGC
GTGACATCGGACTCATCAGGAAGC
TTCTCCATCGGACTCATCAGGAAGC
TCATCGGACTCATCAGGAAGC
ACATCGGACTCATCAGGAAGC
AAAGGCTCATCGGACTCATCAGGAAGC
ACATCGGACTCATCAGGAAGC
TTTATCATCGGACTCATCAGGAAGC
CATCGGACTCATCAGGAAGC
TCATCGGACTCATCAGGAAGC
CCATCGGACTCATCAGGAAGC
CCCATCGGACTCATCAGGAAGC
AGCATCGGACTCATCAGGAAGC
CACATCGGACTCATCAGGAAGC
ATCATCGGACTCATCAGGAAGC
GTGACATCGGACTCATCAGGAAGC
CACACATCGGACTCATCAGGAAGC
ATTGACATCGGACTCATCAGGAAGC
ACATCGGACTCATCAGGAAGC
CCATCGGACTCATCAGGAAGC
CATCGGACTCATCAGGAAGC
CATCGGACTCATCAGGAAGC
ACTTCATCGGACTCATCAGGAAGC
CATTCATCGGACTCATCAGGAAGC
TTCATCGGACTCATCAGGAAGC
TTCTCATCGGACTCATCAGGAAGC
ACATCGGACTCATCAGGAAGC
CAGAACATCGGACTCATCAGGAAGC
AGTCATCGGACTCATCAGGAAGC
GCATCGGACTCATCAGGAAGC
CACATCGGACTCATCAGGAAGC
TCGGACTCATCAGGAAGC
GGCATCGGACTCATCAGGAAGC
AGTCATCGGACTCATCAGGAAGC
GGCATCGGACTCATCAGGAAGC
GAGACATCGGACTCATCAGGAAGC
CCATCGGACTCATCAGGAAGC
GTTCATCGGACTCATCAGGAAGC

AGTGCATCGGACTCATCAGGAAGC
CACATCGGACTCATCAGGAAGC
ACATCGGACTCATCAGGAAGC
CCCATCGGACTCATCAGGAAGC
CACATCGGACTCATCAGGAAGC
ACATCGGACTCATCAGGAAGC
ACACATCGGACTCATCAGGAAGC
GATCATCGGACTCATCAGGAAGC
GCCCATCGGACTCATCAGGAAGC
CCCATCGGACTCATCAGGAAGC
CATCGGACTCATCAGGAAGC
GGGCATCGGACTCATCAGGAAGC
ACATCGGACTCATCAGGAAGC
TTGACATCGGACTCATCAGGAAGC
ATCGGACTCATCAGGAAGC
TTGCATCGGACTCATCAGGAAGC
GAGCATCGGACTCATCAGGAAGC
CATCGGACTCATCAGGAAGC
CTTTCATCGGACTCATCAGGAAGC
AGCCATCGGACTCATCAGGAAGC
TCGGACTCATCAGGAAGC
CAGCATCGGACTCATCAGGAAGC
GCATCGGACTCATCAGGAAGC
TAGGCATCGGACTCATCAGGAAGC
TGTCATCGGACTCATCAGGAAGC
GAGCATCGGACTCATCAGGAAGC
CGCATCGGACTCATCAGGAAGC
GCCCATCGGACTCATCAGGAAGC
AAGCCATCGGACTCATCAGGAAGC
TCATCGGACTCATCAGGAAGC
TCATCGGACTCATCAGGAAGC
TCTTCATCGGACTCATCAGGAAGC
TTCCATCGGACTCATCAGGAAGC
GTTCATCGGACTCATCAGGAAGC
ACATCGGACTCATCAGGAAGC
CCCATCGGACTCATCAGGAAGC
GGTCATCGGACTCATCAGGAAGC
ATCGGACTCATCAGGAAGC
CACCTCATCGGACTCATCAGGAAGC
CCCTCATCGGACTCATCAGGAAGC
CCCATCGGACTCATCAGGAAGC
AAACAGACATCGGACTCATCAGGAAGC
GAACCCATCGGACTCATCAGGAAGC

GCTCATCGGACTCATCAGGAAGC
CTTCCATCGGACTCATCAGGAAGC
TGCATCGGACTCATCAGGAAGC
GCATCGGACTCATCAGGAAGC
TCATCGGACTCATCAGGAAGC
TCCATCGGACTCATCAGGAAGC
GACATCGGACTCATCAGGAAGC
TTCATCGGACTCATCAGGAAGC
GCATCGGACTCATCAGGAAGC
ATCGGACTCATCAGGAAGC
CCCATCGGACTCATCAGGAAGC
TTCCATCGGACTCATCAGGAAGC
ATCGGACTCATCAGGAAGC
CCTTCACATCGGACTCATCAGGAAGC
CATCGGACTCATCAGGAAGC
GGAACATCGGACTCATCAGGAAGC
ACGAGCATCGGACTCATCAGGAAGC
TCATCGGACTCATCAGGAAGC
TTCATCGGACTCATCAGGAAGC
CCTGCATCGGACTCATCAGGAAGC
GCCATCGGACTCATCAGGAAGC
CCATCGGACTCATCAGGAAGC
AACATCGGACTCATCAGGAAGC
GAGCCATCGGACTCATCAGGAAGC
TGGACATCGGACTCATCAGGAAGC
TCATCGGACTCATCAGGAAGC
GGGCATCGGACTCATCAGGAAGC
GTCATCGGACTCATCAGGAAGC
ACATCGGACTCATCAGGAAGC
TCATCGGACTCATCAGGAAGC
GTACATCGGACTCATCAGGAAGC
GCTGCATCGGACTCATCAGGAAGC
CATCATCGGACTCATCAGGAAGC
TCATCGGACTCATCAGGAAGC
TAGTCTCATCGGACTCATCAGGAAGC
GGATCATCGGACTCATCAGGAAGC
CCATCGGACTCATCAGGAAGC
AAGCATCGGACTCATCAGGAAGC
CTTCATCGGACTCATCAGGAAGC
ACATCGGACTCATCAGGAAGC
GCATCGGACTCATCAGGAAGC
ACCATCGGACTCATCAGGAAGC
CATCGGACTCATCAGGAAGC

CACATCGGACTCATCAGGAAGC
CCATCGGACTCATCAGGAAGC
GTGCATCGGACTCATCAGGAAGC
CTGCATCGGACTCATCAGGAAGC
GGAACATCGGACTCATCAGGAAGC
GCACATCGGACTCATCAGGAAGC
GGAACATCGGACTCATCAGGAAGC
GAGCATCGGACTCATCAGGAAGC
AGAGCATCGGACTCATCAGGAAGC
TGGCACATCGGACTCATCAGGAAGC
CACATCGGACTCATCAGGAAGC
TCATCGGACTCATCAGGAAGC
TCCATCGGACTCATCAGGAAGC
CCATCGGACTCATCAGGAAGC
ACCATCGGACTCATCAGGAAGC
CCCATCGGACTCATCAGGAAGC
TGTCATCGGACTCATCAGGAAGC
TCATCGGACTCATCAGGAAGC
CATCGGACTCATCAGGAAGC
AGGCATCGGACTCATCAGGAAGC
ACATCGGACTCATCAGGAAGC
TGTGCATCGGACTCATCAGGAAGC
ACATCGGACTCATCAGGAAGC
ACATCGGACTCATCAGGAAGC
ATCGGACTCATCAGGAAGC
GGGACATCGGACTCATCAGGAAGC
GAACTCATCGGACTCATCAGGAAGC
CATCGGACTCATCAGGAAGC
TCACATCGGACTCATCAGGAAGC
CAGCATCGGACTCATCAGGAAGC
GGCATCGGACTCATCAGGAAGC
GAGACATCGGACTCATCAGGAAGC
ATCATCGGACTCATCAGGAAGC
CTTTCATCGGACTCATCAGGAAGC
TTCATCGGACTCATCAGGAAGC
TCACACATCGGACTCATCAGGAAGC
CAGGCATCATCGGACTCATCAGGAAGC
GTCATCGGACTCATCAGGAAGC
AATGACATCGGACTCATCAGGAAGC
ACATCGGACTCATCAGGAAGC
TGCATCGGACTCATCAGGAAGC
ACATCGGACTCATCAGGAAGC
GCTTATCATCGGACTCATCAGGAAGC

CAGGCATCGGACTCATCAGGAAGC
CCATCGGACTCATCAGGAAGC
AACATCGGACTCATCAGGAAGC
GGCATCGGACTCATCAGGAAGC
GCATCGGACTCATCAGGAAGC
CTTCATCGGACTCATCAGGAAGC
AGGGGACATCGGACTCATCAGGAAGC
TTCCATCGGACTCATCAGGAAGC
AGGCATCGGACTCATCAGGAAGC
AAGCATCGGACTCATCAGGAAGC
CTCTCAACATCGGACTCATCAGGAAGC
CAGTCATCGGACTCATCAGGAAGC
TGTGCCATCGGACTCATCAGGAAGC
CATCGGACTCATCAGGAAGC
AACATCGGACTCATCAGGAAGC
TGCCATCGGACTCATCAGGAAGC
TGTACATCGGACTCATCAGGAAGC
GAAACACATCGGACTCATCAGGAAGC
CTCATCGGACTCATCAGGAAGC
AGCATCGGACTCATCAGGAAGC
GCATCGGACTCATCAGGAAGC
CCAACATCGGACTCATCAGGAAGC
CTCATCGGACTCATCAGGAAGC
CCATCGGACTCATCAGGAAGC
TTCATCGGACTCATCAGGAAGC
CTCATCGGACTCATCAGGAAGC
ATTCATCGGACTCATCAGGAAGC
CATCGGACTCATCAGGAAGC
GATGTCATCGGACTCATCAGGAAGC
GCATCGGACTCATCAGGAAGC
GCACATCGGACTCATCAGGAAGC
AACACATCGGACTCATCAGGAAGC
TCTCTCCATCGGACTCATCAGGAAGC
ACATCGGACTCATCAGGAAGC
GCATCGGACTCATCAGGAAGC
TGTTCATCGGACTCATCAGGAAGC
GGCCATCGGACTCATCAGGAAGC
ATGCCATCGGACTCATCAGGAAGC
TCATCGGACTCATCAGGAAGC
GAGCATCGGACTCATCAGGAAGC
CATCGGACTCATCAGGAAGC
CCATCGGACTCATCAGGAAGC
TGTTCCATCGGACTCATCAGGAAGC

CATCGGACTCATCAGGAAGC
TCACATCGGACTCATCAGGAAGC
CACATCGGACTCATCAGGAAGC
GGTGGCATCGGACTCATCAGGAAGC
TCATCGGACTCATCAGGAAGC
GTGTCCATCGGACTCATCAGGAAGC
TGAGCATCGGACTCATCAGGAAGC
GGTGCCCCATCGGACTCATCAGGAAGC
CCACCATCGGACTCATCAGGAAGC
GTACTGACATCGGACTCATCAGGAAGC
GACATCGGACTCATCAGGAAGC
CCATCGGACTCATCAGGAAGC
TCCATCGGACTCATCAGGAAGC
TCATCGGACTCATCAGGAAGC
GACATCGGACTCATCAGGAAGC
TGTCATCGGACTCATCAGGAAGC
CCATCGGACTCATCAGGAAGC
CTCATCGGACTCATCAGGAAGC
AGCCATCGGACTCATCAGGAAGC
CCCATCGGACTCATCAGGAAGC
GCACATCGGACTCATCAGGAAGC
GGGCATCGGACTCATCAGGAAGC
ATCGGACTCATCAGGAAGC
TCCACATCGGACTCATCAGGAAGC
CATCGGACTCATCAGGAAGC
ACCATCGGACTCATCAGGAAGC
TGTCATCGGACTCATCAGGAAGC
CTGTCATCGGACTCATCAGGAAGC
TCAGACATCGGACTCATCAGGAAGC
GGCCATCGGACTCATCAGGAAGC
CTGCATCGGACTCATCAGGAAGC
AGCGCATCGGACTCATCAGGAAGC
AACATCGGACTCATCAGGAAGC
TCGGACTCATCAGGAAGC
CATCGGACTCATCAGGAAGC
GACACATCGGACTCATCAGGAAGC
GGGCATCGGACTCATCAGGAAGC
TACATCGGACTCATCAGGAAGC
CTCCATCGGACTCATCAGGAAGC
CCATCGGACTCATCAGGAAGC
CATCGGACTCATCAGGAAGC
AGCCATCGGACTCATCAGGAAGC
GCGCTCCATCGGACTCATCAGGAAGC

ACATCGGACTCATCAGGAAGC
CATCATCGGACTCATCAGGAAGC
GACATCGGACTCATCAGGAAGC
CACACATCGGACTCATCAGGAAGC
GCATCGGACTCATCAGGAAGC
GCCCATCGGACTCATCAGGAAGC
TTGCCCATCGGACTCATCAGGAAGC
GGTCATCGGACTCATCAGGAAGC
CTCATCGGACTCATCAGGAAGC
ATCATCATCGGACTCATCAGGAAGC
GTCATCGGACTCATCAGGAAGC
GTTCATCGGACTCATCAGGAAGC
GGCATCGGACTCATCAGGAAGC
AGCACATCGGACTCATCAGGAAGC
ATGGCATCGGACTCATCAGGAAGC
ACCATCGGACTCATCAGGAAGC
TAGCATCGGACTCATCAGGAAGC
CCGTGCATCGGACTCATCAGGAAGC
ACATCGGACTCATCAGGAAGC
AACATCGGACTCATCAGGAAGC
ACATCGGACTCATCAGGAAGC
GGTACATCGGACTCATCAGGAAGC
TCATCGGACTCATCAGGAAGC
ATGCAACATCGGACTCATCAGGAAGC
CACATCGGACTCATCAGGAAGC
AAGCATCGGACTCATCAGGAAGC
ATGCATCGGACTCATCAGGAAGC
CATCGGACTCATCAGGAAGC
GGTACATCGGACTCATCAGGAAGC
CATCGGACTCATCAGGAAGC
TCGGACTCATCAGGAAGC
GTCTCATCGGACTCATCAGGAAGC
CATCGGACTCATCAGGAAGC
CATCGGACTCATCAGGAAGC
GCATCGGACTCATCAGGAAGC
GGCTCATCGGACTCATCAGGAAGC
GCATCGGACTCATCAGGAAGC
ATCGGACTCATCAGGAAGC
GTCCCATCGGACTCATCAGGAAGC
TACATCGGACTCATCAGGAAGC
TAGCATCGGACTCATCAGGAAGC
TCATCGGACTCATCAGGAAGC
CAGCATCGGACTCATCAGGAAGC

GGCACATCGGACTCATCAGGAAGC
CATCGGACTCATCAGGAAGC
TCATCGGACTCATCAGGAAGC
CGCATCGGACTCATCAGGAAGC
TGGGCATCGGACTCATCAGGAAGC
ACCCATCGGACTCATCAGGAAGC
TCATCGGACTCATCAGGAAGC
GCATCGGACTCATCAGGAAGC
GTGCATCGGACTCATCAGGAAGC
GTACATCGGACTCATCAGGAAGC
ATGGCATCGGACTCATCAGGAAGC
AGCATCGGACTCATCAGGAAGC
GGCATCGGACTCATCAGGAAGC
AACATCGGACTCATCAGGAAGC
CATCGGACTCATCAGGAAGC
GCATCGGACTCATCAGGAAGC
GGAGATCATCGGACTCATCAGGAAGC
GGCATCGGACTCATCAGGAAGC
AGGGCATCGGACTCATCAGGAAGC
CCATCGGACTCATCAGGAAGC
AAACATCGGACTCATCAGGAAGC
GCTGACATCGGACTCATCAGGAAGC
TCTGCATCGGACTCATCAGGAAGC
GGCATCGGACTCATCAGGAAGC
TCGGACTCATCAGGAAGC
CCATCGGACTCATCAGGAAGC
TGACATCGGACTCATCAGGAAGC
TTAGTCATCGGACTCATCAGGAAGC
AGGCATCGGACTCATCAGGAAGC
CATCGGACTCATCAGGAAGC
TCCATCGGACTCATCAGGAAGC
ACACAGCATCGGACTCATCAGGAAGC
GACATCGGACTCATCAGGAAGC
AGCATCGGACTCATCAGGAAGC
AACATCGGACTCATCAGGAAGC
ACATCGGACTCATCAGGAAGC
CCACATCGGACTCATCAGGAAGC
ATCGGACTCATCAGGAAGC
AGCATCGGACTCATCAGGAAGC
CACATCGGACTCATCAGGAAGC
AGCATCGGACTCATCAGGAAGC
TCCATCGGACTCATCAGGAAGC
ACATCGGACTCATCAGGAAGC

GATTTCATCGGACTCATCAGGAAGC
ACATCGGACTCATCAGGAAGC
CCCATCGGACTCATCAGGAAGC
AGGTCATCGGACTCATCAGGAAGC
CCCATCGGACTCATCAGGAAGC
TCATCGGACTCATCAGGAAGC
CTCCATCGGACTCATCAGGAAGC
TTTTCATCGGACTCATCAGGAAGC
TCATCGGACTCATCAGGAAGC
AGCCATCGGACTCATCAGGAAGC
ATCCATCGGACTCATCAGGAAGC
AACATCGGACTCATCAGGAAGC
GCATCGGACTCATCAGGAAGC
CCATCGGACTCATCAGGAAGC
TGCATCGGACTCATCAGGAAGC
CTCCATCGGACTCATCAGGAAGC
TCATCATCGGACTCATCAGGAAGC
AGCATCGGACTCATCAGGAAGC
GTGCCATCGGACTCATCAGGAAGC
CTCATCGGACTCATCAGGAAGC
TAATGGTCATCGGACTCATCAGGAAGC
TGTGCATCGGACTCATCAGGAAGC
TCATCGGACTCATCAGGAAGC
CCATCGGACTCATCAGGAAGC
CAATCATCGGACTCATCAGGAAGC
ACATCGGACTCATCAGGAAGC
TGTCATCGGACTCATCAGGAAGC
ATCGGACTCATCAGGAAGC
CCCCCATCGGACTCATCAGGAAGC
CATCGGACTCATCAGGAAGC
CCACATCGGACTCATCAGGAAGC
CCATCGGACTCATCAGGAAGC
GCATCGGACTCATCAGGAAGC
GTGTCATCGGACTCATCAGGAAGC
CCAACATCGGACTCATCAGGAAGC
CTCATCGGACTCATCAGGAAGC
GTCATCGGACTCATCAGGAAGC
CATCGGACTCATCAGGAAGC
ATAACACATCGGACTCATCAGGAAGC
ACCATCGGACTCATCAGGAAGC
AACATCGGACTCATCAGGAAGC
TGCATCGGACTCATCAGGAAGC
AGCATCGGACTCATCAGGAAGC

CCATCGGACTCATCAGGAAGC
ACATCGGACTCATCAGGAAGC
GTCATCGGACTCATCAGGAAGC
TCATCGGACTCATCAGGAAGC
AGCATCATCGGACTCATCAGGAAGC
CCATCGGACTCATCAGGAAGC
GAGATCATCGGACTCATCAGGAAGC
CTCACATCGGACTCATCAGGAAGC
CACATCGGACTCATCAGGAAGC
CATCGGACTCATCAGGAAGC
CATCGGACTCATCAGGAAGC
ATCATCGGACTCATCAGGAAGC
GCCCATCGGACTCATCAGGAAGC
CCCATCGGACTCATCAGGAAGC
ATCGGACTCATCAGGAAGC
ACATCGGACTCATCAGGAAGC
CATTGCATCGGACTCATCAGGAAGC
TGAAGACATCGGACTCATCAGGAAGC
TGTCATCGGACTCATCAGGAAGC
AGACATCGGACTCATCAGGAAGC
CATCGGACTCATCAGGAAGC
ATGCATCGGACTCATCAGGAAGC
TCACCATCGGACTCATCAGGAAGC
TGCATCGGACTCATCAGGAAGC
CATCGGACTCATCAGGAAGC
GGCCATCGGACTCATCAGGAAGC
GTGCATCGGACTCATCAGGAAGC
ATCATCGGACTCATCAGGAAGC
CTGCATCGGACTCATCAGGAAGC
GGTCATCGGACTCATCAGGAAGC
GCATCATCGGACTCATCAGGAAGC
TAGGTCATCGGACTCATCAGGAAGC
CATCATCGGACTCATCAGGAAGC
GACATCGGACTCATCAGGAAGC
ACATCGGACTCATCAGGAAGC
GCATCGGACTCATCAGGAAGC
TCCATCGGACTCATCAGGAAGC
TTCATCGGACTCATCAGGAAGC
GTGCATCGGACTCATCAGGAAGC
ACATCGGACTCATCAGGAAGC
TCATCGGACTCATCAGGAAGC
ACTCATCGGACTCATCAGGAAGC
CCATCGGACTCATCAGGAAGC

AACATCGGACTCATCAGGAAGC
GTCATCGGACTCATCAGGAAGC
TATCATCGGACTCATCAGGAAGC
CCATCGGACTCATCAGGAAGC
AGCCATCGGACTCATCAGGAAGC
GGCATCGGACTCATCAGGAAGC
AACATCGGACTCATCAGGAAGC
ACCCATCGGACTCATCAGGAAGC
TAGCATCGGACTCATCAGGAAGC
TCATCGGACTCATCAGGAAGC
CCCATCGGACTCATCAGGAAGC
ACCATCGGACTCATCAGGAAGC
CCGTCATCGGACTCATCAGGAAGC
ATCATCGGACTCATCAGGAAGC
TGGCATCGGACTCATCAGGAAGC
CCATCGGACTCATCAGGAAGC
CATCGGACTCATCAGGAAGC
ACATCGGACTCATCAGGAAGC
AGGCATCGGACTCATCAGGAAGC
ACATCGGACTCATCAGGAAGC
GTTGCATCGGACTCATCAGGAAGC
TTCATCGGACTCATCAGGAAGC
GAACATCGGACTCATCAGGAAGC
TCCATCGGACTCATCAGGAAGC
GAACATCGGACTCATCAGGAAGC
AACATCGGACTCATCAGGAAGC
CAACATCGGACTCATCAGGAAGC
AACATCGGACTCATCAGGAAGC
GTCATCGGACTCATCAGGAAGC
CATCGGACTCATCAGGAAGC
AGTACATCGGACTCATCAGGAAGC
ATCTACATCGGACTCATCAGGAAGC
TGCATCGGACTCATCAGGAAGC
TCTCATCGGACTCATCAGGAAGC
CATCGGACTCATCAGGAAGC
GTGTCATCGGACTCATCAGGAAGC
ATGCCATCGGACTCATCAGGAAGC
AAAACATCGGACTCATCAGGAAGC
CACATCGGACTCATCAGGAAGC
GACATCGGACTCATCAGGAAGC
AAGCATCGGACTCATCAGGAAGC
TTCTCATCGGACTCATCAGGAAGC
ACCATCGGACTCATCAGGAAGC

CAGTCATCGGACTCATCAGGAAGC
GACATCGGACTCATCAGGAAGC
CTCATCGGACTCATCAGGAAGC
CATCGGACTCATCAGGAAGC
CATCGGACTCATCAGGAAGC
AACATCGGACTCATCAGGAAGC
CTCATCGGACTCATCAGGAAGC
CTGGCATCGGACTCATCAGGAAGC
TACATCGGACTCATCAGGAAGC
GTCATCGGACTCATCAGGAAGC
CTCACATCGGACTCATCAGGAAGC
TCATCGGACTCATCAGGAAGC
GCCATCGGACTCATCAGGAAGC
ACCATCGGACTCATCAGGAAGC
GGACATCGGACTCATCAGGAAGC
CCCATCGGACTCATCAGGAAGC
TCAACATCGGACTCATCAGGAAGC
GGCATCGGACTCATCAGGAAGC
GCCCATCGGACTCATCAGGAAGC
GACATCGGACTCATCAGGAAGC
GCATCGGACTCATCAGGAAGC
ATCGGACTCATCAGGAAGC
AACATCGGACTCATCAGGAAGC
ACATCGGACTCATCAGGAAGC
CATCGGACTCATCAGGAAGC
AACATCGGACTCATCAGGAAGC
TCCATCGGACTCATCAGGAAGC
CCATCGGACTCATCAGGAAGC
TGGCATCGGACTCATCAGGAAGC
AACCACATCGGACTCATCAGGAAGC
GGCATCGGACTCATCAGGAAGC
TGTCATCGGACTCATCAGGAAGC
GCTGCATCGGACTCATCAGGAAGC
TTTACATCGGACTCATCAGGAAGC
AAGACATCGGACTCATCAGGAAGC
AGGCATCGGACTCATCAGGAAGC
CTATCATCGGACTCATCAGGAAGC
ACCATCGGACTCATCAGGAAGC
CATCGGACTCATCAGGAAGC
GTTCATCGGACTCATCAGGAAGC
TCCCATCGGACTCATCAGGAAGC
TGCCATCGGACTCATCAGGAAGC
CCTCATCGGACTCATCAGGAAGC

CCCATCGGACTCATCAGGAAGC
CTCATCGGACTCATCAGGAAGC
ACATCGGACTCATCAGGAAGC
TCATCGGACTCATCAGGAAGC
CATCGGACTCATCAGGAAGC
GGCTCATCGGACTCATCAGGAAGC
ATCGGACTCATCAGGAAGC
TCCCATCGGACTCATCAGGAAGC
TCATCGGACTCATCAGGAAGC
ACCATCGGACTCATCAGGAAGC
CCATCGGACTCATCAGGAAGC
ATCGGACTCATCAGGAAGC
TGGCATCGGACTCATCAGGAAGC
TCTCATCGGACTCATCAGGAAGC
TTCATCGGACTCATCAGGAAGC
ATCGGACTCATCAGGAAGC
GCCATCGGACTCATCAGGAAGC
ACATCGGACTCATCAGGAAGC
TCATCGGACTCATCAGGAAGC
TACATCGGACTCATCAGGAAGC
GAAGCATCGGACTCATCAGGAAGC
ATCGGACTCATCAGGAAGC
TTTCATCGGACTCATCAGGAAGC
CACCGCCATCGGACTCATCAGGAAGC
GAGAGCATCGGACTCATCAGGAAGC
GTGCATCGGACTCATCAGGAAGC
TTTCCCCATCGGACTCATCAGGAAGC
TAGCCATCGGACTCATCAGGAAGC
CGGACTCATCAGGAAGC
TCGGACTCATCAGGAAGC
AGACCATCGGACTCATCAGGAAGC
CCATCGGACTCATCAGGAAGC
CTCTCATCGGACTCATCAGGAAGC
AACATCGGACTCATCAGGAAGC
GTGTTTCATCGGACTCATCAGGAAGC
GTATCATCGGACTCATCAGGAAGC
GACGCATCGGACTCATCAGGAAGC
GTCATCGGACTCATCAGGAAGC
CCATCGGACTCATCAGGAAGC
TCCATCGGACTCATCAGGAAGC
CATCGGACTCATCAGGAAGC
CATCGGACTCATCAGGAAGC
ATCGGACTCATCAGGAAGC

TACATCGGACTCATCAGGAAGC
CATCGGACTCATCAGGAAGC
CATCGGACTCATCAGGAAGC
CTGCCATCGGACTCATCAGGAAGC
ATCGGACTCATCAGGAAGC
TCGGACTCATCAGGAAGC
TCATCGGACTCATCAGGAAGC
TCATCGGACTCATCAGGAAGC
ATGCATCGGACTCATCAGGAAGC
GTCATCGGACTCATCAGGAAGC
AGCCCATCGGACTCATCAGGAAGC
CATCGGACTCATCAGGAAGC
CCCCATCGGACTCATCAGGAAGC
ATCCATCGGACTCATCAGGAAGC
ATTTCATCGGACTCATCAGGAAGC
ATCGGACTCATCAGGAAGC
GTCATCGGACTCATCAGGAAGC
GCATCGGACTCATCAGGAAGC
TACATCGGACTCATCAGGAAGC
GAGGCATCGGACTCATCAGGAAGC
TCATCGGACTCATCAGGAAGC
ACATCGGACTCATCAGGAAGC
CACATCGGACTCATCAGGAAGC
ACCATCGGACTCATCAGGAAGC
GTTCATCGGACTCATCAGGAAGC
ATGTTCATCGGACTCATCAGGAAGC
TGCATCGGACTCATCAGGAAGC
ATACCACATCGGACTCATCAGGAAGC
CCTCATCGGACTCATCAGGAAGC
GCCATCGGACTCATCAGGAAGC
CGCATCGGACTCATCAGGAAGC
ATCGGACTCATCAGGAAGC
CCTGTCATCGGACTCATCAGGAAGC
GCCATCGGACTCATCAGGAAGC
CATCGGACTCATCAGGAAGC
CATCGGACTCATCAGGAAGC
GACATCGGACTCATCAGGAAGC
AGGGAACATCGGACTCATCAGGAAGC
CTGTCATCGGACTCATCAGGAAGC
GTGGCATCGGACTCATCAGGAAGC
GTCATCGGACTCATCAGGAAGC
CATCGGACTCATCAGGAAGC
GACATCGGACTCATCAGGAAGC

TCATCGGACTCATCAGGAAGC
CATCGGACTCATCAGGAAGC
TCATCGGACTCATCAGGAAGC
CACATCGGACTCATCAGGAAGC
CATCGGACTCATCAGGAAGC
CATCGGACTCATCAGGAAGC
AGAACATCGGACTCATCAGGAAGC
AACTCATCGGACTCATCAGGAAGC
GGCATCGGACTCATCAGGAAGC
TTCCTCATCGGACTCATCAGGAAGC
GGAACATCGGACTCATCAGGAAGC
GGCATCGGACTCATCAGGAAGC
GAGGCATCGGACTCATCAGGAAGC
CTCATCGGACTCATCAGGAAGC
AGCCATCGGACTCATCAGGAAGC
TTCATCGGACTCATCAGGAAGC
TACATCGGACTCATCAGGAAGC
CATCGGACTCATCAGGAAGC
CTTCCATCGGACTCATCAGGAAGC
CAGCCATCGGACTCATCAGGAAGC
TTCCATCGGACTCATCAGGAAGC
GTCATCGGACTCATCAGGAAGC
CATCGGACTCATCAGGAAGC
GCATCGGACTCATCAGGAAGC
GGCATCGGACTCATCAGGAAGC
CCATCGGACTCATCAGGAAGC
TCTGCATCGGACTCATCAGGAAGC
ATTCATCGGACTCATCAGGAAGC
CATCGGACTCATCAGGAAGC
TCTCCATCGGACTCATCAGGAAGC
TGCATCGGACTCATCAGGAAGC
GATTCATCGGACTCATCAGGAAGC
ACCATCGGACTCATCAGGAAGC
TCAACATCGGACTCATCAGGAAGC
TCACATCGGACTCATCAGGAAGC
ATCGGACTCATCAGGAAGC
TGGGTCATCGGACTCATCAGGAAGC
TGACATCGGACTCATCAGGAAGC
TGCCATCGGACTCATCAGGAAGC
CATCGGACTCATCAGGAAGC
AGCATCGGACTCATCAGGAAGC
CCCATCGGACTCATCAGGAAGC
TCGGACTCATCAGGAAGC

AAGGCATCGGACTCATCAGGAAGC
ACCATCGGACTCATCAGGAAGC
GCAACATCGGACTCATCAGGAAGC
CAACATCGGACTCATCAGGAAGC
GGCATCGGACTCATCAGGAAGC
CATCGGACTCATCAGGAAGC
CATCGGACTCATCAGGAAGC
TTGCATCGGACTCATCAGGAAGC
TCATCGGACTCATCAGGAAGC
TGCATCGGACTCATCAGGAAGC
GTTCATCGGACTCATCAGGAAGC
AGTCATCGGACTCATCAGGAAGC
ACATCGGACTCATCAGGAAGC
ATCGGACTCATCAGGAAGC
GTCATCGGACTCATCAGGAAGC
TGCATCGGACTCATCAGGAAGC
TTACATCGGACTCATCAGGAAGC
CATCGGACTCATCAGGAAGC
ACATCGGACTCATCAGGAAGC
AACTCATCGGACTCATCAGGAAGC
AACCCCACATCGGACTCATCAGGAAGC
GCTCATCGGACTCATCAGGAAGC
TCATCCATCGGACTCATCAGGAAGC
CCATCGGACTCATCAGGAAGC
CATCGGACTCATCAGGAAGC
ATCATCGGACTCATCAGGAAGC
TCCATCGGACTCATCAGGAAGC
CATCGGACTCATCAGGAAGC
TACATCGGACTCATCAGGAAGC
GATACATCGGACTCATCAGGAAGC
GGCATCGGACTCATCAGGAAGC
CTGACCATCGGACTCATCAGGAAGC
AGACATCGGACTCATCAGGAAGC
TCATCGGACTCATCAGGAAGC
AAACCATCGGACTCATCAGGAAGC
GAAACATCGGACTCATCAGGAAGC
GTGCATCGGACTCATCAGGAAGC
ATAGCATCGGACTCATCAGGAAGC
TTGCATCGGACTCATCAGGAAGC
TCGCATCGGACTCATCAGGAAGC
TCCCCATCGGACTCATCAGGAAGC
GGACCATCGGACTCATCAGGAAGC
CACATCGGACTCATCAGGAAGC

GGCATCGGACTCATCAGGAAGC
GCATCGGACTCATCAGGAAGC
CACATCGGACTCATCAGGAAGC
CACATCGGACTCATCAGGAAGC
CACATCGGACTCATCAGGAAGC
CAACATCGGACTCATCAGGAAGC
CATCGGACTCATCAGGAAGC
GCATCGGACTCATCAGGAAGC
AGACATCGGACTCATCAGGAAGC
GCATCATCGGACTCATCAGGAAGC
GCAACATCGGACTCATCAGGAAGC
CCATCGGACTCATCAGGAAGC
GGAACATCGGACTCATCAGGAAGC
ACATCGGACTCATCAGGAAGC
GGGCCATCGGACTCATCAGGAAGC
GCCATCGGACTCATCAGGAAGC
CCCATCGGACTCATCAGGAAGC
GCCATCGGACTCATCAGGAAGC
AGCATCGGACTCATCAGGAAGC
ACCCATCGGACTCATCAGGAAGC
CCATCGGACTCATCAGGAAGC
GACATCGGACTCATCAGGAAGC
ACATCGGACTCATCAGGAAGC
GGGCATCGGACTCATCAGGAAGC
GATCATCGGACTCATCAGGAAGC
GCATCGGACTCATCAGGAAGC
CACATCGGACTCATCAGGAAGC
ACCATCGGACTCATCAGGAAGC
CATCGGACTCATCAGGAAGC
CAGGCCATCGGACTCATCAGGAAGC
TCATCGGACTCATCAGGAAGC
CATCGGACTCATCAGGAAGC
ATCATCGGACTCATCAGGAAGC
GTCATCGGACTCATCAGGAAGC
GCCATCGGACTCATCAGGAAGC
GCATCGGACTCATCAGGAAGC
GAACATCGGACTCATCAGGAAGC
ACATCGGACTCATCAGGAAGC
ATCGGACTCATCAGGAAGC
GAACATCGGACTCATCAGGAAGC
TGCATCGGACTCATCAGGAAGC
GGCATCGGACTCATCAGGAAGC
ATGTTCATCGGACTCATCAGGAAGC

CATCGGACTCATCAGGAAGC
TACATCGGACTCATCAGGAAGC
ATGCTCATCGGACTCATCAGGAAGC
AAACATCGGACTCATCAGGAAGC
GGCATCGGACTCATCAGGAAGC
GCATCGGACTCATCAGGAAGC
TCGGCATCGGACTCATCAGGAAGC
TCATCGGACTCATCAGGAAGC
ATCGGACTCATCAGGAAGC
CATCGGACTCATCAGGAAGC
CCATCGGACTCATCAGGAAGC
GAATCATCGGACTCATCAGGAAGC
CATCGGACTCATCAGGAAGC
GGCATCGGACTCATCAGGAAGC
ACATCGGACTCATCAGGAAGC
TATAAGTCATCGGACTCATCAGGAAGC
TGGAACATCGGACTCATCAGGAAGC
GCATCGGACTCATCAGGAAGC
ACATCGGACTCATCAGGAAGC
AAACATCGGACTCATCAGGAAGC
CCATCGGACTCATCAGGAAGC
TCATCGGACTCATCAGGAAGC
TCGGACTCATCAGGAAGC
TACATCGGACTCATCAGGAAGC
ATCGGACTCATCAGGAAGC
TGCAGACATCGGACTCATCAGGAAGC
CATCGGACTCATCAGGAAGC
CTCATCGGACTCATCAGGAAGC
ATTCATCGGACTCATCAGGAAGC
TTTCATCGGACTCATCAGGAAGC
GTCATCGGACTCATCAGGAAGC
GCATCGGACTCATCAGGAAGC
ATCGGACTCATCAGGAAGC
GGCATCGGACTCATCAGGAAGC
CGCATCGGACTCATCAGGAAGC
AACATCGGACTCATCAGGAAGC
CACATCGGACTCATCAGGAAGC
GGAACATCGGACTCATCAGGAAGC
CCATCGGACTCATCAGGAAGC
CATCGGACTCATCAGGAAGC
ACACATCGGACTCATCAGGAAGC
TCATCGGACTCATCAGGAAGC
GGACTCATCAGGAAGC

AGCCATCGGACTCATCAGGAAGC
ATCGGACTCATCAGGAAGC
CTCATCGGACTCATCAGGAAGC
ATCATCGGACTCATCAGGAAGC
CTCAACATCGGACTCATCAGGAAGC
ATCGGACTCATCAGGAAGC
ACCATCGGACTCATCAGGAAGC
ATCATCGGACTCATCAGGAAGC
GTGCATCGGACTCATCAGGAAGC
ATTCATCGGACTCATCAGGAAGC
TGCATCGGACTCATCAGGAAGC
CTCATCGGACTCATCAGGAAGC
TTCCATCGGACTCATCAGGAAGC
TCCATCGGACTCATCAGGAAGC
CTTTCTCATCGGACTCATCAGGAAGC
CACATCGGACTCATCAGGAAGC
CCTCATCGGACTCATCAGGAAGC
CTCATCGGACTCATCAGGAAGC
GCCATCGGACTCATCAGGAAGC
CCATCGGACTCATCAGGAAGC
AGACATCGGACTCATCAGGAAGC
CGGACTCATCAGGAAGC
TTCATCGGACTCATCAGGAAGC
TGCATCGGACTCATCAGGAAGC
ATCATCGGACTCATCAGGAAGC
CATCGGACTCATCAGGAAGC
ACATCGGACTCATCAGGAAGC
GGGCCATCGGACTCATCAGGAAGC
AAGCATCGGACTCATCAGGAAGC
GGCACATCGGACTCATCAGGAAGC
CCATCGGACTCATCAGGAAGC
CATCGGACTCATCAGGAAGC
GCATCGGACTCATCAGGAAGC
GCTCATCGGACTCATCAGGAAGC
GCATCGGACTCATCAGGAAGC
TTTCATCGGACTCATCAGGAAGC
GTTCATCGGACTCATCAGGAAGC
TCATCGGACTCATCAGGAAGC
CTGCATCGGACTCATCAGGAAGC
CCATCGGACTCATCAGGAAGC
GGCATCGGACTCATCAGGAAGC
ACCATCGGACTCATCAGGAAGC
ACATCGGACTCATCAGGAAGC

CACATCGGACTCATCAGGAAGC
CCCATCGGACTCATCAGGAAGC
GACATCGGACTCATCAGGAAGC
GAAGAAGCCATCGGACTCATCAGGAAGC
CTTCATCGGACTCATCAGGAAGC
TGCATCGGACTCATCAGGAAGC
TTCATCGGACTCATCAGGAAGC
GCATCGGACTCATCAGGAAGC
CATGTCATCGGACTCATCAGGAAGC
TACCATCGGACTCATCAGGAAGC
ACATCGGACTCATCAGGAAGC
CTCATCGGACTCATCAGGAAGC
TCATCGGACTCATCAGGAAGC
TGCATCGGACTCATCAGGAAGC
CCATCGGACTCATCAGGAAGC
AGACATCGGACTCATCAGGAAGC
TCATCGGACTCATCAGGAAGC
GATTTCATCGGACTCATCAGGAAGC
TCCCATCGGACTCATCAGGAAGC
GGTGACATCGGACTCATCAGGAAGC
AAGCATCGGACTCATCAGGAAGC
AGACATCGGACTCATCAGGAAGC
GATCATCGGACTCATCAGGAAGC
TTTCATCGGACTCATCAGGAAGC
ATCGGACTCATCAGGAAGC
CCACATCGGACTCATCAGGAAGC
TGGGACATCGGACTCATCAGGAAGC
CTCATCGGACTCATCAGGAAGC
GTGTCATCGGACTCATCAGGAAGC
AGGTCATCGGACTCATCAGGAAGC
TCATCGGACTCATCAGGAAGC
ACATCGGACTCATCAGGAAGC
CATCGGACTCATCAGGAAGC
GGTCATCGGACTCATCAGGAAGC
CCATCGGACTCATCAGGAAGC
ACCCATCGGACTCATCAGGAAGC
CATCGGACTCATCAGGAAGC
AATCATCGGACTCATCAGGAAGC
GCATCGGACTCATCAGGAAGC
GTCATCGGACTCATCAGGAAGC
ATCGGACTCATCAGGAAGC
AGAGCATCGGACTCATCAGGAAGC
GTGCCATCGGACTCATCAGGAAGC

AGACATCGGACTCATCAGGAAGC
TCATCGGACTCATCAGGAAGC
GGACATCGGACTCATCAGGAAGC
CCCATCGGACTCATCAGGAAGC
CCATCGGACTCATCAGGAAGC
ATCGGACTCATCAGGAAGC
CCATCGGACTCATCAGGAAGC
GCCCCATCGGACTCATCAGGAAGC
TCCCATCGGACTCATCAGGAAGC
AACATCGGACTCATCAGGAAGC
TCATCGGACTCATCAGGAAGC
GACATCGGACTCATCAGGAAGC
CATCGGACTCATCAGGAAGC
CATCGGACTCATCAGGAAGC
GTCCATCGGACTCATCAGGAAGC
GACATCGGACTCATCAGGAAGC
TCATCGGACTCATCAGGAAGC
TCCATCGGACTCATCAGGAAGC
TGCATCGGACTCATCAGGAAGC
TGACATCGGACTCATCAGGAAGC
CCATCGGACTCATCAGGAAGC
TGCCCATCGGACTCATCAGGAAGC
CATCGGACTCATCAGGAAGC
ATCGGACTCATCAGGAAGC
AACATCGGACTCATCAGGAAGC
TCATCGGACTCATCAGGAAGC
ACATCGGACTCATCAGGAAGC
TCATCGGACTCATCAGGAAGC
ACACATCGGACTCATCAGGAAGC
ACATCGGACTCATCAGGAAGC
CATCGGACTCATCAGGAAGC
CCCCACATCGGACTCATCAGGAAGC
GTACTCATCGGACTCATCAGGAAGC
ACATCGGACTCATCAGGAAGC
GTGGCATCGGACTCATCAGGAAGC
GCATCGGACTCATCAGGAAGC
TCCATCGGACTCATCAGGAAGC
AGACATCGGACTCATCAGGAAGC
CATCGGACTCATCAGGAAGC
GGCATCGGACTCATCAGGAAGC
GGCATCGGACTCATCAGGAAGC
CACCATCGGACTCATCAGGAAGC
TCATCGGACTCATCAGGAAGC

CATCATCGGACTCATCAGGAAGC
TTCATCGGACTCATCAGGAAGC
ACATCGGACTCATCAGGAAGC
ATTCATCGGACTCATCAGGAAGC
ATCGGACTCATCAGGAAGC
TGCCATCGGACTCATCAGGAAGC
GGAGACATCGGACTCATCAGGAAGC
GCCATCGGACTCATCAGGAAGC
ACTACATCGGACTCATCAGGAAGC
TTGCATCGGACTCATCAGGAAGC
CAAATCATCGGACTCATCAGGAAGC
CTCATCGGACTCATCAGGAAGC
GTCATCGGACTCATCAGGAAGC
CACATCGGACTCATCAGGAAGC
GCATCGGACTCATCAGGAAGC
TCCATCGGACTCATCAGGAAGC
ACATCGGACTCATCAGGAAGC
GCATCGGACTCATCAGGAAGC
TCATCGGACTCATCAGGAAGC
ATCATCGGACTCATCAGGAAGC
TTCCTCATCGGACTCATCAGGAAGC
CCATCGGACTCATCAGGAAGC
GCATCGGACTCATCAGGAAGC
GCATCGGACTCATCAGGAAGC
GAACATCGGACTCATCAGGAAGC
TTGCCATCGGACTCATCAGGAAGC
ATGCATCGGACTCATCAGGAAGC
TAACATCGGACTCATCAGGAAGC
TCTTCATCGGACTCATCAGGAAGC
CAAACATCGGACTCATCAGGAAGC
GGACATCGGACTCATCAGGAAGC
CACATCGGACTCATCAGGAAGC
CCACATCGGACTCATCAGGAAGC
TTCTCATCGGACTCATCAGGAAGC
GCATCGGACTCATCAGGAAGC
TGCATCGGACTCATCAGGAAGC
TACATCGGACTCATCAGGAAGC
GACATCGGACTCATCAGGAAGC
GAGCATCGGACTCATCAGGAAGC
CTCATCGGACTCATCAGGAAGC
CATCGGACTCATCAGGAAGC
TGAGAACCATCGGACTCATCAGGAAGC
GGCATCGGACTCATCAGGAAGC

TCCATCGGACTCATCAGGAAGC
CATCGGACTCATCAGGAAGC
GCATCGGACTCATCAGGAAGC
ACATCGGACTCATCAGGAAGC
TGCCATCGGACTCATCAGGAAGC
TCATCGGACTCATCAGGAAGC
CTCATCGGACTCATCAGGAAGC
TCCATCGGACTCATCAGGAAGC
CATCATCGGACTCATCAGGAAGC
CCATCGGACTCATCAGGAAGC
TCATCGGACTCATCAGGAAGC
AACATCGGACTCATCAGGAAGC
CATCGGACTCATCAGGAAGC
TCCATCGGACTCATCAGGAAGC
TCATCGGACTCATCAGGAAGC
GGGGCATCGGACTCATCAGGAAGC
AGCATCGGACTCATCAGGAAGC
CCATCGGACTCATCAGGAAGC
ACATCGGACTCATCAGGAAGC
CACCATCGGACTCATCAGGAAGC
TCATCGGACTCATCAGGAAGC
GCCATCGGACTCATCAGGAAGC
CACATCGGACTCATCAGGAAGC
TCATCGGACTCATCAGGAAGC
ACATCGGACTCATCAGGAAGC
TACATCGGACTCATCAGGAAGC
CATCGGACTCATCAGGAAGC
ACATCGGACTCATCAGGAAGC
GCATCGGACTCATCAGGAAGC
ACATCGGACTCATCAGGAAGC
GTCATCGGACTCATCAGGAAGC
TCGGACTCATCAGGAAGC
CATCGGACTCATCAGGAAGC
TACACATCGGACTCATCAGGAAGC
ACCATCGGACTCATCAGGAAGC
CATCGGACTCATCAGGAAGC
GTCATCGGACTCATCAGGAAGC
CATCGGACTCATCAGGAAGC
TCATCGGACTCATCAGGAAGC
TTCATCGGACTCATCAGGAAGC
CCCATCGGACTCATCAGGAAGC
ACATCGGACTCATCAGGAAGC
TCATCGGACTCATCAGGAAGC

ACCATCGGACTCATCAGGAAGC
TCATCGGACTCATCAGGAAGC
GCTCATCGGACTCATCAGGAAGC
AGAGCATCGGACTCATCAGGAAGC
GAAGACATCGGACTCATCAGGAAGC
CCTCATCGGACTCATCAGGAAGC
TCATCGGACTCATCAGGAAGC
ATCATCGGACTCATCAGGAAGC
CAGAGCATCGGACTCATCAGGAAGC
TACATCGGACTCATCAGGAAGC
GTGTCCATCGGACTCATCAGGAAGC
CCATCGGACTCATCAGGAAGC
TAGCATCGGACTCATCAGGAAGC
AACATCGGACTCATCAGGAAGC
ACGCATCGGACTCATCAGGAAGC
AGCACATCGGACTCATCAGGAAGC
CATAAAGCATCGGACTCATCAGGAAGC
GTGCATCGGACTCATCAGGAAGC
TCCATCGGACTCATCAGGAAGC
CCATCGGACTCATCAGGAAGC
CATCGGACTCATCAGGAAGC
TTTGCATCGGACTCATCAGGAAGC
ACATCGGACTCATCAGGAAGC
ACATCGGACTCATCAGGAAGC
GGCATCGGACTCATCAGGAAGC
AACTGCATCGGACTCATCAGGAAGC
CTAACCATCGGACTCATCAGGAAGC
TAACATCGGACTCATCAGGAAGC
ATTCATCGGACTCATCAGGAAGC
ATCATCGGACTCATCAGGAAGC
CAGGCATCGGACTCATCAGGAAGC
CCACATCGGACTCATCAGGAAGC
AATGCATCGGACTCATCAGGAAGC
GAACATCGGACTCATCAGGAAGC
GAGCATCGGACTCATCAGGAAGC
TGCCATCGGACTCATCAGGAAGC
CACATCGGACTCATCAGGAAGC
GATCATCGGACTCATCAGGAAGC
CCATCGGACTCATCAGGAAGC
GACATCGGACTCATCAGGAAGC
CATCGGACTCATCAGGAAGC
GTCATCGGACTCATCAGGAAGC
ACCATCGGACTCATCAGGAAGC

CAGACATCGGACTCATCAGGAAGC
CCATCGGACTCATCAGGAAGC
ACCATCGGACTCATCAGGAAGC
TCGGACTCATCAGGAAGC
ATCGGACTCATCAGGAAGC
TTTACATCGGACTCATCAGGAAGC
ACAGCATCGGACTCATCAGGAAGC
CATCGGACTCATCAGGAAGC
GCCATCGGACTCATCAGGAAGC
CCTACATCGGACTCATCAGGAAGC
CCCACATCGGACTCATCAGGAAGC
CATCGGACTCATCAGGAAGC
CATCGGACTCATCAGGAAGC
CATCGGACTCATCAGGAAGC
GGTCCATCGGACTCATCAGGAAGC
GCATCGGACTCATCAGGAAGC
TAGCATCGGACTCATCAGGAAGC
TGAGACCATCGGACTCATCAGGAAGC
CATCGGACTCATCAGGAAGC
CTCATCGGACTCATCAGGAAGC
ACATCGGACTCATCAGGAAGC
GAACATCGGACTCATCAGGAAGC
TCATCGGACTCATCAGGAAGC
TCGGACTCATCAGGAAGC
CTGCATCGGACTCATCAGGAAGC
CCTCATCGGACTCATCAGGAAGC
ATCCATCGGACTCATCAGGAAGC
TCTCATCGGACTCATCAGGAAGC
CCATCGGACTCATCAGGAAGC
CATCGGACTCATCAGGAAGC
CATCGGACTCATCAGGAAGC
CAGCACATCGGACTCATCAGGAAGC
CTCTCATCGGACTCATCAGGAAGC
CCATCGGACTCATCAGGAAGC
GTCATCGGACTCATCAGGAAGC
ACATCGGACTCATCAGGAAGC
TCATCGGACTCATCAGGAAGC
ATCATCGGACTCATCAGGAAGC
GCATCGGACTCATCAGGAAGC
CATCGGACTCATCAGGAAGC
CATCGGACTCATCAGGAAGC
CCATCGGACTCATCAGGAAGC
ACATCGGACTCATCAGGAAGC

CCATCGGACTCATCAGGAAGC
GGCATCGGACTCATCAGGAAGC
ATCATCGGACTCATCAGGAAGC
GGACATCGGACTCATCAGGAAGC
CCATCGGACTCATCAGGAAGC
CAACATCGGACTCATCAGGAAGC
ACATCGGACTCATCAGGAAGC
CATCGGACTCATCAGGAAGC
TAAGGCATCGGACTCATCAGGAAGC
CCATCGGACTCATCAGGAAGC
TGTGCATCGGACTCATCAGGAAGC
CCATCGGACTCATCAGGAAGC
TCATCGGACTCATCAGGAAGC
ACATCGGACTCATCAGGAAGC
TCATCGGACTCATCAGGAAGC
AGCATCGGACTCATCAGGAAGC
TACATCGGACTCATCAGGAAGC
CATCGGACTCATCAGGAAGC
CCATCGGACTCATCAGGAAGC
AACATCGGACTCATCAGGAAGC
CCATCGGACTCATCAGGAAGC
TCATCGGACTCATCAGGAAGC
CATCGGACTCATCAGGAAGC
CCATCGGACTCATCAGGAAGC
CTCCATCGGACTCATCAGGAAGC
AACATCGGACTCATCAGGAAGC
CCATCGGACTCATCAGGAAGC
TGGGCATCGGACTCATCAGGAAGC
CCATCGGACTCATCAGGAAGC
GCATCGGACTCATCAGGAAGC
CCCATCGGACTCATCAGGAAGC
TGTCCATCGGACTCATCAGGAAGC
CATCGGACTCATCAGGAAGC
AATGCATCGGACTCATCAGGAAGC
CCATCGGACTCATCAGGAAGC
CCATCGGACTCATCAGGAAGC
CATCGGACTCATCAGGAAGC
CTTCATCGGACTCATCAGGAAGC
TCATCGGACTCATCAGGAAGC
GCATCGGACTCATCAGGAAGC
TGCATCGGACTCATCAGGAAGC
GTCATCGGACTCATCAGGAAGC
ACCATCGGACTCATCAGGAAGC

AGGACATCGGACTCATCAGGAAGC
TCATCGGACTCATCAGGAAGC
AGTACATCGGACTCATCAGGAAGC
CCATCGGACTCATCAGGAAGC
GGGGCCATCGGACTCATCAGGAAGC
CATCGGACTCATCAGGAAGC
TCACATCGGACTCATCAGGAAGC
GCATCGGACTCATCAGGAAGC
CTCTCATCGGACTCATCAGGAAGC
ACATCGGACTCATCAGGAAGC
CATCGGACTCATCAGGAAGC
AGAGAGCATCGGACTCATCAGGAAGC
CACATCGGACTCATCAGGAAGC
CTCATCGGACTCATCAGGAAGC
TCTCATCGGACTCATCAGGAAGC
CATCGGACTCATCAGGAAGC
TGCATCGGACTCATCAGGAAGC
ATCGGACTCATCAGGAAGC
AGCATCGGACTCATCAGGAAGC
ATCATCGGACTCATCAGGAAGC
CCCATCGGACTCATCAGGAAGC
ACATCGGACTCATCAGGAAGC
AACATCGGACTCATCAGGAAGC
GCTCATCGGACTCATCAGGAAGC
AGGCATCGGACTCATCAGGAAGC
GCCATCGGACTCATCAGGAAGC
ATTCATCGGACTCATCAGGAAGC
CAGTCATCGGACTCATCAGGAAGC
AAGGTCATCGGACTCATCAGGAAGC
CATCGGACTCATCAGGAAGC
AACATCGGACTCATCAGGAAGC
CCTGCATCGGACTCATCAGGAAGC
GACATCGGACTCATCAGGAAGC
CGCATCGGACTCATCAGGAAGC
CCATCGGACTCATCAGGAAGC
AACATCGGACTCATCAGGAAGC
TCACATCGGACTCATCAGGAAGC
GTGCATCGGACTCATCAGGAAGC
GCCCATCGGACTCATCAGGAAGC
ATACTCATCGGACTCATCAGGAAGC
GTGTCATCGGACTCATCAGGAAGC
CACATCGGACTCATCAGGAAGC
GCATCGGACTCATCAGGAAGC

TTGCATCGGACTCATCAGGAAGC
TATCATCGGACTCATCAGGAAGC
CCAGCCATCGGACTCATCAGGAAGC
TCATCGGACTCATCAGGAAGC
GGTCATCGGACTCATCAGGAAGC
CATCGGACTCATCAGGAAGC
TTACATCGGACTCATCAGGAAGC
TGCCATCGGACTCATCAGGAAGC
CATCGGACTCATCAGGAAGC
ATCGGACTCATCAGGAAGC
AGGGCATCGGACTCATCAGGAAGC
CCATCGGACTCATCAGGAAGC
GCTTCATCGGACTCATCAGGAAGC
AGCCATCGGACTCATCAGGAAGC
TACATCGGACTCATCAGGAAGC
CCATCGGACTCATCAGGAAGC
GCGGCATCGGACTCATCAGGAAGC
TGCATCGGACTCATCAGGAAGC
AGCCATCGGACTCATCAGGAAGC
GAGGCATCGGACTCATCAGGAAGC
CCCATCGGACTCATCAGGAAGC
ATCGGACTCATCAGGAAGC
ACATCGGACTCATCAGGAAGC
GTCATCGGACTCATCAGGAAGC
GAGCGGCATCGGACTCATCAGGAAGC
AGGCCATCGGACTCATCAGGAAGC
TAACATCGGACTCATCAGGAAGC
TCATCGGACTCATCAGGAAGC
ATACATCGGACTCATCAGGAAGC
TCATCGGACTCATCAGGAAGC
GAACATCGGACTCATCAGGAAGC
TATGCCATCGGACTCATCAGGAAGC
TTACTCACATCGGACTCATCAGGAAGC
ACATCGGACTCATCAGGAAGC
GAACATCGGACTCATCAGGAAGC
AGTCATCGGACTCATCAGGAAGC
GCCCATCGGACTCATCAGGAAGC
TGCATCGGACTCATCAGGAAGC
CCATCGGACTCATCAGGAAGC
TCGGCATCGGACTCATCAGGAAGC
TGCCATCGGACTCATCAGGAAGC
CATCGGACTCATCAGGAAGC
CCATCGGACTCATCAGGAAGC

GTCATCGGACTCATCAGGAAGC
TCATCGGACTCATCAGGAAGC
TCTCATCGGACTCATCAGGAAGC
GCACATCGGACTCATCAGGAAGC
GAAGACATCGGACTCATCAGGAAGC
TCATCGGACTCATCAGGAAGC
CATCGGACTCATCAGGAAGC
GGGCATCGGACTCATCAGGAAGC
CACATCGGACTCATCAGGAAGC
GTGCATCGGACTCATCAGGAAGC
GCAGCATCGGACTCATCAGGAAGC
ACATCATCGGACTCATCAGGAAGC
GACATCGGACTCATCAGGAAGC
ACAGACATCGGACTCATCAGGAAGC
AACATCGGACTCATCAGGAAGC
AGCATCGGACTCATCAGGAAGC
TACATCGGACTCATCAGGAAGC
GGGCATCGGACTCATCAGGAAGC
TCGGACTCATCAGGAAGC
GCATCGGACTCATCAGGAAGC
ATCGGACTCATCAGGAAGC
TTCCCATCGGACTCATCAGGAAGC
ATCGGACTCATCAGGAAGC
CATCGGACTCATCAGGAAGC
TATCATCGGACTCATCAGGAAGC
GACATCGGACTCATCAGGAAGC
CATCGGACTCATCAGGAAGC
GTACATCGGACTCATCAGGAAGC
CACATCGGACTCATCAGGAAGC
GCCATCGGACTCATCAGGAAGC
CTCATCGGACTCATCAGGAAGC
CCATCGGACTCATCAGGAAGC
TCATCGGACTCATCAGGAAGC
GTGTCATCGGACTCATCAGGAAGC
CCATCGGACTCATCAGGAAGC
AGGGCATCGGACTCATCAGGAAGC
TCATCGGACTCATCAGGAAGC
TTCCGTTCATCGGACTCATCAGGAAGC
AGTCATCGGACTCATCAGGAAGC
GTGGCATCGGACTCATCAGGAAGC
TTTCATCGGACTCATCAGGAAGC
CAACATCGGACTCATCAGGAAGC
ACCATCGGACTCATCAGGAAGC

CATCGGACTCATCAGGAAGC
CATCGGACTCATCAGGAAGC
TCATCGGACTCATCAGGAAGC
CCATCGGACTCATCAGGAAGC
TCGGACTCATCAGGAAGC
ACATCGGACTCATCAGGAAGC
GCCTCATCGGACTCATCAGGAAGC
GTGCATCGGACTCATCAGGAAGC
CTCATCGGACTCATCAGGAAGC
GACATCGGACTCATCAGGAAGC
ATGCTTCATCGGACTCATCAGGAAGC
TCATCGGACTCATCAGGAAGC
AAAAACATCGGACTCATCAGGAAGC
GACATCGGACTCATCAGGAAGC
ACCATCGGACTCATCAGGAAGC
CATCGGACTCATCAGGAAGC
TTCTGACATCGGACTCATCAGGAAGC
CCATCGGACTCATCAGGAAGC
ATCATCGGACTCATCAGGAAGC
ACATCGGACTCATCAGGAAGC
CCCATCGGACTCATCAGGAAGC
ACATCGGACTCATCAGGAAGC
AAGAACATCGGACTCATCAGGAAGC
GGCATCGGACTCATCAGGAAGC
TCATCGGACTCATCAGGAAGC
TGCATCGGACTCATCAGGAAGC
CATCGGACTCATCAGGAAGC
CCATCGGACTCATCAGGAAGC
GTTTCATCGGACTCATCAGGAAGC
AACATCGGACTCATCAGGAAGC
ATTACATCGGACTCATCAGGAAGC
GGCATCGGACTCATCAGGAAGC
TGACATCGGACTCATCAGGAAGC
ACATCGGACTCATCAGGAAGC
CATCGGACTCATCAGGAAGC
CATCGGACTCATCAGGAAGC
CATCGGACTCATCAGGAAGC
CATCGGACTCATCAGGAAGC
AGCATCGGACTCATCAGGAAGC
GACATCGGACTCATCAGGAAGC
TGCCCATCGGACTCATCAGGAAGC
GCCATCGGACTCATCAGGAAGC
CATCGGACTCATCAGGAAGC

TAGCATCGGACTCATCAGGAAGC
CCTCATCGGACTCATCAGGAAGC
CACCCATCGGACTCATCAGGAAGC
GTGCATCGGACTCATCAGGAAGC
TGATCATCGGACTCATCAGGAAGC
GACATCGGACTCATCAGGAAGC
GAGCCATCGGACTCATCAGGAAGC
GAAGTCATCGGACTCATCAGGAAGC
TCCCATCGGACTCATCAGGAAGC
ATACATCGGACTCATCAGGAAGC
CATACATCGGACTCATCAGGAAGC
AGCATACATCGGACTCATCAGGAAGC
AGGCATCGGACTCATCAGGAAGC
GACATCGGACTCATCAGGAAGC
CCCATCGGACTCATCAGGAAGC
CCTTCATCGGACTCATCAGGAAGC
CATCGGACTCATCAGGAAGC
AGGCATCGGACTCATCAGGAAGC
TCCATCGGACTCATCAGGAAGC
GGCATCGGACTCATCAGGAAGC
ACTCCCATCGGACTCATCAGGAAGC
CCATCGGACTCATCAGGAAGC
CCCATCGGACTCATCAGGAAGC
ACATCGGACTCATCAGGAAGC
GTGTCATCGGACTCATCAGGAAGC
GGTGTCTCATCGGACTCATCAGGAAGC
AGGCATCGGACTCATCAGGAAGC
AGGCATCGGACTCATCAGGAAGC
TCTCCCATCGGACTCATCAGGAAGC
CCATCGGACTCATCAGGAAGC
GGCACATCGGACTCATCAGGAAGC
TGCATCGGACTCATCAGGAAGC
CCATCGGACTCATCAGGAAGC
TCATCGGACTCATCAGGAAGC
TCATCGGACTCATCAGGAAGC
GCCATCGGACTCATCAGGAAGC
GCATCGGACTCATCAGGAAGC
TAGTCATCGGACTCATCAGGAAGC
TCATCATCGGACTCATCAGGAAGC
GAGACATCGGACTCATCAGGAAGC
ACATCGGACTCATCAGGAAGC
GCTCATCGGACTCATCAGGAAGC
TTTCATCGGACTCATCAGGAAGC

AACGCATCGGACTCATCAGGAAGC
GATGGACATCGGACTCATCAGGAAGC
GTCATCGGACTCATCAGGAAGC
ACATCGGACTCATCAGGAAGC
CATCGGACTCATCAGGAAGC
TCATCGGACTCATCAGGAAGC
AAACATCGGACTCATCAGGAAGC
CATCGGACTCATCAGGAAGC
TGTCATCGGACTCATCAGGAAGC
GATTACACATCGGACTCATCAGGAAGC
ATCGGACTCATCAGGAAGC
CATTCATCGGACTCATCAGGAAGC
TGCATCGGACTCATCAGGAAGC
GCCATCGGACTCATCAGGAAGC
TTTACATCGGACTCATCAGGAAGC
GAGGAACATCGGACTCATCAGGAAGC
TCCCATCGGACTCATCAGGAAGC
CCATCGGACTCATCAGGAAGC
ACATCGGACTCATCAGGAAGC
GTGCCATCGGACTCATCAGGAAGC
ATCATCGGACTCATCAGGAAGC
GTACATCGGACTCATCAGGAAGC
TACAACATCGGACTCATCAGGAAGC
CATCGGACTCATCAGGAAGC
GGCATCGGACTCATCAGGAAGC
CATAACATCGGACTCATCAGGAAGC
AAGGCATCGGACTCATCAGGAAGC
GACATCGGACTCATCAGGAAGC
TCCATCGGACTCATCAGGAAGC
ACCACGCATCGGACTCATCAGGAAGC
CACATCGGACTCATCAGGAAGC
GAGCATCGGACTCATCAGGAAGC
CATCGGACTCATCAGGAAGC
ACACATCGGACTCATCAGGAAGC
TAACATCGGACTCATCAGGAAGC
CACATCGGACTCATCAGGAAGC
GAGATCATCGGACTCATCAGGAAGC
GACATCGGACTCATCAGGAAGC
CTCCATCGGACTCATCAGGAAGC
CCATCGGACTCATCAGGAAGC
AGATCATCGGACTCATCAGGAAGC
TATGACATCGGACTCATCAGGAAGC
GTCATCGGACTCATCAGGAAGC

ACATCGGACTCATCAGGAAGC
CATCGGACTCATCAGGAAGC
GCATCGGACTCATCAGGAAGC
CATCGGACTCATCAGGAAGC
AAACATCGGACTCATCAGGAAGC
GATATGCATCGGACTCATCAGGAAGC
AGCATCGGACTCATCAGGAAGC
CATCGGACTCATCAGGAAGC
CACATCGGACTCATCAGGAAGC
CATCGGACTCATCAGGAAGC
CACATCGGACTCATCAGGAAGC
TACATCGGACTCATCAGGAAGC
GAACATCGGACTCATCAGGAAGC
ATCGGACTCATCAGGAAGC
GCATCGGACTCATCAGGAAGC
CCATCGGACTCATCAGGAAGC
TGTCATCGGACTCATCAGGAAGC
GGCATCGGACTCATCAGGAAGC
TCATCGGACTCATCAGGAAGC
GACCATCGGACTCATCAGGAAGC
CCATCGGACTCATCAGGAAGC
ATCATCGGACTCATCAGGAAGC
CCCATCGGACTCATCAGGAAGC
AACATCGGACTCATCAGGAAGC
AAAACATCGGACTCATCAGGAAGC
ATCCATCGGACTCATCAGGAAGC
AGACATCGGACTCATCAGGAAGC
GCATCGGACTCATCAGGAAGC
CCATCGGACTCATCAGGAAGC
CCATCGGACTCATCAGGAAGC
CATCGGACTCATCAGGAAGC
CGCATCGGACTCATCAGGAAGC
AAAGCATCGGACTCATCAGGAAGC
ACCATCGGACTCATCAGGAAGC
CATCGGACTCATCAGGAAGC
CTGCATCGGACTCATCAGGAAGC
CATCGGACTCATCAGGAAGC
GGTCATCGGACTCATCAGGAAGC
CATCGGACTCATCAGGAAGC
TTGCATCGGACTCATCAGGAAGC
GTGACATCGGACTCATCAGGAAGC
ATCGGACTCATCAGGAAGC
GGCATCGGACTCATCAGGAAGC

TCATCGGACTCATCAGGAAGC
ATCATCGGACTCATCAGGAAGC
CATCGGACTCATCAGGAAGC
GCACATCGGACTCATCAGGAAGC
CCATCGGACTCATCAGGAAGC
GAGGCATCGGACTCATCAGGAAGC
ATTCATCGGACTCATCAGGAAGC
GGTCATCGGACTCATCAGGAAGC
GACATCGGACTCATCAGGAAGC
GCATCGGACTCATCAGGAAGC
GTCCCATCGGACTCATCAGGAAGC
ATACATCGGACTCATCAGGAAGC
ACGCATCGGACTCATCAGGAAGC
TGCATCGGACTCATCAGGAAGC
GGGCATCGGACTCATCAGGAAGC
ACATCGGACTCATCAGGAAGC
GACATCGGACTCATCAGGAAGC
CCATCGGACTCATCAGGAAGC
CCCATCGGACTCATCAGGAAGC
TGTCATCGGACTCATCAGGAAGC
AGCATCGGACTCATCAGGAAGC
CATCGGACTCATCAGGAAGC
CCCATCGGACTCATCAGGAAGC
AGACATCGGACTCATCAGGAAGC
GTCACCATCGGACTCATCAGGAAGC
TCTCATCGGACTCATCAGGAAGC
ATCGGACTCATCAGGAAGC
GCCATCGGACTCATCAGGAAGC
GACATCGGACTCATCAGGAAGC
ACACATCGGACTCATCAGGAAGC
CCATCGGACTCATCAGGAAGC
CATCGGACTCATCAGGAAGC
CATCGGACTCATCAGGAAGC
ATCGGACTCATCAGGAAGC
AGCATCGGACTCATCAGGAAGC
ATCGGACTCATCAGGAAGC
AAACATCGGACTCATCAGGAAGC
AGGCATCGGACTCATCAGGAAGC
TTGTCATCGGACTCATCAGGAAGC
TCATCGGACTCATCAGGAAGC
TTAGCATCGGACTCATCAGGAAGC
ATCGCATCGGACTCATCAGGAAGC
GAGCATCGGACTCATCAGGAAGC

ACATCGGACTCATCAGGAAGC
GCTCATCGGACTCATCAGGAAGC
ACCATCGGACTCATCAGGAAGC
GCATCGGACTCATCAGGAAGC
GGGCATCGGACTCATCAGGAAGC
ACATCGGACTCATCAGGAAGC
GACATCGGACTCATCAGGAAGC
ATCATCGGACTCATCAGGAAGC
TCGGACTCATCAGGAAGC
GAATCATCGGACTCATCAGGAAGC
AATCATCGGACTCATCAGGAAGC
AGACATCGGACTCATCAGGAAGC
AACCCCATCGGACTCATCAGGAAGC
ACATCGGACTCATCAGGAAGC
ATCGGACTCATCAGGAAGC
ACCATCGGACTCATCAGGAAGC
AGCCATCGGACTCATCAGGAAGC
CATCGGACTCATCAGGAAGC
CCATCGGACTCATCAGGAAGC
TCATCGGACTCATCAGGAAGC
ACATCGGACTCATCAGGAAGC
TCATCGGACTCATCAGGAAGC
GAGACCATCGGACTCATCAGGAAGC
GTGTCATCGGACTCATCAGGAAGC
TGGCATCGGACTCATCAGGAAGC
CATCGGACTCATCAGGAAGC
TGGCATCGGACTCATCAGGAAGC
ACCATCGGACTCATCAGGAAGC
AACATCGGACTCATCAGGAAGC
CTCATCGGACTCATCAGGAAGC
CTCATCGGACTCATCAGGAAGC
CCATCGGACTCATCAGGAAGC
GAACACATCGGACTCATCAGGAAGC
CTCATCGGACTCATCAGGAAGC
TACATCGGACTCATCAGGAAGC
GCCATCGGACTCATCAGGAAGC
TCATCGGACTCATCAGGAAGC
AGACATCGGACTCATCAGGAAGC
TGCATCGGACTCATCAGGAAGC
CACATCGGACTCATCAGGAAGC
TCCCATCGGACTCATCAGGAAGC
TAGGCATCGGACTCATCAGGAAGC
ATCATCGGACTCATCAGGAAGC

ACTTCATCGGACTCATCAGGAAGC
CGGCATCGGACTCATCAGGAAGC
GCATCGGACTCATCAGGAAGC
CCACATCGGACTCATCAGGAAGC
CCATCGGACTCATCAGGAAGC
AACCATCGGACTCATCAGGAAGC
CATCGGACTCATCAGGAAGC
CGCATCGGACTCATCAGGAAGC
CCCATCGGACTCATCAGGAAGC
GTACATCGGACTCATCAGGAAGC
GGGTCATCGGACTCATCAGGAAGC
TACATCGGACTCATCAGGAAGC
CATCGGACTCATCAGGAAGC
TCATCGGACTCATCAGGAAGC
ATCGGACTCATCAGGAAGC
GCAGCATCGGACTCATCAGGAAGC
CTTCACATCGGACTCATCAGGAAGC
GCATCGGACTCATCAGGAAGC
GTCATCGGACTCATCAGGAAGC
GGAGATCATCGGACTCATCAGGAAGC
CTCATCGGACTCATCAGGAAGC
CATCGGACTCATCAGGAAGC
AGCATCGGACTCATCAGGAAGC
GAGCATCGGACTCATCAGGAAGC
GATTTCATCGGACTCATCAGGAAGC
CATCGGACTCATCAGGAAGC
TTCATCGGACTCATCAGGAAGC
ACATCGGACTCATCAGGAAGC
TTCATCGGACTCATCAGGAAGC
GCACATCGGACTCATCAGGAAGC
TTCCACATCGGACTCATCAGGAAGC
AGCATCGGACTCATCAGGAAGC
ACCATCGGACTCATCAGGAAGC
CATCGGACTCATCAGGAAGC
ATCATCGGACTCATCAGGAAGC
GGACTCATCAGGAAGC
CCATCGGACTCATCAGGAAGC
GAACCATCGGACTCATCAGGAAGC
TTTCATCGGACTCATCAGGAAGC
GCCATCGGACTCATCAGGAAGC
AAGCCATCGGACTCATCAGGAAGC
CTTCCACATCGGACTCATCAGGAAGC
TCATCGGACTCATCAGGAAGC

CAGCATCGGACTCATCAGGAAGC
GGGTCATCGGACTCATCAGGAAGC
CCCATCGGACTCATCAGGAAGC
GACATCGGACTCATCAGGAAGC
ATCGGACTCATCAGGAAGC
TTCATCGGACTCATCAGGAAGC
AGCACATCGGACTCATCAGGAAGC
ATGGCATCGGACTCATCAGGAAGC
AGATCATCGGACTCATCAGGAAGC
TCATCGGACTCATCAGGAAGC
TCCATCATCGGACTCATCAGGAAGC
TCATCGGACTCATCAGGAAGC
CACATCGGACTCATCAGGAAGC
GTTCATCGGACTCATCAGGAAGC
AGCATCGGACTCATCAGGAAGC
GCATCGGACTCATCAGGAAGC
GCATCGGACTCATCAGGAAGC
TACATCGGACTCATCAGGAAGC
ATCATCGGACTCATCAGGAAGC
CGATTTCATCGGACTCATCAGGAAGC
AGGCCATCGGACTCATCAGGAAGC
GCCTGTCATCGGACTCATCAGGAAGC
CACATCGGACTCATCAGGAAGC
TTGGCATCGGACTCATCAGGAAGC
TATTGCATCGGACTCATCAGGAAGC
GGAAACATCGGACTCATCAGGAAGC
CATCGGACTCATCAGGAAGC
GGCTCACATCGGACTCATCAGGAAGC
ATCATCGGACTCATCAGGAAGC
TTCATCGGACTCATCAGGAAGC
ACATCGGACTCATCAGGAAGC
CCATCGGACTCATCAGGAAGC
GTGTCATCGGACTCATCAGGAAGC
CATGTCATCGGACTCATCAGGAAGC
GGCATCGGACTCATCAGGAAGC
CATCGGACTCATCAGGAAGC
TGCCATCGGACTCATCAGGAAGC
AATCATCGGACTCATCAGGAAGC
CCATCGGACTCATCAGGAAGC
ATCACATCGGACTCATCAGGAAGC
TCGGACTCATCAGGAAGC
TCAACATCGGACTCATCAGGAAGC
ATCTGGCATCGGACTCATCAGGAAGC

GGTGTCATCGGACTCATCAGGAAGC
CATCGGACTCATCAGGAAGC
AGACATCGGACTCATCAGGAAGC
TGCCATCGGACTCATCAGGAAGC
GAGAGCATCGGACTCATCAGGAAGC
ATCGGACTCATCAGGAAGC
AGCAGAGCATCGGACTCATCAGGAAGC
CATCGGACTCATCAGGAAGC
TCCAGCATCGGACTCATCAGGAAGC
GCATCGGACTCATCAGGAAGC
ACTCATCGGACTCATCAGGAAGC
GCATCGGACTCATCAGGAAGC
CTTACATCGGACTCATCAGGAAGC
ATCGGACTCATCAGGAAGC
GGCCATCGGACTCATCAGGAAGC
GCATACATCGGACTCATCAGGAAGC
CTCATCGGACTCATCAGGAAGC
TGGCATCGGACTCATCAGGAAGC
TCATCGGACTCATCAGGAAGC
GAACATCGGACTCATCAGGAAGC
CATCGGACTCATCAGGAAGC
CCACATCGGACTCATCAGGAAGC
TGACATCGGACTCATCAGGAAGC
CATCGGACTCATCAGGAAGC
CATCGGACTCATCAGGAAGC
TGGCATCGGACTCATCAGGAAGC
AGCATCGGACTCATCAGGAAGC
ACCAGCCATCGGACTCATCAGGAAGC
ATCGGACTCATCAGGAAGC
AACCATCGGACTCATCAGGAAGC
ACCATCGGACTCATCAGGAAGC
GCATCGGACTCATCAGGAAGC
GCATCGGACTCATCAGGAAGC
GTTTCATCGGACTCATCAGGAAGC
CATCGGACTCATCAGGAAGC
ATCGGACTCATCAGGAAGC
GTGCATCGGACTCATCAGGAAGC
GCAGCATCGGACTCATCAGGAAGC
GTCATCGGACTCATCAGGAAGC
CATCATCGGACTCATCAGGAAGC
AGACCATCGGACTCATCAGGAAGC
AGCATCGGACTCATCAGGAAGC
GACATCGGACTCATCAGGAAGC

AGGGCATCGGACTCATCAGGAAGC
ATCGGACTCATCAGGAAGC
CACATCGGACTCATCAGGAAGC
AGGGCATCGGACTCATCAGGAAGC
AAGAACATCGGACTCATCAGGAAGC
GCATCGGACTCATCAGGAAGC
AACACATCGGACTCATCAGGAAGC
ATACATCGGACTCATCAGGAAGC
GCCATCGGACTCATCAGGAAGC
AACATCGGACTCATCAGGAAGC
AACATCGGACTCATCAGGAAGC
AGGGCCATCGGACTCATCAGGAAGC
TGTGCATCGGACTCATCAGGAAGC
CATCGGACTCATCAGGAAGC
CATCGGACTCATCAGGAAGC
CATCGGACTCATCAGGAAGC
CTCATCGGACTCATCAGGAAGC
AGTACATCGGACTCATCAGGAAGC
TGGCATCGGACTCATCAGGAAGC
TCCCATCGGACTCATCAGGAAGC
CAGCCATCGGACTCATCAGGAAGC
AAACATCGGACTCATCAGGAAGC
ACATCGGACTCATCAGGAAGC
CTCATCGGACTCATCAGGAAGC
ACATCGGACTCATCAGGAAGC
CATCGGACTCATCAGGAAGC
CTACTGCATCGGACTCATCAGGAAGC
ACATCGGACTCATCAGGAAGC
TAACACATCGGACTCATCAGGAAGC
ATCATCGGACTCATCAGGAAGC
CATCGGACTCATCAGGAAGC
GCATCGGACTCATCAGGAAGC
TCATCGGACTCATCAGGAAGC
GACATCGGACTCATCAGGAAGC
GGCTTCATCGGACTCATCAGGAAGC
GCATCGGACTCATCAGGAAGC
TTGCATCGGACTCATCAGGAAGC
GGTCATCGGACTCATCAGGAAGC
CACCATCGGACTCATCAGGAAGC
CTCATCGGACTCATCAGGAAGC
TCGGACTCATCAGGAAGC
CACATCGGACTCATCAGGAAGC
TCCATCGGACTCATCAGGAAGC

TGCATCGGACTCATCAGGAAGC
GCATCGGACTCATCAGGAAGC
GAGGCATCGGACTCATCAGGAAGC
CAACATCGGACTCATCAGGAAGC
CCACATCGGACTCATCAGGAAGC
GCATCGGACTCATCAGGAAGC
TCATCGGACTCATCAGGAAGC
GCATCGGACTCATCAGGAAGC
ATCATCGGACTCATCAGGAAGC
GTCATCGGACTCATCAGGAAGC
CCATCGGACTCATCAGGAAGC
CCATCGGACTCATCAGGAAGC
ACCCATCGGACTCATCAGGAAGC
CGCTGCATCGGACTCATCAGGAAGC
TGTGCATCGGACTCATCAGGAAGC
GTCATCGGACTCATCAGGAAGC
ATCGGACTCATCAGGAAGC
TAACATCGGACTCATCAGGAAGC
AAACATCGGACTCATCAGGAAGC
AGCCATCGGACTCATCAGGAAGC
CTCATCGGACTCATCAGGAAGC
GTCATCGGACTCATCAGGAAGC
AACCCATCGGACTCATCAGGAAGC
CATCGGACTCATCAGGAAGC
AGCCATCGGACTCATCAGGAAGC
AACATCGGACTCATCAGGAAGC
ATCGGACTCATCAGGAAGC
CTTCATCGGACTCATCAGGAAGC
ACATCGGACTCATCAGGAAGC
CACATCGGACTCATCAGGAAGC
AAACATCGGACTCATCAGGAAGC
GTGCCATCGGACTCATCAGGAAGC
GCCCATCGGACTCATCAGGAAGC
CCCATCGGACTCATCAGGAAGC
CATCATCGGACTCATCAGGAAGC
CCATCGGACTCATCAGGAAGC
AACATCGGACTCATCAGGAAGC
TAACATCGGACTCATCAGGAAGC
ATGGCATCGGACTCATCAGGAAGC
TCATCGGACTCATCAGGAAGC
CATCGGACTCATCAGGAAGC
ACATCGGACTCATCAGGAAGC
TCATCGGACTCATCAGGAAGC

ACCACATCGGACTCATCAGGAAGC
CATCGGACTCATCAGGAAGC
GCCATCGGACTCATCAGGAAGC
AGAGGCATCGGACTCATCAGGAAGC
TGCATCGGACTCATCAGGAAGC
GCATCGGACTCATCAGGAAGC
TACATCGGACTCATCAGGAAGC
GGACATCGGACTCATCAGGAAGC
GGCATCGGACTCATCAGGAAGC
ATCGGACTCATCAGGAAGC
CCATCGGACTCATCAGGAAGC
CCATCGGACTCATCAGGAAGC
ATACATCGGACTCATCAGGAAGC
TTCTCCATCGGACTCATCAGGAAGC
TCATCGGACTCATCAGGAAGC
ACATCGGACTCATCAGGAAGC
ACCATCGGACTCATCAGGAAGC
GCATCGGACTCATCAGGAAGC
GCATCGGACTCATCAGGAAGC
CATCGGACTCATCAGGAAGC
GACATCGGACTCATCAGGAAGC
CCATCGGACTCATCAGGAAGC
GAACTCATCGGACTCATCAGGAAGC
TTCATCGGACTCATCAGGAAGC
ACATCGGACTCATCAGGAAGC
GACATCGGACTCATCAGGAAGC
GCATCGGACTCATCAGGAAGC
CCCCATCGGACTCATCAGGAAGC
TCCATCGGACTCATCAGGAAGC
AGTTCATCGGACTCATCAGGAAGC
ACATCGGACTCATCAGGAAGC
GTCCATCGGACTCATCAGGAAGC
AAGATTCATCGGACTCATCAGGAAGC
AACATCGGACTCATCAGGAAGC
ATCGGACTCATCAGGAAGC
ACCCATCGGACTCATCAGGAAGC
CCCATCGGACTCATCAGGAAGC
ATCATCGGACTCATCAGGAAGC
ACATCGGACTCATCAGGAAGC
ACTCCATCGGACTCATCAGGAAGC
ACATCGGACTCATCAGGAAGC
ATCATCGGACTCATCAGGAAGC
CACATCGGACTCATCAGGAAGC

GCCATCGGACTCATCAGGAAGC
AGCACATCGGACTCATCAGGAAGC
GTCATCGGACTCATCAGGAAGC
TCCATCGGACTCATCAGGAAGC
ACATCGGACTCATCAGGAAGC
AGACATCGGACTCATCAGGAAGC
ACATCGGACTCATCAGGAAGC
CAGTCACATCGGACTCATCAGGAAGC
ACATCGGACTCATCAGGAAGC
GAACATCGGACTCATCAGGAAGC
CAGCATCGGACTCATCAGGAAGC
AGACATCGGACTCATCAGGAAGC
GGCATCGGACTCATCAGGAAGC
CACATCGGACTCATCAGGAAGC
CCATCGGACTCATCAGGAAGC
TACATCGGACTCATCAGGAAGC
TCCCATCGGACTCATCAGGAAGC
ACCCATCGGACTCATCAGGAAGC
TTCATCGGACTCATCAGGAAGC
AGTACATCGGACTCATCAGGAAGC
TCATCGGACTCATCAGGAAGC
CATCGGACTCATCAGGAAGC
ATGCCATCGGACTCATCAGGAAGC
TCATCGGACTCATCAGGAAGC
AACATCGGACTCATCAGGAAGC
CCCATCGGACTCATCAGGAAGC
CATCGGACTCATCAGGAAGC
AGCATCGGACTCATCAGGAAGC
TTCATCGGACTCATCAGGAAGC
CATCGGACTCATCAGGAAGC
CTCATCGGACTCATCAGGAAGC
TTCCCATCGGACTCATCAGGAAGC
GCATCGGACTCATCAGGAAGC
ATCGGACTCATCAGGAAGC
ACTGCCATCGGACTCATCAGGAAGC
GTTCATCGGACTCATCAGGAAGC
CATCGGACTCATCAGGAAGC
CATCGGACTCATCAGGAAGC
CTGCATCGGACTCATCAGGAAGC
GAGCCATCGGACTCATCAGGAAGC
ACCATCGGACTCATCAGGAAGC
CATCGGACTCATCAGGAAGC
TTCCCATCGGACTCATCAGGAAGC

GCATCGGACTCATCAGGAAGC
AAACATCGGACTCATCAGGAAGC
ACATCGGACTCATCAGGAAGC
CCCATCGGACTCATCAGGAAGC
TACATCGGACTCATCAGGAAGC
GCACCATCGGACTCATCAGGAAGC
CAGCATCGGACTCATCAGGAAGC
GGTCATCGGACTCATCAGGAAGC
ACATCGGACTCATCAGGAAGC
TTCATCGGACTCATCAGGAAGC
CATCGGACTCATCAGGAAGC
ATAGCATCGGACTCATCAGGAAGC
TGCATCGGACTCATCAGGAAGC
TCATCGGACTCATCAGGAAGC
GAGCATCGGACTCATCAGGAAGC
TGAGCATCGGACTCATCAGGAAGC
AGTCATCGGACTCATCAGGAAGC
ACACATCGGACTCATCAGGAAGC
GGTCATCGGACTCATCAGGAAGC
CCATCGGACTCATCAGGAAGC
TGCATCGGACTCATCAGGAAGC
AAGACATCGGACTCATCAGGAAGC
AGGCATCGGACTCATCAGGAAGC
AGCCATCGGACTCATCAGGAAGC
CAGCATCGGACTCATCAGGAAGC
GGCATCGGACTCATCAGGAAGC
ACTTCATCGGACTCATCAGGAAGC
ATCGGACTCATCAGGAAGC
GACATCGGACTCATCAGGAAGC
ACATCGGACTCATCAGGAAGC
CACCATCGGACTCATCAGGAAGC
AACATCGGACTCATCAGGAAGC
CCACATCGGACTCATCAGGAAGC
CCATCGGACTCATCAGGAAGC
TGCAAACATCGGACTCATCAGGAAGC
GCCATCGGACTCATCAGGAAGC
CATCGGACTCATCAGGAAGC
TCATCGGACTCATCAGGAAGC
AACATCGGACTCATCAGGAAGC
CCATCGGACTCATCAGGAAGC
GCATCGGACTCATCAGGAAGC
ATACATCGGACTCATCAGGAAGC
TCCATCGGACTCATCAGGAAGC

TCATCGGACTCATCAGGAAGC
TTACATCGGACTCATCAGGAAGC
CATCATCGGACTCATCAGGAAGC
ATCGGACTCATCAGGAAGC
TCATCGGACTCATCAGGAAGC
TCATCGGACTCATCAGGAAGC
ATCGGACTCATCAGGAAGC
ACATCGGACTCATCAGGAAGC
AGCATCGGACTCATCAGGAAGC
ACCATCGGACTCATCAGGAAGC
AGCATCGGACTCATCAGGAAGC
GCCATCGGACTCATCAGGAAGC
GCCATCGGACTCATCAGGAAGC
TGCATCGGACTCATCAGGAAGC
TCATCGGACTCATCAGGAAGC
CACATCGGACTCATCAGGAAGC
CATCGGACTCATCAGGAAGC
CATCGGACTCATCAGGAAGC
GAACATCGGACTCATCAGGAAGC
CATCATCGGACTCATCAGGAAGC
TTCATCGGACTCATCAGGAAGC
ATCGGACTCATCAGGAAGC
ACATCGGACTCATCAGGAAGC
TCATCGGACTCATCAGGAAGC
GCATCGGACTCATCAGGAAGC
CATCGGACTCATCAGGAAGC
ACATCGGACTCATCAGGAAGC
GCATCGGACTCATCAGGAAGC
AAGCATCGGACTCATCAGGAAGC
CACATCGGACTCATCAGGAAGC
TCATCGGACTCATCAGGAAGC
GGGCATCGGACTCATCAGGAAGC
ATCATCGGACTCATCAGGAAGC
ATCGGACTCATCAGGAAGC
GCATCGGACTCATCAGGAAGC
CCATCGGACTCATCAGGAAGC
ACCATCGGACTCATCAGGAAGC
CTCATCGGACTCATCAGGAAGC
GCCATCGGACTCATCAGGAAGC
GGCATCGGACTCATCAGGAAGC
GACATCGGACTCATCAGGAAGC
TCATCGGACTCATCAGGAAGC
TGCATCGGACTCATCAGGAAGC

TGAGACATCGGACTCATCAGGAAGC
CATCGGACTCATCAGGAAGC
CATCGGACTCATCAGGAAGC
CATCACATCGGACTCATCAGGAAGC
CCATCGGACTCATCAGGAAGC
GGCATCGGACTCATCAGGAAGC
ACATCGGACTCATCAGGAAGC
TACATCGGACTCATCAGGAAGC
CCATCGGACTCATCAGGAAGC
ACATCGGACTCATCAGGAAGC
TGCCATCGGACTCATCAGGAAGC
TTACATCGGACTCATCAGGAAGC
CATCGGACTCATCAGGAAGC
AGGCATCGGACTCATCAGGAAGC
TCATCGGACTCATCAGGAAGC
ATCGGACTCATCAGGAAGC
ACCATCGGACTCATCAGGAAGC
GGAACATCGGACTCATCAGGAAGC
GAGCATCGGACTCATCAGGAAGC
TCCCATCGGACTCATCAGGAAGC
CTCCCATCGGACTCATCAGGAAGC
TCACATCGGACTCATCAGGAAGC
CTGCCATCGGACTCATCAGGAAGC
TGGGCATCGGACTCATCAGGAAGC
ACCATCGGACTCATCAGGAAGC
CATCGGACTCATCAGGAAGC
CATCGGACTCATCAGGAAGC
AGCCATCGGACTCATCAGGAAGC
GAGGGCATCGGACTCATCAGGAAGC
CACATCGGACTCATCAGGAAGC
ACATCGGACTCATCAGGAAGC
TATCATCGGACTCATCAGGAAGC
GCCATCGGACTCATCAGGAAGC
GAGACATCGGACTCATCAGGAAGC
TCATCGGACTCATCAGGAAGC
ACATCGGACTCATCAGGAAGC
AGGCATCGGACTCATCAGGAAGC
GCCATCGGACTCATCAGGAAGC
TCATCGGACTCATCAGGAAGC
TGCCATCGGACTCATCAGGAAGC
CACCATCGGACTCATCAGGAAGC
ATACGACATCGGACTCATCAGGAAGC
ACATCGGACTCATCAGGAAGC

CCATCGGACTCATCAGGAAGC
CATCGGACTCATCAGGAAGC
ATCATCGGACTCATCAGGAAGC
TCGGACTCATCAGGAAGC
AGAGGCATCGGACTCATCAGGAAGC
CCATCGGACTCATCAGGAAGC
GTCATCGGACTCATCAGGAAGC
CACATCGGACTCATCAGGAAGC
ACATCGGACTCATCAGGAAGC
CTGCATCGGACTCATCAGGAAGC
GAGCATCGGACTCATCAGGAAGC
ATCGGACTCATCAGGAAGC
CACATCGGACTCATCAGGAAGC
GGCACATCGGACTCATCAGGAAGC
CCTACATCGGACTCATCAGGAAGC
AGTCATCGGACTCATCAGGAAGC
CTTCATCGGACTCATCAGGAAGC
GCATCGGACTCATCAGGAAGC
GTCATCGGACTCATCAGGAAGC
CATCGGACTCATCAGGAAGC
ACATCGGACTCATCAGGAAGC
GCATCGGACTCATCAGGAAGC
CCATCGGACTCATCAGGAAGC
CCATCGGACTCATCAGGAAGC
CCATCGGACTCATCAGGAAGC
CAGCATCGGACTCATCAGGAAGC
AGTCATCGGACTCATCAGGAAGC
CATCGGACTCATCAGGAAGC
AACATCGGACTCATCAGGAAGC
AGACATCGGACTCATCAGGAAGC
TCATCGGACTCATCAGGAAGC
ACATCGGACTCATCAGGAAGC
CTCATCGGACTCATCAGGAAGC
TGCCATCGGACTCATCAGGAAGC
CATCGGACTCATCAGGAAGC
GGACATCGGACTCATCAGGAAGC
TTCATCGGACTCATCAGGAAGC
ATTACATCGGACTCATCAGGAAGC
CTTCATCGGACTCATCAGGAAGC
TCAGCATCGGACTCATCAGGAAGC
AAGCATCGGACTCATCAGGAAGC
CCCATCGGACTCATCAGGAAGC
CTCATCGGACTCATCAGGAAGC

GATCATCGGACTCATCAGGAAGC
GCATCGGACTCATCAGGAAGC
ATCGGACTCATCAGGAAGC
GCCATCGGACTCATCAGGAAGC
CCATCGGACTCATCAGGAAGC
ACTCATCGGACTCATCAGGAAGC
GACATCGGACTCATCAGGAAGC
CTCATCGGACTCATCAGGAAGC
GTTTCATCGGACTCATCAGGAAGC
TCATCGGACTCATCAGGAAGC
TCGGACTCATCAGGAAGC
CATCGGACTCATCAGGAAGC
ACCATCGGACTCATCAGGAAGC
CCCATCGGACTCATCAGGAAGC
GACATCGGACTCATCAGGAAGC
GTCACATCGGACTCATCAGGAAGC
CCATCGGACTCATCAGGAAGC
GTCATCGGACTCATCAGGAAGC
GGTCATCGGACTCATCAGGAAGC
TCATCGGACTCATCAGGAAGC
AACATCGGACTCATCAGGAAGC
GCATCGGACTCATCAGGAAGC
CATCGGACTCATCAGGAAGC
GCATCGGACTCATCAGGAAGC
CATCGGACTCATCAGGAAGC
ACATCGGACTCATCAGGAAGC
CCCATCGGACTCATCAGGAAGC
CACACATCGGACTCATCAGGAAGC
GCCATCGGACTCATCAGGAAGC
TCATCGGACTCATCAGGAAGC
CCATCGGACTCATCAGGAAGC
ATCGGACTCATCAGGAAGC
CTCATCGGACTCATCAGGAAGC
GTCATCGGACTCATCAGGAAGC
CTCATCGGACTCATCAGGAAGC
CCCATCGGACTCATCAGGAAGC
ATGCATCGGACTCATCAGGAAGC
CAACATCGGACTCATCAGGAAGC
TGCATCGGACTCATCAGGAAGC
AGGCATCGGACTCATCAGGAAGC
ATCGGACTCATCAGGAAGC
AGCATCGGACTCATCAGGAAGC
ACCCATCGGACTCATCAGGAAGC

CTTCATCGGACTCATCAGGAAGC
CAGACATCGGACTCATCAGGAAGC
TGTCATCGGACTCATCAGGAAGC
TCACATCGGACTCATCAGGAAGC
CTCATCGGACTCATCAGGAAGC
GGCATCGGACTCATCAGGAAGC
CATCGGACTCATCAGGAAGC
GGCATCGGACTCATCAGGAAGC
CAACCACATCGGACTCATCAGGAAGC
CCATCGGACTCATCAGGAAGC
ATCGGACTCATCAGGAAGC
ATCGGACTCATCAGGAAGC
GCAGTCATCGGACTCATCAGGAAGC
ACATCGGACTCATCAGGAAGC
CACATCGGACTCATCAGGAAGC
CTCATCGGACTCATCAGGAAGC
GAGCATCGGACTCATCAGGAAGC
GGCATCGGACTCATCAGGAAGC
ACATCGGACTCATCAGGAAGC
GCATCGGACTCATCAGGAAGC
GTCATCGGACTCATCAGGAAGC
CATCGGACTCATCAGGAAGC
CATCGGACTCATCAGGAAGC
ATCGGACTCATCAGGAAGC
CCCATCGGACTCATCAGGAAGC
GTCATCGGACTCATCAGGAAGC
GGCATCGGACTCATCAGGAAGC
ACATCGGACTCATCAGGAAGC
ACCATCGGACTCATCAGGAAGC
CTCATCGGACTCATCAGGAAGC
ATCATCGGACTCATCAGGAAGC
CCATCGGACTCATCAGGAAGC
CATCGGACTCATCAGGAAGC
CATCGGACTCATCAGGAAGC
TCATCGGACTCATCAGGAAGC
TCATCGGACTCATCAGGAAGC
CAACCATCGGACTCATCAGGAAGC
CATCGGACTCATCAGGAAGC
CGCATCGGACTCATCAGGAAGC
GCATCGGACTCATCAGGAAGC
CACATCGGACTCATCAGGAAGC
CATCGGACTCATCAGGAAGC
ATCGGACTCATCAGGAAGC

ACCATCGGACTCATCAGGAAGC
CACACATCGGACTCATCAGGAAGC
CTTCATCGGACTCATCAGGAAGC
AACATCGGACTCATCAGGAAGC
TCATCGGACTCATCAGGAAGC
ACTTCATCGGACTCATCAGGAAGC
TCCATCGGACTCATCAGGAAGC
GCACATCGGACTCATCAGGAAGC
GTCCATCGGACTCATCAGGAAGC
GTCTCACATCGGACTCATCAGGAAGC
CCATCGGACTCATCAGGAAGC
CATCATCGGACTCATCAGGAAGC
AACATCGGACTCATCAGGAAGC
AACATCGGACTCATCAGGAAGC
ATCATCGGACTCATCAGGAAGC
ATCGGACTCATCAGGAAGC
TTTCATCGGACTCATCAGGAAGC
TCATCGGACTCATCAGGAAGC
CATCGGACTCATCAGGAAGC
CCATCGGACTCATCAGGAAGC
CATCGGACTCATCAGGAAGC
CATCGGACTCATCAGGAAGC
GCACCATCGGACTCATCAGGAAGC
AGCATCGGACTCATCAGGAAGC
TTCATCGGACTCATCAGGAAGC
CGAGCATCGGACTCATCAGGAAGC
GCAGCCATCGGACTCATCAGGAAGC
TTCATCGGACTCATCAGGAAGC
GACATCGGACTCATCAGGAAGC
GCATCGGACTCATCAGGAAGC
ACATCGGACTCATCAGGAAGC
ATGTCATCGGACTCATCAGGAAGC
CTTAGCATCGGACTCATCAGGAAGC
CAGCATCGGACTCATCAGGAAGC
TAACATCGGACTCATCAGGAAGC
GAAGACATCGGACTCATCAGGAAGC
TGCATCGGACTCATCAGGAAGC
CATCGGACTCATCAGGAAGC
CACATCGGACTCATCAGGAAGC
TCATCGGACTCATCAGGAAGC
TCGATACATCGGACTCATCAGGAAGC
CACATCGGACTCATCAGGAAGC
TCATCGGACTCATCAGGAAGC

GCATCGGACTCATCAGGAAGC
CTGTCCATCGGACTCATCAGGAAGC
GGCTCATCGGACTCATCAGGAAGC
ACATCGGACTCATCAGGAAGC
AAAACATCGGACTCATCAGGAAGC
AACTCACATCGGACTCATCAGGAAGC
ACCAGCATCGGACTCATCAGGAAGC
CCTACATCGGACTCATCAGGAAGC
ATTCATCGGACTCATCAGGAAGC
CATCGGACTCATCAGGAAGC
GTCATCGGACTCATCAGGAAGC
AACCTACATCGGACTCATCAGGAAGC
GTTCATCGGACTCATCAGGAAGC
GTTTCATCGGACTCATCAGGAAGC
GTGACATCGGACTCATCAGGAAGC
ATCCATCGGACTCATCAGGAAGC
CTTCCATCGGACTCATCAGGAAGC
GCATCGGACTCATCAGGAAGC
ATCGGACTCATCAGGAAGC
CACCATCGGACTCATCAGGAAGC
CCCATCGGACTCATCAGGAAGC
CATCGGACTCATCAGGAAGC
GTTCACCATCGGACTCATCAGGAAGC
CATCGGACTCATCAGGAAGC
GGTCATCGGACTCATCAGGAAGC
AGCAGACATCGGACTCATCAGGAAGC
CCATCGGACTCATCAGGAAGC
CTACCATCGGACTCATCAGGAAGC
AACATCGGACTCATCAGGAAGC
TCGGACTCATCAGGAAGC
AAGCATCGGACTCATCAGGAAGC
ACATCGGACTCATCAGGAAGC
ATCGGACTCATCAGGAAGC
TCCATCGGACTCATCAGGAAGC
CATCGGACTCATCAGGAAGC
GCATCGGACTCATCAGGAAGC
AGCATCGGACTCATCAGGAAGC
GGGCATCGGACTCATCAGGAAGC
GGGTGGCATCGGACTCATCAGGAAGC
CATCGGACTCATCAGGAAGC
CTATCATCGGACTCATCAGGAAGC
ACCACTCATCGGACTCATCAGGAAGC
GCATCGGACTCATCAGGAAGC

CCATCGGACTCATCAGGAAGC
GGCATCGGACTCATCAGGAAGC
AGCATCGGACTCATCAGGAAGC
GAACATCGGACTCATCAGGAAGC
ACATCGGACTCATCAGGAAGC
CATCGGACTCATCAGGAAGC
ATCATCGGACTCATCAGGAAGC
TCGGACTCATCAGGAAGC
AACATCGGACTCATCAGGAAGC
AACATCGGACTCATCAGGAAGC
GACATCGGACTCATCAGGAAGC
CATCGGACTCATCAGGAAGC
CAACATCGGACTCATCAGGAAGC
ACACATCGGACTCATCAGGAAGC
GTGTCATCGGACTCATCAGGAAGC
GCCATCGGACTCATCAGGAAGC
TTACATCGGACTCATCAGGAAGC
CATCGGACTCATCAGGAAGC
TCATCGGACTCATCAGGAAGC
GAAACATCGGACTCATCAGGAAGC
ACATCGGACTCATCAGGAAGC
ACAGCATCGGACTCATCAGGAAGC
ACATCGGACTCATCAGGAAGC
GTGTTCATCGGACTCATCAGGAAGC
AAGCATCGGACTCATCAGGAAGC
CCACATCGGACTCATCAGGAAGC
GCTCATCGGACTCATCAGGAAGC
CATCGGACTCATCAGGAAGC
TCATCGGACTCATCAGGAAGC
AGCATCGGACTCATCAGGAAGC
ATGCATCGGACTCATCAGGAAGC
ACATCGGACTCATCAGGAAGC
GACATCGGACTCATCAGGAAGC
TCATCGGACTCATCAGGAAGC
ATCGGACTCATCAGGAAGC
GGCATCGGACTCATCAGGAAGC
GCCATCGGACTCATCAGGAAGC
CATCATCGGACTCATCAGGAAGC
CTCATCGGACTCATCAGGAAGC
CTCCATCGGACTCATCAGGAAGC
GGCACATCGGACTCATCAGGAAGC
GGAAACATCGGACTCATCAGGAAGC
GTCCACATCGGACTCATCAGGAAGC

TATCACATCGGACTCATCAGGAAGC
GTCATCGGACTCATCAGGAAGC
TTACATCGGACTCATCAGGAAGC
TGCATCGGACTCATCAGGAAGC
CGCATCGGACTCATCAGGAAGC
CACATCGGACTCATCAGGAAGC
AGTCGCATCGGACTCATCAGGAAGC
CCATCGGACTCATCAGGAAGC
GACATCGGACTCATCAGGAAGC
CTCACATCGGACTCATCAGGAAGC
AAGCATCGGACTCATCAGGAAGC
AGACATCGGACTCATCAGGAAGC
AATCGCATCGGACTCATCAGGAAGC
GGACATCGGACTCATCAGGAAGC
CCATCGGACTCATCAGGAAGC
GGCCATCGGACTCATCAGGAAGC
CATCGGACTCATCAGGAAGC
CATCGGACTCATCAGGAAGC
GGACATCGGACTCATCAGGAAGC
GATACATCGGACTCATCAGGAAGC
CACATCGGACTCATCAGGAAGC
CACATCGGACTCATCAGGAAGC
AAAATCATCGGACTCATCAGGAAGC
GCATCGGACTCATCAGGAAGC
GTCATCGGACTCATCAGGAAGC
TGTGCATCGGACTCATCAGGAAGC
ATCATCGGACTCATCAGGAAGC
CATCGGACTCATCAGGAAGC
CATCGGACTCATCAGGAAGC
AGATCATCGGACTCATCAGGAAGC
TCATCGGACTCATCAGGAAGC
ATTCATCGGACTCATCAGGAAGC
GGCATCGGACTCATCAGGAAGC
GACACATCGGACTCATCAGGAAGC
GCATCGGACTCATCAGGAAGC
CCATCGGACTCATCAGGAAGC
TGCATCGGACTCATCAGGAAGC
TGCCATCGGACTCATCAGGAAGC
ACATCGGACTCATCAGGAAGC
TCATCGGACTCATCAGGAAGC
AATCCATCGGACTCATCAGGAAGC
GCATCGGACTCATCAGGAAGC
ATCATCGGACTCATCAGGAAGC

CACATCGGACTCATCAGGAAGC
ACCTCATCGGACTCATCAGGAAGC
GTTGCCATCGGACTCATCAGGAAGC
ACAGCATCGGACTCATCAGGAAGC
TGTTCATCGGACTCATCAGGAAGC
CCCATCGGACTCATCAGGAAGC
GTGTCATCGGACTCATCAGGAAGC
GAGACATCGGACTCATCAGGAAGC
CCTCATCGGACTCATCAGGAAGC
CATCGGACTCATCAGGAAGC
CCCATCGGACTCATCAGGAAGC
CCATCGGACTCATCAGGAAGC
ACATCGGACTCATCAGGAAGC
AGAACATCGGACTCATCAGGAAGC
CCTCATCGGACTCATCAGGAAGC
GACATCGGACTCATCAGGAAGC
TCATCGGACTCATCAGGAAGC
CACATCGGACTCATCAGGAAGC
CTCCATCGGACTCATCAGGAAGC
ACACATCGGACTCATCAGGAAGC
TCCATCGGACTCATCAGGAAGC
ACATCGGACTCATCAGGAAGC
GAGCATCGGACTCATCAGGAAGC
CATCGGACTCATCAGGAAGC
CATCGGACTCATCAGGAAGC
TCCATCGGACTCATCAGGAAGC
CCATCGGACTCATCAGGAAGC
CATCGGACTCATCAGGAAGC
AAGGCATCGGACTCATCAGGAAGC
CATCGGACTCATCAGGAAGC
ACATCGGACTCATCAGGAAGC
ATCATCGGACTCATCAGGAAGC
GCTCTCATCGGACTCATCAGGAAGC
TGAGCATCGGACTCATCAGGAAGC
CATCGGACTCATCAGGAAGC
GGAACATCGGACTCATCAGGAAGC
ATCGGACTCATCAGGAAGC
CATCGGACTCATCAGGAAGC
TTCATCGGACTCATCAGGAAGC
CATCGGACTCATCAGGAAGC
ACATCGGACTCATCAGGAAGC
ATCATCGGACTCATCAGGAAGC
ACATCGGACTCATCAGGAAGC

TCCATCGGACTCATCAGGAAGC
GGCATCGGACTCATCAGGAAGC
CATCGGACTCATCAGGAAGC
GTTCATCGGACTCATCAGGAAGC
CACATCGGACTCATCAGGAAGC
ATCGGACTCATCAGGAAGC
GACATCGGACTCATCAGGAAGC
TCCTGACATCGGACTCATCAGGAAGC
TTCTCATCGGACTCATCAGGAAGC
CATCGGACTCATCAGGAAGC
CCCATCGGACTCATCAGGAAGC
GCACATCGGACTCATCAGGAAGC
GAGACAGCATCGGACTCATCAGGAAGC
GACATCGGACTCATCAGGAAGC
TGGGCACATCGGACTCATCAGGAAGC
ATCTCATCGGACTCATCAGGAAGC
AGCATCGGACTCATCAGGAAGC
CATCATCGGACTCATCAGGAAGC
AACATCGGACTCATCAGGAAGC
TGCATCGGACTCATCAGGAAGC
CCTCATCGGACTCATCAGGAAGC
TTCATCGGACTCATCAGGAAGC
CCCATCGGACTCATCAGGAAGC
GTCCATCGGACTCATCAGGAAGC
GGACTCATCAGGAAGC
TCCAACATCGGACTCATCAGGAAGC
TGCCATCGGACTCATCAGGAAGC
AGGCATCGGACTCATCAGGAAGC
CCATCGGACTCATCAGGAAGC
CATCGGACTCATCAGGAAGC
CCATCGGACTCATCAGGAAGC
AGGCATCGGACTCATCAGGAAGC
GCCATCGGACTCATCAGGAAGC
GATAGGCATCGGACTCATCAGGAAGC
CCATCGGACTCATCAGGAAGC
ATGCCATCGGACTCATCAGGAAGC
CCACATCGGACTCATCAGGAAGC
GCCATCGGACTCATCAGGAAGC
CACATCGGACTCATCAGGAAGC
CACATCGGACTCATCAGGAAGC
CATCGGACTCATCAGGAAGC
GTGCCATCGGACTCATCAGGAAGC
AACATCGGACTCATCAGGAAGC

TCATCGGACTCATCAGGAAGC
CTTCATCGGACTCATCAGGAAGC
GCTCATCGGACTCATCAGGAAGC
TCCATCGGACTCATCAGGAAGC
CCATCGGACTCATCAGGAAGC
ATCGGACTCATCAGGAAGC
AACATCGGACTCATCAGGAAGC
ACCATCGGACTCATCAGGAAGC
CATCGGACTCATCAGGAAGC
ACATCGGACTCATCAGGAAGC
CTACATCGGACTCATCAGGAAGC
CATCGGACTCATCAGGAAGC
CATCGGACTCATCAGGAAGC
GCAACATCGGACTCATCAGGAAGC
CATCGGACTCATCAGGAAGC
CATCGGACTCATCAGGAAGC
ACACCATCGGACTCATCAGGAAGC
GTTCATCGGACTCATCAGGAAGC
GCATCGGACTCATCAGGAAGC
TGCCATCGGACTCATCAGGAAGC
CCATCGGACTCATCAGGAAGC
CTCATCGGACTCATCAGGAAGC
TCCCATCGGACTCATCAGGAAGC
GCATCGGACTCATCAGGAAGC
CATCGGACTCATCAGGAAGC
TCATCGGACTCATCAGGAAGC
TCATCGGACTCATCAGGAAGC
ATACATCGGACTCATCAGGAAGC
ATCGGACTCATCAGGAAGC
ACATCGGACTCATCAGGAAGC
GTCATCGGACTCATCAGGAAGC
GGACATCGGACTCATCAGGAAGC
ATGACATCGGACTCATCAGGAAGC
TGGCATCGGACTCATCAGGAAGC
TCATCGGACTCATCAGGAAGC
ACATCGGACTCATCAGGAAGC
AGCCATCGGACTCATCAGGAAGC
CTGTCATCGGACTCATCAGGAAGC
ACATCGGACTCATCAGGAAGC
ACATCGGACTCATCAGGAAGC
GGGCATCGGACTCATCAGGAAGC
AATCATCGGACTCATCAGGAAGC
TCTCATCGGACTCATCAGGAAGC

TGTTCATCGGACTCATCAGGAAGC
CCATCGGACTCATCAGGAAGC
TGCCATCGGACTCATCAGGAAGC
TTCATCGGACTCATCAGGAAGC
AACATCGGACTCATCAGGAAGC
TCCATCGGACTCATCAGGAAGC
CACATCGGACTCATCAGGAAGC
TACACATCGGACTCATCAGGAAGC
TTATCATCGGACTCATCAGGAAGC
GCCATCGGACTCATCAGGAAGC
ATTCATCGGACTCATCAGGAAGC
CATCATCGGACTCATCAGGAAGC
TACATCGGACTCATCAGGAAGC
CACATCGGACTCATCAGGAAGC
GCATCGGACTCATCAGGAAGC
AGCATCGGACTCATCAGGAAGC
ATCGGACTCATCAGGAAGC
TGCATCGGACTCATCAGGAAGC
TGCCATCGGACTCATCAGGAAGC
CACATCGGACTCATCAGGAAGC
CATCGGACTCATCAGGAAGC
AAATCATCGGACTCATCAGGAAGC
GGCCATCGGACTCATCAGGAAGC
ATCATCGGACTCATCAGGAAGC
CACCCATCGGACTCATCAGGAAGC
CCATCGGACTCATCAGGAAGC
AATCACATCGGACTCATCAGGAAGC
ACATCGGACTCATCAGGAAGC
CCATCGGACTCATCAGGAAGC
CATCGGACTCATCAGGAAGC
AGGCATCGGACTCATCAGGAAGC
CACATCGGACTCATCAGGAAGC
CCAACATCGGACTCATCAGGAAGC
CCATCGGACTCATCAGGAAGC
GCTGGACATCGGACTCATCAGGAAGC
GTCCATCGGACTCATCAGGAAGC
ATCATCGGACTCATCAGGAAGC
CCATCGGACTCATCAGGAAGC
ACATCGGACTCATCAGGAAGC
CCACCATCGGACTCATCAGGAAGC
GCATCGGACTCATCAGGAAGC
TCATCGGACTCATCAGGAAGC
GACATCGGACTCATCAGGAAGC

ATCATCGGACTCATCAGGAAGC
TCATCGGACTCATCAGGAAGC
TTTTCATCGGACTCATCAGGAAGC
CCATCGGACTCATCAGGAAGC
CCAGCATCGGACTCATCAGGAAGC
AAACATCGGACTCATCAGGAAGC
GCCATCGGACTCATCAGGAAGC
TCCATCGGACTCATCAGGAAGC
ACCATCGGACTCATCAGGAAGC
CTCATCGGACTCATCAGGAAGC
ACATCGGACTCATCAGGAAGC
TCATCGGACTCATCAGGAAGC
GACATCGGACTCATCAGGAAGC
CTTGCATCGGACTCATCAGGAAGC
CATCGGACTCATCAGGAAGC
TCATCGGACTCATCAGGAAGC
TCATCGGACTCATCAGGAAGC
CCATCGGACTCATCAGGAAGC
GCATCGGACTCATCAGGAAGC
ATCGGACTCATCAGGAAGC
GGGCATCGGACTCATCAGGAAGC
ATCATCGGACTCATCAGGAAGC
GCCATCGGACTCATCAGGAAGC
ACATCGGACTCATCAGGAAGC
CTTCATCGGACTCATCAGGAAGC
GTCATCGGACTCATCAGGAAGC
GTTCATCGGACTCATCAGGAAGC
CATCGGACTCATCAGGAAGC
GCATCGGACTCATCAGGAAGC
CCATCGGACTCATCAGGAAGC
CCATCGGACTCATCAGGAAGC
ACATCGGACTCATCAGGAAGC
TGCATCGGACTCATCAGGAAGC
GGACATCGGACTCATCAGGAAGC
TCATCGGACTCATCAGGAAGC
ACATCGGACTCATCAGGAAGC
GCATCGGACTCATCAGGAAGC
GCATCGGACTCATCAGGAAGC
CATCGGACTCATCAGGAAGC
TGCATCGGACTCATCAGGAAGC
TTCATCGGACTCATCAGGAAGC
ACACATCGGACTCATCAGGAAGC
GGAATGCATCGGACTCATCAGGAAGC

CATCGGACTCATCAGGAAGC
AAGGCATCGGACTCATCAGGAAGC
TCATCGGACTCATCAGGAAGC
AGCATCGGACTCATCAGGAAGC
CATCGGACTCATCAGGAAGC
ATTCATCGGACTCATCAGGAAGC
CCATCGGACTCATCAGGAAGC
ATCCATCGGACTCATCAGGAAGC
GGGACATCGGACTCATCAGGAAGC
GTTCATCGGACTCATCAGGAAGC
TGGCCATCGGACTCATCAGGAAGC
AATCCATCGGACTCATCAGGAAGC
TGGACATCGGACTCATCAGGAAGC
CCATCGGACTCATCAGGAAGC
TCATCGGACTCATCAGGAAGC
CCATCGGACTCATCAGGAAGC
CATCGGACTCATCAGGAAGC
TCATCGGACTCATCAGGAAGC
GCCATCGGACTCATCAGGAAGC
AGACATCGGACTCATCAGGAAGC
GTCATCGGACTCATCAGGAAGC
ATGCATCGGACTCATCAGGAAGC
GTCATCGGACTCATCAGGAAGC
TGCCTCCATCGGACTCATCAGGAAGC
CCTCATCGGACTCATCAGGAAGC
AACATCGGACTCATCAGGAAGC
TTCCCATCGGACTCATCAGGAAGC
TCATCGGACTCATCAGGAAGC
AGCATCGGACTCATCAGGAAGC
TCATCGGACTCATCAGGAAGC
GCATCGGACTCATCAGGAAGC
GCATCGGACTCATCAGGAAGC
CATCGGACTCATCAGGAAGC
AGCATCGGACTCATCAGGAAGC
GATACATCGGACTCATCAGGAAGC
CTCATCGGACTCATCAGGAAGC
CCATCATCGGACTCATCAGGAAGC
CTCATCGGACTCATCAGGAAGC
AGCGCATCGGACTCATCAGGAAGC
ATGAGCATCGGACTCATCAGGAAGC
GTCATCGGACTCATCAGGAAGC
GTCATCGGACTCATCAGGAAGC
AGTCTCATCGGACTCATCAGGAAGC

ACATCGGACTCATCAGGAAGC
TCATCGGACTCATCAGGAAGC
CATCGGACTCATCAGGAAGC
TTACATCGGACTCATCAGGAAGC
TCATCGGACTCATCAGGAAGC
ACACATCGGACTCATCAGGAAGC
GACATCGGACTCATCAGGAAGC
CATCGGACTCATCAGGAAGC
TCATCGGACTCATCAGGAAGC
GCTGGCATCGGACTCATCAGGAAGC
TGTGTCATCGGACTCATCAGGAAGC
GCATCGGACTCATCAGGAAGC
GCCATCGGACTCATCAGGAAGC
GTGTCATCGGACTCATCAGGAAGC
GTGCATCGGACTCATCAGGAAGC
ACTCATCAGGAAGC
CTCATCGGACTCATCAGGAAGC
CCATCGGACTCATCAGGAAGC
CATCGGACTCATCAGGAAGC
CGCATCGGACTCATCAGGAAGC
CATCGGACTCATCAGGAAGC
ATCATCGGACTCATCAGGAAGC
ACTTCATCGGACTCATCAGGAAGC
TTTGCATCGGACTCATCAGGAAGC
TCCCATCGGACTCATCAGGAAGC
CCATCGGACTCATCAGGAAGC
TCACATCGGACTCATCAGGAAGC
CATCGGACTCATCAGGAAGC
GACATCGGACTCATCAGGAAGC
AACCATCGGACTCATCAGGAAGC
ATTCATCGGACTCATCAGGAAGC
GCATCGGACTCATCAGGAAGC
CCCATCGGACTCATCAGGAAGC
GCATCGGACTCATCAGGAAGC
CAATCATCGGACTCATCAGGAAGC
ATCGGACTCATCAGGAAGC
ATCGGACTCATCAGGAAGC
ATTCACATCGGACTCATCAGGAAGC
AGGATCATCGGACTCATCAGGAAGC
ACATCGGACTCATCAGGAAGC
CTCATCGGACTCATCAGGAAGC
ATCGGACTCATCAGGAAGC
CCACATCGGACTCATCAGGAAGC

AGTTTGCCATCGGACTCATCAGGAAGC
CTCATCGGACTCATCAGGAAGC
CATCGGACTCATCAGGAAGC
GCATCGGACTCATCAGGAAGC
ATCGGACTCATCAGGAAGC
GTCCATCGGACTCATCAGGAAGC
GGCATCGGACTCATCAGGAAGC
ACACATCGGACTCATCAGGAAGC
AGGCATCGGACTCATCAGGAAGC
AGACATCGGACTCATCAGGAAGC
GGCCATCGGACTCATCAGGAAGC
CCATCGGACTCATCAGGAAGC
GGATCATCGGACTCATCAGGAAGC
CCATCGGACTCATCAGGAAGC
CCCATCGGACTCATCAGGAAGC
CCATCGGACTCATCAGGAAGC
AGACATCGGACTCATCAGGAAGC
GCATCGGACTCATCAGGAAGC
ATCATCGGACTCATCAGGAAGC
GCCTCATCGGACTCATCAGGAAGC
CACATCGGACTCATCAGGAAGC
GACATCGGACTCATCAGGAAGC
GCATCGGACTCATCAGGAAGC
CTTCATCGGACTCATCAGGAAGC
GCATCGGACTCATCAGGAAGC
TCATCGGACTCATCAGGAAGC
CATCGGACTCATCAGGAAGC
CGCACGCATCGGACTCATCAGGAAGC
AACATCGGACTCATCAGGAAGC
ACATCGGACTCATCAGGAAGC
CTCCATCGGACTCATCAGGAAGC
ATCGGACTCATCAGGAAGC
GGCCATCGGACTCATCAGGAAGC
AGCATCGGACTCATCAGGAAGC
AGACACATCGGACTCATCAGGAAGC
TCATCGGACTCATCAGGAAGC
TGGCATCGGACTCATCAGGAAGC
GATCATCGGACTCATCAGGAAGC
ATCGGACTCATCAGGAAGC
CTCCCATCGGACTCATCAGGAAGC
CATCGGACTCATCAGGAAGC
GCGCATCGGACTCATCAGGAAGC
CAGCATCGGACTCATCAGGAAGC

GTCATCGGACTCATCAGGAAGC
CATCGGACTCATCAGGAAGC
TCGGACTCATCAGGAAGC
GACATCGGACTCATCAGGAAGC
GCATCGGACTCATCAGGAAGC
CATCGGACTCATCAGGAAGC
AGGCATCGGACTCATCAGGAAGC
CATCGGACTCATCAGGAAGC
TGGACATCGGACTCATCAGGAAGC
ATCGGACTCATCAGGAAGC
CCCATCGGACTCATCAGGAAGC
ATTCAACATCGGACTCATCAGGAAGC
TGTCATCGGACTCATCAGGAAGC
ACCCATCGGACTCATCAGGAAGC
CATCGGACTCATCAGGAAGC
CATCGGACTCATCAGGAAGC
CCATCGGACTCATCAGGAAGC
CATCGGACTCATCAGGAAGC
GGGACATCGGACTCATCAGGAAGC
ACATCGGACTCATCAGGAAGC
GGATGCATCGGACTCATCAGGAAGC
ACCATCGGACTCATCAGGAAGC
AAGGGGGCATCGGACTCATCAGGAAGC
CGCATCGGACTCATCAGGAAGC
TCATCGGACTCATCAGGAAGC
TGCATCGGACTCATCAGGAAGC
ATCGGACTCATCAGGAAGC
TCGGACTCATCAGGAAGC
CATCGGACTCATCAGGAAGC
GTGCATCGGACTCATCAGGAAGC
CACATCGGACTCATCAGGAAGC
AGAGCATCGGACTCATCAGGAAGC
ACATCGGACTCATCAGGAAGC
ATCATCGGACTCATCAGGAAGC
AGGGCATCGGACTCATCAGGAAGC
ATTTCACATCGGACTCATCAGGAAGC
GCACACATCGGACTCATCAGGAAGC
ATCGGACTCATCAGGAAGC
TCCATCGGACTCATCAGGAAGC
ACATCGGACTCATCAGGAAGC
GCTCATCGGACTCATCAGGAAGC
CCATCGGACTCATCAGGAAGC
TCATCGGACTCATCAGGAAGC

GACATCGGACTCATCAGGAAGC
ACATCGGACTCATCAGGAAGC
TGCATCGGACTCATCAGGAAGC
TCATCGGACTCATCAGGAAGC
TGACATCGGACTCATCAGGAAGC
GTCATCGGACTCATCAGGAAGC
CCATCGGACTCATCAGGAAGC
GCCATCGGACTCATCAGGAAGC
ACATCGGACTCATCAGGAAGC
TGTCATCGGACTCATCAGGAAGC
CCATCGGACTCATCAGGAAGC
AAACATCGGACTCATCAGGAAGC
CATCGGACTCATCAGGAAGC
TCATCGGACTCATCAGGAAGC
AACATCGGACTCATCAGGAAGC
CCTAACATCGGACTCATCAGGAAGC
CATCGGACTCATCAGGAAGC
CGCATCGGACTCATCAGGAAGC
ACATCGGACTCATCAGGAAGC
GTCACATCGGACTCATCAGGAAGC
TCATCGGACTCATCAGGAAGC
GCATCGGACTCATCAGGAAGC
CCATCGGACTCATCAGGAAGC
TCACATCGGACTCATCAGGAAGC
TCCTCATCGGACTCATCAGGAAGC
CTCCATCGGACTCATCAGGAAGC
ACTGCATCGGACTCATCAGGAAGC
GACATCGGACTCATCAGGAAGC
CCATCGGACTCATCAGGAAGC
CATCGGACTCATCAGGAAGC
GGACATCGGACTCATCAGGAAGC
ACATCGGACTCATCAGGAAGC
AAACATCGGACTCATCAGGAAGC
CCATCGGACTCATCAGGAAGC
GAACACATCGGACTCATCAGGAAGC
GAGAGCATCGGACTCATCAGGAAGC
GACCACATCGGACTCATCAGGAAGC
CCATCGGACTCATCAGGAAGC
CATCGGACTCATCAGGAAGC
AGGCATCGGACTCATCAGGAAGC
GCAGCATCGGACTCATCAGGAAGC
GTAGTCATCGGACTCATCAGGAAGC
AAGCATCGGACTCATCAGGAAGC

AGGCATCGGACTCATCAGGAAGC
ATCGGACTCATCAGGAAGC
ATCATCGGACTCATCAGGAAGC
CCCCCATCGGACTCATCAGGAAGC
GAACATCGGACTCATCAGGAAGC
ACAGCATCGGACTCATCAGGAAGC
TCATCGGACTCATCAGGAAGC
GTCATCGGACTCATCAGGAAGC
TCCATCGGACTCATCAGGAAGC
AGGCCATCGGACTCATCAGGAAGC
CCCATCGGACTCATCAGGAAGC
TTTCATCGGACTCATCAGGAAGC
TGCATCGGACTCATCAGGAAGC
TCCTCATCGGACTCATCAGGAAGC
CTCACATCGGACTCATCAGGAAGC
AAACATCGGACTCATCAGGAAGC
AGTCATCGGACTCATCAGGAAGC
GCATCGGACTCATCAGGAAGC
CTCATCGGACTCATCAGGAAGC
GTGCATCGGACTCATCAGGAAGC
AGGCATCGGACTCATCAGGAAGC
GCATCGGACTCATCAGGAAGC
GAGCATCGGACTCATCAGGAAGC
AGTACATCGGACTCATCAGGAAGC
ATCGGACTCATCAGGAAGC
TCTTCCATCGGACTCATCAGGAAGC
CACATCGGACTCATCAGGAAGC
GCATCGGACTCATCAGGAAGC
GCCATCGGACTCATCAGGAAGC
CTGCATCGGACTCATCAGGAAGC
ACCATCGGACTCATCAGGAAGC
AGATCATCGGACTCATCAGGAAGC
GGGGCATCGGACTCATCAGGAAGC
GAACATCGGACTCATCAGGAAGC
GGGTCATCGGACTCATCAGGAAGC
GCCCATCGGACTCATCAGGAAGC
CACATCGGACTCATCAGGAAGC
CCATCGGACTCATCAGGAAGC
CCATCGGACTCATCAGGAAGC
TCACATCGGACTCATCAGGAAGC
GAGACATCGGACTCATCAGGAAGC
CTGCATCGGACTCATCAGGAAGC
ATCATCGGACTCATCAGGAAGC

TCATCGGACTCATCAGGAAGC
GAGGGCATCGGACTCATCAGGAAGC
GCCATCGGACTCATCAGGAAGC
AGTGTCATCGGACTCATCAGGAAGC
AGCCATCGGACTCATCAGGAAGC
TCATCGGACTCATCAGGAAGC
TCATCGGACTCATCAGGAAGC
ATCATCGGACTCATCAGGAAGC
GACATCGGACTCATCAGGAAGC
AGCATCGGACTCATCAGGAAGC
CATCGGACTCATCAGGAAGC
AGCATCGGACTCATCAGGAAGC
CATCGGACTCATCAGGAAGC
GACATCGGACTCATCAGGAAGC
CACATCGGACTCATCAGGAAGC
ACATCGGACTCATCAGGAAGC
GTACATCGGACTCATCAGGAAGC
TTCATCGGACTCATCAGGAAGC
CATCGGACTCATCAGGAAGC
GTACATCGGACTCATCAGGAAGC
TTCATCGGACTCATCAGGAAGC
GCCATCGGACTCATCAGGAAGC
CTGACATCGGACTCATCAGGAAGC
GGTGCATCGGACTCATCAGGAAGC
CTCATCGGACTCATCAGGAAGC
TGCCATCGGACTCATCAGGAAGC
GCCATCGGACTCATCAGGAAGC
CTCCATCGGACTCATCAGGAAGC
AGGCATCGGACTCATCAGGAAGC
ATCTCTCATCGGACTCATCAGGAAGC
AGCATCGGACTCATCAGGAAGC
GTGCCATCGGACTCATCAGGAAGC
TGCATCGGACTCATCAGGAAGC
CATCGGACTCATCAGGAAGC
ATCGGACTCATCAGGAAGC
GCATCGGACTCATCAGGAAGC
CATTGACATCGGACTCATCAGGAAGC
TCATCGGACTCATCAGGAAGC
CACATCGGACTCATCAGGAAGC
CATCGGACTCATCAGGAAGC
GGACCATCGGACTCATCAGGAAGC
ACATCGGACTCATCAGGAAGC
CTCATCGGACTCATCAGGAAGC

ATCATCGGACTCATCAGGAAGC
GCCCATCGGACTCATCAGGAAGC
GGGGCATCGGACTCATCAGGAAGC
CTCAGCATCGGACTCATCAGGAAGC
ATGCATCGGACTCATCAGGAAGC
CTGTCATCGGACTCATCAGGAAGC
CCCCATCGGACTCATCAGGAAGC
TCTCATCGGACTCATCAGGAAGC
AATCATCGGACTCATCAGGAAGC
CCCTTCATCGGACTCATCAGGAAGC
AGCATCGGACTCATCAGGAAGC
ACTGCCATCGGACTCATCAGGAAGC
GGGCATCGGACTCATCAGGAAGC
CCATCGGACTCATCAGGAAGC
CCGAGCCATCGGACTCATCAGGAAGC
CCATCGGACTCATCAGGAAGC
CTCATCGGACTCATCAGGAAGC
GCATCGGACTCATCAGGAAGC
GGAGCATCGGACTCATCAGGAAGC
TCATCGGACTCATCAGGAAGC
CTCATCGGACTCATCAGGAAGC
CCACATCGGACTCATCAGGAAGC
CCATCGGACTCATCAGGAAGC
ATCGGACTCATCAGGAAGC
TATCATCGGACTCATCAGGAAGC
GGAGCCATCGGACTCATCAGGAAGC
TAACATCGGACTCATCAGGAAGC
CATCGGACTCATCAGGAAGC
GACATCGGACTCATCAGGAAGC
GTCTCATCGGACTCATCAGGAAGC
TCGGACTCATCAGGAAGC
TTCATCGGACTCATCAGGAAGC
CATCGGACTCATCAGGAAGC
TCATCGGACTCATCAGGAAGC
ATCATCGGACTCATCAGGAAGC
TCTCATCGGACTCATCAGGAAGC
GCACATCGGACTCATCAGGAAGC
CTGGCATCGGACTCATCAGGAAGC
ACATCGGACTCATCAGGAAGC
TCATCGGACTCATCAGGAAGC
TCATCGGACTCATCAGGAAGC
TACATCGGACTCATCAGGAAGC
ATCGGACTCATCAGGAAGC

CCATCGGACTCATCAGGAAGC
GCATCGGACTCATCAGGAAGC
CCATCGGACTCATCAGGAAGC
GGCATCGGACTCATCAGGAAGC
ACATCGGACTCATCAGGAAGC
AGCATCGGACTCATCAGGAAGC
TGTGTCATCGGACTCATCAGGAAGC
GTGACCATCGGACTCATCAGGAAGC
ACATCGGACTCATCAGGAAGC
ATGGGTTCATCGGACTCATCAGGAAGC
GCATCGGACTCATCAGGAAGC
AGCATCGGACTCATCAGGAAGC
GGCATCGGACTCATCAGGAAGC
ACATCGGACTCATCAGGAAGC
CCCATCGGACTCATCAGGAAGC
TCATCGGACTCATCAGGAAGC
AGCATCGGACTCATCAGGAAGC
TCCATCGGACTCATCAGGAAGC
GCGCATCGGACTCATCAGGAAGC
TGTCATCGGACTCATCAGGAAGC
AGCATCGGACTCATCAGGAAGC
GCATCGGACTCATCAGGAAGC
AGCCATCGGACTCATCAGGAAGC
ATCATCGGACTCATCAGGAAGC
GCCATCGGACTCATCAGGAAGC
CTGCATCGGACTCATCAGGAAGC
AGGCATCGGACTCATCAGGAAGC
AACATCGGACTCATCAGGAAGC
AGGCACATCGGACTCATCAGGAAGC
TGAGCATCGGACTCATCAGGAAGC
TCCATCGGACTCATCAGGAAGC
TTGCATCGGACTCATCAGGAAGC
TTTCATCGGACTCATCAGGAAGC
TTCATCGGACTCATCAGGAAGC
CGCCATCGGACTCATCAGGAAGC
TGCAGCATCGGACTCATCAGGAAGC
CCATGTCATCGGACTCATCAGGAAGC
TACATCGGACTCATCAGGAAGC
CTCAACATCGGACTCATCAGGAAGC
GTGCATCGGACTCATCAGGAAGC
TGCATCGGACTCATCAGGAAGC
AGGCATCGGACTCATCAGGAAGC
TCATCGGACTCATCAGGAAGC

TTCCATCGGACTCATCAGGAAGC
ACATCGGACTCATCAGGAAGC
CATCGGACTCATCAGGAAGC
TCACATCGGACTCATCAGGAAGC
CATCGGACTCATCAGGAAGC
GCATCGGACTCATCAGGAAGC
CATCGGACTCATCAGGAAGC
GATTCATCGGACTCATCAGGAAGC
GTCATCGGACTCATCAGGAAGC
CCATCGGACTCATCAGGAAGC
TTCCATCGGACTCATCAGGAAGC
TATGTGCATCGGACTCATCAGGAAGC
CCATCGGACTCATCAGGAAGC
GCCATCGGACTCATCAGGAAGC
AACATCGGACTCATCAGGAAGC
CATCGGACTCATCAGGAAGC
GTGTGGCATCGGACTCATCAGGAAGC
TTACATCGGACTCATCAGGAAGC
CCATCGGACTCATCAGGAAGC
TCATCGGACTCATCAGGAAGC
GCATCGGACTCATCAGGAAGC
ACATCGGACTCATCAGGAAGC
TTCATCGGACTCATCAGGAAGC
TTTCATCGGACTCATCAGGAAGC
ACATCGGACTCATCAGGAAGC
CATCGGACTCATCAGGAAGC
ACATCGGACTCATCAGGAAGC
TCATCGGACTCATCAGGAAGC
TTCATCGGACTCATCAGGAAGC
TGCATCGGACTCATCAGGAAGC
ACATCGGACTCATCAGGAAGC
TGACATCGGACTCATCAGGAAGC
TGGGTCATCGGACTCATCAGGAAGC
GTCATCGGACTCATCAGGAAGC
CATCGGACTCATCAGGAAGC
AAAACATCGGACTCATCAGGAAGC
ATCGGACTCATCAGGAAGC
TCCATCGGACTCATCAGGAAGC
AAAGATCATCGGACTCATCAGGAAGC
CATCGGACTCATCAGGAAGC
TGCATCGGACTCATCAGGAAGC
CATCGGACTCATCAGGAAGC
GCATCGGACTCATCAGGAAGC

GAGCATCGGACTCATCAGGAAGC
TCATCGGACTCATCAGGAAGC
TCATCGGACTCATCAGGAAGC
CATCGGACTCATCAGGAAGC
TCCATCGGACTCATCAGGAAGC
GACATCGGACTCATCAGGAAGC
GCATCGGACTCATCAGGAAGC
GTTTCATCGGACTCATCAGGAAGC
CATCGGACTCATCAGGAAGC
CATCGGACTCATCAGGAAGC
TCATCGGACTCATCAGGAAGC
TCCATCGGACTCATCAGGAAGC
CCCATCGGACTCATCAGGAAGC
AGTCCATCGGACTCATCAGGAAGC
TCGGACTCATCAGGAAGC
ACATCGGACTCATCAGGAAGC
TACACATCGGACTCATCAGGAAGC
GTCATCGGACTCATCAGGAAGC
CTCATCGGACTCATCAGGAAGC
TGGCATCGGACTCATCAGGAAGC
AACATCGGACTCATCAGGAAGC
CACATCGGACTCATCAGGAAGC
GTGCATCGGACTCATCAGGAAGC
CCATCGGACTCATCAGGAAGC
TCATCGGACTCATCAGGAAGC
ACACATCGGACTCATCAGGAAGC
ACATCGGACTCATCAGGAAGC
CCACATCGGACTCATCAGGAAGC
GCCATCGGACTCATCAGGAAGC
CCATCGGACTCATCAGGAAGC
CCATCGGACTCATCAGGAAGC
ATCGGACTCATCAGGAAGC
TCATCGGACTCATCAGGAAGC
GTACATCGGACTCATCAGGAAGC
CCTCATCGGACTCATCAGGAAGC
TTCCATCGGACTCATCAGGAAGC
GCTATCCATCGGACTCATCAGGAAGC
TAAACATCGGACTCATCAGGAAGC
CAAACATCGGACTCATCAGGAAGC
ATCGGACTCATCAGGAAGC
GCATCGGACTCATCAGGAAGC
ACCCATCGGACTCATCAGGAAGC
ACATCGGACTCATCAGGAAGC

GTTCATCGGACTCATCAGGAAGC
TGTGCATCGGACTCATCAGGAAGC
AAACATCGGACTCATCAGGAAGC
CACATCGGACTCATCAGGAAGC
GGTCATCGGACTCATCAGGAAGC
AGAGACATCGGACTCATCAGGAAGC
CGGACTCATCAGGAAGC
CATCGGACTCATCAGGAAGC
TCATCGGACTCATCAGGAAGC
TGTCATCGGACTCATCAGGAAGC
ACACCATCGGACTCATCAGGAAGC
GCACATCGGACTCATCAGGAAGC
GCATCGGACTCATCAGGAAGC
CACATCGGACTCATCAGGAAGC
ACCATCGGACTCATCAGGAAGC
ACCCATCGGACTCATCAGGAAGC
GTCACATCGGACTCATCAGGAAGC
CATCGGACTCATCAGGAAGC
TTCATCGGACTCATCAGGAAGC
GGCCATCGGACTCATCAGGAAGC
CTTCATCGGACTCATCAGGAAGC
TCTCCATCGGACTCATCAGGAAGC
ACACATCGGACTCATCAGGAAGC
CATCGGACTCATCAGGAAGC
ATCGGACTCATCAGGAAGC
GTCATCGGACTCATCAGGAAGC
ACACATCGGACTCATCAGGAAGC
GCGCATCGGACTCATCAGGAAGC
TCATCGGACTCATCAGGAAGC
GTCCCATCGGACTCATCAGGAAGC
GATTCATCGGACTCATCAGGAAGC
GTGGAACATCGGACTCATCAGGAAGC
GGCCATCGGACTCATCAGGAAGC
CATCGGACTCATCAGGAAGC
ATCGGACTCATCAGGAAGC
TGAACATCGGACTCATCAGGAAGC
GTCCATCGGACTCATCAGGAAGC
CTCATCGGACTCATCAGGAAGC
TCCATCGGACTCATCAGGAAGC
GAGCATCGGACTCATCAGGAAGC
GAGCCATCGGACTCATCAGGAAGC
GGCATCGGACTCATCAGGAAGC
CTCCATCGGACTCATCAGGAAGC

AGCATCGGACTCATCAGGAAGC
CCATCGGACTCATCAGGAAGC
ATCGGACTCATCAGGAAGC
CATCGGACTCATCAGGAAGC
TCATCGGACTCATCAGGAAGC
GCATCGGACTCATCAGGAAGC
CCATCGGACTCATCAGGAAGC
CCCATCGGACTCATCAGGAAGC
GCCATCGGACTCATCAGGAAGC
CCACATCGGACTCATCAGGAAGC
TCCATCGGACTCATCAGGAAGC
TTGGCATCGGACTCATCAGGAAGC
ATTGGTCATCGGACTCATCAGGAAGC
GACCACATCGGACTCATCAGGAAGC
AACATCGGACTCATCAGGAAGC
GTCATCGGACTCATCAGGAAGC
CCCATCGGACTCATCAGGAAGC
AGCATCGGACTCATCAGGAAGC
GCTGCATCGGACTCATCAGGAAGC
GCATCGGACTCATCAGGAAGC
CTTCATCGGACTCATCAGGAAGC
AACACATCGGACTCATCAGGAAGC
AGCCATCGGACTCATCAGGAAGC
TCATCGGACTCATCAGGAAGC
CCATCGGACTCATCAGGAAGC
GTCTCATCGGACTCATCAGGAAGC
GGCATCGGACTCATCAGGAAGC
TATGCCATCGGACTCATCAGGAAGC
TTGACACATCGGACTCATCAGGAAGC
AGTCATCGGACTCATCAGGAAGC
GCATCGGACTCATCAGGAAGC
GTCATCGGACTCATCAGGAAGC
TACATCGGACTCATCAGGAAGC
GACATCGGACTCATCAGGAAGC
CTCATCGGACTCATCAGGAAGC
ACATCGGACTCATCAGGAAGC
CATCGGACTCATCAGGAAGC
ACATCGGACTCATCAGGAAGC
TACATCGGACTCATCAGGAAGC
CGATCATCGGACTCATCAGGAAGC
GATTCATCGGACTCATCAGGAAGC
TCATCGGACTCATCAGGAAGC
TCATCGGACTCATCAGGAAGC

AACATCGGACTCATCAGGAAGC
TGCATCGGACTCATCAGGAAGC
AGGGCATCGGACTCATCAGGAAGC
CCATCGGACTCATCAGGAAGC
ATCGGACTCATCAGGAAGC
TACATCGGACTCATCAGGAAGC
ATCATCGGACTCATCAGGAAGC
TGTCATCGGACTCATCAGGAAGC
CCATCGGACTCATCAGGAAGC
TCATCGGACTCATCAGGAAGC
ATCGGACTCATCAGGAAGC
GCCATCGGACTCATCAGGAAGC
TTCCATCGGACTCATCAGGAAGC
TGTCATCGGACTCATCAGGAAGC
CATCGGACTCATCAGGAAGC
GAACATCGGACTCATCAGGAAGC
TCGGACTCATCAGGAAGC
ACCCCATCGGACTCATCAGGAAGC
ATCGGACTCATCAGGAAGC
ACCATCGGACTCATCAGGAAGC
AAGCATCGGACTCATCAGGAAGC
TCTCATCGGACTCATCAGGAAGC
TGGCATCGGACTCATCAGGAAGC
ATCGGACTCATCAGGAAGC
TTTCATCGGACTCATCAGGAAGC
GCATCGGACTCATCAGGAAGC
GTACATCGGACTCATCAGGAAGC
GTCTCATCGGACTCATCAGGAAGC
AGCACATCGGACTCATCAGGAAGC
TACACATCGGACTCATCAGGAAGC
AGCATCGGACTCATCAGGAAGC
GCATCGGACTCATCAGGAAGC
TGACATCGGACTCATCAGGAAGC
GACATCGGACTCATCAGGAAGC
CATCGGACTCATCAGGAAGC
TACATCGGACTCATCAGGAAGC
TGCATCGGACTCATCAGGAAGC
CCATCGGACTCATCAGGAAGC
AAGACATCGGACTCATCAGGAAGC
AGTTGCATCGGACTCATCAGGAAGC
ACCATCGGACTCATCAGGAAGC
CTTATCATCGGACTCATCAGGAAGC
CATCGGACTCATCAGGAAGC

TGGCATCGGACTCATCAGGAAGC
CATCGGACTCATCAGGAAGC
GTTCATCGGACTCATCAGGAAGC
TCGGACTCATCAGGAAGC
CACATCGGACTCATCAGGAAGC
CCACATCGGACTCATCAGGAAGC
CTCTTCATCGGACTCATCAGGAAGC
TCATCGGACTCATCAGGAAGC
ATGCCATCGGACTCATCAGGAAGC
CACTGCATCGGACTCATCAGGAAGC
TATTCATCGGACTCATCAGGAAGC
TGTCATCGGACTCATCAGGAAGC
TACATCGGACTCATCAGGAAGC
GTGAGCATCGGACTCATCAGGAAGC
AACACATCGGACTCATCAGGAAGC
ATCGGACTCATCAGGAAGC
ACATCGGACTCATCAGGAAGC
CCATCGGACTCATCAGGAAGC
CATCGGACTCATCAGGAAGC
ACATCGGACTCATCAGGAAGC
CTGGCATCGGACTCATCAGGAAGC
TCATCGGACTCATCAGGAAGC
GACATCGGACTCATCAGGAAGC
CTCATCGGACTCATCAGGAAGC
TCCCATCGGACTCATCAGGAAGC
ATCGGACTCATCAGGAAGC
TCACATCGGACTCATCAGGAAGC
ATCGGACTCATCAGGAAGC
GTCATCGGACTCATCAGGAAGC
AGCCATCGGACTCATCAGGAAGC
GCCATCGGACTCATCAGGAAGC
TCATCGGACTCATCAGGAAGC
TATGACATCGGACTCATCAGGAAGC
TCATCGGACTCATCAGGAAGC
CCATCGGACTCATCAGGAAGC
CTCATCGGACTCATCAGGAAGC
CCATCATCGGACTCATCAGGAAGC
ATCGGACTCATCAGGAAGC
TGTCATCGGACTCATCAGGAAGC
CCATCGGACTCATCAGGAAGC
GGCATCGGACTCATCAGGAAGC
CCATCGGACTCATCAGGAAGC
CACATCGGACTCATCAGGAAGC

GCATCGGACTCATCAGGAAGC
CATCGGACTCATCAGGAAGC
CTCCATCGGACTCATCAGGAAGC
GCACACATCGGACTCATCAGGAAGC
CCATCGGACTCATCAGGAAGC
TCATCGGACTCATCAGGAAGC
TCCATCGGACTCATCAGGAAGC
CCTCATCGGACTCATCAGGAAGC
TACACATCGGACTCATCAGGAAGC
CATCGGACTCATCAGGAAGC
CATCGGACTCATCAGGAAGC
TCATCGGACTCATCAGGAAGC
ATTTCATCGGACTCATCAGGAAGC
TCAAACATCGGACTCATCAGGAAGC
CATCATCGGACTCATCAGGAAGC
CATCGGACTCATCAGGAAGC
GGCATCGGACTCATCAGGAAGC
ACCCCATCGGACTCATCAGGAAGC
CTACATCGGACTCATCAGGAAGC
CCATCGGACTCATCAGGAAGC
CCATCGGACTCATCAGGAAGC
GTGGCATCGGACTCATCAGGAAGC
AACATCGGACTCATCAGGAAGC
CTTTCATCGGACTCATCAGGAAGC
GGACTCATCAGGAAGC
CATCGGACTCATCAGGAAGC
TGGCATCGGACTCATCAGGAAGC
CCCATCGGACTCATCAGGAAGC
TTCATCGGACTCATCAGGAAGC
GACATCGGACTCATCAGGAAGC
ACATCGGACTCATCAGGAAGC
CACCCATCGGACTCATCAGGAAGC
GTCTCATCGGACTCATCAGGAAGC
AGCACATCGGACTCATCAGGAAGC
CCATCGGACTCATCAGGAAGC
TCATCGGACTCATCAGGAAGC
CCCATCGGACTCATCAGGAAGC
AACATCGGACTCATCAGGAAGC
ATCATCGGACTCATCAGGAAGC
TACATCGGACTCATCAGGAAGC
ATCATCGGACTCATCAGGAAGC
CACATCGGACTCATCAGGAAGC
GATGCCATCGGACTCATCAGGAAGC

AACAGACATCGGACTCATCAGGAAGC
GAGAGAGCATCGGACTCATCAGGAAGC
TACATCGGACTCATCAGGAAGC
GCATCGGACTCATCAGGAAGC
TACATCGGACTCATCAGGAAGC
AACATCGGACTCATCAGGAAGC
ATCGGACTCATCAGGAAGC
TCTCTCATCGGACTCATCAGGAAGC
TTCATCGGACTCATCAGGAAGC
GTCATCGGACTCATCAGGAAGC
CATCGGACTCATCAGGAAGC
CCATCGGACTCATCAGGAAGC
CACATCGGACTCATCAGGAAGC
AGGACATCGGACTCATCAGGAAGC
GCATCGGACTCATCAGGAAGC
CCATCGGACTCATCAGGAAGC
TTACATCGGACTCATCAGGAAGC
CCATCGGACTCATCAGGAAGC
ACATCGGACTCATCAGGAAGC
TGCATCGGACTCATCAGGAAGC
ACTCACATCGGACTCATCAGGAAGC
GCCATCGGACTCATCAGGAAGC
CTTGGCATCGGACTCATCAGGAAGC
GTTCATCGGACTCATCAGGAAGC
TCATCGGACTCATCAGGAAGC
CGGACTCATCAGGAAGC
TCATCGGACTCATCAGGAAGC
CATCGGACTCATCAGGAAGC
TCATCGGACTCATCAGGAAGC
TCCATCGGACTCATCAGGAAGC
CATCGGACTCATCAGGAAGC
CTTCATCGGACTCATCAGGAAGC
CATTTCATCGGACTCATCAGGAAGC
CTGCATCGGACTCATCAGGAAGC
CATCATCGGACTCATCAGGAAGC
TCATCGGACTCATCAGGAAGC
CCCATCGGACTCATCAGGAAGC
GTCATCGGACTCATCAGGAAGC
ATCGGACTCATCAGGAAGC
ACATCGGACTCATCAGGAAGC
TCTCATCGGACTCATCAGGAAGC
GACATCGGACTCATCAGGAAGC
ACATCGGACTCATCAGGAAGC

CCATCGGACTCATCAGGAAGC
ATCGGACTCATCAGGAAGC
CGAACTCACATCGGACTCATCAGGAAGC
AACATCGGACTCATCAGGAAGC
GTAGGGGCATCGGACTCATCAGGAAGC
CTAACATCGGACTCATCAGGAAGC
TCATCGGACTCATCAGGAAGC
AACACATCGGACTCATCAGGAAGC
GGACATCGGACTCATCAGGAAGC
GCATCGGACTCATCAGGAAGC
TCGCATCGGACTCATCAGGAAGC
CTATCATCGGACTCATCAGGAAGC
AAGGCATCGGACTCATCAGGAAGC
ACCATCGGACTCATCAGGAAGC
CATCGGACTCATCAGGAAGC
AAACATCGGACTCATCAGGAAGC
AAGTGCATCGGACTCATCAGGAAGC
ATGGTCATCGGACTCATCAGGAAGC
TCATCGGACTCATCAGGAAGC
GCATCGGACTCATCAGGAAGC
TTCATCGGACTCATCAGGAAGC
AAACATCGGACTCATCAGGAAGC
TCCCATCGGACTCATCAGGAAGC
AACGCATCGGACTCATCAGGAAGC
TCATCGGACTCATCAGGAAGC
CCCCATCGGACTCATCAGGAAGC
GCATCGGACTCATCAGGAAGC
TGTCATCGGACTCATCAGGAAGC
CCATCGGACTCATCAGGAAGC
GACATCGGACTCATCAGGAAGC
GCATCGGACTCATCAGGAAGC
TCATCGGACTCATCAGGAAGC
TTGCATCGGACTCATCAGGAAGC
TCCATCGGACTCATCAGGAAGC
AGACATCGGACTCATCAGGAAGC
GATCATCGGACTCATCAGGAAGC
GTTTCCCATCGGACTCATCAGGAAGC
CCATCGGACTCATCAGGAAGC
ATCCATCGGACTCATCAGGAAGC
CCATCGGACTCATCAGGAAGC
CCCATCGGACTCATCAGGAAGC
TCATCGGACTCATCAGGAAGC
CACAACATCGGACTCATCAGGAAGC

GTCCATCGGACTCATCAGGAAGC
TCCATCGGACTCATCAGGAAGC
CACATCGGACTCATCAGGAAGC
TACATCGGACTCATCAGGAAGC
CACCATCGGACTCATCAGGAAGC
AAACATCGGACTCATCAGGAAGC
TGCATCGGACTCATCAGGAAGC
GCATCGGACTCATCAGGAAGC
CCATCGGACTCATCAGGAAGC
TGACATCGGACTCATCAGGAAGC
TTTCATCGGACTCATCAGGAAGC
ACCATCGGACTCATCAGGAAGC
ATACATCGGACTCATCAGGAAGC
CCATCGGACTCATCAGGAAGC
CCCATCGGACTCATCAGGAAGC
CATCGGACTCATCAGGAAGC
ACGCACATCGGACTCATCAGGAAGC
ACCTCATCGGACTCATCAGGAAGC
CCCCATCGGACTCATCAGGAAGC
CCTCCATCGGACTCATCAGGAAGC
AGGCATCGGACTCATCAGGAAGC
AAACATCGGACTCATCAGGAAGC
ATCATCGGACTCATCAGGAAGC
TCATCGGACTCATCAGGAAGC
ACATCGGACTCATCAGGAAGC
AACATCGGACTCATCAGGAAGC
GCATCATCGGACTCATCAGGAAGC
TCATCGGACTCATCAGGAAGC
ACATCGGACTCATCAGGAAGC
TACCATCGGACTCATCAGGAAGC
ACCCATCGGACTCATCAGGAAGC
CATCGGACTCATCAGGAAGC
CATCGGACTCATCAGGAAGC
AGAACATCGGACTCATCAGGAAGC
CATCGGACTCATCAGGAAGC
GCAAACATCGGACTCATCAGGAAGC
TCATCGGACTCATCAGGAAGC
AAACATCGGACTCATCAGGAAGC
GCACATCGGACTCATCAGGAAGC
TTCATCGGACTCATCAGGAAGC
TGACATCGGACTCATCAGGAAGC
GGAAACATCGGACTCATCAGGAAGC
CTCATCGGACTCATCAGGAAGC

TCCCATCGGACTCATCAGGAAGC
CATCGGACTCATCAGGAAGC
ATCGGACTCATCAGGAAGC
CACATCGGACTCATCAGGAAGC
GCCATCGGACTCATCAGGAAGC
AGCCATCGGACTCATCAGGAAGC
GCCCATCGGACTCATCAGGAAGC
TTGAAACATCGGACTCATCAGGAAGC
TCCATCGGACTCATCAGGAAGC
TTCATCGGACTCATCAGGAAGC
TCATCGGACTCATCAGGAAGC
CACTCATCGGACTCATCAGGAAGC
TACATCGGACTCATCAGGAAGC
GCATCGGACTCATCAGGAAGC
CCATCGGACTCATCAGGAAGC
ACATCGGACTCATCAGGAAGC
CCATCGGACTCATCAGGAAGC
ATCATCGGACTCATCAGGAAGC
CATCGGACTCATCAGGAAGC
GACCATCGGACTCATCAGGAAGC
CACATCGGACTCATCAGGAAGC
GTTGCATCGGACTCATCAGGAAGC
CATCGGACTCATCAGGAAGC
TCTCATCGGACTCATCAGGAAGC
TGCATCGGACTCATCAGGAAGC
CATCGGACTCATCAGGAAGC
AGCATCGGACTCATCAGGAAGC
CTCATCGGACTCATCAGGAAGC
TTGCATCGGACTCATCAGGAAGC
CCTCATCGGACTCATCAGGAAGC
CAACATCGGACTCATCAGGAAGC
CATCGGACTCATCAGGAAGC
TCCATCGGACTCATCAGGAAGC
AACATCGGACTCATCAGGAAGC
CATCGGACTCATCAGGAAGC
ACCCCATCGGACTCATCAGGAAGC
ATCGGACTCATCAGGAAGC
CTCATCGGACTCATCAGGAAGC
AACATCGGACTCATCAGGAAGC
ACATCGGACTCATCAGGAAGC
ACAACATCGGACTCATCAGGAAGC
GACATCGGACTCATCAGGAAGC
TACATCGGACTCATCAGGAAGC

AAGGCATCGGACTCATCAGGAAGC
GAAACATCGGACTCATCAGGAAGC
CTGCATCGGACTCATCAGGAAGC
TGTCATCGGACTCATCAGGAAGC
ATCATCGGACTCATCAGGAAGC
AGACATCGGACTCATCAGGAAGC
AAGCATCGGACTCATCAGGAAGC
GCCATCGGACTCATCAGGAAGC
CTCATCGGACTCATCAGGAAGC
TGCTCATCGGACTCATCAGGAAGC
TCCATCGGACTCATCAGGAAGC
GCACATCGGACTCATCAGGAAGC
CCACATCGGACTCATCAGGAAGC
TTCATCGGACTCATCAGGAAGC
CCACATCGGACTCATCAGGAAGC
CCAGCATCGGACTCATCAGGAAGC
TTCATCGGACTCATCAGGAAGC
CCCATCGGACTCATCAGGAAGC
ACACATCGGACTCATCAGGAAGC
TCATCGGACTCATCAGGAAGC
CATCGGACTCATCAGGAAGC
TCGGACTCATCAGGAAGC
TACATCGGACTCATCAGGAAGC
ATACATCGGACTCATCAGGAAGC
TCCATCGGACTCATCAGGAAGC
AGCATCGGACTCATCAGGAAGC
CTCATCGGACTCATCAGGAAGC
GCATCGGACTCATCAGGAAGC
CATCGGACTCATCAGGAAGC
TCATCGGACTCATCAGGAAGC
CTTTCCATCGGACTCATCAGGAAGC
ATCGGACTCATCAGGAAGC
CATCGGACTCATCAGGAAGC
CTATCCATCGGACTCATCAGGAAGC
GCCATCGGACTCATCAGGAAGC
GACATCGGACTCATCAGGAAGC
TCATCGGACTCATCAGGAAGC
GACATCGGACTCATCAGGAAGC
TCTCATCGGACTCATCAGGAAGC
TCATCGGACTCATCAGGAAGC
CAGCATCGGACTCATCAGGAAGC
GTGCATCGGACTCATCAGGAAGC
CTCTCATCGGACTCATCAGGAAGC

TACCATCGGACTCATCAGGAAGC
ACATCGGACTCATCAGGAAGC
ACCACATCGGACTCATCAGGAAGC
GGCATCGGACTCATCAGGAAGC
GGAGCATCGGACTCATCAGGAAGC
GCATCGGACTCATCAGGAAGC
CCCATCGGACTCATCAGGAAGC
GCTACATCGGACTCATCAGGAAGC
ATCATCGGACTCATCAGGAAGC
GCATCGGACTCATCAGGAAGC
CAGACATCGGACTCATCAGGAAGC
ATCATCGGACTCATCAGGAAGC
TCATCGGACTCATCAGGAAGC
CATCGGACTCATCAGGAAGC
AGACATCGGACTCATCAGGAAGC
ACATCGGACTCATCAGGAAGC
CTCATCGGACTCATCAGGAAGC
GCGCATCGGACTCATCAGGAAGC
GAGCATCGGACTCATCAGGAAGC
GTCATCGGACTCATCAGGAAGC
TAACATCGGACTCATCAGGAAGC
AGCATCATCGGACTCATCAGGAAGC
CCCTCATCGGACTCATCAGGAAGC
ACATCGGACTCATCAGGAAGC
GTCCATCGGACTCATCAGGAAGC
ATCATCGGACTCATCAGGAAGC
ACATCGGACTCATCAGGAAGC
ACGACATCGGACTCATCAGGAAGC
AGCATCGGACTCATCAGGAAGC
CATCGGACTCATCAGGAAGC
GGCATCGGACTCATCAGGAAGC
GCCATCGGACTCATCAGGAAGC
GTGACCATCGGACTCATCAGGAAGC
TCCCTCATCGGACTCATCAGGAAGC
AACATCGGACTCATCAGGAAGC
GAGCATCGGACTCATCAGGAAGC
ACATCGGACTCATCAGGAAGC
CGCATCGGACTCATCAGGAAGC
TCCATCGGACTCATCAGGAAGC
GAGCATCGGACTCATCAGGAAGC
ACCCCATCGGACTCATCAGGAAGC
ATCATCGGACTCATCAGGAAGC
ATCCATCGGACTCATCAGGAAGC

CACATCGGACTCATCAGGAAGC
CATCGGACTCATCAGGAAGC
CTGACATCGGACTCATCAGGAAGC
GCCAAACATCGGACTCATCAGGAAGC
GCTACATCGGACTCATCAGGAAGC
ACCCATCGGACTCATCAGGAAGC
CATTCATCGGACTCATCAGGAAGC
TTCATCGGACTCATCAGGAAGC
ATCGGACTCATCAGGAAGC
GAAGACATCGGACTCATCAGGAAGC
GCATCGGACTCATCAGGAAGC
CATCGGACTCATCAGGAAGC
ACATCGGACTCATCAGGAAGC
ATCATCGGACTCATCAGGAAGC
ATCATCGGACTCATCAGGAAGC
TTCATCGGACTCATCAGGAAGC
AGCATCGGACTCATCAGGAAGC
ACCAAGCATCGGACTCATCAGGAAGC
CACCATCGGACTCATCAGGAAGC
TACATCGGACTCATCAGGAAGC
GACATCGGACTCATCAGGAAGC
GTAGTGCATCGGACTCATCAGGAAGC
GCATCGGACTCATCAGGAAGC
CATCGGACTCATCAGGAAGC
TCATCGGACTCATCAGGAAGC
ACCATCGGACTCATCAGGAAGC
CCATCGGACTCATCAGGAAGC
TCCCATCGGACTCATCAGGAAGC
CCATCGGACTCATCAGGAAGC
TGACCATCGGACTCATCAGGAAGC
CCATCGGACTCATCAGGAAGC
GGCATCGGACTCATCAGGAAGC
CATCGGACTCATCAGGAAGC
GAGCCATCGGACTCATCAGGAAGC
TGTCATCGGACTCATCAGGAAGC
GGGCATCGGACTCATCAGGAAGC
ATGACATCGGACTCATCAGGAAGC
AACATCGGACTCATCAGGAAGC
CCCATCGGACTCATCAGGAAGC
CATCGGACTCATCAGGAAGC
AGTTACATCGGACTCATCAGGAAGC
TCGGACTCATCAGGAAGC
CATCGGACTCATCAGGAAGC

TCTTCATCGGACTCATCAGGAAGC
TTCATCGGACTCATCAGGAAGC
CACATCGGACTCATCAGGAAGC
TCATCGGACTCATCAGGAAGC
CCCATCGGACTCATCAGGAAGC
CACATCGGACTCATCAGGAAGC
GCCATCGGACTCATCAGGAAGC
TGCTCATCGGACTCATCAGGAAGC
CCACATCGGACTCATCAGGAAGC
AGACATCGGACTCATCAGGAAGC
GGCCATCGGACTCATCAGGAAGC
AAGCCATCGGACTCATCAGGAAGC
CATCGGACTCATCAGGAAGC
TTCATCGGACTCATCAGGAAGC
TCCATCGGACTCATCAGGAAGC
CTCATCGGACTCATCAGGAAGC
AAAACATCGGACTCATCAGGAAGC
TCATCGGACTCATCAGGAAGC
AATCATCGGACTCATCAGGAAGC
GGACATCGGACTCATCAGGAAGC
TGGCATCGGACTCATCAGGAAGC
ACATCGGACTCATCAGGAAGC
CATCGGACTCATCAGGAAGC
CACATCGGACTCATCAGGAAGC
TCACATCGGACTCATCAGGAAGC
CCCATCGGACTCATCAGGAAGC
CCACATCGGACTCATCAGGAAGC
CATCGGACTCATCAGGAAGC
GCCATCGGACTCATCAGGAAGC
AAGCCATCGGACTCATCAGGAAGC
TCATCGGACTCATCAGGAAGC
ACATCGGACTCATCAGGAAGC
GTACATCGGACTCATCAGGAAGC
TCATCGGACTCATCAGGAAGC
CCCATCGGACTCATCAGGAAGC
GCCATCGGACTCATCAGGAAGC
GTTTCATCGGACTCATCAGGAAGC
AGCATCGGACTCATCAGGAAGC
CATCGGACTCATCAGGAAGC
TTTCATCGGACTCATCAGGAAGC
AGTCATCGGACTCATCAGGAAGC
AGCATCGGACTCATCAGGAAGC
CCATCGGACTCATCAGGAAGC

GGACTCATCAGGAAGC
GCATCGGACTCATCAGGAAGC
TCCATCGGACTCATCAGGAAGC
CACATCGGACTCATCAGGAAGC
GTGTCATCGGACTCATCAGGAAGC
ATGTCATCGGACTCATCAGGAAGC
CCATCGGACTCATCAGGAAGC
TGTCATCGGACTCATCAGGAAGC
CCATCGGACTCATCAGGAAGC
GTGACATCGGACTCATCAGGAAGC
GACATCGGACTCATCAGGAAGC
CATCGGACTCATCAGGAAGC
CTCATCGGACTCATCAGGAAGC
CCATCGGACTCATCAGGAAGC
GTCATCGGACTCATCAGGAAGC
TTCATCGGACTCATCAGGAAGC
GCATCGGACTCATCAGGAAGC
GAAACATCGGACTCATCAGGAAGC
GGCATCGGACTCATCAGGAAGC
ATTCATCGGACTCATCAGGAAGC
CGCATCGGACTCATCAGGAAGC
GGAGCATCGGACTCATCAGGAAGC
CATCGGACTCATCAGGAAGC
CTCACATCGGACTCATCAGGAAGC
TCATCGGACTCATCAGGAAGC
GTAGTTCATCGGACTCATCAGGAAGC
GCATCGGACTCATCAGGAAGC
GTTACATCGGACTCATCAGGAAGC
AGATCATCGGACTCATCAGGAAGC
TCATCGGACTCATCAGGAAGC
TTGCATCGGACTCATCAGGAAGC
TCATCGGACTCATCAGGAAGC
CATCGGACTCATCAGGAAGC
AACGCATCGGACTCATCAGGAAGC
CATCGGACTCATCAGGAAGC
ATCGGACTCATCAGGAAGC
CATCGGACTCATCAGGAAGC
TGGCATCGGACTCATCAGGAAGC
GCATCGGACTCATCAGGAAGC
TTGTCATCGGACTCATCAGGAAGC
AATCACATCGGACTCATCAGGAAGC
TCATCGGACTCATCAGGAAGC
AGTCATCGGACTCATCAGGAAGC

CAGGCATCGGACTCATCAGGAAGC
CGGACTCATCAGGAAGC
CACATCGGACTCATCAGGAAGC
CTCATCGGACTCATCAGGAAGC
TTCATCGGACTCATCAGGAAGC
GCCATCGGACTCATCAGGAAGC
TCGGACTCATCAGGAAGC
CTGCATCGGACTCATCAGGAAGC
CTACATCGGACTCATCAGGAAGC
CTCATCGGACTCATCAGGAAGC
CACATCGGACTCATCAGGAAGC
ATCGGACTCATCAGGAAGC
GGTACATCGGACTCATCAGGAAGC
CCATCGGACTCATCAGGAAGC
TTCCATCGGACTCATCAGGAAGC
ACACATCGGACTCATCAGGAAGC
GGCTCACATCGGACTCATCAGGAAGC
ACATCGGACTCATCAGGAAGC
GGAGACATCGGACTCATCAGGAAGC
CCCATCGGACTCATCAGGAAGC
CTCATCGGACTCATCAGGAAGC
GTCATCGGACTCATCAGGAAGC
TCAACATCGGACTCATCAGGAAGC
GCATCGGACTCATCAGGAAGC
TGGCATCGGACTCATCAGGAAGC
TTCATCGGACTCATCAGGAAGC
ACCATCGGACTCATCAGGAAGC
CATCGGACTCATCAGGAAGC
TCATCGGACTCATCAGGAAGC
CATGACATCGGACTCATCAGGAAGC
ACATCGGACTCATCAGGAAGC
CCATCGGACTCATCAGGAAGC
ATCGGACTCATCAGGAAGC
AAGCATCGGACTCATCAGGAAGC
TGGCATCGGACTCATCAGGAAGC
CGCATCGGACTCATCAGGAAGC
TGCATCGGACTCATCAGGAAGC
GACATCGGACTCATCAGGAAGC
GGTCATCGGACTCATCAGGAAGC
CACATCGGACTCATCAGGAAGC
CATCGGACTCATCAGGAAGC
CACATCGGACTCATCAGGAAGC
TTCATCGGACTCATCAGGAAGC

TTTTCATCGGACTCATCAGGAAGC
TGCATCGGACTCATCAGGAAGC
TGTGCATCGGACTCATCAGGAAGC
TCCCATCGGACTCATCAGGAAGC
CATCGGACTCATCAGGAAGC
CTGCATCGGACTCATCAGGAAGC
TCCCATCGGACTCATCAGGAAGC
TATAACATCGGACTCATCAGGAAGC
TTTTCATCGGACTCATCAGGAAGC
CCAACATCGGACTCATCAGGAAGC
ACATCGGACTCATCAGGAAGC
CCCATCGGACTCATCAGGAAGC
CATCATCGGACTCATCAGGAAGC
TTGACATCGGACTCATCAGGAAGC
TCCATCGGACTCATCAGGAAGC
CCCATCGGACTCATCAGGAAGC
GCCCATCGGACTCATCAGGAAGC
TCATCGGACTCATCAGGAAGC
ACATCGGACTCATCAGGAAGC
CCATCATCGGACTCATCAGGAAGC
GTTCATCGGACTCATCAGGAAGC
AGCCATCGGACTCATCAGGAAGC
ACATCGGACTCATCAGGAAGC
GACATCGGACTCATCAGGAAGC
CTCCTCATCGGACTCATCAGGAAGC
TCATCGGACTCATCAGGAAGC
CTCCCATCGGACTCATCAGGAAGC
CCATCGGACTCATCAGGAAGC
CACATCGGACTCATCAGGAAGC
TACATCGGACTCATCAGGAAGC
TGTGAGACATCGGACTCATCAGGAAGC
CATCGGACTCATCAGGAAGC
CCATCGGACTCATCAGGAAGC
GTCATCGGACTCATCAGGAAGC
TGGACATCGGACTCATCAGGAAGC
CATCGGACTCATCAGGAAGC
CCCACATCGGACTCATCAGGAAGC
GAACCATCGGACTCATCAGGAAGC
GTCATCGGACTCATCAGGAAGC
GTGACATCGGACTCATCAGGAAGC
CACATCGGACTCATCAGGAAGC
GCCATCGGACTCATCAGGAAGC
GTGCATCGGACTCATCAGGAAGC

CTTGCCATCGGACTCATCAGGAAGC
TCCATCGGACTCATCAGGAAGC
TGCGCATCGGACTCATCAGGAAGC
ATCATCGGACTCATCAGGAAGC
TCACATCGGACTCATCAGGAAGC
GGGCATCGGACTCATCAGGAAGC
ATCATCGGACTCATCAGGAAGC
CACCATCGGACTCATCAGGAAGC
CCATCGGACTCATCAGGAAGC
CCATCGGACTCATCAGGAAGC
TACATCGGACTCATCAGGAAGC
ATCGGACTCATCAGGAAGC
ACCCATCGGACTCATCAGGAAGC
TCGGACTCATCAGGAAGC
CACATCGGACTCATCAGGAAGC
TTGGCATCGGACTCATCAGGAAGC
TCCATCGGACTCATCAGGAAGC
GCCATCGGACTCATCAGGAAGC
GCATCGGACTCATCAGGAAGC
CCATCGGACTCATCAGGAAGC
AAGAAGCATCGGACTCATCAGGAAGC
ACACATCGGACTCATCAGGAAGC
CCATCGGACTCATCAGGAAGC
CATCGGACTCATCAGGAAGC
ACATCGGACTCATCAGGAAGC
CATCGGACTCATCAGGAAGC
CCATCGGACTCATCAGGAAGC
CTGACATCGGACTCATCAGGAAGC
CACATCGGACTCATCAGGAAGC
TCCCATCGGACTCATCAGGAAGC
CCATCGGACTCATCAGGAAGC
ATCGGACTCATCAGGAAGC
CATCGGACTCATCAGGAAGC
TGACACATCGGACTCATCAGGAAGC
TCATCGGACTCATCAGGAAGC
TCATCGGACTCATCAGGAAGC
GCATCGGACTCATCAGGAAGC
ATCGGACTCATCAGGAAGC
ATCGGACTCATCAGGAAGC
ACCATCGGACTCATCAGGAAGC
CATCGGACTCATCAGGAAGC
CCATCGGACTCATCAGGAAGC
TCATCGGACTCATCAGGAAGC

GAACATCGGACTCATCAGGAAGC
GACATCGGACTCATCAGGAAGC
ATCATCGGACTCATCAGGAAGC
GATCATCGGACTCATCAGGAAGC
TAGCATCGGACTCATCAGGAAGC
TCGGACTCATCAGGAAGC
ACATCGGACTCATCAGGAAGC
CACATCGGACTCATCAGGAAGC
GCAGCATCGGACTCATCAGGAAGC
TCATCGGACTCATCAGGAAGC
AGCGACATCGGACTCATCAGGAAGC
ACACATCGGACTCATCAGGAAGC
ACATCGGACTCATCAGGAAGC
ATCATCGGACTCATCAGGAAGC
CCCCATCGGACTCATCAGGAAGC
CATCGGACTCATCAGGAAGC
ACATCGGACTCATCAGGAAGC
TGACATCGGACTCATCAGGAAGC
TCATCGGACTCATCAGGAAGC
TCATCGGACTCATCAGGAAGC
TACATCGGACTCATCAGGAAGC
CATCGGACTCATCAGGAAGC
ACATCGGACTCATCAGGAAGC
CTCATCGGACTCATCAGGAAGC
CCCCATCGGACTCATCAGGAAGC
TACATCGGACTCATCAGGAAGC
CACATCGGACTCATCAGGAAGC
TTACATCGGACTCATCAGGAAGC
CACATCGGACTCATCAGGAAGC
AACACATCGGACTCATCAGGAAGC
ATCATCGGACTCATCAGGAAGC
CCATCGGACTCATCAGGAAGC
CCATCGGACTCATCAGGAAGC
ACATCGGACTCATCAGGAAGC
CATCGGACTCATCAGGAAGC
TTTGACATCGGACTCATCAGGAAGC
AATCATCGGACTCATCAGGAAGC
TTGTCCATCGGACTCATCAGGAAGC
ATCATCGGACTCATCAGGAAGC
CTCATCGGACTCATCAGGAAGC
CGCATCGGACTCATCAGGAAGC
ATCGGACTCATCAGGAAGC
GGCTACATCGGACTCATCAGGAAGC

AAGCCATCGGACTCATCAGGAAGC
TCATCGGACTCATCAGGAAGC
ACATCGGACTCATCAGGAAGC
TCATCGGACTCATCAGGAAGC
CATCGGACTCATCAGGAAGC
GGCATCGGACTCATCAGGAAGC
CATCGGACTCATCAGGAAGC
CCCTACATCGGACTCATCAGGAAGC
CTCATCGGACTCATCAGGAAGC
ATCGGACTCATCAGGAAGC
GCATCGGACTCATCAGGAAGC
GACATCGGACTCATCAGGAAGC
CATCGGACTCATCAGGAAGC
GACATCGGACTCATCAGGAAGC
TTCATCGGACTCATCAGGAAGC
ATGACATCGGACTCATCAGGAAGC
TGCCATCGGACTCATCAGGAAGC
ACACATCGGACTCATCAGGAAGC
TCAGCATCGGACTCATCAGGAAGC
CATCGGACTCATCAGGAAGC
ACATCGGACTCATCAGGAAGC
GACCATCGGACTCATCAGGAAGC
GATCATCGGACTCATCAGGAAGC
TACCATCGGACTCATCAGGAAGC
ATCGGACTCATCAGGAAGC
GTACATCGGACTCATCAGGAAGC
GTCATCGGACTCATCAGGAAGC
TTCATCGGACTCATCAGGAAGC
TACCATCGGACTCATCAGGAAGC
CCCATCGGACTCATCAGGAAGC
CATCGGACTCATCAGGAAGC
CATCGGACTCATCAGGAAGC
ACATCGGACTCATCAGGAAGC
GACATCGGACTCATCAGGAAGC
TCATCGGACTCATCAGGAAGC
GGCATCGGACTCATCAGGAAGC
ATCGGACTCATCAGGAAGC
CACATCGGACTCATCAGGAAGC
GACATCGGACTCATCAGGAAGC
CATCGGACTCATCAGGAAGC
GGCATCGGACTCATCAGGAAGC
GGCATCGGACTCATCAGGAAGC
ATCATCGGACTCATCAGGAAGC

GTCCATCGGACTCATCAGGAAGC
TATCATCGGACTCATCAGGAAGC
GTGTGCATCGGACTCATCAGGAAGC
TCCATCGGACTCATCAGGAAGC
AGCCATCGGACTCATCAGGAAGC
GCATCACATCGGACTCATCAGGAAGC
GATCATCGGACTCATCAGGAAGC
AGGTCATCGGACTCATCAGGAAGC
TCATCGGACTCATCAGGAAGC
GTCATCGGACTCATCAGGAAGC
CGGACATCGGACTCATCAGGAAGC
GCTCCATCGGACTCATCAGGAAGC
AGCATCGGACTCATCAGGAAGC
CATCGGACTCATCAGGAAGC
ATGGTCATCGGACTCATCAGGAAGC
TCCACCATCGGACTCATCAGGAAGC
ATACGCATCGGACTCATCAGGAAGC
ATGCCATCGGACTCATCAGGAAGC
TTGCATCGGACTCATCAGGAAGC
AACATCGGACTCATCAGGAAGC
GGACATCGGACTCATCAGGAAGC
TCATCGGACTCATCAGGAAGC
GAACATCGGACTCATCAGGAAGC
TTCATCGGACTCATCAGGAAGC
CCATCGGACTCATCAGGAAGC
TCCATCGGACTCATCAGGAAGC
TCATCGGACTCATCAGGAAGC
ACCATCGGACTCATCAGGAAGC
TACATCGGACTCATCAGGAAGC
CACCATCGGACTCATCAGGAAGC
CTCACATCGGACTCATCAGGAAGC
CATCGGACTCATCAGGAAGC
ATACACATCGGACTCATCAGGAAGC
TGCATCGGACTCATCAGGAAGC
GAACATCGGACTCATCAGGAAGC
ACATCGGACTCATCAGGAAGC
TACTGACATCGGACTCATCAGGAAGC
AGTGCATCGGACTCATCAGGAAGC
TACATCGGACTCATCAGGAAGC
AGTCATCGGACTCATCAGGAAGC
CATCGGACTCATCAGGAAGC
GCCATCGGACTCATCAGGAAGC
TGAGACATCGGACTCATCAGGAAGC

TATCATCGGACTCATCAGGAAGC
TCTCCATCGGACTCATCAGGAAGC
ACATCGGACTCATCAGGAAGC
TACATCGGACTCATCAGGAAGC
TACATCGGACTCATCAGGAAGC
CCATCGGACTCATCAGGAAGC
TCCATCGGACTCATCAGGAAGC
CAGCATCGGACTCATCAGGAAGC
GCATCGGACTCATCAGGAAGC
GCTCCATCGGACTCATCAGGAAGC
GCATCGGACTCATCAGGAAGC
CCTCATCGGACTCATCAGGAAGC
GGACATCGGACTCATCAGGAAGC
ATCGGACTCATCAGGAAGC
TCATCGGACTCATCAGGAAGC
AACATCGGACTCATCAGGAAGC
GCATCGGACTCATCAGGAAGC
TCATCGGACTCATCAGGAAGC
GCATCGGACTCATCAGGAAGC
ATGTCATCGGACTCATCAGGAAGC
AGACATCGGACTCATCAGGAAGC
CACATCGGACTCATCAGGAAGC
CCCCATCGGACTCATCAGGAAGC
CTCCATCGGACTCATCAGGAAGC
ATCGGACTCATCAGGAAGC
ACCCATCGGACTCATCAGGAAGC
CATCGGACTCATCAGGAAGC
TCATCGGACTCATCAGGAAGC
CTCATCGGACTCATCAGGAAGC
GCATCGGACTCATCAGGAAGC
TAACATCGGACTCATCAGGAAGC
TTGACATCGGACTCATCAGGAAGC
GTCATCGGACTCATCAGGAAGC
CACATCGGACTCATCAGGAAGC
ACGGACATCGGACTCATCAGGAAGC
CATCGGACTCATCAGGAAGC
AACATCGGACTCATCAGGAAGC
AGACATCGGACTCATCAGGAAGC
ACATCGGACTCATCAGGAAGC
CTGATCATCGGACTCATCAGGAAGC
CACATCGGACTCATCAGGAAGC
AACATCGGACTCATCAGGAAGC
GCATCGGACTCATCAGGAAGC

CATCGGACTCATCAGGAAGC
ACATCATCGGACTCATCAGGAAGC
ACATCGGACTCATCAGGAAGC
GCGCATCGGACTCATCAGGAAGC
CACATCGGACTCATCAGGAAGC
GGTCATCGGACTCATCAGGAAGC
CCATCGGACTCATCAGGAAGC
CCATCGGACTCATCAGGAAGC
TGTCATCGGACTCATCAGGAAGC
AGGGGCATCGGACTCATCAGGAAGC
GCATCGGACTCATCAGGAAGC
GCCATCGGACTCATCAGGAAGC
TCATCGGACTCATCAGGAAGC
CCATCGGACTCATCAGGAAGC
ACCCATCGGACTCATCAGGAAGC
TTCATCGGACTCATCAGGAAGC
CACATCGGACTCATCAGGAAGC
TGGCATCGGACTCATCAGGAAGC
ATCATCGGACTCATCAGGAAGC
CCCATCGGACTCATCAGGAAGC
TTCATCGGACTCATCAGGAAGC
CCATCGGACTCATCAGGAAGC
ATCGGACTCATCAGGAAGC
GTCTCATCGGACTCATCAGGAAGC
GGGCATCGGACTCATCAGGAAGC
GACATCGGACTCATCAGGAAGC
TCATCGGACTCATCAGGAAGC
TTCATCGGACTCATCAGGAAGC
GTCATCGGACTCATCAGGAAGC
TGTCATCGGACTCATCAGGAAGC
CATCGGACTCATCAGGAAGC
ACATCGGACTCATCAGGAAGC
CCACATCGGACTCATCAGGAAGC
GCATCGGACTCATCAGGAAGC
ACATCGGACTCATCAGGAAGC
GCATCGGACTCATCAGGAAGC
ATCATCGGACTCATCAGGAAGC
CAACATCGGACTCATCAGGAAGC
TCATCGGACTCATCAGGAAGC
GTGTCATCGGACTCATCAGGAAGC
GTCATCGGACTCATCAGGAAGC
AGCATCGGACTCATCAGGAAGC
TCATCGGACTCATCAGGAAGC

CCATCGGACTCATCAGGAAGC
AGACCATCGGACTCATCAGGAAGC
GTACATCGGACTCATCAGGAAGC
GCCATCGGACTCATCAGGAAGC
ATACATCGGACTCATCAGGAAGC
CATCGGACTCATCAGGAAGC
GCCCATCGGACTCATCAGGAAGC
GGGCATCGGACTCATCAGGAAGC
TCACATCGGACTCATCAGGAAGC
CATCGGACTCATCAGGAAGC
CCATCGGACTCATCAGGAAGC
GGACATCGGACTCATCAGGAAGC
CTCCCATCGGACTCATCAGGAAGC
ACATCGGACTCATCAGGAAGC
GATTGACCATCGGACTCATCAGGAAGC
GGTCATCGGACTCATCAGGAAGC
GCATCGGACTCATCAGGAAGC
TGGCATCGGACTCATCAGGAAGC
CATCGGACTCATCAGGAAGC
GCATCGGACTCATCAGGAAGC
GTGCATCGGACTCATCAGGAAGC
GGGACATCGGACTCATCAGGAAGC
GACCATCGGACTCATCAGGAAGC
CATCGGACTCATCAGGAAGC
GGCTCATCGGACTCATCAGGAAGC
ACATCGGACTCATCAGGAAGC
AGTCCATCGGACTCATCAGGAAGC
AGGGCATCGGACTCATCAGGAAGC
GGCCATCGGACTCATCAGGAAGC
GCACATCGGACTCATCAGGAAGC
AGCCATCGGACTCATCAGGAAGC
ACCCATCGGACTCATCAGGAAGC
CACACATCGGACTCATCAGGAAGC
GGCCATCGGACTCATCAGGAAGC
TCCATCGGACTCATCAGGAAGC
CCATCGGACTCATCAGGAAGC
ACATCGGACTCATCAGGAAGC
ATCGGACTCATCAGGAAGC
TCATCGGACTCATCAGGAAGC
CACCACATCGGACTCATCAGGAAGC
CTTCCATCGGACTCATCAGGAAGC
AGCCATCGGACTCATCAGGAAGC
ACATCGGACTCATCAGGAAGC

GTTTTCGCATCGGACTCATCAGGAAGC
TCCATCGGACTCATCAGGAAGC
ATCGGACTCATCAGGAAGC
GTGTCATCGGACTCATCAGGAAGC
GATGCATCGGACTCATCAGGAAGC
ACCATCGGACTCATCAGGAAGC
ATAGACATCGGACTCATCAGGAAGC
TCATCGGACTCATCAGGAAGC
GAGCATCGGACTCATCAGGAAGC
TCTACATCGGACTCATCAGGAAGC
AGACATCGGACTCATCAGGAAGC
GGCATCGGACTCATCAGGAAGC
AACATCGGACTCATCAGGAAGC
GTGCATCGGACTCATCAGGAAGC
CATCGGACTCATCAGGAAGC
CTCCATCGGACTCATCAGGAAGC
ACATTGGCATCGGACTCATCAGGAAGC
CATCCATCGGACTCATCAGGAAGC
ACATCGGACTCATCAGGAAGC
TCATCGGACTCATCAGGAAGC
CAACATCGGACTCATCAGGAAGC
TTCATCGGACTCATCAGGAAGC
TTCCATCGGACTCATCAGGAAGC
CACATCGGACTCATCAGGAAGC
GTCATCGGACTCATCAGGAAGC
CCATCGGACTCATCAGGAAGC
TGCTCATCGGACTCATCAGGAAGC
ACACATCGGACTCATCAGGAAGC
ACATCGGACTCATCAGGAAGC
GCCCATCGGACTCATCAGGAAGC
GAGCATCGGACTCATCAGGAAGC
TGTTCATCGGACTCATCAGGAAGC
ACATCGGACTCATCAGGAAGC
TCAGCATCGGACTCATCAGGAAGC
GCCATCGGACTCATCAGGAAGC
GGGCATCGGACTCATCAGGAAGC
GACATCGGACTCATCAGGAAGC
GCCATCGGACTCATCAGGAAGC
CAGACATCGGACTCATCAGGAAGC
GCCACATCGGACTCATCAGGAAGC
AGACATCGGACTCATCAGGAAGC
ATAGACATCGGACTCATCAGGAAGC
CTCATCGGACTCATCAGGAAGC

GGACTCATCAGGAAGC
CATCGGACTCATCAGGAAGC
GCATCGGACTCATCAGGAAGC
ACACACATCGGACTCATCAGGAAGC
GGCATCGGACTCATCAGGAAGC
TGGTCATCGGACTCATCAGGAAGC
GAACATCGGACTCATCAGGAAGC
GTCATCGGACTCATCAGGAAGC
CCACATCGGACTCATCAGGAAGC
CATCGGACTCATCAGGAAGC
AGTCAGCATCGGACTCATCAGGAAGC
ATCATCGGACTCATCAGGAAGC
TCCATCGGACTCATCAGGAAGC
TCATCGGACTCATCAGGAAGC
CACCATCGGACTCATCAGGAAGC
CTCATCGGACTCATCAGGAAGC
CGACATCGGACTCATCAGGAAGC
TCATCGGACTCATCAGGAAGC
CATCGGACTCATCAGGAAGC
TTCATCGGACTCATCAGGAAGC
TCATCGGACTCATCAGGAAGC
CATCGGACTCATCAGGAAGC
CATCGGACTCATCAGGAAGC
TGACATCGGACTCATCAGGAAGC
CATCGGACTCATCAGGAAGC
CAGCCATCGGACTCATCAGGAAGC
ATCCCATCGGACTCATCAGGAAGC
TTCATCGGACTCATCAGGAAGC
ATTCCATCGGACTCATCAGGAAGC
ATCACATCGGACTCATCAGGAAGC
TCATCGGACTCATCAGGAAGC
TAAATCATCGGACTCATCAGGAAGC
GCATCGGACTCATCAGGAAGC
GGGTCATCGGACTCATCAGGAAGC
TCCATCATCGGACTCATCAGGAAGC
TGTCATCGGACTCATCAGGAAGC
CACATCGGACTCATCAGGAAGC
TTACATCGGACTCATCAGGAAGC
ACCATCGGACTCATCAGGAAGC
CATCGGACTCATCAGGAAGC
ATAGCATCGGACTCATCAGGAAGC
TTCATCGGACTCATCAGGAAGC
TGCCATCGGACTCATCAGGAAGC

GTGCCATCGGACTCATCAGGAAGC
ACATCGGACTCATCAGGAAGC
CATCGGACTCATCAGGAAGC
CACCATCGGACTCATCAGGAAGC
GACATCGGACTCATCAGGAAGC
AACCATCGGACTCATCAGGAAGC
TCATCGGACTCATCAGGAAGC
GAACATCGGACTCATCAGGAAGC
GACATCGGACTCATCAGGAAGC
TCTCCACATCGGACTCATCAGGAAGC
CATCGGACTCATCAGGAAGC
CCATCGGACTCATCAGGAAGC
GTCATCGGACTCATCAGGAAGC
CAGCATCGGACTCATCAGGAAGC
TCATCGGACTCATCAGGAAGC
GCCATCGGACTCATCAGGAAGC
CCATCGGACTCATCAGGAAGC
CATCGGACTCATCAGGAAGC
CCATCGGACTCATCAGGAAGC
AAACATCGGACTCATCAGGAAGC
CATCATCGGACTCATCAGGAAGC
AGGCCATCGGACTCATCAGGAAGC
CTCATCGGACTCATCAGGAAGC
CTCATCGGACTCATCAGGAAGC
CCCCATCGGACTCATCAGGAAGC
CTGATCATCGGACTCATCAGGAAGC
GCCATCGGACTCATCAGGAAGC
CATCGGACTCATCAGGAAGC
GTCTACATCGGACTCATCAGGAAGC
AAACATCGGACTCATCAGGAAGC
ACCATCGGACTCATCAGGAAGC
ATTCATCGGACTCATCAGGAAGC
CATCGGACTCATCAGGAAGC
TACATCGGACTCATCAGGAAGC
GGACATCGGACTCATCAGGAAGC
CCCCATCGGACTCATCAGGAAGC
AGCATCGGACTCATCAGGAAGC
TGTTCATCGGACTCATCAGGAAGC
GTCATCGGACTCATCAGGAAGC
CATCGGACTCATCAGGAAGC
AGGCATCGGACTCATCAGGAAGC
CTGCATCGGACTCATCAGGAAGC
GTCATCGGACTCATCAGGAAGC

ACATCGGACTCATCAGGAAGC
GTTACATCGGACTCATCAGGAAGC
TGCATCGGACTCATCAGGAAGC
CCATCGGACTCATCAGGAAGC
GCCATCGGACTCATCAGGAAGC
GACATCGGACTCATCAGGAAGC
TGCATCGGACTCATCAGGAAGC
TTTCATCGGACTCATCAGGAAGC
ACATCGGACTCATCAGGAAGC
CCATCGGACTCATCAGGAAGC
CATCGGACTCATCAGGAAGC
TCCATCGGACTCATCAGGAAGC
CAATTACCATCGGACTCATCAGGAAGC
ATCATCGGACTCATCAGGAAGC
AACATCGGACTCATCAGGAAGC
AAGCATCGGACTCATCAGGAAGC
TTCATCGGACTCATCAGGAAGC
ATGCATCGGACTCATCAGGAAGC
CACCATCGGACTCATCAGGAAGC
GGCATCGGACTCATCAGGAAGC
GTCATCGGACTCATCAGGAAGC
CCATCGGACTCATCAGGAAGC
GTTACATCGGACTCATCAGGAAGC
TGACATCGGACTCATCAGGAAGC
GCCATCGGACTCATCAGGAAGC
GACATCGGACTCATCAGGAAGC
GTCATCGGACTCATCAGGAAGC
TCCATCGGACTCATCAGGAAGC
GGCCATCGGACTCATCAGGAAGC
AACATCGGACTCATCAGGAAGC
ACCCCATCGGACTCATCAGGAAGC
TCATCGGACTCATCAGGAAGC
AGCATCGGACTCATCAGGAAGC
AAGGCACATCGGACTCATCAGGAAGC
TATGAGCCATCGGACTCATCAGGAAGC
GAACATCGGACTCATCAGGAAGC
GACATCGGACTCATCAGGAAGC
CCACATCGGACTCATCAGGAAGC
CGCATCGGACTCATCAGGAAGC
AGAGACATCGGACTCATCAGGAAGC
CCATCGGACTCATCAGGAAGC
CATCGGACTCATCAGGAAGC
CCATCGGACTCATCAGGAAGC

GCGCATCGGACTCATCAGGAAGC
CATCATCGGACTCATCAGGAAGC
TCATCGGACTCATCAGGAAGC
AAAAGCCATCGGACTCATCAGGAAGC
GGTCATCGGACTCATCAGGAAGC
TCATCGGACTCATCAGGAAGC
ATGCATCGGACTCATCAGGAAGC
ATCGGACTCATCAGGAAGC
GACATCGGACTCATCAGGAAGC
ACATCGGACTCATCAGGAAGC
ATTCATCGGACTCATCAGGAAGC
ACATCGGACTCATCAGGAAGC
GACATCGGACTCATCAGGAAGC
ATCATCGGACTCATCAGGAAGC
GCATCGGACTCATCAGGAAGC
TCATCGGACTCATCAGGAAGC
ACATCGGACTCATCAGGAAGC
CCTCATCGGACTCATCAGGAAGC
ATCGGACTCATCAGGAAGC
TCTCATCGGACTCATCAGGAAGC
CATCGGACTCATCAGGAAGC
TCATCGGACTCATCAGGAAGC
ACTGCATCGGACTCATCAGGAAGC
TCATCGGACTCATCAGGAAGC
CACAACATCGGACTCATCAGGAAGC
AACACATCGGACTCATCAGGAAGC
GACCATCGGACTCATCAGGAAGC
CCATCGGACTCATCAGGAAGC
CAACATCGGACTCATCAGGAAGC
AGAACATCGGACTCATCAGGAAGC
ACATCGGACTCATCAGGAAGC
ACATCGGACTCATCAGGAAGC
TCCATCGGACTCATCAGGAAGC
ATCGGACTCATCAGGAAGC
TCCATCGGACTCATCAGGAAGC
GACATCGGACTCATCAGGAAGC
ACATCGGACTCATCAGGAAGC
CTCATCGGACTCATCAGGAAGC
TCATCGGACTCATCAGGAAGC
GGTCATCGGACTCATCAGGAAGC
AAATGCCATCGGACTCATCAGGAAGC
GGCATCGGACTCATCAGGAAGC
GACATCGGACTCATCAGGAAGC

CATCGGACTCATCAGGAAGC
CCCATCGGACTCATCAGGAAGC
CACTCATCGGACTCATCAGGAAGC
AGTCATCGGACTCATCAGGAAGC
TGACATCGGACTCATCAGGAAGC
TGTCATCGGACTCATCAGGAAGC
TGGCATCGGACTCATCAGGAAGC
GTCATCGGACTCATCAGGAAGC
TCATCGGACTCATCAGGAAGC
GTCATCATCGGACTCATCAGGAAGC
CTCCATCGGACTCATCAGGAAGC
TCTCTCATCGGACTCATCAGGAAGC
CTGCATCGGACTCATCAGGAAGC
CTTCATCGGACTCATCAGGAAGC
GAGCATCGGACTCATCAGGAAGC
CTCATCGGACTCATCAGGAAGC
ACCATCGGACTCATCAGGAAGC
TGCATCGGACTCATCAGGAAGC
AAACATCGGACTCATCAGGAAGC
CATCGGACTCATCAGGAAGC
GTGCATCGGACTCATCAGGAAGC
GCAACATCGGACTCATCAGGAAGC
TCATCGGACTCATCAGGAAGC
GCCATCGGACTCATCAGGAAGC
GGACATCGGACTCATCAGGAAGC
TAGCCATCGGACTCATCAGGAAGC
ACAGCATCGGACTCATCAGGAAGC
AAGTGCATCGGACTCATCAGGAAGC
TCCATCGGACTCATCAGGAAGC
TAGACATCGGACTCATCAGGAAGC
TTACATCGGACTCATCAGGAAGC
GCCATCGGACTCATCAGGAAGC
AAGGCCATCGGACTCATCAGGAAGC
TCTCATCGGACTCATCAGGAAGC
AGCATCGGACTCATCAGGAAGC
ATCTACATCGGACTCATCAGGAAGC
CATCGGACTCATCAGGAAGC
CACATCGGACTCATCAGGAAGC
TAGTCATCGGACTCATCAGGAAGC
AATGCCCATCGGACTCATCAGGAAGC
TCCATCGGACTCATCAGGAAGC
ACTCCATCGGACTCATCAGGAAGC
TCTCATCGGACTCATCAGGAAGC

GTCAGTTCATCGGACTCATCAGGAAGC
TCATCGGACTCATCAGGAAGC
CTCATCGGACTCATCAGGAAGC
CACATCGGACTCATCAGGAAGC
TTCATCGGACTCATCAGGAAGC
ATCCCACATCGGACTCATCAGGAAGC
TCCACATCGGACTCATCAGGAAGC
GCATCGGACTCATCAGGAAGC
TGCATCGGACTCATCAGGAAGC
CCATCGGACTCATCAGGAAGC
AAGGGCATCGGACTCATCAGGAAGC
ATCGGACTCATCAGGAAGC
GCATCGGACTCATCAGGAAGC
TATGCATCGGACTCATCAGGAAGC
CCCATCGGACTCATCAGGAAGC
ACATCGGACTCATCAGGAAGC
CATCGGACTCATCAGGAAGC
GGAATCATCGGACTCATCAGGAAGC
TCATCGGACTCATCAGGAAGC
AAAGCATCGGACTCATCAGGAAGC
ATCGGACTCATCAGGAAGC
GGCATCGGACTCATCAGGAAGC
TCCCCACCATCGGACTCATCAGGAAGC
TTCGTTCATCGGACTCATCAGGAAGC
ATCGGACTCATCAGGAAGC
AAGGCATCGGACTCATCAGGAAGC
GGCCATCGGACTCATCAGGAAGC
ATCGGACTCATCAGGAAGC
CCATCGGACTCATCAGGAAGC
ACATCGGACTCATCAGGAAGC
GCATCGGACTCATCAGGAAGC
ATGCATCGGACTCATCAGGAAGC
CATCGGACTCATCAGGAAGC
TCTGACATCGGACTCATCAGGAAGC
TCCATCGGACTCATCAGGAAGC
TCGGACTCATCAGGAAGC
GCCATCGGACTCATCAGGAAGC
GGCATCGGACTCATCAGGAAGC
CTCCATCGGACTCATCAGGAAGC
TGTCATCGGACTCATCAGGAAGC
TTGCATCGGACTCATCAGGAAGC
CTGGACATCGGACTCATCAGGAAGC
CATCGGACTCATCAGGAAGC

ACATCGGACTCATCAGGAAGC
GACCATCGGACTCATCAGGAAGC
ACATCGGACTCATCAGGAAGC
GGCCATCGGACTCATCAGGAAGC
GCCTCATCGGACTCATCAGGAAGC
CCATCGGACTCATCAGGAAGC
GCCATCGGACTCATCAGGAAGC
CGCATCGGACTCATCAGGAAGC
GCAACATCGGACTCATCAGGAAGC
AGGACATCGGACTCATCAGGAAGC
GGACATCGGACTCATCAGGAAGC
CCATCGGACTCATCAGGAAGC
ATCGGACTCATCAGGAAGC
CACATCGGACTCATCAGGAAGC
TGACATCGGACTCATCAGGAAGC
GGCATCGGACTCATCAGGAAGC
TCAACATCGGACTCATCAGGAAGC
GGTCATCGGACTCATCAGGAAGC
GTTCACATCGGACTCATCAGGAAGC
AGCATCGGACTCATCAGGAAGC
GGCATCGGACTCATCAGGAAGC
TAAAACATCGGACTCATCAGGAAGC
GCTCATCGGACTCATCAGGAAGC
GACATCGGACTCATCAGGAAGC
GGGCCATCGGACTCATCAGGAAGC
AACATCGGACTCATCAGGAAGC
TATGCCATCGGACTCATCAGGAAGC
CCCATCGGACTCATCAGGAAGC
CAAGGCATCGGACTCATCAGGAAGC
CCCATCGGACTCATCAGGAAGC
CATCGGACTCATCAGGAAGC
GATCATCGGACTCATCAGGAAGC
ATCGGACTCATCAGGAAGC
CAACATCGGACTCATCAGGAAGC
CCACTCATCGGACTCATCAGGAAGC
TCATCGGACTCATCAGGAAGC
CGCATCGGACTCATCAGGAAGC
GCACATCGGACTCATCAGGAAGC
TGGCCATCGGACTCATCAGGAAGC
AGCCATCGGACTCATCAGGAAGC
AAGCCCATCGGACTCATCAGGAAGC
ACTGTGCATCGGACTCATCAGGAAGC
GCTCGCATCGGACTCATCAGGAAGC

TCATCGGACTCATCAGGAAGC
ATCATCGGACTCATCAGGAAGC
ACATCGGACTCATCAGGAAGC
ATACACATCGGACTCATCAGGAAGC
ATCGGACTCATCAGGAAGC
GCATCGGACTCATCAGGAAGC
GGTCATCGGACTCATCAGGAAGC
ACGCATCGGACTCATCAGGAAGC
GTCATCGGACTCATCAGGAAGC
ACATCGGACTCATCAGGAAGC
CCATCGGACTCATCAGGAAGC
ATAACTCATCGGACTCATCAGGAAGC
ATCGGACTCATCAGGAAGC
AGCCCATCGGACTCATCAGGAAGC
GCCCATCGGACTCATCAGGAAGC
CTCATCGGACTCATCAGGAAGC
CATCGGACTCATCAGGAAGC
ATTCATCGGACTCATCAGGAAGC
CCATCGGACTCATCAGGAAGC
AGCATCGGACTCATCAGGAAGC
TCATCGGACTCATCAGGAAGC
TTGTTTCATCGGACTCATCAGGAAGC
CTTTCATCGGACTCATCAGGAAGC
ACATCGGACTCATCAGGAAGC
TCCTTGACATCGGACTCATCAGGAAGC
GTACATCGGACTCATCAGGAAGC
ACAGCATCGGACTCATCAGGAAGC
TACATCGGACTCATCAGGAAGC
CATCGGACTCATCAGGAAGC
TTCACATCGGACTCATCAGGAAGC
GCATCGGACTCATCAGGAAGC
GAGGCATCGGACTCATCAGGAAGC
GCCATCGGACTCATCAGGAAGC
ACATCGGACTCATCAGGAAGC
ATCGGACTCATCAGGAAGC
GCATCGGACTCATCAGGAAGC
TCATCGGACTCATCAGGAAGC
TCCATCGGACTCATCAGGAAGC
ATCGGACTCATCAGGAAGC
TCATCGGACTCATCAGGAAGC
GAGCTTCATCGGACTCATCAGGAAGC
ATCGGACTCATCAGGAAGC
TCACATCGGACTCATCAGGAAGC

CTCATCGGACTCATCAGGAAGC
ACCTGCATCGGACTCATCAGGAAGC
AGCATCGGACTCATCAGGAAGC
CCATCGGACTCATCAGGAAGC
TCATCGGACTCATCAGGAAGC
AATCATCGGACTCATCAGGAAGC
GTCCATCGGACTCATCAGGAAGC
AACATCGGACTCATCAGGAAGC
AGCATCGGACTCATCAGGAAGC
ACATCGGACTCATCAGGAAGC
GATCATCGGACTCATCAGGAAGC
CCATCGGACTCATCAGGAAGC
TCTACATCGGACTCATCAGGAAGC
CTACATCGGACTCATCAGGAAGC
TCCATCGGACTCATCAGGAAGC
TCATCGGACTCATCAGGAAGC
CATCGGACTCATCAGGAAGC
TCATCGGACTCATCAGGAAGC
CAGCATCGGACTCATCAGGAAGC
TTCATCGGACTCATCAGGAAGC
AGCCATCGGACTCATCAGGAAGC
TCATCGGACTCATCAGGAAGC
CATCGGACTCATCAGGAAGC
GCTAAACATCGGACTCATCAGGAAGC
AACATCGGACTCATCAGGAAGC
CAACATCGGACTCATCAGGAAGC
AGCATCGGACTCATCAGGAAGC
TCCCATCGGACTCATCAGGAAGC
AATACATCGGACTCATCAGGAAGC
CCATCGGACTCATCAGGAAGC
CTACTCATCGGACTCATCAGGAAGC
GCTAACATCGGACTCATCAGGAAGC
GCCCATCGGACTCATCAGGAAGC
AACACATCGGACTCATCAGGAAGC
GGCTCATCGGACTCATCAGGAAGC
TAACATCGGACTCATCAGGAAGC
ACCATCGGACTCATCAGGAAGC
ACATCGGACTCATCAGGAAGC
AGGCATCGGACTCATCAGGAAGC
AGACATCGGACTCATCAGGAAGC
TTCATCGGACTCATCAGGAAGC
TACATCGGACTCATCAGGAAGC
TACATCGGACTCATCAGGAAGC

CCATCGGACTCATCAGGAAGC
CACATCGGACTCATCAGGAAGC
ACCCCCCATCGGACTCATCAGGAAGC
TTCACATCGGACTCATCAGGAAGC
ACATCGGACTCATCAGGAAGC
CATCGGACTCATCAGGAAGC
TCATCGGACTCATCAGGAAGC
CATCGGACTCATCAGGAAGC
ATCGGACTCATCAGGAAGC
ACACATCGGACTCATCAGGAAGC
AACACATCGGACTCATCAGGAAGC
CCATCGGACTCATCAGGAAGC
TGCCATCGGACTCATCAGGAAGC
CTACATCGGACTCATCAGGAAGC
TCATCGGACTCATCAGGAAGC
TTCATCGGACTCATCAGGAAGC
AGCATCGGACTCATCAGGAAGC
CATCGGACTCATCAGGAAGC
ACTATCATCGGACTCATCAGGAAGC
CATCGGACTCATCAGGAAGC
TCACATCGGACTCATCAGGAAGC
AACATCGGACTCATCAGGAAGC
TCATCGGACTCATCAGGAAGC
AGGCATCGGACTCATCAGGAAGC
TCCATCGGACTCATCAGGAAGC
GTTACATCGGACTCATCAGGAAGC
CTCATCGGACTCATCAGGAAGC
GCTCATCGGACTCATCAGGAAGC
GCATCGGACTCATCAGGAAGC
CCCATCGGACTCATCAGGAAGC
GCATCGGACTCATCAGGAAGC
AACACATCGGACTCATCAGGAAGC
TCGGACTCATCAGGAAGC
CCATCGGACTCATCAGGAAGC
CCATCGGACTCATCAGGAAGC
GAGACATCGGACTCATCAGGAAGC
ACATCGGACTCATCAGGAAGC
ATCGGACTCATCAGGAAGC
CATCGGACTCATCAGGAAGC
CCATCGGACTCATCAGGAAGC
CCCATCGGACTCATCAGGAAGC
ACCAGGGCATCGGACTCATCAGGAAGC
GTCATCGGACTCATCAGGAAGC

CAACATCGGACTCATCAGGAAGC
GATCATCGGACTCATCAGGAAGC
CCACATCGGACTCATCAGGAAGC
CATCGGACTCATCAGGAAGC
TGCATCGGACTCATCAGGAAGC
GACACACATCGGACTCATCAGGAAGC
CATCGGACTCATCAGGAAGC
CCCCATCGGACTCATCAGGAAGC
CATCATCGGACTCATCAGGAAGC
CCTCATCGGACTCATCAGGAAGC
GCTGTCATCGGACTCATCAGGAAGC
CTCATCGGACTCATCAGGAAGC
TCATCGGACTCATCAGGAAGC
AACATCGGACTCATCAGGAAGC
TCATCGGACTCATCAGGAAGC
TTTTCATCGGACTCATCAGGAAGC
GGAACATCGGACTCATCAGGAAGC
CAATCATCGGACTCATCAGGAAGC
AAGCATCGGACTCATCAGGAAGC
TCATCGGACTCATCAGGAAGC
TCCATCGGACTCATCAGGAAGC
CTTCATCGGACTCATCAGGAAGC
GACATCGGACTCATCAGGAAGC
GTCATCGGACTCATCAGGAAGC
GATCATCGGACTCATCAGGAAGC
TCATCGGACTCATCAGGAAGC
ACATCGGACTCATCAGGAAGC
ACATCGGACTCATCAGGAAGC
GCATCGGACTCATCAGGAAGC
GACATCGGACTCATCAGGAAGC
TAAGAACATCGGACTCATCAGGAAGC
ACATCGGACTCATCAGGAAGC
ACATCGGACTCATCAGGAAGC
AGCATCGGACTCATCAGGAAGC
CCCATCGGACTCATCAGGAAGC
CTCATCGGACTCATCAGGAAGC
AACATCGGACTCATCAGGAAGC
ACATCGGACTCATCAGGAAGC
TGACATCGGACTCATCAGGAAGC
GCCATCGGACTCATCAGGAAGC
AACATCGGACTCATCAGGAAGC
GCACATCGGACTCATCAGGAAGC
ACAGGCATCGGACTCATCAGGAAGC

GACATCGGACTCATCAGGAAGC
AATCATCGGACTCATCAGGAAGC
TGCATCGGACTCATCAGGAAGC
ATCATCGGACTCATCAGGAAGC
TCATCGGACTCATCAGGAAGC
CCCATCGGACTCATCAGGAAGC
TCATCGGACTCATCAGGAAGC
ATCATCGGACTCATCAGGAAGC
AAACTCATCGGACTCATCAGGAAGC
TCGGACTCATCAGGAAGC
CGTCATCGGACTCATCAGGAAGC
CATCGGACTCATCAGGAAGC
CACATCGGACTCATCAGGAAGC
AACCATCGGACTCATCAGGAAGC
TCACATCGGACTCATCAGGAAGC
AACATCGGACTCATCAGGAAGC
CTCATCGGACTCATCAGGAAGC
ACCACATCGGACTCATCAGGAAGC
AGATCATCGGACTCATCAGGAAGC
TGACATCGGACTCATCAGGAAGC
CATCGGACTCATCAGGAAGC
GGCATCGGACTCATCAGGAAGC
ATCATCGGACTCATCAGGAAGC
CTGCATCGGACTCATCAGGAAGC
TCATCGGACTCATCAGGAAGC
AACATCGGACTCATCAGGAAGC
TCATCGGACTCATCAGGAAGC
CTCATCGGACTCATCAGGAAGC
ACATCGGACTCATCAGGAAGC
ACATCGGACTCATCAGGAAGC
GACATCGGACTCATCAGGAAGC
GTGCCATCGGACTCATCAGGAAGC
CCATCGGACTCATCAGGAAGC
GTCATCGGACTCATCAGGAAGC
GCCATCGGACTCATCAGGAAGC
CCATCGGACTCATCAGGAAGC
CTCCATCGGACTCATCAGGAAGC
ACATCGGACTCATCAGGAAGC
GGACATCGGACTCATCAGGAAGC
CTACATCGGACTCATCAGGAAGC
ACATCGGACTCATCAGGAAGC
ACATCGGACTCATCAGGAAGC
ATCATCGGACTCATCAGGAAGC

AGCCATCGGACTCATCAGGAAGC
TGGCATCGGACTCATCAGGAAGC
TCTACATCGGACTCATCAGGAAGC
CATCGGACTCATCAGGAAGC
AGACGCATCGGACTCATCAGGAAGC
TCATCGGACTCATCAGGAAGC
CTTCATCGGACTCATCAGGAAGC
CCATCGGACTCATCAGGAAGC
ACACATCGGACTCATCAGGAAGC
CTTCACATCGGACTCATCAGGAAGC
GCATCGGACTCATCAGGAAGC
GCAAACATCGGACTCATCAGGAAGC
AACATCGGACTCATCAGGAAGC
CTCCATCGGACTCATCAGGAAGC
GCCTCATCGGACTCATCAGGAAGC
CTTTCATCGGACTCATCAGGAAGC
CCATCGGACTCATCAGGAAGC
GCCATCGGACTCATCAGGAAGC
ACATCGGACTCATCAGGAAGC
CCATCGGACTCATCAGGAAGC
GTCCACATCGGACTCATCAGGAAGC
CCTGCATCGGACTCATCAGGAAGC
AAACATCGGACTCATCAGGAAGC
AAGTCATCGGACTCATCAGGAAGC
CTCATCGGACTCATCAGGAAGC
AAGCATCGGACTCATCAGGAAGC
CACATCGGACTCATCAGGAAGC
GCAAACATCGGACTCATCAGGAAGC
GCCATCGGACTCATCAGGAAGC
TCATCGGACTCATCAGGAAGC
TCCCCATCGGACTCATCAGGAAGC
AGTCATCGGACTCATCAGGAAGC
GGAGAGCATCGGACTCATCAGGAAGC
ACATCGGACTCATCAGGAAGC
CATTACATCGGACTCATCAGGAAGC
GGTCATCGGACTCATCAGGAAGC
ACATCCATCGGACTCATCAGGAAGC
GGGCATCGGACTCATCAGGAAGC
ACCATCGGACTCATCAGGAAGC
AGACATCGGACTCATCAGGAAGC
ATCGCATCGGACTCATCAGGAAGC
ACATCGGACTCATCAGGAAGC
TGGCATCGGACTCATCAGGAAGC

TCATCGGACTCATCAGGAAGC
CGCATCGGACTCATCAGGAAGC
CATCGGACTCATCAGGAAGC
TCATCGGACTCATCAGGAAGC
TATCATCGGACTCATCAGGAAGC
GTCATCGGACTCATCAGGAAGC
GGACATCGGACTCATCAGGAAGC
GGGCATCGGACTCATCAGGAAGC
CACATCGGACTCATCAGGAAGC
CCATCGGACTCATCAGGAAGC
ACATCGGACTCATCAGGAAGC
CGCATCGGACTCATCAGGAAGC
CATCGGACTCATCAGGAAGC
ATCGGACTCATCAGGAAGC
TTGCATCGGACTCATCAGGAAGC
TAGTTCATCGGACTCATCAGGAAGC
CATCGGACTCATCAGGAAGC
CCCATCGGACTCATCAGGAAGC
TTCTACATCGGACTCATCAGGAAGC
CATCGGACTCATCAGGAAGC
CAGTCATCGGACTCATCAGGAAGC
GAACATCGGACTCATCAGGAAGC
CATCGGACTCATCAGGAAGC
TTGCATCGGACTCATCAGGAAGC
CATCGGACTCATCAGGAAGC
CCATCGGACTCATCAGGAAGC
CACATCGGACTCATCAGGAAGC
GACACATCGGACTCATCAGGAAGC
TCATCGGACTCATCAGGAAGC
GGGCATCGGACTCATCAGGAAGC
TCATCGGACTCATCAGGAAGC
CACCATCGGACTCATCAGGAAGC
AGTCATCGGACTCATCAGGAAGC
TGGCATCGGACTCATCAGGAAGC
TACATCGGACTCATCAGGAAGC
CATCGGACTCATCAGGAAGC
TGTCATCGGACTCATCAGGAAGC
TCATCGGACTCATCAGGAAGC
GCTCATCGGACTCATCAGGAAGC
GCATCGGACTCATCAGGAAGC
CATCGGACTCATCAGGAAGC
GAGACATCGGACTCATCAGGAAGC
TATCATCGGACTCATCAGGAAGC

CCACATCGGACTCATCAGGAAGC
CATCGGACTCATCAGGAAGC
CTGCCATCGGACTCATCAGGAAGC
TCATCGGACTCATCAGGAAGC
AGGCTCATCGGACTCATCAGGAAGC
TCTCATCGGACTCATCAGGAAGC
AATCATCGGACTCATCAGGAAGC
CCATCGGACTCATCAGGAAGC
ATCGGACTCATCAGGAAGC
GGGCATCGGACTCATCAGGAAGC
GCCCACATCGGACTCATCAGGAAGC
TCCATCGGACTCATCAGGAAGC
TCTCATCGGACTCATCAGGAAGC
TACATCGGACTCATCAGGAAGC
CATCGGACTCATCAGGAAGC
AAATCATCGGACTCATCAGGAAGC
GCATCGGACTCATCAGGAAGC
CATCGGACTCATCAGGAAGC
CCATCGGACTCATCAGGAAGC
CTCATCGGACTCATCAGGAAGC
GACATCGGACTCATCAGGAAGC
CATCGGACTCATCAGGAAGC
CACATCGGACTCATCAGGAAGC
AGTCATCGGACTCATCAGGAAGC
ATCGGACTCATCAGGAAGC
CATCGGACTCATCAGGAAGC
CTACATCGGACTCATCAGGAAGC
TACATCGGACTCATCAGGAAGC
GGCATCGGACTCATCAGGAAGC
ACCATCGGACTCATCAGGAAGC
CATCGGACTCATCAGGAAGC
CCATCGGACTCATCAGGAAGC
TAATGACATCGGACTCATCAGGAAGC
CCATCGGACTCATCAGGAAGC
AGGAAACATCGGACTCATCAGGAAGC
GTCACATCGGACTCATCAGGAAGC
GCATCGGACTCATCAGGAAGC
CGCATCGGACTCATCAGGAAGC
CATCGGACTCATCAGGAAGC
GTCATCGGACTCATCAGGAAGC
GCCATCGGACTCATCAGGAAGC
CATCATCGGACTCATCAGGAAGC
AACCATCGGACTCATCAGGAAGC

AGCATCGGACTCATCAGGAAGC
AGTCATCGGACTCATCAGGAAGC
TCCATCGGACTCATCAGGAAGC
GCATCGGACTCATCAGGAAGC
GGACATCGGACTCATCAGGAAGC
GGAACATCGGACTCATCAGGAAGC
AGACATCGGACTCATCAGGAAGC
TCATCGGACTCATCAGGAAGC
ACATCGGACTCATCAGGAAGC
GAGCATCGGACTCATCAGGAAGC
GGCCATCGGACTCATCAGGAAGC
CCTCATCGGACTCATCAGGAAGC
ACATCGGACTCATCAGGAAGC
GGCATCGGACTCATCAGGAAGC
ACATCGGACTCATCAGGAAGC
TACATCGGACTCATCAGGAAGC
ATCATCGGACTCATCAGGAAGC
ACACATCGGACTCATCAGGAAGC
AGAACATCGGACTCATCAGGAAGC
CATCGGACTCATCAGGAAGC
CATCGGACTCATCAGGAAGC
ACACATCGGACTCATCAGGAAGC
GAGACATCGGACTCATCAGGAAGC
AGCATCGGACTCATCAGGAAGC
GAAACATCGGACTCATCAGGAAGC
GTCATCGGACTCATCAGGAAGC
AGCATCGGACTCATCAGGAAGC
TCATCGGACTCATCAGGAAGC
TGCATCGGACTCATCAGGAAGC
GTCCATCGGACTCATCAGGAAGC
GACATCGGACTCATCAGGAAGC
GCCATCGGACTCATCAGGAAGC
CCATCGGACTCATCAGGAAGC
CATCGGACTCATCAGGAAGC
AGACATCGGACTCATCAGGAAGC
CACTCATCGGACTCATCAGGAAGC
GTCATCGGACTCATCAGGAAGC
ACATCGGACTCATCAGGAAGC
TGAAGCATCGGACTCATCAGGAAGC
GCATCGGACTCATCAGGAAGC
AGCATCGGACTCATCAGGAAGC
CTCATCGGACTCATCAGGAAGC
CAGGCATCGGACTCATCAGGAAGC

TGCATCGGACTCATCAGGAAGC
CCATCGGACTCATCAGGAAGC
TCATCGGACTCATCAGGAAGC
GCATCGGACTCATCAGGAAGC
ATTCATCGGACTCATCAGGAAGC
GCAGCATCGGACTCATCAGGAAGC
ACTCATCGGACTCATCAGGAAGC
ACACATCGGACTCATCAGGAAGC
CTTCATCGGACTCATCAGGAAGC
TCCATCGGACTCATCAGGAAGC
GCATCGGACTCATCAGGAAGC
CCCATCGGACTCATCAGGAAGC
TGACATCGGACTCATCAGGAAGC
AACCCCATCGGACTCATCAGGAAGC
GCCATCGGACTCATCAGGAAGC
GTGTCATCGGACTCATCAGGAAGC
CATCGGACTCATCAGGAAGC
CCATCGGACTCATCAGGAAGC
GTCATCGGACTCATCAGGAAGC
CATCATCGGACTCATCAGGAAGC
AGGTCTCATCGGACTCATCAGGAAGC
CATCGGACTCATCAGGAAGC
AGACATCGGACTCATCAGGAAGC
CATCGGACTCATCAGGAAGC
GGTCATCGGACTCATCAGGAAGC
CTCATCGGACTCATCAGGAAGC
GCTCTACATCGGACTCATCAGGAAGC
CATCGGACTCATCAGGAAGC
CCCATCGGACTCATCAGGAAGC
CTCATCGGACTCATCAGGAAGC
AGGCATCGGACTCATCAGGAAGC
GAAACATCGGACTCATCAGGAAGC
GGCATCGGACTCATCAGGAAGC
CGGACTCATCAGGAAGC
AACATCGGACTCATCAGGAAGC
GTCCACATCGGACTCATCAGGAAGC
CATCGGACTCATCAGGAAGC
CCATCGGACTCATCAGGAAGC
ATCGGACTCATCAGGAAGC
ATCGGACTCATCAGGAAGC
TGCATCGGACTCATCAGGAAGC
GGACATCGGACTCATCAGGAAGC
TCCATCGGACTCATCAGGAAGC

CCATCGGACTCATCAGGAAGC
TCACCATCGGACTCATCAGGAAGC
GAGGCATCGGACTCATCAGGAAGC
GGGTCATCGGACTCATCAGGAAGC
AGGCATCGGACTCATCAGGAAGC
CCATCGGACTCATCAGGAAGC
CATCGGACTCATCAGGAAGC
GGCATCGGACTCATCAGGAAGC
AGCATCGGACTCATCAGGAAGC
ACCATCGGACTCATCAGGAAGC
AACAGCATCGGACTCATCAGGAAGC
TCATCGGACTCATCAGGAAGC
GCTCATCGGACTCATCAGGAAGC
GGACATCGGACTCATCAGGAAGC
TGCATCGGACTCATCAGGAAGC
TGTCATCGGACTCATCAGGAAGC
CCATCGGACTCATCAGGAAGC
ACATCGGACTCATCAGGAAGC
ACAACATCGGACTCATCAGGAAGC
GAGCATCGGACTCATCAGGAAGC
GCTCATCGGACTCATCAGGAAGC
AACATCGGACTCATCAGGAAGC
CCATCGGACTCATCAGGAAGC
ATTACATCGGACTCATCAGGAAGC
CTCATCGGACTCATCAGGAAGC
ACACATCGGACTCATCAGGAAGC
TGAACATCGGACTCATCAGGAAGC
AAAAGGCATCGGACTCATCAGGAAGC
ACATCGGACTCATCAGGAAGC
AACATCGGACTCATCAGGAAGC
ACATCGGACTCATCAGGAAGC
AAGTCATCGGACTCATCAGGAAGC
TTCATCGGACTCATCAGGAAGC
ATCGGACTCATCAGGAAGC
AATCATCGGACTCATCAGGAAGC
AGCATCGGACTCATCAGGAAGC
GAATCATCGGACTCATCAGGAAGC
CCCATCGGACTCATCAGGAAGC
ACATCGGACTCATCAGGAAGC
TCATCGGACTCATCAGGAAGC
GCTAAACATCGGACTCATCAGGAAGC
CATCGGACTCATCAGGAAGC
GGTGGCATCGGACTCATCAGGAAGC

CATCGGACTCATCAGGAAGC
ATCATCGGACTCATCAGGAAGC
ACATCGGACTCATCAGGAAGC
TCCATCGGACTCATCAGGAAGC
GGCATCGGACTCATCAGGAAGC
ACACATCGGACTCATCAGGAAGC
GGACATCGGACTCATCAGGAAGC
GGCCATCGGACTCATCAGGAAGC
ATGCATCGGACTCATCAGGAAGC
CTCCATCGGACTCATCAGGAAGC
GAACCTCATCGGACTCATCAGGAAGC
CATCGGACTCATCAGGAAGC
AGACCACATCGGACTCATCAGGAAGC
TTCATCGGACTCATCAGGAAGC
CCATCGGACTCATCAGGAAGC
ATCGGACTCATCAGGAAGC
GCCATCGGACTCATCAGGAAGC
CCATCGGACTCATCAGGAAGC
TGTGTGCATCGGACTCATCAGGAAGC
AGCCCATCGGACTCATCAGGAAGC
GTCATCGGACTCATCAGGAAGC
GTTCATCGGACTCATCAGGAAGC
TAAGCATCGGACTCATCAGGAAGC
TCCACATCGGACTCATCAGGAAGC
TGTCATCGGACTCATCAGGAAGC
TCCCATCGGACTCATCAGGAAGC
ACCATCGGACTCATCAGGAAGC
GCATCGGACTCATCAGGAAGC
CTCATCGGACTCATCAGGAAGC
GCTCATCGGACTCATCAGGAAGC
ACATCGGACTCATCAGGAAGC
TCAGCATCGGACTCATCAGGAAGC
CATCGGACTCATCAGGAAGC
GCATCGGACTCATCAGGAAGC
ACCATCGGACTCATCAGGAAGC
ATCATCGGACTCATCAGGAAGC
CCATCGGACTCATCAGGAAGC
ATCGGACTCATCAGGAAGC
AGCCCCATCGGACTCATCAGGAAGC
GCATCGGACTCATCAGGAAGC
ATACATCGGACTCATCAGGAAGC
AGCCACCATCGGACTCATCAGGAAGC
TACTCATCGGACTCATCAGGAAGC

CATCGGACTCATCAGGAAGC
CCTCATCGGACTCATCAGGAAGC
CCATCGGACTCATCAGGAAGC
TCATCGGACTCATCAGGAAGC
GCCACATCGGACTCATCAGGAAGC
GGGACATCGGACTCATCAGGAAGC
TGCATCGGACTCATCAGGAAGC
CCATCGGACTCATCAGGAAGC
ACGCATCGGACTCATCAGGAAGC
TCATCGGACTCATCAGGAAGC
TGACATCGGACTCATCAGGAAGC
GAGCATCGGACTCATCAGGAAGC
CCATCGGACTCATCAGGAAGC
AACATCGGACTCATCAGGAAGC
GCCATCGGACTCATCAGGAAGC
GGCCATCGGACTCATCAGGAAGC
ACATCGGACTCATCAGGAAGC
CATCGGACTCATCAGGAAGC
CATCGGACTCATCAGGAAGC
CCATCGGACTCATCAGGAAGC
CCTCATCGGACTCATCAGGAAGC
TGCATCGGACTCATCAGGAAGC
ACATCGGACTCATCAGGAAGC
TCATCGGACTCATCAGGAAGC
ACCATCGGACTCATCAGGAAGC
AAAACATCGGACTCATCAGGAAGC
CAACATCGGACTCATCAGGAAGC
TCATCGGACTCATCAGGAAGC
ATCGGACTCATCAGGAAGC
TTTCATCGGACTCATCAGGAAGC
GGCATCGGACTCATCAGGAAGC
TTCATCGGACTCATCAGGAAGC
ACATCGGACTCATCAGGAAGC
ATCGGACTCATCAGGAAGC
TCGGACTCATCAGGAAGC
CACATCGGACTCATCAGGAAGC
CTACATCGGACTCATCAGGAAGC
GGTGCATCGGACTCATCAGGAAGC
TCCCCATCGGACTCATCAGGAAGC
ACATCGGACTCATCAGGAAGC
CATCGGACTCATCAGGAAGC
TCATCGGACTCATCAGGAAGC
TACATCGGACTCATCAGGAAGC

TCATCGGACTCATCAGGAAGC
GTATCATCGGACTCATCAGGAAGC
CTGTCATCGGACTCATCAGGAAGC
GGTCATCGGACTCATCAGGAAGC
TCATCGGACTCATCAGGAAGC
CATCGGACTCATCAGGAAGC
ACCATCGGACTCATCAGGAAGC
TCTGCATCGGACTCATCAGGAAGC
GAGCATCGGACTCATCAGGAAGC
GCTTTCATCGGACTCATCAGGAAGC
CATCGGACTCATCAGGAAGC
GAAGCATCGGACTCATCAGGAAGC
ACATCGGACTCATCAGGAAGC
GCATCGGACTCATCAGGAAGC
ACCATCGGACTCATCAGGAAGC
AGTCATCGGACTCATCAGGAAGC
TGCATCGGACTCATCAGGAAGC
CCCATCGGACTCATCAGGAAGC
TACATCGGACTCATCAGGAAGC
AGCACATCGGACTCATCAGGAAGC
TCATCGGACTCATCAGGAAGC
ACATCGGACTCATCAGGAAGC
CCTCATCGGACTCATCAGGAAGC
TTACATCGGACTCATCAGGAAGC
ATCCATCGGACTCATCAGGAAGC
CATCGGACTCATCAGGAAGC
GAAAACATCGGACTCATCAGGAAGC
TCTCATCGGACTCATCAGGAAGC
ATCGGACTCATCAGGAAGC
GTCCATCGGACTCATCAGGAAGC
CATCGGACTCATCAGGAAGC
ATTCCATCGGACTCATCAGGAAGC
CTACCTCATCGGACTCATCAGGAAGC
ATGCAGCATCGGACTCATCAGGAAGC
CATCGGACTCATCAGGAAGC
TGCTGTGCATCGGACTCATCAGGAAGC
ACCCCATCGGACTCATCAGGAAGC
TCCATCGGACTCATCAGGAAGC
TCGGACTCATCAGGAAGC
CTCATCGGACTCATCAGGAAGC
CATCGGACTCATCAGGAAGC
GGCATCGGACTCATCAGGAAGC
TCACATCGGACTCATCAGGAAGC

CTACCATCGGACTCATCAGGAAGC
ATATTCATCGGACTCATCAGGAAGC
CAGCATCGGACTCATCAGGAAGC
CAGCATCGGACTCATCAGGAAGC
ACATCGGACTCATCAGGAAGC
GCATCATCGGACTCATCAGGAAGC
ACGCATCGGACTCATCAGGAAGC
TGGCATCGGACTCATCAGGAAGC
GGGGGCATCGGACTCATCAGGAAGC
GCATCGGACTCATCAGGAAGC
GCCATCGGACTCATCAGGAAGC
CATCGGACTCATCAGGAAGC
AGCATCGGACTCATCAGGAAGC
CAAATCATCGGACTCATCAGGAAGC
TCATCGGACTCATCAGGAAGC
CAACATCGGACTCATCAGGAAGC
ACATCGGACTCATCAGGAAGC
CAGCATCGGACTCATCAGGAAGC
GTAGACATCGGACTCATCAGGAAGC
TCATCGGACTCATCAGGAAGC
CATCGGACTCATCAGGAAGC
GACATCGGACTCATCAGGAAGC
ATCGCACATCGGACTCATCAGGAAGC
CCATCGGACTCATCAGGAAGC
GACATCGGACTCATCAGGAAGC
TGCATCGGACTCATCAGGAAGC
TCATCGGACTCATCAGGAAGC
GCACATCGGACTCATCAGGAAGC
TAGACATCGGACTCATCAGGAAGC
ACATCGGACTCATCAGGAAGC
TCCATCGGACTCATCAGGAAGC
GCTCATCGGACTCATCAGGAAGC
CGCATCGGACTCATCAGGAAGC
GTACATCGGACTCATCAGGAAGC
ACATCGGACTCATCAGGAAGC
CGCATCGGACTCATCAGGAAGC
GCCCATCGGACTCATCAGGAAGC
TCATCGGACTCATCAGGAAGC
GCCATCGGACTCATCAGGAAGC
CCCATCGGACTCATCAGGAAGC
CGCATCGGACTCATCAGGAAGC
CACATCGGACTCATCAGGAAGC
CATCGGACTCATCAGGAAGC

TTCATCGGACTCATCAGGAAGC
GGTCATCGGACTCATCAGGAAGC
CAACATCGGACTCATCAGGAAGC
GGTCATCGGACTCATCAGGAAGC
CGCATCGGACTCATCAGGAAGC
GCCTCATCGGACTCATCAGGAAGC
ACATCGGACTCATCAGGAAGC
CTACATCGGACTCATCAGGAAGC
CCCATCGGACTCATCAGGAAGC
GCCTCCATCGGACTCATCAGGAAGC
ACCCGCATCGGACTCATCAGGAAGC
CATCGGACTCATCAGGAAGC
TATCATCGGACTCATCAGGAAGC
GCCTTCACATCGGACTCATCAGGAAGC
CACATCGGACTCATCAGGAAGC
GCCATCGGACTCATCAGGAAGC
GGCATCGGACTCATCAGGAAGC
CCCCATCGGACTCATCAGGAAGC
CATCGGACTCATCAGGAAGC
TAGCCATCGGACTCATCAGGAAGC
TGCCATCGGACTCATCAGGAAGC
GGACATCGGACTCATCAGGAAGC
GTAGGCATCGGACTCATCAGGAAGC
ACATCGGACTCATCAGGAAGC
GCCTCATCGGACTCATCAGGAAGC
GGGGCATCGGACTCATCAGGAAGC
CACATCGGACTCATCAGGAAGC
TGAGACATCGGACTCATCAGGAAGC
GCATCGGACTCATCAGGAAGC
TGTCATCGGACTCATCAGGAAGC
AATCATCGGACTCATCAGGAAGC
GACATCGGACTCATCAGGAAGC
CATCGGACTCATCAGGAAGC
CGGACTCATCAGGAAGC
CTGTCATCGGACTCATCAGGAAGC
ACGCATCGGACTCATCAGGAAGC
AGCATCGGACTCATCAGGAAGC
TCATCGGACTCATCAGGAAGC
TCTCATCGGACTCATCAGGAAGC
GCACATCGGACTCATCAGGAAGC
AGACATCGGACTCATCAGGAAGC
CAGCATCGGACTCATCAGGAAGC
CTCCATCGGACTCATCAGGAAGC

TCATCGGACTCATCAGGAAGC
CGCATCGGACTCATCAGGAAGC
CCATCGGACTCATCAGGAAGC
TCATCGGACTCATCAGGAAGC
ATCGGACTCATCAGGAAGC
GCATCGGACTCATCAGGAAGC
ACAGTCATCGGACTCATCAGGAAGC
GAACATCGGACTCATCAGGAAGC
TTCCATCGGACTCATCAGGAAGC
GGGACATCGGACTCATCAGGAAGC
CATCGGACTCATCAGGAAGC
ACATCGGACTCATCAGGAAGC
GTGACATCGGACTCATCAGGAAGC
GCACATCGGACTCATCAGGAAGC
TCATCGGACTCATCAGGAAGC
TATTGCTCATCGGACTCATCAGGAAGC
TGGTCATCGGACTCATCAGGAAGC
TGGACATCGGACTCATCAGGAAGC
ATCGGACTCATCAGGAAGC
AAGCATCGGACTCATCAGGAAGC
AGGCATCGGACTCATCAGGAAGC
GCACATCGGACTCATCAGGAAGC
GCATCGGACTCATCAGGAAGC
GAGCCATCGGACTCATCAGGAAGC
AGACATCGGACTCATCAGGAAGC
TCCCATCGGACTCATCAGGAAGC
CATCGGACTCATCAGGAAGC
CCACATCGGACTCATCAGGAAGC
CTCTCATCGGACTCATCAGGAAGC
CACATCGGACTCATCAGGAAGC
AAACATCGGACTCATCAGGAAGC
GGAGGCATCGGACTCATCAGGAAGC
ACCATCGGACTCATCAGGAAGC
ACATCGGACTCATCAGGAAGC
AGACATCGGACTCATCAGGAAGC
CACATCGGACTCATCAGGAAGC
ACATCGGACTCATCAGGAAGC
CATCATCGGACTCATCAGGAAGC
CTTCATCGGACTCATCAGGAAGC
GCCACATCGGACTCATCAGGAAGC
TCATCGGACTCATCAGGAAGC
TTTCATCGGACTCATCAGGAAGC
GACACATCGGACTCATCAGGAAGC

AAAACATCGGACTCATCAGGAAGC
TCGGACTCATCAGGAAGC
TGCATCGGACTCATCAGGAAGC
TCATCGGACTCATCAGGAAGC
ATCGGACTCATCAGGAAGC
ACATCGGACTCATCAGGAAGC
ACATCGGACTCATCAGGAAGC
CTGTGCGCATCGGACTCATCAGGAAGC
CCCATCGGACTCATCAGGAAGC
GTGCATCGGACTCATCAGGAAGC
CTCCATCGGACTCATCAGGAAGC
CTCATCGGACTCATCAGGAAGC
CATCGGACTCATCAGGAAGC
ACCATCGGACTCATCAGGAAGC
ATCGGACTCATCAGGAAGC
CTACATCGGACTCATCAGGAAGC
CACATCGGACTCATCAGGAAGC
ACATCGGACTCATCAGGAAGC
TGCATCGGACTCATCAGGAAGC
GTCATCGGACTCATCAGGAAGC
CATCGGACTCATCAGGAAGC
GGCATCGGACTCATCAGGAAGC
ATCGGACTCATCAGGAAGC
ATGTCATCGGACTCATCAGGAAGC
ATCGGACTCATCAGGAAGC
CCATCGGACTCATCAGGAAGC
TAACATCGGACTCATCAGGAAGC
ATCGGACTCATCAGGAAGC
AGCATCGGACTCATCAGGAAGC
GGCATCGGACTCATCAGGAAGC
GCTGGCATCGGACTCATCAGGAAGC
ACATCGGACTCATCAGGAAGC
AGTCATCGGACTCATCAGGAAGC
ACCATCGGACTCATCAGGAAGC
TACATCGGACTCATCAGGAAGC
TTCATCGGACTCATCAGGAAGC
CAGCATCGGACTCATCAGGAAGC
CCATCGGACTCATCAGGAAGC
ACATCGGACTCATCAGGAAGC
TTCATCGGACTCATCAGGAAGC
ACCCATCGGACTCATCAGGAAGC
TGGACATCGGACTCATCAGGAAGC
CACATCGGACTCATCAGGAAGC

GGTCATCGGACTCATCAGGAAGC
AGGTCATCGGACTCATCAGGAAGC
AGGTCATCGGACTCATCAGGAAGC
TCATCGGACTCATCAGGAAGC
ACATCGGACTCATCAGGAAGC
TGCATCGGACTCATCAGGAAGC
ATCGGACTCATCAGGAAGC
GGTCATCGGACTCATCAGGAAGC
GTACATCGGACTCATCAGGAAGC
CATCGGACTCATCAGGAAGC
CATCGGACTCATCAGGAAGC
ATCGGACTCATCAGGAAGC
GCGCATCGGACTCATCAGGAAGC
CCCATCGGACTCATCAGGAAGC
CATCATCGGACTCATCAGGAAGC
TTCATCGGACTCATCAGGAAGC
TTCATCGGACTCATCAGGAAGC
GCCATCGGACTCATCAGGAAGC
CTCTCATCGGACTCATCAGGAAGC
AACCTGCATCGGACTCATCAGGAAGC
TCGGACTCATCAGGAAGC
TAGCGTCATCGGACTCATCAGGAAGC
ACACATCGGACTCATCAGGAAGC
CCATCGGACTCATCAGGAAGC
AAAAGCATCGGACTCATCAGGAAGC
TGTTCATCGGACTCATCAGGAAGC
AGTCACATCGGACTCATCAGGAAGC
ACCCACATCGGACTCATCAGGAAGC
ATGAAACATCGGACTCATCAGGAAGC
AGCATCGGACTCATCAGGAAGC
GTTTTCATCGGACTCATCAGGAAGC
GGGCATCGGACTCATCAGGAAGC
CAACATCGGACTCATCAGGAAGC
GTATGCATCGGACTCATCAGGAAGC
CATCGGACTCATCAGGAAGC
GACATCGGACTCATCAGGAAGC
AGGCATCGGACTCATCAGGAAGC
TGCCATCGGACTCATCAGGAAGC
GATCATCGGACTCATCAGGAAGC
CCTCATCGGACTCATCAGGAAGC
AGCACATCGGACTCATCAGGAAGC
GCATCGGACTCATCAGGAAGC
ACATCGGACTCATCAGGAAGC

TGTCATCGGACTCATCAGGAAGC
TCATCGGACTCATCAGGAAGC
GCCCATCGGACTCATCAGGAAGC
ACATCGGACTCATCAGGAAGC
AGGGGCATCGGACTCATCAGGAAGC
TCATCGGACTCATCAGGAAGC
TCACATCGGACTCATCAGGAAGC
TGCATCGGACTCATCAGGAAGC
CCATCGGACTCATCAGGAAGC
GGCACATCGGACTCATCAGGAAGC
ACATACACATCGGACTCATCAGGAAGC
TCATCGGACTCATCAGGAAGC
AGGGGACATCGGACTCATCAGGAAGC
CCACATCGGACTCATCAGGAAGC
GCCCATCGGACTCATCAGGAAGC
GCCATCGGACTCATCAGGAAGC
CACATCGGACTCATCAGGAAGC
AGCATCGGACTCATCAGGAAGC
CACCATCGGACTCATCAGGAAGC
TGGCATCGGACTCATCAGGAAGC
CATCGGACTCATCAGGAAGC
AACAGCATCGGACTCATCAGGAAGC
TGCCCATCGGACTCATCAGGAAGC
ACATCGGACTCATCAGGAAGC
TCATCGGACTCATCAGGAAGC
TCGGACTCATCAGGAAGC
CATCGGACTCATCAGGAAGC
GCATCGGACTCATCAGGAAGC
CAACATCGGACTCATCAGGAAGC
ATCATCGGACTCATCAGGAAGC
ACACATCGGACTCATCAGGAAGC
AGCATCGGACTCATCAGGAAGC
CCATCGGACTCATCAGGAAGC
CCATCGGACTCATCAGGAAGC
GCATCGGACTCATCAGGAAGC
TAACATCGGACTCATCAGGAAGC
GCTACATCGGACTCATCAGGAAGC
GTGCCATCGGACTCATCAGGAAGC
TTTGATCATCGGACTCATCAGGAAGC
TACATCGGACTCATCAGGAAGC
AACCATCGGACTCATCAGGAAGC
GAGCATCGGACTCATCAGGAAGC
ATCCCATCGGACTCATCAGGAAGC

TCTCATCGGACTCATCAGGAAGC
GCATCGGACTCATCAGGAAGC
ACCATCGGACTCATCAGGAAGC
TCAGCATCGGACTCATCAGGAAGC
ACCATCGGACTCATCAGGAAGC
GTCACATCGGACTCATCAGGAAGC
ACCCATCGGACTCATCAGGAAGC
AAGACATCGGACTCATCAGGAAGC
TCCATCGGACTCATCAGGAAGC
CATCGGACTCATCAGGAAGC
CCATCGGACTCATCAGGAAGC
ACATCGGACTCATCAGGAAGC
GCCATCGGACTCATCAGGAAGC
AACATCGGACTCATCAGGAAGC
GGCATCGGACTCATCAGGAAGC
GAACCACATCGGACTCATCAGGAAGC
TCATCGGACTCATCAGGAAGC
ACATCGGACTCATCAGGAAGC
ATCGGACTCATCAGGAAGC
GACATCGGACTCATCAGGAAGC
CATCGGACTCATCAGGAAGC
AGCATCGGACTCATCAGGAAGC
CGCATCGGACTCATCAGGAAGC
ACATCGGACTCATCAGGAAGC
TCACACATCGGACTCATCAGGAAGC
ATCGGACTCATCAGGAAGC
TACATCGGACTCATCAGGAAGC
ATCATCGGACTCATCAGGAAGC
CCTCATCGGACTCATCAGGAAGC
GCCATCGGACTCATCAGGAAGC
AGACATCGGACTCATCAGGAAGC
AAACATCGGACTCATCAGGAAGC
AGGGCATCGGACTCATCAGGAAGC
GAGCATCGGACTCATCAGGAAGC
AACCATCGGACTCATCAGGAAGC
ATCGGACTCATCAGGAAGC
ATCGGACTCATCAGGAAGC
ACATCGGACTCATCAGGAAGC
ATCGGACTCATCAGGAAGC
CCATCGGACTCATCAGGAAGC
TGATCATCGGACTCATCAGGAAGC
CATCGGACTCATCAGGAAGC
CTCATCGGACTCATCAGGAAGC

CCATCGGACTCATCAGGAAGC
ACATCGGACTCATCAGGAAGC
TGGCATCGGACTCATCAGGAAGC
ATCGGACTCATCAGGAAGC
AGTCATCGGACTCATCAGGAAGC
CCATCGGACTCATCAGGAAGC
AGAACCATCGGACTCATCAGGAAGC
GGGCACATCGGACTCATCAGGAAGC
GCATCGGACTCATCAGGAAGC
TTCATCGGACTCATCAGGAAGC
GAGCCATCGGACTCATCAGGAAGC
CATCGGACTCATCAGGAAGC
CGCATCGGACTCATCAGGAAGC
CATCGGACTCATCAGGAAGC
ATCATCGGACTCATCAGGAAGC
ACCATCGGACTCATCAGGAAGC
AGCATCGGACTCATCAGGAAGC
AACATCGGACTCATCAGGAAGC
ACTGCCTCATCGGACTCATCAGGAAGC
TCATCGGACTCATCAGGAAGC
TACATCGGACTCATCAGGAAGC
CCATCGGACTCATCAGGAAGC
ACATCGGACTCATCAGGAAGC
CACATCGGACTCATCAGGAAGC
CTCATCGGACTCATCAGGAAGC
CTGCATCGGACTCATCAGGAAGC
CCATCGGACTCATCAGGAAGC
ATTCATCGGACTCATCAGGAAGC
AAACATCGGACTCATCAGGAAGC
ACATCGGACTCATCAGGAAGC
GCCATCATCGGACTCATCAGGAAGC
TGCATCGGACTCATCAGGAAGC
AGCATCGGACTCATCAGGAAGC
ATCGGACTCATCAGGAAGC
GACATCGGACTCATCAGGAAGC
AAAACATCGGACTCATCAGGAAGC
TGTCATCGGACTCATCAGGAAGC
CTACATCGGACTCATCAGGAAGC
ACATCGGACTCATCAGGAAGC
ATCGGACTCATCAGGAAGC
ACATCGGACTCATCAGGAAGC
GCTCATCGGACTCATCAGGAAGC
GGCATCGGACTCATCAGGAAGC

CAAACATCGGACTCATCAGGAAGC
CAGAAACATCGGACTCATCAGGAAGC
AGCATCGGACTCATCAGGAAGC
CTCCATCGGACTCATCAGGAAGC
CACATCGGACTCATCAGGAAGC
CACATCGGACTCATCAGGAAGC
GCCATCGGACTCATCAGGAAGC
TCATCGGACTCATCAGGAAGC
TCATCGGACTCATCAGGAAGC
TGCATCGGACTCATCAGGAAGC
TGGTTCATCGGACTCATCAGGAAGC
ACGCATCGGACTCATCAGGAAGC
TCACATCGGACTCATCAGGAAGC
CATCGGACTCATCAGGAAGC
TACTCATCGGACTCATCAGGAAGC
CATCGGACTCATCAGGAAGC
TTGCATCGGACTCATCAGGAAGC
GCATCGGACTCATCAGGAAGC
GACATCGGACTCATCAGGAAGC
TGAGCATCGGACTCATCAGGAAGC
AAGGTCATCGGACTCATCAGGAAGC
TGCATCGGACTCATCAGGAAGC
GGCATCGGACTCATCAGGAAGC
ATCGGACTCATCAGGAAGC
CATCGGACTCATCAGGAAGC
CCCACATCGGACTCATCAGGAAGC
AACATCGGACTCATCAGGAAGC
CATCGGACTCATCAGGAAGC
CCATCGGACTCATCAGGAAGC
ACCATCGGACTCATCAGGAAGC
ACACATCGGACTCATCAGGAAGC
AGCATCGGACTCATCAGGAAGC
AAACACATCGGACTCATCAGGAAGC
GAGGCATCGGACTCATCAGGAAGC
TTCATCGGACTCATCAGGAAGC
CATCGGACTCATCAGGAAGC
TTACATCGGACTCATCAGGAAGC
ACATCGGACTCATCAGGAAGC
ATCGGACTCATCAGGAAGC
ATCATCGGACTCATCAGGAAGC
AACATCGGACTCATCAGGAAGC
CTGCATCGGACTCATCAGGAAGC
GTCATCGGACTCATCAGGAAGC

GTACATCGGACTCATCAGGAAGC
TACATCGGACTCATCAGGAAGC
ATCGGACTCATCAGGAAGC
GCAACATCGGACTCATCAGGAAGC
GGTCATCGGACTCATCAGGAAGC
ACCCATCGGACTCATCAGGAAGC
CTCATCGGACTCATCAGGAAGC
AGCATCGGACTCATCAGGAAGC
AGCCATCGGACTCATCAGGAAGC
ACATCGGACTCATCAGGAAGC
TCCATCGGACTCATCAGGAAGC
CCCATCGGACTCATCAGGAAGC
ACAATCATCGGACTCATCAGGAAGC
GGGCCATCGGACTCATCAGGAAGC
CTGACATCGGACTCATCAGGAAGC
ACATCGGACTCATCAGGAAGC
ATCATCGGACTCATCAGGAAGC
ACATCGGACTCATCAGGAAGC
ATGCATCGGACTCATCAGGAAGC
ATATGCATCGGACTCATCAGGAAGC
CATTCATCGGACTCATCAGGAAGC
TCTCATCGGACTCATCAGGAAGC
AGCCATCGGACTCATCAGGAAGC
ACATCGGACTCATCAGGAAGC
GGACATCGGACTCATCAGGAAGC
GCCATCGGACTCATCAGGAAGC
TGTCATCGGACTCATCAGGAAGC
AACATCGGACTCATCAGGAAGC
CACATCGGACTCATCAGGAAGC
TGATCATCGGACTCATCAGGAAGC
ATCATCGGACTCATCAGGAAGC
CTCTCTCATCGGACTCATCAGGAAGC
GCAGGTCATCGGACTCATCAGGAAGC
ACATCGGACTCATCAGGAAGC
TCATCGGACTCATCAGGAAGC
GCATCGGACTCATCAGGAAGC
TAGCCATCGGACTCATCAGGAAGC
TACATCGGACTCATCAGGAAGC
CCCATCGGACTCATCAGGAAGC
GGCATCGGACTCATCAGGAAGC
GTCATCGGACTCATCAGGAAGC
AAACATCGGACTCATCAGGAAGC
ACATCGGACTCATCAGGAAGC

ACCCATCGGACTCATCAGGAAGC
ACATCGGACTCATCAGGAAGC
TACATCGGACTCATCAGGAAGC
GCACATCGGACTCATCAGGAAGC
TGTCATCGGACTCATCAGGAAGC
ATTGTTCATCGGACTCATCAGGAAGC
CATCGGACTCATCAGGAAGC
ACATCGGACTCATCAGGAAGC
CTTACATCGGACTCATCAGGAAGC
GGTCATCGGACTCATCAGGAAGC
CATCGGACTCATCAGGAAGC
TATTCATCGGACTCATCAGGAAGC
CACATCGGACTCATCAGGAAGC
CTCCATCGGACTCATCAGGAAGC
TGCATCGGACTCATCAGGAAGC
TTCATCGGACTCATCAGGAAGC
TCATCGGACTCATCAGGAAGC
AATGCTCATCGGACTCATCAGGAAGC
AACACATCGGACTCATCAGGAAGC
ACATCGGACTCATCAGGAAGC
CATCGGACTCATCAGGAAGC
GCACATCGGACTCATCAGGAAGC
TACATCGGACTCATCAGGAAGC
CTCATCGGACTCATCAGGAAGC
GGGCATCGGACTCATCAGGAAGC
AGAGGCATCGGACTCATCAGGAAGC
CGGACTCATCAGGAAGC
TGACTACATCGGACTCATCAGGAAGC
CATCGGACTCATCAGGAAGC
TGCTACATCGGACTCATCAGGAAGC
GTGCATCGGACTCATCAGGAAGC
ATCATCGGACTCATCAGGAAGC
GCATCGGACTCATCAGGAAGC
GAGAATCATCGGACTCATCAGGAAGC
AGGGCATCGGACTCATCAGGAAGC
CATCGGACTCATCAGGAAGC
GTACATCGGACTCATCAGGAAGC
GGCATCGGACTCATCAGGAAGC
CATCGGACTCATCAGGAAGC
TCATCGGACTCATCAGGAAGC
TGTCATCGGACTCATCAGGAAGC
CCATCGGACTCATCAGGAAGC
TCGGACTCATCAGGAAGC

CACATCGGACTCATCAGGAAGC
CTCATCGGACTCATCAGGAAGC
TTCATCGGACTCATCAGGAAGC
CCCATCGGACTCATCAGGAAGC
ATCATCGGACTCATCAGGAAGC
AAGACATCGGACTCATCAGGAAGC
GCATCGGACTCATCAGGAAGC
TGACATCGGACTCATCAGGAAGC
GGCACATCGGACTCATCAGGAAGC
TCATCGGACTCATCAGGAAGC
TGCATCGGACTCATCAGGAAGC
GGCCATCGGACTCATCAGGAAGC
AGCCATCGGACTCATCAGGAAGC
CTGTCATCGGACTCATCAGGAAGC
AGGCATCGGACTCATCAGGAAGC
CTCCATCGGACTCATCAGGAAGC
CATCGGACTCATCAGGAAGC
GCATCGGACTCATCAGGAAGC
GGCATCGGACTCATCAGGAAGC
ATAGACATCGGACTCATCAGGAAGC
GCATCGGACTCATCAGGAAGC
CTCATCGGACTCATCAGGAAGC
TTCATCGGACTCATCAGGAAGC
GGCATCGGACTCATCAGGAAGC
AGCCATCGGACTCATCAGGAAGC
CATCGGACTCATCAGGAAGC
GTGCATCGGACTCATCAGGAAGC
CATCGGACTCATCAGGAAGC
GTGGCATCGGACTCATCAGGAAGC
CACATCGGACTCATCAGGAAGC
TTACATCGGACTCATCAGGAAGC
TGCATCGGACTCATCAGGAAGC
ACATCGGACTCATCAGGAAGC
GACATCGGACTCATCAGGAAGC
TCATCGGACTCATCAGGAAGC
AGCCATCGGACTCATCAGGAAGC
ACATCGGACTCATCAGGAAGC
GACATCGGACTCATCAGGAAGC
TGCTCATCGGACTCATCAGGAAGC
CCTCATCGGACTCATCAGGAAGC
CCATCGGACTCATCAGGAAGC
GTGGCTCATCGGACTCATCAGGAAGC
CATCGGACTCATCAGGAAGC

TCCATCGGACTCATCAGGAAGC
ACACATCGGACTCATCAGGAAGC
GACATCGGACTCATCAGGAAGC
CAGCATCGGACTCATCAGGAAGC
GGACTCATCAGGAAGC
AACATCGGACTCATCAGGAAGC
AACATCGGACTCATCAGGAAGC
ATCGGACTCATCAGGAAGC
ACATCGGACTCATCAGGAAGC
ACCATCGGACTCATCAGGAAGC
CCATCGGACTCATCAGGAAGC
CTTATCATCGGACTCATCAGGAAGC
CATCGGACTCATCAGGAAGC
ACATCGGACTCATCAGGAAGC
CTCATCGGACTCATCAGGAAGC
CCATCGGACTCATCAGGAAGC
TCGCACATCGGACTCATCAGGAAGC
TCATCGGACTCATCAGGAAGC
AACATCGGACTCATCAGGAAGC
GAACATCGGACTCATCAGGAAGC
TCTCTCATCGGACTCATCAGGAAGC
CATCGGACTCATCAGGAAGC
ACATCGGACTCATCAGGAAGC
CATCGGACTCATCAGGAAGC
ACCATCGGACTCATCAGGAAGC
TGCATCGGACTCATCAGGAAGC
ATCGGACTCATCAGGAAGC
GGCATCGGACTCATCAGGAAGC
TTCATCGGACTCATCAGGAAGC
CATCGGACTCATCAGGAAGC
AGAACATCGGACTCATCAGGAAGC
ACATCGGACTCATCAGGAAGC
CCATCGGACTCATCAGGAAGC
ACCATCGGACTCATCAGGAAGC
ACATCGGACTCATCAGGAAGC
CATCGGACTCATCAGGAAGC
TCATCGGACTCATCAGGAAGC
GACATCGGACTCATCAGGAAGC
AGCATCGGACTCATCAGGAAGC
GTCATCGGACTCATCAGGAAGC
TTGCATCGGACTCATCAGGAAGC
CATCATCGGACTCATCAGGAAGC
GCATCGGACTCATCAGGAAGC

CAGCATCGGACTCATCAGGAAGC
ACAGAGCATCGGACTCATCAGGAAGC
GCCATCGGACTCATCAGGAAGC
GACCATCGGACTCATCAGGAAGC
AGCATCGGACTCATCAGGAAGC
GCATCGGACTCATCAGGAAGC
CGCATCGGACTCATCAGGAAGC
AAACACATCGGACTCATCAGGAAGC
GCATCGGACTCATCAGGAAGC
GGCATCGGACTCATCAGGAAGC
CTGCATCGGACTCATCAGGAAGC
AGCATCGGACTCATCAGGAAGC
CTCAAGCATCGGACTCATCAGGAAGC
GCATCGGACTCATCAGGAAGC
GCATCGGACTCATCAGGAAGC
TCGGACTCATCAGGAAGC
CAGCACATCGGACTCATCAGGAAGC
CATCGGACTCATCAGGAAGC
AACATCGGACTCATCAGGAAGC
ATACATCGGACTCATCAGGAAGC
TTCATCGGACTCATCAGGAAGC
ACATCGGACTCATCAGGAAGC
CGGACTCATCAGGAAGC
ATTTACATCGGACTCATCAGGAAGC
CATCGGACTCATCAGGAAGC
GAACCATCGGACTCATCAGGAAGC
GACCATCGGACTCATCAGGAAGC
ACACCATCGGACTCATCAGGAAGC
TCATCGGACTCATCAGGAAGC
CTCATCGGACTCATCAGGAAGC
CATCGGACTCATCAGGAAGC
AAGACATCGGACTCATCAGGAAGC
GAACCATCGGACTCATCAGGAAGC
AAACATCGGACTCATCAGGAAGC
CTCATCGGACTCATCAGGAAGC
AGCATCGGACTCATCAGGAAGC
CACCCACATCGGACTCATCAGGAAGC
TCATCGGACTCATCAGGAAGC
GGAACATCGGACTCATCAGGAAGC
ACATCGGACTCATCAGGAAGC
TCTCATCGGACTCATCAGGAAGC
ACATCGGACTCATCAGGAAGC
ACATCGGACTCATCAGGAAGC

CCCATCGGACTCATCAGGAAGC
TCATCGGACTCATCAGGAAGC
ATCACCATCGGACTCATCAGGAAGC
TTCATCGGACTCATCAGGAAGC
GACATCGGACTCATCAGGAAGC
CCCATCGGACTCATCAGGAAGC
GTCATCGGACTCATCAGGAAGC
TGCCATCGGACTCATCAGGAAGC
ATCATCGGACTCATCAGGAAGC
TCATCGGACTCATCAGGAAGC
GTCATCGGACTCATCAGGAAGC
ATCATCGGACTCATCAGGAAGC
GTGCCATCGGACTCATCAGGAAGC
TACATCGGACTCATCAGGAAGC
CACATCGGACTCATCAGGAAGC
AGGCATCGGACTCATCAGGAAGC
TCATCGGACTCATCAGGAAGC
TACATCGGACTCATCAGGAAGC
TCATCGGACTCATCAGGAAGC
GACATCGGACTCATCAGGAAGC
TGTGCATCGGACTCATCAGGAAGC
CTCCACATCGGACTCATCAGGAAGC
TCATCGGACTCATCAGGAAGC
AGCATCGGACTCATCAGGAAGC
GCATCGGACTCATCAGGAAGC
GCCATCGGACTCATCAGGAAGC
ACCATCGGACTCATCAGGAAGC
CATCGGACTCATCAGGAAGC
GTGTCATCGGACTCATCAGGAAGC
CCCCCATCGGACTCATCAGGAAGC
ACATCGGACTCATCAGGAAGC
AGTGCATCGGACTCATCAGGAAGC
GACATCGGACTCATCAGGAAGC
GGGACATCGGACTCATCAGGAAGC
TATCATCGGACTCATCAGGAAGC
CCAGGCATCGGACTCATCAGGAAGC
TGAGCATCGGACTCATCAGGAAGC
CCATCGGACTCATCAGGAAGC
CATCGGACTCATCAGGAAGC
AACATCGGACTCATCAGGAAGC
CTTCATCGGACTCATCAGGAAGC
GTCATCGGACTCATCAGGAAGC
ATCGGACTCATCAGGAAGC

TCGGACTCATCAGGAAGC
ATCGGACTCATCAGGAAGC
AGCCATCGGACTCATCAGGAAGC
TGACCATCGGACTCATCAGGAAGC
AGAGGCATCGGACTCATCAGGAAGC
ACCATCGGACTCATCAGGAAGC
TGCCATCGGACTCATCAGGAAGC
CTCATCGGACTCATCAGGAAGC
TGTCATCGGACTCATCAGGAAGC
ACATCGGACTCATCAGGAAGC
CGGACTCATCAGGAAGC
TGCATCGGACTCATCAGGAAGC
GCATCGGACTCATCAGGAAGC
CATCGGACTCATCAGGAAGC
CCATCGGACTCATCAGGAAGC
CCATCGGACTCATCAGGAAGC
TCATCGGACTCATCAGGAAGC
TCATCGGACTCATCAGGAAGC
CCATCGGACTCATCAGGAAGC
TCATCGGACTCATCAGGAAGC
ATCGGACTCATCAGGAAGC
ACATCGGACTCATCAGGAAGC
TACATCGGACTCATCAGGAAGC
CATCGGACTCATCAGGAAGC
GTTCCATCGGACTCATCAGGAAGC
ACATCGGACTCATCAGGAAGC
ACATCGGACTCATCAGGAAGC
GCCATCGGACTCATCAGGAAGC
TCATCGGACTCATCAGGAAGC
GAGCATCGGACTCATCAGGAAGC
TATCATCGGACTCATCAGGAAGC
CATCGGACTCATCAGGAAGC
CCATCGGACTCATCAGGAAGC
CATCGGACTCATCAGGAAGC
ATCATCGGACTCATCAGGAAGC
GTCATCGGACTCATCAGGAAGC
GCCATCGGACTCATCAGGAAGC
GACATCGGACTCATCAGGAAGC
GCCATCGGACTCATCAGGAAGC
CATCGGACTCATCAGGAAGC
CAGCATCGGACTCATCAGGAAGC
GCATCGGACTCATCAGGAAGC
TCATCGGACTCATCAGGAAGC

CATCGGACTCATCAGGAAGC
ACATCGGACTCATCAGGAAGC
CATCGGACTCATCAGGAAGC
CTCCATCGGACTCATCAGGAAGC
CATCGGACTCATCAGGAAGC
AGGACATCGGACTCATCAGGAAGC
GTTCCATCGGACTCATCAGGAAGC
ACATCGGACTCATCAGGAAGC
ATCGGACTCATCAGGAAGC
TGCCATCGGACTCATCAGGAAGC
ATCATCGGACTCATCAGGAAGC
AACATCGGACTCATCAGGAAGC
CTTGCATCGGACTCATCAGGAAGC
AACATCGGACTCATCAGGAAGC
AGCCATCGGACTCATCAGGAAGC
CACCATCGGACTCATCAGGAAGC
TGGTACATCGGACTCATCAGGAAGC
CTCCATCGGACTCATCAGGAAGC
TTGTCATCGGACTCATCAGGAAGC
TCTCTCATCGGACTCATCAGGAAGC
TGGACATCGGACTCATCAGGAAGC
ACATCGGACTCATCAGGAAGC
ACATCGGACTCATCAGGAAGC
GTCCATCGGACTCATCAGGAAGC
GACCATCGGACTCATCAGGAAGC
GTCCATCGGACTCATCAGGAAGC
CACATCGGACTCATCAGGAAGC
CCTCATCGGACTCATCAGGAAGC
CATCGGACTCATCAGGAAGC
CAACATCGGACTCATCAGGAAGC
CCCATCGGACTCATCAGGAAGC
GCCATCATCGGACTCATCAGGAAGC
AAGGCCATCGGACTCATCAGGAAGC
TGCCATCGGACTCATCAGGAAGC
ACATCGGACTCATCAGGAAGC
GCAGCATCGGACTCATCAGGAAGC
ATCCCATCGGACTCATCAGGAAGC
TCCATCGGACTCATCAGGAAGC
GTCAGCATCGGACTCATCAGGAAGC
TCCCATCGGACTCATCAGGAAGC
CATCATCGGACTCATCAGGAAGC
ATCTCATCGGACTCATCAGGAAGC
GCACATCGGACTCATCAGGAAGC

AGGCATCGGACTCATCAGGAAGC
TGTGCATCGGACTCATCAGGAAGC
CCCATCGGACTCATCAGGAAGC
CATCGGACTCATCAGGAAGC
ATCCATCGGACTCATCAGGAAGC
CCACATCGGACTCATCAGGAAGC
GGACATCGGACTCATCAGGAAGC
AACATCGGACTCATCAGGAAGC
AGAGCATCGGACTCATCAGGAAGC
CAGCATCGGACTCATCAGGAAGC
GGACATCGGACTCATCAGGAAGC
TTCATCGGACTCATCAGGAAGC
TCATCGGACTCATCAGGAAGC
CATCGGACTCATCAGGAAGC
CTCTCATCGGACTCATCAGGAAGC
TCTCATCGGACTCATCAGGAAGC
CATCGGACTCATCAGGAAGC
ACCATCGGACTCATCAGGAAGC
GGCACATCGGACTCATCAGGAAGC
ATCCCATCGGACTCATCAGGAAGC
GTCATCGGACTCATCAGGAAGC
AGCCATCGGACTCATCAGGAAGC
TGTCATCGGACTCATCAGGAAGC
ACACATCATCGGACTCATCAGGAAGC
GCATCGGACTCATCAGGAAGC
GAGGGCATCGGACTCATCAGGAAGC
CATTCATCGGACTCATCAGGAAGC
AGGTCATCGGACTCATCAGGAAGC
GTCATCGGACTCATCAGGAAGC
TGCCATCGGACTCATCAGGAAGC
CCATCGGACTCATCAGGAAGC
CCATCGGACTCATCAGGAAGC
ATTTCATCGGACTCATCAGGAAGC
TTCCATCGGACTCATCAGGAAGC
CATACATCGGACTCATCAGGAAGC
CCACATCGGACTCATCAGGAAGC
GGACATCGGACTCATCAGGAAGC
GCCATCGGACTCATCAGGAAGC
CACATCGGACTCATCAGGAAGC
CATCGGACTCATCAGGAAGC
GCATCGGACTCATCAGGAAGC
TCACATCGGACTCATCAGGAAGC
GATGGCACATCGGACTCATCAGGAAGC

ACATCGGACTCATCAGGAAGC
AGCACATCGGACTCATCAGGAAGC
TTCATCGGACTCATCAGGAAGC
CATCGGACTCATCAGGAAGC
CCATCGGACTCATCAGGAAGC
CATCGGACTCATCAGGAAGC
GACATCGGACTCATCAGGAAGC
GTTCATCGGACTCATCAGGAAGC
ACATCGGACTCATCAGGAAGC
ATGCATCGGACTCATCAGGAAGC
GTTCTCCATCGGACTCATCAGGAAGC
TCATCGGACTCATCAGGAAGC
CCATCGGACTCATCAGGAAGC
AGACATCGGACTCATCAGGAAGC
TCCATCGGACTCATCAGGAAGC
TCATCGGACTCATCAGGAAGC
GCATCGGACTCATCAGGAAGC
CACGCATCGGACTCATCAGGAAGC
ACACATCGGACTCATCAGGAAGC
CTCATCGGACTCATCAGGAAGC
CACATCGGACTCATCAGGAAGC
TGATCATCGGACTCATCAGGAAGC
TTGGAGCATCGGACTCATCAGGAAGC
AGAACATCGGACTCATCAGGAAGC
TGCCATCGGACTCATCAGGAAGC
GGTCATCGGACTCATCAGGAAGC
AATCATCGGACTCATCAGGAAGC
ATCGGACTCATCAGGAAGC
CATCGGACTCATCAGGAAGC
ACACATCGGACTCATCAGGAAGC
GAACATCGGACTCATCAGGAAGC
CTTCATCGGACTCATCAGGAAGC
GACATCGGACTCATCAGGAAGC
TACGCATCGGACTCATCAGGAAGC
ATTTCATCGGACTCATCAGGAAGC
GCCATCGGACTCATCAGGAAGC
TGTCATCGGACTCATCAGGAAGC
TTCATCGGACTCATCAGGAAGC
GCATCGGACTCATCAGGAAGC
CCATCGGACTCATCAGGAAGC
TCACATCGGACTCATCAGGAAGC
TCATCGGACTCATCAGGAAGC
CATCGGACTCATCAGGAAGC

CTGCATCGGACTCATCAGGAAGC
CTTACATCGGACTCATCAGGAAGC
ATCGGACTCATCAGGAAGC
GTGCATCGGACTCATCAGGAAGC
GGCCATCGGACTCATCAGGAAGC
AAACATCGGACTCATCAGGAAGC
ATCATCGGACTCATCAGGAAGC
TACATCGGACTCATCAGGAAGC
TTCATCGGACTCATCAGGAAGC
GACTCATCAGGAAGC
CTGCATCGGACTCATCAGGAAGC
GTCATCGGACTCATCAGGAAGC
ACATCGGACTCATCAGGAAGC
TCCATCATCGGACTCATCAGGAAGC
TCACATCGGACTCATCAGGAAGC
GTTCATCGGACTCATCAGGAAGC
GCCATCGGACTCATCAGGAAGC
GCATCGGACTCATCAGGAAGC
TGTCATCGGACTCATCAGGAAGC
GTCCATCGGACTCATCAGGAAGC
CACATCGGACTCATCAGGAAGC
AGCATCGGACTCATCAGGAAGC
TCGGACTCATCAGGAAGC
TCATCGGACTCATCAGGAAGC
AGGTCATCGGACTCATCAGGAAGC
GCCATCGGACTCATCAGGAAGC
CGCATCGGACTCATCAGGAAGC
CATCGGACTCATCAGGAAGC
TCATCGGACTCATCAGGAAGC
CATCGGACTCATCAGGAAGC
GGCATCGGACTCATCAGGAAGC
GGCATCGGACTCATCAGGAAGC
ACCATCGGACTCATCAGGAAGC
ATCATCGGACTCATCAGGAAGC
CACATCGGACTCATCAGGAAGC
ACATCGGACTCATCAGGAAGC
GATGCCATCGGACTCATCAGGAAGC
TCCATCGGACTCATCAGGAAGC
CCATCGGACTCATCAGGAAGC
GCATCGGACTCATCAGGAAGC
CGCATCGGACTCATCAGGAAGC
TGTTTCATCGGACTCATCAGGAAGC
AAGCATCGGACTCATCAGGAAGC

CAAGCATCGGACTCATCAGGAAGC
GGCCATCGGACTCATCAGGAAGC
GTGCCATCGGACTCATCAGGAAGC
ATGGCATCGGACTCATCAGGAAGC
GCACATCGGACTCATCAGGAAGC
CATCGGACTCATCAGGAAGC
ACATCGGACTCATCAGGAAGC
AGACATCGGACTCATCAGGAAGC
TCATCGGACTCATCAGGAAGC
TTTCATCGGACTCATCAGGAAGC
CCATCGGACTCATCAGGAAGC
TCAGCATCGGACTCATCAGGAAGC
TCGGACTCATCAGGAAGC
AGCATCGGACTCATCAGGAAGC
ACATCGGACTCATCAGGAAGC
ACATCGGACTCATCAGGAAGC
GCATCGGACTCATCAGGAAGC
TGGCATCGGACTCATCAGGAAGC
CCCATCGGACTCATCAGGAAGC
CATCGGACTCATCAGGAAGC
GACATCGGACTCATCAGGAAGC
ACATCGGACTCATCAGGAAGC
ATCGGACTCATCAGGAAGC
TGCATCGGACTCATCAGGAAGC
CTCATCGGACTCATCAGGAAGC
GCATCGGACTCATCAGGAAGC
TTCATCGGACTCATCAGGAAGC
AAACATCGGACTCATCAGGAAGC
AAGCATCGGACTCATCAGGAAGC
TGACATCGGACTCATCAGGAAGC
GCCATCGGACTCATCAGGAAGC
ATGCCATCGGACTCATCAGGAAGC
TCGGACTCATCAGGAAGC
ATACATCGGACTCATCAGGAAGC
ATGCATCGGACTCATCAGGAAGC
CCATCGGACTCATCAGGAAGC
AACATCGGACTCATCAGGAAGC
GTGCATCGGACTCATCAGGAAGC
GCCATCGGACTCATCAGGAAGC
TAGCTGCATCGGACTCATCAGGAAGC
ATGGTCATCGGACTCATCAGGAAGC
CCATCGGACTCATCAGGAAGC
TTTCATCGGACTCATCAGGAAGC

CGCATCGGACTCATCAGGAAGC
AACATCGGACTCATCAGGAAGC
AGGGCATCGGACTCATCAGGAAGC
CATCGGACTCATCAGGAAGC
CACATCGGACTCATCAGGAAGC
ACCATCGGACTCATCAGGAAGC
GGCATCGGACTCATCAGGAAGC
AGCCATCGGACTCATCAGGAAGC
CCATCGGACTCATCAGGAAGC
GGGCATCGGACTCATCAGGAAGC
CATCGGACTCATCAGGAAGC
ATGTGTGCATCGGACTCATCAGGAAGC
CACATCGGACTCATCAGGAAGC
GGCATCGGACTCATCAGGAAGC
ACATCGGACTCATCAGGAAGC
CTCCCACATCGGACTCATCAGGAAGC
GTCATCGGACTCATCAGGAAGC
ATCGGACTCATCAGGAAGC
GATCATCGGACTCATCAGGAAGC
CTGTCATCGGACTCATCAGGAAGC
GCTCATCGGACTCATCAGGAAGC
ACTCATCGGACTCATCAGGAAGC
ACAGCATCGGACTCATCAGGAAGC
GGCATCGGACTCATCAGGAAGC
ATCGGACTCATCAGGAAGC
TACTCATCGGACTCATCAGGAAGC
CCATCGGACTCATCAGGAAGC
GTCATCGGACTCATCAGGAAGC
GGCATCGGACTCATCAGGAAGC
CCATCGGACTCATCAGGAAGC
TACCCATCGGACTCATCAGGAAGC
GTCATCGGACTCATCAGGAAGC
TGACATCGGACTCATCAGGAAGC
GCACATCGGACTCATCAGGAAGC
CACATCGGACTCATCAGGAAGC
GGATCATCGGACTCATCAGGAAGC
TTCCATCGGACTCATCAGGAAGC
CATCGGACTCATCAGGAAGC
CCATCGGACTCATCAGGAAGC
ACCATCGGACTCATCAGGAAGC
TAAGCACATCGGACTCATCAGGAAGC
GCCATCGGACTCATCAGGAAGC
CTCCATCGGACTCATCAGGAAGC

TTCATCGGACTCATCAGGAAGC
TGAACATCGGACTCATCAGGAAGC
TACATCGGACTCATCAGGAAGC
TCATCGGACTCATCAGGAAGC
ACATCGGACTCATCAGGAAGC
TCATCGGACTCATCAGGAAGC
CCATCGGACTCATCAGGAAGC
GCCATCGGACTCATCAGGAAGC
TTCATCGGACTCATCAGGAAGC
AACATCGGACTCATCAGGAAGC
ACACACATCGGACTCATCAGGAAGC
GGCATCGGACTCATCAGGAAGC
CATCGGACTCATCAGGAAGC
CCATCGGACTCATCAGGAAGC
CTCATCGGACTCATCAGGAAGC
AGGCATCGGACTCATCAGGAAGC
GTCATCGGACTCATCAGGAAGC
TCCTCATCGGACTCATCAGGAAGC
GGACATCGGACTCATCAGGAAGC
CATCGGACTCATCAGGAAGC
AGCATCGGACTCATCAGGAAGC
TCATCGGACTCATCAGGAAGC
CATCGGACTCATCAGGAAGC
TCATCGGACTCATCAGGAAGC
AACATCGGACTCATCAGGAAGC
CATCGGACTCATCAGGAAGC
ACATCGGACTCATCAGGAAGC
TGCATCGGACTCATCAGGAAGC
CATCGGACTCATCAGGAAGC
CATCGGACTCATCAGGAAGC
AGTCATCGGACTCATCAGGAAGC
ATCATCGGACTCATCAGGAAGC
CATCGGACTCATCAGGAAGC
TATGCATCGGACTCATCAGGAAGC
CCATCGGACTCATCAGGAAGC
AACATCGGACTCATCAGGAAGC
CATCGGACTCATCAGGAAGC
TTCAGCCATCGGACTCATCAGGAAGC
CCCCATCGGACTCATCAGGAAGC
TATCCATCGGACTCATCAGGAAGC
TGACATCGGACTCATCAGGAAGC
CATCGGACTCATCAGGAAGC
TCATCGGACTCATCAGGAAGC

GACATCGGACTCATCAGGAAGC
AAGGTTCATCGGACTCATCAGGAAGC
CAATACATCGGACTCATCAGGAAGC
ACAGACATCGGACTCATCAGGAAGC
GCTTCATCGGACTCATCAGGAAGC
TCTCATCGGACTCATCAGGAAGC
GCATCGGACTCATCAGGAAGC
ACATCGGACTCATCAGGAAGC
TTTCATCGGACTCATCAGGAAGC
TCCTCATCGGACTCATCAGGAAGC
CGGACATCGGACTCATCAGGAAGC
AGGTCATCGGACTCATCAGGAAGC
TGACATCGGACTCATCAGGAAGC
GTCCATCGGACTCATCAGGAAGC
TCATCGGACTCATCAGGAAGC
GAACATCGGACTCATCAGGAAGC
TGAACATCGGACTCATCAGGAAGC
GCATCGGACTCATCAGGAAGC
AATAACATCGGACTCATCAGGAAGC
ACATCGGACTCATCAGGAAGC
CTTCATCGGACTCATCAGGAAGC
CTCATCGGACTCATCAGGAAGC
CATCGGACTCATCAGGAAGC
TTCATCGGACTCATCAGGAAGC
GACATCGGACTCATCAGGAAGC
TTCATCGGACTCATCAGGAAGC
TTTCATCGGACTCATCAGGAAGC
ACTCATCGGACTCATCAGGAAGC
GCCACATCGGACTCATCAGGAAGC
CATCGGACTCATCAGGAAGC
TGCCATCGGACTCATCAGGAAGC
CATCGGACTCATCAGGAAGC
CAACATCGGACTCATCAGGAAGC
TAAACATCGGACTCATCAGGAAGC
GGACATCGGACTCATCAGGAAGC
TGGGCATCGGACTCATCAGGAAGC
ACCCATCGGACTCATCAGGAAGC
GCTCATCGGACTCATCAGGAAGC
GGGCATCGGACTCATCAGGAAGC
ACCTCATCGGACTCATCAGGAAGC
CCATCGGACTCATCAGGAAGC
AGCATCGGACTCATCAGGAAGC
ACATCGGACTCATCAGGAAGC

GGCTCATCGGACTCATCAGGAAGC
TCCATCGGACTCATCAGGAAGC
GCACATCGGACTCATCAGGAAGC
TCATCGGACTCATCAGGAAGC
CTCATCGGACTCATCAGGAAGC
CATCGGACTCATCAGGAAGC
CACTACATCGGACTCATCAGGAAGC
TCGGACTCATCAGGAAGC
CCATCGGACTCATCAGGAAGC
ATCATCGGACTCATCAGGAAGC
GGCATCGGACTCATCAGGAAGC
TGACATCGGACTCATCAGGAAGC
GAGCATCGGACTCATCAGGAAGC
CATCGGACTCATCAGGAAGC
TATCATCGGACTCATCAGGAAGC
CATCGGACTCATCAGGAAGC
TCCATCGGACTCATCAGGAAGC
AGGCCATCGGACTCATCAGGAAGC
TCATCGGACTCATCAGGAAGC
ATCATCGGACTCATCAGGAAGC
ATCGGACTCATCAGGAAGC
GGCATCGGACTCATCAGGAAGC
CATCGGACTCATCAGGAAGC
TGGCATCGGACTCATCAGGAAGC
TACATCGGACTCATCAGGAAGC
TACATCGGACTCATCAGGAAGC
CACATCGGACTCATCAGGAAGC
GTCATCGGACTCATCAGGAAGC
TGACATCGGACTCATCAGGAAGC
TTTCATCGGACTCATCAGGAAGC
ACATCGGACTCATCAGGAAGC
GACATCGGACTCATCAGGAAGC
CATCGGACTCATCAGGAAGC
TCATCGGACTCATCAGGAAGC
AGACATCGGACTCATCAGGAAGC
CATCGGACTCATCAGGAAGC
TGCATCGGACTCATCAGGAAGC
CATCGGACTCATCAGGAAGC
AGCCATCGGACTCATCAGGAAGC
CTTCATCGGACTCATCAGGAAGC
GGAGCCATCGGACTCATCAGGAAGC
TTCATCGGACTCATCAGGAAGC
ACATCGGACTCATCAGGAAGC

GGCATCGGACTCATCAGGAAGC
AACATCGGACTCATCAGGAAGC
ATCGGACTCATCAGGAAGC
CACATCGGACTCATCAGGAAGC
CTTCATCGGACTCATCAGGAAGC
ACATCGGACTCATCAGGAAGC
ACATCGGACTCATCAGGAAGC
AGAGGACATCGGACTCATCAGGAAGC
GTCATCGGACTCATCAGGAAGC
ACATCGGACTCATCAGGAAGC
CATCATCGGACTCATCAGGAAGC
GGCATCGGACTCATCAGGAAGC
ATCGGACTCATCAGGAAGC
TGGCATCGGACTCATCAGGAAGC
TTGCATCGGACTCATCAGGAAGC
TTTGCATCGGACTCATCAGGAAGC
CATCGGACTCATCAGGAAGC
GAGCATCGGACTCATCAGGAAGC
CTCCATCGGACTCATCAGGAAGC
ACATCGGACTCATCAGGAAGC
CATCGGACTCATCAGGAAGC
CATCGGACTCATCAGGAAGC
GATCCCACATCGGACTCATCAGGAAGC
AGCACATCGGACTCATCAGGAAGC
CAGCATCGGACTCATCAGGAAGC
CACCATCGGACTCATCAGGAAGC
GTCATCGGACTCATCAGGAAGC
GGCATCGGACTCATCAGGAAGC
GTCATCGGACTCATCAGGAAGC
CTCATCGGACTCATCAGGAAGC
ATCATCGGACTCATCAGGAAGC
CATCGGACTCATCAGGAAGC
GCATCGGACTCATCAGGAAGC
CATCGGACTCATCAGGAAGC
CCCCTCATCGGACTCATCAGGAAGC
TACATCGGACTCATCAGGAAGC
TTCATCGGACTCATCAGGAAGC
TCATCGGACTCATCAGGAAGC
TCCATCGGACTCATCAGGAAGC
TACTCATCGGACTCATCAGGAAGC
TGCATCGGACTCATCAGGAAGC
GCCATCGGACTCATCAGGAAGC
ACATCGGACTCATCAGGAAGC

CATCGGACTCATCAGGAAGC
CATCGGACTCATCAGGAAGC
CATCGGACTCATCAGGAAGC
CCATCGGACTCATCAGGAAGC
GGTCCATCGGACTCATCAGGAAGC
GACATCGGACTCATCAGGAAGC
TTTCATCGGACTCATCAGGAAGC
GCATCGGACTCATCAGGAAGC
ACAAAACATCGGACTCATCAGGAAGC
CACATCGGACTCATCAGGAAGC
CATCGGACTCATCAGGAAGC
ATGGGGCATCGGACTCATCAGGAAGC
TGCCATCGGACTCATCAGGAAGC
CCCATCGGACTCATCAGGAAGC
TCATCGGACTCATCAGGAAGC
GCATCGGACTCATCAGGAAGC
AAGCCATCGGACTCATCAGGAAGC
CTGTCATCGGACTCATCAGGAAGC
CAGCATCATCGGACTCATCAGGAAGC
ACATCGGACTCATCAGGAAGC
AGCATCGGACTCATCAGGAAGC
CCATCGGACTCATCAGGAAGC
GGACCATCGGACTCATCAGGAAGC
CATCGGACTCATCAGGAAGC
GGCCATCGGACTCATCAGGAAGC
TCATCGGACTCATCAGGAAGC
TCATCGGACTCATCAGGAAGC
GAATTTGCATCGGACTCATCAGGAAGC
TGCATCGGACTCATCAGGAAGC
GTCATCGGACTCATCAGGAAGC
GCTGTCATCGGACTCATCAGGAAGC
GCCATCGGACTCATCAGGAAGC
CAAACATCGGACTCATCAGGAAGC
TCATCGGACTCATCAGGAAGC
TTCATCGGACTCATCAGGAAGC
ACATCGGACTCATCAGGAAGC
AGGCATCGGACTCATCAGGAAGC
CAAAACATCGGACTCATCAGGAAGC
TTCATCGGACTCATCAGGAAGC
TACATCGGACTCATCAGGAAGC
TTCATCGGACTCATCAGGAAGC
GCTACACATCGGACTCATCAGGAAGC
AATCATCGGACTCATCAGGAAGC

GCACATCGGACTCATCAGGAAGC
ATCGGACTCATCAGGAAGC
CCATCGGACTCATCAGGAAGC
AGTCATCGGACTCATCAGGAAGC
GTGGACATCGGACTCATCAGGAAGC
CATCGGACTCATCAGGAAGC
CACATCGGACTCATCAGGAAGC
GTCATCGGACTCATCAGGAAGC
TCCATCGGACTCATCAGGAAGC
CCATCGGACTCATCAGGAAGC
CATACATCGGACTCATCAGGAAGC
TCATCGGACTCATCAGGAAGC
AGCATCGGACTCATCAGGAAGC
CATCGGACTCATCAGGAAGC
AAACATCGGACTCATCAGGAAGC
CATCGGACTCATCAGGAAGC
AGACATCGGACTCATCAGGAAGC
GTCATCGGACTCATCAGGAAGC
ATCGGACTCATCAGGAAGC
CACATCGGACTCATCAGGAAGC
TTATTCATCGGACTCATCAGGAAGC
ACATCGGACTCATCAGGAAGC
TAACCATCGGACTCATCAGGAAGC
TAGCATCGGACTCATCAGGAAGC
CCATCGGACTCATCAGGAAGC
CCATCGGACTCATCAGGAAGC
TTTCATCGGACTCATCAGGAAGC
GTCATCGGACTCATCAGGAAGC
CATCGGACTCATCAGGAAGC
AAAGCATCGGACTCATCAGGAAGC
TTCACATCGGACTCATCAGGAAGC
CATCGGACTCATCAGGAAGC
GCAACATCGGACTCATCAGGAAGC
CATCGGACTCATCAGGAAGC
ATGCCCATCGGACTCATCAGGAAGC
CATCGGACTCATCAGGAAGC
GAGCATCGGACTCATCAGGAAGC
AATCATCGGACTCATCAGGAAGC
GCCATCGGACTCATCAGGAAGC
TGCATCGGACTCATCAGGAAGC
TGCATCGGACTCATCAGGAAGC
GCATCGGACTCATCAGGAAGC
CACATCGGACTCATCAGGAAGC

GTCATCGGACTCATCAGGAAGC
CATCCATCGGACTCATCAGGAAGC
TGTCATCGGACTCATCAGGAAGC
GATCATCGGACTCATCAGGAAGC
TTCATCGGACTCATCAGGAAGC
TCATCGGACTCATCAGGAAGC
CATCGGACTCATCAGGAAGC
ACATCGGACTCATCAGGAAGC
TCATCGGACTCATCAGGAAGC
ATGCCCATCGGACTCATCAGGAAGC
TGAACCATCGGACTCATCAGGAAGC
TGACTACATCGGACTCATCAGGAAGC
GGCATCGGACTCATCAGGAAGC
CATCGGACTCATCAGGAAGC
TCATCGGACTCATCAGGAAGC
GAACATCGGACTCATCAGGAAGC
CATCGGACTCATCAGGAAGC
GCATCGGACTCATCAGGAAGC
ATCGGACTCATCAGGAAGC
TCATCGGACTCATCAGGAAGC
GAGCATCGGACTCATCAGGAAGC
ACACATCGGACTCATCAGGAAGC
TCATCGGACTCATCAGGAAGC
GAGCATCGGACTCATCAGGAAGC
TCATCGGACTCATCAGGAAGC
TGGACATCGGACTCATCAGGAAGC
GTCATCGGACTCATCAGGAAGC
GTCATCGGACTCATCAGGAAGC
TAAACATCGGACTCATCAGGAAGC
TTCCATCGGACTCATCAGGAAGC
TTCCATCGGACTCATCAGGAAGC
ACATCGGACTCATCAGGAAGC
ACATCGGACTCATCAGGAAGC
AGAGCATCGGACTCATCAGGAAGC
TGTCCCATCGGACTCATCAGGAAGC
TCATCGGACTCATCAGGAAGC
ATCATCGGACTCATCAGGAAGC
TGACATCGGACTCATCAGGAAGC
GTGCATCGGACTCATCAGGAAGC
TCATCGGACTCATCAGGAAGC
ATCGGACTCATCAGGAAGC
CCCATCGGACTCATCAGGAAGC
AGACATCGGACTCATCAGGAAGC

GAAACATCGGACTCATCAGGAAGC
TGTGCATCGGACTCATCAGGAAGC
GCATCGGACTCATCAGGAAGC
GCATCGGACTCATCAGGAAGC
CATCGGACTCATCAGGAAGC
ACATCGGACTCATCAGGAAGC
CATCGGACTCATCAGGAAGC
AATGCATCGGACTCATCAGGAAGC
TCATCGGACTCATCAGGAAGC
CATCGGACTCATCAGGAAGC
AAGTCATCGGACTCATCAGGAAGC
CATCGGACTCATCAGGAAGC
CCATCGGACTCATCAGGAAGC
CCATCGGACTCATCAGGAAGC
TGCATCGGACTCATCAGGAAGC
AAACATCGGACTCATCAGGAAGC
AGCATCGGACTCATCAGGAAGC
GCATCGGACTCATCAGGAAGC
AAGCCCATCGGACTCATCAGGAAGC
AGGCATCGGACTCATCAGGAAGC
GTGCATCGGACTCATCAGGAAGC
GCCATCGGACTCATCAGGAAGC
GTCATCGGACTCATCAGGAAGC
GCATCGGACTCATCAGGAAGC
GCATCGGACTCATCAGGAAGC
AACTCCATCGGACTCATCAGGAAGC
GAAAACATCGGACTCATCAGGAAGC
CTCCCATCGGACTCATCAGGAAGC
ACACATCGGACTCATCAGGAAGC
CATCGGACTCATCAGGAAGC
GTTGCCATCGGACTCATCAGGAAGC
ACCATCGGACTCATCAGGAAGC
CTGGCATCGGACTCATCAGGAAGC
TTTCATCGGACTCATCAGGAAGC
CATCGGACTCATCAGGAAGC
AACACATCGGACTCATCAGGAAGC
CCACATCGGACTCATCAGGAAGC
TCCATCGGACTCATCAGGAAGC
CATCGGACTCATCAGGAAGC
TCATCGGACTCATCAGGAAGC
ACATCGGACTCATCAGGAAGC
GGACATCGGACTCATCAGGAAGC
TCATCGGACTCATCAGGAAGC

GACATCGGACTCATCAGGAAGC
TCGGACTCATCAGGAAGC
CTTCATCGGACTCATCAGGAAGC
CCCCATCGGACTCATCAGGAAGC
TTGTCATCGGACTCATCAGGAAGC
CATCGGACTCATCAGGAAGC
GCCATCGGACTCATCAGGAAGC
CCCATCGGACTCATCAGGAAGC
ACACATCGGACTCATCAGGAAGC
GACATCGGACTCATCAGGAAGC
CTACATCGGACTCATCAGGAAGC
GTTACATCGGACTCATCAGGAAGC
GCTCATCGGACTCATCAGGAAGC
CCATCGGACTCATCAGGAAGC
CAATCATCGGACTCATCAGGAAGC
TACATCGGACTCATCAGGAAGC
ACATCGGACTCATCAGGAAGC
CATCGGACTCATCAGGAAGC
GGATCATCGGACTCATCAGGAAGC
CATCGGACTCATCAGGAAGC
TACATCGGACTCATCAGGAAGC
TACATCGGACTCATCAGGAAGC
GTCATCGGACTCATCAGGAAGC
CCGCATCGGACTCATCAGGAAGC
ACACACCCATCGGACTCATCAGGAAGC
GCCATCGGACTCATCAGGAAGC
CATCGGACTCATCAGGAAGC
CTTTCATCGGACTCATCAGGAAGC
TTGTCATCGGACTCATCAGGAAGC
GGACATCGGACTCATCAGGAAGC
TCATCGGACTCATCAGGAAGC
ATCGGACTCATCAGGAAGC
TCATCGGACTCATCAGGAAGC
ATGTCATCGGACTCATCAGGAAGC
AGCATCGGACTCATCAGGAAGC
TGGATCATCGGACTCATCAGGAAGC
TAGGCACATCGGACTCATCAGGAAGC
TGTCATCGGACTCATCAGGAAGC
AGGTCATCGGACTCATCAGGAAGC
GCACACATCGGACTCATCAGGAAGC
GCATCGGACTCATCAGGAAGC
AAGCATCGGACTCATCAGGAAGC
CATCGGACTCATCAGGAAGC

GCCATCGGACTCATCAGGAAGC
TGCATCGGACTCATCAGGAAGC
CATCATCGGACTCATCAGGAAGC
GCCATCGGACTCATCAGGAAGC
TCTCATCGGACTCATCAGGAAGC
CACGCATCGGACTCATCAGGAAGC
GCCATCGGACTCATCAGGAAGC
CATCGGACTCATCAGGAAGC
ACATCGGACTCATCAGGAAGC
TAACATCGGACTCATCAGGAAGC
CGCATCGGACTCATCAGGAAGC
CACCTCATCGGACTCATCAGGAAGC
ACATCGGACTCATCAGGAAGC
ATCACATCGGACTCATCAGGAAGC
CATCGGACTCATCAGGAAGC
CACATCGGACTCATCAGGAAGC
ACATCGGACTCATCAGGAAGC
ACATCGGACTCATCAGGAAGC
TAGCCATCGGACTCATCAGGAAGC
GGTACATCGGACTCATCAGGAAGC
ATCATCGGACTCATCAGGAAGC
CTGTCATCGGACTCATCAGGAAGC
CACATCGGACTCATCAGGAAGC
TGCATCGGACTCATCAGGAAGC
TCATCGGACTCATCAGGAAGC
CCCATCGGACTCATCAGGAAGC
GGCATCGGACTCATCAGGAAGC
ACCATCGGACTCATCAGGAAGC
CATCGGACTCATCAGGAAGC
CCATCGGACTCATCAGGAAGC
GGGTCATCGGACTCATCAGGAAGC
TGGGCATCGGACTCATCAGGAAGC
GCCATCGGACTCATCAGGAAGC
CATCGGACTCATCAGGAAGC
CATCGGACTCATCAGGAAGC
CCATCGGACTCATCAGGAAGC
CATCATCGGACTCATCAGGAAGC
ATCGGACTCATCAGGAAGC
CCATCGGACTCATCAGGAAGC
GCATCGGACTCATCAGGAAGC
GCATCGGACTCATCAGGAAGC
ATCGGACTCATCAGGAAGC
CATCGGACTCATCAGGAAGC

TCATCGGACTCATCAGGAAGC
TCTCTCATCGGACTCATCAGGAAGC
GACATCGGACTCATCAGGAAGC
CACACATCGGACTCATCAGGAAGC
AGCATCGGACTCATCAGGAAGC
TCATCGGACTCATCAGGAAGC
TTGCTCATCGGACTCATCAGGAAGC
CCATCGGACTCATCAGGAAGC
GGGGCATCGGACTCATCAGGAAGC
GCCATCGGACTCATCAGGAAGC
AGCATCGGACTCATCAGGAAGC
CAGCATCGGACTCATCAGGAAGC
CATCATCGGACTCATCAGGAAGC
CATCGGACTCATCAGGAAGC
GCATCGGACTCATCAGGAAGC
GTACATCGGACTCATCAGGAAGC
GGACATCGGACTCATCAGGAAGC
TCTCATCGGACTCATCAGGAAGC
AGTGCATCGGACTCATCAGGAAGC
GGAAACATCGGACTCATCAGGAAGC
AAACATCGGACTCATCAGGAAGC
TCATCGGACTCATCAGGAAGC
AGGGCATCGGACTCATCAGGAAGC
AGCCATCGGACTCATCAGGAAGC
CCATCGGACTCATCAGGAAGC
AGCATCGGACTCATCAGGAAGC
GGCATCGGACTCATCAGGAAGC
CATCGGACTCATCAGGAAGC
GAACATCGGACTCATCAGGAAGC
AGCATCGGACTCATCAGGAAGC
CATCGGACTCATCAGGAAGC
AGACATCGGACTCATCAGGAAGC
ACATCGGACTCATCAGGAAGC
CCATCGGACTCATCAGGAAGC
CTCATCGGACTCATCAGGAAGC
ACATCGGACTCATCAGGAAGC
TCATCGGACTCATCAGGAAGC
CTTCATCGGACTCATCAGGAAGC
AGCATCGGACTCATCAGGAAGC
ACTCATCGGACTCATCAGGAAGC
CACATCGGACTCATCAGGAAGC
GCCATCGGACTCATCAGGAAGC
ACCCATCGGACTCATCAGGAAGC

CCATCGGACTCATCAGGAAGC
CATCGGACTCATCAGGAAGC
TGCGCATCGGACTCATCAGGAAGC
AACATCGGACTCATCAGGAAGC
CATCGGACTCATCAGGAAGC
GATGGCCATCGGACTCATCAGGAAGC
ATCGGACTCATCAGGAAGC
TCCATCGGACTCATCAGGAAGC
GACATCGGACTCATCAGGAAGC
AACATCGGACTCATCAGGAAGC
CCATCGGACTCATCAGGAAGC
TGTCATCGGACTCATCAGGAAGC
AGCATCGGACTCATCAGGAAGC
GCACATCGGACTCATCAGGAAGC
GACATCGGACTCATCAGGAAGC
TGCATCGGACTCATCAGGAAGC
TCATCGGACTCATCAGGAAGC
ACATCGGACTCATCAGGAAGC
ATCGGACTCATCAGGAAGC
CATCGGACTCATCAGGAAGC
TTGCATCGGACTCATCAGGAAGC
GACATCGGACTCATCAGGAAGC
TATGCATCGGACTCATCAGGAAGC
CAAGGCATCGGACTCATCAGGAAGC
CTCCATCGGACTCATCAGGAAGC
CCCATCGGACTCATCAGGAAGC
ATTCATCGGACTCATCAGGAAGC
GTCACATCGGACTCATCAGGAAGC
AACATCGGACTCATCAGGAAGC
TTCATCGGACTCATCAGGAAGC
CCATCGGACTCATCAGGAAGC
TACATCGGACTCATCAGGAAGC
TCATCGGACTCATCAGGAAGC
GTCATCGGACTCATCAGGAAGC
AGCATCGGACTCATCAGGAAGC
ATCGGACTCATCAGGAAGC
AATCATCGGACTCATCAGGAAGC
ACATCGGACTCATCAGGAAGC
CTTCCATCGGACTCATCAGGAAGC
GCCTCATCGGACTCATCAGGAAGC
CATCGGACTCATCAGGAAGC
ACATCGGACTCATCAGGAAGC
ACATCGGACTCATCAGGAAGC

CCCATCGGACTCATCAGGAAGC
CCATCGGACTCATCAGGAAGC
CATCGGACTCATCAGGAAGC
GACATCGGACTCATCAGGAAGC
TGACATCGGACTCATCAGGAAGC
CACCTGCATCGGACTCATCAGGAAGC
TCATCGGACTCATCAGGAAGC
TCATCGGACTCATCAGGAAGC
GTCATCGGACTCATCAGGAAGC
TCATCGGACTCATCAGGAAGC
CATCGGACTCATCAGGAAGC
ACATCGGACTCATCAGGAAGC
CCCATCGGACTCATCAGGAAGC
TTTCTCATCGGACTCATCAGGAAGC
CTCCATCGGACTCATCAGGAAGC
AGCCCCATCGGACTCATCAGGAAGC
AAATCATCGGACTCATCAGGAAGC
GACATCGGACTCATCAGGAAGC
TGTCATCGGACTCATCAGGAAGC
TCATCGGACTCATCAGGAAGC
CATCATCGGACTCATCAGGAAGC
ATCATCGGACTCATCAGGAAGC
GCGCATCGGACTCATCAGGAAGC
GTCATCGGACTCATCAGGAAGC
TCATCGGACTCATCAGGAAGC
GACCATCATCGGACTCATCAGGAAGC
GCCATCGGACTCATCAGGAAGC
AGGACATCGGACTCATCAGGAAGC
GACCATCGGACTCATCAGGAAGC
TCGCTCATCGGACTCATCAGGAAGC
GGCCATCGGACTCATCAGGAAGC
GGTCATCGGACTCATCAGGAAGC
ACCATCGGACTCATCAGGAAGC
AGCCATCGGACTCATCAGGAAGC
TCACATCGGACTCATCAGGAAGC
TAGGCCATCGGACTCATCAGGAAGC
CATCGGACTCATCAGGAAGC
GCCATCGGACTCATCAGGAAGC
GTCATCGGACTCATCAGGAAGC
AAGACATCGGACTCATCAGGAAGC
GTGCATCGGACTCATCAGGAAGC
TTCATCGGACTCATCAGGAAGC
CAGACATCGGACTCATCAGGAAGC

TCATCGGACTCATCAGGAAGC
GTGCCATCGGACTCATCAGGAAGC
ATGCATCGGACTCATCAGGAAGC
TCATCGGACTCATCAGGAAGC
CCCCATCGGACTCATCAGGAAGC
AACATCGGACTCATCAGGAAGC
TGCATCGGACTCATCAGGAAGC
ATCGGACTCATCAGGAAGC
GGCAAACATCGGACTCATCAGGAAGC
CCATCGGACTCATCAGGAAGC
AAGCATCGGACTCATCAGGAAGC
AGCCATCGGACTCATCAGGAAGC
TGGCATCGGACTCATCAGGAAGC
CCATCGGACTCATCAGGAAGC
TCTCATCGGACTCATCAGGAAGC
TCCATCGGACTCATCAGGAAGC
CCATCGGACTCATCAGGAAGC
CCCATCGGACTCATCAGGAAGC
CATCATCGGACTCATCAGGAAGC
GTGGCATCGGACTCATCAGGAAGC
TCATCGGACTCATCAGGAAGC
AAAGCCATCGGACTCATCAGGAAGC
CATCGGACTCATCAGGAAGC
ATCGGACTCATCAGGAAGC
CCATCGGACTCATCAGGAAGC
CACCATCGGACTCATCAGGAAGC
TCATCGGACTCATCAGGAAGC
CAGGCATCGGACTCATCAGGAAGC
ATAACATCGGACTCATCAGGAAGC
AACATCGGACTCATCAGGAAGC
TGCATCGGACTCATCAGGAAGC
CATCGGACTCATCAGGAAGC
AGGTCATCGGACTCATCAGGAAGC
AGCATCGGACTCATCAGGAAGC
CCCATCGGACTCATCAGGAAGC
GCATCGGACTCATCAGGAAGC
TTCATCGGACTCATCAGGAAGC
GAGCATCGGACTCATCAGGAAGC
GGCATCGGACTCATCAGGAAGC
AAGACATCGGACTCATCAGGAAGC
CATCGGACTCATCAGGAAGC
TCATCGGACTCATCAGGAAGC
GGAAACATCGGACTCATCAGGAAGC

CTCATCGGACTCATCAGGAAGC
ACATTCATCGGACTCATCAGGAAGC
CCATCGGACTCATCAGGAAGC
CCATCGGACTCATCAGGAAGC
TCATCGGACTCATCAGGAAGC
CATCGGACTCATCAGGAAGC
CTCATCGGACTCATCAGGAAGC
CGCATCGGACTCATCAGGAAGC
ACATCGGACTCATCAGGAAGC
AGACATCGGACTCATCAGGAAGC
CATCGGACTCATCAGGAAGC
CATCGGACTCATCAGGAAGC
GTGCATCGGACTCATCAGGAAGC
GCATCGGACTCATCAGGAAGC
ACATCGGACTCATCAGGAAGC
TGCAACATCGGACTCATCAGGAAGC
TCAGCATCGGACTCATCAGGAAGC
GCCATCGGACTCATCAGGAAGC
TTCATCGGACTCATCAGGAAGC
ATCGGACTCATCAGGAAGC
GCCCATCGGACTCATCAGGAAGC
GACATCGGACTCATCAGGAAGC
TTTTCATCGGACTCATCAGGAAGC
GCATCGGACTCATCAGGAAGC
TGGCATCGGACTCATCAGGAAGC
ATCATCGGACTCATCAGGAAGC
TGGCCATCGGACTCATCAGGAAGC
GGCATCGGACTCATCAGGAAGC
GGCATCGGACTCATCAGGAAGC
GGCATCGGACTCATCAGGAAGC
CATCGGACTCATCAGGAAGC
AAGACATCGGACTCATCAGGAAGC
AGCCCATCGGACTCATCAGGAAGC
TACCATCGGACTCATCAGGAAGC
CATCATCGGACTCATCAGGAAGC
GTCCTTCATCGGACTCATCAGGAAGC
AGTCATCGGACTCATCAGGAAGC
GCATCGGACTCATCAGGAAGC
GCATCGGACTCATCAGGAAGC
GGCATCGGACTCATCAGGAAGC
GAGCATCGGACTCATCAGGAAGC
CCATCGGACTCATCAGGAAGC
CTCCATCGGACTCATCAGGAAGC

CTCATCGGACTCATCAGGAAGC
GTACATCGGACTCATCAGGAAGC
TGTCATCGGACTCATCAGGAAGC
CATCGGACTCATCAGGAAGC
CCATCGGACTCATCAGGAAGC
GATCATCGGACTCATCAGGAAGC
ATACATCGGACTCATCAGGAAGC
CCCTACATCGGACTCATCAGGAAGC
GGCCATCGGACTCATCAGGAAGC
TCCATCGGACTCATCAGGAAGC
ATACCATCGGACTCATCAGGAAGC
CTCATCGGACTCATCAGGAAGC
TCACATCGGACTCATCAGGAAGC
CACATCGGACTCATCAGGAAGC
CCATCGGACTCATCAGGAAGC
AGGCATCGGACTCATCAGGAAGC
AGTTCATCGGACTCATCAGGAAGC
TGCATCGGACTCATCAGGAAGC
CCACATCGGACTCATCAGGAAGC
GGCATCGGACTCATCAGGAAGC
ACATCGGACTCATCAGGAAGC
GAACATCGGACTCATCAGGAAGC
GAAACATCGGACTCATCAGGAAGC
ATCGGACTCATCAGGAAGC
CCATCGGACTCATCAGGAAGC
AAACATCGGACTCATCAGGAAGC
AGATCATCGGACTCATCAGGAAGC
CCCATCGGACTCATCAGGAAGC
TGCATCGGACTCATCAGGAAGC
CTCCATCGGACTCATCAGGAAGC
CTCATCGGACTCATCAGGAAGC
CCACATCGGACTCATCAGGAAGC
TGTGTCATCGGACTCATCAGGAAGC
GACATCGGACTCATCAGGAAGC
TACATCGGACTCATCAGGAAGC
CATCGGACTCATCAGGAAGC
CCACATCGGACTCATCAGGAAGC
GCATCGGACTCATCAGGAAGC
GTGCCATCGGACTCATCAGGAAGC
TGTCATCGGACTCATCAGGAAGC
CATCGGACTCATCAGGAAGC
GCATCGGACTCATCAGGAAGC
GAGCTGACATCGGACTCATCAGGAAGC

CTCATCGGACTCATCAGGAAGC
CATCGGACTCATCAGGAAGC
GACATCGGACTCATCAGGAAGC
TGCCATCGGACTCATCAGGAAGC
ATCGGACTCATCAGGAAGC
CAGCATCGGACTCATCAGGAAGC
ACATCGGACTCATCAGGAAGC
CTGACATCGGACTCATCAGGAAGC
ATCGGACTCATCAGGAAGC
GGCTCATCGGACTCATCAGGAAGC
ATCGGACTCATCAGGAAGC
GCATCGGACTCATCAGGAAGC
CACCATCGGACTCATCAGGAAGC
TCATCGGACTCATCAGGAAGC
GAACATCGGACTCATCAGGAAGC
ACATCGGACTCATCAGGAAGC
CCCATCGGACTCATCAGGAAGC
GATCATCGGACTCATCAGGAAGC
GCCATCGGACTCATCAGGAAGC
GCATCGGACTCATCAGGAAGC
CTCATCGGACTCATCAGGAAGC
ACATCGGACTCATCAGGAAGC
CCATCGGACTCATCAGGAAGC
CACATCGGACTCATCAGGAAGC
TCCATCGGACTCATCAGGAAGC
ACATCGGACTCATCAGGAAGC
AAGCGCATCGGACTCATCAGGAAGC
AGTCTCATCGGACTCATCAGGAAGC
ACCCATCGGACTCATCAGGAAGC
ATCATCGGACTCATCAGGAAGC
GGTCATCGGACTCATCAGGAAGC
TTCATCGGACTCATCAGGAAGC
GCCATCGGACTCATCAGGAAGC
ACATCGGACTCATCAGGAAGC
ACCATCGGACTCATCAGGAAGC
ATCGGACTCATCAGGAAGC
GCCCACATCGGACTCATCAGGAAGC
CATCGGACTCATCAGGAAGC
CCATCGGACTCATCAGGAAGC
ATCGGACTCATCAGGAAGC
TGTCATCGGACTCATCAGGAAGC
GGACCATCGGACTCATCAGGAAGC
GGTTGCCATCGGACTCATCAGGAAGC

CTTCATCGGACTCATCAGGAAGC
TCATCGGACTCATCAGGAAGC
ATGGAACATCGGACTCATCAGGAAGC
TCCATCGGACTCATCAGGAAGC
GCATCGGACTCATCAGGAAGC
ATCATCGGACTCATCAGGAAGC
ATCACATCGGACTCATCAGGAAGC
ACACATCGGACTCATCAGGAAGC
GGCATCGGACTCATCAGGAAGC
ACCCATCGGACTCATCAGGAAGC
AGTTCATCGGACTCATCAGGAAGC
CATCGGACTCATCAGGAAGC
GCCCCATCGGACTCATCAGGAAGC
CCACATCGGACTCATCAGGAAGC
TGTCATCGGACTCATCAGGAAGC
GGCATCGGACTCATCAGGAAGC
CATCGGACTCATCAGGAAGC
ACCATCGGACTCATCAGGAAGC
CTTCATCGGACTCATCAGGAAGC
CGCATCGGACTCATCAGGAAGC
TCTCATCGGACTCATCAGGAAGC
ACATCGGACTCATCAGGAAGC
GGCATCGGACTCATCAGGAAGC
CCATCGGACTCATCAGGAAGC
CCACATCGGACTCATCAGGAAGC
GTTCATCGGACTCATCAGGAAGC
GTGCCATCGGACTCATCAGGAAGC
GGAACATCGGACTCATCAGGAAGC
ATCATCGGACTCATCAGGAAGC
AGCTTTCATCGGACTCATCAGGAAGC
TCTTCATCGGACTCATCAGGAAGC
GACAACATCGGACTCATCAGGAAGC
CCCATCGGACTCATCAGGAAGC
ACTACATCGGACTCATCAGGAAGC
GACATCGGACTCATCAGGAAGC
CCATCGGACTCATCAGGAAGC
TCATCGGACTCATCAGGAAGC
ATCGGACTCATCAGGAAGC
AATCATCGGACTCATCAGGAAGC
TGCATCGGACTCATCAGGAAGC
GAGTGACATCGGACTCATCAGGAAGC
AATCCATCGGACTCATCAGGAAGC
TCATCGGACTCATCAGGAAGC

AACATCGGACTCATCAGGAAGC
GCATCGGACTCATCAGGAAGC
TGCATCGGACTCATCAGGAAGC
CATCGGACTCATCAGGAAGC
TTCATCGGACTCATCAGGAAGC
ACATCGGACTCATCAGGAAGC
ATCGGACTCATCAGGAAGC
CACATCGGACTCATCAGGAAGC
CTGACATCGGACTCATCAGGAAGC
GCATCGGACTCATCAGGAAGC
GCATCGGACTCATCAGGAAGC
GGATCATCGGACTCATCAGGAAGC
GGCACATCGGACTCATCAGGAAGC
CAATCATCGGACTCATCAGGAAGC
AGGGACATCGGACTCATCAGGAAGC
ACATCGGACTCATCAGGAAGC
TGGCCATCGGACTCATCAGGAAGC
CATCGGACTCATCAGGAAGC
AACATCGGACTCATCAGGAAGC
GGTCATCGGACTCATCAGGAAGC
TCATCGGACTCATCAGGAAGC
GCCATCGGACTCATCAGGAAGC
TACATCGGACTCATCAGGAAGC
GGCATCGGACTCATCAGGAAGC
CACATCGGACTCATCAGGAAGC
CATCGGACTCATCAGGAAGC
CCTCATCGGACTCATCAGGAAGC
TACATCGGACTCATCAGGAAGC
CCATCGGACTCATCAGGAAGC
ATGACATCGGACTCATCAGGAAGC
GTCATCGGACTCATCAGGAAGC
CACATCGGACTCATCAGGAAGC
CTCCATCGGACTCATCAGGAAGC
TTCCTCATCGGACTCATCAGGAAGC
CCTCATCGGACTCATCAGGAAGC
CATCGGACTCATCAGGAAGC
AACATCGGACTCATCAGGAAGC
CCCCATCGGACTCATCAGGAAGC
AACATCGGACTCATCAGGAAGC
ACCCATCGGACTCATCAGGAAGC
ATCGGACTCATCAGGAAGC
CTCCATCGGACTCATCAGGAAGC
AGACATCGGACTCATCAGGAAGC

CAACATCGGACTCATCAGGAAGC
ACATCGGACTCATCAGGAAGC
GGGTCATCGGACTCATCAGGAAGC
CCATCGGACTCATCAGGAAGC
AGCGCATCGGACTCATCAGGAAGC
AACCATCGGACTCATCAGGAAGC
CCCATCGGACTCATCAGGAAGC
GACATCGGACTCATCAGGAAGC
CATCGGACTCATCAGGAAGC
CCATCGGACTCATCAGGAAGC
TCATCGGACTCATCAGGAAGC
AGCTACATCGGACTCATCAGGAAGC
CCATCGGACTCATCAGGAAGC
ACATCGGACTCATCAGGAAGC
TGTATCATCGGACTCATCAGGAAGC
CATCGGACTCATCAGGAAGC
ACATCGGACTCATCAGGAAGC
ACATCGGACTCATCAGGAAGC
GCCATCGGACTCATCAGGAAGC
GACATCGGACTCATCAGGAAGC
CTGGGCATCGGACTCATCAGGAAGC
AAACATCGGACTCATCAGGAAGC
CCATCGGACTCATCAGGAAGC
AAGCACATCGGACTCATCAGGAAGC
CCAACATCGGACTCATCAGGAAGC
AACATCGGACTCATCAGGAAGC
ACCCATCGGACTCATCAGGAAGC
GAGCATCGGACTCATCAGGAAGC
CAGCATCGGACTCATCAGGAAGC
AAACATCGGACTCATCAGGAAGC
CCATCGGACTCATCAGGAAGC
CATCGGACTCATCAGGAAGC
TTCAGCATCGGACTCATCAGGAAGC
AGCATCGGACTCATCAGGAAGC
ATCGGACTCATCAGGAAGC
CATCGGACTCATCAGGAAGC
CCTCATCGGACTCATCAGGAAGC
TCATCGGACTCATCAGGAAGC
ACGCATCGGACTCATCAGGAAGC
CTCATCGGACTCATCAGGAAGC
CAAGCATCGGACTCATCAGGAAGC
ATCATCGGACTCATCAGGAAGC
CATAGTCATCGGACTCATCAGGAAGC

AAGGCATCGGACTCATCAGGAAGC
CATCGGACTCATCAGGAAGC
CACATCGGACTCATCAGGAAGC
GTGCATCGGACTCATCAGGAAGC
TCATCGGACTCATCAGGAAGC
CATCGGACTCATCAGGAAGC
GTCCGCATCGGACTCATCAGGAAGC
CTCATCGGACTCATCAGGAAGC
CATCGGACTCATCAGGAAGC
TGCATCGGACTCATCAGGAAGC
ACCCATCGGACTCATCAGGAAGC
TAGCCATCGGACTCATCAGGAAGC
TCATCGGACTCATCAGGAAGC
CATCGGACTCATCAGGAAGC
GTTCATCGGACTCATCAGGAAGC
TATGTCATCGGACTCATCAGGAAGC
GCATCGGACTCATCAGGAAGC
GGTCATCGGACTCATCAGGAAGC
CCATCGGACTCATCAGGAAGC
GACATCGGACTCATCAGGAAGC
GCTTCATCGGACTCATCAGGAAGC
TACATCGGACTCATCAGGAAGC
CATCGGACTCATCAGGAAGC
ACATCGGACTCATCAGGAAGC
TGACCATCGGACTCATCAGGAAGC
TCATCGGACTCATCAGGAAGC
TGCATCGGACTCATCAGGAAGC
TCATCGGACTCATCAGGAAGC
ACATCGGACTCATCAGGAAGC
CACATCGGACTCATCAGGAAGC
GCCATCGGACTCATCAGGAAGC
CCCATCGGACTCATCAGGAAGC
CTCATCGGACTCATCAGGAAGC
CTCATCGGACTCATCAGGAAGC
TGCCATCGGACTCATCAGGAAGC
CCATCGGACTCATCAGGAAGC
AGGCACATCGGACTCATCAGGAAGC
GCATCGGACTCATCAGGAAGC
AAGCCATCGGACTCATCAGGAAGC
ATCGGACTCATCAGGAAGC
ACCATCGGACTCATCAGGAAGC
GACATCGGACTCATCAGGAAGC
CATCGGACTCATCAGGAAGC

CCCCCATCGGACTCATCAGGAAGC
GCCATCGGACTCATCAGGAAGC
TTTCATCGGACTCATCAGGAAGC
CATCGGACTCATCAGGAAGC
CACATCGGACTCATCAGGAAGC
CCATCGGACTCATCAGGAAGC
TTCATCGGACTCATCAGGAAGC
GTGCATCGGACTCATCAGGAAGC
CTCATCGGACTCATCAGGAAGC
CAACACATCGGACTCATCAGGAAGC
TGCATCGGACTCATCAGGAAGC
AACATCGGACTCATCAGGAAGC
TTCATCGGACTCATCAGGAAGC
TCTCATCGGACTCATCAGGAAGC
TTCATCGGACTCATCAGGAAGC
GATCATCGGACTCATCAGGAAGC
AGCATCGGACTCATCAGGAAGC
CATCATCGGACTCATCAGGAAGC
CACCATCGGACTCATCAGGAAGC
TGCATCGGACTCATCAGGAAGC
TCATCGGACTCATCAGGAAGC
AAGACATCGGACTCATCAGGAAGC
CCATCGGACTCATCAGGAAGC
AGAACATCGGACTCATCAGGAAGC
TCATCGGACTCATCAGGAAGC
ATGCATCGGACTCATCAGGAAGC
CATCGGACTCATCAGGAAGC
ACATCGGACTCATCAGGAAGC
ATCACATCGGACTCATCAGGAAGC
TGGCATCGGACTCATCAGGAAGC
GACATCGGACTCATCAGGAAGC
ACACATCGGACTCATCAGGAAGC
ATCATCGGACTCATCAGGAAGC
TCATCGGACTCATCAGGAAGC
GAACATCGGACTCATCAGGAAGC
AACATCGGACTCATCAGGAAGC
ATCGGACTCATCAGGAAGC
ACATCGGACTCATCAGGAAGC
TGGGCATCGGACTCATCAGGAAGC
AGACAACATCGGACTCATCAGGAAGC
TCATCGGACTCATCAGGAAGC
CATCGGACTCATCAGGAAGC
TCAGTCATCGGACTCATCAGGAAGC

TCACATCGGACTCATCAGGAAGC
ATCGGACTCATCAGGAAGC
ACACATCGGACTCATCAGGAAGC
ACATCGGACTCATCAGGAAGC
GATCATCGGACTCATCAGGAAGC
CACATCGGACTCATCAGGAAGC
ATCGGACTCATCAGGAAGC
TCCATCGGACTCATCAGGAAGC
CATCGGACTCATCAGGAAGC
TCATCGGACTCATCAGGAAGC
CATCGGACTCATCAGGAAGC
AGCATCGGACTCATCAGGAAGC
GACATCGGACTCATCAGGAAGC
CATCGGACTCATCAGGAAGC
GCTCATCGGACTCATCAGGAAGC
CCATCGGACTCATCAGGAAGC
GTCATCGGACTCATCAGGAAGC
GGCATCGGACTCATCAGGAAGC
AGAGGCCATCGGACTCATCAGGAAGC
CTTCATCGGACTCATCAGGAAGC
AGGCATCGGACTCATCAGGAAGC
TCATCGGACTCATCAGGAAGC
GTACATCGGACTCATCAGGAAGC
GCATCGGACTCATCAGGAAGC
AAAACATCGGACTCATCAGGAAGC
GACATCGGACTCATCAGGAAGC
CATCGGACTCATCAGGAAGC
ACACATCGGACTCATCAGGAAGC
ATCATCGGACTCATCAGGAAGC
CACATCGGACTCATCAGGAAGC
CCATCGGACTCATCAGGAAGC
ACCATCGGACTCATCAGGAAGC
GGCATCGGACTCATCAGGAAGC
TGCATCGGACTCATCAGGAAGC
ACATCGGACTCATCAGGAAGC
CATCGGACTCATCAGGAAGC
ATCGGACTCATCAGGAAGC
CCATCGGACTCATCAGGAAGC
TGCATCGGACTCATCAGGAAGC
ACATCGGACTCATCAGGAAGC
GGATCATCGGACTCATCAGGAAGC
TTCATCGGACTCATCAGGAAGC
TGACATCGGACTCATCAGGAAGC

GTTCATCGGACTCATCAGGAAGC
GTGGCATCGGACTCATCAGGAAGC
GTTTCATCGGACTCATCAGGAAGC
ACATCGGACTCATCAGGAAGC
AGGACATCGGACTCATCAGGAAGC
TTCATCGGACTCATCAGGAAGC
TAACATCGGACTCATCAGGAAGC
ACTACATCGGACTCATCAGGAAGC
ACCATCGGACTCATCAGGAAGC
TTCATCGGACTCATCAGGAAGC
CCCATCGGACTCATCAGGAAGC
TCATCGGACTCATCAGGAAGC
AACATCGGACTCATCAGGAAGC
TCATCGGACTCATCAGGAAGC
ATCGGACTCATCAGGAAGC
CACCATCGGACTCATCAGGAAGC
TCACATCGGACTCATCAGGAAGC
TCCCATCGGACTCATCAGGAAGC
TCATCGGACTCATCAGGAAGC
GCATCGGACTCATCAGGAAGC
CCATCGGACTCATCAGGAAGC
CAACATCGGACTCATCAGGAAGC
ACATCGGACTCATCAGGAAGC
TTCATCGGACTCATCAGGAAGC
CATCGGACTCATCAGGAAGC
GCCATCGGACTCATCAGGAAGC
TCCATCGGACTCATCAGGAAGC
TCCCCATCGGACTCATCAGGAAGC
GGCATCGGACTCATCAGGAAGC
GTCCATCGGACTCATCAGGAAGC
CAGTCCATCGGACTCATCAGGAAGC
CCATCGGACTCATCAGGAAGC
TCATCGGACTCATCAGGAAGC
AACATCGGACTCATCAGGAAGC
GCATCGGACTCATCAGGAAGC
TGTGCATCGGACTCATCAGGAAGC
TTCATCGGACTCATCAGGAAGC
CCATCGGACTCATCAGGAAGC
AGCCATCGGACTCATCAGGAAGC
TCTGCATCGGACTCATCAGGAAGC
GTCCATCGGACTCATCAGGAAGC
CCTGACATCGGACTCATCAGGAAGC
TCATCGGACTCATCAGGAAGC

GCGCATCGGACTCATCAGGAAGC
AGGACATCGGACTCATCAGGAAGC
AACATCGGACTCATCAGGAAGC
AACATCGGACTCATCAGGAAGC
ACCCATCGGACTCATCAGGAAGC
GAAACATCGGACTCATCAGGAAGC
CACATCGGACTCATCAGGAAGC
CTTCATCGGACTCATCAGGAAGC
CCCATCGGACTCATCAGGAAGC
AGTGCATCGGACTCATCAGGAAGC
ATCGGACTCATCAGGAAGC
ACATCGGACTCATCAGGAAGC
TCATCGGACTCATCAGGAAGC
CTCCATCGGACTCATCAGGAAGC
GATTCATCGGACTCATCAGGAAGC
AGCCATCGGACTCATCAGGAAGC
TACATCGGACTCATCAGGAAGC
GCCATCGGACTCATCAGGAAGC
GTAAACATCGGACTCATCAGGAAGC
ACATCGGACTCATCAGGAAGC
CCATCGGACTCATCAGGAAGC
TATCATCGGACTCATCAGGAAGC
TGCCATCGGACTCATCAGGAAGC
CATCGGACTCATCAGGAAGC
CATCGGACTCATCAGGAAGC
TCATCGGACTCATCAGGAAGC
TTGCCATCGGACTCATCAGGAAGC
GCCATCGGACTCATCAGGAAGC
TACCATCGGACTCATCAGGAAGC
ACATCGGACTCATCAGGAAGC
TCACATCGGACTCATCAGGAAGC
GTCATCGGACTCATCAGGAAGC
ATTCATCGGACTCATCAGGAAGC
CTTCATCGGACTCATCAGGAAGC
ACATCGGACTCATCAGGAAGC
CTGCATCGGACTCATCAGGAAGC
AGGCATCGGACTCATCAGGAAGC
ATCGGACTCATCAGGAAGC
TCCCATCGGACTCATCAGGAAGC
TCCATCGGACTCATCAGGAAGC
AACATCGGACTCATCAGGAAGC
AACATCGGACTCATCAGGAAGC
GCATCGGACTCATCAGGAAGC

ATTCCACATCGGACTCATCAGGAAGC
AACCATCGGACTCATCAGGAAGC
GAAACATCGGACTCATCAGGAAGC
TTTCATCGGACTCATCAGGAAGC
TCATCGGACTCATCAGGAAGC
TGCATCGGACTCATCAGGAAGC
GTCATCGGACTCATCAGGAAGC
ATCGGACTCATCAGGAAGC
TCGGACTCATCAGGAAGC
CCAGCCATCGGACTCATCAGGAAGC
GGGCATCGGACTCATCAGGAAGC
CACATCGGACTCATCAGGAAGC
TGACATCGGACTCATCAGGAAGC
GTGCATCATCGGACTCATCAGGAAGC
GACATCGGACTCATCAGGAAGC
GCATCGGACTCATCAGGAAGC
CCATCGGACTCATCAGGAAGC
TAACATCGGACTCATCAGGAAGC
CATCGGACTCATCAGGAAGC
TCTTCATCGGACTCATCAGGAAGC
TGCATCGGACTCATCAGGAAGC
GCCATCGGACTCATCAGGAAGC
CCCATCGGACTCATCAGGAAGC
GTTCATCGGACTCATCAGGAAGC
ACATCGGACTCATCAGGAAGC
GGACATCGGACTCATCAGGAAGC
TAAGACATCGGACTCATCAGGAAGC
CCATCGGACTCATCAGGAAGC
TGTACATCGGACTCATCAGGAAGC
TTCATCGGACTCATCAGGAAGC
AGCATCGGACTCATCAGGAAGC
TTCATCGGACTCATCAGGAAGC
TGCATCGGACTCATCAGGAAGC
ACATCGGACTCATCAGGAAGC
GAGCATCGGACTCATCAGGAAGC
TCCATCGGACTCATCAGGAAGC
CTTGACATCGGACTCATCAGGAAGC
GGACATCGGACTCATCAGGAAGC
ACCATCGGACTCATCAGGAAGC
CGGGCATCGGACTCATCAGGAAGC
CTGTCATCGGACTCATCAGGAAGC
ATCGGACTCATCAGGAAGC
TCATCGGACTCATCAGGAAGC

GTCATCGGACTCATCAGGAAGC
GCCCATCGGACTCATCAGGAAGC
GGCATCGGACTCATCAGGAAGC
GTCACATCGGACTCATCAGGAAGC
CTCATCGGACTCATCAGGAAGC
TTTCATCGGACTCATCAGGAAGC
CACATCGGACTCATCAGGAAGC
TGTCATCGGACTCATCAGGAAGC
GCCATCGGACTCATCAGGAAGC
GGCATCGGACTCATCAGGAAGC
AGCATCGGACTCATCAGGAAGC
ACATCGGACTCATCAGGAAGC
CCATCGGACTCATCAGGAAGC
GGTCATCGGACTCATCAGGAAGC
ACATCGGACTCATCAGGAAGC
CTGTCATCGGACTCATCAGGAAGC
TTCATCGGACTCATCAGGAAGC
TCATCGGACTCATCAGGAAGC
TGACATCGGACTCATCAGGAAGC
CCATCGGACTCATCAGGAAGC
AACATCGGACTCATCAGGAAGC
AGACATCGGACTCATCAGGAAGC
TCATCGGACTCATCAGGAAGC
ACATCGGACTCATCAGGAAGC
TCATCGGACTCATCAGGAAGC
TTCATCGGACTCATCAGGAAGC
ATCATCGGACTCATCAGGAAGC
TGCCCATCGGACTCATCAGGAAGC
GCATCGGACTCATCAGGAAGC
CTCCATCGGACTCATCAGGAAGC
TTCATCGGACTCATCAGGAAGC
AGTCAGCATCGGACTCATCAGGAAGC
GGTCATCGGACTCATCAGGAAGC
AGCCATCGGACTCATCAGGAAGC
CATCGGACTCATCAGGAAGC
TCATCATCGGACTCATCAGGAAGC
AAACATCGGACTCATCAGGAAGC
TCATCGGACTCATCAGGAAGC
CCATCGGACTCATCAGGAAGC
ATCATCGGACTCATCAGGAAGC
CACATCGGACTCATCAGGAAGC
GCATCGGACTCATCAGGAAGC
GATACATCGGACTCATCAGGAAGC

TTCATCGGACTCATCAGGAAGC
CCTCATCGGACTCATCAGGAAGC
ACATCGGACTCATCAGGAAGC
ATCATCGGACTCATCAGGAAGC
TCTCATCGGACTCATCAGGAAGC
AGCATCGGACTCATCAGGAAGC
CCTCATCGGACTCATCAGGAAGC
CGCATCGGACTCATCAGGAAGC
GGGCATCGGACTCATCAGGAAGC
ATGCCATCGGACTCATCAGGAAGC
CACCATCGGACTCATCAGGAAGC
AGACACATCGGACTCATCAGGAAGC
CAGCATCGGACTCATCAGGAAGC
GCATCGGACTCATCAGGAAGC
GTACATCGGACTCATCAGGAAGC
ACCATCGGACTCATCAGGAAGC
GAAGCATCGGACTCATCAGGAAGC
CATCATCGGACTCATCAGGAAGC
TCCATCGGACTCATCAGGAAGC
AGCATCGGACTCATCAGGAAGC
GTCCATCGGACTCATCAGGAAGC
CCATCGGACTCATCAGGAAGC
GCATCGGACTCATCAGGAAGC
TCATCGGACTCATCAGGAAGC
ACCATCGGACTCATCAGGAAGC
CTGGTCATCGGACTCATCAGGAAGC
CATCGGACTCATCAGGAAGC
CACATCGGACTCATCAGGAAGC
TCATCGGACTCATCAGGAAGC
AGCATCGGACTCATCAGGAAGC
AAACGCATCGGACTCATCAGGAAGC
ATCGGACTCATCAGGAAGC
CACATCGGACTCATCAGGAAGC
TCATCGGACTCATCAGGAAGC
CCATCGGACTCATCAGGAAGC
GTCATCGGACTCATCAGGAAGC
ATCATCGGACTCATCAGGAAGC
GTCCATCGGACTCATCAGGAAGC
TCATCGGACTCATCAGGAAGC
TGCATCGGACTCATCAGGAAGC
GGACATCGGACTCATCAGGAAGC
GTGTCCATCGGACTCATCAGGAAGC
TTTTCATCGGACTCATCAGGAAGC

GTTCATCGGACTCATCAGGAAGC
CCATCGGACTCATCAGGAAGC
CATCGGACTCATCAGGAAGC
ATCGGACTCATCAGGAAGC
ACCATCGGACTCATCAGGAAGC
CCATCGGACTCATCAGGAAGC
GTCACATCGGACTCATCAGGAAGC
GTTGACATCGGACTCATCAGGAAGC
TGACATCGGACTCATCAGGAAGC
CCATCGGACTCATCAGGAAGC
GGGTCATCGGACTCATCAGGAAGC
AAACATCGGACTCATCAGGAAGC
CCATCGGACTCATCAGGAAGC
TGCATCGGACTCATCAGGAAGC
GCATCGGACTCATCAGGAAGC
CATCGGACTCATCAGGAAGC
TTTCATCGGACTCATCAGGAAGC
ATCATCGGACTCATCAGGAAGC
GCCATCGGACTCATCAGGAAGC
CCATCGGACTCATCAGGAAGC
GGCATCGGACTCATCAGGAAGC
GGCATCGGACTCATCAGGAAGC
TATCACATCGGACTCATCAGGAAGC
TCATCGGACTCATCAGGAAGC
AGCATCGGACTCATCAGGAAGC
CTTCATCGGACTCATCAGGAAGC
GCATCGGACTCATCAGGAAGC
TGCATCGGACTCATCAGGAAGC
TGACATCGGACTCATCAGGAAGC
CCATCGGACTCATCAGGAAGC
GTTTCATCGGACTCATCAGGAAGC
TTTACATCGGACTCATCAGGAAGC
TATCATCGGACTCATCAGGAAGC
GAAGCATCGGACTCATCAGGAAGC
ACATCGGACTCATCAGGAAGC
TCCATCGGACTCATCAGGAAGC
TTCCATCGGACTCATCAGGAAGC
ACCATCGGACTCATCAGGAAGC
AGCAAGCATCGGACTCATCAGGAAGC
CATCGGACTCATCAGGAAGC
GAACATCGGACTCATCAGGAAGC
GGGACATCGGACTCATCAGGAAGC
CATCGGACTCATCAGGAAGC

AGACCATCGGACTCATCAGGAAGC
AACGCATCGGACTCATCAGGAAGC
CCATCGGACTCATCAGGAAGC
CTCATCGGACTCATCAGGAAGC
CATCGGACTCATCAGGAAGC
AGCATCGGACTCATCAGGAAGC
GCCATCGGACTCATCAGGAAGC
TTCATCGGACTCATCAGGAAGC
AACTCATCGGACTCATCAGGAAGC
CTCATCGGACTCATCAGGAAGC
ATCGGACTCATCAGGAAGC
CGTGACATCGGACTCATCAGGAAGC
AAACATCGGACTCATCAGGAAGC
GGAACATCGGACTCATCAGGAAGC
TGCATCGGACTCATCAGGAAGC
TGGCATCGGACTCATCAGGAAGC
AACATCGGACTCATCAGGAAGC
TCATCGGACTCATCAGGAAGC
GTTCATCGGACTCATCAGGAAGC
TCATCGGACTCATCAGGAAGC
CATCATCGGACTCATCAGGAAGC
GCACCATCGGACTCATCAGGAAGC
CCCATCGGACTCATCAGGAAGC
CAGCATCGGACTCATCAGGAAGC
CATCGGACTCATCAGGAAGC
GACATCGGACTCATCAGGAAGC
TTCATCGGACTCATCAGGAAGC
TCATCGGACTCATCAGGAAGC
ATTCATCGGACTCATCAGGAAGC
ACATCGGACTCATCAGGAAGC
AGTTGTCATCGGACTCATCAGGAAGC
GTGTGTCATCGGACTCATCAGGAAGC
AACATCGGACTCATCAGGAAGC
CCATCGGACTCATCAGGAAGC
CATCATCGGACTCATCAGGAAGC
ATCATCGGACTCATCAGGAAGC
TGTGTCATCGGACTCATCAGGAAGC
ACATCGGACTCATCAGGAAGC
AAAGCCATCGGACTCATCAGGAAGC
TGCATCGGACTCATCAGGAAGC
CATCGGACTCATCAGGAAGC
CCCATCGGACTCATCAGGAAGC
AACTCATCGGACTCATCAGGAAGC

CATCGGACTCATCAGGAAGC
TCATCGGACTCATCAGGAAGC
GTCATCGGACTCATCAGGAAGC
GCATCGGACTCATCAGGAAGC
TTTCATCGGACTCATCAGGAAGC
CCATCGGACTCATCAGGAAGC
TCACATCGGACTCATCAGGAAGC
TGACCATCGGACTCATCAGGAAGC
CCATCGGACTCATCAGGAAGC
TCATCGGACTCATCAGGAAGC
AGCATCGGACTCATCAGGAAGC
ATCCCATCGGACTCATCAGGAAGC
TCATCGGACTCATCAGGAAGC
ACACATCGGACTCATCAGGAAGC
AACATCGGACTCATCAGGAAGC
ACATCGGACTCATCAGGAAGC
CCATCGGACTCATCAGGAAGC
TGGTCATCGGACTCATCAGGAAGC
GACATCGGACTCATCAGGAAGC
TCTGGCATCGGACTCATCAGGAAGC
ATCGGACTCATCAGGAAGC
CATCGGACTCATCAGGAAGC
AGAAACATCGGACTCATCAGGAAGC
AGATCATCGGACTCATCAGGAAGC
GGCATCGGACTCATCAGGAAGC
GCCATCGGACTCATCAGGAAGC
TGGCATCGGACTCATCAGGAAGC
ATGTTTACATCGGACTCATCAGGAAGC
CACTACATCGGACTCATCAGGAAGC
GACCATCGGACTCATCAGGAAGC
CATCGGACTCATCAGGAAGC
TCGGACTCATCAGGAAGC
TCATCGGACTCATCAGGAAGC
CTCATCGGACTCATCAGGAAGC
GGCATCGGACTCATCAGGAAGC
CCATCGGACTCATCAGGAAGC
GCATCGGACTCATCAGGAAGC
TTCATCGGACTCATCAGGAAGC
GAACCCATCGGACTCATCAGGAAGC
AAGGCATCGGACTCATCAGGAAGC
TGGCATCGGACTCATCAGGAAGC
CATCGGACTCATCAGGAAGC
TTTCATCGGACTCATCAGGAAGC

CATCGGACTCATCAGGAAGC
CATCGGACTCATCAGGAAGC
CATCGGACTCATCAGGAAGC
GTTCATCGGACTCATCAGGAAGC
GTCATCGGACTCATCAGGAAGC
ATGCATCGGACTCATCAGGAAGC
GCCATCGGACTCATCAGGAAGC
TCGGACTCATCAGGAAGC
TCGGACTCATCAGGAAGC
GTCTCATCGGACTCATCAGGAAGC
TCCATCGGACTCATCAGGAAGC
TTCATCGGACTCATCAGGAAGC
TACATCGGACTCATCAGGAAGC
TGCATCGGACTCATCAGGAAGC
CTACATCGGACTCATCAGGAAGC
GCCATCGGACTCATCAGGAAGC
CTCATCGGACTCATCAGGAAGC
TGGCATCGGACTCATCAGGAAGC
ACATCGGACTCATCAGGAAGC
AACATCGGACTCATCAGGAAGC
CCATCGGACTCATCAGGAAGC
TCGGACTCATCAGGAAGC
GTGTCATCGGACTCATCAGGAAGC
GCTCATCGGACTCATCAGGAAGC
GGGGGCATCGGACTCATCAGGAAGC
GCCATCGGACTCATCAGGAAGC
CCCATCGGACTCATCAGGAAGC
TTCATCGGACTCATCAGGAAGC
ACATCGGACTCATCAGGAAGC
GACATCGGACTCATCAGGAAGC
TCATCGGACTCATCAGGAAGC
CGGACTCATCAGGAAGC
TCTCATCGGACTCATCAGGAAGC
AGTCATCGGACTCATCAGGAAGC
ATCATCGGACTCATCAGGAAGC
GCATCGGACTCATCAGGAAGC
AGACATCGGACTCATCAGGAAGC
TCATCGGACTCATCAGGAAGC
CATCGGACTCATCAGGAAGC
CATCGGACTCATCAGGAAGC
TACATCGGACTCATCAGGAAGC
CAGACATCGGACTCATCAGGAAGC
TCCATCGGACTCATCAGGAAGC

AGAACATCGGACTCATCAGGAAGC
TCGGACTCATCAGGAAGC
GCCATCGGACTCATCAGGAAGC
CATCGGACTCATCAGGAAGC
GACATCGGACTCATCAGGAAGC
GCATCGGACTCATCAGGAAGC
CCATCGGACTCATCAGGAAGC
TCGGACTCATCAGGAAGC
TCTACATCGGACTCATCAGGAAGC
CATCGGACTCATCAGGAAGC
GTCCATCGGACTCATCAGGAAGC
TTCATCGGACTCATCAGGAAGC
GTGCATCGGACTCATCAGGAAGC
CTCCATCGGACTCATCAGGAAGC
TCCATCGGACTCATCAGGAAGC
AACATCGGACTCATCAGGAAGC
TCTATAACATCGGACTCATCAGGAAGC
GCATCGGACTCATCAGGAAGC
ACATCGGACTCATCAGGAAGC
GTGTGTCATCGGACTCATCAGGAAGC
TTTGCATCGGACTCATCAGGAAGC
CCATCGGACTCATCAGGAAGC
CCCATCGGACTCATCAGGAAGC
CACATCGGACTCATCAGGAAGC
CTCCATCGGACTCATCAGGAAGC
ATCGGACTCATCAGGAAGC
GGGGCATCGGACTCATCAGGAAGC
CATCGGACTCATCAGGAAGC
AGAGCATCGGACTCATCAGGAAGC
CCGCATCGGACTCATCAGGAAGC
GGACTCATCAGGAAGC
ACATCGGACTCATCAGGAAGC
GAACATCGGACTCATCAGGAAGC
CTCATCGGACTCATCAGGAAGC
TACATCGGACTCATCAGGAAGC
GGCATCGGACTCATCAGGAAGC
TGTCATCGGACTCATCAGGAAGC
CAGCATCGGACTCATCAGGAAGC
ATGCCATCGGACTCATCAGGAAGC
TTACATCGGACTCATCAGGAAGC
AACCATCGGACTCATCAGGAAGC
GCATCGGACTCATCAGGAAGC
TTCATCGGACTCATCAGGAAGC

GTCATCGGACTCATCAGGAAGC
GCCATCGGACTCATCAGGAAGC
CCATCGGACTCATCAGGAAGC
GCATCGGACTCATCAGGAAGC
ATGGCTCATCGGACTCATCAGGAAGC
CAAACATCGGACTCATCAGGAAGC
CCATCGGACTCATCAGGAAGC
GCCACATCGGACTCATCAGGAAGC
CACATCGGACTCATCAGGAAGC
GTCATCGGACTCATCAGGAAGC
TCATCGGACTCATCAGGAAGC
GTTCATCGGACTCATCAGGAAGC
GAACATCGGACTCATCAGGAAGC
TCATCGGACTCATCAGGAAGC
TCATCGGACTCATCAGGAAGC
GAACTCATCGGACTCATCAGGAAGC
GCACATCGGACTCATCAGGAAGC
CCCATCGGACTCATCAGGAAGC
CAGCATCGGACTCATCAGGAAGC
CCATCGGACTCATCAGGAAGC
GGTCATCGGACTCATCAGGAAGC
GCACATCGGACTCATCAGGAAGC
GAACATCGGACTCATCAGGAAGC
ACTCATCGGACTCATCAGGAAGC
TGGTCATCGGACTCATCAGGAAGC
ACCATCGGACTCATCAGGAAGC
GTGTGTCATCGGACTCATCAGGAAGC
CCATCGGACTCATCAGGAAGC
CCTCATCGGACTCATCAGGAAGC
ATCGGACTCATCAGGAAGC
ATCGGACTCATCAGGAAGC
GACACATCGGACTCATCAGGAAGC
GGCATCGGACTCATCAGGAAGC
CCACATCGGACTCATCAGGAAGC
CCATCGGACTCATCAGGAAGC
CCATCGGACTCATCAGGAAGC
GGCATCGGACTCATCAGGAAGC
TGTGCATCGGACTCATCAGGAAGC
AACATCGGACTCATCAGGAAGC
GCATCGGACTCATCAGGAAGC
TGCATCGGACTCATCAGGAAGC
AGCATCGGACTCATCAGGAAGC
TCATCGGACTCATCAGGAAGC

CCCATCGGACTCATCAGGAAGC
CCATCGGACTCATCAGGAAGC
ACATCGGACTCATCAGGAAGC
ATCGGACTCATCAGGAAGC
GCATCGGACTCATCAGGAAGC
TCATCGGACTCATCAGGAAGC
CATCGGACTCATCAGGAAGC
CATCGGACTCATCAGGAAGC
CGCATCGGACTCATCAGGAAGC
CAGCATCGGACTCATCAGGAAGC
ACATCGGACTCATCAGGAAGC
CATCGGACTCATCAGGAAGC
TGGTCATCGGACTCATCAGGAAGC
GATGGTCATCGGACTCATCAGGAAGC
ACATCGGACTCATCAGGAAGC
GACATCGGACTCATCAGGAAGC
ACATCGGACTCATCAGGAAGC
CATCGGACTCATCAGGAAGC
TTACATCGGACTCATCAGGAAGC
AGCATCGGACTCATCAGGAAGC
TACATCGGACTCATCAGGAAGC
GTAGGGTCATCGGACTCATCAGGAAGC
CATCGGACTCATCAGGAAGC
ATCGGACTCATCAGGAAGC
TGTGTTCATCGGACTCATCAGGAAGC
CTCCATCGGACTCATCAGGAAGC
CATCGGACTCATCAGGAAGC
GACATCGGACTCATCAGGAAGC
GACATCGGACTCATCAGGAAGC
ATCGGACTCATCAGGAAGC
CATCGGACTCATCAGGAAGC
GCATCGGACTCATCAGGAAGC
AACATCGGACTCATCAGGAAGC
ACCATCGGACTCATCAGGAAGC
CTCATCGGACTCATCAGGAAGC
CTCATCGGACTCATCAGGAAGC
CATCGGACTCATCAGGAAGC
CATCGGACTCATCAGGAAGC
TTCATCGGACTCATCAGGAAGC
AGAGACATCGGACTCATCAGGAAGC
ATGTCATCGGACTCATCAGGAAGC
GTCCCATCGGACTCATCAGGAAGC
GCATCGGACTCATCAGGAAGC

ACCATCGGACTCATCAGGAAGC
GTCATCGGACTCATCAGGAAGC
AGACATCGGACTCATCAGGAAGC
TGCATCGGACTCATCAGGAAGC
GGCATCGGACTCATCAGGAAGC
CACATCGGACTCATCAGGAAGC
TGCATCGGACTCATCAGGAAGC
AACATCGGACTCATCAGGAAGC
AAAGACATCGGACTCATCAGGAAGC
TTGTGCATCGGACTCATCAGGAAGC
AACATCGGACTCATCAGGAAGC
TGGCATCGGACTCATCAGGAAGC
CATCGGACTCATCAGGAAGC
AACATCGGACTCATCAGGAAGC
CTCATCGGACTCATCAGGAAGC
GCCATCGGACTCATCAGGAAGC
ATCATCGGACTCATCAGGAAGC
GCCATCGGACTCATCAGGAAGC
GAGCATCGGACTCATCAGGAAGC
GAAACATCGGACTCATCAGGAAGC
GAGCATCGGACTCATCAGGAAGC
CTCAACCATCGGACTCATCAGGAAGC
TCATCGGACTCATCAGGAAGC
AGCTACATCGGACTCATCAGGAAGC
TTCATCGGACTCATCAGGAAGC
CCATCGGACTCATCAGGAAGC
TGCATCGGACTCATCAGGAAGC
CAGCATCGGACTCATCAGGAAGC
AATCATCGGACTCATCAGGAAGC
TGCAACATCGGACTCATCAGGAAGC
AAACATCGGACTCATCAGGAAGC
GTCATCGGACTCATCAGGAAGC
CACATCGGACTCATCAGGAAGC
TCATCGGACTCATCAGGAAGC
TCATCGGACTCATCAGGAAGC
CCCATCGGACTCATCAGGAAGC
CTACGCATCGGACTCATCAGGAAGC
TACATCGGACTCATCAGGAAGC
ACCACATCGGACTCATCAGGAAGC
CCCATCGGACTCATCAGGAAGC
ACATCGGACTCATCAGGAAGC
CATCGGACTCATCAGGAAGC
CCATCGGACTCATCAGGAAGC

TTCATCGGACTCATCAGGAAGC
AGCATCGGACTCATCAGGAAGC
ACATCGGACTCATCAGGAAGC
GTCATCGGACTCATCAGGAAGC
CATCGGACTCATCAGGAAGC
CACATCGGACTCATCAGGAAGC
GGCATCGGACTCATCAGGAAGC
CACATCGGACTCATCAGGAAGC
GTACTCATCGGACTCATCAGGAAGC
ACATCGGACTCATCAGGAAGC
GTCATCGGACTCATCAGGAAGC
ACATCGGACTCATCAGGAAGC
ACATCGGACTCATCAGGAAGC
ACGCATCGGACTCATCAGGAAGC
TTACATCGGACTCATCAGGAAGC
TCTCATCGGACTCATCAGGAAGC
GACTACATCGGACTCATCAGGAAGC
ACATCGGACTCATCAGGAAGC
TACATCGGACTCATCAGGAAGC
AGCATCGGACTCATCAGGAAGC
CATCGGACTCATCAGGAAGC
CCATCGGACTCATCAGGAAGC
TTCATCGGACTCATCAGGAAGC
CATCGGACTCATCAGGAAGC
ATCATCGGACTCATCAGGAAGC
AGGCATCGGACTCATCAGGAAGC
AAGCATCGGACTCATCAGGAAGC
GACCATCGGACTCATCAGGAAGC
CCATCGGACTCATCAGGAAGC
CCATCGGACTCATCAGGAAGC
TCCATCGGACTCATCAGGAAGC
GGCATCGGACTCATCAGGAAGC
TGGCATCGGACTCATCAGGAAGC
GTCATCGGACTCATCAGGAAGC
TCACATCGGACTCATCAGGAAGC
AGCATCGGACTCATCAGGAAGC
CTACATCGGACTCATCAGGAAGC
GGACATCGGACTCATCAGGAAGC
GCCATCGGACTCATCAGGAAGC
ACACATCGGACTCATCAGGAAGC
ATTTCATCGGACTCATCAGGAAGC
GGTTCATCGGACTCATCAGGAAGC
ACATCGGACTCATCAGGAAGC

CGCCATCGGACTCATCAGGAAGC
AGACATCGGACTCATCAGGAAGC
CACATCGGACTCATCAGGAAGC
ATCGGACTCATCAGGAAGC
TACATCGGACTCATCAGGAAGC
ACCATCGGACTCATCAGGAAGC
GCCACATCGGACTCATCAGGAAGC
GCATCGGACTCATCAGGAAGC
TCATCGGACTCATCAGGAAGC
GGATGTCATCGGACTCATCAGGAAGC
CGCATCGGACTCATCAGGAAGC
TCATCGGACTCATCAGGAAGC
TTTCATCGGACTCATCAGGAAGC
ATGTCATCGGACTCATCAGGAAGC
GTGACATCGGACTCATCAGGAAGC
TCATCATCGGACTCATCAGGAAGC
CCATCGGACTCATCAGGAAGC
ATGCATCGGACTCATCAGGAAGC
ACATCGGACTCATCAGGAAGC
GTGCATCGGACTCATCAGGAAGC
AGCCATCGGACTCATCAGGAAGC
TGGCATCGGACTCATCAGGAAGC
ACATCGGACTCATCAGGAAGC
TTGCATCGGACTCATCAGGAAGC
ATCGGACTCATCAGGAAGC
GAAAGCCATCGGACTCATCAGGAAGC
GCATCGGACTCATCAGGAAGC
TTCCATCGGACTCATCAGGAAGC
TCATCGGACTCATCAGGAAGC
CCATCGGACTCATCAGGAAGC
CCACATCGGACTCATCAGGAAGC
TTTTGTCATCGGACTCATCAGGAAGC
GTGCATCGGACTCATCAGGAAGC
ATACATCGGACTCATCAGGAAGC
CCACAACATCGGACTCATCAGGAAGC
TGACATCGGACTCATCAGGAAGC
CACATCGGACTCATCAGGAAGC
CCCATCGGACTCATCAGGAAGC
CCATCGGACTCATCAGGAAGC
TCTCTCATCGGACTCATCAGGAAGC
TTCATCGGACTCATCAGGAAGC
CTGCATCGGACTCATCAGGAAGC
GACATCGGACTCATCAGGAAGC

ACATCGGACTCATCAGGAAGC
ATCATCGGACTCATCAGGAAGC
GATCATCGGACTCATCAGGAAGC
CACCATCGGACTCATCAGGAAGC
TTTCATCGGACTCATCAGGAAGC
GAGCATCGGACTCATCAGGAAGC
CACATCGGACTCATCAGGAAGC
CATCGGACTCATCAGGAAGC
TACATCGGACTCATCAGGAAGC
TGAAGACATCGGACTCATCAGGAAGC
ATCGGACTCATCAGGAAGC
CAACATCGGACTCATCAGGAAGC
GAATCATCGGACTCATCAGGAAGC
CATCGGACTCATCAGGAAGC
GCATCGGACTCATCAGGAAGC
ACCCATCGGACTCATCAGGAAGC
ACATCGGACTCATCAGGAAGC
CTGGCCATCGGACTCATCAGGAAGC
GCCATCGGACTCATCAGGAAGC
TGCATCGGACTCATCAGGAAGC
CACATCGGACTCATCAGGAAGC
GATACATCGGACTCATCAGGAAGC
CATCGGACTCATCAGGAAGC
TGGCATCGGACTCATCAGGAAGC
CATCGGACTCATCAGGAAGC
AGTGTCATCGGACTCATCAGGAAGC
AGCATCGGACTCATCAGGAAGC
CCATCGGACTCATCAGGAAGC
GTACATCGGACTCATCAGGAAGC
GACCATCGGACTCATCAGGAAGC
CATCGGACTCATCAGGAAGC
ACTCATCGGACTCATCAGGAAGC
TACATCGGACTCATCAGGAAGC
ACATCGGACTCATCAGGAAGC
CCATCGGACTCATCAGGAAGC
CATCGGACTCATCAGGAAGC
GAAACATCGGACTCATCAGGAAGC
CCATCGGACTCATCAGGAAGC
AACATCGGACTCATCAGGAAGC
ATCGGACTCATCAGGAAGC
GCCATCGGACTCATCAGGAAGC
GTTCATCGGACTCATCAGGAAGC
AAGCACATCGGACTCATCAGGAAGC

AGGCATCGGACTCATCAGGAAGC
ACATCGGACTCATCAGGAAGC
GCATCGGACTCATCAGGAAGC
TTCCCATCGGACTCATCAGGAAGC
CAGTCATCGGACTCATCAGGAAGC
AAGCATCGGACTCATCAGGAAGC
TAAGCATCGGACTCATCAGGAAGC
CATCGGACTCATCAGGAAGC
AGAGCATCGGACTCATCAGGAAGC
CACCATCGGACTCATCAGGAAGC
ATCGGACTCATCAGGAAGC
ATCATCGGACTCATCAGGAAGC
GATGGACATCGGACTCATCAGGAAGC
AAGCATCGGACTCATCAGGAAGC
GGTCCATCGGACTCATCAGGAAGC
GCCCATCGGACTCATCAGGAAGC
ACCACCCATCGGACTCATCAGGAAGC
ACACATCGGACTCATCAGGAAGC
CATCGGACTCATCAGGAAGC
TTCATCGGACTCATCAGGAAGC
TGGGCATCGGACTCATCAGGAAGC
CTTCATCGGACTCATCAGGAAGC
CCATCGGACTCATCAGGAAGC
GGTCAGCATCGGACTCATCAGGAAGC
CTCCATCGGACTCATCAGGAAGC
GACATCGGACTCATCAGGAAGC
CCATCGGACTCATCAGGAAGC
TCATCGGACTCATCAGGAAGC
AACATCGGACTCATCAGGAAGC
TCACATCGGACTCATCAGGAAGC
TAGTCATCGGACTCATCAGGAAGC
GCTGTCATCGGACTCATCAGGAAGC
AACATCGGACTCATCAGGAAGC
CATCGGACTCATCAGGAAGC
CCATCGGACTCATCAGGAAGC
CATGACATCGGACTCATCAGGAAGC
AACATCGGACTCATCAGGAAGC
CCATCGGACTCATCAGGAAGC
TTCTCATCGGACTCATCAGGAAGC
CATCGGACTCATCAGGAAGC
ATCGGACTCATCAGGAAGC
CATCATCGGACTCATCAGGAAGC
AGCATCGGACTCATCAGGAAGC

ACATCGGACTCATCAGGAAGC
AGGACATCGGACTCATCAGGAAGC
ACATCGGACTCATCAGGAAGC
ACATCGGACTCATCAGGAAGC
ATGCATCGGACTCATCAGGAAGC
ACAGGACATCGGACTCATCAGGAAGC
TCATCGGACTCATCAGGAAGC
TCATCGGACTCATCAGGAAGC
CATCGGACTCATCAGGAAGC
GTGCATCGGACTCATCAGGAAGC
GCCATCGGACTCATCAGGAAGC
TTCCATCGGACTCATCAGGAAGC
AGCCATCGGACTCATCAGGAAGC
TGGCTCATCGGACTCATCAGGAAGC
AACATCGGACTCATCAGGAAGC
CATCGGACTCATCAGGAAGC
GACTCATCAGGAAGC
AGCCATCGGACTCATCAGGAAGC
TCATCGGACTCATCAGGAAGC
GAGCATCGGACTCATCAGGAAGC
TGCTCATCGGACTCATCAGGAAGC
GGCATCGGACTCATCAGGAAGC
GTGCATCGGACTCATCAGGAAGC
ACATCGGACTCATCAGGAAGC
CATCATCGGACTCATCAGGAAGC
AATGCATCGGACTCATCAGGAAGC
CATCGGACTCATCAGGAAGC
AGACATCGGACTCATCAGGAAGC
CCCCATCGGACTCATCAGGAAGC
CATCGGACTCATCAGGAAGC
CATCGGACTCATCAGGAAGC
AACATCGGACTCATCAGGAAGC
TCATCGGACTCATCAGGAAGC
GGCCATCGGACTCATCAGGAAGC
AGCATCGGACTCATCAGGAAGC
GCCATCGGACTCATCAGGAAGC
CCCATCGGACTCATCAGGAAGC
AGCATCGGACTCATCAGGAAGC
GAATCATCGGACTCATCAGGAAGC
CATCGGACTCATCAGGAAGC
GCATCGGACTCATCAGGAAGC
CCCCATCGGACTCATCAGGAAGC
GACATCGGACTCATCAGGAAGC

TGCATCATCGGACTCATCAGGAAGC
CATCGGACTCATCAGGAAGC
CATCGGACTCATCAGGAAGC
CATCGGACTCATCAGGAAGC
CAGCATCGGACTCATCAGGAAGC
ATCAGCATCGGACTCATCAGGAAGC
CCATCGGACTCATCAGGAAGC
TGCATCGGACTCATCAGGAAGC
CCATCGGACTCATCAGGAAGC
ACATCGGACTCATCAGGAAGC
CATCGGACTCATCAGGAAGC
GGGCATCGGACTCATCAGGAAGC
TCATCGGACTCATCAGGAAGC
CATCGGACTCATCAGGAAGC
TGACATCGGACTCATCAGGAAGC
TATGCCATCGGACTCATCAGGAAGC
CAACACATCGGACTCATCAGGAAGC
GAGCCATCGGACTCATCAGGAAGC
GTCATCGGACTCATCAGGAAGC
CATCGGACTCATCAGGAAGC
ACTCATCGGACTCATCAGGAAGC
AGCATCGGACTCATCAGGAAGC
AAGGCATCGGACTCATCAGGAAGC
GTGTGCATCGGACTCATCAGGAAGC
ATCGGACTCATCAGGAAGC
CATCGGACTCATCAGGAAGC
GTAACATCGGACTCATCAGGAAGC
CCACATCGGACTCATCAGGAAGC
CAGTGCATCGGACTCATCAGGAAGC
TTACATCGGACTCATCAGGAAGC
AAAGGCATCGGACTCATCAGGAAGC
GCATCGGACTCATCAGGAAGC
GGCACATCGGACTCATCAGGAAGC
TTGCCATCGGACTCATCAGGAAGC
GTATCTCATCGGACTCATCAGGAAGC
GTGTCATCGGACTCATCAGGAAGC
GCCATCGGACTCATCAGGAAGC
AGCATCATCGGACTCATCAGGAAGC
CGCATCGGACTCATCAGGAAGC
CCCCATCGGACTCATCAGGAAGC
TCATCGGACTCATCAGGAAGC
GGTCATCGGACTCATCAGGAAGC
CATGACATCGGACTCATCAGGAAGC

TTCATCGGACTCATCAGGAAGC
AATCATCGGACTCATCAGGAAGC
CTCCCATCGGACTCATCAGGAAGC
ACATCGGACTCATCAGGAAGC
AAACATCGGACTCATCAGGAAGC
CTCATCGGACTCATCAGGAAGC
ATCGGACTCATCAGGAAGC
AAATGCATCGGACTCATCAGGAAGC
GGACATCGGACTCATCAGGAAGC
GCCATCGGACTCATCAGGAAGC
CCCATCGGACTCATCAGGAAGC
ATCGGACTCATCAGGAAGC
GTTCATCGGACTCATCAGGAAGC
TCCATCGGACTCATCAGGAAGC
TGTCATCGGACTCATCAGGAAGC
CACCATCGGACTCATCAGGAAGC
GCATCGGACTCATCAGGAAGC
ATCATCGGACTCATCAGGAAGC
CATCGGACTCATCAGGAAGC
ATCGGACTCATCAGGAAGC
GAACATCGGACTCATCAGGAAGC
ATCATCGGACTCATCAGGAAGC
AGGGCATCGGACTCATCAGGAAGC
CCTCATCGGACTCATCAGGAAGC
CAGAGGCATCGGACTCATCAGGAAGC
GAACATCGGACTCATCAGGAAGC
GTCATCGGACTCATCAGGAAGC
GGCATCGGACTCATCAGGAAGC
AGTGCATCGGACTCATCAGGAAGC
ATCGGACTCATCAGGAAGC
CCATCGGACTCATCAGGAAGC
ATCATCGGACTCATCAGGAAGC
AAACATCGGACTCATCAGGAAGC
CAACATCGGACTCATCAGGAAGC
CTTCATCGGACTCATCAGGAAGC
ACAATACATCGGACTCATCAGGAAGC
TGGTACATCGGACTCATCAGGAAGC
TAACATCGGACTCATCAGGAAGC
GTTCATCGGACTCATCAGGAAGC
GGTCATCGGACTCATCAGGAAGC
GCCATCGGACTCATCAGGAAGC
GAGCCATCGGACTCATCAGGAAGC
CATCGGACTCATCAGGAAGC

TCAGTCATCGGACTCATCAGGAAGC
CATCGGACTCATCAGGAAGC
GCTCATCGGACTCATCAGGAAGC
AGCACATCGGACTCATCAGGAAGC
CCATCGGACTCATCAGGAAGC
CATCGGACTCATCAGGAAGC
ATCGGACTCATCAGGAAGC
AAACATCGGACTCATCAGGAAGC
TCATCGGACTCATCAGGAAGC
CCATCGGACTCATCAGGAAGC
GTGGCATCGGACTCATCAGGAAGC
GCCATCGGACTCATCAGGAAGC
TTGCATCGGACTCATCAGGAAGC
TCATCGGACTCATCAGGAAGC
TACCATCGGACTCATCAGGAAGC
TAGCCATCGGACTCATCAGGAAGC
TCATCGGACTCATCAGGAAGC
ACATCGGACTCATCAGGAAGC
GTCATCGGACTCATCAGGAAGC
GCCATCGGACTCATCAGGAAGC
GCCCATCGGACTCATCAGGAAGC
GTTGACCATCGGACTCATCAGGAAGC
CTAGTCATCGGACTCATCAGGAAGC
CCCATCGGACTCATCAGGAAGC
CATCGGACTCATCAGGAAGC
TGGCATCGGACTCATCAGGAAGC
GTCATCGGACTCATCAGGAAGC
CATCGGACTCATCAGGAAGC
GCACATCGGACTCATCAGGAAGC
AACCATCGGACTCATCAGGAAGC
TAGTGCATCGGACTCATCAGGAAGC
CATCGGACTCATCAGGAAGC
CTCATCGGACTCATCAGGAAGC
GTTGATCATCGGACTCATCAGGAAGC
GTACACATCGGACTCATCAGGAAGC
ATCATCGGACTCATCAGGAAGC
ACATCGGACTCATCAGGAAGC
ACCATCGGACTCATCAGGAAGC
TTCCATCGGACTCATCAGGAAGC
TCATCGGACTCATCAGGAAGC
ATCGGACTCATCAGGAAGC
GGTCATCGGACTCATCAGGAAGC
CCCATCGGACTCATCAGGAAGC

CATCGGACTCATCAGGAAGC
CCATCGGACTCATCAGGAAGC
CATCGGACTCATCAGGAAGC
GCATCGGACTCATCAGGAAGC
TCATCGGACTCATCAGGAAGC
TGGCATCGGACTCATCAGGAAGC
CGGACTCATCAGGAAGC
CATCGGACTCATCAGGAAGC
AATCATCGGACTCATCAGGAAGC
GGCATCGGACTCATCAGGAAGC
GGTCATCGGACTCATCAGGAAGC
TGCATCGGACTCATCAGGAAGC
GCATCGGACTCATCAGGAAGC
CCCATCGGACTCATCAGGAAGC
CATCGGACTCATCAGGAAGC
ACATCGGACTCATCAGGAAGC
ACATCGGACTCATCAGGAAGC
ACATCGGACTCATCAGGAAGC
AATCATCGGACTCATCAGGAAGC
CCATCGGACTCATCAGGAAGC
GTCATCGGACTCATCAGGAAGC
GAATCATCGGACTCATCAGGAAGC
GCATCGGACTCATCAGGAAGC
TCCATCGGACTCATCAGGAAGC
TCATCGGACTCATCAGGAAGC
CCATCGGACTCATCAGGAAGC
GCTCATCGGACTCATCAGGAAGC
CATCGGACTCATCAGGAAGC
ATCGGACTCATCAGGAAGC
GGCATCGGACTCATCAGGAAGC
AAGCATCGGACTCATCAGGAAGC
ATCGGACTCATCAGGAAGC
TGCATCGGACTCATCAGGAAGC
TGCCATCGGACTCATCAGGAAGC
CATCGGACTCATCAGGAAGC
CATCGGACTCATCAGGAAGC
TGCCATCGGACTCATCAGGAAGC
CATCGGACTCATCAGGAAGC
CATCGGACTCATCAGGAAGC
TTCATCGGACTCATCAGGAAGC
CATCGGACTCATCAGGAAGC
GACATCGGACTCATCAGGAAGC
ACCATCGGACTCATCAGGAAGC

TGACATCGGACTCATCAGGAAGC
TCATCGGACTCATCAGGAAGC
TAGCATCGGACTCATCAGGAAGC
ATCCATCGGACTCATCAGGAAGC
GAATCATCGGACTCATCAGGAAGC
GTCATCGGACTCATCAGGAAGC
GCATCGGACTCATCAGGAAGC
AATCATCGGACTCATCAGGAAGC
GTCATCGGACTCATCAGGAAGC
ATCATCGGACTCATCAGGAAGC
CATCATCGGACTCATCAGGAAGC
CTCATCGGACTCATCAGGAAGC
CATCGGACTCATCAGGAAGC
CATCGGACTCATCAGGAAGC
GAACATCGGACTCATCAGGAAGC
TCCATCGGACTCATCAGGAAGC
GGCATCGGACTCATCAGGAAGC
CACATCGGACTCATCAGGAAGC
AGACACATCGGACTCATCAGGAAGC
CAAGCATCGGACTCATCAGGAAGC
GACCATCGGACTCATCAGGAAGC
ACATCGGACTCATCAGGAAGC
ATCGGACTCATCAGGAAGC
CCATCGGACTCATCAGGAAGC
CTGCATCGGACTCATCAGGAAGC
GACATCGGACTCATCAGGAAGC
TTGTCATCGGACTCATCAGGAAGC
TTCATCGGACTCATCAGGAAGC
CCATCGGACTCATCAGGAAGC
ATTCCCATCGGACTCATCAGGAAGC
GGCATCGGACTCATCAGGAAGC
CACATCGGACTCATCAGGAAGC
CATCGGACTCATCAGGAAGC
CCATCGGACTCATCAGGAAGC
CCATCGGACTCATCAGGAAGC
CATCGGACTCATCAGGAAGC
TACTGCATCGGACTCATCAGGAAGC
GTCATCGGACTCATCAGGAAGC
ATTCATCGGACTCATCAGGAAGC
TCATCGGACTCATCAGGAAGC
CTCATCGGACTCATCAGGAAGC
GACATCGGACTCATCAGGAAGC
CTCATCGGACTCATCAGGAAGC

GCAGTCTCATCGGACTCATCAGGAAGC
TCATCGGACTCATCAGGAAGC
CAAACATCGGACTCATCAGGAAGC
CCATCGGACTCATCAGGAAGC
GAACATCGGACTCATCAGGAAGC
CCCATCGGACTCATCAGGAAGC
CCATCGGACTCATCAGGAAGC
GACATCGGACTCATCAGGAAGC
GCATCGGACTCATCAGGAAGC
GAAGGCATCGGACTCATCAGGAAGC
ACATCGGACTCATCAGGAAGC
TCATCGGACTCATCAGGAAGC
CCATCGGACTCATCAGGAAGC
AACATCGGACTCATCAGGAAGC
ATCGGACTCATCAGGAAGC
GGGACATCGGACTCATCAGGAAGC
CTGACACATCGGACTCATCAGGAAGC
TCATCGGACTCATCAGGAAGC
GCCCCCATCGGACTCATCAGGAAGC
ATCGGACTCATCAGGAAGC
CATCGGACTCATCAGGAAGC
GGGCATCGGACTCATCAGGAAGC
AGACATCGGACTCATCAGGAAGC
GTCATCGGACTCATCAGGAAGC
CAGCATCGGACTCATCAGGAAGC
CATCGGACTCATCAGGAAGC
GTCATCGGACTCATCAGGAAGC
CTGTCATCGGACTCATCAGGAAGC
CATCGGACTCATCAGGAAGC
TCCATCGGACTCATCAGGAAGC
GTGCCATCGGACTCATCAGGAAGC
TCATCGGACTCATCAGGAAGC
TCGGACTCATCAGGAAGC
GCATCGGACTCATCAGGAAGC
ATCGGACTCATCAGGAAGC
TGCATCGGACTCATCAGGAAGC
CTCCATCGGACTCATCAGGAAGC
CATCGGACTCATCAGGAAGC
CGCATCGGACTCATCAGGAAGC
CACATCGGACTCATCAGGAAGC
GTCATCGGACTCATCAGGAAGC
GTCATCGGACTCATCAGGAAGC
CATCGGACTCATCAGGAAGC

GTCATCGGACTCATCAGGAAGC
ACCATGACATCGGACTCATCAGGAAGC
CCCATCGGACTCATCAGGAAGC
ACATCGGACTCATCAGGAAGC
GGGACATCGGACTCATCAGGAAGC
AATCATCGGACTCATCAGGAAGC
CCATCGGACTCATCAGGAAGC
GACATCGGACTCATCAGGAAGC
TGCATCGGACTCATCAGGAAGC
ACATCGGACTCATCAGGAAGC
ACACATCGGACTCATCAGGAAGC
CAACATCGGACTCATCAGGAAGC
TTTCCCATCGGACTCATCAGGAAGC
TGGTCCATCGGACTCATCAGGAAGC
CCATCGGACTCATCAGGAAGC
CTCATCGGACTCATCAGGAAGC
TACATCGGACTCATCAGGAAGC
ATAACATCGGACTCATCAGGAAGC
ACATCGGACTCATCAGGAAGC
CCATCGGACTCATCAGGAAGC
TCATCGGACTCATCAGGAAGC
TGCATCGGACTCATCAGGAAGC
GCATCGGACTCATCAGGAAGC
CATCGGACTCATCAGGAAGC
GGTCATCGGACTCATCAGGAAGC
ATTCATCGGACTCATCAGGAAGC
ATCGGACTCATCAGGAAGC
ACACATCGGACTCATCAGGAAGC
GGCATCGGACTCATCAGGAAGC
TGGCATCGGACTCATCAGGAAGC
CATCGGACTCATCAGGAAGC
GCATCGGACTCATCAGGAAGC
GTGCATCGGACTCATCAGGAAGC
GATGGCATCGGACTCATCAGGAAGC
GCCCATCGGACTCATCAGGAAGC
ACATCGGACTCATCAGGAAGC
CCAACATCGGACTCATCAGGAAGC
TTTGCATCGGACTCATCAGGAAGC
GTCCCATCGGACTCATCAGGAAGC
GAGCATCGGACTCATCAGGAAGC
CATCGGACTCATCAGGAAGC
ACATCGGACTCATCAGGAAGC
CACATCGGACTCATCAGGAAGC

CATCGGACTCATCAGGAAGC
GCATCGGACTCATCAGGAAGC
TCACCACCCATCGGACTCATCAGGAAGC
ATCGGACTCATCAGGAAGC
GACATCGGACTCATCAGGAAGC
ACCATCGGACTCATCAGGAAGC
GCCATCGGACTCATCAGGAAGC
ACCATCGGACTCATCAGGAAGC
CCCCATCGGACTCATCAGGAAGC
CCATCGGACTCATCAGGAAGC
TCATCGGACTCATCAGGAAGC
CATCGGACTCATCAGGAAGC
TTGGCCATCGGACTCATCAGGAAGC
CCATCGGACTCATCAGGAAGC
CATCGGACTCATCAGGAAGC
TCATCGGACTCATCAGGAAGC
CATCGGACTCATCAGGAAGC
ACTCATCGGACTCATCAGGAAGC
GTACATCGGACTCATCAGGAAGC
GGTCATCGGACTCATCAGGAAGC
TGATCATCGGACTCATCAGGAAGC
GCACATCGGACTCATCAGGAAGC
TTCATCGGACTCATCAGGAAGC
ACATCGGACTCATCAGGAAGC
GGCATCGGACTCATCAGGAAGC
ACATCGGACTCATCAGGAAGC
CTCATCGGACTCATCAGGAAGC
AAAAACATCGGACTCATCAGGAAGC
GTCATCGGACTCATCAGGAAGC
GTGTGCATCGGACTCATCAGGAAGC
ACATCGGACTCATCAGGAAGC
CATCGGACTCATCAGGAAGC
ATCGGACTCATCAGGAAGC
GCATCGGACTCATCAGGAAGC
GCCCATCGGACTCATCAGGAAGC
ACAGCATCGGACTCATCAGGAAGC
TGACATCGGACTCATCAGGAAGC
CCATCGGACTCATCAGGAAGC
AACATCGGACTCATCAGGAAGC
GGCATCGGACTCATCAGGAAGC
AACACATCGGACTCATCAGGAAGC
CCATCTCATCGGACTCATCAGGAAGC
TGCCATCGGACTCATCAGGAAGC

[illegible]GAGCATCGGACTCATCAGGAAGC
GCCCATCGGACTCATCAGGAAGC
[illegible]ACATCGGACTCATCAGGAAGC
