## Supplementary material for "Retrospective cell lineage reconstruction in Humans using short tandem repeats": sup file2

| TEID | Sequence |
|---|---|
| 119079 | TATGAGTGTGGAGTCGTTGCGGTACTATGAAGTAAGTATGCATNNNAGATCGGAA |
| 119080 | TATGAGTGTGGAGTCGTTGCAACACAACCTTAAAAAGAGAAAANNNAGATCGGAA |
| 119081 | TATGAGTGTGGAGTCGTTGCTACACTCTGCTAAACTTATTTCANNNAGATCGGAAGA |
| 119082 | TATGAGTGTGGAGTCGTTGCTTTTCAAATCTTCTACAGTTCTGNNNAGATCGGAAGA |
| 119083 | TATGAGTGTGGAGTCGTTGCTGCTAAATAATTTAAATTGCAGCNNNAGATCGGAAG |
| 119084 | TATGAGTGTGGAGTCGTTGCATACATGTTCAATAAGGACAATGNNNAGATCGGAA |
| 119085 | TATGAGTGTGGAGTCGTTGCAATAGTCCAATGAGTGTGTGNNNAGATCGGAAGAG |
| 119086 | TATGAGTGTGGAGTCGTTGCTAGAATCCATTTACAAGGACAGNNNAGATCGGAAG |
| 119087 | TATGAGTGTGGAGTCGTTGCCTTTGTTAGAGCAATACTCAATNNNAGATCGGAAGA |
| 119088 | TATGAGTGTGGAGTCGTTGCCTTGGGATTCAGAATTTTGAGNNNAGATCGGAAGA |
| 119089 | TATGAGTGTGGAGTCGTTGCGTAATCCAGATCCGTAAGCTNNNAGATCGGAAGAG |
| 119090 | TATGAGTGTGGAGTCGTTGCAAGACCTACATGTTCAGAATCNNNAGATCGGAAGAG |
| 119091 | TATGAGTGTGGAGTCGTTGCCATTAGACTTCTTGTTGGAAAGNNNAGATCGGAAG |
| 119092 | TATGAGTGTGGAGTCGTTGCTGTTCTGGATTTAAAGAAATACCNNNAGATCGGAAG |
| 119093 | TATGAGTGTGGAGTCGTTGCGAATGATGATCACACCACTGNNNAGATCGGAAGAG |
| 119094 | TATGAGTGTGGAGTCGTTGCAATCTGGTCTTCATCACCTANNNAGATCGGAAGAGC |
| 119095 | TATGAGTGTGGAGTCGTTGCCCCATGTTGCCTATTTATTTTTTNNNAGATCGGAAGA |
| 119096 | TATGAGTGTGGAGTCGTTGCGGGCAAAAAAAGTTTATTGAAAANNNAGATCGGAA |
| 119097 | TATGAGTGTGGAGTCGTTGCAATAGCTAAATATTGGAAACCACNNNAGATCGGAAG |
| 119098 | TATGAGTGTGGAGTCGTTGCGAGGAACATAACAAACTTTACATNNNAGATCGGAAG |
| 119099 | TATGAGTGTGGAGTCGTTGCCCGAGATACTTGTTAAAAATTTGNNNAGATCGGAAG |
| 119100 | TATGAGTGTGGAGTCGTTGCGAACTTCATCTGCAAGTGAGNNNAGATCGGAAGAG |
| 119101 | TATGAGTGTGGAGTCGTTGCGAGGAGGACTATAAAAGAACAAANNNAGATCGGA |
| 119102 | TATGAGTGTGGAGTCGTTGCTTCTCTATGTTCACCTAACTCTANNNAGATCGGAAGA |
| 119103 | TATGAGTGTGGAGTCGTTGCGATATTCCACCAGAATCGTAGNNNAGATCGGAAGA |
| 119104 | TATGAGTGTGGAGTCGTTGCTCCATGAGAATCTTCCTCAGNNNAGATCGGAAGAGC |
| 119105 | TATGAGTGTGGAGTCGTTGCTGAACCAGCAATAATGTTTTCNNNAGATCGGAAGAG |
| 119106 | TATGAGTGTGGAGTCGTTGCGCATGTACAGAGTGAGTAATGNNNAGATCGGAAGA |
| 119107 | TATGAGTGTGGAGTCGTTGCCTTGTGAAGGAATCTAGCATAANNNAGATCGGAAG |
| 119108 | TATGAGTGTGGAGTCGTTGCTTATTTTTCCTTCCTTTTCTTGCNNNAGATCGGAAGA |
| 119109 | TATGAGTGTGGAGTCGTTGCCAGTAAGTAACGATGCTCTTTANNNAGATCGGAAGA |
| 119110 | TATGAGTGTGGAGTCGTTGCAACAGTTATTTCTAGTGTTGTTGNNNAGATCGGAAG |
| 119111 | TATGAGTGTGGAGTCGTTGCTTTATCAAAAAAGAACTCAACCANNNAGATCGGAAG |
| 119112 | TATGAGTGTGGAGTCGTTGCGAGAAAAGCAAGAGAAGTTAAATNNNAGATCGGA |
| 119113 | TATGAGTGTGGAGTCGTTGCAGTAGCAAGATTATATATGTGCANNNAGATCGGAA |
| 119114 | TATGAGTGTGGAGTCGTTGCGAACGTAAGTGAGGAAAATATAAANNNAGATCGGA |
| 119115 | TATGAGTGTGGAGTCGTTGCGCCTTGGTAACTTTTTCTAATTANNNAGATCGGAAG |
| 119116 | TATGAGTGTGGAGTCGTTGCCAGAATTGGAGTCTCTGTATCNNNAGATCGGAAGA |
| 119117 | TATGAGTGTGGAGTCGTTGCTAATTTCCCCAAAAAAATGAAACNNNAGATCGGAAG |
| 119118 | TATGAGTGTGGAGTCGTTGCTTACTTGTTTGTCCAAAGTTATTNNNAGATCGGAAG |
| 119119 | TATGAGTGTGGAGTCGTTGCCCAAATAGCTAGGACTCCAGNNNAGATCGGAAGAG |
| 119120 | TATGAGTGTGGAGTCGTTGCAATAACAAACCATTTTCATGAGTNNNAGATCGGAAG |

119121 TATGAGTGTGGAGTCGTTGCATTAGCCAACTACCGTTTATAAANNNAGATCGGAAG
119122 TATGAGTGTGGAGTCGTTGCCAACTGACTGAACATGAAAATATNNNAGATCGGAAG
119123 TATGAGTGTGGAGTCGTTGCGTTTTTTTCTGTAGGTAAACTCTNNNAGATCGGAAG
119124 TATGAGTGTGGAGTCGTTGCAGAAGTCTTCTTTTGCTAATGTNNNAGATCGGAAGA
119125 TATGAGTGTGGAGTCGTTGCCTTTTATAGCCTAATGAACACACNNNAGATCGGAAG
119126 TATGAGTGTGGAGTCGTTGCTTGTTTATTCGAGAAAATGATCANNNAGATCGGAAG
119127 TATGAGTGTGGAGTCGTTGCTGACTGAGAAAGACCTTTTAATNNNAGATCGGAAGA
119128 TATGAGTGTGGAGTCGTTGCCAAAGGTTATTCTTAATCCATCCNNNAGATCGGAAG
119129 TATGAGTGTGGAGTCGTTGCACTAATCTGTCTTACTGTTCAATNNNAGATCGGAAGA
119130 TATGAGTGTGGAGTCGTTGCCTTTCAAAAGATCCAGTTCTCTNNNAGATCGGAAGA
119131 TATGAGTGTGGAGTCGTTGCGAGTTATTTGGCAGTAGATTTTTNNNAGATCGGAAG
119132 TATGAGTGTGGAGTCGTTGCAAACACTGATCTATTCACCTATTNNNAGATCGGAAGA
119133 TATGAGTGTGGAGTCGTTGCAAACATATCTACTTCCACGTAAGNNNAGATCGGAAG
119134 TATGAGTGTGGAGTCGTTGCGAGAAGATTTATGAGCCAAGTANNNAGATCGGAAG
119135 TATGAGTGTGGAGTCGTTGCAACTACCTCTACCTCCTCTTNNNAGATCGGAAGAGCA
119136 TATGAGTGTGGAGTCGTTGCTTTGTGAAGAAATTCAGAATGCNNNAGATCGGAAG
119137 TATGAGTGTGGAGTCGTTGCGTACAGAAGACTCAAAAACTTCNNNAGATCGGAAGA
119138 TATGAGTGTGGAGTCGTTGCGAAAAGCTACTATAGTCACAGANNNAGATCGGAAG
119139 TATGAGTGTGGAGTCGTTGCTATACTCACAGTACTCACACTNNNAGATCGGAAGAG
119140 TATGAGTGTGGAGTCGTTGCCTTAATAGGCACATGAGATCTCNNNAGATCGGAAGA
119141 TATGAGTGTGGAGTCGTTGCTGGCTTATTTAGTATCATGTATGNNNAGATCGGAAG
119142 TATGAGTGTGGAGTCGTTGCCCTAACAACAGCAATCAATTAATNNNAGATCGGAAG
119143 TATGAGTGTGGAGTCGTTGCGAGATTTTAGCAGATCCGATATNNNAGATCGGAAG
119144 TATGAGTGTGGAGTCGTTGCGTAAAATGACAAATACTGAAGGTNNNAGATCGGAA
119145 TATGAGTGTGGAGTCGTTGCCTTTTCCTTGTACCCAACAANNNAGATCGGAAGAGC
119146 TATGAGTGTGGAGTCGTTGCGATGCTACTGGACAATCTTTTNNNAGATCGGAAGAG
119147 TATGAGTGTGGAGTCGTTGCATAGTTGAGTGTAACAATCTCATNNNAGATCGGAAG
119149 TATGAGTGTGGAGTCGTTGCTTCAGGAGTTTTTTTATGTTCTTNNNAGATCGGAAG
119150 TATGAGTGTGGAGTCGTTGCCCTTACATGCTTTATCAACTAAGNNNAGATCGGAAG
119151 TATGAGTGTGGAGTCGTTGCAGTTTACAAATGCCTACAAGTTANNNAGATCGGAAG
119152 TATGAGTGTGGAGTCGTTGCCAATGGACTACTTGGAATTTTTANNNAGATCGGAAG
119153 TATGAGTGTGGAGTCGTTGCGTTATGAAGGTAAATGGATTCACNNNAGATCGGAA
119154 TATGAGTGTGGAGTCGTTGCGTTATGAAGGTAAATGGATTCACNNNAGATCGGAA
119155 TATGAGTGTGGAGTCGTTGCATAACTTTCAATTACAGTCCAACNNNAGATCGGAAG
119156 TATGAGTGTGGAGTCGTTGCGTGAATGAAATAGAAAAAATCGTTNNNAGATCGGA
119157 TATGAGTGTGGAGTCGTTGCCAAAGACTAAGTCCACCTAAGNNNAGATCGGAAGA
119158 TATGAGTGTGGAGTCGTTGCGTGTGAGTGCAAAAAAGAAGNNNAGATCGGAAGA
119159 TATGAGTGTGGAGTCGTTGCAATTTTGCTGAAAATCCAGATATNNNAGATCGGAAG
119160 TATGAGTGTGGAGTCGTTGCTGACCTGGTTAATAAATATGCTANNNAGATCGGAAG
119161 TATGAGTGTGGAGTCGTTGCTACATTCAAACTCAATATCCATGNNNAGATCGGAAG
119162 TATGAGTGTGGAGTCGTTGCAAAGAAGTAACCCAAACGTAANNNAGATCGGAAGA
119163 TATGAGTGTGGAGTCGTTGCAAAACTGAGTAGTGTTGTTAAAGNNNAGATCGGAA
119164 TATGAGTGTGGAGTCGTTGCTGTGTAAAAAAAATATGTCCATGNNNAGATCGGAA

119165 TATGAGTGTGGAGTCGTTGCATATAGAATGTATATCTCAGGGCNNNAGATCGGAAG
119166 TATGAGTGTGGAGTCGTTGCAAAATCGAGAAGAGAGGAGANNNAGATCGGAAGA
119167 TATGAGTGTGGAGTCGTTGCAACATGTTAGTGTTAAGGAAAATNNNAGATCGGAA
119168 TATGAGTGTGGAGTCGTTGCATTCCTTATTGACATAGCTTAGCNNNAGATCGGAAG
119169 TATGAGTGTGGAGTCGTTGCCAAATAGTTCTCGCTCAGTGNNNAGATCGGAAGAG
119170 TATGAGTGTGGAGTCGTTGCATTAGGTCCCTTCATTAGAAATCNNNAGATCGGAAG
119171 TATGAGTGTGGAGTCGTTGCATTCTCCTTTAGTCATTCAACAANNNAGATCGGAAGA
119172 TATGAGTGTGGAGTCGTTGCGAAGAAGGAATGAGGCTTAAATANNNAGATCGGAA
119174 TATGAGTGTGGAGTCGTTGCGAAAACACTTTAAACCTCTTTTGNNNAGATCGGAAG
119175 TATGAGTGTGGAGTCGTTGCTTTCTCTTTATAGCAGTCGTTAANNNAGATCGGAAG
119176 TATGAGTGTGGAGTCGTTGCTAAAGGTGTTAGTCAAACAAAACNNNAGATCGGAA
119177 TATGAGTGTGGAGTCGTTGCGGAAAACATAAGTCAAGTATCAANNNAGATCGGAA
119178 TATGAGTGTGGAGTCGTTGCCATAGAAACTGGAAAGTACAGTANNNAGATCGGAA
119179 TATGAGTGTGGAGTCGTTGCCAACAAGATGATTTCTGGAATCNNNAGATCGGAAGA
119180 TATGAGTGTGGAGTCGTTGCCTGTTCACTTTCTCTTACAAGNNNAGATCGGAAGAG
119181 TATGAGTGTGGAGTCGTTGCAAAAGTTTTTAATACACTGCTGANNNAGATCGGAAG
119182 TATGAGTGTGGAGTCGTTGCTGCTTAGGGATTTTCTTATTGATNNNAGATCGGAAG
119183 TATGAGTGTGGAGTCGTTGCTCAAAACAAATAAGTTTTCATCAGNNNAGATCGGAA
119184 TATGAGTGTGGAGTCGTTGCTCACCGATAAGATCATCTATTTTNNNAGATCGGAAG
119185 TATGAGTGTGGAGTCGTTGCTATATGTATGAGAGAGAAACACGNNNAGATCGGAA
119186 TATGAGTGTGGAGTCGTTGCTTTAATTCCTGCAAGAGAAATACNNNAGATCGGAAG
119187 TATGAGTGTGGAGTCGTTGCAAATCCCATACAAGTAAGAGTTANNNAGATCGGAAG
119188 TATGAGTGTGGAGTCGTTGCCTTCTTCTTCATCACTTTCCTTNNNAGATCGGAAGAG
119189 TATGAGTGTGGAGTCGTTGCAGAGAGGTTTTGATTACTCAAATNNNAGATCGGAAG
119190 TATGAGTGTGGAGTCGTTGCCATAATGTCTTTTTCCAACAAACNNNAGATCGGAAGA
119191 TATGAGTGTGGAGTCGTTGCGCAAGTTCATTCTCATAACAAAANNNAGATCGGAAG
119192 TATGAGTGTGGAGTCGTTGCTTTCTCTGGATGCAAAATAAAATNNNAGATCGGAAG
119193 TATGAGTGTGGAGTCGTTGCTTAGTCTTCATTCCCATCCTGNNNAGATCGGAAGAG
119194 TATGAGTGTGGAGTCGTTGCATATCAAGGTTATTGGTAAGTGANNNAGATCGGAA
119195 TATGAGTGTGGAGTCGTTGCTTTTAAATTCCAAGTCATTCCTCNNNAGATCGGAAGA
119196 TATGAGTGTGGAGTCGTTGCCCTCTATTTTGTAAACCTCAGANNNAGATCGGAAGA
119197 TATGAGTGTGGAGTCGTTGCGTTTAAACTACTATGAGAAAAGGANNNAGATCGGA
119198 TATGAGTGTGGAGTCGTTGCCCTTCTTATCTAGTTAGATGCTTNNNAGATCGGAAGA
119199 TATGAGTGTGGAGTCGTTGCATTCTTCACTATGGAGCTCTTTNNNAGATCGGAAGA
119200 TATGAGTGTGGAGTCGTTGCCTCTTAAGGACCCATCTAGATNNNAGATCGGAAGAG
119201 TATGAGTGTGGAGTCGTTGCTACAACCTCAAAGAACGTTATNNNAGATCGGAAGAG
119202 TATGAGTGTGGAGTCGTTGCGAACTAGATTTACTGGAAGACTTNNNAGATCGGAAG
119203 TATGAGTGTGGAGTCGTTGCACCGAGATAGAAAAAAAAGTTTTNNNAGATCGGAA
119204 TATGAGTGTGGAGTCGTTGCCTTTCCTTCTCAACAGAGTGNNNAGATCGGAAGAGC
119205 TATGAGTGTGGAGTCGTTGCGTTTTGACAAACTCAAGCTTANNNAGATCGGAAGAG
119206 TATGAGTGTGGAGTCGTTGCGAAATTCTTTAATAGCAGCTCCNNNAGATCGGAAGA
119207 TATGAGTGTGGAGTCGTTGCTTATAAAGCAAATATTTCCTCCCNNNAGATCGGAAGA
119208 TATGAGTGTGGAGTCGTTGCCTGGTGACCTAAAAAAGAAATAANNNAGATCGGAA

119209 TATGAGTGTGGAGTCGTTGCCTACAAAAGCTTCCACTAAATTCNNNAGATCGGAAG
119210 TATGAGTGTGGAGTCGTTGCGGAATTCCTTATGTGTGAAATTTNNNAGATCGGAAG
119211 TATGAGTGTGGAGTCGTTGCCCTTTCAAATTAGCTTATTTCACNNNAGATCGGAAGA
119212 TATGAGTGTGGAGTCGTTGCTAAAAGAAAAACTTCAGCTGAATNNNAGATCGGAA
119213 TATGAGTGTGGAGTCGTTGCTATCTTCAAAACAACCTGATAGANNNAGATCGGAAG
119214 TATGAGTGTGGAGTCGTTGCCCAGGAATAGTACCATATTCATTNNNAGATCGGAAG
119215 TATGAGTGTGGAGTCGTTGCAAAGTTCTTCAGAAGCATCAANNNAGATCGGAAGA
119216 TATGAGTGTGGAGTCGTTGCCTGAGAGCTTTTGTCTCAAANNNAGATCGGAAGAGC
119217 TATGAGTGTGGAGTCGTTGCCAAAATACATCTATTCCCATCAGNNNAGATCGGAAG
119219 TATGAGTGTGGAGTCGTTGCCTAAATTGGGAAACAGTTGTTAGNNNAGATCGGAA
119220 TATGAGTGTGGAGTCGTTGCGCAGTAAGAGTCATACGAATAANNNAGATCGGAAG
119221 TATGAGTGTGGAGTCGTTGCCTTGGTATTTGTATTCAAGCATCNNNAGATCGGAAG
119222 TATGAGTGTGGAGTCGTTGCCTCAATGGTTTGTGATTTTTTTTNNNAGATCGGAAG
119223 TATGAGTGTGGAGTCGTTGCAATTCCCAGTCTTCATTTCCNNNAGATCGGAAGAGCA
119224 TATGAGTGTGGAGTCGTTGCGCATCATTCACTTTTATCAAATGNNNAGATCGGAAG
119225 TATGAGTGTGGAGTCGTTGCATAGGAACATGAAATAGCAATGANNNAGATCGGAA
119226 TATGAGTGTGGAGTCGTTGCTACCTACTACCATTTTTCAATGTNNNAGATCGGAAGA
119227 TATGAGTGTGGAGTCGTTGCATTAATGCATTCTTCTCCAAATCNNNAGATCGGAAGA
119228 TATGAGTGTGGAGTCGTTGCATGCTATCCTGGACTTTGAGNNNAGATCGGAAGAG
119229 TATGAGTGTGGAGTCGTTGCCAGGCTCTCAAGATAAAAGATANNNAGATCGGAAG
119230 TATGAGTGTGGAGTCGTTGCACAGTGGTAATGTATTTTTGTGNNNAGATCGGAAG
119231 TATGAGTGTGGAGTCGTTGCCTAGGATGCCAACAAAGAAANNNAGATCGGAAGAG
119232 TATGAGTGTGGAGTCGTTGCACATAGGAGAAGACGATCAGNNNAGATCGGAAGA
119233 TATGAGTGTGGAGTCGTTGCGTGATTTCCAAGTGTAACAATAGNNNAGATCGGAA
119234 TATGAGTGTGGAGTCGTTGCAAAGTTGTATTTTGGGTAAAAGANNNAGATCGGAA
119235 TATGAGTGTGGAGTCGTTGCATGAGCCAAATAAGAAGTTCTTNNNAGATCGGAAG
119236 TATGAGTGTGGAGTCGTTGCGAAAAGCTCCTATCAAACAAATANNNAGATCGGAAG
119237 TATGAGTGTGGAGTCGTTGCAGATGAGTAAAAACATCAGCTNNNAGATCGGAAGA
119238 TATGAGTGTGGAGTCGTTGCGAAGTCACATTAATGATGAACTGNNNAGATCGGAA
119239 TATGAGTGTGGAGTCGTTGCGCATCGTCATGAGTATGTACNNNAGATCGGAAGAG
119240 TATGAGTGTGGAGTCGTTGCGCAACTTTATATCAGTGAGAATTNNNAGATCGGAAG
119241 TATGAGTGTGGAGTCGTTGCATTATGAATAAATGAGGAAGGGCNNNAGATCGGAA
119242 TATGAGTGTGGAGTCGTTGCTGAGATTAAGTCCAATCAATCATNNNAGATCGGAAG
119243 TATGAGTGTGGAGTCGTTGCCCAGATCTAAGTAATAAGAAGCANNNAGATCGGAA
119244 TATGAGTGTGGAGTCGTTGCCTCTGTAGTTACATGCAAATAAANNNAGATCGGAAG
119245 TATGAGTGTGGAGTCGTTGCTGCTTATATAGGTTTGATGAGTANNNAGATCGGAA
119246 TATGAGTGTGGAGTCGTTGCTCAAGGAAGAATTGAGCTTATGNNNAGATCGGAAG
119247 TATGAGTGTGGAGTCGTTGCGGAAAGAAAAACTGGAATATAGTNNNAGATCGGAA
119248 TATGAGTGTGGAGTCGTTGCGCTTTAAATTCCTCTCGAAATTNNNAGATCGGAAGA
119249 TATGAGTGTGGAGTCGTTGCTAGCCTCAATAACAAATAAATGCNNNAGATCGGAAG
119250 TATGAGTGTGGAGTCGTTGCCTTCATGTTCAGAAAAACTCCNNNAGATCGGAAGAG
119251 TATGAGTGTGGAGTCGTTGCCACACCTAGTAAGAAAATTTTGTNNNAGATCGGAAG
119252 TATGAGTGTGGAGTCGTTGCCATCAAAACACAGAAACAGGNNNAGATCGGAAGAG

119253 TATGAGTGTGGAGTCGTTGCTAATTGCATTTTAGTTCAAGGGNNNAGATCGGAAGA
119254 TATGAGTGTGGAGTCGTTGCATCTATTAGTACACGAGTAACTCNNNAGATCGGAAG
119255 TATGAGTGTGGAGTCGTTGCAACAGGTCATAGTTTGAACCNNNAGATCGGAAGAG
119256 TATGAGTGTGGAGTCGTTGCGACTGGAGAAAAGGTTTCAGNNNAGATCGGAAGAG
119257 TATGAGTGTGGAGTCGTTGCTGAAAAAATAGGTAGCTTTTTGTNNNAGATCGGAAG
119258 TATGAGTGTGGAGTCGTTGCGTAGTGGAAAGTAATGATCTCTTNNNAGATCGGAAG
119259 TATGAGTGTGGAGTCGTTGCCTGAGGGTCATGATCTATATTCNNNAGATCGGAAGA
119260 TATGAGTGTGGAGTCGTTGCCCAAATACCAAAAACGTTGAGNNNAGATCGGAAGAG
119261 TATGAGTGTGGAGTCGTTGCGATTCCTAACAGACTGGTTGNNNAGATCGGAAGAG
119262 TATGAGTGTGGAGTCGTTGCATTCAGTCTTCTGTTCCCTATNNNAGATCGGAAGAGC
119263 TATGAGTGTGGAGTCGTTGCCTTCTCAAAACATCTTCTTTCCNNNAGATCGGAAGAG
119264 TATGAGTGTGGAGTCGTTGCCCAGTGTATATGCTAAGCAGNNNAGATCGGAAGAG
119265 TATGAGTGTGGAGTCGTTGCAGCTATGAATACAGAGGTATATGNNNAGATCGGAA
119266 TATGAGTGTGGAGTCGTTGCTTCTAAAAGTCATGTTAGACAGTNNNAGATCGGAAG
119267 TATGAGTGTGGAGTCGTTGCTCACACAAAAAAATGTTAAGGATNNNAGATCGGAAG
119268 TATGAGTGTGGAGTCGTTGCGTATTAGTGGAAGACTTTAAGAANNNAGATCGGAA
119269 TATGAGTGTGGAGTCGTTGCATCAAGAGGATAAACTGCATTNNNAGATCGGAAGA
119270 TATGAGTGTGGAGTCGTTGCTGAGATAATAAATTATGGTGTAAGTTNNNAGATCGG
119271 TATGAGTGTGGAGTCGTTGCCCCAGAAAGGTAAGAAAATTAAANNNAGATCGGAA
119272 TATGAGTGTGGAGTCGTTGCTTCTAAAAGTCATGTTAGACAGTNNNAGATCGGAAG
119273 TATGAGTGTGGAGTCGTTGCAATTTTATCACACAACAGTCATCNNNAGATCGGAAGA
119274 TATGAGTGTGGAGTCGTTGCTTGTCTATGCCTCAGATTTTATNNNAGATCGGAAGAG
119275 TATGAGTGTGGAGTCGTTGCAATCTCCTTGTGGTAAATAAGATNNNAGATCGGAAG
119276 TATGAGTGTGGAGTCGTTGCTAATTTAGACCCTGAAAAAGAGANNNAGATCGGAAG
119277 TATGAGTGTGGAGTCGTTGCATGTTTTAGTTCAACAGAATACCNNNAGATCGGAAG
119278 TATGAGTGTGGAGTCGTTGCAACCTAACGGAGTTCTCTATNNNAGATCGGAAGAGC
119279 TATGAGTGTGGAGTCGTTGCAAGGTCAGATAAGGAAAATGAAANNNAGATCGGAA
119280 TATGAGTGTGGAGTCGTTGCATCTACTCCCACAAGATACTTNNNAGATCGGAAGAG
119281 TATGAGTGTGGAGTCGTTGCTGGATTTCTAATTCTTTTCTGTGNNNAGATCGGAAGA
119282 TATGAGTGTGGAGTCGTTGCATGTCAGATTGAAGATATACAGTNNNAGATCGGAAG
119283 TATGAGTGTGGAGTCGTTGCGATAGTGACTATCTCAATAGAGCNNNAGATCGGAAG
119285 TATGAGTGTGGAGTCGTTGCCTGATACTTCTTGAAATACACACNNNAGATCGGAAGA
119286 TATGAGTGTGGAGTCGTTGCCATCCCTGGAGAAAGTAAAGNNNAGATCGGAAGAG
119287 TATGAGTGTGGAGTCGTTGCTATGATTTGTTTGTTTGTTTGTGNNNAGATCGGAAG
119288 TATGAGTGTGGAGTCGTTGCATTTCCACAGAATCTTTTAACAGNNNAGATCGGAAGA
119289 TATGAGTGTGGAGTCGTTGCCTTCATGAAATCAGCATGAGANNNAGATCGGAAGAG
119290 TATGAGTGTGGAGTCGTTGCATCAGAACCTGGTAAATATCTTTNNNAGATCGGAAG
119291 TATGAGTGTGGAGTCGTTGCAATTTTATGGAATCCACACCTNNNAGATCGGAAGAG
119292 TATGAGTGTGGAGTCGTTGCTTTTAGCTTTTTGGAAGACTAAANNNAGATCGGAAG
119293 TATGAGTGTGGAGTCGTTGCCAGTTGCTTTGTACTTAACATCNNNAGATCGGAAGA
119294 TATGAGTGTGGAGTCGTTGCTCCTATTTTATCAGGTATTGTCCNNNAGATCGGAAGA
119295 TATGAGTGTGGAGTCGTTGCAAATTGGATTGAGGTAAATTGTTNNNAGATCGGAA
119296 TATGAGTGTGGAGTCGTTGCACAGCATTATATGGAATACAGATNNNAGATCGGAAG

119297 TATGAGTGTGGAGTCGTTGCAGTTGTTACTCATAGAGGTATTGNNNAGATCGGAAG
119298 TATGAGTGTGGAGTCGTTGCTCCTTTGAATGTTACTTAACCTANNNAGATCGGAAGA
119299 TATGAGTGTGGAGTCGTTGCGAAGTCACTCCTTCTACATTATANNNAGATCGGAAGA
119300 TATGAGTGTGGAGTCGTTGCCAAGTCACAAGAGATACCTAGNNNAGATCGGAAGAG
119301 TATGAGTGTGGAGTCGTTGCCTCGAATTTAAGGTATGGTATTGNNNAGATCGGAAG
119302 TATGAGTGTGGAGTCGTTGCCTTCACTCCTACCACTGAAANNNAGATCGGAAGAGCA
119303 TATGAGTGTGGAGTCGTTGCTGAATACTACCAAACATTCTTTGNNNAGATCGGAAGA
119304 TATGAGTGTGGAGTCGTTGCTAGGAATAGATCCAAACTAGCTANNNAGATCGGAAG
119305 TATGAGTGTGGAGTCGTTGCTAAAGTGCTACTCCCAACTTNNNAGATCGGAAGAGCA
119306 TATGAGTGTGGAGTCGTTGCTATTCGACAAATATGTATGATGCNNNAGATCGGAAG
119307 TATGAGTGTGGAGTCGTTGCAACTGAGGAGGATTGCATTTNNNAGATCGGAAGAG
119308 TATGAGTGTGGAGTCGTTGCTTGGCTTTCTTTCTTTTTCTTTANNNAGATCGGAAGAG
119309 TATGAGTGTGGAGTCGTTGCTTATTGGTCATCATTTCCAGTANNNAGATCGGAAGAG
119310 TATGAGTGTGGAGTCGTTGCGAGAAGGGAGAAAAAGGAAGNNNAGATCGGAAGA
119311 TATGAGTGTGGAGTCGTTGCATTCATTCAGTGTAGGTGTGNNNAGATCGGAAGAG
119312 TATGAGTGTGGAGTCGTTGCTTGTAAGAATTGTTAAGCATTCCNNNAGATCGGAAG
119313 TATGAGTGTGGAGTCGTTGCGAGAAGAACCATATGTAAGTGTNNNAGATCGGAAG
119314 TATGAGTGTGGAGTCGTTGCTTTTTGACCATACCTGAGTATAANNNAGATCGGAAGA
119315 TATGAGTGTGGAGTCGTTGCAGTAAGTCAAAGACGATAAGAATNNNAGATCGGAAG
119316 TATGAGTGTGGAGTCGTTGCTTTAATGATCTTCTGGTCATTTGNNNAGATCGGAAGA
119317 TATGAGTGTGGAGTCGTTGCGATAGTTTGTCTTTCGAATAGTCNNNAGATCGGAAG
119318 TATGAGTGTGGAGTCGTTGCGAAGAAGAAAACAAAGGTAGAACNNNAGATCGGAA
119319 TATGAGTGTGGAGTCGTTGCGATGGGTTCATTATCAAAGATTTNNNAGATCGGAAG
119320 TATGAGTGTGGAGTCGTTGCCTTTTCATCGAAGTATGTGTAAANNNAGATCGGAAG
119321 TATGAGTGTGGAGTCGTTGCAAGGATCTTCTTCAGACAGCNNNAGATCGGAAGAGC
119322 TATGAGTGTGGAGTCGTTGCGAGATTTCAGGTCATTATCAATCNNNAGATCGGAAG
119323 TATGAGTGTGGAGTCGTTGCTTGTATTGTTTCTAAGACAAAGGNNNAGATCGGAAG
119324 TATGAGTGTGGAGTCGTTGCCAAGAAGGAAGGGAAATTAAAGNNNAGATCGGAAG
119325 TATGAGTGTGGAGTCGTTGCCTTTGAGAAGAAAGGTGAGTCNNNAGATCGGAAGA
119326 TATGAGTGTGGAGTCGTTGCCCTGGTTATATGGTGGAAAGNNNAGATCGGAAGAG
119327 TATGAGTGTGGAGTCGTTGCAAAATGGAAAACGTAAACACAANNNAGATCGGAAGA
119328 TATGAGTGTGGAGTCGTTGCTGGTTTCAGAACAGTTTAAAAATNNNAGATCGGAAG
119329 TATGAGTGTGGAGTCGTTGCAGAGAGGGTATTTAGACCAGNNNAGATCGGAAGAG
119330 TATGAGTGTGGAGTCGTTGCTTTGTGGACCTAATTATAAAGGTNNNAGATCGGAAG
119331 TATGAGTGTGGAGTCGTTGCCTCAATTCTCAGGGACTTTAATNNNAGATCGGAAGAG
119332 TATGAGTGTGGAGTCGTTGCTTTTCTGTCCTTGAACTTCTTCNNNAGATCGGAAGAG
119333 TATGAGTGTGGAGTCGTTGCGGTCTGGCTATATATAACACTTTNNNAGATCGGAAGA
119334 TATGAGTGTGGAGTCGTTGCAGTATCAGGGGAAATCTAAATTTNNNAGATCGGAAG
119335 TATGAGTGTGGAGTCGTTGCTTACATTCTTGTCACAGGTCNNNAGATCGGAAGAGCA
119336 TATGAGTGTGGAGTCGTTGCAGAACACTTGGATACCTTAAATANNNAGATCGGAAG
119337 TATGAGTGTGGAGTCGTTGCGTACTGGCTCCTAACGTATTNNNAGATCGGAAGAGC
119338 TATGAGTGTGGAGTCGTTGCCTTTGCAAACTCTAGAGAAAAAANNNAGATCGGAAG
119339 TATGAGTGTGGAGTCGTTGCGTTTTGTGCAAAAGTGTTTGNNNAGATCGGAAGAGC

119340 TATGAGTGTGGAGTCGTTGCTATTTTGGCCATCTGAAAGAANNNAGATCGGAAGA

119341 TATGAGTGTGGAGTCGTTGCGAAGAATTGCAAAAGAAGTTAGANNNAGATCGGAA

119342 TATGAGTGTGGAGTCGTTGCAACAACAAAAAAATTCAACTCTGNNNAGATCGGAAG

119343 TATGAGTGTGGAGTCGTTGCAAGGAAATTCAGTTACAGGAGNNNAGATCGGAAGA

119344 TATGAGTGTGGAGTCGTTGCAATAATCTTGTGCAGGAAGTTNNNAGATCGGAAGA

119345 TATGAGTGTGGAGTCGTTGCGTTTCTCAGTCATGATTCCCNNNAGATCGGAAGAGC

119346 TATGAGTGTGGAGTCGTTGCCTTTCTTGACATCCCTTTAAAAANNNAGATCGGAAGA

119347 TATGAGTGTGGAGTCGTTGCTTACTCCATGCAAAATCTCCNNNAGATCGGAAGAGC

119348 TATGAGTGTGGAGTCGTTGCCATTAACAGGATTCAGACTAAAANNNAGATCGGAAG

119349 TATGAGTGTGGAGTCGTTGCTAAAACCCAGGTAGTTAGATAAGNNNAGATCGGAA

119350 TATGAGTGTGGAGTCGTTGCATTTGATAACCAACACTTAGAACNNNAGATCGGAAG

119351 TATGAGTGTGGAGTCGTTGCCTGCAGGAGTAACTAAGATTATANNNAGATCGGAA

119352 TATGAGTGTGGAGTCGTTGCGTTCTAACCTAATCACCTATCTTNNNAGATCGGAAGA

119353 TATGAGTGTGGAGTCGTTGCTAAAAATACAAAAACTAGCCAGGNNNAGATCGGAA

119354 TATGAGTGTGGAGTCGTTGCATACTGAGAGTTAGAAGTTAAGGNNNAGATCGGAA

119355 TATGAGTGTGGAGTCGTTGCGTAGTGCAGAATCTTTAATTTCANNNAGATCGGAAG

119356 TATGAGTGTGGAGTCGTTGCATTTTGCCAATGCTATTCTATTTNNNAGATCGGAAGA

119357 TATGAGTGTGGAGTCGTTGCAATGTACCACTGAAATTTTTTCTNNNAGATCGGAAG

119358 TATGAGTGTGGAGTCGTTGCAAACTGCTTAATGATGGATTTAGNNNAGATCGGAA

119359 TATGAGTGTGGAGTCGTTGCGAAAAATACCAGCTAGAGAGAANNNAGATCGGAAG

119360 TATGAGTGTGGAGTCGTTGCATCGGCATTTTATATTAAAGTGGNNNAGATCGGAAG

119361 TATGAGTGTGGAGTCGTTGCAAAATGAAAAGAAGGTACTGTACNNNAGATCGGAA

119362 TATGAGTGTGGAGTCGTTGCGGTGATGTTTAAAACAATAAAGCNNNAGATCGGAA

119363 TATGAGTGTGGAGTCGTTGCGGAACACATTAATTTTCTTCTCTNNNAGATCGGAAG

119364 TATGAGTGTGGAGTCGTTGCAAATCTGTAGGATCAACATAGACNNNAGATCGGAA

119365 TATGAGTGTGGAGTCGTTGCTATGAAAGGGTTAAGAATTTGTGNNNAGATCGGAA

119366 TATGAGTGTGGAGTCGTTGCGTGTTCAACAAGAGTTCTTTGNNNAGATCGGAAGA

119367 TATGAGTGTGGAGTCGTTGCGATGATGGCTTCTTCCTAGGNNNAGATCGGAAGAG

119368 TATGAGTGTGGAGTCGTTGCTAGATTGTCTATGGAATTAGCAANNNAGATCGGAA

119369 TATGAGTGTGGAGTCGTTGCTAGAAAAGGAGACACAAAAGAAANNNAGATCGGA

119370 TATGAGTGTGGAGTCGTTGCGGAGGAGACAGCTTCATAAGNNNAGATCGGAAGA

119371 TATGAGTGTGGAGTCGTTGCCCAATCGTCTCTTAAGTTTTTATNNNAGATCGGAAGA

119372 TATGAGTGTGGAGTCGTTGCGAGATGGTCCATTTCATTCATNNNAGATCGGAAGAG

119373 TATGAGTGTGGAGTCGTTGCGTTGCTCTATTACTAAAACTGTTNNNAGATCGGAAG

119374 TATGAGTGTGGAGTCGTTGCAATTTCTTCCTAAAAGCCTACTNNNAGATCGGAAGA

119375 TATGAGTGTGGAGTCGTTGCTAAATGTCATTTGCTGTGAGTATNNNAGATCGGAAG

119376 TATGAGTGTGGAGTCGTTGCCAGATGAACCTGGAATTAAAAAGNNNAGATCGGAA

119377 TATGAGTGTGGAGTCGTTGCTCATCTAAACATTTAAAAGTGCANNNAGATCGGAAG

119378 TATGAGTGTGGAGTCGTTGCCACCTGTTGATCTCTCCTTTNNNAGATCGGAAGAGCA

119379 TATGAGTGTGGAGTCGTTGCTAGTTCTATCCAAATGAGGAAAANNNAGATCGGAA

119380 TATGAGTGTGGAGTCGTTGCTAAGTGATTCTTCCTTGTGTTTANNNAGATCGGAAG

119381 TATGAGTGTGGAGTCGTTGCTCTTGAATTTCAAAAGGATTCTCNNNAGATCGGAAG

119382 TATGAGTGTGGAGTCGTTGCTATAGAAGTTGAAACAAGAAGGANNNAGATCGGAA

119383 TATGAGTGTGGAGTCGTTGCCTTTATGAAGTGTTTCACCAAANNNAGATCGGAAGA
119384 TATGAGTGTGGAGTCGTTGCGTCATTGGTACAGTCTCAAATANNNAGATCGGAAGA
119385 TATGAGTGTGGAGTCGTTGCGGAGGTAAGAATAACAAATCATCNNNAGATCGGAA
119386 TATGAGTGTGGAGTCGTTGCGGAGAATATATCCCCTCATATTCNNNAGATCGGAAG
119387 TATGAGTGTGGAGTCGTTGCCATATCTTCAAACATGTCATCAANNNAGATCGGAAG
119388 TATGAGTGTGGAGTCGTTGCAAAATCACAGTGAAATTGAAATGNNNAGATCGGAA
119389 TATGAGTGTGGAGTCGTTGCTGAAGATAAAGTCAGCCATGNNNAGATCGGAAGAG
119390 TATGAGTGTGGAGTCGTTGCTCATTGTTGTAGGACTCCAGNNNAGATCGGAAGAG
119391 TATGAGTGTGGAGTCGTTGCCATTTTGTCTAACTTTGGAGTGNNNAGATCGGAAGA
119392 TATGAGTGTGGAGTCGTTGCGGAAAGAGGAAGGAGATCTGNNNAGATCGGAAGA
119393 TATGAGTGTGGAGTCGTTGCTTCGATATTCAAAGACTGTTTTGNNNAGATCGGAAG
119394 TATGAGTGTGGAGTCGTTGCCCATGATTCCCTTTCCTTTGNNNAGATCGGAAGAGCA
119395 TATGAGTGTGGAGTCGTTGCTACAAAGGTAAGAGGAGACTGNNNAGATCGGAAG
119396 TATGAGTGTGGAGTCGTTGCCATCTCTTGCCTTCATAGTACNNNAGATCGGAAGAG
119397 TATGAGTGTGGAGTCGTTGCTTGCAAATCAGTCTCTACAGNNNAGATCGGAAGAGC
119398 TATGAGTGTGGAGTCGTTGCCTTGAGATGACAAGATGTTCTNNNAGATCGGAAGA
119399 TATGAGTGTGGAGTCGTTGCATTTTACCATGATATACCCTGAANNNAGATCGGAAG
119400 TATGAGTGTGGAGTCGTTGCTTAGGAAGTTATCAGAAACCAACNNNAGATCGGAAG
119401 TATGAGTGTGGAGTCGTTGCATATTAGTGACGAACATCTTACANNNAGATCGGAAG
119402 TATGAGTGTGGAGTCGTTGCATTCTGTAGATAATGTCTTGGATNNNAGATCGGAAG
119403 TATGAGTGTGGAGTCGTTGCCCAATTCAACTTTTTTCCCAGNNNAGATCGGAAGAGC
119404 TATGAGTGTGGAGTCGTTGCATTCTTTGACTTTTGAGAAGATTNNNAGATCGGAAG
119405 TATGAGTGTGGAGTCGTTGCTAGAAACAAGACAGATGTTAAACNNNAGATCGGAA
119406 TATGAGTGTGGAGTCGTTGCAAAACCCCTATGTACCACTGNNNAGATCGGAAGAGC
119407 TATGAGTGTGGAGTCGTTGCATGACTAGAAGGTCAGATGTNNNAGATCGGAAGAG
119409 TATGAGTGTGGAGTCGTTGCCATAGACATAAAAGGACTCACTNNNAGATCGGAAGA
119410 TATGAGTGTGGAGTCGTTGCATGAAGGTTCTAAGTAGAATTGANNNAGATCGGAA
119411 TATGAGTGTGGAGTCGTTGCTATAAGTTATTATCGCATGCCTANNNAGATCGGAAG
119412 TATGAGTGTGGAGTCGTTGCCTCCACTTCTTTATTAACAACTGNNNAGATCGGAAGA
119413 TATGAGTGTGGAGTCGTTGCCTTCTTTTCCACTTTGCTCTNNNAGATCGGAAGAGCA
119415 TATGAGTGTGGAGTCGTTGCTTGAAAATTGTTGCATTTGTAAANNNAGATCGGAAG
119416 TATGAGTGTGGAGTCGTTGCGGAAGACAGTGTATTCCAAANNNAGATCGGAAGAG
119417 TATGAGTGTGGAGTCGTTGCAACAGATTTTACAGCAAAGATTTNNNAGATCGGAAG
119418 TATGAGTGTGGAGTCGTTGCATAAAATGAGATTCCAGCCAGNNNAGATCGGAAGA
119419 TATGAGTGTGGAGTCGTTGCCAATCCTTTTGAACATACTCATANNNAGATCGGAAGA
119420 TATGAGTGTGGAGTCGTTGCAATTTCCTCCAGCTATTAGTAATNNNAGATCGGAAGA
119421 TATGAGTGTGGAGTCGTTGCCATGTTCTCTGTGTATTATTACANNNAGATCGGAAGA
119422 TATGAGTGTGGAGTCGTTGCGAACTCAAAGACAATATTCTTGTNNNAGATCGGAAG
119423 TATGAGTGTGGAGTCGTTGCGAAACCTAGTCACATTGGAGNNNAGATCGGAAGAG
119424 TATGAGTGTGGAGTCGTTGCCTGACTTACCCTAGTACTTAGTANNNAGATCGGAAG
119425 TATGAGTGTGGAGTCGTTGCTAATACAAACAAACAGCAAGAAANNNAGATCGGAA
119426 TATGAGTGTGGAGTCGTTGCAGATGTTGGTAAGACTTAGTATCNNNAGATCGGAA
119427 TATGAGTGTGGAGTCGTTGCCCTAGGAGAGAATGGGTTTGNNNAGATCGGAAGA

119428 TATGAGTGTGGAGTCGTTGCGTATCCAATGTTTAGCTTTTCANNNAGATCGGAAGA
119429 TATGAGTGTGGAGTCGTTGCACATTAATGTCATCACTGCANNNAGATCGGAAGAGC
119430 TATGAGTGTGGAGTCGTTGCTAAAAAGGGAAACTTGGAAAAACNNNAGATCGGAA
119431 TATGAGTGTGGAGTCGTTGCCTAACCAATAAAACAAGAGATCCNNNAGATCGGAAG
119432 TATGAGTGTGGAGTCGTTGCCATTGATGAGTAGCTTTTCAGNNNAGATCGGAAGA
119433 TATGAGTGTGGAGTCGTTGCCGTTCTGCTTTATTCTTGATAGNNNAGATCGGAAGA
119434 TATGAGTGTGGAGTCGTTGCAGTTTATAATCTGGAATCTCCTCNNNAGATCGGAAG
119435 TATGAGTGTGGAGTCGTTGCACTCCACTGCTTATTTCATCNNNAGATCGGAAGAGCA
119436 TATGAGTGTGGAGTCGTTGCTTAGAATTCATCAGAGAGTTCATNNNAGATCGGAAG
119437 TATGAGTGTGGAGTCGTTGCATGTTTGGAATGTTTTTAAAACCNNNAGATCGGAAG
119438 TATGAGTGTGGAGTCGTTGCAAGGGTTTTTTACATCATAAGACNNNAGATCGGAAG
119439 TATGAGTGTGGAGTCGTTGCCACCTACATTTTGGAAATTGAANNNAGATCGGAAGA
119440 TATGAGTGTGGAGTCGTTGCACCTTCTATTTTAATCTCTCTCCNNNAGATCGGAAGA
119441 TATGAGTGTGGAGTCGTTGCGGATGAAGTAGAATTCCAAAAACNNNAGATCGGAA
119442 TATGAGTGTGGAGTCGTTGCCTATGAAGTCATGGAAGCTCNNNAGATCGGAAGAG
119443 TATGAGTGTGGAGTCGTTGCAATATTTGTAAGTTCGAAATGGGNNNAGATCGGAA
119444 TATGAGTGTGGAGTCGTTGCGTATTCTTGGTTTCTGATGAAATNNNAGATCGGAAG
119445 TATGAGTGTGGAGTCGTTGCGACTGTTTCAAGGAGCTACTNNNAGATCGGAAGAG
119447 TATGAGTGTGGAGTCGTTGCCTCTGGGGTAATATCTAAACTTGNNNAGATCGGAAG
119448 TATGAGTGTGGAGTCGTTGCGAATCTCATCCCCTCTCAACNNNAGATCGGAAGAGCA
119449 TATGAGTGTGGAGTCGTTGCCCTTGTCTTATGTTCCAGTATTNNNAGATCGGAAGAG
119450 TATGAGTGTGGAGTCGTTGCCTCAGCTACACCAAGAGATTNNNAGATCGGAAGAGC
119451 TATGAGTGTGGAGTCGTTGCCTCCTTAAGCCATTCTAAAGTTNNNAGATCGGAAGAG
119452 TATGAGTGTGGAGTCGTTGCGTGGATTCACAAACACAATCNNNAGATCGGAAGAGC
119453 TATGAGTGTGGAGTCGTTGCTTTCTGGAAAGAGAGAGAAATCNNNAGATCGGAAG
119454 TATGAGTGTGGAGTCGTTGCGTTAACTTCAATGGGTTTTCCNNNAGATCGGAAGAG
119455 TATGAGTGTGGAGTCGTTGCCTGTTTCAGGTTTCTTATTGGNNNAGATCGGAAGAG
119456 TATGAGTGTGGAGTCGTTGCGAGCTTAATAAACATGTAGCAAANNNAGATCGGAAG
119457 TATGAGTGTGGAGTCGTTGCTGATTAATGCAGATGGAAAGATANNNAGATCGGAA
119458 TATGAGTGTGGAGTCGTTGCATCTGACAGGTGATATTCAAACNNNAGATCGGAAGA
119459 TATGAGTGTGGAGTCGTTGCATAGGATGCAGATATTGTGATAANNNAGATCGGAA
119460 TATGAGTGTGGAGTCGTTGCATAGAGTGTAGATGCATCAAAATNNNAGATCGGAA
119461 TATGAGTGTGGAGTCGTTGCGGTGAAGATTTTTGGATTCTTTNNNAGATCGGAAGA
119462 TATGAGTGTGGAGTCGTTGCAAAAATGTTTCTGTGTGTACTTANNNAGATCGGAAG
119463 TATGAGTGTGGAGTCGTTGCAGAATATGTACTGTAATAAGGAGANNNAGATCGGA
119464 TATGAGTGTGGAGTCGTTGCAGATTGTATCTCCAAAGGTTAATNNNAGATCGGAAG
119465 TATGAGTGTGGAGTCGTTGCGAAAATAAGCTGGTTTCTATGTANNNAGATCGGAAG
119466 TATGAGTGTGGAGTCGTTGCGTAAGACTCTGTGACAAATCTATNNNAGATCGGAAG
119467 TATGAGTGTGGAGTCGTTGCTATATCATTTGAAGGTAGACACCNNNAGATCGGAAG
119468 TATGAGTGTGGAGTCGTTGCCTGGACTCTATATTGTCAAACTANNNAGATCGGAAG
119469 TATGAGTGTGGAGTCGTTGCAAAACACAGGTACATTATGGTNNNAGATCGGAAGA
119470 TATGAGTGTGGAGTCGTTGCACTGTCTAGAGCTTTTTGATTCNNNAGATCGGAAGA
119471 TATGAGTGTGGAGTCGTTGCGAGCATGTTATACTTCAGAAACNNNAGATCGGAAGA

119472 TATGAGTGTGGAGTCGTTGCTATTCCACTGTTCTTTGTGTAANNNAGATCGGAAGA
119473 TATGAGTGTGGAGTCGTTGCCTCATAGTTCAGAAGAGTTCAGNNNAGATCGGAAGA
119474 TATGAGTGTGGAGTCGTTGCCACAAAATCAACCTTGTCAGNNNAGATCGGAAGAGC
119475 TATGAGTGTGGAGTCGTTGCAATAAGTGCTTAAATGTCTTTGGNNNAGATCGGAAG
119476 TATGAGTGTGGAGTCGTTGCCACTGAAAATTGATGCTTGTANNNAGATCGGAAGAG
119477 TATGAGTGTGGAGTCGTTGCCAGGTCCTTTTCCAAACTTANNNAGATCGGAAGAGC
119478 TATGAGTGTGGAGTCGTTGCATCCCATGAATAGAAAGAGATGNNNAGATCGGAAG
119479 TATGAGTGTGGAGTCGTTGCGAAAGAGAATGATTTTGCTCTTANNNAGATCGGAA
119480 TATGAGTGTGGAGTCGTTGCGTTTTGTTGCTGACACAAATNNNAGATCGGAAGAGC
119481 TATGAGTGTGGAGTCGTTGCCCCTCTTTTATCTATTCCCTAAANNNAGATCGGAAGA
119482 TATGAGTGTGGAGTCGTTGCGACAAGACACAAATCTAAGAACNNNAGATCGGAAG
119483 TATGAGTGTGGAGTCGTTGCGATTTTATGCTTTCTTTACATGGNNNAGATCGGAAG
119484 TATGAGTGTGGAGTCGTTGCTTTTGTCAATTCCAGAATACAAANNNAGATCGGAAG
119485 TATGAGTGTGGAGTCGTTGCAAGTTACTTCCATCCACTTTCNNNAGATCGGAAGAGC
119486 TATGAGTGTGGAGTCGTTGCAATTGCCCAGAGAGTTTCTTNNNAGATCGGAAGAGC
119487 TATGAGTGTGGAGTCGTTGCACATAAAATGTGGTAACATACACNNNAGATCGGAAG
119488 TATGAGTGTGGAGTCGTTGCAATGAATGGCAAATTCTATTTCTNNNAGATCGGAAG
119489 TATGAGTGTGGAGTCGTTGCCAGCATATGGACAAAAAACATNNNAGATCGGAAGA
119490 TATGAGTGTGGAGTCGTTGCTGAAGTAATAGATGAAAGGGTCNNNAGATCGGAAG
119491 TATGAGTGTGGAGTCGTTGCCCAGGTTTTCAGAAAATGAAGNNNAGATCGGAAGA
119492 TATGAGTGTGGAGTCGTTGCTGCTCTCTTCTTCCAAGATGNNNAGATCGGAAGAGC
119493 TATGAGTGTGGAGTCGTTGCAGTGTTCCTTCATTAATTTAAGCNNNAGATCGGAAG
119494 TATGAGTGTGGAGTCGTTGCCTTGTGAAAGACAAATCTGTAAGNNNAGATCGGAA
119495 TATGAGTGTGGAGTCGTTGCCGGCTTTTAAGTGTGATTTTANNNAGATCGGAAGAG
119496 TATGAGTGTGGAGTCGTTGCAGATCATCGAACTCTTAGATATGNNNAGATCGGAAG
119497 TATGAGTGTGGAGTCGTTGCGCAGATTAAAAGACAAAACAAAANNNAGATCGGAA
119498 TATGAGTGTGGAGTCGTTGCCTAAGGTATTCAGTCATCATGGNNNAGATCGGAAGA
119499 TATGAGTGTGGAGTCGTTGCCTCAATAGTTACCAGCCCATNNNAGATCGGAAGAGC
119500 TATGAGTGTGGAGTCGTTGCTTTCCTTGAGCTGTTTTCTGNNNAGATCGGAAGAGC
119501 TATGAGTGTGGAGTCGTTGCTTTCCTTGAGCTGTTTTCTGNNNAGATCGGAAGAGC
119502 TATGAGTGTGGAGTCGTTGCCACATATCATGGTAGTGACGNNNAGATCGGAAGAG
119503 TATGAGTGTGGAGTCGTTGCTGTCTCTAACAGGAGAACTAANNNAGATCGGAAGA
119504 TATGAGTGTGGAGTCGTTGCTTCTCCCCATATTTCACAGATNNNAGATCGGAAGAGC
119505 TATGAGTGTGGAGTCGTTGCCACCATTTTTATAGTCCATGTTTNNNAGATCGGAAGA
119506 TATGAGTGTGGAGTCGTTGCTAAATTCAATACACAGTTCAGTGNNNAGATCGGAAG
119507 TATGAGTGTGGAGTCGTTGCAGCACAGGATCCGAAATTAANNNAGATCGGAAGAG
119508 TATGAGTGTGGAGTCGTTGCTGGTTTTCTTACATAAACATGAGNNNAGATCGGAAG
119509 TATGAGTGTGGAGTCGTTGCTTGGTCACTTTCATTATAAGAAANNNAGATCGGAAG
119510 TATGAGTGTGGAGTCGTTGCCATTCTCCTCATCAAATTTCTCNNNAGATCGGAAGAG
119511 TATGAGTGTGGAGTCGTTGCAACTGACCAAGATGTGTACTATNNNAGATCGGAAGA
119512 TATGAGTGTGGAGTCGTTGCATTTGTAAGAAAATACAGAGCAGNNNAGATCGGAA
119513 TATGAGTGTGGAGTCGTTGCTTTGCTGATTTTCATCTACTTCANNNAGATCGGAAGA
119514 TATGAGTGTGGAGTCGTTGCACTTGGATATCTGCATGAATCNNNAGATCGGAAGAG

119515 TATGAGTGTGGAGTCGTTGCAGACACCGAAGAGAAAAAATTNNNAGATCGGAAGA
119516 TATGAGTGTGGAGTCGTTGCTATCATGAGTGATTCCAACTATGNNNAGATCGGAAG
119517 TATGAGTGTGGAGTCGTTGCTCTTTCCTATGTTTTTCTCCATANNNAGATCGGAAGA
119518 TATGAGTGTGGAGTCGTTGCTCCTAATAACTCATCAAGAAACANNNAGATCGGAAG
119519 TATGAGTGTGGAGTCGTTGCATTATGTGAAAAATAAGATGCGGNNNAGATCGGAA
119521 TATGAGTGTGGAGTCGTTGCTTTGGAATTAAGCTCTTTGTTATNNNAGATCGGAAG
119522 TATGAGTGTGGAGTCGTTGCTCTGCATCTCTTGTTCTTTGNNNAGATCGGAAGAGC
119523 TATGAGTGTGGAGTCGTTGCGATGAGGAGGAAGAGATGATNNNAGATCGGAAGA
119524 TATGAGTGTGGAGTCGTTGCAAGATCAAGAAAATTTAGCTGTTNNNAGATCGGAA
119525 TATGAGTGTGGAGTCGTTGCCTAGTAAGTACGGATAATGGATCNNNAGATCGGAA
119526 TATGAGTGTGGAGTCGTTGCTGCTTCGTAAGGGAAACTATNNNAGATCGGAAGAG
119527 TATGAGTGTGGAGTCGTTGCGGTGAAAGGTTAAAACATATCTTNNNAGATCGGAA
119528 TATGAGTGTGGAGTCGTTGCCAAGCTTGTTGAAGAAGACANNNAGATCGGAAGAG
119529 TATGAGTGTGGAGTCGTTGCCACTGATACTAAGTTCAGAAACTNNNAGATCGGAAG
119530 TATGAGTGTGGAGTCGTTGCTCTAACTTCTCCGAGAACACNNNAGATCGGAAGAGC
119531 TATGAGTGTGGAGTCGTTGCTTCTCACCTTGTCATTAGAAAANNNAGATCGGAAGA
119532 TATGAGTGTGGAGTCGTTGCCATGGTAAAGTTACTAACACAAANNNAGATCGGAAG
119533 TATGAGTGTGGAGTCGTTGCATTTCCCTGAAGTATAAATAAGCNNNAGATCGGAAG
119534 TATGAGTGTGGAGTCGTTGCCATCATTTCTGGAGTCTCAGNNNAGATCGGAAGAGC
119535 TATGAGTGTGGAGTCGTTGCAACAGTCTCCAGATTATAAATCCNNNAGATCGGAAG
119536 TATGAGTGTGGAGTCGTTGCATGTTAGTAATGAGGACAACTAGNNNAGATCGGAA
119537 TATGAGTGTGGAGTCGTTGCCTCTCTGAAGGACATCTACAANNNAGATCGGAAGAG
119538 TATGAGTGTGGAGTCGTTGCCCCAAATTACTACAAAGCATAGNNNAGATCGGAAGA
119539 TATGAGTGTGGAGTCGTTGCTTTTCTGGAATGGAAAATTAAGTNNNAGATCGGAA
119540 TATGAGTGTGGAGTCGTTGCATGATGAAGAAGGAGACCATNNNAGATCGGAAGA
119541 TATGAGTGTGGAGTCGTTGCGGAAGAAATTAATGTTATGCTCANNNAGATCGGAA
119542 TATGAGTGTGGAGTCGTTGCCAACATTTTAGGAACTTTCTAGATNNNAGATCGGAA
119543 TATGAGTGTGGAGTCGTTGCAATTCTCCATCCTTTCCAAAGNNNAGATCGGAAGAG
119544 TATGAGTGTGGAGTCGTTGCCTTCTAAAAGTTCTTCATCTGAANNNAGATCGGAAG
119545 TATGAGTGTGGAGTCGTTGCATTCTTTTCAAAATAGCCTTTGANNNAGATCGGAAG
119546 TATGAGTGTGGAGTCGTTGCTTCAAAGCGTTCTTCACATTNNNAGATCGGAAGAGC
119547 TATGAGTGTGGAGTCGTTGCGTCTATGAAGTGAAAAACAACANNNAGATCGGAAG
119548 TATGAGTGTGGAGTCGTTGCGAGAAGGAGGAAAAAAAACAGNNNAGATCGGAAG
119549 TATGAGTGTGGAGTCGTTGCTTTTCTTACTCCATTCTTGCAANNNAGATCGGAAGAG
119550 TATGAGTGTGGAGTCGTTGCTAGTGACACTGACAACCATCNNNAGATCGGAAGAG
119551 TATGAGTGTGGAGTCGTTGCTATAACTGTGAAGCTTAATTCAGNNNAGATCGGAAG
119552 TATGAGTGTGGAGTCGTTGCTTTGAATGTTCGGTGAAGACNNNAGATCGGAAGAG
119553 TATGAGTGTGGAGTCGTTGCTGGAAGAATAGATACAAGCATAANNNAGATCGGAA
119554 TATGAGTGTGGAGTCGTTGCTGGAAGAATAGATACAAGCATAANNNAGATCGGAA
119555 TATGAGTGTGGAGTCGTTGCATTGGAATGTTTGTTAATCACAANNNAGATCGGAAG
119556 TATGAGTGTGGAGTCGTTGCCTAACTCCAGATAACTCCCTNNNAGATCGGAAGAGC
119557 TATGAGTGTGGAGTCGTTGCGTTCTAGAACTTACCCTTTTTGNNNAGATCGGAAGA
119558 TATGAGTGTGGAGTCGTTGCCAAAGACTTAGAGCAGAAGCNNNAGATCGGAAGAG

119559 TATGAGTGTGGAGTCGTTGCAACCAAATGTTCTTTATCCATCTNNNAGATCGGAAGA
119560 TATGAGTGTGGAGTCGTTGCTAAAACTAGACTCAGTGAAAAGANNNAGATCGGAA
119561 TATGAGTGTGGAGTCGTTGCAATGTCACCTGTATCTGTGANNNAGATCGGAAGAG
119562 TATGAGTGTGGAGTCGTTGCGATGTGACAAGATTGTGGAGNNNAGATCGGAAGA
119563 TATGAGTGTGGAGTCGTTGCCCTCTCCTTCTACCATGTTATNNNAGATCGGAAGAGC
119564 TATGAGTGTGGAGTCGTTGCACCTCCTATGAAAATTGCAGNNNAGATCGGAAGAG
119565 TATGAGTGTGGAGTCGTTGCGTCAAATTTCTCAACTCTGTGNNNAGATCGGAAGAG
119566 TATGAGTGTGGAGTCGTTGCATCTACAACTCTTCCTTCTACTNNNAGATCGGAAGAG
119567 TATGAGTGTGGAGTCGTTGCCTAAATTCCATCATAAGTCGAAANNNAGATCGGAAG
119568 TATGAGTGTGGAGTCGTTGCTAATTGCATTTTAGTTCAAGGGNNNAGATCGGAAGA
119569 TATGAGTGTGGAGTCGTTGCAGGCAAATCATTTTAATCTTGAANNNAGATCGGAAG
119570 TATGAGTGTGGAGTCGTTGCCTATGTACATGTTCCTCCAGNNNAGATCGGAAGAGC
119571 TATGAGTGTGGAGTCGTTGCCAGTACTATTATGAACCACCATANNNAGATCGGAAG
119572 TATGAGTGTGGAGTCGTTGCCACTCTTCTCAAACAAAAAGTCNNNAGATCGGAAGA
119573 TATGAGTGTGGAGTCGTTGCGATAAAACTATGGCGAAGAAAANNNAGATCGGAAG
119574 TATGAGTGTGGAGTCGTTGCGAGAAGTCATCACAACTCATCNNNAGATCGGAAGAG
119575 TATGAGTGTGGAGTCGTTGCTAAGATGGCACCTATATGAGANNNAGATCGGAAGA
119576 TATGAGTGTGGAGTCGTTGCCAAAGTCCCATAGAACACTATGNNNAGATCGGAAGA
119577 TATGAGTGTGGAGTCGTTGCAATCTGAATCTTGTGGTCTGNNNAGATCGGAAGAG
119578 TATGAGTGTGGAGTCGTTGCCATCAGGTGACCATTAAGTTANNNAGATCGGAAGA
119579 TATGAGTGTGGAGTCGTTGCCCTAAAGAAGATCTTATGGAAACNNNAGATCGGAA
119580 TATGAGTGTGGAGTCGTTGCAATACTTGCAATTTTCTTAACGTNNNAGATCGGAAGA
119581 TATGAGTGTGGAGTCGTTGCTTTGGTCGTGTGTACATATTTATNNNAGATCGGAAG
119582 TATGAGTGTGGAGTCGTTGCATGAAAGAAACACAAAAACCTTANNNAGATCGGAA
119583 TATGAGTGTGGAGTCGTTGCAAAGACCAACAGTTCTTTTTGNNNAGATCGGAAGAG
119584 TATGAGTGTGGAGTCGTTGCAAATCTCAGCCGTTTAAAACTTNNNAGATCGGAAGA
119585 TATGAGTGTGGAGTCGTTGCATTTTGGCTACATTAAGCTTTTNNNAGATCGGAAGA
119586 TATGAGTGTGGAGTCGTTGCATTTTGGCTACATTAAGCTTTTNNNAGATCGGAAGA
119587 TATGAGTGTGGAGTCGTTGCAAAAGTCCAGTTTGTATGTTTTGNNNAGATCGGAAG
119588 TATGAGTGTGGAGTCGTTGCGTTAAATTCAAGGTCAATCCAGNNNAGATCGGAAGA
119589 TATGAGTGTGGAGTCGTTGCACCATTGTCACAATGTATATGANNNAGATCGGAAGA
119590 TATGAGTGTGGAGTCGTTGCTCTGGAGAAATAGAAGAAAAGATNNNAGATCGGAA
119591 TATGAGTGTGGAGTCGTTGCGAATCACCTATTACATTTGACTTNNNAGATCGGAAGA
119592 TATGAGTGTGGAGTCGTTGCAATTATTAGCATTGATCTGAGTGNNNAGATCGGAAG
119593 TATGAGTGTGGAGTCGTTGCGGTAAGGTTAAAATGGAAAATCTNNNAGATCGGAA
119594 TATGAGTGTGGAGTCGTTGCCTTTTCTCATACCTTTGACAGANNNAGATCGGAAGA
119595 TATGAGTGTGGAGTCGTTGCCTCAGGTGAAGGACAAAAAGNNNAGATCGGAAGA
119596 TATGAGTGTGGAGTCGTTGCGAAACTTCATAATTAGCACAGATNNNAGATCGGAAG
119597 TATGAGTGTGGAGTCGTTGCTTCTCTTAATAATGCTCACAATGNNNAGATCGGAAGA
119598 TATGAGTGTGGAGTCGTTGCCATGATCGTACTTGTCCTATTATNNNAGATCGGAAGA
119599 TATGAGTGTGGAGTCGTTGCGGAACTCCTGAATAATTTTTTCANNNAGATCGGAAG
119600 TATGAGTGTGGAGTCGTTGCGATAGGACTTGAACAGAAGATTNNNAGATCGGAAG
119601 TATGAGTGTGGAGTCGTTGCAACTGTTGCGGATATACTTTCNNNAGATCGGAAGAG

119602 TATGAGTGTGGAGTCGTTGCGAATGTAGAACGATTTCAGAATGNNNAGATCGGAA
119603 TATGAGTGTGGAGTCGTTGCGAAATATGTGAAAACCAGTAAGTNNNAGATCGGAA
119604 TATGAGTGTGGAGTCGTTGCAATCTTTAACTGCCATGTAAACNNNAGATCGGAAGA
119605 TATGAGTGTGGAGTCGTTGCAAGACATTATTTGGGTGAATAGNNNAGATCGGAA
119606 TATGAGTGTGGAGTCGTTGCGGTTTATCTGGTGGGAATAGNNNAGATCGGAAGAG
119607 TATGAGTGTGGAGTCGTTGCATCCTTTATGTACTAGAAAAGGTNNNAGATCGGAAG
119608 TATGAGTGTGGAGTCGTTGCATGGAGTCATAATATTCTCAGTTNNNAGATCGGAAG
119609 TATGAGTGTGGAGTCGTTGCATAGTGACTATCTCAATAGAACCNNNAGATCGGAAG
119610 TATGAGTGTGGAGTCGTTGCAATTTCTGAGGAGTGGTTTTATANNNAGATCGGAAG
119611 TATGAGTGTGGAGTCGTTGCCAAAGAAGTCCTGGATGATAATNNNAGATCGGAAG
119612 TATGAGTGTGGAGTCGTTGCAGGTGTTAATGAAAAACAAAGAANNNAGATCGGAA
119613 TATGAGTGTGGAGTCGTTGCATTCCTGTATTTCTTTAGTTGTCNNNAGATCGGAAGA
119614 TATGAGTGTGGAGTCGTTGCATCAAAGATATCAACAGAAAACCNNNAGATCGGAAG
119615 TATGAGTGTGGAGTCGTTGCCTTTAGGAATGCTGTGTGTCNNNAGATCGGAAGAG
119616 TATGAGTGTGGAGTCGTTGCGGCTATTTTTCCACTGATTAAATNNNAGATCGGAAG
119617 TATGAGTGTGGAGTCGTTGCCATACATCTTGATTCTTTTCCATNNNAGATCGGAAGA
119618 TATGAGTGTGGAGTCGTTGCTCTCCTTTTATTAATAGGAAGGCNNNAGATCGGAAG
119619 TATGAGTGTGGAGTCGTTGCGACAAGAACACAGAGACAAGNNNAGATCGGAAGAG
119620 TATGAGTGTGGAGTCGTTGCATTTAACTCCTTGGGTTTTTTAGNNNAGATCGGAAG
119621 TATGAGTGTGGAGTCGTTGCGAACAACCCCTTATACTTAATTGNNNAGATCGGAAG
119622 TATGAGTGTGGAGTCGTTGCAACTCCTCATATTCAAATTTGAGNNNAGATCGGAAG
119623 TATGAGTGTGGAGTCGTTGCAGCATCAAAGTGGTAAATAAAATNNNAGATCGGAA
119624 TATGAGTGTGGAGTCGTTGCGTATTGACATCTAATCTGCATTTNNNAGATCGGAAG
119625 TATGAGTGTGGAGTCGTTGCACAAATTAATATCCAGTAGAGAGANNNAGATCGGA
119626 TATGAGTGTGGAGTCGTTGCTAAGCAAATGTCATATTACCTTGNNNAGATCGGAAG
119628 TATGAGTGTGGAGTCGTTGCAATCTCTATGATCGCATCAGTANNNAGATCGGAAGA
119629 TATGAGTGTGGAGTCGTTGCGAGTATGGTGAAGAGGTTAATGNNNAGATCGGAA
119630 TATGAGTGTGGAGTCGTTGCCCGGTATGTAATTTAACATTCAANNNAGATCGGAAG
119631 TATGAGTGTGGAGTCGTTGCCCGGTATGTAATTTAACATTCAANNNAGATCGGAAG
119632 TATGAGTGTGGAGTCGTTGCCCGGTATGTAATTTAACATTCAANNNAGATCGGAAG
119633 TATGAGTGTGGAGTCGTTGCCCGGTATGTAATTTAACATTCAANNNAGATCGGAAG
119634 TATGAGTGTGGAGTCGTTGCCCGGTATGTAATTTAACATTCAANNNAGATCGGAAG
119635 TATGAGTGTGGAGTCGTTGCCCGGTATGTAATTTAACATTCAANNNAGATCGGAAG
119636 TATGAGTGTGGAGTCGTTGCCAACACTTCATAGACATCAATAANNNAGATCGGAAG
119637 TATGAGTGTGGAGTCGTTGCTAGCGACACAAAGTTAAACAANNNAGATCGGAAGA
119638 TATGAGTGTGGAGTCGTTGCATTTTAGCTAAAGATTTACCAGCNNNAGATCGGAAG
119639 TATGAGTGTGGAGTCGTTGCTTTTCCTGTTTAAGCATTCATAANNNAGATCGGAAGA
119640 TATGAGTGTGGAGTCGTTGCCATCATCTGCAAAAACATCCNNNAGATCGGAAGAGC
119641 TATGAGTGTGGAGTCGTTGCAAATTTCCAACTTGTATGTGTTTNNNAGATCGGAAG
119642 TATGAGTGTGGAGTCGTTGCGAAGTAAGTGCATCCACTTGNNNAGATCGGAAGAG
119644 TATGAGTGTGGAGTCGTTGCGGTCTGTCTTAACATTCTTATTGNNNAGATCGGAAG
119645 TATGAGTGTGGAGTCGTTGCCATCAAAATCACAGTGGATGNNNAGATCGGAAGAG
119646 TATGAGTGTGGAGTCGTTGCCTCTTTGTTTCTCATCAACTGNNNAGATCGGAAGAG

119648 TATGAGTGTGGAGTCGTTGCTATCAGGAATGGACTTAGTCTNNNAGATCGGAAGA

119649 TATGAGTGTGGAGTCGTTGCTTGCTTTGGGATCACATTATTTNNNAGATCGGAAGA

119650 TATGAGTGTGGAGTCGTTGCCTAATGGTACAACTGCTGAAANNNAGATCGGAAGA

119651 TATGAGTGTGGAGTCGTTGCACGTCAGAGACTTTGTATAAAGNNNAGATCGGAAG

119652 TATGAGTGTGGAGTCGTTGCAAAACAAAACAAGGGTCACTTNNNAGATCGGAAGA

119653 TATGAGTGTGGAGTCGTTGCCACAGAGGAAGAGAATCTCCNNNAGATCGGAAGAG

119654 TATGAGTGTGGAGTCGTTGCCTTTCTCCTGCTGCTTATTTNNNAGATCGGAAGAGCA

119655 TATGAGTGTGGAGTCGTTGCTTGGTAATCTTTACTTCAAAACCNNNAGATCGGAAG

119656 TATGAGTGTGGAGTCGTTGCCCATCAATAATAGCAAGTCATTTNNNAGATCGGAAG

119657 TATGAGTGTGGAGTCGTTGCCTAATCCAGAACCACTTTGTANNNAGATCGGAAGAG

119658 TATGAGTGTGGAGTCGTTGCGGCTGCAATAAGAGATATTTTAANNNAGATCGGAA

119659 TATGAGTGTGGAGTCGTTGCTATTGTGGAATGAAATTGTGATGNNNAGATCGGAA

119660 TATGAGTGTGGAGTCGTTGCTATTGTGGAATGAAATTGTGATGNNNAGATCGGAA

119661 TATGAGTGTGGAGTCGTTGCTCCATTTCTTCTAAAATAGTCCANNNAGATCGGAAGA

119662 TATGAGTGTGGAGTCGTTGCCCTATTCCCTATAAGAAATACGANNNAGATCGGAAG

119663 TATGAGTGTGGAGTCGTTGCGGAAACATCATGAAGAATAAAGANNNAGATCGGAA

119664 TATGAGTGTGGAGTCGTTGCAGGAGAAAAGAAAACCATTTAAANNNAGATCGGAA

119665 TATGAGTGTGGAGTCGTTGCATCTATAGAACTCTGAACCACTANNNAGATCGGAAG

119666 TATGAGTGTGGAGTCGTTGCTTTTGCTACAGGAAAACAGAAANNNAGATCGGAAG

119667 TATGAGTGTGGAGTCGTTGCCTAATTCAGAATCATTTTGTGGANNNAGATCGGAAG

119668 TATGAGTGTGGAGTCGTTGCAATCATTTGAAGATATTCACCATNNNAGATCGGAAG

119669 TATGAGTGTGGAGTCGTTGCTTCTGCAAAATCATAACTGTTATNNNAGATCGGAAG

119670 TATGAGTGTGGAGTCGTTGCGTTTGCCTTTTTTTCATGTAAATNNNAGATCGGAAGA

119671 TATGAGTGTGGAGTCGTTGCTCAGTACTGGAGATTCATTTGNNNAGATCGGAAGAG

119672 TATGAGTGTGGAGTCGTTGCTAGCTTGACATCCTTGGAAANNNAGATCGGAAGAG

119673 TATGAGTGTGGAGTCGTTGCGAAGAAGAGAGTCAGAGTCANNNAGATCGGAAGA

119674 TATGAGTGTGGAGTCGTTGCCTGGTTGATTTTCTTGATACTTTNNNAGATCGGAAG

119675 TATGAGTGTGGAGTCGTTGCGAATACATCACCACTAGCAANNNAGATCGGAAGAG

119676 TATGAGTGTGGAGTCGTTGCAGACAGTGAAAAAAGATGAAAATNNNAGATCGGAA

119677 TATGAGTGTGGAGTCGTTGCCTGGATCCTGATACAGATAAATGNNNAGATCGGAAG

119678 TATGAGTGTGGAGTCGTTGCGTCATCCTAGAAGGAATGGTNNNAGATCGGAAGAG

119679 TATGAGTGTGGAGTCGTTGCCCTACAGGCTTTAAGTAGATAATNNNAGATCGGAAG

119680 TATGAGTGTGGAGTCGTTGCCAATCTACACAAAATAGAAGCAANNNAGATCGGAAG

119681 TATGAGTGTGGAGTCGTTGCCCTCTCTTTTAATAAACGTTTCTNNNAGATCGGAAGA

119682 TATGAGTGTGGAGTCGTTGCTTCCTCTAGAGTATGTAATCTCTNNNAGATCGGAAG

119683 TATGAGTGTGGAGTCGTTGCGTCATTCTTATTGGTGACGTTNNNAGATCGGAAGAG

119684 TATGAGTGTGGAGTCGTTGCTCTAAGAAAAAATGTAAAAGCCCNNNAGATCGGAAG

119685 TATGAGTGTGGAGTCGTTGCAAGTAAAATATTTAAAAGCAACCANNNAGATCGGAA

119686 TATGAGTGTGGAGTCGTTGCGATAAGGAAGGGTGAGAGTTNNNAGATCGGAAGA

119687 TATGAGTGTGGAGTCGTTGCTATTAATGCTAAGATTTAGTGCCNNNAGATCGGAAG

119688 TATGAGTGTGGAGTCGTTGCTGCAAAGGAAAACAAAAAATAATNNNAGATCGGAA

119689 TATGAGTGTGGAGTCGTTGCGAAACTAATCTAACAGACACAAANNNAGATCGGAAG

119690 TATGAGTGTGGAGTCGTTGCTTGAAAAGAAATCGAAACTGTTANNNAGATCGGAA

119691 TATGAGTGTGGAGTCGTTGCAAAGTCTAAAGTGGTTCTTCTCNNNAGATCGGAAGA
119692 TATGAGTGTGGAGTCGTTGCTCCAAATGCTGCAAAATAATAATNNNAGATCGGAAG
119693 TATGAGTGTGGAGTCGTTGCCATAATTAGTGCCAATATACCACNNNAGATCGGAAG
119694 TATGAGTGTGGAGTCGTTGCAGTATTTTGGGGAATTTTGAATTNNNAGATCGGAA
119695 TATGAGTGTGGAGTCGTTGCATTTATTTTACGACAACAACAACNNNAGATCGGAAG
119696 TATGAGTGTGGAGTCGTTGCATGAGTGATTCCAACTATGTTGNNNAGATCGGAAG
119697 TATGAGTGTGGAGTCGTTGCATCTGACGTACACTTTCCAGNNNAGATCGGAAGAGC
119698 TATGAGTGTGGAGTCGTTGCCATCTGAATCAACAATGTGATAANNNAGATCGGAAG
119699 TATGAGTGTGGAGTCGTTGCATTCCTGTATTTCTTTAGTTGTCNNNAGATCGGAAGA
119700 TATGAGTGTGGAGTCGTTGCATCAAAGATATCAACAGAAAACCNNNAGATCGGAAG
119701 TATGAGTGTGGAGTCGTTGCCTTTAGGAATGCTGTGTGTCNNNAGATCGGAAGAG
119702 TATGAGTGTGGAGTCGTTGCAATCTCGATGAATGTGTAGATATNNNAGATCGGAA
119703 TATGAGTGTGGAGTCGTTGCGGCTATTTTTCCACTGATTAAATNNNAGATCGGAAG
119704 TATGAGTGTGGAGTCGTTGCCATACATCTTGATTCTTTTCCATNNNAGATCGGAAGA
119705 TATGAGTGTGGAGTCGTTGCTCTCCTTTTATTAATAGGAAGGCNNNAGATCGGAAG
119706 TATGAGTGTGGAGTCGTTGCGACAAGAACACAGAGACAAGNNNAGATCGGAAGA
119707 TATGAGTGTGGAGTCGTTGCATTTAACTCCTTGGGTTTTTTAGNNNAGATCGGAAG
119708 TATGAGTGTGGAGTCGTTGCGAACAACCCCTTATACTTAATTGNNNAGATCGGAAG
119709 TATGAGTGTGGAGTCGTTGCAACTCCTCATATTCAAATTTGAGNNNAGATCGGAAG
119710 TATGAGTGTGGAGTCGTTGCAGCATCAAAGTGGTAAATAAAATNNNAGATCGGAA
119711 TATGAGTGTGGAGTCGTTGCGTATTGACATCTAATCTGCATTTNNNAGATCGGAAG
119712 TATGAGTGTGGAGTCGTTGCACAAATTAATATCCAGTAGAGAGANNNAGATCGGA
119713 TATGAGTGTGGAGTCGTTGCTAAGCAAATGTCATATTACCTTGNNNAGATCGGAAG
119715 TATGAGTGTGGAGTCGTTGCAATCTCTATGATCGCATCAGTANNNAGATCGGAAGA
119716 TATGAGTGTGGAGTCGTTGCCATTGGATTTCAAGTAAAACATTNNNAGATCGGAAG
119717 TATGAGTGTGGAGTCGTTGCCCGGTATGTAATTTAACATTCAANNNAGATCGGAAG
119718 TATGAGTGTGGAGTCGTTGCCCGGTATGTAATTTAACATTCAANNNAGATCGGAAG
119719 TATGAGTGTGGAGTCGTTGCCCGGTATGTAATTTAACATTCAANNNAGATCGGAAG
119720 TATGAGTGTGGAGTCGTTGCCCGGTATGTAATTTAACATTCAANNNAGATCGGAAG
119721 TATGAGTGTGGAGTCGTTGCCCGGTATGTAATTTAACATTCAANNNAGATCGGAAG
119722 TATGAGTGTGGAGTCGTTGCCCGGTATGTAATTTAACATTCAANNNAGATCGGAAG
119723 TATGAGTGTGGAGTCGTTGCAATAATTTCCCTATTAGCCAGAANNNAGATCGGAAG
119724 TATGAGTGTGGAGTCGTTGCCTTTATGAAGTGTTTCACCAAANNNAGATCGGAAGA
119725 TATGAGTGTGGAGTCGTTGCCCATCAATAATAGCAAGTCATTTNNNAGATCGGAAG
119726 TATGAGTGTGGAGTCGTTGCCTAGGTTAGTTTGTGTTAACATGNNNAGATCGGAAG
119727 TATGAGTGTGGAGTCGTTGCTCATTAGCATTAAGAGCATAAACNNNAGATCGGAAG
119729 TATGAGTGTGGAGTCGTTGCAGATGTTGACATTTTAATAAGCANNNAGATCGGAAG
119730 TATGAGTGTGGAGTCGTTGCGAAGTGTTCGTTCAGATATTTCNNNAGATCGGAAGA
119731 TATGAGTGTGGAGTCGTTGCTATGTTACAGGATTAGTGGATTTNNNAGATCGGAA
119732 TATGAGTGTGGAGTCGTTGCTTTAAACCAACAATTAAGACAGANNNAGATCGGAAG
119733 TATGAGTGTGGAGTCGTTGCAAAAATGACATTGAATTCTGGACNNNAGATCGGAA
119734 TATGAGTGTGGAGTCGTTGCCTAAAGAAAAAAAAAGTGGTAGCNNNAGATCGGAA
119735 TATGAGTGTGGAGTCGTTGCAAGAGGAACAAAACACCAGTNNNAGATCGGAAGAG

119736 TATGAGTGTGGAGTCGTTGCCCTTCTAGCCATAGTAACATATTNNNAGATCGGAAGA
119737 TATGAGTGTGGAGTCGTTGCGTAACTCTTCATCCGTTGACNNNAGATCGGAAGAGC
119738 TATGAGTGTGGAGTCGTTGCGCCAAGCTATTAAGTTTGTAAATNNNAGATCGGAAG
119739 TATGAGTGTGGAGTCGTTGCTAAGCAGAGAAAGACTGTCAANNNAGATCGGAAGA
119740 TATGAGTGTGGAGTCGTTGCCATACATACTTGACTTCTCATCTNNNAGATCGGAAGA
119741 TATGAGTGTGGAGTCGTTGCCGCCAGCGAATAAATAATAAATANNNAGATCGGAA
119742 TATGAGTGTGGAGTCGTTGCCGCCAGCGAATAAATAATAAATANNNAGATCGGAA
119743 TATGAGTGTGGAGTCGTTGCATTGATTCAGATGAAGAAGATCANNNAGATCGGAA
119744 TATGAGTGTGGAGTCGTTGCTCTCTATAAAAATGGTTCTTTGCNNNAGATCGGAAG
119745 TATGAGTGTGGAGTCGTTGCTTAATAGTTTTTCTCATCATGGCNNNAGATCGGAAG
119746 TATGAGTGTGGAGTCGTTGCCCTTTGGAAAGTAAAACAAAAAANNNAGATCGGAA
119747 TATGAGTGTGGAGTCGTTGCAATGACAATATTAGGACTTCTCANNNAGATCGGAAG
119748 TATGAGTGTGGAGTCGTTGCTTCGAATTACAACTTGTAAAACTNNNAGATCGGAAG
119749 TATGAGTGTGGAGTCGTTGCTAAATACATGTTTGGCATCATCNNNAGATCGGAAGA
119750 TATGAGTGTGGAGTCGTTGCAGAGAAGAAAACAAAGGTAGGNNNAGATCGGAAG
119751 TATGAGTGTGGAGTCGTTGCCAAGATGGTTGATGAAATAATCANNNAGATCGGAA
119752 TATGAGTGTGGAGTCGTTGCTTACAATCTGCAAAGTACAAATTNNNAGATCGGAAG
119753 TATGAGTGTGGAGTCGTTGCTATCATAGAATAATGCAAGTGGANNNAGATCGGAA
119754 TATGAGTGTGGAGTCGTTGCTCTTTAATGATTTGGTCATACTGNNNAGATCGGAAG
119755 TATGAGTGTGGAGTCGTTGCTTTATCACTAAAGAAATGGTTGGNNNAGATCGGAAG
119756 TATGAGTGTGGAGTCGTTGCTCTTTAATGATTTGGTCATACTGNNNAGATCGGAAG
119757 TATGAGTGTGGAGTCGTTGCAAAATATCATCAACTGCATTGAGNNNAGATCGGAAG
119758 TATGAGTGTGGAGTCGTTGCAAAAATGAGTTATGAATACTGCANNNAGATCGGAA
119759 TATGAGTGTGGAGTCGTTGCGACTACCACACCATCATTTAGNNNAGATCGGAAGAG
119760 TATGAGTGTGGAGTCGTTGCTAAAACCAGCTCTGAAAAGAAANNNAGATCGGAAG
119761 TATGAGTGTGGAGTCGTTGCTCTTCAGTATCAGGACAAGGNNNAGATCGGAAGAG
119762 TATGAGTGTGGAGTCGTTGCGATGATCTTCAGTATCAGGACNNNAGATCGGAAGA
119763 TATGAGTGTGGAGTCGTTGCCAACAAGATTCTGAAGAACATGNNNAGATCGGAAG
119764 TATGAGTGTGGAGTCGTTGCTGCTGGTTTGTATTTTAGTTTTANNNAGATCGGAAG
119765 TATGAGTGTGGAGTCGTTGCCAGAGAAACCAAACTAACTTTAGNNNAGATCGGAAG
119766 TATGAGTGTGGAGTCGTTGCGAATTGAGAGAGGAAGAAGAATTNNNAGATCGGA
119767 TATGAGTGTGGAGTCGTTGCAAATTTCCTGTATTCTGAACAAGNNNAGATCGGAAG
119768 TATGAGTGTGGAGTCGTTGCAATTCCAGAAAAGGCTGTCTNNNAGATCGGAAGAG
119769 TATGAGTGTGGAGTCGTTGCTATTTCCACTAAACCTAGTCTTTNNNAGATCGGAAGA
119770 TATGAGTGTGGAGTCGTTGCCCAGGGAAGTATTATACTTACGNNNAGATCGGAAG
119771 TATGAGTGTGGAGTCGTTGCGTTGCTTTTCAAAGTGAAGATNNNAGATCGGAAGA
119772 TATGAGTGTGGAGTCGTTGCGCTGTTTAGATTTATATTGCTGTNNNAGATCGGAAG
119773 TATGAGTGTGGAGTCGTTGCGAGTTTGTCTGGTACAGTAACNNNAGATCGGAAGA
119774 TATGAGTGTGGAGTCGTTGCGAGAAAAAGAGATATGTGATGTTNNNAGATCGGAA
119775 TATGAGTGTGGAGTCGTTGCGAAGGGGTTAACATGACAGTNNNAGATCGGAAGAG
119776 TATGAGTGTGGAGTCGTTGCTTGTTGTCCAGTTGAAAAAAGNNNAGATCGGAAGA
119777 TATGAGTGTGGAGTCGTTGCTATCATACAAATAAGCCAGTCACNNNAGATCGGAAG
119778 TATGAGTGTGGAGTCGTTGCCCATCTGGTTGATCTAAAATTTTNNNAGATCGGAAG

119779 TATGAGTGTGGAGTCGTTGCCTTCAGACCTTGACTTATCTACNNNAGATCGGAAGAG
119780 TATGAGTGTGGAGTCGTTGCCTTTCTGGGAGTTTTCTAAATGNNNAGATCGGAAGA
119781 TATGAGTGTGGAGTCGTTGCCTCAATGGAGAACAGGAAAGNNNAGATCGGAAGAG
119782 TATGAGTGTGGAGTCGTTGCTACATTTGACTCGGAGAAAAATNNNAGATCGGAAGA
119783 TATGAGTGTGGAGTCGTTGCGACAATGTTTTTAAAGCTTGATNNNAGATCGGAAG
119784 TATGAGTGTGGAGTCGTTGCCTCACCTGGTACAAACAATATNNNAGATCGGAAGAG
119785 TATGAGTGTGGAGTCGTTGCTTTCTAAAAGGAGACTCCCTCNNNAGATCGGAAGAG
119786 TATGAGTGTGGAGTCGTTGCGAGTTCTTTGTCTTCAGAAAAGNNNAGATCGGAAGA
119787 TATGAGTGTGGAGTCGTTGCGCTGTTATTATTAAGGTAAGAGCNNNAGATCGGAAG
119788 TATGAGTGTGGAGTCGTTGCTTTGTGATGTGAATACTTTCAAANNNAGATCGGAAG
119789 TATGAGTGTGGAGTCGTTGCCATCCAAGTTCTTCTTACTACTTNNNAGATCGGAAGA
119790 TATGAGTGTGGAGTCGTTGCAAACTGTTATCACACTTAGAAAGNNNAGATCGGAAG
119791 TATGAGTGTGGAGTCGTTGCCATCCAAGTTCTTCTTACTACTTNNNAGATCGGAAGA
119792 TATGAGTGTGGAGTCGTTGCAAAAAGTGCGGAATTTCCATATNNNAGATCGGAAGA
119793 TATGAGTGTGGAGTCGTTGCTGTTCGTAGACATGAAAGAATNNNAGATCGGAAGA
119794 TATGAGTGTGGAGTCGTTGCAAAGTTAAAAACGGAGAGAAAATNNNAGATCGGAA
119795 TATGAGTGTGGAGTCGTTGCAGGTAGGGTGATTTTAATGATATNNNAGATCGGAA
119796 TATGAGTGTGGAGTCGTTGCACTGACTGACTTGGATCTTANNNAGATCGGAAGAG
119797 TATGAGTGTGGAGTCGTTGCGACAGAATTATGTAGGCTTTATCNNNAGATCGGAAG
119798 TATGAGTGTGGAGTCGTTGCGAGTACTGAATGACAGCCATNNNAGATCGGAAGAG
119799 TATGAGTGTGGAGTCGTTGCTAATTTTCATGTAATTGTGTGGTNNNAGATCGGAAG
119800 TATGAGTGTGGAGTCGTTGCCTATCTCTTAGACTTGTTCTGAANNNAGATCGGAAGA
119801 TATGAGTGTGGAGTCGTTGCGATACAATGGTGAGTGAAAATATNNNAGATCGGAA
119802 TATGAGTGTGGAGTCGTTGCTTTAAGAAAGGGAGATTTTAAGCNNNAGATCGGAA
119803 TATGAGTGTGGAGTCGTTGCGTTACCTTACGAAGATATTTCAGNNNAGATCGGAAG
119804 TATGAGTGTGGAGTCGTTGCCTACTACCTTGTATCAACCTCTNNNAGATCGGAAGAG
119805 TATGAGTGTGGAGTCGTTGCCAACTGAAGCCTTCTTTTAAGNNNAGATCGGAAGAG
119806 TATGAGTGTGGAGTCGTTGCACTTAAATTCTGTCTGAACAAAGNNNAGATCGGAAG
119807 TATGAGTGTGGAGTCGTTGCTGCCTCATATTCATGTTAGATAANNNAGATCGGAAG
119808 TATGAGTGTGGAGTCGTTGCTTCTAACTGGAGTGAGATAATACNNNAGATCGGAAG
119809 TATGAGTGTGGAGTCGTTGCCACCTTTAATTTTGTAATGGTCTNNNAGATCGGAAGA
119810 TATGAGTGTGGAGTCGTTGCCTGAAATCTCTTAGGCAATTTCNNNAGATCGGAAGA
119811 TATGAGTGTGGAGTCGTTGCAAGTTGTGATCATGTATCTGTATNNNAGATCGGAAG
119812 TATGAGTGTGGAGTCGTTGCTTTCTGCCTTTTAGTTTTTCCNNNAGATCGGAAGAGC
119813 TATGAGTGTGGAGTCGTTGCTTAGACAGTACCATGTTGATATGNNNAGATCGGAAG
119814 TATGAGTGTGGAGTCGTTGCAGAAAGAACAAAACTCCATCTNNNAGATCGGAAGAG
119815 TATGAGTGTGGAGTCGTTGCCCCTATCTTTTCAACCTAATTTTNNNAGATCGGAAGA
119816 TATGAGTGTGGAGTCGTTGCGTTTCCTTAAAGGCAGTTACNNNAGATCGGAAGAGC
119817 TATGAGTGTGGAGTCGTTGCAAACCCCATCTCTATCTTAAAATNNNAGATCGGAAGA
119818 TATGAGTGTGGAGTCGTTGCCCACTGTTTTTCATTTTTAGTACNNNAGATCGGAAGA
119819 TATGAGTGTGGAGTCGTTGCGGGTAAAAAAATAATTTCCCTTCNNNAGATCGGAAG
119820 TATGAGTGTGGAGTCGTTGCGCAAGGAAAGAGTATTTAAAAACNNNAGATCGGAA
119822 TATGAGTGTGGAGTCGTTGCTTAGTGTCTGGCTTAAATTCCNNNAGATCGGAAGAG

119823 TATGAGTGTGGAGTCGTTGCTTAGTGTCTGGCTTAAATTCCNNNAGATCGGAAGAG
119824 TATGAGTGTGGAGTCGTTGCACTCCACTGCTTATTTCATCNNNAGATCGGAAGAGCA
119825 TATGAGTGTGGAGTCGTTGCGATTTAAATCCCTCGCAAGANNNAGATCGGAAGAGC
119826 TATGAGTGTGGAGTCGTTGCGTATTCTTGGTTTCTGATGAAATNNNAGATCGGAAG
119827 TATGAGTGTGGAGTCGTTGCATACAATAGAAAGTAAACAGCACNNNAGATCGGAAG
119828 TATGAGTGTGGAGTCGTTGCCTATAATCCCAGCACTTTTGNNNAGATCGGAAGAGC
119829 TATGAGTGTGGAGTCGTTGCCTCCTTAAGCCATTCTAAAGTTNNNAGATCGGAAGAG
119830 TATGAGTGTGGAGTCGTTGCCTCCTTAAGCCATTCTAAAGTTNNNAGATCGGAAGAG
119831 TATGAGTGTGGAGTCGTTGCCCATTCTCAACATCATCATAAATNNNAGATCGGAAGA
119832 TATGAGTGTGGAGTCGTTGCGTAACAGATGACAAGATCAAAGNNNAGATCGGAAG
119833 TATGAGTGTGGAGTCGTTGCTTGCCTTTCATGAATACCAGNNNAGATCGGAAGAGCA
119834 TATGAGTGTGGAGTCGTTGCTTGTTTCTTTGCCTATAATTGTTNNNAGATCGGAAGA
119835 TATGAGTGTGGAGTCGTTGCAATCTGAATAACGGCCAAATNNNAGATCGGAAGAGC
119836 TATGAGTGTGGAGTCGTTGCTTGTTTCTTTGCCTATAATTGTTNNNAGATCGGAAGA
119837 TATGAGTGTGGAGTCGTTGCCCAAATAGCTAGGACTCCAGNNNAGATCGGAAGAGC
119838 TATGAGTGTGGAGTCGTTGCGATACTTACTGTAGTTCTCAGAGNNNAGATCGGAAG
119839 TATGAGTGTGGAGTCGTTGCCATTCTCATAAACACTCCCAANNNAGATCGGAAGAGC
119840 TATGAGTGTGGAGTCGTTGCCAGAAGAATCTTCCTCTTTTTCNNNAGATCGGAAGAG
119841 TATGAGTGTGGAGTCGTTGCATTCTGAAGCTTTTGAAAATACTNNNAGATCGGAAGA
119842 TATGAGTGTGGAGTCGTTGCTTTGAGAGATTTCCTTTTTCAAANNNAGATCGGAAGA
119843 TATGAGTGTGGAGTCGTTGCTTTTATTTTTTTGAGACAGAGGCNNNAGATCGGAAG
119844 TATGAGTGTGGAGTCGTTGCAAGGCAGAATCAAAAAATAGAAANNNAGATCGGAA
119845 TATGAGTGTGGAGTCGTTGCCAGATCTGATTGAGAGTTGGNNNAGATCGGAAGAG
119846 TATGAGTGTGGAGTCGTTGCCACCTACTAGTTGTTTATTACCTNNNAGATCGGAAGA
119847 TATGAGTGTGGAGTCGTTGCAAAACAATTTGCACATACCATATNNNAGATCGGAAGA
119848 TATGAGTGTGGAGTCGTTGCGGTCTTCACATTCATTTTCTACNNNAGATCGGAAGAG
119849 TATGAGTGTGGAGTCGTTGCTACTGTTTCTTTGAAACTTAAGGNNNAGATCGGAAGA
119850 TATGAGTGTGGAGTCGTTGCTTTCACATCCTTACATTTCAAAANNNAGATCGGAAGA
119851 TATGAGTGTGGAGTCGTTGCTTTGAGATATTACATGTTCAAGCNNNAGATCGGAAG
119852 TATGAGTGTGGAGTCGTTGCATCCAAACTCTATGAACCTTATTNNNAGATCGGAAGA
119853 TATGAGTGTGGAGTCGTTGCGGAGAAGCCTAAGTAAAAATAAANNNAGATCGGAA
119854 TATGAGTGTGGAGTCGTTGCCTCTACATTCAAAACCTATTGTTNNNAGATCGGAAGA
119855 TATGAGTGTGGAGTCGTTGCAAATCGAAGATTTCTATACCACTNNNAGATCGGAAGA
119856 TATGAGTGTGGAGTCGTTGCATAAAATAGAAAGAATGGCAAGCNNNAGATCGGAA
119857 TATGAGTGTGGAGTCGTTGCAAAATTTATCAACTAAGCACGTANNNAGATCGGAAG
119858 TATGAGTGTGGAGTCGTTGCAAGTCAGTAAGCAACACTTCNNNAGATCGGAAGAGC
119859 TATGAGTGTGGAGTCGTTGCAAACAAACAAATAAAAAGATCTCANNNAGATCGGAA
119860 TATGAGTGTGGAGTCGTTGCCCTACTATGTACCCACAAATAATNNNAGATCGGAAGA
119861 TATGAGTGTGGAGTCGTTGCGAGTGAAATTAGATACATTCACCNNNAGATCGGAAG
119862 TATGAGTGTGGAGTCGTTGCAATTCAAAACCCATTTCCTAATTNNNAGATCGGAAGA
119863 TATGAGTGTGGAGTCGTTGCAGCCTGTATAAAAATAAAAACAGANNNAGATCGGAA
119864 TATGAGTGTGGAGTCGTTGCTCAGAAAGCAAGATTTTAAACTTNNNAGATCGGAAG
119865 TATGAGTGTGGAGTCGTTGCTATGTTGATTTACCTCCCATATTNNNAGATCGGAAGA

119866 TATGAGTGTGGAGTCGTTGCCCAAATAAGGGTAGTAGATTTACNNNAGATCGGAA
119867 TATGAGTGTGGAGTCGTTGCCTGAGAAACAGGGATATGAAAANNNAGATCGGAAG
119868 TATGAGTGTGGAGTCGTTGCTCCGGCTAATTTTTGTATTTTTANNNAGATCGGAAG
119869 TATGAGTGTGGAGTCGTTGCTTTGTAACGAAGTGAAGGAGNNNAGATCGGAAGA
119870 TATGAGTGTGGAGTCGTTGCGATTGGAGTGGTATGATCATAGNNNAGATCGGAAG
119871 TATGAGTGTGGAGTCGTTGCGTCAGCATAACTTGATATTTTGANNNAGATCGGAAG
119872 TATGAGTGTGGAGTCGTTGCATTTTTTTTTTTGCTGTTTTGACNNNAGATCGGAAGA
119873 TATGAGTGTGGAGTCGTTGCGTTAATAACTGCATTGGATTTTCNNNAGATCGGAAG
119874 TATGAGTGTGGAGTCGTTGCCACCACCTTAAATATTTGTCTTTNNNAGATCGGAAGA
119875 TATGAGTGTGGAGTCGTTGCAAGAAGAAGGGGGAAAATTAAGNNNAGATCGGAA
119876 TATGAGTGTGGAGTCGTTGCGTGTTATAGCAGTTTTGCTGNNNAGATCGGAAGAG
119877 TATGAGTGTGGAGTCGTTGCAGTCCAAGGTATTATTCTTATGCNNNAGATCGGAAG
119878 TATGAGTGTGGAGTCGTTGCGAGTGACACAATGGGAAAAGNNNAGATCGGAAGA
119879 TATGAGTGTGGAGTCGTTGCGCCTAAAGCTAGACTATATTTCTNNNAGATCGGAAG
119880 TATGAGTGTGGAGTCGTTGCTTACATGAACTCTATTAGGTGTCNNNAGATCGGAAG
119881 TATGAGTGTGGAGTCGTTGCTCATCTCACTTAATAAGACATGTNNNAGATCGGAAG
119882 TATGAGTGTGGAGTCGTTGCCATGCTAGCAATTAACAATCAANNNAGATCGGAAGA
119883 TATGAGTGTGGAGTCGTTGCTCTGTACTAAGTTTAAAAAGCACNNNAGATCGGAAG
119884 TATGAGTGTGGAGTCGTTGCGGACAAAAACTACAACCATATCNNNAGATCGGAAGA
119885 TATGAGTGTGGAGTCGTTGCCACTAGTACCTCCATGTTATTTANNNAGATCGGAAGA
119886 TATGAGTGTGGAGTCGTTGCCCGTCTCTACTAAAAATACAAAANNNAGATCGGAAG
119887 TATGAGTGTGGAGTCGTTGCTATGTATTCGCCAATTTGTAATTNNNAGATCGGAAG
119888 TATGAGTGTGGAGTCGTTGCGTACTAGGTAATGGTCCATAACNNNAGATCGGAAGA
119889 TATGAGTGTGGAGTCGTTGCCACTGCCTACTCTGAATAAATNNNAGATCGGAAGAG
119890 TATGAGTGTGGAGTCGTTGCACTTTCTTAATAAACTTGCTTGCNNNAGATCGGAAGA
119891 TATGAGTGTGGAGTCGTTGCATAACCAAAGTCACCAATACTTANNNAGATCGGAAG
119892 TATGAGTGTGGAGTCGTTGCATAACCAAAGTCACCAATACTTANNNAGATCGGAAG
119893 TATGAGTGTGGAGTCGTTGCAGAACTAATGAGATCATAATCAGANNNAGATCGGAA
119894 TATGAGTGTGGAGTCGTTGCCTTTAGTAGCCTTCTCTTCATATNNNAGATCGGAAGA
119895 TATGAGTGTGGAGTCGTTGCCTTTAGTAGCCTTCTCTTCATATNNNAGATCGGAAGA
119896 TATGAGTGTGGAGTCGTTGCTTTATCTTTTTTCTGCCATCTTTNNNAGATCGGAAGA
119897 TATGAGTGTGGAGTCGTTGCATGGCCCTTTAACTTATTAAAAANNNAGATCGGAAG
119898 TATGAGTGTGGAGTCGTTGCATTTTACCATGATATACCCTGAANNNAGATCGGAAG
119899 TATGAGTGTGGAGTCGTTGCCGTGAGAATTTAGAGATACACTANNNAGATCGGAA
119900 TATGAGTGTGGAGTCGTTGCAAGGGGATGATTCTAAATCATTNNNAGATCGGAAG
119901 TATGAGTGTGGAGTCGTTGCGAGTTCATAGATAGATGCAAAACNNNAGATCGGAA
119903 TATGAGTGTGGAGTCGTTGCGGCAAGGAAAGAGTATTTAAAAANNNAGATCGGAA
119904 TATGAGTGTGGAGTCGTTGCCAATTTCATTATCTTCTTCACTCTNNNAGATCGGAAG
119905 TATGAGTGTGGAGTCGTTGCTTAGTGTCTGGCTTAAATTCCNNNAGATCGGAAGAG
119906 TATGAGTGTGGAGTCGTTGCTTAGTGTCTGGCTTAAATTCCNNNAGATCGGAAGAG
119907 TATGAGTGTGGAGTCGTTGCATACCATGTTTTGTTTATCCATTNNNAGATCGGAAGA
119908 TATGAGTGTGGAGTCGTTGCTATAATCATTTGCAATGCATACANNNAGATCGGAAG
119909 TATGAGTGTGGAGTCGTTGCGTATTCTTGGTTTCTGATGAAATNNNAGATCGGAAG

119910 TATGAGTGTGGAGTCGTTGCTACTATTGTCTATTAAGTGTGCANNNAGATCGGAAG
119911 TATGAGTGTGGAGTCGTTGCCTTGGGATTCAGAATTTTGAGNNNAGATCGGAAGA
119912 TATGAGTGTGGAGTCGTTGCAATGGGAGAAATCAACAATTCTNNNAGATCGGAAG
119913 TATGAGTGTGGAGTCGTTGCCTCCTTAAGCCATTCTAAAGTTNNNAGATCGGAAGA
119914 TATGAGTGTGGAGTCGTTGCTTATAACTTGATTCTTCCCATTGNNNAGATCGGAAGA
119915 TATGAGTGTGGAGTCGTTGCGTAACAGATGACAAGATCAAAGNNNAGATCGGAAG
119916 TATGAGTGTGGAGTCGTTGCTTGCCTTTCATGAATACCAGNNNAGATCGGAAGAGC
119917 TATGAGTGTGGAGTCGTTGCTTGCCTTTCATGAATACCAGNNNAGATCGGAAGAGC
119918 TATGAGTGTGGAGTCGTTGCAATCTGAATAACGGCCAAATNNNAGATCGGAAGAG
119919 TATGAGTGTGGAGTCGTTGCCATCATTTCTGGAGTCTCAGNNNAGATCGGAAGAGC
119920 TATGAGTGTGGAGTCGTTGCCATCATTTCTGGAGTCTCAGNNNAGATCGGAAGAGC
119922 TATGAGTGTGGAGTCGTTGCCAGAAGAATCTTCCTCTTTTTCNNNAGATCGGAAGA
119923 TATGAGTGTGGAGTCGTTGCCAGAAGAATCTTCCTCTTTTTCNNNAGATCGGAAGA
119924 TATGAGTGTGGAGTCGTTGCCATGAAAACAAGGCAAAATATTTNNNAGATCGGAA
119925 TATGAGTGTGGAGTCGTTGCAGAAAGAGAAAGAAAGAAAGAGANNNAGATCGGA
119926 TATGAGTGTGGAGTCGTTGCTCCTCTAACAATCTTATTCAAGANNNAGATCGGAAG
119927 TATGAGTGTGGAGTCGTTGCAGAAAGAGAAAGAAAGAAAGAGANNNAGATCGGA
119928 TATGAGTGTGGAGTCGTTGCTTTCCAAACTTATTAAGAAGCTTNNNAGATCGGAAG
119929 TATGAGTGTGGAGTCGTTGCCTAATGCTACAAAGGTTACTCTTNNNAGATCGGAAG
119930 TATGAGTGTGGAGTCGTTGCCTTAACCTGAATTTGATTAAAACANNNAGATCGGAA
119931 TATGAGTGTGGAGTCGTTGCATGTTAGTAATGAGGACAACTAGNNNAGATCGGAA
119932 TATGAGTGTGGAGTCGTTGCTTTTCTCTTTCAGGTTAAGCCNNNAGATCGGAAGAG
119933 TATGAGTGTGGAGTCGTTGCCACGTCTGGTTAATTTTTGTATNNNAGATCGGAAGA
119934 TATGAGTGTGGAGTCGTTGCTAAAACCCTGTCTCTACTAAAAGNNNAGATCGGAAG
119935 TATGAGTGTGGAGTCGTTGCCAAAGGAATGACTCATACAATGNNNAGATCGGAAG
119937 TATGAGTGTGGAGTCGTTGCTCTTTTTTGGTTCCATATGAATTNNNAGATCGGAAGA
119938 TATGAGTGTGGAGTCGTTGCAAAGGCTGCAGTTTTATACTNNNAGATCGGAAGAG
119939 TATGAGTGTGGAGTCGTTGCCCTTTTGTGTGTTCTTTTATGANNNAGATCGGAAGA
119940 TATGAGTGTGGAGTCGTTGCCCTTTTGTGTGTTCTTTTATGANNNAGATCGGAAGA
119941 TATGAGTGTGGAGTCGTTGCGTGGAGTTTTTGAGATAAATCAANNNAGATCGGAA
119942 TATGAGTGTGGAGTCGTTGCTTGATTGTTTTCTAAAAGGAGACNNNAGATCGGAAG
119943 TATGAGTGTGGAGTCGTTGCAGAAAAAAGCAAATTAAAACACCNNNAGATCGGAA
119944 TATGAGTGTGGAGTCGTTGCTTCTCTAACTCTGAATCATTCATNNNAGATCGGAAGA
119945 TATGAGTGTGGAGTCGTTGCTTTGTAACGAAGTGAAGGAGNNNAGATCGGAAGAG
119946 TATGAGTGTGGAGTCGTTGCGGATATAGCAGAGAGAGGTTNNNAGATCGGAAGA
119947 TATGAGTGTGGAGTCGTTGCACTGTTAACTACTGTACAACCNNNAGATCGGAAGAG
119948 TATGAGTGTGGAGTCGTTGCTACTGTTAACTACTGTACAACCNNNAGATCGGAAGA
119949 TATGAGTGTGGAGTCGTTGCAAAGGTACTGTTAACTACTGTACNNNAGATCGGAAG
119950 TATGAGTGTGGAGTCGTTGCACCACACAAGACATTTTTCTANNNAGATCGGAAGAG
119951 TATGAGTGTGGAGTCGTTGCACATTGAAGATGCTTTTTTTTTTNNNAGATCGGAAG
119952 TATGAGTGTGGAGTCGTTGCTTTCATTTGTCTAAGGTTTTTCTNNNAGATCGGAAGA
119953 TATGAGTGTGGAGTCGTTGCTTCTGAGAATGACATGAAAATTTNNNAGATCGGAAG
119954 TATGAGTGTGGAGTCGTTGCTACATTTTATAGGTTTGTGCAAANNNAGATCGGAAG

119955 TATGAGTGTGGAGTCGTTGCTATTTATTTTTTGAGACAGGGTCNNNAGATCGGAAG
119956 TATGAGTGTGGAGTCGTTGCTGTAGTTTATTTCAGGAAGACTGNNNAGATCGGAAG
119957 TATGAGTGTGGAGTCGTTGCAATTAACAAATGTTAGGGGAAATNNNAGATCGGAA
119958 TATGAGTGTGGAGTCGTTGCTTCTTCTCTCCAGAACACTAGNNNAGATCGGAAGAG
119959 TATGAGTGTGGAGTCGTTGCAATGGAATACTAAAAAAACCTGCNNNAGATCGGAAG
119960 TATGAGTGTGGAGTCGTTGCTAGAGAGAACACAAAGAAATCATNNNAGATCGGAA
119961 TATGAGTGTGGAGTCGTTGCAAGTTATAGTCATTGGGGTAAAANNNAGATCGGAA
119962 TATGAGTGTGGAGTCGTTGCGGGGGAAAAAAACATCTAATAAANNNAGATCGGAA
119963 TATGAGTGTGGAGTCGTTGCGGGGGAAAAAAACATCTAATAAANNNAGATCGGAA
119964 TATGAGTGTGGAGTCGTTGCCATGTAACAGTGAAGTCAGAGNNNAGATCGGAAGA
119965 TATGAGTGTGGAGTCGTTGCAAAGAATGTTTGTATACAAAGCTNNNAGATCGGAAG
119966 TATGAGTGTGGAGTCGTTGCCTCTGTTTACCTGGTTATGTAANNNAGATCGGAAGA
119967 TATGAGTGTGGAGTCGTTGCCTTGGAAGTTAACTAGCCTTTATNNNAGATCGGAAG
119968 TATGAGTGTGGAGTCGTTGCATTGAGACATGAAGAACAGACNNNAGATCGGAAGA
119969 TATGAGTGTGGAGTCGTTGCTCCATTGTTTCTAAGGTCTTCNNNAGATCGGAAGAG
119970 TATGAGTGTGGAGTCGTTGCATTGAGACATGAAGAACAGACNNNAGATCGGAAGA
119971 TATGAGTGTGGAGTCGTTGCAACCTTTAAACATCAAACTCATTNNNAGATCGGAAGA
119972 TATGAGTGTGGAGTCGTTGCTTGAAGTTTTCCATCAATTAACANNNAGATCGGAAG
119973 TATGAGTGTGGAGTCGTTGCTTGAAGTTTTCCATCAATTAACANNNAGATCGGAAG
119974 TATGAGTGTGGAGTCGTTGCTTGAGAATACATGAGGGAGTTANNNAGATCGGAAG
119975 TATGAGTGTGGAGTCGTTGCTAAAACATATTCTTGTCTCTGTCNNNAGATCGGAAGA
119976 TATGAGTGTGGAGTCGTTGCCTCAAAACCATACTATTACATGGNNNAGATCGGAAG
119977 TATGAGTGTGGAGTCGTTGCGCTCTCATCATAAAATTCAGAAANNNAGATCGGAAG
119978 TATGAGTGTGGAGTCGTTGCTTTTTAAAATGAGATGAAGCCCNNNAGATCGGAAGA
119979 TATGAGTGTGGAGTCGTTGCCATTTATCTGGGATAAATGAGAANNNAGATCGGAA
119980 TATGAGTGTGGAGTCGTTGCGTTGTACTACAAGAACATCAGTNNNAGATCGGAAGA
119981 TATGAGTGTGGAGTCGTTGCTTCTTTCCCTCCACTATGTTATNNNAGATCGGAAGAG
119982 TATGAGTGTGGAGTCGTTGCACATATATTTATCATGTGCAACANNNAGATCGGAAG
119983 TATGAGTGTGGAGTCGTTGCGAGCAATTAAAACAGATTCCTAANNNAGATCGGAAG
119984 TATGAGTGTGGAGTCGTTGCGAAAGTCTGAGAACGGATTCNNNAGATCGGAAGAG
119985 TATGAGTGTGGAGTCGTTGCACAATGCTTAGAAACTAATACACNNNAGATCGGAAG
119986 TATGAGTGTGGAGTCGTTGCAATTATCCTACCTCAGCCTCNNNAGATCGGAAGAGC
119987 TATGAGTGTGGAGTCGTTGCATATATAGTGGAGAGACTCAGAANNNAGATCGGAA
119988 TATGAGTGTGGAGTCGTTGCATATATAGTGGAGAGACTCAGAANNNAGATCGGAA
119989 TATGAGTGTGGAGTCGTTGCTTCCAGAAGTAGAAGACGAGNNNAGATCGGAAGAG
119990 TATGAGTGTGGAGTCGTTGCATTCTTGTTTTTAAACTTAGCCANNNAGATCGGAAGA
119991 TATGAGTGTGGAGTCGTTGCATAAAACCAAAAAACAGGTAAGTNNNAGATCGGAA
119992 TATGAGTGTGGAGTCGTTGCGAAACCATAAATGTAACACTCATNNNAGATCGGAAG
119993 TATGAGTGTGGAGTCGTTGCCTTTAGGCGTTAAAACTGTAAATNNNAGATCGGAAG
119994 TATGAGTGTGGAGTCGTTGCCTGTTTGCATCCCAACATACNNNAGATCGGAAGAGC
119995 TATGAGTGTGGAGTCGTTGCAGAACTCACAGATTCTTTCTTATNNNAGATCGGAAGA
119996 TATGAGTGTGGAGTCGTTGCCGTTAAGTCGTAGCCCTTAANNNAGATCGGAAGAGC
119997 TATGAGTGTGGAGTCGTTGCTACTCAATTTAATTGCAACCTTTNNNAGATCGGAAGA

119998 TATGAGTGTGGAGTCGTTGCGGATACCTATAATTGGATATTTCCNNNAGATCGGAA
119999 TATGAGTGTGGAGTCGTTGCATATCTTAAGGAATAATTGTGGATNNNAGATCGGAA
120000 TATGAGTGTGGAGTCGTTGCATGTGTTTATTTCTATCTATCTATCTNNNAGATCGGA
120001 TATGAGTGTGGAGTCGTTGCGCCCTCTAGTTGAATTCTAAANNNAGATCGGAAGAG
120002 TATGAGTGTGGAGTCGTTGCTATCAGGAATGGACTTAGTCTNNNAGATCGGAAGA
120003 TATGAGTGTGGAGTCGTTGCTATCAGGAATGGACTTAGTCTNNNAGATCGGAAGA
120004 TATGAGTGTGGAGTCGTTGCATTTACTTCTGGATGAGCTCTNNNAGATCGGAAGAG
120005 TATGAGTGTGGAGTCGTTGCCTTCATACAGGGTATTCCATAATNNNAGATCGGAAG
120006 TATGAGTGTGGAGTCGTTGCCTTCATACAGGGTATTCCATAATNNNAGATCGGAAG
120007 TATGAGTGTGGAGTCGTTGCAGAGAAGGTTCACTAACTGTNNNAGATCGGAAGAG
120008 TATGAGTGTGGAGTCGTTGCTTTTGGTCTTGTTTCAATTTCTTNNNAGATCGGAAGA
120009 TATGAGTGTGGAGTCGTTGCTTTTGGTCTTGTTTCAATTTCTTNNNAGATCGGAAGA
120010 TATGAGTGTGGAGTCGTTGCGAGCAATTAAAACAGATTCCTAANNNAGATCGGAAG
120011 TATGAGTGTGGAGTCGTTGCAGAACTCACAGATTCTTTCTTATNNNAGATCGGAAG
120012 TATGAGTGTGGAGTCGTTGCCATGGTAATTAGCACTTTTCCNNNAGATCGGAAGAG
120013 TATGAGTGTGGAGTCGTTGCTGCCTTGTTTTTAAGTAATACAANNNAGATCGGAAG
120014 TATGAGTGTGGAGTCGTTGCAGAGAAGTATTAAGAATGACTCCNNNAGATCGGAA
120015 TATGAGTGTGGAGTCGTTGCACACAATCTGGATAATATTGAGANNNAGATCGGAAG
120016 TATGAGTGTGGAGTCGTTGCTGCCTTGTTTTTAAGTAATACAANNNAGATCGGAAG
120017 TATGAGTGTGGAGTCGTTGCCATTAGTAGCCTGACTGTTAATNNNAGATCGGAAGA
120018 TATGAGTGTGGAGTCGTTGCAGCTAGTTTACTGGCATTATCNNNAGATCGGAAGAG
120019 TATGAGTGTGGAGTCGTTGCAAACTGATTTTTGTTGTTGTAGTNNNAGATCGGAAG
120020 TATGAGTGTGGAGTCGTTGCAAACTGATTTTTGTTGTTGTAGTNNNAGATCGGAAG
120021 TATGAGTGTGGAGTCGTTGCGGCATATCTCAAAAACTGATTTNNNAGATCGGAAGA
120022 TATGAGTGTGGAGTCGTTGCCACCATTCATTTCATTTTGTTTTNNNAGATCGGAAGA
120023 TATGAGTGTGGAGTCGTTGCCACCATTCATTTCATTTTGTTTTNNNAGATCGGAAGA
120024 TATGAGTGTGGAGTCGTTGCCTGTTAAGGTTTCTCTGAGGNNNAGATCGGAAGAG
120025 TATGAGTGTGGAGTCGTTGCGAGAAAGAAGCAATTGTGATGNNNAGATCGGAAG
120026 TATGAGTGTGGAGTCGTTGCCTTAAAAGTGTTGTGTTACTTTGNNNAGATCGGAAG
120027 TATGAGTGTGGAGTCGTTGCACACAATCTGGATAATATTGAGANNNAGATCGGAAG
120028 TATGAGTGTGGAGTCGTTGCATACATGTGTAGATGGATTAGGANNNAGATCGGAA
120029 TATGAGTGTGGAGTCGTTGCTGTATGTCATTCCTGTTCACNNNAGATCGGAAGAGC
120030 TATGAGTGTGGAGTCGTTGCTGTATGTCATTCCTGTTCACNNNAGATCGGAAGAGC
120031 TATGAGTGTGGAGTCGTTGCAAAAATTGGTGGGTTATGCTTNNNAGATCGGAAGA
120032 TATGAGTGTGGAGTCGTTGCATCATGTTTAGATCACAATTAGGNNNAGATCGGAAG
120033 TATGAGTGTGGAGTCGTTGCACATAAAATGTGGTAACATACACNNNAGATCGGAAG
120034 TATGAGTGTGGAGTCGTTGCGTCTATATGCAAAATGAGTTTGANNNAGATCGGAAG
120035 TATGAGTGTGGAGTCGTTGCCTAGTTGATCCAGATCAGTGANNNAGATCGGAAGA
120036 TATGAGTGTGGAGTCGTTGCTCGATTGTCTTCTCTAGTGANNNAGATCGGAAGAGC
120037 TATGAGTGTGGAGTCGTTGCTCGATTGTCTTCTCTAGTGANNNAGATCGGAAGAGC
120038 TATGAGTGTGGAGTCGTTGCCTGAAAGTCGCAAAAGTTCTNNNAGATCGGAAGAG
120039 TATGAGTGTGGAGTCGTTGCCTGAAAGTCGCAAAAGTTCTNNNAGATCGGAAGAG
120040 TATGAGTGTGGAGTCGTTGCGGTACATATATACTGTGGGATACNNNAGATCGGAA

120041 TATGAGTGTGGAGTCGTTGCATAGATATCTGAAGTGTGTGAAGNNNAGATCGGAA
120042 TATGAGTGTGGAGTCGTTGCATAGATATCTGAAGTGTGTGAAGNNNAGATCGGAA
120043 TATGAGTGTGGAGTCGTTGCTTCATGGGATTAGAAGAATTCAANNNAGATCGGAA
120044 TATGAGTGTGGAGTCGTTGCTTCATGGGATTAGAAGAATTCAANNNAGATCGGAA
120045 TATGAGTGTGGAGTCGTTGCTTATTTTGTTTGTTTCATTTGGGNNNAGATCGGAAG
120046 TATGAGTGTGGAGTCGTTGCGAGAATAGGAACCCAGATATGNNNAGATCGGAAGA
120047 TATGAGTGTGGAGTCGTTGCTAGACTGAGCTTAGACAGAGNNNAGATCGGAAGAG
120048 TATGAGTGTGGAGTCGTTGCTTTGGTTTCCATAACCTAAGTATNNNAGATCGGAAG
120049 TATGAGTGTGGAGTCGTTGCATTTCCCTACATGATGTACATTTNNNAGATCGGAAGA
120050 TATGAGTGTGGAGTCGTTGCATTTCCCTACATGATGTACATTTNNNAGATCGGAAGA
120051 TATGAGTGTGGAGTCGTTGCTTTATACCTGATTTTTCCAACACNNNAGATCGGAAGA
120052 TATGAGTGTGGAGTCGTTGCCACCATTTTTATAGTCCATGTTTNNNAGATCGGAAGA
120053 TATGAGTGTGGAGTCGTTGCATTATGTGAGAAGGTGAATGATNNNAGATCGGAAG
120054 TATGAGTGTGGAGTCGTTGCGAAGAGAGCTGTTGAACATGNNNAGATCGGAAGA
120055 TATGAGTGTGGAGTCGTTGCCTGACCATTAGCATCACTTAATNNNAGATCGGAAGA
120056 TATGAGTGTGGAGTCGTTGCCTGACCATTAGCATCACTTAATNNNAGATCGGAAGA
120057 TATGAGTGTGGAGTCGTTGCCTTTTCCTGACCATTAGCATNNNAGATCGGAAGAGC
120058 TATGAGTGTGGAGTCGTTGCCAATGTCAGGATCCAGAAAGNNNAGATCGGAAGAG
120059 TATGAGTGTGGAGTCGTTGCGGATTCTTTAAAGGGGTTGTNNNAGATCGGAAGAG
120060 TATGAGTGTGGAGTCGTTGCGGATTCTTTAAAGGGGTTGTNNNAGATCGGAAGAG
120061 TATGAGTGTGGAGTCGTTGCCTGACTTGTTTTCCACAAACNNNAGATCGGAAGAGC
120062 TATGAGTGTGGAGTCGTTGCCTTTAAGATAAAACACAGGCTTANNNAGATCGGAAG
120063 TATGAGTGTGGAGTCGTTGCGATTTCTGAGGCACATAGTCNNNAGATCGGAAGAG
120064 TATGAGTGTGGAGTCGTTGCGATTTCTGAGGCACATAGTCNNNAGATCGGAAGAG
120065 TATGAGTGTGGAGTCGTTGCAAGGCATATGGAGATGTAGTNNNAGATCGGAAGA
120066 TATGAGTGTGGAGTCGTTGCTTTTCAAGTGAGGTAACCAACNNNAGATCGGAAGA
120067 TATGAGTGTGGAGTCGTTGCTTTTCAAGTGAGGTAACCAACNNNAGATCGGAAGA
120068 TATGAGTGTGGAGTCGTTGCTGCTATGGTAAGATAACCTCTTNNNAGATCGGAAGA
120069 TATGAGTGTGGAGTCGTTGCTGAAAATTAAGTGCTATGGTAAGNNNAGATCGGAA
120070 TATGAGTGTGGAGTCGTTGCTGAAAATTAAGTGCTATGGTAAGNNNAGATCGGAA
120071 TATGAGTGTGGAGTCGTTGCTTACATATCATCTCGTCCAATAANNNAGATCGGAAG
120072 TATGAGTGTGGAGTCGTTGCTTACATATCATCTCGTCCAATAANNNAGATCGGAAG
120073 TATGAGTGTGGAGTCGTTGCGTTTTCCTGCTATTCATGTTTCNNNAGATCGGAAGAG
120074 TATGAGTGTGGAGTCGTTGCGAGAAAGAAGCAATTGTGATGNNNAGATCGGAAG
120075 TATGAGTGTGGAGTCGTTGCATTTTCTTAGCACCATTTCTTAGNNNAGATCGGAAGA
120076 TATGAGTGTGGAGTCGTTGCGCTATTAAAGACACCATTTTCTTNNNAGATCGGAAG
120077 TATGAGTGTGGAGTCGTTGCTCCACATTTTAAAAACTAAGCTTNNNAGATCGGAAG
120078 TATGAGTGTGGAGTCGTTGCTCTTCAAATCCGTATACAATTTGNNNAGATCGGAAG
120079 TATGAGTGTGGAGTCGTTGCCCCACCTTCTGTTTACTTTAGNNNAGATCGGAAGAG
120080 TATGAGTGTGGAGTCGTTGCCTCTTTGTTTCTCATCAACTGNNNAGATCGGAAGAG
120081 TATGAGTGTGGAGTCGTTGCGGCTATTTTTCCACTGATTAAATNNNAGATCGGAAG
120082 TATGAGTGTGGAGTCGTTGCGTTTCAGATAGTGCTGTGTTNNNAGATCGGAAGAG
120083 TATGAGTGTGGAGTCGTTGCGTTGATACTGAAGAATTGATGGNNNAGATCGGAAG

120084 TATGAGTGTGGAGTCGTTGCGTTGATACTGAAGAATTGATGGNNNAGATCGGAAG
120085 TATGAGTGTGGAGTCGTTGCGGAATTTCCAGTGTATTGTTTGNNNAGATCGGAAG
120086 TATGAGTGTGGAGTCGTTGCATTCATTTCAACTTGGTACATTTNNNAGATCGGAAG
120087 TATGAGTGTGGAGTCGTTGCCATGAATGTAATCTTAGCTTTCANNNAGATCGGAAG
120088 TATGAGTGTGGAGTCGTTGCCACATGAGCTTTTGGTAAATGNNNAGATCGGAAGA
120089 TATGAGTGTGGAGTCGTTGCATAAGTTTTTCAGTTTGGGAATCNNNAGATCGGAAG
120090 TATGAGTGTGGAGTCGTTGCGATTCAAGTGCTGTTTCAACNNNAGATCGGAAGAG
120091 TATGAGTGTGGAGTCGTTGCCTTAAAAGTGTTGTGTTACTTTGNNNAGATCGGAAG
120092 TATGAGTGTGGAGTCGTTGCCTGTTTATGAGGCTTATGTTGNNNAGATCGGAAGA
120093 TATGAGTGTGGAGTCGTTGCTTTATTTGTAATTTTTGCTGCTGNNNAGATCGGAAG
120094 TATGAGTGTGGAGTCGTTGCTTTATTTGTAATTTTTGCTGCTGNNNAGATCGGAAG
120095 TATGAGTGTGGAGTCGTTGCGCTTGATTTTGAATACTGATTTTNNNAGATCGGAAG
120096 TATGAGTGTGGAGTCGTTGCGCTTGATTTTGAATACTGATTTTNNNAGATCGGAAG
120097 TATGAGTGTGGAGTCGTTGCTGATTTCATGGTCTGACTATAAGNNNAGATCGGAAG
120098 TATGAGTGTGGAGTCGTTGCTGATTTCATGGTCTGACTATAAGNNNAGATCGGAAG
120099 TATGAGTGTGGAGTCGTTGCAAAAAGTTCAGGATGTGTAGTNNNAGATCGGAAGA
120100 TATGAGTGTGGAGTCGTTGCAAAAAGTTCAGGATGTGTAGTNNNAGATCGGAAGA
120101 TATGAGTGTGGAGTCGTTGCTTGCTTTGGGATCACATTATTTNNNAGATCGGAAGA
120102 TATGAGTGTGGAGTCGTTGCCACCTGTGAAGAAGATCTTGNNNAGATCGGAAGAG
120103 TATGAGTGTGGAGTCGTTGCAAAAACATTTGCATGGTTATGTANNNAGATCGGAAG
120104 TATGAGTGTGGAGTCGTTGCAAAAACATTTGCATGGTTATGTANNNAGATCGGAAG
120105 TATGAGTGTGGAGTCGTTGCTTCTTGCTTCTGTAAGAGATGNNNAGATCGGAAGAG
120106 TATGAGTGTGGAGTCGTTGCCCAGGTATATTCATCTCTAAGAANNNAGATCGGAAG
120107 TATGAGTGTGGAGTCGTTGCACTATGGCAAGACTCAGTTTNNNAGATCGGAAGAG
120108 TATGAGTGTGGAGTCGTTGCTTACCTTTTTCCATGTTTTGTTTNNNAGATCGGAAGA
120109 TATGAGTGTGGAGTCGTTGCTTACCTTTTTCCATGTTTTGTTTNNNAGATCGGAAGA
120110 TATGAGTGTGGAGTCGTTGCCTTATGGTCAAATAACGACTTGNNNAGATCGGAAGA
120112 TATGAGTGTGGAGTCGTTGCACATTTCAACTTTTACTTGCTTNNNAGATCGGAAGAG
120113 TATGAGTGTGGAGTCGTTGCTTTGTCTAGTCAAAACTGTGANNNAGATCGGAAGAG
120114 TATGAGTGTGGAGTCGTTGCGAGTATCTAGTAATTTGGAAGGTNNNAGATCGGAA
120115 TATGAGTGTGGAGTCGTTGCGAGTATCTAGTAATTTGGAAGGTNNNAGATCGGAA
120116 TATGAGTGTGGAGTCGTTGCAAGATGGGAAATCATATTGAGTNNNAGATCGGAAG
120117 TATGAGTGTGGAGTCGTTGCGGAAAATCCCATATCTGAAGATANNNAGATCGGAA
120118 TATGAGTGTGGAGTCGTTGCTTCAGATTTTTATGAGCCTTTACNNNAGATCGGAAG
120119 TATGAGTGTGGAGTCGTTGCTTCAGATTTTTATGAGCCTTTACNNNAGATCGGAAG
120120 TATGAGTGTGGAGTCGTTGCTAACTTGACAAAAAATGTAAGCTNNNAGATCGGAAG
120121 TATGAGTGTGGAGTCGTTGCGAAAAATAAACATGCACGATTATNNNAGATCGGAA
120122 TATGAGTGTGGAGTCGTTGCGAAAAATAAACATGCACGATTATNNNAGATCGGAA
120123 TATGAGTGTGGAGTCGTTGCCTCTCCAGGAGATAATATACAAANNNAGATCGGAAG
120124 TATGAGTGTGGAGTCGTTGCCTATGGTCGTTATTTTCTCATTTNNNAGATCGGAAGA
120125 TATGAGTGTGGAGTCGTTGCCTTAATCCTCAAAACGGTAAATGNNNAGATCGGAAG
120126 TATGAGTGTGGAGTCGTTGCAATAATTTCCCTATTAGCCAGAANNNAGATCGGAAG
120127 TATGAGTGTGGAGTCGTTGCAAAACAAAACAAGGGTCACTTNNNAGATCGGAAGA

120128 TATGAGTGTGGAGTCGTTGCAAATGGAGAAAAGGAAACTTTAGNNNAGATCGGAA
120130 TATGAGTGTGGAGTCGTTGCGGTATGCCTGTATCATATTTTTTNNNAGATCGGAAG
120131 TATGAGTGTGGAGTCGTTGCAACTTTTACATACAGTGAGAAACNNNAGATCGGAAG
120132 TATGAGTGTGGAGTCGTTGCATTGTTGGCATTTTTTTAACAATNNNAGATCGGAAG
120134 TATGAGTGTGGAGTCGTTGCTTTAAACCTTACAACTGAATGACNNNAGATCGGAAG
120135 TATGAGTGTGGAGTCGTTGCTTTTAAAGAGGTAGATTCCCAATNNNAGATCGGAAG
120136 TATGAGTGTGGAGTCGTTGCGTAATTTTGGTCTCATTATGTCANNNAGATCGGAAG
120137 TATGAGTGTGGAGTCGTTGCAAAGCTAACACTTAGTGAATCTANNNAGATCGGAAG
120138 TATGAGTGTGGAGTCGTTGCGCAACCAAAAGATGTATAAAGTTNNNAGATCGGAA
120139 TATGAGTGTGGAGTCGTTGCCAAGGTATACATTTAGGATTTGTNNNAGATCGGAAG
120140 TATGAGTGTGGAGTCGTTGCTGCTATCAAAGGATAAAAAATGANNNAGATCGGAA
120141 TATGAGTGTGGAGTCGTTGCCTCCTTTAAAATAATGGTTCACANNNAGATCGGAAG
120142 TATGAGTGTGGAGTCGTTGCATACACTCAGGAGTATGTATGTANNNAGATCGGAA
120143 TATGAGTGTGGAGTCGTTGCTTATCCATTATCATGAAATGCTGNNNAGATCGGAAG
120145 TATGAGTGTGGAGTCGTTGCAAATGTCATATATACCTAGCACANNNAGATCGGAAG
120146 TATGAGTGTGGAGTCGTTGCGAATATAGACATTTCCCACTCTANNNAGATCGGAAG
120147 TATGAGTGTGGAGTCGTTGCATGCAAATACATATTCGAAGTTTNNNAGATCGGAAG
120148 TATGAGTGTGGAGTCGTTGCAAAGAACACAAATAGGAAATCAANNNAGATCGGAA
120149 TATGAGTGTGGAGTCGTTGCAAAAAGGTGAGCAAACTGATATNNNAGATCGGAAG
120150 TATGAGTGTGGAGTCGTTGCGGAAATAGCCTTAAAAAAAAAACTNNNAGATCGGA
120151 TATGAGTGTGGAGTCGTTGCATTTATCTCATAAGTGGAAATGCNNNAGATCGGAAG
120152 TATGAGTGTGGAGTCGTTGCGAAGCCGAACATTATTTTATGAANNNAGATCGGAAG
120153 TATGAGTGTGGAGTCGTTGCTTCCACTTTCTTTTGAAAAATTTNNNAGATCGGAAGA
120154 TATGAGTGTGGAGTCGTTGCGAGGATTAAAAAGGACATGTAAGNNNAGATCGGA
120155 TATGAGTGTGGAGTCGTTGCTTTAATAAATGAAATTGCTGGGANNNAGATCGGAA
120156 TATGAGTGTGGAGTCGTTGCTTTAATAAATGAAATTGCTGGGANNNAGATCGGAA
120157 TATGAGTGTGGAGTCGTTGCTAACTTTGCAAAATACTTGAGTCNNNAGATCGGAAG
120158 TATGAGTGTGGAGTCGTTGCATTATGCATCGTCCATAACTTTNNNAGATCGGAAGA
120159 TATGAGTGTGGAGTCGTTGCACAGTTGGTATTTTTAATTCTCCNNNAGATCGGAAG
120160 TATGAGTGTGGAGTCGTTGCACAGTTGGTATTTTTAATTCTCCNNNAGATCGGAAG
120161 TATGAGTGTGGAGTCGTTGCACTTACTGTCCATAAAATTTTCTNNNAGATCGGAAGA
120162 TATGAGTGTGGAGTCGTTGCGAATTATATTCTGCCACTTAAGGNNNAGATCGGAAG
120163 TATGAGTGTGGAGTCGTTGCGGCAGCAATGGAAAACTAATNNNAGATCGGAAGAG
120164 TATGAGTGTGGAGTCGTTGCGGCAGCAATGGAAAACTAATNNNAGATCGGAAGAG
120165 TATGAGTGTGGAGTCGTTGCCATACTGTAGGGTAATAAATTTGTNNNAGATCGGAA
120166 TATGAGTGTGGAGTCGTTGCCATACTGTAGGGTAATAAATTTGTNNNAGATCGGAA
120167 TATGAGTGTGGAGTCGTTGCTTCTGGAAAAGCTGGAAAAGNNNAGATCGGAAGAG
120168 TATGAGTGTGGAGTCGTTGCGAAATAAAGTGCACAATAAATGCNNNAGATCGGAA
120169 TATGAGTGTGGAGTCGTTGCGAAATAAAGTGCACAATAAATGCNNNAGATCGGAA
120170 TATGAGTGTGGAGTCGTTGCGCTCCTTATGAGAATCTGATGNNNAGATCGGAAGA
120171 TATGAGTGTGGAGTCGTTGCGCTCCTTATGAGAATCTGATGNNNAGATCGGAAGA
120172 TATGAGTGTGGAGTCGTTGCATTAGATTCTCATAGCAGCACNNNAGATCGGAAGAG
120173 TATGAGTGTGGAGTCGTTGCCTTCATCTGTATTTATAGCCACTNNNAGATCGGAAGA

120174 TATGAGTGTGGAGTCGTTGCCTTCATCTGTATTTATAGCCACTNNNAGATCGGAAGA
120175 TATGAGTGTGGAGTCGTTGCAGTCTCTAGAACAGGGATTCNNNAGATCGGAAGAG
120176 TATGAGTGTGGAGTCGTTGCAGTCTCTAGAACAGGGATTCNNNAGATCGGAAGAG
120177 TATGAGTGTGGAGTCGTTGCAGGTACCACTAGTCAAGGATNNNAGATCGGAAGAG
120178 TATGAGTGTGGAGTCGTTGCATCTTGCAACAGAGAGATTATCNNNAGATCGGAAGA
120179 TATGAGTGTGGAGTCGTTGCATCTTGCAACAGAGAGATTATCNNNAGATCGGAAGA
120180 TATGAGTGTGGAGTCGTTGCGTATTCTTGGTTTCTGATGAAATNNNAGATCGGAAG
120183 TATGAGTGTGGAGTCGTTGCAAGACAGCTGTGTCCTAATCNNNAGATCGGAAGAG
120184 TATGAGTGTGGAGTCGTTGCATTAGAACTATAGCGGCTGGNNNAGATCGGAAGAG
120185 TATGAGTGTGGAGTCGTTGCCAAATTGAAGATCTAATAGAGGCNNNAGATCGGAA
120186 TATGAGTGTGGAGTCGTTGCTTTATTTTTTTGAGACAGAGTCTNNNAGATCGGAAG
120187 TATGAGTGTGGAGTCGTTGCTTTATTTTTTTGAGACAGAGTCTNNNAGATCGGAAG
120188 TATGAGTGTGGAGTCGTTGCAAGAAAAGAGATAAGGTAAAGTTNNNAGATCGGAA
120189 TATGAGTGTGGAGTCGTTGCCTAAACGAATAAAGCAGTTACATNNNAGATCGGAAG
120190 TATGAGTGTGGAGTCGTTGCCTAAACGAATAAAGCAGTTACATNNNAGATCGGAAG
120191 TATGAGTGTGGAGTCGTTGCTATGAGTTCTTTAGCTAAACGAANNNAGATCGGAAG
120192 TATGAGTGTGGAGTCGTTGCTTGATGTAGCTGTGAAAATTAATNNNAGATCGGAAG
120193 TATGAGTGTGGAGTCGTTGCGACATTTGTCAAAAGTTAGTAAANNNAGATCGGAAG
120194 TATGAGTGTGGAGTCGTTGCAACTTTTTTTTGACATGAAAACANNNAGATCGGAAG
120197 TATGAGTGTGGAGTCGTTGCTAATCCTGGTTTTCAAGAGTATTNNNAGATCGGAAG
120198 TATGAGTGTGGAGTCGTTGCAAACCTCTCTGAAAATTAATTGGNNNAGATCGGAAG
120199 TATGAGTGTGGAGTCGTTGCTCAAGTGTTGATGTAGTAATAAATTNNNAGATCGGA
120200 TATGAGTGTGGAGTCGTTGCAAGTCGGTAAGTAAAGAAAGCNNNAGATCGGAAGA
120202 TATGAGTGTGGAGTCGTTGCTTGCATTGATATGATGGATATTGNNNAGATCGGAAG
120203 TATGAGTGTGGAGTCGTTGCAAGAAAAAAAAGAGGTTATCACANNNAGATCGGAA
120204 TATGAGTGTGGAGTCGTTGCCTTCCATCCCTCTCTTCATTNNNAGATCGGAAGAGCA
120205 TATGAGTGTGGAGTCGTTGCCTTCCATCCCTCTCTTCATTNNNAGATCGGAAGAGCA
120206 TATGAGTGTGGAGTCGTTGCACTAAACTCAAGAAGTCAGTATTNNNAGATCGGAAG
120207 TATGAGTGTGGAGTCGTTGCATGGCAGACAGTGAAAAAACNNNAGATCGGAAGAG
120208 TATGAGTGTGGAGTCGTTGCAGAGGTCAGGTACAAATGAANNNAGATCGGAAGAG
120209 TATGAGTGTGGAGTCGTTGCGAACTACATGAGATGTAAGAATGNNNAGATCGGAA
120210 TATGAGTGTGGAGTCGTTGCGAACTACATGAGATGTAAGAATGNNNAGATCGGAA
120211 TATGAGTGTGGAGTCGTTGCGTAATAGGAGCTAGTATCAGAAANNNAGATCGGAA
120212 TATGAGTGTGGAGTCGTTGCGTAATAGGAGCTAGTATCAGAAANNNAGATCGGAA
120213 TATGAGTGTGGAGTCGTTGCATCAGAATTAAGACTAGAGATGGNNNAGATCGGAA
120214 TATGAGTGTGGAGTCGTTGCAGACAAGGAAACTAATGATAGAANNNAGATCGGAA
120215 TATGAGTGTGGAGTCGTTGCAGACAAGGAAACTAATGATAGAANNNAGATCGGAA
120216 TATGAGTGTGGAGTCGTTGCCCATTCTTGAGAGTTCCTTTTTNNNAGATCGGAAGAG
120217 TATGAGTGTGGAGTCGTTGCCCATTCTTGAGAGTTCCTTTTTNNNAGATCGGAAGAG
120218 TATGAGTGTGGAGTCGTTGCTTGAGAGATGAGATGTAAAATGCNNNAGATCGGAA
120219 TATGAGTGTGGAGTCGTTGCCCCATATGCTATCCCTTATATTTNNNAGATCGGAAGA
120220 TATGAGTGTGGAGTCGTTGCCCCATATGCTATCCCTTATATTTNNNAGATCGGAAGA
120221 TATGAGTGTGGAGTCGTTGCTGTATAATAAAACTTCGAAGCTCNNNAGATCGGAAG

120222 TATGAGTGTGGAGTCGTTGCCCTAGTTGTCACCTTTGTATAATNNNAGATCGGAAG
120223 TATGAGTGTGGAGTCGTTGCCCTAGTTGTCACCTTTGTATAATNNNAGATCGGAAG
120224 TATGAGTGTGGAGTCGTTGCTTTATCACCATTTTACAAAGTGGNNNAGATCGGAAG
120225 TATGAGTGTGGAGTCGTTGCGATGCCATAAGAACAAAGATACNNNAGATCGGAAG
120226 TATGAGTGTGGAGTCGTTGCCTAATAGATTACCTTTATCGCCTNNNAGATCGGAAG
120227 TATGAGTGTGGAGTCGTTGCCTAATAGATTACCTTTATCGCCTNNNAGATCGGAAG
120228 TATGAGTGTGGAGTCGTTGCTTTCATAATTTATAAGTGTGGGGNNNAGATCGGAA
120229 TATGAGTGTGGAGTCGTTGCTTTCATAATTTATAAGTGTGGGGNNNAGATCGGAA
120230 TATGAGTGTGGAGTCGTTGCAAAAAGTCCTCCATAAATTATGTNNNAGATCGGAAG
120231 TATGAGTGTGGAGTCGTTGCGCAAAGTTTATTTTCTCTTTCTTNNNAGATCGGAAGA
120232 TATGAGTGTGGAGTCGTTGCCTCTGAATTCATCTTTTCTTCAANNNAGATCGGAAGA
120233 TATGAGTGTGGAGTCGTTGCGAATAAGCATAACTATTGTTCCANNNAGATCGGAAG
120234 TATGAGTGTGGAGTCGTTGCCATGAGTTCATTCAAGTAAAGTANNNAGATCGGAAG
120235 TATGAGTGTGGAGTCGTTGCCATGAGATGCTTTTGAAGATGNNNAGATCGGAAGA
120236 TATGAGTGTGGAGTCGTTGCTTGTACTAAAGACCTGACAACNNNAGATCGGAAGAG
120237 TATGAGTGTGGAGTCGTTGCTTGTACTAAAGACCTGACAACNNNAGATCGGAAGAG
120238 TATGAGTGTGGAGTCGTTGCTGGGAAGAATGGTTTATGTCNNNAGATCGGAAGAG
120239 TATGAGTGTGGAGTCGTTGCCAAAATGGAATTTGTTTTCAGAGNNNAGATCGGAA
120240 TATGAGTGTGGAGTCGTTGCCAAAATGGAATTTGTTTTCAGAGNNNAGATCGGAA
120243 TATGAGTGTGGAGTCGTTGCGAGAACTGTGTGATATCAGAAANNNAGATCGGAAG
120244 TATGAGTGTGGAGTCGTTGCTGTATGTTATTTAACAACAAGCANNNAGATCGGAAG
120245 TATGAGTGTGGAGTCGTTGCTGTATGTTATTTAACAACAAGCANNNAGATCGGAAG
120246 TATGAGTGTGGAGTCGTTGCTTGGAACTTGAATTCTGTTTTAANNNAGATCGGAAG
120247 TATGAGTGTGGAGTCGTTGCTGAATCACTAGCCTCTAAATTATNNNAGATCGGAAG
120248 TATGAGTGTGGAGTCGTTGCTCTTTAATAGAAAACCTTTGGACNNNAGATCGGAAG
120249 TATGAGTGTGGAGTCGTTGCTCTTTAATAGAAAACCTTTGGACNNNAGATCGGAAG
120250 TATGAGTGTGGAGTCGTTGCCTAGTTTGAGAGAAAGTCCTATGNNNAGATCGGAA
120251 TATGAGTGTGGAGTCGTTGCAGAACTCTGATGTTTGTAAAATCNNNAGATCGGAAG
120252 TATGAGTGTGGAGTCGTTGCAGAACTCTGATGTTTGTAAAATCNNNAGATCGGAAG
120253 TATGAGTGTGGAGTCGTTGCGATACAGAAGTAGAGGAAAGTAANNNAGATCGGA
120254 TATGAGTGTGGAGTCGTTGCGCTAGGATCTTTAGAGAAGTTTANNNAGATCGGAA
120255 TATGAGTGTGGAGTCGTTGCGCTAGGATCTTTAGAGAAGTTTANNNAGATCGGAA
120256 TATGAGTGTGGAGTCGTTGCGTTTCCAATGATAGTAGGAAAACNNNAGATCGGAA
120257 TATGAGTGTGGAGTCGTTGCCAAACAGATTGCATATTATCTGANNNAGATCGGAAG
120258 TATGAGTGTGGAGTCGTTGCCAAACAGATTGCATATTATCTGANNNAGATCGGAAG
120259 TATGAGTGTGGAGTCGTTGCAAAATTGCAGAACGAATATAAGTNNNAGATCGGAA
120260 TATGAGTGTGGAGTCGTTGCTAAGTAAAGAATCTGGTCAATTGNNNAGATCGGAA
120261 TATGAGTGTGGAGTCGTTGCAAGAAACATTTGGAAATAGAACANNNAGATCGGAA
120262 TATGAGTGTGGAGTCGTTGCAAGAAACATTTGGAAATAGAACANNNAGATCGGAA
120263 TATGAGTGTGGAGTCGTTGCTAAAAATTATGTTAGACCTGGTCNNNAGATCGGAAG
120264 TATGAGTGTGGAGTCGTTGCTGGAAGAAGAAATGTTTTTAGTTNNNAGATCGGAA
120265 TATGAGTGTGGAGTCGTTGCTGGAAGAAGAAATGTTTTTAGTTNNNAGATCGGAA
120266 TATGAGTGTGGAGTCGTTGCCAATAAAACTTTGCTGCCACNNNAGATCGGAAGAGC

120268 TATGAGTGTGGAGTCGTTGCCAACATCTTTTACATTCTGCAANNNAGATCGGAAGAG
120269 TATGAGTGTGGAGTCGTTGCCACTGCCTACTCTGAATAAATNNNAGATCGGAAGAG
120270 TATGAGTGTGGAGTCGTTGCGGTCAACATCTTTTACATTCTGNNNAGATCGGAAGA
120271 TATGAGTGTGGAGTCGTTGCTTTTCTTTTTCTTTGAGACTGGNNNAGATCGGAAGA
120274 TATGAGTGTGGAGTCGTTGCCTTATTTGACCTCCCTATAACCNNNAGATCGGAAGAG
120275 TATGAGTGTGGAGTCGTTGCATAGAGTGTAGATGCATCAAAATNNNAGATCGGAA
120276 TATGAGTGTGGAGTCGTTGCCTGAACTTGATGAGTCCTCTNNNAGATCGGAAGAGC
120278 TATGAGTGTGGAGTCGTTGCTAAAAGACGAGGGAGATCCTNNNAGATCGGAAGAG
120279 TATGAGTGTGGAGTCGTTGCCTTCCTGAAAACAACGTTCTNNNAGATCGGAAGAGC
120280 TATGAGTGTGGAGTCGTTGCACACACTAGGAAGATACCTCNNNAGATCGGAAGAG
120281 TATGAGTGTGGAGTCGTTGCCACACACTAGGAAGATACCTNNNAGATCGGAAGAG
120282 TATGAGTGTGGAGTCGTTGCATAGTCTGGGCCATACTTTTNNNAGATCGGAAGAGC
120283 TATGAGTGTGGAGTCGTTGCTGTTTCTAGATAGCACAGGANNNAGATCGGAAGAG
120284 TATGAGTGTGGAGTCGTTGCTACAAACGTATGGAATGCTTNNNAGATCGGAAGAG
120285 TATGAGTGTGGAGTCGTTGCTACAAACGTATGGAATGCTTNNNAGATCGGAAGAG
120286 TATGAGTGTGGAGTCGTTGCTCTATTTGTGTGGCAGTTATTNNNAGATCGGAAGAG
120287 TATGAGTGTGGAGTCGTTGCTCTATTTGTGTGGCAGTTATTNNNAGATCGGAAGAG
120288 TATGAGTGTGGAGTCGTTGCAATGTGGTCCTATTTAAGCCNNNAGATCGGAAGAGC
120291 TATGAGTGTGGAGTCGTTGCCTGATCATTTGGCTCTTTTTTTNNNAGATCGGAAGAG
120292 TATGAGTGTGGAGTCGTTGCCTGATCATTTGGCTCTTTTTTTNNNAGATCGGAAGAG
120293 TATGAGTGTGGAGTCGTTGCGTAGCGTATACATTTGTTTTGGNNNAGATCGGAAGA
120294 TATGAGTGTGGAGTCGTTGCGTAGCGTATACATTTGTTTTGGNNNAGATCGGAAGA
120295 TATGAGTGTGGAGTCGTTGCATGACATGTTTGGGTCTTTTATANNNAGATCGGAAG
120296 TATGAGTGTGGAGTCGTTGCATGACATGTTTGGGTCTTTTATANNNAGATCGGAAG
120297 TATGAGTGTGGAGTCGTTGCATACCTGTTTCTTTGGGACATNNNAGATCGGAAGAG
120298 TATGAGTGTGGAGTCGTTGCTCTGTGCAGATAAACAGAAATNNNAGATCGGAAGA
120299 TATGAGTGTGGAGTCGTTGCAGTAAATAGACAGCCTTTCTGNNNAGATCGGAAGAG
120300 TATGAGTGTGGAGTCGTTGCAGTAAATAGACAGCCTTTCTGNNNAGATCGGAAGAG
120301 TATGAGTGTGGAGTCGTTGCTTCACATAAAGGAACATGGTTNNNAGATCGGAAGAG
120302 TATGAGTGTGGAGTCGTTGCTACCAAATTTCAGAACACAAGAANNNAGATCGGAAG
120303 TATGAGTGTGGAGTCGTTGCTACCAAATTTCAGAACACAAGAANNNAGATCGGAAG
120305 TATGAGTGTGGAGTCGTTGCCAGTTCTTTCTTTTTCTCACTATNNNAGATCGGAAGA
120306 TATGAGTGTGGAGTCGTTGCCAGTTCTTTCTTTTTCTCACTATNNNAGATCGGAAGA
120307 TATGAGTGTGGAGTCGTTGCACTCATGTCCCTATTTATGATTTNNNAGATCGGAAGA
120308 TATGAGTGTGGAGTCGTTGCCTCAGGAAAGTTAAGGTATCAANNNAGATCGGAAG
120310 TATGAGTGTGGAGTCGTTGCTTCATTGACTTCAGGTAGATTAANNNAGATCGGAAG
120311 TATGAGTGTGGAGTCGTTGCTGTTTCAAAAAAAATCAAACTGTNNNAGATCGGAAG
120312 TATGAGTGTGGAGTCGTTGCTGTTTCAAAAAAAATCAAACTGTNNNAGATCGGAAG
120313 TATGAGTGTGGAGTCGTTGCCAGACTCTTACCAGCATATATAGNNNAGATCGGAAG
120314 TATGAGTGTGGAGTCGTTGCTCATGCTCTGATTAGATAATCTTNNNAGATCGGAAG
120315 TATGAGTGTGGAGTCGTTGCGTTAACTGAAGATATTCCAACTCNNNAGATCGGAAG
120316 TATGAGTGTGGAGTCGTTGCGTTAACTGAAGATATTCCAACTCNNNAGATCGGAAG
120317 TATGAGTGTGGAGTCGTTGCTGGAGAAAAATGAAGGAAAAAATNNNAGATCGGA

120318 TATGAGTGTGGAGTCGTTGCAAATGTCAGCAAATGTAAAAATTNNNAGATCGGAA
120319 TATGAGTGTGGAGTCGTTGCTTATTCCGTATTACATTTTTGCANNNAGATCGGAAGA
120320 TATGAGTGTGGAGTCGTTGCTTATTCCGTATTACATTTTTGCANNNAGATCGGAAGA
120321 TATGAGTGTGGAGTCGTTGCAAAACACATTCTTTGCAGCTNNNAGATCGGAAGAGC
120322 TATGAGTGTGGAGTCGTTGCGTATACTCAGGGACCATGTCNNNAGATCGGAAGAG
120323 TATGAGTGTGGAGTCGTTGCGGTCTGTCTTAACATTCTTATTGNNNAGATCGGAAG
120324 TATGAGTGTGGAGTCGTTGCGAAACCCTAATTCAATGTTTGAANNNAGATCGGAAG
120325 TATGAGTGTGGAGTCGTTGCTTGTATGTTAATTTCCTAACCAGNNNAGATCGGAAG
120326 TATGAGTGTGGAGTCGTTGCGACTGTTAGTAACTTAGTCTCCNNNAGATCGGAAGA
120327 TATGAGTGTGGAGTCGTTGCGACTGTTAGTAACTTAGTCTCCNNNAGATCGGAAGA
120328 TATGAGTGTGGAGTCGTTGCAAGATAGTCTCCACTTTTTAATGNNNAGATCGGAAG
120329 TATGAGTGTGGAGTCGTTGCAAGATAGTCTCCACTTTTTAATGNNNAGATCGGAAG
120330 TATGAGTGTGGAGTCGTTGCTCGCATAAATATTTTAGGAAGTGNNNAGATCGGAA
120331 TATGAGTGTGGAGTCGTTGCTCGCATAAATATTTTAGGAAGTGNNNAGATCGGAA
120332 TATGAGTGTGGAGTCGTTGCATGATAAAAGTGGCCTTGTTANNNAGATCGGAAGA
120333 TATGAGTGTGGAGTCGTTGCATGATAAAAGTGGCCTTGTTANNNAGATCGGAAGA
120334 TATGAGTGTGGAGTCGTTGCATTGTTGTATCACCAGTTAGTTNNNAGATCGGAAGA
120335 TATGAGTGTGGAGTCGTTGCATTGTTGTATCACCAGTTAGTTNNNAGATCGGAAGA
120336 TATGAGTGTGGAGTCGTTGCTATCAAATATTTAGCATGGATCCNNNAGATCGGAAG
120337 TATGAGTGTGGAGTCGTTGCTATCAAATATTTAGCATGGATCCNNNAGATCGGAAG
120338 TATGAGTGTGGAGTCGTTGCTAATATCTACTTTTGCTCAGCTTNNNAGATCGGAAGA
120339 TATGAGTGTGGAGTCGTTGCACAGTAAATTAATCCTGGTTTTCNNNAGATCGGAAG
120340 TATGAGTGTGGAGTCGTTGCTTGCCTTTCATGAATACCAGNNNAGATCGGAAGAGC
120341 TATGAGTGTGGAGTCGTTGCATTTTCTTAGCACCATTTCTTAGNNNAGATCGGAAGA
120342 TATGAGTGTGGAGTCGTTGCTCCACATTTTAAAAACTAAGCTTNNNAGATCGGAAG
120343 TATGAGTGTGGAGTCGTTGCTTAAGAAAATAGAGGCTTATCCANNNAGATCGGAA
120344 TATGAGTGTGGAGTCGTTGCCATCATTTCTGGAGTCTCAGNNNAGATCGGAAGAGC
120345 TATGAGTGTGGAGTCGTTGCCTCTTTGTTTCTCATCAACTGNNNAGATCGGAAGAG
120346 TATGAGTGTGGAGTCGTTGCTAACATGCGAATGTAGTTCCNNNAGATCGGAAGAG
120347 TATGAGTGTGGAGTCGTTGCAGCTGTAGTGTAATCCAATTATTNNNAGATCGGAAG
120348 TATGAGTGTGGAGTCGTTGCCATCATAAACTAGTTTTTCAGGTNNNAGATCGGAAG
120349 TATGAGTGTGGAGTCGTTGCTAATAAAATGCACATTGAAACCCNNNAGATCGGAAG
120350 TATGAGTGTGGAGTCGTTGCTCTTTTACTGAGAATCTTTTCTGNNNAGATCGGAAGA
120351 TATGAGTGTGGAGTCGTTGCATTCATTTCAACTTGGTACATTTNNNAGATCGGAAGA
120352 TATGAGTGTGGAGTCGTTGCCTTCCAAATTCAGATTCCAGTANNNAGATCGGAAGA
120353 TATGAGTGTGGAGTCGTTGCGATTCAAGTGCTGTTTCAACNNNAGATCGGAAGAG
120354 TATGAGTGTGGAGTCGTTGCAGTCATTATTTGTAATCTGGATGNNNAGATCGGAAG
120355 TATGAGTGTGGAGTCGTTGCCAGAAGAATCTTCCTCTTTTTCNNNAGATCGGAAGA
120356 TATGAGTGTGGAGTCGTTGCATTGAGACATGAAGAACAGACNNNAGATCGGAAGA
120357 TATGAGTGTGGAGTCGTTGCAAAGAATGTTTGTATACAAAGCTNNNAGATCGGAA
120358 TATGAGTGTGGAGTCGTTGCGGTCTAAATGATGAAAGTGAAATNNNAGATCGGAA
120359 TATGAGTGTGGAGTCGTTGCATGATATGTGCTTTCCTGTTTNNNAGATCGGAAGAG
120360 TATGAGTGTGGAGTCGTTGCCCACTATTTTCAATTTATCTGTCTNNNAGATCGGAAG

120361 TATGAGTGTGGAGTCGTTGCCTCAACAGATAGAACATTTTGAANNNAGATCGGAAG
120362 TATGAGTGTGGAGTCGTTGCAAAAACTTTCCCAGTGAAGTATNNNAGATCGGAAGA
120363 TATGAGTGTGGAGTCGTTGCTTCTTGCTTCTGTAAGAGATGNNNAGATCGGAAGAG
120364 TATGAGTGTGGAGTCGTTGCTTTGAGAGATTTCCTTTTTCAAANNNAGATCGGAAG
120365 TATGAGTGTGGAGTCGTTGCGACAGAATTATGTAGGCTTTATCNNNAGATCGGAAG
120366 TATGAGTGTGGAGTCGTTGCACATTTCAACTTTTACTTGCTTNNNAGATCGGAAGAG
120367 TATGAGTGTGGAGTCGTTGCCAATATAATCTCTGTGGCAATCNNNAGATCGGAAGA
120369 TATGAGTGTGGAGTCGTTGCCTTCCTATTGGTTCGTTTTTAGNNNAGATCGGAAGA
120370 TATGAGTGTGGAGTCGTTGCCTAGAATAAAGGCTGAATGTTTANNNAGATCGGAA
120371 TATGAGTGTGGAGTCGTTGCAAAAACTTTCCCAGTGAAGTATNNNAGATCGGAAGA
120372 TATGAGTGTGGAGTCGTTGCCACAGTAAAACATCTGTTAAAAGNNNAGATCGGAAG
120374 TATGAGTGTGGAGTCGTTGCCATGTATGGATTGGTGGATCNNNAGATCGGAAGAG
120375 TATGAGTGTGGAGTCGTTGCAACTTTTGTCTTTCTTATGAACCNNNAGATCGGAAGA
120376 TATGAGTGTGGAGTCGTTGCCATTATACCAAAAAGATACACGANNNAGATCGGAAG
120377 TATGAGTGTGGAGTCGTTGCAAATGGAGAAAAGGAAACTTTAGNNNAGATCGGAA
120378 TATGAGTGTGGAGTCGTTGCTCCTTATCCGTATACTTTGTATCNNNAGATCGGAAGA
120379 TATGAGTGTGGAGTCGTTGCTGATATAAAGAAAACAAGAAGCANNNAGATCGGAA
120380 TATGAGTGTGGAGTCGTTGCATAAGGAAGTAACTCAAAGGAATNNNAGATCGGAA
120381 TATGAGTGTGGAGTCGTTGCCTCATTTGACTATAGAGCTAAGTNNNAGATCGGAAG
120382 TATGAGTGTGGAGTCGTTGCCTCATTCTAGTTCTCTAACAGAGNNNAGATCGGAAG
120383 TATGAGTGTGGAGTCGTTGCATAATATAAACCATGTCAGCAACNNNAGATCGGAAG
120384 TATGAGTGTGGAGTCGTTGCTTACAGGAATTCACCAACATGNNNAGATCGGAAGAG
120385 TATGAGTGTGGAGTCGTTGCGTTGTGAAACCTTCCTTGAANNNAGATCGGAAGAGC
120386 TATGAGTGTGGAGTCGTTGCCAATTCTTTTGGATATGTACTCANNNAGATCGGAAG
120387 TATGAGTGTGGAGTCGTTGCTACTCATCTTTTCCCAGGTATNNNAGATCGGAAGAG
120388 TATGAGTGTGGAGTCGTTGCGGAAGAGACAGGATTTTCTGNNNAGATCGGAAGAG
120389 TATGAGTGTGGAGTCGTTGCCTCTAGCATCTGTAGTTGATGNNNAGATCGGAAGAG
120390 TATGAGTGTGGAGTCGTTGCATAAGTTTTAAACACCATAACGTNNNAGATCGGAAG
120391 TATGAGTGTGGAGTCGTTGCCTTTTTGTTTTTCAGATCCTTCTNNNAGATCGGAAGA
120392 TATGAGTGTGGAGTCGTTGCGAGACCTACATTACACCACTNNNAGATCGGAAGAGC
120393 TATGAGTGTGGAGTCGTTGCACGATTAAGACAAGGAGGATNNNAGATCGGAAGAG
120394 TATGAGTGTGGAGTCGTTGCAGACTTTCTACATATTAAAGAGGTNNNAGATCGGAA
120395 TATGAGTGTGGAGTCGTTGCAAGTTATAGTCATTGGGGTAAAANNNAGATCGGAA
120396 TATGAGTGTGGAGTCGTTGCCTACCATCCTTAGTTTGATAGAGNNNAGATCGGAAG
120397 TATGAGTGTGGAGTCGTTGCTCCCCTCTCCTCTAATTAAAANNNAGATCGGAAGAGC
120398 TATGAGTGTGGAGTCGTTGCGAAAAACTGTAAACTGAGTTCTTNNNAGATCGGAAG
120399 TATGAGTGTGGAGTCGTTGCTGGAAGAGGTATAGTTAAAGATCNNNAGATCGGAA
120400 TATGAGTGTGGAGTCGTTGCGTCAAATTTCTCAACTCTGTGNNNAGATCGGAAGAG
120402 TATGAGTGTGGAGTCGTTGCATTCTTTTGAAGTTAATTCTCCCNNNAGATCGGAAGA
120404 TATGAGTGTGGAGTCGTTGCCCACACTATCTCTTTATCCATAANNNAGATCGGAAGA
120405 TATGAGTGTGGAGTCGTTGCCTCCGTCTCAACAACAAAAANNNAGATCGGAAGAGC
120406 TATGAGTGTGGAGTCGTTGCTGTTTTCTAAGTTTTTTCTCCTTNNNAGATCGGAAGA
120407 TATGAGTGTGGAGTCGTTGCCAAACTATGGCTTACAACATCNNNAGATCGGAAGAG

120408 TATGAGTGTGGAGTCGTTGCCAGAAAGCAATAAGCACCAGNNNAGATCGGAAGAG
120409 TATGAGTGTGGAGTCGTTGCCTGAAGGTTGCCAAAGAAATNNNAGATCGGAAGAG
120410 TATGAGTGTGGAGTCGTTGCGTTTTTTATGTAAGCATCTCCTNNNAGATCGGAAG
120412 TATGAGTGTGGAGTCGTTGCCTACCATATTGGCTTCCTGTNNNAGATCGGAAGAGC
120413 TATGAGTGTGGAGTCGTTGCAAGGAGTTGATAAGGATTTAAGTNNNAGATCGGAA
120414 TATGAGTGTGGAGTCGTTGCAAACTACAATCATAAGTCTCTGANNNAGATCGGAAG
120415 TATGAGTGTGGAGTCGTTGCATTATTCATTGATGGTTCTTACCNNNAGATCGGAAG
120416 TATGAGTGTGGAGTCGTTGCCTGTAGAATTCCCCTCTGTANNNAGATCGGAAGAGC
120419 TATGAGTGTGGAGTCGTTGCTGTTGCTATCCTTTCTTTTATATCNNNAGATCGGAAG
120420 TATGAGTGTGGAGTCGTTGCTTGAGCAAGAAAGAGATGATTNNNAGATCGGAAGA
120421 TATGAGTGTGGAGTCGTTGCTTTCCCACATTGCGTAAATATTNNNAGATCGGAAGA
120422 TATGAGTGTGGAGTCGTTGCATACCTCAATTCCCTTGGTTTNNNAGATCGGAAGAG
120423 TATGAGTGTGGAGTCGTTGCATAGGATTTAGTGTCTTGCTTTANNNAGATCGGAAG
120424 TATGAGTGTGGAGTCGTTGCATTCTTTCCCCACAGTAATAATANNNAGATCGGAAG
120426 TATGAGTGTGGAGTCGTTGCCATGGCCATAAAAATGAAGTAATNNNAGATCGGAA
120427 TATGAGTGTGGAGTCGTTGCCATGGCCATAAAAATGAAGTAATNNNAGATCGGAA
120428 TATGAGTGTGGAGTCGTTGCGAGTTCTTTTGAGAAGACAAAGNNNAGATCGGAAG
120429 TATGAGTGTGGAGTCGTTGCGAGTTCTTTTGAGAAGACAAAGNNNAGATCGGAAG
120430 TATGAGTGTGGAGTCGTTGCTATTATTCCCACTGCCTGATTNNNAGATCGGAAGAG
120431 TATGAGTGTGGAGTCGTTGCACTTTAACCTATGCAATATCTCTNNNAGATCGGAAGA
120432 TATGAGTGTGGAGTCGTTGCGTGAAGTGAATCAATTTGTAATGNNNAGATCGGAA
120433 TATGAGTGTGGAGTCGTTGCAGAAATCTGTCTTGTTATCAGTANNNAGATCGGAAG
120434 TATGAGTGTGGAGTCGTTGCGTGTACAAATGCAGCAATAGNNNAGATCGGAAGAG
120436 TATGAGTGTGGAGTCGTTGCATTAATTTATACTTAGTGGCAGGNNNAGATCGGAAG
120437 TATGAGTGTGGAGTCGTTGCTGTCTCTCTCACATCTCTTTNNNAGATCGGAAGAGCA
120438 TATGAGTGTGGAGTCGTTGCTTTAATGTCCTTTTTACAGTTCCNNNAGATCGGAAGA
120439 TATGAGTGTGGAGTCGTTGCACATTGCAAGATAACTTAAAACTNNNAGATCGGAAG
120440 TATGAGTGTGGAGTCGTTGCAGTACCAAAGTAACAGAGGTAANNNAGATCGGAAG
120441 TATGAGTGTGGAGTCGTTGCGGATTGGTTTATTTGATTATGGANNNAGATCGGAA
120442 TATGAGTGTGGAGTCGTTGCGACAATCTGATCAAACAACATAANNNAGATCGGAAG
120443 TATGAGTGTGGAGTCGTTGCGGCAAGAATTAAAAATCTTTCAGNNNAGATCGGAA
120444 TATGAGTGTGGAGTCGTTGCATACACACATATATGGTCTTGTTNNNAGATCGGAAG
120445 TATGAGTGTGGAGTCGTTGCAATATTACCACAAATTTAGCAGCNNNAGATCGGAAG
120446 TATGAGTGTGGAGTCGTTGCATAGATTCAACCATCCTCAAATANNNAGATCGGAAG
120447 TATGAGTGTGGAGTCGTTGCCATCTGTTTTTCACTTGATCTTANNNAGATCGGAAGA
120448 TATGAGTGTGGAGTCGTTGCCATGTGTATGTAAGTCATAAAGGNNNAGATCGGAA
120449 TATGAGTGTGGAGTCGTTGCTGATTTTTCTTCCCTACAGAAANNNAGATCGGAAGA
120450 TATGAGTGTGGAGTCGTTGCTACTGGTCTGTTAGCAATAAATTNNNAGATCGGAAG
120451 TATGAGTGTGGAGTCGTTGCATGTAAACTAAGAAATGACTGTGNNNAGATCGGAA
120452 TATGAGTGTGGAGTCGTTGCACATACCAAGAGGAGATTTATACNNNAGATCGGAA
120453 TATGAGTGTGGAGTCGTTGCGCTTCCTAATATATGCCTGATCNNNAGATCGGAAGA
120454 TATGAGTGTGGAGTCGTTGCTTGGGATTTTATGTAAGTTCCTNNNAGATCGGAAGA
120455 TATGAGTGTGGAGTCGTTGCAGACATGAGGACTGATTATTTATNNNAGATCGGAA

120456 TATGAGTGTGGAGTCGTTGCTTCATTAATCTTCACAACAACTTNNNAGATCGGAAGA
120457 TATGAGTGTGGAGTCGTTGCAAGAATGAAATAATATCTGTGGCNNNAGATCGGAA
120458 TATGAGTGTGGAGTCGTTGCAAAACAAGCAAAACACTGAAAANNNAGATCGGAAG
120459 TATGAGTGTGGAGTCGTTGCCATAACCGATATGATGAAGCTANNNAGATCGGAAG
120460 TATGAGTGTGGAGTCGTTGCGTACACACAATATATTATCTGCCNNNAGATCGGAAG
120461 TATGAGTGTGGAGTCGTTGCGGTTACTTAGATTTTCAATGGATNNNAGATCGGAAG
120462 TATGAGTGTGGAGTCGTTGCGTACTGTGCTAAACAATTTGATNNNAGATCGGAAGA
120463 TATGAGTGTGGAGTCGTTGCAGAAAATAGAACAGATGAAGCTANNNAGATCGGAA
120464 TATGAGTGTGGAGTCGTTGCGAACTACAAAATGCTCAAGGNNNAGATCGGAAGAG
120466 TATGAGTGTGGAGTCGTTGCGGACTAGATGATTTAGAGCAACNNNAGATCGGAAG
120468 TATGAGTGTGGAGTCGTTGCCATTCCCTCACACTCAATTAGNNNAGATCGGAAGAG
120469 TATGAGTGTGGAGTCGTTGCATAAAACCAAAATGTGTGACACNNNAGATCGGAAG
120470 TATGAGTGTGGAGTCGTTGCCAGACCTTGCTGAATAGGATNNNAGATCGGAAGAG
120471 TATGAGTGTGGAGTCGTTGCGGTGGAGTTATAATGAATCATTCNNNAGATCGGAA
120472 TATGAGTGTGGAGTCGTTGCTTTATTCTGTCCTCTATAACACCNNNAGATCGGAAGA
120473 TATGAGTGTGGAGTCGTTGCCTTCTATAGCCCTAAGAACTTTTNNNAGATCGGAAG
120474 TATGAGTGTGGAGTCGTTGCACATTTATTGAACACCTACTCTANNNAGATCGGAAG
120475 TATGAGTGTGGAGTCGTTGCCTACATTATGGTAGGAGAATGGNNNAGATCGGAAG
120476 TATGAGTGTGGAGTCGTTGCATTTATTTTTTTGGAATGTCACGNNNAGATCGGAAG
120477 TATGAGTGTGGAGTCGTTGCGACAGCAATTTTCGTAAACTTNNNAGATCGGAAGAG
120478 TATGAGTGTGGAGTCGTTGCAAAAGCACAAATCAACAACTGNNNAGATCGGAAGA
120479 TATGAGTGTGGAGTCGTTGCCTGAACAAATGAAATGAACAAAGNNNAGATCGGAA
120480 TATGAGTGTGGAGTCGTTGCTTGTGGAAATGAAAACTTTTAGANNNAGATCGGAA
120481 TATGAGTGTGGAGTCGTTGCGGATGACTATGACCATTAAAAAANNNAGATCGGAA
120482 TATGAGTGTGGAGTCGTTGCCTGTTTCTCAGGCTAATACAANNNAGATCGGAAGAG
120483 TATGAGTGTGGAGTCGTTGCTCAAAAAAGTACAGGAAAAAAGTNNNAGATCGGAA
120484 TATGAGTGTGGAGTCGTTGCGGGTGGAAAAAAAAGAAAAAAAANNNAGATCGGA
120485 TATGAGTGTGGAGTCGTTGCTAGCTGGATTTCACATATGATATNNNAGATCGGAAG
120486 TATGAGTGTGGAGTCGTTGCTTATTTTCTTTGGGCATTGAAANNNAGATCGGAAGA
120487 TATGAGTGTGGAGTCGTTGCGATTCTCTTTGTCTCATTCCTNNNAGATCGGAAGAGC
120488 TATGAGTGTGGAGTCGTTGCAGTAGAAAAAGGATATGAATGGANNNAGATCGGA
120489 TATGAGTGTGGAGTCGTTGCTATGCAAGAGACTTTCTTTAGAGNNNAGATCGGAAG
120490 TATGAGTGTGGAGTCGTTGCCATTCCAATACCCACTGTCTNNNAGATCGGAAGAGC
120491 TATGAGTGTGGAGTCGTTGCATGAAGGCACTAGATATTTCTTANNNAGATCGGAAG
120493 TATGAGTGTGGAGTCGTTGCGATAAACAGCATCTGACTGATNNNAGATCGGAAGA
120494 TATGAGTGTGGAGTCGTTGCTATGAATATGGTTGGTTTAGTGANNNAGATCGGAA
120495 TATGAGTGTGGAGTCGTTGCAATTGATTGATTATAGGGATTGGNNNAGATCGGAA
120496 TATGAGTGTGGAGTCGTTGCAATTGATTGATTATAGGGATTGGNNNAGATCGGAA
120497 TATGAGTGTGGAGTCGTTGCTTAGACTTAGAAGCTAGGATAGANNNAGATCGGAA
120498 TATGAGTGTGGAGTCGTTGCCATCAAGATTGCTAATTGTATCTNNNAGATCGGAAG
120499 TATGAGTGTGGAGTCGTTGCTGTATGCATGATCTAAATGTAGANNNAGATCGGAAG
120500 TATGAGTGTGGAGTCGTTGCCTATTTTTTGCAGCAAGTTGTANNNAGATCGGAAGA
120501 TATGAGTGTGGAGTCGTTGCCTATTTTTTGCAGCAAGTTGTANNNAGATCGGAAGA

120502 TATGAGTGTGGAGTCGTTGCGATTGTTTTAGGACATTTCAAAGNNNAGATCGGAAG
120503 TATGAGTGTGGAGTCGTTGCGATGAAAAAGAGAAGGCTTCTNNNAGATCGGAAGA
120504 TATGAGTGTGGAGTCGTTGCGATGAAAAAGAGAAGGCTTCTNNNAGATCGGAAGA
120505 TATGAGTGTGGAGTCGTTGCCCTTGGTGATCTTCTGAAAGNNNAGATCGGAAGAG
120508 TATGAGTGTGGAGTCGTTGCAGATAAATAAGCACATACACAGANNNAGATCGGAA
120509 TATGAGTGTGGAGTCGTTGCAGAATTATAGAGAGGGCTGGNNNAGATCGGAAGA
120510 TATGAGTGTGGAGTCGTTGCTTCACTGTCCTTGAATTTCTTNNNAGATCGGAAGAG
120511 TATGAGTGTGGAGTCGTTGCTATATGTGGGAAACTAGTTTTTGNNNAGATCGGAA
120512 TATGAGTGTGGAGTCGTTGCCCAGAAGAAACCAAATCTGGNNNAGATCGGAAGAG
120513 TATGAGTGTGGAGTCGTTGCCAACAAACACCTTAGGAATATGNNNAGATCGGAAG
120514 TATGAGTGTGGAGTCGTTGCGTGTGTGAGAGAGTAGAAGTNNNAGATCGGAAGA
120515 TATGAGTGTGGAGTCGTTGCTAGGAAATCTCAGGAACTGTTNNNAGATCGGAAGA
120516 TATGAGTGTGGAGTCGTTGCATGCAGGATTTTTTTTTTAAGGNNNAGATCGGAAG
120517 TATGAGTGTGGAGTCGTTGCTTGAGAGAGAGAGAGAGAGGNNNAGATCGGAAG
120518 TATGAGTGTGGAGTCGTTGCTCTTGTTCTTTATATGGATTCCTNNNAGATCGGAAGA
120519 TATGAGTGTGGAGTCGTTGCGAAGAGGAAGTCCTGATATTTAANNNAGATCGGAA
120520 TATGAGTGTGGAGTCGTTGCCTTGTGCAAAATGAAAATGCNNNAGATCGGAAGAG
120521 TATGAGTGTGGAGTCGTTGCTGAGCAGGATTATCATAAATCTTNNNAGATCGGAAG
120522 TATGAGTGTGGAGTCGTTGCAGTGAATGATAGCTATGGATATCNNNAGATCGGAA
120523 TATGAGTGTGGAGTCGTTGCCTGAACTGCTAGATAAAGACTTNNNAGATCGGAAG
120524 TATGAGTGTGGAGTCGTTGCATAATTACAAACTGTTAGAAGCCNNNAGATCGGAAG
120525 TATGAGTGTGGAGTCGTTGCCTAAATCTGAGGCCTATTCTTCNNNAGATCGGAAGA
120526 TATGAGTGTGGAGTCGTTGCCTCCCAATAAAAGTGACCTTTNNNAGATCGGAAGAG
120527 TATGAGTGTGGAGTCGTTGCCAAAAAAAAAACCCAAAAAGAAANNNAGATCGGAA
120528 TATGAGTGTGGAGTCGTTGCACAAGTTCTTGCTATGTTGCNNNAGATCGGAAGAGC
120529 TATGAGTGTGGAGTCGTTGCCTATCACAATATATCCATTTCCCNNNAGATCGGAAGA
120530 TATGAGTGTGGAGTCGTTGCGATCTCCCATGTTTTTACCCNNNAGATCGGAAGAGC
120531 TATGAGTGTGGAGTCGTTGCGGATAAAGGAAAATACTGCATTTNNNAGATCGGAA
120532 TATGAGTGTGGAGTCGTTGCTTTTCCTTTCTAAATTCGATCTGNNNAGATCGGAAGA
120533 TATGAGTGTGGAGTCGTTGCTAAATATTTTGGCTCTTCAAGTCNNNAGATCGGAAG
120534 TATGAGTGTGGAGTCGTTGCCTGGTCTACTCAAATCACACNNNAGATCGGAAGAGC
120535 TATGAGTGTGGAGTCGTTGCCTGGTGATTTTATAAAACGGTATNNNAGATCGGAAG
120536 TATGAGTGTGGAGTCGTTGCGCACATTTAGCTCTTCAAATATANNNAGATCGGAAG
120537 TATGAGTGTGGAGTCGTTGCGCACATTTAGCTCTTCAAATATANNNAGATCGGAAG
120538 TATGAGTGTGGAGTCGTTGCATCAAAGAGTTGAGGAGAAACNNNAGATCGGAAG
120539 TATGAGTGTGGAGTCGTTGCAAGAAATGCAGTAACTATAGGAANNNAGATCGGAA
120540 TATGAGTGTGGAGTCGTTGCCAAATCAAAGATGGGTTTAGTTTNNNAGATCGGAA
120541 TATGAGTGTGGAGTCGTTGCAAGAATGTCTAATTATGGCATTCNNNAGATCGGAAG
120542 TATGAGTGTGGAGTCGTTGCTTTCTTCAGTACTAAACAAGTTCNNNAGATCGGAAG
120543 TATGAGTGTGGAGTCGTTGCCTTCAGTGCAAAAAAAGTAATTGNNNAGATCGGAA
120544 TATGAGTGTGGAGTCGTTGCTAAGAGTGTCTTCACCATCATTNNNAGATCGGAAGA
120545 TATGAGTGTGGAGTCGTTGCTTGGACATAAATTTATGCAGAAANNNAGATCGGAA
120546 TATGAGTGTGGAGTCGTTGCCACTTTGAGACCATGTAAATATCNNNAGATCGGAAG

120547 TATGAGTGTGGAGTCGTTGCTGAGATTTAGCGAATTTAAAAGGNNNAGATCGGAA
120548 TATGAGTGTGGAGTCGTTGCTTTTTCTCTCCTGTCATACTTTNNNAGATCGGAAGAG
120549 TATGAGTGTGGAGTCGTTGCTTTGAAACAGGATCTCACTCNNNAGATCGGAAGAGC
120550 TATGAGTGTGGAGTCGTTGCCTGCCTTGATTCTGTTTATTTTTNNNAGATCGGAAGA
120551 TATGAGTGTGGAGTCGTTGCTAGGAGAAATATCCACAATTCAGNNNAGATCGGAAG
120552 TATGAGTGTGGAGTCGTTGCTGGAGAATGATGAGTGATAATTANNNAGATCGGAA
120553 TATGAGTGTGGAGTCGTTGCCAAAAAATGTAAAACAAGTGCTANNNAGATCGGAAG
120554 TATGAGTGTGGAGTCGTTGCTTAACTCTAACATTTCCTACCAANNNAGATCGGAAGA
120555 TATGAGTGTGGAGTCGTTGCTTAACTCTAACATTTCCTACCAANNNAGATCGGAAGA
120556 TATGAGTGTGGAGTCGTTGCCAGAAAAGGAAATATTCTCGAAANNNAGATCGGAA
120557 TATGAGTGTGGAGTCGTTGCGACTGAGACAGAAACAAAGANNNAGATCGGAAGAG
120558 TATGAGTGTGGAGTCGTTGCTCAGAGAGTTGTCTAATTAACTTNNNAGATCGGAAG
120559 TATGAGTGTGGAGTCGTTGCATTCAGAAATCTACAGGATGATTNNNAGATCGGAAG
120560 TATGAGTGTGGAGTCGTTGCAATTCTCCTATGAACAGAAAGATNNNAGATCGGAAG
120564 TATGAGTGTGGAGTCGTTGCAATGGTTGATGTTTATTAAGCTTNNNAGATCGGAAG
120565 TATGAGTGTGGAGTCGTTGCCATATGCACCTCACATATACTTNNNAGATCGGAAGAG
120566 TATGAGTGTGGAGTCGTTGCAAGAGAAAAGCAGTCCAATATATNNNAGATCGGAA
120567 TATGAGTGTGGAGTCGTTGCATTGGAGTGATATGTCTACAAACNNNAGATCGGAAG
120568 TATGAGTGTGGAGTCGTTGCTGTGAGCGTGTATAATAGTATATANNNAGATCGGA
120569 TATGAGTGTGGAGTCGTTGCCACCTCAAGATGATAGAGAAGNNNAGATCGGAAGA
120570 TATGAGTGTGGAGTCGTTGCGGTAAGTTTTTCAGGTCATTATTNNNAGATCGGAAG
120571 TATGAGTGTGGAGTCGTTGCCTAACAGGTGTAAGTATGGTACNNNAGATCGGAAG
120572 TATGAGTGTGGAGTCGTTGCAAAAGTTGCTAGAAACGTTTTTANNNAGATCGGAAG
120573 TATGAGTGTGGAGTCGTTGCTTTTTCTAAAGATTTCCAAGCATNNNAGATCGGAAG
120574 TATGAGTGTGGAGTCGTTGCATTAGACCATTTCCTTTGAGATNNNAGATCGGAAGA
120575 TATGAGTGTGGAGTCGTTGCTTTTCTTTGCCTTTGAAAAATTGNNNAGATCGGAAG
120576 TATGAGTGTGGAGTCGTTGCTAAAAATTACTATCCTAGGCCTGNNNAGATCGGAAG
120577 TATGAGTGTGGAGTCGTTGCTTTCAGAGTAAGGTGCTTATATTNNNAGATCGGAAG
120578 TATGAGTGTGGAGTCGTTGCATTTTGCAGTGTATACATATGTCNNNAGATCGGAAG
120579 TATGAGTGTGGAGTCGTTGCGGATGTATGTATTTAACACCATTNNNAGATCGGAAG
120580 TATGAGTGTGGAGTCGTTGCGAGATAACTCCTAGAAAGAAGAANNNAGATCGGAA
120581 TATGAGTGTGGAGTCGTTGCAGTGGCTCTCTCATAAATATTTTNNNAGATCGGAAG
120582 TATGAGTGTGGAGTCGTTGCGTGGAAGAAATGTTACATCTATTNNNAGATCGGAAG
120583 TATGAGTGTGGAGTCGTTGCGGCTTCTAAACAACAGAAATTTANNNAGATCGGAAG
120586 TATGAGTGTGGAGTCGTTGCATTATCTGAGATTGTCTTTCAGTNNNAGATCGGAAG
120587 TATGAGTGTGGAGTCGTTGCTATATAGAATTTTGCCAGAAGGTNNNAGATCGGAAG
120588 TATGAGTGTGGAGTCGTTGCCTCAAAGACAGTTTCAAGATAAGNNNAGATCGGAAG
120589 TATGAGTGTGGAGTCGTTGCAAATCTAGTACTCTTTTGAGCTTNNNAGATCGGAAG
120592 TATGAGTGTGGAGTCGTTGCGGTGATACTATACAACAGTTTTCNNNAGATCGGAAG
120593 TATGAGTGTGGAGTCGTTGCATTTATTGAACCGTTACTGTATCNNNAGATCGGAAG
120594 TATGAGTGTGGAGTCGTTGCATAATAATACCCAAAGAGAAGCTNNNAGATCGGAAG
120595 TATGAGTGTGGAGTCGTTGCATAAAGAAGAAAGAATTAGGTGGNNNAGATCGGA
120596 TATGAGTGTGGAGTCGTTGCGTAGGGTGCCTATATACTTAAAANNNAGATCGGAAG

| | |
|---|---|
| 120597 | TATGAGTGTGGAGTCGTTGCATGAAGGGACATAAATTCCATATNNNAGATCGGAAG |
| 120598 | TATGAGTGTGGAGTCGTTGCAACTTCAATTCCTTGAAATTTGANNNAGATCGGAAG |
| 120599 | TATGAGTGTGGAGTCGTTGCATGTGATATCTCTGGAAAGATACNNNAGATCGGAAG |
| 120600 | TATGAGTGTGGAGTCGTTGCGCCTGTAATCCCAATACTTTNNNAGATCGGAAGAGC |
| 120601 | TATGAGTGTGGAGTCGTTGCTATGGTATGGGAGAATATGTTTTNNNAGATCGGAA |
| 120602 | TATGAGTGTGGAGTCGTTGCAGTAATCAGGTTTTGAGGTTAGNNNAGATCGGAAG |
| 120603 | TATGAGTGTGGAGTCGTTGCGTAGGCATATACATACAGGGTNNNAGATCGGAAGA |
| 120604 | TATGAGTGTGGAGTCGTTGCTTTACATCTATCTACCTAACAGCNNNAGATCGGAAGA |
| 120605 | TATGAGTGTGGAGTCGTTGCATAGATAATTCTCTGCCTTTCTCNNNAGATCGGAAGA |
| 120606 | TATGAGTGTGGAGTCGTTGCGTACATCAATCCATAGCTATGNNNAGATCGGAAGAG |
| 120607 | TATGAGTGTGGAGTCGTTGCCATATAAGTGAAAATGCTGATCANNNAGATCGGAAG |
| 120608 | TATGAGTGTGGAGTCGTTGCAGTTGTATAGCATTAGGTGAAANNNAGATCGGAAG |
| 120609 | TATGAGTGTGGAGTCGTTGCGTAGGGCTAGCTGAATTTTCNNNAGATCGGAAGAG |
| 120610 | TATGAGTGTGGAGTCGTTGCGTAGGGCTAGCTGAATTTTCNNNAGATCGGAAGAG |
| 120611 | TATGAGTGTGGAGTCGTTGCTCCATCTATTGTATTCTTTATCAGNNNAGATCGGAAG |
| 120612 | TATGAGTGTGGAGTCGTTGCATACACACACACCTTCAATTATNNNAGATCGGAAGAG |
| 120613 | TATGAGTGTGGAGTCGTTGCTTTAGCTGTTAAATTCCATATGCNNNAGATCGGAAG |
| 120614 | TATGAGTGTGGAGTCGTTGCCATTTTAAACTGTTGAGTTCTCTNNNAGATCGGAAGA |
| 120615 | TATGAGTGTGGAGTCGTTGCCATTTTAAACTGTTGAGTTCTCTNNNAGATCGGAAGA |
| 120616 | TATGAGTGTGGAGTCGTTGCGTGTCTTCAAATATTGCTTCTAANNNAGATCGGAAG |
| 120617 | TATGAGTGTGGAGTCGTTGCGTGGAAAAAAGAAGGTTATTGTANNNAGATCGGAA |
| 120618 | TATGAGTGTGGAGTCGTTGCAGAGACAGACAGAGAAAGAGNNNAGATCGGAAGA |
| 120619 | TATGAGTGTGGAGTCGTTGCAGAGACAGACAGAGAAAGAGNNNAGATCGGAAGA |
| 120620 | TATGAGTGTGGAGTCGTTGCAAGAATCTTTCCTTGCCTTTNNNAGATCGGAAGAGC |
| 120621 | TATGAGTGTGGAGTCGTTGCTACCCTGACTTTTTATACCTTCTNNNAGATCGGAAGA |
| 120622 | TATGAGTGTGGAGTCGTTGCAAATAACTGTTGAAAATGAGGAGNNNAGATCGGAA |
| 120623 | TATGAGTGTGGAGTCGTTGCGTGTATCACATCCAAAATCCANNNAGATCGGAAGAG |
| 120624 | TATGAGTGTGGAGTCGTTGCTTAAATTGCAGGTTTAAATCCCNNNAGATCGGAAGA |
| 120625 | TATGAGTGTGGAGTCGTTGCATCTAGATGGGTGGAGTAAAGNNNAGATCGGAAGA |
| 120626 | TATGAGTGTGGAGTCGTTGCTTCTCTCTCTCTCTCTCTCTNNNAGATCGGAAGAGCAC |
| 120627 | TATGAGTGTGGAGTCGTTGCTACAAATTGAAAGGAAGATAAACANNNAGATCGGA |
| 120628 | TATGAGTGTGGAGTCGTTGCGCTGCCAATAATAAATTGAGTTANNNAGATCGGAAG |
| 120629 | TATGAGTGTGGAGTCGTTGCAGTATAAAAAAACAATTTGCAGCNNNAGATCGGAAG |
| 120630 | TATGAGTGTGGAGTCGTTGCTCAGACATATTGATGTATACTCANNNAGATCGGAAG |
| 120631 | TATGAGTGTGGAGTCGTTGCCTACTTTTTGCACTGCAGAGNNNAGATCGGAAGAGC |
| 120632 | TATGAGTGTGGAGTCGTTGCCTTCACTGTTTAAGGAGACTTANNNAGATCGGAAGA |
| 120633 | TATGAGTGTGGAGTCGTTGCTTAATTGGCCTCAGATTTTCTNNNAGATCGGAAGAG |
| 120634 | TATGAGTGTGGAGTCGTTGCCAAATCTTTTCTTCTTACCATGANNNAGATCGGAAGA |
| 120635 | TATGAGTGTGGAGTCGTTGCACATGTCTTAGTCACCACTCNNNAGATCGGAAGAGC |
| 120636 | TATGAGTGTGGAGTCGTTGCACATGTCTTAGTCACCACTCNNNAGATCGGAAGAGC |
| 120638 | TATGAGTGTGGAGTCGTTGCCTGGTTTTTGTTATGGTTGTTNNNAGATCGGAAGAG |
| 120639 | TATGAGTGTGGAGTCGTTGCCTCTCTTCTCATTTCACTTCTCNNNAGATCGGAAGAG |
| 120640 | TATGAGTGTGGAGTCGTTGCATTCTAGTTACTGACATGAGTTTNNNAGATCGGAAG |

120641 TATGAGTGTGGAGTCGTTGCATCCTTAAAGACTAACTACCTTGNNNAGATCGGAAG
120642 TATGAGTGTGGAGTCGTTGCAATATCAGAATTCTCTCTACCACNNNAGATCGGAAGA
120643 TATGAGTGTGGAGTCGTTGCAACAATTGCTAAGATTTATTGGGNNNAGATCGGAA
120644 TATGAGTGTGGAGTCGTTGCGTCTGTTCTTGCTCATTAAACNNNAGATCGGAAGAG
120645 TATGAGTGTGGAGTCGTTGCGTCTGTTCTTGCTCATTAAACNNNAGATCGGAAGAG
120647 TATGAGTGTGGAGTCGTTGCGTCTTCAACTTCTAGCAAAATTNNNAGATCGGAAGA
120648 TATGAGTGTGGAGTCGTTGCTCTAAAATATACGCCTAATAGTCANNNAGATCGGAA
120649 TATGAGTGTGGAGTCGTTGCGACAGGTTGTTTCTGATTAAATANNNAGATCGGAAG
120650 TATGAGTGTGGAGTCGTTGCTTCTAAAAGTGACCCTTATGAAANNNAGATCGGAAG
120651 TATGAGTGTGGAGTCGTTGCTTTCTCCTCAATCCAACTTTTATNNNAGATCGGAAGA
120652 TATGAGTGTGGAGTCGTTGCTCTCCTATGGCTCAAAATAACNNNAGATCGGAAGAG
120653 TATGAGTGTGGAGTCGTTGCGTACACACAATATATTATCTGCCNNNAGATCGGAAG
120654 TATGAGTGTGGAGTCGTTGCCAGAACTGTGAGAAAATAAATGTNNNAGATCGGAA
120655 TATGAGTGTGGAGTCGTTGCGAGAAGGCAGCCATCTATAANNNAGATCGGAAGAG
120656 TATGAGTGTGGAGTCGTTGCATGTAAGACTGTGCATTTATAACNNNAGATCGGAAG
120657 TATGAGTGTGGAGTCGTTGCATGTAAGACTGTGCATTTATAACNNNAGATCGGAAG
120658 TATGAGTGTGGAGTCGTTGCCCCCTGTTTAAGAAGGTTAGNNNAGATCGGAAGAG
120659 TATGAGTGTGGAGTCGTTGCTTTCATTTCCTCATCTCTAACATNNNAGATCGGAAGA
120660 TATGAGTGTGGAGTCGTTGCGAGGAAAGGAGAGGTATCAATNNNAGATCGGAAG
120661 TATGAGTGTGGAGTCGTTGCTCATCTGTTTCATAGTCTATTGANNNAGATCGGAAG
120662 TATGAGTGTGGAGTCGTTGCTCATCTGTTTCATAGTCTATTGANNNAGATCGGAAG
120663 TATGAGTGTGGAGTCGTTGCCATCACATCATCCTCTCTAAATNNNAGATCGGAAGAG
120664 TATGAGTGTGGAGTCGTTGCTGTTAAAACACTCATTTTACAACNNNAGATCGGAAG
120665 TATGAGTGTGGAGTCGTTGCGAGAGAATAAGAGCAATTTCTTCNNNAGATCGGAA
120666 TATGAGTGTGGAGTCGTTGCAATTAAGCATACCAAAAGATTGTNNNAGATCGGAAG
120667 TATGAGTGTGGAGTCGTTGCAATACAGTCATATAGTAGCATGGNNNAGATCGGAA
120668 TATGAGTGTGGAGTCGTTGCAATACAGTCATATAGTAGCATGGNNNAGATCGGAA
120669 TATGAGTGTGGAGTCGTTGCAATACAGTCATATAGTAGCATGGNNNAGATCGGAA
120670 TATGAGTGTGGAGTCGTTGCTTGGTAACAGCTCTATATATCAGNNNAGATCGGAAG
120671 TATGAGTGTGGAGTCGTTGCCAATCTCAACAATATTCCACTCNNNAGATCGGAAGAG
120672 TATGAGTGTGGAGTCGTTGCTTAGTTTTGCAATCAGTGATAAANNNAGATCGGAAG
120673 TATGAGTGTGGAGTCGTTGCTGGGAAGAGAATTTTTATTTTGTNNNAGATCGGAA
120674 TATGAGTGTGGAGTCGTTGCACCCTAATCTTGGACTTCTAGNNNAGATCGGAAGAG
120675 TATGAGTGTGGAGTCGTTGCTTGTTTGATTAACTTACTTAGCANNNAGATCGGAAG
120676 TATGAGTGTGGAGTCGTTGCATTCTGTCAAATCTCTTATCAGANNNAGATCGGAAG
120677 TATGAGTGTGGAGTCGTTGCATGCAGGAATAAGAAACTAGTATNNNAGATCGGAA
120678 TATGAGTGTGGAGTCGTTGCCAGATAACTGGGTAGTCTAGTTNNNAGATCGGAAG
120679 TATGAGTGTGGAGTCGTTGCGAAGAAACTACTTCCTGTCTAGNNNAGATCGGAAGA
120680 TATGAGTGTGGAGTCGTTGCAAGAGATAGAGCAATCACATATANNNAGATCGGAA
120681 TATGAGTGTGGAGTCGTTGCGAAAACAGACAACACCAAACNNNAGATCGGAAGAG
120682 TATGAGTGTGGAGTCGTTGCAGTATTTCAGAGTTGATTGACTANNNAGATCGGAAG
120683 TATGAGTGTGGAGTCGTTGCCAAATCAGCCTACAGATCTCNNNAGATCGGAAGAGC
120684 TATGAGTGTGGAGTCGTTGCAATAAAGAGATATGAGCATAGCANNNAGATCGGAA

120685 TATGAGTGTGGAGTCGTTGCAGTATATGTGTTTGTGTTAGGTANNNAGATCGGAAG
120688 TATGAGTGTGGAGTCGTTGCGTAAGTAATACAGTTGTTACTGCNNNAGATCGGAAG
120689 TATGAGTGTGGAGTCGTTGCATTTTTCTTATCCATTCATCTGGNNNAGATCGGAAGA
120690 TATGAGTGTGGAGTCGTTGCCAAACAGATGCAACAAATAATTGNNNAGATCGGAAG
120691 TATGAGTGTGGAGTCGTTGCATACTGCAGAATAATTGGTTCTANNNAGATCGGAAG
120692 TATGAGTGTGGAGTCGTTGCGCTGAACTATTTGAAAGTAAGTTNNNAGATCGGAAG
120693 TATGAGTGTGGAGTCGTTGCGACTCTCAAATAGGAAAAGAAAANNNAGATCGGAAG
120694 TATGAGTGTGGAGTCGTTGCAGATTTTTTATCAATCAGCATGTNNNAGATCGGAAGA
120697 TATGAGTGTGGAGTCGTTGCCATATATGTTGTGGGCATAGAANNNAGATCGGAAGA
120698 TATGAGTGTGGAGTCGTTGCCCAGAGAATTCTAACTGATACATNNNAGATCGGAAG
120699 TATGAGTGTGGAGTCGTTGCTAAACAATGAAAAAGTAGTCAGGNNNAGATCGGAAG
120700 TATGAGTGTGGAGTCGTTGCGAAAATTAAATAAGAGCACATGGNNNAGATCGGAAG
120701 TATGAGTGTGGAGTCGTTGCTTGTCTTCAATTTCTCATCATTTNNNAGATCGGAAGAG
120702 TATGAGTGTGGAGTCGTTGCCTGTTTGGACCTCTTTCTCTNNNAGATCGGAAGAGCA
120704 TATGAGTGTGGAGTCGTTGCTCTTTTACTTTTCTCTTTTCAGGNNNAGATCGGAAGAG
120706 TATGAGTGTGGAGTCGTTGCATTGTGTGTGTTGTGTGTATNNNAGATCGGAAGAGC
120707 TATGAGTGTGGAGTCGTTGCATTGTGTGTGTTGTGTGTATNNNAGATCGGAAGAGC
120709 TATGAGTGTGGAGTCGTTGCCAGAATCACAAAACCAGCTATNNNAGATCGGAAGAG
120710 TATGAGTGTGGAGTCGTTGCATTGTGTCTAGGACAAAACAATANNNAGATCGGAAG
120711 TATGAGTGTGGAGTCGTTGCATTGTGTCTAGGACAAAACAATANNNAGATCGGAAG
120712 TATGAGTGTGGAGTCGTTGCCACTCATTAAGTTTGTTGAATAAANNNAGATCGGAAG
120714 TATGAGTGTGGAGTCGTTGCCCCCTGGAAATGATTTTAAAAANNNAGATCGGAAGA
120715 TATGAGTGTGGAGTCGTTGCACGTTCAGAGAATAAAATTTGAGNNNAGATCGGAAG
120717 TATGAGTGTGGAGTCGTTGCCACAAATACCCTATGATTAGTTGNNNAGATCGGAAGA
120718 TATGAGTGTGGAGTCGTTGCCACAAATACCCTATGATTAGTTGNNNAGATCGGAAGA
120719 TATGAGTGTGGAGTCGTTGCCCAATATATCCATGTAACAACACNNNAGATCGGAAGA
120722 TATGAGTGTGGAGTCGTTGCTGTAGCAATTATTGATTACCTTGNNNAGATCGGAAGA
120723 TATGAGTGTGGAGTCGTTGCCGATTTGTGTATATTGCAATTTTNNNAGATCGGAAGA
120724 TATGAGTGTGGAGTCGTTGCCGTCATTTTAGCATTTCTTTTCNNNAGATCGGAAGAG
120725 TATGAGTGTGGAGTCGTTGCGGTTACTCAAACATTTTTCTACANNNAGATCGGAAGA
120726 TATGAGTGTGGAGTCGTTGCTTTGTGCATCTTGAGTCTATCNNNAGATCGGAAGAGC
120727 TATGAGTGTGGAGTCGTTGCACCTAGAGTAGTAGAGATACATTNNNAGATCGGAAG
120728 TATGAGTGTGGAGTCGTTGCGTTCAAAGCCTAATAGCACTNNNAGATCGGAAGAGC
120729 TATGAGTGTGGAGTCGTTGCACAATAGCATACTGTCTTGATTANNNAGATCGGAAGA
120730 TATGAGTGTGGAGTCGTTGCGGTTGTTATGATTTTTTTCAGAGNNNAGATCGGAAG
120731 TATGAGTGTGGAGTCGTTGCAATTTTGTGATGTGTTAACTTCTNNNAGATCGGAAGA
120733 TATGAGTGTGGAGTCGTTGCTAGTCTGCTTTTGTGTATATGAANNNAGATCGGAAG
120734 TATGAGTGTGGAGTCGTTGCTATACAGGAAATTGCTGTACCNNNAGATCGGAAGAG
120736 TATGAGTGTGGAGTCGTTGCTAAAACCTGTCGTTCAATTTATTNNNAGATCGGAAGA
120737 TATGAGTGTGGAGTCGTTGCCTCAATAGAGGAAGGAGTTCTNNNAGATCGGAAGAG
120738 TATGAGTGTGGAGTCGTTGCAAACTAAATGACATCTCCATTTGNNNAGATCGGAAGA
120739 TATGAGTGTGGAGTCGTTGCGTATCTGCAATCCATAATTTGAANNNAGATCGGAAGA
120740 TATGAGTGTGGAGTCGTTGCCTGTATTGGAGGCTGATGATNNNAGATCGGAAGAG

120741 TATGAGTGTGGAGTCGTTGCTTTCGATAGTTCAGTATTAGTGGNNNAGATCGGAAG
120742 TATGAGTGTGGAGTCGTTGCAAGTATTGGCTGGTATTCTTAGNNNAGATCGGAAGA
120743 TATGAGTGTGGAGTCGTTGCTATTTTTTTCTGGCACACAAATANNNAGATCGGAAGA
120744 TATGAGTGTGGAGTCGTTGCATGCATATATATATGTGTGTGTGNNNAGATCGGAAG
120745 TATGAGTGTGGAGTCGTTGCATTTCTATACACAAGACGTTGANNNAGATCGGAAGA
120746 TATGAGTGTGGAGTCGTTGCGTATGCCTTACATTGGATAATTTNNNAGATCGGAAGA
120747 TATGAGTGTGGAGTCGTTGCAAGGAACACTATTACATGAGTATNNNAGATCGGAAG
120748 TATGAGTGTGGAGTCGTTGCATTAGAAGAAAAACAGGATTTGGNNNAGATCGGAA
120749 TATGAGTGTGGAGTCGTTGCAAATCTCAACTTATTCCTTACCANNNAGATCGGAAGA
120750 TATGAGTGTGGAGTCGTTGCTTCAATTTTCTAGGCAAAATGATNNNAGATCGGAAGA
120751 TATGAGTGTGGAGTCGTTGCTGAGGGAAATTGACCAAAAANNNAGATCGGAAGAG
120752 TATGAGTGTGGAGTCGTTGCTAGCTAAAAATTATTTCAGGGGNNNAGATCGGAAG
120753 TATGAGTGTGGAGTCGTTGCTCTAGGAGAAGATCCAATCCNNNAGATCGGAAGAGC
120754 TATGAGTGTGGAGTCGTTGCTATTTATTGAGGATTCAGAGCTTNNNAGATCGGAAG
120755 TATGAGTGTGGAGTCGTTGCTTATATCACTTCATAGTAGTGCANNNAGATCGGAAGA
120756 TATGAGTGTGGAGTCGTTGCATGCATTTCTGTTTTGATGTTNNNAGATCGGAAGAG
120757 TATGAGTGTGGAGTCGTTGCAGATCTCATGAGAACTCACTATNNNAGATCGGAAGA
120758 TATGAGTGTGGAGTCGTTGCGAGGGAACCACTAGATAAGATNNNAGATCGGAAGA
120759 TATGAGTGTGGAGTCGTTGCGGAGTAGAGAACAAAGCAAAANNNAGATCGGAAGA
120760 TATGAGTGTGGAGTCGTTGCTTACTGTTTTTGAGACAGGATCNNNAGATCGGAAGA
120761 TATGAGTGTGGAGTCGTTGCTTACTGTTTTTGAGACAGGATCNNNAGATCGGAAGA
120762 TATGAGTGTGGAGTCGTTGCCCATAAAATTGGCTACAATGAATNNNAGATCGGAAG
120764 TATGAGTGTGGAGTCGTTGCGCTACACATTTTCCAAACTTTTANNNAGATCGGAAGA
120765 TATGAGTGTGGAGTCGTTGCATATGTGTGTGTGTATGTATGTGNNNAGATCGGAA
120766 TATGAGTGTGGAGTCGTTGCGATTGACAACTATTATCTGAATGTNNNAGATCGGAA
120767 TATGAGTGTGGAGTCGTTGCAAATCAGGAAACAACTCAAAATANNNAGATCGGAAG
120768 TATGAGTGTGGAGTCGTTGCATATAGCAGAGGGAATTTAGTACNNNAGATCGGAA
120769 TATGAGTGTGGAGTCGTTGCATATGGTATTTCCTGAAGAATGTNNNAGATCGGAAG
120770 TATGAGTGTGGAGTCGTTGCCTATGTAGGTAAGAGAGAGAGAANNNAGATCGGAA
120771 TATGAGTGTGGAGTCGTTGCCATGAAATGAATAAAATGCCAAGNNNAGATCGGAA
120772 TATGAGTGTGGAGTCGTTGCATATGTAAATTTGATTGACTGCANNNAGATCGGAAG
120773 TATGAGTGTGGAGTCGTTGCCTTGTAAAACCCTGTCTCAANNNAGATCGGAAGAGCA
120774 TATGAGTGTGGAGTCGTTGCACAGAGTGATTGGAAAAAAATAANNNAGATCGGAA
120775 TATGAGTGTGGAGTCGTTGCTTCAGAGTTATTTGTTTTCTGTCNNNAGATCGGAAGA
120776 TATGAGTGTGGAGTCGTTGCGAAGAATATTGTTTGTTTGCTTGNNNAGATCGGAAG
120777 TATGAGTGTGGAGTCGTTGCCTATATTGAACCCACTTCCTTTNNNAGATCGGAAGAG
120778 TATGAGTGTGGAGTCGTTGCAAACCCACTCCCATGATAATNNNAGATCGGAAGAGCA
120779 TATGAGTGTGGAGTCGTTGCATATGTGCCTACCTATCTATCTNNNAGATCGGAAGAG
120780 TATGAGTGTGGAGTCGTTGCACTTAAGAAACAAAGAACAAAGTNNNAGATCGGAA
120781 TATGAGTGTGGAGTCGTTGCATGATAGGTGGCCATAGAACNNNAGATCGGAAGAG
120782 TATGAGTGTGGAGTCGTTGCCTTAATCCTTCAGAATTTACTGGNNNAGATCGGAAGA
120783 TATGAGTGTGGAGTCGTTGCGCCTATAAGAGACTTGATGTAATNNNAGATCGGAAG
120784 TATGAGTGTGGAGTCGTTGCAATCATACACTCTCCTGGAAANNNAGATCGGAAGAG

120785 TATGAGTGTGGAGTCGTTGCATACTACTTCTTTCATTCCAAGTNNNAGATCGGAAGA
120786 TATGAGTGTGGAGTCGTTGCGACAAAATATAGACATGGACAAANNNAGATCGGAA
120787 TATGAGTGTGGAGTCGTTGCATATGCCACTATATACACACACNNNAGATCGGAAGA
120788 TATGAGTGTGGAGTCGTTGCATATGCCACTATATACACACACNNNAGATCGGAAGA
120789 TATGAGTGTGGAGTCGTTGCCTCTTTGAGGTCCTAGAAATCNNNAGATCGGAAGAG
120790 TATGAGTGTGGAGTCGTTGCGGAAAGAAAAATCTAACGAGAAANNNAGATCGGAA
120791 TATGAGTGTGGAGTCGTTGCGCACAGTATCTAGTGGATATAAANNNAGATCGGAA
120792 TATGAGTGTGGAGTCGTTGCCTGACATAAGAGCCAGAATCNNNAGATCGGAAGAG
120793 TATGAGTGTGGAGTCGTTGCTGACATTTTGGTAGTTATAACAGNNNAGATCGGAAG
120794 TATGAGTGTGGAGTCGTTGCTGACATTTTGGTAGTTATAACAGNNNAGATCGGAAG
120795 TATGAGTGTGGAGTCGTTGCTGACATTTTGGTAGTTATAACAGNNNAGATCGGAAG
120796 TATGAGTGTGGAGTCGTTGCAAACATGTCTCATTTTGATTCTTNNNAGATCGGAAG
120799 TATGAGTGTGGAGTCGTTGCTATTTATATTTGCTTCACCTCTGNNNAGATCGGAAGA
120800 TATGAGTGTGGAGTCGTTGCTATATTTCTTGTCAGAGACAGCNNNAGATCGGAAGA
120801 TATGAGTGTGGAGTCGTTGCCTAGAACAGAGATTTCCTTTGAANNNAGATCGGAAG
120802 TATGAGTGTGGAGTCGTTGCACACTTACTTCTCTGACAATCNNNAGATCGGAAGAG
120803 TATGAGTGTGGAGTCGTTGCCCACATATTCATGGAAACATTTNNNAGATCGGAAGA
120804 TATGAGTGTGGAGTCGTTGCAAGTAGGTGCAGACTAGAAANNNAGATCGGAAGAG
120805 TATGAGTGTGGAGTCGTTGCAGAAATAACACCACACATCTATANNNAGATCGGAAG
120806 TATGAGTGTGGAGTCGTTGCGCTGAAATTATAGCAACATGATANNNAGATCGGAAG
120807 TATGAGTGTGGAGTCGTTGCCTAATCTCCAGAACATGTGAATNNNAGATCGGAAGA
120809 TATGAGTGTGGAGTCGTTGCCATGGAATACCTTTTGTCTAGANNNAGATCGGAAGA
120811 TATGAGTGTGGAGTCGTTGCGTATCAGGTTTACTAGTTCTGTANNNAGATCGGAAG
120812 TATGAGTGTGGAGTCGTTGCCTCTGATCTATGTTCAAACTAGANNNAGATCGGAAG
120813 TATGAGTGTGGAGTCGTTGCACAGATGATAGATAGAATAGCTTNNNAGATCGGAA
120814 TATGAGTGTGGAGTCGTTGCACAGATGATAGATAGAATAGCTTNNNAGATCGGAA
120815 TATGAGTGTGGAGTCGTTGCTTACTCATCAGAATTTGTAATGGNNNAGATCGGAAG
120816 TATGAGTGTGGAGTCGTTGCATTTTCATCATTGGCAACTTTTTNNNAGATCGGAAGA
120817 TATGAGTGTGGAGTCGTTGCTCAAGCAAGTTAGTGTTTATTACNNNAGATCGGAAG
120818 TATGAGTGTGGAGTCGTTGCATTATTACCTTTAGTGCTCATGTNNNAGATCGGAAGA
120819 TATGAGTGTGGAGTCGTTGCATATAATGCATATGCTTTACCAGNNNAGATCGGAAG
120820 TATGAGTGTGGAGTCGTTGCGTATTAGTCTGTTTTCAAGCTGNNNAGATCGGAAGA
120821 TATGAGTGTGGAGTCGTTGCTCTTTATGCAGATACAATGAAACNNNAGATCGGAAG
120822 TATGAGTGTGGAGTCGTTGCAGAGGAAGATGACTTCAGAGNNNAGATCGGAAGA
120823 TATGAGTGTGGAGTCGTTGCGTATATATTTAGCAGCAGGTACTNNNAGATCGGAAG
120824 TATGAGTGTGGAGTCGTTGCTATCTGTTTTTTGTGCCTACATANNNAGATCGGAAGA
120825 TATGAGTGTGGAGTCGTTGCCTATTGCAGCCATTCTACAANNNAGATCGGAAGAGC
120826 TATGAGTGTGGAGTCGTTGCTATTGTGAATAGTGCTCCAATAANNNAGATCGGAAG
120828 TATGAGTGTGGAGTCGTTGCTATATTATCTCTATGGTGACCACNNNAGATCGGAAG
120829 TATGAGTGTGGAGTCGTTGCCATGGGAAGAAAATATAGGTAGANNNAGATCGGAA
120830 TATGAGTGTGGAGTCGTTGCACAATATTGAACTATCCAAAAGTNNNAGATCGGAAG
120831 TATGAGTGTGGAGTCGTTGCACAATATTGAACTATCCAAAAGTNNNAGATCGGAAG
120832 TATGAGTGTGGAGTCGTTGCCTATCTCATGAGTATGGAATCTTNNNAGATCGGAAG

120833 TATGAGTGTGGAGTCGTTGCAAATAAGGATTTAGCATTCATCTNNNAGATCGGAAG
120834 TATGAGTGTGGAGTCGTTGCAAATAGGGGACCATTTTAGTTTANNNAGATCGGAAG
120835 TATGAGTGTGGAGTCGTTGCTTTGAATAAACGGCTAAGATACTNNNAGATCGGAAG
120836 TATGAGTGTGGAGTCGTTGCGTTTTTTTTTCTACCTAGGTCTTNNNAGATCGGAAGA
120837 TATGAGTGTGGAGTCGTTGCAGCAAAACTCAAATCGTTAATATNNNAGATCGGAAG
120838 TATGAGTGTGGAGTCGTTGCTCTTACATATCCAGTTAGACAAGNNNAGATCGGAAG
120839 TATGAGTGTGGAGTCGTTGCGAGTTTCTTTTCCTCAATCTTTTNNNAGATCGGAAGA
120840 TATGAGTGTGGAGTCGTTGCGTAATATTAGAACACCTCATCCTNNNAGATCGGAAG
120842 TATGAGTGTGGAGTCGTTGCGGGTTAGAGATAAACAAGACTATNNNAGATCGGAA
120843 TATGAGTGTGGAGTCGTTGCCTTTGTTTGTCTTGTTCAGTGNNNAGATCGGAAGAG
120844 TATGAGTGTGGAGTCGTTGCTTAAGTTGTGACATTAAGAAGTGNNNAGATCGGAA
120845 TATGAGTGTGGAGTCGTTGCGAACTCTTAATGGCACTGAANNNAGATCGGAAGAG
120846 TATGAGTGTGGAGTCGTTGCAAGTAATTACAAATGAACAAAAGTNNNAGATCGGAA
120847 TATGAGTGTGGAGTCGTTGCATCATTGTTTATCATTCGGTTTTNNNAGATCGGAAGA
120848 TATGAGTGTGGAGTCGTTGCCCTGAAAGAATACAAACCAAAANNNAGATCGGAAGA
120849 TATGAGTGTGGAGTCGTTGCCTTTCTCTTTCTCTCTTGGAGNNNAGATCGGAAGAGC
120850 TATGAGTGTGGAGTCGTTGCTAAAGCTTTCATTGTTTCTCTAGNNNAGATCGGAAGA
120851 TATGAGTGTGGAGTCGTTGCTAAAGCTTTCATTGTTTCTCTAGNNNAGATCGGAAGA
120852 TATGAGTGTGGAGTCGTTGCCTATGCATATCCAATTTCCCTNNNAGATCGGAAGAGC
120853 TATGAGTGTGGAGTCGTTGCGGATTACAGCTACAATTAAGAAGNNNAGATCGGAA
120854 TATGAGTGTGGAGTCGTTGCTTAATATTGTGAATGTGGCTTTTNNNAGATCGGAAG
120855 TATGAGTGTGGAGTCGTTGCATTCAGATATCTTAGGTGACTTTNNNAGATCGGAAG
120856 TATGAGTGTGGAGTCGTTGCGAGAGTCAATGTGTAATTCCANNNAGATCGGAAGA
120857 TATGAGTGTGGAGTCGTTGCCATGAAAGAGCAGATGGTACNNNAGATCGGAAGAG
120858 TATGAGTGTGGAGTCGTTGCACTCAGCAAGAAGAGATTATACNNNAGATCGGAAGA
120859 TATGAGTGTGGAGTCGTTGCACGTACATACTTTATACCAACTTNNNAGATCGGAAGA
120860 TATGAGTGTGGAGTCGTTGCTGTATGATCACAGTTGGATGNNNAGATCGGAAGAG
120861 TATGAGTGTGGAGTCGTTGCCGAGAAGTCAAATAGATTTCAAANNNAGATCGGAAG
120862 TATGAGTGTGGAGTCGTTGCATGCTATCTTGCAGTCATTTTANNNAGATCGGAAGA
120863 TATGAGTGTGGAGTCGTTGCAAATATAGTGAGAACATCAGCTANNNAGATCGGAAG
120864 TATGAGTGTGGAGTCGTTGCTTCATTCAGAACATTTATTAGGCNNNAGATCGGAAG
120865 TATGAGTGTGGAGTCGTTGCTTACTGAGGTATAATCCACATACNNNAGATCGGAAG
120866 TATGAGTGTGGAGTCGTTGCACATATATCCACCATTACAGAATNNNAGATCGGAAG
120868 TATGAGTGTGGAGTCGTTGCCTTTCAAATGAATTTTCCTTCCANNNAGATCGGAAGA
120869 TATGAGTGTGGAGTCGTTGCGTTGAAGTAGAGAAAGTCAAATCNNNAGATCGGAA
120870 TATGAGTGTGGAGTCGTTGCCACTGAAATAGATTCTTCAAGAANNNAGATCGGAAG
120871 TATGAGTGTGGAGTCGTTGCCATCATTATCATTAGCACAAAACNNNAGATCGGAAG
120872 TATGAGTGTGGAGTCGTTGCCTACATATAGGCTAGAACCCTANNNAGATCGGAAGA
120873 TATGAGTGTGGAGTCGTTGCCAACAGCAGAATGACATATTATCNNNAGATCGGAAG
120874 TATGAGTGTGGAGTCGTTGCTGGTAAAATACTGTTTCTAGGAANNNAGATCGGAAG
120875 TATGAGTGTGGAGTCGTTGCAATCAAAACAATAAAGCATGAGANNNAGATCGGAA
120876 TATGAGTGTGGAGTCGTTGCGTATCTTTATTACTGGAAGTGTGNNNAGATCGGAAG
120877 TATGAGTGTGGAGTCGTTGCTGGGGTTGTAATTTATGTTAGATNNNAGATCGGAA

120878 TATGAGTGTGGAGTCGTTGCCTAGCAAATTTATCGTTCATCATNNNAGATCGGAAGA
120879 TATGAGTGTGGAGTCGTTGCAAAGGAGGTCCATTTCTAAGTATNNNAGATCGGAAG
120880 TATGAGTGTGGAGTCGTTGCAGACATAAAGGCATAATAGTACANNNAGATCGGAA
120881 TATGAGTGTGGAGTCGTTGCTGTAAAATGTGTGTGTTCTTTAANNNAGATCGGAAG
120882 TATGAGTGTGGAGTCGTTGCCTATTAATATCCTGAGGAAACCANNNAGATCGGAAG
120883 TATGAGTGTGGAGTCGTTGCTCAGAGAAAGGATTGGTTGGNNNAGATCGGAAGA
120884 TATGAGTGTGGAGTCGTTGCATCTGATAGAGAAGAAGTGATAGNNNAGATCGGAA
120885 TATGAGTGTGGAGTCGTTGCATTGTGAATAGTGCTGAAATAAANNNAGATCGGAA
120886 TATGAGTGTGGAGTCGTTGCGCAAAACAAAAAAACAAAGGATANNNAGATCGGAA
120887 TATGAGTGTGGAGTCGTTGCAGGATATGTTGATATATTCTGCCNNNAGATCGGAAG
120888 TATGAGTGTGGAGTCGTTGCTGTTGCAAAATATGTACCTATTTNNNAGATCGGAAGA
120889 TATGAGTGTGGAGTCGTTGCGAGAAGACAGAGAAAAAAATTCANNNAGATCGGAA
120890 TATGAGTGTGGAGTCGTTGCATAGGAAAAGATCTGAAGAAACANNNAGATCGGAA
120891 TATGAGTGTGGAGTCGTTGCACAATGTAATATTGATGCAACTTNNNAGATCGGAAG
120892 TATGAGTGTGGAGTCGTTGCTATTTTTTTAGATGGAGGCTTACNNNAGATCGGAAG
120893 TATGAGTGTGGAGTCGTTGCTAAGGAGTTGGTCAAATGATTATNNNAGATCGGAA
120894 TATGAGTGTGGAGTCGTTGCTTATTATCTGAATCTTGTCCACANNNAGATCGGAAGA
120895 TATGAGTGTGGAGTCGTTGCTATTTGAGAAGAGAGACGGANNNAGATCGGAAGA
120896 TATGAGTGTGGAGTCGTTGCTGTTGAGATAAGGCTTAGGATANNNAGATCGGAAG
120897 TATGAGTGTGGAGTCGTTGCTAACAATGGTGTTGACCCTATNNNAGATCGGAAGAG
120898 TATGAGTGTGGAGTCGTTGCCACACAATCAAATCTACATAACANNNAGATCGGAAGA
120899 TATGAGTGTGGAGTCGTTGCGTCTTGTTGGTAGAATAAGAAAANNNAGATCGGAA
120900 TATGAGTGTGGAGTCGTTGCACAGTCAAGGTATAGAATAGTTCNNNAGATCGGAA
120901 TATGAGTGTGGAGTCGTTGCAGAAAAATATGGACAACTCTCTANNNAGATCGGAAG
120902 TATGAGTGTGGAGTCGTTGCACGCATACTTATGTAATCATAGANNNAGATCGGAAG
120903 TATGAGTGTGGAGTCGTTGCAGTGTTTATTTTAAAACTCTTGGANNNAGATCGGAA
120904 TATGAGTGTGGAGTCGTTGCCAGCTCAAACCTACATTAATTAANNNAGATCGGAAGA
120905 TATGAGTGTGGAGTCGTTGCGATCACTTTCAATTCTGCCTNNNAGATCGGAAGAGCA
120906 TATGAGTGTGGAGTCGTTGCAAAAATTTAAGTTGCTTTCAGTGNNNAGATCGGAAG
120907 TATGAGTGTGGAGTCGTTGCTGTAATCAAGGTCTCGCTATNNNAGATCGGAAGAGC
120908 TATGAGTGTGGAGTCGTTGCAGGACATTTATTATAGGAATTTGCNNNAGATCGGAA
120909 TATGAGTGTGGAGTCGTTGCTGTGTTTGTTCATTCATTTATTTNNNAGATCGGAAGA
120910 TATGAGTGTGGAGTCGTTGCCTGCATTCTATGTATGACCAATANNNAGATCGGAAGA
120911 TATGAGTGTGGAGTCGTTGCTATGCTTATTCAATAGATCAAGCNNNAGATCGGAAGA
120912 TATGAGTGTGGAGTCGTTGCCCAGAAAGAAATAACACATTACTNNNAGATCGGAAG
120914 TATGAGTGTGGAGTCGTTGCCAGCTTGAATGTTATTGATGTATNNNAGATCGGAAG
120917 TATGAGTGTGGAGTCGTTGCTAGTCTCTCTCAATTTGTGTTTANNNAGATCGGAAGA
120918 TATGAGTGTGGAGTCGTTGCATCAACCTGTATTTTGATTTTGGNNNAGATCGGAAGA
120919 TATGAGTGTGGAGTCGTTGCGTAAGTAAAAAGACACAAACTCTNNNAGATCGGAA
120920 TATGAGTGTGGAGTCGTTGCTCCAACTCTAGTAACAATTCAATNNNAGATCGGAAGA
120921 TATGAGTGTGGAGTCGTTGCTGCTTCAATAAACAAACAAGTATNNNAGATCGGAAG
120922 TATGAGTGTGGAGTCGTTGCCAGCTGTGATATCTATCTTCTCNNNAGATCGGAAGA
120923 TATGAGTGTGGAGTCGTTGCATTTTTCCCAGGCATCTAATATTNNNAGATCGGAAGA

120924 TATGAGTGTGGAGTCGTTGCTATACAGGAAGAGGATAGTATGANNNAGATCGGAA
120925 TATGAGTGTGGAGTCGTTGCATCAACAACACATTAGAAAGATCNNNAGATCGGAAG
120926 TATGAGTGTGGAGTCGTTGCCTGAAAACAACAATAACAAAGACNNNAGATCGGAAG
120927 TATGAGTGTGGAGTCGTTGCATTTGTATTCCTGCAATAACTTCNNNAGATCGGAAGA
120928 TATGAGTGTGGAGTCGTTGCGTTGAGTTTGATTTTGAAAGATGNNNAGATCGGAAG
120929 TATGAGTGTGGAGTCGTTGCTTCCATAGTGCCTTTTGAAAANNNAGATCGGAAGAG
120930 TATGAGTGTGGAGTCGTTGCGAAAATAATCCCTGACTAAAAGTNNNAGATCGGAAG
120931 TATGAGTGTGGAGTCGTTGCCTAAAACCCTTGCTACTATGTAANNNAGATCGGAAGA
120932 TATGAGTGTGGAGTCGTTGCCTAAAACCCTTGCTACTATGTAANNNAGATCGGAAGA
120933 TATGAGTGTGGAGTCGTTGCATGTCCTCAGCTATAGAATGANNNAGATCGGAAGAG
120934 TATGAGTGTGGAGTCGTTGCTCCTCTGTATTTTATTCTGTTCTNNNAGATCGGAAGA
120935 TATGAGTGTGGAGTCGTTGCCAAGGCTGGGTAATTTATGANNNAGATCGGAAGAG
120936 TATGAGTGTGGAGTCGTTGCCGTCTCATTAATCTGAATGTCTANNNAGATCGGAAGA
120937 TATGAGTGTGGAGTCGTTGCGTTTGGATTGTTTGTAGTTCTTANNNAGATCGGAAG
120938 TATGAGTGTGGAGTCGTTGCACAGCACATCAGATTTTGTATTNNNAGATCGGAAGA
120939 TATGAGTGTGGAGTCGTTGCACAGCACATCAGATTTTGTATTNNNAGATCGGAAGA
120940 TATGAGTGTGGAGTCGTTGCTTATTGAAACTAGACTGTTTGTCNNNAGATCGGAAGA
120941 TATGAGTGTGGAGTCGTTGCGTCATAGTGAATGAGTACCTAGNNNAGATCGGAAGA
120942 TATGAGTGTGGAGTCGTTGCTTTGTGGACGTATTTTAAAATCANNNAGATCGGAAG
120943 TATGAGTGTGGAGTCGTTGCCCTCACCTATTCAAATTGCTANNNAGATCGGAAGAG
120944 TATGAGTGTGGAGTCGTTGCCACAGAGATGTTGGGGATAGNNNAGATCGGAAGAG
120945 TATGAGTGTGGAGTCGTTGCCTTCACAGACCTTCCTTTTGNNNAGATCGGAAGAGCA
120946 TATGAGTGTGGAGTCGTTGCTAAGTAGCTTTTGTAGAAGAGAGNNNAGATCGGAAG
120947 TATGAGTGTGGAGTCGTTGCTAAGTAGCTTTTGTAGAAGAGAGNNNAGATCGGAAG
120948 TATGAGTGTGGAGTCGTTGCCAAATTGGGTAAATGTGATCTTTNNNAGATCGGAAG
120949 TATGAGTGTGGAGTCGTTGCCCCATTACATTTATCTAACCTTTNNNAGATCGGAAGA
120950 TATGAGTGTGGAGTCGTTGCTTTGGATAGTATTTTTTTCAGGTNNNAGATCGGAAGA
120951 TATGAGTGTGGAGTCGTTGCGGAGTCTATTGTCAGGATATAAANNNAGATCGGAAG
120952 TATGAGTGTGGAGTCGTTGCTTGCTTTTGTTTGCACTTATTTANNNAGATCGGAAGA
120953 TATGAGTGTGGAGTCGTTGCATTTGTGTTATTCCTTTGGATACNNNAGATCGGAAGA
120954 TATGAGTGTGGAGTCGTTGCTACCTAAAGACATATAGAGCTTCNNNAGATCGGAAG
120955 TATGAGTGTGGAGTCGTTGCAAATCTCATGTCCTTCTCATATTNNNAGATCGGAAGA
120956 TATGAGTGTGGAGTCGTTGCACTTCCCATAAAAGGATTACTTCNNNAGATCGGAAGA
120957 TATGAGTGTGGAGTCGTTGCCATCTGGGTCTAATAGGTCTANNNAGATCGGAAGAG
120958 TATGAGTGTGGAGTCGTTGCTGTAGAAAACAGTTTACAAAGAANNNAGATCGGAAG
120959 TATGAGTGTGGAGTCGTTGCTCATGGTTTTGCTTATCTTCTNNNAGATCGGAAGAG
120960 TATGAGTGTGGAGTCGTTGCTCTATGTTTCAGAGAGAGAGAGNNNAGATCGGAAG
120961 TATGAGTGTGGAGTCGTTGCTATAATAAGTTGCCCCAAACTCNNNAGATCGGAAGA
120962 TATGAGTGTGGAGTCGTTGCAATGAGAATGAGAATGAGAGAGNNNAGATCGGAAG
120963 TATGAGTGTGGAGTCGTTGCAAATAATGAGCTGGTTTATTAGCNNNAGATCGGAAG
120964 TATGAGTGTGGAGTCGTTGCATTGAGAGACTTATCTATTCCAGNNNAGATCGGAAG
120965 TATGAGTGTGGAGTCGTTGCAAAATAACTTCTGTAGCCCAACNNNAGATCGGAAGA
120966 TATGAGTGTGGAGTCGTTGCGCTCCCCTAGAAGTAAAGAANNNAGATCGGAAGAG

120967 TATGAGTGTGGAGTCGTTGCAGATCATTTCTGGAGAACTTTTANNNAGATCGGAAG
120968 TATGAGTGTGGAGTCGTTGCAAAGTGTGACAATGAGAAACTTANNNAGATCGGAA
120969 TATGAGTGTGGAGTCGTTGCTTTCATGGTTTTTATACAGTGTTNNNAGATCGGAAG
120970 TATGAGTGTGGAGTCGTTGCGAAGAAGGTGAACGAGATAATTNNNAGATCGGAA
120971 TATGAGTGTGGAGTCGTTGCCAAATCCATTTGGTCAGAATTNNNAGATCGGAAGAG
120972 TATGAGTGTGGAGTCGTTGCCCTGGAAGAGTGAAAAATCTANNNAGATCGGAAGA
120973 TATGAGTGTGGAGTCGTTGCGGAAAAGGAAATTAAATCTACTCANNNAGATCGGA
120974 TATGAGTGTGGAGTCGTTGCGGAAAAGGAAATTAAATCTACTCANNNAGATCGGA
120975 TATGAGTGTGGAGTCGTTGCGGAAAAGGAAATTAAATCTACTCANNNAGATCGGA
120976 TATGAGTGTGGAGTCGTTGCATAAACCAAGCTTTAGTAGTACANNNAGATCGGAAG
120977 TATGAGTGTGGAGTCGTTGCCTGTCATTGAAGCTATAAATGAGNNNAGATCGGAA
120979 TATGAGTGTGGAGTCGTTGCGTTGGAACTTCATTAAGATCTTTNNNAGATCGGAAG
120980 TATGAGTGTGGAGTCGTTGCATGGCATAATTTGGAAAATACATNNNAGATCGGAA
120981 TATGAGTGTGGAGTCGTTGCCAGACAGGGATACAATTTTAAAANNNAGATCGGAA
120982 TATGAGTGTGGAGTCGTTGCCAGACAGGGATACAATTTTAAAANNNAGATCGGAA
120983 TATGAGTGTGGAGTCGTTGCAAATACAGAAAATCTGGGATTTGNNNAGATCGGAA
120984 TATGAGTGTGGAGTCGTTGCAGAATTTATTTGACTAAGTACGCNNNAGATCGGAAG
120985 TATGAGTGTGGAGTCGTTGCTTTCCATCCAAATCTGATACAANNNAGATCGGAAGA
120986 TATGAGTGTGGAGTCGTTGCTTTCCATCCAAATCTGATACAANNNAGATCGGAAGA
120987 TATGAGTGTGGAGTCGTTGCCCCCCAATAAAATAGGAAATAATNNNAGATCGGAAG
120988 TATGAGTGTGGAGTCGTTGCATAAGTAGATTGAAATGTGTGAGNNNAGATCGGAA
120989 TATGAGTGTGGAGTCGTTGCGGTATAACTGTGAAAAAGACAGNNNAGATCGGAAG
120990 TATGAGTGTGGAGTCGTTGCATCAGTTCTTTTGTCTCAATTTTNNNAGATCGGAAGA
120993 TATGAGTGTGGAGTCGTTGCTGGGGAAAAGTGATTGAATTTANNNAGATCGGAAG
120994 TATGAGTGTGGAGTCGTTGCAATCTATCACTGAAGATTCCTGNNNAGATCGGAAGA
120996 TATGAGTGTGGAGTCGTTGCAAATACTTGGATGTTTGGCTNNNAGATCGGAAGAG
120997 TATGAGTGTGGAGTCGTTGCTTCAATCTTGGGCTCAAATAATNNNAGATCGGAAGA
120998 TATGAGTGTGGAGTCGTTGCGAAGTCAAATAAGTAGTTCTTGGNNNAGATCGGAA
120999 TATGAGTGTGGAGTCGTTGCGGTTAACAAAATCACTCAAAAAANNNAGATCGGAAG
121000 TATGAGTGTGGAGTCGTTGCATAAGGGAAGATAGAAGTTTTGTNNNAGATCGGAA
121001 TATGAGTGTGGAGTCGTTGCCACCCATATAAAACCATTGTACNNNAGATCGGAAGA
121002 TATGAGTGTGGAGTCGTTGCCACCCATATAAAACCATTGTACNNNAGATCGGAAGA
121003 TATGAGTGTGGAGTCGTTGCCACCCATATAAAACCATTGTACNNNAGATCGGAAGA
121004 TATGAGTGTGGAGTCGTTGCGAGGATGAAGAAATTTGGTTAATNNNAGATCGGAA
121007 TATGAGTGTGGAGTCGTTGCTTCCTCTACAAAGTTTTGGAATANNNAGATCGGAAG
121008 TATGAGTGTGGAGTCGTTGCGTGACAGTGAATTGATAAAAAGANNNAGATCGGAA
121010 TATGAGTGTGGAGTCGTTGCAATTAAAACTGAGCATTAGTGAANNNAGATCGGAA
121011 TATGAGTGTGGAGTCGTTGCGGCACTGTAAGTATTTTACTATTNNNAGATCGGAAG
121012 TATGAGTGTGGAGTCGTTGCAGACTTTGTGTTTTAACAGAAAANNNAGATCGGAAG
121013 TATGAGTGTGGAGTCGTTGCCTAACACCAAAAGAGAAAGATAGNNNAGATCGGAA
121014 TATGAGTGTGGAGTCGTTGCGTCTGTAGCTATGTAAATATGGANNNAGATCGGAA
121015 TATGAGTGTGGAGTCGTTGCAGGAAATATGGTAAAAGGAAGAGNNNAGATCGGA
121016 TATGAGTGTGGAGTCGTTGCCAGTGTGTTAAAATGGTTGAATNNNAGATCGGAAG

121017 TATGAGTGTGGAGTCGTTGCGAAGAATAGAATGACAACAGTGNNNAGATCGGAAG
121018 TATGAGTGTGGAGTCGTTGCGGATTTCCATCTACTTTAGAATGNNNAGATCGGAAG
121019 TATGAGTGTGGAGTCGTTGCAATCACAAATGGGAAACAAGTTANNNAGATCGGAA
121020 TATGAGTGTGGAGTCGTTGCGATAGCAGTCAAAAAAATTGAAGNNNAGATCGGAA
121021 TATGAGTGTGGAGTCGTTGCCTGATATTTAGCTATAGTGTTGCNNNAGATCGGAAG
121022 TATGAGTGTGGAGTCGTTGCAGGGAACATATAGGTTTACTTTTNNNAGATCGGAAG
121023 TATGAGTGTGGAGTCGTTGCTGTGAGCTCAAATTTCTTTTAATNNNAGATCGGAAG
121024 TATGAGTGTGGAGTCGTTGCATAAAGAGTTGAATGTGGATTGANNNAGATCGGAA
121025 TATGAGTGTGGAGTCGTTGCGTTGAAAACCTTTTCTACCTTAANNNAGATCGGAAG
121026 TATGAGTGTGGAGTCGTTGCGATTTATGTCACATAAAGGAACANNNAGATCGGAAG
121027 TATGAGTGTGGAGTCGTTGCCCTTACATTCCTACTCCTAAAAANNNAGATCGGAAGA
121028 TATGAGTGTGGAGTCGTTGCGTTCAAAATTCTCTCAAAATTCCNNNAGATCGGAAGA
121029 TATGAGTGTGGAGTCGTTGCTTTGGTGAAATATCTCTGTCTTANNNAGATCGGAAG
121030 TATGAGTGTGGAGTCGTTGCGAAGTTCTTCATTCCTGAGTCNNNAGATCGGAAGAG
121031 TATGAGTGTGGAGTCGTTGCAATTTTTTCCTCTTTGCCAACNNNAGATCGGAAGAGC
121032 TATGAGTGTGGAGTCGTTGCTTTCAATAGATTACAGTTTGTGGNNNAGATCGGAAG
121033 TATGAGTGTGGAGTCGTTGCACTTCTTAATTTTCATGTCTAGCNNNAGATCGGAAGA
121034 TATGAGTGTGGAGTCGTTGCGGGTTTGAAATGTAACATTTACANNNAGATCGGAAG
121035 TATGAGTGTGGAGTCGTTGCAATCTCCTATTGGTTCTATTACCNNNAGATCGGAAGA
121036 TATGAGTGTGGAGTCGTTGCTAAACCTCTGTGTTTTTCAGTTNNNAGATCGGAAGA
121037 TATGAGTGTGGAGTCGTTGCTTAGAAGCAGCAAGTGTAGTANNNAGATCGGAAGA
121038 TATGAGTGTGGAGTCGTTGCCACCGTTTCCCTAAATTATAATTNNNAGATCGGAAGA
121039 TATGAGTGTGGAGTCGTTGCAATGACTATTTCCACTAGCAAANNNAGATCGGAAGA
121040 TATGAGTGTGGAGTCGTTGCAGACCATTATTGTAAGTGAAGTANNNAGATCGGAAG
121041 TATGAGTGTGGAGTCGTTGCTAGAATTATCTGTAAGGAACTCCNNNAGATCGGAAG
121042 TATGAGTGTGGAGTCGTTGCATTTTCTCCCATAGTCATTTTTCNNNAGATCGGAAGA
121043 TATGAGTGTGGAGTCGTTGCAATTTCTCACATTTCATAAAGCANNNAGATCGGAAGA
121044 TATGAGTGTGGAGTCGTTGCTACCCACACATTATATGTATTGTNNNAGATCGGAAGA
121045 TATGAGTGTGGAGTCGTTGCAATTTATTAGGAAGAATTGGCTGNNNAGATCGGAA
121046 TATGAGTGTGGAGTCGTTGCGGATCTAGGTGATTTTATTTTGTANNNAGATCGGAA
121047 TATGAGTGTGGAGTCGTTGCAAATATACAGGTTTAGAGTCTGTNNNAGATCGGAAG
121048 TATGAGTGTGGAGTCGTTGCCACAAACATAGCCAATTACTACNNNAGATCGGAAGA
121049 TATGAGTGTGGAGTCGTTGCGAAATAGAGGTGAGAGACTTTACNNNAGATCGGAA
121050 TATGAGTGTGGAGTCGTTGCAAAGCTCAAGAAAGAAAATTAGTNNNAGATCGGAA
121051 TATGAGTGTGGAGTCGTTGCCACTGCAAAAGGTAAGAATATTANNNAGATCGGAAG
121052 TATGAGTGTGGAGTCGTTGCCCCAAGGGTAAATAATAATTCAGNNNAGATCGGAAG
121053 TATGAGTGTGGAGTCGTTGCCACACAGACACATACATATCATNNNAGATCGGAAGA
121054 TATGAGTGTGGAGTCGTTGCCATTTCTATCAGCTTTCAAATCANNNAGATCGGAAGA
121055 TATGAGTGTGGAGTCGTTGCATTTCCAGCATCTTTTTATAAGANNNAGATCGGAAGA
121056 TATGAGTGTGGAGTCGTTGCATAACCAAGACTGTCTGCTAATNNNAGATCGGAAGA
121057 TATGAGTGTGGAGTCGTTGCTTCTAAGTGTGTATGTATGTAGCNNNAGATCGGAAG
121058 TATGAGTGTGGAGTCGTTGCCTCCTACTATTCCTCCTCTTTNNNAGATCGGAAGAGCA
121059 TATGAGTGTGGAGTCGTTGCTTTCCCAACAGTAGTAGATAAATNNNAGATCGGAAG

121061 TATGAGTGTGGAGTCGTTGCGCTTACTATTTCACTTTATGGAANNNAGATCGGAAG
121062 TATGAGTGTGGAGTCGTTGCGCTTACTATTTCACTTTATGGAANNNAGATCGGAAG
121063 TATGAGTGTGGAGTCGTTGCGGGAGAACTATAACCACATTTNNNAGATCGGAAGA
121064 TATGAGTGTGGAGTCGTTGCGTTACTAGAATCACCCCGACNNNAGATCGGAAGAGC
121065 TATGAGTGTGGAGTCGTTGCCTGTAAGAAAACTACTCAAGGATNNNAGATCGGAAG
121066 TATGAGTGTGGAGTCGTTGCCAAAATCCAGACCCTCTATTATNNNAGATCGGAAGA
121067 TATGAGTGTGGAGTCGTTGCCAAAATCCAGACCCTCTATTATNNNAGATCGGAAGA
121068 TATGAGTGTGGAGTCGTTGCAAAATACCCCAGTATACTCTTACNNNAGATCGGAAG
121069 TATGAGTGTGGAGTCGTTGCAGCTAAAATCTTCCCTTTAGAGNNNAGATCGGAAGA
121070 TATGAGTGTGGAGTCGTTGCCAATTGGAAGTGGATACTGTANNNAGATCGGAAGA
121073 TATGAGTGTGGAGTCGTTGCGTTAAAGGGAAAAGGAGTATAGANNNAGATCGGA
121074 TATGAGTGTGGAGTCGTTGCAAAATAGTCATTCAAGAGGACAGNNNAGATCGGAA
121075 TATGAGTGTGGAGTCGTTGCCAATTTCTGGCTTCCCTTTCNNNAGATCGGAAGAGCA
121076 TATGAGTGTGGAGTCGTTGCAGGATTTAATGCAGTGTGATAGNNNAGATCGGAAG
121077 TATGAGTGTGGAGTCGTTGCATATTCTAGAGAAACTCCTGTACNNNAGATCGGAAG
121078 TATGAGTGTGGAGTCGTTGCTTAGCCCTGAAATAACTGTATTANNNAGATCGGAAG
121079 TATGAGTGTGGAGTCGTTGCACAGTTTAACTTTTTGGAGTTTTNNNAGATCGGAAG
121080 TATGAGTGTGGAGTCGTTGCACAGTTTAACTTTTTGGAGTTTTNNNAGATCGGAAG
121081 TATGAGTGTGGAGTCGTTGCGACACTTTCTAGAGCCTGATNNNAGATCGGAAGAGC
121082 TATGAGTGTGGAGTCGTTGCTCTTGATCAATCTTGAAAGAATGNNNAGATCGGAAG
121083 TATGAGTGTGGAGTCGTTGCAAATAAGGTCAAAAAGGCATGNNNAGATCGGAAGA
121084 TATGAGTGTGGAGTCGTTGCTTTTAAACATGCTCAGTTTTTTTNNNAGATCGGAAGA
121085 TATGAGTGTGGAGTCGTTGCTTTTAAACATGCTCAGTTTTTTTNNNAGATCGGAAGA
121086 TATGAGTGTGGAGTCGTTGCCCTTAAAGAGTTTCAACTGTTTANNNAGATCGGAAG
121087 TATGAGTGTGGAGTCGTTGCGCAATGAAAAGAGGTTTATTACTNNNAGATCGGAA
121088 TATGAGTGTGGAGTCGTTGCCTGCAATACCCAGAGAAAACNNNAGATCGGAAGAG
121089 TATGAGTGTGGAGTCGTTGCCTGTGAATGAGAAAGAAGTAAATNNNAGATCGGAA
121090 TATGAGTGTGGAGTCGTTGCCGAGAAAAAAAAATCAAGGAATANNNAGATCGGAA
121091 TATGAGTGTGGAGTCGTTGCGAAAGAACTTGGCTTCATTTTAANNNAGATCGGAAG
121092 TATGAGTGTGGAGTCGTTGCTTTTCTTGTTATGTTGTCAAAATANNNAGATCGGAA
121093 TATGAGTGTGGAGTCGTTGCGAAAACAAAACACTTGAAAACTTNNNAGATCGGAAG
121094 TATGAGTGTGGAGTCGTTGCGAGGAGACTGCTGAAATTTTANNNAGATCGGAAGA
121095 TATGAGTGTGGAGTCGTTGCGAGATTTATTCTAAAGGAATCGCNNNAGATCGGAA
121096 TATGAGTGTGGAGTCGTTGCAAGAGACATCAAAATAAAACACANNNAGATCGGAA
121097 TATGAGTGTGGAGTCGTTGCCTCAATAATTGCTCTTGGATTAGNNNAGATCGGAAG
121098 TATGAGTGTGGAGTCGTTGCCGTCTTACAAAGATATGGATTTANNNAGATCGGAAG
121099 TATGAGTGTGGAGTCGTTGCGACAAATATAGACACATTCCAAGNNNAGATCGGAAG
121100 TATGAGTGTGGAGTCGTTGCAATATTTGTTGAGCCAATATAGCNNNAGATCGGAAG
121101 TATGAGTGTGGAGTCGTTGCCCATATTGAATACCGTTCAATTTNNNAGATCGGAAG
121103 TATGAGTGTGGAGTCGTTGCATACAATTGTAAATTTCCACAGGNNNAGATCGGAAG
121104 TATGAGTGTGGAGTCGTTGCCAAAATAAGAGCTCTTTATCCAANNNAGATCGGAAG
121105 TATGAGTGTGGAGTCGTTGCTGAGAAGACTAAAAGCTTCTTTANNNAGATCGGAAG
121106 TATGAGTGTGGAGTCGTTGCTGAGAAGACTAAAAGCTTCTTTANNNAGATCGGAAG

121107 TATGAGTGTGGAGTCGTTGCAGAGTAAAGAATATATGGCTGTTNNNAGATCGGAA
121108 TATGAGTGTGGAGTCGTTGCAATTAATTTTGGCTTCCAGTTACNNNAGATCGGAAG
121109 TATGAGTGTGGAGTCGTTGCCATTTCAAAATACGTCAAGTAAANNNAGATCGGAAG
121110 TATGAGTGTGGAGTCGTTGCTTGCAAAAATCCTCAACAAAATANNNAGATCGGAAG
121111 TATGAGTGTGGAGTCGTTGCATAACTTCTTCATTTGAAATCCCNNNAGATCGGAAGA
121112 TATGAGTGTGGAGTCGTTGCTAATGATCTATAAGTGTGCAAAGNNNAGATCGGAA
121113 TATGAGTGTGGAGTCGTTGCGAGCCATTAACTTCTCAATAATTNNNAGATCGGAAG
121114 TATGAGTGTGGAGTCGTTGCTGTTTTGTTTTGTTTTACTAGTCNNNAGATCGGAAGA
121115 TATGAGTGTGGAGTCGTTGCGCCAAACATTTTATTTTTCTGTTNNNAGATCGGAAGA
121116 TATGAGTGTGGAGTCGTTGCCCCTTCCTCTACTTATATTTCTCNNNAGATCGGAAGA
121117 TATGAGTGTGGAGTCGTTGCCCCTTCCTCTACTTATATTTCTCNNNAGATCGGAAGA
121118 TATGAGTGTGGAGTCGTTGCTTCTACATGTATATTTGTGTGTGNNNAGATCGGAAG
121119 TATGAGTGTGGAGTCGTTGCGAGAATCCTAACTAATGCATTTANNNAGATCGGAAG
121120 TATGAGTGTGGAGTCGTTGCTCAAAAGAATAAGTTTGAGAAGGNNNAGATCGGAA
121121 TATGAGTGTGGAGTCGTTGCATAGTAGGACAAATGGAGACATANNNAGATCGGAA
121122 TATGAGTGTGGAGTCGTTGCATGAATTGATTTCAGAAGAACTGNNNAGATCGGAA
121123 TATGAGTGTGGAGTCGTTGCTACTTAAGGGTCAAAGCCAANNNAGATCGGAAGAG
121125 TATGAGTGTGGAGTCGTTGCGAATTAAACACTCAACACTAGACNNNAGATCGGAAG
121126 TATGAGTGTGGAGTCGTTGCATGTACCCTGCAGAGTATAGNNNAGATCGGAAGAG
121127 TATGAGTGTGGAGTCGTTGCTATGTACCCTGCAGAGTATAGNNNAGATCGGAAGA
121128 TATGAGTGTGGAGTCGTTGCTAGGAAGTCACTAATTCATAAGGNNNAGATCGGAA
121129 TATGAGTGTGGAGTCGTTGCCCACTCAAATGTTAATGCAGNNNAGATCGGAAGAGC
121130 TATGAGTGTGGAGTCGTTGCGGATGGAAAAGATCTACAATTTANNNAGATCGGAA
121131 TATGAGTGTGGAGTCGTTGCAACTTTGGGAGACTTTTCTCNNNAGATCGGAAGAGC
121132 TATGAGTGTGGAGTCGTTGCTATGTCCAGCATAAAACCTGNNNAGATCGGAAGAGC
121133 TATGAGTGTGGAGTCGTTGCTCTATGAATATTCCTACTGCATGNNNAGATCGGAAG
121136 TATGAGTGTGGAGTCGTTGCTATGAGAGAGAGAGAAAGAGAGNNNAGATCGGAA
121137 TATGAGTGTGGAGTCGTTGCCACAACCTCAAATTCCATTAGNNNAGATCGGAAGAG
121138 TATGAGTGTGGAGTCGTTGCCAATTTCATCTTTCGTCACCNNNAGATCGGAAGAGCA
121139 TATGAGTGTGGAGTCGTTGCAAATAGCAGATTCAAGACAATCTNNNAGATCGGAAG
121140 TATGAGTGTGGAGTCGTTGCCTTGAATCCAAAGGTTTTTTTTTNNNAGATCGGAAG
121141 TATGAGTGTGGAGTCGTTGCCTTGAATCCAAAGGTTTTTTTTTNNNAGATCGGAAG
121142 TATGAGTGTGGAGTCGTTGCATAGTGAACTGAGTTTCAAAAACNNNAGATCGGAA
121143 TATGAGTGTGGAGTCGTTGCCTTGTAGATCTTCACCCTGGNNNAGATCGGAAGAGC
121144 TATGAGTGTGGAGTCGTTGCATCAAGACGATAGTATTCTAAGCNNNAGATCGGAAG
121145 TATGAGTGTGGAGTCGTTGCTTTTATTGTCAAAGAGTAGGCTANNNAGATCGGAAG
121146 TATGAGTGTGGAGTCGTTGCCAGATGCCATAATGTTTAATGATNNNAGATCGGAAG
121147 TATGAGTGTGGAGTCGTTGCCACATACCCCCTCTTACTTCNNNAGATCGGAAGAGCA
121148 TATGAGTGTGGAGTCGTTGCTTACTTTATACCTTCTACAGCAANNNAGATCGGAAGA
121149 TATGAGTGTGGAGTCGTTGCGAGATATAAAAAACAGGCTTGAANNNAGATCGGAA
121150 TATGAGTGTGGAGTCGTTGCCAAGATAATGGGTACTTCAGATNNNAGATCGGAAG
121151 TATGAGTGTGGAGTCGTTGCTGGTTTACATTTTTAAATGGTTAAACNNNAGATCGG
121152 TATGAGTGTGGAGTCGTTGCTATATGCCTTCTTTCCTCTAATTNNNAGATCGGAAGA

121153 TATGAGTGTGGAGTCGTTGCTTCTAACTCATTTATCCAGATCCNNNAGATCGGAAGA
121154 TATGAGTGTGGAGTCGTTGCCTAACATAACAGATCACTCTGATNNNAGATCGGAAG
121155 TATGAGTGTGGAGTCGTTGCAGCTTCTTGAGTGAATGAAGNNNAGATCGGAAGAG
121156 TATGAGTGTGGAGTCGTTGCCTCACATGATTCTACAAGCTAANNNAGATCGGAAGA
121157 TATGAGTGTGGAGTCGTTGCCACACTTTGCTTTCACTTTATATNNNAGATCGGAAGA
121158 TATGAGTGTGGAGTCGTTGCATGTATATATACACACACACACANNNAGATCGGAAG
121159 TATGAGTGTGGAGTCGTTGCTCATTAGCTAATGACTAAGCTANNNAGATCGGAAGA
121160 TATGAGTGTGGAGTCGTTGCCACACAGAGAAACATAATTCTCNNNAGATCGGAAGA
121161 TATGAGTGTGGAGTCGTTGCATAAACATCTCCAACATAGGATTNNNAGATCGGAAG
121162 TATGAGTGTGGAGTCGTTGCGTGAATAGATTCTCCTATACGTTNNNAGATCGGAAG
121163 TATGAGTGTGGAGTCGTTGCTAGATGTTATTAGGAGTGCATAGNNNAGATCGGAA
121164 TATGAGTGTGGAGTCGTTGCATGGTATGAACATAGCTCACTNNNAGATCGGAAGA
121165 TATGAGTGTGGAGTCGTTGCCAAATCAGCATAACATCTGAACNNNAGATCGGAAGA
121166 TATGAGTGTGGAGTCGTTGCGAACTTATTATGGGCTGAATTCNNNAGATCGGAAGA
121167 TATGAGTGTGGAGTCGTTGCACACATAAAATTAGGACCTTTTGNNNAGATCGGAAG
121168 TATGAGTGTGGAGTCGTTGCATGCACCAAATTAACTTTTCTATNNNAGATCGGAAGA
121169 TATGAGTGTGGAGTCGTTGCGTGATACAAACACATCCCTATNNNAGATCGGAAGAG
121170 TATGAGTGTGGAGTCGTTGCTAGATAATCTTGTTCTCCAATGGNNNAGATCGGAAG
121171 TATGAGTGTGGAGTCGTTGCCTTCTTCTTATCATCCTGGAGNNNAGATCGGAAGAG
121173 TATGAGTGTGGAGTCGTTGCTCAGGAAAAAACACATACAGATANNNAGATCGGAA
121174 TATGAGTGTGGAGTCGTTGCAAAGAAAACTTCAGGTCAATATTNNNAGATCGGAAG
121175 TATGAGTGTGGAGTCGTTGCTTATAGCATCATAGAGGTATTCCNNNAGATCGGAAG
121176 TATGAGTGTGGAGTCGTTGCTGTACATCATCTGTTTAATAAGCNNNAGATCGGAAG
121177 TATGAGTGTGGAGTCGTTGCCTCTCATATCTCTTACAAGGGNNNAGATCGGAAGAG
121178 TATGAGTGTGGAGTCGTTGCCTCTCATATCTCTTACAAGGGNNNAGATCGGAAGAG
121180 TATGAGTGTGGAGTCGTTGCAAATTCCACAAGAGGCAAATAANNNAGATCGGAAG
121181 TATGAGTGTGGAGTCGTTGCCTGAATATGAATTGCAGTCTAACNNNAGATCGGAAG
121182 TATGAGTGTGGAGTCGTTGCCTAATACACATTATGCTGTTCTGNNNAGATCGGAAG
121183 TATGAGTGTGGAGTCGTTGCTCTGTCTCTTTCATAAACATACANNNAGATCGGAAGA
121184 TATGAGTGTGGAGTCGTTGCTCTGTCTCTTTCATAAACATACANNNAGATCGGAAGA
121185 TATGAGTGTGGAGTCGTTGCTCTGTCTCTTTCATAAACATACANNNAGATCGGAAGA
121186 TATGAGTGTGGAGTCGTTGCCTTCTTCTTCTTTTTCTTCTTCTNNNAGATCGGAAGAG
121187 TATGAGTGTGGAGTCGTTGCCTTCTTCCTTCTTCTTCTTTTTCNNNAGATCGGAAGAG
121188 TATGAGTGTGGAGTCGTTGCCAATGTTTAGGACCAAATCTTAANNNAGATCGGAAG
121189 TATGAGTGTGGAGTCGTTGCCTCTTCTTCTTCTTTCTTCTTTCNNNAGATCGGAAGAG
121190 TATGAGTGTGGAGTCGTTGCCTCTTCTTCTTCTTTCTTCTTTCNNNAGATCGGAAGAG
121191 TATGAGTGTGGAGTCGTTGCGAAGGGAACCAATGTATTTTGNNNAGATCGGAAGA
121192 TATGAGTGTGGAGTCGTTGCTACAAGTGACTAGTGGTTCTATNNNAGATCGGAAGA
121193 TATGAGTGTGGAGTCGTTGCCAGAATTACATTACTGGAAAAGTNNNAGATCGGAA
121194 TATGAGTGTGGAGTCGTTGCGTTAAAAAGGCCAAAGATAAAAGNNNAGATCGGAA
121195 TATGAGTGTGGAGTCGTTGCGAGACAAAGAGGACATAAGTGNNNAGATCGGAAG
121196 TATGAGTGTGGAGTCGTTGCGAATAAAGAGAGGAGAGGGANNNAGATCGGAAGA
121197 TATGAGTGTGGAGTCGTTGCGGAGACTGAAGCTTAATTCTTANNNAGATCGGAAGA

121198 TATGAGTGTGGAGTCGTTGCTTGAGATTATGGGCATGAGANNNAGATCGGAAGAG
121199 TATGAGTGTGGAGTCGTTGCGGACATGTTCACAAGCTTAANNNAGATCGGAAGAG
121200 TATGAGTGTGGAGTCGTTGCCAGGTTGAATTCTTTTTCAATGNNNAGATCGGAAG
121201 TATGAGTGTGGAGTCGTTGCTCTTTGTATAGCATTTACCCATANNNAGATCGGAAG
121202 TATGAGTGTGGAGTCGTTGCATCTAATGGAATTCACACACTTNNNAGATCGGAAGA
121203 TATGAGTGTGGAGTCGTTGCTAGAGGTATAAATTTTGTTTGGCNNNAGATCGGAA
121204 TATGAGTGTGGAGTCGTTGCTAGAGGTATAAATTTTGTTTGGCNNNAGATCGGAA
121205 TATGAGTGTGGAGTCGTTGCAGAAAAAGGAACAGAGAGAAAGNNNAGATCGGAA
121206 TATGAGTGTGGAGTCGTTGCCAGAATAGAGCTACAAAAGTAAGNNNAGATCGGAA
121207 TATGAGTGTGGAGTCGTTGCATCTGTAGAGTAGGCATGTATATNNNAGATCGGAA
121208 TATGAGTGTGGAGTCGTTGCATAAGGCTTTTGCTTATCTAATGNNNAGATCGGAAG
121209 TATGAGTGTGGAGTCGTTGCCTTCATGTTGAGAAAGCTTCNNNAGATCGGAAGAG
121210 TATGAGTGTGGAGTCGTTGCAGGCAAAGTTATAGGATTTTTTTNNNAGATCGGAAG
121211 TATGAGTGTGGAGTCGTTGCATAGGAAAAAAGAGAGATGGTAANNNAGATCGGA
121212 TATGAGTGTGGAGTCGTTGCGCAACAGCATATGACAGTATNNNAGATCGGAAGAG
121213 TATGAGTGTGGAGTCGTTGCGCTTGTATAGAAAGAAGAGGGNNNAGATCGGAAG
121214 TATGAGTGTGGAGTCGTTGCGCTTGTATAGAAAGAAGAGGGNNNAGATCGGAAG
121215 TATGAGTGTGGAGTCGTTGCGAGTTCCAGAGAGATAAAGTAATNNNAGATCGGAA
121216 TATGAGTGTGGAGTCGTTGCGAGTTCCAGAGAGATAAAGTAATNNNAGATCGGAA
121217 TATGAGTGTGGAGTCGTTGCTATTAAGAGCTGAAAGGAAAAAGNNNAGATCGGAA
121218 TATGAGTGTGGAGTCGTTGCAAATACTCATGACAATTTACAGGNNNAGATCGGAAG
121219 TATGAGTGTGGAGTCGTTGCGAGTTTTATTATGGTTTTCTGGANNNAGATCGGAAG
121220 TATGAGTGTGGAGTCGTTGCTTTCTATATTAACAGCCCAAATCNNNAGATCGGAAG
121221 TATGAGTGTGGAGTCGTTGCCGGTAACAATGATCTTTTTCATTNNNAGATCGGAAG
121222 TATGAGTGTGGAGTCGTTGCCACTCACACACACATATACAATANNNAGATCGGAAG
121223 TATGAGTGTGGAGTCGTTGCCACTCACACACACATATACAATANNNAGATCGGAAG
121224 TATGAGTGTGGAGTCGTTGCCTTAGGAAATTTTAGCACAGAAGNNNAGATCGGAA
121225 TATGAGTGTGGAGTCGTTGCCTTAGGAAATTTTAGCACAGAAGNNNAGATCGGAA
121226 TATGAGTGTGGAGTCGTTGCTTAAAGGGCGTAACTGAGATTANNNAGATCGGAAG
121227 TATGAGTGTGGAGTCGTTGCGATAAACCTAAAATTAAGTGGGGNNNAGATCGGAA
121228 TATGAGTGTGGAGTCGTTGCTTAAGGAAAAAATGGAAATAGTGCNNNAGATCGGAA
121229 TATGAGTGTGGAGTCGTTGCTGAAAACCAAAGTTAAAACCTATNNNAGATCGGAAG
121230 TATGAGTGTGGAGTCGTTGCCATTGCTTTACAATAGGAATACANNNAGATCGGAAG
121231 TATGAGTGTGGAGTCGTTGCGAGTCTAACTAACTCTGTTGCNNNAGATCGGAAGAG
121232 TATGAGTGTGGAGTCGTTGCGAGTCTAACTAACTCTGTTGCNNNAGATCGGAAGAG
121233 TATGAGTGTGGAGTCGTTGCCAAGCAGAAATATCTAGGGTTAANNNAGATCGGAA
121234 TATGAGTGTGGAGTCGTTGCCAAGCAGAAATATCTAGGGTTAANNNAGATCGGAA
121235 TATGAGTGTGGAGTCGTTGCGTAGCATCAATAACTGTGGAANNNAGATCGGAAGA
121236 TATGAGTGTGGAGTCGTTGCATTCTGTGATCTAGTCTTAGAAGNNNAGATCGGAAG
121237 TATGAGTGTGGAGTCGTTGCATACATACAACAATTTATGACCGNNNAGATCGGAAG
121238 TATGAGTGTGGAGTCGTTGCTAAGATAGCTCAAATTCACAGACNNNAGATCGGAAG
121239 TATGAGTGTGGAGTCGTTGCGAGAGATTGTTGAAATTGATGTTNNNAGATCGGAA
121240 TATGAGTGTGGAGTCGTTGCTGCATTAGGTGACAAATACAANNNAGATCGGAAGA

121241 TATGAGTGTGGAGTCGTTGCATAGTGTTAAAATGAATGATGGCNNNAGATCGGAA
121242 TATGAGTGTGGAGTCGTTGCTGCATTAGGTGACAAATACAANNNAGATCGGAAGA
121243 TATGAGTGTGGAGTCGTTGCAATGTTTTCAGTAGGAACGTTATNNNAGATCGGAAG
121244 TATGAGTGTGGAGTCGTTGCGTTGTTGTTGTTGTTGTTTTGNNNAGATCGGAAGAG
121245 TATGAGTGTGGAGTCGTTGCCCAGAAACATCAAATACTTAACANNNAGATCGGAAG
121247 TATGAGTGTGGAGTCGTTGCTCATTTTTTCTATCCTTTGAAAAANNNAGATCGGAAG
121248 TATGAGTGTGGAGTCGTTGCGAGTATGCTCTCTCTCTTCCNNNAGATCGGAAGAGC
121249 TATGAGTGTGGAGTCGTTGCTTCTAGAGAACTCCTGACAAATANNNAGATCGGAAG
121250 TATGAGTGTGGAGTCGTTGCGGAACCAGTGATCTAGAATATTTNNNAGATCGGAA
121251 TATGAGTGTGGAGTCGTTGCGGAACCAGTGATCTAGAATATTTNNNAGATCGGAA
121252 TATGAGTGTGGAGTCGTTGCTCACAGACTTACAGATTTTAAAGNNNAGATCGGAAG
121253 TATGAGTGTGGAGTCGTTGCATCTGCAGCATTAATCTAGTATANNNAGATCGGAAG
121254 TATGAGTGTGGAGTCGTTGCTATATGTGATAGATTCTCCACTCNNNAGATCGGAAG
121257 TATGAGTGTGGAGTCGTTGCGGTAAAACCAGTATTACAATCATNNNAGATCGGAAG
121258 TATGAGTGTGGAGTCGTTGCGGTAAAACCAGTATTACAATCATNNNAGATCGGAAG
121259 TATGAGTGTGGAGTCGTTGCGGTAAAACCAGTATTACAATCATNNNAGATCGGAAG
121260 TATGAGTGTGGAGTCGTTGCCACTAGGCTATTTTCTAGAGTAANNNAGATCGGAAG
121261 TATGAGTGTGGAGTCGTTGCCACTAGGCTATTTTCTAGAGTAANNNAGATCGGAAG
121262 TATGAGTGTGGAGTCGTTGCGGAAAAGTCCAATTTGCAAGNNNAGATCGGAAGAG
121263 TATGAGTGTGGAGTCGTTGCGGAAAAGTCCAATTTGCAAGNNNAGATCGGAAGAG
121264 TATGAGTGTGGAGTCGTTGCCTTCACAAATGCTGTTAACTATNNNAGATCGGAAGA
121265 TATGAGTGTGGAGTCGTTGCCTCACCTTTGCAAAATATTCTAANNNAGATCGGAAGA
121266 TATGAGTGTGGAGTCGTTGCTTTTTAGGAAAAGAGAAGTTCTGNNNAGATCGGAA
121267 TATGAGTGTGGAGTCGTTGCTTCATGTTGAAACTTAATATCCANNNAGATCGGAAG
121268 TATGAGTGTGGAGTCGTTGCGCAGATAAACAAAATGTGGTATANNNAGATCGGAA
121269 TATGAGTGTGGAGTCGTTGCTCCTCTAGTGTATTAAAATGTGGNNNAGATCGGAAG
121270 TATGAGTGTGGAGTCGTTGCCCAATGCATCAGGATTTTTAAANNNAGATCGGAAGA
121271 TATGAGTGTGGAGTCGTTGCGGTTATTAGAGGTTATTAGCCAANNNAGATCGGAA
121272 TATGAGTGTGGAGTCGTTGCTAATGAAAACTGTCAGTGTATTGNNNAGATCGGAA
121273 TATGAGTGTGGAGTCGTTGCGATCCCATTTTCTCTATACTTCCNNNAGATCGGAAGA
121274 TATGAGTGTGGAGTCGTTGCCCTCTCTCTCTCTCTCTCTCNNNAGATCGGAAGAGCA
121275 TATGAGTGTGGAGTCGTTGCCCTCTCTCTCTCTCTCTCTCNNNAGATCGGAAGAGCA
121276 TATGAGTGTGGAGTCGTTGCGTTAGTGTGTGAAGTTGAGTNNNAGATCGGAAGAG
121277 TATGAGTGTGGAGTCGTTGCTATTAACATCTTACATTGGCATGNNNAGATCGGAAG
121278 TATGAGTGTGGAGTCGTTGCAGTAATAAGAAAGGAGTTCACTTNNNAGATCGGAA
121279 TATGAGTGTGGAGTCGTTGCGAAAGGAAAAAAAAATACATGCANNNAGATCGGAA
121280 TATGAGTGTGGAGTCGTTGCCTGAAAGAGAATCTAATCTCCTTNNNAGATCGGAAG
121281 TATGAGTGTGGAGTCGTTGCCTGAAAGAGAATCTAATCTCCTTNNNAGATCGGAAG
121282 TATGAGTGTGGAGTCGTTGCATGTATATCACAAAGTAACCCTTNNNAGATCGGAAG
121283 TATGAGTGTGGAGTCGTTGCCTTCTTTACGGTCCCATTAGNNNAGATCGGAAGAGC
121284 TATGAGTGTGGAGTCGTTGCACATATATACACAGTGGTAGTTGNNNAGATCGGAA
121285 TATGAGTGTGGAGTCGTTGCATGCATATTGTATTCTGTACCTTNNNAGATCGGAAG
121286 TATGAGTGTGGAGTCGTTGCACATGTAAACAATGTAAATTCTGNNNAGATCGGAAG

121287 TATGAGTGTGGAGTCGTTGCGCCAAAGTAGAAATCAATGATAANNNAGATCGGAA

121288 TATGAGTGTGGAGTCGTTGCGAAAAATATCAAAGGAGGGTTCNNNAGATCGGAAG

121289 TATGAGTGTGGAGTCGTTGCGAAAAATATCAAAGGAGGGTTCNNNAGATCGGAAG

121291 TATGAGTGTGGAGTCGTTGCATTCCCTTAGTCTTGGTATCCNNNAGATCGGAAGAG

121292 TATGAGTGTGGAGTCGTTGCCATGTTGAGTTTTTCCTTCTGNNNAGATCGGAAGAG

121293 TATGAGTGTGGAGTCGTTGCCATGTTGAGTTTTTCCTTCTGNNNAGATCGGAAGAG

121294 TATGAGTGTGGAGTCGTTGCTTTGAGAAAGACAAGAAAAATCANNNAGATCGGAA

121295 TATGAGTGTGGAGTCGTTGCACAAGATAGATATTACCGTTATGANNNAGATCGGAA

121296 TATGAGTGTGGAGTCGTTGCGTTATGAAAGAGTTTGGAAGATTNNNAGATCGGAA

121297 TATGAGTGTGGAGTCGTTGCGTTATGAAAGAGTTTGGAAGATTNNNAGATCGGAA

121298 TATGAGTGTGGAGTCGTTGCGTGAAGTTAGCCATCTTACATNNNAGATCGGAAGAG

121299 TATGAGTGTGGAGTCGTTGCTAAAACATATCCTCTTAGCCATTNNNAGATCGGAAGA

121300 TATGAGTGTGGAGTCGTTGCTTTTTGTAAAGTTGCTAAGATGGNNNAGATCGGAAG

121301 TATGAGTGTGGAGTCGTTGCCCCAAAATTTTCAGTGTTTTTTTNNNAGATCGGAAGA

121302 TATGAGTGTGGAGTCGTTGCTATGGAAACTAGAACATGTCTCNNNAGATCGGAAGA

121303 TATGAGTGTGGAGTCGTTGCAATCAACTACAACCTTTTACAAANNNAGATCGGAAGA

121304 TATGAGTGTGGAGTCGTTGCCTGTATTATAACCATTCCTGAACNNNAGATCGGAAGA

121305 TATGAGTGTGGAGTCGTTGCCATTGACCTGACATTAAAAAAACNNNAGATCGGAAG

121306 TATGAGTGTGGAGTCGTTGCTCTTGGCTATTGAGAATAATACTNNNAGATCGGAAG

121307 TATGAGTGTGGAGTCGTTGCTCTTGGCTATTGAGAATAATACTNNNAGATCGGAAG

121308 TATGAGTGTGGAGTCGTTGCCATTTGAATGATCTATCAGTTGTNNNAGATCGGAAG

121309 TATGAGTGTGGAGTCGTTGCTTAAAGATAAGACTTCCTCTACCNNNAGATCGGAAGA

121310 TATGAGTGTGGAGTCGTTGCTTAAAGATAAGACTTCCTCTACCNNNAGATCGGAAGA

121311 TATGAGTGTGGAGTCGTTGCGGACTGTAAGAAATAAGTCTCTANNNAGATCGGAA

121312 TATGAGTGTGGAGTCGTTGCCAAGAGAGAAGAATAAGAAAGTGNNNAGATCGGAA

121313 TATGAGTGTGGAGTCGTTGCATATGGTAGGTGTTTTTCTTTTGNNNAGATCGGAAG

121314 TATGAGTGTGGAGTCGTTGCTCTTGGCTTGTATATGATTTACTNNNAGATCGGAAGA

121315 TATGAGTGTGGAGTCGTTGCCAGATTAGCTAGATACAGAGTGNNNAGATCGGAAG

121316 TATGAGTGTGGAGTCGTTGCCAAATCCATTTCACAAAAATCAANNNAGATCGGAAGA

121317 TATGAGTGTGGAGTCGTTGCTTTGAATTTTTGTTCTCTCCAAANNNAGATCGGAAGA

121318 TATGAGTGTGGAGTCGTTGCTTTGAATTTTTGTTCTCTCCAAANNNAGATCGGAAGA

121320 TATGAGTGTGGAGTCGTTGCATTTATGAAACAGTCGGGAAATTNNNAGATCGGAA

121321 TATGAGTGTGGAGTCGTTGCTTTTGCAAATAGAGGAAAAGAGNNNAGATCGGAAG

121326 TATGAGTGTGGAGTCGTTGCCTCAGTATCCCATATATCTCCTTNNNAGATCGGAAGA

121327 TATGAGTGTGGAGTCGTTGCCTCAGTATCCCATATATCTCCTTNNNAGATCGGAAGA

121328 TATGAGTGTGGAGTCGTTGCTACTCCCAGTTGTAATCATTTATNNNAGATCGGAAGA

121330 TATGAGTGTGGAGTCGTTGCTTACTCTTGCATTTTCTTTTGCNNNAGATCGGAAGAG

121331 TATGAGTGTGGAGTCGTTGCCTTCTGGAAGAGGTTAACTATTNNNAGATCGGAAGA

121332 TATGAGTGTGGAGTCGTTGCCTTCTGGAAGAGGTTAACTATTNNNAGATCGGAAGA

121333 TATGAGTGTGGAGTCGTTGCTTGAAGATAAGAGGATGTCATTTNNNAGATCGGAA

121334 TATGAGTGTGGAGTCGTTGCTTTCGGCACCAGAATATAAGNNNAGATCGGAAGAG

121336 TATGAGTGTGGAGTCGTTGCGTGTGTATGTAATTGAGAAGGANNNAGATCGGAAG

121337 TATGAGTGTGGAGTCGTTGCTAGAATAGAAATGCTCAGATTCANNNAGATCGGAAG

121338 TATGAGTGTGGAGTCGTTGCGGATACTCAGGTTTAGAATAGAANNNAGATCGGAA
121339 TATGAGTGTGGAGTCGTTGCGATGGGTAGCTTTTTAAATACATNNNAGATCGGAAG
121340 TATGAGTGTGGAGTCGTTGCTAACAGATAAATGGTTTGCAAATNNNAGATCGGAA
121341 TATGAGTGTGGAGTCGTTGCTTGGAGTACAGTGTTGTGATNNNAGATCGGAAGAG
121342 TATGAGTGTGGAGTCGTTGCTATACTAAAATGCCATCTGGTAANNNAGATCGGAAG
121343 TATGAGTGTGGAGTCGTTGCTTTTTCTTGAAGTTATAAAGCCANNNAGATCGGAAG
121344 TATGAGTGTGGAGTCGTTGCGAAGTGAGAAAGAGAGAGAATTNNNAGATCGGAA
121345 TATGAGTGTGGAGTCGTTGCGTTGAAGGAAAACAGCAATTNNNAGATCGGAAGAG
121346 TATGAGTGTGGAGTCGTTGCTAAGTAGGAGAAGAGAACAAGANNNAGATCGGAA
121347 TATGAGTGTGGAGTCGTTGCCTTTTGAGAGAAGAATCAATCACNNNAGATCGGAAG
121348 TATGAGTGTGGAGTCGTTGCTTTCTATGTGTGTATGTATCAGANNNAGATCGGAAG
121349 TATGAGTGTGGAGTCGTTGCTTTCCATTTACTTTAGAGGGAAANNNAGATCGGAAG
121350 TATGAGTGTGGAGTCGTTGCTTTCCATTTACTTTAGAGGGAAANNNAGATCGGAAG
121353 TATGAGTGTGGAGTCGTTGCGCCGAACATATTATTTATCTTAGTNNNAGATCGGAA
121354 TATGAGTGTGGAGTCGTTGCCTGAGTAGTATTCTATGGTATGTNNNAGATCGGAAG
121355 TATGAGTGTGGAGTCGTTGCAGTATTAGACACTCATTCTTTCANNNAGATCGGAAG
121356 TATGAGTGTGGAGTCGTTGCACTGAAAAGTCTTCATTTCTAAGNNNAGATCGGAAG
121357 TATGAGTGTGGAGTCGTTGCTCCCTAGAACCTTAGTCATAATNNNAGATCGGAAGA
121358 TATGAGTGTGGAGTCGTTGCCTTAGGATTACATTGATTGAGTGNNNAGATCGGAA
121359 TATGAGTGTGGAGTCGTTGCCTTAGGATGGAAACTCTCTTAAGNNNAGATCGGAAG
121360 TATGAGTGTGGAGTCGTTGCCTTAGGATGGAAACTCTCTTAAGNNNAGATCGGAAG
121361 TATGAGTGTGGAGTCGTTGCTTTGTTAATTTGAGAAAGATCCTNNNAGATCGGAAG
121362 TATGAGTGTGGAGTCGTTGCTTTTCAATAGAGATGAAGAGTTTNNNAGATCGGAA
121363 TATGAGTGTGGAGTCGTTGCACATGCTTGTATAAGTCAAAATCNNNAGATCGGAAG
121364 TATGAGTGTGGAGTCGTTGCACATGCTTGTATAAGTCAAAATCNNNAGATCGGAAG
121365 TATGAGTGTGGAGTCGTTGCATGTTAATTATCTTGATTGTGGTNNNAGATCGGAAG
121366 TATGAGTGTGGAGTCGTTGCATTACCTATTTTAGGGAACACATNNNAGATCGGAAG
121367 TATGAGTGTGGAGTCGTTGCTGAGTTTCATATTTTTGGTATGTNNNAGATCGGAAG
121368 TATGAGTGTGGAGTCGTTGCATGATTTTATCTTCTTCTGTGTGNNNAGATCGGAAG
121369 TATGAGTGTGGAGTCGTTGCAAGAAAGAAATCCTGTCATTTACNNNAGATCGGAAG
121370 TATGAGTGTGGAGTCGTTGCGTCAAATAGGGAAAGGTTTATTTNNNAGATCGGAA
121371 TATGAGTGTGGAGTCGTTGCGAGTTGAAAAGGTACAATAACTGNNNAGATCGGAA
121372 TATGAGTGTGGAGTCGTTGCATGAAGTTTTAGAGGATCTCTTTNNNAGATCGGAAG
121373 TATGAGTGTGGAGTCGTTGCATATATGTTAAGGTCCTAATCCCNNNAGATCGGAAG
121374 TATGAGTGTGGAGTCGTTGCACATCTCTTATTCTTTGGAAAATNNNAGATCGGAAG
121375 TATGAGTGTGGAGTCGTTGCGAAAACATCTCTTATTCTTTGGANNNAGATCGGAAG
121376 TATGAGTGTGGAGTCGTTGCTCTTCGTTATTTACATCTCTGTTNNNAGATCGGAAGA
121377 TATGAGTGTGGAGTCGTTGCATTTACCTAATCTGCTATTTCCANNNAGATCGGAAGA
121378 TATGAGTGTGGAGTCGTTGCTTTTCACTCAACACTATGATTTTNNNAGATCGGAAGA
121379 TATGAGTGTGGAGTCGTTGCTCTAAGAAAATATCTTTCAGGCTNNNAGATCGGAAG
121380 TATGAGTGTGGAGTCGTTGCCTACACATTAAAGAAACATCAGGNNNAGATCGGAAG
121382 TATGAGTGTGGAGTCGTTGCTATCTCATCACTTTGTGTTTTTTNNNAGATCGGAAGA
121383 TATGAGTGTGGAGTCGTTGCTGTTTTGATATCTCTGTTGAAAANNNAGATCGGAAG

121384 TATGAGTGTGGAGTCGTTGCTGAATTCAAATTACAGGTCAAACNNNAGATCGGAAG
121385 TATGAGTGTGGAGTCGTTGCTATTTCTTCTGGCATTTGATTTCNNNAGATCGGAAGA
121386 TATGAGTGTGGAGTCGTTGCCGTGACTACTTGAAGTAACTAGNNNAGATCGGAAGA
121388 TATGAGTGTGGAGTCGTTGCTAAGCTTCAGACATTAGATGATTNNNAGATCGGAAG
121389 TATGAGTGTGGAGTCGTTGCCCTATGTATAGTGGGGTTTAAAANNNAGATCGGAAG
121390 TATGAGTGTGGAGTCGTTGCGTAAGACATCCAAGTAGAGATATNNNAGATCGGAAG
121391 TATGAGTGTGGAGTCGTTGCGCGTGCGTATTAGATTTTATATANNNAGATCGGAAG
121392 TATGAGTGTGGAGTCGTTGCGCGTGCGTATTAGATTTTATATANNNAGATCGGAAG
121393 TATGAGTGTGGAGTCGTTGCGAGCAAGTCTTTTTGGTTTCNNNAGATCGGAAGAGC
121394 TATGAGTGTGGAGTCGTTGCTTTTCTGGAAACAGTCTCACNNNAGATCGGAAGAGCA
121395 TATGAGTGTGGAGTCGTTGCGATTAAAAGACAGTGTATGCTCNNNAGATCGGAAGA
121396 TATGAGTGTGGAGTCGTTGCTTTGTCCTGGAAATAAATAAACANNNAGATCGGAAG
121397 TATGAGTGTGGAGTCGTTGCTGCTAATATCAGGGGTTTATATANNNAGATCGGAAG
121398 TATGAGTGTGGAGTCGTTGCCGCTACATCTCTATCTATGATTANNNAGATCGGAAGA
121399 TATGAGTGTGGAGTCGTTGCCGCTACATCTCTATCTATGATTANNNAGATCGGAAGA
121400 TATGAGTGTGGAGTCGTTGCCTATTTGATCCTCTTACAGTGAANNNAGATCGGAAGA
121401 TATGAGTGTGGAGTCGTTGCTTTCTTAGCAAATTTCAACTACANNNAGATCGGAAGA
121402 TATGAGTGTGGAGTCGTTGCATAGCACAGTTTACTAATGTGTANNNAGATCGGAAG
121403 TATGAGTGTGGAGTCGTTGCTAGAATGTTTCCAGGTAAGTTCNNNAGATCGGAAGA
121404 TATGAGTGTGGAGTCGTTGCAGTTTTGAAAGGAGAACCTAGNNNAGATCGGAAGA
121405 TATGAGTGTGGAGTCGTTGCGGAGTAGGAAGAACAAAATTTAATNNNAGATCGGA
121406 TATGAGTGTGGAGTCGTTGCATGATAAAGCAAAAAGTAAGGTGNNNAGATCGGAA
121407 TATGAGTGTGGAGTCGTTGCTATTTTAGATTCAGTGGGTACATNNNAGATCGGAAG
121408 TATGAGTGTGGAGTCGTTGCCACTGAATAATGACATGAAGTTCNNNAGATCGGAAG
121409 TATGAGTGTGGAGTCGTTGCGTATCCCATAGAGTAAGCTTTAANNNAGATCGGAAG
121410 TATGAGTGTGGAGTCGTTGCCTGAAAGACAGAAATTGTTTATAAANNNAGATCGGA
121411 TATGAGTGTGGAGTCGTTGCAACTCTGGGTCGTGAAATAGNNNAGATCGGAAGAG
121414 TATGAGTGTGGAGTCGTTGCCTCTTTCTCTCTCTCTCTCTCNNNAGATCGGAAGAGCA
121415 TATGAGTGTGGAGTCGTTGCTCCCTCTAATATTTTTGTAGCTTNNNAGATCGGAAGA
121416 TATGAGTGTGGAGTCGTTGCCACAAAGTATGTATGTGAATGTANNNAGATCGGAAG
121417 TATGAGTGTGGAGTCGTTGCTGAGGCATTTAAAAACAATCTTANNNAGATCGGAAG
121418 TATGAGTGTGGAGTCGTTGCTTCTACAGAATTCATTTGAGAGANNNAGATCGGAAG
121419 TATGAGTGTGGAGTCGTTGCTTTTCTTTTCATGCTTACTTTTATNNNAGATCGGAAGA
121420 TATGAGTGTGGAGTCGTTGCAAGAAAACACACCAAAATGCNNNAGATCGGAAGAGC
121421 TATGAGTGTGGAGTCGTTGCATACATATACATACACACACACANNNAGATCGGAAGA
121422 TATGAGTGTGGAGTCGTTGCCAAGGTTATATCATCCTTTAGTGNNNAGATCGGAAGA
121423 TATGAGTGTGGAGTCGTTGCTCATAGTAATCCTTCTCAAAAGANNNAGATCGGAAGA
121424 TATGAGTGTGGAGTCGTTGCTTAAGACCACAAAGTTACTCTTCNNNAGATCGGAAGA
121425 TATGAGTGTGGAGTCGTTGCCTTCAAGTGTTTAGGGATCTTNNNAGATCGGAAGAG
121426 TATGAGTGTGGAGTCGTTGCCTTCACTTAAAATAAGGCAGAATNNNAGATCGGAAG
121427 TATGAGTGTGGAGTCGTTGCTATGAAGGTTTCTACGCTAATAANNNAGATCGGAAG
121428 TATGAGTGTGGAGTCGTTGCGATATCCCTCTGAGATGTACTNNNAGATCGGAAGAG
121429 TATGAGTGTGGAGTCGTTGCATATACACACACACATCCTATTTNNNAGATCGGAAGA

121431 TATGAGTGTGGAGTCGTTGCCCACATTAAAGAAACAAAAACATNNNAGATCGGAAG
121432 TATGAGTGTGGAGTCGTTGCTATCCTCTGTTACCTTGTTTATTNNNAGATCGGAAGA
121433 TATGAGTGTGGAGTCGTTGCCTAGAGATTTTGACTGAAACTTCNNNAGATCGGAAG
121434 TATGAGTGTGGAGTCGTTGCCTAGAGATTTTGACTGAAACTTCNNNAGATCGGAAG
121435 TATGAGTGTGGAGTCGTTGCGGAGAAAACACTACCACTACNNNAGATCGGAAGAG
121436 TATGAGTGTGGAGTCGTTGCGTATGTGTTTCTAGGAGGAAGNNNAGATCGGAAGA
121437 TATGAGTGTGGAGTCGTTGCGCCGATAGGGAGATAATAATAAANNNAGATCGGAA
121438 TATGAGTGTGGAGTCGTTGCATTTTGGTTTCTAGTTTTCAAGTNNNAGATCGGAAG
121439 TATGAGTGTGGAGTCGTTGCGAGAGAAAGAGAGAATCTAAGTCNNNAGATCGGAA
121440 TATGAGTGTGGAGTCGTTGCCACGTTGAGTAAAATAAGAATGANNNAGATCGGAA
121441 TATGAGTGTGGAGTCGTTGCCAAACGAATTCCTTATCTTAGAANNNAGATCGGAAG
121443 TATGAGTGTGGAGTCGTTGCAATAGAGCCCAGAGAAAGAATTNNNAGATCGGAAG
121444 TATGAGTGTGGAGTCGTTGCGTTTGTGGTCTAGTTAGTAGTATNNNAGATCGGAAG
121445 TATGAGTGTGGAGTCGTTGCTAATGTTTCTATGCTTTTGTTCANNNAGATCGGAAGA
121446 TATGAGTGTGGAGTCGTTGCGGAGTAGTATACTTTATCAGCAANNNAGATCGGAAG
121447 TATGAGTGTGGAGTCGTTGCGAATATCACACGTGTGCTTTNNNAGATCGGAAGAG
121449 TATGAGTGTGGAGTCGTTGCATTGGATCAAGGTCAGTTTCNNNAGATCGGAAGAG
121450 TATGAGTGTGGAGTCGTTGCGAGAGATTATTACCTAACTGCTTNNNAGATCGGAAG
121451 TATGAGTGTGGAGTCGTTGCTATAGAGGAGTGCACCTGTANNNAGATCGGAAGAG
121452 TATGAGTGTGGAGTCGTTGCATCTGTTGGTATTGCAAAAAAAANNNAGATCGGAAG
121453 TATGAGTGTGGAGTCGTTGCGGAGCTTACAGTGATTCTAACNNNAGATCGGAAGAG
121454 TATGAGTGTGGAGTCGTTGCGATTGGAGTGATGCATCTATAANNNAGATCGGAAG
121455 TATGAGTGTGGAGTCGTTGCTGTGTGTCTGTGCATATATATATNNNAGATCGGAAG
121456 TATGAGTGTGGAGTCGTTGCAGGAAAACTGAGTATAAGCATATNNNAGATCGGAA
121457 TATGAGTGTGGAGTCGTTGCAATTTTTCTAAGCAACCTGAATANNNAGATCGGAAG
121458 TATGAGTGTGGAGTCGTTGCTTGGAGATACTTGAATACATCTTNNNAGATCGGAAG
121459 TATGAGTGTGGAGTCGTTGCATAACAGGTATGACTTTCATCTTNNNAGATCGGAAG
121460 TATGAGTGTGGAGTCGTTGCATGAAAAAACAGTGTAGAACATTNNNAGATCGGAA
121461 TATGAGTGTGGAGTCGTTGCATATTTGGGTATTCTCAAAACACNNNAGATCGGAAG
121462 TATGAGTGTGGAGTCGTTGCTTTGTCAGTCTCCAAATCTGNNNAGATCGGAAGAGC
121463 TATGAGTGTGGAGTCGTTGCGGTCTTCACCATCATAATAAAACNNNAGATCGGAAG
121464 TATGAGTGTGGAGTCGTTGCTCTTCCAATCTTTCTATAAGTTTGNNNAGATCGGAAG
121465 TATGAGTGTGGAGTCGTTGCTTGGGGTAGGTTTAAATACTTTTNNNAGATCGGAAG
121466 TATGAGTGTGGAGTCGTTGCATTTAAGGAGAATAACAAATCCCNNNAGATCGGAAG
121467 TATGAGTGTGGAGTCGTTGCTTTACTTTTGAGGTGAATTGAATNNNAGATCGGAAG
121469 TATGAGTGTGGAGTCGTTGCGCTGCATCTTTGTTATTTACTATNNNAGATCGGAAGA
121470 TATGAGTGTGGAGTCGTTGCATAATGAGATGAAGAGACTTTTGNNNAGATCGGAA
121471 TATGAGTGTGGAGTCGTTGCTACTGTAAACTGGAATAAGCATNNNAGATCGGAAGA
121472 TATGAGTGTGGAGTCGTTGCTGCCATTTATTGCAAAGTTATTANNNAGATCGGAAG
121473 TATGAGTGTGGAGTCGTTGCTAAAAGTATGTGTTTGGAACGTNNNAGATCGGAAG
121474 TATGAGTGTGGAGTCGTTGCCTGATTTACAGAGAATCTGAGANNNAGATCGGAAG
121475 TATGAGTGTGGAGTCGTTGCGGAGAAAAGGCATACAAATTTATNNNAGATCGGAA
121476 TATGAGTGTGGAGTCGTTGCTCTTGCTCACAGTAGATATTAAANNNAGATCGGAAG

121477 TATGAGTGTGGAGTCGTTGCCATGTAGATAACAAAGAAACAGANNNAGATCGGAA
121478 TATGAGTGTGGAGTCGTTGCATACTACTCAGCCATAAAAATGANNNAGATCGGAAG
121479 TATGAGTGTGGAGTCGTTGCTAATCAATAGAAGAGTTCAAGGTNNNAGATCGGAA
121480 TATGAGTGTGGAGTCGTTGCGTTTCCCTACACCTAACATATTANNNAGATCGGAAGA
121481 TATGAGTGTGGAGTCGTTGCAAAGACATAAATGGAAACAAAGTNNNAGATCGGAA
121482 TATGAGTGTGGAGTCGTTGCCCTGAGCTAAATGATTAGATGANNNAGATCGGAAG
121483 TATGAGTGTGGAGTCGTTGCCCTGAGCTAAATGATTAGATGANNNAGATCGGAAG
121484 TATGAGTGTGGAGTCGTTGCTCCAGAAATAGGCACTAAATAATNNNAGATCGGAAG
121485 TATGAGTGTGGAGTCGTTGCGCAGCAGTATTTATTTATCTTGANNNAGATCGGAAG
121486 TATGAGTGTGGAGTCGTTGCATGGATATACGCACTGATTAAGNNNAGATCGGAAG
121487 TATGAGTGTGGAGTCGTTGCTGGGTTTCAGTTGAAGTAATTATNNNAGATCGGAAG
121490 TATGAGTGTGGAGTCGTTGCCTTTCTATTCCCTTGATTACCTNNNAGATCGGAAGAG
121491 TATGAGTGTGGAGTCGTTGCAGCGCTGACATTAAATAAATTATNNNAGATCGGAAG
121492 TATGAGTGTGGAGTCGTTGCAGCGCTGACATTAAATAAATTATNNNAGATCGGAAG
121493 TATGAGTGTGGAGTCGTTGCAATATTGAAACCTTTTTGTTCCTNNNAGATCGGAAGA
121494 TATGAGTGTGGAGTCGTTGCTTGCTGTTTCTTCAAGTATACTNNNAGATCGGAAGA
121495 TATGAGTGTGGAGTCGTTGCTTTATTCTGTTTTTGTTTCCTGGNNNAGATCGGAAGA
121496 TATGAGTGTGGAGTCGTTGCTCAAGATTTTGTTTGCACATATANNNAGATCGGAAG
121497 TATGAGTGTGGAGTCGTTGCCAAATGTCAAACAGCAATTATTANNNAGATCGGAAG
121498 TATGAGTGTGGAGTCGTTGCGCCCAATTTTAACTACATATGAGNNNAGATCGGAAG
121499 TATGAGTGTGGAGTCGTTGCTGAGATTTTTTTCTAGCACTTACNNNAGATCGGAAGA
121500 TATGAGTGTGGAGTCGTTGCTTCATACAAAAGCTTTGTACTTTNNNAGATCGGAAGA
121501 TATGAGTGTGGAGTCGTTGCAAAGCACAATCATAGCTACTATANNNAGATCGGAAG
121502 TATGAGTGTGGAGTCGTTGCTGTCTCCTCATTATTTCCTAAATNNNAGATCGGAAGA
121503 TATGAGTGTGGAGTCGTTGCCTTCTTTTCATAGCTAATTTCGTNNNAGATCGGAAGA
121504 TATGAGTGTGGAGTCGTTGCGAGAAAGCATACCCAACATTNNNAGATCGGAAGAG
121505 TATGAGTGTGGAGTCGTTGCGTAATAACCTGTTTTATAGGTCGNNNAGATCGGAAG
121506 TATGAGTGTGGAGTCGTTGCGAAAATCCTAGGGAATGATTCTANNNAGATCGGAA
121507 TATGAGTGTGGAGTCGTTGCGAAAATCCTAGGGAATGATTCTANNNAGATCGGAA
121508 TATGAGTGTGGAGTCGTTGCCCCCATTATCTTCAATCTGTTAANNNAGATCGGAAGA
121509 TATGAGTGTGGAGTCGTTGCTAAAAATGACTTTGCAACAAGATNNNAGATCGGAAG
121510 TATGAGTGTGGAGTCGTTGCTCCAGTGATTAACAAAATCATTTNNNAGATCGGAAG
121511 TATGAGTGTGGAGTCGTTGCGATCTGTTAGATTAGAACCCATANNNAGATCGGAAG
121512 TATGAGTGTGGAGTCGTTGCCAAAACACTTTGATTGACAATGNNNAGATCGGAAGA
121513 TATGAGTGTGGAGTCGTTGCATGGAGAACAGGTACACATTNNNAGATCGGAAGAG
121514 TATGAGTGTGGAGTCGTTGCTTAACCTTAATCACAATTTCTCCNNNAGATCGGAAGA
121515 TATGAGTGTGGAGTCGTTGCGAAGATTAGTCAGAGAGCACNNNAGATCGGAAGAG
121516 TATGAGTGTGGAGTCGTTGCTTATGGCATCTTTCATTCAAATTNNNAGATCGGAAGA
121517 TATGAGTGTGGAGTCGTTGCCCTGCAGAATTATGTAGATTTTTNNNAGATCGGAAG
121518 TATGAGTGTGGAGTCGTTGCGTTGTGTCCAAAATAATCTTCANNNAGATCGGAAGA
121519 TATGAGTGTGGAGTCGTTGCAATACGACACATTGGTATTAGAGNNNAGATCGGAA
121520 TATGAGTGTGGAGTCGTTGCCTTGGAGAACACGATATTGTANNNAGATCGGAAGA
121521 TATGAGTGTGGAGTCGTTGCTGTATGACACTTAAAATGTAACANNNAGATCGGAAG

121522 TATGAGTGTGGAGTCGTTGCGCTACAGAAATTACCAGATAGAANNNAGATCGGAAG
121523 TATGAGTGTGGAGTCGTTGCATGTGTACTGTTTTTAAGAACTCNNNAGATCGGAAG
121524 TATGAGTGTGGAGTCGTTGCATGTGTACTGTTTTTAAGAACTCNNNAGATCGGAAG
121525 TATGAGTGTGGAGTCGTTGCCATTATAGGTCTCTAAAGAACGTNNNAGATCGGAAG
121526 TATGAGTGTGGAGTCGTTGCAGCAACAAAAATCCAGTATTTTNNNAGATCGGAAGA
121527 TATGAGTGTGGAGTCGTTGCAATACACGAAAGTTCCAGAGNNNAGATCGGAAGAG
121528 TATGAGTGTGGAGTCGTTGCCTATCTGGCTGAGATTCAGATATNNNAGATCGGAAG
121529 TATGAGTGTGGAGTCGTTGCTGGATAGATGAGATTAGTGACTANNNAGATCGGAA
121530 TATGAGTGTGGAGTCGTTGCCAAAAATCTGACTCCTCATAGANNNAGATCGGAAGA
121531 TATGAGTGTGGAGTCGTTGCTGCAAATACTCTCAATTTATTCTNNNAGATCGGAAGA
121532 TATGAGTGTGGAGTCGTTGCCTCTTATCCTTCACATCATACTCNNNAGATCGGAAGA
121533 TATGAGTGTGGAGTCGTTGCAACTTGATATTCTTAGCTCGTTTNNNAGATCGGAAGA
121534 TATGAGTGTGGAGTCGTTGCTAAATAACCAGAATTAACATCGGNNNAGATCGGAAG
121535 TATGAGTGTGGAGTCGTTGCAATGACACCTTCCAATTCTAATNNNAGATCGGAAGAG
121536 TATGAGTGTGGAGTCGTTGCAAATAATCTAAGCAGGCTCTATTNNNAGATCGGAAG
121538 TATGAGTGTGGAGTCGTTGCACACCTGTTCTCTACTCATTNNNAGATCGGAAGAGCA
121539 TATGAGTGTGGAGTCGTTGCTATACTGCTCTTTCTCATTGTTANNNAGATCGGAAGA
121540 TATGAGTGTGGAGTCGTTGCTGGTTTCTCTTTCTCTTCTTATANNNAGATCGGAAGA
121541 TATGAGTGTGGAGTCGTTGCAGTATAGAAGAGAAACCAACTAANNNAGATCGGAA
121542 TATGAGTGTGGAGTCGTTGCAAAGTAAAGCCCATTTATTTTCANNNAGATCGGAAG
121543 TATGAGTGTGGAGTCGTTGCGATTTGGTTTTAAGGCTATTTTGNNNAGATCGGAAG
121544 TATGAGTGTGGAGTCGTTGCCATAACTCATTCTTTGGAATTCTNNNAGATCGGAAGA
121545 TATGAGTGTGGAGTCGTTGCTTCTTACCAGGTGTAAAATGTTANNNAGATCGGAAG
121546 TATGAGTGTGGAGTCGTTGCTACTTTGTAGTACCATGAGATACNNNAGATCGGAAG
121547 TATGAGTGTGGAGTCGTTGCCTGTTCCTTTCCTTCAAATCTNNNAGATCGGAAGAGC
121548 TATGAGTGTGGAGTCGTTGCACAACAGAGTTATACTACTTCTCNNNAGATCGGAAG
121549 TATGAGTGTGGAGTCGTTGCGTATTTGGTCTGTTTGGAAATTTNNNAGATCGGAAG
121550 TATGAGTGTGGAGTCGTTGCAAAAAGATAAAACTAGGAAGTGGNNNAGATCGGAA
121551 TATGAGTGTGGAGTCGTTGCCACTGGGATCTAAACTCTATGNNNAGATCGGAAGAG
121552 TATGAGTGTGGAGTCGTTGCCAATTAAACAATTACATAAAGCCANNNAGATCGGAA
121553 TATGAGTGTGGAGTCGTTGCACAGCATAGAGAGTTGAAAAANNNAGATCGGAAGA
121554 TATGAGTGTGGAGTCGTTGCTAAAAACCAACGCTTTTATTCAANNNAGATCGGAAG
121555 TATGAGTGTGGAGTCGTTGCAGGACATAGGACATATTGTATTANNNAGATCGGAA
121556 TATGAGTGTGGAGTCGTTGCTGGCCAATTATAAGATTATTCACNNNAGATCGGAAG
121557 TATGAGTGTGGAGTCGTTGCTAAGGCACTGGTTATATTTTGATNNNAGATCGGAAG
121558 TATGAGTGTGGAGTCGTTGCCTCATTTTCCAAAACAAAACAATNNNAGATCGGAAG
121559 TATGAGTGTGGAGTCGTTGCTCATCTCATGGCTTCCTAATNNNAGATCGGAAGAGC
121560 TATGAGTGTGGAGTCGTTGCGAAGAGACTGTTCTTTCCTTAATNNNAGATCGGAAG
121561 TATGAGTGTGGAGTCGTTGCGTCTCTTACCACTCTTCTCCNNNAGATCGGAAGAGCA
121562 TATGAGTGTGGAGTCGTTGCGTGTGTCTTCTCTTGCTTTCNNNAGATCGGAAGAGC
121564 TATGAGTGTGGAGTCGTTGCGAGCTTATTCTCCTTAAAAATCCNNNAGATCGGAAG
121565 TATGAGTGTGGAGTCGTTGCTATATCTTCTTTTTTTTCTGGCANNNAGATCGGAAGA
121566 TATGAGTGTGGAGTCGTTGCACTGTATTTTCCAGGAATTTTTTNNNAGATCGGAAG

121567 TATGAGTGTGGAGTCGTTGCGCAATTACATTTTCTGCATAATTNNNAGATCGGAAGA
121568 TATGAGTGTGGAGTCGTTGCCTCGGAATGTACAGTTTTATTTANNNAGATCGGAAG
121569 TATGAGTGTGGAGTCGTTGCGTAGTGGTCATGGTAGTCATANNNAGATCGGAAGA
121570 TATGAGTGTGGAGTCGTTGCAATGCAACACTTATAAAGGATTTNNNAGATCGGAAG
121571 TATGAGTGTGGAGTCGTTGCATCAAAGCTGTTCTCAACTTNNNAGATCGGAAGAGC
121572 TATGAGTGTGGAGTCGTTGCTGATCTTGTAACTTTTATACCCTNNNAGATCGGAAGA
121573 TATGAGTGTGGAGTCGTTGCGAGTATCCCGGTGTTTATTATATNNNAGATCGGAAG
121574 TATGAGTGTGGAGTCGTTGCGTACAGGTACCTCATTTCATAGNNNAGATCGGAAGA
121575 TATGAGTGTGGAGTCGTTGCACATATAAGAAGAAAAGAAAGAAAGNNNAGATCGG
121576 TATGAGTGTGGAGTCGTTGCAGGGTAATAACATATAAGAAGAAAANNNAGATCGG
121577 TATGAGTGTGGAGTCGTTGCAAAAAGAAAAGAAATATGAAGGCNNNAGATCGGAA
121578 TATGAGTGTGGAGTCGTTGCCTGAAGTATAATGGCGTGACNNNAGATCGGAAGAG
121579 TATGAGTGTGGAGTCGTTGCCTACCCCTGAACAAAAAAATATTNNNAGATCGGAAGA
121580 TATGAGTGTGGAGTCGTTGCTATCAAATCTGTATAGTTCTGCANNNAGATCGGAAGA
121581 TATGAGTGTGGAGTCGTTGCTATCAAATCTGTATAGTTCTGCANNNAGATCGGAAGA
121582 TATGAGTGTGGAGTCGTTGCGAAGGATTCATGAATTTCTGATCNNNAGATCGGAAG
121584 TATGAGTGTGGAGTCGTTGCGTGCATATACATTTGAGAAGAAANNNAGATCGGAA
121585 TATGAGTGTGGAGTCGTTGCCCAAGTATATGGCAATTTTGTTNNNAGATCGGAAGA
121586 TATGAGTGTGGAGTCGTTGCAATTTCTCTTTCTACTCCCATCNNNAGATCGGAAGAG
121587 TATGAGTGTGGAGTCGTTGCATCTGAGTTTGAGATTGAAGTTNNNAGATCGGAAGA
121588 TATGAGTGTGGAGTCGTTGCCAGTCTGAATATCCAAATCAAATNNNAGATCGGAAG
121589 TATGAGTGTGGAGTCGTTGCGTTTTATCTCTGGGAGAACAGNNNAGATCGGAAGA
121590 TATGAGTGTGGAGTCGTTGCTTAAGGAAGACACAAATAAAACCNNNAGATCGGAA
121591 TATGAGTGTGGAGTCGTTGCTGTCAGGCTAATCTAATCTTTTNNNAGATCGGAAGA
121592 TATGAGTGTGGAGTCGTTGCGGCTATTCCTTGATTATATGCTANNNAGATCGGAAGA
121593 TATGAGTGTGGAGTCGTTGCCTAGACTCTTGAGATGCCTTNNNAGATCGGAAGAGC
121594 TATGAGTGTGGAGTCGTTGCCTAGACTCTTGAGATGCCTTNNNAGATCGGAAGAGC
121595 TATGAGTGTGGAGTCGTTGCAAACAGATGAACAAGACACANNNAGATCGGAAGAG
121596 TATGAGTGTGGAGTCGTTGCAAAAGGATTTGGAGCTATTTCTTNNNAGATCGGAAG
121597 TATGAGTGTGGAGTCGTTGCTTCAGGTTCTAAAAAATATACAGANNNAGATCGGAA
121598 TATGAGTGTGGAGTCGTTGCCGTTTACTGTTCCCTATAAGTATNNNAGATCGGAAGA
121599 TATGAGTGTGGAGTCGTTGCTTAAATCATGTTCTGTGCATACANNNAGATCGGAAGA
121600 TATGAGTGTGGAGTCGTTGCCTAGTCCTCACTCTGTCCTANNNAGATCGGAAGAGCA
121601 TATGAGTGTGGAGTCGTTGCCAACTCTTTCTTCAGGTACTTNNNAGATCGGAAGAG
121602 TATGAGTGTGGAGTCGTTGCGAGAGAGTGTACTCAGTTCATNNNAGATCGGAAGA
121603 TATGAGTGTGGAGTCGTTGCTAATGACCCCTAATATTTAACCANNNAGATCGGAAGA
121604 TATGAGTGTGGAGTCGTTGCGGAAATAGAAACCTAAAAACCTTNNNAGATCGGAAG
121605 TATGAGTGTGGAGTCGTTGCGAAGTTACTGTTAACAACAACAANNNAGATCGGAAG
121606 TATGAGTGTGGAGTCGTTGCCCCCAAAAGTAACCATTTTTATTNNNAGATCGGAAGA
121607 TATGAGTGTGGAGTCGTTGCCAACTATTTGCCCCTAAAGCNNNAGATCGGAAGAGCA
121608 TATGAGTGTGGAGTCGTTGCTGTTTTTAAATAATGAGTCCCAGNNNAGATCGGAAG
121609 TATGAGTGTGGAGTCGTTGCAAACAGGAAGATTTTGTAAAGAANNNAGATCGGAA
121610 TATGAGTGTGGAGTCGTTGCTGAAATTAGGCAATTAACAACATNNNAGATCGGAAG

121611 TATGAGTGTGGAGTCGTTGCTCTGATATCTTCCCTCACATGNNNAGATCGGAAGAG
121612 TATGAGTGTGGAGTCGTTGCCCATCCTACTTCTCCATATTATTNNNAGATCGGAAGA
121613 TATGAGTGTGGAGTCGTTGCTTGAATTTTCAGCCAAGTAATAANNNAGATCGGAAG
121614 TATGAGTGTGGAGTCGTTGCTGACAATGTTATAAGATCTCTGANNNAGATCGGAAG
121615 TATGAGTGTGGAGTCGTTGCTATTTAGGCTAAATGGACACAANNNAGATCGGAAG
121616 TATGAGTGTGGAGTCGTTGCATGACAATTCAACCATATCTAGANNNAGATCGGAAG
121617 TATGAGTGTGGAGTCGTTGCATTATATGTCTGTCCTAAGCTCANNNAGATCGGAAG
121618 TATGAGTGTGGAGTCGTTGCAAAAGTAAACAAGAAACAGATCANNNAGATCGGAA
121619 TATGAGTGTGGAGTCGTTGCAAATAAAATGGAAGAAGGCTTAGNNNAGATCGGAA
121621 TATGAGTGTGGAGTCGTTGCATCTATGAACAAAATCTGCATTTNNNAGATCGGAAG
121622 TATGAGTGTGGAGTCGTTGCGTAATTTCTCTGCCTTTATTCAGNNNAGATCGGAAGA
121623 TATGAGTGTGGAGTCGTTGCAAGAGATAGCCTTGGTAGTTNNNAGATCGGAAGAG
121624 TATGAGTGTGGAGTCGTTGCTAAAAATTAGGTTTTGTGTTGGTNNNAGATCGGAAG
121625 TATGAGTGTGGAGTCGTTGCGAAATAAGTGTTTGGGAGAATGNNNAGATCGGAAG
121626 TATGAGTGTGGAGTCGTTGCATTTCTTTCTACTATTCCAGTTGNNNAGATCGGAAGA
121627 TATGAGTGTGGAGTCGTTGCTATTATCTGGAATAGCTAGTTCCNNNAGATCGGAAG
121628 TATGAGTGTGGAGTCGTTGCCGAAGTTATTGCATCTGTCTNNNAGATCGGAAGAGC
121629 TATGAGTGTGGAGTCGTTGCCTCAAAATGTAGTTGGAAACTCNNNAGATCGGAAGA
121630 TATGAGTGTGGAGTCGTTGCTTTGTGTTTCTTTTTCCTTGAATNNNAGATCGGAAGA
121631 TATGAGTGTGGAGTCGTTGCATATGTTTTCTTGTTTTCTCGTTNNNAGATCGGAAGA
121632 TATGAGTGTGGAGTCGTTGCATAAATCTGCTTCACTGGAAGNNNAGATCGGAAGAG
121633 TATGAGTGTGGAGTCGTTGCAAAGAGGCAGATAATGAAAGTTTNNNAGATCGGAA
121634 TATGAGTGTGGAGTCGTTGCTTCATAATATTCACCACATATGCNNNAGATCGGAAGA
121635 TATGAGTGTGGAGTCGTTGCGTGTAGTTTGGGAGATATATCTCNNNAGATCGGAAG
121636 TATGAGTGTGGAGTCGTTGCAAGTTCTTGGAACAAATGTTGNNNAGATCGGAAGAG
121637 TATGAGTGTGGAGTCGTTGCTCTTTATTTGCTGTGGTAATTTTNNNAGATCGGAAGA
121638 TATGAGTGTGGAGTCGTTGCGAAAGTTGGAATTTAGAGTTAGGNNNAGATCGGAA
121639 TATGAGTGTGGAGTCGTTGCGGGATGTCTAGGGATTTCTANNNAGATCGGAAGAG
121640 TATGAGTGTGGAGTCGTTGCTATTTAAACTGAAAATGTTGGCANNNAGATCGGAAG
121641 TATGAGTGTGGAGTCGTTGCCTCTGGGAGATTAACAGAATAGNNNAGATCGGAAG
121642 TATGAGTGTGGAGTCGTTGCAAAAATTTTCAAAAATGGCAAGANNNAGATCGGAAG
121643 TATGAGTGTGGAGTCGTTGCAGAGGTTAAAACTTGGGTAATAANNNAGATCGGAA
121644 TATGAGTGTGGAGTCGTTGCGCTTAAACTGGAGCATGTTTNNNAGATCGGAAGAG
121645 TATGAGTGTGGAGTCGTTGCTAATGGATTTAAGATGGGGAATANNNAGATCGGAA
121646 TATGAGTGTGGAGTCGTTGCCTCAAAGATATTCTTTCAGCAAANNNAGATCGGAAG
121647 TATGAGTGTGGAGTCGTTGCATTTTCTTTATCCAGTCCATCATNNNAGATCGGAAGA
121648 TATGAGTGTGGAGTCGTTGCAAGATAATTCACTACCAAGAGAANNNAGATCGGAAG
121649 TATGAGTGTGGAGTCGTTGCAGAAAAAAGAAAAGAAAAAAAGGANNNAGATCGG
121651 TATGAGTGTGGAGTCGTTGCGGAATAATGTGAGGAGAAAAATTNNNAGATCGGAA
121652 TATGAGTGTGGAGTCGTTGCTGACAGAGTAAAGTATATGTTCANNNAGATCGGAAG
121653 TATGAGTGTGGAGTCGTTGCACATGGAAGATAAAAGGAGATTTNNNAGATCGGAA
121654 TATGAGTGTGGAGTCGTTGCAGCTATTCGTTAGTATGATATCCNNNAGATCGGAAG
121655 TATGAGTGTGGAGTCGTTGCTACATACCCTTATACCCGTAATANNNAGATCGGAAGA

121656 TATGAGTGTGGAGTCGTTGCTGCTTTTTAAATTTATAGCATTTGANNNAGATCGGAA
121657 TATGAGTGTGGAGTCGTTGCTGCTTTTTAAATTTATAGCATTTGANNNAGATCGGAA
121659 TATGAGTGTGGAGTCGTTGCAAACAATGACCAAAAATGTTCTANNNAGATCGGAAG
121660 TATGAGTGTGGAGTCGTTGCTAAATGAAAAGGAAGAAAAAAGGNNNAGATCGGAA
121661 TATGAGTGTGGAGTCGTTGCGGCAGGAAAAAATATTTTTATCCNNNAGATCGGAAG
121662 TATGAGTGTGGAGTCGTTGCGAACTGGATTTTAAGAATGAGTTNNNAGATCGGAAG
121663 TATGAGTGTGGAGTCGTTGCGTCATGGTGGAAAAATTATTTGANNNAGATCGGAAG
121664 TATGAGTGTGGAGTCGTTGCCAACAGCACTGAAAAGTATTATTNNNAGATCGGAAG
121666 TATGAGTGTGGAGTCGTTGCATCTAAAAATTTGTGGAAGGATCNNNAGATCGGAAG
121667 TATGAGTGTGGAGTCGTTGCATTATTCACTAAAAGGAACCAGTNNNAGATCGGAAG
121668 TATGAGTGTGGAGTCGTTGCTTGTGTCTGTTTTAGTCCAGNNNAGATCGGAAGAGC
121669 TATGAGTGTGGAGTCGTTGCTTGTGTCTGTTTTAGTCCAGNNNAGATCGGAAGAGC
121670 TATGAGTGTGGAGTCGTTGCCTCTTTCCGAAAATCACATATCNNNAGATCGGAAGAG
121671 TATGAGTGTGGAGTCGTTGCTTTAAGAGCTTCACCACAGANNNAGATCGGAAGAGC
121672 TATGAGTGTGGAGTCGTTGCCAAAAATCTATTCCTTTCCTTGANNNAGATCGGAAGA
121673 TATGAGTGTGGAGTCGTTGCAGACTTTTAAAAGAATGAGCAATNNNAGATCGGAAG
121674 TATGAGTGTGGAGTCGTTGCGATACTTTCAGGAAGAGCTGNNNAGATCGGAAGAG
121675 TATGAGTGTGGAGTCGTTGCTATAACTTATTTGCATGCTGTTCNNNAGATCGGAAGA
121676 TATGAGTGTGGAGTCGTTGCGTGCAATGTAAAGACTTACAATANNNAGATCGGAAG
121677 TATGAGTGTGGAGTCGTTGCAATGAAATGAAATTAAGGATGCTNNNAGATCGGAAG
121678 TATGAGTGTGGAGTCGTTGCCAAAACATACACACACACACNNNAGATCGGAAGAGCA
121679 TATGAGTGTGGAGTCGTTGCTACTCCTGGGATTTCAGATTNNNAGATCGGAAGAGC
121680 TATGAGTGTGGAGTCGTTGCTAAGATCATAAATGTAAAACCGCNNNAGATCGGAAG
121681 TATGAGTGTGGAGTCGTTGCCAATCTCCATTGTTCACAGTANNNAGATCGGAAGAG
121683 TATGAGTGTGGAGTCGTTGCGTATGTCTTTTTCCTACTGACTTNNNAGATCGGAAGA
121684 TATGAGTGTGGAGTCGTTGCATAATGAGAAACTAACGGAGAAANNNAGATCGGAA
121686 TATGAGTGTGGAGTCGTTGCTCTGTTTGTGGCATATCTTAAANNNAGATCGGAAGA
121687 TATGAGTGTGGAGTCGTTGCTCTGTTTGTGGCATATCTTAAANNNAGATCGGAAGA
121691 TATGAGTGTGGAGTCGTTGCCTGATTCAGCTTAGAGATTTTTANNNAGATCGGAAGA
121692 TATGAGTGTGGAGTCGTTGCGGTAAAATATGTCAAGCTTTGTANNNAGATCGGAAG
121693 TATGAGTGTGGAGTCGTTGCCATGATAAGCAAAAAAGGGAAGNNNAGATCGGAAG
121694 TATGAGTGTGGAGTCGTTGCTATATGCACACACACACTTTGNNNAGATCGGAAGAG
121695 TATGAGTGTGGAGTCGTTGCAGAAATGAACCTTAAAATCACTCNNNAGATCGGAAG
121696 TATGAGTGTGGAGTCGTTGCAATTGAAGTACTTATTCTTCCCTNNNAGATCGGAAGA
121697 TATGAGTGTGGAGTCGTTGCTTTACTTTAAGTTCCAGGATACANNNAGATCGGAAGA
121698 TATGAGTGTGGAGTCGTTGCCACTCACTGCCAATTATACATANNNAGATCGGAAGAG
121699 TATGAGTGTGGAGTCGTTGCCAATATAATTGCATGCCTGTATCNNNAGATCGGAAGA
121700 TATGAGTGTGGAGTCGTTGCCAATATAATTGCATGCCTGTATCNNNAGATCGGAAGA
121701 TATGAGTGTGGAGTCGTTGCCTTTATAAATGAGAAACAGGGTTNNNAGATCGGAAG
121702 TATGAGTGTGGAGTCGTTGCTTCCTGTATTTAAGTCTACAGAANNNAGATCGGAAGA
121703 TATGAGTGTGGAGTCGTTGCCAAACATTAGGGAAGAAAGATGNNNAGATCGGAAG
121704 TATGAGTGTGGAGTCGTTGCTATGTACTCCCATTCCACCTNNNAGATCGGAAGAGCA
121705 TATGAGTGTGGAGTCGTTGCCTCTGCCATACTAAATGCTGNNNAGATCGGAAGAGCA

121706 TATGAGTGTGGAGTCGTTGCGTGAGATATGTCCAGATATTGATNNNAGATCGGAAG
121707 TATGAGTGTGGAGTCGTTGCCTACCTACTTGATTTCCTCACNNNAGATCGGAAGAGC
121709 TATGAGTGTGGAGTCGTTGCCAAGCTGTTGAATAGATATTGTANNNAGATCGGAAG
121710 TATGAGTGTGGAGTCGTTGCCAAGCTGTTGAATAGATATTGTANNNAGATCGGAAG
121711 TATGAGTGTGGAGTCGTTGCAACAGAATATTCATATAGGGGACNNNAGATCGGAAG
121712 TATGAGTGTGGAGTCGTTGCATTTGAATCTTCTAACTATTGCTNNNAGATCGGAAGA
121713 TATGAGTGTGGAGTCGTTGCCATGCAAGATGAAACATTCCNNNAGATCGGAAGAGC
121714 TATGAGTGTGGAGTCGTTGCCATGCAAGATGAAACATTCCNNNAGATCGGAAGAGC
121715 TATGAGTGTGGAGTCGTTGCACACTGTAGTAAGACTTAAAACTNNNAGATCGGAAG
121716 TATGAGTGTGGAGTCGTTGCTAAAGTAACAGATGTCATCTACANNNAGATCGGAAG
121717 TATGAGTGTGGAGTCGTTGCCTGAAAAGAAGAGTGATATCTGANNNAGATCGGAAG
121719 TATGAGTGTGGAGTCGTTGCTTTGTAGAAACACAGATTGTCATNNNAGATCGGAAG
121720 TATGAGTGTGGAGTCGTTGCCTCAGTATGTTTTACTTGAACTCNNNAGATCGGAAGA
121721 TATGAGTGTGGAGTCGTTGCTGTAAAGTGAGTTCAAATAATCANNNAGATCGGAAG
121722 TATGAGTGTGGAGTCGTTGCGGGAAAAAAAAAAACAACTCTAGNNNAGATCGGAAG
121724 TATGAGTGTGGAGTCGTTGCATGATGGATATTGCATTATTGTCNNNAGATCGGAAG
121725 TATGAGTGTGGAGTCGTTGCAGTTGTTTTTTTTTTTACTGTGGNNNAGATCGGAAGA
121726 TATGAGTGTGGAGTCGTTGCTATTGATATCTGTGCATTTGAAGNNNAGATCGGAAG
121727 TATGAGTGTGGAGTCGTTGCCTCCCCATATTAGGATAATCAATNNNAGATCGGAAGA
121728 TATGAGTGTGGAGTCGTTGCCTGAAACCTAAAAATGAGAAGAANNNAGATCGGAAG
121729 TATGAGTGTGGAGTCGTTGCGTGATATATGCATAACTACTCCTNNNAGATCGGAAGA
121730 TATGAGTGTGGAGTCGTTGCCTAAGTGAGTTTCAAATCTGATTNNNAGATCGGAAG
121731 TATGAGTGTGGAGTCGTTGCTCTGCTATGTTATAGAATCACATNNNAGATCGGAAGA
121732 TATGAGTGTGGAGTCGTTGCGAAAATAGGAAAGAAAAGAGGAANNNAGATCGGAA
121733 TATGAGTGTGGAGTCGTTGCATTATCAAGAGCCAAATCTGTANNNAGATCGGAAGA
121734 TATGAGTGTGGAGTCGTTGCTTGTAAGAGGATAATGCAAAAATNNNAGATCGGAAG
121735 TATGAGTGTGGAGTCGTTGCCCATCATAAAAAAATGTCTCAGAANNNAGATCGGAAG
121736 TATGAGTGTGGAGTCGTTGCGGTAAATGAGAGGGAATACTATGNNNAGATCGGAA
121738 TATGAGTGTGGAGTCGTTGCATCTTCCTAACCTTTCATAGTGNNNAGATCGGAAGAG
121739 TATGAGTGTGGAGTCGTTGCATTTGAAATAGCTATAAACCAGGNNNAGATCGGAAG
121740 TATGAGTGTGGAGTCGTTGCGAACAGAATTAAAGCATTTTCTGNNNAGATCGGAAG
121741 TATGAGTGTGGAGTCGTTGCACGGAATTTATATAGGAAAGTGANNNAGATCGGAAG
121742 TATGAGTGTGGAGTCGTTGCCTGTGTGAGGTAAAAAGAATATTNNNAGATCGGAAG
121743 TATGAGTGTGGAGTCGTTGCGGGGATTGGTAAAAATTAATGTTNNNAGATCGGAAG
121744 TATGAGTGTGGAGTCGTTGCTTGGATCCAGGGATTATAGAANNNAGATCGGAAGAG
121745 TATGAGTGTGGAGTCGTTGCTCTGATAATGTAACAGTTTCTCANNNAGATCGGAAGA
121746 TATGAGTGTGGAGTCGTTGCAGGGATTGTATTCCTTTATTTTGNNNAGATCGGAAGA
121747 TATGAGTGTGGAGTCGTTGCTGTTTCTTTGAGACAGACAGNNNAGATCGGAAGAGC
121748 TATGAGTGTGGAGTCGTTGCAAAGAACACTGTGATTTCCTATNNNAGATCGGAAGA
121749 TATGAGTGTGGAGTCGTTGCTATAACTTCATCACACCTTTACCNNNAGATCGGAAGAG
121750 TATGAGTGTGGAGTCGTTGCAAACTTTTTTGATCTAAGAGGAGNNNAGATCGGAAG
121751 TATGAGTGTGGAGTCGTTGCTCTCAGCATTTTGTTTCTAAAATNNNAGATCGGAAGA
121752 TATGAGTGTGGAGTCGTTGCGTAAGGATAAAGAGTAGTTGTGANNNAGATCGGAA

121753 TATGAGTGTGGAGTCGTTGCATATACACACACACATCCTATTTNNNAGATCGGAAGA
121754 TATGAGTGTGGAGTCGTTGCATATACACACACACATCCTATTTNNNAGATCGGAAGA
121755 TATGAGTGTGGAGTCGTTGCTCTGACAAGAATAATTCAGAAAANNNAGATCGGAAG
121756 TATGAGTGTGGAGTCGTTGCGGGAATTCAGAACATAACAAAAANNNAGATCGGAA
121757 TATGAGTGTGGAGTCGTTGCCAGATACCACACACACAAACNNNAGATCGGAAGAGC
121758 TATGAGTGTGGAGTCGTTGCCTTCATCAGAAACCATCCCTNNNAGATCGGAAGAGC
121759 TATGAGTGTGGAGTCGTTGCGGGATTATGGGATAATTTGTATGNNNAGATCGGAA
121760 TATGAGTGTGGAGTCGTTGCGTATTATCTTGCACTATCACAAGNNNAGATCGGAAG
121761 TATGAGTGTGGAGTCGTTGCCTGTCTATATTAGAACAAGCATGNNNAGATCGGAAG
121762 TATGAGTGTGGAGTCGTTGCACAATCAATTTAGATCAATACGGNNNAGATCGGAAG
121763 TATGAGTGTGGAGTCGTTGCAAATACTAACAAACTGAATGGAGNNNAGATCGGAA
121764 TATGAGTGTGGAGTCGTTGCATACACGAAGATGCACAAACNNNAGATCGGAAGAG
121765 TATGAGTGTGGAGTCGTTGCAAAAGTGATACCATATGTAGAGANNNAGATCGGAA
121766 TATGAGTGTGGAGTCGTTGCAGGTCAAAATAACATGCATTTATNNNAGATCGGAAG
121767 TATGAGTGTGGAGTCGTTGCCACTTTTCTCAGACAAATACTAANNNAGATCGGAAG
121768 TATGAGTGTGGAGTCGTTGCGTTAGACAAATGAACAGTGTTGNNNAGATCGGAAG
121769 TATGAGTGTGGAGTCGTTGCAATTCATGTGTCGGAAACATAATNNNAGATCGGAAG
121770 TATGAGTGTGGAGTCGTTGCGATGAAGAATAGAAACCACACTNNNAGATCGGAAG
121771 TATGAGTGTGGAGTCGTTGCACTACAGAATATTACTCAGCAATNNNAGATCGGAAG
121772 TATGAGTGTGGAGTCGTTGCCTTGTATCATTGTAAGTGTGTGNNNAGATCGGAAGA
121773 TATGAGTGTGGAGTCGTTGCAAAAATAGGCTTTTAAAACTCCANNNAGATCGGAAG
121774 TATGAGTGTGGAGTCGTTGCGCTCTTCCTCTCCCTAAAATNNNAGATCGGAAGAGCA
121775 TATGAGTGTGGAGTCGTTGCTAGATTCCCATTAGTTCTTTCCNNNAGATCGGAAGAG
121776 TATGAGTGTGGAGTCGTTGCGGTATTAAAGTGTGCTGTCTANNNAGATCGGAAGA
121777 TATGAGTGTGGAGTCGTTGCTTTGTAATTCTCTCTGAAGCTATNNNAGATCGGAAG
121778 TATGAGTGTGGAGTCGTTGCGACATTATAAGGAATTGGTTCACNNNAGATCGGAAG
121779 TATGAGTGTGGAGTCGTTGCTTATTCTTCAGTCTTTGAAAAGGNNNAGATCGGAAG
121780 TATGAGTGTGGAGTCGTTGCACAAAGATCCTTGGATACTTATTNNNAGATCGGAAG
121781 TATGAGTGTGGAGTCGTTGCTTCAGAATAATGCTTCAAGATTTNNNAGATCGGAAG
121782 TATGAGTGTGGAGTCGTTGCTCTTCCAGACCAAATAAATATCCNNNAGATCGGAAG
121783 TATGAGTGTGGAGTCGTTGCTCCCATACTACAATAATCAAGAANNNAGATCGGAAG
121784 TATGAGTGTGGAGTCGTTGCCCAAAGAAATGATAAATGTCTGANNNAGATCGGAAG
121785 TATGAGTGTGGAGTCGTTGCCTATATGCACACAAAGTAGAATGNNNAGATCGGAAG
121786 TATGAGTGTGGAGTCGTTGCTATTCTTTGTCATTTCAAGATGCNNNAGATCGGAAG
121787 TATGAGTGTGGAGTCGTTGCTTCTGTAATGCTCTGACAATATANNNAGATCGGAAG
121788 TATGAGTGTGGAGTCGTTGCGGTAATATGAGCTTGAATACAAGNNNAGATCGGAA
121789 TATGAGTGTGGAGTCGTTGCGAACGGGAATGATTAAACAAATNNNAGATCGGAAG
121790 TATGAGTGTGGAGTCGTTGCCCAACATCCTAAAATTATTTCCANNNAGATCGGAAGA
121791 TATGAGTGTGGAGTCGTTGCAAGGTGAGAGGTCCATGATANNNAGATCGGAAGAG
121792 TATGAGTGTGGAGTCGTTGCACCCTATGCATGATAATGAATATNNNAGATCGGAAG
121793 TATGAGTGTGGAGTCGTTGCACCCTATGCATGATAATGAATATNNNAGATCGGAAG
121794 TATGAGTGTGGAGTCGTTGCCATTACCTCCAACTCCATAAGNNNAGATCGGAAGAG
121795 TATGAGTGTGGAGTCGTTGCCTACTTAGTCATAGAAAGGAACANNNAGATCGGAAG

121797 TATGAGTGTGGAGTCGTTGCAATGCTGGGTAGGATCTTTTNNNAGATCGGAAGAG
121798 TATGAGTGTGGAGTCGTTGCGCATATAAGTCTGTGTGTATTTTNNNAGATCGGAAG
121799 TATGAGTGTGGAGTCGTTGCACAGGTGTGAGTTTATCTATTTANNNAGATCGGAAG
121800 TATGAGTGTGGAGTCGTTGCACTCAAAAGTCATTTTTACCAAANNNAGATCGGAAG
121801 TATGAGTGTGGAGTCGTTGCTTTTGGTTTTGGTCCTGAAGNNNAGATCGGAAGAG
121802 TATGAGTGTGGAGTCGTTGCCAGAAAGATAAATACTGCATGATNNNAGATCGGAA
121803 TATGAGTGTGGAGTCGTTGCTATGTGTGTGTGTGTGTGTANNNAGATCGGAAGAG
121804 TATGAGTGTGGAGTCGTTGCACTCACTTAATGAATATCTCAATANNNAGATCGGAA
121805 TATGAGTGTGGAGTCGTTGCGTACAAGTGGTTTTGGGTTANNNAGATCGGAAGAG
121806 TATGAGTGTGGAGTCGTTGCAATCTCCAAATTGTACCTCTCNNNAGATCGGAAGAG
121807 TATGAGTGTGGAGTCGTTGCATGTTAATGGAGTGCAATTAATTNNNAGATCGGAA
121808 TATGAGTGTGGAGTCGTTGCCATTCTCCTTGAAACAAAGATTANNNAGATCGGAAG
121809 TATGAGTGTGGAGTCGTTGCGTCAGTTAACACTTGTTTAATTGNNNAGATCGGAAG
121810 TATGAGTGTGGAGTCGTTGCGCATCTATCTTTTAGGCTTTTCNNNAGATCGGAAGA
121811 TATGAGTGTGGAGTCGTTGCTAAATCGACTTTGCCAAAAATTANNNAGATCGGAAG
121812 TATGAGTGTGGAGTCGTTGCCAAGGAAACATGTAGAATGCNNNAGATCGGAAGAG
121813 TATGAGTGTGGAGTCGTTGCGTCAAGATCTCCTAACTGATACNNNAGATCGGAAGA
121814 TATGAGTGTGGAGTCGTTGCCAAATGTGGTAGATGAGATCATNNNAGATCGGAAG
121815 TATGAGTGTGGAGTCGTTGCCTACGATTCTATCTACTCCAGANNNAGATCGGAAGA
121816 TATGAGTGTGGAGTCGTTGCGCTTTTGTAAAATGACTATCTAACNNNAGATCGGAA
121818 TATGAGTGTGGAGTCGTTGCAAGGAAATAAAAATGAAACGTTGNNNAGATCGGAA
121819 TATGAGTGTGGAGTCGTTGCTACTAATTTACACTCCCACTTACNNNAGATCGGAAGA
121820 TATGAGTGTGGAGTCGTTGCTATAAAAAAGGTGTAAGAGATGGNNNAGATCGGAA
121821 TATGAGTGTGGAGTCGTTGCCATTTCTACTGTGTGTGTCTCNNNAGATCGGAAGAG
121822 TATGAGTGTGGAGTCGTTGCTGAAAGCATTATATGGAATTCAGNNNAGATCGGAA
121823 TATGAGTGTGGAGTCGTTGCTTTCAACTACAGATTTTCAGAAGNNNAGATCGGAAG
121824 TATGAGTGTGGAGTCGTTGCGCTTTGATTCCTTTTTTTTCTTTNNNAGATCGGAAGA
121825 TATGAGTGTGGAGTCGTTGCTGTTTCCCTATTCCGAGTTCNNNAGATCGGAAGAGC
121827 TATGAGTGTGGAGTCGTTGCAATTCTTGCCTTCTCAGAAGNNNAGATCGGAAGAGC
121828 TATGAGTGTGGAGTCGTTGCTCTGACTGGCTTTACTTAATCNNNAGATCGGAAGAG
121829 TATGAGTGTGGAGTCGTTGCCACATATTTCCAGGTCAAGATANNNAGATCGGAAGA
121830 TATGAGTGTGGAGTCGTTGCTTATGAATATGCATTCCCTAAGANNNAGATCGGAAG
121831 TATGAGTGTGGAGTCGTTGCATTGGACTAAGGATAAAAGTTCTNNNAGATCGGAA
121832 TATGAGTGTGGAGTCGTTGCTAAAAATAGAAGAATGAAGCCAGNNNAGATCGGAA
121833 TATGAGTGTGGAGTCGTTGCCATGTATAACTTTTGATTCCCCNNNAGATCGGAAGA
121834 TATGAGTGTGGAGTCGTTGCCATGTATAACTTTTGATTCCCCNNNAGATCGGAAGA
121835 TATGAGTGTGGAGTCGTTGCATTCCTCTTCCTGCAAAATTTNNNAGATCGGAAGAG
121836 TATGAGTGTGGAGTCGTTGCACTGCTTCTGTAATCTCATTTNNNAGATCGGAAGAG
121837 TATGAGTGTGGAGTCGTTGCTTGATATTTCAGTGTGAATTCTTNNNAGATCGGAAG
121838 TATGAGTGTGGAGTCGTTGCGAAAGACAGATGAGATCAAATTGNNNAGATCGGAA
121839 TATGAGTGTGGAGTCGTTGCGGGGGAATAGAATTGCAAATNNNAGATCGGAAGA
121840 TATGAGTGTGGAGTCGTTGCTGTATTCTCATGGATAGGTTTTGNNNAGATCGGAAG
121841 TATGAGTGTGGAGTCGTTGCGAGGTCTCTTCTACACATTTGNNNAGATCGGAAGAG

121842 TATGAGTGTGGAGTCGTTGCATATGGGAAGAGTGCTTGATTNNNAGATCGGAAGA
121843 TATGAGTGTGGAGTCGTTGCATAGAAACAAGACCAAACCATNNNAGATCGGAAGAG
121844 TATGAGTGTGGAGTCGTTGCAATAAAAGTACCACTACCAAAACNNNAGATCGGAAG
121845 TATGAGTGTGGAGTCGTTGCGGAAGGAAATATAGGAGGTTAAGNNNAGATCGGA
121846 TATGAGTGTGGAGTCGTTGCTCTTAGCTAATATGAAACAGTGANNNAGATCGGAAG
121847 TATGAGTGTGGAGTCGTTGCCTTGTTGATAGGTATCACTGGNNNAGATCGGAAGAG
121848 TATGAGTGTGGAGTCGTTGCTAGTCTACCAATGGATGTATTGNNNAGATCGGAAGA
121849 TATGAGTGTGGAGTCGTTGCCAAAGGAAAACTGAAGTCTAATGNNNAGATCGGAA
121850 TATGAGTGTGGAGTCGTTGCAAAGGGTATGCTCATTTTAAATTNNNAGATCGGAAG
121851 TATGAGTGTGGAGTCGTTGCTGTAAAAAGATAGAATTGGTTCCNNNAGATCGGAA
121852 TATGAGTGTGGAGTCGTTGCCGCACATACACAAATATATCTAANNNAGATCGGAAG
121853 TATGAGTGTGGAGTCGTTGCCTATTTTCCAGAATGTTTGTTCANNNAGATCGGAAG
121854 TATGAGTGTGGAGTCGTTGCAATGGGAATGCTTTTTTTTTCTANNNAGATCGGAAG
121855 TATGAGTGTGGAGTCGTTGCTGATATCATAATAATACTGTGCCNNNAGATCGGAAG
121856 TATGAGTGTGGAGTCGTTGCTCAAAAGGAACAAACTTCTCTNNNAGATCGGAAGAG
121857 TATGAGTGTGGAGTCGTTGCGGGGGAATATAGAATTAAAGGAANNNAGATCGGA
121858 TATGAGTGTGGAGTCGTTGCCATAGTATGACTCCTACTTGAATNNNAGATCGGAAG
121859 TATGAGTGTGGAGTCGTTGCATTTCCTACTTCAAAGAAACAAANNNAGATCGGAAG
121860 TATGAGTGTGGAGTCGTTGCCTTATACTAATATGCACTCCGTANNNAGATCGGAAGA
121861 TATGAGTGTGGAGTCGTTGCTAACAATTACCAGGTTTTTGAAANNNAGATCGGAAG
121862 TATGAGTGTGGAGTCGTTGCAGAAAAAGATCTCCAGTATTGGNNNAGATCGGAAG
121863 TATGAGTGTGGAGTCGTTGCATAGTGGAAAGAAATCAATCCTNNNAGATCGGAAG
121864 TATGAGTGTGGAGTCGTTGCATAGTGGAAAGAAATCAATCCTNNNAGATCGGAAG
121865 TATGAGTGTGGAGTCGTTGCAAGTGTGAAGTCATATCTAAACTNNNAGATCGGAAG
121866 TATGAGTGTGGAGTCGTTGCAGAGAAAGAGAGAGAGAGATTTANNNAGATCGGA
121867 TATGAGTGTGGAGTCGTTGCTAGGATTTTCTTTCTGTTAAAGCNNNAGATCGGAAG
121868 TATGAGTGTGGAGTCGTTGCAGAAAATCTTATACTGCTCTGTANNNAGATCGGAAG
121869 TATGAGTGTGGAGTCGTTGCAGTCTTCTTTGATACCCTTTTTANNNAGATCGGAAGA
121870 TATGAGTGTGGAGTCGTTGCCATCATCTCCTCCAGATAAAAGNNNAGATCGGAAGA
121871 TATGAGTGTGGAGTCGTTGCCATGGCTAAATTTGAGAATAGATNNNAGATCGGAAG
121872 TATGAGTGTGGAGTCGTTGCAGAGGGACAGAGAGATATAAAANNNAGATCGGAA
121873 TATGAGTGTGGAGTCGTTGCATATCTAAGATTTTTTTCACGCANNNAGATCGGAAGA
121875 TATGAGTGTGGAGTCGTTGCTAGGAAATTGACCTCTTGTTTTNNNAGATCGGAAGA
121876 TATGAGTGTGGAGTCGTTGCATTTATTGGCCAAAATGGGTNNNAGATCGGAAGAG
121877 TATGAGTGTGGAGTCGTTGCTCTCAATGACTACATTTACTACANNNAGATCGGAAGA
121878 TATGAGTGTGGAGTCGTTGCTTGAATTGTAGCTCCCATAATTNNNAGATCGGAAGA
121879 TATGAGTGTGGAGTCGTTGCGGCCATACTTCTTTTAAAATGAANNNAGATCGGAAG
121880 TATGAGTGTGGAGTCGTTGCGAGGTGAGTTTAGGATATGTGNNNAGATCGGAAGA
121881 TATGAGTGTGGAGTCGTTGCAAATGTGTTAGAACAGACCTAGNNNAGATCGGAAG
121882 TATGAGTGTGGAGTCGTTGCCTTATCAGGACATTGGGTTTANNNAGATCGGAAGAG
121883 TATGAGTGTGGAGTCGTTGCAATTTTTTATTTTCCTGTGAGGANNNAGATCGGAAG
121884 TATGAGTGTGGAGTCGTTGCACTGTAGAATGAAGACAATAACANNNAGATCGGAA
121885 TATGAGTGTGGAGTCGTTGCAGTAAAAATCCTATGGGTTTAGTNNNAGATCGGAAG

121886 TATGAGTGTGGAGTCGTTGCAAGGAGAAAGAGAATTAGAACATNNNAGATCGGAA
121887 TATGAGTGTGGAGTCGTTGCCAAAGAACAAGAGGAGAAATAAANNNAGATCGGAA
121888 TATGAGTGTGGAGTCGTTGCAAGTTAGTTGAGACATTATGACANNNAGATCGGAA
121889 TATGAGTGTGGAGTCGTTGCTCAAATTCTTATTCTGCATACTGNNNAGATCGGAAG
121890 TATGAGTGTGGAGTCGTTGCACCCATGGAATTATATTACAGAANNNAGATCGGAAG
121891 TATGAGTGTGGAGTCGTTGCGTGATAAAGTTGAAAACCACTGNNNAGATCGGAAG
121892 TATGAGTGTGGAGTCGTTGCGGAAAAGAATGAGAAAAAAGGAANNNAGATCGGA
121893 TATGAGTGTGGAGTCGTTGCGACCAACAACAACTATATTAACTNNNAGATCGGAAG
121894 TATGAGTGTGGAGTCGTTGCAAATAAAAGCCCATTTATCTGTCNNNAGATCGGAAG
121895 TATGAGTGTGGAGTCGTTGCACAAAAGTAAATCTGATTGTGTTNNNAGATCGGAAG
121896 TATGAGTGTGGAGTCGTTGCAAGTAAGTTGGAGTTATTTAGGANNNAGATCGGAA
121898 TATGAGTGTGGAGTCGTTGCGGGTCACTAGAAATTCATTATTGNNNAGATCGGAAG
121899 TATGAGTGTGGAGTCGTTGCATGCATGTGTATGTTCCTTTANNNAGATCGGAAGAG
121900 TATGAGTGTGGAGTCGTTGCAAGGGATAATTATATCACACAGGNNNAGATCGGAA
121901 TATGAGTGTGGAGTCGTTGCAGAAAAATGGGACAAGGAAANNNAGATCGGAAGA
121902 TATGAGTGTGGAGTCGTTGCTTTTCATACCTACTCCTACAAATNNNAGATCGGAAGA
121903 TATGAGTGTGGAGTCGTTGCGTAAGGAACTCTCTCTGAAATGNNNAGATCGGAAGA
121904 TATGAGTGTGGAGTCGTTGCATGTATTCCATGCATAAATAGCTNNNAGATCGGAAG
121905 TATGAGTGTGGAGTCGTTGCATCTTGTGTCTCTCCTAATCATNNNAGATCGGAAGAG
121907 TATGAGTGTGGAGTCGTTGCTTCTCTCTCTCTCTCTCTCTNNNAGATCGGAAGAGCAC
121908 TATGAGTGTGGAGTCGTTGCGTGTGAATGTGCTCTACAATNNNAGATCGGAAGAG
121909 TATGAGTGTGGAGTCGTTGCCGAAATAGAAGGAAACATAGAGANNNAGATCGGAA
121910 TATGAGTGTGGAGTCGTTGCTTCTTCCAATAAGTCAACTCTTANNNAGATCGGAAGA
121911 TATGAGTGTGGAGTCGTTGCAAATGTCCAAAGTTCACAAATTNNNAGATCGGAAGA
121912 TATGAGTGTGGAGTCGTTGCACAGGGGATTAAAATAATGTCANNNAGATCGGAAG
121913 TATGAGTGTGGAGTCGTTGCCTCCTATGTCATTTATTATTGGCNNNAGATCGGAAGA
121914 TATGAGTGTGGAGTCGTTGCGTGTGTCTCCTATTTCCTTAGNNNAGATCGGAAGAG
121915 TATGAGTGTGGAGTCGTTGCGTGTAATCCAATCTAAGTTTGAANNNAGATCGGAAG
121916 TATGAGTGTGGAGTCGTTGCGACAGAGGCTAAGCTATTATTAANNNAGATCGGAA
121917 TATGAGTGTGGAGTCGTTGCGACAGAGGCTAAGCTATTATTAANNNAGATCGGAA
121918 TATGAGTGTGGAGTCGTTGCGTTCCGAATATTTCAGGAAATAANNNAGATCGGAAG
121919 TATGAGTGTGGAGTCGTTGCTCATTGTGGTTTTAATATGCATTNNNAGATCGGAAG
121920 TATGAGTGTGGAGTCGTTGCCTAAAGTAGGTCTACTAACAACANNNAGATCGGAAG
121922 TATGAGTGTGGAGTCGTTGCACAGAATTATATATGTAGCTGGGNNNAGATCGGAA
121923 TATGAGTGTGGAGTCGTTGCACAGAATTATATATGTAGCTGGGNNNAGATCGGAA
121924 TATGAGTGTGGAGTCGTTGCATGACTTCAGATGTAGGAATAACNNNAGATCGGAA
121925 TATGAGTGTGGAGTCGTTGCGGGAAGGTTGTGCTATAAAANNNAGATCGGAAGA
121926 TATGAGTGTGGAGTCGTTGCAGAGAATGAAATATTGTGGTCTANNNAGATCGGAA
121927 TATGAGTGTGGAGTCGTTGCAATGTTTGGAGAAGAGGAAATANNNAGATCGGAA
121928 TATGAGTGTGGAGTCGTTGCATCCAATAACCTAGAAGCAAATANNNAGATCGGAAG
121929 TATGAGTGTGGAGTCGTTGCCCCTGTCACTCTCTCTATATATANNNAGATCGGAAGA
121930 TATGAGTGTGGAGTCGTTGCAAATGAAATCAAAACAAAGCATGNNNAGATCGGAA
121931 TATGAGTGTGGAGTCGTTGCCCCTGTCACTCTCTCTATATATANNNAGATCGGAAGA

121932 TATGAGTGTGGAGTCGTTGCCTTGTATTTTGATTTTTCTGCTCNNNAGATCGGAAGA
121933 TATGAGTGTGGAGTCGTTGCTGTGTGTAACATTTTGTATGAAANNNAGATCGGAAG
121934 TATGAGTGTGGAGTCGTTGCGTGGTTACAATACACACTGANNNAGATCGGAAGAG
121935 TATGAGTGTGGAGTCGTTGCCCTTCTATGTGACCAAACACNNNAGATCGGAAGAGC
121936 TATGAGTGTGGAGTCGTTGCGTTGGTTTCTATGTGAGAGACNNNAGATCGGAAGA
121937 TATGAGTGTGGAGTCGTTGCGTTGGTTTCTATGTGAGAGACNNNAGATCGGAAGA
121938 TATGAGTGTGGAGTCGTTGCTCTAAAGTATTTCACCATGTGTANNNAGATCGGAAG
121939 TATGAGTGTGGAGTCGTTGCGAATCAAGACAACAGAAGTAAAANNNAGATCGGAA
121940 TATGAGTGTGGAGTCGTTGCTTATAACAGGGTCATAAGCATAANNNAGATCGGAAG
121941 TATGAGTGTGGAGTCGTTGCATACCCACAAGCTTAGATTTTATNNNAGATCGGAAG
121942 TATGAGTGTGGAGTCGTTGCTTTTAGCTAGTAAATACTGACCANNNAGATCGGAAG
121943 TATGAGTGTGGAGTCGTTGCCACCATTACTTTTCCTTTTGATNNNAGATCGGAAGAG
121944 TATGAGTGTGGAGTCGTTGCCCTCCATGAAGTTTTTTCATATTNNNAGATCGGAAGA
121945 TATGAGTGTGGAGTCGTTGCCAAAACAGAGTATAGTCCTGGNNNAGATCGGAAGA
121946 TATGAGTGTGGAGTCGTTGCCAAAACAGAGTATAGTCCTGGNNNAGATCGGAAGA
121947 TATGAGTGTGGAGTCGTTGCCTTTGTCTGGTTTTGGTATTAGNNNAGATCGGAAGA
121948 TATGAGTGTGGAGTCGTTGCTTGGGAGTATATGAGCCATTNNNAGATCGGAAGAG
121949 TATGAGTGTGGAGTCGTTGCAACAGAACACGTATTAACATCTNNNAGATCGGAAGA
121950 TATGAGTGTGGAGTCGTTGCTGAATTGAAGCAATAATTCTTCANNNAGATCGGAAG
121951 TATGAGTGTGGAGTCGTTGCCTGTGCATGTATGTATGTATATGNNNAGATCGGAAG
121952 TATGAGTGTGGAGTCGTTGCAATGAAACAGGAGATATCACTACNNNAGATCGGAA
121953 TATGAGTGTGGAGTCGTTGCTGTATACTGTCGTATTATTTCCTNNNAGATCGGAAG
121954 TATGAGTGTGGAGTCGTTGCCATCTCATGAAAACCAAAAAAGNNNAGATCGGAAG
121955 TATGAGTGTGGAGTCGTTGCCAGCTTAAACTAGTAGTACCATANNNAGATCGGAAG
121956 TATGAGTGTGGAGTCGTTGCAGTGGTATGAAAATACTGACATTNNNAGATCGGAA
121957 TATGAGTGTGGAGTCGTTGCAATAATGACAATATGTTGAAGGGNNNAGATCGGAA
121958 TATGAGTGTGGAGTCGTTGCACACACCTGAAATTAGCATAATNNNAGATCGGAAGA
121959 TATGAGTGTGGAGTCGTTGCAACTAATGGTCTCTATCTATGAGNNNAGATCGGAAG
121960 TATGAGTGTGGAGTCGTTGCAACTAATGGTCTCTATCTATGAGNNNAGATCGGAAG
121961 TATGAGTGTGGAGTCGTTGCAGCTGAAACATATATTTGATAGCNNNAGATCGGAAG
121962 TATGAGTGTGGAGTCGTTGCGAAGGGATGGTATTAAGATCTACNNNAGATCGGAA
121963 TATGAGTGTGGAGTCGTTGCCATAAATATTCTCTATATAGAAGAGCANNNAGATCG
121964 TATGAGTGTGGAGTCGTTGCACCTATCTTTCCATAAATATTCTCTANNNAGATCGGA
121965 TATGAGTGTGGAGTCGTTGCTGAAATGTTGTTTTTCTCTTTTGNNNAGATCGGAAG
121966 TATGAGTGTGGAGTCGTTGCGTGCATTTATACACACATATGTNNNAGATCGGAAGA
121967 TATGAGTGTGGAGTCGTTGCTTTCTCTTTATCAATACCTTCCANNNAGATCGGAAGA
121968 TATGAGTGTGGAGTCGTTGCTTCAAGGTACTTTATCTGCTAATNNNAGATCGGAAG
121969 TATGAGTGTGGAGTCGTTGCTGTAGCTGAGATAAAATAGATCANNNAGATCGGAA
121970 TATGAGTGTGGAGTCGTTGCCAGTTGGTTGAAAACTTTAAAAANNNAGATCGGAAG
121971 TATGAGTGTGGAGTCGTTGCATTCTTGTCATCTGTAGAATAGGNNNAGATCGGAAG
121972 TATGAGTGTGGAGTCGTTGCTATTTGAGAACTTACCATGCTAGNNNAGATCGGAAG
121973 TATGAGTGTGGAGTCGTTGCAGTCTTGAAAATAAGCTATTTTGANNNAGATCGGAA
121974 TATGAGTGTGGAGTCGTTGCAGTGGTTCATTATTGCTAGACNNNAGATCGGAAGAG

121975 TATGAGTGTGGAGTCGTTGCTAGTGGGAGGAATAAAATAATCANNNAGATCGGAA
121976 TATGAGTGTGGAGTCGTTGCAAAGCCAGAGGTAAACAATTANNNAGATCGGAAGA
121977 TATGAGTGTGGAGTCGTTGCGAGAGGGGGAGGATAAATAGNNNAGATCGGAAGA
121978 TATGAGTGTGGAGTCGTTGCCTTGTTAGAAACATGGTAATGATNNNAGATCGGAA
121979 TATGAGTGTGGAGTCGTTGCTTTTTGGACCATATCATAATTCGNNNAGATCGGAAG
121980 TATGAGTGTGGAGTCGTTGCTTCAAAGTTTTCAAACACCTTAANNNAGATCGGAAG
121981 TATGAGTGTGGAGTCGTTGCAGGAAAAAAAAATTAAGTACTGGANNNAGATCGGA
121982 TATGAGTGTGGAGTCGTTGCCACAACTAAATCCAGGACAGNNNAGATCGGAAGAG
121983 TATGAGTGTGGAGTCGTTGCGAAAGCCCTCTTTTAAGATTAAANNNAGATCGGAAG
121984 TATGAGTGTGGAGTCGTTGCTTTTGTATCATCTGTTTCAAAGGNNNAGATCGGAAG
121985 TATGAGTGTGGAGTCGTTGCTTTTGTATCATCTGTTTCAAAGGNNNAGATCGGAAG
121986 TATGAGTGTGGAGTCGTTGCAAGTATGACCACAATAGAAATTCNNNAGATCGGAAG
121987 TATGAGTGTGGAGTCGTTGCGAGGTATACTCTTTTAAAATGCTNNNAGATCGGAAG
121988 TATGAGTGTGGAGTCGTTGCCCTGAAATATGTTAGAGTAGGAANNNAGATCGGAA
121990 TATGAGTGTGGAGTCGTTGCATTTTTCTGTAAGGGAAATCATANNNAGATCGGAAG
121991 TATGAGTGTGGAGTCGTTGCTGTGATTTCTATTCCATATCAAAGNNNAGATCGGAA
121992 TATGAGTGTGGAGTCGTTGCCGCAAATAACTTTTTAATGATCCNNNAGATCGGAAG
121993 TATGAGTGTGGAGTCGTTGCTTCTTATCCTGACAGCATTATTNNNAGATCGGAAGA
121994 TATGAGTGTGGAGTCGTTGCCAAAGCTGTACAGTTATTGTTCNNNAGATCGGAAGA
121995 TATGAGTGTGGAGTCGTTGCTACATACACATATGTACTGGTGNNNAGATCGGAAGA
121996 TATGAGTGTGGAGTCGTTGCTTAAATTTTTTTGTTTGTGTGCTNNNAGATCGGAAG
121997 TATGAGTGTGGAGTCGTTGCTAGTGGCCTTTAAATTTTTTTGTNNNAGATCGGAAG
121998 TATGAGTGTGGAGTCGTTGCCAATTATGCTTCCAAGATACATTNNNAGATCGGAAG
121999 TATGAGTGTGGAGTCGTTGCATACATAGATGCAATAATGGAACNNNAGATCGGAA
122000 TATGAGTGTGGAGTCGTTGCGAAAGATGTGATGATTCAGTATCNNNAGATCGGAA
122001 TATGAGTGTGGAGTCGTTGCCTTGAGGGTAAAAATTTCATCTTNNNAGATCGGAAG
122002 TATGAGTGTGGAGTCGTTGCCTTTATTGTTTACAGATCCCAGANNNAGATCGGAAG
122003 TATGAGTGTGGAGTCGTTGCTTGTTTATTGGAGGTGTTCATNNNAGATCGGAAGA
122004 TATGAGTGTGGAGTCGTTGCCAGACTACCTTTTTCAGAGTATNNNAGATCGGAAGA
122005 TATGAGTGTGGAGTCGTTGCGAAAATAAGCATTGGTAAGGTTNNNAGATCGGAAG
122006 TATGAGTGTGGAGTCGTTGCATTGTATATACAGCTGGGGATNNNAGATCGGAAGA
122007 TATGAGTGTGGAGTCGTTGCAGAAAACTAGAGACTAGGATAGANNNAGATCGGAA
122008 TATGAGTGTGGAGTCGTTGCTTCTGAGACAAACAAAAATTTTGNNNAGATCGGAAG
122009 TATGAGTGTGGAGTCGTTGCTTGGTTCAGAGATATAAAGAGATNNNAGATCGGAA
122010 TATGAGTGTGGAGTCGTTGCTTGGTTCAGAGATATAAAGAGATNNNAGATCGGAA
122011 TATGAGTGTGGAGTCGTTGCCAGTATTTTCCTTTCATGTTCTTNNNAGATCGGAAGA
122012 TATGAGTGTGGAGTCGTTGCCTCTTATTAAAACCACAGTGAAGNNNAGATCGGAAG
122013 TATGAGTGTGGAGTCGTTGCTTATCTACAATAAAGTAGGCCATNNNAGATCGGAAG
122014 TATGAGTGTGGAGTCGTTGCGGATAAACAATGATGAAAAACACNNNAGATCGGAA
122015 TATGAGTGTGGAGTCGTTGCTACATCAATGTCTTTTTTTTCCCNNNAGATCGGAAGA
122016 TATGAGTGTGGAGTCGTTGCATAATACATATAGCAAGGGATCCNNNAGATCGGAA
122017 TATGAGTGTGGAGTCGTTGCGAATGGACATGAACAACAACNNNAGATCGGAAGAG
122018 TATGAGTGTGGAGTCGTTGCGTCTTATTTGTGGCAACAAGNNNAGATCGGAAGAG

122019 TATGAGTGTGGAGTCGTTGCTCTCTGACTTACCACATTCATNNNAGATCGGAAGAGC
122020 TATGAGTGTGGAGTCGTTGCACAAATATATGTGCATGTGTATCNNNAGATCGGAAG
122021 TATGAGTGTGGAGTCGTTGCTTCTTTAGTTTGCCAACTTTAAANNNAGATCGGAAGA
122023 TATGAGTGTGGAGTCGTTGCCAAATAAAGCTGAGTAAGTGACNNNAGATCGGAAGA
122024 TATGAGTGTGGAGTCGTTGCGAACTCTCATTCATTGATTTAGGNNNAGATCGGAAGA
122025 TATGAGTGTGGAGTCGTTGCGGAAACTCAACTATATGGACATANNNAGATCGGAAG
122026 TATGAGTGTGGAGTCGTTGCTCATTTTCTGGCATTTGTTATGNNNAGATCGGAAGA
122027 TATGAGTGTGGAGTCGTTGCTGAGATAACATATGTCACATTCCNNNAGATCGGAAGA
122028 TATGAGTGTGGAGTCGTTGCAAGGGAAACTTCTTCAGAACNNNAGATCGGAAGAG
122029 TATGAGTGTGGAGTCGTTGCGTGAACAACCTGAATGAATAAATNNNAGATCGGAAG
122030 TATGAGTGTGGAGTCGTTGCAAAAACAGAGGGAAACTAAGAAANNNAGATCGGAA
122031 TATGAGTGTGGAGTCGTTGCCAAACATTTAAGAAAATCTCTGCNNNAGATCGGAAGA
122032 TATGAGTGTGGAGTCGTTGCACTTAGAAAAGCAACAACAATAANNNAGATCGGAAG
122033 TATGAGTGTGGAGTCGTTGCCTGAGGAGAAATTAACTTGAACNNNAGATCGGAAGA
122034 TATGAGTGTGGAGTCGTTGCGGTAAGCTGTGTTATTGATTAAANNNAGATCGGAAG
122035 TATGAGTGTGGAGTCGTTGCTTAAAATCAATCCCAGTGTCATANNNAGATCGGAAGA
122036 TATGAGTGTGGAGTCGTTGCAAGATACACAACTATAAGCTGTCNNNAGATCGGAAG
122037 TATGAGTGTGGAGTCGTTGCAAGATACACAACTATAAGCTGTCNNNAGATCGGAAG
122038 TATGAGTGTGGAGTCGTTGCTTGTTTGTTTTAGAGAGAGACTANNNAGATCGGAAG
122039 TATGAGTGTGGAGTCGTTGCTTGTTTGTTTTAGAGAGAGACTANNNAGATCGGAAG
122040 TATGAGTGTGGAGTCGTTGCCTTTCACCTCTCACTTTGAANNNAGATCGGAAGAGCA
122041 TATGAGTGTGGAGTCGTTGCCCACAAATATCTATTCAACTCATNNNAGATCGGAAGA
122042 TATGAGTGTGGAGTCGTTGCATATGCATGCATACGTATTCATANNNAGATCGGAAGA
122043 TATGAGTGTGGAGTCGTTGCTATTCTCTCCCTTCTCTGAACNNNAGATCGGAAGAGCA
122044 TATGAGTGTGGAGTCGTTGCGTAGGGGTATGAAATCTGATAAANNNAGATCGGAA
122045 TATGAGTGTGGAGTCGTTGCGGGGATATCAGGCCTTATTCNNNAGATCGGAAGAG
122046 TATGAGTGTGGAGTCGTTGCGCTATCAACTACCAACTATTTTTNNNAGATCGGAAGA
122047 TATGAGTGTGGAGTCGTTGCGCTATCAACTACCAACTATTTTTNNNAGATCGGAAGA
122048 TATGAGTGTGGAGTCGTTGCCTCATGCTTATCCTCAGAACNNNAGATCGGAAGAGCA
122049 TATGAGTGTGGAGTCGTTGCGACATCCACAAAGAACAAAGNNNAGATCGGAAGAG
122050 TATGAGTGTGGAGTCGTTGCATTCTCCACAAATGGTAACTTANNNAGATCGGAAGA
122051 TATGAGTGTGGAGTCGTTGCATTCTCCACAAATGGTAACTTANNNAGATCGGAAGA
122052 TATGAGTGTGGAGTCGTTGCATTCTCCACAAATGGTAACTTANNNAGATCGGAAGA
122053 TATGAGTGTGGAGTCGTTGCCAACTCATATCACACATTGATAGNNNAGATCGGAAGA
122054 TATGAGTGTGGAGTCGTTGCATCCATATATGTAGACAGTTAGCNNNAGATCGGAAG
122055 TATGAGTGTGGAGTCGTTGCGACTTTTCCACAAGATGATTTTTNNNAGATCGGAAGA
122056 TATGAGTGTGGAGTCGTTGCCCTAAGAACATTTATTTACGTCTNNNAGATCGGAAGA
122057 TATGAGTGTGGAGTCGTTGCCAAAAAGTATTGTCCTTAATCCANNNAGATCGGAAGA
122058 TATGAGTGTGGAGTCGTTGCCCAAAAATCTGAACCACATGNNNAGATCGGAAGAGC
122059 TATGAGTGTGGAGTCGTTGCCAAATGTGTTCAGACAAAATTTTNNNAGATCGGAAG
122060 TATGAGTGTGGAGTCGTTGCAACTCTTCTCTGGAGACAAATNNNAGATCGGAAGAG
122061 TATGAGTGTGGAGTCGTTGCACAACTTTGTCAACTTTGTAATANNNAGATCGGAAGA
122062 TATGAGTGTGGAGTCGTTGCACAACTTTGTCAACTTTGTAATANNNAGATCGGAAGA

122063 TATGAGTGTGGAGTCGTTGCCAACTTTGCTTATCTTTTTTTGTNNNAGATCGGAAGA
122064 TATGAGTGTGGAGTCGTTGCCAACTTTGCTTATCTTTTTTTGTNNNAGATCGGAAGA
122065 TATGAGTGTGGAGTCGTTGCGAACCTTTGTAAACAAGTGTTATNNNAGATCGGAAG
122066 TATGAGTGTGGAGTCGTTGCTTGACGATCCTATATCAAGTATCNNNAGATCGGAAG
122067 TATGAGTGTGGAGTCGTTGCTCTCTTTATTCCAGGAATAGAAANNNAGATCGGAAG
122068 TATGAGTGTGGAGTCGTTGCAATGTAAAGCCTCTATTTTTGAGNNNAGATCGGAAG
122069 TATGAGTGTGGAGTCGTTGCAATGTAAAGCCTCTATTTTTGAGNNNAGATCGGAAG
122070 TATGAGTGTGGAGTCGTTGCTTTCATAAATCATTAATGAGCCATNNNAGATCGGAA
122071 TATGAGTGTGGAGTCGTTGCTGACAACATTCTTCATCACCNNNAGATCGGAAGAGC
122072 TATGAGTGTGGAGTCGTTGCACACATTGACAATACCATAGATANNNAGATCGGAAG
122074 TATGAGTGTGGAGTCGTTGCGCATTACAATTCTTTAAACCAACNNNAGATCGGAAGA
122075 TATGAGTGTGGAGTCGTTGCATGTAGAACTTATAGTTGCTCTTNNNAGATCGGAAG
122076 TATGAGTGTGGAGTCGTTGCGGAAGAAAAGCAGAGAATATAAANNNAGATCGGAA
122077 TATGAGTGTGGAGTCGTTGCTCCTCCCTTCATTGATATATTTTNNNAGATCGGAAGA
122078 TATGAGTGTGGAGTCGTTGCCCAAATCCCAGAAAACTTTATTANNNAGATCGGAAG
122079 TATGAGTGTGGAGTCGTTGCTTATCCAAATAAGTGTGCTGTATNNNAGATCGGAAG
122080 TATGAGTGTGGAGTCGTTGCCGAAGTCTAAAAGCATTGTCNNNAGATCGGAAGAG
122081 TATGAGTGTGGAGTCGTTGCAATGTGCATCCCTTCTACAANNNAGATCGGAAGAGC
122082 TATGAGTGTGGAGTCGTTGCAGGAAACATGGGAACTTTTATATNNNAGATCGGAA
122083 TATGAGTGTGGAGTCGTTGCTGCAAGAAATAGAAAACATTTTGNNNAGATCGGAA
122084 TATGAGTGTGGAGTCGTTGCTACATTTGAATCCTGACTCTATCNNNAGATCGGAAGA
122085 TATGAGTGTGGAGTCGTTGCGAAGAGTTTTGAAACACTGAACNNNAGATCGGAAG
122086 TATGAGTGTGGAGTCGTTGCCTAAGGTTAGGTTACCAAATGANNNAGATCGGAAG
122087 TATGAGTGTGGAGTCGTTGCTGAATGAATAAGAATGAGAATGCNNNAGATCGGAA
122088 TATGAGTGTGGAGTCGTTGCCTTACAAGCCTGAAAAACAGATANNNAGATCGGAAG
122089 TATGAGTGTGGAGTCGTTGCGAAAATATACCCACAAAGGCTNNNAGATCGGAAGAG
122090 TATGAGTGTGGAGTCGTTGCTTCACCTAGTGAGTCCTTCTNNNAGATCGGAAGAGC
122091 TATGAGTGTGGAGTCGTTGCAATGACTGACATGTATCTTTGATNNNAGATCGGAAG
122092 TATGAGTGTGGAGTCGTTGCAACTTTTGCAACCAATAGATATTNNNAGATCGGAAG
122093 TATGAGTGTGGAGTCGTTGCGTCTTCTGTTATAATGCTAGGTANNNAGATCGGAAG
122094 TATGAGTGTGGAGTCGTTGCTTTTAGACACATTGCAATCAAANNNAGATCGGAAGA
122095 TATGAGTGTGGAGTCGTTGCTAGCAAGAAACTAGAGATTGTCNNNAGATCGGAAG
122096 TATGAGTGTGGAGTCGTTGCCACAGAGGAAGAGGTAACTANNNAGATCGGAAGAG
122097 TATGAGTGTGGAGTCGTTGCTTTTATATTTTAGCAGAGACCTGNNNAGATCGGAAG
122098 TATGAGTGTGGAGTCGTTGCTTTCCCCCTATGTAAATGATATTNNNAGATCGGAAGA
122099 TATGAGTGTGGAGTCGTTGCACAGAACAACAGAGATAGTATACNNNAGATCGGAA
122100 TATGAGTGTGGAGTCGTTGCTTTAAATTTCCCCATCATACTGANNNAGATCGGAAGA
122101 TATGAGTGTGGAGTCGTTGCTACTTGGGGAATGCTTTTTGNNNAGATCGGAAGAG
122102 TATGAGTGTGGAGTCGTTGCTTTAGAAAAGAGTAAAAAGGAGCNNNAGATCGGAA
122103 TATGAGTGTGGAGTCGTTGCTTTCAGTGAAGTATAGAACACTTNNNAGATCGGAAG
122104 TATGAGTGTGGAGTCGTTGCCTAACTGACAGAGTGATTTCANNNAGATCGGAAGAG
122105 TATGAGTGTGGAGTCGTTGCTTAACTTAAAACTGCTCTGAAAANNNAGATCGGAAG
122106 TATGAGTGTGGAGTCGTTGCGAAACCAGTCCAGGATTTTAGNNNAGATCGGAAGA

122107 TATGAGTGTGGAGTCGTTGCGAAACCAGTCCAGGATTTTAGNNNAGATCGGAAGA
122108 TATGAGTGTGGAGTCGTTGCATATACAGAAATTCAGTTGTTCCNNNAGATCGGAAG
122109 TATGAGTGTGGAGTCGTTGCTGAAAACTATCACTGTCTATTCANNNAGATCGGAAG
122110 TATGAGTGTGGAGTCGTTGCTAGGAGGCTATCTTTTTGATCTANNNAGATCGGAAG
122112 TATGAGTGTGGAGTCGTTGCGGATGCAAATGGTACATTTATTNNNAGATCGGAAG
122113 TATGAGTGTGGAGTCGTTGCGCATCAAGAAATTGTTCAGAATANNNAGATCGGAAG
122114 TATGAGTGTGGAGTCGTTGCCTTTCATGCATTTTTCTTTATGGNNNAGATCGGAAGA
122115 TATGAGTGTGGAGTCGTTGCCCAACCTCTAGAACTGTAATAAANNNAGATCGGAAG
122116 TATGAGTGTGGAGTCGTTGCGCATTAGCTAAAATGTATTCAGANNNAGATCGGAAG
122117 TATGAGTGTGGAGTCGTTGCAAGGGTTGTCATCTTCTATATAANNNAGATCGGAAG
122118 TATGAGTGTGGAGTCGTTGCCAGCCTAATATGTCAGTATAGATNNNAGATCGGAAG
122119 TATGAGTGTGGAGTCGTTGCATGTTATCAGAATCTGAACACTANNNAGATCGGAAG
122120 TATGAGTGTGGAGTCGTTGCCTTTGACTGTAAGATCTGTGGNNNAGATCGGAAGA
122121 TATGAGTGTGGAGTCGTTGCTGGAATAAGCCAATATACACATANNNAGATCGGAAG
122124 TATGAGTGTGGAGTCGTTGCCATTCAGCATGTTTACATTTCANNNAGATCGGAAGA
122125 TATGAGTGTGGAGTCGTTGCAAAAGTCATAGCATTCTTGAAACNNNAGATCGGAAG
122127 TATGAGTGTGGAGTCGTTGCATAATGACTTCAGAGAAAAGGAANNNAGATCGGAA
122128 TATGAGTGTGGAGTCGTTGCCTATGCAATACACTTAAAGGATTNNNAGATCGGAAG
122129 TATGAGTGTGGAGTCGTTGCTTTATTTCTTTTCTGAATGGCTTNNNAGATCGGAAGA
122131 TATGAGTGTGGAGTCGTTGCAATCTATTACAGGTAACACCATCNNNAGATCGGAAG
122132 TATGAGTGTGGAGTCGTTGCCATTACACTCCAGAATGTACATNNNAGATCGGAAGA
122133 TATGAGTGTGGAGTCGTTGCTTCTTTGCTGGATAGAAAATAACNNNAGATCGGAAG
122134 TATGAGTGTGGAGTCGTTGCTCATTTTCCTTTATGATCCATTGNNNAGATCGGAAGA
122135 TATGAGTGTGGAGTCGTTGCTCATTTTCCTTTATGATCCATTGNNNAGATCGGAAGA
122136 TATGAGTGTGGAGTCGTTGCCTGTCTCTGTATTGGAACTTTNNNAGATCGGAAGAG
122137 TATGAGTGTGGAGTCGTTGCCTGTCTCTGTATTGGAACTTTNNNAGATCGGAAGAG
122138 TATGAGTGTGGAGTCGTTGCTTGCTACCACTTTTCTGATATTANNNAGATCGGAAGA
122139 TATGAGTGTGGAGTCGTTGCATAGAATAGAACACAAAGACTTTNNNAGATCGGAAG
122140 TATGAGTGTGGAGTCGTTGCCAAGGGGATACTAATCACAAANNNAGATCGGAAGA
122141 TATGAGTGTGGAGTCGTTGCCATGGTCACATAGGTTAATTAACNNNAGATCGGAAG
122142 TATGAGTGTGGAGTCGTTGCCCCACTTCTTAGTTTCAAATTTANNNAGATCGGAAGA
122143 TATGAGTGTGGAGTCGTTGCAGTCTTTGCTGGAATATAAGATANNNAGATCGGAAG
122144 TATGAGTGTGGAGTCGTTGCATCTCTCTCCCTTCTTACAGNNNAGATCGGAAGAGCA
122146 TATGAGTGTGGAGTCGTTGCAAACTGAGTGTTACAAGAAGTAANNNAGATCGGAA
122149 TATGAGTGTGGAGTCGTTGCAAAAAGCTAAACATAGTTACCACNNNAGATCGGAAG
122150 TATGAGTGTGGAGTCGTTGCAGAAGGGAACCACCTAAATTTANNNAGATCGGAAG
122151 TATGAGTGTGGAGTCGTTGCGACTACAATCCAGAATTTCTGANNNAGATCGGAAGA
122152 TATGAGTGTGGAGTCGTTGCTTAATTCTTTTGCATGTGGATAANNNAGATCGGAAG
122153 TATGAGTGTGGAGTCGTTGCTATCATCACCTTCTCCTACCTNNNAGATCGGAAGAGC
122154 TATGAGTGTGGAGTCGTTGCTCTCTCTTTCTTTCTGTTCTATTNNNAGATCGGAAGAG
122155 TATGAGTGTGGAGTCGTTGCGTTTATAAACATATTGCTGTGGTNNNAGATCGGAAG
122156 TATGAGTGTGGAGTCGTTGCTGCATTAAGCTTAAAGAAGAAATNNNAGATCGGAA
122157 TATGAGTGTGGAGTCGTTGCCACTGATTCTTTTCCAATAAGTTNNNAGATCGGAAGA

122158 TATGAGTGTGGAGTCGTTGCGTCTTCTTGGAACACAATTTCNNNAGATCGGAAGAG
122159 TATGAGTGTGGAGTCGTTGCGGTAAGTGGTATCAAATGTATTAGNNNAGATCGGA
122160 TATGAGTGTGGAGTCGTTGCATTCAAGACAATACTGTAAATCCNNNAGATCGGAAG
122161 TATGAGTGTGGAGTCGTTGCAAACACACAGAAAGCAATATAATNNNAGATCGGAA
122162 TATGAGTGTGGAGTCGTTGCGTTTTTACCCTAAGTGATGGGNNNAGATCGGAAGA
122163 TATGAGTGTGGAGTCGTTGCAGAAAATTACAGAAGAACATGAGNNNAGATCGGAA
122164 TATGAGTGTGGAGTCGTTGCAAGAAACTCCAAGTAGATAAGTTNNNAGATCGGAA
122165 TATGAGTGTGGAGTCGTTGCGGGAAAATAAAAAAGCAAATTGANNNAGATCGGAA
122166 TATGAGTGTGGAGTCGTTGCCCTCATGTATATTTTCTTGGAAGNNNAGATCGGAAG
122167 TATGAGTGTGGAGTCGTTGCGAGAGACCTCATGTATATTTTCTNNNAGATCGGAAG
122168 TATGAGTGTGGAGTCGTTGCACATTTCATTCCAAACTTAAACTNNNAGATCGGAAGA
122169 TATGAGTGTGGAGTCGTTGCACTAAGGGTGAGATAAAGAAAGNNNAGATCGGAA
122170 TATGAGTGTGGAGTCGTTGCCTATTCCCTAACTACAAAGGAAGNNNAGATCGGAAG
122171 TATGAGTGTGGAGTCGTTGCTACATTTCTGTTTGTTTTTCTCTNNNAGATCGGAAGA
122172 TATGAGTGTGGAGTCGTTGCTATGAGAATGTCAGACATATCACNNNAGATCGGAAG
122173 TATGAGTGTGGAGTCGTTGCCAATAGTTCACAGGATTAAAGAGNNNAGATCGGAA
122174 TATGAGTGTGGAGTCGTTGCTTTGAGAGAAAGAGCAAGTATTNNNAGATCGGAAG
122175 TATGAGTGTGGAGTCGTTGCGATTTGAAATGCCTATGAGATGNNNAGATCGGAAG
122176 TATGAGTGTGGAGTCGTTGCCATATACTAGACTTCGAATCACTNNNAGATCGGAAG
122177 TATGAGTGTGGAGTCGTTGCTTTCTTGAGAGAGGACTTTTTAGNNNAGATCGGAAG
122178 TATGAGTGTGGAGTCGTTGCGTAGGAATACAGCTAACCATGNNNAGATCGGAAGA
122179 TATGAGTGTGGAGTCGTTGCTATCACTTTCCTTACTGTTAACANNNAGATCGGAAGA
122180 TATGAGTGTGGAGTCGTTGCAGTATAAAAAAACAATTTGCAGCNNNAGATCGGAA
122181 TATGAGTGTGGAGTCGTTGCCTGAGTAAGGAAAGGTGATTTNNNAGATCGGAAGA
122182 TATGAGTGTGGAGTCGTTGCGTGCATACAGTAGCAAATATTTANNNAGATCGGAAG
122183 TATGAGTGTGGAGTCGTTGCACTTTCCTCCTTTTAATACTAGGNNNAGATCGGAAGA
122184 TATGAGTGTGGAGTCGTTGCTTTTTCATTTTTTTTCTCTTGGCNNNAGATCGGAAGA
122185 TATGAGTGTGGAGTCGTTGCGCTCACTTCCAAAGAATAAAGNNNAGATCGGAAGA
122186 TATGAGTGTGGAGTCGTTGCTGTCATCTTTTTACTCAGCTATANNNAGATCGGAAGA
122187 TATGAGTGTGGAGTCGTTGCAGTGTATAAACTGAGTTGTTCTTNNNAGATCGGAAG
122188 TATGAGTGTGGAGTCGTTGCTATATACCAACACTAGGAGATCTNNNAGATCGGAAG
122191 TATGAGTGTGGAGTCGTTGCGGCAATTAAAGTGTTTAGTAAGANNNAGATCGGAA
122192 TATGAGTGTGGAGTCGTTGCGGCAATTAAAGTGTTTAGTAAGANNNAGATCGGAA
122193 TATGAGTGTGGAGTCGTTGCAAGAGAGAGACAGAGAAGAGNNNAGATCGGAAGA
122197 TATGAGTGTGGAGTCGTTGCGCAATTATGCATAACATAAGAGANNNAGATCGGAA
122198 TATGAGTGTGGAGTCGTTGCACAAATAATGTTACATCACACAANNNAGATCGGAAG
122199 TATGAGTGTGGAGTCGTTGCTTAGAGCCAGGGACTTATTCNNNAGATCGGAAGAG
122200 TATGAGTGTGGAGTCGTTGCTTTTAAGAAGTTAAAGTTGCTCCNNNAGATCGGAAG
122201 TATGAGTGTGGAGTCGTTGCGGTGAATCTGGTTTAAAAGTATTNNNAGATCGGAA
122202 TATGAGTGTGGAGTCGTTGCCTTGGTCACTGAAAAATATCTCNNNAGATCGGAAGA
122204 TATGAGTGTGGAGTCGTTGCTATGATTGTTTACCACTTTCTTGNNNAGATCGGAAGA
122205 TATGAGTGTGGAGTCGTTGCTTCTTTTTCTAAAGTTTGGATCCNNNAGATCGGAAGA
122206 TATGAGTGTGGAGTCGTTGCATGCAAACAGTACTATGCTATTANNNAGATCGGAAG

122207 TATGAGTGTGGAGTCGTTGCGATACTATCTGATTCACAGGATTNNNAGATCGGAAG
122209 TATGAGTGTGGAGTCGTTGCACCAGATATTGTACAAGAGTATCNNNAGATCGGAAG
122210 TATGAGTGTGGAGTCGTTGCAACCTTATACTTTCAAAATCTGGNNNAGATCGGAAG
122211 TATGAGTGTGGAGTCGTTGCAAGATTAACACCAAACCATAGTANNNAGATCGGAAG
122213 TATGAGTGTGGAGTCGTTGCTATAAAACATTGTTTGCATGAGANNNAGATCGGAAG
122214 TATGAGTGTGGAGTCGTTGCGACTAGCAAAATCCAGACTAGNNNAGATCGGAAGA
122215 TATGAGTGTGGAGTCGTTGCTCTGCAATTTTATTACTGTGTGNNNAGATCGGAAGA
122216 TATGAGTGTGGAGTCGTTGCTTTGAGAACTATTGTCTTATGGANNNAGATCGGAAG
122217 TATGAGTGTGGAGTCGTTGCGATAGTCTATTGCTTACCAGTAGNNNAGATCGGAAG
122218 TATGAGTGTGGAGTCGTTGCAAAGAAGAGCTTCCTGAATATATNNNAGATCGGAA
122219 TATGAGTGTGGAGTCGTTGCTTCAGTCTTGGATAAAACTAGAANNNAGATCGGAAG
122220 TATGAGTGTGGAGTCGTTGCGTTCCTCAATGAATAGTTGTTAGNNNAGATCGGAAG
122221 TATGAGTGTGGAGTCGTTGCGTCTGGATAACTGAGAATTTTANNNAGATCGGAAG
122222 TATGAGTGTGGAGTCGTTGCTCTTTTTCTTAAAATCAAGGTTGNNNAGATCGGAAG
122223 TATGAGTGTGGAGTCGTTGCAATTTGTCATAAAGAATTGGCTCNNNAGATCGGAAG
122225 TATGAGTGTGGAGTCGTTGCCTTGGACATATTCTGCATAATGNNNAGATCGGAAGA
122226 TATGAGTGTGGAGTCGTTGCCTTGGACATATTCTGCATAATGNNNAGATCGGAAGA
122227 TATGAGTGTGGAGTCGTTGCATTGCAAATTTGATATTTCAGGANNNAGATCGGAAG
122228 TATGAGTGTGGAGTCGTTGCCAATTGAGGTCTCCAAAGACNNNAGATCGGAAGAG
122229 TATGAGTGTGGAGTCGTTGCATGTATATTGGCATGAAACTTTCNNNAGATCGGAAG
122230 TATGAGTGTGGAGTCGTTGCGATAGATACACAGATGCATAGATNNNAGATCGGAA
122231 TATGAGTGTGGAGTCGTTGCGATAGATACACAGATGCATAGATNNNAGATCGGAA
122232 TATGAGTGTGGAGTCGTTGCATTAATCTGCTAGACGTAAACTANNNAGATCGGAAG
122233 TATGAGTGTGGAGTCGTTGCTCCATGAATGAGTACGTACATNNNAGATCGGAAGAG
122234 TATGAGTGTGGAGTCGTTGCAAGTATGTATGTATGTTGGGGNNNAGATCGGAAGA
122235 TATGAGTGTGGAGTCGTTGCGATGAAAGCAGGACTATTTTATCNNNAGATCGGAAG
122236 TATGAGTGTGGAGTCGTTGCAATCAAAGTTTTTGTTGTTGTTTNNNAGATCGGAAG
122237 TATGAGTGTGGAGTCGTTGCAATCAAAGTTTTTGTTGTTGTTTNNNAGATCGGAAG
122238 TATGAGTGTGGAGTCGTTGCATCAATAGTCTAGAAAGTGCTTANNNAGATCGGAAG
122239 TATGAGTGTGGAGTCGTTGCTCATAACCTAGGACTATGTTTTTNNNAGATCGGAAG
122240 TATGAGTGTGGAGTCGTTGCCCTAGAGACTTGAAAGGACTTNNNAGATCGGAAGA
122241 TATGAGTGTGGAGTCGTTGCAAAGGAATTTTCAGTTTCAGATTNNNAGATCGGAAG
122242 TATGAGTGTGGAGTCGTTGCCAGAGCTATAGTGTCTACATTAANNNAGATCGGAAG
122243 TATGAGTGTGGAGTCGTTGCGACTATCTGTTTTCCACTCTTANNNAGATCGGAAGAG
122244 TATGAGTGTGGAGTCGTTGCGTGAAGGCAGTGTATTCTAANNNAGATCGGAAGAG
122245 TATGAGTGTGGAGTCGTTGCGTGAAGGCAGTGTATTCTAANNNAGATCGGAAGAG
122247 TATGAGTGTGGAGTCGTTGCTTTTATAGACAGAAGAAGGAAGANNNAGATCGGAA
122248 TATGAGTGTGGAGTCGTTGCTATAATGGACTACTGCAGCTANNNAGATCGGAAGAG
122249 TATGAGTGTGGAGTCGTTGCAAAATTTTCAAGAGTTCCAAATGNNNAGATCGGAAG
122250 TATGAGTGTGGAGTCGTTGCATGGATGTGTTAATTAACTTGTCNNNAGATCGGAAG
122251 TATGAGTGTGGAGTCGTTGCTATGGACTAAATTATATCCCCACNNNAGATCGGAAG
122252 TATGAGTGTGGAGTCGTTGCTATGGACTAAATTATATCCCCACNNNAGATCGGAAG
122253 TATGAGTGTGGAGTCGTTGCTCTCTAATTCTAGAATGAAGCATNNNAGATCGGAAG

122254 TATGAGTGTGGAGTCGTTGCGTCTGTCTCTTTCTGGACTCNNNAGATCGGAAGAGC
122256 TATGAGTGTGGAGTCGTTGCAATAGTAGGCTTTCATATTAATTTTNNNAGATCGGA
122258 TATGAGTGTGGAGTCGTTGCGGGAAATATGACTGAGAAATATGNNNAGATCGGAA
122259 TATGAGTGTGGAGTCGTTGCGGGAAATATGACTGAGAAATATGNNNAGATCGGAA
122260 TATGAGTGTGGAGTCGTTGCCTAAATCTCAGTAAGAAGAGCANNNAGATCGGAAG
122261 TATGAGTGTGGAGTCGTTGCCTTCAAGTCTTTTTTTTTCACTCNNNAGATCGGAAGA
122262 TATGAGTGTGGAGTCGTTGCAACCAGCTTTTGATTTCATTTATNNNAGATCGGAAGA
122263 TATGAGTGTGGAGTCGTTGCTAATTCTATCTGGCCTTACTTGNNNAGATCGGAAGA
122264 TATGAGTGTGGAGTCGTTGCGATTCTTCCTCTCAAAGACTTNNNAGATCGGAAGAG
122265 TATGAGTGTGGAGTCGTTGCGATTCTTCCTCTCAAAGACTTNNNAGATCGGAAGAG
122266 TATGAGTGTGGAGTCGTTGCGCTATGAGAGTGACTTTTTAAATNNNAGATCGGAAG
122267 TATGAGTGTGGAGTCGTTGCAAATCTTCATTCTCCATAGGAAANNNAGATCGGAAG
122268 TATGAGTGTGGAGTCGTTGCCAATTTCAAGCTACTGATAGTTTNNNAGATCGGAAG
122269 TATGAGTGTGGAGTCGTTGCAGAACTTATTCTGCCAAATACTANNNAGATCGGAAG
122270 TATGAGTGTGGAGTCGTTGCAATATTCTGAGTATTGTTCCCTTNNNAGATCGGAAG
122271 TATGAGTGTGGAGTCGTTGCAATCTCATCTTGTATCTCCCATANNNAGATCGGAAGA
122272 TATGAGTGTGGAGTCGTTGCGCATCGAGAAAAATACTATTTATAGNNNAGATCGGA
122273 TATGAGTGTGGAGTCGTTGCTTATAACGTCACTAATCTCATCANNNAGATCGGAAGA
122274 TATGAGTGTGGAGTCGTTGCCTACATCCTTGTCAATGTACANNNAGATCGGAAGAG
122275 TATGAGTGTGGAGTCGTTGCCTACATCCTTGTCAATGTACANNNAGATCGGAAGAG
122276 TATGAGTGTGGAGTCGTTGCTTTGCCACTATCTCTTTCTTTTNNNAGATCGGAAGAG
122277 TATGAGTGTGGAGTCGTTGCGCATTCAAAATAAAAAGCAGAATNNNAGATCGGAAG
122278 TATGAGTGTGGAGTCGTTGCACCATACTCTTTAGATCTACAAANNNAGATCGGAAGA
122279 TATGAGTGTGGAGTCGTTGCTTGCATCTTTATTGATATTCTCCNNNAGATCGGAAGA
122280 TATGAGTGTGGAGTCGTTGCAGGTGGTAGATATGGTCTGANNNAGATCGGAAGAG
122281 TATGAGTGTGGAGTCGTTGCAATACTTGAACAACTTTTCTTGTNNNAGATCGGAAGA
122282 TATGAGTGTGGAGTCGTTGCTTTAAAGACTGAAAATATGCCAGNNNAGATCGGAAG
122283 TATGAGTGTGGAGTCGTTGCAAAGACAATTTAAGAATGTCTCCNNNAGATCGGAAG
122284 TATGAGTGTGGAGTCGTTGCCTTGTTTTTAGGGCCAGAATNNNAGATCGGAAGAG
122285 TATGAGTGTGGAGTCGTTGCATAATTCCGAGTTCAATACCATNNNAGATCGGAAGA
122286 TATGAGTGTGGAGTCGTTGCATCTTTCTCCCTAAAAACATACTNNNAGATCGGAAGA
122287 TATGAGTGTGGAGTCGTTGCAGGGAAAATTTCATAAAATGGATNNNAGATCGGAA
122288 TATGAGTGTGGAGTCGTTGCTAGGTAGGTAGGTAGGTAGGNNNAGATCGGAAGA
122289 TATGAGTGTGGAGTCGTTGCCTCAGGAGAAGATGACTACTANNNAGATCGGAAGA
122290 TATGAGTGTGGAGTCGTTGCACTGAAGAATGAGCTATTTAAATNNNAGATCGGAAG
122291 TATGAGTGTGGAGTCGTTGCGTATGTATGTCTACACAGAGAGNNNAGATCGGAAG
122292 TATGAGTGTGGAGTCGTTGCTTTACACATTAGAGGGAAATTTGNNNAGATCGGAAG
122293 TATGAGTGTGGAGTCGTTGCGCTGATTAAATTCTTATCCAGAGNNNAGATCGGAAG
122294 TATGAGTGTGGAGTCGTTGCATGTTATATTCTTTCCATGTCAGNNNAGATCGGAAGA
122295 TATGAGTGTGGAGTCGTTGCAAGTTAATCAGACCCTATTCTACNNNAGATCGGAAG
122296 TATGAGTGTGGAGTCGTTGCTAACAGGTAGATTCAATTCTCATNNNAGATCGGAAG
122297 TATGAGTGTGGAGTCGTTGCACAATTTTTAGCCATTCATCTACNNNAGATCGGAAGA
122298 TATGAGTGTGGAGTCGTTGCTACAAAGATTAGAGAGATGACAGNNNAGATCGGAA

122299 TATGAGTGTGGAGTCGTTGCTACAAAGATTAGAGAGATGACAGNNNAGATCGGAA
122300 TATGAGTGTGGAGTCGTTGCTTTTCACTTCTGATTTTGCTTATNNNAGATCGGAAGA
122301 TATGAGTGTGGAGTCGTTGCAATGAGAGGCCTTAATATTTAGANNNAGATCGGAA
122302 TATGAGTGTGGAGTCGTTGCCTTTCATTGCTACTTTACTGATTNNNAGATCGGAAGA
122303 TATGAGTGTGGAGTCGTTGCCACTCTTACTAGCTCTGAGANNNAGATCGGAAGAGC
122304 TATGAGTGTGGAGTCGTTGCGAGATTGGATAGAATGCATAGTNNNAGATCGGAAG
122305 TATGAGTGTGGAGTCGTTGCGCAAAGAATTAGGTAGACATTATNNNAGATCGGAA
122306 TATGAGTGTGGAGTCGTTGCGAGTACAGAGGTTAGCATTATTANNNAGATCGGAA
122307 TATGAGTGTGGAGTCGTTGCCAGAACCTATGAGATTCTACTAANNNAGATCGGAAG
122309 TATGAGTGTGGAGTCGTTGCAAAGTTTTCAGAATTTGTCTCTTNNNAGATCGGAAG
122310 TATGAGTGTGGAGTCGTTGCGTTTTGAGGGAGAAAGAAGANNNAGATCGGAAGA
122311 TATGAGTGTGGAGTCGTTGCGAAGGACATTCTACAAAACAACNNNAGATCGGAAG
122312 TATGAGTGTGGAGTCGTTGCGAGTACTAGTGCTTTCATTTTCNNNAGATCGGAAGA
122313 TATGAGTGTGGAGTCGTTGCAAAGAAGGAGTATGAGAAAGGANNNAGATCGGAA
122314 TATGAGTGTGGAGTCGTTGCGCATTTTCATCTTGGACTTCNNNAGATCGGAAGAGC
122315 TATGAGTGTGGAGTCGTTGCGGCATTTTCATCTTGGACTTNNNAGATCGGAAGAGC
122316 TATGAGTGTGGAGTCGTTGCGCCCAGACATTGATATTAAAAANNNAGATCGGAAGA
122317 TATGAGTGTGGAGTCGTTGCAAATGCTCAGAAATGCCTATNNNAGATCGGAAGAG
122318 TATGAGTGTGGAGTCGTTGCTAGACCTGAAATTCTAGAAAGAANNNAGATCGGAA
122319 TATGAGTGTGGAGTCGTTGCATATACGTGTATGCCCGAATNNNAGATCGGAAGAG
122320 TATGAGTGTGGAGTCGTTGCAATTGAGCCAAAGAGTTCAAGNNNAGATCGGAAGA
122321 TATGAGTGTGGAGTCGTTGCAGTAACTCAAGAAATGCCTTCNNNAGATCGGAAGAG
122322 TATGAGTGTGGAGTCGTTGCAATTCCTTGTACTACATCTATGGNNNAGATCGGAAG
122323 TATGAGTGTGGAGTCGTTGCATAGTAATGGAAAGAAGGAAGTTNNNAGATCGGAA
122324 TATGAGTGTGGAGTCGTTGCAGAAGCTTAAAAAGTTTTTTTCCNNNAGATCGGAAG
122325 TATGAGTGTGGAGTCGTTGCAGAAGCTTAAAAAGTTTTTTTCCNNNAGATCGGAAG
122326 TATGAGTGTGGAGTCGTTGCTGAGTGCTTATGATTTAAAATGANNNAGATCGGAAG
122327 TATGAGTGTGGAGTCGTTGCTTTCCTTAAGTCTAGTTCATGAGNNNAGATCGGAAG
122328 TATGAGTGTGGAGTCGTTGCCTTTACATCAGTCTTAATCTTGCNNNAGATCGGAAGA
122329 TATGAGTGTGGAGTCGTTGCTTGGTTACTAGAAAGTTCATAGANNNAGATCGGAAG
122330 TATGAGTGTGGAGTCGTTGCGACACTGTGGCTAAGAAAACNNNAGATCGGAAGAG
122331 TATGAGTGTGGAGTCGTTGCTTTGTTTATTTGTTTCTGTTTCTNNNAGATCGGAAGA
122332 TATGAGTGTGGAGTCGTTGCCAACAAAACCTAGGGTAGATATNNNAGATCGGAAG
122333 TATGAGTGTGGAGTCGTTGCATCTTGTTGAACTTCCTACATAANNNAGATCGGAAGA
122334 TATGAGTGTGGAGTCGTTGCTAAATGAGTGGACAGTTTATTTGNNNAGATCGGAA
122336 TATGAGTGTGGAGTCGTTGCAATATCCTCAAAATTAGCTTGTGNNNAGATCGGAAG
122337 TATGAGTGTGGAGTCGTTGCAAAGGACATATGTGAGATCAAGNNNAGATCGGAAG
122338 TATGAGTGTGGAGTCGTTGCAAAGGACATATGTGAGATCAAGNNNAGATCGGAAG
122339 TATGAGTGTGGAGTCGTTGCAAAATGAAACATCTGAGGGAANNNAGATCGGAAGA
122340 TATGAGTGTGGAGTCGTTGCATTTCGTTGTTGACATACAATTNNNAGATCGGAAGA
122341 TATGAGTGTGGAGTCGTTGCGAAATATTCTGGCGGGAATANNNAGATCGGAAGAG
122342 TATGAGTGTGGAGTCGTTGCAAAACACATTTGAGCTTTCTAATNNNAGATCGGAAG
122343 TATGAGTGTGGAGTCGTTGCAAACTGAACATTTTGTACTTCCNNNAGATCGGAAGA

122344 TATGAGTGTGGAGTCGTTGCTTTTCAAAAGAAGGAAAGAACTTNNNAGATCGGAAG
122345 TATGAGTGTGGAGTCGTTGCAAATGTGTAGAAAGCATACATACNNNAGATCGGAAG
122346 TATGAGTGTGGAGTCGTTGCAAACTGTCCAGAATTTGAGTTNNNAGATCGGAAGAG
122349 TATGAGTGTGGAGTCGTTGCGTTTGAGAAGCAGAATTCTTGNNNAGATCGGAAGAG
122350 TATGAGTGTGGAGTCGTTGCCAGAACCTGTGAATGTATTATTTNNNAGATCGGAAG
122351 TATGAGTGTGGAGTCGTTGCAACAAAATGTACTGCTTAAGTCNNNAGATCGGAAGA
122352 TATGAGTGTGGAGTCGTTGCGAAAATGACTGAGAGAAGTCANNNAGATCGGAAGA
122353 TATGAGTGTGGAGTCGTTGCCACACAAAGTAAAAGTCCTAAAANNNAGATCGGAAG
122354 TATGAGTGTGGAGTCGTTGCGATACTGTTGATACTAGTCTGTTNNNAGATCGGAAG
122355 TATGAGTGTGGAGTCGTTGCCAGCAAGTTCCAATAGCTTANNNAGATCGGAAGAGC
122356 TATGAGTGTGGAGTCGTTGCTACAAGATTACCTTAACCTTTGANNNAGATCGGAAGA
122357 TATGAGTGTGGAGTCGTTGCACTATTTTACTGAGACATAGCATNNNAGATCGGAAGA
122358 TATGAGTGTGGAGTCGTTGCATCCAGACCTCATGATGTATAANNNAGATCGGAAGA
122359 TATGAGTGTGGAGTCGTTGCTTCTATCTTTATTTCAGCCTTTGNNNAGATCGGAAGAG
122360 TATGAGTGTGGAGTCGTTGCGACAGAATCTTACTTTGTTGCNNNAGATCGGAAGAG
122361 TATGAGTGTGGAGTCGTTGCACACACATACATACACACACNNNAGATCGGAAGAGCA
122362 TATGAGTGTGGAGTCGTTGCCCAAGTAGTCTTTTTCCATTTAGNNNAGATCGGAAGA
122363 TATGAGTGTGGAGTCGTTGCAACATCTTGACCTTTGTCTATACNNNAGATCGGAAGA
122364 TATGAGTGTGGAGTCGTTGCCACAGAACACCTAGTAAATATTTNNNAGATCGGAAGA
122366 TATGAGTGTGGAGTCGTTGCAAACAACTAATGTGAATTTCAGTNNNAGATCGGAAG
122367 TATGAGTGTGGAGTCGTTGCATTTCTAACTTCTACTTGGAAGANNNAGATCGGAAGA
122368 TATGAGTGTGGAGTCGTTGCAGAGATTGGTCAGATCCTTACNNNAGATCGGAAGAG
122370 TATGAGTGTGGAGTCGTTGCGATAGAATGTGGACAATCATTGNNNAGATCGGAAGA
122371 TATGAGTGTGGAGTCGTTGCGGTATGCTTGGGTTTATCATNNNAGATCGGAAGAGC
122372 TATGAGTGTGGAGTCGTTGCATCAGAGAAAGATGTTACATGTANNNAGATCGGAAG
122373 TATGAGTGTGGAGTCGTTGCAGTCTTTCCCAAAAAAAAAAAAAANNNAGATCGGAAG
122374 TATGAGTGTGGAGTCGTTGCATGCTGTTAATAAAGACATACACNNNAGATCGGAAG
122375 TATGAGTGTGGAGTCGTTGCCCAGGTAGAGAATATGGAATAAANNNAGATCGGAA
122379 TATGAGTGTGGAGTCGTTGCCATGAAGACACAAGGAGAAGNNNAGATCGGAAGAG
122380 TATGAGTGTGGAGTCGTTGCACACTAGAACCTGGAAAGTGNNNAGATCGGAAGAG
122381 TATGAGTGTGGAGTCGTTGCAGAGAACTTTGCCTTCTAAAGNNNAGATCGGAAGAG
122382 TATGAGTGTGGAGTCGTTGCCAAGCTAAAAATTGGAATGACCNNNAGATCGGAAGA
122383 TATGAGTGTGGAGTCGTTGCATCACCATGACTGAAACATTNNNAGATCGGAAGAGC
122384 TATGAGTGTGGAGTCGTTGCGAATGATATAAGAAAGTCACGTCNNNAGATCGGAAG
122385 TATGAGTGTGGAGTCGTTGCGTTTCTCTCTGTAATTTGTGTGNNNAGATCGGAAGAG
122386 TATGAGTGTGGAGTCGTTGCACCTAAGAAATAACTTATCTTGGANNNAGATCGGAAG
122387 TATGAGTGTGGAGTCGTTGCATTGAAAAACACCTGAGAATAAANNNAGATCGGAAG
122388 TATGAGTGTGGAGTCGTTGCATTGCACTGAATCAAAAGTTTAGNNNAGATCGGAAG
122389 TATGAGTGTGGAGTCGTTGCGTCCACGTAAGGTTAAATAAATANNNAGATCGGAAG
122390 TATGAGTGTGGAGTCGTTGCTGCATTCATGTATTTATGTAAACANNNAGATCGGAAG
122391 TATGAGTGTGGAGTCGTTGCATCATTTGCCTGAAAGACCTANNNAGATCGGAAGAG
122392 TATGAGTGTGGAGTCGTTGCTTTCCTCCTCTATTACTTTCTTTNNNAGATCGGAAGAG
122393 TATGAGTGTGGAGTCGTTGCAAAAATGCCTATGACACAAAAANNNAGATCGGAAGA

122394 TATGAGTGTGGAGTCGTTGCAGAAAATAAGGGTTGTCTTTTTTNNNAGATCGGAAG
122395 TATGAGTGTGGAGTCGTTGCTGTATTCATCCACTTAGGAATAANNNAGATCGGAAG
122396 TATGAGTGTGGAGTCGTTGCTTTTTCTGTGCTGCTTTTATTTNNNAGATCGGAAGAG
122397 TATGAGTGTGGAGTCGTTGCTAGGTGAGATATGCTATCTAACTNNNAGATCGGAAG
122398 TATGAGTGTGGAGTCGTTGCTAGGTGAGATATGCTATCTAACTNNNAGATCGGAAG
122399 TATGAGTGTGGAGTCGTTGCCAGTTCTAGAATTGGTCAAGANNNAGATCGGAAGA
122400 TATGAGTGTGGAGTCGTTGCAATCTAGAAACACAGGCATTTTANNNAGATCGGAAG
122401 TATGAGTGTGGAGTCGTTGCTTGCTTTCAATCTTTAGACCCNNNAGATCGGAAGAG
122402 TATGAGTGTGGAGTCGTTGCCCTCATTTTAACGAATACATCTGNNNAGATCGGAAG
122405 TATGAGTGTGGAGTCGTTGCAGCTTCTCTTCTTAGTCATTTAGNNNAGATCGGAAG
122406 TATGAGTGTGGAGTCGTTGCCACAATCCATAGTATCTGTATTGNNNAGATCGGAAG
122407 TATGAGTGTGGAGTCGTTGCGGAAATATGAAGTCATGAATACCNNNAGATCGGAA
122408 TATGAGTGTGGAGTCGTTGCTATGTAATGAAGCCTGCATAAANNNAGATCGGAAG
122409 TATGAGTGTGGAGTCGTTGCTAGTTAGCATGGGGAGATCTNNNAGATCGGAAGAG
122410 TATGAGTGTGGAGTCGTTGCCATCAAAAGGAAATTGTTGATTTNNNAGATCGGAAG
122411 TATGAGTGTGGAGTCGTTGCACCTTAATATATATGGCTAGTGGNNNAGATCGGAAG
122412 TATGAGTGTGGAGTCGTTGCAATTTATTCCAACGAAGGCTAANNNAGATCGGAAGA
122414 TATGAGTGTGGAGTCGTTGCAGTTGACTGAAATAAAAGATAGCNNNAGATCGGAA
122415 TATGAGTGTGGAGTCGTTGCTTTCTGTCATCCATCTTTCTATTNNNAGATCGGAAGA
122416 TATGAGTGTGGAGTCGTTGCTCCAGAATCAAAACAGTCTGNNNAGATCGGAAGAG
122417 TATGAGTGTGGAGTCGTTGCGAAGGAAAGCAGAATAAATTAGTNNNAGATCGGAA
122418 TATGAGTGTGGAGTCGTTGCAATGGAGTTGCTGTGTTTAATNNNAGATCGGAAGA
122419 TATGAGTGTGGAGTCGTTGCTCTTTCTGCTATCATTTAGAAGANNNAGATCGGAAG
122420 TATGAGTGTGGAGTCGTTGCCTGACTGATTGAATGATTCTTTCNNNAGATCGGAAG
122421 TATGAGTGTGGAGTCGTTGCAGGTACTCTACAGGAATTTTTATNNNAGATCGGAAG
122422 TATGAGTGTGGAGTCGTTGCTAATAATTATCTGAAGCTTGGACNNNAGATCGGAAG
122423 TATGAGTGTGGAGTCGTTGCACCTAATCTAAGAAACACTACAANNNAGATCGGAAG
122424 TATGAGTGTGGAGTCGTTGCTAATATTGTTTGTACAAGCTGACNNNAGATCGGAAG
122425 TATGAGTGTGGAGTCGTTGCACAACAGTTTTTCTGAAACTTTANNNAGATCGGAAG
122426 TATGAGTGTGGAGTCGTTGCTCACTCACTCACTCATACAGNNNAGATCGGAAGAGC
122427 TATGAGTGTGGAGTCGTTGCTAATTGCATATAAAAACCCAAGTNNNAGATCGGAAG
122428 TATGAGTGTGGAGTCGTTGCTGCATTTTAGATGAGACCTAAANNNAGATCGGAAGA
122429 TATGAGTGTGGAGTCGTTGCTGCATTTTAGATGAGACCTAAANNNAGATCGGAAGA
122430 TATGAGTGTGGAGTCGTTGCAACACAAAGACACAGATACATATNNNAGATCGGAAG
122431 TATGAGTGTGGAGTCGTTGCTGATATATAAGAAACTCAGAGCANNNAGATCGGAA
122432 TATGAGTGTGGAGTCGTTGCATTTGAACTCTGTGATAGAATTGNNNAGATCGGAAG
122433 TATGAGTGTGGAGTCGTTGCAAATGGCATTTCTAGATTTAGTGNNNAGATCGGAAG
122434 TATGAGTGTGGAGTCGTTGCGGGCAATTTATTTTCAAAAATTAAANNNAGATCGGA
122435 TATGAGTGTGGAGTCGTTGCAATGCCTATATACATGGGTATTCNNNAGATCGGAAG
122436 TATGAGTGTGGAGTCGTTGCGAGTTATTTACAAAGAGATGCAGNNNAGATCGGAA
122437 TATGAGTGTGGAGTCGTTGCACTCTGAATATCTCATCTTATGGNNNAGATCGGAAG
122438 TATGAGTGTGGAGTCGTTGCCATTGTTTCAAAGTCAATAATGTNNNAGATCGGAAG
122439 TATGAGTGTGGAGTCGTTGCTAGGAAGAATCATTCTGCTTTTNNNAGATCGGAAGA

122440 TATGAGTGTGGAGTCGTTGCACTATAGAGCCCAATTACTTTAGNNNAGATCGGAAG
122441 TATGAGTGTGGAGTCGTTGCTATAAAGAAGTATCTGAGACTGGNNNAGATCGGAA
122442 TATGAGTGTGGAGTCGTTGCCATTTGGAAACTTACATAATTGCNNNAGATCGGAAG
122443 TATGAGTGTGGAGTCGTTGCTGTGTTATATACATGACATACCANNNAGATCGGAAG
122444 TATGAGTGTGGAGTCGTTGCCAAAATGAAAACCTCTGTGTAGNNNAGATCGGAAGA
122445 TATGAGTGTGGAGTCGTTGCGGAGAGTAGGAATGATTGTACNNNAGATCGGAAGA
122446 TATGAGTGTGGAGTCGTTGCTCTCACCTGTATTCTATAGAAGANNNAGATCGGAAG
122447 TATGAGTGTGGAGTCGTTGCCATACATATATATGTGTGTGTGTNNNAGATCGGAAG
122448 TATGAGTGTGGAGTCGTTGCCTTGAAGGTTTATTTTACCCTTTNNNAGATCGGAAGA
122449 TATGAGTGTGGAGTCGTTGCCTATGAGTCTGCTTTGAAATTTANNNAGATCGGAAG
122450 TATGAGTGTGGAGTCGTTGCTACATACACACTATTTATAGCCCNNNAGATCGGAAGA
122451 TATGAGTGTGGAGTCGTTGCCACATACACATACCCCTATATATNNNAGATCGGAAGA
122452 TATGAGTGTGGAGTCGTTGCCACATACACATACCCCTATATATNNNAGATCGGAAGA
122453 TATGAGTGTGGAGTCGTTGCTTTGGTGAGGTAACTAGGTTTANNNAGATCGGAAG
122454 TATGAGTGTGGAGTCGTTGCGAATAAGAGGAGACAATTCAACNNNAGATCGGAAG
122455 TATGAGTGTGGAGTCGTTGCAACAGAGAATGTGAAGGTATGNNNAGATCGGAAGA
122456 TATGAGTGTGGAGTCGTTGCGGGTGGTCCATAAAATACAANNNAGATCGGAAGAG
122457 TATGAGTGTGGAGTCGTTGCAGCAAATTGTCAATGTGTAATACNNNAGATCGGAAG
122458 TATGAGTGTGGAGTCGTTGCGACTATAAAAAGAAATCACGACANNNAGATCGGAA
122459 TATGAGTGTGGAGTCGTTGCGCTGACATTCTAGAAATGAAATANNNAGATCGGAA
122460 TATGAGTGTGGAGTCGTTGCTCTTTTCTAAATTGTACATTTCTGNNNAGATCGGAAG
122461 TATGAGTGTGGAGTCGTTGCAGGTACTAAGCTAGTATTAGGAANNNAGATCGGAA
122462 TATGAGTGTGGAGTCGTTGCAAAAACTGTTTTTCTGAGTCAGANNNAGATCGGAAG
122463 TATGAGTGTGGAGTCGTTGCTCCCCTCCAAAGTAAATACTNNNAGATCGGAAGAGC
122464 TATGAGTGTGGAGTCGTTGCTACATATAGAGTAGTCCTGCATANNNAGATCGGAAG
122465 TATGAGTGTGGAGTCGTTGCGGATAGATTCTAGATTTTAGGCTNNNAGATCGGAA
122466 TATGAGTGTGGAGTCGTTGCCTATATCCAGTCCCCTAGTTTTNNNAGATCGGAAGAG
122467 TATGAGTGTGGAGTCGTTGCAATAAAATTGCAACCTTAGAGGNNNAGATCGGAAGA
122468 TATGAGTGTGGAGTCGTTGCTTTAAATTTGGTAATTATGGCCANNNAGATCGGAAG
122469 TATGAGTGTGGAGTCGTTGCCTGTATCTTTCACTCAGTTTTTTNNNAGATCGGAAGA
122470 TATGAGTGTGGAGTCGTTGCGTGTAACCTTTTACAGCATTTTNNNAGATCGGAAGA
122471 TATGAGTGTGGAGTCGTTGCTATCTAGGGAATACACCTATCAGNNNAGATCGGAAG
122472 TATGAGTGTGGAGTCGTTGCAATGTGTCTTAGAGGATAATGGNNNAGATCGGAAG
122473 TATGAGTGTGGAGTCGTTGCCAGGGTTTTCTTTAAAGTGATTANNNAGATCGGAAG
122474 TATGAGTGTGGAGTCGTTGCCCTTATCATGTAACATAAGATGTNNNAGATCGGAAG
122475 TATGAGTGTGGAGTCGTTGCGTTTTGGGAAACTATATGCTAAANNNAGATCGGAA
122476 TATGAGTGTGGAGTCGTTGCGGTGAATGGAAACTATTAACATTNNNAGATCGGAA
122477 TATGAGTGTGGAGTCGTTGCTATCCTTATTTGGAAATAGGGCNNNAGATCGGAAGA
122478 TATGAGTGTGGAGTCGTTGCATTTAGGTTGATTCCAGATCTTCNNNAGATCGGAAG
122479 TATGAGTGTGGAGTCGTTGCACTCTTTACATGAACTTAAAACCNNNAGATCGGAAGA
122480 TATGAGTGTGGAGTCGTTGCGGGTGATGGAAAACTAAATATTTNNNAGATCGGAA
122481 TATGAGTGTGGAGTCGTTGCTATGGAAAGTTGATAAGTTTGGTNNNAGATCGGAA
122482 TATGAGTGTGGAGTCGTTGCCACAAGGACAGAAAATGTTTTNNNAGATCGGAAGA

122485 TATGAGTGTGGAGTCGTTGCATTAGAAAGGGCTTCATTCATANNNAGATCGGAAG
122486 TATGAGTGTGGAGTCGTTGCAATGGTCTCAACAAATCAGACNNNAGATCGGAAGA
122487 TATGAGTGTGGAGTCGTTGCTTCTGTACAGAAAAAAATAGCAANNNAGATCGGAA
122488 TATGAGTGTGGAGTCGTTGCGAGAGATACAGAGAGAGATAATGNNNAGATCGG
122489 TATGAGTGTGGAGTCGTTGCAGATACATACAGAGAGAGATACANNNAGATCGGA
122490 TATGAGTGTGGAGTCGTTGCCTGATGAAAGACAATAATGACAANNNAGATCGGA
122491 TATGAGTGTGGAGTCGTTGCTTTATCTGAATTCCAAGACTTGNNNAGATCGGAAG
122492 TATGAGTGTGGAGTCGTTGCTATACATGCTATGTAATGTGGTGNNNAGATCGGAA
122493 TATGAGTGTGGAGTCGTTGCCCTGTAATCATCAGTGACTATTTNNNAGATCGGAAG
122494 TATGAGTGTGGAGTCGTTGCTTTACTTTCCAATCAAAATTCCTNNNAGATCGGAAGA
122495 TATGAGTGTGGAGTCGTTGCGAATCACTGTCAAAAGTCAATTANNNAGATCGGAAG
122496 TATGAGTGTGGAGTCGTTGCTCCTCTGAGGAATTTGTATTTANNNAGATCGGAAG
122497 TATGAGTGTGGAGTCGTTGCTTTCTCAAGATAAGATAACCCANNNAGATCGGAAG
122498 TATGAGTGTGGAGTCGTTGCGTATATGCGTTTATATACATCTGGNNNAGATCGGA
122499 TATGAGTGTGGAGTCGTTGCTACAAAAATGACTACAAAACACTNNNAGATCGGAAG
122500 TATGAGTGTGGAGTCGTTGCTTGTTTATGTTTAAGTTCCACAANNNAGATCGGAAG
122501 TATGAGTGTGGAGTCGTTGCAATCTCAGTAAACCAAAATCCAANNNAGATCGGAAG
122502 TATGAGTGTGGAGTCGTTGCTTAGGGTAAGGAAAACAACAATANNNAGATCGGAA
122503 TATGAGTGTGGAGTCGTTGCTATTTTTCTTTAGGCAAATACCCNNNAGATCGGAAGA
122504 TATGAGTGTGGAGTCGTTGCTATTTTTCCTGACCTCCCTAATTNNNAGATCGGAAGA
122505 TATGAGTGTGGAGTCGTTGCGACAGAAGGGTAATAGCATAGNNNAGATCGGAAG
122506 TATGAGTGTGGAGTCGTTGCGATCTTGTCAGTATCAACAATAANNNAGATCGGAAG
122507 TATGAGTGTGGAGTCGTTGCAGGAAAAACAGAATTAGGAAAACNNNAGATCGGAA
122508 TATGAGTGTGGAGTCGTTGCACAGTTTGGAATGAATTCTCATNNNAGATCGGAAGA
122509 TATGAGTGTGGAGTCGTTGCCAGATAAACAAAATCTACCTTGTNNNAGATCGGAAG
122510 TATGAGTGTGGAGTCGTTGCCAGATTCTCCTGGAAGTTTGNNNAGATCGGAAGAG
122511 TATGAGTGTGGAGTCGTTGCAATAATTACTACCTACCTTCTCANNNAGATCGGAAGA
122512 TATGAGTGTGGAGTCGTTGCGAAGTAGGACTGTAGATGTTGNNNAGATCGGAAGA
122513 TATGAGTGTGGAGTCGTTGCCTGAACAGAGGTGTGAGAAANNNAGATCGGAAGA
122514 TATGAGTGTGGAGTCGTTGCCTGATTCAGCACTGAGAAATNNNAGATCGGAAGAG
122515 TATGAGTGTGGAGTCGTTGCTGAAAATGGTTCTTTCTTTATCCNNNAGATCGGAAG
122516 TATGAGTGTGGAGTCGTTGCTACTTTACAGTCTGAAGATTTCTNNNAGATCGGAAG
122517 TATGAGTGTGGAGTCGTTGCGTACTCTTTCAAGTCCCTAAGNNNAGATCGGAAGAG
122520 TATGAGTGTGGAGTCGTTGCCACTTATTCTGTTTCATTGGTCNNNAGATCGGAAGA
122521 TATGAGTGTGGAGTCGTTGCTGTGTATGTGTATGTGTGTATATNNNAGATCGGAA
122522 TATGAGTGTGGAGTCGTTGCGCATGAAATTGACTTTGACATNNNAGATCGGAAGA
122523 TATGAGTGTGGAGTCGTTGCCCCTCATTACTCTTCATTTATGANNNAGATCGGAAGA
122524 TATGAGTGTGGAGTCGTTGCCCAGGGAAATAAAAGAAATAGAGNNNAGATCGGAA
122525 TATGAGTGTGGAGTCGTTGCGATGGTATTTTCACATTCTTCATNNNAGATCGGAAG
122526 TATGAGTGTGGAGTCGTTGCATATTTATTAGCTGACTTCCCAGNNNAGATCGGAAG
122527 TATGAGTGTGGAGTCGTTGCAATATTTATTAGCTGACTTCCCANNNAGATCGGAAG
122528 TATGAGTGTGGAGTCGTTGCAAATGCATTTTATGTGGAAACTANNNAGATCGGAAG
122529 TATGAGTGTGGAGTCGTTGCCTAATCTTGCAGAGTTTTGTAGNNNAGATCGGAAGA

122530 TATGAGTGTGGAGTCGTTGCGAAACTGCAAAAACCCAAAGNNNAGATCGGAAGAG
122531 TATGAGTGTGGAGTCGTTGCTCATTAGAAATGCTTCCTAGATTNNNAGATCGGAAG
122532 TATGAGTGTGGAGTCGTTGCCTCAGAATGAAGAGTTTTGAATTNNNAGATCGGAAG
122533 TATGAGTGTGGAGTCGTTGCAAGAGGGAAAGCCTTTAAAAGNNNAGATCGGAAGA
122534 TATGAGTGTGGAGTCGTTGCTTATAAATGAAGAACTTCAGAGCNNNAGATCGGAAG
122535 TATGAGTGTGGAGTCGTTGCAAATAAATCCTGAGATAAGTTGCNNNAGATCGGAAG
122536 TATGAGTGTGGAGTCGTTGCCATACAGGATCATTTTGTGATAANNNAGATCGGAAG
122537 TATGAGTGTGGAGTCGTTGCGTACCATAGTTATGTGAGATGTCNNNAGATCGGAAG
122538 TATGAGTGTGGAGTCGTTGCAAATAATCTCAGTCTTTTTCCTGNNNAGATCGGAAGA
122539 TATGAGTGTGGAGTCGTTGCGGTTTGATTAGTAGAAAAAAGGANNNAGATCGGAA
122540 TATGAGTGTGGAGTCGTTGCAGTAGTTTTAGAGTTTTATGCTGNNNAGATCGGAAG
122541 TATGAGTGTGGAGTCGTTGCCTGCACAGAAGAATCATCTGNNNAGATCGGAAGAG
122542 TATGAGTGTGGAGTCGTTGCGTTTCCTCCTCTTTTTTTTTTTTNNNAGATCGGAAGAG
122543 TATGAGTGTGGAGTCGTTGCAAGGTTGAAAACATAACATTACANNNAGATCGGAAG
122544 TATGAGTGTGGAGTCGTTGCCTCCAAATATTTTGGCTGAAAGNNNAGATCGGAAGA
122545 TATGAGTGTGGAGTCGTTGCGAATTAGACGTGATAGGGATGNNNAGATCGGAAGA
122546 TATGAGTGTGGAGTCGTTGCCTTAGCTGACCATTCTTTTTCNNNAGATCGGAAGAGC
122547 TATGAGTGTGGAGTCGTTGCCCTGGACTAATTTGTAAAGTCTANNNAGATCGGAAG
122548 TATGAGTGTGGAGTCGTTGCATCACACTTTATTTACTCATTGANNNAGATCGGAAGA
122549 TATGAGTGTGGAGTCGTTGCAGAAAAAAATAAGAGGATAGAGCANNNAGATCGGAA
122550 TATGAGTGTGGAGTCGTTGCTGACATAACTGTGTTGTAGTTATNNNAGATCGGAAG
122551 TATGAGTGTGGAGTCGTTGCTATAAGTGGTGATGGAGAGGNNNAGATCGGAAGA
122552 TATGAGTGTGGAGTCGTTGCATAGATACAGAGGTAAACAAGACNNNAGATCGGAA
122553 TATGAGTGTGGAGTCGTTGCAATCTTATGCTGTGTAGGTCTTNNNAGATCGGAAGA
122554 TATGAGTGTGGAGTCGTTGCCACACAAACTTTCACAATCTTATNNNAGATCGGAAGA
122555 TATGAGTGTGGAGTCGTTGCGTACTAGAGCCTAGAACATTGNNNAGATCGGAAGA
122556 TATGAGTGTGGAGTCGTTGCCCCAAATCTCAAATTTGAATCANNNAGATCGGAAGA
122557 TATGAGTGTGGAGTCGTTGCTTTACTAGAATAAAGGCCAAGAGNNNAGATCGGAA
122558 TATGAGTGTGGAGTCGTTGCCTACCACCTACACTAGATAGAANNNAGATCGGAAGA
122559 TATGAGTGTGGAGTCGTTGCCACTGACTAAAGACATTTCATCNNNAGATCGGAAGA
122560 TATGAGTGTGGAGTCGTTGCTAAAACACAATTTGAAGCAGAANNNAGATCGGAAG
122561 TATGAGTGTGGAGTCGTTGCTATGCATATATAGAGAGAGTGTGNNNAGATCGGAA
122562 TATGAGTGTGGAGTCGTTGCAGATTCTCTGAAATTTTAAAATTCCNNNAGATCGGA
122563 TATGAGTGTGGAGTCGTTGCAGATTCTCTGAAATTTTAAAATTCCNNNAGATCGGA
122564 TATGAGTGTGGAGTCGTTGCATTTATTCTCTCTAGGTCACTTTNNNAGATCGGAAGA
122565 TATGAGTGTGGAGTCGTTGCAAAGAATAAGGACATGTTAGCTANNNAGATCGGAA
122566 TATGAGTGTGGAGTCGTTGCCTCTAATGACCTCATTTAACTTTNNNAGATCGGAAGA
122567 TATGAGTGTGGAGTCGTTGCAAAGTCAGTGAAGTTTTCTTTAGNNNAGATCGGAAG
122568 TATGAGTGTGGAGTCGTTGCGTGCATTCTTTTTGTTTTGTTTNNNAGATCGGAAGAG
122569 TATGAGTGTGGAGTCGTTGCTACAAAACATGGGTGAAATAAATNNNAGATCGGAA
122570 TATGAGTGTGGAGTCGTTGCTATTATATTTACAGCCAGTGTCTNNNAGATCGGAAG
122571 TATGAGTGTGGAGTCGTTGCGCTGTATCCTCATATAGAACTAGNNNAGATCGGAAG
122572 TATGAGTGTGGAGTCGTTGCCGCCTTTATATTTAAATACCTTCTNNNAGATCGGAAG

122573 TATGAGTGTGGAGTCGTTGCTAAAGTTTATCAAGGACCACTTANNNAGATCGGAAG
122574 TATGAGTGTGGAGTCGTTGCGTACAAGGCCTCTAATTTTTAATNNNAGATCGGAAG
122575 TATGAGTGTGGAGTCGTTGCGTACAAGGCCTCTAATTTTTAATNNNAGATCGGAAG
122576 TATGAGTGTGGAGTCGTTGCTGAAAGGTTATCATCTAGTACAGNNNAGATCGGAAG
122577 TATGAGTGTGGAGTCGTTGCCTACCATGAAAAAAATCCTCAATNNNAGATCGGAAG
122578 TATGAGTGTGGAGTCGTTGCAACAGGCTAAAATATACAGTTTCNNNAGATCGGAAG
122579 TATGAGTGTGGAGTCGTTGCTTTGTATAATGCTTTCATTCCTTNNNAGATCGGAAGA
122580 TATGAGTGTGGAGTCGTTGCTTACTGTATTCCTGATTTTGATGNNNAGATCGGAAG
122581 TATGAGTGTGGAGTCGTTGCATGTTTGATTGAAAGAATGCTTTNNNAGATCGGAAG
122582 TATGAGTGTGGAGTCGTTGCGTGAATAATTTCCCAAATCAGAANNNAGATCGGAAG
122583 TATGAGTGTGGAGTCGTTGCCATCAATACTATGCTTTTGATCANNNAGATCGGAAG
122584 TATGAGTGTGGAGTCGTTGCGACAAAAAGCTTTTTTTCCATTANNNAGATCGGAAG
122585 TATGAGTGTGGAGTCGTTGCTATTTGAAAGCCATTGATTTAGTNNNAGATCGGAAG
122586 TATGAGTGTGGAGTCGTTGCAAAAGTACAGCCACTCTATTTTANNNAGATCGGAAG
122587 TATGAGTGTGGAGTCGTTGCTGTGTATGTGTATGTATGTGTATNNNAGATCGGAAG
122588 TATGAGTGTGGAGTCGTTGCATGGCCAAAACATATATCTTTACNNNAGATCGGAAG
122589 TATGAGTGTGGAGTCGTTGCTATTGTGCTCTTAGATCTATTCANNNAGATCGGAAG
122590 TATGAGTGTGGAGTCGTTGCACATACAGTTTGACATAGTACTANNNAGATCGGAAG
122591 TATGAGTGTGGAGTCGTTGCAAAAAGTAGCTCAAGAAAAGTTTNNNAGATCGGAA
122592 TATGAGTGTGGAGTCGTTGCGTAGCTATGTCAGACATATACTCNNNAGATCGGAAG
122593 TATGAGTGTGGAGTCGTTGCCTATAAAACGGACACATCTTTCNNNAGATCGGAAGA
122595 TATGAGTGTGGAGTCGTTGCAAATTAGCTGAGACATGGTGNNNAGATCGGAAGAG
122596 TATGAGTGTGGAGTCGTTGCCTAGTTGTATGACTTCACTCTCNNNAGATCGGAAGA
122597 TATGAGTGTGGAGTCGTTGCATATCTCCTGATATTCTGTGTCTNNNAGATCGGAAGA
122598 TATGAGTGTGGAGTCGTTGCGAGAGATACAATACAATATCTCCTNNNAGATCGGAA
122599 TATGAGTGTGGAGTCGTTGCAAGCGTATAAATAAGCGTATAAGNNNAGATCGGAA
122600 TATGAGTGTGGAGTCGTTGCAAGCGTATAAATAAGCGTATAAGNNNAGATCGGAA
122601 TATGAGTGTGGAGTCGTTGCATTTGAAGATTTTGGTAATGTGTNNNAGATCGGAAG
122602 TATGAGTGTGGAGTCGTTGCATTTTCATAACTTTTTGGAGCTGNNNAGATCGGAAG
122603 TATGAGTGTGGAGTCGTTGCGAATCCAAATGTTAGGAAACTTTNNNAGATCGGAAG
122604 TATGAGTGTGGAGTCGTTGCGAATCCAAATGTTAGGAAACTTTNNNAGATCGGAAG
122605 TATGAGTGTGGAGTCGTTGCGTATTGGTCTTTCGTTCATGTANNNAGATCGGAAGA
122606 TATGAGTGTGGAGTCGTTGCTACACTCTATTCTCCCATGTTAANNNAGATCGGAAGA
122607 TATGAGTGTGGAGTCGTTGCCAATCAACAGCAAATCAGAACNNNAGATCGGAAGAG
122608 TATGAGTGTGGAGTCGTTGCGTCTAAGATCCTTTTGTTGTTTGNNNAGATCGGAAG
122609 TATGAGTGTGGAGTCGTTGCTATGTATTTTTGAGACAGGGTATNNNAGATCGGAAG
122610 TATGAGTGTGGAGTCGTTGCATGCCTAACATTAAACATCTATGNNNAGATCGGAAG
122611 TATGAGTGTGGAGTCGTTGCAATTACAGGGTATTTTCAAACAANNNAGATCGGAAG
122612 TATGAGTGTGGAGTCGTTGCAATTTAAGCATCTCCTTAGAATGNNNAGATCGGAAG
122613 TATGAGTGTGGAGTCGTTGCATAACCATGTAAACATTAGGACANNNAGATCGGAAG
122614 TATGAGTGTGGAGTCGTTGCGAACAGAGCTCATAAACATTTAANNNAGATCGGAAG
122615 TATGAGTGTGGAGTCGTTGCATCCTTAATCCTTAGACCATTTTNNNAGATCGGAAGA
122616 TATGAGTGTGGAGTCGTTGCTCTTTTTTTGTGACAAGGTTTTNNNAGATCGGAAGA

122617 TATGAGTGTGGAGTCGTTGCTAAATCTGAGTACTCTGCTTTTTNNNAGATCGGAAG
122618 TATGAGTGTGGAGTCGTTGCAATGTTAGCAAGATGGAAGTTANNNAGATCGGAAG
122619 TATGAGTGTGGAGTCGTTGCTATAGGGAGAAGACAGAGGANNNAGATCGGAAGA
122620 TATGAGTGTGGAGTCGTTGCCTTTTGTGAGGGCTTAAAATCNNNAGATCGGAAGAG
122621 TATGAGTGTGGAGTCGTTGCATATCTTTGTATTCCCATTACCANNNAGATCGGAAGA
122622 TATGAGTGTGGAGTCGTTGCACATGTATATGATCCCATTTAGANNNAGATCGGAAG
122623 TATGAGTGTGGAGTCGTTGCTATGTTTCCCTCCCATAAATACNNNAGATCGGAAGAG
122624 TATGAGTGTGGAGTCGTTGCCATTGGGATTTTGATAATGCATANNNAGATCGGAAG
122625 TATGAGTGTGGAGTCGTTGCTAAAGGGGATATTTATCACAAGTNNNAGATCGGAAG
122626 TATGAGTGTGGAGTCGTTGCTAAAACTGGGTAACAGAATATCTNNNAGATCGGAAG
122627 TATGAGTGTGGAGTCGTTGCTATATTGTCATTATCTGTGGACCNNNAGATCGGAAG
122628 TATGAGTGTGGAGTCGTTGCGGCAAGATAAATGCTTGAATATANNNAGATCGGAA
122629 TATGAGTGTGGAGTCGTTGCGCACATATCCCTTATACTCATATNNNAGATCGGAAGA
122630 TATGAGTGTGGAGTCGTTGCTGAAGATAATGATACACATGTCTNNNAGATCGGAAG
122632 TATGAGTGTGGAGTCGTTGCATACTAAAACCTTATTCTGTGTGNNNAGATCGGAAG
122633 TATGAGTGTGGAGTCGTTGCGAATTCAGATAAACAGGAGAAACNNNAGATCGGAA
122634 TATGAGTGTGGAGTCGTTGCATTTCTTATAGTGAATGTCCTCANNNAGATCGGAAG
122635 TATGAGTGTGGAGTCGTTGCTGAAAATAAGCACTTTCAAATCTNNNAGATCGGAAG
122636 TATGAGTGTGGAGTCGTTGCGATTCCAGAATGTGCATATAAGNNNAGATCGGAAG
122637 TATGAGTGTGGAGTCGTTGCTAGATAAGATAGAGGAGTGTTCCNNNAGATCGGAA
122638 TATGAGTGTGGAGTCGTTGCCTAATGGCATAAACAAGAAGAACNNNAGATCGGAA
122640 TATGAGTGTGGAGTCGTTGCCTAATATAAGCCTCTTGGGTAGNNNAGATCGGAAGA
122641 TATGAGTGTGGAGTCGTTGCAAAATCATGGTATCAGGAATGTANNNAGATCGGAA
122642 TATGAGTGTGGAGTCGTTGCTTACTTTAGTAGTATACACAGCCNNNAGATCGGAAG
122643 TATGAGTGTGGAGTCGTTGCGTTTTGGAATATGGAATTAGGAANNNAGATCGGAA
122644 TATGAGTGTGGAGTCGTTGCTTAGATGGAATCTCACTCTGTTNNNAGATCGGAAGA
122645 TATGAGTGTGGAGTCGTTGCCTTCCATGATAAACCTGATTGANNNAGATCGGAAGA
122646 TATGAGTGTGGAGTCGTTGCACTGATAAAGCTGCAGAAAANNNAGATCGGAAGAG
122647 TATGAGTGTGGAGTCGTTGCCTTGAGTTTTGATTTGCTGATTNNNAGATCGGAAGA
122648 TATGAGTGTGGAGTCGTTGCTATCAGAGAGATACAGAAGAGAANNNAGATCGGAA
122649 TATGAGTGTGGAGTCGTTGCATAGCTTATCAACCAAACTTCTGNNNAGATCGGAAG
122650 TATGAGTGTGGAGTCGTTGCATAGCTTATCAACCAAACTTCTGNNNAGATCGGAAG
122651 TATGAGTGTGGAGTCGTTGCAAAGAAGAAAGGACAGTAAGATANNNAGATCGGA
122653 TATGAGTGTGGAGTCGTTGCTCACTACTCAGTATTAACTTCAANNNAGATCGGAAGA
122654 TATGAGTGTGGAGTCGTTGCGGTGGACTATAATTAAATCCAATTNNNAGATCGGAA
122655 TATGAGTGTGGAGTCGTTGCTATAAATAGCAGTTGTGGAAGTANNNAGATCGGAA
122656 TATGAGTGTGGAGTCGTTGCAAATCTGAATCTTTTCTGTGATGNNNAGATCGGAAG
122657 TATGAGTGTGGAGTCGTTGCTCTACTAGGAATGTTCTTTCTTTNNNAGATCGGAAGA
122658 TATGAGTGTGGAGTCGTTGCGTATGTGTGTATCTGTGTGTNNNAGATCGGAAGAG
122659 TATGAGTGTGGAGTCGTTGCGCTAATATTATAATGGTGGAACANNNAGATCGGAA
122660 TATGAGTGTGGAGTCGTTGCGGTGGCATCAACTAAGATATATANNNAGATCGGAA
122661 TATGAGTGTGGAGTCGTTGCGTTTTGCACATATTCATCTCTATNNNAGATCGGAAGA
122662 TATGAGTGTGGAGTCGTTGCAGAAAGAAGAATTAGAAAAGAAACNNNAGATCGG

122663 TATGAGTGTGGAGTCGTTGCCATTTTTCTACCTTGCATACATCNNNAGATCGGAAGA
122664 TATGAGTGTGGAGTCGTTGCGACTGAATCACTAAAGTACATTGNNNAGATCGGAAG
122665 TATGAGTGTGGAGTCGTTGCGGCCTTACTAAAACTCAGAATNNNAGATCGGAAGAG
122666 TATGAGTGTGGAGTCGTTGCAATGGTGATTGCATAAGGTTATANNNAGATCGGAA
122667 TATGAGTGTGGAGTCGTTGCAGATTTATTAAGCAAGTTCACTGNNNAGATCGGAAG
122668 TATGAGTGTGGAGTCGTTGCAGACAAAAATTCACCTTTTATGGNNNAGATCGGAAG
122670 TATGAGTGTGGAGTCGTTGCTTACCCTGAAAAAAAAAAACGANNNAGATCGGAAG
122671 TATGAGTGTGGAGTCGTTGCCGTTCCTTTGAATACCTAAAAANNNAGATCGGAAGA
122672 TATGAGTGTGGAGTCGTTGCGAACCCTAAAAGAGAAGAGAATANNNAGATCGGAA
122673 TATGAGTGTGGAGTCGTTGCTTTTTATATAAAGAAGCAAAGGCNNNAGATCGGAA
122674 TATGAGTGTGGAGTCGTTGCCTACCTATAATGTATCACGATGTNNNAGATCGGAAG
122675 TATGAGTGTGGAGTCGTTGCAAGAGATATGAACTGAGGACTATNNNAGATCGGAA
122676 TATGAGTGTGGAGTCGTTGCATAATATACACAGCATACCTCAGNNNAGATCGGAAG
122677 TATGAGTGTGGAGTCGTTGCGCTCAGACTTTTACATTATCCTANNNAGATCGGAAGA
122679 TATGAGTGTGGAGTCGTTGCGTTCAAAATTCTCTCAAAATTCCNNNAGATCGGAAGA
122680 TATGAGTGTGGAGTCGTTGCCTGAAATCTGACATATCTTCCTTNNNAGATCGGAAGA
122681 TATGAGTGTGGAGTCGTTGCCATCTGTTGTATTTTCTTGAGTGNNNAGATCGGAAG
122682 TATGAGTGTGGAGTCGTTGCCTAATGAATTGTTGGTTCTTTTCNNNAGATCGGAAG
122683 TATGAGTGTGGAGTCGTTGCCACTCCTAAAGTATTAACATGTCNNNAGATCGGAAG
122684 TATGAGTGTGGAGTCGTTGCGGAATTAAGACAGAATGCTTTAANNNAGATCGGAA
122685 TATGAGTGTGGAGTCGTTGCACAGAGACAGAAATTTGAGTAATNNNAGATCGGAA
122686 TATGAGTGTGGAGTCGTTGCCTTGGGCAAAGAAAGCTAATNNNAGATCGGAAGAG
122687 TATGAGTGTGGAGTCGTTGCCTTAGTAAGTGAGAGAATAGCTANNNAGATCGGAA
122689 TATGAGTGTGGAGTCGTTGCCATTAAAATGCTTCGATCAGCNNNAGATCGGAAGAG
122690 TATGAGTGTGGAGTCGTTGCGTCCATAAATAGTGGCTGTACNNNAGATCGGAAGA
122691 TATGAGTGTGGAGTCGTTGCGAATTGAAAACTAAATGACAGTGNNNAGATCGGAA
122692 TATGAGTGTGGAGTCGTTGCGTTAGAGGTTTTTTGCTTTCTTNNNAGATCGGAAGA
122693 TATGAGTGTGGAGTCGTTGCCCTCCTAGTTCTCTATTTACAAANNNAGATCGGAAGA
122694 TATGAGTGTGGAGTCGTTGCCCTCCTAGTTCTCTATTTACAAANNNAGATCGGAAGA
122695 TATGAGTGTGGAGTCGTTGCGGATAGGAGTAAGAGGTAATGANNNAGATCGGAA
122696 TATGAGTGTGGAGTCGTTGCCAAGAAGCATGGCTAATGATNNNAGATCGGAAGAG
122697 TATGAGTGTGGAGTCGTTGCTCTCTCTGTATGACTGGTTTANNNAGATCGGAAGAG
122698 TATGAGTGTGGAGTCGTTGCCTAAAGGCTTCTTGGTTGTGNNNAGATCGGAAGAG
122699 TATGAGTGTGGAGTCGTTGCAGTTATATATGTATCGAGTGGGANNNAGATCGGAA
122700 TATGAGTGTGGAGTCGTTGCAGTCATAGCATTTCTAAAATTCCNNNAGATCGGAAG
122701 TATGAGTGTGGAGTCGTTGCGTTTGGAGATTGAGTGTTATAAANNNAGATCGGAA
122702 TATGAGTGTGGAGTCGTTGCTACACTTGGAACTCTAACGTTANNNAGATCGGAAGA
122703 TATGAGTGTGGAGTCGTTGCCTCCTATATTGACATTTTAGCTGNNNAGATCGGAAG
122704 TATGAGTGTGGAGTCGTTGCTATATAATCTATGTCCTGGACCTNNNAGATCGGAAG
122705 TATGAGTGTGGAGTCGTTGCGATTCAGAAAATTTTATAGCCACNNNAGATCGGAAG
122706 TATGAGTGTGGAGTCGTTGCTTTGCCTTCTTTTTTTTTTATCANNNAGATCGGAAGA
122707 TATGAGTGTGGAGTCGTTGCCTTCATGTCTTTCTATCCCCNNNAGATCGGAAGAGCA
122708 TATGAGTGTGGAGTCGTTGCTATTCCTCCTCTATTAAACACAGNNNAGATCGGAAGA

122709 TATGAGTGTGGAGTCGTTGCCTGGAAGAGATTCTCCTTTGNNNAGATCGGAAGAG
122710 TATGAGTGTGGAGTCGTTGCTAATTCTTGAGTCACTTTGAGTANNNAGATCGGAAG
122711 TATGAGTGTGGAGTCGTTGCATGGTGTTCTGTTCAGAGTTNNNAGATCGGAAGAG
122712 TATGAGTGTGGAGTCGTTGCGAAAGAGTAGGGTCAGTTCTNNNAGATCGGAAGAG
122713 TATGAGTGTGGAGTCGTTGCACAGAAAGAACTTTTTGTAATGANNNAGATCGGAA
122714 TATGAGTGTGGAGTCGTTGCGGATATAGAAGGCAATGTCTTNNNAGATCGGAAGA
122716 TATGAGTGTGGAGTCGTTGCCAAAGAGGTTGAGTATCTTATCANNNAGATCGGAA
122717 TATGAGTGTGGAGTCGTTGCACTAAAAGCAGTAAATATTCCTGNNNAGATCGGAAG
122718 TATGAGTGTGGAGTCGTTGCACTAAAAGCAGTAAATATTCCTGNNNAGATCGGAAG
122719 TATGAGTGTGGAGTCGTTGCTGTCTTTTCTTTGCATATGTTTTNNNAGATCGGAAGA
122720 TATGAGTGTGGAGTCGTTGCTGTCTTTTCTTTGCATATGTTTTNNNAGATCGGAAGA
122721 TATGAGTGTGGAGTCGTTGCTAAAAGACACATGGTAGATTCATNNNAGATCGGAA
122722 TATGAGTGTGGAGTCGTTGCCTGGAGTAAGGTGACATAATCNNNAGATCGGAAGA
122723 TATGAGTGTGGAGTCGTTGCTCTAGTAATTTTCATACGCTTATGNNNAGATCGGAA
122724 TATGAGTGTGGAGTCGTTGCGGTTTTAACCGATTACCTGANNNAGATCGGAAGAG
122725 TATGAGTGTGGAGTCGTTGCGAGTTGTCTATTCTAAGCTAGATNNNAGATCGGAAG
122726 TATGAGTGTGGAGTCGTTGCGAGTCTCACTCTGTTGTTCANNNAGATCGGAAGAGC
122727 TATGAGTGTGGAGTCGTTGCCTGGTTACAGTGTTTGTTATTTANNNAGATCGGAAG
122728 TATGAGTGTGGAGTCGTTGCGAAGTAGTGGAATAGGAGAAAGNNNAGATCGGAA
122729 TATGAGTGTGGAGTCGTTGCATTATCCCATTTCAGAGAAGAGNNNAGATCGGAAGA
122730 TATGAGTGTGGAGTCGTTGCAACAACTTTATATGTGTGTGTACNNNAGATCGGAAG
122731 TATGAGTGTGGAGTCGTTGCTAATTTTTGCGAGCAATTTTTTCNNNAGATCGGAAGA
122732 TATGAGTGTGGAGTCGTTGCTTCAGCTCAAAAGATATGTATTTNNNAGATCGGAAG
122733 TATGAGTGTGGAGTCGTTGCATTAATCCCTAAATGCAACATAGNNNAGATCGGAAG
122735 TATGAGTGTGGAGTCGTTGCAAACACAAATATTTCTCTCTCTCNNNAGATCGGAAGA
122736 TATGAGTGTGGAGTCGTTGCCACTGTCTCAAGAGTATTAAAAGNNNAGATCGGAAG
122737 TATGAGTGTGGAGTCGTTGCAGAAAGAGAAATTCCTACTGTTANNNAGATCGGAA
122738 TATGAGTGTGGAGTCGTTGCTCTATGTATGTATTTCCTGAGGNNNAGATCGGAAG
122739 TATGAGTGTGGAGTCGTTGCCATTATTCACCCATTCCTCTTNNNAGATCGGAAGAGC
122740 TATGAGTGTGGAGTCGTTGCTTTCATGTAAAAAAAATCTCCCCNNNAGATCGGAAGA
122741 TATGAGTGTGGAGTCGTTGCCTGGACTGATGGAAAGTAAGNNNAGATCGGAAGAG
122742 TATGAGTGTGGAGTCGTTGCAATTTGAAAATAATGAGGTCACCNNNAGATCGGAA
122743 TATGAGTGTGGAGTCGTTGCCTTTTTCTCTCTCTCTCTCTCTNNNAGATCGGAAGAGC
122744 TATGAGTGTGGAGTCGTTGCATTTGCAAATAGAGGTAGTTTTTNNNAGATCGGAAG
122745 TATGAGTGTGGAGTCGTTGCGCAAGTTTACAATATTCTTGTTGNNNAGATCGGAAG
122746 TATGAGTGTGGAGTCGTTGCTATGCCAAGTCCAGTTTTTATNNNAGATCGGAAGAG
122748 TATGAGTGTGGAGTCGTTGCCATGCTTCAAATTAATCAAGAACNNNAGATCGGAAG
122749 TATGAGTGTGGAGTCGTTGCATGGCTTCATTCAACACTTANNNAGATCGGAAGAGC
122750 TATGAGTGTGGAGTCGTTGCGGTATAAGGTGTTGACTACATAANNNAGATCGGAA
122751 TATGAGTGTGGAGTCGTTGCTTCCTTTACTCCATTACCATAATNNNAGATCGGAAGA
122752 TATGAGTGTGGAGTCGTTGCCAGTGACTTAGAAATAGAGTCAANNNAGATCGGAA
122753 TATGAGTGTGGAGTCGTTGCCAGTGACTTAGAAATAGAGTCAANNNAGATCGGAA
122754 TATGAGTGTGGAGTCGTTGCGCACTTGTAATCTCAGTTACTTNNNAGATCGGAAGA

122755 TATGAGTGTGGAGTCGTTGCAGGTAGAGAGGTAGAGAGAGNNNAGATCGGAAGA
122756 TATGAGTGTGGAGTCGTTGCGTTAAAGCACGGAATTATCTTACNNNAGATCGGAAG
122757 TATGAGTGTGGAGTCGTTGCGATAGTAGAAGCTTCCTTCAGNNNAGATCGGAAGAG
122758 TATGAGTGTGGAGTCGTTGCTCAAATTTAAAGACCTTCTCATGNNNAGATCGGAAGA
122759 TATGAGTGTGGAGTCGTTGCATTTCTTCCTCCAGATTCTTGNNNAGATCGGAAGAGC
122761 TATGAGTGTGGAGTCGTTGCTGAAGGGAATACAAAGGTATGNNNAGATCGGAAGA
122762 TATGAGTGTGGAGTCGTTGCGAATACAGTGATGCAATCTTGNNNAGATCGGAAGA
122763 TATGAGTGTGGAGTCGTTGCGTGATCATTCTGTAGTTTTCTTTNNNAGATCGGAAGA
122764 TATGAGTGTGGAGTCGTTGCAAGTCATCCCGTTCTACTATNNNAGATCGGAAGAGCA
122765 TATGAGTGTGGAGTCGTTGCCTCTTGTTTTACTTGTTCCTTATNNNAGATCGGAAGA
122766 TATGAGTGTGGAGTCGTTGCCTCTTGTTTTACTTGTTCCTTATNNNAGATCGGAAGA
122767 TATGAGTGTGGAGTCGTTGCAGGTGATGGCTTAATCATATTTANNNAGATCGGAAG
122768 TATGAGTGTGGAGTCGTTGCTTTCCTACTGAAACTATTCCAAANNNAGATCGGAAGA
122769 TATGAGTGTGGAGTCGTTGCAAATGGTAGTTAATTTGCTTCCNNNAGATCGGAAGA
122770 TATGAGTGTGGAGTCGTTGCAAATGGTAGTTAATTTGCTTCCNNNAGATCGGAAGA
122772 TATGAGTGTGGAGTCGTTGCTTGAGAGTTGTGGTTATGTTAGNNNAGATCGGAAGA
122773 TATGAGTGTGGAGTCGTTGCCTCATAAGACTTTCTAATGGCTANNNAGATCGGAAGA
122774 TATGAGTGTGGAGTCGTTGCGATCCTTACACCTTGACCTCNNNAGATCGGAAGAGCA
122775 TATGAGTGTGGAGTCGTTGCGAGTAAACTTTGTGAACATGAANNNAGATCGGAAGA
122776 TATGAGTGTGGAGTCGTTGCCTCTGATTTTTCAGGTTTTTCTNNNAGATCGGAAGAG
122777 TATGAGTGTGGAGTCGTTGCCTTTTCAAACTGAAAGCTCTAAANNNAGATCGGAAGA
122778 TATGAGTGTGGAGTCGTTGCTTATATGGTGTCTGCATATTGAGNNNAGATCGGAAG
122779 TATGAGTGTGGAGTCGTTGCCATACCTGTGGATTTTTCCATNNNAGATCGGAAGAG
122780 TATGAGTGTGGAGTCGTTGCTAGAGTCTGCACTTTTATCCTATNNNAGATCGGAAGA
122781 TATGAGTGTGGAGTCGTTGCCAAGATCCTCATCACTTCTATTNNNAGATCGGAAGAG
122782 TATGAGTGTGGAGTCGTTGCGAGAAAAAGACAAACATGATAGANNNAGATCGGAA
122783 TATGAGTGTGGAGTCGTTGCAAACATAAAAACAAACAAAACCCNNNAGATCGGAAG
122784 TATGAGTGTGGAGTCGTTGCTGTATGTGTGCTAAAAACATAAANNNAGATCGGAAG
122785 TATGAGTGTGGAGTCGTTGCACAAAACAATACCTTTCCTTCTANNNAGATCGGAAGA
122786 TATGAGTGTGGAGTCGTTGCGAGAAAATCTCTGGCCTCTTNNNAGATCGGAAGAGC
122787 TATGAGTGTGGAGTCGTTGCATCTAATTTCTAGAAGGAGTAGCNNNAGATCGGAAG
122788 TATGAGTGTGGAGTCGTTGCACCTGAAATGTATATTAGAGCTTNNNAGATCGGAAG
122790 TATGAGTGTGGAGTCGTTGCCTGTTTCCAAAGTCAATAGTGNNNAGATCGGAAGAG
122791 TATGAGTGTGGAGTCGTTGCCTTCTCACTTTACACTAGTCATANNNAGATCGGAAGA
122792 TATGAGTGTGGAGTCGTTGCGAAAGAAAACAAGGAAATGACTTNNNAGATCGGAA
122793 TATGAGTGTGGAGTCGTTGCTAGAATCTACGATTCACGATTTTNNNAGATCGGAAGA
122794 TATGAGTGTGGAGTCGTTGCTCTCCTTTGAATCTTTAGCTCNNNAGATCGGAAGAGC
122795 TATGAGTGTGGAGTCGTTGCAAGTAAATGTGTTGATTTTGTCTNNNAGATCGGAAG
122796 TATGAGTGTGGAGTCGTTGCGGAGAAATACTGGATATATGACANNNAGATCGGAA
122797 TATGAGTGTGGAGTCGTTGCTTCCATGTGTAATGCCTAATATANNNAGATCGGAAGA
122798 TATGAGTGTGGAGTCGTTGCATGAATCTAATTTCAGGAACACTNNNAGATCGGAAG
122799 TATGAGTGTGGAGTCGTTGCACAGCTTCTTATACAAATAAACTNNNAGATCGGAAGA
122801 TATGAGTGTGGAGTCGTTGCCTTTCAGTTTAGAACAACCTAAGNNNAGATCGGAAGA

122802 TATGAGTGTGGAGTCGTTGCGCTAGAATAGAAATGCAAAATGANNNAGATCGGAA
122803 TATGAGTGTGGAGTCGTTGCTTATTCTATTACCATAGCCTTCANNNAGATCGGAAGA
122804 TATGAGTGTGGAGTCGTTGCACTCATTGTTATCTTATTTCTGCNNNAGATCGGAAGA
122805 TATGAGTGTGGAGTCGTTGCACGTGTGTATATGTCTATGAAANNNAGATCGGAAG
122806 TATGAGTGTGGAGTCGTTGCTACCTGTAAGTATTTTGCTTTACNNNAGATCGGAAG
122807 TATGAGTGTGGAGTCGTTGCCTGAAGAGCAACCAAAATAATATNNNAGATCGGAA
122808 TATGAGTGTGGAGTCGTTGCGGCTCAAACAATTATGTAGTTTNNNAGATCGGAAGA
122809 TATGAGTGTGGAGTCGTTGCAAGATTGCTTGTTTTTAGGTAANNNAGATCGGAAG
122810 TATGAGTGTGGAGTCGTTGCGAGATAAATCCCTCACTCTTAATNNNAGATCGGAAG
122811 TATGAGTGTGGAGTCGTTGCGTTTATAAAGAAGGATCCTGAATNNNAGATCGGAA
122812 TATGAGTGTGGAGTCGTTGCAAACATTCTTTGAACTAATGAGGNNNAGATCGGAAG
122813 TATGAGTGTGGAGTCGTTGCAAGTCAAGGTCACTATTCCTANNNAGATCGGAAGAG
122814 TATGAGTGTGGAGTCGTTGCTACAGTGTTATAAGGCAATTTTTNNNAGATCGGAAG
122815 TATGAGTGTGGAGTCGTTGCCTTGGAAGTTATTGGTGTATATGNNNAGATCGGAA
122816 TATGAGTGTGGAGTCGTTGCTTTGCAAATAAACAAATTGAGACNNNAGATCGGAAG
122817 TATGAGTGTGGAGTCGTTGCAAAGACACTGTAAGAAGTAAAACNNNAGATCGGAA
122818 TATGAGTGTGGAGTCGTTGCAGGTATGTATGACTTCTGTTTTTNNNAGATCGGAAG
122819 TATGAGTGTGGAGTCGTTGCGAACAACCAAAGAGAGAAAATAANNNAGATCGGAA
122820 TATGAGTGTGGAGTCGTTGCCATGAAACACCTAAACATCAAGNNNAGATCGGAAGA
122821 TATGAGTGTGGAGTCGTTGCCATGTATGTATTAGAGGATATGCNNNAGATCGGAA
122822 TATGAGTGTGGAGTCGTTGCGTGTGCTTATGAATAACTCATTANNNAGATCGGAAG
122823 TATGAGTGTGGAGTCGTTGCAGAAAGGTAAACAAATCATTCAANNNAGATCGGAA
122824 TATGAGTGTGGAGTCGTTGCAAGTTCACAAAATATAAGGGATTNNNAGATCGGAA
122825 TATGAGTGTGGAGTCGTTGCTATCAAAGATTGGGAATGGAGNNNAGATCGGAAGA
122826 TATGAGTGTGGAGTCGTTGCTAGACAGACAGGAATATGAATTTNNNAGATCGGAA
122827 TATGAGTGTGGAGTCGTTGCTACAGCAAACATCTTACTTAAAGNNNAGATCGGAAG
122828 TATGAGTGTGGAGTCGTTGCGTATCCAAAATTTTTTATGCTCTNNNAGATCGGAAGA
122829 TATGAGTGTGGAGTCGTTGCGGATTTATTGATAAACACCATGTNNNAGATCGGAAG
122830 TATGAGTGTGGAGTCGTTGCGTCATTTCATTCTCAACTTACTAANNNAGATCGGAAG
122831 TATGAGTGTGGAGTCGTTGCCTGGTATAAAGAGTACTAGAGTTNNNAGATCGGAA
122832 TATGAGTGTGGAGTCGTTGCGGAAAAGATACAACAGGAAATAANNNAGATCGGAA
122833 TATGAGTGTGGAGTCGTTGCACAGATAATAAACTAGATGACCANNNAGATCGGAA
122834 TATGAGTGTGGAGTCGTTGCCCTGTACTAGAACAAAAATGTAANNNAGATCGGAAG
122835 TATGAGTGTGGAGTCGTTGCAAATGCCTCATGAGAATGATAATNNNAGATCGGAA
122836 TATGAGTGTGGAGTCGTTGCAAAAGTGTGAGCAATTATTCTAANNNAGATCGGAA
122837 TATGAGTGTGGAGTCGTTGCTTTATTTGACACCGACATAAAATNNNAGATCGGAAG
122839 TATGAGTGTGGAGTCGTTGCAAGAAATAGAAATGGGCTTGGNNNAGATCGGAAGA
122840 TATGAGTGTGGAGTCGTTGCTGCAGATAGAGAAGTTAAATAGATNNNAGATCGGA
122841 TATGAGTGTGGAGTCGTTGCTGCAGATAGAGAAGTTAAATAGATNNNAGATCGGA
122842 TATGAGTGTGGAGTCGTTGCAGAATCCAGCTCTTCATAGTNNNAGATCGGAAGAGC
122843 TATGAGTGTGGAGTCGTTGCTGGGATTTAAAAATTTGTCTTCANNNAGATCGGAAG
122844 TATGAGTGTGGAGTCGTTGCAGAGCAGAACAATTTAAAACTTTNNNAGATCGGAAG
122845 TATGAGTGTGGAGTCGTTGCCAAGGAAACTAGCTTTTTAGAAGNNNAGATCGGAA

122846 TATGAGTGTGGAGTCGTTGCCGCTTTCTTTTCCTAGGATTANNNAGATCGGAAGAG
122847 TATGAGTGTGGAGTCGTTGCAACTATGCCAGGAAGTAAAAATANNNAGATCGGAA
122848 TATGAGTGTGGAGTCGTTGCTACTTCTTAAGTGAATGAGTGATNNNAGATCGGAAG
122849 TATGAGTGTGGAGTCGTTGCCGTGTCTCTCCTATTCTTATAAGNNNAGATCGGAAGA
122850 TATGAGTGTGGAGTCGTTGCAGAGAACATATGAAAAACACAATNNNAGATCGGAA
122851 TATGAGTGTGGAGTCGTTGCTCCAGTAAGCCTTTTTAAAAAATNNNAGATCGGAAG
122852 TATGAGTGTGGAGTCGTTGCACTAGATTAAGAATGAGTTAGGCNNNAGATCGGAA
122853 TATGAGTGTGGAGTCGTTGCGTTCCTTCTTTACAGCATAATATNNNAGATCGGAAGA
122854 TATGAGTGTGGAGTCGTTGCTGGTCAGAACACTAAACTATTAANNNAGATCGGAAG
122855 TATGAGTGTGGAGTCGTTGCCAGGGAAAAGGAAAAAAAAAAAANNNAGATCGGA
122856 TATGAGTGTGGAGTCGTTGCGGTATGAAATAGAAGGGAATGATNNNAGATCGGA
122857 TATGAGTGTGGAGTCGTTGCTAGTCTGAGTCATTTCCTAAATTNNNAGATCGGAAG
122858 TATGAGTGTGGAGTCGTTGCTCTCAAAATTAGTTCAGAAAAGGNNNAGATCGGAAG
122859 TATGAGTGTGGAGTCGTTGCCGATGTCAAATAACTAGTTGATTNNNAGATCGGAAG
122860 TATGAGTGTGGAGTCGTTGCATATTTTACTGAATGTGGGACATNNNAGATCGGAAG
122861 TATGAGTGTGGAGTCGTTGCTTAAAAATGGCTGGTAAAACTTANNNAGATCGGAAG
122862 TATGAGTGTGGAGTCGTTGCACATTTCACAGGAAATTACATTTNNNAGATCGGAAG
122863 TATGAGTGTGGAGTCGTTGCATATATGTGTGTTTATGCAATGCNNNAGATCGGAAG
122864 TATGAGTGTGGAGTCGTTGCAAAACAGCAGATAATTGAAGAAANNNAGATCGGAA
122866 TATGAGTGTGGAGTCGTTGCGATATTCTTCACTTGTGAGTTACNNNAGATCGGAAG
122867 TATGAGTGTGGAGTCGTTGCATGACCTGTTCGCTTATATTATANNNAGATCGGAAG
122868 TATGAGTGTGGAGTCGTTGCGAAGTTAGCAAATCCAAAATCTNNNAGATCGGAAGA
122869 TATGAGTGTGGAGTCGTTGCTGAACAAAAAATTTTACTGTAGCNNNAGATCGGAAG
122870 TATGAGTGTGGAGTCGTTGCTAAATAAAATGTGTTGTCCATGTNNNAGATCGGAAG
122871 TATGAGTGTGGAGTCGTTGCGCATGACTGTAAGAAAGTAAAATNNNAGATCGGAA
122872 TATGAGTGTGGAGTCGTTGCCCTAGGTTGAATAACAAACAACNNNAGATCGGAAGA
122873 TATGAGTGTGGAGTCGTTGCGTTTGGTAAAAGCAGAACTGNNNAGATCGGAAGAG
122874 TATGAGTGTGGAGTCGTTGCGCAGTAATAGAAATCGAACATAGNNNAGATCGGAA
122875 TATGAGTGTGGAGTCGTTGCTTTTCTTTGCAAAAGTTTAATCCNNNAGATCGGAAGA
122876 TATGAGTGTGGAGTCGTTGCCAGAAATAGAGTGGAAACAGGNNNAGATCGGAAG
122877 TATGAGTGTGGAGTCGTTGCTCATTGTTTTTACTGCACATTATNNNAGATCGGAAGA
122878 TATGAGTGTGGAGTCGTTGCTGTCTCTTTCAGGTTATTTCATANNNAGATCGGAAGA
122879 TATGAGTGTGGAGTCGTTGCCTTCTCCAAATACAATATCTTGGNNNAGATCGGAAGA
122880 TATGAGTGTGGAGTCGTTGCTAATAGCAGACAGGTAACATTAANNNAGATCGGAA
122881 TATGAGTGTGGAGTCGTTGCGTGTGATGTGTGTATATTTACAANNNAGATCGGAA
122882 TATGAGTGTGGAGTCGTTGCCTGTAACAAAGCCAATTAATGTANNNAGATCGGAAG
122883 TATGAGTGTGGAGTCGTTGCCTAACACAGTTTAAATGCTTTTCNNNAGATCGGAAG
122884 TATGAGTGTGGAGTCGTTGCTGTAATTAGAAAGGAAAGTAGGANNNAGATCGGAA
122886 TATGAGTGTGGAGTCGTTGCGTTTTGTGAAAGGTAAGTAGATTNNNAGATCGGAA
122887 TATGAGTGTGGAGTCGTTGCCATGCACAAAAGAAACATTTGNNNAGATCGGAAGA
122888 TATGAGTGTGGAGTCGTTGCCACAGATTTGTCTTATACACAATNNNAGATCGGAAG
122889 TATGAGTGTGGAGTCGTTGCCACAGATTTGTCTTATACACAATNNNAGATCGGAAG
122890 TATGAGTGTGGAGTCGTTGCTAGAGTGGAAATTAAAATTCCACNNNAGATCGGAAG

122891 TATGAGTGTGGAGTCGTTGCGTTGGTGATATTTTTCTGTAGAANNNAGATCGGAAG
122892 TATGAGTGTGGAGTCGTTGCGAGGAGGGAAAGGAATTTATATTNNNAGATCGGAA
122893 TATGAGTGTGGAGTCGTTGCCTGAGGCTTCTATTTATTTTACCNNNAGATCGGAAGA
122894 TATGAGTGTGGAGTCGTTGCTATTCAGTATTTTCCTGTCTGAGNNNAGATCGGAAG
122895 TATGAGTGTGGAGTCGTTGCCTCCAAAATGCAAGTCACAANNNAGATCGGAAGAG
122896 TATGAGTGTGGAGTCGTTGCTTCTACATGTATATTTGTGTGTGNNNAGATCGGAAG
122897 TATGAGTGTGGAGTCGTTGCCCTCAAAGAAACAGAAATCTATGNNNAGATCGGAAG
122898 TATGAGTGTGGAGTCGTTGCTTGAGTGTTTAAGCATTTTAAGTNNNAGATCGGAAG
122899 TATGAGTGTGGAGTCGTTGCAAATAACACAAGACATCAATGTTNNNAGATCGGAAG
122900 TATGAGTGTGGAGTCGTTGCGGTTAGACAGGATAAAAGTTTGNNNAGATCGGAAG
122901 TATGAGTGTGGAGTCGTTGCGTTTCTCTGAATAACTCTTATGCNNNAGATCGGAAG
122902 TATGAGTGTGGAGTCGTTGCAATTTGAGAAACAAGGTTCTAAGNNNAGATCGGAA
122903 TATGAGTGTGGAGTCGTTGCACACATATATTGCTTATGAGGTANNNAGATCGGAAG
122904 TATGAGTGTGGAGTCGTTGCACACATATATTGCTTATGAGGTANNNAGATCGGAAG
122905 TATGAGTGTGGAGTCGTTGCTATCTTTCTCCTTTCGATCATACNNNAGATCGGAAGA
122906 TATGAGTGTGGAGTCGTTGCTATCTTTCTCCTTTCGATCATACNNNAGATCGGAAGA
122907 TATGAGTGTGGAGTCGTTGCCATCTTGCTTATCTTTGATTTCTNNNAGATCGGAAGA
122909 TATGAGTGTGGAGTCGTTGCGTTTAACTATAGTCAGTGGTGTNNNAGATCGGAAG
122910 TATGAGTGTGGAGTCGTTGCCAAATATAGCAAACACACACANNNAGATCGGAAGAG
122911 TATGAGTGTGGAGTCGTTGCTTATTCCTTATGGTTCCTTATGTNNNAGATCGGAAGA
122912 TATGAGTGTGGAGTCGTTGCTATGTGGTGCATATAGATTTGTNNNAGATCGGAAG
122913 TATGAGTGTGGAGTCGTTGCATATGTAATATAGCTTGTGCATCNNNAGATCGGAAG
122914 TATGAGTGTGGAGTCGTTGCGTGATTCCCACTTCCAATTCNNNAGATCGGAAGAGC
122915 TATGAGTGTGGAGTCGTTGCTTACCTACTCATTTATTGATGGGNNNAGATCGGAAG
122916 TATGAGTGTGGAGTCGTTGCCTCCAGAGAGCAATATTTCTGNNNAGATCGGAAGAG
122917 TATGAGTGTGGAGTCGTTGCGTAGGCTTTTCTGAATATCAAAGNNNAGATCGGAAG
122918 TATGAGTGTGGAGTCGTTGCACACACATACATAAACATCTACANNNAGATCGGAAG
122919 TATGAGTGTGGAGTCGTTGCACACACATACATAAACATCTACANNNAGATCGGAAG
122920 TATGAGTGTGGAGTCGTTGCACACACATACATAAACATCTACANNNAGATCGGAAG
122921 TATGAGTGTGGAGTCGTTGCTGTATGTATATCACAGTTTCCTTNNNAGATCGGAAG
122922 TATGAGTGTGGAGTCGTTGCTTTATAAATTTTGGGAAGTGGTTNNNAGATCGGAA
122923 TATGAGTGTGGAGTCGTTGCGAGATTAGCTCACATGATTATAGNNNAGATCGGAA
122926 TATGAGTGTGGAGTCGTTGCTTCATGACTTGAAACAAACATTTNNNAGATCGGAAG
122927 TATGAGTGTGGAGTCGTTGCTAATTTATTCTTTGAGCTCCTCTNNNAGATCGGAAGA
122928 TATGAGTGTGGAGTCGTTGCCTGGCATTTATCTCAGTTTTAAANNNAGATCGGAAG
122929 TATGAGTGTGGAGTCGTTGCATAATGGCTGTTGAATTTCATCNNNAGATCGGAAGA
122930 TATGAGTGTGGAGTCGTTGCCTTTACAGAGGAGAGGCTTCNNNAGATCGGAAGAG
122931 TATGAGTGTGGAGTCGTTGCGTTCAAGTGATTCTCTTGTCTNNNAGATCGGAAGAG
122932 TATGAGTGTGGAGTCGTTGCAAAAACATGTTGAGCAAAGTAANNNAGATCGGAAG
122933 TATGAGTGTGGAGTCGTTGCTTCACTAATGGCTGAATGATTNNNAGATCGGAAGAG
122934 TATGAGTGTGGAGTCGTTGCGGTTCACCTAAATGCATTTATATNNNAGATCGGAAG
122935 TATGAGTGTGGAGTCGTTGCATAAAGCAAATGTTAGCAGTATCNNNAGATCGGAA
122936 TATGAGTGTGGAGTCGTTGCATGACAAGTACAATGTTTTTCTTNNNAGATCGGAAG

122937 TATGAGTGTGGAGTCGTTGCACTGGATAATTCTGTGAATTTTCNNNAGATCGGAAG
122938 TATGAGTGTGGAGTCGTTGCTATTTTGCGAAAGGATTTCAAGNNNAGATCGGAAG
122939 TATGAGTGTGGAGTCGTTGCGAAATACCACTTTTCCTAAAGAGNNNAGATCGGAAG
122940 TATGAGTGTGGAGTCGTTGCACATGCATTACTCCATATAGTTANNNAGATCGGAAG
122941 TATGAGTGTGGAGTCGTTGCACATGCATTACTCCATATAGTTANNNAGATCGGAAG
122942 TATGAGTGTGGAGTCGTTGCACATGCATTACTCCATATAGTTANNNAGATCGGAAG
122943 TATGAGTGTGGAGTCGTTGCCTGAAGGATATAGGTCAATAGAGNNNAGATCGGAA
122944 TATGAGTGTGGAGTCGTTGCCACATACATAGGTTTTTGTTCATNNNAGATCGGAAG
122945 TATGAGTGTGGAGTCGTTGCCATACGTTTATTGACTGAACTAANNNAGATCGGAAG
122946 TATGAGTGTGGAGTCGTTGCGTAATAACCCATGTACATACAACNNNAGATCGGAAG
122947 TATGAGTGTGGAGTCGTTGCGCCTTCTTAAAAATACTCAGAATNNNAGATCGGAAG
122948 TATGAGTGTGGAGTCGTTGCATAAGTGTGTCTAACTTAGAAGCNNNAGATCGGAA
122949 TATGAGTGTGGAGTCGTTGCGGCTGAAGAGAGACTTTATCNNNAGATCGGAAGAG
122950 TATGAGTGTGGAGTCGTTGCCTTGTTAGATTGTTGAGATCCTNNNAGATCGGAAGA
122951 TATGAGTGTGGAGTCGTTGCCTAATGTGTGTATGTGCCTGNNNAGATCGGAAGAG
122952 TATGAGTGTGGAGTCGTTGCAAAATAAGAACTTTCAAAAGCTGNNNAGATCGGAA
122953 TATGAGTGTGGAGTCGTTGCATTAGACAAAGTCTTTGGTCTTNNNAGATCGGAAGA
122954 TATGAGTGTGGAGTCGTTGCAGTTAGCTACACATGACCATNNNAGATCGGAAGAG
122955 TATGAGTGTGGAGTCGTTGCCTTTTTGTTTATGGCTATCAAAGNNNAGATCGGAAG
122956 TATGAGTGTGGAGTCGTTGCCTTATGTGAATGCCTTTTATTCTNNNAGATCGGAAG
122957 TATGAGTGTGGAGTCGTTGCTAGGAAAAAAGCATACAAAGTTTNNNAGATCGGAA
122959 TATGAGTGTGGAGTCGTTGCCATGTACGCCATAGAAAAGANNNAGATCGGAAGAG
122960 TATGAGTGTGGAGTCGTTGCCTGTGTACTTTTGAGAAAATTCTNNNAGATCGGAAG
122961 TATGAGTGTGGAGTCGTTGCGGTCTTCATATCTCTATTTGGTTNNNAGATCGGAAG
122962 TATGAGTGTGGAGTCGTTGCCTTAAGGTCAGGAGTTCAAGNNNAGATCGGAAGAG
122963 TATGAGTGTGGAGTCGTTGCCCCCATACTATGCCTTTACTNNNAGATCGGAAGAGCA
122964 TATGAGTGTGGAGTCGTTGCACAACTGAGAATAAGAGAATTCTNNNAGATCGGAA
122965 TATGAGTGTGGAGTCGTTGCTTACAGCCTATTCACCACATANNNAGATCGGAAGAG
122966 TATGAGTGTGGAGTCGTTGCCCATAAGATTATAGTGTAGCTGANNNAGATCGGAA
122967 TATGAGTGTGGAGTCGTTGCGTTATTCACCACATAATGACATTNNNAGATCGGAAG
122968 TATGAGTGTGGAGTCGTTGCTTTTAATCATGCTTTTACCTACCNNNAGATCGGAAGA
122969 TATGAGTGTGGAGTCGTTGCTTTTTGCATAAGAGTTTCTGTAANNNAGATCGGAAG
122970 TATGAGTGTGGAGTCGTTGCTCATTTTCTTTATGACTTTGTTCNNNAGATCGGAAGA
122971 TATGAGTGTGGAGTCGTTGCCTTTTTTTGAGTCTCCAAGTCNNNAGATCGGAAGAG
122972 TATGAGTGTGGAGTCGTTGCATAAGAAAACACTCTGGCTTTTANNNAGATCGGAAG
122974 TATGAGTGTGGAGTCGTTGCTTCAAATATGTGAAAAGTCGTTTNNNAGATCGGAAG
122975 TATGAGTGTGGAGTCGTTGCAAAGAAAGAGGTTGTTTATCCAGNNNAGATCGGAA
122976 TATGAGTGTGGAGTCGTTGCCCTTTTTGGCTTTTGAATTTAATNNNAGATCGGAAGA
122977 TATGAGTGTGGAGTCGTTGCATTACCCACATGAATTTGATATGNNNAGATCGGAAG
122978 TATGAGTGTGGAGTCGTTGCCCACAACTTTTCATAGAAACTTTNNNAGATCGGAAGA
122979 TATGAGTGTGGAGTCGTTGCTTTTTCTTTTGTTTTGTCACAAANNNAGATCGGAAGA
122980 TATGAGTGTGGAGTCGTTGCTTTTTCTTTTGTTTTGTCACAAANNNAGATCGGAAGA
122981 TATGAGTGTGGAGTCGTTGCTTACTTTGGTCCATTTATTTCAGNNNAGATCGGAAGA

122982 TATGAGTGTGGAGTCGTTGCGTGAAAAGCTCTCTAACAAGATANNNAGATCGGAAG
122983 TATGAGTGTGGAGTCGTTGCTTATCCAATTCTAAACACTTCTGNNNAGATCGGAAGA
122984 TATGAGTGTGGAGTCGTTGCGAGATGCAAAAATCCTCAATAANNNAGATCGGAAGA
122985 TATGAGTGTGGAGTCGTTGCCACTCTGGTAACTATTGCAGNNNAGATCGGAAGAGC
122986 TATGAGTGTGGAGTCGTTGCAAGTTAGAGACTGAAACACAGNNNAGATCGGAAGA
122987 TATGAGTGTGGAGTCGTTGCCTAAAAACCTTTTCCTTCCTTTTNNNAGATCGGAAGAC
122988 TATGAGTGTGGAGTCGTTGCTATAGATGCTTCCTGATGATTCTNNNAGATCGGAAGA
122989 TATGAGTGTGGAGTCGTTGCTGGTGTAATTTAGGAGAAGTTCNNNAGATCGGAAGA
122990 TATGAGTGTGGAGTCGTTGCTTCAAATAAAAATCTAGGCCTTGNNNAGATCGGAAG
122991 TATGAGTGTGGAGTCGTTGCGAACCATTTGAAAGTAAAGAAGTNNNAGATCGGAAG
122992 TATGAGTGTGGAGTCGTTGCCAATACATAGCAATGATAGGTAAANNNAGATCGGAA
122993 TATGAGTGTGGAGTCGTTGCGATCTTGGAATGCCTATTCTATANNNAGATCGGAAGA
122994 TATGAGTGTGGAGTCGTTGCGATCTTGGAATGCCTATTCTATANNNAGATCGGAAGA
122995 TATGAGTGTGGAGTCGTTGCGGCAAATATAAAGCTTACCTATTNNNAGATCGGAAG
122996 TATGAGTGTGGAGTCGTTGCAGTATCTGTTATATTTCTGTGCTNNNAGATCGGAAGA
122997 TATGAGTGTGGAGTCGTTGCAGATCTAATTGGTTCATCATGTANNNAGATCGGAAG
122998 TATGAGTGTGGAGTCGTTGCTCTAGGGAGTTTTGAAATTATGANNNAGATCGGAAG
122999 TATGAGTGTGGAGTCGTTGCGTTGATGCTTTTTTAATTGTCTCNNNAGATCGGAAGA
123000 TATGAGTGTGGAGTCGTTGCAATTCTTACAAGAACCCTAGGNNNAGATCGGAAGAG
123001 TATGAGTGTGGAGTCGTTGCAGTGTGAGAGTAGGAGAAAANNNAGATCGGAAGAG
123002 TATGAGTGTGGAGTCGTTGCGCACATTAAAAAAAAAAAAACCCNNNAGATCGGAAG
123003 TATGAGTGTGGAGTCGTTGCTTCATTTCTTTCTATTTCTCCCTNNNAGATCGGAAGAG
123004 TATGAGTGTGGAGTCGTTGCGTGGTATGTTTTAATTTCTTCCTNNNAGATCGGAAGA
123005 TATGAGTGTGGAGTCGTTGCGATAAATCTAGATTCCCAATGAGNNNAGATCGGAAG
123006 TATGAGTGTGGAGTCGTTGCTGCAATAATTGTTTTAAATAACCANNNAGATCGGAAG
123007 TATGAGTGTGGAGTCGTTGCGTTGTCTGATTGCTCAACTGNNNAGATCGGAAGAGC
123008 TATGAGTGTGGAGTCGTTGCAAGATCCAGAGAAAAAGAGATAGNNNAGATCGGAA
123009 TATGAGTGTGGAGTCGTTGCTTTACTATTTTTGCTTTCCAGCNNNAGATCGGAAGAG
123010 TATGAGTGTGGAGTCGTTGCTCAGACATCATACACACAGGNNNAGATCGGAAGAGC
123011 TATGAGTGTGGAGTCGTTGCTTTGATACTTTGGCATCTTATTCNNNAGATCGGAAGA
123012 TATGAGTGTGGAGTCGTTGCTACATATATGGCTATGTGAGATGNNNAGATCGGAAG
123013 TATGAGTGTGGAGTCGTTGCGCACAATGAAATTCATACCTATATNNNAGATCGGAAG
123016 TATGAGTGTGGAGTCGTTGCCTATGAGCTGAACATTGAATTTTNNNAGATCGGAAG
123017 TATGAGTGTGGAGTCGTTGCTGCTAAAGCTTTTGAGATTTTAANNNAGATCGGAAG
123018 TATGAGTGTGGAGTCGTTGCCACATACATAGGTTTTTGTTCATNNNAGATCGGAAGA
123019 TATGAGTGTGGAGTCGTTGCTTTTATTTTGTCATACCCATCTCNNNAGATCGGAAGAG
123020 TATGAGTGTGGAGTCGTTGCGAGAGACTGCAGATACTGAGNNNAGATCGGAAGAG
123021 TATGAGTGTGGAGTCGTTGCACATTTTCTTTGTGGAACATTATNNNAGATCGGAAGA
123022 TATGAGTGTGGAGTCGTTGCGAAACCCCTTATAAAAGTGAAATNNNAGATCGGAAG
123023 TATGAGTGTGGAGTCGTTGCGCACACCTAAACTATAAGAAATATNNNAGATCGGAAG
123024 TATGAGTGTGGAGTCGTTGCCTTTCCTGGAAACCAAAATTATANNNAGATCGGAAGA
123025 TATGAGTGTGGAGTCGTTGCTATATATGTATATTTGGCCTGGGNNNAGATCGGAAG
123026 TATGAGTGTGGAGTCGTTGCGAGAGAAATATCTGTCAGTGTAANNNAGATCGGAAG

123027 TATGAGTGTGGAGTCGTTGCGCCATAATTCTTCCTCATACTTANNNAGATCGGAAGA
123028 TATGAGTGTGGAGTCGTTGCGGCTCTGCTATTCTATTTTTAAGNNNAGATCGGAAG
123029 TATGAGTGTGGAGTCGTTGCAACACCAGTACTATTGGATTAGNNNAGATCGGAAG
123030 TATGAGTGTGGAGTCGTTGCAACACCAGTACTATTGGATTAGNNNAGATCGGAAG
123031 TATGAGTGTGGAGTCGTTGCAACACCAGTACTATTGGATTAGNNNAGATCGGAAG
123032 TATGAGTGTGGAGTCGTTGCGCACTCCTACATGTATACATATGNNNAGATCGGAAG
123033 TATGAGTGTGGAGTCGTTGCATCTCAGTCATTTCAAGTTTGANNNAGATCGGAAGA
123034 TATGAGTGTGGAGTCGTTGCTTCTGTTTTAGACAAAATCGAAANNNAGATCGGAAG
123035 TATGAGTGTGGAGTCGTTGCGGTGACTAAGGATTATGAATGTNNNAGATCGGAAG
123036 TATGAGTGTGGAGTCGTTGCAAATTCTGCTGTAATTTCAACTTNNNAGATCGGAAG
123037 TATGAGTGTGGAGTCGTTGCCTTCTATTTCCTACATTGTCTCTNNNAGATCGGAAGA
123038 TATGAGTGTGGAGTCGTTGCGTTGGTTTTTTTGTTGTTGTTNNNAGATCGGAAGAG
123039 TATGAGTGTGGAGTCGTTGCCATTCAACCTTGGATTGAAAATANNNAGATCGGAAG
123040 TATGAGTGTGGAGTCGTTGCCTAAAAGGAACTAAGGAAAAACANNNAGATCGGAA
123041 TATGAGTGTGGAGTCGTTGCAGTTTCATCATTTACTTGGAAGTNNNAGATCGGAAG
123042 TATGAGTGTGGAGTCGTTGCAGTTTCATCATTTACTTGGAAGTNNNAGATCGGAAG
123043 TATGAGTGTGGAGTCGTTGCAAAAGAATCATGATGTCTTTTGTNNNAGATCGGAAG
123044 TATGAGTGTGGAGTCGTTGCCTGACACAGGAGATATTTTTTTANNNAGATCGGAAG
123045 TATGAGTGTGGAGTCGTTGCCCACAGAGCCACCATATTAANNNAGATCGGAAGAGC
123046 TATGAGTGTGGAGTCGTTGCGGTCTAGTGCTAGTATTTATGTGNNNAGATCGGAAG
123047 TATGAGTGTGGAGTCGTTGCCTCACAGGTACAAGTCAATAATNNNAGATCGGAAGA
123048 TATGAGTGTGGAGTCGTTGCCATCCTTTGAGATTATAGTTTCGNNNAGATCGGAAG
123049 TATGAGTGTGGAGTCGTTGCATTTATACCTCCCTAATGACTACNNNAGATCGGAAGA
123050 TATGAGTGTGGAGTCGTTGCATAGGAACATGATCTAAATGGTTNNNAGATCGGAA
123051 TATGAGTGTGGAGTCGTTGCTAGAGAAGGTTTGTTTTGCATNNNAGATCGGAAGA
123052 TATGAGTGTGGAGTCGTTGCGATGTAGAGAAGGTTTGTTTTGNNNAGATCGGAAG
123053 TATGAGTGTGGAGTCGTTGCCAATATCTTGACCCAAACTTTCNNNAGATCGGAAGA
123054 TATGAGTGTGGAGTCGTTGCTATAGTGTCAAAGCATTGTAAACNNNAGATCGGAAG
123056 TATGAGTGTGGAGTCGTTGCCATTGAGTACTGGTAGTGGGNNNAGATCGGAAGAG
123057 TATGAGTGTGGAGTCGTTGCGGTTGTCAGGTATTTTTATCTTTNNNAGATCGGAAG
123058 TATGAGTGTGGAGTCGTTGCTGACCAATAGTTATTCCAATTTCNNNAGATCGGAAG
123059 TATGAGTGTGGAGTCGTTGCAAGGAGACTACAATGAGAATTTTNNNAGATCGGAA
123060 TATGAGTGTGGAGTCGTTGCCTGTGCTAATTTCAATAGTACTTNNNAGATCGGAAG
123061 TATGAGTGTGGAGTCGTTGCGTCACATAGTTTGTAGAATGTTCNNNAGATCGGAAG
123062 TATGAGTGTGGAGTCGTTGCACATTATACTGGGTTAATGACCNNNAGATCGGAAGA
123063 TATGAGTGTGGAGTCGTTGCCCCATAAATACTGCCATAATGANNNAGATCGGAAGA
123064 TATGAGTGTGGAGTCGTTGCCCCATAAATACTGCCATAATGANNNAGATCGGAAGA
123065 TATGAGTGTGGAGTCGTTGCACAGTGTTCTTTATTGGTTTTAANNNAGATCGGAAG
123066 TATGAGTGTGGAGTCGTTGCAGAAATGTTTATTGCAGCTTTATNNNAGATCGGAAG
123067 TATGAGTGTGGAGTCGTTGCATATAAAATAAGGAATTGGCTCGNNNAGATCGGAA
123068 TATGAGTGTGGAGTCGTTGCTATGTGTGCTATTGTCAACTTANNNAGATCGGAAGA
123069 TATGAGTGTGGAGTCGTTGCGCTGGCACAATATGATAATAATANNNAGATCGGAAG
123070 TATGAGTGTGGAGTCGTTGCATTTCTAACATGCCCATGATATANNNAGATCGGAAG

123071 TATGAGTGTGGAGTCGTTGCAACACAGATACAAAAATGTTGATNNNAGATCGGAA
123072 TATGAGTGTGGAGTCGTTGCCTGCACTCATTTTTTAGCTTTANNNAGATCGGAAGAG
123074 TATGAGTGTGGAGTCGTTGCCTCATTTGGTTATTTTCCTGGNNNAGATCGGAAGAG
123075 TATGAGTGTGGAGTCGTTGCCATCATAATACCAAACACTTGTTNNNAGATCGGAAG
123076 TATGAGTGTGGAGTCGTTGCCTATGCATGTACTTGTGTATGTNNNAGATCGGAAGA
123077 TATGAGTGTGGAGTCGTTGCCTCTGTCTTTGCCTTAATAAATTNNNAGATCGGAAGA
123078 TATGAGTGTGGAGTCGTTGCAATTATGAACAAGAGGGAGTATTNNNAGATCGGAA
123079 TATGAGTGTGGAGTCGTTGCAGAACATATTTTTCTTGGAAACANNNAGATCGGAAG
123080 TATGAGTGTGGAGTCGTTGCGGACAATTAGGTAAAAAAAGAGANNNAGATCGGA
123081 TATGAGTGTGGAGTCGTTGCTCTGTGTAGACATGGTCTATATANNNAGATCGGAAG
123082 TATGAGTGTGGAGTCGTTGCGTACTTGCATTGGTAGTGATNNNAGATCGGAAGAG
123083 TATGAGTGTGGAGTCGTTGCTCTACTAAAAATAAAAACTTAGCCANNNAGATCGGA
123084 TATGAGTGTGGAGTCGTTGCCCAGAGATTAAACCAGGAAGNNNAGATCGGAAGAG
123085 TATGAGTGTGGAGTCGTTGCGCTTTTACATTTATTGAAGCATTNNNAGATCGGAAG
123086 TATGAGTGTGGAGTCGTTGCAAAGACATCTCTAATGAGGTCANNNAGATCGGAAG
123087 TATGAGTGTGGAGTCGTTGCAGAAGTGAAGTTGAGAATAGAATNNNAGATCGGA
123088 TATGAGTGTGGAGTCGTTGCGATCAACTGAAGAAAATCTAACANNNAGATCGGAA
123089 TATGAGTGTGGAGTCGTTGCCTTTTGGTATATTATGGCATACCNNNAGATCGGAAG
123090 TATGAGTGTGGAGTCGTTGCCTTCAAAATACACAAAGAAGACANNNAGATCGGAAG
123091 TATGAGTGTGGAGTCGTTGCCTTCAAAATACACAAAGAAGACANNNAGATCGGAAG
123092 TATGAGTGTGGAGTCGTTGCAGTCACACACATGGTGATAGNNNAGATCGGAAGAG
123093 TATGAGTGTGGAGTCGTTGCAGTCACACACATGGTGATAGNNNAGATCGGAAGAG
123094 TATGAGTGTGGAGTCGTTGCGTGTTTTATCCATTTACCTAGTCNNNAGATCGGAAGA
123095 TATGAGTGTGGAGTCGTTGCGATCTAAAGAAACACTGCAATTNNNAGATCGGAAGA
123096 TATGAGTGTGGAGTCGTTGCCTGTTGAATTCACTCTGTGTNNNAGATCGGAAGAGC
123097 TATGAGTGTGGAGTCGTTGCCATCTTGTCCTAAAAATAAGGAGNNNAGATCGGAAG
123098 TATGAGTGTGGAGTCGTTGCATAGTATCCAATACACACACAGNNNAGATCGGAAGA
123099 TATGAGTGTGGAGTCGTTGCTATAGTATCCAATACACACACAGNNNAGATCGGAAG
123100 TATGAGTGTGGAGTCGTTGCGGCAAAGAAATATCAGAGTACTANNNAGATCGGAA
123101 TATGAGTGTGGAGTCGTTGCTTCCTATGAATTTGTCAAAATGTNNNAGATCGGAAG
123102 TATGAGTGTGGAGTCGTTGCTAATGACTGACTTTCATATTTGCNNNAGATCGGAAG
123103 TATGAGTGTGGAGTCGTTGCAGACATAATGACTGACTTTCATANNNAGATCGGAAG
123104 TATGAGTGTGGAGTCGTTGCTATCTTGTATCATCTACCTACCTNNNAGATCGGAAGA
123105 TATGAGTGTGGAGTCGTTGCGGATAACCCAGACATATATGTATNNNAGATCGGAAG
123106 TATGAGTGTGGAGTCGTTGCCTTTCTGGCTACTTGTTTTATTTNNNAGATCGGAAGA
123107 TATGAGTGTGGAGTCGTTGCACAAATACTTGAGGTTTCTATCANNNAGATCGGAAG
123108 TATGAGTGTGGAGTCGTTGCCTGGGCAACAAGTTTAATTGNNNAGATCGGAAGAG
123109 TATGAGTGTGGAGTCGTTGCCATAGAAGTATGTTTACAAAGCCNNNAGATCGGAAG
123110 TATGAGTGTGGAGTCGTTGCAGAGAGTTTTCATGCATTATAACNNNAGATCGGAAG
123111 TATGAGTGTGGAGTCGTTGCGCACAGTATCTAGTGGATATAAANNNAGATCGGAA
123112 TATGAGTGTGGAGTCGTTGCCTTAATTAGTGGAGCAAATGAAANNNAGATCGGAA
123113 TATGAGTGTGGAGTCGTTGCAGAGATCTTCCTCTCCTTATAANNNAGATCGGAAGA
123114 TATGAGTGTGGAGTCGTTGCGTACAGATGTGTGCTAGTTAATNNNAGATCGGAAG

123115 TATGAGTGTGGAGTCGTTGCTTATCTGTACTTTTGCTTTTCATNNNAGATCGGAAGA
123116 TATGAGTGTGGAGTCGTTGCGGTTTTTGTCATGATTTCTGGNNNAGATCGGAAGAG
123117 TATGAGTGTGGAGTCGTTGCTCTCAGTTTTGCTCTGTTTCNNNAGATCGGAAGAGCA
123118 TATGAGTGTGGAGTCGTTGCACTTAAAATGCTATACTCAGAGANNNAGATCGGAAG
123119 TATGAGTGTGGAGTCGTTGCTTTATTCCAAATACTGAACTCAGNNNAGATCGGAAG
123120 TATGAGTGTGGAGTCGTTGCAGACAAGAGAAATTGGAGTTAANNNAGATCGGAAG
123121 TATGAGTGTGGAGTCGTTGCCAGCAGAAGAGATAGTCATAAATNNNAGATCGGAA
123122 TATGAGTGTGGAGTCGTTGCAGGTTTCTATTTTCCTTGGATAANNNAGATCGGAAG
123123 TATGAGTGTGGAGTCGTTGCCAATCTTTATGAAGAAGAACGAANNNAGATCGGAA
123125 TATGAGTGTGGAGTCGTTGCCATGGGTCTCTAGCTTCTAGNNNAGATCGGAAGAGC
123126 TATGAGTGTGGAGTCGTTGCCATGGGTCTCTAGCTTCTAGNNNAGATCGGAAGAGC
123127 TATGAGTGTGGAGTCGTTGCTCTCATATGTAATGAATCCCTAGNNNAGATCGGAAG
123128 TATGAGTGTGGAGTCGTTGCGTATGTATGTTGAGACAGAGAANNNAGATCGGAAG
123129 TATGAGTGTGGAGTCGTTGCGTATGTATGTTGAGACAGAGAANNNAGATCGGAAG
123130 TATGAGTGTGGAGTCGTTGCGAATGCCTTACATATCATTCTATNNNAGATCGGAAG
123131 TATGAGTGTGGAGTCGTTGCCTAGGTTCAAGTGATTCTCCNNNAGATCGGAAGAGC
123132 TATGAGTGTGGAGTCGTTGCACTTTAGACATAAAGGGTGATAANNNAGATCGGAA
123133 TATGAGTGTGGAGTCGTTGCACTTTAGACATAAAGGGTGATAANNNAGATCGGAA
123134 TATGAGTGTGGAGTCGTTGCAAGAAACTACAACAGAGTTTGTANNNAGATCGGAA
123135 TATGAGTGTGGAGTCGTTGCGATTTTTCATGACCTCAAGTAACNNNAGATCGGAAG
123136 TATGAGTGTGGAGTCGTTGCCATCATACTCATCTTCAAAGAAANNNAGATCGGAAG
123137 TATGAGTGTGGAGTCGTTGCTACTCACATGCTATGAATTTGANNNAGATCGGAAGA
123138 TATGAGTGTGGAGTCGTTGCTGGTACAATCTTAACTCACTGNNNAGATCGGAAGAG
123139 TATGAGTGTGGAGTCGTTGCCATGTCACCTGGAAATCTTGNNNAGATCGGAAGAGC
123140 TATGAGTGTGGAGTCGTTGCTTGCTAGCTCAATAACTTTCTATNNNAGATCGGAAGA
123141 TATGAGTGTGGAGTCGTTGCAAAATACAGGACAAGGATATACANNNAGATCGGAA
123142 TATGAGTGTGGAGTCGTTGCCTACTAAAGTGAGGGAACTAAGNNNAGATCGGAAG
123143 TATGAGTGTGGAGTCGTTGCTTTATAAAGTCATCTGTTGTTGGNNNAGATCGGAAG
123145 TATGAGTGTGGAGTCGTTGCATTCAGTGAGTATGGAGAGAANNNAGATCGGAAGA
123146 TATGAGTGTGGAGTCGTTGCAACTAGAGTTTTCAAAGTACAAGNNNAGATCGGAA
123147 TATGAGTGTGGAGTCGTTGCTTTCCCTGAATAGGACACATNNNAGATCGGAAGAGC
123148 TATGAGTGTGGAGTCGTTGCCTGACAACAGGTACCATTTATTNNNAGATCGGAAGA
123149 TATGAGTGTGGAGTCGTTGCCTGTGTGCCTACCATATTACNNNAGATCGGAAGAGC
123150 TATGAGTGTGGAGTCGTTGCGACTGATTGAAAGCCATTATAACNNNAGATCGGAAG
123151 TATGAGTGTGGAGTCGTTGCGACAGAATTTAGCAGAAAGAACNNNAGATCGGAAG
123152 TATGAGTGTGGAGTCGTTGCAACACAAACAGATCCATCTTTTANNNAGATCGGAAG
123153 TATGAGTGTGGAGTCGTTGCGATCATTCGTGACCCTTTATANNNAGATCGGAAGAG
123154 TATGAGTGTGGAGTCGTTGCCAATCCAAAGTAGAAGTAAATGTNNNAGATCGGAA
123155 TATGAGTGTGGAGTCGTTGCATCTAGCAAGTTCTTATTCTGGNNNAGATCGGAAGA
123156 TATGAGTGTGGAGTCGTTGCATTATAACCATGGCAACCTTTANNNAGATCGGAAGA
123157 TATGAGTGTGGAGTCGTTGCCCATGTATATGCATAAAAGAACANNNAGATCGGAAG
123158 TATGAGTGTGGAGTCGTTGCATAGTGCAATATATGAAGGCTTANNNAGATCGGAA
123159 TATGAGTGTGGAGTCGTTGCTAAAAGATTCCTCTTGGGAGATANNNAGATCGGAA

123160 TATGAGTGTGGAGTCGTTGCGACATGCATCTCAATGAATAGANNNAGATCGGAAGA
123161 TATGAGTGTGGAGTCGTTGCGAGGGTGAAGTTGTTTTTTTTNNNAGATCGGAAGA
123162 TATGAGTGTGGAGTCGTTGCTTTCATATATCATGATGCTTCACNNNAGATCGGAAGA
123163 TATGAGTGTGGAGTCGTTGCCTAAGATCATAGTGAAGTGAGGNNNAGATCGGAAG
123164 TATGAGTGTGGAGTCGTTGCCATCACTGCATTGTAGAAAAANNNAGATCGGAAGAG
123165 TATGAGTGTGGAGTCGTTGCAAATAGCATTTCTCTACATTTGGNNNAGATCGGAAG
123166 TATGAGTGTGGAGTCGTTGCTACGATCTCATCTTAGCTAACTANNNAGATCGGAAGA
123167 TATGAGTGTGGAGTCGTTGCCCTGTTTCATCTTTAAACTCATTNNNAGATCGGAAGA
123168 TATGAGTGTGGAGTCGTTGCATTATTTGCACTGTCATGTTTATNNNAGATCGGAAGA
123169 TATGAGTGTGGAGTCGTTGCTAAAACCTCTGCCTTTTGTTNNNAGATCGGAAGAGC
123170 TATGAGTGTGGAGTCGTTGCATAGGAAATGAAACACCATTGTTNNNAGATCGGAAG
123171 TATGAGTGTGGAGTCGTTGCGATAGACAGAGAGATAGAGAATGNNNAGATCGGA
123172 TATGAGTGTGGAGTCGTTGCGATCTCAGAAGAAACCTAACTTTNNNAGATCGGAAG
123173 TATGAGTGTGGAGTCGTTGCAAACACAACTGAGATCTAAAAATNNNAGATCGGAAG
123174 TATGAGTGTGGAGTCGTTGCCTTTCCTTCTTTCTTTCTTTCTTNNNAGATCGGAAGAG
123175 TATGAGTGTGGAGTCGTTGCATAGATAACAGAAGAACATCTGGNNNAGATCGGAA
123177 TATGAGTGTGGAGTCGTTGCTACCAGTGGATCTAGATGTACTANNNAGATCGGAAG
123178 TATGAGTGTGGAGTCGTTGCTACCAGTGGATCTAGATGTACTANNNAGATCGGAAG
123179 TATGAGTGTGGAGTCGTTGCTACCAGTGGATCTAGATGTACTANNNAGATCGGAAG
123180 TATGAGTGTGGAGTCGTTGCTAATCCTGACAAACATAGTCTTCNNNAGATCGGAAG
123181 TATGAGTGTGGAGTCGTTGCCAGAATAGCTGACAACAAATGNNNAGATCGGAAGA
123182 TATGAGTGTGGAGTCGTTGCTCCTATAACACTACCCTAGTTATNNNAGATCGGAAGA
123183 TATGAGTGTGGAGTCGTTGCAATAATTGGAAGACTTTTGATGTNNNAGATCGGAAG
123184 TATGAGTGTGGAGTCGTTGCGCTGAGATAATTTTATGTTAGACANNNAGATCGGAA
123185 TATGAGTGTGGAGTCGTTGCTCTGTTCATTTGTTTGGTCTANNNAGATCGGAAGAG
123186 TATGAGTGTGGAGTCGTTGCTCACTATAGGATTCTCAAAATGTNNNAGATCGGAAG
123187 TATGAGTGTGGAGTCGTTGCGGGGTGTTTATGTATGTCTTCNNNAGATCGGAAGA
123188 TATGAGTGTGGAGTCGTTGCTAGGAAGCAGGTGACATATGNNNAGATCGGAAGAG
123189 TATGAGTGTGGAGTCGTTGCTCTCCATATCTATCTGTATCTCANNNAGATCGGAAGA
123190 TATGAGTGTGGAGTCGTTGCTAAAATGGTTGATATGGGAAATGNNNAGATCGGAA
123191 TATGAGTGTGGAGTCGTTGCTTTACGTTTCTCTCTTACTCTCNNNAGATCGGAAGAG
123192 TATGAGTGTGGAGTCGTTGCTTTCTGCACCTGTCTATAGTANNNAGATCGGAAGAG
123193 TATGAGTGTGGAGTCGTTGCATATCTTAGTGCATATGTGTATGNNNAGATCGGAAG
123194 TATGAGTGTGGAGTCGTTGCTCCTAAAGGAACAAAAAGGGNNNAGATCGGAAGAG
123195 TATGAGTGTGGAGTCGTTGCCTGAGTGTTCTGGATTTAAATACNNNAGATCGGAAG
123196 TATGAGTGTGGAGTCGTTGCCTCTCAAACTCAAGTGATCTGNNNAGATCGGAAGAG
123197 TATGAGTGTGGAGTCGTTGCTTATGAATACATTTAGGGTGGATNNNAGATCGGAA
123198 TATGAGTGTGGAGTCGTTGCCCACCTAAGTAATAGCTATGTTNNNAGATCGGAAGA
123199 TATGAGTGTGGAGTCGTTGCATAATTCTGGGGTTATGTTTACANNNAGATCGGAAG
123200 TATGAGTGTGGAGTCGTTGCGAATGGAACATTGAGACTTTTGNNNAGATCGGAAG
123201 TATGAGTGTGGAGTCGTTGCGTGGAAAGTAAATCCGGAGANNNAGATCGGAAGA
123202 TATGAGTGTGGAGTCGTTGCAAATGGAGATACTTCAGCTATAGNNNAGATCGGAA
123203 TATGAGTGTGGAGTCGTTGCATAGTAAAGTTGGAGTAGTTGTTNNNAGATCGGAA

123204 TATGAGTGTGGAGTCGTTGCTTTGTGATATGTAAATCCACAAGNNNAGATCGGAAG
123205 TATGAGTGTGGAGTCGTTGCGTGTAGAAGACTGACATGAGNNNAGATCGGAAGAG
123206 TATGAGTGTGGAGTCGTTGCATTTTGGTAACCATATGTACAACNNNAGATCGGAAGA
123207 TATGAGTGTGGAGTCGTTGCGCCTGTATCTCAGAGAACTTNNNAGATCGGAAGAGC
123208 TATGAGTGTGGAGTCGTTGCGCCTATATTAGTCTGTTTTGTTTNNNAGATCGGAAGA
123209 TATGAGTGTGGAGTCGTTGCTAAACTGTCAATTTGTTTCTTCTNNNAGATCGGAAGA
123210 TATGAGTGTGGAGTCGTTGCCTACTGATGCAAAAGGAGGANNNAGATCGGAAGAG
123211 TATGAGTGTGGAGTCGTTGCAAAATCTACCATTTCTACCAGATNNNAGATCGGAAGA
123212 TATGAGTGTGGAGTCGTTGCATAAGCACTCCATACAGAGAATNNNAGATCGGAAGA
123213 TATGAGTGTGGAGTCGTTGCAGCTTAATTATTACTCAGTTGTGNNNAGATCGGAAGA
123214 TATGAGTGTGGAGTCGTTGCTATCTGCATCATACACTTTTTACNNNAGATCGGAAGA
123215 TATGAGTGTGGAGTCGTTGCATTTGAGATAGGGCTACATTTTNNNAGATCGGAAGA
123216 TATGAGTGTGGAGTCGTTGCTACAGTTTTGCCTACAATTTCANNNAGATCGGAAGAG
123217 TATGAGTGTGGAGTCGTTGCGGGACATAGCAAAATGGTAATNNNAGATCGGAAGA
123218 TATGAGTGTGGAGTCGTTGCGATGATTTTCTGGTTTTGTTTAGNNNAGATCGGAAG
123219 TATGAGTGTGGAGTCGTTGCTGTCATGGGTAGTCAATTTTATANNNAGATCGGAAG
123220 TATGAGTGTGGAGTCGTTGCATATGTAAATATGGAGTACCAGCNNNAGATCGGAAG
123221 TATGAGTGTGGAGTCGTTGCATATGTAAATATGGAGTACCAGCNNNAGATCGGAAG
123222 TATGAGTGTGGAGTCGTTGCAGTTGTGTTGATTTAATGAATGANNNAGATCGGAAG
123223 TATGAGTGTGGAGTCGTTGCGGGGTTGTTATCTCTTATTTTTTNNNAGATCGGAAGA
123224 TATGAGTGTGGAGTCGTTGCCTCCAGAGAGACTACATCTGNNNAGATCGGAAGAGC
123225 TATGAGTGTGGAGTCGTTGCATATACGCACAATATATCTGGATNNNAGATCGGAAG
123226 TATGAGTGTGGAGTCGTTGCGGTCTCACTATGTTATTCAGGNNNAGATCGGAAGAG
123227 TATGAGTGTGGAGTCGTTGCCTCTCAGTACCATACCATAATTCNNNAGATCGGAAGA
123228 TATGAGTGTGGAGTCGTTGCAACAGAGTAGGTAAAGAATCTTTNNNAGATCGGAAG
123229 TATGAGTGTGGAGTCGTTGCTTTTACATTCTCCTCTACTTTCTNNNAGATCGGAAGAG
123230 TATGAGTGTGGAGTCGTTGCATATACCTGGACATCTGTACTATNNNAGATCGGAAGA
123231 TATGAGTGTGGAGTCGTTGCTAAAGACTTTCTCAAACAAACAANNNAGATCGGAAGA
123232 TATGAGTGTGGAGTCGTTGCCTTTCTATTCTGGGAGATTTTCTNNNAGATCGGAAGA
123234 TATGAGTGTGGAGTCGTTGCCTTGGATTTAAGAGAACGTCTNNNAGATCGGAAGAG
123235 TATGAGTGTGGAGTCGTTGCTTCTGGAAGATATTGTGTAGAATNNNAGATCGGAAG
123236 TATGAGTGTGGAGTCGTTGCCTGTTACTCTTCTTAGACACCNNNAGATCGGAAGAGC
123237 TATGAGTGTGGAGTCGTTGCTACATATGGTCTAAGCTGAACATNNNAGATCGGAAG
123238 TATGAGTGTGGAGTCGTTGCGTCTGACCTTTACTTTCTCTCNNNAGATCGGAAGAGC
123240 TATGAGTGTGGAGTCGTTGCAGATAGATAGATATCACAATTTCTTNNNAGATCGGAA
123241 TATGAGTGTGGAGTCGTTGCGCCATAAAATCAGACTAATCCTANNNAGATCGGAAGA
123242 TATGAGTGTGGAGTCGTTGCGTCTATCTAGCTCTACAACTTACNNNAGATCGGAAGA
123243 TATGAGTGTGGAGTCGTTGCTTTTGATAACTTTCCTTTCCAAGNNNAGATCGGAAGA
123244 TATGAGTGTGGAGTCGTTGCGGTGCTACACTTATTTTTGTTTTNNNAGATCGGAAGA
123245 TATGAGTGTGGAGTCGTTGCGTGAGAGAGAGAGACATTATAATNNNAGATCGGAA
123246 TATGAGTGTGGAGTCGTTGCTTTCCCTTATATGTCATCTCAAANNNAGATCGGAAGA
123247 TATGAGTGTGGAGTCGTTGCGTGTAACTCCCAAATCTACATATNNNAGATCGGAAGA
123248 TATGAGTGTGGAGTCGTTGCGTGTAACTCCCAAATCTACATATNNNAGATCGGAAGA

123249 TATGAGTGTGGAGTCGTTGCTTTGTTTCATATTCCCTATGTTCNNNAGATCGGAAGA
123250 TATGAGTGTGGAGTCGTTGCAACACCACCAATTTCATTATAACNNNAGATCGGAAGA
123251 TATGAGTGTGGAGTCGTTGCTATAAGGGATTGATTCATGTGATNNNAGATCGGAAG
123252 TATGAGTGTGGAGTCGTTGCTATAAGGGATTGATTCATGTGATNNNAGATCGGAAG
123253 TATGAGTGTGGAGTCGTTGCTGTATCTAGACTACAGTAAGTCANNNAGATCGGAAG
123254 TATGAGTGTGGAGTCGTTGCTAACTTCCATTATGTTAGCAATGNNNAGATCGGAAG
123255 TATGAGTGTGGAGTCGTTGCGTAGAGATAAGGAAGAGAGAGTANNNAGATCGGA
123256 TATGAGTGTGGAGTCGTTGCTATATACACCACACTTTATCCACNNNAGATCGGAAGA
123257 TATGAGTGTGGAGTCGTTGCTGTATAAAAGCAACCAGTATCTANNNAGATCGGAAG
123258 TATGAGTGTGGAGTCGTTGCCTTTTTCCATTGTTTTATTGCTCNNNAGATCGGAAGA
123259 TATGAGTGTGGAGTCGTTGCTATAGTCAAAATACAGAACAGGTNNNAGATCGGAAG
123260 TATGAGTGTGGAGTCGTTGCCATATCTTCAGAGGTCATCATTNNNAGATCGGAAGA
123261 TATGAGTGTGGAGTCGTTGCCTACTAAGGAAGAATCCACTTCNNNAGATCGGAAGA
123262 TATGAGTGTGGAGTCGTTGCATTATGTTGTGCATAGATACTTGNNNAGATCGGAAG
123263 TATGAGTGTGGAGTCGTTGCGAGTTTGCCCATTAGTACATNNNAGATCGGAAGAG
123264 TATGAGTGTGGAGTCGTTGCGAGTTTGCCCATTAGTACATNNNAGATCGGAAGAG
123265 TATGAGTGTGGAGTCGTTGCAAAATGGGTAGATTTTATTGCATNNNAGATCGGAAG
123266 TATGAGTGTGGAGTCGTTGCATAACCACAGCATTGTACATAANNNAGATCGGAAGA
123267 TATGAGTGTGGAGTCGTTGCATAACCACAGCATTGTACATAANNNAGATCGGAAGA
123268 TATGAGTGTGGAGTCGTTGCTAATGATTTATGAGATCACACCTNNNAGATCGGAAG
123269 TATGAGTGTGGAGTCGTTGCATTCATGCACATGTACACACNNNAGATCGGAAGAGC
123270 TATGAGTGTGGAGTCGTTGCTAACCCTCCTCTTTTCTAATTCNNNAGATCGGAAGAG
123271 TATGAGTGTGGAGTCGTTGCAATGAAATTTGTCAATGTCTTTGNNNAGATCGGAAG
123272 TATGAGTGTGGAGTCGTTGCTCTATTGCTCCTCTAGACTGNNNAGATCGGAAGAGC
123273 TATGAGTGTGGAGTCGTTGCCCCCTCTCCTCTAAATAATTATTNNNAGATCGGAAGA
123274 TATGAGTGTGGAGTCGTTGCTGTGTCCTCTTATAATGACATAANNNAGATCGGAAG
123275 TATGAGTGTGGAGTCGTTGCCTTTGACGGTTGTAGATATTTTTNNNAGATCGGAAG
123276 TATGAGTGTGGAGTCGTTGCTATGTAAAAGACTAAGATCCTGGNNNAGATCGGAAG
123277 TATGAGTGTGGAGTCGTTGCTATGTAAAAGACTAAGATCCTGGNNNAGATCGGAAG
123278 TATGAGTGTGGAGTCGTTGCTTTAACTGCCTTTTTGATATCTCNNNAGATCGGAAGA
123279 TATGAGTGTGGAGTCGTTGCCACTTTTCAATTCCTTGAATTTGNNNAGATCGGAAGA
123280 TATGAGTGTGGAGTCGTTGCCAAGAATTCATAGGACATGTTACNNNAGATCGGAAG
123283 TATGAGTGTGGAGTCGTTGCTATATGTAGGAAACTGTCTCTTCNNNAGATCGGAAG
123284 TATGAGTGTGGAGTCGTTGCCTTATGAAATGAGAAGCTGAATGNNNAGATCGGAA
123285 TATGAGTGTGGAGTCGTTGCTGTTTCCTTTACATTCCCTGNNNAGATCGGAAGAGCA
123286 TATGAGTGTGGAGTCGTTGCATTCTCAGGTTTATTCATGAATGNNNAGATCGGAAG
123287 TATGAGTGTGGAGTCGTTGCTTCTCTCCCAAATATCCTTCTAGNNNAGATCGGAAGA
123288 TATGAGTGTGGAGTCGTTGCTACATCCAGTAAGAAATTCACATNNNAGATCGGAAG
123289 TATGAGTGTGGAGTCGTTGCTACATCCAGTAAGAAATTCACATNNNAGATCGGAAG
123290 TATGAGTGTGGAGTCGTTGCCTCTCAATCTCTCTCTCTCTCNNNAGATCGGAAGAGC
123291 TATGAGTGTGGAGTCGTTGCTTTAACAGTGAAAATGGTGATACNNNAGATCGGAA
123292 TATGAGTGTGGAGTCGTTGCAGAGAGTTTTCATGCATTATAACNNNAGATCGGAAG
123293 TATGAGTGTGGAGTCGTTGCATCTTTTTTTTGAGACGAAGTTTNNNAGATCGGAAG

123294 TATGAGTGTGGAGTCGTTGCATGTGTGTTGATGACAATGATTANNNAGATCGGAA
123295 TATGAGTGTGGAGTCGTTGCCTCTTGGGTTTATCATGTTTTCNNNAGATCGGAAGA
123296 TATGAGTGTGGAGTCGTTGCCTCTTGGGTTTATCATGTTTTCNNNAGATCGGAAGA
123297 TATGAGTGTGGAGTCGTTGCCTCTTGGGTTTATCATGTTTTCNNNAGATCGGAAGA
123298 TATGAGTGTGGAGTCGTTGCTTCCCTGAAAAACTAGTAATACTNNNAGATCGGAAG
123299 TATGAGTGTGGAGTCGTTGCTCTAGTGATAAATCGGGAAATTTNNNAGATCGGAA
123300 TATGAGTGTGGAGTCGTTGCTGCTCTATTTCATTAGTTTATGTCNNNAGATCGGAAG
123302 TATGAGTGTGGAGTCGTTGCCAGCTTTATTTCTTTTCTTTTCTNNNAGATCGGAAGA
123303 TATGAGTGTGGAGTCGTTGCCCTCATCCAATATTATGCTCTTANNNAGATCGGAAGA
123304 TATGAGTGTGGAGTCGTTGCGTACACTCAGGAGACTTTCTNNNAGATCGGAAGAG
123306 TATGAGTGTGGAGTCGTTGCTTAAAAATGATCACATCTCATGCNNNAGATCGGAAG
123307 TATGAGTGTGGAGTCGTTGCTTTATTTGTTTTGCAGATGGTCNNNAGATCGGAAGA
123308 TATGAGTGTGGAGTCGTTGCAGACTTATGCCTTATTTTGGATANNNAGATCGGAAG
123309 TATGAGTGTGGAGTCGTTGCAGACTTATGCCTTATTTTGGATANNNAGATCGGAAG
123310 TATGAGTGTGGAGTCGTTGCTGAACTAGATAGAACACATTACTNNNAGATCGGAAG
123311 TATGAGTGTGGAGTCGTTGCCATGATTCTAGAAGCCTCAGNNNAGATCGGAAGAG
123312 TATGAGTGTGGAGTCGTTGCTGTTATAGTCACACATAGTAAGTNNNAGATCGGAAG
123313 TATGAGTGTGGAGTCGTTGCTTATTACCTATTTCGGAGAAGTTNNNAGATCGGAAG
123314 TATGAGTGTGGAGTCGTTGCTATCTCTTTGGGGAAATGGAGNNNAGATCGGAAGA
123315 TATGAGTGTGGAGTCGTTGCACCAGTAACTATATTCCCAAATTNNNAGATCGGAAGA
123316 TATGAGTGTGGAGTCGTTGCTCTAAGGAACTAACGGTTTTATTNNNAGATCGGAAG
123317 TATGAGTGTGGAGTCGTTGCTGTACATTAGGTCTCTAAAACATNNNAGATCGGAAG
123318 TATGAGTGTGGAGTCGTTGCGTGAGTAGATATGTAACCATACANNNAGATCGGAA
123319 TATGAGTGTGGAGTCGTTGCTAACTCTTGTCCATCTAAAAGTANNNAGATCGGAAG
123320 TATGAGTGTGGAGTCGTTGCAAAACACCAAGGGTTAATTTATTNNNAGATCGGAAG
123321 TATGAGTGTGGAGTCGTTGCAAATTCAGCAGTTTGAAAACATANNNAGATCGGAAG
123322 TATGAGTGTGGAGTCGTTGCGTCCTCTGTTTATTTAGTCAGTTNNNAGATCGGAAG
123323 TATGAGTGTGGAGTCGTTGCACACATATTCTAACAGTTCATTGNNNAGATCGGAAG
123324 TATGAGTGTGGAGTCGTTGCCACATTTACCTGGTTTGGTANNNAGATCGGAAGAGC
123325 TATGAGTGTGGAGTCGTTGCGCTAGAAAATAAAAAGAACTTGCNNNAGATCGGAA
123326 TATGAGTGTGGAGTCGTTGCCACTGAATCCATCTCTCTCCNNNAGATCGGAAGAGCA
123327 TATGAGTGTGGAGTCGTTGCCTATACCTTCGTTTCCTGTACNNNAGATCGGAAGAG
123328 TATGAGTGTGGAGTCGTTGCCTCTTTCTCTTTTCTTTTCTTCANNNAGATCGGAAGAG
123329 TATGAGTGTGGAGTCGTTGCAAAAGACACCAAAATTAACAGATNNNAGATCGGAA
123330 TATGAGTGTGGAGTCGTTGCGTCTCTCTATTAGTACTGATACANNNAGATCGGAAG
123331 TATGAGTGTGGAGTCGTTGCTAACTCGACTACCTGTTAGGNNNAGATCGGAAGAG
123332 TATGAGTGTGGAGTCGTTGCTATCTCATTTTCTGCATTACCTANNNAGATCGGAAGA
123334 TATGAGTGTGGAGTCGTTGCTGGTATTTACTGGAATTACAGAANNNAGATCGGAA
123335 TATGAGTGTGGAGTCGTTGCATTTTAGAGATACCGAGATATGGNNNAGATCGGAA
123336 TATGAGTGTGGAGTCGTTGCCTGTAATTTAGCTCTATGTATACAGNNNAGATCGGA
123337 TATGAGTGTGGAGTCGTTGCACATTGCTTATATTTGTTAGATTGNNNAGATCGGAA
123338 TATGAGTGTGGAGTCGTTGCAAGAGATCAGACAACAATTAGGNNNAGATCGGAAG
123339 TATGAGTGTGGAGTCGTTGCTCTCTATGCACTATGTCTATGTANNNAGATCGGAAG

123340 TATGAGTGTGGAGTCGTTGCAGCATAATTCAGAAAGCAATTAANNNAGATCGGAAG
123341 TATGAGTGTGGAGTCGTTGCCTGGAATTCTCTTCCTCTTTCNNNAGATCGGAAGAGC
123342 TATGAGTGTGGAGTCGTTGCAATGAAGGAAATAGCATCAACTANNNAGATCGGAA
123344 TATGAGTGTGGAGTCGTTGCAAATTACTTCATTGTAAGCCTTTNNNAGATCGGAAGA
123347 TATGAGTGTGGAGTCGTTGCGCAATTTCTCTTCTAGTTTTTCTNNNAGATCGGAAGA
123348 TATGAGTGTGGAGTCGTTGCATATATGTAAGTTCCAGTGAGGANNNAGATCGGAA
123349 TATGAGTGTGGAGTCGTTGCCTGGACTGTACTCTCAAATGNNNAGATCGGAAGAGC
123350 TATGAGTGTGGAGTCGTTGCCATGACAAGATACTCCCTTTTNNNAGATCGGAAGAG
123351 TATGAGTGTGGAGTCGTTGCACAAAAAGTACTTTGAAAGGAATNNNAGATCGGAA
123354 TATGAGTGTGGAGTCGTTGCATCTTTTTCTCTTCTCTCTGACNNNAGATCGGAAGAG
123355 TATGAGTGTGGAGTCGTTGCATGAGTACCCTTCAAAGAATAATNNNAGATCGGAAG
123356 TATGAGTGTGGAGTCGTTGCGTGTGGAAGGTTGATATACTTNNNAGATCGGAAGA
123357 TATGAGTGTGGAGTCGTTGCACTGAGACCAATAGTGTATATTTNNNAGATCGGAAG
123358 TATGAGTGTGGAGTCGTTGCAAGGGGGAAATAAAGGAATTTNNNAGATCGGAAGA
123359 TATGAGTGTGGAGTCGTTGCGTCAAGTCTTTAGAGAGATTAGGNNNAGATCGGAA
123360 TATGAGTGTGGAGTCGTTGCTAAACCCCTTGGATTTTATTCTTNNNAGATCGGAAGA
123361 TATGAGTGTGGAGTCGTTGCTTATACAGGTTGAGTATCTCTTGNNNAGATCGGAAG
123362 TATGAGTGTGGAGTCGTTGCGGAGAGAGAGAGTGAATAAAAGNNNAGATCGGAA
123363 TATGAGTGTGGAGTCGTTGCAGAATTAAATTCCCTTAAGAGGANNNAGATCGGAAG
123364 TATGAGTGTGGAGTCGTTGCTCATGTTTAGAAAAGTTCAAACTNNNAGATCGGAAG
123365 TATGAGTGTGGAGTCGTTGCCTACTTATTTCCCCAAGGTCNNNAGATCGGAAGAGCA
123366 TATGAGTGTGGAGTCGTTGCTATATTGTCATTATCTGTGGACCNNNAGATCGGAAGA
123368 TATGAGTGTGGAGTCGTTGCTAGTTAAGGAAAAATGCATTTCTNNNAGATCGGAAG
123373 TATGAGTGTGGAGTCGTTGCGATAAAAGAAAATTCCTGACCATNNNAGATCGGAAG
123374 TATGAGTGTGGAGTCGTTGCAACAGACATATTTCTAAGTAGCTNNNAGATCGGAAG
123375 TATGAGTGTGGAGTCGTTGCGTTGATGCTTTTTTAATTGTCTCNNNAGATCGGAAGA
123376 TATGAGTGTGGAGTCGTTGCTCTGAAGTTCAAAAGCACTGNNNAGATCGGAAGAG
123377 TATGAGTGTGGAGTCGTTGCTCTGAAGTTCAAAAGCACTGNNNAGATCGGAAGAG
123378 TATGAGTGTGGAGTCGTTGCAAAATAAAGGCAATATCAAGAGGNNNAGATCGGAA
123379 TATGAGTGTGGAGTCGTTGCTCTACACAATACAGTTTGAAAGANNNAGATCGGAAG
123380 TATGAGTGTGGAGTCGTTGCCTCCTTGAATTGGATACCATAATNNNAGATCGGAAGA
123381 TATGAGTGTGGAGTCGTTGCTGCACTACCACTATAATATATCGNNNAGATCGGAAGA
123382 TATGAGTGTGGAGTCGTTGCGATAGTTCATTGAGTTGTTTCTTNNNAGATCGGAAGA
123383 TATGAGTGTGGAGTCGTTGCGAAAAATTGTTGGGTTTTTTTCTNNNAGATCGGAAG
123384 TATGAGTGTGGAGTCGTTGCCTAGAGTGAGGTCTCCTAATGNNNAGATCGGAAGA
123385 TATGAGTGTGGAGTCGTTGCCACTTTTGTCTTGGAATTCCNNNAGATCGGAAGAGCA
123386 TATGAGTGTGGAGTCGTTGCTGACATTTGCAGATATTATTTGTNNNAGATCGGAAGA
123387 TATGAGTGTGGAGTCGTTGCGAACTGAGACCTTCTACCAANNNAGATCGGAAGAGC
123388 TATGAGTGTGGAGTCGTTGCATTGATCATAATAGCAGAGACTTNNNAGATCGGAAG
123389 TATGAGTGTGGAGTCGTTGCGTATCATAATTTGTAAACCCCAANNNAGATCGGAAGA
123390 TATGAGTGTGGAGTCGTTGCCACATATGAAGATGAGAGAGAGNNNAGATCGGAAG
123391 TATGAGTGTGGAGTCGTTGCAATAAAAAGAAATCTGCACCTAGNNNAGATCGGAAG
123392 TATGAGTGTGGAGTCGTTGCTTGTTCTTTATAGTCATCTTGATATNNNAGATCGGAA

123393 TATGAGTGTGGAGTCGTTGCGAGATTACAGGAATGAGCTACNNNAGATCGGAAGA
123394 TATGAGTGTGGAGTCGTTGCCTACAGAAAATGTCTTTTTTCCCNNNAGATCGGAAGA
123395 TATGAGTGTGGAGTCGTTGCCTAACTGTCCTGGTCTGAAGNNNAGATCGGAAGAGC
123396 TATGAGTGTGGAGTCGTTGCTATTTGTAATTCTGTAACCTGGANNNAGATCGGAAGA
123397 TATGAGTGTGGAGTCGTTGCCTTTTTTTTTTTTCGGAGACAGNNNAGATCGGAAGAC
123398 TATGAGTGTGGAGTCGTTGCAGATTCACAAAACCCATATAACTNNNAGATCGGAAGA
123399 TATGAGTGTGGAGTCGTTGCCATTATGCATCATCTAGAATCAANNNAGATCGGAAGA
123400 TATGAGTGTGGAGTCGTTGCCTCAAAGGATATATGCATTTGATNNNAGATCGGAAG
123401 TATGAGTGTGGAGTCGTTGCACGATGTCTATCAATAAGTATTTGNNNAGATCGGAAC
123402 TATGAGTGTGGAGTCGTTGCAAACTGTCTTTATTCACAAATGANNNAGATCGGAAGA
123403 TATGAGTGTGGAGTCGTTGCTACTTACTATGTGTCAAGCATATNNNAGATCGGAAGA
123404 TATGAGTGTGGAGTCGTTGCTTATATACGTATACACACACACANNNAGATCGGAAGA
123405 TATGAGTGTGGAGTCGTTGCTAGGATGAAGTTTGGAAAAGAANNNAGATCGGAAG
123406 TATGAGTGTGGAGTCGTTGCCCATCAATTCATTAGGAGTTTTANNNAGATCGGAAGA
123407 TATGAGTGTGGAGTCGTTGCTAAGGAAGTGTTACATGTCTAAANNNAGATCGGAAG
123408 TATGAGTGTGGAGTCGTTGCTTCCCTTATATCTAGTAAGTCACNNNAGATCGGAAGA
123409 TATGAGTGTGGAGTCGTTGCCAGGAAGGGGAAGATTATGGNNNAGATCGGAAGAC
123410 TATGAGTGTGGAGTCGTTGCTACTCGTGTGTGTATGTGTCNNNAGATCGGAAGAGC
123411 TATGAGTGTGGAGTCGTTGCAAAGTTGCTCATTTTCTTGAAGNNNAGATCGGAAGA
123412 TATGAGTGTGGAGTCGTTGCTTTTGATAATACCTTTTGTGGTTNNNAGATCGGAAGA
123413 TATGAGTGTGGAGTCGTTGCACTGGTTAAAGTTTATAACTAGAGNNNAGATCGGAA
123414 TATGAGTGTGGAGTCGTTGCGAGACAAACATAGTATGTCAGGNNNAGATCGGAAG
123415 TATGAGTGTGGAGTCGTTGCGTAAATAAATAAGACGTGCCATTNNNAGATCGGAAG
123416 TATGAGTGTGGAGTCGTTGCATTTCTTCCCTATATCTTTCCTTNNNAGATCGGAAGAG
123417 TATGAGTGTGGAGTCGTTGCCATTTTTTGTCTTTGTTGATTGTNNNAGATCGGAAGA
123418 TATGAGTGTGGAGTCGTTGCGTTACCTTTTGGTTTTCACTAGNNNAGATCGGAAGAC
123419 TATGAGTGTGGAGTCGTTGCCAACTCCCTTTGTCTACTTTNNNAGATCGGAAGAGCA
123420 TATGAGTGTGGAGTCGTTGCGTGGTCAATATTTAGAGAAGGATNNNAGATCGGAAC
123421 TATGAGTGTGGAGTCGTTGCCCCCAACATTTTTAATTTCTGTTNNNAGATCGGAAGAC
123422 TATGAGTGTGGAGTCGTTGCCAATGACAAGGAGAATAATAGTCNNNAGATCGGAAC
123423 TATGAGTGTGGAGTCGTTGCCTTCTAAAATAGCCTCTATTCCTNNNAGATCGGAAGA
123424 TATGAGTGTGGAGTCGTTGCCCCCATTTAATTCTTCAAAAAATNNNAGATCGGAAGA
123425 TATGAGTGTGGAGTCGTTGCCCCCATTTAATTCTTCAAAAAATNNNAGATCGGAAGA
123426 TATGAGTGTGGAGTCGTTGCGAAATCATATCCGCCTCTTTTNNNAGATCGGAAGAGC
123427 TATGAGTGTGGAGTCGTTGCCATAACAATAAAACTTGCCATTTNNNAGATCGGAAGA
123428 TATGAGTGTGGAGTCGTTGCCATAACAATAAAACTTGCCATTTNNNAGATCGGAAGA
123429 TATGAGTGTGGAGTCGTTGCTACTTCTGTATTCATAGCTTTCTNNNAGATCGGAAGA
123430 TATGAGTGTGGAGTCGTTGCACCTGAGGTATACATGTTATTAANNNAGATCGGAAG
123431 TATGAGTGTGGAGTCGTTGCATAGATATACACACACACAATGTNNNAGATCGGAAG
123432 TATGAGTGTGGAGTCGTTGCGAGATTTCCTTTTTCTTTCTCCNNNAGATCGGAAGAG
123434 TATGAGTGTGGAGTCGTTGCGTTGAATTTCATCAAATGCTTTTNNNAGATCGGAAGA
123435 TATGAGTGTGGAGTCGTTGCATAAACAGGCTGTGTACCTANNNAGATCGGAAGAGC
123436 TATGAGTGTGGAGTCGTTGCCACATTATAGCATTTTACTACCCNNNAGATCGGAAGA

123437 TATGAGTGTGGAGTCGTTGCCTTACAGGAACTCTTGTGATTTNNNAGATCGGAAGA
123438 TATGAGTGTGGAGTCGTTGCATTATGATCATCGTCATCCATGNNNAGATCGGAAGA
123439 TATGAGTGTGGAGTCGTTGCGATTTCAGTGTCAATTCATTCTCNNNAGATCGGAAG
123440 TATGAGTGTGGAGTCGTTGCAAAAACATGAAAGGAAAAGGTAANNNAGATCGGAA
123441 TATGAGTGTGGAGTCGTTGCGTGTAAATAAGTGACCAGATGTNNNAGATCGGAAG
123442 TATGAGTGTGGAGTCGTTGCACATGTACGTATAGTTTTGTTTTNNNAGATCGGAAG
123443 TATGAGTGTGGAGTCGTTGCTTAATCCATAACAAAGAGAAGGGNNNAGATCGGAA
123444 TATGAGTGTGGAGTCGTTGCTTTGTTTTAGAGATAAGTCCTCANNNAGATCGGAAG
123445 TATGAGTGTGGAGTCGTTGCTGAAATAATAATCTTGTTCAGCTNNNAGATCGGAAG
123446 TATGAGTGTGGAGTCGTTGCGGGAGCCTTCTGTATTTATAATNNNAGATCGGAAGA
123447 TATGAGTGTGGAGTCGTTGCACTCTTCACTGACTGTATCANNNAGATCGGAAGAGC
123448 TATGAGTGTGGAGTCGTTGCGTTTCTTCTCCTTCTCCTTCNNNAGATCGGAAGAGCA
123449 TATGAGTGTGGAGTCGTTGCCAGCTTCTGAATCCTAAAATTTANNNAGATCGGAAG
123450 TATGAGTGTGGAGTCGTTGCGATCATTCTCATGACCAAGATAANNNAGATCGGAAG
123451 TATGAGTGTGGAGTCGTTGCCAGATTAGGGCTTTCCATTGNNNAGATCGGAAGAG
123452 TATGAGTGTGGAGTCGTTGCCAAGGACAGGACAGATTAGGNNNAGATCGGAAGA
123455 TATGAGTGTGGAGTCGTTGCTTTCTATTTCTTCACACATACACNNNAGATCGGAAGA
123456 TATGAGTGTGGAGTCGTTGCTTATAAACGTTCTGCCATTTAAANNNAGATCGGAAG
123457 TATGAGTGTGGAGTCGTTGCTACATTTTCTTTATCCACTCGTTNNNAGATCGGAAGA
123458 TATGAGTGTGGAGTCGTTGCTTAGCTCTAGTAGATTTCTTGTGNNNAGATCGGAAG
123459 TATGAGTGTGGAGTCGTTGCTTGCAATTTTTACAAAGTGAAAGNNNAGATCGGAAG
123460 TATGAGTGTGGAGTCGTTGCCTATATCCACATACTTTCAGAGANNNAGATCGGAAG
123461 TATGAGTGTGGAGTCGTTGCAAAAATGTTGCTACAATTAGTGANNNAGATCGGAAG
123462 TATGAGTGTGGAGTCGTTGCATTATGAAATTTCATGTCCATCGNNNAGATCGGAAG
123463 TATGAGTGTGGAGTCGTTGCTTTGTTTCCTTCCTGTGTAAAATNNNAGATCGGAAGA
123464 TATGAGTGTGGAGTCGTTGCTTTGTTTCCTTCCTGTGTAAAATNNNAGATCGGAAGA
123465 TATGAGTGTGGAGTCGTTGCCAACAAATCTAAGACTAGAACTGNNNAGATCGGAAG
123466 TATGAGTGTGGAGTCGTTGCGCACTGAAATTCTTGAGTTTTTANNNAGATCGGAAG
123467 TATGAGTGTGGAGTCGTTGCTATATTACTTCCACATCATCCTCNNNAGATCGGAAGA
123468 TATGAGTGTGGAGTCGTTGCCACACAAATTTTTTCACTCATATNNNAGATCGGAAGA
123469 TATGAGTGTGGAGTCGTTGCGTAATTGCCCATAAAAGTGAAANNNAGATCGGAAG
123470 TATGAGTGTGGAGTCGTTGCTAATGGGATACTATGTTACAACANNNAGATCGGAAG
123471 TATGAGTGTGGAGTCGTTGCAATCTTAGGCATTCACTTATTTGNNNAGATCGGAAG
123472 TATGAGTGTGGAGTCGTTGCCCAAAATCATATTCAGAACAAGTNNNAGATCGGAAG
123473 TATGAGTGTGGAGTCGTTGCGAAAAGTTATGGAGTTTGTCTGNNNAGATCGGAAG
123474 TATGAGTGTGGAGTCGTTGCAGAATTATGAGACACTGATTAGANNNAGATCGGAA
123475 TATGAGTGTGGAGTCGTTGCTATATAGCATTGCTGGAGATAAANNNAGATCGGAA
123476 TATGAGTGTGGAGTCGTTGCACTAATGTGTTAAGGTAGACTTTNNNAGATCGGAAG
123477 TATGAGTGTGGAGTCGTTGCAAATGAGAGATGTTTTGGATATCNNNAGATCGGAA
123479 TATGAGTGTGGAGTCGTTGCCTTTGGAAAACTATTCTACTGTCNNNAGATCGGAAG
123480 TATGAGTGTGGAGTCGTTGCTCAAAAAAAATCCTAGTATCGAGNNNAGATCGGAAG
123482 TATGAGTGTGGAGTCGTTGCTATGTGTGTGTGTGTGTATATATNNNAGATCGGAAG
123483 TATGAGTGTGGAGTCGTTGCACACACGTGTATATATATGTGTNNNAGATCGGAAGA

123484 TATGAGTGTGGAGTCGTTGCAGTTACTTTTGGTACAATTCATTNNNAGATCGGAAG
123485 TATGAGTGTGGAGTCGTTGCGGAACTCCCATATACTTAATAAGANNNAGATCGGAA
123486 TATGAGTGTGGAGTCGTTGCGCTAAAGCTTTACTCTCTTTTTNNNAGATCGGAAGAG
123487 TATGAGTGTGGAGTCGTTGCTTTTGAAAAAAGTGTTTTGTTGTNNNAGATCGGAAG
123488 TATGAGTGTGGAGTCGTTGCTATGTTCTGATAAACCCATCATANNNAGATCGGAAG
123489 TATGAGTGTGGAGTCGTTGCATAAGTCACTGAAATGATTTTGTNNNAGATCGGAAG
123490 TATGAGTGTGGAGTCGTTGCAGTTCTTCACTTTTCAGGAATNNNAGATCGGAAGAG
123491 TATGAGTGTGGAGTCGTTGCATGGAACACCTAATTAAAAACTCNNNAGATCGGAAG
123492 TATGAGTGTGGAGTCGTTGCGCCCAAAAAAGGAAAAATTTTAANNNAGATCGGAA
123493 TATGAGTGTGGAGTCGTTGCGCCCAAAAAAGGAAAAATTTTAANNNAGATCGGAA
123494 TATGAGTGTGGAGTCGTTGCATATGCATATTAGAAGCGTAAAGNNNAGATCGGAA
123495 TATGAGTGTGGAGTCGTTGCTGACAATCTATATTGCTTTTAGCNNNAGATCGGAAG
123496 TATGAGTGTGGAGTCGTTGCTTCTATGGTACTAGGTCAAGAANNNAGATCGGAAG
123497 TATGAGTGTGGAGTCGTTGCCTGAATTGATATGGATTCCCANNNAGATCGGAAGAG
123498 TATGAGTGTGGAGTCGTTGCTGTGTTTGTGATTTCTTTTTCTANNNAGATCGGAAGA
123499 TATGAGTGTGGAGTCGTTGCAATTTATTTCGCTAGCTGTATTTNNNAGATCGGAAG
123500 TATGAGTGTGGAGTCGTTGCTTATGAGGTCAATTCCTTCTTAANNNAGATCGGAAG
123501 TATGAGTGTGGAGTCGTTGCTTATGAGGTCAATTCCTTCTTAANNNAGATCGGAAG
123502 TATGAGTGTGGAGTCGTTGCTATCTAAACTTTTTTCCTGTTCCNNNAGATCGGAAGA
123503 TATGAGTGTGGAGTCGTTGCTCTTGCTATAGAATAAACTGTCTNNNAGATCGGAAG
123504 TATGAGTGTGGAGTCGTTGCTAGTACAGCTATTCTATGTGAAGNNNAGATCGGAAG
123505 TATGAGTGTGGAGTCGTTGCGAGAGTTTCTATCATATGTCCTGNNNAGATCGGAAG
123506 TATGAGTGTGGAGTCGTTGCGAGAGAAAAGCGTATACATTTTTNNNAGATCGGAA
123507 TATGAGTGTGGAGTCGTTGCGTTATGGTGATTGATTCAGTAATNNNAGATCGGAA
123508 TATGAGTGTGGAGTCGTTGCGTGTGTATTTGTATGTGTTCTTANNNAGATCGGAAG
123509 TATGAGTGTGGAGTCGTTGCTGATTTCCCACAACTAAGTTTNNNAGATCGGAAGAG
123510 TATGAGTGTGGAGTCGTTGCATATCCTCAGTAGATTCATTTCANNNAGATCGGAAG
123511 TATGAGTGTGGAGTCGTTGCGCCATTATTCCTAAAACCATATTNNNAGATCGGAAGA
123512 TATGAGTGTGGAGTCGTTGCAGATCAGTATTGAATTTCACTTCNNNAGATCGGAAG
123513 TATGAGTGTGGAGTCGTTGCTACTTAGTGGTGTCATAGTACTTNNNAGATCGGAAG
123514 TATGAGTGTGGAGTCGTTGCTACTTAGTGGTGTCATAGTACTTNNNAGATCGGAAG
123515 TATGAGTGTGGAGTCGTTGCCAATCATAAACAGATAGGAAACANNNAGATCGGAA
123516 TATGAGTGTGGAGTCGTTGCAGAGAGAAGATATACAAACAACCNNNAGATCGGAA
123517 TATGAGTGTGGAGTCGTTGCCATAATGCAACATCCTAGAAAGNNNAGATCGGAAGA
123518 TATGAGTGTGGAGTCGTTGCGCTTCAACAGGATAGTTTCTAATNNNAGATCGGAAG
123519 TATGAGTGTGGAGTCGTTGCGAAAGAATGGATGACTTGTTAGNNNAGATCGGAAG
123520 TATGAGTGTGGAGTCGTTGCCAAGGCTGGTAAGAATAATTTTCNNNAGATCGGAA
123522 TATGAGTGTGGAGTCGTTGCTATTTTTATTCATTCCTAGGCCTNNNAGATCGGAAGA
123523 TATGAGTGTGGAGTCGTTGCTATTTTTATTCATTCCTAGGCCTNNNAGATCGGAAGA
123524 TATGAGTGTGGAGTCGTTGCCTGATTTTCTAGTTTTGCTGATTNNNAGATCGGAAG
123525 TATGAGTGTGGAGTCGTTGCCAAAATGAAAACCTGGTAAACTANNNAGATCGGAA
123526 TATGAGTGTGGAGTCGTTGCAGAGATGAGAGAGAGAAAGAGNNNAGATCGGAAG
123527 TATGAGTGTGGAGTCGTTGCCTTTTCATCCTTCTAATCAATGGNNNAGATCGGAAGA

123528 TATGAGTGTGGAGTCGTTGCAGAGATGAGAGAGAGAAAGAGNNNAGATCGGAAG
123529 TATGAGTGTGGAGTCGTTGCTGTTCTCCAGAAGAAATGTAATNNNAGATCGGAAG
123530 TATGAGTGTGGAGTCGTTGCGTGAACATGACTCATCTTGGNNNAGATCGGAAGAG
123531 TATGAGTGTGGAGTCGTTGCTTTTTCACTTTTCCAAGGTTTATNNNAGATCGGAAGA
123532 TATGAGTGTGGAGTCGTTGCTACTTTTTGTTTTGATTCAGTGANNNAGATCGGAAG
123533 TATGAGTGTGGAGTCGTTGCAACATTTAGATCTATTTCTGCACNNNAGATCGGAAG
123534 TATGAGTGTGGAGTCGTTGCCCAATATAAGCCAAGGTAATAACNNNAGATCGGAAG
123535 TATGAGTGTGGAGTCGTTGCATAGAATTGTCTTTAAGTTCTGGNNNAGATCGGAAG
123536 TATGAGTGTGGAGTCGTTGCAAAGATGTAAGAGGATTCTAGCNNNAGATCGGAAG
123537 TATGAGTGTGGAGTCGTTGCTTACCAGAAATTCAAAGCAGTNNNAGATCGGAAGA
123540 TATGAGTGTGGAGTCGTTGCAAGTTTTTCTGTTCACAAATCANNNAGATCGGAAGA
123542 TATGAGTGTGGAGTCGTTGCACATTTTATCTGGTTATGTTCAGNNNAGATCGGAAG
123543 TATGAGTGTGGAGTCGTTGCTTGATCTAGAGAGAATGAGACTANNNAGATCGGAA
123544 TATGAGTGTGGAGTCGTTGCAATTTTCACTTCAGGTAGATAGTNNNAGATCGGAAG
123545 TATGAGTGTGGAGTCGTTGCGGGTAATAACATAGGATGTAGTTNNNAGATCGGAA
123546 TATGAGTGTGGAGTCGTTGCAAAAGTAGAGAACCAGAAACTTTNNNAGATCGGAA
123547 TATGAGTGTGGAGTCGTTGCGCATGTTTACTGGAAAAGTAANNNAGATCGGAAGA
123548 TATGAGTGTGGAGTCGTTGCGTGCAAATACACAAAACTAAAAANNNAGATCGGAA
123549 TATGAGTGTGGAGTCGTTGCAAAGTTTTAGTCTTAATTCCTCANNNAGATCGGAAG
123550 TATGAGTGTGGAGTCGTTGCCCATAAAAGTGAAGGTCATACTANNNAGATCGGAA
123551 TATGAGTGTGGAGTCGTTGCCCATATTCAGCAGAAATCTAGANNNAGATCGGAAGA
123552 TATGAGTGTGGAGTCGTTGCCAGAGATAGTTTTACGTTTTTCTNNNAGATCGGAAG
123553 TATGAGTGTGGAGTCGTTGCATGCTGATCCAAAGTAATAATTGNNNAGATCGGAAG
123554 TATGAGTGTGGAGTCGTTGCTATGATATTGTAAGATGTCACCANNNAGATCGGAAG
123555 TATGAGTGTGGAGTCGTTGCTTTGTATATCAGGTGAGCTATATNNNAGATCGGAAG
123556 TATGAGTGTGGAGTCGTTGCTTACAAAATAGGATGGTGCATTTNNNAGATCGGAA
123557 TATGAGTGTGGAGTCGTTGCTAGTACAGGTGGCATACTATTTNNNAGATCGGAAG
123558 TATGAGTGTGGAGTCGTTGCAGAATAATTACTGCTTTTCCTCTNNNAGATCGGAAG
123559 TATGAGTGTGGAGTCGTTGCCATCAAGAAAAGATGACCTCANNNAGATCGGAAGA
123560 TATGAGTGTGGAGTCGTTGCGTTTATAACATAGAAACGCTTCTNNNAGATCGGAAG
123561 TATGAGTGTGGAGTCGTTGCAGGACTAATTCCCTTGCTAGNNNAGATCGGAAGAG
123562 TATGAGTGTGGAGTCGTTGCGATAATAACCTTTCTATTCCAGCNNNAGATCGGAAG
123563 TATGAGTGTGGAGTCGTTGCAAGGATATGGAGGAGAAACANNNAGATCGGAAGA
123564 TATGAGTGTGGAGTCGTTGCATAACCAAAAATGGGTTAAGTTCNNNAGATCGGAA
123565 TATGAGTGTGGAGTCGTTGCAAGATGTATTAAGGAGAAATTGCNNNAGATCGGAA
123566 TATGAGTGTGGAGTCGTTGCCAGATCATAAAGTTTTTAGGCTTNNNAGATCGGAAG
123567 TATGAGTGTGGAGTCGTTGCAACAGTAATGCAAAAATAAAACGNNNAGATCGGAA
123568 TATGAGTGTGGAGTCGTTGCCTCAAAAAGGTAAAGTCAAAGAANNNAGATCGGAA
123569 TATGAGTGTGGAGTCGTTGCGTATCTTTATTACTGGAAGTGTGNNNAGATCGGAAG
123570 TATGAGTGTGGAGTCGTTGCTTTGTTTTAGCCATCTTTGTATTNNNAGATCGGAAGA
123571 TATGAGTGTGGAGTCGTTGCACCCAGAATTTAGACAGAATACNNNAGATCGGAAGA
123572 TATGAGTGTGGAGTCGTTGCCCGTTTACACTCAATATAATTGTNNNAGATCGGAAG
123573 TATGAGTGTGGAGTCGTTGCATATCTCTATATTCAGGCAGAACNNNAGATCGGAAG

123574 TATGAGTGTGGAGTCGTTGCGCACATAGTAGGTATTCAATAAANNNAGATCGGAA
123575 TATGAGTGTGGAGTCGTTGCGTTCACAATACAATACCCATAAANNNAGATCGGAAG
123576 TATGAGTGTGGAGTCGTTGCTTGCTACTTCTGTAGGTCACNNNAGATCGGAAGAGC
123577 TATGAGTGTGGAGTCGTTGCAAGTTATGGAATAAACTGAAGTGNNNAGATCGGAA
123578 TATGAGTGTGGAGTCGTTGCTAGCACATTTTCTTTATCCATTCNNNAGATCGGAAGA
123579 TATGAGTGTGGAGTCGTTGCGCCAACATATTACTTTAACAGTTNNNAGATCGGAAG
123580 TATGAGTGTGGAGTCGTTGCTTGTTTTGTTTTTTTGAGGACTGNNNAGATCGGAAG
123581 TATGAGTGTGGAGTCGTTGCGGTGAAACATGAAGAAAAATAAGNNNAGATCGGAA
123582 TATGAGTGTGGAGTCGTTGCATAATCAGGAAAAGTTGGAAATGNNNAGATCGGAA
123583 TATGAGTGTGGAGTCGTTGCTCATAAAGAGACAAAAAGAGAGANNNAGATCGGAA
123584 TATGAGTGTGGAGTCGTTGCTCATAAAGAGACAAAAAGAGAGANNNAGATCGGAA
123585 TATGAGTGTGGAGTCGTTGCATTTAAAATCTCCACCTTCTCTGNNNAGATCGGAAGA
123586 TATGAGTGTGGAGTCGTTGCAGCTCATTTTTGATGTGTTTTATNNNAGATCGGAAG
123587 TATGAGTGTGGAGTCGTTGCTATTTTCACACTGAAACTACCTANNNAGATCGGAAG
123588 TATGAGTGTGGAGTCGTTGCGAATAGTGCTGCAAATAAACAANNNAGATCGGAAG
123589 TATGAGTGTGGAGTCGTTGCTGCTATGAACTCAATCTCTATTANNNAGATCGGAAG
123590 TATGAGTGTGGAGTCGTTGCTTTGTAAAGGAGAAAGGAAAAGANNNAGATCGGA
123591 TATGAGTGTGGAGTCGTTGCCACATAAAATACACTAACAGACANNNAGATCGGAAG
123592 TATGAGTGTGGAGTCGTTGCATGCCAAACTTTAGATTGTTTAANNNAGATCGGAAG
123593 TATGAGTGTGGAGTCGTTGCCAAACCTGAAAAATCAAGAAGTANNNAGATCGGAA
123594 TATGAGTGTGGAGTCGTTGCCAAACCTGAAAAATCAAGAAGTANNNAGATCGGAA
123595 TATGAGTGTGGAGTCGTTGCTTCTTGAAACACTAAGGATAAGANNNAGATCGGAA
123597 TATGAGTGTGGAGTCGTTGCAACATTATGCATAGTGAAAGAAGNNNAGATCGGAA
123598 TATGAGTGTGGAGTCGTTGCCATAGTTATAAACCTGAGAGACTNNNAGATCGGAAG
123599 TATGAGTGTGGAGTCGTTGCTATTTGTAAAGCATTCTCTATGGNNNAGATCGGAAG
123601 TATGAGTGTGGAGTCGTTGCTTAAATTTTTAATTCAGGGCACTNNNAGATCGGAAG
123602 TATGAGTGTGGAGTCGTTGCTAGCCCAATTTAATACAACTACTNNNAGATCGGAAG
123603 TATGAGTGTGGAGTCGTTGCCCAACTTTATATGCCAAGTATTTNNNAGATCGGAAG
123604 TATGAGTGTGGAGTCGTTGCGCAGAAGTAATATACATTCTTGTNNNAGATCGGAAG
123605 TATGAGTGTGGAGTCGTTGCCAAAAAAGGCAAAATGGTGANNNAGATCGGAAGA
123606 TATGAGTGTGGAGTCGTTGCCTGCTTTCAATTCTCTATACCTANNNAGATCGGAAGA
123607 TATGAGTGTGGAGTCGTTGCGTGCACATAATTGTTTTTATTCANNNAGATCGGAAG
123608 TATGAGTGTGGAGTCGTTGCATTCCTCCTCATAGTTTATTCATNNNAGATCGGAAGA
123610 TATGAGTGTGGAGTCGTTGCACTTTGCTATTATAGACTTCACANNNAGATCGGAAG
123611 TATGAGTGTGGAGTCGTTGCACTGTTATTGCTTTCATAGAAACNNNAGATCGGAAG
123612 TATGAGTGTGGAGTCGTTGCCGTGGAATTAACTGGTCATATANNNAGATCGGAAG
123613 TATGAGTGTGGAGTCGTTGCGTTTTTAGAAAAGCCAAAGAAAANNNAGATCGGAA
123615 TATGAGTGTGGAGTCGTTGCAGACTTATATAATCGTGGACAAANNNAGATCGGAA
123616 TATGAGTGTGGAGTCGTTGCGAATGGAATAAATCTAGCAATGANNNAGATCGGAA
123617 TATGAGTGTGGAGTCGTTGCGGTTTATTATAATGAACTGGCTCNNNAGATCGGAAG
123618 TATGAGTGTGGAGTCGTTGCGTTTTTAGAAAAGGTGTTTGATCNNNAGATCGGAA
123619 TATGAGTGTGGAGTCGTTGCGTTTTTAGAAAAGGTGTTTGATCNNNAGATCGGAA
123620 TATGAGTGTGGAGTCGTTGCGTGTTAGGGCAATAAGATTATTTNNNAGATCGGAA

123621 TATGAGTGTGGAGTCGTTGCGTGTTAGGGCAATAAGATTATTTNNNAGATCGGAA
123622 TATGAGTGTGGAGTCGTTGCTGTCTATGTATATATGTGTGCACNNNAGATCGGAAG
123623 TATGAGTGTGGAGTCGTTGCATTTCTACTTGCTCTCTATGACNNNAGATCGGAAGAG
123624 TATGAGTGTGGAGTCGTTGCTAATCTTCAGGAAAATTCCCAAANNNAGATCGGAAG
123625 TATGAGTGTGGAGTCGTTGCTTTACTGGAAGGGAATTGATAATNNNAGATCGGAA
123626 TATGAGTGTGGAGTCGTTGCTATGCGTGTATGTGTGTATATATNNNAGATCGGAAG
123627 TATGAGTGTGGAGTCGTTGCATATAGTATCACTAGTTCTTGGCNNNAGATCGGAAG
123628 TATGAGTGTGGAGTCGTTGCCCTGAGAGAATGGAGTTTTGNNNAGATCGGAAGAG
123630 TATGAGTGTGGAGTCGTTGCCTCTGAACCATATGAGAGTAAGNNNAGATCGGAAG
123631 TATGAGTGTGGAGTCGTTGCATTTTCTGGAGTTCAGGTTCNNNAGATCGGAAGAG
123632 TATGAGTGTGGAGTCGTTGCTATATTCATTTTCTGGAGTTCAGNNNAGATCGGAAG
123633 TATGAGTGTGGAGTCGTTGCCTAGTAACTCAGAATGTGACAATNNNAGATCGGAAG
123634 TATGAGTGTGGAGTCGTTGCATATACACACACATTCACATATGNNNAGATCGGAAG
123635 TATGAGTGTGGAGTCGTTGCAATCTGCTAAATTACTCATGACTNNNAGATCGGAAG
123636 TATGAGTGTGGAGTCGTTGCACTTCACGTATCAATCCATACNNNAGATCGGAAGAG
123637 TATGAGTGTGGAGTCGTTGCTACTCATGGTTGTTCAATAATTGNNNAGATCGGAAG
123638 TATGAGTGTGGAGTCGTTGCTGAAATTTCTGATACCCTTATGANNNAGATCGGAAG
123639 TATGAGTGTGGAGTCGTTGCATGGTTTCTAGAACAGGAAAAGNNNAGATCGGAAG
123640 TATGAGTGTGGAGTCGTTGCTAGATTGGGAGAAGAGAAAAAAANNNAGATCGGA
123641 TATGAGTGTGGAGTCGTTGCTGATGAGGACATGCTTTTATATANNNAGATCGGAAG
123642 TATGAGTGTGGAGTCGTTGCGAGAGAGTGAAACTAAAATGTTCNNNAGATCGGAA
123643 TATGAGTGTGGAGTCGTTGCATTTCACTTTTGTTGCCCAANNNAGATCGGAAGAGC
123644 TATGAGTGTGGAGTCGTTGCTAGCTTTATTCATTCCTTTCATGNNNAGATCGGAAGA
123645 TATGAGTGTGGAGTCGTTGCCCATTACCAAGTTATTCACATTTNNNAGATCGGAAGA
123647 TATGAGTGTGGAGTCGTTGCCTTCCCAGGAGATAAAATTCTATNNNAGATCGGAAG
123648 TATGAGTGTGGAGTCGTTGCACAGTAGTGTGTTTAAAATCATANNNAGATCGGAAG
123649 TATGAGTGTGGAGTCGTTGCACAGTAGTGTGTTTAAAATCATANNNAGATCGGAAG
123650 TATGAGTGTGGAGTCGTTGCATTTTCATAAAATAGCATCATGATNNNAGATCGGAA
123651 TATGAGTGTGGAGTCGTTGCCATAGTGAAGCTTGCTTTATTATNNNAGATCGGAAG
123652 TATGAGTGTGGAGTCGTTGCTAAATGTTTATTGGCCCCAAGNNNAGATCGGAAGAG
123653 TATGAGTGTGGAGTCGTTGCGAGATGGAGTTTCATTCACTCNNNAGATCGGAAGAG
123654 TATGAGTGTGGAGTCGTTGCATGATCAGATACTGATGAATGTTNNNAGATCGGAAG
123655 TATGAGTGTGGAGTCGTTGCATTTATCCAACTGTTTACTGGANNNAGATCGGAAGA
123656 TATGAGTGTGGAGTCGTTGCACCTTTGCATATTTAGAATTCAANNNAGATCGGAAG
123657 TATGAGTGTGGAGTCGTTGCAAATGGGGGTTTTCACTCTTNNNAGATCGGAAGAG
123658 TATGAGTGTGGAGTCGTTGCTTACTATAATCCCTAACTTGTCCNNNAGATCGGAAGA
123659 TATGAGTGTGGAGTCGTTGCTAATACAGTATTGGCCTACAAATNNNAGATCGGAAG
123660 TATGAGTGTGGAGTCGTTGCCAGGATCACTGCTACTTTTTNNNAGATCGGAAGAGC
123661 TATGAGTGTGGAGTCGTTGCCAGATTGAGACAGTTCAATTTAANNNAGATCGGAAG
123662 TATGAGTGTGGAGTCGTTGCATGGATGTATATCAACTTGTCAANNNAGATCGGAAG
123663 TATGAGTGTGGAGTCGTTGCAAACATGACCTTAGGAATAAGCNNNAGATCGGAAG
123664 TATGAGTGTGGAGTCGTTGCTTTTAAACATGTTTTCATTTCCCNNNAGATCGGAAGA
123665 TATGAGTGTGGAGTCGTTGCATCTGTAATTCCAGAAATCCAGNNNAGATCGGAAGA

123666 TATGAGTGTGGAGTCGTTGCCAGTATCCATTATCCATTGAAAANNNAGATCGGAAG
123667 TATGAGTGTGGAGTCGTTGCCTGAAGTCCCTGATATTAAATTGNNNAGATCGGAAG
123668 TATGAGTGTGGAGTCGTTGCTATATATGCATGCATATGTCTGTNNNAGATCGGAAG
123670 TATGAGTGTGGAGTCGTTGCATGACAGTTTACAGAAAATATGGNNNAGATCGGAA
123671 TATGAGTGTGGAGTCGTTGCCTTCAATATCAACATGCCTTAGNNNAGATCGGAAGA
123674 TATGAGTGTGGAGTCGTTGCTAATATACACATAGGCTTCCTTTNNNAGATCGGAAG
123675 TATGAGTGTGGAGTCGTTGCAAGTGCTGTAAGAATAAAAGAAGNNNAGATCGGAA
123676 TATGAGTGTGGAGTCGTTGCATTACATTATGGTAAAGGGATCANNNAGATCGGAA
123677 TATGAGTGTGGAGTCGTTGCTGGATAATTCTTTTTAAAGGGTTNNNAGATCGGAAG
123678 TATGAGTGTGGAGTCGTTGCTCTCTGAGTTTTGTATAGTCTTCNNNAGATCGGAAG
123679 TATGAGTGTGGAGTCGTTGCTACATATACACACCATGAGGTNNNAGATCGGAAGAG
123680 TATGAGTGTGGAGTCGTTGCAAAATCTCTTCATCTTCTCGAANNNAGATCGGAAGA
123681 TATGAGTGTGGAGTCGTTGCCTGATAAACATCCCACAGTGNNNAGATCGGAAGAG
123682 TATGAGTGTGGAGTCGTTGCGTTGAAGACTCTCTAGAACTTTANNNAGATCGGAAG
123683 TATGAGTGTGGAGTCGTTGCCTATTCCTTCCCATTTTCCTTNNNAGATCGGAAGAGC
123684 TATGAGTGTGGAGTCGTTGCCTCCAGTATAGCATGATAAATCTNNNAGATCGGAAG
123685 TATGAGTGTGGAGTCGTTGCCTCCAGTATAGCATGATAAATCTNNNAGATCGGAAG
123686 TATGAGTGTGGAGTCGTTGCTCAAAATCTCAATTTCCATCTTTNNNAGATCGGAAGA
123688 TATGAGTGTGGAGTCGTTGCACATGTATATATATGTGTGTGTGNNNAGATCGGAAG
123689 TATGAGTGTGGAGTCGTTGCGATCTACTCTGTTAGCAAATTTCNNNAGATCGGAAG
123690 TATGAGTGTGGAGTCGTTGCAAATGTATTGAAGACTTGAACAGNNNAGATCGGAA
123691 TATGAGTGTGGAGTCGTTGCAACCTCTTATTATCATTTTGCTCNNNAGATCGGAAGA
123692 TATGAGTGTGGAGTCGTTGCCCATAAAAATAGCAAAGAAGACTNNNAGATCGGAA
123693 TATGAGTGTGGAGTCGTTGCCTCTAGACACTAAGCACAATATTNNNAGATCGGAAG
123694 TATGAGTGTGGAGTCGTTGCCTTACAAGGACACTAGTCAATAGNNNAGATCGGAAG
123695 TATGAGTGTGGAGTCGTTGCGAAGGAAGACTGGCTAAATTNNNAGATCGGAAGAG
123696 TATGAGTGTGGAGTCGTTGCTGAAAAGAACATTTTGAGAAGAANNNAGATCGGAA
123697 TATGAGTGTGGAGTCGTTGCAAACATTCCTTCCTCCAATTCNNNAGATCGGAAGAGC
123698 TATGAGTGTGGAGTCGTTGCATGAATTATAGGGTTAGGAAGTCNNNAGATCGGAA
123699 TATGAGTGTGGAGTCGTTGCCAAGGAATATGATACACTATCCTNNNAGATCGGAAG
123700 TATGAGTGTGGAGTCGTTGCGAATACCAGAGCTATCACTTTCNNNAGATCGGAAGA
123701 TATGAGTGTGGAGTCGTTGCGAATACCAGAGCTATCACTTTCNNNAGATCGGAAGA
123702 TATGAGTGTGGAGTCGTTGCCTAATATTACCTGCTCTTTGAGANNNAGATCGGAAG
123703 TATGAGTGTGGAGTCGTTGCTTTTTAGCATTTTTCTGAATGTGNNNAGATCGGAAG
123704 TATGAGTGTGGAGTCGTTGCATACACAGATACATATACACACANNNAGATCGGAAG
123705 TATGAGTGTGGAGTCGTTGCTTTCCCATTAAAGTAGTTTCTTCNNNAGATCGGAAGA
123706 TATGAGTGTGGAGTCGTTGCCGAAAATGAAGAAAAATGCAATANNNAGATCGGAA
123707 TATGAGTGTGGAGTCGTTGCGTAAGTGAGAAAAGATGTGAAAANNNAGATCGGA
123708 TATGAGTGTGGAGTCGTTGCTAATAATAAAGGATACAGCATGCNNNAGATCGGAA
123709 TATGAGTGTGGAGTCGTTGCTTATCACTGCAGTCTTAATTACCNNNAGATCGGAAGA
123710 TATGAGTGTGGAGTCGTTGCTCAGTCTCCAGAGTAACTTGNNNAGATCGGAAGAGC
123711 TATGAGTGTGGAGTCGTTGCAGATGAAATCTCACTCTGTTGNNNAGATCGGAAGAG
123712 TATGAGTGTGGAGTCGTTGCTTTTCAAAGCCTATCTAAAGATGNNNAGATCGGAAG

123713 TATGAGTGTGGAGTCGTTGCTAGGAATAAGATGTTTTAGGAGGNNNAGATCGGAA
123714 TATGAGTGTGGAGTCGTTGCGCACATAGTAAGTGATCGATAAANNNAGATCGGAA
123715 TATGAGTGTGGAGTCGTTGCGTCATAAAGTTGTTCTCCAAATANNNAGATCGGAAG
123716 TATGAGTGTGGAGTCGTTGCGGTTGAAACAGAAATGATGTATCNNNAGATCGGAA
123717 TATGAGTGTGGAGTCGTTGCCTAAGATCCAGAAGAGACAGANNNAGATCGGAAGA
123718 TATGAGTGTGGAGTCGTTGCGTGAATTGAAATGAATGCCTTANNNAGATCGGAAG
123719 TATGAGTGTGGAGTCGTTGCATTTGTGTCTCTTCTTCACTGNNNAGATCGGAAGAG
123720 TATGAGTGTGGAGTCGTTGCGCAGTTTGAGATATTTCAACAAANNNAGATCGGAAG
123722 TATGAGTGTGGAGTCGTTGCGGTCTTTGTATAAGATGAAATGGNNNAGATCGGAA
123724 TATGAGTGTGGAGTCGTTGCTTTTGTTTCAGTTTCAGGATTTNNNAGATCGGAAGA
123725 TATGAGTGTGGAGTCGTTGCTGATTACTTTGATGTTTCTTACCNNNAGATCGGAAGA
123726 TATGAGTGTGGAGTCGTTGCAACATTTTGTGATTTTGATTTGTNNNAGATCGGAAG
123727 TATGAGTGTGGAGTCGTTGCAACATTTTGTGATTTTGATTTGTNNNAGATCGGAAG
123728 TATGAGTGTGGAGTCGTTGCGCACTTATATGTCCATAATATGCNNNAGATCGGAAG
123729 TATGAGTGTGGAGTCGTTGCCTGAGAAAAGATAAGGATGTCTTNNNAGATCGGAA
123730 TATGAGTGTGGAGTCGTTGCGACATTAGTCCATTTCATTAGTCNNNAGATCGGAAG
123731 TATGAGTGTGGAGTCGTTGCTCTTATTGATTTACTGATGTCTGNNNAGATCGGAAG
123732 TATGAGTGTGGAGTCGTTGCCCCGTCTAAAATATTTCAATGTANNNAGATCGGAAG
123733 TATGAGTGTGGAGTCGTTGCCATTTATTGAATAGTCCCTTCTTNNNAGATCGGAAGA
123734 TATGAGTGTGGAGTCGTTGCTCTTCTATCCAGGTATCCAAGNNNAGATCGGAAGAG
123735 TATGAGTGTGGAGTCGTTGCTATTCTGGCAGTTTTTTAAAGTTNNNAGATCGGAAG
123736 TATGAGTGTGGAGTCGTTGCGGACCATTCTTATAAGATTTTGGNNNAGATCGGAAG
123737 TATGAGTGTGGAGTCGTTGCCTCTCATTTCTGTCCATTAGGNNNAGATCGGAAGAG
123739 TATGAGTGTGGAGTCGTTGCCAGATAAGTGACACAATTCACNNNAGATCGGAAGAG
123740 TATGAGTGTGGAGTCGTTGCATGTATCCTATCTGATGACCTAANNNAGATCGGAAG
123741 TATGAGTGTGGAGTCGTTGCATGTATCCTATCTGATGACCTAANNNAGATCGGAAG
123742 TATGAGTGTGGAGTCGTTGCCCATCCATCCTATAGATAACAATNNNAGATCGGAAG
123743 TATGAGTGTGGAGTCGTTGCCAAAAATAGACTCTTCAAGGAAANNNAGATCGGAAG
123744 TATGAGTGTGGAGTCGTTGCCTAGAACAAATGGGTCAAATTTTNNNAGATCGGAAG
123745 TATGAGTGTGGAGTCGTTGCCTAGAACAAATGGGTCAAATTTTNNNAGATCGGAAG
123746 TATGAGTGTGGAGTCGTTGCTCACAGAAGTTTCTTCTAGATTANNNAGATCGGAAG
123747 TATGAGTGTGGAGTCGTTGCATGTATATATGTGGTAAGTCCTCNNNAGATCGGAAG
123748 TATGAGTGTGGAGTCGTTGCTTATTTTCTTTTGTTTCCACATGNNNAGATCGGAAGA
123749 TATGAGTGTGGAGTCGTTGCATTGACTATTTATTCTTCTTGGCNNNAGATCGGAAGA
123750 TATGAGTGTGGAGTCGTTGCCTTCAAAAGTTAAAGGTGAGTATNNNAGATCGGAA
123751 TATGAGTGTGGAGTCGTTGCTTTATACGTAGACCATATGAACANNNAGATCGGAAG
123752 TATGAGTGTGGAGTCGTTGCAGCACAATGAAAGAAAAGTAATCNNNAGATCGGAA
123753 TATGAGTGTGGAGTCGTTGCTAAGAAAAAGTCACTCCTTGTATNNNAGATCGGAAG
123754 TATGAGTGTGGAGTCGTTGCACGTTCATTCATTAATTCAATCANNNAGATCGGAAGA
123755 TATGAGTGTGGAGTCGTTGCCACATCGGCCTATCAAATAGNNNAGATCGGAAGAG
123756 TATGAGTGTGGAGTCGTTGCGCTTTAGGCTATAATATGAGTAGANNNAGATCGGA
123757 TATGAGTGTGGAGTCGTTGCCTTTGAAAAGGGCAATAGACTNNNAGATCGGAAGA
123758 TATGAGTGTGGAGTCGTTGCACAAACTTCCAACTCATCTTCNNNAGATCGGAAGAG

123759 TATGAGTGTGGAGTCGTTGCATGTTGGATTCATTTATCTTGAGNNNAGATCGGAAG
123760 TATGAGTGTGGAGTCGTTGCTATACATATTTTCCTGCATGATGNNNAGATCGGAAG
123761 TATGAGTGTGGAGTCGTTGCTTCTTATTCACACACTTCATCTANNNAGATCGGAAGA
123762 TATGAGTGTGGAGTCGTTGCTAGAGACACGGTCTTACTATCNNNAGATCGGAAGAG
123763 TATGAGTGTGGAGTCGTTGCTGTTCTTGACTTCCATATAAATGNNNAGATCGGAAG
123764 TATGAGTGTGGAGTCGTTGCAAAGAGATACTCAGTTTTATCCTNNNAGATCGGAAG
123765 TATGAGTGTGGAGTCGTTGCGGAAATTTGTTTGAATAGACAATNNNAGATCGGAA
123766 TATGAGTGTGGAGTCGTTGCTATTGGCTGTTCTTATAGTAGAGNNNAGATCGGAAG
123767 TATGAGTGTGGAGTCGTTGCCCATCCTAGTCCAATCATGGNNNAGATCGGAAGAGC
123768 TATGAGTGTGGAGTCGTTGCGTAAGGATTCTTTGCTTCCANNNAGATCGGAAGAGC
123769 TATGAGTGTGGAGTCGTTGCACTCAGGAAGTTTTGATATAGAANNNAGATCGGAA
123770 TATGAGTGTGGAGTCGTTGCTACATTTATGCAAATACATGTCCNNNAGATCGGAAG
123771 TATGAGTGTGGAGTCGTTGCGCCAAAGATGTATTCACAAATATNNNAGATCGGAAG
123772 TATGAGTGTGGAGTCGTTGCTAAAAATGGATTACCAGATGAGANNNAGATCGGAA
123773 TATGAGTGTGGAGTCGTTGCTGAATGGTCAGTAATAAGTCAAGNNNAGATCGGAA
123774 TATGAGTGTGGAGTCGTTGCATAATGTGTGATTTCAAATGTCANNNAGATCGGAAG
123775 TATGAGTGTGGAGTCGTTGCACCTTGTCAGTTTCTAAAAAAAANNNAGATCGGAAG
123776 TATGAGTGTGGAGTCGTTGCCTGAGTTTGCTTTATTGAATACTNNNAGATCGGAAG
123777 TATGAGTGTGGAGTCGTTGCATTAGTTATGGAATTTGGTGGAANNNAGATCGGAA
123781 TATGAGTGTGGAGTCGTTGCGGAGATTATTCCATGCAAATTATNNNAGATCGGAAG
123782 TATGAGTGTGGAGTCGTTGCGCGTTTTAAGAACAATAATCTTCNNNAGATCGGAAG
123783 TATGAGTGTGGAGTCGTTGCGCGTTTTAAGAACAATAATCTTCNNNAGATCGGAAG
123784 TATGAGTGTGGAGTCGTTGCCTAAGATATTCAACTAAGGAAGCNNNAGATCGGAA
123785 TATGAGTGTGGAGTCGTTGCGAAAATGCATTCTTACCAATGTANNNAGATCGGAAG
123786 TATGAGTGTGGAGTCGTTGCCAGTAATATAAGGTTGAGCCATANNNAGATCGGAA
123787 TATGAGTGTGGAGTCGTTGCTTTCACACATAGTACTTTCCAAANNNAGATCGGAAGA
123788 TATGAGTGTGGAGTCGTTGCTTTCACACATAGTACTTTCCAAANNNAGATCGGAAGA
123790 TATGAGTGTGGAGTCGTTGCCAGAGGAGAAATTTTTCAGTAATNNNAGATCGGAA
123791 TATGAGTGTGGAGTCGTTGCCTATGGCTTAAGATCCAAAACANNNAGATCGGAAGA
123792 TATGAGTGTGGAGTCGTTGCATACAGAAGAGTGGATATGTCTANNNAGATCGGAA
123793 TATGAGTGTGGAGTCGTTGCAGAGAAGAGTTCAAAAAATAAAGTNNNAGATCGGA
123794 TATGAGTGTGGAGTCGTTGCCCAGAAGAACTATAAAGAGATGANNNAGATCGGAA
123795 TATGAGTGTGGAGTCGTTGCTTTCTCCAACCCTAATTAGTTAANNNAGATCGGAAGA
123796 TATGAGTGTGGAGTCGTTGCGGCAGAAAAAAAATGTAAAAGTTNNNAGATCGGAA
123797 TATGAGTGTGGAGTCGTTGCTTTTCTTCATAGCTACTATCACTNNNAGATCGGAAGA
123798 TATGAGTGTGGAGTCGTTGCGAGGAACCCTAGACCTTTTCNNNAGATCGGAAGAGC
123799 TATGAGTGTGGAGTCGTTGCGAGATGAAGTTTTGTTCTTGTTNNNAGATCGGAAGA
123800 TATGAGTGTGGAGTCGTTGCTTATTAGAGCTAAAGAGGTAAGGNNNAGATCGGAA
123801 TATGAGTGTGGAGTCGTTGCCTTTATGTACATGGAACTTTTTCNNNAGATCGGAAGA
123802 TATGAGTGTGGAGTCGTTGCAGGAGTTTGGGATACAAATATTANNNAGATCGGAA
123803 TATGAGTGTGGAGTCGTTGCTACCAGAAGAAAAATTGCTAAAANNNAGATCGGAA
123804 TATGAGTGTGGAGTCGTTGCTACTTGTGCAGGTTTCATATATNNNAGATCGGAAGA
123805 TATGAGTGTGGAGTCGTTGCTTGTTGTTGTTTGTTTTGTTTTANNNAGATCGGAAGA

123806 TATGAGTGTGGAGTCGTTGCGTAGGCATCATCATTGACTGNNNAGATCGGAAGAG
123807 TATGAGTGTGGAGTCGTTGCGATAAATTCCCCAGGATAAAATNNNAGATCGGAAG
123808 TATGAGTGTGGAGTCGTTGCAAGTGAAGTGGCATGATCTTNNNAGATCGGAAGAG
123809 TATGAGTGTGGAGTCGTTGCGAACAAAGTAGGTGTATAACAAANNNAGATCGGAA
123810 TATGAGTGTGGAGTCGTTGCGTTTGTCCTTAGAGTTTTGATAGNNNAGATCGGAAG
123811 TATGAGTGTGGAGTCGTTGCAAGTCAACCAGGAATATAGATACNNNAGATCGGAA
123812 TATGAGTGTGGAGTCGTTGCATTCATTAAAAATGTCTGGCCNNNAGATCGGAAGAG
123813 TATGAGTGTGGAGTCGTTGCTTCTACCTAGAAAAAGACACATTNNNAGATCGGAAG
123814 TATGAGTGTGGAGTCGTTGCCACAACCTCTGAATAATAGAGTANNNAGATCGGAAG
123815 TATGAGTGTGGAGTCGTTGCTTATCCTTATATTCATCCTAGGCNNNAGATCGGAAGA
123816 TATGAGTGTGGAGTCGTTGCGAGTTCTCTATTTTCTAGGTCTTNNNAGATCGGAAG
123817 TATGAGTGTGGAGTCGTTGCGAGAGAAGGAGAGAGAGAAANNNAGATCGGAAG
123818 TATGAGTGTGGAGTCGTTGCAACAGAACTTCATTTTACAATGANNNAGATCGGAAG
123819 TATGAGTGTGGAGTCGTTGCCAACAATCCTTAAAATCAACATGNNNAGATCGGAAG
123820 TATGAGTGTGGAGTCGTTGCCTCTTCCTTTGAAAAATGGTAGNNNAGATCGGAAGA
123821 TATGAGTGTGGAGTCGTTGCGGGGAGATTTATTTCAAGAAATTNNNAGATCGGAA
123822 TATGAGTGTGGAGTCGTTGCAAGCATAGTTATTAGTCATTTCGNNNAGATCGGAAG
123823 TATGAGTGTGGAGTCGTTGCATGTGTGTCTATGTGTCCAGNNNAGATCGGAAGAG
123824 TATGAGTGTGGAGTCGTTGCACACCTACATCTTGGTTTATAATNNNAGATCGGAAG
123825 TATGAGTGTGGAGTCGTTGCTTCAGATCTTCCCTTTGAGANNNAGATCGGAAGAGC
123826 TATGAGTGTGGAGTCGTTGCCTTTTTCTTTGGGTCTTCATTTNNNAGATCGGAAGAG
123827 TATGAGTGTGGAGTCGTTGCTGGTACAGATTATGTAACCTAAANNNAGATCGGAAG
123828 TATGAGTGTGGAGTCGTTGCCATGAAGAAGAACAAGATCATTGNNNAGATCGGAA
123829 TATGAGTGTGGAGTCGTTGCGCTTGATAGATTTTCTGCTATACNNNAGATCGGAAG
123830 TATGAGTGTGGAGTCGTTGCTTTGAGGAGTAACTGAACTATAGNNNAGATCGGAA
123831 TATGAGTGTGGAGTCGTTGCAAGTCTCTAACCAGCTTTATAATNNNAGATCGGAAG
123832 TATGAGTGTGGAGTCGTTGCCCTTTAATTACTCATTCACCAAANNNAGATCGGAAGA
123833 TATGAGTGTGGAGTCGTTGCATTTCTGGAAACTGAAATGATTTNNNAGATCGGAAG
123834 TATGAGTGTGGAGTCGTTGCAGCATATCAATGGAGATCTATATNNNAGATCGGAAG
123835 TATGAGTGTGGAGTCGTTGCTCATCCTTAAATAATTCAGCATGNNNAGATCGGAAG
123836 TATGAGTGTGGAGTCGTTGCCTGGAATAACTTGTTTTGAATCTNNNAGATCGGAAG
123837 TATGAGTGTGGAGTCGTTGCGTCAGGAGGTAAAGATGAAATNNNAGATCGGAAGA
123838 TATGAGTGTGGAGTCGTTGCGTCAGGAGGTAAAGATGAAATNNNAGATCGGAAGA
123839 TATGAGTGTGGAGTCGTTGCGCCTAACAGTCTAATCACTTAATNNNAGATCGGAAG
123840 TATGAGTGTGGAGTCGTTGCCATAATGCTTCACTGTCATAAGNNNAGATCGGAAGA
123841 TATGAGTGTGGAGTCGTTGCTCATTTTATTAATTCCTCTGGGANNNAGATCGGAAG
123842 TATGAGTGTGGAGTCGTTGCATTTCTGCTATAAAACATAAGCANNNAGATCGGAAG
123843 TATGAGTGTGGAGTCGTTGCTTTTGAGTTTACTATCCTGTCTTNNNAGATCGGAAGA
123844 TATGAGTGTGGAGTCGTTGCTCTTTACTAAATCAAGATCTGGTNNNAGATCGGAAG
123845 TATGAGTGTGGAGTCGTTGCGTAAGTTTTCAAGGCTGTCTTNNNAGATCGGAAGAG
123846 TATGAGTGTGGAGTCGTTGCGTGTCTTTGTGTGTATGTTTGNNNAGATCGGAAGAG
123847 TATGAGTGTGGAGTCGTTGCAGTATTTGTATTTTTGTGACTGGNNNAGATCGGAAG
123848 TATGAGTGTGGAGTCGTTGCGCTTTTTTTCTGGAAAGACAANNNAGATCGGAAGAG

123849 TATGAGTGTGGAGTCGTTGCCTGAGATCACTTTCTCCTTTTTNNNAGATCGGAAGAG
123850 TATGAGTGTGGAGTCGTTGCAATTACAAATTTTCAGACATTGANNNAGATCGGAAG
123851 TATGAGTGTGGAGTCGTTGCAATACATGTCAGAAAAACGAAATNNNAGATCGGAA
123852 TATGAGTGTGGAGTCGTTGCATATTTTCTCTATATAGCCAGGCNNNAGATCGGAAG
123853 TATGAGTGTGGAGTCGTTGCACTTACTTCACTGTTATTAAAGCNNNAGATCGGAAG
123854 TATGAGTGTGGAGTCGTTGCTATTTTGTTTACACAATTGGTCTNNNAGATCGGAAG
123855 TATGAGTGTGGAGTCGTTGCTATAGGCTAAATCACTACAACAANNNAGATCGGAAG
123856 TATGAGTGTGGAGTCGTTGCCTTCCTCCATCCCTTTATTTTNNNAGATCGGAAGAGC
123857 TATGAGTGTGGAGTCGTTGCGATTGTACCTGTTTCCTTACTTANNNAGATCGGAAG
123858 TATGAGTGTGGAGTCGTTGCGGCTAAAACTAGTGTAAATACAGNNNAGATCGGAA
123859 TATGAGTGTGGAGTCGTTGCTATTTTTAGAGACAAATCTCCCTNNNAGATCGGAAG
123860 TATGAGTGTGGAGTCGTTGCAGTTTCACCTCAATTTCCAAGNNNAGATCGGAAGAG
123861 TATGAGTGTGGAGTCGTTGCGCAGGCAATGTTAAAATGTATATNNNAGATCGGAA
123862 TATGAGTGTGGAGTCGTTGCCTGAGTCTCTATCTACAAATACANNNAGATCGGAAG
123863 TATGAGTGTGGAGTCGTTGCCTAACAGTATCTTCAATGCTTTCNNNAGATCGGAAG
123864 TATGAGTGTGGAGTCGTTGCTTAGACTGCAGAATCACTTTATANNNAGATCGGAAG
123865 TATGAGTGTGGAGTCGTTGCTCTAAACAATTTATTTCAACCCTNNNAGATCGGAAGA
123866 TATGAGTGTGGAGTCGTTGCTGTGTATTTCATCTATTCTGAAGNNNAGATCGGAAG
123867 TATGAGTGTGGAGTCGTTGCCTTAATCCCCTTAGGCTTTTCNNNAGATCGGAAGAGC
123868 TATGAGTGTGGAGTCGTTGCTTGTTAATCATAAGCACAAAGTTNNNAGATCGGAAG
123869 TATGAGTGTGGAGTCGTTGCGGTGAATTCTAACCAACATTTTANNNAGATCGGAAG
123870 TATGAGTGTGGAGTCGTTGCGGTATTGAGTCAACTATTTTCTCNNNAGATCGGAAG
123871 TATGAGTGTGGAGTCGTTGCTTAATTTTGCTATACGGTATGAGNNNAGATCGGAAG
123872 TATGAGTGTGGAGTCGTTGCTGAGTATTTAATACAAATCAGAGCNNNAGATCGGAA
123873 TATGAGTGTGGAGTCGTTGCTGAGTATTTAATACAAATCAGAGCNNNAGATCGGAA
123875 TATGAGTGTGGAGTCGTTGCCACTTCACCTATAAAATACTTGCNNNAGATCGGAAGA
123876 TATGAGTGTGGAGTCGTTGCTGCACCTACAAAACCATAATATANNNAGATCGGAAG
123877 TATGAGTGTGGAGTCGTTGCAAAAAGGACTAAAAACCAAGATTNNNAGATCGGAA
123878 TATGAGTGTGGAGTCGTTGCGAAACTGTGGTTTATGTGTGNNNAGATCGGAAGAG
123879 TATGAGTGTGGAGTCGTTGCCATTATGGTTGGATGTAATGATTNNNAGATCGGAAG
123880 TATGAGTGTGGAGTCGTTGCCAAATTCTTGTGAGTTTATTGTGNNNAGATCGGAAG
123881 TATGAGTGTGGAGTCGTTGCTTTTTGCATTGAATGACTTTTAANNNAGATCGGAAG
123882 TATGAGTGTGGAGTCGTTGCTTTACTTCACAGGATTTTGTTTTNNNAGATCGGAAGA
123884 TATGAGTGTGGAGTCGTTGCTCAAATAAGAGCATTTAGATGTGNNNAGATCGGAA
123885 TATGAGTGTGGAGTCGTTGCTCAAATAAGAGCATTTAGATGTGNNNAGATCGGAA
123886 TATGAGTGTGGAGTCGTTGCCCATGGGTTCATAAACAAAAANNNAGATCGGAAGA
123887 TATGAGTGTGGAGTCGTTGCGTATGCAGCATTCTACATAAAATNNNAGATCGGAAG
123888 TATGAGTGTGGAGTCGTTGCTGGCTAAAAGACATTCTTTAAAANNNAGATCGGAAG
123889 TATGAGTGTGGAGTCGTTGCGTGTCACAAGTGAGATCAAANNNAGATCGGAAGAG
123890 TATGAGTGTGGAGTCGTTGCTTAGGTTTCTATTATCTTCTCCCNNNAGATCGGAAGA
123891 TATGAGTGTGGAGTCGTTGCCCACTAAAATTTTTGTTGTCTTCNNNAGATCGGAAGA
123892 TATGAGTGTGGAGTCGTTGCGTTTTCTGTATGTGACACTACTANNNAGATCGGAAG
123893 TATGAGTGTGGAGTCGTTGCGGCATACTTGTTGAAAAAAATTTNNNAGATCGGAAG

123894 TATGAGTGTGGAGTCGTTGCATGATAATTTCCTGCCAAATCTNNNAGATCGGAAGA
123895 TATGAGTGTGGAGTCGTTGCTTTGTAATGGAAAGAATGTTGAANNNAGATCGGAA
123896 TATGAGTGTGGAGTCGTTGCTTTGTAATGGAAAGAATGTTGAANNNAGATCGGAA
123898 TATGAGTGTGGAGTCGTTGCCAACAAATAGCAGCAATTTAATTNNNAGATCGGAAG
123899 TATGAGTGTGGAGTCGTTGCCCTAAGTTTAAGCTATAGCAAATNNNAGATCGGAAG
123900 TATGAGTGTGGAGTCGTTGCGGAATCTCACCATCACCTAGNNNAGATCGGAAGAGC
123901 TATGAGTGTGGAGTCGTTGCGGGTAAAGGATATGTTACTTTTGNNNAGATCGGAA
123902 TATGAGTGTGGAGTCGTTGCGTCTTTAGATCCTTTCCTCTGNNNAGATCGGAAGAG
123903 TATGAGTGTGGAGTCGTTGCACTAGAAGAAACTAACATGAAGGNNNAGATCGGAA
123904 TATGAGTGTGGAGTCGTTGCTATAGTCTGGAAAAGAAAAGAGANNNAGATCGGAA
123905 TATGAGTGTGGAGTCGTTGCAGAATGATATATTAACTCCTGGGNNNAGATCGGAA
123906 TATGAGTGTGGAGTCGTTGCAAATGTGTTCAGCAACAATATATNNNAGATCGGAAG
123907 TATGAGTGTGGAGTCGTTGCCTAGTTATTTTAAAGAGTCTGGCNNNAGATCGGAAG
123908 TATGAGTGTGGAGTCGTTGCCCTAGAACTTTTACTTGCCANNNAGATCGGAAGAGC
123909 TATGAGTGTGGAGTCGTTGCGAGGTACTGGAGAGCTATACNNNAGATCGGAAGAG
123910 TATGAGTGTGGAGTCGTTGCTTTAAAAGCTCCAAGTCCATTNNNAGATCGGAAGAG
123911 TATGAGTGTGGAGTCGTTGCGGGTTTCCTACATCTAAGATTATNNNAGATCGGAAG
123914 TATGAGTGTGGAGTCGTTGCTTCTCCTAGTTTCCTGTAGAANNNAGATCGGAAGAG
123915 TATGAGTGTGGAGTCGTTGCAGAATGCGCTAGTTCTATTTAATNNNAGATCGGAAG
123916 TATGAGTGTGGAGTCGTTGCCAGTTTCAAGCAATTCTCCTNNNAGATCGGAAGAGC
123917 TATGAGTGTGGAGTCGTTGCGAAAAATATTGTTCCACTCTCTTNNNAGATCGGAAG
123918 TATGAGTGTGGAGTCGTTGCACTAACAATAAGTAAAAGATTTGAGNNNAGATCGG
123919 TATGAGTGTGGAGTCGTTGCTTGTTTTGTTTTTCTTTACTCTGNNNAGATCGGAAGA
123920 TATGAGTGTGGAGTCGTTGCCCACTGAACTTTATTTTTACTGANNNAGATCGGAAG
123921 TATGAGTGTGGAGTCGTTGCCTTTCAACTAAGACCTTCCTTAANNNAGATCGGAAGA
123922 TATGAGTGTGGAGTCGTTGCCTTAGGAACTATGACGTCTAATANNNAGATCGGAAG
123923 TATGAGTGTGGAGTCGTTGCAATGACTACTCTGTCTCTAGATANNNAGATCGGAAG
123924 TATGAGTGTGGAGTCGTTGCAAGAATCCAGGAGAAAAATAAGGNNNAGATCGGA
123925 TATGAGTGTGGAGTCGTTGCCCCAGTAGTATAAAAAACTCAACNNNAGATCGGAAG
123926 TATGAGTGTGGAGTCGTTGCAAGATTACTTTAGCTAACATGTGNNNAGATCGGAAG
123927 TATGAGTGTGGAGTCGTTGCTAATGAACTGATTTCCTCATCTANNNAGATCGGAAG
123929 TATGAGTGTGGAGTCGTTGCATAATAAGGAAAGAGAAGACAGGNNNAGATCGGA
123930 TATGAGTGTGGAGTCGTTGCCCAATCAATTTACTTTAGCTGTANNNAGATCGGAAG
123932 TATGAGTGTGGAGTCGTTGCTTAAATATCATGCATGGACATTTNNNAGATCGGAAG
123933 TATGAGTGTGGAGTCGTTGCCTATTATAGCTCACTGCAGCNNNAGATCGGAAGAGC
123934 TATGAGTGTGGAGTCGTTGCCTTTAAAGTGCTAAAACGAAAAGNNNAGATCGGAA
123935 TATGAGTGTGGAGTCGTTGCATCAACAATTTCAAACAACAAAGNNNAGATCGGAAG
123936 TATGAGTGTGGAGTCGTTGCATACATAATTCACTGACCTTTCTNNNAGATCGGAAGA
123937 TATGAGTGTGGAGTCGTTGCTCCATTATTTCTGTCTTCCTATTNNNAGATCGGAAGA
123938 TATGAGTGTGGAGTCGTTGCATCAAATAAACTCCACATGAAACNNNAGATCGGAAG
123939 TATGAGTGTGGAGTCGTTGCATCAAATAAACTCCACATGAAACNNNAGATCGGAAG
123940 TATGAGTGTGGAGTCGTTGCGTTTTACTTTTACCCCACATTTANNNAGATCGGAAGA
123941 TATGAGTGTGGAGTCGTTGCCTAGGGCCTCTTAGTAAAATCNNNAGATCGGAAGAG

123942 TATGAGTGTGGAGTCGTTGCACTCATACTTACAATTCCTACTGNNNAGATCGGAAGA
123943 TATGAGTGTGGAGTCGTTGCTTGTGTTAGTAAAACAATTCAGTNNNAGATCGGAAG
123944 TATGAGTGTGGAGTCGTTGCAGAAAACCTATTTACAATTCCTGNNNAGATCGGAAGA
123945 TATGAGTGTGGAGTCGTTGCTCAGCCTAACTCAAGCATAANNNAGATCGGAAGAGC
123946 TATGAGTGTGGAGTCGTTGCTCTATGCATATGCCATCTATATTNNNAGATCGGAAGA
123947 TATGAGTGTGGAGTCGTTGCTACTCCATCCTCATACTAGAAGNNNAGATCGGAAGAG
123948 TATGAGTGTGGAGTCGTTGCTCTGTGGATTTGGATTTCATTNNNAGATCGGAAGAG
123949 TATGAGTGTGGAGTCGTTGCTACCCCTGAGATACTGTATAAANNNAGATCGGAAGA
123950 TATGAGTGTGGAGTCGTTGCTACCCCTGAGATACTGTATAAANNNAGATCGGAAGA
123951 TATGAGTGTGGAGTCGTTGCACATAATTCTACCAAAAAGACACNNNAGATCGGAAGA
123952 TATGAGTGTGGAGTCGTTGCACATAATTCTACCAAAAAGACACNNNAGATCGGAAGA
123953 TATGAGTGTGGAGTCGTTGCATCAGTCAAACTCATTATTAGGTNNNAGATCGGAAGA
123954 TATGAGTGTGGAGTCGTTGCATTTTGATGCTAATTTTATCCCCNNNAGATCGGAAGA
123955 TATGAGTGTGGAGTCGTTGCCTATAGCAAACTGCATACTTTAGNNNAGATCGGAAGA
123956 TATGAGTGTGGAGTCGTTGCCTATAGCAAACTGCATACTTTAGNNNAGATCGGAAGA
123957 TATGAGTGTGGAGTCGTTGCCAAATGGGACATAATTGAAGTGNNNAGATCGGAAG
123958 TATGAGTGTGGAGTCGTTGCGACATTTCATTTAAGGTAGAAGANNNAGATCGGAAG
123959 TATGAGTGTGGAGTCGTTGCGACATTTCATTTAAGGTAGAAGANNNAGATCGGAAG
123960 TATGAGTGTGGAGTCGTTGCCAAAGCATACTCTGTCCTTTNNNAGATCGGAAGAGCA
123961 TATGAGTGTGGAGTCGTTGCAGAGTAGATTGAGGACATTTAATNNNAGATCGGAAG
123962 TATGAGTGTGGAGTCGTTGCCTCATCTTCATAAGGACCTTGNNNAGATCGGAAGAGC
123964 TATGAGTGTGGAGTCGTTGCCTTTTGCTTTCTTTCTCTTTCTNNNAGATCGGAAGAGC
123965 TATGAGTGTGGAGTCGTTGCTTTATGTGTGCCAAGAATAAAGNNNAGATCGGAAGA
123966 TATGAGTGTGGAGTCGTTGCTTGGTACTAAGATAAACTCTACGNNNAGATCGGAAG
123967 TATGAGTGTGGAGTCGTTGCTTGGTACTAAGATAAACTCTACGNNNAGATCGGAAG
123968 TATGAGTGTGGAGTCGTTGCCATCTGTTATCTTTTGAAGTACTNNNAGATCGGAAGA
123969 TATGAGTGTGGAGTCGTTGCTCTTACGTAGAGCTTTTACATATNNNAGATCGGAAGA
123970 TATGAGTGTGGAGTCGTTGCTCTTACGTAGAGCTTTTACATATNNNAGATCGGAAGA
123971 TATGAGTGTGGAGTCGTTGCCTAATGAATGACAGCATCTTTCNNNAGATCGGAAGAG
123972 TATGAGTGTGGAGTCGTTGCACAGAATGAGATAGCCATTTTTNNNAGATCGGAAGA
123973 TATGAGTGTGGAGTCGTTGCAGTTCTGTGTAATTTTATCACACNNNAGATCGGAAGA
123974 TATGAGTGTGGAGTCGTTGCGGAAAGATTCTTGTTCTAACTTTNNNAGATCGGAAGA
123975 TATGAGTGTGGAGTCGTTGCACTTGGATTTTCACTAATGTTTTNNNAGATCGGAAGA
123976 TATGAGTGTGGAGTCGTTGCGCCATGCTTTTAATAGATAGTTTNNNAGATCGGAAGA
123977 TATGAGTGTGGAGTCGTTGCATTTCTCCCACTTCTTCATCTANNNAGATCGGAAGAGC
123978 TATGAGTGTGGAGTCGTTGCCAATAAGTAACATGCAACAGATNNNAGATCGGAAGA
123980 TATGAGTGTGGAGTCGTTGCGCCCACAAAAATGTTTTCAANNNAGATCGGAAGAGC
123981 TATGAGTGTGGAGTCGTTGCTACAGATTCAAGCATATCAAATCNNNAGATCGGAAGA
123982 TATGAGTGTGGAGTCGTTGCCAATACAGATTCAAGCATATCAANNNAGATCGGAAG
123983 TATGAGTGTGGAGTCGTTGCCTCTCATCCAGAACCAATATACNNNAGATCGGAAGAG
123984 TATGAGTGTGGAGTCGTTGCGGACAACTCCTTGTAAATATCTANNNAGATCGGAAGA
123985 TATGAGTGTGGAGTCGTTGCTATTACGCATTTCTGCTTTTTTNNNAGATCGGAAGAG
123986 TATGAGTGTGGAGTCGTTGCTATTACGCATTTCTGCTTTTTTNNNAGATCGGAAGAG

123987 TATGAGTGTGGAGTCGTTGCCGTTTTAAAAAATCAGCCTAAATNNNAGATCGGAAG
123988 TATGAGTGTGGAGTCGTTGCCATCTCAAATTTAGAATTGGGTTNNNAGATCGGAAG
123990 TATGAGTGTGGAGTCGTTGCGAACTCTTTCTCAGGTTCATTNNNAGATCGGAAGAG
123991 TATGAGTGTGGAGTCGTTGCATATATTTTTGAGGTCTGCATACNNNAGATCGGAAG
123993 TATGAGTGTGGAGTCGTTGCGGGATGATTCAGTATCTATTGTANNNAGATCGGAAG
123994 TATGAGTGTGGAGTCGTTGCGTATTATTCTGTACACAATCCATNNNAGATCGGAAG
123995 TATGAGTGTGGAGTCGTTGCAAGATAGAGAGAGACAATATGAGNNNAGATCGGA
123996 TATGAGTGTGGAGTCGTTGCACATAGGGGAATGAATGTATAAANNNAGATCGGAA
123997 TATGAGTGTGGAGTCGTTGCAATATGAAAATAGTGGAATGCTGNNNAGATCGGAA
123998 TATGAGTGTGGAGTCGTTGCAATATGAAAATAGTGGAATGCTGNNNAGATCGGAA
123999 TATGAGTGTGGAGTCGTTGCTAGAAAGAAAGGCAAAGAGTATTNNNAGATCGGAA
124000 TATGAGTGTGGAGTCGTTGCTAGAAAGAAAGGCAAAGAGTATTNNNAGATCGGAA
124001 TATGAGTGTGGAGTCGTTGCATACTAATAAGGTGTCAGTCAAGNNNAGATCGGAA
124002 TATGAGTGTGGAGTCGTTGCCTTGAGAAAGGCATTATAGGATNNNAGATCGGAAG
124003 TATGAGTGTGGAGTCGTTGCCTTGAGAAAGGCATTATAGGATNNNAGATCGGAAG
124004 TATGAGTGTGGAGTCGTTGCTGAAGTAATTTGGATGTTAAGTTNNNAGATCGGAA
124005 TATGAGTGTGGAGTCGTTGCCTAAAAAAAAAATCTTTTGCGTGNNNAGATCGGAAG
124006 TATGAGTGTGGAGTCGTTGCGAATGCTCAAGTCTCTGATATANNNAGATCGGAAGA
124008 TATGAGTGTGGAGTCGTTGCATCTATCTATCTCTCTCTCTCTCNNNAGATCGGAAGAG
124009 TATGAGTGTGGAGTCGTTGCGCACAAATGGGAGATTATTTAANNNAGATCGGAA
124010 TATGAGTGTGGAGTCGTTGCCGTAATAAACATGACAACCTAAANNNAGATCGGAAG
124011 TATGAGTGTGGAGTCGTTGCCGTAATAAACATGACAACCTAAANNNAGATCGGAAG
124012 TATGAGTGTGGAGTCGTTGCAAATATTGGAGACAGCTTCTAAGNNNAGATCGGAA
124013 TATGAGTGTGGAGTCGTTGCCTTTTCCATACTTATCACTTCACNNNAGATCGGAAGA
124014 TATGAGTGTGGAGTCGTTGCCTGAGTGAGGTATGGGAATGNNNAGATCGGAAGA
124015 TATGAGTGTGGAGTCGTTGCCACTTATGTTTTTCTCACATCTTNNNAGATCGGAAGA
124016 TATGAGTGTGGAGTCGTTGCTAATATTTTCAAGATTCACAGCCNNNAGATCGGAAG
124017 TATGAGTGTGGAGTCGTTGCGGCTTTCTTAATTAACCTGAAAANNNAGATCGGAAG
124018 TATGAGTGTGGAGTCGTTGCACTTCAGTTAGATATAAGTCACCNNNAGATCGGAAG
124019 TATGAGTGTGGAGTCGTTGCGTCTCATCAGGCTATATTCAAANNNAGATCGGAAGA
124020 TATGAGTGTGGAGTCGTTGCTTTTTGTATCATCTCTGGAAAAGNNNAGATCGGAAG
124021 TATGAGTGTGGAGTCGTTGCTCCATTTAAAATCTCTTTTCACANNNAGATCGGAAGA
124022 TATGAGTGTGGAGTCGTTGCCAGGTTTATCAGCCCAAAAGNNNAGATCGGAAGAG
124023 TATGAGTGTGGAGTCGTTGCGGTCATGGAAAAGGTAAATTTAANNNAGATCGGAA
124024 TATGAGTGTGGAGTCGTTGCCTTCAGTATTAGATGGGCTATATNNNAGATCGGAAG
124025 TATGAGTGTGGAGTCGTTGCCATGAAAAGGTAGTTTTGACTTTNNNAGATCGGAAG
124026 TATGAGTGTGGAGTCGTTGCCAACATATCTTTCTAGTCCAGAANNNAGATCGGAAG
124027 TATGAGTGTGGAGTCGTTGCTAAAGGTTCATTCTTCCTTACTTNNNAGATCGGAAGA
124028 TATGAGTGTGGAGTCGTTGCAAAACCAAGATAAAGGGACTCNNNAGATCGGAAGA
124030 TATGAGTGTGGAGTCGTTGCTATTTCTCCAACTATAAGAGTGCNNNAGATCGGAAG
124031 TATGAGTGTGGAGTCGTTGCCAGAGTGGGTATATAGAGCANNNAGATCGGAAGAG
124032 TATGAGTGTGGAGTCGTTGCGGCATAGACTATGTAATTTTATGTNNNAGATCGGAA
124033 TATGAGTGTGGAGTCGTTGCCTTGAGCTCTTCTTTTTAACTTTNNNAGATCGGAAGA

124034 TATGAGTGTGGAGTCGTTGCGCAAAATTCAACAGTTGATAAAANNNAGATCGGAAG
124035 TATGAGTGTGGAGTCGTTGCGAATAGGCAATGGAATCTCTCNNNAGATCGGAAGA
124036 TATGAGTGTGGAGTCGTTGCAATTTGAACAGGAGAGAGTATAANNNAGATCGGAA
124037 TATGAGTGTGGAGTCGTTGCGGAAACACAAAGAGCTAATAATANNNAGATCGGAA
124038 TATGAGTGTGGAGTCGTTGCGAATGAATGAAGAGAGACAAAACNNNAGATCGGAA
124040 TATGAGTGTGGAGTCGTTGCAATGAATTTTCCATTTTTTGTCCNNNAGATCGGAAGA
124041 TATGAGTGTGGAGTCGTTGCAAAAACAAACAAAAAAAAACACCNNNAGATCGGAAG
124042 TATGAGTGTGGAGTCGTTGCTTTACTACTACAGAATCATCTGGNNNAGATCGGAAG
124043 TATGAGTGTGGAGTCGTTGCGGCATTGGTGAAGTAATATATTTNNNAGATCGGAAG
124044 TATGAGTGTGGAGTCGTTGCGAAAGTAGTCAACATCTCTTACTNNNAGATCGGAAG
124045 TATGAGTGTGGAGTCGTTGCTAGCCAAACAGATGTATTTCTAANNNAGATCGGAAG
124046 TATGAGTGTGGAGTCGTTGCTCTTTTAGGCATATAAAACTCAANNNAGATCGGAAG
124047 TATGAGTGTGGAGTCGTTGCAATTCAACCACCATCTACCTNNNAGATCGGAAGAGCA
124048 TATGAGTGTGGAGTCGTTGCATTGAAACAAAGACTACCATTTTNNNAGATCGGAAG
124049 TATGAGTGTGGAGTCGTTGCCATTTCTAAGTTAAAGCACTTGANNNAGATCGGAAG
124050 TATGAGTGTGGAGTCGTTGCAAGGTATCATGAAAAGAACAAAANNNAGATCGGAA
124051 TATGAGTGTGGAGTCGTTGCGTAAGGTCAAACTTCTGTTAATTNNNAGATCGGAAG
124052 TATGAGTGTGGAGTCGTTGCTAAGTGAGCAACCTCTACAANNNAGATCGGAAGAG
124053 TATGAGTGTGGAGTCGTTGCTAAGTGAGCAACCTCTACAANNNAGATCGGAAGAG
124054 TATGAGTGTGGAGTCGTTGCGGGCAATAAAACTATAACATCTTNNNAGATCGGAAG
124055 TATGAGTGTGGAGTCGTTGCCATTTTTTGCTTCCCTAGATTACNNNAGATCGGAAGA
124056 TATGAGTGTGGAGTCGTTGCAAGCTTTCCTTCTTTATATCACNNNAGATCGGAAGA
124057 TATGAGTGTGGAGTCGTTGCTTCCACGTATATTCTTATACAGTNNNAGATCGGAAGA
124058 TATGAGTGTGGAGTCGTTGCTTTTACTTTACACAGTAGTGAACNNNAGATCGGAAG
124059 TATGAGTGTGGAGTCGTTGCAGTGTTTTTTTAATATAGCAGCANNNAGATCGGAAG
124060 TATGAGTGTGGAGTCGTTGCAGAAATGTTTATGCTTTTTGATGNNNAGATCGGAAG
124061 TATGAGTGTGGAGTCGTTGCAAGTCAGAAAGGGTCTTATATACNNNAGATCGGAAG
124062 TATGAGTGTGGAGTCGTTGCGCAATTCTATAAATTCGAAAGTGNNNAGATCGGAAG
124063 TATGAGTGTGGAGTCGTTGCGTATCCTGGGCATTTAAAATATTNNNAGATCGGAAG
124064 TATGAGTGTGGAGTCGTTGCCATCAACATCAAATCTATCAAGANNNAGATCGGAAG
124065 TATGAGTGTGGAGTCGTTGCGTGAGAATGTAAATTGGTACAACNNNAGATCGGAA
124066 TATGAGTGTGGAGTCGTTGCGAGAACCTTGTGAGCGATATNNNAGATCGGAAGAG
124067 TATGAGTGTGGAGTCGTTGCCATCATATTGCCTATCAATTTCTNNNAGATCGGAAGA
124068 TATGAGTGTGGAGTCGTTGCACAGATTCATCTTGTAATTGAGTNNNAGATCGGAAG
124069 TATGAGTGTGGAGTCGTTGCCTTAGTAGTTCCCCTTTTCAGNNNAGATCGGAAGAG
124070 TATGAGTGTGGAGTCGTTGCAATTTCTATTGACCTGTCTTCAANNNAGATCGGAAGA
124071 TATGAGTGTGGAGTCGTTGCAGAATAAATCAGGGAATGAGAGNNNAGATCGGAAG
124072 TATGAGTGTGGAGTCGTTGCATCAAAACAACCTATAATGCTTCNNNAGATCGGAAG
124073 TATGAGTGTGGAGTCGTTGCGTGAAAGAGAAAGAGAGAGAGNNNAGATCGGAAG
124074 TATGAGTGTGGAGTCGTTGCGCATAGAAACAGTATTTAAAGGGNNNAGATCGGAA
124075 TATGAGTGTGGAGTCGTTGCTAAATTGAAACAGAAAATCACCCNNNAGATCGGAAG
124076 TATGAGTGTGGAGTCGTTGCGAAACTGGCTTCAAGTAAAATAANNNAGATCGGAAG
124077 TATGAGTGTGGAGTCGTTGCCTATACAGGACAAGAGGTCANNNAGATCGGAAGAG

124078 TATGAGTGTGGAGTCGTTGCGCTTCAGGTATTAGAAATACCANNNAGATCGGAAGA
124079 TATGAGTGTGGAGTCGTTGCTTTACTTATCTCCTTAGATCTGCNNNAGATCGGAAGA
124080 TATGAGTGTGGAGTCGTTGCTTTTCAAATAAGAAAATGGAAGCNNNAGATCGGAAG
124081 TATGAGTGTGGAGTCGTTGCTTTTTCCAAAGAATCTGTCTCTNNNAGATCGGAAGA
124082 TATGAGTGTGGAGTCGTTGCTAATACAGTAGAAGAAACAGGAGNNNAGATCGGAA
124083 TATGAGTGTGGAGTCGTTGCGGAAATAGAAATGGTAGTTGAAGNNNAGATCGGA
124084 TATGAGTGTGGAGTCGTTGCATGTGGAAATCTTAGAGTACTGNNNAGATCGGAAG
124085 TATGAGTGTGGAGTCGTTGCGAACACAATTGCTATATCCATACNNNAGATCGGAAG
124086 TATGAGTGTGGAGTCGTTGCGCTGATGTAATGTCAATTCATAANNNAGATCGGAAG
124087 TATGAGTGTGGAGTCGTTGCAATCGTAACGAGGTTAAATGATANNNAGATCGGAA
124088 TATGAGTGTGGAGTCGTTGCCTATCGCCAATATGTGACTGNNNAGATCGGAAGAG
124089 TATGAGTGTGGAGTCGTTGCCTAGAGAGAGAGAGAGAGATTATNNNAGATCGGA
124091 TATGAGTGTGGAGTCGTTGCTTTAGCCAAGAACTGTACAGNNNAGATCGGAAGAG
124092 TATGAGTGTGGAGTCGTTGCTTCAAGGGAATTTAAGGGATGNNNAGATCGGAAGA
124093 TATGAGTGTGGAGTCGTTGCCTGATTCAGTTCCCACTAAATNNNAGATCGGAAGAG
124094 TATGAGTGTGGAGTCGTTGCCCTTCTCAATGTTTTTCTTCTTTNNNAGATCGGAAGA
124095 TATGAGTGTGGAGTCGTTGCGCATATCTGTCTTGAGAAGTTNNNAGATCGGAAGA
124096 TATGAGTGTGGAGTCGTTGCTTCTCATTTTCTGGCACTATAANNNAGATCGGAAGA
124097 TATGAGTGTGGAGTCGTTGCGAAATCTAATTACAAGTGCTCATNNNAGATCGGAAG
124098 TATGAGTGTGGAGTCGTTGCTAAATCTTTGAATTGCAATCTCTNNNAGATCGGAAG
124099 TATGAGTGTGGAGTCGTTGCATTTTCTGAAAGATTGAAGCAATNNNAGATCGGAAG
124100 TATGAGTGTGGAGTCGTTGCTAATCTCTGTTTTGAATACTTGGNNNAGATCGGAAG
124101 TATGAGTGTGGAGTCGTTGCCATACACACTACTGAGGTATTTTNNNAGATCGGAAG
124102 TATGAGTGTGGAGTCGTTGCCATACACACTACTGAGGTATTTTNNNAGATCGGAAG
124103 TATGAGTGTGGAGTCGTTGCTATAAATAGGAAAGGAGAAGGTCNNNAGATCGGAA
124104 TATGAGTGTGGAGTCGTTGCACTTTGGTTTGAATTTTGTATGANNNAGATCGGAAG
124105 TATGAGTGTGGAGTCGTTGCATATAGATGCTCCATGAATAACANNNAGATCGGAAG
124106 TATGAGTGTGGAGTCGTTGCTAAAACAAAAACAGCTCAGAAAANNNAGATCGGAA
124107 TATGAGTGTGGAGTCGTTGCGTCACAAAGGACTGTACTTGNNNAGATCGGAAGAG
124108 TATGAGTGTGGAGTCGTTGCATTTAGTGTTACATTTGGAGTACNNNAGATCGGAAG
124109 TATGAGTGTGGAGTCGTTGCGAGGTAACATGATAATAGAGGAANNNAGATCGGAA
124110 TATGAGTGTGGAGTCGTTGCTCATAATCCTGCAGGTATAGANNNAGATCGGAAGA
124111 TATGAGTGTGGAGTCGTTGCAGAGAAATCACTCATAATCCTGNNNAGATCGGAAGA
124112 TATGAGTGTGGAGTCGTTGCAAGACAAAGAAATTATGAGAAGCNNNAGATCGGAA
124113 TATGAGTGTGGAGTCGTTGCGATGAAACTGGGGAAAGTTTNNNAGATCGGAAGA
124114 TATGAGTGTGGAGTCGTTGCGGTGAAATTAAATCACTCTGTAANNNAGATCGGAAG
124115 TATGAGTGTGGAGTCGTTGCTTTAGCAATAAGGCTTATTTCTGNNNAGATCGGAAG
124116 TATGAGTGTGGAGTCGTTGCGTATAAGAACTTTGCAACACTATNNNAGATCGGAAG
124117 TATGAGTGTGGAGTCGTTGCGGTTTATTTTTCTCAGAATTGGTNNNAGATCGGAAG
124118 TATGAGTGTGGAGTCGTTGCTGCATTTCTATATGTGTAAATACCNNNAGATCGGAA
124119 TATGAGTGTGGAGTCGTTGCCCAAATGCAATCTTCCTATAAATNNNAGATCGGAAG
124120 TATGAGTGTGGAGTCGTTGCCCAAATGCAATCTTCCTATAAATNNNAGATCGGAAG
124121 TATGAGTGTGGAGTCGTTGCACATAATCTACACAGAGATAGTTNNNAGATCGGAAG

124122 TATGAGTGTGGAGTCGTTGCTGGATTAGAGACTAAGTAGGTATNNNAGATCGGAA
124123 TATGAGTGTGGAGTCGTTGCAATGTTTTTTAGTAAGTTCCCAGNNNAGATCGGAAG
124124 TATGAGTGTGGAGTCGTTGCAAAATTCACTTTATTTCTGAGCTNNNAGATCGGAAG
124125 TATGAGTGTGGAGTCGTTGCTTGGAACCTCTCAATTTTTTTATNNNAGATCGGAAGA
124126 TATGAGTGTGGAGTCGTTGCCAGGAGAAAATCTTCTGATAGTNNNAGATCGGAAG
124127 TATGAGTGTGGAGTCGTTGCCATTGCATATACAATTTGGGTAANNNAGATCGGAAG
124128 TATGAGTGTGGAGTCGTTGCAGAGATAACTGTACAAAAATGTCNNNAGATCGGAA
124129 TATGAGTGTGGAGTCGTTGCCACCAAAACAGGAATCAAAATNNNAGATCGGAAGA
124130 TATGAGTGTGGAGTCGTTGCGTCTTACTAGTTACAAACAAGAANNNAGATCGGAAG
124131 TATGAGTGTGGAGTCGTTGCACAATAGTGTAAGGACATAAGAANNNAGATCGGAA
124132 TATGAGTGTGGAGTCGTTGCGCTATGTCATGAATGCATAAAATNNNAGATCGGAAG
124133 TATGAGTGTGGAGTCGTTGCGTCTGCAAATTCTATGAATTCTANNNAGATCGGAAG
124134 TATGAGTGTGGAGTCGTTGCAGATAGACAGAAAGAAAGCAAGNNNAGATCGGAA
124135 TATGAGTGTGGAGTCGTTGCTTGTTTTGTTTTGTTTTGTTTTGNNNAGATCGGAAGA
124136 TATGAGTGTGGAGTCGTTGCACATAATTTGGGGATATAACTGANNNAGATCGGAA
124137 TATGAGTGTGGAGTCGTTGCGTGTTCTCTGAGAAAAGGAGNNNAGATCGGAAGAG
124138 TATGAGTGTGGAGTCGTTGCATTTTAAATGAGGGAAGTTCACANNNAGATCGGAA
124139 TATGAGTGTGGAGTCGTTGCCTTCGGTCATAGGATCAAGTNNNAGATCGGAAGAG
124140 TATGAGTGTGGAGTCGTTGCGATTTCCATGCAAATTTATCCANNNAGATCGGAAGA
124141 TATGAGTGTGGAGTCGTTGCCATCATTTCATTTTATAACCCCANNNAGATCGGAAGA
124142 TATGAGTGTGGAGTCGTTGCCAGAAAAGGTGGCATTTATTTNNNAGATCGGAAGA
124143 TATGAGTGTGGAGTCGTTGCTTAGTGGAAATAAGAATGGGTTANNNAGATCGGAA
124144 TATGAGTGTGGAGTCGTTGCGCATGCATTTCAAAACTTAATAGNNNAGATCGGAAG
124145 TATGAGTGTGGAGTCGTTGCGAAAAGGTTTTGACTGTGAGNNNAGATCGGAAGAG
124147 TATGAGTGTGGAGTCGTTGCATTCTGCTGATTTTAGTTTCTTCNNNAGATCGGAAGA
124148 TATGAGTGTGGAGTCGTTGCATTCTGCTGATTTTAGTTTCTTCNNNAGATCGGAAGA
124149 TATGAGTGTGGAGTCGTTGCGAATGAAACTCTGCAACCAANNNAGATCGGAAGAG
124150 TATGAGTGTGGAGTCGTTGCAGATAAGGTCTCACTGTCACNNNAGATCGGAAGAG
124151 TATGAGTGTGGAGTCGTTGCCAAGAAGCAAACAAGTCTTCNNNAGATCGGAAGAG
124152 TATGAGTGTGGAGTCGTTGCATTCTTTAAGATAGCCTTTTGAGNNNAGATCGGAAG
124153 TATGAGTGTGGAGTCGTTGCTTGAGGATATGTTTGTGTAAATGNNNAGATCGGAA
124154 TATGAGTGTGGAGTCGTTGCTGTACATCACCTAAGTTCTAAATNNNAGATCGGAAG
124155 TATGAGTGTGGAGTCGTTGCGTATATAGGTCCAGTAAAGTCATNNNAGATCGGAA
124156 TATGAGTGTGGAGTCGTTGCCTTGTGGTGTTTACAATCTATTTNNNAGATCGGAAG
124157 TATGAGTGTGGAGTCGTTGCTCTTGCAAGAATTTATGTGTTAGNNNAGATCGGAAG
124158 TATGAGTGTGGAGTCGTTGCCTCAAACCCCAGAAAGAGATNNNAGATCGGAAGAG
124159 TATGAGTGTGGAGTCGTTGCAAAAGACACTTTTTTTGTACCTANNNAGATCGGAAG
124160 TATGAGTGTGGAGTCGTTGCATGTTGTGTAGGTATAGAAGAAANNNAGATCGGAA
124161 TATGAGTGTGGAGTCGTTGCAGTAGTTTGATAGGAATAGCATTNNNAGATCGGAA
124162 TATGAGTGTGGAGTCGTTGCAGAAAAGAGAAAAACAAGAAGTTNNNAGATCGGAA
124163 TATGAGTGTGGAGTCGTTGCGGCCATAAGGAGATAATTTACTNNNAGATCGGAAG
124164 TATGAGTGTGGAGTCGTTGCGAGAGTAAATGCAGGTCACTNNNAGATCGGAAGAG
124165 TATGAGTGTGGAGTCGTTGCGCTTATCCACAATCACAGATNNNAGATCGGAAGAGC

124166 TATGAGTGTGGAGTCGTTGCGCTTGTGATGCATTTTTCTATNNNAGATCGGAAGAG
124167 TATGAGTGTGGAGTCGTTGCAGATACATTATTCTGGAAGTGAANNNAGATCGGAA
124168 TATGAGTGTGGAGTCGTTGCACAAAATACATGAAACATACAGTNNNAGATCGGAAG
124169 TATGAGTGTGGAGTCGTTGCTTATAATGCTTTATAGTTTACAAGCNNNAGATCGGA
124170 TATGAGTGTGGAGTCGTTGCCTATGCAAAGATATAATCCTCACNNNAGATCGGAAG
124171 TATGAGTGTGGAGTCGTTGCCACACTTAGTTCTATGCAATATTNNNAGATCGGAAG
124173 TATGAGTGTGGAGTCGTTGCTTTTTGCTTGATTTTCTCCAAGNNNAGATCGGAAGAG
124174 TATGAGTGTGGAGTCGTTGCTGGAAATGGCTTGCTATTATTNNNAGATCGGAAGA
124175 TATGAGTGTGGAGTCGTTGCCTCAGTTTCCTCATATGTATCATNNNAGATCGGAAGA
124176 TATGAGTGTGGAGTCGTTGCTTTTGATTTGATTTTTGTCTGTGNNNAGATCGGAAG
124177 TATGAGTGTGGAGTCGTTGCCATCGTATAAGCAGTAATTTTGANNNAGATCGGAAG
124178 TATGAGTGTGGAGTCGTTGCTAGATACTGCATAAAAATTTCCCNNNAGATCGGAAG
124179 TATGAGTGTGGAGTCGTTGCCATCTAACCCACTACAACACNNNAGATCGGAAGAGC
124180 TATGAGTGTGGAGTCGTTGCGTGGAAGTATTTATGCATGTAATNNNAGATCGGAA
124181 TATGAGTGTGGAGTCGTTGCGGGAGAGAGAAATTAGGAATATTNNNAGATCGGA
124182 TATGAGTGTGGAGTCGTTGCTAAATAAGATTCAGTATGCATGCNNNAGATCGGAAG
124183 TATGAGTGTGGAGTCGTTGCCTTATTGGTGGACCTAGTATATCNNNAGATCGGAAG
124184 TATGAGTGTGGAGTCGTTGCAATTCTCTGCAGGTAAAATAGATNNNAGATCGGAAG
124185 TATGAGTGTGGAGTCGTTGCAATACGCAGATGATACAGAAATNNNAGATCGGAAG
124186 TATGAGTGTGGAGTCGTTGCTAATCAAATTATGTGGATGTTGCNNNAGATCGGAAG
124188 TATGAGTGTGGAGTCGTTGCAAGAATAGGATGTGATAGCAAANNNAGATCGGAAG
124189 TATGAGTGTGGAGTCGTTGCCTCTAGTCTTGAGATAGTAACACNNNAGATCGGAAG
124190 TATGAGTGTGGAGTCGTTGCATGAGGAAGAGATCTATAAACTGNNNAGATCGGAA
124191 TATGAGTGTGGAGTCGTTGCCTTATATGTTGAGCAGACTTTTCNNNAGATCGGAAG
124192 TATGAGTGTGGAGTCGTTGCGCTAACAACTAAATACGTAAAGANNNAGATCGGAA
124193 TATGAGTGTGGAGTCGTTGCTAAAATCCACTAATTGTAGTCCANNNAGATCGGAAG
124194 TATGAGTGTGGAGTCGTTGCTGCAAAAATCCTCAATAAAATACNNNAGATCGGAAG
124195 TATGAGTGTGGAGTCGTTGCTATATTTTTCTTGTATGAAGGCTNNNAGATCGGAAG
124196 TATGAGTGTGGAGTCGTTGCCAATAGAAATCTACACTGGGGNNNAGATCGGAAGA
124197 TATGAGTGTGGAGTCGTTGCTGTATTTTATAGCTTACCTCACTNNNAGATCGGAAGA
124198 TATGAGTGTGGAGTCGTTGCGATTTTCTCTTTCAATCTTGCTANNNAGATCGGAAGA
124199 TATGAGTGTGGAGTCGTTGCAGTAAGAATTATCACCAAAATGGNNNAGATCGGAA
124200 TATGAGTGTGGAGTCGTTGCAGTAAGAATTATCACCAAAATGGNNNAGATCGGAA
124201 TATGAGTGTGGAGTCGTTGCTCTCCTTCATAGCCAAATGTANNNAGATCGGAAGAG
124202 TATGAGTGTGGAGTCGTTGCAAAAAGGTTATCATGTTTTGTCTNNNAGATCGGAAG
124203 TATGAGTGTGGAGTCGTTGCACCCTAAAGTATCTTTTGAAAACNNNAGATCGGAAG
124204 TATGAGTGTGGAGTCGTTGCAAAAACAAGAAAACCCACAAAATNNNAGATCGGAA
124205 TATGAGTGTGGAGTCGTTGCAATTATTTGTCAGTGAAGAGTTGNNNAGATCGGAA
124206 TATGAGTGTGGAGTCGTTGCCCCTACTAGGATATAAGCTCTATNNNAGATCGGAAG
124207 TATGAGTGTGGAGTCGTTGCCACATACAACACACACATTTGNNNAGATCGGAAGAG
124208 TATGAGTGTGGAGTCGTTGCAGATACTTGTATTAGGGTTCTGNNNAGATCGGAAG
124209 TATGAGTGTGGAGTCGTTGCTTTGTATACATTTTCAGTTGACCNNNAGATCGGAAG
124210 TATGAGTGTGGAGTCGTTGCTATTCTCAGTAGCTGCATATATCNNNAGATCGGAAG

124211 TATGAGTGTGGAGTCGTTGCACAGAATGAGATAGCCATTTTTNNNAGATCGGAAGA
124212 TATGAGTGTGGAGTCGTTGCTCTATGAATCATGAAGTTCATCTNNNAGATCGGAAG
124213 TATGAGTGTGGAGTCGTTGCTATTATTCCTGGCTGCAATAAAANNNAGATCGGAAG
124215 TATGAGTGTGGAGTCGTTGCAGCTTACCATATTTCTAATGTCTNNNAGATCGGAAGA
124216 TATGAGTGTGGAGTCGTTGCATGGTATCACACAGGAATTATTANNNAGATCGGAAG
124217 TATGAGTGTGGAGTCGTTGCATTTGGACTGCGATTTATTTCNNNAGATCGGAAGAG
124218 TATGAGTGTGGAGTCGTTGCTAAAGAAATGTTCCCTTTAGACTNNNAGATCGGAAG
124219 TATGAGTGTGGAGTCGTTGCCATCACTTTACAGATTGTCATAGNNNAGATCGGAAG
124220 TATGAGTGTGGAGTCGTTGCGTGGGGTTGTTTTATCTTTTGNNNAGATCGGAAGAG
124221 TATGAGTGTGGAGTCGTTGCGTTATTCCCATTTGACAGATTGNNNAGATCGGAAGA
124222 TATGAGTGTGGAGTCGTTGCGGAAACTTGCCTCTTCTTTANNNAGATCGGAAGAGC
124223 TATGAGTGTGGAGTCGTTGCGTTGCTGGAAAATATCTTACTAGNNNAGATCGGAAG
124224 TATGAGTGTGGAGTCGTTGCTCTTATTTTGTCTCTGTTTCATCNNNAGATCGGAAGA
124225 TATGAGTGTGGAGTCGTTGCTTCATTTGCTCTGATCTTTTATCNNNAGATCGGAAGA
124226 TATGAGTGTGGAGTCGTTGCTTTGTTAGATAGTTTTTGACCAGNNNAGATCGGAAG
124227 TATGAGTGTGGAGTCGTTGCATTATTCTTTGGACAAATGGTTANNNAGATCGGAAG
124230 TATGAGTGTGGAGTCGTTGCAAGAGAGAGAGAGAGAGAGAGNNNAGATCGGAAG
124231 TATGAGTGTGGAGTCGTTGCCTGAAAATAGAGAAATGGAAACANNNAGATCGGAA
124232 TATGAGTGTGGAGTCGTTGCCTATTGTAAATGGTTTTGCATTCNNNAGATCGGAAG
124233 TATGAGTGTGGAGTCGTTGCTTTTCTTATTAGAGATGAGGTCTNNNAGATCGGAAG
124234 TATGAGTGTGGAGTCGTTGCTAAGTCAACTTAGAATCTCCTCTNNNAGATCGGAAGA
124236 TATGAGTGTGGAGTCGTTGCGGTGCAACAAATAAAAATATGAGNNNAGATCGGAA
124237 TATGAGTGTGGAGTCGTTGCAAAGAGAGAGAATATAGAAGCAGNNNAGATCGGAA
124238 TATGAGTGTGGAGTCGTTGCCACACAATAATGGTGGATGTNNNAGATCGGAAGAG
124239 TATGAGTGTGGAGTCGTTGCCTTCAGCCTTTATTGTAGCTNNNAGATCGGAAGAGC
124240 TATGAGTGTGGAGTCGTTGCCTCCTTCAGCCTTTATTGTAGNNNAGATCGGAAGAG
124241 TATGAGTGTGGAGTCGTTGCGCCATGGTTACTTTTACTATTTANNNAGATCGGAAGA
124242 TATGAGTGTGGAGTCGTTGCAAACAAGTCAGTTTTAGTTTCAANNNAGATCGGAAG
124243 TATGAGTGTGGAGTCGTTGCAATCATGTTTTCAGGAGAATGTANNNAGATCGGAAG
124244 TATGAGTGTGGAGTCGTTGCCACCAATAAATGAACTGAGTTTNNNAGATCGGAAGA
124245 TATGAGTGTGGAGTCGTTGCATGTGATAAATTCATTGACAGAANNNAGATCGGAAG
124246 TATGAGTGTGGAGTCGTTGCACTTTTCAAACTAAAAAATCCCANNNAGATCGGAAGA
124247 TATGAGTGTGGAGTCGTTGCTCTCCTTCATAGCCAAATGTANNNAGATCGGAAGAG
124248 TATGAGTGTGGAGTCGTTGCTTTCCTCTAGGTTTTCTAACTTGNNNAGATCGGAAGA
124251 TATGAGTGTGGAGTCGTTGCGTAGACCTTATCTTCCTGGGNNNAGATCGGAAGAGC
124252 TATGAGTGTGGAGTCGTTGCACCCCTCTCTTACCTATAAATATNNNAGATCGGAAGA
124253 TATGAGTGTGGAGTCGTTGCGTCTCAGCACTTAAAATTGTATTNNNAGATCGGAAG
124254 TATGAGTGTGGAGTCGTTGCAAGTCTTGATAAACAGATCCTTNNNAGATCGGAAGA
124255 TATGAGTGTGGAGTCGTTGCTTATGAACAGCTTATACTTTGTGNNNAGATCGGAAG
124256 TATGAGTGTGGAGTCGTTGCTTATGAACAGCTTATACTTTGTGNNNAGATCGGAAG
124257 TATGAGTGTGGAGTCGTTGCCTTTGGGGAGCTATTTGTAGNNNAGATCGGAAGAG
124258 TATGAGTGTGGAGTCGTTGCCTTAGAACCCAACTGTAGTTCNNNAGATCGGAAGAG
124259 TATGAGTGTGGAGTCGTTGCAAGCAAAGAAATGAATTTTTTCTNNNAGATCGGAAG

124260 TATGAGTGTGGAGTCGTTGCCACTTTGATGGTTTGAAGATAGNNNAGATCGGAAG
124261 TATGAGTGTGGAGTCGTTGCGGTTCTCCTTTCATAAATATTCANNNAGATCGGAAGA
124263 TATGAGTGTGGAGTCGTTGCTAATGACACATTGCAACTTATTTNNNAGATCGGAAG
124264 TATGAGTGTGGAGTCGTTGCTTCCTATAATCTCCAGGTGTAGNNNAGATCGGAAGA
124265 TATGAGTGTGGAGTCGTTGCCTCCAAATCTCAAGTTAAAATGTNNNAGATCGGAAG
124266 TATGAGTGTGGAGTCGTTGCTATTTCTTAGACTGTAGAATGCANNNAGATCGGAAG
124267 TATGAGTGTGGAGTCGTTGCCTGATATAGTTGGTCCAGAATGNNNAGATCGGAAG
124268 TATGAGTGTGGAGTCGTTGCAAAAGCAGTCACTAAAAATATCTNNNAGATCGGAAG
124269 TATGAGTGTGGAGTCGTTGCAGAAAGAAAGATGAAGGAAAAAGNNNAGATCGGA
124270 TATGAGTGTGGAGTCGTTGCAACAGAGAAGTTACTGTATCTAGNNNAGATCGGAA
124271 TATGAGTGTGGAGTCGTTGCAAAGGTGTTATGGTTGGAAATANNNAGATCGGAAG
124272 TATGAGTGTGGAGTCGTTGCATAAGTAGATTGAAATGTGTGAGNNNAGATCGGAA
124273 TATGAGTGTGGAGTCGTTGCTAGAAATGGAGATGGTAGTTATGNNNAGATCGGAA
124274 TATGAGTGTGGAGTCGTTGCCATTGATTGCCTTATTATTCTCANNNAGATCGGAAGA
124275 TATGAGTGTGGAGTCGTTGCATGAAGATGTGAATGATCAATTTNNNAGATCGGAA
124276 TATGAGTGTGGAGTCGTTGCCACCAAAAAAAAAAAAGACAGAGNNNAGATCGGAA
124277 TATGAGTGTGGAGTCGTTGCTATACCAATGGAAAATACTCTCANNNAGATCGGAAG
124278 TATGAGTGTGGAGTCGTTGCGTACACATGTATGGCACTTANNNAGATCGGAAGAG
124279 TATGAGTGTGGAGTCGTTGCATTATCTCATCTCTGGCCACNNNAGATCGGAAGAGC
124280 TATGAGTGTGGAGTCGTTGCCTAAAAAGTGCCATGGAAAATNNNAGATCGGAAGA
124281 TATGAGTGTGGAGTCGTTGCATTTTATCCAGCTTTTCTACTTGNNNAGATCGGAAGA
124282 TATGAGTGTGGAGTCGTTGCCAGATTACCAAATCAGATTCAATNNNAGATCGGAAG
124283 TATGAGTGTGGAGTCGTTGCATTATACATGCATGCATATTTGTNNNAGATCGGAAG
124284 TATGAGTGTGGAGTCGTTGCAAGAGCATTACTATGGTAATTGTNNNAGATCGGAAG
124285 TATGAGTGTGGAGTCGTTGCTGGACATTGTGTTTATATTTCTGNNNAGATCGGAAG
124286 TATGAGTGTGGAGTCGTTGCCTAAATCTCTAGAGCTGCAACNNNAGATCGGAAGAG
124287 TATGAGTGTGGAGTCGTTGCAACCTCCACTTCAGTGATTCNNNAGATCGGAAGAGC
124288 TATGAGTGTGGAGTCGTTGCAATTTAAAAGCTTGAGACAATCANNNAGATCGGAAG
124289 TATGAGTGTGGAGTCGTTGCTAACACCAGTTTTCCTTTGTNNNAGATCGGAAGAGC
124290 TATGAGTGTGGAGTCGTTGCCTATGCCCTCATCTTAACTAATNNNAGATCGGAAGAG
124291 TATGAGTGTGGAGTCGTTGCTATTTCACTCACATAAGGAAATGNNNAGATCGGAAG
124292 TATGAGTGTGGAGTCGTTGCCACTACTAAAAGTATACCTGGATNNNAGATCGGAAG
124293 TATGAGTGTGGAGTCGTTGCGCCTTGTAGAATCATTTATGAAANNNAGATCGGAAG
124294 TATGAGTGTGGAGTCGTTGCTATAGATATCCAGTAGGTTCTGTNNNAGATCGGAAG
124295 TATGAGTGTGGAGTCGTTGCATCATAATTCTATTTGGCCAAAGNNNAGATCGGAAG
124296 TATGAGTGTGGAGTCGTTGCTAGAATTTACTTTTGGTGTTGTCNNNAGATCGGAAG
124297 TATGAGTGTGGAGTCGTTGCCCACTCTAGACCTTCTTTGANNNAGATCGGAAGAGC
124298 TATGAGTGTGGAGTCGTTGCGAGGAAAGAAGTAAGCTATTTTGNNNAGATCGGAA
124299 TATGAGTGTGGAGTCGTTGCTGCCATAACAAAATACCACANNNAGATCGGAAGAGC
124300 TATGAGTGTGGAGTCGTTGCCATCTATCATCTTCTCCACATTTNNNAGATCGGAAGA
124302 TATGAGTGTGGAGTCGTTGCGAGTGAGAATGAATATGAGAATGNNNAGATCGGA
124303 TATGAGTGTGGAGTCGTTGCCTTGAAGGCAATGTATAATCAGNNNAGATCGGAAG
124304 TATGAGTGTGGAGTCGTTGCAGCTATAGAACATATGGGAAAATNNNAGATCGGAA

124305 TATGAGTGTGGAGTCGTTGCCTCATTCCAAATATTGTCAGTTTNNNAGATCGGAAG
124306 TATGAGTGTGGAGTCGTTGCAATTAACTGGCCTTAGAATATGANNNAGATCGGAAG
124307 TATGAGTGTGGAGTCGTTGCTTAGTGGTTGTTATGAGAGTTCNNNAGATCGGAAG
124308 TATGAGTGTGGAGTCGTTGCGCATCAGCTTTAAATTAAAACAANNNAGATCGGAAG
124309 TATGAGTGTGGAGTCGTTGCGCAGTTTGTGTTTTTCATTAAATNNNAGATCGGAAG
124310 TATGAGTGTGGAGTCGTTGCCTCAGAAATTAATGAGGACAATGNNNAGATCGGAA
124311 TATGAGTGTGGAGTCGTTGCTAATTGATATGTTACCCATGAGTNNNAGATCGGAAG
124312 TATGAGTGTGGAGTCGTTGCGTGTTTTGTCTGAGAAGTCTNNNAGATCGGAAGAG
124313 TATGAGTGTGGAGTCGTTGCTCTAAACATCAAAGTCAGTATCCNNNAGATCGGAAG
124314 TATGAGTGTGGAGTCGTTGCCAGATTCTGCTGAAATCATAATTNNNAGATCGGAAG
124315 TATGAGTGTGGAGTCGTTGCTTTGTTTGTTTTAGAGATGGGNNNAGATCGGAAGA
124316 TATGAGTGTGGAGTCGTTGCTAGATTCACTAAAAATGAGGCANNNAGATCGGAAG
124317 TATGAGTGTGGAGTCGTTGCTGGGGAGAAACTATTAGTTTTAANNNAGATCGGAA
124318 TATGAGTGTGGAGTCGTTGCTTTTGTAAGCTTCTTTCACTTAANNNAGATCGGAAGA
124319 TATGAGTGTGGAGTCGTTGCCTGTTTTGTATCATTGTGTCATNNNAGATCGGAAGA
124320 TATGAGTGTGGAGTCGTTGCCTGTGGTAATTTGCAAGATATTANNNAGATCGGAAG
124321 TATGAGTGTGGAGTCGTTGCATGAAGAATCAACTGAGGTAATANNNAGATCGGAA
124322 TATGAGTGTGGAGTCGTTGCCATAAAACTCAACTCCCAATAAANNNAGATCGGAAG
124323 TATGAGTGTGGAGTCGTTGCGATATTTTCTCATGTGGCATATTNNNAGATCGGAAG
124324 TATGAGTGTGGAGTCGTTGCAGAAGCTAATCACTATGAGTTTANNNAGATCGGAAG
124325 TATGAGTGTGGAGTCGTTGCATGCTTATTCTCTTTTACTTTCCNNNAGATCGGAAGA
124326 TATGAGTGTGGAGTCGTTGCATATAGGGGAATTATTTGTCACGNNNAGATCGGAA
124327 TATGAGTGTGGAGTCGTTGCATATAGGGGAATTATTTGTCACGNNNAGATCGGAA
124330 TATGAGTGTGGAGTCGTTGCTCAGGTGATCTAACTTGTTAAAANNNAGATCGGAAG
124331 TATGAGTGTGGAGTCGTTGCGTAAAGATCCACATGCAAAATNNNAGATCGGAAGA
124332 TATGAGTGTGGAGTCGTTGCTGTTTCTTCTCCAAATTATCAAANNNAGATCGGAAGA
124333 TATGAGTGTGGAGTCGTTGCTTTAAGTGTACTATAACATCCTGNNNAGATCGGAAG
124334 TATGAGTGTGGAGTCGTTGCACTTCCTGTCTTTCATATTATCTNNNAGATCGGAAGA
124335 TATGAGTGTGGAGTCGTTGCAGTCAGAATGGCTATTACTAAAANNNAGATCGGAA
124336 TATGAGTGTGGAGTCGTTGCTATAAACATCAACACATTCCATCNNNAGATCGGAAGA
124337 TATGAGTGTGGAGTCGTTGCCCATGTGAAGATGAAAATAGAGANNNAGATCGGAA
124339 TATGAGTGTGGAGTCGTTGCATAATAACAGAGCAAAGGCAATANNNAGATCGGAA
124340 TATGAGTGTGGAGTCGTTGCATGTGTATGTTTGAAGACTAAAGNNNAGATCGGAA
124341 TATGAGTGTGGAGTCGTTGCACCATTTTTATTGCATAATTCAGNNNAGATCGGAAG
124342 TATGAGTGTGGAGTCGTTGCCTAAGAGAAAAGAACAATCTTCANNNAGATCGGAA
124343 TATGAGTGTGGAGTCGTTGCTAATAACTACCACTCATTAGCACNNNAGATCGGAAG
124344 TATGAGTGTGGAGTCGTTGCCTCACCAACTACATGGATATTTNNNAGATCGGAAGA
124345 TATGAGTGTGGAGTCGTTGCACCATCATTTACATACTTTCAAANNNAGATCGGAAGA
124346 TATGAGTGTGGAGTCGTTGCCTTCCTTCCTTCCTTTTTTTTGNNNAGATCGGAAGAG
124347 TATGAGTGTGGAGTCGTTGCATAACATGTAAGCTGTATTGTACNNNAGATCGGAAG
124348 TATGAGTGTGGAGTCGTTGCATATAAGGAAGCATCTAATCTGGNNNAGATCGGAA
124349 TATGAGTGTGGAGTCGTTGCAAATGGAAATGCTTGGTTATAATNNNAGATCGGAA
124350 TATGAGTGTGGAGTCGTTGCAATCTTGGTAGGTCTAGAATATGNNNAGATCGGAA

124351 TATGAGTGTGGAGTCGTTGCCATACATCCCATCATTAGAGTCNNNAGATCGGAAGA
124352 TATGAGTGTGGAGTCGTTGCCTGTGGTTTGTTTTCATTTATTTNNNAGATCGGAAGA
124353 TATGAGTGTGGAGTCGTTGCTAAATAATGCTTCAGTGAACATGNNNAGATCGGAAG
124354 TATGAGTGTGGAGTCGTTGCACTAAGTCCTTTTAAAACAACACNNNAGATCGGAAG
124355 TATGAGTGTGGAGTCGTTGCGAAGAAATTATCCAGAAAGAGTCNNNAGATCGGAA
124356 TATGAGTGTGGAGTCGTTGCTCTTGGTATTGTTAAGATTTAGGNNNAGATCGGAAG
124357 TATGAGTGTGGAGTCGTTGCATCATTTTCTAAAACCAGTCTTCNNNAGATCGGAAGA
124358 TATGAGTGTGGAGTCGTTGCGGTGATGATAGCTACGATTTCNNNAGATCGGAAGA
124359 TATGAGTGTGGAGTCGTTGCGGATAAAACTACTAAAGCTACTTTNNNAGATCGGAA
124360 TATGAGTGTGGAGTCGTTGCGTGTTCTGAAAGGATAAAACTACNNNAGATCGGAA
124361 TATGAGTGTGGAGTCGTTGCAAGAACCGTATTTATTTTCATCTNNNAGATCGGAAG
124362 TATGAGTGTGGAGTCGTTGCATACATATGAAACCCAAACAAAANNNAGATCGGAAG
124363 TATGAGTGTGGAGTCGTTGCAAAAATGATTCTGGTGCTCTTTNNNAGATCGGAAGA
124365 TATGAGTGTGGAGTCGTTGCTTTCATATATCATGATGCTTCACNNNAGATCGGAAGA
124366 TATGAGTGTGGAGTCGTTGCCCTCACAATCATATTTGGTATTTNNNAGATCGGAAGA
124367 TATGAGTGTGGAGTCGTTGCCTTTTCCCCTTTGGTTTTTATCNNNAGATCGGAAGAG
124368 TATGAGTGTGGAGTCGTTGCTGAAGTGAAATTGAAGGTAAAATNNNAGATCGGAA
124369 TATGAGTGTGGAGTCGTTGCAATATTCTATTCTCCTCCACAAANNNAGATCGGAAGA
124370 TATGAGTGTGGAGTCGTTGCTCATGCCTGGATATAACTATTATATNNNAGATCGGA
124371 TATGAGTGTGGAGTCGTTGCCATTCACAGTGACAAATTCTCNNNAGATCGGAAGAG
124372 TATGAGTGTGGAGTCGTTGCTATGTATATACACACACAGATGGNNNAGATCGGAAG
124373 TATGAGTGTGGAGTCGTTGCCTGTTCATCATTTTTTTTTCCTGNNNAGATCGGAAGA
124374 TATGAGTGTGGAGTCGTTGCTTTCTTCAACATTTACATCAAGTNNNAGATCGGAAGA
124375 TATGAGTGTGGAGTCGTTGCTATTCTCTCCTCGTCTATTTTTTNNNAGATCGGAAGA
124376 TATGAGTGTGGAGTCGTTGCCCTGGGATAACTATATGTCTTTTNNNAGATCGGAAG
124377 TATGAGTGTGGAGTCGTTGCGTGACTTTGGAATAAACATTGATNNNAGATCGGAAG
124378 TATGAGTGTGGAGTCGTTGCTTTGGACAAACTGTTACCTATTANNNAGATCGGAAG
124379 TATGAGTGTGGAGTCGTTGCCACTAAAATGTTTACAGTGGTTANNNAGATCGGAAG
124380 TATGAGTGTGGAGTCGTTGCTACTATAAGGAATTGGTTCACATNNNAGATCGGAAG
124382 TATGAGTGTGGAGTCGTTGCACCTTGAAAATGAATAAAAAGGANNNAGATCGGAA
124383 TATGAGTGTGGAGTCGTTGCGGAGCAAATGTAGTAAAAACTTANNNAGATCGGAA
124384 TATGAGTGTGGAGTCGTTGCTATATTTTTTCCCTTGGTTTCTGNNNAGATCGGAAGA
124385 TATGAGTGTGGAGTCGTTGCCAAGCATTTAGTTTTTAACCAAANNNAGATCGGAAG
124386 TATGAGTGTGGAGTCGTTGCTAGCTTCCACTAATTAACTACAANNNAGATCGGAAGA
124387 TATGAGTGTGGAGTCGTTGCAGCATTTATCAAAATGACTGATTNNNAGATCGGAAG
124388 TATGAGTGTGGAGTCGTTGCGAATATCAATGTCAGGATGGAGNNNAGATCGGAAG
124389 TATGAGTGTGGAGTCGTTGCGGAAATGTAAATTTTACTGTCTGNNNAGATCGGAAG
124390 TATGAGTGTGGAGTCGTTGCTATATATCTGTCTCTCTCTCTCCNNNAGATCGGAAGA
124391 TATGAGTGTGGAGTCGTTGCCGTTTTTCTTAAAAGGCAATAACNNNAGATCGGAAG
124392 TATGAGTGTGGAGTCGTTGCTAGAAACATGGAATTATCTCACCNNNAGATCGGAAG
124393 TATGAGTGTGGAGTCGTTGCAACCAAAAGGATAGAAGAGTATTNNNAGATCGGAA
124394 TATGAGTGTGGAGTCGTTGCATGTATTTATTAGGTATAGCCCCNNNAGATCGGAAG
124395 TATGAGTGTGGAGTCGTTGCATGTATTTATTAGGTATAGCCCCNNNAGATCGGAAG

124396 TATGAGTGTGGAGTCGTTGCTATATTAGATTGTGGTGGTCATTNNNAGATCGGAAG
124398 TATGAGTGTGGAGTCGTTGCGAATAAAATAAATATCCGTGAGCNNNAGATCGGAAG
124399 TATGAGTGTGGAGTCGTTGCACAAATGACTTAAATAACAGAGANNNAGATCGGAAG
124400 TATGAGTGTGGAGTCGTTGCGTATCAGGCACAGTACAAATATNNNAGATCGGAAGA
124401 TATGAGTGTGGAGTCGTTGCCAGGAATGTGTATGTATCAAAAANNNAGATCGGAAG
124402 TATGAGTGTGGAGTCGTTGCGGAATGTAAAAGAAACAGTAGAANNNAGATCGGAA
124403 TATGAGTGTGGAGTCGTTGCCTCCTGAAGTTTTGGAATGTANNNAGATCGGAAGAG
124405 TATGAGTGTGGAGTCGTTGCTATATAGAGAGAGAGAGAGAGAGNNNAGATCGGA
124407 TATGAGTGTGGAGTCGTTGCATTTCATTTGCAAAATAATCTGCNNNAGATCGGAAGA
124408 TATGAGTGTGGAGTCGTTGCGTATAAAAACATGGGGTGATAAGNNNAGATCGGAA
124409 TATGAGTGTGGAGTCGTTGCAAAACTACATGGAAATGGAAAAGNNNAGATCGGAA
124410 TATGAGTGTGGAGTCGTTGCGTACTTCACATCAGAGTTAAAAGNNNAGATCGGAAG
124412 TATGAGTGTGGAGTCGTTGCACAATCTAAAAGTCAACTAAAGGNNNAGATCGGAAG
124413 TATGAGTGTGGAGTCGTTGCTAAAATTTTGGAGGGTATCAATGNNNAGATCGGAAG
124414 TATGAGTGTGGAGTCGTTGCTTGATAAGAAGCTTTAATTGACTNNNAGATCGGAAG
124415 TATGAGTGTGGAGTCGTTGCTATAGCAGTAATTTGTAACCAGANNNAGATCGGAAG
124416 TATGAGTGTGGAGTCGTTGCTATAGCAGTAATTTGTAACCAGANNNAGATCGGAAG
124419 TATGAGTGTGGAGTCGTTGCCCCTAACTCAAATTTAGAACTACNNNAGATCGGAAGA
124420 TATGAGTGTGGAGTCGTTGCGTATTTCCTTGCTATCTCAGATNNNAGATCGGAAGAG
124421 TATGAGTGTGGAGTCGTTGCTTCTGAAGACATATTATAGACCTNNNAGATCGGAAGA
124422 TATGAGTGTGGAGTCGTTGCGTGGCTTATGTCTTTAATCTCTNNNAGATCGGAAGAG
124423 TATGAGTGTGGAGTCGTTGCAAGTAGTCCTGTGGATATGTNNNAGATCGGAAGAGC
124424 TATGAGTGTGGAGTCGTTGCGACTTGCTGTATTGGTTTTTTNNNAGATCGGAAGAG
124425 TATGAGTGTGGAGTCGTTGCCCTCTGCCTCACTCTTATAANNNAGATCGGAAGAGCA
124426 TATGAGTGTGGAGTCGTTGCCTGGGAAGGTTATGTTTCTTATANNNAGATCGGAAG
124427 TATGAGTGTGGAGTCGTTGCGAAGACCTAAGGAAAGTAATACTNNNAGATCGGAAG
124428 TATGAGTGTGGAGTCGTTGCGGAAGAATTAGGAGAATTTCTTTNNNAGATCGGAAG
124429 TATGAGTGTGGAGTCGTTGCTCCTCTCTGTTCTCTTTTTTTTNNNAGATCGGAAGAGC
124430 TATGAGTGTGGAGTCGTTGCTTTTTGACAACCATGTAGAATTTNNNAGATCGGAAGA
124431 TATGAGTGTGGAGTCGTTGCTTGTTTGTTTGTTTGTTTTTTTGNNNAGATCGGAAGA
124432 TATGAGTGTGGAGTCGTTGCCTTTTTTTTTTTTTTGACAGGGTNNNAGATCGGAAGA
124433 TATGAGTGTGGAGTCGTTGCTAAAAGGTAAAGACTATATGGCCNNNAGATCGGAAG
124434 TATGAGTGTGGAGTCGTTGCAGATTGTTTTATGAAAGAAAGTCNNNAGATCGGAAG
124435 TATGAGTGTGGAGTCGTTGCATAAAGCTGCAATAACTATCTTGNNNAGATCGGAAG
124436 TATGAGTGTGGAGTCGTTGCTGCTCATAAGAAATTCAAAATCTNNNAGATCGGAAGA
124437 TATGAGTGTGGAGTCGTTGCATCTCATACATTGCTGATAAAAGNNNAGATCGGAAG
124438 TATGAGTGTGGAGTCGTTGCGTGTCTACTACCTAGTGTAAAGNNNAGATCGGAAGA
124439 TATGAGTGTGGAGTCGTTGCCTTGGATGGAAGGAAAACAANNNAGATCGGAAGAG
124440 TATGAGTGTGGAGTCGTTGCAGGTGTATTCCTCAAATAGATTCNNNAGATCGGAAG
124441 TATGAGTGTGGAGTCGTTGCGGGGCATGTATATATGAAATATGNNNAGATCGGAAG
124442 TATGAGTGTGGAGTCGTTGCACAGAGAGAGAGAGATTTATTATNNNAGATCGGAAG
124443 TATGAGTGTGGAGTCGTTGCATAAATATATACACACACAGGCTNNNAGATCGGAAG
124444 TATGAGTGTGGAGTCGTTGCTGCATTAGGTGACAAATACAANNNAGATCGGAAGAG

124445 TATGAGTGTGGAGTCGTTGCGAATAATTTCTCATCTCAGTAGCNNNAGATCGGAAG
124446 TATGAGTGTGGAGTCGTTGCCAATACAACCTCATCATAACTAGNNNAGATCGGAAG
124447 TATGAGTGTGGAGTCGTTGCCATGTGACAATCATCTTATCTTTNNNAGATCGGAAGA
124448 TATGAGTGTGGAGTCGTTGCATACACACACACACATTCTAAGNNNAGATCGGAAGA
124449 TATGAGTGTGGAGTCGTTGCAATTCATGTATCTACAACATTGCNNNAGATCGGAAG
124450 TATGAGTGTGGAGTCGTTGCCCTCATCTCAAATCCAGAATTATNNNAGATCGGAAGA
124451 TATGAGTGTGGAGTCGTTGCGCTTCTAAATATCTCAAGCATTANNNAGATCGGAAG
124452 TATGAGTGTGGAGTCGTTGCCTTTCATGTCTTTGTGTAATGATNNNAGATCGGAAG
124453 TATGAGTGTGGAGTCGTTGCAAAGATATTCTTGGGAAAAAAGTNNNAGATCGGAA
124454 TATGAGTGTGGAGTCGTTGCATTAGAGGGTAGTTTGAAAAATCNNNAGATCGGAA
124455 TATGAGTGTGGAGTCGTTGCTGTAGATACGTGAGATAAGTTTTNNNAGATCGGAA
124456 TATGAGTGTGGAGTCGTTGCAATTCAATGGGAATGTTTATTGTNNNAGATCGGAAG
124457 TATGAGTGTGGAGTCGTTGCTTATTTAGTGTCTCCCAGTCATANNNAGATCGGAAG
124458 TATGAGTGTGGAGTCGTTGCCCAGGAAACCTATACTGTTAATANNNAGATCGGAAG
124459 TATGAGTGTGGAGTCGTTGCTTTCTAGATGTTCCAGATGTAATNNNAGATCGGAAG
124460 TATGAGTGTGGAGTCGTTGCCAAGTCATTTCTATCCATAGTTGNNNAGATCGGAAG
124461 TATGAGTGTGGAGTCGTTGCATAGAACGTAGAAAGTGGAAAATNNNAGATCGGAA
124462 TATGAGTGTGGAGTCGTTGCGTGAGAAGACTTGAGATCTACNNNAGATCGGAAGA
124463 TATGAGTGTGGAGTCGTTGCAAGTCTTAAATAATTCCAGCATCNNNAGATCGGAAG
124464 TATGAGTGTGGAGTCGTTGCCTATCTTTGCTGATCTCCAGNNNAGATCGGAAGAGC
124465 TATGAGTGTGGAGTCGTTGCCTATCTTAGAATCATCTGAGCTTNNNAGATCGGAAG
124466 TATGAGTGTGGAGTCGTTGCCTATCTTAGAATCATCTGAGCTTNNNAGATCGGAAG
124467 TATGAGTGTGGAGTCGTTGCGGAGTTCGTTGTCTCATTAGNNNAGATCGGAAGAG
124468 TATGAGTGTGGAGTCGTTGCTTTGAAAAAAATAGGATGTGTGTNNNAGATCGGAA
124469 TATGAGTGTGGAGTCGTTGCTGGATATATTGGAAATCTACTGANNNAGATCGGAA
124470 TATGAGTGTGGAGTCGTTGCGAGTACAGTAGTCAAGAATCTCNNNAGATCGGAAG
124471 TATGAGTGTGGAGTCGTTGCGTTGCCTTCTCTTTTTATCTTCNNNAGATCGGAAGAG
124472 TATGAGTGTGGAGTCGTTGCTTTATGATTGATAACACCTCAAGNNNAGATCGGAAG
124473 TATGAGTGTGGAGTCGTTGCTTTATGATTGATAACACCTCAAGNNNAGATCGGAAG
124474 TATGAGTGTGGAGTCGTTGCAAAACACAAGATGTATATGGGATNNNAGATCGGAA
124475 TATGAGTGTGGAGTCGTTGCATCCAGAATTTCTAGATAGTAGGNNNAGATCGGAA
124476 TATGAGTGTGGAGTCGTTGCGTGATTATTACACATTGCATTCNNNAGATCGGAAGA
124477 TATGAGTGTGGAGTCGTTGCCAATTGAAAGTAAACCCCTTAAANNNAGATCGGAAG
124478 TATGAGTGTGGAGTCGTTGCCTTATTATCTACTGTCCTCTGAGNNNAGATCGGAAGA
124479 TATGAGTGTGGAGTCGTTGCTATGAAAACGGTATCTATATGCTNNNAGATCGGAAG
124480 TATGAGTGTGGAGTCGTTGCGGCAAATATGTGTTTGAAAATTANNNAGATCGGAA
124482 TATGAGTGTGGAGTCGTTGCTTCTTTTAATGAGAAAAAGCAACNNNAGATCGGAAG
124483 TATGAGTGTGGAGTCGTTGCGTAGGTAACAAATGCTAGTTTGNNNAGATCGGAAG
124484 TATGAGTGTGGAGTCGTTGCTCGCTCTGTTATATGTGTATATANNNAGATCGGAAG
124485 TATGAGTGTGGAGTCGTTGCTCGCTCTGTTATATGTGTATATANNNAGATCGGAAG
124486 TATGAGTGTGGAGTCGTTGCCAGGAATCCACCTTCTAGAGNNNAGATCGGAAGAG
124487 TATGAGTGTGGAGTCGTTGCCCTTGGTTTCTTAAAGAGTTAATNNNAGATCGGAAG
124488 TATGAGTGTGGAGTCGTTGCGGTGCAATTATAGCTCACTGNNNAGATCGGAAGAG

124489 TATGAGTGTGGAGTCGTTGCCTGTGTATTCAATTCATCTACACNNNAGATCGGAAGA
124490 TATGAGTGTGGAGTCGTTGCGACCTGACTCCTAGTTCATANNNAGATCGGAAGAGCA
124491 TATGAGTGTGGAGTCGTTGCGTGTATGGTATGGTATGTTACTANNNAGATCGGAA
124492 TATGAGTGTGGAGTCGTTGCCTGAAGACAACTAACTCTTTTTTNNNAGATCGGAAGA
124493 TATGAGTGTGGAGTCGTTGCTTATGGGTTTTGGGTTTTTTTTNNNAGATCGGAAGA
124494 TATGAGTGTGGAGTCGTTGCTCAATATCAAACCTGCAAAAATANNNAGATCGGAAGA
124495 TATGAGTGTGGAGTCGTTGCAGAGAAAGAGAGAGGAGATGNNNAGATCGGAAGA
124496 TATGAGTGTGGAGTCGTTGCTATGAGAGAGAGAAATGCTAACNNNAGATCGGAAG
124497 TATGAGTGTGGAGTCGTTGCTATGCCTGACATATTCCTCAANNNAGATCGGAAGAG
124498 TATGAGTGTGGAGTCGTTGCGAGTTTTAGGTTTCCTACTTCTANNNAGATCGGAAGA
124499 TATGAGTGTGGAGTCGTTGCTAAGTTCAAGAGAGAGAGAAAAGNNNAGATCGGAA
124500 TATGAGTGTGGAGTCGTTGCTTAAGTTTAAATCAAGCCAATGGNNNAGATCGGAAG
124501 TATGAGTGTGGAGTCGTTGCGTTTTGTTTTGAGACAGAATCTNNNAGATCGGAAGA
124502 TATGAGTGTGGAGTCGTTGCACATTTCCTGAAAACTGTTTAAANNNAGATCGGAAGA
124503 TATGAGTGTGGAGTCGTTGCTCATTACTGGATTTCTGTGATATNNNAGATCGGAAGA
124504 TATGAGTGTGGAGTCGTTGCATCCATGAGGTTCTAATAGGAANNNAGATCGGAAGA
124505 TATGAGTGTGGAGTCGTTGCAACATTAAAGACGCTGTTATAAANNNAGATCGGAAG
124508 TATGAGTGTGGAGTCGTTGCCTTTGCTGTCTGAGATTGTCNNNAGATCGGAAGAGC
124509 TATGAGTGTGGAGTCGTTGCGCTATTTCCCAATGAGTATTTTANNNAGATCGGAAGA
124510 TATGAGTGTGGAGTCGTTGCAAAGATTTGGAGAATGTGGATATNNNAGATCGGAA
124511 TATGAGTGTGGAGTCGTTGCCACTCTCATATACTCTTGGTAGNNNAGATCGGAAGA
124513 TATGAGTGTGGAGTCGTTGCGCCCAAAAGTTGTTTATTTTTTTNNNAGATCGGAAGA
124514 TATGAGTGTGGAGTCGTTGCAACTTCAATTTTTTTCAAATGCCNNNAGATCGGAAGA
124515 TATGAGTGTGGAGTCGTTGCATACAGAGAGAAGACATCCATNNNAGATCGGAAGA
124516 TATGAGTGTGGAGTCGTTGCTAGCTCCAGATAAGAGGATCTNNNAGATCGGAAGA
124517 TATGAGTGTGGAGTCGTTGCATCATGGAAAACAGTTTACTTAGNNNAGATCGGAAG
124518 TATGAGTGTGGAGTCGTTGCACCTATCAGTAAAATCTACTTGGNNNAGATCGGAAGA
124519 TATGAGTGTGGAGTCGTTGCCACTCTCTCTCAAAGATACATATNNNAGATCGGAAGA
124520 TATGAGTGTGGAGTCGTTGCTGTAAAGATTTTCTGTGTATTGGNNNAGATCGGAAG
124521 TATGAGTGTGGAGTCGTTGCTATTAGGATGACTTGATCCAGANNNAGATCGGAAGA
124522 TATGAGTGTGGAGTCGTTGCTAGAAATATTGAGACCTGAATGTNNNAGATCGGAA
124523 TATGAGTGTGGAGTCGTTGCGCTGAAATAATACCTTAGATTTCTNNNAGATCGGAA
124524 TATGAGTGTGGAGTCGTTGCCTCAGGAATGGAAAACTAAATAGNNNAGATCGGAA
124525 TATGAGTGTGGAGTCGTTGCGAAGATGGAATGACTGTAGAGNNNAGATCGGAAGA
124526 TATGAGTGTGGAGTCGTTGCCATATGGCTGTGTTATAGCTCNNNAGATCGGAAGAG
124527 TATGAGTGTGGAGTCGTTGCAATTTTACCAAGAGAAATCAAGGNNNAGATCGGAA
124528 TATGAGTGTGGAGTCGTTGCCTTGCTAGAGATGCCTTTATATNNNAGATCGGAAGA
124529 TATGAGTGTGGAGTCGTTGCGCAATTTCCTGGTTATAAAATCANNNAGATCGGAAGA
124530 TATGAGTGTGGAGTCGTTGCAATTCTTATTTGGAAAGTACGACNNNAGATCGGAAG
124531 TATGAGTGTGGAGTCGTTGCCTGAAAAATTTAAGTAACATCTGGNNNAGATCGGAA
124532 TATGAGTGTGGAGTCGTTGCCTTTTAAACAGTCAAATTTCCATNNNAGATCGGAAGA
124533 TATGAGTGTGGAGTCGTTGCAAGTCAGGTCTTTATTGAAATTGNNNAGATCGGAAG
124534 TATGAGTGTGGAGTCGTTGCAATTACTTATTCAGCCACAAATTNNNAGATCGGAAGA

124536 TATGAGTGTGGAGTCGTTGCTTTTCCTTAGCTAAAAATTGACANNNAGATCGGAAG
124537 TATGAGTGTGGAGTCGTTGCCTTTTCATCATGCTCTTTACAGNNNAGATCGGAAGAG
124538 TATGAGTGTGGAGTCGTTGCTTCACTACACTGTATTGTCTTTNNNAGATCGGAAGAG
124539 TATGAGTGTGGAGTCGTTGCGAGGGATGAATTGAACACAGNNNAGATCGGAAGA
124540 TATGAGTGTGGAGTCGTTGCCCTGTTTCATCTTTAAACTCATTNNNAGATCGGAAGA
124541 TATGAGTGTGGAGTCGTTGCTATCTTTACATATCAGGACAGAGNNNAGATCGGAAG
124542 TATGAGTGTGGAGTCGTTGCTCTCTGTGTACCTGTATCATTNNNAGATCGGAAGAG
124543 TATGAGTGTGGAGTCGTTGCACATACTAAAATTTCAGCACAATNNNAGATCGGAAG
124544 TATGAGTGTGGAGTCGTTGCCAACTAAATAACTGAAGAAGGAANNNAGATCGGAA
124545 TATGAGTGTGGAGTCGTTGCTTGCCTTAACTTCCATTTTTTATNNNAGATCGGAAGA
124546 TATGAGTGTGGAGTCGTTGCCCTGTATATCCAGAAATTAGAGANNNAGATCGGAAG
124547 TATGAGTGTGGAGTCGTTGCTGTTTGGACAGAGATTTCTTTTANNNAGATCGGAAG
124548 TATGAGTGTGGAGTCGTTGCAAGTCATTTCAAAATACAGAGTCNNNAGATCGGAAG
124549 TATGAGTGTGGAGTCGTTGCGTAATCAAGTCACACACATTACNNNAGATCGGAAGA
124550 TATGAGTGTGGAGTCGTTGCGGGATTTTTTGTATAGGCAATTANNNAGATCGGAA
124551 TATGAGTGTGGAGTCGTTGCAGTACCTACTTCAGAAGTTTTAANNNAGATCGGAAG
124552 TATGAGTGTGGAGTCGTTGCTAAGGTTGAGACAGGAATGATTANNNAGATCGGAA
124553 TATGAGTGTGGAGTCGTTGCCTCTTTTCTCAGGGTCAGAGNNNAGATCGGAAGAG
124554 TATGAGTGTGGAGTCGTTGCTCAAACTAGCTTTAATACCATTGNNNAGATCGGAAG
124555 TATGAGTGTGGAGTCGTTGCGTCTTACATAAGCAGAGATAATGNNNAGATCGGAA
124556 TATGAGTGTGGAGTCGTTGCTTACCAAGATATTTGACTGTTTGNNNAGATCGGAAG
124557 TATGAGTGTGGAGTCGTTGCCAAAGAGAAGGAAAAAAGAAAAGNNNAGATCGGA
124558 TATGAGTGTGGAGTCGTTGCCAAAGAGAAGGAAAAAAGAAAAGNNNAGATCGGA
124559 TATGAGTGTGGAGTCGTTGCAACAAAATTAAACTCAGAGACATNNNAGATCGGAAG
124560 TATGAGTGTGGAGTCGTTGCTTGATTTAGCACATGATAATCATNNNAGATCGGAAG
124561 TATGAGTGTGGAGTCGTTGCATACATTTATTATGGGAATTGGCNNNAGATCGGAAG
124562 TATGAGTGTGGAGTCGTTGCATGAGGAAGATGTCATTTTCTATNNNAGATCGGAAG
124563 TATGAGTGTGGAGTCGTTGCGGACATTATGCTGAAAGAAATAGNNNAGATCGGAA
124564 TATGAGTGTGGAGTCGTTGCAAAAACAGCCATATTATGACCANNNAGATCGGAAGA
124565 TATGAGTGTGGAGTCGTTGCCCTCCAGCTCATTTTTAATATTTNNNAGATCGGAAGA
124566 TATGAGTGTGGAGTCGTTGCATAATCACTCATGTAACACACTNNNAGATCGGAAGA
124567 TATGAGTGTGGAGTCGTTGCACATTCCTGTCTACCACTTTNNNAGATCGGAAGAGCA
124568 TATGAGTGTGGAGTCGTTGCACAGACTGATAAAATCTACTGTTNNNAGATCGGAAG
124570 TATGAGTGTGGAGTCGTTGCCTGTCAATGTTAGCTTCCAGNNNAGATCGGAAGAGC
124571 TATGAGTGTGGAGTCGTTGCGAAACCACCCAGTTGTAAAGNNNAGATCGGAAGAG
124572 TATGAGTGTGGAGTCGTTGCGAAACCACCCAGTTGTAAAGNNNAGATCGGAAGAG
124574 TATGAGTGTGGAGTCGTTGCAATGACTATTTCCACTAGCAAANNNAGATCGGAAGA
124575 TATGAGTGTGGAGTCGTTGCCAAGTACACTAGAAATGTTGATANNNAGATCGGAA
124576 TATGAGTGTGGAGTCGTTGCGTATCCTAGCTGAAAGAAGAGNNNAGATCGGAAGA
124578 TATGAGTGTGGAGTCGTTGCCCCAAAGAATTACTCCTAGAAANNNAGATCGGAAGA
124579 TATGAGTGTGGAGTCGTTGCTATTTTCATGACTTTGAACTAGCNNNAGATCGGAAG
124580 TATGAGTGTGGAGTCGTTGCATATCTTAGTGCATATGTGTATGNNNAGATCGGAAG
124581 TATGAGTGTGGAGTCGTTGCGCTGTTTGAGGAAAAAAAAATAANNNAGATCGGAA

124582 TATGAGTGTGGAGTCGTTGCAATAGTTTCTTACTTCTCAAGGANNNAGATCGGAAG
124583 TATGAGTGTGGAGTCGTTGCCATACACGTATTTGTGAATGCNNNAGATCGGAAGAG
124585 TATGAGTGTGGAGTCGTTGCGCTGTCTACTTCATAGGATTATTNNNAGATCGGAAG
124586 TATGAGTGTGGAGTCGTTGCAGTTACTTTTTTCCCCAATATGNNNAGATCGGAAGA
124587 TATGAGTGTGGAGTCGTTGCTATTATTTGGTGAGACTTTCCTTNNNAGATCGGAAG
124588 TATGAGTGTGGAGTCGTTGCTCAGAGCTAAATTGTTTTTCTTTNNNAGATCGGAAG
124589 TATGAGTGTGGAGTCGTTGCTAAAAGAGATCCAGGCAATATTTNNNAGATCGGAA
124590 TATGAGTGTGGAGTCGTTGCAAAATAGTAACTCTGACTACAGANNNAGATCGGAAG
124591 TATGAGTGTGGAGTCGTTGCTCAGGCATATGGAGTAGATTNNNAGATCGGAAGAG
124592 TATGAGTGTGGAGTCGTTGCACTAATGGCAATCTGAATAAAATNNNAGATCGGAAG
124593 TATGAGTGTGGAGTCGTTGCTTTCTGGACCTCTTTTAAAAATTNNNAGATCGGAAG
124594 TATGAGTGTGGAGTCGTTGCAAACAAGAAGTAAAATCACAACTNNNAGATCGGAA
124595 TATGAGTGTGGAGTCGTTGCGATGCTGTAATTCCTAGTCATATNNNAGATCGGAAG
124596 TATGAGTGTGGAGTCGTTGCGATGCTGTAATTCCTAGTCATATNNNAGATCGGAAG
124597 TATGAGTGTGGAGTCGTTGCACCTTTAAAAATCTCAAAACTCANNNAGATCGGAAG
124598 TATGAGTGTGGAGTCGTTGCGTTTACAACTCTGCCTTAGCNNNAGATCGGAAGAGC
124599 TATGAGTGTGGAGTCGTTGCCCATACCCACTTCTCAAAATATNNNAGATCGGAAGAG
124600 TATGAGTGTGGAGTCGTTGCTTGGACTAAAAGGGAAATTTTAGNNNAGATCGGAA
124601 TATGAGTGTGGAGTCGTTGCCTTTTCAAATGCAGCATTAATACNNNAGATCGGAAG
124602 TATGAGTGTGGAGTCGTTGCCGTGATTTAATCTCTAGGAATTCNNNAGATCGGAAG
124603 TATGAGTGTGGAGTCGTTGCGCTGCTCTATTAAGTATGTAATAGNNNAGATCGGAA
124604 TATGAGTGTGGAGTCGTTGCAGTAATTTGATTAGGATATGCCTNNNAGATCGGAAG
124605 TATGAGTGTGGAGTCGTTGCGCAAAAGATCAAAATTCTGTTTTNNNAGATCGGAAG
124606 TATGAGTGTGGAGTCGTTGCTATAGGTTTGCTTTATGATCTTCNNNAGATCGGAAG
124607 TATGAGTGTGGAGTCGTTGCATACTGATTTAGATTTCCAGTCTNNNAGATCGGAAG
124608 TATGAGTGTGGAGTCGTTGCTCAATGTCATGATGTCATTTATGNNNAGATCGGAAG
124609 TATGAGTGTGGAGTCGTTGCATTCATTTGGCCAACATGTANNNAGATCGGAAGAGC
124610 TATGAGTGTGGAGTCGTTGCCACTTTTTCAAGCCTCTGTTNNNAGATCGGAAGAGC
124611 TATGAGTGTGGAGTCGTTGCATTCTGAATAGTTGCTTACCTTTNNNAGATCGGAAG
124613 TATGAGTGTGGAGTCGTTGCCTTCATACTTCCTGTCAATGANNNAGATCGGAAGAG
124614 TATGAGTGTGGAGTCGTTGCTGTATCCTTTGAAATAGCCTTTANNNAGATCGGAAG
124615 TATGAGTGTGGAGTCGTTGCTGTATCCTTTGAAATAGCCTTTANNNAGATCGGAAG
124616 TATGAGTGTGGAGTCGTTGCAAACCCTCTTGATTATGTCTTTTNNNAGATCGGAAGA
124617 TATGAGTGTGGAGTCGTTGCTAATCTCAGCTCACTGCAATNNNAGATCGGAAGAGC
124618 TATGAGTGTGGAGTCGTTGCTGGATGACTAATATTTGAACTCTNNNAGATCGGAAG
124619 TATGAGTGTGGAGTCGTTGCACCTTGATATCTGTTAAACAAACNNNAGATCGGAAG
124620 TATGAGTGTGGAGTCGTTGCATAACCTCCTTCCTTGTTTATTNNNAGATCGGAAGAG
124621 TATGAGTGTGGAGTCGTTGCAAGAAGTTATTGGGTAGAGAATTNNNAGATCGGAA
124622 TATGAGTGTGGAGTCGTTGCTTTAGTCTACACCTGTGAAAGNNNAGATCGGAAGAG
124623 TATGAGTGTGGAGTCGTTGCAATTGGTATAATCATTACTGCAGNNNAGATCGGAAG
124624 TATGAGTGTGGAGTCGTTGCATCTGGATATGATAGACAGAAAGNNNAGATCGGAA
124625 TATGAGTGTGGAGTCGTTGCATTTTTTTCCTCTGTCTTTGTCNNNAGATCGGAAGAG
124626 TATGAGTGTGGAGTCGTTGCATTTTTCAGATTGTCTCACTATCNNNAGATCGGAAGA

124628 TATGAGTGTGGAGTCGTTGCCAAAGACCTTGTCAAAACATCNNNAGATCGGAAGAG
124629 TATGAGTGTGGAGTCGTTGCTACTTTGATTTCCCAATTACTTGNNNAGATCGGAAGA
124630 TATGAGTGTGGAGTCGTTGCATTTATCACATGCACATCTCTTANNNAGATCGGAAGA
124631 TATGAGTGTGGAGTCGTTGCTGTCTGTCTTTTCTTAACCTATTNNNAGATCGGAAGA
124632 TATGAGTGTGGAGTCGTTGCGTCTGTTTTAGTTGCCATTTAGNNNAGATCGGAAGA
124633 TATGAGTGTGGAGTCGTTGCTAATCCTTACAAGTCCTAGAGTTNNNAGATCGGAAG
124634 TATGAGTGTGGAGTCGTTGCTTTGTCTAGATTAGCTGTCCTNNNAGATCGGAAGAG
124636 TATGAGTGTGGAGTCGTTGCCAATACCCCCAGGAATTCATNNNAGATCGGAAGAGC
124637 TATGAGTGTGGAGTCGTTGCAAAAGTTGGAAGCTTCAGATNNNAGATCGGAAGAG
124638 TATGAGTGTGGAGTCGTTGCGGTAATTCTAACCCTACAAAATAGNNNAGATCGGAA
124639 TATGAGTGTGGAGTCGTTGCTTTTCTCCCCATATTCTATTTCTNNNAGATCGGAAGAG
124640 TATGAGTGTGGAGTCGTTGCGGAACCAGTGATCTAGAATATTTNNNAGATCGGAAG
124641 TATGAGTGTGGAGTCGTTGCATACTTATTGAGCATCTATGTGTNNNAGATCGGAAG
124642 TATGAGTGTGGAGTCGTTGCAACTATATTTGGGCTGAAATTTTNNNAGATCGGAAG
124643 TATGAGTGTGGAGTCGTTGCTGCTTCAATAAACAAACAAGTATNNNAGATCGGAAG
124644 TATGAGTGTGGAGTCGTTGCTCAATAAGCTCTAGCCACTTNNNAGATCGGAAGAGC
124645 TATGAGTGTGGAGTCGTTGCCATAAACAGCAAAGAGACAAGNNNAGATCGGAAGA
124646 TATGAGTGTGGAGTCGTTGCTGAATTTTCTGGGTTTCTTAAAANNNAGATCGGAAG
124648 TATGAGTGTGGAGTCGTTGCTTCCTAAACATCAATGTATGGTANNNAGATCGGAAG
124649 TATGAGTGTGGAGTCGTTGCTCAAAATTACCAAACACTGTAAANNNAGATCGGAAG
124650 TATGAGTGTGGAGTCGTTGCTGTTCATTCATAGTCACACCNNNAGATCGGAAGAGC
124651 TATGAGTGTGGAGTCGTTGCAATAGGACTACTAAGAATCTTGGNNNAGATCGGAA
124652 TATGAGTGTGGAGTCGTTGCTATGTGAGCAAGTGTGTATTAATNNNAGATCGGAA
124653 TATGAGTGTGGAGTCGTTGCTCAAGTGTCATTTCAGTTTTTAANNNAGATCGGAAG
124654 TATGAGTGTGGAGTCGTTGCGTCATTTGATGGGGTCTTTGNNNAGATCGGAAGAG
124655 TATGAGTGTGGAGTCGTTGCGAAAAGAGACATATGAAAGCAGNNNAGATCGGAAG
124656 TATGAGTGTGGAGTCGTTGCGGAAGTGTTAACAACTTTCTAGNNNAGATCGGAAGA
124657 TATGAGTGTGGAGTCGTTGCAGTATGAAGTGTCAACTTGATTNNNAGATCGGAAGA
124658 TATGAGTGTGGAGTCGTTGCGACACAGTCATGTTGGATTANNNAGATCGGAAGAG
124659 TATGAGTGTGGAGTCGTTGCCATATATACACACACACAGTTTTNNNAGATCGGAAGA
124660 TATGAGTGTGGAGTCGTTGCTTCTAGGCTACTAAGTGTCAATNNNAGATCGGAAGA
124661 TATGAGTGTGGAGTCGTTGCAAATGTGGGACAATATTCAAAATNNNAGATCGGAA
124662 TATGAGTGTGGAGTCGTTGCGAGTCATTTAACATCCAGAACTNNNAGATCGGAAGA
124663 TATGAGTGTGGAGTCGTTGCAGTTTTAGAAATGACTTTGACATNNNAGATCGGAAG
124664 TATGAGTGTGGAGTCGTTGCAATTATTTTAATCACTTCACAATGTNNNAGATCGGAA
124665 TATGAGTGTGGAGTCGTTGCTTCTAAGTGTGTATGTATGTAGCNNNAGATCGGAAG
124666 TATGAGTGTGGAGTCGTTGCTTTCCCAACAGTAGTAGATAAATNNNAGATCGGAAG
124667 TATGAGTGTGGAGTCGTTGCGGCAGTGTTTGTAATTTTTTTTTNNNAGATCGGAAG
124668 TATGAGTGTGGAGTCGTTGCCAAGTAAAATCAAAACCAAAAGGNNNAGATCGGAA
124669 TATGAGTGTGGAGTCGTTGCATAATGTATATAAGATACTTTGCACNNNAGATCGGA
124670 TATGAGTGTGGAGTCGTTGCGTAAAAATGGGGATGTCATTTCNNNAGATCGGAAG
124671 TATGAGTGTGGAGTCGTTGCTTCAAACACTGAGGTAAATCTGNNNAGATCGGAAGA
124672 TATGAGTGTGGAGTCGTTGCTTCAAACACTGAGGTAAATCTGNNNAGATCGGAAGA

124673 TATGAGTGTGGAGTCGTTGCAGATGTTCATTCCTACCTAATGNNNAGATCGGAAGA
124674 TATGAGTGTGGAGTCGTTGCACAGTTCATCCTGCTACTTCNNNAGATCGGAAGAGC
124675 TATGAGTGTGGAGTCGTTGCTTCATTTTGCTTCTGTATTTTTGNNNAGATCGGAAGA
124676 TATGAGTGTGGAGTCGTTGCCCTTTAGATCTAATTACCCCTATNNNAGATCGGAAGA
124677 TATGAGTGTGGAGTCGTTGCCACCATTTTTTTTTGTTATCCAANNNAGATCGGAAGA
124678 TATGAGTGTGGAGTCGTTGCACTTTTTTTTCTTTAAGTTTGGGNNNAGATCGGAAG
124679 TATGAGTGTGGAGTCGTTGCTTAGACATGTGACATTAGCTTTNNNAGATCGGAAGA
124680 TATGAGTGTGGAGTCGTTGCTTCAAGGAAGAGGAGATGTTNNNAGATCGGAAGA
124681 TATGAGTGTGGAGTCGTTGCACTATTTGTGTCTCTGTATCGNNNAGATCGGAAGAG
124682 TATGAGTGTGGAGTCGTTGCGAAAACCTTGGGCACTAAATNNNAGATCGGAAGAG
124683 TATGAGTGTGGAGTCGTTGCTATTTTTATGAATGGTTACCACCNNNAGATCGGAAG
124684 TATGAGTGTGGAGTCGTTGCATAGTGCATGTATAGCATTTACANNNAGATCGGAAG
124685 TATGAGTGTGGAGTCGTTGCGATTCTTTTCTCCATATTGCATTNNNAGATCGGAAGA
124686 TATGAGTGTGGAGTCGTTGCCAGGCAATGAGAAAAAAAGGNNNAGATCGGAAGA
124687 TATGAGTGTGGAGTCGTTGCCATGACATTCTCTTTAGTTAGTGNNNAGATCGGAAG
124688 TATGAGTGTGGAGTCGTTGCTCTTCCTTATTCTTTGTTTTTCCNNNAGATCGGAAGA
124689 TATGAGTGTGGAGTCGTTGCTCTCCTTCTTTGCAATACATAATNNNAGATCGGAAGA
124690 TATGAGTGTGGAGTCGTTGCAAAAAGTGAAAGCAGTACCAATNNNAGATCGGAAG
124691 TATGAGTGTGGAGTCGTTGCATAACTTTCTTTATCAAAAGCCCNNNAGATCGGAAGA
124692 TATGAGTGTGGAGTCGTTGCCAGTAATCCATGGGGTTTAGNNNAGATCGGAAGAG
124693 TATGAGTGTGGAGTCGTTGCACTGATACAGAAATATCAAGTGGNNNAGATCGGAA
124694 TATGAGTGTGGAGTCGTTGCGGTCTTCATCTAGAGATCATAAGNNNAGATCGGAAG
124695 TATGAGTGTGGAGTCGTTGCATGCTATTTTCCTGCTTTTTTATNNNAGATCGGAAGA
124696 TATGAGTGTGGAGTCGTTGCGGTGTAAAAGCAATATACATTCANNNAGATCGGAA
124697 TATGAGTGTGGAGTCGTTGCGCTATCCCTTTCAAAAATAGAATNNNAGATCGGAAG
124698 TATGAGTGTGGAGTCGTTGCTCAGTAAATACAGATCTTTCTCTNNNAGATCGGAAG
124699 TATGAGTGTGGAGTCGTTGCTCTAAATGGGAAATGTTACAGTNNNAGATCGGAAG
124701 TATGAGTGTGGAGTCGTTGCGAGAGCATGTTTAAATTGAAATGNNNAGATCGGAA
124702 TATGAGTGTGGAGTCGTTGCTTCTTTTTCTTTCCCTACCCTATNNNAGATCGGAAGAG
124703 TATGAGTGTGGAGTCGTTGCCACTATTCATCTAATTACCTCCTNNNAGATCGGAAGA
124704 TATGAGTGTGGAGTCGTTGCATATCTCTTTTCAGAAACTCAGANNNAGATCGGAAG
124705 TATGAGTGTGGAGTCGTTGCGGACAGAGGTTATATTAGCTTAGNNNAGATCGGAA
124706 TATGAGTGTGGAGTCGTTGCGTACTATGTAAGTGACAGAGAATNNNAGATCGGAA
124707 TATGAGTGTGGAGTCGTTGCCAGGCTTTTTTGAAAGAGTAGNNNAGATCGGAAGA
124708 TATGAGTGTGGAGTCGTTGCAAATTCATGCATGGTTCAATTNNNAGATCGGAAGAG
124709 TATGAGTGTGGAGTCGTTGCAAATTCATGCATGGTTCAATTNNNAGATCGGAAGAG
124710 TATGAGTGTGGAGTCGTTGCATCTACCTATCTATCTATCTGCANNNAGATCGGAAGA
124711 TATGAGTGTGGAGTCGTTGCGTATATTGGGTACTCAGTACTTCNNNAGATCGGAAG
124712 TATGAGTGTGGAGTCGTTGCTGAAGAAAGGTTCACAAATTAATNNNAGATCGGAA
124713 TATGAGTGTGGAGTCGTTGCCACAAAAGCATAGAAGATTGTNNNAGATCGGAAGA
124714 TATGAGTGTGGAGTCGTTGCAGACTTTTCAGTAGATACCTTACNNNAGATCGGAAG
124715 TATGAGTGTGGAGTCGTTGCATTACACCTTGTGAATTTATAGGNNNAGATCGGAAG
124716 TATGAGTGTGGAGTCGTTGCCTAGGGTTCATTTTGTGTTTATCNNNAGATCGGAAG

124717 TATGAGTGTGGAGTCGTTGCCAATGAACTCTCTAAGGGAAGNNNAGATCGGAAGA
124718 TATGAGTGTGGAGTCGTTGCACATTTAATAGAATCCCTCCATANNNAGATCGGAAG
124719 TATGAGTGTGGAGTCGTTGCTATTTATTCGTTTAGTGCCTTTTNNNAGATCGGAAGA
124720 TATGAGTGTGGAGTCGTTGCATTCTGTGATCTAGTCTTAGAAGNNNAGATCGGAAG
124721 TATGAGTGTGGAGTCGTTGCTGGATTATGTGTATGTTAGCATANNNAGATCGGAA
124722 TATGAGTGTGGAGTCGTTGCGAGAATGACAGTTCTACCATTNNNAGATCGGAAGA
124723 TATGAGTGTGGAGTCGTTGCTGCCTTAGTCAATATAAGTCAATNNNAGATCGGAAG
124724 TATGAGTGTGGAGTCGTTGCATACTTATAACAGTCCCAGTGTTNNNAGATCGGAAG
124725 TATGAGTGTGGAGTCGTTGCTTCTACCTCCAAATTATAACCTCNNNAGATCGGAAGA
124726 TATGAGTGTGGAGTCGTTGCGGTACAAACACTGTTCACTGNNNAGATCGGAAGAG
124727 TATGAGTGTGGAGTCGTTGCGGTACAAACACTGTTCACTGNNNAGATCGGAAGAG
124728 TATGAGTGTGGAGTCGTTGCCCTGTTCTAAATCCACATATCTNNNAGATCGGAAGAG
124729 TATGAGTGTGGAGTCGTTGCCCATATGTTTCCTAAGTGACTATNNNAGATCGGAAG
124730 TATGAGTGTGGAGTCGTTGCAAAATGTGGATATGAACAAATGANNNAGATCGGAA
124731 TATGAGTGTGGAGTCGTTGCTAAAGACAAACCAATCAACCANNNAGATCGGAAGAG
124732 TATGAGTGTGGAGTCGTTGCATTCACAGTGTATGTTTTTTCTTNNNAGATCGGAAGA
124733 TATGAGTGTGGAGTCGTTGCGTATTTGCCCCAATCAGATGNNNAGATCGGAAGAG
124734 TATGAGTGTGGAGTCGTTGCTCATGTGTATTATATAGAGAGAGTNNNAGATCGGA
124735 TATGAGTGTGGAGTCGTTGCAGTAATTTATAAGAGCATGGAGTNNNAGATCGGAA
124737 TATGAGTGTGGAGTCGTTGCTCCATTTCAAGAAAGAAATTCATNNNAGATCGGAAG
124738 TATGAGTGTGGAGTCGTTGCGGTGAGAGAGAGAAGAGAAGNNNAGATCGGAAG
124739 TATGAGTGTGGAGTCGTTGCTTTGGAAAGTTTGTTTCTTTTTTNNNAGATCGGAAG
124741 TATGAGTGTGGAGTCGTTGCCAGTTCTAATTAAAGAGACTTTGTNNNAGATCGGAA
124742 TATGAGTGTGGAGTCGTTGCGATTTCAATGCAAAAAGACATCNNNAGATCGGAAGA
124743 TATGAGTGTGGAGTCGTTGCGTTTTGTGTTTTCTTGAATGAAANNNAGATCGGAAG
124744 TATGAGTGTGGAGTCGTTGCTTATTCCCCATCTAGTTAGAATGNNNAGATCGGAAG
124745 TATGAGTGTGGAGTCGTTGCAACTATAGCATGTTGGTACTACNNNAGATCGGAAGA
124747 TATGAGTGTGGAGTCGTTGCTAAAATTGGTATAAAAGGTGACCNNNAGATCGGAA
124748 TATGAGTGTGGAGTCGTTGCCAAATCTTTCTAATAGCAGCATTNNNAGATCGGAAG
124749 TATGAGTGTGGAGTCGTTGCTACTTCACAAACCTAACCAATTANNNAGATCGGAAGA
124750 TATGAGTGTGGAGTCGTTGCCTGGCTTCTTTCAATATAAACATNNNAGATCGGAAG
124751 TATGAGTGTGGAGTCGTTGCGATTACAAAACCTCTCAATACTGNNNAGATCGGAAG
124752 TATGAGTGTGGAGTCGTTGCGTGTGTACACAGTATTCATATTGNNNAGATCGGAAG
124753 TATGAGTGTGGAGTCGTTGCGTTAACTGTAACAATGCTATACANNNAGATCGGAAG
124754 TATGAGTGTGGAGTCGTTGCTTTTGGTAGGGAATCATTCTTTTNNNAGATCGGAAG
124755 TATGAGTGTGGAGTCGTTGCATTTCTTTTCTTTGGGTAGATACNNNAGATCGGAAG
124756 TATGAGTGTGGAGTCGTTGCGAAATCTGGTTGTCTATGGATANNNAGATCGGAAG
124757 TATGAGTGTGGAGTCGTTGCATCAATACAGAATGTAGAGCATNNNAGATCGGAAG
124758 TATGAGTGTGGAGTCGTTGCCTCCAGAGAGACTACATCTGNNNAGATCGGAAGAG
124759 TATGAGTGTGGAGTCGTTGCTTGTGTTGTAAAATGGATTTTTGNNNAGATCGGAAG
124760 TATGAGTGTGGAGTCGTTGCACTGTTATGTTGGTAATTGTTAGNNNAGATCGGAAG
124761 TATGAGTGTGGAGTCGTTGCTTTCAAAATCCAGACCTTGTNNNAGATCGGAAGAGC
124762 TATGAGTGTGGAGTCGTTGCTCAATAACAGAATGTATTGAAACANNNAGATCGGAA

124763 TATGAGTGTGGAGTCGTTGCTTTCAATGTTTTCAAGGGAATTANNNAGATCGGAAG
124764 TATGAGTGTGGAGTCGTTGCTTTAGTCTCTCTAAAAGTGGGATNNNAGATCGGAAG
124765 TATGAGTGTGGAGTCGTTGCGTGTTTTCAAACTTCCCTATTATNNNAGATCGGAAGA
124766 TATGAGTGTGGAGTCGTTGCAATTTCATTCCTTTTTTATGGCTNNNAGATCGGAAGA
124767 TATGAGTGTGGAGTCGTTGCAATAAGAAAGCAACAGTAAGTTTNNNAGATCGGAA
124768 TATGAGTGTGGAGTCGTTGCAATGGACATTGAAAGGGAAGNNNAGATCGGAAGA
124769 TATGAGTGTGGAGTCGTTGCTGGTTAAGACTATCATCTTTACANNNAGATCGGAAG
124770 TATGAGTGTGGAGTCGTTGCTACATCTGCAAAGACATTATTACNNNAGATCGGAAG
124771 TATGAGTGTGGAGTCGTTGCAGTCATTGTATGATGATTCCATNNNAGATCGGAAGA
124772 TATGAGTGTGGAGTCGTTGCCATGATCTAGGTTCACTGTAACNNNAGATCGGAAGA
124773 TATGAGTGTGGAGTCGTTGCCAAGTATAAAAAGAAACTGCAACNNNAGATCGGAA
124774 TATGAGTGTGGAGTCGTTGCATTGAGCAACAAAATAAACAATGNNNAGATCGGAA
124775 TATGAGTGTGGAGTCGTTGCTGGAATAAATTGAGCAACAAAATNNNAGATCGGAA
124776 TATGAGTGTGGAGTCGTTGCATTGTACAATAGTAACAGGATGTNNNAGATCGGAA
124778 TATGAGTGTGGAGTCGTTGCAAGATCAGAGAAAAATTCCCTTNNNAGATCGGAAGA
124779 TATGAGTGTGGAGTCGTTGCTAGCTTTAGATTTCAGTACATACANNNAGATCGGAA
124780 TATGAGTGTGGAGTCGTTGCTGGTATTCTAGTTAGGTTATTGTNNNAGATCGGAAG
124781 TATGAGTGTGGAGTCGTTGCTGGTATTCTAGTTAGGTTATTGTNNNAGATCGGAAG
124782 TATGAGTGTGGAGTCGTTGCTGAATTTAGCCAGAGATTAGTAANNNAGATCGGAA
124783 TATGAGTGTGGAGTCGTTGCCTGTTTCTAGGAAACAAATACAANNNAGATCGGAAG
124784 TATGAGTGTGGAGTCGTTGCATAGTGAATGTGACTGTAATGTANNNAGATCGGAA
124785 TATGAGTGTGGAGTCGTTGCGCGATCTTTTAAAGGACATATTTNNNAGATCGGAAG
124786 TATGAGTGTGGAGTCGTTGCATATATGTTTCCACCACTCTATCNNNAGATCGGAAGA
124787 TATGAGTGTGGAGTCGTTGCTTCTTACAAAATTCATTCTGTGTNNNAGATCGGAAG
124790 TATGAGTGTGGAGTCGTTGCCAAAAAATAAATTCCCTGGAAAGNNNAGATCGGAA
124791 TATGAGTGTGGAGTCGTTGCCTAGAACAGGATAATTACCCAAGNNNAGATCGGAA
124793 TATGAGTGTGGAGTCGTTGCTGAATCCTGTCTATCATATTAGCNNNAGATCGGAAG
124794 TATGAGTGTGGAGTCGTTGCAATTCATAATCCCTGAAGTAGGNNNAGATCGGAAGA
124795 TATGAGTGTGGAGTCGTTGCTAGTACATGCATATATGTAAGCTNNNAGATCGGAAG
124798 TATGAGTGTGGAGTCGTTGCAGTCTGCAAACCAGAAAGATNNNAGATCGGAAGAG
124799 TATGAGTGTGGAGTCGTTGCAAATTTCTTGTCTTGTACTGTGNNNAGATCGGAAGA
124800 TATGAGTGTGGAGTCGTTGCCTAAAGTCACTGTAAATCCAAAGNNNAGATCGGAAG
124801 TATGAGTGTGGAGTCGTTGCTATTCCTTCCTCATTACTTTGAANNNAGATCGGAAGA
124802 TATGAGTGTGGAGTCGTTGCATTTTGGTTATTTCTTGCCTTTTNNNAGATCGGAAGA
124803 TATGAGTGTGGAGTCGTTGCCAAGATTAAGCGTGGAATAGTANNNAGATCGGAAG
124804 TATGAGTGTGGAGTCGTTGCTTTGTTTAAAAAATGTTGGCTTTNNNAGATCGGAAG
124805 TATGAGTGTGGAGTCGTTGCAGGAACAAAGAAATAAAAGAGCNNNAGATCGGAA
124808 TATGAGTGTGGAGTCGTTGCTTCTCTCTCACATCTCACACNNNAGATCGGAAGAGCA
124809 TATGAGTGTGGAGTCGTTGCAAGAGGAACATGTTAAATGACATNNNAGATCGGAA
124810 TATGAGTGTGGAGTCGTTGCTAAACATGAAGGCACAGATCNNNAGATCGGAAGAG
124811 TATGAGTGTGGAGTCGTTGCCTACCTCCATTCACTGAATGNNNAGATCGGAAGAGC
124812 TATGAGTGTGGAGTCGTTGCTCCTTCTTGTTTCATTGCTTNNNAGATCGGAAGAGCA
124813 TATGAGTGTGGAGTCGTTGCCTTTCTTGTACAAAGACCTTAAANNNAGATCGGAAG

124814 TATGAGTGTGGAGTCGTTGCGTTTCTTGTTGAGGGAAGATNNNAGATCGGAAGAG

124815 TATGAGTGTGGAGTCGTTGCGTTGTTGTTTTTGAGACAGANNNAGATCGGAAGAG

124817 TATGAGTGTGGAGTCGTTGCAAATTGAGGATCTGATATCTGATNNNAGATCGGAAG

124818 TATGAGTGTGGAGTCGTTGCGGAATTTCCCTCAAGTTGTANNNAGATCGGAAGAGC

124819 TATGAGTGTGGAGTCGTTGCTATCACATCTGGGTAATTATACCNNNAGATCGGAAGA

124820 TATGAGTGTGGAGTCGTTGCGGAATATTAATTGTGATTGGTCANNNAGATCGGAAG

124821 TATGAGTGTGGAGTCGTTGCAGAAGATCCATTGTTAGAGAAATNNNAGATCGGAAG

124822 TATGAGTGTGGAGTCGTTGCCGTAGTTTTACAATTTGAAGAGANNNAGATCGGAAG

124823 TATGAGTGTGGAGTCGTTGCTGTCATGCAGTTGTATTATTTGNNNAGATCGGAAGA

124824 TATGAGTGTGGAGTCGTTGCATAGTTTTGCTTCCTTTTTTCTTNNNAGATCGGAAGA

124827 TATGAGTGTGGAGTCGTTGCAAGATTAACACCAAACCATAGTANNNAGATCGGAAG

124828 TATGAGTGTGGAGTCGTTGCATTACAATTCAAGATGGGATTTGNNNAGATCGGAAG

124829 TATGAGTGTGGAGTCGTTGCCAAAAATGTCTCCAGATTGAGNNNAGATCGGAAGAG

124830 TATGAGTGTGGAGTCGTTGCCACCCCAAGATATAAATCAATTTNNNAGATCGGAAGA

124831 TATGAGTGTGGAGTCGTTGCTTTTTCTCCGTATTCTTTGTTTTNNNAGATCGGAAGA

124832 TATGAGTGTGGAGTCGTTGCTTTTTGTCGTGGAGATTATAAAGNNNAGATCGGAAG

124833 TATGAGTGTGGAGTCGTTGCCAAAAGAGAATGTGACCATATCNNNAGATCGGAAGA

124834 TATGAGTGTGGAGTCGTTGCCCAAGATTCTGACAAATTAAAACNNNAGATCGGAAG

124835 TATGAGTGTGGAGTCGTTGCCTGGTTGTCATACTTTTAACTTTNNNAGATCGGAAGA

124836 TATGAGTGTGGAGTCGTTGCCAAGATTATTGCATGGAAACATANNNAGATCGGAAG

124837 TATGAGTGTGGAGTCGTTGCCAAGATTATTGCATGGAAACATANNNAGATCGGAAG

124838 TATGAGTGTGGAGTCGTTGCGAGCAAGACAGTTATTTTAACTTNNNAGATCGGAAG

124839 TATGAGTGTGGAGTCGTTGCGGAAGGGGAATTAAAACATATGNNNAGATCGGAAG

124840 TATGAGTGTGGAGTCGTTGCCCCAAGAATAGATGTGAGGANNNAGATCGGAAGAG

124841 TATGAGTGTGGAGTCGTTGCGCAAAGACAGATCAAGAAATATTNNNAGATCGGAAG

124842 TATGAGTGTGGAGTCGTTGCCAAAGCTAAATATAGATCGGAAGNNNAGATCGGAAG

124843 TATGAGTGTGGAGTCGTTGCAATTTGTCTCTTTTGAGTTTCAGNNNAGATCGGAAGA

124844 TATGAGTGTGGAGTCGTTGCTACTTGTTAACCTGTCACATTTANNNAGATCGGAAGA

124845 TATGAGTGTGGAGTCGTTGCAGAACAGAAAAACATTTGTCTCNNNAGATCGGAAGA

124846 TATGAGTGTGGAGTCGTTGCTTCCTCCAAGAGCTGAATATNNNAGATCGGAAGAGC

124847 TATGAGTGTGGAGTCGTTGCTCACTGAAAAACATAAAAGAAGANNNAGATCGGAAG

124848 TATGAGTGTGGAGTCGTTGCGAAAGAAGATGAAGTCTCGAATANNNAGATCGGAA

124849 TATGAGTGTGGAGTCGTTGCAGAGTAGGAAACAAGAAGAAATANNNAGATCGGAA

124850 TATGAGTGTGGAGTCGTTGCTTACCTTACATACTTACTGCTTTNNNAGATCGGAAGA

124851 TATGAGTGTGGAGTCGTTGCTGCAAATTAAAGTGATAATGAGANNNAGATCGGAAG

124852 TATGAGTGTGGAGTCGTTGCGTGTGTTTTATTTTCTGTATGGANNNAGATCGGAAG

124853 TATGAGTGTGGAGTCGTTGCTAGAATGCCTTTGGGAAATAAATNNNAGATCGGAAG

124854 TATGAGTGTGGAGTCGTTGCTATTTGATTCCCCCAAATATACTNNNAGATCGGAAGA

124855 TATGAGTGTGGAGTCGTTGCCTCCAACTCTCTTCTCTCTATNNNAGATCGGAAGAGCA

124856 TATGAGTGTGGAGTCGTTGCAATACATATGAAAATGCAAAGGANNNAGATCGGAAG

124857 TATGAGTGTGGAGTCGTTGCGAAAAAGGAGCACTTTGTATAATNNNAGATCGGAAG

124859 TATGAGTGTGGAGTCGTTGCGAGTTTAGGAAATGTGTGTGTNNNAGATCGGAAGA

124860 TATGAGTGTGGAGTCGTTGCTTTGTGCTGAGTTTAGGAAGNNNAGATCGGAAGAG

124861 TATGAGTGTGGAGTCGTTGCCATTGCTACACAAATTTTTCTTCNNNAGATCGGAAGA
124862 TATGAGTGTGGAGTCGTTGCAAAATCCTGGTAGGAAGAAATTTNNNAGATCGGAA
124863 TATGAGTGTGGAGTCGTTGCATACTTTTTATAACCTACAGGCANNNAGATCGGAAG
124864 TATGAGTGTGGAGTCGTTGCAATATGTGAGAATTATGGGAACTNNNAGATCGGAA
124865 TATGAGTGTGGAGTCGTTGCGCAAATAGGAAGATATCTGGAANNNAGATCGGAAG
124866 TATGAGTGTGGAGTCGTTGCTTATTCCATATCAACCTCTAAGTNNNAGATCGGAAGA
124867 TATGAGTGTGGAGTCGTTGCTTCTGTATCTGTATACCTAGGATNNNAGATCGGAAG
124868 TATGAGTGTGGAGTCGTTGCATCTTGAAGAATAACATCAAAGCNNNAGATCGGAAG
124869 TATGAGTGTGGAGTCGTTGCTCTAAAACTTACATGAAAATGCANNNAGATCGGAAG
124870 TATGAGTGTGGAGTCGTTGCGGAAGAAGAAAACTATGAGAAAANNNAGATCGGA
124872 TATGAGTGTGGAGTCGTTGCCTTTCTCTTCATCCTATCTACAANNNAGATCGGAAGA
124873 TATGAGTGTGGAGTCGTTGCAAAGGATAAGAATAGACTAGTGGNNNAGATCGGA
124874 TATGAGTGTGGAGTCGTTGCTATGATTGTGACTTCTAGTTCTTNNNAGATCGGAAG
124875 TATGAGTGTGGAGTCGTTGCCTAGGATTTTTTTTTTCTTTCCCNNNAGATCGGAAGA
124876 TATGAGTGTGGAGTCGTTGCAATGGTAAGAGTTGGAAGAATTANNNAGATCGGAA
124877 TATGAGTGTGGAGTCGTTGCTTTACTTAACTGACAGGTTGTATNNNAGATCGGAAG
124878 TATGAGTGTGGAGTCGTTGCTTTAGTAGCCTTGAGAAAGTTATNNNAGATCGGAAG
124879 TATGAGTGTGGAGTCGTTGCATAGCAAGTTTGTTTTGAATGTNNNAGATCGGAAG
124880 TATGAGTGTGGAGTCGTTGCATAGCAAGTTTGTTTTGAATGTNNNAGATCGGAAG
124881 TATGAGTGTGGAGTCGTTGCTTCCCATATTATGTCCATAGCTNNNAGATCGGAAGA
124884 TATGAGTGTGGAGTCGTTGCGAATTCTGAAGAGTTTAAGAGAANNNAGATCGGAA
124885 TATGAGTGTGGAGTCGTTGCGAAATATAAAATGGTGTAGCCANNNAGATCGGAAG
124886 TATGAGTGTGGAGTCGTTGCGAGAAAGGAGTGAAGATTAAGTNNNAGATCGGAA
124887 TATGAGTGTGGAGTCGTTGCAATGTGAAGAAATAACTCACAAANNNAGATCGGAA
124888 TATGAGTGTGGAGTCGTTGCGTCTGAAAATGGAGTTTCCGNNNAGATCGGAAGAG
124889 TATGAGTGTGGAGTCGTTGCCAGTAACAGAATCCAAAGAAAAGNNNAGATCGGAA
124890 TATGAGTGTGGAGTCGTTGCATGAATGTACAAAAGCATACATTNNNAGATCGGAAG
124891 TATGAGTGTGGAGTCGTTGCAAGGCCTAACTTTGAAAAATAAANNNAGATCGGAA
124892 TATGAGTGTGGAGTCGTTGCCTCATTTCGTAGTGAGATCTCNNNAGATCGGAAGAG
124893 TATGAGTGTGGAGTCGTTGCATATTTACACACACTCTTGAAACNNNAGATCGGAAG
124895 TATGAGTGTGGAGTCGTTGCGATAAATTCCCCCAGGATAAAATNNNAGATCGGAAG
124896 TATGAGTGTGGAGTCGTTGCTAAAATAAGGCTGAGACCACTNNNAGATCGGAAGA
124897 TATGAGTGTGGAGTCGTTGCCTATGGACTCAATACCTCCTANNNAGATCGGAAGAG
124898 TATGAGTGTGGAGTCGTTGCCAGCATTTCTAAGAAGACTAATCNNNAGATCGGAAG
124899 TATGAGTGTGGAGTCGTTGCTAATACAACGTGCTTCTTTTTAANNNAGATCGGAAG
124900 TATGAGTGTGGAGTCGTTGCTAATACAACGTGCTTCTTTTTAANNNAGATCGGAAG
124901 TATGAGTGTGGAGTCGTTGCTTGATATAAATCAAAGCAGGTTGNNNAGATCGGAA
124902 TATGAGTGTGGAGTCGTTGCAGTATGGCAATATATTATCCTCANNNAGATCGGAAG
124903 TATGAGTGTGGAGTCGTTGCTTTTGTACTTATCATCTTCCTGANNNAGATCGGAAGA
124904 TATGAGTGTGGAGTCGTTGCTCATTTATATACACAAATGTCCTNNNAGATCGGAAGA
124905 TATGAGTGTGGAGTCGTTGCCCATCATTCACCTTATTCATTTTNNNAGATCGGAAGA
124906 TATGAGTGTGGAGTCGTTGCGTGTGATGTAGAAAAGGGTATTANNNAGATCGGAA
124907 TATGAGTGTGGAGTCGTTGCCATATTTTTATTTCTTTTGGGCANNNAGATCGGAAGA

124908 TATGAGTGTGGAGTCGTTGCATGAGGACAAAGAAAACAGAATANNNAGATCGGAA
124909 TATGAGTGTGGAGTCGTTGCATCTTTTTGTTCTCTTCATAGTGNNNAGATCGGAAGA
124910 TATGAGTGTGGAGTCGTTGCACACACAGAGAAACAGATAGANNNAGATCGGAAGA
124911 TATGAGTGTGGAGTCGTTGCTATGGAGTAGGAATGTGAGATTANNNAGATCGGAA
124912 TATGAGTGTGGAGTCGTTGCAGAAACAGATGAAAACAATGTATNNNAGATCGGAA
124913 TATGAGTGTGGAGTCGTTGCGGTTCGGTATTATTTTCTTCAATNNNAGATCGGAAG
124915 TATGAGTGTGGAGTCGTTGCCTCAATGATCCAAATAACTATGGNNNAGATCGGAAG
124918 TATGAGTGTGGAGTCGTTGCTACACATATACTAGATACCAACANNNAGATCGGAAG
124919 TATGAGTGTGGAGTCGTTGCTTATCTTACACTTCTAGAGGTTGNNNAGATCGGAAG
124920 TATGAGTGTGGAGTCGTTGCTGTAATTTTTATCTCTGTTCCTCNNNAGATCGGAAGA
124921 TATGAGTGTGGAGTCGTTGCCCCTCAAATTCCTATGGTAAANNNAGATCGGAAGAG
124922 TATGAGTGTGGAGTCGTTGCCTACATTATCTCCATTCTATGCTNNNAGATCGGAAGA
124923 TATGAGTGTGGAGTCGTTGCGAGGTGATGAAATACAAAGAATANNNAGATCGGAA
124924 TATGAGTGTGGAGTCGTTGCTTCTACTACATGAAAGAGTTTCTNNNAGATCGGAAG
124925 TATGAGTGTGGAGTCGTTGCAATAGTATCTTCAGGTACTAGGANNNAGATCGGAAG
124926 TATGAGTGTGGAGTCGTTGCTATCTTAAGCAATTTGATGTGAGNNNAGATCGGAAG
124927 TATGAGTGTGGAGTCGTTGCCTTCTGAAATTTACATGAACTGANNNAGATCGGAAG
124928 TATGAGTGTGGAGTCGTTGCAGACAGTATCTCACCATATTTCNNNAGATCGGAAGA
124929 TATGAGTGTGGAGTCGTTGCATATAAAGTAGATGGTCCTGTTGNNNAGATCGGAA
124930 TATGAGTGTGGAGTCGTTGCCTCAGATTGTAGAGCAAAATATCNNNAGATCGGAAG
124931 TATGAGTGTGGAGTCGTTGCTCCGTGTGTATTATGGGATGNNNAGATCGGAAGAG
124932 TATGAGTGTGGAGTCGTTGCTTCCACAGAGAAAAATCAGGNNNAGATCGGAAGAG
124933 TATGAGTGTGGAGTCGTTGCATTAACACTTAATTCCACAGGTTNNNAGATCGGAAG
124934 TATGAGTGTGGAGTCGTTGCAAATTCTCTAGAAGAAAGAGTGTNNNAGATCGGAA
124935 TATGAGTGTGGAGTCGTTGCTAGAATCCATAAATTAGTGCAGANNNAGATCGGAAG
124936 TATGAGTGTGGAGTCGTTGCGTATCTGGTCAGGGGAAATCNNNAGATCGGAAGAG
124937 TATGAGTGTGGAGTCGTTGCATAGTTTCACCAAAACAAATAGGNNNAGATCGGAAG
124939 TATGAGTGTGGAGTCGTTGCCCAAGACAGAAAAAATACAACTTNNNAGATCGGAAG
124940 TATGAGTGTGGAGTCGTTGCCAGAAACCACAGTTTTTTTAAANNNAGATCGGAAG
124941 TATGAGTGTGGAGTCGTTGCCAGACAGAATAATTAGCTAATCGNNNAGATCGGAAG
124942 TATGAGTGTGGAGTCGTTGCTATACACACAGAGAGAGAGAATANNNAGATCGGAA
124943 TATGAGTGTGGAGTCGTTGCATGTATGTATGGAGAAAGGAATTNNNAGATCGGAA
124944 TATGAGTGTGGAGTCGTTGCGAACATAGATTCCTTCTCCAATNNNAGATCGGAAGA
124945 TATGAGTGTGGAGTCGTTGCAAACCAGTGACTGAAATATTATGNNNAGATCGGAAG
124946 TATGAGTGTGGAGTCGTTGCCTGAGTGATAATGCTGTGAGNNNAGATCGGAAGAG
124947 TATGAGTGTGGAGTCGTTGCGAATTTGCAATTGGTGATGAANNNAGATCGGAAGA
124948 TATGAGTGTGGAGTCGTTGCAGTTTTAGGTGTGACAATAAAAGNNNAGATCGGAA
124949 TATGAGTGTGGAGTCGTTGCGGATAAATTACCCCCAATTATTTNNNAGATCGGAAG
124950 TATGAGTGTGGAGTCGTTGCTAGGCTAATCAATTATGTGCAANNNAGATCGGAAGA
124951 TATGAGTGTGGAGTCGTTGCATAATCAGCTTATGTTGGTCTATNNNAGATCGGAAG
124952 TATGAGTGTGGAGTCGTTGCGAGATTGTGAAAACATATTCCTTNNNAGATCGGAAG
124953 TATGAGTGTGGAGTCGTTGCTAGATGAAAAAGCAGATACAAGANNNAGATCGGAA
124954 TATGAGTGTGGAGTCGTTGCAATCATATATAAAACCATGTGGCNNNAGATCGGAAG

124955 TATGAGTGTGGAGTCGTTGCATGTGGGACTATATCTTCAAAAANNNAGATCGGAAG
124956 TATGAGTGTGGAGTCGTTGCAAGCAATGACTAATTCTTGAAAGNNNAGATCGGAAG
124957 TATGAGTGTGGAGTCGTTGCCCAGAATCTTTCTTCACTTTTTNNNAGATCGGAAGAG
124958 TATGAGTGTGGAGTCGTTGCTAAGGAAACCCTCAAGAAAAATANNNAGATCGGAA
124959 TATGAGTGTGGAGTCGTTGCCACATTTTTTCAAGAGACAGGNNNAGATCGGAAGAG
124960 TATGAGTGTGGAGTCGTTGCAGGAATTCCTTCTGTTTCACNNNAGATCGGAAGAGC
124961 TATGAGTGTGGAGTCGTTGCCTATGACTTGTACTTTTTCTTGANNNAGATCGGAAGA
124962 TATGAGTGTGGAGTCGTTGCCAGAATAGTCAAAACAATCTTGANNNAGATCGGAAG
124963 TATGAGTGTGGAGTCGTTGCTTATCCCAAACATAACACAAAAANNNAGATCGGAAG
124964 TATGAGTGTGGAGTCGTTGCATGGGTAAAAGTAAGCACATTNNNAGATCGGAAGA
124965 TATGAGTGTGGAGTCGTTGCGAAGAAGAAGAAGAAGAAACATTNNNAGATCGGA
124966 TATGAGTGTGGAGTCGTTGCCAATCAGTCAATGTTTAGTCAAGNNNAGATCGGAAG
124967 TATGAGTGTGGAGTCGTTGCTTGTTGTTGTTATTTACCCCTNNNAGATCGGAAGAG
124968 TATGAGTGTGGAGTCGTTGCCTGCTCTCAAATATTGGATTAAANNNAGATCGGAAG
124969 TATGAGTGTGGAGTCGTTGCTCTATGAAGATTAGTCACCTTTTNNNAGATCGGAAG
124970 TATGAGTGTGGAGTCGTTGCACATCAAACATATTACACCTTTCNNNAGATCGGAAGA
124971 TATGAGTGTGGAGTCGTTGCGATAGACTATGAGATCCACATTTNNNAGATCGGAAG
124972 TATGAGTGTGGAGTCGTTGCCTGGGTTAAAAATGTATGCATATNNNAGATCGGAAG
124973 TATGAGTGTGGAGTCGTTGCAGAAACCTAAGTTACTCTTCATTNNNAGATCGGAAG
124974 TATGAGTGTGGAGTCGTTGCTTCTTTATGGTTTATCTCCTCATNNNAGATCGGAAGA
124975 TATGAGTGTGGAGTCGTTGCATTAGGACTCTATTTTTAGGACCNNNAGATCGGAAG
124976 TATGAGTGTGGAGTCGTTGCTAAACCTTACCTGAATTCTGATCNNNAGATCGGAAGA
124977 TATGAGTGTGGAGTCGTTGCTGTTCATTTTTTAGAACTGTCTTNNNAGATCGGAAGA
124978 TATGAGTGTGGAGTCGTTGCGCTGAGACCCATGATAATATAAANNNAGATCGGAAG
124979 TATGAGTGTGGAGTCGTTGCACATCAAACATATTACACCTTTCNNNAGATCGGAAGA
124980 TATGAGTGTGGAGTCGTTGCTTGTTATCAGGAACATCCCANNNAGATCGGAAGAGC
124981 TATGAGTGTGGAGTCGTTGCTCATAAATCTGTGCATTTTCTAGNNNAGATCGGAAG
124982 TATGAGTGTGGAGTCGTTGCTCTTCTCTTTATAAGATTTCCTACTNNNAGATCGGAA
124983 TATGAGTGTGGAGTCGTTGCCATCTTGGGGTTGTTATCAGNNNAGATCGGAAGAG
124984 TATGAGTGTGGAGTCGTTGCTACATGAAGGTCTCCAGATTTANNNAGATCGGAAGA
124985 TATGAGTGTGGAGTCGTTGCAGTTACCAAGAAAGAAAATTGTTNNNAGATCGGAA
124986 TATGAGTGTGGAGTCGTTGCTCTGTTAATGCTATGTAGTTACTNNNAGATCGGAAG
124987 TATGAGTGTGGAGTCGTTGCGCTAAGATTTCTGATTGACTTTANNNAGATCGGAAG
124988 TATGAGTGTGGAGTCGTTGCAGATAGATAGATAGATAGAGAGAGANNNAGATCG
124989 TATGAGTGTGGAGTCGTTGCTAGGGGATTGTATACTGTCTTNNNAGATCGGAAGA
124990 TATGAGTGTGGAGTCGTTGCAAATAAATAGTCAAAGTTTGGGGNNNAGATCGGAA
124991 TATGAGTGTGGAGTCGTTGCCCATACTCTGTCTACCCTACNNNAGATCGGAAGAGCA
124992 TATGAGTGTGGAGTCGTTGCAATATACATTTTTTTCCGCACCNNNAGATCGGAAGAG
124993 TATGAGTGTGGAGTCGTTGCCAACAGCAGATTTTTTTTTTCTTNNNAGATCGGAAGA
124994 TATGAGTGTGGAGTCGTTGCGCTACCTCAGTATCTTGATATAANNNAGATCGGAAG
124995 TATGAGTGTGGAGTCGTTGCTAATAGTCATATTTCCACTTGGTNNNAGATCGGAAG
124996 TATGAGTGTGGAGTCGTTGCGGTGTACATTTATGTTGTGTTTANNNAGATCGGAAG
124997 TATGAGTGTGGAGTCGTTGCATCAATATTTTCCATGGAATTGTNNNAGATCGGAAG

124998 TATGAGTGTGGAGTCGTTGCCAATAGGTACATTTTACAGACAGNNNAGATCGGAAG
124999 TATGAGTGTGGAGTCGTTGCAATGTAAAATGACACAGCCATTANNNAGATCGGAAG
125000 TATGAGTGTGGAGTCGTTGCTAGCAGAATTGTGGTAAATTTTTNNNAGATCGGAAG
125001 TATGAGTGTGGAGTCGTTGCCTCTTTCTACAGTCTCCCTCNNNAGATCGGAAGAGCA
125002 TATGAGTGTGGAGTCGTTGCATATATCCCTTTCAATGGACTTTNNNAGATCGGAAGA
125003 TATGAGTGTGGAGTCGTTGCGAAAGAGAAGGACAGAAAGAGNNNAGATCGGAAG
125004 TATGAGTGTGGAGTCGTTGCGCTGAACACTTGTAAGAATAAATNNNAGATCGGAAG
125005 TATGAGTGTGGAGTCGTTGCATATCCTGGAGTTTTAGGTTGNNNAGATCGGAAGAG
125007 TATGAGTGTGGAGTCGTTGCGATAGACAGAGAGATAGAGAATGNNNAGATCGGAA
125008 TATGAGTGTGGAGTCGTTGCCTGAACCTCCAATATTTTTCAAGNNNAGATCGGAAGA
125010 TATGAGTGTGGAGTCGTTGCATTAGACATGTAGAGACAGAGANNNAGATCGGAAG
125011 TATGAGTGTGGAGTCGTTGCAGTGTATGAAAAAAACAAAAACANNNAGATCGGAA
125012 TATGAGTGTGGAGTCGTTGCCATGTATCTGAAACATATGATCTNNNAGATCGGAAG
125013 TATGAGTGTGGAGTCGTTGCAAATTTCTGTCTAAAGGATCACTNNNAGATCGGAAG
125014 TATGAGTGTGGAGTCGTTGCTGATCCTAAAAGCAATACAAAAANNNAGATCGGAAG
125015 TATGAGTGTGGAGTCGTTGCGAGCTGAAGAAAATTTTTTCAAANNNAGATCGGAAG
125017 TATGAGTGTGGAGTCGTTGCAAGACAGGAGAAAGTAATCAATANNNAGATCGGAA
125018 TATGAGTGTGGAGTCGTTGCATATAAAAGAAACTGAGCAAACANNNAGATCGGAA
125019 TATGAGTGTGGAGTCGTTGCATAAAATCAGTTCTGTGGCTATANNNAGATCGGAAG
125020 TATGAGTGTGGAGTCGTTGCGGGGAGAAAATATTCTGTACTATNNNAGATCGGAA
125021 TATGAGTGTGGAGTCGTTGCGTTTTCTTGAATTGTAGCTCCNNNAGATCGGAAGAG
125022 TATGAGTGTGGAGTCGTTGCAGCTATTTTCCCCTAATCATATTNNNAGATCGGAAGA
125023 TATGAGTGTGGAGTCGTTGCGATTTTATTCCTTCCCTTTTTCANNNAGATCGGAAGA
125024 TATGAGTGTGGAGTCGTTGCATTTTGGGGGTTACTATCTTTAGNNNAGATCGGAAG
125025 TATGAGTGTGGAGTCGTTGCATTTTGGGGGTTACTATCTTTAGNNNAGATCGGAAG
125026 TATGAGTGTGGAGTCGTTGCTTCAGAATGTATGTGTGTGTGNNNAGATCGGAAGA
125027 TATGAGTGTGGAGTCGTTGCTAAACATAGTAATACTGACCCTCNNNAGATCGGAAG
125028 TATGAGTGTGGAGTCGTTGCCTTATACAGGTCCATTGAAGGNNNAGATCGGAAGAG
125029 TATGAGTGTGGAGTCGTTGCGTGTGTTCCAAGACTATATTCTNNNAGATCGGAAGA
125030 TATGAGTGTGGAGTCGTTGCCAAAATGAACAAATAAACCGATTNNNAGATCGGAAG
125031 TATGAGTGTGGAGTCGTTGCTTTGAAATGCATTTCTCACTCNNNAGATCGGAAGAG
125032 TATGAGTGTGGAGTCGTTGCTGCTTCATTTCTCTGTAATAAAGNNNAGATCGGAAG
125033 TATGAGTGTGGAGTCGTTGCTTCATATCAGAATGAGACAGAGNNNAGATCGGAAG
125034 TATGAGTGTGGAGTCGTTGCCCAAATATGTTGTTTTCTGACAANNNAGATCGGAAG
125035 TATGAGTGTGGAGTCGTTGCACACACATATCCAAACAACANNNAGATCGGAAGAGC
125036 TATGAGTGTGGAGTCGTTGCCTTTTGGGCTCTCTATAACAANNNAGATCGGAAGAG
125037 TATGAGTGTGGAGTCGTTGCAACTTCCTTGAAAAATGTTGAANNNAGATCGGAAGA
125038 TATGAGTGTGGAGTCGTTGCGTTTGTCTGCATTATATGTGTAGNNNAGATCGGAAG
125039 TATGAGTGTGGAGTCGTTGCTACTTGCATTTAAATGAAGCCTANNNAGATCGGAAG
125040 TATGAGTGTGGAGTCGTTGCCAATCTACCATCTCAAGCTATAGNNNAGATCGGAAG
125041 TATGAGTGTGGAGTCGTTGCATCTTTGCTTCTTAGGAAAAAAANNNAGATCGGAAG
125042 TATGAGTGTGGAGTCGTTGCATATCCAATCTCACCCTTTTTATNNNAGATCGGAAGA
125043 TATGAGTGTGGAGTCGTTGCCTGCTAGATAACATGGATTATTCNNNAGATCGGAAG

125044 TATGAGTGTGGAGTCGTTGCGTCAGTGATCATTTAGTCATTTCNNNAGATCGGAAG
125045 TATGAGTGTGGAGTCGTTGCGGTTCCTGAAGATTTATTTTCTTNNNAGATCGGAAG
125046 TATGAGTGTGGAGTCGTTGCCACCAAAAAAAAAAAAATGCTAANNNAGATCGGAAG
125047 TATGAGTGTGGAGTCGTTGCATATTTGCTTTGTCCATTTATCCNNNAGATCGGAAGA
125048 TATGAGTGTGGAGTCGTTGCAGAAACCTGGAAATACACATTNNNAGATCGGAAGAG
125049 TATGAGTGTGGAGTCGTTGCCATTATCCTTAGCAAACTAACGNNNAGATCGGAAGA
125050 TATGAGTGTGGAGTCGTTGCAAACCTTCAGAGAGAGACTTTTNNNAGATCGGAAGA
125051 TATGAGTGTGGAGTCGTTGCCTTCTTCTGTACCAAATCACTNNNAGATCGGAAGAGC
125052 TATGAGTGTGGAGTCGTTGCTTATAACAGGTCTGAAACAAAAGNNNAGATCGGAAG
125053 TATGAGTGTGGAGTCGTTGCAGATTCAAGTGATTCTTCTCCNNNAGATCGGAAGAG
125054 TATGAGTGTGGAGTCGTTGCGTCATTGTGTGTGATGTGATNNNAGATCGGAAGAG
125055 TATGAGTGTGGAGTCGTTGCTTTGTAATCAATACAGGTTTGACNNNAGATCGGAAG
125057 TATGAGTGTGGAGTCGTTGCCTTTAACTTCAAGTTCACTGTCNNNAGATCGGAAGAG
125058 TATGAGTGTGGAGTCGTTGCAGCTGTATTCATTCCCACTTNNNAGATCGGAAGAGC
125059 TATGAGTGTGGAGTCGTTGCTGTATGAAATCACATCAGTTAGANNNAGATCGGAAG
125060 TATGAGTGTGGAGTCGTTGCTATATCTTTGGAAAAATTTGGGGNNNAGATCGGAAG
125061 TATGAGTGTGGAGTCGTTGCTGAGGTTTCAGAGAATGTATATTNNNAGATCGGAAG
125062 TATGAGTGTGGAGTCGTTGCGAAATGACAGTCACTCTTTATTTNNNAGATCGGAAG
125063 TATGAGTGTGGAGTCGTTGCCTCAGGAAATATCCATCCTTCNNNAGATCGGAAGAG
125064 TATGAGTGTGGAGTCGTTGCTTTTCCTGACCGTCTTTTTGNNNAGATCGGAAGAGC
125065 TATGAGTGTGGAGTCGTTGCACAGCTATTATAGAAAAACAGTCNNNAGATCGGAAG
125066 TATGAGTGTGGAGTCGTTGCTGAGTTTATACTTCATTTTTGGGNNNAGATCGGAAG
125067 TATGAGTGTGGAGTCGTTGCGGAAAAGAGAATAGATTGGAAAANNNAGATCGGA
125068 TATGAGTGTGGAGTCGTTGCGAAAGAGAGAGAGAGAGAGATANNNAGATCGGAA
125069 TATGAGTGTGGAGTCGTTGCAATGGAGAGACTTAATAGTTGACNNNAGATCGGAA
125070 TATGAGTGTGGAGTCGTTGCAATCGTATTAGAAAGATTGAGGANNNAGATCGGAA
125071 TATGAGTGTGGAGTCGTTGCTGTCAATAGCCATCTTTGTTANNNAGATCGGAAGAG
125072 TATGAGTGTGGAGTCGTTGCCATGATGGATTTCTCTAGACTAANNNAGATCGGAAG
125073 TATGAGTGTGGAGTCGTTGCAAGGTTGCAAATATTTCACAATANNNAGATCGGAAG
125074 TATGAGTGTGGAGTCGTTGCTGCTAACAGTGTTTACCTTGNNNAGATCGGAAGAGC
125075 TATGAGTGTGGAGTCGTTGCTTATATGCATGTAAGTTAGCATGNNNAGATCGGAAG
125076 TATGAGTGTGGAGTCGTTGCCACACATATTCAAGCAAAAAAATNNNAGATCGGAAG
125077 TATGAGTGTGGAGTCGTTGCGATTGTGATTGGAATCTTTTTTTNNNAGATCGGAAG
125078 TATGAGTGTGGAGTCGTTGCTTCTAGTCTTTTTCTTTGTCTTCNNNAGATCGGAAGA
125079 TATGAGTGTGGAGTCGTTGCAGATTCTAATTTGTCAAATGTGGNNNAGATCGGAAG
125082 TATGAGTGTGGAGTCGTTGCTTCACATCCTTGGAGATTATAAGNNNAGATCGGAAG
125083 TATGAGTGTGGAGTCGTTGCCTCTTCTCTATGCTCTTACCTNNNAGATCGGAAGAGC
125084 TATGAGTGTGGAGTCGTTGCTGTTGTTTTCCCTTTATTTCAAANNNAGATCGGAAGA
125085 TATGAGTGTGGAGTCGTTGCTTCAAGCAGAGAACTTAAAGAANNNAGATCGGAAG
125086 TATGAGTGTGGAGTCGTTGCGTGTACACATATGAAAGGGGNNNAGATCGGAAGAG
125087 TATGAGTGTGGAGTCGTTGCGTGTACACATATGAAAGGGGNNNAGATCGGAAGAG
125088 TATGAGTGTGGAGTCGTTGCTATGATGTCAGTATTTATGAGGTNNNAGATCGGAAG
125089 TATGAGTGTGGAGTCGTTGCACACACATATATATACACACACANNNAGATCGGAAG

125091 TATGAGTGTGGAGTCGTTGCCTTGGAACCTTCAGAGAATTAANNNAGATCGGAAGA
125092 TATGAGTGTGGAGTCGTTGCAAATTCAATACCACAGAAAGTAANNNAGATCGGAAG
125093 TATGAGTGTGGAGTCGTTGCGCAGAATGGGATATAGCATATATNNNAGATCGGAA
125094 TATGAGTGTGGAGTCGTTGCTTTCTTTATCCTTTCATCTGTTGNNNAGATCGGAAGA
125095 TATGAGTGTGGAGTCGTTGCAGAGGAGAGAGAAAGAGTATATANNNAGATCGGA
125096 TATGAGTGTGGAGTCGTTGCTAGAGACCTACATTCTAAGGCNNNAGATCGGAAGAG
125097 TATGAGTGTGGAGTCGTTGCATTCTTGAAGCTACAGGTTTATANNNAGATCGGAAG
125098 TATGAGTGTGGAGTCGTTGCTCAACCTTTAATGTACATACAAANNNAGATCGGAAG
125099 TATGAGTGTGGAGTCGTTGCTACAAAACAAGGGAAATTAGGTANNNAGATCGGAA
125100 TATGAGTGTGGAGTCGTTGCACAGAGACAGAGAAATTGAAATANNNAGATCGGAA
125101 TATGAGTGTGGAGTCGTTGCACAGAGACAGAGAAATTGAAATANNNAGATCGGAA
125102 TATGAGTGTGGAGTCGTTGCTATTCTCTTCACCACAATGAATNNNAGATCGGAAGAG
125103 TATGAGTGTGGAGTCGTTGCTTAAAGGGAAGACCAAGTATTATNNNAGATCGGAA
125104 TATGAGTGTGGAGTCGTTGCGAGTACGTTGGCATGATTACNNNAGATCGGAAGAG
125105 TATGAGTGTGGAGTCGTTGCATATGATTTTAGACAGACACGTNNNAGATCGGAAGA
125106 TATGAGTGTGGAGTCGTTGCATATGATTTTAGACAGACACGTNNNAGATCGGAAGA
125107 TATGAGTGTGGAGTCGTTGCCATTACTTGTAACTCCTCTTTTTNNNAGATCGGAAGA
125108 TATGAGTGTGGAGTCGTTGCTAGGACTGACAACCTTATTATGNNNAGATCGGAAGA
125109 TATGAGTGTGGAGTCGTTGCATTATCTTTATGGGTAGAAAGCANNNAGATCGGAAG
125110 TATGAGTGTGGAGTCGTTGCGTAGAAAGAATGGACATATATGCNNNAGATCGGAA
125111 TATGAGTGTGGAGTCGTTGCGATTTTGGGAGATAAGACTTTTGNNNAGATCGGAA
125112 TATGAGTGTGGAGTCGTTGCTTAGAAGTTATTTTGGGGTCATANNNAGATCGGAAG
125113 TATGAGTGTGGAGTCGTTGCGAGAAAGCTGAAAATCAAAAACNNNAGATCGGAAG
125114 TATGAGTGTGGAGTCGTTGCGACAAAGTATTTAATGAGCCATCNNNAGATCGGAAG
125115 TATGAGTGTGGAGTCGTTGCTACACACATTCTTTTATAGAGGANNNAGATCGGAAG
125116 TATGAGTGTGGAGTCGTTGCTATGCCTGACACATAAGCATNNNAGATCGGAAGAGC
125117 TATGAGTGTGGAGTCGTTGCCTACTTCCATGACTTCTGAAGNNNAGATCGGAAGAG
125118 TATGAGTGTGGAGTCGTTGCCTACTTCCATGACTTCTGAAGNNNAGATCGGAAGAG
125119 TATGAGTGTGGAGTCGTTGCCAAAGATAGGATTTTATCAGGTCNNNAGATCGGAAG
125120 TATGAGTGTGGAGTCGTTGCTAATGAGAGACATTTTAGAAGCTNNNAGATCGGAAG
125122 TATGAGTGTGGAGTCGTTGCATTTTTAGTAGAGATGGGGTTTNNNAGATCGGAAG
125123 TATGAGTGTGGAGTCGTTGCTTAGATTTTGAAGGTAGTGAGAANNNAGATCGGAA
125124 TATGAGTGTGGAGTCGTTGCATGAATGCCATGATTTCTCANNNAGATCGGAAGAGC
125125 TATGAGTGTGGAGTCGTTGCAACTGTCTCTCTAATAATTGGTTNNNAGATCGGAAG
125126 TATGAGTGTGGAGTCGTTGCGAAAAGATGATTGCATGCAAANNNAGATCGGAAGA
125127 TATGAGTGTGGAGTCGTTGCGAAAAGATGATTGCATGCAAANNNAGATCGGAAGA
125128 TATGAGTGTGGAGTCGTTGCGACTATTACCTCCTACTTGATGNNNAGATCGGAAGA
125129 TATGAGTGTGGAGTCGTTGCAGATGACTGTATTCCACATTATTNNNAGATCGGAAG
125130 TATGAGTGTGGAGTCGTTGCCTGGAAGTTTCTTACTGAAAGANNNAGATCGGAAGA
125131 TATGAGTGTGGAGTCGTTGCTGAAATCAGTTCTTTTTGTGATANNNAGATCGGAAG
125132 TATGAGTGTGGAGTCGTTGCTGAAATCAGTTCTTTTTGTGATANNNAGATCGGAAG
125133 TATGAGTGTGGAGTCGTTGCTGAAATCAGTTCTTTTTGTGATANNNAGATCGGAAG
125134 TATGAGTGTGGAGTCGTTGCTTTTAAAATGGAAAAATTCAGGCNNNAGATCGGAA

125135 TATGAGTGTGGAGTCGTTGCGGGGAGTAATTATTGGCTATTAANNNAGATCGGAA
125136 TATGAGTGTGGAGTCGTTGCCATTTGGTAATGTCTGGAGACNNNAGATCGGAAGA
125137 TATGAGTGTGGAGTCGTTGCGATGGCAACAAGGAGAATAGNNNAGATCGGAAGA
125138 TATGAGTGTGGAGTCGTTGCGCAACTCTAATTTGGGAATTAAANNNAGATCGGAAG
125139 TATGAGTGTGGAGTCGTTGCGTTTGTTTTCATGTGGTTTTGNNNAGATCGGAAGAG
125140 TATGAGTGTGGAGTCGTTGCAATGCTTAGAAGAATTAAGAAGCNNNAGATCGGAA
125141 TATGAGTGTGGAGTCGTTGCTTACTTTCTGCCAAATAGTATCTNNNAGATCGGAAGA
125142 TATGAGTGTGGAGTCGTTGCAAAGAGAAATAATTCACCTTTCCNNNAGATCGGAAG
125143 TATGAGTGTGGAGTCGTTGCCTTCATTCTTTTCTCAAATCTCTNNNAGATCGGAAGA
125144 TATGAGTGTGGAGTCGTTGCTTTCTCTTCTCTCTTCTTTCTTTNNNAGATCGGAAGAG
125145 TATGAGTGTGGAGTCGTTGCTTTCTCTTCTCTCTTCTTTCTTTNNNAGATCGGAAGAG
125146 TATGAGTGTGGAGTCGTTGCCATCTTTCTCTGTGATAAACAAGNNNAGATCGGAAG
125147 TATGAGTGTGGAGTCGTTGCCATTACCAGAAGTTGAAATCTTNNNAGATCGGAAGA
125148 TATGAGTGTGGAGTCGTTGCCAGCATAAACACACATATTCTATNNNAGATCGGAAG
125149 TATGAGTGTGGAGTCGTTGCTGGTATTTTTAATTTTAAGCCTGANNNAGATCGGAA
125150 TATGAGTGTGGAGTCGTTGCCCTTTGTGGAGACATAATATTTTNNNAGATCGGAAG
125151 TATGAGTGTGGAGTCGTTGCAGTACCACACTTTCATCCATNNNAGATCGGAAGAGC
125152 TATGAGTGTGGAGTCGTTGCTGAAAATCTTTAAAATGCAACAGNNNAGATCGGAAG
125153 TATGAGTGTGGAGTCGTTGCTTTTTATGCATTCCTGTCCTCNNNAGATCGGAAGAGC
125154 TATGAGTGTGGAGTCGTTGCAGAAGGCAGACAGGTTTATTNNNAGATCGGAAGAG
125155 TATGAGTGTGGAGTCGTTGCTTTTGCGTATTTTAACAGAAGATNNNAGATCGGAAG
125156 TATGAGTGTGGAGTCGTTGCCTAGGAATACATCTAACCAAGAGNNNAGATCGGAA
125157 TATGAGTGTGGAGTCGTTGCGCATTTAACAAGTTTTGATACAGNNNAGATCGGAAG
125158 TATGAGTGTGGAGTCGTTGCCTACTTACAGGAAATGTATGAGANNNAGATCGGAA
125159 TATGAGTGTGGAGTCGTTGCTAAGATCACCACAGATCCTATTNNNAGATCGGAAGA
125160 TATGAGTGTGGAGTCGTTGCAAACCAGAGAAATCAATGTGTNNNAGATCGGAAGA
125162 TATGAGTGTGGAGTCGTTGCCCAACTTGACCTATTTTATATGGNNNAGATCGGAAG
125163 TATGAGTGTGGAGTCGTTGCAAGTCTTTCTTTAGACAACAAAANNNAGATCGGAAG
125164 TATGAGTGTGGAGTCGTTGCGCATATTTTATGGATAAAAACTGGNNNAGATCGGAA
125165 TATGAGTGTGGAGTCGTTGCGAAATTATGTTAAGTTGGACACTNNNAGATCGGAA
125166 TATGAGTGTGGAGTCGTTGCTTCACTTCAGGGAATCATTTTTANNNAGATCGGAAG
125167 TATGAGTGTGGAGTCGTTGCCTTCAGTGAATCCAATCTGATATNNNAGATCGGAAG
125168 TATGAGTGTGGAGTCGTTGCGATAAAAGGTGCAATCTGATAAANNNAGATCGGAA
125169 TATGAGTGTGGAGTCGTTGCAAATTATGTTACAAAGCTACAGCNNNAGATCGGAAG
125170 TATGAGTGTGGAGTCGTTGCCAGAAATAGATGAGGAGAAACCNNNAGATCGGAAG
125171 TATGAGTGTGGAGTCGTTGCTGCAAATCATTTCACAATGTATANNNAGATCGGAAG
125172 TATGAGTGTGGAGTCGTTGCGAGCACTTCTTTTTCCTTTTATNNNAGATCGGAAGAG
125173 TATGAGTGTGGAGTCGTTGCTATTTTAGCAATGCATTTCACTTNNNAGATCGGAAGA
125174 TATGAGTGTGGAGTCGTTGCTATCACAGAGAGTAAGGTAAGTNNNAGATCGGAAG
125175 TATGAGTGTGGAGTCGTTGCCCTTTTTTTTTTGACAAAGTCTNNNAGATCGGAAGA
125176 TATGAGTGTGGAGTCGTTGCCCCTTCTTTTCCATTCTTATAAGNNNAGATCGGAAGA
125177 TATGAGTGTGGAGTCGTTGCATGGCTCCTTTTGTAGAATATATNNNAGATCGGAAG
125178 TATGAGTGTGGAGTCGTTGCAAATGTTCCAAAGCATCTTAAATNNNAGATCGGAAG

125179 TATGAGTGTGGAGTCGTTGCATGAAAACTATTGGCTATAAGGANNNAGATCGGAA
125180 TATGAGTGTGGAGTCGTTGCATGAAAACTATTGGCTATAAGGANNNAGATCGGAA
125181 TATGAGTGTGGAGTCGTTGCTTCTGTTGTAAAAGATCTGTCTANNNAGATCGGAAG
125182 TATGAGTGTGGAGTCGTTGCATTGAAATTGTCTGCAATACTTTNNNAGATCGGAAG
125183 TATGAGTGTGGAGTCGTTGCCAACTTAACATATTGCTTTAGGTNNNAGATCGGAAG
125184 TATGAGTGTGGAGTCGTTGCCAGAACCTGAGTATACCAAAANNNAGATCGGAAGA
125185 TATGAGTGTGGAGTCGTTGCCCAATAACAATCACTTCCATAATNNNAGATCGGAAGA
125186 TATGAGTGTGGAGTCGTTGCTGAAATGCACATTTACTTGATATNNNAGATCGGAAG
125187 TATGAGTGTGGAGTCGTTGCAACATATCAGCAGCTTAAAATCNNNAGATCGGAAGA
125188 TATGAGTGTGGAGTCGTTGCAAAGGTCCTTGACTAACAAAANNNAGATCGGAAGA
125189 TATGAGTGTGGAGTCGTTGCACATCATTTTCTTTCTTCAACGNNNAGATCGGAAGAG
125190 TATGAGTGTGGAGTCGTTGCGTCTCTCTAGTTTCCAGTTGNNNAGATCGGAAGAGC
125191 TATGAGTGTGGAGTCGTTGCGGAGTTTATTAGGAGAATTGACTNNNAGATCGGAA
125192 TATGAGTGTGGAGTCGTTGCTAGGGTTTTCTACATATAAGGTCNNNAGATCGGAAG
125193 TATGAGTGTGGAGTCGTTGCTATCTCCTAAGAATGAGGACATTNNNAGATCGGAAG
125194 TATGAGTGTGGAGTCGTTGCCTTCACTTTTTAATAACATCACGNNNAGATCGGAAGA
125196 TATGAGTGTGGAGTCGTTGCTAAAGGGTAATGTCAATTTCAAGNNNAGATCGGAA
125197 TATGAGTGTGGAGTCGTTGCTTTTTCTTGGTCAGTCTTCAANNNAGATCGGAAGAG
125198 TATGAGTGTGGAGTCGTTGCAAGTAGATGAACAAGAGGTTTTANNNAGATCGGAA
125199 TATGAGTGTGGAGTCGTTGCTGCAGAATTTTTACTGATCAATTNNNAGATCGGAAG
125200 TATGAGTGTGGAGTCGTTGCGCATCATTCTAGACTTTTCTTCNNNAGATCGGAAGAG
125201 TATGAGTGTGGAGTCGTTGCCTACAGATTAATGTTGCCTCTNNNAGATCGGAAGAG
125202 TATGAGTGTGGAGTCGTTGCATGCTATTTGTCAAGAGTAAAGANNNAGATCGGAA
125203 TATGAGTGTGGAGTCGTTGCTACAAGAAGTCAATCAGATATCCNNNAGATCGGAAG
125204 TATGAGTGTGGAGTCGTTGCATGAAAGCTTCCCCTAGAAGNNNAGATCGGAAGAG
125205 TATGAGTGTGGAGTCGTTGCATGAAAGCTTCCCCTAGAAGNNNAGATCGGAAGAG
125206 TATGAGTGTGGAGTCGTTGCCATTATCTTTACTCCTAACCCTCNNNAGATCGGAAGA
125207 TATGAGTGTGGAGTCGTTGCGATAGATGATAAAGTTGAGAAGCNNNAGATCGGAA
125208 TATGAGTGTGGAGTCGTTGCCATAGACACTGTCAGATGTACNNNAGATCGGAAGAG
125209 TATGAGTGTGGAGTCGTTGCAAAATAAAAAGTAGAAGTTGGGGNNNAGATCGGAA
125210 TATGAGTGTGGAGTCGTTGCAACAAATATTGAGTGAATCAGACNNNAGATCGGAA
125211 TATGAGTGTGGAGTCGTTGCTCAAAGGACATTCCATCTAATATNNNAGATCGGAAG
125212 TATGAGTGTGGAGTCGTTGCCAGAACAATGAGCTACATAAATTNNNAGATCGGAAG
125213 TATGAGTGTGGAGTCGTTGCGGTATAATGGGGAAGGAAGGNNNAGATCGGAAGA
125214 TATGAGTGTGGAGTCGTTGCCTGACTTTGCTATTGTGAATAGNNNAGATCGGAAGA
125215 TATGAGTGTGGAGTCGTTGCTTGGCTTTCTTCATTTTCATTTANNNAGATCGGAAGA
125216 TATGAGTGTGGAGTCGTTGCCAAAGGCTGACTTTTACATTAANNNAGATCGGAAGA
125217 TATGAGTGTGGAGTCGTTGCTGAATGAATCTAAACTATTTTGCTNNNAGATCGGAA
125218 TATGAGTGTGGAGTCGTTGCTATTCTATTTTGCTAAGCCCTAANNNAGATCGGAAGA
125219 TATGAGTGTGGAGTCGTTGCCAATTCATTAGGGCATTTTTAAANNNAGATCGGAAG
125220 TATGAGTGTGGAGTCGTTGCTTATACTTCACTTGGAACCATATNNNAGATCGGAAGA
125222 TATGAGTGTGGAGTCGTTGCGCAAAAAATGTTACTGTTCTCTTNNNAGATCGGAAG
125223 TATGAGTGTGGAGTCGTTGCGTGAGCATACTTCTTTCATTTCNNNAGATCGGAAGA

125224 TATGAGTGTGGAGTCGTTGCCATATCACAAGGATGTACACCNNNAGATCGGAAGAG
125225 TATGAGTGTGGAGTCGTTGCCTAGTATTTGCATACAACCTATGNNNAGATCGGAAG
125226 TATGAGTGTGGAGTCGTTGCTAAAATTGATTGTGATGATGGTTNNNAGATCGGAA
125227 TATGAGTGTGGAGTCGTTGCCACCTTAGAAAAGTTTTCTGTAANNNAGATCGGAAG
125228 TATGAGTGTGGAGTCGTTGCTTTAGAAAAAAAAAAGCACAGTCNNNAGATCGGAA
125229 TATGAGTGTGGAGTCGTTGCCTACTTCTATCCCAAGAGACTNNNAGATCGGAAGAG
125230 TATGAGTGTGGAGTCGTTGCAGGAAAAAATACATCTACTCCATNNNAGATCGGAAG
125231 TATGAGTGTGGAGTCGTTGCAGGAAAAAATACATCTACTCCATNNNAGATCGGAAG
125232 TATGAGTGTGGAGTCGTTGCCTAGGAAAAAATACATCTACTCCNNNAGATCGGAAG
125233 TATGAGTGTGGAGTCGTTGCCTCATGCCTGTAGATACAAATTNNNAGATCGGAAGA
125234 TATGAGTGTGGAGTCGTTGCATTAAAAATTCTGATCACAGTCTNNNAGATCGGAAG
125236 TATGAGTGTGGAGTCGTTGCTTATATAGCCCAGAGATTAGCTANNNAGATCGGAAG
125237 TATGAGTGTGGAGTCGTTGCCCAACATCTAATACACAGAATACNNNAGATCGGAAG
125238 TATGAGTGTGGAGTCGTTGCCTATTTTTCTCACTTGGCAATATNNNAGATCGGAAGA
125239 TATGAGTGTGGAGTCGTTGCGATATTGTTTCTAATATCTCGGGNNNAGATCGGAAG
125240 TATGAGTGTGGAGTCGTTGCATAATAAATGATTGTGCTACTGGNNNAGATCGGAA
125242 TATGAGTGTGGAGTCGTTGCGAATTTTCAGAGCTCTCTACTATNNNAGATCGGAAG
125243 TATGAGTGTGGAGTCGTTGCCAACCTTGTATATGCATTGAATANNNAGATCGGAAG
125244 TATGAGTGTGGAGTCGTTGCACTCAGAATTCACATAGAATACANNNAGATCGGAAG
125245 TATGAGTGTGGAGTCGTTGCCGCTACTCTTAGAATCAACTCNNNAGATCGGAAGAG
125246 TATGAGTGTGGAGTCGTTGCGGAGAATCAGGAGAATTCAATNNNAGATCGGAAGA
125247 TATGAGTGTGGAGTCGTTGCTCAATATTAAGTTTTGGTAGGGANNNAGATCGGAA
125248 TATGAGTGTGGAGTCGTTGCCACATTTATGGATTACGTTTACTNNNAGATCGGAAG
125249 TATGAGTGTGGAGTCGTTGCGGAAGTGGATTAGAAAGTCAGNNNAGATCGGAAG
125250 TATGAGTGTGGAGTCGTTGCCATATGCACCACTTATTTAAGTCNNNAGATCGGAAG
125251 TATGAGTGTGGAGTCGTTGCGTTGCTTTAGATAACCATATGAGNNNAGATCGGAAG
125252 TATGAGTGTGGAGTCGTTGCCTCTGAGTCTTAAAGGAGAGANNNAGATCGGAAGA
125253 TATGAGTGTGGAGTCGTTGCGCAAGCTGTTTAGAAATTTATCTNNNAGATCGGAAG
125254 TATGAGTGTGGAGTCGTTGCAGTGTGACCATAATTCTTAAGTANNNAGATCGGAAG
125255 TATGAGTGTGGAGTCGTTGCGGTAGAATCAGAGGAGGTAGNNNAGATCGGAAGA
125256 TATGAGTGTGGAGTCGTTGCGAACAAACACAAATAAATGCATTNNNAGATCGGAAG
125257 TATGAGTGTGGAGTCGTTGCCTACTGAGTACCTTGAACAATCNNNAGATCGGAAGA
125258 TATGAGTGTGGAGTCGTTGCCTACTGAGTACCTTGAACAATCNNNAGATCGGAAGA
125259 TATGAGTGTGGAGTCGTTGCCACAAATTCTGTACCTTTATCATNNNAGATCGGAAGA
125260 TATGAGTGTGGAGTCGTTGCTTGTTAAAGACTTACAGAACAAGNNNAGATCGGAA
125261 TATGAGTGTGGAGTCGTTGCCCTGGAACTAAGTGAAAACANNNAGATCGGAAGAG
125262 TATGAGTGTGGAGTCGTTGCTATTTGAGGATGAGACCATTATTNNNAGATCGGAAG
125263 TATGAGTGTGGAGTCGTTGCACTGTGTGGTAAAGATAAAAAAGNNNAGATCGGAA
125264 TATGAGTGTGGAGTCGTTGCTCCAATATTGCATCATACAAAATNNNAGATCGGAAG
125265 TATGAGTGTGGAGTCGTTGCCAAGCAAGGATTGTAAAATGTANNNAGATCGGAAG
125266 TATGAGTGTGGAGTCGTTGCCGATTATCAAGTAAAAACGTGAANNNAGATCGGAA
125267 TATGAGTGTGGAGTCGTTGCCCTTTGTTCTCCAAGTTTAAAANNNAGATCGGAAGA
125268 TATGAGTGTGGAGTCGTTGCGAAGTTCAACTCCCTAACAGNNNAGATCGGAAGAG

125269 TATGAGTGTGGAGTCGTTGCTGTCTCCCTAAGATGTATAAAAGNNNAGATCGGAAG
125270 TATGAGTGTGGAGTCGTTGCCCTCTACGATTTTAGGAATCTAANNNAGATCGGAAG
125271 TATGAGTGTGGAGTCGTTGCCCAAAGATTTACGAAATACCTTANNNAGATCGGAAG
125272 TATGAGTGTGGAGTCGTTGCAGAACAAAATAGTAGATGCCATANNNAGATCGGAA
125273 TATGAGTGTGGAGTCGTTGCCAATGATTGGTCTAAGGGTTNNNAGATCGGAAGAG
125274 TATGAGTGTGGAGTCGTTGCAAGAGAGTAGATTTGGATTTCTTNNNAGATCGGAA
125275 TATGAGTGTGGAGTCGTTGCCAGGGAAAGAGGGAGAAATANNNAGATCGGAAGA
125276 TATGAGTGTGGAGTCGTTGCCAGGGAAAGAGGGAGAAATANNNAGATCGGAAGA
125277 TATGAGTGTGGAGTCGTTGCCTTCCATTAGCATATAGGAAAATNNNAGATCGGAAG
125278 TATGAGTGTGGAGTCGTTGCCTTTTTTGGAAGTAAGCAAGAGNNNAGATCGGAAG
125279 TATGAGTGTGGAGTCGTTGCAGAAATTCTCATACAGTAACTGANNNAGATCGGAAG
125280 TATGAGTGTGGAGTCGTTGCCACTAAATTAAGGTTCAGCATAANNNAGATCGGAAG
125281 TATGAGTGTGGAGTCGTTGCTCATCGAGTACTTTTTTCCCNNNAGATCGGAAGAGC
125282 TATGAGTGTGGAGTCGTTGCAGTCTCATTTCTCTGTACTTTTCNNNAGATCGGAAGA
125283 TATGAGTGTGGAGTCGTTGCCTTCATCTTTCTTTTCTGTGTTANNNAGATCGGAAGA
125285 TATGAGTGTGGAGTCGTTGCCTTCTCAGCTTCTGCATAAGNNNAGATCGGAAGAGC
125286 TATGAGTGTGGAGTCGTTGCCAACCAAATGCGACTTTTATTNNNAGATCGGAAGAG
125287 TATGAGTGTGGAGTCGTTGCATAAATGGAATTGTGTTCTTGATNNNAGATCGGAAG
125288 TATGAGTGTGGAGTCGTTGCAGTCTTGAGATAAAACTTTGTTCNNNAGATCGGAAG
125290 TATGAGTGTGGAGTCGTTGCACTTTTAGCTAATAGACCAAAGTNNNAGATCGGAAG
125291 TATGAGTGTGGAGTCGTTGCGAATAGTTTGTTCAGTAATGTGANNNAGATCGGAA
125292 TATGAGTGTGGAGTCGTTGCGACTTCTAGTACTACGTTGAATANNNAGATCGGAAG
125293 TATGAGTGTGGAGTCGTTGCATGAAAGGGAATTTATTAGGGAANNNAGATCGGAA
125294 TATGAGTGTGGAGTCGTTGCATGAAAGGGAATTTATTAGGGAANNNAGATCGGAA
125295 TATGAGTGTGGAGTCGTTGCTATAAGTCATATGGTGAAGCCTANNNAGATCGGAA
125296 TATGAGTGTGGAGTCGTTGCACTTTATTCCCCTATCAGAATTTNNNAGATCGGAAGA
125297 TATGAGTGTGGAGTCGTTGCGATGCAATCATACCTCACTGNNNAGATCGGAAGAGC
125298 TATGAGTGTGGAGTCGTTGCCGTCCTTTATTTGAATACATTGANNNAGATCGGAAG
125299 TATGAGTGTGGAGTCGTTGCTAAAATCCAGCTTGTTTTATGAGNNNAGATCGGAAG
125300 TATGAGTGTGGAGTCGTTGCAGTAACAGGATTATCATCTACAANNNAGATCGGAAG
125301 TATGAGTGTGGAGTCGTTGCGTAGATTGGTCAAGAAAGAGTATNNNAGATCGGAA
125302 TATGAGTGTGGAGTCGTTGCCATACTCCTCTTCCTTTTTCTCNNNAGATCGGAAGAG
125304 TATGAGTGTGGAGTCGTTGCGACAAGGATCTCCAGATATATAGNNNAGATCGGAA
125305 TATGAGTGTGGAGTCGTTGCGTTCCTTATTTGACTTGCTAATTNNNAGATCGGAAGA
125306 TATGAGTGTGGAGTCGTTGCACTAGCTTTTTGATGCTAAATGNNNAGATCGGAAGA
125307 TATGAGTGTGGAGTCGTTGCTAGTGAATCTAGGTAAAAGGATGNNNAGATCGGAA
125308 TATGAGTGTGGAGTCGTTGCATAATAAGACTTACTGCTGCCNNNAGATCGGAAGAG
125309 TATGAGTGTGGAGTCGTTGCAGACATTCATACATCTTAGTACCNNNAGATCGGAAG
125310 TATGAGTGTGGAGTCGTTGCGTTTATGTTACTCTGGGTATTGNNNAGATCGGAAGA
125311 TATGAGTGTGGAGTCGTTGCCTAGTAAAGGACAACAGGAAAGNNNAGATCGGAAG
125312 TATGAGTGTGGAGTCGTTGCGGGTACCATACAAATATACTTCANNNAGATCGGAAG
125313 TATGAGTGTGGAGTCGTTGCGCTAAGAAGGAAGCTTATCTTNNNAGATCGGAAGA
125314 TATGAGTGTGGAGTCGTTGCATATCATGCTTTGAAGGAGTAATNNNAGATCGGAA

125315 TATGAGTGTGGAGTCGTTGCCAGGAATATATCTGGATACTCATNNNAGATCGGAAG
125316 TATGAGTGTGGAGTCGTTGCGGAAATTGCTCAGTACATAAAATNNNAGATCGGAAG
125317 TATGAGTGTGGAGTCGTTGCATTTTGTTGGCTGTGAAATCNNNAGATCGGAAGAGC
125318 TATGAGTGTGGAGTCGTTGCCACTATTCTTTCTTTACCTCTCNNNAGATCGGAAGAG
125319 TATGAGTGTGGAGTCGTTGCGACTTCTTGTTTCCTTTGAATACNNNAGATCGGAAGA
125320 TATGAGTGTGGAGTCGTTGCTTAACATTTTCACATCACTTTCTNNNAGATCGGAAGA
125321 TATGAGTGTGGAGTCGTTGCACAGAAACAAACTTCAATCATTANNNAGATCGGAAGA
125322 TATGAGTGTGGAGTCGTTGCAGGACCTGAGAAAAGAGAAATANNNAGATCGGAAG
125323 TATGAGTGTGGAGTCGTTGCGACTCTCTGAGTACATTCATCNNNAGATCGGAAGAG
125324 TATGAGTGTGGAGTCGTTGCTGTGTACAATCTCCAAATCGNNNAGATCGGAAGAGC
125325 TATGAGTGTGGAGTCGTTGCGTTTGTGACAATAAGGAAGAAANNNAGATCGGAAG
125326 TATGAGTGTGGAGTCGTTGCCTTCTATAAGGACTCTTGTGATTNNNAGATCGGAAGA
125327 TATGAGTGTGGAGTCGTTGCCCATTTATCTGGAGCTTCTAGNNNAGATCGGAAGAG
125328 TATGAGTGTGGAGTCGTTGCGGATTTCTAATTCATGTTCGTTTNNNAGATCGGAAGA
125329 TATGAGTGTGGAGTCGTTGCGAGACAATAGAAGGTAGTAGTAANNNAGATCGGAA
125330 TATGAGTGTGGAGTCGTTGCGAGACAATAGAAGGTAGTAGTAANNNAGATCGGAA
125331 TATGAGTGTGGAGTCGTTGCCCATAAGAAATAAATAAGCAGCTNNNAGATCGGAAG
125332 TATGAGTGTGGAGTCGTTGCACTCTTAGGCAAATATTCATGTANNNAGATCGGAAGA
125333 TATGAGTGTGGAGTCGTTGCGTCTCTAACACCAGGAGTTCNNNAGATCGGAAGAGC
125334 TATGAGTGTGGAGTCGTTGCCTGAGAGATGGTCATGTATTCNNNAGATCGGAAGAC
125335 TATGAGTGTGGAGTCGTTGCAATTCAGTTCATGTATCTTTGGNNNAGATCGGAAGA
125336 TATGAGTGTGGAGTCGTTGCCTACCTGTTTTGTGTTTTCATCNNNAGATCGGAAGAG
125337 TATGAGTGTGGAGTCGTTGCGACATACACTCCAGGAGAAGNNNAGATCGGAAGAG
125338 TATGAGTGTGGAGTCGTTGCGACATACACTCCAGGAGAAGNNNAGATCGGAAGAG
125339 TATGAGTGTGGAGTCGTTGCTGGATAGTCCAAACGATTTCNNNAGATCGGAAGAGC
125340 TATGAGTGTGGAGTCGTTGCTCATCATAATAAAAGATGACCCANNNAGATCGGAAG
125341 TATGAGTGTGGAGTCGTTGCGATAGTGTTCCCTTTGATAGAANNNAGATCGGAAGA
125342 TATGAGTGTGGAGTCGTTGCCCTGGTACTATTCTAGGAATTTTNNNAGATCGGAAGA
125343 TATGAGTGTGGAGTCGTTGCGCTTGTCTTAACAGTATAACTGNNNAGATCGGAAGA
125344 TATGAGTGTGGAGTCGTTGCTAATGCTTTTTCTTCTAATCCTGNNNAGATCGGAAGA
125345 TATGAGTGTGGAGTCGTTGCAGTTTTTTTTAGCAGTTTACTCTNNNAGATCGGAAGA
125346 TATGAGTGTGGAGTCGTTGCGTACCATAGCCTAAAGAAATAAANNNAGATCGGAAG
125347 TATGAGTGTGGAGTCGTTGCCTGTCTCTCTCTGTCTCTTTNNNAGATCGGAAGAGCAC
125348 TATGAGTGTGGAGTCGTTGCCTTCTTAGTGTGGTTTTAGACTNNNAGATCGGAAGAC
125349 TATGAGTGTGGAGTCGTTGCACCTCATGAACATCTCAAATAANNNAGATCGGAAGAC
125350 TATGAGTGTGGAGTCGTTGCAGAACTGCTTTATAATTTTTCAGTNNNAGATCGGAAG
125351 TATGAGTGTGGAGTCGTTGCAAATAACCAAATCACAGGAATTTNNNAGATCGGAAG
125352 TATGAGTGTGGAGTCGTTGCAAATAACCAAATCACAGGAATTTNNNAGATCGGAAG
125353 TATGAGTGTGGAGTCGTTGCTATTACGAAAAATATCACAGGGANNNAGATCGGAAG
125354 TATGAGTGTGGAGTCGTTGCAAAACATTCAAATCCCTTTTTCTNNNAGATCGGAAGA
125355 TATGAGTGTGGAGTCGTTGCCATCATGATTTTAAGTTTCCTGANNNAGATCGGAAGA
125356 TATGAGTGTGGAGTCGTTGCTTTATAGCTGCTTATATTGCAAGNNNAGATCGGAAGA
125357 TATGAGTGTGGAGTCGTTGCCTTTGACTAGGTAGCTTTGAGNNNAGATCGGAAGAC

125358 TATGAGTGTGGAGTCGTTGCCCTTATTCCACCTTTTTGTTTTANNNAGATCGGAAGA
125359 TATGAGTGTGGAGTCGTTGCGAAATTCTCACAGAAACATACAGNNNAGATCGGAAG
125360 TATGAGTGTGGAGTCGTTGCACAGAGGATGTATTAACTTTACANNNAGATCGGAAG
125361 TATGAGTGTGGAGTCGTTGCCCTGTATCCTCATAGTTTAAAGTNNNAGATCGGAAG
125362 TATGAGTGTGGAGTCGTTGCTTCAGTCTAGTTTTCAGTATTGTNNNAGATCGGAAG
125363 TATGAGTGTGGAGTCGTTGCTAATTATCTTCACCTTTCTCTCANNNAGATCGGAAGA
125364 TATGAGTGTGGAGTCGTTGCCTCATCTTGTGCTTTCTCTGNNNAGATCGGAAGAGC
125365 TATGAGTGTGGAGTCGTTGCATTTCAATGAAAGTGCTAAAGAANNNAGATCGGAA
125366 TATGAGTGTGGAGTCGTTGCAAACATAGGCATGTAGGTATAGNNNAGATCGGAAG
125367 TATGAGTGTGGAGTCGTTGCTTACACATACTTCATTGTGAAAGNNNAGATCGGAAG
125368 TATGAGTGTGGAGTCGTTGCTTGTTACTGTTGATTTTCTGCNNNAGATCGGAAGAG
125369 TATGAGTGTGGAGTCGTTGCTACAACAGAAATCTATTTCCTCANNNAGATCGGAAG
125370 TATGAGTGTGGAGTCGTTGCGGAAAAGTTCTGGAAGTATATTGNNNAGATCGGAA
125373 TATGAGTGTGGAGTCGTTGCGCTTTCAAAAAAACAAAACACTANNNAGATCGGAAG
125374 TATGAGTGTGGAGTCGTTGCAGGTGAGACTTTAGTAAAGAAAGNNNAGATCGGAA
125376 TATGAGTGTGGAGTCGTTGCACTGAATTCTGAACTTTAGTTTCNNNAGATCGGAAG
125378 TATGAGTGTGGAGTCGTTGCGGAAGCAGTTAAATCTGAATTTTNNNAGATCGGAA
125379 TATGAGTGTGGAGTCGTTGCATTCAAGGTTTGGAATTAGGTANNNAGATCGGAAG
125380 TATGAGTGTGGAGTCGTTGCGGGAAATCTTGTGAATTTCATTTNNNAGATCGGAAG
125381 TATGAGTGTGGAGTCGTTGCACCAAATACATTGTTGCTATTATNNNAGATCGGAAG
125382 TATGAGTGTGGAGTCGTTGCGCCTTTTTTTTGTTTGTTACCNNNAGATCGGAAGAGC
125383 TATGAGTGTGGAGTCGTTGCTCTAGTTGTATTCAGAGTTAGTCNNNAGATCGGAAG
125384 TATGAGTGTGGAGTCGTTGCAGTGTAGTAGCAGATAAAAAGAANNNAGATCGGAA
125385 TATGAGTGTGGAGTCGTTGCCCCATCTTGGATTAGACAAATNNNAGATCGGAAGAG
125386 TATGAGTGTGGAGTCGTTGCATCTGGAGGATATTAAGTGAAATNNNAGATCGGAA
125387 TATGAGTGTGGAGTCGTTGCCCTCATCTATTGCTATTTTCAATNNNAGATCGGAAGA
125388 TATGAGTGTGGAGTCGTTGCCCATCTCTACCAAAAATACAAAANNNAGATCGGAAG
125389 TATGAGTGTGGAGTCGTTGCGAGAAATTGGCTTACATGATTATNNNAGATCGGAA
125390 TATGAGTGTGGAGTCGTTGCGATTTTGGAGGAACATCTTTAAANNNAGATCGGAA
125391 TATGAGTGTGGAGTCGTTGCTGGTCCTCCACAATTCTTACNNNAGATCGGAAGAGC
125392 TATGAGTGTGGAGTCGTTGCTATCACATTTGATCCTTCTATCANNNAGATCGGAAGA
125393 TATGAGTGTGGAGTCGTTGCGTTATCTTGAAGAAGAGCATTCNNNAGATCGGAAG
125394 TATGAGTGTGGAGTCGTTGCCCTTTTCTGAGGATAAATTCAAANNNAGATCGGAAG
125395 TATGAGTGTGGAGTCGTTGCAAAGGAAACAGCAGATATTAGAANNNAGATCGGAA
125396 TATGAGTGTGGAGTCGTTGCTATATGCCTGTGTCTATGGTGNNNAGATCGGAAGA
125397 TATGAGTGTGGAGTCGTTGCGCCATGCTTATACTTTGTTATTTNNNAGATCGGAAGA
125398 TATGAGTGTGGAGTCGTTGCATAGGAATTCAGGACCTATTAACNNNAGATCGGAAG
125399 TATGAGTGTGGAGTCGTTGCCACATCTTTATGGGTCAATAGGNNNAGATCGGAAGA
125400 TATGAGTGTGGAGTCGTTGCGTAAAAATATAGGGTACTGGAGTNNNAGATCGGAA
125401 TATGAGTGTGGAGTCGTTGCTCCATAAATAGTTTGCAGAAAAANNNAGATCGGAAG
125402 TATGAGTGTGGAGTCGTTGCGGCAAATTTTTTTCCCTAAAATANNNAGATCGGAAG
125403 TATGAGTGTGGAGTCGTTGCTTAACTTGATCACCTACAAAGACNNNAGATCGGAAG
125404 TATGAGTGTGGAGTCGTTGCCCTAGAAGTCTCTCTTGAAATAANNNAGATCGGAAG

125405 TATGAGTGTGGAGTCGTTGCGGAATTTTTTTAAGTCACATTGTNNNAGATCGGAAG
125406 TATGAGTGTGGAGTCGTTGCGAGTTAGGGAGCTCTTTCAGNNNAGATCGGAAGAG
125407 TATGAGTGTGGAGTCGTTGCCACCAACGTTTGAAAATATTTATNNNAGATCGGAAG
125408 TATGAGTGTGGAGTCGTTGCTATTATTCATGTTTGAGTCAGGANNNAGATCGGAAG
125409 TATGAGTGTGGAGTCGTTGCTTTCTAGAGAACGGCACATTNNNAGATCGGAAGAG
125410 TATGAGTGTGGAGTCGTTGCGTTCGTGTATATCTCAGATTCTANNNAGATCGGAAG
125411 TATGAGTGTGGAGTCGTTGCAAGCTAGAGAAAGGAAATATACTNNNAGATCGGAA
125412 TATGAGTGTGGAGTCGTTGCATATATGCAATATGATGTGTCTGNNNAGATCGGAAG
125413 TATGAGTGTGGAGTCGTTGCATGAGCTATTAGAACATTTCTGANNNAGATCGGAAG
125414 TATGAGTGTGGAGTCGTTGCCTCCTAACTTATCCTCTTCCTNNNAGATCGGAAGAGC
125415 TATGAGTGTGGAGTCGTTGCCTTTCTAAATCCTCTGGAAAAAANNNAGATCGGAAG
125416 TATGAGTGTGGAGTCGTTGCCTTTCTAAATCCTCTGGAAAAAANNNAGATCGGAAG
125417 TATGAGTGTGGAGTCGTTGCATAAGAATTTCAGTTTCCCTACANNNAGATCGGAAG
125418 TATGAGTGTGGAGTCGTTGCCTTCCTAGCCAGAAAGAAGANNNAGATCGGAAGAG
125419 TATGAGTGTGGAGTCGTTGCAAAACAAAGTTGAGAATCTCTTTNNNAGATCGGAAG
125420 TATGAGTGTGGAGTCGTTGCCATTCCTTGATATTACTCCTCCNNNAGATCGGAAGAG
125421 TATGAGTGTGGAGTCGTTGCTGACGGATATGTTAATTAACTTGNNNAGATCGGAAG
125422 TATGAGTGTGGAGTCGTTGCAATAACCAACCCTTTAAGACATANNNAGATCGGAAG
125423 TATGAGTGTGGAGTCGTTGCCACCTCCATTTATATTTCATCTGNNNAGATCGGAAGA
125424 TATGAGTGTGGAGTCGTTGCAGAAAGAGAAAGAACATTCATTCNNNAGATCGGAA
125425 TATGAGTGTGGAGTCGTTGCGACGGAAATTCTTTTTCTTTTTTNNNAGATCGGAAGA
125426 TATGAGTGTGGAGTCGTTGCGACGGAAATTCTTTTTCTTTTTTNNNAGATCGGAAGA
125427 TATGAGTGTGGAGTCGTTGCTTGTTTTTTTTCACTGCTCTTANNNAGATCGGAAGAG
125428 TATGAGTGTGGAGTCGTTGCTAACGTTAATTTGAGTACTCTCTNNNAGATCGGAAG
125429 TATGAGTGTGGAGTCGTTGCTTATTCTTCTTCATATCCCATCCNNNAGATCGGAAGA
125430 TATGAGTGTGGAGTCGTTGCCTACTAACATGAGTATTTTGCATNNNAGATCGGAAG
125431 TATGAGTGTGGAGTCGTTGCAAAATGAATCTGAACTCCGTAANNNAGATCGGAAGA
125432 TATGAGTGTGGAGTCGTTGCGGAAAAATTACGCTAAAACAAAANNNAGATCGGAA
125433 TATGAGTGTGGAGTCGTTGCCTAGAAGCCACTCAGATCTCNNNAGATCGGAAGAGC
125434 TATGAGTGTGGAGTCGTTGCTATTTAGATGGATGGATAGATGGNNNAGATCGGAA
125435 TATGAGTGTGGAGTCGTTGCTAAAAGAAGTGGTTCTCATAAACNNNAGATCGGAA
125436 TATGAGTGTGGAGTCGTTGCTTATTTGCTATTGTGTATGACTGNNNAGATCGGAAG
125437 TATGAGTGTGGAGTCGTTGCTTTGCAAATTAAAGTAGAGACAANNNAGATCGGAA
125438 TATGAGTGTGGAGTCGTTGCATCAACAGGATACTTACATAGCNNNAGATCGGAAGA
125439 TATGAGTGTGGAGTCGTTGCGGGCTTAATAGATATTCCTTACANNNAGATCGGAAG
125440 TATGAGTGTGGAGTCGTTGCGGAAACAAAGGGAGAATAAATTANNNAGATCGGAA
125441 TATGAGTGTGGAGTCGTTGCGAATATCAAGGCTTAACTTTGATNNNAGATCGGAAG
125442 TATGAGTGTGGAGTCGTTGCGAATATCAAGGCTTAACTTTGATNNNAGATCGGAAG
125443 TATGAGTGTGGAGTCGTTGCCTTCAGGATAACCAAATATGGAANNNAGATCGGAA
125444 TATGAGTGTGGAGTCGTTGCCTACCTTTCTTTCTACTCCCTNNNAGATCGGAAGAGC
125445 TATGAGTGTGGAGTCGTTGCCTTTAAGTTTTAGGGTACATGTGNNNAGATCGGAAG
125446 TATGAGTGTGGAGTCGTTGCGTACATAGTTGATGGTTTTTCTANNNAGATCGGAAG
125449 TATGAGTGTGGAGTCGTTGCCTCTTTTTCTTTTTCTTTTCTTTTCTTNNNAGATCGGAA

125450 TATGAGTGTGGAGTCGTTGCGTTCAATTTGTGGTCAACTAATANNNAGATCGGAAG
125451 TATGAGTGTGGAGTCGTTGCAAAACCTCTGCCAAGTTATCNNNAGATCGGAAGAGC
125452 TATGAGTGTGGAGTCGTTGCGAAAGTCAGGACTATTTTATGACNNNAGATCGGAA
125453 TATGAGTGTGGAGTCGTTGCCACTTTAGACTCCTTCTTACCNNNAGATCGGAAGAGC
125454 TATGAGTGTGGAGTCGTTGCGCTAAAAGTTTCAGGGAAAATATNNNAGATCGGAA
125455 TATGAGTGTGGAGTCGTTGCTCTTTGAAACTTCATCAAATACANNNAGATCGGAAG
125456 TATGAGTGTGGAGTCGTTGCACACTCATGGAGAAGTTTTTTNNNAGATCGGAAGAG
125457 TATGAGTGTGGAGTCGTTGCAGACAGTAAAATAAGCTCTATGTNNNAGATCGGAA
125458 TATGAGTGTGGAGTCGTTGCAGAAGTTAGCTATTCTCATTCTTNNNAGATCGGAAG
125459 TATGAGTGTGGAGTCGTTGCAGAAGTTAGCTATTCTCATTCTTNNNAGATCGGAAG
125460 TATGAGTGTGGAGTCGTTGCAGTTTTCCTAAAGAATGTAATACCNNNAGATCGGAA
125461 TATGAGTGTGGAGTCGTTGCATATTCTGGCCATAATTGCAANNNAGATCGGAAGAG
125462 TATGAGTGTGGAGTCGTTGCCTATGCTATCCCCCTTATCANNNAGATCGGAAGAGCA
125463 TATGAGTGTGGAGTCGTTGCTCTTGTAATCATCATTCATTGTGNNNAGATCGGAAG
125464 TATGAGTGTGGAGTCGTTGCATAGGGAAGTGATAGATTACTGANNNAGATCGGAA
125466 TATGAGTGTGGAGTCGTTGCGTGTAAGTAGGGATGAATAGTATNNNAGATCGGAA
125467 TATGAGTGTGGAGTCGTTGCTTCCAGGGATTTTTCTTTTTTTTNNNAGATCGGAAGA
125468 TATGAGTGTGGAGTCGTTGCTTACATAGCACATACAATACAGANNNAGATCGGAAG
125469 TATGAGTGTGGAGTCGTTGCTATCTGAATGTAAGAGTTACAGGNNNAGATCGGAA
125470 TATGAGTGTGGAGTCGTTGCGAAGATGGAGCAGATATCAACNNNAGATCGGAAGA
125471 TATGAGTGTGGAGTCGTTGCCTATAATGAGATTTGCAACTCTCNNNAGATCGGAAG
125472 TATGAGTGTGGAGTCGTTGCTCTTGAACTCACCATTTTACTAANNNAGATCGGAAGA
125473 TATGAGTGTGGAGTCGTTGCACTATTGTAGTACAATGTTGGTANNNAGATCGGAAG
125474 TATGAGTGTGGAGTCGTTGCGAGAGAAGAAAATTAGAGAGGATNNNAGATCGGA
125475 TATGAGTGTGGAGTCGTTGCATCAGCTATATAGAGAGAGACATNNNAGATCGGAA
125476 TATGAGTGTGGAGTCGTTGCTTTAATTGGTCAAACTATGTGTTNNNAGATCGGAAG
125477 TATGAGTGTGGAGTCGTTGCTATATGCCTGTGTCTATGGTGNNNAGATCGGAAGA
125478 TATGAGTGTGGAGTCGTTGCTAACTTATGTAAAAGTTCACACCNNNAGATCGGAAG
125479 TATGAGTGTGGAGTCGTTGCTTTGAACTGAGGACTTTTCGNNNAGATCGGAAGAG
125480 TATGAGTGTGGAGTCGTTGCCTTGTAGCAAGGTTAGCAATNNNAGATCGGAAGAG
125481 TATGAGTGTGGAGTCGTTGCAGACAATTCTGTACATTTTGTTTNNNAGATCGGAAG
125482 TATGAGTGTGGAGTCGTTGCCCACTGCAGTTAATATACAAATTNNNAGATCGGAAG
125484 TATGAGTGTGGAGTCGTTGCTTATCCAGCCTTTTATTTGAATGNNNAGATCGGAAG
125485 TATGAGTGTGGAGTCGTTGCGTTTTGGACTGATAAAATCCAANNNAGATCGGAAGA
125486 TATGAGTGTGGAGTCGTTGCGTTTTGTTGTCTTGTTTTTCTAANNNAGATCGGAAGA
125487 TATGAGTGTGGAGTCGTTGCAAGTAGCCAAAGAGTTAGTAGANNNAGATCGGAAG
125488 TATGAGTGTGGAGTCGTTGCGAGTTTGTCTACATATCCTTTTCNNNAGATCGGAAGA
125491 TATGAGTGTGGAGTCGTTGCCACCAGATGACAGAGTGAATNNNAGATCGGAAGAG
125492 TATGAGTGTGGAGTCGTTGCAAACACTGGTCCAGACTAATNNNAGATCGGAAGAG
125493 TATGAGTGTGGAGTCGTTGCTTATAAGTATCTGCAGTTCTCAGNNNAGATCGGAAG
125494 TATGAGTGTGGAGTCGTTGCTATAGATCTTACAAAGAGCACAANNNAGATCGGAAG
125495 TATGAGTGTGGAGTCGTTGCTAGTCCTGTAACATTTTTTTCACNNNAGATCGGAAGA
125496 TATGAGTGTGGAGTCGTTGCTAGATGGGAAGAAATGACATTATNNNAGATCGGAA

125497 TATGAGTGTGGAGTCGTTGCTGATGATTCTAATAATTGTCTGCNNNAGATCGGAAG
125498 TATGAGTGTGGAGTCGTTGCCTACCCAAAATCTTTTACACAGNNNAGATCGGAAGA
125499 TATGAGTGTGGAGTCGTTGCGAATGAATGGGTACAAGAAACNNNAGATCGGAAGA
125500 TATGAGTGTGGAGTCGTTGCGAGGGAGAAATAAAAAAAATGGANNNAGATCGGA
125501 TATGAGTGTGGAGTCGTTGCGACTGTGATGGTTAGTTTTATGNNNAGATCGGAAG
125502 TATGAGTGTGGAGTCGTTGCATAAAGACTGGTACAGAGACATANNNAGATCGGAA
125503 TATGAGTGTGGAGTCGTTGCGCAACTGGGATTCAATATAATAANNNAGATCGGAAG
125504 TATGAGTGTGGAGTCGTTGCTTAGACCACAGAAATAAAAGGAANNNAGATCGGAA
125505 TATGAGTGTGGAGTCGTTGCTGGGAGGAAATCCAAGTATCNNNAGATCGGAAGAG
125506 TATGAGTGTGGAGTCGTTGCTGGGAGGAAATCCAAGTATCNNNAGATCGGAAGAG
125508 TATGAGTGTGGAGTCGTTGCGAATCATGAAATGTTCTACTCTGNNNAGATCGGAAG
125509 TATGAGTGTGGAGTCGTTGCTCAGATTTGTTTCATAAATTCCTNNNAGATCGGAAGA
125510 TATGAGTGTGGAGTCGTTGCAAGTCAGATTACAAGTGATTTTCNNNAGATCGGAAG
125512 TATGAGTGTGGAGTCGTTGCCTGGGAATCTTTACTAACTGTATNNNAGATCGGAAG
125513 TATGAGTGTGGAGTCGTTGCTCAAGTGGAAATAGGATATGAAANNNAGATCGGAA
125514 TATGAGTGTGGAGTCGTTGCTCAAGTGGAAATAGGATATGAAANNNAGATCGGAA
125516 TATGAGTGTGGAGTCGTTGCATTTTGAACAAAACGAATACTTGNNNAGATCGGAAG
125517 TATGAGTGTGGAGTCGTTGCCTTTTTCACTTCTTAATTGCTTGNNNAGATCGGAAGA
125518 TATGAGTGTGGAGTCGTTGCGATGTGAAAATGCTCAAAAAAATNNNAGATCGGAA
125519 TATGAGTGTGGAGTCGTTGCATTTTTTTCCATACCCAGTGAAANNNAGATCGGAAGA
125520 TATGAGTGTGGAGTCGTTGCATTTTTTTCCATACCCAGTGAAANNNAGATCGGAAGA
125521 TATGAGTGTGGAGTCGTTGCATTCTCCCACTGATATGTATTACNNNAGATCGGAAGA
125522 TATGAGTGTGGAGTCGTTGCTAAATGTATTTGCTAAACCCATCNNNAGATCGGAAG
125523 TATGAGTGTGGAGTCGTTGCTTAAACAGCATCACTTCTGATNNNAGATCGGAAGAG
125524 TATGAGTGTGGAGTCGTTGCGTGATTTGACCATACTCTGTCNNNAGATCGGAAGAG
125525 TATGAGTGTGGAGTCGTTGCCAGAGACTGGATGAATAAAATTGNNNAGATCGGAA
125526 TATGAGTGTGGAGTCGTTGCGGAGAGATTGAAACTTTTCTAAANNNAGATCGGAAG
125527 TATGAGTGTGGAGTCGTTGCTATGGTGGGAAATATTGGAATTANNNAGATCGGAA
125528 TATGAGTGTGGAGTCGTTGCGTATTGCAGATGCTGATAGATTNNNAGATCGGAAG
125529 TATGAGTGTGGAGTCGTTGCGTATTGCAGATGCTGATAGATTNNNAGATCGGAAG
125530 TATGAGTGTGGAGTCGTTGCATACTTTGATTTTCATTGCTTCANNNAGATCGGAAGA
125531 TATGAGTGTGGAGTCGTTGCATACTTTGATTTTCATTGCTTCANNNAGATCGGAAGA
125532 TATGAGTGTGGAGTCGTTGCATACTTTGATTTTCATTGCTTCANNNAGATCGGAAGA
125533 TATGAGTGTGGAGTCGTTGCACCTATCATGGTAGAATAAAGTCNNNAGATCGGAAG
125534 TATGAGTGTGGAGTCGTTGCTAGAACAAAGCTATCTGTGAAATNNNAGATCGGAAG
125535 TATGAGTGTGGAGTCGTTGCATGAATGGATTTTAAATCCTAGAAANNNAGATCGGA
125536 TATGAGTGTGGAGTCGTTGCAGGTTCTTCTTATTACACATTTTNNNAGATCGGAAGA
125537 TATGAGTGTGGAGTCGTTGCTTTCTAAATTTTCCACATGAACTNNNAGATCGGAAGA
125538 TATGAGTGTGGAGTCGTTGCTTTTCTTAGAGGCCATCTTTTTANNNAGATCGGAAGA
125539 TATGAGTGTGGAGTCGTTGCCACAACAAAAAGCCATTTACATNNNAGATCGGAAGA
125540 TATGAGTGTGGAGTCGTTGCTTTGATAGTGTGTTTGAAAGCNNNAGATCGGAAGA
125541 TATGAGTGTGGAGTCGTTGCAATGGTAAAGCAATTACAAGTTTNNNAGATCGGAAG
125542 TATGAGTGTGGAGTCGTTGCGCAGATATTGTTGTTATAGAGTCNNNAGATCGGAAG

125543 TATGAGTGTGGAGTCGTTGCCTTTTCAGGATAAGCAAGTTAACNNNAGATCGGAAG
125544 TATGAGTGTGGAGTCGTTGCCCATTTAGAAGTTGCATATCTTANNNAGATCGGAAG
125545 TATGAGTGTGGAGTCGTTGCCTTGAACTGATGGATAATGAATGNNNAGATCGGAA
125546 TATGAGTGTGGAGTCGTTGCTACTGTAGCATATAGGGTGAAATNNNAGATCGGAA
125547 TATGAGTGTGGAGTCGTTGCGGAGACACATTAGGTTACATAANNNAGATCGGAAG
125548 TATGAGTGTGGAGTCGTTGCTCAAAAAAAAGACCGATAAGCNNNAGATCGGAAGA
125549 TATGAGTGTGGAGTCGTTGCCTCTCTCAAACCTAACCTAAAGNNNAGATCGGAAGA
125551 TATGAGTGTGGAGTCGTTGCCATTCTATTACCATTGCAATGTANNNAGATCGGAAG
125553 TATGAGTGTGGAGTCGTTGCAAAGGAAGGATAGAAGATCATGNNNAGATCGGAA
125554 TATGAGTGTGGAGTCGTTGCAGAGAATCAGAGACCAGATTATANNNAGATCGGAA
125555 TATGAGTGTGGAGTCGTTGCTCATTTACAAGATGAGTAGAAATGNNNAGATCGGA
125556 TATGAGTGTGGAGTCGTTGCAATGGGTTTTCACTTGGTAAATNNNAGATCGGAAG
125557 TATGAGTGTGGAGTCGTTGCTAACTTGTTATGTCTAAACAGGANNNAGATCGGAAG
125559 TATGAGTGTGGAGTCGTTGCCCATTGTGAGTGGAATTTATTTNNNAGATCGGAAGA
125560 TATGAGTGTGGAGTCGTTGCTATACAGAGGGAGTGAGGATNNNAGATCGGAAGA
125561 TATGAGTGTGGAGTCGTTGCACCAAAATCTTAGTTGAAATCTTNNNAGATCGGAAG
125562 TATGAGTGTGGAGTCGTTGCGAAGACAGATCTCAACTGAGNNNAGATCGGAAGAG
125563 TATGAGTGTGGAGTCGTTGCTCTGAGACAGATCTAAATCAATTNNNAGATCGGAAG
125564 TATGAGTGTGGAGTCGTTGCGAATTGAGGGAAGAATTTAAAGTNNNAGATCGGAA
125565 TATGAGTGTGGAGTCGTTGCGAGAAAATGTGTTCTGTGTATTANNNAGATCGGAA
125566 TATGAGTGTGGAGTCGTTGCCTTCTAACTGACTCTGACATTNNNAGATCGGAAGAG
125567 TATGAGTGTGGAGTCGTTGCGAATCATGTTTTCCTATGCATAANNNAGATCGGAAG
125568 TATGAGTGTGGAGTCGTTGCATCCATGTCTTATTTCCAGTTTNNNAGATCGGAAGAG
125569 TATGAGTGTGGAGTCGTTGCAACATAAAGCTGGTGAAATATTTNNNAGATCGGAAG
125570 TATGAGTGTGGAGTCGTTGCATTAGTCCTTTCTCCTTATTCTCNNNAGATCGGAAGA
125571 TATGAGTGTGGAGTCGTTGCCTTCACTTCCTTTCCTACTTNNNAGATCGGAAGAGCA
125572 TATGAGTGTGGAGTCGTTGCCAATTTGTGCTATGAATTTGTCNNNAGATCGGAAGA
125573 TATGAGTGTGGAGTCGTTGCAAATTGTTTACATATTGAATGGCNNNAGATCGGAAG
125574 TATGAGTGTGGAGTCGTTGCATGACTAGTTATTTCCAGAACATNNNAGATCGGAAG
125575 TATGAGTGTGGAGTCGTTGCGAAGACATAGAGAAAGTCCTAAANNNAGATCGGAA
125577 TATGAGTGTGGAGTCGTTGCTACAGCATTATCAAATAGGCAGNNNAGATCGGAAGA
125578 TATGAGTGTGGAGTCGTTGCTACAGCATTATCAAATAGGCAGNNNAGATCGGAAGA
125579 TATGAGTGTGGAGTCGTTGCATTGAATAGTGTGTTTCCTTGTANNNAGATCGGAAG
125580 TATGAGTGTGGAGTCGTTGCATTGAATAGTGTGTTTCCTTGTANNNAGATCGGAAG
125581 TATGAGTGTGGAGTCGTTGCTACTACTCAGCCATAAAAAAAAGNNNAGATCGGAAG
125582 TATGAGTGTGGAGTCGTTGCGCATACATGCACATATACATATGNNNAGATCGGAAG
125583 TATGAGTGTGGAGTCGTTGCGTTACTGAGGTTTGGTTTACANNNAGATCGGAAGA
125584 TATGAGTGTGGAGTCGTTGCACGAATCTCTCTCTAGTTTTAAANNNAGATCGGAAGA
125585 TATGAGTGTGGAGTCGTTGCACGAATCTCTCTCTAGTTTTAAANNNAGATCGGAAGA
125587 TATGAGTGTGGAGTCGTTGCTTCTAAGGAGTGGAGGTAAATANNNAGATCGGAAG
125588 TATGAGTGTGGAGTCGTTGCTACCTTGTAAAAGTACCAAAAGANNNAGATCGGAAG
125589 TATGAGTGTGGAGTCGTTGCGAATTTTAGCTATCGTTGGTACNNNAGATCGGAAGA
125590 TATGAGTGTGGAGTCGTTGCAAAATACATTAAGCTACACATCCNNNAGATCGGAAG

125591 TATGAGTGTGGAGTCGTTGCTACCTTTTTTGTGTGGAGACNNNAGATCGGAAGAG
125592 TATGAGTGTGGAGTCGTTGCAGGGTTATGAGAATGTTGTAAANNNAGATCGGAAG
125594 TATGAGTGTGGAGTCGTTGCTCTTTGACTCTTAATAGATGAGGNNNAGATCGGAAG
125595 TATGAGTGTGGAGTCGTTGCGAGATAAGGTAAATGTGAGCANNNAGATCGGAAG
125596 TATGAGTGTGGAGTCGTTGCTCTAGAAAACCCTGACTAATATGNNNAGATCGGAAG
125597 TATGAGTGTGGAGTCGTTGCATGTTGTAAAGTACATTTCTTCCNNNAGATCGGAAG
125598 TATGAGTGTGGAGTCGTTGCAATCAGGATAACAGTATTTCAAGNNNAGATCGGAAG
125599 TATGAGTGTGGAGTCGTTGCAAAGAGAATAGAACTACCTGAAGNNNAGATCGGAA
125600 TATGAGTGTGGAGTCGTTGCGTCCTTTGATCTGTTCTTAGAGNNNAGATCGGAAGA
125601 TATGAGTGTGGAGTCGTTGCCAAAAGTGAAATAGCCTTGCNNNAGATCGGAAGAG
125602 TATGAGTGTGGAGTCGTTGCATTTTCGAAGTAATGTCTAAGACNNNAGATCGGAAG
125603 TATGAGTGTGGAGTCGTTGCGTTCTGGTGTCTCTTCTTATAANNNAGATCGGAAGA
125604 TATGAGTGTGGAGTCGTTGCAATTGGGTTCTTTATGTTCTTTTNNNAGATCGGAAG
125605 TATGAGTGTGGAGTCGTTGCGGAAATCTCATCATTTGCAATAANNNAGATCGGAAG
125606 TATGAGTGTGGAGTCGTTGCGCAATTATAGTAGGAGTGTAGATNNNAGATCGGAA
125607 TATGAGTGTGGAGTCGTTGCAGAAAACAAACCTACAATTTTCTNNNAGATCGGAAG
125608 TATGAGTGTGGAGTCGTTGCCCGTATGTTAAGATAAGAAGAGANNNAGATCGGAA
125609 TATGAGTGTGGAGTCGTTGCGTTATGGATGAAGTAATGGAATCNNNAGATCGGAA
125610 TATGAGTGTGGAGTCGTTGCTACATTAATGAACCTCATTGTTCNNNAGATCGGAAG
125611 TATGAGTGTGGAGTCGTTGCGTATTGGCTCACAAGATTATAAANNNAGATCGGAAG
125612 TATGAGTGTGGAGTCGTTGCATAGAATTCTCTCTGAAGGAGTNNNAGATCGGAAG
125613 TATGAGTGTGGAGTCGTTGCCTATCATTAGAGATTCAGTGGACNNNAGATCGGAAG
125614 TATGAGTGTGGAGTCGTTGCCTTTCTGGAATAACTTTAAGCTCNNNAGATCGGAAG
125615 TATGAGTGTGGAGTCGTTGCATAATTCTGGTTCCTTTGTATTGNNNAGATCGGAAG
125616 TATGAGTGTGGAGTCGTTGCCTTTCTAACTTTCTGTCACTACANNNAGATCGGAAGA
125617 TATGAGTGTGGAGTCGTTGCTGTGTGTACTTTCTAACTTTCTNNNAGATCGGAAGAG
125618 TATGAGTGTGGAGTCGTTGCGAGTCTTGTCTAAGAATTCAATTNNNAGATCGGAAG
125619 TATGAGTGTGGAGTCGTTGCTGTATATTAGGTATGTGTGTGTTNNNAGATCGGAAG
125620 TATGAGTGTGGAGTCGTTGCATTTTCTAATTGGGTGGTAAAACNNNAGATCGGAAG
125621 TATGAGTGTGGAGTCGTTGCGCTTTAAGTGTCTTTTCATTACTNNNAGATCGGAAGA
125622 TATGAGTGTGGAGTCGTTGCTATAAAAACCTTTTTGCGGCNNNAGATCGGAAGAGC
125623 TATGAGTGTGGAGTCGTTGCTTTTGTATTCTTTTTGGGTTTTGNNNAGATCGGAAG
125624 TATGAGTGTGGAGTCGTTGCTATAGCTTACCTAGATAATCTGCNNNAGATCGGAAG
125625 TATGAGTGTGGAGTCGTTGCGGATTAATAGCCATTAACTAAGGNNNAGATCGGAAG
125627 TATGAGTGTGGAGTCGTTGCTATTTTTCCAATCCTTTATGGTCNNNAGATCGGAAGA
125628 TATGAGTGTGGAGTCGTTGCTGATAATGATAGCTACATGTCTTNNNAGATCGGAAG
125629 TATGAGTGTGGAGTCGTTGCTGTGCAGTAGGAAAATAATAATGNNNAGATCGGAA
125630 TATGAGTGTGGAGTCGTTGCTATAATAAGGGAGAAAAAGTGGGNNNAGATCGGA
125631 TATGAGTGTGGAGTCGTTGCTAAATAGATTTTGCCCATTTTCCNNNAGATCGGAAGA
125632 TATGAGTGTGGAGTCGTTGCCCAAATTTTCTTTATCCATCCATNNNAGATCGGAAGA
125633 TATGAGTGTGGAGTCGTTGCCTCATTCCTAGGCTTTCAAATANNNAGATCGGAAGA
125634 TATGAGTGTGGAGTCGTTGCCAGAATTATCTCTCTAAGTGTGANNNAGATCGGAAG
125635 TATGAGTGTGGAGTCGTTGCTCTCTAAGGAGGAATAACATCTANNNAGATCGGAAG

125636 TATGAGTGTGGAGTCGTTGCCATTTTCTTTATCCACTTGTAGGNNNAGATCGGAAGA

125637 TATGAGTGTGGAGTCGTTGCGATGCAAATATGTAATTAGACCCNNNAGATCGGAAG

125638 TATGAGTGTGGAGTCGTTGCCTATGACTTCTGCCATTAATAAGNNNAGATCGGAAG

125639 TATGAGTGTGGAGTCGTTGCAATCTTTTCCCTCACTTATTTACNNNAGATCGGAAGA

125640 TATGAGTGTGGAGTCGTTGCTTCTGTTACACTCAAAAATTTGANNNAGATCGGAAG

125641 TATGAGTGTGGAGTCGTTGCTTCTGTTACACTCAAAAATTTGANNNAGATCGGAAG

125642 TATGAGTGTGGAGTCGTTGCATTCTACTGGTTCTGTTTCTCNNNAGATCGGAAGAG

125643 TATGAGTGTGGAGTCGTTGCATTCTACTGGTTCTGTTTCTCNNNAGATCGGAAGAG

125644 TATGAGTGTGGAGTCGTTGCCCAGCTTTGAAAAGAAACATGNNNAGATCGGAAGA

125645 TATGAGTGTGGAGTCGTTGCATTTGGTCTAAATACTTTTCACANNNAGATCGGAAG

125646 TATGAGTGTGGAGTCGTTGCGATTTGTCCTAATACAACTGTGNNNAGATCGGAAGA

125647 TATGAGTGTGGAGTCGTTGCGAAGAGAAATTGATAAACCAGTTNNNAGATCGGAA

125648 TATGAGTGTGGAGTCGTTGCCTCTAATAGTCTTCCAATAGAGGNNNAGATCGGAAG

125649 TATGAGTGTGGAGTCGTTGCAACGTCATTTACTCAGATAATTGNNNAGATCGGAAG

125650 TATGAGTGTGGAGTCGTTGCCTTTGTCTCTGGATTTCTCTGNNNAGATCGGAAGAG

125651 TATGAGTGTGGAGTCGTTGCGCTCCCAAACTTTGATTTATTTANNNAGATCGGAAG

125653 TATGAGTGTGGAGTCGTTGCAGTGGGTGTTATTGCATTTAANNNAGATCGGAAGA

125654 TATGAGTGTGGAGTCGTTGCGCTGTCTTCTAACAAAGATAATTNNNAGATCGGAAG

125655 TATGAGTGTGGAGTCGTTGCGCTGTCTTCTAACAAAGATAATTNNNAGATCGGAAG

125656 TATGAGTGTGGAGTCGTTGCTATCAATAACTCGGTAATGAGAANNNAGATCGGAA

125657 TATGAGTGTGGAGTCGTTGCTCAAAGGATTGGAATAAAATCAGNNNAGATCGGAA

125658 TATGAGTGTGGAGTCGTTGCTACTTTACGTTCTGGGATACATNNNAGATCGGAAGA

125659 TATGAGTGTGGAGTCGTTGCCATATTCACTGCAGCATTATTCNNNAGATCGGAAGA

125661 TATGAGTGTGGAGTCGTTGCATATATGTGTTTTCACAAACTGGNNNAGATCGGAAG

125662 TATGAGTGTGGAGTCGTTGCAATCCTTATGGTCACAAGATAACNNNAGATCGGAAG

125663 TATGAGTGTGGAGTCGTTGCTTTAGAATCAAAATAGAGAGGGGNNNAGATCGGAA

125664 TATGAGTGTGGAGTCGTTGCCATTTTCTTTATTCCTTCTGCANNNAGATCGGAAGAG

125665 TATGAGTGTGGAGTCGTTGCTTAATTTATGGCTCTTTTCTCTGNNNAGATCGGAAGA

125666 TATGAGTGTGGAGTCGTTGCTTGAAAAAGCGAAATAGATCAGNNNAGATCGGAAG

125667 TATGAGTGTGGAGTCGTTGCCCTTTTCTTATTTTGGCTTATCTNNNAGATCGGAAGA

125668 TATGAGTGTGGAGTCGTTGCTTTACTCCCTTCCATGATGATNNNAGATCGGAAGAG

125669 TATGAGTGTGGAGTCGTTGCGAACAAAATTTGAAGCTAGATTCNNNAGATCGGAA

125670 TATGAGTGTGGAGTCGTTGCGAACAAAATTTGAAGCTAGATTCNNNAGATCGGAA

125672 TATGAGTGTGGAGTCGTTGCCCACGATCAATTGTTCAAAGNNNAGATCGGAAGAG

125673 TATGAGTGTGGAGTCGTTGCTTCCTTCAAAAATTTTTTTCTGTNNNAGATCGGAAGA

125674 TATGAGTGTGGAGTCGTTGCTTCCTTCAAAAATTTTTTTCTGTNNNAGATCGGAAGA

125675 TATGAGTGTGGAGTCGTTGCTTCCTTCAAAAATTTTTTTCTGTNNNAGATCGGAAGA

125676 TATGAGTGTGGAGTCGTTGCTGTAAATGAGGTCATAATTCATGNNNAGATCGGAA

125677 TATGAGTGTGGAGTCGTTGCCTCTTGAAAACACTAGGTAAATCNNNAGATCGGAAG

125678 TATGAGTGTGGAGTCGTTGCCTATTGTTATGTGAAACATCCTCNNNAGATCGGAAG

125679 TATGAGTGTGGAGTCGTTGCCAAACACTTTGGAGTTATGTCNNNAGATCGGAAGAG

125680 TATGAGTGTGGAGTCGTTGCGTGAGACCTTTGGACTTTATNNNAGATCGGAAGAG

125681 TATGAGTGTGGAGTCGTTGCCTATGGAAGAACCAGGAACANNNAGATCGGAAGAG

125682 TATGAGTGTGGAGTCGTTGCCCTCTTTGGAAGGTATAAAGATANNNAGATCGGAAG
125684 TATGAGTGTGGAGTCGTTGCGTATTCCCACAACTGAATTTAATNNNAGATCGGAAG
125685 TATGAGTGTGGAGTCGTTGCATTTCATAAACAGAGATTGGTTCNNNAGATCGGAAG
125686 TATGAGTGTGGAGTCGTTGCGTTGACTGTTGTTAGTAACTTAANNNAGATCGGAAG
125687 TATGAGTGTGGAGTCGTTGCGATGATTCTGTTGTTCTCTAAACNNNAGATCGGAAG
125688 TATGAGTGTGGAGTCGTTGCAGGAAGAAAAATAACAAAGCTAGNNNAGATCGGAA
125689 TATGAGTGTGGAGTCGTTGCCCTATTATTTGCTATTCTCCTTGNNNAGATCGGAAGA
125690 TATGAGTGTGGAGTCGTTGCCTGTTTTGTTTGTTTGTTTTAGANNNAGATCGGAAG
125691 TATGAGTGTGGAGTCGTTGCGAGAACTCTGAACAATACAATGNNNAGATCGGAAG
125692 TATGAGTGTGGAGTCGTTGCATTGACCAGATAGTCCTCTACNNNAGATCGGAAGAG
125693 TATGAGTGTGGAGTCGTTGCATGAGGTAGATATCATTTCTTCCNNNAGATCGGAAG
125694 TATGAGTGTGGAGTCGTTGCTTGTTTGTTTGCTTATTTAGAGANNNAGATCGGAAG
125695 TATGAGTGTGGAGTCGTTGCCCAATTAACAATCTGTGGATTAANNNAGATCGGAAG
125696 TATGAGTGTGGAGTCGTTGCCTAAAACGCACATACACTTAGNNNAGATCGGAAGAG
125697 TATGAGTGTGGAGTCGTTGCTGGAATACTATTCAGTCTGAAAANNNAGATCGGAAG
125698 TATGAGTGTGGAGTCGTTGCCTTTTTTGTTGTTGTTGTTGAGNNNAGATCGGAAGA
125699 TATGAGTGTGGAGTCGTTGCCGTTTATATTATCTCCATCTCCANNNAGATCGGAAGA
125700 TATGAGTGTGGAGTCGTTGCATCTGTCTTGATCCTGAACTNNNAGATCGGAAGAGC
125701 TATGAGTGTGGAGTCGTTGCGAATGAAGGAATGAATGAAGACNNNAGATCGGAAG
125702 TATGAGTGTGGAGTCGTTGCCAAATTCTGGTTAAAAACACTGNNNAGATCGGAAGA
125703 TATGAGTGTGGAGTCGTTGCGTTTTTTTGAGATAGGGTCTTGNNNAGATCGGAAG
125704 TATGAGTGTGGAGTCGTTGCTAGATTTCTGGATTGATTTTCTCNNNAGATCGGAAG
125705 TATGAGTGTGGAGTCGTTGCTTTTCATTAGACCTCATCTTATGNNNAGATCGGAAGA
125707 TATGAGTGTGGAGTCGTTGCCACATTAAAGGCAGTTAATCAANNNAGATCGGAAGA
125708 TATGAGTGTGGAGTCGTTGCCACAGGCATAATGATGAATTAAANNNAGATCGGAAG
125709 TATGAGTGTGGAGTCGTTGCTACAAAAACATTTTGGCTACTATNNNAGATCGGAAG
125710 TATGAGTGTGGAGTCGTTGCATCTTGACTTCACAACAGATTANNNAGATCGGAAGA
125711 TATGAGTGTGGAGTCGTTGCGTAAATTCCTAGGGTGGAAAAAANNNAGATCGGAAG
125712 TATGAGTGTGGAGTCGTTGCATATACAGGCTTTCAGTCTATTTNNNAGATCGGAAG
125713 TATGAGTGTGGAGTCGTTGCTTGGCAACATTTCTCAAATAATANNNAGATCGGAAG
125714 TATGAGTGTGGAGTCGTTGCACTAATTTTGTAACTTTAGAGCCNNNAGATCGGAAG
125715 TATGAGTGTGGAGTCGTTGCTTTCTAAGTGAATACAATTCCACNNNAGATCGGAAG
125716 TATGAGTGTGGAGTCGTTGCTTTATTTTTTGAGATGGAGTCTGNNNAGATCGGAAG
125717 TATGAGTGTGGAGTCGTTGCGTTTTTCAATTTGGATACCCTTTNNNAGATCGGAAGA
125718 TATGAGTGTGGAGTCGTTGCGGTAAGTAAGCACTGTAAAAATGNNNAGATCGGAA
125719 TATGAGTGTGGAGTCGTTGCATTATTCAGAGGACATGATAGTCNNNAGATCGGAAG
125720 TATGAGTGTGGAGTCGTTGCATATATACACAAACACCTCAGTTNNNAGATCGGAAG
125722 TATGAGTGTGGAGTCGTTGCTGGAAGATCCAGAATTTATCTTTNNNAGATCGGAAG
125723 TATGAGTGTGGAGTCGTTGCATGAATTTAAGCATGTGGATTACNNNAGATCGGAAG
125725 TATGAGTGTGGAGTCGTTGCATTGATACTGATCTCTCCAATTCNNNAGATCGGAAGA
125726 TATGAGTGTGGAGTCGTTGCTTCTGCCATTTTATTTTTCCTAGNNNAGATCGGAAGA
125727 TATGAGTGTGGAGTCGTTGCTTTGATGATACTACACACTACACNNNAGATCGGAAG
125728 TATGAGTGTGGAGTCGTTGCCATGTTTGGACCATGAGTACNNNAGATCGGAAGAG

125729 TATGAGTGTGGAGTCGTTGCGGTATATGTTTGCAATTCAATTCNNNAGATCGGAAG
125730 TATGAGTGTGGAGTCGTTGCAAGTATGCATTTTATTTTCACAGNNNAGATCGGAAG
125731 TATGAGTGTGGAGTCGTTGCGCTGCCATTTTAATACAAAATAANNNAGATCGGAAG
125732 TATGAGTGTGGAGTCGTTGCATGTTTTAGGAGCATAATATGGTNNNAGATCGGAA
125733 TATGAGTGTGGAGTCGTTGCTGCCATGAAATAATTTAGATGAANNNAGATCGGAA
125734 TATGAGTGTGGAGTCGTTGCGTTACATTTTATTGGGCTGTAAGNNNAGATCGGAAG
125735 TATGAGTGTGGAGTCGTTGCCCTCTTGTTAAAAAACATTGAGANNNAGATCGGAAG
125736 TATGAGTGTGGAGTCGTTGCGTACGATCATAGCTCATTGTAGNNNAGATCGGAAGA
125737 TATGAGTGTGGAGTCGTTGCGTAGGAAAATATCACTGTCCTAANNNAGATCGGAAG
125738 TATGAGTGTGGAGTCGTTGCCATATGAGTTCCATGATAGTTTTNNNAGATCGGAAG
125739 TATGAGTGTGGAGTCGTTGCACTTTATTACTTTCTGGCACTANNNAGATCGGAAGAG
125740 TATGAGTGTGGAGTCGTTGCTGTGAATGCATTTGTTTCTGNNNAGATCGGAAGAG
125741 TATGAGTGTGGAGTCGTTGCCATGAGTTCATCTTTAGTTAGTGNNNAGATCGGAAG
125742 TATGAGTGTGGAGTCGTTGCTATGCATGAGAAATTTCACATTTNNNAGATCGGAAG
125743 TATGAGTGTGGAGTCGTTGCCTACCTTGTCCAGTTTCATATATNNNAGATCGGAAGA
125744 TATGAGTGTGGAGTCGTTGCGAACACTTCTATCATGAGACTAANNNAGATCGGAAG
125745 TATGAGTGTGGAGTCGTTGCCATACATATACACACAGAATCAGNNNAGATCGGAAG
125746 TATGAGTGTGGAGTCGTTGCATTTTGAGAGGGGAAGGAAGNNNAGATCGGAAGA
125747 TATGAGTGTGGAGTCGTTGCAAAGGAGGGGAGTTCATTTTNNNAGATCGGAAGAG
125748 TATGAGTGTGGAGTCGTTGCGACGAAAGTTTCATGAAATACTANNNAGATCGGAAG
125749 TATGAGTGTGGAGTCGTTGCGAAGATGAAGGAAATCACAATTNNNAGATCGGAAG
125750 TATGAGTGTGGAGTCGTTGCTCCTTCTATATGCTATCAAAACTNNNAGATCGGAAGA
125754 TATGAGTGTGGAGTCGTTGCCACTCAGTGCTTAACATAGTANNNAGATCGGAAGAG
125755 TATGAGTGTGGAGTCGTTGCGCAGACACTGAGATTTTAAAAANNNAGATCGGAAG
125756 TATGAGTGTGGAGTCGTTGCATAGATAGATAAAGTTGGCCATTNNNAGATCGGAA
125757 TATGAGTGTGGAGTCGTTGCATGCTTGGAGATGATTTTTATACNNNAGATCGGAAG
125758 TATGAGTGTGGAGTCGTTGCCAGGACTTTAGGATTTATTGTAGNNNAGATCGGAA
125759 TATGAGTGTGGAGTCGTTGCTTAAATGATCACAGTTAATTGGTNNNAGATCGGAAG
125760 TATGAGTGTGGAGTCGTTGCATACAGGAAACGGAACTTTTAANNNAGATCGGAAG
125761 TATGAGTGTGGAGTCGTTGCCTGAATGTGAGATGAAAGTTTGNNNAGATCGGAAG
125762 TATGAGTGTGGAGTCGTTGCTCTGATTTTGTTGTAATTTTGGTNNNAGATCGGAAG
125763 TATGAGTGTGGAGTCGTTGCTCTGATTTTGTTGTAATTTTGGTNNNAGATCGGAAG
125764 TATGAGTGTGGAGTCGTTGCTTTAGTCATTAAATGGAAGTGTCNNNAGATCGGAAG
125765 TATGAGTGTGGAGTCGTTGCCAGAATGTGTGTATTTTTGTGTNNNAGATCGGAAGA
125766 TATGAGTGTGGAGTCGTTGCCAGAATGTGTGTATTTTTGTGTNNNAGATCGGAAGA
125767 TATGAGTGTGGAGTCGTTGCACACATATTAACTGTGAGAAAACNNNAGATCGGAAG
125768 TATGAGTGTGGAGTCGTTGCGGAGAGACCACATATTTAATACANNNAGATCGGAA
125769 TATGAGTGTGGAGTCGTTGCGGAGAGACCACATATTTAATACANNNAGATCGGAA
125770 TATGAGTGTGGAGTCGTTGCGGAGAGACCACATATTTAATACANNNAGATCGGAA
125771 TATGAGTGTGGAGTCGTTGCTAATTGTTTTATCTGTTCTCAGCNNNAGATCGGAAGA
125772 TATGAGTGTGGAGTCGTTGCAAAAAGCAAAACAAATAAACGAANNNAGATCGGAA
125773 TATGAGTGTGGAGTCGTTGCGCATTATTTATAAAGAGACGTGTNNNAGATCGGAA
125774 TATGAGTGTGGAGTCGTTGCAATGCTGAAACCATATGAGATANNNAGATCGGAAG

125775 TATGAGTGTGGAGTCGTTGCGCAGAGAAACTGATTTATTTTTGNNNAGATCGGAAG
125776 TATGAGTGTGGAGTCGTTGCTGTGTGTCCTGAGTATTTATTTANNNAGATCGGAAG
125777 TATGAGTGTGGAGTCGTTGCGCTACCAAAATCTTTAGAAAACTNNNAGATCGGAAG
125778 TATGAGTGTGGAGTCGTTGCCCATGTATGATATGACACTTACANNNAGATCGGAAG
125779 TATGAGTGTGGAGTCGTTGCTCATGTATGAATATGTCTTCCCNNNAGATCGGAAGA
125780 TATGAGTGTGGAGTCGTTGCTATTTTTTTTTCAAAAGCCAACTNNNAGATCGGAAGA
125781 TATGAGTGTGGAGTCGTTGCAAGATACATAAGCCTATATGTGCNNNAGATCGGAAG
125782 TATGAGTGTGGAGTCGTTGCAAGATACATAAGCCTATATGTGCNNNAGATCGGAAG
125783 TATGAGTGTGGAGTCGTTGCAAGATACATAAGCCTATATGTGCNNNAGATCGGAAG
125784 TATGAGTGTGGAGTCGTTGCCATAACCAAATGTTCCTGAATATNNNAGATCGGAAG
125785 TATGAGTGTGGAGTCGTTGCATGACTTTATAATCACACTGTTTNNNAGATCGGAAG
125786 TATGAGTGTGGAGTCGTTGCAACTGAAAAATTCAACTCACAATNNNAGATCGGAAG
125787 TATGAGTGTGGAGTCGTTGCCTATCAGAAATTTTCACCTCAATNNNAGATCGGAAG
125788 TATGAGTGTGGAGTCGTTGCGATAGGTAAAGATATGACTGGTGNNNAGATCGGAA
125789 TATGAGTGTGGAGTCGTTGCTTTGCTTCCTAAAATGTAACTGNNNAGATCGGAAGA
125790 TATGAGTGTGGAGTCGTTGCATTTCTTGACTTGCATAGAAGTNNNAGATCGGAAGA
125791 TATGAGTGTGGAGTCGTTGCTGCAATTCCTAAATGTACTTTACNNNAGATCGGAAG
125792 TATGAGTGTGGAGTCGTTGCGAAATGAAAAACTGAAGAAATAGGNNNAGATCGG
125793 TATGAGTGTGGAGTCGTTGCCAAATCCTACAAAACAATGTTTCNNNAGATCGGAAG
125794 TATGAGTGTGGAGTCGTTGCATAAGGACTTCAGACTGAGGNNNAGATCGGAAGAG
125795 TATGAGTGTGGAGTCGTTGCTCAGTTCTGACTTTTTTTTACATNNNAGATCGGAAGA
125796 TATGAGTGTGGAGTCGTTGCTACAATGCGAGAGATAAGAGTANNNAGATCGGAAG
125797 TATGAGTGTGGAGTCGTTGCAGAGAAATTTGAAGGGTTACTATNNNAGATCGGAA
125798 TATGAGTGTGGAGTCGTTGCTTTGACTTTATAGTGGAGTAGCNNNAGATCGGAAG
125799 TATGAGTGTGGAGTCGTTGCCTACATCTGGCTATGATAAAGTANNNAGATCGGAAG
125800 TATGAGTGTGGAGTCGTTGCTCTGTGAAACTTTTGGACATTNNNAGATCGGAAGAG
125801 TATGAGTGTGGAGTCGTTGCGGAATAAGTACAATATCCAGAAAANNNAGATCGGA
125802 TATGAGTGTGGAGTCGTTGCGGAATAAGTACAATATCCAGAAAANNNAGATCGGA
125803 TATGAGTGTGGAGTCGTTGCTATATTTACTTATTTGGTCGGTGNNNAGATCGGAAG
125804 TATGAGTGTGGAGTCGTTGCCTCGAGGATTTTACATTTACAAGNNNAGATCGGAAG
125805 TATGAGTGTGGAGTCGTTGCAGATGACTGTATTCCACATTATTNNNAGATCGGAAG
125806 TATGAGTGTGGAGTCGTTGCGTGTATAAAAATAGCTGCTTCATNNNAGATCGGAAG
125807 TATGAGTGTGGAGTCGTTGCTATAGCAATGAAAATTGTGAGTTNNNAGATCGGAA
125808 TATGAGTGTGGAGTCGTTGCTTTGAAGAACTTTTTTCATTGTTNNNAGATCGGAAG
125809 TATGAGTGTGGAGTCGTTGCTTTTCTTACTTAATGTAGCTGTGNNNAGATCGGAAG
125810 TATGAGTGTGGAGTCGTTGCTATTGTGCTTACTATGTATGAGGNNNAGATCGGAAG
125811 TATGAGTGTGGAGTCGTTGCGAATATAAGATGATGGTCCCATANNNAGATCGGAA
125812 TATGAGTGTGGAGTCGTTGCATTTATTTTCATGTGTGATTGCTNNNAGATCGGAAG
125813 TATGAGTGTGGAGTCGTTGCAAATTTTTCTGTTTCTTCCATCCNNNAGATCGGAAGA
125814 TATGAGTGTGGAGTCGTTGCAAGGATTCACACTAAATAGCTAANNNAGATCGGAAG
125815 TATGAGTGTGGAGTCGTTGCATAATGGCATAAGATTCTGTGATNNNAGATCGGAA
125816 TATGAGTGTGGAGTCGTTGCCTACTCCTTTAGAATCATACTGANNNAGATCGGAAG
125817 TATGAGTGTGGAGTCGTTGCATCTTCCCTCCTTTCTCTTTTNNNAGATCGGAAGAGC

125818 TATGAGTGTGGAGTCGTTGCAGTCTTTTCTGGAATAAAGAATTNNNAGATCGGAAG
125819 TATGAGTGTGGAGTCGTTGCGGTCTTTCATAAAGGAATGATGNNNAGATCGGAAG
125820 TATGAGTGTGGAGTCGTTGCCGTATCAATTAAGAAGTCCAATTNNNAGATCGGAAG
125821 TATGAGTGTGGAGTCGTTGCTTATATGATATGGCATTTTGTGTNNNAGATCGGAAG
125822 TATGAGTGTGGAGTCGTTGCAACAAGAATTCACATATGATACCNNNAGATCGGAAG
125823 TATGAGTGTGGAGTCGTTGCTTCATTCATTTTTGTTCAACAAANNNAGATCGGAAGA
125824 TATGAGTGTGGAGTCGTTGCAAAAATGAATCCTAGAACAATGANNNAGATCGGAA
125826 TATGAGTGTGGAGTCGTTGCGAAGTCACAGGAAATAATTTAGTNNNAGATCGGAA
125827 TATGAGTGTGGAGTCGTTGCGTACAATCTCTTAGCTTCCTTTNNNAGATCGGAAGAG
125828 TATGAGTGTGGAGTCGTTGCCTGATCCATGGAAAACACTAANNNAGATCGGAAGAG
125829 TATGAGTGTGGAGTCGTTGCCAGAGGTATGACTGTTTACATCNNNAGATCGGAAGA
125830 TATGAGTGTGGAGTCGTTGCTTGCAAGAAATTATAAAGTCCATNNNAGATCGGAAG
125831 TATGAGTGTGGAGTCGTTGCCAGAATTACCAGAAACAGTAGTANNNAGATCGGAA
125832 TATGAGTGTGGAGTCGTTGCTGCTTAGAAGAGTGTAAATCCNNNAGATCGGAAGA
125833 TATGAGTGTGGAGTCGTTGCTCTGTGTAAAAATAACACTTTGTNNNAGATCGGAAG
125834 TATGAGTGTGGAGTCGTTGCTTTCTTGCATAATGAACAACATNNNAGATCGGAAGA
125836 TATGAGTGTGGAGTCGTTGCAGGTAAAAAGTATAGTACCTGATNNNAGATCGGAA
125837 TATGAGTGTGGAGTCGTTGCTGCCTGATTTTAATGCATTTATANNNAGATCGGAAG
125838 TATGAGTGTGGAGTCGTTGCGTTAATTATTTTTGAGACGGAGTNNNAGATCGGAA
125839 TATGAGTGTGGAGTCGTTGCCTTTAGGATTTGATTGTGACAGNNNAGATCGGAAGA
125840 TATGAGTGTGGAGTCGTTGCCAAATCTGTCTGCTCAATTATTTNNNAGATCGGAAGA
125841 TATGAGTGTGGAGTCGTTGCATTACATACATTACTTATATGCCTNNNAGATCGGAAG
125842 TATGAGTGTGGAGTCGTTGCCTATCTCTATCTGCAATCATTTGNNNAGATCGGAAGA
125843 TATGAGTGTGGAGTCGTTGCTAAGGAGAAACTGAGTTTGTTATNNNAGATCGGAA
125844 TATGAGTGTGGAGTCGTTGCGTCTTCTATTTCAGGCATCAGNNNAGATCGGAAGAG
125845 TATGAGTGTGGAGTCGTTGCACCTCTGAATACAAAACAATTTTNNNAGATCGGAAG
125846 TATGAGTGTGGAGTCGTTGCTTGCATCTATACTCTTAGGAAAANNNAGATCGGAAG
125847 TATGAGTGTGGAGTCGTTGCCTGAGAGAAAAATTTCACTGTTANNNAGATCGGAAG
125848 TATGAGTGTGGAGTCGTTGCTTAATCAGGATTAAAGCTTTTCCNNNAGATCGGAAG
125849 TATGAGTGTGGAGTCGTTGCCTAAGAAGGCTTTAGTCAAAGANNNAGATCGGAAG
125850 TATGAGTGTGGAGTCGTTGCCTAAGAAGGCTTTAGTCAAAGANNNAGATCGGAAG
125851 TATGAGTGTGGAGTCGTTGCTATGAAAGACAAGATACTCAAGTNNNAGATCGGAA
125852 TATGAGTGTGGAGTCGTTGCAGTTAGTGTATTTTTCTAGGACTNNNAGATCGGAAG
125853 TATGAGTGTGGAGTCGTTGCTTAAATTCATTCCAAAATGCAACNNNAGATCGGAAG
125854 TATGAGTGTGGAGTCGTTGCCTTCAGGGCTTCTAATTACAGNNNAGATCGGAAGAG
125855 TATGAGTGTGGAGTCGTTGCTTGATGTCTCAAATTATGGACTANNNAGATCGGAAG
125856 TATGAGTGTGGAGTCGTTGCACTTGATACTTTAAGATATCTGAAANNNAGATCGGA
125858 TATGAGTGTGGAGTCGTTGCGAAAGAAAGAAAAGAGACTTCACNNNAGATCGGAA
125859 TATGAGTGTGGAGTCGTTGCGTGGGGTTGTTTTATCTTTTGNNNAGATCGGAAGAG
125860 TATGAGTGTGGAGTCGTTGCTGTCAGAGGTTTTGTCATTTNNNAGATCGGAAGAGC
125861 TATGAGTGTGGAGTCGTTGCGGATCCTATATATACCTGGTAGANNNAGATCGGAAG
125862 TATGAGTGTGGAGTCGTTGCTATAATATGTGCGTGTGTGTATANNNAGATCGGAA
125863 TATGAGTGTGGAGTCGTTGCTTTCTGATAAAAACACCTTCTTCNNNAGATCGGAAGA

125864 TATGAGTGTGGAGTCGTTGCTGTTCTTAAAACAAACAGCAATANNNAGATCGGAAG
125865 TATGAGTGTGGAGTCGTTGCCAAAAAAAATCACTTCCAAATGANNNAGATCGGAAG
125866 TATGAGTGTGGAGTCGTTGCTCTTTCTACTTCTAACCTTTCATNNNAGATCGGAAGAG
125867 TATGAGTGTGGAGTCGTTGCAACATTTCTACTACTGTCTTTCTNNNAGATCGGAAGA
125868 TATGAGTGTGGAGTCGTTGCGCTGCTCTTTTTTTTTGTAATTTNNNAGATCGGAAGA
125869 TATGAGTGTGGAGTCGTTGCATATAGTAAGTGATCTTTGGGATNNNAGATCGGAAG
125870 TATGAGTGTGGAGTCGTTGCTAAGTTGGATTCCTAGGTATTTTNNNAGATCGGAAG
125871 TATGAGTGTGGAGTCGTTGCATTGGACTTATATCTGTAGGAAGNNNAGATCGGAAG
125872 TATGAGTGTGGAGTCGTTGCTGCTAGAATTAGAAAATATGGGANNNAGATCGGAA
125873 TATGAGTGTGGAGTCGTTGCCAATAAATTCATCCCATACTCAANNNAGATCGGAAGA
125874 TATGAGTGTGGAGTCGTTGCCCTTGGTTTATGCTGAAAAATNNNAGATCGGAAGAG
125875 TATGAGTGTGGAGTCGTTGCCCTTGGTTTATGCTGAAAAATNNNAGATCGGAAGAG
125876 TATGAGTGTGGAGTCGTTGCATTTATACATACCAAGTCTCAGTNNNAGATCGGAAGA
125877 TATGAGTGTGGAGTCGTTGCTACCAGTGTTCATTTTCTAGTATNNNAGATCGGAAGA
125879 TATGAGTGTGGAGTCGTTGCTTCCTCAGTCATACACATTATAANNNAGATCGGAAGA
125880 TATGAGTGTGGAGTCGTTGCATATTCTAACATTTCAAGTGAGCNNNAGATCGGAAGA
125881 TATGAGTGTGGAGTCGTTGCAGTACTTGACTTTATGCGATTNNNAGATCGGAAGAG
125882 TATGAGTGTGGAGTCGTTGCTATAAAGGGTCTCCTAAAATGTCNNNAGATCGGAAG
125883 TATGAGTGTGGAGTCGTTGCGTTTTTTTCCCCTTTATCTCTTCNNNAGATCGGAAGAG
125884 TATGAGTGTGGAGTCGTTGCATCATGCTCATATAATGTTTTGGNNNAGATCGGAAGA
125885 TATGAGTGTGGAGTCGTTGCCAACACAAAGAAGTAAGAGAATGNNNAGATCGGAA
125886 TATGAGTGTGGAGTCGTTGCAATGAAGACTAAACCTTGGAGNNNAGATCGGAAGA
125887 TATGAGTGTGGAGTCGTTGCCTTTCTTTCAATGTTTTTTGTCTNNNAGATCGGAAGA
125888 TATGAGTGTGGAGTCGTTGCAGCATGGTTTCTTTGTAATTTATNNNAGATCGGAAGA
125890 TATGAGTGTGGAGTCGTTGCTCTCTGAATCAAAAGAAGTCATANNNAGATCGGAAG
125891 TATGAGTGTGGAGTCGTTGCTTCTCTTTCTCTTACTGTGTTATNNNAGATCGGAAGA
125892 TATGAGTGTGGAGTCGTTGCCTCTCCTTTTGGGGTATTTCNNNAGATCGGAAGAGCA
125893 TATGAGTGTGGAGTCGTTGCTAAGATACCCCATTATTATTGCANNNAGATCGGAAGA
125894 TATGAGTGTGGAGTCGTTGCCTTCACCTAGTGTAATCTTCTTANNNAGATCGGAAGA
125895 TATGAGTGTGGAGTCGTTGCCATGTATGAATCTGATGATTTCANNNAGATCGGAAG
125896 TATGAGTGTGGAGTCGTTGCGAGCTAAAGTCAGACTGTTATAANNNAGATCGGAAG
125898 TATGAGTGTGGAGTCGTTGCTAATACTTGTACAGCTAGGATTTNNNAGATCGGAAG
125901 TATGAGTGTGGAGTCGTTGCAATAGAGAAAGAGAGAGTTGTAGNNNAGATCGGAA
125902 TATGAGTGTGGAGTCGTTGCTCTAGATGAATGTTCCTTATAGGNNNAGATCGGAAG
125903 TATGAGTGTGGAGTCGTTGCCAAGAGAGCACAGTTTAAATTTNNNAGATCGGAAGA
125904 TATGAGTGTGGAGTCGTTGCATTTTTGATCGTCTGATGATTTCNNNAGATCGGAAGA
125905 TATGAGTGTGGAGTCGTTGCAGTTAAATTTACTGTCTCAGAGANNNAGATCGGAAG
125906 TATGAGTGTGGAGTCGTTGCCAGAATTTAGCATGCAATATGTNNNAGATCGGAAGA
125907 TATGAGTGTGGAGTCGTTGCAATTCTGATTTCAATGTCAGTGNNNAGATCGGAAGA
125908 TATGAGTGTGGAGTCGTTGCAATTCTGATTTCAATGTCAGTGNNNAGATCGGAAGA
125909 TATGAGTGTGGAGTCGTTGCTTTATCCAAAGCCTCCTTAGATANNNAGATCGGAAGA
125910 TATGAGTGTGGAGTCGTTGCATAAGTCAACACACACATATAACNNNAGATCGGAAGA
125911 TATGAGTGTGGAGTCGTTGCAAAAGCAGCCTTTGTGAATTNNNAGATCGGAAGAG

125912 TATGAGTGTGGAGTCGTTGCGAGAATGCTCCCATAGATCCNNNAGATCGGAAGAG
125913 TATGAGTGTGGAGTCGTTGCGCAGATGTATAAGGACATTTTTTNNNAGATCGGAAG
125914 TATGAGTGTGGAGTCGTTGCTAAATGACTCACCAAAGATACATNNNAGATCGGAAG
125915 TATGAGTGTGGAGTCGTTGCGAACTACCAGGCTTTGTTACNNNAGATCGGAAGAG
125916 TATGAGTGTGGAGTCGTTGCTCCAGTGTTAGTTTTTTTTTTCNNNAGATCGGAAGA
125917 TATGAGTGTGGAGTCGTTGCAAACACTCAGTATTCTGTGTTANNNAGATCGGAAGA
125918 TATGAGTGTGGAGTCGTTGCCCGTAGTCTGTTTAGTTTAACTANNNAGATCGGAAG
125919 TATGAGTGTGGAGTCGTTGCCAGTACTAAAAGATTAGGACCTTNNNAGATCGGAAG
125920 TATGAGTGTGGAGTCGTTGCGCACACGTTTTTTATTCTAATCNNNAGATCGGAAGA
125921 TATGAGTGTGGAGTCGTTGCATTTTTCTTTACTGTTTGACTGTNNNAGATCGGAAGA
125922 TATGAGTGTGGAGTCGTTGCACGTAGGTTTCTAAACTATCTTCNNNAGATCGGAAG
125923 TATGAGTGTGGAGTCGTTGCGACAAGAGATTAGGTGTTCTCNNNAGATCGGAAGA
125925 TATGAGTGTGGAGTCGTTGCGCTAGGTTTTTCGGTTTTTAATANNNAGATCGGAAG
125926 TATGAGTGTGGAGTCGTTGCCTCAATTGTGTAAGAGGAAAATCNNNAGATCGGAA
125927 TATGAGTGTGGAGTCGTTGCGTGAGAAACAGAAAAAAAATTACANNNAGATCGGA
125928 TATGAGTGTGGAGTCGTTGCGAAAGGAATATAAGGTAGTGGAANNNAGATCGGA
125929 TATGAGTGTGGAGTCGTTGCTGTTTTGTTTTGTTTTGTTTTTGNNNAGATCGGAAGA
125930 TATGAGTGTGGAGTCGTTGCTGAAGGCCTAGAAACAATAATATNNNAGATCGGAA
125931 TATGAGTGTGGAGTCGTTGCAACAATTGAAGATGCAATTCTTGNNNAGATCGGAAG
125932 TATGAGTGTGGAGTCGTTGCAGTGAGAGAAAGAGAAGAATTTTNNNAGATCGGAA
125933 TATGAGTGTGGAGTCGTTGCCTTCTGCTGCTACAGATAAANNNAGATCGGAAGAGC
125934 TATGAGTGTGGAGTCGTTGCGAAAGAAGGAAGGAAGGAAAANNNAGATCGGAAG
125935 TATGAGTGTGGAGTCGTTGCTTTGTCTGGAGAAAGTTCATTGNNNAGATCGGAAG
125936 TATGAGTGTGGAGTCGTTGCCCAGTGAACTGATATTTAGAAGTNNNAGATCGGAAG
125937 TATGAGTGTGGAGTCGTTGCCCAGTGAACTGATATTTAGAAGTNNNAGATCGGAAG
125938 TATGAGTGTGGAGTCGTTGCTTTCTTCCCAATGATTCCTATTANNNAGATCGGAAGA
125939 TATGAGTGTGGAGTCGTTGCATTTTACTTCATCTCCTTTGTTGNNNAGATCGGAAGA
125940 TATGAGTGTGGAGTCGTTGCAGAAAGATACAAAATTGCATGATNNNAGATCGGAA
125941 TATGAGTGTGGAGTCGTTGCTGTTTTTCAAAGTACCTAAAACANNNAGATCGGAAG
125942 TATGAGTGTGGAGTCGTTGCGTACAGGTTGAGTGTGTATCNNNAGATCGGAAGAG
125943 TATGAGTGTGGAGTCGTTGCAAATGCAGTAGAGAAGGACTNNNAGATCGGAAGAG
125944 TATGAGTGTGGAGTCGTTGCCTAGAATGTTTTTGCTTTTCTTGNNNAGATCGGAAG
125945 TATGAGTGTGGAGTCGTTGCCATATTTCATGAAAACGCAGATNNNAGATCGGAAGA
125946 TATGAGTGTGGAGTCGTTGCGTATTCAGAAGTCTTATCATCCTNNNAGATCGGAAG
125947 TATGAGTGTGGAGTCGTTGCGGAGTGATTTTCACCAAAATATTNNNAGATCGGAAG
125948 TATGAGTGTGGAGTCGTTGCATTTGTAATGCTGAAGAGTATTCNNNAGATCGGAAG
125949 TATGAGTGTGGAGTCGTTGCAAATTTCCAGCATACAATACAATNNNAGATCGGAAG
125950 TATGAGTGTGGAGTCGTTGCCTTTTTTCTTCTATTCCTCCAAANNNAGATCGGAAGA
125951 TATGAGTGTGGAGTCGTTGCTATGATCTACCCTGTGCTTACNNNAGATCGGAAGAG
125952 TATGAGTGTGGAGTCGTTGCTGTGTTTTTTCAAAGTTTACCTTNNNAGATCGGAAGA
125953 TATGAGTGTGGAGTCGTTGCCTTGATTACCTCCACCTGTTNNNAGATCGGAAGAGCA
125954 TATGAGTGTGGAGTCGTTGCATGAAATGCATGGCTTTATTTATNNNAGATCGGAAG
125955 TATGAGTGTGGAGTCGTTGCGTGATAACAGTAAGGATAGAGAANNNAGATCGGA

125956 TATGAGTGTGGAGTCGTTGCCCTGTGATGAAGTAAGTAAGGNNNAGATCGGAAGA
125957 TATGAGTGTGGAGTCGTTGCTGTGAATAATCTAAATGAGCATGNNNAGATCGGAA
125958 TATGAGTGTGGAGTCGTTGCACAAACTGGATGAAGTGTAAANNNAGATCGGAAGA
125959 TATGAGTGTGGAGTCGTTGCTGATTTATGTATTTGTTTTCCGTNNNAGATCGGAAG
125960 TATGAGTGTGGAGTCGTTGCAAGGGGATGTGAGTAAAGATANNNAGATCGGAAG
125961 TATGAGTGTGGAGTCGTTGCACTTTTCAAATTCTCATGAGATCNNNAGATCGGAAG
125962 TATGAGTGTGGAGTCGTTGCCTTCAAAACAGCTACTATTACCNNNAGATCGGAAGA
125963 TATGAGTGTGGAGTCGTTGCCTAGCTATCTTGGGAATCTATTTNNNAGATCGGAAG
125964 TATGAGTGTGGAGTCGTTGCCTTAGCAGAAGGCAGAATAGNNNAGATCGGAAGAG
125965 TATGAGTGTGGAGTCGTTGCTGCATGATATTTTTATGAAACCANNNAGATCGGAAG
125966 TATGAGTGTGGAGTCGTTGCGACATTTTGCATGTTAAAATCATNNNAGATCGGAAG
125967 TATGAGTGTGGAGTCGTTGCGAGAACTAATGGGTGAGAAGNNNAGATCGGAAGA
125968 TATGAGTGTGGAGTCGTTGCGTAACAATGGTGAAACTTGAATANNNAGATCGGAA
125970 TATGAGTGTGGAGTCGTTGCTACTAAATTGAAAGTAGGTACCANNNAGATCGGAA
125971 TATGAGTGTGGAGTCGTTGCCAGAGAGGTTAGGAATTGGGNNNAGATCGGAAGA
125972 TATGAGTGTGGAGTCGTTGCGCAGTCAGGATTTTATTTTTAGANNNAGATCGGAAG
125973 TATGAGTGTGGAGTCGTTGCGAAACAAGCTATGGTGAGATNNNAGATCGGAAGAG
125974 TATGAGTGTGGAGTCGTTGCGTGGTAATGCTAAAACTCAGANNNAGATCGGAAGA
125975 TATGAGTGTGGAGTCGTTGCAAAGAGAGATACCAGGAAAAGTANNNAGATCGGAA
125976 TATGAGTGTGGAGTCGTTGCAATTGAATAAATGAATGTGACCANNNAGATCGGAA
125979 TATGAGTGTGGAGTCGTTGCTATCAACAAAAGTTTTAACCCAGNNNAGATCGGAAG
125980 TATGAGTGTGGAGTCGTTGCTAGTGGAAATATGGTAGAATGTTNNNAGATCGGAA
125981 TATGAGTGTGGAGTCGTTGCTATTTGATAAACAATGCTGAGACNNNAGATCGGAAG
125982 TATGAGTGTGGAGTCGTTGCTATTTGATAAACAATGCTGAGACNNNAGATCGGAAG
125983 TATGAGTGTGGAGTCGTTGCGACTTTTCCTCTTAACATATCTCNNNAGATCGGAAGA
125984 TATGAGTGTGGAGTCGTTGCCAGGCTTTGCTCATTAGTTGNNNAGATCGGAAGAGC
125985 TATGAGTGTGGAGTCGTTGCGAGAGGCATACAGACCTATGNNNAGATCGGAAGAG
125986 TATGAGTGTGGAGTCGTTGCCTACATTATCTGGCATGGAGNNNAGATCGGAAGAG
125987 TATGAGTGTGGAGTCGTTGCCAGGTACATAGTTGTAGGATACNNNAGATCGGAAG
125988 TATGAGTGTGGAGTCGTTGCTATCTGAATGTAAGAGTTACAGGNNNAGATCGGAA
125989 TATGAGTGTGGAGTCGTTGCAAGAAATATGAAGAGATGTTTGGNNNAGATCGGAA
125990 TATGAGTGTGGAGTCGTTGCGAGAAGGCTGACAAATTAGTNNNAGATCGGAAGAG
125991 TATGAGTGTGGAGTCGTTGCGCCTTTTCCATGTTATTTATTTTCNNNAGATCGGAAG
125992 TATGAGTGTGGAGTCGTTGCTATGCCTTTTCTTCCTCTATATTNNNAGATCGGAAGA
125993 TATGAGTGTGGAGTCGTTGCCAAGTTTAAAAACCCACAAAAATNNNAGATCGGAAG
125994 TATGAGTGTGGAGTCGTTGCTTTGTATAATGTATCCTGAAGGTNNNAGATCGGAAG
125995 TATGAGTGTGGAGTCGTTGCTTTCTTCAAGTTACTGATTTCAGNNNAGATCGGAAG
125996 TATGAGTGTGGAGTCGTTGCTAAATACACATCCCATCTGAAANNNAGATCGGAAGA
125997 TATGAGTGTGGAGTCGTTGCAATATCTGGGTGAAAAAACTATCNNNAGATCGGAA
125998 TATGAGTGTGGAGTCGTTGCTCTTGAACATCAATAAGGAAATGNNNAGATCGGAA
125999 TATGAGTGTGGAGTCGTTGCAATTTCTCATCTTCCAAATTTCANNNAGATCGGAAGA
126000 TATGAGTGTGGAGTCGTTGCAAGTAAAAAAAATGTAGGGAAGCNNNAGATCGGAA
126001 TATGAGTGTGGAGTCGTTGCTGCCATGCTTTTTATTATAGAAANNNAGATCGGAAG

126002 TATGAGTGTGGAGTCGTTGCCACAATTTCCTTTTGTTCAAAATNNNAGATCGGAAGA
126003 TATGAGTGTGGAGTCGTTGCCACAATTTCCTTTTGTTCAAAATNNNAGATCGGAAGA
126004 TATGAGTGTGGAGTCGTTGCTTTGTTCAGTGTGAAACAAATAGNNNAGATCGGAAG
126005 TATGAGTGTGGAGTCGTTGCTTTGTTCAGTGTGAAACAAATAGNNNAGATCGGAAG
126006 TATGAGTGTGGAGTCGTTGCCAGAATTTCTCACATTGACATTANNNAGATCGGAAG
126007 TATGAGTGTGGAGTCGTTGCAGTGAACATTTAAAACAAAGTCTNNNAGATCGGAAG
126008 TATGAGTGTGGAGTCGTTGCAAGACATAAGAATTGAACAGATGNNNAGATCGGAA
126009 TATGAGTGTGGAGTCGTTGCAAATGTAGCTTAGGGGAATAAAGNNNAGATCGGAA
126011 TATGAGTGTGGAGTCGTTGCCTGGAACTAGACTATTCTTACATNNNAGATCGGAAG
126012 TATGAGTGTGGAGTCGTTGCTCCCTAATAATTGCAACATATCTNNNAGATCGGAAGA
126013 TATGAGTGTGGAGTCGTTGCTCTCTAGAAAGATACTCAATTGGNNNAGATCGGAAG
126014 TATGAGTGTGGAGTCGTTGCTCTTTTTCTAACTCTGAAAATGTNNNAGATCGGAAGA
126015 TATGAGTGTGGAGTCGTTGCATACTGTTCCTTCTGTCTAAACNNNAGATCGGAAGAG
126016 TATGAGTGTGGAGTCGTTGCCACAGGTTCTTTTCCTTTATTATNNNAGATCGGAAGA
126017 TATGAGTGTGGAGTCGTTGCCGCCCTATATATTTTGGTCTTNNNAGATCGGAAGAG
126018 TATGAGTGTGGAGTCGTTGCGGTATGTCTTTCTTTTTCCTTAANNNAGATCGGAAGA
126019 TATGAGTGTGGAGTCGTTGCCTTTTGGTCTACATGGTACTCNNNAGATCGGAAGAG
126020 TATGAGTGTGGAGTCGTTGCAAAGTACAAGCGTTGTGATTANNNAGATCGGAAGA
126021 TATGAGTGTGGAGTCGTTGCATATACAAATGCATCGTATATCCNNNAGATCGGAAG
126022 TATGAGTGTGGAGTCGTTGCGTATGGATGATTCTGTTTTTCTTNNNAGATCGGAAG
126023 TATGAGTGTGGAGTCGTTGCTTGTTAATTTTTGCCAATTATGCNNNAGATCGGAAG
126024 TATGAGTGTGGAGTCGTTGCGCTTTCAAAAGTACAACTTAATGNNNAGATCGGAAG
126025 TATGAGTGTGGAGTCGTTGCAACAACACAGACACAAATAATAANNNAGATCGGAAG
126026 TATGAGTGTGGAGTCGTTGCCTCCCAAAGTGCTGAATTTANNNAGATCGGAAGAGC
126027 TATGAGTGTGGAGTCGTTGCATACAGAAAGAAAGAAGAAAGGANNNAGATCGGA
126028 TATGAGTGTGGAGTCGTTGCTCCTAGAAAAACAAATTCAAACANNNAGATCGGAAG
126029 TATGAGTGTGGAGTCGTTGCGTTCCAATTTACACAATGGAGNNNAGATCGGAAGAG
126030 TATGAGTGTGGAGTCGTTGCAATTCTTTTGTGATGTTCTTTCTNNNAGATCGGAAGA
126031 TATGAGTGTGGAGTCGTTGCCTAGGATGCAAGTAGTTTTAGTTNNNAGATCGGAAG
126032 TATGAGTGTGGAGTCGTTGCAAGGTTTTATCAAGATAATGCTGNNNAGATCGGAAG
126033 TATGAGTGTGGAGTCGTTGCACTTCCTTACCTATTTGTATTCCNNNAGATCGGAAGA
126034 TATGAGTGTGGAGTCGTTGCATATGTTTAAATGGAGCAAAACANNNAGATCGGAAG
126035 TATGAGTGTGGAGTCGTTGCAAATTTTTCCCTTGCCATGANNNAGATCGGAAGAGC
126036 TATGAGTGTGGAGTCGTTGCGGAGACTAGAATTGAATCTCTAANNNAGATCGGAAG
126037 TATGAGTGTGGAGTCGTTGCCAGCTATCTTTTTTTTTCCCTAANNNAGATCGGAAGA
126038 TATGAGTGTGGAGTCGTTGCTTTAGGTGTCCAGACATTGANNNAGATCGGAAGAG
126039 TATGAGTGTGGAGTCGTTGCTGTTTCTTTGTTTCTTATTACCANNNAGATCGGAAGA
126040 TATGAGTGTGGAGTCGTTGCTTATCTTCAAGGTCTCTAGTAACNNNAGATCGGAAG
126041 TATGAGTGTGGAGTCGTTGCCTGTGGCTAAATATGAATTAGACNNNAGATCGGAAG
126042 TATGAGTGTGGAGTCGTTGCCATGAAAACATAGTGAGGAATCNNNAGATCGGAAG
126044 TATGAGTGTGGAGTCGTTGCTAGGCAGATGACTAGAATCAAANNNAGATCGGAAG
126045 TATGAGTGTGGAGTCGTTGCCATGAGCAAAATCACTCACTNNNAGATCGGAAGAGC
126046 TATGAGTGTGGAGTCGTTGCAACTTTAATTCCTTATGCATGAGNNNAGATCGGAAG

126047 TATGAGTGTGGAGTCGTTGCCTCGTGTTTTTAGTTTAGATTGANNNAGATCGGAAG
126048 TATGAGTGTGGAGTCGTTGCCAGGCTGAGTATGATTATTTTTGNNNAGATCGGAAG
126049 TATGAGTGTGGAGTCGTTGCAAACAGAACAAAACGAAAACTANNNAGATCGGAAG
126051 TATGAGTGTGGAGTCGTTGCTATACAGTTTTACATCATGTAGCNNNAGATCGGAAG
126052 TATGAGTGTGGAGTCGTTGCAAATTTAGCCCATAAATTGTTCANNNAGATCGGAAG
126053 TATGAGTGTGGAGTCGTTGCGCCTAAAAATATTTACTCCTCACNNNAGATCGGAAGA
126054 TATGAGTGTGGAGTCGTTGCGCAGCACCATTTATTGATATTTTNNNAGATCGGAAG
126055 TATGAGTGTGGAGTCGTTGCTTTATAGGTGGAATCACTACCTANNNAGATCGGAAG
126056 TATGAGTGTGGAGTCGTTGCAAAGTGAATAAGAGGTGGTATAANNNAGATCGGAA
126057 TATGAGTGTGGAGTCGTTGCCTCTACTCCTCATCTGTGTGNNNAGATCGGAAGAGC
126058 TATGAGTGTGGAGTCGTTGCGTCTGAAAATAAGGCTTTTTTTGNNNAGATCGGAAG
126059 TATGAGTGTGGAGTCGTTGCCTATACCTAATTCAAATACTGCCNNNAGATCGGAAGA
126060 TATGAGTGTGGAGTCGTTGCTATAGAGAGAGAGAGAGAGAGAGNNNAGATCGGA
126061 TATGAGTGTGGAGTCGTTGCTTTGTGAATGTATTTACCTAGTGNNNAGATCGGAAG
126062 TATGAGTGTGGAGTCGTTGCCTAAAGGCAGGTTAATAGAAGAANNNAGATCGGAA
126063 TATGAGTGTGGAGTCGTTGCTGTCCATTTTTTTCAAAAGTTTTNNNAGATCGGAAGA
126064 TATGAGTGTGGAGTCGTTGCTGTCCATTTTTTTCAAAAGTTTTNNNAGATCGGAAGA
126065 TATGAGTGTGGAGTCGTTGCCCATGATATGTGGGAATTAAGANNNAGATCGGAAG
126066 TATGAGTGTGGAGTCGTTGCCAAGTCTCAAGATTTTTTGTTTGNNNAGATCGGAAG
126068 TATGAGTGTGGAGTCGTTGCAAAATGAAATGGGGTTAAATGAANNNAGATCGGAA
126069 TATGAGTGTGGAGTCGTTGCATATTCCTTAGAAATTTTGGAGGNNNAGATCGGAAG
126070 TATGAGTGTGGAGTCGTTGCATATTCCTTAGAAATTTTGGAGGNNNAGATCGGAAG
126071 TATGAGTGTGGAGTCGTTGCGATTCCTCTTAAAGATAAGCAAGNNNAGATCGGAAG
126072 TATGAGTGTGGAGTCGTTGCTTTTTGGATTTTTTGTAAAGCTCNNNAGATCGGAAG
126073 TATGAGTGTGGAGTCGTTGCCAGAAAGAGGAAAAAAGAGGTNNNAGATCGGAAG
126074 TATGAGTGTGGAGTCGTTGCCAGAAAGAGGAAAAAAGAGGTNNNAGATCGGAAG
126075 TATGAGTGTGGAGTCGTTGCCTTGAAGAATCTGGAGAGTTAATNNNAGATCGGAA
126076 TATGAGTGTGGAGTCGTTGCAGAACAAGGAATGAAGAAGTTANNNAGATCGGAAG
126077 TATGAGTGTGGAGTCGTTGCCACTTTTTGGTTAATAGCTTCTANNNAGATCGGAAGA
126078 TATGAGTGTGGAGTCGTTGCTACCCATTTCCTTGAATATTAGGNNNAGATCGGAAG
126079 TATGAGTGTGGAGTCGTTGCTGGTTAAATTATCTGGAGTCTTTNNNAGATCGGAAG
126080 TATGAGTGTGGAGTCGTTGCGAATCAAAACTGAAAGGACAATANNNAGATCGGAA
126081 TATGAGTGTGGAGTCGTTGCCTTTTTAGTCTGTATCTCCAGGNNNAGATCGGAAGA
126082 TATGAGTGTGGAGTCGTTGCGAGAGAAAGAGAAAGAAAGAGATNNNAGATCGGA
126083 TATGAGTGTGGAGTCGTTGCAATGGACTCTTTAAAAAATAGCTNNNAGATCGGAAG
126084 TATGAGTGTGGAGTCGTTGCTTATACCACTATCTTGTTCGTAGNNNAGATCGGAAG
126085 TATGAGTGTGGAGTCGTTGCTATTAGTCCGTTCTCATGCTNNNAGATCGGAAGAGC
126086 TATGAGTGTGGAGTCGTTGCTATTAGTCCGTTCTCATGCTNNNAGATCGGAAGAGC
126087 TATGAGTGTGGAGTCGTTGCCAAATGCCTTTTATTTCTTGTCTNNNAGATCGGAAGA
126089 TATGAGTGTGGAGTCGTTGCTAATCTTAGCCTTTTCTAATCGANNNAGATCGGAAGA
126090 TATGAGTGTGGAGTCGTTGCCCATTTAGACTCCATTCATTTATNNNAGATCGGAAGA
126091 TATGAGTGTGGAGTCGTTGCTTGATCATTTATACAGGATACCTNNNAGATCGGAAG
126092 TATGAGTGTGGAGTCGTTGCTTGATCATTTATACAGGATACCTNNNAGATCGGAAG

126093 TATGAGTGTGGAGTCGTTGCGTAGCATCAATAACTGTGGAANNNAGATCGGAAGA
126094 TATGAGTGTGGAGTCGTTGCTGCTTCCTGTGGTTTTTTTTNNNAGATCGGAAGAGC
126095 TATGAGTGTGGAGTCGTTGCCAAACCTTGTTTAGGGTCTTNNNAGATCGGAAGAGC
126096 TATGAGTGTGGAGTCGTTGCGTTCCATATTTTTGCAATTGAGANNNAGATCGGAAG
126097 TATGAGTGTGGAGTCGTTGCGCTTGTATGTCTGTGTTTTATTNNNAGATCGGAAGA
126098 TATGAGTGTGGAGTCGTTGCATTTAAGCACTAAGATACCATCTNNNAGATCGGAAG
126099 TATGAGTGTGGAGTCGTTGCTATCATAAACACCTCTTCAATCTNNNAGATCGGAAGA
126100 TATGAGTGTGGAGTCGTTGCAGAATTTGTTCAGACAGACTTNNNAGATCGGAAGA
126101 TATGAGTGTGGAGTCGTTGCCCTATGTATATACCGAGTGTGANNNAGATCGGAAGA
126102 TATGAGTGTGGAGTCGTTGCCTAGCAATGAATTAAGACATACCNNNAGATCGGAAG
126103 TATGAGTGTGGAGTCGTTGCATGGTATGAGAAAATGATCTTCTNNNAGATCGGAA
126104 TATGAGTGTGGAGTCGTTGCAAGAAGCTAATGAGAATGGAATANNNAGATCGGAA
126105 TATGAGTGTGGAGTCGTTGCTCAATAGTTTTGTTGTAGATGTGNNNAGATCGGAAG
126106 TATGAGTGTGGAGTCGTTGCCCTCACATAACTTTAACTCAAATNNNAGATCGGAAGA
126107 TATGAGTGTGGAGTCGTTGCAAAGACTTATCGGTTGATTTTTANNNAGATCGGAAG
126108 TATGAGTGTGGAGTCGTTGCCACAACAGAGATTCACATACATANNNAGATCGGAAG
126109 TATGAGTGTGGAGTCGTTGCGGTAAAGGACAGGAATATAGAAANNNAGATCGGA
126110 TATGAGTGTGGAGTCGTTGCTCTCATATATTGCTCTTCACAATNNNAGATCGGAAGA
126111 TATGAGTGTGGAGTCGTTGCGAATTGTCTTGACACTGATAAAANNNAGATCGGAAG
126112 TATGAGTGTGGAGTCGTTGCTGAGTTTATGTATGTATATGGGTNNNAGATCGGAA
126113 TATGAGTGTGGAGTCGTTGCTCTATATCTATGTGTGCCTATCTNNNAGATCGGAAGA
126114 TATGAGTGTGGAGTCGTTGCTCTATATCTATGTGTGCCTATCTNNNAGATCGGAAGA
126115 TATGAGTGTGGAGTCGTTGCTGTAAGTTATACCTGAATGTACCNNNAGATCGGAAG
126116 TATGAGTGTGGAGTCGTTGCAGTAAGATTTCTCCTGATCAGTANNNAGATCGGAAG
126117 TATGAGTGTGGAGTCGTTGCGAACATGATGTCTCAAGTTAATTNNNAGATCGGAAG
126118 TATGAGTGTGGAGTCGTTGCCTTTATTCCACAAGTAGGTTTGNNNAGATCGGAAGA
126119 TATGAGTGTGGAGTCGTTGCACTTTTGTAATAGAGACACTTCANNNAGATCGGAAG
126120 TATGAGTGTGGAGTCGTTGCAACGACAATCTAATAGACTAACANNNAGATCGGAAG
126121 TATGAGTGTGGAGTCGTTGCACCATGTTCTAATGCATTATTTTNNNAGATCGGAAGA
126122 TATGAGTGTGGAGTCGTTGCTTTTATGATCTTTTCCTTCTGTGNNNAGATCGGAAGA
126123 TATGAGTGTGGAGTCGTTGCAATCTTACAATCAGCAATTTCTCNNNAGATCGGAAGA
126124 TATGAGTGTGGAGTCGTTGCCCCTGTAAAACACATTACTAAAGNNNAGATCGGAAG
126125 TATGAGTGTGGAGTCGTTGCAAATGAGAATTTTCTGTTTGACTNNNAGATCGGAAG
126126 TATGAGTGTGGAGTCGTTGCCTCTGTTGAATCTGATTGTAAAANNNAGATCGGAAG
126127 TATGAGTGTGGAGTCGTTGCATATTTTCCTGTCATTCACATTCNNNAGATCGGAAGA
126128 TATGAGTGTGGAGTCGTTGCGACAGATGAGCTACTACTTAAATNNNAGATCGGAAG
126129 TATGAGTGTGGAGTCGTTGCCACATGTTTATCCAAAATTTGACNNNAGATCGGAAG
126131 TATGAGTGTGGAGTCGTTGCTGCATTTCTATTCTTTGTTTTCTNNNAGATCGGAAGA
126132 TATGAGTGTGGAGTCGTTGCAGTAATGACAAAACTTCCAAAATNNNAGATCGGAAG
126133 TATGAGTGTGGAGTCGTTGCGAAAGAAAGAGAGTAGGAGAAANNNAGATCGGAA
126134 TATGAGTGTGGAGTCGTTGCGAATTCAAAGTGAAAACATTGTTNNNAGATCGGAA
126135 TATGAGTGTGGAGTCGTTGCTTACATTTAAAACCCTAAATCCGNNNAGATCGGAAG
126136 TATGAGTGTGGAGTCGTTGCTTACATTTAAAACCCTAAATCCGNNNAGATCGGAAG

126137 TATGAGTGTGGAGTCGTTGCATGAGTTCAGTTTTATACCTGTTNNNAGATCGGAAG
126138 TATGAGTGTGGAGTCGTTGCATTTACACTGCTCCACTAATCNNNAGATCGGAAGAG
126139 TATGAGTGTGGAGTCGTTGCTAAGTAAGTAAAATGCAGTGATGNNNAGATCGGAA
126140 TATGAGTGTGGAGTCGTTGCCCTGTTGTATAAGAAATCCTTTGNNNAGATCGGAAG
126141 TATGAGTGTGGAGTCGTTGCAGCTTATTCAGATGATACTAACANNNAGATCGGAAG
126142 TATGAGTGTGGAGTCGTTGCAGAAGATTAATTGTTGAAGTCTTNNNAGATCGGAA
126143 TATGAGTGTGGAGTCGTTGCGGGATGATCATATGACAGTAATANNNAGATCGGAA
126144 TATGAGTGTGGAGTCGTTGCCGGAAGACTTGAATATAAAATGTNNNAGATCGGAA
126145 TATGAGTGTGGAGTCGTTGCTCAAGATGACTTACTAGTAGTATCNNNAGATCGGAA
126147 TATGAGTGTGGAGTCGTTGCTGAATTATCACTATACAGTACCGNNNAGATCGGAAG
126148 TATGAGTGTGGAGTCGTTGCAAAATCTTTGCCCTTACAATATTNNNAGATCGGAAG
126149 TATGAGTGTGGAGTCGTTGCCACTCACATACACATATACATGTNNNAGATCGGAAG
126150 TATGAGTGTGGAGTCGTTGCGTGATAATGAAAGAGGAAGAATCNNNAGATCGGA
126151 TATGAGTGTGGAGTCGTTGCGGGATGATAATCATACACAGTATNNNAGATCGGAA
126152 TATGAGTGTGGAGTCGTTGCGTTCTGTGCCTATAAACAGAGNNNAGATCGGAAGA
126153 TATGAGTGTGGAGTCGTTGCACAACAAAATATTAACAAGCCATNNNAGATCGGAAG
126154 TATGAGTGTGGAGTCGTTGCTTATTCTAGATCCTTGCTGATTTNNNAGATCGGAAG
126155 TATGAGTGTGGAGTCGTTGCGAAGTAATGGTGTACTTTCATTCNNNAGATCGGAAG
126156 TATGAGTGTGGAGTCGTTGCATAACAGAATGGTCAACCTTTTANNNAGATCGGAAG
126157 TATGAGTGTGGAGTCGTTGCAAATTTGGGGATTATAAAGCATTNNNAGATCGGAA
126158 TATGAGTGTGGAGTCGTTGCTAACAATTTAGAAGCATCAAACGNNNAGATCGGAAG
126159 TATGAGTGTGGAGTCGTTGCACAACATTCAGATACATATTATCANNNAGATCGGAA
126160 TATGAGTGTGGAGTCGTTGCAAACTGAAAAATCAAGAAACAACNNNAGATCGGAA
126161 TATGAGTGTGGAGTCGTTGCTTCACTTAAAAATATGTCATGGGNNNAGATCGGAAG
126162 TATGAGTGTGGAGTCGTTGCAACTTTATCCTGTGAGTGATTTANNNAGATCGGAAG
126163 TATGAGTGTGGAGTCGTTGCGGGCATTTTAAATAAAAAGGATCNNNAGATCGGAA
126164 TATGAGTGTGGAGTCGTTGCGTGATTAGGCAGAAATAAATTCANNNAGATCGGAA
126165 TATGAGTGTGGAGTCGTTGCAAAGGGAATTACAAATGCAAAANNNAGATCGGAAG
126166 TATGAGTGTGGAGTCGTTGCCATTTGGCTAAAGAGATTTTGATNNNAGATCGGAAG
126167 TATGAGTGTGGAGTCGTTGCCCTAGAATAGAGCAGATGTTTAANNNAGATCGGAA
126168 TATGAGTGTGGAGTCGTTGCCGTCTGCACACATATACATATATNNNAGATCGGAAG
126169 TATGAGTGTGGAGTCGTTGCACATGACATAAAATTTACCATTTTNNNAGATCGGAA
126170 TATGAGTGTGGAGTCGTTGCCTTCATAACACATAGAAGGAATGNNNAGATCGGAA
126171 TATGAGTGTGGAGTCGTTGCCTTCATAACACATAGAAGGAATGNNNAGATCGGAA
126172 TATGAGTGTGGAGTCGTTGCTGTTAATTAACTGTACTATGTGCNNNAGATCGGAAG
126173 TATGAGTGTGGAGTCGTTGCAGTAGTATATGGACAAAACAACTNNNAGATCGGAA
126175 TATGAGTGTGGAGTCGTTGCTCCACTCCTCTTTTGATTAAAANNNAGATCGGAAGAG
126176 TATGAGTGTGGAGTCGTTGCCACAAAAAGAATCTTCCTAAAGTNNNAGATCGGAAG
126177 TATGAGTGTGGAGTCGTTGCGTCACAAAAAGAATCTTCCTAAANNNAGATCGGAAG
126178 TATGAGTGTGGAGTCGTTGCCTCTCAAGAATTTTCTTCCCCNNNAGATCGGAAGAGC
126179 TATGAGTGTGGAGTCGTTGCCTCCTGAGTAACTGGGATTANNNAGATCGGAAGAG
126180 TATGAGTGTGGAGTCGTTGCAAGCATATATGGCAAAAAGATAGNNNAGATCGGAA
126181 TATGAGTGTGGAGTCGTTGCTTCTAGAGAACTCCTGACAAATANNNAGATCGGAAG

126182 TATGAGTGTGGAGTCGTTGCCTGTCTTCCTTTCTTTTCTCTNNNAGATCGGAAGAGC
126183 TATGAGTGTGGAGTCGTTGCCTGTCTTCCTTTCTTTTCTCTNNNAGATCGGAAGAGC
126184 TATGAGTGTGGAGTCGTTGCACGTTGAATATATTGATGTAAGANNNAGATCGGAA
126185 TATGAGTGTGGAGTCGTTGCAGCCCACAAAATAATATCTTATCNNNAGATCGGAAG
126186 TATGAGTGTGGAGTCGTTGCAGATAGGGACAGACAAAATATTTNNNAGATCGGAA
126187 TATGAGTGTGGAGTCGTTGCATAGAGATTTTTCAGTCTGAAGANNNAGATCGGAA
126188 TATGAGTGTGGAGTCGTTGCGAAAATAGCTTCATGTTTGACATNNNAGATCGGAAG
126189 TATGAGTGTGGAGTCGTTGCACAGTAATTATCAAGAACTTTGGNNNAGATCGGAAG
126190 TATGAGTGTGGAGTCGTTGCGAAAAAGCAAATATTTTCAGGACNNNAGATCGGAA
126191 TATGAGTGTGGAGTCGTTGCGCAAATATAAATAAGTCTAGGGCNNNAGATCGGAA
126192 TATGAGTGTGGAGTCGTTGCTCTTGAGTATTATTTTTCCCAAANNNAGATCGGAAGA
126193 TATGAGTGTGGAGTCGTTGCATCTGTGTTGTTAAATGCCANNNAGATCGGAAGAG
126195 TATGAGTGTGGAGTCGTTGCTTAACACAAAAGAAAACAGAGAANNNAGATCGGAA
126196 TATGAGTGTGGAGTCGTTGCTCCATGGTGTGTATATATATCACNNNAGATCGGAAG
126197 TATGAGTGTGGAGTCGTTGCATATATAATGCATCCTGAATGGANNNAGATCGGAAG
126198 TATGAGTGTGGAGTCGTTGCTGTTTTACTCTGTTTTGTAAGAGNNNAGATCGGAAG
126199 TATGAGTGTGGAGTCGTTGCACACAGAAATAAATCTTTTCCTTNNNAGATCGGAAG
126200 TATGAGTGTGGAGTCGTTGCAAAGAAGAAAGTGACCCAGTANNNAGATCGGAAGA
126201 TATGAGTGTGGAGTCGTTGCTTTCCATCTTCTGAAATTTGGTTNNNAGATCGGAAGA
126202 TATGAGTGTGGAGTCGTTGCAATTATCTCTGGCAGAATTTGATNNNAGATCGGAAG
126203 TATGAGTGTGGAGTCGTTGCACTGTATATTGAAATCATTGTCANNNAGATCGGAAG
126204 TATGAGTGTGGAGTCGTTGCAAATCAAAGAAAGATCAGCTTAGNNNAGATCGGAA
126205 TATGAGTGTGGAGTCGTTGCTAGATATTTGAAACGGGTAAAGANNNAGATCGGAA
126206 TATGAGTGTGGAGTCGTTGCATCTGCAGCATTAATCTAGTATANNNAGATCGGAAG
126207 TATGAGTGTGGAGTCGTTGCTTAAGCCCCTAAGTTTATTGTATNNNAGATCGGAAG
126208 TATGAGTGTGGAGTCGTTGCTTTCATGGGGAAATGATTTTTCNNNAGATCGGAAGA
126209 TATGAGTGTGGAGTCGTTGCCAGCCATAAAAAGGAATGAAATANNNAGATCGGAA
126210 TATGAGTGTGGAGTCGTTGCATGGTATGAGCAAGACCTTANNNAGATCGGAAGAG
126211 TATGAGTGTGGAGTCGTTGCGTGATTTTTTGTGTTAAATTCCANNNAGATCGGAAG
126212 TATGAGTGTGGAGTCGTTGCGTATCCATACAATTATGTATCGCNNNAGATCGGAAG
126213 TATGAGTGTGGAGTCGTTGCGGATAAAGAACAGGTGGTAAATANNNAGATCGGA
126214 TATGAGTGTGGAGTCGTTGCATATATTTGAGAGCTATCAGCATNNNAGATCGGAAG
126215 TATGAGTGTGGAGTCGTTGCATTATTTATTTGGAGCTGGATTGNNNAGATCGGAAG
126217 TATGAGTGTGGAGTCGTTGCATCTTTCTTGGAGATAGACCANNNAGATCGGAAGAG
126218 TATGAGTGTGGAGTCGTTGCATAAAAGGAGAGACTGAGAATGNNNAGATCGGAA
126219 TATGAGTGTGGAGTCGTTGCATAAAAGGAGAGACTGAGAATGNNNAGATCGGAA
126220 TATGAGTGTGGAGTCGTTGCATGGTGCGTGTATATATATATGGNNNAGATCGGAA
126223 TATGAGTGTGGAGTCGTTGCTAAAAGACCTTTCTGATAGATGTNNNAGATCGGAAG
126224 TATGAGTGTGGAGTCGTTGCTTTCTATGTGTGTATGTATCAGANNNAGATCGGAAG
126225 TATGAGTGTGGAGTCGTTGCCCAATGAAATCATTAGGAACTAANNNAGATCGGAAG
126226 TATGAGTGTGGAGTCGTTGCCCTATTGGAAAGCTTCTTTTTCNNNAGATCGGAAGA
126227 TATGAGTGTGGAGTCGTTGCGTGTATTTCTTGCTATTGCTTGNNNAGATCGGAAGA
126228 TATGAGTGTGGAGTCGTTGCAAGTTACTTGAGTCTAATGTCTTNNNAGATCGGAAG

126229 TATGAGTGTGGAGTCGTTGCCAATAAGAAACAATGGAAGGAAANNNAGATCGGAA
126230 TATGAGTGTGGAGTCGTTGCAACTAGAACACTTTTCTCATTTCNNNAGATCGGAAGA
126231 TATGAGTGTGGAGTCGTTGCGTTGACCAGATATCTGAAATGTNNNAGATCGGAAG
126232 TATGAGTGTGGAGTCGTTGCATCATTTTTTTGTTTTTGAGACGNNNAGATCGGAAG
126233 TATGAGTGTGGAGTCGTTGCTCTTAGTGATTAGTTTCTAGTGANNNAGATCGGAAG
126235 TATGAGTGTGGAGTCGTTGCACAACATATTTTCATCTTTGGAANNNAGATCGGAAG
126236 TATGAGTGTGGAGTCGTTGCGTTTTTTAATTCAACTTCCTTCGNNNAGATCGGAAGA
126237 TATGAGTGTGGAGTCGTTGCGTAAACAGAGAGATTTGTTCTAANNNAGATCGGAA
126238 TATGAGTGTGGAGTCGTTGCATGGATCTAATAATCACATGCTANNNAGATCGGAAG
126239 TATGAGTGTGGAGTCGTTGCTATTGTGTTAAATGTTACAACCTNNNAGATCGGAAG
126240 TATGAGTGTGGAGTCGTTGCGGAGTGAACACAGCTTAAAANNNAGATCGGAAGAG
126241 TATGAGTGTGGAGTCGTTGCGGAGTGAACACAGCTTAAAANNNAGATCGGAAGAG
126242 TATGAGTGTGGAGTCGTTGCTGAATTTCTTTGGATGGTTTTGNNNAGATCGGAAGA
126243 TATGAGTGTGGAGTCGTTGCATTACTACTGGACCTAAGAAATGNNNAGATCGGAAG
126245 TATGAGTGTGGAGTCGTTGCTTATACTGTGATAAACTGTTGGTNNNAGATCGGAAG
126246 TATGAGTGTGGAGTCGTTGCCTTTCTGTCTTCTGCTGTTCNNNAGATCGGAAGAGCA
126247 TATGAGTGTGGAGTCGTTGCATTTTGGTTATTTCTTGCCTTTTNNNAGATCGGAAGA
126250 TATGAGTGTGGAGTCGTTGCTATAATTTCAACTTGCTTTTCCANNNAGATCGGAAGA
126251 TATGAGTGTGGAGTCGTTGCAAAGAATAGGAACTAATAGAGGGNNNAGATCGGAA
126252 TATGAGTGTGGAGTCGTTGCAGATTGATTCACAGTTGTCATNNNAGATCGGAAGAG
126253 TATGAGTGTGGAGTCGTTGCAGATTGATTCACAGTTGTCATNNNAGATCGGAAGAG
126254 TATGAGTGTGGAGTCGTTGCGTAGGCTTATGCAACTTATTTCNNNAGATCGGAAGA
126255 TATGAGTGTGGAGTCGTTGCAAGAAACCTCTTTTGAGAACTTNNNAGATCGGAAGA
126256 TATGAGTGTGGAGTCGTTGCATATACTTTGCTCCATAACATCTNNNAGATCGGAAGA
126257 TATGAGTGTGGAGTCGTTGCATATACTTTGCTCCATAACATCTNNNAGATCGGAAGA
126258 TATGAGTGTGGAGTCGTTGCTAGGAATTCTCTGTACTTTTGAANNNAGATCGGAAG
126259 TATGAGTGTGGAGTCGTTGCCAAACTTTACTGAGAGCTATAATNNNAGATCGGAAG
126260 TATGAGTGTGGAGTCGTTGCCCTTGATTTCTTTTTCTTGTTTANNNAGATCGGAAGA
126261 TATGAGTGTGGAGTCGTTGCAAATTGGGCTATATGATCTTCTANNNAGATCGGAAG
126262 TATGAGTGTGGAGTCGTTGCTAAAAGTTTGAAATTTCTCCCTCNNNAGATCGGAAG
126263 TATGAGTGTGGAGTCGTTGCGATTTAGTTTTAAATCCTGACCCNNNAGATCGGAAG
126264 TATGAGTGTGGAGTCGTTGCAAACTGAAAGGATCCTTAAGTTTNNNAGATCGGAAG
126265 TATGAGTGTGGAGTCGTTGCCATGTTCCTATAGTTTTAACCTCNNNAGATCGGAAGA
126266 TATGAGTGTGGAGTCGTTGCTTTTTGGAAAAATTCTTGGAGATNNNAGATCGGAAG
126267 TATGAGTGTGGAGTCGTTGCACTAATTTAACAAGTGCAACTTANNNAGATCGGAAG
126269 TATGAGTGTGGAGTCGTTGCGTTTCAGCTATAGTAAATAATGCTNNNAGATCGGAA
126270 TATGAGTGTGGAGTCGTTGCAATCTCTGAAAGAGCTATTTGTANNNAGATCGGAAG
126271 TATGAGTGTGGAGTCGTTGCATACAACAAAGAATCCTCAAATCNNNAGATCGGAAG
126272 TATGAGTGTGGAGTCGTTGCTTGAATAGGTTACCATTTTCTTGNNNAGATCGGAAG
126273 TATGAGTGTGGAGTCGTTGCCCAATTCTGCTTTGTTTTTATTTNNNAGATCGGAAGA
126274 TATGAGTGTGGAGTCGTTGCTTACAACTTTATTTGAGAAAGCANNNAGATCGGAAG
126275 TATGAGTGTGGAGTCGTTGCAAAATAGTTTATTTGGGGGAAAANNNAGATCGGAA
126276 TATGAGTGTGGAGTCGTTGCCAGATAGCTTCAATTATCAAGTTNNNAGATCGGAAG

126277 TATGAGTGTGGAGTCGTTGCGCCATAATGTAAAAAACAAACAANNNAGATCGGAAG
126278 TATGAGTGTGGAGTCGTTGCATAAATCTGTAGGAAGAAACAGTNNNAGATCGGAA
126279 TATGAGTGTGGAGTCGTTGCTATTGTTCAGGCATAAAGAAGANNNAGATCGGAAG
126280 TATGAGTGTGGAGTCGTTGCTGGGATAATCCTAAGTAGATACTNNNAGATCGGAAG
126281 TATGAGTGTGGAGTCGTTGCTAGTAAAGGCTTTAAACCATAGANNNAGATCGGAAG
126282 TATGAGTGTGGAGTCGTTGCAGCTCCATATAATTGGTACATAGNNNAGATCGGAAG
126283 TATGAGTGTGGAGTCGTTGCGCTTCTTACTTACATTTACCAAANNNAGATCGGAAGA
126284 TATGAGTGTGGAGTCGTTGCACAAGTATTTTTTGAGTGTCTACNNNAGATCGGAAG
126285 TATGAGTGTGGAGTCGTTGCCTAGGGAAAACAATGAAAGAATTNNNAGATCGGAA
126286 TATGAGTGTGGAGTCGTTGCGGAAATTTGCTTTTGAAAGATAGNNNAGATCGGAA
126287 TATGAGTGTGGAGTCGTTGCGTGTCTATAAATTCTAGCCCTGNNNAGATCGGAAGA
126288 TATGAGTGTGGAGTCGTTGCCTGACTTTGTTCCTTTCCTCNNNAGATCGGAAGAGCA
126289 TATGAGTGTGGAGTCGTTGCATATACTCTGCAGTGGTATCAAANNNAGATCGGAAG
126290 TATGAGTGTGGAGTCGTTGCCTGTATCTAGGGATGCTAAATAANNNAGATCGGAAG
126291 TATGAGTGTGGAGTCGTTGCTGTATACATTTTTGAGAATGTGCNNNAGATCGGAAG
126292 TATGAGTGTGGAGTCGTTGCAAGAAACAATAAAAACACAAAGGNNNAGATCGGAA
126293 TATGAGTGTGGAGTCGTTGCAATTTTTCTGTGAACCTAAAACTNNNAGATCGGAAG
126294 TATGAGTGTGGAGTCGTTGCATTCATACATACATACAAGTCTGNNNAGATCGGAAG
126295 TATGAGTGTGGAGTCGTTGCCAGAATCATGAGCTAATGAGANNNAGATCGGAAGA
126297 TATGAGTGTGGAGTCGTTGCCCCAATAATCTGAGCTTTATGANNNAGATCGGAAGA
126298 TATGAGTGTGGAGTCGTTGCGTCTTCTATTCTTCAGCTGTATNNNAGATCGGAAGAG
126299 TATGAGTGTGGAGTCGTTGCTTGTTATTTGGGGAGAGTGTNNNAGATCGGAAGAG
126300 TATGAGTGTGGAGTCGTTGCTTTGTCCAAGATCAATTATGTACNNNAGATCGGAAG
126301 TATGAGTGTGGAGTCGTTGCGAGAAGATAACTCTGATCAAAACNNNAGATCGGAAG
126302 TATGAGTGTGGAGTCGTTGCAACTGTTGCCTATTTTTAGTTACNNNAGATCGGAAG
126303 TATGAGTGTGGAGTCGTTGCCATATGTGCCTGTGTATTATAGANNNAGATCGGAAG
126304 TATGAGTGTGGAGTCGTTGCTAAAAACCCAGTTAAGAGGATAANNNAGATCGGAA
126305 TATGAGTGTGGAGTCGTTGCTAGAAACTTTGATGACTTTTTGTNNNAGATCGGAAG
126306 TATGAGTGTGGAGTCGTTGCAACAAAACCTCATTATTCAGTTTNNNAGATCGGAAG
126307 TATGAGTGTGGAGTCGTTGCCTTATAGGGATGAATGAGCAAANNNAGATCGGAAG
126308 TATGAGTGTGGAGTCGTTGCCTTATAGGGATGAATGAGCAAANNNAGATCGGAAG
126309 TATGAGTGTGGAGTCGTTGCTTCATTATGCCATTCTTCTTTAGNNNAGATCGGAAGA
126310 TATGAGTGTGGAGTCGTTGCGTGGCCAGTCATAGAATTCTNNNAGATCGGAAGAG
126313 TATGAGTGTGGAGTCGTTGCTTCCTTAACTTTTTATCTTTCCTNNNAGATCGGAAGAG
126314 TATGAGTGTGGAGTCGTTGCCCATTCACTCCCTTGAAAAANNNAGATCGGAAGAGC
126315 TATGAGTGTGGAGTCGTTGCAACTCATCAGGAAAAGTTCTTAGNNNAGATCGGAAG
126316 TATGAGTGTGGAGTCGTTGCCACAGAAATACACATGAAAAATGNNNAGATCGGAAG
126317 TATGAGTGTGGAGTCGTTGCTACATTGTGAATATTTTGCTTGTNNNAGATCGGAAG
126318 TATGAGTGTGGAGTCGTTGCAACCTCATGTTGAAATTTGATCNNNAGATCGGAAGA
126319 TATGAGTGTGGAGTCGTTGCACCTGATCCTAAGATATTGTTAGNNNAGATCGGAAG
126320 TATGAGTGTGGAGTCGTTGCGACCTGATCCTAAGATATTGTTANNNAGATCGGAAG
126321 TATGAGTGTGGAGTCGTTGCCAAAATGGAAGGAATGAACTTATNNNAGATCGGAA
126322 TATGAGTGTGGAGTCGTTGCGAAATTTTGGGGAGGTTTAATATNNNAGATCGGAA

126323 TATGAGTGTGGAGTCGTTGCCTAGCATTTTAGACACATGACTNNNAGATCGGAAGA
126324 TATGAGTGTGGAGTCGTTGCATAAATGTATGTGTTACGGCTTANNNAGATCGGAAG
126325 TATGAGTGTGGAGTCGTTGCATTCTATCAAGTTGCAAATGTCNNNAGATCGGAAGA
126326 TATGAGTGTGGAGTCGTTGCCTCTTTGTTTCCTTTCTTAGAAANNNAGATCGGAAGA
126327 TATGAGTGTGGAGTCGTTGCTATCCCCTCCTGGCTATTATNNNAGATCGGAAGAGCA
126328 TATGAGTGTGGAGTCGTTGCCAGATTTGTCAGCAATTTTTTTCNNNAGATCGGAAGA
126329 TATGAGTGTGGAGTCGTTGCAGAGTAGCTTAAACAATAGAAGTNNNAGATCGGAA
126330 TATGAGTGTGGAGTCGTTGCTATGATACGAAATACAAACGTTCNNNAGATCGGAAG
126331 TATGAGTGTGGAGTCGTTGCTGAAGTAATATATGTGTGTAGGGNNNAGATCGGAA
126332 TATGAGTGTGGAGTCGTTGCTAAAAATTGAAGGGACATCATCCNNNAGATCGGAAG
126333 TATGAGTGTGGAGTCGTTGCTCATTGTAAATGTTACAAAATGTNNNAGATCGGAAG
126334 TATGAGTGTGGAGTCGTTGCGATATTATCTTACAGGCCAAGTANNNAGATCGGAAG
126337 TATGAGTGTGGAGTCGTTGCAAAATATGGTACCTTGTCATTTGNNNAGATCGGAAG
126338 TATGAGTGTGGAGTCGTTGCGCCAATACAATTTCATCTCAATANNNAGATCGGAAGA
126339 TATGAGTGTGGAGTCGTTGCTAAACAAGAAGCCAATACAATTTNNNAGATCGGAAG
126340 TATGAGTGTGGAGTCGTTGCCTTTTACCAGTAGATGGATTTTGNNNAGATCGGAAG
126341 TATGAGTGTGGAGTCGTTGCCTTTTACCAGTAGATGGATTTTGNNNAGATCGGAAG
126342 TATGAGTGTGGAGTCGTTGCAAGATTGCCTGAAAAGAGAGNNNAGATCGGAAGAG
126343 TATGAGTGTGGAGTCGTTGCGAGGGATATCTAGATAAACTTGGNNNAGATCGGAA
126344 TATGAGTGTGGAGTCGTTGCGTGATGCACATTCTAAATTAGGNNNAGATCGGAAGA
126345 TATGAGTGTGGAGTCGTTGCTTTTCTTTACGTCTATGATGGTANNNAGATCGGAAGA
126346 TATGAGTGTGGAGTCGTTGCATATTGGGGGTGGATATGTGNNNAGATCGGAAGAG
126347 TATGAGTGTGGAGTCGTTGCAAAATAATAGCAGGCATGGTGNNNAGATCGGAAGA
126348 TATGAGTGTGGAGTCGTTGCCTATCACTGTAGAGAACCAATATNNNAGATCGGAAG
126349 TATGAGTGTGGAGTCGTTGCAAGAGTAGCATTGAAATTTGATGNNNAGATCGGAA
126350 TATGAGTGTGGAGTCGTTGCAGAAATGAGGTCTCACTATGTNNNAGATCGGAAGA
126351 TATGAGTGTGGAGTCGTTGCTTTACAGACATTTTTACAGAGCNNNAGATCGGAAGA
126352 TATGAGTGTGGAGTCGTTGCTTCCTGTTTTTTGGAAAAAATTCNNNAGATCGGAAGA
126353 TATGAGTGTGGAGTCGTTGCTATTTGTTTCCTAATGCAGATTCNNNAGATCGGAAGA
126354 TATGAGTGTGGAGTCGTTGCATCAACTCTCATGTGAAATATTCNNNAGATCGGAAGA
126355 TATGAGTGTGGAGTCGTTGCGCCCTGAATATTAGGCAAAAANNNAGATCGGAAGA
126356 TATGAGTGTGGAGTCGTTGCTAATAAGGGAAAGAGCTAGTCTANNNAGATCGGAA
126357 TATGAGTGTGGAGTCGTTGCGTTCAAACTATCAGGACTAGAGNNNAGATCGGAAGA
126358 TATGAGTGTGGAGTCGTTGCGATCACTCCTTTCTTCACAGNNNAGATCGGAAGAGCA
126359 TATGAGTGTGGAGTCGTTGCCGTTACTATAGAGACCTCTTTCNNNAGATCGGAAGA
126360 TATGAGTGTGGAGTCGTTGCCTGGTGCAGAATTAGTTAATTATNNNAGATCGGAAG
126361 TATGAGTGTGGAGTCGTTGCCAACTATTTGCCCCTAAAGCNNNAGATCGGAAGAGC
126362 TATGAGTGTGGAGTCGTTGCGTTATGAAAGAGTTTGGAAGATTNNNAGATCGGAA
126363 TATGAGTGTGGAGTCGTTGCGAAACTTGGTTGGTAAGAATTTNNNAGATCGGAAG
126364 TATGAGTGTGGAGTCGTTGCATTGGTCCATTTTACAGAGAGNNNAGATCGGAAGA
126365 TATGAGTGTGGAGTCGTTGCGAGAGTCTGAATCAGTTAGAAANNNAGATCGGAAG
126366 TATGAGTGTGGAGTCGTTGCTCTCCTCATAACTTTCCTTAAAANNNAGATCGGAAGA
126367 TATGAGTGTGGAGTCGTTGCAAGCATTTTCTTGTTCACTAAATNNNAGATCGGAAGA

126368 TATGAGTGTGGAGTCGTTGCTTTCATCTTTTTTCTGTGTTTTGNNNAGATCGGAAGA
126369 TATGAGTGTGGAGTCGTTGCCATCTATAAAATTTTAAAAATTGCCCNNNAGATCGGA
126370 TATGAGTGTGGAGTCGTTGCATTATGTATCATAGCCCAACATTNNNAGATCGGAAG
126371 TATGAGTGTGGAGTCGTTGCAGGTTTGACTCTGATTAGATAAANNNAGATCGGAA
126373 TATGAGTGTGGAGTCGTTGCGAAGACGGAAAGAAACTAAATTANNNAGATCGGAA
126374 TATGAGTGTGGAGTCGTTGCTTAAACTATACTATGTAGCAGGCNNNAGATCGGAAG
126375 TATGAGTGTGGAGTCGTTGCCTAGAAAATGCAAGATAATCCAGNNNAGATCGGAA
126376 TATGAGTGTGGAGTCGTTGCTGTAAGTGGAATCATAAGTATGTNNNAGATCGGAA
126377 TATGAGTGTGGAGTCGTTGCCATTTCGTAAGGTTTTATGTACANNNAGATCGGAAG
126378 TATGAGTGTGGAGTCGTTGCATACATACACACATATACACACANNNAGATCGGAAG
126379 TATGAGTGTGGAGTCGTTGCCTCTAGTCGAACTATTTTCAAATNNNAGATCGGAAG
126380 TATGAGTGTGGAGTCGTTGCCAGAGGAGAAAGGAACTATTAGNNNAGATCGGAA
126381 TATGAGTGTGGAGTCGTTGCCAGAGGAGAAAGGAACTATTAGNNNAGATCGGAA
126382 TATGAGTGTGGAGTCGTTGCTGTATGTGAGTTTATGTTGTTTTNNNAGATCGGAAG
126383 TATGAGTGTGGAGTCGTTGCCTACGTGAAGTGTATTTCTTTGNNNAGATCGGAAGA
126384 TATGAGTGTGGAGTCGTTGCGATGTCTGCACTATTACTGTCNNNAGATCGGAAGAG
126385 TATGAGTGTGGAGTCGTTGCCACCACTATTTTCTTTTCTTTTTTNNNAGATCGGAAGA
126386 TATGAGTGTGGAGTCGTTGCTAGACATAGAGAAAGTCTGAAAGNNNAGATCGGAA
126387 TATGAGTGTGGAGTCGTTGCTATTTCCTATTTTCCTCCTCACNNNAGATCGGAAGAG
126388 TATGAGTGTGGAGTCGTTGCGCAGTGATACAGAGAATTTTTTANNNAGATCGGAAG
126389 TATGAGTGTGGAGTCGTTGCTGATAGTTTTGCAACATTGTAATNNNAGATCGGAAG
126390 TATGAGTGTGGAGTCGTTGCACCCTATCTCTGTTAAAAAAAGANNNAGATCGGAAG
126391 TATGAGTGTGGAGTCGTTGCAAAGTGGTGCAAGACTATATATTNNNAGATCGGAA
126392 TATGAGTGTGGAGTCGTTGCGCTTTTGGTTTCTATCTATCTTTNNNAGATCGGAAGA
126393 TATGAGTGTGGAGTCGTTGCTTCCATCGCTATGAAGAAAATTNNNAGATCGGAAGA
126394 TATGAGTGTGGAGTCGTTGCTCAGCTGACTAGTAAGAAAAATNNNAGATCGGAAG
126395 TATGAGTGTGGAGTCGTTGCGACTATCATTCTACTGTTTACCTNNNAGATCGGAAGA
126396 TATGAGTGTGGAGTCGTTGCTTTCATGTGACTATAATGCAAAANNNAGATCGGAAG
126397 TATGAGTGTGGAGTCGTTGCCGATAGAAAAGAAAACCTCATTTNNNAGATCGGAAG
126398 TATGAGTGTGGAGTCGTTGCTTTCCTAGTTTACTTGTGGGNNNAGATCGGAAGAGC
126399 TATGAGTGTGGAGTCGTTGCACCTATTATATATGTGAGAGAGTTNNNAGATCGGAA
126400 TATGAGTGTGGAGTCGTTGCGAGTGGGATACTTTTGTTTTAGNNNAGATCGGAAG
126401 TATGAGTGTGGAGTCGTTGCCAAGATATCCTAACATAACCCTTNNNAGATCGGAAG
126402 TATGAGTGTGGAGTCGTTGCTTGAGGATTTGAGACAGTAGTAANNNAGATCGGAA
126403 TATGAGTGTGGAGTCGTTGCAGTAAGGAAGCGTTTTTAATTAANNNAGATCGGAA
126404 TATGAGTGTGGAGTCGTTGCTGTGGTTTTTTGCTTACATGNNNAGATCGGAAGAGC
126405 TATGAGTGTGGAGTCGTTGCATGTTGGTATATGGTAAGTACTCNNNAGATCGGAA
126406 TATGAGTGTGGAGTCGTTGCAACAACATTCTTGTATCTGAGANNNAGATCGGAAGA
126407 TATGAGTGTGGAGTCGTTGCCACCTAGATCCTAAGTTAACTTANNNAGATCGGAAG
126408 TATGAGTGTGGAGTCGTTGCAATCATATATAAAACCATGTGGCNNNAGATCGGAAG
126409 TATGAGTGTGGAGTCGTTGCAATCATATATAAAACCATGTGGCNNNAGATCGGAAG
126410 TATGAGTGTGGAGTCGTTGCCCTAAAATCAAGGGCATCTTNNNAGATCGGAAGAGC
126411 TATGAGTGTGGAGTCGTTGCTACACATCATACCCTACTCATAANNNAGATCGGAAGA

126412 TATGAGTGTGGAGTCGTTGCAAAGAGAAAGTAAAAAGAATCCCNNNAGATCGGAA
126413 TATGAGTGTGGAGTCGTTGCAGAGAGAAGAAAAGAGAAAGTAANNNAGATCGGA
126414 TATGAGTGTGGAGTCGTTGCCAATCTGAATTCCTATGTGGANNNAGATCGGAAGAG
126415 TATGAGTGTGGAGTCGTTGCAAATTTAACAAACAGCAAGAAAGNNNAGATCGGAA
126416 TATGAGTGTGGAGTCGTTGCGTATGTTTGTGTGTGCATATATANNNAGATCGGAAG
126417 TATGAGTGTGGAGTCGTTGCAAATAACAGGAATTTGTTTCTCANNNAGATCGGAAG
126419 TATGAGTGTGGAGTCGTTGCATCGTCCATAACAGTAAAGATTANNNAGATCGGAAG
126420 TATGAGTGTGGAGTCGTTGCGAAAACTGATTTGGAAACTACTANNNAGATCGGAAG
126421 TATGAGTGTGGAGTCGTTGCTAAATAAGGAATAGGGTGAGAGGNNNAGATCGGA
126422 TATGAGTGTGGAGTCGTTGCTAGATTCCTTTCTGACTCTGAANNNAGATCGGAAGA
126423 TATGAGTGTGGAGTCGTTGCGGGAGGTGAAGAGCATATATNNNAGATCGGAAGA
126424 TATGAGTGTGGAGTCGTTGCTAATCACGTATTCACTTTGAATCNNNAGATCGGAAGA
126425 TATGAGTGTGGAGTCGTTGCTATCCAATGGCAGAAAATTAATCNNNAGATCGGAAG
126426 TATGAGTGTGGAGTCGTTGCATTCATATTACTCTCTAGGGTCANNNAGATCGGAAGA
126427 TATGAGTGTGGAGTCGTTGCAAATGTTTGATAGGATACTTTGCNNNAGATCGGAAG
126429 TATGAGTGTGGAGTCGTTGCGCTGCCTAGATTAAAAATAAGATNNNAGATCGGAAG
126430 TATGAGTGTGGAGTCGTTGCTTTTTGGGAGAAATAACACAAAGNNNAGATCGGAA
126431 TATGAGTGTGGAGTCGTTGCTCAGTCTCTCTCAAAATACTTCNNNAGATCGGAAGAG
126432 TATGAGTGTGGAGTCGTTGCCAAGTCCATCTCATCTTTCTGNNNAGATCGGAAGAG
126433 TATGAGTGTGGAGTCGTTGCCATCCTCAGTACAGTTTTAGAGNNNAGATCGGAAGA
126434 TATGAGTGTGGAGTCGTTGCTATCAAGATCCTAAAACAGCAAANNNAGATCGGAAG
126435 TATGAGTGTGGAGTCGTTGCGTTTTGCTCCTAAGAATTATTCTNNNAGATCGGAAGA
126436 TATGAGTGTGGAGTCGTTGCTAACATTCACACATAAACACACNNNAGATCGGAAGA
126437 TATGAGTGTGGAGTCGTTGCAAAAGTAACCCAAACCACAAATNNNAGATCGGAAGA
126438 TATGAGTGTGGAGTCGTTGCAGAGACATTTGCTTTCCATTANNNAGATCGGAAGAG
126439 TATGAGTGTGGAGTCGTTGCAGAGATCAGAAGCCTTCCTANNNAGATCGGAAGAG
126440 TATGAGTGTGGAGTCGTTGCTAGGTCTAACCAGCTCTCTANNNAGATCGGAAGAGC
126441 TATGAGTGTGGAGTCGTTGCTTTTGCAATAACGTGTGTGTNNNAGATCGGAAGAG
126442 TATGAGTGTGGAGTCGTTGCCACCTCCATTCTTGTTATTAAAGNNNAGATCGGAAGA
126443 TATGAGTGTGGAGTCGTTGCCCTAGAAGACTTGGGAATTTTNNNAGATCGGAAGAG
126444 TATGAGTGTGGAGTCGTTGCAATGGAAAGTATCTTGTGCTTTANNNAGATCGGAAG
126445 TATGAGTGTGGAGTCGTTGCGGAGAGGATAGTATCACATTATCNNNAGATCGGAA
126446 TATGAGTGTGGAGTCGTTGCTACACATTCTGAGATAAGACCTANNNAGATCGGAAG
126447 TATGAGTGTGGAGTCGTTGCACTAATATATCTTTGAGGCCATTNNNAGATCGGAAGA
126448 TATGAGTGTGGAGTCGTTGCGTACAGTACAGTGATATTTTGAGNNNAGATCGGAAG
126449 TATGAGTGTGGAGTCGTTGCAAACAGTTCCTCAATAAGTGGNNNAGATCGGAAGAG
126450 TATGAGTGTGGAGTCGTTGCCAGAGCAAGTCAGATTCTTGNNNAGATCGGAAGAG
126451 TATGAGTGTGGAGTCGTTGCGAAACAGTATTTGTAGTTGACATNNNAGATCGGAAG
126452 TATGAGTGTGGAGTCGTTGCCAGGCAGCTAATCTTAAAATAAANNNAGATCGGAAG
126453 TATGAGTGTGGAGTCGTTGCATTATGAAATTTCATGTCCATCGNNNAGATCGGAAGA
126454 TATGAGTGTGGAGTCGTTGCTACTTCAAATCGATAAATGTGTCNNNAGATCGGAAGA
126455 TATGAGTGTGGAGTCGTTGCTATGTATTGCTTGAAATACTTGCNNNAGATCGGAAGA
126456 TATGAGTGTGGAGTCGTTGCTATCGTTATTTCATCAGCTTCTTNNNAGATCGGAAGA

126458 TATGAGTGTGGAGTCGTTGCTAGCCAGGAAATATATATTGGTGNNNAGATCGGAA
126459 TATGAGTGTGGAGTCGTTGCCTAGCTATCTTGGGAATCTATTTNNNAGATCGGAAG
126460 TATGAGTGTGGAGTCGTTGCTTTTGAGAGGGCAACTAATTANNNAGATCGGAAGA
126461 TATGAGTGTGGAGTCGTTGCCTATGAGTCTGCTTTGAAATTTANNNAGATCGGAAG
126462 TATGAGTGTGGAGTCGTTGCAAAACTCAAAAAGCCAGAGTANNNAGATCGGAAGA
126463 TATGAGTGTGGAGTCGTTGCTGAAAAATCAGTACGATTAACTGNNNAGATCGGAAG
126464 TATGAGTGTGGAGTCGTTGCGTGTATATTTGCACACAATATATGNNNAGATCGGAA
126465 TATGAGTGTGGAGTCGTTGCTTTAAGTTAAGCACCTTTTAACCNNNAGATCGGAAG
126466 TATGAGTGTGGAGTCGTTGCCTAATAATGCCTAAGCCTCAGNNNAGATCGGAAGAG
126467 TATGAGTGTGGAGTCGTTGCAGCTATAAAGAAGAACCAGATAGNNNAGATCGGAA
126468 TATGAGTGTGGAGTCGTTGCTATAGCAGGACAGTGACTTCNNNAGATCGGAAGAG
126469 TATGAGTGTGGAGTCGTTGCTTTGCTTAGGCTATTGACCTANNNAGATCGGAAGAG
126470 TATGAGTGTGGAGTCGTTGCCCACCACGAAAAATACAAAATNNNAGATCGGAAGAG
126471 TATGAGTGTGGAGTCGTTGCTATAAAATACACACACAGACACANNNAGATCGGAAG
126472 TATGAGTGTGGAGTCGTTGCTTCCATTTCCTCAACTATAACATNNNAGATCGGAAGA
126473 TATGAGTGTGGAGTCGTTGCCACAGTAAACTATATGTACGTTCNNNAGATCGGAAG
126474 TATGAGTGTGGAGTCGTTGCAATAGTGCTGTGGTAAATGTATNNNAGATCGGAAG
126475 TATGAGTGTGGAGTCGTTGCAAATGGAGTTTCCCTATTTTTCNNNAGATCGGAAGA
126476 TATGAGTGTGGAGTCGTTGCGCAGGGGAAATACTATAGATAATNNNAGATCGGAA
126477 TATGAGTGTGGAGTCGTTGCAAAAGCATGAGAAAAACTCTAAANNNAGATCGGAA
126478 TATGAGTGTGGAGTCGTTGCCTCTAAAGTAGTGCACATAGAAANNNAGATCGGAAG
126479 TATGAGTGTGGAGTCGTTGCCATGATTAAGGCTCACTGTAGNNNAGATCGGAAGA
126480 TATGAGTGTGGAGTCGTTGCCAGGACAAATCTATTTTGGATAGNNNAGATCGGAAG
126481 TATGAGTGTGGAGTCGTTGCCTAAATTCTCTGCATGGTAAAAGNNNAGATCGGAAG
126482 TATGAGTGTGGAGTCGTTGCGGCATCTATAAAATTCCCATTAGNNNAGATCGGAAG
126483 TATGAGTGTGGAGTCGTTGCAAGAGTAAGGGATTTGAATATGTNNNAGATCGGAA
126484 TATGAGTGTGGAGTCGTTGCCACTACATACTATGCGGAAGNNNAGATCGGAAGAG
126485 TATGAGTGTGGAGTCGTTGCCTGTCTTTTTCCAGTTCTTAGANNNAGATCGGAAGAG
126486 TATGAGTGTGGAGTCGTTGCTAGGATTTTGTAATTCCAAAGTCNNNAGATCGGAAG
126487 TATGAGTGTGGAGTCGTTGCTATGACTGGAGAATCAACTTTATNNNAGATCGGAAG
126488 TATGAGTGTGGAGTCGTTGCTAGACACTGAGTAAATACTTGTTNNNAGATCGGAAG
126489 TATGAGTGTGGAGTCGTTGCTGGAAAACACAATTTGGAAACNNNAGATCGGAAGA
126490 TATGAGTGTGGAGTCGTTGCCTGAGGAATCACTCAAGACANNNAGATCGGAAGAG
126491 TATGAGTGTGGAGTCGTTGCGGTCATTACCCTCTAAATTATATGNNNAGATCGGAA
126492 TATGAGTGTGGAGTCGTTGCAAATCAGACACCTCTCAAAAGNNNAGATCGGAAGAG
126493 TATGAGTGTGGAGTCGTTGCGAAGAATAAAGTGGAAGTTCTTTNNNAGATCGGAA
126494 TATGAGTGTGGAGTCGTTGCGAAGAATAAAGTGGAAGTTCTTTNNNAGATCGGAA
126495 TATGAGTGTGGAGTCGTTGCCTCCTGAACTATCTAAGAGTAAGNNNAGATCGGAAG
126496 TATGAGTGTGGAGTCGTTGCATGTATACAGTTACATGTTTGAGNNNAGATCGGAAG
126497 TATGAGTGTGGAGTCGTTGCCTCAGGGAAGAGATGATACTNNNAGATCGGAAGAG
126498 TATGAGTGTGGAGTCGTTGCAGAACCTTAGAGGAAGAAGAGNNNAGATCGGAAG
126499 TATGAGTGTGGAGTCGTTGCTGAAAAAAAGAGACAAATACTGTNNNAGATCGGAA
126500 TATGAGTGTGGAGTCGTTGCCCCTAGCCTGTAGTATTTCTNNNAGATCGGAAGAGC

126501 TATGAGTGTGGAGTCGTTGCCCTTTAGCTATCAGTCCATTGNNNAGATCGGAAGAG
126502 TATGAGTGTGGAGTCGTTGCTACATGAGGGATGAGAAAATTAANNNAGATCGGAA
126503 TATGAGTGTGGAGTCGTTGCAGACTATGATTGGTGTTACAAATNNNAGATCGGAAG
126505 TATGAGTGTGGAGTCGTTGCAACAGAAACAAAAGTCCAACTNNNAGATCGGAAGAG
126506 TATGAGTGTGGAGTCGTTGCCATCATTGCATCAATTAATGAAGNNNAGATCGGAAG
126507 TATGAGTGTGGAGTCGTTGCCAGTCTAAATCATTTTGTTGGATNNNAGATCGGAAG
126508 TATGAGTGTGGAGTCGTTGCACTGAGAGGTTATTAAATGATCANNNAGATCGGAAG
126509 TATGAGTGTGGAGTCGTTGCTTTCTGGGTCTAATGGATAATCNNNAGATCGGAAGA
126510 TATGAGTGTGGAGTCGTTGCTCAGCATTCTTTCATGATTTTTNNNAGATCGGAAGAG
126511 TATGAGTGTGGAGTCGTTGCCCTTCACTTCTCAAACTGAANNNAGATCGGAAGAGC
126512 TATGAGTGTGGAGTCGTTGCTAAATATAGTGTGACAGGTTCAANNNAGATCGGAAG
126515 TATGAGTGTGGAGTCGTTGCTGTCTAACTAGCTCTTACAATAGNNNAGATCGGAAG
126516 TATGAGTGTGGAGTCGTTGCTGTCTAACTAGCTCTTACAATAGNNNAGATCGGAAG
126517 TATGAGTGTGGAGTCGTTGCATTCCCAGATATTCTGATTGAATNNNAGATCGGAAG
126518 TATGAGTGTGGAGTCGTTGCATTTTGCCTCATATTTTGGAATANNNAGATCGGAAG
126519 TATGAGTGTGGAGTCGTTGCTATTTTCCTAAGTTCATGTTCAGNNNAGATCGGAAG
126520 TATGAGTGTGGAGTCGTTGCGGCTTTAGGTGAAAAGTAAAATANNNAGATCGGAA
126522 TATGAGTGTGGAGTCGTTGCCCCAAGGTACTAAAATATAAGGANNNAGATCGGAAG
126524 TATGAGTGTGGAGTCGTTGCTGCACACTTCTAATATCCAATATNNNAGATCGGAAGA
126525 TATGAGTGTGGAGTCGTTGCCTCTTTCTCCTACTCTACAAAATNNNAGATCGGAAGA
126526 TATGAGTGTGGAGTCGTTGCGGCAGTTATTAAATCTTTACACTNNNAGATCGGAAG
126527 TATGAGTGTGGAGTCGTTGCAGTTTTCCTATATACTGTAGGGTNNNAGATCGGAAG
126528 TATGAGTGTGGAGTCGTTGCAGCTATTTGATAAATCCTTACAGNNNAGATCGGAAG
126529 TATGAGTGTGGAGTCGTTGCGTCATTTCCTCCAGTTTTTTAGNNNAGATCGGAAGAG
126530 TATGAGTGTGGAGTCGTTGCTTATGACAGACACATTGCTAAATNNNAGATCGGAAG
126531 TATGAGTGTGGAGTCGTTGCCCCTTTATCTCAAAATTGGATTTNNNAGATCGGAAGA
126532 TATGAGTGTGGAGTCGTTGCCAAATTCCTATGTTGAAACCTAANNNAGATCGGAAG
126533 TATGAGTGTGGAGTCGTTGCCTCAGTGAGGATATATTGAGTANNNAGATCGGAAG
126534 TATGAGTGTGGAGTCGTTGCCATGATTAAGCCTCTATCATCTANNNAGATCGGAAG
126535 TATGAGTGTGGAGTCGTTGCTAGGCTTTCAAATTCTTGATAACNNNAGATCGGAAG
126536 TATGAGTGTGGAGTCGTTGCATACAAACATGTTTGGTAGAAACNNNAGATCGGAAG
126537 TATGAGTGTGGAGTCGTTGCCTTACACAAATATAAGGTTGAGGNNNAGATCGGAAG
126538 TATGAGTGTGGAGTCGTTGCCAATATTTGTTGGTGTTATTTCTNNNAGATCGGAAG
126539 TATGAGTGTGGAGTCGTTGCTCTCCTAACAATACCGTTTATTTNNNAGATCGGAAGA
126540 TATGAGTGTGGAGTCGTTGCTCATCTGCACATATTTTTGTTTANNNAGATCGGAAGA
126541 TATGAGTGTGGAGTCGTTGCTGAAGTATTTGACGATGATTTTTNNNAGATCGGAAG
126542 TATGAGTGTGGAGTCGTTGCTAATTTTGCCAAGTAAAGACATTNNNAGATCGGAAG
126543 TATGAGTGTGGAGTCGTTGCGCCATTTGTAAGAACTGTAACNNNAGATCGGAAGAG
126544 TATGAGTGTGGAGTCGTTGCTTATTTCAATGGATAAATGCCAANNNAGATCGGAAG
126545 TATGAGTGTGGAGTCGTTGCGAAAATTTTATACACACACACACNNNAGATCGGAAG
126546 TATGAGTGTGGAGTCGTTGCAAATGTACACTTTTTGGAAACTANNNAGATCGGAAG
126547 TATGAGTGTGGAGTCGTTGCGAATGATGACCTGTGAACTCNNNAGATCGGAAGAG
126549 TATGAGTGTGGAGTCGTTGCCAACTGGTTGAAAGAGTTATTATNNNAGATCGGAAG

126550 TATGAGTGTGGAGTCGTTGCACATAAAAATACAGAGACTTTCGNNNAGATCGGAAG
126551 TATGAGTGTGGAGTCGTTGCAAAACAACTATTGGTTAGTGATCNNNAGATCGGAAG
126552 TATGAGTGTGGAGTCGTTGCTGCTAAAATAGATTCCACTAAGANNNAGATCGGAAG
126553 TATGAGTGTGGAGTCGTTGCGGGGTTTTGTATTTTGAAGATTNNNAGATCGGAAG
126554 TATGAGTGTGGAGTCGTTGCCTACTGATTGGCCCATTTTANNNAGATCGGAAGAGC
126555 TATGAGTGTGGAGTCGTTGCAGTAGAGAAATAACGAGATGAAANNNAGATCGGA
126556 TATGAGTGTGGAGTCGTTGCAGTAGAGAAATAACGAGATGAAANNNAGATCGGA
126557 TATGAGTGTGGAGTCGTTGCTCAGAAGAAGACAGAAAGATGNNNAGATCGGAAG
126558 TATGAGTGTGGAGTCGTTGCTTGTGGTAAAATACAAATGATGTNNNAGATCGGAA
126559 TATGAGTGTGGAGTCGTTGCCACGCGAAAAACATTTAATTAAANNNAGATCGGAAG
126560 TATGAGTGTGGAGTCGTTGCTATTTCAGTGAAACAGGAAGATANNNAGATCGGAA
126561 TATGAGTGTGGAGTCGTTGCAATCAATCTTAGTGTTTTGAACCNNNAGATCGGAAG
126562 TATGAGTGTGGAGTCGTTGCGAACGGTTTGATCAGGTTAGNNNAGATCGGAAGAG
126563 TATGAGTGTGGAGTCGTTGCTAAAAACACATTTTAGCAAAGCANNNAGATCGGAAG
126564 TATGAGTGTGGAGTCGTTGCTATGTTCTCTCTAATTGTCCAATNNNAGATCGGAAGA
126565 TATGAGTGTGGAGTCGTTGCCATTAAAGAGAGCTCCTAAATTTNNNAGATCGGAAG
126566 TATGAGTGTGGAGTCGTTGCCTAGTTTGTAATCTCCATGAGGNNNAGATCGGAAGA
126567 TATGAGTGTGGAGTCGTTGCGTAATGGAATCTTAGTATGTTGCNNNAGATCGGAA
126568 TATGAGTGTGGAGTCGTTGCCCAAAAGCTTGGAAAGTATTATTNNNAGATCGGAAG
126569 TATGAGTGTGGAGTCGTTGCAATTCACTCTAATCTGACCACNNNAGATCGGAAGAG
126570 TATGAGTGTGGAGTCGTTGCATAGGAACAGCTCCAGTCTANNNAGATCGGAAGAG
126571 TATGAGTGTGGAGTCGTTGCCTTCCTGGTATCACAAGAGGNNNAGATCGGAAGAG
126572 TATGAGTGTGGAGTCGTTGCGAAGTTTGAGTAGGTAAACAAAGNNNAGATCGGAA
126573 TATGAGTGTGGAGTCGTTGCTCACTAACAATGTATCAATCAAANNNAGATCGGAAG
126574 TATGAGTGTGGAGTCGTTGCAAAGTTCTAAAATTGATTGTGGTNNNAGATCGGAA
126575 TATGAGTGTGGAGTCGTTGCTGTTTAATACTTCATTCAAAGCTNNNAGATCGGAAG
126576 TATGAGTGTGGAGTCGTTGCAGGAGATCTGATGCTTTTATAGNNNAGATCGGAAG
126578 TATGAGTGTGGAGTCGTTGCTTCCTAATTAGCAATTCATGTTCNNNAGATCGGAAGA
126579 TATGAGTGTGGAGTCGTTGCCTTCACTGTTTAAGGAGACTTANNNAGATCGGAAGA
126580 TATGAGTGTGGAGTCGTTGCATTTGAAAGCTAAAACAACAATGNNNAGATCGGAA
126581 TATGAGTGTGGAGTCGTTGCTTATTTCACATGTTCTACCCTTANNNAGATCGGAAGA
126582 TATGAGTGTGGAGTCGTTGCAATAGGTACTCATTGTGTTATCANNNAGATCGGAAG
126583 TATGAGTGTGGAGTCGTTGCGGGAAAATCTTCAAGAAAAAAATNNNAGATCGGAA
126584 TATGAGTGTGGAGTCGTTGCTGAGATACCACAAAATAGTAAGTNNNAGATCGGAA
126585 TATGAGTGTGGAGTCGTTGCAGCAAAGGAAATACTGTAATTTCNNNAGATCGGAA
126586 TATGAGTGTGGAGTCGTTGCGAATGACTGGAAGTAGGTATTGNNNAGATCGGAAG
126587 TATGAGTGTGGAGTCGTTGCAATGTAACTGTATTTGGAGACAANNNAGATCGGAA
126588 TATGAGTGTGGAGTCGTTGCTCAGAAGCTCTTTAAAGTCTTCNNNAGATCGGAAGA
126589 TATGAGTGTGGAGTCGTTGCAAAGAAAAACAGATAAATTGGGTNNNAGATCGGAA
126590 TATGAGTGTGGAGTCGTTGCGGTAAGTTTTTCTTAGGTCATTTNNNAGATCGGAAG
126591 TATGAGTGTGGAGTCGTTGCGGTTCTTTTAGGTATTAATTTCCANNNAGATCGGAA
126592 TATGAGTGTGGAGTCGTTGCTCCTACTAAAAAATGTCTCTCTGNNNAGATCGGAAG
126594 TATGAGTGTGGAGTCGTTGCTTAGCAAACACAGATTATAAAGCNNNAGATCGGAAG

126596 TATGAGTGTGGAGTCGTTGCATCAGACTCAGGAAATTAACATNNNAGATCGGAAGA

126597 TATGAGTGTGGAGTCGTTGCCTCACCCTCTTTTCTTTTCTTNNNAGATCGGAAGAGCA

126598 TATGAGTGTGGAGTCGTTGCGGAATAGTGCTGTGATAAAAATANNNAGATCGGAA

126599 TATGAGTGTGGAGTCGTTGCTAACTAGTAGAATATGGCAAAGGNNNAGATCGGAA

126600 TATGAGTGTGGAGTCGTTGCGAATTGTTGTCATTGAATTTGAANNNAGATCGGAAG

126601 TATGAGTGTGGAGTCGTTGCTCTGCCTAGGTATTCTTTAAAATNNNAGATCGGAAGA

126602 TATGAGTGTGGAGTCGTTGCATTTCTTTTTGAGATGGAATTTCNNNAGATCGGAAGA

126603 TATGAGTGTGGAGTCGTTGCGCTTTCTGACAGTATAATTGTAGNNNAGATCGGAAG

126604 TATGAGTGTGGAGTCGTTGCTATTTATTGAGGATTCAGAGCTTNNNAGATCGGAAG

126606 TATGAGTGTGGAGTCGTTGCCACAGGGTAAATCTGTAAGATTNNNAGATCGGAAGA

126607 TATGAGTGTGGAGTCGTTGCCTCTTCCTTTTCTGTTTTCTAAANNNAGATCGGAAGA

126608 TATGAGTGTGGAGTCGTTGCCAGCACTATTCATAATAGGAAAGNNNAGATCGGAAG

126609 TATGAGTGTGGAGTCGTTGCTAGAAGCAGAGGGATATTTCTNNNAGATCGGAAGA

126610 TATGAGTGTGGAGTCGTTGCATTTTGTTCCAGCTGTGTTTANNNAGATCGGAAGAG

126611 TATGAGTGTGGAGTCGTTGCGGGGGATAAAAATGTTCTAAAATNNNAGATCGGAA

126612 TATGAGTGTGGAGTCGTTGCCTTCCTTGCACATAGTTGAGNNNAGATCGGAAGAGC

126613 TATGAGTGTGGAGTCGTTGCATGTGTAAAGAACTTTTACTGAGNNNAGATCGGAAG

126614 TATGAGTGTGGAGTCGTTGCAACAACTTTGGGGTATGATCNNNAGATCGGAAGAG

126615 TATGAGTGTGGAGTCGTTGCGGTTTGTTACATAGGTATATGTGNNNAGATCGGAAG

126616 TATGAGTGTGGAGTCGTTGCAGTTCATTATATCAATCATCCGTNNNAGATCGGAAGA

126617 TATGAGTGTGGAGTCGTTGCAATGGCTGTTAAATAATAAAAGGNNNAGATCGGAA

126618 TATGAGTGTGGAGTCGTTGCAAAGGATAAGGTTTTCACAGTANNNAGATCGGAAGA

126619 TATGAGTGTGGAGTCGTTGCATATCTAATGTCTGAAAACAAGCNNNAGATCGGAAG

126620 TATGAGTGTGGAGTCGTTGCTTTTCTGAAGAGACTGTTTACTANNNAGATCGGAAGA

126621 TATGAGTGTGGAGTCGTTGCCCCCACTAGTAGATTATAAGTTCNNNAGATCGGAAGA

126623 TATGAGTGTGGAGTCGTTGCATAACTCACAGGCTAAAGTAATANNNAGATCGGAAG

126624 TATGAGTGTGGAGTCGTTGCGGCATATTGATAGCTTGAAATTANNNAGATCGGAAG

126625 TATGAGTGTGGAGTCGTTGCTCTTTCTTTCTTTTTTTTTTTTTGGNNNAGATCGGAAG

126626 TATGAGTGTGGAGTCGTTGCGTTTTCTTGAGGCTCAAAAATNNNAGATCGGAAGAG

126627 TATGAGTGTGGAGTCGTTGCGCCTAGGCTTTTCAAAGTTAANNNAGATCGGAAGAG

126628 TATGAGTGTGGAGTCGTTGCGGAAACAAAGAATACCTTTTACANNNAGATCGGAAG

126629 TATGAGTGTGGAGTCGTTGCCTTTATCAAAGGCATCCATTTGNNNAGATCGGAAGA

126630 TATGAGTGTGGAGTCGTTGCAGAGATATACATTAGAACGTTCANNNAGATCGGAAG

126631 TATGAGTGTGGAGTCGTTGCGGGCACTAAATATTCAATGATAANNNAGATCGGAAG

126632 TATGAGTGTGGAGTCGTTGCGTACTTCTTCATCCCTATAGTTCNNNAGATCGGAAGA

126633 TATGAGTGTGGAGTCGTTGCGAATTTACTCATGTAATTGTGGANNNAGATCGGAAG

126634 TATGAGTGTGGAGTCGTTGCGAATTTACTCATGTAATTGTGGANNNAGATCGGAAG

126635 TATGAGTGTGGAGTCGTTGCTCTTTGTCCATTTTGTATTGTTANNNAGATCGGAAGA

126636 TATGAGTGTGGAGTCGTTGCATAACCTTAGCATAAAGAACCTTNNNAGATCGGAAGA

126637 TATGAGTGTGGAGTCGTTGCGATAAATGTGCAGGGAAAAATAGNNNAGATCGGAA

126638 TATGAGTGTGGAGTCGTTGCTTTGGTGATGAGTATTTTCAAAANNNAGATCGGAAG

126639 TATGAGTGTGGAGTCGTTGCGGTCATATTGGTATGTGATTAACNNNAGATCGGAAG

126640 TATGAGTGTGGAGTCGTTGCGGTCATATTGGTATGTGATTAACNNNAGATCGGAAG

126641 TATGAGTGTGGAGTCGTTGCCATCCTGAATCCTTGGTATCNNNAGATCGGAAGAGC
126642 TATGAGTGTGGAGTCGTTGCGCTTGTACTCCCAATACTTTNNNAGATCGGAAGAGC
126643 TATGAGTGTGGAGTCGTTGCATTCTTGATGAAAATAGTCTGAGNNNAGATCGGAAG
126645 TATGAGTGTGGAGTCGTTGCGGGGCATGTATATGTGTATAAANNNAGATCGGAAG
126646 TATGAGTGTGGAGTCGTTGCATGAGAAATAGTTGACCTGATCNNNAGATCGGAAG
126647 TATGAGTGTGGAGTCGTTGCCACTGAGTTTTAGGAATTTCTTGNNNAGATCGGAAG
126648 TATGAGTGTGGAGTCGTTGCGGCTTAGATTATGCATAACATTTNNNAGATCGGAAG
126650 TATGAGTGTGGAGTCGTTGCATAGTTAGTTAGGTGGATAGTCTNNNAGATCGGAA
126651 TATGAGTGTGGAGTCGTTGCAGATAGTAAGACAAGGCATTTTANNNAGATCGGAA
126652 TATGAGTGTGGAGTCGTTGCACTCTTCAGCAATTTATCTCCNNNAGATCGGAAGAG
126653 TATGAGTGTGGAGTCGTTGCGAGGACTCTTCAGCAATTTATNNNAGATCGGAAGAG
126654 TATGAGTGTGGAGTCGTTGCGTTATGGAAGGTTTTTCTTTTGANNNAGATCGGAAG
126655 TATGAGTGTGGAGTCGTTGCTTGAGGAAGTGCTTGATTTTANNNAGATCGGAAGA
126656 TATGAGTGTGGAGTCGTTGCATGTTTTCAATAAATTGCTCTTGNNNAGATCGGAAG
126657 TATGAGTGTGGAGTCGTTGCCTACAGTCAGCTCCAGAATCNNNAGATCGGAAGAGC
126658 TATGAGTGTGGAGTCGTTGCGTGAAAGAGGAGAAATAGAAAAGNNNAGATCGGA
126659 TATGAGTGTGGAGTCGTTGCAAGAGAGTGTATGTGTGTTTGNNNAGATCGGAAGA
126660 TATGAGTGTGGAGTCGTTGCCTCCTATGGTTCTATTTCTCTGNNNAGATCGGAAGAG
126661 TATGAGTGTGGAGTCGTTGCTGCTTTAATAACTTCAAAAGTGTNNNAGATCGGAAG
126662 TATGAGTGTGGAGTCGTTGCAACATGTTATACGTAAAGTACACNNNAGATCGGAAG
126663 TATGAGTGTGGAGTCGTTGCGTAATTTGAACAAGGTTAGTGTCNNNAGATCGGAA
126664 TATGAGTGTGGAGTCGTTGCGCTAAGGGTGTAAAGAAAGATAANNNAGATCGGAA
126665 TATGAGTGTGGAGTCGTTGCGTGTTAGTCAATTTGATTGTAGTNNNAGATCGGAAG
126666 TATGAGTGTGGAGTCGTTGCGGTACTATCCAATCCTCTATTACNNNAGATCGGAAGA
126667 TATGAGTGTGGAGTCGTTGCAAAAGTAGAGAAGAGAAAAAACGNNNAGATCGGA
126668 TATGAGTGTGGAGTCGTTGCCCAATATGTGTATCTCTTCTCAANNNAGATCGGAAGA
126669 TATGAGTGTGGAGTCGTTGCCTGTGTACTGAGCTTAACTTTNNNAGATCGGAAGAG
126670 TATGAGTGTGGAGTCGTTGCCTCCTGAACTATCTAAGAGTAAGNNNAGATCGGAAG
126671 TATGAGTGTGGAGTCGTTGCAATGGGATGATCTTGTCTCANNNAGATCGGAAGAG
126672 TATGAGTGTGGAGTCGTTGCTGTGTAGACATTCCACTCAANNNAGATCGGAAGAGC
126673 TATGAGTGTGGAGTCGTTGCATATTGTTACTAATATCCAGGGGNNNAGATCGGAAG
126674 TATGAGTGTGGAGTCGTTGCCTTTCTGACTAGGTTCTCACNNNAGATCGGAAGAGC
126675 TATGAGTGTGGAGTCGTTGCGAATTTGGGCTGAGTTGTATNNNAGATCGGAAGAG
126676 TATGAGTGTGGAGTCGTTGCACATACACACATACACAGACNNNAGATCGGAAGAGC
126677 TATGAGTGTGGAGTCGTTGCCATCATCTCCCTGGTATTTTCNNNAGATCGGAAGAG
126679 TATGAGTGTGGAGTCGTTGCGAATCTTGGGGAAAACTAAATGNNNAGATCGGAAG
126680 TATGAGTGTGGAGTCGTTGCCTAGACTGATGTGGAGGTAGNNNAGATCGGAAGA
126681 TATGAGTGTGGAGTCGTTGCCTAACTACCAATACTTTGCATTTNNNAGATCGGAAGA
126682 TATGAGTGTGGAGTCGTTGCTAGACCTGAGTGTATATCTGATANNNAGATCGGAAG
126683 TATGAGTGTGGAGTCGTTGCTACTCACTCATCTGCCTATCNNNAGATCGGAAGAGCA
126684 TATGAGTGTGGAGTCGTTGCAATTTTAACAAATTGCACTTGAGNNNAGATCGGAAG
126685 TATGAGTGTGGAGTCGTTGCGATCTCTCGGCTCATTGTAANNNAGATCGGAAGAGC
126686 TATGAGTGTGGAGTCGTTGCGTCACGCTCCATAAATTATATTTNNNAGATCGGAAG

126687 TATGAGTGTGGAGTCGTTGCTCCTGGATTCAAGAGATTCTNNNAGATCGGAAGAGC

126688 TATGAGTGTGGAGTCGTTGCCAGCATCAAAGCAATTAAAAATANNNAGATCGGAAG

126689 TATGAGTGTGGAGTCGTTGCTATCTCACAGGGTTAAAACTTTANNNAGATCGGAAG

126690 TATGAGTGTGGAGTCGTTGCTGTAGGGAAAATCTCATATTTCANNNAGATCGGAAG

126691 TATGAGTGTGGAGTCGTTGCAAGCATAATGCCAATTTTATCTTNNNAGATCGGAAG

126692 TATGAGTGTGGAGTCGTTGCGTATTCTGTTAGACTTGGGCNNNAGATCGGAAGAG

126693 TATGAGTGTGGAGTCGTTGCGGTACCATTCCTACTAAAACTATNNNAGATCGGAAG

126694 TATGAGTGTGGAGTCGTTGCGCTAAAAAAAAAAACCTAAGAGGNNNAGATCGGAA

126695 TATGAGTGTGGAGTCGTTGCCAGAAGAAAAATAGGAAACAGAANNNAGATCGGAA

126696 TATGAGTGTGGAGTCGTTGCGAACTGAAAAGCACTATCTAAATNNNAGATCGGAAG

126697 TATGAGTGTGGAGTCGTTGCCATTATGGGTATGTTAAATGCTGNNNAGATCGGAA

126698 TATGAGTGTGGAGTCGTTGCAGACAGTCAGCCACTAAAATNNNAGATCGGAAGAG

126699 TATGAGTGTGGAGTCGTTGCGTACCTCTGATGTTGATTTCCNNNAGATCGGAAGAG

126700 TATGAGTGTGGAGTCGTTGCGACTTTAAAATAGGGACATGATCNNNAGATCGGAA

126701 TATGAGTGTGGAGTCGTTGCGACTCAAATTATGAAACTTGGAGNNNAGATCGGAA

126702 TATGAGTGTGGAGTCGTTGCAAACAATTCTCAGAGAACCGNNNAGATCGGAAGAG

126703 TATGAGTGTGGAGTCGTTGCATGTAAGATTAGAGGAGCTCANNNAGATCGGAAGA

126704 TATGAGTGTGGAGTCGTTGCATTTGAGAGTAAGCTGACTATTTNNNAGATCGGAAG

126705 TATGAGTGTGGAGTCGTTGCCTCATTATCGGCAATTTATACTCNNNAGATCGGAAGA

126706 TATGAGTGTGGAGTCGTTGCAAAAATAACAGAGGGAAAGAAAGNNNAGATCGGA

126707 TATGAGTGTGGAGTCGTTGCTAAATGATGCTAACCCTCTATACNNNAGATCGGAAG

126708 TATGAGTGTGGAGTCGTTGCTATTTAAATTGCTCCTCTGTCTNNNAGATCGGAAGAG

126709 TATGAGTGTGGAGTCGTTGCGAGTTTAGAGTAGGAGAAAACANNNAGATCGGAA

126710 TATGAGTGTGGAGTCGTTGCGTAGAGGTCATCTTGAAGAAAATNNNAGATCGGAA

126711 TATGAGTGTGGAGTCGTTGCCATCCTGGATTCAAGTGATGNNNAGATCGGAAGAG

126712 TATGAGTGTGGAGTCGTTGCTTGGTAAAAATGCAGACTCCNNNAGATCGGAAGAG

126713 TATGAGTGTGGAGTCGTTGCCTTAAGGAATAATGATGAGCAAGNNNAGATCGGAA

126714 TATGAGTGTGGAGTCGTTGCTTCAAACTTTAAATGCACCTTTANNNAGATCGGAAGA

126715 TATGAGTGTGGAGTCGTTGCTTAGATTTGAGGCAGTATGCNNNAGATCGGAAGAG

126716 TATGAGTGTGGAGTCGTTGCTTAGATTTGAGGCAGTATGCNNNAGATCGGAAGAG

126718 TATGAGTGTGGAGTCGTTGCCTTTGGCTTCCCATAAATTTTATNNNAGATCGGAAGA

126719 TATGAGTGTGGAGTCGTTGCCTTAGGATGATGTGTGTCCANNNAGATCGGAAGAG

126720 TATGAGTGTGGAGTCGTTGCATGTACCTCTGGAAAAATTCTTANNNAGATCGGAAG

126721 TATGAGTGTGGAGTCGTTGCTTTCTCTTTCTTTCTCTTCCTTNNNAGATCGGAAGAGC

126722 TATGAGTGTGGAGTCGTTGCTACAATGATAGTGGTTCTGCTNNNAGATCGGAAGA

126723 TATGAGTGTGGAGTCGTTGCGGTTCAACCAATTCCCTTACNNNAGATCGGAAGAGC

126724 TATGAGTGTGGAGTCGTTGCATTCTAGAAAAATCCTTCACTCTNNNAGATCGGAAGA

126725 TATGAGTGTGGAGTCGTTGCCCTTAATATTTTTGTAAGCCCATNNNAGATCGGAAGA

126726 TATGAGTGTGGAGTCGTTGCTCAAGTTTAACTGGTGTCTTATANNNAGATCGGAAG

126727 TATGAGTGTGGAGTCGTTGCATCTGAAATTCAAGTTTAACTGGNNNAGATCGGAAG

126728 TATGAGTGTGGAGTCGTTGCAATACAGCACTCCAGAATTGNNNAGATCGGAAGAG

126729 TATGAGTGTGGAGTCGTTGCTTAAACCTCTGCTCTTAAACTCNNNAGATCGGAAGAG

126730 TATGAGTGTGGAGTCGTTGCGGATCGTTCCTTATACTTCTTGNNNAGATCGGAAGA

126731 TATGAGTGTGGAGTCGTTGCATAAAGAAATGGAGTTTGTTACCNNNAGATCGGAA
126732 TATGAGTGTGGAGTCGTTGCCATACGTTCTAGATGTCTATGAANNNAGATCGGAAG
126733 TATGAGTGTGGAGTCGTTGCGTCATTACAGGAAGCTTAAAATTNNNAGATCGGAAG
126734 TATGAGTGTGGAGTCGTTGCGGGCATAATATTATTTGAAGCTTNNNAGATCGGAAG
126735 TATGAGTGTGGAGTCGTTGCGTCTCACTCTATCAGCTATCANNNAGATCGGAAGAG
126736 TATGAGTGTGGAGTCGTTGCACTTTAATAATGCTCACTACCTANNNAGATCGGAAGA
126738 TATGAGTGTGGAGTCGTTGCTTTCTTATCTGTGGGATTTCATNNNAGATCGGAAGA
126739 TATGAGTGTGGAGTCGTTGCTGGCTTATGTTAATTTTGTATCCNNNAGATCGGAAG
126740 TATGAGTGTGGAGTCGTTGCAAGAAACAGGTTTGGCTTATNNNAGATCGGAAGAG
126741 TATGAGTGTGGAGTCGTTGCAAGTATTAGTAAGCAGCAACANNNAGATCGGAAGA
126742 TATGAGTGTGGAGTCGTTGCTTGTAGTGGTAGACAATTTCTTTNNNAGATCGGAAG
126743 TATGAGTGTGGAGTCGTTGCTTTGCTTATTGATTTGCTTATTGNNNAGATCGGAAG
126744 TATGAGTGTGGAGTCGTTGCCTTTTACAAAATAGTTCTGGGGNNNAGATCGGAAGA
126745 TATGAGTGTGGAGTCGTTGCCCCTTTTACAAAATAGTTCTGGNNNAGATCGGAAGA
126746 TATGAGTGTGGAGTCGTTGCGTTCATAAGCTGCTCCTATGNNNAGATCGGAAGAGC
126747 TATGAGTGTGGAGTCGTTGCCTTTCATTGAAGACCCTCAATNNNAGATCGGAAGAG
126748 TATGAGTGTGGAGTCGTTGCGCATCAGAATTTTTTGGTAAATTNNNAGATCGGAAG
126749 TATGAGTGTGGAGTCGTTGCCACAAGATTCTGCTCTTAATTCNNNAGATCGGAAGA
126750 TATGAGTGTGGAGTCGTTGCGATTTTCCCTAGAAATATTGGTGNNNAGATCGGAAG
126751 TATGAGTGTGGAGTCGTTGCATATTTGCATAACTCAGTAGACANNNAGATCGGAAG
126752 TATGAGTGTGGAGTCGTTGCCAATGTGCTCAGTTATTACTTATNNNAGATCGGAAG
126754 TATGAGTGTGGAGTCGTTGCATTCAGTGAAGCCTTAAATAAACNNNAGATCGGAAG
126755 TATGAGTGTGGAGTCGTTGCATATAACAGTTATCTGGAGGAACNNNAGATCGGAA
126756 TATGAGTGTGGAGTCGTTGCGTTTGTTACTATTCAAGGACAATNNNAGATCGGAAG
126757 TATGAGTGTGGAGTCGTTGCTTCAAGCAATTTTCATGCCTNNNAGATCGGAAGAGC
126758 TATGAGTGTGGAGTCGTTGCGGATTTATTAGTTAGGGTTCATGNNNAGATCGGAA
126759 TATGAGTGTGGAGTCGTTGCGATGCCTATAAATGTTGGAATTCNNNAGATCGGAAG
126760 TATGAGTGTGGAGTCGTTGCAAGCTATAGGACCATGTTATAGANNNAGATCGGAA
126761 TATGAGTGTGGAGTCGTTGCGAGGTTATTTCTGTGATTTGATTNNNAGATCGGAAG
126762 TATGAGTGTGGAGTCGTTGCGAGACAGGGTTTTACTTTGTNNNAGATCGGAAGAG
126763 TATGAGTGTGGAGTCGTTGCGACAGGGTTAATATCTGTTTTTANNNAGATCGGAAG
126764 TATGAGTGTGGAGTCGTTGCGTACAATCATGGTTCTCTGTAGNNNAGATCGGAAGA
126765 TATGAGTGTGGAGTCGTTGCCTCTGCCTTATCCTGGTTATNNNAGATCGGAAGAGC
126766 TATGAGTGTGGAGTCGTTGCTTAAGGGTAGAACAGGAATTAAANNNAGATCGGAA
126767 TATGAGTGTGGAGTCGTTGCTCATCACTCTGTATCTCTCTCNNNAGATCGGAAGAGC
126769 TATGAGTGTGGAGTCGTTGCTGTCACTTTAATCATGTTCTTAGNNNAGATCGGAAG
126770 TATGAGTGTGGAGTCGTTGCATATCTTGGCAGAAAATAAAACGNNNAGATCGGAA
126771 TATGAGTGTGGAGTCGTTGCACAAATATCTTGGCAGAAAATAANNNAGATCGGAA
126772 TATGAGTGTGGAGTCGTTGCGTCAAGTTTAGGTTCCACACNNNAGATCGGAAGAG
126773 TATGAGTGTGGAGTCGTTGCACAATAGATAAAATAATGCGTGANNNAGATCGGAA
126774 TATGAGTGTGGAGTCGTTGCCCTTGACTTTCTGGATTCCANNNAGATCGGAAGAGC
126775 TATGAGTGTGGAGTCGTTGCAGTGTTATCTAAGTGTGTTGTACNNNAGATCGGAAG
126776 TATGAGTGTGGAGTCGTTGCATGTTGTTCTCAGAAGTTCTCNNNAGATCGGAAGAG

126777 TATGAGTGTGGAGTCGTTGCTACCTTATTGAAGAATTTCTCCANNNAGATCGGAAG
126778 TATGAGTGTGGAGTCGTTGCAAATTATCACATCTTATGCCTTGNNNAGATCGGAAG
126779 TATGAGTGTGGAGTCGTTGCGAAGAGAGAGGAAGGGAAAGNNNAGATCGGAAG
126780 TATGAGTGTGGAGTCGTTGCCATATAGGAATAATGCCTAGGGNNNAGATCGGAAG
126781 TATGAGTGTGGAGTCGTTGCCATATAGGAATAATGCCTAGGGNNNAGATCGGAAG
126782 TATGAGTGTGGAGTCGTTGCTTACTGGTTATCTTTATGAGAGGNNNAGATCGGAAG
126783 TATGAGTGTGGAGTCGTTGCTTACTGGTTATCTTTATGAGAGGNNNAGATCGGAAG
126785 TATGAGTGTGGAGTCGTTGCATACATGTTGGCCAGTATATTTNNNAGATCGGAAGA
126786 TATGAGTGTGGAGTCGTTGCAACATTTATTGGGCAATACTATGNNNAGATCGGAAG
126787 TATGAGTGTGGAGTCGTTGCTATTTATTTGATTTCCCTGAGCTNNNAGATCGGAAG
126788 TATGAGTGTGGAGTCGTTGCTAATGATAAAAGACAGTCCTTTGNNNAGATCGGAAG
126789 TATGAGTGTGGAGTCGTTGCTAATGATAAAAGACAGTCCTTTGNNNAGATCGGAAG
126790 TATGAGTGTGGAGTCGTTGCGAAAGTTGTCTACATAGAAATGGNNNAGATCGGAA
126791 TATGAGTGTGGAGTCGTTGCGACAGAAAAAAGAGTTACAAGATNNNAGATCGGAA
126792 TATGAGTGTGGAGTCGTTGCAAAGAGAAGTTAATGGAAGAGATNNNAGATCGGA
126793 TATGAGTGTGGAGTCGTTGCTGTACATAAGTATTGCACTCTATNNNAGATCGGAAG
126794 TATGAGTGTGGAGTCGTTGCATAAATTCATAACCATGCTGAAGNNNAGATCGGAAG
126795 TATGAGTGTGGAGTCGTTGCGACAAGCTGCACAGATATTCNNNAGATCGGAAGAG
126796 TATGAGTGTGGAGTCGTTGCTTCAGAGGAACTATTAACACATTNNNAGATCGGAAG
126797 TATGAGTGTGGAGTCGTTGCCTTCTTAAGTTTTTACTTGCTCANNNAGATCGGAAGA
126798 TATGAGTGTGGAGTCGTTGCCTTGTTATTTACTAGCTGTGTGNNNAGATCGGAAGA
126799 TATGAGTGTGGAGTCGTTGCTATCTGCATGGGATAAATACTTTNNNAGATCGGAAG
126800 TATGAGTGTGGAGTCGTTGCTGAAACTCCTTCTCTACTAAAAANNNAGATCGGAAG
126801 TATGAGTGTGGAGTCGTTGCTAAAGCATCATGGGCATAGANNNAGATCGGAAGAG
126802 TATGAGTGTGGAGTCGTTGCCTTGTAACTTGCTTTGACTAATANNNAGATCGGAAG
126805 TATGAGTGTGGAGTCGTTGCTCTTTTTTCTTTTTTCTCAGACANNNAGATCGGAAGA
126806 TATGAGTGTGGAGTCGTTGCCCCTTTATCTCAAAATTGGATTTNNNAGATCGGAAG
126807 TATGAGTGTGGAGTCGTTGCCCCTTTATCTCAAAATTGGATTTNNNAGATCGGAAG
126808 TATGAGTGTGGAGTCGTTGCGAGTAAAGTGGTAAAAACAGATTNNNAGATCGGAA
126809 TATGAGTGTGGAGTCGTTGCCTTGTTTCTGAATTGTATCCTCNNNAGATCGGAAGA
126810 TATGAGTGTGGAGTCGTTGCGGCTTTCCAAAATGTCATATAANNNAGATCGGAAGA
126811 TATGAGTGTGGAGTCGTTGCCAAAACTTTGCTCCAAAAATGNNNAGATCGGAAGAG
126812 TATGAGTGTGGAGTCGTTGCTAATCTGAGTGTTATCCATGGAANNNAGATCGGAAG
126813 TATGAGTGTGGAGTCGTTGCCACCAGGTATGCTTTAAAATCNNNAGATCGGAAGAG
126814 TATGAGTGTGGAGTCGTTGCCAGAATCCTGTTTATTCTCTCTANNNAGATCGGAAGA
126815 TATGAGTGTGGAGTCGTTGCTTAACTAAGCATCTACTATGTGTNNNAGATCGGAAG
126816 TATGAGTGTGGAGTCGTTGCATTTACATGCATTGCTCAATTATNNNAGATCGGAAG
126817 TATGAGTGTGGAGTCGTTGCATGTGTGTTTCCCTAAGTTTATANNNAGATCGGAAG
126818 TATGAGTGTGGAGTCGTTGCTCCATTAAATTAGAGCTTTAAGTNNNAGATCGGAAG
126819 TATGAGTGTGGAGTCGTTGCAAATTCTCTTTCAATGTGCAAATNNNAGATCGGAAG
126820 TATGAGTGTGGAGTCGTTGCAGAATAAACACTAGACCATTAATTTNNNAGATCGGA
126821 TATGAGTGTGGAGTCGTTGCGTGCATGTTTACTTTTCATTTAANNNAGATCGGAAG
126822 TATGAGTGTGGAGTCGTTGCTGTGTGTTTAAAGATAGGTCTTNNNAGATCGGAAG

126823 TATGAGTGTGGAGTCGTTGCGTGATTTTGAAGTCATTAGATCANNNAGATCGGAAG
126824 TATGAGTGTGGAGTCGTTGCATTTGACATCTGATTAACCTTTCNNNAGATCGGAAGA
126825 TATGAGTGTGGAGTCGTTGCCTTTTAGACTTCAGTGTGATTGNNNAGATCGGAAGA
126826 TATGAGTGTGGAGTCGTTGCTTAATAACCAAGTGAGTCACAAANNNAGATCGGAAG
126827 TATGAGTGTGGAGTCGTTGCCAACATTATGGTACATTTTCCTANNNAGATCGGAAGA
126828 TATGAGTGTGGAGTCGTTGCTGTGTCTATTGTCAGTATTAGTTNNNAGATCGGAAGA
126829 TATGAGTGTGGAGTCGTTGCCATGCCCTTCCTTTTATTATTCNNNAGATCGGAAGAG
126830 TATGAGTGTGGAGTCGTTGCATACATTTTCCTGAGTTTACCTANNNAGATCGGAAGA
126831 TATGAGTGTGGAGTCGTTGCAGATGAAGTCTCACTCTCTCNNNAGATCGGAAGAGCA
126832 TATGAGTGTGGAGTCGTTGCGTATATACTTTCATATATGCTGGTNNNAGATCGGAA
126833 TATGAGTGTGGAGTCGTTGCGAAAAATTTGTCTTCAGTGTTTCNNNAGATCGGAAGA
126834 TATGAGTGTGGAGTCGTTGCGTTTCTTTTAGAAGTAGGAGAGGNNNAGATCGGAA
126835 TATGAGTGTGGAGTCGTTGCGTTTCTTTTAGAAGTAGGAGAGGNNNAGATCGGAA
126836 TATGAGTGTGGAGTCGTTGCGTTTCTTTTAGAAGTAGGAGAGGNNNAGATCGGAA
126837 TATGAGTGTGGAGTCGTTGCTGTTTTAATTCCTTTAAGCCTACNNNAGATCGGAAGA
126838 TATGAGTGTGGAGTCGTTGCGAGAGAGGTTTTACATTCCAGNNNAGATCGGAAGA
126839 TATGAGTGTGGAGTCGTTGCTTGCACTACTCAAACATATAGAANNNAGATCGGAAGA
126840 TATGAGTGTGGAGTCGTTGCCCTGCCATCAGCTTTTTATTANNNAGATCGGAAGAG
126841 TATGAGTGTGGAGTCGTTGCCACAGTTTTTAGTCAACATTTCTNNNAGATCGGAAGA
126842 TATGAGTGTGGAGTCGTTGCTACTTGCCATTTTGACACTTAGNNNAGATCGGAAGA
126843 TATGAGTGTGGAGTCGTTGCCATGGATGGAAAAACAAAAAAGNNNAGATCGGAAG
126844 TATGAGTGTGGAGTCGTTGCTATATACACGAGAAACCTGTTAGNNNAGATCGGAAG
126845 TATGAGTGTGGAGTCGTTGCTTTATACTGATTCAGTTTGAGAGNNNAGATCGGAAG
126846 TATGAGTGTGGAGTCGTTGCTATATGCAAATACACATGGAAAGNNNAGATCGGAA
126847 TATGAGTGTGGAGTCGTTGCAAAAGAAGTATACCATTTGTCTCNNNAGATCGGAAG
126848 TATGAGTGTGGAGTCGTTGCGAATAGCTGGGACTACAGACNNNAGATCGGAAGAG
126849 TATGAGTGTGGAGTCGTTGCGAGGATGGAAGAAACAAGGANNNAGATCGGAAGA
126850 TATGAGTGTGGAGTCGTTGCAAGAATCATTCCATTTCTAGACANNNAGATCGGAAGA
126851 TATGAGTGTGGAGTCGTTGCATCCAACAATGGTTTAAAATTCTNNNAGATCGGAAGA
126852 TATGAGTGTGGAGTCGTTGCAAAACAAAACATGGTTTTTATCCNNNAGATCGGAAGA
126853 TATGAGTGTGGAGTCGTTGCCTACAAAACTACTGGAATTACAGNNNAGATCGGAAG
126854 TATGAGTGTGGAGTCGTTGCGACCCCAAGATACAGAAAATTNNNAGATCGGAAGA
126855 TATGAGTGTGGAGTCGTTGCTTAGATGAAATCCTGCTGCTNNNAGATCGGAAGAG
126856 TATGAGTGTGGAGTCGTTGCTATACACCAGATTTGGATTATGTNNNAGATCGGAAG
126857 TATGAGTGTGGAGTCGTTGCTCTTTCCACAAACTAAGAAGAGNNNAGATCGGAAGA
126858 TATGAGTGTGGAGTCGTTGCTCTAAACAAATATACTCATCCTGANNNAGATCGGAAG
126859 TATGAGTGTGGAGTCGTTGCGGAAATGTGGATAACTAAGATTTNNNAGATCGGAA
126860 TATGAGTGTGGAGTCGTTGCGGAAATGTGGATAACTAAGATTTNNNAGATCGGAA
126861 TATGAGTGTGGAGTCGTTGCCCTTCTTAGTTCCTTTTTGTTATNNNAGATCGGAAGA
126862 TATGAGTGTGGAGTCGTTGCCCATTTCCCAAACTAAGTTTTATNNNAGATCGGAAGA
126863 TATGAGTGTGGAGTCGTTGCGTTCAAGCGATACTCCTATCNNNAGATCGGAAGAGCA
126864 TATGAGTGTGGAGTCGTTGCACACATGTTTTCTACCTTTATTTNNNAGATCGGAAGA
126865 TATGAGTGTGGAGTCGTTGCCATGGTCTTAGCTAACTGTAANNNAGATCGGAAGAG

126866 TATGAGTGTGGAGTCGTTGCCTCATATGTAACAGGTGAAAAAANNNAGATCGGAAG
126867 TATGAGTGTGGAGTCGTTGCCCTTCTGGAGATCAAACATATNNNAGATCGGAAGAG
126868 TATGAGTGTGGAGTCGTTGCAACAGAAAATCCATCTCATACTANNNAGATCGGAAGA
126869 TATGAGTGTGGAGTCGTTGCTTTCAAACAACTCCTGATTAAATNNNAGATCGGAAGA
126870 TATGAGTGTGGAGTCGTTGCCTCTGGAGAACACTGATAATAATNNNAGATCGGAAG
126872 TATGAGTGTGGAGTCGTTGCAATCTCTATGGATACCTAAGACANNNAGATCGGAAG
126873 TATGAGTGTGGAGTCGTTGCGGGCTATATCGTTTGTTAAATTTNNNAGATCGGAAG
126874 TATGAGTGTGGAGTCGTTGCGCAAGATTTGGATGTCTTTTGNNNAGATCGGAAGAG
126875 TATGAGTGTGGAGTCGTTGCGAGGCATGAGAATCTTGAATNNNAGATCGGAAGAG
126876 TATGAGTGTGGAGTCGTTGCGAGAAATAGAAATAACAGAGTGGNNNAGATCGGAA
126877 TATGAGTGTGGAGTCGTTGCTTTCACTATGTTGCCCATACNNNAGATCGGAAGAGCA
126878 TATGAGTGTGGAGTCGTTGCGAGAGCTGTCTCAATCAATCNNNAGATCGGAAGAGC
126879 TATGAGTGTGGAGTCGTTGCTAAACTGTTTTAATGTATGGACCNNNAGATCGGAAG
126880 TATGAGTGTGGAGTCGTTGCGCTTCTCATTAGGTCTATTATCTNNNAGATCGGAAGA
126881 TATGAGTGTGGAGTCGTTGCAAGAAACTTAAAATGACAACAGTNNNAGATCGGAAG
126882 TATGAGTGTGGAGTCGTTGCAGTTTAAAATTTACTTTTGGGTTNNNAGATCGGAAGA
126883 TATGAGTGTGGAGTCGTTGCTTGCTCATATACACTTGGAATAANNNAGATCGGAAGA
126884 TATGAGTGTGGAGTCGTTGCTAAACTCATCTAGCATTCACTAGNNNAGATCGGAAGA
126885 TATGAGTGTGGAGTCGTTGCGAAAGCTTGAAATGTAAGACAATNNNAGATCGGAAG
126886 TATGAGTGTGGAGTCGTTGCAACACCAAATGAAGAAAATAGTCNNNAGATCGGAAG
126887 TATGAGTGTGGAGTCGTTGCCCTTCAAAGAAGTACTTAAGTTANNNAGATCGGAAG
126888 TATGAGTGTGGAGTCGTTGCGATTCAATTCAAGGTTAATTCCTNNNAGATCGGAAGA
126889 TATGAGTGTGGAGTCGTTGCCAAAGTTTTTCCAGAAACAGGNNNAGATCGGAAGAG
126890 TATGAGTGTGGAGTCGTTGCAATACAGAGTATGAACTTTCTCANNNAGATCGGAAG
126891 TATGAGTGTGGAGTCGTTGCGAAAGCTAAACCAGTAAAAGCNNNAGATCGGAAGAG
126892 TATGAGTGTGGAGTCGTTGCCCCGTGTAGAATGTTTTATAGTANNNAGATCGGAAG
126893 TATGAGTGTGGAGTCGTTGCCCCGTGTAGAATGTTTTATAGTANNNAGATCGGAAG
126894 TATGAGTGTGGAGTCGTTGCTATCCGTAATACTTATAATCCCTNNNAGATCGGAAGA
126895 TATGAGTGTGGAGTCGTTGCGAGCTGAAGTCAGAAAAATGNNNAGATCGGAAGAG
126896 TATGAGTGTGGAGTCGTTGCTTGGATAGGGAGACATGATAANNNAGATCGGAAGA
126897 TATGAGTGTGGAGTCGTTGCTATCCGTAATACTTATAATCCCTNNNAGATCGGAAGA
126898 TATGAGTGTGGAGTCGTTGCTCTCTGTCTTTGCTTATGTGNNNAGATCGGAAGAGCA
126899 TATGAGTGTGGAGTCGTTGCGAATTTCCTGTTATTAGACTCCANNNAGATCGGAAGA
126900 TATGAGTGTGGAGTCGTTGCTATATCAATTCAATGTGTAGTGCNNNAGATCGGAAG
126901 TATGAGTGTGGAGTCGTTGCCAAAATCTAAACTCCATTTCTGANNNAGATCGGAAGA
126902 TATGAGTGTGGAGTCGTTGCTTATCAGAAACTTCATAAGGAAGNNNAGATCGGAAG
126903 TATGAGTGTGGAGTCGTTGCGTAACTTTGGTCAAAAGAGTAAGNNNAGATCGGAAG
126904 TATGAGTGTGGAGTCGTTGCATTATGGGTGAGTTTTTAAATGGNNNAGATCGGAAG
126905 TATGAGTGTGGAGTCGTTGCGGAGCAATTTCATGGTATCTATANNNAGATCGGAAG
126906 TATGAGTGTGGAGTCGTTGCTTTTTACATACGTATACATGTGCNNNAGATCGGAAGA
126907 TATGAGTGTGGAGTCGTTGCGTTCTGGCCTAACTTATACAGNNNAGATCGGAAGAG
126908 TATGAGTGTGGAGTCGTTGCGAACTTTGTATATTTTTCCACCTNNNAGATCGGAAGA
126909 TATGAGTGTGGAGTCGTTGCCATATACCTGTTATGACATTGTCNNNAGATCGGAAGA

126910 TATGAGTGTGGAGTCGTTGCACACAAACATATACCTGTTATGANNNAGATCGGAAG
126911 TATGAGTGTGGAGTCGTTGCTTTTCTTAGAACAAATGTTGACCNNNAGATCGGAAG
126912 TATGAGTGTGGAGTCGTTGCTTTTCTTAGAACAAATGTTGACCNNNAGATCGGAAG
126913 TATGAGTGTGGAGTCGTTGCCCAGTGTTAGGACAATTACATNNNAGATCGGAAGA
126914 TATGAGTGTGGAGTCGTTGCGAATTTCCCTTTCCATCCTTTNNNAGATCGGAAGAGC
126915 TATGAGTGTGGAGTCGTTGCTTTTAGACTTTGCCAAATCAATANNNAGATCGGAAG
126916 TATGAGTGTGGAGTCGTTGCCTCTCATTCTCTCAACATAGTTANNNAGATCGGAAGA
126917 TATGAGTGTGGAGTCGTTGCGCATCTCTAAGAATAAGAACATCNNNAGATCGGAAG
126918 TATGAGTGTGGAGTCGTTGCAAAGCTACATGTCCTGTCTTNNNAGATCGGAAGAGC
126919 TATGAGTGTGGAGTCGTTGCTTTGTGTTTTGCTAAACTATGTANNNAGATCGGAAG
126920 TATGAGTGTGGAGTCGTTGCTCAGAAATATACACAAGTTCAGANNNAGATCGGAAG
126921 TATGAGTGTGGAGTCGTTGCCAGCTCTAGTGGTTCTCTACNNNAGATCGGAAGAGC
126922 TATGAGTGTGGAGTCGTTGCGATAAGGAACCCAGAACTAAGNNNAGATCGGAAGA
126923 TATGAGTGTGGAGTCGTTGCAACAACTGGAATTATCACATTGNNNAGATCGGAAGA
126924 TATGAGTGTGGAGTCGTTGCAGACATGTACAAAAGTATCTCCNNNAGATCGGAAGA
126925 TATGAGTGTGGAGTCGTTGCTTAACATAATTGCTGGAGTTTAGNNNAGATCGGAAG
126926 TATGAGTGTGGAGTCGTTGCATACTAAGAACAGCCCTCAAANNNAGATCGGAAGAG
126927 TATGAGTGTGGAGTCGTTGCCATGGAATACTTAGCCATAAAAANNNAGATCGGAAG
126928 TATGAGTGTGGAGTCGTTGCCATCATACCTGGAATTAGGAGNNNAGATCGGAAGA
126930 TATGAGTGTGGAGTCGTTGCAGAATAAAATCATGCAAAGGAAANNNAGATCGGAA
126931 TATGAGTGTGGAGTCGTTGCGGAGATGGGACAGGAATTTANNNAGATCGGAAGA
126932 TATGAGTGTGGAGTCGTTGCACCTTGACTGATATAAATTTTGGNNNAGATCGGAAG
126933 TATGAGTGTGGAGTCGTTGCACCTTGACTGATATAAATTTTGGNNNAGATCGGAAG
126934 TATGAGTGTGGAGTCGTTGCCTCACACTGAGATATATGAAAGANNNAGATCGGAAG
126935 TATGAGTGTGGAGTCGTTGCAAGTAATTCTCCTGCCTCAGNNNAGATCGGAAGAGC
126936 TATGAGTGTGGAGTCGTTGCATGAACGTTAACATTTCCTAAAANNNAGATCGGAAG
126937 TATGAGTGTGGAGTCGTTGCGCAAAAAAAAACTTTAGATTTCTGNNNAGATCGGAA
126938 TATGAGTGTGGAGTCGTTGCAGTACATACACACACATACACNNNAGATCGGAAGAG
126940 TATGAGTGTGGAGTCGTTGCTGAGTGAACTATTTGTATGTGTANNNAGATCGGAAG
126941 TATGAGTGTGGAGTCGTTGCGCACACAGAATTGAACATATTATNNNAGATCGGAAG
126942 TATGAGTGTGGAGTCGTTGCGGAACTATGGCTGGTATTTTTANNNAGATCGGAAGA
126943 TATGAGTGTGGAGTCGTTGCCATCAGAGGTTGAACAGAGANNNAGATCGGAAGAG
126944 TATGAGTGTGGAGTCGTTGCTATTAATTGATAAAGGCACCACTNNNAGATCGGAAG
126945 TATGAGTGTGGAGTCGTTGCTAGAATAACATCCAAGCTCCTTNNNAGATCGGAAGA
126946 TATGAGTGTGGAGTCGTTGCAAAGACATATTTAGTAGCCAAGANNNAGATCGGAAG
126947 TATGAGTGTGGAGTCGTTGCAATGAAATGATCAAAGATTCTGGNNNAGATCGGAA
126948 TATGAGTGTGGAGTCGTTGCTATTCTAAACTGGAGAGAATACCNNNAGATCGGAAG
126949 TATGAGTGTGGAGTCGTTGCCTGTTCTTACTGTGTTATGTTATNNNAGATCGGAAGA
126950 TATGAGTGTGGAGTCGTTGCAAAATAAATAATGGCAGCACATTNNNAGATCGGAAG
126953 TATGAGTGTGGAGTCGTTGCTAACTGAAGGTGGAATTGAAATNNNAGATCGGAAG
126955 TATGAGTGTGGAGTCGTTGCACAAACTTTGGAAACTAAAAGATNNNAGATCGGAAG
126956 TATGAGTGTGGAGTCGTTGCTGTAAAAACCATAAAGCTATCACNNNAGATCGGAAG
126957 TATGAGTGTGGAGTCGTTGCTTTCTTCTATTTTTTGGTCTCTGNNNAGATCGGAAGA

126958 TATGAGTGTGGAGTCGTTGCACCATAATTAGGATAATGAGCATNNNAGATCGGAAG
126959 TATGAGTGTGGAGTCGTTGCCACATTTATGTCAACACTTTCANNNAGATCGGAAGAG
126960 TATGAGTGTGGAGTCGTTGCTGCTTTCCTTTTACTTACAAAATNNNAGATCGGAAGA
126961 TATGAGTGTGGAGTCGTTGCAATAAAGGATTTGTGAGACTAGTNNNAGATCGGAA
126962 TATGAGTGTGGAGTCGTTGCAACACCAAATAATGTTGCTATAGNNNAGATCGGAAG
126963 TATGAGTGTGGAGTCGTTGCCTCTTGATCAGTATCATTTCCANNNAGATCGGAAGAG
126964 TATGAGTGTGGAGTCGTTGCAACTATGCTTTCTAAATCTCTCTNNNAGATCGGAAGA
126965 TATGAGTGTGGAGTCGTTGCACCTTTCACATGAAACTTCTTNNNAGATCGGAAGAG
126966 TATGAGTGTGGAGTCGTTGCCTATTGACAGATCAGGATACAGNNNAGATCGGAAG
126967 TATGAGTGTGGAGTCGTTGCTATATGTGTGTGTGTGTGTGNNNAGATCGGAAGAG
126968 TATGAGTGTGGAGTCGTTGCAATCTTAGAAGTGAGGTTAGACTNNNAGATCGGAA
126969 TATGAGTGTGGAGTCGTTGCAAAGACTTAGGTTTCTTAAGCTANNNAGATCGGAAG
126970 TATGAGTGTGGAGTCGTTGCCCTTTAGATCTAATTACCCCTATNNNAGATCGGAAGA
126971 TATGAGTGTGGAGTCGTTGCCCTTTAGATCTAATTACCCCTATNNNAGATCGGAAGA
126972 TATGAGTGTGGAGTCGTTGCCACAAAGTAAAGACTACCAAATTNNNAGATCGGAAG
126973 TATGAGTGTGGAGTCGTTGCTTTTCTCAGTGAGATTGTTTTACNNNAGATCGGAAG
126974 TATGAGTGTGGAGTCGTTGCTTGAAAATAACAGGATAGGTGATNNNAGATCGGAA
126975 TATGAGTGTGGAGTCGTTGCCAACACACTGAAGCATATAATCNNNAGATCGGAAGA
126976 TATGAGTGTGGAGTCGTTGCAGCCTTGATTTATCATTTTGAAANNNAGATCGGAAG
126977 TATGAGTGTGGAGTCGTTGCATACATATTAGCTTCCTGTGACNNNAGATCGGAAGA
126978 TATGAGTGTGGAGTCGTTGCCATTACGAACTGTTTTTTTTTTTNNNAGATCGGAAG
126979 TATGAGTGTGGAGTCGTTGCAAAATGAAGGAGAGAAAGACAGNNNAGATCGGAA
126980 TATGAGTGTGGAGTCGTTGCACACTCTAACAACTGTTAAAATCNNNAGATCGGAAG
126981 TATGAGTGTGGAGTCGTTGCCTAGTGCTTCAACCAGATCANNNAGATCGGAAGAGC
126982 TATGAGTGTGGAGTCGTTGCCATATGCTACTTACTCTGGAACNNNAGATCGGAAGA
126983 TATGAGTGTGGAGTCGTTGCGAATAGAGAGCCCAGAAATAAGNNNAGATCGGAAG
126984 TATGAGTGTGGAGTCGTTGCATATGCTATTGGTGCCTGATANNNAGATCGGAAGAG
126985 TATGAGTGTGGAGTCGTTGCAAAACAACTCCTCTAAAAAAGTCNNNAGATCGGAAG
126986 TATGAGTGTGGAGTCGTTGCGTGTTGTCCAATAGAATATTCTGNNNAGATCGGAAG
126987 TATGAGTGTGGAGTCGTTGCTTGCCTTTAATGTCGATGAATNNNAGATCGGAAGAG
126988 TATGAGTGTGGAGTCGTTGCATTTTTAGTAGAGATGGGGTTTCNNNAGATCGGAA
126989 TATGAGTGTGGAGTCGTTGCCTTAAAGCAACAGAAATGTATTGNNNAGATCGGAAG
126990 TATGAGTGTGGAGTCGTTGCCTTAGTGAGAGGAAGTTTCTAAANNNAGATCGGAA
126991 TATGAGTGTGGAGTCGTTGCCTATGCTATTTCACACTTCTCTNNNAGATCGGAAGAG
126992 TATGAGTGTGGAGTCGTTGCCTATGCTATTTCACACTTCTCTNNNAGATCGGAAGAG
126993 TATGAGTGTGGAGTCGTTGCAACATAGACGTAAAAATCGTTAGNNNAGATCGGAA
126994 TATGAGTGTGGAGTCGTTGCAAACATCTGGGAACTTATACTCNNNAGATCGGAAGA
126996 TATGAGTGTGGAGTCGTTGCGATCATTTGTTCCTTCCACTTANNNAGATCGGAAGAG
126997 TATGAGTGTGGAGTCGTTGCTTTATGAAGCGTTTCTCCAGNNNAGATCGGAAGAGC
126998 TATGAGTGTGGAGTCGTTGCAATAATGTGTTTTGCATAGTGTTNNNAGATCGGAAG
126999 TATGAGTGTGGAGTCGTTGCAACTCAATCTTCTTTTTCTGAATNNNAGATCGGAAGA
127001 TATGAGTGTGGAGTCGTTGCAAGGAATCCTACAAAATCTCATNNNAGATCGGAAGA
127002 TATGAGTGTGGAGTCGTTGCATGCATTCTTCAAATTTTCTACANNNAGATCGGAAGA

127003 TATGAGTGTGGAGTCGTTGCACTTTGCTTACACTAGAGAATAANNNAGATCGGAAG
127004 TATGAGTGTGGAGTCGTTGCATGTGTATATACTGAGGTTGGGNNNAGATCGGAAG
127006 TATGAGTGTGGAGTCGTTGCCAAACAGAGAATCATACAGCTATNNNAGATCGGAAG
127007 TATGAGTGTGGAGTCGTTGCCAACCACAAGGATTACATTTATTNNNAGATCGGAAG
127009 TATGAGTGTGGAGTCGTTGCACTTGAGAACATTATTTATATGCANNNAGATCGGAA
127010 TATGAGTGTGGAGTCGTTGCGGGTCTATCTAGATAATTATCCCNNNAGATCGGAAG
127012 TATGAGTGTGGAGTCGTTGCAGGCTAATATACCTGATCTCTATNNNAGATCGGAAG
127014 TATGAGTGTGGAGTCGTTGCAGACACTCTTCTGAACATTTTATNNNAGATCGGAAG
127016 TATGAGTGTGGAGTCGTTGCAAATCTTGGTCTCAACTCAAGNNNAGATCGGAAGAG
127017 TATGAGTGTGGAGTCGTTGCCCTTCATAATGCCTGATTTCTNNNAGATCGGAAGAG
127018 TATGAGTGTGGAGTCGTTGCAGTGTTTTTCTGAACAGAAATAANNNAGATCGGAAG
127019 TATGAGTGTGGAGTCGTTGCAGATTCTGTCAATAGATAGAGTGNNNAGATCGGAA
127020 TATGAGTGTGGAGTCGTTGCAAATGTAACCTTACTGACTTACTNNNAGATCGGAAG
127021 TATGAGTGTGGAGTCGTTGCTAAATGATGATGAAAACCTGTTTNNNAGATCGGAAG
127022 TATGAGTGTGGAGTCGTTGCGAAATACCACTTTTCCTAAAGAGNNNAGATCGGAAG
127023 TATGAGTGTGGAGTCGTTGCTAGACAGTCATACACCATATAACNNNAGATCGGAAG
127024 TATGAGTGTGGAGTCGTTGCTAGACAGTCATACACCATATAACNNNAGATCGGAAG
127025 TATGAGTGTGGAGTCGTTGCCTTAGAGCTCACCTTTGACTNNNAGATCGGAAGAGC
127026 TATGAGTGTGGAGTCGTTGCCTAAGCACTGGAAAGGTAAGNNNAGATCGGAAGAG
127027 TATGAGTGTGGAGTCGTTGCATACAGGGTTTCTCCATGTTNNNAGATCGGAAGAGC
127028 TATGAGTGTGGAGTCGTTGCTGTCATTATTTCTGAGCATAGTNNNAGATCGGAAGA
127029 TATGAGTGTGGAGTCGTTGCTGAAAGAATCTCATATATGCTGTNNNAGATCGGAAG
127030 TATGAGTGTGGAGTCGTTGCGAGAAAAATCCTTTGCTATTTTGNNNAGATCGGAAG
127031 TATGAGTGTGGAGTCGTTGCCCCATAGACCTTACATTCTATTTNNNAGATCGGAAGA
127032 TATGAGTGTGGAGTCGTTGCCACTTAGTATAGCGATTTCATTCNNNAGATCGGAAG
127033 TATGAGTGTGGAGTCGTTGCCCCACCTCTTATAGTCATATTACNNNAGATCGGAAGA
127034 TATGAGTGTGGAGTCGTTGCTGGCTATAGTGGCTATAGTAGNNNAGATCGGAAGA
127035 TATGAGTGTGGAGTCGTTGCGCCCACAAAAAATATATGTGAATNNNAGATCGGAAG
127036 TATGAGTGTGGAGTCGTTGCCCTCAGTTGATCTTTAAAAACAANNNAGATCGGAAG
127037 TATGAGTGTGGAGTCGTTGCCATTCAACAATGTATGATTTGTCNNNAGATCGGAAG
127039 TATGAGTGTGGAGTCGTTGCCCAGTGGAGGATCTTTATTAGNNNAGATCGGAAGA
127040 TATGAGTGTGGAGTCGTTGCAATAAGAAAAAGTCTCAAGATCCNNNAGATCGGAA
127041 TATGAGTGTGGAGTCGTTGCATAATAGCATTTTGGTCAATGACNNNAGATCGGAAG
127042 TATGAGTGTGGAGTCGTTGCGAAAAGGAAAACCCAAGAAACNNNAGATCGGAAGA
127043 TATGAGTGTGGAGTCGTTGCCACATGATTAAACCTCACTAAATNNNAGATCGGAAG
127044 TATGAGTGTGGAGTCGTTGCACACATTTTTTTTTTTGAGATGANNNAGATCGGAAG
127045 TATGAGTGTGGAGTCGTTGCTTTGAATAATACATCAAACCTGGNNNAGATCGGAAG
127046 TATGAGTGTGGAGTCGTTGCCTCCTCCATGTCTGCTTATTNNNAGATCGGAAGAGCA
127047 TATGAGTGTGGAGTCGTTGCGAGTCTGCTTAATAGGAAACANNNAGATCGGAAGA
127048 TATGAGTGTGGAGTCGTTGCTCATCTCTGCTACATAAAGTGNNNAGATCGGAAGAG
127049 TATGAGTGTGGAGTCGTTGCCTAAGTTTATGAACATGCAGTCNNNAGATCGGAAGA
127050 TATGAGTGTGGAGTCGTTGCTAAATATTGACCAGCAAAAAAGANNNAGATCGGAA
127051 TATGAGTGTGGAGTCGTTGCTTTTACTGCAGTGAAAAAATGAANNNAGATCGGAA

127052 TATGAGTGTGGAGTCGTTGCAGCATATAAGAGTTTTGTTCTTCNNNAGATCGGAAG
127053 TATGAGTGTGGAGTCGTTGCCTTGAAGTGTGAACCAAAATGNNNAGATCGGAAGA
127054 TATGAGTGTGGAGTCGTTGCGATCTCACTCTGTCACCTAGNNNAGATCGGAAGAGC
127055 TATGAGTGTGGAGTCGTTGCAAAGAGAAAGAGAGAAAGAAAACNNNAGATCGGA
127057 TATGAGTGTGGAGTCGTTGCGTGTTTTCCCTCTGTTGAATNNNAGATCGGAAGAGC
127058 TATGAGTGTGGAGTCGTTGCTGCTTTCCTTTTACTTACAAAATNNNAGATCGGAAGA
127059 TATGAGTGTGGAGTCGTTGCAAAACAATGGTGTACAAAATCANNNAGATCGGAAG
127060 TATGAGTGTGGAGTCGTTGCTTTAGACTTATTTGAGGACAAGANNNAGATCGGAAG
127061 TATGAGTGTGGAGTCGTTGCCAGAATACACATATTTGACAAGTNNNAGATCGGAAG
127062 TATGAGTGTGGAGTCGTTGCGGACAGAGGTTATATTAGCTTAGNNNAGATCGGAA
127063 TATGAGTGTGGAGTCGTTGCTATGGACACGAAAGGTACTTTNNNAGATCGGAAGA
127064 TATGAGTGTGGAGTCGTTGCTGTATGTTTATGTGACCTAAAACNNNAGATCGGAAG
127065 TATGAGTGTGGAGTCGTTGCAGATACACTGCAAGAATGACNNNAGATCGGAAGAG
127066 TATGAGTGTGGAGTCGTTGCCCTCTTATCCCATAGTTAACATTNNNAGATCGGAAGA
127067 TATGAGTGTGGAGTCGTTGCGTCTTCTCTGTCCAACTACTNNNAGATCGGAAGAGCA
127068 TATGAGTGTGGAGTCGTTGCATACTTAGCTAGGACTTAGGGNNNAGATCGGAAGA
127069 TATGAGTGTGGAGTCGTTGCTTTTAAAAGGAAAGAATGTCTCTNNNAGATCGGAAG
127070 TATGAGTGTGGAGTCGTTGCATAAGTGAGAACATGCCATATATNNNAGATCGGAAG
127071 TATGAGTGTGGAGTCGTTGCCGGGACTGAATTTATTGTTTTATNNNAGATCGGAAG
127072 TATGAGTGTGGAGTCGTTGCCATTGATAAGCATTGATAGTACTNNNAGATCGGAAG
127073 TATGAGTGTGGAGTCGTTGCTCTGAGTTTTTTTCCTCTAGATTNNNAGATCGGAAGA
127074 TATGAGTGTGGAGTCGTTGCCTCTCAGGCATAGCTTTTAAANNNAGATCGGAAGAG
127075 TATGAGTGTGGAGTCGTTGCCAAGAGCAGACATTGATAAATACNNNAGATCGGAAG
127076 TATGAGTGTGGAGTCGTTGCGAAGAACTAGGAATTGATGAGTNNNAGATCGGAAG
127077 TATGAGTGTGGAGTCGTTGCCAGACTTCCTTGAGAAAGTACNNNAGATCGGAAGAG
127078 TATGAGTGTGGAGTCGTTGCCAATAATGTAAATCAGGATGTGGNNNAGATCGGAA
127079 TATGAGTGTGGAGTCGTTGCAGGATTCAAAGCAAAATGAAAATNNNAGATCGGAA
127081 TATGAGTGTGGAGTCGTTGCGTTGTGAGCTGTGTATTTAGANNNAGATCGGAAGA
127082 TATGAGTGTGGAGTCGTTGCTTATCCTAAAAATCAAATGGGAGNNNAGATCGGAAG
127083 TATGAGTGTGGAGTCGTTGCAATGCATGCTCTAGAAATCCNNNAGATCGGAAGAGC
127084 TATGAGTGTGGAGTCGTTGCTGAATAGAAAACTGGTGAAATACNNNAGATCGGAA
127085 TATGAGTGTGGAGTCGTTGCGGAGAGAGAAACAAACAAAATATNNNAGATCGGAA
127086 TATGAGTGTGGAGTCGTTGCAGTACAAAAACTGTCACTTTGANNNAGATCGGAAGA
127087 TATGAGTGTGGAGTCGTTGCGTTGAGAGACAAATTGCACTNNNAGATCGGAAGAG
127089 TATGAGTGTGGAGTCGTTGCTAGGTCATAAAAATATGTGTGGTNNNAGATCGGAA
127090 TATGAGTGTGGAGTCGTTGCTAGGTCATAAAAATATGTGTGGTNNNAGATCGGAA
127091 TATGAGTGTGGAGTCGTTGCTTAAGGTTAAGTAAAGCAGCAAANNNAGATCGGAA
127092 TATGAGTGTGGAGTCGTTGCTTTTGCATAATCACTTGGATTTANNNAGATCGGAAG
127093 TATGAGTGTGGAGTCGTTGCCCATCACTACAATCAGGATAAGNNNAGATCGGAAGA
127094 TATGAGTGTGGAGTCGTTGCGAGTAATTCACATTGAAACCTTANNNAGATCGGAAG
127095 TATGAGTGTGGAGTCGTTGCATCTAAAAGAAAGTGCTTACATGNNNAGATCGGAAG
127096 TATGAGTGTGGAGTCGTTGCTTTTCCTGAGTTGAATTAGAATTNNNAGATCGGAAG
127098 TATGAGTGTGGAGTCGTTGCTAAAACCCTTACTTCTTTTCTCTNNNAGATCGGAAGA

127099 TATGAGTGTGGAGTCGTTGCGTTCCACAAAAAGTTTCTGGNNNAGATCGGAAGAG
127100 TATGAGTGTGGAGTCGTTGCACCCTAAGGGATATATGGTAAANNNAGATCGGAAG
127101 TATGAGTGTGGAGTCGTTGCTTTTAATCTTCCTAATGTTGGAGNNNAGATCGGAAG
127102 TATGAGTGTGGAGTCGTTGCAAGTTACCACAGGAAAAATTTAGNNNAGATCGGAA
127103 TATGAGTGTGGAGTCGTTGCATAAATTGACCTGTTCTTCAACTNNNAGATCGGAAG
127104 TATGAGTGTGGAGTCGTTGCCACTTAATGTCATTGATAGGTTCNNNAGATCGGAAG
127105 TATGAGTGTGGAGTCGTTGCCACTTAATGTCATTGATAGGTTCNNNAGATCGGAAG
127106 TATGAGTGTGGAGTCGTTGCGCTCACTAAATATTATTTGTCCCNNNAGATCGGAAG
127107 TATGAGTGTGGAGTCGTTGCTTTTGCTAGCTGGAGATTTGNNNAGATCGGAAGAG
127108 TATGAGTGTGGAGTCGTTGCTTGCTTCTGGAACTTCATATATANNNAGATCGGAAG
127109 TATGAGTGTGGAGTCGTTGCATCTTCAGTACTCTCTCCTATAANNNAGATCGGAAGA
127110 TATGAGTGTGGAGTCGTTGCGCACTTACAGAGTCTCTCTCNNNAGATCGGAAGAGC
127111 TATGAGTGTGGAGTCGTTGCTTTGCTAGCAAAAAAAAAATCTGNNNAGATCGGAAG
127112 TATGAGTGTGGAGTCGTTGCAAGGAGTTGGTGAAATATTCCNNNAGATCGGAAGA
127113 TATGAGTGTGGAGTCGTTGCAATTTCAAATTCCTTGTACTTGTNNNAGATCGGAAGA
127114 TATGAGTGTGGAGTCGTTGCGTTCTCACAGTACTGAGTTATTTNNNAGATCGGAAG
127115 TATGAGTGTGGAGTCGTTGCACACACTATAATACTGGCATAAGNNNAGATCGGAAG
127116 TATGAGTGTGGAGTCGTTGCACTCTCTTTTAAAAAGATATGCAANNNAGATCGGAA
127117 TATGAGTGTGGAGTCGTTGCTAAAAGGTCAGAAACTCAAGATANNNAGATCGGAA
127118 TATGAGTGTGGAGTCGTTGCCAAGACCAAAAAAAAGCTTTTNNNAGATCGGAAGA
127119 TATGAGTGTGGAGTCGTTGCCAAGACCAAAAAAAAGCTTTTNNNAGATCGGAAGA
127120 TATGAGTGTGGAGTCGTTGCATCTTTTCCTTCTTTTTCCTTTTNNNAGATCGGAAGAG
127121 TATGAGTGTGGAGTCGTTGCAGCAAACAAAATTCAAAATCAAANNNAGATCGGAAG
127122 TATGAGTGTGGAGTCGTTGCGAAAAACTGTTTCTTGAATTCTGNNNAGATCGGAAG
127123 TATGAGTGTGGAGTCGTTGCATTTAAAAGCGTACAATTCAGGNNNAGATCGGAAGA
127125 TATGAGTGTGGAGTCGTTGCGAAGAGAAATAGCATCCCAATNNNAGATCGGAAGA
127126 TATGAGTGTGGAGTCGTTGCAACCTAGGAAAACTTGTTCTCNNNAGATCGGAAGAG
127127 TATGAGTGTGGAGTCGTTGCAAAATACATGCTCAAGAAAGGTANNNAGATCGGAA
127128 TATGAGTGTGGAGTCGTTGCAGTGTGGAGTTATACAGAAATTANNNAGATCGGAA
127129 TATGAGTGTGGAGTCGTTGCAGTTGAACACAGTAAATTATAGGNNNAGATCGGAA
127130 TATGAGTGTGGAGTCGTTGCAAAATTAATTAATAGGCTAGGCCNNNAGATCGGAA
127131 TATGAGTGTGGAGTCGTTGCGCTTAAATGATTCTTCTGTCTTGNNNAGATCGGAAG
127132 TATGAGTGTGGAGTCGTTGCAAAATTAATTAATAGGCTAGGCCNNNAGATCGGAA
127133 TATGAGTGTGGAGTCGTTGCTTATTCTTGAGTAGATTACTGGTNNNAGATCGGAAG
127134 TATGAGTGTGGAGTCGTTGCGAATAGATCCCAAAGACTTTTACNNNAGATCGGAAG
127135 TATGAGTGTGGAGTCGTTGCCTTCATCTCATTTGGAAGTTTATNNNAGATCGGAAG
127137 TATGAGTGTGGAGTCGTTGCGGCCAAGACAATTTAAGAAATATNNNAGATCGGAA
127138 TATGAGTGTGGAGTCGTTGCGAGTGGAGTATGGAAATTATACTNNNAGATCGGAA
127139 TATGAGTGTGGAGTCGTTGCGTTATGCGAATAAATAAGACACTNNNAGATCGGAA
127140 TATGAGTGTGGAGTCGTTGCCTTGAATTTGATGGGACATAAAANNNAGATCGGAA
127141 TATGAGTGTGGAGTCGTTGCGAAAAGGCACAGAGATAAAAATTNNNAGATCGGAA
127142 TATGAGTGTGGAGTCGTTGCGACAGAGAGAGAGAGATTAGTNNNAGATCGGAAG
127143 TATGAGTGTGGAGTCGTTGCCTTTTATCCAAAGTGGGATCTNNNAGATCGGAAGAG

127144 TATGAGTGTGGAGTCGTTGCTGTCTTATCTGAGATACATTTCANNNAGATCGGAAG
127145 TATGAGTGTGGAGTCGTTGCAAGACTTTCTTGATGAAGGAAGNNNAGATCGGAAG
127146 TATGAGTGTGGAGTCGTTGCGAATCTCACATGCATCCTTTANNNAGATCGGAAGAG
127147 TATGAGTGTGGAGTCGTTGCCAAAAAACAGATGCTTCTCTGNNNAGATCGGAAGAG
127148 TATGAGTGTGGAGTCGTTGCACCATGAGTAGCTGTAACTANNNAGATCGGAAGAG
127149 TATGAGTGTGGAGTCGTTGCATTTCCAGATCTTTCTTTTCATTNNNAGATCGGAAGA
127150 TATGAGTGTGGAGTCGTTGCCTTGAAAATGCTCTAGTTCTATGNNNAGATCGGAAG
127151 TATGAGTGTGGAGTCGTTGCTTCCAATATAGCCTATGTTTTCTNNNAGATCGGAAGA
127152 TATGAGTGTGGAGTCGTTGCGTTTGAATGTTGGTTCCATGNNNAGATCGGAAGAG
127153 TATGAGTGTGGAGTCGTTGCATTGCTGACTTTGATAACACTATNNNAGATCGGAAG
127154 TATGAGTGTGGAGTCGTTGCTAAAGACAGGAACAAATAAAGGANNNAGATCGGAA
127155 TATGAGTGTGGAGTCGTTGCCAGAGTAGCTGGGATAACAGNNNAGATCGGAAGAG
127156 TATGAGTGTGGAGTCGTTGCCTAGTGAGCTGTAAGTTTCTTNNNAGATCGGAAGAG
127158 TATGAGTGTGGAGTCGTTGCACTAGAAAGAGCTACTTGAATAGNNNAGATCGGAA
127159 TATGAGTGTGGAGTCGTTGCAGCCTTCAAATACTTTTCTTTTNNNAGATCGGAAGA
127160 TATGAGTGTGGAGTCGTTGCTTATTTTCTGGTCAAATGAAAGGNNNAGATCGGAAG
127161 TATGAGTGTGGAGTCGTTGCGACAAACTTCCATCAGTCATNNNAGATCGGAAGAGC
127162 TATGAGTGTGGAGTCGTTGCCGAGAAAAAAAGACAAAAATACANNNAGATCGGAA
127163 TATGAGTGTGGAGTCGTTGCTATCATTGCCTGCTTTTTTTTTTNNNAGATCGGAAGA
127164 TATGAGTGTGGAGTCGTTGCAAGGAGAAAGTACAGGGAAANNNAGATCGGAAGA
127165 TATGAGTGTGGAGTCGTTGCCCAACTTTATATGCCAAGTATTTNNNAGATCGGAAGA
127167 TATGAGTGTGGAGTCGTTGCTATTACTATATTTGTGCCACAGTNNNAGATCGGAAG
127168 TATGAGTGTGGAGTCGTTGCCTAAAGTCACTGTAAATCCAAAGNNNAGATCGGAAG
127169 TATGAGTGTGGAGTCGTTGCTACAGGTTCTCTAGTGTGTATNNNAGATCGGAAGAG
127170 TATGAGTGTGGAGTCGTTGCCTCAAATCTCATTTCTGTTGTTNNNAGATCGGAAGAG
127171 TATGAGTGTGGAGTCGTTGCAGGTGTCTATGTAAACTGATTTANNNAGATCGGAAG
127172 TATGAGTGTGGAGTCGTTGCAGTTCAGTACTTGGTTCATAATANNNAGATCGGAAG
127173 TATGAGTGTGGAGTCGTTGCGATAGAGCCATTAACAAGGATANNNAGATCGGAAG
127174 TATGAGTGTGGAGTCGTTGCGTTGTAGACAATTGTTCACATACNNNAGATCGGAAG
127175 TATGAGTGTGGAGTCGTTGCAGAAAGACAGACAATAGAAACTTNNNAGATCGGAA
127176 TATGAGTGTGGAGTCGTTGCAAACTTAGAAAAAGAAACACACANNNAGATCGGAA
127177 TATGAGTGTGGAGTCGTTGCCAGTAGAAAGTGTAAGCTAGTATNNNAGATCGGAA
127178 TATGAGTGTGGAGTCGTTGCAAATCAAGCTTCTAATCCAGAANNNAGATCGGAAGA
127179 TATGAGTGTGGAGTCGTTGCTAGCCAACGATAATTATCTCAAANNNAGATCGGAAG
127180 TATGAGTGTGGAGTCGTTGCGAACAAAGAAATAAGGAAGGTAGNNNAGATCGGA
127182 TATGAGTGTGGAGTCGTTGCATACATAGGTTCATACAATGCATNNNAGATCGGAAG
127183 TATGAGTGTGGAGTCGTTGCCTCTCTATGCATTTCTTCCATANNNAGATCGGAAGAG
127184 TATGAGTGTGGAGTCGTTGCTATTTCCAAACACACCTTGCNNNAGATCGGAAGAGC
127185 TATGAGTGTGGAGTCGTTGCCTGGGCATATGAAACTCTAGNNNAGATCGGAAGAG
127186 TATGAGTGTGGAGTCGTTGCCGTATATACTGCAAACACAAATNNNAGATCGGAAGA
127188 TATGAGTGTGGAGTCGTTGCCTCTTCATCTAGGATGCTTTCNNNAGATCGGAAGAG
127189 TATGAGTGTGGAGTCGTTGCAATTGAAAGGAAAACCCTAACTTNNNAGATCGGAAG
127190 TATGAGTGTGGAGTCGTTGCGGAGAAGCCTACTTTTTTGTNNNAGATCGGAAGAG

127191 TATGAGTGTGGAGTCGTTGCAGATTGAGGCAGGAGAATAANNNAGATCGGAAGA
127192 TATGAGTGTGGAGTCGTTGCTATCATAGGGACTTTGAAATTGANNNAGATCGGAAG
127193 TATGAGTGTGGAGTCGTTGCGCCCTGGACTGTAATTTAATANNNAGATCGGAAGAG
127194 TATGAGTGTGGAGTCGTTGCAGAATGAATTGGGAAATAGTTTCNNNAGATCGGAA
127195 TATGAGTGTGGAGTCGTTGCTGAATTTTTTCCCACTAATTTGANNNAGATCGGAAG
127196 TATGAGTGTGGAGTCGTTGCATGCTGGAGTTTTATTATTGTTTNNNAGATCGGAAG
127197 TATGAGTGTGGAGTCGTTGCGTAGTCGAAAATTCATGCATAANNNAGATCGGAAG
127198 TATGAGTGTGGAGTCGTTGCATAGCTCTTATGGAGGTTACATTNNNAGATCGGAAG
127199 TATGAGTGTGGAGTCGTTGCAGAAGCTAATCACTATGAGTTTANNNAGATCGGAAG
127200 TATGAGTGTGGAGTCGTTGCGAGGCAACTCAATTTCTAATTTANNNAGATCGGAAG
127201 TATGAGTGTGGAGTCGTTGCCACAAAAGGAGAGTACAAATATCNNNAGATCGGAA
127202 TATGAGTGTGGAGTCGTTGCTGAAGTGTACAATATTATACTGGTNNNAGATCGGAA
127203 TATGAGTGTGGAGTCGTTGCACTGTATATATGTTCCAGTGTGNNNAGATCGGAAGA
127204 TATGAGTGTGGAGTCGTTGCTATGCTCCTTTTCTTTCTAGAAGNNNAGATCGGAAG
127205 TATGAGTGTGGAGTCGTTGCTACATGCATAGAGAATGTGTAATNNNAGATCGGAA
127206 TATGAGTGTGGAGTCGTTGCAAGTTTACCTTTGCCTGTATTTNNNAGATCGGAAGA
127207 TATGAGTGTGGAGTCGTTGCACACTTGTAGATTTTTATGACACNNNAGATCGGAAG
127208 TATGAGTGTGGAGTCGTTGCAAATAGAGTTGCACAGAGAAAANNNAGATCGGAAG
127209 TATGAGTGTGGAGTCGTTGCCATGAACCTAGAGAATCTGATCNNNAGATCGGAAGA
127210 TATGAGTGTGGAGTCGTTGCGTTTTTAGAAAAGCCAAAGAAAANNNAGATCGGAA
127211 TATGAGTGTGGAGTCGTTGCCTCCAATTCATGAGATCAGAATNNNAGATCGGAAGA
127212 TATGAGTGTGGAGTCGTTGCGAAGAGGAGACTCAGTTTTATATNNNAGATCGGAA
127213 TATGAGTGTGGAGTCGTTGCAGAGTTTATGTTCTTATTTGCAGNNNAGATCGGAAG
127214 TATGAGTGTGGAGTCGTTGCCATGAACCTAGAGAATCTGATCNNNAGATCGGAAGA
127215 TATGAGTGTGGAGTCGTTGCATTTTATGGTGAAAACAATGAGTNNNAGATCGGAA
127216 TATGAGTGTGGAGTCGTTGCCTATAATCTCAGCTACTCAGAAGNNNAGATCGGAAG
127217 TATGAGTGTGGAGTCGTTGCCTACATACTTATTTTCCCTCTCTNNNAGATCGGAAGA
127218 TATGAGTGTGGAGTCGTTGCTGTCTATACCTGATTTAGCATTTNNNAGATCGGAAG
127219 TATGAGTGTGGAGTCGTTGCAGACAAGGAAGGAAGGTAAATANNNAGATCGGAA
127220 TATGAGTGTGGAGTCGTTGCCTTCTGGGTCTCATAATTTTATGNNNAGATCGGAAG
127222 TATGAGTGTGGAGTCGTTGCTTCCTTATGTGCTTAGGTAACNNNAGATCGGAAGAG
127223 TATGAGTGTGGAGTCGTTGCCATTTTTGCTTATGTGTTTCAGNNNAGATCGGAAGA
127224 TATGAGTGTGGAGTCGTTGCAAGAAAGAAAGAAAAAACTCAGANNNAGATCGGAA
127225 TATGAGTGTGGAGTCGTTGCTAGGCCTAAAGAGCAGGTATNNNAGATCGGAAGAG
127226 TATGAGTGTGGAGTCGTTGCTTATCAGCATCTTTGGTATTTACNNNAGATCGGAAG
127227 TATGAGTGTGGAGTCGTTGCAACAATCCTGTGAGATATTCTTANNNAGATCGGAAG
127228 TATGAGTGTGGAGTCGTTGCTATCATCAAATGAGCAGAACAATNNNAGATCGGAAG
127229 TATGAGTGTGGAGTCGTTGCCATTATGGGATGATCCATCAANNNAGATCGGAAGA
127230 TATGAGTGTGGAGTCGTTGCTTTTAAGATTTTTAAAGGTGCACNNNAGATCGGAAG
127231 TATGAGTGTGGAGTCGTTGCCAAAGAAAAGAGAGGAAGAAAGNNNAGATCGGAA
127232 TATGAGTGTGGAGTCGTTGCCAAAGAAAAGAGAGGAAGAAAGNNNAGATCGGAA
127233 TATGAGTGTGGAGTCGTTGCAAACAGCAATCTTATACTGTCTANNNAGATCGGAAG
127234 TATGAGTGTGGAGTCGTTGCATTATCCTCATTTTAGAGATGGGNNNAGATCGGAAG

127235 TATGAGTGTGGAGTCGTTGCAAATTATAGGAAGAAAGTGCAATNNNAGATCGGAA
127236 TATGAGTGTGGAGTCGTTGCCTTGCCTAAGTTATCTAACCTCNNNAGATCGGAAGA
127237 TATGAGTGTGGAGTCGTTGCTATTTGTTCTGTGATTGTAGAACNNNAGATCGGAAG
127238 TATGAGTGTGGAGTCGTTGCGATTTTCTTTGGGGAATCTGANNNAGATCGGAAGA
127239 TATGAGTGTGGAGTCGTTGCCTTATCTTCAGATGTAGTAGGAGNNNAGATCGGAA
127240 TATGAGTGTGGAGTCGTTGCCCAGGGCTATACTTTTTATTTTTNNNAGATCGGAAG
127241 TATGAGTGTGGAGTCGTTGCAATATCTCTGTTTCAGGTAGATTNNNAGATCGGAAG
127242 TATGAGTGTGGAGTCGTTGCAACCATAATAACCAGTCTGTATTNNNAGATCGGAAG
127244 TATGAGTGTGGAGTCGTTGCCGAAAATCAAAGAAAAAGAGGANNNAGATCGGAA
127245 TATGAGTGTGGAGTCGTTGCCCATACCAGTACTACAGATTATGNNNAGATCGGAAG
127246 TATGAGTGTGGAGTCGTTGCAATTAGGTGTGATGGTTTATTTTNNNAGATCGGAAG
127247 TATGAGTGTGGAGTCGTTGCCTGAACTTGAGAAAAGCAAAANNNAGATCGGAAGA
127248 TATGAGTGTGGAGTCGTTGCGTACTAAGAACAAAGTGTGATTTNNNAGATCGGAA
127249 TATGAGTGTGGAGTCGTTGCCTGATTCTGGATACAATACAAAGNNNAGATCGGAAG
127251 TATGAGTGTGGAGTCGTTGCTTAATGTCTTTGATGGGTTCTTNNNAGATCGGAAGA
127252 TATGAGTGTGGAGTCGTTGCCTAAAAACCTAGAACCTTTTCAGNNNAGATCGGAAG
127253 TATGAGTGTGGAGTCGTTGCATTATACTCTCTACCTCCATGAGNNNAGATCGGAAG
127254 TATGAGTGTGGAGTCGTTGCTACTTAGCAAATACAAAAACCAANNNAGATCGGAAG
127255 TATGAGTGTGGAGTCGTTGCCAGAAAGTTTCACACATATGATTNNNAGATCGGAAG
127256 TATGAGTGTGGAGTCGTTGCCTTTTACCTCTACATCTTTTTCCNNNAGATCGGAAGA
127258 TATGAGTGTGGAGTCGTTGCATGCAAGAAATGTTTATACCTTTNNNAGATCGGAAG
127259 TATGAGTGTGGAGTCGTTGCCAGAGAAGATTCCCGAATATTCNNNAGATCGGAAG
127260 TATGAGTGTGGAGTCGTTGCCTTGGATTTCTCTCTTCTTCTTNNNAGATCGGAAGAG
127261 TATGAGTGTGGAGTCGTTGCTTTTCTTAAAAATCTGACTGTCTNNNAGATCGGAAG
127262 TATGAGTGTGGAGTCGTTGCCTACACATATCAGATGCTGTATTNNNAGATCGGAAG
127263 TATGAGTGTGGAGTCGTTGCCTAGTTCTAAATGATTCCCCTTTNNNAGATCGGAAG
127264 TATGAGTGTGGAGTCGTTGCTCTGAGTTCAAGCAATTCTCNNNAGATCGGAAGAGC
127265 TATGAGTGTGGAGTCGTTGCCAGAGAAGTAAAAGGCCTACNNNAGATCGGAAGAG
127266 TATGAGTGTGGAGTCGTTGCTAAAAGACCAACTGATTGTCTTTNNNAGATCGGAAG
127267 TATGAGTGTGGAGTCGTTGCGTTAGAGGAAATAAGTCTTACACNNNAGATCGGAA
127268 TATGAGTGTGGAGTCGTTGCGTGGTTAGAGGAAATAAGTCTTANNNAGATCGGAA
127269 TATGAGTGTGGAGTCGTTGCTTTTAGAGCTCTTCCTTATCTTTNNNAGATCGGAAGA
127270 TATGAGTGTGGAGTCGTTGCCACTGCAGAAAGAAAGAAAANNNAGATCGGAAGAG
127271 TATGAGTGTGGAGTCGTTGCACTATTTTTGGAGCTAACCTAAGNNNAGATCGGAAG
127272 TATGAGTGTGGAGTCGTTGCTTTGTTTGTTTTTTGAGACAGANNNAGATCGGAAGA
127273 TATGAGTGTGGAGTCGTTGCTAAACATGGGTAAAAATTCAGCNNNAGATCGGAAG
127274 TATGAGTGTGGAGTCGTTGCATAACTTGTGTGTGTATTTCAATNNNAGATCGGAAG
127275 TATGAGTGTGGAGTCGTTGCATGTAAGCATAGGTTAAACATGGNNNAGATCGGAA
127276 TATGAGTGTGGAGTCGTTGCCTCTACTATGTGTGGGTTTTTNNNAGATCGGAAGAG
127278 TATGAGTGTGGAGTCGTTGCAAATGGTTTGAGTAATCATCAGANNNAGATCGGAA
127279 TATGAGTGTGGAGTCGTTGCTAAGTCTAACCCTAAAGGAGAGNNNAGATCGGAAG
127280 TATGAGTGTGGAGTCGTTGCTTATACATTTAGGTGCTAATTGGNNNAGATCGGAAG
127281 TATGAGTGTGGAGTCGTTGCATTTTGATTTTTCCAGCTACAAGNNNAGATCGGAAG

127282 TATGAGTGTGGAGTCGTTGCTAGAGAGAGAGAGAAGATCTTGNNNAGATCGGAA
127284 TATGAGTGTGGAGTCGTTGCTCTAAAAGTTGTAAAAGTTTGCTNNNAGATCGGAAG
127285 TATGAGTGTGGAGTCGTTGCATCTTTCCCATTAAGAGAAACANNNAGATCGGAAGA
127286 TATGAGTGTGGAGTCGTTGCAAACATCCTTTTAGAAGAGTGANNNAGATCGGAAG
127287 TATGAGTGTGGAGTCGTTGCAATAACTGATAAAGGCCAAAAAANNNAGATCGGAA
127288 TATGAGTGTGGAGTCGTTGCTACTAAGTGGTACCTAAGACCNNNAGATCGGAAGAG
127290 TATGAGTGTGGAGTCGTTGCGTATGATAGCTGAGTAAAGATGANNNAGATCGGAA
127291 TATGAGTGTGGAGTCGTTGCACTTAACCAAATTTCAATTTCCANNNAGATCGGAAGA
127292 TATGAGTGTGGAGTCGTTGCTATGTAACTTTTACTCTGTGTGANNNAGATCGGAAG
127293 TATGAGTGTGGAGTCGTTGCCTCAGACAAGTCTTCCATCTNNNAGATCGGAAGAGC
127294 TATGAGTGTGGAGTCGTTGCCATTACTGCGACCTTATCACNNNAGATCGGAAGAGC
127295 TATGAGTGTGGAGTCGTTGCCAACCAGCAAAAGAAAAAAATTANNNAGATCGGAA
127296 TATGAGTGTGGAGTCGTTGCTATCTGGGACCTATTTCAGTCNNNAGATCGGAAGAG
127297 TATGAGTGTGGAGTCGTTGCGAGTGCTGTAGTAGGATAATAGNNNAGATCGGAAG
127300 TATGAGTGTGGAGTCGTTGCTCCTCTTATAAGGAAACTAATCCNNNAGATCGGAAG
127301 TATGAGTGTGGAGTCGTTGCCTTCAGCAGTTTATATACAACAANNNAGATCGGAAG
127302 TATGAGTGTGGAGTCGTTGCCTTCAGCAGTTTATATACAACAANNNAGATCGGAAG
127303 TATGAGTGTGGAGTCGTTGCGTAATAAACCTATCATGAGTTGGNNNAGATCGGAAG
127304 TATGAGTGTGGAGTCGTTGCTGTAGGGATTTAAAATGACAAAGNNNAGATCGGAA
127305 TATGAGTGTGGAGTCGTTGCGTGTTTTTGATCATCCTAACTTANNNAGATCGGAAGA
127306 TATGAGTGTGGAGTCGTTGCGACTACAGTGGTTTGAACTTNNNAGATCGGAAGAG
127307 TATGAGTGTGGAGTCGTTGCGTCAGATCTAAAGCCAATACANNNAGATCGGAAGAG
127308 TATGAGTGTGGAGTCGTTGCATCAAAGTAGCTAACAATCCTATNNNAGATCGGAAG
127309 TATGAGTGTGGAGTCGTTGCGTGTATGTCTGTATGTTATTTGGNNNAGATCGGAAG
127310 TATGAGTGTGGAGTCGTTGCCTCAGGAAATAAAGGCTACATNNNAGATCGGAAGA
127311 TATGAGTGTGGAGTCGTTGCTACTATTGTCTGTTTTTGAAAGGNNNAGATCGGAAG
127312 TATGAGTGTGGAGTCGTTGCCCCTTTTAGCAAAGAATTTAAACNNNAGATCGGAAG
127313 TATGAGTGTGGAGTCGTTGCGATTTTATCCACATAAGAATCCCNNNAGATCGGAAG
127314 TATGAGTGTGGAGTCGTTGCGATTTTATCCACATAAGAATCCCNNNAGATCGGAAG
127315 TATGAGTGTGGAGTCGTTGCGTAAAACATGCCTGATTGAAANNNAGATCGGAAGA
127317 TATGAGTGTGGAGTCGTTGCTATGTGTGATTGTGGCATAAATANNNAGATCGGAA
127318 TATGAGTGTGGAGTCGTTGCCATTTAGCTATTCTGAACCATCNNNAGATCGGAAGA
127319 TATGAGTGTGGAGTCGTTGCGATCTGACTGTACTAAGAATGTTNNNAGATCGGAAG
127321 TATGAGTGTGGAGTCGTTGCTATTATTGTTGAACTGTTTGGAGNNNAGATCGGAAG
127322 TATGAGTGTGGAGTCGTTGCACAAATGTCTGCAGAAAGATANNNAGATCGGAAGA
127323 TATGAGTGTGGAGTCGTTGCCCATGTAAGGACATAATGAGAAGNNNAGATCGGAA
127324 TATGAGTGTGGAGTCGTTGCCCATGTAAGGACATAATGAGAAGNNNAGATCGGAA
127325 TATGAGTGTGGAGTCGTTGCATATATACACACACACAGTATCANNNAGATCGGAAG
127326 TATGAGTGTGGAGTCGTTGCCATGATCATAGCTCACTGTAGNNNAGATCGGAAGAG
127327 TATGAGTGTGGAGTCGTTGCCTTACCAACAGAAGCTTGATNNNAGATCGGAAGAGC
127328 TATGAGTGTGGAGTCGTTGCTAAACCTTGTAAGACGGATATTCNNNAGATCGGAAG
127329 TATGAGTGTGGAGTCGTTGCAAATATTCTTTCTGCTATTCACCNNNAGATCGGAAGA
127330 TATGAGTGTGGAGTCGTTGCTTCCTTATCTACAACATCTGTTTNNNAGATCGGAAGA

127331 TATGAGTGTGGAGTCGTTGCGAAATTGTATTCCTCCTAACAATNNNAGATCGGAAG
127332 TATGAGTGTGGAGTCGTTGCGAAATTGTATTCCTCCTAACAATNNNAGATCGGAAG
127333 TATGAGTGTGGAGTCGTTGCTCATTCTACAGATGAGAAAAATGNNNAGATCGGAA
127334 TATGAGTGTGGAGTCGTTGCTTTACCTTCAAAGTTAACTGTACNNNAGATCGGAAG
127335 TATGAGTGTGGAGTCGTTGCAAGAAAAGAGAAAGGAAAGAAAGNNNAGATCGGA
127336 TATGAGTGTGGAGTCGTTGCCATGTGAGGATATAGCAACTGNNNAGATCGGAAGA
127337 TATGAGTGTGGAGTCGTTGCCATGTGAGGATATAGCAACTGNNNAGATCGGAAGA
127338 TATGAGTGTGGAGTCGTTGCCATGTGAGGATATAGCAACTGNNNAGATCGGAAGA
127339 TATGAGTGTGGAGTCGTTGCATACAGAATTGGGAGGGGTANNNAGATCGGAAGA
127340 TATGAGTGTGGAGTCGTTGCACCTAGTAGGAGACTAAATTGATNNNAGATCGGAA
127341 TATGAGTGTGGAGTCGTTGCCATTTCTTTGGAGAACTTTAACTNNNAGATCGGAAG
127342 TATGAGTGTGGAGTCGTTGCCATTTCTTTGGAGAACTTTAACTNNNAGATCGGAAG
127343 TATGAGTGTGGAGTCGTTGCGAACCAATGTGGGAAGTATATATNNNAGATCGGAA
127347 TATGAGTGTGGAGTCGTTGCATTACAGATGTGTACTACAATGTNNNAGATCGGAAG
127348 TATGAGTGTGGAGTCGTTGCTACATCGTCGCTATTTTTTTGNNNAGATCGGAAGAG
127349 TATGAGTGTGGAGTCGTTGCTTTCTAGCTCTATCCATAAAGAGNNNAGATCGGAAG
127351 TATGAGTGTGGAGTCGTTGCCTTGGAATTGCAGATCTCTAGNNNAGATCGGAAGAG
127352 TATGAGTGTGGAGTCGTTGCTCAGACTTGAGTTTGTTTGCNNNAGATCGGAAGAGC
127353 TATGAGTGTGGAGTCGTTGCTCTCTCTGACATTTTACTTCTCNNNAGATCGGAAGAG
127354 TATGAGTGTGGAGTCGTTGCTGAACCATTTTTTTTCTACAGAANNNAGATCGGAAG
127355 TATGAGTGTGGAGTCGTTGCTGAACCATTTTTTTTCTACAGAANNNAGATCGGAAG
127356 TATGAGTGTGGAGTCGTTGCTGAACCATTTTTTTTCTACAGAANNNAGATCGGAAG
127357 TATGAGTGTGGAGTCGTTGCAAAAATAGTTTTGTTCAACACACNNNAGATCGGAAG
127358 TATGAGTGTGGAGTCGTTGCGGGAATTTTAACAGCTACAATTANNNAGATCGGAAG
127359 TATGAGTGTGGAGTCGTTGCTCTTTCTCCTGTCTCTCTACNNNAGATCGGAAGAGCA
127360 TATGAGTGTGGAGTCGTTGCTCTGTTCTAGAATGAAAACCCNNNAGATCGGAAGAG
127361 TATGAGTGTGGAGTCGTTGCAAGTAACCTAAAATCACTGACTCNNNAGATCGGAAG
127362 TATGAGTGTGGAGTCGTTGCGTTGAGAGATATTCTTCACCTGNNNAGATCGGAAGA
127363 TATGAGTGTGGAGTCGTTGCGTTGAGAGATATTCTTCACCTGNNNAGATCGGAAGA
127364 TATGAGTGTGGAGTCGTTGCTATATACTACCAAACATTAGGGGNNNAGATCGGAAG
127367 TATGAGTGTGGAGTCGTTGCATATTTTGTAGTCATGCTCAATCNNNAGATCGGAAG
127368 TATGAGTGTGGAGTCGTTGCTTGGAAAATGAAATCACGTTTTNNNAGATCGGAAGA
127369 TATGAGTGTGGAGTCGTTGCTTCATGAAAGACAATTTTTCCATNNNAGATCGGAAG
127370 TATGAGTGTGGAGTCGTTGCCCTGACTGTATCTGTAATATCATNNNAGATCGGAAG
127371 TATGAGTGTGGAGTCGTTGCATTTTCTTTGAATTGAGCAAAACNNNAGATCGGAAG
127372 TATGAGTGTGGAGTCGTTGCGGAAACAAAGTTCACAAAATAACNNNAGATCGGAA
127373 TATGAGTGTGGAGTCGTTGCATTTTGCACTGTAATAACTCAATNNNAGATCGGAAG
127375 TATGAGTGTGGAGTCGTTGCTACTAAGGGGATCTAGAATCAATNNNAGATCGGAA
127376 TATGAGTGTGGAGTCGTTGCAGTGTCTGAACTTAGGAGAANNNAGATCGGAAGAG
127378 TATGAGTGTGGAGTCGTTGCCTCCCTAAAGAAAAATCTTGTTTNNNAGATCGGAAG
127379 TATGAGTGTGGAGTCGTTGCTTTAGGATTTGTTTTCTTTTGGTNNNAGATCGGAAG
127380 TATGAGTGTGGAGTCGTTGCCCCAAAAAATTTTAGAGAAGTCTNNNAGATCGGAAG
127381 TATGAGTGTGGAGTCGTTGCAAGAAATATGAAGAGATGTTTGGNNNAGATCGGAA

127382 TATGAGTGTGGAGTCGTTGCAGTTAAATTTGACTTGAACAGTTNNNAGATCGGAAG
127384 TATGAGTGTGGAGTCGTTGCGTATGTGATATGGGATAACCTCNNNAGATCGGAAGA
127386 TATGAGTGTGGAGTCGTTGCCTTTTACTGCAATACAGGTATCNNNAGATCGGAAGA
127387 TATGAGTGTGGAGTCGTTGCCTAAATAATTGGACAGCTATCTCNNNAGATCGGAAG
127388 TATGAGTGTGGAGTCGTTGCTTATTTGATCAGCCAGGTAATAANNNAGATCGGAAG
127389 TATGAGTGTGGAGTCGTTGCAGTATCCTAGTTGATCCTCTGNNNAGATCGGAAGAG
127390 TATGAGTGTGGAGTCGTTGCTCATGATTCTTATCCACCAATAANNNAGATCGGAAGA
127391 TATGAGTGTGGAGTCGTTGCAAAACAGCATAGGAAAAACTTCNNNAGATCGGAAGA
127392 TATGAGTGTGGAGTCGTTGCTATGAGAATGAGACAACAACAANNNAGATCGGAAG
127393 TATGAGTGTGGAGTCGTTGCAATCTAGAGGAAGAAACATTTCANNNAGATCGGAA
127394 TATGAGTGTGGAGTCGTTGCATCAAAATCAAAACACAGTACTTNNNAGATCGGAAG
127395 TATGAGTGTGGAGTCGTTGCCTGTATGGTGCTATAATTAACACNNNAGATCGGAAG
127396 TATGAGTGTGGAGTCGTTGCCTTATTATGTCACAATCTCACAGNNNAGATCGGAAGA
127397 TATGAGTGTGGAGTCGTTGCTCCAAATACAATGTTTTCAAAGANNNAGATCGGAAG
127398 TATGAGTGTGGAGTCGTTGCTACAATGGAGTTACATTCAGATANNNAGATCGGAAG
127399 TATGAGTGTGGAGTCGTTGCAAATCTTCCATTTGATTAGGAGANNNAGATCGGAAG
127400 TATGAGTGTGGAGTCGTTGCTTGTAGTAACTCGTATCAAGAGNNNAGATCGGAAGA
127401 TATGAGTGTGGAGTCGTTGCGAGTTATAGCTGACTTCTTTTTTNNNAGATCGGAAGA
127402 TATGAGTGTGGAGTCGTTGCGCTTCTAAATATTGTTTTCCACTNNNAGATCGGAAGA
127403 TATGAGTGTGGAGTCGTTGCGTATATCAAAGAGATATCTGCACNNNAGATCGGAAG
127404 TATGAGTGTGGAGTCGTTGCTCTTATAATATGTCTCATGGTGTNNNAGATCGGAAGA
127405 TATGAGTGTGGAGTCGTTGCTATATATGTGTGTATGTGTGTGTNNNAGATCGGAA
127406 TATGAGTGTGGAGTCGTTGCCTCAGGTTCAAGCAATTCTCNNNAGATCGGAAGAGC
127407 TATGAGTGTGGAGTCGTTGCTATCATCACCATAACCATTTTTGNNNAGATCGGAAGA
127408 TATGAGTGTGGAGTCGTTGCTATATATGTGTGTGTGTGTGTATNNNAGATCGGAA
127409 TATGAGTGTGGAGTCGTTGCTACTTCTGAAACTTCTTTTTCTCNNNAGATCGGAAGA
127410 TATGAGTGTGGAGTCGTTGCGAAAGGGGGACAGTACATTTNNNAGATCGGAAGA
127411 TATGAGTGTGGAGTCGTTGCATTTAAAGGAAGTGTGTTTTGGNNNAGATCGGAAG
127412 TATGAGTGTGGAGTCGTTGCCTTCTCTTTCTGGGGTATAGANNNAGATCGGAAGAG
127413 TATGAGTGTGGAGTCGTTGCACTGATCTCTTGAGTATTCATTTNNNAGATCGGAAGA
127414 TATGAGTGTGGAGTCGTTGCTGGTGGTATTTACACAAATCTATNNNAGATCGGAAG
127415 TATGAGTGTGGAGTCGTTGCATTGCTTGTTTTTTTAAGTTCTGNNNAGATCGGAAGA
127416 TATGAGTGTGGAGTCGTTGCCTGAGAGTGAAGAATGCTTTANNNAGATCGGAAGA
127417 TATGAGTGTGGAGTCGTTGCAAAGTACCCAAGGAGTATCTATNNNAGATCGGAAGA
127418 TATGAGTGTGGAGTCGTTGCTTTACCCTTTGTTTTGAATCAGNNNAGATCGGAAGA
127419 TATGAGTGTGGAGTCGTTGCTTTCACCAAATATTTTACACCTTNNNAGATCGGAAGA
127421 TATGAGTGTGGAGTCGTTGCGATAGTCCTTACTTAACAGATCANNNAGATCGGAAG
127422 TATGAGTGTGGAGTCGTTGCCATATGGGATACAATGAAAAGACNNNAGATCGGAA
127423 TATGAGTGTGGAGTCGTTGCCTTTCCCACTTTTGCAATTGNNNAGATCGGAAGAGCA
127424 TATGAGTGTGGAGTCGTTGCGCAGTTTTATCATGTGTAAATTCNNNAGATCGGAAG
127425 TATGAGTGTGGAGTCGTTGCACCTGTAAGATATACAACAGTTTNNNAGATCGGAAG
127426 TATGAGTGTGGAGTCGTTGCTCCCAGGTTACTAAATGTCATNNNAGATCGGAAGAG
127427 TATGAGTGTGGAGTCGTTGCCATCTCACATTCCTCACTACNNNAGATCGGAAGAGCA

127428 TATGAGTGTGGAGTCGTTGCGGAGTGAGAATATTTGTTTAGTGNNNAGATCGGAA
127429 TATGAGTGTGGAGTCGTTGCCATCAATGAACACTAAGTTGTATNNNAGATCGGAAG
127430 TATGAGTGTGGAGTCGTTGCCACATAGAGATAGTACATAGCATNNNAGATCGGAAG
127431 TATGAGTGTGGAGTCGTTGCGTTCTTCCTCCACACAATATCNNNAGATCGGAAGAGC
127432 TATGAGTGTGGAGTCGTTGCCTAATTGCTAGGCTTTTGTAGNNNAGATCGGAAGAG
127434 TATGAGTGTGGAGTCGTTGCAATCAATTCTCCCTGTGCTTANNNAGATCGGAAGAG
127435 TATGAGTGTGGAGTCGTTGCACTAGTACTAAAATCTGTGTCTGNNNAGATCGGAAG
127436 TATGAGTGTGGAGTCGTTGCAAGACCCTGTCTCAAAAAAAANNNAGATCGGAAGAG
127437 TATGAGTGTGGAGTCGTTGCCTGTGGGTAAACATCCTTAGNNNAGATCGGAAGAG
127438 TATGAGTGTGGAGTCGTTGCCTTTGTATGGCAGATGTAATTAGNNNAGATCGGAAG
127439 TATGAGTGTGGAGTCGTTGCCTATTGTGAAGAGTGCTGTAATANNNAGATCGGAAG
127440 TATGAGTGTGGAGTCGTTGCAATAGTACTGCACTAAACATGATNNNAGATCGGAAG
127443 TATGAGTGTGGAGTCGTTGCTATTCTGAGAAATGGGTATGATTNNNAGATCGGAAG
127444 TATGAGTGTGGAGTCGTTGCCCAAGAAATCAAAGGGAGAANNNAGATCGGAAGAG
127445 TATGAGTGTGGAGTCGTTGCGTAGAATGAGTTGTGGTAGAATNNNAGATCGGAAG
127446 TATGAGTGTGGAGTCGTTGCTGTGTATATATGCATGTATATGTGNNNAGATCGGAA
127447 TATGAGTGTGGAGTCGTTGCGTTTACTTAGAGGAATCCAAATGNNNAGATCGGAAG
127448 TATGAGTGTGGAGTCGTTGCTAATTCCCCTATGCCTTCTCNNNAGATCGGAAGAGCA
127449 TATGAGTGTGGAGTCGTTGCGCTTTTAAAATCAGAAAACAGTCNNNAGATCGGAAG
127450 TATGAGTGTGGAGTCGTTGCCTGGAGGTAACACTGAAATGNNNAGATCGGAAGAG
127451 TATGAGTGTGGAGTCGTTGCGATAAATATGCTTTACAGAACGANNNAGATCGGAAG
127453 TATGAGTGTGGAGTCGTTGCTTTGGGAATATTTGCATTATACTNNNAGATCGGAAG
127454 TATGAGTGTGGAGTCGTTGCATCTCACATTGTCTTTATTCATCNNNAGATCGGAAGA
127455 TATGAGTGTGGAGTCGTTGCAAGACACATCAAAAGAAGAAAATNNNAGATCGGAA
127456 TATGAGTGTGGAGTCGTTGCGCATGTAATGATGGAATGATATTNNNAGATCGGAA
127457 TATGAGTGTGGAGTCGTTGCCTGATGAATATTCACAAGCATAANNNAGATCGGAAG
127458 TATGAGTGTGGAGTCGTTGCCATTCTTCGTTAGCCACTTTANNNAGATCGGAAGAG
127459 TATGAGTGTGGAGTCGTTGCTATTTTGTTGTTGTTGTTGTAGANNNAGATCGGAAG
127460 TATGAGTGTGGAGTCGTTGCAATGGATGAATATTTTGTTCTGANNNAGATCGGAAG
127462 TATGAGTGTGGAGTCGTTGCGTGTCACTTTATTCATACCTTTANNNAGATCGGAAGA
127463 TATGAGTGTGGAGTCGTTGCATAATCACTCCTTATTTGGGAATNNNAGATCGGAAG
127464 TATGAGTGTGGAGTCGTTGCACTTAACCTCTCTGAATTTATGTNNNAGATCGGAAGA
127465 TATGAGTGTGGAGTCGTTGCTAGACTGGAAACTTAGTTCTAAGNNNAGATCGGAAG
127466 TATGAGTGTGGAGTCGTTGCAAGTCATACGGAAAACATTTTCNNNAGATCGGAAGA
127467 TATGAGTGTGGAGTCGTTGCTATTTAGGTACAGGGATTAGAGGNNNAGATCGGAA
127468 TATGAGTGTGGAGTCGTTGCACAGAAAGACCAAAAGCTACNNNAGATCGGAAGAG
127469 TATGAGTGTGGAGTCGTTGCACAGAAAGACCAAAAGCTACNNNAGATCGGAAGAG
127470 TATGAGTGTGGAGTCGTTGCATACCATAATAGCAAAGACATCTNNNAGATCGGAAG
127471 TATGAGTGTGGAGTCGTTGCTTTATTCTACGAGACCTGAATTANNNAGATCGGAAG
127472 TATGAGTGTGGAGTCGTTGCTAGAAGTATGGTGTTACTTTTGANNNAGATCGGAAG
127473 TATGAGTGTGGAGTCGTTGCATTTCAAAACACTGAAAAGAAGGNNNAGATCGGAA
127474 TATGAGTGTGGAGTCGTTGCGTAGATAGTGGAGTTGGAATAATNNNAGATCGGAA
127475 TATGAGTGTGGAGTCGTTGCACAGATTGAACATAGGTATGTATNNNAGATCGGAA

127476 TATGAGTGTGGAGTCGTTGCCTGTTTTGCTTACTGATGTTTTNNNAGATCGGAAGA
127477 TATGAGTGTGGAGTCGTTGCGTACCAAGTAGGAATAGCTAAGNNNAGATCGGAAG
127478 TATGAGTGTGGAGTCGTTGCGTACCAAGTAGGAATAGCTAAGNNNAGATCGGAAG
127479 TATGAGTGTGGAGTCGTTGCTACCTAATGTTATTTGGTCTCTTNNNAGATCGGAAG
127480 TATGAGTGTGGAGTCGTTGCTCTCAACGATTATTATATTTCTGGNNNAGATCGGAA
127481 TATGAGTGTGGAGTCGTTGCTATGACTGTGCTTGAAGATATTANNNAGATCGGAAG
127482 TATGAGTGTGGAGTCGTTGCGGGACTTCAAGTTTTTATGAATANNNAGATCGGAAG
127484 TATGAGTGTGGAGTCGTTGCGAACTCTATACCTACCCTGAGNNNAGATCGGAAGAG
127485 TATGAGTGTGGAGTCGTTGCGCATACACGTAATATATATGAGGNNNAGATCGGAA
127487 TATGAGTGTGGAGTCGTTGCAAGAGAGAGAGAGAGTTCTGTNNNAGATCGGAAG
127488 TATGAGTGTGGAGTCGTTGCCTCATCAGAACATGTTTCAATAGNNNAGATCGGAAG
127489 TATGAGTGTGGAGTCGTTGCACATGTACATTCATACCAGAATANNNAGATCGGAAG
127490 TATGAGTGTGGAGTCGTTGCATGTGTTTATGTGAGTGTATCTTNNNAGATCGGAAG
127491 TATGAGTGTGGAGTCGTTGCTCTCTATTGACTCTTACTCCAANNNAGATCGGAAGAG
127492 TATGAGTGTGGAGTCGTTGCATCAAACCCTAAACAAAACAGANNNAGATCGGAAGA
127493 TATGAGTGTGGAGTCGTTGCATCAAACCCTAAACAAAACAGANNNAGATCGGAAGA
127494 TATGAGTGTGGAGTCGTTGCTTTAGTATGTGTGTGTCTATGTANNNAGATCGGAAG
127495 TATGAGTGTGGAGTCGTTGCAGAAGTCAAAAATACAGTATTGGNNNAGATCGGAA
127496 TATGAGTGTGGAGTCGTTGCTGAAAGGAAAAATGTTGACATTANNNAGATCGGAA
127497 TATGAGTGTGGAGTCGTTGCAAGAGAGAGAATGAATAACAAGTNNNAGATCGGAA
127498 TATGAGTGTGGAGTCGTTGCGAAAACCTGGCTTATTCTCTTNNNAGATCGGAAGAG
127499 TATGAGTGTGGAGTCGTTGCGTGAGATATGTCCAGATATTGATNNNAGATCGGAA
127500 TATGAGTGTGGAGTCGTTGCAAGTTAATTGTAAGTGTTCCTTCNNNAGATCGGAAG
127501 TATGAGTGTGGAGTCGTTGCAACTCCAACCAAGTTGATTCNNNAGATCGGAAGAGC
127502 TATGAGTGTGGAGTCGTTGCTCAACAAATATTTGCTCTCTTGNNNAGATCGGAAGA
127503 TATGAGTGTGGAGTCGTTGCTTTTCTTTTTCTCCTGGTGAAANNNAGATCGGAAGA
127504 TATGAGTGTGGAGTCGTTGCTAGACTAATATGGATGAACATGGNNNAGATCGGAA
127505 TATGAGTGTGGAGTCGTTGCGTGAAAATTTCAAACAATAGCAANNNAGATCGGAA
127506 TATGAGTGTGGAGTCGTTGCATTAAGCCTGCAAAGAATTAAAANNNAGATCGGAA
127507 TATGAGTGTGGAGTCGTTGCTTCAAATGCCATTAGTTACTATTNNNAGATCGGAAG
127508 TATGAGTGTGGAGTCGTTGCTAGAAAGATCTCTACAAGGAGAANNNAGATCGGAA
127509 TATGAGTGTGGAGTCGTTGCTTCCCTTTATGGAAACCTGTNNNAGATCGGAAGAGC
127510 TATGAGTGTGGAGTCGTTGCTTCTTTTTCTTCTACTTTTCCTCNNNAGATCGGAAGAG
127511 TATGAGTGTGGAGTCGTTGCTATACACTTCAGGATTCCACATNNNAGATCGGAAGA
127512 TATGAGTGTGGAGTCGTTGCAGTAAGGACTTCCAAAAACTTATNNNAGATCGGAAG
127513 TATGAGTGTGGAGTCGTTGCTTTACATTCACCCAGTATTTCTNNNAGATCGGAAGAG
127514 TATGAGTGTGGAGTCGTTGCCTGTCTGTCTGTCTTTGAATANNNAGATCGGAAGAG
127515 TATGAGTGTGGAGTCGTTGCCTCAGGTAATCATTAGCATTTTTNNNAGATCGGAAG
127516 TATGAGTGTGGAGTCGTTGCAAACATCCTCAGACCCATATCNNNAGATCGGAAGAG
127517 TATGAGTGTGGAGTCGTTGCATAAAAGAAGTGTAGCAGTGTATNNNAGATCGGAA
127518 TATGAGTGTGGAGTCGTTGCTATTTGCTTTATATATCTGGGTGNNNAGATCGGAAG
127519 TATGAGTGTGGAGTCGTTGCTAATATTTCCCCTTCCTCTGANNNAGATCGGAAGAGC
127520 TATGAGTGTGGAGTCGTTGCAACTACTAAATCCACTTCTCTACNNNAGATCGGAAGA

127521 TATGAGTGTGGAGTCGTTGCTTAGAATATACTGAACCTATGCANNNAGATCGGAAG
127522 TATGAGTGTGGAGTCGTTGCAGTACCTGTTAAACTTATTCTCANNNAGATCGGAAG
127523 TATGAGTGTGGAGTCGTTGCGTTAGATTCATTAGGTCTAAAGCNNNAGATCGGAAG
127526 TATGAGTGTGGAGTCGTTGCGATCAACATGCAGTATACAAATANNNAGATCGGAAG
127527 TATGAGTGTGGAGTCGTTGCTTTCTTTCATTGTACCCTAACTTNNNAGATCGGAAGA
127528 TATGAGTGTGGAGTCGTTGCATAGTATTGTCAAAAGCCATTTTNNNAGATCGGAAG
127529 TATGAGTGTGGAGTCGTTGCTATCAGTATAACAAAGGCAGAAANNNAGATCGGAA
127530 TATGAGTGTGGAGTCGTTGCCTACAGTATCTGGAACATATGTCNNNAGATCGGAAG
127531 TATGAGTGTGGAGTCGTTGCGTGTAGTAATTGCATATAACCTCNNNAGATCGGAAG
127532 TATGAGTGTGGAGTCGTTGCAACAATGAATTCTAAGTGTGTTGNNNAGATCGGAA
127534 TATGAGTGTGGAGTCGTTGCCTGAAGCCTCTAAGATATATCTTNNNAGATCGGAAG
127535 TATGAGTGTGGAGTCGTTGCCCTGAAAAGGCTATTATCCATNNNAGATCGGAAGAG
127537 TATGAGTGTGGAGTCGTTGCAGTCTGCTACCAAATTTATCCNNNAGATCGGAAGAG
127538 TATGAGTGTGGAGTCGTTGCCATCAAAGACTACCTAAAAACAGNNNAGATCGGAAG
127539 TATGAGTGTGGAGTCGTTGCTGTAAAATAAGATGTGTTGACTTNNNAGATCGGAA
127540 TATGAGTGTGGAGTCGTTGCTGTAAGGAGACCCATACTTTNNNAGATCGGAAGAG
127541 TATGAGTGTGGAGTCGTTGCTACAGACAAGAGAAATGAAATACNNNAGATCGGAA
127542 TATGAGTGTGGAGTCGTTGCGTAGTAGAGATGGGGTTTCANNNAGATCGGAAGA
127543 TATGAGTGTGGAGTCGTTGCAACACCTTAAACATAACTCATTCNNNAGATCGGAAGA
127544 TATGAGTGTGGAGTCGTTGCATGATGAAAGGTCAATTATCAGANNNAGATCGGAA
127545 TATGAGTGTGGAGTCGTTGCTTTGGGATGAAGGGACAAATNNNAGATCGGAAGA
127546 TATGAGTGTGGAGTCGTTGCCATATGTTCACAAAAGCTCAANNNAGATCGGAAGAG
127547 TATGAGTGTGGAGTCGTTGCAAAAATTCTATGAAAAGCACTGANNNAGATCGGAA
127548 TATGAGTGTGGAGTCGTTGCTTACATAATAGTGGCATTTTAGCNNNAGATCGGAAG
127549 TATGAGTGTGGAGTCGTTGCTTACATAATAGTGGCATTTTAGCNNNAGATCGGAAG
127550 TATGAGTGTGGAGTCGTTGCGATGTTGTGTCTCTTTTTACAGNNNAGATCGGAAGA
127552 TATGAGTGTGGAGTCGTTGCAGAGAGATTATAAAGTAACAGCANNNAGATCGGAA
127553 TATGAGTGTGGAGTCGTTGCCCCTTTATTTCTCTTCTTTTCAGNNNAGATCGGAAGA
127554 TATGAGTGTGGAGTCGTTGCGAGAAGAGCCATTATACACAATANNNAGATCGGAA
127555 TATGAGTGTGGAGTCGTTGCCCTTTTTATAAGGAGACAGTCTANNNAGATCGGAAG
127556 TATGAGTGTGGAGTCGTTGCGTACACCACAATTCAGATGANNNAGATCGGAAGAG
127557 TATGAGTGTGGAGTCGTTGCCTCAATATTCCCTGCTTCTAATNNNAGATCGGAAGAG
127558 TATGAGTGTGGAGTCGTTGCCTGCCTGGTTAATTTTAATGTTTNNNAGATCGGAAG
127559 TATGAGTGTGGAGTCGTTGCAGATAGATTTACTGTTTTTGTGTNNNAGATCGGAAG
127560 TATGAGTGTGGAGTCGTTGCCCGTTGGATATAATCTGAATTTANNNAGATCGGAAG
127561 TATGAGTGTGGAGTCGTTGCAGAAGGAAGCATTGCTAATATAGNNNAGATCGGAA
127562 TATGAGTGTGGAGTCGTTGCTTTTACTCCCCAGAATAATCTTCNNNAGATCGGAAGA
127563 TATGAGTGTGGAGTCGTTGCTATTCCATAGAATATGTGCAGGNNNAGATCGGAAG
127564 TATGAGTGTGGAGTCGTTGCATTTTTAGTTATAGTCAACCTGCNNNAGATCGGAAG
127565 TATGAGTGTGGAGTCGTTGCAAATTTTCATGTTGTTCTCCATANNNAGATCGGAAGA
127566 TATGAGTGTGGAGTCGTTGCATTGCATCACATTATATATCACTNNNAGATCGGAAGA
127567 TATGAGTGTGGAGTCGTTGCATTGACCATGTTCTGTGAAGNNNAGATCGGAAGAG
127568 TATGAGTGTGGAGTCGTTGCTGTATTATTGTTTTCATCATCATTNNNAGATCGGAAG

127569 TATGAGTGTGGAGTCGTTGCGAAACCCATATATAAGTCAGCTANNNAGATCGGAAG
127570 TATGAGTGTGGAGTCGTTGCTATATATTACACACACAGACACANNNAGATCGGAAG
127571 TATGAGTGTGGAGTCGTTGCATTTTCCCAAGGATTACAACATNNNAGATCGGAAGA
127572 TATGAGTGTGGAGTCGTTGCGTTTCAATATGTAGCCTTATAGCNNNAGATCGGAAG
127573 TATGAGTGTGGAGTCGTTGCCTTACTCTCTAAGGTTGTTTACTNNNAGATCGGAAGA
127574 TATGAGTGTGGAGTCGTTGCCAGGGCAAATGAAAATAGTTATANNNAGATCGGAA
127575 TATGAGTGTGGAGTCGTTGCTTGGTATCTATACAAACTCTTGTNNNAGATCGGAAG
127576 TATGAGTGTGGAGTCGTTGCGATGTGCATAACTTTAAACTCAANNNAGATCGGAAG
127577 TATGAGTGTGGAGTCGTTGCTGGAACATAAACTGAAGTCAATANNNAGATCGGAA
127578 TATGAGTGTGGAGTCGTTGCCTGATTACATTTTACATTGCTGANNNAGATCGGAAG
127579 TATGAGTGTGGAGTCGTTGCATTTAAAGTGGGTAGTGAATTTGNNNAGATCGGAA
127580 TATGAGTGTGGAGTCGTTGCCACTTAGAATCTGAAGCTCAATNNNAGATCGGAAGA
127581 TATGAGTGTGGAGTCGTTGCATTTATTTTCTTTCTCAGCCATGNNNAGATCGGAAGA
127582 TATGAGTGTGGAGTCGTTGCTACTGAAAACTGGCTTAGATTTTNNNAGATCGGAAG
127583 TATGAGTGTGGAGTCGTTGCAGTGTAAAGTGATATCTCATTGTNNNAGATCGGAAG
127584 TATGAGTGTGGAGTCGTTGCCAGTTTTCTTCTTTGATCTTCTTNNNAGATCGGAAGA
127585 TATGAGTGTGGAGTCGTTGCGTCTTCAGTAAAGTACATCCATANNNAGATCGGAAG
127587 TATGAGTGTGGAGTCGTTGCTACATGTGTGTATATATGTGTGNNNAGATCGGAAGA
127589 TATGAGTGTGGAGTCGTTGCCCAAAGTGGTGCCTAATAATNNNAGATCGGAAGAG
127590 TATGAGTGTGGAGTCGTTGCCCAAAGTGGTGCCTAATAATNNNAGATCGGAAGAG
127591 TATGAGTGTGGAGTCGTTGCATGAAAGACAGGGTTTGTAATNNNAGATCGGAAGA
127592 TATGAGTGTGGAGTCGTTGCATGCTAGATTAAATTACTGGTCTNNNAGATCGGAAG
127593 TATGAGTGTGGAGTCGTTGCGAGTACAATGGCACAATCTTNNNAGATCGGAAGAG
127594 TATGAGTGTGGAGTCGTTGCTTTATTTTTTGAGAAACAGGTCTNNNAGATCGGAAG
127595 TATGAGTGTGGAGTCGTTGCTCCTTCACTAGAACACACTGNNNAGATCGGAAGAGC
127596 TATGAGTGTGGAGTCGTTGCGTGTTTCTATGTGTGTATGTATCNNNAGATCGGAAG
127597 TATGAGTGTGGAGTCGTTGCCTTGTATTTTTGTATTTGTCGTGNNNAGATCGGAAG
127598 TATGAGTGTGGAGTCGTTGCCTTGTATTTTTGTATTTGTCGTGNNNAGATCGGAAG
127599 TATGAGTGTGGAGTCGTTGCTACATTCTATTACTCTGTGTGCNNNAGATCGGAAGA
127600 TATGAGTGTGGAGTCGTTGCGGTACAAACTTTCTGAAAGATAGNNNAGATCGGAA
127601 TATGAGTGTGGAGTCGTTGCCTGAAAAATGCTTGCTATCATTNNNAGATCGGAAGA
127602 TATGAGTGTGGAGTCGTTGCGAAAGCAGTCAAGTACAATAAGNNNAGATCGGAAG
127603 TATGAGTGTGGAGTCGTTGCCCCTACCTAGAATGTTTACATTANNNAGATCGGAAG
127604 TATGAGTGTGGAGTCGTTGCGTGTGTTTTGTCACTTTGTTNNNAGATCGGAAGAGC
127605 TATGAGTGTGGAGTCGTTGCTAACTGCAATCTCTTACTTCTGNNNAGATCGGAAGA
127606 TATGAGTGTGGAGTCGTTGCTTCAGTCTTTCCCAGTCTTTNNNAGATCGGAAGAGCA
127607 TATGAGTGTGGAGTCGTTGCCTAAAAGTCCAGACTAGAAATGTNNNAGATCGGAA
127608 TATGAGTGTGGAGTCGTTGCTAAATACACCAGATGGAAAGACNNNAGATCGGAAG
127609 TATGAGTGTGGAGTCGTTGCTTATATGTCAAGGTTTGTTCAAGNNNAGATCGGAAG
127610 TATGAGTGTGGAGTCGTTGCGTGTTTATCCAGTTTCATTCATANNNAGATCGGAAG
127611 TATGAGTGTGGAGTCGTTGCAAATGCAAAATTAAAACAACTGGNNNAGATCGGAA
127612 TATGAGTGTGGAGTCGTTGCGACTGTCACTAACCTCTCTTNNNAGATCGGAAGAGC
127613 TATGAGTGTGGAGTCGTTGCACTTTGTCAAAAATAATCAGGGNNNAGATCGGAAG

127614 TATGAGTGTGGAGTCGTTGCTGGGACTTTGTCAAAAATAATCNNNAGATCGGAAGA
127616 TATGAGTGTGGAGTCGTTGCAGCTAAAAAATGTATAATCGTCTNNNAGATCGGAAG
127617 TATGAGTGTGGAGTCGTTGCGGAACTCTACTAAGTAATCGATANNNAGATCGGAAG
127618 TATGAGTGTGGAGTCGTTGCCTAGTACAGTAAATTAATTGCCCNNNAGATCGGAAGA
127619 TATGAGTGTGGAGTCGTTGCTTCCTTATCTACAACATCTGTTTNNNAGATCGGAAGA
127620 TATGAGTGTGGAGTCGTTGCCCTTTCTTATCCCCCTACTTNNNAGATCGGAAGAGCA
127621 TATGAGTGTGGAGTCGTTGCCAAGGCAATTTATCCCTAAATATNNNAGATCGGAAGA
127622 TATGAGTGTGGAGTCGTTGCCATCATCAAAAGAGTAAAAGAGTNNNAGATCGGAA
127625 TATGAGTGTGGAGTCGTTGCATGATTTCACGTCTCACATCNNNAGATCGGAAGAGCA
127626 TATGAGTGTGGAGTCGTTGCAATACAAATAGAAGTCAAGGCTANNNAGATCGGAA
127627 TATGAGTGTGGAGTCGTTGCTATACCTAACCAAGAACTTGAAANNNAGATCGGAAG
127628 TATGAGTGTGGAGTCGTTGCTACATTTCTTTGCTTCTCCAGNNNAGATCGGAAGAG
127629 TATGAGTGTGGAGTCGTTGCTATTGGCTCATGTAATTATGGAGNNNAGATCGGAA
127630 TATGAGTGTGGAGTCGTTGCTAACACCCAGCATCCTATAANNNAGATCGGAAGAGCA
127631 TATGAGTGTGGAGTCGTTGCTTTCTTGTACACACATACCTGNNNAGATCGGAAGAG
127632 TATGAGTGTGGAGTCGTTGCCTTCCTATTTGCTATTCCCTCNNNAGATCGGAAGAGCA
127633 TATGAGTGTGGAGTCGTTGCCATTTGAGTTCAGAAGTATGAGNNNAGATCGGAAGA
127634 TATGAGTGTGGAGTCGTTGCTAGTGGCATTCATTGTTTCANNNAGATCGGAAGAG
127635 TATGAGTGTGGAGTCGTTGCCTATGGAAAGAGAGAGGGAGNNNAGATCGGAAGA
127636 TATGAGTGTGGAGTCGTTGCTTATATACTTACATGCTCACACANNNAGATCGGAAGA
127637 TATGAGTGTGGAGTCGTTGCCTCTTTTCTCAGTTGTAGTTCTNNNAGATCGGAAGAG
127638 TATGAGTGTGGAGTCGTTGCTATGTACACATCTTATTGGTTCTNNNAGATCGGAAGA
127639 TATGAGTGTGGAGTCGTTGCATAAAGGATAGGAAGTCATTTCANNNAGATCGGAA
127640 TATGAGTGTGGAGTCGTTGCAGTTCTTTTGATATACAGTACTCNNNAGATCGGAAGA
127641 TATGAGTGTGGAGTCGTTGCGTAACAACATTTTAGTCAACAATNNNAGATCGGAAGA
127642 TATGAGTGTGGAGTCGTTGCGTAACAACATTTTAGTCAACAATNNNAGATCGGAAGA
127643 TATGAGTGTGGAGTCGTTGCAAAGACAGAGAGAGAGGAAGNNNAGATCGGAAGA
127644 TATGAGTGTGGAGTCGTTGCAAAATAAGTCTTGGACTGAAGTANNNAGATCGGAA
127645 TATGAGTGTGGAGTCGTTGCTAATTGTGAGAAAGAAGTAATGGNNNAGATCGGAA
127646 TATGAGTGTGGAGTCGTTGCGTCAAGGTAGAATAAATATCTGGNNNAGATCGGAA
127647 TATGAGTGTGGAGTCGTTGCTTCATTTTGCAATCGACATGNNNAGATCGGAAGAGCA
127648 TATGAGTGTGGAGTCGTTGCCCTACAAACTATCAACATTAACCNNNAGATCGGAAGA
127649 TATGAGTGTGGAGTCGTTGCAATTAAATAAATAGGCCCGGATANNNAGATCGGAA
127650 TATGAGTGTGGAGTCGTTGCTTTTTCCTATCTTCATTTGCTAGNNNAGATCGGAAGA
127651 TATGAGTGTGGAGTCGTTGCGCATTCTCTGTGGATTTAATTATNNNAGATCGGAAGA
127652 TATGAGTGTGGAGTCGTTGCATGTGTGTGTGTATTTAGATAGANNNAGATCGGAA
127653 TATGAGTGTGGAGTCGTTGCTCTCATAGTTCAGAGATTCAAATNNNAGATCGGAAGA
127654 TATGAGTGTGGAGTCGTTGCAAAATAGGAAGACATCAATGCAANNNAGATCGGAA
127655 TATGAGTGTGGAGTCGTTGCAAGCATTTGATAAAATTCGGTATNNNAGATCGGAAG
127656 TATGAGTGTGGAGTCGTTGCACTCTCATAGGACTGACTTATATNNNAGATCGGAAGA
127657 TATGAGTGTGGAGTCGTTGCGAGTGAAATTATTCATGGAGAATNNNAGATCGGAA
127658 TATGAGTGTGGAGTCGTTGCCCTATCTTCCAGAATATTGCTAANNNAGATCGGAAGA
127659 TATGAGTGTGGAGTCGTTGCAGAGATGAGAGAGAGAAAGAGNNNAGATCGGAAG

127660 TATGAGTGTGGAGTCGTTGCGAAATCAACCAGCAGAGATGNNNAGATCGGAAGAG
127661 TATGAGTGTGGAGTCGTTGCATGTTTTGATGCATCTGATTTACNNNAGATCGGAAG
127662 TATGAGTGTGGAGTCGTTGCGTTCCCTAAACATTAATAGAAGANNNAGATCGGAAG
127663 TATGAGTGTGGAGTCGTTGCTGACTTTCATTTAAACTTGACTTNNNAGATCGGAAGA
127664 TATGAGTGTGGAGTCGTTGCAGATAGTCAAATAAATGGGGTAANNNAGATCGGAA
127665 TATGAGTGTGGAGTCGTTGCGCTGGAAATGTTGTAAATTACAANNNAGATCGGAA
127666 TATGAGTGTGGAGTCGTTGCAAGGCAGTTTTAATTTTATGACANNNAGATCGGAAG
127668 TATGAGTGTGGAGTCGTTGCTTCAACTTTTGTTTTAGATTTGGNNNAGATCGGAAG
127669 TATGAGTGTGGAGTCGTTGCGGTGTATATATTTGTAGGGTACANNNAGATCGGAA
127670 TATGAGTGTGGAGTCGTTGCGGAATCACTAATTTTCCCATTCNNNAGATCGGAAGA
127671 TATGAGTGTGGAGTCGTTGCGGAATCACTAATTTTCCCATTCNNNAGATCGGAAGA
127672 TATGAGTGTGGAGTCGTTGCAAATAGTTTTTGAGCAATTACCANNNAGATCGGAAG
127673 TATGAGTGTGGAGTCGTTGCAAAGAAAGAGGTTGTTTATCCAGNNNAGATCGGAA
127674 TATGAGTGTGGAGTCGTTGCCTCTACTTGTTCTTCCAGTTTTNNNAGATCGGAAGAG
127675 TATGAGTGTGGAGTCGTTGCATAAGCAGATTCCAAGATTTTCTNNNAGATCGGAAG
127676 TATGAGTGTGGAGTCGTTGCGTTTGTGTTTTACTCCTACAAAGNNNAGATCGGAAG
127677 TATGAGTGTGGAGTCGTTGCACATATGTTTTCATCCACAGTTNNNAGATCGGAAGA
127678 TATGAGTGTGGAGTCGTTGCTTCTGGGTTTTTTCATTGTAATTNNNAGATCGGAAG
127680 TATGAGTGTGGAGTCGTTGCTATGGCTTTTCTTCCTGATTTTNNNAGATCGGAAGAG
127681 TATGAGTGTGGAGTCGTTGCAATATCGACCAAAAAAATCCATTNNNAGATCGGAAG
127682 TATGAGTGTGGAGTCGTTGCTTATAGGATGAGTTAGGGAGAATNNNAGATCGGAA
127683 TATGAGTGTGGAGTCGTTGCACATGTCTGGATTATTCTTTAATNNNAGATCGGAAG
127684 TATGAGTGTGGAGTCGTTGCCATTTAGTGGTCAAAGTTACAGNNNAGATCGGAAGA
127685 TATGAGTGTGGAGTCGTTGCGTATTGTCACTGTTTTTTGTTTTNNNAGATCGGAAGA
127686 TATGAGTGTGGAGTCGTTGCATAACAAATAAGCCTTCAATCAGNNNAGATCGGAAG
127687 TATGAGTGTGGAGTCGTTGCATAGATCATATACATTTGAGGGGNNNAGATCGGAA
127688 TATGAGTGTGGAGTCGTTGCTGCACAACTCTAAAAATTCTAAANNNAGATCGGAAG
127689 TATGAGTGTGGAGTCGTTGCGACTGAGACAGAAACAAAGANNNAGATCGGAAGAG
127690 TATGAGTGTGGAGTCGTTGCCTCTTTCCTCATAATTTGAAACTNNNAGATCGGAAGA
127691 TATGAGTGTGGAGTCGTTGCCAAGAACTTCTAACTCTATTCCTNNNAGATCGGAAGA
127692 TATGAGTGTGGAGTCGTTGCCCTTTCTTATCCCCCTACTTNNNAGATCGGAAGAGCA
127693 TATGAGTGTGGAGTCGTTGCCAGTCTGTTTTTGTCTCTACTNNNAGATCGGAAGAG
127694 TATGAGTGTGGAGTCGTTGCGATGGTTATAAAGAAGCGTCANNNAGATCGGAAGA
127695 TATGAGTGTGGAGTCGTTGCCTGCTCTCTAACTCAGTTTCNNNAGATCGGAAGAGCA
127697 TATGAGTGTGGAGTCGTTGCTGAGATATCCAGTTAGTACTTTCNNNAGATCGGAAG
127698 TATGAGTGTGGAGTCGTTGCCACAATTCCCTTAAATGACAATTNNNAGATCGGAAGA
127699 TATGAGTGTGGAGTCGTTGCCAAGGCCAAGAGTCTAGAATNNNAGATCGGAAGAG
127700 TATGAGTGTGGAGTCGTTGCAAAACTTGAATACTTGTGACATGNNNAGATCGGAAG
127701 TATGAGTGTGGAGTCGTTGCAAAACTTGAATACTTGTGACATGNNNAGATCGGAAG
127702 TATGAGTGTGGAGTCGTTGCTGGCAGAAAAATTCTTTAAAAACNNNAGATCGGAAG
127703 TATGAGTGTGGAGTCGTTGCCAGGTTCAAGTGATTCTCCTNNNAGATCGGAAGAGC
127705 TATGAGTGTGGAGTCGTTGCATGTTCAAAATCTTCTCATGTTTNNNAGATCGGAAGA
127706 TATGAGTGTGGAGTCGTTGCAAGATAAAATGAGGGGGAGGNNNAGATCGGAAGA

127707 TATGAGTGTGGAGTCGTTGCGTAATGATGTTGGAGATCATTTTNNNAGATCGGAAG
127708 TATGAGTGTGGAGTCGTTGCGCAGAATGTAAAGATATGGTTGNNNAGATCGGAAG
127709 TATGAGTGTGGAGTCGTTGCTCAGTTTCTTCATACATGAAATGNNNAGATCGGAAG
127710 TATGAGTGTGGAGTCGTTGCTTTGTTTCATTTTCTTGGTATGTNNNAGATCGGAAGA
127711 TATGAGTGTGGAGTCGTTGCGATGGCACTAAAGAGATACTATANNNAGATCGGAA
127712 TATGAGTGTGGAGTCGTTGCCAAGAGGTATACATAGAAGACTTNNNAGATCGGAA
127713 TATGAGTGTGGAGTCGTTGCGGAAAAAGACAAACATCAAAATTNNNAGATCGGAA
127714 TATGAGTGTGGAGTCGTTGCCCTAGTGAGATTTGAGCATCNNNAGATCGGAAGAG
127716 TATGAGTGTGGAGTCGTTGCGGGAGACATTTGGATATAAAATGNNNAGATCGGAA
127717 TATGAGTGTGGAGTCGTTGCAGTTTTTGTTATGATTTGATGGTNNNAGATCGGAAG
127718 TATGAGTGTGGAGTCGTTGCTCACATATGCATACATCTCTGNNNAGATCGGAAGAG
127719 TATGAGTGTGGAGTCGTTGCCTACTAATTCTGTTTCCCTAGAANNNAGATCGGAAGA
127720 TATGAGTGTGGAGTCGTTGCTTCTTTTACGGTAAAATGAGATCNNNAGATCGGAAG
127721 TATGAGTGTGGAGTCGTTGCTATTCATTCTAGGTATCCAATGTNNNAGATCGGAAG
127722 TATGAGTGTGGAGTCGTTGCTGTTTACTTGTTTTAGAGTGTTGNNNAGATCGGAAG
127723 TATGAGTGTGGAGTCGTTGCAATCTTCAACTTTGCCAAATAAANNNAGATCGGAAG
127724 TATGAGTGTGGAGTCGTTGCGTCCTCAAGATTTTCGGAACNNNAGATCGGAAGAGC
127725 TATGAGTGTGGAGTCGTTGCCACTTTCCTAAATTCTGGTTAAANNNAGATCGGAAGA
127726 TATGAGTGTGGAGTCGTTGCTAAACATTTTAGTACACCAAGTGNNNAGATCGGAAG
127727 TATGAGTGTGGAGTCGTTGCATTTCTTTTCTCCCTCACATCNNNAGATCGGAAGAGC
127728 TATGAGTGTGGAGTCGTTGCAAAAATTTTCCTCAGTCTTCTTCNNNAGATCGGAAGA
127729 TATGAGTGTGGAGTCGTTGCCTTATGACAGTTTGCTTAGAATGNNNAGATCGGAAG
127730 TATGAGTGTGGAGTCGTTGCCAATTGCCTGATGTTGAAATTNNNAGATCGGAAGAG
127731 TATGAGTGTGGAGTCGTTGCATTGCTAGTCTGCTACAGTCNNNAGATCGGAAGAGC
127732 TATGAGTGTGGAGTCGTTGCCCAAATGTTCCTTTACTCTTTATNNNAGATCGGAAGA
127733 TATGAGTGTGGAGTCGTTGCCACCTGTGTAAGGGATATTATAANNNAGATCGGAAG
127734 TATGAGTGTGGAGTCGTTGCATTCTCTTTCTGTGCCACATNNNAGATCGGAAGAGCA
127735 TATGAGTGTGGAGTCGTTGCATTTACTGTGCATTCTCCAGNNNAGATCGGAAGAGC
127736 TATGAGTGTGGAGTCGTTGCAGAATTACTATCTCCCCAAATTTNNNAGATCGGAAGA
127737 TATGAGTGTGGAGTCGTTGCTTTTTCTGATTATCCAAGAACTGNNNAGATCGGAAG
127738 TATGAGTGTGGAGTCGTTGCCAACACTATACAAACATCTTCAGNNNAGATCGGAAG
127739 TATGAGTGTGGAGTCGTTGCCCTTGGATAAAGCATATCTTTAGNNNAGATCGGAAG
127740 TATGAGTGTGGAGTCGTTGCCCTTGGATAAAGCATATCTTTAGNNNAGATCGGAAG
127741 TATGAGTGTGGAGTCGTTGCTTTATCATTTTGCATTCTGATGANNNAGATCGGAAGA
127742 TATGAGTGTGGAGTCGTTGCAGTAATAAAAGTGATACCATATGTNNNAGATCGGAA
127743 TATGAGTGTGGAGTCGTTGCTATTGTGAATTTCCTGAATTAGCNNNAGATCGGAAG
127744 TATGAGTGTGGAGTCGTTGCAATAAAATCATGTATTTTGCAGCNNNAGATCGGAAG
127745 TATGAGTGTGGAGTCGTTGCAATAATAGTTGGAGCATCAGTTNNNAGATCGGAAG
127746 TATGAGTGTGGAGTCGTTGCAAACTATAGCAGAATGTCACTAGNNNAGATCGGAAG
127747 TATGAGTGTGGAGTCGTTGCATGGGAATATATTGGCATTTCTNNNAGATCGGAAGA
127748 TATGAGTGTGGAGTCGTTGCCTGATAGTCCTTGCCTCTAANNNAGATCGGAAGAGC
127749 TATGAGTGTGGAGTCGTTGCGTTGGTTTTTGTTTTTCCTTTTTNNNAGATCGGAAGA
127750 TATGAGTGTGGAGTCGTTGCGAGACTAAGAACCTAAAACCTCNNNAGATCGGAAGA

127751 TATGAGTGTGGAGTCGTTGCACATTTAGTAATCTGTTCTAAGATNNNAGATCGGAA
127752 TATGAGTGTGGAGTCGTTGCGACTGTACTATGTCATTATCATTNNNAGATCGGAAG
127753 TATGAGTGTGGAGTCGTTGCTTAAGGTGGCAAATGTTGTCNNNAGATCGGAAGAG
127754 TATGAGTGTGGAGTCGTTGCTCTTGGACTTTGGAAACAATTNNNAGATCGGAAGAG
127755 TATGAGTGTGGAGTCGTTGCACATGCAGTATTTATTTTTCCATNNNAGATCGGAAGA
127756 TATGAGTGTGGAGTCGTTGCCTCTATTAGTCTTAGAGACAGGTNNNAGATCGGAAG
127757 TATGAGTGTGGAGTCGTTGCACCACTAATTGACTTTCTATCTTNNNAGATCGGAAGA
127758 TATGAGTGTGGAGTCGTTGCAAACTGTGACTATTAGCTGTCNNNAGATCGGAAGAG
127761 TATGAGTGTGGAGTCGTTGCGGTTTCCGTCTCTATATAATGTANNNAGATCGGAAG
127762 TATGAGTGTGGAGTCGTTGCTTATTTTTTGGCTTCTGTGTTTANNNAGATCGGAAGA
127763 TATGAGTGTGGAGTCGTTGCTTTATAGACTCAATCCTTCTCTGNNNAGATCGGAAGA
127764 TATGAGTGTGGAGTCGTTGCAATTCTAAACCTTGTTTCTACCANNNAGATCGGAAGA
127765 TATGAGTGTGGAGTCGTTGCGTTTGAAGAACAATGCAAGTANNNAGATCGGAAGA
127766 TATGAGTGTGGAGTCGTTGCCTATCTCAAGAAGAACTCTCCTNNNAGATCGGAAGA
127767 TATGAGTGTGGAGTCGTTGCCTTTGAATTATTTTCTGGGTCTCNNNAGATCGGAAGA
127768 TATGAGTGTGGAGTCGTTGCGTTACAAAAGACTGCAGAGGNNNAGATCGGAAGAG
127769 TATGAGTGTGGAGTCGTTGCTGAATGAGTTTTCTTTACATTCANNNAGATCGGAAG
127770 TATGAGTGTGGAGTCGTTGCTAAAAGAAATGGTTTCACAAAGANNNAGATCGGAA
127771 TATGAGTGTGGAGTCGTTGCAACTGCTACATTTTCTAGAGTAANNNAGATCGGAAG
127772 TATGAGTGTGGAGTCGTTGCTCTCCTGTTATTATTTCAATAAGTCNNNAGATCGGAA
127773 TATGAGTGTGGAGTCGTTGCCAGATAAGGGGGGATTACTATANNNAGATCGGAAG
127774 TATGAGTGTGGAGTCGTTGCCGTATACATACATAGTTAGCATCNNNAGATCGGAAG
127775 TATGAGTGTGGAGTCGTTGCTCTTACTTGCAAGTATGTTTTGNNNAGATCGGAAGA
127776 TATGAGTGTGGAGTCGTTGCTATACATATGGCATGTTAGAAGTNNNAGATCGGAAG
127777 TATGAGTGTGGAGTCGTTGCAAGGATCATGTGAATGGGATNNNAGATCGGAAGAG
127778 TATGAGTGTGGAGTCGTTGCTGCCTAATACCATAAAGTAAAGANNNAGATCGGAAG
127779 TATGAGTGTGGAGTCGTTGCAGTTGTTAGAAGCAAAGCATATNNNAGATCGGAAG
127780 TATGAGTGTGGAGTCGTTGCATAGTGAATAAGTCTCACAAGATNNNAGATCGGAAG
127781 TATGAGTGTGGAGTCGTTGCTATGTTAATTAGCTCAATTGTGGNNNAGATCGGAAG
127782 TATGAGTGTGGAGTCGTTGCTTTTATTTAAATGCCTGTCATCCNNNAGATCGGAAGA
127783 TATGAGTGTGGAGTCGTTGCTAGTATACTTCCTGGGAAACTATNNNAGATCGGAAG
127784 TATGAGTGTGGAGTCGTTGCGTTTCACCAAATATCTGGGCNNNAGATCGGAAGAGC
127785 TATGAGTGTGGAGTCGTTGCCAATTTATCAAGACAGGGAAATTNNNAGATCGGAAG
127786 TATGAGTGTGGAGTCGTTGCATACACACACACTCACATACTNNNAGATCGGAAGAG
127788 TATGAGTGTGGAGTCGTTGCGAGTTTTGATGGGTACAGAANNNAGATCGGAAGAG
127789 TATGAGTGTGGAGTCGTTGCGCATACACATGTATATACACAGANNNAGATCGGAAG
127790 TATGAGTGTGGAGTCGTTGCTTGTGGCTACATATGAATTTTAGNNNAGATCGGAAG
127791 TATGAGTGTGGAGTCGTTGCTCTCAATATGACTTACTTTTGGANNNAGATCGGAAG
127792 TATGAGTGTGGAGTCGTTGCTGTGTAGATTTACATTGTGTAGANNNAGATCGGAAG
127793 TATGAGTGTGGAGTCGTTGCTTATGTTGTAGTCAAGGACAGNNNAGATCGGAAGA
127794 TATGAGTGTGGAGTCGTTGCTCCAATTTTCTAGCACTATAAGANNNAGATCGGAAG
127795 TATGAGTGTGGAGTCGTTGCCTTAATCACCTTGATATGGTTTAGNNNAGATCGGAA
127796 TATGAGTGTGGAGTCGTTGCCTTGTTCTTAGAGCTCCCAANNNAGATCGGAAGAGC

127797 TATGAGTGTGGAGTCGTTGCTTGCCATCATTGATTCATAGAANNNAGATCGGAAGA

127798 TATGAGTGTGGAGTCGTTGCGAGTTTCCTTGGGATTTGAGNNNAGATCGGAAGAG

127799 TATGAGTGTGGAGTCGTTGCAACTTTCTCATAGTACTCAATGTNNNAGATCGGAAGA

127800 TATGAGTGTGGAGTCGTTGCAACCAGTATATTGATCCAGATTTNNNAGATCGGAAG

127801 TATGAGTGTGGAGTCGTTGCCAATGCACTTTACAGATATCACNNNAGATCGGAAGA

127802 TATGAGTGTGGAGTCGTTGCGGCAATATCTATCCAATTGTAAANNNAGATCGGAAG

127803 TATGAGTGTGGAGTCGTTGCGTGAGACAGAGAGTGATAAAAGNNNAGATCGGAA

127804 TATGAGTGTGGAGTCGTTGCGACATTTGGATTGTGTATTTTTCNNNAGATCGGAAG

127805 TATGAGTGTGGAGTCGTTGCCATATTAGGTACAATGTACACTGNNNAGATCGGAAG

127806 TATGAGTGTGGAGTCGTTGCCTTAATTGGTCTGTTAGGTACTNNNAGATCGGAAGA

127809 TATGAGTGTGGAGTCGTTGCCTGGAATTAGGGGAGAAAACNNNAGATCGGAAGA

127810 TATGAGTGTGGAGTCGTTGCGAAGAGATCAGCATTTGAATTGNNNAGATCGGAAG

127811 TATGAGTGTGGAGTCGTTGCGATCTCACTCTATCTCCCAGNNNAGATCGGAAGAGC

127812 TATGAGTGTGGAGTCGTTGCAACAGGAATGTAAAGATTATTCANNNAGATCGGAA

127813 TATGAGTGTGGAGTCGTTGCTAGTGCCTAAAGAAATTCTTCTANNNAGATCGGAAG

127815 TATGAGTGTGGAGTCGTTGCAAGCAATTTCTATTTCAGTTTCANNNAGATCGGAAGA

127817 TATGAGTGTGGAGTCGTTGCGTTGTACTAAAGAGAACATTTGTNNNAGATCGGAA

127818 TATGAGTGTGGAGTCGTTGCGGCTCCTCTGTTTATTCTTTNNNAGATCGGAAGAGC

127819 TATGAGTGTGGAGTCGTTGCTTTATGCACCCACCATTTATTNNNAGATCGGAAGAG

127821 TATGAGTGTGGAGTCGTTGCCTAATACTATACAACAAACAGAGGNNNAGATCGGAA

127822 TATGAGTGTGGAGTCGTTGCTCTCTTCCTCTTTCTTGATAAATNNNAGATCGGAAGA

127825 TATGAGTGTGGAGTCGTTGCCGAGAGGAATTTTCAGAGAAANNNAGATCGGAAGA

127826 TATGAGTGTGGAGTCGTTGCTTCCTTCCTTCTTTTCTTTTTTTNNNAGATCGGAAGAG

127827 TATGAGTGTGGAGTCGTTGCGTACAAGGTAACCTTTGTTAAGNNNAGATCGGAAGA

127828 TATGAGTGTGGAGTCGTTGCCAATCAAGAATGCATTATCTGGNNNAGATCGGAAGA

127829 TATGAGTGTGGAGTCGTTGCTAAAAACAATCGATACACACATTNNNAGATCGGAAG

127830 TATGAGTGTGGAGTCGTTGCTAGCCAGGAAATATATATTGGTGNNNAGATCGGAA

127831 TATGAGTGTGGAGTCGTTGCAACTCTGGGTCGTGAAATAGNNNAGATCGGAAGAG

127832 TATGAGTGTGGAGTCGTTGCAATTTATTTTTGAGATGGAGTCCNNNAGATCGGAAG

127833 TATGAGTGTGGAGTCGTTGCATAAATTACATTCCCATCAACAANNNAGATCGGAAGA

127834 TATGAGTGTGGAGTCGTTGCTATGCACTTTTCTATAGATGTGTNNNAGATCGGAAG

127835 TATGAGTGTGGAGTCGTTGCTGGTCAATTATAGGGCATATAAANNNAGATCGGAA

127836 TATGAGTGTGGAGTCGTTGCTTGCCAAATATTCTTTTACTGAANNNAGATCGGAAGA

127837 TATGAGTGTGGAGTCGTTGCCCTTAATAGATAACCACATTCATNNNAGATCGGAAGA

127838 TATGAGTGTGGAGTCGTTGCTAAAGCAAACATTGACAGAATAANNNAGATCGGAA

127839 TATGAGTGTGGAGTCGTTGCTTCACTGTAATCTTAAACTCGTNNNAGATCGGAAGA

127840 TATGAGTGTGGAGTCGTTGCTTTTCAGGAACCATTGAAATTTANNNAGATCGGAAG

127841 TATGAGTGTGGAGTCGTTGCCTTTTCAATGTGGTGTCTAGTNNNAGATCGGAAGAG

127842 TATGAGTGTGGAGTCGTTGCGCTATTAAGAGCATGCTACTCNNNAGATCGGAAGAG

127843 TATGAGTGTGGAGTCGTTGCCCATCAGAGGAATAAAAGAAAATNNNAGATCGGAA

127844 TATGAGTGTGGAGTCGTTGCGAGTGAAATGGTGTGATCTTNNNAGATCGGAAGAG

127845 TATGAGTGTGGAGTCGTTGCCCTGTCTCTACAAAATACAAAAANNNAGATCGGAAG

127846 TATGAGTGTGGAGTCGTTGCTTATGAATAGAAGAATGGCATGANNNAGATCGGAA

127847 TATGAGTGTGGAGTCGTTGCTTCTCAAATCCTAACCTTCCTATNNNAGATCGGAAGA
127848 TATGAGTGTGGAGTCGTTGCCATAGGCCTTATCAGGAATGNNNAGATCGGAAGAG
127849 TATGAGTGTGGAGTCGTTGCCATTCAATGTCTTTCAAGAAGAGNNNAGATCGGAAG
127850 TATGAGTGTGGAGTCGTTGCCTCTATGTCTTTTCATTGTATGGNNNAGATCGGAAG
127851 TATGAGTGTGGAGTCGTTGCAATATAGACTTTGGTACCAAGAGNNNAGATCGGAA
127852 TATGAGTGTGGAGTCGTTGCAATATAGACTTTGGTACCAAGAGNNNAGATCGGAA
127853 TATGAGTGTGGAGTCGTTGCATCCAGATTCATTGTAAAATACCNNNAGATCGGAAG
127854 TATGAGTGTGGAGTCGTTGCAATTCTATTTCAAGTTAAGCCTCNNNAGATCGGAAG
127855 TATGAGTGTGGAGTCGTTGCTATGTCTCTACATGTTTCTGTTTNNNAGATCGGAAGA
127856 TATGAGTGTGGAGTCGTTGCCTACTTAGTCATAGAAAGGAACANNNAGATCGGAA
127857 TATGAGTGTGGAGTCGTTGCTTACCCTGAAAAAAAAAAACGANNNAGATCGGAAG
127858 TATGAGTGTGGAGTCGTTGCTTTCTTTTTTTTTTTTTTGCGACNNNAGATCGGAAGA
127859 TATGAGTGTGGAGTCGTTGCAATCATTTGGTGTATCAAAATGTNNNAGATCGGAAG
127860 TATGAGTGTGGAGTCGTTGCTAGAACAGATCACAAAAAAAAGGNNNAGATCGGAA
127862 TATGAGTGTGGAGTCGTTGCACTATATGATATTCTGGATTGGANNNAGATCGGAAG
127863 TATGAGTGTGGAGTCGTTGCGTGTATACAGACGCAATTATATTNNNAGATCGGAAG
127865 TATGAGTGTGGAGTCGTTGCGAATGAATTTTCCACTGTCTATTNNNAGATCGGAAG
127866 TATGAGTGTGGAGTCGTTGCTTCAATACCACTTGCATTTTTATNNNAGATCGGAAGA
127867 TATGAGTGTGGAGTCGTTGCTTTTCAATCCAGTGAGTACTATGNNNAGATCGGAAG
127868 TATGAGTGTGGAGTCGTTGCGCCAACAGAGTGATTACATCNNNAGATCGGAAGAG
127869 TATGAGTGTGGAGTCGTTGCTTTGTGAATAAGAGCTATTAGGANNNAGATCGGAA
127870 TATGAGTGTGGAGTCGTTGCACACATAAGAATTGCACATATTTNNNAGATCGGAAG
127871 TATGAGTGTGGAGTCGTTGCGTTGATTTCTAGAGAAGTGCTTNNNAGATCGGAAG
127873 TATGAGTGTGGAGTCGTTGCAGCAAAAACCCTAGTAGTATATTNNNAGATCGGAAG
127874 TATGAGTGTGGAGTCGTTGCTATTTTTTTCATGAGATAAGCCCNNNAGATCGGAAG
127875 TATGAGTGTGGAGTCGTTGCCATAGGTCTCAATATTGACTGATNNNAGATCGGAAG
127876 TATGAGTGTGGAGTCGTTGCTTGATAACAGTGAGACTTCTTCNNNAGATCGGAAGA
127877 TATGAGTGTGGAGTCGTTGCGAATTACAGATAAGGGAATGACANNNAGATCGGAA
127878 TATGAGTGTGGAGTCGTTGCAGATGTTCAGTTTATTTCAAAAGANNNAGATCGGAA
127879 TATGAGTGTGGAGTCGTTGCCTTTAGTGGAAAGAGCTAGACNNNAGATCGGAAGA
127880 TATGAGTGTGGAGTCGTTGCAACCACTTGAGAAAACAATTTTTNNNAGATCGGAAG
127881 TATGAGTGTGGAGTCGTTGCCTGAAGTATAGTGTTGCAATCANNNAGATCGGAAG
127882 TATGAGTGTGGAGTCGTTGCGCATGCTTCAATTCTTCATTTNNNAGATCGGAAGAG
127883 TATGAGTGTGGAGTCGTTGCTAACTAACATTGGTCTGATTCTANNNAGATCGGAAG
127884 TATGAGTGTGGAGTCGTTGCGATTGTCTTTTCCTACCAGGNNNAGATCGGAAGAGC
127885 TATGAGTGTGGAGTCGTTGCCAAAATTCAAAAACAAAAGCAACNNNAGATCGGAAG
127886 TATGAGTGTGGAGTCGTTGCTATATAAAGTCCATACCAGGCCNNNAGATCGGAAGA
127887 TATGAGTGTGGAGTCGTTGCAGTACATGTATTAGTCTGTTCTCNNNAGATCGGAAG
127888 TATGAGTGTGGAGTCGTTGCTATGAGACAGAGAGAGAGAGAANNNAGATCGGAA
127889 TATGAGTGTGGAGTCGTTGCAGCACATAAATCAATTACAGAATANNNAGATCGGAA
127890 TATGAGTGTGGAGTCGTTGCAACAAACCTGTGTTTTTTTTTTANNNAGATCGGAAGA
127891 TATGAGTGTGGAGTCGTTGCGAATCATCTTTAGGGTTTTCTACNNNAGATCGGAAG
127892 TATGAGTGTGGAGTCGTTGCGATCACAAGGTCAAGAGATAGNNNAGATCGGAAGA

127893 TATGAGTGTGGAGTCGTTGCGACATTTTGTACCGGATTTTATANNNAGATCGGAAG
127894 TATGAGTGTGGAGTCGTTGCATTTGCAACTGGTATTAGAGACNNNAGATCGGAAGA
127895 TATGAGTGTGGAGTCGTTGCCCATCAAGTGAATCATAAAACAANNNAGATCGGAAG
127896 TATGAGTGTGGAGTCGTTGCGCATGCACTATCCCTATAATTTANNNAGATCGGAAGA
127897 TATGAGTGTGGAGTCGTTGCAAGAATGACTTGAACTAATCTCTNNNAGATCGGAAG
127899 TATGAGTGTGGAGTCGTTGCCCAAAATACACAAAGAATTCCTANNNAGATCGGAAG
127900 TATGAGTGTGGAGTCGTTGCCTCTCTTCATCTGTCTCTCTTNNNAGATCGGAAGAGCA
127901 TATGAGTGTGGAGTCGTTGCTATACCTTTGTTTATCCCATTCTNNNAGATCGGAAGA
127903 TATGAGTGTGGAGTCGTTGCCTTCTTTCTTCTGAAACATACTCNNNAGATCGGAAGA
127904 TATGAGTGTGGAGTCGTTGCCATTTGCAGATCTTCTTTACAGNNNAGATCGGAAGA
127905 TATGAGTGTGGAGTCGTTGCCACATACATACATACATACATACANNNAGATCGGAAG
127907 TATGAGTGTGGAGTCGTTGCTTTCTTCAGCAATGTTTAAGTTANNNAGATCGGAAGA
127908 TATGAGTGTGGAGTCGTTGCGTCTTTTCTTCAGCAATGTTTANNNAGATCGGAAGAG
127909 TATGAGTGTGGAGTCGTTGCAAAACCACACATGATGAAATTATNNNAGATCGGAAG
127910 TATGAGTGTGGAGTCGTTGCATTCTCTTCTTTTTATCTCTGGTNNNAGATCGGAAGA
127911 TATGAGTGTGGAGTCGTTGCCTCTGCTGAACTATTTCTGCNNNAGATCGGAAGAGCA
127912 TATGAGTGTGGAGTCGTTGCTATATACAGAGAGAGAGAGAGAGNNNAGATCGGA
127913 TATGAGTGTGGAGTCGTTGCACTGAGAGTCTAACAGAAATATCNNNAGATCGGAAG
127914 TATGAGTGTGGAGTCGTTGCTTAAGTCACTTACCCAAGGTTANNNAGATCGGAAGA
127915 TATGAGTGTGGAGTCGTTGCATATCAAATGGGTATCAAGGAAANNNAGATCGGAA
127916 TATGAGTGTGGAGTCGTTGCCTCTTTACTCCTATTCTTCAAGANNNAGATCGGAAGA
127917 TATGAGTGTGGAGTCGTTGCATCACACTCTTGCTTCATTTNNNAGATCGGAAGAGCA
127918 TATGAGTGTGGAGTCGTTGCACAAGAGAGACCATTTAAAATAAANNNAGATCGGA
127919 TATGAGTGTGGAGTCGTTGCGTTCTCCAACTTAAAACACTTGNNNAGATCGGAAGA
127920 TATGAGTGTGGAGTCGTTGCCCTGCATCCTTTTCTTTTTTTANNNAGATCGGAAGAG
127921 TATGAGTGTGGAGTCGTTGCCCACACAAATAAAAAAAGACTACNNNAGATCGGAAG
127922 TATGAGTGTGGAGTCGTTGCACTAATACAATCTCCTATCAACCNNNAGATCGGAAGA
127923 TATGAGTGTGGAGTCGTTGCTTTCTCTGAGCATTAGTAGTTCNNNAGATCGGAAGA
127924 TATGAGTGTGGAGTCGTTGCCACTCCAATATCCCTGACTCNNNAGATCGGAAGAGCA
127925 TATGAGTGTGGAGTCGTTGCATCTTTTATCCAAAATGCTTAGGNNNAGATCGGAAG
127926 TATGAGTGTGGAGTCGTTGCGAATATAGGATATGTACCAGTGTNNNAGATCGGAA
127927 TATGAGTGTGGAGTCGTTGCTGCAAAAATAAATGAATGATTGTNNNAGATCGGAA
127928 TATGAGTGTGGAGTCGTTGCAAACAAACAAACAAACAAACAAANNNAGATCGGAA
127929 TATGAGTGTGGAGTCGTTGCCTCCCTTTCTTATAACATATTGCNNNAGATCGGAAGA
127930 TATGAGTGTGGAGTCGTTGCAATAACCATTTATCACATTTTGCNNNAGATCGGAAGA
127931 TATGAGTGTGGAGTCGTTGCCTTCCTAATTAAAAAGGCTGAGNNNAGATCGGAAGA
127932 TATGAGTGTGGAGTCGTTGCATGTCTCAGTCTCACAGAATNNNAGATCGGAAGAGC
127933 TATGAGTGTGGAGTCGTTGCCTCCTCAAAATTAACACTTGATANNNAGATCGGAAG
127934 TATGAGTGTGGAGTCGTTGCCCAAAGTACTGGGATTACAGNNNAGATCGGAAGAG
127935 TATGAGTGTGGAGTCGTTGCCATACTAGGCAATCATGTTCTTNNNAGATCGGAAGA
127936 TATGAGTGTGGAGTCGTTGCGTCTGCTGATGAGGAAATAGNNNAGATCGGAAGAG
127937 TATGAGTGTGGAGTCGTTGCATAGAAAGAGGGAAATCAACTGNNNAGATCGGAAG
127938 TATGAGTGTGGAGTCGTTGCCTTTGATCTCAGACTTCTAGTCNNNAGATCGGAAGA

127939 TATGAGTGTGGAGTCGTTGCTATACAGACTAACAGATACACACNNNAGATCGGAAG
127940 TATGAGTGTGGAGTCGTTGCCCAAAAACAGGAAAGAAAAAATANNNAGATCGGAA
127941 TATGAGTGTGGAGTCGTTGCTATATAAAGTCACCCGAAATACANNNAGATCGGAAG
127942 TATGAGTGTGGAGTCGTTGCTATATAAAGTCACCCGAAATACANNNAGATCGGAAG
127943 TATGAGTGTGGAGTCGTTGCATCTTTCCGAGGTACAGAAGNNNAGATCGGAAGAG
127944 TATGAGTGTGGAGTCGTTGCTCTTTTTTTTTCTTTTTTGTCTGNNNAGATCGGAAGA
127945 TATGAGTGTGGAGTCGTTGCAAGACAGATCAATAGTGATTCATNNNAGATCGGAAG
127946 TATGAGTGTGGAGTCGTTGCTAAAAGTATAGCGAGGTCAATTTNNNAGATCGGAA
127947 TATGAGTGTGGAGTCGTTGCTCATTTAATTCTCTGTCTCCTACNNNAGATCGGAAGA
127948 TATGAGTGTGGAGTCGTTGCCCAGACTGGAATCTATACTATAGNNNAGATCGGAAG
127949 TATGAGTGTGGAGTCGTTGCCAAATAGGCTATGAAGACAAGTNNNAGATCGGAAG
127950 TATGAGTGTGGAGTCGTTGCTGTCTTGATTTTCTTTAGTTCAGNNNAGATCGGAAGA
127951 TATGAGTGTGGAGTCGTTGCTGGAAAATTAAGGTTGTAGAACTNNNAGATCGGAA
127953 TATGAGTGTGGAGTCGTTGCAATAATTAGCTCAAATCTCCAACNNNAGATCGGAAGA
127954 TATGAGTGTGGAGTCGTTGCTGTTTTCTAATTAATTCAGCACCNNNAGATCGGAAGA
127955 TATGAGTGTGGAGTCGTTGCTTCCCTCTCTCCTTCACTTANNNAGATCGGAAGAGCA
127956 TATGAGTGTGGAGTCGTTGCTTGTCCATATTGGAGGTTACTNNNAGATCGGAAGAG
127957 TATGAGTGTGGAGTCGTTGCTAACAATGTTTTCTGAAAGGTCNNNAGATCGGAAGA
127958 TATGAGTGTGGAGTCGTTGCTATACACATACCCAAATACACATNNNAGATCGGAAGA
127959 TATGAGTGTGGAGTCGTTGCCAAACCTAAAAATCAGAAGTACANNNAGATCGGAAG
127960 TATGAGTGTGGAGTCGTTGCATTTTTTCTTTTTAGCATGCATTNNNAGATCGGAAGA
127961 TATGAGTGTGGAGTCGTTGCGAGAATTATCCTGGAACTTGTNNNAGATCGGAAGAG
127962 TATGAGTGTGGAGTCGTTGCCTGTATTTCCTTGATATTGTGCNNNAGATCGGAAGA
127963 TATGAGTGTGGAGTCGTTGCAAATGCTATAACTTAGCTTTTGANNNAGATCGGAAG
127965 TATGAGTGTGGAGTCGTTGCTTAGTCTTTTTCTGAACGATTTGNNNAGATCGGAAGA
127967 TATGAGTGTGGAGTCGTTGCAGGCTTTCAGTAAAAATGTAAAANNNAGATCGGAA
127968 TATGAGTGTGGAGTCGTTGCTGATAATCCATTGTGAGAAATTCNNNAGATCGGAAG
127969 TATGAGTGTGGAGTCGTTGCTTAGCCATCTAATCTCATTCAATNNNAGATCGGAAGA
127970 TATGAGTGTGGAGTCGTTGCCCAGCATTGATATAAATACAGTCNNNAGATCGGAAG
127971 TATGAGTGTGGAGTCGTTGCAAAACAAAAAACCTCGGAGANNNAGATCGGAAGAG
127972 TATGAGTGTGGAGTCGTTGCTCTTTCAACACAAATTTATTGTGNNNAGATCGGAAGA
127973 TATGAGTGTGGAGTCGTTGCAATTCTGTCCTGAAGAATATGTNNNAGATCGGAAGA
127975 TATGAGTGTGGAGTCGTTGCTTCTCAACCCTCTCAACAAANNNAGATCGGAAGAGCA
127976 TATGAGTGTGGAGTCGTTGCATAAAGAAGCAGCTTCCTTTTNNNAGATCGGAAGAG
127977 TATGAGTGTGGAGTCGTTGCCCTTGCATCATAGTTATTTATGTNNNAGATCGGAAGA
127978 TATGAGTGTGGAGTCGTTGCGAACACATGTTAATATTGATGTGNNNAGATCGGAAG
127979 TATGAGTGTGGAGTCGTTGCTACAAATGACATATTGCTCAAATNNNAGATCGGAAG
127980 TATGAGTGTGGAGTCGTTGCCCGTCTAGTAGAAATAGAAGTAANNNAGATCGGAA
127981 TATGAGTGTGGAGTCGTTGCAGGGGTCACAATTTATTCTAATANNNAGATCGGAAG
127982 TATGAGTGTGGAGTCGTTGCCTATTTTGGCTTTACCCTTTTCNNNAGATCGGAAGAG
127983 TATGAGTGTGGAGTCGTTGCCGGAGAAGGGAAGGTTATAAANNNAGATCGGAAG
127984 TATGAGTGTGGAGTCGTTGCCTTGAGTTTTGATTTGCTGATTNNNAGATCGGAAGA
127985 TATGAGTGTGGAGTCGTTGCTAGATTTACTGGTTTCTACTTCCNNNAGATCGGAAGA

127986 TATGAGTGTGGAGTCGTTGCAGTTTTATCCTTTTGCTACTTTCNNNAGATCGGAAGA

127987 TATGAGTGTGGAGTCGTTGCGAAATTATAAAGTGTGTGTGTGTNNNAGATCGGAA

127988 TATGAGTGTGGAGTCGTTGCCTCCTTCTCTATCATCTCTTTCNNNAGATCGGAAGAG

127990 TATGAGTGTGGAGTCGTTGCAACAGAAGTACAAAATGGAAATTNNNAGATCGGAA

127991 TATGAGTGTGGAGTCGTTGCGGAGTAAGCATATGGTATAAGTANNNAGATCGGAA

127994 TATGAGTGTGGAGTCGTTGCGATAAACAGGTAACTTCAAGTAGNNNAGATCGGAA

127995 TATGAGTGTGGAGTCGTTGCTGAAGAGAACATCTGTAAAATTGNNNAGATCGGAA

127996 TATGAGTGTGGAGTCGTTGCCCCTCAGAAATCACATAGTTATNNNAGATCGGAAGA

127997 TATGAGTGTGGAGTCGTTGCACCCTTATTTTCCCAGTTTCNNNAGATCGGAAGAGCA

127998 TATGAGTGTGGAGTCGTTGCTTGAAATTTTCAAGAGCTTCAGNNNAGATCGGAAGA

127999 TATGAGTGTGGAGTCGTTGCTATGTCCAAATTTCCTCTTCTTNNNAGATCGGAAGAG

128000 TATGAGTGTGGAGTCGTTGCTATGTCCAAATTTCCTCTTCTTNNNAGATCGGAAGAG

128001 TATGAGTGTGGAGTCGTTGCGAGAAGGGTGTGTTTGTAAGNNNAGATCGGAAGA

128002 TATGAGTGTGGAGTCGTTGCGTAATTGCTAATTCAGGTTGTAANNNAGATCGGAAG

128003 TATGAGTGTGGAGTCGTTGCAGAGAATAACTGTTATCCACTGNNNAGATCGGAAGA

128004 TATGAGTGTGGAGTCGTTGCAATTCCTGACCTCAAGTGATNNNAGATCGGAAGAGC

128005 TATGAGTGTGGAGTCGTTGCCACACATAAATGCAGACATATTANNNAGATCGGAAG

128006 TATGAGTGTGGAGTCGTTGCCCTTTTGAGACTTCTTTTCTACNNNAGATCGGAAGAG

128007 TATGAGTGTGGAGTCGTTGCGGTGGATTTACTTTAAACAGTTTNNNAGATCGGAAG

128008 TATGAGTGTGGAGTCGTTGCTTTTGGCATCTTTAAAATGAAGANNNAGATCGGAAG

128009 TATGAGTGTGGAGTCGTTGCGTTTGTCCTTAGAGTTTTGATAGNNNAGATCGGAAG

128010 TATGAGTGTGGAGTCGTTGCGTAATTCATGCGCATTTTTTTCNNNAGATCGGAAGA

128011 TATGAGTGTGGAGTCGTTGCAATTCCAAAAAACTGCATAGTTNNNAGATCGGAAGA

128012 TATGAGTGTGGAGTCGTTGCATTAAGACATTTTTTGGCCATTANNNAGATCGGAAG

128013 TATGAGTGTGGAGTCGTTGCGGAGATATATAAAGACAGCACTTNNNAGATCGGAA

128014 TATGAGTGTGGAGTCGTTGCATCAGAGACAAATTTACACAATCNNNAGATCGGAAG

128015 TATGAGTGTGGAGTCGTTGCGACACCAATAAGAGTTAATGTTTNNNAGATCGGAAG

128016 TATGAGTGTGGAGTCGTTGCTAGGAGTTGTCAAATATTTTGTGNNNAGATCGGAAG

128017 TATGAGTGTGGAGTCGTTGCGCTATGCTATAATGTTTTCTGATNNNAGATCGGAAG

128018 TATGAGTGTGGAGTCGTTGCCTATGTGGCATTAAAAGTAAGACNNNAGATCGGAAG

128019 TATGAGTGTGGAGTCGTTGCCCAGATATGTTCAATTCATTCTCNNNAGATCGGAAGA

128020 TATGAGTGTGGAGTCGTTGCATGTTGGTCAAGAGAATAACAANNNAGATCGGAAG

128021 TATGAGTGTGGAGTCGTTGCATTTGTTTATGTTGTACAAAGGGNNNAGATCGGAAG

128022 TATGAGTGTGGAGTCGTTGCCCTGAAGAAAATTTTAGGAAAGANNNAGATCGGAA

128023 TATGAGTGTGGAGTCGTTGCCACCTTACTTTCATAATTTGTGTNNNAGATCGGAAGA

128024 TATGAGTGTGGAGTCGTTGCAATTGCTGTTGTGTGAAGTCNNNAGATCGGAAGAG

128025 TATGAGTGTGGAGTCGTTGCCTTTTGAACTCCAGGACTATACNNNAGATCGGAAGA

128027 TATGAGTGTGGAGTCGTTGCATTTCCTCTTCTTTCCCTTTTTTNNNAGATCGGAAGAG

128028 TATGAGTGTGGAGTCGTTGCGCATATATCCCTAATGTAATCCTNNNAGATCGGAAGA

128029 TATGAGTGTGGAGTCGTTGCATTCAGGGAAAGATGGAGAANNNAGATCGGAAGA

128030 TATGAGTGTGGAGTCGTTGCATTCAGGGAAAGATGGAGAANNNAGATCGGAAGA

128031 TATGAGTGTGGAGTCGTTGCATTCAGGGAAAGATGGAGAANNNAGATCGGAAGA

128032 TATGAGTGTGGAGTCGTTGCCAGTATTTGTTGAATGCAATTCTNNNAGATCGGAAG

128033 TATGAGTGTGGAGTCGTTGCATCTTTAAGAGGTAGTGATTTCANNNAGATCGGAAG
128034 TATGAGTGTGGAGTCGTTGCCATTCCATCTCCTTCTACTTATTNNNAGATCGGAAGA
128035 TATGAGTGTGGAGTCGTTGCAAGTACGTACCAGAGTTCTAANNNAGATCGGAAGA
128036 TATGAGTGTGGAGTCGTTGCCTTCCACTAACTTCAAAGAATCNNNAGATCGGAAGA
128037 TATGAGTGTGGAGTCGTTGCACAGAAAGGAAAGAAGTACATTANNNAGATCGGAA
128038 TATGAGTGTGGAGTCGTTGCCATGACTGGACTTTGATTATTTTNNNAGATCGGAAG
128039 TATGAGTGTGGAGTCGTTGCTTTATGTTTGTATTGCCTGTAAANNNAGATCGGAAG
128040 TATGAGTGTGGAGTCGTTGCAAGAAAGTTCTAAAGCTCTCTATNNNAGATCGGAAG
128041 TATGAGTGTGGAGTCGTTGCAAGCATTCCTTTGTTCTATCTTNNNAGATCGGAAGAG
128042 TATGAGTGTGGAGTCGTTGCCCACATTTTCTTTATTCACTCATNNNAGATCGGAAGA
128044 TATGAGTGTGGAGTCGTTGCGTCTCTCTCTCTTCTGGATCNNNAGATCGGAAGAGC
128045 TATGAGTGTGGAGTCGTTGCTTTGATCCTCTCCTCATATGATANNNAGATCGGAAGA
128046 TATGAGTGTGGAGTCGTTGCTTTATTTGACTTTACCCCCTAAANNNAGATCGGAAGA
128047 TATGAGTGTGGAGTCGTTGCGACATTCTGAAAAGTCTTATGAANNNAGATCGGAAG
128049 TATGAGTGTGGAGTCGTTGCTATTTGTTTTCTATTGCTGCATTNNNAGATCGGAAGA
128050 TATGAGTGTGGAGTCGTTGCTCCTTTTCTTGTCTAATTAAAACANNNAGATCGGAAG
128051 TATGAGTGTGGAGTCGTTGCTTTATTTGTCTAAGCCACTATTGNNNAGATCGGAAG
128052 TATGAGTGTGGAGTCGTTGCTTTATTTGTCTAAGCCACTATTGNNNAGATCGGAAG
128053 TATGAGTGTGGAGTCGTTGCGTTCCAAGTATTAACGTACATGNNNAGATCGGAAGA
128054 TATGAGTGTGGAGTCGTTGCTCATTATTGGTATGAACAATTGGNNNAGATCGGAAG
128055 TATGAGTGTGGAGTCGTTGCACTACTCATCCATAAAAAGGAATNNNAGATCGGAAG
128056 TATGAGTGTGGAGTCGTTGCAGAATTTCCAATGAATTTTGACANNNAGATCGGAAG
128057 TATGAGTGTGGAGTCGTTGCAAGATTCAATACCTAGGAGATTGNNNAGATCGGAA
128058 TATGAGTGTGGAGTCGTTGCGAGAGTATTAGTAAACTCCTGTANNNAGATCGGAA
128059 TATGAGTGTGGAGTCGTTGCCTCTCTGGTATTTGCTGAATATANNNAGATCGGAAG
128060 TATGAGTGTGGAGTCGTTGCAGTGAAGTGGTACAATCATGNNNAGATCGGAAGAG
128061 TATGAGTGTGGAGTCGTTGCTTCAAGAATGAGGATCATGATTTNNNAGATCGGAA
128062 TATGAGTGTGGAGTCGTTGCAACCTTGACTAATACAACAATTTNNNAGATCGGAAG
128063 TATGAGTGTGGAGTCGTTGCGAATCTGTGATTAGTCAGAGTAANNNAGATCGGAA
128064 TATGAGTGTGGAGTCGTTGCGGATCACTTTCCATACTGTATTANNNAGATCGGAAG
128065 TATGAGTGTGGAGTCGTTGCTACAAGTATATGCTGCTTTTCTANNNAGATCGGAAG
128066 TATGAGTGTGGAGTCGTTGCCTCTTCAGGGTGACATCTAGNNNAGATCGGAAGAG
128067 TATGAGTGTGGAGTCGTTGCTACTTTCTGCATTATTTACTGAGNNNAGATCGGAAG
128068 TATGAGTGTGGAGTCGTTGCTGCATTCCTACATATATATGTGTANNNAGATCGGAA
128069 TATGAGTGTGGAGTCGTTGCTGTGTATGTGGTCTATATTGTAGNNNAGATCGGAA
128070 TATGAGTGTGGAGTCGTTGCCCCATAGGATTTGAGAATGAGNNNAGATCGGAAGA
128071 TATGAGTGTGGAGTCGTTGCCACATGTTTTGAAGGGAATGNNNAGATCGGAAGAG
128072 TATGAGTGTGGAGTCGTTGCTCTGTTTCATTTATTCCACAAATNNNAGATCGGAAGA
128073 TATGAGTGTGGAGTCGTTGCGTGTCTCCCATTCTCTCTTCNNNAGATCGGAAGAGCA
128074 TATGAGTGTGGAGTCGTTGCGTCATGATGCTATTAGATACTGANNNAGATCGGAAG
128076 TATGAGTGTGGAGTCGTTGCATCACATTTTCTTTATCCATTCANNNAGATCGGAAGA
128077 TATGAGTGTGGAGTCGTTGCGTGTATGTCTCTGTGTGTCTNNNAGATCGGAAGAG
128078 TATGAGTGTGGAGTCGTTGCCCCTTTTTTTTTTTTTTGACAGANNNAGATCGGAAGA

128079 TATGAGTGTGGAGTCGTTGCCTCAGAACACATAAGTATTTTCANNNAGATCGGAAG
128080 TATGAGTGTGGAGTCGTTGCGTTATAAAGCCATTCACATTTCTNNNAGATCGGAAG
128082 TATGAGTGTGGAGTCGTTGCAGAGATTTTTTTTGAGGAGCTNNNAGATCGGAAGA
128083 TATGAGTGTGGAGTCGTTGCTCCTTGGTGATTCTAGATGTNNNAGATCGGAAGAG
128084 TATGAGTGTGGAGTCGTTGCTCCTTGGTGATTCTAGATGTNNNAGATCGGAAGAG
128085 TATGAGTGTGGAGTCGTTGCGATTCTAGATGTGAGTGAAGTGNNNAGATCGGAAG
128086 TATGAGTGTGGAGTCGTTGCACTGCATTTTGTTCCTTTATTTNNNAGATCGGAAGAG
128087 TATGAGTGTGGAGTCGTTGCTTTACATCTTTACATGTCATTGGNNNAGATCGGAAG
128088 TATGAGTGTGGAGTCGTTGCGACAAAAGAATACACATACACATNNNAGATCGGAAG
128089 TATGAGTGTGGAGTCGTTGCGCATTTTCTCTAAAATATTTGCCNNNAGATCGGAAG
128090 TATGAGTGTGGAGTCGTTGCAAAAGTTCCAGTCTCAGTCTNNNAGATCGGAAGAG
128091 TATGAGTGTGGAGTCGTTGCCACATATACACCACATCTCTTTNNNAGATCGGAAGAG
128092 TATGAGTGTGGAGTCGTTGCGCAGAAAAGCCAAACTATTTATANNNAGATCGGAAG
128093 TATGAGTGTGGAGTCGTTGCAACTGCTACATTTTCTAGAGTAANNNAGATCGGAAG
128094 TATGAGTGTGGAGTCGTTGCATGCTTTGATCTATTCTGTACATNNNAGATCGGAAG
128095 TATGAGTGTGGAGTCGTTGCATGCTTTGATCTATTCTGTACATNNNAGATCGGAAG
128096 TATGAGTGTGGAGTCGTTGCTTTTTGTTTTTTTTGTCACTCTCNNNAGATCGGAAGA
128097 TATGAGTGTGGAGTCGTTGCAGACAGGATCTCATTCTGTTNNNAGATCGGAAGAG
128098 TATGAGTGTGGAGTCGTTGCCCTCATAATAGTTTGCAAAAGAANNNAGATCGGAAG
128099 TATGAGTGTGGAGTCGTTGCATACAGAATCACAAAATCTTCTGNNNAGATCGGAAG
128100 TATGAGTGTGGAGTCGTTGCCAGCATATCAATGGAGATCTATANNNAGATCGGAAG
128101 TATGAGTGTGGAGTCGTTGCCAAATAGAGAAGCTCATAAACAGNNNAGATCGGAA
128102 TATGAGTGTGGAGTCGTTGCTTAGCTGGATTTTTGTTAAGTTTNNNAGATCGGAAG
128103 TATGAGTGTGGAGTCGTTGCAATCAATGGGAGAAGAGAGTAANNNAGATCGGAA
128104 TATGAGTGTGGAGTCGTTGCTGTGTTTTGTTTTGTTTTTGAGNNNAGATCGGAAGA
128105 TATGAGTGTGGAGTCGTTGCGGAGTATTTGAAGCTTTATGATTNNNAGATCGGAA
128106 TATGAGTGTGGAGTCGTTGCTTATAAAGAATGGGGTCTTTGAGNNNAGATCGGAA
128107 TATGAGTGTGGAGTCGTTGCTAGTTCTTACCTACAGTTTCTCTNNNAGATCGGAAGA
128109 TATGAGTGTGGAGTCGTTGCACTTTCTGAGTGCATAACACNNNAGATCGGAAGAG
128110 TATGAGTGTGGAGTCGTTGCGTGTATGTGTCAGTGTTGTANNNAGATCGGAAGAG
128111 TATGAGTGTGGAGTCGTTGCACAAAAGTTTCTGGAAAAAATACNNNAGATCGGAA
128112 TATGAGTGTGGAGTCGTTGCAAAGGGTAATGCATTTCTTTAAANNNAGATCGGAAG
128113 TATGAGTGTGGAGTCGTTGCTAAATGTGTAAGAGGTAGGAAATNNNAGATCGGAA
128114 TATGAGTGTGGAGTCGTTGCATGTGAGGATATAACAAAGAAGANNNAGATCGGAA
128115 TATGAGTGTGGAGTCGTTGCATGTGAGGATATAACAAAGAAGANNNAGATCGGAA
128116 TATGAGTGTGGAGTCGTTGCCTAAACCATGCCTATTTTACAAANNNAGATCGGAAG
128117 TATGAGTGTGGAGTCGTTGCCGAGAATGAAGTGCTACTATGNNNAGATCGGAAGA
128120 TATGAGTGTGGAGTCGTTGCTCTTCTTAGACTACTTTTTCCATNNNAGATCGGAAGA
128121 TATGAGTGTGGAGTCGTTGCAAACAAGAAATCCCACAGAAGNNNAGATCGGAAGA
128122 TATGAGTGTGGAGTCGTTGCAAACAAGAAATCCCACAGAAGNNNAGATCGGAAGA
128124 TATGAGTGTGGAGTCGTTGCTATGATTATGTCTGAGAAGTCTCNNNAGATCGGAAG
128125 TATGAGTGTGGAGTCGTTGCGTTAATTTTTGCTTGTGTCATAGNNNAGATCGGAAG
128126 TATGAGTGTGGAGTCGTTGCCTCACTTTTGCTTACCTTCTNNNAGATCGGAAGAGCA

128127 TATGAGTGTGGAGTCGTTGCCTCACTTTTGCTTACCTTCTNNNAGATCGGAAGAGCA
128128 TATGAGTGTGGAGTCGTTGCAAGCATACAAAGAAAACATAACANNNAGATCGGAAG
128129 TATGAGTGTGGAGTCGTTGCGTTTTATTGTGCTTTGAGTGANNNAGATCGGAAGAG
128130 TATGAGTGTGGAGTCGTTGCATGTGAGTGTTTAGAAAGATACANNNAGATCGGAA
128131 TATGAGTGTGGAGTCGTTGCGAAGCTAGAAAAACAAGACTTAANNNAGATCGGAA
128132 TATGAGTGTGGAGTCGTTGCACATTTTAATTGATAATGGCTCCNNNAGATCGGAAG
128135 TATGAGTGTGGAGTCGTTGCCAAACAAAAATCAAGCTCTAAAANNNAGATCGGAAG
128136 TATGAGTGTGGAGTCGTTGCCTAGGAAGCATTCATGGTACNNNAGATCGGAAGAG
128137 TATGAGTGTGGAGTCGTTGCTTTGTATAATGTCTGTTTTACCANNNAGATCGGAAG
128138 TATGAGTGTGGAGTCGTTGCTTTGTATAATGTCTGTTTTACCANNNAGATCGGAAG
128139 TATGAGTGTGGAGTCGTTGCATGAGAATTAAATGCAATGACATNNNAGATCGGAAG
128140 TATGAGTGTGGAGTCGTTGCAGCAACTATTAAATAATGGGTCTNNNAGATCGGAAG
128141 TATGAGTGTGGAGTCGTTGCCTGATAGGATCGGTGTGTACNNNAGATCGGAAGAG
128142 TATGAGTGTGGAGTCGTTGCTAGTGTGGAGCCTTAAGAGANNNAGATCGGAAGAG
128143 TATGAGTGTGGAGTCGTTGCAATAGAGAAAGAGAGAGTTGTAGNNNAGATCGGA
128144 TATGAGTGTGGAGTCGTTGCATTGAGATGTGAGAATTAAGCAANNNAGATCGGAA
128145 TATGAGTGTGGAGTCGTTGCATTGATTGTAGAGCTTAATGTGANNNAGATCGGAAG
128146 TATGAGTGTGGAGTCGTTGCTTCTGTAAGCAAATGACAGAGNNNAGATCGGAAGA
128147 TATGAGTGTGGAGTCGTTGCCATGCTAAAAAGTCAAGACTTAANNNAGATCGGAAG
128148 TATGAGTGTGGAGTCGTTGCCATAGAAGAGGGTTGGACTANNNAGATCGGAAGAG
128149 TATGAGTGTGGAGTCGTTGCTTCCATAATAAAAATGGTATCGGNNNAGATCGGAAG
128150 TATGAGTGTGGAGTCGTTGCAATACACATTAATCTCAAGCACANNNAGATCGGAAG
128151 TATGAGTGTGGAGTCGTTGCATCCAGTAAGATTATAATGGACCNNNAGATCGGAAG
128152 TATGAGTGTGGAGTCGTTGCCTGAACTTCATGCCAAATTCNNNAGATCGGAAGAGC
128153 TATGAGTGTGGAGTCGTTGCCAGTCAGAGAAATCTACAGCNNNAGATCGGAAGAG
128154 TATGAGTGTGGAGTCGTTGCATCAGTACTGACTTTGTCAAATANNNAGATCGGAAG
128155 TATGAGTGTGGAGTCGTTGCATCTCTATCTGCACAGTTATAAANNNAGATCGGAAG
128156 TATGAGTGTGGAGTCGTTGCAGTAAATACGGCCAGAATTTAGNNNAGATCGGAAG
128157 TATGAGTGTGGAGTCGTTGCTTTTCAATGGACCAGCTATTTNNNAGATCGGAAGAG
128158 TATGAGTGTGGAGTCGTTGCAGTTCTTTCTTAGTAATTCTAGCTNNNAGATCGGAAG
128159 TATGAGTGTGGAGTCGTTGCGATAACAACCGTAAAAATCTCATNNNAGATCGGAAG
128160 TATGAGTGTGGAGTCGTTGCCTGGAAGTTCTTTTTGTAATCTTNNNAGATCGGAAG
128161 TATGAGTGTGGAGTCGTTGCATGTATACCAGATAAAAGACCTCNNNAGATCGGAAG
128162 TATGAGTGTGGAGTCGTTGCGGATTGAGGTGAAATTAGAAAATNNNAGATCGGAA
128163 TATGAGTGTGGAGTCGTTGCAAAGTTCACCTCCTGTAGTAANNNAGATCGGAAGAG
128164 TATGAGTGTGGAGTCGTTGCATCATCACAAAACATTTGTATCANNNAGATCGGAAG
128166 TATGAGTGTGGAGTCGTTGCCATGTTAAAATGCAGGTCATAATNNNAGATCGGAAG
128167 TATGAGTGTGGAGTCGTTGCTTTCATTTCTTATTTGCATATGCNNNAGATCGGAAGA
128169 TATGAGTGTGGAGTCGTTGCCAGCATCTTCATTTTGACATTANNNAGATCGGAAGA
128170 TATGAGTGTGGAGTCGTTGCAGACAGTTCTCAGAATTAATCATNNNAGATCGGAAG
128171 TATGAGTGTGGAGTCGTTGCGATGTATTGAAAATAATACCCCCNNNAGATCGGAAG
128172 TATGAGTGTGGAGTCGTTGCAGATTGTGTGAGAGAGAGATTNNNAGATCGGAAGA
128173 TATGAGTGTGGAGTCGTTGCTTGGTAAAATAATATTTGCAGCTNNNAGATCGGAAG

128174 TATGAGTGTGGAGTCGTTGCCCTAGAAAGAATGAGTTGAGTNNNAGATCGGAAGA
128175 TATGAGTGTGGAGTCGTTGCCCTAGAAAGAATGAGTTGAGTNNNAGATCGGAAGA
128176 TATGAGTGTGGAGTCGTTGCAATAAGATGCATTCGACAAAAATNNNAGATCGGAA
128177 TATGAGTGTGGAGTCGTTGCAATAAGATGCATTCGACAAAAATNNNAGATCGGAA
128178 TATGAGTGTGGAGTCGTTGCGAGAACAGGAAAGAAGAGAATNNNAGATCGGAAG
128179 TATGAGTGTGGAGTCGTTGCCTAATTCTTTTTCCTCTCCCTGNNNAGATCGGAAGAG
128180 TATGAGTGTGGAGTCGTTGCCTAATTCTTTTTCCTCTCCCTGNNNAGATCGGAAGAG
128181 TATGAGTGTGGAGTCGTTGCTATTTGTATGCATGCATGTGTATNNNAGATCGGAAG
128182 TATGAGTGTGGAGTCGTTGCAGGGAGAGAGAGAAGGAAAANNNAGATCGGAAG
128183 TATGAGTGTGGAGTCGTTGCACCTTTGTAACTAATCATTATAGCNNNAGATCGGAA
128184 TATGAGTGTGGAGTCGTTGCCTTTCTTTGGTACTACCCATTNNNAGATCGGAAGAG
128185 TATGAGTGTGGAGTCGTTGCTATTATGGAGAATTTCAAGCATGNNNAGATCGGAA
128186 TATGAGTGTGGAGTCGTTGCCTGTAAAGCTATTCCTCACCNNNAGATCGGAAGAGC
128187 TATGAGTGTGGAGTCGTTGCCTGTAAAGCTATTCCTCACCNNNAGATCGGAAGAGC
128188 TATGAGTGTGGAGTCGTTGCTACACTCATAATACATAGTGAGGNNNAGATCGGAAG
128189 TATGAGTGTGGAGTCGTTGCATTTTACTACACAGAAACAAACTNNNAGATCGGAAG
128190 TATGAGTGTGGAGTCGTTGCTGAATTTAGGGTTACTGATAACANNNAGATCGGAA
128191 TATGAGTGTGGAGTCGTTGCTTTGGTCTATGCAGTACATAAAANNNAGATCGGAAG
128192 TATGAGTGTGGAGTCGTTGCATCTCTGCAATTTTCTTAAAACANNNAGATCGGAAGA
128193 TATGAGTGTGGAGTCGTTGCAAAGAAGAAAAATCTGCCATTTANNNAGATCGGAA
128194 TATGAGTGTGGAGTCGTTGCAGAGGGACAGAGAGATATAAAANNNAGATCGGAA
128195 TATGAGTGTGGAGTCGTTGCCATACTTGGAATAGTCGATCCNNNAGATCGGAAGAG
128196 TATGAGTGTGGAGTCGTTGCGCTCATATGCATAAGTAGTTTTTNNNAGATCGGAAG
128197 TATGAGTGTGGAGTCGTTGCTATCTCCTTCTTTCCTGTATTACNNNAGATCGGAAGA
128198 TATGAGTGTGGAGTCGTTGCGGGAAATGAAAGACACAGAGNNNAGATCGGAAGA
128199 TATGAGTGTGGAGTCGTTGCAAGGTTAGGGAACATAGTCATANNNAGATCGGAAG
128200 TATGAGTGTGGAGTCGTTGCCCTCTGTGTGAACTAGAAAAANNNAGATCGGAAGA
128201 TATGAGTGTGGAGTCGTTGCCAATGATGGAAACTGGGAAGNNNAGATCGGAAGA
128203 TATGAGTGTGGAGTCGTTGCCTGATGTAGAAGATGGTGTTGNNNAGATCGGAAGA
128204 TATGAGTGTGGAGTCGTTGCTACAATCAAACAAACCATTTTTGNNNAGATCGGAAG
128205 TATGAGTGTGGAGTCGTTGCTATACAGTAGAGGCAGCAAAANNNAGATCGGAAGA
128206 TATGAGTGTGGAGTCGTTGCTATACCTTGGTAACAAGTTGTATNNNAGATCGGAAG
128207 TATGAGTGTGGAGTCGTTGCATATCCAGGATTTATTTCCAGAGNNNAGATCGGAAG
128208 TATGAGTGTGGAGTCGTTGCCTCATAGAAATGTACCTCTTTTTNNNAGATCGGAAGA
128209 TATGAGTGTGGAGTCGTTGCTGTTAAAATAATTGGAAGAGCTCNNNAGATCGGAA
128210 TATGAGTGTGGAGTCGTTGCTTTGCTATTCAATCACCCTTCNNNAGATCGGAAGAGC
128211 TATGAGTGTGGAGTCGTTGCATGCATATATACACATACACACTNNNAGATCGGAAG
128212 TATGAGTGTGGAGTCGTTGCAAAAACTAGAACCAGAACCAGNNNAGATCGGAAGA
128213 TATGAGTGTGGAGTCGTTGCATAGAAAGAATGTTAGTCCCTTTNNNAGATCGGAAG
128214 TATGAGTGTGGAGTCGTTGCTAGAATTCATCTTTTGACCCTACNNNAGATCGGAAGA
128215 TATGAGTGTGGAGTCGTTGCAAATTAAAATGCATATTTCAGCCNNNAGATCGGAAG
128216 TATGAGTGTGGAGTCGTTGCTATATGACCTTGGGCAAATACNNNAGATCGGAAGAG
128217 TATGAGTGTGGAGTCGTTGCTCTTTTCTTATATTCTCTCTGGGNNNAGATCGGAAGA

128218 TATGAGTGTGGAGTCGTTGCGAGGAGCTCTACTTTGAAAATNNNAGATCGGAAGA
128219 TATGAGTGTGGAGTCGTTGCCAAAAGATATTGAAGCACAAATTNNNAGATCGGAAG
128220 TATGAGTGTGGAGTCGTTGCCAAAAGATATTGAAGCACAAATTNNNAGATCGGAAG
128221 TATGAGTGTGGAGTCGTTGCTGTAATGCACAGATTTATTTCTGNNNAGATCGGAAG
128222 TATGAGTGTGGAGTCGTTGCTTCCGAAGTTCTTATATTTCTGANNNAGATCGGAAG
128223 TATGAGTGTGGAGTCGTTGCATTTTCCACACTCTAACTGTCNNNAGATCGGAAGAGC
128224 TATGAGTGTGGAGTCGTTGCAATACTTTCATTCATTCTGCTTTNNNAGATCGGAAGA
128225 TATGAGTGTGGAGTCGTTGCGATAGCTTTAGAGTTGATTTGTTNNNAGATCGGAAG
128226 TATGAGTGTGGAGTCGTTGCAGCTAGGAAGTGTAAATATATGAANNNAGATCGGA
128227 TATGAGTGTGGAGTCGTTGCTTTGGTGGGAAATACTATAATCTNNNAGATCGGAAG
128228 TATGAGTGTGGAGTCGTTGCTGCAAGTATAATTAACTCCATTTNNNAGATCGGAAG
128229 TATGAGTGTGGAGTCGTTGCGCTGGAACAGAAATAACTTATAGNNNAGATCGGAA
128230 TATGAGTGTGGAGTCGTTGCCTTTCTTATTTGGGTGCTCCNNNAGATCGGAAGAGC
128231 TATGAGTGTGGAGTCGTTGCAATGAGAGAGAGGAACAGAANNNAGATCGGAAGA
128232 TATGAGTGTGGAGTCGTTGCATAGTCTAAAGAGATTTGGATCANNNAGATCGGAA
128235 TATGAGTGTGGAGTCGTTGCCTAGCATCCTAATCTCCAACNNNAGATCGGAAGAGC
128236 TATGAGTGTGGAGTCGTTGCACTATTTCATATACATGAGTGCCNNNAGATCGGAAG
128237 TATGAGTGTGGAGTCGTTGCGTGAGCTGAATTTAGTTATAATCANNNAGATCGGAA
128238 TATGAGTGTGGAGTCGTTGCTGTGTATATATATATGTATGTCTGGANNNAGATCGG
128239 TATGAGTGTGGAGTCGTTGCATTCTCTCCTATCTCAGTAAAAGNNNAGATCGGAAG
128241 TATGAGTGTGGAGTCGTTGCTATACTCCAATACATACACATCCNNNAGATCGGAAGA
128242 TATGAGTGTGGAGTCGTTGCCTCTGTCTCTCTCTCTCTCTNNNAGATCGGAAGAGCA
128243 TATGAGTGTGGAGTCGTTGCACAAAACGGTAATGTAAACATTANNNAGATCGGAAG
128244 TATGAGTGTGGAGTCGTTGCGTTTGTTTTTGTATTTGTCTCGNNNAGATCGGAAGA
128245 TATGAGTGTGGAGTCGTTGCATTCATTCTCCATAGCATTCTAANNNAGATCGGAAGA
128246 TATGAGTGTGGAGTCGTTGCAAATTTCAGTTAGCATAGTTGTGNNNAGATCGGAAG
128247 TATGAGTGTGGAGTCGTTGCGTCATACAGGCTAAACTGTAANNNAGATCGGAAGA
128248 TATGAGTGTGGAGTCGTTGCAAATAGCATTTAGAATTTCGGAANNNAGATCGGAAG
128249 TATGAGTGTGGAGTCGTTGCGATACAGTCTCACTTTGTGGNNNAGATCGGAAGAG
128250 TATGAGTGTGGAGTCGTTGCAATAGCAATACACAGGCTTATATNNNAGATCGGAAG
128252 TATGAGTGTGGAGTCGTTGCTTTTAGAAATGAAAGAGTGAGTGNNNAGATCGGAA
128253 TATGAGTGTGGAGTCGTTGCGGTAAATTCAGCTAGAGGATAATNNNAGATCGGAA
128254 TATGAGTGTGGAGTCGTTGCGTTGAGTTGTAGGAGTTCTTATANNNAGATCGGAA
128255 TATGAGTGTGGAGTCGTTGCTAAGTTGATCCTCCTATCTCAGNNNAGATCGGAAGA
128256 TATGAGTGTGGAGTCGTTGCCAGTCCTTGGAATAAGATCAGNNNAGATCGGAAGA
128257 TATGAGTGTGGAGTCGTTGCAATATGCTCAGCAGAGGTAGNNNAGATCGGAAGAG
128258 TATGAGTGTGGAGTCGTTGCAAATAAAAGCCCATTTATCTGTCNNNAGATCGGAAG
128259 TATGAGTGTGGAGTCGTTGCCTTGAGTTACTGTAGCAGAATNNNAGATCGGAAGA
128260 TATGAGTGTGGAGTCGTTGCAGAGACTGCCATTATAATGTATTNNNAGATCGGAAG
128261 TATGAGTGTGGAGTCGTTGCCAAATCAGACCTTCCTGTAGNNNAGATCGGAAGAGC
128262 TATGAGTGTGGAGTCGTTGCTTTCCAAAGATCTCTAGATAAGGNNNAGATCGGAAG
128263 TATGAGTGTGGAGTCGTTGCACCTAGTATTTTATTTTTAGTGCCNNNAGATCGGAAG
128264 TATGAGTGTGGAGTCGTTGCGGGTATATGTGTGTGTGTATATANNNAGATCGGAA

128265 TATGAGTGTGGAGTCGTTGCAAGAAACTCTGTCCTTACCTATNNNAGATCGGAAGA
128266 TATGAGTGTGGAGTCGTTGCCCTCTACATGTAAATGAACAATCNNNAGATCGGAAG
128267 TATGAGTGTGGAGTCGTTGCCCTCTACATGTAAATGAACAATCNNNAGATCGGAAG
128268 TATGAGTGTGGAGTCGTTGCGAATGTGCTTCTAAATTATTTGGNNNAGATCGGAAG
128269 TATGAGTGTGGAGTCGTTGCCATCTCAGCCTTTCAAGTAGNNNAGATCGGAAGAGC
128270 TATGAGTGTGGAGTCGTTGCACACATGTGTACACGTTTTCNNNAGATCGGAAGAGC
128271 TATGAGTGTGGAGTCGTTGCTCCCAGAATCTTTTTTAAACAAANNNAGATCGGAAG
128272 TATGAGTGTGGAGTCGTTGCTATAATTTGACACTTTGGAAGACNNNAGATCGGAAG
128273 TATGAGTGTGGAGTCGTTGCGATTTCAATAGTGTGGATGATTCNNNAGATCGGAAG
128274 TATGAGTGTGGAGTCGTTGCTAGCTATGCATACAGAAATTGAANNNAGATCGGAAG
128275 TATGAGTGTGGAGTCGTTGCAAGGATGAGGATGATAAGCTNNNAGATCGGAAGA
128277 TATGAGTGTGGAGTCGTTGCAGAAGTGTTTTAGGATCCTTAATNNNAGATCGGAAG
128278 TATGAGTGTGGAGTCGTTGCGGTGCTCTTTAAATGATATTGATNNNAGATCGGAAG
128279 TATGAGTGTGGAGTCGTTGCGAATAAGGTTTATCCCAAGAATGNNNAGATCGGAAG
128280 TATGAGTGTGGAGTCGTTGCTGGTATTTTGGTTTTTCAGAATANNNAGATCGGAAG
128281 TATGAGTGTGGAGTCGTTGCCTTTATATTTTGCCAGGACCTNNNAGATCGGAAGAG
128282 TATGAGTGTGGAGTCGTTGCTCCTGTATACTATTAATTCTGCANNNAGATCGGAAGA
128283 TATGAGTGTGGAGTCGTTGCTCCTGTATACTATTAATTCTGCANNNAGATCGGAAGA
128284 TATGAGTGTGGAGTCGTTGCAGAATACCTCTAGAGTTTAGAGANNNAGATCGGAAG
128285 TATGAGTGTGGAGTCGTTGCAGAATACCTCTAGAGTTTAGAGANNNAGATCGGAAG
128288 TATGAGTGTGGAGTCGTTGCTTTGAAAGACAATCCCTCTCTNNNAGATCGGAAGAG
128289 TATGAGTGTGGAGTCGTTGCAGGCCCCCTATAATTTATTATTTNNNAGATCGGAAGA
128290 TATGAGTGTGGAGTCGTTGCACAGAAAAACATCAAAATCTTCANNNAGATCGGAAG
128291 TATGAGTGTGGAGTCGTTGCACAGAAAAACATCAAAATCTTCANNNAGATCGGAAG
128292 TATGAGTGTGGAGTCGTTGCTGTTCACTTCAGTATGCAAANNNAGATCGGAAGAGC
128293 TATGAGTGTGGAGTCGTTGCATAGTGAATGAGTTCTCTTGAAANNNAGATCGGAAG
128294 TATGAGTGTGGAGTCGTTGCATTTATATAAAATCCCCCTTCCANNNAGATCGGAAGA
128295 TATGAGTGTGGAGTCGTTGCATGCTAAGACACCTATGAATATTNNNAGATCGGAAG
128296 TATGAGTGTGGAGTCGTTGCGCTATAAACTGAGATTTCACAAANNNAGATCGGAAG
128297 TATGAGTGTGGAGTCGTTGCCCATTATCGTATTCACTTCTTTGNNNAGATCGGAAGA
128298 TATGAGTGTGGAGTCGTTGCGTACATACACATACATACATCCCNNNAGATCGGAAGA
128299 TATGAGTGTGGAGTCGTTGCTTAGGTTTCTATTATCTTCTCCCNNNAGATCGGAAGA
128300 TATGAGTGTGGAGTCGTTGCCTGGAGATGTGTTTTATACATTCNNNAGATCGGAAG
128301 TATGAGTGTGGAGTCGTTGCACTGACTTGTCATACTATTCTTANNNAGATCGGAAGA
128302 TATGAGTGTGGAGTCGTTGCCAGAATGGACATAAGCAAATCNNNAGATCGGAAGA
128303 TATGAGTGTGGAGTCGTTGCAATGTTCTAAGTCAGAAGGTAATNNNAGATCGGAAG
128304 TATGAGTGTGGAGTCGTTGCCTTGCTCATGTATTAGGAATTATNNNAGATCGGAAG
128305 TATGAGTGTGGAGTCGTTGCTTACCAGACTATCTATCTTTGACNNNAGATCGGAAGA
128306 TATGAGTGTGGAGTCGTTGCCCACTCTAAGAAGATAAGATTTGNNNAGATCGGAAG
128307 TATGAGTGTGGAGTCGTTGCATAGGAGCAGTATCAGAGGAANNNAGATCGGAAGA
128308 TATGAGTGTGGAGTCGTTGCCTATGGTGAGACAAAAAAGAAATNNNAGATCGGAA
128309 TATGAGTGTGGAGTCGTTGCCTATGGTGAGACAAAAAAGAAATNNNAGATCGGAA
128310 TATGAGTGTGGAGTCGTTGCCTATGGTGAGACAAAAAAGAAATNNNAGATCGGAA

128311 TATGAGTGTGGAGTCGTTGCCCAAGTAGCTGGGATTATAGNNNAGATCGGAAGAG
128312 TATGAGTGTGGAGTCGTTGCGATAAATGCAACCTGTTCTAACNNNAGATCGGAAGA
128313 TATGAGTGTGGAGTCGTTGCTTAAGAAGAAAGACAGTGATGAANNNAGATCGGAA
128314 TATGAGTGTGGAGTCGTTGCTAATGCAAGAAAACACTACTATGNNNAGATCGGAAG
128315 TATGAGTGTGGAGTCGTTGCATATTTTACCCAGGCAATTTACTNNNAGATCGGAAGA
128316 TATGAGTGTGGAGTCGTTGCCTAGAATTAAGCTGAAATTCACANNNAGATCGGAAG
128317 TATGAGTGTGGAGTCGTTGCCTAGAATTAAGCTGAAATTCACANNNAGATCGGAAG
128318 TATGAGTGTGGAGTCGTTGCTAGTACTTATTTCTGGCATGTANNNAGATCGGAAGA
128319 TATGAGTGTGGAGTCGTTGCTTTTCCCTTGCTATTTCATCANNNAGATCGGAAGAGCA
128320 TATGAGTGTGGAGTCGTTGCCTATGCAATTTTTGGAGGATATTNNNAGATCGGAAG
128321 TATGAGTGTGGAGTCGTTGCCTAGAGAGTGAAACAGATTTAGANNNAGATCGGAA
128322 TATGAGTGTGGAGTCGTTGCACCAAATATTTACAGAAGAATCANNNAGATCGGAAG
128323 TATGAGTGTGGAGTCGTTGCGTATCAGGCTATTTATCTTTTCCNNNAGATCGGAAGA
128324 TATGAGTGTGGAGTCGTTGCAGGTTAACAATTAAGTGTTTTCANNNAGATCGGAAG
128325 TATGAGTGTGGAGTCGTTGCCAAGAGAGACAGAAAGTAGAGNNNAGATCGGAAGA
128327 TATGAGTGTGGAGTCGTTGCCATGGACAGGGATTAATAGATAANNNAGATCGGAA
128328 TATGAGTGTGGAGTCGTTGCGTTAGTCATTACACTTACATACGNNNAGATCGGAAGA
128329 TATGAGTGTGGAGTCGTTGCTGAAGAGAGCTATAAAAAGAAGANNNAGATCGGAA
128330 TATGAGTGTGGAGTCGTTGCGTTACTGGTTTATTGGGAAAATTNNNAGATCGGAAG
128331 TATGAGTGTGGAGTCGTTGCACAGACTACAATAAAACAGAAATAANNNAGATCGGA
128332 TATGAGTGTGGAGTCGTTGCTACTTAGAACACAAAGAAAAAGCNNNAGATCGGAAG
128333 TATGAGTGTGGAGTCGTTGCGAACCTTCCTCATAGAGTTTTNNNAGATCGGAAGAG
128334 TATGAGTGTGGAGTCGTTGCGATAAGATAACAGTGGTGAATTTNNNAGATCGGAA
128335 TATGAGTGTGGAGTCGTTGCATATTCTCCATATAGTCTTCCCANNNAGATCGGAAGA
128336 TATGAGTGTGGAGTCGTTGCATGAAACCCTCAGAGAGTAGNNNAGATCGGAAGAG
128337 TATGAGTGTGGAGTCGTTGCCTCACTCTGTTACCTAGAGTNNNAGATCGGAAGAGCA
128338 TATGAGTGTGGAGTCGTTGCCTAAAAGATGACCGGTATGTTNNNAGATCGGAAGA
128339 TATGAGTGTGGAGTCGTTGCGAGATTACAGGTAGGGAGAGNNNAGATCGGAAGA
128340 TATGAGTGTGGAGTCGTTGCAGAAACAGAGACAGAGTGAGNNNAGATCGGAAGA
128341 TATGAGTGTGGAGTCGTTGCAGAAACAGAGACAGAGTGAGNNNAGATCGGAAGA
128342 TATGAGTGTGGAGTCGTTGCCTCACTTAGGATCCAAAATTGTANNNAGATCGGAAG
128343 TATGAGTGTGGAGTCGTTGCCATGTGTGAGTGAAGAGAGANNNAGATCGGAAGA
128344 TATGAGTGTGGAGTCGTTGCATAGGGCAGATTCATCAAATATTNNNAGATCGGAAG
128345 TATGAGTGTGGAGTCGTTGCATCTCTCTCTCTCTCTTTTTTTTNNNAGATCGGAAGAG
128346 TATGAGTGTGGAGTCGTTGCTTGTTTTGTTTTGTTTTTTTCCANNNAGATCGGAAGA
128347 TATGAGTGTGGAGTCGTTGCCTTTTACTCTCTTTCTGTCTCTTNNNAGATCGGAAGAG
128348 TATGAGTGTGGAGTCGTTGCCTTTTACTCTCTTTCTGTCTCTTNNNAGATCGGAAGAG
128349 TATGAGTGTGGAGTCGTTGCCTATACTGTTGACACAAAAGAAANNNAGATCGGAAG
128350 TATGAGTGTGGAGTCGTTGCCAGAAACCTCATAATCTTTGTTANNNAGATCGGAAGA
128351 TATGAGTGTGGAGTCGTTGCTACAATAGTACAGGAGGGAAGNNNAGATCGGAAGA
128352 TATGAGTGTGGAGTCGTTGCATACATATTTTATATATGTGTGTGTGTNNNAGATCGG
128353 TATGAGTGTGGAGTCGTTGCGAAAGCTGCAAATTGATATTTGNNNAGATCGGAAGA
128354 TATGAGTGTGGAGTCGTTGCCACATCCTATCATTTTTGTTTCTNNNAGATCGGAAGA

128355 TATGAGTGTGGAGTCGTTGCCAGCATTGATAAAAAGGAAAATGNNNAGATCGGAA
128356 TATGAGTGTGGAGTCGTTGCATGCTCATGGTATATTTTCCCNNNAGATCGGAAGAG
128357 TATGAGTGTGGAGTCGTTGCGTCTTCCTGATCTTAAAGGAANNNAGATCGGAAGAG
128358 TATGAGTGTGGAGTCGTTGCCTAGAGCATCCCAGTAATAAAANNNAGATCGGAAGA
128359 TATGAGTGTGGAGTCGTTGCTGAATATAGTGAATGTATTTGCANNNAGATCGGAAG
128360 TATGAGTGTGGAGTCGTTGCTAAAGTGCCAACAGATAGAAGNNNAGATCGGAAGA
128362 TATGAGTGTGGAGTCGTTGCTGTGTCAGGAGAATGAGTAGNNNAGATCGGAAGAG
128363 TATGAGTGTGGAGTCGTTGCGTGTGTCAGAAGAATGAGTAGNNNAGATCGGAAGA
128364 TATGAGTGTGGAGTCGTTGCCTTTCAGTCATAACTTTTCAGATNNNAGATCGGAAGA
128365 TATGAGTGTGGAGTCGTTGCTACATACATATAAACTCACAGCANNNAGATCGGAAG
128366 TATGAGTGTGGAGTCGTTGCGACTAAAGAGCCTAAATTCTCATNNNAGATCGGAAG
128367 TATGAGTGTGGAGTCGTTGCCTACTGGGTACAATATTCACTATNNNAGATCGGAAG
128368 TATGAGTGTGGAGTCGTTGCCTTGTCCTTAACATTGTCAATGNNNAGATCGGAAGA
128369 TATGAGTGTGGAGTCGTTGCGGTATGTCTATGTCTGAGTTTGNNNAGATCGGAAGA
128370 TATGAGTGTGGAGTCGTTGCCCTCAAATCACCAAAACAACNNNAGATCGGAAGAGC
128371 TATGAGTGTGGAGTCGTTGCCATATATTCCCTACTTCACTGATNNNAGATCGGAAGA
128372 TATGAGTGTGGAGTCGTTGCAGTTATACAAGATGCTAACATTGNNNAGATCGGAAG
128373 TATGAGTGTGGAGTCGTTGCGCTATATTCTCCAAGGTGAAANNNAGATCGGAAGAG
128374 TATGAGTGTGGAGTCGTTGCGTTCAAGAATTTCTCCAGTAACNNNAGATCGGAAGA
128375 TATGAGTGTGGAGTCGTTGCACATTCATAGTAAACAATTCACANNNAGATCGGAAG
128376 TATGAGTGTGGAGTCGTTGCTATATGTTGGCTGTTGTGAATANNNAGATCGGAAGA
128377 TATGAGTGTGGAGTCGTTGCAAAATAAAATGCTTCCCCAATAANNNAGATCGGAAG
128378 TATGAGTGTGGAGTCGTTGCGTTATAGGAACCCATCCAATTNNNAGATCGGAAGAG
128380 TATGAGTGTGGAGTCGTTGCGAAGCACCAAATTGGAGAAANNNAGATCGGAAGAG
128381 TATGAGTGTGGAGTCGTTGCCTTCCCTTTTTTTTTTTTTTTGTNNNAGATCGGAAGAG
128382 TATGAGTGTGGAGTCGTTGCTTCTAAGGTAGAGCTGACTTTATNNNAGATCGGAAG
128383 TATGAGTGTGGAGTCGTTGCCTGATAACAACAGGTTTTGTGNNNAGATCGGAAGAG
128384 TATGAGTGTGGAGTCGTTGCAAAGAAAGAAGGTGAAGAAGAANNNAGATCGGAA
128385 TATGAGTGTGGAGTCGTTGCCCCTGTCACTCTCTCTATATATANNNAGATCGGAAGA
128386 TATGAGTGTGGAGTCGTTGCGGATCAAAGTGTTTACTTCATATNNNAGATCGGAAG
128387 TATGAGTGTGGAGTCGTTGCTTTTGAATTTGTATCCCTAAACCNNNAGATCGGAAGA
128388 TATGAGTGTGGAGTCGTTGCGGCTTGTAGGTTATTTATTACTAANNNAGATCGGAA
128389 TATGAGTGTGGAGTCGTTGCGTAGGGGTATGAAATCTGATAAANNNAGATCGGAA
128390 TATGAGTGTGGAGTCGTTGCGGCTATTACCTCTATAGAACTTCNNNAGATCGGAAG
128391 TATGAGTGTGGAGTCGTTGCGAAAGCTGCAAATTGATATTTGNNNAGATCGGAAG
128392 TATGAGTGTGGAGTCGTTGCTTTTTCTGATTATCCAAGAACTGNNNAGATCGGAAG
128393 TATGAGTGTGGAGTCGTTGCATCATTTCTTTCTGTTATTGCTCNNNAGATCGGAAGA
128394 TATGAGTGTGGAGTCGTTGCGTAGACATTCACAACTTCTTTGNNNAGATCGGAAGA
128395 TATGAGTGTGGAGTCGTTGCCAGAGACACTGATTTTTTCATTNNNAGATCGGAAGA
128396 TATGAGTGTGGAGTCGTTGCCAGAGACACTGATTTTTTCATTNNNAGATCGGAAGA
128397 TATGAGTGTGGAGTCGTTGCCAAATTTGTACTGGGAAAAATTTNNNAGATCGGAAG
128398 TATGAGTGTGGAGTCGTTGCGGAAAAGGCACAAAAGTTTTNNNAGATCGGAAGAG
128399 TATGAGTGTGGAGTCGTTGCCTTTCCAATTTCCAGACATTCNNNAGATCGGAAGAG

128400 TATGAGTGTGGAGTCGTTGCCTTTCGAAAACCAAGGAGAANNNAGATCGGAAGAG
128401 TATGAGTGTGGAGTCGTTGCGCACAGAATAAGCACTGAATAANNNAGATCGGAAG
128402 TATGAGTGTGGAGTCGTTGCCTGGATATGCTATACATATGACANNNAGATCGGAAG
128403 TATGAGTGTGGAGTCGTTGCCCCATAGGAAAATAATGTTCAAANNNAGATCGGAAG
128404 TATGAGTGTGGAGTCGTTGCTTCATCAATACATTTGTGATTGANNNAGATCGGAAG
128405 TATGAGTGTGGAGTCGTTGCGAGTGATGTAGTGATGTGATCNNNAGATCGGAAGA
128406 TATGAGTGTGGAGTCGTTGCGTTTGAATGCTTGACTAATCTATNNNAGATCGGAAG
128407 TATGAGTGTGGAGTCGTTGCGCAAAATTTCCACTAAATTAAGAANNNAGATCGGAA
128408 TATGAGTGTGGAGTCGTTGCATACCATTGCAGTTTGAATAAAANNNAGATCGGAAG
128409 TATGAGTGTGGAGTCGTTGCATACCATTGCAGTTTGAATAAAANNNAGATCGGAAG
128410 TATGAGTGTGGAGTCGTTGCCTCTATTATGAGACTGTAGTCAANNNAGATCGGAAG
128411 TATGAGTGTGGAGTCGTTGCTATATGGCTTTAAATGGGTTACTNNNAGATCGGAAG
128412 TATGAGTGTGGAGTCGTTGCTTTGTAAATTCAGCACAATATGANNNAGATCGGAAG
128413 TATGAGTGTGGAGTCGTTGCGATAGATTCATCATCCAAAATGGNNNAGATCGGAAG
128414 TATGAGTGTGGAGTCGTTGCATTAGGAAACTGTAAAAGACACTNNNAGATCGGAAG
128415 TATGAGTGTGGAGTCGTTGCCTGGCAACTCAGTAATAAAAAGNNNAGATCGGAAG
128416 TATGAGTGTGGAGTCGTTGCTATCTAAGGACAGAGGGTAATATNNNAGATCGGAA
128417 TATGAGTGTGGAGTCGTTGCAGGAATGGAATACTTTAATCACTNNNAGATCGGAAG
128418 TATGAGTGTGGAGTCGTTGCTATTATTATTGAGTGCAGGTACTNNNAGATCGGAAG
128419 TATGAGTGTGGAGTCGTTGCAACTGTGGAAACAAGTTTAAAANNNAGATCGGAAG
128420 TATGAGTGTGGAGTCGTTGCGTATTTAAATTCACATACAGGCTNNNAGATCGGAAG
128421 TATGAGTGTGGAGTCGTTGCTTTAGAGAATTGGCTCATATCATNNNAGATCGGAAG
128422 TATGAGTGTGGAGTCGTTGCATCCAGTGAACAAGTTGAGATNNNAGATCGGAAGA
128424 TATGAGTGTGGAGTCGTTGCATAACTTCTTCATTTGAAATCCCNNNAGATCGGAAGA
128425 TATGAGTGTGGAGTCGTTGCCTTTGTTTCTGGTGATCTTTGNNNAGATCGGAAGAG
128426 TATGAGTGTGGAGTCGTTGCTGCATTATATCAAATGGCTAAAANNNAGATCGGAAG
128427 TATGAGTGTGGAGTCGTTGCTAAGTCACTTGCTAAACACTTNNNAGATCGGAAGAG
128428 TATGAGTGTGGAGTCGTTGCATATTATGTTTTGGGGTTTCTCNNNAGATCGGAAG
128429 TATGAGTGTGGAGTCGTTGCGTGGAAATACAAGGGATTTTTATNNNAGATCGGAA
128430 TATGAGTGTGGAGTCGTTGCACACTCTTTTTCTGGCTTTTANNNAGATCGGAAGAG
128431 TATGAGTGTGGAGTCGTTGCACACTCTTTTTCTGGCTTTTANNNAGATCGGAAGAG
128432 TATGAGTGTGGAGTCGTTGCAAAAGTATGTCACATTTTAGTGGNNNAGATCGGAAG
128433 TATGAGTGTGGAGTCGTTGCGATTTTTTAGGCCAAATTTTCAANNNAGATCGGAAG
128434 TATGAGTGTGGAGTCGTTGCATTCTAGTCAATCAACTCTATCCNNNAGATCGGAAGA
128435 TATGAGTGTGGAGTCGTTGCAGATGAAGTCTCATTCTGTCANNNAGATCGGAAGAG
128436 TATGAGTGTGGAGTCGTTGCTCTAACCATGTTGTCTAGGATATNNNAGATCGGAAG
128437 TATGAGTGTGGAGTCGTTGCTTATTTACCTAAATGTCCAGAGCNNNAGATCGGAAG
128438 TATGAGTGTGGAGTCGTTGCCAAAATTTACGAGAGAATTTCAGNNNAGATCGGAAG
128440 TATGAGTGTGGAGTCGTTGCAGATGGAAAATAGGCTTGTAGNNNAGATCGGAAGA
128441 TATGAGTGTGGAGTCGTTGCGAGTAATAATAGAGAGTAGCGAGNNNAGATCGGAA
128442 TATGAGTGTGGAGTCGTTGCGAGTAAACAACTTTTTCCCAATTNNNAGATCGGAAG
128443 TATGAGTGTGGAGTCGTTGCGTTGTTTATATGGTAGACACTTTNNNAGATCGGAAG
128444 TATGAGTGTGGAGTCGTTGCATAAAGATAGAGGAATGCTATGANNNAGATCGGAA

128446 TATGAGTGTGGAGTCGTTGCCTTGAGGGATAACAAGAAAAATANNNAGATCGGAA
128447 TATGAGTGTGGAGTCGTTGCCTGTCTGAAAAACATGGAATTTNNNAGATCGGAAGA
128448 TATGAGTGTGGAGTCGTTGCAGATTGGCTTAATATGAACTAGANNNAGATCGGAAG
128449 TATGAGTGTGGAGTCGTTGCAGTTATGCTCTTTCTCCCAGNNNAGATCGGAAGAGC
128450 TATGAGTGTGGAGTCGTTGCATGGCATAGTATTTGCATATAACNNNAGATCGGAAG
128451 TATGAGTGTGGAGTCGTTGCATGGCATAGTATTTGCATATAACNNNAGATCGGAAG
128453 TATGAGTGTGGAGTCGTTGCATAGTAAGTATCTTCGGAATGAANNNAGATCGGAAG
128454 TATGAGTGTGGAGTCGTTGCATAGTAAGTATCTTCGGAATGAANNNAGATCGGAAG
128455 TATGAGTGTGGAGTCGTTGCACCAACCACAAATCAAAAATATTNNNAGATCGGAAG
128456 TATGAGTGTGGAGTCGTTGCAAATGAAAGTCATAATAATGGCANNNAGATCGGAA
128457 TATGAGTGTGGAGTCGTTGCATTCACAAGTATTCCAAAGTAGTNNNAGATCGGAAG
128458 TATGAGTGTGGAGTCGTTGCATAATATACACAGCATACCTCAGNNNAGATCGGAAG
128459 TATGAGTGTGGAGTCGTTGCCAGTCTGTTTTTGTCTCTACTNNNAGATCGGAAGAG
128460 TATGAGTGTGGAGTCGTTGCCATTGATGGCTATCTTAAACATTNNNAGATCGGAAG
128461 TATGAGTGTGGAGTCGTTGCATTCCATCTCCCAAATTCAAATNNNAGATCGGAAGAG
128462 TATGAGTGTGGAGTCGTTGCGTAGGTTGATTGATTGATTGATTNNNAGATCGGAA
128463 TATGAGTGTGGAGTCGTTGCCTGTAGCTTACAATAAAATAGCANNNAGATCGGAAG
128464 TATGAGTGTGGAGTCGTTGCTCAACTTTGGCAGAAAATATAAANNNAGATCGGAAG
128465 TATGAGTGTGGAGTCGTTGCCCACTTTCTTTTCGTCTTTGNNNAGATCGGAAGAGCA
128466 TATGAGTGTGGAGTCGTTGCTTGAGGATTTGAGACAGTAGTAANNNAGATCGGAA
128467 TATGAGTGTGGAGTCGTTGCTGTATATATATGTGTGTGCGTATNNNAGATCGGAAG
128468 TATGAGTGTGGAGTCGTTGCGTTCTATCACAATTTTGGCTAAGNNNAGATCGGAAG
128469 TATGAGTGTGGAGTCGTTGCGTTCTATCACAATTTTGGCTAAGNNNAGATCGGAAG
128470 TATGAGTGTGGAGTCGTTGCCTCTGTTTTCCTAATGTGTACANNNAGATCGGAAGAG
128471 TATGAGTGTGGAGTCGTTGCATAGTGTCACTGTCTTTTCTTTTNNNAGATCGGAAGA
128472 TATGAGTGTGGAGTCGTTGCAACATGATATACATTTCTGGAGANNNAGATCGGAAG
128473 TATGAGTGTGGAGTCGTTGCTATTACACTGTGTCAGACCTNNNAGATCGGAAGAGC
128474 TATGAGTGTGGAGTCGTTGCAAGATATATAGGACAGTTGTTGGNNNAGATCGGAA
128475 TATGAGTGTGGAGTCGTTGCAAGATATATAGGACAGTTGTTGGNNNAGATCGGAA
128476 TATGAGTGTGGAGTCGTTGCGAGGAGAGCATACTGAAATGNNNAGATCGGAAGAG
128477 TATGAGTGTGGAGTCGTTGCGTTCAAGTTATTCTCCTGTCTNNNAGATCGGAAGAG
128478 TATGAGTGTGGAGTCGTTGCTCCTTTTTCTGAATAAGTGAATGNNNAGATCGGAAG
128479 TATGAGTGTGGAGTCGTTGCCTTACAAATGCAAATCAAAGTTTNNNAGATCGGAAG
128480 TATGAGTGTGGAGTCGTTGCGCCTAAAATCAGCAATCTTCNNNAGATCGGAAGAGC
128481 TATGAGTGTGGAGTCGTTGCCTTTTAGTCTATAGCCGTCTTCNNNAGATCGGAAGAG
128482 TATGAGTGTGGAGTCGTTGCAGACTGTGTAGACAAAATATGAGNNNAGATCGGAA
128483 TATGAGTGTGGAGTCGTTGCGTTAAATCACCAGATGTCTAAAANNNAGATCGGAAG
128484 TATGAGTGTGGAGTCGTTGCTAGAAGACCTATATGTTAGAGCTNNNAGATCGGAAG
128485 TATGAGTGTGGAGTCGTTGCCTAGAGTGCAGTGGGATAACNNNAGATCGGAAGAG
128486 TATGAGTGTGGAGTCGTTGCATATGGGCTCTCACTTTGATATNNNAGATCGGAAGA
128489 TATGAGTGTGGAGTCGTTGCATAGAACATTGAGCCTAAAAAAGNNNAGATCGGAA
128490 TATGAGTGTGGAGTCGTTGCATTCTAGGTAAATGTCAGATTCCNNNAGATCGGAAG
128491 TATGAGTGTGGAGTCGTTGCCAATTATTTCTGCCATGCTCNNNAGATCGGAAGAGC

128494 TATGAGTGTGGAGTCGTTGCTACATAATCACAAACACTGGATTNNNAGATCGGAAG
128495 TATGAGTGTGGAGTCGTTGCTACATAATCACAAACACTGGATTNNNAGATCGGAAG
128497 TATGAGTGTGGAGTCGTTGCCTGCACAAAGGAAAAAAATCTNNNAGATCGGAAGA
128498 TATGAGTGTGGAGTCGTTGCCAAACAGATGCTTAAAGAGTTGNNNAGATCGGAAG
128499 TATGAGTGTGGAGTCGTTGCCCTAATGATCAGTAATATTCAGCNNNAGATCGGAAG
128500 TATGAGTGTGGAGTCGTTGCAGTACAATGTTCTGGGATTGNNNAGATCGGAAGAG
128501 TATGAGTGTGGAGTCGTTGCCATAATCCAGTGAGACAGAACNNNAGATCGGAAGA
128502 TATGAGTGTGGAGTCGTTGCGAAATAATCTATCATGGGCCATANNNAGATCGGAAG
128503 TATGAGTGTGGAGTCGTTGCAGATGCCTGAAACTCAAAATANNNAGATCGGAAGAG
128504 TATGAGTGTGGAGTCGTTGCTCTTTGACATTCCACTTTTTAAANNNAGATCGGAAGA
128505 TATGAGTGTGGAGTCGTTGCGCAAGCACACTTTTATATACATANNNAGATCGGAAG
128506 TATGAGTGTGGAGTCGTTGCCAACTCAGGAGAAAGATAAAAATNNNAGATCGGAA
128507 TATGAGTGTGGAGTCGTTGCATAGACATATGAGGAAGAGATTTNNNAGATCGGAA
128508 TATGAGTGTGGAGTCGTTGCATTCACTCTCAACAAGTCTAAAANNNAGATCGGAAG
128509 TATGAGTGTGGAGTCGTTGCTATGCCCTAAGTCTTCTAATTTTNNNAGATCGGAAGA
128510 TATGAGTGTGGAGTCGTTGCTTTACTAAGTGACAACTAAAAGCNNNAGATCGGAAG
128511 TATGAGTGTGGAGTCGTTGCTATTCATTGTACTGCTAAACCANNNAGATCGGAAGA
128512 TATGAGTGTGGAGTCGTTGCCATCTAAACATTTACAACACTCTNNNAGATCGGAAGA
128513 TATGAGTGTGGAGTCGTTGCGTAGTCCACCTTTCCTTATTCNNNAGATCGGAAGAGC
128514 TATGAGTGTGGAGTCGTTGCGTTGCTTTGTCTATGGGAATNNNAGATCGGAAGAG
128515 TATGAGTGTGGAGTCGTTGCATAATTACTGTGAAGAGTAGGAGNNNAGATCGGAA
128516 TATGAGTGTGGAGTCGTTGCATAATTACTGTGAAGAGTAGGAGNNNAGATCGGAA
128517 TATGAGTGTGGAGTCGTTGCGTATCTCTCTCTCTGCCATATNNNAGATCGGAAGAGC
128518 TATGAGTGTGGAGTCGTTGCACTGTCCTTTTTGTAAGTAAAAANNNAGATCGGAAG
128519 TATGAGTGTGGAGTCGTTGCGAAATAAGGAGAGCCATTGTTNNNAGATCGGAAGA
128520 TATGAGTGTGGAGTCGTTGCAGTATGACATGTCTGGTAATAAGNNNAGATCGGAA
128521 TATGAGTGTGGAGTCGTTGCCCTGTCCACTTTGTATATAAAATNNNAGATCGGAAG
128522 TATGAGTGTGGAGTCGTTGCGTCTGATTATCTAGCCTTGTCNNNAGATCGGAAGAG
128523 TATGAGTGTGGAGTCGTTGCGAAATGGGATTTCTCTCTTGTNNNAGATCGGAAGAG
128524 TATGAGTGTGGAGTCGTTGCCACAAAAAGAGATCTTCTGAGNNNAGATCGGAAGA
128525 TATGAGTGTGGAGTCGTTGCATAAACATTTCTGGACAGAGGNNNAGATCGGAAGA
128526 TATGAGTGTGGAGTCGTTGCATTGAATTTGATTTCTGGATTCANNNAGATCGGAAG
128528 TATGAGTGTGGAGTCGTTGCGCAACTGGAATGGAGATTAGNNNAGATCGGAAGAG
128529 TATGAGTGTGGAGTCGTTGCAATAGTGACAATAGGTGGAAATANNNAGATCGGAA
128530 TATGAGTGTGGAGTCGTTGCATAAGGAAGAAGAAAAGAGAAGANNNAGATCGGA
128531 TATGAGTGTGGAGTCGTTGCCAAAAACTATGTTGAATAGGAGTNNNAGATCGGAA
128532 TATGAGTGTGGAGTCGTTGCAAAATACACGTTACCCATAAAATNNNAGATCGGAAG
128533 TATGAGTGTGGAGTCGTTGCTGAGCTGCAATAGTTTTCTTANNNAGATCGGAAGAG
128534 TATGAGTGTGGAGTCGTTGCTTCCATTACTGCTACTGATATTGNNNAGATCGGAAG
128535 TATGAGTGTGGAGTCGTTGCGCTACTGTTGTTTTTAAGTTCTANNNAGATCGGAAG
128536 TATGAGTGTGGAGTCGTTGCATGACCACATTTACATATTGAAGNNNAGATCGGAAG
128537 TATGAGTGTGGAGTCGTTGCAAAAAGACTGCTAACCAAGAATTNNNAGATCGGAAG
128538 TATGAGTGTGGAGTCGTTGCCTTTCTCTTTCTCTCTTTCTTTCNNNAGATCGGAAGAG

128539 TATGAGTGTGGAGTCGTTGCGCAAGAGGAAAAATATACCTTATNNNAGATCGGAA
128541 TATGAGTGTGGAGTCGTTGCCATGCCCAGCTAATTTTTAATTNNNAGATCGGAAGA
128542 TATGAGTGTGGAGTCGTTGCACTATCTCTCAACCTCAGAGNNNAGATCGGAAGAGC
128543 TATGAGTGTGGAGTCGTTGCCATTGATACTTGCCAATTTTTATNNNAGATCGGAAG
128544 TATGAGTGTGGAGTCGTTGCCTATTTGCTTGTTCAACTCATNNNAGATCGGAAGAG
128545 TATGAGTGTGGAGTCGTTGCAGGACAACATTCAAATTCATAAANNNAGATCGGAAG
128546 TATGAGTGTGGAGTCGTTGCCCTTTTGACTCCAAGACTTGNNNAGATCGGAAGAGC
128547 TATGAGTGTGGAGTCGTTGCAGAATTTCTTTGAAGAGTTTCTCNNNAGATCGGAAG
128548 TATGAGTGTGGAGTCGTTGCAGAATTTCTTTGAAGAGTTTCTCNNNAGATCGGAAG
128549 TATGAGTGTGGAGTCGTTGCGAAGAACAGAGGGAAAGAAAANNNAGATCGGAAG
128550 TATGAGTGTGGAGTCGTTGCATACAGCTATATGGGATTTTTCTNNNAGATCGGAAG
128551 TATGAGTGTGGAGTCGTTGCTCTTATATGCTACAGTGTGTATTNNNAGATCGGAAG
128552 TATGAGTGTGGAGTCGTTGCACTCAATTAGAAGTCATTGTTTGNNNAGATCGGAAG
128553 TATGAGTGTGGAGTCGTTGCTATTTTCTAAGAGCTTATCCAGTNNNAGATCGGAAG
128554 TATGAGTGTGGAGTCGTTGCCAAACCTGGAAGAATGAATTACNNNAGATCGGAAG
128555 TATGAGTGTGGAGTCGTTGCAGGAAATATAACCTGATAACCTGNNNAGATCGGAA
128556 TATGAGTGTGGAGTCGTTGCCTAGCTCATCAGATTCCACTNNNAGATCGGAAGAGC
128557 TATGAGTGTGGAGTCGTTGCGAACTTAGCCAACAATTAAGAATNNNAGATCGGAAG
128558 TATGAGTGTGGAGTCGTTGCAAGGAACTAGAAAATCCTACAATNNNAGATCGGAA
128559 TATGAGTGTGGAGTCGTTGCATTTTCCCTTGATTTTCCTTTCNNNAGATCGGAAGAG
128560 TATGAGTGTGGAGTCGTTGCATTTTCCCTTGATTTTCCTTTCNNNAGATCGGAAGAG
128561 TATGAGTGTGGAGTCGTTGCTATAACCATTTCAGGCTAAAGATNNNAGATCGGAAG
128562 TATGAGTGTGGAGTCGTTGCCTGTACTTTCTTACTTTCTGGAANNNAGATCGGAAGA
128563 TATGAGTGTGGAGTCGTTGCCCAAAGTGCTGGGATTATAGNNNAGATCGGAAGAG
128564 TATGAGTGTGGAGTCGTTGCAGAAACAGAAATCCCTTCATATTNNNAGATCGGAAG
128565 TATGAGTGTGGAGTCGTTGCAACAGATAACTCTAGGGTCATATNNNAGATCGGAAG
128566 TATGAGTGTGGAGTCGTTGCATATTGTATATTGCTGCTTTGTTNNNAGATCGGAAG
128568 TATGAGTGTGGAGTCGTTGCAGTGAACATTTAAAACAAAGTCTNNNAGATCGGAAG
128569 TATGAGTGTGGAGTCGTTGCTATATACGAATAGGGAGCTATGANNNAGATCGGAA
128570 TATGAGTGTGGAGTCGTTGCGCTTTTCTCTTAGAACCTTAAAANNNAGATCGGAAGA
128571 TATGAGTGTGGAGTCGTTGCTAAGGATGGGGTAACTATTCTAGNNNAGATCGGAA
128572 TATGAGTGTGGAGTCGTTGCGTTTTTCAGAAATGAGGTCTCNNNAGATCGGAAGAG
128573 TATGAGTGTGGAGTCGTTGCATTGGAATTTTTGAGTTATGACANNNAGATCGGAAG
128574 TATGAGTGTGGAGTCGTTGCATAGCCATTCATTTTGAAAAGTCNNNAGATCGGAAG
128575 TATGAGTGTGGAGTCGTTGCGTATGCGTGTGTATGTGTATNNNAGATCGGAAGAG
128576 TATGAGTGTGGAGTCGTTGCGTTCTAATAAATTTCAGTCTGCANNNAGATCGGAAG
128577 TATGAGTGTGGAGTCGTTGCGAAACTGATTAGAAAATCCCAACNNNAGATCGGAAG
128578 TATGAGTGTGGAGTCGTTGCGAAACTGATTAGAAAATCCCAACNNNAGATCGGAAG
128579 TATGAGTGTGGAGTCGTTGCTGTAGCAATCTCTCTCTCTATATNNNAGATCGGAAGA
128580 TATGAGTGTGGAGTCGTTGCTCTGCATTAATTTTGGAAAGTTANNNAGATCGGAAG
128581 TATGAGTGTGGAGTCGTTGCCAAGGTCTAGCTCTGTTACCNNNAGATCGGAAGAGC
128582 TATGAGTGTGGAGTCGTTGCCAGGAATAATTTTGCTAGCAATANNNAGATCGGAAG
128583 TATGAGTGTGGAGTCGTTGCCATGTATGACTGTGAGTGTGNNNAGATCGGAAGAG

128584 TATGAGTGTGGAGTCGTTGCTCCTTTGTCTCAAAAAAAAGTTANNNAGATCGGAAG
128585 TATGAGTGTGGAGTCGTTGCCCAGGGATTTGTGTATTTTAAATNNNAGATCGGAAG
128586 TATGAGTGTGGAGTCGTTGCACACAGACATTTATCTACATCTANNNAGATCGGAAG
128587 TATGAGTGTGGAGTCGTTGCACTGAGGATCAAAATATTTTGANNNAGATCGGAAG
128588 TATGAGTGTGGAGTCGTTGCAGGGATAGGACTTATTTATTTCTTNNNAGATCGGAA
128589 TATGAGTGTGGAGTCGTTGCTCTTTTTCTAACTCTGAAAATGTNNNAGATCGGAAG
128590 TATGAGTGTGGAGTCGTTGCATGCAGATTTGTAGAAGAATTTTNNNAGATCGGAAG
128591 TATGAGTGTGGAGTCGTTGCGGGCCAATTCTTTATACATATTTNNNAGATCGGAAG
128593 TATGAGTGTGGAGTCGTTGCAAGTGCCATGGGAATTATAATNNNAGATCGGAAGA
128595 TATGAGTGTGGAGTCGTTGCCTCTTTACTAAGTGCCCAATAANNNAGATCGGAAGA
128596 TATGAGTGTGGAGTCGTTGCACACTTAAAAGTCTGAAACATTTNNNAGATCGGAAG
128597 TATGAGTGTGGAGTCGTTGCCCTTTCTATAACAGTGCAATAAGNNNAGATCGGAAG
128598 TATGAGTGTGGAGTCGTTGCCCTTTCTATAACAGTGCAATAAGNNNAGATCGGAAG
128599 TATGAGTGTGGAGTCGTTGCGACAGAAAAGCAGTAAATCTTACNNNAGATCGGAA
128600 TATGAGTGTGGAGTCGTTGCGAAAACCAAACGAATACTTTTTNNNAGATCGGAAG
128601 TATGAGTGTGGAGTCGTTGCTATACACACAGACATACTGAAGNNNAGATCGGAAG
128602 TATGAGTGTGGAGTCGTTGCTGCAATGATGGAAATACTTTTTNNNAGATCGGAAGA
128603 TATGAGTGTGGAGTCGTTGCCCAAGATCAGTTGACTATAAGTANNNAGATCGGAAG
128604 TATGAGTGTGGAGTCGTTGCGGTTCAAGCGATTATCCTACNNNAGATCGGAAGAGC
128605 TATGAGTGTGGAGTCGTTGCCTCTATCACAGGCTACAGTGNNNAGATCGGAAGAGC
128606 TATGAGTGTGGAGTCGTTGCATATTTTCCTGTCTTTCCTTAGTNNNAGATCGGAAGA
128607 TATGAGTGTGGAGTCGTTGCGATAGTGCCAGATAAGAAATCTTNNNAGATCGGAA
128608 TATGAGTGTGGAGTCGTTGCGTTTTGCCTTCTTAGATTATTCANNNAGATCGGAAGA
128609 TATGAGTGTGGAGTCGTTGCTTATATTTTTACCCCTAAGCCATNNNAGATCGGAAGA
128610 TATGAGTGTGGAGTCGTTGCCCACTTTATCTTTCTTTATAGCCNNNAGATCGGAAGA
128611 TATGAGTGTGGAGTCGTTGCAGAAAGAGAGAGAGAGAGAAGNNNAGATCGGAAG
128612 TATGAGTGTGGAGTCGTTGCGGCTATTACAAATAATGTTGCTANNNAGATCGGAAG
128613 TATGAGTGTGGAGTCGTTGCTTGTATTTTTTTAGTAGAGACGGNNNAGATCGGAAG
128614 TATGAGTGTGGAGTCGTTGCTTTCTGGTCTCTTTTATAGCATANNNAGATCGGAAGA
128615 TATGAGTGTGGAGTCGTTGCGCTAAGATGACAAAATGGAATAANNNAGATCGGAA
128616 TATGAGTGTGGAGTCGTTGCCTCATGATTTTGAGAAAGAATGANNNAGATCGGAA
128618 TATGAGTGTGGAGTCGTTGCGTATATGTCCATTTGTCAACATTNNNAGATCGGAAG
128619 TATGAGTGTGGAGTCGTTGCAATGGATTTATTATTCTGGGTCANNNAGATCGGAAG
128620 TATGAGTGTGGAGTCGTTGCTTTAGGAAACAAAGCCCTTTNNNAGATCGGAAGAGC
128621 TATGAGTGTGGAGTCGTTGCTTTATTCACATTGAGACCCATGNNNAGATCGGAAGA
128622 TATGAGTGTGGAGTCGTTGCTTTATTCACATTGAGACCCATNNNAGATCGGAAGAG
128623 TATGAGTGTGGAGTCGTTGCCATGGATAGATGTGTGTGTATNNNAGATCGGAAGA
128624 TATGAGTGTGGAGTCGTTGCAGTATGTGCGATATCCAAATATANNNAGATCGGAAG
128625 TATGAGTGTGGAGTCGTTGCTATAAAAACTGTGCAGTAGTTACNNNAGATCGGAAG
128626 TATGAGTGTGGAGTCGTTGCGATGGATAAAAACTCAGTTGAATNNNAGATCGGAA
128627 TATGAGTGTGGAGTCGTTGCATAGGACAAGCTTAGCTGTGNNNAGATCGGAAGAG
128628 TATGAGTGTGGAGTCGTTGCTAATGTGCTAGAAAGAATGTCTANNNAGATCGGAA
128629 TATGAGTGTGGAGTCGTTGCCCTGAGTTCTTATTTTTTCTTCANNNAGATCGGAAGA

128630 TATGAGTGTGGAGTCGTTGCCAAATGACAGCCATTATTTCTATNNNAGATCGGAAG
128631 TATGAGTGTGGAGTCGTTGCCCTGTTTAAGGTACAGTTAAAAGNNNAGATCGGAA
128632 TATGAGTGTGGAGTCGTTGCCATTACAACATTCCTCCCTGNNNAGATCGGAAGAGC
128633 TATGAGTGTGGAGTCGTTGCCTCAGTTTACTTAGGTGCAGNNNAGATCGGAAGAG
128634 TATGAGTGTGGAGTCGTTGCCCCAGTTAGGAGTAAAATGACNNNAGATCGGAAGA
128635 TATGAGTGTGGAGTCGTTGCATGCAGATAATTGTTTTTAGCAANNNAGATCGGAAG
128636 TATGAGTGTGGAGTCGTTGCCTCTTGGTAAAAATGATCATCACNNNAGATCGGAAG
128637 TATGAGTGTGGAGTCGTTGCGCTTGTACCTAATTTTGTATTCTNNNAGATCGGAAGA
128638 TATGAGTGTGGAGTCGTTGCCGTATTTTGATATAGAGTCTCACNNNAGATCGGAAG
128639 TATGAGTGTGGAGTCGTTGCCTAAAAATCTGTATTAGGGATGCNNNAGATCGGAA
128640 TATGAGTGTGGAGTCGTTGCGTTCTCTATTGATATGGTTAGGANNNAGATCGGAAG
128641 TATGAGTGTGGAGTCGTTGCTTTACTCTGGAATGTAAACAGATNNNAGATCGGAAG
128642 TATGAGTGTGGAGTCGTTGCGGGAAAATTGTAATTATTTTGGGNNNAGATCGGAA
128643 TATGAGTGTGGAGTCGTTGCTATTCCTATAACCTAGTGTTTCCNNNAGATCGGAAGA
128644 TATGAGTGTGGAGTCGTTGCCTGGAGAAGCTAGGACTATAGNNNAGATCGGAAGA
128645 TATGAGTGTGGAGTCGTTGCTTATTTATGCTTGAAAAAGGACCNNNAGATCGGAAG
128646 TATGAGTGTGGAGTCGTTGCAAGAAAGAGAAAAAGAAAGAGGNNNAGATCGGAA
128647 TATGAGTGTGGAGTCGTTGCCCTTCTATTCTACTTTTCAAACCNNNAGATCGGAAGA
128648 TATGAGTGTGGAGTCGTTGCCCTTCTATTCTACTTTTCAAACCNNNAGATCGGAAGA
128649 TATGAGTGTGGAGTCGTTGCACACATATACATTTATTCCATCCNNNAGATCGGAAGA
128650 TATGAGTGTGGAGTCGTTGCTATTTGTCAGATTGCTAGGTAATNNNAGATCGGAAG
128651 TATGAGTGTGGAGTCGTTGCATCAAGTAATGCTGATGAAAATTNNNAGATCGGAA
128652 TATGAGTGTGGAGTCGTTGCAGCTTTTCTTTTTTTCTCCTTCNNNAGATCGGAAGAG
128653 TATGAGTGTGGAGTCGTTGCGACCCAGATATTTCTAGTCATTTNNNAGATCGGAAG
128654 TATGAGTGTGGAGTCGTTGCTTATACCACACATCTTACTCTACNNNAGATCGGAAGA
128655 TATGAGTGTGGAGTCGTTGCGGTTATAAGCATACATCTGAATGNNNAGATCGGAA
128656 TATGAGTGTGGAGTCGTTGCCACATATGCTTTGTCTTTTTTTTNNNAGATCGGAAGA
128657 TATGAGTGTGGAGTCGTTGCTGCTCAATTTACTTCTTTGTTTANNNAGATCGGAAGA
128658 TATGAGTGTGGAGTCGTTGCGCTTGTTAAATGCTTTTGATTATNNNAGATCGGAAG
128659 TATGAGTGTGGAGTCGTTGCACACCTTTAGTTCAATTCTTTAGNNNAGATCGGAAGA
128660 TATGAGTGTGGAGTCGTTGCGAATCTCCAGAGTGATAAAAATCNNNAGATCGGAA
128661 TATGAGTGTGGAGTCGTTGCCATGAGTGGAAAGCTTTATTTAANNNAGATCGGAA
128662 TATGAGTGTGGAGTCGTTGCGATCTCAATCTCCTGACCTCNNNAGATCGGAAGAGC
128663 TATGAGTGTGGAGTCGTTGCGTCAAGATCACAATTCAGATTTANNNAGATCGGAAG
128664 TATGAGTGTGGAGTCGTTGCAGAAAAGAAAATCTTCAAGATGTNNNAGATCGGAA
128665 TATGAGTGTGGAGTCGTTGCAGTCAAATACAGAATTGGAAAAANNNAGATCGGAA
128666 TATGAGTGTGGAGTCGTTGCTTTTGTTGATTTTTTGCATGAATNNNAGATCGGAAG
128667 TATGAGTGTGGAGTCGTTGCTCCCTATTATATCATCTGTGGTANNNAGATCGGAAG
128668 TATGAGTGTGGAGTCGTTGCGAGACTAGTGAAGACAATAGTAGNNNAGATCGGAA
128669 TATGAGTGTGGAGTCGTTGCTTTCCATAGAAAAGGTATAATGTNNNAGATCGGAAG
128670 TATGAGTGTGGAGTCGTTGCTTTCTTTCTACAACTGCATGANNNAGATCGGAAGAG
128671 TATGAGTGTGGAGTCGTTGCTTTGAATTGATGAATACCTGATGNNNAGATCGGAAG
128672 TATGAGTGTGGAGTCGTTGCATAATGATATCCAATGGGTCTTTNNNAGATCGGAAG

128673 TATGAGTGTGGAGTCGTTGCATTAGGGGTCTTATAGTTAGAAANNNAGATCGGAA(
128674 TATGAGTGTGGAGTCGTTGCTTCCACCATTTAAAAAAAAATCTNNNAGATCGGAAGA
128675 TATGAGTGTGGAGTCGTTGCCCAGGTTCAAGCTATTCTCTNNNAGATCGGAAGAGC
128676 TATGAGTGTGGAGTCGTTGCCGACTTTTCTATTGTCTAGAATAGNNNAGATCGGAA(
128677 TATGAGTGTGGAGTCGTTGCCAACAATGATAGACTGGATTAAGNNNAGATCGGAA(
128678 TATGAGTGTGGAGTCGTTGCCTACAACTCCATGTATTATTGTGNNNAGATCGGAAGA
128679 TATGAGTGTGGAGTCGTTGCGACAAAGAGGGAAAGAATGTAANNNAGATCGGAA(
128680 TATGAGTGTGGAGTCGTTGCTTTTGGAGATTTAGTCTGACTATNNNAGATCGGAAG
128681 TATGAGTGTGGAGTCGTTGCTTGTTGTAACAAAAATTTGATCANNNAGATCGGAAG
128682 TATGAGTGTGGAGTCGTTGCACAACAGGTTCTTTTTATTTTGANNNAGATCGGAAGA
128683 TATGAGTGTGGAGTCGTTGCCTTCTACCAGTAGTAAGACCANNNAGATCGGAAGAG
128684 TATGAGTGTGGAGTCGTTGCTTTGGACTAAATTTTCATTGGAANNNAGATCGGAAG
128685 TATGAGTGTGGAGTCGTTGCCATTACTTTCATTCTCAGAGAAGNNNAGATCGGAAGA
128686 TATGAGTGTGGAGTCGTTGCTTAGCTTTTAAAATGCCATTTTGNNNAGATCGGAAGA
128687 TATGAGTGTGGAGTCGTTGCGAATTTGCTGCATAGTCTTGNNNAGATCGGAAGAG(
128688 TATGAGTGTGGAGTCGTTGCGTTTTGATACCAGTTAGTTGTTTNNNAGATCGGAAG.
128689 TATGAGTGTGGAGTCGTTGCGTAGAATGGTGACTGACATATAANNNAGATCGGAA(
128690 TATGAGTGTGGAGTCGTTGCTTTACCATAAAACGAAGTAGGANNNAGATCGGAAGA
128691 TATGAGTGTGGAGTCGTTGCCTTCTGTCCTGGCATTATTAGNNNAGATCGGAAGAG
128692 TATGAGTGTGGAGTCGTTGCGTTTCTACAGGATACTTCACTAGNNNAGATCGGAAG.
128693 TATGAGTGTGGAGTCGTTGCGCTTATTGCCTTCTTTTGTATAANNNAGATCGGAAGA
128695 TATGAGTGTGGAGTCGTTGCGCATGTATAAAACATATGCCTTNNNAGATCGGAAGA
128696 TATGAGTGTGGAGTCGTTGCGTGTGTATGTGTGTCTAGTTGNNNAGATCGGAAGA(
128697 TATGAGTGTGGAGTCGTTGCGATAGAAAAATTATAAAAGCAGCCNNNAGATCGGAA
128698 TATGAGTGTGGAGTCGTTGCGGATAGAAAAAATAGGTTAGAGCNNNAGATCGGAA
128699 TATGAGTGTGGAGTCGTTGCTTTTGTTTGCTAATGAGATGAGNNNAGATCGGAAGA
128700 TATGAGTGTGGAGTCGTTGCTGTATTCTCTCTAATAGGCTAGANNNAGATCGGAAG.
128701 TATGAGTGTGGAGTCGTTGCCTTTCATTGAACTCTATTGTGAANNNAGATCGGAAGA
128702 TATGAGTGTGGAGTCGTTGCTCTGTTTACAGCTATTCTTACTTNNNAGATCGGAAGA
128704 TATGAGTGTGGAGTCGTTGCCGCTTTCTAAAGTACTATGTTTTNNNAGATCGGAAGA
128706 TATGAGTGTGGAGTCGTTGCCATGTTTTGGATTTACTTCTCAANNNAGATCGGAAGA
128707 TATGAGTGTGGAGTCGTTGCGCACTCTCAGATTTACATTTTATNNNAGATCGGAAGA
128708 TATGAGTGTGGAGTCGTTGCCAAATCTTTCTAATAGCAGCATTNNNAGATCGGAAGA
128710 TATGAGTGTGGAGTCGTTGCGAGTGAAATCTACTCATTCATTCNNNAGATCGGAAGA
128711 TATGAGTGTGGAGTCGTTGCTTACAGGAGGCCTCTTTTTTANNNAGATCGGAAGAG
128712 TATGAGTGTGGAGTCGTTGCAATGTTCATGTATGTGCATAGTANNNAGATCGGAAG
128713 TATGAGTGTGGAGTCGTTGCTGATTATAGAGGCTAAGAAGTTCNNNAGATCGGAA(
128714 TATGAGTGTGGAGTCGTTGCTTCTGTCTTTCTTTCTTTTCTTTNNNAGATCGGAAGAG
128715 TATGAGTGTGGAGTCGTTGCTAGCTAACTAAGAAAAGACTTCANNNAGATCGGAAG
128716 TATGAGTGTGGAGTCGTTGCTCTTTTTTTTTTTTTGAGATGGCNNNAGATCGGAAGA
128717 TATGAGTGTGGAGTCGTTGCTGTGTTGAAAAGCTTTCTAACNNNAGATCGGAAGAG
128718 TATGAGTGTGGAGTCGTTGCGAAAAGAACATGTTCCTGTAAANNNAGATCGGAAGA
128719 TATGAGTGTGGAGTCGTTGCCAACGAGAAAACTTGTTCTGNNNAGATCGGAAGAG(

128720 TATGAGTGTGGAGTCGTTGCCTATTCTCTGTGGATGTCACNNNAGATCGGAAGAGC
128721 TATGAGTGTGGAGTCGTTGCGTTTCTCTTTTCTTTTGGAAACTNNNAGATCGGAAGA
128722 TATGAGTGTGGAGTCGTTGCGCCTTTAAAATTTCTATAACCCANNNAGATCGGAAGA
128724 TATGAGTGTGGAGTCGTTGCTTTCTTTTTTTTTCTGAGACAGANNNAGATCGGAAGA
128725 TATGAGTGTGGAGTCGTTGCAATCACATTTATTCATCTATGCCNNNAGATCGGAAGA
128726 TATGAGTGTGGAGTCGTTGCAATCACATTTATTCATCTATGCCNNNAGATCGGAAGA
128727 TATGAGTGTGGAGTCGTTGCCTTTATGGTTCTAACTTTGAGAGNNNAGATCGGAAG
128728 TATGAGTGTGGAGTCGTTGCCACTTGCTAGAGAATATTCATTTNNNAGATCGGAAGA
128729 TATGAGTGTGGAGTCGTTGCTGTCCATTTTTTAAAAGCTTAGANNNAGATCGGAAGA
128731 TATGAGTGTGGAGTCGTTGCTATATGATGGAATACTACTCAGCNNNAGATCGGAAG
128732 TATGAGTGTGGAGTCGTTGCTTAGGCTGAGGTGAAGAAATANNNAGATCGGAAGA
128733 TATGAGTGTGGAGTCGTTGCTGTCATACTACTACCATCACTATNNNAGATCGGAAGA
128734 TATGAGTGTGGAGTCGTTGCCTTAAGATTAAAAGACGCTCAGNNNAGATCGGAAGA
128735 TATGAGTGTGGAGTCGTTGCTTTGAGGGGGTTATCTATTTTAANNNAGATCGGAAG
128736 TATGAGTGTGGAGTCGTTGCTATCAGGTCTAAACAAGAACTTCNNNAGATCGGAAG
128737 TATGAGTGTGGAGTCGTTGCCTAATTTACACTCACTGAAGTTCNNNAGATCGGAAGA
128738 TATGAGTGTGGAGTCGTTGCAATAACTGAAGTTCTTCCTTGANNNAGATCGGAAGA
128739 TATGAGTGTGGAGTCGTTGCGGACATCATTAATCAACTAAACANNNAGATCGGAAG
128740 TATGAGTGTGGAGTCGTTGCAGCATATTGTTCATCAGTTTTAGNNNAGATCGGAAGA
128741 TATGAGTGTGGAGTCGTTGCCTGACAGCTGGAAATACATCNNNAGATCGGAAGAGC
128742 TATGAGTGTGGAGTCGTTGCTATATCACCCCATATCAAAAGAGNNNAGATCGGAAGA
128744 TATGAGTGTGGAGTCGTTGCAGTTTTATTTTCTTCTCTCTCCANNNAGATCGGAAGA
128745 TATGAGTGTGGAGTCGTTGCCTGGTTTGAAATTCAAGAAGTCNNNAGATCGGAAGA
128746 TATGAGTGTGGAGTCGTTGCGAAATTTCCAAAACATTGTGATCNNNAGATCGGAAG
128747 TATGAGTGTGGAGTCGTTGCGTGTGTCTCCTTATACATACATANNNAGATCGGAAGA
128748 TATGAGTGTGGAGTCGTTGCCTAAATGCTGAAGGTCATACANNNAGATCGGAAGAG
128749 TATGAGTGTGGAGTCGTTGCTTAGTTCTGCATGTCAACATTTANNNAGATCGGAAGA
128750 TATGAGTGTGGAGTCGTTGCTTTGTTTTATTTTTGAGACAAGCNNNAGATCGGAAGA
128751 TATGAGTGTGGAGTCGTTGCATTATCTCCCCATCTCAAGATNNNAGATCGGAAGAGC
128752 TATGAGTGTGGAGTCGTTGCGTGATGCCATCATGTTTAATTANNNAGATCGGAAGA
128753 TATGAGTGTGGAGTCGTTGCGAGAGATGACCTTGACAATAGNNNAGATCGGAAGA
128754 TATGAGTGTGGAGTCGTTGCCTAATATTATACCTAGCCCGAGNNNAGATCGGAAGA
128755 TATGAGTGTGGAGTCGTTGCCTGAAAGCTATTGTACATACAAANNNAGATCGGAAG
128756 TATGAGTGTGGAGTCGTTGCCAATAGCTTAGAAAGTGAAGATTNNNAGATCGGAAG
128757 TATGAGTGTGGAGTCGTTGCAATGTCTATATTCCCACACCTANNNAGATCGGAAGAG
128758 TATGAGTGTGGAGTCGTTGCCATCCTCCTTAGAGAAGTACATNNNAGATCGGAAGA
128759 TATGAGTGTGGAGTCGTTGCTGCTAGTCCAAAATCTATCAATANNNAGATCGGAAGA
128760 TATGAGTGTGGAGTCGTTGCAAGAATTTGTTGTTTGATGAGATNNNAGATCGGAAG
128761 TATGAGTGTGGAGTCGTTGCCATCAAGTTCTTTGTTCGTTAGNNNAGATCGGAAGA
128764 TATGAGTGTGGAGTCGTTGCCATCTGGTGAGGCTAATAGANNNAGATCGGAAGAG
128765 TATGAGTGTGGAGTCGTTGCATTTTGAATGAAGAATGAAGACANNNAGATCGGAAG
128766 TATGAGTGTGGAGTCGTTGCCTCAAAGTCTACAAGATTGAACNNNAGATCGGAAGA
128767 TATGAGTGTGGAGTCGTTGCTATTTATCCACTCATTGATTGACNNNAGATCGGAAGA

128768 TATGAGTGTGGAGTCGTTGCCCAATAGGAATATAATATAAGCCAANNNAGATCGGA
128769 TATGAGTGTGGAGTCGTTGCCCAAATGAGAGACCAAATTAATANNNAGATCGGAAG
128770 TATGAGTGTGGAGTCGTTGCCACTGAAACCTAAAACTGAACNNNAGATCGGAAGAG
128771 TATGAGTGTGGAGTCGTTGCGACCACCCTGTACAAAAAAGNNNAGATCGGAAGAG
128772 TATGAGTGTGGAGTCGTTGCGATTAGAATTCAGAAGAGAGCCNNNAGATCGGAAG
128773 TATGAGTGTGGAGTCGTTGCCTGTCTCTATGTCTCTCTATCTANNNAGATCGGAAGA
128776 TATGAGTGTGGAGTCGTTGCAAAACAATACCATCTGACTAAACNNNAGATCGGAAG
128777 TATGAGTGTGGAGTCGTTGCAGATTGTGTACTCTTTTTCTTTTNNNAGATCGGAAGA
128778 TATGAGTGTGGAGTCGTTGCCTAAATAATGGTGAGTTGATGAGNNNAGATCGGAA
128779 TATGAGTGTGGAGTCGTTGCTCCTAATAAGAATTCTACATAGCTNNNAGATCGGAAG
128780 TATGAGTGTGGAGTCGTTGCCCACATTTTGTTTAGATGTAAGTNNNAGATCGGAAG
128781 TATGAGTGTGGAGTCGTTGCTGAAACACACAAAGCTGAAGNNNAGATCGGAAGAG
128782 TATGAGTGTGGAGTCGTTGCCATTTTCTTTATCCACTCTTTGANNNAGATCGGAAGA
128783 TATGAGTGTGGAGTCGTTGCCTTTCAGTATGTCTTCAAATTCANNNAGATCGGAAGA
128784 TATGAGTGTGGAGTCGTTGCTATTATATGGAATTGTTGCAACTNNNAGATCGGAAG
128786 TATGAGTGTGGAGTCGTTGCGAAATACACATGAAAAATGCAATNNNAGATCGGAAG
128787 TATGAGTGTGGAGTCGTTGCTGTATCTGAAACAGTATCCAAGNNNAGATCGGAAGA
128788 TATGAGTGTGGAGTCGTTGCGATCCTTAAATTCCAAACAAAACNNNAGATCGGAAG
128789 TATGAGTGTGGAGTCGTTGCAAGTGTAATTTTTCACAGTTGTANNNAGATCGGAAG
128790 TATGAGTGTGGAGTCGTTGCCTCTTTAAGGCTCAGTGTTTATANNNAGATCGGAAG
128791 TATGAGTGTGGAGTCGTTGCTGGTGTATGATTTAATTACTTGCNNNAGATCGGAAG
128792 TATGAGTGTGGAGTCGTTGCGAGAGTGAGTGATTTTTATTGTTNNNAGATCGGAAG
128793 TATGAGTGTGGAGTCGTTGCCTTTATGTAAAGGGAGAAGAAATNNNAGATCGGAA
128794 TATGAGTGTGGAGTCGTTGCCTTCATCTCTTTTTATGACACCNNNAGATCGGAAGAG
128795 TATGAGTGTGGAGTCGTTGCCTTATGGCAGAACTGTAAAAAGNNNAGATCGGAAG
128796 TATGAGTGTGGAGTCGTTGCCTCAAGCAGAAAGAGCTATTTNNNAGATCGGAAGAG
128798 TATGAGTGTGGAGTCGTTGCAAATAGGGACTAAATGTTCTTCTNNNAGATCGGAAG
128799 TATGAGTGTGGAGTCGTTGCGATAAAAATTCAAGGCGTTCTCNNNAGATCGGAAGA
128801 TATGAGTGTGGAGTCGTTGCCAGTGACCTTTGTTTCTATCTNNNAGATCGGAAGAG
128802 TATGAGTGTGGAGTCGTTGCAGCAATGAGATAAAAAACCTAACNNNAGATCGGAAG
128803 TATGAGTGTGGAGTCGTTGCAGCAATGAGATAAAAAACCTAACNNNAGATCGGAAG
128804 TATGAGTGTGGAGTCGTTGCCCTTTGTCACCACTATCTTAGNNNAGATCGGAAGAG
128805 TATGAGTGTGGAGTCGTTGCGCAGCCCTATCTATAATGTATAANNNAGATCGGAAG
128806 TATGAGTGTGGAGTCGTTGCCTCACACTTCTCTCTGTCTCNNNAGATCGGAAGAGCA
128807 TATGAGTGTGGAGTCGTTGCAAAGAAAATATCCCAGGACAAANNNAGATCGGAAG
128808 TATGAGTGTGGAGTCGTTGCCAGCTTCTATCCATCTTTCACNNNAGATCGGAAGAGC
128809 TATGAGTGTGGAGTCGTTGCTATTATGTGGTGAGAGACGATNNNAGATCGGAAGA
128810 TATGAGTGTGGAGTCGTTGCGAAATGCCTAGAATAACTAATGGNNNAGATCGGAA
128811 TATGAGTGTGGAGTCGTTGCTCACAACTCATAGACATCTATACNNNAGATCGGAAGA
128812 TATGAGTGTGGAGTCGTTGCTAAGTCTATTAATCACAGGAAGGNNNAGATCGGAA
128813 TATGAGTGTGGAGTCGTTGCTATCATCCCAAGGCTAGAATGNNNAGATCGGAAGAG
128814 TATGAGTGTGGAGTCGTTGCTTCTAGAGTTAGTTCTTCACCTNNNAGATCGGAAGA
128816 TATGAGTGTGGAGTCGTTGCTTACATCAGTAGAAATGGCAANNNAGATCGGAAGA

128817 TATGAGTGTGGAGTCGTTGCTTACATCAGTAGAAATGGCAANNNAGATCGGAAGA
128818 TATGAGTGTGGAGTCGTTGCTTACATCAGTAGAAATGGCAANNNAGATCGGAAGA
128819 TATGAGTGTGGAGTCGTTGCCCACAAGTTTACTTCTTTATCTCNNNAGATCGGAAGA
128820 TATGAGTGTGGAGTCGTTGCCATAGGTATATTATGGAGCAGAANNNAGATCGGAA
128821 TATGAGTGTGGAGTCGTTGCTTTTTCATGAATTGGCTTATGTANNNAGATCGGAAG
128822 TATGAGTGTGGAGTCGTTGCTCCGTGTGTATTATGGGATGNNNAGATCGGAAGAG
128823 TATGAGTGTGGAGTCGTTGCGAAAACCAACTCCTCTTTTGNNNAGATCGGAAGAGC
128824 TATGAGTGTGGAGTCGTTGCCAGGTAATTCTAATGTGTGTGTNNNAGATCGGAAGA
128825 TATGAGTGTGGAGTCGTTGCAGTCTTCACTAGCTTCCTATTTNNNAGATCGGAAGAG
128826 TATGAGTGTGGAGTCGTTGCAACTCACAGAACTCTTTAAGGNNNAGATCGGAAGAG
128827 TATGAGTGTGGAGTCGTTGCTGGTCTACATAGAGCTAATTAAGNNNAGATCGGAAG
128828 TATGAGTGTGGAGTCGTTGCACTCTGACCAATATACCTAGAATNNNAGATCGGAAG
128829 TATGAGTGTGGAGTCGTTGCTCTGAACTCATGGTATACCTAATNNNAGATCGGAAG
128830 TATGAGTGTGGAGTCGTTGCCACATTTTCTTTACCCACTAATTNNNAGATCGGAAGA
128831 TATGAGTGTGGAGTCGTTGCAATCTGTTTTATCTGTTTCTTCCNNNAGATCGGAAGA
128832 TATGAGTGTGGAGTCGTTGCCATATGTGCCTGGACATTTTNNNAGATCGGAAGAGC
128833 TATGAGTGTGGAGTCGTTGCCAAATCTTGAAATAAGGGAAAGTNNNAGATCGGAA
128834 TATGAGTGTGGAGTCGTTGCCCTCCATCCTATTTTCCAAGNNNAGATCGGAAGAGCA
128835 TATGAGTGTGGAGTCGTTGCGTTCCCATCAGATAAATTTCATCNNNAGATCGGAAGA
128836 TATGAGTGTGGAGTCGTTGCTTACATGGTAAGAAAAAAGGTTTNNNAGATCGGAA
128837 TATGAGTGTGGAGTCGTTGCCAAGAATTACTCTTATTTTCCCANNNAGATCGGAAGA
128838 TATGAGTGTGGAGTCGTTGCTTAGAGAACTAATGGCTGATTTTNNNAGATCGGAAG
128839 TATGAGTGTGGAGTCGTTGCAATTTCCCATAATGTAAAATGGTNNNAGATCGGAAG
128840 TATGAGTGTGGAGTCGTTGCATTTTACCTCTCTTTTAGTTGTGNNNAGATCGGAAGA
128841 TATGAGTGTGGAGTCGTTGCATTTGTTGGAATCCTTGAATATGNNNAGATCGGAAG
128842 TATGAGTGTGGAGTCGTTGCCTAGGTTCAAGCGATTCTTCNNNAGATCGGAAGAGC
128844 TATGAGTGTGGAGTCGTTGCCTTCTAAATTCCACCGTAAAAAGNNNAGATCGGAAG
128845 TATGAGTGTGGAGTCGTTGCGGATTCCCTCTTTTTTTATTGTTNNNAGATCGGAAGA
128846 TATGAGTGTGGAGTCGTTGCTAGTTAGTAAGAATCTCAAGAGCNNNAGATCGGAA
128847 TATGAGTGTGGAGTCGTTGCTTACAGGAGGCCTCTTTTTTANNNAGATCGGAAGAG
128848 TATGAGTGTGGAGTCGTTGCTTTCTCTTTAAGTCTCAGTATCGNNNAGATCGGAAGA
128849 TATGAGTGTGGAGTCGTTGCAATCCTATTGTATTAGAACCCTGNNNAGATCGGAAG
128850 TATGAGTGTGGAGTCGTTGCAATCCTATTGTATTAGAACCCTGNNNAGATCGGAAG
128851 TATGAGTGTGGAGTCGTTGCAATCCTATTGTATTAGAACCCTGNNNAGATCGGAAG
128852 TATGAGTGTGGAGTCGTTGCGACCTCTGCTTTTACTAAAATTANNNAGATCGGAAGA
128853 TATGAGTGTGGAGTCGTTGCGGATTAGGAAAAACGACATAATTNNNAGATCGGAA
128854 TATGAGTGTGGAGTCGTTGCCCTTTTGATGCATATAATGTAATNNNAGATCGGAAG
128855 TATGAGTGTGGAGTCGTTGCGTAGGTGGAATAAAGAGAGATTCNNNAGATCGGAA
128856 TATGAGTGTGGAGTCGTTGCATATGTATCATACAGAAGGAGTGNNNAGATCGGAA
128857 TATGAGTGTGGAGTCGTTGCTCTTTTGTTTCTACCTCAGTCNNNAGATCGGAAGAGC
128858 TATGAGTGTGGAGTCGTTGCCATACCTTATCACCCAATTTTCNNNAGATCGGAAGAG
128859 TATGAGTGTGGAGTCGTTGCCTCCCTTAAGATGTCCTCTTNNNAGATCGGAAGAGCA
128860 TATGAGTGTGGAGTCGTTGCCCCAGGTTAGTTTCAAACTCNNNAGATCGGAAGAGC

128861 TATGAGTGTGGAGTCGTTGCAACTTTTGCCATAAAAATGAAAGNNNAGATCGGAAG
128862 TATGAGTGTGGAGTCGTTGCCTGGCATATTCAGATAATTTTCTNNNAGATCGGAAG
128863 TATGAGTGTGGAGTCGTTGCAACTGGGGCTACTTTGAAATNNNAGATCGGAAGAG
128864 TATGAGTGTGGAGTCGTTGCAGTTTTGCTATTATCACCCAGNNNAGATCGGAAGAG
128865 TATGAGTGTGGAGTCGTTGCTATCTTCCTTCCTTTCAAGATACNNNAGATCGGAAGA
128866 TATGAGTGTGGAGTCGTTGCGTTACAGTCCCTCAATACAAATANNNAGATCGGAAG
128867 TATGAGTGTGGAGTCGTTGCCTCCCATAGTTGACAGAATGNNNAGATCGGAAGAG
128868 TATGAGTGTGGAGTCGTTGCTTCATGCGTGATTTGATGAGNNNAGATCGGAAGAG
128869 TATGAGTGTGGAGTCGTTGCCTTTGTTTCCCTCTTCCTTCNNNAGATCGGAAGAGCA
128872 TATGAGTGTGGAGTCGTTGCCAATGGAGAAAGAGTAATTCACNNNAGATCGGAAG
128873 TATGAGTGTGGAGTCGTTGCCATTGTTTTCCCAAATGATTTCNNNAGATCGGAAGAG
128874 TATGAGTGTGGAGTCGTTGCGAAGAGAAATAGCATCCCAATNNNAGATCGGAAGA
128878 TATGAGTGTGGAGTCGTTGCATAAATTACTGTTGTTCTCTTGGNNNAGATCGGAAG
128879 TATGAGTGTGGAGTCGTTGCATAAATTACTGTTGTTCTCTTGGNNNAGATCGGAAG
128880 TATGAGTGTGGAGTCGTTGCTATAGGGAGTTAGTTGTCTTTAGNNNAGATCGGAA
128881 TATGAGTGTGGAGTCGTTGCCTCAGAAGGAGAGAAAACAACNNNAGATCGGAAGA
128882 TATGAGTGTGGAGTCGTTGCGAATTGCCTCTAAATTTAGGAAGNNNAGATCGGAAG
128883 TATGAGTGTGGAGTCGTTGCACTATGGCATGAATGGATATACNNNAGATCGGAAG
128884 TATGAGTGTGGAGTCGTTGCATATGAGAGAAAGCTGTCTTGNNNAGATCGGAAGA
128886 TATGAGTGTGGAGTCGTTGCCATCTGTACTGAACATGTACANNNAGATCGGAAGAG
128887 TATGAGTGTGGAGTCGTTGCCTCTGCAGTGATCTGTTTACNNNAGATCGGAAGAGC
128890 TATGAGTGTGGAGTCGTTGCAATTTGCTTTCCAACATTTATTGNNNAGATCGGAAGA
128891 TATGAGTGTGGAGTCGTTGCAAGCTGCCAGTCTTCTTATTNNNAGATCGGAAGAGC
128892 TATGAGTGTGGAGTCGTTGCGTTGAGTTTAGGGTTAAATCAAANNNAGATCGGAA
128893 TATGAGTGTGGAGTCGTTGCTAAGCTGTCAATTGTCTGAAAANNNAGATCGGAAGA
128894 TATGAGTGTGGAGTCGTTGCTAGTCCCAGGTTACTTTACAANNNAGATCGGAAGAG
128895 TATGAGTGTGGAGTCGTTGCAAATACAAAAAAATAGCCTAGCANNNAGATCGGAAG
128896 TATGAGTGTGGAGTCGTTGCAAAAAGAATTCTAAGGCTAGCCNNNAGATCGGAAG
128897 TATGAGTGTGGAGTCGTTGCCTTTGCACCTGTGTTTCTTTNNNAGATCGGAAGAGCA
128898 TATGAGTGTGGAGTCGTTGCATAACCAAAAGAAACAGTTAACCNNNAGATCGGAAG
128899 TATGAGTGTGGAGTCGTTGCCTTTCTCCAAAAAAGGGAATAGNNNAGATCGGAAG
128900 TATGAGTGTGGAGTCGTTGCATGGAAATAGAATCTTCAGAGAANNNAGATCGGAA
128901 TATGAGTGTGGAGTCGTTGCATGGAAATAGAATCTTCAGAGAANNNAGATCGGAA
128902 TATGAGTGTGGAGTCGTTGCCTGGAGTGAAGATTACAGCTNNNAGATCGGAAGAG
128903 TATGAGTGTGGAGTCGTTGCCCTGATATAAAGTGGTGTAGTATNNNAGATCGGAA
128904 TATGAGTGTGGAGTCGTTGCCTTTCCACAGGGTGTATTGANNNAGATCGGAAGAG
128905 TATGAGTGTGGAGTCGTTGCCTGATGTAGGAAGATTTCTTGANNNAGATCGGAAG
128906 TATGAGTGTGGAGTCGTTGCAGTTGGTTCATTAGAATAAAAGANNNAGATCGGAA
128907 TATGAGTGTGGAGTCGTTGCAGATAATGAAAGGGAAACCATTANNNAGATCGGAA
128908 TATGAGTGTGGAGTCGTTGCGAAAATTCTTGCTCATCTTTTTGNNNAGATCGGAAG
128909 TATGAGTGTGGAGTCGTTGCGTGTTCTCTGAGAAAAGGAGNNNAGATCGGAAGAG
128910 TATGAGTGTGGAGTCGTTGCCCTAAGTTTAGCTCCACTATATANNNAGATCGGAAG
128911 TATGAGTGTGGAGTCGTTGCGAAGCATAGCCATTGATAAACNNNAGATCGGAAGA

128912 TATGAGTGTGGAGTCGTTGCCACTTTCACCTCCCACTATTNNNAGATCGGAAGAGCA
128913 TATGAGTGTGGAGTCGTTGCCTGATTTTTGTACATTGGTTTTGNNNAGATCGGAAG
128914 TATGAGTGTGGAGTCGTTGCGCTGTCCAAAACAATAGAAATTTNNNAGATCGGAAG
128915 TATGAGTGTGGAGTCGTTGCAGAACTTTGTAAATGTTTTCCAANNNAGATCGGAAG
128916 TATGAGTGTGGAGTCGTTGCATGTTTACTCAGAACACCTGNNNAGATCGGAAGAGC
128917 TATGAGTGTGGAGTCGTTGCTTTTATTTTTTGAGATGGCATCTNNNAGATCGGAAG
128918 TATGAGTGTGGAGTCGTTGCAAGGTAAGGCATAAAATTCAAATNNNAGATCGGAA
128919 TATGAGTGTGGAGTCGTTGCCTTTCTTTCTTAGACAGGGTCNNNAGATCGGAAGAG
128921 TATGAGTGTGGAGTCGTTGCACTTTTTTTTTCTTTTTTCAGACNNNAGATCGGAAG
128922 TATGAGTGTGGAGTCGTTGCCTGGCTTCATCGTATGTATTNNNAGATCGGAAGAGC
128923 TATGAGTGTGGAGTCGTTGCTAGTAGGTCCTCAATAAACAATTNNNAGATCGGAAG
128924 TATGAGTGTGGAGTCGTTGCCTTTGGAGCTCATTTTCATGNNNAGATCGGAAGAGC
128925 TATGAGTGTGGAGTCGTTGCATGCTTGTAGATTACAGCTGNNNAGATCGGAAGAG
128926 TATGAGTGTGGAGTCGTTGCGCAATTTTATATAGTGATGATGGANNNAGATCGGA
128927 TATGAGTGTGGAGTCGTTGCACTCTAAAGATTCACCATAAGATNNNAGATCGGAAG
128928 TATGAGTGTGGAGTCGTTGCTGGGAAAAAATAACATTAGCTTTNNNAGATCGGAA
128929 TATGAGTGTGGAGTCGTTGCCTAACTCTTAGGGAAACAACCNNNAGATCGGAAGAG
128930 TATGAGTGTGGAGTCGTTGCTTTATTCTATTATCTGGCCCCANNNAGATCGGAAGAG
128931 TATGAGTGTGGAGTCGTTGCATACCCTCTTGAAGACCAATTATNNNAGATCGGAAG
128932 TATGAGTGTGGAGTCGTTGCTTGGTCAGGTCAGCTTTTTANNNAGATCGGAAGAGC
128933 TATGAGTGTGGAGTCGTTGCATTTTTAAGACTGTTTCATAGGTNNNAGATCGGAAG
128934 TATGAGTGTGGAGTCGTTGCTATATGCTGCTATATGTATTGGGNNNAGATCGGAAG
128935 TATGAGTGTGGAGTCGTTGCCCTAAGCATCTATATCCTGAAGNNNAGATCGGAAGA
128936 TATGAGTGTGGAGTCGTTGCCAATCTCAGGTCCAAGATACNNNAGATCGGAAGAGC
128937 TATGAGTGTGGAGTCGTTGCGATCTACTTGTCTATTCCCATAANNNAGATCGGAAGA
128938 TATGAGTGTGGAGTCGTTGCCAACAAATGTTTGCTGAATAAATNNNAGATCGGAAG
128939 TATGAGTGTGGAGTCGTTGCGATAGCTAAATTCTTGGCACTANNNAGATCGGAAGA
128940 TATGAGTGTGGAGTCGTTGCGTGCTGATTTAAAAACCAGAATNNNAGATCGGAAG
128941 TATGAGTGTGGAGTCGTTGCAATTATCTTCACTTCTTTCTCCTNNNAGATCGGAAGA
128942 TATGAGTGTGGAGTCGTTGCCAGCTACAAGTAACTCCAAGNNNAGATCGGAAGAG
128943 TATGAGTGTGGAGTCGTTGCGAAGAAGCATCCACATTTCANNNAGATCGGAAGAG
128945 TATGAGTGTGGAGTCGTTGCCTTATGAGGACACTAATCCTATCNNNAGATCGGAAG
128946 TATGAGTGTGGAGTCGTTGCCCTATATCAGCAAATAGAAAAGGNNNAGATCGGAA
128947 TATGAGTGTGGAGTCGTTGCACTACATTTCTTATGCAATTCTCNNNAGATCGGAAGA
128948 TATGAGTGTGGAGTCGTTGCAGGCAGATTCAAAGTTATAATTGNNNAGATCGGAA
128949 TATGAGTGTGGAGTCGTTGCCACACTACATACACATACATTGNNNAGATCGGAAGA
128950 TATGAGTGTGGAGTCGTTGCCCACTTGAAGAAAATGATGTGNNNAGATCGGAAGA
128951 TATGAGTGTGGAGTCGTTGCCTCCATCCTCTTTCTCTTTGNNNAGATCGGAAGAGCA
128952 TATGAGTGTGGAGTCGTTGCAGATCAGTTGGCTACATTTATATNNNAGATCGGAAG
128953 TATGAGTGTGGAGTCGTTGCTCTCAAAAAACTACAAGGATTTTNNNAGATCGGAAG
128954 TATGAGTGTGGAGTCGTTGCAAAAGATCAGGTTCAAAAAGAGNNNAGATCGGAAG
128955 TATGAGTGTGGAGTCGTTGCTAGGATTGAGATCTGTTCCANNNAGATCGGAAGAG
128956 TATGAGTGTGGAGTCGTTGCCATATTGAGCAGCTGATCTCNNNAGATCGGAAGAGC

128957 TATGAGTGTGGAGTCGTTGCGGAAGAAAAGCAGAGAATATAAANNNAGATCGGAA
128958 TATGAGTGTGGAGTCGTTGCTTGTGACTTATTAGTGTTCTTTGNNNAGATCGGAAG
128959 TATGAGTGTGGAGTCGTTGCCTCTCTTGCACTTCATTCTCNNNAGATCGGAAGAGCA
128960 TATGAGTGTGGAGTCGTTGCCTGGCTTTTTTCTTTTACTTTTTNNNAGATCGGAAGA
128961 TATGAGTGTGGAGTCGTTGCAATTACTGAAAAGCATTTTTGACNNNAGATCGGAAG
128962 TATGAGTGTGGAGTCGTTGCATGAGTCTTGAACCAATGAATNNNAGATCGGAAGAG
128963 TATGAGTGTGGAGTCGTTGCACATATTTTTTCAATTTCCTCCCNNNAGATCGGAAGA
128964 TATGAGTGTGGAGTCGTTGCCCTATACCAAAGGTGTGTATATANNNAGATCGGAAG
128966 TATGAGTGTGGAGTCGTTGCTGTAGAATGAATTTGACAGAAAGNNNAGATCGGAA
128967 TATGAGTGTGGAGTCGTTGCCGTTACTCATTTTGAGAACTTTNNNAGATCGGAAGA
128968 TATGAGTGTGGAGTCGTTGCGAATACATTCCAGAGGAGACTNNNAGATCGGAAGA
128969 TATGAGTGTGGAGTCGTTGCCAGTCTCCTTGTCTGTAAAATNNNAGATCGGAAGAG
128970 TATGAGTGTGGAGTCGTTGCCCTTGGGATATATTTAGATAAAGCNNNAGATCGGAA
128971 TATGAGTGTGGAGTCGTTGCCCTTTCTCAGATACTAAAGTGTNNNAGATCGGAAGA
128972 TATGAGTGTGGAGTCGTTGCTTTCCTAACACCTTTCTTTATCTNNNAGATCGGAAGAG
128973 TATGAGTGTGGAGTCGTTGCAAGAAGTTTATCGGAAGGTAGNNNAGATCGGAAGA
128974 TATGAGTGTGGAGTCGTTGCATTCCTCTTTCATTTTCTCAATGNNNAGATCGGAAGA
128975 TATGAGTGTGGAGTCGTTGCCCTGACATCTTTGCCAATATTANNNAGATCGGAAGAG
128976 TATGAGTGTGGAGTCGTTGCGGACTGAATTGTTTAGAAAAAACNNNAGATCGGAA
128977 TATGAGTGTGGAGTCGTTGCAGCAATTTTTTTTCAAGTAAGTAANNNAGATCGGAA
128978 TATGAGTGTGGAGTCGTTGCCTTTCTTTCTCTTTCATTGAGATNNNAGATCGGAAGA
128979 TATGAGTGTGGAGTCGTTGCTATGAAAATCATACCCAAAAACCNNNAGATCGGAAG
128980 TATGAGTGTGGAGTCGTTGCTATAGAAGTCAAAGAAGTCAAGTNNNAGATCGGAA
128981 TATGAGTGTGGAGTCGTTGCGATAGATAACAAAAGCCATGATTNNNAGATCGGAAG
128982 TATGAGTGTGGAGTCGTTGCATCTTTAAACAGAGGGCTAAATNNNAGATCGGAAGA
128983 TATGAGTGTGGAGTCGTTGCATTCATGTATACCTATGGTAACCNNNAGATCGGAAG
128984 TATGAGTGTGGAGTCGTTGCTTTCTTTCTTCTTTCTTTCTCTCNNNAGATCGGAAGAG
128985 TATGAGTGTGGAGTCGTTGCGATTCGTCTAGAAATGGATTTGNNNAGATCGGAAGA
128986 TATGAGTGTGGAGTCGTTGCGATTTTGATTCGTCTAGAAATGGNNNAGATCGGAAG
128987 TATGAGTGTGGAGTCGTTGCGAATCTCCACCTAGATTTCACNNNAGATCGGAAGAG
128988 TATGAGTGTGGAGTCGTTGCCATGACTCAGATGTGTATAAAACNNNAGATCGGAAG
128989 TATGAGTGTGGAGTCGTTGCGAGATTATTGTTTCCCTACATGANNNAGATCGGAAG
128990 TATGAGTGTGGAGTCGTTGCCTGCCTTTTGTGAAATTATTAAANNNAGATCGGAAG
128991 TATGAGTGTGGAGTCGTTGCGTATACAGCCATTGAAAAAGAAGNNNAGATCGGAA
128992 TATGAGTGTGGAGTCGTTGCAAACACAGAAACATACCTACANNNAGATCGGAAGAG
128993 TATGAGTGTGGAGTCGTTGCTAAAGGACCAGGTAATAAATGTTNNNAGATCGGAA
128994 TATGAGTGTGGAGTCGTTGCTTTTCATTCTCTTGCTCTCACNNNAGATCGGAAGAGC
128995 TATGAGTGTGGAGTCGTTGCCTCTGTTTTCCAAATCAATCTGNNNAGATCGGAAGAG
128996 TATGAGTGTGGAGTCGTTGCGTATGTATGTATGTGAGACAGAGNNNAGATCGGAA
128997 TATGAGTGTGGAGTCGTTGCACTACTTGCATATTTTAACATAATTNNNAGATCGGAA
128998 TATGAGTGTGGAGTCGTTGCAACTCAAGCCAACTTACAGANNNAGATCGGAAGAGC
128999 TATGAGTGTGGAGTCGTTGCCTCTCTCAGAGATTCCCATGNNNAGATCGGAAGAGC
129000 TATGAGTGTGGAGTCGTTGCCTCTTTCTCATTCTGTCTCTCNNNAGATCGGAAGAGC

129001 TATGAGTGTGGAGTCGTTGCTTCTGATTTTAGTACTGAAAGTCNNNAGATCGGAAG
129002 TATGAGTGTGGAGTCGTTGCCATGAAGAAGAACAAGATCATTGNNNAGATCGGAA
129003 TATGAGTGTGGAGTCGTTGCGGACATGTTACTCTTTCATTATTNNNAGATCGGAAG
129004 TATGAGTGTGGAGTCGTTGCCAGCTGATATTTCTTTTTCTTTTNNNAGATCGGAAGA
129005 TATGAGTGTGGAGTCGTTGCCTGTGGTATGAATGATAAAGTTCNNNAGATCGGAA
129006 TATGAGTGTGGAGTCGTTGCAGGAAAACGTCTCATTTATACCNNNAGATCGGAAGA
129008 TATGAGTGTGGAGTCGTTGCCTCAACTGCTATTCTTTTTCTAGNNNAGATCGGAAGA
129009 TATGAGTGTGGAGTCGTTGCTAAATATCAGACAATGAGTAGGGNNNAGATCGGAA
129010 TATGAGTGTGGAGTCGTTGCCATGTAATCTGCATCTCTGTCNNNAGATCGGAAGAG
129011 TATGAGTGTGGAGTCGTTGCAAAGAGACCAAAACCTAAAAAGANNNAGATCGGAA
129012 TATGAGTGTGGAGTCGTTGCAGACCTCAGCTACAAAATATGNNNAGATCGGAAGAG
129013 TATGAGTGTGGAGTCGTTGCACATTTAGGTTCCTTTTCATTTTNNNAGATCGGAAGA
129014 TATGAGTGTGGAGTCGTTGCATTAGAATGAGTTAAACATCACCNNNAGATCGGAAG
129015 TATGAGTGTGGAGTCGTTGCGGCTGTATAAGAGAAGGTATTCNNNAGATCGGAAG
129016 TATGAGTGTGGAGTCGTTGCGGCTGTATAAGAGAAGGTATTCNNNAGATCGGAAG
129017 TATGAGTGTGGAGTCGTTGCGGGCTGTATAAGAGAAGGTANNNAGATCGGAAGA
129018 TATGAGTGTGGAGTCGTTGCTTTGTGAATAAGAGCTATTAGGANNNAGATCGGAA
129019 TATGAGTGTGGAGTCGTTGCTATGTACACACACAGAAGTCANNNAGATCGGAAGAG
129021 TATGAGTGTGGAGTCGTTGCATTACCAGCAAACCTTCAAANNNAGATCGGAAGAGC
129022 TATGAGTGTGGAGTCGTTGCTGGTCAGAGTTATTTAATTTACCNNNAGATCGGAAG
129023 TATGAGTGTGGAGTCGTTGCTGTCAGCTTACTTCTTATTTTTCNNNAGATCGGAAGA
129024 TATGAGTGTGGAGTCGTTGCAAAATGATTTCCAAAACTGAGCNNNAGATCGGAAGA
129025 TATGAGTGTGGAGTCGTTGCAAATAAGTACAGCAATCAGAAAGNNNAGATCGGAA
129026 TATGAGTGTGGAGTCGTTGCGTCAAACCTGTTAGAAAAATCTGNNNAGATCGGAAG
129027 TATGAGTGTGGAGTCGTTGCAGGATTATTACAAAAGGGATTCTNNNAGATCGGAA
129028 TATGAGTGTGGAGTCGTTGCGAGATACCAGTTTTGGAATTAACNNNAGATCGGAAG
129029 TATGAGTGTGGAGTCGTTGCGAGATACCAGTTTTGGAATTAACNNNAGATCGGAAG
129030 TATGAGTGTGGAGTCGTTGCTACTCTTGTTATCCTCCTATTTCNNNAGATCGGAAGA
129031 TATGAGTGTGGAGTCGTTGCTACTCTTGTTATCCTCCTATTTCNNNAGATCGGAAGA
129032 TATGAGTGTGGAGTCGTTGCAGATGTCTACAAGGAAATTCTCNNNAGATCGGAAGA
129033 TATGAGTGTGGAGTCGTTGCGTGACTATACATGAAGTTACTCTNNNAGATCGGAAG
129034 TATGAGTGTGGAGTCGTTGCACAAGTTGAATAATAGGGACTAANNNAGATCGGAA
129036 TATGAGTGTGGAGTCGTTGCTTAGTTCAGAGACTAAAGGTAAGNNNAGATCGGAA
129037 TATGAGTGTGGAGTCGTTGCAAGATTCCTAACTAGTTCATCATNNNAGATCGGAAG
129038 TATGAGTGTGGAGTCGTTGCCGAATCATGTACCAGTTTTATTTNNNAGATCGGAAG
129039 TATGAGTGTGGAGTCGTTGCTATACTTAGGATTTCTGGTGGTANNNAGATCGGAAG
129040 TATGAGTGTGGAGTCGTTGCACGAATGGTAATGAAATGAAAATNNNAGATCGGAA
129041 TATGAGTGTGGAGTCGTTGCCATCAAAAGGCAAAACACTTNNNAGATCGGAAGAG
129042 TATGAGTGTGGAGTCGTTGCTAACAAATGCTTACCAAGAAGTTNNNAGATCGGAAG
129043 TATGAGTGTGGAGTCGTTGCATGACTTCTAAACCCATTTCCNNNAGATCGGAAGAG
129044 TATGAGTGTGGAGTCGTTGCGAAAGTGGAAAATGGGTAGAGNNNAGATCGGAAG
129045 TATGAGTGTGGAGTCGTTGCCATAGAGAAAGAGCAAAAGTAAANNNAGATCGGAA
129046 TATGAGTGTGGAGTCGTTGCACACCATCTATAATTGCTTTTTAANNNAGATCGGAAG

129047 TATGAGTGTGGAGTCGTTGCTTGGCTTTGATTAAATGTGAAATNNNAGATCGGAAG
129048 TATGAGTGTGGAGTCGTTGCGAAGCTCTAGGAAAGAATCTTANNNAGATCGGAAG
129049 TATGAGTGTGGAGTCGTTGCATGTTTCTGTGTGTATGTGTCNNNAGATCGGAAGAG
129050 TATGAGTGTGGAGTCGTTGCGTGTGTCTATTTGAGATGTGTNNNAGATCGGAAGA
129051 TATGAGTGTGGAGTCGTTGCCCAAGCTAATCAATTGTCTTAGNNNAGATCGGAAGA
129052 TATGAGTGTGGAGTCGTTGCATATAAAGGTGACAGAAAAACAGNNNAGATCGGAA
129053 TATGAGTGTGGAGTCGTTGCATTCACTAAACACTTTTGAGTGNNNAGATCGGAAGA
129054 TATGAGTGTGGAGTCGTTGCAATAGCAAAACAGACACATTTAANNNAGATCGGAAG
129055 TATGAGTGTGGAGTCGTTGCTTGACGACTATTTCATTTCTCCNNNAGATCGGAAGAG
129056 TATGAGTGTGGAGTCGTTGCGTGGATTAAATAGCTTTCAAAGANNNAGATCGGAA
129057 TATGAGTGTGGAGTCGTTGCGAATTCTGACTCAATCTTTTTCTNNNAGATCGGAAGA
129058 TATGAGTGTGGAGTCGTTGCGTTCACAACAGTAATACAAGTAANNNAGATCGGAAG
129059 TATGAGTGTGGAGTCGTTGCTCTCTTCTTCACATCTACTTCTANNNAGATCGGAAGA
129060 TATGAGTGTGGAGTCGTTGCCTTCATCATTTTTTATGGTGTCTNNNAGATCGGAAGA
129061 TATGAGTGTGGAGTCGTTGCATATTTGAAAACACTAATCAGCANNNAGATCGGAAG
129062 TATGAGTGTGGAGTCGTTGCGTTACCTAGGATGATTAGCTGNNNAGATCGGAAGA
129063 TATGAGTGTGGAGTCGTTGCAAGGAAAAAGATTTTAATGGACTNNNAGATCGGAA
129064 TATGAGTGTGGAGTCGTTGCAGTATAAAATCTCATGTCCATCTNNNAGATCGGAAG
129065 TATGAGTGTGGAGTCGTTGCCCATGATTGTAAGTTTTCTGAGNNNAGATCGGAAGA
129066 TATGAGTGTGGAGTCGTTGCGGGTCATTCTTCATAAATTACAANNNAGATCGGAAG
129067 TATGAGTGTGGAGTCGTTGCAACAAGTAAAAATTTTTGGAACCNNNAGATCGGAAG
129068 TATGAGTGTGGAGTCGTTGCTTACAACTGACACTAAATGAAAGNNNAGATCGGAAG
129069 TATGAGTGTGGAGTCGTTGCTCATTATGTTCTGTTTTGCATAANNNAGATCGGAAG
129070 TATGAGTGTGGAGTCGTTGCATTGGATTCTCAGAACTTGTTANNNAGATCGGAAGA
129071 TATGAGTGTGGAGTCGTTGCGCTGATTCATTTCAGATGTTTAANNNAGATCGGAAG
129072 TATGAGTGTGGAGTCGTTGCAGACAAAATCTGGCTTATAGCNNNAGATCGGAAGAG
129073 TATGAGTGTGGAGTCGTTGCATAGGTCTTCACCATCATAATAANNNAGATCGGAAG
129074 TATGAGTGTGGAGTCGTTGCACTGATCTATCATCTATCTACCTNNNAGATCGGAAGA
129075 TATGAGTGTGGAGTCGTTGCACTGTCAGTTATAGTAATTCTGCNNNAGATCGGAAG
129076 TATGAGTGTGGAGTCGTTGCTATTCTGCAAGGATAATTGAATGNNNAGATCGGAAG
129078 TATGAGTGTGGAGTCGTTGCTATTCGAGGGTGAGTCTTTCNNNAGATCGGAAGAG
129079 TATGAGTGTGGAGTCGTTGCGGGTATTCAAATATTCAGTTTCANNNAGATCGGAAG
129081 TATGAGTGTGGAGTCGTTGCCAGAGAAATATTTTGTGAAAGGANNNAGATCGGAA
129082 TATGAGTGTGGAGTCGTTGCTCAATGAGATAGGAGACATTTAGNNNAGATCGGAA
129083 TATGAGTGTGGAGTCGTTGCAATTCTAATGAGGTAGGTAACACNNNAGATCGGAA
129084 TATGAGTGTGGAGTCGTTGCTTTGTAACTGACATGATCTGATCNNNAGATCGGAAG
129085 TATGAGTGTGGAGTCGTTGCATTTCCCCACTTCATATTGATCNNNAGATCGGAAGAG
129086 TATGAGTGTGGAGTCGTTGCCACTTGAAATTTTTTACTCCCTCNNNAGATCGGAAGA
129087 TATGAGTGTGGAGTCGTTGCCACTTGAAATTTTTTACTCCCTCNNNAGATCGGAAGA
129088 TATGAGTGTGGAGTCGTTGCCATGTGTGTCAATAATTCTAGGNNNAGATCGGAAGA
129089 TATGAGTGTGGAGTCGTTGCCCTTTTGAGACTTCTTTTCTACNNNAGATCGGAAGAG
129090 TATGAGTGTGGAGTCGTTGCCTCTCAACTTTACATAATGAGATNNNAGATCGGAAG
129091 TATGAGTGTGGAGTCGTTGCTGTGTGTATCTATACTGTATATCTNNNAGATCGGAA

129092 TATGAGTGTGGAGTCGTTGCTATCAGAATGCATACCACATAATNNNAGATCGGAAG
129093 TATGAGTGTGGAGTCGTTGCTTTGTGTAATGATTAGTGATGTTNNNAGATCGGAAG
129096 TATGAGTGTGGAGTCGTTGCGCATACCTTCAATTGCTATAATTNNNAGATCGGAAG
129097 TATGAGTGTGGAGTCGTTGCCTTCTCAGTTCAGCTACCTTNNNAGATCGGAAGAGC
129098 TATGAGTGTGGAGTCGTTGCAGAATACCTACTTGAAGGAGTTNNNAGATCGGAAG
129099 TATGAGTGTGGAGTCGTTGCTCTCTTTCTTTGTCTCTCTTTTANNNAGATCGGAAGA
129100 TATGAGTGTGGAGTCGTTGCTCTGATTTGAGGAATGTAAACTANNNAGATCGGAAG
129101 TATGAGTGTGGAGTCGTTGCAGTTTGAGAATCCTTGACTTATANNNAGATCGGAAG
129103 TATGAGTGTGGAGTCGTTGCGTAGTTTGAGGGAAAGACCTNNNAGATCGGAAGAG
129104 TATGAGTGTGGAGTCGTTGCCTCTCCAACTAAGAAACTGGNNNAGATCGGAAGAG
129105 TATGAGTGTGGAGTCGTTGCCTCTCCAACTAAGAAACTGGNNNAGATCGGAAGAG
129106 TATGAGTGTGGAGTCGTTGCATGATTGTAAGTTTCCTGAGGNNNAGATCGGAAGA
129108 TATGAGTGTGGAGTCGTTGCGAATCTTTTCCTTCCTTTGAACNNNAGATCGGAAGAG
129109 TATGAGTGTGGAGTCGTTGCTAATTCACAAACACAAGGGTGNNNAGATCGGAAGA
129110 TATGAGTGTGGAGTCGTTGCTTTCTCTTTCTCTCTTTCTGTTTNNNAGATCGGAAGAG
129111 TATGAGTGTGGAGTCGTTGCGTTTCCTCAGCTTCTCTTTCNNNAGATCGGAAGAGCA
129112 TATGAGTGTGGAGTCGTTGCCTCAGATCTCATCTCTTATGACNNNAGATCGGAAGAG
129113 TATGAGTGTGGAGTCGTTGCAAAAATAAACTGTTGTGAGATCANNNAGATCGGAAG
129114 TATGAGTGTGGAGTCGTTGCCTATAATGCTATACTGCCTCTTTNNNAGATCGGAAGA
129115 TATGAGTGTGGAGTCGTTGCCATACACATACATACATGTTTGGNNNAGATCGGAAG
129116 TATGAGTGTGGAGTCGTTGCATAGGTCTTCACCATCATAATAANNNAGATCGGAAG
129117 TATGAGTGTGGAGTCGTTGCCCTGTATACTGCTTCTAGAAATANNNAGATCGGAAG
129118 TATGAGTGTGGAGTCGTTGCGACAAATAATCACAGCAAATAGTNNNAGATCGGAAG
129119 TATGAGTGTGGAGTCGTTGCAGATACATAGGTAGATAGGTAGANNNAGATCGGAA
129122 TATGAGTGTGGAGTCGTTGCAATCACATAAACTAGCAAAGATCNNNAGATCGGAAG
129123 TATGAGTGTGGAGTCGTTGCTAAATTTTGTGTATATCTTCCCGNNNAGATCGGAAG
129124 TATGAGTGTGGAGTCGTTGCCCAATTGTGAAATAAAGGGAATTNNNAGATCGGAA
129125 TATGAGTGTGGAGTCGTTGCCATACAATTTACATGGCTGAATANNNAGATCGGAAG
129127 TATGAGTGTGGAGTCGTTGCGCCTAGTATGGTCTTCTCTANNNAGATCGGAAGAGC
129128 TATGAGTGTGGAGTCGTTGCAGAGAAGAAAAGAGAAACCAAGNNNAGATCGGAA
129129 TATGAGTGTGGAGTCGTTGCTGAATAATAAGCATCATAGCATGNNNAGATCGGAA
129130 TATGAGTGTGGAGTCGTTGCTCTTTTAGGGACTTAGGGATAANNNAGATCGGAAG
129131 TATGAGTGTGGAGTCGTTGCGTTTTGTATCTACCCAGCAAANNNAGATCGGAAGAG
129132 TATGAGTGTGGAGTCGTTGCTTTGTTTTATGAACAAACCAGAGNNNAGATCGGAAG
129134 TATGAGTGTGGAGTCGTTGCCGAAGTTCATATATGGAAATCCTNNNAGATCGGAAG
129135 TATGAGTGTGGAGTCGTTGCTCAGTTCTGTATTTTACTGAGTGNNNAGATCGGAAG
129136 TATGAGTGTGGAGTCGTTGCAACGTACAAAACAAGGGAAATNNNAGATCGGAAGA
129137 TATGAGTGTGGAGTCGTTGCAACAGTAGTTCATCTAAGACTGNNNAGATCGGAAGA
129138 TATGAGTGTGGAGTCGTTGCTCTCTCTGGAAATTCAACTATAANNNAGATCGGAAG
129139 TATGAGTGTGGAGTCGTTGCAAATAAGCAAGGAAAAAAAACAGNNNAGATCGGAA
129140 TATGAGTGTGGAGTCGTTGCTTTGGTGGTTTAATGTTCTATTTNNNAGATCGGAAG
129141 TATGAGTGTGGAGTCGTTGCTATATCCCTCACACATCTAGTAANNNAGATCGGAAGA
129142 TATGAGTGTGGAGTCGTTGCGACTCAGAGTAACTTTTCTTTATNNNAGATCGGAAG

129143 TATGAGTGTGGAGTCGTTGCATCTAGATAATTCTCACTCCAACNNNAGATCGGAAGA
129144 TATGAGTGTGGAGTCGTTGCGATTATTGTAGCTCTGTGGTATNNNAGATCGGAAGA
129145 TATGAGTGTGGAGTCGTTGCCTTTGTCTCTATGAATTTCCAANNNAGATCGGAAGA
129146 TATGAGTGTGGAGTCGTTGCGTAGTATTGCTTGATCCTTGTANNNAGATCGGAAGA
129147 TATGAGTGTGGAGTCGTTGCCACCTGTAATCCTATTGCTTTNNNAGATCGGAAGAGC
129148 TATGAGTGTGGAGTCGTTGCTGGCTTGTCTATTTAGTCATGNNNAGATCGGAAGAG
129149 TATGAGTGTGGAGTCGTTGCTAATTTCTTCTATAAGTCAGCCANNNAGATCGGAAGA
129150 TATGAGTGTGGAGTCGTTGCTCTGTTTTTGCATGTTTACATAGNNNAGATCGGAAGA
129151 TATGAGTGTGGAGTCGTTGCCTTAGGACATTGGATTGGAGNNNAGATCGGAAGAG
129152 TATGAGTGTGGAGTCGTTGCATCATGCATTCACTTATAAACTTNNNAGATCGGAAGA
129153 TATGAGTGTGGAGTCGTTGCAAATTTCTTGAGGTTTAGAAGTCNNNAGATCGGAAG
129154 TATGAGTGTGGAGTCGTTGCACATTCAAAATGCAAATTTTCATNNNAGATCGGAAGA
129155 TATGAGTGTGGAGTCGTTGCCATAGCAACAATGAACAATGGNNNAGATCGGAAGAG
129156 TATGAGTGTGGAGTCGTTGCACTTTACACATTCCATTATAAGCNNNAGATCGGAAGA
129157 TATGAGTGTGGAGTCGTTGCAATTTTTACAAAAAAAAAGGCCTNNNAGATCGGAAG
129158 TATGAGTGTGGAGTCGTTGCATATGAGGCATTTTATATGCTTTNNNAGATCGGAAGA
129159 TATGAGTGTGGAGTCGTTGCCCTTATTTAGGCATCCTAACATNNNAGATCGGAAGAG
129160 TATGAGTGTGGAGTCGTTGCGATTTCTGCTTTAAGGAAAACTNNNAGATCGGAAGA
129161 TATGAGTGTGGAGTCGTTGCGTTTTTCTACCAGTTACATTCTTNNNAGATCGGAAGA
129162 TATGAGTGTGGAGTCGTTGCGTTTTTCTACCAGTTACATTCTTNNNAGATCGGAAGA
129163 TATGAGTGTGGAGTCGTTGCGTTTTTCTACCAGTTACATTCTTNNNAGATCGGAAGA
129164 TATGAGTGTGGAGTCGTTGCCAATCCCCTTCTCATAATTATTGNNNAGATCGGAAGA
129165 TATGAGTGTGGAGTCGTTGCGGGCAGAACTTTCTATTTATTAANNNAGATCGGAAG
129166 TATGAGTGTGGAGTCGTTGCATGTATCCTTCCTTCTTTTCTTCNNNAGATCGGAAGAG
129167 TATGAGTGTGGAGTCGTTGCATGTATCCTTCCTTCTTTTCTTCNNNAGATCGGAAGAG
129168 TATGAGTGTGGAGTCGTTGCTAATTATAAACTAGTGCCATTCGNNNAGATCGGAAGA
129169 TATGAGTGTGGAGTCGTTGCCAAAACCCCCTTTCTACTAAANNNAGATCGGAAGAGC
129170 TATGAGTGTGGAGTCGTTGCACAATCAATTTCAACTATGTGAANNNAGATCGGAAGA
129171 TATGAGTGTGGAGTCGTTGCAAAAGAGAAACAGACTGTCATNNNAGATCGGAAGAG
129172 TATGAGTGTGGAGTCGTTGCGTGATAGTTTGGGACCCTATNNNAGATCGGAAGAGC
129173 TATGAGTGTGGAGTCGTTGCCTTTCCAAAGATTCTCAAAAGTNNNAGATCGGAAGAG
129174 TATGAGTGTGGAGTCGTTGCACTTGATGATCTTGTTTAGTTTCNNNAGATCGGAAGA
129175 TATGAGTGTGGAGTCGTTGCCTTTTCCTTTATGCTAGGGTATTNNNAGATCGGAAGA
129176 TATGAGTGTGGAGTCGTTGCCTGGCAACTCAGTAATAAAAAGNNNAGATCGGAAGA
129177 TATGAGTGTGGAGTCGTTGCCTACACTGTAACGTGAGTTTTATNNNAGATCGGAAGA
129178 TATGAGTGTGGAGTCGTTGCGAGAGTTTCCATGAACAGAAANNNAGATCGGAAGAG
129179 TATGAGTGTGGAGTCGTTGCCAGTGGTCCCATAAGATTATAATNNNAGATCGGAAG
129180 TATGAGTGTGGAGTCGTTGCTTGGTCTCTCTTACTAATGAAATNNNAGATCGGAAGA
129181 TATGAGTGTGGAGTCGTTGCGGCCAAAGGATTATGAAAATATTNNNAGATCGGAAG
129182 TATGAGTGTGGAGTCGTTGCTTTTAGAATAAGTTGCACATGAANNNAGATCGGAAG
129183 TATGAGTGTGGAGTCGTTGCCTACCCTGGTTTCTATAAGTATGNNNAGATCGGAAGA
129184 TATGAGTGTGGAGTCGTTGCTTCACAGAATTTTTATGCACTAANNNAGATCGGAAGA
129186 TATGAGTGTGGAGTCGTTGCATGATTTTCATCCCTTCTCTGNNNAGATCGGAAGAGC

129187 TATGAGTGTGGAGTCGTTGCAGATGTATTTTTGAACGTAGAAGNNNAGATCGGAA
129188 TATGAGTGTGGAGTCGTTGCCTATTTGCTGTTATTTGTTGCANNNAGATCGGAAGA
129189 TATGAGTGTGGAGTCGTTGCGTACTTGTCATTAATTTCGGAAANNNAGATCGGAAG
129190 TATGAGTGTGGAGTCGTTGCCCTCTAAAGTCTCTGGAAATTTTNNNAGATCGGAAG
129191 TATGAGTGTGGAGTCGTTGCTTAGTAAATCTATTTGGCTTGTGNNNAGATCGGAAG
129192 TATGAGTGTGGAGTCGTTGCTTTCCAGAAAACAAAACAGAAAANNNAGATCGGAA
129193 TATGAGTGTGGAGTCGTTGCTAGTTTTACTACTTCTTTCCACTNNNAGATCGGAAGA
129195 TATGAGTGTGGAGTCGTTGCGAAATGGAGGAAGGAGAAAGNNNAGATCGGAAG
129196 TATGAGTGTGGAGTCGTTGCATTCTTCATACAGATTCCTAACANNNAGATCGGAAG
129197 TATGAGTGTGGAGTCGTTGCATTCTTCATACAGATTCCTAACANNNAGATCGGAAG
129198 TATGAGTGTGGAGTCGTTGCATTCTTCATACAGATTCCTAACANNNAGATCGGAAG
129199 TATGAGTGTGGAGTCGTTGCAATCAGAGGAGAAGCAATTAGNNNAGATCGGAAG
129200 TATGAGTGTGGAGTCGTTGCCCTCTCCTTTTCTGTTTTTTATTNNNAGATCGGAAGA
129201 TATGAGTGTGGAGTCGTTGCCCTCTCCTTTTCTGTTTTTTATTNNNAGATCGGAAGA
129202 TATGAGTGTGGAGTCGTTGCCCAAAGTGCTAGGATTACAGNNNAGATCGGAAGAG
129203 TATGAGTGTGGAGTCGTTGCTTTTATGATCTTTTCCTTCTGTGNNNAGATCGGAAGA
129204 TATGAGTGTGGAGTCGTTGCTTTTCAAGGATATTGGTTATTCANNNAGATCGGAAG
129205 TATGAGTGTGGAGTCGTTGCGACTTATATGTTTGCCTGTTAATNNNAGATCGGAAG
129206 TATGAGTGTGGAGTCGTTGCTCCACCTAGGAAAACTAAGGNNNAGATCGGAAGAG
129207 TATGAGTGTGGAGTCGTTGCTCCACCTAGGAAAACTAAGGNNNAGATCGGAAGAG
129208 TATGAGTGTGGAGTCGTTGCTTTCTCTCTCACTTACTTTTACANNNAGATCGGAAGA
129209 TATGAGTGTGGAGTCGTTGCGGCTCAAAATCTGATATTGTATCNNNAGATCGGAAG
129210 TATGAGTGTGGAGTCGTTGCGGCTCAAAATCTGATATTGTATCNNNAGATCGGAAG
129211 TATGAGTGTGGAGTCGTTGCACTTCTAGTTTACTTTTATCATCCNNNAGATCGGAAG
129212 TATGAGTGTGGAGTCGTTGCATGATGCATAAAAAGACATTGAGNNNAGATCGGAA
129213 TATGAGTGTGGAGTCGTTGCAAGGTGATAGGGTAAGACTGNNNAGATCGGAAGA
129214 TATGAGTGTGGAGTCGTTGCCACCATCATAACTCACTACAGNNNAGATCGGAAGAG
129215 TATGAGTGTGGAGTCGTTGCACATTTTTTAAACAGAAAGCTTGNNNAGATCGGAAG
129216 TATGAGTGTGGAGTCGTTGCATTTTAGTTTGGAGTGTTACATGNNNAGATCGGAAG
129217 TATGAGTGTGGAGTCGTTGCCTGACTTTGCAAAAGAAATAAAANNNAGATCGGAA
129218 TATGAGTGTGGAGTCGTTGCCTGACCTAACTAACTCTATCTTGNNNAGATCGGAAG
129219 TATGAGTGTGGAGTCGTTGCGGGTTTTTTTTTGTTTTTGTTTTNNNAGATCGGAAG
129220 TATGAGTGTGGAGTCGTTGCTTGATAAATAAACATTCAGGCAGNNNAGATCGGAA
129221 TATGAGTGTGGAGTCGTTGCAAATGAAATGTTGCAATAAGTTGNNNAGATCGGAA
129222 TATGAGTGTGGAGTCGTTGCTATTGATGAGTCTCTCAAATCATNNNAGATCGGAAG
129223 TATGAGTGTGGAGTCGTTGCGCTGGGAGGTAATAAAAGATAGNNNAGATCGGAA
129224 TATGAGTGTGGAGTCGTTGCCCATCTGAGTAAAAGTAATTCTTAANNNAGATCGGA
129225 TATGAGTGTGGAGTCGTTGCTTTTGCATTGTTTACAGGATAATNNNAGATCGGAAG
129226 TATGAGTGTGGAGTCGTTGCACTTCTTCTTCCTATAATTTCCANNNAGATCGGAAGA
129227 TATGAGTGTGGAGTCGTTGCGAGAGAGGAGAGAGAGATATATANNNAGATCGGA
129228 TATGAGTGTGGAGTCGTTGCTGATGATTGTTATCATTTCCAAANNNAGATCGGAAG
129229 TATGAGTGTGGAGTCGTTGCGATTGCATTTTTTTCCTCTTTGNNNAGATCGGAAGAG
129230 TATGAGTGTGGAGTCGTTGCCAAAGATGGAGAGTAAATACAAGNNNAGATCGGAA

129231 TATGAGTGTGGAGTCGTTGCATTTACAAAGCAGTTACCAAATTNNNAGATCGGAAG
129232 TATGAGTGTGGAGTCGTTGCGACACTTATTATAAGGAATTGGCNNNAGATCGGAAG
129233 TATGAGTGTGGAGTCGTTGCTCAATGTAACACTGCTATCAANNNAGATCGGAAGAG
129234 TATGAGTGTGGAGTCGTTGCCTTTTTGGAGGGGTAAGGAANNNAGATCGGAAGAG
129235 TATGAGTGTGGAGTCGTTGCGAATGGCATTTTGATTGTCTATNNNAGATCGGAAGA
129236 TATGAGTGTGGAGTCGTTGCGTGTAGTTTAGAATGAAGTGAAANNNAGATCGGAA
129237 TATGAGTGTGGAGTCGTTGCCTGGATACACATAGGTATACATTNNNAGATCGGAAG
129238 TATGAGTGTGGAGTCGTTGCATTTTTTCTTTTTAGCATGCATTNNNAGATCGGAAGA
129239 TATGAGTGTGGAGTCGTTGCCACATTAGTAAGAGTGATCTACTNNNAGATCGGAAG
129240 TATGAGTGTGGAGTCGTTGCCATCTGTTTATGCTTTTGGAATANNNAGATCGGAAG
129242 TATGAGTGTGGAGTCGTTGCATGATGTCATCAATGTATTGGATNNNAGATCGGAAG
129243 TATGAGTGTGGAGTCGTTGCTTTCTCAAGGATCCTTTCTCATNNNAGATCGGAAGAG
129244 TATGAGTGTGGAGTCGTTGCCACACTTGCATTTTCTCAAGNNNAGATCGGAAGAGC
129245 TATGAGTGTGGAGTCGTTGCTTTCCATGGAAAAATGCCAANNNAGATCGGAAGAG
129246 TATGAGTGTGGAGTCGTTGCTCATCCAGAAAACAATTAACAAANNNAGATCGGAAG
129247 TATGAGTGTGGAGTCGTTGCATTGGGTTAGATTGTTGACATANNNAGATCGGAAG
129248 TATGAGTGTGGAGTCGTTGCAAAGAGAAAGCAAAAGGAATAAANNNAGATCGGA
129249 TATGAGTGTGGAGTCGTTGCCTTGCTTTTCAGAAAGTCACNNNAGATCGGAAGAGC
129250 TATGAGTGTGGAGTCGTTGCCAGAAGATGCCTAATTTCTATCTNNNAGATCGGAAG
129251 TATGAGTGTGGAGTCGTTGCGAATAATACCCACAGCAGAATANNNAGATCGGAAG
129253 TATGAGTGTGGAGTCGTTGCTAAAAAACTCCTCTACTTGTACTNNNAGATCGGAAG
129254 TATGAGTGTGGAGTCGTTGCTAAGGAATTCAGACAAGACAGNNNAGATCGGAAGA
129255 TATGAGTGTGGAGTCGTTGCGAAAGAGAATTCCAGGTAAAAAGNNNAGATCGGAA
129256 TATGAGTGTGGAGTCGTTGCACACTAAGACAGTAATTTTTTCCNNNAGATCGGAAG
129257 TATGAGTGTGGAGTCGTTGCCTGAATCACCACTAATTATACCTNNNAGATCGGAAG
129258 TATGAGTGTGGAGTCGTTGCCTGAATCACCACTAATTATACCTNNNAGATCGGAAG
129259 TATGAGTGTGGAGTCGTTGCGAAAGAAAATCTACAGCTATCACNNNAGATCGGAAG
129260 TATGAGTGTGGAGTCGTTGCGCTTTTTTCTTGCAAGGAATTNNNAGATCGGAAGAG
129261 TATGAGTGTGGAGTCGTTGCTGTCAACAGGATGATTTCTATAANNNAGATCGGAAG
129262 TATGAGTGTGGAGTCGTTGCCACCACATTATCTCACTAGATCNNNAGATCGGAAGA
129263 TATGAGTGTGGAGTCGTTGCGCCAAGTAAAGTAGAAAGTAGTNNNAGATCGGAAG
129264 TATGAGTGTGGAGTCGTTGCATATAAAGAACTGTGACAGATCANNNAGATCGGAA
129265 TATGAGTGTGGAGTCGTTGCAGATTCATAGTAAATTCTCCTGTNNNAGATCGGAAG
129266 TATGAGTGTGGAGTCGTTGCCTCAATCTTGAAGCTTCTTATAGNNNAGATCGGAAG
129267 TATGAGTGTGGAGTCGTTGCCTCTAATAACTTTCCATCCTCAGNNNAGATCGGAAGA
129268 TATGAGTGTGGAGTCGTTGCGGGTTATAAATGTGGTTGCANNNAGATCGGAAGAG
129269 TATGAGTGTGGAGTCGTTGCGAGAACTATGCAATGGAGTTTNNNAGATCGGAAGA
129270 TATGAGTGTGGAGTCGTTGCTTTTAGAGTAAATTGTCACACAGNNNAGATCGGAAG
129271 TATGAGTGTGGAGTCGTTGCCCCAAAATGTGTATATGCTAATANNNAGATCGGAAG
129272 TATGAGTGTGGAGTCGTTGCTCAAGCAAAGATGAAAAATGAGNNNAGATCGGAAG
129273 TATGAGTGTGGAGTCGTTGCGGTAACTAATCATTAAACCCACTNNNAGATCGGAAG
129274 TATGAGTGTGGAGTCGTTGCGTGAAACTACTATGATCTTCCATNNNAGATCGGAAG
129275 TATGAGTGTGGAGTCGTTGCTTTACAAGTTTATGTCTGGGTTNNNAGATCGGAAGA

129276 TATGAGTGTGGAGTCGTTGCTTTACAAGTTTATGTCTGGGTTNNNAGATCGGAAGA
129277 TATGAGTGTGGAGTCGTTGCGTTTCCATATCTATCAAAAACCANNNAGATCGGAAG
129278 TATGAGTGTGGAGTCGTTGCTATATAACAGCACCTTTCTGAGANNNAGATCGGAAG
129279 TATGAGTGTGGAGTCGTTGCTTTCTGTTGTAGACTGAATAATGNNNAGATCGGAAG
129280 TATGAGTGTGGAGTCGTTGCTTTCTGTTGTAGACTGAATAATGNNNAGATCGGAAG
129281 TATGAGTGTGGAGTCGTTGCAATAGTGTTGGAACTGAATTAGANNNAGATCGGAA
129282 TATGAGTGTGGAGTCGTTGCCTTGGAAGTTTCTCTAGTATTTGNNNAGATCGGAAG
129283 TATGAGTGTGGAGTCGTTGCTTTCTATAATAGCTATCCTCCAGNNNAGATCGGAAG
129284 TATGAGTGTGGAGTCGTTGCGCTCTAAAATCCTAAGACAAATTNNNAGATCGGAAG
129285 TATGAGTGTGGAGTCGTTGCGAAAGAGAGAGAGAAAGCAGNNNAGATCGGAAGA
129287 TATGAGTGTGGAGTCGTTGCGAAAGTACTGTCATCTTTGTATTNNNAGATCGGAAG
129288 TATGAGTGTGGAGTCGTTGCTTTATCTGGGAGGTCAGAAAATANNNAGATCGGAA
129289 TATGAGTGTGGAGTCGTTGCTTCACAAATTAAGGGAATAATGGNNNAGATCGGAA
129290 TATGAGTGTGGAGTCGTTGCTTCACAAATTAAGGGAATAATGGNNNAGATCGGAA
129291 TATGAGTGTGGAGTCGTTGCAAGAACAGGAACTAACATTTTTTNNNAGATCGGAAG
129292 TATGAGTGTGGAGTCGTTGCTGATGCAAGAGATTAGTATAATCTNNNAGATCGGA
129293 TATGAGTGTGGAGTCGTTGCGAAAACTCACCATTTGTCAAGNNNAGATCGGAAGAG
129294 TATGAGTGTGGAGTCGTTGCATCAAGATGTATTATGAGGGATTNNNAGATCGGAA
129295 TATGAGTGTGGAGTCGTTGCTCTTCTCATTTCTTTCTTTTTTTNNNAGATCGGAAGAG
129296 TATGAGTGTGGAGTCGTTGCTTAAAGCACCTCATCTCTCTTNNNAGATCGGAAGAG
129297 TATGAGTGTGGAGTCGTTGCTCAAGTCTTTTCATTTAAAACAANNNAGATCGGAAG
129298 TATGAGTGTGGAGTCGTTGCACTGCCAATATGAATAGAAAAAANNNAGATCGGAA
129299 TATGAGTGTGGAGTCGTTGCCCTTGTGAGAGACATTCAACNNNAGATCGGAAGAG
129300 TATGAGTGTGGAGTCGTTGCACTCACTTACTAAAGGATTAACTNNNAGATCGGAAG
129301 TATGAGTGTGGAGTCGTTGCGTCTATCAAGAGGGAAAAAAATANNNAGATCGGAA
129302 TATGAGTGTGGAGTCGTTGCAGAGTTTTCTATTTTTTTTGTGTNNNAGATCGGAAG
129303 TATGAGTGTGGAGTCGTTGCCTCAAGAGAAACAAATATTTTAAGGNNNAGATCGG
129304 TATGAGTGTGGAGTCGTTGCTAATGAACATATTAGGTCAGGAGNNNAGATCGGAA
129305 TATGAGTGTGGAGTCGTTGCATGTATGAGTGTTTAGTTCTAAANNNAGATCGGAAG
129306 TATGAGTGTGGAGTCGTTGCCTACATATAGGCTAGAACCCTANNNAGATCGGAAGA
129307 TATGAGTGTGGAGTCGTTGCGTCTTACTGATGGACCTCTANNNAGATCGGAAGAGC
129308 TATGAGTGTGGAGTCGTTGCGCGGAAACTGTAAAAAAGAATATNNNAGATCGGAA
129309 TATGAGTGTGGAGTCGTTGCTCAGAAAGAAGACTATTGAGAAGNNNAGATCGGAA
129310 TATGAGTGTGGAGTCGTTGCGAAGAAGACAAGTGTGACTATNNNAGATCGGAAGA
129311 TATGAGTGTGGAGTCGTTGCTGAATCTATAACAGCATTTTGAGNNNAGATCGGAAG
129312 TATGAGTGTGGAGTCGTTGCTATAGAAAGGACAAAATAGGACANNNAGATCGGAA
129313 TATGAGTGTGGAGTCGTTGCTACCACAGAAGTATTTTAAAAGTNNNAGATCGGAAG
129314 TATGAGTGTGGAGTCGTTGCCTTTATCCAACTTCAGCAATAGNNNAGATCGGAAGA
129315 TATGAGTGTGGAGTCGTTGCCCTCTGCAGCAGATAATAAATNNNAGATCGGAAGAG
129316 TATGAGTGTGGAGTCGTTGCCTATGGGGTTCTTAGAAAGAAANNNAGATCGGAAG
129317 TATGAGTGTGGAGTCGTTGCGCTTTTCAGAAGAGAAGAGATNNNAGATCGGAAGA
129318 TATGAGTGTGGAGTCGTTGCCTTTATCCATTTCCTCTTTTGTGNNNAGATCGGAAGA
129319 TATGAGTGTGGAGTCGTTGCTAGAGAAGAAGGGGATCTTATAGNNNAGATCGGAA

129320 TATGAGTGTGGAGTCGTTGCACATTTCTCAGTAGTATCTTAGGNNNAGATCGGAAG
129321 TATGAGTGTGGAGTCGTTGCTTTGTTTCCTTTTGATGCATATANNNAGATCGGAAGA
129322 TATGAGTGTGGAGTCGTTGCCACTACCTCTTGCAATATTTTAANNNAGATCGGAAGA
129323 TATGAGTGTGGAGTCGTTGCGAACACTGAGAAAGATAAAAGACNNNAGATCGGAA
129324 TATGAGTGTGGAGTCGTTGCCGAAGGATACACAAATAGATTGTNNNAGATCGGAA
129325 TATGAGTGTGGAGTCGTTGCAACAAAACAATAACAATCATTACANNNAGATCGGAA
129326 TATGAGTGTGGAGTCGTTGCCCTTGACTACTTTTCTTTCTGANNNAGATCGGAAGAG
129327 TATGAGTGTGGAGTCGTTGCTTTATACCAAGCATTGAGGAATANNNAGATCGGAAG
129329 TATGAGTGTGGAGTCGTTGCTAGTACTCAGAACATCTTTCAACNNNAGATCGGAAG
129330 TATGAGTGTGGAGTCGTTGCTTTTTGTTTTTGTTTTGAAGGAGNNNAGATCGGAAG
129331 TATGAGTGTGGAGTCGTTGCAGAAAATTGAGAGATGACAATTGNNNAGATCGGAA
129332 TATGAGTGTGGAGTCGTTGCAAGAGACAAGGAGAAAGGAAATNNNAGATCGGAA
129333 TATGAGTGTGGAGTCGTTGCAACACAGAATATTTCTAGATGGANNNAGATCGGAAG
129334 TATGAGTGTGGAGTCGTTGCGAAGGAGAAGAGACAGATTTTNNNAGATCGGAAG
129335 TATGAGTGTGGAGTCGTTGCGAGAGGGATATAGCATTTGTTANNNAGATCGGAAG
129336 TATGAGTGTGGAGTCGTTGCCAAAGTTCAGAAGTCAATGCNNNAGATCGGAAGAG
129337 TATGAGTGTGGAGTCGTTGCGGAAGTAATGATTAATGCTCATTNNNAGATCGGAAG
129338 TATGAGTGTGGAGTCGTTGCACATGCCTACATTCTACATTTNNNAGATCGGAAGAG
129339 TATGAGTGTGGAGTCGTTGCAGTTTGAGAATACAGCAAAAAANNNAGATCGGAAG
129340 TATGAGTGTGGAGTCGTTGCCTAATTATCTTTCTGACAAAGGGNNNAGATCGGAAG
129341 TATGAGTGTGGAGTCGTTGCCAAACTCTGATTGGAAAATCCNNNAGATCGGAAGAG
129343 TATGAGTGTGGAGTCGTTGCTATTTCTTTTAAGGAATTGGCTCNNNAGATCGGAAG
129344 TATGAGTGTGGAGTCGTTGCGAATTTGCTAGCCCTGAATGNNNAGATCGGAAGAG
129345 TATGAGTGTGGAGTCGTTGCCTGATTCAGCATGTTTAGTCANNNAGATCGGAAGAG
129346 TATGAGTGTGGAGTCGTTGCTTTTTATTTTTGATCCCTAGGACNNNAGATCGGAAGA
129347 TATGAGTGTGGAGTCGTTGCAAAGAAAATAGCTGAAAAGAACCNNNAGATCGGAA
129348 TATGAGTGTGGAGTCGTTGCCATCTACACCAATGTTTACAAATNNNAGATCGGAAGA
129349 TATGAGTGTGGAGTCGTTGCAAAAGACGAGGATGACCTTTANNNAGATCGGAAGA
129350 TATGAGTGTGGAGTCGTTGCCTTTTCTCAGTTCTTTCATTACTNNNAGATCGGAAGA
129351 TATGAGTGTGGAGTCGTTGCGGCCAAAAATTCTTTGAAAGAANNNAGATCGGAAG
129352 TATGAGTGTGGAGTCGTTGCTCTTTCTTTCTTTCTTTCTTTCTNNNAGATCGGAAGAG
129355 TATGAGTGTGGAGTCGTTGCTTATTTTTCGTTATTTGGAGGTTNNNAGATCGGAAG
129356 TATGAGTGTGGAGTCGTTGCAACAGACTGTAATCTCTTTTCTANNNAGATCGGAAGA
129357 TATGAGTGTGGAGTCGTTGCGAGCTGCGTTTATAATTAGTATTNNNAGATCGGAAG
129358 TATGAGTGTGGAGTCGTTGCGCTGTGTTTTTTTTCTGTTTTAANNNAGATCGGAAGA
129359 TATGAGTGTGGAGTCGTTGCTACTTAAAGCCTTAGCCATTTANNNAGATCGGAAGA
129360 TATGAGTGTGGAGTCGTTGCAAAAAGCCAATTAGTTCAATAGTNNNAGATCGGAAG
129361 TATGAGTGTGGAGTCGTTGCAAGTAGGAAAAGGTATGGATTAANNNAGATCGGAA
129362 TATGAGTGTGGAGTCGTTGCTTTTGTGTGGTGAGATAAGGNNNAGATCGGAAGAG
129363 TATGAGTGTGGAGTCGTTGCAGATCCTCAGAAGAAAGAGATTNNNAGATCGGAAG
129364 TATGAGTGTGGAGTCGTTGCTAATGACAGAGTATATGCGTATTNNNAGATCGGAAG
129365 TATGAGTGTGGAGTCGTTGCTAATGACAGAGTATATGCGTATTNNNAGATCGGAAG
129366 TATGAGTGTGGAGTCGTTGCAAGATAAATAAACGAAGGGAAAANNNAGATCGGAA

129367 TATGAGTGTGGAGTCGTTGCTTGATACCAGGGAAGAATGAGNNNAGATCGGAAGA
129368 TATGAGTGTGGAGTCGTTGCCGAATGTTTTCTTTAATAGGCATNNNAGATCGGAAGA
129371 TATGAGTGTGGAGTCGTTGCAGAGGTACCAAAAACCTTTAGNNNAGATCGGAAGA
129372 TATGAGTGTGGAGTCGTTGCTCATTGGTATAGATAAAGAGTGGNNNAGATCGGAA
129373 TATGAGTGTGGAGTCGTTGCTACACACATACGTATATTCAGATNNNAGATCGGAAGA
129374 TATGAGTGTGGAGTCGTTGCGTTCTAAGTGCTTCTTGAATATGNNNAGATCGGAAG
129375 TATGAGTGTGGAGTCGTTGCCTTACATGACATAGCTGGATTTANNNAGATCGGAAG
129376 TATGAGTGTGGAGTCGTTGCGGTTATTTCCACCTTTTGAATAANNNAGATCGGAAGA
129377 TATGAGTGTGGAGTCGTTGCGTAAAATAGCCAAATGTTGAGTANNNAGATCGGAA
129378 TATGAGTGTGGAGTCGTTGCAAATCCTATACTGCAACAAATACNNNAGATCGGAAGA
129379 TATGAGTGTGGAGTCGTTGCATAGAAAGGAAATACTCATGACANNNAGATCGGAA
129380 TATGAGTGTGGAGTCGTTGCAGCATTTTTTTCAGAGCTTAAGNNNAGATCGGAAGA
129381 TATGAGTGTGGAGTCGTTGCTATGTACATGTGTATATACAGGGNNNAGATCGGAA
129383 TATGAGTGTGGAGTCGTTGCGTGGTGGTCATAATAGTTTAAAANNNAGATCGGAA
129384 TATGAGTGTGGAGTCGTTGCCATTTGAGTTAAGTCTTGAAGGNNNAGATCGGAAGA
129385 TATGAGTGTGGAGTCGTTGCCCTGAATGTTATTGGTATATGGANNNAGATCGGAAG
129386 TATGAGTGTGGAGTCGTTGCCGTAGGCTTTAGATTTTTTTTATTNNNAGATCGGAAG
129387 TATGAGTGTGGAGTCGTTGCTCTAAAAGAGAATGAATCTGAGTNNNAGATCGGAA
129388 TATGAGTGTGGAGTCGTTGCAAAAGAAGAAAAATCTGAGTTCGNNNAGATCGGAA
129389 TATGAGTGTGGAGTCGTTGCCAAACAGGAAAAGAAAAATACTGNNNAGATCGGAA
129390 TATGAGTGTGGAGTCGTTGCAACAGTCCTCTTTTTTTAAAAAANNNAGATCGGAAGA
129391 TATGAGTGTGGAGTCGTTGCGTTTGTTTAGAAATGACGTCTCNNNAGATCGGAAGA
129392 TATGAGTGTGGAGTCGTTGCAAAAATCATATATTGAACACCTGNNNAGATCGGAAG
129393 TATGAGTGTGGAGTCGTTGCTCCTTCCTTCCTTTTTTTGATNNNAGATCGGAAGAGCA
129394 TATGAGTGTGGAGTCGTTGCTCATCTCATGGGAATAAAAGATANNNAGATCGGAAG
129395 TATGAGTGTGGAGTCGTTGCAAGAGCTGATTGTTAAATATTCCNNNAGATCGGAAG
129396 TATGAGTGTGGAGTCGTTGCTCTATAGATCTTAACCAAAGTGGNNNAGATCGGAAG
129397 TATGAGTGTGGAGTCGTTGCTTTTTATATTCTGGCTGGATTTGNNNAGATCGGAAGA
129398 TATGAGTGTGGAGTCGTTGCTTTTTATATTCTGGCTGGATTTGNNNAGATCGGAAGA
129399 TATGAGTGTGGAGTCGTTGCGAGAAGTTTCAGTGTTGTAAACNNNAGATCGGAAGA
129400 TATGAGTGTGGAGTCGTTGCACATATGACATTCACAGAAAAAGNNNAGATCGGAAG
129401 TATGAGTGTGGAGTCGTTGCCATACATATGACATTCACAGAAANNNAGATCGGAAG
129402 TATGAGTGTGGAGTCGTTGCACATATGCTAAATGAGTTATTCCNNNAGATCGGAAGA
129403 TATGAGTGTGGAGTCGTTGCTTTTACCACTATAATGCATTTGTNNNAGATCGGAAGA
129404 TATGAGTGTGGAGTCGTTGCCAGAGCATATGATGACTGATAANNNAGATCGGAAGA
129405 TATGAGTGTGGAGTCGTTGCTATAGTACACTGCTTTATACCTGNNNAGATCGGAAGA
129407 TATGAGTGTGGAGTCGTTGCTGCTTTTATTGTCATTGTGTTTANNNAGATCGGAAGA
129408 TATGAGTGTGGAGTCGTTGCTTTAGTAGAACCTTCTGATCCTCNNNAGATCGGAAGA
129409 TATGAGTGTGGAGTCGTTGCTTGATGCAGCTTTTATTCTTAAANNNAGATCGGAAGA
129410 TATGAGTGTGGAGTCGTTGCCCCTTCACTAACAGAATTTAAAGNNNAGATCGGAAGA
129411 TATGAGTGTGGAGTCGTTGCGAGTTACAGATTTGAAAGAGTTTNNNAGATCGGAA
129412 TATGAGTGTGGAGTCGTTGCCTCTCTTTCTCTCTCTCTCTTATNNNAGATCGGAAGAG
129413 TATGAGTGTGGAGTCGTTGCGGGATTTTTATTGGGATTATGTTNNNAGATCGGAAG

129414 TATGAGTGTGGAGTCGTTGCTATTTGTTTCTAGGACTGTACCTNNNAGATCGGAAG
129415 TATGAGTGTGGAGTCGTTGCTAAAATGAAATTCCTTGCTGATGNNNAGATCGGAAG
129416 TATGAGTGTGGAGTCGTTGCATTTTTGTTTTACTGAATTGCTCNNNAGATCGGAAG
129417 TATGAGTGTGGAGTCGTTGCCCCTAAAATGTTTACATTGAAGTNNNAGATCGGAAG
129418 TATGAGTGTGGAGTCGTTGCGTACATTTGTTACATTCGATCAANNNAGATCGGAAG
129419 TATGAGTGTGGAGTCGTTGCAAGAAAGTTCTAAAGCTCTCTATNNNAGATCGGAAG
129420 TATGAGTGTGGAGTCGTTGCTGCTTGAATTTGATCCATAATAGNNNAGATCGGAAG
129421 TATGAGTGTGGAGTCGTTGCCAGAATATCCACAGAATTTTACANNNAGATCGGAAG
129422 TATGAGTGTGGAGTCGTTGCCAATTATCTTCCAATCACATCTTNNNAGATCGGAAGA
129423 TATGAGTGTGGAGTCGTTGCGTACTCTCTTTCATGATGACAATNNNAGATCGGAAG
129424 TATGAGTGTGGAGTCGTTGCCAAATGTTCTCTCTTTACTGTATNNNAGATCGGAAGA
129425 TATGAGTGTGGAGTCGTTGCAATACGTTTTCCATTATTTTCCANNNAGATCGGAAGA
129426 TATGAGTGTGGAGTCGTTGCGCACACACTCATTATTATTAGAGNNNAGATCGGAAG
129427 TATGAGTGTGGAGTCGTTGCTACAAGTTTATCAGAATGGTAGGNNNAGATCGGAA
129428 TATGAGTGTGGAGTCGTTGCGATCTAGGTGCTCCAACAATNNNAGATCGGAAGAG
129429 TATGAGTGTGGAGTCGTTGCTTCCATTGCAAATATATTAGCTCNNNAGATCGGAAG
129430 TATGAGTGTGGAGTCGTTGCAATTTAAATGTTTGGAAATTCGGNNNAGATCGGAAG
129431 TATGAGTGTGGAGTCGTTGCTAACATTGATCACTCATGAAGAANNNAGATCGGAAG
129432 TATGAGTGTGGAGTCGTTGCTTGTTGAATCTCCTCATAGAAAANNNAGATCGGAAG
129433 TATGAGTGTGGAGTCGTTGCGACAAAAACGTCAGTTATAGAAGNNNAGATCGGAA
129434 TATGAGTGTGGAGTCGTTGCGGTTTTGTGAGAATGTTTTTATGNNNAGATCGGAAG
129435 TATGAGTGTGGAGTCGTTGCGTTTCCATTATCCTCAAGTTATTNNNAGATCGGAAGA
129436 TATGAGTGTGGAGTCGTTGCGTTTCCATTATCCTCAAGTTATTNNNAGATCGGAAGA
129437 TATGAGTGTGGAGTCGTTGCGTGACATTCAAATTCTGGATAATNNNAGATCGGAAG
129438 TATGAGTGTGGAGTCGTTGCAGTCTTCTGTGTTTTGTCTCNNNAGATCGGAAGAGC
129439 TATGAGTGTGGAGTCGTTGCGAGAATATTACCCTGGATTATGTNNNAGATCGGAAG
129440 TATGAGTGTGGAGTCGTTGCTAATGAAAACATATTGGCTTAGCNNNAGATCGGAAG
129441 TATGAGTGTGGAGTCGTTGCCTTATAACTATGTGCAATTGAGGNNNAGATCGGAAG
129442 TATGAGTGTGGAGTCGTTGCAGAAGGTATTTACTAGTAATGGGNNNAGATCGGAA
129443 TATGAGTGTGGAGTCGTTGCGACAGCTTGATGTGAACAAGNNNAGATCGGAAGAG
129444 TATGAGTGTGGAGTCGTTGCAGGGAAATGGAGTTTAGAAACNNNAGATCGGAAG
129445 TATGAGTGTGGAGTCGTTGCCCTTGGGGTATTCTTAATTAAACNNNAGATCGGAAG
129446 TATGAGTGTGGAGTCGTTGCCATCTATGGAAAACAAAGCCNNNAGATCGGAAGAG
129447 TATGAGTGTGGAGTCGTTGCAGTCATTTCAGAAAGTTGATTTTNNNAGATCGGAAG
129448 TATGAGTGTGGAGTCGTTGCGGATAATTCCTCCCAAGAAAGNNNAGATCGGAAGA
129449 TATGAGTGTGGAGTCGTTGCACTTTAGCCAAAGATGTATCCNNNAGATCGGAAGAG
129450 TATGAGTGTGGAGTCGTTGCCAACTGATATCTGAAGTGTAGGNNNAGATCGGAAG
129451 TATGAGTGTGGAGTCGTTGCGCAAATAAACAGCAAACAAATAANNNAGATCGGAA
129452 TATGAGTGTGGAGTCGTTGCTATAAATCAGCACACTTTTCAAANNNAGATCGGAAG
129453 TATGAGTGTGGAGTCGTTGCCTATGAGAGGGATTGAGAATAAANNNAGATCGGAA
129454 TATGAGTGTGGAGTCGTTGCGCAAATAAACAGCAAACAAATAANNNAGATCGGAA
129457 TATGAGTGTGGAGTCGTTGCCATTTCTTTTCTGTTTGTTGAAANNNAGATCGGAAGA
129458 TATGAGTGTGGAGTCGTTGCCCTTCAGGTAATTCTGGATTCNNNAGATCGGAAGAG

129459 TATGAGTGTGGAGTCGTTGCTAGTTTTTCAAGGACCTCAATCNNNAGATCGGAAGA
129460 TATGAGTGTGGAGTCGTTGCAAACTTAGAGAAAAATAAGCCAGNNNAGATCGGAA
129461 TATGAGTGTGGAGTCGTTGCGAAGAGGATACTACAGCAGAANNNAGATCGGAAGA
129462 TATGAGTGTGGAGTCGTTGCTTCCACATTTTCAGACAGAATNNNAGATCGGAAGAG
129463 TATGAGTGTGGAGTCGTTGCGTCTGTCTGAAGGTCATTTTTANNNAGATCGGAAGA
129464 TATGAGTGTGGAGTCGTTGCCATTAGGATAATTCTGCATGTTANNNAGATCGGAAG
129465 TATGAGTGTGGAGTCGTTGCGTAATTTGGCAAGCTTATTTTTTNNNAGATCGGAAG
129466 TATGAGTGTGGAGTCGTTGCTGCCTATATATTGCAACAGAANNNAGATCGGAAGAG
129467 TATGAGTGTGGAGTCGTTGCGTTACAGTGTTTGTTATAATGCTNNNAGATCGGAAG
129468 TATGAGTGTGGAGTCGTTGCCCCTATTCCTTTCTATTTTTCCTNNNAGATCGGAAGAG
129469 TATGAGTGTGGAGTCGTTGCCTATACAGCCACAAAGAATTTGNNNAGATCGGAAGA
129470 TATGAGTGTGGAGTCGTTGCATGAATGAAAAATAGTCGATGAGNNNAGATCGGAA
129471 TATGAGTGTGGAGTCGTTGCCAGAACAATCTCTTCTCTTAATTNNNAGATCGGAAGA
129472 TATGAGTGTGGAGTCGTTGCATCAATGGAAATGAAGGTTGANNNAGATCGGAAGA
129473 TATGAGTGTGGAGTCGTTGCTATAGCACATGATCTCTCTTTCNNNAGATCGGAAGAG
129474 TATGAGTGTGGAGTCGTTGCAAGGTTTGTTATTCTCTGTTGATNNNAGATCGGAAG
129475 TATGAGTGTGGAGTCGTTGCTCACTCACAATAATGAATAGACANNNAGATCGGAAG
129476 TATGAGTGTGGAGTCGTTGCGAAACACTGTAGCAGGAGATNNNAGATCGGAAGAG
129477 TATGAGTGTGGAGTCGTTGCGCCCTTTTTATATTTGGTCTTTANNNAGATCGGAAGA
129478 TATGAGTGTGGAGTCGTTGCATTTCAAGGATGCTCTGACTNNNAGATCGGAAGAGC
129479 TATGAGTGTGGAGTCGTTGCTTTTCTAATATAGAAGAGGAGCCNNNAGATCGGAAG
129480 TATGAGTGTGGAGTCGTTGCCATTCCTCTCTACCTTATTCATANNNAGATCGGAAGA
129481 TATGAGTGTGGAGTCGTTGCTCTTTTTTTCTCTCTCTCTCTCNNNAGATCGGAAGAGC
129482 TATGAGTGTGGAGTCGTTGCTCACATTCACCTTTGTATCCNNNAGATCGGAAGAGCA
129483 TATGAGTGTGGAGTCGTTGCTAACATTATTCCCAGTTCCACNNNAGATCGGAAGAGC
129484 TATGAGTGTGGAGTCGTTGCCGTGTGTGTATGTATGTGTATATNNNAGATCGGAAG
129485 TATGAGTGTGGAGTCGTTGCATACTATCAGGAGATGAGGAATANNNAGATCGGAA
129486 TATGAGTGTGGAGTCGTTGCTCAATCAATCTTTTTAAGTGAAATCNNNAGATCGGAA
129487 TATGAGTGTGGAGTCGTTGCCTCTACTTACTCCTTTCTGATTTNNNAGATCGGAAGA
129488 TATGAGTGTGGAGTCGTTGCTTATATAAAAGCAGTAACACAGTNNNAGATCGGAAG
129489 TATGAGTGTGGAGTCGTTGCTTGTGGTAAAATACAAATGATGTNNNAGATCGGAA
129490 TATGAGTGTGGAGTCGTTGCGTAGGACTTTGATTGTTAAAACTNNNAGATCGGAAG
129491 TATGAGTGTGGAGTCGTTGCAAACTTTGTCTTAGAAACCTTGTNNNAGATCGGAAG
129492 TATGAGTGTGGAGTCGTTGCATCCATCATGATTGTAAGTTTCNNNAGATCGGAAG
129493 TATGAGTGTGGAGTCGTTGCCTTTTAATATGTTGGCAGATTTGNNNAGATCGGAAG
129494 TATGAGTGTGGAGTCGTTGCGTGTCATTTTCCTTTTTCTTTTTNNNAGATCGGAAGA
129495 TATGAGTGTGGAGTCGTTGCGTATTTATAGACATGGCAGGTCNNNAGATCGGAAG
129496 TATGAGTGTGGAGTCGTTGCGTATTTATAGACATGGCAGGTCNNNAGATCGGAAG
129497 TATGAGTGTGGAGTCGTTGCTAGCAGACACTTACAAATGATAANNNAGATCGGAAG
129498 TATGAGTGTGGAGTCGTTGCGCATTTGAGAACACTTTGTTNNNAGATCGGAAGAGC
129499 TATGAGTGTGGAGTCGTTGCAAGATGTTTGACATTTAGAACTGNNNAGATCGGAAG
129500 TATGAGTGTGGAGTCGTTGCCTCCTGGATTCAAATAATCCTCNNNAGATCGGAAGA
129501 TATGAGTGTGGAGTCGTTGCTGTTTTGATAACTTCCTTTCTTTNNNAGATCGGAAGA

129502 TATGAGTGTGGAGTCGTTGCTTGGTAGGATATGTTTTTCATCNNNAGATCGGAAG
129503 TATGAGTGTGGAGTCGTTGCGTGTCTGTTTCTATTTGTCTTCNNNAGATCGGAAGAG
129504 TATGAGTGTGGAGTCGTTGCTAGAGTACAATGGTGTGATCTTNNNAGATCGGAAG
129505 TATGAGTGTGGAGTCGTTGCCAGGTAAACGTGCTATTCTGNNNAGATCGGAAGAG
129506 TATGAGTGTGGAGTCGTTGCCAGGTAAACGTGCTATTCTGNNNAGATCGGAAGAG
129507 TATGAGTGTGGAGTCGTTGCCAGGTAAACGTGCTATTCTGNNNAGATCGGAAGAG
129508 TATGAGTGTGGAGTCGTTGCAATAGATATTTTCACCCCCTCTTNNNAGATCGGAAGA
129509 TATGAGTGTGGAGTCGTTGCCATACTGCAAGAACAATATTTCANNNAGATCGGAAG
129510 TATGAGTGTGGAGTCGTTGCGATTTCCTTGAATCTCACAATTCNNNAGATCGGAAGA
129511 TATGAGTGTGGAGTCGTTGCTAAAGCACATGTTGTTCAATATTNNNAGATCGGAAG
129512 TATGAGTGTGGAGTCGTTGCGTCCTTTCTCTCTTATTTTAGAGNNNAGATCGGAAGA
129513 TATGAGTGTGGAGTCGTTGCTAAAAGCATCATTCCCAGAGNNNAGATCGGAAGAG
129514 TATGAGTGTGGAGTCGTTGCAGTCTGTGATGTATGATTAGAAANNNAGATCGGAA
129515 TATGAGTGTGGAGTCGTTGCAGTCTGTGATGTATGATTAGAAANNNAGATCGGAA
129516 TATGAGTGTGGAGTCGTTGCATCATAGTTGCTCAACATCATANNNAGATCGGAAGA
129517 TATGAGTGTGGAGTCGTTGCGTTGAAGTGGATTAGACCAGNNNAGATCGGAAGAG
129518 TATGAGTGTGGAGTCGTTGCAGAAAGTGAAACAAGATGACTNNNAGATCGGAAGA
129519 TATGAGTGTGGAGTCGTTGCACAGAAGATGTGTAGTCTGTNNNAGATCGGAAGAG
129520 TATGAGTGTGGAGTCGTTGCTATAAAGCACCTGACATGAACNNNAGATCGGAAGAG
129521 TATGAGTGTGGAGTCGTTGCTTCTGTGGTCTTTGTATCTAACNNNAGATCGGAAGA
129522 TATGAGTGTGGAGTCGTTGCCTCCTCTCTCTTCTTGTCTCNNNAGATCGGAAGAGCA
129523 TATGAGTGTGGAGTCGTTGCGTGCCTATGTATAACATTTTCTTNNNAGATCGGAAGA
129524 TATGAGTGTGGAGTCGTTGCATTGAATCATGTTTTAGCCATTTNNNAGATCGGAAG
129525 TATGAGTGTGGAGTCGTTGCATCTTTTATAATACTGTTTGGCCNNNAGATCGGAAGA
129526 TATGAGTGTGGAGTCGTTGCCAGAACTCTTGTTTGTTGAATTNNNAGATCGGAAGA
129527 TATGAGTGTGGAGTCGTTGCCAGTTTTAGCTAGCATGATAAAANNNAGATCGGAAG
129528 TATGAGTGTGGAGTCGTTGCGAATTTCTCCATAGCAGTTAAATNNNAGATCGGAAG
129529 TATGAGTGTGGAGTCGTTGCCATCATTGCATCAATTAATGAAGNNNAGATCGGAAG
129530 TATGAGTGTGGAGTCGTTGCGATATCTTCGTGGGTATATATCCNNNAGATCGGAAG
129531 TATGAGTGTGGAGTCGTTGCTTCTTGAAGGATTTATCCCTTTNNNAGATCGGAAGAG
129532 TATGAGTGTGGAGTCGTTGCGTTTCTTATGTCAGGTTAATCTGNNNAGATCGGAAG
129533 TATGAGTGTGGAGTCGTTGCAGTTTTATTCATCATTTTCCCTTNNNAGATCGGAAGA
129534 TATGAGTGTGGAGTCGTTGCCTATTTTGGCTTTACCCTTTTCNNNAGATCGGAAGAG
129535 TATGAGTGTGGAGTCGTTGCCTATGGACTAAATGTGTCTCCNNNAGATCGGAAGAG
129536 TATGAGTGTGGAGTCGTTGCTAATACTGGTGAAACCTATCAACNNNAGATCGGAAG
129537 TATGAGTGTGGAGTCGTTGCAGCAATAAAAAATACAATCACACNNNAGATCGGAAG
129538 TATGAGTGTGGAGTCGTTGCTTCATTCATGTCATTTTGTATGTNNNAGATCGGAAGA
129539 TATGAGTGTGGAGTCGTTGCAAGTCATACGGAAAACATTTTCNNNAGATCGGAAGA
129540 TATGAGTGTGGAGTCGTTGCACTATCCTCCTTGATCTAATTTCNNNAGATCGGAAGA
129541 TATGAGTGTGGAGTCGTTGCAAAACAGCTTCCATGAAGTATNNNAGATCGGAAGAG
129542 TATGAGTGTGGAGTCGTTGCTAGAATTACTAAACGTTCTAGGCNNNAGATCGGAAG
129543 TATGAGTGTGGAGTCGTTGCGTCTTCCTTATTCTGGATAATGTNNNAGATCGGAAG
129544 TATGAGTGTGGAGTCGTTGCATCTAGAGATTTCACTAGAACAGNNNAGATCGGAAG

129545 TATGAGTGTGGAGTCGTTGCTAATGACTGAGGAAAGGAATTACNNNAGATCGGAA
129546 TATGAGTGTGGAGTCGTTGCTAGGTGGTATGACTGTAAAGTAANNNAGATCGGAA
129547 TATGAGTGTGGAGTCGTTGCCATGCCTGTAAAGTGTTAGCNNNAGATCGGAAGAG
129548 TATGAGTGTGGAGTCGTTGCTTAGCTTCTCCAACCTCTAGNNNAGATCGGAAGAGC
129549 TATGAGTGTGGAGTCGTTGCCAGTAGTGTGTAATCTTTTCAAANNNAGATCGGAAG
129550 TATGAGTGTGGAGTCGTTGCTAGAACTATACTACACTGTTCAANNNAGATCGGAAG
129551 TATGAGTGTGGAGTCGTTGCTTCAAATTATGGTCTACCCTTTNNNAGATCGGAAGAG
129552 TATGAGTGTGGAGTCGTTGCTATGATACCTTCCACCATGTTATNNNAGATCGGAAGA
129553 TATGAGTGTGGAGTCGTTGCTTTCTGTACCTGGTTTATTCTATNNNAGATCGGAAGA
129554 TATGAGTGTGGAGTCGTTGCATAGGGTAAAATGCTAAAAACAGNNNAGATCGGAA
129555 TATGAGTGTGGAGTCGTTGCATCTTAGCATTAGCATTCCTATCNNNAGATCGGAAGA
129556 TATGAGTGTGGAGTCGTTGCTTTGACATGTCTTTTAAGTCTTTNNNAGATCGGAAGA
129557 TATGAGTGTGGAGTCGTTGCCTTTGTATGATTTCTCTAGAGCANNNAGATCGGAAG
129558 TATGAGTGTGGAGTCGTTGCAAATTTCCTTTATCCATTCATCCNNNAGATCGGAAGA
129559 TATGAGTGTGGAGTCGTTGCCAAAACACATAGGTTTACCAAGNNNAGATCGGAAGA
129560 TATGAGTGTGGAGTCGTTGCATCTTTGCAATGGTTGTATATTCNNNAGATCGGAAG
129561 TATGAGTGTGGAGTCGTTGCATGACTTCATTATTGTGAGTAGTNNNAGATCGGAAG
129562 TATGAGTGTGGAGTCGTTGCATGACTTCATTATTGTGAGTAGTNNNAGATCGGAAG
129563 TATGAGTGTGGAGTCGTTGCAGAAAAATACAACTTGCTAACACNNNAGATCGGAAG
129564 TATGAGTGTGGAGTCGTTGCTGTATATCACAGTTTCCTTATCCNNNAGATCGGAAGA
129565 TATGAGTGTGGAGTCGTTGCGTTCATTTCCTGACTTTTTTTTTNNNAGATCGGAAGA
129566 TATGAGTGTGGAGTCGTTGCGTTCATTTCCTGACTTTTTTTTTNNNAGATCGGAAGA
129567 TATGAGTGTGGAGTCGTTGCCCTAGCCAAATGTTTTATGAATANNNAGATCGGAAG
129568 TATGAGTGTGGAGTCGTTGCTGTATATGCTGTTTCTTCTATCTNNNAGATCGGAAGA
129569 TATGAGTGTGGAGTCGTTGCATACACAAATACAGATACACACANNNAGATCGGAAG
129570 TATGAGTGTGGAGTCGTTGCCTTCTTTGCTAATAGACATGTACNNNAGATCGGAAG
129571 TATGAGTGTGGAGTCGTTGCGCACACATATATATATATGGAAAGGNNNAGATCGG
129573 TATGAGTGTGGAGTCGTTGCCTATTGAGCTAGCAGTAAAGGNNNAGATCGGAAGA
129574 TATGAGTGTGGAGTCGTTGCGTATGCATGGATCATGACATNNNAGATCGGAAGAG
129575 TATGAGTGTGGAGTCGTTGCTTAGTCCCAGTAAACTTTGATTNNNAGATCGGAAGA
129576 TATGAGTGTGGAGTCGTTGCAATCATTTGGAATCATTGTCTTTNNNAGATCGGAAG
129577 TATGAGTGTGGAGTCGTTGCAAAAACATTAGACAAGGGAAACNNNAGATCGGAAG
129578 TATGAGTGTGGAGTCGTTGCGCAAAATGGCAAGATACTTTATTNNNAGATCGGAAG
129580 TATGAGTGTGGAGTCGTTGCTCTTTTTTGAATCTTGTTTCTGANNNAGATCGGAAGA
129581 TATGAGTGTGGAGTCGTTGCTATTGTTTGGTTGCTTGTTTTATNNNAGATCGGAAG
129582 TATGAGTGTGGAGTCGTTGCAACATGTGGTCAATATATGTCATNNNAGATCGGAAG
129583 TATGAGTGTGGAGTCGTTGCGCGAATCAACTGCATTAATATATNNNAGATCGGAAG
129584 TATGAGTGTGGAGTCGTTGCAGGGAATTTTAGCTTGAACTTNNNAGATCGGAAGA
129585 TATGAGTGTGGAGTCGTTGCAGGGAATTTTAGCTTGAACTTNNNAGATCGGAAGA
129586 TATGAGTGTGGAGTCGTTGCTTTTTATTATAGCCAGAGATGGANNNAGATCGGAAG
129587 TATGAGTGTGGAGTCGTTGCATAAAAATCTGCTACTGGGAATNNNAGATCGGAAG
129588 TATGAGTGTGGAGTCGTTGCCTGCTGATATCCTGTTTTTGANNNAGATCGGAAGAG
129589 TATGAGTGTGGAGTCGTTGCGTTATCGCAGTTGTTATTTTAGANNNAGATCGGAAG

129590 TATGAGTGTGGAGTCGTTGCTTTATACACAAATATACACGCACNNNAGATCGGAAG
129591 TATGAGTGTGGAGTCGTTGCACAAAGCAGAAGTACTATGATATNNNAGATCGGAA
129592 TATGAGTGTGGAGTCGTTGCAAAATGAGCTCTTACCACAAANNNAGATCGGAAGAG
129593 TATGAGTGTGGAGTCGTTGCCAGGGTCAAATAGTTATTGAGTNNNAGATCGGAAG
129594 TATGAGTGTGGAGTCGTTGCGAATAGATAGCCTCTTAGGATACNNNAGATCGGAA
129595 TATGAGTGTGGAGTCGTTGCTGAGAAATGACAGTGTAGTATTANNNAGATCGGAA
129596 TATGAGTGTGGAGTCGTTGCTTCTTTCCTATTCCCCTCTCNNNAGATCGGAAGAGCA
129597 TATGAGTGTGGAGTCGTTGCGATACAGATATAAAGATAGGGCCNNNAGATCGGAA
129598 TATGAGTGTGGAGTCGTTGCCCATAATATTTGCCTTGTATTCANNNAGATCGGAAG
129599 TATGAGTGTGGAGTCGTTGCAGATTCATTACAAAATTTGGCTTNNNAGATCGGAAG
129600 TATGAGTGTGGAGTCGTTGCAAAACCACATACAAGGATAAATGNNNAGATCGGAA
129601 TATGAGTGTGGAGTCGTTGCGAGGATTAGGGATTGTAGATATANNNAGATCGGAA
129602 TATGAGTGTGGAGTCGTTGCGTGATAAATGGCAGACTACATATNNNAGATCGGAA
129603 TATGAGTGTGGAGTCGTTGCGCAGAATCTTACCTACTGATAGNNNAGATCGGAAGA
129604 TATGAGTGTGGAGTCGTTGCATACAGACATTGGCTGAGTTNNNAGATCGGAAGAG
129605 TATGAGTGTGGAGTCGTTGCGTTGACATGTCTTGAGGTTATNNNAGATCGGAAGA
129606 TATGAGTGTGGAGTCGTTGCCTACTGTCAAGTGGTAAACAANNNAGATCGGAAGA
129607 TATGAGTGTGGAGTCGTTGCCTTTTACTCCTGTAACTGCATNNNAGATCGGAAGAG
129608 TATGAGTGTGGAGTCGTTGCTTTATCCTCACTTCCTAAATTCANNNAGATCGGAAGA
129609 TATGAGTGTGGAGTCGTTGCTAGAAGGTTAAGGTGCTTTTAATNNNAGATCGGAA
129610 TATGAGTGTGGAGTCGTTGCATTTTCTTTGTTGGTCCATTGNNNAGATCGGAAGAG
129611 TATGAGTGTGGAGTCGTTGCGAGAACAGACCTATGCAGTTNNNAGATCGGAAGAG
129612 TATGAGTGTGGAGTCGTTGCGCTACCACATTATACAAAACATANNNAGATCGGAAG
129613 TATGAGTGTGGAGTCGTTGCATATGAGAAGGGATTTATTAGGGNNNAGATCGGAA
129614 TATGAGTGTGGAGTCGTTGCCTCTAAAGAACCCTAACCATTAANNNAGATCGGAAG
129615 TATGAGTGTGGAGTCGTTGCCTCTAAAGAACCCTAACCATTAANNNAGATCGGAAG
129616 TATGAGTGTGGAGTCGTTGCTATTGTTGCAGAACCATTTGANNNAGATCGGAAGAG
129617 TATGAGTGTGGAGTCGTTGCTTGAGACAGTTATTTTCTATCGANNNAGATCGGAAG
129618 TATGAGTGTGGAGTCGTTGCACAACCAGATTACCAAAAAAAGNNNAGATCGGAAGA
129619 TATGAGTGTGGAGTCGTTGCTATAAAATCACATGAGGCAATTCNNNAGATCGGAAG
129620 TATGAGTGTGGAGTCGTTGCTGGGTAAATATACAAAAGGGTTANNNAGATCGGAA
129621 TATGAGTGTGGAGTCGTTGCATGAACTGAGAAAGAAAAAATGANNNAGATCGGAA
129622 TATGAGTGTGGAGTCGTTGCCAAAGAATAAAGGGAGCTTTCNNNAGATCGGAAGA
129623 TATGAGTGTGGAGTCGTTGCCAGCCTATATATGCAATTATGAANNNAGATCGGAAG
129624 TATGAGTGTGGAGTCGTTGCGATTTCAAAGATTCCAGAAGAGNNNAGATCGGAAG
129625 TATGAGTGTGGAGTCGTTGCAAGCAATCTTTCCACCTAAGNNNAGATCGGAAGAGC
129626 TATGAGTGTGGAGTCGTTGCGAGAGGAAAACTAGAAGCTATAANNNAGATCGGAA
129627 TATGAGTGTGGAGTCGTTGCAGAGACTGAGAGATGGAGAGNNNAGATCGGAAGA
129628 TATGAGTGTGGAGTCGTTGCCTTCACTTTAGTTCCTTTCTAGANNNAGATCGGAAGA
129629 TATGAGTGTGGAGTCGTTGCCTATTCTAGGCAGAACATTAGATNNNAGATCGGAAG
129630 TATGAGTGTGGAGTCGTTGCTTCTAAGTAAACTAACTTGGGAANNNAGATCGGAAG
129631 TATGAGTGTGGAGTCGTTGCTCAAATGTTCAGATGCTTAGATNNNAGATCGGAAGA
129632 TATGAGTGTGGAGTCGTTGCCTTGCTTTTTTTTTAGAGACAGNNNAGATCGGAAG

129633 TATGAGTGTGGAGTCGTTGCGCAGATTACATAGAAAGATAACTNNNAGATCGGAA
129634 TATGAGTGTGGAGTCGTTGCTCTCCAAATCTGAAGTCATATATNNNAGATCGGAAG
129635 TATGAGTGTGGAGTCGTTGCTATACAGACATGTTCAATGAAACNNNAGATCGGAAG
129636 TATGAGTGTGGAGTCGTTGCTCTTTTTTTCTCTCTCTCTCNNNAGATCGGAAGAGC
129638 TATGAGTGTGGAGTCGTTGCTAAGGACCTTTTAGCTTCTTGNNNAGATCGGAAGAG
129639 TATGAGTGTGGAGTCGTTGCATGATTGGGTTTCTTGGACTNNNAGATCGGAAGAG
129641 TATGAGTGTGGAGTCGTTGCAAACATGTCTCATTATCTTCAACNNNAGATCGGAAGA
129643 TATGAGTGTGGAGTCGTTGCAATTGATATCCTATCCCCTTTGNNNAGATCGGAAGA
129644 TATGAGTGTGGAGTCGTTGCCACGTGATATGTATATGAAGTTGNNNAGATCGGAA
129645 TATGAGTGTGGAGTCGTTGCCATGTGTCTAATTCATGATTCTGNNNAGATCGGAAG
129646 TATGAGTGTGGAGTCGTTGCTGTATTCATCATGTCATCTATCANNNAGATCGGAAGA
129647 TATGAGTGTGGAGTCGTTGCAATAAGCACTATAGTCTGGAAATNNNAGATCGGAA
129648 TATGAGTGTGGAGTCGTTGCTTTTGTGTGGTGAGATAAGGNNNAGATCGGAAGAG
129649 TATGAGTGTGGAGTCGTTGCTTAAATACTACCGTGTGTCTAAGNNNAGATCGGAAG
129650 TATGAGTGTGGAGTCGTTGCATGTCCTTTGACTTAAATGAAAANNNAGATCGGAAG
129651 TATGAGTGTGGAGTCGTTGCGAAGTGCTGTCTTTTATTCATTNNNAGATCGGAAGA
129652 TATGAGTGTGGAGTCGTTGCGTGTTTCTCATTCTCCTTCTATNNNAGATCGGAAGAG
129653 TATGAGTGTGGAGTCGTTGCATCAGAATTCAAATAATGCTCTGNNNAGATCGGAAG
129654 TATGAGTGTGGAGTCGTTGCCACTAACATGGAAACAGAAAANNNAGATCGGAAGA
129655 TATGAGTGTGGAGTCGTTGCGACAAACTTTCTGATGGTTAAATNNNAGATCGGAAG
129656 TATGAGTGTGGAGTCGTTGCCAAAGTAAATGATCTGGTTCATTNNNAGATCGGAAG
129657 TATGAGTGTGGAGTCGTTGCAGCCCTCTGTCTTGTAAATANNNAGATCGGAAGAGC
129658 TATGAGTGTGGAGTCGTTGCTATTTCTTCTCTAAGGTCAACAANNNAGATCGGAAG
129659 TATGAGTGTGGAGTCGTTGCTATTCTAAGTGAAGTAACTCAGGNNNAGATCGGAAG
129660 TATGAGTGTGGAGTCGTTGCTCTGTCCTGCTTTCTTCTATNNNAGATCGGAAGAGCA
129661 TATGAGTGTGGAGTCGTTGCAGAAATAACACCACACATCTATANNNAGATCGGAAG
129663 TATGAGTGTGGAGTCGTTGCTGGTGGAAATCTAGATTTAGTCNNNAGATCGGAAG
129664 TATGAGTGTGGAGTCGTTGCTAATTTTGCTAAGTGTTTACCATNNNAGATCGGAAG
129665 TATGAGTGTGGAGTCGTTGCGTGAACGCTTTTTTGTAAGGNNNAGATCGGAAGAG
129666 TATGAGTGTGGAGTCGTTGCTAATTTCTGGTACACAAGTGTTANNNAGATCGGAAG
129667 TATGAGTGTGGAGTCGTTGCTCATGGAATGATATTTGTCCTATNNNAGATCGGAAG
129668 TATGAGTGTGGAGTCGTTGCTATGAGATACTTCAGCTGGTAATNNNAGATCGGAAG
129669 TATGAGTGTGGAGTCGTTGCACAAAGGATTTTTAAATACATGGANNNAGATCGGAA
129670 TATGAGTGTGGAGTCGTTGCTGATATTTTGCTTTTTATCACCANNNAGATCGGAAGA
129671 TATGAGTGTGGAGTCGTTGCAAATACTATATTCCTGCTGAGAGNNNAGATCGGAAG
129672 TATGAGTGTGGAGTCGTTGCAATCATAGCTTACTGTAACCTCNNNAGATCGGAAGA
129674 TATGAGTGTGGAGTCGTTGCGCTAACAACTAAATACGTAAAGANNNAGATCGGAA
129675 TATGAGTGTGGAGTCGTTGCGTGTGTAATTTTTGTCTTTCAAANNNAGATCGGAAG
129676 TATGAGTGTGGAGTCGTTGCCACATGGAGTTTTCTGGTAANNNAGATCGGAAGAG
129677 TATGAGTGTGGAGTCGTTGCGGCCCTGAAATAGATATGTAATNNNAGATCGGAAG
129678 TATGAGTGTGGAGTCGTTGCTGTCATTATTACCTATTTGGAGGNNNAGATCGGAAG
129679 TATGAGTGTGGAGTCGTTGCAATTCGTTGTTCTATTTAGTGCNNNAGATCGGAAGA
129680 TATGAGTGTGGAGTCGTTGCTCTTTGTCATTCAGGCTAGGNNNAGATCGGAAGAG

129681 TATGAGTGTGGAGTCGTTGCATCTGGATGCATATTGTATTTGANNNAGATCGGAAG
129682 TATGAGTGTGGAGTCGTTGCCTGTATGACTGGGTTTCAAATANNNAGATCGGAAGA
129683 TATGAGTGTGGAGTCGTTGCGGATCATGAGGTCGAGATAGNNNAGATCGGAAGA
129684 TATGAGTGTGGAGTCGTTGCCTTCAGCTAATTGCAACCTCNNNAGATCGGAAGAGC
129685 TATGAGTGTGGAGTCGTTGCCTTCAGCTAATTGCAACCTCNNNAGATCGGAAGAGC
129686 TATGAGTGTGGAGTCGTTGCTTCAGGAAATTTATCCCAAATTTNNNAGATCGGAAG
129687 TATGAGTGTGGAGTCGTTGCATTGTTCAGCCAATATTTTGTAANNNAGATCGGAAG
129688 TATGAGTGTGGAGTCGTTGCTTGAAGTCACTTCTATTCACTATNNNAGATCGGAAGA
129689 TATGAGTGTGGAGTCGTTGCGTAATTGTTTCAGGTCCATTTTANNNAGATCGGAAG
129690 TATGAGTGTGGAGTCGTTGCAATGATTAAAGAAAAGCTCTGAGNNNAGATCGGAA
129692 TATGAGTGTGGAGTCGTTGCGTATCTAGCATGATTTTAGGGTANNNAGATCGGAAG
129695 TATGAGTGTGGAGTCGTTGCAAAGAAAGGCTTAGACAATCATNNNAGATCGGAAG
129696 TATGAGTGTGGAGTCGTTGCCTCTTTCTCTTCTTATAAGACCANNNAGATCGGAAGA
129697 TATGAGTGTGGAGTCGTTGCGAAAGAGAGAGAACAGATGATGNNNAGATCGGAA
129698 TATGAGTGTGGAGTCGTTGCGAAAGAGAGAGAACAGATGATGNNNAGATCGGAA
129699 TATGAGTGTGGAGTCGTTGCAATCTCTCTCTTTTTCTCTCAATNNNAGATCGGAAGA
129700 TATGAGTGTGGAGTCGTTGCTAGCTTTGATTTGACTTAAAGTTNNNAGATCGGAAG
129701 TATGAGTGTGGAGTCGTTGCTGATCTCTCCCTACTTTATTTTTNNNAGATCGGAAGA
129703 TATGAGTGTGGAGTCGTTGCGATAAGACTTTATTTCCCCTTCTNNNAGATCGGAAGA
129704 TATGAGTGTGGAGTCGTTGCAAGAGTGCTAGTCTGGTATTNNNAGATCGGAAGAG
129706 TATGAGTGTGGAGTCGTTGCATGAATGTAAACTGGAGGATATCNNNAGATCGGAA
129707 TATGAGTGTGGAGTCGTTGCATTAAATAAATGACTCACCCCANNNAGATCGGAAGA
129708 TATGAGTGTGGAGTCGTTGCTGGATAATCCAGGATAATCTCCNNNAGATCGGAAGA
129709 TATGAGTGTGGAGTCGTTGCGAAAAAAACAAAAACACATGAACNNNAGATCGGAA
129710 TATGAGTGTGGAGTCGTTGCACTATTATTCCCTTCATTGTACANNNAGATCGGAAGA
129711 TATGAGTGTGGAGTCGTTGCATACAGGCAGATCTAAAGTAAATNNNAGATCGGAA
129712 TATGAGTGTGGAGTCGTTGCAGCAGAATTTTGAGCTAAATAAANNNAGATCGGAA
129713 TATGAGTGTGGAGTCGTTGCAGAGTAGAATCATTGTATGTTCTNNNAGATCGGAAG
129714 TATGAGTGTGGAGTCGTTGCAGGTCACTGGGTATATACATATANNNAGATCGGAA
129715 TATGAGTGTGGAGTCGTTGCATGAAAATTCTGTGAAAGAGTTTNNNAGATCGGAA
129716 TATGAGTGTGGAGTCGTTGCCATCTCTTTGGCATAAAAAAAAANNNAGATCGGAAG
129717 TATGAGTGTGGAGTCGTTGCTTGATACAAAAAGGAAGCCTTNNNAGATCGGAAGA
129718 TATGAGTGTGGAGTCGTTGCAGACAGGTAAAGTGGATATATAANNNAGATCGGAA
129719 TATGAGTGTGGAGTCGTTGCAGGCATGTAAGTAACATTTTTTTNNNAGATCGGAAG
129720 TATGAGTGTGGAGTCGTTGCCTGATCCATGCTGATCTATTTNNNAGATCGGAAGAG
129721 TATGAGTGTGGAGTCGTTGCTTTTATCACATGTGAAGTTTCATNNNAGATCGGAAG
129722 TATGAGTGTGGAGTCGTTGCACTATGGATACAGTGATGATTAANNNAGATCGGAA
129723 TATGAGTGTGGAGTCGTTGCGGAATCAAGTGAAAGGTATTTAANNNAGATCGGAA
129724 TATGAGTGTGGAGTCGTTGCTTCTTTCTCTTTCTCTCTTTCTCNNNAGATCGGAAGAG
129725 TATGAGTGTGGAGTCGTTGCAGAATGCCATGTAAGATGAAGNNNAGATCGGAAGA
129726 TATGAGTGTGGAGTCGTTGCTCCTTTATCTAAATAAGAGCTGTNNNAGATCGGAAG
129727 TATGAGTGTGGAGTCGTTGCATTACTGAGGCAGAGAAGTTATANNNAGATCGGAA
129728 TATGAGTGTGGAGTCGTTGCATTACTGAGGCAGAGAAGTTATANNNAGATCGGAA

129729 TATGAGTGTGGAGTCGTTGCCATTTTCACTACTCGTATTCAACNNNAGATCGGAAGA
129730 TATGAGTGTGGAGTCGTTGCTGATTATAAAGGTCAGACAGATNNNAGATCGGAA
129731 TATGAGTGTGGAGTCGTTGCCAGCTTTTCAGAATATCGTATTTNNNAGATCGGAAG
129732 TATGAGTGTGGAGTCGTTGCGAGAGAATTTACCAGAAACCANNNAGATCGGAAGA
129733 TATGAGTGTGGAGTCGTTGCCTTTACCTTCTACCACAATTGTANNNAGATCGGAAGA
129734 TATGAGTGTGGAGTCGTTGCGCCAAATAAAATAGAGACGTTTTNNNAGATCGGAA
129735 TATGAGTGTGGAGTCGTTGCCCCCTTTCACATTTGTTTTATATNNNAGATCGGAAGA
129736 TATGAGTGTGGAGTCGTTGCAGACAGACAGAAAGACACATANNNAGATCGGAAGA
129737 TATGAGTGTGGAGTCGTTGCCTTTCCTTCAACTAAGTATGAAANNNAGATCGGAAG
129738 TATGAGTGTGGAGTCGTTGCAAACAGTAAAGAGAGAAGGGTAANNNAGATCGGA
129739 TATGAGTGTGGAGTCGTTGCCTCTCTTTCTCTCTCATCATTTANNNAGATCGGAAGA
129740 TATGAGTGTGGAGTCGTTGCTGTGTAGAGAGATAGAGATAGAGNNNAGATCGGA
129741 TATGAGTGTGGAGTCGTTGCGTAAAATAGCCAAATGTTGAGTANNNAGATCGGAA
129742 TATGAGTGTGGAGTCGTTGCCTCTTTCTCTTTCTCTCTTCATTNNNAGATCGGAAGAG
129743 TATGAGTGTGGAGTCGTTGCGGTTATTTCCACCTTTTGAATAANNNAGATCGGAAG
129744 TATGAGTGTGGAGTCGTTGCACATATTACGTGATTTAGTCTTTNNNAGATCGGAAG
129745 TATGAGTGTGGAGTCGTTGCTTTCTTACATTTCAGACAAAACCNNNAGATCGGAAGA
129746 TATGAGTGTGGAGTCGTTGCAGGGAGGTAGAGATTATGTATATNNNAGATCGGAA
129747 TATGAGTGTGGAGTCGTTGCGTAATTTGGGTGAAGGGTATATNNNAGATCGGAAG
129748 TATGAGTGTGGAGTCGTTGCATTTCTGCCCTACTCATTTACNNNAGATCGGAAGAGC
129749 TATGAGTGTGGAGTCGTTGCGGAGAGCCTCTATATTATACCTANNNAGATCGGAAG
129750 TATGAGTGTGGAGTCGTTGCGAAATCTGAGTTCCATTTAGATCNNNAGATCGGAAG
129751 TATGAGTGTGGAGTCGTTGCTCTCTTTCATATCTTTATGAGAGANNNAGATCGGAA
129752 TATGAGTGTGGAGTCGTTGCTTCTCTCTCTTTCTCTCTCTCNNNAGATCGGAAGAGCA
129753 TATGAGTGTGGAGTCGTTGCCTTGAAAATAAGTTGTCAACATGNNNAGATCGGAAG
129754 TATGAGTGTGGAGTCGTTGCGACTTTGTCTCCTAAAGAAAAGNNNAGATCGGAAGA
129755 TATGAGTGTGGAGTCGTTGCTATTTTCTTCGTTTCACTTTTGANNNAGATCGGAAGA
129756 TATGAGTGTGGAGTCGTTGCGTCAGCTCTTTTTCTTCTATTATTNNNAGATCGGAAG
129758 TATGAGTGTGGAGTCGTTGCCTTTATCCCTGATGTAATGTCTNNNAGATCGGAAGA
129759 TATGAGTGTGGAGTCGTTGCCTCTTCTCTGACCTCTAGTCNNNAGATCGGAAGAGCA
129760 TATGAGTGTGGAGTCGTTGCGTAGCAATTTCCTTATGCCATNNNAGATCGGAAGAG
129761 TATGAGTGTGGAGTCGTTGCGAAGTTTTCTGCATTGTTGTTNNNAGATCGGAAGAG
129762 TATGAGTGTGGAGTCGTTGCCCTTGGATAAAAATGTTTACCTTNNNAGATCGGAAG
129763 TATGAGTGTGGAGTCGTTGCAAGAAAAAAGTAGTATAGGGGAANNNAGATCGGA
129764 TATGAGTGTGGAGTCGTTGCTACCTAACTAACAGACTGTACTCNNNAGATCGGAAG
129765 TATGAGTGTGGAGTCGTTGCCTTTTTCTCCTACAAGACAATTGNNNAGATCGGAAGA
129768 TATGAGTGTGGAGTCGTTGCAGTACAAGTACATAGAGTTTTGTNNNAGATCGGAA
129769 TATGAGTGTGGAGTCGTTGCCTGTTACTTTAGAGACAAATCTGNNNAGATCGGAAG
129773 TATGAGTGTGGAGTCGTTGCGTGCAAGAGTTTCCATATATATGNNNAGATCGGAAG
129774 TATGAGTGTGGAGTCGTTGCTCAGTGTCTTAGCTGATAATTATNNNAGATCGGAAG
129775 TATGAGTGTGGAGTCGTTGCGAGTTCTATTCAATCCAACCCNNNAGATCGGAAGAG
129776 TATGAGTGTGGAGTCGTTGCATCCAGAATATGTAAAGAACTCTNNNAGATCGGAAG
129777 TATGAGTGTGGAGTCGTTGCCGAGAAGTCAAATAGATTTCAAANNNAGATCGGAA

129778 TATGAGTGTGGAGTCGTTGCAATTTGAGAAAAAGAGAGAAAGGNNNAGATCGGA
129779 TATGAGTGTGGAGTCGTTGCAATTTGAGAAAAAGAGAGAAAGGNNNAGATCGGA
129780 TATGAGTGTGGAGTCGTTGCCAACACTTTATAATTCAGGCAAANNNAGATCGGAAG
129782 TATGAGTGTGGAGTCGTTGCTATATAGTGACCATCCTTCAAAANNNAGATCGGAAG
129783 TATGAGTGTGGAGTCGTTGCAGATAAGGAAAAAATAAAGCTGGNNNAGATCGGAA
129784 TATGAGTGTGGAGTCGTTGCCTCTTTCTCTCTCTTTCTCTCTNNNAGATCGGAAGAGC
129785 TATGAGTGTGGAGTCGTTGCCACAAGAATGAGCTTAGAAGTNNNAGATCGGAAGA
129786 TATGAGTGTGGAGTCGTTGCGATACTGTTCCTCAAGTTACTATNNNAGATCGGAAG
129787 TATGAGTGTGGAGTCGTTGCATGAGTAAACCTTTAATGCATTTNNNAGATCGGAAG
129788 TATGAGTGTGGAGTCGTTGCTGTGTGTATATATATGTATGTGTGNNNAGATCGGAA
129789 TATGAGTGTGGAGTCGTTGCTGTGTGTATATATATGTATGTGTGNNNAGATCGGAA
129790 TATGAGTGTGGAGTCGTTGCCTGGCTAATTCTTACAAAAATGTNNNAGATCGGAAG
129791 TATGAGTGTGGAGTCGTTGCTAGAATATTCTTTCTGGTTTCCTNNNAGATCGGAAGA
129794 TATGAGTGTGGAGTCGTTGCTCAGGAGGTAAACATGATTTAGNNNAGATCGGAAG
129795 TATGAGTGTGGAGTCGTTGCTACATGCATATATATGTGAGTGTNNNAGATCGGAAG
129796 TATGAGTGTGGAGTCGTTGCCAAAATGTTCACAATAAAATCCCNNNAGATCGGAAG
129797 TATGAGTGTGGAGTCGTTGCGTTTTTATCACACTAGTCAACTCNNNAGATCGGAAGA
129798 TATGAGTGTGGAGTCGTTGCGAAGGATTACAAATTTGAACTCTNNNAGATCGGAAG
129799 TATGAGTGTGGAGTCGTTGCATTAATAGCAATGAAAGGAATCANNNAGATCGGAA
129800 TATGAGTGTGGAGTCGTTGCGAAGGGACTTTAGAAAAATGTACNNNAGATCGGAA
129801 TATGAGTGTGGAGTCGTTGCTACATTATTTAAGCATGGCAATTNNNAGATCGGAAG
129802 TATGAGTGTGGAGTCGTTGCGTTTGCTTTTTAGGAGTAATGAANNNAGATCGGAAG
129803 TATGAGTGTGGAGTCGTTGCAAATTACAATCATAATCTTCCCCNNNAGATCGGAAGA
129804 TATGAGTGTGGAGTCGTTGCGTATAGATTGATTGATTGAGGTGNNNAGATCGGAA
129805 TATGAGTGTGGAGTCGTTGCTATATTTCAGGAATTGTCTTCCANNNAGATCGGAAG
129807 TATGAGTGTGGAGTCGTTGCAAGTTACTCGTTACTTCATGTATNNNAGATCGGAAG
129808 TATGAGTGTGGAGTCGTTGCAGATATTTGGACACAATGTTTTTNNNAGATCGGAAG
129809 TATGAGTGTGGAGTCGTTGCAAAAGGTACATTATCCCTCATATNNNAGATCGGAAG
129810 TATGAGTGTGGAGTCGTTGCGCAATCTGTAAAGAAATAGAGTCNNNAGATCGGAA
129811 TATGAGTGTGGAGTCGTTGCGTGAAAAAGCTCTATAATGAGAANNNAGATCGGAA
129812 TATGAGTGTGGAGTCGTTGCCTATTATTATTCCCACTTTGCAGNNNAGATCGGAAGA
129813 TATGAGTGTGGAGTCGTTGCGTGATTTATGTGGGAAGTGATNNNAGATCGGAAGA
129814 TATGAGTGTGGAGTCGTTGCAAAAGAAGAAAAGTGTGTAAGTTNNNAGATCGGAA
129815 TATGAGTGTGGAGTCGTTGCAAAGTCTGCTCACTTTCAATTANNNAGATCGGAAGA
129816 TATGAGTGTGGAGTCGTTGCGAGTACTTTGAATTATCAAGACCNNNAGATCGGAAG
129817 TATGAGTGTGGAGTCGTTGCAAAGTAAGGTCTATAAAGCTAGGNNNAGATCGGAA
129818 TATGAGTGTGGAGTCGTTGCAAAGTAAGGTCTATAAAGCTAGGNNNAGATCGGAA
129819 TATGAGTGTGGAGTCGTTGCAGTCAAAGTCTAAGTCTAGTTAGNNNAGATCGGAAG
129820 TATGAGTGTGGAGTCGTTGCCAAGTCAAAGTCTAAGTCTAGTTNNNAGATCGGAAG
129821 TATGAGTGTGGAGTCGTTGCCCAGAACTGTCAGAAATAAATTTNNNAGATCGGAAG
129822 TATGAGTGTGGAGTCGTTGCTTCCACTCTGATCTTGATTATTTNNNAGATCGGAAGA
129823 TATGAGTGTGGAGTCGTTGCCTTTGATGGTAAGGATGGGANNNAGATCGGAAGAG
129824 TATGAGTGTGGAGTCGTTGCAGATAACAAATCCAATGACAAAGNNNAGATCGGAA

129825 TATGAGTGTGGAGTCGTTGCTTTCTTTCCATTTATCAGTAAGCNNNAGATCGGAAGA
129826 TATGAGTGTGGAGTCGTTGCCTTCTGACATCAAGGATAGTAATNNNAGATCGGAAG
129827 TATGAGTGTGGAGTCGTTGCATCTTATCTTTGTTTCTCTCTGTNNNAGATCGGAAGA
129828 TATGAGTGTGGAGTCGTTGCAATGCATGGTCTTATAACTAAACNNNAGATCGGAAG
129829 TATGAGTGTGGAGTCGTTGCGGGAAGGAGATTTTTTTTTTTTNNNAGATCGGAAG
129830 TATGAGTGTGGAGTCGTTGCCCTCACTTTACACTTCAGTAGNNNAGATCGGAAGAG
129831 TATGAGTGTGGAGTCGTTGCAATAAATAGAGAGCTACCTGAAANNNAGATCGGAA
129832 TATGAGTGTGGAGTCGTTGCCAGCAGTTTGAGTTTGAAAGNNNAGATCGGAAGAG
129833 TATGAGTGTGGAGTCGTTGCTTTTGCTCTTGTTGAGAATGGNNNAGATCGGAAGA
129834 TATGAGTGTGGAGTCGTTGCTTTTGCTCTTGTTGAGAATGGNNNAGATCGGAAGA
129835 TATGAGTGTGGAGTCGTTGCTTGTAATTACTTTTTGAGACAGGNNNAGATCGGAAG
129836 TATGAGTGTGGAGTCGTTGCGGCCAATCTTATTCTATCTACATNNNAGATCGGAAG
129837 TATGAGTGTGGAGTCGTTGCCCTGTAACCTCAAGATGTAATAANNNAGATCGGAAG
129838 TATGAGTGTGGAGTCGTTGCTATTTTTTTCATGAGATAAGCCCNNNAGATCGGAAG
129839 TATGAGTGTGGAGTCGTTGCATATTCTAGAGAAACTCCTGTACNNNAGATCGGAAG
129840 TATGAGTGTGGAGTCGTTGCATTCTAGGTAAATGTCAGATTCCNNNAGATCGGAAG
129842 TATGAGTGTGGAGTCGTTGCGACAGACAGAGACAGAGATANNNAGATCGGAAGA
129843 TATGAGTGTGGAGTCGTTGCGATGAGATACAGATAGATGAGAGNNNAGATCGGA
129844 TATGAGTGTGGAGTCGTTGCTTTGAATGATAAACTTTCCTCTCNNNAGATCGGAAG
129845 TATGAGTGTGGAGTCGTTGCTGTTGATCATTTGTTTGCATATTNNNAGATCGGAAG
129846 TATGAGTGTGGAGTCGTTGCATTGCTTCTTATTTTACCCTTTTNNNAGATCGGAAGA
129847 TATGAGTGTGGAGTCGTTGCCAAAAAGTAAGAAAATGGACTTGNNNAGATCGGAA
129849 TATGAGTGTGGAGTCGTTGCCTGGTTGAGTTCAAAAATTTTTTNNNAGATCGGAAG
129850 TATGAGTGTGGAGTCGTTGCCAAACTGTGGGAATCTTGAGNNNAGATCGGAAGAG
129851 TATGAGTGTGGAGTCGTTGCCAAACTGTGGGAATCTTGAGNNNAGATCGGAAGAG
129852 TATGAGTGTGGAGTCGTTGCATAAAGGGTGTTAAAACAAAACTNNNAGATCGGAA
129853 TATGAGTGTGGAGTCGTTGCGTGTAGATTTCTTCCTGTAGATTNNNAGATCGGAAG
129854 TATGAGTGTGGAGTCGTTGCAAAGAAGAAAAAAGGAAGTCAGTNNNAGATCGGA
129855 TATGAGTGTGGAGTCGTTGCAAAGAAGAAAAAAGGAAGTCAGTNNNAGATCGGA
129856 TATGAGTGTGGAGTCGTTGCATACAGATGTAGAAAGACTGATCNNNAGATCGGAA
129857 TATGAGTGTGGAGTCGTTGCGAATGTTTCTTGAAATGTCAAAGNNNAGATCGGAA
129859 TATGAGTGTGGAGTCGTTGCCTAATGATCCATGGACATAACANNNAGATCGGAAGA
129860 TATGAGTGTGGAGTCGTTGCTAATTCATGCGCATTTTTTTCATNNNAGATCGGAAGA
129861 TATGAGTGTGGAGTCGTTGCCCACAGCAACACAAAATTAGNNNAGATCGGAAGAG
129862 TATGAGTGTGGAGTCGTTGCCCACTATAGGACTTCACTATTATNNNAGATCGGAAG
129863 TATGAGTGTGGAGTCGTTGCTTAACTCTAATAGTGGTGCTTTANNNAGATCGGAAG
129864 TATGAGTGTGGAGTCGTTGCTGAGGAGCAGAATTAACAATTTANNNAGATCGGAA
129865 TATGAGTGTGGAGTCGTTGCTGAGGAGCAGAATTAACAATTTANNNAGATCGGAA
129866 TATGAGTGTGGAGTCGTTGCTCTTTCTCTCTCTCTCTATAANNNAGATCGGAAGA
129867 TATGAGTGTGGAGTCGTTGCGGGTTTTTATAGTACCATTTCTCNNNAGATCGGAAG
129868 TATGAGTGTGGAGTCGTTGCATCTCTAAAGTATCAACCATAGANNNAGATCGGAAG
129869 TATGAGTGTGGAGTCGTTGCGGAATTATCAGTATAGCACAGTANNNAGATCGGAA
129870 TATGAGTGTGGAGTCGTTGCAGAAGGCAAGCTAAGAAGAGNNNAGATCGGAAGA

129871 TATGAGTGTGGAGTCGTTGCTCTCATTTATTGATCCAAAAGTGNNNAGATCGGAAG
129872 TATGAGTGTGGAGTCGTTGCTAAGCAATTGTAAAAGGTAGGTANNNAGATCGGAA
129873 TATGAGTGTGGAGTCGTTGCATCACATAACTCTGCATCTTCTANNNAGATCGGAAGA
129874 TATGAGTGTGGAGTCGTTGCCTATTTCCTACCCGATCTCANNNAGATCGGAAGAGCA
129875 TATGAGTGTGGAGTCGTTGCAAACTTGTACTCCATCCTCATANNNAGATCGGAAGAG
129876 TATGAGTGTGGAGTCGTTGCGTGTTCATTTTGGTCCATTGNNNAGATCGGAAGAGC
129877 TATGAGTGTGGAGTCGTTGCAGCTGCAGTTAAAAAAAAAATTCNNNAGATCGGAAG
129878 TATGAGTGTGGAGTCGTTGCAGCTGCAGTTAAAAAAAAAATTCNNNAGATCGGAAG
129879 TATGAGTGTGGAGTCGTTGCCTCATTATTCTCTTTTACCAAGTNNNAGATCGGAAGA
129880 TATGAGTGTGGAGTCGTTGCATCTCTCAAGTACTTTTGCTTTTNNNAGATCGGAAGA
129881 TATGAGTGTGGAGTCGTTGCCTAATTGTCTCCTACAGTAACATNNNAGATCGGAAG
129882 TATGAGTGTGGAGTCGTTGCTACTCGATAAAGATGGTCAACNNNAGATCGGAAGAG
129884 TATGAGTGTGGAGTCGTTGCGTTTCATTTATCTCTATATCCCCNNNAGATCGGAAGA
129885 TATGAGTGTGGAGTCGTTGCATCTCTAATAACCAACTACAGACNNNAGATCGGAAG
129886 TATGAGTGTGGAGTCGTTGCAAATGTTGAAAAGAATATGGAGANNNAGATCGGAA
129887 TATGAGTGTGGAGTCGTTGCAAATGTTGAAAAGAATATGGAGANNNAGATCGGAA
129888 TATGAGTGTGGAGTCGTTGCAAATGTTGAAAAGAATATGGAGANNNAGATCGGAA
129890 TATGAGTGTGGAGTCGTTGCTTTACCAGCCAATCACATTTNNNAGATCGGAAGAGC
129891 TATGAGTGTGGAGTCGTTGCCCAGTATTCAAGTACAGGAAANNNAGATCGGAAGA
129892 TATGAGTGTGGAGTCGTTGCGTAATTTTGGCATGATTGTTATGNNNAGATCGGAAG
129893 TATGAGTGTGGAGTCGTTGCGTGGCATAATTCTTATTAGTCTTNNNAGATCGGAAG
129894 TATGAGTGTGGAGTCGTTGCAAATTCTAGCTATCTTGACTTCTNNNAGATCGGAAGA
129895 TATGAGTGTGGAGTCGTTGCTATGGAGATACCTTATTTCTGACNNNAGATCGGAAG
129896 TATGAGTGTGGAGTCGTTGCAAGATTTGTTAGGAATTGTGTTGNNNAGATCGGAA
129897 TATGAGTGTGGAGTCGTTGCCACAATATAAACAACTGGTGAATNNNAGATCGGAAG
129898 TATGAGTGTGGAGTCGTTGCCCAACATATCTGAGTGCAATANNNAGATCGGAAGAG
129899 TATGAGTGTGGAGTCGTTGCTTTCATGTGACTATAATGCAAAANNNAGATCGGAAG
129900 TATGAGTGTGGAGTCGTTGCATGCATGTTCAACTTTTATCTAGNNNAGATCGGAAG
129901 TATGAGTGTGGAGTCGTTGCAAACATGTAACACAAGGAGAGNNNAGATCGGAAGA
129902 TATGAGTGTGGAGTCGTTGCATATAGGTAGCATCTACATGGTTNNNAGATCGGAAG
129903 TATGAGTGTGGAGTCGTTGCGGTAAAAGCAAGCTGAGATGNNNAGATCGGAAGAG
129904 TATGAGTGTGGAGTCGTTGCGAATAAATACAGAAAATCCTGGGNNNAGATCGGAA
129905 TATGAGTGTGGAGTCGTTGCCTACCTAGGGAAGTTTTCTTTAGNNNAGATCGGAAG
129906 TATGAGTGTGGAGTCGTTGCCTTATCTGTAAAATGGGGGTAGNNNAGATCGGAAG
129907 TATGAGTGTGGAGTCGTTGCGTGCTGATTTGAATAAAGAGTAANNNAGATCGGAA
129908 TATGAGTGTGGAGTCGTTGCTTACTAGAAGCTACACAATTTTGNNNAGATCGGAAG
129909 TATGAGTGTGGAGTCGTTGCGTGTAAAGCTGCTCTAAAAAAANNNAGATCGGAAG
129910 TATGAGTGTGGAGTCGTTGCCAAGAATTACAATTGCCATTTGNNNAGATCGGAAGA
129911 TATGAGTGTGGAGTCGTTGCAAAATCTGTCTGTAGTTCTGTAGNNNAGATCGGAAG
129912 TATGAGTGTGGAGTCGTTGCAAAATCTGTCTGTAGTTCTGTAGNNNAGATCGGAAG
129913 TATGAGTGTGGAGTCGTTGCCTCAGAAATCTGCAATGGTTNNNAGATCGGAAGAG
129914 TATGAGTGTGGAGTCGTTGCCTCTACTTGGACACATTGAGNNNAGATCGGAAGAG
129915 TATGAGTGTGGAGTCGTTGCACTATTCAGCCATAAAAAATTGANNNAGATCGGAAG

129916 TATGAGTGTGGAGTCGTTGCGGATACAAGTGATAGCTTTGANNNAGATCGGAAGA
129917 TATGAGTGTGGAGTCGTTGCCACAGATCTTATCTCCATTTTCTNNNAGATCGGAAGA
129918 TATGAGTGTGGAGTCGTTGCTGCTTAAAAAGGAAAAATAAAGCNNNAGATCGGAA
129919 TATGAGTGTGGAGTCGTTGCGCTTGAGTGTTATTCATGTATAGNNNAGATCGGAAG
129920 TATGAGTGTGGAGTCGTTGCGTTTCGTTGAGGTTTTAAAGAGNNNAGATCGGAAG
129921 TATGAGTGTGGAGTCGTTGCCTCTCTTTCTCTCTCAACTTTATNNNAGATCGGAAGAG
129922 TATGAGTGTGGAGTCGTTGCCTCTCTTTGAGATAGTCAGATAGNNNAGATCGGAAG
129924 TATGAGTGTGGAGTCGTTGCTGTGTTTCTTGTTTTCCTTTTAANNNAGATCGGAAGA
129925 TATGAGTGTGGAGTCGTTGCCCTATCCTGTCTTTTCTATGAATNNNAGATCGGAAGA
129926 TATGAGTGTGGAGTCGTTGCTTTGGAATTTTCAAGTTTTCCTANNNAGATCGGAAG
129927 TATGAGTGTGGAGTCGTTGCAAAGAAATAGGACCAAGCAACNNNAGATCGGAAGA
129928 TATGAGTGTGGAGTCGTTGCCAAACAAGATGCTGACATAATACNNNAGATCGGAAG
129929 TATGAGTGTGGAGTCGTTGCCAAGGAACTAGCTTATGTGATTNNNAGATCGGAAG
129930 TATGAGTGTGGAGTCGTTGCACACAAAGAAATAACTAGAACAGNNNAGATCGGAA
129931 TATGAGTGTGGAGTCGTTGCACAGCATGTTTAAAAATACGTATNNNAGATCGGAAG
129932 TATGAGTGTGGAGTCGTTGCGCACATTGTAAGCATTTATTTCNNNAGATCGGAAGA
129933 TATGAGTGTGGAGTCGTTGCTGTGTTGAATGTAAAATTTTCAGNNNAGATCGGAAG
129934 TATGAGTGTGGAGTCGTTGCGGTGAGGTGATGTACTAAATAANNNAGATCGGAAG
129936 TATGAGTGTGGAGTCGTTGCATTATAAGAAATTGGCTCACATGNNNAGATCGGAAG
129937 TATGAGTGTGGAGTCGTTGCCCATACCTCTCTGAGTATTTAGNNNAGATCGGAAGA
129938 TATGAGTGTGGAGTCGTTGCACTTTTGGGAGCTTTAAAATATCNNNAGATCGGAAG
129939 TATGAGTGTGGAGTCGTTGCACATTATGTCTTCAATTTAGCAGNNNAGATCGGAAG
129940 TATGAGTGTGGAGTCGTTGCTGAATCTGAATTTTCACGATTAANNNAGATCGGAAG
129941 TATGAGTGTGGAGTCGTTGCACATATTTTGTGCACATGTAANNNAGATCGGAAGAG
129943 TATGAGTGTGGAGTCGTTGCAAAGAAATTCCATAGCATGTAAGNNNAGATCGGAA
129944 TATGAGTGTGGAGTCGTTGCATAGGGAAAATGGAATTCACTTNNNAGATCGGAAG
129945 TATGAGTGTGGAGTCGTTGCGCTCAAACTCAACTAGAAGTTANNNAGATCGGAAGA
129946 TATGAGTGTGGAGTCGTTGCTGTTGTTGAATTCTCTTTGTAATNNNAGATCGGAAG
129947 TATGAGTGTGGAGTCGTTGCAGAGAAGTATCAGAACTGAAAATNNNAGATCGGAA
129948 TATGAGTGTGGAGTCGTTGCAGAGAAGTATCAGAACTGAAAATNNNAGATCGGAA
129949 TATGAGTGTGGAGTCGTTGCTAGACAGTCTTTGGGTAACANNNAGATCGGAAGAG
129950 TATGAGTGTGGAGTCGTTGCCTCACAAAAGTTCTCCTAAAATTNNNAGATCGGAAG
129951 TATGAGTGTGGAGTCGTTGCCACACCTTTAATTTCTTATACAGTNNNAGATCGGAAG
129952 TATGAGTGTGGAGTCGTTGCACCTACACACCTTTAATTTCTTANNNAGATCGGAAGA
129953 TATGAGTGTGGAGTCGTTGCTATACACATCAAAATATCGCATTNNNAGATCGGAAG
129954 TATGAGTGTGGAGTCGTTGCATAAGCTCACAAATGTATCAGANNNAGATCGGAAGA
129955 TATGAGTGTGGAGTCGTTGCTTGGAAGACAAGTTGTATCTACNNNAGATCGGAAG
129956 TATGAGTGTGGAGTCGTTGCAAGCATCACTTGAGTTCTTATTANNNAGATCGGAAG
129957 TATGAGTGTGGAGTCGTTGCATGATTAGTTGATTTGAAACACANNNAGATCGGAAG
129958 TATGAGTGTGGAGTCGTTGCAGAATATGCAGGTAGATATGTTTNNNAGATCGGAA
129959 TATGAGTGTGGAGTCGTTGCAGAATATGCAGGTAGATATGTTTNNNAGATCGGAA
129960 TATGAGTGTGGAGTCGTTGCAATGTAATGTCTTCAGTGACTATNNNAGATCGGAAG
129961 TATGAGTGTGGAGTCGTTGCGAAAGAAGTTGGGAGAATTAAGNNNAGATCGGAA

129962 TATGAGTGTGGAGTCGTTGCATTAATCTTTTTCATAGCGCTCNNNAGATCGGAAGA
129963 TATGAGTGTGGAGTCGTTGCATTAATCTTTTTCATAGCGCTCNNNAGATCGGAAGA
129964 TATGAGTGTGGAGTCGTTGCCTAGAAAAAGATTCTGGCGANNNAGATCGGAAGAG
129965 TATGAGTGTGGAGTCGTTGCCTAGGTCTTCTTCAGCTTTCNNNAGATCGGAAGAGC
129966 TATGAGTGTGGAGTCGTTGCGTAAAGTACCCTTTTGAGAAAAANNNAGATCGGAAG
129967 TATGAGTGTGGAGTCGTTGCTTTACAACTATTCTTGCACCTTNNNAGATCGGAAGAG
129968 TATGAGTGTGGAGTCGTTGCGAGGATGCTAGCTACACTACNNNAGATCGGAAGAG
129969 TATGAGTGTGGAGTCGTTGCTAGAGAAGAAGTTTTATTCCTGTNNNAGATCGGAAG
129970 TATGAGTGTGGAGTCGTTGCGAGAACCAGCTATTTATGAAAAANNNAGATCGGAA
129971 TATGAGTGTGGAGTCGTTGCTAGCAGTTAGAATAGAGAAAACTNNNAGATCGGAA
129972 TATGAGTGTGGAGTCGTTGCTTTTGCTCTGGTGTGTGATANNNAGATCGGAAGAG
129973 TATGAGTGTGGAGTCGTTGCCTCTGAGGAAAAAAAGAAACAANNNAGATCGGAAG
129974 TATGAGTGTGGAGTCGTTGCTGGATGTCACAGATGAATAATATNNNAGATCGGAA
129975 TATGAGTGTGGAGTCGTTGCTTTATAGTGGAGAGTTTTGGAATNNNAGATCGGAA
129976 TATGAGTGTGGAGTCGTTGCAGAAAATGAGTAAAGATGGGGNNNAGATCGGAAG
129977 TATGAGTGTGGAGTCGTTGCAGAGACAGAAAATGAGTAAAGATNNNAGATCGGA
129978 TATGAGTGTGGAGTCGTTGCAGAGACAGAAAATGAGTAAAGATNNNAGATCGGA
129979 TATGAGTGTGGAGTCGTTGCTTAAAATTTACCTTTTCCCTTGGNNNAGATCGGAAGA
129980 TATGAGTGTGGAGTCGTTGCTTAAAATTTACCTTTTCCCTTGGNNNAGATCGGAAGA
129981 TATGAGTGTGGAGTCGTTGCTATCTCCTAATGCAGAAATGACNNNAGATCGGAAGA
129982 TATGAGTGTGGAGTCGTTGCTTTACAATGGTCCTTCTACTGNNNAGATCGGAAGAG
129983 TATGAGTGTGGAGTCGTTGCGAAGATGGAGGAAGAGATTAAANNNAGATCGGAA
129984 TATGAGTGTGGAGTCGTTGCGGTCGATTTTTCTAGAGGTACNNNAGATCGGAAGA
129985 TATGAGTGTGGAGTCGTTGCCGCCATGATTGTATAAGTTTCNNNAGATCGGAAGAG
129986 TATGAGTGTGGAGTCGTTGCAATCCAAGACAACACTACTAAGNNNAGATCGGAAGA
129987 TATGAGTGTGGAGTCGTTGCCTGTTATCACTGGTTCTAGGNNNAGATCGGAAGAGC
129988 TATGAGTGTGGAGTCGTTGCTTAGTTCTAACAGTTTTTAGTGCNNNAGATCGGAAG
129989 TATGAGTGTGGAGTCGTTGCATGTCAACACATAAGGATACTAANNNAGATCGGAAG
129990 TATGAGTGTGGAGTCGTTGCGCAATGTCAGTAGGTATTTAATANNNAGATCGGAAG
129991 TATGAGTGTGGAGTCGTTGCGCAATGTCAGTAGGTATTTAATANNNAGATCGGAAG
129992 TATGAGTGTGGAGTCGTTGCAGGTTCTTAGTTCAAAAAAGTACNNNAGATCGGAAG
129993 TATGAGTGTGGAGTCGTTGCAAGAGATTTCAGGGATCAATTATNNNAGATCGGAA
129994 TATGAGTGTGGAGTCGTTGCAAGTGAATGATTCTTTGTGATAGNNNAGATCGGAA
129995 TATGAGTGTGGAGTCGTTGCCTTTCTCTTCCTCTATCTATGAGNNNAGATCGGAAGA
129996 TATGAGTGTGGAGTCGTTGCCTTTCTCTTCCTCTATCTATGAGNNNAGATCGGAAGA
129997 TATGAGTGTGGAGTCGTTGCCAGGTAGGGGAAGAGTAAATTANNNAGATCGGAA
129998 TATGAGTGTGGAGTCGTTGCCAGGTAGGGGAAGAGTAAATTANNNAGATCGGAA
129999 TATGAGTGTGGAGTCGTTGCATCTCCTAGTTAAGACTGTACTTNNNAGATCGGAAG
130000 TATGAGTGTGGAGTCGTTGCCTCTAGATATGATTGGTTTCCTTNNNAGATCGGAAG
130001 TATGAGTGTGGAGTCGTTGCGCCTCCATAAAAGGAAATAAAATNNNAGATCGGAAG
130002 TATGAGTGTGGAGTCGTTGCGCCTCCATAAAAGGAAATAAAATNNNAGATCGGAAG
130003 TATGAGTGTGGAGTCGTTGCAAAGAGACACAGGAATCAACNNNAGATCGGAAGAG
130004 TATGAGTGTGGAGTCGTTGCGACTAATGTGTGGTTTGCAGNNNAGATCGGAAGAG

130005 TATGAGTGTGGAGTCGTTGCTTTAAATTCGCATTTGTTTAGTCNNNAGATCGGAAG
130006 TATGAGTGTGGAGTCGTTGCTCTTCTTTTTAAATTCGCATTTGNNNAGATCGGAAGA
130007 TATGAGTGTGGAGTCGTTGCTGAATAAAGTAGAGTTGTTTTGANNNAGATCGGAA
130008 TATGAGTGTGGAGTCGTTGCTTATTGGTAGGTGGTTAGGTAATNNNAGATCGGAA
130009 TATGAGTGTGGAGTCGTTGCGAAACCATTCATTAGTCACATGNNNAGATCGGAAGA
130010 TATGAGTGTGGAGTCGTTGCGAAACCATTCATTAGTCACATGNNNAGATCGGAAGA
130011 TATGAGTGTGGAGTCGTTGCGTCTCTCTCTCTCTCTCTCTNNNAGATCGGAAGAGCA
130012 TATGAGTGTGGAGTCGTTGCTGTGAGTAGAAACCATAAACTAANNNAGATCGGAA
130014 TATGAGTGTGGAGTCGTTGCGGTTAGCCTATTAAAATGAAAAANNNAGATCGGAA
130015 TATGAGTGTGGAGTCGTTGCAGAAGGAAGAAATGATACTTGATNNNAGATCGGAA
130016 TATGAGTGTGGAGTCGTTGCGAAAGTTGAATTTTCTATGGGATNNNAGATCGGAA
130017 TATGAGTGTGGAGTCGTTGCATAAAGTCTCACTGTGCCTCNNNAGATCGGAAGAGC
130018 TATGAGTGTGGAGTCGTTGCCTCAATAGTACCCCCTACCTNNNAGATCGGAAGAGC
130023 TATGAGTGTGGAGTCGTTGCAAATTACATTACCAGCTGTCTNNNAGATCGGAAGAG
130024 TATGAGTGTGGAGTCGTTGCTTTTGATGCTAAAGTCTGTTTCNNNAGATCGGAAGA
130025 TATGAGTGTGGAGTCGTTGCAGGTAAACATAAGTTAAAACCACNNNAGATCGGAA
130026 TATGAGTGTGGAGTCGTTGCAAACTAGTGTATATGTCCTTCAANNNAGATCGGAAG
130027 TATGAGTGTGGAGTCGTTGCCTGTAGGAGAAGATATTTTGAGANNNAGATCGGAA
130028 TATGAGTGTGGAGTCGTTGCTTAGGAAACTAATCTAACATGGCNNNAGATCGGAAG
130029 TATGAGTGTGGAGTCGTTGCACATTACTAGAGAATATTAGGGCNNNAGATCGGAA
130030 TATGAGTGTGGAGTCGTTGCGGCTCGTTAAATGTTCTATATTANNNAGATCGGAAG
130031 TATGAGTGTGGAGTCGTTGCCATTTAGCCCTTCATCATTTAAANNNAGATCGGAAGA
130032 TATGAGTGTGGAGTCGTTGCCAATAAATGCTTCTCTCTACCTANNNAGATCGGAAGA
130033 TATGAGTGTGGAGTCGTTGCGCTATCAACATGCATATGAAAGNNNAGATCGGAAG
130034 TATGAGTGTGGAGTCGTTGCAATTTAAATGTTTGGAAATTCGGNNNAGATCGGAA
130035 TATGAGTGTGGAGTCGTTGCGATTTCCAACTCATGAACATAATNNNAGATCGGAAG
130036 TATGAGTGTGGAGTCGTTGCTCGTGTATGTTTCCTTATCTTTNNNAGATCGGAAGAG
130037 TATGAGTGTGGAGTCGTTGCTCGTGTATGTTTCCTTATCTTTNNNAGATCGGAAGAG
130038 TATGAGTGTGGAGTCGTTGCAGATATTTACTGCAAGAAATTGGNNNAGATCGGAA
130039 TATGAGTGTGGAGTCGTTGCGTTAGAGGTTTTTTGCTTTCTTNNNAGATCGGAAGA
130040 TATGAGTGTGGAGTCGTTGCTGGGCATATTTAAATTTCTTATGTNNNAGATCGGAA
130041 TATGAGTGTGGAGTCGTTGCGTTAGAGGTTTTTTGCTTTCTTNNNAGATCGGAAGA
130042 TATGAGTGTGGAGTCGTTGCGAACATGGAGACAATGAAATTATNNNAGATCGGAA
130043 TATGAGTGTGGAGTCGTTGCGTTAAGTCTTGAGGTATAGAGTTNNNAGATCGGAA
130044 TATGAGTGTGGAGTCGTTGCGCAGCTATATGTTGTAGTATAGANNNAGATCGGAA
130045 TATGAGTGTGGAGTCGTTGCGTGACATTTATGATCATGTGTATNNNAGATCGGAAG
130046 TATGAGTGTGGAGTCGTTGCCACAGTCTTTAGCTCCAAAGNNNAGATCGGAAGAGC
130047 TATGAGTGTGGAGTCGTTGCGTTACCTGTTCAAAGCAAACNNNAGATCGGAAGAGC
130048 TATGAGTGTGGAGTCGTTGCAGTAATTTCTCACATCTAAATGCNNNAGATCGGAAG
130049 TATGAGTGTGGAGTCGTTGCAGTAATTTCTCACATCTAAATGCNNNAGATCGGAAG
130050 TATGAGTGTGGAGTCGTTGCCTGAGTATCTGGGACTACAGNNNAGATCGGAAGAG
130051 TATGAGTGTGGAGTCGTTGCTTCTTCTTCCTAAAATGACATTGNNNAGATCGGAAGA
130052 TATGAGTGTGGAGTCGTTGCAATCAAGAATGCTAGTAATCTGTNNNAGATCGGAAG

130053 TATGAGTGTGGAGTCGTTGCAATCAAGAATGCTAGTAATCTGTNNNAGATCGGAAG
130054 TATGAGTGTGGAGTCGTTGCTACTAATAAACGAATTGAGGCATNNNAGATCGGAA
130055 TATGAGTGTGGAGTCGTTGCGTACCATCATGTATAAAAGGGTTNNNAGATCGGAA
130056 TATGAGTGTGGAGTCGTTGCCACTAGGAGACCTAAGTATTTTTNNNAGATCGGAAG
130058 TATGAGTGTGGAGTCGTTGCGAGAGAAACAGGTAGAAATAGAGNNNAGATCGGA
130059 TATGAGTGTGGAGTCGTTGCATGATTTGGAGAACAAGCAATNNNAGATCGGAAGA
130060 TATGAGTGTGGAGTCGTTGCAAACTACTCATTGAGATTACTCANNNAGATCGGAAG
130061 TATGAGTGTGGAGTCGTTGCTATGTGTATATGTGCAGACATAGNNNAGATCGGAA
130062 TATGAGTGTGGAGTCGTTGCCTAAAATGCAACGTAGGGAANNNAGATCGGAAGAG
130063 TATGAGTGTGGAGTCGTTGCTGTTTAGAGACAAGGAAGGGNNNAGATCGGAAGA
130064 TATGAGTGTGGAGTCGTTGCCATAAATTGTGTATGTCTCCAATNNNAGATCGGAAG
130065 TATGAGTGTGGAGTCGTTGCCATACAATGAAATTATCTGAGGCNNNAGATCGGAAG
130066 TATGAGTGTGGAGTCGTTGCCCTTTTCCAAGATGAGTAAAAAANNNAGATCGGAAG
130067 TATGAGTGTGGAGTCGTTGCGAATCAGTAACAGAGGTTGGNNNAGATCGGAAGAG
130068 TATGAGTGTGGAGTCGTTGCAAGAGGTATGAGGGAACTTTNNNAGATCGGAAGAG
130069 TATGAGTGTGGAGTCGTTGCATACATGTATACACACACACATTNNNAGATCGGAAG
130071 TATGAGTGTGGAGTCGTTGCAGTCTGAAGTTTGTAGGGTANNNAGATCGGAAGAG
130072 TATGAGTGTGGAGTCGTTGCAACTATTAAGGGTCTTTCGTATTNNNAGATCGGAAG
130073 TATGAGTGTGGAGTCGTTGCCATTCTTTCTCATTCACATACTCNNNAGATCGGAAGA
130074 TATGAGTGTGGAGTCGTTGCCATTCTTTCTCATTCACATACTCNNNAGATCGGAAGA
130075 TATGAGTGTGGAGTCGTTGCCCATACATTCCTGTGAACTTANNNAGATCGGAAGAG
130076 TATGAGTGTGGAGTCGTTGCTAGAAGGAGAGAGAGATGAATTANNNAGATCGGA
130077 TATGAGTGTGGAGTCGTTGCTAGAAGGAGAGAGAGATGAATTANNNAGATCGGA
130078 TATGAGTGTGGAGTCGTTGCTTATCTCCTGTACTATGATCTGTNNNAGATCGGAAGA
130079 TATGAGTGTGGAGTCGTTGCGACTGAAGACTGGAAGTGAANNNAGATCGGAAGA
130080 TATGAGTGTGGAGTCGTTGCAGAATAACCCTCAGATTTACTAANNNAGATCGGAAG
130081 TATGAGTGTGGAGTCGTTGCCTTATTGCATCATTCATGGATACNNNAGATCGGAAG
130082 TATGAGTGTGGAGTCGTTGCTGGTATTTACTGGAATTACAGAANNNAGATCGGAAG
130083 TATGAGTGTGGAGTCGTTGCTTGATTAGTGCTTTATCTATCCTNNNAGATCGGAAGA
130084 TATGAGTGTGGAGTCGTTGCTTGATTAGTGCTTTATCTATCCTNNNAGATCGGAAGA
130085 TATGAGTGTGGAGTCGTTGCCCCAGTGGGAAATACATATAGNNNAGATCGGAAGA
130086 TATGAGTGTGGAGTCGTTGCGTTTTGAATTGAAATAATTGAACAANNNAGATCGGA
130087 TATGAGTGTGGAGTCGTTGCCAATTGGACGAATAAAAAACAAANNNAGATCGGAA
130088 TATGAGTGTGGAGTCGTTGCGATAGGTTAAGTTACTTACCTGTNNNAGATCGGAAG
130089 TATGAGTGTGGAGTCGTTGCTCGACTGAGACTACTTATTTAAANNNAGATCGGAAG
130090 TATGAGTGTGGAGTCGTTGCTATCCCATTTCATTGAAAAAATGNNNAGATCGGAAG
130091 TATGAGTGTGGAGTCGTTGCAAGCTCTCAACAGAATATTAAACNNNAGATCGGAAG
130092 TATGAGTGTGGAGTCGTTGCCCAGCAAGTGTCACTATATATAANNNAGATCGGAAG
130093 TATGAGTGTGGAGTCGTTGCGAGAGCACTCATAAAGGAAATNNNAGATCGGAAGA
130094 TATGAGTGTGGAGTCGTTGCACTTTATGTCATACTGTCCTTTANNNAGATCGGAAGA
130096 TATGAGTGTGGAGTCGTTGCATATCTTGGCTATTGCGAATAATNNNAGATCGGAAG
130099 TATGAGTGTGGAGTCGTTGCTACATAGCAACCACATTTTTAAANNNAGATCGGAAG
130100 TATGAGTGTGGAGTCGTTGCCCACTTTCCTCAGGTATAAATTNNNAGATCGGAAGA

130101 TATGAGTGTGGAGTCGTTGCGTTGTTAACTGTTTCTGTTTAGTNNNAGATCGGAAG
130102 TATGAGTGTGGAGTCGTTGCATTAGTACCAGTGTTAATTTCCTNNNAGATCGGAAG
130103 TATGAGTGTGGAGTCGTTGCGTTTTTATGGTGATCCAGCTNNNAGATCGGAAGAG
130104 TATGAGTGTGGAGTCGTTGCATATATTCCCTCCCTCCTAATATNNNAGATCGGAAGA
130105 TATGAGTGTGGAGTCGTTGCCACTATGAGGATTACCTAGAAAGNNNAGATCGGAAG
130106 TATGAGTGTGGAGTCGTTGCCTTTGAGAGAGAGACAGAGANNNAGATCGGAAGA
130107 TATGAGTGTGGAGTCGTTGCAATATTTCCCTTCACAGTATGTANNNAGATCGGAAG
130108 TATGAGTGTGGAGTCGTTGCTTTTCTTGACCAGATATGCATATNNNAGATCGGAAG
130109 TATGAGTGTGGAGTCGTTGCAACATAAAGATGAAAAGCGTAATNNNAGATCGGAA
130110 TATGAGTGTGGAGTCGTTGCCTGTCAATACTGTCAACTTTTCNNNAGATCGGAAGA
130111 TATGAGTGTGGAGTCGTTGCCTTAGGAATATATCCAACCAAGANNNAGATCGGAAG
130112 TATGAGTGTGGAGTCGTTGCAATTACCTGTCATGATGGAAATNNNAGATCGGAAGA
130113 TATGAGTGTGGAGTCGTTGCCCAACCATACTATGTCATTTTATNNNAGATCGGAAGA
130114 TATGAGTGTGGAGTCGTTGCAATGACTTCAGCAGACTGTANNNAGATCGGAAGAG
130115 TATGAGTGTGGAGTCGTTGCAGATATGCAAATCAGATCCTATGNNNAGATCGGAAG
130116 TATGAGTGTGGAGTCGTTGCCAAGTGTATATCCCTGAGGTNNNAGATCGGAAGAG
130117 TATGAGTGTGGAGTCGTTGCCAAACACTTTAAATGGTTGAATTNNNAGATCGGAAG
130119 TATGAGTGTGGAGTCGTTGCCGCTCTGGAGATAACTAACTNNNAGATCGGAAGAG
130120 TATGAGTGTGGAGTCGTTGCCGCTCTGGAGATAACTAACTNNNAGATCGGAAGAG
130121 TATGAGTGTGGAGTCGTTGCTTCAATGAATACTTTTGAGCTACNNNAGATCGGAAG
130122 TATGAGTGTGGAGTCGTTGCGAAAGGATAGAAATCTGCATAGANNNAGATCGGAA
130123 TATGAGTGTGGAGTCGTTGCATAAGGTGTATGATCCCGAGNNNAGATCGGAAGAG
130124 TATGAGTGTGGAGTCGTTGCGTTGAAATGTTAATTGTGGAATCNNNAGATCGGAA
130125 TATGAGTGTGGAGTCGTTGCGACTAAAGATGTATCACTGGTAANNNAGATCGGAA
130126 TATGAGTGTGGAGTCGTTGCTTTGAACTCCCTACTGTAGATTNNNAGATCGGAAGA
130127 TATGAGTGTGGAGTCGTTGCATCTAATACAGCTAAATGTCCTCNNNAGATCGGAAG
130128 TATGAGTGTGGAGTCGTTGCGAAAAGGTAATTAAATGCATGACNNNAGATCGGAA
130129 TATGAGTGTGGAGTCGTTGCGAAAAGGTAATTAAATGCATGACNNNAGATCGGAA
130130 TATGAGTGTGGAGTCGTTGCATGTGTTTCTTTGACAGGTAANNNAGATCGGAAGA
130131 TATGAGTGTGGAGTCGTTGCCTTACATCATGGCCAGAGTANNNAGATCGGAAGAG
130132 TATGAGTGTGGAGTCGTTGCCTAGAGTAAGCACTGTAAGATTNNNAGATCGGAAG
130133 TATGAGTGTGGAGTCGTTGCTGGAGACACACATAAGAAACNNNAGATCGGAAGAG
130134 TATGAGTGTGGAGTCGTTGCTAAGCAAAGAATAGAACAAAAGCNNNAGATCGGAA
130135 TATGAGTGTGGAGTCGTTGCAATAAATTGCCCACAGAATTAATNNNAGATCGGAAG
130137 TATGAGTGTGGAGTCGTTGCGTCCTTATTTAGTAGGTCAATCCNNNAGATCGGAAG
130138 TATGAGTGTGGAGTCGTTGCGTGGAAAGTAAATCCGGAGANNNAGATCGGAAGA
130139 TATGAGTGTGGAGTCGTTGCTTATACCATGTGCTATTTATTGCNNNAGATCGGAAG
130141 TATGAGTGTGGAGTCGTTGCGTGTATTTTTAGCTTCCTGATTTNNNAGATCGGAAG
130142 TATGAGTGTGGAGTCGTTGCACACACATATAGCTACTCTATATNNNAGATCGGAAG
130143 TATGAGTGTGGAGTCGTTGCCATGCTCTTCTTTTGGTAATAATNNNAGATCGGAAG
130144 TATGAGTGTGGAGTCGTTGCTTTTAACTCCATTTTAGACCCTANNNAGATCGGAAGA
130145 TATGAGTGTGGAGTCGTTGCTAGGAGTTATAGCTACGTAAAAGNNNAGATCGGAA
130146 TATGAGTGTGGAGTCGTTGCTACAACTATTTTCTCATTGCTTCNNNAGATCGGAAGA

130147 TATGAGTGTGGAGTCGTTGCATAATAACAAAAAGGAAATGGCANNNAGATCGGAA
130148 TATGAGTGTGGAGTCGTTGCCATTGAGAAGAAGGATGAAGAANNNAGATCGGAAG
130149 TATGAGTGTGGAGTCGTTGCTCCCATTAGATGATATCATAAGCNNNAGATCGGAAG
130150 TATGAGTGTGGAGTCGTTGCCTATTCTACTCAATGGAGACTAGNNNAGATCGGAAG
130153 TATGAGTGTGGAGTCGTTGCTCTTTCTTTCTTTCTTTCTTTCTNNNAGATCGGAAGAG
130154 TATGAGTGTGGAGTCGTTGCAATAGATAAGAAGAAGCTACCTGNNNAGATCGGAA
130155 TATGAGTGTGGAGTCGTTGCCAGCGTTCTTTCATTTTTTAAANNNAGATCGGAAGA
130156 TATGAGTGTGGAGTCGTTGCGTTACTCACACATTTTTGTATCTNNNAGATCGGAAGA
130157 TATGAGTGTGGAGTCGTTGCAAAAGAGAAAGAGTATGTGAATGNNNAGATCGGAA
130158 TATGAGTGTGGAGTCGTTGCGGGCCTCAGATATTATCTATAAGNNNAGATCGGAAG
130159 TATGAGTGTGGAGTCGTTGCATTGAAAGTAAATGATTTGGGTTNNNAGATCGGAA
130160 TATGAGTGTGGAGTCGTTGCTAGATCCCATATCTTTGCTATTGNNNAGATCGGAAGA
130161 TATGAGTGTGGAGTCGTTGCTAGATCCCATATCTTTGCTATTGNNNAGATCGGAAGA
130162 TATGAGTGTGGAGTCGTTGCATTCCTATGTTGAAATTGTAACCNNNAGATCGGAAG
130163 TATGAGTGTGGAGTCGTTGCTAATTGATGCCTGTTAAGATACANNNAGATCGGAAG
130164 TATGAGTGTGGAGTCGTTGCTAATTGATGCCTGTTAAGATACANNNAGATCGGAAG
130165 TATGAGTGTGGAGTCGTTGCCCAAAGTAATTATAGCATGCAGNNNAGATCGGAAGA
130166 TATGAGTGTGGAGTCGTTGCGATAGTTCATTGAGTTGTTTCTTNNNAGATCGGAAG
130167 TATGAGTGTGGAGTCGTTGCTAAGTTTTATGATAGTGGCTACCNNNAGATCGGAAG
130168 TATGAGTGTGGAGTCGTTGCCACTCAATAGTCTCTTCTGAATNNNAGATCGGAAGAG
130169 TATGAGTGTGGAGTCGTTGCCTATGAAAGTAAATCCTGCAAAGNNNAGATCGGAAG
130170 TATGAGTGTGGAGTCGTTGCGTTACTTGGACTAGCTGAAGNNNAGATCGGAAGAG
130171 TATGAGTGTGGAGTCGTTGCTTTTTACTAAGAAAAACTGGTGGNNNAGATCGGAAG
130172 TATGAGTGTGGAGTCGTTGCATTACTGGTTCAAGGCATTTTTANNNAGATCGGAAG
130173 TATGAGTGTGGAGTCGTTGCAGTCTCATGGTTATTTGTTTTTCNNNAGATCGGAAGA
130174 TATGAGTGTGGAGTCGTTGCCATTACAGTGTGGCTTTTTCNNNAGATCGGAAGAGC
130176 TATGAGTGTGGAGTCGTTGCACAACAAGAAATCCTACTTTCTANNNAGATCGGAAGA
130178 TATGAGTGTGGAGTCGTTGCATATTCTTATTACTCTCTCCTGCNNNAGATCGGAAGA
130179 TATGAGTGTGGAGTCGTTGCTCTTTCTCTCTCTCTCTCTCNNNAGATCGGAAGAGCAC
130182 TATGAGTGTGGAGTCGTTGCTAGCTCACTGTAACCTTGAANNNAGATCGGAAGAGC
130183 TATGAGTGTGGAGTCGTTGCGAATTTCATCAGCATTTGAAGTANNNAGATCGGAAG
130184 TATGAGTGTGGAGTCGTTGCTCAATGATAGGTGAAGTATTAGANNNAGATCGGAA
130185 TATGAGTGTGGAGTCGTTGCTTTAGCACACTTGTCTATATTCTNNNAGATCGGAAGA
130186 TATGAGTGTGGAGTCGTTGCCATTAACGCACAGAATAGTTCNNNAGATCGGAAGAG
130187 TATGAGTGTGGAGTCGTTGCGACCCAGATAAAGAAAAAATTTTTNNNAGATCGGAA
130188 TATGAGTGTGGAGTCGTTGCGACCCAGATAAAGAAAAAATTTTTNNNAGATCGGAA
130189 TATGAGTGTGGAGTCGTTGCGGGGAAATAATTTAGATTGAGTTNNNAGATCGGAA
130190 TATGAGTGTGGAGTCGTTGCAATTGGCTTATGTGACTATACANNNAGATCGGAAGA
130191 TATGAGTGTGGAGTCGTTGCTGTGTCTGTCTATAATTGGTAAANNNAGATCGGAAG
130192 TATGAGTGTGGAGTCGTTGCTTTACGTTAGAGGGTACTGAATANNNAGATCGGAA
130193 TATGAGTGTGGAGTCGTTGCCGAACTTCTCATGATTATATCTCNNNAGATCGGAAGA
130194 TATGAGTGTGGAGTCGTTGCCTCAGGAAACACAATGTAAATGNNNAGATCGGAAGA
130195 TATGAGTGTGGAGTCGTTGCGAAAGGCAAAAATGCATTTATAANNNAGATCGGAA

130197 TATGAGTGTGGAGTCGTTGCCACTTTTTCCTTCTTTTCTGGNNNAGATCGGAAGAG
130198 TATGAGTGTGGAGTCGTTGCAATCCTAGCACCATTCATATAATNNNAGATCGGAAG
130199 TATGAGTGTGGAGTCGTTGCAAACTTTATTTACAACTGAACCANNNAGATCGGAAG
130200 TATGAGTGTGGAGTCGTTGCAGGTGGTCGAGTATCTCATANNNAGATCGGAAGAG
130201 TATGAGTGTGGAGTCGTTGCAACATAAAATACTTTCTTAGGCANNNAGATCGGAAG
130202 TATGAGTGTGGAGTCGTTGCACATTTTAAAATCCATTCATCCANNNAGATCGGAAGA
130203 TATGAGTGTGGAGTCGTTGCCCCCAACATTATAACAAAAGACNNNAGATCGGAAGA
130204 TATGAGTGTGGAGTCGTTGCCACAGCTGGATTAGATAACAGNNNAGATCGGAAGA
130205 TATGAGTGTGGAGTCGTTGCTATGTTGTGTTTTGGTTTGTATGNNNAGATCGGAAG
130206 TATGAGTGTGGAGTCGTTGCCTTCCTAGTTTCCTTCTCTAAAGNNNAGATCGGAAGA
130207 TATGAGTGTGGAGTCGTTGCTTAAGATACAGAACACTTCCATCNNNAGATCGGAAG
130208 TATGAGTGTGGAGTCGTTGCTTAAGATACAGAACACTTCCATCNNNAGATCGGAAG
130209 TATGAGTGTGGAGTCGTTGCTAAAATAATTTAAGTGGCGACAANNNAGATCGGAA
130210 TATGAGTGTGGAGTCGTTGCTTTCTTTAGTATGACACTCTCATNNNAGATCGGAAGA
130211 TATGAGTGTGGAGTCGTTGCCTGATTCAAATGCTAATCTCTTCNNNAGATCGGAAGA
130213 TATGAGTGTGGAGTCGTTGCACTTATATCACCATCTATCAGTGNNNAGATCGGAAG
130215 TATGAGTGTGGAGTCGTTGCACAAAGAATAGGTCACTTGAGNNNAGATCGGAAGA
130216 TATGAGTGTGGAGTCGTTGCTGCAATTTGACAAATAATATTGANNNAGATCGGAAG
130217 TATGAGTGTGGAGTCGTTGCGATGTCTGAGAAATCTCTAAACTNNNAGATCGGAAG
130218 TATGAGTGTGGAGTCGTTGCCAACTTGATCTTGGACTTCCNNNAGATCGGAAGAGC
130219 TATGAGTGTGGAGTCGTTGCTTAGGAAAAATTAGGAATTCCACNNNAGATCGGAA
130220 TATGAGTGTGGAGTCGTTGCCTCATCTTGTATTTTTCCAGTTCNNNAGATCGGAAGA
130221 TATGAGTGTGGAGTCGTTGCTATGTTTAATTGTTTTCCTCTGGNNNAGATCGGAAG
130222 TATGAGTGTGGAGTCGTTGCTAACCAACTCCATTTTACCTTTANNNAGATCGGAAGA
130223 TATGAGTGTGGAGTCGTTGCCAGCAAAATTACTAGTTTGGATTNNNAGATCGGAAG
130225 TATGAGTGTGGAGTCGTTGCACATAATAAGTCTGGAGGTAGANNNAGATCGGAAG
130226 TATGAGTGTGGAGTCGTTGCGTCATCTTCAGTCTGAACTAGNNNAGATCGGAAGAG
130227 TATGAGTGTGGAGTCGTTGCCTAAACCTAAACCTTTCCCTCNNNAGATCGGAAGAGC
130228 TATGAGTGTGGAGTCGTTGCTCTAACAAGGCAGAAAATTCTANNNAGATCGGAAG
130229 TATGAGTGTGGAGTCGTTGCAGGAGAATAAGCAAAGGTATTTANNNAGATCGGAA
130230 TATGAGTGTGGAGTCGTTGCTTCATTGCACCATTTTCTATAATNNNAGATCGGAAGA
130231 TATGAGTGTGGAGTCGTTGCAGGAGAAGTAATCATCAATATGGNNNAGATCGGAA
130232 TATGAGTGTGGAGTCGTTGCACAATACATACATAGACATATGGANNNAGATCGGAA
130233 TATGAGTGTGGAGTCGTTGCCTCTCACAATACATACATAGACANNNAGATCGGAAG
130234 TATGAGTGTGGAGTCGTTGCAGAGATTTGTAAGATTACCCAAANNNAGATCGGAA
130235 TATGAGTGTGGAGTCGTTGCTGTACATGAAAACATTGTATTGTNNNAGATCGGAAG
130236 TATGAGTGTGGAGTCGTTGCTCATTATGTTCTGTTTTGCATAANNNAGATCGGAAG
130237 TATGAGTGTGGAGTCGTTGCGACTAACACTGTCTTCTATCATANNNAGATCGGAAG
130238 TATGAGTGTGGAGTCGTTGCTCTGGACATATAACTACTCAAAANNNAGATCGGAAG
130239 TATGAGTGTGGAGTCGTTGCATTTGAATATGACCTAAAGGATGNNNAGATCGGAA
130240 TATGAGTGTGGAGTCGTTGCGAACGTCACATTTTCAATTTTAGNNNAGATCGGAAG
130241 TATGAGTGTGGAGTCGTTGCGGCATCTTTAGTTTTCTTTACTANNNAGATCGGAAGA
130242 TATGAGTGTGGAGTCGTTGCGGCATCTTTAGTTTTCTTTACTANNNAGATCGGAAGA

130243 TATGAGTGTGGAGTCGTTGCCAAGCACATATGAGTAAAATGTANNNAGATCGGAA
130244 TATGAGTGTGGAGTCGTTGCTAGAAAGGGAAGGTTTAGATATGNNNAGATCGGA
130245 TATGAGTGTGGAGTCGTTGCTTTAGAATTTCACTGCCAGGNNNAGATCGGAAGAG
130246 TATGAGTGTGGAGTCGTTGCTACAGGGAAAGAAAATGATCATANNNAGATCGGAA
130247 TATGAGTGTGGAGTCGTTGCCACATGCATATATCTGTGTATACNNNAGATCGGAAG
130248 TATGAGTGTGGAGTCGTTGCCAAGTCACTTTCCAAGAGAGNNNAGATCGGAAGAG
130249 TATGAGTGTGGAGTCGTTGCTATCCATTTATCTGTCAAGAAACNNNAGATCGGAAG
130250 TATGAGTGTGGAGTCGTTGCCTTGAGGAGGTCTTTTTAGTTTANNNAGATCGGAAG
130251 TATGAGTGTGGAGTCGTTGCATCTTAACAAAAAGAGATGGTTGNNNAGATCGGAA
130252 TATGAGTGTGGAGTCGTTGCACTGACCTGTGTGTAATAGGNNNAGATCGGAAGAG
130253 TATGAGTGTGGAGTCGTTGCCACAACAATGTCCATTCTGTNNNAGATCGGAAGAGC
130254 TATGAGTGTGGAGTCGTTGCCATTACCTGAGAACTTGATAATGNNNAGATCGGAAG
130255 TATGAGTGTGGAGTCGTTGCATGGGTCATTTTCTAGAATTAGTNNNAGATCGGAAG
130256 TATGAGTGTGGAGTCGTTGCAATGAGTTATTTACAGAGTAGGANNNAGATCGGAA
130257 TATGAGTGTGGAGTCGTTGCACATTACATGGAGCAAGGAANNNAGATCGGAAGAG
130258 TATGAGTGTGGAGTCGTTGCTTCATCCAAGGCCTTTATTAAATNNNAGATCGGAAG
130259 TATGAGTGTGGAGTCGTTGCTTCATCCAAGGCCTTTATTAAATNNNAGATCGGAAG
130260 TATGAGTGTGGAGTCGTTGCGTAAACGATAGACTTACCCAGNNNAGATCGGAAGA
130262 TATGAGTGTGGAGTCGTTGCTGATCATAATAAACGCATTAACANNNAGATCGGAAG
130263 TATGAGTGTGGAGTCGTTGCGAAATGTGTGGAGAATGATAATTNNNAGATCGGAA
130264 TATGAGTGTGGAGTCGTTGCCCCAGATTCTTTTTTTCTTTTTTNNNAGATCGGAAGA
130265 TATGAGTGTGGAGTCGTTGCGAGAGCTATAACTCAAGGTGNNNAGATCGGAAGAG
130266 TATGAGTGTGGAGTCGTTGCTGAACAGCTCAATAAAGAAAAATNNNAGATCGGAA
130268 TATGAGTGTGGAGTCGTTGCGCCTTCTTATTTGGTTTCTTATANNNAGATCGGAAGA
130269 TATGAGTGTGGAGTCGTTGCTTCTTGCAGCAATAAAAAAAAAANNNAGATCGGAAG
130270 TATGAGTGTGGAGTCGTTGCATCAATGGAAATGAAGGTTGANNNAGATCGGAAGA
130271 TATGAGTGTGGAGTCGTTGCCATGGGTAAAAAGATTGAGATAANNNAGATCGGAA
130272 TATGAGTGTGGAGTCGTTGCTAAATGGTTGTCAATTATGATGGNNNAGATCGGAA
130273 TATGAGTGTGGAGTCGTTGCTAAATGGTTGTCAATTATGATGGNNNAGATCGGAA
130274 TATGAGTGTGGAGTCGTTGCTAAATGGTTGTCAATTATGATGGNNNAGATCGGAA
130275 TATGAGTGTGGAGTCGTTGCCACATGTAAAATACACTAACACTNNNAGATCGGAAG
130276 TATGAGTGTGGAGTCGTTGCATATAAAACAGATGTGCTTGAAANNNAGATCGGAA
130277 TATGAGTGTGGAGTCGTTGCGATTTGTGGAGAAGGAAGAGNNNAGATCGGAAGA
130279 TATGAGTGTGGAGTCGTTGCAGTACAGACATTTCTATTACACANNNAGATCGGAAG
130280 TATGAGTGTGGAGTCGTTGCCTCACACATAAATGCACATGNNNAGATCGGAAGAGC
130281 TATGAGTGTGGAGTCGTTGCAAGTTGGGAAGTTAAGTAAAAAANNNAGATCGGAA
130282 TATGAGTGTGGAGTCGTTGCGGAGAATATTTACGTAGTAAAGCNNNAGATCGGAA
130283 TATGAGTGTGGAGTCGTTGCAGCAGATGTGAACTGAAATAGNNNAGATCGGAAGA
130284 TATGAGTGTGGAGTCGTTGCAATTATTATAAGAATTGGCTCAAGNNNAGATCGGAA
130285 TATGAGTGTGGAGTCGTTGCTCACTCACAATAATGAATAGACANNNAGATCGGAAG
130286 TATGAGTGTGGAGTCGTTGCTAAAAATCTTGAGGGTTGAATACNNNAGATCGGAA
130287 TATGAGTGTGGAGTCGTTGCCTAAGTAGTTGGTTTGAAAAAGTNNNAGATCGGAA
130288 TATGAGTGTGGAGTCGTTGCCCTCCATTAGTTCTATGAACTTANNNAGATCGGAAGA

130289 TATGAGTGTGGAGTCGTTGCCACTTTCCATTATCTAGTTTCATNNNAGATCGGAAGA
130290 TATGAGTGTGGAGTCGTTGCGCCAAAATACTATGTTAATAGGTNNNAGATCGGAAG
130291 TATGAGTGTGGAGTCGTTGCGTCCTCTCTACGTTTAAGTAATANNNAGATCGGAAG
130292 TATGAGTGTGGAGTCGTTGCATTATTTTAAGTGAAACAGCTCANNNAGATCGGAAG
130293 TATGAGTGTGGAGTCGTTGCATATTTTAAGGAATTGTCTCACGNNNAGATCGGAAG
130294 TATGAGTGTGGAGTCGTTGCTAGACTGGAAACTTAGTTCTAAGNNNAGATCGGAAG
130295 TATGAGTGTGGAGTCGTTGCACAACTCTTTTTAAAATCTTCCTNNNAGATCGGAAGA
130296 TATGAGTGTGGAGTCGTTGCCTGAAGGAACAAGGCATAGANNNAGATCGGAAGAG
130297 TATGAGTGTGGAGTCGTTGCTTTGCATGAAGAATTTTTTACTCNNNAGATCGGAAG
130298 TATGAGTGTGGAGTCGTTGCGAGGCAGAGAGTATCTTTAAATANNNAGATCGGAA
130299 TATGAGTGTGGAGTCGTTGCCTCTTATATAAAAGAGGTCTGCANNNAGATCGGAAG
130301 TATGAGTGTGGAGTCGTTGCTACTTTGTCCAGAAATAAGTCAANNNAGATCGGAAG
130302 TATGAGTGTGGAGTCGTTGCCATGTGAGGAAACAACAAGANNNAGATCGGAAGAG
130303 TATGAGTGTGGAGTCGTTGCCTCTGGGCATCGAGAATAATNNNAGATCGGAAGAG
130304 TATGAGTGTGGAGTCGTTGCCACCTGCTTTGATTTACTTATGNNNAGATCGGAAGAG
130305 TATGAGTGTGGAGTCGTTGCACTCGAAGGAAAGATTTGAGNNNAGATCGGAAGAG
130306 TATGAGTGTGGAGTCGTTGCTTCAAGTCCAGTCATTCTCTCNNNAGATCGGAAGAG
130307 TATGAGTGTGGAGTCGTTGCAGGATAATGTCAGCCTAATTTTANNNAGATCGGAAG
130308 TATGAGTGTGGAGTCGTTGCTGTTCTACATCCCTCTATCTATCNNNAGATCGGAAGA
130309 TATGAGTGTGGAGTCGTTGCCATGAAACATTCCTAAGGATTTGNNNAGATCGGAAG
130310 TATGAGTGTGGAGTCGTTGCCACTTCTGAAAGCATGATTATAANNNAGATCGGAAG
130311 TATGAGTGTGGAGTCGTTGCTTTGGAAACATTGAGCACTATATNNNAGATCGGAAG
130312 TATGAGTGTGGAGTCGTTGCCACAAAATAAGGAGTGGGTGNNNAGATCGGAAGAG
130313 TATGAGTGTGGAGTCGTTGCCTAATCGCAAGTGGATAGATTTNNNAGATCGGAAGA
130314 TATGAGTGTGGAGTCGTTGCAGATGCTTATTCCATGCTTAGNNNAGATCGGAAGAG
130315 TATGAGTGTGGAGTCGTTGCATTATTACAAGTTAAGTGGTGTCNNNAGATCGGAAG
130316 TATGAGTGTGGAGTCGTTGCTGCCTTTTATTTCTTCTTCTTTTNNNAGATCGGAAGAG
130317 TATGAGTGTGGAGTCGTTGCGTAAAAAGTACAACAGGAAATTGNNNAGATCGGAA
130319 TATGAGTGTGGAGTCGTTGCACATTTTTCACCTCTTTCTTATTNNNAGATCGGAAGAG
130320 TATGAGTGTGGAGTCGTTGCCACATATATAGGTTTACTTGTGCNNNAGATCGGAAG
130321 TATGAGTGTGGAGTCGTTGCCTGTAGTTTCCCTCTCCTTCNNNAGATCGGAAGAGCA
130322 TATGAGTGTGGAGTCGTTGCAATTTGGCTTACATAATTATGGGNNNAGATCGGAAG
130323 TATGAGTGTGGAGTCGTTGCAATTTGGCTTACATAATTATGGGNNNAGATCGGAAG
130324 TATGAGTGTGGAGTCGTTGCTGCAGAGAAAATAAAATCTCAAANNNAGATCGGAAG
130325 TATGAGTGTGGAGTCGTTGCCATAACTCTTGTCATTTCCTAAGNNNAGATCGGAAGA
130326 TATGAGTGTGGAGTCGTTGCTATAAGGAATTATCTTACGCAGTNNNAGATCGGAAG
130327 TATGAGTGTGGAGTCGTTGCTATAAGGAATTATCTTACGCAGTNNNAGATCGGAAG
130329 TATGAGTGTGGAGTCGTTGCCTTGGCTTTAGAAGTAGAGAGNNNAGATCGGAAGA
130331 TATGAGTGTGGAGTCGTTGCGTAATAATCCAGGATGATCTGATNNNAGATCGGAAG
130332 TATGAGTGTGGAGTCGTTGCGTTTCTTCTATATTGCAGGAAATNNNAGATCGGAAG
130333 TATGAGTGTGGAGTCGTTGCGAGCTCAGTATTATCTGTCAGNNNAGATCGGAAGAG
130334 TATGAGTGTGGAGTCGTTGCGAGTGAAAAGACTCATGGAGNNNAGATCGGAAGAG
130335 TATGAGTGTGGAGTCGTTGCATATAGAGAGAGAGAGAGAGAGANNNAGATCGGA

130337 TATGAGTGTGGAGTCGTTGCGTATGCATCTCTATAGCTAGCNNNAGATCGGAAGAG
130338 TATGAGTGTGGAGTCGTTGCGTATGCATCTCTATAGCTAGCNNNAGATCGGAAGAG
130339 TATGAGTGTGGAGTCGTTGCACACTTCCTTCTTTTTCTTTTTNNNAGATCGGAAGAG
130340 TATGAGTGTGGAGTCGTTGCAATCCTGTCTCTGTCATTTATTANNNAGATCGGAAGA
130341 TATGAGTGTGGAGTCGTTGCGGTCACAAATTCAACTACCTANNNAGATCGGAAGAG
130342 TATGAGTGTGGAGTCGTTGCGAAAAGAGCCAGCAAATGTANNNAGATCGGAAGAG
130343 TATGAGTGTGGAGTCGTTGCCACAGAATAATCAAACATCAAGANNNAGATCGGAAG
130344 TATGAGTGTGGAGTCGTTGCAATGAGGTTGTGAAAGAAATAACNNNAGATCGGAA
130345 TATGAGTGTGGAGTCGTTGCTATATAAATACACAGACACACACNNNAGATCGGAAG
130346 TATGAGTGTGGAGTCGTTGCCATTGATGCATTGTTCACTATATNNNAGATCGGAAG
130347 TATGAGTGTGGAGTCGTTGCATGAATTGTAGTGTATGTGAATGNNNAGATCGGAA
130348 TATGAGTGTGGAGTCGTTGCTTCGTAACCTTTTTATTCAGAGANNNAGATCGGAAG
130349 TATGAGTGTGGAGTCGTTGCTTCGTAACCTTTTTATTCAGAGANNNAGATCGGAAG
130350 TATGAGTGTGGAGTCGTTGCTATACATAACCCTTCAACAGTTTNNNAGATCGGAAGA
130351 TATGAGTGTGGAGTCGTTGCGAAAGAATATATAGGCCACTGGNNNAGATCGGAAG
130352 TATGAGTGTGGAGTCGTTGCATCTATTTTGTGTGTGTGTCTTANNNAGATCGGAAG
130353 TATGAGTGTGGAGTCGTTGCAATGTTAAGAACTGTTGAATCTTNNNAGATCGGAAG
130354 TATGAGTGTGGAGTCGTTGCAGTATTGTGACCAAGAAAAAAAANNNAGATCGGAA
130355 TATGAGTGTGGAGTCGTTGCAACTTTGAAAATAAAACTGATGCNNNAGATCGGAAG
130357 TATGAGTGTGGAGTCGTTGCTAATATTGCAACAAATATGGGAGNNNAGATCGGAA
130358 TATGAGTGTGGAGTCGTTGCATTTTATAATTCCAACACCTGTGNNNAGATCGGAAGA
130359 TATGAGTGTGGAGTCGTTGCCATATATATAATCTCCAGGCCAGNNNAGATCGGAAG
130360 TATGAGTGTGGAGTCGTTGCCATATATATAATCTCCAGGCCAGNNNAGATCGGAAG
130362 TATGAGTGTGGAGTCGTTGCTGTGTGTAGATAGATATGAATGTNNNAGATCGGAA
130363 TATGAGTGTGGAGTCGTTGCCACATATCACCATAAATGACTTTNNNAGATCGGAAGA
130364 TATGAGTGTGGAGTCGTTGCCAGACTTGATAGTTGTTGAAATGNNNAGATCGGAA
130365 TATGAGTGTGGAGTCGTTGCCCAGTCTCTAATTTGAAAACATANNNAGATCGGAAG
130367 TATGAGTGTGGAGTCGTTGCAGAAAGAAAGATAAGGTCTTAGGNNNAGATCGGAA
130368 TATGAGTGTGGAGTCGTTGCTATAAGCATAGGTACTGGTACTANNNAGATCGGAA
130370 TATGAGTGTGGAGTCGTTGCGCCTTTAAATTTGTTTCTTCTAAANNNAGATCGGAAG
130371 TATGAGTGTGGAGTCGTTGCGGGTTCTTTTATTTCCTTGATAANNNAGATCGGAAG
130372 TATGAGTGTGGAGTCGTTGCTACATCATCTTGCACTCCAGNNNAGATCGGAAGAGC
130373 TATGAGTGTGGAGTCGTTGCCAAAGAGTACAGTGTGATGANNNAGATCGGAAGAG
130374 TATGAGTGTGGAGTCGTTGCTATACACCACATGCACATATAATNNNAGATCGGAAG
130375 TATGAGTGTGGAGTCGTTGCACTTCTAAATGTAAAGAGAGGGNNNAGATCGGAAG
130376 TATGAGTGTGGAGTCGTTGCATTCATTCAATCTCAAGTGAGANNNAGATCGGAAGA
130377 TATGAGTGTGGAGTCGTTGCAATCAAAACCACAATAAGATACCNNNAGATCGGAAG
130378 TATGAGTGTGGAGTCGTTGCGGGTACTTCCTAATCTCAATTAANNNAGATCGGAAG
130379 TATGAGTGTGGAGTCGTTGCTAAAAAATGACACAGAAGGAAAANNNAGATCGGAA
130380 TATGAGTGTGGAGTCGTTGCAATAGCAATTTCAATGCAACATNNNAGATCGGAAGA
130381 TATGAGTGTGGAGTCGTTGCGCATAGAAAACTTTTGACAAAATNNNAGATCGGAAG
130382 TATGAGTGTGGAGTCGTTGCATTTATTAACATGGCAATCTGAGNNNAGATCGGAAG
130383 TATGAGTGTGGAGTCGTTGCACTAAGCAGGAAAATAAAAAGAGNNNAGATCGGAA

130384 TATGAGTGTGGAGTCGTTGCGGAGATAACAGTACATGTGTATNNNAGATCGGAAG
130385 TATGAGTGTGGAGTCGTTGCAAAGGCTCATTTTTTTAATCATCNNNAGATCGGAAG
130386 TATGAGTGTGGAGTCGTTGCAAAGGCTCATTTTTTTAATCATCNNNAGATCGGAAG
130387 TATGAGTGTGGAGTCGTTGCAAAATAAATTGGGAAGGATTCTGNNNAGATCGGAA
130388 TATGAGTGTGGAGTCGTTGCATAGATTCAACCAACCTCAGATNNNAGATCGGAAGA
130389 TATGAGTGTGGAGTCGTTGCCACATTTGGAGTTGAATCCTANNNAGATCGGAAGAG
130390 TATGAGTGTGGAGTCGTTGCGGTGGTTTTTGTATTTTTCTTTTNNNAGATCGGAAG
130391 TATGAGTGTGGAGTCGTTGCTTTGGAACAAATTCTAATATGCANNNAGATCGGAAG
130392 TATGAGTGTGGAGTCGTTGCGAAAGGAAATTGCTTGACTTCNNNAGATCGGAAGA
130393 TATGAGTGTGGAGTCGTTGCGCCAAAAGCATAAAATTTTTTTGNNNAGATCGGAAG
130394 TATGAGTGTGGAGTCGTTGCAGAATAAAGTGAGAATCTGGAAGNNNAGATCGGAA
130395 TATGAGTGTGGAGTCGTTGCCTGAAGATACATCAAAAGTGAAGNNNAGATCGGAA
130396 TATGAGTGTGGAGTCGTTGCTAGTGGAGGGTCATAAAAATAAGNNNAGATCGGAA
130397 TATGAGTGTGGAGTCGTTGCTTTTGCTGCTATAATATAAACGTNNNAGATCGGAAG
130398 TATGAGTGTGGAGTCGTTGCTATAATGTAGAGTTCTCTGTTCCNNNAGATCGGAAG
130399 TATGAGTGTGGAGTCGTTGCGTGCTATGAAAAACTAAAAATGGNNNAGATCGGAA
130400 TATGAGTGTGGAGTCGTTGCATATAAATAAGCATGGTTTCTGCNNNAGATCGGAAG
130401 TATGAGTGTGGAGTCGTTGCTATAGATGTATATTCCCTGATGCNNNAGATCGGAAG
130402 TATGAGTGTGGAGTCGTTGCGGATAAATATTTCCAGAGGTGGNNNAGATCGGAAG
130403 TATGAGTGTGGAGTCGTTGCTAGATAAGAAAACAGATGTTCAGNNNAGATCGGAA
130405 TATGAGTGTGGAGTCGTTGCAGAAAATACTGTAAGAAAGCAAGNNNAGATCGGAA
130406 TATGAGTGTGGAGTCGTTGCCTAACAAAGTGAGGGTTTTTTTNNNAGATCGGAAGA
130407 TATGAGTGTGGAGTCGTTGCCTTCCAAGCACTGATTTATTTAGNNNAGATCGGAAG
130408 TATGAGTGTGGAGTCGTTGCCTTCCAAGCACTGATTTATTTAGNNNAGATCGGAAG
130409 TATGAGTGTGGAGTCGTTGCATTTTAGGAAGGCAAGATGTAANNNAGATCGGAAG
130410 TATGAGTGTGGAGTCGTTGCGATCTCCAAAATGTTTTCAGTATNNNAGATCGGAAG
130411 TATGAGTGTGGAGTCGTTGCCTGCAATTTGGATATTTGAAAAANNNAGATCGGAAG
130412 TATGAGTGTGGAGTCGTTGCCTACTTTACCTGGGTTAGAACNNNAGATCGGAAGAG
130413 TATGAGTGTGGAGTCGTTGCCAGTACCAAAACCTAGAGAAAGNNNAGATCGGAAG
130414 TATGAGTGTGGAGTCGTTGCAAGTGAAGTGTTATAGAGTGATTNNNAGATCGGAA
130415 TATGAGTGTGGAGTCGTTGCGGGACTGAAATAAAAATATTCCANNNAGATCGGAA
130416 TATGAGTGTGGAGTCGTTGCAGATAGTTTGCAGACGAAAGNNNAGATCGGAAGAG
130417 TATGAGTGTGGAGTCGTTGCATTTCTTGTAGTTGTTGTACTCTNNNAGATCGGAAG
130419 TATGAGTGTGGAGTCGTTGCGTTAGAAAATGAGGCCTCTGNNNAGATCGGAAGAG
130420 TATGAGTGTGGAGTCGTTGCGCTAATTTTAGATGGGGTTTTGNNNAGATCGGAAG
130421 TATGAGTGTGGAGTCGTTGCGTAGAAGCAGTGAAATTAGTTAGNNNAGATCGGAA
130422 TATGAGTGTGGAGTCGTTGCCCAAAATCCATGTGTTGAAGNNNAGATCGGAAGAG
130423 TATGAGTGTGGAGTCGTTGCCCCCTTATCTCTCTGATATTGTNNNAGATCGGAAGAG
130424 TATGAGTGTGGAGTCGTTGCCCCCCCTTATCTCTCTGATATTNNNAGATCGGAAGAGC
130425 TATGAGTGTGGAGTCGTTGCATTTCTTCCTCTTCAATTATCTGNNNAGATCGGAAGA
130426 TATGAGTGTGGAGTCGTTGCAAGAGATTTCAGGGATCAATTATNNNAGATCGGAA
130427 TATGAGTGTGGAGTCGTTGCGTATTTATAGACATGGCAGGTCNNNAGATCGGAAG
130428 TATGAGTGTGGAGTCGTTGCGGTTCTGTCTTGTTTATAGTACTNNNAGATCGGAAG

130429 TATGAGTGTGGAGTCGTTGCCCCAAAGGAAAATAAATCATTCTNNNAGATCGGAAG
130430 TATGAGTGTGGAGTCGTTGCTATAGTTATCTATTGCTGCCTACNNNAGATCGGAAG
130431 TATGAGTGTGGAGTCGTTGCAGATTCTATGAATTTCAGGTGAGNNNAGATCGGAA
130433 TATGAGTGTGGAGTCGTTGCAGAGTTTAAACAAAATCAGTTTCNNNAGATCGGAAG
130434 TATGAGTGTGGAGTCGTTGCGCAGCAGACTTTATTCTTTATAANNNAGATCGGAAG
130435 TATGAGTGTGGAGTCGTTGCTTTTCCGATTATCTGATCATAGTNNNAGATCGGAAG
130436 TATGAGTGTGGAGTCGTTGCAAAGTTACAATCTTCTGTTTGAANNNAGATCGGAAG
130437 TATGAGTGTGGAGTCGTTGCGCTATATCTACATATCTGCACATNNNAGATCGGAAG
130438 TATGAGTGTGGAGTCGTTGCAATCTGGACTTTAGAGAGAAATTNNNAGATCGGAA
130439 TATGAGTGTGGAGTCGTTGCGAGTCCAAGTACATAGTTCATATNNNAGATCGGAAG
130440 TATGAGTGTGGAGTCGTTGCATACTACTCAGCCATAAAAAAGANNNAGATCGGAAG
130441 TATGAGTGTGGAGTCGTTGCATACTACTCAGCCATAAAAAAGANNNAGATCGGAAG
130442 TATGAGTGTGGAGTCGTTGCGATCTCAGTTCACTGCAATCNNNAGATCGGAAGAGC
130445 TATGAGTGTGGAGTCGTTGCCTCCAAAGCTAAATCCATGTATNNNAGATCGGAAGA
130446 TATGAGTGTGGAGTCGTTGCATTATAACTGTGGATAAGAGCTCNNNAGATCGGAA
130447 TATGAGTGTGGAGTCGTTGCGACTATAAGGCTCTGATAAAGATNNNAGATCGGAA
130449 TATGAGTGTGGAGTCGTTGCGCTGAAATCAAGAAACATGTTANNNAGATCGGAAG
130450 TATGAGTGTGGAGTCGTTGCTTAGAAAGAATGGTGGATTCATNNNAGATCGGAAG
130451 TATGAGTGTGGAGTCGTTGCAATGAAAGAGAACCCAATTTATCNNNAGATCGGAAG
130453 TATGAGTGTGGAGTCGTTGCTCTTATTTGTTCATCTCTGTAACNNNAGATCGGAAGA
130454 TATGAGTGTGGAGTCGTTGCATTAGAGAAACCCCTGTCTCNNNAGATCGGAAGAGC
130455 TATGAGTGTGGAGTCGTTGCAGACAGAGAGGATCCATAATATANNNAGATCGGAA
130456 TATGAGTGTGGAGTCGTTGCGAATGAGACCTAGAGAGAGTATNNNAGATCGGAAG
130457 TATGAGTGTGGAGTCGTTGCATATATGGTAGTGTTGTTTCAGANNNAGATCGGAA
130458 TATGAGTGTGGAGTCGTTGCGGGATTTTCTTTTTAGAATCACANNNAGATCGGAAG
130459 TATGAGTGTGGAGTCGTTGCAAACTGATACAGGAAACAATGAANNNAGATCGGAA
130461 TATGAGTGTGGAGTCGTTGCTCAATTAAAAAAAATGCAGTGTGNNNAGATCGGAA
130462 TATGAGTGTGGAGTCGTTGCGTTGTGTTTACTCTGAAGTTTCNNNAGATCGGAAGA
130463 TATGAGTGTGGAGTCGTTGCTTTAAGTTACAAAGCTGCTTTATNNNAGATCGGAAG
130464 TATGAGTGTGGAGTCGTTGCCAGAGTAGAGAATAAAGTACACANNNAGATCGGAA
130465 TATGAGTGTGGAGTCGTTGCTTGTTTTCTCGTTTCCCTTTNNNAGATCGGAAGAGCA
130466 TATGAGTGTGGAGTCGTTGCTTTTACAAAGGCTTATAATCTGGNNNAGATCGGAAG
130467 TATGAGTGTGGAGTCGTTGCCTTGAAGGAAGGAGTAAGTAAAANNNAGATCGGAA
130468 TATGAGTGTGGAGTCGTTGCGCTTCAGCTTAAATTGTTTTATTNNNAGATCGGAAG
130469 TATGAGTGTGGAGTCGTTGCGATTTTATAGGGATTGCATTGAANNNAGATCGGAA
130470 TATGAGTGTGGAGTCGTTGCTTAGGGAACATAAGGAGGATCNNNAGATCGGAAGA
130471 TATGAGTGTGGAGTCGTTGCATTCACGGTTTTAACTTGTTAATNNNAGATCGGAAG
130472 TATGAGTGTGGAGTCGTTGCCAATAAGGAGCTCCATGAATNNNAGATCGGAAGAG
130473 TATGAGTGTGGAGTCGTTGCCAATAAGGAGCTCCATGAATNNNAGATCGGAAGAG
130474 TATGAGTGTGGAGTCGTTGCTGTCTTGAATTATTAACCCATTGNNNAGATCGGAAG
130475 TATGAGTGTGGAGTCGTTGCGAGAAACCCTAACCTGTATTAATNNNAGATCGGAAG
130476 TATGAGTGTGGAGTCGTTGCTCCCAAGATTTTTTTGTATGTTANNNAGATCGGAAG
130477 TATGAGTGTGGAGTCGTTGCATTCAGCTTATAAAAATGAACCTNNNAGATCGGAAG

130479 TATGAGTGTGGAGTCGTTGCTATTCAAATATTTTTTGGCCTCCNNNAGATCGGAAGA
130480 TATGAGTGTGGAGTCGTTGCCTTAGAAAAGATGAAAAGGGAAGNNNAGATCGGAA
130481 TATGAGTGTGGAGTCGTTGCAATATCCTGAGCTATGCTGATNNNAGATCGGAAGAG
130482 TATGAGTGTGGAGTCGTTGCTTCAGTAAACATAAGTGGTAGTGNNNAGATCGGAA
130483 TATGAGTGTGGAGTCGTTGCGTATGCATAAGTTACTCTTGTTTNNNAGATCGGAAG
130484 TATGAGTGTGGAGTCGTTGCGTATGCATAAGTTACTCTTGTTTNNNAGATCGGAAG
130485 TATGAGTGTGGAGTCGTTGCCAGATGTCTGCTTTTTGATATTANNNAGATCGGAAG
130486 TATGAGTGTGGAGTCGTTGCAGTACATATATCCCCACAATCTNNNAGATCGGAAGA
130487 TATGAGTGTGGAGTCGTTGCTACAATATGTAAGTATGGTCACTNNNAGATCGGAAG
130489 TATGAGTGTGGAGTCGTTGCACTTAGGACATTTATCTACTGTTNNNAGATCGGAAG
130490 TATGAGTGTGGAGTCGTTGCACTTAGGACATTTATCTACTGTTNNNAGATCGGAAG
130491 TATGAGTGTGGAGTCGTTGCTCACAGGTATCTATTGAGTACTANNNAGATCGGAAG
130492 TATGAGTGTGGAGTCGTTGCTACTTACTATGTGTCAAGCATATNNNAGATCGGAAG
130493 TATGAGTGTGGAGTCGTTGCGAGTGAAATAATGTCTTTTGGATNNNAGATCGGAA
130497 TATGAGTGTGGAGTCGTTGCGGATGAAGTGTTATCAAGAGAANNNAGATCGGAAG
130498 TATGAGTGTGGAGTCGTTGCTTTGTTATGTGGAGGTTTTTCTANNNAGATCGGAAG
130499 TATGAGTGTGGAGTCGTTGCAAGAGAGAAAGGAAAGGAGGNNNAGATCGGAAGA
130500 TATGAGTGTGGAGTCGTTGCTATTCTCTAAGTGAAGTAACTCANNNAGATCGGAAG
130501 TATGAGTGTGGAGTCGTTGCTTGTATTATTATGGCATGTTTGCNNNAGATCGGAAG
130502 TATGAGTGTGGAGTCGTTGCATTCATTTCTTTAGTGCATTTCTNNNAGATCGGAAGA
130503 TATGAGTGTGGAGTCGTTGCAAAGGCAGTACACATGGTATANNNAGATCGGAAGA
130504 TATGAGTGTGGAGTCGTTGCTCTGTCTCTCTCTCTTTCTATATNNNAGATCGGAAGA
130505 TATGAGTGTGGAGTCGTTGCGTTTAGGAAGATACTTTCCATCTNNNAGATCGGAAG
130506 TATGAGTGTGGAGTCGTTGCCTTCATTTAGAATTCTCACTCAGNNNAGATCGGAAGA
130508 TATGAGTGTGGAGTCGTTGCCCAGTAGTTTGTTAATCCCTTNNNAGATCGGAAGAG
130509 TATGAGTGTGGAGTCGTTGCGGAGAACCCTTACTAATACATTNNNAGATCGGAAGA
130510 TATGAGTGTGGAGTCGTTGCCAGACAGACAGAGATTTATTTTGNNNAGATCGGAA
130511 TATGAGTGTGGAGTCGTTGCGAGATTGTAGTCAAGCTGTTTNNNAGATCGGAAGA
130512 TATGAGTGTGGAGTCGTTGCGAGATTGTAGTCAAGCTGTTTNNNAGATCGGAAGA
130513 TATGAGTGTGGAGTCGTTGCCAAATGAAAATCTCTGCTCAGNNNAGATCGGAAGAG
130514 TATGAGTGTGGAGTCGTTGCGATGCATGTTATGAGGAGTGNNNAGATCGGAAGA
130515 TATGAGTGTGGAGTCGTTGCCTTGTTCCCAGACAAAAAACNNNAGATCGGAAGAGC
130516 TATGAGTGTGGAGTCGTTGCGGTTTACCATTGTTTAAAACTTTTNNNAGATCGGAA
130517 TATGAGTGTGGAGTCGTTGCCTGTCTCTCTCTGTCTCTTTNNNAGATCGGAAGAGCA
130518 TATGAGTGTGGAGTCGTTGCCTGTCTCTCTCTGTCTCTTTNNNAGATCGGAAGAGCA
130519 TATGAGTGTGGAGTCGTTGCCCAATTCACTCATTTTAAGTGTANNNAGATCGGAAG
130520 TATGAGTGTGGAGTCGTTGCTAAAATAAGCAGGTACATGACTANNNAGATCGGAA
130522 TATGAGTGTGGAGTCGTTGCTAATAGGACTATTTTCCCACAAANNNAGATCGGAAG
130523 TATGAGTGTGGAGTCGTTGCTTTCTAAAGAAAGTCAAGATGTCNNNAGATCGGAAG
130524 TATGAGTGTGGAGTCGTTGCATAATCACAGTGGTAATAAAGGTNNNAGATCGGAA
130525 TATGAGTGTGGAGTCGTTGCTTTTTATTATTGGCATTTCCTCANNNAGATCGGAAGA
130526 TATGAGTGTGGAGTCGTTGCGGATATAGTCATCCCTTAGTATCNNNAGATCGGAAG
130527 TATGAGTGTGGAGTCGTTGCACAGTTTTCTGATTCTTTGAATTNNNAGATCGGAAGA

130528 TATGAGTGTGGAGTCGTTGCTTTACCTCCTGTCATGAATAAAANNNAGATCGGAAGA
130529 TATGAGTGTGGAGTCGTTGCGTTCTTCGCATATTCATTTCTAANNNAGATCGGAAGA
130530 TATGAGTGTGGAGTCGTTGCGAATCCTGTATGTTTGGGTCNNNAGATCGGAAGAGC
130531 TATGAGTGTGGAGTCGTTGCGAAAAGTTTTGGACTGAGAATAANNNAGATCGGAAG
130532 TATGAGTGTGGAGTCGTTGCTTTCCTCTTCCATATAATGACAANNNAGATCGGAAGA
130533 TATGAGTGTGGAGTCGTTGCGACCTTCCAACTAAACTTTTTATNNNAGATCGGAAGA
130534 TATGAGTGTGGAGTCGTTGCTAAATTAGGAGAAACACATTGTCNNNAGATCGGAAG
130535 TATGAGTGTGGAGTCGTTGCTTGCAATGACTATATTCAGTGTANNNAGATCGGAAG
130536 TATGAGTGTGGAGTCGTTGCCTGACTTTATTCAAATTGCTCTANNNAGATCGGAAGA
130537 TATGAGTGTGGAGTCGTTGCCTGCAATGGGCCAAATAATANNNAGATCGGAAGAGC
130538 TATGAGTGTGGAGTCGTTGCCTCCCTAGCATATTCCAAAATANNNAGATCGGAAGAG
130539 TATGAGTGTGGAGTCGTTGCTATACAGAAAATTGAAAAGAGGTNNNAGATCGGAAG
130540 TATGAGTGTGGAGTCGTTGCGATTTTGGTTATTTCTTGTCCTCNNNAGATCGGAAGA
130541 TATGAGTGTGGAGTCGTTGCGAGAAAAGCACAGTTTTAATTTCNNNAGATCGGAAG
130542 TATGAGTGTGGAGTCGTTGCACCTCTATAAAGCAAAGAAAAATANNNAGATCGGAA
130543 TATGAGTGTGGAGTCGTTGCACCTCTATAAAGCAAAGAAAAATANNNAGATCGGAA
130545 TATGAGTGTGGAGTCGTTGCTCTGATTGTTCCTTTTCCATATANNNAGATCGGAAGA
130546 TATGAGTGTGGAGTCGTTGCTAATCTGAAAGAAAATAGCTCCTNNNAGATCGGAAG
130547 TATGAGTGTGGAGTCGTTGCGAGTGTTGTCCTCATACACANNNAGATCGGAAGAGC
130548 TATGAGTGTGGAGTCGTTGCTTTCTTTTTTAGACAGAGTTTCGNNNAGATCGGAAGA
130549 TATGAGTGTGGAGTCGTTGCATGTTCTGATGCAGGAATAAGNNNAGATCGGAAGAG
130550 TATGAGTGTGGAGTCGTTGCCTGAATAACATCAGTTCTTATTGNNNAGATCGGAAGA
130551 TATGAGTGTGGAGTCGTTGCATTTATTTCACTGAATAACATCAGNNNAGATCGGAAG
130552 TATGAGTGTGGAGTCGTTGCCCTGGTTCTGATATTTCTACTATNNNAGATCGGAAGA
130553 TATGAGTGTGGAGTCGTTGCAAAAATTATCTAATCAGCTCGGTNNNAGATCGGAAG
130554 TATGAGTGTGGAGTCGTTGCCAACTAAGATTGTGTGTATGTGNNNAGATCGGAAGA
130555 TATGAGTGTGGAGTCGTTGCATGGGATGTAAGATTCACATTAANNNAGATCGGAAG
130556 TATGAGTGTGGAGTCGTTGCAACGGTATTACTCATAAACTAGANNNAGATCGGAAG
130557 TATGAGTGTGGAGTCGTTGCACCTAGTTTCAAAAATTACAACANNNAGATCGGAAGA
130558 TATGAGTGTGGAGTCGTTGCGCCCTAGGTAAATACCAATTTNNNAGATCGGAAGAG
130559 TATGAGTGTGGAGTCGTTGCTCTAATAGTAGATGCTGGAGATANNNAGATCGGAAG
130560 TATGAGTGTGGAGTCGTTGCAAATACTCTTTCCTCTATATGCANNNAGATCGGAAGA
130561 TATGAGTGTGGAGTCGTTGCCTTCTAAAAATTCTCTCTTGAGCNNNAGATCGGAAGA
130562 TATGAGTGTGGAGTCGTTGCACAAATTCCTATAATTCCAACTGNNNAGATCGGAAGA
130563 TATGAGTGTGGAGTCGTTGCTTATACCCTTCAGACCAATAATGNNNAGATCGGAAGA
130564 TATGAGTGTGGAGTCGTTGCCCAAGATTTCTCTCTAGGAAATANNNAGATCGGAAGA
130565 TATGAGTGTGGAGTCGTTGCCACGTTTTCTATACTTGAAGAAANNNAGATCGGAAGA
130566 TATGAGTGTGGAGTCGTTGCCACGTTTTCTATACTTGAAGAAANNNAGATCGGAAGA
130567 TATGAGTGTGGAGTCGTTGCGTGCTGGGTAATATTTTATAACANNNAGATCGGAAG
130568 TATGAGTGTGGAGTCGTTGCATGTATACATCTTTTAGCATCCTNNNAGATCGGAAGA
130569 TATGAGTGTGGAGTCGTTGCTATGCCCAATTCTTAGAAATAGTNNNAGATCGGAAGA
130570 TATGAGTGTGGAGTCGTTGCTATGCCCAATTCTTAGAAATAGTNNNAGATCGGAAGA
130571 TATGAGTGTGGAGTCGTTGCGATTTGTCCTAATACAACTGTGNNNAGATCGGAAGA

130573 TATGAGTGTGGAGTCGTTGCGGTCTCACTATATCACCCAANNNAGATCGGAAGAGC
130574 TATGAGTGTGGAGTCGTTGCCAGTATCAAAAGATCAAGCATTTNNNAGATCGGAAG
130575 TATGAGTGTGGAGTCGTTGCTAAAACATACTACAAAGCTACAGNNNAGATCGGAAG
130576 TATGAGTGTGGAGTCGTTGCTATCTTGTATGTCCTGAAATTTGNNNAGATCGGAAG
130577 TATGAGTGTGGAGTCGTTGCCTGGTTTCATGAAGAAGTTAAATNNNAGATCGGAAG
130579 TATGAGTGTGGAGTCGTTGCAGTGAATGAAGATTGAGAATCTANNNAGATCGGAA
130580 TATGAGTGTGGAGTCGTTGCCCTAGGACCAAACTGTCTATNNNAGATCGGAAGAGC
130582 TATGAGTGTGGAGTCGTTGCGAAGAATGTTTGGTGTCAGTNNNAGATCGGAAGAG
130584 TATGAGTGTGGAGTCGTTGCGAGACTGAGAGAGAGAGATGNNNAGATCGGAAGA
130585 TATGAGTGTGGAGTCGTTGCAGAGAGATGGAGACTGAGAGNNNAGATCGGAAGA
130586 TATGAGTGTGGAGTCGTTGCCTTCTGGTAAAAAGGGTAGAANNNAGATCGGAAGA
130587 TATGAGTGTGGAGTCGTTGCCATTCTCCTTGAAACAAAGATTANNNAGATCGGAAG
130588 TATGAGTGTGGAGTCGTTGCGGATAAAAGGAAGAATCCATCTNNNAGATCGGAAG
130589 TATGAGTGTGGAGTCGTTGCGAGAGACTGAGTTTGAAGACNNNAGATCGGAAGAG
130590 TATGAGTGTGGAGTCGTTGCGAGAGACTGAGTTTGAAGACNNNAGATCGGAAGAG
130591 TATGAGTGTGGAGTCGTTGCATAAGTTGCTCAGTCCCTATNNNAGATCGGAAGAGC
130592 TATGAGTGTGGAGTCGTTGCGTGATTTTGAACACCTTTCTATANNNAGATCGGAAG
130593 TATGAGTGTGGAGTCGTTGCTTACACTTGAGAGAGGAAAAATGNNNAGATCGGAA
130594 TATGAGTGTGGAGTCGTTGCGAGGGAGGATGTTCATATCTNNNAGATCGGAAGAG
130595 TATGAGTGTGGAGTCGTTGCCATTTGCCTAATCTCTCTGTCNNNAGATCGGAAGAG
130596 TATGAGTGTGGAGTCGTTGCCTTCCAACTACAGGAAAATCCNNNAGATCGGAAGAG
130598 TATGAGTGTGGAGTCGTTGCGAATTCAGAAGTAGCATTTGAGNNNAGATCGGAAG
130601 TATGAGTGTGGAGTCGTTGCGCATTATCGGTTTTTGGATATATNNNAGATCGGAAG
130602 TATGAGTGTGGAGTCGTTGCTCAAAATTAAGAATGCTGTGTTANNNAGATCGGAAG
130603 TATGAGTGTGGAGTCGTTGCCAACTACAAGAAGAAAAGAGTTANNNAGATCGGAA
130604 TATGAGTGTGGAGTCGTTGCGCTAGAGGATTCCTATTTACATANNNAGATCGGAAG
130605 TATGAGTGTGGAGTCGTTGCTTATGTACACAGCAAACATTCTANNNAGATCGGAAG
130606 TATGAGTGTGGAGTCGTTGCTTATGTACACAGCAAACATTCTANNNAGATCGGAAG
130607 TATGAGTGTGGAGTCGTTGCAAGTTTACTTTTTCTAGCTACCTNNNAGATCGGAAGA
130608 TATGAGTGTGGAGTCGTTGCCTTTGAGATAAAAGTCTATGGTGNNNAGATCGGAA
130609 TATGAGTGTGGAGTCGTTGCCAGTGTATGTGTTTGTCTTACNNNAGATCGGAAGAG
130610 TATGAGTGTGGAGTCGTTGCAAGGTAGTAGAAATGAAAGTGTANNNAGATCGGAA
130611 TATGAGTGTGGAGTCGTTGCTGTATCTACTTAAGGATGTTCAGNNNAGATCGGAAG
130612 TATGAGTGTGGAGTCGTTGCAGTCACATTGAACTTGATAAGATNNNAGATCGGAAG
130613 TATGAGTGTGGAGTCGTTGCGTATGTGTGTGTGTGTAGTTTNNNAGATCGGAAGA
130614 TATGAGTGTGGAGTCGTTGCGATTGGGAGTTCACATAAAAATTNNNAGATCGGAA
130615 TATGAGTGTGGAGTCGTTGCGTTTCAATCACCTTTAAGTTCTANNNAGATCGGAAGA
130616 TATGAGTGTGGAGTCGTTGCTGATTTAATTTGGTAGCAAGATGNNNAGATCGGAA
130617 TATGAGTGTGGAGTCGTTGCCTTCGTTTATATTTGGGATACAANNNAGATCGGAAG
130618 TATGAGTGTGGAGTCGTTGCAAAGAGAGAGAAGGAGAGAAANNNAGATCGGAAG
130619 TATGAGTGTGGAGTCGTTGCAAGAGATAGTTCTAGAGGTAGTCNNNAGATCGGAA
130620 TATGAGTGTGGAGTCGTTGCCTGTTGATAAGGACATAGAGAATNNNAGATCGGAA
130621 TATGAGTGTGGAGTCGTTGCGGTGTAGAAATGAATTCACTTTTNNNAGATCGGAAG

130622 TATGAGTGTGGAGTCGTTGCGGTGTAGAAATGAATTCACTTTTNNNAGATCGGAAG

130623 TATGAGTGTGGAGTCGTTGCAGGTGATTGATTATGTCTGTAGNNNAGATCGGAAG

130624 TATGAGTGTGGAGTCGTTGCTATTATTCCACTCACATGAAGTANNNAGATCGGAAG

130625 TATGAGTGTGGAGTCGTTGCATAATGTCTTCAGGGTTCATCNNNAGATCGGAAGAG

130626 TATGAGTGTGGAGTCGTTGCCAACTCACAAAAGACAGATTAATNNNAGATCGGAAG

130627 TATGAGTGTGGAGTCGTTGCCTACATTTTTCTGGGTGAGAGNNNAGATCGGAAGA

130628 TATGAGTGTGGAGTCGTTGCTCAACTAGCCTAAAAGAAAGATANNNAGATCGGAA

130629 TATGAGTGTGGAGTCGTTGCATGAAAAGGTTCTTGAGGATTNNNAGATCGGAAGA

130630 TATGAGTGTGGAGTCGTTGCTGATAGAGCAAATAGGATAAAATGNNNAGATCGGA

130631 TATGAGTGTGGAGTCGTTGCTGAGACCTCTATATATCATCATTNNNAGATCGGAAG

130632 TATGAGTGTGGAGTCGTTGCCATTGATAAAAACAAGTTTGAGGNNNAGATCGGAA

130633 TATGAGTGTGGAGTCGTTGCTGTATTAAATCTTCCTTGAAACANNNAGATCGGAAG

130634 TATGAGTGTGGAGTCGTTGCCTTTCCCCAGCAAATTTCTCNNNAGATCGGAAGAGCA

130636 TATGAGTGTGGAGTCGTTGCTATATAAACACACACAAATGTGGNNNAGATCGGAAG

130637 TATGAGTGTGGAGTCGTTGCTTCTAAGCAGTTCCTTCTGAGNNNAGATCGGAAGAG

130638 TATGAGTGTGGAGTCGTTGCACAATTAGTTAAGGAAAAATCCTNNNAGATCGGAAG

130639 TATGAGTGTGGAGTCGTTGCACCTGATCAAAATAACATGTTTTNNNAGATCGGAAG

130640 TATGAGTGTGGAGTCGTTGCTTGTTTCAATAGGAGAAAAGAAGNNNAGATCGGAA

130641 TATGAGTGTGGAGTCGTTGCAAGAGTAAGGGATTTGAATATGTNNNAGATCGGAA

130642 TATGAGTGTGGAGTCGTTGCCAAGAACCAATAAACTATTGACTNNNAGATCGGAAG

130643 TATGAGTGTGGAGTCGTTGCCTTATACCATTGTTTCTCAAACTNNNAGATCGGAAGA

130644 TATGAGTGTGGAGTCGTTGCAATAATAAAGATCTACTCAGGGTNNNAGATCGGAA

130645 TATGAGTGTGGAGTCGTTGCCTTTATTCAGTGGTTTGTGTTNNNAGATCGGAAGAG

130646 TATGAGTGTGGAGTCGTTGCTGAGTTGATTTGGTAATTTATCCNNNAGATCGGAAG

130647 TATGAGTGTGGAGTCGTTGCAATGCAATGGCTTGATCATANNNAGATCGGAAGAG

130648 TATGAGTGTGGAGTCGTTGCAATTTTAGAATTTCTTCTCCTGGNNNAGATCGGAAG

130649 TATGAGTGTGGAGTCGTTGCCTGTTAAGTTTCCTCTTTTGATANNNAGATCGGAAGA

130650 TATGAGTGTGGAGTCGTTGCCAATTTCCTTCCTTCTGTTTAANNNAGATCGGAAGAG

130651 TATGAGTGTGGAGTCGTTGCTTGTCATTTTAACTGGAGTGTTNNNAGATCGGAAGA

130652 TATGAGTGTGGAGTCGTTGCTTTAATAAAAGAGAAACCTTGGGNNNAGATCGGAA

130654 TATGAGTGTGGAGTCGTTGCTTCTCTCTCTCTCTCTTTCTTCNNNAGATCGGAAGAGC

130655 TATGAGTGTGGAGTCGTTGCGATCACTCCTTTCTTCACAGNNNAGATCGGAAGAGC

130656 TATGAGTGTGGAGTCGTTGCTCTGATAAGAAACTAATACCTGANNNAGATCGGAAG

130657 TATGAGTGTGGAGTCGTTGCTAATGATCTATAAGTGTGCAAAGNNNAGATCGGAA

130659 TATGAGTGTGGAGTCGTTGCATACTTATACTTCAGATGACGTTNNNAGATCGGAAG

130660 TATGAGTGTGGAGTCGTTGCCCCTGAAATTTCAAATTCCTGNNNAGATCGGAAGAG

130661 TATGAGTGTGGAGTCGTTGCTCTTTTTTCTTAGAGATAGGTCTNNNAGATCGGAAG

130662 TATGAGTGTGGAGTCGTTGCTCTGTCTGAAATAAGGGATTTATNNNAGATCGGAAG

130663 TATGAGTGTGGAGTCGTTGCCAGGAAGATGAAAGGTACAGNNNAGATCGGAAGA

130664 TATGAGTGTGGAGTCGTTGCAATCCCCCTTTTAGTTCTTAGTANNNAGATCGGAAGA

130665 TATGAGTGTGGAGTCGTTGCCTTCTAGTAACTTTCCCATAGAANNNAGATCGGAAG

130666 TATGAGTGTGGAGTCGTTGCCTTCTAGTAACTTTCCCATAGAANNNAGATCGGAAG

130667 TATGAGTGTGGAGTCGTTGCGAGTGGGATACTTTTGTTTTAGNNNAGATCGGAAG

130668 TATGAGTGTGGAGTCGTTGCGGAGTAGGTATATCTTCAGTATGNNNAGATCGGAA
130669 TATGAGTGTGGAGTCGTTGCTTTTAAGGCATCTAAACATTCTGNNNAGATCGGAAG
130670 TATGAGTGTGGAGTCGTTGCCAGGAAAAATATAAAAGAGCTGANNNAGATCGGAA
130671 TATGAGTGTGGAGTCGTTGCTCTGAAGAACCTTGACTAATAAANNNAGATCGGAAG
130672 TATGAGTGTGGAGTCGTTGCCACTCACCCATACTAGGATCNNNAGATCGGAAGAGC
130673 TATGAGTGTGGAGTCGTTGCCTTACAGATGTTTGAAGGCATNNNAGATCGGAAGA
130674 TATGAGTGTGGAGTCGTTGCCTATCATCCTGTGGACATTGNNNAGATCGGAAGAGC
130675 TATGAGTGTGGAGTCGTTGCCTAGAAAAGGGTACTACACAGNNNAGATCGGAAGA
130676 TATGAGTGTGGAGTCGTTGCTATGGTTCATCCTAGGTTACANNNAGATCGGAAGAG
130677 TATGAGTGTGGAGTCGTTGCCATCTACGAAATGAGGTAATTTTNNNAGATCGGAAG
130678 TATGAGTGTGGAGTCGTTGCAATTTCTCGATATTTCCCTCTCNNNAGATCGGAAGAG
130679 TATGAGTGTGGAGTCGTTGCTTTCTTACAAAGTTAGTGTGTACNNNAGATCGGAAG
130681 TATGAGTGTGGAGTCGTTGCCTTACAACCTCATGTAACCTTNNNAGATCGGAAGAG
130682 TATGAGTGTGGAGTCGTTGCGGATAATTAACTAGCATGGAAGANNNAGATCGGAA
130683 TATGAGTGTGGAGTCGTTGCCAGGTAGAAAGAATAGTTGGGNNNAGATCGGAAG
130684 TATGAGTGTGGAGTCGTTGCCAGGTAGAAAGAATAGTTGGGNNNAGATCGGAAG
130685 TATGAGTGTGGAGTCGTTGCAAATAATGGCTGAAGGTTTCTAANNNAGATCGGAA
130686 TATGAGTGTGGAGTCGTTGCAAGCATTCACTCTTGTCGTANNNAGATCGGAAGAGC
130687 TATGAGTGTGGAGTCGTTGCTGTTATTAGATAAAATTGCTGGGNNNAGATCGGAA
130688 TATGAGTGTGGAGTCGTTGCGAGGATAAAGAGGAATATCCTGNNNAGATCGGAAG
130689 TATGAGTGTGGAGTCGTTGCGAGGATAAAGAGGAATATCCTGNNNAGATCGGAAG
130692 TATGAGTGTGGAGTCGTTGCAATCTCCTAGTACTTCCTGAATNNNAGATCGGAAGA
130694 TATGAGTGTGGAGTCGTTGCCAAAGTCTCATTGTTATTCTGACNNNAGATCGGAAG
130695 TATGAGTGTGGAGTCGTTGCATGGAGGAAATATGCAGATAAATNNNAGATCGGAA
130696 TATGAGTGTGGAGTCGTTGCTCTCCCTCATCTCAGTATAATTANNNAGATCGGAAGA
130697 TATGAGTGTGGAGTCGTTGCTATTTTCATAAATAGCCAGGAGANNNAGATCGGAAG
130698 TATGAGTGTGGAGTCGTTGCGAAGAGTGGTGTACATATGTAGNNNAGATCGGAAG
130699 TATGAGTGTGGAGTCGTTGCTGAAAACCATTAGTTTATACTGGNNNAGATCGGAAG
130700 TATGAGTGTGGAGTCGTTGCCAAGTAAGCAATCAGTAAAAATGNNNAGATCGGAA
130701 TATGAGTGTGGAGTCGTTGCATATTTTAGATGGACGTCACTTANNNAGATCGGAAG
130704 TATGAGTGTGGAGTCGTTGCTTTGACTTGAATTCACTTAGGATNNNAGATCGGAAG
130705 TATGAGTGTGGAGTCGTTGCGAATTCTAGGTAGGGTTCCTANNNAGATCGGAAGA
130706 TATGAGTGTGGAGTCGTTGCGATCAATGCAGGTCCAATAGNNNAGATCGGAAGAG
130707 TATGAGTGTGGAGTCGTTGCGTTAGCATTTTCAAACACATTCNNNAGATCGGAAGA
130708 TATGAGTGTGGAGTCGTTGCTTTCCCTGAGAGGATTTCAGNNNAGATCGGAAGAG
130709 TATGAGTGTGGAGTCGTTGCATTGATTCACACAATTATGAAGANNNAGATCGGAAG
130710 TATGAGTGTGGAGTCGTTGCAACATTGTGTGAAAAATTGATTCNNNAGATCGGAAG
130711 TATGAGTGTGGAGTCGTTGCGTGTAGAGTATTTTTTCTCTCTGNNNAGATCGGAAG
130712 TATGAGTGTGGAGTCGTTGCGTAGACTGTGTATCTATTTGACTNNNAGATCGGAAG
130713 TATGAGTGTGGAGTCGTTGCGAGATTAAGGAAATTTCAGAAGGNNNAGATCGGAA
130714 TATGAGTGTGGAGTCGTTGCATCTCAATATCATGGTTGTGATANNNAGATCGGAAG
130715 TATGAGTGTGGAGTCGTTGCGGTTCCTGTAATAGAAGTAGTGNNNAGATCGGAAG
130716 TATGAGTGTGGAGTCGTTGCCAAGAGAGAAGAGAAAAGAGAGNNNAGATCGGAA

130718 TATGAGTGTGGAGTCGTTGCTACTGATATAAGCACGCATATAANNNAGATCGGAAG
130720 TATGAGTGTGGAGTCGTTGCTTATTAGGGTAAGCTTCACATAANNNAGATCGGAAG
130721 TATGAGTGTGGAGTCGTTGCGGTGAAGGGAATACTAGTTATTANNNAGATCGGAA
130722 TATGAGTGTGGAGTCGTTGCTAAGTAAAATTGGTGAGCAAATTNNNAGATCGGAA
130723 TATGAGTGTGGAGTCGTTGCTACTACTTTTGTACAGTTTTCCTNNNAGATCGGAAGA
130724 TATGAGTGTGGAGTCGTTGCTCTTGAAATGTCTTAATTTCTCCNNNAGATCGGAAGA
130725 TATGAGTGTGGAGTCGTTGCCCTTTTGAAGTATCAGATAGTCTNNNAGATCGGAAG
130726 TATGAGTGTGGAGTCGTTGCAAATTCTTCAAGGCTATGTCTAANNNAGATCGGAAG
130727 TATGAGTGTGGAGTCGTTGCAATTGACTTATGCAGTTGTTGNNNAGATCGGAAGAG
130728 TATGAGTGTGGAGTCGTTGCGAACTCCACAAATGACTCTTANNNAGATCGGAAGAG
130729 TATGAGTGTGGAGTCGTTGCCTTTGAGGATCTGTTCTCAGNNNAGATCGGAAGAG
130730 TATGAGTGTGGAGTCGTTGCTAGGGAACATTTTTGTTTTGTTANNNAGATCGGAAG
130731 TATGAGTGTGGAGTCGTTGCAACTCATCAGGAAAAGTTCTTAGNNNAGATCGGAAG
130732 TATGAGTGTGGAGTCGTTGCAATGCTACTATCACTCATGAATTNNNAGATCGGAAG
130733 TATGAGTGTGGAGTCGTTGCTTCAAATATAGAGCAATGCACATNNNAGATCGGAAG
130734 TATGAGTGTGGAGTCGTTGCGAAAAAAAAACACTCAACAAAGTNNNAGATCGGAA
130735 TATGAGTGTGGAGTCGTTGCTGGTAAATATTATGTCATGTTCTNNNAGATCGGAAG
130736 TATGAGTGTGGAGTCGTTGCCCACAGTGAAATACAGTTATAACNNNAGATCGGAAG
130738 TATGAGTGTGGAGTCGTTGCTTCTCCATTAACTTCAACTCTTNNNAGATCGGAAGAG
130739 TATGAGTGTGGAGTCGTTGCGAGAGACAGAGAGAGATCTAGNNNAGATCGGAAG
130740 TATGAGTGTGGAGTCGTTGCTATACGGGCATTAAAAAAAATCANNNAGATCGGAA
130741 TATGAGTGTGGAGTCGTTGCGTGTCTCTCAGTATCTCCTTNNNAGATCGGAAGAGC
130742 TATGAGTGTGGAGTCGTTGCAAGCAAATATGTATTAATACACATTNNNAGATCGGA
130743 TATGAGTGTGGAGTCGTTGCGAACCAGTTATTTATTTCATTGACNNNAGATCGGAA
130744 TATGAGTGTGGAGTCGTTGCTTTTCTTTCTTTCTTTCTCCTGTNNNAGATCGGAAGAG
130745 TATGAGTGTGGAGTCGTTGCGACCTCTTAGAACCCTCTTTNNNAGATCGGAAGAGC
130746 TATGAGTGTGGAGTCGTTGCTTCTACAGGCCAGAAAAAAANNNAGATCGGAAGAG
130747 TATGAGTGTGGAGTCGTTGCATATAGCTCAGATCTCTCCAAANNNAGATCGGAAGA
130748 TATGAGTGTGGAGTCGTTGCAACTGGCATCTTTTTGTAAAAAANNNAGATCGGAAG
130749 TATGAGTGTGGAGTCGTTGCATCTATCTAGTGGTGAGGAAAGNNNAGATCGGAAG
130750 TATGAGTGTGGAGTCGTTGCACACTCTTATCTATCTATCTATCTNNNAGATCGGAAG
130751 TATGAGTGTGGAGTCGTTGCTATCACCTTGCTTCATAATACTTNNNAGATCGGAAGA
130752 TATGAGTGTGGAGTCGTTGCTAGCAATAGCAAGAGTGGATATNNNAGATCGGAAG
130753 TATGAGTGTGGAGTCGTTGCGACTTACACACAAATGATTCTACNNNAGATCGGAAG
130754 TATGAGTGTGGAGTCGTTGCTCCATTATTTAGCAGTAATGTCTNNNAGATCGGAAG
130755 TATGAGTGTGGAGTCGTTGCAAGACATAAAATCAAAGAAAAGCNNNAGATCGGAA
130756 TATGAGTGTGGAGTCGTTGCATTCTGGAAAGATTGCTTTACTCNNNAGATCGGAAG
130757 TATGAGTGTGGAGTCGTTGCTTGATCTTCTACATTCATAGTGANNNAGATCGGAAG
130758 TATGAGTGTGGAGTCGTTGCGTAGGTTTGAGGGGTTATATACNNNAGATCGGAAG
130759 TATGAGTGTGGAGTCGTTGCATTTCGCTTATAACCAAAGATACNNNAGATCGGAAG
130760 TATGAGTGTGGAGTCGTTGCATTTTTCTGCCTAAAATCCAAGNNNAGATCGGAAGA
130763 TATGAGTGTGGAGTCGTTGCAAGGAATCAAAGTCTGACAACNNNAGATCGGAAGA
130764 TATGAGTGTGGAGTCGTTGCGTTTGTATGCACGTATGTATAAANNNAGATCGGAAG

130765 TATGAGTGTGGAGTCGTTGCGATATAGTCCTTATGACCTCTGNNNAGATCGGAAGA
130766 TATGAGTGTGGAGTCGTTGCCTGATTTGTTTATACCTTCTTCTANNNAGATCGGAAG
130768 TATGAGTGTGGAGTCGTTGCAGATTTATTATGGGAATTGGCATNNNAGATCGGAA
130769 TATGAGTGTGGAGTCGTTGCTAGAGTCCTATCTAGAGCCTAGNNNAGATCGGAAGA
130770 TATGAGTGTGGAGTCGTTGCGTGCAAGATACCTAAGAATAAAGNNNAGATCGGAA
130771 TATGAGTGTGGAGTCGTTGCGTGCTGAAATATAATTCCTTCAANNNAGATCGGAAG
130772 TATGAGTGTGGAGTCGTTGCGAAGTAAAATAAAGCAGGATGATNNNAGATCGGAA
130773 TATGAGTGTGGAGTCGTTGCTCTTATCCTTATCTTCCCTCATANNNAGATCGGAAGA
130774 TATGAGTGTGGAGTCGTTGCCAAACGCCTCAACTTTCTAGNNNAGATCGGAAGAGC
130775 TATGAGTGTGGAGTCGTTGCATTATTTATGCTCTTCTCCAGACNNNAGATCGGAAGA
130776 TATGAGTGTGGAGTCGTTGCCCAGGAAATAAACCTAGAAAATAGNNNAGATCGGA
130777 TATGAGTGTGGAGTCGTTGCTTGAGTTGGTAATAAAATAAGCCNNNAGATCGGAA
130778 TATGAGTGTGGAGTCGTTGCCCAAATCATATTTTCTTGAATCCNNNAGATCGGAAGA
130779 TATGAGTGTGGAGTCGTTGCCATCCCTTTACTCACCATATATTNNNAGATCGGAAGA
130780 TATGAGTGTGGAGTCGTTGCGATGTGTTTACCTATAATTTGCTNNNAGATCGGAAG
130781 TATGAGTGTGGAGTCGTTGCATAGATATACACACACACAATGTNNNAGATCGGAAG
130782 TATGAGTGTGGAGTCGTTGCAAGTATCCAAAACAAAAAGACATNNNAGATCGGAA
130783 TATGAGTGTGGAGTCGTTGCCCACTCCTAAAAAGTCAAAAATANNNAGATCGGAAG
130784 TATGAGTGTGGAGTCGTTGCGTAGAGAGGGGAATTAAAATAGTNNNAGATCGGA
130785 TATGAGTGTGGAGTCGTTGCTAGCACTGTTATCTCTCAGTCNNNAGATCGGAAGAG
130786 TATGAGTGTGGAGTCGTTGCGTATCTATATTTGGGGCAGAGNNNAGATCGGAAGA
130787 TATGAGTGTGGAGTCGTTGCGAGAGATGTATCTATATTTGGGGNNNAGATCGGAA
130788 TATGAGTGTGGAGTCGTTGCGAGAGATGTATCTATATTTGGGGNNNAGATCGGAA
130789 TATGAGTGTGGAGTCGTTGCGTCTGTTCAGTCAATCCTAGNNNAGATCGGAAGAGC
130790 TATGAGTGTGGAGTCGTTGCTTGGTTTTTTGAATGTTAGTTTGNNNAGATCGGAAG
130791 TATGAGTGTGGAGTCGTTGCAATTTGAAAAAATTTCAGGCATTNNNAGATCGGAAG
130792 TATGAGTGTGGAGTCGTTGCACAAAAGGTCATCTATGAAAAATNNNAGATCGGAA
130793 TATGAGTGTGGAGTCGTTGCATAAGAGCTGAGAAAAGTCATTTNNNAGATCGGAA
130794 TATGAGTGTGGAGTCGTTGCACATACATACACACACATATGTANNNAGATCGGAAG
130795 TATGAGTGTGGAGTCGTTGCGATTTCAAGTGACAAAAATGGTNNNAGATCGGAAG
130796 TATGAGTGTGGAGTCGTTGCGTGGCTATACTATGTTCTACTTANNNAGATCGGAAG
130798 TATGAGTGTGGAGTCGTTGCCTTTTCTTTAGCAGACTTCATTANNNAGATCGGAAGA
130799 TATGAGTGTGGAGTCGTTGCAAGTGTTATGGGTTAAATTGTGANNNAGATCGGAA
130800 TATGAGTGTGGAGTCGTTGCTTTACCACTAAATACTCAAGTGANNNAGATCGGAAG
130801 TATGAGTGTGGAGTCGTTGCCTTAACACCCAGATCATTTCTANNNAGATCGGAAGA
130802 TATGAGTGTGGAGTCGTTGCGAGTGGAGTATGGAAATTATACTNNNAGATCGGAA
130804 TATGAGTGTGGAGTCGTTGCCAACACAGGGGAGATACATTNNNAGATCGGAAGAG
130805 TATGAGTGTGGAGTCGTTGCGTGTATGCTGTCAATAAACAATNNNAGATCGGAAGA
130806 TATGAGTGTGGAGTCGTTGCAAGTTAAGTTAGCAATCTGGAGNNNAGATCGGAAG
130807 TATGAGTGTGGAGTCGTTGCGATGGAGGCAAGTTAAGTTAGNNNAGATCGGAAGA
130808 TATGAGTGTGGAGTCGTTGCTGTCTCTACAAAAAAGACAAAAANNNAGATCGGAAG
130809 TATGAGTGTGGAGTCGTTGCAAACAGAGAGATTGAGAAAGAGNNNAGATCGGAA
130810 TATGAGTGTGGAGTCGTTGCAATGAAAGGGTACATTGAAGATNNNAGATCGGAAG

130812 TATGAGTGTGGAGTCGTTGCCTTATATCTTTCTTCTTGACTCGNNNAGATCGGAAGA
130813 TATGAGTGTGGAGTCGTTGCAATTGAAAGGAAAACCCTAACTTNNNAGATCGGAAG
130814 TATGAGTGTGGAGTCGTTGCCCATACAATATGGTAATCATGAGNNNAGATCGGAAG
130815 TATGAGTGTGGAGTCGTTGCATGGGGGATTTAGAATCTTATTTNNNAGATCGGAA
130816 TATGAGTGTGGAGTCGTTGCACAGAAAAGGAGAGATAGAAAAANNNAGATCGGA
130817 TATGAGTGTGGAGTCGTTGCAAGGAAGATATGATAGAGGACATNNNAGATCGGA
130818 TATGAGTGTGGAGTCGTTGCTAGATTGTGGTAATCATTTCACANNNAGATCGGAAG
130819 TATGAGTGTGGAGTCGTTGCCTCTCTGTCTCTCTCTCTCTNNNAGATCGGAAGAGCA
130821 TATGAGTGTGGAGTCGTTGCATTATTGGCAATGATTACGAAAGNNNAGATCGGAA
130822 TATGAGTGTGGAGTCGTTGCCATGAACCTAGAGAATCTGATCNNNAGATCGGAAG
130823 TATGAGTGTGGAGTCGTTGCTCGTTTATTTGTGGCTAAATAAANNNAGATCGGAAG
130824 TATGAGTGTGGAGTCGTTGCACTGAAACTAATAGGCTTTGATNNNAGATCGGAAG
130825 TATGAGTGTGGAGTCGTTGCAATTGAAACTATGTCTAGACTGTNNNAGATCGGAAG
130826 TATGAGTGTGGAGTCGTTGCCATATGTGTTTACATGGCACNNNAGATCGGAAGAG
130827 TATGAGTGTGGAGTCGTTGCCTGAAACACCATTCTTTTCTTCNNNAGATCGGAAGAG
130828 TATGAGTGTGGAGTCGTTGCGAATCTCTGGAGGGAATAAAAGNNNAGATCGGAAG
130829 TATGAGTGTGGAGTCGTTGCGCTGTTAACAATATGATGATCAANNNAGATCGGAAG
130830 TATGAGTGTGGAGTCGTTGCTAGAGGAAGAGTTACTGGGANNNAGATCGGAAGA
130831 TATGAGTGTGGAGTCGTTGCTAAATGTCAGCTCTTATGTTACANNNAGATCGGAAG
130832 TATGAGTGTGGAGTCGTTGCAGCATTTTATAAGGAAGCTACTANNNAGATCGGAAG
130833 TATGAGTGTGGAGTCGTTGCAAAACATAACAGGTGGTGATATTNNNAGATCGGAA
130834 TATGAGTGTGGAGTCGTTGCTCATATACATATGCAAACATTGGNNNAGATCGGAAG
130835 TATGAGTGTGGAGTCGTTGCCTGGATGGAATATGATGAACTCNNNAGATCGGAAG
130836 TATGAGTGTGGAGTCGTTGCAAAACAAGGAAGAAAAGAGATTTNNNAGATCGGAA
130837 TATGAGTGTGGAGTCGTTGCTGTATGCCTTTAATTTTTATCCANNNAGATCGGAAGA
130838 TATGAGTGTGGAGTCGTTGCAGAGAGAAAGAGAGAGTGAGNNNAGATCGGAAG
130839 TATGAGTGTGGAGTCGTTGCTATATGATGGAATACTACTCAGCNNNAGATCGGAAG
130840 TATGAGTGTGGAGTCGTTGCAGAGATAGAAAGAGAAGGACAGNNNAGATCGGAA
130842 TATGAGTGTGGAGTCGTTGCCAGGGTATCTATCACCTAGGNNNAGATCGGAAGAG
130843 TATGAGTGTGGAGTCGTTGCTATGTAGTTGACCTTTGAACAATNNNAGATCGGAAG
130844 TATGAGTGTGGAGTCGTTGCTATGTAGTTGACCTTTGAACAATNNNAGATCGGAAG
130845 TATGAGTGTGGAGTCGTTGCTGAAGATATGTTCACATCTATGTNNNAGATCGGAAG
130846 TATGAGTGTGGAGTCGTTGCGTGAGTATTCAATAATTGGGTACNNNAGATCGGAA
130847 TATGAGTGTGGAGTCGTTGCTTTCCCTTCTGGTATCTCTTNNNAGATCGGAAGAGCA
130848 TATGAGTGTGGAGTCGTTGCGTCTTGTATAACTTAATCTCTGAANNNAGATCGGAA
130849 TATGAGTGTGGAGTCGTTGCTTCTTGAGCTATGGTTTCATTNNNAGATCGGAAGAG
130850 TATGAGTGTGGAGTCGTTGCGTGTTCATCCACCAATCTAGNNNAGATCGGAAGAGC
130851 TATGAGTGTGGAGTCGTTGCGTGTTTCTGAAGTTCTCTGANNNAGATCGGAAGAG
130852 TATGAGTGTGGAGTCGTTGCAGCTGAAAATGAGAAGCTATATANNNAGATCGGAA
130854 TATGAGTGTGGAGTCGTTGCGGTTTCAACACTCTGGATATAANNNAGATCGGAAGA
130855 TATGAGTGTGGAGTCGTTGCGATGATGTTTATCGGCAGAANNNAGATCGGAAGAG
130856 TATGAGTGTGGAGTCGTTGCACTTTTTTAGTGTGTATATTTGTGNNNAGATCGGAA
130857 TATGAGTGTGGAGTCGTTGCCATCTTGTCTTTCTATTCTCATGNNNAGATCGGAAGA

130859 TATGAGTGTGGAGTCGTTGCATACTTTCCCCAGTAAAAGTAATNNNAGATCGGAAG
130860 TATGAGTGTGGAGTCGTTGCCTCATGTATCCTGGTCTAAAATNNNAGATCGGAAGA
130861 TATGAGTGTGGAGTCGTTGCTCTTATTGGTCAAGAGTAAAAGANNNAGATCGGAAG
130862 TATGAGTGTGGAGTCGTTGCGTCAGCAAAATTTTTAAGACATTNNNAGATCGGAAG
130863 TATGAGTGTGGAGTCGTTGCGGTTTCTATTTCTGACTCTCATNNNAGATCGGAAGAG
130864 TATGAGTGTGGAGTCGTTGCTATTTTTTATTGTGGCAAAACACNNNAGATCGGAAG
130865 TATGAGTGTGGAGTCGTTGCCACATAGAAGACCACAACATNNNAGATCGGAAGAG
130866 TATGAGTGTGGAGTCGTTGCGACAGAAAGTAAGACTCAATTAGNNNAGATCGGAA
130867 TATGAGTGTGGAGTCGTTGCAGAAAAGCAACATGTTGAAAANNNAGATCGGAAGA
130868 TATGAGTGTGGAGTCGTTGCCTCCCAAATTCTAAGGTAAGTTNNNAGATCGGAAGA
130869 TATGAGTGTGGAGTCGTTGCAATCCCGACGTTATGAGAATNNNAGATCGGAAGAG
130871 TATGAGTGTGGAGTCGTTGCTGGAAGGCAACATATAATATACTNNNAGATCGGAAG
130872 TATGAGTGTGGAGTCGTTGCATAGCTTCAAGTTGTAGAAAGGNNNAGATCGGAAG
130874 TATGAGTGTGGAGTCGTTGCATCTGAAGACAGCTTACATTTNNNAGATCGGAAGAG
130875 TATGAGTGTGGAGTCGTTGCGTGTTGTGTGCTAGAGAAAGNNNAGATCGGAAGAG
130877 TATGAGTGTGGAGTCGTTGCCACAGAAAGAGATGACATGGNNNAGATCGGAAGAG
130878 TATGAGTGTGGAGTCGTTGCCCCAATATATCCATTTTCCCTAGNNNAGATCGGAAGA
130879 TATGAGTGTGGAGTCGTTGCTGGGTTTACTTATGCATAAATTTNNNAGATCGGAAG
130880 TATGAGTGTGGAGTCGTTGCCATCTAGGTTTTTAGCTATTTGTNNNAGATCGGAAG
130881 TATGAGTGTGGAGTCGTTGCTTATATTGCTTGGGATTTGCTAANNNAGATCGGAAG
130882 TATGAGTGTGGAGTCGTTGCGTGATGTCAAATATGGTATGTTTNNNAGATCGGAAG
130883 TATGAGTGTGGAGTCGTTGCTTTTTGTTTGTTTGTTAGTTTGTNNNAGATCGGAAGA
130884 TATGAGTGTGGAGTCGTTGCAGTTTTTCTTTTTTGGTTAGAGANNNAGATCGGAAG
130885 TATGAGTGTGGAGTCGTTGCATATACATGTACAGAAGTCAGCTNNNAGATCGGAAG
130886 TATGAGTGTGGAGTCGTTGCGAACTGTGAGGATCCATTTATANNNAGATCGGAAGA
130887 TATGAGTGTGGAGTCGTTGCGCCTGGTTTTAGAGGATTAGNNNAGATCGGAAGAG
130888 TATGAGTGTGGAGTCGTTGCTTGTGTGTTTACAGAATCTCTNNNAGATCGGAAGAG
130889 TATGAGTGTGGAGTCGTTGCCCAGTTCAATAGACAGGATGNNNAGATCGGAAGAG
130890 TATGAGTGTGGAGTCGTTGCGGAGTAGTGCACAAATATGTATANNNAGATCGGAA
130891 TATGAGTGTGGAGTCGTTGCGGAGTAGTGCACAAATATGTATANNNAGATCGGAA
130892 TATGAGTGTGGAGTCGTTGCTTGGAAGTAATAACCCTCTAAAGNNNAGATCGGAAG
130893 TATGAGTGTGGAGTCGTTGCGAGAAATATTTTTTCTTGCCAAGNNNAGATCGGAAG
130894 TATGAGTGTGGAGTCGTTGCGAAGAATAGCTTAAGACCACTCNNNAGATCGGAAGA
130895 TATGAGTGTGGAGTCGTTGCGGAATTTGGGAAGAAAATAAAAGNNNAGATCGGAA
130896 TATGAGTGTGGAGTCGTTGCCACATCTGTGAGAAATATTCAAANNNAGATCGGAAG
130897 TATGAGTGTGGAGTCGTTGCTAACTTTGAACAGTTATCGTTGNNNAGATCGGAAGA
130898 TATGAGTGTGGAGTCGTTGCAGGAGTACTGGACTATAGTTCNNNAGATCGGAAGA
130899 TATGAGTGTGGAGTCGTTGCAAAGTAGGAACTGCTAGAAAAANNNAGATCGGAAG
130900 TATGAGTGTGGAGTCGTTGCCAGAGATGGACATATATGGATTTNNNAGATCGGAA
130901 TATGAGTGTGGAGTCGTTGCTTTGCTGTTATGTATACTTTGAGNNNAGATCGGAAG
130902 TATGAGTGTGGAGTCGTTGCCTCAGTTGTTTTATTCAGCTATGNNNAGATCGGAAG
130903 TATGAGTGTGGAGTCGTTGCGATAGACTCTGTGACTTCAACNNNAGATCGGAAGAG
130904 TATGAGTGTGGAGTCGTTGCCATGAAGAGAATGATCAACTTTANNNAGATCGGAAG

130906 TATGAGTGTGGAGTCGTTGCTGTTTGTATATTTAGTGTTGGAGNNNAGATCGGAAG
130907 TATGAGTGTGGAGTCGTTGCATTTTCTCTATTCCTTCCATTCANNNAGATCGGAAGAG
130908 TATGAGTGTGGAGTCGTTGCGTCTATAATAAGGAAGGGTTTCCNNNAGATCGGAAG
130909 TATGAGTGTGGAGTCGTTGCCCAAATGGCAAAATAAAGATTTANNNAGATCGGAAG
130910 TATGAGTGTGGAGTCGTTGCAATCAGAGACATGATATTTTAGACNNNAGATCGGA
130911 TATGAGTGTGGAGTCGTTGCGAATCTTGTGCTGGAAAAGGNNNAGATCGGAAGAG
130913 TATGAGTGTGGAGTCGTTGCGCTAACAGGATCTTAATTTCTTNNNAGATCGGAAGA
130914 TATGAGTGTGGAGTCGTTGCAGTAAGTGAAAAGAACCATTTTTNNNAGATCGGAAG
130915 TATGAGTGTGGAGTCGTTGCTCAGATTTTTATACACGTTTTGGNNNAGATCGGAAGA
130916 TATGAGTGTGGAGTCGTTGCGGCGTTTATGATAGATTTCTAAANNNAGATCGGAAG
130917 TATGAGTGTGGAGTCGTTGCAGAACATGAGTATCCATATTGAGNNNAGATCGGAAG
130918 TATGAGTGTGGAGTCGTTGCAACAGAATGGTGATTATGGAGNNNAGATCGGAAGA
130920 TATGAGTGTGGAGTCGTTGCGTAGATGACACCACTATTCTCNNNAGATCGGAAGAG
130921 TATGAGTGTGGAGTCGTTGCTAAGTACTTCTAGGTCACCAGNNNAGATCGGAAGAG
130922 TATGAGTGTGGAGTCGTTGCGATTTTCTGAGTCACAATACAATNNNAGATCGGAAGA
130923 TATGAGTGTGGAGTCGTTGCTATAGAACTGTAATACACACACANNNAGATCGGAAG
130924 TATGAGTGTGGAGTCGTTGCCCCATTACCAAAAAATAAAGGAANNNAGATCGGAAG
130925 TATGAGTGTGGAGTCGTTGCCATTGATACAATGAGATGCATAANNNAGATCGGAAG
130927 TATGAGTGTGGAGTCGTTGCTTCTTTTTCTTTTTCAGAGTCTCNNNAGATCGGAAGA
130928 TATGAGTGTGGAGTCGTTGCAAAATAATAGGAAAGAGGACAGTNNNAGATCGGAA
130929 TATGAGTGTGGAGTCGTTGCAATTACTCACCCATTAGGATACTNNNAGATCGGAAGA
130930 TATGAGTGTGGAGTCGTTGCTGTCCATATTTTCAATTGTGAAANNNAGATCGGAAGA
130933 TATGAGTGTGGAGTCGTTGCCTCTCTATTGTTTTCCTGTTTTANNNAGATCGGAAGA
130934 TATGAGTGTGGAGTCGTTGCATATGTAGGTAAACACAGACATANNNAGATCGGAAG
130935 TATGAGTGTGGAGTCGTTGCGCTTGAAAGGAATTATTTTTTGTNNNAGATCGGAAG
130936 TATGAGTGTGGAGTCGTTGCACTGCAGTAGTGTAATCATGNNNAGATCGGAAGAG
130938 TATGAGTGTGGAGTCGTTGCCTTTGTTTGACTCTCTTAGGATNNNAGATCGGAAGA
130939 TATGAGTGTGGAGTCGTTGCACTGATCAAGAAACAAGACTATANNNAGATCGGAAG
130940 TATGAGTGTGGAGTCGTTGCGAGATGAAAATTATCATGTCCTGNNNAGATCGGAAG
130941 TATGAGTGTGGAGTCGTTGCCAACAATCTTAGAAAAGATGTGGNNNAGATCGGAA
130942 TATGAGTGTGGAGTCGTTGCCATAAGAGTACTAATCCCATCATNNNAGATCGGAAGA
130943 TATGAGTGTGGAGTCGTTGCTAGAGTATGATCACCTTTGTTCNNNAGATCGGAAGA
130944 TATGAGTGTGGAGTCGTTGCCATACATTTATACAACTCTGCTGNNNAGATCGGAAGA
130945 TATGAGTGTGGAGTCGTTGCCTTCAGTTTATTCTATCACTGTGNNNAGATCGGAAGA
130946 TATGAGTGTGGAGTCGTTGCATAATGGTACTTTGTAATCCAAGNNNAGATCGGAAG
130947 TATGAGTGTGGAGTCGTTGCATAATGGTACTTTGTAATCCAAGNNNAGATCGGAAG
130949 TATGAGTGTGGAGTCGTTGCTCATTCTCCCCTCTTTATTTATCNNNAGATCGGAAGAG
130950 TATGAGTGTGGAGTCGTTGCGGTGGAGATGGAGATAATTATTTNNNAGATCGGAA
130952 TATGAGTGTGGAGTCGTTGCTTGGGGTGTATAGATGGAATNNNAGATCGGAAGAG
130953 TATGAGTGTGGAGTCGTTGCAAACCTCAAGATAAATACATCTGNNNAGATCGGAAG
130954 TATGAGTGTGGAGTCGTTGCCATGTTTTTGCTATTGTGAATAGNNNAGATCGGAAG
130956 TATGAGTGTGGAGTCGTTGCCAGGTATAGGATGGTTACATTTNNNAGATCGGAAGA
130957 TATGAGTGTGGAGTCGTTGCGGTTTAAGTCATTAGAAGGAAAANNNAGATCGGAA

130958 TATGAGTGTGGAGTCGTTGCTATTAACTCAAGAAGGATGAAGANNNAGATCGGAA
130959 TATGAGTGTGGAGTCGTTGCTTATTCCATTCAGTGCCAATAANNNAGATCGGAAGA
130960 TATGAGTGTGGAGTCGTTGCGAAAATTGGAAATCTGGATAGAANNNAGATCGGAA
130961 TATGAGTGTGGAGTCGTTGCAGGTATATTCAATTGACTATGCTNNNAGATCGGAAG
130962 TATGAGTGTGGAGTCGTTGCAACAAATACTGCCTACCAATTNNNAGATCGGAAGAG
130963 TATGAGTGTGGAGTCGTTGCCATCTTTGGAATAATGCAATCTANNNAGATCGGAAG
130964 TATGAGTGTGGAGTCGTTGCCAGCAGGAGGTTTTAAAATATAANNNAGATCGGAA
130965 TATGAGTGTGGAGTCGTTGCCTACCTTACTTTAGGGAAACAAANNNAGATCGGAAG
130966 TATGAGTGTGGAGTCGTTGCCAGGAACATACTTAACAAATGTTNNNAGATCGGAAG
130967 TATGAGTGTGGAGTCGTTGCATATTATGCTGAGTGTAATAAGCNNNAGATCGGAA
130968 TATGAGTGTGGAGTCGTTGCATGTCACCCTATCATAAAGTTTTNNNAGATCGGAAG
130969 TATGAGTGTGGAGTCGTTGCGGGAAGAACTCTCTAGAGAANNNAGATCGGAAGAG
130970 TATGAGTGTGGAGTCGTTGCGGGAAGAACTCTCTAGAGAANNNAGATCGGAAGAG
130971 TATGAGTGTGGAGTCGTTGCTGGCTCTATATGTATATTGATGTNNNAGATCGGAAG
130972 TATGAGTGTGGAGTCGTTGCCTAATTACTTTCATTTGCTCTCANNNAGATCGGAAGA
130973 TATGAGTGTGGAGTCGTTGCTGAAATGTTTCTTTCTCTAGAGANNNAGATCGGAAG
130974 TATGAGTGTGGAGTCGTTGCGAGGCAGAATCATTATCATTATTNNNAGATCGGAAG
130976 TATGAGTGTGGAGTCGTTGCTGTAAACTGGATGAGGTTTCNNNAGATCGGAAGAG
130977 TATGAGTGTGGAGTCGTTGCGTCCTTTCTCTTTAATGACATACNNNAGATCGGAAGA
130978 TATGAGTGTGGAGTCGTTGCAATAGAAAGAGGCATAGATAAGGNNNAGATCGGAA
130979 TATGAGTGTGGAGTCGTTGCATCTTTAAATCATCTCAGCTGANNNAGATCGGAAGA
130980 TATGAGTGTGGAGTCGTTGCAGTGTCTGATTTATGTTGGGNNNAGATCGGAAGAG
130981 TATGAGTGTGGAGTCGTTGCGCCATATATAGGAGTATTCCAAANNNAGATCGGAAG
130982 TATGAGTGTGGAGTCGTTGCTAAATTTGAAATTTCAGAGCTGTNNNAGATCGGAAG
130983 TATGAGTGTGGAGTCGTTGCACATAAAAGTGTCCATTTACAATNNNAGATCGGAAG
130985 TATGAGTGTGGAGTCGTTGCCAACATTTTAGAGAGAAACATCCNNNAGATCGGAAG
130987 TATGAGTGTGGAGTCGTTGCTATTTTTCATTCCTCATTTGACCNNNAGATCGGAAGA
130988 TATGAGTGTGGAGTCGTTGCTAGAAATGTATGTTGATTGAACANNNAGATCGGAA
130989 TATGAGTGTGGAGTCGTTGCCAAGACTTTTGCAAATACTTAACNNNAGATCGGAAG
130990 TATGAGTGTGGAGTCGTTGCGAGTATGCTCAATTTCCAGTNNNAGATCGGAAGAGC
130991 TATGAGTGTGGAGTCGTTGCGTATTCTCTTCACTTCATTTCTTNNNAGATCGGAAGA
130992 TATGAGTGTGGAGTCGTTGCCAATAATTGTCACTGGTCATTTNNNAGATCGGAAGA
130993 TATGAGTGTGGAGTCGTTGCCTTCCACTGAAAAGTCTGTTNNNAGATCGGAAGAGC
130994 TATGAGTGTGGAGTCGTTGCCTCAATGTTTTTATTGCTTGTACNNNAGATCGGAAGA
130995 TATGAGTGTGGAGTCGTTGCGGCTCCCATTAATCATAATTAAGNNNAGATCGGAAG
130996 TATGAGTGTGGAGTCGTTGCAGACTGGAGCTTTAGTGTATTNNNAGATCGGAAGA
130997 TATGAGTGTGGAGTCGTTGCACTAATATGCAAATGCTATATGTNNNAGATCGGAAG
130998 TATGAGTGTGGAGTCGTTGCCCATCTGCTCTTTTTATACATTTNNNAGATCGGAAGA
131000 TATGAGTGTGGAGTCGTTGCCAGTCTGTAAGCTTGAAAAACNNNAGATCGGAAGA
131001 TATGAGTGTGGAGTCGTTGCATAGCCTTCACCTCTCTTAAAANNNAGATCGGAAGA
131002 TATGAGTGTGGAGTCGTTGCTGAAATTATACTACAGTTTTGCCNNNAGATCGGAAG
131003 TATGAGTGTGGAGTCGTTGCTCCTAGTTCCTGAAAGTTCTANNNAGATCGGAAGAG
131004 TATGAGTGTGGAGTCGTTGCCATAATTTGGGGTTAACATAGGNNNAGATCGGAAG

131005 TATGAGTGTGGAGTCGTTGCGGCAAACCTGGTTTATTAATAAANNNAGATCGGAAG
131006 TATGAGTGTGGAGTCGTTGCTTTATCCCAAAGAAAACAAAGAANNNAGATCGGAAG
131007 TATGAGTGTGGAGTCGTTGCTTTAGAGGGACAAGAATAGCAANNNAGATCGGAAG
131008 TATGAGTGTGGAGTCGTTGCGTGTCCTCTGTATTATGGTTGNNNAGATCGGAAGAG
131009 TATGAGTGTGGAGTCGTTGCGAGATATAGCATCCAAGGAGTNNNAGATCGGAAGA
131010 TATGAGTGTGGAGTCGTTGCGGTCACAGCTAGTATTTATAATGNNNAGATCGGAAG
131011 TATGAGTGTGGAGTCGTTGCAATAGTGAACTGACAATAGCTTNNNAGATCGGAAG
131012 TATGAGTGTGGAGTCGTTGCCATTAAAAAAGGGTAGGAAAGACNNNAGATCGGAA
131013 TATGAGTGTGGAGTCGTTGCCTTCAACAGTTCCCTCAAAGNNNAGATCGGAAGAGC
131014 TATGAGTGTGGAGTCGTTGCCAAGATATAGGAGAGGAGAAACNNNAGATCGGAA
131015 TATGAGTGTGGAGTCGTTGCCAAGATATAGGAGAGGAGAAACNNNAGATCGGAA
131016 TATGAGTGTGGAGTCGTTGCGAAGGATGTTTTTGACCCTGNNNAGATCGGAAGAG
131017 TATGAGTGTGGAGTCGTTGCGAAGGATGTTTTTGACCCTGNNNAGATCGGAAGAG
131021 TATGAGTGTGGAGTCGTTGCCACCCAAATATTATACTAAGCCTNNNAGATCGGAAG
131022 TATGAGTGTGGAGTCGTTGCAGAACAACGTTTTATCTTTTTTTNNNAGATCGGAAGA
131023 TATGAGTGTGGAGTCGTTGCACGTGCTTAGTAGTATCTCTAANNNAGATCGGAAGA
131024 TATGAGTGTGGAGTCGTTGCACACATTTTTGCAACATACATANNNAGATCGGAAGA
131025 TATGAGTGTGGAGTCGTTGCTTCCAACAATACCAAGTAAGTTCNNNAGATCGGAAG
131026 TATGAGTGTGGAGTCGTTGCATTAGTACCATGTTCAGAGATGNNNAGATCGGAAG
131027 TATGAGTGTGGAGTCGTTGCGAACGTAACTATATTTTAGAGGCNNNAGATCGGAAG
131028 TATGAGTGTGGAGTCGTTGCTACTACTTTAGATTTCACCAACTNNNAGATCGGAAGA
131029 TATGAGTGTGGAGTCGTTGCATTTTTTGTAGAGATGGGGTTTTNNNAGATCGGAAG
131031 TATGAGTGTGGAGTCGTTGCTGTATATATATATATGTATGTGTGTGTNNNAGATCG
131032 TATGAGTGTGGAGTCGTTGCAGAATAAATTTTAATCACCCTGGNNNAGATCGGAAG
131033 TATGAGTGTGGAGTCGTTGCGTTAGTCTCATTAGCCACAATANNNAGATCGGAAGA
131034 TATGAGTGTGGAGTCGTTGCGTTAGTCTCATTAGCCACAATANNNAGATCGGAAGA
131035 TATGAGTGTGGAGTCGTTGCCATGGATCCTTTACATCAATAAANNNAGATCGGAAG
131036 TATGAGTGTGGAGTCGTTGCCATGGATCCTTTACATCAATAAANNNAGATCGGAAG
131037 TATGAGTGTGGAGTCGTTGCGAGTGTTGAGATCAGCAATGNNNAGATCGGAAGAG
131038 TATGAGTGTGGAGTCGTTGCTTGACAGAGAGAGAAAAAAAAAGNNNAGATCGGA
131039 TATGAGTGTGGAGTCGTTGCCTTTCACTCTGAACAGGTAGNNNAGATCGGAAGAGC
131040 TATGAGTGTGGAGTCGTTGCGCAAAATGCATATTATAAGAGACNNNAGATCGGAA
131041 TATGAGTGTGGAGTCGTTGCAAGGCAAAACAGAATAGGTAGNNNAGATCGGAAGA
131042 TATGAGTGTGGAGTCGTTGCTTTTGCTTCCAACTCTTTAACNNNAGATCGGAAGAGC
131043 TATGAGTGTGGAGTCGTTGCTAATTGAGACAAACAATGACAACNNNAGATCGGAA
131045 TATGAGTGTGGAGTCGTTGCGTATTTACTCCAGACAGCTAGNNNAGATCGGAAGAG
131046 TATGAGTGTGGAGTCGTTGCCCTGTGTACAAGGAAATTAATTANNNAGATCGGAAG
131047 TATGAGTGTGGAGTCGTTGCTGGTGAAGAGTACAGATTCTANNNAGATCGGAAGA
131049 TATGAGTGTGGAGTCGTTGCAACAACAAAAAATTATTTCCTCANNNAGATCGGAAG
131050 TATGAGTGTGGAGTCGTTGCTCAAACATAAAGAACTGAAGAACNNNAGATCGGAA
131051 TATGAGTGTGGAGTCGTTGCCTGAGGAAAAGGACAGAATGNNNAGATCGGAAGA
131052 TATGAGTGTGGAGTCGTTGCCATTTCTCCAAGGCTAGAATGNNNAGATCGGAAGAG
131053 TATGAGTGTGGAGTCGTTGCTACGCACGTTATAGAAATATTTGNNNAGATCGGAAG

131054 TATGAGTGTGGAGTCGTTGCTACGCACGTTATAGAAATATTTGNNNAGATCGGAAG
131055 TATGAGTGTGGAGTCGTTGCCCTTTATCCCTTCTCTATTTCTANNNAGATCGGAAGAG
131056 TATGAGTGTGGAGTCGTTGCACAGAAAAAGAGAGACAGAAAGNNNAGATCGGAA
131057 TATGAGTGTGGAGTCGTTGCCTCAAAGTGCTGAGATTACANNNAGATCGGAAGAG
131058 TATGAGTGTGGAGTCGTTGCTATTTTCCTTTATTTCTCTCCCTNNNAGATCGGAAGAG
131059 TATGAGTGTGGAGTCGTTGCTCATTCATATTAATGTGAGAAGCNNNAGATCGGAAG
131060 TATGAGTGTGGAGTCGTTGCGTAGCTATCTGGTCATATTTCTANNNAGATCGGAAG
131061 TATGAGTGTGGAGTCGTTGCATTCACACTGTCCAAAGAATAGNNNAGATCGGAAGA
131062 TATGAGTGTGGAGTCGTTGCGGAATAAATTACAATACTCCAACANNNAGATCGGAA
131063 TATGAGTGTGGAGTCGTTGCAAGCAAATGTCCAGTTTTATTAANNNAGATCGGAAG
131064 TATGAGTGTGGAGTCGTTGCGTCATAACCTAACTGGGCATNNNAGATCGGAAGAG
131065 TATGAGTGTGGAGTCGTTGCTGTTGAAGGCTATTCATTATAGTNNNAGATCGGAAG
131066 TATGAGTGTGGAGTCGTTGCCTATGTTTGTGTAAGTTTGTGTANNNAGATCGGAAG
131067 TATGAGTGTGGAGTCGTTGCGTATGTGATTACTAGTGTCAGATNNNAGATCGGAAG
131068 TATGAGTGTGGAGTCGTTGCGGTTTGTTTTTTTTGTTTTGTTTNNNAGATCGGAAG
131069 TATGAGTGTGGAGTCGTTGCTCTTCATCCACTTCATATAACAANNNAGATCGGAAGA
131070 TATGAGTGTGGAGTCGTTGCAGCAACAATAAGAAAATCTAGTGNNNAGATCGGAA
131071 TATGAGTGTGGAGTCGTTGCAGGTATTTAGATATTTGGTGACTNNNAGATCGGAAG
131072 TATGAGTGTGGAGTCGTTGCGTTGGTAAATTTAATGGAGACATNNNAGATCGGAA
131073 TATGAGTGTGGAGTCGTTGCAATTGACAAGTGCATTTTTTTTTNNNAGATCGGAAG
131074 TATGAGTGTGGAGTCGTTGCATCTGATATACTGAAGGATAGGANNNAGATCGGAA
131075 TATGAGTGTGGAGTCGTTGCCATAGGGTCAGAAAAGACACNNNAGATCGGAAGAG
131076 TATGAGTGTGGAGTCGTTGCCACATTCTGGAAGTGATGAGNNNAGATCGGAAGAG
131077 TATGAGTGTGGAGTCGTTGCTAATCAGGCAGAATAAATTTTCCNNNAGATCGGAAG
131078 TATGAGTGTGGAGTCGTTGCTTGATCAGATTCTAGGACAATAANNNAGATCGGAAG
131079 TATGAGTGTGGAGTCGTTGCTTAACTTAGCTACTTTGGCATTNNNAGATCGGAAGA
131080 TATGAGTGTGGAGTCGTTGCCAGAGCATAGGTGTACTTAAATNNNAGATCGGAAG
131081 TATGAGTGTGGAGTCGTTGCAGTCTTAAATATCCTTGGAAGAGNNNAGATCGGAAG
131082 TATGAGTGTGGAGTCGTTGCATTTTTAGTGGCTGCATATTTTTNNNAGATCGGAAG
131083 TATGAGTGTGGAGTCGTTGCTTAAGCCAAGAAGTTTGCTTTNNNAGATCGGAAGAG
131084 TATGAGTGTGGAGTCGTTGCGGACTCACATAGGATTTGTCNNNAGATCGGAAGAG
131085 TATGAGTGTGGAGTCGTTGCGAGATGGAGTAATTGACTACATANNNAGATCGGAA
131086 TATGAGTGTGGAGTCGTTGCATCGTTTATCTGGGGAAATACNNNAGATCGGAAGAG
131087 TATGAGTGTGGAGTCGTTGCAGTGGGTAAATAAATACTACAGTNNNAGATCGGAA
131088 TATGAGTGTGGAGTCGTTGCGTCATCAATTTCATCTGGGANNNAGATCGGAAGAGC
131089 TATGAGTGTGGAGTCGTTGCCTCTCCCCTACCTTATAAGTAAANNNAGATCGGAAGA
131092 TATGAGTGTGGAGTCGTTGCTGCGTAATAAAGCACATTAATATNNNAGATCGGAAG
131093 TATGAGTGTGGAGTCGTTGCGAACCTGAAGTAATGCTTTTATTNNNAGATCGGAAG
131094 TATGAGTGTGGAGTCGTTGCGAACCTGAAGTAATGCTTTTATTNNNAGATCGGAAG
131095 TATGAGTGTGGAGTCGTTGCCTCATGGTTTTACTTCCCTATATNNNAGATCGGAAGA
131096 TATGAGTGTGGAGTCGTTGCAGTGGCTAGAAGTACTAATACTANNNAGATCGGAAG
131097 TATGAGTGTGGAGTCGTTGCAAAAATGTAGAAGTGATTTTGGANNNAGATCGGAA
131098 TATGAGTGTGGAGTCGTTGCATGATTTTCTTCTATTTCTCCATNNNAGATCGGAAGA

131099 TATGAGTGTGGAGTCGTTGCTAAATCCTGTAATTTTGGACATCNNNAGATCGGAAG
131100 TATGAGTGTGGAGTCGTTGCTCTCAGTCTATTACCCAGTTTNNNAGATCGGAAGAG
131101 TATGAGTGTGGAGTCGTTGCAGTGGTTTCTAATTCATGTTTTTNNNAGATCGGAAG
131102 TATGAGTGTGGAGTCGTTGCACTAATACAGCACAGTTATGAAGNNNAGATCGGAA
131103 TATGAGTGTGGAGTCGTTGCAAACACAGGTAAACTTTATTGACNNNAGATCGGAAG
131104 TATGAGTGTGGAGTCGTTGCAAACACAGGTAAACTTTATTGACNNNAGATCGGAAG
131105 TATGAGTGTGGAGTCGTTGCCATGAGTGGAAAGCTTTATTTAANNNAGATCGGAA
131106 TATGAGTGTGGAGTCGTTGCTGTATGTGTGATTGGCATATATANNNAGATCGGAA
131107 TATGAGTGTGGAGTCGTTGCTCTAAGTTCAAACTCCAACTATANNNAGATCGGAAG
131108 TATGAGTGTGGAGTCGTTGCAAACAAAATTCCTCCAAAACTATNNNAGATCGGAAG
131110 TATGAGTGTGGAGTCGTTGCTGGCTAGTTAAATATTGTCAGATNNNAGATCGGAAG
131111 TATGAGTGTGGAGTCGTTGCTTAATAAGGATCCCCTAGGTAATNNNAGATCGGAAG
131112 TATGAGTGTGGAGTCGTTGCGCTAGAGGTTACAGATCTTTCNNNAGATCGGAAGA
131113 TATGAGTGTGGAGTCGTTGCGATTCTGGCTATGGATATAACTANNNAGATCGGAAG
131114 TATGAGTGTGGAGTCGTTGCGATTCTGGCTATGGATATAACTANNNAGATCGGAAG
131115 TATGAGTGTGGAGTCGTTGCGATTAGGGTGGGAAAAACATNNNAGATCGGAAGA
131116 TATGAGTGTGGAGTCGTTGCTTGTTGGTGATGTATACTTTTGNNNAGATCGGAAG
131117 TATGAGTGTGGAGTCGTTGCCTAATTTCCCCAAGTTGTTTAGNNNAGATCGGAAGA
131118 TATGAGTGTGGAGTCGTTGCCTAATTTCCCCAAGTTGTTTAGNNNAGATCGGAAGA
131119 TATGAGTGTGGAGTCGTTGCGAGAAAGAGCTCAATATTCAATCNNNAGATCGGAAG
131120 TATGAGTGTGGAGTCGTTGCTAAACATGGGTAAAAATTCAGCNNNAGATCGGAAG
131121 TATGAGTGTGGAGTCGTTGCCAAGCAGACAATAGAAGAAAACNNNAGATCGGAAG
131122 TATGAGTGTGGAGTCGTTGCCCTCTACTTCTTTTTCAAAAGTCNNNAGATCGGAAGA
131124 TATGAGTGTGGAGTCGTTGCCTTATGCTAGAAAAGTAGTGAAGNNNAGATCGGAA
131125 TATGAGTGTGGAGTCGTTGCAGTGAGAGAGAGAAGGTAAAGNNNAGATCGGAAG
131126 TATGAGTGTGGAGTCGTTGCACAGAAACTTCTCTAGCTGGNNNAGATCGGAAGAG
131127 TATGAGTGTGGAGTCGTTGCGCTGTAGGTAGGAGAAGAAANNNAGATCGGAAGA
131128 TATGAGTGTGGAGTCGTTGCGAGTGGTAGGTAGGTAGTTTNNNAGATCGGAAGA
131130 TATGAGTGTGGAGTCGTTGCAATAGCAGAATGGCATGGATANNNAGATCGGAAGA
131131 TATGAGTGTGGAGTCGTTGCAATAGCAGAATGGCATGGATANNNAGATCGGAAGA
131132 TATGAGTGTGGAGTCGTTGCCATACCTTATGTATTGGGTTTTGNNNAGATCGGAAG
131133 TATGAGTGTGGAGTCGTTGCTGATTGTTATTATGCAAGATGTTNNNAGATCGGAAG
131134 TATGAGTGTGGAGTCGTTGCGTTTAAACAGAGCAGAGTAGATNNNAGATCGGAAG
131135 TATGAGTGTGGAGTCGTTGCGTTAATATTGAGTGTCAACTTGANNNAGATCGGAAG
131136 TATGAGTGTGGAGTCGTTGCAGGTAACATCTAAAACTGAAAAANNNAGATCGGAA
131137 TATGAGTGTGGAGTCGTTGCATGTTATGGTGGTAAGTAGATAGNNNAGATCGGAA
131138 TATGAGTGTGGAGTCGTTGCTCTGTGTATCAACTTGTATACTCNNNAGATCGGAAG
131139 TATGAGTGTGGAGTCGTTGCTCTGTGTATCAACTTGTATACTCNNNAGATCGGAAG
131140 TATGAGTGTGGAGTCGTTGCTGTATTCAGCAACTAGTTTTAAGNNNAGATCGGAAG
131141 TATGAGTGTGGAGTCGTTGCTATACCAAAAGCAAATAACATCGNNNAGATCGGAAG
131142 TATGAGTGTGGAGTCGTTGCTAACAGTGAACAGACAGTTGNNNAGATCGGAAGAG
131143 TATGAGTGTGGAGTCGTTGCTTGATAATACTTCAGCTAAAACCNNNAGATCGGAAG
131144 TATGAGTGTGGAGTCGTTGCCTCTCTGACTTAGGAAACAAATNNNAGATCGGAAGA

131145 TATGAGTGTGGAGTCGTTGCAATAGCACAATCATAGCTCCNNNAGATCGGAAGAGC
131146 TATGAGTGTGGAGTCGTTGCATAAAGCCTATAATTGTACCCATNNNAGATCGGAAG
131147 TATGAGTGTGGAGTCGTTGCATTATTTGAGAGTCTCTGTTACCNNNAGATCGGAAG
131148 TATGAGTGTGGAGTCGTTGCGGAATGAACTGTTTTAGATAAGCNNNAGATCGGAA
131149 TATGAGTGTGGAGTCGTTGCAACATATTTATTTTCCTCTGTCTNNNAGATCGGAAGA
131150 TATGAGTGTGGAGTCGTTGCCTCTTCCACTCTTATATCTTCTTNNNAGATCGGAAGA
131151 TATGAGTGTGGAGTCGTTGCATAAATCTTCATCTCAGTTGTCTNNNAGATCGGAAGA
131152 TATGAGTGTGGAGTCGTTGCCGAAGTCCTAGATGTTAGAATTANNNAGATCGGAA
131153 TATGAGTGTGGAGTCGTTGCCTCATTTTCTACCACTGACTGNNNAGATCGGAAGAG
131154 TATGAGTGTGGAGTCGTTGCAAATGACATTTCTTTCTCTCTCTNNNAGATCGGAAGA
131155 TATGAGTGTGGAGTCGTTGCGTGTATGAGGTGATACATGTATANNNAGATCGGAA
131156 TATGAGTGTGGAGTCGTTGCATTATTCTTTCTGCCTAGAGCNNNAGATCGGAAGAG
131157 TATGAGTGTGGAGTCGTTGCTAGACAGTCTTGCTATGCTANNNAGATCGGAAGAGC
131158 TATGAGTGTGGAGTCGTTGCAAAGAGAGAGAGAGAGAAACANNNAGATCGGAAG
131159 TATGAGTGTGGAGTCGTTGCATTATATAAGAGTGACATCAGCCNNNAGATCGGAAG
131160 TATGAGTGTGGAGTCGTTGCTGTATATCTCTTGGAAAAGCTTANNNAGATCGGAAG
131161 TATGAGTGTGGAGTCGTTGCACTGAGAGTAAAATCTAAGTAGCNNNAGATCGGAA
131162 TATGAGTGTGGAGTCGTTGCGAGCAGAGCTATCACATATGNNNAGATCGGAAGAG
131163 TATGAGTGTGGAGTCGTTGCATGATCTTTCAAGATGTAACAAGNNNAGATCGGAAG
131164 TATGAGTGTGGAGTCGTTGCTTCCTTCAGAATGTAAGCATTNNNAGATCGGAAGAG
131165 TATGAGTGTGGAGTCGTTGCCACTCCAGTTACCATGTTAGNNNAGATCGGAAGAGC
131166 TATGAGTGTGGAGTCGTTGCAAATGCAGAGCTTTTCATTTCNNNAGATCGGAAGAG
131167 TATGAGTGTGGAGTCGTTGCCTGGTGATTTTTTTTTGTCTTTTNNNAGATCGGAAGA
131168 TATGAGTGTGGAGTCGTTGCCTATTTCAGAGTATCAGCAGTTNNNAGATCGGAAGA
131169 TATGAGTGTGGAGTCGTTGCTGTTTACTAATTCCAACAAAAGGNNNAGATCGGAAG
131170 TATGAGTGTGGAGTCGTTGCTAGTATCAGTGCTTGAGATATTTNNNAGATCGGAAG
131171 TATGAGTGTGGAGTCGTTGCGAATGATACATGGAATATTGCTTNNNAGATCGGAA
131172 TATGAGTGTGGAGTCGTTGCGCATGATGTTTTGAATAATTAGCNNNAGATCGGAAG
131173 TATGAGTGTGGAGTCGTTGCAAAACAAAAACAAAACATACACANNNAGATCGGAAG
131175 TATGAGTGTGGAGTCGTTGCTCAAATCAAGTATATATCCCCAANNNAGATCGGAAG
131176 TATGAGTGTGGAGTCGTTGCTATGAATTCAGGGTGTATGTGNNNAGATCGGAAGA
131178 TATGAGTGTGGAGTCGTTGCGGAGTAAATGAAGTCAGTAAGANNNAGATCGGAA
131179 TATGAGTGTGGAGTCGTTGCTTCAATATTCATATGCTGCTTTTNNNAGATCGGAAGA
131180 TATGAGTGTGGAGTCGTTGCTTCAATATTCATATGCTGCTTTTNNNAGATCGGAAGA
131181 TATGAGTGTGGAGTCGTTGCCAGTAGTTAATGTCAGTGTTAACNNNAGATCGGAAG
131182 TATGAGTGTGGAGTCGTTGCTAGAAATGGGGACTTACTAGATNNNAGATCGGAAG
131183 TATGAGTGTGGAGTCGTTGCTTGGAGAAACATTACAGAATTTTNNNAGATCGGAAG
131184 TATGAGTGTGGAGTCGTTGCATAAGATGAGAAAGAGTCCCTTTNNNAGATCGGAA
131185 TATGAGTGTGGAGTCGTTGCAGATTTTTATGGGTGACAGAAGNNNAGATCGGAAG
131186 TATGAGTGTGGAGTCGTTGCAGCATTTGTCTTCTAAAAAAAGANNNAGATCGGAAG
131187 TATGAGTGTGGAGTCGTTGCAATATCCCTGACAAACATAGAGNNNAGATCGGAAGA
131189 TATGAGTGTGGAGTCGTTGCCTGATTAACATGTTGCTTTACTNNNAGATCGGAAGA
131190 TATGAGTGTGGAGTCGTTGCCTGTTTCCTCTGATGTCAAANNNAGATCGGAAGAGC

131191 TATGAGTGTGGAGTCGTTGCTCCTCTTTAGAATAATAAGCTGGNNNAGATCGGAAG
131192 TATGAGTGTGGAGTCGTTGCCAGAAGGAAACATAGTCCAGNNNAGATCGGAAGAG
131193 TATGAGTGTGGAGTCGTTGCTACTGTGTCTATTTGTTTTCAAANNNAGATCGGAAG
131194 TATGAGTGTGGAGTCGTTGCTACTGTGTCTATTTGTTTTCAAANNNAGATCGGAAG
131195 TATGAGTGTGGAGTCGTTGCCTTGTATTTCAATATTGGTGTCANNNAGATCGGAAG
131196 TATGAGTGTGGAGTCGTTGCCTTTGAAGGAATTCGTATAGTAGNNNAGATCGGAA
131197 TATGAGTGTGGAGTCGTTGCCTCACATCTTGGGAGTTACTANNNAGATCGGAAGAG
131198 TATGAGTGTGGAGTCGTTGCGTAGAAAATTCAGAAAGGTCTTGNNNAGATCGGAA
131199 TATGAGTGTGGAGTCGTTGCCATTACATTGAATGTCAGAACTANNNAGATCGGAAG
131201 TATGAGTGTGGAGTCGTTGCTTCTGTGAGATCTTAATGAGATGNNNAGATCGGAA
131202 TATGAGTGTGGAGTCGTTGCTTCTGTGAGATCTTAATGAGATGNNNAGATCGGAA
131203 TATGAGTGTGGAGTCGTTGCCAATTGTGAGTTGTACTACAATANNNAGATCGGAAG
131204 TATGAGTGTGGAGTCGTTGCTAATGGCACTTTGAAATTTACAANNNAGATCGGAAG
131205 TATGAGTGTGGAGTCGTTGCGAGTAAAGAATTATGTCCACAAANNNAGATCGGAA
131206 TATGAGTGTGGAGTCGTTGCAAGCTTGAAATTTTCCATAATTTNNNAGATCGGAAG
131207 TATGAGTGTGGAGTCGTTGCACATGTGAATAATAAATGCTTCTNNNAGATCGGAAG
131208 TATGAGTGTGGAGTCGTTGCGATACCAAACTCCTGTCCTCNNNAGATCGGAAGAGC
131209 TATGAGTGTGGAGTCGTTGCATAAGAATTACAGAGGACCTTTTNNNAGATCGGAAG
131210 TATGAGTGTGGAGTCGTTGCATAAGAATTACAGAGGACCTTTTNNNAGATCGGAAG
131211 TATGAGTGTGGAGTCGTTGCGAAGTCATCATTCAATAGACATTNNNAGATCGGAAG
131212 TATGAGTGTGGAGTCGTTGCAAGCTATGTACATAATAGGGAAGNNNAGATCGGAA
131213 TATGAGTGTGGAGTCGTTGCCTGTTATGATACTACCACTGGNNNAGATCGGAAGAG
131214 TATGAGTGTGGAGTCGTTGCTTGAATGTTTTGTTTTTTGGAAGNNNAGATCGGAAG
131215 TATGAGTGTGGAGTCGTTGCCTTTGTTGGCTTCTTATCTATACNNNAGATCGGAAGA
131216 TATGAGTGTGGAGTCGTTGCGTAGCTTCTCATTTCGTCTGNNNAGATCGGAAGAGC
131217 TATGAGTGTGGAGTCGTTGCGGAAGTTTAGTTTTGATTTGGAANNNAGATCGGAA
131218 TATGAGTGTGGAGTCGTTGCCACACATGACTTCCATATACTCNNNAGATCGGAAGA
131219 TATGAGTGTGGAGTCGTTGCTCAATTATTTTATGTTGTGTAAACANNNAGATCGGA
131220 TATGAGTGTGGAGTCGTTGCTGAAGTTTCAGTTATTCACTCTANNNAGATCGGAAG
131221 TATGAGTGTGGAGTCGTTGCCACAAACACACATATTTTAGAGTNNNAGATCGGAAG
131222 TATGAGTGTGGAGTCGTTGCGGTAATCAGCATTTGTAACTACNNNAGATCGGAAGA
131223 TATGAGTGTGGAGTCGTTGCAATGATTCAACTATCAAGCTTAGNNNAGATCGGAAG
131224 TATGAGTGTGGAGTCGTTGCATACAGAGAGAAAAGAGAGAAAGNNNAGATCGGA
131225 TATGAGTGTGGAGTCGTTGCGATGAATCCTTTTTCTTTTTTGGNNNAGATCGGAAG
131226 TATGAGTGTGGAGTCGTTGCCCTATTTTCATTTGATTTCTCCANNNAGATCGGAAGA
131227 TATGAGTGTGGAGTCGTTGCCCTATTTTCATTTGATTTCTCCANNNAGATCGGAAGA
131229 TATGAGTGTGGAGTCGTTGCATCTATTAGTGTCATCACAGAAGNNNAGATCGGAAG
131230 TATGAGTGTGGAGTCGTTGCTGTTTTAATAAAAAGAGGTCTGTNNNAGATCGGAA
131231 TATGAGTGTGGAGTCGTTGCTATTATGCTTAATGGCAATTGAANNNAGATCGGAAG
131232 TATGAGTGTGGAGTCGTTGCTACATGTTTTTCACACTAAATCCNNNAGATCGGAAGA
131233 TATGAGTGTGGAGTCGTTGCATTATATGCAGCCAGATTTCATNNNAGATCGGAAGA
131234 TATGAGTGTGGAGTCGTTGCTGCTCCTAATAGCTACTTTTAATNNNAGATCGGAAGA
131235 TATGAGTGTGGAGTCGTTGCCTCTTGAAGTCCTTTGATTTATCNNNAGATCGGAAGA

131236 TATGAGTGTGGAGTCGTTGCAAAAATCCTGAACAAGAGAAAAANNNAGATCGGAA
131237 TATGAGTGTGGAGTCGTTGCCAGACTTTTGGTGTAATAGTTTCNNNAGATCGGAAG
131238 TATGAGTGTGGAGTCGTTGCAATCACTGTGGCATTTGTATANNNAGATCGGAAGAG
131239 TATGAGTGTGGAGTCGTTGCTATGTTAGCTTGTGAAGTAAACTNNNAGATCGGAAG
131240 TATGAGTGTGGAGTCGTTGCGAACAATTTGAGCCATTTAAAAANNNAGATCGGAAG
131241 TATGAGTGTGGAGTCGTTGCCATCATCTGCCCTCAAAATGNNNAGATCGGAAGAGC
131242 TATGAGTGTGGAGTCGTTGCATTATGGTAGATGGGATAAGTTTNNNAGATCGGAA
131243 TATGAGTGTGGAGTCGTTGCAAATATTTGGCAGATAAATCAGCNNNAGATCGGAAG
131244 TATGAGTGTGGAGTCGTTGCAATTCACTCTAATCTGACCACNNNAGATCGGAAGAG
131245 TATGAGTGTGGAGTCGTTGCCATCTTCTCTCCCACAAATGNNNAGATCGGAAGAGC
131246 TATGAGTGTGGAGTCGTTGCCCTGTGCATAAACTACATACANNNAGATCGGAAGAG
131247 TATGAGTGTGGAGTCGTTGCCTAGAAAGTCTGAAGTCAAGAANNNAGATCGGAAG
131248 TATGAGTGTGGAGTCGTTGCTACTCCAACTTACTACAAGGATNNNAGATCGGAAGA
131249 TATGAGTGTGGAGTCGTTGCCCACAAGAAGCTGATCATTANNNAGATCGGAAGAG
131250 TATGAGTGTGGAGTCGTTGCATTACAGGCATGATCCATTTTNNNAGATCGGAAGAG
131251 TATGAGTGTGGAGTCGTTGCATATTGACTTCACTCTCCATTTCNNNAGATCGGAAGA
131252 TATGAGTGTGGAGTCGTTGCTTGTTTTTAATGTGCAGTGATANNNAGATCGGAAG
131253 TATGAGTGTGGAGTCGTTGCGAGACTATTTCAGAAGGTTACATNNNAGATCGGAAG
131254 TATGAGTGTGGAGTCGTTGCGCACTAAATCAAATGTTAAAGTGNNNAGATCGGAAG
131255 TATGAGTGTGGAGTCGTTGCTCCCATACATAAATTTGACATTTNNNAGATCGGAAGA
131256 TATGAGTGTGGAGTCGTTGCTTTCTATAATAGCTATCCTCCAGNNNAGATCGGAAGA
131257 TATGAGTGTGGAGTCGTTGCCCTACTTTCCACTGAATTTATTCNNNAGATCGGAAGA
131258 TATGAGTGTGGAGTCGTTGCCAGGTAAACGTGCTATTCTGNNNAGATCGGAAGAG
131259 TATGAGTGTGGAGTCGTTGCCTTAGAATTAAGGAGGAAAGCTNNNAGATCGGAAG
131260 TATGAGTGTGGAGTCGTTGCCTTAGAATTAAGGAGGAAAGCTNNNAGATCGGAAG
131261 TATGAGTGTGGAGTCGTTGCCCTAAAAAAGTTTTTGCTAACTACNNNAGATCGGAAG
131262 TATGAGTGTGGAGTCGTTGCACATTTACTCAGAAGGCTTACNNNAGATCGGAAGAG
131263 TATGAGTGTGGAGTCGTTGCCTCCAATCTCTACTGTAATAAGANNNAGATCGGAAGA
131264 TATGAGTGTGGAGTCGTTGCCTCCAATCTCTACTGTAATAAGANNNAGATCGGAAGA
131265 TATGAGTGTGGAGTCGTTGCGGATATAAAAAAACGATCAACCTNNNAGATCGGAAG
131267 TATGAGTGTGGAGTCGTTGCCATCATGGACATTATGTGGTNNNAGATCGGAAGAG
131268 TATGAGTGTGGAGTCGTTGCTGAAAGATAAACATAATGTGTCANNNAGATCGGAAG
131269 TATGAGTGTGGAGTCGTTGCAGAGAGTCTTGAAAGATAAACATNNNAGATCGGAA
131270 TATGAGTGTGGAGTCGTTGCCTTTATAACGTGATCCATACATCNNNAGATCGGAAGA
131271 TATGAGTGTGGAGTCGTTGCTTAGGAAAATAAGACCTAGAGTGNNNAGATCGGAA
131272 TATGAGTGTGGAGTCGTTGCCATCTTTCACCATGATGAGATNNNAGATCGGAAGAG
131273 TATGAGTGTGGAGTCGTTGCGTGAAATGATTATCAGACTATGTNNNAGATCGGAAG
131274 TATGAGTGTGGAGTCGTTGCCATTTGTGAAGAAAAACTCATTTNNNAGATCGGAAG
131275 TATGAGTGTGGAGTCGTTGCCCAAGAACACTGTCTCTTAGNNNAGATCGGAAGAGC
131276 TATGAGTGTGGAGTCGTTGCGTGTATATGCACATTCACATGNNNAGATCGGAAGAG
131277 TATGAGTGTGGAGTCGTTGCGTGTATATGCACATTCACATGNNNAGATCGGAAGAG
131278 TATGAGTGTGGAGTCGTTGCGTTTTTGGTAGTGAGTGTTGNNNAGATCGGAAGAG
131279 TATGAGTGTGGAGTCGTTGCGCAGAATGAGGATAAGATAGTTANNNAGATCGGAA

131280 TATGAGTGTGGAGTCGTTGCAAATAAGTTTTACGGCAAATAGANNNAGATCGGAAG
131281 TATGAGTGTGGAGTCGTTGCTTATAGCTTCTCACAATTCCTTTNNNAGATCGGAAGA
131282 TATGAGTGTGGAGTCGTTGCAAATGATGTTGTTATATTCCTCGNNNAGATCGGAAG
131283 TATGAGTGTGGAGTCGTTGCTATTATGTAGTAAAATGCCAGGANNNAGATCGGAAG
131284 TATGAGTGTGGAGTCGTTGCCTGCATGAATACACTCGAATNNNAGATCGGAAGAGC
131285 TATGAGTGTGGAGTCGTTGCTCATGATGTTAGTAGAATCGTTANNNAGATCGGAAG
131286 TATGAGTGTGGAGTCGTTGCGAAATTAATGCATGTGGAAAATCNNNAGATCGGAAG
131287 TATGAGTGTGGAGTCGTTGCTACCAATTTTCAGAATAAGAAGTNNNAGATCGGAAG
131288 TATGAGTGTGGAGTCGTTGCCACTAATTTCAGATAACTCTTGGNNNAGATCGGAAG
131289 TATGAGTGTGGAGTCGTTGCGTGGTATTTCAAAAAGGTTTTTCNNNAGATCGGAAG
131290 TATGAGTGTGGAGTCGTTGCCTATATATTTTCCCCTGAGTTCANNNAGATCGGAAGA
131291 TATGAGTGTGGAGTCGTTGCTTATGAAAGCTTTGATCTTTTGTNNNAGATCGGAAG
131292 TATGAGTGTGGAGTCGTTGCAAAACTGCTCTAAAAAGTCTATTNNNAGATCGGAAG
131293 TATGAGTGTGGAGTCGTTGCGTATGTTTACAGGAGTTGTTTAGNNNAGATCGGAAG
131294 TATGAGTGTGGAGTCGTTGCTAGTATACTTCCTGGGAAACTATNNNAGATCGGAAG
131295 TATGAGTGTGGAGTCGTTGCGTAATGTAGCCAATCAGTAAAATNNNAGATCGGAAG
131296 TATGAGTGTGGAGTCGTTGCCTAATAGGTGATCTGTTTTGTGNNNAGATCGGAAGA
131297 TATGAGTGTGGAGTCGTTGCAAATGGGGTAATGCCTTCATNNNAGATCGGAAGAG
131298 TATGAGTGTGGAGTCGTTGCAGATCTCTGGATAGTAAGACTAANNNAGATCGGAAG
131299 TATGAGTGTGGAGTCGTTGCCAACAGAAACATGAACCAATAATNNNAGATCGGAAG
131300 TATGAGTGTGGAGTCGTTGCACTTTTGCATGGAGAAATTTTTNNNAGATCGGAAGA
131301 TATGAGTGTGGAGTCGTTGCGGGTATTGAGAAGATAAGTGTTANNNAGATCGGAA
131302 TATGAGTGTGGAGTCGTTGCTCTGAAAAACCCTTACTAATACANNNAGATCGGAAG
131303 TATGAGTGTGGAGTCGTTGCCACATACACACACACAAATATATNNNAGATCGGAAG
131304 TATGAGTGTGGAGTCGTTGCATATAGAGAGAGTGAAATAGGGANNNAGATCGGA
131305 TATGAGTGTGGAGTCGTTGCTATTGGAATTATCAGACAAGGAANNNAGATCGGAAG
131306 TATGAGTGTGGAGTCGTTGCCACATCTTTTATTCACTGACATANNNAGATCGGAAGA
131307 TATGAGTGTGGAGTCGTTGCTAAACAGAGCCTCAAAGATGNNNAGATCGGAAGAG
131308 TATGAGTGTGGAGTCGTTGCCTCTTCTTAGTCTACCCTGTATANNNAGATCGGAAGA
131309 TATGAGTGTGGAGTCGTTGCTTTTATTCTTCTTAACCTGGCTANNNAGATCGGAAGA
131310 TATGAGTGTGGAGTCGTTGCCCAGAAAGATTGCATAGATTTATATNNNAGATCGGA
131311 TATGAGTGTGGAGTCGTTGCTAGGCATAGATTCCTCTATCTTNNNAGATCGGAAGA
131312 TATGAGTGTGGAGTCGTTGCCTATAAGGAATTACCCAAGACTGNNNAGATCGGAAG
131313 TATGAGTGTGGAGTCGTTGCGAGATGTTATGAAGGAGCTGNNNAGATCGGAAGAG
131314 TATGAGTGTGGAGTCGTTGCACTACTCATCCATAAAAAGGAATNNNAGATCGGAAG
131315 TATGAGTGTGGAGTCGTTGCCCAGGATGCCAATTTATTAATTTNNNAGATCGGAAG
131316 TATGAGTGTGGAGTCGTTGCTTTGTACTGATACATCTGACATANNNAGATCGGAAG
131317 TATGAGTGTGGAGTCGTTGCTACATGAAATAAAGATGTGTTGTNNNAGATCGGAAG
131318 TATGAGTGTGGAGTCGTTGCATAGGTAAAAATGGATGCCTTNNNAGATCGGAAGA
131319 TATGAGTGTGGAGTCGTTGCATATTTAGTTGCACAAACATCTCNNNAGATCGGAAG
131320 TATGAGTGTGGAGTCGTTGCTATGCTTATTTTGTAGGTAAGGANNNAGATCGGAAG
131321 TATGAGTGTGGAGTCGTTGCATAGGAAAAAGGGAAAAGTAGGNNNAGATCGGAA
131322 TATGAGTGTGGAGTCGTTGCTATGATTTCCCCATGTGATAAANNNAGATCGGAAGA

131324 TATGAGTGTGGAGTCGTTGCACACTCATTTTCCCTCTCTCNNNAGATCGGAAGAGCA
131325 TATGAGTGTGGAGTCGTTGCCCCATAAGATTATAATGGAGCTNNNAGATCGGAAGA
131326 TATGAGTGTGGAGTCGTTGCTAAAAAGTGCTTGGAGGAAAATNNNAGATCGGAAG
131327 TATGAGTGTGGAGTCGTTGCGGAAGCCATGTTTATTTTAGAATNNNAGATCGGAAG
131328 TATGAGTGTGGAGTCGTTGCTATAAAGCCAAAAGAAAAGAGAGNNNAGATCGGAA
131329 TATGAGTGTGGAGTCGTTGCTATATGACTTTTCTGAGGGTATCNNNAGATCGGAAG
131330 TATGAGTGTGGAGTCGTTGCCAATGTCTTAACATATGTTCACCNNNAGATCGGAAGA
131333 TATGAGTGTGGAGTCGTTGCATTTTTGCCCTTTACCATTTATTNNNAGATCGGAAGA
131334 TATGAGTGTGGAGTCGTTGCCTCATAGTGGATGGCTATAAGNNNAGATCGGAAGA
131335 TATGAGTGTGGAGTCGTTGCAGATTGTATTTTCCAAAGATGACNNNAGATCGGAAG
131336 TATGAGTGTGGAGTCGTTGCTTTTTAGTTGATGGACATTTAGGNNNAGATCGGAAG
131337 TATGAGTGTGGAGTCGTTGCTGTAACTCCTTAAACCTGTTTATNNNAGATCGGAAGA
131338 TATGAGTGTGGAGTCGTTGCCTCACTGCAACCTTGAATTCNNNAGATCGGAAGAGCA
131339 TATGAGTGTGGAGTCGTTGCTAATACCTCAGAGAAAAAAAGTGNNNAGATCGGAA
131340 TATGAGTGTGGAGTCGTTGCATATACACATGTACATACACACANNNAGATCGGAAG
131341 TATGAGTGTGGAGTCGTTGCTATGTGTGTGGTTTCAAATTTCNNNAGATCGGAAGA
131343 TATGAGTGTGGAGTCGTTGCGTTCATGTAGTCCTTTCAATACNNNAGATCGGAAGA
131344 TATGAGTGTGGAGTCGTTGCTGTTTATAATAGGGCCAGAATTANNNAGATCGGAAG
131345 TATGAGTGTGGAGTCGTTGCGGTTATCTTTCAGTTTAGTTCTGNNNAGATCGGAAG
131346 TATGAGTGTGGAGTCGTTGCGAAAAACTGCTCAATTCTCTAGNNNAGATCGGAAGA
131347 TATGAGTGTGGAGTCGTTGCCTTCCTGTTACTTCTCCATTATNNNAGATCGGAAGAG
131348 TATGAGTGTGGAGTCGTTGCCTTCCTGTTACTTCTCCATTATNNNAGATCGGAAGAG
131349 TATGAGTGTGGAGTCGTTGCAGCAAAATTCCAAGTTTATAACTNNNAGATCGGAAG
131350 TATGAGTGTGGAGTCGTTGCGTCTCTCTCTCTCTCTCTCTCTNNNAGATCGGAAGAGCA
131351 TATGAGTGTGGAGTCGTTGCTTCATTCAATTATCCTATTTGGCNNNAGATCGGAAGA
131353 TATGAGTGTGGAGTCGTTGCCCACAAAGGATTCTTAATAATCANNNAGATCGGAAG
131354 TATGAGTGTGGAGTCGTTGCTATAAACATAAAGAGTGTGTGTGNNNAGATCGGAA
131356 TATGAGTGTGGAGTCGTTGCATCATGTCTAGGAATAATCATGANNNAGATCGGAAG
131357 TATGAGTGTGGAGTCGTTGCGAATGACAGGGTCTCATCATNNNAGATCGGAAGAG
131358 TATGAGTGTGGAGTCGTTGCCTGGAGTTTTATTTCCTGGATAANNNAGATCGGAAG
131359 TATGAGTGTGGAGTCGTTGCGTCTGACATATAACAACTTTCAANNNAGATCGGAAG
131361 TATGAGTGTGGAGTCGTTGCTTATTCATGTAACCAAACACTACNNNAGATCGGAAGA
131362 TATGAGTGTGGAGTCGTTGCAAGAGAACCTCTAAGGACAAANNNAGATCGGAAGA
131363 TATGAGTGTGGAGTCGTTGCTTTATTTTAGGAAATTGGCTCATNNNAGATCGGAAG
131364 TATGAGTGTGGAGTCGTTGCAACTGGAGAACATTATGTTATTGNNNAGATCGGAAG
131365 TATGAGTGTGGAGTCGTTGCCTCTCATGTAGGTAAACTGTGNNNAGATCGGAAGAG
131366 TATGAGTGTGGAGTCGTTGCAAGTCATGATTGTTCTGAAGTTNNNAGATCGGAAGA
131367 TATGAGTGTGGAGTCGTTGCTGTTCAACCCTATTATTAACAAANNNAGATCGGAAGA
131369 TATGAGTGTGGAGTCGTTGCCCTGTTACCCGTTTCATTTANNNAGATCGGAAGAGCA
131370 TATGAGTGTGGAGTCGTTGCCTCCATGTATAGATATCTCCAACNNNAGATCGGAAGA
131371 TATGAGTGTGGAGTCGTTGCCTTCAAGTGATTTTATACCATCCNNNAGATCGGAAGA
131372 TATGAGTGTGGAGTCGTTGCCCTTTTAAATCGAGGTCATTAAANNNAGATCGGAAG
131373 TATGAGTGTGGAGTCGTTGCTACACATTTATTTATGCACAGTGNNNAGATCGGAAG

131374 TATGAGTGTGGAGTCGTTGCGAGATAAGGAACAAAAGTGTTTTNNNAGATCGGAA
131375 TATGAGTGTGGAGTCGTTGCGAGATAAGGAACAAAAGTGTTTTNNNAGATCGGAA
131376 TATGAGTGTGGAGTCGTTGCATCTTGCATCACTCAGAATAGNNNAGATCGGAAGAG
131377 TATGAGTGTGGAGTCGTTGCACATCTTAGAGTAGTCTTCCTATNNNAGATCGGAAG
131378 TATGAGTGTGGAGTCGTTGCGGCATGTAGAGATAAAGAAGTNNNAGATCGGAAG
131379 TATGAGTGTGGAGTCGTTGCCTTTCAGAGAAGAGACAAGAANNNAGATCGGAAGA
131380 TATGAGTGTGGAGTCGTTGCGACAGAGAGAGAGACAGATATANNNAGATCGGAAG
131381 TATGAGTGTGGAGTCGTTGCGATGGAAGATGTACAGAATGTNNNAGATCGGAAGA
131383 TATGAGTGTGGAGTCGTTGCCTCTGCATTCTATTCTTGAAGANNNAGATCGGAAGA
131384 TATGAGTGTGGAGTCGTTGCGACAGTCTCTAGTTATTTAAGCTNNNAGATCGGAAG
131385 TATGAGTGTGGAGTCGTTGCTATTTCTTCTCTAAGGTCAACAANNNAGATCGGAAG
131386 TATGAGTGTGGAGTCGTTGCCAAAGTCCTGGAGTCAATACNNNAGATCGGAAGAG
131387 TATGAGTGTGGAGTCGTTGCAAAGAAAATGTTTCAAGCAGATANNNAGATCGGAA
131388 TATGAGTGTGGAGTCGTTGCTTGGGATTTTCTACATAGACAATNNNAGATCGGAAG
131389 TATGAGTGTGGAGTCGTTGCAGAAGAAGCAGAGTTATATGGANNNAGATCGGAAG
131390 TATGAGTGTGGAGTCGTTGCTAGAAGTTTAAACCAAAGGTTCTNNNAGATCGGAAG
131391 TATGAGTGTGGAGTCGTTGCGCTATGACATATTTCATTTCCCNNNAGATCGGAAGAG
131392 TATGAGTGTGGAGTCGTTGCTCTCATTTATTTACTCACCACAGNNNAGATCGGAAGA
131393 TATGAGTGTGGAGTCGTTGCTCTCATTTATTTACTCACCACAGNNNAGATCGGAAGA
131394 TATGAGTGTGGAGTCGTTGCCTGGAGGTTTACATTTTGTAAAGNNNAGATCGGAAG
131395 TATGAGTGTGGAGTCGTTGCAAATACTCCCTTGGGTAACTCNNNAGATCGGAAGAG
131397 TATGAGTGTGGAGTCGTTGCTTTCTTTCTCCCTCTGTGATNNNAGATCGGAAGAGCA
131398 TATGAGTGTGGAGTCGTTGCTCTCTTTCTCTCTTTCTCTCTTNNNAGATCGGAAGAGC
131399 TATGAGTGTGGAGTCGTTGCTTGACCCTTTCTAAGATTCTGNNNAGATCGGAAGAG
131400 TATGAGTGTGGAGTCGTTGCGAGATGCTGATATATGTCCTATANNNAGATCGGAAG
131401 TATGAGTGTGGAGTCGTTGCCAGAAAAAAGATCTCCATGTTTTNNNAGATCGGAAG
131402 TATGAGTGTGGAGTCGTTGCCAATATTCTTCTCCTCTTTCAGANNNAGATCGGAAGA
131403 TATGAGTGTGGAGTCGTTGCCAAACTCCACCATAAAAGTAAATNNNAGATCGGAAG
131404 TATGAGTGTGGAGTCGTTGCATACACATTTTGTCAGAATACAGNNNAGATCGGAAG
131405 TATGAGTGTGGAGTCGTTGCCCAAAAAGATCCTGAAACAAANNNAGATCGGAAGAG
131406 TATGAGTGTGGAGTCGTTGCGAGGAGTCTTTCTTTTTCTATTGNNNAGATCGGAAG
131407 TATGAGTGTGGAGTCGTTGCAAAGCAGAAGGTGATTTAGAANNNAGATCGGAAGA
131408 TATGAGTGTGGAGTCGTTGCGGGTGAACAGTCTATCATATTTANNNAGATCGGAAG
131409 TATGAGTGTGGAGTCGTTGCAGATGGAATCTTGCTCTGTCNNNAGATCGGAAGAG
131410 TATGAGTGTGGAGTCGTTGCGTTGAGGACATTGCGATTTTNNNAGATCGGAAGAG
131412 TATGAGTGTGGAGTCGTTGCAAGAGGGTTCTAAGAGATCTTANNNAGATCGGAAG
131413 TATGAGTGTGGAGTCGTTGCGAAAATGTGTGTTAGACCAAAANNNAGATCGGAAG
131414 TATGAGTGTGGAGTCGTTGCAAGTAAAGGGTTTCTAAAACTTGNNNAGATCGGAA
131415 TATGAGTGTGGAGTCGTTGCGAAGGGAATTATGTTATTTGTGANNNAGATCGGAA
131416 TATGAGTGTGGAGTCGTTGCGTGTAGAGGTAAAATGGCAAANNNAGATCGGAAGA
131417 TATGAGTGTGGAGTCGTTGCTTGTATCAGAGAACGGAAAAAAANNNAGATCGGAAG
131418 TATGAGTGTGGAGTCGTTGCTTAGGGTTTGCTATTTTGAATACNNNAGATCGGAAG
131419 TATGAGTGTGGAGTCGTTGCATTACTGGTTCAAGGCATTTTTANNNAGATCGGAAG

131420 TATGAGTGTGGAGTCGTTGCATCAATAGAGCTTGAGAATAGAGNNNAGATCGGAA
131421 TATGAGTGTGGAGTCGTTGCTCTCTTTATCCACTCATTGATTGNNNAGATCGGAAGA
131422 TATGAGTGTGGAGTCGTTGCAATGTTTATAATGCCTAAAAGCCNNNAGATCGGAAG
131423 TATGAGTGTGGAGTCGTTGCAGGATGTAATAATATATGGCCTCNNNAGATCGGAA
131424 TATGAGTGTGGAGTCGTTGCACATGATTTTGCAATGATTTTAGNNNAGATCGGAAG
131425 TATGAGTGTGGAGTCGTTGCGGTTGGAAAGAACTGATACTATNNNAGATCGGAAG
131426 TATGAGTGTGGAGTCGTTGCGGTTGGAAAGAACTGATACTATNNNAGATCGGAAG
131427 TATGAGTGTGGAGTCGTTGCCTCTCTATTATTCCCCCTTAAATNNNAGATCGGAAGA
131428 TATGAGTGTGGAGTCGTTGCCATGCTCATGATTCACTGAGNNNAGATCGGAAGAG
131429 TATGAGTGTGGAGTCGTTGCTAGTGTGCCTTAATAACTGATAGNNNAGATCGGAAG
131430 TATGAGTGTGGAGTCGTTGCATATTGAAGGTTGACAATTCCAANNNAGATCGGAAG
131431 TATGAGTGTGGAGTCGTTGCATATTGAAGGTTGACAATTCCAANNNAGATCGGAAG
131432 TATGAGTGTGGAGTCGTTGCTTTCACTCTAGTCAGAAAAGACNNNAGATCGGAAGA
131433 TATGAGTGTGGAGTCGTTGCGCATTGTTTAATTGTAATAGTGCNNNAGATCGGAAG
131434 TATGAGTGTGGAGTCGTTGCTGTTTGGTACCTCAATTTTTTTANNNAGATCGGAAG
131435 TATGAGTGTGGAGTCGTTGCTCCTGACTTCATATAATGAGTTTNNNAGATCGGAAG
131436 TATGAGTGTGGAGTCGTTGCATTTTGATGGCCACTCTATATTNNNAGATCGGAAGA
131437 TATGAGTGTGGAGTCGTTGCTTGAATTGCACTATATCTAAGGTNNNAGATCGGAAG
131438 TATGAGTGTGGAGTCGTTGCAATGACACTAGAAACTGGTATCNNNAGATCGGAAG
131439 TATGAGTGTGGAGTCGTTGCATATAATGCAAGCAATTTGGTAANNNAGATCGGAA
131440 TATGAGTGTGGAGTCGTTGCCACATAATGAGAAGATTTGACATNNNAGATCGGAA
131441 TATGAGTGTGGAGTCGTTGCGTCCAGAAGAATGATTTGAAATANNNAGATCGGAA
131442 TATGAGTGTGGAGTCGTTGCGTAACAAATTAACAAACCCATTCNNNAGATCGGAAG
131443 TATGAGTGTGGAGTCGTTGCATGAAAAACATTTTCCAAAACAANNNAGATCGGAAG
131444 TATGAGTGTGGAGTCGTTGCATAGAGACAGAGAGAAACAGAGNNNAGATCGGAA
131445 TATGAGTGTGGAGTCGTTGCATAGAGACAGAGAGAAACAGAGNNNAGATCGGAA
131446 TATGAGTGTGGAGTCGTTGCTTTCACGATTTGGTGGTTTATNNNAGATCGGAAGAG
131447 TATGAGTGTGGAGTCGTTGCCTATGCCCTCATCTTAACTAATNNNAGATCGGAAGAG
131448 TATGAGTGTGGAGTCGTTGCTTATACTTTAAATGGATGGCATGNNNAGATCGGAAG
131449 TATGAGTGTGGAGTCGTTGCGCACCCATATATAATATTTTTTGCNNNAGATCGGAAG
131450 TATGAGTGTGGAGTCGTTGCCCATTCCTATGTAGTCATCTACNNNAGATCGGAAGA
131451 TATGAGTGTGGAGTCGTTGCCTTTTCTTCTGATCTCCTAAGTNNNAGATCGGAAGAG
131452 TATGAGTGTGGAGTCGTTGCTATTATTTTTTTCTAACCGGCCNNNAGATCGGAAGAG
131453 TATGAGTGTGGAGTCGTTGCGCTATTTTTTAGCTGTGATTTTGNNNAGATCGGAAG
131455 TATGAGTGTGGAGTCGTTGCGTGTATTCATCTATTCGTAATGGNNNAGATCGGAAG
131456 TATGAGTGTGGAGTCGTTGCGTTTATCAAACCCATATTCCTGNNNAGATCGGAAGA
131457 TATGAGTGTGGAGTCGTTGCTATGGAATCTTCTGGATACAAATNNNAGATCGGAAG
131458 TATGAGTGTGGAGTCGTTGCTTAAATCACCAGATCACGTTTNNNAGATCGGAAGAG
131459 TATGAGTGTGGAGTCGTTGCTCATTAAATGTATTAGCTTCTGCNNNAGATCGGAAG
131460 TATGAGTGTGGAGTCGTTGCATACAGCCAACTTTTCTCATGNNNAGATCGGAAGAG
131461 TATGAGTGTGGAGTCGTTGCTTTGAAAATATTGTCTGCTCTCTNNNAGATCGGAAG
131462 TATGAGTGTGGAGTCGTTGCGTATGTGGATACAGGTAGGTNNNAGATCGGAAGA
131463 TATGAGTGTGGAGTCGTTGCCGATAGTAATTATGGGTGGGNNNAGATCGGAAGA

131464 TATGAGTGTGGAGTCGTTGCGAAGAGTAAAGACATGCATCTNNNAGATCGGAAGA
131465 TATGAGTGTGGAGTCGTTGCGAAGAGTAAAGACATGCATCTNNNAGATCGGAAGA
131466 TATGAGTGTGGAGTCGTTGCGATACAGAATCTCTCTTTGTCANNNAGATCGGAAGA
131467 TATGAGTGTGGAGTCGTTGCATTCAGTGGTTATAGGATGAATTNNNAGATCGGAA
131468 TATGAGTGTGGAGTCGTTGCACTGGGAGTTTATGAATATGTTANNNAGATCGGAA
131469 TATGAGTGTGGAGTCGTTGCTTCAAAGTTTATTTCTCCTTGTGNNNAGATCGGAAG
131470 TATGAGTGTGGAGTCGTTGCTGTTCATATTATCCTGACTTCAANNNAGATCGGAAG
131471 TATGAGTGTGGAGTCGTTGCTTGTTGACAAACTGTTAAAAATCNNNAGATCGGAAG
131472 TATGAGTGTGGAGTCGTTGCTTGTTGACAAACTGTTAAAAATCNNNAGATCGGAAG
131473 TATGAGTGTGGAGTCGTTGCTTGTTGACAAACTGTTAAAAATCNNNAGATCGGAAG
131474 TATGAGTGTGGAGTCGTTGCAAGCTTGAATTTCTTACGTATTTNNNAGATCGGAAG
131475 TATGAGTGTGGAGTCGTTGCAGGAAAAACAGAATTAGGAAAACNNNAGATCGGAA
131476 TATGAGTGTGGAGTCGTTGCTTTCTCAACATCCCTTTCCNNNAGATCGGAAGAGCA
131478 TATGAGTGTGGAGTCGTTGCAATCTCAGAGATTTGTTTCCTTNNNAGATCGGAAGA
131479 TATGAGTGTGGAGTCGTTGCGGATGTATTTATTGTCTATCTGCNNNAGATCGGAAG
131480 TATGAGTGTGGAGTCGTTGCATATTTGGCAAATGAATCAAATGNNNAGATCGGAA
131481 TATGAGTGTGGAGTCGTTGCATCATCATCAACAAACGTTTATTNNNAGATCGGAAG
131482 TATGAGTGTGGAGTCGTTGCATCATCATCAACAAACGTTTATTNNNAGATCGGAAG
131484 TATGAGTGTGGAGTCGTTGCTATTTGTGTTTTTCTGCACTATTNNNAGATCGGAAGA
131485 TATGAGTGTGGAGTCGTTGCAAAGAGAATAATGCACACTGATNNNAGATCGGAAG
131486 TATGAGTGTGGAGTCGTTGCTTCTAAACCAAGGAGAATATCTGNNNAGATCGGAAG
131487 TATGAGTGTGGAGTCGTTGCAGAGATTTATTATGAGAAACTGGNNNAGATCGGAA
131488 TATGAGTGTGGAGTCGTTGCAATTTTACCATTGCTTATGTGTANNNAGATCGGAAG
131489 TATGAGTGTGGAGTCGTTGCAAGGAATAGAAATCTTCTTAGCTNNNAGATCGGAAG
131490 TATGAGTGTGGAGTCGTTGCGAAATTTGACACCCACTTAAGNNNAGATCGGAAGAG
131491 TATGAGTGTGGAGTCGTTGCGTTTTTTTCCCCTTTATCTCTTCNNNAGATCGGAAGAG
131492 TATGAGTGTGGAGTCGTTGCGAAGATACAATGAGAGGATGGNNNAGATCGGAAG
131494 TATGAGTGTGGAGTCGTTGCAAAGACAGAGAGAGAGACAGNNNAGATCGGAAGA
131496 TATGAGTGTGGAGTCGTTGCTGTCCTCAAAAAATAAAACAATGNNNAGATCGGAAG
131497 TATGAGTGTGGAGTCGTTGCGAGAGTGTCCTCAAAAAATAAAANNNAGATCGGAA
131498 TATGAGTGTGGAGTCGTTGCTCCACTGTCTGTATATAATATCTGNNNAGATCGGAA
131500 TATGAGTGTGGAGTCGTTGCAACTAGTATATAGACAAGCTTCCNNNAGATCGGAAG
131501 TATGAGTGTGGAGTCGTTGCTCTATCTTTCTATCTATCTATCTATCTNNNAGATCGGA
131502 TATGAGTGTGGAGTCGTTGCCTTGATTTTTTCCTCTAATGGAANNNAGATCGGAAG
131503 TATGAGTGTGGAGTCGTTGCCTGTAGAGAGGACATTTTGAGNNNAGATCGGAAGA
131504 TATGAGTGTGGAGTCGTTGCTTGAAACCAGGAGAACTTCTNNNAGATCGGAAGAG
131505 TATGAGTGTGGAGTCGTTGCTATGGTGTATTGGATATATGACCNNNAGATCGGAA
131506 TATGAGTGTGGAGTCGTTGCAAAATGGCTTACAGAGAGAAATNNNAGATCGGAAG
131507 TATGAGTGTGGAGTCGTTGCAATTTATCTTCCTACAATGTCCTNNNAGATCGGAAGA
131508 TATGAGTGTGGAGTCGTTGCAATTTATCTTCCTACAATGTCCTNNNAGATCGGAAGA
131509 TATGAGTGTGGAGTCGTTGCAATTTATCTTCCTACAATGTCCTNNNAGATCGGAAGA
131510 TATGAGTGTGGAGTCGTTGCATTAAAAGACAGATTGATACAGCNNNAGATCGGAA
131511 TATGAGTGTGGAGTCGTTGCATGATGAATTGACTTACAGGTATNNNAGATCGGAA

131512 TATGAGTGTGGAGTCGTTGCTACACTCAACAAATATATTTGCANNNAGATCGGAAGA
131513 TATGAGTGTGGAGTCGTTGCTTGAGAATCCCTATAAGTAACATNNNAGATCGGAAG
131514 TATGAGTGTGGAGTCGTTGCGCTTACTCACTTCAGATTTTGNNNAGATCGGAAGAG
131516 TATGAGTGTGGAGTCGTTGCAAAAAGTTCTCACAGAATACAAANNNAGATCGGAAG
131517 TATGAGTGTGGAGTCGTTGCTTCTCTCTCTCTCTCTCTCTNNNAGATCGGAAGAGCAC
131518 TATGAGTGTGGAGTCGTTGCGTCAGAATTGGGTTGTCTTCNNNAGATCGGAAGAG
131519 TATGAGTGTGGAGTCGTTGCGTTACACAGAAATGCTCATAATANNNAGATCGGAAG
131520 TATGAGTGTGGAGTCGTTGCGAACAGAGTGACACAAGTACNNNAGATCGGAAGAG
131521 TATGAGTGTGGAGTCGTTGCGAACTACTGGAATCACAAATCANNNAGATCGGAAGA
131522 TATGAGTGTGGAGTCGTTGCAAAAGAGTGTGTAGAGAGTCATANNNAGATCGGAA
131523 TATGAGTGTGGAGTCGTTGCTTTGGCATAGTCCTCCATTTNNNAGATCGGAAGAGCA
131524 TATGAGTGTGGAGTCGTTGCGACTTCACTGGGTTAGAACTNNNAGATCGGAAGAG
131525 TATGAGTGTGGAGTCGTTGCCTTCAGGATCTTACACAAGACNNNAGATCGGAAGAG
131526 TATGAGTGTGGAGTCGTTGCAAATTTTATGCTAGAAGCCAAGNNNAGATCGGAAGA
131528 TATGAGTGTGGAGTCGTTGCGCATATTTTGGAATCCTACACNNNAGATCGGAAGAG
131529 TATGAGTGTGGAGTCGTTGCGATTAAAAAGACCATCTTAGTGGNNNAGATCGGAAG
131530 TATGAGTGTGGAGTCGTTGCTTGTCTGTAGTTTGCATAAAATTNNNAGATCGGAAGA
131531 TATGAGTGTGGAGTCGTTGCATATGTTTTGGAATCTTCACAACNNNAGATCGGAAGA
131532 TATGAGTGTGGAGTCGTTGCCTTCTCATTTTCCTGGAATGTNNNAGATCGGAAGAG
131533 TATGAGTGTGGAGTCGTTGCCTCAAACTCCTCTACTAAACTCNNNAGATCGGAAGAG
131534 TATGAGTGTGGAGTCGTTGCGAATGCTAAAGTTTCCTAACATTNNNAGATCGGAAGA
131535 TATGAGTGTGGAGTCGTTGCGACACATTTTCAAGAAATAATGCNNNAGATCGGAAG
131536 TATGAGTGTGGAGTCGTTGCTTCTTCTTTGCTTCACTTATCATNNNAGATCGGAAGAG
131537 TATGAGTGTGGAGTCGTTGCAAGTAATATGGTATGGGATCTTGNNNAGATCGGAAG
131538 TATGAGTGTGGAGTCGTTGCGCCTTCCAAAATATACTTAAGACNNNAGATCGGAAGA
131539 TATGAGTGTGGAGTCGTTGCATCCATTCCAGATGAAACTATATNNNAGATCGGAAGA
131540 TATGAGTGTGGAGTCGTTGCATGGAAAATTTAACGAATTCTGANNNAGATCGGAAG
131541 TATGAGTGTGGAGTCGTTGCCAAATCCTGATCAGTACTGATNNNAGATCGGAAGAG
131542 TATGAGTGTGGAGTCGTTGCCAAATCCTGATCAGTACTGATNNNAGATCGGAAGAG
131543 TATGAGTGTGGAGTCGTTGCTTAATCCTCACCACATTATATGANNNAGATCGGAAGA
131544 TATGAGTGTGGAGTCGTTGCAGATGAAAAACCAAAAGAATGAANNNAGATCGGAA
131545 TATGAGTGTGGAGTCGTTGCAATTCGTTGTTCTATTTAGTGCNNNAGATCGGAAGAG
131546 TATGAGTGTGGAGTCGTTGCCTTCCACTGTTCAAACTATATTCNNNAGATCGGAAGA
131547 TATGAGTGTGGAGTCGTTGCACCAAGACAATAATTTTTACTGTNNNAGATCGGAAGA
131548 TATGAGTGTGGAGTCGTTGCGGAAAAGTGAAGAGGGTATAGNNNAGATCGGAAG
131549 TATGAGTGTGGAGTCGTTGCTTCTGACAGCAAAACTCTAATNNNAGATCGGAAGAG
131550 TATGAGTGTGGAGTCGTTGCCATTTCTGACTAAACACTGAACNNNAGATCGGAAGAG
131551 TATGAGTGTGGAGTCGTTGCCCTTTAGCTATCAGTCCATTGNNNAGATCGGAAGAG
131552 TATGAGTGTGGAGTCGTTGCCCTTTAGTCACTTTATGTTAACANNNAGATCGGAAGA
131553 TATGAGTGTGGAGTCGTTGCGGTCTGGTTATATACATTGATGANNNAGATCGGAAG
131554 TATGAGTGTGGAGTCGTTGCGGTCTGGTTATATACATTGATGANNNAGATCGGAAG
131555 TATGAGTGTGGAGTCGTTGCTATACAATTGTGAGATTGTGTGANNNAGATCGGAAG
131556 TATGAGTGTGGAGTCGTTGCTAATCAGTCTCCTTTTTATCAGTNNNAGATCGGAAGA

131557 TATGAGTGTGGAGTCGTTGCCTATTGTATCCCCTAGAACAATTNNNAGATCGGAAG
131558 TATGAGTGTGGAGTCGTTGCGTGTGATATCTTGGCTCACTNNNAGATCGGAAGAG
131559 TATGAGTGTGGAGTCGTTGCATATGTGAGAATTTTGCTCAATCNNNAGATCGGAAG
131560 TATGAGTGTGGAGTCGTTGCGTGGCTTCTATTTCTAAAATGTANNNAGATCGGAAG
131561 TATGAGTGTGGAGTCGTTGCTGATAGTGGAAAATTGCAAATTANNNAGATCGGAA
131562 TATGAGTGTGGAGTCGTTGCTGATAGTGGAAAATTGCAAATTANNNAGATCGGAA
131563 TATGAGTGTGGAGTCGTTGCACAGAGAGAGAGAGAGACAGNNNAGATCGGAAGA
131564 TATGAGTGTGGAGTCGTTGCAAATCAAGCAAAGAGAACAATATNNNAGATCGGAA
131565 TATGAGTGTGGAGTCGTTGCATGTCTAGCTGGGTTTTCTTNNNAGATCGGAAGAGC
131566 TATGAGTGTGGAGTCGTTGCGCCTGTCTTAATACATAAGTCTTNNNAGATCGGAAG
131567 TATGAGTGTGGAGTCGTTGCATACATACATTGTGTATTCATGTNNNAGATCGGAAG
131568 TATGAGTGTGGAGTCGTTGCAACTTATTTTCTTTTCAAAACTCTTNNNAGATCGGAA
131570 TATGAGTGTGGAGTCGTTGCTCTGTTTTTTAATTCCATTTCCANNNAGATCGGAAGA
131571 TATGAGTGTGGAGTCGTTGCCAAATTGTATTTCTGCTCCAGNNNAGATCGGAAGAG
131572 TATGAGTGTGGAGTCGTTGCCCTCTAGACCTGTGAAATAATAANNNAGATCGGAAG
131573 TATGAGTGTGGAGTCGTTGCCGTCATAGTGGCTTAAAATAATTNNNAGATCGGAAG
131574 TATGAGTGTGGAGTCGTTGCCGTCATAGTGGCTTAAAATAATTNNNAGATCGGAAG
131575 TATGAGTGTGGAGTCGTTGCTAGCTGTTCTCCCCAAATACNNNAGATCGGAAGAGC
131576 TATGAGTGTGGAGTCGTTGCTGACTTACAGAGTGTTTATTTTGNNNAGATCGGAAG
131577 TATGAGTGTGGAGTCGTTGCAGATAAAATCTGGCTTTGGGNNNAGATCGGAAGAG
131579 TATGAGTGTGGAGTCGTTGCAGGAAGATGTAGAAGAAAACAANNNAGATCGGAA
131580 TATGAGTGTGGAGTCGTTGCAAAATGCAGTTCATATAACAACANNNAGATCGGAAG
131581 TATGAGTGTGGAGTCGTTGCAAAATATCTGAATATGATGCTGTNNNAGATCGGAAG
131582 TATGAGTGTGGAGTCGTTGCCCCCATGCTAAAAAATAGCANNNAGATCGGAAGAGC
131583 TATGAGTGTGGAGTCGTTGCCCTGAATTGAATAACCAAAGAAANNNAGATCGGAA
131584 TATGAGTGTGGAGTCGTTGCGTAATGTCAGTGGTCCTTTCNNNAGATCGGAAGAGC
131585 TATGAGTGTGGAGTCGTTGCTATCTAACAAGTTTCCAGTTGATNNNAGATCGGAAG
131586 TATGAGTGTGGAGTCGTTGCGACTTGAAGTCTATATACAGGAGNNNAGATCGGAA
131587 TATGAGTGTGGAGTCGTTGCAGCAAAAATGGATTTTCTGTAGNNNAGATCGGAAG
131588 TATGAGTGTGGAGTCGTTGCATCATTTGTTATATGCAGTTTGGNNNAGATCGGAAG
131589 TATGAGTGTGGAGTCGTTGCACTCTCTAGGAATCTAGTCAATANNNAGATCGGAAG
131590 TATGAGTGTGGAGTCGTTGCCTGTATCTCTGGAGAACTCTAANNNAGATCGGAAGA
131591 TATGAGTGTGGAGTCGTTGCGATACAGTTTAGTTCATGGAGGNNNAGATCGGAAG
131592 TATGAGTGTGGAGTCGTTGCCTCTGCTTGTCCATCTATCTNNNAGATCGGAAGAGCA
131593 TATGAGTGTGGAGTCGTTGCGGCCTTACAAGAGAATATGTGNNNAGATCGGAAGA
131594 TATGAGTGTGGAGTCGTTGCGTGAAATTTAAAATCCCTCATTGNNNAGATCGGAAG
131595 TATGAGTGTGGAGTCGTTGCCCTGTATATCCAGAAATTAGAGANNNAGATCGGAAG
131596 TATGAGTGTGGAGTCGTTGCAAAAAGCGAAGTCATCTTTCTNNNAGATCGGAAGAG
131597 TATGAGTGTGGAGTCGTTGCGAAATGGGTTACTTTTAGATTGANNNAGATCGGAA
131598 TATGAGTGTGGAGTCGTTGCCAACTCTGATTTTGGTATCCTNNNAGATCGGAAGAG
131599 TATGAGTGTGGAGTCGTTGCGCACTATATATTCTCTCGAGTATNNNAGATCGGAAG
131600 TATGAGTGTGGAGTCGTTGCCTCTTGTATCTCTTCGTCTTCNNNAGATCGGAAGAGC
131601 TATGAGTGTGGAGTCGTTGCGGTAGCTCACACTTGTAATCNNNAGATCGGAAGAGC

131602 TATGAGTGTGGAGTCGTTGCTAACATATGCAGTATATGATCCCNNNAGATCGGAAG
131603 TATGAGTGTGGAGTCGTTGCGAGTACTACTCAGCCATAAAAANNNAGATCGGAAGA
131604 TATGAGTGTGGAGTCGTTGCGAGATCCTGTATTGATGAAAAATNNNAGATCGGAAG
131605 TATGAGTGTGGAGTCGTTGCAAACCGTTATTATAAAGTGTTGGNNNAGATCGGAAG
131606 TATGAGTGTGGAGTCGTTGCAAAAAGTAATTCAAAGAGGACTCNNNAGATCGGAA
131607 TATGAGTGTGGAGTCGTTGCCTAAGGAGCTATAAGATTCCTTTNNNAGATCGGAAG
131608 TATGAGTGTGGAGTCGTTGCCTTTTCAATGTGGTGTCTAGTNNNAGATCGGAAGAG
131609 TATGAGTGTGGAGTCGTTGCTAGAGTAATACAGGGAAAGGATNNNAGATCGGAAG
131610 TATGAGTGTGGAGTCGTTGCCTCCATATATACACAGAGAGATTNNNAGATCGGAAG
131611 TATGAGTGTGGAGTCGTTGCTTCAAGACTGTTTTTTTTTCTGNNNAGATCGGAAGA
131612 TATGAGTGTGGAGTCGTTGCAAATAGTTCAGTTACATACCTCANNNAGATCGGAAG
131613 TATGAGTGTGGAGTCGTTGCAAGTTATGTTTGCCTTTCCTNNNAGATCGGAAGAGC
131614 TATGAGTGTGGAGTCGTTGCCATCTATAATCCCAATCCCTGNNNAGATCGGAAGAG
131615 TATGAGTGTGGAGTCGTTGCGATACAGGATGAGTAGAAGTTAGNNNAGATCGGAA
131617 TATGAGTGTGGAGTCGTTGCCAGGAGTTCTCTGAATTTTCTNNNAGATCGGAAGAG
131618 TATGAGTGTGGAGTCGTTGCCTAACTTGATTGTGGTAATCATTNNNAGATCGGAAG
131619 TATGAGTGTGGAGTCGTTGCGGAGTGTCAGAAATAATAGATTGNNNAGATCGGAA
131620 TATGAGTGTGGAGTCGTTGCTTACCAAATTTAAAACATCTGCTNNNAGATCGGAAGA
131621 TATGAGTGTGGAGTCGTTGCATTTGAGCAACATCATAAAGTAGNNNAGATCGGAAG
131622 TATGAGTGTGGAGTCGTTGCATAGATTGTTCCTAAATCCCTACNNNAGATCGGAAGA
131623 TATGAGTGTGGAGTCGTTGCAAGAGAGAGAAATCATAAAACACNNNAGATCGGAA
131624 TATGAGTGTGGAGTCGTTGCCCTTCTCATGTAAGACTATATCANNNAGATCGGAAGA
131625 TATGAGTGTGGAGTCGTTGCGATATTTTTGTGATCATAGGGACNNNAGATCGGAAG
131626 TATGAGTGTGGAGTCGTTGCGATATTTTTGTGATCATAGGGACNNNAGATCGGAAG
131627 TATGAGTGTGGAGTCGTTGCCAGATAAATGGAGGAAAAACACNNNAGATCGGAAG
131628 TATGAGTGTGGAGTCGTTGCTACATCTTGGTTTCTAAATACCANNNAGATCGGAAGA
131629 TATGAGTGTGGAGTCGTTGCGTTTATTCCATGCAAATAGAAACNNNAGATCGGAAG
131631 TATGAGTGTGGAGTCGTTGCGAGTACAGTGACTATTAACAGATNNNAGATCGGAAG
131632 TATGAGTGTGGAGTCGTTGCTGACTAAAATATTCAGTCAGAGTNNNAGATCGGAAG
131633 TATGAGTGTGGAGTCGTTGCGGGGAAAACAAAGACTAATAAATNNNAGATCGGAA
131634 TATGAGTGTGGAGTCGTTGCCATTCATTCATCATTGTCACTTNNNAGATCGGAAGAG
131635 TATGAGTGTGGAGTCGTTGCAGAACAGAAAAGGTGTAAGATGNNNAGATCGGAAG
131637 TATGAGTGTGGAGTCGTTGCGGATAGTTGAAGAATGAAAAACANNNAGATCGGAA
131638 TATGAGTGTGGAGTCGTTGCGATTAAGAAAGCCAAGAAATAGCNNNAGATCGGAA
131639 TATGAGTGTGGAGTCGTTGCCTGATTCAGCACTGAGAAATNNNAGATCGGAAGAG
131640 TATGAGTGTGGAGTCGTTGCATTTCCATCTCTTCCATGAAAAANNNAGATCGGAAGA
131641 TATGAGTGTGGAGTCGTTGCGCAGATCAAAGAGAGAAAATAAANNNAGATCGGAA
131642 TATGAGTGTGGAGTCGTTGCTATAATCATCCACAAAGAGTGATNNNAGATCGGAAG
131643 TATGAGTGTGGAGTCGTTGCTATAATCATCCACAAAGAGTGATNNNAGATCGGAAG
131644 TATGAGTGTGGAGTCGTTGCGCTGCAGATAACTATAGGATTTANNNAGATCGGAAG
131645 TATGAGTGTGGAGTCGTTGCTAAACTGAATAGGATTGCACANNNAGATCGGAAGA
131646 TATGAGTGTGGAGTCGTTGCCTTCCTTAGAACACTCCTATAAANNNAGATCGGAAGA
131647 TATGAGTGTGGAGTCGTTGCGCTGAGATAGAAGTCATCATATANNNAGATCGGAA

131649 TATGAGTGTGGAGTCGTTGCTAGATATGTTGGCTCTTCTTATTNNNAGATCGGAAG
131650 TATGAGTGTGGAGTCGTTGCCTGGTAGTAGCTGACTTTATGNNNAGATCGGAAGA
131651 TATGAGTGTGGAGTCGTTGCACTTGAATTAAAAAAAATGCTACANNNAGATCGGAA
131654 TATGAGTGTGGAGTCGTTGCATGAATTTCCACCCCTACTTNNNAGATCGGAAGAGC
131655 TATGAGTGTGGAGTCGTTGCCATTTTCTTTATCCACTCTTTGANNNAGATCGGAAGA
131656 TATGAGTGTGGAGTCGTTGCGTTCTGGTTATTTGAACTTTCTGNNNAGATCGGAAG
131657 TATGAGTGTGGAGTCGTTGCTATATATGTAATGTGTGTGTGTGNNNAGATCGGAA
131658 TATGAGTGTGGAGTCGTTGCCTTCCTGCTAAACAGATGTCNNNAGATCGGAAGAGC
131659 TATGAGTGTGGAGTCGTTGCCTAGACTTTGTCAATTCTTCACNNNAGATCGGAAGA
131660 TATGAGTGTGGAGTCGTTGCCACACCCTTTTCTTACAGTATNNNAGATCGGAAGAG
131663 TATGAGTGTGGAGTCGTTGCATACATACATTGTGTATTCATGTNNNAGATCGGAAG
131664 TATGAGTGTGGAGTCGTTGCAGCCTCAAATAAATGCTTTTTAANNNAGATCGGAAG
131665 TATGAGTGTGGAGTCGTTGCCAGTACTGAATAAAGAAACAGACNNNAGATCGGAA
131666 TATGAGTGTGGAGTCGTTGCCCAGGAATTCATGACTACAGNNNAGATCGGAAGAG
131667 TATGAGTGTGGAGTCGTTGCATTTTCCTTCTCATGAACCTTNNNAGATCGGAAGAGC
131668 TATGAGTGTGGAGTCGTTGCAGCTATGCAATTTAAAATTGATTATNNNAGATCGGA
131669 TATGAGTGTGGAGTCGTTGCCTGTGTATCTTATGATTTGAACANNNAGATCGGAAG
131671 TATGAGTGTGGAGTCGTTGCGTGATTGTGCTGGATCATACNNNAGATCGGAAGAG
131672 TATGAGTGTGGAGTCGTTGCAAATTTTACTTTTATAACCCCCCNNNAGATCGGAAGA
131673 TATGAGTGTGGAGTCGTTGCCATTGTAATGTAATTGGTGTCTCNNNAGATCGGAAG
131674 TATGAGTGTGGAGTCGTTGCCTCGACAAAGACTAGGAAGTNNNAGATCGGAAGAG
131676 TATGAGTGTGGAGTCGTTGCGTAGAATAACTGGAAAATGCTTTNNNAGATCGGAA
131677 TATGAGTGTGGAGTCGTTGCGATAGCTGGGTCAACAAAATANNNAGATCGGAAGA
131678 TATGAGTGTGGAGTCGTTGCATAAGACAGAATATTCACAGCTANNNAGATCGGAAG
131679 TATGAGTGTGGAGTCGTTGCGTTGATAAAATTATTGAGCTGGANNNAGATCGGAA
131680 TATGAGTGTGGAGTCGTTGCGGCTTAAAAAATGTCAGAAAAATNNNAGATCGGAA
131681 TATGAGTGTGGAGTCGTTGCACCCTAAACAAACAAAAAAAATCNNNAGATCGGAAG
131682 TATGAGTGTGGAGTCGTTGCTGTAAATTTATTTTCTCTGCAGTNNNAGATCGGAAG
131683 TATGAGTGTGGAGTCGTTGCACAATAAGAACTACAAAAGATGGNNNAGATCGGAA
131684 TATGAGTGTGGAGTCGTTGCCACATGTAAAATTTTATTTTTTGGGNNNAGATCGGA
131685 TATGAGTGTGGAGTCGTTGCCAACAGCAGATTGAAAATATTTGNNNAGATCGGAA
131686 TATGAGTGTGGAGTCGTTGCATATTTTGTTAATATATGTGTGTGTNNNAGATCGGA
131687 TATGAGTGTGGAGTCGTTGCTTCACCATCTGCATTCTTAAATANNNAGATCGGAAGA
131688 TATGAGTGTGGAGTCGTTGCTGGGAAGTAAAAATCTTCAAGATNNNAGATCGGAA
131689 TATGAGTGTGGAGTCGTTGCTGGACTGTTTATTTTGCTTATTTNNNAGATCGGAAGA
131690 TATGAGTGTGGAGTCGTTGCTGAATAAGAAACAATGTAGATCGNNNAGATCGGAA
131691 TATGAGTGTGGAGTCGTTGCGTTATACCTAGCAGGGTGAGNNNAGATCGGAAGAG
131692 TATGAGTGTGGAGTCGTTGCAGAAACTTGGAGTTCTTTCAGNNNAGATCGGAAGA
131693 TATGAGTGTGGAGTCGTTGCCCATTTACAACACCTAGAGTGNNNAGATCGGAAGAG
131694 TATGAGTGTGGAGTCGTTGCACAAATGTTTCACCATGTTGNNNAGATCGGAAGAGC
131695 TATGAGTGTGGAGTCGTTGCTCTTTTTAACAGAGTATTGGTACNNNAGATCGGAAG
131696 TATGAGTGTGGAGTCGTTGCAAGCTTTGAGAGTATTGGATATGNNNAGATCGGAA
131697 TATGAGTGTGGAGTCGTTGCCCATTTCTCTTTCTCCCAAGNNNAGATCGGAAGAGCA

131698 TATGAGTGTGGAGTCGTTGCTAGAAAATCTACACAAGTCAAGANNNAGATCGGAAG
131699 TATGAGTGTGGAGTCGTTGCCAACATACTGGTTTCTAAATTGANNNAGATCGGAAG
131700 TATGAGTGTGGAGTCGTTGCCAAAAGATGACTGTCTTAGAAATNNNAGATCGGAAG
131701 TATGAGTGTGGAGTCGTTGCCTACATTCAAAGCCTGATCANNNAGATCGGAAGAGC
131702 TATGAGTGTGGAGTCGTTGCGGTGGAGATTCTAGACAAAGNNNAGATCGGAAGAG
131703 TATGAGTGTGGAGTCGTTGCTTTCTTTAAATTTAACCAAGGGCNNNAGATCGGAAG
131705 TATGAGTGTGGAGTCGTTGCACCAGTAATTCAAAGACTAAATGNNNAGATCGGAAG
131706 TATGAGTGTGGAGTCGTTGCTAACTGAAAAGGATGGATAAGATNNNAGATCGGAA
131707 TATGAGTGTGGAGTCGTTGCAAGCAAAGGTTATTGATTTTCTANNNAGATCGGAAG
131708 TATGAGTGTGGAGTCGTTGCGTGAATGTCGATGCAAAAATNNNAGATCGGAAGAG
131709 TATGAGTGTGGAGTCGTTGCTGTTTTTTTAACTACTACTGGTTNNNAGATCGGAAGA
131710 TATGAGTGTGGAGTCGTTGCCTTACAATAGCATCATGAAGTGNNNAGATCGGAAGA
131711 TATGAGTGTGGAGTCGTTGCGTTGCATAAGATATGTTTCACANNNAGATCGGAAGA
131712 TATGAGTGTGGAGTCGTTGCGTAGCTTAATCCTATCCCTTGNNNAGATCGGAAGAG
131713 TATGAGTGTGGAGTCGTTGCTGAGAAATCATTGACAAGATATTTNNNAGATCGGAA
131714 TATGAGTGTGGAGTCGTTGCATAGACTAAGATGACAACTACAGNNNAGATCGGAAG
131715 TATGAGTGTGGAGTCGTTGCTAACACAGACTGTCTTTCCTNNNAGATCGGAAGAGC
131716 TATGAGTGTGGAGTCGTTGCAAAAGAAACGGAAAATAGGAACNNNAGATCGGAAG
131717 TATGAGTGTGGAGTCGTTGCCTTAAGAAACTTATGCATCTCTGNNNAGATCGGAAG
131718 TATGAGTGTGGAGTCGTTGCTAGCTGTATAACAAAAAGGAAACNNNAGATCGGAA
131719 TATGAGTGTGGAGTCGTTGCGTCTCCAATCCTAGCACTTTNNNAGATCGGAAGAGC
131720 TATGAGTGTGGAGTCGTTGCGAAAATTCTGCAATCTGTAGAGNNNAGATCGGAAG
131721 TATGAGTGTGGAGTCGTTGCCAAAGGAAAGAAGAATCTCTTCNNNAGATCGGAAG
131722 TATGAGTGTGGAGTCGTTGCGTTCTGGTTCCATATGAATTTTANNNAGATCGGAAG
131723 TATGAGTGTGGAGTCGTTGCATACCATTTCAAAGAAACACAATNNNAGATCGGAAG
131724 TATGAGTGTGGAGTCGTTGCTTCATGAAATCACATATGTACTGNNNAGATCGGAAG
131725 TATGAGTGTGGAGTCGTTGCAGAATGATTAATTAGCAAGCAAGNNNAGATCGGAA
131726 TATGAGTGTGGAGTCGTTGCCTTTCACCCTTGATGGAATGNNNAGATCGGAAGAGC
131727 TATGAGTGTGGAGTCGTTGCGTCCAGGGCATTATATAAACANNNAGATCGGAAGAG
131728 TATGAGTGTGGAGTCGTTGCGCTTAAAGAAAACAATATTTTGCNNNAGATCGGAAG
131729 TATGAGTGTGGAGTCGTTGCCCTCCTGTATGAAAAAAACTTAANNNAGATCGGAAG
131730 TATGAGTGTGGAGTCGTTGCTCCAATAAATTAAATACTGATGCANNNAGATCGGAA
131731 TATGAGTGTGGAGTCGTTGCATAAAATCAGCTAGACCACTGNNNAGATCGGAAGAG
131733 TATGAGTGTGGAGTCGTTGCGTTAGATTTATTGATGAACGCTGNNNAGATCGGAAG
131734 TATGAGTGTGGAGTCGTTGCATGTATATATTTGAGGGTGTTTTNNNAGATCGGAAG
131735 TATGAGTGTGGAGTCGTTGCTATATGTAAGACAGAGTCTCACTNNNAGATCGGAAG
131736 TATGAGTGTGGAGTCGTTGCGGGAGAGAAATGATATAATTCAAGNNNAGATCGGA
131737 TATGAGTGTGGAGTCGTTGCCCATGTCTAGATTTGTTTCTTTGNNNAGATCGGAAGA
131738 TATGAGTGTGGAGTCGTTGCCAAGAACAAGAAGAAGCTTATTTNNNAGATCGGAAG
131740 TATGAGTGTGGAGTCGTTGCTAGATGGAACTGGAGACTATTATNNNAGATCGGAA
131741 TATGAGTGTGGAGTCGTTGCCCTCCTGTATGAAAAAAACTTAANNNAGATCGGAAG
131742 TATGAGTGTGGAGTCGTTGCATATTTTTTTGAGACGGAGTTTCNNNAGATCGGAAG
131743 TATGAGTGTGGAGTCGTTGCCATTATCTGAGAGGGAGTTTAGNNNAGATCGGAAG

131744 TATGAGTGTGGAGTCGTTGCTGTCATTATCTTCAACATCCTTANNNAGATCGGAAGA
131745 TATGAGTGTGGAGTCGTTGCATCTGTAATCATTAGGCTTGTATNNNAGATCGGAAGA
131746 TATGAGTGTGGAGTCGTTGCATACATTGTAAGGAAAGATGGAANNNAGATCGGAA
131747 TATGAGTGTGGAGTCGTTGCCATTATGTCTCTCAGGTCTTCNNNAGATCGGAAGAGC
131748 TATGAGTGTGGAGTCGTTGCTTACAAGTGAAAAATGCATTTAANNNAGATCGGAAG
131749 TATGAGTGTGGAGTCGTTGCAGGTTTTCTCTTTTTTCTAATCTNNNAGATCGGAAGAG
131750 TATGAGTGTGGAGTCGTTGCATTTTAAGACAAGGCCTCTCNNNAGATCGGAAGAGC
131752 TATGAGTGTGGAGTCGTTGCAAAGCTATGTAAAAGTTACTCTGNNNAGATCGGAAG
131753 TATGAGTGTGGAGTCGTTGCGAAATCCAATTTGAGGTCAACNNNAGATCGGAAGAG
131754 TATGAGTGTGGAGTCGTTGCTCTACCTTTTATTCTTCCATTCTNNNAGATCGGAAGAG
131755 TATGAGTGTGGAGTCGTTGCTTTTGAAAGAGGAAAGTAAGAAGNNNAGATCGGAA
131756 TATGAGTGTGGAGTCGTTGCATGGACAGGTCTGATATTTTTATNNNAGATCGGAAG
131757 TATGAGTGTGGAGTCGTTGCCTTCTAAAGGGAGAGAGCAGNNNAGATCGGAAGAG
131758 TATGAGTGTGGAGTCGTTGCTACTAAAATTACCAGTTTTCCCANNNAGATCGGAAGA
131759 TATGAGTGTGGAGTCGTTGCAAAACTTGTTTTTAGCACTCAANNNAGATCGGAAGAG
131760 TATGAGTGTGGAGTCGTTGCTGTTATTTTGCATTTTAACATCCNNNAGATCGGAAGA
131761 TATGAGTGTGGAGTCGTTGCGCTCTGATTTTCTTACATTTAGTNNNAGATCGGAAGA
131764 TATGAGTGTGGAGTCGTTGCCATGAAAACTTCAAAGAGAAAATNNNAGATCGGAAG
131765 TATGAGTGTGGAGTCGTTGCCCCTTTCATTCTTATTCAACAATNNNAGATCGGAAGAG
131766 TATGAGTGTGGAGTCGTTGCATTGTTTTTGGGTTTTTTTTGTTNNNAGATCGGAAGA
131767 TATGAGTGTGGAGTCGTTGCAAGACTTTACCAGAAATGTGAANNNAGATCGGAAGA
131768 TATGAGTGTGGAGTCGTTGCGAAATGTGCCACAAGTCTTTNNNAGATCGGAAGAGC
131769 TATGAGTGTGGAGTCGTTGCCTAAGTTAGTAACTGCTGAACTTNNNAGATCGGAAG
131770 TATGAGTGTGGAGTCGTTGCCCAACATAAAATTACAAACTCCTNNNAGATCGGAAGA
131771 TATGAGTGTGGAGTCGTTGCTAACCAGTATTTTTGCTATTTCCNNNAGATCGGAAGA
131772 TATGAGTGTGGAGTCGTTGCAATTCTTGAACAAAAGCCTTATTNNNAGATCGGAAGA
131774 TATGAGTGTGGAGTCGTTGCCAACTACAATCAAGATACAGAACNNNAGATCGGAAG
131775 TATGAGTGTGGAGTCGTTGCTGTTACGAGGATAAAGATTTTTGNNNAGATCGGAAG
131776 TATGAGTGTGGAGTCGTTGCGATATTCAGGACGCTGAGATNNNAGATCGGAAGAG
131777 TATGAGTGTGGAGTCGTTGCCTGTCTCTGAATTCCAAAATCTNNNAGATCGGAAGAG
131778 TATGAGTGTGGAGTCGTTGCGTTTTGGGAAATATATGTGCTATNNNAGATCGGAAG
131779 TATGAGTGTGGAGTCGTTGCGCTTAGATTTAGAAAATCATAGGTNNNAGATCGGAA
131780 TATGAGTGTGGAGTCGTTGCATAACCCTTAAATCACAGTACATNNNAGATCGGAAGA
131781 TATGAGTGTGGAGTCGTTGCTATAAGTACAAAGGCATGCAATANNNAGATCGGAAG
131782 TATGAGTGTGGAGTCGTTGCAAAAATCCTCAACAAAATACCAGNNNAGATCGGAAG
131783 TATGAGTGTGGAGTCGTTGCTCTTTCTCACCCTTAAATATCCNNNAGATCGGAAGAG
131784 TATGAGTGTGGAGTCGTTGCCCATTCATTTTGACTATTCCATANNNAGATCGGAAGA
131785 TATGAGTGTGGAGTCGTTGCCCATTCATTTTGACTATTCCATANNNAGATCGGAAGA
131786 TATGAGTGTGGAGTCGTTGCCTTGGGGATTATAAAACAAAGAANNNAGATCGGAAG
131787 TATGAGTGTGGAGTCGTTGCAAGTCAGTTGGTTAATTGAGAANNNAGATCGGAAGA
131788 TATGAGTGTGGAGTCGTTGCCCCGTTTCCTAACTTAATTTTCNNNAGATCGGAAGAG
131789 TATGAGTGTGGAGTCGTTGCATGAGACAAGAGTTTTGCTCNNNAGATCGGAAGAGC
131790 TATGAGTGTGGAGTCGTTGCATAATACTTCTAGCATAGTGACCNNNAGATCGGAAGA

131791 TATGAGTGTGGAGTCGTTGCATTGCAATGAAATTGATGCTTTANNNAGATCGGAAG
131792 TATGAGTGTGGAGTCGTTGCGAAATCAATAGCTCAAAGAGATANNNAGATCGGAA
131793 TATGAGTGTGGAGTCGTTGCACTGGAGACTATTATTCTAAGTGNNNAGATCGGAAG
131794 TATGAGTGTGGAGTCGTTGCGTAGGTTAGTTAAGATGAGCTCNNNAGATCGGAAG
131795 TATGAGTGTGGAGTCGTTGCAAACAAATTAGCAAAATTTCTGGNNNAGATCGGAAG
131796 TATGAGTGTGGAGTCGTTGCTAATCCCAGTGACACAGAAANNNAGATCGGAAGAG
131797 TATGAGTGTGGAGTCGTTGCATAGTAGAATGGATAATTGTGGTNNNAGATCGGAA
131799 TATGAGTGTGGAGTCGTTGCCACATTTTCTGTTAACCTTTCTNNNAGATCGGAAGAG
131801 TATGAGTGTGGAGTCGTTGCCCTTGTTAATCCCTTACAAATCNNNAGATCGGAAGAG
131802 TATGAGTGTGGAGTCGTTGCATGTCAGCAATAAGTAATACATGNNNAGATCGGAAG
131803 TATGAGTGTGGAGTCGTTGCATTTTTTTTTTGCATTTCTGGAANNNAGATCGGAAGA
131804 TATGAGTGTGGAGTCGTTGCTGACATTTAAATAAAGTGTGGAGNNNAGATCGGAA
131805 TATGAGTGTGGAGTCGTTGCGAGACTGGAGACTGTTATTCTANNNAGATCGGAAGA
131806 TATGAGTGTGGAGTCGTTGCGACTTATAATGGAAAGCCGANNNAGATCGGAAGAG
131807 TATGAGTGTGGAGTCGTTGCATACCCATTTTTCCAGTATAGATNNNAGATCGGAAGA
131809 TATGAGTGTGGAGTCGTTGCTCTCCTTGGATGATATCTTTTGNNNAGATCGGAAGA
131810 TATGAGTGTGGAGTCGTTGCAGTGTAGGGATAGATTAAGGAGNNNAGATCGGAA
131811 TATGAGTGTGGAGTCGTTGCCTGAAATCATCTGCTTAAAAAAANNNAGATCGGAAG
131812 TATGAGTGTGGAGTCGTTGCAGAAGGAGTCTCCCTATATTGNNNAGATCGGAAGA
131813 TATGAGTGTGGAGTCGTTGCAAGATGGCAAGAGTAAATTTTTTNNNAGATCGGAA
131814 TATGAGTGTGGAGTCGTTGCTTCATTCTGTGCAGCAATTANNNAGATCGGAAGAGC
131815 TATGAGTGTGGAGTCGTTGCTAATCAATTGTTGGGATACTCANNNAGATCGGAAGA
131816 TATGAGTGTGGAGTCGTTGCCCTGACATTTTTACCGTATTGNNNAGATCGGAAGAG
131817 TATGAGTGTGGAGTCGTTGCCCTGCATTTTGTATAGATGTAATNNNAGATCGGAAG
131818 TATGAGTGTGGAGTCGTTGCAAGAGAGAGAGACAAATTTATGANNNAGATCGGAA
131819 TATGAGTGTGGAGTCGTTGCTCAAGATGGATTAAAGACTTAAATNNNAGATCGGAA
131820 TATGAGTGTGGAGTCGTTGCGGAATACTACTCAGCAATAAAAANNNAGATCGGAAG
131821 TATGAGTGTGGAGTCGTTGCAATCTATTTGGATCTCTACAGTGNNNAGATCGGAAG
131822 TATGAGTGTGGAGTCGTTGCCCTTTCTACATTTGGATGTTGNNNAGATCGGAAGAG
131823 TATGAGTGTGGAGTCGTTGCTATTCAGCTTTAAAAAAGAAGGGNNNAGATCGGAA
131824 TATGAGTGTGGAGTCGTTGCCTCTCTTCTTCAATCTCCTTCNNNAGATCGGAAGAGC
131825 TATGAGTGTGGAGTCGTTGCCATGTTGAATTGTAATCCTCAATNNNAGATCGGAAG
131826 TATGAGTGTGGAGTCGTTGCGGAGGGTATAACTTTTCTAAATGNNNAGATCGGAA
131827 TATGAGTGTGGAGTCGTTGCAAAGAACTCTTTGTTATGAGGGNNNAGATCGGAAG
131828 TATGAGTGTGGAGTCGTTGCATTTGGATGCTCTTTACTTTTTTNNNAGATCGGAAGA
131829 TATGAGTGTGGAGTCGTTGCCTTGAGCTTAGGAGTCAAGANNNAGATCGGAAGAG
131830 TATGAGTGTGGAGTCGTTGCTAAAACAAAAAGATAACGGACTCNNNAGATCGGAA
131831 TATGAGTGTGGAGTCGTTGCCATGCTATAGCTTGGATATTTGNNNAGATCGGAAGA
131832 TATGAGTGTGGAGTCGTTGCAAGTAATGGCTTCACTGTAAANNNAGATCGGAAGA
131833 TATGAGTGTGGAGTCGTTGCAGGAAAACATTCTCCTTAGAAANNNAGATCGGAAGA
131834 TATGAGTGTGGAGTCGTTGCTTTGAACAATTCAGGTATTAGAGNNNAGATCGGAAG
131835 TATGAGTGTGGAGTCGTTGCATAGACTACCTCTCTTCCTATTTNNNAGATCGGAAGA
131837 TATGAGTGTGGAGTCGTTGCTTACTCTTCCAATCTAACCATTCNNNAGATCGGAAGA

131838 TATGAGTGTGGAGTCGTTGCCTTACAAGGATTCCCAATTTAAANNNAGATCGGAAG
131839 TATGAGTGTGGAGTCGTTGCTATCTTGGCTATTGTAAATAGCGNNNAGATCGGAAG
131840 TATGAGTGTGGAGTCGTTGCATCCACATCAAGTAAAGAAGAANNNAGATCGGAAG
131841 TATGAGTGTGGAGTCGTTGCACAACAGATACTATTGTTTTTCTNNNAGATCGGAAG
131842 TATGAGTGTGGAGTCGTTGCGAGTCTAAGTACCATGTTCAATNNNAGATCGGAAGA
131843 TATGAGTGTGGAGTCGTTGCACATACATGCAAATTTGGATTCNNNAGATCGGAAGA
131844 TATGAGTGTGGAGTCGTTGCATTATTAGAAAGTGACTTCCCAANNNAGATCGGAAG
131845 TATGAGTGTGGAGTCGTTGCCTCTGTCAGATTGGTAGTCTNNNAGATCGGAAGAG
131846 TATGAGTGTGGAGTCGTTGCGAGTCATCTGATATATAACATTCTGNNNAGATCGGA
131847 TATGAGTGTGGAGTCGTTGCCACTGTCTCTTTAAAAGAAATGANNNAGATCGGAAG
131848 TATGAGTGTGGAGTCGTTGCATTAAAAGCTGACACAATTACAANNNAGATCGGAAG
131849 TATGAGTGTGGAGTCGTTGCAAATGTACCTTCCCATCCAGNNNAGATCGGAAGAGC
131850 TATGAGTGTGGAGTCGTTGCATTTCCCCTTTTCATCTACATATNNNAGATCGGAAGA
131851 TATGAGTGTGGAGTCGTTGCTTTGTTAAAAACATACTTTTGGGNNNAGATCGGAAG
131852 TATGAGTGTGGAGTCGTTGCTTTTTCTTTTTTTGAGACAGAGTNNNAGATCGGAAG
131853 TATGAGTGTGGAGTCGTTGCTATTCTTAGTTCTTCCTTTGCTTNNNAGATCGGAAGA
131854 TATGAGTGTGGAGTCGTTGCACATAAATCTGAAAAATGAACCTNNNAGATCGGAAG
131855 TATGAGTGTGGAGTCGTTGCTTTAAATGGTTCTATAGCGATAGNNNAGATCGGAAG
131856 TATGAGTGTGGAGTCGTTGCTAAGTCCTCATCTGATCTTTTTTNNNAGATCGGAAGA
131857 TATGAGTGTGGAGTCGTTGCTATTTTGTCACAATTCTAGAGGANNNAGATCGGAAG
131858 TATGAGTGTGGAGTCGTTGCTATTCCCTATAGATAAGCACTTGNNNAGATCGGAAG
131859 TATGAGTGTGGAGTCGTTGCACAGAGGAGAGAGAAAAAATATANNNAGATCGGA
131860 TATGAGTGTGGAGTCGTTGCATTTTATTTTTTTGAGACAGGGANNNAGATCGGAAG
131861 TATGAGTGTGGAGTCGTTGCGCTGAACGATTTTAAATTGAATTNNNAGATCGGAAG
131862 TATGAGTGTGGAGTCGTTGCTTTGGGGTTTATGACTCGTANNNAGATCGGAAGAG
131863 TATGAGTGTGGAGTCGTTGCGATCAAGATTCTCATGAAGTTTTNNNAGATCGGAAG
131864 TATGAGTGTGGAGTCGTTGCGAGCTCTGATGCTATAGCTATNNNAGATCGGAAGA
131865 TATGAGTGTGGAGTCGTTGCCATAGAGAAAGGCCTTAGAATAANNNAGATCGGAA
131867 TATGAGTGTGGAGTCGTTGCGTGTCTCAGTCTTTCTCTCCNNNAGATCGGAAGAGC
131868 TATGAGTGTGGAGTCGTTGCATAGAATCTCACTCTGTCACCNNNAGATCGGAAGAG
131869 TATGAGTGTGGAGTCGTTGCGATCATCAATATCACTGTCTTCANNNAGATCGGAAG
131870 TATGAGTGTGGAGTCGTTGCATACAAACATATATGTGTGTGTGNNNAGATCGGAA
131871 TATGAGTGTGGAGTCGTTGCAATTTAACAGCAGACCTTTAAATNNNAGATCGGAAG
131872 TATGAGTGTGGAGTCGTTGCTTTGTTTAGATCTGGCAACTTTNNNAGATCGGAAGA
131873 TATGAGTGTGGAGTCGTTGCACTCTAGCACAGTTAATCTTTATNNNAGATCGGAAG
131874 TATGAGTGTGGAGTCGTTGCCAAAACATATCAAACTCTGAAAAANNNAGATCGGAA
131875 TATGAGTGTGGAGTCGTTGCTAAATGAGATGTTTTGATAGAGGNNNAGATCGGAA
131876 TATGAGTGTGGAGTCGTTGCTAGATGAGACTGGAGACTATTATNNNAGATCGGAA
131877 TATGAGTGTGGAGTCGTTGCATCTTTCTAAAAGAATAGGAACGNNNAGATCGGAA
131878 TATGAGTGTGGAGTCGTTGCGCATGAGATACTATTTTCTACACNNNAGATCGGAAG
131879 TATGAGTGTGGAGTCGTTGCCTTCTCTTGTGTAACCAAAATTANNNAGATCGGAAG
131880 TATGAGTGTGGAGTCGTTGCATATATCAATTCATCAACACAGGNNNAGATCGGAAG
131881 TATGAGTGTGGAGTCGTTGCTTTTCTGTTTGTCTCTTCTGTATNNNAGATCGGAAGA

131882 TATGAGTGTGGAGTCGTTGCTAATGGATGTGACTGCTATAAAGNNNAGATCGGAA
131883 TATGAGTGTGGAGTCGTTGCATAAAATGTACTTCTTACCATGGNNNAGATCGGAAG
131884 TATGAGTGTGGAGTCGTTGCTTTCTAATTTCCAGTCGAAAGATNNNAGATCGGAAG
131885 TATGAGTGTGGAGTCGTTGCTATTGGTTATAGAGTCTTAGCACNNNAGATCGGAAG
131886 TATGAGTGTGGAGTCGTTGCATATTGTAGAGATAGGGTTTCACNNNAGATCGGAA
131887 TATGAGTGTGGAGTCGTTGCCAGTTTTCTCTACAGAGGAATGNNNAGATCGGAAGA
131888 TATGAGTGTGGAGTCGTTGCTGTATGTGTATATACGTGTGTATNNNAGATCGGAAG
131889 TATGAGTGTGGAGTCGTTGCCAGAGACAACATGAGTGAAGNNNAGATCGGAAGA
131890 TATGAGTGTGGAGTCGTTGCAGAGAAAGAGAAAAATGGAGAATNNNAGATCGGA
131891 TATGAGTGTGGAGTCGTTGCCAAACATACATTACAGGAACTATNNNAGATCGGAAG
131892 TATGAGTGTGGAGTCGTTGCCTTAAAGTGTTGGGAATTTTTTGNNNAGATCGGAAG
131893 TATGAGTGTGGAGTCGTTGCTTGACATCACCACAAAACAANNNAGATCGGAAGAGC
131894 TATGAGTGTGGAGTCGTTGCACACTTAACAAACATATTTTGCTNNNAGATCGGAAG
131895 TATGAGTGTGGAGTCGTTGCGGTCAGTCCCATAATAAAACTTNNNAGATCGGAAGA
131896 TATGAGTGTGGAGTCGTTGCTTTTGTTTGTTTGTTTGTTTGTTNNNAGATCGGAAG
131897 TATGAGTGTGGAGTCGTTGCGGAGGGCTATAAACAAGAAAANNNAGATCGGAAG
131898 TATGAGTGTGGAGTCGTTGCAAGCAGAGTAAAAATTCTAACACNNNAGATCGGAA
131899 TATGAGTGTGGAGTCGTTGCCCTGTAATCCCAGCATTTTGNNNAGATCGGAAGAGC
131900 TATGAGTGTGGAGTCGTTGCAAAAGCAGAGTCAAAATTCTAACNNNAGATCGGAA
131901 TATGAGTGTGGAGTCGTTGCGAGATACAGCTCTACAGAATACNNNAGATCGGAAG
131902 TATGAGTGTGGAGTCGTTGCTATTCATCAAAATGGCTCAATCANNNAGATCGGAAG
131903 TATGAGTGTGGAGTCGTTGCTTTCATCTCATTTCAAGCAGAANNNAGATCGGAAGA
131904 TATGAGTGTGGAGTCGTTGCTTGTAGAATGTCTCACATTATGGNNNAGATCGGAAG
131905 TATGAGTGTGGAGTCGTTGCTTTCCTAAGTGATATTGTTCAACNNNAGATCGGAAG
131906 TATGAGTGTGGAGTCGTTGCATACAGTGAAGTAGGACAGAGNNNAGATCGGAAG
131907 TATGAGTGTGGAGTCGTTGCCATCGCTTCTTTCCATGTATNNNAGATCGGAAGAGC
131908 TATGAGTGTGGAGTCGTTGCAAGGATGAGAAGACCAAAAAATANNNAGATCGGAA
131909 TATGAGTGTGGAGTCGTTGCTATAGTAAGCAGCACATAGTAAGNNNAGATCGGAA
131910 TATGAGTGTGGAGTCGTTGCTAGTCATATTAGCATTTTCTCGTNNNAGATCGGAAG
131911 TATGAGTGTGGAGTCGTTGCTTAAGTGACTATTCTACTAAGGCNNNAGATCGGAAG
131912 TATGAGTGTGGAGTCGTTGCTTAGAACTTTATCAAAGAACACCNNNAGATCGGAAG
131913 TATGAGTGTGGAGTCGTTGCAAAATGAAAAATACCTACCTTCCNNNAGATCGGAAG
131914 TATGAGTGTGGAGTCGTTGCAATTAGAACTGGACAAGAATGAANNNAGATCGGAA
131915 TATGAGTGTGGAGTCGTTGCGGGTTCAAACGATTCTCATANNNAGATCGGAAGAG
131916 TATGAGTGTGGAGTCGTTGCAAAGCATGTAAAGCATCTAAATANNNAGATCGGAA
131917 TATGAGTGTGGAGTCGTTGCTCCCATCCTAGTCCAATCATNNNAGATCGGAAGAGC
131918 TATGAGTGTGGAGTCGTTGCAATGTATCCTTTTTCTGAAGAACNNNAGATCGGAAG
131919 TATGAGTGTGGAGTCGTTGCGGCGTAATTTTTCATTTTCAAANNNAGATCGGAAGA
131921 TATGAGTGTGGAGTCGTTGCCCTACACTTGCTAATTAAACTTTNNNAGATCGGAAGA
131922 TATGAGTGTGGAGTCGTTGCCTAAGAATTACTAAGCTTGGAGTNNNAGATCGGAAG
131923 TATGAGTGTGGAGTCGTTGCCTGAGTTACAGCACATTATTTCNNNAGATCGGAAGA
131924 TATGAGTGTGGAGTCGTTGCTACACTATTATGGCAAAATCATGNNNAGATCGGAAG
131925 TATGAGTGTGGAGTCGTTGCGAGTCATTTCATTCTCCATTGNNNAGATCGGAAGAG

131926 TATGAGTGTGGAGTCGTTGCGTTCCTCTTAAGAATTCTGACTANNNAGATCGGAAG
131927 TATGAGTGTGGAGTCGTTGCAAAATGATGAGATGTGATTTCTGNNNAGATCGGAA
131928 TATGAGTGTGGAGTCGTTGCTCCCATCCTAGTCCAATCATNNNAGATCGGAAGAGC
131929 TATGAGTGTGGAGTCGTTGCCGTTTAAACTGAAAATAAGAACCNNNAGATCGGAAG
131930 TATGAGTGTGGAGTCGTTGCCCTTCTATGAATGTAGACTTTTNNNAGATCGGAAG
131931 TATGAGTGTGGAGTCGTTGCTGCAGGTATAGAAAATTATTCCNNNAGATCGGAAG
131933 TATGAGTGTGGAGTCGTTGCAATATTTATTGAGGGCTTACTCANNNAGATCGGAAG
131934 TATGAGTGTGGAGTCGTTGCTGGAAGTCCAAAATAGAAGAAANNNAGATCGGAAG
131935 TATGAGTGTGGAGTCGTTGCTTCTTATATTTTCAGAGATGGGANNNAGATCGGAAG
131936 TATGAGTGTGGAGTCGTTGCTTCCTAGAATTATTGATAGAGCANNNAGATCGGAAG
131937 TATGAGTGTGGAGTCGTTGCCATCATAGAAGACTGATAGACAANNNAGATCGGAA
131938 TATGAGTGTGGAGTCGTTGCAGCTCTCAGCAGTAAACTTANNNAGATCGGAAGAG
131939 TATGAGTGTGGAGTCGTTGCTAATTTAGCACATTGTTTGAAAGNNNAGATCGGAAG
131940 TATGAGTGTGGAGTCGTTGCAGCATGAATAACAAAACGATTAANNNAGATCGGAA
131941 TATGAGTGTGGAGTCGTTGCCTGAAAAAGGGCTTAAGAAAGNNNAGATCGGAAGA
131942 TATGAGTGTGGAGTCGTTGCGAGAGAAAGAGAGAGAGAGAGNNNAGATCGGAA
131943 TATGAGTGTGGAGTCGTTGCCATTTGAACTGAGTTTGGAAANNNAGATCGGAAGA
131944 TATGAGTGTGGAGTCGTTGCCTTCAACTTTTATTACCTTCTGTNNNAGATCGGAAGA
131945 TATGAGTGTGGAGTCGTTGCATTCCCATGTAGTATCCAAATAANNNAGATCGGAAG
131946 TATGAGTGTGGAGTCGTTGCGTCTCCATGGCAAATTTATTATANNNAGATCGGAAG
131947 TATGAGTGTGGAGTCGTTGCCTAAAAATTTCAGTGTCTCACTTNNNAGATCGGAAGA
131948 TATGAGTGTGGAGTCGTTGCTTTGTTATCTCACACTAGCTATTNNNAGATCGGAAGA
131949 TATGAGTGTGGAGTCGTTGCATAACCAAGCCTTAACTATACTTNNNAGATCGGAAGA
131950 TATGAGTGTGGAGTCGTTGCTGTAGAAAAAATGGAAAATGTCTNNNAGATCGGAA
131952 TATGAGTGTGGAGTCGTTGCCAATGCCTCTTTAGTTTTTTATTNNNAGATCGGAAGA
131953 TATGAGTGTGGAGTCGTTGCCTTTGGGAAATCACATGGTANNNAGATCGGAAGAG
131954 TATGAGTGTGGAGTCGTTGCAAACAACTCACAGAACAACAANNNAGATCGGAAGAG
131955 TATGAGTGTGGAGTCGTTGCTATATATTTTGAGACACGGTCTTNNNAGATCGGAAG
131956 TATGAGTGTGGAGTCGTTGCTGGATGGAATTGAAGACTATTATNNNAGATCGGAA
131957 TATGAGTGTGGAGTCGTTGCTGAATACCTATATGATTTGCAAGNNNAGATCGGAAG
131958 TATGAGTGTGGAGTCGTTGCACTCAGACATAAAAAGAAATGAANNNAGATCGGAA
131959 TATGAGTGTGGAGTCGTTGCTTTAGAGGTGATAAGAACAGTCNNNAGATCGGAAG
131960 TATGAGTGTGGAGTCGTTGCATGTGGTAAACTGTTAACATTTTNNNAGATCGGAAG
131961 TATGAGTGTGGAGTCGTTGCAAAGTCCTCATCTTTGACTATTANNNAGATCGGAAGA
131962 TATGAGTGTGGAGTCGTTGCCCCGAAAGTATTGGAATTCANNNAGATCGGAAGAG
131963 TATGAGTGTGGAGTCGTTGCTGGTCTGTGTATAGTTTTCATTANNNAGATCGGAAG
131964 TATGAGTGTGGAGTCGTTGCGGTATTGGAATTATCAAACAGAANNNAGATCGGAA
131965 TATGAGTGTGGAGTCGTTGCAAATATTCCAGATGGTTTTGCTNNNAGATCGGAAGA
131966 TATGAGTGTGGAGTCGTTGCGTTTTGGTTTAGGGAAAAGTAANNNAGATCGGAAG
131967 TATGAGTGTGGAGTCGTTGCTCTTTCTTTCTTTTTTTGAGACGNNNAGATCGGAAGA
131968 TATGAGTGTGGAGTCGTTGCGAAATGGAATCCATCCTTAATTCNNNAGATCGGAAG
131970 TATGAGTGTGGAGTCGTTGCAATACCTAAAAAGGAACAAAAGGNNNAGATCGGAA
131971 TATGAGTGTGGAGTCGTTGCGGTGGTTCAGAATATCCAACNNNAGATCGGAAGAG

131972 TATGAGTGTGGAGTCGTTGCCATTCAAAAAAGGGAAGCTGNNNAGATCGGAAGAG
131973 TATGAGTGTGGAGTCGTTGCCTCACTACCACATATCTCTCANNNAGATCGGAAGAGC
131974 TATGAGTGTGGAGTCGTTGCTACATACAGAAGTGGGATGGNNNAGATCGGAAGAG
131975 TATGAGTGTGGAGTCGTTGCCTACATATGATTTCAACTTAGCTNNNAGATCGGAAGA
131976 TATGAGTGTGGAGTCGTTGCTTTTTAGATATAGAGTCTTGCTCNNNAGATCGGAAGA
131977 TATGAGTGTGGAGTCGTTGCATTAGAGAGCCAAAGAAATAATGNNNAGATCGGAA
131978 TATGAGTGTGGAGTCGTTGCAAAATATTAATTTGGCAGATCCANNNAGATCGGAAG
131979 TATGAGTGTGGAGTCGTTGCCTTGAAACTTTCACAACACTCNNNAGATCGGAAGAG
131980 TATGAGTGTGGAGTCGTTGCCATGTACAAGAATGCTCTTTATANNNAGATCGGAAG
131981 TATGAGTGTGGAGTCGTTGCGTTTTATTTTTCCTTCAATGCTTNNNAGATCGGAAGA
131982 TATGAGTGTGGAGTCGTTGCAGATAGGAACATGGTGATTTATANNNAGATCGGAA
131983 TATGAGTGTGGAGTCGTTGCTCTAGTACTAGCTATTCAGGAAGNNNAGATCGGAAG
131986 TATGAGTGTGGAGTCGTTGCCATAATGATGTGGTCAACAATGNNNAGATCGGAAGA
131988 TATGAGTGTGGAGTCGTTGCATACCTCATCTCCATCACACNNNAGATCGGAAGAGCA
131989 TATGAGTGTGGAGTCGTTGCAGAAAGGAAAACTCCATTTTATCNNNAGATCGGAAG
131990 TATGAGTGTGGAGTCGTTGCGTTAATATTCATTGTTGTTGGTGNNNAGATCGGAAG
131991 TATGAGTGTGGAGTCGTTGCTAAACTTCCTCAACTTGATTTTTNNNAGATCGGAAGA
131992 TATGAGTGTGGAGTCGTTGCTTTCTCTGGTGAACCCTAATATANNNAGATCGGAAGA
131993 TATGAGTGTGGAGTCGTTGCCCATAAAAAAGAATGTGATCATGNNNAGATCGGAA
131994 TATGAGTGTGGAGTCGTTGCTTGCAGGTTCAAGTAATTCTCNNNAGATCGGAAGAG
131995 TATGAGTGTGGAGTCGTTGCTCTCCTGAGCTATATAACTTAGANNNAGATCGGAAGA
131996 TATGAGTGTGGAGTCGTTGCTCAGAAAATCGTCCTAACAATNNNAGATCGGAAGAG
131998 TATGAGTGTGGAGTCGTTGCGTTTCTCTCAGGTCTAACATTANNNAGATCGGAAGA
131999 TATGAGTGTGGAGTCGTTGCGGTACTAGCCACTCTTAGAGNNNAGATCGGAAGAG
132000 TATGAGTGTGGAGTCGTTGCCCCGTTTCATTAACTTATTGTTNNNAGATCGGAAGAG
132001 TATGAGTGTGGAGTCGTTGCCTGAGATCAGGATTTCAAGATCNNNAGATCGGAAGA
132002 TATGAGTGTGGAGTCGTTGCAAAGACTAAGAACTGCTACTTTANNNAGATCGGAAG
132003 TATGAGTGTGGAGTCGTTGCGCAATTGATTATCTTGGTAAGTANNNAGATCGGAAG
132004 TATGAGTGTGGAGTCGTTGCTTAACTATAGTCATCATGTTGCTNNNAGATCGGAAGA
132005 TATGAGTGTGGAGTCGTTGCTCATTTTAAATGTAGAAGGAAGGNNNAGATCGGAA
132006 TATGAGTGTGGAGTCGTTGCAGAAAAAGGATCTTGTTATGTTGNNNAGATCGGAA
132007 TATGAGTGTGGAGTCGTTGCACATATTTTCAAGTAGAGGAGACNNNAGATCGGAA
132008 TATGAGTGTGGAGTCGTTGCATTTTACAACTAAACAACCAAGTNNNAGATCGGAAGA
132009 TATGAGTGTGGAGTCGTTGCGAAGCCAATATTAGTACAACAATNNNAGATCGGAAG
132010 TATGAGTGTGGAGTCGTTGCCTCAGAAGAGTAATGTGACAGNNNAGATCGGAAGA
132011 TATGAGTGTGGAGTCGTTGCTGTTTTCCATAGTTGTACTAGTTNNNAGATCGGAAGA
132012 TATGAGTGTGGAGTCGTTGCGGAGTCAGAAAAGTCTTTACTNNNAGATCGGAAGA
132013 TATGAGTGTGGAGTCGTTGCAAAGCAAACAAGCACATAATTGNNNAGATCGGAAGA
132014 TATGAGTGTGGAGTCGTTGCATGTTAAGCTTCTATTTGGGATANNNAGATCGGAAG
132015 TATGAGTGTGGAGTCGTTGCGTGACTCTAATTACTGTTTGTTTNNNAGATCGGAAGA
132016 TATGAGTGTGGAGTCGTTGCCTAATACAAGCACCTTTCAATCNNNAGATCGGAAGA
132017 TATGAGTGTGGAGTCGTTGCTTATTTTCATTCCACACATCCTANNNAGATCGGAAGA
132018 TATGAGTGTGGAGTCGTTGCAGAACTTACAGCCTATAAGAAAGNNNAGATCGGAA

132019 TATGAGTGTGGAGTCGTTGCCTGAGGAAGAAAAAATTCAGTCNNNAGATCGGAAG
132020 TATGAGTGTGGAGTCGTTGCTCTTTCTTGAAAATATTACTGGANNNAGATCGGAAG
132021 TATGAGTGTGGAGTCGTTGCTGTGGTAGATATGAATTTACTCANNNAGATCGGAAG
132022 TATGAGTGTGGAGTCGTTGCAATTTATTATGAGAAATCTGCCCNNNAGATCGGAAG
132023 TATGAGTGTGGAGTCGTTGCCTTAAACCTTCTTCCATATGGANNNAGATCGGAAGA
132024 TATGAGTGTGGAGTCGTTGCACTTTTGTTTTCTTGTTTTGTTTNNNAGATCGGAAGA
132025 TATGAGTGTGGAGTCGTTGCACAACAAAATACTTCAGCTTTATNNNAGATCGGAAG
132026 TATGAGTGTGGAGTCGTTGCACAAGCTCAAATTTTAAATGATTNNNAGATCGGAAG
132027 TATGAGTGTGGAGTCGTTGCAGAAAGGAAAAACATAGCTTTACNNNAGATCGGAA
132028 TATGAGTGTGGAGTCGTTGCCAAGAGTTGTTTTTTTGAAAAAANNNAGATCGGAAG
132029 TATGAGTGTGGAGTCGTTGCACTATTGTGAAAACAACTTCTTTNNNAGATCGGAAG
132030 TATGAGTGTGGAGTCGTTGCAAGAGAGAGAGAGACAGAGANNNAGATCGGAAGA
132031 TATGAGTGTGGAGTCGTTGCCAATCTCAGCTCACTACAACNNNAGATCGGAAGAGC
132032 TATGAGTGTGGAGTCGTTGCGTTTTTCTGATACCATCATCTTTNNNAGATCGGAAGA
132033 TATGAGTGTGGAGTCGTTGCAATGAGGACTAGAAGGAAAATAANNNAGATCGGAA
132034 TATGAGTGTGGAGTCGTTGCTAGCTGTCCTCATTTTTTTTTTTNNNAGATCGGAAGA
132035 TATGAGTGTGGAGTCGTTGCGTTTGAGTCATTTTCATCTTTTGNNNAGATCGGAAG
132036 TATGAGTGTGGAGTCGTTGCCCTATTTTTATTTGTGTCCTCTTNNNAGATCGGAAGA
132037 TATGAGTGTGGAGTCGTTGCAGTAGGAGCCTCTAGTTTACNNNAGATCGGAAGAG
132038 TATGAGTGTGGAGTCGTTGCTTTTTAGGAAAAAACAAAATGGTNNNAGATCGGAA
132039 TATGAGTGTGGAGTCGTTGCGATTAAGTTTCAGAGAACGAAAGNNNAGATCGGAA
132040 TATGAGTGTGGAGTCGTTGCTAATAGGACTACTCTGAGAGTTANNNAGATCGGAA
132041 TATGAGTGTGGAGTCGTTGCCAAGCAATCCTCTTACTTCANNNAGATCGGAAGAGC
132042 TATGAGTGTGGAGTCGTTGCGTTAAAATGCTGTCTCTGATTTANNNAGATCGGAAG
132043 TATGAGTGTGGAGTCGTTGCAATGGATGGCTTTTAGAAAAAGNNNAGATCGGAAG
132045 TATGAGTGTGGAGTCGTTGCTTGTTTTGTTTTTGTTTTTTTGAGNNNAGATCGGAAG
132046 TATGAGTGTGGAGTCGTTGCCATTAATTACCAAAGACTCTGACNNNAGATCGGAAG
132047 TATGAGTGTGGAGTCGTTGCCTCAGATATGCCTGTCTTTTTANNNAGATCGGAAGA
132048 TATGAGTGTGGAGTCGTTGCCATTAGAAGAGCTGTAAGAGTAGNNNAGATCGGAA
132049 TATGAGTGTGGAGTCGTTGCGAGAGAAGAGACAGATTGATTTNNNAGATCGGAAG
132050 TATGAGTGTGGAGTCGTTGCCTTCTGGCTTTTTTTTTTGTTTANNNAGATCGGAAGA
132051 TATGAGTGTGGAGTCGTTGCAGAGAACCCTGACTAATATAGATNNNAGATCGGAA
132052 TATGAGTGTGGAGTCGTTGCGCAGTTGTCAGAGTTCTTACNNNAGATCGGAAGAG
132053 TATGAGTGTGGAGTCGTTGCAAGGGAGAGAAAGAAAGAAAAGNNNAGATCGGAA
132054 TATGAGTGTGGAGTCGTTGCTCCCTACCATCATACAACTATTANNNAGATCGGAAGA
132055 TATGAGTGTGGAGTCGTTGCCAACCTCTACCTTATGGATTCNNNAGATCGGAAGAG
132056 TATGAGTGTGGAGTCGTTGCGAGAACTCTGACTAATATACTACANNNAGATCGGAA
132057 TATGAGTGTGGAGTCGTTGCCATTTCAAAAATTGTAGCACATTANNNAGATCGGAA
132058 TATGAGTGTGGAGTCGTTGCGAATGGACATAATCACATCAATANNNAGATCGGAAG
132059 TATGAGTGTGGAGTCGTTGCAATTTGTTAATATCACCACAGCNNNAGATCGGAAGA
132060 TATGAGTGTGGAGTCGTTGCATATGAATAGGCTTGTCATACATNNNAGATCGGAAG
132061 TATGAGTGTGGAGTCGTTGCATTATCTCCTAGATGTTTGGTTTNNNAGATCGGAAG
132062 TATGAGTGTGGAGTCGTTGCTCTCCCATATCCTTGATTATTTTNNNAGATCGGAAGA

132063 TATGAGTGTGGAGTCGTTGCAATATCCATCACATAGATAACCCNNNAGATCGGAAG
132064 TATGAGTGTGGAGTCGTTGCATCAACCATGTCAATATTAACTGNNNAGATCGGAAG
132066 TATGAGTGTGGAGTCGTTGCCTCCTTTGAAGAGGTTCTTAGNNNAGATCGGAAGAG
132067 TATGAGTGTGGAGTCGTTGCGATAATTTCCTGATTCCAGTTTCNNNAGATCGGAAG
132068 TATGAGTGTGGAGTCGTTGCTTCTGATAAGTGAGATCAAGTTANNNAGATCGGAA
132069 TATGAGTGTGGAGTCGTTGCCCTTCTGTTCTAATGGTCACNNNAGATCGGAAGAGC
132070 TATGAGTGTGGAGTCGTTGCAAAGAATTTGAACAAAAACTGACNNNAGATCGGAA
132071 TATGAGTGTGGAGTCGTTGCATTTTGTTTGTTTGTTTGTTTGTNNNAGATCGGAAG
132072 TATGAGTGTGGAGTCGTTGCAAGTTAAAATAACCAATTTGGGTNNNAGATCGGAA
132073 TATGAGTGTGGAGTCGTTGCCACAAAGTCTTTCTTTACAAAACNNNAGATCGGAAG
132074 TATGAGTGTGGAGTCGTTGCATAAACCCTTCTCTTCTCCANNNAGATCGGAAGAGCA
132075 TATGAGTGTGGAGTCGTTGCAGAGTGAATATCATACTTAATGGTNNNAGATCGGA
132076 TATGAGTGTGGAGTCGTTGCTTTTTAAATGCTCTGGTATCCAANNNAGATCGGAAG
132077 TATGAGTGTGGAGTCGTTGCGAGAGAGCAAGCTTCTATATATTNNNAGATCGGAA
132078 TATGAGTGTGGAGTCGTTGCCCGTATTAGTGATTTTTGACTTTNNNAGATCGGAAG
132079 TATGAGTGTGGAGTCGTTGCTCTATGCCACATTTTCTTTATTCNNNAGATCGGAAGA
132080 TATGAGTGTGGAGTCGTTGCCTGAGGTCACCTAAAAAAAATTTNNNAGATCGGAAG
132081 TATGAGTGTGGAGTCGTTGCAGATTCTAGTTCTCAAATGAGTTNNNAGATCGGAAG
132082 TATGAGTGTGGAGTCGTTGCACATTAATGTGCATTCTGGCNNNAGATCGGAAGAGC
132083 TATGAGTGTGGAGTCGTTGCGAGAGATCTGTGCTTCATTTANNNAGATCGGAAGA
132084 TATGAGTGTGGAGTCGTTGCGTTCCCTATATAAAATGGCAAAANNNAGATCGGAAG
132085 TATGAGTGTGGAGTCGTTGCCATGGATCTCTAAAGTTGTCTNNNAGATCGGAAGAG
132086 TATGAGTGTGGAGTCGTTGCTGAGGAAACTTACCTTTAAAAAATNNNAGATCGGAA
132087 TATGAGTGTGGAGTCGTTGCGGAACTAATTAGAAATGGTGATGNNNAGATCGGAA
132089 TATGAGTGTGGAGTCGTTGCATCCATGTTTAGGAAAATTTCAANNNAGATCGGAAG
132090 TATGAGTGTGGAGTCGTTGCCAATAGTTATCACGAAATCCAAGNNNAGATCGGAAG
132091 TATGAGTGTGGAGTCGTTGCATAGATTGTATACCAATGACCAANNNAGATCGGAAG
132092 TATGAGTGTGGAGTCGTTGCCAAATCAAAACTGAGCAGAAGNNNAGATCGGAAGA
132093 TATGAGTGTGGAGTCGTTGCTTGTAAATCACTCTTTACTGTTGNNNAGATCGGAAG
132094 TATGAGTGTGGAGTCGTTGCGTGTAAGTAATCAAATCCATCAANNNAGATCGGAAG
132095 TATGAGTGTGGAGTCGTTGCAAAGACAAGATAATGTTCAACTTNNNAGATCGGAAG
132096 TATGAGTGTGGAGTCGTTGCTTGTGGGGACATAATTTTATTTTNNNAGATCGGAAG
132097 TATGAGTGTGGAGTCGTTGCTACCTAAGTTGATTACATGTCTTNNNAGATCGGAAG
132098 TATGAGTGTGGAGTCGTTGCTATATTCTTTTTGAGACAGAATCTNNNAGATCGGAA
132099 TATGAGTGTGGAGTCGTTGCCCTTGCACTTTTGTGTTATGNNNAGATCGGAAGAGC
132100 TATGAGTGTGGAGTCGTTGCAAAGAAACAGAGACAGCAATTTNNNAGATCGGAAG
132101 TATGAGTGTGGAGTCGTTGCAAATTCAAAGACTTTGCAAGAANNNAGATCGGAAG
132102 TATGAGTGTGGAGTCGTTGCGCTATACAAGTCTGGAATTTTAGNNNAGATCGGAAG
132103 TATGAGTGTGGAGTCGTTGCCTAAAAGGGCTCTCAGTAATTNNNAGATCGGAAGA
132104 TATGAGTGTGGAGTCGTTGCACAGTTTCTTTATCCATTCTTTGNNNAGATCGGAAGA
132105 TATGAGTGTGGAGTCGTTGCAATGTAGACTGACTGTACTTTTTNNNAGATCGGAAG
132106 TATGAGTGTGGAGTCGTTGCTTTTTAAATGAGGTGGAGTCTTCNNNAGATCGGAAG
132107 TATGAGTGTGGAGTCGTTGCAGATGAAACTTGAGGACATTATANNNAGATCGGAA

132108 TATGAGTGTGGAGTCGTTGCTTGTGTTTAATATAGAGACAGGGNNNAGATCGGAA
132109 TATGAGTGTGGAGTCGTTGCACAGAGTCTCAATCTACTGCNNNAGATCGGAAGAGC
132110 TATGAGTGTGGAGTCGTTGCCATGTTCGAGCTATTCTCCTNNNAGATCGGAAGAGC
132111 TATGAGTGTGGAGTCGTTGCGAGAGAGGGAAAAGGTATATTAGNNNAGATCGGA
132112 TATGAGTGTGGAGTCGTTGCAATTTCTGTTCTAAATTACTCCGNNNAGATCGGAAGA
132113 TATGAGTGTGGAGTCGTTGCTAAAAGAAAAAAAATTTGGCCANNNAGATCGGAA
132114 TATGAGTGTGGAGTCGTTGCTACACACACACATACAGTTAGNNNAGATCGGAAGAG
132115 TATGAGTGTGGAGTCGTTGCAAGCTTAGAGATTAAGAATAGCANNNAGATCGGAA
132116 TATGAGTGTGGAGTCGTTGCATAAGACTCTCTAAAGCATAGAGNNNAGATCGGAA
132117 TATGAGTGTGGAGTCGTTGCTAATTACTAAGTTTTCCTGCTTCNNNAGATCGGAAGA
132118 TATGAGTGTGGAGTCGTTGCCAGTACCTCATCTTCTTATAGTTNNNAGATCGGAAGA
132119 TATGAGTGTGGAGTCGTTGCCATGATCATGGCTCACTATAGNNNAGATCGGAAGAG
132120 TATGAGTGTGGAGTCGTTGCCTTGCCTGTTTGAGACTTTTNNNAGATCGGAAGAGC
132121 TATGAGTGTGGAGTCGTTGCATGGTAAATCATTTTCTCCTACTNNNAGATCGGAAGA
132122 TATGAGTGTGGAGTCGTTGCCTTGTAATCCCAGCTTCTTGNNNAGATCGGAAGAGC
132123 TATGAGTGTGGAGTCGTTGCTACTTCTTCAAATACTTCTAGGGNNNAGATCGGAAG
132124 TATGAGTGTGGAGTCGTTGCTGGTAATTCTGTCAATTTACTTTNNNAGATCGGAAG
132125 TATGAGTGTGGAGTCGTTGCTTTCTCTTAGAATTGCCTTTGTNNNAGATCGGAAGAG
132126 TATGAGTGTGGAGTCGTTGCAATTTCCTCACTATTTCTTACCTNNNAGATCGGAAGA
132127 TATGAGTGTGGAGTCGTTGCAACAAAAAAACAAAAAAAACCTCNNNAGATCGGAAG
132128 TATGAGTGTGGAGTCGTTGCGTAGGCAGAACTCTTTTATCANNNAGATCGGAAGAG
132129 TATGAGTGTGGAGTCGTTGCATTACTTGATTAATGTCTCTCCCNNNAGATCGGAAGA
132130 TATGAGTGTGGAGTCGTTGCAACTTACATTGTAATGGTGATGANNNAGATCGGAAG
132131 TATGAGTGTGGAGTCGTTGCCCCTCTGTTTTTTAGAAATTCTANNNAGATCGGAAGA
132132 TATGAGTGTGGAGTCGTTGCTCCTTTTAAATTATTTCCTTAGGANNNAGATCGGAAG
132133 TATGAGTGTGGAGTCGTTGCCTGGGATCCCTTGAGATTAANNNAGATCGGAAGAG
132134 TATGAGTGTGGAGTCGTTGCCTCTATGGCACATTGGATTCNNNAGATCGGAAGAGC
132135 TATGAGTGTGGAGTCGTTGCATATATTACTAGTCCATTGCTGCNNNAGATCGGAAG
132136 TATGAGTGTGGAGTCGTTGCAATTGTGTGTAGTTTATTGTCTGNNNAGATCGGAAG
132137 TATGAGTGTGGAGTCGTTGCAATACAATGAAATATGAGTCAGCNNNAGATCGGAA
132138 TATGAGTGTGGAGTCGTTGCCTCTTAAAGCTGTAGACTATGTCNNNAGATCGGAAG
132139 TATGAGTGTGGAGTCGTTGCTTTTGTATACACTGAAACTAAGCNNNAGATCGGAAG
132140 TATGAGTGTGGAGTCGTTGCATCAAGAATTAAGTGTATTCTGGNNNAGATCGGAA
132141 TATGAGTGTGGAGTCGTTGCGTATCTCACTGTTTCCTGAAANNNAGATCGGAAGAG
132142 TATGAGTGTGGAGTCGTTGCGCATACTAATAGGTCTTTTTAGCNNNAGATCGGAAG
132143 TATGAGTGTGGAGTCGTTGCCCATAATACCATTTTGTACTTCCNNNAGATCGGAAGA
132144 TATGAGTGTGGAGTCGTTGCAAAGGACCAAATATGTGATTATGNNNAGATCGGAA
132145 TATGAGTGTGGAGTCGTTGCACTAGAAATAATAATTTGTCACCTNNNAGATCGGAA
132146 TATGAGTGTGGAGTCGTTGCCCCATTAAAATTCACATACCTTTNNNAGATCGGAAGA
132147 TATGAGTGTGGAGTCGTTGCTATTACTCAGCCTTAAAAAAGGANNNAGATCGGAAG
132148 TATGAGTGTGGAGTCGTTGCTTATGATCTCTCTTAGCAACTTTNNNAGATCGGAAGA
132149 TATGAGTGTGGAGTCGTTGCAACATATGTATTGAAACTTCACTNNNAGATCGGAAG
132150 TATGAGTGTGGAGTCGTTGCAATTTAAGTGAAAATAAGTGGGGNNNAGATCGGAA

132151 TATGAGTGTGGAGTCGTTGCTGCTAAGAAATTATACTCAAGGTNNNAGATCGGAAG
132152 TATGAGTGTGGAGTCGTTGCAAGATTGAGACACTTAATGGCNNNAGATCGGAAGA
132153 TATGAGTGTGGAGTCGTTGCCTATCCATTTGTATCCTCAATTATNNNAGATCGGAAG
132154 TATGAGTGTGGAGTCGTTGCCTTTATACTAGGGATTTAGGGTGNNNAGATCGGAA
132155 TATGAGTGTGGAGTCGTTGCTGATAGTGGGTTAAGTTCTTTCNNNAGATCGGAAG
132156 TATGAGTGTGGAGTCGTTGCAAAATTATAGGGAGGGAGATAACNNNAGATCGGA
132157 TATGAGTGTGGAGTCGTTGCCACAGCATCTACCAAATTTTATTNNNAGATCGGAAG
132158 TATGAGTGTGGAGTCGTTGCTCCTATATTTTTTCTGGGGAACNNNAGATCGGAAGA
132159 TATGAGTGTGGAGTCGTTGCTTAGTGACTGGTTGGAATTTTNNNAGATCGGAAGA
132160 TATGAGTGTGGAGTCGTTGCGATATCTGCTGTTAGTGTTTTGNNNAGATCGGAAGA
132161 TATGAGTGTGGAGTCGTTGCTGTAACAGTTAAGAGAATCTACTNNNAGATCGGAAG
132162 TATGAGTGTGGAGTCGTTGCAAATGTTCCTGAAGATCTCTAGNNNAGATCGGAAGA
132164 TATGAGTGTGGAGTCGTTGCCATTTTCTTTAAAGACCTCCTTTNNNAGATCGGAAGA
132165 TATGAGTGTGGAGTCGTTGCACCTTACATAAGTCACCTAATACNNNAGATCGGAAG
132166 TATGAGTGTGGAGTCGTTGCATCAGTTATGGGTATGTATTACANNNAGATCGGAAG
132167 TATGAGTGTGGAGTCGTTGCCTGAGACAGGTGGATTATCTNNNAGATCGGAAGAG
132168 TATGAGTGTGGAGTCGTTGCCCAAGTCCATTATTATCAGGAGNNNAGATCGGAAGA
132169 TATGAGTGTGGAGTCGTTGCACATCTAGCATTAAACCTAATCTNNNAGATCGGAAG
132171 TATGAGTGTGGAGTCGTTGCCATATTTTGATAGGTGCTTTTCTNNNAGATCGGAAG
132173 TATGAGTGTGGAGTCGTTGCGAAAAACAATGACGATATAAAGCNNNAGATCGGAA
132174 TATGAGTGTGGAGTCGTTGCCTCCCTGCAAAAATCTAATAATTNNNAGATCGGAAG
132175 TATGAGTGTGGAGTCGTTGCCAATAATCAATCTGCGAAATGANNNAGATCGGAAG
132176 TATGAGTGTGGAGTCGTTGCTATGTACACTAGTTGAAATGTGANNNAGATCGGAAG
132177 TATGAGTGTGGAGTCGTTGCCAGCCAAAATATTTTATAGGCTTNNNAGATCGGAAG
132178 TATGAGTGTGGAGTCGTTGCGGCATGATCATATCTCACTATAGNNNAGATCGGAAG
132179 TATGAGTGTGGAGTCGTTGCGTGGATACTGAAATCTATGGGNNNAGATCGGAAGA
132180 TATGAGTGTGGAGTCGTTGCGAACTTATCCTTAAATTTGTCCANNNAGATCGGAAG
132181 TATGAGTGTGGAGTCGTTGCGTCAGATCTCTCCTTGAATATGNNNAGATCGGAAGA
132182 TATGAGTGTGGAGTCGTTGCTCTGTCACTAATTTTGGAAATTTNNNAGATCGGAAG
132183 TATGAGTGTGGAGTCGTTGCAAATAGAGATTCCAGAAAACAGANNNAGATCGGAA
132184 TATGAGTGTGGAGTCGTTGCCTGTGGTTCTAGCTACTCAANNNAGATCGGAAGAGC
132185 TATGAGTGTGGAGTCGTTGCCTTTTTCTTGGTCTCAGTAGTANNNAGATCGGAAGA
132186 TATGAGTGTGGAGTCGTTGCACTAACTAATATCTCTGATAAGCANNNAGATCGGAA
132187 TATGAGTGTGGAGTCGTTGCTTCTCTAACACTTGGTAGTAAAANNNAGATCGGAAG
132188 TATGAGTGTGGAGTCGTTGCAATGATTTCTAGACCTTTGTTTCNNNAGATCGGAAG
132189 TATGAGTGTGGAGTCGTTGCATTTTAGATGGAGTTTTGCTCNNNAGATCGGAAGAG
132190 TATGAGTGTGGAGTCGTTGCTGGTTTGTGAAAAATGAAGAATANNNAGATCGGAA
132191 TATGAGTGTGGAGTCGTTGCTCACATTTTAAAAAGTCCATTCANNNAGATCGGAAG
132192 TATGAGTGTGGAGTCGTTGCAAGACAGAGAACCAATTATCGNNNAGATCGGAAGA
132193 TATGAGTGTGGAGTCGTTGCCAGGGAGGAATAACTGACTTNNNAGATCGGAAGAG
132194 TATGAGTGTGGAGTCGTTGCCAAGCTATTCTAAGAACAGTTTCNNNAGATCGGAAG
132195 TATGAGTGTGGAGTCGTTGCAGTCAAGATACAGAACAATTCTNNNAGATCGGAAGA
132196 TATGAGTGTGGAGTCGTTGCATGCTTCAGAATACGTAAAAAATNNNAGATCGGAAG

132197 TATGAGTGTGGAGTCGTTGCAGTGGTATTTCCTTAACTAACAANNNAGATCGGAAG
132198 TATGAGTGTGGAGTCGTTGCCGCTATTGATTGATAAACATTTGNNNAGATCGGAAG
132199 TATGAGTGTGGAGTCGTTGCATTTCTGCTAGATCACAATTCTNNNAGATCGGAAGAG
132200 TATGAGTGTGGAGTCGTTGCGAAGAAAGTTGTACATTCTGAAANNNAGATCGGAA
132201 TATGAGTGTGGAGTCGTTGCAAATAAGACTTTTTAGGAAGGTGNNNAGATCGGAA
132202 TATGAGTGTGGAGTCGTTGCTTATAAGCGGCCCTATAGAGNNNAGATCGGAAGAG
132203 TATGAGTGTGGAGTCGTTGCGCTTAAACAACAGAAATGTATTGNNNAGATCGGAAG
132204 TATGAGTGTGGAGTCGTTGCCACATTCTTTTTAATGGCTTTATNNNAGATCGGAAGA
132205 TATGAGTGTGGAGTCGTTGCGATATAAGCTCTCTCTACATACANNNAGATCGGAAG
132206 TATGAGTGTGGAGTCGTTGCGAGAAATGAAGTTCTTTAACAGTNNNAGATCGGAA
132207 TATGAGTGTGGAGTCGTTGCCTTTTTATAACCCAGATTCCTGNNNAGATCGGAAGAG
132208 TATGAGTGTGGAGTCGTTGCAAATATGCAAGGAAATAAACCAANNNAGATCGGAA
132209 TATGAGTGTGGAGTCGTTGCATCATCACCCTTGAATGTTTCNNNAGATCGGAAGAG
132210 TATGAGTGTGGAGTCGTTGCGAGAAGGAAAATGATATGAAGACNNNAGATCGGA
132211 TATGAGTGTGGAGTCGTTGCTTATTTAGTTATTTTGAGGCAGGNNNAGATCGGAAG
132212 TATGAGTGTGGAGTCGTTGCGGTATGATCTCAGTTCACTGNNNAGATCGGAAGAG
132213 TATGAGTGTGGAGTCGTTGCGTTATAAACCCTTTTTCCTTCCNNNAGATCGGAAGAG
132214 TATGAGTGTGGAGTCGTTGCTAAGATAATGAAAGGGTGCTTAGNNNAGATCGGAA
132215 TATGAGTGTGGAGTCGTTGCATGTTAATGTTCACATTCCATTTNNNAGATCGGAAGA
132216 TATGAGTGTGGAGTCGTTGCTGTATATAAAAAGAGGTTTTGCCNNNAGATCGGAAG
132217 TATGAGTGTGGAGTCGTTGCCCTATTATTGTAAAACTGCTGAANNNAGATCGGAAG
132218 TATGAGTGTGGAGTCGTTGCCTATTTAGACTCTGTTTTGGTTCNNNAGATCGGAAG
132219 TATGAGTGTGGAGTCGTTGCAATTAACAAATTTGACTGGAGAANNNAGATCGGAA
132220 TATGAGTGTGGAGTCGTTGCGTTTTCGAAACAGAGTCTTGNNNAGATCGGAAGAG
132221 TATGAGTGTGGAGTCGTTGCAATGTTGGCTCACTGTAATCNNNAGATCGGAAGAGC
132222 TATGAGTGTGGAGTCGTTGCATTCTCCGTTGAAACAGATCNNNAGATCGGAAGAGC
132223 TATGAGTGTGGAGTCGTTGCATCAGAAGTGGAAAATTCATTTTNNNAGATCGGAAG
132224 TATGAGTGTGGAGTCGTTGCCTCCCATTTATTGTTGTAACAGNNNAGATCGGAAGA
132225 TATGAGTGTGGAGTCGTTGCATTGGAGACGATTATTCTGAGTANNNAGATCGGAA
132226 TATGAGTGTGGAGTCGTTGCGTTAAAAATGCGTATGCTCTATANNNAGATCGGAAG
132227 TATGAGTGTGGAGTCGTTGCAGGAAATCTAGTGCTAAATACAANNNAGATCGGAA
132229 TATGAGTGTGGAGTCGTTGCTCTACTAACATGTTTAGCAAATGNNNAGATCGGAAG
132230 TATGAGTGTGGAGTCGTTGCCAAATAGTACACCAGCTCATATNNNAGATCGGAAGA
132232 TATGAGTGTGGAGTCGTTGCGATTTATTTGAGTCCTTGAGAAANNNAGATCGGAAG
132233 TATGAGTGTGGAGTCGTTGCCAGTAGAAAGGCATTTTTCATCNNNAGATCGGAAGA
132234 TATGAGTGTGGAGTCGTTGCGCTAATAAAATCATTTCCTGGTTNNNAGATCGGAAG
132236 TATGAGTGTGGAGTCGTTGCTTACATTTAACAAGTGAGGAAACNNNAGATCGGAA
132237 TATGAGTGTGGAGTCGTTGCGGCATTTGTATATGTAGATATGGNNNAGATCGGAA
132238 TATGAGTGTGGAGTCGTTGCGATACAGTAAACTCTGATACAGANNNAGATCGGAA
132239 TATGAGTGTGGAGTCGTTGCTTTCATGGAAGAAGTAACCATTNNNAGATCGGAAGA
132240 TATGAGTGTGGAGTCGTTGCTGAATACATCAAGCACATTTTTCNNNAGATCGGAAG
132241 TATGAGTGTGGAGTCGTTGCGGATGGAATTAGAGACTATCATTNNNAGATCGGAA
132242 TATGAGTGTGGAGTCGTTGCTAGAATCAGAGGTTGTAAAAGAANNNAGATCGGAA

132243 TATGAGTGTGGAGTCGTTGCTAATCCTTACAGTTCCATTTCCNNNAGATCGGAAGAG
132244 TATGAGTGTGGAGTCGTTGCCACACGATTGAGGTCCTAATNNNAGATCGGAAGAGC
132245 TATGAGTGTGGAGTCGTTGCATGAATCAACTCTATTGTGGTCNNNAGATCGGAAGA
132246 TATGAGTGTGGAGTCGTTGCTTAGCAAAGTAGTTTTAAAAATCANNNAGATCGGAA
132247 TATGAGTGTGGAGTCGTTGCGTCTTGTTGCTCTATTTCTATTTNNNAGATCGGAAGA
132248 TATGAGTGTGGAGTCGTTGCCACAAATAAAAGTTCAAGAGCTNNNAGATCGGAAGA
132249 TATGAGTGTGGAGTCGTTGCATGATTTCCTCCATTCTATGACNNNAGATCGGAAGAG
132250 TATGAGTGTGGAGTCGTTGCAGAATTATCAAGTTGTGAATACCNNNAGATCGGAAG
132251 TATGAGTGTGGAGTCGTTGCAAGTTGACTCACCTTTAAGATATNNNAGATCGGAAGA
132252 TATGAGTGTGGAGTCGTTGCTTGGAACCTACATTACCATTTATNNNAGATCGGAAGA
132253 TATGAGTGTGGAGTCGTTGCTGCTTAATCAAGAGTTTATTTCCNNNAGATCGGAAGA
132254 TATGAGTGTGGAGTCGTTGCTTATTATGAGGAATTAGCTCAGGNNNAGATCGGAAG
132255 TATGAGTGTGGAGTCGTTGCCCGTTCTAGAGATGATTACTGNNNAGATCGGAAGAG
132257 TATGAGTGTGGAGTCGTTGCAAAGAACGCTGGTCTAGTTTNNNAGATCGGAAGAGC
132258 TATGAGTGTGGAGTCGTTGCTTGTTGTTGTTTTTAGAGATGANNNAGATCGGAAG
132259 TATGAGTGTGGAGTCGTTGCGTTAATGTTAATGACAGACTCAANNNAGATCGGAAG
132260 TATGAGTGTGGAGTCGTTGCTCTGTTTAACTTTGAAATGCTTANNNAGATCGGAAGA
132261 TATGAGTGTGGAGTCGTTGCTTTAACAAAGAGAAGCAAAACANNNAGATCGGAAGA
132262 TATGAGTGTGGAGTCGTTGCTAAGTAGGACATTTCTCTGTAAGNNNAGATCGGAAG
132263 TATGAGTGTGGAGTCGTTGCGTTTTGAAACATTTGGGATTTTANNNAGATCGGAAG
132264 TATGAGTGTGGAGTCGTTGCCATGGGATGTGAAAAAACAAAANNNAGATCGGAAG
132265 TATGAGTGTGGAGTCGTTGCACAAATAGAAACATTTTCAGAGGNNNAGATCGGAAG
132266 TATGAGTGTGGAGTCGTTGCAAAAGAATTTACCCTGGAAATAGNNNAGATCGGAAG
132267 TATGAGTGTGGAGTCGTTGCTAAAGGGAGACACTACAAATTAANNNAGATCGGAAG
132268 TATGAGTGTGGAGTCGTTGCATCTGAAGTAAGTAGGATTCCTNNNAGATCGGAAGA
132269 TATGAGTGTGGAGTCGTTGCTATATATGAAGCCTGTTGTGATTNNNAGATCGGAAG
132270 TATGAGTGTGGAGTCGTTGCCACACTGTCTTTAAAGTTATGACNNNAGATCGGAAGA
132271 TATGAGTGTGGAGTCGTTGCGAAAAAAAAAAAGCCTTGACAGNNNAGATCGGAAG
132272 TATGAGTGTGGAGTCGTTGCGTAGTGACTAACAGGATCTCCNNNAGATCGGAAGAG
132273 TATGAGTGTGGAGTCGTTGCCAAGCCAAGGTGATAATAAAATANNNAGATCGGAAG
132274 TATGAGTGTGGAGTCGTTGCTTGCAATCATACATCTTGTTAATNNNAGATCGGAAGA
132275 TATGAGTGTGGAGTCGTTGCATTAATCCAAGGTTGACATTTTTNNNAGATCGGAAGA
132276 TATGAGTGTGGAGTCGTTGCCCCAGAAAAGATTTAGAAAGTATNNNAGATCGGAAG
132277 TATGAGTGTGGAGTCGTTGCTTTCATTACAACAGAGTATTAGCNNNAGATCGGAAGA
132279 TATGAGTGTGGAGTCGTTGCTTTCCAAAATGTTGTTTTCATTGNNNAGATCGGAAGA
132280 TATGAGTGTGGAGTCGTTGCCAGTAAGTACTTGTCAAATGAAANNNAGATCGGAAG
132281 TATGAGTGTGGAGTCGTTGCCTTATCTAGGCATCATTTGAAAANNNAGATCGGAAGA
132282 TATGAGTGTGGAGTCGTTGCTAACTTTCACTGACTACTTGTTCNNNAGATCGGAAGA
132283 TATGAGTGTGGAGTCGTTGCATATATTTTGAGATGCAGTCTTGNNNAGATCGGAAG
132284 TATGAGTGTGGAGTCGTTGCTTTAATACGGAGTCTCGCTTNNNAGATCGGAAGAGC
132285 TATGAGTGTGGAGTCGTTGCGACTCTACCTTTCTTGTCATTTNNNAGATCGGAAGAG
132286 TATGAGTGTGGAGTCGTTGCGACACAAGTTCTTTCTAGGTTNNNAGATCGGAAGAG
132287 TATGAGTGTGGAGTCGTTGCCTCCTAGGTTTATTAATGTGGATNNNAGATCGGAAG

132288 TATGAGTGTGGAGTCGTTGCATTTGTGAATTTGCCTAGTTTATNNNAGATCGGAAG
132289 TATGAGTGTGGAGTCGTTGCTCCAACTGCTTCTAAGTTTTATANNNAGATCGGAAGA
132290 TATGAGTGTGGAGTCGTTGCCTCTTACACAAAGATCTTTCATCNNNAGATCGGAAGA
132291 TATGAGTGTGGAGTCGTTGCCTCTTTTTTTCCATTAGAAACTGNNNAGATCGGAAGA
132292 TATGAGTGTGGAGTCGTTGCATGTAAAGCATTATCTTGGTTAGNNNAGATCGGAAG
132293 TATGAGTGTGGAGTCGTTGCCTATTACGATTAGCACGACTATANNNAGATCGGAAG
132294 TATGAGTGTGGAGTCGTTGCACAAAAGAAAAAGGAAATTGGATNNNAGATCGGAA
132295 TATGAGTGTGGAGTCGTTGCGCTAGAGACCTAAAGAAAAGTTANNNAGATCGGAA
132296 TATGAGTGTGGAGTCGTTGCTGGGGTGTTTTAATTTTCTTTAANNNAGATCGGAAG
132297 TATGAGTGTGGAGTCGTTGCAGAGAATTTAGAATAAAGGAATAGTNNNAGATCGG
132298 TATGAGTGTGGAGTCGTTGCAATTCCTGGGCTCAAATGATNNNAGATCGGAAGAG
132299 TATGAGTGTGGAGTCGTTGCACAGGTACTAAAACATGAATGAANNNAGATCGGAA
132300 TATGAGTGTGGAGTCGTTGCAGATGAGGATATAGGAGAAAAACNNNAGATCGGAA
132301 TATGAGTGTGGAGTCGTTGCCTTAGCAATTAGGCAAACTAAAANNNAGATCGGAAG
132302 TATGAGTGTGGAGTCGTTGCGATAAATTGTCCTTGACCTCTNNNAGATCGGAAGAG
132303 TATGAGTGTGGAGTCGTTGCGTTCTTATGGAGTGATCTGGNNNAGATCGGAAGAG
132304 TATGAGTGTGGAGTCGTTGCCTTTGGATCTGTCAGTAATTTCNNNAGATCGGAAGA
132305 TATGAGTGTGGAGTCGTTGCCAGACTATGCACTAGGAACTNNNAGATCGGAAGAG
132306 TATGAGTGTGGAGTCGTTGCCAGACTATGCACTAGGAACTNNNAGATCGGAAGAG
132307 TATGAGTGTGGAGTCGTTGCCTTTCCCTCTGTTCCTTTAAGNNNAGATCGGAAGAGC
132308 TATGAGTGTGGAGTCGTTGCTATAGTTAAACACAGACAAGAAGNNNAGATCGGAA
132309 TATGAGTGTGGAGTCGTTGCGAAAATCCTGTTGCATAAAAATGNNNAGATCGGAAG
132310 TATGAGTGTGGAGTCGTTGCAAATATGAAACAACTGATAACCANNNAGATCGGAAG
132311 TATGAGTGTGGAGTCGTTGCACCTTTAAAGTGTAGTAATTCCANNNAGATCGGAAG
132312 TATGAGTGTGGAGTCGTTGCGCTTTCAATTACAGATTCAATTTNNNAGATCGGAAGA
132313 TATGAGTGTGGAGTCGTTGCATGTTCATAGCAACATTATTTGTNNNAGATCGGAAG
132314 TATGAGTGTGGAGTCGTTGCCCAATTTTAAGTTCTGGCAATNNNAGATCGGAAGAG
132315 TATGAGTGTGGAGTCGTTGCATGTGTGTATACAAAGGAGAAANNNAGATCGGAAG
132316 TATGAGTGTGGAGTCGTTGCTCAACAATATCCAAATAAGCATTNNNAGATCGGAAG
132317 TATGAGTGTGGAGTCGTTGCTTCTAGAAGATTAGAAAGCTCATNNNAGATCGGAAG
132318 TATGAGTGTGGAGTCGTTGCTTCAGTAACAGATGACTATTTCANNNAGATCGGAAG
132319 TATGAGTGTGGAGTCGTTGCCAAATTCTCTGCATTGTAAAGANNNAGATCGGAAGA
132320 TATGAGTGTGGAGTCGTTGCCAACTGACTTTGAAATGTATGATNNNAGATCGGAAG
132321 TATGAGTGTGGAGTCGTTGCAATTTCAGTCACAGAAAGACATANNNAGATCGGAAG
132322 TATGAGTGTGGAGTCGTTGCACTGATGCTAGGTATGAAAATATNNNAGATCGGAA
132323 TATGAGTGTGGAGTCGTTGCTTTGTCAGGTTCTAGATTATAGGNNNAGATCGGAAG
132324 TATGAGTGTGGAGTCGTTGCGACATTAAAAGGGAAAGTTTCTGNNNAGATCGGAA
132326 TATGAGTGTGGAGTCGTTGCTATTTTTTGAGACAGAGTTTCACNNNAGATCGGAAG
132327 TATGAGTGTGGAGTCGTTGCGACGTATTTGAAAAACTTCATTGNNNAGATCGGAAG
132328 TATGAGTGTGGAGTCGTTGCTATTTTTTGAGACAGAGTTTCACNNNAGATCGGAAG
132329 TATGAGTGTGGAGTCGTTGCTCTAATAGGCTTACTAACAGAACNNNAGATCGGAAG
132330 TATGAGTGTGGAGTCGTTGCGGGATAAATGTGCAGAACATNNNAGATCGGAAGAG
132331 TATGAGTGTGGAGTCGTTGCCCAGATTACTTTGGTTTTGAATNNNAGATCGGAAGA

132332 TATGAGTGTGGAGTCGTTGCTCCTTTGAAAGTTCTATGCTTTNNNAGATCGGAAGAG
132333 TATGAGTGTGGAGTCGTTGCTACTTTTAGTTTTTAGAGCAGTNNNAGATCGGAAGA
132335 TATGAGTGTGGAGTCGTTGCTTAATGGAAATTGTAGCTGATTCNNNAGATCGGAAG
132336 TATGAGTGTGGAGTCGTTGCGAACTGAGAACAGAAATAACATTNNNAGATCGGAAG
132337 TATGAGTGTGGAGTCGTTGCGATATGATCTTGGCTCACTGNNNAGATCGGAAGAGC
132338 TATGAGTGTGGAGTCGTTGCCTTCACGGTAGAGAACAATGNNNAGATCGGAAGAGC
132339 TATGAGTGTGGAGTCGTTGCAATCCCATTTCACAGATGAAGNNNAGATCGGAAGAG
132340 TATGAGTGTGGAGTCGTTGCTGCAGAAATAAATGAATTAATGAATNNNAGATCGGA
132341 TATGAGTGTGGAGTCGTTGCAACAAAGAGGAACACTAATTTACNNNAGATCGGAAG
132342 TATGAGTGTGGAGTCGTTGCTATTTTTTGAGACAGAGTTTCACNNNAGATCGGAAGA
132343 TATGAGTGTGGAGTCGTTGCTTAAAGATAGATTTTAGGCTGGANNNAGATCGGAAG
132345 TATGAGTGTGGAGTCGTTGCCAACAGCCTTTAAAAATAGGTACNNNAGATCGGAAG
132346 TATGAGTGTGGAGTCGTTGCTAATGCAGTCATGAGATAATACANNNAGATCGGAAG
132347 TATGAGTGTGGAGTCGTTGCTGTATTTTGGGGTGTATATATATATNNNAGATCGGA
132348 TATGAGTGTGGAGTCGTTGCGTCATCATTGGTTGTAGTTATTTNNNAGATCGGAAGA
132349 TATGAGTGTGGAGTCGTTGCATACTTGGAAAACCTAAAATGTGNNNAGATCGGAAG
132350 TATGAGTGTGGAGTCGTTGCGAAAACTAGTAACTGTGTAACACNNNAGATCGGAAG
132351 TATGAGTGTGGAGTCGTTGCAAAAATTAGAATAGGGGTTCCAANNNAGATCGGAAG
132352 TATGAGTGTGGAGTCGTTGCGATTTTGTTCTCAGCTTGAATATNNNAGATCGGAAGA
132353 TATGAGTGTGGAGTCGTTGCAGACAGAGAGAGCAAATAAAAANNNAGATCGGAAG
132355 TATGAGTGTGGAGTCGTTGCCATTATTTCACTCATTTGAGGTANNNAGATCGGAAGA
132356 TATGAGTGTGGAGTCGTTGCTTTATTATTCAACCATGTGCTAGNNNAGATCGGAAGA
132357 TATGAGTGTGGAGTCGTTGCGGATTTAACTTTTCTAAGCACTANNNAGATCGGAAGA
132358 TATGAGTGTGGAGTCGTTGCCACCAGAAACAAATACAGAATATNNNAGATCGGAAG
132359 TATGAGTGTGGAGTCGTTGCAGTCATAATTTCATAGTGCAGANNNAGATCGGAAGA
132360 TATGAGTGTGGAGTCGTTGCGTTTCCTGCATTCACTTAATTANNNAGATCGGAAGAG
132361 TATGAGTGTGGAGTCGTTGCATTTTATAATGACAGAGTGAGGANNNAGATCGGAAG
132362 TATGAGTGTGGAGTCGTTGCCAACCACCAAAAATAAAAAAGGNNNAGATCGGAAGA
132363 TATGAGTGTGGAGTCGTTGCGAATCTCATTCTATCAGCCAGNNNAGATCGGAAGAG
132364 TATGAGTGTGGAGTCGTTGCTGAGGTCTTTTTTCCATTTTTAANNNAGATCGGAAGA
132365 TATGAGTGTGGAGTCGTTGCACAGTGTTTTCAAAAATTGTTTTNNNAGATCGGAAGA
132366 TATGAGTGTGGAGTCGTTGCTATTTGGATAAACAAATTAGGGCNNNAGATCGGAAG
132367 TATGAGTGTGGAGTCGTTGCTACACAGTTGATGAGACATAAATNNNAGATCGGAAG
132368 TATGAGTGTGGAGTCGTTGCCATATAGAACCCTTACAACCTAANNNAGATCGGAAGA
132369 TATGAGTGTGGAGTCGTTGCTACCAGGAATGATGATTCAATTANNNAGATCGGAAG
132370 TATGAGTGTGGAGTCGTTGCGCTCTTAATTCTATCAAATTCCTTNNNAGATCGGAAGA
132371 TATGAGTGTGGAGTCGTTGCATAAAAGATCCTTAACAATGCAGNNNAGATCGGAAG
132372 TATGAGTGTGGAGTCGTTGCGTAAATGGATGGGAAAATACATTNNNAGATCGGAAG
132373 TATGAGTGTGGAGTCGTTGCGAGCATAGCTTCTTACTAGATCNNNAGATCGGAAGA
132374 TATGAGTGTGGAGTCGTTGCGTCATCTGTTACAATGGGAATNNNAGATCGGAAGAG
132375 TATGAGTGTGGAGTCGTTGCAGAGTTGTGTTCTACCTCTAGNNNAGATCGGAAGAG
132376 TATGAGTGTGGAGTCGTTGCCTTTTAAATGCTGGAACCAAGNNNAGATCGGAAGAG
132377 TATGAGTGTGGAGTCGTTGCCTGTATGAGATTGGAGACTATTANNNAGATCGGAAG

132378 TATGAGTGTGGAGTCGTTGCATTTATTGAAAGGAATTGGCTTANNNAGATCGGAAG
132379 TATGAGTGTGGAGTCGTTGCGTAATATAGATTTTGGAACCAGGNNNAGATCGGAA[illegible]
132380 TATGAGTGTGGAGTCGTTGCACAGTCAAAAATCCATGTATAACNNNAGATCGGAAG

AGCACACGTCTGAACTCTTTCCCTACACGACGCTCTTCCGATCTNNNCTGATCAATGAGGAGTCTGA
AGCACACGTCTGAACTCTTTCCCTACACGACGCTCTTCCGATCTNNNTTACCTCTCTCTCAACATTAAT
GCACACGTCTGAACTCTTTCCCTACACGACGCTCTTCCGATCTNNNCCCATTATGAATACAAAACCAAA
GCACACGTCTGAACTCTTTCCCTACACGACGCTCTTCCGATCTNNNGTTGGGAAAATTATCACTATGA
AGCACACGTCTGAACTCTTTCCCTACACGACGCTCTTCCGATCTNNNCCAAAAGAGTAAAATCTCAAT
AGCACACGTCTGAACTCTTTCCCTACACGACGCTCTTCCGATCTNNNCTTTTTCCTTTGGGTTTCCTCAT
CACACGTCTGAACTCTTTCCCTACACGACGCTCTTCCGATCTNNNCTCTCTCTATCTTGCCTCTTCATCG
GCACACGTCTGAACTCTTTCCCTACACGACGCTCTTCCGATCTNNNGACTTTTTCACAGAAAACAAAA
GCACACGTCTGAACTCTTTCCCTACACGACGCTCTTCCGATCTNNNTTTGCTCCAAGCTATTAAAATAG
CACACGTCTGAACTCTTTCCCTACACGACGCTCTTCCGATCTNNNTAGAGTGTAGATGCATCAAAATC
ACACGTCTGAACTCTTTCCCTACACGACGCTCTTCCGATCTNNNATCTCCCAAAGGATATGCTCCATCG
CACACGTCTGAACTCTTTCCCTACACGACGCTCTTCCGATCTNNNGCATCAGATAGAAGGTAATGTAT
GCACACGTCTGAACTCTTTCCCTACACGACGCTCTTCCGATCTNNNCCTCTTACTCATTCATCATCTAAC
AGCACACGTCTGAACTCTTTCCCTACACGACGCTCTTCCGATCTNNNTTAGAAAACAAAAGCCAATGT
ACACGTCTGAACTCTTTCCCTACACGACGCTCTTCCGATCTNNNCCAGTATGACAAGGTATGATAAGC
CACGTCTGAACTCTTTCCCTACACGACGCTCTTCCGATCTNNNGAGTTAGATATGGAATTGGGTAGC
GCACACGTCTGAACTCTTTCCCTACACGACGCTCTTCCGATCTNNNCTGGAATCTTTACAGGAGAACC
GAGCACACGTCTGAACTCTTTCCCTACACGACGCTCTTCCGATCTNNNTCTTCAGAACTTTCAGAGAAA
AGCACACGTCTGAACTCTTTCCCTACACGACGCTCTTCCGATCTNNNCTGTCACATTTCATAGAAACTG
AGCACACGTCTGAACTCTTTCCCTACACGACGCTCTTCCGATCTNNNTATTTGAAAACGTTACCTGAA
AGCACACGTCTGAACTCTTTCCCTACACGACGCTCTTCCGATCTNNNCTCTTTCTATTCTCTGAGTGAT
ACACGTCTGAACTCTTTCCCTACACGACGCTCTTCCGATCTNNNCATCGAGGTTCACAGTTATGCATC
GAGCACACGTCTGAACTCTTTCCCTACACGACGCTCTTCCGATCTNNNTTAGGCTTTACAAATCATATC
GCACACGTCTGAACTCTTTCCCTACACGACGCTCTTCCGATCTNNNTACATCACTGAAGTTCATATCAT
CACACGTCTGAACTCTTTCCCTACACGACGCTCTTCCGATCTNNNGATTAAATGGACTTTACCTTCAGC
ACACGTCTGAACTCTTTCCCTACACGACGCTCTTCCGATCTNNNTGTTTTTCTTCTAAAGGCTTTTCAT
CACACGTCTGAACTCTTTCCCTACACGACGCTCTTCCGATCTNNNCACATCAAGCATGACAGATGCATC
GCACACGTCTGAACTCTTTCCCTACACGACGCTCTTCCGATCTNNNGTTATTTAGATCACCTTTTGGCC
AGCACACGTCTGAACTCTTTCCCTACACGACGCTCTTCCGATCTNNNCATTTAGATCCAGATACCGAAT
CACACGTCTGAACTCTTTCCCTACACGACGCTCTTCCGATCTNNNTTTTTCATAAGGTTATAGCTCAGC
GCACACGTCTGAACTCTTTCCCTACACGACGCTCTTCCGATCTNNNTTCTGTATCCAGTGTAACAAAAA
AGCACACGTCTGAACTCTTTCCCTACACGACGCTCTTCCGATCTNNNGGATCATTCCCATATAGAAGA
AGCACACGTCTGAACTCTTTCCCTACACGACGCTCTTCCGATCTNNNACATCTTTATGGTATAACTGTG
GAGCACACGTCTGAACTCTTTCCCTACACGACGCTCTTCCGATCTNNNGATTATTTGCTCGATACTGA
GAGCACACGTCTGAACTCTTTCCCTACACGACGCTCTTCCGATCTNNNTCACTGGTTTCATTTTGATTAA
AGAGCACACGTCTGAACTCTTTCCCTACACGACGCTCTTCCGATCTNNNCTGGTTGTGGATACTGAGA
AGCACACGTCTGAACTCTTTCCCTACACGACGCTCTTCCGATCTNNNATACATTTGTCTTAGGTTTGGA
CACACGTCTGAACTCTTTCCCTACACGACGCTCTTCCGATCTNNNGTGATTCTCAGCTTAAGCATCATC
AGCACACGTCTGAACTCTTTCCCTACACGACGCTCTTCCGATCTNNNTAAATGGAGAAATCTTCTTCTG
AGCACACGTCTGAACTCTTTCCCTACACGACGCTCTTCCGATCTNNNCTGGGTTCAGAATCTGTATACC
ACACGTCTGAACTCTTTCCCTACACGACGCTCTTCCGATCTNNNCTGAAGAAATAAAGTTCTTTTTTTT
AGCACACGTCTGAACTCTTTCCCTACACGACGCTCTTCCGATCTNNNGTATGGAATATAGCAGAGATG

AGCACACGTCTGAACTCTTTCCCTACACGACGCTCTTCCGATCTNNNAAAGTATGTGTGTTTTCTTACA
AGCACACGTCTGAACTCTTTCCCTACACGACGCTCTTCCGATCTNNNGTAATCTGAGGTAAAATGATG
AGCACACGTCTGAACTCTTTCCCTACACGACGCTCTTCCGATCTNNNTAATTGATGATCATCTGATTGA
GCACACGTCTGAACTCTTTCCCTACACGACGCTCTTCCGATCTNNNGCCTGTGTTCTTCTTTGTAGCAT
AGCACACGTCTGAACTCTTTCCCTACACGACGCTCTTCCGATCTNNNCGAGAATACTTTATTGCTATGT
AGCACACGTCTGAACTCTTTCCCTACACGACGCTCTTCCGATCTNNNGCATATTGTCCAGTCTCATTAC
GCACACGTCTGAACTCTTTCCCTACACGACGCTCTTCCGATCTNNNTATTGCTGAGTCCTCTGATTACA
GCACACGTCTGAACTCTTTCCCTACACGACGCTCTTCCGATCTNNNGAAAGTGCAGGCTATATAACTT
GCACACGTCTGAACTCTTTCCCTACACGACGCTCTTCCGATCTNNNTATTCTTCAGGGTTATTGAATCA
CACACGTCTGAACTCTTTCCCTACACGACGCTCTTCCGATCTNNNGTGGACTTTAGAAAATGTTAAGT
AGCACACGTCTGAACTCTTTCCCTACACGACGCTCTTCCGATCTNNNTTCTTCTTCCTTGATAGCCTCAT
GCACACGTCTGAACTCTTTCCCTACACGACGCTCTTCCGATCTNNNTGTAATTTATTGGCAGATCTCTA
AGCACACGTCTGAACTCTTTCCCTACACGACGCTCTTCCGATCTNNNTGCAGTTATAACTCATATCAAG
AGCACACGTCTGAACTCTTTCCCTACACGACGCTCTTCCGATCTNNNGGATTCGGCAGTTTTTATTAG
CACGTCTGAACTCTTTCCCTACACGACGCTCTTCCGATCTNNNAGGCTTCATTTCTGTCATTTCATCGGA
AGCACACGTCTGAACTCTTTCCCTACACGACGCTCTTCCGATCTNNNCTTAAAATCTCCAAATCAGAGA
GCACACGTCTGAACTCTTTCCCTACACGACGCTCTTCCGATCTNNNTTTACCACTAGAAACTAACAGTT
AGCACACGTCTGAACTCTTTCCCTACACGACGCTCTTCCGATCTNNNATACCTGCAAGATCTTCCTTCAT
ACACGTCTGAACTCTTTCCCTACACGACGCTCTTCCGATCTNNNCTTCTTTATGATGTCGCATTATGCA
GCACACGTCTGAACTCTTTCCCTACACGACGCTCTTCCGATCTNNNGTGAAAATAAAGAAGATTGTCC
AGCACACGTCTGAACTCTTTCCCTACACGACGCTCTTCCGATCTNNNCTGATGTTTTCACAGTGAATTT
AGCACACGTCTGAACTCTTTCCCTACACGACGCTCTTCCGATCTNNNGTATCCTTCAGAACCTTTGGCAT
AGCACACGTCTGAACTCTTTCCCTACACGACGCTCTTCCGATCTNNNCATCCCCTATGTAGAATCTCAT
GAGCACACGTCTGAACTCTTTCCCTACACGACGCTCTTCCGATCTNNNGGTGGTGCTTATTTTAAAATT
CACGTCTGAACTCTTTCCCTACACGACGCTCTTCCGATCTNNNTAGTCATAATGGCTGTCAAAACATC
CACACGTCTGAACTCTTTCCCTACACGACGCTCTTCCGATCTNNNCAAGTTCTACAATTACCACAAGCA
AGCACACGTCTGAACTCTTTCCCTACACGACGCTCTTCCGATCTNNNCTCATGAATAAATGCCTGTTGC
GCACACGTCTGAACTCTTTCCCTACACGACGCTCTTCCGATCTNNNTTTGTTGGATTTTTCACAAAA
AGCACACGTCTGAACTCTTTCCCTACACGACGCTCTTCCGATCTNNNGCTGTAGCAGTTTCTAAAAATA
AGCACACGTCTGAACTCTTTCCCTACACGACGCTCTTCCGATCTNNNCTGGATCTAAAGATTATGCAG
AGCACACGTCTGAACTCTTTCCCTACACGACGCTCTTCCGATCTNNNTGTTCACTCATTAAATTTTGTCT
GAGCACACGTCTGAACTCTTTCCCTACACGACGCTCTTCCGATCTNNNGAGATGTCAGGAGACTTTTA
GAGCACACGTCTGAACTCTTTCCCTACACGACGCTCTTCCGATCTNNNGAGATGTCAGGAGACTTTTA
AGCACACGTCTGAACTCTTTCCCTACACGACGCTCTTCCGATCTNNNACCCCCTCGACTATAGTTAGCAT
AGAGCACACGTCTGAACTCTTTCCCTACACGACGCTCTTCCGATCTNNNGAGAAAACCAAGATTGCTT
CACACGTCTGAACTCTTTCCCTACACGACGCTCTTCCGATCTNNNTTCCTACGAAGGTCATGAAACAT
GCACACGTCTGAACTCTTTCCCTACACGACGCTCTTCCGATCTNNNTATTTTATCTTTAGGGCTTAGGT
AGCACACGTCTGAACTCTTTCCCTACACGACGCTCTTCCGATCTNNNTGGACTAAAATTTCTAACAAAC
AGCACACGTCTGAACTCTTTCCCTACACGACGCTCTTCCGATCTNNNTAATTACAGCAACCAGTTTTAT
AGCACACGTCTGAACTCTTTCCCTACACGACGCTCTTCCGATCTNNNAAAATAGAAAACAATGAATGG
GCACACGTCTGAACTCTTTCCCTACACGACGCTCTTCCGATCTNNNTCTTTCTGTGTCGAATTTTATAGC
GAGCACACGTCTGAACTCTTTCCCTACACGACGCTCTTCCGATCTNNNCCAAATGTTTTTATGATCTCT
AGCACACGTCTGAACTCTTTCCCTACACGACGCTCTTCCGATCTNNNTATGTTGAATTCATTTTCAGG

AGCACACGTCTGAACTCTTTCCCTACACGACGCTCTTCCGATCTNNNTTGACTTTTTAATGACATGAG
GCACACGTCTGAACTCTTTCCCTACACGACGCTCTTCCGATCTNNNCTCTACTTGATGGAGTTCTTAAT
AGCACACGTCTGAACTCTTTCCCTACACGACGCTCTTCCGATCTNNNGATTTGGTTTATCCCGAAATA
GCACACGTCTGAACTCTTTCCCTACACGACGCTCTTCCGATCTNNNATCCCATTTTTTTTGTGGTAATA
ACACGTCTGAACTCTTTCCCTACACGACGCTCTTCCGATCTNNNGAAGAGAATTTGAGAGAAAAGAG
GCACACGTCTGAACTCTTTCCCTACACGACGCTCTTCCGATCTNNNAGTTTATTGTGAAAACATGATG
GCACACGTCTGAACTCTTTCCCTACACGACGCTCTTCCGATCTNNNTGATATAAAGGCAAATCTTCTTG
GAGCACACGTCTGAACTCTTTCCCTACACGACGCTCTTCCGATCTNNNTACTCTCATTGTAGGTCCAA
GCACACGTCTGAACTCTTTCCCTACACGACGCTCTTCCGATCTNNNTCAATTTGAAAATACTCATCGA
GCACACGTCTGAACTCTTTCCCTACACGACGCTCTTCCGATCTNNNAAGTATAAGAAAGCTCTCCATT
AGCACACGTCTGAACTCTTTCCCTACACGACGCTCTTCCGATCTNNNCAGATTCATGAGATCGTACAT
AGCACACGTCTGAACTCTTTCCCTACACGACGCTCTTCCGATCTNNNCCTTTTCTTCCAATACTATTGA
AGCACACGTCTGAACTCTTTCCCTACACGACGCTCTTCCGATCTNNNTCCAAGCTTTCAATAGTAAAT
GCACACGTCTGAACTCTTTCCCTACACGACGCTCTTCCGATCTNNNTTACCAACCTCATTTCTTTACTTC
ACACGTCTGAACTCTTTCCCTACACGACGCTCTTCCGATCTNNNTGAACAGGTTAAAGACGAAACATC
AGCACACGTCTGAACTCTTTCCCTACACGACGCTCTTCCGATCTNNNTATCAGAGAAGGGTTTCAGAA
AGCACACGTCTGAACTCTTTCCCTACACGACGCTCTTCCGATCTNNNCATTCTGATACTAGTCCTATTCC
AGCACACGTCTGAACTCTTTCCCTACACGACGCTCTTCCGATCTNNNTGAGAGTATTTACCTTTTGAT
GCACACGTCTGAACTCTTTCCCTACACGACGCTCTTCCGATCTNNNTAAGAGTGTAGTATGAATGAT
GAGCACACGTCTGAACTCTTTCCCTACACGACGCTCTTCCGATCTNNNAGTTTGAGTCATTATCCTATT
AGCACACGTCTGAACTCTTTCCCTACACGACGCTCTTCCGATCTNNNTTTTTTTTTTTTTTAACAGGCAG
AGCACACGTCTGAACTCTTTCCCTACACGACGCTCTTCCGATCTNNNAAATTCATCTTAAATCAGGAA
ACACGTCTGAACTCTTTCCCTACACGACGCTCTTCCGATCTNNNCTTCTTCTCCATTGTCTCCTCATCGG
AGCACACGTCTGAACTCTTTCCCTACACGACGCTCTTCCGATCTNNNAAAATGTCTTCTATACCCCAGA
GCACACGTCTGAACTCTTTCCCTACACGACGCTCTTCCGATCTNNNATATGAAAAAGTACTGGTAGAC
AGCACACGTCTGAACTCTTTCCCTACACGACGCTCTTCCGATCTNNNCATCAGTTTGCATCATGGAGCA
AGCACACGTCTGAACTCTTTCCCTACACGACGCTCTTCCGATCTNNNATTTCGGGAATCTTGAATGTAT
ACACGTCTGAACTCTTTCCCTACACGACGCTCTTCCGATCTNNNCTCCACCCTAGATCACTTTGCATCG
AGCACACGTCTGAACTCTTTCCCTACACGACGCTCTTCCGATCTNNNATATTTGTGTATCTCCAATAG
GCACACGTCTGAACTCTTTCCCTACACGACGCTCTTCCGATCTNNNCTGTTTGGCTCAAAATTAGATGC
CACACGTCTGAACTCTTTCCCTACACGACGCTCTTCCGATCTNNNAGATATAGTTCAGGCAAACTAGC
GAGCACACGTCTGAACTCTTTCCCTACACGACGCTCTTCCGATCTNNNTATTCATCTGCTCTCCCTTTT
GCACACGTCTGAACTCTTTCCCTACACGACGCTCTTCCGATCTNNNTCATTGTAGCTATAGACTATTG
CACACGTCTGAACTCTTTCCCTACACGACGCTCTTCCGATCTNNNGCTATACTTTATGACACTAAGGA
CACACGTCTGAACTCTTTCCCTACACGACGCTCTTCCGATCTNNNCATTTTTTATTGTTTCCGGAATGC
CACACGTCTGAACTCTTTCCCTACACGACGCTCTTCCGATCTNNNTTAATGTAAACGGTTATCTTTCCC
AGCACACGTCTGAACTCTTTCCCTACACGACGCTCTTCCGATCTNNNAAGACGTAGATTTTTCTGGAA
AGCACACGTCTGAACTCTTTCCCTACACGACGCTCTTCCGATCTNNNAGAGAGGAATAAAGACAGA
ACACGTCTGAACTCTTTCCCTACACGACGCTCTTCCGATCTNNNAGAGGTCACACTAGTACCTTCATCG
CACACGTCTGAACTCTTTCCCTACACGACGCTCTTCCGATCTNNNTTCCTCTTTATCTCAGATATGTGCA
GCACACGTCTGAACTCTTTCCCTACACGACGCTCTTCCGATCTNNNATTAATACCAGGATGCAACTGCA
GCACACGTCTGAACTCTTTCCCTACACGACGCTCTTCCGATCTNNNCTTCTTTTCCAAGTCATTCATTAC
AGCACACGTCTGAACTCTTTCCCTACACGACGCTCTTCCGATCTNNNCATACATTTCTGTGATGGTCT

AGCACACGTCTGAACTCTTTCCCTACACGACGCTCTTCCGATCTNNNGTTGATTTATACCCTGATAGAGT
AGCACACGTCTGAACTCTTTCCCTACACGACGCTCTTCCGATCTNNNACTAAAATAACCACTGTCAACA
GCACACGTCTGAACTCTTTCCCTACACGACGCTCTTCCGATCTNNNCAACTTATCAAGCATGTTTGGCA
GAGCACACGTCTGAACTCTTTCCCTACACGACGCTCTTCCGATCTNNNGAGCTTACCTATTATCCTCAGC
AGCACACGTCTGAACTCTTTCCCTACACGACGCTCTTCCGATCTNNNATTCAGATGATTGGACTATTCA
AGCACACGTCTGAACTCTTTCCCTACACGACGCTCTTCCGATCTNNNATTATTTGCTTCATTTCTCTGTTC
GCACACGTCTGAACTCTTTCCCTACACGACGCTCTTCCGATCTNNNGGCAATATTTTAACCATACTCAACA
CACACGTCTGAACTCTTTCCCTACACGACGCTCTTCCGATCTNNNATTGTGTGTCTCATGATGTTCATCG
AGCACACGTCTGAACTCTTTCCCTACACGACGCTCTTCCGATCTNNNCATTAGAAAACATCAGCACTCCA
GAGCACACGTCTGAACTCTTTCCCTACACGACGCTCTTCCGATCTNNNCAGCTGGTTGTATATCCAATAT
AGCACACGTCTGAACTCTTTCCCTACACGACGCTCTTCCGATCTNNNTCCATGAACATTGGATTTATCTA
AGCACACGTCTGAACTCTTTCCCTACACGACGCTCTTCCGATCTNNNTAGTGAAATCTCTCAGATTGATT
AGCACACGTCTGAACTCTTTCCCTACACGACGCTCTTCCGATCTNNNCTTTTTCTCCCTCTCAGATGCAT
ACACGTCTGAACTCTTTCCCTACACGACGCTCTTCCGATCTNNNAGTTATGTTAGGTATACCCGACATCG
AGCACACGTCTGAACTCTTTCCCTACACGACGCTCTTCCGATCTNNNGAAGACAAACTCTATGACATTTT
GAGCACACGTCTGAACTCTTTCCCTACACGACGCTCTTCCGATCTNNNATCTTGGACGTAGATAAGTGT
GCACACGTCTGAACTCTTTCCCTACACGACGCTCTTCCGATCTNNNTTTATTGTTACGAAGGACTTTTC
AGCACACGTCTGAACTCTTTCCCTACACGACGCTCTTCCGATCTNNNCAATATCCTTCAAAAACTTCAGAC
CACACGTCTGAACTCTTTCCCTACACGACGCTCTTCCGATCTNNNAATGGTGCTGAAATGAAGATCATC
AGCACACGTCTGAACTCTTTCCCTACACGACGCTCTTCCGATCTNNNTTGTTTTCTTCTTTCCCATTTAAC
AGCACACGTCTGAACTCTTTCCCTACACGACGCTCTTCCGATCTNNNAGTCTAAAGTTAGATCTGTCTTG
CACACGTCTGAACTCTTTCCCTACACGACGCTCTTCCGATCTNNNGTGAAAAACAAATCATCATCATCCA
GCACACGTCTGAACTCTTTCCCTACACGACGCTCTTCCGATCTNNNTAAGTTGATGGAGGAAGTAAATA
GAGCACACGTCTGAACTCTTTCCCTACACGACGCTCTTCCGATCTNNNTCTTTATCTCATAGTTTTTGCAC
GAGCACACGTCTGAACTCTTTCCCTACACGACGCTCTTCCGATCTNNNACTAATAGAAATAGTCACTTG
AGCACACGTCTGAACTCTTTCCCTACACGACGCTCTTCCGATCTNNNCTTCGAACTTCTACTTTAACAAAC
AGCACACGTCTGAACTCTTTCCCTACACGACGCTCTTCCGATCTNNNGTTTTTCAGGTAAATGGTCTTA
GCACACGTCTGAACTCTTTCCCTACACGACGCTCTTCCGATCTNNNACCTTGAAGAAAATCAGTTTCCAT
GAGCACACGTCTGAACTCTTTCCCTACACGACGCTCTTCCGATCTNNNGAATGTAGAGGATGAGGAAA
CACACGTCTGAACTCTTTCCCTACACGACGCTCTTCCGATCTNNNCTATCTCTCAACATGTGCTGCATCGC
AGCACACGTCTGAACTCTTTCCCTACACGACGCTCTTCCGATCTNNNAATGAAGGAGGAGAGTTCATC
AGAGCACACGTCTGAACTCTTTCCCTACACGACGCTCTTCCGATCTNNNACAGTGAGGCATTTTGTAAA
AGCACACGTCTGAACTCTTTCCCTACACGACGCTCTTCCGATCTNNNTACCATCTCAACAAGTCAATTCA
GAGCACACGTCTGAACTCTTTCCCTACACGACGCTCTTCCGATCTNNNGAATAAGCAGATCCATGGAGC
AGCACACGTCTGAACTCTTTCCCTACACGACGCTCTTCCGATCTNNNTAAGACTTCCTTTTACTTTCTCTC
GAGCACACGTCTGAACTCTTTCCCTACACGACGCTCTTCCGATCTNNNTATACAGTTCAGGTACTTTGTC
AGCACACGTCTGAACTCTTTCCCTACACGACGCTCTTCCGATCTNNNCTGAGGATAAGGATGACTTAAC
AGAGCACACGTCTGAACTCTTTCCCTACACGACGCTCTTCCGATCTNNNCAAATTGGAGTGGATATGAT
GCACACGTCTGAACTCTTTCCCTACACGACGCTCTTCCGATCTNNNGCTCAGAAAGTTTTGATATGTTAC
AGCACACGTCTGAACTCTTTCCCTACACGACGCTCTTCCGATCTNNNGTCCATACTTACCTAATTTTGAG
CACACGTCTGAACTCTTTCCCTACACGACGCTCTTCCGATCTNNNTGAGAGAGAAATGCATCCATCATC
AGCACACGTCTGAACTCTTTCCCTACACGACGCTCTTCCGATCTNNNGGGTTTTACAAATATAAGTCAC
CACACGTCTGAACTCTTTCCCTACACGACGCTCTTCCGATCTNNNGAAATTAATTAGAAGAGTTGCCAC

GCACACGTCTGAACTCTTTCCCTACACGACGCTCTTCCGATCTNNNATACTCTTTCTGACTGATGACTA
AGCACACGTCTGAACTCTTTCCCTACACGACGCTCTTCCGATCTNNNTCCCCCATATAAGTTCTTAATTT
CACACGTCTGAACTCTTTCCCTACACGACGCTCTTCCGATCTNNNTCTTTTGAGTAAAGTTTTCTTGTCA
CACACGTCTGAACTCTTTCCCTACACGACGCTCTTCCGATCTNNNCTTTGGTGGCTTATCTGATGCATC
AGCACACGTCTGAACTCTTTCCCTACACGACGCTCTTCCGATCTNNNCGTTACGAAATATATCCTGTTT
AGCACACGTCTGAACTCTTTCCCTACACGACGCTCTTCCGATCTNNNGTGACCTCGATGCTAAATAGC
GCACACGTCTGAACTCTTTCCCTACACGACGCTCTTCCGATCTNNNACTTATGGGTGAATCATGTTAT
CACACGTCTGAACTCTTTCCCTACACGACGCTCTTCCGATCTNNNTCGACTATAGCTGTTATGTTAATC
CACACGTCTGAACTCTTTCCCTACACGACGCTCTTCCGATCTNNNCTTACAAAGTAGTAGAAGGCATCA
ACACGTCTGAACTCTTTCCCTACACGACGCTCTTCCGATCTNNNGAAGAAGACATTGTTGAAGTGCAT
CACACGTCTGAACTCTTTCCCTACACGACGCTCTTCCGATCTNNNATAATGAGTACAAGTAAGATGGT
CACACGTCTGAACTCTTTCCCTACACGACGCTCTTCCGATCTNNNGAATATGGCAAGCATTTTGGCATC
GAGCACACGTCTGAACTCTTTCCCTACACGACGCTCTTCCGATCTNNNCATGTCAACCTTTTTCTTTTCT
AGCACACGTCTGAACTCTTTCCCTACACGACGCTCTTCCGATCTNNNGCTGTTTTAACTTTATTGTGAT
AGCACACGTCTGAACTCTTTCCCTACACGACGCTCTTCCGATCTNNNACGTGGTGAAATTAAACAAA
GAGCACACGTCTGAACTCTTTCCCTACACGACGCTCTTCCGATCTNNNGAACAATAATGGAAATGTTA
GCACACGTCTGAACTCTTTCCCTACACGACGCTCTTCCGATCTNNNATTATACTTTAATGACCTCCCCCA
AAGAGCACACGTCTGAACTCTTTCCCTACACGACGCTCTTCCGATCTNNNTTTGGCATTGGTCTGACT
GAGCACACGTCTGAACTCTTTCCCTACACGACGCTCTTCCGATCTNNNTCAAGATTCTTGCTCAATTTA
AGCACACGTCTGAACTCTTTCCCTACACGACGCTCTTCCGATCTNNNGCTGTTTTAACTTTATTGTGAT
GCACACGTCTGAACTCTTTCCCTACACGACGCTCTTCCGATCTNNNGTTATGTGGCCATCTGTTATCAT
GCACACGTCTGAACTCTTTCCCTACACGACGCTCTTCCGATCTNNNGGACAAAAAATCTTAAGTGTACT
AGCACACGTCTGAACTCTTTCCCTACACGACGCTCTTCCGATCTNNNTATTTTGGAAAAACAAACGGA
AGCACACGTCTGAACTCTTTCCCTACACGACGCTCTTCCGATCTNNNGGAGATTTTCCTATTTCAGAG
AGCACACGTCTGAACTCTTTCCCTACACGACGCTCTTCCGATCTNNNCAGATCTTCAACTTACTTTCTTT
ACACGTCTGAACTCTTTCCCTACACGACGCTCTTCCGATCTNNNAGTGAGATCAAGAGGAAGACCATC
GAGCACACGTCTGAACTCTTTCCCTACACGACGCTCTTCCGATCTNNNGGTTGTCAAAAGGGAAGTA
CACACGTCTGAACTCTTTCCCTACACGACGCTCTTCCGATCTNNNATTATTCTGAGTTTCTTTCCTTGCAT
GCACACGTCTGAACTCTTTCCCTACACGACGCTCTTCCGATCTNNNTGCAATTATCTTAACAAGTCTAA
AGCACACGTCTGAACTCTTTCCCTACACGACGCTCTTCCGATCTNNNAAAGATCAATCTGAATGTGTT
AGCACACGTCTGAACTCTTTCCCTACACGACGCTCTTCCGATCTNNNAGAGGATTGTCCCATGATAAA
AGCACACGTCTGAACTCTTTCCCTACACGACGCTCTTCCGATCTNNNTCACATTTTTTTGAACAATTCAC
CACACGTCTGAACTCTTTCCCTACACGACGCTCTTCCGATCTNNNATGTTGGAGACATACCAGACCATC
AGCACACGTCTGAACTCTTTCCCTACACGACGCTCTTCCGATCTNNNTTTAAAATATCAAGAGATCGTC
AGCACACGTCTGAACTCTTTCCCTACACGACGCTCTTCCGATCTNNNAGTATACATGATGGAAAAGAG
GCACACGTCTGAACTCTTTCCCTACACGACGCTCTTCCGATCTNNNGATGAAACGTATGCAGATGGCA
AGCACACGTCTGAACTCTTTCCCTACACGACGCTCTTCCGATCTNNNATTGACCTTCTGTTGTCTATGC
CACACGTCTGAACTCTTTCCCTACACGACGCTCTTCCGATCTNNNTTTGGTTTTATTTTTGTCCTACTCA
AGCACACGTCTGAACTCTTTCCCTACACGACGCTCTTCCGATCTNNNCTTCTACGACTGTGTCATAGCA
GCACACGTCTGAACTCTTTCCCTACACGACGCTCTTCCGATCTNNNCCTTAAAATAGAATGTGGACCTA
GCACACGTCTGAACTCTTTCCCTACACGACGCTCTTCCGATCTNNNTCTGATTTACATAGTTTTAAGGC
AGCACACGTCTGAACTCTTTCCCTACACGACGCTCTTCCGATCTNNNAATTTATCATTTTATCCTGCCA
AGCACACGTCTGAACTCTTTCCCTACACGACGCTCTTCCGATCTNNNAAGTATATCTGTTTTTGAAGC

AGCACACGTCTGAACTCTTTCCCTACACGACGCTCTTCCGATCTNNNAAGTTACGAGTCTCAGAAATT

GCACACGTCTGAACTCTTTCCCTACACGACGCTCTTCCGATCTNNNTGTGTACAAGTTTCACATTTTAC

GCACACGTCTGAACTCTTTCCCTACACGACGCTCTTCCGATCTNNNTAGAATGACAGCCTTTTTCAGC

CACACGTCTGAACTCTTTCCCTACACGACGCTCTTCCGATCTNNNCTATTACCTTCTGATGACTTCCCA

AGCACACGTCTGAACTCTTTCCCTACACGACGCTCTTCCGATCTNNNAAGACCAATACATCTTTATCCA

CACGTCTGAACTCTTTCCCTACACGACGCTCTTCCGATCTNNNCAACTTCTACAACTTCAGGAGCATCG

GCACACGTCTGAACTCTTTCCCTACACGACGCTCTTCCGATCTNNNTTTGCTTTTCCATTAGATCAGC

AGCACACGTCTGAACTCTTTCCCTACACGACGCTCTTCCGATCTNNNCCTTTTCTAAAGGATGAAAAA

CACGTCTGAACTCTTTCCCTACACGACGCTCTTCCGATCTNNNAATCCTGGTCCAGTAAAGAGCATCG

AGCACACGTCTGAACTCTTTCCCTACACGACGCTCTTCCGATCTNNNCGTGTATGATGAAATGAGTAA

CACACGTCTGAACTCTTTCCCTACACGACGCTCTTCCGATCTNNNCCTCCTAAGGGATTGATTGGCATC

CACACGTCTGAACTCTTTCCCTACACGACGCTCTTCCGATCTNNNGCAGCTTATAGTGATTTAGGATA

CACACGTCTGAACTCTTTCCCTACACGACGCTCTTCCGATCTNNNCTTATGTACCTCAATATCATCCTC

GCACACGTCTGAACTCTTTCCCTACACGACGCTCTTCCGATCTNNNATATTCATTGCACCAAAATGTAT

ACACGTCTGAACTCTTTCCCTACACGACGCTCTTCCGATCTNNNCAGGCATAGTCAAACAATAGATCA

AGCACACGTCTGAACTCTTTCCCTACACGACGCTCTTCCGATCTNNNGGAGTTTTCAATAAGCTCTTT

AGCACACGTCTGAACTCTTTCCCTACACGACGCTCTTCCGATCTNNNATATACCCAACTACTCTGCATAT

GCACACGTCTGAACTCTTTCCCTACACGACGCTCTTCCGATCTNNNTAAATATCAGCTCTGGAATAAC

GAGCACACGTCTGAACTCTTTCCCTACACGACGCTCTTCCGATCTNNNGAATTTTACGACCAGAATTAT

GCACACGTCTGAACTCTTTCCCTACACGACGCTCTTCCGATCTNNNATTTGAAAAATTGCTTCAATCT

AGCACACGTCTGAACTCTTTCCCTACACGACGCTCTTCCGATCTNNNTCTATCTTATCTGCTATCCCATA

GAGCACACGTCTGAACTCTTTCCCTACACGACGCTCTTCCGATCTNNNGACACTTATTTTTTCTTTGTA

AGCACACGTCTGAACTCTTTCCCTACACGACGCTCTTCCGATCTNNNGTGAAGGTTTCTGAGAAAGT

AGCACACGTCTGAACTCTTTCCCTACACGACGCTCTTCCGATCTNNNTTTTCTTTTAGTTGGTCGAATT

ACACGTCTGAACTCTTTCCCTACACGACGCTCTTCCGATCTNNNTTGAACATCAGAGAGGCAATCAT

AGCACACGTCTGAACTCTTTCCCTACACGACGCTCTTCCGATCTNNNCTAAATCCCAACAATATCTCCTT

AGCACACGTCTGAACTCTTTCCCTACACGACGCTCTTCCGATCTNNNACAAAAGTGTTGAATATTGTG

GAGCACACGTCTGAACTCTTTCCCTACACGACGCTCTTCCGATCTNNNGATATGTGGATGATGATTTT

GCACACGTCTGAACTCTTTCCCTACACGACGCTCTTCCGATCTNNNCATCCTGGATGACTATGACTCAT

CACACGTCTGAACTCTTTCCCTACACGACGCTCTTCCGATCTNNNTGATGTGAACAGCATATTCGCATC

AGCACACGTCTGAACTCTTTCCCTACACGACGCTCTTCCGATCTNNNTTGTCAACAAGCGAAGAAAAC

AGCACACGTCTGAACTCTTTCCCTACACGACGCTCTTCCGATCTNNNATCAAAATCCTACCTTTACTCTT

CACACGTCTGAACTCTTTCCCTACACGACGCTCTTCCGATCTNNNCACCTTCTTCTGTGAAACTGCATC

AGCACACGTCTGAACTCTTTCCCTACACGACGCTCTTCCGATCTNNNGCCTTATTTAAGCTAAATATGC

CACACGTCTGAACTCTTTCCCTACACGACGCTCTTCCGATCTNNNGTGAAATTAACTGTTAGGAGAA

CACACGTCTGAACTCTTTCCCTACACGACGCTCTTCCGATCTNNNAAGGAAGTCTTGAATTTGGCCATC

AGCACACGTCTGAACTCTTTCCCTACACGACGCTCTTCCGATCTNNNACTGTTTATGAAGATGAAACTT

GAGCACACGTCTGAACTCTTTCCCTACACGACGCTCTTCCGATCTNNNTCCATTTACTGAATTTATGGA

ACACGTCTGAACTCTTTCCCTACACGACGCTCTTCCGATCTNNNGCGTACTTTGTCTTCTTGTAACATCG

AGCACACGTCTGAACTCTTTCCCTACACGACGCTCTTCCGATCTNNNATCTTTATACCTATGCTGGAAA

ACACGTCTGAACTCTTTCCCTACACGACGCTCTTCCGATCTNNNAGTAACATATCCCGGATGAGCATC

AGCACACGTCTGAACTCTTTCCCTACACGACGCTCTTCCGATCTNNNTCAGGATTATCCAACTTTACAG

ACACGTCTGAACTCTTTCCCTACACGACGCTCTTCCGATCTNNNGAGACCACATGAATACACACCATC

CACACGTCTGAACTCTTTCCCTACACGACGCTCTTCCGATCTNNNTTACAGTTGTCGGATTTGTCCATC
GAGCACACGTCTGAACTCTTTCCCTACACGACGCTCTTCCGATCTNNNCTTTCTGTTCTTCGTTCATTCC
AGCACACGTCTGAACTCTTTCCCTACACGACGCTCTTCCGATCTNNNTTTTTAGGTGGCCTTTTCATCA
GCACACGTCTGAACTCTTTCCCTACACGACGCTCTTCCGATCTNNNTGTTGAAGAAATTCTTGAGAAC
CACACGTCTGAACTCTTTCCCTACACGACGCTCTTCCGATCTNNNCTTCCTTTACTTGATGCTCTTCATC
CACGTCTGAACTCTTTCCCTACACGACGCTCTTCCGATCTNNNGATGTTCCTCAGTCCAAAACCATCG
GCACACGTCTGAACTCTTTCCCTACACGACGCTCTTCCGATCTNNNTAACTATATTCGGTCCTTCATAC
CACGTCTGAACTCTTTCCCTACACGACGCTCTTCCGATCTNNNTAAAAAATAAGGCTATTGGATGCCA
AGCACACGTCTGAACTCTTTCCCTACACGACGCTCTTCCGATCTNNNCGGCAAAAAGGTTATATTTAT
AGCACACGTCTGAACTCTTTCCCTACACGACGCTCTTCCGATCTNNNAATTGAACCTTTAAGTATGAG
AGCACACGTCTGAACTCTTTCCCTACACGACGCTCTTCCGATCTNNNTGTTAACAACATGAACTATCTT
AGCACACGTCTGAACTCTTTCCCTACACGACGCTCTTCCGATCTNNNTTCACCTACTGTAATCTAGTTT
GCACACGTCTGAACTCTTTCCCTACACGACGCTCTTCCGATCTNNNACAGACATCCATGAAATACAATA
AGCACACGTCTGAACTCTTTCCCTACACGACGCTCTTCCGATCTNNNCAAACAGTTGGAGGAAAGAG
GAGCACACGTCTGAACTCTTTCCCTACACGACGCTCTTCCGATCTNNNCCTTTTTTCATACTCATGCTA
AGCACACGTCTGAACTCTTTCCCTACACGACGCTCTTCCGATCTNNNTAGTGGCTTAGATAAAAAATC
GCACACGTCTGAACTCTTTCCCTACACGACGCTCTTCCGATCTNNNGTTAACATTCATTCTCCTTTGAC
GCACACGTCTGAACTCTTTCCCTACACGACGCTCTTCCGATCTNNNAAGAATTTCCAGAGAGTTGTAT
AGCACACGTCTGAACTCTTTCCCTACACGACGCTCTTCCGATCTNNNGAGAAAGCAAAGGAATTCTT
AGCACACGTCTGAACTCTTTCCCTACACGACGCTCTTCCGATCTNNNAAATTCTTCATCATATGTTTGC
AGCACACGTCTGAACTCTTTCCCTACACGACGCTCTTCCGATCTNNNTACTGATAGATAGATAGGGTC
GAGCACACGTCTGAACTCTTTCCCTACACGACGCTCTTCCGATCTNNNTAATTGTGTTTCTGTATAGGT
AGCACACGTCTGAACTCTTTCCCTACACGACGCTCTTCCGATCTNNNCTCTGCTATGATCAGGGTATC
GCACACGTCTGAACTCTTTCCCTACACGACGCTCTTCCGATCTNNNATTAACAATCGGGGAAAAATAA
AGCACACGTCTGAACTCTTTCCCTACACGACGCTCTTCCGATCTNNNTAAATCTAGATATCAGCTGAT
GAGCACACGTCTGAACTCTTTCCCTACACGACGCTCTTCCGATCTNNNTCTCTACCAATATTAAGCCTT
CACACGTCTGAACTCTTTCCCTACACGACGCTCTTCCGATCTNNNAGAGGAATTCTTAGACAGATAAC
ACACGTCTGAACTCTTTCCCTACACGACGCTCTTCCGATCTNNNCCACCATCATCATAATCACCCATCG
AGCACACGTCTGAACTCTTTCCCTACACGACGCTCTTCCGATCTNNNATGCTTTTTAACGTTTTCTTTT
GAGCACACGTCTGAACTCTTTCCCTACACGACGCTCTTCCGATCTNNNATAACATTAAGCCAATTTTCT
CACACGTCTGAACTCTTTCCCTACACGACGCTCTTCCGATCTNNNTTGATAGTACTCTGTCATTGTCCA
GCACACGTCTGAACTCTTTCCCTACACGACGCTCTTCCGATCTNNNATCTCCATTTATTGCTTCTGTTAC
CACACGTCTGAACTCTTTCCCTACACGACGCTCTTCCGATCTNNNTTTTTTGTTTTAAGATAGCTTGCC
GCACACGTCTGAACTCTTTCCCTACACGACGCTCTTCCGATCTNNNATCCTTAAAGCTTCTGATTGTTT
CACACGTCTGAACTCTTTCCCTACACGACGCTCTTCCGATCTNNNTATCAAGGACTTCTTTTAAGTGAC
AGCACACGTCTGAACTCTTTCCCTACACGACGCTCTTCCGATCTNNNTTACAGATCATCAGTATTTGG
AGCACACGTCTGAACTCTTTCCCTACACGACGCTCTTCCGATCTNNNAGCAATTACCATTTTTTTTTG
AGCACACGTCTGAACTCTTTCCCTACACGACGCTCTTCCGATCTNNNCATTTTAATTCAGTTCCGTTTAC
CACGTCTGAACTCTTTCCCTACACGACGCTCTTCCGATCTNNNCGTATAATTTTTTGGGAAGTGAACA
AGCACACGTCTGAACTCTTTCCCTACACGACGCTCTTCCGATCTNNNCTTCAAGTTTTCCATGATCTTT
GCACACGTCTGAACTCTTTCCCTACACGACGCTCTTCCGATCTNNNCTTGCAGGGATTTCATATTAAA
AGCACACGTCTGAACTCTTTCCCTACACGACGCTCTTCCGATCTNNNCTACTTCTTCCACTTCCTCTACAT
GAGCACACGTCTGAACTCTTTCCCTACACGACGCTCTTCCGATCTNNNTTTCTACCTTATTTTACTTGCA

GCACACGTCTGAACTCTTTCCCTACACGACGCTCTTCCGATCTNNNATTCTCTAAAATAACCAAAGCAG
AGCACACGTCTGAACTCTTTCCCTACACGACGCTCTTCCGATCTNNNATAAATCATCTAAGCCATCCTCCA
GAGCACACGTCTGAACTCTTTCCCTACACGACGCTCTTCCGATCTNNNTATTTTCTTTTTCAGTTCTTTCG
AGCACACGTCTGAACTCTTTCCCTACACGACGCTCTTCCGATCTNNNAAGTGATAGAAATCGATCAGTT
AGCACACGTCTGAACTCTTTCCCTACACGACGCTCTTCCGATCTNNNGTCAGTGTATGTAGTAACCATAT
GAGCACACGTCTGAACTCTTTCCCTACACGACGCTCTTCCGATCTNNNGATTCTTCAATAGACTGGAAG
CACACGTCTGAACTCTTTCCCTACACGACGCTCTTCCGATCTNNNGCCGTTAGGAGTATTTGAATCATC
CACACGTCTGAACTCTTTCCCTACACGACGCTCTTCCGATCTNNNATTGATCAGGATTTATTCACAGACA
GCACACGTCTGAACTCTTTCCCTACACGACGCTCTTCCGATCTNNNAACTTGTTAAAATTCACAAAGTG
GCACACGTCTGAACTCTTTCCCTACACGACGCTCTTCCGATCTNNNATTACCTTTCTGCTCAGGAGCATG
AGCACACGTCTGAACTCTTTCCCTACACGACGCTCTTCCGATCTNNNTTTTTTTGTTGGGATTTCAGATA
ACACGTCTGAACTCTTTCCCTACACGACGCTCTTCCGATCTNNNCTGTTGTTAAACTGTCTTCACCATCGG
AGCACACGTCTGAACTCTTTCCCTACACGACGCTCTTCCGATCTNNNAACTGCTATCTACTGCTTAACCAT
CACACGTCTGAACTCTTTCCCTACACGACGCTCTTCCGATCTNNNTGTTTACAGGGAATGTAGTTTCCAT
ACACGTCTGAACTCTTTCCCTACACGACGCTCTTCCGATCTNNNTGTTTACTGTAAGACAAAAAACACA
GCACACGTCTGAACTCTTTCCCTACACGACGCTCTTCCGATCTNNNTATTACTATGAGTCAAGCCACTAC
AGCACACGTCTGAACTCTTTCCCTACACGACGCTCTTCCGATCTNNNATGAACACACTCTTTAATTTCATG
GAGCACACGTCTGAACTCTTTCCCTACACGACGCTCTTCCGATCTNNNCAAACAATCATGTAATAGACAG
AGCACACGTCTGAACTCTTTCCCTACACGACGCTCTTCCGATCTNNNAAACATTCATGAAACATTTGTTG
AGCACACGTCTGAACTCTTTCCCTACACGACGCTCTTCCGATCTNNNTTTATGACAGAACCAATTTCAA
CACACGTCTGAACTCTTTCCCTACACGACGCTCTTCCGATCTNNNTGAAGAAGATGTGGATAACTACCA
AGCACACGTCTGAACTCTTTCCCTACACGACGCTCTTCCGATCTNNNTGAAATAAAAATCTACAATGGG
GAGCACACGTCTGAACTCTTTCCCTACACGACGCTCTTCCGATCTNNNGAAATATTAAGAAGTCTCCCTT
ACACGTCTGAACTCTTTCCCTACACGACGCTCTTCCGATCTNNNCAATCCATGTTCATAAAGAGGAACA
CACACGTCTGAACTCTTTCCCTACACGACGCTCTTCCGATCTNNNTCTTCCTACCCAGATCCTAACATCG
AGCACACGTCTGAACTCTTTCCCTACACGACGCTCTTCCGATCTNNNAATCTTATGAATGTCAACAATGT
GAGCACACGTCTGAACTCTTTCCCTACACGACGCTCTTCCGATCTNNNAAGTTTTGTTTAAAAAATGTG
AGCACACGTCTGAACTCTTTCCCTACACGACGCTCTTCCGATCTNNNCTCAGCATATTTTTTCATCTCTTC
GCACACGTCTGAACTCTTTCCCTACACGACGCTCTTCCGATCTNNNTTCTTCTTCACTTCTGTTATAGTCA
CACGTCTGAACTCTTTCCCTACACGACGCTCTTCCGATCTNNNTTATAGATGCGAATCCTTTCCCATCGG
AGCACACGTCTGAACTCTTTCCCTACACGACGCTCTTCCGATCTNNNATGAATCTTTCTATAATTGCAG
CACACGTCTGAACTCTTTCCCTACACGACGCTCTTCCGATCTNNNCTGATAGATTCCAGAGAGATTGCA
AGCACACGTCTGAACTCTTTCCCTACACGACGCTCTTCCGATCTNNNGTAAGTGCACATTCTAAGAACT
GCACACGTCTGAACTCTTTCCCTACACGACGCTCTTCCGATCTNNNGAGATGACAAAAATGAGACACCA
AGCACACGTCTGAACTCTTTCCCTACACGACGCTCTTCCGATCTNNNGACATTAAAAGACCTACCAATTG
AGCACACGTCTGAACTCTTTCCCTACACGACGCTCTTCCGATCTNNNTAGGCTGTGTAGGTAAAATTTA
AGCACACGTCTGAACTCTTTCCCTACACGACGCTCTTCCGATCTNNNATGGGATTGAAAACATAGACTT
AGCACACGTCTGAACTCTTTCCCTACACGACGCTCTTCCGATCTNNNTAAACTGTGTCTTTTGTAGTTTG
CACACGTCTGAACTCTTTCCCTACACGACGCTCTTCCGATCTNNNAATTCTTTTGGTTTTTGTCTAGGCA
AGCACACGTCTGAACTCTTTCCCTACACGACGCTCTTCCGATCTNNNTTAGTACTGGATCTAACTCATTG
GAGCACACGTCTGAACTCTTTCCCTACACGACGCTCTTCCGATCTNNNGTATCTATGCTTTCTAGACAGA
GAGCACACGTCTGAACTCTTTCCCTACACGACGCTCTTCCGATCTNNNTTAATTTTGGTTCCACAGAATC
GCACACGTCTGAACTCTTTCCCTACACGACGCTCTTCCGATCTNNNGAGATATGTCTAGGTCAGGCCAT

GCACACGTCTGAACTCTTTCCCTACACGACGCTCTTCCGATCTNNNCCCAGAAACTTAAATATCTGAATC
ACACGTCTGAACTCTTTCCCTACACGACGCTCTTCCGATCTNNNTTACCTAGGTAATTTTCCATACCCATC
GAGCACACGTCTGAACTCTTTCCCTACACGACGCTCTTCCGATCTNNNCTGTTTTAATCTGAAAAGAGC
AGCACACGTCTGAACTCTTTCCCTACACGACGCTCTTCCGATCTNNNTTCTGACATCTCTTCTTTTCTTTC
AGCACACGTCTGAACTCTTTCCCTACACGACGCTCTTCCGATCTNNNATTGATTTCACTTATTACCTTGGC
GCACACGTCTGAACTCTTTCCCTACACGACGCTCTTCCGATCTNNNAATAGCAGAATTCCTAGTCAAAAC
AGCACACGTCTGAACTCTTTCCCTACACGACGCTCTTCCGATCTNNNTTAATGTACAGAGTTTTTGGATC
ACACGTCTGAACTCTTTCCCTACACGACGCTCTTCCGATCTNNNAAATGCTAGTTGCTTGTTTGCATCGG
AGCACACGTCTGAACTCTTTCCCTACACGACGCTCTTCCGATCTNNNTGAGAAAACCTATGAATGTAG
AGCACACGTCTGAACTCTTTCCCTACACGACGCTCTTCCGATCTNNNGTTCTCACCCATATATTGATTTT
AGCACACGTCTGAACTCTTTCCCTACACGACGCTCTTCCGATCTNNNTTTTGCAAACCATCAATATCTAA
GCACACGTCTGAACTCTTTCCCTACACGACGCTCTTCCGATCTNNNGTTGCATATTCTTCATTTCATTGC
GCACACGTCTGAACTCTTTCCCTACACGACGCTCTTCCGATCTNNNAATCATCTTTGTTGATTTCATCACA
GAGCACACGTCTGAACTCTTTCCCTACACGACGCTCTTCCGATCTNNNCTGTTTTTGGAGTCTTTTCTTA
CACACGTCTGAACTCTTTCCCTACACGACGCTCTTCCGATCTNNNTGCACCATTTTCTCTATCAGCATCGC
GAGCACACGTCTGAACTCTTTCCCTACACGACGCTCTTCCGATCTNNNAATGAGCTGAGTTGCATTTCC
AGCACACGTCTGAACTCTTTCCCTACACGACGCTCTTCCGATCTNNNCTCATCTCTTCTCTTTTCCTCCAT
CACACGTCTGAACTCTTTCCCTACACGACGCTCTTCCGATCTNNNGGAAGTTTTCTGAAGATCAAAGCA
AGCACACGTCTGAACTCTTTCCCTACACGACGCTCTTCCGATCTNNNTTAAATATGAGATTTACCAAGC
ACACGTCTGAACTCTTTCCCTACACGACGCTCTTCCGATCTNNNAGGAAGAATGCGATGTAAAATCATC
GCACACGTCTGAACTCTTTCCCTACACGACGCTCTTCCGATCTNNNCTTAAAGAATGAGAACTTCGACCA
CACACGTCTGAACTCTTTCCCTACACGACGCTCTTCCGATCTNNNCTTTCAAACATGAGGAGATGCATC
GCACACGTCTGAACTCTTTCCCTACACGACGCTCTTCCGATCTNNNATCAAAGTTCTGAGACTTAATCTC
CACACGTCTGAACTCTTTCCCTACACGACGCTCTTCCGATCTNNNCTGATTTTCTGATCCAGCACCATCGC
AGCACACGTCTGAACTCTTTCCCTACACGACGCTCTTCCGATCTNNNGGAGAATGACAGAGATATCAA
CACACGTCTGAACTCTTTCCCTACACGACGCTCTTCCGATCTNNNTATAACATTGAAGAACAAGAAGCC
CACACGTCTGAACTCTTTCCCTACACGACGCTCTTCCGATCTNNNTTGTAAATTCAACTCATTCTTTTCCA
GAGCACACGTCTGAACTCTTTCCCTACACGACGCTCTTCCGATCTNNNAACAAGAAGGTACAGAACACC
GAGCACACGTCTGAACTCTTTCCCTACACGACGCTCTTCCGATCTNNNTTTACCTGGAACTCCTGATTCA
AGCACACGTCTGAACTCTTTCCCTACACGACGCTCTTCCGATCTNNNATCATAGATGTTTTCACCTCTACC
GAGCACACGTCTGAACTCTTTCCCTACACGACGCTCTTCCGATCTNNNCTTTCTTGGCTTTAAGTAAGTA
GAGCACACGTCTGAACTCTTTCCCTACACGACGCTCTTCCGATCTNNNCTCCAATAAGAGCATCCATGCA
AGCACACGTCTGAACTCTTTCCCTACACGACGCTCTTCCGATCTNNNATACTTTTACTTCTTTACTGGGTC
AGCACACGTCTGAACTCTTTCCCTACACGACGCTCTTCCGATCTNNNTACTAAATCAAACTTCTCGACTT
AGAGCACACGTCTGAACTCTTTCCCTACACGACGCTCTTCCGATCTNNNGATGTGTAAAGAATATACAC
AGCACACGTCTGAACTCTTTCCCTACACGACGCTCTTCCGATCTNNNCATCAACACCTTCTGTAGAGCA
GAGCACACGTCTGAACTCTTTCCCTACACGACGCTCTTCCGATCTNNNGAGTCTTTGCTTTTAACAAAAT
AGCACACGTCTGAACTCTTTCCCTACACGACGCTCTTCCGATCTNNNCTAACCTTTTGTGTCTTGTATGC
AGCACACGTCTGAACTCTTTCCCTACACGACGCTCTTCCGATCTNNNCACATCCCAACTTGAATAAAAG
AGCACACGTCTGAACTCTTTCCCTACACGACGCTCTTCCGATCTNNNTAGCTAGTGATGTTTTTTTGTTC
GCACACGTCTGAACTCTTTCCCTACACGACGCTCTTCCGATCTNNNATTATATCAGAATAGCCTCTAGCC
GCACACGTCTGAACTCTTTCCCTACACGACGCTCTTCCGATCTNNNCTTTTCATTCGTTTCACTTGACATC
AGCACACGTCTGAACTCTTTCCCTACACGACGCTCTTCCGATCTNNNAAACTCAACTCTGTAATAAGAAA

CACACGTCTGAACTCTTTCCCTACACGACGCTCTTCCGATCTNNNTAGAATGAGGAAGAACCACTCAT

GCACACGTCTGAACTCTTTCCCTACACGACGCTCTTCCGATCTNNNCTTAGATCATAGCCATGAAAGA

ACACGTCTGAACTCTTTCCCTACACGACGCTCTTCCGATCTNNNTAGAAAAAGTTAAACCAAACAGGC

AGCACACGTCTGAACTCTTTCCCTACACGACGCTCTTCCGATCTNNNACTAACATTTCTTACCTGGTTA

CACACGTCTGAACTCTTTCCCTACACGACGCTCTTCCGATCTNNNCTTACCTGAGAGAGAGGTTCCAT

CACGTCTGAACTCTTTCCCTACACGACGCTCTTCCGATCTNNNATCCCATCCAGATTGAAAATGCATC

AGCACACGTCTGAACTCTTTCCCTACACGACGCTCTTCCGATCTNNNATGTTATTGTCACAATAAGCAT

AGCACACGTCTGAACTCTTTCCCTACACGACGCTCTTCCGATCTNNNATGTAGTTGCCAAATATCCTT

ACACGTCTGAACTCTTTCCCTACACGACGCTCTTCCGATCTNNNAAAGTCCTGTAGCTCATAGTCATCG

CACACGTCTGAACTCTTTCCCTACACGACGCTCTTCCGATCTNNNTCAAAATCACTGATTTTCTTCTTC

GCACACGTCTGAACTCTTTCCCTACACGACGCTCTTCCGATCTNNNTTGTTTTCTTTAACTAGCATGTG

GCACACGTCTGAACTCTTTCCCTACACGACGCTCTTCCGATCTNNNAAGTCCAGACTGAATAATACTA

AGCACACGTCTGAACTCTTTCCCTACACGACGCTCTTCCGATCTNNNTATGTTTGTTTATGTGCTGTTT

ACACGTCTGAACTCTTTCCCTACACGACGCTCTTCCGATCTNNNTCATATCATTAGGTACACAGAAACA

ACACGTCTGAACTCTTTCCCTACACGACGCTCTTCCGATCTNNNTAGAACTTACATAAGGATTGCCCAT

AGCACACGTCTGAACTCTTTCCCTACACGACGCTCTTCCGATCTNNNTGTTGCAATGTATTCTGATAT

AGCACACGTCTGAACTCTTTCCCTACACGACGCTCTTCCGATCTNNNCATTTTGGAATCAATATGTCTC

CACACGTCTGAACTCTTTCCCTACACGACGCTCTTCCGATCTNNNGCATCACATTCCTCAAGAAGCAT

AGCACACGTCTGAACTCTTTCCCTACACGACGCTCTTCCGATCTNNNATTATTTGTTGTTTGATCACTC

CACACGTCTGAACTCTTTCCCTACACGACGCTCTTCCGATCTNNNTGCTTCTAATTCACCTTGTGCATC

CACGTCTGAACTCTTTCCCTACACGACGCTCTTCCGATCTNNNGTGTGTTGCCATAACAAAATACATC

GCACACGTCTGAACTCTTTCCCTACACGACGCTCTTCCGATCTNNNTACCAAATGTAAAAATAACTGG

AGCACACGTCTGAACTCTTTCCCTACACGACGCTCTTCCGATCTNNNATAATGATAGTATTCCATTGC

CACACGTCTGAACTCTTTCCCTACACGACGCTCTTCCGATCTNNNGATGTTAACACAAATGAAGTCCC

AGCACACGTCTGAACTCTTTCCCTACACGACGCTCTTCCGATCTNNNCTTTTTCAGTACATCCACTTAA

GAGCACACGTCTGAACTCTTTCCCTACACGACGCTCTTCCGATCTNNNTTTTACATACCTCATCCTCATT

GCACACGTCTGAACTCTTTCCCTACACGACGCTCTTCCGATCTNNNGTCTCTTGAATCCTCATCAGCAT

ACACGTCTGAACTCTTTCCCTACACGACGCTCTTCCGATCTNNNGTTCTTCCACTGCTTACTACCATCGG

ACACGTCTGAACTCTTTCCCTACACGACGCTCTTCCGATCTNNNGTGGCTTTCACTGTAAATTTGCATC

ACACGTCTGAACTCTTTCCCTACACGACGCTCTTCCGATCTNNNGTGGCTTTCACTGTAAATTTGCATC

CACACGTCTGAACTCTTTCCCTACACGACGCTCTTCCGATCTNNNGGGTATAATTATGATCTCAGACAC

CACACGTCTGAACTCTTTCCCTACACGACGCTCTTCCGATCTNNNTTCTCCTTCATAGGCTATTTTAGC

ACACGTCTGAACTCTTTCCCTACACGACGCTCTTCCGATCTNNNCTAGTGAAAAAAAGAAAAGAAGC

GCACACGTCTGAACTCTTTCCCTACACGACGCTCTTCCGATCTNNNAAGCATCTTTGATAACCCATTAC

AGCACACGTCTGAACTCTTTCCCTACACGACGCTCTTCCGATCTNNNATTAATGTGGATGATGAGATT

CACACGTCTGAACTCTTTCCCTACACGACGCTCTTCCGATCTNNNTTAAACAGGACTCACAAACGCATC

AGCACACGTCTGAACTCTTTCCCTACACGACGCTCTTCCGATCTNNNAACAATTATGTGTGGATAAGA

AGCACACGTCTGAACTCTTTCCCTACACGACGCTCTTCCGATCTNNNCCAAACTCAAGACAATTACTTAT

CACACGTCTGAACTCTTTCCCTACACGACGCTCTTCCGATCTNNNGAAAGCTGTTCAGAAACATCCATC

GCACACGTCTGAACTCTTTCCCTACACGACGCTCTTCCGATCTNNNGGAACTGCAACTATCAACACCAT

GAGCACACGTCTGAACTCTTTCCCTACACGACGCTCTTCCGATCTNNNATGATAAAAACCAAGATGTT

GCACACGTCTGAACTCTTTCCCTACACGACGCTCTTCCGATCTNNNCTGTTTTTCAGGTTCATTCTGCAT

CACACGTCTGAACTCTTTCCCTACACGACGCTCTTCCGATCTNNNAGAAAGAAAAGTCAAAGTGTAA

GCACACGTCTGAACTCTTTCCCTACACGACGCTCTTCCGATCTNNNTCTGAACTTTTCCACTTTAGAGCA
AGCACACGTCTGAACTCTTTCCCTACACGACGCTCTTCCGATCTNNNCAGGATAATACACATCACAGTA
GCACACGTCTGAACTCTTTCCCTACACGACGCTCTTCCGATCTNNNGTAAAGGTGCCTCTTAAAGAACA
AGCACACGTCTGAACTCTTTCCCTACACGACGCTCTTCCGATCTNNNATTAACCTGATTCACTGCTCCAT
GAGCACACGTCTGAACTCTTTCCCTACACGACGCTCTTCCGATCTNNNTCTTTTTTTCTTTTAGGTTCTG
AGCACACGTCTGAACTCTTTCCCTACACGACGCTCTTCCGATCTNNNATAGACACAATTCTATCCTAAGC
ACACGTCTGAACTCTTTCCCTACACGACGCTCTTCCGATCTNNNTAGAAAATCCAGGCCCATAACATCGG
GCACACGTCTGAACTCTTTCCCTACACGACGCTCTTCCGATCTNNNCATCCATCTCCTTTGCTTTGCATCC
GAGCACACGTCTGAACTCTTTCCCTACACGACGCTCTTCCGATCTNNNACTTTTATTTTTGTCTAGGATG
GAGCACACGTCTGAACTCTTTCCCTACACGACGCTCTTCCGATCTNNNGCAATTTCTTTTCTAATTAGTG
CACACGTCTGAACTCTTTCCCTACACGACGCTCTTCCGATCTNNNCTAGAAATGTGGTTCTCGTGCATCC
GAGCACACGTCTGAACTCTTTCCCTACACGACGCTCTTCCGATCTNNNCTTAATGTGAGAGAGACAAAA
CACACGTCTGAACTCTTTCCCTACACGACGCTCTTCCGATCTNNNGACAGAAGGAGAAGATGGAGCAT
AGCACACGTCTGAACTCTTTCCCTACACGACGCTCTTCCGATCTNNNGAAGAGATTGATCATGTAATTC
ACACGTCTGAACTCTTTCCCTACACGACGCTCTTCCGATCTNNNCACATGTGTATGTCCTGTCTCATCGG
GCACACGTCTGAACTCTTTCCCTACACGACGCTCTTCCGATCTNNNTCTATTACATGCTTAGGGAAATTC
GAGCACACGTCTGAACTCTTTCCCTACACGACGCTCTTCCGATCTNNNAATATTGATCTTGCCTTCTTTTG
AGCACACGTCTGAACTCTTTCCCTACACGACGCTCTTCCGATCTNNNTACTAGACAGTCCTCAAACATAC
ACACGTCTGAACTCTTTCCCTACACGACGCTCTTCCGATCTNNNGAAGATACTTACTGTAGTTCTCACAT
AGCACACGTCTGAACTCTTTCCCTACACGACGCTCTTCCGATCTNNNGAAAGCTTTGCAAGAGTATGCA
GAGCACACGTCTGAACTCTTTCCCTACACGACGCTCTTCCGATCTNNNGGTGATTGTATTCTGTGTACT
CACACGTCTGAACTCTTTCCCTACACGACGCTCTTCCGATCTNNNGATTCCACTGACTCAGAAAGCATC
GCACACGTCTGAACTCTTTCCCTACACGACGCTCTTCCGATCTNNNCAGAATTCACCCTTCATACACCAT
GAGCACACGTCTGAACTCTTTCCCTACACGACGCTCTTCCGATCTNNNAATATGTTTTTTTTCCTTCCCC
GCACACGTCTGAACTCTTTCCCTACACGACGCTCTTCCGATCTNNNGTTGTAGAGGATGGGGTTTACAT
GAGCACACGTCTGAACTCTTTCCCTACACGACGCTCTTCCGATCTNNNGTTTAGAACAGAAACTTAGCT
GAGCACACGTCTGAACTCTTTCCCTACACGACGCTCTTCCGATCTNNNCAATATTGTGAGAGAAGAAG
CACACGTCTGAACTCTTTCCCTACACGACGCTCTTCCGATCTNNNAAGATAAACAATAATTGGGTTCCCA
AGCACACGTCTGAACTCTTTCCCTACACGACGCTCTTCCGATCTNNNAATTCAAGCCATGGAAATTAAG
AGCACACGTCTGAACTCTTTCCCTACACGACGCTCTTCCGATCTNNNCTGTTGAATGGTTAACTGTAAAT
ACACGTCTGAACTCTTTCCCTACACGACGCTCTTCCGATCTNNNCTTCTCTCTCCATGATTACTGCATCGG
AGCACACGTCTGAACTCTTTCCCTACACGACGCTCTTCCGATCTNNNCTCTTCTAAATCAGCCTTTTTCCA
AGCACACGTCTGAACTCTTTCCCTACACGACGCTCTTCCGATCTNNNGATGATGGTAACCAGGTACATC
CACACGTCTGAACTCTTTCCCTACACGACGCTCTTCCGATCTNNNTTTGAATTTATGGTATTCTGTGCCA
CACACGTCTGAACTCTTTCCCTACACGACGCTCTTCCGATCTNNNACAAAAGCATCAGAATTATCTTTCAT
AGCACACGTCTGAACTCTTTCCCTACACGACGCTCTTCCGATCTNNNAGACACTAAAACCAATTTTGAA
CACACGTCTGAACTCTTTCCCTACACGACGCTCTTCCGATCTNNNCAACTTCATACCGTCTTCTTCATCGC
GAGCACACGTCTGAACTCTTTCCCTACACGACGCTCTTCCGATCTNNNTAAGTGATCATATTTACCACCA
GAGCACACGTCTGAACTCTTTCCCTACACGACGCTCTTCCGATCTNNNTAAGTGATCATATTTACCACCA
AGCACACGTCTGAACTCTTTCCCTACACGACGCTCTTCCGATCTNNNTGAAATACCACCTAATCTTTCTT
ACACGTCTGAACTCTTTCCCTACACGACGCTCTTCCGATCTNNNATCCAAGAATATGCCCTTTCCATCGG
GCACACGTCTGAACTCTTTCCCTACACGACGCTCTTCCGATCTNNNCTGACACATACAATCAGTTTTTACA
CACACGTCTGAACTCTTTCCCTACACGACGCTCTTCCGATCTNNNCGTCATTGCTTTTGGACTATCATCG

AGCACACGTCTGAACTCTTTCCCTACACGACGCTCTTCCGATCTNNNAAATATACACAGGGCAGAGATA
GAGCACACGTCTGAACTCTTTCCCTACACGACGCTCTTCCGATCTNNNCTTCATTAGTTCAACAGCTTGC
CACACGTCTGAACTCTTTCCCTACACGACGCTCTTCCGATCTNNNATTCAGGATTTCAATGACTTTTACAT
GCACACGTCTGAACTCTTTCCCTACACGACGCTCTTCCGATCTNNNCTTGTATTTTTCAGGCTATTTTCCA
ACACGTCTGAACTCTTTCCCTACACGACGCTCTTCCGATCTNNNATGTTATAGTCCTTTGCATCCCATCG
CACACGTCTGAACTCTTTCCCTACACGACGCTCTTCCGATCTNNNCTATGAAATACAGTGACAGTGTCAT
CACACGTCTGAACTCTTTCCCTACACGACGCTCTTCCGATCTNNNGAAACGTCTAGATGCTCAACCATCC
CACACGTCTGAACTCTTTCCCTACACGACGCTCTTCCGATCTNNNAGGCAATACAATTCCTTTAATTTCA
AGCACACGTCTGAACTCTTTCCCTACACGACGCTCTTCCGATCTNNNATAATCTACAATTATCCAGGCTT
AGCACACGTCTGAACTCTTTCCCTACACGACGCTCTTCCGATCTNNNATACTCTTTCTGACTGATGACTAC
AGCACACGTCTGAACTCTTTCCCTACACGACGCTCTTCCGATCTNNNTTAAAGCAAATACATTGTTCTG
ACACGTCTGAACTCTTTCCCTACACGACGCTCTTCCGATCTNNNTAACTTTCAGCTGTTAGGATTCCATC
AGCACACGTCTGAACTCTTTCCCTACACGACGCTCTTCCGATCTNNNAACATCAGAAAATACCATTTCAT
GCACACGTCTGAACTCTTTCCCTACACGACGCTCTTCCGATCTNNNTGTTGGATTTTATTTGCATTTTCCA
AGCACACGTCTGAACTCTTTCCCTACACGACGCTCTTCCGATCTNNNAGCCAATATGACTTATTTACAA
GCACACGTCTGAACTCTTTCCCTACACGACGCTCTTCCGATCTNNNATAGACTCCAATCGCATTCTTCATC
GCACACGTCTGAACTCTTTCCCTACACGACGCTCTTCCGATCTNNNGGTCTTGACACCATAGATTATGCA
AGCACACGTCTGAACTCTTTCCCTACACGACGCTCTTCCGATCTNNNGTAGGGCATATTTTACATCTTTG
CACACGTCTGAACTCTTTCCCTACACGACGCTCTTCCGATCTNNNATGTGATTTGTTTGAATTCAGATCA
GCACACGTCTGAACTCTTTCCCTACACGACGCTCTTCCGATCTNNNTTTCAGCTCCTATATCATTTTTGCA
GAGCACACGTCTGAACTCTTTCCCTACACGACGCTCTTCCGATCTNNNACAATCCTCTTCATGTACTTGCA
AGCACACGTCTGAACTCTTTCCCTACACGACGCTCTTCCGATCTNNNCAAGAAGCATTTTTAAAGCATTT
AGCACACGTCTGAACTCTTTCCCTACACGACGCTCTTCCGATCTNNNAGCAAGCTATCAAAAGAAATTT
GAGCACACGTCTGAACTCTTTCCCTACACGACGCTCTTCCGATCTNNNAAAGATACACATAGCCAGAAA
CACACGTCTGAACTCTTTCCCTACACGACGCTCTTCCGATCTNNNCCCTCATTAACTTAATGGAATTGCA
GCACACGTCTGAACTCTTTCCCTACACGACGCTCTTCCGATCTNNNAATGACTACAAAAAACTGTCAATG
GCACACGTCTGAACTCTTTCCCTACACGACGCTCTTCCGATCTNNNGATTTACTGAATACATCCATGACC
GCACACGTCTGAACTCTTTCCCTACACGACGCTCTTCCGATCTNNNGATTTACTGAATACATCCATGACC
AGCACACGTCTGAACTCTTTCCCTACACGACGCTCTTCCGATCTNNNGAAAGCAAATGTGTAACCATCC
AGCACACGTCTGAACTCTTTCCCTACACGACGCTCTTCCGATCTNNNAAATGAGAACTTTACCCACAATC
GCACACGTCTGAACTCTTTCCCTACACGACGCTCTTCCGATCTNNNGAGTAAGTCTATCATCAATTTGG
AGAGCACACGTCTGAACTCTTTCCCTACACGACGCTCTTCCGATCTNNNGGGTAGAAAAAATGAATACT
AGCACACGTCTGAACTCTTTCCCTACACGACGCTCTTCCGATCTNNNCAGATTCATCTAACATTGCAATA
GAGCACACGTCTGAACTCTTTCCCTACACGACGCTCTTCCGATCTNNNGTTCCATAACCTGTTTATAACAA
GAGCACACGTCTGAACTCTTTCCCTACACGACGCTCTTCCGATCTNNNCCCCTCTTAGAAGTTTTTTAAA
GCACACGTCTGAACTCTTTCCCTACACGACGCTCTTCCGATCTNNNAGGAACTATGACAGTGTAAAATA
GCACACGTCTGAACTCTTTCCCTACACGACGCTCTTCCGATCTNNNAAAGCTTTGAGAATGTTCTTAAAC
AGCACACGTCTGAACTCTTTCCCTACACGACGCTCTTCCGATCTNNNTTCTTTCAGTATGTTACAGAAG
AGCACACGTCTGAACTCTTTCCCTACACGACGCTCTTCCGATCTNNNTCTTGGAAGTAACAGAACTTAAT
AGCACACGTCTGAACTCTTTCCCTACACGACGCTCTTCCGATCTNNNCATCCACACATTGTATTTTCTTAC
AGCACACGTCTGAACTCTTTCCCTACACGACGCTCTTCCGATCTNNNCGTAGTAACTTTTATTCGTGAAA
AGCACACGTCTGAACTCTTTCCCTACACGACGCTCTTCCGATCTNNNAAATCAAAGATGTTCGGATTCC
CACACGTCTGAACTCTTTCCCTACACGACGCTCTTCCGATCTNNNCTGAATAGCGGATAGTTGTCCATC

GAGCACACGTCTGAACTCTTTCCCTACACGACGCTCTTCCGATCTNNNGGATTGTAGGAATTCTGACTT
GAGCACACGTCTGAACTCTTTCCCTACACGACGCTCTTCCGATCTNNNTTAGCTCACTTAATGACGTATC
GCACACGTCTGAACTCTTTCCCTACACGACGCTCTTCCGATCTNNNGATCAGGCTTATTTTATTGAACTC
GAGCACACGTCTGAACTCTTTCCCTACACGACGCTCTTCCGATCTNNNGTATTTTCTTAGCTTTCCTGTA
GCACACGTCTGAACTCTTTCCCTACACGACGCTCTTCCGATCTNNNAATTCTTGGTCTTCTGTATCTTTCA
AGCACACGTCTGAACTCTTTCCCTACACGACGCTCTTCCGATCTNNNAGAAATGAAGTGGAAATATGA
AGCACACGTCTGAACTCTTTCCCTACACGACGCTCTTCCGATCTNNNAATGTTATAGACAGCTTTTTCTC
AGCACACGTCTGAACTCTTTCCCTACACGACGCTCTTCCGATCTNNNCGAGAATTGTTCCATGATAAAT
GAGCACACGTCTGAACTCTTTCCCTACACGACGCTCTTCCGATCTNNNTAGTATTAGACACTGATGGAT
AGCACACGTCTGAACTCTTTCCCTACACGACGCTCTTCCGATCTNNNTACTGCAATGAAAAATATTCCTC
GAGCACACGTCTGAACTCTTTCCCTACACGACGCTCTTCCGATCTNNNAGTAAAAGATTTGGGATGTT
AGCACACGTCTGAACTCTTTCCCTACACGACGCTCTTCCGATCTNNNCCAGTGGGATATTATCTAACTTT
GAGCACACGTCTGAACTCTTTCCCTACACGACGCTCTTCCGATCTNNNGTAATGAAGTTCTTCTAGTTCA
CACACGTCTGAACTCTTTCCCTACACGACGCTCTTCCGATCTNNNATAACCTTCCTAGTACCTGAGCATC
AGCACACGTCTGAACTCTTTCCCTACACGACGCTCTTCCGATCTNNNACACCTCAGATATATTTCTTCATC
GCACACGTCTGAACTCTTTCCCTACACGACGCTCTTCCGATCTNNNAGTTTCTTCTTGATTTTTTTGGAC
AGCACACGTCTGAACTCTTTCCCTACACGACGCTCTTCCGATCTNNNTATAAGTAAAAGTGTGGGCATA
GCACACGTCTGAACTCTTTCCCTACACGACGCTCTTCCGATCTNNNCAGGGACCTTACCTTATACACATC
AGCACACGTCTGAACTCTTTCCCTACACGACGCTCTTCCGATCTNNNTACCAGTGAATTTTTGCAATAAT
AGCACACGTCTGAACTCTTTCCCTACACGACGCTCTTCCGATCTNNNACAGCAAGTATTTTAAGTTTTGT
AGCACACGTCTGAACTCTTTCCCTACACGACGCTCTTCCGATCTNNNCTAATGTTGGCAAAAAATTGAAT
GAGCACACGTCTGAACTCTTTCCCTACACGACGCTCTTCCGATCTNNNATCTATTTTCTCAACTGCCATAT
AGCACACGTCTGAACTCTTTCCCTACACGACGCTCTTCCGATCTNNNATTCACATCATCTTTGTAGGTAT
AGAGCACACGTCTGAACTCTTTCCCTACACGACGCTCTTCCGATCTNNNTATACAAACAAAGCTTCAACA
AGCACACGTCTGAACTCTTTCCCTACACGACGCTCTTCCGATCTNNNCCTGGTAATTCTCATGAAAAATA
GCACACGTCTGAACTCTTTCCCTACACGACGCTCTTCCGATCTNNNCTTTAACCAGCTATAAACAAGAG
GAGCACACGTCTGAACTCTTTCCCTACACGACGCTCTTCCGATCTNNNCTGTCCACAAATACCATGATAT
AGCACACGTCTGAACTCTTTCCCTACACGACGCTCTTCCGATCTNNNATATTACATTTCAGAGTTCACCT
AGCACACGTCTGAACTCTTTCCCTACACGACGCTCTTCCGATCTNNNATATTACATTTCAGAGTTCACCT
AGCACACGTCTGAACTCTTTCCCTACACGACGCTCTTCCGATCTNNNATATTACATTTCAGAGTTCACCT
AGCACACGTCTGAACTCTTTCCCTACACGACGCTCTTCCGATCTNNNATATTACATTTCAGAGTTCACCT
AGCACACGTCTGAACTCTTTCCCTACACGACGCTCTTCCGATCTNNNATATTACATTTCAGAGTTCACCT
AGCACACGTCTGAACTCTTTCCCTACACGACGCTCTTCCGATCTNNNATATTACATTTCAGAGTTCACCT
AGCACACGTCTGAACTCTTTCCCTACACGACGCTCTTCCGATCTNNNATCAACTGTTACAGAAATGAAA
GCACACGTCTGAACTCTTTCCCTACACGACGCTCTTCCGATCTNNNTATACTGACGTCTCCAACATCATC
AGCACACGTCTGAACTCTTTCCCTACACGACGCTCTTCCGATCTNNNACTTAAAGTGGAAATTACAAAG
AGCACACGTCTGAACTCTTTCCCTACACGACGCTCTTCCGATCTNNNCTATATTTTCCAAAATCCCTACCC
ACACGTCTGAACTCTTTCCCTACACGACGCTCTTCCGATCTNNNGTTCAAATTCTGGTTGAAAGATCATC
AGCACACGTCTGAACTCTTTCCCTACACGACGCTCTTCCGATCTNNNAAGCCATTATCTGCAAAAATATC
CACACGTCTGAACTCTTTCCCTACACGACGCTCTTCCGATCTNNNAACCTCATTCTGTTTTGTTCTCATCG
AGCACACGTCTGAACTCTTTCCCTACACGACGCTCTTCCGATCTNNNAAGATTTAATGACCTCAGGTTTC
CACACGTCTGAACTCTTTCCCTACACGACGCTCTTCCGATCTNNNTGATTGGTTATTCAACAGATATGCA
CACACGTCTGAACTCTTTCCCTACACGACGCTCTTCCGATCTNNNCTCATTACGTTTTAGATGGGATTCA

GCACACGTCTGAACTCTTTCCCTACACGACGCTCTTCCGATCTNNNACAGGGTATTCCATAATAACTTTC
GCACACGTCTGAACTCTTTCCCTACACGACGCTCTTCCGATCTNNNGTTATGTATCAAGTACCTTTGCCA
GCACACGTCTGAACTCTTTCCCTACACGACGCTCTTCCGATCTNNNTGAATGATGTTCACCATTTTTATCA
AGCACACGTCTGAACTCTTTCCCTACACGACGCTCTTCCGATCTNNNCTTGGCTTCTAGTTTCCTTTCATC
GCACACGTCTGAACTCTTTCCCTACACGACGCTCTTCCGATCTNNNTGAGAATCACAATCACTGCACATC
CACACGTCTGAACTCTTTCCCTACACGACGCTCTTCCGATCTNNNAGCCTGGTTTTTAAATGGGCATC
CACGTCTGAACTCTTTCCCTACACGACGCTCTTCCGATCTNNNTATTTGGATGACAGAAACACTTTCATC
AGCACACGTCTGAACTCTTTCCCTACACGACGCTCTTCCGATCTNNNTTGCAAGTAATTACAATGATGAT
AGCACACGTCTGAACTCTTTCCCTACACGACGCTCTTCCGATCTNNNCAGGATTCTTACAGAAAACAAG
CACACGTCTGAACTCTTTCCCTACACGACGCTCTTCCGATCTNNNAGCTTTAAAGTACTGTAGATGTGC
GAGCACACGTCTGAACTCTTTCCCTACACGACGCTCTTCCGATCTNNNATGTTCACATGTGGCTTAAACA
GAGCACACGTCTGAACTCTTTCCCTACACGACGCTCTTCCGATCTNNNTGAAACCATTCCAGTGTAAAA
GAGCACACGTCTGAACTCTTTCCCTACACGACGCTCTTCCGATCTNNNTGAAACCATTCCAGTGTAAAA
AGCACACGTCTGAACTCTTTCCCTACACGACGCTCTTCCGATCTNNNACATCTCCATTCTTCTCTTTAACA
AGCACACGTCTGAACTCTTTCCCTACACGACGCTCTTCCGATCTNNNCAGCTCCATTTCTTCTAAAATAG
GAGCACACGTCTGAACTCTTTCCCTACACGACGCTCTTCCGATCTNNNTGTAGTGTTTCAATATAATGC
GAGCACACGTCTGAACTCTTTCCCTACACGACGCTCTTCCGATCTNNNCGTTTCCTTTAACCACATAATT
AGCACACGTCTGAACTCTTTCCCTACACGACGCTCTTCCGATCTNNNCATCCCCTTAAATTGGATTAAAA
AGCACACGTCTGAACTCTTTCCCTACACGACGCTCTTCCGATCTNNNTTATAAGCAGTGCCAAAAATAA
AGCACACGTCTGAACTCTTTCCCTACACGACGCTCTTCCGATCTNNNGTATTAACCTTATGTGTGACAT
AGCACACGTCTGAACTCTTTCCCTACACGACGCTCTTCCGATCTNNNCAAGTAGTAATTGATGGAGAAA
AGCACACGTCTGAACTCTTTCCCTACACGACGCTCTTCCGATCTNNNAATAAATGTGATTTGCCTTCTAG
AGCACACGTCTGAACTCTTTCCCTACACGACGCTCTTCCGATCTNNNGATATCTTCCCAGCAAAATAATC
GCACACGTCTGAACTCTTTCCCTACACGACGCTCTTCCGATCTNNNCTCATATCAAAAAGGCCCTGCATC
CACACGTCTGAACTCTTTCCCTACACGACGCTCTTCCGATCTNNNGGTGTTTGAAGAGTTTTGTCCATCC
GCACACGTCTGAACTCTTTCCCTACACGACGCTCTTCCGATCTNNNCAAAATCAAGAAGCTGGAAGCAT
AGCACACGTCTGAACTCTTTCCCTACACGACGCTCTTCCGATCTNNNACAATCTATTCTGTGAATCATTTC
CACACGTCTGAACTCTTTCCCTACACGACGCTCTTCCGATCTNNNCATAAAGGGTGTGATCTAGAGCAT
AGAGCACACGTCTGAACTCTTTCCCTACACGACGCTCTTCCGATCTNNNTGATGAGATTATGAGTGATT
GAGCACACGTCTGAACTCTTTCCCTACACGACGCTCTTCCGATCTNNNCATTATCCTGTCTATAGATCCAC
CACACGTCTGAACTCTTTCCCTACACGACGCTCTTCCGATCTNNNATATCAACGTAGGGTATTACACCA
AGCACACGTCTGAACTCTTTCCCTACACGACGCTCTTCCGATCTNNNGAGACTTCCTTGTGTCATAATA
GAGCACACGTCTGAACTCTTTCCCTACACGACGCTCTTCCGATCTNNNTGTCTTTTCTTGTCATTAATAGC
GCACACGTCTGAACTCTTTCCCTACACGACGCTCTTCCGATCTNNNTGATGTAGAAAGTTAAAGTGTTC
AGCACACGTCTGAACTCTTTCCCTACACGACGCTCTTCCGATCTNNNGATTTTAAGTGTCACAGGTAAA
CACACGTCTGAACTCTTTCCCTACACGACGCTCTTCCGATCTNNNTACTGTACAAATGCATAACCTTTCA
GAGCACACGTCTGAACTCTTTCCCTACACGACGCTCTTCCGATCTNNNCAATACCCCATCTCTATTTAAGT
AGAGCACACGTCTGAACTCTTTCCCTACACGACGCTCTTCCGATCTNNNCATCTCCTTTCTTCATCACCCAT
GCACACGTCTGAACTCTTTCCCTACACGACGCTCTTCCGATCTNNNGAATTTCTGAATGATGAAAACAA
AGCACACGTCTGAACTCTTTCCCTACACGACGCTCTTCCGATCTNNNGTCACACAATAGCCTAAATATTC
GAGCACACGTCTGAACTCTTTCCCTACACGACGCTCTTCCGATCTNNNAATACATACACTGTTCTTTTGT
GAGCACACGTCTGAACTCTTTCCCTACACGACGCTCTTCCGATCTNNNCTCGTTTTCCAATCATTTTATTC
GAGCACACGTCTGAACTCTTTCCCTACACGACGCTCTTCCGATCTNNNGATTCCACTAATGACTGACATC

GCACACGTCTGAACTCTTTCCCTACACGACGCTCTTCCGATCTNNNCTCAACAATAGATGGAAATTTCA
AGCACACGTCTGAACTCTTTCCCTACACGACGCTCTTCCGATCTNNNACTTCTTGAAATATCTACTCTGT
AGCACACGTCTGAACTCTTTCCCTACACGACGCTCTTCCGATCTNNNAGTCCAACCAACATATTTTAAAT
AGCACACGTCTGAACTCTTTCCCTACACGACGCTCTTCCGATCTNNNGCATTCATCATATTTTCTATTT
AGCACACGTCTGAACTCTTTCCCTACACGACGCTCTTCCGATCTNNNCAACTAACCTCACTTTCTAGAAA
GCACACGTCTGAACTCTTTCCCTACACGACGCTCTTCCGATCTNNNAGGATAATACACATCACAGTAA
ACACGTCTGAACTCTTTCCCTACACGACGCTCTTCCGATCTNNNAACTGTAGCCCTCTGTAGTACATCG
AGCACACGTCTGAACTCTTTCCCTACACGACGCTCTTCCGATCTNNNAATACACCTTATCACACAATGA
GCACACGTCTGAACTCTTTCCCTACACGACGCTCTTCCGATCTNNNCCAGTGGGATATTATCTAACTTT
AGCACACGTCTGAACTCTTTCCCTACACGACGCTCTTCCGATCTNNNGTAATGAAGTTCTTCTAGTTCA
CACACGTCTGAACTCTTTCCCTACACGACGCTCTTCCGATCTNNNATAACCTTCCTAGTACCTGAGCATC
AGCACACGTCTGAACTCTTTCCCTACACGACGCTCTTCCGATCTNNNTTATTTACTTTCCCTCAGAAGA
AGCACACGTCTGAACTCTTTCCCTACACGACGCTCTTCCGATCTNNNACACCTCAGATATATTTCTTCAT
GCACACGTCTGAACTCTTTCCCTACACGACGCTCTTCCGATCTNNNAGTTTCTTCTTGATTTTTTTGGAC
AGCACACGTCTGAACTCTTTCCCTACACGACGCTCTTCCGATCTNNNTATAAGTAAAAGTGTGGGCAT
CACACGTCTGAACTCTTTCCCTACACGACGCTCTTCCGATCTNNNCAGGGACCTTACCTTATACACATC
AGCACACGTCTGAACTCTTTCCCTACACGACGCTCTTCCGATCTNNNTACCAGTGAATTTTTGCAATAAT
AGCACACGTCTGAACTCTTTCCCTACACGACGCTCTTCCGATCTNNNACAGCAAGTATTTTAAGTTTTG
AGCACACGTCTGAACTCTTTCCCTACACGACGCTCTTCCGATCTNNNCTAATGTTGGCAAAAATTGAAT
GAGCACACGTCTGAACTCTTTCCCTACACGACGCTCTTCCGATCTNNNATCTATTTTCTCAACTGCCATA
AGCACACGTCTGAACTCTTTCCCTACACGACGCTCTTCCGATCTNNNATTCACATCATCTTTGTAGGTAT
GAGCACACGTCTGAACTCTTTCCCTACACGACGCTCTTCCGATCTNNNTATACAAACAAAGCTTCAAC
AGCACACGTCTGAACTCTTTCCCTACACGACGCTCTTCCGATCTNNNCCTGGTAATTCTCATGAAAAAT
GCACACGTCTGAACTCTTTCCCTACACGACGCTCTTCCGATCTNNNCTTTAACCAGCTATAAACAAGAG
AGCACACGTCTGAACTCTTTCCCTACACGACGCTCTTCCGATCTNNNATTGTAATAGAATGGAAGGAA
AGCACACGTCTGAACTCTTTCCCTACACGACGCTCTTCCGATCTNNNATATTACATTTCAGAGTTCACCT
AGCACACGTCTGAACTCTTTCCCTACACGACGCTCTTCCGATCTNNNATATTACATTTCAGAGTTCACCT
AGCACACGTCTGAACTCTTTCCCTACACGACGCTCTTCCGATCTNNNATATTACATTTCAGAGTTCACCT
AGCACACGTCTGAACTCTTTCCCTACACGACGCTCTTCCGATCTNNNATATTACATTTCAGAGTTCACCT
AGCACACGTCTGAACTCTTTCCCTACACGACGCTCTTCCGATCTNNNATATTACATTTCAGAGTTCACCT
AGCACACGTCTGAACTCTTTCCCTACACGACGCTCTTCCGATCTNNNATATTACATTTCAGAGTTCACCT
AGCACACGTCTGAACTCTTTCCCTACACGACGCTCTTCCGATCTNNNGATGAAATTTTAGAGAAAAGG
GCACACGTCTGAACTCTTTCCCTACACGACGCTCTTCCGATCTNNNATTCTCTAAAATAACCAAAGCAG
AGCACACGTCTGAACTCTTTCCCTACACGACGCTCTTCCGATCTNNNCAGGATTCTTACAGAAAACAAG
AGCACACGTCTGAACTCTTTCCCTACACGACGCTCTTCCGATCTNNNTTAAAGCTATACGACTAAGTCT
AGCACACGTCTGAACTCTTTCCCTACACGACGCTCTTCCGATCTNNNAAATTAAAATCTCTTACCTTCC
AGCACACGTCTGAACTCTTTCCCTACACGACGCTCTTCCGATCTNNNCTGGAGCAAGATTATCTAATA
GCACACGTCTGAACTCTTTCCCTACACGACGCTCTTCCGATCTNNNAGATATTCCTCATATGTTACCAT
AGCACACGTCTGAACTCTTTCCCTACACGACGCTCTTCCGATCTNNNCCTTAAAAGGGAGTAACAATT
AGCACACGTCTGAACTCTTTCCCTACACGACGCTCTTCCGATCTNNNGTTCCTTTTGAGAACAATTCTA
AGCACACGTCTGAACTCTTTCCCTACACGACGCTCTTCCGATCTNNNGCACTGTGATGTATTATTTTCA
GAGCACACGTCTGAACTCTTTCCCTACACGACGCTCTTCCGATCTNNNAAAACGAAAATAAGGAGAG
CACACGTCTGAACTCTTTCCCTACACGACGCTCTTCCGATCTNNNATGAGTTCTTAAATTCTCAGTCTC

AGCACACGTCTGAACTCTTTCCCTACACGACGCTCTTCCGATCTNNNCTCTATTCCTACCACTCATTAAG

ACACGTCTGAACTCTTTCCCTACACGACGCTCTTCCGATCTNNNCTCCTGAAATATCAACCCATGCATCG

AGCACACGTCTGAACTCTTTCCCTACACGACGCTCTTCCGATCTNNNAAAATTTAGGATGGTTTTCTT

GCACACGTCTGAACTCTTTCCCTACACGACGCTCTTCCGATCTNNNGACCACGTTTAGATTCTGAGCAT

GCACACGTCTGAACTCTTTCCCTACACGACGCTCTTCCGATCTNNNAAAGCCATCCACTATAAAATTT

GAGCACACGTCTGAACTCTTTCCCTACACGACGCTCTTCCGATCTNNNGTGATTATACCAACGTAGGG

GAGCACACGTCTGAACTCTTTCCCTACACGACGCTCTTCCGATCTNNNGTGATTATACCAACGTAGGG

GAGCACACGTCTGAACTCTTTCCCTACACGACGCTCTTCCGATCTNNNATGATTACTTAGATGCTATCGA

AGCACACGTCTGAACTCTTTCCCTACACGACGCTCTTCCGATCTNNNTTACTTTTGGAATTTTTTGCAAA

AGCACACGTCTGAACTCTTTCCCTACACGACGCTCTTCCGATCTNNNCATTTTGGACAAGGGATATTTA

GAGCACACGTCTGAACTCTTTCCCTACACGACGCTCTTCCGATCTNNNTAATACACATGAACCATTTTTG

AGCACACGTCTGAACTCTTTCCCTACACGACGCTCTTCCGATCTNNNGTTGAAGAAAAGAAAATTGTT

AGCACACGTCTGAACTCTTTCCCTACACGACGCTCTTCCGATCTNNNTCTTCAAATATACTTTGCTTACG

GCACACGTCTGAACTCTTTCCCTACACGACGCTCTTCCGATCTNNNTTCTGAGAGATGTATTTTCTTTT

AGCACACGTCTGAACTCTTTCCCTACACGACGCTCTTCCGATCTNNNATGTAAAATGTTTGCTTAGCAT

GAGCACACGTCTGAACTCTTTCCCTACACGACGCTCTTCCGATCTNNNTGATATGAAAATTGAATAGC

AGCACACGTCTGAACTCTTTCCCTACACGACGCTCTTCCGATCTNNNTAGTAGTACTACCAGTATAGGA

GAGCACACGTCTGAACTCTTTCCCTACACGACGCTCTTCCGATCTNNNCAAAAAAAGGAGTACTATCAT

AGCACACGTCTGAACTCTTTCCCTACACGACGCTCTTCCGATCTNNNCTCATCTTCTAGCTCCTTCTTCAT

GAGCACACGTCTGAACTCTTTCCCTACACGACGCTCTTCCGATCTNNNGCTTGAATTTATGCACATTCTT

AGCACACGTCTGAACTCTTTCCCTACACGACGCTCTTCCGATCTNNNCTCATCTTCTAGCTCCTTCTTCAT

AGCACACGTCTGAACTCTTTCCCTACACGACGCTCTTCCGATCTNNNCCAGTTTCATGCTTTTATTTCAT

GAGCACACGTCTGAACTCTTTCCCTACACGACGCTCTTCCGATCTNNNCTCTTCTTTTGACTCTTTTTCTT

CACACGTCTGAACTCTTTCCCTACACGACGCTCTTCCGATCTNNNAACAGTTCATTGTTTCTTTTTTCCAT

AGCACACGTCTGAACTCTTTCCCTACACGACGCTCTTCCGATCTNNNCTTTTGATGATCTTCAGTATCAG

CACACGTCTGAACTCTTTCCCTACACGACGCTCTTCCGATCTNNNAGAATGAGCTGTCACCATAGCATC

GCACACGTCTGAACTCTTTCCCTACACGACGCTCTTCCGATCTNNNGAGAATGAGCTGTCACCATACAT

AGCACACGTCTGAACTCTTTCCCTACACGACGCTCTTCCGATCTNNNATTCTCCTTAAAATTCATACCAG

AGCACACGTCTGAACTCTTTCCCTACACGACGCTCTTCCGATCTNNNATACAAATTCTTTATTTCCTCCC

GAGCACACGTCTGAACTCTTTCCCTACACGACGCTCTTCCGATCTNNNGGCCTTTTTTTTCTATTATGAGA

AGAGCACACGTCTGAACTCTTTCCCTACACGACGCTCTTCCGATCTNNNCTACATCAGTATCATCACTTG

AGCACACGTCTGAACTCTTTCCCTACACGACGCTCTTCCGATCTNNNGAAAATGATTCTGGGGTCTATT

CACACGTCTGAACTCTTTCCCTACACGACGCTCTTCCGATCTNNNGTGTTCCTAGCAAAACTGTGCATC

GCACACGTCTGAACTCTTTCCCTACACGACGCTCTTCCGATCTNNNACTCTTACTTAGGGAAAGAAAAT

AGCACACGTCTGAACTCTTTCCCTACACGACGCTCTTCCGATCTNNNCTGAAATCACTTGTGATCTTGCA

GCACACGTCTGAACTCTTTCCCTACACGACGCTCTTCCGATCTNNNTTTCCAGGGAAGCAGTTTAACAT

AGCACACGTCTGAACTCTTTCCCTACACGACGCTCTTCCGATCTNNNTTTTAGGATATTTTTGAGTGTC

GCACACGTCTGAACTCTTTCCCTACACGACGCTCTTCCGATCTNNNATGAATGAATTTGAGACGGAACA

AGAGCACACGTCTGAACTCTTTCCCTACACGACGCTCTTCCGATCTNNNTTTTTAAACTTCTCATATGGT

GCACACGTCTGAACTCTTTCCCTACACGACGCTCTTCCGATCTNNNACTGTATAGAAGGTGGTCTCCAT

GCACACGTCTGAACTCTTTCCCTACACGACGCTCTTCCGATCTNNNTGAAAACAAACTTGAATAAACTG

AGCACACGTCTGAACTCTTTCCCTACACGACGCTCTTCCGATCTNNNCTGTAAATACAAAGCTTTCAAA

AGCACACGTCTGAACTCTTTCCCTACACGACGCTCTTCCGATCTNNNAAAAAATAAGGCAGAATCACAT

CACACGTCTGAACTCTTTCCCTACACGACGCTCTTCCGATCTNNNCCATGTTTACATTCCCTTTATTTCA
GCACACGTCTGAACTCTTTCCCTACACGACGCTCTTCCGATCTNNNTATTGTTCTACAAAGGCAGAAA
CACACGTCTGAACTCTTTCCCTACACGACGCTCTTCCGATCTNNNGGCTGATCTTGACAAACATGCAT
GCACACGTCTGAACTCTTTCCCTACACGACGCTCTTCCGATCTNNNATATCTTGAGCTCAGGAGTTCA
AGCACACGTCTGAACTCTTTCCCTACACGACGCTCTTCCGATCTNNNAAATATCTCAAAAAGTCAGTG
CACACGTCTGAACTCTTTCCCTACACGACGCTCTTCCGATCTNNNCTGTTCTTGGTTTCACAGATCATC
CACACGTCTGAACTCTTTCCCTACACGACGCTCTTCCGATCTNNNTTGGTTTGTAACGAAGTGAACAT
GCACACGTCTGAACTCTTTCCCTACACGACGCTCTTCCGATCTNNNGTGTCATTTTTCATTCCCTTAAC
AGCACACGTCTGAACTCTTTCCCTACACGACGCTCTTCCGATCTNNNAGATGGTTTTAGTATTCATCA
AGCACACGTCTGAACTCTTTCCCTACACGACGCTCTTCCGATCTNNNCAATCTCCTGAAGACAGATGC
GCACACGTCTGAACTCTTTCCCTACACGACGCTCTTCCGATCTNNNTCATAAAGGAATCCATTGGCCAT
AGCACACGTCTGAACTCTTTCCCTACACGACGCTCTTCCGATCTNNNTTTTGAGGTGTAGTACTTCATT
GCACACGTCTGAACTCTTTCCCTACACGACGCTCTTCCGATCTNNNTCATAAAGGAATCCATTGGCCAT
GCACACGTCTGAACTCTTTCCCTACACGACGCTCTTCCGATCTNNNAGGAGAAAATCTGAGAATAAG
CACACGTCTGAACTCTTTCCCTACACGACGCTCTTCCGATCTNNNCATAGGTCTTTCATCCCTTAATAC
GAGCACACGTCTGAACTCTTTCCCTACACGACGCTCTTCCGATCTNNNTTACATTTTGAAAGACTGGT
AGCACACGTCTGAACTCTTTCCCTACACGACGCTCTTCCGATCTNNNGTAATTACCCTTCAGCTCTTA
ACACGTCTGAACTCTTTCCCTACACGACGCTCTTCCGATCTNNNTTCTGGTAAGCACATTTAATAGACA
AGCACACGTCTGAACTCTTTCCCTACACGACGCTCTTCCGATCTNNNTCATCATCTTGATCTTCAAATT
CACACGTCTGAACTCTTTCCCTACACGACGCTCTTCCGATCTNNNAAGCTGCTGTATTTTAGTGAGCAT
AGCACACGTCTGAACTCTTTCCCTACACGACGCTCTTCCGATCTNNNAACTTTCTTTCTAACACTGTTTT
GCACACGTCTGAACTCTTTCCCTACACGACGCTCTTCCGATCTNNNGAGATACTGGCTCATCATACTA
GAGCACACGTCTGAACTCTTTCCCTACACGACGCTCTTCCGATCTNNNCCTCACATTATATTGATAGAG
AGCACACGTCTGAACTCTTTCCCTACACGACGCTCTTCCGATCTNNNTATTTTGCAGCTCATAGACAA
AGCACACGTCTGAACTCTTTCCCTACACGACGCTCTTCCGATCTNNNTACACATCCTAAATTTGGTAGG
CACACGTCTGAACTCTTTCCCTACACGACGCTCTTCCGATCTNNNCTACTGCCTTTTTTTTTTTTTCTCAT
CACACGTCTGAACTCTTTCCCTACACGACGCTCTTCCGATCTNNNTTGTATTTCTAGCAGAGACGCATC
AGCACACGTCTGAACTCTTTCCCTACACGACGCTCTTCCGATCTNNNAAAACTTACTTACTAGGCACTA
GCACACGTCTGAACTCTTTCCCTACACGACGCTCTTCCGATCTNNNCCTTTGGTCATAACATTTTCATT
AGCACACGTCTGAACTCTTTCCCTACACGACGCTCTTCCGATCTNNNGTACTGATTTTCTTTCTTTTGG
GCACACGTCTGAACTCTTTCCCTACACGACGCTCTTCCGATCTNNNTTTTGTGTGTCAGCATAATAATT
CACACGTCTGAACTCTTTCCCTACACGACGCTCTTCCGATCTNNNTTAATTAGCTTAGAATCCCTGAA
AGCACACGTCTGAACTCTTTCCCTACACGACGCTCTTCCGATCTNNNTATGACAAAAAGAAGGGTGTT
ACACGTCTGAACTCTTTCCCTACACGACGCTCTTCCGATCTNNNAGTAATTAGAACGGAAGGGAACAT
AGCACACGTCTGAACTCTTTCCCTACACGACGCTCTTCCGATCTNNNGCAGAAAGAATTAACAGACAT
CACACGTCTGAACTCTTTCCCTACACGACGCTCTTCCGATCTNNNCTAAAAATACAAAAATTAGCTGG
CACACGTCTGAACTCTTTCCCTACACGACGCTCTTCCGATCTNNNCAATTAACCTAAGCTTGATGATG
ACACGTCTGAACTCTTTCCCTACACGACGCTCTTCCGATCTNNNATGAATGTTCCATCAAAGGGCATC
GCACACGTCTGAACTCTTTCCCTACACGACGCTCTTCCGATCTNNNCTGTAAAAATGACCTCCCTTCCA
GCACACGTCTGAACTCTTTCCCTACACGACGCTCTTCCGATCTNNNCTTCACTCTTATCTGTTACGTAAC
AGCACACGTCTGAACTCTTTCCCTACACGACGCTCTTCCGATCTNNNAACTGTTGATACTCATTCAATT
GAGCACACGTCTGAACTCTTTCCCTACACGACGCTCTTCCGATCTNNNCTGATATGACCCTTCGTTTTC
CACACGTCTGAACTCTTTCCCTACACGACGCTCTTCCGATCTNNNCTAGAGCTTTTCCTTTTAAGAGAC

CACACGTCTGAACTCTTTCCCTACACGACGCTCTTCCGATCTNNNCTAGAGCTTTTCCTTTTAAGAGAC
CACGTCTGAACTCTTTCCCTACACGACGCTCTTCCGATCTNNNAAATGCTAGTTGCTTGTTTGCATCGG
ACACGTCTGAACTCTTTCCCTACACGACGCTCTTCCGATCTNNNCTGATTGAGAAGACTCTGTACCAT
AGCACACGTCTGAACTCTTTCCCTACACGACGCTCTTCCGATCTNNNCTCATCTCTTCTCTTTTCCTCCAT
AGCACACGTCTGAACTCTTTCCCTACACGACGCTCTTCCGATCTNNNTATCATCAATGGCTGCTAAAA
ACACGTCTGAACTCTTTCCCTACACGACGCTCTTCCGATCTNNNAGACTCTATCAAGACAGTAAAAAC
CACACGTCTGAACTCTTTCCCTACACGACGCTCTTCCGATCTNNNATCAAAGTTCTGAGACTTAATCT
CACACGTCTGAACTCTTTCCCTACACGACGCTCTTCCGATCTNNNATCAAAGTTCTGAGACTTAATCT
GCACACGTCTGAACTCTTTCCCTACACGACGCTCTTCCGATCTNNNGTACTTGCTTTTTGTGGTTGCAT
AGCACACGTCTGAACTCTTTCCCTACACGACGCTCTTCCGATCTNNNCACCAGGGAATCTGAATAACC
ACACGTCTGAACTCTTTCCCTACACGACGCTCTTCCGATCTNNNATATATGTGCCATCTTTCTACATCAT
GCACACGTCTGAACTCTTTCCCTACACGACGCTCTTCCGATCTNNNAATTTTGATTCTGTTACCTTGTT
ACACGTCTGAACTCTTTCCCTACACGACGCTCTTCCGATCTNNNCTTGGTAATGATCCTACAGATCAT
GCACACGTCTGAACTCTTTCCCTACACGACGCTCTTCCGATCTNNNAATTTTGATTCTGTTACCTTGTT
ACACGTCTGAACTCTTTCCCTACACGACGCTCTTCCGATCTNNNCTGAAGAAATAAAGTTCTTTTTTT
AGCACACGTCTGAACTCTTTCCCTACACGACGCTCTTCCGATCTNNNTAGAGTCCTAAATCTCGAACA
ACACGTCTGAACTCTTTCCCTACACGACGCTCTTCCGATCTNNNATGCAGTCTCATAAACACTCCATCG
CACACGTCTGAACTCTTTCCCTACACGACGCTCTTCCGATCTNNNTTTCAAATAAAACTTGTACTGGG
AGCACACGTCTGAACTCTTTCCCTACACGACGCTCTTCCGATCTNNNTTTTAACTTACGGTTGATGAA
GCACACGTCTGAACTCTTTCCCTACACGACGCTCTTCCGATCTNNNTCACTTAGCTTTTCCCTAAAATA
AGCACACGTCTGAACTCTTTCCCTACACGACGCTCTTCCGATCTNNNAAATCTTTCTTTTAAAATGCTG
GAGCACACGTCTGAACTCTTTCCCTACACGACGCTCTTCCGATCTNNNTTAACATTTGATGGTTTGAA
CACACGTCTGAACTCTTTCCCTACACGACGCTCTTCCGATCTNNNAATTAACTAGTTCCATGCATTTTC
GCACACGTCTGAACTCTTTCCCTACACGACGCTCTTCCGATCTNNNGTATTTTGGTGTTTGTAACCATA
GCACACGTCTGAACTCTTTCCCTACACGACGCTCTTCCGATCTNNNTTATAGAGGAACGTAATAGTC
CACACGTCTGAACTCTTTCCCTACACGACGCTCTTCCGATCTNNNCTCAAATTCTCTTGCTTTTCTATCA
AGCACACGTCTGAACTCTTTCCCTACACGACGCTCTTCCGATCTNNNTAATCAAAGTCAGCAATTTATG
GCACACGTCTGAACTCTTTCCCTACACGACGCTCTTCCGATCTNNNATAATCCAATCACTTCTTTCTCTC
AGCACACGTCTGAACTCTTTCCCTACACGACGCTCTTCCGATCTNNNCCAGCTCTATTCTTATAAACTTC
GCACACGTCTGAACTCTTTCCCTACACGACGCTCTTCCGATCTNNNCTGACATATCATTGTTACCAAAG
GAGCACACGTCTGAACTCTTTCCCTACACGACGCTCTTCCGATCTNNNCTCTGTATGGAATCCACATAC
GCACACGTCTGAACTCTTTCCCTACACGACGCTCTTCCGATCTNNNGTCTTCATTCTTATCATCTTCACC
GCACACGTCTGAACTCTTTCCCTACACGACGCTCTTCCGATCTNNNATATCCAATTTCATATACTGGC
GAGCACACGTCTGAACTCTTTCCCTACACGACGCTCTTCCGATCTNNNTATTCTGTTCTTATCCATGAG
AGCACACGTCTGAACTCTTTCCCTACACGACGCTCTTCCGATCTNNNGTTAAGATCTGGGAATCGACC
ACACGTCTGAACTCTTTCCCTACACGACGCTCTTCCGATCTNNNCTCCGTCTCAAAAAAAACAAAACAT
GAGCACACGTCTGAACTCTTTCCCTACACGACGCTCTTCCGATCTNNNGTATATTGCATACCTGATCCA
GCACACGTCTGAACTCTTTCCCTACACGACGCTCTTCCGATCTNNNATAAGTATATTAAAGAGACAGC
AGCACACGTCTGAACTCTTTCCCTACACGACGCTCTTCCGATCTNNNCATAATATAAGTCAGCAGGTA
GCACACGTCTGAACTCTTTCCCTACACGACGCTCTTCCGATCTNNNCTGAGCTAACACATCATTGACAT
GAGCACACGTCTGAACTCTTTCCCTACACGACGCTCTTCCGATCTNNNTATAATTCTTCCTTGTCCTTT
AGCACACGTCTGAACTCTTTCCCTACACGACGCTCTTCCGATCTNNNCTAAGATGCAGTCCACAATGC
GCACACGTCTGAACTCTTTCCCTACACGACGCTCTTCCGATCTNNNGGAGACTCCTTTCAATTGACCAT

AGCACACGTCTGAACTCTTTCCCTACACGACGCTCTTCCGATCTNNNATCTAACAGATTTCCTAAAAT
AGCACACGTCTGAACTCTTTCCCTACACGACGCTCTTCCGATCTNNNGTCAGTTGTTTGCTGATTTTAT
GCACACGTCTGAACTCTTTCCCTACACGACGCTCTTCCGATCTNNNTTTTGAAATGGAGTCTCACTTTC
CACACGTCTGAACTCTTTCCCTACACGACGCTCTTCCGATCTNNNTTCATAATGGTGCTTTCAAACCAT
AGCACACGTCTGAACTCTTTCCCTACACGACGCTCTTCCGATCTNNNAAGTTTACACCATTGCACTCCA
AGCACACGTCTGAACTCTTTCCCTACACGACGCTCTTCCGATCTNNNTTTTTAGAATATGTTATGCTGG
GCACACGTCTGAACTCTTTCCCTACACGACGCTCTTCCGATCTNNNATTAAGCATTAACTCATCAACAA
AGCACACGTCTGAACTCTTTCCCTACACGACGCTCTTCCGATCTNNNCTTCTATCTCTATTGGGCTTTTA
GCACACGTCTGAACTCTTTCCCTACACGACGCTCTTCCGATCTNNNTTAATTTGATCACATCTGATTCC
GAGCACACGTCTGAACTCTTTCCCTACACGACGCTCTTCCGATCTNNNAGAACTACATTCTATAGGAA
ACACGTCTGAACTCTTTCCCTACACGACGCTCTTCCGATCTNNNGACAAAGCAAAGGTATGAAAGCA
AGCACACGTCTGAACTCTTTCCCTACACGACGCTCTTCCGATCTNNNGGTGTTTGATTACATGAATAA
CACACGTCTGAACTCTTTCCCTACACGACGCTCTTCCGATCTNNNGAATTTCAGGTATTAAAGGAAC
GCACACGTCTGAACTCTTTCCCTACACGACGCTCTTCCGATCTNNNCCTAACAGTGAGAAGATTAAG
AGCACACGTCTGAACTCTTTCCCTACACGACGCTCTTCCGATCTNNNTTTTATTTCTTCTTTCACAACCC
GCACACGTCTGAACTCTTTCCCTACACGACGCTCTTCCGATCTNNNCTTTCCTGATCTAATTGTTCATA
GCACACGTCTGAACTCTTTCCCTACACGACGCTCTTCCGATCTNNNACATCATTTTAGCCCTTACTCCAT
AGCACACGTCTGAACTCTTTCCCTACACGACGCTCTTCCGATCTNNNGAACATCAATAACAGAAAGAC
GCACACGTCTGAACTCTTTCCCTACACGACGCTCTTCCGATCTNNNTATAAAACCATCAGATCTCATGA
GCACACGTCTGAACTCTTTCCCTACACGACGCTCTTCCGATCTNNNTATTGTTCTACCTTTCAGATGAA
GCACACGTCTGAACTCTTTCCCTACACGACGCTCTTCCGATCTNNNTAGACACTTCACAAAAAGTAGA
AGCACACGTCTGAACTCTTTCCCTACACGACGCTCTTCCGATCTNNNAGGATATTCTTCCTGCTTAATG
GCACACGTCTGAACTCTTTCCCTACACGACGCTCTTCCGATCTNNNGAATGACTGAAGTATCAACATT
CACACGTCTGAACTCTTTCCCTACACGACGCTCTTCCGATCTNNNTGAGTTTATTTTGAAGGTTAACTC
GCACACGTCTGAACTCTTTCCCTACACGACGCTCTTCCGATCTNNNTTACCCTACATGGTCTAAAAAG
GCACACGTCTGAACTCTTTCCCTACACGACGCTCTTCCGATCTNNNGCTTAACGGAACTTATAAAACA
GCACACGTCTGAACTCTTTCCCTACACGACGCTCTTCCGATCTNNNGCTTAACGGAACTTATAAAACA
GAGCACACGTCTGAACTCTTTCCCTACACGACGCTCTTCCGATCTNNNCTGGTGGGTTTTATCTTTTT
GCACACGTCTGAACTCTTTCCCTACACGACGCTCTTCCGATCTNNNATAATGAGGTTTTTAACTTGTG
GCACACGTCTGAACTCTTTCCCTACACGACGCTCTTCCGATCTNNNATAATGAGGTTTTTAACTTGTG
CACACGTCTGAACTCTTTCCCTACACGACGCTCTTCCGATCTNNNGGGGATCTGTACCATTTTAATAT
GCACACGTCTGAACTCTTTCCCTACACGACGCTCTTCCGATCTNNNCAATTTTATCTTGAAACCATTCG
GCACACGTCTGAACTCTTTCCCTACACGACGCTCTTCCGATCTNNNATGAACACACTCTTTAATTTCAT
AGCACACGTCTGAACTCTTTCCCTACACGACGCTCTTCCGATCTNNNGAGAAAATGAGGAAGCAAA
GCACACGTCTGAACTCTTTCCCTACACGACGCTCTTCCGATCTNNNAAACTTACAGTCATAGCAGAAG
AGCACACGTCTGAACTCTTTCCCTACACGACGCTCTTCCGATCTNNNGTATTATGATCAGGACTCACA
GAGCACACGTCTGAACTCTTTCCCTACACGACGCTCTTCCGATCTNNNGAATCTGATATGACCCTTCG
GCACACGTCTGAACTCTTTCCCTACACGACGCTCTTCCGATCTNNNTCCTTAGTGTCTGGCTTAAATCA
CACACGTCTGAACTCTTTCCCTACACGACGCTCTTCCGATCTNNNCTAGAGCTTTTCCTTTTAAGAGAC
CACACGTCTGAACTCTTTCCCTACACGACGCTCTTCCGATCTNNNCTAGAGCTTTTCCTTTTAAGAGAC
GCACACGTCTGAACTCTTTCCCTACACGACGCTCTTCCGATCTNNNGTCTGTGAAATCATACAGTATTT
GCACACGTCTGAACTCTTTCCCTACACGACGCTCTTCCGATCTNNNAACTAAAAAAATCAAGAAACTA
AGCACACGTCTGAACTCTTTCCCTACACGACGCTCTTCCGATCTNNNCTCATCTCTTCTCTTTTCCTCCAT

AGCACACGTCTGAACTCTTTCCCTACACGACGCTCTTCCGATCTNNNTACTTTCACAACTGTAAGCATA
CACACGTCTGAACTCTTTCCCTACACGACGCTCTTCCGATCTNNNTAGAGTGTAGATGCATCAAAAT
GCACACGTCTGAACTCTTTCCCTACACGACGCTCTTCCGATCTNNNTTATTCTATTGCCCAGAAATAG
CACACGTCTGAACTCTTTCCCTACACGACGCTCTTCCGATCTNNNATCAAAGTTCTGAGACTTAATCT
GCACACGTCTGAACTCTTTCCCTACACGACGCTCTTCCGATCTNNNCTTAGGCTGGAGTTTACAATAA
AGCACACGTCTGAACTCTTTCCCTACACGACGCTCTTCCGATCTNNNCACCAGGGAATCTGAATAACC
ACACGTCTGAACTCTTTCCCTACACGACGCTCTTCCGATCTNNNATATATGTGCCATCTTTCTACATCAT
ACACGTCTGAACTCTTTCCCTACACGACGCTCTTCCGATCTNNNATATATGTGCCATCTTTCTACATCAT
ACACGTCTGAACTCTTTCCCTACACGACGCTCTTCCGATCTNNNCTTGGTAATGATCCTACAGATCAT
ACACGTCTGAACTCTTTCCCTACACGACGCTCTTCCGATCTNNNGAAGATACTTACTGTAGTTCTCACA
ACACGTCTGAACTCTTTCCCTACACGACGCTCTTCCGATCTNNNGAAGATACTTACTGTAGTTCTCACA
CACACGTCTGAACTCTTTCCCTACACGACGCTCTTCCGATCTNNNTTTCAAATAAAACTTGTACTGGG
CACACGTCTGAACTCTTTCCCTACACGACGCTCTTCCGATCTNNNTTTCAAATAAAACTTGTACTGGG
AGCACACGTCTGAACTCTTTCCCTACACGACGCTCTTCCGATCTNNNAAATGTTAATAAGTCCCTTTTG
AGAGCACACGTCTGAACTCTTTCCCTACACGACGCTCTTCCGATCTNNNACCTTGTCAAAAAAAAAAA
GCACACGTCTGAACTCTTTCCCTACACGACGCTCTTCCGATCTNNNATAGAGAACATTTCCAGAATAC
AGAGCACACGTCTGAACTCTTTCCCTACACGACGCTCTTCCGATCTNNNACCTTGTCAAAAAAAAAAA
GCACACGTCTGAACTCTTTCCCTACACGACGCTCTTCCGATCTNNNTGTGCATTTTAATTTCAAAAAG
GCACACGTCTGAACTCTTTCCCTACACGACGCTCTTCCGATCTNNNGAAATTATCTCCTGCTCATAGT
AGCACACGTCTGAACTCTTTCCCTACACGACGCTCTTCCGATCTNNNAGTAACATTTAAACCTGTAAA
GAGCACACGTCTGAACTCTTTCCCTACACGACGCTCTTCCGATCTNNNGGTGATTGTATTCTGTGTACT
ACACGTCTGAACTCTTTCCCTACACGACGCTCTTCCGATCTNNNGCTGACTCAACTGGAAATTTCATC
GCACACGTCTGAACTCTTTCCCTACACGACGCTCTTCCGATCTNNNATTTTTTGAGACAGTCTCGCCAT
GCACACGTCTGAACTCTTTCCCTACACGACGCTCTTCCGATCTNNNACACCCAGCTAATTTTTTAAAA
GCACACGTCTGAACTCTTTCCCTACACGACGCTCTTCCGATCTNNNATATACATCAAACACCTACTTCT
GCACACGTCTGAACTCTTTCCCTACACGACGCTCTTCCGATCTNNNATTGGTCTATATCTTTGTTTTGG
ACACGTCTGAACTCTTTCCCTACACGACGCTCTTCCGATCTNNNTAAAAAAACATACAAAACGTACTG
GCACACGTCTGAACTCTTTCCCTACACGACGCTCTTCCGATCTNNNGTTTCTCACTTTAGCCAATTCCAT
CACACGTCTGAACTCTTTCCCTACACGACGCTCTTCCGATCTNNNGTTTCTCACTTTAGCCAATTCCAT
GAGCACACGTCTGAACTCTTTCCCTACACGACGCTCTTCCGATCTNNNCATATGGTTATCTTGGAATAT
AGCACACGTCTGAACTCTTTCCCTACACGACGCTCTTCCGATCTNNNCTCTTCCCCTTGGTTTGTAACA
GAGCACACGTCTGAACTCTTTCCCTACACGACGCTCTTCCGATCTNNNTCCCCTTCAAACATATACATTA
GCACACGTCTGAACTCTTTCCCTACACGACGCTCTTCCGATCTNNNCAAAGGGGTAAAAAAAAAAAG
CACACGTCTGAACTCTTTCCCTACACGACGCTCTTCCGATCTNNNTTCATAATGGTGCTTTCAAACCAT
GCACACGTCTGAACTCTTTCCCTACACGACGCTCTTCCGATCTNNNTTGAAAATTCTTAAGCTCATCGC
CACACGTCTGAACTCTTTCCCTACACGACGCTCTTCCGATCTNNNCACAAGACATTTTTCTACAGTATCA
GCACACGTCTGAACTCTTTCCCTACACGACGCTCTTCCGATCTNNNCACAAGACATTTTTCTACAGTATC
AGCACACGTCTGAACTCTTTCCCTACACGACGCTCTTCCGATCTNNNACCACACAAGACATTTTTCTAC
ACACGTCTGAACTCTTTCCCTACACGACGCTCTTCCGATCTNNNTGATCAGATAGGAGCACAAGCAT
GCACACGTCTGAACTCTTTCCCTACACGACGCTCTTCCGATCTNNNACCTGTATTAAGAATCTGTATT
GCACACGTCTGAACTCTTTCCCTACACGACGCTCTTCCGATCTNNNGTATGTAGCTTTCCTTTTTCTTA
AGCACACGTCTGAACTCTTTCCCTACACGACGCTCTTCCGATCTNNNAATTTGTAATGATTTTTCTGTG
AGCACACGTCTGAACTCTTTCCCTACACGACGCTCTTCCGATCTNNNGATCAACTCTGAAACATACATA

AGCACACGTCTGAACTCTTTCCCTACACGACGCTCTTCCGATCTNNNATTTTGTATTTTTAGTAGAGAC

AGCACACGTCTGAACTCTTTCCCTACACGACGCTCTTCCGATCTNNNCCTTTATTCCTTCTCTTCTTTTC

GAGCACACGTCTGAACTCTTTCCCTACACGACGCTCTTCCGATCTNNNCTGCTGTTTTACTAAGAACGA

ACACGTCTGAACTCTTTCCCTACACGACGCTCTTCCGATCTNNNCAAGCATGTTCCAGTATTTGCATCG

AGCACACGTCTGAACTCTTTCCCTACACGACGCTCTTCCGATCTNNNCAATCTGGAGAAGAATTTTGA

GAGCACACGTCTGAACTCTTTCCCTACACGACGCTCTTCCGATCTNNNGGATAGTCAGTCTTCCATATT

GAGCACACGTCTGAACTCTTTCCCTACACGACGCTCTTCCGATCTNNNTATCAAGATAGTACAGAGAA

GAGCACACGTCTGAACTCTTTCCCTACACGACGCTCTTCCGATCTNNNAAAGAATGTTTGTATACAAA

GAGCACACGTCTGAACTCTTTCCCTACACGACGCTCTTCCGATCTNNNAAAGAATGTTTGTATACAAA

GCACACGTCTGAACTCTTTCCCTACACGACGCTCTTCCGATCTNNNAGCTCAGAATGTAATCTTCTTTC

AGCACACGTCTGAACTCTTTCCCTACACGACGCTCTTCCGATCTNNNTAAGAAATTGGTGGCTAGAA

GCACACGTCTGAACTCTTTCCCTACACGACGCTCTTCCGATCTNNNGAGTGGACTGGTAAAACTTTCA

AGCACACGTCTGAACTCTTTCCCTACACGACGCTCTTCCGATCTNNNGCACTGGTAAAATAAATACTC

GCACACGTCTGAACTCTTTCCCTACACGACGCTCTTCCGATCTNNNATCATACTGCTAAGAATTCAAA

ACACGTCTGAACTCTTTCCCTACACGACGCTCTTCCGATCTNNNTAAAAATGTGGTTTCTTTAGTCCCA

GCACACGTCTGAACTCTTTCCCTACACGACGCTCTTCCGATCTNNNATCATACTGCTAAGAATTCAAA

GCACACGTCTGAACTCTTTCCCTACACGACGCTCTTCCGATCTNNNAGGTAATGCATATTAACAACTT

GCACACGTCTGAACTCTTTCCCTACACGACGCTCTTCCGATCTNNNTAGAAACAATACGTATTTGCTT

GCACACGTCTGAACTCTTTCCCTACACGACGCTCTTCCGATCTNNNTAGAAACAATACGTATTTGCTT

AGCACACGTCTGAACTCTTTCCCTACACGACGCTCTTCCGATCTNNNTCTGAAAGTGGGAAAGGAAT

GCACACGTCTGAACTCTTTCCCTACACGACGCTCTTCCGATCTNNNAAGTAAAAGTTTCAAAGGTACT

GCACACGTCTGAACTCTTTCCCTACACGACGCTCTTCCGATCTNNNGCATATGGATGATACTCTAAG

AGCACACGTCTGAACTCTTTCCCTACACGACGCTCTTCCGATCTNNNACGGAAGTATTATATAGCACTT

GCACACGTCTGAACTCTTTCCCTACACGACGCTCTTCCGATCTNNNTTTGACAATGTTCTTAGTAACTG

AGCACACGTCTGAACTCTTTCCCTACACGACGCTCTTCCGATCTNNNTAGTAGTTAATGGTATGGAT

GCACACGTCTGAACTCTTTCCCTACACGACGCTCTTCCGATCTNNNTTTATAAAACCGTCAGACCCCAT

CACACGTCTGAACTCTTTCCCTACACGACGCTCTTCCGATCTNNNAGTGTAGATAGATGAAACCTTTAC

GCACACGTCTGAACTCTTTCCCTACACGACGCTCTTCCGATCTNNNCTGACATTATCTGCTAAGAGAG

AGCACACGTCTGAACTCTTTCCCTACACGACGCTCTTCCGATCTNNNTGATTCCACTTTATTAAAAAGC

CACACGTCTGAACTCTTTCCCTACACGACGCTCTTCCGATCTNNNTTAGAGTAGATTATATACGACCA

AGCACACGTCTGAACTCTTTCCCTACACGACGCTCTTCCGATCTNNNAAAGCTATTTTGTGATTGTTTT

CACGTCTGAACTCTTTCCCTACACGACGCTCTTCCGATCTNNNCACGCCTGGCTATTATTATTATCATC

GAGCACACGTCTGAACTCTTTCCCTACACGACGCTCTTCCGATCTNNNCCTTATCACTACTTCCAGAAG

GAGCACACGTCTGAACTCTTTCCCTACACGACGCTCTTCCGATCTNNNCCTTATCACTACTTCCAGAAG

CACACGTCTGAACTCTTTCCCTACACGACGCTCTTCCGATCTNNNGAAATATCTGTCTTATCCTCTGTC

GCACACGTCTGAACTCTTTCCCTACACGACGCTCTTCCGATCTNNNCTAGTGAGGTTAATGATCCTTC

GAGCACACGTCTGAACTCTTTCCCTACACGACGCTCTTCCGATCTNNNCACTCAGCTGATTTACCTAAG

AGCACACGTCTGAACTCTTTCCCTACACGACGCTCTTCCGATCTNNNGACTATCTAAGGCAAGCATGC

AGCACACGTCTGAACTCTTTCCCTACACGACGCTCTTCCGATCTNNNCTTTCAAAACATTCCTTTTCTTC

ACACGTCTGAACTCTTTCCCTACACGACGCTCTTCCGATCTNNNATCTTCTGTGAAATGTTTAGGACAT

GCACACGTCTGAACTCTTTCCCTACACGACGCTCTTCCGATCTNNNAGTAATACAATGTTCTTGTCATG

ACACGTCTGAACTCTTTCCCTACACGACGCTCTTCCGATCTNNNGTAAAGAAACACACGGTCTGCATC

GCACACGTCTGAACTCTTTCCCTACACGACGCTCTTCCGATCTNNNTCAGGTATGCTTCCATTTAATAG

GAGCACACGTCTGAACTCTTTCCCTACACGACGCTCTTCCGATCTNNNAAAGGCAAGTAAACAATCTTG
AGAGCACACGTCTGAACTCTTTCCCTACACGACGCTCTTCCGATCTNNNGTTCTATCCTGTTCCATCAGCA
AGAGCACACGTCTGAACTCTTTCCCTACACGACGCTCTTCCGATCTNNNATGGTATCAGGAATGGACTT
CACACGTCTGAACTCTTTCCCTACACGACGCTCTTCCGATCTNNNTCCATTCTGGAGCTTTGTAGCATCC
GCACACGTCTGAACTCTTTCCCTACACGACGCTCTTCCGATCTNNNACAGGGTATTCCATAATAACTTTC
GCACACGTCTGAACTCTTTCCCTACACGACGCTCTTCCGATCTNNNACAGGGTATTCCATAATAACTTTC
CACACGTCTGAACTCTTTCCCTACACGACGCTCTTCCGATCTNNNTTTTATTCCTCCATTTGCAAACATC
AGCACACGTCTGAACTCTTTCCCTACACGACGCTCTTCCGATCTNNNTGTTTCAATTTCTTGATCTGAAG
AGCACACGTCTGAACTCTTTCCCTACACGACGCTCTTCCGATCTNNNTGTTTCAATTTCTTGATCTGAAG
CACACGTCTGAACTCTTTCCCTACACGACGCTCTTCCGATCTNNNATTCTTTCTTATCACTCAAATCGCAT
GCACACGTCTGAACTCTTTCCCTACACGACGCTCTTCCGATCTNNNTGAAATCTTTAACTGCATTTTCTC
GCACACGTCTGAACTCTTTCCCTACACGACGCTCTTCCGATCTNNNTGAAATCTTTAACTGCATTTTCTC
GAGCACACGTCTGAACTCTTTCCCTACACGACGCTCTTCCGATCTNNNTGATTCCACTTTATTAAAAAGC
AGCACACGTCTGAACTCTTTCCCTACACGACGCTCTTCCGATCTNNNAGTAATACAATGTTCTTGTCATG
CACACGTCTGAACTCTTTCCCTACACGACGCTCTTCCGATCTNNNTGGAAACACAATCTGGATAATATC
AGCACACGTCTGAACTCTTTCCCTACACGACGCTCTTCCGATCTNNNCATTTATGTGAGATGTGGTTTT
GAGCACACGTCTGAACTCTTTCCCTACACGACGCTCTTCCGATCTNNNTAATTTCTAAGGCAAGAATCTT
GAGCACACGTCTGAACTCTTTCCCTACACGACGCTCTTCCGATCTNNNCTCAGAGTTAGAGGTCTGTGC
AGCACACGTCTGAACTCTTTCCCTACACGACGCTCTTCCGATCTNNNCATTTATGTGAGATGTGGTTTT
GCACACGTCTGAACTCTTTCCCTACACGACGCTCTTCCGATCTNNNTACTGGCATTATCAGCATCATCAT
CACACGTCTGAACTCTTTCCCTACACGACGCTCTTCCGATCTNNNAACCTGAACAGAATTCTTCACCATC
AGCACACGTCTGAACTCTTTCCCTACACGACGCTCTTCCGATCTNNNTTTTGTTTTAAATACCGTTCAGA
AGCACACGTCTGAACTCTTTCCCTACACGACGCTCTTCCGATCTNNNTTTTGTTTTAAATACCGTTCAGA
GCACACGTCTGAACTCTTTCCCTACACGACGCTCTTCCGATCTNNNAAGCACCATTCATTTCATTTTCAT
GCACACGTCTGAACTCTTTCCCTACACGACGCTCTTCCGATCTNNNAGTGCTTTTTCCTTCTGAAGCATC
GCACACGTCTGAACTCTTTCCCTACACGACGCTCTTCCGATCTNNNAGTGCTTTTTCCTTCTGAAGCATC
CACACGTCTGAACTCTTTCCCTACACGACGCTCTTCCGATCTNNNATAAAGGAATGAGTCATCTGTTTCA
AGCACACGTCTGAACTCTTTCCCTACACGACGCTCTTCCGATCTNNNTTTACCCTCTATTTTCACTTCCCAT
AGCACACGTCTGAACTCTTTCCCTACACGACGCTCTTCCGATCTNNNCAGTCTGTATGTCATTCCTGCAT
GAGCACACGTCTGAACTCTTTCCCTACACGACGCTCTTCCGATCTNNNCTCAGAGTTAGAGGTCTGTGC
GAGCACACGTCTGAACTCTTTCCCTACACGACGCTCTTCCGATCTNNNTTATGCTTGAGGTGTTCTGAC
ACACGTCTGAACTCTTTCCCTACACGACGCTCTTCCGATCTNNNCTCTTTGTTTCTCATCAACTGCATCGG
ACACGTCTGAACTCTTTCCCTACACGACGCTCTTCCGATCTNNNCTCTTTGTTTCTCATCAACTGCATCGG
GCACACGTCTGAACTCTTTCCCTACACGACGCTCTTCCGATCTNNNAATCCATCCTGGTTTCAGATACAT
AGCACACGTCTGAACTCTTTCCCTACACGACGCTCTTCCGATCTNNNAAACACAGTAGCTTCTCTTCCAT
GAGCACACGTCTGAACTCTTTCCCTACACGACGCTCTTCCGATCTNNNTGTTGCAATGTATTCTGATATA
GAGCACACGTCTGAACTCTTTCCCTACACGACGCTCTTCCGATCTNNNCATGAACAACCAAAAGAGAAG
GCACACGTCTGAACTCTTTCCCTACACGACGCTCTTCCGATCTNNNAAATTCTCGATTGTCTTCTCTAGCA
ACACGTCTGAACTCTTTCCCTACACGACGCTCTTCCGATCTNNNGAAAAGAACTCAGTACCTGAATCAT
ACACGTCTGAACTCTTTCCCTACACGACGCTCTTCCGATCTNNNGAAAAGAACTCAGTACCTGAATCAT
CACACGTCTGAACTCTTTCCCTACACGACGCTCTTCCGATCTNNNGATCCAGAAGCAGAATAAGAGCAT
CACACGTCTGAACTCTTTCCCTACACGACGCTCTTCCGATCTNNNGATCCAGAAGCAGAATAAGAGCAT
GAGCACACGTCTGAACTCTTTCCCTACACGACGCTCTTCCGATCTNNNCACAATGGAATATCATTCAACT

GAGCACACGTCTGAACTCTTTCCCTACACGACGCTCTTCCGATCTNNNCATTGCATTGATATGATGGAT
GAGCACACGTCTGAACTCTTTCCCTACACGACGCTCTTCCGATCTNNNCATTGCATTGATATGATGGAT
GAGCACACGTCTGAACTCTTTCCCTACACGACGCTCTTCCGATCTNNNTTGGATACCTATACAGATCCAT
GAGCACACGTCTGAACTCTTTCCCTACACGACGCTCTTCCGATCTNNNTTGGATACCTATACAGATCCAT
AGCACACGTCTGAACTCTTTCCCTACACGACGCTCTTCCGATCTNNNATATTCTTCATCTCAAAAGACAC
AGCACACGTCTGAACTCTTTCCCTACACGACGCTCTTCCGATCTNNNTCCATAACCTAAGTATTTAGATG
CACACGTCTGAACTCTTTCCCTACACGACGCTCTTCCGATCTNNNCTACATGATGTACATTTGGTCTACA
AGCACACGTCTGAACTCTTTCCCTACACGACGCTCTTCCGATCTNNNCTCAGATTGGGTGGTAATCTAC
AGCACACGTCTGAACTCTTTCCCTACACGACGCTCTTCCGATCTNNNTAGTTATGGATTATGTGAGAAG
AGCACACGTCTGAACTCTTTCCCTACACGACGCTCTTCCGATCTNNNTAGTTATGGATTATGTGAGAAG
GCACACGTCTGAACTCTTTCCCTACACGACGCTCTTCCGATCTNNNTGTTATTTATTTTACTTGCATGTG
AGCACACGTCTGAACTCTTTCCCTACACGACGCTCTTCCGATCTNNNAAGCATCTTTGATAACCCATTACA
AGCACACGTCTGAACTCTTTCCCTACACGACGCTCTTCCGATCTNNNGCAGTTCTATCATGGTTAAGAC
GCACACGTCTGAACTCTTTCCCTACACGACGCTCTTCCGATCTNNNGACCATTAGCATCACTTAATTTGCA
GCACACGTCTGAACTCTTTCCCTACACGACGCTCTTCCGATCTNNNGAGAATTGACCCATGAGTTGCAT
GCACACGTCTGAACTCTTTCCCTACACGACGCTCTTCCGATCTNNNGAGAATTGACCCATGAGTTGCAT
ACACGTCTGAACTCTTTCCCTACACGACGCTCTTCCGATCTNNNCTCAATGAGAATTGACCCATCATCGG
CACACGTCTGAACTCTTTCCCTACACGACGCTCTTCCGATCTNNNATGCTCTTATGCTGGTAAAAGCAT
CACACGTCTGAACTCTTTCCCTACACGACGCTCTTCCGATCTNNNATACTTTTCACACTCTTCCTCCATCG
CACACGTCTGAACTCTTTCCCTACACGACGCTCTTCCGATCTNNNATACTTTTCACACTCTTCCTCCATCG
ACACGTCTGAACTCTTTCCCTACACGACGCTCTTCCGATCTNNNATTTTCCTGTCTTCATGAATGACATC
AGCACACGTCTGAACTCTTTCCCTACACGACGCTCTTCCGATCTNNNATGGTTGAATGAATGGTTAAA
CACACGTCTGAACTCTTTCCCTACACGACGCTCTTCCGATCTNNNAAGGCATATGGAGATGTAGTTACA
CACACGTCTGAACTCTTTCCCTACACGACGCTCTTCCGATCTNNNAAGGCATATGGAGATGTAGTTACA
GCACACGTCTGAACTCTTTCCCTACACGACGCTCTTCCGATCTNNNTCATGAATACTGTGACCTTTCCATC
GCACACGTCTGAACTCTTTCCCTACACGACGCTCTTCCGATCTNNNTGAAAATTAAGTGCTATGGTAAG
GCACACGTCTGAACTCTTTCCCTACACGACGCTCTTCCGATCTNNNTGAAAATTAAGTGCTATGGTAAG
GCACACGTCTGAACTCTTTCCCTACACGACGCTCTTCCGATCTNNNTATCATCTCGTCCAATAATAGAAC
GAGCACACGTCTGAACTCTTTCCCTACACGACGCTCTTCCGATCTNNNTTTACATATCATCTCGTCCAAT
GAGCACACGTCTGAACTCTTTCCCTACACGACGCTCTTCCGATCTNNNTTTACATATCATCTCGTCCAAT
AGCACACGTCTGAACTCTTTCCCTACACGACGCTCTTCCGATCTNNNCAGATGGTATTTTAAGATTCCTG
AGCACACGTCTGAACTCTTTCCCTACACGACGCTCTTCCGATCTNNNCAGATGGTATTTTAAGATTCCTG
GCACACGTCTGAACTCTTTCCCTACACGACGCTCTTCCGATCTNNNAATGAAAAGGTACCCCTATTATTC
AGCACACGTCTGAACTCTTTCCCTACACGACGCTCTTCCGATCTNNNTTTACCCTCTATTTTCACTTCCCAT
AGCACACGTCTGAACTCTTTCCCTACACGACGCTCTTCCGATCTNNNAAAAACTAAGCTTGTGTAAGAAT
AGCACACGTCTGAACTCTTTCCCTACACGACGCTCTTCCGATCTNNNGTCCACATTTTAAAAACTAAGCT
AGCACACGTCTGAACTCTTTCCCTACACGACGCTCTTCCGATCTNNNGCTTCTAAAACACCATTTAAAAT
AGCACACGTCTGAACTCTTTCCCTACACGACGCTCTTCCGATCTNNNCATCTTTACATTTTTTTGCAGTTC
CACACGTCTGAACTCTTTCCCTACACGACGCTCTTCCGATCTNNNTTTTCTGTCGCTTTAATCTCATTCATC
CACACGTCTGAACTCTTTCCCTACACGACGCTCTTCCGATCTNNNCTCATTACGTTTTAGATGGGATTCA
AGCACACGTCTGAACTCTTTCCCTACACGACGCTCTTCCGATCTNNNACACCTCAGATATATTTCTTCATC
CACACGTCTGAACTCTTTCCCTACACGACGCTCTTCCGATCTNNNGAATTTCCAGTGTATTGTTTGGCAT
AGCACACGTCTGAACTCTTTCCCTACACGACGCTCTTCCGATCTNNNATTCATTTCAACTTGGTACATTT

AGCACACGTCTGAACTCTTTCCCTACACGACGCTCTTCCGATCTNNNATTCATTTCAACTTGGTACATTT
GCACACGTCTGAACTCTTTCCCTACACGACGCTCTTCCGATCTNNNTTTTTGGAACTGGAGATGTTCA
GCACACGTCTGAACTCTTTCCCTACACGACGCTCTTCCGATCTNNNTTTGTTGAAAATGAAATCTAGT
AGCACACGTCTGAACTCTTTCCCTACACGACGCTCTTCCGATCTNNNAACAGGTTAATTCCCAATTTAA
CACACGTCTGAACTCTTTCCCTACACGACGCTCTTCCGATCTNNNCAAAAGGTGTGAGTTTGAAAAT
AGCACACGTCTGAACTCTTTCCCTACACGACGCTCTTCCGATCTNNNTTTTGAGGGAGATGTGAACTC
ACACGTCTGAACTCTTTCCCTACACGACGCTCTTCCGATCTNNNTTTATTTGTAATTTTTGCTGCTGCA
AGCACACGTCTGAACTCTTTCCCTACACGACGCTCTTCCGATCTNNNCAGTCTGTATGTCATTCCTGCA
CACACGTCTGAACTCTTTCCCTACACGACGCTCTTCCGATCTNNNTTCTCTTTGCTGATCATTGTTCAT
GCACACGTCTGAACTCTTTCCCTACACGACGCTCTTCCGATCTNNNAACTCGCTTGATTTTGAATACCA
GCACACGTCTGAACTCTTTCCCTACACGACGCTCTTCCGATCTNNNAACTCGCTTGATTTTGAATACCA
AGCACACGTCTGAACTCTTTCCCTACACGACGCTCTTCCGATCTNNNGAAACTAGGTGTGTATTGTTT
AGCACACGTCTGAACTCTTTCCCTACACGACGCTCTTCCGATCTNNNGAAACTAGGTGTGTATTGTTT
AGCACACGTCTGAACTCTTTCCCTACACGACGCTCTTCCGATCTNNNTATTTGGAAAATATTGCATGT
AGCACACGTCTGAACTCTTTCCCTACACGACGCTCTTCCGATCTNNNTATTTGGAAAATATTGCATGT
GCACACGTCTGAACTCTTTCCCTACACGACGCTCTTCCGATCTNNNTTTATATCCCTGTAGAACTGAAG
GCACACGTCTGAACTCTTTCCCTACACGACGCTCTTCCGATCTNNNTTTATATCCCTGTAGAACTGAAG
GCACACGTCTGAACTCTTTCCCTACACGACGCTCTTCCGATCTNNNGTTATGTATCAAGTACCTTTGCC
CACACGTCTGAACTCTTTCCCTACACGACGCTCTTCCGATCTNNNGGAAAAACATTTGCATGGTTATCA
AGCACACGTCTGAACTCTTTCCCTACACGACGCTCTTCCGATCTNNNCTATGGCAAGACTCAGTTTGC
AGCACACGTCTGAACTCTTTCCCTACACGACGCTCTTCCGATCTNNNCTATGGCAAGACTCAGTTTGC
CACACGTCTGAACTCTTTCCCTACACGACGCTCTTCCGATCTNNNCTTAGTTACCTTTTTCCATGTTTCA
AGCACACGTCTGAACTCTTTCCCTACACGACGCTCTTCCGATCTNNNACAGAAACAGTGATAAAACAA
ACACGTCTGAACTCTTTCCCTACACGACGCTCTTCCGATCTNNNCTCTTTTGCTGCACATTATCCATCG
GCACACGTCTGAACTCTTTCCCTACACGACGCTCTTCCGATCTNNNTTCTGTGATTTGAGAGTAAGAG
GCACACGTCTGAACTCTTTCCCTACACGACGCTCTTCCGATCTNNNTTCTGTGATTTGAGAGTAAGAG
GCACACGTCTGAACTCTTTCCCTACACGACGCTCTTCCGATCTNNNGTCTTTTCTGAAGAAGTTGTTTC
CACACGTCTGAACTCTTTCCCTACACGACGCTCTTCCGATCTNNNAAAATTATTGAGAACAGAAGCAG
CACACGTCTGAACTCTTTCCCTACACGACGCTCTTCCGATCTNNNGAGTATCTAGTAATTTGGAAGGT
GAGCACACGTCTGAACTCTTTCCCTACACGACGCTCTTCCGATCTNNNCCCATATCTGAAGATAAATTT
GAGCACACGTCTGAACTCTTTCCCTACACGACGCTCTTCCGATCTNNNCCCATATCTGAAGATAAATTT
AGCACACGTCTGAACTCTTTCCCTACACGACGCTCTTCCGATCTNNNTTTTTATGAGCCTTTACAAATT
AGCACACGTCTGAACTCTTTCCCTACACGACGCTCTTCCGATCTNNNTAATTCTTTTGGAAAAACTGC
GCACACGTCTGAACTCTTTCCCTACACGACGCTCTTCCGATCTNNNCATTCAAATTTTACAGGATCTGA
GCACACGTCTGAACTCTTTCCCTACACGACGCTCTTCCGATCTNNNCATTCAAATTTTACAGGATCTGA
AGCACACGTCTGAACTCTTTCCCTACACGACGCTCTTCCGATCTNNNGCGAATTGATCTAAAACAGAT
AGCACACGTCTGAACTCTTTCCCTACACGACGCTCTTCCGATCTNNNCTATGGTCGTTATTTTCTCATT
AGCACACGTCTGAACTCTTTCCCTACACGACGCTCTTCCGATCTNNNCTATGGTCGTTATTTTCTCATT
AGCACACGTCTGAACTCTTTCCCTACACGACGCTCTTCCGATCTNNNTAATTTCCCTATTAGCCAGAAA
GCACACGTCTGAACTCTTTCCCTACACGACGCTCTTCCGATCTNNNTAAGAGTGTCAGCATTATCATTC
AGCACACGTCTGAACTCTTTCCCTACACGACGCTCTTCCGATCTNNNAACAAAAGTCACTGTTAATTCT
AGCACACGTCTGAACTCTTTCCCTACACGACGCTCTTCCGATCTNNNGATGAAATTTTAGAGAAAAGG
CACACGTCTGAACTCTTTCCCTACACGACGCTCTTCCGATCTNNNTGAGAATCACAATCACTGCACAT

GAGCACACGTCTGAACTCTTTCCCTACACGACGCTCTTCCGATCTNNNATTCATTTTATTGTGGTATGG
AGCACACGTCTGAACTCTTTCCCTACACGACGCTCTTCCGATCTNNNTGCTATTGCACACTTAGTATGC
AGCACACGTCTGAACTCTTTCCCTACACGACGCTCTTCCGATCTNNNTCCACCAGCAAAAAGATTAGC
AGCACACGTCTGAACTCTTTCCCTACACGACGCTCTTCCGATCTNNNCTTATTTTAAGAAATTGCCACAC
AGCACACGTCTGAACTCTTTCCCTACACGACGCTCTTCCGATCTNNNTCTTTACTTTCTTTTTAACAGGC
AGCACACGTCTGAACTCTTTCCCTACACGACGCTCTTCCGATCTNNNAATTCTTTCAATTTTTCCAGAG
AGCACACGTCTGAACTCTTTCCCTACACGACGCTCTTCCGATCTNNNATTCTGTCTGGACTCTACTTTCA
AGCACACGTCTGAACTCTTTCCCTACACGACGCTCTTCCGATCTNNNTTCATGACTTGTATTTGATGAT
GAGCACACGTCTGAACTCTTTCCCTACACGACGCTCTTCCGATCTNNNTCTATGTATTGTAATTCTAGCG
AGCACACGTCTGAACTCTTTCCCTACACGACGCTCTTCCGATCTNNNTCATAAAGTTGTGATCTAAAT
GAGCACACGTCTGAACTCTTTCCCTACACGACGCTCTTCCGATCTNNNTCTCCAATAAGTGTAGATTTT
AGCACACGTCTGAACTCTTTCCCTACACGACGCTCTTCCGATCTNNNCAGTCATTATCTCATCAACATTT
GAGCACACGTCTGAACTCTTTCCCTACACGACGCTCTTCCGATCTNNNCTGGAAACTATCAAACACAAC
AGCACACGTCTGAACTCTTTCCCTACACGACGCTCTTCCGATCTNNNCAACTGTATCTTTTGCATAAAA
AGCACACGTCTGAACTCTTTCCCTACACGACGCTCTTCCGATCTNNNATGAAGTTTTAGTGTGGTCTA
AGCACACGTCTGAACTCTTTCCCTACACGACGCTCTTCCGATCTNNNGCAATTTACGTAGATGATTTTC
AGCACACGTCTGAACTCTTTCCCTACACGACGCTCTTCCGATCTNNNGTATTTTTCGTATTTCTTGCAA
GAGCACACGTCTGAACTCTTTCCCTACACGACGCTCTTCCGATCTNNNTGTAAACTCACTCTAGAAAAT
AGCACACGTCTGAACTCTTTCCCTACACGACGCTCTTCCGATCTNNNAAATATCTTACCTGCAGTAATC
AGAGCACACGTCTGAACTCTTTCCCTACACGACGCTCTTCCGATCTNNNATTATTTGGGACATATTGAT
AGCACACGTCTGAACTCTTTCCCTACACGACGCTCTTCCGATCTNNNTCAATGCAACTTGTTTTATGTA
AGCACACGTCTGAACTCTTTCCCTACACGACGCTCTTCCGATCTNNNATGTTGCTTTTCTGATTATTTC
GCACACGTCTGAACTCTTTCCCTACACGACGCTCTTCCGATCTNNNCTGGTGTGAGTAATTTTTTTCTA
AGAGCACACGTCTGAACTCTTTCCCTACACGACGCTCTTCCGATCTNNNCACATTTAATCCATAACTCCT
GAGCACACGTCTGAACTCTTTCCCTACACGACGCTCTTCCGATCTNNNTTTAATGAAGAAGAGAAGTC
GAGCACACGTCTGAACTCTTTCCCTACACGACGCTCTTCCGATCTNNNTTTAATGAAGAAGAGAAGTC
AGCACACGTCTGAACTCTTTCCCTACACGACGCTCTTCCGATCTNNNACAGTTGGTATTTTTAATTCTC
GCACACGTCTGAACTCTTTCCCTACACGACGCTCTTCCGATCTNNNTTACTGGAGAGTTATTTTGTACA
AGCACACGTCTGAACTCTTTCCCTACACGACGCTCTTCCGATCTNNNTAGAAAAGTTAGGAAGACAGA
AGCACACGTCTGAACTCTTTCCCTACACGACGCTCTTCCGATCTNNNTAGAAAAGTTAGGAAGACAGA
GCACACGTCTGAACTCTTTCCCTACACGACGCTCTTCCGATCTNNNTTTAGTTCAATGGATAGATTCCT
AGCACACGTCTGAACTCTTTCCCTACACGACGCTCTTCCGATCTNNNGGATAAACTGGTTGCATAATA
CACACGTCTGAACTCTTTCCCTACACGACGCTCTTCCGATCTNNNTTCTGGAAAAGCTGGAAAAGCAT
CACACGTCTGAACTCTTTCCCTACACGACGCTCTTCCGATCTNNNTTCTGGAAAAGCTGGAAAAGCAT
GAGCACACGTCTGAACTCTTTCCCTACACGACGCTCTTCCGATCTNNNCATGGAAAAATTGTGTTCCG
GAGCACACGTCTGAACTCTTTCCCTACACGACGCTCTTCCGATCTNNNCATGGAAAAATTGTGTTCCG
CACACGTCTGAACTCTTTCCCTACACGACGCTCTTCCGATCTNNNTGTAATAATAGTAGAAATAAAGT
GAGCACACGTCTGAACTCTTTCCCTACACGACGCTCTTCCGATCTNNNCTGATGATCTGTCACTGTCTC
GAGCACACGTCTGAACTCTTTCCCTACACGACGCTCTTCCGATCTNNNCTGATGATCTGTCACTGTCTC
GCACACGTCTGAACTCTTTCCCTACACGACGCTCTTCCGATCTNNNCTTCATCTGTATTTATAGCCACTC
GCACACGTCTGAACTCTTTCCCTACACGACGCTCTTCCGATCTNNNCTTCATCTGTATTTATAGCCACTC
CACACGTCTGAACTCTTTCCCTACACGACGCTCTTCCGATCTNNNGTCCATGGCCTATTAGGAAGCATC
GCACACGTCTGAACTCTTTCCCTACACGACGCTCTTCCGATCTNNNAAGGAGGTACCACTAGTCAACA

GCACACGTCTGAACTCTTTCCCTACACGACGCTCTTCCGATCTNNNAAGGAGGTACCACTAGTCAACA
ACACGTCTGAACTCTTTCCCTACACGACGCTCTTCCGATCTNNNAGAGATTATCCTGGATTGTCTCAT
ACACGTCTGAACTCTTTCCCTACACGACGCTCTTCCGATCTNNNAGAGATTATCCTGGATTGTCTCAT
CACACGTCTGAACTCTTTCCCTACACGACGCTCTTCCGATCTNNNCTGGCTGAGTTAAGGATCTTCATC
GCACACGTCTGAACTCTTTCCCTACACGACGCTCTTCCGATCTNNNGTATTTTTAGTAGAGACAGGGT
GCACACGTCTGAACTCTTTCCCTACACGACGCTCTTCCGATCTNNNGTATTTTTAGTAGAGACAGGGT
AGCACACGTCTGAACTCTTTCCCTACACGACGCTCTTCCGATCTNNNCTCATCTCTTCTCTTTTCCTCCAT
ACACGTCTGAACTCTTTCCCTACACGACGCTCTTCCGATCTNNNCTTCCTGCTTTAGAATCCCTCATCG
CACACGTCTGAACTCTTTCCCTACACGACGCTCTTCCGATCTNNNTATGGTTTGAATCATTTTTGTGTC
AGCACACGTCTGAACTCTTTCCCTACACGACGCTCTTCCGATCTNNNTGAATGAGGATATCTTCATAG
GCACACGTCTGAACTCTTTCCCTACACGACGCTCTTCCGATCTNNNGAACAGAATTACAGGAAGTTT
GCACACGTCTGAACTCTTTCCCTACACGACGCTCTTCCGATCTNNNGAACAGAATTACAGGAAGTTT
GAGCACACGTCTGAACTCTTTCCCTACACGACGCTCTTCCGATCTNNNTGGACTTTTCTATGAGTTCTT
AGCACACGTCTGAACTCTTTCCCTACACGACGCTCTTCCGATCTNNNAAAAGAAGAAATTCTTTCCAG
AGCACACGTCTGAACTCTTTCCCTACACGACGCTCTTCCGATCTNNNAAAAGAAGAAATTCTTTCCAG
AGCACACGTCTGAACTCTTTCCCTACACGACGCTCTTCCGATCTNNNCACAAAGCGTTTTAAAAAAGA
AGCACACGTCTGAACTCTTTCCCTACACGACGCTCTTCCGATCTNNNCGGATGTAAATTTCGAGTTTT
AGCACACGTCTGAACTCTTTCCCTACACGACGCTCTTCCGATCTNNNCTGACGACTCTTTTTCAGATCA
GCACACGTCTGAACTCTTTCCCTACACGACGCTCTTCCGATCTNNNTGATAGGAATGTTTGATAGTG
AGCACACGTCTGAACTCTTTCCCTACACGACGCTCTTCCGATCTNNNTTTGATCACAATGGGAAAATTA
AGCACACGTCTGAACTCTTTCCCTACACGACGCTCTTCCGATCTNNNGAAACATCACACATCTAAGATT
AGAGCACACGTCTGAACTCTTTCCCTACACGACGCTCTTCCGATCTNNNGTAAAGGAGTTAGTAGAA
GCACACGTCTGAACTCTTTCCCTACACGACGCTCTTCCGATCTNNNATTTCCTTGCCAACTAGTAAAAA
AGCACACGTCTGAACTCTTTCCCTACACGACGCTCTTCCGATCTNNNCTGGTAAAGTGTATGGATGG
GAGCACACGTCTGAACTCTTTCCCTACACGACGCTCTTCCGATCTNNNGAAAACAAAATGCTGAAGAA
ACGTCTGAACTCTTTCCCTACACGACGCTCTTCCGATCTNNNTCTTTACCCTCTAAGCTGAACATCGGA
ACGTCTGAACTCTTTCCCTACACGACGCTCTTCCGATCTNNNTCTTTACCCTCTAAGCTGAACATCGGA
AGCACACGTCTGAACTCTTTCCCTACACGACGCTCTTCCGATCTNNNGGATTCCATATTGTTCATAACA
CACACGTCTGAACTCTTTCCCTACACGACGCTCTTCCGATCTNNNCTAGAATGAAGCTGGAGAATGC
CACACGTCTGAACTCTTTCCCTACACGACGCTCTTCCGATCTNNNTAGTATCAGAAAAGCTTAAGGCC
GAGCACACGTCTGAACTCTTTCCCTACACGACGCTCTTCCGATCTNNNATCAGAATTAAGACTAGAGA
GAGCACACGTCTGAACTCTTTCCCTACACGACGCTCTTCCGATCTNNNATCAGAATTAAGACTAGAGA
GAGCACACGTCTGAACTCTTTCCCTACACGACGCTCTTCCGATCTNNNAGACAAGGAAACTAATGATA
GAGCACACGTCTGAACTCTTTCCCTACACGACGCTCTTCCGATCTNNNAGACAAGGAAACTAATGATA
GAGCACACGTCTGAACTCTTTCCCTACACGACGCTCTTCCGATCTNNNTTAGCTTTAGTCACTGAGAA
GAGCACACGTCTGAACTCTTTCCCTACACGACGCTCTTCCGATCTNNNACTTGAGAGATGAGATGTA
GAGCACACGTCTGAACTCTTTCCCTACACGACGCTCTTCCGATCTNNNACTTGAGAGATGAGATGTA
CACACGTCTGAACTCTTTCCCTACACGACGCTCTTCCGATCTNNNCCATATGCTATCCCTTATATTTCA
CACACGTCTGAACTCTTTCCCTACACGACGCTCTTCCGATCTNNNCCATATGCTATCCCTTATATTTCA
GAGCACACGTCTGAACTCTTTCCCTACACGACGCTCTTCCGATCTNNNCACAGTCCTTCCCATTATTAC
GCACACGTCTGAACTCTTTCCCTACACGACGCTCTTCCGATCTNNNACCTAGTTGTCACCTTTGTATAA
GCACACGTCTGAACTCTTTCCCTACACGACGCTCTTCCGATCTNNNACCTAGTTGTCACCTTTGTATAA
AGCACACGTCTGAACTCTTTCCCTACACGACGCTCTTCCGATCTNNNAAGGTTTACAAAATAATTTTTC

GCACACGTCTGAACTCTTTCCCTACACGACGCTCTTCCGATCTNNNATACCTATATTGTGCTTTAAGGT
GCACACGTCTGAACTCTTTCCCTACACGACGCTCTTCCGATCTNNNATACCTATATTGTGCTTTAAGGT
AGCACACGTCTGAACTCTTTCCCTACACGACGCTCTTCCGATCTNNNCCATAAGAACAAAGATACATGT
AGCACACGTCTGAACTCTTTCCCTACACGACGCTCTTCCGATCTNNNGCAATGTAAGAGATAATTATTC
GCACACGTCTGAACTCTTTCCCTACACGACGCTCTTCCGATCTNNNAGTCCTCCATAAATTATGTAACC
GCACACGTCTGAACTCTTTCCCTACACGACGCTCTTCCGATCTNNNAGTCCTCCATAAATTATGTAACC
AGCACACGTCTGAACTCTTTCCCTACACGACGCTCTTCCGATCTNNNCAGAAGGTTTTGCTATATGTA
AGCACACGTCTGAACTCTTTCCCTACACGACGCTCTTCCGATCTNNNCAGAAGGTTTTGCTATATGTA
AGCACACGTCTGAACTCTTTCCCTACACGACGCTCTTCCGATCTNNNCTGGGAGTTGTAAAAATCTTTT
GCACACGTCTGAACTCTTTCCCTACACGACGCTCTTCCGATCTNNNCTTATTGAAGCTCTGTCCTCCATC
GCACACGTCTGAACTCTTTCCCTACACGACGCTCTTCCGATCTNNNCTATGTTTTGTGGAAAGAGAAG
AGCACACGTCTGAACTCTTTCCCTACACGACGCTCTTCCGATCTNNNGGAAAAAATTACTGTCTTCTAC
AGCACACGTCTGAACTCTTTCCCTACACGACGCTCTTCCGATCTNNNTCTCTAGAGTCAGAATCCTGCA
GCACACGTCTGAACTCTTTCCCTACACGACGCTCTTCCGATCTNNNGAATGGTTTATGTCAACAAAATG
CACACGTCTGAACTCTTTCCCTACACGACGCTCTTCCGATCTNNNTTCTGAAACTGAAGAGTCAAACAT
CACACGTCTGAACTCTTTCCCTACACGACGCTCTTCCGATCTNNNTTCTGAAACTGAAGAGTCAAACAT
CACACGTCTGAACTCTTTCCCTACACGACGCTCTTCCGATCTNNNGTTAATGAATCCGGTAGAAGATC
AGCACACGTCTGAACTCTTTCCCTACACGACGCTCTTCCGATCTNNNTGGTATTGCCCTATAATAATTC
AGCACACGTCTGAACTCTTTCCCTACACGACGCTCTTCCGATCTNNNTGGTATTGCCCTATAATAATTC
AGCACACGTCTGAACTCTTTCCCTACACGACGCTCTTCCGATCTNNNTGTTAACACAAGATTTTTGTAT
AGCACACGTCTGAACTCTTTCCCTACACGACGCTCTTCCGATCTNNNCTCTAAATTATCCAGACTGAAG
AGCACACGTCTGAACTCTTTCCCTACACGACGCTCTTCCGATCTNNNCTCTAAATTATCCAGACTGAAG
AGCACACGTCTGAACTCTTTCCCTACACGACGCTCTTCCGATCTNNNCTTTGCCTGGGTATAGAGTACA
AGCACACGTCTGAACTCTTTCCCTACACGACGCTCTTCCGATCTNNNTTAACTCAGAGAAGTTTCCATTC
AGCACACGTCTGAACTCTTTCCCTACACGACGCTCTTCCGATCTNNNTAGTGATTCAGATAGTAGTTGT
AGCACACGTCTGAACTCTTTCCCTACACGACGCTCTTCCGATCTNNNTAGTGATTCAGATAGTAGTTGT
AGCACACGTCTGAACTCTTTCCCTACACGACGCTCTTCCGATCTNNNGAGAACTCTGATGTTTGTAAA
AGCACACGTCTGAACTCTTTCCCTACACGACGCTCTTCCGATCTNNNTTAGAGAAGTTTAAGAGGCAA
AGCACACGTCTGAACTCTTTCCCTACACGACGCTCTTCCGATCTNNNTTAGAGAAGTTTAAGAGGCAA
GAGCACACGTCTGAACTCTTTCCCTACACGACGCTCTTCCGATCTNNNTTTCCAATGATAGTAGGAAA
AGCACACGTCTGAACTCTTTCCCTACACGACGCTCTTCCGATCTNNNCACAAACAGATTGCATATTAT
AGCACACGTCTGAACTCTTTCCCTACACGACGCTCTTCCGATCTNNNCACAAACAGATTGCATATTAT
AGCACACGTCTGAACTCTTTCCCTACACGACGCTCTTCCGATCTNNNGAATATAAGTTATGGGCTAG
AGCACACGTCTGAACTCTTTCCCTACACGACGCTCTTCCGATCTNNNCAGCACTTTAAGTAAAGAATCT
AGCACACGTCTGAACTCTTTCCCTACACGACGCTCTTCCGATCTNNNCAGCACTTTAAGTAAAGAATCT
AGCACACGTCTGAACTCTTTCCCTACACGACGCTCTTCCGATCTNNNTAAAAATTATGTTAGACCTGG
AGCACACGTCTGAACTCTTTCCCTACACGACGCTCTTCCGATCTNNNAAAACTAAATCAACTGAAACA
GAGCACACGTCTGAACTCTTTCCCTACACGACGCTCTTCCGATCTNNNAAGAAATGTTTTTAGTTATGG
GAGCACACGTCTGAACTCTTTCCCTACACGACGCTCTTCCGATCTNNNAAGAAATGTTTTTAGTTATGG
AGCACACGTCTGAACTCTTTCCCTACACGACGCTCTTCCGATCTNNNAGAGAAAAAAAAAGAATCTAG
GAGCACACGTCTGAACTCTTTCCCTACACGACGCTCTTCCGATCTNNNATAGAATTCAGAATCAACCAA
GAGCACACGTCTGAACTCTTTCCCTACACGACGCTCTTCCGATCTNNNATAGAATTCAGAATCAACCAA
ACACGTCTGAACTCTTTCCCTACACGACGCTCTTCCGATCTNNNCTGGGATTACAATTGTGAGCCATCG

CACACGTCTGAACTCTTTCCCTACACGACGCTCTTCCGATCTNNNCTTTTGCATGTTTTGTAGAGATCA

CACACGTCTGAACTCTTTCCCTACACGACGCTCTTCCGATCTNNNTGAGTTTATTTTGAAGGTTAACTC

GCACACGTCTGAACTCTTTCCCTACACGACGCTCTTCCGATCTNNNCAACTTTTGCATGTTTTGTAGACA

CACACGTCTGAACTCTTTCCCTACACGACGCTCTTCCGATCTNNNGTGAACCTTAGTACCTAAAAGGC

CACACGTCTGAACTCTTTCCCTACACGACGCTCTTCCGATCTNNNCAGAAACACTTTTCGACATAGCAT

GAGCACACGTCTGAACTCTTTCCCTACACGACGCTCTTCCGATCTNNNCTCCAATAAGAGCATCCATGC

ACACGTCTGAACTCTTTCCCTACACGACGCTCTTCCGATCTNNNCTAAAAGTTTTGAGCTTCTCAACAT

CACACGTCTGAACTCTTTCCCTACACGACGCTCTTCCGATCTNNNCTTTTTGCCATGGGATTTCCATC

ACACGTCTGAACTCTTTCCCTACACGACGCTCTTCCGATCTNNNAAACATTTTCAGACCTATGGAAACA

ACACGTCTGAACTCTTTCCCTACACGACGCTCTTCCGATCTNNNTACCACAGTTTCTTCTGTAAATCAT

ACACGTCTGAACTCTTTCCCTACACGACGCTCTTCCGATCTNNNGTACCACAGTTTCTTCTGTAACATC

ACACGTCTGAACTCTTTCCCTACACGACGCTCTTCCGATCTNNNGATGTAGTTACTGTGGAGTCTCAT

CACACGTCTGAACTCTTTCCCTACACGACGCTCTTCCGATCTNNNTCACAAAACATCCTGAAAATTCCAT

ACACGTCTGAACTCTTTCCCTACACGACGCTCTTCCGATCTNNNTATAACACATCTACTGAAAAAGCC

ACACGTCTGAACTCTTTCCCTACACGACGCTCTTCCGATCTNNNTATAACACATCTACTGAAAAAGCC

CACACGTCTGAACTCTTTCCCTACACGACGCTCTTCCGATCTNNNAACTGTTGGTTATTTGTGTTATTC

CACACGTCTGAACTCTTTCCCTACACGACGCTCTTCCGATCTNNNAACTGTTGGTTATTTGTGTTATTC

ACACGTCTGAACTCTTTCCCTACACGACGCTCTTCCGATCTNNNGTTCTAGGGGAAATGAATGTGCAT

CACACGTCTGAACTCTTTCCCTACACGACGCTCTTCCGATCTNNNGTTTGACAATAATCTCACAAAACC

CACACGTCTGAACTCTTTCCCTACACGACGCTCTTCCGATCTNNNGTTTGACAATAATCTCACAAAACC

GCACACGTCTGAACTCTTTCCCTACACGACGCTCTTCCGATCTNNNCGTCCTGATGATTTTTATTTTTG

GCACACGTCTGAACTCTTTCCCTACACGACGCTCTTCCGATCTNNNCGTCCTGATGATTTTTATTTTTG

AGCACACGTCTGAACTCTTTCCCTACACGACGCTCTTCCGATCTNNNATTAGAAAGGAAATGAATGGT

AGCACACGTCTGAACTCTTTCCCTACACGACGCTCTTCCGATCTNNNATTAGAAAGGAAATGAATGGT

CACACGTCTGAACTCTTTCCCTACACGACGCTCTTCCGATCTNNNACAATCAGTGAGTAAATAGACAG

GCACACGTCTGAACTCTTTCCCTACACGACGCTCTTCCGATCTNNNCTTTCTCCTGTTTATGGGTTTCAT

CACACGTCTGAACTCTTTCCCTACACGACGCTCTTCCGATCTNNNCTTCACATAAAGGAACATGGTTC

CACACGTCTGAACTCTTTCCCTACACGACGCTCTTCCGATCTNNNCTTCACATAAAGGAACATGGTTC

GCACACGTCTGAACTCTTTCCCTACACGACGCTCTTCCGATCTNNNATATCTCCCTTTTTTTCTCTCTTCAT

AGCACACGTCTGAACTCTTTCCCTACACGACGCTCTTCCGATCTNNNCTATCTGAAAATACATTTCCCT

AGCACACGTCTGAACTCTTTCCCTACACGACGCTCTTCCGATCTNNNCTATCTGAAAATACATTTCCCT

GCACACGTCTGAACTCTTTCCCTACACGACGCTCTTCCGATCTNNNGAAAGTTAAGGTATCAATGAGA

GCACACGTCTGAACTCTTTCCCTACACGACGCTCTTCCGATCTNNNGAAAGTTAAGGTATCAATGAGA

GCACACGTCTGAACTCTTTCCCTACACGACGCTCTTCCGATCTNNNCTTCAGGTAGATTAAGGTATCTT

AGCACACGTCTGAACTCTTTCCCTACACGACGCTCTTCCGATCTNNNAAAATCAAACTGTAGTTGTTTT

AGCACACGTCTGAACTCTTTCCCTACACGACGCTCTTCCGATCTNNNCAGACTCTTACCAGCATATATA

AGCACACGTCTGAACTCTTTCCCTACACGACGCTCTTCCGATCTNNNTCATGCTCTGATTAGATAATCT

AGCACACGTCTGAACTCTTTCCCTACACGACGCTCTTCCGATCTNNNTCATGCTCTGATTAGATAATCT

AGCACACGTCTGAACTCTTTCCCTACACGACGCTCTTCCGATCTNNNTAACTGAAGATATTCCAACTCA

AGCACACGTCTGAACTCTTTCCCTACACGACGCTCTTCCGATCTNNNCCCAAATTAAACGTAAATGTTTT

AGCACACGTCTGAACTCTTTCCCTACACGACGCTCTTCCGATCTNNNTTAATTTCTGAAAAGCACCATT

AGCACACGTCTGAACTCTTTCCCTACACGACGCTCTTCCGATCTNNNTTAATTTCTGAAAAGCACCATT

GAGCACACGTCTGAACTCTTTCCCTACACGACGCTCTTCCGATCTNNNTTGCAGCTAAATTACCATAA

AGCACACGTCTGAACTCTTTCCCTACACGACGCTCTTCCGATCTNNNGTTCTTATCTAGAGAAGAGCT
GCACACGTCTGAACTCTTTCCCTACACGACGCTCTTCCGATCTNNNGTAAGAGTTAAAGAGTAAAGC
GCACACGTCTGAACTCTTTCCCTACACGACGCTCTTCCGATCTNNNGTAAGAGTTAAAGAGTAAAGC
ACACGTCTGAACTCTTTCCCTACACGACGCTCTTCCGATCTNNNCTACGATCAGTCCTACCAATACATCG
ACACGTCTGAACTCTTTCCCTACACGACGCTCTTCCGATCTNNNCATCACTGTCTTCACAAACCCATCG
GCACACGTCTGAACTCTTTCCCTACACGACGCTCTTCCGATCTNNNAAGATTTAATGACCTCAGGTTT
AGCACACGTCTGAACTCTTTCCCTACACGACGCTCTTCCGATCTNNNCATTGAAAATGAAGGAAAACA
GCACACGTCTGAACTCTTTCCCTACACGACGCTCTTCCGATCTNNNTTATTTGATCATTTGTTTTTGGC
GCACACGTCTGAACTCTTTCCCTACACGACGCTCTTCCGATCTNNNCAAAGACAGAACTTAACCTCTCA
GCACACGTCTGAACTCTTTCCCTACACGACGCTCTTCCGATCTNNNCAAAGACAGAACTTAACCTCTCA
GCACACGTCTGAACTCTTTCCCTACACGACGCTCTTCCGATCTNNNGGAACTTCCAAATAATGTGTTT
GCACACGTCTGAACTCTTTCCCTACACGACGCTCTTCCGATCTNNNGGAACTTCCAAATAATGTGTTT
AGCACACGTCTGAACTCTTTCCCTACACGACGCTCTTCCGATCTNNNAGTTTTGCATGATAAAAGTGG
AGCACACGTCTGAACTCTTTCCCTACACGACGCTCTTCCGATCTNNNAGTTTTGCATGATAAAAGTGG
CACACGTCTGAACTCTTTCCCTACACGACGCTCTTCCGATCTNNNGTTTAAATTCTCCACAGATTGTAC
CACACGTCTGAACTCTTTCCCTACACGACGCTCTTCCGATCTNNNGTTTAAATTCTCCACAGATTGTAC
GCACACGTCTGAACTCTTTCCCTACACGACGCTCTTCCGATCTNNNATTAATCCTTCACAAAGTCTTTGC
GCACACGTCTGAACTCTTTCCCTACACGACGCTCTTCCGATCTNNNATTAATCCTTCACAAAGTCTTTGC
AGCACACGTCTGAACTCTTTCCCTACACGACGCTCTTCCGATCTNNNCGTAACAATAATAGTTCCTAAC
AGCACACGTCTGAACTCTTTCCCTACACGACGCTCTTCCGATCTNNNCGTAACAATAATAGTTCCTAAC
GCACACGTCTGAACTCTTTCCCTACACGACGCTCTTCCGATCTNNNAAGGTAAAAGAAATCATTGAGT
GCACACGTCTGAACTCTTTCCCTACACGACGCTCTTCCGATCTNNNGACTGATGTTTGATCACAATGC
ACACGTCTGAACTCTTTCCCTACACGACGCTCTTCCGATCTNNNATATATGTGCCATCTTTCTACATCAT
GCACACGTCTGAACTCTTTCCCTACACGACGCTCTTCCGATCTNNNAAAAACTAAGCTTGTGTAAGAA
GCACACGTCTGAACTCTTTCCCTACACGACGCTCTTCCGATCTNNNGCTTCTAAAACACCATTTAAAAT
AGCACACGTCTGAACTCTTTCCCTACACGACGCTCTTCCGATCTNNNTCCTACCTTTCCCATTTATAAAT
ACACGTCTGAACTCTTTCCCTACACGACGCTCTTCCGATCTNNNGAAGATACTTACTGTAGTTCTCACA
ACACGTCTGAACTCTTTCCCTACACGACGCTCTTCCGATCTNNNCTCATTACGTTTTAGATGGGATTCA
ACACGTCTGAACTCTTTCCCTACACGACGCTCTTCCGATCTNNNTGCATATCATTTATATGGTATATCA
AGCACACGTCTGAACTCTTTCCCTACACGACGCTCTTCCGATCTNNNCTTTAGGGAGGCTTTAAAAA
GCACACGTCTGAACTCTTTCCCTACACGACGCTCTTCCGATCTNNNGGCAATTAGTTGAAAGAGTTA
AGCACACGTCTGAACTCTTTCCCTACACGACGCTCTTCCGATCTNNNACACCCTTGCTGATCTTAATAT
GCACACGTCTGAACTCTTTCCCTACACGACGCTCTTCCGATCTNNNTTTCCTTAATATTGGCATGAATT
GCACACGTCTGAACTCTTTCCCTACACGACGCTCTTCCGATCTNNNTTTGTTGAAAATGAAATCTAGT
CACACGTCTGAACTCTTTCCCTACACGACGCTCTTCCGATCTNNNCTGAAGGAGTTATCTCTCTTTAA
ACACGTCTGAACTCTTTCCCTACACGACGCTCTTCCGATCTNNNTTTATTTGTAATTTTTGCTGCTGCAT
AGCACACGTCTGAACTCTTTCCCTACACGACGCTCTTCCGATCTNNNTAGTTTCAGAATTGAATTGCA
CACACGTCTGAACTCTTTCCCTACACGACGCTCTTCCGATCTNNNTTTCAAATAAAACTTGTACTGGG
GCACACGTCTGAACTCTTTCCCTACACGACGCTCTTCCGATCTNNNATCATACTGCTAAGAATTCAAA
AGCACACGTCTGAACTCTTTCCCTACACGACGCTCTTCCGATCTNNNTAAGAAATTGGTGGCTAGAA
GAGCACACGTCTGAACTCTTTCCCTACACGACGCTCTTCCGATCTNNNGAATGAATCTGAGTTTATCT
CACACGTCTGAACTCTTTCCCTACACGACGCTCTTCCGATCTNNNGCAGGTTTGTAGAGTGATATATA
GCACACGTCTGAACTCTTTCCCTACACGACGCTCTTCCGATCTNNNAATCATTAATTATTGCACAATG

AGCACACGTCTGAACTCTTTCCCTACACGACGCTCTTCCGATCTNNNCTTGTACTAAAGACCTGACAAC
GCACACGTCTGAACTCTTTCCCTACACGACGCTCTTCCGATCTNNNAGGACAATGAACTCTTTAGAATT
CACACGTCTGAACTCTTTCCCTACACGACGCTCTTCCGATCTNNNCTTAGTTACCTTTTTCCATGTTTCAT
AGCACACGTCTGAACTCTTTCCCTACACGACGCTCTTCCGATCTNNNTCACTTAGCTTTTCCCTAAAATA
AGCACACGTCTGAACTCTTTCCCTACACGACGCTCTTCCGATCTNNNTCATCATCTTGATCTTCAAATTC
CACACGTCTGAACTCTTTCCCTACACGACGCTCTTCCGATCTNNNAAAATTATTGAGAACAGAAGCAG
GCACACGTCTGAACTCTTTCCCTACACGACGCTCTTCCGATCTNNNAATTGGAAATTCAGAAGTAGAA
CACACGTCTGAACTCTTTCCCTACACGACGCTCTTCCGATCTNNNCTATTGATTGGACAGCTTGAATC
AGCACACGTCTGAACTCTTTCCCTACACGACGCTCTTCCGATCTNNNTATGAAGATGACACTGGTAAT
GCACACGTCTGAACTCTTTCCCTACACGACGCTCTTCCGATCTNNNAGGACAATGAACTCTTTAGAATT
AGCACACGTCTGAACTCTTTCCCTACACGACGCTCTTCCGATCTNNNAAATCATCTCCAAAAAAAGTCT
CACACGTCTGAACTCTTTCCCTACACGACGCTCTTCCGATCTNNNGGTTCATGGGATTAGAAGAATTC
GCACACGTCTGAACTCTTTCCCTACACGACGCTCTTCCGATCTNNNTAGTAAACACTCTATCAATTGGT
AGCACACGTCTGAACTCTTTCCCTACACGACGCTCTTCCGATCTNNNTAATTTTAGGAGTAGGATTGA
GAGCACACGTCTGAACTCTTTCCCTACACGACGCTCTTCCGATCTNNNATTCATTTTATTGTGGTATGG
GCACACGTCTGAACTCTTTCCCTACACGACGCTCTTCCGATCTNNNATTTAGATCATCTTAAAGGCTGT
GAGCACACGTCTGAACTCTTTCCCTACACGACGCTCTTCCGATCTNNNAGATTAGTTGAGACACTAGT
GAGCACACGTCTGAACTCTTTCCCTACACGACGCTCTTCCGATCTNNNCTTCTACTAATGAATGGTAGT
AGCACACGTCTGAACTCTTTCCCTACACGACGCTCTTCCGATCTNNNTGGTGGTAACTTGATGTATAA
AGCACACGTCTGAACTCTTTCCCTACACGACGCTCTTCCGATCTNNNATCTGGAGGCAGGTATAAAAC
AGCACACGTCTGAACTCTTTCCCTACACGACGCTCTTCCGATCTNNNACATGCTTATAAAAATGTCCAT
CACACGTCTGAACTCTTTCCCTACACGACGCTCTTCCGATCTNNNTTTTCTTTTTTTCAGACAGAGTCA
ACACGTCTGAACTCTTTCCCTACACGACGCTCTTCCGATCTNNNTACATCAAGAGGTCTCCTTGCATCG
AGCACACGTCTGAACTCTTTCCCTACACGACGCTCTTCCGATCTNNNTTCCATTGTATGTCTATATCACA
ACACGTCTGAACTCTTTCCCTACACGACGCTCTTCCGATCTNNNTACTTTTTTCTTTGCATTTGTTCCAT
CACACGTCTGAACTCTTTCCCTACACGACGCTCTTCCGATCTNNNAACAGGGAGAAGAAAGACAGCAT
CACACGTCTGAACTCTTTCCCTACACGACGCTCTTCCGATCTNNNAAAATCACAGCATTGTAAGAAAT
AGCACACGTCTGAACTCTTTCCCTACACGACGCTCTTCCGATCTNNNGTACTTTTGTGATTCTGATGTAT
GCACACGTCTGAACTCTTTCCCTACACGACGCTCTTCCGATCTNNNAGTATCCTTCTTTCATAATGAATG
ACACGTCTGAACTCTTTCCCTACACGACGCTCTTCCGATCTNNNTTCACCCCTCAAAAATATTGTAACAT
CACACGTCTGAACTCTTTCCCTACACGACGCTCTTCCGATCTNNNTTTTCTTAATAGCTGAGTTCATGC
GAGCACACGTCTGAACTCTTTCCCTACACGACGCTCTTCCGATCTNNNTTCCTCATTCCTTATTCCATGC
GAGCACACGTCTGAACTCTTTCCCTACACGACGCTCTTCCGATCTNNNTATCAAGATAGTACAGAGAA
AGCACACGTCTGAACTCTTTCCCTACACGACGCTCTTCCGATCTNNNTGTGATCTTTGATGTGTCTATT
ACACGTCTGAACTCTTTCCCTACACGACGCTCTTCCGATCTNNNGTACCTAAATCTGGATCACTCCATC
AGCACACGTCTGAACTCTTTCCCTACACGACGCTCTTCCGATCTNNNTACATATTCATGGGATACATCA
GAGCACACGTCTGAACTCTTTCCCTACACGACGCTCTTCCGATCTNNNTAAAAATTCAACCATATCTGG
CACACGTCTGAACTCTTTCCCTACACGACGCTCTTCCGATCTNNNGAAACGTCTAGATGCTCAACCATC
GCACACGTCTGAACTCTTTCCCTACACGACGCTCTTCCGATCTNNNTATGACGTTGTCAATATACAGTA
GCACACGTCTGAACTCTTTCCCTACACGACGCTCTTCCGATCTNNNACCATGCCTAGTTGATTTTTAAA
ACACGTCTGAACTCTTTCCCTACACGACGCTCTTCCGATCTNNNAAAAATACAAAAATGAGCCGGCAT
GCACACGTCTGAACTCTTTCCCTACACGACGCTCTTCCGATCTNNNGTAAAAACTCTGGGGAAATAAA
CACACGTCTGAACTCTTTCCCTACACGACGCTCTTCCGATCTNNNTTTAGACTTTGTACCTGTTTTACCA

CACACGTCTGAACTCTTTCCCTACACGACGCTCTTCCGATCTNNNAAAATAACATTGTCTTTGAGCCCA
CACACGTCTGAACTCTTTCCCTACACGACGCTCTTCCGATCTNNNATCTGTCAATTTAAGCTGGTTCAT
GCACACGTCTGAACTCTTTCCCTACACGACGCTCTTCCGATCTNNNATCTCCCTTTTGTTTTACAAGTC
ACACGTCTGAACTCTTTCCCTACACGACGCTCTTCCGATCTNNNAATGAACATGTACCTGCACTCATCG
GAGCACACGTCTGAACTCTTTCCCTACACGACGCTCTTCCGATCTNNNGTGTTCTCTACTTATTTCTCCT
AGCACACGTCTGAACTCTTTCCCTACACGACGCTCTTCCGATCTNNNGGCTTTTAGTTAATTTTGTTTG
GCACACGTCTGAACTCTTTCCCTACACGACGCTCTTCCGATCTNNNTTCAGTTTATCAATTGCTAACAA
ACACGTCTGAACTCTTTCCCTACACGACGCTCTTCCGATCTNNNCATTTGTTAAGAGAAACAATTGCCA
AGCACACGTCTGAACTCTTTCCCTACACGACGCTCTTCCGATCTNNNTAGTTCTTCAGTTCCAGACTATA
GCACACGTCTGAACTCTTTCCCTACACGACGCTCTTCCGATCTNNNTAGAAATGCAGATACCATTCAA
GCACACGTCTGAACTCTTTCCCTACACGACGCTCTTCCGATCTNNNGAAATTGTTTTCTGTTTTCAAGTC
ACACGTCTGAACTCTTTCCCTACACGACGCTCTTCCGATCTNNNCTTAAACAAATCAGGATAGCTTTCA
AGCACACGTCTGAACTCTTTCCCTACACGACGCTCTTCCGATCTNNNGAATTACTCCTGTCATTTTGTT
GCACACGTCTGAACTCTTTCCCTACACGACGCTCTTCCGATCTNNNCTCTCTCTCTGTTATCCTCGCATC
AGCACACGTCTGAACTCTTTCCCTACACGACGCTCTTCCGATCTNNNTTGCAATTGTAAAAATATGGA
AGCACACGTCTGAACTCTTTCCCTACACGACGCTCTTCCGATCTNNNTTGCAATTGTAAAAATATGGA
AGCACACGTCTGAACTCTTTCCCTACACGACGCTCTTCCGATCTNNNCTCAAAAAAAAAAAAGGGCTT
AGCACACGTCTGAACTCTTTCCCTACACGACGCTCTTCCGATCTNNNCTCAAAAAAAAAAAAGGGCTT
ACACGTCTGAACTCTTTCCCTACACGACGCTCTTCCGATCTNNNTTTGACACAAACCTCAAAATTATCA
GCACACGTCTGAACTCTTTCCCTACACGACGCTCTTCCGATCTNNNGTGGGCCAATAAATAAAAAAAT
AGCACACGTCTGAACTCTTTCCCTACACGACGCTCTTCCGATCTNNNTCACAAGAAATGCAGAATTTT
AGCACACGTCTGAACTCTTTCCCTACACGACGCTCTTCCGATCTNNNCTTAAAGCTAATTTAGTTGTCC
CACACGTCTGAACTCTTTCCCTACACGACGCTCTTCCGATCTNNNGATTTGGACTGAGAAAGTAGAGC
AGCACACGTCTGAACTCTTTCCCTACACGACGCTCTTCCGATCTNNNTTTTGGTGTATCTCCTTTTAGA
CACGTCTGAACTCTTTCCCTACACGACGCTCTTCCGATCTNNNTAACTTCTGAGACTGTGTTATAGCAT
GCACACGTCTGAACTCTTTCCCTACACGACGCTCTTCCGATCTNNNATATTTTATCACCGAATGCTTTTC
AGCACACGTCTGAACTCTTTCCCTACACGACGCTCTTCCGATCTNNNGTTTCTTTTCTTTCTTTGGTTTG
AGCACACGTCTGAACTCTTTCCCTACACGACGCTCTTCCGATCTNNNTATCCTATTGGTCTATATGTTG
AGCACACGTCTGAACTCTTTCCCTACACGACGCTCTTCCGATCTNNNGGTTATAATACCTAACAGTAG
AGCACACGTCTGAACTCTTTCCCTACACGACGCTCTTCCGATCTNNNAGAAAATGCTCAAACACATGC
AGCACACGTCTGAACTCTTTCCCTACACGACGCTCTTCCGATCTNNNTAATGTCTTGGTTATTCTCAGT
AGCACACGTCTGAACTCTTTCCCTACACGACGCTCTTCCGATCTNNNCTGGAATATAATCTGTCTGATG
AGCACACGTCTGAACTCTTTCCCTACACGACGCTCTTCCGATCTNNNGTAAAGTATAAGAAGTAGTCC
AGCACACGTCTGAACTCTTTCCCTACACGACGCTCTTCCGATCTNNNGATGATGCATTATGAGAGAAT
GCACACGTCTGAACTCTTTCCCTACACGACGCTCTTCCGATCTNNNAAGGATGAGATATTGTATGTCT
AGCACACGTCTGAACTCTTTCCCTACACGACGCTCTTCCGATCTNNNGTGCAAATGAACTAATATATC
GCACACGTCTGAACTCTTTCCCTACACGACGCTCTTCCGATCTNNNAGTTCCATTGTGTTAATTTCTGCA
AGCACACGTCTGAACTCTTTCCCTACACGACGCTCTTCCGATCTNNNAAGAAGAAAATAATATGCACT
AGCACACGTCTGAACTCTTTCCCTACACGACGCTCTTCCGATCTNNNAATGGAAAAGAGGAAAGTA
AGCACACGTCTGAACTCTTTCCCTACACGACGCTCTTCCGATCTNNNGTGTCTCTTAAAATTATACCC
GCACACGTCTGAACTCTTTCCCTACACGACGCTCTTCCGATCTNNNGAAATGGACAGTGTTCACTGCAT
GCACACGTCTGAACTCTTTCCCTACACGACGCTCTTCCGATCTNNNAGGATTCAAAAATATACGTTTCT
AGCACACGTCTGAACTCTTTCCCTACACGACGCTCTTCCGATCTNNNCACTTCTAAATTGGGAAAAGA

GCACACGTCTGAACTCTTTCCCTACACGACGCTCTTCCGATCTNNNTACAATATGTAATAGCACAACAG
GAGCACACGTCTGAACTCTTTCCCTACACGACGCTCTTCCGATCTNNNCAATAGCAAAGATATGGAATC
AGCACACGTCTGAACTCTTTCCCTACACGACGCTCTTCCGATCTNNNCTTCTGTCTTCACACCAATCCATC
AGCACACGTCTGAACTCTTTCCCTACACGACGCTCTTCCGATCTNNNTATAATGAGGAATGGAGTTGTT
AGCACACGTCTGAACTCTTTCCCTACACGACGCTCTTCCGATCTNNNAATTCCCCTAAAGATACCAAAAT
AGCACACGTCTGAACTCTTTCCCTACACGACGCTCTTCCGATCTNNNAATATTTTCCCACTAATTACGAC
AGCACACGTCTGAACTCTTTCCCTACACGACGCTCTTCCGATCTNNNGAGGAATATGATTTCACTGGATC
GAGCACACGTCTGAACTCTTTCCCTACACGACGCTCTTCCGATCTNNNTAAAATGCAACATATTTCCTCT
CACACGTCTGAACTCTTTCCCTACACGACGCTCTTCCGATCTNNNTTTTTTTTCAAATGTTCCCAATCCAT
AGCACACGTCTGAACTCTTTCCCTACACGACGCTCTTCCGATCTNNNGCATGATTATTCAAATATCCCAT
CACACGTCTGAACTCTTTCCCTACACGACGCTCTTCCGATCTNNNCTCCATCAGAAATTAAAAGATCACA
AGCACACGTCTGAACTCTTTCCCTACACGACGCTCTTCCGATCTNNNATATATATATAGTGTGGCCATGC
CACACGTCTGAACTCTTTCCCTACACGACGCTCTTCCGATCTNNNAACTATCTCTAAGGCTGAGATACAT
GAGCACACGTCTGAACTCTTTCCCTACACGACGCTCTTCCGATCTNNNTGCAACTTAATATTAACTATGC
GCACACGTCTGAACTCTTTCCCTACACGACGCTCTTCCGATCTNNNGGTTGGTATCCAAGAAATATTAC
AGCACACGTCTGAACTCTTTCCCTACACGACGCTCTTCCGATCTNNNGATTATTTTTCTTTCTTTCCTTCAC
AGCACACGTCTGAACTCTTTCCCTACACGACGCTCTTCCGATCTNNNACAGAATATGCAAACAATAAATC
AGCACACGTCTGAACTCTTTCCCTACACGACGCTCTTCCGATCTNNNTTATCAGACCCAATTAGCATAAT
AGCACACGTCTGAACTCTTTCCCTACACGACGCTCTTCCGATCTNNNGTCTTACTAAAGCATCTAACGTA
CACACGTCTGAACTCTTTCCCTACACGACGCTCTTCCGATCTNNNGTGTAATCAAGGGTAGTTCTGCAT
GCACACGTCTGAACTCTTTCCCTACACGACGCTCTTCCGATCTNNNCGTGCACACATATACATATACATC
GAGCACACGTCTGAACTCTTTCCCTACACGACGCTCTTCCGATCTNNNAATTTTGACTGTGCTCTTAATA
GAGCACACGTCTGAACTCTTTCCCTACACGACGCTCTTCCGATCTNNNATTAACACTAGCCATTGATTTT
GAGCACACGTCTGAACTCTTTCCCTACACGACGCTCTTCCGATCTNNNGTGATGGTATGCATGCATATA
CACACGTCTGAACTCTTTCCCTACACGACGCTCTTCCGATCTNNNCTTGGAAAGGAAGGAAAGGACAT
GAGCACACGTCTGAACTCTTTCCCTACACGACGCTCTTCCGATCTNNNGGAAATGTTTCATATCTTGAT
AGAGCACACGTCTGAACTCTTTCCCTACACGACGCTCTTCCGATCTNNNTAATTTTATACAAGTGGGTT
AGCACACGTCTGAACTCTTTCCCTACACGACGCTCTTCCGATCTNNNGTTCAGTTAGCTAGAGTTTTTT
GCACACGTCTGAACTCTTTCCCTACACGACGCTCTTCCGATCTNNNAATGGGAAAGATATTCAAACGTA
ACACGTCTGAACTCTTTCCCTACACGACGCTCTTCCGATCTNNNCCTAAACTGTAATGTGACAAAAGCA
AGAGCACACGTCTGAACTCTTTCCCTACACGACGCTCTTCCGATCTNNNCAAATTGCAGAGTAATGAGA
GAGCACACGTCTGAACTCTTTCCCTACACGACGCTCTTCCGATCTNNNGTTGGAAATGCTAACACTATGC
ACACGTCTGAACTCTTTCCCTACACGACGCTCTTCCGATCTNNNAAAAGGATTTCTTGCAAAACAAACAT
AGCACACGTCTGAACTCTTTCCCTACACGACGCTCTTCCGATCTNNNCTTTCTCTCTCTCTTTCTCTCA
GCACACGTCTGAACTCTTTCCCTACACGACGCTCTTCCGATCTNNNCTCTTAACCTCTTTTAGCTAGATCA
GAGCACACGTCTGAACTCTTTCCCTACACGACGCTCTTCCGATCTNNNATTTAGTTCTGTTTCGTTTCAT
GAGCACACGTCTGAACTCTTTCCCTACACGACGCTCTTCCGATCTNNNGCAGAAGAAAATCTGTGTATA
GAGCACACGTCTGAACTCTTTCCCTACACGACGCTCTTCCGATCTNNNGCAGAAGAAAATCTGTGTATA
GAGCACACGTCTGAACTCTTTCCCTACACGACGCTCTTCCGATCTNNNATGATTTCATTCTTTACTATGG
AGCACACGTCTGAACTCTTTCCCTACACGACGCTCTTCCGATCTNNNTCCATATTCTTTTTTTAAGCAGG
GAGCACACGTCTGAACTCTTTCCCTACACGACGCTCTTCCGATCTNNNCTTAGACGGAAGGTATTTATG
GCACACGTCTGAACTCTTTCCCTACACGACGCTCTTCCGATCTNNNTCAAAAAGTTTTTGATTTTGGAG
GCACACGTCTGAACTCTTTCCCTACACGACGCTCTTCCGATCTNNNTCAAAAAGTTTTTGATTTTGGAG

AGCACACGTCTGAACTCTTTCCCTACACGACGCTCTTCCGATCTNNNCTTTGCATCCAAGACTTTTATC
GCACACGTCTGAACTCTTTCCCTACACGACGCTCTTCCGATCTNNNAGGAAAATAGAAAAAGAAAGC
GCACACGTCTGAACTCTTTCCCTACACGACGCTCTTCCGATCTNNNAGGAAAATAGAAAAAGAAAGC
ACACGTCTGAACTCTTTCCCTACACGACGCTCTTCCGATCTNNNAAAATCACAAAAAGAACTGAACAC
AGCACACGTCTGAACTCTTTCCCTACACGACGCTCTTCCGATCTNNNGGTAAATGGAAGAATTCTTG
CACACGTCTGAACTCTTTCCCTACACGACGCTCTTCCGATCTNNNCATGTCAATATAGAATCAACTGG
ACACGTCTGAACTCTTTCCCTACACGACGCTCTTCCGATCTNNNGAGATTGTGACTTCAAAATATCAC
AGCACACGTCTGAACTCTTTCCCTACACGACGCTCTTCCGATCTNNNGAATTAACATTACCTGATGTC
CACACGTCTGAACTCTTTCCCTACACGACGCTCTTCCGATCTNNNAGCCTTTTGTCCACTGTTATCATCG
GCACACGTCTGAACTCTTTCCCTACACGACGCTCTTCCGATCTNNNATAGTTTTTAGAGGGACATTGA
GCACACGTCTGAACTCTTTCCCTACACGACGCTCTTCCGATCTNNNATTTCACCTTGCATGATACACATC
CACACGTCTGAACTCTTTCCCTACACGACGCTCTTCCGATCTNNNTGGATTCTTATTCATACAGGTTTC
AGCACACGTCTGAACTCTTTCCCTACACGACGCTCTTCCGATCTNNNCTCCTACACTCTCAGATTTTATA
GCACACGTCTGAACTCTTTCCCTACACGACGCTCTTCCGATCTNNNGTTACCATTATACCAAAGCCTCA
GCACACGTCTGAACTCTTTCCCTACACGACGCTCTTCCGATCTNNNTTTGTATTTTTAATAGAGGTGA
AGCACACGTCTGAACTCTTTCCCTACACGACGCTCTTCCGATCTNNNAAGATAACTGTCCTTCAACTA
ACACGTCTGAACTCTTTCCCTACACGACGCTCTTCCGATCTNNNTTGAAGTTTTCCCATCTGTACCATC
AGCACACGTCTGAACTCTTTCCCTACACGACGCTCTTCCGATCTNNNTTAACCTTATTCTCACTGACTTT
AGCACACGTCTGAACTCTTTCCCTACACGACGCTCTTCCGATCTNNNAAAATAAATAGCCTCCATAAG
GCACACGTCTGAACTCTTTCCCTACACGACGCTCTTCCGATCTNNNTCCTTTTGTGAATCAAAGAAATT
AGCACACGTCTGAACTCTTTCCCTACACGACGCTCTTCCGATCTNNNATCTCTACCTGATGGAACTTAA
CACACGTCTGAACTCTTTCCCTACACGACGCTCTTCCGATCTNNNGAACAGACTTTGAAACTAGTTAG
CACACGTCTGAACTCTTTCCCTACACGACGCTCTTCCGATCTNNNCAGGACACTACCAGAATATATAGC
AGCACACGTCTGAACTCTTTCCCTACACGACGCTCTTCCGATCTNNNTACAAGGAAGCTTCAATACAT
ACACGTCTGAACTCTTTCCCTACACGACGCTCTTCCGATCTNNNCTTTGACCATTGAGTATGATGCAT
GCACACGTCTGAACTCTTTCCCTACACGACGCTCTTCCGATCTNNNTATAGTCCTACACAATCTGTTACC
CACGTCTGAACTCTTTCCCTACACGACGCTCTTCCGATCTNNNTTTATTACAAACTAGCGTCTTCCATC
AGCACACGTCTGAACTCTTTCCCTACACGACGCTCTTCCGATCTNNNAAATAAAGAGAAGAAAAAAC
GCACACGTCTGAACTCTTTCCCTACACGACGCTCTTCCGATCTNNNTGATGAATAAATGGGTTTTCATT
AGCACACGTCTGAACTCTTTCCCTACACGACGCTCTTCCGATCTNNNAGGCTCTTAGCAATTACATATA
ACACGTCTGAACTCTTTCCCTACACGACGCTCTTCCGATCTNNNAAACCCTACAATACACAGGACATCG
AGCACACGTCTGAACTCTTTCCCTACACGACGCTCTTCCGATCTNNNTCATCAACATCCATGTATCAGC
AGCACACGTCTGAACTCTTTCCCTACACGACGCTCTTCCGATCTNNNAACTCTATGCTATTAGTGCTCCA
AGCACACGTCTGAACTCTTTCCCTACACGACGCTCTTCCGATCTNNNAACTCTATGCTATTAGTGCTCCA
GCACACGTCTGAACTCTTTCCCTACACGACGCTCTTCCGATCTNNNAAGATAGAAGAGAACAGATTG
GAGCACACGTCTGAACTCTTTCCCTACACGACGCTCTTCCGATCTNNNAAAGAACATGCAATTGATTC
AGCACACGTCTGAACTCTTTCCCTACACGACGCTCTTCCGATCTNNNAGTTTTCTCATTTATATTGGCT
AGCACACGTCTGAACTCTTTCCCTACACGACGCTCTTCCGATCTNNNAGTAAGAATCAAGGAAGAGA
AGCACACGTCTGAACTCTTTCCCTACACGACGCTCTTCCGATCTNNNTTCTTACAAGTTTTTGAAATTGC
AGCACACGTCTGAACTCTTTCCCTACACGACGCTCTTCCGATCTNNNATATAATGACACAGGTAGGTT
CACACGTCTGAACTCTTTCCCTACACGACGCTCTTCCGATCTNNNTTGATCTCACAATCTTTCTCTTTCA
AGCACACGTCTGAACTCTTTCCCTACACGACGCTCTTCCGATCTNNNATAGAGAGCTTTGTACTTACT
AGCACACGTCTGAACTCTTTCCCTACACGACGCTCTTCCGATCTNNNTTGATGATAAAATCAAGGTGT

GAGCACACGTCTGAACTCTTTCCCTACACGACGCTCTTCCGATCTNNNTAGAATTCCAAAAATAATGCT
CACACGTCTGAACTCTTTCCCTACACGACGCTCTTCCGATCTNNNTTAGAACTAAACACTAGTGCTAACA
ACACGTCTGAACTCTTTCCCTACACGACGCTCTTCCGATCTNNNCCACCTCATAGTATAGTTGTATTCAT
GCACACGTCTGAACTCTTTCCCTACACGACGCTCTTCCGATCTNNNGTGTTCATAATCCTTCTTGTTTAC
GAGCACACGTCTGAACTCTTTCCCTACACGACGCTCTTCCGATCTNNNTTGTTAGACTTTACGTATCTGA
GAGCACACGTCTGAACTCTTTCCCTACACGACGCTCTTCCGATCTNNNCAAGGTTCAACATAGTTTCTT
GAGCACACGTCTGAACTCTTTCCCTACACGACGCTCTTCCGATCTNNNAGGAAGATTATAACGTAAAGT
GCACACGTCTGAACTCTTTCCCTACACGACGCTCTTCCGATCTNNNCCTAATATGTTGTTTGGATATCC
GCACACGTCTGAACTCTTTCCCTACACGACGCTCTTCCGATCTNNNCCTAATATGTTGTTTGGATATCC
GAGCACACGTCTGAACTCTTTCCCTACACGACGCTCTTCCGATCTNNNGATATACGATTTCTGTTAGGA
GCACACGTCTGAACTCTTTCCCTACACGACGCTCTTCCGATCTNNNCTTCATGTTACGAGATGCTATCAT
AGCACACGTCTGAACTCTTTCCCTACACGACGCTCTTCCGATCTNNNAGAATTGGTTGTTATTTTTGAA
GAGCACACGTCTGAACTCTTTCCCTACACGACGCTCTTCCGATCTNNNCGTTTATGAAAACTTCAATTAG
AGCACACGTCTGAACTCTTTCCCTACACGACGCTCTTCCGATCTNNNGCAAAATGATGAGAAACACAT
AGCACACGTCTGAACTCTTTCCCTACACGACGCTCTTCCGATCTNNNAGTAGCTACACAAATCATCTAA
GCACACGTCTGAACTCTTTCCCTACACGACGCTCTTCCGATCTNNNATAATGCAATATCCCATCATACACA
GAGCACACGTCTGAACTCTTTCCCTACACGACGCTCTTCCGATCTNNNCTGAAATACATTTACTAGATG
GAGCACACGTCTGAACTCTTTCCCTACACGACGCTCTTCCGATCTNNNTGTAATTAGTTAAGATGAGGT
AGAGCACACGTCTGAACTCTTTCCCTACACGACGCTCTTCCGATCTNNNTTCATTCTTGAAGTCAGTGA
GCACACGTCTGAACTCTTTCCCTACACGACGCTCTTCCGATCTNNNTCCATTGGTCAACTTGGATACAT
AGCACACGTCTGAACTCTTTCCCTACACGACGCTCTTCCGATCTNNNGTCTTTTCAACTTTTCAAAAACA
AGCACACGTCTGAACTCTTTCCCTACACGACGCTCTTCCGATCTNNNGTGGTTGCATTATTTTACATTT
GAGCACACGTCTGAACTCTTTCCCTACACGACGCTCTTCCGATCTNNNCAGTTCTTCATTGCTTTGATAC
AGCACACGTCTGAACTCTTTCCCTACACGACGCTCTTCCGATCTNNNAGCCTACAACTTAAAGTATAACA
GCACACGTCTGAACTCTTTCCCTACACGACGCTCTTCCGATCTNNNTTAATATAATCCCTGTCCTGAACC
AGCACACGTCTGAACTCTTTCCCTACACGACGCTCTTCCGATCTNNNAACACTCTTTTTTTACAAATTGT
AGCACACGTCTGAACTCTTTCCCTACACGACGCTCTTCCGATCTNNNGAAGAGACATAGTTCTATGTT
AGCACACGTCTGAACTCTTTCCCTACACGACGCTCTTCCGATCTNNNGTATATAACTATTGTGTGTGT
AGCACACGTCTGAACTCTTTCCCTACACGACGCTCTTCCGATCTNNNATGTATATTTGAACCTTGCTAA
AGCACACGTCTGAACTCTTTCCCTACACGACGCTCTTCCGATCTNNNTAGACAGAGTACTAACTTCTAC
GAGCACACGTCTGAACTCTTTCCCTACACGACGCTCTTCCGATCTNNNACTAATGTTCTACAACACAGA
AGCACACGTCTGAACTCTTTCCCTACACGACGCTCTTCCGATCTNNNGGGCTATCTATGAATATAAATG
GAGCACACGTCTGAACTCTTTCCCTACACGACGCTCTTCCGATCTNNNCTTAGAGGTCATCTAATCTAGT
AGCACACGTCTGAACTCTTTCCCTACACGACGCTCTTCCGATCTNNNACTGTTAAGGAATTTTTGTATG
AGCACACGTCTGAACTCTTTCCCTACACGACGCTCTTCCGATCTNNNTATAAAAATCACTTAGCACAACA
GAGCACACGTCTGAACTCTTTCCCTACACGACGCTCTTCCGATCTNNNAAAGGTAAAGCTTTTAACAAA
GAGCACACGTCTGAACTCTTTCCCTACACGACGCTCTTCCGATCTNNNAAAAGGCAGTTGTATTCACACA
AGCACACGTCTGAACTCTTTCCCTACACGACGCTCTTCCGATCTNNNGTTTTTCTCATTTGTAGAATGGT
AGCACACGTCTGAACTCTTTCCCTACACGACGCTCTTCCGATCTNNNGAAAGCACAATAACATATTACC
AGCACACGTCTGAACTCTTTCCCTACACGACGCTCTTCCGATCTNNNAGTAAAATCTTGCTATTCAGAAA
GAGCACACGTCTGAACTCTTTCCCTACACGACGCTCTTCCGATCTNNNGAACGTTGAAGAATAAAGTTG
AGAGCACACGTCTGAACTCTTTCCCTACACGACGCTCTTCCGATCTNNNTGAATTAATTGGTTTCTAGG
GAGCACACGTCTGAACTCTTTCCCTACACGACGCTCTTCCGATCTNNNTTTTCTGTCTAGATAATCTGTC

AGCACACGTCTGAACTCTTTCCCTACACGACGCTCTTCCGATCTNNNTTCTTATTAAGGACAACAGAA
GCACACGTCTGAACTCTTTCCCTACACGACGCTCTTCCGATCTNNNAGAAAGAATGGAGGGGTTAG
AGCACACGTCTGAACTCTTTCCCTACACGACGCTCTTCCGATCTNNNAGATTCCAAACATTGTTAAAT
CACGTCTGAACTCTTTCCCTACACGACGCTCTTCCGATCTNNNTAAGAAATTTACTGGCACTGTACAT
AGCACACGTCTGAACTCTTTCCCTACACGACGCTCTTCCGATCTNNNAGTAGTAGTAGTAGTAGTAG
AGCACACGTCTGAACTCTTTCCCTACACGACGCTCTTCCGATCTNNNTTACACGAGCCAATATATTGAA
GCACACGTCTGAACTCTTTCCCTACACGACGCTCTTCCGATCTNNNGAGTTGTCTCTATATGGATGTG
GCACACGTCTGAACTCTTTCCCTACACGACGCTCTTCCGATCTNNNATCTAGTAGTAGTTGATTTTGC
GCACACGTCTGAACTCTTTCCCTACACGACGCTCTTCCGATCTNNNAGAACCAAGACTTGTATTCTAG
CACACGTCTGAACTCTTTCCCTACACGACGCTCTTCCGATCTNNNGTTTATGACTATGCTCTCAAACCA
AGCACACGTCTGAACTCTTTCCCTACACGACGCTCTTCCGATCTNNNTTACTATCTATTGAGATGGGG
AGCACACGTCTGAACTCTTTCCCTACACGACGCTCTTCCGATCTNNNTAATTAATCTCATGGGATTTCT
ACACGTCTGAACTCTTTCCCTACACGACGCTCTTCCGATCTNNNCCTTGTCTCAAAAAAAAAAAAATT
ACACGTCTGAACTCTTTCCCTACACGACGCTCTTCCGATCTNNNCCTTGTCTCAAAAAAAAAAAAATT
AGCACACGTCTGAACTCTTTCCCTACACGACGCTCTTCCGATCTNNNCGATATCCTGTTCATTTTATTCA
CACACGTCTGAACTCTTTCCCTACACGACGCTCTTCCGATCTNNNAAAGTGAATGATGACAAAGTATT
GCACACGTCTGAACTCTTTCCCTACACGACGCTCTTCCGATCTNNNTTTTGGAGAGAGAGAGAGAAA
GCACACGTCTGAACTCTTTCCCTACACGACGCTCTTCCGATCTNNNGGCTGAAATTTCTCCTTTTGCAT
GCACACGTCTGAACTCTTTCCCTACACGACGCTCTTCCGATCTNNNGGCTGAAATTTCTCCTTTTGCAT
GCACACGTCTGAACTCTTTCCCTACACGACGCTCTTCCGATCTNNNTCAAGCTCTTTACTTTGATTTTT
GAGCACACGTCTGAACTCTTTCCCTACACGACGCTCTTCCGATCTNNNCAGGTAATTTTTATCAGGAT
GCACACGTCTGAACTCTTTCCCTACACGACGCTCTTCCGATCTNNNAAATTGACAAACCTATACCCAAA
GCACACGTCTGAACTCTTTCCCTACACGACGCTCTTCCGATCTNNNAAATTGACAAACCTATACCCAAA
CACGTCTGAACTCTTTCCCTACACGACGCTCTTCCGATCTNNNCCTTCTTTTAATATTACAGTGCTCAT
CACACGTCTGAACTCTTTCCCTACACGACGCTCTTCCGATCTNNNCAATTTTCTCTCTGGACCTTTACA
GAGCACACGTCTGAACTCTTTCCCTACACGACGCTCTTCCGATCTNNNTGCCTAAGCATAGCTACTATC
CACACGTCTGAACTCTTTCCCTACACGACGCTCTTCCGATCTNNNATATTTTAGAAAATGCTCTGTGGC
GCACACGTCTGAACTCTTTCCCTACACGACGCTCTTCCGATCTNNNAGAAATGATACTGGCATTATAA
GCACACGTCTGAACTCTTTCCCTACACGACGCTCTTCCGATCTNNNTACTACAGGTAGTAAAACACAC
ACGTCTGAACTCTTTCCCTACACGACGCTCTTCCGATCTNNNTTTTCCAAATAAGGTCATAGTCACATC
GAGCACACGTCTGAACTCTTTCCCTACACGACGCTCTTCCGATCTNNNTATGTTAAACCTCAAAGTTC
AGCACACGTCTGAACTCTTTCCCTACACGACGCTCTTCCGATCTNNNGTTCATTCCTTTTGTTTGTTTT
AGCACACGTCTGAACTCTTTCCCTACACGACGCTCTTCCGATCTNNNTACATGATCAAACACAGATGT
AGCACACGTCTGAACTCTTTCCCTACACGACGCTCTTCCGATCTNNNCTTGATTCTGCTACACTTTATT
ACACGTCTGAACTCTTTCCCTACACGACGCTCTTCCGATCTNNNCCCATAGCCTGAAACATCTACATCG
GCACACGTCTGAACTCTTTCCCTACACGACGCTCTTCCGATCTNNNCAATTACCACCCAATCTGAGCAT
ACACGTCTGAACTCTTTCCCTACACGACGCTCTTCCGATCTNNNGAGTTTTTGTTTCAGGAAATTTCC
GCACACGTCTGAACTCTTTCCCTACACGACGCTCTTCCGATCTNNNGCTATGAGATACAGATTAGTCT
CACGTCTGAACTCTTTCCCTACACGACGCTCTTCCGATCTNNNCCCAACAAAGATTCCTTCATCATCGG
CACGTCTGAACTCTTTCCCTACACGACGCTCTTCCGATCTNNNCCCAACAAAGATTCCTTCATCATCGG
CACACGTCTGAACTCTTTCCCTACACGACGCTCTTCCGATCTNNNTAAACAGTTCCACTTTTAAAACCC
ACACGTCTGAACTCTTTCCCTACACGACGCTCTTCCGATCTNNNGCTGGTAAAATATTTCCATTTTGCA
AGCACACGTCTGAACTCTTTCCCTACACGACGCTCTTCCGATCTNNNTCTAGAACAATGTTGAATAGA

AGCACACGTCTGAACTCTTTCCCTACACGACGCTCTTCCGATCTNNNGATTCAAAATAAGGACTGATTC
AGCACACGTCTGAACTCTTTCCCTACACGACGCTCTTCCGATCTNNNTAAGATTACAAGACTAGCAATGT
GAGCACACGTCTGAACTCTTTCCCTACACGACGCTCTTCCGATCTNNNATCTTATATCCAGGCCTTTTACC
CACACGTCTGAACTCTTTCCCTACACGACGCTCTTCCGATCTNNNCGAGCTATAGATAACTCAGTAAACA
CACACGTCTGAACTCTTTCCCTACACGACGCTCTTCCGATCTNNNCGAGCTATAGATAACTCAGTAAACA
GCACACGTCTGAACTCTTTCCCTACACGACGCTCTTCCGATCTNNNTAATATATGTGAAATCGCCCTCCA
GAGCACACGTCTGAACTCTTTCCCTACACGACGCTCTTCCGATCTNNNATTTAGTATTCATGGGGGAT
GAGCACACGTCTGAACTCTTTCCCTACACGACGCTCTTCCGATCTNNNTTTAACTCTCAAGAAATAGGTG
AGCACACGTCTGAACTCTTTCCCTACACGACGCTCTTCCGATCTNNNGAATATCTGCTCTGATCACTCCA
GCACACGTCTGAACTCTTTCCCTACACGACGCTCTTCCGATCTNNNGTATTTGACACACAGTAATTGTAC
CACACGTCTGAACTCTTTCCCTACACGACGCTCTTCCGATCTNNNCTTTTAGCAAATGATCTCATCTCCAT
AGCACACGTCTGAACTCTTTCCCTACACGACGCTCTTCCGATCTNNNAATTCCCCTAAAGATACCAAAAT
GAGCACACGTCTGAACTCTTTCCCTACACGACGCTCTTCCGATCTNNNTCTCTCTCTCTCTCTCTAGCA
CACACGTCTGAACTCTTTCCCTACACGACGCTCTTCCGATCTNNNTAATTAGGTTTATGTGAGGTTCTC
AGCACACGTCTGAACTCTTTCCCTACACGACGCTCTTCCGATCTNNNACAATTACAAACATTTCTGGAA
AGCACACGTCTGAACTCTTTCCCTACACGACGCTCTTCCGATCTNNNACAATTACAAACATTTCTGGAA
CACACGTCTGAACTCTTTCCCTACACGACGCTCTTCCGATCTNNNGAACTAGGTATCATTTAAGCTTCCA
GCACACGTCTGAACTCTTTCCCTACACGACGCTCTTCCGATCTNNNCTTTTTATTTTATAACAGCAGCACA
AGCACACGTCTGAACTCTTTCCCTACACGACGCTCTTCCGATCTNNNCAGAGATAGGACATAGAGGTG
AGCACACGTCTGAACTCTTTCCCTACACGACGCTCTTCCGATCTNNNGGAAAATCAAATCTCTCTCTCTA
AGCACACGTCTGAACTCTTTCCCTACACGACGCTCTTCCGATCTNNNGGAAAATCAAATCTCTCTCTCTA
CACACGTCTGAACTCTTTCCCTACACGACGCTCTTCCGATCTNNNCATCAAGCTGATTCTGAAATCCATC
AGCACACGTCTGAACTCTTTCCCTACACGACGCTCTTCCGATCTNNNCTTATTCTTTGATGCTAGCTTGCA
GAGCACACGTCTGAACTCTTTCCCTACACGACGCTCTTCCGATCTNNNCTTGAATTTCTGTCCTGATAAC
GAGCACACGTCTGAACTCTTTCCCTACACGACGCTCTTCCGATCTNNNTTTCTCATCTTAAAAAGGGGAT
GAGCACACGTCTGAACTCTTTCCCTACACGACGCTCTTCCGATCTNNNCAGAATTTCTATTTGTTTCGGT
GAGCACACGTCTGAACTCTTTCCCTACACGACGCTCTTCCGATCTNNNCAGAATTTCTATTTGTTTCGGT
GAGCACACGTCTGAACTCTTTCCCTACACGACGCTCTTCCGATCTNNNCAGAATTTCTATTTGTTTCGGT
AGCACACGTCTGAACTCTTTCCCTACACGACGCTCTTCCGATCTNNNTCCCTCATCATCCTCTTTTTCATC
GCACACGTCTGAACTCTTTCCCTACACGACGCTCTTCCGATCTNNNCTTGTCCTCAAAAGCATGTCCATCC
AGCACACGTCTGAACTCTTTCCCTACACGACGCTCTTCCGATCTNNNCCTTTCTTTTTTTCTATGGAAAC
GAGCACACGTCTGAACTCTTTCCCTACACGACGCTCTTCCGATCTNNNTTCGCTTTGCATTCAATATTATA
CACACGTCTGAACTCTTTCCCTACACGACGCTCTTCCGATCTNNNTGTACCCTTATAAGAAGCAATATCA
AGCACACGTCTGAACTCTTTCCCTACACGACGCTCTTCCGATCTNNNCAAGCATCAAACACATAGAATAC
AGCACACGTCTGAACTCTTTCCCTACACGACGCTCTTCCGATCTNNNTGACTAAGAAGTTTTTGTTGTTC
GAGCACACGTCTGAACTCTTTCCCTACACGACGCTCTTCCGATCTNNNAGTATAGGAGATAGAAAAAA
AGCACACGTCTGAACTCTTTCCCTACACGACGCTCTTCCGATCTNNNATTTAGGACAGAAGCAAGATC
AGCACACGTCTGAACTCTTTCCCTACACGACGCTCTTCCGATCTNNNGTTCTAGAAATCAAAAGCCATAT
GAGCACACGTCTGAACTCTTTCCCTACACGACGCTCTTCCGATCTNNNACTAGGTTAAATCAAAACTGA
CACACGTCTGAACTCTTTCCCTACACGACGCTCTTCCGATCTNNNACAGCTGAAAATAAGACAAAATAC
GAGCACACGTCTGAACTCTTTCCCTACACGACGCTCTTCCGATCTNNNCATGTAAGCAAAGCATATTGAC
ACACGTCTGAACTCTTTCCCTACACGACGCTCTTCCGATCTNNNACTGGCTATCATTTTAAGGATCCATC
GAGCACACGTCTGAACTCTTTCCCTACACGACGCTCTTCCGATCTNNNGCTACTTACCTTATTTGAACAA

GAGCACACGTCTGAACTCTTTCCCTACACGACGCTCTTCCGATCTNNNCAGAAGAAATAGTGTAGGCAT
GAGCACACGTCTGAACTCTTTCCCTACACGACGCTCTTCCGATCTNNNCAATGACAAAATCTCCTAATGA
GCACACGTCTGAACTCTTTCCCTACACGACGCTCTTCCGATCTNNNTACTTCACTTAACATAGTGTCCCA
GAGCACACGTCTGAACTCTTTCCCTACACGACGCTCTTCCGATCTNNNCCTAAGCATATTATGTACTTGA
AGCACACGTCTGAACTCTTTCCCTACACGACGCTCTTCCGATCTNNNAGCATAAAAGATCTCAAAATGA
GAGCACACGTCTGAACTCTTTCCCTACACGACGCTCTTCCGATCTNNNGCTTATATTGGAAACTTCATTG
GAGCACACGTCTGAACTCTTTCCCTACACGACGCTCTTCCGATCTNNNAAACCCTTTAACTAATAAGCCT
AGCACACGTCTGAACTCTTTCCCTACACGACGCTCTTCCGATCTNNNTCTTTGCTCAAAAATGCATATAA
AGCACACGTCTGAACTCTTTCCCTACACGACGCTCTTCCGATCTNNNTTGATTTGTAAAATGGGATATG
AGCACACGTCTGAACTCTTTCCCTACACGACGCTCTTCCGATCTNNNGCACCAAGTTCTTACAATAAAT
GAGCACACGTCTGAACTCTTTCCCTACACGACGCTCTTCCGATCTNNNTGAGTACATATACATACATAC
GAGCACACGTCTGAACTCTTTCCCTACACGACGCTCTTCCGATCTNNNATTGGCATATCACAAAACTTTA
GCACACGTCTGAACTCTTTCCCTACACGACGCTCTTCCGATCTNNNTCTTCAGAGAAATAGAACCAATA
ACACGTCTGAACTCTTTCCCTACACGACGCTCTTCCGATCTNNNCTCTCCTCTCTCCTTTCTTCCATCGGAC
GCACACGTCTGAACTCTTTCCCTACACGACGCTCTTCCGATCTNNNTCTTTCCAGTGTAAATGTTTATGC
CACACGTCTGAACTCTTTCCCTACACGACGCTCTTCCGATCTNNNGGTATGTATGTGTATGTTTATGGC
CACACGTCTGAACTCTTTCCCTACACGACGCTCTTCCGATCTNNNGGTATGTATGTGTATGTTTATGGC
GCACACGTCTGAACTCTTTCCCTACACGACGCTCTTCCGATCTNNNGAGGATTCACGTTAAGTTTTAAGC
GAGCACACGTCTGAACTCTTTCCCTACACGACGCTCTTCCGATCTNNNCATAATCATAAGAGCCAATTCT
GAGCACACGTCTGAACTCTTTCCCTACACGACGCTCTTCCGATCTNNNCATAATCATAAGAGCCAATTCT
GAGCACACGTCTGAACTCTTTCCCTACACGACGCTCTTCCGATCTNNNCTGTTTCCTTAAGTTGAAATCA
AGCACACGTCTGAACTCTTTCCCTACACGACGCTCTTCCGATCTNNNCTGACTAGCAAAGGAAACAGCA
GAGCACACGTCTGAACTCTTTCCCTACACGACGCTCTTCCGATCTNNNAAATAGAGACTATGCTGAGAA
AGCACACGTCTGAACTCTTTCCCTACACGACGCTCTTCCGATCTNNNAATGAGTGTTATTTGATCCACTA
AGCACACGTCTGAACTCTTTCCCTACACGACGCTCTTCCGATCTNNNAATGAGTGTTATTTGATCCACTA
AGCACACGTCTGAACTCTTTCCCTACACGACGCTCTTCCGATCTNNNCCATCTTAAATTTTTGTATATGG
AGCACACGTCTGAACTCTTTCCCTACACGACGCTCTTCCGATCTNNNATTTTAAACTCATGGTGTCTAAC
AGCACACGTCTGAACTCTTTCCCTACACGACGCTCTTCCGATCTNNNTGTTATTCCATTTTTGGCATTAT
CACACGTCTGAACTCTTTCCCTACACGACGCTCTTCCGATCTNNNTTTTAAGGCTGAATAGTATTCCTCA
AGCACACGTCTGAACTCTTTCCCTACACGACGCTCTTCCGATCTNNNATGTATAACCCTCAAGAAGATAC
CACACGTCTGAACTCTTTCCCTACACGACGCTCTTCCGATCTNNNGATCTCTCCTTTCTCTTCTATTCCATG
GAGCACACGTCTGAACTCTTTCCCTACACGACGCTCTTCCGATCTNNNACTGTTATAAACCATTAAAACT
CACACGTCTGAACTCTTTCCCTACACGACGCTCTTCCGATCTNNNTACAGATTTCAGACTTGACAGCATC
AGCACACGTCTGAACTCTTTCCCTACACGACGCTCTTCCGATCTNNNAATTGCCTTTGAAACATTTTTCC
AGCACACGTCTGAACTCTTTCCCTACACGACGCTCTTCCGATCTNNNATTGACTACAACACAATTTTACA
AGCACACGTCTGAACTCTTTCCCTACACGACGCTCTTCCGATCTNNNTCAAAAAATCAGAAAAGTAGCA
AGCACACGTCTGAACTCTTTCCCTACACGACGCTCTTCCGATCTNNNGAGAGATAGTACAGCAAAGAT
GCACACGTCTGAACTCTTTCCCTACACGACGCTCTTCCGATCTNNNCTGTATCCCGATGTATAACATAGC
AGCACACGTCTGAACTCTTTCCCTACACGACGCTCTTCCGATCTNNNAAACCTGTTATGTTATTGTACAT
GCACACGTCTGAACTCTTTCCCTACACGACGCTCTTCCGATCTNNNAACTGTGATGAAAACATACATAC
AGCACACGTCTGAACTCTTTCCCTACACGACGCTCTTCCGATCTNNNCATGAGAAGTTCCCTAAAAGAA
AGCACACGTCTGAACTCTTTCCCTACACGACGCTCTTCCGATCTNNNGACAAAGACACCAATTATGATA
CACACGTCTGAACTCTTTCCCTACACGACGCTCTTCCGATCTNNNACAGGGTAATTTTATCATCTGTACA

AGCACACGTCTGAACTCTTTCCCTACACGACGCTCTTCCGATCTNNNAATTTTCAGTGTGTAAGTCTTT
GCACACGTCTGAACTCTTTCCCTACACGACGCTCTTCCGATCTNNNTTATTTTTGTGACCCTGCATTAC
GCACACGTCTGAACTCTTTCCCTACACGACGCTCTTCCGATCTNNNTTTAATCGTATTTTTCTCCTCAC
AGCACACGTCTGAACTCTTTCCCTACACGACGCTCTTCCGATCTNNNTTCAGTTCAGATACTATACTCT
GCACACGTCTGAACTCTTTCCCTACACGACGCTCTTCCGATCTNNNATCAGAGTATTCCATTTTATTGC
AGCACACGTCTGAACTCTTTCCCTACACGACGCTCTTCCGATCTNNNCCAAGAAGAGAGAGAAGTCTC
AGCACACGTCTGAACTCTTTCCCTACACGACGCTCTTCCGATCTNNNTACTGTTGTCATAAGGAGTAA
GAGCACACGTCTGAACTCTTTCCCTACACGACGCTCTTCCGATCTNNNGAGACAAAGGTCAAGAAAT
GCACACGTCTGAACTCTTTCCCTACACGACGCTCTTCCGATCTNNNAATTGATGTTATAATGGGTATG
AGCACACGTCTGAACTCTTTCCCTACACGACGCTCTTCCGATCTNNNCAGAATTAGCTACCAGTTTTTA
CACACGTCTGAACTCTTTCCCTACACGACGCTCTTCCGATCTNNNGTTAGAAAACCATCTAGACAGAC
AGCACACGTCTGAACTCTTTCCCTACACGACGCTCTTCCGATCTNNNAAAAGTATCTGATCATGTTCA
CACACGTCTGAACTCTTTCCCTACACGACGCTCTTCCGATCTNNNGCAACTTAACAACACAAATTTATCA
AGCACACGTCTGAACTCTTTCCCTACACGACGCTCTTCCGATCTNNNGGTGTGTTGAACATACTTTTT
GCACACGTCTGAACTCTTTCCCTACACGACGCTCTTCCGATCTNNNATTTATCTCCTCTCTTAGCATTC
CACACGTCTGAACTCTTTCCCTACACGACGCTCTTCCGATCTNNNGAAAAATCTCTTCAGTGTTTGACA
GCACACGTCTGAACTCTTTCCCTACACGACGCTCTTCCGATCTNNNGAAGATTACAATCATGACAGAA
GCACACGTCTGAACTCTTTCCCTACACGACGCTCTTCCGATCTNNNATGACAGATGACAATGAAATAA
GCACACGTCTGAACTCTTTCCCTACACGACGCTCTTCCGATCTNNNTTTTTTAGACATAAGGGGGCCA
GCACACGTCTGAACTCTTTCCCTACACGACGCTCTTCCGATCTNNNATTTGTAATTTGGCTAAATCAGT
GCACACGTCTGAACTCTTTCCCTACACGACGCTCTTCCGATCTNNNATTTGTAATTTGGCTAAATCAGT
AGCACACGTCTGAACTCTTTCCCTACACGACGCTCTTCCGATCTNNNAACTGAAGATAATACTGCAAC
GCACACGTCTGAACTCTTTCCCTACACGACGCTCTTCCGATCTNNNTGGCTAATAACAATAGACTCATT
AGCACACGTCTGAACTCTTTCCCTACACGACGCTCTTCCGATCTNNNCCTACACCTCACCCTAAAAATC
GAGCACACGTCTGAACTCTTTCCCTACACGACGCTCTTCCGATCTNNNATGATGAATTCAAGGTAACA
AGCACACGTCTGAACTCTTTCCCTACACGACGCTCTTCCGATCTNNNTAAACGTGAAGGATTTGCATA
AGCACACGTCTGAACTCTTTCCCTACACGACGCTCTTCCGATCTNNNTCATTACACAAGACAGATTCT
AGCACACGTCTGAACTCTTTCCCTACACGACGCTCTTCCGATCTNNNGATGCATTTCTCAGAACATAT
GAGCACACGTCTGAACTCTTTCCCTACACGACGCTCTTCCGATCTNNNAATTCTGTGAGTGGGTCAA
AGCACACGTCTGAACTCTTTCCCTACACGACGCTCTTCCGATCTNNNCATTCAGCTCTAATTAATGGA
AGCACACGTCTGAACTCTTTCCCTACACGACGCTCTTCCGATCTNNNGTAAGTCTAGAGATTGAGTGT
CACGTCTGAACTCTTTCCCTACACGACGCTCTTCCGATCTNNNTGCCATAAAGCAGAGATATGCATCG
GAGCACACGTCTGAACTCTTTCCCTACACGACGCTCTTCCGATCTNNNTAAGCAATGGGAAGAAAAT
GCACACGTCTGAACTCTTTCCCTACACGACGCTCTTCCGATCTNNNATGTAGTGACCAATTAGATCTG
AGCACACGTCTGAACTCTTTCCCTACACGACGCTCTTCCGATCTNNNTTCCTGGAGAAGTTATATTTA
CACACGTCTGAACTCTTTCCCTACACGACGCTCTTCCGATCTNNNATTACCAACAAAGGAAGCAGCAT
CACGTCTGAACTCTTTCCCTACACGACGCTCTTCCGATCTNNNCTTTTTGCTGTATTATCCCATGCATC
CACACGTCTGAACTCTTTCCCTACACGACGCTCTTCCGATCTNNNTGGAAACTAATATTTAGAGACCA
AGCACACGTCTGAACTCTTTCCCTACACGACGCTCTTCCGATCTNNNTACCCATCTATCTATTTAGCAT
CACACGTCTGAACTCTTTCCCTACACGACGCTCTTCCGATCTNNNATCTATCTAATCTAATCTATCTGCC
GCACACGTCTGAACTCTTTCCCTACACGACGCTCTTCCGATCTNNNCCTCACACTACAGATTAAAAGT
AGCACACGTCTGAACTCTTTCCCTACACGACGCTCTTCCGATCTNNNAAATAACTAGACTGGGAATA
CACACGTCTGAACTCTTTCCCTACACGACGCTCTTCCGATCTNNNGTTTCAAGCACAAAATTGGGCAT

GCACACGTCTGAACTCTTTCCCTACACGACGCTCTTCCGATCTNNNTTAATTTTGTAACCTCCTACCTTCA
GAGCACACGTCTGAACTCTTTCCCTACACGACGCTCTTCCGATCTNNNATATTTAGGAGTAGAAGGGT
GCACACGTCTGAACTCTTTCCCTACACGACGCTCTTCCGATCTNNNGAAGCAGTAGAAACATTTACTGC
GCACACGTCTGAACTCTTTCCCTACACGACGCTCTTCCGATCTNNNGAAGCAGTAGAAACATTTACTGC
CACACGTCTGAACTCTTTCCCTACACGACGCTCTTCCGATCTNNNTATGAAACAATGTTACCAAGGATC
AGAGCACACGTCTGAACTCTTTCCCTACACGACGCTCTTCCGATCTNNNATAGATACTAAATCAGTAGCA
GAGCACACGTCTGAACTCTTTCCCTACACGACGCTCTTCCGATCTNNNTTAAATTGTGTACCTCTGCTAA
CACACGTCTGAACTCTTTCCCTACACGACGCTCTTCCGATCTNNNAGGTCTGTTGAACTCTAAAATATCA
GAGCACACGTCTGAACTCTTTCCCTACACGACGCTCTTCCGATCTNNNAGATAGATTAGCTAGACAAAC
GAGCACACGTCTGAACTCTTTCCCTACACGACGCTCTTCCGATCTNNNAGATAGATTAGCTAGACAAAC
GAGCACACGTCTGAACTCTTTCCCTACACGACGCTCTTCCGATCTNNNAGATAGATTAGCTAGACAAAC
AGCACACGTCTGAACTCTTTCCCTACACGACGCTCTTCCGATCTNNNCTTTGAATAACTTGTTCTTTCCTC
GCACACGTCTGAACTCTTTCCCTACACGACGCTCTTCCGATCTNNNAGGTCTCACATGTACCAATTCATC
GCACACGTCTGAACTCTTTCCCTACACGACGCTCTTCCGATCTNNNTAGAGCAGGTAAAATTGATCTAC
AGCACACGTCTGAACTCTTTCCCTACACGACGCTCTTCCGATCTNNNGTGGAATAAGGAAGAGATAAC
CACACGTCTGAACTCTTTCCCTACACGACGCTCTTCCGATCTNNNTTCTGCTTTCATTCAATTCTAACCATC
GCACACGTCTGAACTCTTTCCCTACACGACGCTCTTCCGATCTNNNGGATCATGAACAAAAATTTTTCAC
GCACACGTCTGAACTCTTTCCCTACACGACGCTCTTCCGATCTNNNTTGTAATTTCTCATGAATTGGTCCA
AGCACACGTCTGAACTCTTTCCCTACACGACGCTCTTCCGATCTNNNCACACATATATATACACACACAT
GAGCACACGTCTGAACTCTTTCCCTACACGACGCTCTTCCGATCTNNNAGAAAAAAAGAAACAGAAACA
GCACACGTCTGAACTCTTTCCCTACACGACGCTCTTCCGATCTNNNTCTGGTAAAAATCTATAAACAGA
GCACACGTCTGAACTCTTTCCCTACACGACGCTCTTCCGATCTNNNATAGACACACGTAGACACACCAT
AGCACACGTCTGAACTCTTTCCCTACACGACGCTCTTCCGATCTNNNTAAATGTTTTGTAAGTACTGCA
AGCACACGTCTGAACTCTTTCCCTACACGACGCTCTTCCGATCTNNNTACATACCTACACACATTCATAC
GAGCACACGTCTGAACTCTTTCCCTACACGACGCTCTTCCGATCTNNNAGTCTTTTAAATCATATGGTTT
GAGCACACGTCTGAACTCTTTCCCTACACGACGCTCTTCCGATCTNNNAGTCTTTTAAATCATATGGTTT
AGCACACGTCTGAACTCTTTCCCTACACGACGCTCTTCCGATCTNNNCTTAAGACATGCCTGAATTTCCA
AGCACACGTCTGAACTCTTTCCCTACACGACGCTCTTCCGATCTNNNAGACATTATGTAAGTGAATCTGT
AGCACACGTCTGAACTCTTTCCCTACACGACGCTCTTCCGATCTNNNTTAATCATAACACTATCATGCTG
AGCACACGTCTGAACTCTTTCCCTACACGACGCTCTTCCGATCTNNNTATAGTGAAGCAAGTTACCATAT
AGCACACGTCTGAACTCTTTCCCTACACGACGCTCTTCCGATCTNNNGTTGAGAAATTAGCAGATGTAA
GCACACGTCTGAACTCTTTCCCTACACGACGCTCTTCCGATCTNNNATAGAAATGGTCAAGAGCTATAC
AGCACACGTCTGAACTCTTTCCCTACACGACGCTCTTCCGATCTNNNAATATAGAGGCTTACATTGAGA
GCACACGTCTGAACTCTTTCCCTACACGACGCTCTTCCGATCTNNNGCAAATGTGTTTTCTGAACTCATC
GAGCACACGTCTGAACTCTTTCCCTACACGACGCTCTTCCGATCTNNNCTTTTGTATTACAGAAACCAGA
AGCACACGTCTGAACTCTTTCCCTACACGACGCTCTTCCGATCTNNNATTTTGAGCCATGTGATCATATC
ACACGTCTGAACTCTTTCCCTACACGACGCTCTTCCGATCTNNNGCTATGAAGTTTCTGAGAAAATTCA
AGCACACGTCTGAACTCTTTCCCTACACGACGCTCTTCCGATCTNNNTTTTATGGTTGCATAATATTCCA
AGCACACGTCTGAACTCTTTCCCTACACGACGCTCTTCCGATCTNNNGGTTGTTGTTGTTACTGTTTATA
AGAGCACACGTCTGAACTCTTTCCCTACACGACGCTCTTCCGATCTNNNAAGGTTTGTACTCACTGAAAT
AGCACACGTCTGAACTCTTTCCCTACACGACGCTCTTCCGATCTNNNGAATGAAGCTAAATTTGAAATC
AGCACACGTCTGAACTCTTTCCCTACACGACGCTCTTCCGATCTNNNGAATGAAGCTAAATTTGAAATC
AGCACACGTCTGAACTCTTTCCCTACACGACGCTCTTCCGATCTNNNATTGTTAGTTTTGCTTAACTGTT

AGCACACGTCTGAACTCTTTCCCTACACGACGCTCTTCCGATCTNNNAATTAGCTAAAGAACTCAAAA
AGCACACGTCTGAACTCTTTCCCTACACGACGCTCTTCCGATCTNNNGGGATGAAATTGGAAGATTA
AGCACACGTCTGAACTCTTTCCCTACACGACGCTCTTCCGATCTNNNGTCAAAAAATGAGGCAAATAT
GCACACGTCTGAACTCTTTCCCTACACGACGCTCTTCCGATCTNNNTCTTCCCTATCTAATATTTCACCC
AGCACACGTCTGAACTCTTTCCCTACACGACGCTCTTCCGATCTNNNGTTTTTCTGCTTTTTGTTAACTC
AGCACACGTCTGAACTCTTTCCCTACACGACGCTCTTCCGATCTNNNTTTAGACATACATGAGACACAA
GCACACGTCTGAACTCTTTCCCTACACGACGCTCTTCCGATCTNNNTTTTGCACTGACGATTATCATCAT
GCACACGTCTGAACTCTTTCCCTACACGACGCTCTTCCGATCTNNNCAAGTCAAGGATCTGTTTATAA
GAGCACACGTCTGAACTCTTTCCCTACACGACGCTCTTCCGATCTNNNATTAAGAGTCACATAGTCACT
CACACGTCTGAACTCTTTCCCTACACGACGCTCTTCCGATCTNNNCTAACTAAAGCCTAAATTCTCACCA
AGCACACGTCTGAACTCTTTCCCTACACGACGCTCTTCCGATCTNNNTAACCTTGTCGAAAATCAATT
ACACGTCTGAACTCTTTCCCTACACGACGCTCTTCCGATCTNNNGAGTGAGTTAATCCAATAAATGTC
GAGCACACGTCTGAACTCTTTCCCTACACGACGCTCTTCCGATCTNNNGATTTTGAATATAGTGAGCC
GCACACGTCTGAACTCTTTCCCTACACGACGCTCTTCCGATCTNNNTTTAAATCCTAAGACAAGAGAG
GCACACGTCTGAACTCTTTCCCTACACGACGCTCTTCCGATCTNNNAGTGTATATAACATGATACCCTT
ACACGTCTGAACTCTTTCCCTACACGACGCTCTTCCGATCTNNNCTGTGCCTACTCAAATTCTAGCATC
GCACACGTCTGAACTCTTTCCCTACACGACGCTCTTCCGATCTNNNATGGGAATTTGACCTTTAAAAA
GCACACGTCTGAACTCTTTCCCTACACGACGCTCTTCCGATCTNNNATGGGAATTTGACCTTTAAAAA
ACACGTCTGAACTCTTTCCCTACACGACGCTCTTCCGATCTNNNCTTAAGAAGAGAGCTTGAACTCAT
AGCACACGTCTGAACTCTTTCCCTACACGACGCTCTTCCGATCTNNNTGTATTTCTTACTCTTTGGATG
AGCACACGTCTGAACTCTTTCCCTACACGACGCTCTTCCGATCTNNNCTTCAGGAAAATTGTCATTTTT
AGCACACGTCTGAACTCTTTCCCTACACGACGCTCTTCCGATCTNNNTTAGAAAAAAAGCCTGAGTATT
CACACGTCTGAACTCTTTCCCTACACGACGCTCTTCCGATCTNNNGTATTAGAGTTCCTCAGAGAAAC
CACACGTCTGAACTCTTTCCCTACACGACGCTCTTCCGATCTNNNGCCCAAATGACTTTTAGTTTATTC
GCACACGTCTGAACTCTTTCCCTACACGACGCTCTTCCGATCTNNNATATTATACATAACCCAATATCG
GCACACGTCTGAACTCTTTCCCTACACGACGCTCTTCCGATCTNNNTTTTTAATCTAGGTCTCTCAGAG
CACACGTCTGAACTCTTTCCCTACACGACGCTCTTCCGATCTNNNCGAGGAGGCTTAAAAACATACAT
AGCACACGTCTGAACTCTTTCCCTACACGACGCTCTTCCGATCTNNNAACCCTAATCTGTTATTCCATT
CACACGTCTGAACTCTTTCCCTACACGACGCTCTTCCGATCTNNNGACTACTTTGTTCACTTGTTCCAT
AGCACACGTCTGAACTCTTTCCCTACACGACGCTCTTCCGATCTNNNATATCCTTCAGAAAAATCACTC
GCACACGTCTGAACTCTTTCCCTACACGACGCTCTTCCGATCTNNNTTACTGGACAGCACTAATCTAT
AGCACACGTCTGAACTCTTTCCCTACACGACGCTCTTCCGATCTNNNTAAGTGTTCGTTTAAGATAGA
GCACACGTCTGAACTCTTTCCCTACACGACGCTCTTCCGATCTNNNGTGATGAATCTATATTGAACAC
GCACACGTCTGAACTCTTTCCCTACACGACGCTCTTCCGATCTNNNTTTGCTCTGGTTTTAGTAATTT
GAGCACACGTCTGAACTCTTTCCCTACACGACGCTCTTCCGATCTNNNACATATCTTTGAGTATGCACA
AGCACACGTCTGAACTCTTTCCCTACACGACGCTCTTCCGATCTNNNAAATTGTCTTCTAACTTTGAGA
GCACACGTCTGAACTCTTTCCCTACACGACGCTCTTCCGATCTNNNAATCTGTTTGTGAAATTTCAGT
GCACACGTCTGAACTCTTTCCCTACACGACGCTCTTCCGATCTNNNTTTAATATTAAAGTCAGGCAGT
AGCACACGTCTGAACTCTTTCCCTACACGACGCTCTTCCGATCTNNNATAGGGTACATTGACTGAAAT
AGCACACGTCTGAACTCTTTCCCTACACGACGCTCTTCCGATCTNNNTAAGAGGACTCATGTGAAATA
AGCACACGTCTGAACTCTTTCCCTACACGACGCTCTTCCGATCTNNNACTGTAGAAGGGAGAAATTT
AGCACACGTCTGAACTCTTTCCCTACACGACGCTCTTCCGATCTNNNGATGAGAACTGAAGACTCATA
AGCACACGTCTGAACTCTTTCCCTACACGACGCTCTTCCGATCTNNNAAAATCAATCCTTAGATCAGT

GCACACGTCTGAACTCTTTCCCTACACGACGCTCTTCCGATCTNNNGCACAAATTGAACATGTTTATA

AGCACACGTCTGAACTCTTTCCCTACACGACGCTCTTCCGATCTNNNCATTTCCTCCAATCCAGGAACA

GAGCACACGTCTGAACTCTTTCCCTACACGACGCTCTTCCGATCTNNNCACATCATTTTCTCTTATGTCA

AGCACACGTCTGAACTCTTTCCCTACACGACGCTCTTCCGATCTNNNCTCATGTCGACAAAAAGCTAC

AGCACACGTCTGAACTCTTTCCCTACACGACGCTCTTCCGATCTNNNTCAGGATATTAAAGGCTTTGA

CACACGTCTGAACTCTTTCCCTACACGACGCTCTTCCGATCTNNNGGGGAGGGTAGTCATAAAAATA

GAGCACACGTCTGAACTCTTTCCCTACACGACGCTCTTCCGATCTNNNGACATTTTCAGTCTTCTAAAC

AGCACACGTCTGAACTCTTTCCCTACACGACGCTCTTCCGATCTNNNCATGACAGGATTTTCTTACTTT

GAGCACACGTCTGAACTCTTTCCCTACACGACGCTCTTCCGATCTNNNTTTCCATTAACAGAAATCTAA

AGCACACGTCTGAACTCTTTCCCTACACGACGCTCTTCCGATCTNNNTAACAAACAGAGCTGGAAATC

GCACACGTCTGAACTCTTTCCCTACACGACGCTCTTCCGATCTNNNCTGCAATGAATTTCAGATAAAA

GAGCACACGTCTGAACTCTTTCCCTACACGACGCTCTTCCGATCTNNNAATTTATATGGTCTTGAGTC

GAGCACACGTCTGAACTCTTTCCCTACACGACGCTCTTCCGATCTNNNTAGCTGTTTAAAAGAATTGG

AGCACACGTCTGAACTCTTTCCCTACACGACGCTCTTCCGATCTNNNATAAAACCAAACTGATAGCTA

AGCACACGTCTGAACTCTTTCCCTACACGACGCTCTTCCGATCTNNNGTAAGCCTAGATCATGCTACTC

AGCACACGTCTGAACTCTTTCCCTACACGACGCTCTTCCGATCTNNNAGATTTAGTAAATATTAGGA

GCACACGTCTGAACTCTTTCCCTACACGACGCTCTTCCGATCTNNNTAATGTTAGAAATTACTCCCACA

CACACGTCTGAACTCTTTCCCTACACGACGCTCTTCCGATCTNNNTTATGCCTTGTAGATAAAAAAGC

AGCACACGTCTGAACTCTTTCCCTACACGACGCTCTTCCGATCTNNNACAAGTTTCAGCTCATACTAAT

CACACGTCTGAACTCTTTCCCTACACGACGCTCTTCCGATCTNNNCTTTTTAACTCAGAGGAAAGAAA

GCACACGTCTGAACTCTTTCCCTACACGACGCTCTTCCGATCTNNNCTTAAGGGAGTGAAAAACTTG

GAGCACACGTCTGAACTCTTTCCCTACACGACGCTCTTCCGATCTNNNTTTTTTGATTTCTTAGTTGCAT

AGCACACGTCTGAACTCTTTCCCTACACGACGCTCTTCCGATCTNNNAATCTACCAATCTTATTCAGAG

AGCACACGTCTGAACTCTTTCCCTACACGACGCTCTTCCGATCTNNNCATCTGCCTAGAATTTCTCTGC

AGCACACGTCTGAACTCTTTCCCTACACGACGCTCTTCCGATCTNNNGTGAAACTTGAATTTTAAAGC

GAGCACACGTCTGAACTCTTTCCCTACACGACGCTCTTCCGATCTNNNGTTTGTGGTGTATACATGTAT

GCACACGTCTGAACTCTTTCCCTACACGACGCTCTTCCGATCTNNNTCCATTCATCCATCAAATATACA

CACGTCTGAACTCTTTCCCTACACGACGCTCTTCCGATCTNNNCGGTTACTGAACATGTGTTTATCATC

AGCACACGTCTGAACTCTTTCCCTACACGACGCTCTTCCGATCTNNNTAAGAGTCATTCTTATTGAAG

ACACGTCTGAACTCTTTCCCTACACGACGCTCTTCCGATCTNNNATTTCAACCTCAACTTGCTGCATCG

GAGCACACGTCTGAACTCTTTCCCTACACGACGCTCTTCCGATCTNNNTATTTTATGAGGCAATTGAT

GCACACGTCTGAACTCTTTCCCTACACGACGCTCTTCCGATCTNNNATCCATTTCTTCTAGGTTTTCTAC

GCACACGTCTGAACTCTTTCCCTACACGACGCTCTTCCGATCTNNNACTAACTCCTTTTGAAGAAGAT

AGCACACGTCTGAACTCTTTCCCTACACGACGCTCTTCCGATCTNNNGGCTTGATGTCATTAAACATTT

AGCACACGTCTGAACTCTTTCCCTACACGACGCTCTTCCGATCTNNNAAACAGGTTTAATCCAAAGAA

AGCACACGTCTGAACTCTTTCCCTACACGACGCTCTTCCGATCTNNNGATTTCTTTCAGTAGTGTTTTG

GCACACGTCTGAACTCTTTCCCTACACGACGCTCTTCCGATCTNNNCCTCTAGAATTGTGGAAAATAA

GCACACGTCTGAACTCTTTCCCTACACGACGCTCTTCCGATCTNNNTCAGATCTCTCTCTGTTTGTACA

AGCACACGTCTGAACTCTTTCCCTACACGACGCTCTTCCGATCTNNNTAGTATTTGTTAATGAGGCTG

GCACACGTCTGAACTCTTTCCCTACACGACGCTCTTCCGATCTNNNGTGAAAGAAAATGACACATTTT

AGCACACGTCTGAACTCTTTCCCTACACGACGCTCTTCCGATCTNNNTTTTACTGCTATGTAGTATTCC

CACACGTCTGAACTCTTTCCCTACACGACGCTCTTCCGATCTNNNCCTTAGAATAAGGAAGATCTGCC

GCACACGTCTGAACTCTTTCCCTACACGACGCTCTTCCGATCTNNNAGGATGTTACAGCTTTCTTATAT

GAGCACACGTCTGAACTCTTTCCCTACACGACGCTCTTCCGATCTNNNTGCTATTTTAAAACAAGTAG
AGCACACGTCTGAACTCTTTCCCTACACGACGCTCTTCCGATCTNNNCATCCAAAGACACCAATATTAC
AGCACACGTCTGAACTCTTTCCCTACACGACGCTCTTCCGATCTNNNCCCATTTATTACAACGAACTTA
GCACACGTCTGAACTCTTTCCCTACACGACGCTCTTCCGATCTNNNCAAAATTGCTGATTTGTATTCAT
AGCACACGTCTGAACTCTTTCCCTACACGACGCTCTTCCGATCTNNNCAAAGCAATGTGAGAGGAAT
CACACGTCTGAACTCTTTCCCTACACGACGCTCTTCCGATCTNNNGGTTGTTTTAATTACTATGGCTTC
AGCACACGTCTGAACTCTTTCCCTACACGACGCTCTTCCGATCTNNNTCCATATTGGGAAAGTTTCTAA
AGCACACGTCTGAACTCTTTCCCTACACGACGCTCTTCCGATCTNNNACAAACAAAGAAAAAGATAGA
AGCACACGTCTGAACTCTTTCCCTACACGACGCTCTTCCGATCTNNNACAAACAAAGAAAAAGATAGA
CACACGTCTGAACTCTTTCCCTACACGACGCTCTTCCGATCTNNNGGTAAACAGAGTTACAAAGAAAA
GCACACGTCTGAACTCTTTCCCTACACGACGCTCTTCCGATCTNNNTGTTTATTGTTTTGCTTATGGAT
CACACGTCTGAACTCTTTCCCTACACGACGCTCTTCCGATCTNNNCTACCTTCATTTTTACCTAGCACAT
AGCACACGTCTGAACTCTTTCCCTACACGACGCTCTTCCGATCTNNNCCTACAAATACATACACATACAG
AGCACACGTCTGAACTCTTTCCCTACACGACGCTCTTCCGATCTNNNATAATCCCTCTGTATTGTTTGT
GCACACGTCTGAACTCTTTCCCTACACGACGCTCTTCCGATCTNNNCTCCATGAACAGTAAGTGTACCA
GCACACGTCTGAACTCTTTCCCTACACGACGCTCTTCCGATCTNNNCTCCATGAACAGTAAGTGTACCA
AGCACACGTCTGAACTCTTTCCCTACACGACGCTCTTCCGATCTNNNGTTTGTTTGTTTGAATTAGGAT
AGCACACGTCTGAACTCTTTCCCTACACGACGCTCTTCCGATCTNNNATCAGGTACAGTCATAGTGTAT
AGCACACGTCTGAACTCTTTCCCTACACGACGCTCTTCCGATCTNNNCAAATGACTTTTATGGTTGAAT
ACACGTCTGAACTCTTTCCCTACACGACGCTCTTCCGATCTNNNAGGAATTGAAAGAGGAAGTATAG
CACACGTCTGAACTCTTTCCCTACACGACGCTCTTCCGATCTNNNATTCTTTGGCAGTGATTTTAAAG
CACGTCTGAACTCTTTCCCTACACGACGCTCTTCCGATCTNNNATCTCTTCTTCATTGCTGTTTCATCGG
GAGCACACGTCTGAACTCTTTCCCTACACGACGCTCTTCCGATCTNNNACATTCTACTTCCAGTTGTTTC
GAGCACACGTCTGAACTCTTTCCCTACACGACGCTCTTCCGATCTNNNACATTCTACTTCCAGTTGTTTC
AGCACACGTCTGAACTCTTTCCCTACACGACGCTCTTCCGATCTNNNTTAGAATGTCATCTAATGCCTT
GCACACGTCTGAACTCTTTCCCTACACGACGCTCTTCCGATCTNNNACGTTACTTTCAACTAATATGTTC
AGCACACGTCTGAACTCTTTCCCTACACGACGCTCTTCCGATCTNNNTTATTTGTCATGGTTATTCAGT
GAGCACACGTCTGAACTCTTTCCCTACACGACGCTCTTCCGATCTNNNCTGGAGGTATGAAACAGTAC
GCACACGTCTGAACTCTTTCCCTACACGACGCTCTTCCGATCTNNNGCATTTCCAAAAGTGAAAATTTT
AGCACACGTCTGAACTCTTTCCCTACACGACGCTCTTCCGATCTNNNCTAGAATCGAACGCAATTTAAG
AGCACACGTCTGAACTCTTTCCCTACACGACGCTCTTCCGATCTNNNCCAGTGATAAGGATATAAGTC
GCACACGTCTGAACTCTTTCCCTACACGACGCTCTTCCGATCTNNNATGAAAAATCTATCCCCATGATT
AGCACACGTCTGAACTCTTTCCCTACACGACGCTCTTCCGATCTNNNGAAAAGAAGAAAGAACAACAA
CACACGTCTGAACTCTTTCCCTACACGACGCTCTTCCGATCTNNNAAGAGATTACTGAGCATGAAGC
GAGCACACGTCTGAACTCTTTCCCTACACGACGCTCTTCCGATCTNNNTTAGAGTTACTTTAAGAGGA
ACACGTCTGAACTCTTTCCCTACACGACGCTCTTCCGATCTNNNAGGTATGTAAGGCAAATACATATC
AGCACACGTCTGAACTCTTTCCCTACACGACGCTCTTCCGATCTNNNGGAATTAAGATTTCAACGTAT
GCACACGTCTGAACTCTTTCCCTACACGACGCTCTTCCGATCTNNNTAGCCAGATGAAATACTTTTAGA
GAGCACACGTCTGAACTCTTTCCCTACACGACGCTCTTCCGATCTNNNTAAGTGCTATTACTGAGATAC
AGCACACGTCTGAACTCTTTCCCTACACGACGCTCTTCCGATCTNNNCAAAAGAGATTCCTTTCATCTA
AGCACACGTCTGAACTCTTTCCCTACACGACGCTCTTCCGATCTNNNCAAGATTCTGATATGTAAGGA
GCACACGTCTGAACTCTTTCCCTACACGACGCTCTTCCGATCTNNNCTTGACTGTTTACATGTGAATTC
ACACGTCTGAACTCTTTCCCTACACGACGCTCTTCCGATCTNNNTAGAGATATGGATATGGGTAGAG

AGCACACGTCTGAACTCTTTCCCTACACGACGCTCTTCCGATCTNNNTTCCTAGTACCTCTGTTCATTCA
GAGCACACGTCTGAACTCTTTCCCTACACGACGCTCTTCCGATCTNNNGTATGTGTGTATGTTTGTGTA
AGCACACGTCTGAACTCTTTCCCTACACGACGCTCTTCCGATCTNNNTAAGCTGAATTTCTTAAGGAAA
GAGCACACGTCTGAACTCTTTCCCTACACGACGCTCTTCCGATCTNNNAATTTTCATTGACACCTTTATCA
CACACGTCTGAACTCTTTCCCTACACGACGCTCTTCCGATCTNNNCATGGAGCTTTTTTGTTAGAGCAT
GCACACGTCTGAACTCTTTCCCTACACGACGCTCTTCCGATCTNNNTTCCTGAACACATAAGTACTTATC
AGAGCACACGTCTGAACTCTTTCCCTACACGACGCTCTTCCGATCTNNNGTTATATACTCCTCAAAACAC
AGAGCACACGTCTGAACTCTTTCCCTACACGACGCTCTTCCGATCTNNNGTTATATACTCCTCAAAACAC
AGAGCACACGTCTGAACTCTTTCCCTACACGACGCTCTTCCGATCTNNNGTTATATACTCCTCAAAACAC
AGCACACGTCTGAACTCTTTCCCTACACGACGCTCTTCCGATCTNNNGGGTGGTTGTAGTAGTAGTTC
GAGCACACGTCTGAACTCTTTCCCTACACGACGCTCTTCCGATCTNNNTTTCAGGATTATCTTTGCTAAT
AGCACACGTCTGAACTCTTTCCCTACACGACGCTCTTCCGATCTNNNTTTGTCATTAGTTACACTGAATC
GAGCACACGTCTGAACTCTTTCCCTACACGACGCTCTTCCGATCTNNNTACAAAGAAGAGGATTATCAG
GAGCACACGTCTGAACTCTTTCCCTACACGACGCTCTTCCGATCTNNNCCTAACTCTTATCTCTCTCTGTC
GAGCACACGTCTGAACTCTTTCCCTACACGACGCTCTTCCGATCTNNNCCTAACTCTTATCTCTCTCTGTC
GAGCACACGTCTGAACTCTTTCCCTACACGACGCTCTTCCGATCTNNNTGTCCTTCATTTCTTTTATTTCA
AGCACACGTCTGAACTCTTTCCCTACACGACGCTCTTCCGATCTNNNAGATTACCTATATGAAGTGCTT
GCACACGTCTGAACTCTTTCCCTACACGACGCTCTTCCGATCTNNNTGAAAATAACTGTCTTAGTGAGA
GCACACGTCTGAACTCTTTCCCTACACGACGCTCTTCCGATCTNNNTGAAAATAACTGTCTTAGTGAGA
AGCACACGTCTGAACTCTTTCCCTACACGACGCTCTTCCGATCTNNNCTTGACTATTAGAGGGACAAC
GAGCACACGTCTGAACTCTTTCCCTACACGACGCTCTTCCGATCTNNNTATTAGTGGTTGCTATTATCA
AGCACACGTCTGAACTCTTTCCCTACACGACGCTCTTCCGATCTNNNCTCATTCTAGTTCTCTAACAGAC
GCACACGTCTGAACTCTTTCCCTACACGACGCTCTTCCGATCTNNNAGCCTCATTTACTTATTTAAATGT
AGCACACGTCTGAACTCTTTCCCTACACGACGCTCTTCCGATCTNNNTGTAATACTCTGTTATACAGCA
GCACACGTCTGAACTCTTTCCCTACACGACGCTCTTCCGATCTNNNAAGAGCAAAGTAATAGAATACA
CACACGTCTGAACTCTTTCCCTACACGACGCTCTTCCGATCTNNNAAATATTTTAACAGCAAAATTCCCA
GCACACGTCTGAACTCTTTCCCTACACGACGCTCTTCCGATCTNNNCCCAAACACTTACCTATTTAGTAC
GAGCACACGTCTGAACTCTTTCCCTACACGACGCTCTTCCGATCTNNNGTTTCTATGTGCATTGTGAAG
GAGCACACGTCTGAACTCTTTCCCTACACGACGCTCTTCCGATCTNNNCACTTTAAGATGACTTACCAAA
GAGCACACGTCTGAACTCTTTCCCTACACGACGCTCTTCCGATCTNNNATTCTTCTTTCCTATTGGAGAT
GCACACGTCTGAACTCTTTCCCTACACGACGCTCTTCCGATCTNNNAGACAGGAAATTGTATGTGTATA
GCACACGTCTGAACTCTTTCCCTACACGACGCTCTTCCGATCTNNNAGACAGGAAATTGTATGTGTATA
GCACACGTCTGAACTCTTTCCCTACACGACGCTCTTCCGATCTNNNAGACAGGAAATTGTATGTGTATA
GAGCACACGTCTGAACTCTTTCCCTACACGACGCTCTTCCGATCTNNNATATATGGGAGCTAAAAGAG
AGCACACGTCTGAACTCTTTCCCTACACGACGCTCTTCCGATCTNNNAGAATTTTTCCATCTATGTTCAC
GAGCACACGTCTGAACTCTTTCCCTACACGACGCTCTTCCGATCTNNNTGCTGTAAAAAATAAGAAGA
GAGCACACGTCTGAACTCTTTCCCTACACGACGCTCTTCCGATCTNNNAACATGATTATGATGTCTAATC
AGCACACGTCTGAACTCTTTCCCTACACGACGCTCTTCCGATCTNNNTCTGAGATGATTGATATAGCTT
GAGCACACGTCTGAACTCTTTCCCTACACGACGCTCTTCCGATCTNNNTTAAACTAAGAATACTCCACCA
GAGCACACGTCTGAACTCTTTCCCTACACGACGCTCTTCCGATCTNNNCATGGTCACATAGGTTAATTA
GAGCACACGTCTGAACTCTTTCCCTACACGACGCTCTTCCGATCTNNNAAGACATATAGCTTATTTTTGC
AGAGCACACGTCTGAACTCTTTCCCTACACGACGCTCTTCCGATCTNNNACAATTAGGCTTGCTATTTTT
AGCACACGTCTGAACTCTTTCCCTACACGACGCTCTTCCGATCTNNNTAAACTTTAAATAACGAGGCAA

AGCACACGTCTGAACTCTTTCCCTACACGACGCTCTTCCGATCTNNNTGCAAAGAAATAAACTCAAAG
AGCACACGTCTGAACTCTTTCCCTACACGACGCTCTTCCGATCTNNNATGATAACAATGGAATAAACT
GAGCACACGTCTGAACTCTTTCCCTACACGACGCTCTTCCGATCTNNNCCATCATTCAGGAAACTAACA
GAGCACACGTCTGAACTCTTTCCCTACACGACGCTCTTCCGATCTNNNAACATTTATTGCACAAAGTAA
AGCACACGTCTGAACTCTTTCCCTACACGACGCTCTTCCGATCTNNNGAACCAGAGAAGTGGATAAA
AGCACACGTCTGAACTCTTTCCCTACACGACGCTCTTCCGATCTNNNGTCAACTTTTGGTTAAGTCTCT
GCACACGTCTGAACTCTTTCCCTACACGACGCTCTTCCGATCTNNNTAGGTTTTCTCTCTTACTTGCCA
GAGCACACGTCTGAACTCTTTCCCTACACGACGCTCTTCCGATCTNNNTGTTTTTAAACCATTGACATG
GCACACGTCTGAACTCTTTCCCTACACGACGCTCTTCCGATCTNNNAGTCAAAAGAAAAAAAAAGAT
AGCACACGTCTGAACTCTTTCCCTACACGACGCTCTTCCGATCTNNNACTGTAATTGAACGTGAATTA
GCACACGTCTGAACTCTTTCCCTACACGACGCTCTTCCGATCTNNNTTACCTTTGAGCTATTGTACAAC
GCACACGTCTGAACTCTTTCCCTACACGACGCTCTTCCGATCTNNNAATGGTTGGCTGAAAAATTATA
AGCACACGTCTGAACTCTTTCCCTACACGACGCTCTTCCGATCTNNNTTTTATTTTGGTCATTCTGATGT
CACACGTCTGAACTCTTTCCCTACACGACGCTCTTCCGATCTNNNCTGGATTAGAGTAAGGGGAGCA
ACACGTCTGAACTCTTTCCCTACACGACGCTCTTCCGATCTNNNGAGATACAGATCTGATGAGAAGCA
AGCACACGTCTGAACTCTTTCCCTACACGACGCTCTTCCGATCTNNNATAGTGGCATTAGGTACAAAT
GCACACGTCTGAACTCTTTCCCTACACGACGCTCTTCCGATCTNNNTTGGAGAAAAATGAATCAAGTT
AGCACACGTCTGAACTCTTTCCCTACACGACGCTCTTCCGATCTNNNTTAAACTAACAATGGGTACAG
GCACACGTCTGAACTCTTTCCCTACACGACGCTCTTCCGATCTNNNTAGGATTCAGACTGAATTACAT
GCACACGTCTGAACTCTTTCCCTACACGACGCTCTTCCGATCTNNNTTGGTAACCCTATCTTTCATTTAC
GCACACGTCTGAACTCTTTCCCTACACGACGCTCTTCCGATCTNNNCACTTGATTAATTTGTTTGAACC
GCACACGTCTGAACTCTTTCCCTACACGACGCTCTTCCGATCTNNNTTGGTTGTAAAAATTACTCATG
GCACACGTCTGAACTCTTTCCCTACACGACGCTCTTCCGATCTNNNCAATCTACATGAAGGAAACAAA
AGCACACGTCTGAACTCTTTCCCTACACGACGCTCTTCCGATCTNNNATCAATGAGTGGATAAAGAA
AGCACACGTCTGAACTCTTTCCCTACACGACGCTCTTCCGATCTNNNAAATAAAGGAAGCTAAGAGA
GCACACGTCTGAACTCTTTCCCTACACGACGCTCTTCCGATCTNNNCTAGTACAGTTTTTGCTTTAGAG
GCACACGTCTGAACTCTTTCCCTACACGACGCTCTTCCGATCTNNNGTTCCTGAATCTCTTTTGATAAG
GCACACGTCTGAACTCTTTCCCTACACGACGCTCTTCCGATCTNNNGAATCTGGAAAGAAACGTCTAT
AGCACACGTCTGAACTCTTTCCCTACACGACGCTCTTCCGATCTNNNATTAGAGGATGCATTAGTTA
GAGCACACGTCTGAACTCTTTCCCTACACGACGCTCTTCCGATCTNNNGGAAGTAAATGGGTTAAAC
AGCACACGTCTGAACTCTTTCCCTACACGACGCTCTTCCGATCTNNNTTTGAATTTTCACCCCTAAATA
GCACACGTCTGAACTCTTTCCCTACACGACGCTCTTCCGATCTNNNAGATTTGAGGGCTTTACTATGCA
GAGCACACGTCTGAACTCTTTCCCTACACGACGCTCTTCCGATCTNNNCAACCGAGATACTAAGACTA
AGCACACGTCTGAACTCTTTCCCTACACGACGCTCTTCCGATCTNNNTTAGTTCTCTGCTCTAGATCA
AGCACACGTCTGAACTCTTTCCCTACACGACGCTCTTCCGATCTNNNATTCAAACCCAAATCAAAGGA
AGCACACGTCTGAACTCTTTCCCTACACGACGCTCTTCCGATCTNNNACTGTGTTTGATGATAAGAAA
GCACACGTCTGAACTCTTTCCCTACACGACGCTCTTCCGATCTNNNCTAAACCAGAGAACTGATGAGC
GCACACGTCTGAACTCTTTCCCTACACGACGCTCTTCCGATCTNNNCTATTCTAGATTTCGTTTTCCACC
GCACACGTCTGAACTCTTTCCCTACACGACGCTCTTCCGATCTNNNATTCTGTTTTGAACCACTTTTAA
GCACACGTCTGAACTCTTTCCCTACACGACGCTCTTCCGATCTNNNGCTTACAGAATCTCTAACATACA
AGCACACGTCTGAACTCTTTCCCTACACGACGCTCTTCCGATCTNNNCGTTTAGTTATGTTTTCTAAGC
ACACGTCTGAACTCTTTCCCTACACGACGCTCTTCCGATCTNNNAAGACACTAGCCTCATTCATCATCGG
AGCACACGTCTGAACTCTTTCCCTACACGACGCTCTTCCGATCTNNNATATGTGTATTTACAAAATGCC

AGCACACGTCTGAACTCTTTCCCTACACGACGCTCTTCCGATCTNNNCACTTCTCTTTTGATTCATAGTA
AGCACACGTCTGAACTCTTTCCCTACACGACGCTCTTCCGATCTNNNCACTTCTCTTTTGATTCATAGTA
GCACACGTCTGAACTCTTTCCCTACACGACGCTCTTCCGATCTNNNTGACTATTTGTGCTTGCATCCATC
ACACGTCTGAACTCTTTCCCTACACGACGCTCTTCCGATCTNNNCATATCATCCCTTCCATGATTATCAT
GAGCACACGTCTGAACTCTTTCCCTACACGACGCTCTTCCGATCTNNNGTTTGGAGAAGTCACAGTAG
GCACACGTCTGAACTCTTTCCCTACACGACGCTCTTCCGATCTNNNTTAGGAAAGCTAAGGGATAAAT
GCACACGTCTGAACTCTTTCCCTACACGACGCTCTTCCGATCTNNNTTAGGAAAGCTAAGGGATAAAT
AGCACACGTCTGAACTCTTTCCCTACACGACGCTCTTCCGATCTNNNTGTATAATGTCAGGCAAAAATA
GCACACGTCTGAACTCTTTCCCTACACGACGCTCTTCCGATCTNNNTTCTCTTTCAAAATCCAACTCTCA
GCACACGTCTGAACTCTTTCCCTACACGACGCTCTTCCGATCTNNNAACGTGCAAAATATATTCCAATA
AGAGCACACGTCTGAACTCTTTCCCTACACGACGCTCTTCCGATCTNNNTTGCCTTATTGGATTAGGAA
GAGCACACGTCTGAACTCTTTCCCTACACGACGCTCTTCCGATCTNNNGCAGATAATTTGACAGGAAG
CACGTCTGAACTCTTTCCCTACACGACGCTCTTCCGATCTNNNGATTGAGGAGAAATGAAGGAGCAT
AGCACACGTCTGAACTCTTTCCCTACACGACGCTCTTCCGATCTNNNCTTTCTGCCTTTTCATTTTAAAC
AGCACACGTCTGAACTCTTTCCCTACACGACGCTCTTCCGATCTNNNAGTAGAAAGTTGGAAAACTAG
AGCACACGTCTGAACTCTTTCCCTACACGACGCTCTTCCGATCTNNNAAGGCTTTGCTTTGGTTAATAT
AGCACACGTCTGAACTCTTTCCCTACACGACGCTCTTCCGATCTNNNGCATGGTAAATATATTTGGGA
AGCACACGTCTGAACTCTTTCCCTACACGACGCTCTTCCGATCTNNNGCATGGTAAATATATTTGGGA
CACACGTCTGAACTCTTTCCCTACACGACGCTCTTCCGATCTNNNTATCTCCAAAGATTCTCCTATTCCAT
AGCACACGTCTGAACTCTTTCCCTACACGACGCTCTTCCGATCTNNNTAATATTAGGAAAAGCTGAGA
GCACACGTCTGAACTCTTTCCCTACACGACGCTCTTCCGATCTNNNATAGTAAGTACAAAGATCTTGA
GCACACGTCTGAACTCTTTCCCTACACGACGCTCTTCCGATCTNNNACATGCTAATATATGTACACTGA
GCACACGTCTGAACTCTTTCCCTACACGACGCTCTTCCGATCTNNNACATGCTAATATATGTACACTGA
AGCACACGTCTGAACTCTTTCCCTACACGACGCTCTTCCGATCTNNNATGTACCCACAATTCTAAGAAT
GAGCACACGTCTGAACTCTTTCCCTACACGACGCTCTTCCGATCTNNNCTTTAGTTAGTGTCACACAAA
CACACGTCTGAACTCTTTCCCTACACGACGCTCTTCCGATCTNNNCCAGCCCTATATCTTGGATACATCG
GAGCACACGTCTGAACTCTTTCCCTACACGACGCTCTTCCGATCTNNNAGACTTGTTTGCTTTCAGTCC
GAGCACACGTCTGAACTCTTTCCCTACACGACGCTCTTCCGATCTNNNTGAAATAAAGTCAAATTACC
AGCACACGTCTGAACTCTTTCCCTACACGACGCTCTTCCGATCTNNNTGCTCTTAACTAACCAAATAAT
GAGCACACGTCTGAACTCTTTCCCTACACGACGCTCTTCCGATCTNNNCAAAACAGACATACTTGTTAC
GAGCACACGTCTGAACTCTTTCCCTACACGACGCTCTTCCGATCTNNNAGGTTTCTAATTTTTTGGCTTA
GCACACGTCTGAACTCTTTCCCTACACGACGCTCTTCCGATCTNNNTGACCTCATGAAGTTTATATTCT
GAGCACACGTCTGAACTCTTTCCCTACACGACGCTCTTCCGATCTNNNAAATTTGTATTAGGGTTCTCC
GAGCACACGTCTGAACTCTTTCCCTACACGACGCTCTTCCGATCTNNNTCTCTATTATAATTTTGTTCCT
AGCACACGTCTGAACTCTTTCCCTACACGACGCTCTTCCGATCTNNNACTAAAAATTCTTTATACGAAT
AGCACACGTCTGAACTCTTTCCCTACACGACGCTCTTCCGATCTNNNAGAAAGTCCTCAGTCATTTTTC
GAGCACACGTCTGAACTCTTTCCCTACACGACGCTCTTCCGATCTNNNAACTTCTCAGCTTCCATAACCA
AGCACACGTCTGAACTCTTTCCCTACACGACGCTCTTCCGATCTNNNAACTCCTACAAAAAACAGAAA
AGCACACGTCTGAACTCTTTCCCTACACGACGCTCTTCCGATCTNNNTCCCATGAATATGACTTATATCC
AGCACACGTCTGAACTCTTTCCCTACACGACGCTCTTCCGATCTNNNCCTTACTGTTCTTGTGATAGTG
AGCACACGTCTGAACTCTTTCCCTACACGACGCTCTTCCGATCTNNNTTTAGTGTCAAATAGTAAACCT
GAGCACACGTCTGAACTCTTTCCCTACACGACGCTCTTCCGATCTNNNTCTTTCTAATCATTACCAGCAT
GAGCACACGTCTGAACTCTTTCCCTACACGACGCTCTTCCGATCTNNNTCTTTCTAATCATTACCAGCAT

GAGCACACGTCTGAACTCTTTCCCTACACGACGCTCTTCCGATCTNNNTACACAATAGAACTGTGCAAT
AGCACACGTCTGAACTCTTTCCCTACACGACGCTCTTCCGATCTNNNCATTGCTCTCTATCTATGTATCTC
AGCACACGTCTGAACTCTTTCCCTACACGACGCTCTTCCGATCTNNNATTTAATCTCTCGTAGATACAGC
AGCACACGTCTGAACTCTTTCCCTACACGACGCTCTTCCGATCTNNNGTACTATTCCAAAAAATTGAGG
AGCACACGTCTGAACTCTTTCCCTACACGACGCTCTTCCGATCTNNNCGAATGAATTATTTAAGTTCTGC
GAGCACACGTCTGAACTCTTTCCCTACACGACGCTCTTCCGATCTNNNTACACGTTTGTCATTTTTTTACT
AGCACACGTCTGAACTCTTTCCCTACACGACGCTCTTCCGATCTNNNTACCAAAAATCACTGTATTGAAC
AGCACACGTCTGAACTCTTTCCCTACACGACGCTCTTCCGATCTNNNGTTCAGAAAAAAAGTGTGTAAA
AGCACACGTCTGAACTCTTTCCCTACACGACGCTCTTCCGATCTNNNAACTTTATATGTCTTTACCCAGTC
GCACACGTCTGAACTCTTTCCCTACACGACGCTCTTCCGATCTNNNGAGAATATATATGTGTAAGGGGC
GCACACGTCTGAACTCTTTCCCTACACGACGCTCTTCCGATCTNNNGAGAATATATATGTGTAAGGGGC
AGCACACGTCTGAACTCTTTCCCTACACGACGCTCTTCCGATCTNNNTTTTGGTGAAAAGAATTAGGA
AGCACACGTCTGAACTCTTTCCCTACACGACGCTCTTCCGATCTNNNGATCTTTTAAGCTTCCACAATCC
GAGCACACGTCTGAACTCTTTCCCTACACGACGCTCTTCCGATCTNNNATTTCAAGGTAAAAAAGCATA
GAGCACACGTCTGAACTCTTTCCCTACACGACGCTCTTCCGATCTNNNAATGAACAGAACCTCTCCTCCA
GAGCACACGTCTGAACTCTTTCCCTACACGACGCTCTTCCGATCTNNNAGTATTTGCACTAGTTTTTGTA
CACACGTCTGAACTCTTTCCCTACACGACGCTCTTCCGATCTNNNCTCTCACTCACTTTCATCTGCATCGG
AGCACACGTCTGAACTCTTTCCCTACACGACGCTCTTCCGATCTNNNTGTAACTCTTACAAATACCTTCC
CACACGTCTGAACTCTTTCCCTACACGACGCTCTTCCGATCTNNNATTTTTTCTCTCACATCTCAAAGCAT
GCACACGTCTGAACTCTTTCCCTACACGACGCTCTTCCGATCTNNNATTTTTTCTCTCACATCTCAAAGCA
GAGCACACGTCTGAACTCTTTCCCTACACGACGCTCTTCCGATCTNNNATTTCCATTTCTGAATGAAAGA
CACACGTCTGAACTCTTTCCCTACACGACGCTCTTCCGATCTNNNGACTGAGAGCAGAATAGAAGCATC
GAGCACACGTCTGAACTCTTTCCCTACACGACGCTCTTCCGATCTNNNTTAATTTTTCCATGGGATGTAT
ACACGTCTGAACTCTTTCCCTACACGACGCTCTTCCGATCTNNNGTTAGAACATTTTGAAGGAAGGCA
CACACGTCTGAACTCTTTCCCTACACGACGCTCTTCCGATCTNNNGCTAGATGTTATTAGCACTCCCATCC
AGCACACGTCTGAACTCTTTCCCTACACGACGCTCTTCCGATCTNNNTCTCTCTCTCTCTCTGTATATCA
GAGCACACGTCTGAACTCTTTCCCTACACGACGCTCTTCCGATCTNNNCATGAAGAAGAAAACAACTA
CACACGTCTGAACTCTTTCCCTACACGACGCTCTTCCGATCTNNNTGAAACCAAAACTATAGGACTAACA
ACACGTCTGAACTCTTTCCCTACACGACGCTCTTCCGATCTNNNTGGTAAAAAAAAAAATCTAGGACACA
GAGCACACGTCTGAACTCTTTCCCTACACGACGCTCTTCCGATCTNNNTTTCACCGCTCAATTTTTCTAAC
AGCACACGTCTGAACTCTTTCCCTACACGACGCTCTTCCGATCTNNNTCTATACAATAAGAATGCAGGG
AGCACACGTCTGAACTCTTTCCCTACACGACGCTCTTCCGATCTNNNTCTATACAATAAGAATGCAGGG
GAGCACACGTCTGAACTCTTTCCCTACACGACGCTCTTCCGATCTNNNGCAGCAAAATTTTCATATCAAA
ACACGTCTGAACTCTTTCCCTACACGACGCTCTTCCGATCTNNNAGAAAGTCTTCAAACTACAAGTCATC
GAGCACACGTCTGAACTCTTTCCCTACACGACGCTCTTCCGATCTNNNACTGAATGGAAATTTGGTTTTA
GAGCACACGTCTGAACTCTTTCCCTACACGACGCTCTTCCGATCTNNNCTAGGGAATGTGGTACTATTCT
AGCACACGTCTGAACTCTTTCCCTACACGACGCTCTTCCGATCTNNNATAACAATTATGCCTTAGTCCTC
CACGTCTGAACTCTTTCCCTACACGACGCTCTTCCGATCTNNNGGTGACAGAGTGAAATATACACATCC
AGCACACGTCTGAACTCTTTCCCTACACGACGCTCTTCCGATCTNNNTGTTGGGATTACAAGCATAAGCA
GAGCACACGTCTGAACTCTTTCCCTACACGACGCTCTTCCGATCTNNNCCAATTTACAACTATCTACACA
AGCACACGTCTGAACTCTTTCCCTACACGACGCTCTTCCGATCTNNNCAGAAATATACCTTGGCACACAT
AAGAGCACACGTCTGAACTCTTTCCCTACACGACGCTCTTCCGATCTNNNCTGTCTTTCTCTTTGCCTCTC
GCACACGTCTGAACTCTTTCCCTACACGACGCTCTTCCGATCTNNNCTCTCCCCTACAGAATAGAAACAT

GCACACGTCTGAACTCTTTCCCTACACGACGCTCTTCCGATCTNNNGAAGCTGAAGAAAAAGTATGTG
AGCACACGTCTGAACTCTTTCCCTACACGACGCTCTTCCGATCTNNNGCATTCCAAAGATTTCTATTGTT
CACACGTCTGAACTCTTTCCCTACACGACGCTCTTCCGATCTNNNGTTCTGAGTAATTTCCATTTCATCA
GCACACGTCTGAACTCTTTCCCTACACGACGCTCTTCCGATCTNNNTGGGACAAACAATTTACAAATTA
GCACACGTCTGAACTCTTTCCCTACACGACGCTCTTCCGATCTNNNTGTAGGTCACAGATATTTATCAA
AGCACACGTCTGAACTCTTTCCCTACACGACGCTCTTCCGATCTNNNAGGTAAAATCACTTGTTACAAA
GCACACGTCTGAACTCTTTCCCTACACGACGCTCTTCCGATCTNNNGAGACTGGTTGAATGTTGAACA
GCACACGTCTGAACTCTTTCCCTACACGACGCTCTTCCGATCTNNNCTCTTCTCTTAGGAACTCCTTCATG
AGCACACGTCTGAACTCTTTCCCTACACGACGCTCTTCCGATCTNNNAAAAAAAAAAAAAAAAAGAGAC
AGCACACGTCTGAACTCTTTCCCTACACGACGCTCTTCCGATCTNNNCTATGAGTCCATAAACTCTCATT
GAGCACACGTCTGAACTCTTTCCCTACACGACGCTCTTCCGATCTNNNTGTATTTGAGAACACATTTACA
GCACACGTCTGAACTCTTTCCCTACACGACGCTCTTCCGATCTNNNTATAGAATAAGGATGTACAGCAA
GCACACGTCTGAACTCTTTCCCTACACGACGCTCTTCCGATCTNNNTCTATTTCTCTCTATTACTCTTCCC
AGCACACGTCTGAACTCTTTCCCTACACGACGCTCTTCCGATCTNNNCTATGTATCCCAATGGACATTTCA
AGCACACGTCTGAACTCTTTCCCTACACGACGCTCTTCCGATCTNNNTTTTCTTTCTTTCTCCATTACTGC
AGCACACGTCTGAACTCTTTCCCTACACGACGCTCTTCCGATCTNNNCAGTCAGATTTCTGTCTTCTGCAT
CACACGTCTGAACTCTTTCCCTACACGACGCTCTTCCGATCTNNNAGATAATATCCAGAAGAATTTTCTG
AGCACACGTCTGAACTCTTTCCCTACACGACGCTCTTCCGATCTNNNAGTATATCTATTTGGCCCTTTAG
CACACGTCTGAACTCTTTCCCTACACGACGCTCTTCCGATCTNNNTACACATGGTGAATTAACTGATCAT
GAGCACACGTCTGAACTCTTTCCCTACACGACGCTCTTCCGATCTNNNATATGTATATCTGCTGACACAT
GAGCACACGTCTGAACTCTTTCCCTACACGACGCTCTTCCGATCTNNNAACTATTCCAAAAATTTGAGGA
AGCACACGTCTGAACTCTTTCCCTACACGACGCTCTTCCGATCTNNNCATAATTCAGTCCATAACACTTT
AGCACACGTCTGAACTCTTTCCCTACACGACGCTCTTCCGATCTNNNTTCACTGTAAAGATGTCATTCTA
CACACGTCTGAACTCTTTCCCTACACGACGCTCTTCCGATCTNNNGAAGTCCAAGATCAAGGTATAGCA
CACACGTCTGAACTCTTTCCCTACACGACGCTCTTCCGATCTNNNGAAGTCCAAGATCAAGGTATAGCA
AGCACACGTCTGAACTCTTTCCCTACACGACGCTCTTCCGATCTNNNAGCAAGACTCCATCTCAAAACAT
AGCACACGTCTGAACTCTTTCCCTACACGACGCTCTTCCGATCTNNNATTTGGCTCTATTAACCACACCA
AGCACACGTCTGAACTCTTTCCCTACACGACGCTCTTCCGATCTNNNTGGAATTTCAAATTCTTTCTACG
AGCACACGTCTGAACTCTTTCCCTACACGACGCTCTTCCGATCTNNNTTCTTCTTCTTCTTCTTCTTCTTCAT
AGCACACGTCTGAACTCTTTCCCTACACGACGCTCTTCCGATCTNNNTTCTTCTTCTTCTTCTTCTTCTTCAT
AGCACACGTCTGAACTCTTTCCCTACACGACGCTCTTCCGATCTNNNTTCTTCTTCTTCTTCTTCTTCTTCAT
CACACGTCTGAACTCTTTCCCTACACGACGCTCTTCCGATCTNNNGCAAAGACCTATAAGATTAACACC
CACACGTCTGAACTCTTTCCCTACACGACGCTCTTCCGATCTNNNTTCTATTGCAAAGACCTATAAGACA
AGCACACGTCTGAACTCTTTCCCTACACGACGCTCTTCCGATCTNNNGTATAGCAAGGCATAGTATGA
CACACGTCTGAACTCTTTCCCTACACGACGCTCTTCCGATCTNNNTTTTTTGTTTTGCCTTTGTTTTCATG
CACACGTCTGAACTCTTTCCCTACACGACGCTCTTCCGATCTNNNTTTTTTGTTTTGCCTTTGTTTTCATG
GCACACGTCTGAACTCTTTCCCTACACGACGCTCTTCCGATCTNNNGTTTTACATCATTACTTCAACCCCA
AGCACACGTCTGAACTCTTTCCCTACACGACGCTCTTCCGATCTNNNTTTGTGTGTGAGTCACTTATTCA
GAGCACACGTCTGAACTCTTTCCCTACACGACGCTCTTCCGATCTNNNGCATATGATTGGAAATCTAAG
GAGCACACGTCTGAACTCTTTCCCTACACGACGCTCTTCCGATCTNNNGTTCATGATACAGTTCAAAGA
AGCACACGTCTGAACTCTTTCCCTACACGACGCTCTTCCGATCTNNNACAGAGTGAGATCCAGTTATAT
AGCACACGTCTGAACTCTTTCCCTACACGACGCTCTTCCGATCTNNNGCATAATACTATATGTGCATGTG
AGCACACGTCTGAACTCTTTCCCTACACGACGCTCTTCCGATCTNNNTTCCTTCTAATCTTACAGGCTACA

GCACACGTCTGAACTCTTTCCCTACACGACGCTCTTCCGATCTNNNGCCCAGCTAAGTTTTATCTTCATCG
CACACGTCTGAACTCTTTCCCTACACGACGCTCTTCCGATCTNNNAGCTTTTCCTTTCTGATAAACTACAT
AGCACACGTCTGAACTCTTTCCCTACACGACGCTCTTCCGATCTNNNGACATTAAGGATGCCAATATAA
AGCACACGTCTGAACTCTTTCCCTACACGACGCTCTTCCGATCTNNNACACATTTTTTCAGTCCTTAAATC
GCACACGTCTGAACTCTTTCCCTACACGACGCTCTTCCGATCTNNNTGAATAGAAAAGGAAGAGAGAT
GAGCACACGTCTGAACTCTTTCCCTACACGACGCTCTTCCGATCTNNNTGAAGAAAGAATTGGATTAAA
GAGCACACGTCTGAACTCTTTCCCTACACGACGCTCTTCCGATCTNNNTGAAGAAAGAATTGGATTAAA
GAGCACACGTCTGAACTCTTTCCCTACACGACGCTCTTCCGATCTNNNATAAGTTGTTATGATCTTTCCA
GAGCACACGTCTGAACTCTTTCCCTACACGACGCTCTTCCGATCTNNNATACAAAGAATACAACAGGA
GAGCACACGTCTGAACTCTTTCCCTACACGACGCTCTTCCGATCTNNNGTACAGGAGATATATAAGACA
AGCACACGTCTGAACTCTTTCCCTACACGACGCTCTTCCGATCTNNNGAAGGTCATCATATTTACTCATC
CACACGTCTGAACTCTTTCCCTACACGACGCTCTTCCGATCTNNNCTTTAAGTCCCCAACCCTTACATCGG
GAGCACACGTCTGAACTCTTTCCCTACACGACGCTCTTCCGATCTNNNGATCTAAGGAAAAACTATTCGA
AGAGCACACGTCTGAACTCTTTCCCTACACGACGCTCTTCCGATCTNNNTAAGGGTCTAAGATTGTAGT
CACACGTCTGAACTCTTTCCCTACACGACGCTCTTCCGATCTNNNAATTAGGAAGAAGGTAGAGACCA
AGCACACGTCTGAACTCTTTCCCTACACGACGCTCTTCCGATCTNNNACTAGTGATTCTGGGGAAGACA
AGCACACGTCTGAACTCTTTCCCTACACGACGCTCTTCCGATCTNNNACTAGTGATTCTGGGGAAGACA
GAGCACACGTCTGAACTCTTTCCCTACACGACGCTCTTCCGATCTNNNTCTCTGCATTTTACATGTGTTA
GAGCACACGTCTGAACTCTTTCCCTACACGACGCTCTTCCGATCTNNNTCTCTGCATTTTACATGTGTTA
AGAGCACACGTCTGAACTCTTTCCCTACACGACGCTCTTCCGATCTNNNGATCTGACTCCTTGAATGTAT
AGCACACGTCTGAACTCTTTCCCTACACGACGCTCTTCCGATCTNNNTGTGATATAAAGTCACTACATC
AGCACACGTCTGAACTCTTTCCCTACACGACGCTCTTCCGATCTNNNATTATTCTGTCCACTACATGTAC
AGCACACGTCTGAACTCTTTCCCTACACGACGCTCTTCCGATCTNNNAATTATTCAACATCACTGCTATCC
AGCACACGTCTGAACTCTTTCCCTACACGACGCTCTTCCGATCTNNNCTAGGTGCAACATCTGAAATACA
AGCACACGTCTGAACTCTTTCCCTACACGACGCTCTTCCGATCTNNNGAAATAATTCACCCATAGAAGCC
AGCACACGTCTGAACTCTTTCCCTACACGACGCTCTTCCGATCTNNNGAAATAATTCACCCATAGAAGCC
GAGCACACGTCTGAACTCTTTCCCTACACGACGCTCTTCCGATCTNNNTATACAAAAGAGGAAAAGAG
GAGCACACGTCTGAACTCTTTCCCTACACGACGCTCTTCCGATCTNNNTATACAAAAGAGGAAAAGAG
AGCACACGTCTGAACTCTTTCCCTACACGACGCTCTTCCGATCTNNNATGAGCTAGGTCTTCAAACTCA
GAGCACACGTCTGAACTCTTTCCCTACACGACGCTCTTCCGATCTNNNACTAAAACATTGCAGAATACA
GAGCACACGTCTGAACTCTTTCCCTACACGACGCTCTTCCGATCTNNNCAAATAAGAAAGAGAAAAAT
AGCACACGTCTGAACTCTTTCCCTACACGACGCTCTTCCGATCTNNNAAGATTCAAGAAGTACAGTAA
AGCACACGTCTGAACTCTTTCCCTACACGACGCTCTTCCGATCTNNNCAGTAGAGAAGGTAAAAATTT
CACACGTCTGAACTCTTTCCCTACACGACGCTCTTCCGATCTNNNGAAGAATCCAGGTCTTTTAATTTCA
CACACGTCTGAACTCTTTCCCTACACGACGCTCTTCCGATCTNNNGAAGAATCCAGGTCTTTTAATTTCA
GAGCACACGTCTGAACTCTTTCCCTACACGACGCTCTTCCGATCTNNNTATTCCCTTGTGCTTGTATGCA
GAGCACACGTCTGAACTCTTTCCCTACACGACGCTCTTCCGATCTNNNTATTCCCTTGTGCTTGTATGCA
GCACACGTCTGAACTCTTTCCCTACACGACGCTCTTCCGATCTNNNATTATGTGCTTATGAGAGATAGG
AGCACACGTCTGAACTCTTTCCCTACACGACGCTCTTCCGATCTNNNCTTATAACATGGCAACTGATTT
AGCACACGTCTGAACTCTTTCCCTACACGACGCTCTTCCGATCTNNNGTGATTTTTCTTTATGAACAT
AGCACACGTCTGAACTCTTTCCCTACACGACGCTCTTCCGATCTNNNTGTAGAAAGAGAGAGAGAGA
GAGCACACGTCTGAACTCTTTCCCTACACGACGCTCTTCCGATCTNNNGATCTCTGTTATGCCCTTTAAC
GCACACGTCTGAACTCTTTCCCTACACGACGCTCTTCCGATCTNNNTTATAGTGTTCTCTCTTTCTCAGCA

GAGCACACGTCTGAACTCTTTCCCTACACGACGCTCTTCCGATCTNNNCATCTGGAATTTCTCGTACAA
GCACACGTCTGAACTCTTTCCCTACACGACGCTCTTCCGATCTNNNTTATAGTGTTCTCTCTTTCTCAGCA
GAGCACACGTCTGAACTCTTTCCCTACACGACGCTCTTCCGATCTNNNCATGAATGTTCATCAAGGATAT
GCACACGTCTGAACTCTTTCCCTACACGACGCTCTTCCGATCTNNNGGTAAATAAGATTTTTGATGTGC
AGCACACGTCTGAACTCTTTCCCTACACGACGCTCTTCCGATCTNNNATTCATATAGATGCACATTCTCT
AGCACACGTCTGAACTCTTTCCCTACACGACGCTCTTCCGATCTNNNAATATGCAGGATTTTATTCCTT
ACACGTCTGAACTCTTTCCCTACACGACGCTCTTCCGATCTNNNGGATGGTGAAACTTATGAAGTCATC
AGCACACGTCTGAACTCTTTCCCTACACGACGCTCTTCCGATCTNNNCCTCCACAATAATGAAAACTAAT
GAGCACACGTCTGAACTCTTTCCCTACACGACGCTCTTCCGATCTNNNATTTAACGTAGTAGGTCTAAT
GAGCACACGTCTGAACTCTTTCCCTACACGACGCTCTTCCGATCTNNNATTTAACGTAGTAGGTCTAAT
AGCACACGTCTGAACTCTTTCCCTACACGACGCTCTTCCGATCTNNNTTACACATACACTCTCAAATAGA
AGCACACGTCTGAACTCTTTCCCTACACGACGCTCTTCCGATCTNNNAATAGAGGGAAATAAACACACA
AGCACACGTCTGAACTCTTTCCCTACACGACGCTCTTCCGATCTNNNGTTGTATAAATTGGTCTGGATT
AGCACACGTCTGAACTCTTTCCCTACACGACGCTCTTCCGATCTNNNGTTTTTCACTTAATACAGCCTTC
AGCACACGTCTGAACTCTTTCCCTACACGACGCTCTTCCGATCTNNNGTTTTTCACTTAATACAGCCTTC
AGCACACGTCTGAACTCTTTCCCTACACGACGCTCTTCCGATCTNNNGTTTTTCACTTAATACAGCCTTC
AGCACACGTCTGAACTCTTTCCCTACACGACGCTCTTCCGATCTNNNAATTTCTGAGAAAAATTCCCAA
AGCACACGTCTGAACTCTTTCCCTACACGACGCTCTTCCGATCTNNNAATTTCTGAGAAAAATTCCCAA
CACACGTCTGAACTCTTTCCCTACACGACGCTCTTCCGATCTNNNATGAAAATAATGTGTAGTGTGTG
CACACGTCTGAACTCTTTCCCTACACGACGCTCTTCCGATCTNNNATGAAAATAATGTGTAGTGTGTG
GCACACGTCTGAACTCTTTCCCTACACGACGCTCTTCCGATCTNNNATAGTCAATAATAAGGGGAGAT
AGCACACGTCTGAACTCTTTCCCTACACGACGCTCTTCCGATCTNNNCTTAGTTTTTTTTCTGTTCCACTCA
GAGCACACGTCTGAACTCTTTCCCTACACGACGCTCTTCCGATCTNNNTTGTAGAACCCAAATACAAAA
AGCACACGTCTGAACTCTTTCCCTACACGACGCTCTTCCGATCTNNNTGCTTGGAAGTGAAAAAAATAA
GAGCACACGTCTGAACTCTTTCCCTACACGACGCTCTTCCGATCTNNNACCAAACATTTAAAACAGTTTA
AGCACACGTCTGAACTCTTTCCCTACACGACGCTCTTCCGATCTNNNATGTATGAGTGGATATTGGTC
GCACACGTCTGAACTCTTTCCCTACACGACGCTCTTCCGATCTNNNTTTAAAAATTACTGATGCCTAGG
GAGCACACGTCTGAACTCTTTCCCTACACGACGCTCTTCCGATCTNNNCACAAAAAGCTAATAGGTTAC
GAGCACACGTCTGAACTCTTTCCCTACACGACGCTCTTCCGATCTNNNTCTTCGACAAAACCATAAAAAA
GCACACGTCTGAACTCTTTCCCTACACGACGCTCTTCCGATCTNNNACTTTTCCTAGAGATGGACACAT
CACGTCTGAACTCTTTCCCTACACGACGCTCTTCCGATCTNNNCAAGAAATAGGGATAAAAACCATCAT
CACGTCTGAACTCTTTCCCTACACGACGCTCTTCCGATCTNNNCAAGAAATAGGGATAAAAACCATCAT
GCACACGTCTGAACTCTTTCCCTACACGACGCTCTTCCGATCTNNNATATATATACACACACCCTGAGACA
AGCACACGTCTGAACTCTTTCCCTACACGACGCTCTTCCGATCTNNNGCTTAATTTAGTCATTTTTGGTG
GAGCACACGTCTGAACTCTTTCCCTACACGACGCTCTTCCGATCTNNNCACCTCATGTCTAATGAATAAT
GAGCACACGTCTGAACTCTTTCCCTACACGACGCTCTTCCGATCTNNNCTTCTAAAATGTTGTTGTACT
AGCACACGTCTGAACTCTTTCCCTACACGACGCTCTTCCGATCTNNNTATTGCAAGAAAAAAGAGAAA
AGCACACGTCTGAACTCTTTCCCTACACGACGCTCTTCCGATCTNNNTATTGCAAGAAAAAAGAGAAA
AGCACACGTCTGAACTCTTTCCCTACACGACGCTCTTCCGATCTNNNTTTTGTTTCTCTTAGAAGGAAT
ACACGTCTGAACTCTTTCCCTACACGACGCTCTTCCGATCTNNNTTTATTTCTTTGCTCAGATTGTTCATC
GAGCACACGTCTGAACTCTTTCCCTACACGACGCTCTTCCGATCTNNNCTTAGGAATGAATGTGGTACT
AGCACACGTCTGAACTCTTTCCCTACACGACGCTCTTCCGATCTNNNCAAATGCATATACATATGTGTGA
AGCACACGTCTGAACTCTTTCCCTACACGACGCTCTTCCGATCTNNNATGAATTTACCTTGTAGACTTC

GAGCACACGTCTGAACTCTTTCCCTACACGACGCTCTTCCGATCTNNNAAAGGAGGTTTGAGAAAGAA
AGCACACGTCTGAACTCTTTCCCTACACGACGCTCTTCCGATCTNNNAGAAATTTACATCTGTCCAGTG
AGCACACGTCTGAACTCTTTCCCTACACGACGCTCTTCCGATCTNNNAGAAATTTACATCTGTCCAGTG
CACACGTCTGAACTCTTTCCCTACACGACGCTCTTCCGATCTNNNTATCAGTGTGTGATATTCCTTTACA
CACACGTCTGAACTCTTTCCCTACACGACGCTCTTCCGATCTNNNATGTGATTCAATGCATGAGTCATC
CACACGTCTGAACTCTTTCCCTACACGACGCTCTTCCGATCTNNNATGTGATTCAATGCATGAGTCATC
GAGCACACGTCTGAACTCTTTCCCTACACGACGCTCTTCCGATCTNNNACATAACAGAAACACACAATT
AGAGCACACGTCTGAACTCTTTCCCTACACGACGCTCTTCCGATCTNNNGTTTGCTTTTTAAATTAATCC
GAGCACACGTCTGAACTCTTTCCCTACACGACGCTCTTCCGATCTNNNGGTATATGTATGTATGTGGA
GAGCACACGTCTGAACTCTTTCCCTACACGACGCTCTTCCGATCTNNNGGTATATGTATGTATGTGGA
GCACACGTCTGAACTCTTTCCCTACACGACGCTCTTCCGATCTNNNCCTAATTTCAACATTGTATCTCACA
AGCACACGTCTGAACTCTTTCCCTACACGACGCTCTTCCGATCTNNNCAGATAAGTTACATATGCAAGTC
GAGCACACGTCTGAACTCTTTCCCTACACGACGCTCTTCCGATCTNNNGAACTCCATTTCTCAAGTACAT
AGCACACGTCTGAACTCTTTCCCTACACGACGCTCTTCCGATCTNNNATATTTGATACGAAAAGATGCTC
AGCACACGTCTGAACTCTTTCCCTACACGACGCTCTTCCGATCTNNNATGATTTTCATAGAAAAGCTTGT
AGCACACGTCTGAACTCTTTCCCTACACGACGCTCTTCCGATCTNNNTTAGCTGCTGTACTAATAACATA
AGCACACGTCTGAACTCTTTCCCTACACGACGCTCTTCCGATCTNNNAATTGCTATCTTTTTAACAAGCA
AGCACACGTCTGAACTCTTTCCCTACACGACGCTCTTCCGATCTNNNCCTTTGATTTAAAACAGGAGTT
AGCACACGTCTGAACTCTTTCCCTACACGACGCTCTTCCGATCTNNNCAGAGTTTCTGTAATTTTATGG
AGCACACGTCTGAACTCTTTCCCTACACGACGCTCTTCCGATCTNNNCAGAGTTTCTGTAATTTTATGG
AGCACACGTCTGAACTCTTTCCCTACACGACGCTCTTCCGATCTNNNTATATTTCCGTGTCATGTCATTC
AGCACACGTCTGAACTCTTTCCCTACACGACGCTCTTCCGATCTNNNTAAAAACCCCTAAAAACAAAACA
AGCACACGTCTGAACTCTTTCCCTACACGACGCTCTTCCGATCTNNNTAAAAACCCCTAAAAACAAAACA
GAGCACACGTCTGAACTCTTTCCCTACACGACGCTCTTCCGATCTNNNGGGCCTAATGGAAGATATTTA
AGAGCACACGTCTGAACTCTTTCCCTACACGACGCTCTTCCGATCTNNNTAAAATAAGGCAAACCATAG
AGCACACGTCTGAACTCTTTCCCTACACGACGCTCTTCCGATCTNNNCGAACCTACATATGTACATACA
AGCACACGTCTGAACTCTTTCCCTACACGACGCTCTTCCGATCTNNNCTTCCTCACTGTCAAACTTTCATC
AGCACACGTCTGAACTCTTTCCCTACACGACGCTCTTCCGATCTNNNCTTGCTTTTATTCTCTTATCTGCC
AGCACACGTCTGAACTCTTTCCCTACACGACGCTCTTCCGATCTNNNCTTATGCTGGAGGTTTCATTCAT
AGCACACGTCTGAACTCTTTCCCTACACGACGCTCTTCCGATCTNNNCTTTGCAATGTTACAGATATGTA
AGCACACGTCTGAACTCTTTCCCTACACGACGCTCTTCCGATCTNNNCTTTGCAATGTTACAGATATGTA
GAGCACACGTCTGAACTCTTTCCCTACACGACGCTCTTCCGATCTNNNGCTCTTGATTCTGACAATCTAT
AGCACACGTCTGAACTCTTTCCCTACACGACGCTCTTCCGATCTNNNAAATTCAGAATACTTAGAAGAT
GCACACGTCTGAACTCTTTCCCTACACGACGCTCTTCCGATCTNNNTATATTACTTCCTCTTAGAAGCCC
GCACACGTCTGAACTCTTTCCCTACACGACGCTCTTCCGATCTNNNTATATTACTTCCTCTTAGAAGCCC
AGCACACGTCTGAACTCTTTCCCTACACGACGCTCTTCCGATCTNNNTTTCTACTATGCCTGGCTATCATC
CACACGTCTGAACTCTTTCCCTACACGACGCTCTTCCGATCTNNNTGTCTTACCCGATAGATTTATACCA
AGCACACGTCTGAACTCTTTCCCTACACGACGCTCTTCCGATCTNNNAAAGAACCTCAATGATTCTCAAC
AGCACACGTCTGAACTCTTTCCCTACACGACGCTCTTCCGATCTNNNAAAGAACCTCAATGATTCTCAAC
GAGCACACGTCTGAACTCTTTCCCTACACGACGCTCTTCCGATCTNNNTACTTGTTCATTGTGAAGGAG
CACACGTCTGAACTCTTTCCCTACACGACGCTCTTCCGATCTNNNTAGGAGATTCTCTGGGTCTTCATCC
GAGCACACGTCTGAACTCTTTCCCTACACGACGCTCTTCCGATCTNNNAAGATGCTTTGATTATTACATG
GAGCACACGTCTGAACTCTTTCCCTACACGACGCTCTTCCGATCTNNNAGTTAATCCTTAGTTGATGAGA

GAGCACACGTCTGAACTCTTTCCCTACACGACGCTCTTCCGATCTNNNCAGGATTGTTTCATACAGTTA
GAGCACACGTCTGAACTCTTTCCCTACACGACGCTCTTCCGATCTNNNATTGCCTACTCATTAAAGATTTG
GAGCACACGTCTGAACTCTTTCCCTACACGACGCTCTTCCGATCTNNNACCCATTTTTAAATTAGGTTAT
CACACGTCTGAACTCTTTCCCTACACGACGCTCTTCCGATCTNNNCTGTCAAGAAAGAAAGAAAGAAA
AGCACACGTCTGAACTCTTTCCCTACACGACGCTCTTCCGATCTNNNTAATCATATGCTAGTAGACTGT
AGCACACGTCTGAACTCTTTCCCTACACGACGCTCTTCCGATCTNNNATGTCTTCCTTTTTCCTTTTAACC
GAGCACACGTCTGAACTCTTTCCCTACACGACGCTCTTCCGATCTNNNGTCCAGTCATAAGGATCTTTA
CACACGTCTGAACTCTTTCCCTACACGACGCTCTTCCGATCTNNNAGAGCTACAACTCCAATTTAGCATG
GAGCACACGTCTGAACTCTTTCCCTACACGACGCTCTTCCGATCTNNNCTCACTCAATGTGGTCATAGCA
GAGCACACGTCTGAACTCTTTCCCTACACGACGCTCTTCCGATCTNNNTATTGCATGACAGCTTTCTTACA
AGCACACGTCTGAACTCTTTCCCTACACGACGCTCTTCCGATCTNNNTGAATGTTACATATGTTCTCTT
AGCACACGTCTGAACTCTTTCCCTACACGACGCTCTTCCGATCTNNNCTCTCTCTCTCTGTCTCTCATC
AGCACACGTCTGAACTCTTTCCCTACACGACGCTCTTCCGATCTNNNCTCTCTCTCTCTGTCTCTCATC
GAGCACACGTCTGAACTCTTTCCCTACACGACGCTCTTCCGATCTNNNTCATCAACATAGTAACATGTGA
GAGCACACGTCTGAACTCTTTCCCTACACGACGCTCTTCCGATCTNNNAAGAACACGTGTAAGACATAT
AGCACACGTCTGAACTCTTTCCCTACACGACGCTCTTCCGATCTNNNTATTACCTGATGGCATCTTATAC
AGCACACGTCTGAACTCTTTCCCTACACGACGCTCTTCCGATCTNNNAAGTCTTGCAAGTTATCTTTCTA
GCACACGTCTGAACTCTTTCCCTACACGACGCTCTTCCGATCTNNNCATCCACATTCAAGTATGGCCATC
GAGCACACGTCTGAACTCTTTCCCTACACGACGCTCTTCCGATCTNNNGAATCACCAAGAACCATAAACC
GAGCACACGTCTGAACTCTTTCCCTACACGACGCTCTTCCGATCTNNNAAACTCTCTCTCTCTCTCTTTCAT
GAGCACACGTCTGAACTCTTTCCCTACACGACGCTCTTCCGATCTNNNAAACTCTCTCTCTCTCTCTTTCAT
AGCACACGTCTGAACTCTTTCCCTACACGACGCTCTTCCGATCTNNNGAATAAAAAGGTAGTTGGGTA
GAGCACACGTCTGAACTCTTTCCCTACACGACGCTCTTCCGATCTNNNGTCTAATTCTTCTTCTGGAGAG
AGCACACGTCTGAACTCTTTCCCTACACGACGCTCTTCCGATCTNNNGCATTTTTAAAAAACAAAAGTG
AGCACACGTCTGAACTCTTTCCCTACACGACGCTCTTCCGATCTNNNGCATTTTTAAAAAACAAAAGTG
AGCACACGTCTGAACTCTTTCCCTACACGACGCTCTTCCGATCTNNNCATGGTGACTATAGGTAACAAT
AGCACACGTCTGAACTCTTTCCCTACACGACGCTCTTCCGATCTNNNTTTAATTCCAACTTCTTTAGTCCC
AGCACACGTCTGAACTCTTTCCCTACACGACGCTCTTCCGATCTNNNCAGTCTTATGTGCTATCAGTATA
AGCACACGTCTGAACTCTTTCCCTACACGACGCTCTTCCGATCTNNNTCTGAAGAGATAAATTTGATGG
AGCACACGTCTGAACTCTTTCCCTACACGACGCTCTTCCGATCTNNNAAAACATGAAAACAACCTAAGT
GAGCACACGTCTGAACTCTTTCCCTACACGACGCTCTTCCGATCTNNNTGGGTTGATATTTTATACTGTG
GAGCACACGTCTGAACTCTTTCCCTACACGACGCTCTTCCGATCTNNNAAAAAGCAGGTGATAAAACA
AGCACACGTCTGAACTCTTTCCCTACACGACGCTCTTCCGATCTNNNTTTTTTTTTTACAAAATGCCAGC
AGCACACGTCTGAACTCTTTCCCTACACGACGCTCTTCCGATCTNNNAATTCTCATTCATTTTTCATGGG
AGCACACGTCTGAACTCTTTCCCTACACGACGCTCTTCCGATCTNNNACTTATTGAGTAGACTGTAATTC
AGCACACGTCTGAACTCTTTCCCTACACGACGCTCTTCCGATCTNNNCAGTACTTATTGAGTAGACTGT
GCACACGTCTGAACTCTTTCCCTACACGACGCTCTTCCGATCTNNNCATTAATTAATCCATCTCACTGGC
GCACACGTCTGAACTCTTTCCCTACACGACGCTCTTCCGATCTNNNATTCTTCACTATCTTGTTAGCTCCA
GCACACGTCTGAACTCTTTCCCTACACGACGCTCTTCCGATCTNNNTTACAGTTCCATCACATATGTATC
AGCACACGTCTGAACTCTTTCCCTACACGACGCTCTTCCGATCTNNNTAATTCAATAAACACACACACAT
GAGCACACGTCTGAACTCTTTCCCTACACGACGCTCTTCCGATCTNNNTACTAGCTATATGACCAGTACA
GCACACGTCTGAACTCTTTCCCTACACGACGCTCTTCCGATCTNNNAAAATTGCAAAATGAACTTACAG
AGCACACGTCTGAACTCTTTCCCTACACGACGCTCTTCCGATCTNNNCTTATCCTGGTATTATTGTGCTC

AGCACACGTCTGAACTCTTTCCCTACACGACGCTCTTCCGATCTNNNAACTACTCTAACCCATATCTCTT
GCACACGTCTGAACTCTTTCCCTACACGACGCTCTTCCGATCTNNNTATGTTTTAATTTTTGTTGCGTT
GCACACGTCTGAACTCTTTCCCTACACGACGCTCTTCCGATCTNNNCAGAGATAAATAAGCTTAGCAA
AGCACACGTCTGAACTCTTTCCCTACACGACGCTCTTCCGATCTNNNTTCTTGCAATAAATCTCATAAG
AGCACACGTCTGAACTCTTTCCCTACACGACGCTCTTCCGATCTNNNATAAATTTCTTTCAAACCCTGA
AGCACACGTCTGAACTCTTTCCCTACACGACGCTCTTCCGATCTNNNCTAGATTTGCTATTTGATGAC
AGCACACGTCTGAACTCTTTCCCTACACGACGCTCTTCCGATCTNNNGTGTAAAAATACCTCATTTCTG
AGCACACGTCTGAACTCTTTCCCTACACGACGCTCTTCCGATCTNNNGTGTAAAAATACCTCATTTCTG
ACACGTCTGAACTCTTTCCCTACACGACGCTCTTCCGATCTNNNTGTTATCTTTCTGTCTCTATCTCCAT
ACACGTCTGAACTCTTTCCCTACACGACGCTCTTCCGATCTNNNGAATCTTCATCTGAAAGCCGCATCG
GCACACGTCTGAACTCTTTCCCTACACGACGCTCTTCCGATCTNNNAGTTAAACAATGCTATGTGAAT
AGCACACGTCTGAACTCTTTCCCTACACGACGCTCTTCCGATCTNNNCAGAGAATTAAATGCTAGAGA
AGCACACGTCTGAACTCTTTCCCTACACGACGCTCTTCCGATCTNNNGCAGCTAAATGAACTTATGAA
GCACACGTCTGAACTCTTTCCCTACACGACGCTCTTCCGATCTNNNAGAAAAGGAAGATGTGATGAT
GCACACGTCTGAACTCTTTCCCTACACGACGCTCTTCCGATCTNNNAGAAAAGGAAGATGTGATGAT
GCACACGTCTGAACTCTTTCCCTACACGACGCTCTTCCGATCTNNNACATATCATGAATTTGGAATGA
GCACACGTCTGAACTCTTTCCCTACACGACGCTCTTCCGATCTNNNGTACATACACATTGTGAAATGA
AGCACACGTCTGAACTCTTTCCCTACACGACGCTCTTCCGATCTNNNCTTAGTGATGAATATGTGTAT
GCACACGTCTGAACTCTTTCCCTACACGACGCTCTTCCGATCTNNNTATTCACTTTAAAAATGGTTCGT
GCACACGTCTGAACTCTTTCCCTACACGACGCTCTTCCGATCTNNNCTATCATCTTGTTTGGTGTGTCA
AGAGCACACGTCTGAACTCTTTCCCTACACGACGCTCTTCCGATCTNNNGGAGGAGAGTGATAAAAA
GAGCACACGTCTGAACTCTTTCCCTACACGACGCTCTTCCGATCTNNNTACTATGATTACAGAAGAGA
AGCACACGTCTGAACTCTTTCCCTACACGACGCTCTTCCGATCTNNNTCTTTTCTTCTGTCTTTCTGTAT
AGCACACGTCTGAACTCTTTCCCTACACGACGCTCTTCCGATCTNNNTCATTCATGCTTAATTCAGTAT
AGCACACGTCTGAACTCTTTCCCTACACGACGCTCTTCCGATCTNNNATGTTTGCACAATGATGAAAT
AGAGCACACGTCTGAACTCTTTCCCTACACGACGCTCTTCCGATCTNNNTTTTTTGCCCAACTGAATTT
CACACGTCTGAACTCTTTCCCTACACGACGCTCTTCCGATCTNNNCATTTCCGCCTATCCTATTTCCATC
CACGTCTGAACTCTTTCCCTACACGACGCTCTTCCGATCTNNNAGTAACTAAGGTAAAATGGGATTCA
GCACACGTCTGAACTCTTTCCCTACACGACGCTCTTCCGATCTNNNAACAATATCTCACCATACTTGAG
AGCACACGTCTGAACTCTTTCCCTACACGACGCTCTTCCGATCTNNNAACATCAAATCTATCACCAAAT
AGCACACGTCTGAACTCTTTCCCTACACGACGCTCTTCCGATCTNNNAAAACTACCTCTTTACAGGAAA
AGCACACGTCTGAACTCTTTCCCTACACGACGCTCTTCCGATCTNNNACAAAATCTTGAAGAAGAGAT
GCACACGTCTGAACTCTTTCCCTACACGACGCTCTTCCGATCTNNNCTACCTGGAAGAAAACACACCA
CACACGTCTGAACTCTTTCCCTACACGACGCTCTTCCGATCTNNNGAAAGAGATGCATACAAACATTTC
GCACACGTCTGAACTCTTTCCCTACACGACGCTCTTCCGATCTNNNAGTGGGAATCTAAATTTATTCC
AGCACACGTCTGAACTCTTTCCCTACACGACGCTCTTCCGATCTNNNTATTCCTTTTTGAGTGAAAACG
AGCACACGTCTGAACTCTTTCCCTACACGACGCTCTTCCGATCTNNNCAAGATTACAAACAAACTCCTTT
AGCACACGTCTGAACTCTTTCCCTACACGACGCTCTTCCGATCTNNNTATTTAGATTCTTGCATAACAG
CACACGTCTGAACTCTTTCCCTACACGACGCTCTTCCGATCTNNNCTAACGCCTTATTTACTCTGTCATC
AGCACACGTCTGAACTCTTTCCCTACACGACGCTCTTCCGATCTNNNACATTAGTAAATGATTGCAAA
AGCACACGTCTGAACTCTTTCCCTACACGACGCTCTTCCGATCTNNNTTTGTTTGACTAACTTCAGATA
CACACGTCTGAACTCTTTCCCTACACGACGCTCTTCCGATCTNNNAGGGTTGGATATTTTGAATTTTTC
GCACACGTCTGAACTCTTTCCCTACACGACGCTCTTCCGATCTNNNGGGGATTATTGGGATTATAATC

AGCACACGTCTGAACTCTTTCCCTACACGACGCTCTTCCGATCTNNNCACAGTTATACTAAATGCAGAA
GCACACGTCTGAACTCTTTCCCTACACGACGCTCTTCCGATCTNNNCTATGGCATTAGTAGAATGAATT
AGCACACGTCTGAACTCTTTCCCTACACGACGCTCTTCCGATCTNNNGAAGTCACTTTGCTTATGTTGC
AGCACACGTCTGAACTCTTTCCCTACACGACGCTCTTCCGATCTNNNGAAGTCACTTTGCTTATGTTGC
CACACGTCTGAACTCTTTCCCTACACGACGCTCTTCCGATCTNNNCTTTCCTCCAGAATCATGAATACATG
GCACACGTCTGAACTCTTTCCCTACACGACGCTCTTCCGATCTNNNTGCTTTGAAGGATAAATAAAGA
GAGCACACGTCTGAACTCTTTCCCTACACGACGCTCTTCCGATCTNNNGTAAAAGAAATGGGCAAGTG
AGCACACGTCTGAACTCTTTCCCTACACGACGCTCTTCCGATCTNNNAGGACATAAATGATCAAGCTTC
AGAGCACACGTCTGAACTCTTTCCCTACACGACGCTCTTCCGATCTNNNCAAACAAAAAGAAGTCAATG
GAGCACACGTCTGAACTCTTTCCCTACACGACGCTCTTCCGATCTNNNGGGAGAAAAAACTCTCAATTT
AGCACACGTCTGAACTCTTTCCCTACACGACGCTCTTCCGATCTNNNAAACCAGACTATGATAAGGTAT
AGCACACGTCTGAACTCTTTCCCTACACGACGCTCTTCCGATCTNNNACTAATCATGAATTATACGGGA
GAGCACACGTCTGAACTCTTTCCCTACACGACGCTCTTCCGATCTNNNTGGTGGTTTCACAAATCTATAT
AGCACACGTCTGAACTCTTTCCCTACACGACGCTCTTCCGATCTNNNTGGACCTATACTATCTTTCTAGAG
GAGCACACGTCTGAACTCTTTCCCTACACGACGCTCTTCCGATCTNNNGGACTATAGCGAATTCATTAAA
CACACGTCTGAACTCTTTCCCTACACGACGCTCTTCCGATCTNNNCCTCTCTTCATTCCTCTAATGCATCGG
CACACGTCTGAACTCTTTCCCTACACGACGCTCTTCCGATCTNNNTATGATGTATGTTGGCTATGGCATG
AGCACACGTCTGAACTCTTTCCCTACACGACGCTCTTCCGATCTNNNAAAAAACCAAAACCTTACAAAA
CACACGTCTGAACTCTTTCCCTACACGACGCTCTTCCGATCTNNNGAACATACGATTTATCTTGGAAGC
GAGCACACGTCTGAACTCTTTCCCTACACGACGCTCTTCCGATCTNNNAGTTAAAGAGAATTCACATTTT
GCACACGTCTGAACTCTTTCCCTACACGACGCTCTTCCGATCTNNNTTAGTTTCTGTGTCTCTCTATCTCA
AGCACACGTCTGAACTCTTTCCCTACACGACGCTCTTCCGATCTNNNGATAAGTTTCTACTGGATTAGG
AGCACACGTCTGAACTCTTTCCCTACACGACGCTCTTCCGATCTNNNCAAGAGCAGATTTACACTGATA
GAGCACACGTCTGAACTCTTTCCCTACACGACGCTCTTCCGATCTNNNAAATAAACATAGGAACTCACA
AGCACACGTCTGAACTCTTTCCCTACACGACGCTCTTCCGATCTNNNTAAACATAAACTTTGGAAGGTT
AGCACACGTCTGAACTCTTTCCCTACACGACGCTCTTCCGATCTNNNGAAATTGCTTCCATAAGTCTTTA
AGCACACGTCTGAACTCTTTCCCTACACGACGCTCTTCCGATCTNNNTCTGAGATACCTTAGATGAAGA
GAGCACACGTCTGAACTCTTTCCCTACACGACGCTCTTCCGATCTNNNTCTGATGACTTTTAGACTCTAA
AGCACACGTCTGAACTCTTTCCCTACACGACGCTCTTCCGATCTNNNCTTTGAAACTCTCACAAAATTGC
ACACGTCTGAACTCTTTCCCTACACGACGCTCTTCCGATCTNNNGAGAATCATTTGAACCCAGGCATCG
AGCACACGTCTGAACTCTTTCCCTACACGACGCTCTTCCGATCTNNNTCAGAATTTATTTATTGGGGGA
AGCACACGTCTGAACTCTTTCCCTACACGACGCTCTTCCGATCTNNNCTTAGGAAATACACACTGAGAT
GAGCACACGTCTGAACTCTTTCCCTACACGACGCTCTTCCGATCTNNNGTGATTTTTTGTTTGTTTTGGT
GAGCACACGTCTGAACTCTTTCCCTACACGACGCTCTTCCGATCTNNNTCCCTAAAGTTATAAATTCCTGT
AGCACACGTCTGAACTCTTTCCCTACACGACGCTCTTCCGATCTNNNCATGAGTATTGAAGCACTATTC
AGCACACGTCTGAACTCTTTCCCTACACGACGCTCTTCCGATCTNNNACTATTAAAACTATCACAAAGCA
GAGCACACGTCTGAACTCTTTCCCTACACGACGCTCTTCCGATCTNNNTTTTAAAAAAGACCTCATATCC
AGCACACGTCTGAACTCTTTCCCTACACGACGCTCTTCCGATCTNNNGCCTAAATGATCCCAGTATGCAT
AGCACACGTCTGAACTCTTTCCCTACACGACGCTCTTCCGATCTNNNACAAAGGTTAACAATTTAGAAT
AGCACACGTCTGAACTCTTTCCCTACACGACGCTCTTCCGATCTNNNGGGATGTTAGGTTCAATATCTT
AGCACACGTCTGAACTCTTTCCCTACACGACGCTCTTCCGATCTNNNACTGGCAATAAGAAATAAATTA
GAGCACACGTCTGAACTCTTTCCCTACACGACGCTCTTCCGATCTNNNTAAAAAACAAGAAACATACAC
AGCACACGTCTGAACTCTTTCCCTACACGACGCTCTTCCGATCTNNNATTGAGCTATCTCTAATTTCCAT

GAGCACACGTCTGAACTCTTTCCCTACACGACGCTCTTCCGATCTNNNATAAAACACCTATCTGCTATAC
AGCACACGTCTGAACTCTTTCCCTACACGACGCTCTTCCGATCTNNNGGCTGTTGTAACAAATTACTAT
GAGCACACGTCTGAACTCTTTCCCTACACGACGCTCTTCCGATCTNNNACAATCTTTTAAAATGCAAATC
AGCACACGTCTGAACTCTTTCCCTACACGACGCTCTTCCGATCTNNNACACTGGTATGTTTTTCCTATAAC
GAGCACACGTCTGAACTCTTTCCCTACACGACGCTCTTCCGATCTNNNTTTAAGCATATCTGAAACCAA
AGCACACGTCTGAACTCTTTCCCTACACGACGCTCTTCCGATCTNNNTGACTCTTTCAACTTATCCTTATC
AGCACACGTCTGAACTCTTTCCCTACACGACGCTCTTCCGATCTNNNTGACTCTTTCAACTTATCCTTATC
GAGCACACGTCTGAACTCTTTCCCTACACGACGCTCTTCCGATCTNNNAGATTTAATGCAATTCCAAATC
AGCACACGTCTGAACTCTTTCCCTACACGACGCTCTTCCGATCTNNNATTTAAGTCTGCAGAGGTTACC
AGCACACGTCTGAACTCTTTCCCTACACGACGCTCTTCCGATCTNNNCACTTTAGAATTACAAACCACAT
GAGCACACGTCTGAACTCTTTCCCTACACGACGCTCTTCCGATCTNNNACTATAAATCTTTCCATACACCT
CACACGTCTGAACTCTTTCCCTACACGACGCTCTTCCGATCTNNNATGTTGATTTTTCTCCCTAGATACA
AGCACACGTCTGAACTCTTTCCCTACACGACGCTCTTCCGATCTNNNGTATAGAACTCCAATTTTTTGCT
AGCACACGTCTGAACTCTTTCCCTACACGACGCTCTTCCGATCTNNNGTATAGAACTCCAATTTTTTGCT
AGCACACGTCTGAACTCTTTCCCTACACGACGCTCTTCCGATCTNNNCTTTGCTTCATAATTTGTTCTTTC
GCACACGTCTGAACTCTTTCCCTACACGACGCTCTTCCGATCTNNNAGTATAAATACAAAAACCTCCCAC
AGCACACGTCTGAACTCTTTCCCTACACGACGCTCTTCCGATCTNNNCTCATTTCAAACATGAAAGACTTC
AGCACACGTCTGAACTCTTTCCCTACACGACGCTCTTCCGATCTNNNAATTTATCATCTGTCTGATAGCT
AGCACACGTCTGAACTCTTTCCCTACACGACGCTCTTCCGATCTNNNATATGTGATATCAGTCATTGCT
AGCACACGTCTGAACTCTTTCCCTACACGACGCTCTTCCGATCTNNNAGCAATGGTTTATACTAAGGAT
AGCACACGTCTGAACTCTTTCCCTACACGACGCTCTTCCGATCTNNNGTTGATCTTTTTACCTACTCTTGC
AGCACACGTCTGAACTCTTTCCCTACACGACGCTCTTCCGATCTNNNATCTTTTGTGTATCTATGTCAGA
AGCACACGTCTGAACTCTTTCCCTACACGACGCTCTTCCGATCTNNNATATACTAGAGATAGAATAGTC
GCACACGTCTGAACTCTTTCCCTACACGACGCTCTTCCGATCTNNNATATTTTATGGATGGATATGCTT
GCACACGTCTGAACTCTTTCCCTACACGACGCTCTTCCGATCTNNNAGCCTCTTAGGAAAATCACTCATC
CACACGTCTGAACTCTTTCCCTACACGACGCTCTTCCGATCTNNNACAAACTAGAAAGATCAGAGAAAC
AGCACACGTCTGAACTCTTTCCCTACACGACGCTCTTCCGATCTNNNCATCTGACAGGCTTTAGATTCC
GAGCACACGTCTGAACTCTTTCCCTACACGACGCTCTTCCGATCTNNNAATAATGGTCTCTCTCTCTCTTC
GAGCACACGTCTGAACTCTTTCCCTACACGACGCTCTTCCGATCTNNNAATAATGGTCTCTCTCTCTCTTC
GCACACGTCTGAACTCTTTCCCTACACGACGCTCTTCCGATCTNNNGAAATGTTTAACTATACCTTCCCC
GAGCACACGTCTGAACTCTTTCCCTACACGACGCTCTTCCGATCTNNNAATATGCAGGTAAAAGTTCTTT
AGCACACGTCTGAACTCTTTCCCTACACGACGCTCTTCCGATCTNNNGAAACTGTAGTACCAGATCAGC
AGCACACGTCTGAACTCTTTCCCTACACGACGCTCTTCCGATCTNNNGAACACACACACAGTCATATCA
AGCACACGTCTGAACTCTTTCCCTACACGACGCTCTTCCGATCTNNNGAACGAAAAGAGAAGTTGAAAC
CACACGTCTGAACTCTTTCCCTACACGACGCTCTTCCGATCTNNNAAAAGGAATCCTCCCCATTATACAT
GCACACGTCTGAACTCTTTCCCTACACGACGCTCTTCCGATCTNNNTTTATCTATACTAACCCAACCTCCA
GCACACGTCTGAACTCTTTCCCTACACGACGCTCTTCCGATCTNNNGAAGGGCAAGGACTGATATACAT
AGCACACGTCTGAACTCTTTCCCTACACGACGCTCTTCCGATCTNNNTAAATAACTGATTGAGTTTCTGC
AGCACACGTCTGAACTCTTTCCCTACACGACGCTCTTCCGATCTNNNCCTATTTTTTTCCAGTTCTCTTTC
GCACACGTCTGAACTCTTTCCCTACACGACGCTCTTCCGATCTNNNGAAGAGAGATCCTCAATACTGCA
GAGCACACGTCTGAACTCTTTCCCTACACGACGCTCTTCCGATCTNNNAAAGATCATATATTCCATACCC
GCACACGTCTGAACTCTTTCCCTACACGACGCTCTTCCGATCTNNNTTTACAAGGTTTCTAGTCTAGGCA
AGCACACGTCTGAACTCTTTCCCTACACGACGCTCTTCCGATCTNNNCAAAATCAAGGCTATTACTTGG

GAGCACACGTCTGAACTCTTTCCCTACACGACGCTCTTCCGATCTNNNAACAAGTAGTAGATTGTACTG
AGCACACGTCTGAACTCTTTCCCTACACGACGCTCTTCCGATCTNNNGTCCTAAATTCTGTTTGTATTCA
AGCACACGTCTGAACTCTTTCCCTACACGACGCTCTTCCGATCTNNNGTCCTAAATTCTGTTTGTATTCA
AGCACACGTCTGAACTCTTTCCCTACACGACGCTCTTCCGATCTNNNGTGTGTGTGTATTTTAAAAT
GCACACGTCTGAACTCTTTCCCTACACGACGCTCTTCCGATCTNNNTCTCTTTTGAAGTAACTTTACCTC
CACACGTCTGAACTCTTTCCCTACACGACGCTCTTCCGATCTNNNGGCCAATAACGTATTCAATTATGCA
AGCACACGTCTGAACTCTTTCCCTACACGACGCTCTTCCGATCTNNNGCTGGGAGATGAATACATTTCA
GAGCACACGTCTGAACTCTTTCCCTACACGACGCTCTTCCGATCTNNNACATATGTTATCCAAAGGAGA
GCACACGTCTGAACTCTTTCCCTACACGACGCTCTTCCGATCTNNNTAGTCCATCCTAAGTATATGCTAC
AGCACACGTCTGAACTCTTTCCCTACACGACGCTCTTCCGATCTNNNACATTAATTACATTCCCCATTTCCA
GCACACGTCTGAACTCTTTCCCTACACGACGCTCTTCCGATCTNNNCAATCTTAGTATCAGAGACACCCA
AGCACACGTCTGAACTCTTTCCCTACACGACGCTCTTCCGATCTNNNGCAAACAAAACTCTCTGATATCC
GAGCACACGTCTGAACTCTTTCCCTACACGACGCTCTTCCGATCTNNNCCTTTTATATTGTTTCTAGACTC
GCACACGTCTGAACTCTTTCCCTACACGACGCTCTTCCGATCTNNNGGTAATCTGTCAGTTTCTTAAAAC
AGCACACGTCTGAACTCTTTCCCTACACGACGCTCTTCCGATCTNNNTTTATAGTTTCCAGCTTAATGTC
ACACGTCTGAACTCTTTCCCTACACGACGCTCTTCCGATCTNNNCTTTCTCACTTAACAAGGCACATCGGA
GCACACGTCTGAACTCTTTCCCTACACGACGCTCTTCCGATCTNNNTATTTACATGGCCTTTCAGGCAT
GCACACGTCTGAACTCTTTCCCTACACGACGCTCTTCCGATCTNNNAGATCAATATGCCTGGAGATTCA
GAGCACACGTCTGAACTCTTTCCCTACACGACGCTCTTCCGATCTNNNAGTGAGCCAAAGATATTTTAA
AGCACACGTCTGAACTCTTTCCCTACACGACGCTCTTCCGATCTNNNTCCCAAGTCTCTTATTTTTATCAC
AGCACACGTCTGAACTCTTTCCCTACACGACGCTCTTCCGATCTNNNGTAATGAGAGAAAAATTAGCC
AGCACACGTCTGAACTCTTTCCCTACACGACGCTCTTCCGATCTNNNTTCTGAGTAAATCAAATAGCAAC
AGCACACGTCTGAACTCTTTCCCTACACGACGCTCTTCCGATCTNNNGGTTCTGTATTTCCTTTCCTACC
AGCACACGTCTGAACTCTTTCCCTACACGACGCTCTTCCGATCTNNNGAGTAACTGAATCTTGCTGAAC
ACACGTCTGAACTCTTTCCCTACACGACGCTCTTCCGATCTNNNCTAGTTTGATATGGCTGGAGCATCG
AGCACACGTCTGAACTCTTTCCCTACACGACGCTCTTCCGATCTNNNCTGAGCCTACTCTTTTTATAGTA
AGCACACGTCTGAACTCTTTCCCTACACGACGCTCTTCCGATCTNNNGTAGGGAGAAAATGGGTTTTA
AGAGCACACGTCTGAACTCTTTCCCTACACGACGCTCTTCCGATCTNNNATAGATAAAAATGTCATCCA
GCACACGTCTGAACTCTTTCCCTACACGACGCTCTTCCGATCTNNNGAAAGCTGAGTCTTAGAGATGCA
GAGCACACGTCTGAACTCTTTCCCTACACGACGCTCTTCCGATCTNNNGGATTACTATAGTAAGTAGA
AGCACACGTCTGAACTCTTTCCCTACACGACGCTCTTCCGATCTNNNTTCAAAGAGTTTGCAGTCTAGCA
AGCACACGTCTGAACTCTTTCCCTACACGACGCTCTTCCGATCTNNNCCCGATTTTTTTTCTTCTTATTCA
GAGCACACGTCTGAACTCTTTCCCTACACGACGCTCTTCCGATCTNNNCTAAACTATCCACAGGACACCA
AGCACACGTCTGAACTCTTTCCCTACACGACGCTCTTCCGATCTNNNATTTAGTTACATAGTTGCCAAC
AGCACACGTCTGAACTCTTTCCCTACACGACGCTCTTCCGATCTNNNTAAATCTGCTTTGGATTGAGTT
AGCACACGTCTGAACTCTTTCCCTACACGACGCTCTTCCGATCTNNNGATTCATTAACCGAAATTTTGAG
ACACGTCTGAACTCTTTCCCTACACGACGCTCTTCCGATCTNNNAAAAAAATCTGTCTTCCTCTTTGCATC
AGCACACGTCTGAACTCTTTCCCTACACGACGCTCTTCCGATCTNNNAATTTGATTTTTGTAGATGGTG
CACGTCTGAACTCTTTCCCTACACGACGCTCTTCCGATCTNNNAAGTTCTTACAATAGTCTGCAGCATCC
ACACGTCTGAACTCTTTCCCTACACGACGCTCTTCCGATCTNNNGTATGAACAGAACTCAAATAAGGCA
AGCACACGTCTGAACTCTTTCCCTACACGACGCTCTTCCGATCTNNNGAATTCCAGCTGATAGGGAACA
GCACACGTCTGAACTCTTTCCCTACACGACGCTCTTCCGATCTNNNCTGTTTTATTTTCTTGCTCAAACA
AGCACACGTCTGAACTCTTTCCCTACACGACGCTCTTCCGATCTNNNATTTGGGCATATTTCTTCTTTAG

GCACACGTCTGAACTCTTTCCCTACACGACGCTCTTCCGATCTNNNAACAATTTCAATAAGTGATACG
AGCACACGTCTGAACTCTTTCCCTACACGACGCTCTTCCGATCTNNNATAACAAATAAATGGATGCTG
GCACACGTCTGAACTCTTTCCCTACACGACGCTCTTCCGATCTNNNAAGAGTTTAGTAAAGGAAAGG
AGCACACGTCTGAACTCTTTCCCTACACGACGCTCTTCCGATCTNNNTGATGAAAAAATATAGATCCA
CACGTCTGAACTCTTTCCCTACACGACGCTCTTCCGATCTNNNGAAAAAAAAAGAAGGCGTCTCATC
GCACACGTCTGAACTCTTTCCCTACACGACGCTCTTCCGATCTNNNGTAACCTCTATTAACTTGTCAAC
AGCACACGTCTGAACTCTTTCCCTACACGACGCTCTTCCGATCTNNNAAGATCCAAAGCAGAAAATGT
GCACACGTCTGAACTCTTTCCCTACACGACGCTCTTCCGATCTNNNAAATCCCAAATAAAACGAACTC
AAGAGCACACGTCTGAACTCTTTCCCTACACGACGCTCTTCCGATCTNNNATTTATTCACCTAACTAGT
AAGAGCACACGTCTGAACTCTTTCCCTACACGACGCTCTTCCGATCTNNNGCCTGTATTTATTCACCT
GAGCACACGTCTGAACTCTTTCCCTACACGACGCTCTTCCGATCTNNNAACCAGATTTACAGATGAGA
CACACGTCTGAACTCTTTCCCTACACGACGCTCTTCCGATCTNNNAGACTTTAGGACTAAAACTTACGC
GCACACGTCTGAACTCTTTCCCTACACGACGCTCTTCCGATCTNNNTGTATCAGAATAATCCTGTTTC
GCACACGTCTGAACTCTTTCCCTACACGACGCTCTTCCGATCTNNNTAGCAAACATTTAACTTGTTTT
GCACACGTCTGAACTCTTTCCCTACACGACGCTCTTCCGATCTNNNTAGCAAACATTTAACTTGTTTT
AGCACACGTCTGAACTCTTTCCCTACACGACGCTCTTCCGATCTNNNAGCACTTTGTTAATCATTTGAT
AGCACACGTCTGAACTCTTTCCCTACACGACGCTCTTCCGATCTNNNAGAATGAAAAAACTGCAGAA
GCACACGTCTGAACTCTTTCCCTACACGACGCTCTTCCGATCTNNNACTACCTGTAGTAAGAACCTAG
ACACGTCTGAACTCTTTCCCTACACGACGCTCTTCCGATCTNNNCATATAACCCAATGCTACTAGATCA
GCACACGTCTGAACTCTTTCCCTACACGACGCTCTTCCGATCTNNNATTTTTATTTGCAAAATCTAGCA
AGCACACGTCTGAACTCTTTCCCTACACGACGCTCTTCCGATCTNNNCTCTCAAGCCATTTCAGAAATC
CACACGTCTGAACTCTTTCCCTACACGACGCTCTTCCGATCTNNNCCAAAGTGCTGGGATTATAGCAT
AGCACACGTCTGAACTCTTTCCCTACACGACGCTCTTCCGATCTNNNGAGAAAGGCCTTTTCCATACC
CACACGTCTGAACTCTTTCCCTACACGACGCTCTTCCGATCTNNNAACTTGAGAAAATTTGAAAGCTA
GCACACGTCTGAACTCTTTCCCTACACGACGCTCTTCCGATCTNNNAGAATACAGGAAAATGGCTGC
ACACGTCTGAACTCTTTCCCTACACGACGCTCTTCCGATCTNNNATTTAGGCTTTGCAATAGCTAACAT
ACACGTCTGAACTCTTTCCCTACACGACGCTCTTCCGATCTNNNATTTAGGCTTTGCAATAGCTAACAT
CACACGTCTGAACTCTTTCCCTACACGACGCTCTTCCGATCTNNNCACCTTGATTTTAAACTTCTGATCA
AGCACACGTCTGAACTCTTTCCCTACACGACGCTCTTCCGATCTNNNTAAAAATCAGCCCCTGAGTTA
GAGCACACGTCTGAACTCTTTCCCTACACGACGCTCTTCCGATCTNNNTTTAGCACAGTTAAACATTCA
GCACACGTCTGAACTCTTTCCCTACACGACGCTCTTCCGATCTNNNCAAGAAGTTCCAAGCTATTAAA
GCACACGTCTGAACTCTTTCCCTACACGACGCTCTTCCGATCTNNNCTCTCCAAGTAGAGAACTAAAG
CACGTCTGAACTCTTTCCCTACACGACGCTCTTCCGATCTNNNTCAACCTGGAAGTCTAAAAAATACAT
ACACGTCTGAACTCTTTCCCTACACGACGCTCTTCCGATCTNNNTTCCAAACTGTGTTTTTAAATTGCA
CACACGTCTGAACTCTTTCCCTACACGACGCTCTTCCGATCTNNNAATGTGAGAACACTAACATCTCC
GCACACGTCTGAACTCTTTCCCTACACGACGCTCTTCCGATCTNNNGTATTTTTCAAATTGCCTACAAG
AGCACACGTCTGAACTCTTTCCCTACACGACGCTCTTCCGATCTNNNTAAGAACTGCTGAATTACTAA
AGCACACGTCTGAACTCTTTCCCTACACGACGCTCTTCCGATCTNNNCATGGTGATTTTCTTGAGAAT
GCACACGTCTGAACTCTTTCCCTACACGACGCTCTTCCGATCTNNNTTAAAATATTTGCAATTCCCTCT
ACACGTCTGAACTCTTTCCCTACACGACGCTCTTCCGATCTNNNGATCTTTAAGGACAAGCTGAGCAT
AGCACACGTCTGAACTCTTTCCCTACACGACGCTCTTCCGATCTNNNCTGTCTAGAAGTCTGTGCTCCA
GAGCACACGTCTGAACTCTTTCCCTACACGACGCTCTTCCGATCTNNNTACTGAACTTTACCAATTATC
AGCACACGTCTGAACTCTTTCCCTACACGACGCTCTTCCGATCTNNNAAAGGTTTTAGAGTTTCTTTG

ACACGTCTGAACTCTTTCCCTACACGACGCTCTTCCGATCTNNNGCCAATTTCTGAGTTTTTAAAGACA
CACACGTCTGAACTCTTTCCCTACACGACGCTCTTCCGATCTNNNGTGTCATATTTCACTTTCACTAAC
AGCACACGTCTGAACTCTTTCCCTACACGACGCTCTTCCGATCTNNNCTCTCAGTTGATAAATCCCACC
AGCACACGTCTGAACTCTTTCCCTACACGACGCTCTTCCGATCTNNNGACTAAGACAGGTGTAAATTT
GCACACGTCTGAACTCTTTCCCTACACGACGCTCTTCCGATCTNNNTCTAAAAGGTTCTCTTGCTATTA
AGCACACGTCTGAACTCTTTCCCTACACGACGCTCTTCCGATCTNNNTTAAGGATTCCTTTTTTCACTCT
GCACACGTCTGAACTCTTTCCCTACACGACGCTCTTCCGATCTNNNTATGACCAAGCAGAGATATTTT
AGCACACGTCTGAACTCTTTCCCTACACGACGCTCTTCCGATCTNNNGAAATATTAAAACCCAAAAAC
GAGCACACGTCTGAACTCTTTCCCTACACGACGCTCTTCCGATCTNNNGATTCACCTGACTTAGTGAC
GCACACGTCTGAACTCTTTCCCTACACGACGCTCTTCCGATCTNNNTATAGAAAGCCTGTGTTCATTT
GCACACGTCTGAACTCTTTCCCTACACGACGCTCTTCCGATCTNNNTACATGTTTTATCATCTAGAGCC
CACACGTCTGAACTCTTTCCCTACACGACGCTCTTCCGATCTNNNGTCAAAGTTGAACTTGTATAAGG
AGCACACGTCTGAACTCTTTCCCTACACGACGCTCTTCCGATCTNNNTAATTTTGTTTGCTCCCTCTTCA
AGCACACGTCTGAACTCTTTCCCTACACGACGCTCTTCCGATCTNNNTAGGGAATTCATTAATCATGA
GCACACGTCTGAACTCTTTCCCTACACGACGCTCTTCCGATCTNNNCAATGTCCTTGACCTACAATATC
AGCACACGTCTGAACTCTTTCCCTACACGACGCTCTTCCGATCTNNNTTTTTTACGGTGGAATCTTAAC
ACACGTCTGAACTCTTTCCCTACACGACGCTCTTCCGATCTNNNACTTATTTCTCATTCATTTCTTCTCAT
GCACACGTCTGAACTCTTTCCCTACACGACGCTCTTCCGATCTNNNCCATAACAGAAAATGGCAAAAC
GCACACGTCTGAACTCTTTCCCTACACGACGCTCTTCCGATCTNNNTGTACTTTAGAAGGCTAGCTCAT
GCACACGTCTGAACTCTTTCCCTACACGACGCTCTTCCGATCTNNNTTGCCTAGGTCACCATAATTCAT
CACACGTCTGAACTCTTTCCCTACACGACGCTCTTCCGATCTNNNTTTCCATTAAACCAGAACTCTATC
GAGCACACGTCTGAACTCTTTCCCTACACGACGCTCTTCCGATCTNNNCTCCTAAGGAATGATTGTTCT
GCACACGTCTGAACTCTTTCCCTACACGACGCTCTTCCGATCTNNNTATACATGAGTCATTGTTTTTGG
AGCACACGTCTGAACTCTTTCCCTACACGACGCTCTTCCGATCTNNNTGTATTTTGACACACAAATCTG
CACACGTCTGAACTCTTTCCCTACACGACGCTCTTCCGATCTNNNAGACTTTAGCCTTTGTGTTCCATC
GCACACGTCTGAACTCTTTCCCTACACGACGCTCTTCCGATCTNNNTACTTTTAAAGAGAAGGTTTCC
GAGCACACGTCTGAACTCTTTCCCTACACGACGCTCTTCCGATCTNNNGTCTTCTGGTCTTGTGTTTAT
CACACGTCTGAACTCTTTCCCTACACGACGCTCTTCCGATCTNNNTTTATTCAAGGGATAGATATGGG
AGCACACGTCTGAACTCTTTCCCTACACGACGCTCTTCCGATCTNNNTCTTCTAAGTGTATGACGTATT
AGCACACGTCTGAACTCTTTCCCTACACGACGCTCTTCCGATCTNNNTCTAAAACCTTCCTGTTAGTTTC
AGCACACGTCTGAACTCTTTCCCTACACGACGCTCTTCCGATCTNNNCTTAGATCCCCTTTAGAACTAG
GAGCACACGTCTGAACTCTTTCCCTACACGACGCTCTTCCGATCTNNNATACTCAAAGAGTTATCCCAA
ACACGTCTGAACTCTTTCCCTACACGACGCTCTTCCGATCTNNNGAAGAATCTGGAAATCTGGATTCA
GAGCACACGTCTGAACTCTTTCCCTACACGACGCTCTTCCGATCTNNNCCATATGAAGAGAAATGTAG
AGCACACGTCTGAACTCTTTCCCTACACGACGCTCTTCCGATCTNNNTTGAACATATTTGAGGTTTACA
GCACACGTCTGAACTCTTTCCCTACACGACGCTCTTCCGATCTNNNCATGATCTCGTTCCTTTTTATGCA
AGCACACGTCTGAACTCTTTCCCTACACGACGCTCTTCCGATCTNNNCTTTATAACAGCCAGAGTGCC
AAGAGCACACGTCTGAACTCTTTCCCTACACGACGCTCTTCCGATCTNNNATCTGGCTTGTAATTTAAA
GAGCACACGTCTGAACTCTTTCCCTACACGACGCTCTTCCGATCTNNNAGTATAGCAATTTCAAAAAG
AGCACACGTCTGAACTCTTTCCCTACACGACGCTCTTCCGATCTNNNGTATCAGCAGTGTGAATCATC
GAGCACACGTCTGAACTCTTTCCCTACACGACGCTCTTCCGATCTNNNTTCGAGATGAAGAACTTTAG
AGCACACGTCTGAACTCTTTCCCTACACGACGCTCTTCCGATCTNNNGAAGCTAGTTTTGTTCTAAAGG
GCACACGTCTGAACTCTTTCCCTACACGACGCTCTTCCGATCTNNNAGCACAAATAATATAGATGTTG

GAGCACACGTCTGAACTCTTTCCCTACACGACGCTCTTCCGATCTNNNTCCAATTATATTTGCAATGAA
GAGCACACGTCTGAACTCTTTCCCTACACGACGCTCTTCCGATCTNNNTCCAATTATATTTGCAATGAA
AGCACACGTCTGAACTCTTTCCCTACACGACGCTCTTCCGATCTNNNGGACTACAAAAAATATCAACA
AGAGCACACGTCTGAACTCTTTCCCTACACGACGCTCTTCCGATCTNNNCACTAATGTGTAAGGAGAT
AGCACACGTCTGAACTCTTTCCCTACACGACGCTCTTCCGATCTNNNATAACATGGTAACTCAAATAC
GAGCACACGTCTGAACTCTTTCCCTACACGACGCTCTTCCGATCTNNNATTTATAATAAATCTCTGCCTG
GAGCACACGTCTGAACTCTTTCCCTACACGACGCTCTTCCGATCTNNNTACTCAAAGAATATCTGTGAA
AGCACACGTCTGAACTCTTTCCCTACACGACGCTCTTCCGATCTNNNATGTTCCTTTTCAATTATGAAC
GAGCACACGTCTGAACTCTTTCCCTACACGACGCTCTTCCGATCTNNNGATAAAGTGATTTGGAAAAC
AGCACACGTCTGAACTCTTTCCCTACACGACGCTCTTCCGATCTNNNTGTTATACATGTATGTGTGTG
ACACGTCTGAACTCTTTCCCTACACGACGCTCTTCCGATCTNNNCCCTATGTCTCAATACTGTATTTCAT
ACACGTCTGAACTCTTTCCCTACACGACGCTCTTCCGATCTNNNCCCTATGTCTCAATACTGTATTTCAT
CACACGTCTGAACTCTTTCCCTACACGACGCTCTTCCGATCTNNNTTCTTTCATTTGTTTTTCCTTCTCAT
ACACGTCTGAACTCTTTCCCTACACGACGCTCTTCCGATCTNNNATAAACACCATGTGATGAGTATCAT
GCACACGTCTGAACTCTTTCCCTACACGACGCTCTTCCGATCTNNNTTTAATTTAGGCTTCTTTCTGAC
GAGCACACGTCTGAACTCTTTCCCTACACGACGCTCTTCCGATCTNNNCATCAGGAGAGTATTTTGAG
CACACGTCTGAACTCTTTCCCTACACGACGCTCTTCCGATCTNNNGCTCTACTGATTCAGTAAAATTCC
AGCACACGTCTGAACTCTTTCCCTACACGACGCTCTTCCGATCTNNNGAATAAAAATGTAGTCCTTCAG
AGCACACGTCTGAACTCTTTCCCTACACGACGCTCTTCCGATCTNNNGCAAAAGAAGTATGAATTTTC
GAGCACACGTCTGAACTCTTTCCCTACACGACGCTCTTCCGATCTNNNAAGTTATCAACACCATTTACA
ACACGTCTGAACTCTTTCCCTACACGACGCTCTTCCGATCTNNNCTGTAGTGAGCTATGATCATGCATC
ACACGTCTGAACTCTTTCCCTACACGACGCTCTTCCGATCTNNNCTGCTCAAAATGACTGATGGCATCG
AGCACACGTCTGAACTCTTTCCCTACACGACGCTCTTCCGATCTNNNAGAACTGGATGAAGCAATCTC
CACACGTCTGAACTCTTTCCCTACACGACGCTCTTCCGATCTNNNGCCTACATAAGAACTATTTCTATGC
GCACACGTCTGAACTCTTTCCCTACACGACGCTCTTCCGATCTNNNATTTTCCTTTACATCTGGAGATA
GAGCACACGTCTGAACTCTTTCCCTACACGACGCTCTTCCGATCTNNNATTACAAAGAGGTAATGGTG
GCACACGTCTGAACTCTTTCCCTACACGACGCTCTTCCGATCTNNNTCAAAGGGTCTTTAGAAGTTTG
GCACACGTCTGAACTCTTTCCCTACACGACGCTCTTCCGATCTNNNTCAAAGGGTCTTTAGAAGTTTG
AGCACACGTCTGAACTCTTTCCCTACACGACGCTCTTCCGATCTNNNGCTACTAAGTAAACTTTGCTTT
AGCACACGTCTGAACTCTTTCCCTACACGACGCTCTTCCGATCTNNNGAGACCTACTTCTTTGTATGTA
AGCACACGTCTGAACTCTTTCCCTACACGACGCTCTTCCGATCTNNNAAATACAAAAAATTAGTCAGG
CACACGTCTGAACTCTTTCCCTACACGACGCTCTTCCGATCTNNNCCTCTTGGAAGTTCTGTCATCATCG
AGCACACGTCTGAACTCTTTCCCTACACGACGCTCTTCCGATCTNNNGAAGACTAAAAATTCACAGAT
GCACACGTCTGAACTCTTTCCCTACACGACGCTCTTCCGATCTNNNCCTGTCAAACTCTCTCTATATAG
AGCACACGTCTGAACTCTTTCCCTACACGACGCTCTTCCGATCTNNNTCAATAGTCACACACACATATAT
CACACGTCTGAACTCTTTCCCTACACGACGCTCTTCCGATCTNNNATTCAGAAGAACATGATGGGCAT
AGCACACGTCTGAACTCTTTCCCTACACGACGCTCTTCCGATCTNNNCTGGGAATCTTGGAATACATCC
AGCACACGTCTGAACTCTTTCCCTACACGACGCTCTTCCGATCTNNNCTGGGAATCTTGGAATACATCC
AGCACACGTCTGAACTCTTTCCCTACACGACGCTCTTCCGATCTNNNAAAGATAGAGAACATTTGTA
AGCACACGTCTGAACTCTTTCCCTACACGACGCTCTTCCGATCTNNNTAGGATCACTATGTTTTGCTAA
AGCACACGTCTGAACTCTTTCCCTACACGACGCTCTTCCGATCTNNNAATGTTGCCTTTTCCAAAACTA
CACGTCTGAACTCTTTCCCTACACGACGCTCTTCCGATCTNNNGTCTCAGCTATGTAGACCTTCATCGG
ACACGTCTGAACTCTTTCCCTACACGACGCTCTTCCGATCTNNNTTTGATCTCCCAACCCTTATACATCG

GAGCACACGTCTGAACTCTTTCCCTACACGACGCTCTTCCGATCTNNNAACTATTCTTTCTTTTTCTCCC
ACACGTCTGAACTCTTTCCCTACACGACGCTCTTCCGATCTNNNTTGTGAAAACAGATAAAAGAAGGC
GAGCACACGTCTGAACTCTTTCCCTACACGACGCTCTTCCGATCTNNNGCGTGTGTGTGTATATATGTA
GAGCACACGTCTGAACTCTTTCCCTACACGACGCTCTTCCGATCTNNNGCGTGTGTGTGTATATATGTA
GAGCACACGTCTGAACTCTTTCCCTACACGACGCTCTTCCGATCTNNNAAAATGACATTGTTGAGTATA
AGCACACGTCTGAACTCTTTCCCTACACGACGCTCTTCCGATCTNNNGTGTCTATGCTTTGATATGTAAA
CACACGTCTGAACTCTTTCCCTACACGACGCTCTTCCGATCTNNNGAATTTTCTGTGAACTTGAGATTCA
CACACGTCTGAACTCTTTCCCTACACGACGCTCTTCCGATCTNNNGAATTTTCTGTGAACTTGAGATTCA
AGCACACGTCTGAACTCTTTCCCTACACGACGCTCTTCCGATCTNNNCCTTGATATATTGTTTTTCCCTT
AGCACACGTCTGAACTCTTTCCCTACACGACGCTCTTCCGATCTNNNATGGAGTAAGCTGATTAGGAC
GAGCACACGTCTGAACTCTTTCCCTACACGACGCTCTTCCGATCTNNNAGTATCTGCAAAGTATCCATA
AGCACACGTCTGAACTCTTTCCCTACACGACGCTCTTCCGATCTNNNACATAATACTAATGGAATCTCC
AGCACACGTCTGAACTCTTTCCCTACACGACGCTCTTCCGATCTNNNATTGTTTGTGAGGTGTTTTGCAT
GAGCACACGTCTGAACTCTTTCCCTACACGACGCTCTTCCGATCTNNNTCATGACGATACTCAGTAAATA
GAGCACACGTCTGAACTCTTTCCCTACACGACGCTCTTCCGATCTNNNAAATAAAAATCCTTGCCTTTCT
AGCACACGTCTGAACTCTTTCCCTACACGACGCTCTTCCGATCTNNNACATATTCTCAACCCTATTCTAC
AGCACACGTCTGAACTCTTTCCCTACACGACGCTCTTCCGATCTNNNTTGTAAAAGGAAATAGAACAGA
AGCACACGTCTGAACTCTTTCCCTACACGACGCTCTTCCGATCTNNNATTTGTAGACCTTTATTTCTTGC
AGCACACGTCTGAACTCTTTCCCTACACGACGCTCTTCCGATCTNNNAGTTGAATAAATGCTTTTATTTG
GAGCACACGTCTGAACTCTTTCCCTACACGACGCTCTTCCGATCTNNNCATGAAAGCAATAAATAGAGT
AGCACACGTCTGAACTCTTTCCCTACACGACGCTCTTCCGATCTNNNTAGAGTAGGATATGTTCTCTCA
AGCACACGTCTGAACTCTTTCCCTACACGACGCTCTTCCGATCTNNNGAACTTACTGGCTGAATGTAAT
AGCACACGTCTGAACTCTTTCCCTACACGACGCTCTTCCGATCTNNNAGAGATCTGGCTAAAATGAATA
AGAGCACACGTCTGAACTCTTTCCCTACACGACGCTCTTCCGATCTNNNATCCAAGAGATATTATCCTC
AGCACACGTCTGAACTCTTTCCCTACACGACGCTCTTCCGATCTNNNGAGATTAGTGCCTATATAAGAA
GAGCACACGTCTGAACTCTTTCCCTACACGACGCTCTTCCGATCTNNNCTGTTCACTAAGAGTAGGATA
AGCACACGTCTGAACTCTTTCCCTACACGACGCTCTTCCGATCTNNNTTCGTTAAGAAAACAGAATGAA
AGAGCACACGTCTGAACTCTTTCCCTACACGACGCTCTTCCGATCTNNNCTCCTAATGAGACACTTGTCC
GCACACGTCTGAACTCTTTCCCTACACGACGCTCTTCCGATCTNNNGGGATGAATGAATAAACCTTGAT
GAGCACACGTCTGAACTCTTTCCCTACACGACGCTCTTCCGATCTNNNAAGTCTTCTTTAGCTATGATCT
GAGCACACGTCTGAACTCTTTCCCTACACGACGCTCTTCCGATCTNNNTACAGCTAAGCCTAATCAAAGC
GAGCACACGTCTGAACTCTTTCCCTACACGACGCTCTTCCGATCTNNNGCCCATGGTCATTTAGTAAAT
GAGCACACGTCTGAACTCTTTCCCTACACGACGCTCTTCCGATCTNNNCTCTCTTTCTGTATCTTTCTCTC
GAGCACACGTCTGAACTCTTTCCCTACACGACGCTCTTCCGATCTNNNTATGACTAGATCTGTTCTTTCA
GCACACGTCTGAACTCTTTCCCTACACGACGCTCTTCCGATCTNNNAACATGGGTTTATTTATGAGAAC
AGCACACGTCTGAACTCTTTCCCTACACGACGCTCTTCCGATCTNNNGATAAAACTGCAGGAGTTCTCA
AGCACACGTCTGAACTCTTTCCCTACACGACGCTCTTCCGATCTNNNTAACTACATCTTGATTTCAACAC
CACACGTCTGAACTCTTTCCCTACACGACGCTCTTCCGATCTNNNAGCTAGAGTACATAACTTTAAACTC
GCACACGTCTGAACTCTTTCCCTACACGACGCTCTTCCGATCTNNNAACATTTAATTTTAGTGGATCTG
GCACACGTCTGAACTCTTTCCCTACACGACGCTCTTCCGATCTNNNCTTTGTTTCTGGAGACCTAATCAT
GAGCACACGTCTGAACTCTTTCCCTACACGACGCTCTTCCGATCTNNNCAAAACCTATTTCTTTCCGTCCA
AGCACACGTCTGAACTCTTTCCCTACACGACGCTCTTCCGATCTNNNAGCATAGAAATCAAGCTAAAAT
AGAGCACACGTCTGAACTCTTTCCCTACACGACGCTCTTCCGATCTNNNATTTTTCAGTTGAAACGTACT

GCACACGTCTGAACTCTTTCCCTACACGACGCTCTTCCGATCTNNNGGGGATTATTGGGATTATAATC

GCACACGTCTGAACTCTTTCCCTACACGACGCTCTTCCGATCTNNNGGGGATTATTGGGATTATAATC

AGCACACGTCTGAACTCTTTCCCTACACGACGCTCTTCCGATCTNNNTTCTCTTTACCTCTCAAAATCTT

GAGCACACGTCTGAACTCTTTCCCTACACGACGCTCTTCCGATCTNNNCCCTAAAAAAAAATTGACAAT

ACACGTCTGAACTCTTTCCCTACACGACGCTCTTCCGATCTNNNTACAGATACAAACATGGGGACATC

CACGTCTGAACTCTTTCCCTACACGACGCTCTTCCGATCTNNNGACCTCTCTCTCTCATTCCATCGGA

GAGCACACGTCTGAACTCTTTCCCTACACGACGCTCTTCCGATCTNNNCTTTAATTGATTTGACTAGTG

AGCACACGTCTGAACTCTTTCCCTACACGACGCTCTTCCGATCTNNNTTTCAGATTTGCTATCCTCTAGC

AGCACACGTCTGAACTCTTTCCCTACACGACGCTCTTCCGATCTNNNGATTTCTCCCATTTTATTGTTCT

AGCACACGTCTGAACTCTTTCCCTACACGACGCTCTTCCGATCTNNNGTTTTTCTCTTAAATTTGACTG

AGCACACGTCTGAACTCTTTCCCTACACGACGCTCTTCCGATCTNNNCTAACTCATTCTTCAAAAACAG

ACACGTCTGAACTCTTTCCCTACACGACGCTCTTCCGATCTNNNCTTTCATCCACCTTCAACTCCATCGG

GAGCACACGTCTGAACTCTTTCCCTACACGACGCTCTTCCGATCTNNNTTTATTGCATGCATATTGAAA

AGCACACGTCTGAACTCTTTCCCTACACGACGCTCTTCCGATCTNNNAAGAAAAAACAAGAGAGGTT

GCACACGTCTGAACTCTTTCCCTACACGACGCTCTTCCGATCTNNNACATATAATAGGATCCATGCAT

AGCACACGTCTGAACTCTTTCCCTACACGACGCTCTTCCGATCTNNNCTGGAGTTTCAGTAGAATTTCA

AGCACACGTCTGAACTCTTTCCCTACACGACGCTCTTCCGATCTNNNTCAATATTTTGTTCTTCTTGCTT

GCACACGTCTGAACTCTTTCCCTACACGACGCTCTTCCGATCTNNNTCTCAAAACAAAAACAAAAACA

AGCACACGTCTGAACTCTTTCCCTACACGACGCTCTTCCGATCTNNNTACACAAATGTTTACAGCAATT

GCACACGTCTGAACTCTTTCCCTACACGACGCTCTTCCGATCTNNNCATAGTCCAAGTGAATGTTTAT

AGCACACGTCTGAACTCTTTCCCTACACGACGCTCTTCCGATCTNNNCTTTAAAATACTGTCAGTCCT

CACGTCTGAACTCTTTCCCTACACGACGCTCTTCCGATCTNNNGAGAATGAAATCTACATGCTTAGCA

CACACGTCTGAACTCTTTCCCTACACGACGCTCTTCCGATCTNNNAAATAAATGTCTTTGTGATTGGT

CACACGTCTGAACTCTTTCCCTACACGACGCTCTTCCGATCTNNNTTGTCTGCTATATTCGCTATAATC

GCACACGTCTGAACTCTTTCCCTACACGACGCTCTTCCGATCTNNNTTTCAACAACTGAAACTGATAA

AGCACACGTCTGAACTCTTTCCCTACACGACGCTCTTCCGATCTNNNTCTCAAAAGTTCCAGTTATCTA

AGCACACGTCTGAACTCTTTCCCTACACGACGCTCTTCCGATCTNNNTTTCTCCACTTTCTTTCTTAGAA

AGCACACGTCTGAACTCTTTCCCTACACGACGCTCTTCCGATCTNNNCCATGTATTTAGAGGTTTTTGT

AGCACACGTCTGAACTCTTTCCCTACACGACGCTCTTCCGATCTNNNTTTCGTTTGCTATATTTGCTTTA

GCACACGTCTGAACTCTTTCCCTACACGACGCTCTTCCGATCTNNNAATTGGTGGTCTTCTATTCTGC

GCACACGTCTGAACTCTTTCCCTACACGACGCTCTTCCGATCTNNNATTATAATCTTATGGGACCACT

AGCACACGTCTGAACTCTTTCCCTACACGACGCTCTTCCGATCTNNNGGGATGTAATCTGATATTTTT

AGCACACGTCTGAACTCTTTCCCTACACGACGCTCTTCCGATCTNNNAAGGTTAAGAAAGCAGTATCT

GCACACGTCTGAACTCTTTCCCTACACGACGCTCTTCCGATCTNNNTTACTCAAATTTTTCCAGCTTTC

GCACACGTCTGAACTCTTTCCCTACACGACGCTCTTCCGATCTNNNATTTCAATTCTCTTTTTGACTCA

GAGCACACGTCTGAACTCTTTCCCTACACGACGCTCTTCCGATCTNNNAGAAAAAGCTTATTGAAGTA

AGCACACGTCTGAACTCTTTCCCTACACGACGCTCTTCCGATCTNNNGAAAAATTAAACTCTTAAGCCC

GCACACGTCTGAACTCTTTCCCTACACGACGCTCTTCCGATCTNNNAGGATTTGGCAAAATTACCTAC

CACACGTCTGAACTCTTTCCCTACACGACGCTCTTCCGATCTNNNGAAAACTGAGGGATTCACACCAT

AGCACACGTCTGAACTCTTTCCCTACACGACGCTCTTCCGATCTNNNCAATTCCAGTTAATGTCCTGTA

AGCACACGTCTGAACTCTTTCCCTACACGACGCTCTTCCGATCTNNNCAATTCCAGTTAATGTCCTGTA

ACACGTCTGAACTCTTTCCCTACACGACGCTCTTCCGATCTNNNAGATATTTGAAGGCTGAATGATAC

AGCACACGTCTGAACTCTTTCCCTACACGACGCTCTTCCGATCTNNNAGTGGAATAAAATGTGAGAT

CACACGTCTGAACTCTTTCCCTACACGACGCTCTTCCGATCTNNNCAGATTGCATTTTTTCTTTTTCTCAT
AGCACACGTCTGAACTCTTTCCCTACACGACGCTCTTCCGATCTNNNTTCTACACAACAAGGTCAGACA
AGCACACGTCTGAACTCTTTCCCTACACGACGCTCTTCCGATCTNNNTGTCAATGCTAATTATTTTGCA
AGCACACGTCTGAACTCTTTCCCTACACGACGCTCTTCCGATCTNNNCAAAAGAGCTTGAATACTTACA
CACACGTCTGAACTCTTTCCCTACACGACGCTCTTCCGATCTNNNCCAAACATCTACTTTAGAAGAAGCA
GAGCACACGTCTGAACTCTTTCCCTACACGACGCTCTTCCGATCTNNNTTCTTATGAGCCTTACAAGTTA
CACACGTCTGAACTCTTTCCCTACACGACGCTCTTCCGATCTNNNAAAACGAGGATTTGGAATAAAAT
GAGCACACGTCTGAACTCTTTCCCTACACGACGCTCTTCCGATCTNNNGAGGGTCAGTTATAAAAACTT
CACACGTCTGAACTCTTTCCCTACACGACGCTCTTCCGATCTNNNCCTGGCACCATCTTCATATACATCG
CACACGTCTGAACTCTTTCCCTACACGACGCTCTTCCGATCTNNNTATTTTCCTTCATTCATGAGAGTCAT
GAGCACACGTCTGAACTCTTTCCCTACACGACGCTCTTCCGATCTNNNTTCTAGATTCAAATCTGTTGTT
AGCACACGTCTGAACTCTTTCCCTACACGACGCTCTTCCGATCTNNNACTAAAATCAAACTTACCGATTC
AGCACACGTCTGAACTCTTTCCCTACACGACGCTCTTCCGATCTNNNTTCTTGTTGGCTTCTCTCTATCAT
GCACACGTCTGAACTCTTTCCCTACACGACGCTCTTCCGATCTNNNATAATTTCTTTGAGCACTATGGTC
AGCACACGTCTGAACTCTTTCCCTACACGACGCTCTTCCGATCTNNNAAAACTTGAATAGAACAAAACA
CACACGTCTGAACTCTTTCCCTACACGACGCTCTTCCGATCTNNNCCTAAATTTTAGCTTTGATGACTCA
GCACACGTCTGAACTCTTTCCCTACACGACGCTCTTCCGATCTNNNGAAACAGGACATTTTTCTAATCA
AGCACACGTCTGAACTCTTTCCCTACACGACGCTCTTCCGATCTNNNCATCATAGATAATCCACAAAGT
GCACACGTCTGAACTCTTTCCCTACACGACGCTCTTCCGATCTNNNTGCAAATATGATCCAATTTTACAC
GAGCACACGTCTGAACTCTTTCCCTACACGACGCTCTTCCGATCTNNNGCTCTCTCAAAAAAACCTTATT
GAGCACACGTCTGAACTCTTTCCCTACACGACGCTCTTCCGATCTNNNAAATGACAGAAATCACATTCA
GCACACGTCTGAACTCTTTCCCTACACGACGCTCTTCCGATCTNNNATAATACTGCCTTGATCTTTCTTC
AGAGCACACGTCTGAACTCTTTCCCTACACGACGCTCTTCCGATCTNNNACCAAGGGTTTTTTTTCTTTA
CACACGTCTGAACTCTTTCCCTACACGACGCTCTTCCGATCTNNNGAGATAGAGCTCGTCTAGTGCATC
GAGCACACGTCTGAACTCTTTCCCTACACGACGCTCTTCCGATCTNNNACTGAGGGATGACTATCTATA
AGCACACGTCTGAACTCTTTCCCTACACGACGCTCTTCCGATCTNNNCTATTTTAAAAAGCACATCAAC
GCACACGTCTGAACTCTTTCCCTACACGACGCTCTTCCGATCTNNNGACTTACTGTCAACATTTTCTTACA
ACACGTCTGAACTCTTTCCCTACACGACGCTCTTCCGATCTNNNAGTAAACATGCAGATTCTCAGCATCG
ACACGTCTGAACTCTTTCCCTACACGACGCTCTTCCGATCTNNNATAATAAATCTCTCTCTCCTTCCCATC
CACACGTCTGAACTCTTTCCCTACACGACGCTCTTCCGATCTNNNTTCAAAAAGAAGGTCAAGATACAC
GCACACGTCTGAACTCTTTCCCTACACGACGCTCTTCCGATCTNNNATCTTAGTTAACGCTAGTTTCACA
AGCACACGTCTGAACTCTTTCCCTACACGACGCTCTTCCGATCTNNNAAAATGGCAGAAAACAAAAATA
GAGCACACGTCTGAACTCTTTCCCTACACGACGCTCTTCCGATCTNNNTTATCAACAAGCTTTATTTAGC
GAGCACACGTCTGAACTCTTTCCCTACACGACGCTCTTCCGATCTNNNGAGAACTCCTTCTTTTGAGAG
GCACACGTCTGAACTCTTTCCCTACACGACGCTCTTCCGATCTNNNACCTTGCCTCAAAATAAATAAATCA
GCACACGTCTGAACTCTTTCCCTACACGACGCTCTTCCGATCTNNNACCTTGCCTCAAAATAAATAAATCA
CACACGTCTGAACTCTTTCCCTACACGACGCTCTTCCGATCTNNNGTATGAAGACTAACAGTGAATTGCA
CACACGTCTGAACTCTTTCCCTACACGACGCTCTTCCGATCTNNNATTAGAGAGAGAGAGAGAGAGTGCA
AGCACACGTCTGAACTCTTTCCCTACACGACGCTCTTCCGATCTNNNTCAATGCTTCTGTTTCCTATTATC
GAGCACACGTCTGAACTCTTTCCCTACACGACGCTCTTCCGATCTNNNTTCATGTGTGTGTATCTCTAT
GCACACGTCTGAACTCTTTCCCTACACGACGCTCTTCCGATCTNNNAATATTTACTCCAAGACTTTCCCCA
AGCACACGTCTGAACTCTTTCCCTACACGACGCTCTTCCGATCTNNNTTAAAAACTCAGAAGTGAAGA
CACACGTCTGAACTCTTTCCCTACACGACGCTCTTCCGATCTNNNACTTTATGTCATTTTCAAGTCTTCAT

GCACACGTCTGAACTCTTTCCCTACACGACGCTCTTCCGATCTNNNTATAAATAAAGTCACTGATCATG

CACACGTCTGAACTCTTTCCCTACACGACGCTCTTCCGATCTNNNATGATTATTATTTGATGGTGGTG

AGCACACGTCTGAACTCTTTCCCTACACGACGCTCTTCCGATCTNNNTATTCATCTGCCTTTCATTTTTC

GAGCACACGTCTGAACTCTTTCCCTACACGACGCTCTTCCGATCTNNNGAAAGGAACTGAGTTATTG

AGCACACGTCTGAACTCTTTCCCTACACGACGCTCTTCCGATCTNNNAAACTAAGACAAGTACCTAGT

CACACGTCTGAACTCTTTCCCTACACGACGCTCTTCCGATCTNNNCTTTATGGCTTTAGACAGTATTG

GCACACGTCTGAACTCTTTCCCTACACGACGCTCTTCCGATCTNNNAAATATATTTTTGAGATGCCTCC

GAGCACACGTCTGAACTCTTTCCCTACACGACGCTCTTCCGATCTNNNGAACCTCTCTGGAAAAAAATA

AGCACACGTCTGAACTCTTTCCCTACACGACGCTCTTCCGATCTNNNGTGCTCCATTTTATAGATATAT

AGCACACGTCTGAACTCTTTCCCTACACGACGCTCTTCCGATCTNNNCACACATATCCCCATAACAGCA

AGCACACGTCTGAACTCTTTCCCTACACGACGCTCTTCCGATCTNNNGATAAATTCAACACTTGTCCTA

GCACACGTCTGAACTCTTTCCCTACACGACGCTCTTCCGATCTNNNTGATGAAGAAGTAAAATCTCT

AGCACACGTCTGAACTCTTTCCCTACACGACGCTCTTCCGATCTNNNAGAAACTACTTTGAAAAATAG

AGCACACGTCTGAACTCTTTCCCTACACGACGCTCTTCCGATCTNNNCATAATCATGTAAGTCAATTCC

CACACGTCTGAACTCTTTCCCTACACGACGCTCTTCCGATCTNNNTCTGAAAGATACAAAGAGCAGCA

GAGCACACGTCTGAACTCTTTCCCTACACGACGCTCTTCCGATCTNNNATGTTAAAATCCTCTTACAC

AGCACACGTCTGAACTCTTTCCCTACACGACGCTCTTCCGATCTNNNTCAGATAAACTATAGGAAATG

AGCACACGTCTGAACTCTTTCCCTACACGACGCTCTTCCGATCTNNNTAGCTGAATATTTTAAAGGTG

GCACACGTCTGAACTCTTTCCCTACACGACGCTCTTCCGATCTNNNAGAACATTCTTATATGGGTCTT

AGCACACGTCTGAACTCTTTCCCTACACGACGCTCTTCCGATCTNNNAAGTAACCTACTTACGAGAAA

AGCACACGTCTGAACTCTTTCCCTACACGACGCTCTTCCGATCTNNNATATAATACCTGTACAGTACAC

AGCACACGTCTGAACTCTTTCCCTACACGACGCTCTTCCGATCTNNNAGGAAGAATTTTACAAAATCTC

GCACACGTCTGAACTCTTTCCCTACACGACGCTCTTCCGATCTNNNAGGAAGAATTTTACAAAATCTC

AGCACACGTCTGAACTCTTTCCCTACACGACGCTCTTCCGATCTNNNAGTTAATACCATTCTAGTGAG

GAGCACACGTCTGAACTCTTTCCCTACACGACGCTCTTCCGATCTNNNTTTGTGGATAGCAATAAATT

AGCACACGTCTGAACTCTTTCCCTACACGACGCTCTTCCGATCTNNNTGGGTCTTTAAGTTAAAAAATT

AGCACACGTCTGAACTCTTTCCCTACACGACGCTCTTCCGATCTNNNCTTATTTGGGTCTTAGCATAAA

GCACACGTCTGAACTCTTTCCCTACACGACGCTCTTCCGATCTNNNTCTCCTTGAAACTCTGTAAAATA

CACACGTCTGAACTCTTTCCCTACACGACGCTCTTCCGATCTNNNTCATTTAGAAGACTGCCAAAATC

AGCACACGTCTGAACTCTTTCCCTACACGACGCTCTTCCGATCTNNNATACACACACATATATATGCT

AGCACACGTCTGAACTCTTTCCCTACACGACGCTCTTCCGATCTNNNTGTTTTTAGGAAATAGATGTT

GCACACGTCTGAACTCTTTCCCTACACGACGCTCTTCCGATCTNNNTATCACTTTCACACTGTCATAAA

GCACACGTCTGAACTCTTTCCCTACACGACGCTCTTCCGATCTNNNCTACAATGGGGGAATACTTTTCA

ACACGTCTGAACTCTTTCCCTACACGACGCTCTTCCGATCTNNNGTTTCTGTGGCTCCATCTATCATCG

GCACACGTCTGAACTCTTTCCCTACACGACGCTCTTCCGATCTNNNGAATGCATTACACTAAGTATTG

CACACGTCTGAACTCTTTCCCTACACGACGCTCTTCCGATCTNNNTCTCTCTCTTTCTTTTGTCTGCATC

AGCACACGTCTGAACTCTTTCCCTACACGACGCTCTTCCGATCTNNNTGAAAAAAATGTTCTGATCACT

GCACACGTCTGAACTCTTTCCCTACACGACGCTCTTCCGATCTNNNATAATATGAGGTATCTGACACT

AGCACACGTCTGAACTCTTTCCCTACACGACGCTCTTCCGATCTNNNGTTGAGGTTGAGACGTATTATA

CACACGTCTGAACTCTTTCCCTACACGACGCTCTTCCGATCTNNNTTGCAGCAACATAACTCTAATCAT

AGCACACGTCTGAACTCTTTCCCTACACGACGCTCTTCCGATCTNNNTTTTTGAAGAAAGAAATACTGG

AGCACACGTCTGAACTCTTTCCCTACACGACGCTCTTCCGATCTNNNTGAAAGAGAAAAATGTAATG

AGCACACGTCTGAACTCTTTCCCTACACGACGCTCTTCCGATCTNNNATCTATGAATTATGCATTGCTT

AGAGCACACGTCTGAACTCTTTCCCTACACGACGCTCTTCCGATCTNNNCAAATAATGATTGAAAATCCC
AGAGCACACGTCTGAACTCTTTCCCTACACGACGCTCTTCCGATCTNNNAGAATGTCTACCGTGTTACCC
GAGCACACGTCTGAACTCTTTCCCTACACGACGCTCTTCCGATCTNNNACAGTCAAGACATATATCATTT
AGCACACGTCTGAACTCTTTCCCTACACGACGCTCTTCCGATCTNNNTACAGTATCATCTTTAAACCCTTC
AGCACACGTCTGAACTCTTTCCCTACACGACGCTCTTCCGATCTNNNTTTCATATTCATTCCTAAAGGTC
AGCACACGTCTGAACTCTTTCCCTACACGACGCTCTTCCGATCTNNNGAAGCCCTTGTTTTATCTCTCAT
AGAGCACACGTCTGAACTCTTTCCCTACACGACGCTCTTCCGATCTNNNGTATTCATTGCTTGATTGGA
AGCACACGTCTGAACTCTTTCCCTACACGACGCTCTTCCGATCTNNNCTAAACCTCATTTCTCACATCTCA
AGCACACGTCTGAACTCTTTCCCTACACGACGCTCTTCCGATCTNNNAGTGAGTACTCTTTGTATATGTC
GAGCACACGTCTGAACTCTTTCCCTACACGACGCTCTTCCGATCTNNNAAGTCTTCAACATTTCTTATTGC
GAGCACACGTCTGAACTCTTTCCCTACACGACGCTCTTCCGATCTNNNTGCAACAACAGTTTATACAAA
GAGCACACGTCTGAACTCTTTCCCTACACGACGCTCTTCCGATCTNNNATGTTTTACAAAAGCTTTTCTTC
CACACGTCTGAACTCTTTCCCTACACGACGCTCTTCCGATCTNNNATTACCTGTCAACAATATAATGCCA
GAGCACACGTCTGAACTCTTTCCCTACACGACGCTCTTCCGATCTNNNGATGTTATGTGAATGCTAGTT
GCACACGTCTGAACTCTTTCCCTACACGACGCTCTTCCGATCTNNNGGATGGGATTTTCCTGAAAACAT
GCACACGTCTGAACTCTTTCCCTACACGACGCTCTTCCGATCTNNNAACAGATTAGGCCACTATCTACAT
AGCACACGTCTGAACTCTTTCCCTACACGACGCTCTTCCGATCTNNNTTTACTTTTCAGGTAAATAAGTC
AGCACACGTCTGAACTCTTTCCCTACACGACGCTCTTCCGATCTNNNAATCCTGTTTCACCTACATATAT
GCACACGTCTGAACTCTTTCCCTACACGACGCTCTTCCGATCTNNNTAATCTCTCTCTCTCTTTCTCCAT
CACGTCTGAACTCTTTCCCTACACGACGCTCTTCCGATCTNNNAAACTTGTACCTCCTTGTTTCCATCGGA
CACACGTCTGAACTCTTTCCCTACACGACGCTCTTCCGATCTNNNTATGTCTGCATTGCTCCTAGCATCG
AGAGCACACGTCTGAACTCTTTCCCTACACGACGCTCTTCCGATCTNNNCTTTCTGACATTTAATCCACTC
AGCACACGTCTGAACTCTTTCCCTACACGACGCTCTTCCGATCTNNNTCTGTTACATACCAGAACATAAA
GCACACGTCTGAACTCTTTCCCTACACGACGCTCTTCCGATCTNNNTTTGCTAGAGGTATCTTAGATC
AGCACACGTCTGAACTCTTTCCCTACACGACGCTCTTCCGATCTNNNCAACCTGAGTTTAGTAGAGTGC
AGCACACGTCTGAACTCTTTCCCTACACGACGCTCTTCCGATCTNNNACATACATACACTTATACACACAC
CACACGTCTGAACTCTTTCCCTACACGACGCTCTTCCGATCTNNNCAATATTGGGACCTATGCATTACAT
AGCACACGTCTGAACTCTTTCCCTACACGACGCTCTTCCGATCTNNNCAAAAATAGAGAACCCATTGTC
GAGCACACGTCTGAACTCTTTCCCTACACGACGCTCTTCCGATCTNNNATTCTTTTCTTTCTATCCATCCA
GAGCACACGTCTGAACTCTTTCCCTACACGACGCTCTTCCGATCTNNNATTCTTTTCTTTCTATCCATCCA
GAGCACACGTCTGAACTCTTTCCCTACACGACGCTCTTCCGATCTNNNGCAAACCATTTTTAAAACCTTA
AGCACACGTCTGAACTCTTTCCCTACACGACGCTCTTCCGATCTNNNAATCTTAGCTCTTTATTCATTGC
AGCACACGTCTGAACTCTTTCCCTACACGACGCTCTTCCGATCTNNNATTACATAACTGCTTTTACCATT
GAGCACACGTCTGAACTCTTTCCCTACACGACGCTCTTCCGATCTNNNATCCTCTTTCTATTGCATCATGC
GAGCACACGTCTGAACTCTTTCCCTACACGACGCTCTTCCGATCTNNNATCCTCTTTCTATTGCATCATGC
GAGCACACGTCTGAACTCTTTCCCTACACGACGCTCTTCCGATCTNNNGGGCATTTAATTTAATGTCTA
GCACACGTCTGAACTCTTTCCCTACACGACGCTCTTCCGATCTNNNCCAATAAATCCATAAAGCTAACCCA
GAGCACACGTCTGAACTCTTTCCCTACACGACGCTCTTCCGATCTNNNCATACAAATGTGCTACTTGCCA
GAGCACACGTCTGAACTCTTTCCCTACACGACGCTCTTCCGATCTNNNCCAAAAGATCAAAACAAAACAA
AGCACACGTCTGAACTCTTTCCCTACACGACGCTCTTCCGATCTNNNAGATCAGCTACATCATTATTAG
GCACACGTCTGAACTCTTTCCCTACACGACGCTCTTCCGATCTNNNATATCCAGGGCCATTTCTATCATC
GAGCACACGTCTGAACTCTTTCCCTACACGACGCTCTTCCGATCTNNNTAAGTTGTAAATATCTTTCTTC
GCACACGTCTGAACTCTTTCCCTACACGACGCTCTTCCGATCTNNNATATCCAGGGCCATTTCTATCATC

GCACACGTCTGAACTCTTTCCCTACACGACGCTCTTCCGATCTNNNTATCCTGTGATATCTGACTGTAT
AGCACACGTCTGAACTCTTTCCCTACACGACGCTCTTCCGATCTNNNTTTAAAATGGCTTTCTTACATC
CACACGTCTGAACTCTTTCCCTACACGACGCTCTTCCGATCTNNNAGCAGACATTTAAATATCTGTTGC
ACACGTCTGAACTCTTTCCCTACACGACGCTCTTCCGATCTNNNTATGAAACAGCCTATTGTGTCCATC
GCACACGTCTGAACTCTTTCCCTACACGACGCTCTTCCGATCTNNNTATATACATACCTCAAAACCCAGC
GCACACGTCTGAACTCTTTCCCTACACGACGCTCTTCCGATCTNNNTATATACATACCTCAAAACCCAGC
AGCACACGTCTGAACTCTTTCCCTACACGACGCTCTTCCGATCTNNNTGCACATTTCTTATCTTAAAACT
GAGCACACGTCTGAACTCTTTCCCTACACGACGCTCTTCCGATCTNNNCCGTCTATATACTACACAGTA
AGCACACGTCTGAACTCTTTCCCTACACGACGCTCTTCCGATCTNNNCAGAACTCAACTAAAGCTAAA
AGCACACGTCTGAACTCTTTCCCTACACGACGCTCTTCCGATCTNNNGAAGAAAATGGTGAGCTCAGC
AGCACACGTCTGAACTCTTTCCCTACACGACGCTCTTCCGATCTNNNACTGAGAAATTTTGAAGTCAAT
CACACGTCTGAACTCTTTCCCTACACGACGCTCTTCCGATCTNNNATTCAAACCTTGACATCTAGATCAT
GCACACGTCTGAACTCTTTCCCTACACGACGCTCTTCCGATCTNNNAAGTTTAGGAAAGTTTCTTTCAG
GCACACGTCTGAACTCTTTCCCTACACGACGCTCTTCCGATCTNNNGAAAGAAAATCAAGGATCCATG
GCACACGTCTGAACTCTTTCCCTACACGACGCTCTTCCGATCTNNNGAAAGAAAATCAAGGATCCATG
GCACACGTCTGAACTCTTTCCCTACACGACGCTCTTCCGATCTNNNTGATCTTTGTAATGTATTGTTGA
CACACGTCTGAACTCTTTCCCTACACGACGCTCTTCCGATCTNNNATTTCATGAATCACATTATTCCCCA
GCACACGTCTGAACTCTTTCCCTACACGACGCTCTTCCGATCTNNNATGAAAATTAGAAACTGTCCTTG
AGCACACGTCTGAACTCTTTCCCTACACGACGCTCTTCCGATCTNNNCCCTACAACATATTTTGTCTATC
AGCACACGTCTGAACTCTTTCCCTACACGACGCTCTTCCGATCTNNNCTTCCATGCCTCCTTAGTAACAT
AGCACACGTCTGAACTCTTTCCCTACACGACGCTCTTCCGATCTNNNATACAAATAAGAGGCAAAAT
GCACACGTCTGAACTCTTTCCCTACACGACGCTCTTCCGATCTNNNATTTCCTTTGGACAATTACATAC
GCACACGTCTGAACTCTTTCCCTACACGACGCTCTTCCGATCTNNNAAATGTGTTAGTGCCTAAGTCC
AGCACACGTCTGAACTCTTTCCCTACACGACGCTCTTCCGATCTNNNAAATATATTTCTTCCCCAACAAG
AGCACACGTCTGAACTCTTTCCCTACACGACGCTCTTCCGATCTNNNGGGAAGACAAATAATTGAAA
GAGCACACGTCTGAACTCTTTCCCTACACGACGCTCTTCCGATCTNNNCCTTAGAGAAATAATGGGTT
GCACACGTCTGAACTCTTTCCCTACACGACGCTCTTCCGATCTNNNATGAGACTTGTTCCCTGTAACAT
AGCACACGTCTGAACTCTTTCCCTACACGACGCTCTTCCGATCTNNNCTCAAAAATATAATGCCCATTT
AGCACACGTCTGAACTCTTTCCCTACACGACGCTCTTCCGATCTNNNCTCAAAAATATAATGCCCATTT
AGCACACGTCTGAACTCTTTCCCTACACGACGCTCTTCCGATCTNNNACACTATCTTTTGTAGATGAAG
GAGCACACGTCTGAACTCTTTCCCTACACGACGCTCTTCCGATCTNNNCAACAAACACGTAAATAATTT
AAGAGCACACGTCTGAACTCTTTCCCTACACGACGCTCTTCCGATCTNNNGAAAATAACCAGCCCAAA
GAGCACACGTCTGAACTCTTTCCCTACACGACGCTCTTCCGATCTNNNGGCCAAAGGAAAATAACCA
GCACACGTCTGAACTCTTTCCCTACACGACGCTCTTCCGATCTNNNTCAAACTCTAAGAGTGTCATGC
GCACACGTCTGAACTCTTTCCCTACACGACGCTCTTCCGATCTNNNACACTGTCTTTATGTGTATGTAT
CACACGTCTGAACTCTTTCCCTACACGACGCTCTTCCGATCTNNNAATAAACTGCAAGCCCTTTATTCA
GCACACGTCTGAACTCTTTCCCTACACGACGCTCTTCCGATCTNNNTAGTGTTCAACATAAACTAATG
AGCACACGTCTGAACTCTTTCCCTACACGACGCTCTTCCGATCTNNNTGAACAGGCAATTATAAATTT
AGCACACGTCTGAACTCTTTCCCTACACGACGCTCTTCCGATCTNNNAAAAACTTCCCCAACAGATTCA
AGCACACGTCTGAACTCTTTCCCTACACGACGCTCTTCCGATCTNNNTCACTATAAGTGGATCCTTCTT
AGCACACGTCTGAACTCTTTCCCTACACGACGCTCTTCCGATCTNNNTATGATATCTGGAGTTTGGAT
GAGCACACGTCTGAACTCTTTCCCTACACGACGCTCTTCCGATCTNNNACATGTACTCTCTGAATCTAA
CACACGTCTGAACTCTTTCCCTACACGACGCTCTTCCGATCTNNNAGAGATAATGACTAGGAGGTGC

GAGCACACGTCTGAACTCTTTCCCTACACGACGCTCTTCCGATCTNNNATCATAACCTCATTGATATGCA
GCACACGTCTGAACTCTTTCCCTACACGACGCTCTTCCGATCTNNNTTAATGCTTCTAACTAACATGCCA
AGCACACGTCTGAACTCTTTCCCTACACGACGCTCTTCCGATCTNNNCATGATATAGGCTGTCACTTGCA
GAGCACACGTCTGAACTCTTTCCCTACACGACGCTCTTCCGATCTNNNTGGAAGATGAAGTTTGATATG
AGCACACGTCTGAACTCTTTCCCTACACGACGCTCTTCCGATCTNNNAGTACTATCTATGTTTTGAGTAC
AGCACACGTCTGAACTCTTTCCCTACACGACGCTCTTCCGATCTNNNATATAGGAATATGATCCCAACAA
AGAGCACACGTCTGAACTCTTTCCCTACACGACGCTCTTCCGATCTNNNAAACTGACAAACATGAAACA
CACACGTCTGAACTCTTTCCCTACACGACGCTCTTCCGATCTNNNAGTGAATCTTCTTTTGGATAGAACA
AGCACACGTCTGAACTCTTTCCCTACACGACGCTCTTCCGATCTNNNCTCTCTCTCTCTCTCTATATAC
AGCACACGTCTGAACTCTTTCCCTACACGACGCTCTTCCGATCTNNNAAGGAGTTCAAGACTAAACATA
AGCACACGTCTGAACTCTTTCCCTACACGACGCTCTTCCGATCTNNNAAGGAGTTCAAGACTAAACATA
GAGCACACGTCTGAACTCTTTCCCTACACGACGCTCTTCCGATCTNNNGCCAGGTATACAAGTACATATA
AGCACACGTCTGAACTCTTTCCCTACACGACGCTCTTCCGATCTNNNCATACCACATTCATAGCTCAATA
GAGCACACGTCTGAACTCTTTCCCTACACGACGCTCTTCCGATCTNNNTACCCTTAATGATCCTATAGAC
AGCACACGTCTGAACTCTTTCCCTACACGACGCTCTTCCGATCTNNNGGAACTTGAGATGATATATAC
GAGCACACGTCTGAACTCTTTCCCTACACGACGCTCTTCCGATCTNNNAAGGAGACAAATGAAAAGAA
AGCACACGTCTGAACTCTTTCCCTACACGACGCTCTTCCGATCTNNNTGAAAAAGCAAGAAAATGTATC
GCACACGTCTGAACTCTTTCCCTACACGACGCTCTTCCGATCTNNNATTTACTCAAACAAAGCCTGATAC
GCACACGTCTGAACTCTTTCCCTACACGACGCTCTTCCGATCTNNNACTTTCTTCTTTCCTGACTTCCATC
AGCACACGTCTGAACTCTTTCCCTACACGACGCTCTTCCGATCTNNNATTAGAAAATATGTCTTTTGGGC
AGCACACGTCTGAACTCTTTCCCTACACGACGCTCTTCCGATCTNNNATTTCTCTTTCCCTTTCTCTCCATC
AGCACACGTCTGAACTCTTTCCCTACACGACGCTCTTCCGATCTNNNCAGTTGATTTCTCTTTCCCTTCAT
AGCACACGTCTGAACTCTTTCCCTACACGACGCTCTTCCGATCTNNNCAGAGTGAGCTAGAAATACGCA
GAGCACACGTCTGAACTCTTTCCCTACACGACGCTCTTCCGATCTNNNGTAATGATAGAACACGGGTTC
GAGCACACGTCTGAACTCTTTCCCTACACGACGCTCTTCCGATCTNNNATATTTGACGTGGAATACTAG
AGCACACGTCTGAACTCTTTCCCTACACGACGCTCTTCCGATCTNNNCTCTTCAGCACAGATACTCTCAT
AGCACACGTCTGAACTCTTTCCCTACACGACGCTCTTCCGATCTNNNGATCATCTCCCTGTGAAGTTCAT
GCACACGTCTGAACTCTTTCCCTACACGACGCTCTTCCGATCTNNNTCTTATGAGAGTTGTGGTTTCCAT
GCACACGTCTGAACTCTTTCCCTACACGACGCTCTTCCGATCTNNNCTTAGACTCAGATTGGGATAGCA
AGCACACGTCTGAACTCTTTCCCTACACGACGCTCTTCCGATCTNNNACTTAAAAAATGCTGAACATCT
GCACACGTCTGAACTCTTTCCCTACACGACGCTCTTCCGATCTNNNCTTTTAAACTTTTTGTCCAAAGTCA
AGAGCACACGTCTGAACTCTTTCCCTACACGACGCTCTTCCGATCTNNNGAAGGTATCTTAAAATGTGA
AGCACACGTCTGAACTCTTTCCCTACACGACGCTCTTCCGATCTNNNCAACTAAGAAGAGAATAGAGT
GAGCACACGTCTGAACTCTTTCCCTACACGACGCTCTTCCGATCTNNNTATGTGTCAGCCAATTTCTTAT
GAGCACACGTCTGAACTCTTTCCCTACACGACGCTCTTCCGATCTNNNTATGTGTCAGCCAATTTCTTAT
GCACACGTCTGAACTCTTTCCCTACACGACGCTCTTCCGATCTNNNCAGGAAGACTATATCCAATCAAT
AGCACACGTCTGAACTCTTTCCCTACACGACGCTCTTCCGATCTNNNAAAAATATAAACCTCATGTCTG
AGCACACGTCTGAACTCTTTCCCTACACGACGCTCTTCCGATCTNNNTAAGGTTACCAATCAATTAATG
GAGCACACGTCTGAACTCTTTCCCTACACGACGCTCTTCCGATCTNNNTGAATGAATGTCACAGTATAG
GCACACGTCTGAACTCTTTCCCTACACGACGCTCTTCCGATCTNNNAGGTGTGAAAAAAGACTGTTTCA
GAGCACACGTCTGAACTCTTTCCCTACACGACGCTCTTCCGATCTNNNAGAAAATATTAAAGTGCATTG
CACACGTCTGAACTCTTTCCCTACACGACGCTCTTCCGATCTNNNGACATCTTTCACAATCGGATCATCG
CACACGTCTGAACTCTTTCCCTACACGACGCTCTTCCGATCTNNNAGTTTTACAGCAAATACTACCTCAT

ACACGTCTGAACTCTTTCCCTACACGACGCTCTTCCGATCTNNNGTGTGAGAATGGACTAATACAACA
AGCACACGTCTGAACTCTTTCCCTACACGACGCTCTTCCGATCTNNNGAAAAAAAAGATACTCACACA
GCACACGTCTGAACTCTTTCCCTACACGACGCTCTTCCGATCTNNNATTTATGGTTACAGATACGATC
AGCACACGTCTGAACTCTTTCCCTACACGACGCTCTTCCGATCTNNNCTGCCTTAAAAATGAGAGTAA
AGCACACGTCTGAACTCTTTCCCTACACGACGCTCTTCCGATCTNNNAAATCTATGTAGCTTTGAGTTT
AGCACACGTCTGAACTCTTTCCCTACACGACGCTCTTCCGATCTNNNGCCAATATGAAATGATACAAA
CACACGTCTGAACTCTTTCCCTACACGACGCTCTTCCGATCTNNNTGTTTAAGGTGTAAATTGGTTAT
AGCACACGTCTGAACTCTTTCCCTACACGACGCTCTTCCGATCTNNNTCCTTTACCGACTAATTAACTT
ACACGTCTGAACTCTTTCCCTACACGACGCTCTTCCGATCTNNNTAAGACTTATTGTTAGAAACCGAC
AGCACACGTCTGAACTCTTTCCCTACACGACGCTCTTCCGATCTNNNCAAACTCCACCTATACACTACC
GAGCACACGTCTGAACTCTTTCCCTACACGACGCTCTTCCGATCTNNNGGGTTAGAGTTTAGAGCAT
AGCACACGTCTGAACTCTTTCCCTACACGACGCTCTTCCGATCTNNNAAGAACCAAGTTGTAAATTTCT
AGCACACGTCTGAACTCTTTCCCTACACGACGCTCTTCCGATCTNNNAAACAATCACAACAGTAACTTT
GCACACGTCTGAACTCTTTCCCTACACGACGCTCTTCCGATCTNNNGTCTCTTATTGTCTTTCCATTAC
AGCACACGTCTGAACTCTTTCCCTACACGACGCTCTTCCGATCTNNNTCAGAATAGGGAATTAAACT
GCACACGTCTGAACTCTTTCCCTACACGACGCTCTTCCGATCTNNNAAAACAAAAAAAAAACACCCAA
AGCACACGTCTGAACTCTTTCCCTACACGACGCTCTTCCGATCTNNNGAAATAGGGAAGGAAGAATT
AGCACACGTCTGAACTCTTTCCCTACACGACGCTCTTCCGATCTNNNGAAATAGGGAAGGAAGAATT
AGCACACGTCTGAACTCTTTCCCTACACGACGCTCTTCCGATCTNNNAGAATCACCATCTCAAATATGT
AGCACACGTCTGAACTCTTTCCCTACACGACGCTCTTCCGATCTNNNAGAATCACCATCTCAAATATGT
CACGTCTGAACTCTTTCCCTACACGACGCTCTTCCGATCTNNNGCCTGTATATATCATTTAAAGCACAT
GCACACGTCTGAACTCTTTCCCTACACGACGCTCTTCCGATCTNNNCATAGAAAGACCCCATCTCTACA
AGCACACGTCTGAACTCTTTCCCTACACGACGCTCTTCCGATCTNNNCTTTCTGAACTCATACATTGAA
ACACGTCTGAACTCTTTCCCTACACGACGCTCTTCCGATCTNNNGCTTAAAACACACCCTATCTTTACAT
GAGCACACGTCTGAACTCTTTCCCTACACGACGCTCTTCCGATCTNNNCCAAGAGGTAGCATTATGTA
ACACGTCTGAACTCTTTCCCTACACGACGCTCTTCCGATCTNNNTTTCAAAGACTTGACAGAGGCAT
GCACACGTCTGAACTCTTTCCCTACACGACGCTCTTCCGATCTNNNTCTACTGGATTTCCTTTCTTTTAC
GCACACGTCTGAACTCTTTCCCTACACGACGCTCTTCCGATCTNNNTCTACTGGATTTCCTTTCTTTTAC
CACGTCTGAACTCTTTCCCTACACGACGCTCTTCCGATCTNNNTATGATTTCTGAGACTAGGTTAGCA
CACACGTCTGAACTCTTTCCCTACACGACGCTCTTCCGATCTNNNATCGAGGTAGAGTTAGATTCTTAC
CACACGTCTGAACTCTTTCCCTACACGACGCTCTTCCGATCTNNNTATGATTGTTTTGATGTGTTCAT
CACACGTCTGAACTCTTTCCCTACACGACGCTCTTCCGATCTNNNTATGATTGTTTTGATGTGTTCAT
CACACGTCTGAACTCTTTCCCTACACGACGCTCTTCCGATCTNNNTATGATTGTTTTGATGTGTTCAT
GCACACGTCTGAACTCTTTCCCTACACGACGCTCTTCCGATCTNNNAAAGTAAGAATGTAAAGTTGA
AGCACACGTCTGAACTCTTTCCCTACACGACGCTCTTCCGATCTNNNGATAGATAGATAGAGATATG
GCACACGTCTGAACTCTTTCCCTACACGACGCTCTTCCGATCTNNNCTCAATTAATTTTGCTTTTGGAG
GCACACGTCTGAACTCTTTCCCTACACGACGCTCTTCCGATCTNNNACATAGACAATATCATGCTAGA
GCACACGTCTGAACTCTTTCCCTACACGACGCTCTTCCGATCTNNNTACTCTGTATGTGTATATGTTG
ACACGTCTGAACTCTTTCCCTACACGACGCTCTTCCGATCTNNNTGTGGGTATATTTGTTTTATACGCA
AGCACACGTCTGAACTCTTTCCCTACACGACGCTCTTCCGATCTNNNCAGGATCCAATTTGTACTTTAA
CACACGTCTGAACTCTTTCCCTACACGACGCTCTTCCGATCTNNNACTAATACAGCAGATTTAGGAAT
GCACACGTCTGAACTCTTTCCCTACACGACGCTCTTCCGATCTNNNTACCTGTTTAATAGAATTGACC
GCACACGTCTGAACTCTTTCCCTACACGACGCTCTTCCGATCTNNNTACCTGTTTAATAGAATTGACC

GCACACGTCTGAACTCTTTCCCTACACGACGCTCTTCCGATCTNNNTGCTTCTATTTTATTTATTTGGGT

GCACACGTCTGAACTCTTTCCCTACACGACGCTCTTCCGATCTNNNTGCTTCTATTTTATTTATTTGGGT

AGCACACGTCTGAACTCTTTCCCTACACGACGCTCTTCCGATCTNNNCAAATTTGAAGTGGAATCTAAC

AGCACACGTCTGAACTCTTTCCCTACACGACGCTCTTCCGATCTNNNAAACACTGAAGGAAAAAGAAG

AGCACACGTCTGAACTCTTTCCCTACACGACGCTCTTCCGATCTNNNTTCTCTCTCTGTTTCTCTCTCAT

AGCACACGTCTGAACTCTTTCCCTACACGACGCTCTTCCGATCTNNNCTGGAATGAATTTGTGAAGGC

AGCACACGTCTGAACTCTTTCCCTACACGACGCTCTTCCGATCTNNNCTGGAATGAATTTGTGAAGGC

GAGCACACGTCTGAACTCTTTCCCTACACGACGCTCTTCCGATCTNNNATTCACACTATGACAGCCAACA

ACACGTCTGAACTCTTTCCCTACACGACGCTCTTCCGATCTNNNATGAAAGCCTAGGTCTCAAGCATCG

AGCACACGTCTGAACTCTTTCCCTACACGACGCTCTTCCGATCTNNNTGCTTTGTTTGTATGTTAGATT

AGCACACGTCTGAACTCTTTCCCTACACGACGCTCTTCCGATCTNNNTACACACACAGCATAATTTATTT

AGCACACGTCTGAACTCTTTCCCTACACGACGCTCTTCCGATCTNNNTGTATTCCTGAATTTGCTTATTT

AGAGCACACGTCTGAACTCTTTCCCTACACGACGCTCTTCCGATCTNNNCCTCCCTAATTTTATTCTTGTT

GCACACGTCTGAACTCTTTCCCTACACGACGCTCTTCCGATCTNNNTCTTCACTCACATTTTAACAAAACA

AGCACACGTCTGAACTCTTTCCCTACACGACGCTCTTCCGATCTNNNCAAAGTGCTGGGATTATAGGCA

AGCACACGTCTGAACTCTTTCCCTACACGACGCTCTTCCGATCTNNNTTATTTCTGCTAAGAACAAGTT

CACACGTCTGAACTCTTTCCCTACACGACGCTCTTCCGATCTNNNTAACAATACTTCAGAAAGCAAAGCA

ACACGTCTGAACTCTTTCCCTACACGACGCTCTTCCGATCTNNNCAGATAGATGTTTTTGGTAAGTGCA

GAGCACACGTCTGAACTCTTTCCCTACACGACGCTCTTCCGATCTNNNAAGTCTCCATCATCCATGATAC

GAGCACACGTCTGAACTCTTTCCCTACACGACGCTCTTCCGATCTNNNTTTTAAGATTTTCTCCTCATCAC

AGCACACGTCTGAACTCTTTCCCTACACGACGCTCTTCCGATCTNNNCTTCCTTCACTGAGAAAAAATAA

AGCACACGTCTGAACTCTTTCCCTACACGACGCTCTTCCGATCTNNNGTGTGTGATTCTGAAATTGAGC

AGCACACGTCTGAACTCTTTCCCTACACGACGCTCTTCCGATCTNNNATGGGTTATGTTTAGCTTCAAA

GAGCACACGTCTGAACTCTTTCCCTACACGACGCTCTTCCGATCTNNNCTTCTGCTATCTCTCTCTTGCA

GAGCACACGTCTGAACTCTTTCCCTACACGACGCTCTTCCGATCTNNNTTCCACTGTGATATTTTGCTTCA

GCACACGTCTGAACTCTTTCCCTACACGACGCTCTTCCGATCTNNNGTATGAAAAGCTCATTGAGTGCA

ACACGTCTGAACTCTTTCCCTACACGACGCTCTTCCGATCTNNNATTTTTAAATGCGGTCGTTACCATCG

AGCACACGTCTGAACTCTTTCCCTACACGACGCTCTTCCGATCTNNNAATTTCCAACCATTCCTTCTTCAT

AGCACACGTCTGAACTCTTTCCCTACACGACGCTCTTCCGATCTNNNACATACTAATACATATCCTGACC

AGCACACGTCTGAACTCTTTCCCTACACGACGCTCTTCCGATCTNNNTGTATGGTGTTATGATACATAC

GCACACGTCTGAACTCTTTCCCTACACGACGCTCTTCCGATCTNNNAATACTCAAAGAGTTAGTGATGT

AGCACACGTCTGAACTCTTTCCCTACACGACGCTCTTCCGATCTNNNATTCATGGCAGCTAATTTCTTTA

GCACACGTCTGAACTCTTTCCCTACACGACGCTCTTCCGATCTNNNCTAAGGTCTCCTAAATAGTAAGTC

AGCACACGTCTGAACTCTTTCCCTACACGACGCTCTTCCGATCTNNNGTGATAAACAGATAGACAAAG

AGCACACGTCTGAACTCTTTCCCTACACGACGCTCTTCCGATCTNNNGAAAACTTTTTAAAGCCACTTTA

GAGCACACGTCTGAACTCTTTCCCTACACGACGCTCTTCCGATCTNNNCAGTCACTAAACCAAGATTTG

AGCACACGTCTGAACTCTTTCCCTACACGACGCTCTTCCGATCTNNNGACAACAATCAGGATAGAAAAA

CACACGTCTGAACTCTTTCCCTACACGACGCTCTTCCGATCTNNNGAGTTTGAGGCTATAGTGAGCATC

AGAGCACACGTCTGAACTCTTTCCCTACACGACGCTCTTCCGATCTNNNGTTATGTACAGCAATTAGTGT

AGCACACGTCTGAACTCTTTCCCTACACGACGCTCTTCCGATCTNNNGAATGCATCTCTTTGTATAACTT

GCACACGTCTGAACTCTTTCCCTACACGACGCTCTTCCGATCTNNNAGAAAGAGGGCCTTAGTTTTACA

AGCACACGTCTGAACTCTTTCCCTACACGACGCTCTTCCGATCTNNNCTAAAATAACGTGGATTTCAGG

GCACACGTCTGAACTCTTTCCCTACACGACGCTCTTCCGATCTNNNAGATATAACTAGAGGCCGAGCAT

GCACACGTCTGAACTCTTTCCCTACACGACGCTCTTCCGATCTNNNAGATATAACTAGAGGCCGAGCAT
AGCACACGTCTGAACTCTTTCCCTACACGACGCTCTTCCGATCTNNNGATCTATTGATCTATTTTGCGTT
AGCACACGTCTGAACTCTTTCCCTACACGACGCTCTTCCGATCTNNNGCTAATGACTTACAGATCAAATT
AGCACACGTCTGAACTCTTTCCCTACACGACGCTCTTCCGATCTNNNGAACGGTTGGTTATCTAAATAA
AGCACACGTCTGAACTCTTTCCCTACACGACGCTCTTCCGATCTNNNCATATGACTTCTTAGACCTACAG
GAGCACACGTCTGAACTCTTTCCCTACACGACGCTCTTCCGATCTNNNGCACTTTCCTTATTCTGTTTTTA
AGCACACGTCTGAACTCTTTCCCTACACGACGCTCTTCCGATCTNNNTACACTCACATGCCTTGATATCAT
AGCACACGTCTGAACTCTTTCCCTACACGACGCTCTTCCGATCTNNNCTGTGATCTTAGTGTCCTTATA
GAGCACACGTCTGAACTCTTTCCCTACACGACGCTCTTCCGATCTNNNCCTGTATGAAAAACAAAAACAA
AGCACACGTCTGAACTCTTTCCCTACACGACGCTCTTCCGATCTNNNTAGAATTCTTGTATCAGTTCTC
AGCACACGTCTGAACTCTTTCCCTACACGACGCTCTTCCGATCTNNNTTTTACAATTAGATTAACAGGG
AGCACACGTCTGAACTCTTTCCCTACACGACGCTCTTCCGATCTNNNGCATGAAATCTACCATTTTCTAT
GCACACGTCTGAACTCTTTCCCTACACGACGCTCTTCCGATCTNNNAAAAGAAAGAAAAAGTGTTCAA
GAGCACACGTCTGAACTCTTTCCCTACACGACGCTCTTCCGATCTNNNAAGAGATACAAAGGTAGGAA
GCACACGTCTGAACTCTTTCCCTACACGACGCTCTTCCGATCTNNNTTTTTTAACCAGAATGTTTCAGAC
AGCACACGTCTGAACTCTTTCCCTACACGACGCTCTTCCGATCTNNNAGGTTCATAGATATGTGTTAT
GAGCACACGTCTGAACTCTTTCCCTACACGACGCTCTTCCGATCTNNNTATAATGGAGACTACCAGAG
AGCACACGTCTGAACTCTTTCCCTACACGACGCTCTTCCGATCTNNNATTGAGATAAAACAATCCATCT
AGCACACGTCTGAACTCTTTCCCTACACGACGCTCTTCCGATCTNNNTAGAAATTTCAAAAATAGTGCCA
AGCACACGTCTGAACTCTTTCCCTACACGACGCTCTTCCGATCTNNNTATAAGTTTGTATTCTTCCACCA
GCACACGTCTGAACTCTTTCCCTACACGACGCTCTTCCGATCTNNNTGTACATGTTTGACTGTAATTGCA
AGCACACGTCTGAACTCTTTCCCTACACGACGCTCTTCCGATCTNNNTTCAAAGCCACTAAATTCTCTTA
GCACACGTCTGAACTCTTTCCCTACACGACGCTCTTCCGATCTNNNGAAGGGGAGAGTGATAGATTT
GCACACGTCTGAACTCTTTCCCTACACGACGCTCTTCCGATCTNNNGAAGGGGAGAGTGATAGATTT
CACACGTCTGAACTCTTTCCCTACACGACGCTCTTCCGATCTNNNGTTAGGTTTCAATAAGCTTTACACA
CACACGTCTGAACTCTTTCCCTACACGACGCTCTTCCGATCTNNNGTTAGGTTTCAATAAGCTTTACACA
AGCACACGTCTGAACTCTTTCCCTACACGACGCTCTTCCGATCTNNNAACAGATTTACAAATTCCAGAAA
GAGCACACGTCTGAACTCTTTCCCTACACGACGCTCTTCCGATCTNNNTAAAAATGTTACCTAGACAGT
GCACACGTCTGAACTCTTTCCCTACACGACGCTCTTCCGATCTNNNGTATGCATGCTTGTGTATTTATA
AGCACACGTCTGAACTCTTTCCCTACACGACGCTCTTCCGATCTNNNTGGGCTTAATCCTGATAATAAA
GCACACGTCTGAACTCTTTCCCTACACGACGCTCTTCCGATCTNNNTTACCAAATCTAAACTCTGTTTCC
GAGCACACGTCTGAACTCTTTCCCTACACGACGCTCTTCCGATCTNNNATCCATTCAATACACACTTATT
CACGTCTGAACTCTTTCCCTACACGACGCTCTTCCGATCTNNNAAAAAGAAAAATTAGGTAGTCCCAT
GAGCACACGTCTGAACTCTTTCCCTACACGACGCTCTTCCGATCTNNNTGGTACACTATTCTAATTACT
AGCACACGTCTGAACTCTTTCCCTACACGACGCTCTTCCGATCTNNNAATTAGTTTCCTTTGTCATCCTA
AGCACACGTCTGAACTCTTTCCCTACACGACGCTCTTCCGATCTNNNAGAGAAAACATCTAGACAATAC
GCACACGTCTGAACTCTTTCCCTACACGACGCTCTTCCGATCTNNNGTAATTGTATGCACTTTGAGTAT
AGCACACGTCTGAACTCTTTCCCTACACGACGCTCTTCCGATCTNNNGAGTTTTGTAGTTTTAGCACTA
ACACGTCTGAACTCTTTCCCTACACGACGCTCTTCCGATCTNNNTATACCTAAATTGGTTAACAAAGGCA
GCACACGTCTGAACTCTTTCCCTACACGACGCTCTTCCGATCTNNNAACAGTGTTATAGACTAGTAACCC
AGCACACGTCTGAACTCTTTCCCTACACGACGCTCTTCCGATCTNNNTGTAAAGGTCTTACACATTTTA
GAGCACACGTCTGAACTCTTTCCCTACACGACGCTCTTCCGATCTNNNTAATTGGAATCCTGGATAAAA
AGCACACGTCTGAACTCTTTCCCTACACGACGCTCTTCCGATCTNNNGCATTACAAAATATAGCTACACA

CACACGTCTGAACTCTTTCCCTACACGACGCTCTTCCGATCTNNNTCATAATACTACTATCTGAGAGTAT

AGAGCACACGTCTGAACTCTTTCCCTACACGACGCTCTTCCGATCTNNNTACATGCTGAAGTACTTAGA

AGCACACGTCTGAACTCTTTCCCTACACGACGCTCTTCCGATCTNNNCCTAACATTTGTGGAAAATTTC

GAGCACACGTCTGAACTCTTTCCCTACACGACGCTCTTCCGATCTNNNACAAAGAAGGACATCAAAAA

GCACACGTCTGAACTCTTTCCCTACACGACGCTCTTCCGATCTNNNAATACCATATTTTTGTTCCTTCACA

GAGCACACGTCTGAACTCTTTCCCTACACGACGCTCTTCCGATCTNNNCAATCTCTCATTTATAGGCCC

GAGCACACGTCTGAACTCTTTCCCTACACGACGCTCTTCCGATCTNNNCTAATGGAAGTGAACAAATG

GAGCACACGTCTGAACTCTTTCCCTACACGACGCTCTTCCGATCTNNNACATGTTCCATCATATTCTAT

AGCACACGTCTGAACTCTTTCCCTACACGACGCTCTTCCGATCTNNNCTCATATAACTCATAGGAAGTG

AGCACACGTCTGAACTCTTTCCCTACACGACGCTCTTCCGATCTNNNAGAGACTCATATAACTCATAGG

GCACACGTCTGAACTCTTTCCCTACACGACGCTCTTCCGATCTNNNTGTATGTATGAATGTGTATGGA

GAGCACACGTCTGAACTCTTTCCCTACACGACGCTCTTCCGATCTNNNTATATGAGTGTGTGTGTTACA

AGCACACGTCTGAACTCTTTCCCTACACGACGCTCTTCCGATCTNNNTTGATGCTTTTCCTCATGTTCAT

GCACACGTCTGAACTCTTTCCCTACACGACGCTCTTCCGATCTNNNCTCAATAAACCCCCCTTTATAACA

GAGCACACGTCTGAACTCTTTCCCTACACGACGCTCTTCCGATCTNNNAGATTTTATTAGTTAAGCTCTC

GAGCACACGTCTGAACTCTTTCCCTACACGACGCTCTTCCGATCTNNNAATATAGGGTGGAATAAAAC

AGCACACGTCTGAACTCTTTCCCTACACGACGCTCTTCCGATCTNNNGAGTGAGAAATCTAAGGAGA

AGCACACGTCTGAACTCTTTCCCTACACGACGCTCTTCCGATCTNNNTTAAAAGAGTTAACAAAAGAA

AGCACACGTCTGAACTCTTTCCCTACACGACGCTCTTCCGATCTNNNAGGATTTAAGGGTACTAATTTC

GAGCACACGTCTGAACTCTTTCCCTACACGACGCTCTTCCGATCTNNNTTTTTTTCTCCAAGTATTTTCCA

GCACACGTCTGAACTCTTTCCCTACACGACGCTCTTCCGATCTNNNTAAAATCAATGTGCAAAAATCAC

GCACACGTCTGAACTCTTTCCCTACACGACGCTCTTCCGATCTNNNCTTAGCCAGAAAATAACAAGAC

GAGCACACGTCTGAACTCTTTCCCTACACGACGCTCTTCCGATCTNNNTACATGATCAAACACAGATGT

GCACACGTCTGAACTCTTTCCCTACACGACGCTCTTCCGATCTNNNAATGAATCAGTGTGAACTTTTC

GAGCACACGTCTGAACTCTTTCCCTACACGACGCTCTTCCGATCTNNNATAATCTTTTCTCCCTGAAATA

GCACACGTCTGAACTCTTTCCCTACACGACGCTCTTCCGATCTNNNATATTTCTTGTCTTCCAAAGTTG

GCACACGTCTGAACTCTTTCCCTACACGACGCTCTTCCGATCTNNNTTATTATACCTTGTAATGTTGCCC

GCACACGTCTGAACTCTTTCCCTACACGACGCTCTTCCGATCTNNNTATATAGAAACTTCCACCTCCACAT

GCACACGTCTGAACTCTTTCCCTACACGACGCTCTTCCGATCTNNNGCATTATTCACAATAGCCAATAT

AGCACACGTCTGAACTCTTTCCCTACACGACGCTCTTCCGATCTNNNGATTTTCTAAAGCAAGTCTGA

AGCACACGTCTGAACTCTTTCCCTACACGACGCTCTTCCGATCTNNNCTTCTCCTTTTCTGTGTCTTTCAT

GAGCACACGTCTGAACTCTTTCCCTACACGACGCTCTTCCGATCTNNNGTCAATCCATTTAATTCGACC

GAGCACACGTCTGAACTCTTTCCCTACACGACGCTCTTCCGATCTNNNGTCAATCCATTTAATTCGACC

GCACACGTCTGAACTCTTTCCCTACACGACGCTCTTCCGATCTNNNCTTTTAGAAGGTAGCTGACATT

GAGCACACGTCTGAACTCTTTCCCTACACGACGCTCTTCCGATCTNNNGCTTCAACATATGACTTTGAG

AGCACACGTCTGAACTCTTTCCCTACACGACGCTCTTCCGATCTNNNCCTAAGGGAATAATCAGAACAT

CACACGTCTGAACTCTTTCCCTACACGACGCTCTTCCGATCTNNNAATAGATTACTGGTTGTTAGGAAC

AGCACACGTCTGAACTCTTTCCCTACACGACGCTCTTCCGATCTNNNGCTCTCCTGTAGTTTTTCATATA

GAGCACACGTCTGAACTCTTTCCCTACACGACGCTCTTCCGATCTNNNTTAGGAATAAAAGGGCTTGAT

GCACACGTCTGAACTCTTTCCCTACACGACGCTCTTCCGATCTNNNCAAACACTGATAACTTACTAGTG

AGCACACGTCTGAACTCTTTCCCTACACGACGCTCTTCCGATCTNNNAAATCTATAATCCGAGTGTTCTT

AGCACACGTCTGAACTCTTTCCCTACACGACGCTCTTCCGATCTNNNACTCATTGTAGAAAATTTGGAA

AGCACACGTCTGAACTCTTTCCCTACACGACGCTCTTCCGATCTNNNTTCTAGTGACCACAATTATGAA

AGCACACGTCTGAACTCTTTCCCTACACGACGCTCTTCCGATCTNNNTGCTAACATAATTAACATAAGC
AGCACACGTCTGAACTCTTTCCCTACACGACGCTCTTCCGATCTNNNACACACACATATATGTCTCTTG
AGCACACGTCTGAACTCTTTCCCTACACGACGCTCTTCCGATCTNNNCATTCACAGTATTACACCAAGAC
AGCACACGTCTGAACTCTTTCCCTACACGACGCTCTTCCGATCTNNNTATTTTAGTTCTGTTAGCACT
AGCACACGTCTGAACTCTTTCCCTACACGACGCTCTTCCGATCTNNNAATATCAACACAAAAGCCATTT
GCACACGTCTGAACTCTTTCCCTACACGACGCTCTTCCGATCTNNNTTTACCAGTAATGCAAATCAAAT
GCACACGTCTGAACTCTTTCCCTACACGACGCTCTTCCGATCTNNNTGTCACATGTCAACATACTATAT
AGCACACGTCTGAACTCTTTCCCTACACGACGCTCTTCCGATCTNNNCACTTAGGAATTTGTTAAAATG
AGCACACGTCTGAACTCTTTCCCTACACGACGCTCTTCCGATCTNNNTGAATCTTTTCTTATGGACAGT
AGCACACGTCTGAACTCTTTCCCTACACGACGCTCTTCCGATCTNNNTCCATGAATGTTCTATTCACAT
AGCACACGTCTGAACTCTTTCCCTACACGACGCTCTTCCGATCTNNNATCTGTCTACTGTTGTTTACTT
AGCACACGTCTGAACTCTTTCCCTACACGACGCTCTTCCGATCTNNNTAGATCATTGTTTTCTTAATGC
AGCACACGTCTGAACTCTTTCCCTACACGACGCTCTTCCGATCTNNNAGTATGTCTGGATAGCTGAGC
AGCACACGTCTGAACTCTTTCCCTACACGACGCTCTTCCGATCTNNNTTCTCTTTTATACAAAGTTGCTC
AGCACACGTCTGAACTCTTTCCCTACACGACGCTCTTCCGATCTNNNGACTAGAATATCTAATGGGCA
GCACACGTCTGAACTCTTTCCCTACACGACGCTCTTCCGATCTNNNACTCCATTTGAAAGATGTTTTAC
GCACACGTCTGAACTCTTTCCCTACACGACGCTCTTCCGATCTNNNACTCCATTTGAAAGATGTTTTAC
AGCACACGTCTGAACTCTTTCCCTACACGACGCTCTTCCGATCTNNNGTTTATTCAAAGACCCATTTAG
CACACGTCTGAACTCTTTCCCTACACGACGCTCTTCCGATCTNNNTTTTATGCCTGGCCATAAAGCATCG
AGCACACGTCTGAACTCTTTCCCTACACGACGCTCTTCCGATCTNNNTTATGTAACAAAAAGTCTGGAT
GAGCACACGTCTGAACTCTTTCCCTACACGACGCTCTTCCGATCTNNNATAGATATGGATGGAGAGAC
GAGCACACGTCTGAACTCTTTCCCTACACGACGCTCTTCCGATCTNNNATAGATATGGATGGAGAGAC
AGCACACGTCTGAACTCTTTCCCTACACGACGCTCTTCCGATCTNNNTATGTCTTCTTTTCCCTGTATTT
GCACACGTCTGAACTCTTTCCCTACACGACGCTCTTCCGATCTNNNTTTAGATGCAAGAGGCTTATAGC
GCACACGTCTGAACTCTTTCCCTACACGACGCTCTTCCGATCTNNNGTAAGGCAGGCTAAATATGATC
AGCACACGTCTGAACTCTTTCCCTACACGACGCTCTTCCGATCTNNNTGATGGTAAAGAGTGATTGA
AGCACACGTCTGAACTCTTTCCCTACACGACGCTCTTCCGATCTNNNATTATACTAGGAAACAAGGCTT
AGCACACGTCTGAACTCTTTCCCTACACGACGCTCTTCCGATCTNNNATTATACTAGGAAACAAGGCTT
AGCACACGTCTGAACTCTTTCCCTACACGACGCTCTTCCGATCTNNNAAGAAGCAAGCAAATACTTTA
AGCACACGTCTGAACTCTTTCCCTACACGACGCTCTTCCGATCTNNNGCATGGAGTTATTAAATTTCGT
GCACACGTCTGAACTCTTTCCCTACACGACGCTCTTCCGATCTNNNAAATCATAATCAAAACCTGTCTCC
AGCACACGTCTGAACTCTTTCCCTACACGACGCTCTTCCGATCTNNNGATGCTGAGAAGAGTCTATAT
AGCACACGTCTGAACTCTTTCCCTACACGACGCTCTTCCGATCTNNNTTTTATTTTGCTTCTCTTGTCTT
CACACGTCTGAACTCTTTCCCTACACGACGCTCTTCCGATCTNNNTTAAATATTTTCACCTACATGCTCA
CACACGTCTGAACTCTTTCCCTACACGACGCTCTTCCGATCTNNNTATCAATTACCTCTTCATGGTTACA
CACACGTCTGAACTCTTTCCCTACACGACGCTCTTCCGATCTNNNTATCAATTACCTCTTCATGGTTACA
GAGCACACGTCTGAACTCTTTCCCTACACGACGCTCTTCCGATCTNNNCTAGTAGTATGCAACTGAGA
GCACACGTCTGAACTCTTTCCCTACACGACGCTCTTCCGATCTNNNCAAAATTGCCTGATTGAGAACCAT
AGCACACGTCTGAACTCTTTCCCTACACGACGCTCTTCCGATCTNNNAACAATTTCACCCAATTTAAAA
AGCACACGTCTGAACTCTTTCCCTACACGACGCTCTTCCGATCTNNNCTGAGGACCTATATACAATGT
AGCACACGTCTGAACTCTTTCCCTACACGACGCTCTTCCGATCTNNNCTTCAGAAGATTGTGATCCAAT
AGCACACGTCTGAACTCTTTCCCTACACGACGCTCTTCCGATCTNNNCTTCAGAAGATTGTGATCCAAT
AGCACACGTCTGAACTCTTTCCCTACACGACGCTCTTCCGATCTNNNATTTTGAAGTAGTTTCATCTTC

ACACGTCTGAACTCTTTCCCTACACGACGCTCTTCCGATCTNNNAGCATAGTTGATTCTGTGTGCATCG
AGAGCACACGTCTGAACTCTTTCCCTACACGACGCTCTTCCGATCTNNNTGGAATTCTCTCTCTCTCTAT
GAGCACACGTCTGAACTCTTTCCCTACACGACGCTCTTCCGATCTNNNATTCCTCCTTCTATATTTTGAG
GAGCACACGTCTGAACTCTTTCCCTACACGACGCTCTTCCGATCTNNNATTCCTCCTTCTATATTTTGAG
AGCACACGTCTGAACTCTTTCCCTACACGACGCTCTTCCGATCTNNNCAACAAATCAATAAAAGCACTA
GCACACGTCTGAACTCTTTCCCTACACGACGCTCTTCCGATCTNNNCTTTTATGGTAAACAGTTCCTTC
AGCACACGTCTGAACTCTTTCCCTACACGACGCTCTTCCGATCTNNNTATTATGCTCTTAAGATCTGTGG
GCACACGTCTGAACTCTTTCCCTACACGACGCTCTTCCGATCTNNNAATTTTTGCAGAGTGTTTCAGCAT
CACACGTCTGAACTCTTTCCCTACACGACGCTCTTCCGATCTNNNAATTCATCAGACTTGCCTTCCATCG
CACACGTCTGAACTCTTTCCCTACACGACGCTCTTCCGATCTNNNAATTCATCAGACTTGCCTTCCATCG
AGCACACGTCTGAACTCTTTCCCTACACGACGCTCTTCCGATCTNNNAAACTTTCTGGAGAACTTGTAT
AGCACACGTCTGAACTCTTTCCCTACACGACGCTCTTCCGATCTNNNAACCAGATTTACTATATAGCCAT
AGCACACGTCTGAACTCTTTCCCTACACGACGCTCTTCCGATCTNNNTTGAGCAAATGCATACATTATA
AGCACACGTCTGAACTCTTTCCCTACACGACGCTCTTCCGATCTNNNAGAAATAAAAGCTGTTATTGG
AGCACACGTCTGAACTCTTTCCCTACACGACGCTCTTCCGATCTNNNCAGCATTTTTTCCATATACTTCTC
GCACACGTCTGAACTCTTTCCCTACACGACGCTCTTCCGATCTNNNAAATATGATTGCTTTCCAAGTAC
AGAGCACACGTCTGAACTCTTTCCCTACACGACGCTCTTCCGATCTNNNGCACTAGTGAAGTCCTTAA
AGCACACGTCTGAACTCTTTCCCTACACGACGCTCTTCCGATCTNNNTCTAGAAGATCAATATCAGAAG
CACACGTCTGAACTCTTTCCCTACACGACGCTCTTCCGATCTNNNTTACATATATATGAGAGCAGAGCC
CACACGTCTGAACTCTTTCCCTACACGACGCTCTTCCGATCTNNNTTACATATATATGAGAGCAGAGCC
CACACGTCTGAACTCTTTCCCTACACGACGCTCTTCCGATCTNNNCTCTTCCTCTCTCCATATATACAT
GAGCACACGTCTGAACTCTTTCCCTACACGACGCTCTTCCGATCTNNNTTTATGAAAAAAAAAAACCCC
AGCACACGTCTGAACTCTTTCCCTACACGACGCTCTTCCGATCTNNNATTTAGACAGGATATGCACTTA
GCACACGTCTGAACTCTTTCCCTACACGACGCTCTTCCGATCTNNNTTCCTCAAACTAGATAATCTCAG
GCACACGTCTGAACTCTTTCCCTACACGACGCTCTTCCGATCTNNNGAAGTGATCTAATGGTGTCCCAT
AGCACACGTCTGAACTCTTTCCCTACACGACGCTCTTCCGATCTNNNTAGATTGGTTCAGTTTTTCTGTA
GAGCACACGTCTGAACTCTTTCCCTACACGACGCTCTTCCGATCTNNNCTTGTTAATAAGCTTTCTTCTA
AGCACACGTCTGAACTCTTTCCCTACACGACGCTCTTCCGATCTNNNTTAAAGAAACATTTTGATTTGG
CACACGTCTGAACTCTTTCCCTACACGACGCTCTTCCGATCTNNNCCTGTTCTCCTTTTTTCTTATTCCATC
GCACACGTCTGAACTCTTTCCCTACACGACGCTCTTCCGATCTNNNTATTGTCAATTAAATTCCACCTCC
GCACACGTCTGAACTCTTTCCCTACACGACGCTCTTCCGATCTNNNATAGGCTCTCTAGTCATGATAAA
GAGCACACGTCTGAACTCTTTCCCTACACGACGCTCTTCCGATCTNNNTCACATATTTTACCTGAAGTG
GCACACGTCTGAACTCTTTCCCTACACGACGCTCTTCCGATCTNNNCATTGCCAAGTCTCTTAGAACAT
GCACACGTCTGAACTCTTTCCCTACACGACGCTCTTCCGATCTNNNTCTCCATCTTTGGTCTGTTCCATC
GAGCACACGTCTGAACTCTTTCCCTACACGACGCTCTTCCGATCTNNNAAGACAATAAAATGTGTGGT
AGCACACGTCTGAACTCTTTCCCTACACGACGCTCTTCCGATCTNNNGTTCCCAACAATTGCATAAGCA
GAGCACACGTCTGAACTCTTTCCCTACACGACGCTCTTCCGATCTNNNCTTTGGGAGAAATCAATAAGA
AGCACACGTCTGAACTCTTTCCCTACACGACGCTCTTCCGATCTNNNCAGGGAAAATTTTGTAAGTTA
AGCACACGTCTGAACTCTTTCCCTACACGACGCTCTTCCGATCTNNNGTACCATTTGAATATCTTGTGAT
AGCACACGTCTGAACTCTTTCCCTACACGACGCTCTTCCGATCTNNNACAATAATGGGTTTTAATCCCTA
AGCACACGTCTGAACTCTTTCCCTACACGACGCTCTTCCGATCTNNNTATTTATACAGAAACAAAAACG
GCACACGTCTGAACTCTTTCCCTACACGACGCTCTTCCGATCTNNNTCATATACATCATTGCAAATAGC
GAGCACACGTCTGAACTCTTTCCCTACACGACGCTCTTCCGATCTNNNAAAACATGCATTTTAGAATTG

GAGCACACGTCTGAACTCTTTCCCTACACGACGCTCTTCCGATCTNNNAAAACATGCATTTTAGAATTG
GCACACGTCTGAACTCTTTCCCTACACGACGCTCTTCCGATCTNNNGTTGTATGTTTCCAGGAATTTATG
GAGCACACGTCTGAACTCTTTCCCTACACGACGCTCTTCCGATCTNNNTTTACAGCTTTTTACATGTCAA
GCACACGTCTGAACTCTTTCCCTACACGACGCTCTTCCGATCTNNNTTTTCTCTATGTTTTAGGGCCCAT
ACACGTCTGAACTCTTTCCCTACACGACGCTCTTCCGATCTNNNATCATCCTACTATGCTTTGCCATCGG
AGCACACGTCTGAACTCTTTCCCTACACGACGCTCTTCCGATCTNNNCATCGTAGTCTATCAAACAAAT
GAGCACACGTCTGAACTCTTTCCCTACACGACGCTCTTCCGATCTNNNCTAGCTTAAGAGAGAAAAGAA
GAGCACACGTCTGAACTCTTTCCCTACACGACGCTCTTCCGATCTNNNTGTGTGAATCAAGTATATTTG
AGCACACGTCTGAACTCTTTCCCTACACGACGCTCTTCCGATCTNNNCTAAACCTAGACCTGGAATATT
AGCACACGTCTGAACTCTTTCCCTACACGACGCTCTTCCGATCTNNNTTCTCTCTCTTTTTTCACTCTATCA
GCACACGTCTGAACTCTTTCCCTACACGACGCTCTTCCGATCTNNNCTTTGATGAGTACAACGTAGGCA
AGCACACGTCTGAACTCTTTCCCTACACGACGCTCTTCCGATCTNNNTAATATGATGTGATAAGAATGG
GCACACGTCTGAACTCTTTCCCTACACGACGCTCTTCCGATCTNNNGAAATTGAGAGAATTCTTTCCATG
GAGCACACGTCTGAACTCTTTCCCTACACGACGCTCTTCCGATCTNNNAAATAGAAGAAAGGAGACAA
ACACGTCTGAACTCTTTCCCTACACGACGCTCTTCCGATCTNNNCTTATAAAAAGAGACAAGAGCTCCA
ACACGTCTGAACTCTTTCCCTACACGACGCTCTTCCGATCTNNNCTCTTATAAAAAGAGACAAGAGCCA
AGCACACGTCTGAACTCTTTCCCTACACGACGCTCTTCCGATCTNNNTATATACATGCACTCACACACCAT
CACACGTCTGAACTCTTTCCCTACACGACGCTCTTCCGATCTNNNGATACTGTACACATAGAACTCCCAT
GAGCACACGTCTGAACTCTTTCCCTACACGACGCTCTTCCGATCTNNNATGATTTGAGAAAGCTAATGA
CACACGTCTGAACTCTTTCCCTACACGACGCTCTTCCGATCTNNNCCCTAAATCAGAATTTTTAACCACAT
GCACACGTCTGAACTCTTTCCCTACACGACGCTCTTCCGATCTNNNGATGGAACCCTCTACAAAAAATA
GCACACGTCTGAACTCTTTCCCTACACGACGCTCTTCCGATCTNNNATGGATCTAAAAAGGCTAAGTATG
AGCACACGTCTGAACTCTTTCCCTACACGACGCTCTTCCGATCTNNNTTTCTTGTCTTTTCTGAGAAAAA
AGAGCACACGTCTGAACTCTTTCCCTACACGACGCTCTTCCGATCTNNNATGGATGTGATATCTATTCTG
AGCACACGTCTGAACTCTTTCCCTACACGACGCTCTTCCGATCTNNNTTTTTTGTAAAGTGGATCCTCTT
AGCACACGTCTGAACTCTTTCCCTACACGACGCTCTTCCGATCTNNNTTTTTTGTAAAGTGGATCCTCTT
GAGCACACGTCTGAACTCTTTCCCTACACGACGCTCTTCCGATCTNNNATTGTGATTCTGACTGTAGTGG
AGCACACGTCTGAACTCTTTCCCTACACGACGCTCTTCCGATCTNNNCTTATTGAAGGGAAAACACAAA
AGCACACGTCTGAACTCTTTCCCTACACGACGCTCTTCCGATCTNNNTGCTGCAATTTCATGTAATATAG
GAGCACACGTCTGAACTCTTTCCCTACACGACGCTCTTCCGATCTNNNGAAACTTTGAGATTACTCTTCA
CACACGTCTGAACTCTTTCCCTACACGACGCTCTTCCGATCTNNNAAATGTCTATTCTTTTAAGCCACCA
GCACACGTCTGAACTCTTTCCCTACACGACGCTCTTCCGATCTNNNTCCCTAATGAATCTAACTAATCAC
AGCACACGTCTGAACTCTTTCCCTACACGACGCTCTTCCGATCTNNNAGTCACTTTTTGTTAACAGAATA
AGCACACGTCTGAACTCTTTCCCTACACGACGCTCTTCCGATCTNNNCTATACCATGAAAAATGCTGAGG
GAGCACACGTCTGAACTCTTTCCCTACACGACGCTCTTCCGATCTNNNAGGAAGAAAACAAACCAAATA
AGCACACGTCTGAACTCTTTCCCTACACGACGCTCTTCCGATCTNNNGCACTCTATACTCAAAATAACTT
AGCACACGTCTGAACTCTTTCCCTACACGACGCTCTTCCGATCTNNNCTTTCTTTTCTTTCTTGCCTTCCA
AGCACACGTCTGAACTCTTTCCCTACACGACGCTCTTCCGATCTNNNCTTTCTTTTCTTTCTTGCCTTCCA
AGCACACGTCTGAACTCTTTCCCTACACGACGCTCTTCCGATCTNNNTTAAATATTGGCATGTGTTAAGC
GCACACGTCTGAACTCTTTCCCTACACGACGCTCTTCCGATCTNNNAAATTTTGTGTAATGGATAGACG
CACACGTCTGAACTCTTTCCCTACACGACGCTCTTCCGATCTNNNCTGAAACGTGTGTATTTTTTGCATG
AGCACACGTCTGAACTCTTTCCCTACACGACGCTCTTCCGATCTNNNTTATTAGATTTCAGGTCCTTTCT
GCACACGTCTGAACTCTTTCCCTACACGACGCTCTTCCGATCTNNNTAGATAACAACCTTATTTGTCCTCA

GAGCACACGTCTGAACTCTTTCCCTACACGACGCTCTTCCGATCTNNNATCAATTACATAAACCAGGAA
GAGCACACGTCTGAACTCTTTCCCTACACGACGCTCTTCCGATCTNNNAGTTGAAGTATCAGCTTTATC
GCACACGTCTGAACTCTTTCCCTACACGACGCTCTTCCGATCTNNNGCCATTCCTCTATATGTATACATC
GCACACGTCTGAACTCTTTCCCTACACGACGCTCTTCCGATCTNNNCTCCCCTGATAAACCATCTATCAT
AGCACACGTCTGAACTCTTTCCCTACACGACGCTCTTCCGATCTNNNGAACATTATCATTGGTTCATCA
AGCACACGTCTGAACTCTTTCCCTACACGACGCTCTTCCGATCTNNNTAGATGGTAAATCATGAGACTT
AGCACACGTCTGAACTCTTTCCCTACACGACGCTCTTCCGATCTNNNGAATTTAAGGTAGTGCTCACAA
GAGCACACGTCTGAACTCTTTCCCTACACGACGCTCTTCCGATCTNNNTAAGAGAATGTACTGGTTTTA
AGCACACGTCTGAACTCTTTCCCTACACGACGCTCTTCCGATCTNNNAACAATAGCCAAGCAATAAATA
CACACGTCTGAACTCTTTCCCTACACGACGCTCTTCCGATCTNNNACCTACTTAACTCTGATTCATTTCAT
AGCACACGTCTGAACTCTTTCCCTACACGACGCTCTTCCGATCTNNNGGTTTTTTTGTTTTGTTTTGTT
AGCACACGTCTGAACTCTTTCCCTACACGACGCTCTTCCGATCTNNNAGCAAGTAGTGTTTTTTACTTTT
GCACACGTCTGAACTCTTTCCCTACACGACGCTCTTCCGATCTNNNGAAGCAGGAGAATCACTTAAGC
GCACACGTCTGAACTCTTTCCCTACACGACGCTCTTCCGATCTNNNTTTGCTGAATAAAAGTCATCATT
CACACGTCTGAACTCTTTCCCTACACGACGCTCTTCCGATCTNNNAAAGATGCAGATTCAAATTTAGG
ACACGTCTGAACTCTTTCCCTACACGACGCTCTTCCGATCTNNNCTTAAAAAGTCAGCTTCAAGTGCAT
AGCACACGTCTGAACTCTTTCCCTACACGACGCTCTTCCGATCTNNNACATGAACATTTTTTATTTCTGC
AGCACACGTCTGAACTCTTTCCCTACACGACGCTCTTCCGATCTNNNCCACACAGTTTTTGGAATATATA
AGCACACGTCTGAACTCTTTCCCTACACGACGCTCTTCCGATCTNNNTTTCTCTCTTTTGTTAGTCTTTC
AGCACACGTCTGAACTCTTTCCCTACACGACGCTCTTCCGATCTNNNTAGGTCTTCCTTCACTTCTTTCA
AGCACACGTCTGAACTCTTTCCCTACACGACGCTCTTCCGATCTNNNCATAATGAGACATGTCCATTTC
GCACACGTCTGAACTCTTTCCCTACACGACGCTCTTCCGATCTNNNTAAGTTTATTCTCCATAGAGTAGT
AGCACACGTCTGAACTCTTTCCCTACACGACGCTCTTCCGATCTNNNATGTATCCTCACTACCCACTCAT
CACACGTCTGAACTCTTTCCCTACACGACGCTCTTCCGATCTNNNTGGGGTTCCTATTTATCTATTTACAT
GAGCACACGTCTGAACTCTTTCCCTACACGACGCTCTTCCGATCTNNNTTTGAGTCCTAATTCCCTTGAT
AGCACACGTCTGAACTCTTTCCCTACACGACGCTCTTCCGATCTNNNTCCTAATAAATTAATCAACATA
AGCACACGTCTGAACTCTTTCCCTACACGACGCTCTTCCGATCTNNNTACTTATCTTGTTCAAATTTCCC
GAGCACACGTCTGAACTCTTTCCCTACACGACGCTCTTCCGATCTNNNAATTGAGAATATGGTGAGGA
GCACACGTCTGAACTCTTTCCCTACACGACGCTCTTCCGATCTNNNATAGGTCCTTGAAGAGCTTATTA
CACACGTCTGAACTCTTTCCCTACACGACGCTCTTCCGATCTNNNATGCTATTCATGTTCATGTCTTACA
GCACACGTCTGAACTCTTTCCCTACACGACGCTCTTCCGATCTNNNCATTTCATTCTTTTTTGCTTTTGCAT
AGCACACGTCTGAACTCTTTCCCTACACGACGCTCTTCCGATCTNNNATATACCCATTTCCAACTAACAT
ACACGTCTGAACTCTTTCCCTACACGACGCTCTTCCGATCTNNNAAAAATAACTCACAAGTCTCTTGCAT
GAGCACACGTCTGAACTCTTTCCCTACACGACGCTCTTCCGATCTNNNTTGGAGTGTGTATATGAAATA
GCACACGTCTGAACTCTTTCCCTACACGACGCTCTTCCGATCTNNNCTTACATCAGTTGTTTGTTTGATC
GAGCACACGTCTGAACTCTTTCCCTACACGACGCTCTTCCGATCTNNNTATCCATCCATGACTTATGAT
GAGCACACGTCTGAACTCTTTCCCTACACGACGCTCTTCCGATCTNNNGCGATTTATTTTTTTTCTGACA
GAGCACACGTCTGAACTCTTTCCCTACACGACGCTCTTCCGATCTNNNATCTTCCATTGGTGTATCTCTCA
GAGCACACGTCTGAACTCTTTCCCTACACGACGCTCTTCCGATCTNNNTGACCTCTAACTAATATCTCTCT
GAGCACACGTCTGAACTCTTTCCCTACACGACGCTCTTCCGATCTNNNTTCTTGCGATGTAAGAAATTA
CACACGTCTGAACTCTTTCCCTACACGACGCTCTTCCGATCTNNNCTTCCAAAAACTAGTTCAGATTGCA
GCACACGTCTGAACTCTTTCCCTACACGACGCTCTTCCGATCTNNNACATACTGAATAGTCACAGATTTC
AGCACACGTCTGAACTCTTTCCCTACACGACGCTCTTCCGATCTNNNTCATTTTTATTATTAGCTGCATCT

AGCACACGTCTGAACTCTTTCCCTACACGACGCTCTTCCGATCTNNNGTTAGGATAGAATGGAAGGT
AGCACACGTCTGAACTCTTTCCCTACACGACGCTCTTCCGATCTNNNGACTGATTTAAATCAAACCTTT
CACACGTCTGAACTCTTTCCCTACACGACGCTCTTCCGATCTNNNCAGGCTTTTATATCTTGGATTTTC
AGCACACGTCTGAACTCTTTCCCTACACGACGCTCTTCCGATCTNNNTTGAATGAGGGAAGAGAAAA
AGCACACGTCTGAACTCTTTCCCTACACGACGCTCTTCCGATCTNNNTTGAATGAGGGAAGAGAAAA
GCACACGTCTGAACTCTTTCCCTACACGACGCTCTTCCGATCTNNNCTCACTTTTCTCTACCTAACATACA
AGCACACGTCTGAACTCTTTCCCTACACGACGCTCTTCCGATCTNNNATAATACATGCATACATACCAT
ACACGTCTGAACTCTTTCCCTACACGACGCTCTTCCGATCTNNNCATTTGAGAGAGATCTTCCACCATC
GCACACGTCTGAACTCTTTCCCTACACGACGCTCTTCCGATCTNNNCTCATTGCTTGAAGAATCATGC
GCACACGTCTGAACTCTTTCCCTACACGACGCTCTTCCGATCTNNNGATGGAAATTGACAGCTATAAA
AGCACACGTCTGAACTCTTTCCCTACACGACGCTCTTCCGATCTNNNCAATATAGTGAGACCCTGTTAT
GAGCACACGTCTGAACTCTTTCCCTACACGACGCTCTTCCGATCTNNNTTCTAAAGATGTAATTTCTCCC
GCACACGTCTGAACTCTTTCCCTACACGACGCTCTTCCGATCTNNNTTATAGCCAAAGACCAAAATAA
CACACGTCTGAACTCTTTCCCTACACGACGCTCTTCCGATCTNNNTCCAAAATAAATATGCTAAAACCT
AGCACACGTCTGAACTCTTTCCCTACACGACGCTCTTCCGATCTNNNTGGTTTGGAGTAGTTTTATCT
AGCACACGTCTGAACTCTTTCCCTACACGACGCTCTTCCGATCTNNNACATCTCTTATACTCTTGTTCAT
GCACACGTCTGAACTCTTTCCCTACACGACGCTCTTCCGATCTNNNCAAAAATATAGAAGAAACCACC
GAGCACACGTCTGAACTCTTTCCCTACACGACGCTCTTCCGATCTNNNAAACAAAAACAAATACTGTCA
GCACACGTCTGAACTCTTTCCCTACACGACGCTCTTCCGATCTNNNTTTATACCTGTATGTCTGTTTTGC
ACACGTCTGAACTCTTTCCCTACACGACGCTCTTCCGATCTNNNAGCCCAAAGAGGAGAATAAACATC
GAGCACACGTCTGAACTCTTTCCCTACACGACGCTCTTCCGATCTNNNACGGTAATAGCTATGTGACT
GCACACGTCTGAACTCTTTCCCTACACGACGCTCTTCCGATCTNNNGTTTTCAGATAATCAGAACATGT
AGCACACGTCTGAACTCTTTCCCTACACGACGCTCTTCCGATCTNNNTTTTACATCTATGTTCATCAGG
AGCACACGTCTGAACTCTTTCCCTACACGACGCTCTTCCGATCTNNNTTCATTCCTTTTGTCTTTCTTTC
AGCACACGTCTGAACTCTTTCCCTACACGACGCTCTTCCGATCTNNNAAAAAACCAAACAGTAAACAA
AGCACACGTCTGAACTCTTTCCCTACACGACGCTCTTCCGATCTNNNCAGAACAGTAAATGAGATAAT
AGCACACGTCTGAACTCTTTCCCTACACGACGCTCTTCCGATCTNNNCTTTTATTCTAACTTCTAGCTGG
AGCACACGTCTGAACTCTTTCCCTACACGACGCTCTTCCGATCTNNNGTAATGTATGCTTTGGAAATA
AGCACACGTCTGAACTCTTTCCCTACACGACGCTCTTCCGATCTNNNAAAACCTGGATATCAGATTGTA
ACACGTCTGAACTCTTTCCCTACACGACGCTCTTCCGATCTNNNTTATCTCTCATATCTCATTGCCCATCG
AGCACACGTCTGAACTCTTTCCCTACACGACGCTCTTCCGATCTNNNAAGTCTCAAAATGAGTAAGAG
GCACACGTCTGAACTCTTTCCCTACACGACGCTCTTCCGATCTNNNAATCTCTGTCTTTCTCTATCTCTC
GCACACGTCTGAACTCTTTCCCTACACGACGCTCTTCCGATCTNNNAATCTCTGTCTTTCTCTATCTCTC
AGCACACGTCTGAACTCTTTCCCTACACGACGCTCTTCCGATCTNNNTTGTTATAGTTCTGCCTATGAA
GAGCACACGTCTGAACTCTTTCCCTACACGACGCTCTTCCGATCTNNNAAACTTATTGAAAGTATCACC
AGCACACGTCTGAACTCTTTCCCTACACGACGCTCTTCCGATCTNNNTGTTTACTTAAAAAGAAAGAG
AGCACACGTCTGAACTCTTTCCCTACACGACGCTCTTCCGATCTNNNTGTGTTTTGTTTCTATATTTGG
AGAGCACACGTCTGAACTCTTTCCCTACACGACGCTCTTCCGATCTNNNCCTTGGCTGTTTCACATATA
AGCACACGTCTGAACTCTTTCCCTACACGACGCTCTTCCGATCTNNNACTGGATACTCAGTATAAAACA
GAGCACACGTCTGAACTCTTTCCCTACACGACGCTCTTCCGATCTNNNAAATGAAAAGTGTCAAGATA
AGCACACGTCTGAACTCTTTCCCTACACGACGCTCTTCCGATCTNNNCTTGTTTGTTGTTAAAACTTGG
AGCACACGTCTGAACTCTTTCCCTACACGACGCTCTTCCGATCTNNNTCACTTATATTTTATCTTTCTCCC
GCACACGTCTGAACTCTTTCCCTACACGACGCTCTTCCGATCTNNNGAGTCTACTACAGCAATCTATAA

AGCACACGTCTGAACTCTTTCCCTACACGACGCTCTTCCGATCTNNNAGATCAATACCAGGCTAAATAT
GAGCACACGTCTGAACTCTTTCCCTACACGACGCTCTTCCGATCTNNNTATCACAGGACAGTTTGATTA
AGCACACGTCTGAACTCTTTCCCTACACGACGCTCTTCCGATCTNNNAATATGCATACCAGACAGAATA
AGCACACGTCTGAACTCTTTCCCTACACGACGCTCTTCCGATCTNNNTAGGATGCAAAAGTAACTTTCT
AGCACACGTCTGAACTCTTTCCCTACACGACGCTCTTCCGATCTNNNGCAATGTTTTCTAGTAAGTATGT
AGCACACGTCTGAACTCTTTCCCTACACGACGCTCTTCCGATCTNNNTATGCTACTATGAAATCAGTGTT
AGCACACGTCTGAACTCTTTCCCTACACGACGCTCTTCCGATCTNNNAAAGAGTACATGTATGTGTGTA
AGCACACGTCTGAACTCTTTCCCTACACGACGCTCTTCCGATCTNNNAAGAAGAACATATCAGGCAAT
AGCACACGTCTGAACTCTTTCCCTACACGACGCTCTTCCGATCTNNNGCGTGTAAATAATCATATTTCCT
AGCACACGTCTGAACTCTTTCCCTACACGACGCTCTTCCGATCTNNNTGAGTGGGATAGGTTGTAATC
AGCACACGTCTGAACTCTTTCCCTACACGACGCTCTTCCGATCTNNNGTTTACTGTATCATCTTTAACCTC
AGCACACGTCTGAACTCTTTCCCTACACGACGCTCTTCCGATCTNNNTGCTAATAAGTAGGAGCTAAATT
AGCACACGTCTGAACTCTTTCCCTACACGACGCTCTTCCGATCTNNNTGCTAATAAGTAGGAGCTAAATT
AGCACACGTCTGAACTCTTTCCCTACACGACGCTCTTCCGATCTNNNCTTGCTACACAACATATATATGC
AGCACACGTCTGAACTCTTTCCCTACACGACGCTCTTCCGATCTNNNAAAATAAAGAATTAACAGCCCT
AGCACACGTCTGAACTCTTTCCCTACACGACGCTCTTCCGATCTNNNTTTGTGGAAGTGGAGAGATTCA
CACACGTCTGAACTCTTTCCCTACACGACGCTCTTCCGATCTNNNAAGCTTGTCATAAATGCCTGCATCC
GAGCACACGTCTGAACTCTTTCCCTACACGACGCTCTTCCGATCTNNNTGATATACATAGCGCAAGTATT
GAGCACACGTCTGAACTCTTTCCCTACACGACGCTCTTCCGATCTNNNTGTGAGGTATAATATACAAAA
GAGCACACGTCTGAACTCTTTCCCTACACGACGCTCTTCCGATCTNNNTGAAAATTCTCAGGGCTTATTC
AGCACACGTCTGAACTCTTTCCCTACACGACGCTCTTCCGATCTNNNTTTGTTAATAACCCCAATAAGTC
GAGCACACGTCTGAACTCTTTCCCTACACGACGCTCTTCCGATCTNNNCAAGACTTTCCTTTGTAATTGT
AGCACACGTCTGAACTCTTTCCCTACACGACGCTCTTCCGATCTNNNGTAAACAGGCCTGAAAAATGC
ACACGTCTGAACTCTTTCCCTACACGACGCTCTTCCGATCTNNNTAAATATAGGGACAACAGGTATGCA
AGCACACGTCTGAACTCTTTCCCTACACGACGCTCTTCCGATCTNNNCAATCCCCTACAATAAATCTCTC
GAGCACACGTCTGAACTCTTTCCCTACACGACGCTCTTCCGATCTNNNCTCTTCCTACCTGAAAAAGTGC
GCACACGTCTGAACTCTTTCCCTACACGACGCTCTTCCGATCTNNNAAAATTCATCATCCCAACTAATGCA
AGCACACGTCTGAACTCTTTCCCTACACGACGCTCTTCCGATCTNNNCTTGTTAATGGTGGTACATAAA
AGCACACGTCTGAACTCTTTCCCTACACGACGCTCTTCCGATCTNNNATCATCCATAGTGTGTAATGCC
GCACACGTCTGAACTCTTTCCCTACACGACGCTCTTCCGATCTNNNCTTTCTCTCATTCTTTCTTGTTTCA
GCACACGTCTGAACTCTTTCCCTACACGACGCTCTTCCGATCTNNNTTTAGGTAATATTACAGGGCAAT
AGCACACGTCTGAACTCTTTCCCTACACGACGCTCTTCCGATCTNNNAATAATCAGTTACAGGTAGAAA
AGCACACGTCTGAACTCTTTCCCTACACGACGCTCTTCCGATCTNNNAAGGGAATTTTTTAGATTATAC
AGCACACGTCTGAACTCTTTCCCTACACGACGCTCTTCCGATCTNNNTTGGGAAACTATCTTCTATTCTT
AGCACACGTCTGAACTCTTTCCCTACACGACGCTCTTCCGATCTNNNCTTGCTGAAGATAAGGGAAATA
GAGCACACGTCTGAACTCTTTCCCTACACGACGCTCTTCCGATCTNNNCAAAGTGTAACCTTAGCAATAC
GAGCACACGTCTGAACTCTTTCCCTACACGACGCTCTTCCGATCTNNNGAAATGAACTTGGAATGCTTA
AGCACACGTCTGAACTCTTTCCCTACACGACGCTCTTCCGATCTNNNACTTAGCTTTATCTTTCATGAGG
AGCACACGTCTGAACTCTTTCCCTACACGACGCTCTTCCGATCTNNNCTTTTTTATGGCTGCATAGTATT
AGCACACGTCTGAACTCTTTCCCTACACGACGCTCTTCCGATCTNNNAAGTACAGCAGACATTATCTTCC
GAGCACACGTCTGAACTCTTTCCCTACACGACGCTCTTCCGATCTNNNGCTACTTCAGAATTATCAGAA
GAGCACACGTCTGAACTCTTTCCCTACACGACGCTCTTCCGATCTNNNTACTTCTCTACCTTCATCTCATA
GCACACGTCTGAACTCTTTCCCTACACGACGCTCTTCCGATCTNNNGAATCTTGTGTTTGTTAATCACTC

GCACACGTCTGAACTCTTTCCCTACACGACGCTCTTCCGATCTNNNATGCTTTAACGTAGAATCAGAT
CACACGTCTGAACTCTTTCCCTACACGACGCTCTTCCGATCTNNNATCTTTTTGAGCATGTAATTCTC
AGCACACGTCTGAACTCTTTCCCTACACGACGCTCTTCCGATCTNNNCTTTGTCAGATAAAGAAGTGA
GAGCACACGTCTGAACTCTTTCCCTACACGACGCTCTTCCGATCTNNNATGGACATGATTTGAAAAC
GAGCACACGTCTGAACTCTTTCCCTACACGACGCTCTTCCGATCTNNNTAATACATTTCCATGGACATG
AGCACACGTCTGAACTCTTTCCCTACACGACGCTCTTCCGATCTNNNTAGTGAAGTGGTCTCTAATCA
GCACACGTCTGAACTCTTTCCCTACACGACGCTCTTCCGATCTNNNTATACATCAAGATCAATTCTGA
AGCACACGTCTGAACTCTTTCCCTACACGACGCTCTTCCGATCTNNNCCAATAATGTATGTTTCTGTCT
GCACACGTCTGAACTCTTTCCCTACACGACGCTCTTCCGATCTNNNGACAGAACCTGCTTCCTTATCAT
GCACACGTCTGAACTCTTTCCCTACACGACGCTCTTCCGATCTNNNCCTTGTTAAACTCTAGTTCTAGT
AGCACACGTCTGAACTCTTTCCCTACACGACGCTCTTCCGATCTNNNTTGAGTTAATTTTTATGTGTG
GCACACGTCTGAACTCTTTCCCTACACGACGCTCTTCCGATCTNNNTATTGTGACATAAATTTGACAG
GCACACGTCTGAACTCTTTCCCTACACGACGCTCTTCCGATCTNNNATAGGCATTTCTCAAAATACAT
GAGCACACGTCTGAACTCTTTCCCTACACGACGCTCTTCCGATCTNNNTTTCATGAGAAATAGTATCCA
AGCACACGTCTGAACTCTTTCCCTACACGACGCTCTTCCGATCTNNNTATCCATGAATAACGATCAAG
GCACACGTCTGAACTCTTTCCCTACACGACGCTCTTCCGATCTNNNTAGAAAAGTGGTAGCTTACAAT
AGCACACGTCTGAACTCTTTCCCTACACGACGCTCTTCCGATCTNNNGTTAAGGTCTAGTCAAGAAGA
GAGCACACGTCTGAACTCTTTCCCTACACGACGCTCTTCCGATCTNNNGTTTGTATTGAGTATTTGCTA
GCACACGTCTGAACTCTTTCCCTACACGACGCTCTTCCGATCTNNNGTTGATAGACCCTTAGATTGGC
GCACACGTCTGAACTCTTTCCCTACACGACGCTCTTCCGATCTNNNACTAAAGCTTACTGAGATACATC
GCACACGTCTGAACTCTTTCCCTACACGACGCTCTTCCGATCTNNNGATTACAATTCGTGATGAGATT
AGCACACGTCTGAACTCTTTCCCTACACGACGCTCTTCCGATCTNNNAAAGAGACAGAGAGAGAGAA
GAGCACACGTCTGAACTCTTTCCCTACACGACGCTCTTCCGATCTNNNCCTTTTTTTTCTGAAGAATT
GCACACGTCTGAACTCTTTCCCTACACGACGCTCTTCCGATCTNNNTTTCAAGATATGTGAACTCTGA
AGCACACGTCTGAACTCTTTCCCTACACGACGCTCTTCCGATCTNNNCATCTTATCCCTACACTCCTACA
ACACGTCTGAACTCTTTCCCTACACGACGCTCTTCCGATCTNNNAGTGGAGGGATAATTCAGAGCAT
GCACACGTCTGAACTCTTTCCCTACACGACGCTCTTCCGATCTNNNAAAAATGGCAAGAAATTCCTAC
GCACACGTCTGAACTCTTTCCCTACACGACGCTCTTCCGATCTNNNGGGAAGAAGCTTGTATTTTATT
CACACGTCTGAACTCTTTCCCTACACGACGCTCTTCCGATCTNNNTGACAAAGAATGAGGAATAGTC
ACACGTCTGAACTCTTTCCCTACACGACGCTCTTCCGATCTNNNCTCTCCTAACTCTGACTTGGCATCG
GCACACGTCTGAACTCTTTCCCTACACGACGCTCTTCCGATCTNNNTTAGTAAGTCATTCGATGCAAC
GCACACGTCTGAACTCTTTCCCTACACGACGCTCTTCCGATCTNNNTTTTTCTCTGCATATGTATTCAT
CACACGTCTGAACTCTTTCCCTACACGACGCTCTTCCGATCTNNNATTAGCCTAAATAAATTCATGGAC
CACACGTCTGAACTCTTTCCCTACACGACGCTCTTCCGATCTNNNTTTTGTGCCTTTGTTGAAAATCAT
AGCACACGTCTGAACTCTTTCCCTACACGACGCTCTTCCGATCTNNNCACTATTGGCTGAAAAGACTA
CACACGTCTGAACTCTTTCCCTACACGACGCTCTTCCGATCTNNNCTCTCTTAGCATATCCTGAGTCAT
GCACACGTCTGAACTCTTTCCCTACACGACGCTCTTCCGATCTNNNATATCTTATCTAGAATCCCAAGC
GAGCACACGTCTGAACTCTTTCCCTACACGACGCTCTTCCGATCTNNNTAACATCTCAAACTATCAAAC
GCACACGTCTGAACTCTTTCCCTACACGACGCTCTTCCGATCTNNNGCTTGTCTTTTATGACTGATATG
GCACACGTCTGAACTCTTTCCCTACACGACGCTCTTCCGATCTNNNAAGTATAGCACTAGAACAAAA
GCACACGTCTGAACTCTTTCCCTACACGACGCTCTTCCGATCTNNNGTGATAAGTATAGCACTAGAAC
AGCACACGTCTGAACTCTTTCCCTACACGACGCTCTTCCGATCTNNNAAATAAAACAGCTATTTCCCAT
GCACACGTCTGAACTCTTTCCCTACACGACGCTCTTCCGATCTNNNAATAGATTCCTCAAGCAAGAGC

CACACGTCTGAACTCTTTCCCTACACGACGCTCTTCCGATCTNNNAATTTCGGAGGAGAGAAGAAACA
AGCACACGTCTGAACTCTTTCCCTACACGACGCTCTTCCGATCTNNNTTAGTTGTGTCTATACTTGTAG
AGCACACGTCTGAACTCTTTCCCTACACGACGCTCTTCCGATCTNNNGGCATAGATAGGTAATTTCAA
GCACACGTCTGAACTCTTTCCCTACACGACGCTCTTCCGATCTNNNCTATTTCAGAAAGACTTCCAAG
AGCACACGTCTGAACTCTTTCCCTACACGACGCTCTTCCGATCTNNNTAGTTAATGATGCAAACACTG
AGCACACGTCTGAACTCTTTCCCTACACGACGCTCTTCCGATCTNNNAGAGAAATTTAGCCAAGGAT
AGCACACGTCTGAACTCTTTCCCTACACGACGCTCTTCCGATCTNNNCATTTTGATTGTTCATTTGGTA
AGCACACGTCTGAACTCTTTCCCTACACGACGCTCTTCCGATCTNNNCGCATGTGATAAAACTGTATA
GCACACGTCTGAACTCTTTCCCTACACGACGCTCTTCCGATCTNNNATTGTGGAGTTTTTGACATTAT
GAGCACACGTCTGAACTCTTTCCCTACACGACGCTCTTCCGATCTNNNTAAGAAATTCTTACCAGCCA
AGCACACGTCTGAACTCTTTCCCTACACGACGCTCTTCCGATCTNNNTGTGTGTGTGTGTATATATAT
CACACGTCTGAACTCTTTCCCTACACGACGCTCTTCCGATCTNNNAATAGAGAAGTAGCTGCAGATCAT
CACACGTCTGAACTCTTTCCCTACACGACGCTCTTCCGATCTNNNATCAGTGTTTATCAAGGATATTG
AGCACACGTCTGAACTCTTTCCCTACACGACGCTCTTCCGATCTNNNAAGAAGACAGTTTAAATCCTA
GCACACGTCTGAACTCTTTCCCTACACGACGCTCTTCCGATCTNNNCTCTTCATCACAAGCTATAGTGC
GCACACGTCTGAACTCTTTCCCTACACGACGCTCTTCCGATCTNNNTCATGTTCATATGGTCTTATAG
ACACGTCTGAACTCTTTCCCTACACGACGCTCTTCCGATCTNNNCAGAGAACAGATATCAAAAACAAC
AGCACACGTCTGAACTCTTTCCCTACACGACGCTCTTCCGATCTNNNTTGGGTAATATAATATGGTGA
GCACACGTCTGAACTCTTTCCCTACACGACGCTCTTCCGATCTNNNCATTATTTGTTCCTTTTCATGG
GAGCACACGTCTGAACTCTTTCCCTACACGACGCTCTTCCGATCTNNNCTACTAAAAGACAACTGGGT
AGCACACGTCTGAACTCTTTCCCTACACGACGCTCTTCCGATCTNNNCCCAAAGCTACATATTTGATAA
GCACACGTCTGAACTCTTTCCCTACACGACGCTCTTCCGATCTNNNGTTCCTTTGTATGAGTGCATATC
GAGCACACGTCTGAACTCTTTCCCTACACGACGCTCTTCCGATCTNNNTTATCTTACTCTTTTTGCTACC
GCACACGTCTGAACTCTTTCCCTACACGACGCTCTTCCGATCTNNNAATTGTCTCCTTGTACAAATTTT
GCACACGTCTGAACTCTTTCCCTACACGACGCTCTTCCGATCTNNNTTTGGTAGCAAAGAAATTGTCC
GCACACGTCTGAACTCTTTCCCTACACGACGCTCTTCCGATCTNNNAAAAATACGTATTTGTTTTGGTG
CACACGTCTGAACTCTTTCCCTACACGACGCTCTTCCGATCTNNNAACAACTAGGCCTATCCTTCCAT
GAGCACACGTCTGAACTCTTTCCCTACACGACGCTCTTCCGATCTNNNAATGATTACCATTTACTGAAC
GCACACGTCTGAACTCTTTCCCTACACGACGCTCTTCCGATCTNNNAATAAAACATTCTTGTGGTGCCA
GCACACGTCTGAACTCTTTCCCTACACGACGCTCTTCCGATCTNNNGAATTCCACAATGTTTTTGTAAC
AGCACACGTCTGAACTCTTTCCCTACACGACGCTCTTCCGATCTNNNTGACTGAGAAAGATTAACTTAG
GAGCACACGTCTGAACTCTTTCCCTACACGACGCTCTTCCGATCTNNNGTGAGTGAATGGATAAAGA
GAGCACACGTCTGAACTCTTTCCCTACACGACGCTCTTCCGATCTNNNTATAGTACCTCTGTCTATCTG
GAGCACACGTCTGAACTCTTTCCCTACACGACGCTCTTCCGATCTNNNTATAGTACCTCTGTCTATCTG
GCACACGTCTGAACTCTTTCCCTACACGACGCTCTTCCGATCTNNNATTAGATTTTCCTCTTTTCCTCTC
GAGCACACGTCTGAACTCTTTCCCTACACGACGCTCTTCCGATCTNNNCACAAAAGTAACTCCATTCAT
GCACACGTCTGAACTCTTTCCCTACACGACGCTCTTCCGATCTNNNTTTCTTCTCTCTCTGTCTTTACCAT
AGCACACGTCTGAACTCTTTCCCTACACGACGCTCTTCCGATCTNNNTACCATTTTTCCTTCATAACATG
CACACGTCTGAACTCTTTCCCTACACGACGCTCTTCCGATCTNNNAAACTTTATTTAGCCACATCACCA
GAGCACACGTCTGAACTCTTTCCCTACACGACGCTCTTCCGATCTNNNAATTGAAAACTCAATCCCTTTT
AGCACACGTCTGAACTCTTTCCCTACACGACGCTCTTCCGATCTNNNGTACCAAAAAAACAAAACAAAG
AGCACACGTCTGAACTCTTTCCCTACACGACGCTCTTCCGATCTNNNAACTAAAAAGCAGTGAATACA
AGCACACGTCTGAACTCTTTCCCTACACGACGCTCTTCCGATCTNNNTATATATCAAAACCAGAAACGG

AGCACACGTCTGAACTCTTTCCCTACACGACGCTCTTCCGATCTNNNGTAAGACGGAAATAATCAATCT

AGCACACGTCTGAACTCTTTCCCTACACGACGCTCTTCCGATCTNNNAGAAAAAACTTACATACAAAGC

AGCACACGTCTGAACTCTTTCCCTACACGACGCTCTTCCGATCTNNNAGAAAAAACTTACATACAAAGC

GAGCACACGTCTGAACTCTTTCCCTACACGACGCTCTTCCGATCTNNNACCAAATTATATACAACACCAA

AGCACACGTCTGAACTCTTTCCCTACACGACGCTCTTCCGATCTNNNTTTAATTTTTACGTGGCAAAATC

AGCACACGTCTGAACTCTTTCCCTACACGACGCTCTTCCGATCTNNNTAGCTGGATATAGATAAAAGC

GCACACGTCTGAACTCTTTCCCTACACGACGCTCTTCCGATCTNNNATCTTTAGCTGAACTAAGGAAAA

AGCACACGTCTGAACTCTTTCCCTACACGACGCTCTTCCGATCTNNNAAAGTGCTTCTTGAGAGATATT

GAGCACACGTCTGAACTCTTTCCCTACACGACGCTCTTCCGATCTNNNGGCCAGATTGAAATCTATATTC

AGCACACGTCTGAACTCTTTCCCTACACGACGCTCTTCCGATCTNNNTTGCTTACTAACTCACTTCTATC

AGCACACGTCTGAACTCTTTCCCTACACGACGCTCTTCCGATCTNNNGAAATTGCTCTCTCTAGATTTAC

AGCACACGTCTGAACTCTTTCCCTACACGACGCTCTTCCGATCTNNNATGATGACCTATGACTCCTTTCA

AGCACACGTCTGAACTCTTTCCCTACACGACGCTCTTCCGATCTNNNATGTCAGAGTACATCACTTATA

AGCACACGTCTGAACTCTTTCCCTACACGACGCTCTTCCGATCTNNNTTATCACCATATTATACCTCAGG

GAGCACACGTCTGAACTCTTTCCCTACACGACGCTCTTCCGATCTNNNGGCTTCTAGAGGTTTTATAAAT

AGCACACGTCTGAACTCTTTCCCTACACGACGCTCTTCCGATCTNNNTTACCTAAGTATACACCCTTTGC

AGCACACGTCTGAACTCTTTCCCTACACGACGCTCTTCCGATCTNNNACTTATAATTTGGTATGAATGAT

AGCACACGTCTGAACTCTTTCCCTACACGACGCTCTTCCGATCTNNNATTCAACTTTGTCACTTTTCTTAC

GAGCACACGTCTGAACTCTTTCCCTACACGACGCTCTTCCGATCTNNNTCATCAGAACATGAAATAAAT

AGCACACGTCTGAACTCTTTCCCTACACGACGCTCTTCCGATCTNNNATAATATAAATTTGAGCATGGC

GCACACGTCTGAACTCTTTCCCTACACGACGCTCTTCCGATCTNNNGTTAGGATGGGATGGAGATATA

CACACGTCTGAACTCTTTCCCTACACGACGCTCTTCCGATCTNNNTCAAAAAAATGGTGATTAGGGCA

GCACACGTCTGAACTCTTTCCCTACACGACGCTCTTCCGATCTNNNCGCTCACATTTCTTCTTATAATTCA

AGCACACGTCTGAACTCTTTCCCTACACGACGCTCTTCCGATCTNNNTTAGATTTTAGTATCCACGGAAG

GAGCACACGTCTGAACTCTTTCCCTACACGACGCTCTTCCGATCTNNNGTGAGTAGCCTTAGATTTTAG

GAGCACACGTCTGAACTCTTTCCCTACACGACGCTCTTCCGATCTNNNTTTCTTATTTAAAGCACAGATC

GAGCACACGTCTGAACTCTTTCCCTACACGACGCTCTTCCGATCTNNNTTTCTTATTTAAAGCACAGATC

GAGCACACGTCTGAACTCTTTCCCTACACGACGCTCTTCCGATCTNNNAGCTTATATTTACTGAGCTCTT

AGCACACGTCTGAACTCTTTCCCTACACGACGCTCTTCCGATCTNNNGAGGAAGGGTTGCATTTTAGCA

GAGCACACGTCTGAACTCTTTCCCTACACGACGCTCTTCCGATCTNNNGAGATGATTCATGATTTTGTCT

GAGCACACGTCTGAACTCTTTCCCTACACGACGCTCTTCCGATCTNNNGAGATGATTCATGATTTTGTCT

GCACACGTCTGAACTCTTTCCCTACACGACGCTCTTCCGATCTNNNTATGAAGGGCATCTTGAAAATCA

GCACACGTCTGAACTCTTTCCCTACACGACGCTCTTCCGATCTNNNGAATGAACTAAGCTAACATATCA

GCACACGTCTGAACTCTTTCCCTACACGACGCTCTTCCGATCTNNNAAAGACAAAATAAGCAACTGCCAT

AGCACACGTCTGAACTCTTTCCCTACACGACGCTCTTCCGATCTNNNCAAAATGGAAAATGCATCTTAT

GAGCACACGTCTGAACTCTTTCCCTACACGACGCTCTTCCGATCTNNNAACCCTTTGGAAAATCACTATT

AGCACACGTCTGAACTCTTTCCCTACACGACGCTCTTCCGATCTNNNATGTAACATCATCTTTAAAGTGC

AGCACACGTCTGAACTCTTTCCCTACACGACGCTCTTCCGATCTNNNTCTAAGAAAAGGAGGATTTGT

AGCACACGTCTGAACTCTTTCCCTACACGACGCTCTTCCGATCTNNNGGTTCTATATGGTCAGAGAAAT

GAGCACACGTCTGAACTCTTTCCCTACACGACGCTCTTCCGATCTNNNCAGAGCTTCAAATTATCTCTATC

GAGCACACGTCTGAACTCTTTCCCTACACGACGCTCTTCCGATCTNNNCAAAGAAAGGATTTACAAAGA

GCACACGTCTGAACTCTTTCCCTACACGACGCTCTTCCGATCTNNNATTAGATAAGTGGCCAGAACTAC

GCACACGTCTGAACTCTTTCCCTACACGACGCTCTTCCGATCTNNNCTGTATGTTTGCAACAGTTTTCAT

GCACACGTCTGAACTCTTTCCCTACACGACGCTCTTCCGATCTNNNAATTCATGCTATCAATCCTCAAC

AGCACACGTCTGAACTCTTTCCCTACACGACGCTCTTCCGATCTNNNAACATGGCAACTTTGTAAGAC

GCACACGTCTGAACTCTTTCCCTACACGACGCTCTTCCGATCTNNNATAACCATTGTGTTATGTTGTTT

CACACGTCTGAACTCTTTCCCTACACGACGCTCTTCCGATCTNNNTTATATGCCTTTGACTAGTAAGTC

GCACACGTCTGAACTCTTTCCCTACACGACGCTCTTCCGATCTNNNTTCGGTAATTTGGAGCTATATAT

AGCACACGTCTGAACTCTTTCCCTACACGACGCTCTTCCGATCTNNNAATTGCAGAAAAACATGTATG

CACACGTCTGAACTCTTTCCCTACACGACGCTCTTCCGATCTNNNCAAAAGAAACACAAAATTCTCAAC

AGCACACGTCTGAACTCTTTCCCTACACGACGCTCTTCCGATCTNNNTGGATAATGAAAATGTGTTGT

AGCACACGTCTGAACTCTTTCCCTACACGACGCTCTTCCGATCTNNNGATCACATGCATATTCAAAAG

AGCACACGTCTGAACTCTTTCCCTACACGACGCTCTTCCGATCTNNNCTTCACTAAAAAGTAAAACAT

GCACACGTCTGAACTCTTTCCCTACACGACGCTCTTCCGATCTNNNCAAGATCCATTTCTTCCCAACAT

AGCACACGTCTGAACTCTTTCCCTACACGACGCTCTTCCGATCTNNNTTACAGGCACACATATATGTT

GCACACGTCTGAACTCTTTCCCTACACGACGCTCTTCCGATCTNNNAAAGAAAACAAAGCAAACTAAA

AGCACACGTCTGAACTCTTTCCCTACACGACGCTCTTCCGATCTNNNACATTTAAAAAGTACTCAGAT

GCACACGTCTGAACTCTTTCCCTACACGACGCTCTTCCGATCTNNNTGAATAATCTTCCAAAGTGATT

GAGCACACGTCTGAACTCTTTCCCTACACGACGCTCTTCCGATCTNNNATTGAAGGTATAGGAGAGA

GCACACGTCTGAACTCTTTCCCTACACGACGCTCTTCCGATCTNNNATGAAACAGAATAGAAAATGC

AGCACACGTCTGAACTCTTTCCCTACACGACGCTCTTCCGATCTNNNCAAAGAAGTAAGACATTTCTTG

GCACACGTCTGAACTCTTTCCCTACACGACGCTCTTCCGATCTNNNTATCACTGATTACTGAACTCATT

GAGCACACGTCTGAACTCTTTCCCTACACGACGCTCTTCCGATCTNNNCACCACACCCTGATTTATTTCA

AGCACACGTCTGAACTCTTTCCCTACACGACGCTCTTCCGATCTNNNGAGGTGTCTCTTAATCACAGC

GCACACGTCTGAACTCTTTCCCTACACGACGCTCTTCCGATCTNNNGATCTGTGGATCCTAAGGAGCA

AGCACACGTCTGAACTCTTTCCCTACACGACGCTCTTCCGATCTNNNAATAAAATGGAAAACAAATG

GCACACGTCTGAACTCTTTCCCTACACGACGCTCTTCCGATCTNNNTGTCATAAGGGAAGAAAAGAA

GAGCACACGTCTGAACTCTTTCCCTACACGACGCTCTTCCGATCTNNNTCCTATTCCTCTTATTTCCTTTT

GCACACGTCTGAACTCTTTCCCTACACGACGCTCTTCCGATCTNNNCTAAAAACTCGTTGCCAAATCCA

GCACACGTCTGAACTCTTTCCCTACACGACGCTCTTCCGATCTNNNATATGTCAGTCAAGTCACCACAT

CACACGTCTGAACTCTTTCCCTACACGACGCTCTTCCGATCTNNNTATTCCTCCATATATATGCCTTTCA

GCACACGTCTGAACTCTTTCCCTACACGACGCTCTTCCGATCTNNNCAACAAGAACAAAACTTCGTCAT

GAGCACACGTCTGAACTCTTTCCCTACACGACGCTCTTCCGATCTNNNATCTTCAGTAAAGTTTACCAA

GCACACGTCTGAACTCTTTCCCTACACGACGCTCTTCCGATCTNNNTAGTAGAACTATCATCATTCCAC

GCACACGTCTGAACTCTTTCCCTACACGACGCTCTTCCGATCTNNNTAGTAGAACTATCATCATTCCAC

GAGCACACGTCTGAACTCTTTCCCTACACGACGCTCTTCCGATCTNNNTAAACAATGAATAAGCCTTC

GCACACGTCTGAACTCTTTCCCTACACGACGCTCTTCCGATCTNNNGCATGCTTTTTGTTTTATTTTGA

GAGCACACGTCTGAACTCTTTCCCTACACGACGCTCTTCCGATCTNNNAATACCATCATTAGCCACAAT

AGCACACGTCTGAACTCTTTCCCTACACGACGCTCTTCCGATCTNNNTCTTTTTTCTACCTTTTCCTGAT

AGCACACGTCTGAACTCTTTCCCTACACGACGCTCTTCCGATCTNNNTTCCGATAAAATAAAAGTTCCT

GCACACGTCTGAACTCTTTCCCTACACGACGCTCTTCCGATCTNNNAGAACCTTGGAAATTATATGAA

CACACGTCTGAACTCTTTCCCTACACGACGCTCTTCCGATCTNNNGTGAAACAATACAGCTTTCAGCAT

AGCACACGTCTGAACTCTTTCCCTACACGACGCTCTTCCGATCTNNNGCAAGATACACAGAACTAAAA

AGCACACGTCTGAACTCTTTCCCTACACGACGCTCTTCCGATCTNNNACTTTGTTTTCAGCTTCATAAG

GCACACGTCTGAACTCTTTCCCTACACGACGCTCTTCCGATCTNNNCTCCCCACTCTCATTTACTCCATC

AGAGCACACGTCTGAACTCTTTCCCTACACGACGCTCTTCCGATCTNNNCCTTACCATTGTATAATGAC

GCACACGTCTGAACTCTTTCCCTACACGACGCTCTTCCGATCTNNNACTAGTCTATCTGTTTTCAGTTTC
AGCACACGTCTGAACTCTTTCCCTACACGACGCTCTTCCGATCTNNNAAAATCATGTTATTGGGGAAA
CACACGTCTGAACTCTTTCCCTACACGACGCTCTTCCGATCTNNNACTTCACGATATGATTTTCAATTC
GAGCACACGTCTGAACTCTTTCCCTACACGACGCTCTTCCGATCTNNNGTGTATCACCTATGTATCTTG
AGCACACGTCTGAACTCTTTCCCTACACGACGCTCTTCCGATCTNNNGGTCATTACATGCAATCAATTA
AGCACACGTCTGAACTCTTTCCCTACACGACGCTCTTCCGATCTNNNCAGTTAGTTTTAGAAATCTGAC
AGCACACGTCTGAACTCTTTCCCTACACGACGCTCTTCCGATCTNNNGAGACTTCTTTCTTCATTCAAAT
GCACACGTCTGAACTCTTTCCCTACACGACGCTCTTCCGATCTNNNCTTCTGATGCTTTTCGATTTTACA
GAGCACACGTCTGAACTCTTTCCCTACACGACGCTCTTCCGATCTNNNATAGAAGAAATCACTATGCA
AGCACACGTCTGAACTCTTTCCCTACACGACGCTCTTCCGATCTNNNTGATCAGTTATTATTTACCAG
AGCACACGTCTGAACTCTTTCCCTACACGACGCTCTTCCGATCTNNNCATTGCAATTCTATTTTGTGTAC
GAGCACACGTCTGAACTCTTTCCCTACACGACGCTCTTCCGATCTNNNGAGTTGTAGCAAATCTAAAA
AGCACACGTCTGAACTCTTTCCCTACACGACGCTCTTCCGATCTNNNACACTCCTAAAGAAAAAAACTG
GCACACGTCTGAACTCTTTCCCTACACGACGCTCTTCCGATCTNNNCTACAGTAGTATTTTGAGTTTG
GCACACGTCTGAACTCTTTCCCTACACGACGCTCTTCCGATCTNNNAATGGTTGGCTGAAAAATTATA
GCACACGTCTGAACTCTTTCCCTACACGACGCTCTTCCGATCTNNNCTCATGTTCCTATGTGCATAATC
AGCACACGTCTGAACTCTTTCCCTACACGACGCTCTTCCGATCTNNNGTAAGAATTTGCATTTATGTGA
AGCACACGTCTGAACTCTTTCCCTACACGACGCTCTTCCGATCTNNNGAATAAATTAGGGGTGAGACA
AGCACACGTCTGAACTCTTTCCCTACACGACGCTCTTCCGATCTNNNGTTAGCTCTAATCTCTTGTTTT
GAGCACACGTCTGAACTCTTTCCCTACACGACGCTCTTCCGATCTNNNAAAATTGCAAGGTTCCTATAA
GAGCACACGTCTGAACTCTTTCCCTACACGACGCTCTTCCGATCTNNNTTCGTTAAAATGAAGTCATAC
CACACGTCTGAACTCTTTCCCTACACGACGCTCTTCCGATCTNNNAGAGATGGAGCTAATTTTAGTGC
GAGCACACGTCTGAACTCTTTCCCTACACGACGCTCTTCCGATCTNNNAGATAGAAACAAAAGAAAAG
CACACGTCTGAACTCTTTCCCTACACGACGCTCTTCCGATCTNNNTATTGTTTTGTGTTCCCCTATTACA
GCACACGTCTGAACTCTTTCCCTACACGACGCTCTTCCGATCTNNNATACTCACAAAGCATACAGAGCA
GAGCACACGTCTGAACTCTTTCCCTACACGACGCTCTTCCGATCTNNNAGAGTACTTGGCAAATCTTAT
GCACACGTCTGAACTCTTTCCCTACACGACGCTCTTCCGATCTNNNGTTTAGATGAGGTATGTGACTC
GCACACGTCTGAACTCTTTCCCTACACGACGCTCTTCCGATCTNNNAAAATTTTCCTTCAATTCCCATCC
GCACACGTCTGAACTCTTTCCCTACACGACGCTCTTCCGATCTNNNAAAATTTTCCTTCAATTCCCATCC
GAGCACACGTCTGAACTCTTTCCCTACACGACGCTCTTCCGATCTNNNATAGTGTCTTTGTTAAATTAC
CACACGTCTGAACTCTTTCCCTACACGACGCTCTTCCGATCTNNNGCTCTATGATTCTCAAACATTTACA
CACACGTCTGAACTCTTTCCCTACACGACGCTCTTCCGATCTNNNTCATGTGACTGCAGTAAGATCATC
ACACGTCTGAACTCTTTCCCTACACGACGCTCTTCCGATCTNNNACCTACTGAAACGTCTTTTACCATC
GAGCACACGTCTGAACTCTTTCCCTACACGACGCTCTTCCGATCTNNNACTACGTGTTGAAATCTATAG
AGCACACGTCTGAACTCTTTCCCTACACGACGCTCTTCCGATCTNNNCTTGATTCTTCTACCCTTCTCCAT
GAGCACACGTCTGAACTCTTTCCCTACACGACGCTCTTCCGATCTNNNTGAGGAGACATTTAAGTATT
GCACACGTCTGAACTCTTTCCCTACACGACGCTCTTCCGATCTNNNGTTTTGACAGCAGTAAACACCAT
AGCACACGTCTGAACTCTTTCCCTACACGACGCTCTTCCGATCTNNNTGGTATTTATTTTCTTTTTGCCTC
AGCACACGTCTGAACTCTTTCCCTACACGACGCTCTTCCGATCTNNNGTAACCTTCAGCTATTTCATTCC
AGCACACGTCTGAACTCTTTCCCTACACGACGCTCTTCCGATCTNNNTGAGAATGATCTAGATAAACC
CACACGTCTGAACTCTTTCCCTACACGACGCTCTTCCGATCTNNNAATCTCTCTCTCTCTCTCTGCATCG
CACGTCTGAACTCTTTCCCTACACGACGCTCTTCCGATCTNNNCTTTGGCTCTGTTTCTTTCTCATCGGA
GCACACGTCTGAACTCTTTCCCTACACGACGCTCTTCCGATCTNNNAGGATCAACTCATAATGATTTCT

CACACGTCTGAACTCTTTCCCTACACGACGCTCTTCCGATCTNNNCACCTATATGGATATGCACATGCAT

AGCACACGTCTGAACTCTTTCCCTACACGACGCTCTTCCGATCTNNNCAAAGTAAATGTTTGCAAAACT

CACACGTCTGAACTCTTTCCCTACACGACGCTCTTCCGATCTNNNTTTTCTCTGAGTACTCTGGTTTCAT

GCACACGTCTGAACTCTTTCCCTACACGACGCTCTTCCGATCTNNNCTTCTTCTTTCATGGGTTCTTACAT

GAGCACACGTCTGAACTCTTTCCCTACACGACGCTCTTCCGATCTNNNACTTTATTACGTCTTCTAGAGA

GCACACGTCTGAACTCTTTCCCTACACGACGCTCTTCCGATCTNNNCCTATTCAGCATTACTTGTCTCAT

GAGCACACGTCTGAACTCTTTCCCTACACGACGCTCTTCCGATCTNNNCTCTCTCTCTCTTTCTCTTACA

AGCACACGTCTGAACTCTTTCCCTACACGACGCTCTTCCGATCTNNNAAAGAACACATCCAAAGAGAT

AGCACACGTCTGAACTCTTTCCCTACACGACGCTCTTCCGATCTNNNAAAGAACACATCCAAAGAGAT

GCACACGTCTGAACTCTTTCCCTACACGACGCTCTTCCGATCTNNNATTGAGAATAGAGTTAGAAAG

GCACACGTCTGAACTCTTTCCCTACACGACGCTCTTCCGATCTNNNATTGAGAATAGAGTTAGAAAG

GAGCACACGTCTGAACTCTTTCCCTACACGACGCTCTTCCGATCTNNNTCTAAAATTGGTTTCTCCCTAG

GCACACGTCTGAACTCTTTCCCTACACGACGCTCTTCCGATCTNNNTTTAAACTCATTTTCTCCTGTAGCA

GAGCACACGTCTGAACTCTTTCCCTACACGACGCTCTTCCGATCTNNNAAGAAATACATTTTGAAAGCT

CACACGTCTGAACTCTTTCCCTACACGACGCTCTTCCGATCTNNNTTTTATGTTTTCTTTCCATGGAACAT

AGCACACGTCTGAACTCTTTCCCTACACGACGCTCTTCCGATCTNNNATTACTCACATCTAGGAGAAGA

ACACGTCTGAACTCTTTCCCTACACGACGCTCTTCCGATCTNNNTGATAGAAAGTCACTACTCCTCATC

AGCACACGTCTGAACTCTTTCCCTACACGACGCTCTTCCGATCTNNNTAATATATATGGCATTGTTGGA

GAGCACACGTCTGAACTCTTTCCCTACACGACGCTCTTCCGATCTNNNCCAATAACTTCCAAACAATGAT

AGCACACGTCTGAACTCTTTCCCTACACGACGCTCTTCCGATCTNNNACTACTACTGCAATATATTCACA

AGCACACGTCTGAACTCTTTCCCTACACGACGCTCTTCCGATCTNNNCACTGAAGTATTTAAGGGTAAA

AGCACACGTCTGAACTCTTTCCCTACACGACGCTCTTCCGATCTNNNTAATTAAGATGTGCTCAGCATAT

AGCACACGTCTGAACTCTTTCCCTACACGACGCTCTTCCGATCTNNNCTGGATGCAGAGAAAATTCTCA

AGCACACGTCTGAACTCTTTCCCTACACGACGCTCTTCCGATCTNNNATTAAACCCAAAATGAGTATAC

GCACACGTCTGAACTCTTTCCCTACACGACGCTCTTCCGATCTNNNTGAGAGAGAAAACACAAATATT

GAGCACACGTCTGAACTCTTTCCCTACACGACGCTCTTCCGATCTNNNTTATAAATTACTACCGTGACTG

GAGCACACGTCTGAACTCTTTCCCTACACGACGCTCTTCCGATCTNNNTATAGAAATAACCATGACTAG

AGCACACGTCTGAACTCTTTCCCTACACGACGCTCTTCCGATCTNNNGTCTGGAATTTAGAGAGTTAGA

ACACGTCTGAACTCTTTCCCTACACGACGCTCTTCCGATCTNNNGGAATAACTCCCTCCTCTTCCATCGG

AGCACACGTCTGAACTCTTTCCCTACACGACGCTCTTCCGATCTNNNCCTTTCTGAACAACATTAACTTCA

GCACACGTCTGAACTCTTTCCCTACACGACGCTCTTCCGATCTNNNACAAGTCTGTATCATAGAACTGCA

GAGCACACGTCTGAACTCTTTCCCTACACGACGCTCTTCCGATCTNNNCTAGGTATGCTAATTGCCATTA

ACACGTCTGAACTCTTTCCCTACACGACGCTCTTCCGATCTNNNTCACTAAAATTCAGAGACTTGTCAT

GAGCACACGTCTGAACTCTTTCCCTACACGACGCTCTTCCGATCTNNNTTGTATCCTACAACCTTATTGAA

AGCACACGTCTGAACTCTTTCCCTACACGACGCTCTTCCGATCTNNNAAATATTCATTGCTGTGAGATT

CACACGTCTGAACTCTTTCCCTACACGACGCTCTTCCGATCTNNNTCCCAGCAAAACAATACATATATCA

AGCACACGTCTGAACTCTTTCCCTACACGACGCTCTTCCGATCTNNNTAAAATAGGAATGGATGGATA

ACACGTCTGAACTCTTTCCCTACACGACGCTCTTCCGATCTNNNTGCTCAGTTAACTAAATTGTAAGCAT

GAGCACACGTCTGAACTCTTTCCCTACACGACGCTCTTCCGATCTNNNTATACAATTAACAAATGGCAA

GCACACGTCTGAACTCTTTCCCTACACGACGCTCTTCCGATCTNNNAAACAACTGTGTATAGCATTAAA

GAGCACACGTCTGAACTCTTTCCCTACACGACGCTCTTCCGATCTNNNGAGGTGTATCTGTCAGTTTG

GAGCACACGTCTGAACTCTTTCCCTACACGACGCTCTTCCGATCTNNNGAGGTGTATCTGTCAGTTTG

GCACACGTCTGAACTCTTTCCCTACACGACGCTCTTCCGATCTNNNGTGTAATCCCAGCATTTTGGCAT

GCACACGTCTGAACTCTTTCCCTACACGACGCTCTTCCGATCTNNNATGGACATGTGAACATTTTAAA
AGCACACGTCTGAACTCTTTCCCTACACGACGCTCTTCCGATCTNNNTCTTTTCCCTTGTCTCCTTACAT
CACACGTCTGAACTCTTTCCCTACACGACGCTCTTCCGATCTNNNGGTTACAGAATTTGTAGTACCTC
GCACACGTCTGAACTCTTTCCCTACACGACGCTCTTCCGATCTNNNCTAACTGCTTACCTGAATTTGCA
ACACGTCTGAACTCTTTCCCTACACGACGCTCTTCCGATCTNNNCCTCTCAAAAGATTTATTTCTCCCAT
GCACACGTCTGAACTCTTTCCCTACACGACGCTCTTCCGATCTNNNCCAAGCTCAATCTGATCTTTCAT
CACACGTCTGAACTCTTTCCCTACACGACGCTCTTCCGATCTNNNTTGTTGTTGTTCTTCTTCCAT
GCACACGTCTGAACTCTTTCCCTACACGACGCTCTTCCGATCTNNNTGTGATTTCACCTATGTCATATA
CACGTCTGAACTCTTTCCCTACACGACGCTCTTCCGATCTNNNATATCACAATGCATCCTATACCATC
CACACGTCTGAACTCTTTCCCTACACGACGCTCTTCCGATCTNNNTATAACATTTGGTATTTTTGCCCC
CACACGTCTGAACTCTTTCCCTACACGACGCTCTTCCGATCTNNNTATAACATTTGGTATTTTTGCCCC
AGCACACGTCTGAACTCTTTCCCTACACGACGCTCTTCCGATCTNNNAAGTTATGCACTGAATTATTCT
GCACACGTCTGAACTCTTTCCCTACACGACGCTCTTCCGATCTNNNAATAACCAAAATTAGAGCTGAA
GCACACGTCTGAACTCTTTCCCTACACGACGCTCTTCCGATCTNNNCATTCTGCCAATTACAAAATATG
GCACACGTCTGAACTCTTTCCCTACACGACGCTCTTCCGATCTNNNCATTCTGCCAATTACAAAATATG
AGCACACGTCTGAACTCTTTCCCTACACGACGCTCTTCCGATCTNNNCATCCTAGGTCAATTCATATTCT
AGCACACGTCTGAACTCTTTCCCTACACGACGCTCTTCCGATCTNNNATCTTCAATGACTTTTTCCATGC
CACGTCTGAACTCTTTCCCTACACGACGCTCTTCCGATCTNNNTAAGTCTCCACACTTCATGGCATCGG
AGCACACGTCTGAACTCTTTCCCTACACGACGCTCTTCCGATCTNNNTGTCATAATTTGTTAAGAAGAG
CACACGTCTGAACTCTTTCCCTACACGACGCTCTTCCGATCTNNNATTTTGGGTTTTACAAGATTTCTC
AGCACACGTCTGAACTCTTTCCCTACACGACGCTCTTCCGATCTNNNAAAAAATTAAATTGGACACTTG
AGCACACGTCTGAACTCTTTCCCTACACGACGCTCTTCCGATCTNNNGAAATAAGGAACAAATGCATA
ACACGTCTGAACTCTTTCCCTACACGACGCTCTTCCGATCTNNNCTAATCTGGCCTCTAGAGTTTCATC
GCACACGTCTGAACTCTTTCCCTACACGACGCTCTTCCGATCTNNNCTCTATATTTGACTGAGTGTACT
CACACGTCTGAACTCTTTCCCTACACGACGCTCTTCCGATCTNNNGCATACATATAGTGAAAATGTGT
GAGCACACGTCTGAACTCTTTCCCTACACGACGCTCTTCCGATCTNNNTTTTGCATTTGGAGAACATA
AGCACACGTCTGAACTCTTTCCCTACACGACGCTCTTCCGATCTNNNTTTTGGAAAGTTTTCAGAGTA
AGCACACGTCTGAACTCTTTCCCTACACGACGCTCTTCCGATCTNNNGGAGGTATTATATTGGATATT
GCACACGTCTGAACTCTTTCCCTACACGACGCTCTTCCGATCTNNNATTATTTACAATTAGGACCTTG
ACACGTCTGAACTCTTTCCCTACACGACGCTCTTCCGATCTNNNCTTTCCTCTTTCCACTACTGCATCGG
AGCACACGTCTGAACTCTTTCCCTACACGACGCTCTTCCGATCTNNNGTTTTAGCTATAATTGGGTAA
AGCACACGTCTGAACTCTTTCCCTACACGACGCTCTTCCGATCTNNNTATAAAATACAGGACAAGGTC
CACACGTCTGAACTCTTTCCCTACACGACGCTCTTCCGATCTNNNTAATGGCAAATGACAATTCTATCC
GCACACGTCTGAACTCTTTCCCTACACGACGCTCTTCCGATCTNNNCTCCTGAAATGCATACATTGCAT
GAGCACACGTCTGAACTCTTTCCCTACACGACGCTCTTCCGATCTNNNTTATGTGATCTGTGATAAGA
AGCACACGTCTGAACTCTTTCCCTACACGACGCTCTTCCGATCTNNNGAGGTTACATGTTTGTTTATTT
ACACGTCTGAACTCTTTCCCTACACGACGCTCTTCCGATCTNNNCCCTGCTTTTATTTTTATTTCTCCATC
AGCACACGTCTGAACTCTTTCCCTACACGACGCTCTTCCGATCTNNNAGGAGATGACAAATCTCTAAT
GAGCACACGTCTGAACTCTTTCCCTACACGACGCTCTTCCGATCTNNNATTTTTGATATATGGAAGCTC
AGCACACGTCTGAACTCTTTCCCTACACGACGCTCTTCCGATCTNNNTTATTGTGCATTTGAATTATCCA
AGCACACGTCTGAACTCTTTCCCTACACGACGCTCTTCCGATCTNNNTTTTCTAAACCATCTCAGAGAA
AGCACACGTCTGAACTCTTTCCCTACACGACGCTCTTCCGATCTNNNCTAACCCTCTATTAGCACATTAG
AGCACACGTCTGAACTCTTTCCCTACACGACGCTCTTCCGATCTNNNAGAAACCAACAACCCATAATCC

GAGCACACGTCTGAACTCTTTCCCTACACGACGCTCTTCCGATCTNNNACCTTGAGATATTTCCTAACA
GCACACGTCTGAACTCTTTCCCTACACGACGCTCTTCCGATCTNNNTTAGTTGATCAACAAGTTGTTAG
GCACACGTCTGAACTCTTTCCCTACACGACGCTCTTCCGATCTNNNGCTGATGTATTTTTCTTGGTAAA
GCACACGTCTGAACTCTTTCCCTACACGACGCTCTTCCGATCTNNNTATTTGCCACATAGAGAAAAA
GCACACGTCTGAACTCTTTCCCTACACGACGCTCTTCCGATCTNNNTATCATTTCTTGCATATGTTACG
AGCACACGTCTGAACTCTTTCCCTACACGACGCTCTTCCGATCTNNNATACTTCTGGGTAAATGATGT
GCACACGTCTGAACTCTTTCCCTACACGACGCTCTTCCGATCTNNNGGTTACATCCCAATAAATGTATC
AGCACACGTCTGAACTCTTTCCCTACACGACGCTCTTCCGATCTNNNCCAGGAAGTATGTTTGGTAGC
GCACACGTCTGAACTCTTTCCCTACACGACGCTCTTCCGATCTNNNAAGAGAGAGAATTGTGTTTCT
AGCACACGTCTGAACTCTTTCCCTACACGACGCTCTTCCGATCTNNNATATTGTTTGAGTGTACATTC
AGCACACGTCTGAACTCTTTCCCTACACGACGCTCTTCCGATCTNNNAGCATGAGAATAGACTAATAC
CACACGTCTGAACTCTTTCCCTACACGACGCTCTTCCGATCTNNNAAGAAGTCCTTTTTCAACAAAGCA
AGCACACGTCTGAACTCTTTCCCTACACGACGCTCTTCCGATCTNNNAATGTATCCTTCATTAAAGACA
AGCACACGTCTGAACTCTTTCCCTACACGACGCTCTTCCGATCTNNNATAGATCTTGTTGTAGGTCTT
AGCACACGTCTGAACTCTTTCCCTACACGACGCTCTTCCGATCTNNNAACAGCTAATTGGTATAGAAG
GAGCACACGTCTGAACTCTTTCCCTACACGACGCTCTTCCGATCTNNNTATAATTAAAGATGGATGGG
AGCACACGTCTGAACTCTTTCCCTACACGACGCTCTTCCGATCTNNNTATGATCACTTTGGAATAATGT
GAGCACACGTCTGAACTCTTTCCCTACACGACGCTCTTCCGATCTNNNATCTTATAGTGCAGAAAACA
GCACACGTCTGAACTCTTTCCCTACACGACGCTCTTCCGATCTNNNCAGGTATATATGTGGGTATACA
AGCACACGTCTGAACTCTTTCCCTACACGACGCTCTTCCGATCTNNNATTATACTTTAGTAGTCACTG
AGCACACGTCTGAACTCTTTCCCTACACGACGCTCTTCCGATCTNNNTTCTTGCTAAACTTCATTTCAAT
AGCACACGTCTGAACTCTTTCCCTACACGACGCTCTTCCGATCTNNNTAGCATTTTAAGACCTGACTA
AGCACACGTCTGAACTCTTTCCCTACACGACGCTCTTCCGATCTNNNAGTCATCTCTTATCAGACATTA
GCACACGTCTGAACTCTTTCCCTACACGACGCTCTTCCGATCTNNNTAAGTTACAACTCTACACAAAAG
AGCACACGTCTGAACTCTTTCCCTACACGACGCTCTTCCGATCTNNNCATCTTCAATAAACCATGTCTC
AGCACACGTCTGAACTCTTTCCCTACACGACGCTCTTCCGATCTNNNCTAGCAAACTCAGCAAATTAAA
GCACACGTCTGAACTCTTTCCCTACACGACGCTCTTCCGATCTNNNACGTATGGAATGTATTTGTATC
AGCACACGTCTGAACTCTTTCCCTACACGACGCTCTTCCGATCTNNNGTCTCAAACTGTCTTAGAAAAC
AGCACACGTCTGAACTCTTTCCCTACACGACGCTCTTCCGATCTNNNATTAGAGACTAAAAGCTTCAAT
AGCACACGTCTGAACTCTTTCCCTACACGACGCTCTTCCGATCTNNNATATATTTCCTGCTAAAGTCTC
GAGCACACGTCTGAACTCTTTCCCTACACGACGCTCTTCCGATCTNNNGTAGAGCACCTCACAATATG
AGCACACGTCTGAACTCTTTCCCTACACGACGCTCTTCCGATCTNNNAAATCTGGTCTTAGAAATCCC
AGCACACGTCTGAACTCTTTCCCTACACGACGCTCTTCCGATCTNNNTATAATCCAACTCTGCTTCTAA
AGCACACGTCTGAACTCTTTCCCTACACGACGCTCTTCCGATCTNNNGGCACATATAATATCTCCAAG
AGCACACGTCTGAACTCTTTCCCTACACGACGCTCTTCCGATCTNNNCCAGGAGAGGTAGAATATAA
GCACACGTCTGAACTCTTTCCCTACACGACGCTCTTCCGATCTNNNTTCTTGGTTTTCTTATTAGAGG
GCACACGTCTGAACTCTTTCCCTACACGACGCTCTTCCGATCTNNNCAGTGTAAGGTTTAAATAAGG
AGAGCACACGTCTGAACTCTTTCCCTACACGACGCTCTTCCGATCTNNNCTCTCCTACCTCCTAATCATA
AGAGCACACGTCTGAACTCTTTCCCTACACGACGCTCTTCCGATCTNNNCTCTCCTACCTCCTAATCATA
ACACGTCTGAACTCTTTCCCTACACGACGCTCTTCCGATCTNNNAAAACTCTGCAAAGATAGATTTTCA
AGCACACGTCTGAACTCTTTCCCTACACGACGCTCTTCCGATCTNNNCTGATCAGTAGTGGGATCATC
AGCACACGTCTGAACTCTTTCCCTACACGACGCTCTTCCGATCTNNNTTAAAATGAGTTAGGCAGGTA
AGCACACGTCTGAACTCTTTCCCTACACGACGCTCTTCCGATCTNNNAAAAATCATTAAAAGGAACA

ACACGTCTGAACTCTTTCCCTACACGACGCTCTTCCGATCTNNNCTACCAAGCCATCATTGTTGCATCG
AGCACACGTCTGAACTCTTTCCCTACACGACGCTCTTCCGATCTNNNTATATCTATGTAACAAACCTG
AGCACACGTCTGAACTCTTTCCCTACACGACGCTCTTCCGATCTNNNCTAATAACAAGGCTTTAGACTT
GCACACGTCTGAACTCTTTCCCTACACGACGCTCTTCCGATCTNNNCATAATTAGCTCCTGGTGAGCA
AGCACACGTCTGAACTCTTTCCCTACACGACGCTCTTCCGATCTNNNGGGTATCCAGAAATTAACAA
AGCACACGTCTGAACTCTTTCCCTACACGACGCTCTTCCGATCTNNNTATAAAAACAAGAGCAAGAAG
GAGCACACGTCTGAACTCTTTCCCTACACGACGCTCTTCCGATCTNNNGGTCTTTCCTCATCTCAATATA
GCACACGTCTGAACTCTTTCCCTACACGACGCTCTTCCGATCTNNNGTTGAGATTACAGGTGTGAGC
AGCACACGTCTGAACTCTTTCCCTACACGACGCTCTTCCGATCTNNNCAGATTACTCTTCTCAGTACAT
GAGCACACGTCTGAACTCTTTCCCTACACGACGCTCTTCCGATCTNNNAGCAGACTAAATACTTGATA
GAGCACACGTCTGAACTCTTTCCCTACACGACGCTCTTCCGATCTNNNCTTAAGTTATTAAACTGAAG
GCACACGTCTGAACTCTTTCCCTACACGACGCTCTTCCGATCTNNNTGCAGAAATTTCAAAAATATGT
AGCACACGTCTGAACTCTTTCCCTACACGACGCTCTTCCGATCTNNNAATTTACTGTTTTTGTCCCA
AGCACACGTCTGAACTCTTTCCCTACACGACGCTCTTCCGATCTNNNTGCATAGTATATGTATTTTGTC
AGCACACGTCTGAACTCTTTCCCTACACGACGCTCTTCCGATCTNNNTGGTTTAGGATGTCACTAAAG
AGCACACGTCTGAACTCTTTCCCTACACGACGCTCTTCCGATCTNNNATCTAGAATGAAAGAGAGTA
GCACACGTCTGAACTCTTTCCCTACACGACGCTCTTCCGATCTNNNATAGATTTACTCCTTAGACACC
AGCACACGTCTGAACTCTTTCCCTACACGACGCTCTTCCGATCTNNNGAACGTCAGTTATACCCTTTCC
GAGCACACGTCTGAACTCTTTCCCTACACGACGCTCTTCCGATCTNNNTGTACCTGCTTATGTTTGTAA
GCACACGTCTGAACTCTTTCCCTACACGACGCTCTTCCGATCTNNNGATTAAATTGAGTAGCCAAGT
GCACACGTCTGAACTCTTTCCCTACACGACGCTCTTCCGATCTNNNAGCTAGTAGGAATGAGCTAGC
GCACACGTCTGAACTCTTTCCCTACACGACGCTCTTCCGATCTNNNCTGGGAAATAGCATTGTATTAG
AGCACACGTCTGAACTCTTTCCCTACACGACGCTCTTCCGATCTNNNAGCAATCTCATAAGATGTATT
AGCACACGTCTGAACTCTTTCCCTACACGACGCTCTTCCGATCTNNNTAATTTTTCTATAGGGTGCGA
GAGCACACGTCTGAACTCTTTCCCTACACGACGCTCTTCCGATCTNNNCTAACATTAAAGACAAAAGG
GCACACGTCTGAACTCTTTCCCTACACGACGCTCTTCCGATCTNNNCATAGTCTATGATTTTCCCATTC
CACACGTCTGAACTCTTTCCCTACACGACGCTCTTCCGATCTNNNTTATCTGCAGATATGCTAAACATC
GAGCACACGTCTGAACTCTTTCCCTACACGACGCTCTTCCGATCTNNNTACGGATCATCTTTGGTTATT
GCACACGTCTGAACTCTTTCCCTACACGACGCTCTTCCGATCTNNNGGTTTAGGAATACATCACTGAT
GCACACGTCTGAACTCTTTCCCTACACGACGCTCTTCCGATCTNNNCTTCTATACCTATCTGAAACCTT
GCACACGTCTGAACTCTTTCCCTACACGACGCTCTTCCGATCTNNNGTTATCATGACTGTCTAGAGCC
GCACACGTCTGAACTCTTTCCCTACACGACGCTCTTCCGATCTNNNCTCTCCCCTATGAAGAGTATATA
GCACACGTCTGAACTCTTTCCCTACACGACGCTCTTCCGATCTNNNGCCAAGTGTACATGAACATACA
GAGCACACGTCTGAACTCTTTCCCTACACGACGCTCTTCCGATCTNNNACAGGTTGATTTGTTATTTT
AGCACACGTCTGAACTCTTTCCCTACACGACGCTCTTCCGATCTNNNATGTATGTATATGAGTATGCA
AGCACACGTCTGAACTCTTTCCCTACACGACGCTCTTCCGATCTNNNAACCCCTTAGTAGTGATAAAA
GCACACGTCTGAACTCTTTCCCTACACGACGCTCTTCCGATCTNNNTGTTATGTGTATACATGTGTGT
GAGCACACGTCTGAACTCTTTCCCTACACGACGCTCTTCCGATCTNNNGCCTTTATCCTGAAATATATG
GAGCACACGTCTGAACTCTTTCCCTACACGACGCTCTTCCGATCTNNNTTTATGTGTAATGTGGCATTA
CACACGTCTGAACTCTTTCCCTACACGACGCTCTTCCGATCTNNNCACAATACAGAGAGGTGAAAAT
GCACACGTCTGAACTCTTTCCCTACACGACGCTCTTCCGATCTNNNTTTTAGCATTACTAGAGGCTTA
GCACACGTCTGAACTCTTTCCCTACACGACGCTCTTCCGATCTNNNTTTTAGCATTACTAGAGGCTTA
AGCACACGTCTGAACTCTTTCCCTACACGACGCTCTTCCGATCTNNNAGTGTATGTGTATGTTCAAAG

AGCACACGTCTGAACTCTTTCCCTACACGACGCTCTTCCGATCTNNNTTTGAGGATTTTGGATTTGTA
GAGCACACGTCTGAACTCTTTCCCTACACGACGCTCTTCCGATCTNNNTTGAATTCTTCCTTTGTGTTG
GCACACGTCTGAACTCTTTCCCTACACGACGCTCTTCCGATCTNNNGGGTTTCTTTTTTTCTACTTTTTC
GCACACGTCTGAACTCTTTCCCTACACGACGCTCTTCCGATCTNNNTTATGCAATAACACAATAAGGA
ACACGTCTGAACTCTTTCCCTACACGACGCTCTTCCGATCTNNNAAAATTTTGTTTAAACTCCTTGCCA
AGCACACGTCTGAACTCTTTCCCTACACGACGCTCTTCCGATCTNNNTTTTGGTGAAAAGAATTAGGA
AGCACACGTCTGAACTCTTTCCCTACACGACGCTCTTCCGATCTNNNGTAAAATCTGGAAAAGTTGTC
AGCACACGTCTGAACTCTTTCCCTACACGACGCTCTTCCGATCTNNNTCATTGAAAAGGAATACAATA
AGCACACGTCTGAACTCTTTCCCTACACGACGCTCTTCCGATCTNNNATTACCATAGGTAGCTTTTACA
AGCACACGTCTGAACTCTTTCCCTACACGACGCTCTTCCGATCTNNNCCGATACATAGACATAGGGAC
GCACACGTCTGAACTCTTTCCCTACACGACGCTCTTCCGATCTNNNCTGCAGATCTTAGGACTTATAA
AGCACACGTCTGAACTCTTTCCCTACACGACGCTCTTCCGATCTNNNAGTAAGTCAGATAATTAGAA
AGCACACGTCTGAACTCTTTCCCTACACGACGCTCTTCCGATCTNNNAGTCTATGGTTATCAGAGTAG
AGCACACGTCTGAACTCTTTCCCTACACGACGCTCTTCCGATCTNNNAGTCTATGGTTATCAGAGTAG
CACACGTCTGAACTCTTTCCCTACACGACGCTCTTCCGATCTNNNATAACACAAAACACACATATACC
CACACGTCTGAACTCTTTCCCTACACGACGCTCTTCCGATCTNNNATAACACAAAACACACATATACC
GCACACGTCTGAACTCTTTCCCTACACGACGCTCTTCCGATCTNNNTCACTAACCACTCTATTTAACATC
GCACACGTCTGAACTCTTTCCCTACACGACGCTCTTCCGATCTNNNCTCTAACAATAAGTCCTTGCTAA
CACACGTCTGAACTCTTTCCCTACACGACGCTCTTCCGATCTNNNCTCATCTCTCTTCCTCTCTTCATCG
GCACACGTCTGAACTCTTTCCCTACACGACGCTCTTCCGATCTNNNTGATTACAAAAATGAGAACAAC
GCACACGTCTGAACTCTTTCCCTACACGACGCTCTTCCGATCTNNNCGAGTGTGTTGTATATTCATAA
AGCACACGTCTGAACTCTTTCCCTACACGACGCTCTTCCGATCTNNNTACAGTAGAGAAACAAAGTGT
CACGTCTGAACTCTTTCCCTACACGACGCTCTTCCGATCTNNNAAGATTCAGTCATGTATGGATACCA
GCACACGTCTGAACTCTTTCCCTACACGACGCTCTTCCGATCTNNNAGTTACTTCACTTAGAACAGTA
CACACGTCTGAACTCTTTCCCTACACGACGCTCTTCCGATCTNNNCCTTTATTCTTCATGTCTATTGGCA
AGCACACGTCTGAACTCTTTCCCTACACGACGCTCTTCCGATCTNNNAGTGAGACCATCTCTACATACC
GCACACGTCTGAACTCTTTCCCTACACGACGCTCTTCCGATCTNNNTAAATAAAGAAGTGAATCTCCC
GCACACGTCTGAACTCTTTCCCTACACGACGCTCTTCCGATCTNNNTAAATAAAGAAGTGAATCTCCC
GCACACGTCTGAACTCTTTCCCTACACGACGCTCTTCCGATCTNNNTAAATAAAGAAGTGAATCTCCC
GCACACGTCTGAACTCTTTCCCTACACGACGCTCTTCCGATCTNNNGCTGAGTAGTATTCCATCATAT
AGCACACGTCTGAACTCTTTCCCTACACGACGCTCTTCCGATCTNNNCCTTTACTTCAAATTGTACTCA
AGCACACGTCTGAACTCTTTCCCTACACGACGCTCTTCCGATCTNNNTATATTAGTCAACCTTCTCCAG
GCACACGTCTGAACTCTTTCCCTACACGACGCTCTTCCGATCTNNNTATAAGGTTCTGCTAAAAGGA
GCACACGTCTGAACTCTTTCCCTACACGACGCTCTTCCGATCTNNNATGGTTACTGTTGATGTAGAAA
GCACACGTCTGAACTCTTTCCCTACACGACGCTCTTCCGATCTNNNAGAACATTTGATCCATAATAAC
GCACACGTCTGAACTCTTTCCCTACACGACGCTCTTCCGATCTNNNTCATAGGAGATCACATCCATTCA
CACACGTCTGAACTCTTTCCCTACACGACGCTCTTCCGATCTNNNTGTATATATATGTGTGTGTGTATG
CACACGTCTGAACTCTTTCCCTACACGACGCTCTTCCGATCTNNNGATTCTCCTGGTTCTTTTTTGCATC
AGCACACGTCTGAACTCTTTCCCTACACGACGCTCTTCCGATCTNNNAAGGAAATGGTTTGTTCTCTA
CACACGTCTGAACTCTTTCCCTACACGACGCTCTTCCGATCTNNNTGGTTTTTTGTAAAAACAACAAAC
GCACACGTCTGAACTCTTTCCCTACACGACGCTCTTCCGATCTNNNGCTTTCTCTTTCTTTTTCTCTTCA
AGCACACGTCTGAACTCTTTCCCTACACGACGCTCTTCCGATCTNNNAGTGGATTTCTTAGGATTTAC
GCACACGTCTGAACTCTTTCCCTACACGACGCTCTTCCGATCTNNNGAAATACTAAGTTGACTGCAAT

AGCACACGTCTGAACTCTTTCCCTACACGACGCTCTTCCGATCTNNNGGAAATCCTCACTAAAAAGAAA
AGCACACGTCTGAACTCTTTCCCTACACGACGCTCTTCCGATCTNNNTTTACACTGTCTCAAAGAGATAT
AGCACACGTCTGAACTCTTTCCCTACACGACGCTCTTCCGATCTNNNTGCCTATTTTTTTCACTCTTTTA
AGCACACGTCTGAACTCTTTCCCTACACGACGCTCTTCCGATCTNNNTATTCATCATGTAGAACACAAT
AGCACACGTCTGAACTCTTTCCCTACACGACGCTCTTCCGATCTNNNTATTCATCATGTAGAACACAAT
AGCACACGTCTGAACTCTTTCCCTACACGACGCTCTTCCGATCTNNNTATTCATCATGTAGAACACAAT
GAGCACACGTCTGAACTCTTTCCCTACACGACGCTCTTCCGATCTNNNAATCAAAAGAAATTGCTAGG
AGCACACGTCTGAACTCTTTCCCTACACGACGCTCTTCCGATCTNNNTAAAGATTGTACCTTATGGAAG
AGCACACGTCTGAACTCTTTCCCTACACGACGCTCTTCCGATCTNNNTTTTTAAAGTGTTCAAGAGACC
AGCACACGTCTGAACTCTTTCCCTACACGACGCTCTTCCGATCTNNNGTATTAGAGAAAGGTGGTAGA
AGCACACGTCTGAACTCTTTCCCTACACGACGCTCTTCCGATCTNNNGCTAAAAAGTAACTGTTTTCAT
GAGCACACGTCTGAACTCTTTCCCTACACGACGCTCTTCCGATCTNNNCAGAAATCAAGACATAAATTT
CACACGTCTGAACTCTTTCCCTACACGACGCTCTTCCGATCTNNNTAAGTGCTCAAAAATTATCCATCCA
GCACACGTCTGAACTCTTTCCCTACACGACGCTCTTCCGATCTNNNCACAATCGTTCAGACTAACTCAT
CACACGTCTGAACTCTTTCCCTACACGACGCTCTTCCGATCTNNNCTCTCAAAATGGTCGAATTATGCAT
GAGCACACGTCTGAACTCTTTCCCTACACGACGCTCTTCCGATCTNNNGATGAAGTGAAGACAGACAG
GCACACGTCTGAACTCTTTCCCTACACGACGCTCTTCCGATCTNNNAAAAGAGGAAAAAAGGACACTA
CACACGTCTGAACTCTTTCCCTACACGACGCTCTTCCGATCTNNNCTCTCCTGCTTCCAGTATTTCATCGG
AGCACACGTCTGAACTCTTTCCCTACACGACGCTCTTCCGATCTNNNATTCTTTTTGTATTTGTAGGCA
AGCACACGTCTGAACTCTTTCCCTACACGACGCTCTTCCGATCTNNNTGCATTTATTAGTTCTAACAGTG
GAGCACACGTCTGAACTCTTTCCCTACACGACGCTCTTCCGATCTNNNCTGGGTTGCATATAAATACAT
CACACGTCTGAACTCTTTCCCTACACGACGCTCTTCCGATCTNNNTGTTGAGATGGAATTTATAAACCC
AGCACACGTCTGAACTCTTTCCCTACACGACGCTCTTCCGATCTNNNGAGAGTGGCTTAATATGTACC
AGCACACGTCTGAACTCTTTCCCTACACGACGCTCTTCCGATCTNNNGACCTAAACATGGAAATGTAAA
CACACGTCTGAACTCTTTCCCTACACGACGCTCTTCCGATCTNNNAAAAATCTAAACAATAAAGCCATCA
ACACGTCTGAACTCTTTCCCTACACGACGCTCTTCCGATCTNNNATGTAAGTTTTTAAAAGCTCCCCATC
GAGCACACGTCTGAACTCTTTCCCTACACGACGCTCTTCCGATCTNNNTCTTTCATATGAGCCTGTTATT
CACACGTCTGAACTCTTTCCCTACACGACGCTCTTCCGATCTNNNTCACTAACTGAAACATCATAATGCA
GAGCACACGTCTGAACTCTTTCCCTACACGACGCTCTTCCGATCTNNNGAATACATTTCTTCTGGGATA
AGCACACGTCTGAACTCTTTCCCTACACGACGCTCTTCCGATCTNNNCACCAATTTCAAAAGTAGAAAA
GCACACGTCTGAACTCTTTCCCTACACGACGCTCTTCCGATCTNNNCTGACATTCCACAATAATGAATT
AGCACACGTCTGAACTCTTTCCCTACACGACGCTCTTCCGATCTNNNAAATGGCTGTAAGTTATTCAAT
GCACACGTCTGAACTCTTTCCCTACACGACGCTCTTCCGATCTNNNCTCATGTCATTACACCATTTGCAT
CACACGTCTGAACTCTTTCCCTACACGACGCTCTTCCGATCTNNNGAATGTTGTGAATGTTGGATGCAT
AGCACACGTCTGAACTCTTTCCCTACACGACGCTCTTCCGATCTNNNCCATCAAAAAAAAGAAAGAAA
AGCACACGTCTGAACTCTTTCCCTACACGACGCTCTTCCGATCTNNNGAAAGACAAGGGATAAGAGA
GAGCACACGTCTGAACTCTTTCCCTACACGACGCTCTTCCGATCTNNNGAAACTCCATCTCAAAAAAAA
AGCACACGTCTGAACTCTTTCCCTACACGACGCTCTTCCGATCTNNNTTTGGAATTTTGCTAAGCTAAAT
AGCACACGTCTGAACTCTTTCCCTACACGACGCTCTTCCGATCTNNNAAATAACCTTCACCATTCTTAAC
AGCACACGTCTGAACTCTTTCCCTACACGACGCTCTTCCGATCTNNNGCTTGAATTCATAACCTAGTTTT
GCACACGTCTGAACTCTTTCCCTACACGACGCTCTTCCGATCTNNNTACTCTTCTCTGTGTGTATTCTCA
GCACACGTCTGAACTCTTTCCCTACACGACGCTCTTCCGATCTNNNTACTCTTCTCTGTGTGTATTCTCA
AGCACACGTCTGAACTCTTTCCCTACACGACGCTCTTCCGATCTNNNTTAAATAAATTTGAGAGACAGC

GAGCACACGTCTGAACTCTTTCCCTACACGACGCTCTTCCGATCTNNNATCAGGGTAATTTTTTATTGC
AGCACACGTCTGAACTCTTTCCCTACACGACGCTCTTCCGATCTNNNAAGTAACAACAACAGAAACAATA
AGCACACGTCTGAACTCTTTCCCTACACGACGCTCTTCCGATCTNNNCTTCAAAAAAAATTGAGGATGA
CACACGTCTGAACTCTTTCCCTACACGACGCTCTTCCGATCTNNNATGATTTGCATCTCTAAACACTCATC
GCACACGTCTGAACTCTTTCCCTACACGACGCTCTTCCGATCTNNNATTCCTGAATTGCAATATTTTCTC
GCACACGTCTGAACTCTTTCCCTACACGACGCTCTTCCGATCTNNNTCAAAGACTAGAAATGATGAAAC
AGCACACGTCTGAACTCTTTCCCTACACGACGCTCTTCCGATCTNNNCACTATTCAACTCCAGGTAATGC
AGCACACGTCTGAACTCTTTCCCTACACGACGCTCTTCCGATCTNNNAACCCTTTACATATACATGTGTA
AGCACACGTCTGAACTCTTTCCCTACACGACGCTCTTCCGATCTNNNCTGATTCAAACAGATGCCAGCA
GAGCACACGTCTGAACTCTTTCCCTACACGACGCTCTTCCGATCTNNNTAGTAGAAAGAATTCCTATGT
AGAGCACACGTCTGAACTCTTTCCCTACACGACGCTCTTCCGATCTNNNCAAGACCCTGTCATAGAAAG
AGCACACGTCTGAACTCTTTCCCTACACGACGCTCTTCCGATCTNNNAAAATGGGTAGAGATGTAACA
AGCACACGTCTGAACTCTTTCCCTACACGACGCTCTTCCGATCTNNNAAAATGGGTAGAGATGTAACA
AGCACACGTCTGAACTCTTTCCCTACACGACGCTCTTCCGATCTNNNTTTCTATCATATTTAGTGATGCA
AGCACACGTCTGAACTCTTTCCCTACACGACGCTCTTCCGATCTNNNGTATAATATCTGGAAGGTACTG
AGCACACGTCTGAACTCTTTCCCTACACGACGCTCTTCCGATCTNNNTGTAACTAGAAAACAAAGTACT
GAGCACACGTCTGAACTCTTTCCCTACACGACGCTCTTCCGATCTNNNCGATACACAGAAGAGAAAATA
AGCACACGTCTGAACTCTTTCCCTACACGACGCTCTTCCGATCTNNNCAAAACAAAACAAAAAACAACTC
GCACACGTCTGAACTCTTTCCCTACACGACGCTCTTCCGATCTNNNAATTATTTGTGGAAAGAAGGAAG
GCACACGTCTGAACTCTTTCCCTACACGACGCTCTTCCGATCTNNNTTTACATTTGAAGGAATGGAAAC
GAGCACACGTCTGAACTCTTTCCCTACACGACGCTCTTCCGATCTNNNCTAGGCCAAAAAAAAAAAAAG
GCACACGTCTGAACTCTTTCCCTACACGACGCTCTTCCGATCTNNNGAAAACAATAAAATAAGGGCTCTC
AGCACACGTCTGAACTCTTTCCCTACACGACGCTCTTCCGATCTNNNACTCTTGTCATTCATTATGCTATC
GAGCACACGTCTGAACTCTTTCCCTACACGACGCTCTTCCGATCTNNNCCTATTATTGAAGGCATATTTG
GAGCACACGTCTGAACTCTTTCCCTACACGACGCTCTTCCGATCTNNNCCCTAGAAAATACTTTTTTTTC
CACACGTCTGAACTCTTTCCCTACACGACGCTCTTCCGATCTNNNTGAGTGACAAAGGTAGAATTAGCA
AGAGCACACGTCTGAACTCTTTCCCTACACGACGCTCTTCCGATCTNNNTACAAACACACTCAAAGTAAA
CACACGTCTGAACTCTTTCCCTACACGACGCTCTTCCGATCTNNNTGTCCATATCCTTAATGTAGATCCA
CACACGTCTGAACTCTTTCCCTACACGACGCTCTTCCGATCTNNNAGAGATCCTTGTGTGTCTTCCATCG
AGCACACGTCTGAACTCTTTCCCTACACGACGCTCTTCCGATCTNNNCCAATTTTGGGTATGTCTTTATTC
GAGCACACGTCTGAACTCTTTCCCTACACGACGCTCTTCCGATCTNNNGATGAATAAGTCTAGAGCTCT
GAGCACACGTCTGAACTCTTTCCCTACACGACGCTCTTCCGATCTNNNCTCTCTGGCTATGTGTATGTAT
AGCACACGTCTGAACTCTTTCCCTACACGACGCTCTTCCGATCTNNNCAACTTCGTATAGTTACCAAAAC
AGCACACGTCTGAACTCTTTCCCTACACGACGCTCTTCCGATCTNNNATGAAGTAGAGTATAAGCAGT
AGCACACGTCTGAACTCTTTCCCTACACGACGCTCTTCCGATCTNNNTAAAGATTGTACCTTATGGAAG
GCACACGTCTGAACTCTTTCCCTACACGACGCTCTTCCGATCTNNNACTGTTTTTGTATTATGAAAATC
GCACACGTCTGAACTCTTTCCCTACACGACGCTCTTCCGATCTNNNCCTTTCCTAACTTCATTCATAGTCA
AGCACACGTCTGAACTCTTTCCCTACACGACGCTCTTCCGATCTNNNTATTTTTTCCATTCCTGAGTGAC
GAGCACACGTCTGAACTCTTTCCCTACACGACGCTCTTCCGATCTNNNGATTACTTTTGGGAGATTGAT
GAGCACACGTCTGAACTCTTTCCCTACACGACGCTCTTCCGATCTNNNCAAGAGTACTTGGAGAAGTC
AGCACACGTCTGAACTCTTTCCCTACACGACGCTCTTCCGATCTNNNTAGACCATTTACCTCCTAGTCCA
GAGCACACGTCTGAACTCTTTCCCTACACGACGCTCTTCCGATCTNNNATATTCTCTCTGTGTTGTCTTAT
GAGCACACGTCTGAACTCTTTCCCTACACGACGCTCTTCCGATCTNNNTTCCATATATTGTTCCTAAGTG

GCACACGTCTGAACTCTTTCCCTACACGACGCTCTTCCGATCTNNNAAAAACAAATTTGCTTGTCATCC
AGCACACGTCTGAACTCTTTCCCTACACGACGCTCTTCCGATCTNNNCAGAAAGTCCAAAAATCTATGA
GCACACGTCTGAACTCTTTCCCTACACGACGCTCTTCCGATCTNNNCTTTATTCTCTCTTTCTCTGTCTC
GCACACGTCTGAACTCTTTCCCTACACGACGCTCTTCCGATCTNNNCTTTATTCTCTCTTTCTCTGTCTC
GCACACGTCTGAACTCTTTCCCTACACGACGCTCTTCCGATCTNNNCTTTATTCTCTCTTTCTCTGTCTC
AGCACACGTCTGAACTCTTTCCCTACACGACGCTCTTCCGATCTNNNGTCTTAGGATTTTCATTCATGTT
GCACACGTCTGAACTCTTTCCCTACACGACGCTCTTCCGATCTNNNTAACCTCAAATCATCCAGAAAGC
AGCACACGTCTGAACTCTTTCCCTACACGACGCTCTTCCGATCTNNNATTCTGTGTGGATGGATGAAT
AGCACACGTCTGAACTCTTTCCCTACACGACGCTCTTCCGATCTNNNCAATGACAATTAGAGGGAGA
GCACACGTCTGAACTCTTTCCCTACACGACGCTCTTCCGATCTNNNCTATACAGATCATTTATTGGTG
GCACACGTCTGAACTCTTTCCCTACACGACGCTCTTCCGATCTNNNTTTTAATTTCTAACTTGCTCTGGC
CACACGTCTGAACTCTTTCCCTACACGACGCTCTTCCGATCTNNNGAGAATATAGAGAAATTGGGTCA
AGCACACGTCTGAACTCTTTCCCTACACGACGCTCTTCCGATCTNNNATACACTGTGAGCATTCAAATC
GAGCACACGTCTGAACTCTTTCCCTACACGACGCTCTTCCGATCTNNNCTTTTTTGAAACATGCACATA
AGCACACGTCTGAACTCTTTCCCTACACGACGCTCTTCCGATCTNNNTCTGCCAGTAATGTAAATCTCC
AGCACACGTCTGAACTCTTTCCCTACACGACGCTCTTCCGATCTNNNTCTGCCAGTAATGTAAATCTCC
AGCACACGTCTGAACTCTTTCCCTACACGACGCTCTTCCGATCTNNNACTGAGGAATTTTACACTGAT
AGCACACGTCTGAACTCTTTCCCTACACGACGCTCTTCCGATCTNNNAATAACTTTGGGAAGCTAACA
ACACGTCTGAACTCTTTCCCTACACGACGCTCTTCCGATCTNNNGGGAAAGACATTTACACACTTACAT
AGCACACGTCTGAACTCTTTCCCTACACGACGCTCTTCCGATCTNNNGTTCAGATAGAAATTGGCATT
GCACACGTCTGAACTCTTTCCCTACACGACGCTCTTCCGATCTNNNAGTAAAACTAACCTACTATGACA
AGCACACGTCTGAACTCTTTCCCTACACGACGCTCTTCCGATCTNNNGACTTTCTTCTTTTTCTGTAACT
GCACACGTCTGAACTCTTTCCCTACACGACGCTCTTCCGATCTNNNAGCAATATATGAAAACTTCAGTT
GAGCACACGTCTGAACTCTTTCCCTACACGACGCTCTTCCGATCTNNNTTATGCACATTTTGTAAGACA
GCACACGTCTGAACTCTTTCCCTACACGACGCTCTTCCGATCTNNNGGTTTTATCTTCTGTGTGTATGC
AGCACACGTCTGAACTCTTTCCCTACACGACGCTCTTCCGATCTNNNTGGTTTTATCTTCTGTGTGTAT
CACACGTCTGAACTCTTTCCCTACACGACGCTCTTCCGATCTNNNGAAAGCAGCTGAAATTTTAAAAA
AGCACACGTCTGAACTCTTTCCCTACACGACGCTCTTCCGATCTNNNGATGTGAAAGGTATTAAAACT
CACACGTCTGAACTCTTTCCCTACACGACGCTCTTCCGATCTNNNTCAGTAAAGTGAAGACCTGGCAT
AGCACACGTCTGAACTCTTTCCCTACACGACGCTCTTCCGATCTNNNATTTAATTTTTTTTCTGCCCAAG
AGCACACGTCTGAACTCTTTCCCTACACGACGCTCTTCCGATCTNNNCAGTAGTACACACACATATAGA
GAGCACACGTCTGAACTCTTTCCCTACACGACGCTCTTCCGATCTNNNCTGTAAGGTTACTGTGTTAG
AGCACACGTCTGAACTCTTTCCCTACACGACGCTCTTCCGATCTNNNTTAGGTTTTTCTCTAGTTACTCC
AGCACACGTCTGAACTCTTTCCCTACACGACGCTCTTCCGATCTNNNTTATTGCATCTAGTCATCATAG
GCACACGTCTGAACTCTTTCCCTACACGACGCTCTTCCGATCTNNNGAATGAACAAAAGGAAAAGAA
GCACACGTCTGAACTCTTTCCCTACACGACGCTCTTCCGATCTNNNATTCTTGTAATTCTTTTTGTCCCC
GCACACGTCTGAACTCTTTCCCTACACGACGCTCTTCCGATCTNNNATTCTTGTAATTCTTTTTGTCCCC
AGCACACGTCTGAACTCTTTCCCTACACGACGCTCTTCCGATCTNNNGTGACAGTACTTTTATAACATC
AGCACACGTCTGAACTCTTTCCCTACACGACGCTCTTCCGATCTNNNTATAGCTTCATAAAAGGAAGT
GAGCACACGTCTGAACTCTTTCCCTACACGACGCTCTTCCGATCTNNNGATGTACATGTCAGTTACAGT
GCACACGTCTGAACTCTTTCCCTACACGACGCTCTTCCGATCTNNNGTTTTTAGGATGATATGCGTTTT
AGCACACGTCTGAACTCTTTCCCTACACGACGCTCTTCCGATCTNNNGTTCCACCATTTACCAGTTATC
AGCACACGTCTGAACTCTTTCCCTACACGACGCTCTTCCGATCTNNNAAATCAGCTTCTCTTTGTTTAGC

AGCACACGTCTGAACTCTTTCCCTACACGACGCTCTTCCGATCTNNNTAGGTTGAAAAACTTTAGGA
CACACGTCTGAACTCTTTCCCTACACGACGCTCTTCCGATCTNNNAGACAAGGTTTGTGTTTGTGCAT
ACACGTCTGAACTCTTTCCCTACACGACGCTCTTCCGATCTNNNCTTTGAAAACAAATCCCATTACCA
GCACACGTCTGAACTCTTTCCCTACACGACGCTCTTCCGATCTNNNTTCCAAAGATTTAAAAAAGCAA
GCACACGTCTGAACTCTTTCCCTACACGACGCTCTTCCGATCTNNNCATTTACCTCAGCAAGTATTTAT
GCACACGTCTGAACTCTTTCCCTACACGACGCTCTTCCGATCTNNNCTTTCTGAAATGGGGGTATAGC
GAGCACACGTCTGAACTCTTTCCCTACACGACGCTCTTCCGATCTNNNAGATTTTTCAAAAACTACCAA
AGCACACGTCTGAACTCTTTCCCTACACGACGCTCTTCCGATCTNNNTTCATCTTCCTTGTCTTCAAACA
GAGCACACGTCTGAACTCTTTCCCTACACGACGCTCTTCCGATCTNNNGGAAGGAAGTGTAATAGAA
AGCACACGTCTGAACTCTTTCCCTACACGACGCTCTTCCGATCTNNNGTGTTGTTCAGTAATCAAACA
ACACGTCTGAACTCTTTCCCTACACGACGCTCTTCCGATCTNNNGGGTTTCTCAAATATATTTCCTGCA
GAGCACACGTCTGAACTCTTTCCCTACACGACGCTCTTCCGATCTNNNTATTGACAACATTAAGTGAG
CACACGTCTGAACTCTTTCCCTACACGACGCTCTTCCGATCTNNNCAGCATTTAAATTCTTGTCTTTCCA
GCACACGTCTGAACTCTTTCCCTACACGACGCTCTTCCGATCTNNNAATGTTTGAACCATTTATCCTTT
GCACACGTCTGAACTCTTTCCCTACACGACGCTCTTCCGATCTNNNTTGATCACTGCTATAGAGAAAA
GAGCACACGTCTGAACTCTTTCCCTACACGACGCTCTTCCGATCTNNNGTTTGCATTCAAAATTAGAT
AGCACACGTCTGAACTCTTTCCCTACACGACGCTCTTCCGATCTNNNAAACCTCTTTAAATTCCTTTTG
GCACACGTCTGAACTCTTTCCCTACACGACGCTCTTCCGATCTNNNCAATCATAGATCCTTCCTGTTTC
AGCACACGTCTGAACTCTTTCCCTACACGACGCTCTTCCGATCTNNNTTTCTTCACTGGACACTATAAT
AGCACACGTCTGAACTCTTTCCCTACACGACGCTCTTCCGATCTNNNTTTCTTCACTGGACACTATAAT
CACACGTCTGAACTCTTTCCCTACACGACGCTCTTCCGATCTNNNCACAATTACTCTGTGGACTTCATCG
CACACGTCTGAACTCTTTCCCTACACGACGCTCTTCCGATCTNNNCACAATTACTCTGTGGACTTCATCG
GCACACGTCTGAACTCTTTCCCTACACGACGCTCTTCCGATCTNNNTGGAATATCTTTTTCCATCTCTT
GCACACGTCTGAACTCTTTCCCTACACGACGCTCTTCCGATCTNNNATGTACAAACCTAGCATATTTT
ACACGTCTGAACTCTTTCCCTACACGACGCTCTTCCGATCTNNNGTGTGTGTAGTTCCCTATCATCG
AGCACACGTCTGAACTCTTTCCCTACACGACGCTCTTCCGATCTNNNAAAGAGCTCTAGAATTTCTGA
GCACACGTCTGAACTCTTTCCCTACACGACGCTCTTCCGATCTNNNTATGGTAAAAGGCTAAGTGTTT
AGCACACGTCTGAACTCTTTCCCTACACGACGCTCTTCCGATCTNNNTATGGTAAAAGGCTAAGTGTT
GAGCACACGTCTGAACTCTTTCCCTACACGACGCTCTTCCGATCTNNNGGATGAGTACTATAAAACAG
GCACACGTCTGAACTCTTTCCCTACACGACGCTCTTCCGATCTNNNACAGAAGTACTTGAGTTTCTTG
GCACACGTCTGAACTCTTTCCCTACACGACGCTCTTCCGATCTNNNTATTCTTCAAACTCAAGGGAAA
AGCACACGTCTGAACTCTTTCCCTACACGACGCTCTTCCGATCTNNNGCATCTATTCTTCAAACTCAAG
GCACACGTCTGAACTCTTTCCCTACACGACGCTCTTCCGATCTNNNCATCTATCTAGCCATCAACTAATC
AGCACACGTCTGAACTCTTTCCCTACACGACGCTCTTCCGATCTNNNAAAAAGGTTTCTTAAAGTTCT
GCACACGTCTGAACTCTTTCCCTACACGACGCTCTTCCGATCTNNNTATCTTTAAAAAATGGGCTTCAA
AGCACACGTCTGAACTCTTTCCCTACACGACGCTCTTCCGATCTNNNGTTGTAAATATAAGGCAGTTC
ACACGTCTGAACTCTTTCCCTACACGACGCTCTTCCGATCTNNNTTCTCTAAAGAAGGAATGGTTCCA
AGCACACGTCTGAACTCTTTCCCTACACGACGCTCTTCCGATCTNNNTTAACATTGACTCATACCTTGA
AGCACACGTCTGAACTCTTTCCCTACACGACGCTCTTCCGATCTNNNAATAAAGAACCAGTTATTTCCT
AGCACACGTCTGAACTCTTTCCCTACACGACGCTCTTCCGATCTNNNTTAAATTGTGTACCTCTGCTAA
AGCACACGTCTGAACTCTTTCCCTACACGACGCTCTTCCGATCTNNNGCATGACTTTGTCAAACTATA
CACACGTCTGAACTCTTTCCCTACACGACGCTCTTCCGATCTNNNTTCTTGCTATGTCTCACATGCATC
GCACACGTCTGAACTCTTTCCCTACACGACGCTCTTCCGATCTNNNCTGAATCAAATATTTGGTTGGT

GCACACGTCTGAACTCTTTCCCTACACGACGCTCTTCCGATCTNNNGTTTAGTCTAGTGACAGAAAAT
CACACGTCTGAACTCTTTCCCTACACGACGCTCTTCCGATCTNNNCTATCCATTTGAATGATTCGATTC
CACGTCTGAACTCTTTCCCTACACGACGCTCTTCCGATCTNNNATAGGTGAAAAAAACTAATCAGGCA
AGCACACGTCTGAACTCTTTCCCTACACGACGCTCTTCCGATCTNNNTAATATAAGAATGACAACGGA
GCACACGTCTGAACTCTTTCCCTACACGACGCTCTTCCGATCTNNNATCTCTATTTGCTGCTTCTATTA
AGCACACGTCTGAACTCTTTCCCTACACGACGCTCTTCCGATCTNNNTAACATGTCTAACAGATGAGT
GAGCACACGTCTGAACTCTTTCCCTACACGACGCTCTTCCGATCTNNNTTTTCTTTTCCATTTCTCTGTA
GCACACGTCTGAACTCTTTCCCTACACGACGCTCTTCCGATCTNNNTGAGACATAGAGAGATTATCA
GAGCACACGTCTGAACTCTTTCCCTACACGACGCTCTTCCGATCTNNNTTCTCTCCAAATTGATCTATAG
ACACGTCTGAACTCTTTCCCTACACGACGCTCTTCCGATCTNNNCATGGAACTGACAAGTTAGCCATC
ACACGTCTGAACTCTTTCCCTACACGACGCTCTTCCGATCTNNNCATGGAACTGACAAGTTAGCCATC
GCACACGTCTGAACTCTTTCCCTACACGACGCTCTTCCGATCTNNNGTTCATAGCGACTTTATTCATA
AGCACACGTCTGAACTCTTTCCCTACACGACGCTCTTCCGATCTNNNTATGAAAAGAACGTGAGAGG
AGCACACGTCTGAACTCTTTCCCTACACGACGCTCTTCCGATCTNNNTATGAAAAGAACGTGAGAGG
GCACACGTCTGAACTCTTTCCCTACACGACGCTCTTCCGATCTNNNTCCTTCCATTTTATTTTATTGGG
ACACGTCTGAACTCTTTCCCTACACGACGCTCTTCCGATCTNNNCTTCTCTAAGATACCATTCTTCACAT
GAGCACACGTCTGAACTCTTTCCCTACACGACGCTCTTCCGATCTNNNACAGGTGTTTTCATAAGAATT
GAGCACACGTCTGAACTCTTTCCCTACACGACGCTCTTCCGATCTNNNACAGGTGTTTTCATAAGAATT
GAGCACACGTCTGAACTCTTTCCCTACACGACGCTCTTCCGATCTNNNGAGGAAGGAGAATTGTTTGA
GCACACGTCTGAACTCTTTCCCTACACGACGCTCTTCCGATCTNNNATGTCCAATGCCTCTATAGTAC
GCACACGTCTGAACTCTTTCCCTACACGACGCTCTTCCGATCTNNNTCACAAACATATGCTACCTACCA
GCACACGTCTGAACTCTTTCCCTACACGACGCTCTTCCGATCTNNNTCAAAAGCTCAAAGAATTTTAGA
CACACGTCTGAACTCTTTCCCTACACGACGCTCTTCCGATCTNNNCTCTATCTTTCCCTCTATTTTGTCAT
ACACGTCTGAACTCTTTCCCTACACGACGCTCTTCCGATCTNNNATGGAGTTACAGTTGCAGTTACAT
GCACACGTCTGAACTCTTTCCCTACACGACGCTCTTCCGATCTNNNGATGTGTTTTTGTCCATACACCA
GAGCACACGTCTGAACTCTTTCCCTACACGACGCTCTTCCGATCTNNNAAAAAGACGCATAAGTAAAG
AGCACACGTCTGAACTCTTTCCCTACACGACGCTCTTCCGATCTNNNAAACGATATATCAGAGAGTAC
AGCACACGTCTGAACTCTTTCCCTACACGACGCTCTTCCGATCTNNNCACTTAAAATAATGTCCTACAG
GCACACGTCTGAACTCTTTCCCTACACGACGCTCTTCCGATCTNNNTTGCAGTGCCTAATTGAAATCA
AGCACACGTCTGAACTCTTTCCCTACACGACGCTCTTCCGATCTNNNAATAAAACAAAATAGCAACAA
ACACGTCTGAACTCTTTCCCTACACGACGCTCTTCCGATCTNNNATATACACACACACATATATATAATT
GCACACGTCTGAACTCTTTCCCTACACGACGCTCTTCCGATCTNNNGTTTGTGTATGTGTTTGAGAAA
ACACGTCTGAACTCTTTCCCTACACGACGCTCTTCCGATCTNNNAGTTACTTGAGCTTTGGAGGCATCG
AGCACACGTCTGAACTCTTTCCCTACACGACGCTCTTCCGATCTNNNTTCCAATCATTGTGACAACTTA
AGCACACGTCTGAACTCTTTCCCTACACGACGCTCTTCCGATCTNNNAGTAAAGTCAACATTCTGTTTT
GCACACGTCTGAACTCTTTCCCTACACGACGCTCTTCCGATCTNNNGATATATTGGTGGATGCCAGC
CACACGTCTGAACTCTTTCCCTACACGACGCTCTTCCGATCTNNNTCTCCTTAAGATAAACTATACAGTC
GAGCACACGTCTGAACTCTTTCCCTACACGACGCTCTTCCGATCTNNNTGAAGTTCTTCAATATCCTTGT
GCACACGTCTGAACTCTTTCCCTACACGACGCTCTTCCGATCTNNNACTGAACAGTAATTTCATTTTGA
GCACACGTCTGAACTCTTTCCCTACACGACGCTCTTCCGATCTNNNTGTCATTGACAACACTTTTTAAAC
AGCACACGTCTGAACTCTTTCCCTACACGACGCTCTTCCGATCTNNNTTATATGTACTCACTTGAACAG
GAGCACACGTCTGAACTCTTTCCCTACACGACGCTCTTCCGATCTNNNCAAAATTCTGAGAGTTATGAG
AGCACACGTCTGAACTCTTTCCCTACACGACGCTCTTCCGATCTNNNAGTCTCAATATTGGAGTTTTCT

GCACACGTCTGAACTCTTTCCCTACACGACGCTCTTCCGATCTNNNTAACCTTTTCTCTCCTGTCATCAT
CACACGTCTGAACTCTTTCCCTACACGACGCTCTTCCGATCTNNNTCTGTCTATCTTATCAATGCTTTC
GCACACGTCTGAACTCTTTCCCTACACGACGCTCTTCCGATCTNNNTTAGATTCAAAATTGTTGACACA
AGCACACGTCTGAACTCTTTCCCTACACGACGCTCTTCCGATCTNNNCATAGTTTATTCTGGGTGACTA
CACACGTCTGAACTCTTTCCCTACACGACGCTCTTCCGATCTNNNGTGAGATAAATGGTTCATGTAAG
AGCACACGTCTGAACTCTTTCCCTACACGACGCTCTTCCGATCTNNNTATTTCTATTCTTCTTCTCTGGG
GCACACGTCTGAACTCTTTCCCTACACGACGCTCTTCCGATCTNNNCTCTCTTCATCTTCACATGGCAT
GCACACGTCTGAACTCTTTCCCTACACGACGCTCTTCCGATCTNNNGATCAAATGTTTCCATCAGAAGC
GCACACGTCTGAACTCTTTCCCTACACGACGCTCTTCCGATCTNNNAAACAGTATGGAAGTTCCTAAA
CACGTCTGAACTCTTTCCCTACACGACGCTCTTCCGATCTNNNCTAGGATGGAACCACTGTACCATCG
AGCACACGTCTGAACTCTTTCCCTACACGACGCTCTTCCGATCTNNNATAATTCACAACTTCAAATTGG
GAGCACACGTCTGAACTCTTTCCCTACACGACGCTCTTCCGATCTNNNACAGCTCTTAGTTTCTTATTG
AGCACACGTCTGAACTCTTTCCCTACACGACGCTCTTCCGATCTNNNCTTATAAGAAGGCAAAGAAAC
AGCACACGTCTGAACTCTTTCCCTACACGACGCTCTTCCGATCTNNNCAAACCTTTGAACAAATAAATG
CACACGTCTGAACTCTTTCCCTACACGACGCTCTTCCGATCTNNNGTTACAGCAGTTCTTTATACAATC
GAGCACACGTCTGAACTCTTTCCCTACACGACGCTCTTCCGATCTNNNGGGAAAGATACAATTCAATG
AGCACACGTCTGAACTCTTTCCCTACACGACGCTCTTCCGATCTNNNATAGTCTGAAATAAAGCTCTT
AGCACACGTCTGAACTCTTTCCCTACACGACGCTCTTCCGATCTNNNATAGTCTGAAATAAAGCTCTT
AGCACACGTCTGAACTCTTTCCCTACACGACGCTCTTCCGATCTNNNATAGTCTGAAATAAAGCTCTT
AGCACACGTCTGAACTCTTTCCCTACACGACGCTCTTCCGATCTNNNCTCTTATCCTAGACACTAGAGTT
GCACACGTCTGAACTCTTTCCCTACACGACGCTCTTCCGATCTNNNCTTATTAAAGAATGCAGCAGTTC
GCACACGTCTGAACTCTTTCCCTACACGACGCTCTTCCGATCTNNNATATTCATAGGATTTGTTTCCCT
AGCACACGTCTGAACTCTTTCCCTACACGACGCTCTTCCGATCTNNNGCCGTATGTAAATATTCTGAA
GAGCACACGTCTGAACTCTTTCCCTACACGACGCTCTTCCGATCTNNNAGCAAGAGCATCCATGTATA
CACACGTCTGAACTCTTTCCCTACACGACGCTCTTCCGATCTNNNGAACTTTTCTTATTGTCCTACAGCA
AGCACACGTCTGAACTCTTTCCCTACACGACGCTCTTCCGATCTNNNGCTTTTACAAAATTACGCAATA
CACACGTCTGAACTCTTTCCCTACACGACGCTCTTCCGATCTNNNCCAACTTTTCCTTTCAAATACTACA
CACACGTCTGAACTCTTTCCCTACACGACGCTCTTCCGATCTNNNTCTCACGATACTCGACTTTTATAC
GCACACGTCTGAACTCTTTCCCTACACGACGCTCTTCCGATCTNNNCAGAGCAGATTGTCGATATTACA
GAGCACACGTCTGAACTCTTTCCCTACACGACGCTCTTCCGATCTNNNAAAATTCAGGTTGCTTTTTTT
CACACGTCTGAACTCTTTCCCTACACGACGCTCTTCCGATCTNNNAATATCTTTTTTAGACTGTTCTCTC
CACACGTCTGAACTCTTTCCCTACACGACGCTCTTCCGATCTNNNAAAGATAAAACCAACACCTGACAT
AGCACACGTCTGAACTCTTTCCCTACACGACGCTCTTCCGATCTNNNCTCATCATATACTCAAGAGAGA
CACACGTCTGAACTCTTTCCCTACACGACGCTCTTCCGATCTNNNCCTGTTCTTATAACTGTATGTTCCA
AGCACACGTCTGAACTCTTTCCCTACACGACGCTCTTCCGATCTNNNTTTTGACTTCTGCGTCTTATATT
CACACGTCTGAACTCTTTCCCTACACGACGCTCTTCCGATCTNNNGAGTAATCCTTACCTGGTATTAGC
GAGCACACGTCTGAACTCTTTCCCTACACGACGCTCTTCCGATCTNNNATTTTGTAAAACACATAGTTG
GCACACGTCTGAACTCTTTCCCTACACGACGCTCTTCCGATCTNNNGTTATCTTGGGTCTTGGATACCA
AGCACACGTCTGAACTCTTTCCCTACACGACGCTCTTCCGATCTNNNTCCTATCACTCCTATTATCTGAA
AGCACACGTCTGAACTCTTTCCCTACACGACGCTCTTCCGATCTNNNATTAGCAAATGACATTCTCATG
CACACGTCTGAACTCTTTCCCTACACGACGCTCTTCCGATCTNNNAAGCAGAGGAAGAAGGAAAGCA
GAGCACACGTCTGAACTCTTTCCCTACACGACGCTCTTCCGATCTNNNGATTATGAGATTTTTGAGCCT
GAGCACACGTCTGAACTCTTTCCCTACACGACGCTCTTCCGATCTNNNCATACTTTGAAAGAGGTCTTC

AGCACACGTCTGAACTCTTTCCCTACACGACGCTCTTCCGATCTNNNCTCTCTCTATCTCTCTCTCTAAA
CACACGTCTGAACTCTTTCCCTACACGACGCTCTTCCGATCTNNNGTGTGTCAATAAAAGGCTCTCAT
AGCACACGTCTGAACTCTTTCCCTACACGACGCTCTTCCGATCTNNNGAAGCAGAGACAGATATATTT
ACACGTCTGAACTCTTTCCCTACACGACGCTCTTCCGATCTNNNATATCCTCACGTCTCAAATGCATCG
GCACACGTCTGAACTCTTTCCCTACACGACGCTCTTCCGATCTNNNATCAACCAATAGAACAAAACAG
GCACACGTCTGAACTCTTTCCCTACACGACGCTCTTCCGATCTNNNTCTCTTAAGTACTGCAAATAAGA
CACACGTCTGAACTCTTTCCCTACACGACGCTCTTCCGATCTNNNCTTCTCTGAGAACCAGTCTGCATC
GCACACGTCTGAACTCTTTCCCTACACGACGCTCTTCCGATCTNNNTGGAGACAAAGTAAGATAAAC
GCACACGTCTGAACTCTTTCCCTACACGACGCTCTTCCGATCTNNNAGAAATGAAAGTGTCCTATCAG
AGCACACGTCTGAACTCTTTCCCTACACGACGCTCTTCCGATCTNNNCACTGAGCTTATTTAGAACTTA
GCACACGTCTGAACTCTTTCCCTACACGACGCTCTTCCGATCTNNNTAAACACACAAGACATGTATCTA
GCACACGTCTGAACTCTTTCCCTACACGACGCTCTTCCGATCTNNNCACACACATACCTGTATATACAT
CACACGTCTGAACTCTTTCCCTACACGACGCTCTTCCGATCTNNNCAGAGGAGTTTTTGTTTTTGATA
GCACACGTCTGAACTCTTTCCCTACACGACGCTCTTCCGATCTNNNCATCTCTAAGTCAGCAGACCCAT
AGCACACGTCTGAACTCTTTCCCTACACGACGCTCTTCCGATCTNNNGTAAGTTTTCCAGTTTATTGAG
AGCACACGTCTGAACTCTTTCCCTACACGACGCTCTTCCGATCTNNNGGTTACAAAAACAAGTACTAG
AGCACACGTCTGAACTCTTTCCCTACACGACGCTCTTCCGATCTNNNCAGGGTCTTAGGATAAATAG
AGCACACGTCTGAACTCTTTCCCTACACGACGCTCTTCCGATCTNNNCAGGGTCTTAGGATAAATAG
AGCACACGTCTGAACTCTTTCCCTACACGACGCTCTTCCGATCTNNNGTTATAACCATTTTGCAAATG
GCACACGTCTGAACTCTTTCCCTACACGACGCTCTTCCGATCTNNNGAAACCTGTATTCAAAAAGAAA
ACACGTCTGAACTCTTTCCCTACACGACGCTCTTCCGATCTNNNGTACCAATACAGGTCTGAAAGCAT
AGCACACGTCTGAACTCTTTCCCTACACGACGCTCTTCCGATCTNNNTAAGTTTGATTTTCAATGTGAA
CACACGTCTGAACTCTTTCCCTACACGACGCTCTTCCGATCTNNNATATCCAATAAAGAACATGCATCC
GCACACGTCTGAACTCTTTCCCTACACGACGCTCTTCCGATCTNNNTTGAAATAGACACAGGTGTAGC
AGCACACGTCTGAACTCTTTCCCTACACGACGCTCTTCCGATCTNNNTAAATGATTATTATTCCCCCAC
CACACGTCTGAACTCTTTCCCTACACGACGCTCTTCCGATCTNNNATGCATTACTAAATAACTTGCAAC
GCACACGTCTGAACTCTTTCCCTACACGACGCTCTTCCGATCTNNNAACAAGCTTAAAACAATTGAG
AGCACACGTCTGAACTCTTTCCCTACACGACGCTCTTCCGATCTNNNGTGCAGTGAAATATTTAGTTTT
GCACACGTCTGAACTCTTTCCCTACACGACGCTCTTCCGATCTNNNATGCATTTGTGTGAAGAAACTA
CACACGTCTGAACTCTTTCCCTACACGACGCTCTTCCGATCTNNNTACTCCTCTCCCCATAGAAGCATC
AGCACACGTCTGAACTCTTTCCCTACACGACGCTCTTCCGATCTNNNTATTCTGCAAGTACTGGTTTA
ACACGTCTGAACTCTTTCCCTACACGACGCTCTTCCGATCTNNNTAGAATTCTAGGTTTCATTCTGACA
AGCACACGTCTGAACTCTTTCCCTACACGACGCTCTTCCGATCTNNNATGTTTTTGAATGTATGAGCA
ACACGTCTGAACTCTTTCCCTACACGACGCTCTTCCGATCTNNNCAGAGATTGAAACTGACAATACCA
GAGCACACGTCTGAACTCTTTCCCTACACGACGCTCTTCCGATCTNNNAAGGTCATTATTTTGTTCCT
AGCACACGTCTGAACTCTTTCCCTACACGACGCTCTTCCGATCTNNNATAGAATGAGGGGTTAGGAG
GCACACGTCTGAACTCTTTCCCTACACGACGCTCTTCCGATCTNNNTTACTGTATACACATGCATGTAT
GCACACGTCTGAACTCTTTCCCTACACGACGCTCTTCCGATCTNNNCTTTTGCCTTTTTTGCTGATTATC
GCACACGTCTGAACTCTTTCCCTACACGACGCTCTTCCGATCTNNNAGTTTGTAAATTACATACTGCA
GAGCACACGTCTGAACTCTTTCCCTACACGACGCTCTTCCGATCTNNNAGACAATTACCAGGTCATTA
GCACACGTCTGAACTCTTTCCCTACACGACGCTCTTCCGATCTNNNTTTGAGATTTACTTCTTGCAGC
GCACACGTCTGAACTCTTTCCCTACACGACGCTCTTCCGATCTNNNGGGTTGATCTTTAGGAGATTA
GCACACGTCTGAACTCTTTCCCTACACGACGCTCTTCCGATCTNNNGGGTTGATCTTTAGGAGATTA

GCACACGTCTGAACTCTTTCCCTACACGACGCTCTTCCGATCTNNNATACACACACTACACACCTTACAT
GCACACGTCTGAACTCTTTCCCTACACGACGCTCTTCCGATCTNNNCCCTCTTAGAATACAAATGTAG
AGCACACGTCTGAACTCTTTCCCTACACGACGCTCTTCCGATCTNNNATATTAAGCTTGTATCTTGCA
AGCACACGTCTGAACTCTTTCCCTACACGACGCTCTTCCGATCTNNNATATTAAGCTTGTATCTTGCA
AGCACACGTCTGAACTCTTTCCCTACACGACGCTCTTCCGATCTNNNATAATGGTTTCGATTTGTGTA
AGCACACGTCTGAACTCTTTCCCTACACGACGCTCTTCCGATCTNNNCAATCTTTCTCAATGTGATGAT
AGAGCACACGTCTGAACTCTTTCCCTACACGACGCTCTTCCGATCTNNNCTTAGGCTGAGAAGTTTGA
GCACACGTCTGAACTCTTTCCCTACACGACGCTCTTCCGATCTNNNACTTCACTTAGAATAATGGTCTC
AGCACACGTCTGAACTCTTTCCCTACACGACGCTCTTCCGATCTNNNTTGAAAGTGCAGTTAAATTCT
GCACACGTCTGAACTCTTTCCCTACACGACGCTCTTCCGATCTNNNATATAGATTTATGGGTCATGACA
AGCACACGTCTGAACTCTTTCCCTACACGACGCTCTTCCGATCTNNNGTTCAGATTTCTTCAGTTTTAC
GCACACGTCTGAACTCTTTCCCTACACGACGCTCTTCCGATCTNNNGTAGTAATGTGAGGCTGGAGCA
GCACACGTCTGAACTCTTTCCCTACACGACGCTCTTCCGATCTNNNGCCCTATGTATTACAGTCTATTT
AGCACACGTCTGAACTCTTTCCCTACACGACGCTCTTCCGATCTNNNGAATATGCACATTTGTTACACT
ACACGTCTGAACTCTTTCCCTACACGACGCTCTTCCGATCTNNNATGAAAGACATTGTTCATTGCCATC
ACACGTCTGAACTCTTTCCCTACACGACGCTCTTCCGATCTNNNATGAAAGACATTGTTCATTGCCATC
AGCACACGTCTGAACTCTTTCCCTACACGACGCTCTTCCGATCTNNNCATGAGGGAAACTTTTCATAA
GCACACGTCTGAACTCTTTCCCTACACGACGCTCTTCCGATCTNNNATCATTGAAACAGAACAAATCA
GCACACGTCTGAACTCTTTCCCTACACGACGCTCTTCCGATCTNNNATCATTGAAACAGAACAAATCA
AGCACACGTCTGAACTCTTTCCCTACACGACGCTCTTCCGATCTNNNTTATATTCCCCAAGCATCTGAA
ACACGTCTGAACTCTTTCCCTACACGACGCTCTTCCGATCTNNNAGAAATATAAAGGGGTAAAATGAC
CACACGTCTGAACTCTTTCCCTACACGACGCTCTTCCGATCTNNNCATTAAGTTAAGCCCATTATTCCCA
AGCACACGTCTGAACTCTTTCCCTACACGACGCTCTTCCGATCTNNNTTATCTCTCTAGGAAGTACTGC
ACACGTCTGAACTCTTTCCCTACACGACGCTCTTCCGATCTNNNATTTTAAAAATCATCCATGCCTCAT
GCACACGTCTGAACTCTTTCCCTACACGACGCTCTTCCGATCTNNNAGAAAGTCGTTTTTACCTTCTCAT
AGCACACGTCTGAACTCTTTCCCTACACGACGCTCTTCCGATCTNNNGTTCTGAAGATTGGATCAGGC
AGCACACGTCTGAACTCTTTCCCTACACGACGCTCTTCCGATCTNNNTATATTGAGATTCTTTGTGGCT
AGCACACGTCTGAACTCTTTCCCTACACGACGCTCTTCCGATCTNNNATATTCACCAGTATATTTCCTC
AGCACACGTCTGAACTCTTTCCCTACACGACGCTCTTCCGATCTNNNATATTCACCAGTATATTTCCTC
GCACACGTCTGAACTCTTTCCCTACACGACGCTCTTCCGATCTNNNTTCTATACTTCTTTGTTGATTGCC
GCACACGTCTGAACTCTTTCCCTACACGACGCTCTTCCGATCTNNNTAACTTCCCATCAATGATATACT
AGCACACGTCTGAACTCTTTCCCTACACGACGCTCTTCCGATCTNNNCTGAAGTGAACATATGTAGAC
AGCACACGTCTGAACTCTTTCCCTACACGACGCTCTTCCGATCTNNNTTTGGAACCATCAAGGGAATC
GAGCACACGTCTGAACTCTTTCCCTACACGACGCTCTTCCGATCTNNNAATCAGTTGATGAGGATTAG
CACGTCTGAACTCTTTCCCTACACGACGCTCTTCCGATCTNNNAGACCTGAAGGAAACTATCACATCG
AGCACACGTCTGAACTCTTTCCCTACACGACGCTCTTCCGATCTNNNCGTTCCTTTGTAGTGTTTATAC
GCACACGTCTGAACTCTTTCCCTACACGACGCTCTTCCGATCTNNNTTGGTATCAGAGAAATTAAGTC
AGCACACGTCTGAACTCTTTCCCTACACGACGCTCTTCCGATCTNNNATTCTCAAAAGCAACTCTCTCCA
AGCACACGTCTGAACTCTTTCCCTACACGACGCTCTTCCGATCTNNNATTCTCAAAAGCAACTCTCTCCA
ACACGTCTGAACTCTTTCCCTACACGACGCTCTTCCGATCTNNNAAATGATTTTCCCAAGTTATTCACAT
AGCACACGTCTGAACTCTTTCCCTACACGACGCTCTTCCGATCTNNNTTCTGCCTTTAGACTGAACTAC
AGCACACGTCTGAACTCTTTCCCTACACGACGCTCTTCCGATCTNNNAATAAAGAACCAGTTATTTCCT
AGCACACGTCTGAACTCTTTCCCTACACGACGCTCTTCCGATCTNNNTAGACTTAATTGTTTAGAGCAG

GAGCACACGTCTGAACTCTTTCCCTACACGACGCTCTTCCGATCTNNNATCTTCTTTTGTGATTTTGTCAT
GCACACGTCTGAACTCTTTCCCTACACGACGCTCTTCCGATCTNNNCAACAAATTTTAATTCTCTTCCCCA
GCACACGTCTGAACTCTTTCCCTACACGACGCTCTTCCGATCTNNNCAACAAATTTTAATTCTCTTCCCCA
GCACACGTCTGAACTCTTTCCCTACACGACGCTCTTCCGATCTNNNCAACAAATTTTAATTCTCTTCCCCA
AGCACACGTCTGAACTCTTTCCCTACACGACGCTCTTCCGATCTNNNCATATGAATACCAGGATAGCTT
GAGCACACGTCTGAACTCTTTCCCTACACGACGCTCTTCCGATCTNNNGGAAATACGTGATTAATATGC
AGCACACGTCTGAACTCTTTCCCTACACGACGCTCTTCCGATCTNNNAAAATGTTGGGATTACAAGTA
GCACACGTCTGAACTCTTTCCCTACACGACGCTCTTCCGATCTNNNAAGAAGACTTTTGTGGATGATCA
GCACACGTCTGAACTCTTTCCCTACACGACGCTCTTCCGATCTNNNCTTTCAAACTCTTCTCACTCCCATC
CACACGTCTGAACTCTTTCCCTACACGACGCTCTTCCGATCTNNNTTCTAATGTGCATTAATTTGTCACAT
AGCACACGTCTGAACTCTTTCCCTACACGACGCTCTTCCGATCTNNNAGCACACATATATATGTATACGT
GCACACGTCTGAACTCTTTCCCTACACGACGCTCTTCCGATCTNNNCTGTCTTATGGTACTGAAAGAAC
AGCACACGTCTGAACTCTTTCCCTACACGACGCTCTTCCGATCTNNNGAATTCCATTACTCATTTGAGA
AGCACACGTCTGAACTCTTTCCCTACACGACGCTCTTCCGATCTNNNGAATTCCATTACTCATTTGAGA
AGCACACGTCTGAACTCTTTCCCTACACGACGCTCTTCCGATCTNNNAGTAAGGGGCAGATTATAAAA
CACACGTCTGAACTCTTTCCCTACACGACGCTCTTCCGATCTNNNTTGTTCATATTTCCTCAAAAACACAT
AGCACACGTCTGAACTCTTTCCCTACACGACGCTCTTCCGATCTNNNATCTATGAATACACATGTGATC
AGCACACGTCTGAACTCTTTCCCTACACGACGCTCTTCCGATCTNNNGCCTTTTTTTTCTTCAGATATTC
GCACACGTCTGAACTCTTTCCCTACACGACGCTCTTCCGATCTNNNACAGAAGAACAGGTTTCCTTCAT
AGCACACGTCTGAACTCTTTCCCTACACGACGCTCTTCCGATCTNNNAGTCTGTTACATTTTTTCAAACTC
AGCACACGTCTGAACTCTTTCCCTACACGACGCTCTTCCGATCTNNNGGAAAGCATCTTCTTAGTAAGA
AGCACACGTCTGAACTCTTTCCCTACACGACGCTCTTCCGATCTNNNAATTAAAATATCCATCACCTCAC
GAGCACACGTCTGAACTCTTTCCCTACACGACGCTCTTCCGATCTNNNTAGAAGAATGTCCTTATTCTCA
AGCACACGTCTGAACTCTTTCCCTACACGACGCTCTTCCGATCTNNNTAAAAAAACCCAAAACCACTTTA
AGCACACGTCTGAACTCTTTCCCTACACGACGCTCTTCCGATCTNNNTTAAGATACAAATGATGATGTC
GAGCACACGTCTGAACTCTTTCCCTACACGACGCTCTTCCGATCTNNNGAGAAACTTGAGTGATCTGCC
AGCACACGTCTGAACTCTTTCCCTACACGACGCTCTTCCGATCTNNNTTTCTCTTTCTGTGAATGAGGCA
AGCACACGTCTGAACTCTTTCCCTACACGACGCTCTTCCGATCTNNNCTCCAAATAAGAAAGAAAACAG
ACACGTCTGAACTCTTTCCCTACACGACGCTCTTCCGATCTNNNTCCTCTCTCTTATAAGAACTCTACATC
GAGCACACGTCTGAACTCTTTCCCTACACGACGCTCTTCCGATCTNNNAATTGGACTTCAGAAGTAAAC
CACGTCTGAACTCTTTCCCTACACGACGCTCTTCCGATCTNNNCTCTTCTGATTCTCTTATTCTTTCCATC
CACACGTCTGAACTCTTTCCCTACACGACGCTCTTCCGATCTNNNCAATCACTATTTGACAGAAAAGGCA
CACACGTCTGAACTCTTTCCCTACACGACGCTCTTCCGATCTNNNACTAAAACTTGACCTTTAAGAAGC
GAGCACACGTCTGAACTCTTTCCCTACACGACGCTCTTCCGATCTNNNATCACTGTCTACTTTTTTCCAAA
AGCACACGTCTGAACTCTTTCCCTACACGACGCTCTTCCGATCTNNNGCATAACATATGCAAAGTATCAT
CACACGTCTGAACTCTTTCCCTACACGACGCTCTTCCGATCTNNNCAAAATGAAGAAAGGGTAAGAAAC
GCACACGTCTGAACTCTTTCCCTACACGACGCTCTTCCGATCTNNNGGAGTTGGTAAGTCTAAAAAATC
GAGCACACGTCTGAACTCTTTCCCTACACGACGCTCTTCCGATCTNNNGATGTTCTTTTTATTTACCCCA
GAGCACACGTCTGAACTCTTTCCCTACACGACGCTCTTCCGATCTNNNTAAAATAAGTTAGATTAGCCC
AGAGCACACGTCTGAACTCTTTCCCTACACGACGCTCTTCCGATCTNNNCATCCCTGTATGTATGTATGT
GAGCACACGTCTGAACTCTTTCCCTACACGACGCTCTTCCGATCTNNNTCATGGATATTTATCTCAGAGA
AGCACACGTCTGAACTCTTTCCCTACACGACGCTCTTCCGATCTNNNAAAATTAGATATGTGGTGAAG
AGCACACGTCTGAACTCTTTCCCTACACGACGCTCTTCCGATCTNNNCTGTGTACTTAGCATGTGTGCAT

AGCACACGTCTGAACTCTTTCCCTACACGACGCTCTTCCGATCTNNNAATGATGGAAATCAACTAATG
ACACGTCTGAACTCTTTCCCTACACGACGCTCTTCCGATCTNNNTTGTATTTGAGATACTTTCAGGACA
AGCACACGTCTGAACTCTTTCCCTACACGACGCTCTTCCGATCTNNNCACCTTGTCATAAATCTCTTTC
GCACACGTCTGAACTCTTTCCCTACACGACGCTCTTCCGATCTNNNTATTATCATATACCTAGCCTCCC
GCACACGTCTGAACTCTTTCCCTACACGACGCTCTTCCGATCTNNNTATTTTAGTTTTACACCTCAGGTC
AGCACACGTCTGAACTCTTTCCCTACACGACGCTCTTCCGATCTNNNTCAATATATTGATAATGAAGC
ACACGTCTGAACTCTTTCCCTACACGACGCTCTTCCGATCTNNNGACTTAGCCTTCATATTCATGTCAT
ACACGTCTGAACTCTTTCCCTACACGACGCTCTTCCGATCTNNNTTTATTTGTTCAAATTGTTCCTGCA
AGCACACGTCTGAACTCTTTCCCTACACGACGCTCTTCCGATCTNNNACATAGGGAATGAACTAAAA
ACACGTCTGAACTCTTTCCCTACACGACGCTCTTCCGATCTNNNCATAGAGGAGACAGATGAAAGCA
AGCACACGTCTGAACTCTTTCCCTACACGACGCTCTTCCGATCTNNNGATTTAGGGCAACAGATGAAT
GCACACGTCTGAACTCTTTCCCTACACGACGCTCTTCCGATCTNNNAGGAAATATTTATCCTGTTGATT
AGCACACGTCTGAACTCTTTCCCTACACGACGCTCTTCCGATCTNNNAATAGACAAAATTTCACATGG
GCACACGTCTGAACTCTTTCCCTACACGACGCTCTTCCGATCTNNNGTGAAACTTAAAAATTCTAAGG
AGCACACGTCTGAACTCTTTCCCTACACGACGCTCTTCCGATCTNNNAACTTGATATCTGCATACAAA
GCACACGTCTGAACTCTTTCCCTACACGACGCTCTTCCGATCTNNNAAATACAACCAAGAATACAAGA
AGCACACGTCTGAACTCTTTCCCTACACGACGCTCTTCCGATCTNNNCTGAGTAAATCCATATTCCACT
GAGCACACGTCTGAACTCTTTCCCTACACGACGCTCTTCCGATCTNNNCTGTCTTGTATTGATATCAAA
AGCACACGTCTGAACTCTTTCCCTACACGACGCTCTTCCGATCTNNNTTGGACTGACATTGTAAGTAG
AGCACACGTCTGAACTCTTTCCCTACACGACGCTCTTCCGATCTNNNTATGCAATAACGATCTTTTAAC
CACGTCTGAACTCTTTCCCTACACGACGCTCTTCCGATCTNNNCCTCCTTCTTGTTTTGTAGATCATCG
GCACACGTCTGAACTCTTTCCCTACACGACGCTCTTCCGATCTNNNCAAGATCCATTTCTTCCCAACAT
AGCACACGTCTGAACTCTTTCCCTACACGACGCTCTTCCGATCTNNNCAACACATTTGCAAGTTTTTCC
AGCACACGTCTGAACTCTTTCCCTACACGACGCTCTTCCGATCTNNNAAAATTGAGGTTGTATAATGC
AGCACACGTCTGAACTCTTTCCCTACACGACGCTCTTCCGATCTNNNTATAAAGGAAAGAGGGTAAA
GCACACGTCTGAACTCTTTCCCTACACGACGCTCTTCCGATCTNNNCAAAACAAAACAAAAAACAACT
ACACGTCTGAACTCTTTCCCTACACGACGCTCTTCCGATCTNNNTTGAGGAGTTTTCATTAGGATTCA
ACACGTCTGAACTCTTTCCCTACACGACGCTCTTCCGATCTNNNTTGAGGAGTTTTCATTAGGATTCA
GAGCACACGTCTGAACTCTTTCCCTACACGACGCTCTTCCGATCTNNNTGAATTGAAGTTTACCTCCTA
AGCACACGTCTGAACTCTTTCCCTACACGACGCTCTTCCGATCTNNNTATACAAACGTACACTGTGATC
GCACACGTCTGAACTCTTTCCCTACACGACGCTCTTCCGATCTNNNGTCAGAAATTTGAATTACTTCC
GCACACGTCTGAACTCTTTCCCTACACGACGCTCTTCCGATCTNNNATTTTGGTTGTATTATCTTCTGG
AGCACACGTCTGAACTCTTTCCCTACACGACGCTCTTCCGATCTNNNAAAGTCTTCTGAAATTTGTTAC
AGCACACGTCTGAACTCTTTCCCTACACGACGCTCTTCCGATCTNNNCATTCAAGTCTCAGTTTTCAAT
CACACGTCTGAACTCTTTCCCTACACGACGCTCTTCCGATCTNNNGGAAACACATCAAGAGAAGACA
CACGTCTGAACTCTTTCCCTACACGACGCTCTTCCGATCTNNNATTTTTATAGCTATGCAAAGTGGCA
AGCACACGTCTGAACTCTTTCCCTACACGACGCTCTTCCGATCTNNNTGATTTTATTCTGAAGGAGTAC
ACACGTCTGAACTCTTTCCCTACACGACGCTCTTCCGATCTNNNTCTTTTGTGCTTTCTCTCTATATCAT
AGCACACGTCTGAACTCTTTCCCTACACGACGCTCTTCCGATCTNNNCAGAGTAGAGACAAAATATTT
GCACACGTCTGAACTCTTTCCCTACACGACGCTCTTCCGATCTNNNATCTCTCAATGATAGGCCTATC
AGCACACGTCTGAACTCTTTCCCTACACGACGCTCTTCCGATCTNNNCAGAGATCTAGGTCTTTATTT
AGCACACGTCTGAACTCTTTCCCTACACGACGCTCTTCCGATCTNNNAGAAAGATATGAAAGAATAA
GAGCACACGTCTGAACTCTTTCCCTACACGACGCTCTTCCGATCTNNNTGAGACCATGTCCCTAATATA

GCACACGTCTGAACTCTTTCCCTACACGACGCTCTTCCGATCTNNNGTGTGTGTGTGTGTAATATATA
GCACACGTCTGAACTCTTTCCCTACACGACGCTCTTCCGATCTNNNATAACATTTTGTATACTACGCAG
CACACGTCTGAACTCTTTCCCTACACGACGCTCTTCCGATCTNNNGATGTGTTTAGTCTCACCCACATCG
AGCACACGTCTGAACTCTTTCCCTACACGACGCTCTTCCGATCTNNNTGCCTATTTCTTTTCTTTGATTT
CACACGTCTGAACTCTTTCCCTACACGACGCTCTTCCGATCTNNNGAATGTAATGAAGGTGACATGC
AGCACACGTCTGAACTCTTTCCCTACACGACGCTCTTCCGATCTNNNGTTTGTTTTCCAAATTTTACACT
AGCACACGTCTGAACTCTTTCCCTACACGACGCTCTTCCGATCTNNNTAGAAATACTCATTGTTTCTTG
AGCACACGTCTGAACTCTTTCCCTACACGACGCTCTTCCGATCTNNNATTTAGGCTTTTTCTAGTTTTG
GAGCACACGTCTGAACTCTTTCCCTACACGACGCTCTTCCGATCTNNNTATACACACACAGATATGTAC
AGCACACGTCTGAACTCTTTCCCTACACGACGCTCTTCCGATCTNNNTGATATCTACTTTCACCATTTCT
AGCACACGTCTGAACTCTTTCCCTACACGACGCTCTTCCGATCTNNNTTCCCTTCTAATAAGACAAAAG
AGCACACGTCTGAACTCTTTCCCTACACGACGCTCTTCCGATCTNNNAATAACATGGAAAATGCCATTA
AGCACACGTCTGAACTCTTTCCCTACACGACGCTCTTCCGATCTNNNACACACACATATATATATACAC
AGCACACGTCTGAACTCTTTCCCTACACGACGCTCTTCCGATCTNNNAAAGTACTATATGAGCAGTGTT
AGCACACGTCTGAACTCTTTCCCTACACGACGCTCTTCCGATCTNNNTTTTGAGTACTGCTGATAAGT
GCACACGTCTGAACTCTTTCCCTACACGACGCTCTTCCGATCTNNNGCATTTATGTGTTTCTCTTTGTA
GCACACGTCTGAACTCTTTCCCTACACGACGCTCTTCCGATCTNNNAATAGATAAAAGGGGCTGTTCC
CACACGTCTGAACTCTTTCCCTACACGACGCTCTTCCGATCTNNNGTGTTCTAATATGTGTCCGTTTCAT
GCACACGTCTGAACTCTTTCCCTACACGACGCTCTTCCGATCTNNNTGGAGTACAGAAATAGGTCCCA
AGCACACGTCTGAACTCTTTCCCTACACGACGCTCTTCCGATCTNNNAGAAAAATAAAGAGCAAAAAT
GAGCACACGTCTGAACTCTTTCCCTACACGACGCTCTTCCGATCTNNNGCACATTCAGAAAATTATAC
AGCACACGTCTGAACTCTTTCCCTACACGACGCTCTTCCGATCTNNNGGTAGCAAAGAGTTCTTATAA
AGCACACGTCTGAACTCTTTCCCTACACGACGCTCTTCCGATCTNNNAATGTTTTTACCCTGTCTTAAA
CACACGTCTGAACTCTTTCCCTACACGACGCTCTTCCGATCTNNNAGTATTGTCTCATTCTTACTCTTCA
GCACACGTCTGAACTCTTTCCCTACACGACGCTCTTCCGATCTNNNTGAGTTTGTAGATTCTTTCTTTT
CACACGTCTGAACTCTTTCCCTACACGACGCTCTTCCGATCTNNNGCAGCCTTCTTAAATAAATAAGA
CACGTCTGAACTCTTTCCCTACACGACGCTCTTCCGATCTNNNCTCACTGTTCCTAATTCTCAGCATCGG
GAGCACACGTCTGAACTCTTTCCCTACACGACGCTCTTCCGATCTNNNGAGTAGCTCATATTACACCAA
GCACACGTCTGAACTCTTTCCCTACACGACGCTCTTCCGATCTNNNTCTTACAAAAAAACTGTTGTAGT
AGCACACGTCTGAACTCTTTCCCTACACGACGCTCTTCCGATCTNNNGAGGAGAAAAAAGAAAAG
GCACACGTCTGAACTCTTTCCCTACACGACGCTCTTCCGATCTNNNGAAACACGAATCAATTTTGAAA
GCACACGTCTGAACTCTTTCCCTACACGACGCTCTTCCGATCTNNNAGCTGTGTATTTAGAAAGTCTC
GCACACGTCTGAACTCTTTCCCTACACGACGCTCTTCCGATCTNNNAGCTGTGTATTTAGAAAGTCTC
CACACGTCTGAACTCTTTCCCTACACGACGCTCTTCCGATCTNNNCCAATGAAAAGTTTAGATTTCCTCA
AGCACACGTCTGAACTCTTTCCCTACACGACGCTCTTCCGATCTNNNTACTTTCTCACCCAGATGTTCAT
AGCACACGTCTGAACTCTTTCCCTACACGACGCTCTTCCGATCTNNNTACTTTCTCACCCAGATGTTCAT
GCACACGTCTGAACTCTTTCCCTACACGACGCTCTTCCGATCTNNNGTAAGGGCTCATAAATATTTGTT
AGCACACGTCTGAACTCTTTCCCTACACGACGCTCTTCCGATCTNNNCTTTGGAAGCATCAAAAATTAT
AGCACACGTCTGAACTCTTTCCCTACACGACGCTCTTCCGATCTNNNGATGGTTGGATGAAGCTAATA
CACACGTCTGAACTCTTTCCCTACACGACGCTCTTCCGATCTNNNAAATAAAACTGTAATGCACTCTCC
AGCACACGTCTGAACTCTTTCCCTACACGACGCTCTTCCGATCTNNNAGGTTCCACAAAAAGTTAAAAA
CACACGTCTGAACTCTTTCCCTACACGACGCTCTTCCGATCTNNNCTTGTTCCCACAGAAACTAAACATC
GCACACGTCTGAACTCTTTCCCTACACGACGCTCTTCCGATCTNNNATATGATTTTTTCAGCATGGTTG

GCACACGTCTGAACTCTTTCCCTACACGACGCTCTTCCGATCTNNNGAACTCCTGACCTCATGATCCATC
GCACACGTCTGAACTCTTTCCCTACACGACGCTCTTCCGATCTNNNAAACTCTCTTCTGGTTATTGTTCA
AGCACACGTCTGAACTCTTTCCCTACACGACGCTCTTCCGATCTNNNCTCTGGTTTTTCTTCATTCATAC
AGAGCACACGTCTGAACTCTTTCCCTACACGACGCTCTTCCGATCTNNNTATGGCATCATTACATAACTT
AGCACACGTCTGAACTCTTTCCCTACACGACGCTCTTCCGATCTNNNGTCCTGTGAAGCTTTTATTTTCC
AGCACACGTCTGAACTCTTTCCCTACACGACGCTCTTCCGATCTNNNACTTGGTTGAAAGTCTATTTTG
GAGCACACGTCTGAACTCTTTCCCTACACGACGCTCTTCCGATCTNNNAGCAGAGAAGCCATGTAAATA
AGCACACGTCTGAACTCTTTCCCTACACGACGCTCTTCCGATCTNNNCATCCATAAGGGTCATTTCACCA
AGCACACGTCTGAACTCTTTCCCTACACGACGCTCTTCCGATCTNNNAAGAGAAGGAATTTCATTGGA
AGCACACGTCTGAACTCTTTCCCTACACGACGCTCTTCCGATCTNNNAGTTGTCATTCATATCTGTAGGC
ACACGTCTGAACTCTTTCCCTACACGACGCTCTTCCGATCTNNNAATTTCCTTCATGCTCACATAACATCG
CACGTCTGAACTCTTTCCCTACACGACGCTCTTCCGATCTNNNGATATCAGGGTACTAGCACTCATCGG
AGCACACGTCTGAACTCTTTCCCTACACGACGCTCTTCCGATCTNNNACTGTCAAAATAAAAAAGGACA
AGCACACGTCTGAACTCTTTCCCTACACGACGCTCTTCCGATCTNNNTTACTCCTATTATAAAACAGGG
CACACGTCTGAACTCTTTCCCTACACGACGCTCTTCCGATCTNNNGAAGAGTTGTCCTTTCTCTCCATCG
GCACACGTCTGAACTCTTTCCCTACACGACGCTCTTCCGATCTNNNCTACTGGACATGGAAGAGTTCAT
GCACACGTCTGAACTCTTTCCCTACACGACGCTCTTCCGATCTNNNTAATTAGATATGACTTGAGCTCA
AGCACACGTCTGAACTCTTTCCCTACACGACGCTCTTCCGATCTNNNCACTCGAATCAAATACTGAAATA
GCACACGTCTGAACTCTTTCCCTACACGACGCTCTTCCGATCTNNNGGAGGTACTTCACTTAGAATAAT
AGCACACGTCTGAACTCTTTCCCTACACGACGCTCTTCCGATCTNNNGCTATGATAAATGGATTTACTC
GAGCACACGTCTGAACTCTTTCCCTACACGACGCTCTTCCGATCTNNNCAATGAGAGGAATATATGTAG
AGCACACGTCTGAACTCTTTCCCTACACGACGCTCTTCCGATCTNNNTGAGGATATTCAGATGATAAGA
GAGCACACGTCTGAACTCTTTCCCTACACGACGCTCTTCCGATCTNNNGGATTTCAAAATTATTTAGTA
AGCACACGTCTGAACTCTTTCCCTACACGACGCTCTTCCGATCTNNNCTTCTTTCATCTCTCTCATTCTCA
AGCACACGTCTGAACTCTTTCCCTACACGACGCTCTTCCGATCTNNNCAGAAAGTATTAAGGAGTGAG
AGCACACGTCTGAACTCTTTCCCTACACGACGCTCTTCCGATCTNNNCAGAAAGTATTAAGGAGTGAG
GAGCACACGTCTGAACTCTTTCCCTACACGACGCTCTTCCGATCTNNNTTATATATTTATTGAAGGCTCG
AGCACACGTCTGAACTCTTTCCCTACACGACGCTCTTCCGATCTNNNGTCTTTTGTGAAAAGCAATTCT
GCACACGTCTGAACTCTTTCCCTACACGACGCTCTTCCGATCTNNNGTTGGTGTGTTTTTAGGTATTTA
GCACACGTCTGAACTCTTTCCCTACACGACGCTCTTCCGATCTNNNAGGATGTCAAAGGTAAGTAAAA
AGCACACGTCTGAACTCTTTCCCTACACGACGCTCTTCCGATCTNNNATAAAAATCAGTGGTGTATTTG
GAGCACACGTCTGAACTCTTTCCCTACACGACGCTCTTCCGATCTNNNGTTTCTTTTTGCTTAATTGCATC
AGCACACGTCTGAACTCTTTCCCTACACGACGCTCTTCCGATCTNNNGAAATTACAGTCTTGGTCAGAT
AGCACACGTCTGAACTCTTTCCCTACACGACGCTCTTCCGATCTNNNTTAGTTATGGTAATAGGCATAC
AGCACACGTCTGAACTCTTTCCCTACACGACGCTCTTCCGATCTNNNTATTTCTCCTATCTGCATGTTTA
GAGCACACGTCTGAACTCTTTCCCTACACGACGCTCTTCCGATCTNNNCCACAAAGAATTATTTTCAAC
GAGCACACGTCTGAACTCTTTCCCTACACGACGCTCTTCCGATCTNNNGGAAAAATATCACATGCAAAT
GAGCACACGTCTGAACTCTTTCCCTACACGACGCTCTTCCGATCTNNNATAGTAGATTTGAATAGGGAG
GAGCACACGTCTGAACTCTTTCCCTACACGACGCTCTTCCGATCTNNNTGGAAAAATTATCTTCCATGA
AGCACACGTCTGAACTCTTTCCCTACACGACGCTCTTCCGATCTNNNCAGGTACCCAAAAAATTTAAAA
GAGCACACGTCTGAACTCTTTCCCTACACGACGCTCTTCCGATCTNNNATGATTAAAGGCTTATTTGAG
GAGCACACGTCTGAACTCTTTCCCTACACGACGCTCTTCCGATCTNNNCCTACCCCTGAATTTAGTAATG
AGCACACGTCTGAACTCTTTCCCTACACGACGCTCTTCCGATCTNNNTAAGTTAGATTCTGTAGGACCTA

AGCACACGTCTGAACTCTTTCCCTACACGACGCTCTTCCGATCTNNNGACTGTGTATGTATATATGCAT
GAGCACACGTCTGAACTCTTTCCCTACACGACGCTCTTCCGATCTNNNTTCCTTTTTGAAAATTTGCACT
GCACACGTCTGAACTCTTTCCCTACACGACGCTCTTCCGATCTNNNCTGTTCACTGACACAATATATACCA
AGCACACGTCTGAACTCTTTCCCTACACGACGCTCTTCCGATCTNNNGAAATGACTTGGTCAATTTTCT
AGCACACGTCTGAACTCTTTCCCTACACGACGCTCTTCCGATCTNNNTTCTGAATATTTTCAATCCATGG
AGCACACGTCTGAACTCTTTCCCTACACGACGCTCTTCCGATCTNNNACTGACTATTTCTCAAGATACA
CACACGTCTGAACTCTTTCCCTACACGACGCTCTTCCGATCTNNNATTTAGAGTTCTAATCAAGGTGAC
AGCACACGTCTGAACTCTTTCCCTACACGACGCTCTTCCGATCTNNNATCATGTCATGTTATTTCTCAA
GAGCACACGTCTGAACTCTTTCCCTACACGACGCTCTTCCGATCTNNNATACACACACACACATATGTAT
GAGCACACGTCTGAACTCTTTCCCTACACGACGCTCTTCCGATCTNNNATACACACACACACATATGTAT
GAGCACACGTCTGAACTCTTTCCCTACACGACGCTCTTCCGATCTNNNTATATGTAGATACCAAAGGA
AGCACACGTCTGAACTCTTTCCCTACACGACGCTCTTCCGATCTNNNACTTTCCTTTATGAGTGAAAGTA
AGCACACGTCTGAACTCTTTCCCTACACGACGCTCTTCCGATCTNNNCCAGATCCCCTTATAACATAAAA
GCACACGTCTGAACTCTTTCCCTACACGACGCTCTTCCGATCTNNNAGACACTAAAGCTGTCTAAAATCA
AGCACACGTCTGAACTCTTTCCCTACACGACGCTCTTCCGATCTNNNTTTTCTTCCCGAATATTTCAGTTC
AGCACACGTCTGAACTCTTTCCCTACACGACGCTCTTCCGATCTNNNTATCTTCTCGTTTTATGGTTTTC
AGCACACGTCTGAACTCTTTCCCTACACGACGCTCTTCCGATCTNNNCTATAGGTACCAAGTTCTTTCCC
AGCACACGTCTGAACTCTTTCCCTACACGACGCTCTTCCGATCTNNNCTATAGGTACCAAGTTCTTTCCC
GCACACGTCTGAACTCTTTCCCTACACGACGCTCTTCCGATCTNNNGCTAAATCAAAGAAGATGCATAA
AGCACACGTCTGAACTCTTTCCCTACACGACGCTCTTCCGATCTNNNGCGTTTCTCATAGGATAATTAA
GAGCACACGTCTGAACTCTTTCCCTACACGACGCTCTTCCGATCTNNNGCAATATTCTTTTCTCTGTTTTG
AGCACACGTCTGAACTCTTTCCCTACACGACGCTCTTCCGATCTNNNTGTAGTCCTATCTACATACAAT
GAGCACACGTCTGAACTCTTTCCCTACACGACGCTCTTCCGATCTNNNTAGTACGTGATTGTCTATTTTT
GAGCACACGTCTGAACTCTTTCCCTACACGACGCTCTTCCGATCTNNNCATTATTCTAGGAGCACATTCT
AGCACACGTCTGAACTCTTTCCCTACACGACGCTCTTCCGATCTNNNCCTTGGAGTTAAAAGAAGAAT
CACACGTCTGAACTCTTTCCCTACACGACGCTCTTCCGATCTNNNATTCCTAATGATCTGATAAGCTTCA
AGCACACGTCTGAACTCTTTCCCTACACGACGCTCTTCCGATCTNNNTATAGCTGGTGACTATCATAAA
AGCACACGTCTGAACTCTTTCCCTACACGACGCTCTTCCGATCTNNNTAAAACTGCCTTATCCTTTGTAT
AGCACACGTCTGAACTCTTTCCCTACACGACGCTCTTCCGATCTNNNGATCATATCATGACTGAGCTCCA
AGCACACGTCTGAACTCTTTCCCTACACGACGCTCTTCCGATCTNNNATTCATCACAATCTACAGGATA
AGCACACGTCTGAACTCTTTCCCTACACGACGCTCTTCCGATCTNNNATTCATCACAATCTACAGGATA
GAGCACACGTCTGAACTCTTTCCCTACACGACGCTCTTCCGATCTNNNTTTTTTGTTATGTGAATTTCAC
GAGCACACGTCTGAACTCTTTCCCTACACGACGCTCTTCCGATCTNNNTTCATTCATCAGACATTTTCTT
AGCACACGTCTGAACTCTTTCCCTACACGACGCTCTTCCGATCTNNNCTAAATTGAGAAGCTAAATCAG
AGCACACGTCTGAACTCTTTCCCTACACGACGCTCTTCCGATCTNNNAAAGATACGGTTACATAGTAA
AGCACACGTCTGAACTCTTTCCCTACACGACGCTCTTCCGATCTNNNCTGAAGAAAACAAAAACAATC
GAGCACACGTCTGAACTCTTTCCCTACACGACGCTCTTCCGATCTNNNTGAATTGGTTATGCAATCTTTT
GCACACGTCTGAACTCTTTCCCTACACGACGCTCTTCCGATCTNNNACATTTCTATTTTGTCAAAATCAC
GCACACGTCTGAACTCTTTCCCTACACGACGCTCTTCCGATCTNNNACATTTCTATTTTGTCAAAATCAC
AGCACACGTCTGAACTCTTTCCCTACACGACGCTCTTCCGATCTNNNTTCAATCTAAGTGTCCAACTCCA
GAGCACACGTCTGAACTCTTTCCCTACACGACGCTCTTCCGATCTNNNGTCCTTTGGATAGAGAAATCA
GAGCACACGTCTGAACTCTTTCCCTACACGACGCTCTTCCGATCTNNNCATTCTCACATATGGGAGGAA
AGCACACGTCTGAACTCTTTCCCTACACGACGCTCTTCCGATCTNNNACCAGTTGTTTTTTCATAGTTA

AGCACACGTCTGAACTCTTTCCCTACACGACGCTCTTCCGATCTNNNCATTCTCACATATGGGAGGAA
GCACACGTCTGAACTCTTTCCCTACACGACGCTCTTCCGATCTNNNGAAAAAAAAAGAGGAAGGGA
ACACGTCTGAACTCTTTCCCTACACGACGCTCTTCCGATCTNNNGGACTTCCTACTGGAGATAATACA
GCACACGTCTGAACTCTTTCCCTACACGACGCTCTTCCGATCTNNNGCATAGCATCTATCATAGAACTA
GCACACGTCTGAACTCTTTCCCTACACGACGCTCTTCCGATCTNNNTGGAAAATACGATTGAAGAGA
GCACACGTCTGAACTCTTTCCCTACACGACGCTCTTCCGATCTNNNCATTTGGATATTATCTTGGTGT
AGCACACGTCTGAACTCTTTCCCTACACGACGCTCTTCCGATCTNNNGGGATTTGGCATAATAATATG
AGCACACGTCTGAACTCTTTCCCTACACGACGCTCTTCCGATCTNNNCTTACTGTGTGCTAAATAATTG
AGCACACGTCTGAACTCTTTCCCTACACGACGCTCTTCCGATCTNNNAGTAGAAAAAAAAAACTCAAG
CACACGTCTGAACTCTTTCCCTACACGACGCTCTTCCGATCTNNNGAAATACAAAGGTGAAGTAGGA
CACACGTCTGAACTCTTTCCCTACACGACGCTCTTCCGATCTNNNTATTGGTTTCTTCCTCTAGTTCCA
GCACACGTCTGAACTCTTTCCCTACACGACGCTCTTCCGATCTNNNCTGGAGTTTGAATTTCCAATCC
AGCACACGTCTGAACTCTTTCCCTACACGACGCTCTTCCGATCTNNNCAAATCTCTGCAATATTGAAG
AGCACACGTCTGAACTCTTTCCCTACACGACGCTCTTCCGATCTNNNCACATTGTTTCTTACTCAACATA
GAGCACACGTCTGAACTCTTTCCCTACACGACGCTCTTCCGATCTNNNAAAGAGAACAGTAGAAAAC
AGCACACGTCTGAACTCTTTCCCTACACGACGCTCTTCCGATCTNNNAAGCCTATAAAGTTGCTTTAT
CACACGTCTGAACTCTTTCCCTACACGACGCTCTTCCGATCTNNNTTTGAAATGAGCCAAAAGATTTA
AGCACACGTCTGAACTCTTTCCCTACACGACGCTCTTCCGATCTNNNAAATAAAAGTCAATGTGAAG
GCACACGTCTGAACTCTTTCCCTACACGACGCTCTTCCGATCTNNNCTTACAAAATCCTGGTTCTAATG
AGCACACGTCTGAACTCTTTCCCTACACGACGCTCTTCCGATCTNNNAAGAAAGACTTTACCTAACTC
GCACACGTCTGAACTCTTTCCCTACACGACGCTCTTCCGATCTNNNAGATGTTTCTTATTGGGAACAC
GCACACGTCTGAACTCTTTCCCTACACGACGCTCTTCCGATCTNNNGTAACAGTAGATTTTGTATCCT
AGCACACGTCTGAACTCTTTCCCTACACGACGCTCTTCCGATCTNNNCAAATACACTGCTGAAATCAA
AGCACACGTCTGAACTCTTTCCCTACACGACGCTCTTCCGATCTNNNGTGAAACTACACATCTAAATGT
AGCACACGTCTGAACTCTTTCCCTACACGACGCTCTTCCGATCTNNNTGTGTAATGGCTCTATTATATG
AGCACACGTCTGAACTCTTTCCCTACACGACGCTCTTCCGATCTNNNACCTGTGATGATTAGTATTAG
GCACACGTCTGAACTCTTTCCCTACACGACGCTCTTCCGATCTNNNCTTAAAATCTCTCTTCTCACATG
GCACACGTCTGAACTCTTTCCCTACACGACGCTCTTCCGATCTNNNAATGGTTCTCAGAAAAGTCTAA
CACACGTCTGAACTCTTTCCCTACACGACGCTCTTCCGATCTNNNATTACAGAGGAACCTGAAGTCAT
AGCACACGTCTGAACTCTTTCCCTACACGACGCTCTTCCGATCTNNNAATGACAGCTAATATGTATTG
ACACGTCTGAACTCTTTCCCTACACGACGCTCTTCCGATCTNNNGGATAACCTCCTAATTTTATTGCCA
GCACACGTCTGAACTCTTTCCCTACACGACGCTCTTCCGATCTNNNATAAAACCAAGCCAAATCAAAA
CACACGTCTGAACTCTTTCCCTACACGACGCTCTTCCGATCTNNNCAAAGCATAAAGAGAGAAGAAG
AGCACACGTCTGAACTCTTTCCCTACACGACGCTCTTCCGATCTNNNTATATATACATAGGTGTCTGT
GAGCACACGTCTGAACTCTTTCCCTACACGACGCTCTTCCGATCTNNNAAAAAATGTTTAAAGGCTAT
AGCACACGTCTGAACTCTTTCCCTACACGACGCTCTTCCGATCTNNNTTTCCTTCGATTCTTTGATGTAT
AGCACACGTCTGAACTCTTTCCCTACACGACGCTCTTCCGATCTNNNGAGAAAAAACTTGTACTTACC
AGCACACGTCTGAACTCTTTCCCTACACGACGCTCTTCCGATCTNNNTTTCATTATAACCATCATGCA
AGCACACGTCTGAACTCTTTCCCTACACGACGCTCTTCCGATCTNNNGATGAGAACTGAAGACTCATA
GCACACGTCTGAACTCTTTCCCTACACGACGCTCTTCCGATCTNNNCTTACTCAAAACTCCAATAAGCC
GCACACGTCTGAACTCTTTCCCTACACGACGCTCTTCCGATCTNNNGTCCAAATAGAATAAAGAGCAT
GCACACGTCTGAACTCTTTCCCTACACGACGCTCTTCCGATCTNNNTTCTTCACTTTCAGTCTATATGT
AGCACACGTCTGAACTCTTTCCCTACACGACGCTCTTCCGATCTNNNCATCAATCTTTGAAGAACTATG

GAGCACACGTCTGAACTCTTTCCCTACACGACGCTCTTCCGATCTNNNTAAGACAACTGTGTGACCTAT
AGCACACGTCTGAACTCTTTCCCTACACGACGCTCTTCCGATCTNNNCTTGGGGTTCGATATAGATATA
ACACGTCTGAACTCTTTCCCTACACGACGCTCTTCCGATCTNNNCCTTTACAGTTTTTACTCTTGAACAT
GAGCACACGTCTGAACTCTTTCCCTACACGACGCTCTTCCGATCTNNNTTACCTCCCAAATTTAAAATCA
GCACACGTCTGAACTCTTTCCCTACACGACGCTCTTCCGATCTNNNTTAATATAATGTCCTCCGAGTTT
AGCACACGTCTGAACTCTTTCCCTACACGACGCTCTTCCGATCTNNNAAACAGAAGCAATTTAAAGTTT
AGCACACGTCTGAACTCTTTCCCTACACGACGCTCTTCCGATCTNNNCTCACAGCTCTGATCAATTTCAT
GAGCACACGTCTGAACTCTTTCCCTACACGACGCTCTTCCGATCTNNNTCTGCTTCCTACATTAAAA
GAGCACACGTCTGAACTCTTTCCCTACACGACGCTCTTCCGATCTNNNCATAGTCCCATAGTCTTAGCT
GAGCACACGTCTGAACTCTTTCCCTACACGACGCTCTTCCGATCTNNNAAATGTTAAGGGACATTGAA
GAGCACACGTCTGAACTCTTTCCCTACACGACGCTCTTCCGATCTNNNAAATGTTAAGGGACATTGAA
GCACACGTCTGAACTCTTTCCCTACACGACGCTCTTCCGATCTNNNATAGATTTTCTGCCTAACTGATC
AGCACACGTCTGAACTCTTTCCCTACACGACGCTCTTCCGATCTNNNTTAAACTTTGTTCTGTTGAATCA
GCACACGTCTGAACTCTTTCCCTACACGACGCTCTTCCGATCTNNNATCTCTTAGAGCAGGAAATAAT
AGCACACGTCTGAACTCTTTCCCTACACGACGCTCTTCCGATCTNNNTTATGGCTGAATAATATCCCAT
AGCACACGTCTGAACTCTTTCCCTACACGACGCTCTTCCGATCTNNNTCTTGATTTATCACTTTGATGTT
AGAGCACACGTCTGAACTCTTTCCCTACACGACGCTCTTCCGATCTNNNGTCTGATGGAGGTGATTTT
AGCACACGTCTGAACTCTTTCCCTACACGACGCTCTTCCGATCTNNNATTGAGAGAGAACATCCCTAT
AGCACACGTCTGAACTCTTTCCCTACACGACGCTCTTCCGATCTNNNGAATTTGAGTGATTTGAGTGA
GAGCACACGTCTGAACTCTTTCCCTACACGACGCTCTTCCGATCTNNNTAACAATTCTGGAATGATAAG
GAGCACACGTCTGAACTCTTTCCCTACACGACGCTCTTCCGATCTNNNTAACAATTCTGGAATGATAAG
GAGCACACGTCTGAACTCTTTCCCTACACGACGCTCTTCCGATCTNNNTGGAAATGTTACATTAGTTAA
GAGCACACGTCTGAACTCTTTCCCTACACGACGCTCTTCCGATCTNNNTTATACCTAGTAAAGCAGTTG
GAGCACACGTCTGAACTCTTTCCCTACACGACGCTCTTCCGATCTNNNGTGGTATCTGTTTCATCTACAG
AGCACACGTCTGAACTCTTTCCCTACACGACGCTCTTCCGATCTNNNACATGGAAAAGTAGAAGGAA
AGCACACGTCTGAACTCTTTCCCTACACGACGCTCTTCCGATCTNNNTCAATCTTTCTCTCAACTTATTC
AGCACACGTCTGAACTCTTTCCCTACACGACGCTCTTCCGATCTNNNCTGCTAGTACTACTACTACTACT
AGCACACGTCTGAACTCTTTCCCTACACGACGCTCTTCCGATCTNNNATAACAACAAAGAAGCTTTCAT
AGCACACGTCTGAACTCTTTCCCTACACGACGCTCTTCCGATCTNNNATCATTCCCAATAACTCACTTAT
GCACACGTCTGAACTCTTTCCCTACACGACGCTCTTCCGATCTNNNATAGTCAGGAGAGAACTCTTCCA
GCACACGTCTGAACTCTTTCCCTACACGACGCTCTTCCGATCTNNNGTAAGTGATCAGTTACCAAATTT
AGCACACGTCTGAACTCTTTCCCTACACGACGCTCTTCCGATCTNNNGAAAACAAACTTGAGTATAGA
GCACACGTCTGAACTCTTTCCCTACACGACGCTCTTCCGATCTNNNAGTGGTTGAAAAGACAATTATT
AGCACACGTCTGAACTCTTTCCCTACACGACGCTCTTCCGATCTNNNAGATGGAGCTGGTATTTCAACA
AGCACACGTCTGAACTCTTTCCCTACACGACGCTCTTCCGATCTNNNACTTGACTATATAGCATTCACTT
AGCACACGTCTGAACTCTTTCCCTACACGACGCTCTTCCGATCTNNNGGAAGGATAAATTATTTGTTCT
GAGCACACGTCTGAACTCTTTCCCTACACGACGCTCTTCCGATCTNNNCAGGTTACATAATTTGCTCAT
GAGCACACGTCTGAACTCTTTCCCTACACGACGCTCTTCCGATCTNNNTGATATATAACCTAGAGGGAT
GAGCACACGTCTGAACTCTTTCCCTACACGACGCTCTTCCGATCTNNNCTGCTCTTAATCGAGATTTTT
AGCACACGTCTGAACTCTTTCCCTACACGACGCTCTTCCGATCTNNNTAGCCATCTACATCCTGATTCA
GAGCACACGTCTGAACTCTTTCCCTACACGACGCTCTTCCGATCTNNNCTCTATTCCTCTCTGTCTCTCCA
GAGCACACGTCTGAACTCTTTCCCTACACGACGCTCTTCCGATCTNNNCTCTATTCCTCTCTGTCTCTCCA
GAGCACACGTCTGAACTCTTTCCCTACACGACGCTCTTCCGATCTNNNGGTAACATAATTCAACTGCAT

GAGCACACGTCTGAACTCTTTCCCTACACGACGCTCTTCCGATCTNNNGGTAACATAATTCAACTGCATT
AGCACACGTCTGAACTCTTTCCCTACACGACGCTCTTCCGATCTNNNAAATTTAGCTCCTCAAAAGAAA
GCACACGTCTGAACTCTTTCCCTACACGACGCTCTTCCGATCTNNNCTTTGTAGTTCCTACCCAATTTCAT
AGCACACGTCTGAACTCTTTCCCTACACGACGCTCTTCCGATCTNNNTTTTAAAAAAAGTCTTCTTCCCA
GAGCACACGTCTGAACTCTTTCCCTACACGACGCTCTTCCGATCTNNNGACATAAAATCATCAGATGAG
GAGCACACGTCTGAACTCTTTCCCTACACGACGCTCTTCCGATCTNNNTATCTGTCTTAATTCATGAGTG
AGCACACGTCTGAACTCTTTCCCTACACGACGCTCTTCCGATCTNNNTTATACCTTTTCTGAGCTGTAGC
CACACGTCTGAACTCTTTCCCTACACGACGCTCTTCCGATCTNNNTAACTGTCTTTTAAAATTGCTGTCA
AGCACACGTCTGAACTCTTTCCCTACACGACGCTCTTCCGATCTNNNGTGTGACTTTTACTGAAAAAAA
CACACGTCTGAACTCTTTCCCTACACGACGCTCTTCCGATCTNNNTATTTTTGACAAAGGGCAATTTACA
AGCACACGTCTGAACTCTTTCCCTACACGACGCTCTTCCGATCTNNNTATGCAATATTTTTGACAAAGG
GAGCACACGTCTGAACTCTTTCCCTACACGACGCTCTTCCGATCTNNNTAGTAGGGGTTTGGAATTTTT
AGCACACGTCTGAACTCTTTCCCTACACGACGCTCTTCCGATCTNNNCTAAGTCCTTCCTCTAACTTAACC
AGCACACGTCTGAACTCTTTCCCTACACGACGCTCTTCCGATCTNNNTTCCCTGTCCCAAATACTTACATC
CACACGTCTGAACTCTTTCCCTACACGACGCTCTTCCGATCTNNNCTACTTTGCTTACACACCCTCATCGG
AGCACACGTCTGAACTCTTTCCCTACACGACGCTCTTCCGATCTNNNTAAATAAAAGAGTGCCTTGATT
AGCACACGTCTGAACTCTTTCCCTACACGACGCTCTTCCGATCTNNNGAAATATCGATTGCTTCTATTCC
AGCACACGTCTGAACTCTTTCCCTACACGACGCTCTTCCGATCTNNNCACAAACAAACAAAAAAAAATA
AGAGCACACGTCTGAACTCTTTCCCTACACGACGCTCTTCCGATCTNNNCTTTGATTCACTTTACAGGA
GAGCACACGTCTGAACTCTTTCCCTACACGACGCTCTTCCGATCTNNNTAGATTTTTCTCTTTCTTTCTGG
GAGCACACGTCTGAACTCTTTCCCTACACGACGCTCTTCCGATCTNNNTTTCAGATAAAAAGATCCAAC
ACACGTCTGAACTCTTTCCCTACACGACGCTCTTCCGATCTNNNACAATTCAGCCAAATTCTTTGCATCGC
GCACACGTCTGAACTCTTTCCCTACACGACGCTCTTCCGATCTNNNGTCTCTACTTCTGCTTCTTGCATCC
GCACACGTCTGAACTCTTTCCCTACACGACGCTCTTCCGATCTNNNTGCCAATCTAAAATTTATTAGGG
AGCACACGTCTGAACTCTTTCCCTACACGACGCTCTTCCGATCTNNNAAAATTCTCAACATGCCAAAACA
GAGCACACGTCTGAACTCTTTCCCTACACGACGCTCTTCCGATCTNNNTTGCTGTTTCCAAAGATAAAGC
GAGCACACGTCTGAACTCTTTCCCTACACGACGCTCTTCCGATCTNNNTTGCTGTTTCCAAAGATAAAGC
GAGCACACGTCTGAACTCTTTCCCTACACGACGCTCTTCCGATCTNNNGCAGTATAAACAGTGAAGACT
AGCACACGTCTGAACTCTTTCCCTACACGACGCTCTTCCGATCTNNNACATAAATACAAAGAAGAGAAG
GCACACGTCTGAACTCTTTCCCTACACGACGCTCTTCCGATCTNNNCTGAAGTCCTTTCTCATTGTCATCG
GCACACGTCTGAACTCTTTCCCTACACGACGCTCTTCCGATCTNNNTTTTGTCTCCCACAATCCTTCATCG
GAGCACACGTCTGAACTCTTTCCCTACACGACGCTCTTCCGATCTNNNCTAAATGTCAGAACTTAGATGT
GCACACGTCTGAACTCTTTCCCTACACGACGCTCTTCCGATCTNNNGGTGGTATATTTAAATGATCACC
AGCACACGTCTGAACTCTTTCCCTACACGACGCTCTTCCGATCTNNNAGGTTTCTTTTATTTTAACCTGT
CACACGTCTGAACTCTTTCCCTACACGACGCTCTTCCGATCTNNNTAATCATTGGTCCTCTTCGATTACAT
GCACACGTCTGAACTCTTTCCCTACACGACGCTCTTCCGATCTNNNGCAACTTTACGTAAACAATTTACC
AGCACACGTCTGAACTCTTTCCCTACACGACGCTCTTCCGATCTNNNTTTGTTATTTCAGTTGTCTAGTC
ACACGTCTGAACTCTTTCCCTACACGACGCTCTTCCGATCTNNNTTAATCCATAGTTGTGTACTGGCATC
GAGCACACGTCTGAACTCTTTCCCTACACGACGCTCTTCCGATCTNNNTGGAGTTTACCAATATTTGAAA
AGCACACGTCTGAACTCTTTCCCTACACGACGCTCTTCCGATCTNNNCTATTCTCTCTCTCTGTCTCCA
AGCACACGTCTGAACTCTTTCCCTACACGACGCTCTTCCGATCTNNNCTTTCCATTTTGCTGTAAATCTCA
GCACACGTCTGAACTCTTTCCCTACACGACGCTCTTCCGATCTNNNCTCTCTCACTTTCTCAATATTTGCA
GCACACGTCTGAACTCTTTCCCTACACGACGCTCTTCCGATCTNNNTGTAAGTATTATGTATGAAGGG

AGCACACGTCTGAACTCTTTCCCTACACGACGCTCTTCCGATCTNNNGTGAAAATCTCATTCATGTGTA
AGCACACGTCTGAACTCTTTCCCTACACGACGCTCTTCCGATCTNNNTATATATGCATGCATATGTCTG
AGCACACGTCTGAACTCTTTCCCTACACGACGCTCTTCCGATCTNNNCATCTTTAAAGATAGGATCAAT
GAGCACACGTCTGAACTCTTTCCCTACACGACGCTCTTCCGATCTNNNACACAAAAACCTATATATACC
GCACACGTCTGAACTCTTTCCCTACACGACGCTCTTCCGATCTNNNAAGACCCTGTCTGAAAAAAATAC
AGCACACGTCTGAACTCTTTCCCTACACGACGCTCTTCCGATCTNNNTGTGCTGAATATAGCTAGAATT
AGAGCACACGTCTGAACTCTTTCCCTACACGACGCTCTTCCGATCTNNNACTTAAGTGGTTAAAAAAAA
GAGCACACGTCTGAACTCTTTCCCTACACGACGCTCTTCCGATCTNNNAAATGGATGGAGGAAGATCT
AGCACACGTCTGAACTCTTTCCCTACACGACGCTCTTCCGATCTNNNGGAAAATGGGTATATGACAA
AGCACACGTCTGAACTCTTTCCCTACACGACGCTCTTCCGATCTNNNGCTATTCTATTTTCAAGGAGAA
GCACACGTCTGAACTCTTTCCCTACACGACGCTCTTCCGATCTNNNACATCTTTGTTTGAATTTCTGATC
GCACACGTCTGAACTCTTTCCCTACACGACGCTCTTCCGATCTNNNCACCCTTTAGGAAGTATATTCCCA
CACACGTCTGAACTCTTTCCCTACACGACGCTCTTCCGATCTNNNACATTCATCAGTGTCTGGAGCATCG
AGCACACGTCTGAACTCTTTCCCTACACGACGCTCTTCCGATCTNNNCTTAGACCTCAACACTAGATGC
ACACGTCTGAACTCTTTCCCTACACGACGCTCTTCCGATCTNNNAAATCTTTTAAGCAGTTCTCATCCAT
AGCACACGTCTGAACTCTTTCCCTACACGACGCTCTTCCGATCTNNNGTATCTAACATTGAAAGAGAA
AGCACACGTCTGAACTCTTTCCCTACACGACGCTCTTCCGATCTNNNGTATCTAACATTGAAAGAGAA
GCACACGTCTGAACTCTTTCCCTACACGACGCTCTTCCGATCTNNNCAGTGTTTTCAGTTACATACATA
GAGCACACGTCTGAACTCTTTCCCTACACGACGCTCTTCCGATCTNNNTGCAAGAAAATCAGACTAATA
AGCACACGTCTGAACTCTTTCCCTACACGACGCTCTTCCGATCTNNNTTATCTGATATAGGTGTACGTT
GAGCACACGTCTGAACTCTTTCCCTACACGACGCTCTTCCGATCTNNNACCCAGAAGAAATAAGTAAAT
GCACACGTCTGAACTCTTTCCCTACACGACGCTCTTCCGATCTNNNCTTAATGTATACATGCACACATA
GAGCACACGTCTGAACTCTTTCCCTACACGACGCTCTTCCGATCTNNNCCTAGAGGCTTTATTTTTGAAT
AGCACACGTCTGAACTCTTTCCCTACACGACGCTCTTCCGATCTNNNTTCCTTCATTAGTTGGTTTCACA
GAGCACACGTCTGAACTCTTTCCCTACACGACGCTCTTCCGATCTNNNCTTGTCTTGTGTCTTGTTCCCAT
GCACACGTCTGAACTCTTTCCCTACACGACGCTCTTCCGATCTNNNTTGTGAAACGTGAAAATCACCAT
GAGCACACGTCTGAACTCTTTCCCTACACGACGCTCTTCCGATCTNNNATTGAAGAAAATGTCAGTTA
ACACGTCTGAACTCTTTCCCTACACGACGCTCTTCCGATCTNNNCCAATCTTAGAGATAGTGACATACA
GAGCACACGTCTGAACTCTTTCCCTACACGACGCTCTTCCGATCTNNNCTAACTCATACACAGTATCAT
AGCACACGTCTGAACTCTTTCCCTACACGACGCTCTTCCGATCTNNNTGACTTTCTGAATGTTTCCTTC
GCACACGTCTGAACTCTTTCCCTACACGACGCTCTTCCGATCTNNNTCTTCTAGGCTATGAGATAGAG
GCACACGTCTGAACTCTTTCCCTACACGACGCTCTTCCGATCTNNNTCTTCTAGGCTATGAGATAGAG
AGCACACGTCTGAACTCTTTCCCTACACGACGCTCTTCCGATCTNNNAATACTGGCCTATAGATAAGGT
AGCACACGTCTGAACTCTTTCCCTACACGACGCTCTTCCGATCTNNNGTAAATATTTTCCTACTCAGGT
AGCACACGTCTGAACTCTTTCCCTACACGACGCTCTTCCGATCTNNNGCATATGCAAGTGTTTAATATG
GCACACGTCTGAACTCTTTCCCTACACGACGCTCTTCCGATCTNNNTTCTATAAGGTTTTTCTTTGGTT
GAGCACACGTCTGAACTCTTTCCCTACACGACGCTCTTCCGATCTNNNATCAACATTTTAACAGCTGTG
AGAGCACACGTCTGAACTCTTTCCCTACACGACGCTCTTCCGATCTNNNATAATACATATCTCTGTGAG
GAGCACACGTCTGAACTCTTTCCCTACACGACGCTCTTCCGATCTNNNTATAAGTTTATGCAAAATGAC
AGCACACGTCTGAACTCTTTCCCTACACGACGCTCTTCCGATCTNNNTTTGTTGTTGAAAGAATGACAT
CACACGTCTGAACTCTTTCCCTACACGACGCTCTTCCGATCTNNNCCTTCTTAGTTATTCCCTCTTACATC
GCACACGTCTGAACTCTTTCCCTACACGACGCTCTTCCGATCTNNNTTTTTTTTTTGTAGAGATGGAGT
AGCACACGTCTGAACTCTTTCCCTACACGACGCTCTTCCGATCTNNNTAAAAAACCTCCTCCATACAAAT

GAGCACACGTCTGAACTCTTTCCCTACACGACGCTCTTCCGATCTNNNTTATTTGAGAAACCATGCAA
AGCACACGTCTGAACTCTTTCCCTACACGACGCTCTTCCGATCTNNNCATAGCTAGTGATATCAGACT
AGCACACGTCTGAACTCTTTCCCTACACGACGCTCTTCCGATCTNNNAGTCTTTTCTTTACAAAGGTTTT
AGCACACGTCTGAACTCTTTCCCTACACGACGCTCTTCCGATCTNNNCAAGGAAAATAATTGGAGAT
GCACACGTCTGAACTCTTTCCCTACACGACGCTCTTCCGATCTNNNAACTAAGTACTTAGACCAACATT
AGCACACGTCTGAACTCTTTCCCTACACGACGCTCTTCCGATCTNNNGAAGGGTCCTTAGAAATTATCT
ACACGTCTGAACTCTTTCCCTACACGACGCTCTTCCGATCTNNNACATTTGATTTGAGTGGAAAATAC
AGCACACGTCTGAACTCTTTCCCTACACGACGCTCTTCCGATCTNNNGCAACATAAATTTATTTTTCAC
AGCACACGTCTGAACTCTTTCCCTACACGACGCTCTTCCGATCTNNNAATTGTCATTGTAATACCCTT
GCACACGTCTGAACTCTTTCCCTACACGACGCTCTTCCGATCTNNNAAGAGTTAAGGTTGCTGAAGCA
GCACACGTCTGAACTCTTTCCCTACACGACGCTCTTCCGATCTNNNTAAATAATTACTCCAATCTGGCT
AGCACACGTCTGAACTCTTTCCCTACACGACGCTCTTCCGATCTNNNTCTCAGTCTCTCAATATCTCTAT
AGCACACGTCTGAACTCTTTCCCTACACGACGCTCTTCCGATCTNNNTCTCAGTCTCTCAATATCTCTAT
AGCACACGTCTGAACTCTTTCCCTACACGACGCTCTTCCGATCTNNNTTTGATGCTAATTACCATTACAG
AGCACACGTCTGAACTCTTTCCCTACACGACGCTCTTCCGATCTNNNATACTATCACTGAGAAGACAT
GCACACGTCTGAACTCTTTCCCTACACGACGCTCTTCCGATCTNNNCTAGGAATCCGTGAATACATTC
AGCACACGTCTGAACTCTTTCCCTACACGACGCTCTTCCGATCTNNNCTGAACAACTGAATAAAACCCC
AGCACACGTCTGAACTCTTTCCCTACACGACGCTCTTCCGATCTNNNACTCTAAAGTTCTTCAGAATCTT
GCACACGTCTGAACTCTTTCCCTACACGACGCTCTTCCGATCTNNNAAGTTTCTGATTCATCTTGAATT
CACACGTCTGAACTCTTTCCCTACACGACGCTCTTCCGATCTNNNTCAATGAGTTGATCTTCGATGCAT
AGCACACGTCTGAACTCTTTCCCTACACGACGCTCTTCCGATCTNNNAACTTAAAAACTTCAGAGTTCT
AGCACACGTCTGAACTCTTTCCCTACACGACGCTCTTCCGATCTNNNCCAAAGAGAAAACAAGTTGAT
ACACGTCTGAACTCTTTCCCTACACGACGCTCTTCCGATCTNNNGTCAATCAGGCTATTTTATTTCCCA
CACACGTCTGAACTCTTTCCCTACACGACGCTCTTCCGATCTNNNCCTTACATTGTAGGAAATCGATA
AGCACACGTCTGAACTCTTTCCCTACACGACGCTCTTCCGATCTNNNCTTGTCCTCATTTTTTATTAGCT
AGCACACGTCTGAACTCTTTCCCTACACGACGCTCTTCCGATCTNNNCTTGTCCTCATTTTTTATTAGCT
GCACACGTCTGAACTCTTTCCCTACACGACGCTCTTCCGATCTNNNCCATTAATGGCAGTTGTTTCCAT
AGCACACGTCTGAACTCTTTCCCTACACGACGCTCTTCCGATCTNNNCTGATCTTCTATAAGGATGTC
AGCACACGTCTGAACTCTTTCCCTACACGACGCTCTTCCGATCTNNNTTAATGAGGCAATGTTTTAGA
AGCACACGTCTGAACTCTTTCCCTACACGACGCTCTTCCGATCTNNNTTAATGAGGCAATGTTTTAGA
AGCACACGTCTGAACTCTTTCCCTACACGACGCTCTTCCGATCTNNNTCCTTCTAAGGTTTGTTTTTACT
AGCACACGTCTGAACTCTTTCCCTACACGACGCTCTTCCGATCTNNNAAGATTGACAATAACCTAAAT
GCACACGTCTGAACTCTTTCCCTACACGACGCTCTTCCGATCTNNNATTATTGACACTGTTTTTCAGAA
GCACACGTCTGAACTCTTTCCCTACACGACGCTCTTCCGATCTNNNTGCACTATTTAGTTATTTTCCTG
AGCACACGTCTGAACTCTTTCCCTACACGACGCTCTTCCGATCTNNNTAAAAGCTATCAGAAACTTTG
AGCACACGTCTGAACTCTTTCCCTACACGACGCTCTTCCGATCTNNNTTCACCATCTGACCTTAATTATT
AGCACACGTCTGAACTCTTTCCCTACACGACGCTCTTCCGATCTNNNACAGTTAGTAAATAAGTTAT
AGCACACGTCTGAACTCTTTCCCTACACGACGCTCTTCCGATCTNNNCAAAGACCTCAGGAGAATATTT
GCACACGTCTGAACTCTTTCCCTACACGACGCTCTTCCGATCTNNNGTATGTGAACCCTGTCTCTTCAT
ACACGTCTGAACTCTTTCCCTACACGACGCTCTTCCGATCTNNNATAAATAAAGCACCCAAGAAAAAC
GAGCACACGTCTGAACTCTTTCCCTACACGACGCTCTTCCGATCTNNNATATTACCAATCCTCATATGT
GCACACGTCTGAACTCTTTCCCTACACGACGCTCTTCCGATCTNNNTATTACTACTAATTCTCTGGGTG
ACACGTCTGAACTCTTTCCCTACACGACGCTCTTCCGATCTNNNCCTTCAGCTATAGTTTGTGTCCATC

AGCACACGTCTGAACTCTTTCCCTACACGACGCTCTTCCGATCTNNNTAAAAGCTGCATTTTCAATACT

GCACACGTCTGAACTCTTTCCCTACACGACGCTCTTCCGATCTNNNCTCCTTCAACCTCTGTTTTATTC

GCACACGTCTGAACTCTTTCCCTACACGACGCTCTTCCGATCTNNNCATTGAATTTCTTTAGCTCCTCCA

CACACGTCTGAACTCTTTCCCTACACGACGCTCTTCCGATCTNNNGAGTAGAGTGGCATGATCATCAT

GCACACGTCTGAACTCTTTCCCTACACGACGCTCTTCCGATCTNNNATTCCATCAACCAGAAGATTCC

AGCACACGTCTGAACTCTTTCCCTACACGACGCTCTTCCGATCTNNNAGGTTAGAACATGGAAATATC

AGCACACGTCTGAACTCTTTCCCTACACGACGCTCTTCCGATCTNNNGTATGCAAGGATATCATTCAT

AGCACACGTCTGAACTCTTTCCCTACACGACGCTCTTCCGATCTNNNGTACCTTAGTCATAATGTGGT

ACACGTCTGAACTCTTTCCCTACACGACGCTCTTCCGATCTNNNAGATCCTGTTAGTCACTACAGCATC

ACACGTCTGAACTCTTTCCCTACACGACGCTCTTCCGATCTNNNAAACTAATTTTGTGTCTTTCCCCATC

AGCACACGTCTGAACTCTTTCCCTACACGACGCTCTTCCGATCTNNNACAGGTTCAGTTATATTACCA

GCACACGTCTGAACTCTTTCCCTACACGACGCTCTTCCGATCTNNNGGAATGATGAAAAAAACACTT

AGCACACGTCTGAACTCTTTCCCTACACGACGCTCTTCCGATCTNNNCTAAAGTGATGTATTGTTGGT

GAGCACACGTCTGAACTCTTTCCCTACACGACGCTCTTCCGATCTNNNCTCCTTTTAAGATAACATTGA

GAGCACACGTCTGAACTCTTTCCCTACACGACGCTCTTCCGATCTNNNCTTGGATGGAAGGAACAAAG

AGCACACGTCTGAACTCTTTCCCTACACGACGCTCTTCCGATCTNNNTATGTGTATATGTGTACTCTGT

AGCACACGTCTGAACTCTTTCCCTACACGACGCTCTTCCGATCTNNNTCTTTTTCAAAATTGTTTCAACT

GCACACGTCTGAACTCTTTCCCTACACGACGCTCTTCCGATCTNNNAAATACAGATTAAGAGGGACA

GAGCACACGTCTGAACTCTTTCCCTACACGACGCTCTTCCGATCTNNNCATGAGAAGCTTTCAGAAAA

AGCACACGTCTGAACTCTTTCCCTACACGACGCTCTTCCGATCTNNNATTAGAAGGAAGGAAGTTTT

AGCACACGTCTGAACTCTTTCCCTACACGACGCTCTTCCGATCTNNNGTTTAGGGAGAAATACAGGTA

AGCACACGTCTGAACTCTTTCCCTACACGACGCTCTTCCGATCTNNNGTTTAGGGAGAAATACAGGTA

AGCACACGTCTGAACTCTTTCCCTACACGACGCTCTTCCGATCTNNNTATTAACTAAAATGCCCCATCT

AGCACACGTCTGAACTCTTTCCCTACACGACGCTCTTCCGATCTNNNCTGAGTTGTAGCCTCTATATAA

AGCACACGTCTGAACTCTTTCCCTACACGACGCTCTTCCGATCTNNNGACTCCAACATATCTTTCACAT

GCACACGTCTGAACTCTTTCCCTACACGACGCTCTTCCGATCTNNNGAAAAACAAAATCAACCCAAAG

GCACACGTCTGAACTCTTTCCCTACACGACGCTCTTCCGATCTNNNGAAAAACAAAATCAACCCAAAG

AGCACACGTCTGAACTCTTTCCCTACACGACGCTCTTCCGATCTNNNCAGATTCTCAAATCCTGTAATT

GCACACGTCTGAACTCTTTCCCTACACGACGCTCTTCCGATCTNNNGAAATAAAAGAGCCCTATTTAC

GAGCACACGTCTGAACTCTTTCCCTACACGACGCTCTTCCGATCTNNNATTTGCAAGTTAAGATAGAC

AGAGCACACGTCTGAACTCTTTCCCTACACGACGCTCTTCCGATCTNNNGTGGAAGGTAAATATATTG

GAGCACACGTCTGAACTCTTTCCCTACACGACGCTCTTCCGATCTNNNAACAAATTAGACCAAATGATT

GCACACGTCTGAACTCTTTCCCTACACGACGCTCTTCCGATCTNNNACAAAATATACAGCAAAAAGTG

GAGCACACGTCTGAACTCTTTCCCTACACGACGCTCTTCCGATCTNNNGTAGCTTGTGTTACTATTTCT

GCACACGTCTGAACTCTTTCCCTACACGACGCTCTTCCGATCTNNNTAATCTATACCTTTATGGGAGTG

ACACGTCTGAACTCTTTCCCTACACGACGCTCTTCCGATCTNNNTGGAAGTTTAGTGTGGCTAGCATC

GCACACGTCTGAACTCTTTCCCTACACGACGCTCTTCCGATCTNNNCTGTTGATCCTTCTAACAGTGCA

GAGCACACGTCTGAACTCTTTCCCTACACGACGCTCTTCCGATCTNNNGGAAAGGCATTACATAATGA

GCACACGTCTGAACTCTTTCCCTACACGACGCTCTTCCGATCTNNNATAAGCAAATAGGGCAGATAA

AGCACACGTCTGAACTCTTTCCCTACACGACGCTCTTCCGATCTNNNAATTGGATCTCAGTTTCCATA

AGCACACGTCTGAACTCTTTCCCTACACGACGCTCTTCCGATCTNNNGGGAAGATTGAAACTAAATG

GCACACGTCTGAACTCTTTCCCTACACGACGCTCTTCCGATCTNNNTATGAAGAAAGTTTTATGCACAT

GCACACGTCTGAACTCTTTCCCTACACGACGCTCTTCCGATCTNNNCTCCAGAAATAAACCAGGAGCA

CACACGTCTGAACTCTTTCCCTACACGACGCTCTTCCGATCTNNNGTGCATCTCAGTTCTAAAGATCATC
AGCACACGTCTGAACTCTTTCCCTACACGACGCTCTTCCGATCTNNNAAATATACAGCAAACAACATTG
CACACGTCTGAACTCTTTCCCTACACGACGCTCTTCCGATCTNNNAAATATATCTTTCCTGCTCTCTTCAT
GAGCACACGTCTGAACTCTTTCCCTACACGACGCTCTTCCGATCTNNNCTCTTGTTTATTTATTACTCCCA
AGCACACGTCTGAACTCTTTCCCTACACGACGCTCTTCCGATCTNNNTTTAAGATTTTGCAATCATCTCT
GAGCACACGTCTGAACTCTTTCCCTACACGACGCTCTTCCGATCTNNNTGGACATATAATAAAACAGCA
CACACGTCTGAACTCTTTCCCTACACGACGCTCTTCCGATCTNNNTTAGTGTACAGGAACTACTCTCCAT
AGCACACGTCTGAACTCTTTCCCTACACGACGCTCTTCCGATCTNNNCTAACTGGAGATAAAGGATTTC
AGCACACGTCTGAACTCTTTCCCTACACGACGCTCTTCCGATCTNNNTAAACTCATGTTAGTAAAGCCA
AGCACACGTCTGAACTCTTTCCCTACACGACGCTCTTCCGATCTNNNAAAATGAGAATTTGTAACTCACA
AGCACACGTCTGAACTCTTTCCCTACACGACGCTCTTCCGATCTNNNTTCTTATAATGAATGTGAAAGG
AGCACACGTCTGAACTCTTTCCCTACACGACGCTCTTCCGATCTNNNCATCTTCCCTCCATATCTATTCCAT
AGCACACGTCTGAACTCTTTCCCTACACGACGCTCTTCCGATCTNNNAGACTGAACAGAGACCATTCCA
AGCACACGTCTGAACTCTTTCCCTACACGACGCTCTTCCGATCTNNNTAAGTTTAACCAGTTATCTCTGT
GCACACGTCTGAACTCTTTCCCTACACGACGCTCTTCCGATCTNNNGATCAGTGGTCATGAACTATCCA
GAGCACACGTCTGAACTCTTTCCCTACACGACGCTCTTCCGATCTNNNCTACAGTCTATGACATTTGCAC
AGCACACGTCTGAACTCTTTCCCTACACGACGCTCTTCCGATCTNNNTTAATATCTTGGTTGATCACAA
CACACGTCTGAACTCTTTCCCTACACGACGCTCTTCCGATCTNNNCAAGAACATGGAGATACAGTCCAT
AGCACACGTCTGAACTCTTTCCCTACACGACGCTCTTCCGATCTNNNAAGGTACTATATTGTTGCCTAAA
ACACGTCTGAACTCTTTCCCTACACGACGCTCTTCCGATCTNNNATTATGGCTGAATAATAGTCCATCAT
CACACGTCTGAACTCTTTCCCTACACGACGCTCTTCCGATCTNNNTACCATTAAATCCTACTTCCTTTCAT
GAGCACACGTCTGAACTCTTTCCCTACACGACGCTCTTCCGATCTNNNATAGGAGCCTGAGAAAAACTC
GAGCACACGTCTGAACTCTTTCCCTACACGACGCTCTTCCGATCTNNNGTGACAGATTGGATAAAGAA
AGCACACGTCTGAACTCTTTCCCTACACGACGCTCTTCCGATCTNNNTACCTAGGGTAAACTTTAATCAC
GAGCACACGTCTGAACTCTTTCCCTACACGACGCTCTTCCGATCTNNNCCTGTTACCCTTCCTTTACATAC
AGCACACGTCTGAACTCTTTCCCTACACGACGCTCTTCCGATCTNNNTTAGAGAGAACCATATTGAGAT
GCACACGTCTGAACTCTTTCCCTACACGACGCTCTTCCGATCTNNNTGAAAAATTGGCATAAGAAATTC
AGCACACGTCTGAACTCTTTCCCTACACGACGCTCTTCCGATCTNNNCAAAACTTGTCTTTCTCTGTCCA
GAGCACACGTCTGAACTCTTTCCCTACACGACGCTCTTCCGATCTNNNCTCTAGGACTTCCTTTATCTCTC
AGCACACGTCTGAACTCTTTCCCTACACGACGCTCTTCCGATCTNNNATGTTCATATTCCTTCTACATGAC
AGCACACGTCTGAACTCTTTCCCTACACGACGCTCTTCCGATCTNNNCATTTTCAGTCAAGTTCTAATGA
AGCACACGTCTGAACTCTTTCCCTACACGACGCTCTTCCGATCTNNNTAGAATAGACCAGACCCTTGCAT
AGCACACGTCTGAACTCTTTCCCTACACGACGCTCTTCCGATCTNNNTAGAATAGACCAGACCCTTGCAT
AGCACACGTCTGAACTCTTTCCCTACACGACGCTCTTCCGATCTNNNAAAAGAAAAAATAGCTGGACAC
GCACACGTCTGAACTCTTTCCCTACACGACGCTCTTCCGATCTNNNAAGTCAACTAAATCCACAAAATTC
AGCACACGTCTGAACTCTTTCCCTACACGACGCTCTTCCGATCTNNNTCTATACTATACTGGTCTGACCCA
AGCACACGTCTGAACTCTTTCCCTACACGACGCTCTTCCGATCTNNNGTGTGCATATGTATATATACAC
AGCACACGTCTGAACTCTTTCCCTACACGACGCTCTTCCGATCTNNNCATTTTTTTTCTTCTCTCTTCCTCA
AGCACACGTCTGAACTCTTTCCCTACACGACGCTCTTCCGATCTNNNTAAAACATGTCATATTTCCCTCTC
GCACACGTCTGAACTCTTTCCCTACACGACGCTCTTCCGATCTNNNACAGGTAGGAGAAGAACTAAGCA
GCACACGTCTGAACTCTTTCCCTACACGACGCTCTTCCGATCTNNNTTTTAAGATGCCATTCTAAAAGCC
AGCACACGTCTGAACTCTTTCCCTACACGACGCTCTTCCGATCTNNNACAATCAAACATACAATAATGG
GCACACGTCTGAACTCTTTCCCTACACGACGCTCTTCCGATCTNNNCTTTGGTCTCACATACATTATTTCA

CACACGTCTGAACTCTTTCCCTACACGACGCTCTTCCGATCTNNNATGTATATGCATATGTGGTATGT
AGCACACGTCTGAACTCTTTCCCTACACGACGCTCTTCCGATCTNNNTCGATAGGAATAAAGTTCAAG
AGCACACGTCTGAACTCTTTCCCTACACGACGCTCTTCCGATCTNNNTAGGGTTCCAAATAATTCAAT
AGCACACGTCTGAACTCTTTCCCTACACGACGCTCTTCCGATCTNNNTAATGAACATCAGTGGAGAAA
AGCACACGTCTGAACTCTTTCCCTACACGACGCTCTTCCGATCTNNNAGAATTTGAGAATCTGAATTTC
AGCACACGTCTGAACTCTTTCCCTACACGACGCTCTTCCGATCTNNNGATTGCCTTTGAAAATGAATAA
AGCACACGTCTGAACTCTTTCCCTACACGACGCTCTTCCGATCTNNNTTGGAAAATCAGTAGTAATCA
CACGTCTGAACTCTTTCCCTACACGACGCTCTTCCGATCTNNNTGATCCCTTTACCATTATGTAATCAT
GCACACGTCTGAACTCTTTCCCTACACGACGCTCTTCCGATCTNNNGGTATAAGAAAAGGCCATAATT
AGCACACGTCTGAACTCTTTCCCTACACGACGCTCTTCCGATCTNNNCCTGGTGTATTTTCAAAATTC
GCACACGTCTGAACTCTTTCCCTACACGACGCTCTTCCGATCTNNNTATACACACCCTTTATTCTTTCT
CACACGTCTGAACTCTTTCCCTACACGACGCTCTTCCGATCTNNNAATGTGAGAAAGGACTAATACAA
AGCACACGTCTGAACTCTTTCCCTACACGACGCTCTTCCGATCTNNNTACAGAGAATAATTCAGAATG
AGCACACGTCTGAACTCTTTCCCTACACGACGCTCTTCCGATCTNNNTTCATTGAATGAACAGTTACAA
GCACACGTCTGAACTCTTTCCCTACACGACGCTCTTCCGATCTNNNAATACACATCACCTTAGTTTTC
AGCACACGTCTGAACTCTTTCCCTACACGACGCTCTTCCGATCTNNNGAAGTCAAACATTTTCTCCTTT
GCACACGTCTGAACTCTTTCCCTACACGACGCTCTTCCGATCTNNNTCTTTCCATTCTACTCTTTTTCTCA
AGCACACGTCTGAACTCTTTCCCTACACGACGCTCTTCCGATCTNNNTTTGGATTTCTGAAATTTCACC
ACACGTCTGAACTCTTTCCCTACACGACGCTCTTCCGATCTNNNAGAGAAGAGAGAGGAGAGAAAC
AGCACACGTCTGAACTCTTTCCCTACACGACGCTCTTCCGATCTNNNCCTTATTTTTTGTTGTGGTTTTT
AGCACACGTCTGAACTCTTTCCCTACACGACGCTCTTCCGATCTNNNATAAAAAGCCTAAAGAGAGCT
AGCACACGTCTGAACTCTTTCCCTACACGACGCTCTTCCGATCTNNNTAATACCTTTATAGCTACACACA
AGCACACGTCTGAACTCTTTCCCTACACGACGCTCTTCCGATCTNNNCTGGATCATCTGTATTTACTTC
GAGCACACGTCTGAACTCTTTCCCTACACGACGCTCTTCCGATCTNNNTCTAATTTTGTTTTGGTATCT
GAGCACACGTCTGAACTCTTTCCCTACACGACGCTCTTCCGATCTNNNTCTAATTTTGTTTTGGTATCT
GCACACGTCTGAACTCTTTCCCTACACGACGCTCTTCCGATCTNNNCTAGAGTAAGCGTTTCTGTCCA
AGCACACGTCTGAACTCTTTCCCTACACGACGCTCTTCCGATCTNNNTATGACCTTTGTATTGGAAAAG
AGCACACGTCTGAACTCTTTCCCTACACGACGCTCTTCCGATCTNNNCCTATGGGATTACAAAAGGG
CACACGTCTGAACTCTTTCCCTACACGACGCTCTTCCGATCTNNNGGCATTCTTCTTTTTTCATAAACCA
AGCACACGTCTGAACTCTTTCCCTACACGACGCTCTTCCGATCTNNNACTGCATTTTATTTTGCATTAA
AGCACACGTCTGAACTCTTTCCCTACACGACGCTCTTCCGATCTNNNAATTTAATTCTAAGTTTGCCAG
AGCACACGTCTGAACTCTTTCCCTACACGACGCTCTTCCGATCTNNNTAAGAGAAAGAAATGACATAG
GCACACGTCTGAACTCTTTCCCTACACGACGCTCTTCCGATCTNNNTGTAAAAATCAACCTTCAAGAA
AGCACACGTCTGAACTCTTTCCCTACACGACGCTCTTCCGATCTNNNTACACTGTACTTGGTAAATAC
AGCACACGTCTGAACTCTTTCCCTACACGACGCTCTTCCGATCTNNNTACACTGTACTTGGTAAATAC
GCACACGTCTGAACTCTTTCCCTACACGACGCTCTTCCGATCTNNNATGATTTTAAACTCAAACTCACA
AGCACACGTCTGAACTCTTTCCCTACACGACGCTCTTCCGATCTNNNATTTACAGCTTCATCCTTAAAA
AGCACACGTCTGAACTCTTTCCCTACACGACGCTCTTCCGATCTNNNGTTGAGTAAACAATAAAATGA
CACACGTCTGAACTCTTTCCCTACACGACGCTCTTCCGATCTNNNTATGTGAAACCATCATAACCAACA
GCACACGTCTGAACTCTTTCCCTACACGACGCTCTTCCGATCTNNNGGAATATTTAAAGTTCCTTCAGG
GCACACGTCTGAACTCTTTCCCTACACGACGCTCTTCCGATCTNNNTTTGTTTCAGAACTTACCATTT
AGCACACGTCTGAACTCTTTCCCTACACGACGCTCTTCCGATCTNNNCTACTTCAAAATGTTACTGTCA
AGCACACGTCTGAACTCTTTCCCTACACGACGCTCTTCCGATCTNNNAACACTTTTATGCTAATAGGA

GCACACGTCTGAACTCTTTCCCTACACGACGCTCTTCCGATCTNNNTGTATACAAAGTGTCAGTGTAAG

GAGCACACGTCTGAACTCTTTCCCTACACGACGCTCTTCCGATCTNNNTAATACATTGTTGTGTCCAAAA

GAGCACACGTCTGAACTCTTTCCCTACACGACGCTCTTCCGATCTNNNTAATACATTGTTGTGTCCAAAA

AGCACACGTCTGAACTCTTTCCCTACACGACGCTCTTCCGATCTNNNAGAGCATTTTCATTACTTTCAAA

AGCACACGTCTGAACTCTTTCCCTACACGACGCTCTTCCGATCTNNNTGTAGCAGAAACAATGGTAGC

ACACGTCTGAACTCTTTCCCTACACGACGCTCTTCCGATCTNNNGATGATACTGATGAGGGAGTCATC

GAGCACACGTCTGAACTCTTTCCCTACACGACGCTCTTCCGATCTNNNCTGTATATTTACCATATGGGCT

CACACGTCTGAACTCTTTCCCTACACGACGCTCTTCCGATCTNNNAATGTGGTTTATTTAGAATGACCCA

GAGCACACGTCTGAACTCTTTCCCTACACGACGCTCTTCCGATCTNNNGCTTTATGTTACTACATTCTTG

AGAGCACACGTCTGAACTCTTTCCCTACACGACGCTCTTCCGATCTNNNTAAAAGAAAGGTAGAATGAC

GAGCACACGTCTGAACTCTTTCCCTACACGACGCTCTTCCGATCTNNNCTAGACATTTCCTCTAGTTGTA

AGCACACGTCTGAACTCTTTCCCTACACGACGCTCTTCCGATCTNNNTTTATTAGTTGAGAAGACTGTC

AGCACACGTCTGAACTCTTTCCCTACACGACGCTCTTCCGATCTNNNAAACATTTAGAGCTGACTGAAT

ACACGTCTGAACTCTTTCCCTACACGACGCTCTTCCGATCTNNNGTGACTATTCTCTGTCTTATCTACATC

GCACACGTCTGAACTCTTTCCCTACACGACGCTCTTCCGATCTNNNCAGTTCTTTCAGTAAACTTCATGCA

CACACGTCTGAACTCTTTCCCTACACGACGCTCTTCCGATCTNNNTAATAAAGCAGTGGTCAATATGAC

AGCACACGTCTGAACTCTTTCCCTACACGACGCTCTTCCGATCTNNNTTTAGTGTATAGAGAAGCAAC

CACACGTCTGAACTCTTTCCCTACACGACGCTCTTCCGATCTNNNGAAGAGGTTATTGATACTCTCATCA

AGCACACGTCTGAACTCTTTCCCTACACGACGCTCTTCCGATCTNNNGTCTTGGTCTACGTAAAAATTC

ACACGTCTGAACTCTTTCCCTACACGACGCTCTTCCGATCTNNNCCTTCTCTTTTCTCTTCTTTCTCATCGG

AGCACACGTCTGAACTCTTTCCCTACACGACGCTCTTCCGATCTNNNCAAGTAGCTGAGATTACAGGCA

AAGAGCACACGTCTGAACTCTTTCCCTACACGACGCTCTTCCGATCTNNNGAGACACAGGATTAAATAA

GCACACGTCTGAACTCTTTCCCTACACGACGCTCTTCCGATCTNNNGAGAAGATTTAGTTTCAATGCTA

AGCACACGTCTGAACTCTTTCCCTACACGACGCTCTTCCGATCTNNNTCTTTAGCCTAATCTTTTCCATAC

AGCACACGTCTGAACTCTTTCCCTACACGACGCTCTTCCGATCTNNNAAGGAGGTTTTGGTTGTTAGCA

AGCACACGTCTGAACTCTTTCCCTACACGACGCTCTTCCGATCTNNNTAACTGATGTGGACTACTGTTC

AGCACACGTCTGAACTCTTTCCCTACACGACGCTCTTCCGATCTNNNTATTCGTTGATGTCCCTATTGCA

AGAGCACACGTCTGAACTCTTTCCCTACACGACGCTCTTCCGATCTNNNCATATCATTCTGCTCCAGTATC

AGCACACGTCTGAACTCTTTCCCTACACGACGCTCTTCCGATCTNNNTACATTTGTATCATTCTGCATTT

AGCACACGTCTGAACTCTTTCCCTACACGACGCTCTTCCGATCTNNNAATGCGAATTGTATTCTAACTA

AGCACACGTCTGAACTCTTTCCCTACACGACGCTCTTCCGATCTNNNTTTGTTTTTCTCTAAAAGAAGGA

AGAGCACACGTCTGAACTCTTTCCCTACACGACGCTCTTCCGATCTNNNAGAAAGAGACAGAGTTGAC

AGCACACGTCTGAACTCTTTCCCTACACGACGCTCTTCCGATCTNNNCTCCCAAAGTGCTAGGATTACAT

AGCACACGTCTGAACTCTTTCCCTACACGACGCTCTTCCGATCTNNNCAATTTACAGGAAATACAGGAA

ACACGTCTGAACTCTTTCCCTACACGACGCTCTTCCGATCTNNNGACATTTTGGTTGCTTTCAACATCGC

GAGCACACGTCTGAACTCTTTCCCTACACGACGCTCTTCCGATCTNNNCTCTATACCCACAATGAAGATG

AGCACACGTCTGAACTCTTTCCCTACACGACGCTCTTCCGATCTNNNGTAGTTGATTATGAAAAAGGT

AGCACACGTCTGAACTCTTTCCCTACACGACGCTCTTCCGATCTNNNCTTGAATGTTATTACAGGAACAC

GCACACGTCTGAACTCTTTCCCTACACGACGCTCTTCCGATCTNNNATCATTATTGCTAGGGTTAGTTAC

AGCACACGTCTGAACTCTTTCCCTACACGACGCTCTTCCGATCTNNNCTTTGAAAAAGAGAACAACAA

AGCACACGTCTGAACTCTTTCCCTACACGACGCTCTTCCGATCTNNNCTTTGAAAAAGAGAACAACAA

GCACACGTCTGAACTCTTTCCCTACACGACGCTCTTCCGATCTNNNAATAAATGTGTGACTTGGATAAC

GCACACGTCTGAACTCTTTCCCTACACGACGCTCTTCCGATCTNNNACATCCTTTTCCCTTGTATATGCAT

GCACACGTCTGAACTCTTTCCCTACACGACGCTCTTCCGATCTNNNATACAGAGTAAAAAAACCCAAT
AGCACACGTCTGAACTCTTTCCCTACACGACGCTCTTCCGATCTNNNATAATTTAAGGTTTCCTGTCAC
AGCACACGTCTGAACTCTTTCCCTACACGACGCTCTTCCGATCTNNNCCAAATATTCCAGTGTATTCAAT
ACACGTCTGAACTCTTTCCCTACACGACGCTCTTCCGATCTNNNTTTCCCATTCTCAAAAGTGGCATCG
GCACACGTCTGAACTCTTTCCCTACACGACGCTCTTCCGATCTNNNACTTCATGAAATTATTTTTGCA
GCACACGTCTGAACTCTTTCCCTACACGACGCTCTTCCGATCTNNNCTTCAAAAGAATAGTTACACACA
CACACGTCTGAACTCTTTCCCTACACGACGCTCTTCCGATCTNNNAAATATGTGAGGATATGAGAGA
GCACACGTCTGAACTCTTTCCCTACACGACGCTCTTCCGATCTNNNTAATGTAATGTATCCCAAAGAAC
GCACACGTCTGAACTCTTTCCCTACACGACGCTCTTCCGATCTNNNTAATGTAATGTATCCCAAAGAAC
AGCACACGTCTGAACTCTTTCCCTACACGACGCTCTTCCGATCTNNNTCTCAAAGAACTTAAAACAAAC
AGCACACGTCTGAACTCTTTCCCTACACGACGCTCTTCCGATCTNNNTCTCAAAGAACTTAAAACAAAC
AGCACACGTCTGAACTCTTTCCCTACACGACGCTCTTCCGATCTNNNATCTGAATTTTAGCCTGACTAT
GCACACGTCTGAACTCTTTCCCTACACGACGCTCTTCCGATCTNNNTGTTCCTCAATTTTAGAAAGATG
AGCACACGTCTGAACTCTTTCCCTACACGACGCTCTTCCGATCTNNNGTTTCTGTCTCCCTCTCTACCAT
AGCACACGTCTGAACTCTTTCCCTACACGACGCTCTTCCGATCTNNNGTTTCTGTCTCCCTCTCTACCAT
AGCACACGTCTGAACTCTTTCCCTACACGACGCTCTTCCGATCTNNNTTTTTATTTTGAAAGGTTGAAG
GAGCACACGTCTGAACTCTTTCCCTACACGACGCTCTTCCGATCTNNNATGCCATAGTTTACTGTATGT
GAGCACACGTCTGAACTCTTTCCCTACACGACGCTCTTCCGATCTNNNATGCCATAGTTTACTGTATGT
ACACGTCTGAACTCTTTCCCTACACGACGCTCTTCCGATCTNNNCTAGTTCACCTATTGAGTATGTTCAT
GAGCACACGTCTGAACTCTTTCCCTACACGACGCTCTTCCGATCTNNNAAATAGAACCATTAGATGTCC
CACACGTCTGAACTCTTTCCCTACACGACGCTCTTCCGATCTNNNGAAAATTCAGCCTCTATGAATATC
CACACGTCTGAACTCTTTCCCTACACGACGCTCTTCCGATCTNNNATTATAAGAGCATACCCACATAGC
AGCACACGTCTGAACTCTTTCCCTACACGACGCTCTTCCGATCTNNNTCTTTTTCATAAACAAAACGTG
AGCACACGTCTGAACTCTTTCCCTACACGACGCTCTTCCGATCTNNNAAGATGTTTTAGACTTGAAAG
AGCACACGTCTGAACTCTTTCCCTACACGACGCTCTTCCGATCTNNNAAGATGTTTTAGACTTGAAAG
AGCACACGTCTGAACTCTTTCCCTACACGACGCTCTTCCGATCTNNNCACTGTATCCAAAAAAAAAAAG
AGCACACGTCTGAACTCTTTCCCTACACGACGCTCTTCCGATCTNNNGACTTCTCAGCCTTCATAATTCA
AGCACACGTCTGAACTCTTTCCCTACACGACGCTCTTCCGATCTNNNGACTTCTCAGCCTTCATAATTCA
GCACACGTCTGAACTCTTTCCCTACACGACGCTCTTCCGATCTNNNTTTGAATTTTGGAAATGACAAAG
GCACACGTCTGAACTCTTTCCCTACACGACGCTCTTCCGATCTNNNAGACATAAAATGAAGATTGAAG
AGCACACGTCTGAACTCTTTCCCTACACGACGCTCTTCCGATCTNNNCCCAATAGTAACATTTTGCAAA
AGCACACGTCTGAACTCTTTCCCTACACGACGCTCTTCCGATCTNNNTCTTCATTTTTCCTGAAATATGT
AGCACACGTCTGAACTCTTTCCCTACACGACGCTCTTCCGATCTNNNCATTGGATTTGCAATTTATTCAA
AGCACACGTCTGAACTCTTTCCCTACACGACGCTCTTCCGATCTNNNGCAAGTAGAGTTTTTGTTTTTT
CACACGTCTGAACTCTTTCCCTACACGACGCTCTTCCGATCTNNNCTTTCCAAATCCTTATCTTTTCTCAT
AGCACACGTCTGAACTCTTTCCCTACACGACGCTCTTCCGATCTNNNAAAACTCAGGCTGGACAATACA
ACACGTCTGAACTCTTTCCCTACACGACGCTCTTCCGATCTNNNATAAAACTCACTCAAAGTAACACCA
AGCACACGTCTGAACTCTTTCCCTACACGACGCTCTTCCGATCTNNNTAATGCAGAAAATAGGGCAAT
AGCACACGTCTGAACTCTTTCCCTACACGACGCTCTTCCGATCTNNNAAAAGTTAATGCAGAAAATAG
GCACACGTCTGAACTCTTTCCCTACACGACGCTCTTCCGATCTNNNTTATTGTTTCTAGGCCCTTTCCAT
AGCACACGTCTGAACTCTTTCCCTACACGACGCTCTTCCGATCTNNNCTTTGAAAAGCTCCTAGATTA
CACACGTCTGAACTCTTTCCCTACACGACGCTCTTCCGATCTNNNTATAGCTATCTGGTGATTTTAGG
CACACGTCTGAACTCTTTCCCTACACGACGCTCTTCCGATCTNNNTATAGCTATCTGGTGATTTTAGG

AGCACACGTCTGAACTCTTTCCCTACACGACGCTCTTCCGATCTNNNACCCTGTCTCATATACGTATATA
AGCACACGTCTGAACTCTTTCCCTACACGACGCTCTTCCGATCTNNNCCATCTATATTCAGCAATCTGTA
CACACGTCTGAACTCTTTCCCTACACGACGCTCTTCCGATCTNNNTTTGCAGAAGTGTAAAGTCCCATC
AGCACACGTCTGAACTCTTTCCCTACACGACGCTCTTCCGATCTNNNAAAAATGCCAGTGATTCCTTAC
GAGCACACGTCTGAACTCTTTCCCTACACGACGCTCTTCCGATCTNNNGTTAAACATGACTCAATTTCCA
AGCACACGTCTGAACTCTTTCCCTACACGACGCTCTTCCGATCTNNNATGCTTTTATCTCTACATCAAAC
AGAGCACACGTCTGAACTCTTTCCCTACACGACGCTCTTCCGATCTNNNTTCAGATGATGCAAATATTT
GAGCACACGTCTGAACTCTTTCCCTACACGACGCTCTTCCGATCTNNNGTAGGTACTGATGCCATACT
GAGCACACGTCTGAACTCTTTCCCTACACGACGCTCTTCCGATCTNNNATTTTTCCCTTTATATAACTGC
GAGCACACGTCTGAACTCTTTCCCTACACGACGCTCTTCCGATCTNNNATTTTTCCCTTTATATAACTGC
AGAGCACACGTCTGAACTCTTTCCCTACACGACGCTCTTCCGATCTNNNTGACACAAAATTGCTCTTATT
AGAGCACACGTCTGAACTCTTTCCCTACACGACGCTCTTCCGATCTNNNTGACACAAAATTGCTCTTATT
GAGCACACGTCTGAACTCTTTCCCTACACGACGCTCTTCCGATCTNNNATCAATTGACTGTTAGTTCTAG
AGCACACGTCTGAACTCTTTCCCTACACGACGCTCTTCCGATCTNNNTTGAAAAGACTAGAAATACAGT
AGCACACGTCTGAACTCTTTCCCTACACGACGCTCTTCCGATCTNNNTTGAAAAGACTAGAAATACAGT
GAGCACACGTCTGAACTCTTTCCCTACACGACGCTCTTCCGATCTNNNTGATAAGAATGAGTCTGGTAA
AGCACACGTCTGAACTCTTTCCCTACACGACGCTCTTCCGATCTNNNCTATGCAATAAAAGCCCATAAA
AGCACACGTCTGAACTCTTTCCCTACACGACGCTCTTCCGATCTNNNGTTTTCAATAATTGACTTTCAGT
GCACACGTCTGAACTCTTTCCCTACACGACGCTCTTCCGATCTNNNGCTCAGGTTTCTATATCCTTATACA
GAGCACACGTCTGAACTCTTTCCCTACACGACGCTCTTCCGATCTNNNAGAAAGACATATACTCTAAGT
AGCACACGTCTGAACTCTTTCCCTACACGACGCTCTTCCGATCTNNNTATATACACTTCTTTCTGTTCGT
AGCACACGTCTGAACTCTTTCCCTACACGACGCTCTTCCGATCTNNNTATATACACTTCTTTCTGTTCGT
GAGCACACGTCTGAACTCTTTCCCTACACGACGCTCTTCCGATCTNNNCATAAGAAACACACAGAACAT
GCACACGTCTGAACTCTTTCCCTACACGACGCTCTTCCGATCTNNNCAGTTCTGTTTGAAAATTTGTGCA
GCACACGTCTGAACTCTTTCCCTACACGACGCTCTTCCGATCTNNNGGAAGAGAAAAGGTCACAATCA
GCACACGTCTGAACTCTTTCCCTACACGACGCTCTTCCGATCTNNNCTCCAGGTACATCATTCATATTAC
AGCACACGTCTGAACTCTTTCCCTACACGACGCTCTTCCGATCTNNNCAAGAATTTTACAATTAGAAAA
AGCACACGTCTGAACTCTTTCCCTACACGACGCTCTTCCGATCTNNNTTCTTATCTGGAAAAATGAAGA
AGCACACGTCTGAACTCTTTCCCTACACGACGCTCTTCCGATCTNNNCATTTAGCACCTCTCCTATGCAT
GCACACGTCTGAACTCTTTCCCTACACGACGCTCTTCCGATCTNNNAAAACTGTCATACATTTCAATCAC
AGCACACGTCTGAACTCTTTCCCTACACGACGCTCTTCCGATCTNNNCTCTCTCTTCTTTTCTGTCTTTCA
GCACACGTCTGAACTCTTTCCCTACACGACGCTCTTCCGATCTNNNGAACAAGTAACTGCTTTTTATTG
CACACGTCTGAACTCTTTCCCTACACGACGCTCTTCCGATCTNNNAGACATCCAAAACACTTGATACATC
GAGCACACGTCTGAACTCTTTCCCTACACGACGCTCTTCCGATCTNNNCATTCCTTGACAAATACAAAT
AGCACACGTCTGAACTCTTTCCCTACACGACGCTCTTCCGATCTNNNCCAACATTACACAAATAATATG
GAGCACACGTCTGAACTCTTTCCCTACACGACGCTCTTCCGATCTNNNCTTGTGCATATAATGGCCACCA
AGCACACGTCTGAACTCTTTCCCTACACGACGCTCTTCCGATCTNNNATTCCCCAAATGTTAACACTATCA
AGCACACGTCTGAACTCTTTCCCTACACGACGCTCTTCCGATCTNNNAGAGAATGATGTGATTCAACACA
GCACACGTCTGAACTCTTTCCCTACACGACGCTCTTCCGATCTNNNAAGAGTGTGTGAGAAATTTCTT
AGCACACGTCTGAACTCTTTCCCTACACGACGCTCTTCCGATCTNNNATCATACTTAATGCAGAGAAAC
GCACACGTCTGAACTCTTTCCCTACACGACGCTCTTCCGATCTNNNGTCAAAGCTGCATTTTTAGCCATC
GAGCACACGTCTGAACTCTTTCCCTACACGACGCTCTTCCGATCTNNNAATAATTGTGAATATTTTGG
GCACACGTCTGAACTCTTTCCCTACACGACGCTCTTCCGATCTNNNTGGCTAAGGTATAAAAGTTGTA

GAGCACACGTCTGAACTCTTTCCCTACACGACGCTCTTCCGATCTNNNAACGCAGTATTTTTAAATGTT

GCACACGTCTGAACTCTTTCCCTACACGACGCTCTTCCGATCTNNNAGCATTTTTTCTGATCACTAAACA

GAGCACACGTCTGAACTCTTTCCCTACACGACGCTCTTCCGATCTNNNTGGATGCTTTGTTTTATTTGG

GAGCACACGTCTGAACTCTTTCCCTACACGACGCTCTTCCGATCTNNNTAAAATCCTAATCAGAATCCCA

AGAGCACACGTCTGAACTCTTTCCCTACACGACGCTCTTCCGATCTNNNACACTCTCTCTCTCTCTACA

AGCACACGTCTGAACTCTTTCCCTACACGACGCTCTTCCGATCTNNNTTATTTGTTCCTAAAACTTAGCAC

GAGCACACGTCTGAACTCTTTCCCTACACGACGCTCTTCCGATCTNNNTTCTAGGAGAAGAAAGAAAGC

AGCACACGTCTGAACTCTTTCCCTACACGACGCTCTTCCGATCTNNNTGAAAACTTTAATTACATGGACC

GAGCACACGTCTGAACTCTTTCCCTACACGACGCTCTTCCGATCTNNNCTTTATATGGCAATACAGTTC

AGCACACGTCTGAACTCTTTCCCTACACGACGCTCTTCCGATCTNNNGTGAGTGTTCAGTACATGTAGC

AGCACACGTCTGAACTCTTTCCCTACACGACGCTCTTCCGATCTNNNAATGGCAAAGAAAAAGAAAAA

AGCACACGTCTGAACTCTTTCCCTACACGACGCTCTTCCGATCTNNNTTGATAATCCCCAAGTAGTTATT

ACACGTCTGAACTCTTTCCCTACACGACGCTCTTCCGATCTNNNGCATTATCCTTCACCACAAACATCGGA

AGCACACGTCTGAACTCTTTCCCTACACGACGCTCTTCCGATCTNNNATTATTCCATATTGTTCACACAG

AGCACACGTCTGAACTCTTTCCCTACACGACGCTCTTCCGATCTNNNATTACTTGACTCCAGGAGTTCAT

GAGCACACGTCTGAACTCTTTCCCTACACGACGCTCTTCCGATCTNNNTTTCTAGGCTTTCTGGGATAC

AGCACACGTCTGAACTCTTTCCCTACACGACGCTCTTCCGATCTNNNCAAATAGTCATTAGTAAACACC

CACACGTCTGAACTCTTTCCCTACACGACGCTCTTCCGATCTNNNTCTACCTAATTATGTGATTAGTGTCA

CACACGTCTGAACTCTTTCCCTACACGACGCTCTTCCGATCTNNNTCTACCTAATTATGTGATTAGTGTCA

AGCACACGTCTGAACTCTTTCCCTACACGACGCTCTTCCGATCTNNNGGAAATTAGACAACTAGACAA

GCACACGTCTGAACTCTTTCCCTACACGACGCTCTTCCGATCTNNNAAACAGAAGTGTAGGAGATCTT

GCACACGTCTGAACTCTTTCCCTACACGACGCTCTTCCGATCTNNNATCTCTCAAGTCAGTGTATTTGCA

AGCACACGTCTGAACTCTTTCCCTACACGACGCTCTTCCGATCTNNNTGTTTACTTATTTTTAGGGCAAA

AGCACACGTCTGAACTCTTTCCCTACACGACGCTCTTCCGATCTNNNAAACTTGTTCCTAACTTTGAAAA

AGCACACGTCTGAACTCTTTCCCTACACGACGCTCTTCCGATCTNNNTTTGACATAAGGAGAAACTAAT

AGCACACGTCTGAACTCTTTCCCTACACGACGCTCTTCCGATCTNNNTCACAAATCCCTTCTTCTATTAA

GAGCACACGTCTGAACTCTTTCCCTACACGACGCTCTTCCGATCTNNNGCACAAGAACAGAGAAATATT

GAGCACACGTCTGAACTCTTTCCCTACACGACGCTCTTCCGATCTNNNGTACTTTTGAGTTTTCTCTTCTT

AGCACACGTCTGAACTCTTTCCCTACACGACGCTCTTCCGATCTNNNCTTGTATGTCAATTCTAATGGC

AGCACACGTCTGAACTCTTTCCCTACACGACGCTCTTCCGATCTNNNAATTGGAATTAAAACACCATCAT

GAGCACACGTCTGAACTCTTTCCCTACACGACGCTCTTCCGATCTNNNTTCATCCACGTTGATATAAATC

CACACGTCTGAACTCTTTCCCTACACGACGCTCTTCCGATCTNNNAATTCATTTGGATGCCTTTTCCATC

GCACACGTCTGAACTCTTTCCCTACACGACGCTCTTCCGATCTNNNGTTATCCCAGGTCCAATTTTCATC

AGCACACGTCTGAACTCTTTCCCTACACGACGCTCTTCCGATCTNNNCTCACTTTCTCTGGTTTCTAGCA

CACACGTCTGAACTCTTTCCCTACACGACGCTCTTCCGATCTNNNAATCAGATATGCGTTTGTCTCCATC

AGCACACGTCTGAACTCTTTCCCTACACGACGCTCTTCCGATCTNNNATTTCCTTTGGTTTGAATATGAG

GAGCACACGTCTGAACTCTTTCCCTACACGACGCTCTTCCGATCTNNNCTAGTTTCCACTGCATAAATAA

AGCACACGTCTGAACTCTTTCCCTACACGACGCTCTTCCGATCTNNNCTTACCTTCATCTTGCTAATATGC

GAGCACACGTCTGAACTCTTTCCCTACACGACGCTCTTCCGATCTNNNACTTCAGATTCTCAATACTGTG

GAGCACACGTCTGAACTCTTTCCCTACACGACGCTCTTCCGATCTNNNATAAGGCATACATGGAAATAC

AGCACACGTCTGAACTCTTTCCCTACACGACGCTCTTCCGATCTNNNCTGTTTCTCCTAACATCCTGCAT

GAGCACACGTCTGAACTCTTTCCCTACACGACGCTCTTCCGATCTNNNAAGGTTGATTAATTTTTGGAC

CACACGTCTGAACTCTTTCCCTACACGACGCTCTTCCGATCTNNNATATAAGGTTTTGTTATGCCTCTCA

AGCACACGTCTGAACTCTTTCCCTACACGACGCTCTTCCGATCTNNNAAAGAATCCTCAGAACTAGAATT
GCACACGTCTGAACTCTTTCCCTACACGACGCTCTTCCGATCTNNNAAAAGTTCACCACCTTCTTTTTAC
GAGCACACGTCTGAACTCTTTCCCTACACGACGCTCTTCCGATCTNNNTTGAAGACAGAATAAGACCCT
GCACACGTCTGAACTCTTTCCCTACACGACGCTCTTCCGATCTNNNCAGAAATCTGAATACTTGCATATC
AGAGCACACGTCTGAACTCTTTCCCTACACGACGCTCTTCCGATCTNNNTAATAGTTACCATCTATCAAG
AGAGCACACGTCTGAACTCTTTCCCTACACGACGCTCTTCCGATCTNNNTGTATGTAAATTTCTAGGATC
AGCACACGTCTGAACTCTTTCCCTACACGACGCTCTTCCGATCTNNNAAGCTTTGATCCCAGAAAATATA
AGCACACGTCTGAACTCTTTCCCTACACGACGCTCTTCCGATCTNNNAGAGATTGGAGTTAGAAAAAA
AGCACACGTCTGAACTCTTTCCCTACACGACGCTCTTCCGATCTNNNCATTATCTGCCTTGTCTTTTTCCA
GAGCACACGTCTGAACTCTTTCCCTACACGACGCTCTTCCGATCTNNNAGTGACTATAGATGATACCAA
CACACGTCTGAACTCTTTCCCTACACGACGCTCTTCCGATCTNNNGTTAGCCATACCTTCTTATACTACAT
AGAGCACACGTCTGAACTCTTTCCCTACACGACGCTCTTCCGATCTNNNTTAAAAGTTGGGCACAAATG
CACACGTCTGAACTCTTTCCCTACACGACGCTCTTCCGATCTNNNGACCTAGAAATCTTCAACTTGACAT
AGCACACGTCTGAACTCTTTCCCTACACGACGCTCTTCCGATCTNNNGAAGAAAGGAAGAAAGAAAGT
CACACGTCTGAACTCTTTCCCTACACGACGCTCTTCCGATCTNNNGAGGAACTTTGAGAAGAACTCCAT
GCACACGTCTGAACTCTTTCCCTACACGACGCTCTTCCGATCTNNNTTTTTGGAATCTATGTTCATCAGC
GCACACGTCTGAACTCTTTCCCTACACGACGCTCTTCCGATCTNNNTAAAGATTCAGAGAAGGAAAAGA
GCACACGTCTGAACTCTTTCCCTACACGACGCTCTTCCGATCTNNNCTATTTCGTTGGTTGAATTCTTCAT
AGCACACGTCTGAACTCTTTCCCTACACGACGCTCTTCCGATCTNNNGTGTGACAAATTTGAATAACCT
AGCACACGTCTGAACTCTTTCCCTACACGACGCTCTTCCGATCTNNNAGTATTTTCAGGTGTTGAAGAT
GAGCACACGTCTGAACTCTTTCCCTACACGACGCTCTTCCGATCTNNNTACTCAAAAAGCCATTCCTATAT
AGCACACGTCTGAACTCTTTCCCTACACGACGCTCTTCCGATCTNNNAATTATTTTTAGTGCAACAGACT
AGCACACGTCTGAACTCTTTCCCTACACGACGCTCTTCCGATCTNNNATAAGGTCTAGATTTAGCCATA
AGCACACGTCTGAACTCTTTCCCTACACGACGCTCTTCCGATCTNNNATAAGGTCTAGATTTAGCCATA
AGAGCACACGTCTGAACTCTTTCCCTACACGACGCTCTTCCGATCTNNNTTTAATGGAGAGTGCTGTCT
AGCACACGTCTGAACTCTTTCCCTACACGACGCTCTTCCGATCTNNNTTCTAATCACATACATCCTACTC
AGCACACGTCTGAACTCTTTCCCTACACGACGCTCTTCCGATCTNNNAATACCCTATTCTCTATTTCTGC
GAGCACACGTCTGAACTCTTTCCCTACACGACGCTCTTCCGATCTNNNGAGAGCTAGAACAAGATTGAA
CACACGTCTGAACTCTTTCCCTACACGACGCTCTTCCGATCTNNNGTAGTATTGTTAGTGCAGTTCAACA
AGCACACGTCTGAACTCTTTCCCTACACGACGCTCTTCCGATCTNNNTCCACTAACAAAAGTTCATTATC
AGAGCACACGTCTGAACTCTTTCCCTACACGACGCTCTTCCGATCTNNNCATGAAATATGCAACTTTCAA
GCACACGTCTGAACTCTTTCCCTACACGACGCTCTTCCGATCTNNNGCAGTCATTGATGAAAAATATTCC
AGCACACGTCTGAACTCTTTCCCTACACGACGCTCTTCCGATCTNNNCATTATGCAGTCATTGATGAAAA
GAGCACACGTCTGAACTCTTTCCCTACACGACGCTCTTCCGATCTNNNAATTTTTCCTCTTCCTTTTAGTC
GCACACGTCTGAACTCTTTCCCTACACGACGCTCTTCCGATCTNNNTGTTTTGCATTTTCAAGAGGCATC
GAGCACACGTCTGAACTCTTTCCCTACACGACGCTCTTCCGATCTNNNATTTTACAGAGGCTACAACTGC
AGCACACGTCTGAACTCTTTCCCTACACGACGCTCTTCCGATCTNNNTTTTGGCAGTATAATTTTTTAGC
AGCACACGTCTGAACTCTTTCCCTACACGACGCTCTTCCGATCTNNNTTCCTCTGTTGTCTCTTTTATTAC
AGCACACGTCTGAACTCTTTCCCTACACGACGCTCTTCCGATCTNNNCACCATGTCTTCCTTTGTTATCAT
GAGCACACGTCTGAACTCTTTCCCTACACGACGCTCTTCCGATCTNNNATATCTCCTGCTCACTACATATA
AGCACACGTCTGAACTCTTTCCCTACACGACGCTCTTCCGATCTNNNTGAAGCTATCTGGTTTATTCTTA
AGCACACGTCTGAACTCTTTCCCTACACGACGCTCTTCCGATCTNNNTGAAGCTATCTGGTTTATTCTTA
AGCACACGTCTGAACTCTTTCCCTACACGACGCTCTTCCGATCTNNNATCTCTTTGTATTGATGAGGAA

GAGCACACGTCTGAACTCTTTCCCTACACGACGCTCTTCCGATCTNNNTCAAGATCATTTCAAAATGTG
AGCACACGTCTGAACTCTTTCCCTACACGACGCTCTTCCGATCTNNNCAGTCCTTCTCAAACTCTTATGC
AGCACACGTCTGAACTCTTTCCCTACACGACGCTCTTCCGATCTNNNTTGATAGATTAGATTTTGGGGA
AGCACACGTCTGAACTCTTTCCCTACACGACGCTCTTCCGATCTNNNCCTATGAAAACTTGCTTTTATGA
AGCACACGTCTGAACTCTTTCCCTACACGACGCTCTTCCGATCTNNNCAAAAAAGAGAATGATCATGAA
AGCACACGTCTGAACTCTTTCCCTACACGACGCTCTTCCGATCTNNNAGTCAGTGTGTGTTATATACAT
GAGCACACGTCTGAACTCTTTCCCTACACGACGCTCTTCCGATCTNNNCTGAGACTGCTTAATTAACATA
GCACACGTCTGAACTCTTTCCCTACACGACGCTCTTCCGATCTNNNGTCATGTTTACACATATCTACTGCA
AGCACACGTCTGAACTCTTTCCCTACACGACGCTCTTCCGATCTNNNGATGAAGAAAGTTCCGTTATAA
GAGCACACGTCTGAACTCTTTCCCTACACGACGCTCTTCCGATCTNNNTTAAGTTTCTTGTGAGAATCT
GAGCACACGTCTGAACTCTTTCCCTACACGACGCTCTTCCGATCTNNNATTAAAAAAAAAATGATGGTG
AGCACACGTCTGAACTCTTTCCCTACACGACGCTCTTCCGATCTNNNGTGTATAAATGAGGATGCATAA
GAGCACACGTCTGAACTCTTTCCCTACACGACGCTCTTCCGATCTNNNCAGATTCTATTCCTGGGGAGC
AGCACACGTCTGAACTCTTTCCCTACACGACGCTCTTCCGATCTNNNAGCAATTCCAGTCCTAGATATAC
GAGCACACGTCTGAACTCTTTCCCTACACGACGCTCTTCCGATCTNNNTCTGGTATAGTTCTTTTCTGTT
GCACACGTCTGAACTCTTTCCCTACACGACGCTCTTCCGATCTNNNGCCTCACTGTATTTTAGAAATATCA
GAGCACACGTCTGAACTCTTTCCCTACACGACGCTCTTCCGATCTNNNCAGACATATCTGCTTTATTCCT
CACACGTCTGAACTCTTTCCCTACACGACGCTCTTCCGATCTNNNCTTGTTCATCATCCACACTGCATCG
GCACACGTCTGAACTCTTTCCCTACACGACGCTCTTCCGATCTNNNATTTACTTACTTCTCCCAAAGTTCA
GCACACGTCTGAACTCTTTCCCTACACGACGCTCTTCCGATCTNNNACCTCTGCTCCAAAATATAATAAC
GCACACGTCTGAACTCTTTCCCTACACGACGCTCTTCCGATCTNNNAGAATATGGAGGATTATCTGCTC
GAGCACACGTCTGAACTCTTTCCCTACACGACGCTCTTCCGATCTNNNTTATATTCACAAGAACATAGC
AGCACACGTCTGAACTCTTTCCCTACACGACGCTCTTCCGATCTNNNAACTGGAGAAGTATGAATTGA
GCACACGTCTGAACTCTTTCCCTACACGACGCTCTTCCGATCTNNNCATAGAAAAACTGGGAAGAGACA
AGCACACGTCTGAACTCTTTCCCTACACGACGCTCTTCCGATCTNNNTACAGTTCATACAATATCCAAGA
AGCACACGTCTGAACTCTTTCCCTACACGACGCTCTTCCGATCTNNNTACAGTTCATACAATATCCAAGA
CACACGTCTGAACTCTTTCCCTACACGACGCTCTTCCGATCTNNNGTGACTTGGGTCCATTTTATCATCG
CACACGTCTGAACTCTTTCCCTACACGACGCTCTTCCGATCTNNNAGATGGGGTTTCTCCATATTCATCG
CACACGTCTGAACTCTTTCCCTACACGACGCTCTTCCGATCTNNNGTCCTATTGCCTGACTTTAATCATC
AGCACACGTCTGAACTCTTTCCCTACACGACGCTCTTCCGATCTNNNATTTTATAATCATGTTGTGCTCA
GAGCACACGTCTGAACTCTTTCCCTACACGACGCTCTTCCGATCTNNNTTTTCACATAAATGAGTTGGA
AGCACACGTCTGAACTCTTTCCCTACACGACGCTCTTCCGATCTNNNTGTTTGTTGTTTTTGTTATTTGA
GAGCACACGTCTGAACTCTTTCCCTACACGACGCTCTTCCGATCTNNNAACTTGCACAGAAGAATCTAAA
AGCACACGTCTGAACTCTTTCCCTACACGACGCTCTTCCGATCTNNNTCAGGTGAAATCATATGAGATT
AGCACACGTCTGAACTCTTTCCCTACACGACGCTCTTCCGATCTNNNCAATTTAATCTACCACATTTTGC
CACACGTCTGAACTCTTTCCCTACACGACGCTCTTCCGATCTNNNAGAAGGAAATGTAGCCTACTCATC
AGCACACGTCTGAACTCTTTCCCTACACGACGCTCTTCCGATCTNNNAAGGAACATATCTGATCATAACA
GAGCACACGTCTGAACTCTTTCCCTACACGACGCTCTTCCGATCTNNNTAAAATATGTGCTACTGTTAC
GAGCACACGTCTGAACTCTTTCCCTACACGACGCTCTTCCGATCTNNNGTATAGCTTGAAGTTGGATAA
AGAGCACACGTCTGAACTCTTTCCCTACACGACGCTCTTCCGATCTNNNAGCAGATAACTAAATAATCA
AGCACACGTCTGAACTCTTTCCCTACACGACGCTCTTCCGATCTNNNTTATATGCCCTGAAAATAGCCCA
CACACGTCTGAACTCTTTCCCTACACGACGCTCTTCCGATCTNNNTCTCTGCTGTTCTTCTAAGACATCG
ACACGTCTGAACTCTTTCCCTACACGACGCTCTTCCGATCTNNNCCCTCACAATATCTATCTGATGCATC

CACACGTCTGAACTCTTTCCCTACACGACGCTCTTCCGATCTNNNGACTAATACACCATTTTTCTTCACA
GAGCACACGTCTGAACTCTTTCCCTACACGACGCTCTTCCGATCTNNNGAATTGTCTTGAATCATTCAG
AGCACACGTCTGAACTCTTTCCCTACACGACGCTCTTCCGATCTNNNTCATAAAATGGAAGATTAGAG
AGAGCACACGTCTGAACTCTTTCCCTACACGACGCTCTTCCGATCTNNNCATATGGCATGTGTGTCTAG
AGCACACGTCTGAACTCTTTCCCTACACGACGCTCTTCCGATCTNNNATGCAATAATAAGAAAATGGC
AGCACACGTCTGAACTCTTTCCCTACACGACGCTCTTCCGATCTNNNAATATCAAAACTGGAAAACTGA
GCACACGTCTGAACTCTTTCCCTACACGACGCTCTTCCGATCTNNNAACTGAATTCTTGATCTCCTTCCAT
GCACACGTCTGAACTCTTTCCCTACACGACGCTCTTCCGATCTNNNAAAAAAATTCAATTGTGCTGTCC
GCACACGTCTGAACTCTTTCCCTACACGACGCTCTTCCGATCTNNNATCCGTAAGACACAGTGAAGCA
AGCACACGTCTGAACTCTTTCCCTACACGACGCTCTTCCGATCTNNNCAGAGTTTGAGGTCTTAAATTT
AGCACACGTCTGAACTCTTTCCCTACACGACGCTCTTCCGATCTNNNATTTTCAGCCTTATTCTTTTCAT
AGCACACGTCTGAACTCTTTCCCTACACGACGCTCTTCCGATCTNNNACATCCTAACTGACTGTATTAT
ACACGTCTGAACTCTTTCCCTACACGACGCTCTTCCGATCTNNNATAAACTGGAGACAACAAAAGTACA
AGCACACGTCTGAACTCTTTCCCTACACGACGCTCTTCCGATCTNNNTAAGCTCATACTTCCTAGAGA
GAGCACACGTCTGAACTCTTTCCCTACACGACGCTCTTCCGATCTNNNGAAGGACAAATAAAAATTC
AGCACACGTCTGAACTCTTTCCCTACACGACGCTCTTCCGATCTNNNGGTTTCTAAAGGAAAAAAGTC
AGCACACGTCTGAACTCTTTCCCTACACGACGCTCTTCCGATCTNNNGGGAAAAAACAATGTATGACA
AGCACACGTCTGAACTCTTTCCCTACACGACGCTCTTCCGATCTNNNCTTCTCAGATGAAGTGTTTTTC
AGCACACGTCTGAACTCTTTCCCTACACGACGCTCTTCCGATCTNNNCAATCATATCAGGAAGCAAAA
AGCACACGTCTGAACTCTTTCCCTACACGACGCTCTTCCGATCTNNNGGATTTACATCAAAGAAATCA
AGCACACGTCTGAACTCTTTCCCTACACGACGCTCTTCCGATCTNNNTACTTTCCTTAGCCATTTAAAA
AGCACACGTCTGAACTCTTTCCCTACACGACGCTCTTCCGATCTNNNGGCACTCTGTAGATACAATATA
GAGCACACGTCTGAACTCTTTCCCTACACGACGCTCTTCCGATCTNNNGATCAGGGAAAGTATAGGT
AGCACACGTCTGAACTCTTTCCCTACACGACGCTCTTCCGATCTNNNAATTCTTTTGTTTTACTGCTAGT
AGCACACGTCTGAACTCTTTCCCTACACGACGCTCTTCCGATCTNNNTGATAATGTTACATGGAAAAA
AGCACACGTCTGAACTCTTTCCCTACACGACGCTCTTCCGATCTNNNCTATACATGGGTATACAATAGC
AGCACACGTCTGAACTCTTTCCCTACACGACGCTCTTCCGATCTNNNAAAAAAATTGAGGATGAGGT
AGCACACGTCTGAACTCTTTCCCTACACGACGCTCTTCCGATCTNNNTAATTCAACAAAAGCAATTCCA
GCACACGTCTGAACTCTTTCCCTACACGACGCTCTTCCGATCTNNNGTGTGTATATTTCTACTCAGCAA
GCACACGTCTGAACTCTTTCCCTACACGACGCTCTTCCGATCTNNNTACTCTTCCTTTCTCTCTCAATAC
GCACACGTCTGAACTCTTTCCCTACACGACGCTCTTCCGATCTNNNTATCATCTATTTCTACCCTACCCC
AGCACACGTCTGAACTCTTTCCCTACACGACGCTCTTCCGATCTNNNGGTATTGACCAATTAAGTTAT
AGCACACGTCTGAACTCTTTCCCTACACGACGCTCTTCCGATCTNNNGGTATTGACCAATTAAGTTAT
CACACGTCTGAACTCTTTCCCTACACGACGCTCTTCCGATCTNNNAAAAATTAATCCCCATTCAAAGAC
AGCACACGTCTGAACTCTTTCCCTACACGACGCTCTTCCGATCTNNNCCTGATTATATCTTTGCCATTTT
AGCACACGTCTGAACTCTTTCCCTACACGACGCTCTTCCGATCTNNNAAAAAAAAAAAAACCCAACAA
AGCACACGTCTGAACTCTTTCCCTACACGACGCTCTTCCGATCTNNNCTACAAGGAAGGCTGAGAAT
AGCACACGTCTGAACTCTTTCCCTACACGACGCTCTTCCGATCTNNNCAGAAATCCATCTTCTAGTTCT
AGCACACGTCTGAACTCTTTCCCTACACGACGCTCTTCCGATCTNNNCAATGTCTTTTCTACCTATTTCA
CACACGTCTGAACTCTTTCCCTACACGACGCTCTTCCGATCTNNNCTACCCCTCAAACACACATACATCG
GCACACGTCTGAACTCTTTCCCTACACGACGCTCTTCCGATCTNNNCGTATGAATCTAAGAAAATCA
GCACACGTCTGAACTCTTTCCCTACACGACGCTCTTCCGATCTNNNAGATGCTCAATTATCATCCTATT
GCACACGTCTGAACTCTTTCCCTACACGACGCTCTTCCGATCTNNNAGTTATTTGAATCAGTCCTTTTC

GCACACGTCTGAACTCTTTCCCTACACGACGCTCTTCCGATCTNNNAGACATAAAATGAAGATTGAAG
AGCACACGTCTGAACTCTTTCCCTACACGACGCTCTTCCGATCTNNNAATTCCTTTTTCAAATATTCCCC
AGCACACGTCTGAACTCTTTCCCTACACGACGCTCTTCCGATCTNNNTCTATTTCAAGATTATGGGTTC
GCACACGTCTGAACTCTTTCCCTACACGACGCTCTTCCGATCTNNNTTACTAGCAAAGAATTCATCCTA
AGCACACGTCTGAACTCTTTCCCTACACGACGCTCTTCCGATCTNNNGTGACAGCTAATGATAAATTT
CACACGTCTGAACTCTTTCCCTACACGACGCTCTTCCGATCTNNNTTGTCTAAGACAATATGTGGTCCA
AGCACACGTCTGAACTCTTTCCCTACACGACGCTCTTCCGATCTNNNGCGTGTACATGTATGGTATAT
AGCACACGTCTGAACTCTTTCCCTACACGACGCTCTTCCGATCTNNNGTTAGTTTAGGGCTTACCAAGC
CACACGTCTGAACTCTTTCCCTACACGACGCTCTTCCGATCTNNNTTTCAGGATTAGAAAATTCAGCTC
GCACACGTCTGAACTCTTTCCCTACACGACGCTCTTCCGATCTNNNCTTCTTACTAACGGTGAATAGTA
ACACGTCTGAACTCTTTCCCTACACGACGCTCTTCCGATCTNNNCAATTCAGAAGTTTCACCTTCCATCG
AGCACACGTCTGAACTCTTTCCCTACACGACGCTCTTCCGATCTNNNTGTCTGTATTTTCTATAGAAGG
GCACACGTCTGAACTCTTTCCCTACACGACGCTCTTCCGATCTNNNGTCCTCAGTTCATTTATAAAACAC
GCACACGTCTGAACTCTTTCCCTACACGACGCTCTTCCGATCTNNNGAGATTCTTTATCAGAGCAGATC
AGCACACGTCTGAACTCTTTCCCTACACGACGCTCTTCCGATCTNNNTCTTGAGGATTTTCTTTCAATTT
AGCACACGTCTGAACTCTTTCCCTACACGACGCTCTTCCGATCTNNNAATAATCTGAATTCCTTCTTCCT
GAGCACACGTCTGAACTCTTTCCCTACACGACGCTCTTCCGATCTNNNCTAGTTCAAAGACTCAAAATG
GAGCACACGTCTGAACTCTTTCCCTACACGACGCTCTTCCGATCTNNNGCAGATAATACCTAAACTTCT
AGCACACGTCTGAACTCTTTCCCTACACGACGCTCTTCCGATCTNNNTTTCTTTCAACAGTGTTTTGTAG
AGCACACGTCTGAACTCTTTCCCTACACGACGCTCTTCCGATCTNNNGTTGTTTTGGAGACTCAATGC
GCACACGTCTGAACTCTTTCCCTACACGACGCTCTTCCGATCTNNNTAGGAGACAGTCATTGACAACA
GAGCACACGTCTGAACTCTTTCCCTACACGACGCTCTTCCGATCTNNNTTAAGATGCTGTACAGATGC
AGAGCACACGTCTGAACTCTTTCCCTACACGACGCTCTTCCGATCTNNNAAATGTTTATTATGGCACTT
CACACGTCTGAACTCTTTCCCTACACGACGCTCTTCCGATCTNNNATACAGCCCTGTTCTACAACCATCG
ACACGTCTGAACTCTTTCCCTACACGACGCTCTTCCGATCTNNNAAAAAAAGTATTCCTCCACTTACCAT
CACACGTCTGAACTCTTTCCCTACACGACGCTCTTCCGATCTNNNAAAAAAAAAGTATTCCTCCACTTCA
GCACACGTCTGAACTCTTTCCCTACACGACGCTCTTCCGATCTNNNCCTCAAAAGAAGAAAAAAAAAG
AGCACACGTCTGAACTCTTTCCCTACACGACGCTCTTCCGATCTNNNGATTAGGATGATGACCAAGAA
AGCACACGTCTGAACTCTTTCCCTACACGACGCTCTTCCGATCTNNNCATGCTCACAGAACATAATTG
GCACACGTCTGAACTCTTTCCCTACACGACGCTCTTCCGATCTNNNCCTTTACTTTGAGGATATTGAAG
AGCACACGTCTGAACTCTTTCCCTACACGACGCTCTTCCGATCTNNNTAACACTAAATTCTTCCAATGA
GCACACGTCTGAACTCTTTCCCTACACGACGCTCTTCCGATCTNNNAAAAAATAAAAGAATCACCCAC
CACACGTCTGAACTCTTTCCCTACACGACGCTCTTCCGATCTNNNAAAAATTAATCCCCATTCAAAGAC
GCACACGTCTGAACTCTTTCCCTACACGACGCTCTTCCGATCTNNNCAAAAAACAAACAAACAAACAA
ACACGTCTGAACTCTTTCCCTACACGACGCTCTTCCGATCTNNNCTCTTACCAACAACAACCACCATCGG
GCACACGTCTGAACTCTTTCCCTACACGACGCTCTTCCGATCTNNNACTGTATGTGTTCTAATATCTGA
AGCACACGTCTGAACTCTTTCCCTACACGACGCTCTTCCGATCTNNNTTCCTCTCTCTCTCTTTATATC
GCACACGTCTGAACTCTTTCCCTACACGACGCTCTTCCGATCTNNNATTTTAATATGTAGCCAGGATCA
AGCACACGTCTGAACTCTTTCCCTACACGACGCTCTTCCGATCTNNNAGATGGTTGAATGAATTTAAG
AGCACACGTCTGAACTCTTTCCCTACACGACGCTCTTCCGATCTNNNAGATGGTTGAATGAATTTAAG
CACACGTCTGAACTCTTTCCCTACACGACGCTCTTCCGATCTNNNTGTGTATTAGCATTAGTCCATTAC
CACACGTCTGAACTCTTTCCCTACACGACGCTCTTCCGATCTNNNAAATGATTACTTCCTACCAGTTCCA
AGCACACGTCTGAACTCTTTCCCTACACGACGCTCTTCCGATCTNNNTGTGTGTAAGTATGACTAAAA

AGCACACGTCTGAACTCTTTCCCTACACGACGCTCTTCCGATCTNNNATCACAATTAGAGGAAAGGAG
GCACACGTCTGAACTCTTTCCCTACACGACGCTCTTCCGATCTNNNCTTCACCTTTGACATCAGAACA
AGCACACGTCTGAACTCTTTCCCTACACGACGCTCTTCCGATCTNNNCTAGGTCATCTGTGTACCTCCAT
GCACACGTCTGAACTCTTTCCCTACACGACGCTCTTCCGATCTNNNGGTCTAGTTCAAGATACTGAAT
AGCACACGTCTGAACTCTTTCCCTACACGACGCTCTTCCGATCTNNNTATCTATATGTGCCCTCAATAA
AGCACACGTCTGAACTCTTTCCCTACACGACGCTCTTCCGATCTNNNTGTTTATTACAAGGTACATCAT
AGCACACGTCTGAACTCTTTCCCTACACGACGCTCTTCCGATCTNNNTGTGACATCAATATTCATGCTA
AGCACACGTCTGAACTCTTTCCCTACACGACGCTCTTCCGATCTNNNTATGTATATATGTGCCTTTCTG
AGAGCACACGTCTGAACTCTTTCCCTACACGACGCTCTTCCGATCTNNNTGAATTGACTTTGATATAA
AGCACACGTCTGAACTCTTTCCCTACACGACGCTCTTCCGATCTNNNCTAAATGGAAAGCCTATCTCA
AGCACACGTCTGAACTCTTTCCCTACACGACGCTCTTCCGATCTNNNTCTGTGTGATAATTTGAAAAT
GAGCACACGTCTGAACTCTTTCCCTACACGACGCTCTTCCGATCTNNNTATTAGTGGTTGCTATTATCA
GAGCACACGTCTGAACTCTTTCCCTACACGACGCTCTTCCGATCTNNNTGTCAAAAGTCCTATTACAAT
GCACACGTCTGAACTCTTTCCCTACACGACGCTCTTCCGATCTNNNATTGAAACAAATATCCTTGCACC
AGCACACGTCTGAACTCTTTCCCTACACGACGCTCTTCCGATCTNNNTATAAATATATACTTTGCCCAG
GAGCACACGTCTGAACTCTTTCCCTACACGACGCTCTTCCGATCTNNNTATTTGAATTTGTAGACCCAG
AGCACACGTCTGAACTCTTTCCCTACACGACGCTCTTCCGATCTNNNGGTGCATATATATGTGCCTATA
ACACGTCTGAACTCTTTCCCTACACGACGCTCTTCCGATCTNNNATTCTAAAGGTGGTGAGTAAAATC
ACACGTCTGAACTCTTTCCCTACACGACGCTCTTCCGATCTNNNCCAACATTATAGCAAAAGACTGCAT
GCACACGTCTGAACTCTTTCCCTACACGACGCTCTTCCGATCTNNNTATGTTAGTGATTAAGCTTCTCA
GCACACGTCTGAACTCTTTCCCTACACGACGCTCTTCCGATCTNNNCAAGGAGAAAAAAATGCAGAA
AGCACACGTCTGAACTCTTTCCCTACACGACGCTCTTCCGATCTNNNAACCAGCATCTCTGTTTATATAT
AGCACACGTCTGAACTCTTTCCCTACACGACGCTCTTCCGATCTNNNTTCCTAAAAGTCCTCCTCTATCC
AGCACACGTCTGAACTCTTTCCCTACACGACGCTCTTCCGATCTNNNTCATCATTAGCAAATCGAAAA
AGCACACGTCTGAACTCTTTCCCTACACGACGCTCTTCCGATCTNNNGGTTTTTTGCAAATGCTTATGC
CACACGTCTGAACTCTTTCCCTACACGACGCTCTTCCGATCTNNNAAAAATGCATACCTTCTTTGTACC
ACACGTCTGAACTCTTTCCCTACACGACGCTCTTCCGATCTNNNAATATTTGCATGCCTACTAAATGCAT
AGCACACGTCTGAACTCTTTCCCTACACGACGCTCTTCCGATCTNNNCAAACCCACTTATTCTTTTAGG
ACACGTCTGAACTCTTTCCCTACACGACGCTCTTCCGATCTNNNTAAATTACCAGAGAAGGACTGTACA
CACACGTCTGAACTCTTTCCCTACACGACGCTCTTCCGATCTNNNAATATTTGCCTTCATCTTCAAATCA
AGCACACGTCTGAACTCTTTCCCTACACGACGCTCTTCCGATCTNNNATGATGGTAAGGATGGAGTT
AGCACACGTCTGAACTCTTTCCCTACACGACGCTCTTCCGATCTNNNTCAGAAATAGTTTTTGTGTAA
AGCACACGTCTGAACTCTTTCCCTACACGACGCTCTTCCGATCTNNNTTCTCAGGAGAGTAAAATACA
AGCACACGTCTGAACTCTTTCCCTACACGACGCTCTTCCGATCTNNNCTACTCTACAGATTTCAGATTT
AGCACACGTCTGAACTCTTTCCCTACACGACGCTCTTCCGATCTNNNCTGATAGAGAACATTGTAGGA
AGCACACGTCTGAACTCTTTCCCTACACGACGCTCTTCCGATCTNNNAAGAAATGGAAATGGTACTGA
ACACGTCTGAACTCTTTCCCTACACGACGCTCTTCCGATCTNNNAAACGACAAACAGATGAAGCCATCG
GAGCACACGTCTGAACTCTTTCCCTACACGACGCTCTTCCGATCTNNNAAATACCAAAGCTCAAGAAA
ACACGTCTGAACTCTTTCCCTACACGACGCTCTTCCGATCTNNNCCAAAGACAATTCAACTCAAACATC
GCACACGTCTGAACTCTTTCCCTACACGACGCTCTTCCGATCTNNNCTTTGGTTCTCAAGTAATGAACC
AGAGCACACGTCTGAACTCTTTCCCTACACGACGCTCTTCCGATCTNNNAGATGTTAAGAATCTTGATT
AGCACACGTCTGAACTCTTTCCCTACACGACGCTCTTCCGATCTNNNGGTTTTCTTGATAATCGATCTG
GAGCACACGTCTGAACTCTTTCCCTACACGACGCTCTTCCGATCTNNNGAAAATATTATTTTTCCCCATG

GCACACGTCTGAACTCTTTCCCTACACGACGCTCTTCCGATCTNNNGGGGGTAATTTTGTAAAAGAAT

AGCACACGTCTGAACTCTTTCCCTACACGACGCTCTTCCGATCTNNNTAAAGGCACTGAGAAATCTTAT

GCACACGTCTGAACTCTTTCCCTACACGACGCTCTTCCGATCTNNNAATATTTTACAATGAGCATGTG

AGCACACGTCTGAACTCTTTCCCTACACGACGCTCTTCCGATCTNNNACAAGAAGAAACCATGTGAAC

GCACACGTCTGAACTCTTTCCCTACACGACGCTCTTCCGATCTNNNCATAATCTAAAAGAGGGGAAG

AGCACACGTCTGAACTCTTTCCCTACACGACGCTCTTCCGATCTNNNATAACTGAGATGAAACAGAA

AGCACACGTCTGAACTCTTTCCCTACACGACGCTCTTCCGATCTNNNATGTATTACTGCGTGATGTTAT

ACACGTCTGAACTCTTTCCCTACACGACGCTCTTCCGATCTNNNAAATTACGAGAGATCTTTACTCTCA

GCACACGTCTGAACTCTTTCCCTACACGACGCTCTTCCGATCTNNNCGTCTTTCTCACTTTTACATTTA

AGCACACGTCTGAACTCTTTCCCTACACGACGCTCTTCCGATCTNNNAATAGAAATAAAACAACCAAC

CACACGTCTGAACTCTTTCCCTACACGACGCTCTTCCGATCTNNNGTAATCATAGCTCACTGCAGCAT

GCACACGTCTGAACTCTTTCCCTACACGACGCTCTTCCGATCTNNNTCTTTATGAGAAAGACCTTCTG

AGCACACGTCTGAACTCTTTCCCTACACGACGCTCTTCCGATCTNNNAGACCTTGTCTCTCAAAAAAA

GCACACGTCTGAACTCTTTCCCTACACGACGCTCTTCCGATCTNNNCTTTACTTTTTTAGTCCAACCATC

CACACGTCTGAACTCTTTCCCTACACGACGCTCTTCCGATCTNNNTTCTAAATGTTATGGTTGGCTCA

AGCACACGTCTGAACTCTTTCCCTACACGACGCTCTTCCGATCTNNNCATCCCATTACAAAAAAAACTT

AGCACACGTCTGAACTCTTTCCCTACACGACGCTCTTCCGATCTNNNTTCAGATCCAAGAAGAGAAT

GCACACGTCTGAACTCTTTCCCTACACGACGCTCTTCCGATCTNNNCTCCAGAAGACAAAAAATGAG

GCACACGTCTGAACTCTTTCCCTACACGACGCTCTTCCGATCTNNNCTTACATTTTTGAGTAGGGAAA

AGCACACGTCTGAACTCTTTCCCTACACGACGCTCTTCCGATCTNNNGTAAGTCCATCGTAAGTTGAG

CACACGTCTGAACTCTTTCCCTACACGACGCTCTTCCGATCTNNNTAACCACTACCCTAATGAGATTAC

AGCACACGTCTGAACTCTTTCCCTACACGACGCTCTTCCGATCTNNNTCTCTCTCTCTCTCTCTATAT

AGCACACGTCTGAACTCTTTCCCTACACGACGCTCTTCCGATCTNNNTCTCTCTCTCTCTCTCTATAT

AGCACACGTCTGAACTCTTTCCCTACACGACGCTCTTCCGATCTNNNTAACTTACTTCTGTGTAGTGTA

CACACGTCTGAACTCTTTCCCTACACGACGCTCTTCCGATCTNNNATTTTTAAAAGATGCCATGAGAG

GCACACGTCTGAACTCTTTCCCTACACGACGCTCTTCCGATCTNNNAGTATTAGAAAGATTTCATTCCT

GCACACGTCTGAACTCTTTCCCTACACGACGCTCTTCCGATCTNNNAAGTTTGAACATATGCATATAC

CACACGTCTGAACTCTTTCCCTACACGACGCTCTTCCGATCTNNNATTTACTTCCATATCACATTCCTC

AGCACACGTCTGAACTCTTTCCCTACACGACGCTCTTCCGATCTNNNGACCCTCAGGAAAAAAAAAAT

GCACACGTCTGAACTCTTTCCCTACACGACGCTCTTCCGATCTNNNCATGTATTTGTGCATGTGAGCA

GAGCACACGTCTGAACTCTTTCCCTACACGACGCTCTTCCGATCTNNNGGTCATGCAGATTTAATTAG

AGCACACGTCTGAACTCTTTCCCTACACGACGCTCTTCCGATCTNNNAAAGTAAAACCCAATATGTTC

AGCACACGTCTGAACTCTTTCCCTACACGACGCTCTTCCGATCTNNNTCACCTGGATATTTAATAACA

GCACACGTCTGAACTCTTTCCCTACACGACGCTCTTCCGATCTNNNTAACTACAGCACATCCAAATCCA

AGCACACGTCTGAACTCTTTCCCTACACGACGCTCTTCCGATCTNNNAGCTAATATGAACAGAAAAAC

GCACACGTCTGAACTCTTTCCCTACACGACGCTCTTCCGATCTNNNGTTGCTAACTACACATTTCTTTA

CACACGTCTGAACTCTTTCCCTACACGACGCTCTTCCGATCTNNNAGAAAAATAGGTGGCATTTTATT

GCACACGTCTGAACTCTTTCCCTACACGACGCTCTTCCGATCTNNNAGGCTGTCTGAAAAAATAAAAA

ACACGTCTGAACTCTTTCCCTACACGACGCTCTTCCGATCTNNNTTATTGAGTTTGGAATGCAATACC

AGCACACGTCTGAACTCTTTCCCTACACGACGCTCTTCCGATCTNNNCTATTATTGGTTTGGTGTATGT

AGCACACGTCTGAACTCTTTCCCTACACGACGCTCTTCCGATCTNNNAAAGATCTCATACAGACAACT

AGCACACGTCTGAACTCTTTCCCTACACGACGCTCTTCCGATCTNNNAATATATCAAGTAGGTGTTTT

AGCACACGTCTGAACTCTTTCCCTACACGACGCTCTTCCGATCTNNNCCAGTTTGTGGTAATTTGTTA

GCACACGTCTGAACTCTTTCCCTACACGACGCTCTTCCGATCTNNNGATATTCATGGTTATGTGAATGG
AGCACACGTCTGAACTCTTTCCCTACACGACGCTCTTCCGATCTNNNGGTTAAATCACTGATGCTTTATA
GAGCACACGTCTGAACTCTTTCCCTACACGACGCTCTTCCGATCTNNNTCTTTATTTTTTTAGGCTGAACA
AGCACACGTCTGAACTCTTTCCCTACACGACGCTCTTCCGATCTNNNCCTCAGAAATATGTCTTTTACTG
GAGCACACGTCTGAACTCTTTCCCTACACGACGCTCTTCCGATCTNNNTGTCCATTTAGTATAAAGTGT
GAGCACACGTCTGAACTCTTTCCCTACACGACGCTCTTCCGATCTNNNGCAAAACATCTCAAATTGTTTA
AGCACACGTCTGAACTCTTTCCCTACACGACGCTCTTCCGATCTNNNTTCATACATTTTTATTTGGTGGTC
GCACACGTCTGAACTCTTTCCCTACACGACGCTCTTCCGATCTNNNAATGATCAAAGTACATTGGAAAG
GAGCACACGTCTGAACTCTTTCCCTACACGACGCTCTTCCGATCTNNNAGTTGGCAAATGGTATAAAA
GAGCACACGTCTGAACTCTTTCCCTACACGACGCTCTTCCGATCTNNNGTATAAGTAGAAAAGCAGTTC
AGCACACGTCTGAACTCTTTCCCTACACGACGCTCTTCCGATCTNNNATTTGAGGATTCTATGGATTCA
AGCACACGTCTGAACTCTTTCCCTACACGACGCTCTTCCGATCTNNNGAACCATTCTGATGTCCTTCCAT
GCACACGTCTGAACTCTTTCCCTACACGACGCTCTTCCGATCTNNNCAAAGGTATTTGGTCTCTATCTCA
AGCACACGTCTGAACTCTTTCCCTACACGACGCTCTTCCGATCTNNNTTAGATTCAAAATTGTTGACACA
AGCACACGTCTGAACTCTTTCCCTACACGACGCTCTTCCGATCTNNNTATCAAGGAAATAAAAGAGGAC
CACACGTCTGAACTCTTTCCCTACACGACGCTCTTCCGATCTNNNATTTTCTTTTTTGTTTGCTTTCTCAT
GAGCACACGTCTGAACTCTTTCCCTACACGACGCTCTTCCGATCTNNNACTATAAGGAAGAAATCAAG
GCACACGTCTGAACTCTTTCCCTACACGACGCTCTTCCGATCTNNNCACAAAATGATATGGGGTTATGC
AGAGCACACGTCTGAACTCTTTCCCTACACGACGCTCTTCCGATCTNNNGGAATGAATATTATCTATGT
CACACGTCTGAACTCTTTCCCTACACGACGCTCTTCCGATCTNNNTTTTGTGATAATCCGTGAGTATACA
GAGCACACGTCTGAACTCTTTCCCTACACGACGCTCTTCCGATCTNNNACTAAGTGATAGGAATTCTTCA
GCACACGTCTGAACTCTTTCCCTACACGACGCTCTTCCGATCTNNNATATATAGGCAAGGATGAAAACA
AGCACACGTCTGAACTCTTTCCCTACACGACGCTCTTCCGATCTNNNGAATAGACCTCCTTTCCTCTCATC
GCACACGTCTGAACTCTTTCCCTACACGACGCTCTTCCGATCTNNNTTTGTTCAGTGATATTAGCATTTC
AGCACACGTCTGAACTCTTTCCCTACACGACGCTCTTCCGATCTNNNAAGAAGGTGGTAAAGAACAGC
GAGCACACGTCTGAACTCTTTCCCTACACGACGCTCTTCCGATCTNNNTCCCTTTCTCTCTCCTCTATCATC
AGCACACGTCTGAACTCTTTCCCTACACGACGCTCTTCCGATCTNNNTAACAAAATGTTAGTCTTCTCTC
AGCACACGTCTGAACTCTTTCCCTACACGACGCTCTTCCGATCTNNNGACTAATTCTGCATGTAAGAAA
AGCACACGTCTGAACTCTTTCCCTACACGACGCTCTTCCGATCTNNNTTAAAAGATTATCATGCTTCCAC
GAGCACACGTCTGAACTCTTTCCCTACACGACGCTCTTCCGATCTNNNGGAAACAGATCAACTATTTCA
GAGCACACGTCTGAACTCTTTCCCTACACGACGCTCTTCCGATCTNNNATTCTGTCTCAAAAAGAAACA
GCACACGTCTGAACTCTTTCCCTACACGACGCTCTTCCGATCTNNNGCTTATCCTAAAGTAAAGGAAAC
AGCACACGTCTGAACTCTTTCCCTACACGACGCTCTTCCGATCTNNNTCCTTAGAGTAATGTTCTTAAAC
AGCACACGTCTGAACTCTTTCCCTACACGACGCTCTTCCGATCTNNNTTTATTTCTGCTACTATCCATGTC
AGCACACGTCTGAACTCTTTCCCTACACGACGCTCTTCCGATCTNNNATGTGATACGTATATGTATGCA
AGCACACGTCTGAACTCTTTCCCTACACGACGCTCTTCCGATCTNNNCAAAACAGTCTCTTCTCAGATCA
GAGCACACGTCTGAACTCTTTCCCTACACGACGCTCTTCCGATCTNNNATAATTTTATGTTTGACTTGGC
GCACACGTCTGAACTCTTTCCCTACACGACGCTCTTCCGATCTNNNGGTTTCCTTATAAAATCCTATGGC
AGCACACGTCTGAACTCTTTCCCTACACGACGCTCTTCCGATCTNNNTAGTGCTATAAAATCCAAAATG
AGCACACGTCTGAACTCTTTCCCTACACGACGCTCTTCCGATCTNNNTGTTTACCTATTTCTTCATCGTA
GAGCACACGTCTGAACTCTTTCCCTACACGACGCTCTTCCGATCTNNNTCTTAATACTATCACACTGGTT
AGCACACGTCTGAACTCTTTCCCTACACGACGCTCTTCCGATCTNNNCTATTTTACAAGGATCTCTCTGT
AGCACACGTCTGAACTCTTTCCCTACACGACGCTCTTCCGATCTNNNCTATTTTACAAGGATCTCTCTGT

AGCACACGTCTGAACTCTTTCCCTACACGACGCTCTTCCGATCTNNNGTTCTGAACCCTAATATATAGC
GAGCACACGTCTGAACTCTTTCCCTACACGACGCTCTTCCGATCTNNNCAAAACCATGTCTCAAAAAAT
GAGCACACGTCTGAACTCTTTCCCTACACGACGCTCTTCCGATCTNNNTCAGAATTATATAACCCAACA
AGCACACGTCTGAACTCTTTCCCTACACGACGCTCTTCCGATCTNNNATCAGCAAACAGATACAAACTT
GAGCACACGTCTGAACTCTTTCCCTACACGACGCTCTTCCGATCTNNNTTTCTGTGGTTTATGGATTTC
GAGCACACGTCTGAACTCTTTCCCTACACGACGCTCTTCCGATCTNNNAATCAATTTTATACTGATGA
GCACACGTCTGAACTCTTTCCCTACACGACGCTCTTCCGATCTNNNAGAGAAATAGAACCAATAGGAT
AGAGCACACGTCTGAACTCTTTCCCTACACGACGCTCTTCCGATCTNNNGCCTTAAAATGATCTTCAAG
AGCACACGTCTGAACTCTTTCCCTACACGACGCTCTTCCGATCTNNNTCTTGAGAACATACATTTTGTTT
GAGCACACGTCTGAACTCTTTCCCTACACGACGCTCTTCCGATCTNNNAAACAAACCAACCATATATGT
GAGCACACGTCTGAACTCTTTCCCTACACGACGCTCTTCCGATCTNNNGCTTGTTTTGAGGGTATTTAT
AGCACACGTCTGAACTCTTTCCCTACACGACGCTCTTCCGATCTNNNCTCTATCACACACATACACACC
AGCACACGTCTGAACTCTTTCCCTACACGACGCTCTTCCGATCTNNNATTATTTACATCAGAGCAATTC
GAGCACACGTCTGAACTCTTTCCCTACACGACGCTCTTCCGATCTNNNTATACTAAGAAAAAAGCCTGC
AGCACACGTCTGAACTCTTTCCCTACACGACGCTCTTCCGATCTNNNGAATCATTACCTTATATCTGCT
AGCACACGTCTGAACTCTTTCCCTACACGACGCTCTTCCGATCTNNNGATGTGTAAGAATACAAAATG
AGCACACGTCTGAACTCTTTCCCTACACGACGCTCTTCCGATCTNNNGATGTGTAAGAATACAAAATG
AGCACACGTCTGAACTCTTTCCCTACACGACGCTCTTCCGATCTNNNGTTTGTTTTTGTTTTAGTGAG
GCACACGTCTGAACTCTTTCCCTACACGACGCTCTTCCGATCTNNNTAATGGATTAGAGTTGGAGAAA
AGCACACGTCTGAACTCTTTCCCTACACGACGCTCTTCCGATCTNNNCTTAGTTATTTTGGTTGGAATG
GCACACGTCTGAACTCTTTCCCTACACGACGCTCTTCCGATCTNNNGGAGGAGAAGAAATGAAGACC
CACACGTCTGAACTCTTTCCCTACACGACGCTCTTCCGATCTNNNGTAAATGTGAGTTTTCTTGCTATC
CACACGTCTGAACTCTTTCCCTACACGACGCTCTTCCGATCTNNNGAAAACAATCTGTTGACTAACACA
CACGTCTGAACTCTTTCCCTACACGACGCTCTTCCGATCTNNNATCTGTTTCTTGACTGTTCTGCATCG
AGCACACGTCTGAACTCTTTCCCTACACGACGCTCTTCCGATCTNNNGGGATTATTTGCAGAAATAAG
GAGCACACGTCTGAACTCTTTCCCTACACGACGCTCTTCCGATCTNNNCTTGAAGAAAGAGTTAATCG
GAGCACACGTCTGAACTCTTTCCCTACACGACGCTCTTCCGATCTNNNTATCTGACAAGATGAAAGAA
CACACGTCTGAACTCTTTCCCTACACGACGCTCTTCCGATCTNNNGCTAAATGGTAGTAGAGCTGCATC
AGCACACGTCTGAACTCTTTCCCTACACGACGCTCTTCCGATCTNNNTTTCTATTTCTGACTTTGTGT
GCACACGTCTGAACTCTTTCCCTACACGACGCTCTTCCGATCTNNNTCTTGAACTCCTGATCTCAGCAT
GCACACGTCTGAACTCTTTCCCTACACGACGCTCTTCCGATCTNNNTTTTTAATCAGTCTTGATACGG
GAGCACACGTCTGAACTCTTTCCCTACACGACGCTCTTCCGATCTNNNAAAAGCTTTGTTTAATCTCTA
GAGCACACGTCTGAACTCTTTCCCTACACGACGCTCTTCCGATCTNNNATATGACCATTTCAGGCTAAA
AGCACACGTCTGAACTCTTTCCCTACACGACGCTCTTCCGATCTNNNTATTTCCATCCACCCTTTTATTC
AGCACACGTCTGAACTCTTTCCCTACACGACGCTCTTCCGATCTNNNATTAGTCATCTGTAGGACTATC
AGCACACGTCTGAACTCTTTCCCTACACGACGCTCTTCCGATCTNNNGGAAATCAATCTCTTCAACAAA
GCACACGTCTGAACTCTTTCCCTACACGACGCTCTTCCGATCTNNNAATGGGATCATATATATGCTTG
GCACACGTCTGAACTCTTTCCCTACACGACGCTCTTCCGATCTNNNGAAAGTTAACCTTCTCAGAAACC
AGCACACGTCTGAACTCTTTCCCTACACGACGCTCTTCCGATCTNNNGTTAGGAACAGGTGTGAATGC
GAGCACACGTCTGAACTCTTTCCCTACACGACGCTCTTCCGATCTNNNAAGTACTTTACTATTAACGCT
GAGCACACGTCTGAACTCTTTCCCTACACGACGCTCTTCCGATCTNNNAACTCCATCTCAAAAAAGAAA
AGCACACGTCTGAACTCTTTCCCTACACGACGCTCTTCCGATCTNNNTATTCTGTAGTCCCTTCATTGC
GCACACGTCTGAACTCTTTCCCTACACGACGCTCTTCCGATCTNNNTTATAGTGTTCTCTCTTTCTCAGC

AGCACACGTCTGAACTCTTTCCCTACACGACGCTCTTCCGATCTNNNAAATCTTTCTCTGTTCCATTTTCC
AGCACACGTCTGAACTCTTTCCCTACACGACGCTCTTCCGATCTNNNCACAGTTTTAGTCTTCACATGCA
AGCACACGTCTGAACTCTTTCCCTACACGACGCTCTTCCGATCTNNNAGAATGAATGTCCTGTTAATAGA
GCACACGTCTGAACTCTTTCCCTACACGACGCTCTTCCGATCTNNNAAAGAATGACAGAAGGAACAAC
AGCACACGTCTGAACTCTTTCCCTACACGACGCTCTTCCGATCTNNNTGTCCTGAATGTGTATACTTGCA
AGCACACGTCTGAACTCTTTCCCTACACGACGCTCTTCCGATCTNNNAAAAGAAAAAGAATTGAGACCC
AGCACACGTCTGAACTCTTTCCCTACACGACGCTCTTCCGATCTNNNATGAAATCATATTTACACCAAGC
AGCACACGTCTGAACTCTTTCCCTACACGACGCTCTTCCGATCTNNNTTATGTGATAAAAACTTTGCTGA
GAGCACACGTCTGAACTCTTTCCCTACACGACGCTCTTCCGATCTNNNAGCTAATAGAAAACTTCCAAA
GAGCACACGTCTGAACTCTTTCCCTACACGACGCTCTTCCGATCTNNNGTAAAAATATAGACCTCTGCA
GAGCACACGTCTGAACTCTTTCCCTACACGACGCTCTTCCGATCTNNNTATAACAAGATCTAGCTCAAAC
GAGCACACGTCTGAACTCTTTCCCTACACGACGCTCTTCCGATCTNNNCTAAATAAGTACTGTCCCTGTG
AGCACACGTCTGAACTCTTTCCCTACACGACGCTCTTCCGATCTNNNTGTAATCAGAAAGACACTCCTAC
AGCACACGTCTGAACTCTTTCCCTACACGACGCTCTTCCGATCTNNNCCTTTCTTCCTAAATTTGGAGGC
AGCACACGTCTGAACTCTTTCCCTACACGACGCTCTTCCGATCTNNNTAAGTTGGGTTTTTTTCCTCTA
AGCACACGTCTGAACTCTTTCCCTACACGACGCTCTTCCGATCTNNNCTACATCTAACTACATTGAGACA
AGAGCACACGTCTGAACTCTTTCCCTACACGACGCTCTTCCGATCTNNNCAGGATATGTGAGTGAAGAT
GCACACGTCTGAACTCTTTCCCTACACGACGCTCTTCCGATCTNNNATTCGAGGTATACAAAGTGATAT
AGCACACGTCTGAACTCTTTCCCTACACGACGCTCTTCCGATCTNNNTTAGGATGTCAGTATATGCATT
ACACGTCTGAACTCTTTCCCTACACGACGCTCTTCCGATCTNNNAAGTGTCTCACTAAATGGATCCATCC
AGCACACGTCTGAACTCTTTCCCTACACGACGCTCTTCCGATCTNNNAAAGAATGGGGGCTCTATATTT
AGCACACGTCTGAACTCTTTCCCTACACGACGCTCTTCCGATCTNNNAAAGAATGGGGGCTCTATATTT
CACACGTCTGAACTCTTTCCCTACACGACGCTCTTCCGATCTNNNTCTTCTTCTGTAATTCCTGTATTCAT
GAGCACACGTCTGAACTCTTTCCCTACACGACGCTCTTCCGATCTNNNATAGGAACAGGAAAGAGTAA
GAGCACACGTCTGAACTCTTTCCCTACACGACGCTCTTCCGATCTNNNAATGAAGTCTTTCAGTTGTATT
AGCACACGTCTGAACTCTTTCCCTACACGACGCTCTTCCGATCTNNNCTCGACTTGAACTTCTCAAATCA
CACACGTCTGAACTCTTTCCCTACACGACGCTCTTCCGATCTNNNTCACAGTAAACTATCAATAAGGTCA
AGCACACGTCTGAACTCTTTCCCTACACGACGCTCTTCCGATCTNNNAGTGAAAAGTACAATCTTTTGA
AGCACACGTCTGAACTCTTTCCCTACACGACGCTCTTCCGATCTNNNAGTGAAAAGTACAATCTTTTGA
GAGCACACGTCTGAACTCTTTCCCTACACGACGCTCTTCCGATCTNNNTATTTGTAGAATCCCCCAAATA
GAGCACACGTCTGAACTCTTTCCCTACACGACGCTCTTCCGATCTNNNTAAGAGTAAGCATGCTAACTTC
GCACACGTCTGAACTCTTTCCCTACACGACGCTCTTCCGATCTNNNGGTGAGCATAGTCAATAATAATT
AGCACACGTCTGAACTCTTTCCCTACACGACGCTCTTCCGATCTNNNATTATTGAATGCTTCACATACAT
AGCACACGTCTGAACTCTTTCCCTACACGACGCTCTTCCGATCTNNNGAAGATTCTTCTTTTTAGTCTGA
AGCACACGTCTGAACTCTTTCCCTACACGACGCTCTTCCGATCTNNNGATATATTCAAAAGCTTGGTGT
GAGCACACGTCTGAACTCTTTCCCTACACGACGCTCTTCCGATCTNNNTATGTGACTAATATGTTTGCTC
AGCACACGTCTGAACTCTTTCCCTACACGACGCTCTTCCGATCTNNNATACGCAAAGTTACACGAATCA
AGCACACGTCTGAACTCTTTCCCTACACGACGCTCTTCCGATCTNNNGAGTAAGGGAATGGTATAGAA
AGCACACGTCTGAACTCTTTCCCTACACGACGCTCTTCCGATCTNNNAATAAGCATATAGTCTCTTCTAC
AGCACACGTCTGAACTCTTTCCCTACACGACGCTCTTCCGATCTNNNAATAAGCATATAGTCTCTTCTAC
CACACGTCTGAACTCTTTCCCTACACGACGCTCTTCCGATCTNNNAAATAACAGAATTTCGCAGATGCAT
AGCACACGTCTGAACTCTTTCCCTACACGACGCTCTTCCGATCTNNNGATACTGCTTGACAACATACACA
CACACGTCTGAACTCTTTCCCTACACGACGCTCTTCCGATCTNNNGAAAAAAAAAGCCTCCCCTACATCG

GCACACGTCTGAACTCTTTCCCTACACGACGCTCTTCCGATCTNNNTTCTTCTTTTTCTCCTTCTCTATCA
ACACGTCTGAACTCTTTCCCTACACGACGCTCTTCCGATCTNNNGATTATCCATTCAAAAAACAGTGCAT
GAGCACACGTCTGAACTCTTTCCCTACACGACGCTCTTCCGATCTNNNATAATTACACCCATAATTTGCT
AGCACACGTCTGAACTCTTTCCCTACACGACGCTCTTCCGATCTNNNATTTTCATACACGTGGTAAAAT
GCACACGTCTGAACTCTTTCCCTACACGACGCTCTTCCGATCTNNNCACCAATACATAATATTGTCAGTC
AGCACACGTCTGAACTCTTTCCCTACACGACGCTCTTCCGATCTNNNATCACTTTTCTCAAATCTGATTC
GCACACGTCTGAACTCTTTCCCTACACGACGCTCTTCCGATCTNNNGAATCTGGAAGTCAAAGTCCCAT
AGCACACGTCTGAACTCTTTCCCTACACGACGCTCTTCCGATCTNNNTTTGTATTAAAATTGTAAGCTC
CACACGTCTGAACTCTTTCCCTACACGACGCTCTTCCGATCTNNNCTTTTTAAATTATTGATCCGCCCAT
AGCACACGTCTGAACTCTTTCCCTACACGACGCTCTTCCGATCTNNNTTCCAGTTTTTATAAACAGTACG
AGAGCACACGTCTGAACTCTTTCCCTACACGACGCTCTTCCGATCTNNNTTGAGATGAATCACTGAAGT
GAGCACACGTCTGAACTCTTTCCCTACACGACGCTCTTCCGATCTNNNAGGAATATTCTTAGTTCCCATT
GCACACGTCTGAACTCTTTCCCTACACGACGCTCTTCCGATCTNNNCTTTAGGAGCAATGTTTCACTTCA
AGCACACGTCTGAACTCTTTCCCTACACGACGCTCTTCCGATCTNNNAAAGAAGACAGTTTGGAACACA
AGCACACGTCTGAACTCTTTCCCTACACGACGCTCTTCCGATCTNNNCTTTTCTTTACCCTTTTTCCTACA
AGCACACGTCTGAACTCTTTCCCTACACGACGCTCTTCCGATCTNNNCTTTTCTCTCTCTTTCTTTCTCCA
GAGCACACGTCTGAACTCTTTCCCTACACGACGCTCTTCCGATCTNNNCTCTTGTTTGGGGGTAAAAGC
ACACGTCTGAACTCTTTCCCTACACGACGCTCTTCCGATCTNNNTTTATGTGAAATTAGTTTTCCTAGTC
AGCACACGTCTGAACTCTTTCCCTACACGACGCTCTTCCGATCTNNNTTTGTTACCCCAACTCAAGACAT
GAGCACACGTCTGAACTCTTTCCCTACACGACGCTCTTCCGATCTNNNGAAAGAATGAGATTCCGTTTC
GCACACGTCTGAACTCTTTCCCTACACGACGCTCTTCCGATCTNNNAAAACAAACATACAATATCCACCC
AGCACACGTCTGAACTCTTTCCCTACACGACGCTCTTCCGATCTNNNGGCACTGTTTAAGCAATAATCAT
GCACACGTCTGAACTCTTTCCCTACACGACGCTCTTCCGATCTNNNACAGTTTTGATGTAAGTGAAAAT
GCACACGTCTGAACTCTTTCCCTACACGACGCTCTTCCGATCTNNNTAATTAGGTTTAGATGAGGTCAT
GCACACGTCTGAACTCTTTCCCTACACGACGCTCTTCCGATCTNNNCTCCCCATCCCTTACTTATCCATCG
GAGCACACGTCTGAACTCTTTCCCTACACGACGCTCTTCCGATCTNNNTTGTCAAGATTCCTTTAAATGA
AGCACACGTCTGAACTCTTTCCCTACACGACGCTCTTCCGATCTNNNTTCAGTGTGTAAAGCTAAGAAC
GCACACGTCTGAACTCTTTCCCTACACGACGCTCTTCCGATCTNNNAAAATAAACGTTATGGTTGAAG
GAGCACACGTCTGAACTCTTTCCCTACACGACGCTCTTCCGATCTNNNGAATGAGAGCTTTCTTAACTAA
AGCACACGTCTGAACTCTTTCCCTACACGACGCTCTTCCGATCTNNNCCAGTTATGTGATCAATTTTAGA
GAGCACACGTCTGAACTCTTTCCCTACACGACGCTCTTCCGATCTNNNAACAAAGTAGAAAAGGTAACA
GAGCACACGTCTGAACTCTTTCCCTACACGACGCTCTTCCGATCTNNNTAGATTATGTGATTTTTGTTGC
GAGCACACGTCTGAACTCTTTCCCTACACGACGCTCTTCCGATCTNNNAATGAGTGGATAAAGAAAAT
AGCACACGTCTGAACTCTTTCCCTACACGACGCTCTTCCGATCTNNNGTTTTGATGAGCTATGATTGTG
GCACACGTCTGAACTCTTTCCCTACACGACGCTCTTCCGATCTNNNAGGATTTTTCTCAGTCATTTTGCAT
GAGCACACGTCTGAACTCTTTCCCTACACGACGCTCTTCCGATCTNNNATCAAGCAATCTGATAATGGA
GCACACGTCTGAACTCTTTCCCTACACGACGCTCTTCCGATCTNNNGAAGATAAATAATGTTTCCCTCCC
AGCACACGTCTGAACTCTTTCCCTACACGACGCTCTTCCGATCTNNNCAAATCTACTCAGGGATAAAAC
AGCACACGTCTGAACTCTTTCCCTACACGACGCTCTTCCGATCTNNNTTGATTAGATAGATCTGGAAAC
GAGCACACGTCTGAACTCTTTCCCTACACGACGCTCTTCCGATCTNNNCTCAAAGAACAAAATGCAATA
GCACACGTCTGAACTCTTTCCCTACACGACGCTCTTCCGATCTNNNAGAAATAGCAGAAAATGTGAAT
GAGCACACGTCTGAACTCTTTCCCTACACGACGCTCTTCCGATCTNNNATTATTCAGCTAGAAAATGCAA
AGCACACGTCTGAACTCTTTCCCTACACGACGCTCTTCCGATCTNNNTTTCTAGCATTGATTGAGGAATA

AGCACACGTCTGAACTCTTTCCCTACACGACGCTCTTCCGATCTNNNGTTCATAATATGGTCACTTCTTC
GCACACGTCTGAACTCTTTCCCTACACGACGCTCTTCCGATCTNNNACAAATAGGTAGTAGGTAGGTAG
GCACACGTCTGAACTCTTTCCCTACACGACGCTCTTCCGATCTNNNTGTGATCATGTGAAGTTATTTTC
GCACACGTCTGAACTCTTTCCCTACACGACGCTCTTCCGATCTNNNCGTTCACAATATTTCTCGATTTACA
GCACACGTCTGAACTCTTTCCCTACACGACGCTCTTCCGATCTNNNGATCAAATGTTTCCATCAGAAGC
AGCACACGTCTGAACTCTTTCCCTACACGACGCTCTTCCGATCTNNNTAAGAACTGAAAGACAACATCT
CACACGTCTGAACTCTTTCCCTACACGACGCTCTTCCGATCTNNNATTGAATTTTAATTCCCTTTGGTCAT
AGCACACGTCTGAACTCTTTCCCTACACGACGCTCTTCCGATCTNNNATGTGATAGAACTAGAAACATG
GAGCACACGTCTGAACTCTTTCCCTACACGACGCTCTTCCGATCTNNNCAGAATGGTTTGTATAGCAAC
GCACACGTCTGAACTCTTTCCCTACACGACGCTCTTCCGATCTNNNATAGCTCTCCTACTTATCTACTTCA
GAGCACACGTCTGAACTCTTTCCCTACACGACGCTCTTCCGATCTNNNACTCAGAAATAAACCCACATAT
AGCACACGTCTGAACTCTTTCCCTACACGACGCTCTTCCGATCTNNNACTTGGTGACTTTTCTAAACTAC
AGCACACGTCTGAACTCTTTCCCTACACGACGCTCTTCCGATCTNNNCAAACAAAAAACAGAAGCTAAA
GCACACGTCTGAACTCTTTCCCTACACGACGCTCTTCCGATCTNNNTAGGCTTAATAGAGTTATTGTGA
GAGCACACGTCTGAACTCTTTCCCTACACGACGCTCTTCCGATCTNNNCACCAATTGTACATTGTGAATA
AGCACACGTCTGAACTCTTTCCCTACACGACGCTCTTCCGATCTNNNCTTGCTAAGTGTATGACTGTTCA
GAGCACACGTCTGAACTCTTTCCCTACACGACGCTCTTCCGATCTNNNTTTGGAACTTTAGATCATGA
CACACGTCTGAACTCTTTCCCTACACGACGCTCTTCCGATCTNNNTCTCTACTCTGTCACTCTCTCATCGGA
AGCACACGTCTGAACTCTTTCCCTACACGACGCTCTTCCGATCTNNNCTCATGAGACCCTTTTATATACA
GAGCACACGTCTGAACTCTTTCCCTACACGACGCTCTTCCGATCTNNNCGATTTGTATTTGTTGATTTTG
AGCACACGTCTGAACTCTTTCCCTACACGACGCTCTTCCGATCTNNNTAAAGTTGGGATCTGTAAAGTT
AGAGCACACGTCTGAACTCTTTCCCTACACGACGCTCTTCCGATCTNNNAGGCAAACTATGATTCATAT
AGAGCACACGTCTGAACTCTTTCCCTACACGACGCTCTTCCGATCTNNNAGGCAAACTATGATTCATAT
GAGCACACGTCTGAACTCTTTCCCTACACGACGCTCTTCCGATCTNNNTTGTAAATGAGATCACATAGCT
AGCACACGTCTGAACTCTTTCCCTACACGACGCTCTTCCGATCTNNNTCATATGTGCATAGTCATTGAA
GAGCACACGTCTGAACTCTTTCCCTACACGACGCTCTTCCGATCTNNNAATCTCCAAAGGTAGTTCTTGC
GAGCACACGTCTGAACTCTTTCCCTACACGACGCTCTTCCGATCTNNNCTATTGAAAGTTGAACTGAACT
GAGCACACGTCTGAACTCTTTCCCTACACGACGCTCTTCCGATCTNNNGGTGAATGAATGGATAAAGA
AGCACACGTCTGAACTCTTTCCCTACACGACGCTCTTCCGATCTNNNAGACACTCTTGAACTTCCAACATG
GCACACGTCTGAACTCTTTCCCTACACGACGCTCTTCCGATCTNNNACTACCATTTTTGTTTCAGAAACC
GCACACGTCTGAACTCTTTCCCTACACGACGCTCTTCCGATCTNNNCAAGTACATAGTTTCAGCATATAC
ACACGTCTGAACTCTTTCCCTACACGACGCTCTTCCGATCTNNNGCCTTGAACTTAGAATGATTTATCATG
AGCACACGTCTGAACTCTTTCCCTACACGACGCTCTTCCGATCTNNNTTGGAAGTATTTTTGAATGTTG
ACACGTCTGAACTCTTTCCCTACACGACGCTCTTCCGATCTNNNAAAAATGCCAACTTGAAGAATCATC
CACACGTCTGAACTCTTTCCCTACACGACGCTCTTCCGATCTNNNTTATTTCCTCACTGTGGTAAATCATG
CACACGTCTGAACTCTTTCCCTACACGACGCTCTTCCGATCTNNNTTATTTCCTCACTGTGGTAAATCATG
GCACACGTCTGAACTCTTTCCCTACACGACGCTCTTCCGATCTNNNCAATCTACATGAAGGAAACAAAC
GAGCACACGTCTGAACTCTTTCCCTACACGACGCTCTTCCGATCTNNNAAAAACGACAATCAAAACAAA
GCACACGTCTGAACTCTTTCCCTACACGACGCTCTTCCGATCTNNNGTAGCAAGTATTCCCTTCTACCAT
GCACACGTCTGAACTCTTTCCCTACACGACGCTCTTCCGATCTNNNGCATAAATGTAAGAGGAAAAGA
AGCACACGTCTGAACTCTTTCCCTACACGACGCTCTTCCGATCTNNNAAAAGGTAGGATTTATTTTCCC
AGCACACGTCTGAACTCTTTCCCTACACGACGCTCTTCCGATCTNNNCTCATCATATACTCAAGAGAGA
GAGCACACGTCTGAACTCTTTCCCTACACGACGCTCTTCCGATCTNNNCATGTAAACAATCAGTGAAAA

AGCACACGTCTGAACTCTTTCCCTACACGACGCTCTTCCGATCTNNNCATGGGTTTTTGTTCTTTTTTA
CACACGTCTGAACTCTTTCCCTACACGACGCTCTTCCGATCTNNNTAAATTATATAGCACATGCCTTGC
AGCACACGTCTGAACTCTTTCCCTACACGACGCTCTTCCGATCTNNNATATGTCCCATCAATACGTAATT
GCACACGTCTGAACTCTTTCCCTACACGACGCTCTTCCGATCTNNNCCAATATTGTCTTCTAAAAGCTT
AGCACACGTCTGAACTCTTTCCCTACACGACGCTCTTCCGATCTNNNCAGTTTCTCTTTCTCTCTATCTTC
AGCACACGTCTGAACTCTTTCCCTACACGACGCTCTTCCGATCTNNNTTTCAGAAAGAAAGAAGCATAT
AGCACACGTCTGAACTCTTTCCCTACACGACGCTCTTCCGATCTNNNCATTAGAAAAATCAATTCCATG
AGCACACGTCTGAACTCTTTCCCTACACGACGCTCTTCCGATCTNNNCCATCTCTCCAAATAAAAGTAT
CACACGTCTGAACTCTTTCCCTACACGACGCTCTTCCGATCTNNNTTATCCCCATTTTTAGACAAAGCA
AGCACACGTCTGAACTCTTTCCCTACACGACGCTCTTCCGATCTNNNCAGAATCAGGACTAAATGAAA
GCACACGTCTGAACTCTTTCCCTACACGACGCTCTTCCGATCTNNNGTATTTGAAGGGTATACAGAG
AGCACACGTCTGAACTCTTTCCCTACACGACGCTCTTCCGATCTNNNCTTCAAATTGCTTCGTTGTAAC
AGCACACGTCTGAACTCTTTCCCTACACGACGCTCTTCCGATCTNNNCAGCCTTTCACAAAGACATCCAT
AGCACACGTCTGAACTCTTTCCCTACACGACGCTCTTCCGATCTNNNCAGCCTTTCACAAAGACATCCAT
AGCACACGTCTGAACTCTTTCCCTACACGACGCTCTTCCGATCTNNNGGGTGAGAGTATACAAATACTT
ACACGTCTGAACTCTTTCCCTACACGACGCTCTTCCGATCTNNNACTCTGAAGTCTCTATTTTGTTACAT
CACACGTCTGAACTCTTTCCCTACACGACGCTCTTCCGATCTNNNTCATAGAAGAATGTTACCACTGCA
GAGCACACGTCTGAACTCTTTCCCTACACGACGCTCTTCCGATCTNNNAAAAACATATTTGGTGCAACT
AGCACACGTCTGAACTCTTTCCCTACACGACGCTCTTCCGATCTNNNTCAACAAAATAAAAGGTAACCA
AGCACACGTCTGAACTCTTTCCCTACACGACGCTCTTCCGATCTNNNTATTGGTTTACATTTTGGAAAG
GAGCACACGTCTGAACTCTTTCCCTACACGACGCTCTTCCGATCTNNNCAGTGGTCATAAACTTATCAA
AGCACACGTCTGAACTCTTTCCCTACACGACGCTCTTCCGATCTNNNGAAGTCTACTGTCATTCTTATC
AGCACACGTCTGAACTCTTTCCCTACACGACGCTCTTCCGATCTNNNAAGGACAGAGATGTTAAGCTA
AGCACACGTCTGAACTCTTTCCCTACACGACGCTCTTCCGATCTNNNGAAAACTGCTATTCAGACTTAA
AGCACACGTCTGAACTCTTTCCCTACACGACGCTCTTCCGATCTNNNTACTTCTCACACTAAGTTTCAAT
AGCACACGTCTGAACTCTTTCCCTACACGACGCTCTTCCGATCTNNNGTTGATGTCATCTAATTCTTCAG
ACACGTCTGAACTCTTTCCCTACACGACGCTCTTCCGATCTNNNCTAGAGTGTGATTTGATACCAGCAT
CACGTCTGAACTCTTTCCCTACACGACGCTCTTCCGATCTNNNTATTGCCCATAGGATAAAGTTGCATC
AGCACACGTCTGAACTCTTTCCCTACACGACGCTCTTCCGATCTNNNTTAATCTGGATAGCCTGTTTTAT
CACACGTCTGAACTCTTTCCCTACACGACGCTCTTCCGATCTNNNTTCACTCCTCAGGGTTATTTTCATC
AGCACACGTCTGAACTCTTTCCCTACACGACGCTCTTCCGATCTNNNAAGACCTTGCCATTTATATGTAC
AGCACACGTCTGAACTCTTTCCCTACACGACGCTCTTCCGATCTNNNAAGACCTTGCCATTTATATGTAC
GCACACGTCTGAACTCTTTCCCTACACGACGCTCTTCCGATCTNNNTGACCTTTGTATAATCTCCATTTC
ACACGTCTGAACTCTTTCCCTACACGACGCTCTTCCGATCTNNNCATGCTACCTTTGGAGTTTCCATCGG
AGCACACGTCTGAACTCTTTCCCTACACGACGCTCTTCCGATCTNNNTCACTTATTATATAGCTTGAAG
AGCACACGTCTGAACTCTTTCCCTACACGACGCTCTTCCGATCTNNNGTAACAGTTTCCTCTCTCTTTCAT
CACACGTCTGAACTCTTTCCCTACACGACGCTCTTCCGATCTNNNTTTTTATCCAAATATCCACTTGGCA
GAGCACACGTCTGAACTCTTTCCCTACACGACGCTCTTCCGATCTNNNTGTACTAGTTTCCTATCACTAC
CACACGTCTGAACTCTTTCCCTACACGACGCTCTTCCGATCTNNNCAATATGCGTATGAAAAATCAAG
AGCACACGTCTGAACTCTTTCCCTACACGACGCTCTTCCGATCTNNNCACCAATCTTATGCTAATTTTTG
GAGCACACGTCTGAACTCTTTCCCTACACGACGCTCTTCCGATCTNNNCTATCTGTGAATGGGAGTAT
CACACGTCTGAACTCTTTCCCTACACGACGCTCTTCCGATCTNNNCTTAGAAATTGGATGAGGATTTTC
GCACACGTCTGAACTCTTTCCCTACACGACGCTCTTCCGATCTNNNCATTATGGGATTACGTTCTTTTT

CACACGTCTGAACTCTTTCCCTACACGACGCTCTTCCGATCTNNNGATAAATGGGCTTGAAATTAAAC
GCACACGTCTGAACTCTTTCCCTACACGACGCTCTTCCGATCTNNNTGTCTTTCCATTACCAGTTTATAC
GCACACGTCTGAACTCTTTCCCTACACGACGCTCTTCCGATCTNNNCTCAAAATAGTATTTTGCTGACT
GCACACGTCTGAACTCTTTCCCTACACGACGCTCTTCCGATCTNNNGATTGTAAAGTTGTAAGACCATT
GCACACGTCTGAACTCTTTCCCTACACGACGCTCTTCCGATCTNNNTAGAAAGAAAAATAAGCCCAAT
AGCACACGTCTGAACTCTTTCCCTACACGACGCTCTTCCGATCTNNNATTTTCATGTATTCTGACACAAC
CACACGTCTGAACTCTTTCCCTACACGACGCTCTTCCGATCTNNNCAGTGATGAAAAACAACATAGAT
ACACGTCTGAACTCTTTCCCTACACGACGCTCTTCCGATCTNNNAATATCCTTCTGTCTCTCTGTCATCG
CACACGTCTGAACTCTTTCCCTACACGACGCTCTTCCGATCTNNNCTGTGTTTGTGTATCTGTGTCATC
GAGCACACGTCTGAACTCTTTCCCTACACGACGCTCTTCCGATCTNNNGTATTTTTATTTCTGGGGAAC
CACACGTCTGAACTCTTTCCCTACACGACGCTCTTCCGATCTNNNAGCACACATATATCTTCTGTTATC
AGCACACGTCTGAACTCTTTCCCTACACGACGCTCTTCCGATCTNNNATTTAACGTAGTAGGTCTAAT
AGCACACGTCTGAACTCTTTCCCTACACGACGCTCTTCCGATCTNNNCATAAACACACATATATGCTCAT
AGCACACGTCTGAACTCTTTCCCTACACGACGCTCTTCCGATCTNNNTTTCATTACCAAAGAAGATTCT
AGCACACGTCTGAACTCTTTCCCTACACGACGCTCTTCCGATCTNNNTTTTACTGCTATGTAGTATTCC
ACACGTCTGAACTCTTTCCCTACACGACGCTCTTCCGATCTNNNAATTCTTCAACATTGAACATGGCAT
GCACACGTCTGAACTCTTTCCCTACACGACGCTCTTCCGATCTNNNATTTCTATCAGAGTCCATTGTTT
AGCACACGTCTGAACTCTTTCCCTACACGACGCTCTTCCGATCTNNNAGATTTAGTCATTCTACACAAC
AGCACACGTCTGAACTCTTTCCCTACACGACGCTCTTCCGATCTNNNAAAGGATGGTAAGGATGAAA
AGCACACGTCTGAACTCTTTCCCTACACGACGCTCTTCCGATCTNNNTTACTATATTACCATGTATGCCC
ACACGTCTGAACTCTTTCCCTACACGACGCTCTTCCGATCTNNNCAGGAAATTGATGTTGGTATAAAC
AGCACACGTCTGAACTCTTTCCCTACACGACGCTCTTCCGATCTNNNTCAAATTATTCAAATTTCCATG
AGCACACGTCTGAACTCTTTCCCTACACGACGCTCTTCCGATCTNNNTTATATTCTGATAATTGCATG
AGCACACGTCTGAACTCTTTCCCTACACGACGCTCTTCCGATCTNNNCAGAAAAAAAAAAACCAGTTG
CACACGTCTGAACTCTTTCCCTACACGACGCTCTTCCGATCTNNNGCAAAAGAGATACAGGGAGACAT
AGCACACGTCTGAACTCTTTCCCTACACGACGCTCTTCCGATCTNNNCTCTTCCTACTTAGTGCATTCC
AGCACACGTCTGAACTCTTTCCCTACACGACGCTCTTCCGATCTNNNGGATGAGAAAAAGAAAACATA
AGCACACGTCTGAACTCTTTCCCTACACGACGCTCTTCCGATCTNNNGTATTTGCAACTTCTTTTCAAAT
CACACGTCTGAACTCTTTCCCTACACGACGCTCTTCCGATCTNNNCAAATCTTCCATGTTCCTCGCATCG
AGCACACGTCTGAACTCTTTCCCTACACGACGCTCTTCCGATCTNNNAGTAGAGAATGGATTTGGAAT
GCACACGTCTGAACTCTTTCCCTACACGACGCTCTTCCGATCTNNNTTATAGTGTGGATACCCAGTTAT
AGCACACGTCTGAACTCTTTCCCTACACGACGCTCTTCCGATCTNNNACCCTTGAGAAATATACACTA
GCACACGTCTGAACTCTTTCCCTACACGACGCTCTTCCGATCTNNNCAGATAGAAAACGCTGATATTAT
AGCACACGTCTGAACTCTTTCCCTACACGACGCTCTTCCGATCTNNNAGTTATTTTTAGGAGGAGATG
GAGCACACGTCTGAACTCTTTCCCTACACGACGCTCTTCCGATCTNNNAAGATAGTCGAGGAGAGGA
AGCACACGTCTGAACTCTTTCCCTACACGACGCTCTTCCGATCTNNNCGTTTAGTTATGTTTTCTAAGC
AGCACACGTCTGAACTCTTTCCCTACACGACGCTCTTCCGATCTNNNATATGTGTATTTACAAAATGCC
AGCACACGTCTGAACTCTTTCCCTACACGACGCTCTTCCGATCTNNNAGTGAAGTTAGATCTTAAAGG
AGCACACGTCTGAACTCTTTCCCTACACGACGCTCTTCCGATCTNNNATTTTTCTCCTAGTCTCATTTG
AGAGCACACGTCTGAACTCTTTCCCTACACGACGCTCTTCCGATCTNNNCGAAAATAGCAATCTTCAGT
AGCACACGTCTGAACTCTTTCCCTACACGACGCTCTTCCGATCTNNNTGAATTAGAACCAGCAAAAATT
GCACACGTCTGAACTCTTTCCCTACACGACGCTCTTCCGATCTNNNGCACTTGATGTTTGTTAAATTTA
GCACACGTCTGAACTCTTTCCCTACACGACGCTCTTCCGATCTNNNGCACTTGATGTTTGTTAAATTTA

GCACACGTCTGAACTCTTTCCCTACACGACGCTCTTCCGATCTNNNATACTTCATCAGATTTTCAAGGAC
ACACGTCTGAACTCTTTCCCTACACGACGCTCTTCCGATCTNNNTATGTCTTGCCCTACTATGGCATCGG
GCACACGTCTGAACTCTTTCCCTACACGACGCTCTTCCGATCTNNNGTGTTCATAGAAAATGATCCTAC
GCACACGTCTGAACTCTTTCCCTACACGACGCTCTTCCGATCTNNNTGAGGAAATAAGAAAGTTTTAC
GCACACGTCTGAACTCTTTCCCTACACGACGCTCTTCCGATCTNNNAAATGACCCTCATACCTCTTTCATC
AGCACACGTCTGAACTCTTTCCCTACACGACGCTCTTCCGATCTNNNCACACTAGACAGTTGGTTTTTCA
GCACACGTCTGAACTCTTTCCCTACACGACGCTCTTCCGATCTNNNGCAGAATCATAAGAAACAGTAAA
GCACACGTCTGAACTCTTTCCCTACACGACGCTCTTCCGATCTNNNCAGATCTCTTTTCATCAGTTATCCA
CACACGTCTGAACTCTTTCCCTACACGACGCTCTTCCGATCTNNNGTTGTAAACTCAACTCGGAGCATC
CACACGTCTGAACTCTTTCCCTACACGACGCTCTTCCGATCTNNNAATTGATGAATGTGATTTACTGTCA
AGCACACGTCTGAACTCTTTCCCTACACGACGCTCTTCCGATCTNNNCACTCAAAATCATAACCAGTTTC
AGCACACGTCTGAACTCTTTCCCTACACGACGCTCTTCCGATCTNNNTGGCTGATTTTAGAATTAAAAA
GCACACGTCTGAACTCTTTCCCTACACGACGCTCTTCCGATCTNNNGCTGAAGGTAAATTCTAAAAAGT
GCACACGTCTGAACTCTTTCCCTACACGACGCTCTTCCGATCTNNNTATGTCATTCTTTCTGGACATGCA
AGCACACGTCTGAACTCTTTCCCTACACGACGCTCTTCCGATCTNNNTGATTATTTGGTTTTTTGAAGG
GCACACGTCTGAACTCTTTCCCTACACGACGCTCTTCCGATCTNNNGCCTCTATTTTTTATTACCATGTCA
GCACACGTCTGAACTCTTTCCCTACACGACGCTCTTCCGATCTNNNAGGTAAGAGAACAATTGGAATT
AGCACACGTCTGAACTCTTTCCCTACACGACGCTCTTCCGATCTNNNGAAGTAAAGGCCACAGTAAGC
AGCACACGTCTGAACTCTTTCCCTACACGACGCTCTTCCGATCTNNNTATGTTAGAAGATCTGTTGATGA
CACACGTCTGAACTCTTTCCCTACACGACGCTCTTCCGATCTNNNATAAATATTTTGCCAGGAAGTTTCA
GAGCACACGTCTGAACTCTTTCCCTACACGACGCTCTTCCGATCTNNNTTCCTAGGGGAAGATATCTGC
GAGCACACGTCTGAACTCTTTCCCTACACGACGCTCTTCCGATCTNNNGATTTTCACCCTAGTACTTCATC
GCACACGTCTGAACTCTTTCCCTACACGACGCTCTTCCGATCTNNNTAAAGATCACAATAGAAATGCTG
GAGCACACGTCTGAACTCTTTCCCTACACGACGCTCTTCCGATCTNNNATCTTGATAACACAATTATCTG
AGCACACGTCTGAACTCTTTCCCTACACGACGCTCTTCCGATCTNNNAAACATTCTGAAATTTGGCTTCA
AGCACACGTCTGAACTCTTTCCCTACACGACGCTCTTCCGATCTNNNATGGGTAAAATGATATTTCACAC
AGCACACGTCTGAACTCTTTCCCTACACGACGCTCTTCCGATCTNNNGTAAAGAAAAGAAGATTGGCTT
GAGCACACGTCTGAACTCTTTCCCTACACGACGCTCTTCCGATCTNNNTAACCCAGTAACCATAAGAAA
GCACACGTCTGAACTCTTTCCCTACACGACGCTCTTCCGATCTNNNAAGAGAACATTGTCCAAAGATAAC
GCACACGTCTGAACTCTTTCCCTACACGACGCTCTTCCGATCTNNNAATGTCCTTTTGATTCCCTACCATC
AGCACACGTCTGAACTCTTTCCCTACACGACGCTCTTCCGATCTNNNAGATCCTGTCTCAAAAAAATAAA
GAGCACACGTCTGAACTCTTTCCCTACACGACGCTCTTCCGATCTNNNTTGTTTCATCAATAAAGACTGA
GAGCACACGTCTGAACTCTTTCCCTACACGACGCTCTTCCGATCTNNNTTAGAACACAGTATAAGAGTT
GCACACGTCTGAACTCTTTCCCTACACGACGCTCTTCCGATCTNNNCAACTACTTCTACTCAATAGGATC
CACACGTCTGAACTCTTTCCCTACACGACGCTCTTCCGATCTNNNGGGGACAATACTGATAGTATTCCA
CACACGTCTGAACTCTTTCCCTACACGACGCTCTTCCGATCTNNNGGGGACAATACTGATAGTATTCCA
GCACACGTCTGAACTCTTTCCCTACACGACGCTCTTCCGATCTNNNTAGTATCCTCCAGGTATATTCATC
AGCACACGTCTGAACTCTTTCCCTACACGACGCTCTTCCGATCTNNNTCATGAGTTAATATAAACACTG
GAGCACACGTCTGAACTCTTTCCCTACACGACGCTCTTCCGATCTNNNAAAAGCATGGAAGATTTTGTG
GCACACGTCTGAACTCTTTCCCTACACGACGCTCTTCCGATCTNNNAGCAATGGGTAAAGAATAAAAA
AGCACACGTCTGAACTCTTTCCCTACACGACGCTCTTCCGATCTNNNAGATTTAACCCAAAGAAGTCTA
AGCACACGTCTGAACTCTTTCCCTACACGACGCTCTTCCGATCTNNNTCAGTTTATTTGAACAAAGTGA
AGCACACGTCTGAACTCTTTCCCTACACGACGCTCTTCCGATCTNNNATTAATGAGGATGAACAATTGT

GCACACGTCTGAACTCTTTCCCTACACGACGCTCTTCCGATCTNNNAACTTGAAATTCTGTTTTACCAAC
AGCACACGTCTGAACTCTTTCCCTACACGACGCTCTTCCGATCTNNNATTCAGTGTATGATCTAAAGACA
AGCACACGTCTGAACTCTTTCCCTACACGACGCTCTTCCGATCTNNNGTTTTTGGACAGTTAGCAATATC
AGCACACGTCTGAACTCTTTCCCTACACGACGCTCTTCCGATCTNNNCTTATAACATGGCAACTGATTT
GAGCACACGTCTGAACTCTTTCCCTACACGACGCTCTTCCGATCTNNNTGATTTATCCTGAAGTGAAAAT
GCACACGTCTGAACTCTTTCCCTACACGACGCTCTTCCGATCTNNNTAACTTAGCTGTGTTATTCCAGCA
AGCACACGTCTGAACTCTTTCCCTACACGACGCTCTTCCGATCTNNNTTAGCTCTCCATTTTTCCTGCATC
AGCACACGTCTGAACTCTTTCCCTACACGACGCTCTTCCGATCTNNNGTAGACAGGAATGTTTTACTTG
GCACACGTCTGAACTCTTTCCCTACACGACGCTCTTCCGATCTNNNTAGAATTCACTGCCTCAGTAAACA
CACACGTCTGAACTCTTTCCCTACACGACGCTCTTCCGATCTNNNTTTGCTTATATGCATTGAAAATGCA
CACACGTCTGAACTCTTTCCCTACACGACGCTCTTCCGATCTNNNTTTGCTTATATGCATTGAAAATGCA
GCACACGTCTGAACTCTTTCCCTACACGACGCTCTTCCGATCTNNNAAAGAAAAAGTATCATAGGTTCCC
AGCACACGTCTGAACTCTTTCCCTACACGACGCTCTTCCGATCTNNNGCCTTATTACCAACTAAAGATTT
GAGCACACGTCTGAACTCTTTCCCTACACGACGCTCTTCCGATCTNNNTATCTGGTTTCAAATCTCAAAT
GCACACGTCTGAACTCTTTCCCTACACGACGCTCTTCCGATCTNNNCCTGACTTCTAATAACATTGATGCA
AGCACACGTCTGAACTCTTTCCCTACACGACGCTCTTCCGATCTNNNGACTCCAAAGTACTATAGATTCA
CACACGTCTGAACTCTTTCCCTACACGACGCTCTTCCGATCTNNNAAGCTTCCTTCAACAGAGAGCATCG
AGAGCACACGTCTGAACTCTTTCCCTACACGACGCTCTTCCGATCTNNNGTCATTGACCCAAATATTGTT
GAGCACACGTCTGAACTCTTTCCCTACACGACGCTCTTCCGATCTNNNTTTTAACATAGCATTTTTGCT
AGCACACGTCTGAACTCTTTCCCTACACGACGCTCTTCCGATCTNNNATGCTACCAAGAACTATGTTATT
AGCACACGTCTGAACTCTTTCCCTACACGACGCTCTTCCGATCTNNNAAGTTTCAGGAATCAGAGTTCCA
AGCACACGTCTGAACTCTTTCCCTACACGACGCTCTTCCGATCTNNNAAATAAAAATGAAGAGCAAAAC
GAGCACACGTCTGAACTCTTTCCCTACACGACGCTCTTCCGATCTNNNCATTTGCAAAATTCAAGTACA
AGCACACGTCTGAACTCTTTCCCTACACGACGCTCTTCCGATCTNNNAGAAGAAATGACTCAGAAGAAA
AGCACACGTCTGAACTCTTTCCCTACACGACGCTCTTCCGATCTNNNCTTTTCTGAAACCAAGGTATTTA
AGCACACGTCTGAACTCTTTCCCTACACGACGCTCTTCCGATCTNNNATCTGCACATTCAAATTATCTGC
AGCACACGTCTGAACTCTTTCCCTACACGACGCTCTTCCGATCTNNNCCTGTTTTTGCAAATAAAGTTTTC
GAGCACACGTCTGAACTCTTTCCCTACACGACGCTCTTCCGATCTNNNTTATTACAATAAGTCTCTCCCTC
AGCACACGTCTGAACTCTTTCCCTACACGACGCTCTTCCGATCTNNNTTCTTACTGAATCCTCACATAAAC
AGCACACGTCTGAACTCTTTCCCTACACGACGCTCTTCCGATCTNNNTAATATCCTTTTAGCCAAATCCTC
AGCACACGTCTGAACTCTTTCCCTACACGACGCTCTTCCGATCTNNNATTTTACTACACTTGATCTTAGCC
AGCACACGTCTGAACTCTTTCCCTACACGACGCTCTTCCGATCTNNNCCAGTTTGTAATAAAGCATGAA
AGCACACGTCTGAACTCTTTCCCTACACGACGCTCTTCCGATCTNNNTATATTAGTCAGGATTCTCCAG
AGCACACGTCTGAACTCTTTCCCTACACGACGCTCTTCCGATCTNNNATTCCCCTATAATCAAATTTCCTC
AGCACACGTCTGAACTCTTTCCCTACACGACGCTCTTCCGATCTNNNATTTTTATTTCCTCTTGCTAACCC
AGCACACGTCTGAACTCTTTCCCTACACGACGCTCTTCCGATCTNNNCATTTTCTTTATCCAATCTACCAC
AGCACACGTCTGAACTCTTTCCCTACACGACGCTCTTCCGATCTNNNAAGACTCCGTCTAAAAAATAAAC
AGCACACGTCTGAACTCTTTCCCTACACGACGCTCTTCCGATCTNNNTTATTGTGGATAAAGTTGAAGT
CACACGTCTGAACTCTTTCCCTACACGACGCTCTTCCGATCTNNNGTACCAATACAGGTCTGAAAGCATC
GAGCACACGTCTGAACTCTTTCCCTACACGACGCTCTTCCGATCTNNNCTTTGGAGTAATCTATGTATG
GAGCACACGTCTGAACTCTTTCCCTACACGACGCTCTTCCGATCTNNNTTCCTCTACCAATGAATCATAAA
ACACGTCTGAACTCTTTCCCTACACGACGCTCTTCCGATCTNNNTACTTTCAAAATCCAGATCTTTGCATC
AGAGCACACGTCTGAACTCTTTCCCTACACGACGCTCTTCCGATCTNNNGTGATAAGTTAAATAGAAGA

AGCACACGTCTGAACTCTTTCCCTACACGACGCTCTTCCGATCTNNNAACTATCTGTGAAAGAGATTA
AGCACACGTCTGAACTCTTTCCCTACACGACGCTCTTCCGATCTNNNGAATGCCATTTCATTACTCTTT
GCACACGTCTGAACTCTTTCCCTACACGACGCTCTTCCGATCTNNNTTCTGAGAAAGGTAAAATGGAT
GCACACGTCTGAACTCTTTCCCTACACGACGCTCTTCCGATCTNNNTCTACAGATTTGGACATTTCATA
AGCACACGTCTGAACTCTTTCCCTACACGACGCTCTTCCGATCTNNNAAGGGGATTATACAAGAGAT
CACACGTCTGAACTCTTTCCCTACACGACGCTCTTCCGATCTNNNCTTGTTGGGAAATGTTTTACTAA
AGCACACGTCTGAACTCTTTCCCTACACGACGCTCTTCCGATCTNNNATTATAGGTCCTTTTTACTTCCA
AGCACACGTCTGAACTCTTTCCCTACACGACGCTCTTCCGATCTNNNATGTTTCTCTCTCTGTCTCTCCAT
GCACACGTCTGAACTCTTTCCCTACACGACGCTCTTCCGATCTNNNAATCTCTGCTTCTGTCTTTTCCAT
GCACACGTCTGAACTCTTTCCCTACACGACGCTCTTCCGATCTNNNCCTACAAACCTAAATTCCACTATC
AGCACACGTCTGAACTCTTTCCCTACACGACGCTCTTCCGATCTNNNGTATGAGATGTTGAATGTTG
AGCACACGTCTGAACTCTTTCCCTACACGACGCTCTTCCGATCTNNNGAATGTGCTAAAACATACATA
AGCACACGTCTGAACTCTTTCCCTACACGACGCTCTTCCGATCTNNNGAAAATAAGAATGTGCTAAA
AGCACACGTCTGAACTCTTTCCCTACACGACGCTCTTCCGATCTNNNACACGAATCTAACATGAAATA
GCACACGTCTGAACTCTTTCCCTACACGACGCTCTTCCGATCTNNNCAGTGTAGACAAATCTAAGAGA
AGCACACGTCTGAACTCTTTCCCTACACGACGCTCTTCCGATCTNNNTCTATTTGGTACATAGAATCT
AGCACACGTCTGAACTCTTTCCCTACACGACGCTCTTCCGATCTNNNGAGAAAGAATGGATCTCCTTT
AGCACACGTCTGAACTCTTTCCCTACACGACGCTCTTCCGATCTNNNGAGAAAGAATGGATCTCCTTT
AGCACACGTCTGAACTCTTTCCCTACACGACGCTCTTCCGATCTNNNAAAAAATCTATACACACACAT
AGCACACGTCTGAACTCTTTCCCTACACGACGCTCTTCCGATCTNNNCTAATGCTTATGACCTTGCTAC
AGCACACGTCTGAACTCTTTCCCTACACGACGCTCTTCCGATCTNNNGGCAATTTGTTTAGATGTAAA
AGCACACGTCTGAACTCTTTCCCTACACGACGCTCTTCCGATCTNNNTACTCTTGCTAGTCCATTTTCCA
GCACACGTCTGAACTCTTTCCCTACACGACGCTCTTCCGATCTNNNTGATTCTTTTATATGAAGCTGTG
GCACACGTCTGAACTCTTTCCCTACACGACGCTCTTCCGATCTNNNTTACAACGATCATTAATAAGCTC
AGCACACGTCTGAACTCTTTCCCTACACGACGCTCTTCCGATCTNNNCCAGTCTGTGCCTATTTTTATA
AGCACACGTCTGAACTCTTTCCCTACACGACGCTCTTCCGATCTNNNATCATCTCCAGCTCCATTATCA
AGCACACGTCTGAACTCTTTCCCTACACGACGCTCTTCCGATCTNNNTATTTATAAATCACAGCCAGCA
GCACACGTCTGAACTCTTTCCCTACACGACGCTCTTCCGATCTNNNAAGACATTTAGCACTTTGAAGA
AGCACACGTCTGAACTCTTTCCCTACACGACGCTCTTCCGATCTNNNACTTTAATTTTTAAAGGATGG
CACACGTCTGAACTCTTTCCCTACACGACGCTCTTCCGATCTNNNTGTTCTTCTAAGAAGGGGAACATC
CACACGTCTGAACTCTTTCCCTACACGACGCTCTTCCGATCTNNNAAAGTGTACAATCTGTAATCCAT
AGCACACGTCTGAACTCTTTCCCTACACGACGCTCTTCCGATCTNNNTTCAGAAAAAGACACAAAACA
GCACACGTCTGAACTCTTTCCCTACACGACGCTCTTCCGATCTNNNCTCAAATAACATGCTTATCCAAG
GCACACGTCTGAACTCTTTCCCTACACGACGCTCTTCCGATCTNNNTTTATGATTTTTTTGGAAAGCC
AGCACACGTCTGAACTCTTTCCCTACACGACGCTCTTCCGATCTNNNCTACTGTAATTCTTTATAGCTT
AGCACACGTCTGAACTCTTTCCCTACACGACGCTCTTCCGATCTNNNGAAATGCACTTGTATGACATA
AGCACACGTCTGAACTCTTTCCCTACACGACGCTCTTCCGATCTNNNCAGACAAAAACAAAACAAAAC
CACGTCTGAACTCTTTCCCTACACGACGCTCTTCCGATCTNNNTCTAAGAGGTGATTAGGTCAACATC
AGCACACGTCTGAACTCTTTCCCTACACGACGCTCTTCCGATCTNNNATGTGATGGAATAGGAGGA
CACACGTCTGAACTCTTTCCCTACACGACGCTCTTCCGATCTNNNAAGTTAAAATGAGGTAATTAGGG
CACGTCTGAACTCTTTCCCTACACGACGCTCTTCCGATCTNNNCTATTTTTGTACATGGGAAGTACCA
CACGTCTGAACTCTTTCCCTACACGACGCTCTTCCGATCTNNNAATACAAAATACCTTGTAAGCTGCAT
GCACACGTCTGAACTCTTTCCCTACACGACGCTCTTCCGATCTNNNGCAGGAAGAAAGAAAGAAATT

CACACGTCTGAACTCTTTCCCTACACGACGCTCTTCCGATCTNNNAAGCATAATACAGTAAAACTTCTC
CACACGTCTGAACTCTTTCCCTACACGACGCTCTTCCGATCTNNNGATGATCAAGTTGTGAGAGATT
AGCACACGTCTGAACTCTTTCCCTACACGACGCTCTTCCGATCTNNNTTTTTCTTTTAAACCAGCAAT
ACACGTCTGAACTCTTTCCCTACACGACGCTCTTCCGATCTNNNGCCTGAAAATAAAGAGGAAATACC
GCACACGTCTGAACTCTTTCCCTACACGACGCTCTTCCGATCTNNNTCACCTTACTTTTGTTTTTATGT
AGCACACGTCTGAACTCTTTCCCTACACGACGCTCTTCCGATCTNNNTGTGTGTGTATATATGCTCAT
AGCACACGTCTGAACTCTTTCCCTACACGACGCTCTTCCGATCTNNNAAAGACAGAGACAGAGACAT
AGCACACGTCTGAACTCTTTCCCTACACGACGCTCTTCCGATCTNNNGGAACTTTTTAGATTGATGCA
GCACACGTCTGAACTCTTTCCCTACACGACGCTCTTCCGATCTNNNTATCAAACCCAGAGAATTATGAT
CACACGTCTGAACTCTTTCCCTACACGACGCTCTTCCGATCTNNNTATTGTGTATAGTCCTTGTGTTC
AGCACACGTCTGAACTCTTTCCCTACACGACGCTCTTCCGATCTNNNTATTTTTAGTTCTGTTAGCACT
AGCACACGTCTGAACTCTTTCCCTACACGACGCTCTTCCGATCTNNNGCAGGAGAAACTACCATTTAT
CACACGTCTGAACTCTTTCCCTACACGACGCTCTTCCGATCTNNNATTTTAAGCTGGGTAATTCTTTG
GCACACGTCTGAACTCTTTCCCTACACGACGCTCTTCCGATCTNNNCTCAAAGGTAATTTTTTTCATCC
CACACGTCTGAACTCTTTCCCTACACGACGCTCTTCCGATCTNNNTATAAATTATTAAGACCCAGGCT
AGCACACGTCTGAACTCTTTCCCTACACGACGCTCTTCCGATCTNNNCAGGGATTCTGATTTTATTTC
GCACACGTCTGAACTCTTTCCCTACACGACGCTCTTCCGATCTNNNCACTGACTCTGATATCTTCTGCA
AGCACACGTCTGAACTCTTTCCCTACACGACGCTCTTCCGATCTNNNGAACTAGCAGACAAATGTAAA
GCACACGTCTGAACTCTTTCCCTACACGACGCTCTTCCGATCTNNNTGGTTTTGGTATGATTGAAATA
AGCACACGTCTGAACTCTTTCCCTACACGACGCTCTTCCGATCTNNNACAACAATACTTTAAAGTCCTT
AGCACACGTCTGAACTCTTTCCCTACACGACGCTCTTCCGATCTNNNACAACAATACTTTAAAGTCCTT
AGCACACGTCTGAACTCTTTCCCTACACGACGCTCTTCCGATCTNNNAAAGTCCTCTTTTAAACCATTG
AGCACACGTCTGAACTCTTTCCCTACACGACGCTCTTCCGATCTNNNTCTTCCTTTTTTAGTTCAATTCC
CACACGTCTGAACTCTTTCCCTACACGACGCTCTTCCGATCTNNNGTATGCAAATACAGGTCTGTCAT
AGCACACGTCTGAACTCTTTCCCTACACGACGCTCTTCCGATCTNNNATGTGACTGTTTTCTAAAATC
AGCACACGTCTGAACTCTTTCCCTACACGACGCTCTTCCGATCTNNNGCCCAAATAAATGTAAATACA
GCACACGTCTGAACTCTTTCCCTACACGACGCTCTTCCGATCTNNNTTGTACACATACTACATCCAAGC
GCACACGTCTGAACTCTTTCCCTACACGACGCTCTTCCGATCTNNNACTCCAATTTCCTTGACATAATA
GCACACGTCTGAACTCTTTCCCTACACGACGCTCTTCCGATCTNNNGACACAATTCAGTCCATCATACA
ACACGTCTGAACTCTTTCCCTACACGACGCTCTTCCGATCTNNNCTCTTACCACTTTGACAAGGCATCG
AGCACACGTCTGAACTCTTTCCCTACACGACGCTCTTCCGATCTNNNAGAAAATCAATAGTATACCCC
GAGCACACGTCTGAACTCTTTCCCTACACGACGCTCTTCCGATCTNNNATTCAGAATCACTAGAGAAA
GAGCACACGTCTGAACTCTTTCCCTACACGACGCTCTTCCGATCTNNNGTGAATGTTGAAAAGATAT
GCACACGTCTGAACTCTTTCCCTACACGACGCTCTTCCGATCTNNNGTTCATAAGTCATGTTTGTATCA
AGCACACGTCTGAACTCTTTCCCTACACGACGCTCTTCCGATCTNNNCAGGAAATCCAATAAAAGAA
AGCACACGTCTGAACTCTTTCCCTACACGACGCTCTTCCGATCTNNNAGGGTAAATGTGATAAATAAA
AGCACACGTCTGAACTCTTTCCCTACACGACGCTCTTCCGATCTNNNTTTAGTGTGTTTCTCCTTCATA
GCACACGTCTGAACTCTTTCCCTACACGACGCTCTTCCGATCTNNNCCCTTTGGAGAAACTGAAACCAT
CACGTCTGAACTCTTTCCCTACACGACGCTCTTCCGATCTNNNCACTTCATGAGGATACATATCTACAT
AGCACACGTCTGAACTCTTTCCCTACACGACGCTCTTCCGATCTNNNCATAACGTTCATGAATTGAAA
AGCACACGTCTGAACTCTTTCCCTACACGACGCTCTTCCGATCTNNNACAATACTCTTTTTAGTAAATG
GCACACGTCTGAACTCTTTCCCTACACGACGCTCTTCCGATCTNNNGGACCTATTTCCTCTTTTCTCCAT
CACACGTCTGAACTCTTTCCCTACACGACGCTCTTCCGATCTNNNTAATAGGTTCCCTTTCATGATAGC

GCACACGTCTGAACTCTTTCCCTACACGACGCTCTTCCGATCTNNNAAGAAGAAATACGAAAAAAAC
GAGCACACGTCTGAACTCTTTCCCTACACGACGCTCTTCCGATCTNNNCTTTAGAACATGCCTTTATTCT
AGCACACGTCTGAACTCTTTCCCTACACGACGCTCTTCCGATCTNNNCATATAATTGACCAGCATTAACT
GAGCACACGTCTGAACTCTTTCCCTACACGACGCTCTTCCGATCTNNNCAAGAAGTATGAGTGAAGTG
AGCACACGTCTGAACTCTTTCCCTACACGACGCTCTTCCGATCTNNNAATAGGTTTGGGAGGTTGAG
GCACACGTCTGAACTCTTTCCCTACACGACGCTCTTCCGATCTNNNTGTGAGATAATTGTCTATAGCTT
AGCACACGTCTGAACTCTTTCCCTACACGACGCTCTTCCGATCTNNNACTTTGGAATAGTATTTCTTTGT
AGCACACGTCTGAACTCTTTCCCTACACGACGCTCTTCCGATCTNNNCAAACTGATGTGAAAAAAACT
AGCACACGTCTGAACTCTTTCCCTACACGACGCTCTTCCGATCTNNNTGAATTTAAAACTACTGCTACA
AGAGCACACGTCTGAACTCTTTCCCTACACGACGCTCTTCCGATCTNNNGTTCATTAGTTTGGAAAAGA
GCACACGTCTGAACTCTTTCCCTACACGACGCTCTTCCGATCTNNNTTTCTAACAAGCTCCTGGAACATC
AGAGCACACGTCTGAACTCTTTCCCTACACGACGCTCTTCCGATCTNNNCACTCCTTATAAGAATCTAAT
AGCACACGTCTGAACTCTTTCCCTACACGACGCTCTTCCGATCTNNNTAAAAATTTTTAGCTTGACAGG
GCACACGTCTGAACTCTTTCCCTACACGACGCTCTTCCGATCTNNNTGGACATCATTACCTGTTATTTCA
GAGCACACGTCTGAACTCTTTCCCTACACGACGCTCTTCCGATCTNNNAAAATATTGAGTGAAAAAGG
AGCACACGTCTGAACTCTTTCCCTACACGACGCTCTTCCGATCTNNNTCACTTGTATTTCTCTAAGAACT
AGCACACGTCTGAACTCTTTCCCTACACGACGCTCTTCCGATCTNNNAAAGGGTAGAGATGGATAAT
AGCACACGTCTGAACTCTTTCCCTACACGACGCTCTTCCGATCTNNNCCAACTTGAAAAAAGAGAAAAT
AGCACACGTCTGAACTCTTTCCCTACACGACGCTCTTCCGATCTNNNCCAACTTGAAAAAAGAGAAAAT
GCACACGTCTGAACTCTTTCCCTACACGACGCTCTTCCGATCTNNNTTGTATATATAAGGAGCACATGG
GAGCACACGTCTGAACTCTTTCCCTACACGACGCTCTTCCGATCTNNNGAATGAGAGGGAGAGAGAA
AGCACACGTCTGAACTCTTTCCCTACACGACGCTCTTCCGATCTNNNCTTTCAATTAAAAGGAAACAC
GAGCACACGTCTGAACTCTTTCCCTACACGACGCTCTTCCGATCTNNNCAAATATTTTTGAGCCAAGTT
GAGCACACGTCTGAACTCTTTCCCTACACGACGCTCTTCCGATCTNNNTAAATTGATAATAACCAGTCC
CACACGTCTGAACTCTTTCCCTACACGACGCTCTTCCGATCTNNNTAGGAACAAGCAGGATTCATATCA
GAGCACACGTCTGAACTCTTTCCCTACACGACGCTCTTCCGATCTNNNCAAATTTGAAAAGATGACTGT
AGCACACGTCTGAACTCTTTCCCTACACGACGCTCTTCCGATCTNNNTAAAGACAAATGGATTCTTAA
GAGCACACGTCTGAACTCTTTCCCTACACGACGCTCTTCCGATCTNNNATAACTCTATTAGGAGGAGTG
CACACGTCTGAACTCTTTCCCTACACGACGCTCTTCCGATCTNNNCTTATAGTTTTGGTCTTCTCTTGCA
AGCACACGTCTGAACTCTTTCCCTACACGACGCTCTTCCGATCTNNNAGCTAAGCATTTTTGAAACATTA
AGCACACGTCTGAACTCTTTCCCTACACGACGCTCTTCCGATCTNNNAAATATACAGCAAACAACATTG
GCACACGTCTGAACTCTTTCCCTACACGACGCTCTTCCGATCTNNNTCAAAAATTGTCCAGCAATAATA
CACACGTCTGAACTCTTTCCCTACACGACGCTCTTCCGATCTNNNAAAGATGTCTGTCTTTTTTGAATCA
AGCACACGTCTGAACTCTTTCCCTACACGACGCTCTTCCGATCTNNNTAGATTTATTTTACTGCCTCCT
AGCACACGTCTGAACTCTTTCCCTACACGACGCTCTTCCGATCTNNNGTCTTACTCACATAGAATACAGA
AGCACACGTCTGAACTCTTTCCCTACACGACGCTCTTCCGATCTNNNGTCTTACTCACATAGAATACAGA
GAGCACACGTCTGAACTCTTTCCCTACACGACGCTCTTCCGATCTNNNTTTATACCCTTTCAATTCATTGA
AGCACACGTCTGAACTCTTTCCCTACACGACGCTCTTCCGATCTNNNATTACTTATCATGGGGTACATA
GCACACGTCTGAACTCTTTCCCTACACGACGCTCTTCCGATCTNNNGTAAGAAAATTACAATGGTCACT
AGCACACGTCTGAACTCTTTCCCTACACGACGCTCTTCCGATCTNNNATTATGGAAATGATGTATGACA
GCACACGTCTGAACTCTTTCCCTACACGACGCTCTTCCGATCTNNNAGTTCTATATTATAAATGGAGGG
GAGCACACGTCTGAACTCTTTCCCTACACGACGCTCTTCCGATCTNNNTCTTTGAGAGTTTTTTTCTTTC
AGCACACGTCTGAACTCTTTCCCTACACGACGCTCTTCCGATCTNNNTTACCATGAATTTACCACAATTTC

GAGCACACGTCTGAACTCTTTCCCTACACGACGCTCTTCCGATCTNNNAATTAAGTAGGCTTTTGGTT
GCACACGTCTGAACTCTTTCCCTACACGACGCTCTTCCGATCTNNNAGCTGTTCGTTTTCTTATTATTG
GCACACGTCTGAACTCTTTCCCTACACGACGCTCTTCCGATCTNNNCAATGCTTCTTAGTCATTGTCCA
GAGCACACGTCTGAACTCTTTCCCTACACGACGCTCTTCCGATCTNNNTTCTTTAGGAGAAATTCAAC
AGCACACGTCTGAACTCTTTCCCTACACGACGCTCTTCCGATCTNNNTGTTTAACCTGAATATGTTAG
GCACACGTCTGAACTCTTTCCCTACACGACGCTCTTCCGATCTNNNATAGTAGTTTTAGGTTATCGC
AGCACACGTCTGAACTCTTTCCCTACACGACGCTCTTCCGATCTNNNATTGTTTACTGAAGAAATTCTG
GCACACGTCTGAACTCTTTCCCTACACGACGCTCTTCCGATCTNNNCTATATATGGATTTCAGAGGCC
GCACACGTCTGAACTCTTTCCCTACACGACGCTCTTCCGATCTNNNAAATTGTTAAGTCAACATGGTA
GCACACGTCTGAACTCTTTCCCTACACGACGCTCTTCCGATCTNNNTGTGAAAAAGAGAGAGAAAGA
CACACGTCTGAACTCTTTCCCTACACGACGCTCTTCCGATCTNNNTGTTTGCTTTTCTGAATCATATTCA
GCACACGTCTGAACTCTTTCCCTACACGACGCTCTTCCGATCTNNNATAATTTTGCAAAGATGGTTTCA
GAGCACACGTCTGAACTCTTTCCCTACACGACGCTCTTCCGATCTNNNGTAGGCTTTTCCATCAGTAA
GCACACGTCTGAACTCTTTCCCTACACGACGCTCTTCCGATCTNNNCTCAATGTGTAGGTCTCTAAAA
AGCACACGTCTGAACTCTTTCCCTACACGACGCTCTTCCGATCTNNNCTCAGAAATGTCTCTAGATTA
AGCACACGTCTGAACTCTTTCCCTACACGACGCTCTTCCGATCTNNNACACCTGTTTCCTAAATGGACA
AGCACACGTCTGAACTCTTTCCCTACACGACGCTCTTCCGATCTNNNTTCTTTATAAAACAAAGGGCTT
CACACGTCTGAACTCTTTCCCTACACGACGCTCTTCCGATCTNNNCCTTTCTTCCAGTATTTCTATTGC
AGCACACGTCTGAACTCTTTCCCTACACGACGCTCTTCCGATCTNNNAAGAGGATTGCTTGAGTCTA
AGCACACGTCTGAACTCTTTCCCTACACGACGCTCTTCCGATCTNNNAAAAGCTATAAAGGTCTGGA
CACACGTCTGAACTCTTTCCCTACACGACGCTCTTCCGATCTNNNAAAAGGTCTGCAATATTTTGAGCA
CACACGTCTGAACTCTTTCCCTACACGACGCTCTTCCGATCTNNNCAACCCCTCACTTCTTATTGCATCG
GCACACGTCTGAACTCTTTCCCTACACGACGCTCTTCCGATCTNNNTATTTATTGTCTGTTATTTGCCG
AGCACACGTCTGAACTCTTTCCCTACACGACGCTCTTCCGATCTNNNGAGATGAAGGGAAGAAGTA
AGCACACGTCTGAACTCTTTCCCTACACGACGCTCTTCCGATCTNNNAGACTTATTATCAGTAGCAAC
CACACGTCTGAACTCTTTCCCTACACGACGCTCTTCCGATCTNNNCAGTCTGTATATACTTGATAGGG
AGCACACGTCTGAACTCTTTCCCTACACGACGCTCTTCCGATCTNNNTAAGACGAATCTAGGTCATCT
AGCACACGTCTGAACTCTTTCCCTACACGACGCTCTTCCGATCTNNNTTCAGGAGGCTTCTTATACATA
GCACACGTCTGAACTCTTTCCCTACACGACGCTCTTCCGATCTNNNTCTTGTTTCTCCTTTCTTTAGTA
AGCACACGTCTGAACTCTTTCCCTACACGACGCTCTTCCGATCTNNNAATACATTTTCCAGAGTTTCAA
GAGCACACGTCTGAACTCTTTCCCTACACGACGCTCTTCCGATCTNNNTAGTTGTATCTAACTTCTAGG
GAGCACACGTCTGAACTCTTTCCCTACACGACGCTCTTCCGATCTNNNTGGTAAATATTTACACTTCTC
CACACGTCTGAACTCTTTCCCTACACGACGCTCTTCCGATCTNNNCCTGAAGAGAAAAGATCCAATCA
AGCACACGTCTGAACTCTTTCCCTACACGACGCTCTTCCGATCTNNNCTAAATATAGGGGGAAAAAA
CACACGTCTGAACTCTTTCCCTACACGACGCTCTTCCGATCTNNNTTCCTTGAGATCCCAGATAAACAT
GCACACGTCTGAACTCTTTCCCTACACGACGCTCTTCCGATCTNNNACTACTCTCATTATCCCCAGCATC
AGCACACGTCTGAACTCTTTCCCTACACGACGCTCTTCCGATCTNNNCTATTGATTCTTTTCCTCTGGA
GCACACGTCTGAACTCTTTCCCTACACGACGCTCTTCCGATCTNNNCTCATATCAAAGAAGAGTGAG
GCACACGTCTGAACTCTTTCCCTACACGACGCTCTTCCGATCTNNNAATGTAAAGACTGGTATCTGTA
AGCACACGTCTGAACTCTTTCCCTACACGACGCTCTTCCGATCTNNNTTGCATCACTTAACTAAATTTG
AGCACACGTCTGAACTCTTTCCCTACACGACGCTCTTCCGATCTNNNAAAAAAATTCTCTCTCCTCATT
GAGCACACGTCTGAACTCTTTCCCTACACGACGCTCTTCCGATCTNNNGAGTGGAGATTGCATTGTTG
AGCACACGTCTGAACTCTTTCCCTACACGACGCTCTTCCGATCTNNNTAGGAGAATACCATAGTGGCC

AGCACACGTCTGAACTCTTTCCCTACACGACGCTCTTCCGATCTNNNATTTGTGTTAAAATAGCATGT
AGCACACGTCTGAACTCTTTCCCTACACGACGCTCTTCCGATCTNNNTGTGATAGTTTCTAGAAGCAA
CACACGTCTGAACTCTTTCCCTACACGACGCTCTTCCGATCTNNNAATGTAATCAATGAAGACTTTGG
AGCACACGTCTGAACTCTTTCCCTACACGACGCTCTTCCGATCTNNNAGTGATAATAAAATCTCAGC
CACACGTCTGAACTCTTTCCCTACACGACGCTCTTCCGATCTNNNAGAAATAGAGAGACCAAGAAAT
CACGTCTGAACTCTTTCCCTACACGACGCTCTTCCGATCTNNNAGTATGTTCCAACCTATAATCCCAT
GCACACGTCTGAACTCTTTCCCTACACGACGCTCTTCCGATCTNNNAAAATTTCTGGTCTGATATTTC
AGCACACGTCTGAACTCTTTCCCTACACGACGCTCTTCCGATCTNNNGTTGTGTGTGTGTAGATTCCC
GCACACGTCTGAACTCTTTCCCTACACGACGCTCTTCCGATCTNNNGAAACCACTTTAAGGATTAGTT
GCACACGTCTGAACTCTTTCCCTACACGACGCTCTTCCGATCTNNNCATCTTACAAGAAATGCTAAAGG
GAGCACACGTCTGAACTCTTTCCCTACACGACGCTCTTCCGATCTNNNTTATCCAGACAAAGATAATT
AGCACACGTCTGAACTCTTTCCCTACACGACGCTCTTCCGATCTNNNGAAGCAGATTTATCGTGACTA
ACACGTCTGAACTCTTTCCCTACACGACGCTCTTCCGATCTNNNCTCTTCTATAAAATGGGGATGATC
AGCACACGTCTGAACTCTTTCCCTACACGACGCTCTTCCGATCTNNNGAAAAAAAAATACAGAGAGG
GCACACGTCTGAACTCTTTCCCTACACGACGCTCTTCCGATCTNNNTTAGTACTTAGTCAAACATCTG
GCACACGTCTGAACTCTTTCCCTACACGACGCTCTTCCGATCTNNNTCCCATGATTTTTATAGAATCCT
AGCACACGTCTGAACTCTTTCCCTACACGACGCTCTTCCGATCTNNNTTCATTATACAAAGGCTTTGAT
AGCACACGTCTGAACTCTTTCCCTACACGACGCTCTTCCGATCTNNNAGTATGAATGCACCAAAAATA
GCACACGTCTGAACTCTTTCCCTACACGACGCTCTTCCGATCTNNNTCCAAATAATTAACATTTCTGCT
GCACACGTCTGAACTCTTTCCCTACACGACGCTCTTCCGATCTNNNCCAGGTTCATATTTGATAAATCC
AGCACACGTCTGAACTCTTTCCCTACACGACGCTCTTCCGATCTNNNATCTACCGGTTAAACAACAAAC
GCACACGTCTGAACTCTTTCCCTACACGACGCTCTTCCGATCTNNNTATCCTGAGGAAATATTGCAGT
GCACACGTCTGAACTCTTTCCCTACACGACGCTCTTCCGATCTNNNTCCAAATATTTTTTATCTGTGGT
AGCACACGTCTGAACTCTTTCCCTACACGACGCTCTTCCGATCTNNNAATATTTCTTATCCCTCATCCA
GCACACGTCTGAACTCTTTCCCTACACGACGCTCTTCCGATCTNNNTCCCATGATTTTTATAGAATCCT
ACACGTCTGAACTCTTTCCCTACACGACGCTCTTCCGATCTNNNATCTCTAACAATTCTGTTTCCACATC
GCACACGTCTGAACTCTTTCCCTACACGACGCTCTTCCGATCTNNNCTTAGCACAAGATAAAAACAGA
AGCACACGTCTGAACTCTTTCCCTACACGACGCTCTTCCGATCTNNNCATTTAGCTTCCTTCTTGAAAT
ACACGTCTGAACTCTTTCCCTACACGACGCTCTTCCGATCTNNNAAGGGATCTCTAACAATTCTAAGC
GCACACGTCTGAACTCTTTCCCTACACGACGCTCTTCCGATCTNNNAAAATCCTATCATTTTCTTCTGG
AGCACACGTCTGAACTCTTTCCCTACACGACGCTCTTCCGATCTNNNAGAAGGAAGAGAAGAAGGT
GCACACGTCTGAACTCTTTCCCTACACGACGCTCTTCCGATCTNNNTGTATATGTAGCTAAAGGTAC
AGCACACGTCTGAACTCTTTCCCTACACGACGCTCTTCCGATCTNNNAACTTAGCCAATTTACCTCTTAT
AAGAGCACACGTCTGAACTCTTTCCCTACACGACGCTCTTCCGATCTNNNAGAGAAACAATAGCGGA
CACACGTCTGAACTCTTTCCCTACACGACGCTCTTCCGATCTNNNGAAATAGTCCGGTAACTCATCCA
GAGCACACGTCTGAACTCTTTCCCTACACGACGCTCTTCCGATCTNNNTTTTCTCTCCCTCAGTATATCT
CACGTCTGAACTCTTTCCCTACACGACGCTCTTCCGATCTNNNAATCTTGCAAAGCAGGTATTATACAT
CACACGTCTGAACTCTTTCCCTACACGACGCTCTTCCGATCTNNNCTGAGACCATTTTTAGTTGTAAGC
GCACACGTCTGAACTCTTTCCCTACACGACGCTCTTCCGATCTNNNAGCAAGAGAGAAATGATATAC
GCACACGTCTGAACTCTTTCCCTACACGACGCTCTTCCGATCTNNNTTAGAAAACACTGTAAATCAGA
GCACACGTCTGAACTCTTTCCCTACACGACGCTCTTCCGATCTNNNGATAGAGGACTTTCTCTTTCCC
AGCACACGTCTGAACTCTTTCCCTACACGACGCTCTTCCGATCTNNNACAAAAAAATGAATAAAAAGC
GCACACGTCTGAACTCTTTCCCTACACGACGCTCTTCCGATCTNNNAATGTCTTGCAAGTTCATTATT

AGCACACGTCTGAACTCTTTCCCTACACGACGCTCTTCCGATCTNNNGCAACAAGATTTATTGCTAAC

AGCACACGTCTGAACTCTTTCCCTACACGACGCTCTTCCGATCTNNNAAAAGAAAACCATAAGATACC

AGCACACGTCTGAACTCTTTCCCTACACGACGCTCTTCCGATCTNNNAGTTGATACTAAGAAAGCCAT

CACGTCTGAACTCTTTCCCTACACGACGCTCTTCCGATCTNNNAGATGTATCATAAAACCTCAGAACAT

GCACACGTCTGAACTCTTTCCCTACACGACGCTCTTCCGATCTNNNTTCTATTTGGATTCCTTTCCTATC

AGCACACGTCTGAACTCTTTCCCTACACGACGCTCTTCCGATCTNNNCTGTGAAGTTAATTCCATGTG

AGCACACGTCTGAACTCTTTCCCTACACGACGCTCTTCCGATCTNNNCCAAAAGAAAAACAAACAAAC

CACACGTCTGAACTCTTTCCCTACACGACGCTCTTCCGATCTNNNAGAAAGTTCTGATCTGTAATGAC

GAGCACACGTCTGAACTCTTTCCCTACACGACGCTCTTCCGATCTNNNACAGCTCTTAGTTTCTTATTG

GCACACGTCTGAACTCTTTCCCTACACGACGCTCTTCCGATCTNNNTTCTATTTCACCTATCCTATGTCC

AGCACACGTCTGAACTCTTTCCCTACACGACGCTCTTCCGATCTNNNGAGAGAAAAGGGAGATAAAA

GAGCACACGTCTGAACTCTTTCCCTACACGACGCTCTTCCGATCTNNNACAAAGGTCCTTTAGTTTACA

AGCACACGTCTGAACTCTTTCCCTACACGACGCTCTTCCGATCTNNNCTTATTTTGTCAGGCAGTTTAT

AGCACACGTCTGAACTCTTTCCCTACACGACGCTCTTCCGATCTNNNTTATGAAAATATGCCTGTTGTA

AGCACACGTCTGAACTCTTTCCCTACACGACGCTCTTCCGATCTNNNACACTTCAAATTTAGTCGAAA

AGCACACGTCTGAACTCTTTCCCTACACGACGCTCTTCCGATCTNNNGTTAGTTCTTTTTTTTCAGGT

GAGCACACGTCTGAACTCTTTCCCTACACGACGCTCTTCCGATCTNNNAAAAAAACTAACCTTAAGGG

GAGCACACGTCTGAACTCTTTCCCTACACGACGCTCTTCCGATCTNNNTAATAGAAACCCTTTCAAAAG

AGCACACGTCTGAACTCTTTCCCTACACGACGCTCTTCCGATCTNNNCATTAGTTTGGTCAGGTTTTC

GAGCACACGTCTGAACTCTTTCCCTACACGACGCTCTTCCGATCTNNNATAAGGATTTTGTGAGCATTA

CACACGTCTGAACTCTTTCCCTACACGACGCTCTTCCGATCTNNNCTTGTAAAGTCTCTACAGTGTGCA

GCACACGTCTGAACTCTTTCCCTACACGACGCTCTTCCGATCTNNNCAAAAAAGGTAATACATCGGTA

GCACACGTCTGAACTCTTTCCCTACACGACGCTCTTCCGATCTNNNAGGATAAATGAATTCTACAGACA

AGCACACGTCTGAACTCTTTCCCTACACGACGCTCTTCCGATCTNNNACTTTTTCTGGTTTACCATATTC

AGCACACGTCTGAACTCTTTCCCTACACGACGCTCTTCCGATCTNNNACTTTTTCTGGTTTACCATATTC

GCACACGTCTGAACTCTTTCCCTACACGACGCTCTTCCGATCTNNNGATTTTGATGGTTTAGAACTAGT

AGCACACGTCTGAACTCTTTCCCTACACGACGCTCTTCCGATCTNNNGAATTCTTATCTTCACTTGTCAG

GCACACGTCTGAACTCTTTCCCTACACGACGCTCTTCCGATCTNNNTCTCATATAACTGAACAATGGAA

GCACACGTCTGAACTCTTTCCCTACACGACGCTCTTCCGATCTNNNGGCTGGCTTAAAAGATAAAATA

AGCACACGTCTGAACTCTTTCCCTACACGACGCTCTTCCGATCTNNNACTCTAAAGATGCAGTAATTC

ACACGTCTGAACTCTTTCCCTACACGACGCTCTTCCGATCTNNNTTATGTACAGATGATGGATACTCC

AGCACACGTCTGAACTCTTTCCCTACACGACGCTCTTCCGATCTNNNACAAAGAGTGAACAATGAATA

AGCACACGTCTGAACTCTTTCCCTACACGACGCTCTTCCGATCTNNNTGTAAAGAAAGTAAACTAAAG

AGCACACGTCTGAACTCTTTCCCTACACGACGCTCTTCCGATCTNNNTGTGTGTGTATATATGTATG

ACACGTCTGAACTCTTTCCCTACACGACGCTCTTCCGATCTNNNGTCAAATAACCACAGAAGAATTACA

CACACGTCTGAACTCTTTCCCTACACGACGCTCTTCCGATCTNNNGCGACTTTCCTTTTATAGTGTCATC

GCACACGTCTGAACTCTTTCCCTACACGACGCTCTTCCGATCTNNNATTAGGATCTGAGTTCAGCCCAT

AGCACACGTCTGAACTCTTTCCCTACACGACGCTCTTCCGATCTNNNCTTTGGATGATAATGTGTCAA

AGCACACGTCTGAACTCTTTCCCTACACGACGCTCTTCCGATCTNNNTTGAATTTGTCTGGTTTTTTTA

GCACACGTCTGAACTCTTTCCCTACACGACGCTCTTCCGATCTNNNATATTAGTCAAAGTTCTCTGGA

AGCACACGTCTGAACTCTTTCCCTACACGACGCTCTTCCGATCTNNNTCTACTTTGAGTTAATTTTTGT

GCACACGTCTGAACTCTTTCCCTACACGACGCTCTTCCGATCTNNNAAAATAAAACCAACAACTGTGTC

AGCACACGTCTGAACTCTTTCCCTACACGACGCTCTTCCGATCTNNNTCTAATCAAGACAAATGGCTA

AGCACACGTCTGAACTCTTTCCCTACACGACGCTCTTCCGATCTNNNATAATATCTAATGGGATTGGCT

AGCACACGTCTGAACTCTTTCCCTACACGACGCTCTTCCGATCTNNNCAATGAACTACAAAGCAAGATA

GAGCACACGTCTGAACTCTTTCCCTACACGACGCTCTTCCGATCTNNNGATACACTGATTTTCTCCTCCCA

GCACACGTCTGAACTCTTTCCCTACACGACGCTCTTCCGATCTNNNCTCTAGTTCTATCCATGTTGATGC

GCACACGTCTGAACTCTTTCCCTACACGACGCTCTTCCGATCTNNNCATTCTAAGAAATGTGATCTTCAC

GCACACGTCTGAACTCTTTCCCTACACGACGCTCTTCCGATCTNNNAATCCATCTGGTAGAAAAGTAAT

AGCACACGTCTGAACTCTTTCCCTACACGACGCTCTTCCGATCTNNNAACAAGACACAGACATCTTCCAT

CACACGTCTGAACTCTTTCCCTACACGACGCTCTTCCGATCTNNNTTACTAATTTCCTGCTTAAGAGACAT

GAGCACACGTCTGAACTCTTTCCCTACACGACGCTCTTCCGATCTNNNACTTTTTCCAAAAAATGATGT

CACACGTCTGAACTCTTTCCCTACACGACGCTCTTCCGATCTNNNAAACGCACATACATTTAGATATGCA

CACACGTCTGAACTCTTTCCCTACACGACGCTCTTCCGATCTNNNATTCAAAGGATTGGTCTCTATCCAT

AGCACACGTCTGAACTCTTTCCCTACACGACGCTCTTCCGATCTNNNACGTCACAATAATAAAGTGAAA

GCACACGTCTGAACTCTTTCCCTACACGACGCTCTTCCGATCTNNNTCTGTTCATTGATTTTTCTGATTCA

ACACGTCTGAACTCTTTCCCTACACGACGCTCTTCCGATCTNNNGTATACATGTGCCATGTTGGCATCGC

GAGCACACGTCTGAACTCTTTCCCTACACGACGCTCTTCCGATCTNNNCTGATCAGATTATCCAGTGTTT

GAGCACACGTCTGAACTCTTTCCCTACACGACGCTCTTCCGATCTNNNATTCCTGATTGTTTTCAAGATT

GAGCACACGTCTGAACTCTTTCCCTACACGACGCTCTTCCGATCTNNNTAACTAGATCCTATTAGACAAC

AGCACACGTCTGAACTCTTTCCCTACACGACGCTCTTCCGATCTNNNTGCAGAAAATGTAAATAAACAG

CACACGTCTGAACTCTTTCCCTACACGACGCTCTTCCGATCTNNNCTCATTAATGTAAGCCATCAATCCA

ACACGTCTGAACTCTTTCCCTACACGACGCTCTTCCGATCTNNNCATTACTCTCTCATCACTCTTTTCATCG

GAGCACACGTCTGAACTCTTTCCCTACACGACGCTCTTCCGATCTNNNAGATATTATCTCGACCCAATTA

AGCACACGTCTGAACTCTTTCCCTACACGACGCTCTTCCGATCTNNNTTTTCTACAAGACTGTTCTTTTG

AGAGCACACGTCTGAACTCTTTCCCTACACGACGCTCTTCCGATCTNNNGTGTTAGGAAGAGAGAGAC

AGAGCACACGTCTGAACTCTTTCCCTACACGACGCTCTTCCGATCTNNNATACACATTGCCTGTAACATC

GAGCACACGTCTGAACTCTTTCCCTACACGACGCTCTTCCGATCTNNNATGTGTATGTATGCATGCATC

GAGCACACGTCTGAACTCTTTCCCTACACGACGCTCTTCCGATCTNNNATCTCTTATTCTGTCACTCATA

CACACGTCTGAACTCTTTCCCTACACGACGCTCTTCCGATCTNNNAATTAAAGCCTTCTCACTACTCCATC

AGCACACGTCTGAACTCTTTCCCTACACGACGCTCTTCCGATCTNNNACATATGTGCGTATATATATCCA

AGCACACGTCTGAACTCTTTCCCTACACGACGCTCTTCCGATCTNNNACAAAACCAATATACTATACATC

CACACGTCTGAACTCTTTCCCTACACGACGCTCTTCCGATCTNNNGAAAATAAAGGAAGGGTGGTCATC

GAGCACACGTCTGAACTCTTTCCCTACACGACGCTCTTCCGATCTNNNGAGGAAATAAAAGTCTTGTCA

AGCACACGTCTGAACTCTTTCCCTACACGACGCTCTTCCGATCTNNNTCTCTGCAGAAAAAAAAAATTA

AGCACACGTCTGAACTCTTTCCCTACACGACGCTCTTCCGATCTNNNTGATTTTCTTACTCTTCCATCTTC

GCACACGTCTGAACTCTTTCCCTACACGACGCTCTTCCGATCTNNNGTGTAAATGTAAGCTTTTCAAAG

GAGCACACGTCTGAACTCTTTCCCTACACGACGCTCTTCCGATCTNNNCTTATTCTTGGAAACCTGAAAA

AGCACACGTCTGAACTCTTTCCCTACACGACGCTCTTCCGATCTNNNATAAAACCAGTGTATAATCAAC

ACACGTCTGAACTCTTTCCCTACACGACGCTCTTCCGATCTNNNATCTCCAGATTCACACCCTACATCGG

GCACACGTCTGAACTCTTTCCCTACACGACGCTCTTCCGATCTNNNGTGAATAATATGAGAGCAGTTG

AGCACACGTCTGAACTCTTTCCCTACACGACGCTCTTCCGATCTNNNCAATGAAGAAAGAATGGAGAA

GCACACGTCTGAACTCTTTCCCTACACGACGCTCTTCCGATCTNNNGGAAGAAAAAGAAAGATAGTGT

GCACACGTCTGAACTCTTTCCCTACACGACGCTCTTCCGATCTNNNGGAAGAAAAAGAAAGATAGTGT

GAGCACACGTCTGAACTCTTTCCCTACACGACGCTCTTCCGATCTNNNTAGCTTTTATTTCCATTTCTTCC

AGCACACGTCTGAACTCTTTCCCTACACGACGCTCTTCCGATCTNNNCTGCTTTGTAGATTTCAGTTTCC

GCACACGTCTGAACTCTTTCCCTACACGACGCTCTTCCGATCTNNNAGTTAAAGAGTCAAAACAATGA
AGCACACGTCTGAACTCTTTCCCTACACGACGCTCTTCCGATCTNNNGAGGAAACTTGCAATTGTATT
GAGCACACGTCTGAACTCTTTCCCTACACGACGCTCTTCCGATCTNNNCTCAGAATACCTTACTTGATA
GCACACGTCTGAACTCTTTCCCTACACGACGCTCTTCCGATCTNNNCCTGGCTTATTTCACTTAATATAC
GAGCACACGTCTGAACTCTTTCCCTACACGACGCTCTTCCGATCTNNNATGCTGGAAAGTTTATTGC
CACACGTCTGAACTCTTTCCCTACACGACGCTCTTCCGATCTNNNCATCCCTTTAGTTTGGTTTCATCA
AGCACACGTCTGAACTCTTTCCCTACACGACGCTCTTCCGATCTNNNTCCTCTGCCTACTTTTAATATTT
GCACACGTCTGAACTCTTTCCCTACACGACGCTCTTCCGATCTNNNCACTCCAAATTTTTATCTCTGTA
GAGCACACGTCTGAACTCTTTCCCTACACGACGCTCTTCCGATCTNNNAATTCCCCTTGAAATTGTGTC
GAGCACACGTCTGAACTCTTTCCCTACACGACGCTCTTCCGATCTNNNCCTTCATAAAAGCAAAATACA
GAGCACACGTCTGAACTCTTTCCCTACACGACGCTCTTCCGATCTNNNCCTTCATAAAAGCAAAATACA
CACACGTCTGAACTCTTTCCCTACACGACGCTCTTCCGATCTNNNCTCTCTGTGTGTGTATGTGTCATC
AGCACACGTCTGAACTCTTTCCCTACACGACGCTCTTCCGATCTNNNAGATGGATGAGGGAAACAA
CACACGTCTGAACTCTTTCCCTACACGACGCTCTTCCGATCTNNNAGGCTTTTCATAAATTGGTCTAAC
GCACACGTCTGAACTCTTTCCCTACACGACGCTCTTCCGATCTNNNCATACACAATTTTGCAAATAACA
GCACACGTCTGAACTCTTTCCCTACACGACGCTCTTCCGATCTNNNCATACACAATTTTGCAAATAACA
GCACACGTCTGAACTCTTTCCCTACACGACGCTCTTCCGATCTNNNCATCTCTACTTTATGTCAGTTTGC
GCACACGTCTGAACTCTTTCCCTACACGACGCTCTTCCGATCTNNNATTTTACAGTGACCATGAAAGA
AGCACACGTCTGAACTCTTTCCCTACACGACGCTCTTCCGATCTNNNTCACATTTTTGTCAATTCTACA
GAGCACACGTCTGAACTCTTTCCCTACACGACGCTCTTCCGATCTNNNAGTAGAAAATTTTTCCCAAAT
AGCACACGTCTGAACTCTTTCCCTACACGACGCTCTTCCGATCTNNNCATTTATGAGCATTTTACTCTG
AGCACACGTCTGAACTCTTTCCCTACACGACGCTCTTCCGATCTNNNATAAAACAAAGCAGTTGGTA
AGCACACGTCTGAACTCTTTCCCTACACGACGCTCTTCCGATCTNNNACTCTTGTACATGTAGAATCTA
AGCACACGTCTGAACTCTTTCCCTACACGACGCTCTTCCGATCTNNNCTTTTCATATGTGAGTGACAA
AGCACACGTCTGAACTCTTTCCCTACACGACGCTCTTCCGATCTNNNTATTGGGAAACTATTATTCTGC
ACACGTCTGAACTCTTTCCCTACACGACGCTCTTCCGATCTNNNTGTGTGTATGTATGTATGTATGTC
CACACGTCTGAACTCTTTCCCTACACGACGCTCTTCCGATCTNNNCCTTATAGGCTCCATAAAATAGTC
CACACGTCTGAACTCTTTCCCTACACGACGCTCTTCCGATCTNNNCCTTATAGGCTCCATAAAATAGTC
AGCACACGTCTGAACTCTTTCCCTACACGACGCTCTTCCGATCTNNNTTTAGAAGTGGGCACATATT
AGCACACGTCTGAACTCTTTCCCTACACGACGCTCTTCCGATCTNNNTTGAGATAGACCTATACCTAG
GCACACGTCTGAACTCTTTCCCTACACGACGCTCTTCCGATCTNNNATCTCAGCTCATTACAACTCTCA
GAGCACACGTCTGAACTCTTTCCCTACACGACGCTCTTCCGATCTNNNTTACTCTATGTCAGGCACTAG
ACACGTCTGAACTCTTTCCCTACACGACGCTCTTCCGATCTNNNTACCAAAAGTTCTGAAAACTTGCAT
GCACACGTCTGAACTCTTTCCCTACACGACGCTCTTCCGATCTNNNGTAGGTAGCTAGTCAGGTATA
GCACACGTCTGAACTCTTTCCCTACACGACGCTCTTCCGATCTNNNTAAGATAGAATCTAAGCAGTGA
GCACACGTCTGAACTCTTTCCCTACACGACGCTCTTCCGATCTNNNTAAGATAGAATCTAAGCAGTGA
CACACGTCTGAACTCTTTCCCTACACGACGCTCTTCCGATCTNNNCATGAAGTAGTTTCCAAATTTGA
GCACACGTCTGAACTCTTTCCCTACACGACGCTCTTCCGATCTNNNAGTACTAGTGATATTGATTTTA
GCACACGTCTGAACTCTTTCCCTACACGACGCTCTTCCGATCTNNNCCAACTTTCATATAACTAGGTCT
AGCACACGTCTGAACTCTTTCCCTACACGACGCTCTTCCGATCTNNNATATTTCTCTCCCTCTCTCTTTCA
AGCACACGTCTGAACTCTTTCCCTACACGACGCTCTTCCGATCTNNNATATTTCTCTCCCTCTCTCTTTCA
AGCACACGTCTGAACTCTTTCCCTACACGACGCTCTTCCGATCTNNNATATTTCTCTCCCTCTCTCTTTCA
AGCACACGTCTGAACTCTTTCCCTACACGACGCTCTTCCGATCTNNNCTTTCTCCCCAGTTGAGAATCA

GAGCACACGTCTGAACTCTTTCCCTACACGACGCTCTTCCGATCTNNNACTTTTTATTATGTGTGTAGG
GCACACGTCTGAACTCTTTCCCTACACGACGCTCTTCCGATCTNNNACAGCTCTAAATGAATCCATTCAT
GCACACGTCTGAACTCTTTCCCTACACGACGCTCTTCCGATCTNNNAAAACTGACCTATACATGCATCAT
GAGCACACGTCTGAACTCTTTCCCTACACGACGCTCTTCCGATCTNNNTTTTCCTTTCTCTAGGGACTCAT
CACACGTCTGAACTCTTTCCCTACACGACGCTCTTCCGATCTNNNGTTTATCCTGTTTGGGGAACCATC
GAGCACACGTCTGAACTCTTTCCCTACACGACGCTCTTCCGATCTNNNTCAGATTTTAGTGTAAGAGG
AGCACACGTCTGAACTCTTTCCCTACACGACGCTCTTCCGATCTNNNAGTCATTGTCAAGTTTATGTAAC
AGCACACGTCTGAACTCTTTCCCTACACGACGCTCTTCCGATCTNNNGAAAGAATGTGTGAAAGAGTG
GCACACGTCTGAACTCTTTCCCTACACGACGCTCTTCCGATCTNNNAAAATAAAAAAAGGAGATGCAG
CACACGTCTGAACTCTTTCCCTACACGACGCTCTTCCGATCTNNNTGACATCACATTTGAGTTTTGCATC
CACACGTCTGAACTCTTTCCCTACACGACGCTCTTCCGATCTNNNTGACATCACATTTGAGTTTTGCATC
AGCACACGTCTGAACTCTTTCCCTACACGACGCTCTTCCGATCTNNNTTCAGTATAAATGGTTGGCTAA
GCACACGTCTGAACTCTTTCCCTACACGACGCTCTTCCGATCTNNNTTAGTTTCTGCCCTCTTTTGCATC
AGCACACGTCTGAACTCTTTCCCTACACGACGCTCTTCCGATCTNNNGAGATATAGGGTTTGGTACTAT
GAGCACACGTCTGAACTCTTTCCCTACACGACGCTCTTCCGATCTNNNTTTTAGACGGTACATTGTAGC
AGCACACGTCTGAACTCTTTCCCTACACGACGCTCTTCCGATCTNNNCCCTCTGAAAAACATGCTTACAT
ACACGTCTGAACTCTTTCCCTACACGACGCTCTTCCGATCTNNNATTGACTCACAATTCAGCATCATCGG
GAGCACACGTCTGAACTCTTTCCCTACACGACGCTCTTCCGATCTNNNAACCCCAACAAGATACAGAGCA
ACACGTCTGAACTCTTTCCCTACACGACGCTCTTCCGATCTNNNTATTTATCTTGCATAGGCTATCCCAT
CACACGTCTGAACTCTTTCCCTACACGACGCTCTTCCGATCTNNNCTATCATAAGAGCCTGGGACCATC
AGCACACGTCTGAACTCTTTCCCTACACGACGCTCTTCCGATCTNNNGAAGGAGGAAATTTTATTTAG
GAGCACACGTCTGAACTCTTTCCCTACACGACGCTCTTCCGATCTNNNTCAGGACAAAATCAACATAAAA
AGCACACGTCTGAACTCTTTCCCTACACGACGCTCTTCCGATCTNNNTAAATGCAAGTTTGAAATCTCT
GAGCACACGTCTGAACTCTTTCCCTACACGACGCTCTTCCGATCTNNNATAATTTTCTCCTTTCCTGTTTC
GCACACGTCTGAACTCTTTCCCTACACGACGCTCTTCCGATCTNNNAAAAAAATCGATAAGTCTCTAGC
GCACACGTCTGAACTCTTTCCCTACACGACGCTCTTCCGATCTNNNACTGAAAGGTTATGTGCTATTTA
AGCACACGTCTGAACTCTTTCCCTACACGACGCTCTTCCGATCTNNNCTACATAATCAGGGCTTTATGTC
AGCACACGTCTGAACTCTTTCCCTACACGACGCTCTTCCGATCTNNNCTTTGAATGCCATACATCATCCA
AGAGCACACGTCTGAACTCTTTCCCTACACGACGCTCTTCCGATCTNNNTATAGACATTATATGCAGAG
GAGCACACGTCTGAACTCTTTCCCTACACGACGCTCTTCCGATCTNNNCTGTCATCTAGTATACATTCCAT
AGCACACGTCTGAACTCTTTCCCTACACGACGCTCTTCCGATCTNNNTGGCAAATCCTTACATTAAATAG
AGCACACGTCTGAACTCTTTCCCTACACGACGCTCTTCCGATCTNNNGATATAAGGGTCAAGGGAATG
GAGCACACGTCTGAACTCTTTCCCTACACGACGCTCTTCCGATCTNNNGGGTGAAAAATCTTTTCAATA
AGCACACGTCTGAACTCTTTCCCTACACGACGCTCTTCCGATCTNNNAAAAAAAGCCCTAAAATTGGTA
GAGCACACGTCTGAACTCTTTCCCTACACGACGCTCTTCCGATCTNNNCAAGGAGAATTCCAAAAAGTA
AGCACACGTCTGAACTCTTTCCCTACACGACGCTCTTCCGATCTNNNTGTATACTTGAAATTTGCTAAGA
CACACGTCTGAACTCTTTCCCTACACGACGCTCTTCCGATCTNNNTGGTTTCAAAATCTGTATTTCAGCA
AGCACACGTCTGAACTCTTTCCCTACACGACGCTCTTCCGATCTNNNAATTTACCCACTATAGACTTCAC
AGCACACGTCTGAACTCTTTCCCTACACGACGCTCTTCCGATCTNNNACAACTTGATCATGTATCTTAAT
GCACACGTCTGAACTCTTTCCCTACACGACGCTCTTCCGATCTNNNTTGATTCAGTAAAATAGGTATGC
GCACACGTCTGAACTCTTTCCCTACACGACGCTCTTCCGATCTNNNCATCACATAAAAAAATCAAACCGC
AGCACACGTCTGAACTCTTTCCCTACACGACGCTCTTCCGATCTNNNAAAGCATCTACAATTTTAGTTGC
AGCACACGTCTGAACTCTTTCCCTACACGACGCTCTTCCGATCTNNNTACTTTAGGGAATTTCACACTTT

AGCACACGTCTGAACTCTTTCCCTACACGACGCTCTTCCGATCTNNNCTAAGATTTCATAGCCCAAAA

AGCACACGTCTGAACTCTTTCCCTACACGACGCTCTTCCGATCTNNNCTAAGATTTCATAGCCCAAAA

AGCACACGTCTGAACTCTTTCCCTACACGACGCTCTTCCGATCTNNNGGAAACAGGAAAATAATTTGG

GCACACGTCTGAACTCTTTCCCTACACGACGCTCTTCCGATCTNNNATACTCCCAGTTTTTAGAAGA

GCACACGTCTGAACTCTTTCCCTACACGACGCTCTTCCGATCTNNNTAGTCAGCGAAGGTAACATTAT

CACACGTCTGAACTCTTTCCCTACACGACGCTCTTCCGATCTNNNAAAGTTCTTGGAAATACATACAC

GCACACGTCTGAACTCTTTCCCTACACGACGCTCTTCCGATCTNNNCTAAAATCTGATCTTGGACAAA

AGCACACGTCTGAACTCTTTCCCTACACGACGCTCTTCCGATCTNNNATGAAATAGTACAAGTACAGG

GCACACGTCTGAACTCTTTCCCTACACGACGCTCTTCCGATCTNNNCATTCATTCTTACATGTATGTCTC

CACACGTCTGAACTCTTTCCCTACACGACGCTCTTCCGATCTNNNTATTATCAGTTAGGGGGAGAAC

CACACGTCTGAACTCTTTCCCTACACGACGCTCTTCCGATCTNNNGAAAGGGTTCTTCTGACTTCCATC

CACGTCTGAACTCTTTCCCTACACGACGCTCTTCCGATCTNNNGATCAGTTTACTCAAAGAAGAATCA

AGCACACGTCTGAACTCTTTCCCTACACGACGCTCTTCCGATCTNNNTGTTTCTCTCTAACTGTATTAG

AGCACACGTCTGAACTCTTTCCCTACACGACGCTCTTCCGATCTNNNTAGATTATGCACTGCTATTGAT

AGCACACGTCTGAACTCTTTCCCTACACGACGCTCTTCCGATCTNNNTGCCACATTTTCATTTTGTATAT

GCACACGTCTGAACTCTTTCCCTACACGACGCTCTTCCGATCTNNNTCTCTTTCAGTGTTTGATTGACA

AGCACACGTCTGAACTCTTTCCCTACACGACGCTCTTCCGATCTNNNGAGTTTCAAGAAGAGTACAA

ACACGTCTGAACTCTTTCCCTACACGACGCTCTTCCGATCTNNNGAGGCGATTTTCAGAACTTACATC

GAGCACACGTCTGAACTCTTTCCCTACACGACGCTCTTCCGATCTNNNTATAGGAAATGGTTTTACTTC

GCACACGTCTGAACTCTTTCCCTACACGACGCTCTTCCGATCTNNNTGAAGAGAAGTAATCTACTCTT

CACACGTCTGAACTCTTTCCCTACACGACGCTCTTCCGATCTNNNATTCTAGGAATCTACAGACAAATC

CACACGTCTGAACTCTTTCCCTACACGACGCTCTTCCGATCTNNNTAAACTCCACAAACACATACAACAT

AGCACACGTCTGAACTCTTTCCCTACACGACGCTCTTCCGATCTNNNATAATCAAGTCAGAAGCTAAT

AGCACACGTCTGAACTCTTTCCCTACACGACGCTCTTCCGATCTNNNTTTTCTAAGTACTATTGAACCT

ACACGTCTGAACTCTTTCCCTACACGACGCTCTTCCGATCTNNNACTGAGGAATACTTTTTAAAGCC

ACACGTCTGAACTCTTTCCCTACACGACGCTCTTCCGATCTNNNACTGAGGAATACTTTTTAAAGCC

CACACGTCTGAACTCTTTCCCTACACGACGCTCTTCCGATCTNNNCTTAGAAAGAACTCCATTAGAGA

GAGCACACGTCTGAACTCTTTCCCTACACGACGCTCTTCCGATCTNNNTAACTGGCATGTATTGATAA

CACACGTCTGAACTCTTTCCCTACACGACGCTCTTCCGATCTNNNCTCTCACATGCACACTATCTCATCG

GAGCACACGTCTGAACTCTTTCCCTACACGACGCTCTTCCGATCTNNNTGTCTTTTATTAGATATTCAA

AGCACACGTCTGAACTCTTTCCCTACACGACGCTCTTCCGATCTNNNTTATGCAACTTTTCTTTCAAAA

GCACACGTCTGAACTCTTTCCCTACACGACGCTCTTCCGATCTNNNGGTTGATCCCAGAATTTCAACA

AGCACACGTCTGAACTCTTTCCCTACACGACGCTCTTCCGATCTNNNCTAATAAAAGGATGAGCCCAA

GCACACGTCTGAACTCTTTCCCTACACGACGCTCTTCCGATCTNNNCTCCATTTTGCCCATAGATGCAT

GCACACGTCTGAACTCTTTCCCTACACGACGCTCTTCCGATCTNNNTCTTTTTTATGGTAGCATAGGAT

GCACACGTCTGAACTCTTTCCCTACACGACGCTCTTCCGATCTNNNATATTTCACCGAAATACATCATG

GCACACGTCTGAACTCTTTCCCTACACGACGCTCTTCCGATCTNNNAGATTCTGAGAGGTTAAGGAAC

AGCACACGTCTGAACTCTTTCCCTACACGACGCTCTTCCGATCTNNNTCCTGAAAAGTAAAAATCTAA

GCACACGTCTGAACTCTTTCCCTACACGACGCTCTTCCGATCTNNNTCTATTGAATACATTCTGCTTTG

AGCACACGTCTGAACTCTTTCCCTACACGACGCTCTTCCGATCTNNNTTGTTAATAACAGTACCACAGT

GCACACGTCTGAACTCTTTCCCTACACGACGCTCTTCCGATCTNNNCATCTTTGTTATATATCTGCAGG

GCACACGTCTGAACTCTTTCCCTACACGACGCTCTTCCGATCTNNNCATTCAACAGGTGATTGTTTAT

CACACGTCTGAACTCTTTCCCTACACGACGCTCTTCCGATCTNNNTGAATCACCTATTAATTAAGGGG

GCACACGTCTGAACTCTTTCCCTACACGACGCTCTTCCGATCTNNNTCCACAATACAGAGAGTAATATCC
AGCACACGTCTGAACTCTTTCCCTACACGACGCTCTTCCGATCTNNNTAGCAAAATGACAGAAGTAAAT
GAGCACACGTCTGAACTCTTTCCCTACACGACGCTCTTCCGATCTNNNGAATAAGGATGGTAGCTAAA
AGCACACGTCTGAACTCTTTCCCTACACGACGCTCTTCCGATCTNNNAGATATTAGACGCATAATTTG
GAGCACACGTCTGAACTCTTTCCCTACACGACGCTCTTCCGATCTNNNAATGGATTAAAATCACTCAGG
CACACGTCTGAACTCTTTCCCTACACGACGCTCTTCCGATCTNNNAGAAAGATCTCACCCATTTTATACA
AGCACACGTCTGAACTCTTTCCCTACACGACGCTCTTCCGATCTNNNTCAGAGAAATCTTATTTTGTGT
AGCACACGTCTGAACTCTTTCCCTACACGACGCTCTTCCGATCTNNNTCAGAGAAATCTTATTTTGTGT
AGCACACGTCTGAACTCTTTCCCTACACGACGCTCTTCCGATCTNNNGACCTTTCAGAGAAATCTTATTT
GCACACGTCTGAACTCTTTCCCTACACGACGCTCTTCCGATCTNNNTTCACATATTCACTCTAATCCAGCA
AGCACACGTCTGAACTCTTTCCCTACACGACGCTCTTCCGATCTNNNACCCTGTCTCAAAAAAAAATTAT
AGCACACGTCTGAACTCTTTCCCTACACGACGCTCTTCCGATCTNNNAAAACAGACACATAAACACATA
AGCACACGTCTGAACTCTTTCCCTACACGACGCTCTTCCGATCTNNNATGGAGAATCTCAGATCAGTCA
AGCACACGTCTGAACTCTTTCCCTACACGACGCTCTTCCGATCTNNNACAAATAATTCATAACGTTAGCA
AGCACACGTCTGAACTCTTTCCCTACACGACGCTCTTCCGATCTNNNGGCCAAAATGATGATATTACTA
GAGCACACGTCTGAACTCTTTCCCTACACGACGCTCTTCCGATCTNNNTATGGCATTTCATTTTCAAAAA
AGCACACGTCTGAACTCTTTCCCTACACGACGCTCTTCCGATCTNNNGGCCTCAAATCTTTATCTACATA
AGCACACGTCTGAACTCTTTCCCTACACGACGCTCTTCCGATCTNNNCTGATATTTGTTCCAAAAAGCT
AGCACACGTCTGAACTCTTTCCCTACACGACGCTCTTCCGATCTNNNGTTATTGTTGCCCTTACTAATTT
CACACGTCTGAACTCTTTCCCTACACGACGCTCTTCCGATCTNNNCATGTACATATTTGCACACAGCATC
AGCACACGTCTGAACTCTTTCCCTACACGACGCTCTTCCGATCTNNNAGTATCAGTTTTTACTGTCTCTCC
GAGCACACGTCTGAACTCTTTCCCTACACGACGCTCTTCCGATCTNNNAGATGAGAAAATTGAGGTGC
AGCACACGTCTGAACTCTTTCCCTACACGACGCTCTTCCGATCTNNNAATTGGCCTAATTTCAATATTGT
AGCACACGTCTGAACTCTTTCCCTACACGACGCTCTTCCGATCTNNNATAAATATCATTGGGGGTAAG
AGCACACGTCTGAACTCTTTCCCTACACGACGCTCTTCCGATCTNNNACTAGATGGACGCATATGTACA
GAGCACACGTCTGAACTCTTTCCCTACACGACGCTCTTCCGATCTNNNTTTAACCATAGGCAAAATTTCA
GCACACGTCTGAACTCTTTCCCTACACGACGCTCTTCCGATCTNNNCTTCTCTCTCTCTCTCTGCATCG
AGCACACGTCTGAACTCTTTCCCTACACGACGCTCTTCCGATCTNNNAGTCTGTTATGACCTTTTCTATT
AGCACACGTCTGAACTCTTTCCCTACACGACGCTCTTCCGATCTNNNTAATCAGATCATGAGGGTAGG
GCACACGTCTGAACTCTTTCCCTACACGACGCTCTTCCGATCTNNNGCAATGAGACATAGTAAACATA
GAGCACACGTCTGAACTCTTTCCCTACACGACGCTCTTCCGATCTNNNTAAATTTCTTATCTAGCAGTTG
GCACACGTCTGAACTCTTTCCCTACACGACGCTCTTCCGATCTNNNCACTCACATTACTTCTGATGACAT
GCACACGTCTGAACTCTTTCCCTACACGACGCTCTTCCGATCTNNNCACTCACATTACTTCTGATGACAT
AGCACACGTCTGAACTCTTTCCCTACACGACGCTCTTCCGATCTNNNTCAAATCTTTTGCCCATATTTTAC
GAGCACACGTCTGAACTCTTTCCCTACACGACGCTCTTCCGATCTNNNTTCGATATATCTTCCACACATA
CACACGTCTGAACTCTTTCCCTACACGACGCTCTTCCGATCTNNNGAATTTTATTGGCTCAGAATTCTCA
GAGCACACGTCTGAACTCTTTCCCTACACGACGCTCTTCCGATCTNNNGTAGAATGTCATTATCTTTGTC
GAGCACACGTCTGAACTCTTTCCCTACACGACGCTCTTCCGATCTNNNAGTATACAAAATACAGGCATA
AGCACACGTCTGAACTCTTTCCCTACACGACGCTCTTCCGATCTNNNACCTAGGTTCATAAATTTGTACT
AGCACACGTCTGAACTCTTTCCCTACACGACGCTCTTCCGATCTNNNGTAATTTCTGTCTCCCTTCCCAT
GAGCACACGTCTGAACTCTTTCCCTACACGACGCTCTTCCGATCTNNNCCGTCAAGTGTTGAATAAATG
GCACACGTCTGAACTCTTTCCCTACACGACGCTCTTCCGATCTNNNTCATTTTTACCACCTGTAAGATACA
CACACGTCTGAACTCTTTCCCTACACGACGCTCTTCCGATCTNNNGAAACCCTGTCCTACAAACTCATCG

AGCACACGTCTGAACTCTTTCCCTACACGACGCTCTTCCGATCTNNNCAATTGTCAACCAGAAAAATTT
GCACACGTCTGAACTCTTTCCCTACACGACGCTCTTCCGATCTNNNAGGTGGTTAGATGTTCAAAAT
AGCACACGTCTGAACTCTTTCCCTACACGACGCTCTTCCGATCTNNNAACTGTATTAGCAAAAAAGTC
AGCACACGTCTGAACTCTTTCCCTACACGACGCTCTTCCGATCTNNNTAATAGTGACTGCAAATCAAA
CACACGTCTGAACTCTTTCCCTACACGACGCTCTTCCGATCTNNNATCCTTTGTTCTAGGAGAGGCATC
AGCACACGTCTGAACTCTTTCCCTACACGACGCTCTTCCGATCTNNNCCTGTTTATCTTTGTATTTCTC
GCACACGTCTGAACTCTTTCCCTACACGACGCTCTTCCGATCTNNNAAGTTATCTCCATGAAGTATGCC
GCACACGTCTGAACTCTTTCCCTACACGACGCTCTTCCGATCTNNNAAGTTATCTCCATGAAGTATGCC
AGCACACGTCTGAACTCTTTCCCTACACGACGCTCTTCCGATCTNNNTGGAGTCAAACAAATCTTAATT
GCACACGTCTGAACTCTTTCCCTACACGACGCTCTTCCGATCTNNNTAAATAATTAGTGGATTGGAT
AGCACACGTCTGAACTCTTTCCCTACACGACGCTCTTCCGATCTNNNCATAGTTAGATAACCCAAGAG
AGCACACGTCTGAACTCTTTCCCTACACGACGCTCTTCCGATCTNNNGAAAAGAACCTACGTTGAAAT
CACGTCTGAACTCTTTCCCTACACGACGCTCTTCCGATCTNNNTTGATAATATATCCATCATCCCGCAT
GCACACGTCTGAACTCTTTCCCTACACGACGCTCTTCCGATCTNNNTTCTAAACTGTAAGCCTATAAGG
CACACGTCTGAACTCTTTCCCTACACGACGCTCTTCCGATCTNNNTTCTTCAAATAGTCTTTCAGTTCC
CACGTCTGAACTCTTTCCCTACACGACGCTCTTCCGATCTNNNCAAAGAGCACAATCATTCTTCCATCG
CACACGTCTGAACTCTTTCCCTACACGACGCTCTTCCGATCTNNNATTTAAATGGGCATGTGAAACCAT
AGCACACGTCTGAACTCTTTCCCTACACGACGCTCTTCCGATCTNNNTGTTTTTCCATTTACTTGTGTT
AGCACACGTCTGAACTCTTTCCCTACACGACGCTCTTCCGATCTNNNTACTTTGTCCTTGATATCCTTAG
AGCACACGTCTGAACTCTTTCCCTACACGACGCTCTTCCGATCTNNNTCAATATGGTTATGGGATTAT
AGCACACGTCTGAACTCTTTCCCTACACGACGCTCTTCCGATCTNNNTAAACTGCATAGAGTTGAATG
AGCACACGTCTGAACTCTTTCCCTACACGACGCTCTTCCGATCTNNNCTCCTGCAACTTTAATGAATTC
GAGCACACGTCTGAACTCTTTCCCTACACGACGCTCTTCCGATCTNNNACACCCCTACACATATTTTTA
GAGCACACGTCTGAACTCTTTCCCTACACGACGCTCTTCCGATCTNNNACACCCCTACACATATTTTTA
AGCACACGTCTGAACTCTTTCCCTACACGACGCTCTTCCGATCTNNNCACATTTCATTTACCAAAGAAT
GCACACGTCTGAACTCTTTCCCTACACGACGCTCTTCCGATCTNNNTTGGAACACATATAAACCAGATT
ACACGTCTGAACTCTTTCCCTACACGACGCTCTTCCGATCTNNNTATAATCTGTGGGCAAGAAAAACA
GCACACGTCTGAACTCTTTCCCTACACGACGCTCTTCCGATCTNNNTCATCTTCTCTTTCTTGCTATAA
AGCACACGTCTGAACTCTTTCCCTACACGACGCTCTTCCGATCTNNNGACTTTTCATTTCAAGTCTACTT
AGCACACGTCTGAACTCTTTCCCTACACGACGCTCTTCCGATCTNNNTTCTTTGATTACACATATATGC
GAGCACACGTCTGAACTCTTTCCCTACACGACGCTCTTCCGATCTNNNGTAGGAGAAACAGATAACA
ACACGTCTGAACTCTTTCCCTACACGACGCTCTTCCGATCTNNNTATGTTTTCTTCCCTGAACTAATCAT
AGCACACGTCTGAACTCTTTCCCTACACGACGCTCTTCCGATCTNNNAGAATGTATGTTGGGTGAAA
GCACACGTCTGAACTCTTTCCCTACACGACGCTCTTCCGATCTNNNAATTGAGACATGAGACAATTAT
GCACACGTCTGAACTCTTTCCCTACACGACGCTCTTCCGATCTNNNATTGAACACCTTACAAGTTTTTA
GAGCACACGTCTGAACTCTTTCCCTACACGACGCTCTTCCGATCTNNNCACCTTACTTTCAAATAGTTC
CACACGTCTGAACTCTTTCCCTACACGACGCTCTTCCGATCTNNNATACGTTTATTTATACACCTGGTC
GCACACGTCTGAACTCTTTCCCTACACGACGCTCTTCCGATCTNNNTATTAACTAAACCTCTGTGAGTT
GCACACGTCTGAACTCTTTCCCTACACGACGCTCTTCCGATCTNNNACTCATAGGTTCTAGAAATAAG
AGCACACGTCTGAACTCTTTCCCTACACGACGCTCTTCCGATCTNNNCACCATATGACAGCAATTTAAT
AGCACACGTCTGAACTCTTTCCCTACACGACGCTCTTCCGATCTNNNTTGCAAATAAAGTCGTATAACA
CACACGTCTGAACTCTTTCCCTACACGACGCTCTTCCGATCTNNNAGAACACAAACTTTTTGACAATA
AGCACACGTCTGAACTCTTTCCCTACACGACGCTCTTCCGATCTNNNGAAAGGAATAGAACATGTCT

AGCACACGTCTGAACTCTTTCCCTACACGACGCTCTTCCGATCTNNNCTCTACCTATTCTTAGAGTGAC
AGCACACGTCTGAACTCTTTCCCTACACGACGCTCTTCCGATCTNNNGGGTGATTTTAATTTGGAGA
ACACGTCTGAACTCTTTCCCTACACGACGCTCTTCCGATCTNNNTTATTTCTGGTCAGAGATATTGTCA
CACACGTCTGAACTCTTTCCCTACACGACGCTCTTCCGATCTNNNCTCAGTTTATTTTCTCTCTTCTGCA
GCACACGTCTGAACTCTTTCCCTACACGACGCTCTTCCGATCTNNNCTATTTAGAAAATGGTCTTGGT
GCACACGTCTGAACTCTTTCCCTACACGACGCTCTTCCGATCTNNNTCAGATGATAATGTTTGACTACT
AGCACACGTCTGAACTCTTTCCCTACACGACGCTCTTCCGATCTNNNCATCTTAACTATAGCTTCAGGC
AGCACACGTCTGAACTCTTTCCCTACACGACGCTCTTCCGATCTNNNGTTGCAGTAAGTTGAGATTG
CACACGTCTGAACTCTTTCCCTACACGACGCTCTTCCGATCTNNNCCTACCTTCTATTTTTCTCCTGCATC
ACACGTCTGAACTCTTTCCCTACACGACGCTCTTCCGATCTNNNAGCTTCAAGTGAAAAAGAGATCAT
AGCACACGTCTGAACTCTTTCCCTACACGACGCTCTTCCGATCTNNNCAAAGTATTCAATTTCCCCATA
AGCACACGTCTGAACTCTTTCCCTACACGACGCTCTTCCGATCTNNNACATAGCATCTTAACACCTTTTC
CACACGTCTGAACTCTTTCCCTACACGACGCTCTTCCGATCTNNNCTAAGTACTATGAATCTCTGGCCA
AGCACACGTCTGAACTCTTTCCCTACACGACGCTCTTCCGATCTNNNAAAAAGGAAAGACAAGAAAAA
GAGCACACGTCTGAACTCTTTCCCTACACGACGCTCTTCCGATCTNNNATAAGATATTTGAAAGAGA
GAGCACACGTCTGAACTCTTTCCCTACACGACGCTCTTCCGATCTNNNATAAGATATTTGAAAGAGA
AGCACACGTCTGAACTCTTTCCCTACACGACGCTCTTCCGATCTNNNGTAAAACATGTTCACAAGGTC
AGCACACGTCTGAACTCTTTCCCTACACGACGCTCTTCCGATCTNNNTTTATATTGATGACATTGGTTG
ACACGTCTGAACTCTTTCCCTACACGACGCTCTTCCGATCTNNNAAAAAATAAATGCAAGCACTTTGC
GCACACGTCTGAACTCTTTCCCTACACGACGCTCTTCCGATCTNNNAGGAAAAGAAAAAAGAGTGAAT
AGCACACGTCTGAACTCTTTCCCTACACGACGCTCTTCCGATCTNNNAGCATTGATACAGACAAAATAA
CACACGTCTGAACTCTTTCCCTACACGACGCTCTTCCGATCTNNNCATACACACAGAAGAAAATATGT
CACACGTCTGAACTCTTTCCCTACACGACGCTCTTCCGATCTNNNATTTTAATCTTCAAGTTGTGGAGC
CACACGTCTGAACTCTTTCCCTACACGACGCTCTTCCGATCTNNNATTTTAATCTTCAAGTTGTGGAGC
ACACGTCTGAACTCTTTCCCTACACGACGCTCTTCCGATCTNNNCAAGAAATTGTTACTAAGAGCCCA
AGCACACGTCTGAACTCTTTCCCTACACGACGCTCTTCCGATCTNNNACAATGTGCATATTTAAACTGT
GCACACGTCTGAACTCTTTCCCTACACGACGCTCTTCCGATCTNNNACACAGTTCTGTTCCAAATATCA
AGCACACGTCTGAACTCTTTCCCTACACGACGCTCTTCCGATCTNNNCTGGAAATAAAAGTGTGGTAA
GCACACGTCTGAACTCTTTCCCTACACGACGCTCTTCCGATCTNNNGCTTCTCCTCATTCTACAATCCAT
GCACACGTCTGAACTCTTTCCCTACACGACGCTCTTCCGATCTNNNTTTAATAAGCCAGCTTAGAAAAC
GCACACGTCTGAACTCTTTCCCTACACGACGCTCTTCCGATCTNNNCCACAAAACTTGAATTTGTAAAC
AGCACACGTCTGAACTCTTTCCCTACACGACGCTCTTCCGATCTNNNTATAATTCTGGGTGTTCCTAA
CACGTCTGAACTCTTTCCCTACACGACGCTCTTCCGATCTNNNATGGATTCATGCCAATTATAAAGCAT
GCACACGTCTGAACTCTTTCCCTACACGACGCTCTTCCGATCTNNNATCTTCACAGATTATGCTGATGC
GCACACGTCTGAACTCTTTCCCTACACGACGCTCTTCCGATCTNNNAAATAAGGTTTTATGACTGTAGC
AGCACACGTCTGAACTCTTTCCCTACACGACGCTCTTCCGATCTNNNCTGACTTTCTCATGTAAAAACA
AGCACACGTCTGAACTCTTTCCCTACACGACGCTCTTCCGATCTNNNTGAACCAAATTGATATCTCTTT
AGCACACGTCTGAACTCTTTCCCTACACGACGCTCTTCCGATCTNNNTGAACCAAATTGATATCTCTTT
AGCACACGTCTGAACTCTTTCCCTACACGACGCTCTTCCGATCTNNNTCTGTGATATGTGTATATTCCT
GCACACGTCTGAACTCTTTCCCTACACGACGCTCTTCCGATCTNNNCTCGTTCCAGTCTGTTAGAGCAT
AGCACACGTCTGAACTCTTTCCCTACACGACGCTCTTCCGATCTNNNCTTGCTATTCTCATGATAGTGA
AGCACACGTCTGAACTCTTTCCCTACACGACGCTCTTCCGATCTNNNAATGCTTAGCTGTTTTTTAGAA
GCACACGTCTGAACTCTTTCCCTACACGACGCTCTTCCGATCTNNNTGCAAATACTGAGAGTAAATAGT

CACACGTCTGAACTCTTTCCCTACACGACGCTCTTCCGATCTNNNTTTTAGTCTGAGTCTGAAGGACA
AGCACACGTCTGAACTCTTTCCCTACACGACGCTCTTCCGATCTNNNGTAAATGTTTTCTTCAAAAGC
AGCACACGTCTGAACTCTTTCCCTACACGACGCTCTTCCGATCTNNNGAAATGAGGAATGTCATTGTT
GCACACGTCTGAACTCTTTCCCTACACGACGCTCTTCCGATCTNNNAACTTTTTAGAACCTGTTTCTC
GCACACGTCTGAACTCTTTCCCTACACGACGCTCTTCCGATCTNNNTTGCCATACTAGGATTATCTCTA
CACACGTCTGAACTCTTTCCCTACACGACGCTCTTCCGATCTNNNAAACACAAAACTGAAAATTAGTG
CACGTCTGAACTCTTTCCCTACACGACGCTCTTCCGATCTNNNTCAACATTTATTAACTGGACTTCCAT
AGCACACGTCTGAACTCTTTCCCTACACGACGCTCTTCCGATCTNNNTATCCGTTTCTAATAATCCAAG
AGCACACGTCTGAACTCTTTCCCTACACGACGCTCTTCCGATCTNNNGTATGAGCATGTGTATATGTC
AGCACACGTCTGAACTCTTTCCCTACACGACGCTCTTCCGATCTNNNTTATTACTACCTCTTGGAAGAT
ACACGTCTGAACTCTTTCCCTACACGACGCTCTTCCGATCTNNNTTTTCCTCTCTTCTATCTTTCCCATC
GCACACGTCTGAACTCTTTCCCTACACGACGCTCTTCCGATCTNNNAATATGTATGAAACAGAAAGC
GAGCACACGTCTGAACTCTTTCCCTACACGACGCTCTTCCGATCTNNNACAGAAAGTGTCAATAATAG
AGCACACGTCTGAACTCTTTCCCTACACGACGCTCTTCCGATCTNNNCCTTCTAGATTCCAAGATTTT
GAGCACACGTCTGAACTCTTTCCCTACACGACGCTCTTCCGATCTNNNGGGTTCTTTCATTGAACTTTT
GCACACGTCTGAACTCTTTCCCTACACGACGCTCTTCCGATCTNNNAACATGGCATATAATAAACATT
AGCACACGTCTGAACTCTTTCCCTACACGACGCTCTTCCGATCTNNNTGACACACCATTTCTGAAAGC
GCACACGTCTGAACTCTTTCCCTACACGACGCTCTTCCGATCTNNNTTTCTAGAAAACTCACACATAA
AGCACACGTCTGAACTCTTTCCCTACACGACGCTCTTCCGATCTNNNCATCTAGACCTCGGATTCTCCA
GCACACGTCTGAACTCTTTCCCTACACGACGCTCTTCCGATCTNNNGCAAGTACCTTACAATAACAAA
ACACGTCTGAACTCTTTCCCTACACGACGCTCTTCCGATCTNNNCTTAATGAGCATATTCCTTTCCCATC
AGCACACGTCTGAACTCTTTCCCTACACGACGCTCTTCCGATCTNNNAATTTTGTCTTTTAATTCCCCAA
GAGCACACGTCTGAACTCTTTCCCTACACGACGCTCTTCCGATCTNNNCAGGTAAATCTGAGAAATGT
CACACGTCTGAACTCTTTCCCTACACGACGCTCTTCCGATCTNNNGAAGACCCTTACTTACGATGCAT
AGCACACGTCTGAACTCTTTCCCTACACGACGCTCTTCCGATCTNNNATTCGTAATAGTCAGATATG
GCACACGTCTGAACTCTTTCCCTACACGACGCTCTTCCGATCTNNNCTGGATTTTTTAATACAGAGGT
GCACACGTCTGAACTCTTTCCCTACACGACGCTCTTCCGATCTNNNTTGTTCAATTTAAATCTTGAGG
AGCACACGTCTGAACTCTTTCCCTACACGACGCTCTTCCGATCTNNNTAAAGCTTATACAAGGAAACA
AGCACACGTCTGAACTCTTTCCCTACACGACGCTCTTCCGATCTNNNATCTTTCTTTACAGTGACAGTC
CACGTCTGAACTCTTTCCCTACACGACGCTCTTCCGATCTNNNAAATCAATCCCTCTCCTCACCATCGG
GCACACGTCTGAACTCTTTCCCTACACGACGCTCTTCCGATCTNNNATTTGTTTTATTCTCTTGGATCCC
GCACACGTCTGAACTCTTTCCCTACACGACGCTCTTCCGATCTNNNAACACTTGGTACATAACATGTA
AGCACACGTCTGAACTCTTTCCCTACACGACGCTCTTCCGATCTNNNACTGTAATTCATTTAACCAATC
GAGCACACGTCTGAACTCTTTCCCTACACGACGCTCTTCCGATCTNNNCCTTTCATTCTGTTTTGTAG
CACACGTCTGAACTCTTTCCCTACACGACGCTCTTCCGATCTNNNCTTCATCCTCAAACCTTGTTTCAT
GCACACGTCTGAACTCTTTCCCTACACGACGCTCTTCCGATCTNNNCTATCATTTCAATGTAACACCAA
AGCACACGTCTGAACTCTTTCCCTACACGACGCTCTTCCGATCTNNNAGTCTTCAATGTCAGAACTAC
GCACACGTCTGAACTCTTTCCCTACACGACGCTCTTCCGATCTNNNATATCTAGTTCTTCACATTGGAG
GAGCACACGTCTGAACTCTTTCCCTACACGACGCTCTTCCGATCTNNNGTAAAAATGTACTGAAAGAA
AGCACACGTCTGAACTCTTTCCCTACACGACGCTCTTCCGATCTNNNATGTACATTACCACAGTTATAC
GCACACGTCTGAACTCTTTCCCTACACGACGCTCTTCCGATCTNNNAGTACTATGAAATTCAGAATAA
GCACACGTCTGAACTCTTTCCCTACACGACGCTCTTCCGATCTNNNCTTTATCTTCACACAGCTTCTCA
AGCACACGTCTGAACTCTTTCCCTACACGACGCTCTTCCGATCTNNNGAAAGGAAAGTATTGCTATTC

AGCACACGTCTGAACTCTTTCCCTACACGACGCTCTTCCGATCTNNNTGTTTGGGTTTATTTTTTCTTCT
CACACGTCTGAACTCTTTCCCTACACGACGCTCTTCCGATCTNNNCATGAGACTAAGGAAGCTCCCATC
AGCACACGTCTGAACTCTTTCCCTACACGACGCTCTTCCGATCTNNNAAGGAAGAAAAAAAACCAAAA
AGCACACGTCTGAACTCTTTCCCTACACGACGCTCTTCCGATCTNNNTTAACAGGATTAAACTCATCTC
CACACGTCTGAACTCTTTCCCTACACGACGCTCTTCCGATCTNNNCTTTGTCCCTTGCAAAACATCATCGC
AGCACACGTCTGAACTCTTTCCCTACACGACGCTCTTCCGATCTNNNTAGTTTTAACTAGTTCCATGAAC
GAGCACACGTCTGAACTCTTTCCCTACACGACGCTCTTCCGATCTNNNCAGGAATTTGAAGTTACAGTC
AGCACACGTCTGAACTCTTTCCCTACACGACGCTCTTCCGATCTNNNGTGATACTGAAATCTAAGAGA
AGCACACGTCTGAACTCTTTCCCTACACGACGCTCTTCCGATCTNNNATCTCTAAAACTCATTTGCAATC
ACACGTCTGAACTCTTTCCCTACACGACGCTCTTCCGATCTNNNTTTCTGAAATCTTCTAATTGCTCCATC
AGCACACGTCTGAACTCTTTCCCTACACGACGCTCTTCCGATCTNNNATTATTCTTTCCCATTGTCCTTTC
AGCACACGTCTGAACTCTTTCCCTACACGACGCTCTTCCGATCTNNNATTATTCTTTCCCATTGTCCTTTC
AGCACACGTCTGAACTCTTTCCCTACACGACGCTCTTCCGATCTNNNAATAAACTTCCTTTTTTCTTGGTC
CACACGTCTGAACTCTTTCCCTACACGACGCTCTTCCGATCTNNNCTAACACACCCAACCTTATAACATCC
AGCACACGTCTGAACTCTTTCCCTACACGACGCTCTTCCGATCTNNNAGTAAATTCCCTAAGAAGAACA
CACACGTCTGAACTCTTTCCCTACACGACGCTCTTCCGATCTNNNTTATTCAGATTTCAGCGGTAATACA
AGCACACGTCTGAACTCTTTCCCTACACGACGCTCTTCCGATCTNNNAACTCTGGTTATAACAGTGTAT
AGCACACGTCTGAACTCTTTCCCTACACGACGCTCTTCCGATCTNNNATTCACATTTATACAGAGCCAAA
GCACACGTCTGAACTCTTTCCCTACACGACGCTCTTCCGATCTNNNCTTAGCCTTTTGATACTCTTACTCA
GAGCACACGTCTGAACTCTTTCCCTACACGACGCTCTTCCGATCTNNNCATAGGAATGAAACCTCAAAT
GCACACGTCTGAACTCTTTCCCTACACGACGCTCTTCCGATCTNNNGGATAAAGCAGGACACTAAGCA
GCACACGTCTGAACTCTTTCCCTACACGACGCTCTTCCGATCTNNNGGATAAAGCAGGACACTAAGCA
CACACGTCTGAACTCTTTCCCTACACGACGCTCTTCCGATCTNNNTGTCAATATTACTAACCTCTTGTCA
AGCACACGTCTGAACTCTTTCCCTACACGACGCTCTTCCGATCTNNNATATCTATTTTGCATTTCCTCTCC
GCACACGTCTGAACTCTTTCCCTACACGACGCTCTTCCGATCTNNNTACTTAACATTCAGGATCTCATTCA
AGCACACGTCTGAACTCTTTCCCTACACGACGCTCTTCCGATCTNNNAAACCTATTCTCCAAATAAAGTG
GCACACGTCTGAACTCTTTCCCTACACGACGCTCTTCCGATCTNNNTGTAAAGCCTTCTCTGATTGCAT
GAGCACACGTCTGAACTCTTTCCCTACACGACGCTCTTCCGATCTNNNATAAAGATTACAACATTTTCAC
ACACGTCTGAACTCTTTCCCTACACGACGCTCTTCCGATCTNNNAGACTTTTGTGTGAACATAATTTCAT
GAGCACACGTCTGAACTCTTTCCCTACACGACGCTCTTCCGATCTNNNCTTGTCTGATTTGCTCATCTCA
AGCACACGTCTGAACTCTTTCCCTACACGACGCTCTTCCGATCTNNNTTTTTGTAATGAAGAGACAAA
AGCACACGTCTGAACTCTTTCCCTACACGACGCTCTTCCGATCTNNNTATCTGATAAACCACCAGAATG
AGCACACGTCTGAACTCTTTCCCTACACGACGCTCTTCCGATCTNNNGACTGTCCCTTTTCTCTACTCAT
GCACACGTCTGAACTCTTTCCCTACACGACGCTCTTCCGATCTNNNCACCATTTTGCAACAATTAAATAC
AGCACACGTCTGAACTCTTTCCCTACACGACGCTCTTCCGATCTNNNTCCCTTTTGAACTTGTGATATAT
GAGCACACGTCTGAACTCTTTCCCTACACGACGCTCTTCCGATCTNNNAGTTATTAGACAAATCCCGTT
AGCACACGTCTGAACTCTTTCCCTACACGACGCTCTTCCGATCTNNNTACATAGTGTCCATTATCAGTG
AGCACACGTCTGAACTCTTTCCCTACACGACGCTCTTCCGATCTNNNTACATAGTGTCCATTATCAGTG
AGCACACGTCTGAACTCTTTCCCTACACGACGCTCTTCCGATCTNNNCTTGTTATATGAGCAATAAACC
ACACGTCTGAACTCTTTCCCTACACGACGCTCTTCCGATCTNNNTTGTACGTTCCCAGATTCTTCATCGGA
AGCACACGTCTGAACTCTTTCCCTACACGACGCTCTTCCGATCTNNNTTATAGCATGTTCACATCTCTCC
AGCACACGTCTGAACTCTTTCCCTACACGACGCTCTTCCGATCTNNNGTTCTATTCTTGCTGTTCATGCA
GAGCACACGTCTGAACTCTTTCCCTACACGACGCTCTTCCGATCTNNNCTCTGCCTCTTCTCCATAACCA

AGCACACGTCTGAACTCTTTCCCTACACGACGCTCTTCCGATCTNNNGATTTTGACACTTACTCAGAAT
ACACGTCTGAACTCTTTCCCTACACGACGCTCTTCCGATCTNNNAAGTGAAGTGTGGTAGTCAGCATC
AGCACACGTCTGAACTCTTTCCCTACACGACGCTCTTCCGATCTNNNTTTATGCATCTCAGAACAAAA
ACACGTCTGAACTCTTTCCCTACACGACGCTCTTCCGATCTNNNTCTCTCAGAGCCTTTATTTTTTACAT
AGCACACGTCTGAACTCTTTCCCTACACGACGCTCTTCCGATCTNNNGAAAGAAGTGACATTTTTGTC
GCACACGTCTGAACTCTTTCCCTACACGACGCTCTTCCGATCTNNNTGTCATTTTCTATTTTGATTCCA
CACACGTCTGAACTCTTTCCCTACACGACGCTCTTCCGATCTNNNTAATACCGTCTCCCAATGTATACAT
AGCACACGTCTGAACTCTTTCCCTACACGACGCTCTTCCGATCTNNNAAAACTGTTTCTTAAAACTGA
GCACACGTCTGAACTCTTTCCCTACACGACGCTCTTCCGATCTNNNCATTGTAGGGAAATACTCTAAG
GCACACGTCTGAACTCTTTCCCTACACGACGCTCTTCCGATCTNNNCATTGTAGGGAAATACTCTAAG
AGCACACGTCTGAACTCTTTCCCTACACGACGCTCTTCCGATCTNNNTAATCTAGTATCTAGCATGCA
CACACGTCTGAACTCTTTCCCTACACGACGCTCTTCCGATCTNNNTCTTTCTCTTTTTTCTCTTTCTCCAT
CACGTCTGAACTCTTTCCCTACACGACGCTCTTCCGATCTNNNTATGAGCACTGTGACTTTGACATCGG
GCACACGTCTGAACTCTTTCCCTACACGACGCTCTTCCGATCTNNNCATTTGTGTACATATGCTCTATG
GAGCACACGTCTGAACTCTTTCCCTACACGACGCTCTTCCGATCTNNNTATATAAGAATGTGGCAAAA
GAGCACACGTCTGAACTCTTTCCCTACACGACGCTCTTCCGATCTNNNCTGAGACATTGAAATTATTT
GCACACGTCTGAACTCTTTCCCTACACGACGCTCTTCCGATCTNNNAACATTTCTTCTCTTTTTCCTTGC
AGCACACGTCTGAACTCTTTCCCTACACGACGCTCTTCCGATCTNNNCCATACAACAGATGCAAAATAT
AGCACACGTCTGAACTCTTTCCCTACACGACGCTCTTCCGATCTNNNATTACACATACAAACATACAC
GCACACGTCTGAACTCTTTCCCTACACGACGCTCTTCCGATCTNNNGTGTAATGAAAGTTAAGATGA
GCACACGTCTGAACTCTTTCCCTACACGACGCTCTTCCGATCTNNNAATGAAAGCATTGCTCAGTCCA
GCACACGTCTGAACTCTTTCCCTACACGACGCTCTTCCGATCTNNNGGAAGCAGAAATCTGTTATATG
AGCACACGTCTGAACTCTTTCCCTACACGACGCTCTTCCGATCTNNNATGAACAAAAAGAATGAGGA
GAGCACACGTCTGAACTCTTTCCCTACACGACGCTCTTCCGATCTNNNCAACAACAAAAAAAGTCAAT
AGCACACGTCTGAACTCTTTCCCTACACGACGCTCTTCCGATCTNNNACATTTCTGTTCTTTTAAACCA
AGCACACGTCTGAACTCTTTCCCTACACGACGCTCTTCCGATCTNNNTAAATCTGAACTGTGACCAAG
CACACGTCTGAACTCTTTCCCTACACGACGCTCTTCCGATCTNNNCTTCATCCTCAAACCTTGTTTCAT
GCACACGTCTGAACTCTTTCCCTACACGACGCTCTTCCGATCTNNNAAGTAAGCTATACTTGTACTCT
ACACGTCTGAACTCTTTCCCTACACGACGCTCTTCCGATCTNNNCTTATGCTCCAGATGAAATAAAAC
CACACGTCTGAACTCTTTCCCTACACGACGCTCTTCCGATCTNNNAGTAAATGAGAGGAAGTGATGCA
GCACACGTCTGAACTCTTTCCCTACACGACGCTCTTCCGATCTNNNACCAACCTCAGTGAAAAATCCA
GCACACGTCTGAACTCTTTCCCTACACGACGCTCTTCCGATCTNNNCATTTTACAACATTTGCTACAG
GCACACGTCTGAACTCTTTCCCTACACGACGCTCTTCCGATCTNNNTTATCTGAATGTGATTATCATG
GCACACGTCTGAACTCTTTCCCTACACGACGCTCTTCCGATCTNNNAGATCAACTTGAATTTATACTG
GCACACGTCTGAACTCTTTCCCTACACGACGCTCTTCCGATCTNNNCTTAAGAAAGGGTCTTTGGTCC
AGCACACGTCTGAACTCTTTCCCTACACGACGCTCTTCCGATCTNNNTTCAATGACAATGACAGTTTTA
GCACACGTCTGAACTCTTTCCCTACACGACGCTCTTCCGATCTNNNATATTATGTCTTCATTTGGGTGA
CACACGTCTGAACTCTTTCCCTACACGACGCTCTTCCGATCTNNNTATTCTCTGCAGTGTCCTTTCATCG
ACACGTCTGAACTCTTTCCCTACACGACGCTCTTCCGATCTNNNTAATTTCCACAGCATAAATTACCCAT
AGCACACGTCTGAACTCTTTCCCTACACGACGCTCTTCCGATCTNNNGGCATAAGAAACACATTCATTC
AGCACACGTCTGAACTCTTTCCCTACACGACGCTCTTCCGATCTNNNCTCTTATACTGAACATCGTAGA
GCACACGTCTGAACTCTTTCCCTACACGACGCTCTTCCGATCTNNNTAACATCTTGCATAATCAAAGTG
GAGCACACGTCTGAACTCTTTCCCTACACGACGCTCTTCCGATCTNNNAAAATGCCTAAAGTAAACCTA

AGCACACGTCTGAACTCTTTCCCTACACGACGCTCTTCCGATCTNNNCAAAGTCATCTAGAATATCCAG
CACACGTCTGAACTCTTTCCCTACACGACGCTCTTCCGATCTNNNTATACTACATTTAGAACAGCAGT
GCACACGTCTGAACTCTTTCCCTACACGACGCTCTTCCGATCTNNNTTAAATATGCATAAAACATGGC
AGAGCACACGTCTGAACTCTTTCCCTACACGACGCTCTTCCGATCTNNNGATCATTAGATTTGACAACC
AGCACACGTCTGAACTCTTTCCCTACACGACGCTCTTCCGATCTNNNTATTTACTGCCATAAGTGCAA
GAGCACACGTCTGAACTCTTTCCCTACACGACGCTCTTCCGATCTNNNTTCGGGAGATAAATGTAATC
AGCACACGTCTGAACTCTTTCCCTACACGACGCTCTTCCGATCTNNNAGTTACCATTTTAAGGTGATT
GAGCACACGTCTGAACTCTTTCCCTACACGACGCTCTTCCGATCTNNNGGACTTGGTTTATTTCTACAA
CACACGTCTGAACTCTTTCCCTACACGACGCTCTTCCGATCTNNNGTCTCTCTCTCTCTCTTGCATCG
CACACGTCTGAACTCTTTCCCTACACGACGCTCTTCCGATCTNNNGTCTCTCTCTCTCTCTTGCATCG
AGCACACGTCTGAACTCTTTCCCTACACGACGCTCTTCCGATCTNNNTTACAATAAATTTCTTCACACGT
GCACACGTCTGAACTCTTTCCCTACACGACGCTCTTCCGATCTNNNAAGATTCTTTCATGCTTACTTTG
AGCACACGTCTGAACTCTTTCCCTACACGACGCTCTTCCGATCTNNNGTTGAAATTTGACCCCTAAATA
AGCACACGTCTGAACTCTTTCCCTACACGACGCTCTTCCGATCTNNNCACACACTAATCTCCTAGAAGC
GAGCACACGTCTGAACTCTTTCCCTACACGACGCTCTTCCGATCTNNNGAAGGTAGACAGGAAAATA
GAGCACACGTCTGAACTCTTTCCCTACACGACGCTCTTCCGATCTNNNGAAGGTAGACAGGAAAATA
AGCACACGTCTGAACTCTTTCCCTACACGACGCTCTTCCGATCTNNNCTCCCTGCTTCTAATATATACAT
GCACACGTCTGAACTCTTTCCCTACACGACGCTCTTCCGATCTNNNTCCATTACTTATATTCAAAGCTGC
AGCACACGTCTGAACTCTTTCCCTACACGACGCTCTTCCGATCTNNNTGAAACTATTCCAAACAATTG
GCACACGTCTGAACTCTTTCCCTACACGACGCTCTTCCGATCTNNNATATATACACAACACAGATCAC
GCACACGTCTGAACTCTTTCCCTACACGACGCTCTTCCGATCTNNNATATATACACAACACAGATCAC
GCACACGTCTGAACTCTTTCCCTACACGACGCTCTTCCGATCTNNNATTCTTATGTTTTACATGATGGG
GCACACGTCTGAACTCTTTCCCTACACGACGCTCTTCCGATCTNNNGTTTCCAAGCTATAAGTAAAGA
CACACGTCTGAACTCTTTCCCTACACGACGCTCTTCCGATCTNNNTAGAAAATACATGCACAGCACATC
CACACGTCTGAACTCTTTCCCTACACGACGCTCTTCCGATCTNNNTAAACAGAGTTCAATCATAGAGG
GAGCACACGTCTGAACTCTTTCCCTACACGACGCTCTTCCGATCTNNNAATTATTCTGGAAGTGATGG
AGCACACGTCTGAACTCTTTCCCTACACGACGCTCTTCCGATCTNNNGTTGCCTTTCATCTCTTTAACC
GAGCACACGTCTGAACTCTTTCCCTACACGACGCTCTTCCGATCTNNNGAAAACTGGCAAAAAGGTAA
GCACACGTCTGAACTCTTTCCCTACACGACGCTCTTCCGATCTNNNGAGAACCAGAACCTAGAGAGC
GCACACGTCTGAACTCTTTCCCTACACGACGCTCTTCCGATCTNNNGAGAACCAGAACCTAGAGAGC
GCACACGTCTGAACTCTTTCCCTACACGACGCTCTTCCGATCTNNNTATTTAAAGGTTAAACTTGTGG
GCACACGTCTGAACTCTTTCCCTACACGACGCTCTTCCGATCTNNNTATTTAAAGGTTAAACTTGTGG
GCACACGTCTGAACTCTTTCCCTACACGACGCTCTTCCGATCTNNNTATTTAAAGGTTAAACTTGTGG
AGCACACGTCTGAACTCTTTCCCTACACGACGCTCTTCCGATCTNNNAAAGGGAATAACATCAAAAAC
AGCACACGTCTGAACTCTTTCCCTACACGACGCTCTTCCGATCTNNNTTAAGCCTTTCTTTTATTCAAC
AGAGCACACGTCTGAACTCTTTCCCTACACGACGCTCTTCCGATCTNNNCTCCAGTAGTGCCCTTTATC
GCACACGTCTGAACTCTTTCCCTACACGACGCTCTTCCGATCTNNNAGTCTGATATAGAACTTGTTAG
GCACACGTCTGAACTCTTTCCCTACACGACGCTCTTCCGATCTNNNCTGTAAGGATGTGAGTGAATAT
GCACACGTCTGAACTCTTTCCCTACACGACGCTCTTCCGATCTNNNTAGAACTTACAATCTAGTTGGA
GCACACGTCTGAACTCTTTCCCTACACGACGCTCTTCCGATCTNNNAACTTTTCCTCCTTCCTTTTTCATC
CACACGTCTGAACTCTTTCCCTACACGACGCTCTTCCGATCTNNNGAAATTTCACAAAGCTCTTACTCA
AGCACACGTCTGAACTCTTTCCCTACACGACGCTCTTCCGATCTNNNGAAGACTGAAATCAATAGATA
AGCACACGTCTGAACTCTTTCCCTACACGACGCTCTTCCGATCTNNNTACTTATCAGGATAACAGTTC

AGCACACGTCTGAACTCTTTCCCTACACGACGCTCTTCCGATCTNNNTGACTATTTTTACAACTCAGA
AGCACACGTCTGAACTCTTTCCCTACACGACGCTCTTCCGATCTNNNATTTAGGATGTGCACCTTGCA
GAGCACACGTCTGAACTCTTTCCCTACACGACGCTCTTCCGATCTNNNTTGACAGGAAGAGACCAAAT
GAGCACACGTCTGAACTCTTTCCCTACACGACGCTCTTCCGATCTNNNCCGGCCTAAGATTTATTTAT
AGCACACGTCTGAACTCTTTCCCTACACGACGCTCTTCCGATCTNNNATCACCTGAGCAGTTTCTACCAT
GCACACGTCTGAACTCTTTCCCTACACGACGCTCTTCCGATCTNNNTTACAGCTACTCAGGAAACTCAT
GCACACGTCTGAACTCTTTCCCTACACGACGCTCTTCCGATCTNNNCTAATAAATGTTTCAAGCTGCTAC
AGCACACGTCTGAACTCTTTCCCTACACGACGCTCTTCCGATCTNNNGAAACTAAACTGAATGGCATAA
GAGCACACGTCTGAACTCTTTCCCTACACGACGCTCTTCCGATCTNNNGTGGTCCCACTAATAGGATGC
GAGCACACGTCTGAACTCTTTCCCTACACGACGCTCTTCCGATCTNNNAAAAACACATGATACAAAAA
AGAGCACACGTCTGAACTCTTTCCCTACACGACGCTCTTCCGATCTNNNGAATAGAAACCCTACAAGTA
AGCACACGTCTGAACTCTTTCCCTACACGACGCTCTTCCGATCTNNNTTATAAGAGCAACATCTTAGACA
AGCACACGTCTGAACTCTTTCCCTACACGACGCTCTTCCGATCTNNNTTGTAGATCACTAAGGTTTTTT
AGCACACGTCTGAACTCTTTCCCTACACGACGCTCTTCCGATCTNNNATAGTTGATTTTGTTTGTTCTGT
GCACACGTCTGAACTCTTTCCCTACACGACGCTCTTCCGATCTNNNCTCTTGAATCCTTAATAGGGTTA
AGCACACGTCTGAACTCTTTCCCTACACGACGCTCTTCCGATCTNNNATAGCTAATTGAGAAGCTTTAG
CACACGTCTGAACTCTTTCCCTACACGACGCTCTTCCGATCTNNNCAGAGCGGAATGATTTTTTTAAC
AGCACACGTCTGAACTCTTTCCCTACACGACGCTCTTCCGATCTNNNACATTTTATTATTTTTTTACGGC
AGAGCACACGTCTGAACTCTTTCCCTACACGACGCTCTTCCGATCTNNNATATAACCGATATCTGTTCAC
GAGCACACGTCTGAACTCTTTCCCTACACGACGCTCTTCCGATCTNNNTATTTGAGATCCACACAAACTT
CACACGTCTGAACTCTTTCCCTACACGACGCTCTTCCGATCTNNNAATTCACTGTGGCTATAGAAGCAT
AGCACACGTCTGAACTCTTTCCCTACACGACGCTCTTCCGATCTNNNAGTGTGATTATGCAAAGAGAG
GCACACGTCTGAACTCTTTCCCTACACGACGCTCTTCCGATCTNNNTGATAGTTTTGTTTGTTCTTTCCCA
GAGCACACGTCTGAACTCTTTCCCTACACGACGCTCTTCCGATCTNNNCTGGACATTTTTTTGAGATCTA
GCACACGTCTGAACTCTTTCCCTACACGACGCTCTTCCGATCTNNNCATGGGAGATAGTGTCATTACCA
CACGTCTGAACTCTTTCCCTACACGACGCTCTTCCGATCTNNNCTCAATTCCTCATTCTCTGGCATCGGA
GCACACGTCTGAACTCTTTCCCTACACGACGCTCTTCCGATCTNNNTTCTTACCAGAAAGTCTTTTGGCA
AGCACACGTCTGAACTCTTTCCCTACACGACGCTCTTCCGATCTNNNCCACTAAATGAAACAAGTCAAT
AGCACACGTCTGAACTCTTTCCCTACACGACGCTCTTCCGATCTNNNCATTTAGAAAACTCTTCTTCAGA
GAGCACACGTCTGAACTCTTTCCCTACACGACGCTCTTCCGATCTNNNTATGATGAGTAGAATTATGG
AGCACACGTCTGAACTCTTTCCCTACACGACGCTCTTCCGATCTNNNTTTACTGAGAGCTTCATACAAAA
AGCACACGTCTGAACTCTTTCCCTACACGACGCTCTTCCGATCTNNNTTTACTGAGAGCTTCATACAAAA
AGCACACGTCTGAACTCTTTCCCTACACGACGCTCTTCCGATCTNNNTTCTGGAAATTCTCTTATTCTCT
AGCACACGTCTGAACTCTTTCCCTACACGACGCTCTTCCGATCTNNNTTCTGGAAATTCTCTTATTCTCT
AGCACACGTCTGAACTCTTTCCCTACACGACGCTCTTCCGATCTNNNCATATTTATAGCAGCAAAATTC
AGCACACGTCTGAACTCTTTCCCTACACGACGCTCTTCCGATCTNNNGTTTATAATTCAAAAGACTTGG
GCACACGTCTGAACTCTTTCCCTACACGACGCTCTTCCGATCTNNNCCTCTATTTGTTTGAGTGTATAGC
AGCACACGTCTGAACTCTTTCCCTACACGACGCTCTTCCGATCTNNNTACTAAACTGCAGTATTTCTCAC
AGCACACGTCTGAACTCTTTCCCTACACGACGCTCTTCCGATCTNNNTACTAAACTGCAGTATTTCTCAC
AGCACACGTCTGAACTCTTTCCCTACACGACGCTCTTCCGATCTNNNCATTAGAAGGAAGAAACAAGT
GAGCACACGTCTGAACTCTTTCCCTACACGACGCTCTTCCGATCTNNNGGAGATATTTGGTGAAAAATC
AGCACACGTCTGAACTCTTTCCCTACACGACGCTCTTCCGATCTNNNAGTATTCACTAGCTAATTCTTGG
AGCACACGTCTGAACTCTTTCCCTACACGACGCTCTTCCGATCTNNNATACATAATGGAAGTTACACAA

CACACGTCTGAACTCTTTCCCTACACGACGCTCTTCCGATCTNNNATTCTACAATTCTATTCATGGGTCAT
AGCACACGTCTGAACTCTTTCCCTACACGACGCTCTTCCGATCTNNNCTTAAAATCTTGCAATGTACACT
AGCACACGTCTGAACTCTTTCCCTACACGACGCTCTTCCGATCTNNNTAATCTCAGCCAATCCTTACTCA
AGCACACGTCTGAACTCTTTCCCTACACGACGCTCTTCCGATCTNNNGTATTCATCCTTCATCCATCATCA
AGCACACGTCTGAACTCTTTCCCTACACGACGCTCTTCCGATCTNNNTGTGTGTGTGTGTATATATA
AGCACACGTCTGAACTCTTTCCCTACACGACGCTCTTCCGATCTNNNTGAAGCAGAATAGAATAAAACA
GAGCACACGTCTGAACTCTTTCCCTACACGACGCTCTTCCGATCTNNNTTCCTAGTGTTCTTTTATGTTTC
GAGCACACGTCTGAACTCTTTCCCTACACGACGCTCTTCCGATCTNNNATGGTATACTAAGTACTGGAC
GCACACGTCTGAACTCTTTCCCTACACGACGCTCTTCCGATCTNNNTGTTTCTGGCAGATAAAACACAT
CACACGTCTGAACTCTTTCCCTACACGACGCTCTTCCGATCTNNNAAGGTTTATTTCAGATGCCAACATC
AGCACACGTCTGAACTCTTTCCCTACACGACGCTCTTCCGATCTNNNAAATTCACCCTCTATTCACTATT
GCACACGTCTGAACTCTTTCCCTACACGACGCTCTTCCGATCTNNNCAACAAAACAGATTACATCCTGCA
AGCACACGTCTGAACTCTTTCCCTACACGACGCTCTTCCGATCTNNNACTGTTTTCATCAGTTTTCATGCA
AGCACACGTCTGAACTCTTTCCCTACACGACGCTCTTCCGATCTNNNAAGTCAAGATATAGAATCAACC
GAGCACACGTCTGAACTCTTTCCCTACACGACGCTCTTCCGATCTNNNTCCACATATAATTGACTTCATG
AGCACACGTCTGAACTCTTTCCCTACACGACGCTCTTCCGATCTNNNTCTTAGAAAACCACTGTTACATA
GAGCACACGTCTGAACTCTTTCCCTACACGACGCTCTTCCGATCTNNNCAGGAATTCACATGAGAGAC
GAGCACACGTCTGAACTCTTTCCCTACACGACGCTCTTCCGATCTNNNGACACACATATGCACATGTAT
AGCACACGTCTGAACTCTTTCCCTACACGACGCTCTTCCGATCTNNNAGCCTCCATAATCTCATGAGCAT
GAGCACACGTCTGAACTCTTTCCCTACACGACGCTCTTCCGATCTNNNACAATCATTTTACATTGTGTTA
AGCACACGTCTGAACTCTTTCCCTACACGACGCTCTTCCGATCTNNNCAGTTCTCTTTACTTTCCAATTTC
GAGCACACGTCTGAACTCTTTCCCTACACGACGCTCTTCCGATCTNNNTGTGAAAAAAAAAAGGAGTAT
AGCACACGTCTGAACTCTTTCCCTACACGACGCTCTTCCGATCTNNNTATTCAAATTGTTTTAGCTCTGG
AGCACACGTCTGAACTCTTTCCCTACACGACGCTCTTCCGATCTNNNTGTGTGTGTATGTACATATA
GCACACGTCTGAACTCTTTCCCTACACGACGCTCTTCCGATCTNNNTTTCCTCGATTTTTTCAACTTTCAT
GCACACGTCTGAACTCTTTCCCTACACGACGCTCTTCCGATCTNNNTTTATCTTGTTTCCTCGATTTTTCAT
AGCACACGTCTGAACTCTTTCCCTACACGACGCTCTTCCGATCTNNNAACCTGATCAAAAATTTAGGTA
GAGCACACGTCTGAACTCTTTCCCTACACGACGCTCTTCCGATCTNNNTGTTTCTACCAAGATGATGTAT
GAGCACACGTCTGAACTCTTTCCCTACACGACGCTCTTCCGATCTNNNTAACCCCCCAATATACATTTCTA
AGCACACGTCTGAACTCTTTCCCTACACGACGCTCTTCCGATCTNNNTATCCAACATATATGTGTGTGTA
CACACGTCTGAACTCTTTCCCTACACGACGCTCTTCCGATCTNNNATGGCACGAATAAGATTCTTTCATC
AGCACACGTCTGAACTCTTTCCCTACACGACGCTCTTCCGATCTNNNATGTATTTTTATTTCTCTGGCTG
AGCACACGTCTGAACTCTTTCCCTACACGACGCTCTTCCGATCTNNNAGGTATTTCAAATGGAAGTATC
GAGCACACGTCTGAACTCTTTCCCTACACGACGCTCTTCCGATCTNNNTGCTATGAAAGAAATATTCCCA
GCACACGTCTGAACTCTTTCCCTACACGACGCTCTTCCGATCTNNNCTCAAAGAAATAATGTACCCAAA
AGCACACGTCTGAACTCTTTCCCTACACGACGCTCTTCCGATCTNNNTTTGGAAGACAAAAGAAAATG
GAGCACACGTCTGAACTCTTTCCCTACACGACGCTCTTCCGATCTNNNAGTGAAGACTTGATTCTATTG
AGAGCACACGTCTGAACTCTTTCCCTACACGACGCTCTTCCGATCTNNNAACTACTGCTCTAGTATGGA
AGCACACGTCTGAACTCTTTCCCTACACGACGCTCTTCCGATCTNNNTTGCCAGAATTTATTTTCAGTTAC
GCACACGTCTGAACTCTTTCCCTACACGACGCTCTTCCGATCTNNNTCCTTTTTTGTCTTTTCACTTTACA
GCACACGTCTGAACTCTTTCCCTACACGACGCTCTTCCGATCTNNNACTAGGGAGATACATTTTTAGAC
AGCACACGTCTGAACTCTTTCCCTACACGACGCTCTTCCGATCTNNNATACTTTCATTTGAGAAGGAGA
GAGCACACGTCTGAACTCTTTCCCTACACGACGCTCTTCCGATCTNNNGATGACCCAAAAGAAAAAAAA

AGCACACGTCTGAACTCTTTCCCTACACGACGCTCTTCCGATCTNNNTTTACTTAGAATAATGGTCTCCA

GAGCACACGTCTGAACTCTTTCCCTACACGACGCTCTTCCGATCTNNNAAAACAGCATACAATTTTTTTC

AGCACACGTCTGAACTCTTTCCCTACACGACGCTCTTCCGATCTNNNGTTACTTGAAAGTGTGTAGCAC

GCACACGTCTGAACTCTTTCCCTACACGACGCTCTTCCGATCTNNNAGTTACTTGTTCTTCAGGTAAGCA

AGCACACGTCTGAACTCTTTCCCTACACGACGCTCTTCCGATCTNNNGAGGCTACTTTTTCTTTCTCTTCA

AGCACACGTCTGAACTCTTTCCCTACACGACGCTCTTCCGATCTNNNGAGGCTACTTTTTCTTTCTCTTCA

CACACGTCTGAACTCTTTCCCTACACGACGCTCTTCCGATCTNNNTACAACACATAAACCAATTCTTTCAT

CACACGTCTGAACTCTTTCCCTACACGACGCTCTTCCGATCTNNNTACAACACATAAACCAATTCTTTCAT

GCACACGTCTGAACTCTTTCCCTACACGACGCTCTTCCGATCTNNNACAGAGAAAAACATTTCACTATTC

AGCACACGTCTGAACTCTTTCCCTACACGACGCTCTTCCGATCTNNNGTGTACGTATGTATACCCATACC

GCACACGTCTGAACTCTTTCCCTACACGACGCTCTTCCGATCTNNNGTTTCTATCTCAAACTGTGTAAGC

GAGCACACGTCTGAACTCTTTCCCTACACGACGCTCTTCCGATCTNNNGTTATTCACAGTCTATTTCTGC

AGCACACGTCTGAACTCTTTCCCTACACGACGCTCTTCCGATCTNNNCAACTTATAGATGGGAGATCTT

AGCACACGTCTGAACTCTTTCCCTACACGACGCTCTTCCGATCTNNNCTTCGACTATTTTGGGGTAAAT

CACACGTCTGAACTCTTTCCCTACACGACGCTCTTCCGATCTNNNTCCTAATATAAATGTAGCACACTCA

AGCACACGTCTGAACTCTTTCCCTACACGACGCTCTTCCGATCTNNNTAGATAATTTAGGAGTGGGAAC

GCACACGTCTGAACTCTTTCCCTACACGACGCTCTTCCGATCTNNNCACCTGCATCTAATTCTGAGCATC

AGCACACGTCTGAACTCTTTCCCTACACGACGCTCTTCCGATCTNNNTACATATTGATTTTGTATCCTGC

AGCACACGTCTGAACTCTTTCCCTACACGACGCTCTTCCGATCTNNNTACATATTGATTTTGTATCCTGC

GAGCACACGTCTGAACTCTTTCCCTACACGACGCTCTTCCGATCTNNNGGAACTTTTCACTATACATCCTA

GAGCACACGTCTGAACTCTTTCCCTACACGACGCTCTTCCGATCTNNNCATACTTTCTTGTCAAGCTATA

GCACACGTCTGAACTCTTTCCCTACACGACGCTCTTCCGATCTNNNTGTGTGTGTATATATATGTGTGT

GCACACGTCTGAACTCTTTCCCTACACGACGCTCTTCCGATCTNNNCAACTTCTGGAGATAAAGATAGA

AGCACACGTCTGAACTCTTTCCCTACACGACGCTCTTCCGATCTNNNATCATTCTGCCTGGGAAATTCA

AGCACACGTCTGAACTCTTTCCCTACACGACGCTCTTCCGATCTNNNCTACAATGATCTAAACGAGAAA

AGAGCACACGTCTGAACTCTTTCCCTACACGACGCTCTTCCGATCTNNNATACAAGATGTTACAAGTTTC

CACACGTCTGAACTCTTTCCCTACACGACGCTCTTCCGATCTNNNATAGTGTGAAACTGTCTTTCCCATC

AGCACACGTCTGAACTCTTTCCCTACACGACGCTCTTCCGATCTNNNTGTCATCATCATTCTTATCTGTAC

AGCACACGTCTGAACTCTTTCCCTACACGACGCTCTTCCGATCTNNNCTACAATCATTACCATCATCTTC

GCACACGTCTGAACTCTTTCCCTACACGACGCTCTTCCGATCTNNNCTTTAAGTTGGAGTATGTAACAG

CACACGTCTGAACTCTTTCCCTACACGACGCTCTTCCGATCTNNNTAGGCTGTACTTCACAATATAAACA

GAGCACACGTCTGAACTCTTTCCCTACACGACGCTCTTCCGATCTNNNTGGAAAGAGATACAAGAATTA

GAGCACACGTCTGAACTCTTTCCCTACACGACGCTCTTCCGATCTNNNTGGAAAGAGATACAAGAATTA

CACACGTCTGAACTCTTTCCCTACACGACGCTCTTCCGATCTNNNTTCTATAGCAAAGGAGGTCTTTCAT

GCACACGTCTGAACTCTTTCCCTACACGACGCTCTTCCGATCTNNNTTCAAAAACAGATTCCATTGTTAC

GCACACGTCTGAACTCTTTCCCTACACGACGCTCTTCCGATCTNNNTTCAAAAACAGATTCCATTGTTAC

GCACACGTCTGAACTCTTTCCCTACACGACGCTCTTCCGATCTNNNTTCAAAAACAGATTCCATTGTTAC

GAGCACACGTCTGAACTCTTTCCCTACACGACGCTCTTCCGATCTNNNAAGTAAATAACGTCTAATCCCA

AGCACACGTCTGAACTCTTTCCCTACACGACGCTCTTCCGATCTNNNACTGTAATACCCAGCATATTTTA

AGCACACGTCTGAACTCTTTCCCTACACGACGCTCTTCCGATCTNNNGTATAATTCCATAGGATGACTG

GCACACGTCTGAACTCTTTCCCTACACGACGCTCTTCCGATCTNNNAGAGACAGTTTCCCAATAATGCAT

CACACGTCTGAACTCTTTCCCTACACGACGCTCTTCCGATCTNNNGGGGACTTTTAAAAACAAATAGAC

CACACGTCTGAACTCTTTCCCTACACGACGCTCTTCCGATCTNNNTGTGTTTCTTATTCATAGTACATGC

AGCACACGTCTGAACTCTTTCCCTACACGACGCTCTTCCGATCTNNNGTTTATATGCTTATATTCGTAC
GCACACGTCTGAACTCTTTCCCTACACGACGCTCTTCCGATCTNNNCTACACAGAAGTTATCACTTTTC
AGCACACGTCTGAACTCTTTCCCTACACGACGCTCTTCCGATCTNNNACTCATTAATTAAAGAAAGAA
AGCACACGTCTGAACTCTTTCCCTACACGACGCTCTTCCGATCTNNNAATGACTTCTGATTGAGCTAT
GCACACGTCTGAACTCTTTCCCTACACGACGCTCTTCCGATCTNNNTCTACATTTTAAAAACACAGTCT
GAGCACACGTCTGAACTCTTTCCCTACACGACGCTCTTCCGATCTNNNTGACATTCTGCAGTTTTAATA
GCACACGTCTGAACTCTTTCCCTACACGACGCTCTTCCGATCTNNNTTGTAATGTGGTAAAGTTACTTG
GCACACGTCTGAACTCTTTCCCTACACGACGCTCTTCCGATCTNNNATTTCCCCATCAAATGACAACAT
GCACACGTCTGAACTCTTTCCCTACACGACGCTCTTCCGATCTNNNCAGATTTTTGACTAAGTAGGAA
CACACGTCTGAACTCTTTCCCTACACGACGCTCTTCCGATCTNNNTAAAATCTAAATTTCCCAAGCAGC
AGCACACGTCTGAACTCTTTCCCTACACGACGCTCTTCCGATCTNNNTCATCTGTAATGGTCTTATTAT
AGCACACGTCTGAACTCTTTCCCTACACGACGCTCTTCCGATCTNNNTTCTTTCTATTGTGGAAAATCT
AGCACACGTCTGAACTCTTTCCCTACACGACGCTCTTCCGATCTNNNTTTTTTCAGAGGACCTCATAGC
CACACGTCTGAACTCTTTCCCTACACGACGCTCTTCCGATCTNNNCAAATGAAACACTGTTTTCAACCA
AGCACACGTCTGAACTCTTTCCCTACACGACGCTCTTCCGATCTNNNTTCCTTGTAGAACTATTCGTAA
GCACACGTCTGAACTCTTTCCCTACACGACGCTCTTCCGATCTNNNCAAATATTTATCCTTTGAGGGTG
GCACACGTCTGAACTCTTTCCCTACACGACGCTCTTCCGATCTNNNCAGAATTTCATTTCTTTTATGGCC
CACGTCTGAACTCTTTCCCTACACGACGCTCTTCCGATCTNNNATAGAATTGAACTTTAAGGGGTCAT
AGCACACGTCTGAACTCTTTCCCTACACGACGCTCTTCCGATCTNNNAGAAAGTATCATTCATGTGTG
GCACACGTCTGAACTCTTTCCCTACACGACGCTCTTCCGATCTNNNGTGTTTTTTTTCCTTCCTTTTTC
GCACACGTCTGAACTCTTTCCCTACACGACGCTCTTCCGATCTNNNCACCTGGCCTTTATTCTTAGCAT
GCACACGTCTGAACTCTTTCCCTACACGACGCTCTTCCGATCTNNNCAGTGGTGACCATTATTAATAA
GCACACGTCTGAACTCTTTCCCTACACGACGCTCTTCCGATCTNNNCTCGAGTTTAATACAGAAAGAA
GCACACGTCTGAACTCTTTCCCTACACGACGCTCTTCCGATCTNNNTACCTAACCATCTTTTCTTGATAC
AGCACACGTCTGAACTCTTTCCCTACACGACGCTCTTCCGATCTNNNAACTTTAACAGTAGTGAATGT
GCACACGTCTGAACTCTTTCCCTACACGACGCTCTTCCGATCTNNNAGCAAAACATCTCTAAACTATT
CACACGTCTGAACTCTTTCCCTACACGACGCTCTTCCGATCTNNNGTGCATCTTTTTTTTCTTTCTTCA
AGCACACGTCTGAACTCTTTCCCTACACGACGCTCTTCCGATCTNNNAATAACAAGCAGAAATTTTGT
GCACACGTCTGAACTCTTTCCCTACACGACGCTCTTCCGATCTNNNGAAAAATGTCAGGCTTAGAAT
GCACACGTCTGAACTCTTTCCCTACACGACGCTCTTCCGATCTNNNCCTGTATTTGATCCCATATATT
GCACACGTCTGAACTCTTTCCCTACACGACGCTCTTCCGATCTNNNTGCTTGAAATTGAATAGAAATC
GCACACGTCTGAACTCTTTCCCTACACGACGCTCTTCCGATCTNNNATAAACAATAATGCACTCTCTTC
AGCACACGTCTGAACTCTTTCCCTACACGACGCTCTTCCGATCTNNNTAAGGTATGTCATTTGAATAC
GCACACGTCTGAACTCTTTCCCTACACGACGCTCTTCCGATCTNNNGTATCTGCAACTTTACTCAATT
GAGCACACGTCTGAACTCTTTCCCTACACGACGCTCTTCCGATCTNNNTATGTTATGAGGAACTGGGA
AGCACACGTCTGAACTCTTTCCCTACACGACGCTCTTCCGATCTNNNATCATTTATCATCAACATTGA
GCACACGTCTGAACTCTTTCCCTACACGACGCTCTTCCGATCTNNNTATTTCACTTAGCATAATGTCCT
AGCACACGTCTGAACTCTTTCCCTACACGACGCTCTTCCGATCTNNNAAAGGATTCATGGATTCCTGC
AGCACACGTCTGAACTCTTTCCCTACACGACGCTCTTCCGATCTNNNCTGAGCCAATATGTCTTTATTT
GCACACGTCTGAACTCTTTCCCTACACGACGCTCTTCCGATCTNNNACAGAGCTGGTAATTATTCATA
GCACACGTCTGAACTCTTTCCCTACACGACGCTCTTCCGATCTNNNCTCATGGTTCCTAAACTAGAATA
GCACACGTCTGAACTCTTTCCCTACACGACGCTCTTCCGATCTNNNGAATTAAACACCTGGTAGTGAC
ACACGTCTGAACTCTTTCCCTACACGACGCTCTTCCGATCTNNNCAGGAATAAATGAAGAAACTCAAC

AGCACACGTCTGAACTCTTTCCCTACACGACGCTCTTCCGATCTNNNTAGAGATTATTACAGTCTCTCC
AGCACACGTCTGAACTCTTTCCCTACACGACGCTCTTCCGATCTNNNTAGATAGATAGATGACCAAAA
AGCACACGTCTGAACTCTTTCCCTACACGACGCTCTTCCGATCTNNNCCAAAATGCAATTGTCATAATT
AGCACACGTCTGAACTCTTTCCCTACACGACGCTCTTCCGATCTNNNAAGTTTTAAAGAGGAGAGAG
AGCACACGTCTGAACTCTTTCCCTACACGACGCTCTTCCGATCTNNNAAGAAAAATGAATGCATTACA
AGCACACGTCTGAACTCTTTCCCTACACGACGCTCTTCCGATCTNNNTGTTTAGATACAGCTTATTAG
AGCACACGTCTGAACTCTTTCCCTACACGACGCTCTTCCGATCTNNNCATCCTAGCACATATACTACAT
GCACACGTCTGAACTCTTTCCCTACACGACGCTCTTCCGATCTNNNTTAACCATTGAAATGATGAATG
AGCACACGTCTGAACTCTTTCCCTACACGACGCTCTTCCGATCTNNNCAACTTGGCAATTTGTATATTT
AGCACACGTCTGAACTCTTTCCCTACACGACGCTCTTCCGATCTNNNATTTGAAATCAGCTATTGTGAT
CACACGTCTGAACTCTTTCCCTACACGACGCTCTTCCGATCTNNNATTTTTTGTGTTGCTCAAATTGCA
ACACGTCTGAACTCTTTCCCTACACGACGCTCTTCCGATCTNNNGTATGTCTGTGGTCTGATTTTCAT
AGCACACGTCTGAACTCTTTCCCTACACGACGCTCTTCCGATCTNNNTTGAACGAAGAGGCTAGAAA
AGCACACGTCTGAACTCTTTCCCTACACGACGCTCTTCCGATCTNNNGAATTATCTTGTTCCCTTTGTA
GCACACGTCTGAACTCTTTCCCTACACGACGCTCTTCCGATCTNNNAGCAATAATCCTTTCCATTAGGC
AGCACACGTCTGAACTCTTTCCCTACACGACGCTCTTCCGATCTNNNTTAAGACCTTATCTCCATTGAT
AGCACACGTCTGAACTCTTTCCCTACACGACGCTCTTCCGATCTNNNAGTATTTTTGTAATTGGGGAT
GCACACGTCTGAACTCTTTCCCTACACGACGCTCTTCCGATCTNNNAAGAGGTATTAGAAAACCCAGA
CACACGTCTGAACTCTTTCCCTACACGACGCTCTTCCGATCTNNNCCTTAAAGATGTCTAGAACCTGCA
AGCACACGTCTGAACTCTTTCCCTACACGACGCTCTTCCGATCTNNNCACCTTAAACTAGTACTTTGAA
AGCACACGTCTGAACTCTTTCCCTACACGACGCTCTTCCGATCTNNNTAGAATGGGTTTTCCTCATGCA
GCACACGTCTGAACTCTTTCCCTACACGACGCTCTTCCGATCTNNNAAAACACAAACTAACTTTTCAAC
CACACGTCTGAACTCTTTCCCTACACGACGCTCTTCCGATCTNNNTAAACAAATTCTGGATAGTCAGA
GCACACGTCTGAACTCTTTCCCTACACGACGCTCTTCCGATCTNNNCAGGCTTTATTGTTTTTCATATT
AGCACACGTCTGAACTCTTTCCCTACACGACGCTCTTCCGATCTNNNTTTGCTTCATATATCTGGTAA
AGCACACGTCTGAACTCTTTCCCTACACGACGCTCTTCCGATCTNNNAAGAAGGATTTTTCTTATGTT
AGCACACGTCTGAACTCTTTCCCTACACGACGCTCTTCCGATCTNNNCCTAAATGTGTTCTAAGTATT
AGCACACGTCTGAACTCTTTCCCTACACGACGCTCTTCCGATCTNNNCTGTTTTTCAAGCAGGAAATT
GCACACGTCTGAACTCTTTCCCTACACGACGCTCTTCCGATCTNNNGAGAGTGTTTCTTCTTTTTTCTA
AGCACACGTCTGAACTCTTTCCCTACACGACGCTCTTCCGATCTNNNTCAAGAGGAAGAAATAGAAC
GCACACGTCTGAACTCTTTCCCTACACGACGCTCTTCCGATCTNNNCTACTGTGAAAAATAAGAGTCT
GCACACGTCTGAACTCTTTCCCTACACGACGCTCTTCCGATCTNNNCTACTGTGAAAAATAAGAGTCT
AGCACACGTCTGAACTCTTTCCCTACACGACGCTCTTCCGATCTNNNCCTTCAGTATCTTCAAGGTTTC
GCACACGTCTGAACTCTTTCCCTACACGACGCTCTTCCGATCTNNNTCAAATAATTGCTAGGATTTTGT
GCACACGTCTGAACTCTTTCCCTACACGACGCTCTTCCGATCTNNNTCAAATAATTGCTAGGATTTTGT
AGCACACGTCTGAACTCTTTCCCTACACGACGCTCTTCCGATCTNNNGGTACACTGCTGACAATATGC
AGCACACGTCTGAACTCTTTCCCTACACGACGCTCTTCCGATCTNNNAATCTCATTCATGATCATAGGT
AGCACACGTCTGAACTCTTTCCCTACACGACGCTCTTCCGATCTNNNAATCTCATTCATGATCATAGGT
AGCACACGTCTGAACTCTTTCCCTACACGACGCTCTTCCGATCTNNNAATCTCATTCATGATCATAGGT
GCACACGTCTGAACTCTTTCCCTACACGACGCTCTTCCGATCTNNNATTATTCCCAAGGGTATATAAG
GAGCACACGTCTGAACTCTTTCCCTACACGACGCTCTTCCGATCTNNNAGTTTCTCATTTCTATCTGAG
AGCACACGTCTGAACTCTTTCCCTACACGACGCTCTTCCGATCTNNNTCTTTTTTAGAGACAAATCCAT
GCACACGTCTGAACTCTTTCCCTACACGACGCTCTTCCGATCTNNNCTCTTTTTTGACAACTGGTACCA

AGCACACGTCTGAACTCTTTCCCTACACGACGCTCTTCCGATCTNNNCTGACCTCTTTTATGTAGCTA
AGCACACGTCTGAACTCTTTCCCTACACGACGCTCTTCCGATCTNNNTATTTCATGAATCCTTTCCATTC
AGCACACGTCTGAACTCTTTCCCTACACGACGCTCTTCCGATCTNNNAGATACTTACAAGAGACAAGA
AGCACACGTCTGAACTCTTTCCCTACACGACGCTCTTCCGATCTNNNGACAGACACATTACAGCTATGC
GCACACGTCTGAACTCTTTCCCTACACGACGCTCTTCCGATCTNNNTTTATTCATCCACAAGAACTACC
GCACACGTCTGAACTCTTTCCCTACACGACGCTCTTCCGATCTNNNTATATTCACATGGGTCAGTCTAA
AGCACACGTCTGAACTCTTTCCCTACACGACGCTCTTCCGATCTNNNTCAGTAGATTCAAAGATACAA
AGCACACGTCTGAACTCTTTCCCTACACGACGCTCTTCCGATCTNNNTCAGTAGATTCAAAGATACAA
AGCACACGTCTGAACTCTTTCCCTACACGACGCTCTTCCGATCTNNNTCAGTAGATTCAAAGATACAA
AGCACACGTCTGAACTCTTTCCCTACACGACGCTCTTCCGATCTNNNGCTACATAAAATAAAGCTACAC
GCACACGTCTGAACTCTTTCCCTACACGACGCTCTTCCGATCTNNNTATTGCAAGATTATTTCAGCATT
AGCACACGTCTGAACTCTTTCCCTACACGACGCTCTTCCGATCTNNNAAAGTTTACCATAAAGTTCCAA
GCACACGTCTGAACTCTTTCCCTACACGACGCTCTTCCGATCTNNNCCAAGGATTTAGCTATTACAAAT
GAGCACACGTCTGAACTCTTTCCCTACACGACGCTCTTCCGATCTNNNATTAACCAATCCAATCAAAAA
GCACACGTCTGAACTCTTTCCCTACACGACGCTCTTCCGATCTNNNTTAAATCTCCTGGTGGTGAAACA
GCACACGTCTGAACTCTTTCCCTACACGACGCTCTTCCGATCTNNNTCTCCAGGATATCTATGTGTATA
GCACACGTCTGAACTCTTTCCCTACACGACGCTCTTCCGATCTNNNCATTACTTGTTATACCCTCTCTAC
AGAGCACACGTCTGAACTCTTTCCCTACACGACGCTCTTCCGATCTNNNTCACTAGAAGTCAAATGGT
GCACACGTCTGAACTCTTTCCCTACACGACGCTCTTCCGATCTNNNCTAGTCCAGACATTATACAACTA
CACACGTCTGAACTCTTTCCCTACACGACGCTCTTCCGATCTNNNCACACATAAATATACACGATTGTC
GCACACGTCTGAACTCTTTCCCTACACGACGCTCTTCCGATCTNNNGGTCAACTAAAGTTTCTACAGTA
AGCACACGTCTGAACTCTTTCCCTACACGACGCTCTTCCGATCTNNNAACGTTAGCACTGGATAACTC
GAGCACACGTCTGAACTCTTTCCCTACACGACGCTCTTCCGATCTNNNTTTTATACAACCATTCACCTT
GCACACGTCTGAACTCTTTCCCTACACGACGCTCTTCCGATCTNNNCGGTTGATTCACTTTGATATATT
AGCACACGTCTGAACTCTTTCCCTACACGACGCTCTTCCGATCTNNNCTCAATGAACTTCTCTGTATCT
CACACGTCTGAACTCTTTCCCTACACGACGCTCTTCCGATCTNNNACAGACTTTCTCTCTTTAGAAGCA
GAGCACACGTCTGAACTCTTTCCCTACACGACGCTCTTCCGATCTNNNTTTCAGATAGCCTCAAGAGA
GAGCACACGTCTGAACTCTTTCCCTACACGACGCTCTTCCGATCTNNNTTTCAGATAGCCTCAAGAGA
AGCACACGTCTGAACTCTTTCCCTACACGACGCTCTTCCGATCTNNNAGATTTACTTATTTGGTCATTG
AGCACACGTCTGAACTCTTTCCCTACACGACGCTCTTCCGATCTNNNTCCTTGGTATGTCTTTTTGTTC
GCACACGTCTGAACTCTTTCCCTACACGACGCTCTTCCGATCTNNNAGTACTAGTGATATTGATTTA
AGCACACGTCTGAACTCTTTCCCTACACGACGCTCTTCCGATCTNNNGAATAAAAGTTATGTGTTCCA
GAGCACACGTCTGAACTCTTTCCCTACACGACGCTCTTCCGATCTNNNAGTATAGCCTCATGAAGTTTT
GCACACGTCTGAACTCTTTCCCTACACGACGCTCTTCCGATCTNNNAGTGATGTAGAATATACTGCAT
GCACACGTCTGAACTCTTTCCCTACACGACGCTCTTCCGATCTNNNTTCTTTATTGAAGTGTATGTGT
AGCACACGTCTGAACTCTTTCCCTACACGACGCTCTTCCGATCTNNNTACTCAAAAAGCCACTTTAATT
GAGCACACGTCTGAACTCTTTCCCTACACGACGCTCTTCCGATCTNNNCCAACATAATCCTGTGTTTTCC
GCACACGTCTGAACTCTTTCCCTACACGACGCTCTTCCGATCTNNNTTATACATGGATTTTTGACTGT
GCACACGTCTGAACTCTTTCCCTACACGACGCTCTTCCGATCTNNNTATTTAATTTCATGTGCTGTAGG
AGCACACGTCTGAACTCTTTCCCTACACGACGCTCTTCCGATCTNNNTGTAAAACTGACAATCATAAT
GAGCACACGTCTGAACTCTTTCCCTACACGACGCTCTTCCGATCTNNNAAAACAAAAGCTCACCTATTG
GCACACGTCTGAACTCTTTCCCTACACGACGCTCTTCCGATCTNNNTTTCACCAGTTTTATGTACATTC
CACGTCTGAACTCTTTCCCTACACGACGCTCTTCCGATCTNNNTCTATTTCTTTTCTTGGAGACTCATC

AGCACACGTCTGAACTCTTTCCCTACACGACGCTCTTCCGATCTNNNTTCAAACAATGTCAGGTAATTA
AGCACACGTCTGAACTCTTTCCCTACACGACGCTCTTCCGATCTNNNTGAATTACCCATCAAACACAGCA
AGCACACGTCTGAACTCTTTCCCTACACGACGCTCTTCCGATCTNNNGCTATTATCAATACCTCATTGCC
AGCACACGTCTGAACTCTTTCCCTACACGACGCTCTTCCGATCTNNNAAAAGAAACAAAGTTTAGCTTT
AGCACACGTCTGAACTCTTTCCCTACACGACGCTCTTCCGATCTNNNGCAAGACTTTGTCTCAAAAATA
GCACACGTCTGAACTCTTTCCCTACACGACGCTCTTCCGATCTNNNCTTTTAAATGTGAGAAGGTGAA
GAGCACACGTCTGAACTCTTTCCCTACACGACGCTCTTCCGATCTNNNAACTTTTCCTTAGAACTTCAAT
GAGCACACGTCTGAACTCTTTCCCTACACGACGCTCTTCCGATCTNNNATTAATTTCCTGACAGGTTTTA
CACACGTCTGAACTCTTTCCCTACACGACGCTCTTCCGATCTNNNCCAATATGACCTCATCTTAACTACA
CACACGTCTGAACTCTTTCCCTACACGACGCTCTTCCGATCTNNNTCCTTCCCTTGTTTCCTAATCATCG
GCACACGTCTGAACTCTTTCCCTACACGACGCTCTTCCGATCTNNNCAGTTAGTGTTGTTTTGGTTTCA
AGCACACGTCTGAACTCTTTCCCTACACGACGCTCTTCCGATCTNNNTGTCCGTGTGTATAAATATAGA
GAGCACACGTCTGAACTCTTTCCCTACACGACGCTCTTCCGATCTNNNGAAATGCATTACAAATATTCT
GCACACGTCTGAACTCTTTCCCTACACGACGCTCTTCCGATCTNNNAACATGAATCCTCTCTTATCAGCA
AGCACACGTCTGAACTCTTTCCCTACACGACGCTCTTCCGATCTNNNAATTTTCACGTATCTATCACATG
GCACACGTCTGAACTCTTTCCCTACACGACGCTCTTCCGATCTNNNTGTTTCTACCTTAGTACACTATTCA
GAGCACACGTCTGAACTCTTTCCCTACACGACGCTCTTCCGATCTNNNGGAAGTGATTTTTTCCTAATT
AGCACACGTCTGAACTCTTTCCCTACACGACGCTCTTCCGATCTNNNAGTTTAATAGACAGAAGAAAG
GAGCACACGTCTGAACTCTTTCCCTACACGACGCTCTTCCGATCTNNNGTTTTTTCTCAGTTTGCTTCTCA
AGCACACGTCTGAACTCTTTCCCTACACGACGCTCTTCCGATCTNNNATCTAACAAAACGTAGAGGAAT
GCACACGTCTGAACTCTTTCCCTACACGACGCTCTTCCGATCTNNNCTATGACTTACCATGGAGTTAG
AGCACACGTCTGAACTCTTTCCCTACACGACGCTCTTCCGATCTNNNTAGTGAAAGAAAGATGGAAA
GCACACGTCTGAACTCTTTCCCTACACGACGCTCTTCCGATCTNNNATAGTTGGTCTATGTGAAGAGT
GAGCACACGTCTGAACTCTTTCCCTACACGACGCTCTTCCGATCTNNNGAGTTGAATTGAATTAGAAG
CACACGTCTGAACTCTTTCCCTACACGACGCTCTTCCGATCTNNNATCTCTTTGGCCATTCATCTCATCG
AGCACACGTCTGAACTCTTTCCCTACACGACGCTCTTCCGATCTNNNAGAGTCAAAAAGAAAATACAC
AGCACACGTCTGAACTCTTTCCCTACACGACGCTCTTCCGATCTNNNTATGGCATATCATTTTTGAAGT
GAGCACACGTCTGAACTCTTTCCCTACACGACGCTCTTCCGATCTNNNTTTTTTTGCCTGTATAGTTTCTT
AGCACACGTCTGAACTCTTTCCCTACACGACGCTCTTCCGATCTNNNGTCATAAATTGAATATGAGACC
AGCACACGTCTGAACTCTTTCCCTACACGACGCTCTTCCGATCTNNNTCCTCACATCTCTAAGGAAGCAT
AGCACACGTCTGAACTCTTTCCCTACACGACGCTCTTCCGATCTNNNTCCTCACATCTCTAAGGAAGCAT
GAGCACACGTCTGAACTCTTTCCCTACACGACGCTCTTCCGATCTNNNGTTTAGGATGCCAATTTAAAC
AGCACACGTCTGAACTCTTTCCCTACACGACGCTCTTCCGATCTNNNTTTCATTTATTTGCCTTCTCTGCA
AGCACACGTCTGAACTCTTTCCCTACACGACGCTCTTCCGATCTNNNAAGAGACAGCATATCAAGAAAA
CACACGTCTGAACTCTTTCCCTACACGACGCTCTTCCGATCTNNNATTAGATGTTCATGTTGGGAAAT
AGCACACGTCTGAACTCTTTCCCTACACGACGCTCTTCCGATCTNNNTACTGACATAGAAATTGATTCC
AGAGCACACGTCTGAACTCTTTCCCTACACGACGCTCTTCCGATCTNNNGCAAGACCTCATCTCATCTA
GAGCACACGTCTGAACTCTTTCCCTACACGACGCTCTTCCGATCTNNNTTTTCCTCTTCCTCTAGTTTTT
GCACACGTCTGAACTCTTTCCCTACACGACGCTCTTCCGATCTNNNTTTCAGGATTAGAAAATTCAGCT
CACACGTCTGAACTCTTTCCCTACACGACGCTCTTCCGATCTNNNTTATAAAACAGCTTGCAGAATTCCA
GAGCACACGTCTGAACTCTTTCCCTACACGACGCTCTTCCGATCTNNNAATTGCATGTAATCAGCTAAAT
GAGCACACGTCTGAACTCTTTCCCTACACGACGCTCTTCCGATCTNNNTGTATCCAATACAACATAGAAA
GCACACGTCTGAACTCTTTCCCTACACGACGCTCTTCCGATCTNNNAGACTCCGTCTCAAAACAAACAT

AGCACACGTCTGAACTCTTTCCCTACACGACGCTCTTCCGATCTNNNTTTAGGAATCTTTTGAGGAACA
AGCACACGTCTGAACTCTTTCCCTACACGACGCTCTTCCGATCTNNNTGTAATCCCAGCTACTTAGGCA
GCACACGTCTGAACTCTTTCCCTACACGACGCTCTTCCGATCTNNNACATATTTTGACTTGTTTTCTCACA
GCACACGTCTGAACTCTTTCCCTACACGACGCTCTTCCGATCTNNNCACATCATTTCCTTGCACATCATCC
GCACACGTCTGAACTCTTTCCCTACACGACGCTCTTCCGATCTNNNCACTCATCTCAGTTAAGAATTTTC
GAGCACACGTCTGAACTCTTTCCCTACACGACGCTCTTCCGATCTNNNTGAAGAGATTTAAACTGAACT
AGCACACGTCTGAACTCTTTCCCTACACGACGCTCTTCCGATCTNNNTAAAATATGTGATTGTGTTCGA
GAGCACACGTCTGAACTCTTTCCCTACACGACGCTCTTCCGATCTNNNGTTGTATTGAATGATCTGCATT
GAGCACACGTCTGAACTCTTTCCCTACACGACGCTCTTCCGATCTNNNATAAACATATCAAAGCGTGTT
AGCACACGTCTGAACTCTTTCCCTACACGACGCTCTTCCGATCTNNNCAACTGGTTTCTCCCATAATCATC
GCACACGTCTGAACTCTTTCCCTACACGACGCTCTTCCGATCTNNNTTAATAATTCTGCCTCTTGAATCCA
GCACACGTCTGAACTCTTTCCCTACACGACGCTCTTCCGATCTNNNTTAATAATTCTGCCTCTTGAATCCA
AGCACACGTCTGAACTCTTTCCCTACACGACGCTCTTCCGATCTNNNAAGATATTCAACATCATTGTTCA
AGCACACGTCTGAACTCTTTCCCTACACGACGCTCTTCCGATCTNNNAATTATGTACTTTGATTGTGGTC
AGCACACGTCTGAACTCTTTCCCTACACGACGCTCTTCCGATCTNNNACATAGAAGTTCATCTTTAAGCT
AGCACACGTCTGAACTCTTTCCCTACACGACGCTCTTCCGATCTNNNAATCACAAGAAAAATCTCTCTAT
CACACGTCTGAACTCTTTCCCTACACGACGCTCTTCCGATCTNNNAGAGTATTACCAACTGAATTACTCA
AGCACACGTCTGAACTCTTTCCCTACACGACGCTCTTCCGATCTNNNACTTTTGGACCTATCTAGTCTCA
GCACACGTCTGAACTCTTTCCCTACACGACGCTCTTCCGATCTNNNTAAGTGGAATTGATGGGAAATAT
AGCACACGTCTGAACTCTTTCCCTACACGACGCTCTTCCGATCTNNNCCTGGAATAAATACAAAGTTGA
GAGCACACGTCTGAACTCTTTCCCTACACGACGCTCTTCCGATCTNNNTTGAAAGCAGATTGTTCTTTC
GCACACGTCTGAACTCTTTCCCTACACGACGCTCTTCCGATCTNNNGGGAGGGGAATCATTAATATTC
GCACACGTCTGAACTCTTTCCCTACACGACGCTCTTCCGATCTNNNTAGTTGGAGTCTTTTAAGGTTTCA
AGCACACGTCTGAACTCTTTCCCTACACGACGCTCTTCCGATCTNNNAGAATAATGTCAAATCTCAAAG
AGCACACGTCTGAACTCTTTCCCTACACGACGCTCTTCCGATCTNNNTTTGTTAATGGGAGTGTCTTCC
GCACACGTCTGAACTCTTTCCCTACACGACGCTCTTCCGATCTNNNAGGTGATCAAATTGTATAAAGGA
ACACGTCTGAACTCTTTCCCTACACGACGCTCTTCCGATCTNNNAATTTTAAAATGCAAACTTGGAGCAT
AGCACACGTCTGAACTCTTTCCCTACACGACGCTCTTCCGATCTNNNATATTCTCATCCAAATTCTCCTCC
AGCACACGTCTGAACTCTTTCCCTACACGACGCTCTTCCGATCTNNNACAAATATGTCCCTTAAATCTTGC
AGCACACGTCTGAACTCTTTCCCTACACGACGCTCTTCCGATCTNNNCATATCAGTGAAAAGATAAGCT
GAGCACACGTCTGAACTCTTTCCCTACACGACGCTCTTCCGATCTNNNTTTCTACCCTTTCTAGATACTCT
AGCACACGTCTGAACTCTTTCCCTACACGACGCTCTTCCGATCTNNNGAATGTCATTCAACTTTCTTCAC
AGAGCACACGTCTGAACTCTTTCCCTACACGACGCTCTTCCGATCTNNNAGAGTTCAGATAACCTAAAC
AGCACACGTCTGAACTCTTTCCCTACACGACGCTCTTCCGATCTNNNCTATGCAGACATGAAATATCAA
AGCACACGTCTGAACTCTTTCCCTACACGACGCTCTTCCGATCTNNNATTATAGAATGGGGGACAGATT
AGCACACGTCTGAACTCTTTCCCTACACGACGCTCTTCCGATCTNNNACAAGAACAACAAAACTTATTCT
AGCACACGTCTGAACTCTTTCCCTACACGACGCTCTTCCGATCTNNNTGGGTTATAAGGAATACAAGA
AGCACACGTCTGAACTCTTTCCCTACACGACGCTCTTCCGATCTNNNTAGAAGTTATTATCTGGTTGTGT
GCACACGTCTGAACTCTTTCCCTACACGACGCTCTTCCGATCTNNNCTCTCTCTCTCTCTTTCTCCATCC
GCACACGTCTGAACTCTTTCCCTACACGACGCTCTTCCGATCTNNNCTCTCTCTCTCTCTTTCTCCATCC
AGCACACGTCTGAACTCTTTCCCTACACGACGCTCTTCCGATCTNNNAGTGTATGTACTTTATCAGAACA
AGCACACGTCTGAACTCTTTCCCTACACGACGCTCTTCCGATCTNNNCTAGATCTACACAAATGCATTCC
CACACGTCTGAACTCTTTCCCTACACGACGCTCTTCCGATCTNNNCTCACTTTTCCTTACATTGAATAGCA

ACACGTCTGAACTCTTTCCCTACACGACGCTCTTCCGATCTNNNTATATTCCAGTCATGGTAGACTACA
AGCACACGTCTGAACTCTTTCCCTACACGACGCTCTTCCGATCTNNNAACAATCCACATATAAATAGG
AGCACACGTCTGAACTCTTTCCCTACACGACGCTCTTCCGATCTNNNCTCTGAAATAGTAGCTAACA
ACACGTCTGAACTCTTTCCCTACACGACGCTCTTCCGATCTNNNGTTTCAAAAAAAAAGGAAGTTGAC
GCACACGTCTGAACTCTTTCCCTACACGACGCTCTTCCGATCTNNNCATCTTTCCTTCTTTGTATTCATC
GCACACGTCTGAACTCTTTCCCTACACGACGCTCTTCCGATCTNNNCTTCCCTTCAGATATAACACTGCA
AGCACACGTCTGAACTCTTTCCCTACACGACGCTCTTCCGATCTNNNAGATTTGTTAACTGCAATTTGA
AGCACACGTCTGAACTCTTTCCCTACACGACGCTCTTCCGATCTNNNTTTCTAGTGAGGTCTCTCTTTC
GCACACGTCTGAACTCTTTCCCTACACGACGCTCTTCCGATCTNNNATATTATCATGCTTCTCAAGTTG
GCACACGTCTGAACTCTTTCCCTACACGACGCTCTTCCGATCTNNNTAGAAAACAGCCTTCCAATATAT
AGCACACGTCTGAACTCTTTCCCTACACGACGCTCTTCCGATCTNNNGAATGCAGATGAACATCTATTG
GCACACGTCTGAACTCTTTCCCTACACGACGCTCTTCCGATCTNNNCATTACTGTGGGGAATATAGGC
AGCACACGTCTGAACTCTTTCCCTACACGACGCTCTTCCGATCTNNNTAACTATTGCTCTTCTCTGCTAC
AGCACACGTCTGAACTCTTTCCCTACACGACGCTCTTCCGATCTNNNAATAATTCAAGTTAGTGACTG
AGAGCACACGTCTGAACTCTTTCCCTACACGACGCTCTTCCGATCTNNNTTTATAAAGACAAGCAAA
AGAGCACACGTCTGAACTCTTTCCCTACACGACGCTCTTCCGATCTNNNATGTATACATTGAAGGAGC
GCACACGTCTGAACTCTTTCCCTACACGACGCTCTTCCGATCTNNNAAGAGAATGATTATGGTTGATC
AGCACACGTCTGAACTCTTTCCCTACACGACGCTCTTCCGATCTNNNTAAGGATTGGAAATATTGGT
AGCACACGTCTGAACTCTTTCCCTACACGACGCTCTTCCGATCTNNNTGTAAGACACAGCCTTTATGC
GAGCACACGTCTGAACTCTTTCCCTACACGACGCTCTTCCGATCTNNNAATGGGAGACATATTTGCTT
ACACGTCTGAACTCTTTCCCTACACGACGCTCTTCCGATCTNNNGGATTTATGAAGATGGGTGGCATC
AGCACACGTCTGAACTCTTTCCCTACACGACGCTCTTCCGATCTNNNTATTTTGTAGCTTAAAGGTCTT
AGCACACGTCTGAACTCTTTCCCTACACGACGCTCTTCCGATCTNNNTAAATAAGGAATCTTTGACTGG
AGCACACGTCTGAACTCTTTCCCTACACGACGCTCTTCCGATCTNNNATTTCTATGTCACAGGTACCTA
AGCACACGTCTGAACTCTTTCCCTACACGACGCTCTTCCGATCTNNNATTTCTATGTCACAGGTACCTA
GCACACGTCTGAACTCTTTCCCTACACGACGCTCTTCCGATCTNNNCTACATATGTGGGTGTCAATATC
GCACACGTCTGAACTCTTTCCCTACACGACGCTCTTCCGATCTNNNATATGATGCTTAAATTGGTGTTT
GAGCACACGTCTGAACTCTTTCCCTACACGACGCTCTTCCGATCTNNNGCTGAAGCTGAAAAGATATT
AGCACACGTCTGAACTCTTTCCCTACACGACGCTCTTCCGATCTNNNTGAATGTGAAGAATGTGTGTC
CACACGTCTGAACTCTTTCCCTACACGACGCTCTTCCGATCTNNNGATTAGATGCTCAGAAATACTTGC
CACACGTCTGAACTCTTTCCCTACACGACGCTCTTCCGATCTNNNCCAGAACAATTTTTCTCCTTAATC
AGCACACGTCTGAACTCTTTCCCTACACGACGCTCTTCCGATCTNNNAATCACCCAGTCTCATTTCTCAT
GCACACGTCTGAACTCTTTCCCTACACGACGCTCTTCCGATCTNNNCAATTCAACAGTACAGTACAAA
AGCACACGTCTGAACTCTTTCCCTACACGACGCTCTTCCGATCTNNNCAAAATAAGTTGTCTTGTCTCTT
AGCACACGTCTGAACTCTTTCCCTACACGACGCTCTTCCGATCTNNNTTGCAGAATACCTTGGAAATT
AGCACACGTCTGAACTCTTTCCCTACACGACGCTCTTCCGATCTNNNTTATTTTTGTCTCCCTTTTCTT
AGCACACGTCTGAACTCTTTCCCTACACGACGCTCTTCCGATCTNNNTAGTGAAATGATTTCTACAGTC
CACACGTCTGAACTCTTTCCCTACACGACGCTCTTCCGATCTNNNATCCCTCAATGTCATTAAAGTATC
ACACGTCTGAACTCTTTCCCTACACGACGCTCTTCCGATCTNNNGCAAGGCCATTACAAAAGTACATC
GCACACGTCTGAACTCTTTCCCTACACGACGCTCTTCCGATCTNNNAAGTATTTCCTGTGGAGAATAA
CACGTCTGAACTCTTTCCCTACACGACGCTCTTCCGATCTNNNTTAGATCCTTTTTTTCCTTTCACCATC
AGCACACGTCTGAACTCTTTCCCTACACGACGCTCTTCCGATCTNNNATAGTATGTTTTGGTATCCCTG
GAGCACACGTCTGAACTCTTTCCCTACACGACGCTCTTCCGATCTNNNTATGTTGTAGAAAGGAACCT

GCACACGTCTGAACTCTTTCCCTACACGACGCTCTTCCGATCTNNNTTTCTCACACACACATACTGCATC
GAGCACACGTCTGAACTCTTTCCCTACACGACGCTCTTCCGATCTNNNTTCATTCTTTTTTATGGCTGAA
GCACACGTCTGAACTCTTTCCCTACACGACGCTCTTCCGATCTNNNCTTTAAAGACTGAAAAGCATTTC
AGCACACGTCTGAACTCTTTCCCTACACGACGCTCTTCCGATCTNNNCAATTAGTGCAATCATATACACT
AGCACACGTCTGAACTCTTTCCCTACACGACGCTCTTCCGATCTNNNTGCTACCCTAATAGAAACTTATT
AGCACACGTCTGAACTCTTTCCCTACACGACGCTCTTCCGATCTNNNCAATTGTTCCAAATACGTAATTT
GCACACGTCTGAACTCTTTCCCTACACGACGCTCTTCCGATCTNNNGAAAAGTAAGAGCAAAATTCCTA
AGCACACGTCTGAACTCTTTCCCTACACGACGCTCTTCCGATCTNNNGTGGATATCAACATAGTAAAG
CACACGTCTGAACTCTTTCCCTACACGACGCTCTTCCGATCTNNNAATATCAAACTGGAGAGACCTTA
AGCACACGTCTGAACTCTTTCCCTACACGACGCTCTTCCGATCTNNNAAAGCTTCCCAACAAAAATGCA
AGCACACGTCTGAACTCTTTCCCTACACGACGCTCTTCCGATCTNNNTTTACAACAGTTGAGGGTTTTT
GCACACGTCTGAACTCTTTCCCTACACGACGCTCTTCCGATCTNNNTAGAAAAGCAAATGTTTACAGG
GAGCACACGTCTGAACTCTTTCCCTACACGACGCTCTTCCGATCTNNNATTCTGAATACCTTGTGATAG
GAGCACACGTCTGAACTCTTTCCCTACACGACGCTCTTCCGATCTNNNTCTAACATGCTTGAAAAAGAA
GCACACGTCTGAACTCTTTCCCTACACGACGCTCTTCCGATCTNNNGATCCATTCAGGTTGATTTTGCA
AGCACACGTCTGAACTCTTTCCCTACACGACGCTCTTCCGATCTNNNAACTCATTTGAATGCTCTGTCA
CACACGTCTGAACTCTTTCCCTACACGACGCTCTTCCGATCTNNNTAATAAATGATTCTGAATCCAGG
GCACACGTCTGAACTCTTTCCCTACACGACGCTCTTCCGATCTNNNGCTCTAAAGGAGTTTATTACAAT
GAGCACACGTCTGAACTCTTTCCCTACACGACGCTCTTCCGATCTNNNCCAAAAGTCCTATTCCCTTGC
GAGCACACGTCTGAACTCTTTCCCTACACGACGCTCTTCCGATCTNNNATTTGTTACCAATCTCTTTCATT
AGCACACGTCTGAACTCTTTCCCTACACGACGCTCTTCCGATCTNNNACATATTACTGACTTTTTGGAG
GAGCACACGTCTGAACTCTTTCCCTACACGACGCTCTTCCGATCTNNNATTAATGTGTTAGGAGGAAA
AGCACACGTCTGAACTCTTTCCCTACACGACGCTCTTCCGATCTNNNATAAAAATAGGCCAATGGAAA
AGCACACGTCTGAACTCTTTCCCTACACGACGCTCTTCCGATCTNNNATAAAAATAGGCCAATGGAAA
GCACACGTCTGAACTCTTTCCCTACACGACGCTCTTCCGATCTNNNGATAAGATTCCATGAGTTAGGG
ACACGTCTGAACTCTTTCCCTACACGACGCTCTTCCGATCTNNNCAGGTTTCTAACTCAGTAGGTCATC
CACACGTCTGAACTCTTTCCCTACACGACGCTCTTCCGATCTNNNCTTAGAAGAAGAGGATTTGAGAG
ACACGTCTGAACTCTTTCCCTACACGACGCTCTTCCGATCTNNNGTCAGAGTCAAACATAGCTAGCAT
AGCACACGTCTGAACTCTTTCCCTACACGACGCTCTTCCGATCTNNNAACTTCTTTTCACTTTCTGTATC
GAGCACACGTCTGAACTCTTTCCCTACACGACGCTCTTCCGATCTNNNATTACACATACAAACATACAC
GAGCACACGTCTGAACTCTTTCCCTACACGACGCTCTTCCGATCTNNNTAGCTTTCTCTCAATCTCTCTC
CACACGTCTGAACTCTTTCCCTACACGACGCTCTTCCGATCTNNNTGATATTTGACTTATTCCCTCTGC
AGCACACGTCTGAACTCTTTCCCTACACGACGCTCTTCCGATCTNNNGACTGCCATCATTTTCTTTATAG
GCACACGTCTGAACTCTTTCCCTACACGACGCTCTTCCGATCTNNNGAACTCACTTATTAGTTATGGGA
AGCACACGTCTGAACTCTTTCCCTACACGACGCTCTTCCGATCTNNNAGCTCCAAAGTATGTCAAAGC
AGCACACGTCTGAACTCTTTCCCTACACGACGCTCTTCCGATCTNNNTGTCAATTTCTTATGTATCCTTC
AGCACACGTCTGAACTCTTTCCCTACACGACGCTCTTCCGATCTNNNGTGTTAGAGCTGAATAATATTC
GCACACGTCTGAACTCTTTCCCTACACGACGCTCTTCCGATCTNNNCATTTCTATAAAACTCACTGAGTC
GAGCACACGTCTGAACTCTTTCCCTACACGACGCTCTTCCGATCTNNNTGTTTTCTTTTCTTTTTTACTGC
GAGCACACGTCTGAACTCTTTCCCTACACGACGCTCTTCCGATCTNNNGAAGAGTAATGCATCATATGA
GCACACGTCTGAACTCTTTCCCTACACGACGCTCTTCCGATCTNNNCTGGTTCTAACTAAAGATTATGC
GAGCACACGTCTGAACTCTTTCCCTACACGACGCTCTTCCGATCTNNNTTCTTGTCTTATATGCATGAG
AGCACACGTCTGAACTCTTTCCCTACACGACGCTCTTCCGATCTNNNCCTCAAAGAAAGGAAGATGGC

AGCACACGTCTGAACTCTTTCCCTACACGACGCTCTTCCGATCTNNNCTATTTCCTTTCTTTCTCCCTCCAT

AGCACACGTCTGAACTCTTTCCCTACACGACGCTCTTCCGATCTNNNCTATTTCCTTTCTTTCTCCCTCCAT

GAGCACACGTCTGAACTCTTTCCCTACACGACGCTCTTCCGATCTNNNACTTACTGGATGTAATACTCTT

GAGCACACGTCTGAACTCTTTCCCTACACGACGCTCTTCCGATCTNNNACTTACTGGATGTAATACTCTT

AGCACACGTCTGAACTCTTTCCCTACACGACGCTCTTCCGATCTNNNAATGAAATCAAACCTCAAAGATT

GAGCACACGTCTGAACTCTTTCCCTACACGACGCTCTTCCGATCTNNNGAAGAAAGAGAGTTAGGGAA

GAGCACACGTCTGAACTCTTTCCCTACACGACGCTCTTCCGATCTNNNTAAAAGGTGCCTGAACTTTAT

AGAGCACACGTCTGAACTCTTTCCCTACACGACGCTCTTCCGATCTNNNGGAAAGAGAAAGATTGTTTT

AGCACACGTCTGAACTCTTTCCCTACACGACGCTCTTCCGATCTNNNTATGTAACATATGCCAAAATGA

AGCACACGTCTGAACTCTTTCCCTACACGACGCTCTTCCGATCTNNNCTTTTGCCAATAAAAGTTTCTCTC

AGCACACGTCTGAACTCTTTCCCTACACGACGCTCTTCCGATCTNNNGATAGTCTCAGATTTCTGTGAT

AGCACACGTCTGAACTCTTTCCCTACACGACGCTCTTCCGATCTNNNTTGTTCAGTGAAACCTCTAAATA

GCACACGTCTGAACTCTTTCCCTACACGACGCTCTTCCGATCTNNNCATATCTGGCCCCTTATTTCCATCG

GCACACGTCTGAACTCTTTCCCTACACGACGCTCTTCCGATCTNNNTAATGAGAACAGAGGTTTCACCA

CACACGTCTGAACTCTTTCCCTACACGACGCTCTTCCGATCTNNNGAGTACAAGAAGGAAAAGGGCAT

AGCACACGTCTGAACTCTTTCCCTACACGACGCTCTTCCGATCTNNNCCTTTCTTCCCTCAATTTCTCATCG

CACACGTCTGAACTCTTTCCCTACACGACGCTCTTCCGATCTNNNTTTGGGACTTAGGTACAGATACAT

GCACACGTCTGAACTCTTTCCCTACACGACGCTCTTCCGATCTNNNCAAGATTTCATTACACTGCTCCAT

AGCACACGTCTGAACTCTTTCCCTACACGACGCTCTTCCGATCTNNNTGGTAAATGGTGAAAATTTAGA

AGCACACGTCTGAACTCTTTCCCTACACGACGCTCTTCCGATCTNNNAAAGAATTTTTGTCAACACCATA

AGCACACGTCTGAACTCTTTCCCTACACGACGCTCTTCCGATCTNNNTAATGAGTAGAAGGAACAAGTT

AGCACACGTCTGAACTCTTTCCCTACACGACGCTCTTCCGATCTNNNCACTAGGAGATCATGAAGAAA

GAGCACACGTCTGAACTCTTTCCCTACACGACGCTCTTCCGATCTNNNTATTTCAGATGAGTGAATAAG

ACACGTCTGAACTCTTTCCCTACACGACGCTCTTCCGATCTNNNCTTTTGAGTAGCTGAGACTACCATC

AGAGCACACGTCTGAACTCTTTCCCTACACGACGCTCTTCCGATCTNNNGTCACTAGAAGGAAGTAAG

AGCACACGTCTGAACTCTTTCCCTACACGACGCTCTTCCGATCTNNNAAAAAAATATGTGTTTGTGAGT

GCACACGTCTGAACTCTTTCCCTACACGACGCTCTTCCGATCTNNNGAATTACAATTCCACATGAGATTC

AGCACACGTCTGAACTCTTTCCCTACACGACGCTCTTCCGATCTNNNATAAGATAAACCTGGTTTTCAGA

GAGCACACGTCTGAACTCTTTCCCTACACGACGCTCTTCCGATCTNNNTATCTAAGCTACAACACCTAAT

GAGCACACGTCTGAACTCTTTCCCTACACGACGCTCTTCCGATCTNNNGATTCAGTTTCCCAGTATTTTG

GCACACGTCTGAACTCTTTCCCTACACGACGCTCTTCCGATCTNNNATGATATAATCAAGCCCATTCATC

GAGCACACGTCTGAACTCTTTCCCTACACGACGCTCTTCCGATCTNNNATTCATGCAGAATCAAAATGA

ACACGTCTGAACTCTTTCCCTACACGACGCTCTTCCGATCTNNNGTTTGACAGTGTACTCAATATTGCAT

GAGCACACGTCTGAACTCTTTCCCTACACGACGCTCTTCCGATCTNNNGGAGTTTACATTCCATTTTATG

GCACACGTCTGAACTCTTTCCCTACACGACGCTCTTCCGATCTNNNATGTTTGTAGCTTAATTATCAGCC

CACACGTCTGAACTCTTTCCCTACACGACGCTCTTCCGATCTNNNAATTCCCAATATAGCAACAGGCATC

GCACACGTCTGAACTCTTTCCCTACACGACGCTCTTCCGATCTNNNGGTTTGTATCCATGCATTTATATC

AGCACACGTCTGAACTCTTTCCCTACACGACGCTCTTCCGATCTNNNCTGTGAGATTATATGCAAGATT

GAGCACACGTCTGAACTCTTTCCCTACACGACGCTCTTCCGATCTNNNTAGCTTACTTCCTTCTTTTTAC

AGCACACGTCTGAACTCTTTCCCTACACGACGCTCTTCCGATCTNNNAATTAATCTGCTTTTGTGAGTTC

AGCACACGTCTGAACTCTTTCCCTACACGACGCTCTTCCGATCTNNNGTCTTGAATTCACTGGGTTTCAT

ACACGTCTGAACTCTTTCCCTACACGACGCTCTTCCGATCTNNNAAAGGTTGACAGGAATAATTTTGCA

AGCACACGTCTGAACTCTTTCCCTACACGACGCTCTTCCGATCTNNNTCTTTTTTAATCTTACGGGTTTTC

AGCACACGTCTGAACTCTTTCCCTACACGACGCTCTTCCGATCTNNNAGAGAATTGCTTGAGTTAAGAT
GAGCACACGTCTGAACTCTTTCCCTACACGACGCTCTTCCGATCTNNNAGTCCAAATGAAATCCTTAGTT
AGCACACGTCTGAACTCTTTCCCTACACGACGCTCTTCCGATCTNNNGCAGTATAAATCACTGTTACTCT
AGCACACGTCTGAACTCTTTCCCTACACGACGCTCTTCCGATCTNNNAAAATGAGGAAACTTCATGTTC
AGCACACGTCTGAACTCTTTCCCTACACGACGCTCTTCCGATCTNNNCTTTTCTCAAATACAACCTGAAA
AGCACACGTCTGAACTCTTTCCCTACACGACGCTCTTCCGATCTNNNGCTGAAAAAGTATGGACTCTATG
AGCACACGTCTGAACTCTTTCCCTACACGACGCTCTTCCGATCTNNNTAAGGCCTTACTTTTAAAAAACA
AGCACACGTCTGAACTCTTTCCCTACACGACGCTCTTCCGATCTNNNATTCTAATAAAATGTGCATCTG
AGAGCACACGTCTGAACTCTTTCCCTACACGACGCTCTTCCGATCTNNNTGGTGGATATTTAAATTAAT
ACACGTCTGAACTCTTTCCCTACACGACGCTCTTCCGATCTNNNACTCCGTCTCAAAAAAAAAAAGCATG
AGCACACGTCTGAACTCTTTCCCTACACGACGCTCTTCCGATCTNNNCCAGAACTCTTTTTGTATCTAAT
AGCACACGTCTGAACTCTTTCCCTACACGACGCTCTTCCGATCTNNNCTTCCAGTCACATAGAGTAATCC
AAGAGCACACGTCTGAACTCTTTCCCTACACGACGCTCTTCCGATCTNNNACACATATCTATCTGTATCTA
AGCACACGTCTGAACTCTTTCCCTACACGACGCTCTTCCGATCTNNNGTGTTTTTTTCTCATTTCCAAAT
GAGCACACGTCTGAACTCTTTCCCTACACGACGCTCTTCCGATCTNNNCAGTCTCTTAGTCATGAGAAT
AGCACACGTCTGAACTCTTTCCCTACACGACGCTCTTCCGATCTNNNGGTAAAAAAGAAACAGAAAAAC
AGCACACGTCTGAACTCTTTCCCTACACGACGCTCTTCCGATCTNNNGGTAAAAAAGAAACAGAAAAAC
AGCACACGTCTGAACTCTTTCCCTACACGACGCTCTTCCGATCTNNNTCAGGAAACTTACAATCAAGGC
AGCACACGTCTGAACTCTTTCCCTACACGACGCTCTTCCGATCTNNNAGGTTATATATACACACATGCAT
AGAGCACACGTCTGAACTCTTTCCCTACACGACGCTCTTCCGATCTNNNTATACATACACATGGAAAGAG
GAGCACACGTCTGAACTCTTTCCCTACACGACGCTCTTCCGATCTNNNTTTATTTTTCACTCCAGTAACTG
GAGCACACGTCTGAACTCTTTCCCTACACGACGCTCTTCCGATCTNNNTTTATTTTTCACTCCAGTAACTG
GAGCACACGTCTGAACTCTTTCCCTACACGACGCTCTTCCGATCTNNNGTTGTTTATCTGAACAGTGTAA
AGCACACGTCTGAACTCTTTCCCTACACGACGCTCTTCCGATCTNNNCTGTATGTGAAGTGTGGATGCA
AGCACACGTCTGAACTCTTTCCCTACACGACGCTCTTCCGATCTNNNGACCTTCAGTTACTATCCTTTCCA
AGCACACGTCTGAACTCTTTCCCTACACGACGCTCTTCCGATCTNNNGACCTTCAGTTACTATCCTTTCCA
GAGCACACGTCTGAACTCTTTCCCTACACGACGCTCTTCCGATCTNNNAACATGGCTACTTCCAGTAATG
GAGCACACGTCTGAACTCTTTCCCTACACGACGCTCTTCCGATCTNNNTCTTTAGTTATGATGTGGTATG
AGCACACGTCTGAACTCTTTCCCTACACGACGCTCTTCCGATCTNNNTACAAAAAATGGTCATCTGAAAA
AGCACACGTCTGAACTCTTTCCCTACACGACGCTCTTCCGATCTNNNCCAGGTTTTGAGATACAATATTA
AGCACACGTCTGAACTCTTTCCCTACACGACGCTCTTCCGATCTNNNTCACTATGACGTCATAACAATAT
GAGCACACGTCTGAACTCTTTCCCTACACGACGCTCTTCCGATCTNNNGGATTACAATGTTGATACTGA
GCACACGTCTGAACTCTTTCCCTACACGACGCTCTTCCGATCTNNNGTGTTCTTCTTTACACTTGATACCA
AGAGCACACGTCTGAACTCTTTCCCTACACGACGCTCTTCCGATCTNNNCTAATAATAGTGTTCATGTG
AGCACACGTCTGAACTCTTTCCCTACACGACGCTCTTCCGATCTNNNCATCTTGTATATTGCAAAGCTA
AGCACACGTCTGAACTCTTTCCCTACACGACGCTCTTCCGATCTNNNGTAAGGGTGAGTCAACATATCC
ACACGTCTGAACTCTTTCCCTACACGACGCTCTTCCGATCTNNNTTCCAAAGCTAGGTTATAAAATGCAT
ACACGTCTGAACTCTTTCCCTACACGACGCTCTTCCGATCTNNNTTCCAAAGCTAGGTTATAAAATGCAT
GCACACGTCTGAACTCTTTCCCTACACGACGCTCTTCCGATCTNNNAATAGGAGTTAGAAACCAGGAA
AGCACACGTCTGAACTCTTTCCCTACACGACGCTCTTCCGATCTNNNAAGTAGTAGTAAACAAGTTCTG
GCACACGTCTGAACTCTTTCCCTACACGACGCTCTTCCGATCTNNNTACAATTGGGATTTCAGTTATG
AGCACACGTCTGAACTCTTTCCCTACACGACGCTCTTCCGATCTNNNCAGCTTTAGGGTTTTTTAGAAT
AGCACACGTCTGAACTCTTTCCCTACACGACGCTCTTCCGATCTNNNCAGCTTTAGGGTTTTTTAGAAT

GCACACGTCTGAACTCTTTCCCTACACGACGCTCTTCCGATCTNNNATTATGTGCTTATGAGAGATAG
ACACGTCTGAACTCTTTCCCTACACGACGCTCTTCCGATCTNNNAAATGGTAGGGCTCTTGATTCATCG
ACACGTCTGAACTCTTTCCCTACACGACGCTCTTCCGATCTNNNATTTATATTTGAGCTTTTGTGGTCA
AGCACACGTCTGAACTCTTTCCCTACACGACGCTCTTCCGATCTNNNGTACAATTCAAGGTGAGATTT
GCACACGTCTGAACTCTTTCCCTACACGACGCTCTTCCGATCTNNNGCGGAGATTTGATTAAAATAGA
AGCACACGTCTGAACTCTTTCCCTACACGACGCTCTTCCGATCTNNNAAGGACACAAACAGAAAATAT
GCACACGTCTGAACTCTTTCCCTACACGACGCTCTTCCGATCTNNNTTATTATTTCTGGAAAATCTGCA
CACACGTCTGAACTCTTTCCCTACACGACGCTCTTCCGATCTNNNTTTGTTTCAAAAAAGGCAAATAC
GCACACGTCTGAACTCTTTCCCTACACGACGCTCTTCCGATCTNNNTATCAAAGAGAATTTACACTGG
AGCACACGTCTGAACTCTTTCCCTACACGACGCTCTTCCGATCTNNNAAAGTATTTTATATTCCAGTGC
AGCACACGTCTGAACTCTTTCCCTACACGACGCTCTTCCGATCTNNNATGTGTCATGAAGAATTAAGA
GAGCACACGTCTGAACTCTTTCCCTACACGACGCTCTTCCGATCTNNNCTTAACTCTCACAGCAATTCT
AGCACACGTCTGAACTCTTTCCCTACACGACGCTCTTCCGATCTNNNAAGACAATCTTGAGATTTTGA
GCACACGTCTGAACTCTTTCCCTACACGACGCTCTTCCGATCTNNNTAGTTGAAGGTTTAGGTCAAAT
AGCACACGTCTGAACTCTTTCCCTACACGACGCTCTTCCGATCTNNNATTTAGTTGAGGAATATCGGA
AGCACACGTCTGAACTCTTTCCCTACACGACGCTCTTCCGATCTNNNTTATAAATACAGAGAGACCCT
GAGCACACGTCTGAACTCTTTCCCTACACGACGCTCTTCCGATCTNNNTGTTGGTTTGTTTTTTCAAA
GCACACGTCTGAACTCTTTCCCTACACGACGCTCTTCCGATCTNNNAATGTACAAGGAAATTAAACCA
AGCACACGTCTGAACTCTTTCCCTACACGACGCTCTTCCGATCTNNNTCTGATTTCCTGGTATCTTTTA
AGCACACGTCTGAACTCTTTCCCTACACGACGCTCTTCCGATCTNNNCATATCTCCTGGTTGTATTAG
GCACACGTCTGAACTCTTTCCCTACACGACGCTCTTCCGATCTNNNTAGATGAGTGGAACCATTATAA
GCACACGTCTGAACTCTTTCCCTACACGACGCTCTTCCGATCTNNNTAGATGAGTGGAACCATTATAA
AGCACACGTCTGAACTCTTTCCCTACACGACGCTCTTCCGATCTNNNATTTTTAGGGTAACTTTTTG
AGCACACGTCTGAACTCTTTCCCTACACGACGCTCTTCCGATCTNNNCAGTGGTTCTTGATGTAAAAT
AGCACACGTCTGAACTCTTTCCCTACACGACGCTCTTCCGATCTNNNCTTCTTATGTTTTTAAGGCTGA
GCACACGTCTGAACTCTTTCCCTACACGACGCTCTTCCGATCTNNNAGCAAAAAACAACAAATATTACA
AGCACACGTCTGAACTCTTTCCCTACACGACGCTCTTCCGATCTNNNGTGGTAGATCTCTTTGAAAATT
AGCACACGTCTGAACTCTTTCCCTACACGACGCTCTTCCGATCTNNNATTTACTGCATAATAATCAGAG
GCACACGTCTGAACTCTTTCCCTACACGACGCTCTTCCGATCTNNNGCTGTCTTTATTTAAGTGAAGA
GCACACGTCTGAACTCTTTCCCTACACGACGCTCTTCCGATCTNNNAAAATGCCTTGAGTTGACAAAC
GCACACGTCTGAACTCTTTCCCTACACGACGCTCTTCCGATCTNNNTATCCCTTTGAGATCCTCTTATA
AGCACACGTCTGAACTCTTTCCCTACACGACGCTCTTCCGATCTNNNGTTCCAACATTTTCCCATATAAT
AGCACACGTCTGAACTCTTTCCCTACACGACGCTCTTCCGATCTNNNCTACTATATTCACCTTGGTCTTG
AGCACACGTCTGAACTCTTTCCCTACACGACGCTCTTCCGATCTNNNGCAGAAATTAATAGGAAGCAT
GCACACGTCTGAACTCTTTCCCTACACGACGCTCTTCCGATCTNNNCTGTATACCATATGGGAATTTCT
AGCACACGTCTGAACTCTTTCCCTACACGACGCTCTTCCGATCTNNNAATTTAGGTATCCAAATGAGG
AGCACACGTCTGAACTCTTTCCCTACACGACGCTCTTCCGATCTNNNAGGAAAAAGAATAGTCATGAG
GCACACGTCTGAACTCTTTCCCTACACGACGCTCTTCCGATCTNNNCATACTTACGGATACATTTTTGTC
AGCACACGTCTGAACTCTTTCCCTACACGACGCTCTTCCGATCTNNNACAAAAAGAATAAAAAGGTG
GAGCACACGTCTGAACTCTTTCCCTACACGACGCTCTTCCGATCTNNNATTTAGGGAAATTACTCATG
AGCACACGTCTGAACTCTTTCCCTACACGACGCTCTTCCGATCTNNNTCCAGAATATGATGAAAAAAT
GCACACGTCTGAACTCTTTCCCTACACGACGCTCTTCCGATCTNNNTAATCATTATGTTTTCTGAGACG
GCACACGTCTGAACTCTTTCCCTACACGACGCTCTTCCGATCTNNNTAATCATTATGTTTTCTGAGACG

AGCACACGTCTGAACTCTTTCCCTACACGACGCTCTTCCGATCTNNNGATACATGGTAAAAACAGAGA
CACACGTCTGAACTCTTTCCCTACACGACGCTCTTCCGATCTNNNGTCTTGGAATACATACATATACACC
GAGCACACGTCTGAACTCTTTCCCTACACGACGCTCTTCCGATCTNNNCATTATACAGCTAATCCAATCT
AGCACACGTCTGAACTCTTTCCCTACACGACGCTCTTCCGATCTNNNTTTGCTTTTCAAGGTTACTTTA
AGCACACGTCTGAACTCTTTCCCTACACGACGCTCTTCCGATCTNNNATCTTTATTGCATTGAGGTTAA
GAGCACACGTCTGAACTCTTTCCCTACACGACGCTCTTCCGATCTNNNATGAAAACTGAAAAGGTTCTA
GAGCACACGTCTGAACTCTTTCCCTACACGACGCTCTTCCGATCTNNNCTCTTGGAATGAACTAAGTAA
GAGCACACGTCTGAACTCTTTCCCTACACGACGCTCTTCCGATCTNNNTGAAATTTTCATTTGGATGTTT
GAGCACACGTCTGAACTCTTTCCCTACACGACGCTCTTCCGATCTNNNCTAGAAAGCAAGAAGTAAAC
AGCACACGTCTGAACTCTTTCCCTACACGACGCTCTTCCGATCTNNNATAAAGCAGAATACTTCATTGT
AGCACACGTCTGAACTCTTTCCCTACACGACGCTCTTCCGATCTNNNAGAGCAAAGTATAGAAAGGTTT
AGCACACGTCTGAACTCTTTCCCTACACGACGCTCTTCCGATCTNNNGCCAAGACCAGCATTTATTAACA
AGAGCACACGTCTGAACTCTTTCCCTACACGACGCTCTTCCGATCTNNNACAAACTATACTCAGACTATG
GAGCACACGTCTGAACTCTTTCCCTACACGACGCTCTTCCGATCTNNNTACACAAATGACATGGATAAT
GCACACGTCTGAACTCTTTCCCTACACGACGCTCTTCCGATCTNNNCACACACACAAAACAAGATGCATC
AGCACACGTCTGAACTCTTTCCCTACACGACGCTCTTCCGATCTNNNAGTTACTTTTGTTTTCTAATTCT
AGCACACGTCTGAACTCTTTCCCTACACGACGCTCTTCCGATCTNNNACACTTAGTGAATGTTCTATGGC
AGCACACGTCTGAACTCTTTCCCTACACGACGCTCTTCCGATCTNNNCTCCAGAGGAATATAACAATAC
AGCACACGTCTGAACTCTTTCCCTACACGACGCTCTTCCGATCTNNNTGGAATGAGTAGGATCATGAT
GAGCACACGTCTGAACTCTTTCCCTACACGACGCTCTTCCGATCTNNNTTCTAATCTACTCCCAAAATCAA
GAGCACACGTCTGAACTCTTTCCCTACACGACGCTCTTCCGATCTNNNCACTACAAAAATCTGAGAACAT
GAGCACACGTCTGAACTCTTTCCCTACACGACGCTCTTCCGATCTNNNTCAGTAAAAACCAGTAAGGAA
GAGCACACGTCTGAACTCTTTCCCTACACGACGCTCTTCCGATCTNNNGAGAGAAAGGAAGTGTGTAT
AGCACACGTCTGAACTCTTTCCCTACACGACGCTCTTCCGATCTNNNTTCTCCCAGATATTTTTCTATGG
AGCACACGTCTGAACTCTTTCCCTACACGACGCTCTTCCGATCTNNNCAGAAATGATAGTTCTATGTCA
GAGCACACGTCTGAACTCTTTCCCTACACGACGCTCTTCCGATCTNNNTCAGAGTGAGAGTTTATTATA
GAGCACACGTCTGAACTCTTTCCCTACACGACGCTCTTCCGATCTNNNGAGTAAAATGTGTGTTTAGAA
AGCACACGTCTGAACTCTTTCCCTACACGACGCTCTTCCGATCTNNNTATTTATAATGAACACCAACTG
GAGCACACGTCTGAACTCTTTCCCTACACGACGCTCTTCCGATCTNNNCAGTTCTTAATTTGCTCATTTTC
GAGCACACGTCTGAACTCTTTCCCTACACGACGCTCTTCCGATCTNNNACTTTGATGCTTTCTTTAATTCA
AGCACACGTCTGAACTCTTTCCCTACACGACGCTCTTCCGATCTNNNTGAGCTAAACATCACATCAAATT
GAGCACACGTCTGAACTCTTTCCCTACACGACGCTCTTCCGATCTNNNAGTGAAAATCCTTGTCTTAATT
GAGCACACGTCTGAACTCTTTCCCTACACGACGCTCTTCCGATCTNNNATATGGCTTCTGATATTTGAGA
GAGCACACGTCTGAACTCTTTCCCTACACGACGCTCTTCCGATCTNNNATATGGCTTCTGATATTTGAGA
AGCACACGTCTGAACTCTTTCCCTACACGACGCTCTTCCGATCTNNNCATTGTGTTGTTTCATTTAATGT
GAGCACACGTCTGAACTCTTTCCCTACACGACGCTCTTCCGATCTNNNGTGAGATTGATAATTGACCAA
GCACACGTCTGAACTCTTTCCCTACACGACGCTCTTCCGATCTNNNGCACCTGACTGTATATATACATACA
AGCACACGTCTGAACTCTTTCCCTACACGACGCTCTTCCGATCTNNNGAATGTCATGATGACCAGACCA
AGCACACGTCTGAACTCTTTCCCTACACGACGCTCTTCCGATCTNNNATGAATGTCATGATGACCAGCA
ACACGTCTGAACTCTTTCCCTACACGACGCTCTTCCGATCTNNNGGTAATTAATTACTCATTTGGCTCAT
CACACGTCTGAACTCTTTCCCTACACGACGCTCTTCCGATCTNNNCGTATTTTTCCATATTCTTTTCCCATC
GAGCACACGTCTGAACTCTTTCCCTACACGACGCTCTTCCGATCTNNNCTTGTTTTTAGGAAATCTGTA
AGCACACGTCTGAACTCTTTCCCTACACGACGCTCTTCCGATCTNNNCCTCCACAATAATGAAAACTAAT

ACACGTCTGAACTCTTTCCCTACACGACGCTCTTCCGATCTNNNATTTCTATAGATATCTTCCACCACATC
ACACGTCTGAACTCTTTCCCTACACGACGCTCTTCCGATCTNNNATTTCTATAGATATCTTCCACCACATC
GAGCACACGTCTGAACTCTTTCCCTACACGACGCTCTTCCGATCTNNNACCCTTATGAATATAGATGCAA
AGCACACGTCTGAACTCTTTCCCTACACGACGCTCTTCCGATCTNNNCTCACTGAGTAAAGAATAACATC
GAGCACACGTCTGAACTCTTTCCCTACACGACGCTCTTCCGATCTNNNGATAGGCAATAGAGATTTTTC
GAGCACACGTCTGAACTCTTTCCCTACACGACGCTCTTCCGATCTNNNACTGGAGAGAAAACTAAAGTA
AGCACACGTCTGAACTCTTTCCCTACACGACGCTCTTCCGATCTNNNAAATCCAGCATTTTTAGTATCAC
GAGCACACGTCTGAACTCTTTCCCTACACGACGCTCTTCCGATCTNNNGAATGGATAGTAGGATCAAGC
GAGCACACGTCTGAACTCTTTCCCTACACGACGCTCTTCCGATCTNNNAGAAATAGTTCAACTTAACAG
GAGCACACGTCTGAACTCTTTCCCTACACGACGCTCTTCCGATCTNNNTGGACTTTATTCCAAAAGCTAT
AGCACACGTCTGAACTCTTTCCCTACACGACGCTCTTCCGATCTNNNCTCCCTAATTTTCCAAAAGTATAC
CACACGTCTGAACTCTTTCCCTACACGACGCTCTTCCGATCTNNNATAAATGGGACCCTATGACTCATCG
GAGCACACGTCTGAACTCTTTCCCTACACGACGCTCTTCCGATCTNNNAACTCTCTCTCTCTAAATATACA
AGCACACGTCTGAACTCTTTCCCTACACGACGCTCTTCCGATCTNNNTTACTTCGCTTAGAATAATGGTA
GAGCACACGTCTGAACTCTTTCCCTACACGACGCTCTTCCGATCTNNNTATATATGTGTGTGTGCATATC
AGCACACGTCTGAACTCTTTCCCTACACGACGCTCTTCCGATCTNNNAAAGGCAAATTCTAGTCAATTT
AGCACACGTCTGAACTCTTTCCCTACACGACGCTCTTCCGATCTNNNAATCTATGTCATTCATACACACAC
AGCACACGTCTGAACTCTTTCCCTACACGACGCTCTTCCGATCTNNNGTGAACGTGACAAAAATCTTTCA
AGCACACGTCTGAACTCTTTCCCTACACGACGCTCTTCCGATCTNNNAACCTTCTTCTTGTCTAAAACTAC
AGCACACGTCTGAACTCTTTCCCTACACGACGCTCTTCCGATCTNNNGGTTCTCCAAAAAAATAAAACA
AGCACACGTCTGAACTCTTTCCCTACACGACGCTCTTCCGATCTNNNTAACCATAATGCTACCTAGAGA
GAGCACACGTCTGAACTCTTTCCCTACACGACGCTCTTCCGATCTNNNCAAGAGAATGATTCTTTTAGT
GAGCACACGTCTGAACTCTTTCCCTACACGACGCTCTTCCGATCTNNNGTATAGTGCTTTGCTCATTAT
AGCACACGTCTGAACTCTTTCCCTACACGACGCTCTTCCGATCTNNNAATAGAGGGAAATAAACACACA
AGCACACGTCTGAACTCTTTCCCTACACGACGCTCTTCCGATCTNNNGGATTAGTGTCCTTACAAGACCA
AGCACACGTCTGAACTCTTTCCCTACACGACGCTCTTCCGATCTNNNATGTAAATTGTTTCAACTAGTGG
GAGCACACGTCTGAACTCTTTCCCTACACGACGCTCTTCCGATCTNNNAAAAAATATGGAACCTAACCA
CACACGTCTGAACTCTTTCCCTACACGACGCTCTTCCGATCTNNNCTCAAGAATGTCACAAAACCCATCG
AGCACACGTCTGAACTCTTTCCCTACACGACGCTCTTCCGATCTNNNAACAACCCTTTAAAAATGTGTAT
AGCACACGTCTGAACTCTTTCCCTACACGACGCTCTTCCGATCTNNNTAGGATTTCATCTACTAATGCAA
AGAGCACACGTCTGAACTCTTTCCCTACACGACGCTCTTCCGATCTNNNCTCCTAGAAACTTATTTTTGG
AGCACACGTCTGAACTCTTTCCCTACACGACGCTCTTCCGATCTNNNTTGATCGCATTTTTGGATATAT
GAGCACACGTCTGAACTCTTTCCCTACACGACGCTCTTCCGATCTNNNGCAGAAAATGAGACATTAAAA
GCACACGTCTGAACTCTTTCCCTACACGACGCTCTTCCGATCTNNNATTTTCTTAAAACTGACAATCCTCA
GAGCACACGTCTGAACTCTTTCCCTACACGACGCTCTTCCGATCTNNNTCTAAATTTAGAGTTGATATGC
GAGCACACGTCTGAACTCTTTCCCTACACGACGCTCTTCCGATCTNNNTCTAAATTTAGAGTTGATATGC
GAGCACACGTCTGAACTCTTTCCCTACACGACGCTCTTCCGATCTNNNGAAACATATGGAATCAACCTA
AGCACACGTCTGAACTCTTTCCCTACACGACGCTCTTCCGATCTNNNTCCCATAAAAATGATTACAAGA
AGCACACGTCTGAACTCTTTCCCTACACGACGCTCTTCCGATCTNNNTGAATGTTACATATGTTCTCTCT
GAGCACACGTCTGAACTCTTTCCCTACACGACGCTCTTCCGATCTNNNAATTCTGGAAACACAATATCAA
GCACACGTCTGAACTCTTTCCCTACACGACGCTCTTCCGATCTNNNGTTTGTTTTTGCTTCTAGTTTTCCA
GCACACGTCTGAACTCTTTCCCTACACGACGCTCTTCCGATCTNNNTAGAATAGAAATGATCAGAGTGC
AGCACACGTCTGAACTCTTTCCCTACACGACGCTCTTCCGATCTNNNGGAAACATGTGAACCTATTTAT

GAGCACACGTCTGAACTCTTTCCCTACACGACGCTCTTCCGATCTNNNCATTGTAGGTAATCTCCAAG
GCACACGTCTGAACTCTTTCCCTACACGACGCTCTTCCGATCTNNNCAGATACTGTGTCTGTATGACC
GCACACGTCTGAACTCTTTCCCTACACGACGCTCTTCCGATCTNNNCTTAATGTGCTATTTGGTGATA
GCACACGTCTGAACTCTTTCCCTACACGACGCTCTTCCGATCTNNNTACAATCAAGAAATCTCAAAGA
AGCACACGTCTGAACTCTTTCCCTACACGACGCTCTTCCGATCTNNNGTTTAGCCTTTCTAATTGACAT
GCACACGTCTGAACTCTTTCCCTACACGACGCTCTTCCGATCTNNNTTGTGTGTATTTTACGTGTATT
GCACACGTCTGAACTCTTTCCCTACACGACGCTCTTCCGATCTNNNTCTATCTAAATTCGCTAGTTGTT
AGCACACGTCTGAACTCTTTCCCTACACGACGCTCTTCCGATCTNNNAAATAGTCCCACTCACATAAA
AGCACACGTCTGAACTCTTTCCCTACACGACGCTCTTCCGATCTNNNAGTATTCCAGTATTATTCTGC
GCACACGTCTGAACTCTTTCCCTACACGACGCTCTTCCGATCTNNNTAGAGACAAAGCAATTCTATGT
CACACGTCTGAACTCTTTCCCTACACGACGCTCTTCCGATCTNNNTAGTTTAAGTGCAGGTTGTATTTC
CACACGTCTGAACTCTTTCCCTACACGACGCTCTTCCGATCTNNNTAGTTTAAGTGCAGGTTGTATTTC
GCACACGTCTGAACTCTTTCCCTACACGACGCTCTTCCGATCTNNNCCATATACGTACATATATATGGC
AGCACACGTCTGAACTCTTTCCCTACACGACGCTCTTCCGATCTNNNACACACACATACACATATATGT
AGCACACGTCTGAACTCTTTCCCTACACGACGCTCTTCCGATCTNNNGGGAAAAAGTAAAGGTATAA
CACGTCTGAACTCTTTCCCTACACGACGCTCTTCCGATCTNNNCTTTGAATTGCTCCTCAAAGCATCG
GCACACGTCTGAACTCTTTCCCTACACGACGCTCTTCCGATCTNNNTTTATGATTTTTTTGGAAAGCC
GCACACGTCTGAACTCTTTCCCTACACGACGCTCTTCCGATCTNNNCTTTGAGTGATAAAGATGTTCAT
GAGCACACGTCTGAACTCTTTCCCTACACGACGCTCTTCCGATCTNNNAGTGAACTAAGTCAAAAGA
CACACGTCTGAACTCTTTCCCTACACGACGCTCTTCCGATCTNNNCCAGAGTCTAATCTAGTTTTCATC
CACACGTCTGAACTCTTTCCCTACACGACGCTCTTCCGATCTNNNCCAGAGTCTAATCTAGTTTTCATC
GCACACGTCTGAACTCTTTCCCTACACGACGCTCTTCCGATCTNNNTATCCTGAATTTGGTACTTATCA
GCACACGTCTGAACTCTTTCCCTACACGACGCTCTTCCGATCTNNNTTCAAAGTAAGAGAACCAAGAA
GCACACGTCTGAACTCTTTCCCTACACGACGCTCTTCCGATCTNNNAAAGCCTTAGGTCTTCATAAATA
GCACACGTCTGAACTCTTTCCCTACACGACGCTCTTCCGATCTNNNAAAGCCTTAGGTCTTCATAAATA
AGCACACGTCTGAACTCTTTCCCTACACGACGCTCTTCCGATCTNNNTGTAACAAACATGCCATAATAA
AGCACACGTCTGAACTCTTTCCCTACACGACGCTCTTCCGATCTNNNCAGAAGATGTTAGTACTCTCA
GCACACGTCTGAACTCTTTCCCTACACGACGCTCTTCCGATCTNNNTCATTGATTAGTTCTAACAGGAT
AGCACACGTCTGAACTCTTTCCCTACACGACGCTCTTCCGATCTNNNCCTATTTGTGGTACTAATCTGT
GCACACGTCTGAACTCTTTCCCTACACGACGCTCTTCCGATCTNNNATCTCTTGATTGTTATTTGACA
GCACACGTCTGAACTCTTTCCCTACACGACGCTCTTCCGATCTNNNGTAGAGTTAAGTATCTGGATT
AGCACACGTCTGAACTCTTTCCCTACACGACGCTCTTCCGATCTNNNGACATAACTCATGAAACACTTC
GCACACGTCTGAACTCTTTCCCTACACGACGCTCTTCCGATCTNNNACTTTTATCTACTAGTTTCCACA
AGCACACGTCTGAACTCTTTCCCTACACGACGCTCTTCCGATCTNNNAAAGTAGCAGATTTAAAGATT
AGCACACGTCTGAACTCTTTCCCTACACGACGCTCTTCCGATCTNNNAACTACAGATCTTGGAACTTGC
GAGCACACGTCTGAACTCTTTCCCTACACGACGCTCTTCCGATCTNNNCTAGATTTTGCTTATCCATTC
AGCACACGTCTGAACTCTTTCCCTACACGACGCTCTTCCGATCTNNNCTATTAATACACTTTCTCTCCCT
AGCACACGTCTGAACTCTTTCCCTACACGACGCTCTTCCGATCTNNNGATCTCTCTAGACAGATGGACC
GCACACGTCTGAACTCTTTCCCTACACGACGCTCTTCCGATCTNNNATTGCGAAACATTGGTTATAAA
GCACACGTCTGAACTCTTTCCCTACACGACGCTCTTCCGATCTNNNTATTCATTTCTTAGTCAGACTGT
AGCACACGTCTGAACTCTTTCCCTACACGACGCTCTTCCGATCTNNNGGGAATAAACAAGTGCATTAA
GAGCACACGTCTGAACTCTTTCCCTACACGACGCTCTTCCGATCTNNNCTTCCAAAAGCACAAAACACC
AGCACACGTCTGAACTCTTTCCCTACACGACGCTCTTCCGATCTNNNCAACACATAAACATGCACAATA

AGCACACGTCTGAACTCTTTCCCTACACGACGCTCTTCCGATCTNNNCCAATAATAGTGCAGTGTATT
GAGCACACGTCTGAACTCTTTCCCTACACGACGCTCTTCCGATCTNNNGTTTTCTAATTTATTGCAAGC
AGCACACGTCTGAACTCTTTCCCTACACGACGCTCTTCCGATCTNNNCTATTGGTAGTTAAGTTTTTGG
AGCACACGTCTGAACTCTTTCCCTACACGACGCTCTTCCGATCTNNNTTTAGCCATTACATCACTGTAT
AGCACACGTCTGAACTCTTTCCCTACACGACGCTCTTCCGATCTNNNTCTTATGTAGGCTAAAAGATT
AGCACACGTCTGAACTCTTTCCCTACACGACGCTCTTCCGATCTNNNCTTGATTCTCTCTGAACTTTTAT
GCACACGTCTGAACTCTTTCCCTACACGACGCTCTTCCGATCTNNNTAAATCATAGCAGACTAGTTAG
AGCACACGTCTGAACTCTTTCCCTACACGACGCTCTTCCGATCTNNNTTTTATTTTGCTTTGTATTTGG
GAGCACACGTCTGAACTCTTTCCCTACACGACGCTCTTCCGATCTNNNTGTAGAAAGTCAACATAAAG
GAGCACACGTCTGAACTCTTTCCCTACACGACGCTCTTCCGATCTNNNATTTGATGAGATGTAGAATC
GCACACGTCTGAACTCTTTCCCTACACGACGCTCTTCCGATCTNNNGATAAATATCATCTATACACTGA
CACGTCTGAACTCTTTCCCTACACGACGCTCTTCCGATCTNNNCTTCTTTCACCTTTTCTGTCATACATCG
AGCACACGTCTGAACTCTTTCCCTACACGACGCTCTTCCGATCTNNNGGGAGACAGATTTTTTCTTTTT
AGCACACGTCTGAACTCTTTCCCTACACGACGCTCTTCCGATCTNNNAAAGGATGGAGAATAAAAAG
AGCACACGTCTGAACTCTTTCCCTACACGACGCTCTTCCGATCTNNNATTGATGGGTTCATAGATGAA
GAGCACACGTCTGAACTCTTTCCCTACACGACGCTCTTCCGATCTNNNACAAATGATTAGGAAACAGT
AGCACACGTCTGAACTCTTTCCCTACACGACGCTCTTCCGATCTNNNCAATGCACATATATCAGTTGTA
AGCACACGTCTGAACTCTTTCCCTACACGACGCTCTTCCGATCTNNNCTTTATGCTGTATTATGACTTG
GCACACGTCTGAACTCTTTCCCTACACGACGCTCTTCCGATCTNNNTGGGTGAAAACCAATGTATATA
GCACACGTCTGAACTCTTTCCCTACACGACGCTCTTCCGATCTNNNTATACACACATACATACACCTAC
CACACGTCTGAACTCTTTCCCTACACGACGCTCTTCCGATCTNNNTTTTCATTATTTTACACCCTGGCA
CACACGTCTGAACTCTTTCCCTACACGACGCTCTTCCGATCTNNNCATTCCTATTTCCAGAGAGTTCCAT
AGCACACGTCTGAACTCTTTCCCTACACGACGCTCTTCCGATCTNNNACCAATCTATGTGCTAAATACT
AGCACACGTCTGAACTCTTTCCCTACACGACGCTCTTCCGATCTNNNCAAAACAAAACAAACAAAAAA
AGCACACGTCTGAACTCTTTCCCTACACGACGCTCTTCCGATCTNNNAAAACTTAGAGCCCTTGAAGCA
AGCACACGTCTGAACTCTTTCCCTACACGACGCTCTTCCGATCTNNNTTACCCAGAGTATTAGAGTAG
GAGCACACGTCTGAACTCTTTCCCTACACGACGCTCTTCCGATCTNNNCTTCAGAGCTTGAGTATTTAA
AGCACACGTCTGAACTCTTTCCCTACACGACGCTCTTCCGATCTNNNCCAAATCTTCTGTCTGTATTTAC
AGCACACGTCTGAACTCTTTCCCTACACGACGCTCTTCCGATCTNNNCCTCCCATTATATACTTTTTCCTC
AGCACACGTCTGAACTCTTTCCCTACACGACGCTCTTCCGATCTNNNAAATGTAAATATCAAGGGTGG
AGCACACGTCTGAACTCTTTCCCTACACGACGCTCTTCCGATCTNNNAAATGTAAATATCAAGGGTGG
GCACACGTCTGAACTCTTTCCCTACACGACGCTCTTCCGATCTNNNTCTTTCATTTTCACCAGTTATTTC
CACACGTCTGAACTCTTTCCCTACACGACGCTCTTCCGATCTNNNTAATACAAACATGTGGGACCCATC
CACACGTCTGAACTCTTTCCCTACACGACGCTCTTCCGATCTNNNGCCTTTCTCATATATATACACACC
ACACGTCTGAACTCTTTCCCTACACGACGCTCTTCCGATCTNNNCAGACACTGAACTAGCTTCACATCG
AGCACACGTCTGAACTCTTTCCCTACACGACGCTCTTCCGATCTNNNAAGGAAAACATGTAAAACTGA
AGCACACGTCTGAACTCTTTCCCTACACGACGCTCTTCCGATCTNNNCACCAGTCTATATTCTAAGAA
AGCACACGTCTGAACTCTTTCCCTACACGACGCTCTTCCGATCTNNNATGATCTAGCTGTTTCTTTCTC
GCACACGTCTGAACTCTTTCCCTACACGACGCTCTTCCGATCTNNNATTGCATGTAAAAATAGTGTTTC
AGCACACGTCTGAACTCTTTCCCTACACGACGCTCTTCCGATCTNNNGGAATACTAGAACTACATGGT
AGCACACGTCTGAACTCTTTCCCTACACGACGCTCTTCCGATCTNNNCTGGAATACTAGAACTACATG
GAGCACACGTCTGAACTCTTTCCCTACACGACGCTCTTCCGATCTNNNACCATAAACAATAAACTTTCT
GAGCACACGTCTGAACTCTTTCCCTACACGACGCTCTTCCGATCTNNNCTGTGTCAAAATATCCTATGG

GCACACGTCTGAACTCTTTCCCTACACGACGCTCTTCCGATCTNNNCTGTTCCTTATATCTTTCATTCCCA
AGCACACGTCTGAACTCTTTCCCTACACGACGCTCTTCCGATCTNNNGTTTGTTTTTTGTTGTTGTTTT
GCACACGTCTGAACTCTTTCCCTACACGACGCTCTTCCGATCTNNNTAATTCCTGTACCAAAGAATACT
GCACACGTCTGAACTCTTTCCCTACACGACGCTCTTCCGATCTNNNCCTAAAGTAAACAACCTTGAATA
CACGTCTGAACTCTTTCCCTACACGACGCTCTTCCGATCTNNNCATTTCTACAACCCAGAGACCATCGG
GCACACGTCTGAACTCTTTCCCTACACGACGCTCTTCCGATCTNNNCTCAATTATGAAATTTTTCCATC
GAGCACACGTCTGAACTCTTTCCCTACACGACGCTCTTCCGATCTNNNCTTCTCTGCTGGTAAGTTTTCA
AGCACACGTCTGAACTCTTTCCCTACACGACGCTCTTCCGATCTNNNTACAAGTTTCAAAAGTACAAA
GAGCACACGTCTGAACTCTTTCCCTACACGACGCTCTTCCGATCTNNNGCAAGTTACAGACTCAATTAT
AGCACACGTCTGAACTCTTTCCCTACACGACGCTCTTCCGATCTNNNTTCTTGCTCTGTCACTCTTTCAT
AGCACACGTCTGAACTCTTTCCCTACACGACGCTCTTCCGATCTNNNAGAGAGACATTCTTTATAGACT
AGCACACGTCTGAACTCTTTCCCTACACGACGCTCTTCCGATCTNNNATAAGTAATCAGAAAGAAGAC
AGCACACGTCTGAACTCTTTCCCTACACGACGCTCTTCCGATCTNNNACTATTATATGCAACAGTAAGG
GCACACGTCTGAACTCTTTCCCTACACGACGCTCTTCCGATCTNNNACTTTTCTCTTTTGACAGTTTTAC
AGCACACGTCTGAACTCTTTCCCTACACGACGCTCTTCCGATCTNNNTTGTCTTACTTTTCTCTTTTGAC
AGCACACGTCTGAACTCTTTCCCTACACGACGCTCTTCCGATCTNNNAATTCAGGATTAATTGGGTAT
AGCACACGTCTGAACTCTTTCCCTACACGACGCTCTTCCGATCTNNNAATTCAGGATTAATTGGGTAT
CACACGTCTGAACTCTTTCCCTACACGACGCTCTTCCGATCTNNNTCAAACCACTGGAGTTCTAGCATC
GAGCACACGTCTGAACTCTTTCCCTACACGACGCTCTTCCGATCTNNNATTCTGGGTCAATCGAGATT
GCACACGTCTGAACTCTTTCCCTACACGACGCTCTTCCGATCTNNNAGCTTCAGAGAAGATCTAGGCA
GCACACGTCTGAACTCTTTCCCTACACGACGCTCTTCCGATCTNNNTAAATTTTGTCTGTTGCTAAACA
GCACACGTCTGAACTCTTTCCCTACACGACGCTCTTCCGATCTNNNGGGATTTTAAGGGATTCTTGAAC
GCACACGTCTGAACTCTTTCCCTACACGACGCTCTTCCGATCTNNNCTGTAATCCCAACACTTTGTCAT
AGCACACGTCTGAACTCTTTCCCTACACGACGCTCTTCCGATCTNNNACACTAGTATTCTAACCGAAAT
GAGCACACGTCTGAACTCTTTCCCTACACGACGCTCTTCCGATCTNNNGTCACATCAACTGTTGTATAT
GCACACGTCTGAACTCTTTCCCTACACGACGCTCTTCCGATCTNNNTATCATAGTTCACTGCAGCCCATC
GCACACGTCTGAACTCTTTCCCTACACGACGCTCTTCCGATCTNNNTGTTAACTGGAAAGAAGTACTA
GCACACGTCTGAACTCTTTCCCTACACGACGCTCTTCCGATCTNNNTTTGAAATGAAAACCTGTACAT
GCACACGTCTGAACTCTTTCCCTACACGACGCTCTTCCGATCTNNNCTGTTTTAGAAGCTTGTATCTT
GCACACGTCTGAACTCTTTCCCTACACGACGCTCTTCCGATCTNNNTTAGCAAAATCTCATCTGTGTAT
CACACGTCTGAACTCTTTCCCTACACGACGCTCTTCCGATCTNNNCTTTTGCTCCTACCATATTCCCATC
GAGCACACGTCTGAACTCTTTCCCTACACGACGCTCTTCCGATCTNNNGTGTGTGTTTTTGCAATATAA
GCACACGTCTGAACTCTTTCCCTACACGACGCTCTTCCGATCTNNNATATAAGAAGAGAAAGATGGC
CACGTCTGAACTCTTTCCCTACACGACGCTCTTCCGATCTNNNTTTATCATCGCTCTACACATTTCATCG
CACACGTCTGAACTCTTTCCCTACACGACGCTCTTCCGATCTNNNTAATTAATAGCTCATCAGGGTAG
AGCACACGTCTGAACTCTTTCCCTACACGACGCTCTTCCGATCTNNNCCATCCAGAAGTAGAAATAAA
ACACGTCTGAACTCTTTCCCTACACGACGCTCTTCCGATCTNNNGATCTTTAAGGACAAGCTGAGCATC
GAGCACACGTCTGAACTCTTTCCCTACACGACGCTCTTCCGATCTNNNGGTATATGTATGTATGTGGA
AGCACACGTCTGAACTCTTTCCCTACACGACGCTCTTCCGATCTNNNCCTAACACTTAGTCTCACAAATC
CACACGTCTGAACTCTTTCCCTACACGACGCTCTTCCGATCTNNNTAGTTCAAAAAACCAAAAAAACC
AGCACACGTCTGAACTCTTTCCCTACACGACGCTCTTCCGATCTNNNTTTCATCTATTTACCCTTTTGC
GCACACGTCTGAACTCTTTCCCTACACGACGCTCTTCCGATCTNNNGTAGCATAATCAAAATCATGTGT
GCACACGTCTGAACTCTTTCCCTACACGACGCTCTTCCGATCTNNNAATCCTTATGTGAAAGCAAGAT

GCACACGTCTGAACTCTTTCCCTACACGACGCTCTTCCGATCTNNNGTAAGAACCATCCAGTATATGTA

AGAGCACACGTCTGAACTCTTTCCCTACACGACGCTCTTCCGATCTNNNTAAAAAATAAATTGAGCTT

GCACACGTCTGAACTCTTTCCCTACACGACGCTCTTCCGATCTNNNTTACACCCCTAAACTTACATTT

AGCACACGTCTGAACTCTTTCCCTACACGACGCTCTTCCGATCTNNNCTTTTTCATCATCTTCAGCACC

GAGCACACGTCTGAACTCTTTCCCTACACGACGCTCTTCCGATCTNNNGTGAATTTTGGTAAATGTCC

AGCACACGTCTGAACTCTTTCCCTACACGACGCTCTTCCGATCTNNNTAACTTCTCTTTTTGTTATTGCA

AGCACACGTCTGAACTCTTTCCCTACACGACGCTCTTCCGATCTNNNCTTTAACCATTAGGGAAATTC

AGCACACGTCTGAACTCTTTCCCTACACGACGCTCTTCCGATCTNNNATCATCTTGCAAATCAAACTAT

AGCACACGTCTGAACTCTTTCCCTACACGACGCTCTTCCGATCTNNNCTCTGTATATGCACCTTTCATAC

GCACACGTCTGAACTCTTTCCCTACACGACGCTCTTCCGATCTNNNCTATTTTCCTCCACCTATTGAAT

GCACACGTCTGAACTCTTTCCCTACACGACGCTCTTCCGATCTNNNTATTTGTTTTGGTAACTCACACC

AGCACACGTCTGAACTCTTTCCCTACACGACGCTCTTCCGATCTNNNGATGGTTTTAGAAACACACAT

AGCACACGTCTGAACTCTTTCCCTACACGACGCTCTTCCGATCTNNNGATGGTTTTAGAAACACACAT

AGCACACGTCTGAACTCTTTCCCTACACGACGCTCTTCCGATCTNNNGGCACTATATCATCATGTTAAA

GCACACGTCTGAACTCTTTCCCTACACGACGCTCTTCCGATCTNNNCTATGAGAATCTAACTAATGCCT

CACACGTCTGAACTCTTTCCCTACACGACGCTCTTCCGATCTNNNTTTACTCCCTTGGTAAAAATAGAC

GCACACGTCTGAACTCTTTCCCTACACGACGCTCTTCCGATCTNNNGGTCTAGTTCAAGTTCCTTGCAT

GAGCACACGTCTGAACTCTTTCCCTACACGACGCTCTTCCGATCTNNNAAGTTGCCAAACAAAATGTA

ACACGTCTGAACTCTTTCCCTACACGACGCTCTTCCGATCTNNNAAAAAGCTTGACTTCCATGATTAC

AGCACACGTCTGAACTCTTTCCCTACACGACGCTCTTCCGATCTNNNGCTATTTCTGGCGAAATAATA

AGCACACGTCTGAACTCTTTCCCTACACGACGCTCTTCCGATCTNNNACAAAAACAAAAACAAAAAAC

AGCACACGTCTGAACTCTTTCCCTACACGACGCTCTTCCGATCTNNNAAATAACATATTTTGGACAAG

AGCACACGTCTGAACTCTTTCCCTACACGACGCTCTTCCGATCTNNNTTTCAAACTGCCATAATAAAC

GCACACGTCTGAACTCTTTCCCTACACGACGCTCTTCCGATCTNNNACGGAAGTCATATATTATCTACA

GCACACGTCTGAACTCTTTCCCTACACGACGCTCTTCCGATCTNNNACTAAACTACAAACCTTATTTGG

GCACACGTCTGAACTCTTTCCCTACACGACGCTCTTCCGATCTNNNGGATGAAGTCAGTGAAGTTTA

GCACACGTCTGAACTCTTTCCCTACACGACGCTCTTCCGATCTNNNCTATGAAGCATGAGAACTTATT

AGCACACGTCTGAACTCTTTCCCTACACGACGCTCTTCCGATCTNNNATACTATAGTGCATCCTTGATT

AGCACACGTCTGAACTCTTTCCCTACACGACGCTCTTCCGATCTNNNTTGAACTTGAGAGAGATGATT

ACACGTCTGAACTCTTTCCCTACACGACGCTCTTCCGATCTNNNAATCTAAAGAGAGAATAACCAGGC

GAGCACACGTCTGAACTCTTTCCCTACACGACGCTCTTCCGATCTNNNCTCTCTCTCTCTCTCTCTAC

GCACACGTCTGAACTCTTTCCCTACACGACGCTCTTCCGATCTNNNTTGTCTCTCTCCTAAACTTATTC

GCACACGTCTGAACTCTTTCCCTACACGACGCTCTTCCGATCTNNNAGCCAGAATTTTCTCATAGAAT

GAGCACACGTCTGAACTCTTTCCCTACACGACGCTCTTCCGATCTNNNATAAATAAAGCATCTACGTG

AGCACACGTCTGAACTCTTTCCCTACACGACGCTCTTCCGATCTNNNCTTTGGTAAGAATTTTGAGCT

ACACGTCTGAACTCTTTCCCTACACGACGCTCTTCCGATCTNNNCTGAGATGGAAGGATCATTTGCAT

AGCACACGTCTGAACTCTTTCCCTACACGACGCTCTTCCGATCTNNNGTTTGCTTAGTGGATGAATAA

GCACACGTCTGAACTCTTTCCCTACACGACGCTCTTCCGATCTNNNGGAGGAGAGAAATAGAATTTG

GCACACGTCTGAACTCTTTCCCTACACGACGCTCTTCCGATCTNNNATAAACATATAGTGTCTGTGTG

AGCACACGTCTGAACTCTTTCCCTACACGACGCTCTTCCGATCTNNNTAGGAGAATACCATAGTGGCC

AGCACACGTCTGAACTCTTTCCCTACACGACGCTCTTCCGATCTNNNTAGGAGAATACCATAGTGGCC

ACACGTCTGAACTCTTTCCCTACACGACGCTCTTCCGATCTNNNCTATGAAAACAATCCTTACTGGCAT

GCACACGTCTGAACTCTTTCCCTACACGACGCTCTTCCGATCTNNNCCACCTAAAAGTCATTATTATAC

GAGCACACGTCTGAACTCTTTCCCTACACGACGCTCTTCCGATCTNNNTTCTTAAAGTTCTGCCTTACCC
AGAGCACACGTCTGAACTCTTTCCCTACACGACGCTCTTCCGATCTNNNAAGACATCAGCTTCTTAAAG
CACACGTCTGAACTCTTTCCCTACACGACGCTCTTCCGATCTNNNAGAAGGTTGCTATCTTTATGGCAT
AGCACACGTCTGAACTCTTTCCCTACACGACGCTCTTCCGATCTNNNTATATATGTATGTTTGTGTGT
AGCACACGTCTGAACTCTTTCCCTACACGACGCTCTTCCGATCTNNNGTTTTAAGGAGGAAAGCAATA
AGCACACGTCTGAACTCTTTCCCTACACGACGCTCTTCCGATCTNNNTGATATCTGTGAAATTTTTCTG
AGCACACGTCTGAACTCTTTCCCTACACGACGCTCTTCCGATCTNNNAAGTAATTACAGTAACTCATGG
AGCACACGTCTGAACTCTTTCCCTACACGACGCTCTTCCGATCTNNNCTGAAAATATTTAGTGAACAC
AGAGCACACGTCTGAACTCTTTCCCTACACGACGCTCTTCCGATCTNNNCCTCAAAAATAAAAGAGTTT
GCACACGTCTGAACTCTTTCCCTACACGACGCTCTTCCGATCTNNNATGATCATGCCACTGTACTACATC
GCACACGTCTGAACTCTTTCCCTACACGACGCTCTTCCGATCTNNNAGTAACCTCTTTAGCTAATGTCCA
AGCACACGTCTGAACTCTTTCCCTACACGACGCTCTTCCGATCTNNNAAAATACATCTATACTGCTTTCC
AGCACACGTCTGAACTCTTTCCCTACACGACGCTCTTCCGATCTNNNAAGAACTCACTCTCAATGAAAT
AGCACACGTCTGAACTCTTTCCCTACACGACGCTCTTCCGATCTNNNATACTCTCTTTTTGAATCTGTTG
AGCACACGTCTGAACTCTTTCCCTACACGACGCTCTTCCGATCTNNNGTGAAAGTGTCTACTGAGATG
AGCACACGTCTGAACTCTTTCCCTACACGACGCTCTTCCGATCTNNNAAGTATAAGAAGGCTTACCAT
GAGCACACGTCTGAACTCTTTCCCTACACGACGCTCTTCCGATCTNNNCTTCCAAAATGCTGGGATTAC
CACACGTCTGAACTCTTTCCCTACACGACGCTCTTCCGATCTNNNATGTCTTCCATTCTATTTTCTTGCAT
CACACGTCTGAACTCTTTCCCTACACGACGCTCTTCCGATCTNNNAGTCTATACCCAACATACCAGCATC
GCACACGTCTGAACTCTTTCCCTACACGACGCTCTTCCGATCTNNNTGTTTATTTTTGACTGTGATGAT
AGCACACGTCTGAACTCTTTCCCTACACGACGCTCTTCCGATCTNNNATGAATGAATAAGAATGAATG
AGCACACGTCTGAACTCTTTCCCTACACGACGCTCTTCCGATCTNNNGACTGAGACTATTTTTTAGTGT
GCACACGTCTGAACTCTTTCCCTACACGACGCTCTTCCGATCTNNNAAATTCCACACTCACTCTATTTTCA
GCACACGTCTGAACTCTTTCCCTACACGACGCTCTTCCGATCTNNNGCTCATTTTAGAACATGACACCA
CACACGTCTGAACTCTTTCCCTACACGACGCTCTTCCGATCTNNNTACTGAATGCATATTTGTTGAAAC
CACACGTCTGAACTCTTTCCCTACACGACGCTCTTCCGATCTNNNTCTTTGTATCTTTCATAAACACGCAT
ACACGTCTGAACTCTTTCCCTACACGACGCTCTTCCGATCTNNNGAGCAGTCATTTTCGAAATTTCATC
CACACGTCTGAACTCTTTCCCTACACGACGCTCTTCCGATCTNNNGAGAGTTCTCCCAAGCTATTACATC
GCACACGTCTGAACTCTTTCCCTACACGACGCTCTTCCGATCTNNNTACTTAAGGTTCTACCCCTTTTCA
GCACACGTCTGAACTCTTTCCCTACACGACGCTCTTCCGATCTNNNAAACTTGTATGAGAAGGTGAGC
AGCACACGTCTGAACTCTTTCCCTACACGACGCTCTTCCGATCTNNNAGGGAGCTGATTCTAATGAAT
GAGCACACGTCTGAACTCTTTCCCTACACGACGCTCTTCCGATCTNNNAAATCTAGCAGAAAAACAAG
AGCACACGTCTGAACTCTTTCCCTACACGACGCTCTTCCGATCTNNNTGACCTCATATAGCCTTAATTA
AGCACACGTCTGAACTCTTTCCCTACACGACGCTCTTCCGATCTNNNCATATGAGCCAATACCTTATTTT
GAGCACACGTCTGAACTCTTTCCCTACACGACGCTCTTCCGATCTNNNTTGTTTGGGTGGATTTAGATA
GCACACGTCTGAACTCTTTCCCTACACGACGCTCTTCCGATCTNNNATAGCAAAATCCATAATCAACAG
CACACGTCTGAACTCTTTCCCTACACGACGCTCTTCCGATCTNNNGATCTGGTTTATAAGATAGACCAC
AGCACACGTCTGAACTCTTTCCCTACACGACGCTCTTCCGATCTNNNGGATAGGTAACATCTTTCCTTA
AGCACACGTCTGAACTCTTTCCCTACACGACGCTCTTCCGATCTNNNGAAGTTTAAGTTTACCTTCAGT
AGCACACGTCTGAACTCTTTCCCTACACGACGCTCTTCCGATCTNNNCTTCTTTCATCTCTCTCATTCTCA
AGCACACGTCTGAACTCTTTCCCTACACGACGCTCTTCCGATCTNNNAAGGCAGGATAATCACTTGAA
AGCACACGTCTGAACTCTTTCCCTACACGACGCTCTTCCGATCTNNNAAAAATATTGCAAAGTCATTCA
GCACACGTCTGAACTCTTTCCCTACACGACGCTCTTCCGATCTNNNGTTAGTCCTATCTGCTGTTTCCAT

GAGCACACGTCTGAACTCTTTCCCTACACGACGCTCTTCCGATCTNNNTTATTTGATTGGTAGGAAAA

AGCACACGTCTGAACTCTTTCCCTACACGACGCTCTTCCGATCTNNNGTGGATATCAACATAGTAAAG

GCACACGTCTGAACTCTTTCCCTACACGACGCTCTTCCGATCTNNNGCAGGTTTGTTACATATGTATAC

AGCACACGTCTGAACTCTTTCCCTACACGACGCTCTTCCGATCTNNNTGAGTGGGATAGGTTGTAATC

GCACACGTCTGAACTCTTTCCCTACACGACGCTCTTCCGATCTNNNAAAAACGAACCTAAAACAATAAC

GAGCACACGTCTGAACTCTTTCCCTACACGACGCTCTTCCGATCTNNNTTAATAAAATTGCCAGTAAGC

GAGCACACGTCTGAACTCTTTCCCTACACGACGCTCTTCCGATCTNNNTAATTCTAGGAGAGTTTTTG

AGCACACGTCTGAACTCTTTCCCTACACGACGCTCTTCCGATCTNNNAGGTAAAATGAGGGAAAAGA

CACACGTCTGAACTCTTTCCCTACACGACGCTCTTCCGATCTNNNAAGGCTTCATCCTGCATTTACATC

GAGCACACGTCTGAACTCTTTCCCTACACGACGCTCTTCCGATCTNNNGTAACAAGATATTCCTACCTC

CACACGTCTGAACTCTTTCCCTACACGACGCTCTTCCGATCTNNNTTTTTTTTTTTTCTAGTGAGCACCA

CACACGTCTGAACTCTTTCCCTACACGACGCTCTTCCGATCTNNNCATGGACCTAACTGGAATATCCAT

GCACACGTCTGAACTCTTTCCCTACACGACGCTCTTCCGATCTNNNTATAAACTTGAGGTAGGCCACAT

AGCACACGTCTGAACTCTTTCCCTACACGACGCTCTTCCGATCTNNNCTTACCCAAATTCCCTAAAATTG

GCACACGTCTGAACTCTTTCCCTACACGACGCTCTTCCGATCTNNNAGAATAGCACAGAAAGAAGAG

AGCACACGTCTGAACTCTTTCCCTACACGACGCTCTTCCGATCTNNNGTAATAACTACATAACTGCATC

AGCACACGTCTGAACTCTTTCCCTACACGACGCTCTTCCGATCTNNNAAAGACGTGATTCCATTCTTTT

GCACACGTCTGAACTCTTTCCCTACACGACGCTCTTCCGATCTNNNAGTTTTTTTTTTAGAGACAAGGT

GAGCACACGTCTGAACTCTTTCCCTACACGACGCTCTTCCGATCTNNNTGTGATTTCTCTCCTAGTTTT

GAGCACACGTCTGAACTCTTTCCCTACACGACGCTCTTCCGATCTNNNTCAAAGGTATTTGTTGAGTTT

GAGCACACGTCTGAACTCTTTCCCTACACGACGCTCTTCCGATCTNNNCAAAATGGAGTTCAAAGAATT

GCACACGTCTGAACTCTTTCCCTACACGACGCTCTTCCGATCTNNNGAAATAAATGCCTCTTCTCATCC

GAGCACACGTCTGAACTCTTTCCCTACACGACGCTCTTCCGATCTNNNTTTCTAGTCCCTATGCTGTATC

AGCACACGTCTGAACTCTTTCCCTACACGACGCTCTTCCGATCTNNNGAGGCTAAAAAATACCTGATA

AGCACACGTCTGAACTCTTTCCCTACACGACGCTCTTCCGATCTNNNCGTCCCTTATCTTATAAACAAA

GAGCACACGTCTGAACTCTTTCCCTACACGACGCTCTTCCGATCTNNNACATTAAAAATGTCATGATG

CACACGTCTGAACTCTTTCCCTACACGACGCTCTTCCGATCTNNNACCAAACATGTCCTTAAAGGCATC

GCACACGTCTGAACTCTTTCCCTACACGACGCTCTTCCGATCTNNNTATATCATCTGAAAAAAGGACA

AGCACACGTCTGAACTCTTTCCCTACACGACGCTCTTCCGATCTNNNGGAAATCTTGTAAGCATCATT

AGCACACGTCTGAACTCTTTCCCTACACGACGCTCTTCCGATCTNNNCCTGTTTCATACCACTTTATTT

AGCACACGTCTGAACTCTTTCCCTACACGACGCTCTTCCGATCTNNNACTGAAACAAGTCTATATGCT

GCACACGTCTGAACTCTTTCCCTACACGACGCTCTTCCGATCTNNNATCAAATAAAAATGGAGCTATCC

CACACGTCTGAACTCTTTCCCTACACGACGCTCTTCCGATCTNNNGAAGCTGCCCTGAAATTTAGCATC

GAGCACACGTCTGAACTCTTTCCCTACACGACGCTCTTCCGATCTNNNTATAGAGAATACACATGAGA

GCACACGTCTGAACTCTTTCCCTACACGACGCTCTTCCGATCTNNNGGAATATGTGTTTATTTCAAGCTC

GAGCACACGTCTGAACTCTTTCCCTACACGACGCTCTTCCGATCTNNNTACAGTATTTAGAAACCAAGT

GAGCACACGTCTGAACTCTTTCCCTACACGACGCTCTTCCGATCTNNNTACAGTATTTAGAAACCAAGT

AGCACACGTCTGAACTCTTTCCCTACACGACGCTCTTCCGATCTNNNAATTCAAGGTAAGATTTGGGT

GAGCACACGTCTGAACTCTTTCCCTACACGACGCTCTTCCGATCTNNNGAAGTTTTGGTAGTTAAGAG

CACACGTCTGAACTCTTTCCCTACACGACGCTCTTCCGATCTNNNTCAAGGAAATGTATACACATAGA

AGCACACGTCTGAACTCTTTCCCTACACGACGCTCTTCCGATCTNNNTGTAAAATAGTCATTCTTTGCTT

GAGCACACGTCTGAACTCTTTCCCTACACGACGCTCTTCCGATCTNNNCACACTTATTTATTACTCCTG

ACACGTCTGAACTCTTTCCCTACACGACGCTCTTCCGATCTNNNGAAATTGAATTAGCGTTGGCCATCG

CACACGTCTGAACTCTTTCCCTACACGACGCTCTTCCGATCTNNNGAGTCTACCTGAGTTTGTATTCCAT
GAGCACACGTCTGAACTCTTTCCCTACACGACGCTCTTCCGATCTNNNGTAACATTCTGCTACAGGTTA
GAGCACACGTCTGAACTCTTTCCCTACACGACGCTCTTCCGATCTNNNTAAATAACTAGGCACAGTAAA
GCACACGTCTGAACTCTTTCCCTACACGACGCTCTTCCGATCTNNNCACCTTTGATAGAAGCTTTTATTCA
AGCACACGTCTGAACTCTTTCCCTACACGACGCTCTTCCGATCTNNNGCATTTATTATGTGACACAGTA
AGCACACGTCTGAACTCTTTCCCTACACGACGCTCTTCCGATCTNNNTGCTTCATTTTTCTATTTTCAGA
GAGCACACGTCTGAACTCTTTCCCTACACGACGCTCTTCCGATCTNNNGATATACAGAGACACAACACA
AGCACACGTCTGAACTCTTTCCCTACACGACGCTCTTCCGATCTNNNCAAGACAAAACAAGAAAAAACA
GCACACGTCTGAACTCTTTCCCTACACGACGCTCTTCCGATCTNNNTAAGAAGGAAATAAAAGCGAATT
ACACGTCTGAACTCTTTCCCTACACGACGCTCTTCCGATCTNNNAAAGCTGAAGGAAGACGAATACATC
GAGCACACGTCTGAACTCTTTCCCTACACGACGCTCTTCCGATCTNNNGTGAGTATTAAGTAATCATGC
AGCACACGTCTGAACTCTTTCCCTACACGACGCTCTTCCGATCTNNNAGAACTTCAGATCACCAAATAG
AGCACACGTCTGAACTCTTTCCCTACACGACGCTCTTCCGATCTNNNAGAACTTCAGATCACCAAATAG
AGCACACGTCTGAACTCTTTCCCTACACGACGCTCTTCCGATCTNNNAAGGGCTAAAGTAAAAATTATC
AGCACACGTCTGAACTCTTTCCCTACACGACGCTCTTCCGATCTNNNAACCAAAATAATATCATGCTTCA
AGCACACGTCTGAACTCTTTCCCTACACGACGCTCTTCCGATCTNNNATAGGACAGAATCATACAAAAG
GAGCACACGTCTGAACTCTTTCCCTACACGACGCTCTTCCGATCTNNNGGACTATATCCTGACTTTAGA
GAGCACACGTCTGAACTCTTTCCCTACACGACGCTCTTCCGATCTNNNTCCAAATGATGAAACTAACTAC
AGCACACGTCTGAACTCTTTCCCTACACGACGCTCTTCCGATCTNNNAATTAAATCTGCCAGTGTTTGCA
GCACACGTCTGAACTCTTTCCCTACACGACGCTCTTCCGATCTNNNCTCATCACCTTGTCTGACATCATCC
AGCACACGTCTGAACTCTTTCCCTACACGACGCTCTTCCGATCTNNNCAAGCTAGTTACTTCCTAGATAC
AGCACACGTCTGAACTCTTTCCCTACACGACGCTCTTCCGATCTNNNTTAAGACCTTGTATTAAATGCTC
AGCACACGTCTGAACTCTTTCCCTACACGACGCTCTTCCGATCTNNNCATCAGCATAATCACTGAATTCC
GCACACGTCTGAACTCTTTCCCTACACGACGCTCTTCCGATCTNNNGAGACACATCATGTTTTAGTTGCA
AGCACACGTCTGAACTCTTTCCCTACACGACGCTCTTCCGATCTNNNGTAAGGGCCCAGATTATTACCA
AGCACACGTCTGAACTCTTTCCCTACACGACGCTCTTCCGATCTNNNTCTATTATTTGATGCAGTGACAT
AGCACACGTCTGAACTCTTTCCCTACACGACGCTCTTCCGATCTNNNGTCAAAATGGCTCATCTAAATTC
AGCACACGTCTGAACTCTTTCCCTACACGACGCTCTTCCGATCTNNNTATCTCCTTTCATCTTCCCCCATC
AGCACACGTCTGAACTCTTTCCCTACACGACGCTCTTCCGATCTNNNGCAATGATGTAATTTTTTTCTCTC
AGCACACGTCTGAACTCTTTCCCTACACGACGCTCTTCCGATCTNNNGAGCTCTGTTATGTATTTTTAGT
GAGCACACGTCTGAACTCTTTCCCTACACGACGCTCTTCCGATCTNNNATGGATGTGATTCACTTTTTAA
GAGCACACGTCTGAACTCTTTCCCTACACGACGCTCTTCCGATCTNNNCTCTGAAAGTCTAAATTACAAC
AGCACACGTCTGAACTCTTTCCCTACACGACGCTCTTCCGATCTNNNTATAGTAGAGAATGTGCTTCAA
GCACACGTCTGAACTCTTTCCCTACACGACGCTCTTCCGATCTNNNTGGATACATACACACAAAAGGCA
AGCACACGTCTGAACTCTTTCCCTACACGACGCTCTTCCGATCTNNNTTCTCTGCATAAAAAGGATATCT
AGCACACGTCTGAACTCTTTCCCTACACGACGCTCTTCCGATCTNNNTCTGATCACTAATATCTCCCTAT
AGCACACGTCTGAACTCTTTCCCTACACGACGCTCTTCCGATCTNNNCTTTCTCATTAGTTGTCCCATAC
GCACACGTCTGAACTCTTTCCCTACACGACGCTCTTCCGATCTNNNGACAATAAGGGAATAAAAGCTGC
AGCACACGTCTGAACTCTTTCCCTACACGACGCTCTTCCGATCTNNNGAGAAAACAAATGAAGCAAAT
AGCACACGTCTGAACTCTTTCCCTACACGACGCTCTTCCGATCTNNNCTGAATAGGAAAAAAATGAAGG
AGCACACGTCTGAACTCTTTCCCTACACGACGCTCTTCCGATCTNNNAAAAATCTGAAACAGCTATAGA
CACACGTCTGAACTCTTTCCCTACACGACGCTCTTCCGATCTNNNCAAACATAACTGGGTAGATTCTCAT
GAGCACACGTCTGAACTCTTTCCCTACACGACGCTCTTCCGATCTNNNTATACATTTTAGGGAGACATG

AGCACACGTCTGAACTCTTTCCCTACACGACGCTCTTCCGATCTNNNATAAATTACAGGTTGAGTATC

AGCACACGTCTGAACTCTTTCCCTACACGACGCTCTTCCGATCTNNNCATATTTCTACATTCCAAGGAA

AGCACACGTCTGAACTCTTTCCCTACACGACGCTCTTCCGATCTNNNTGGAGTACTTTTTCATCTCTTT

GCACACGTCTGAACTCTTTCCCTACACGACGCTCTTCCGATCTNNNAGGAGAATAGCTTGAACCTGC

CACGTCTGAACTCTTTCCCTACACGACGCTCTTCCGATCTNNNCAGTAAGATTTACTGCAAAGAGCAT

GAGCACACGTCTGAACTCTTTCCCTACACGACGCTCTTCCGATCTNNNTATTTGTCACTTTTTCTGTCTT

GAGCACACGTCTGAACTCTTTCCCTACACGACGCTCTTCCGATCTNNNTATTTGTCACTTTTTCTGTCTT

GCACACGTCTGAACTCTTTCCCTACACGACGCTCTTCCGATCTNNNCTCTTTATAACAGTGTGAAAAC

AGCACACGTCTGAACTCTTTCCCTACACGACGCTCTTCCGATCTNNNTGTTTTTCCTGTGGATATAGA

AGCACACGTCTGAACTCTTTCCCTACACGACGCTCTTCCGATCTNNNTTGTAGTTGGAGGTAGGAAT

AGCACACGTCTGAACTCTTTCCCTACACGACGCTCTTCCGATCTNNNTTGTAGTGGTTGTTGTTAGA

GCACACGTCTGAACTCTTTCCCTACACGACGCTCTTCCGATCTNNNATTAAAAACAATCATATGCACA

CACACGTCTGAACTCTTTCCCTACACGACGCTCTTCCGATCTNNNCATCATCCCATTCATCTACCCATCG

AGCACACGTCTGAACTCTTTCCCTACACGACGCTCTTCCGATCTNNNCTGTGGCTTTTTCTATGTATAT

GCACACGTCTGAACTCTTTCCCTACACGACGCTCTTCCGATCTNNNGACTCTGTATCGAAAAAAAACTT

AGCACACGTCTGAACTCTTTCCCTACACGACGCTCTTCCGATCTNNNTCATAACTTAGAAGTGGAGAA

GCACACGTCTGAACTCTTTCCCTACACGACGCTCTTCCGATCTNNNAGGACATGGCTCTATTTTAAGC

AGCACACGTCTGAACTCTTTCCCTACACGACGCTCTTCCGATCTNNNAAGCATTTCATGATTTCAGGT

AGCACACGTCTGAACTCTTTCCCTACACGACGCTCTTCCGATCTNNNCTCAATTTTCCATCCATCCTACA

ACACGTCTGAACTCTTTCCCTACACGACGCTCTTCCGATCTNNNCAGTTCCATGCTACATCATTCATCG

ACACGTCTGAACTCTTTCCCTACACGACGCTCTTCCGATCTNNNAACTACTACAATAAAACATTGGGC

ACACGTCTGAACTCTTTCCCTACACGACGCTCTTCCGATCTNNNTGGTACAATATGAGGCTGAACATC

GAGCACACGTCTGAACTCTTTCCCTACACGACGCTCTTCCGATCTNNNGGCACACCAGGAAATTATAT

GCACACGTCTGAACTCTTTCCCTACACGACGCTCTTCCGATCTNNNATATCCAACTTTCTTGCCTAGCA

AGCACACGTCTGAACTCTTTCCCTACACGACGCTCTTCCGATCTNNNAAATGCCACATATTGTATAAC

GCACACGTCTGAACTCTTTCCCTACACGACGCTCTTCCGATCTNNNATTGATCTATGTAATCACCGAG

GCACACGTCTGAACTCTTTCCCTACACGACGCTCTTCCGATCTNNNCAAATCTCATCTTGAATTGTAGT

GCACACGTCTGAACTCTTTCCCTACACGACGCTCTTCCGATCTNNNAATCAGATGATAAACTCCTTGAT

GCACACGTCTGAACTCTTTCCCTACACGACGCTCTTCCGATCTNNNCAATTACCACCCAATCTGAGCAT

AGCACACGTCTGAACTCTTTCCCTACACGACGCTCTTCCGATCTNNNTATTCATTGAGAGAGAAAGA

GCACACGTCTGAACTCTTTCCCTACACGACGCTCTTCCGATCTNNNAACATTGCAAACACAAAGTCCAT

AGCACACGTCTGAACTCTTTCCCTACACGACGCTCTTCCGATCTNNNACATAAGTCTAGGAGTTTGAA

GAGCACACGTCTGAACTCTTTCCCTACACGACGCTCTTCCGATCTNNNGTCGTGACCAAGTATAAGAT

AGCACACGTCTGAACTCTTTCCCTACACGACGCTCTTCCGATCTNNNTAAGGATGACAAGTGATATTT

AGCACACGTCTGAACTCTTTCCCTACACGACGCTCTTCCGATCTNNNTAATCTAACGGAGGAGTGAG

AGCACACGTCTGAACTCTTTCCCTACACGACGCTCTTCCGATCTNNNTTGATTCAGTCTGTGATCATTC

AGCACACGTCTGAACTCTTTCCCTACACGACGCTCTTCCGATCTNNNATAAAAGAGGAAATAAAAGG

GCACACGTCTGAACTCTTTCCCTACACGACGCTCTTCCGATCTNNNAATATGGAAAACAGGACAAAAT

GAGCACACGTCTGAACTCTTTCCCTACACGACGCTCTTCCGATCTNNNCTTCTCAAAAAAATGTGTCTC

AGCACACGTCTGAACTCTTTCCCTACACGACGCTCTTCCGATCTNNNTTGTGGCTGGTACAAGTTTAC

AGCACACGTCTGAACTCTTTCCCTACACGACGCTCTTCCGATCTNNNATATGTAAAACCCCCAAAAAT

GCACACGTCTGAACTCTTTCCCTACACGACGCTCTTCCGATCTNNNATTTTGTAGCCCATGGAGATAT

AGCACACGTCTGAACTCTTTCCCTACACGACGCTCTTCCGATCTNNNTTTATTTGTGGAGATTCTTTG

GCACACGTCTGAACTCTTTCCCTACACGACGCTCTTCCGATCTNNNCAAAGAAGTTCCAAGTACAATA
CACGTCTGAACTCTTTCCCTACACGACGCTCTTCCGATCTNNNATATTTAGGCTTTGCAGGACCATCG
GAGCACACGTCTGAACTCTTTCCCTACACGACGCTCTTCCGATCTNNNTCTTTTTATGGCAGAGTAGT
GAGCACACGTCTGAACTCTTTCCCTACACGACGCTCTTCCGATCTNNNTTGTACACAAGAATCACCTGC
AGCACACGTCTGAACTCTTTCCCTACACGACGCTCTTCCGATCTNNNATATAAAACATGTTTTACAGG
AGCACACGTCTGAACTCTTTCCCTACACGACGCTCTTCCGATCTNNNGGCCTGGTTAGAATATTTTTAA
AGCACACGTCTGAACTCTTTCCCTACACGACGCTCTTCCGATCTNNNTCAACATAAATTCATATTTGCCT
AGCACACGTCTGAACTCTTTCCCTACACGACGCTCTTCCGATCTNNNAAGCTGCATATCAGAGTCTAC
AGCACACGTCTGAACTCTTTCCCTACACGACGCTCTTCCGATCTNNNGGTGTGTTGAACATACTTTTA
AGCACACGTCTGAACTCTTTCCCTACACGACGCTCTTCCGATCTNNNTAAGGTGAACTAAAGACTATTC
CACACGTCTGAACTCTTTCCCTACACGACGCTCTTCCGATCTNNNGTCACCTAAATCTTTAACTTCAAC
AGCACACGTCTGAACTCTTTCCCTACACGACGCTCTTCCGATCTNNNTGTTTCTGAAGTAAAAGTTGA
GCACACGTCTGAACTCTTTCCCTACACGACGCTCTTCCGATCTNNNGCTTATTTTAATAGGTGTGTGAA
CACACGTCTGAACTCTTTCCCTACACGACGCTCTTCCGATCTNNNGAATAATTCTTGCATTCCTTAGTCA
GAGCACACGTCTGAACTCTTTCCCTACACGACGCTCTTCCGATCTNNNAATAGTGGTTTCCTGGAACT
ACACGTCTGAACTCTTTCCCTACACGACGCTCTTCCGATCTNNNTTAAGCCTTGTGGTCTCTTTACATC
AGCACACGTCTGAACTCTTTCCCTACACGACGCTCTTCCGATCTNNNAAACAAACAAACAAACAAAAT
ACACGTCTGAACTCTTTCCCTACACGACGCTCTTCCGATCTNNNTAATGCATTGCTTCTGTAATAAAC
AGCACACGTCTGAACTCTTTCCCTACACGACGCTCTTCCGATCTNNNTAAGGAAAATAGCTCTAGTAC
AGCACACGTCTGAACTCTTTCCCTACACGACGCTCTTCCGATCTNNNGTGTTTGGTTACATGTATAAGT
GAGCACACGTCTGAACTCTTTCCCTACACGACGCTCTTCCGATCTNNNTATACATGATTGCTACAATCT
AGCACACGTCTGAACTCTTTCCCTACACGACGCTCTTCCGATCTNNNTCACACTTATGAATATCACAGA
AGCACACGTCTGAACTCTTTCCCTACACGACGCTCTTCCGATCTNNNTAATTCTTGTTTTTCTAGGAGC
AGCACACGTCTGAACTCTTTCCCTACACGACGCTCTTCCGATCTNNNCACTGTCTAATAGGGTTGTAAT
AGCACACGTCTGAACTCTTTCCCTACACGACGCTCTTCCGATCTNNNATACACAAACACAAATATGAGA
AGCACACGTCTGAACTCTTTCCCTACACGACGCTCTTCCGATCTNNNACCTTGAAAGTCTGTTTTATTT
AGCACACGTCTGAACTCTTTCCCTACACGACGCTCTTCCGATCTNNNACATAAACTTATGGTTAAACA
AGCACACGTCTGAACTCTTTCCCTACACGACGCTCTTCCGATCTNNNATCTCATTTGTTGTTTTGATTT
CACACGTCTGAACTCTTTCCCTACACGACGCTCTTCCGATCTNNNTATGTTACCATTTGTGGAGAGCAT
CACACGTCTGAACTCTTTCCCTACACGACGCTCTTCCGATCTNNNTTTTTCCTTCTTGCCTTCTTCATCG
AGCACACGTCTGAACTCTTTCCCTACACGACGCTCTTCCGATCTNNNTTATTATTGGAGAAGCTCTACT
GCACACGTCTGAACTCTTTCCCTACACGACGCTCTTCCGATCTNNNTTATCAATGAGAAAACAGTTGCC
AGCACACGTCTGAACTCTTTCCCTACACGACGCTCTTCCGATCTNNNGAAAATTGAGTGGTATGAAC
AGCACACGTCTGAACTCTTTCCCTACACGACGCTCTTCCGATCTNNNTAAATCTCTCTTTCTCTCCATAC
GCACACGTCTGAACTCTTTCCCTACACGACGCTCTTCCGATCTNNNATTCTAGAGAGACAGTTATCAA
AGCACACGTCTGAACTCTTTCCCTACACGACGCTCTTCCGATCTNNNTTTGTGATATTCAACATCTAGG
AGCACACGTCTGAACTCTTTCCCTACACGACGCTCTTCCGATCTNNNTTTGTGATATTCAACATCTAGG
GCACACGTCTGAACTCTTTCCCTACACGACGCTCTTCCGATCTNNNATTTTATCAATAACTCCACCATG
AGCACACGTCTGAACTCTTTCCCTACACGACGCTCTTCCGATCTNNNGAGAAGAGAGATGAGACTTT
GAGCACACGTCTGAACTCTTTCCCTACACGACGCTCTTCCGATCTNNNGGAAACAAAAAGAGCTTGA
AGCACACGTCTGAACTCTTTCCCTACACGACGCTCTTCCGATCTNNNCTCATACAATGCAAAAGATACT
AGCACACGTCTGAACTCTTTCCCTACACGACGCTCTTCCGATCTNNNTATTTTTCTGAAGCATAGTCA
AGCACACGTCTGAACTCTTTCCCTACACGACGCTCTTCCGATCTNNNTATTTTTCTGAAGCATAGTCA

ACACGTCTGAACTCTTTCCCTACACGACGCTCTTCCGATCTNNNTGTAAAATAAAACTGCCATCTTTCAT
ACACGTCTGAACTCTTTCCCTACACGACGCTCTTCCGATCTNNNGAATTGCTTGAACCCAGAAGCATCGG
GAGCACACGTCTGAACTCTTTCCCTACACGACGCTCTTCCGATCTNNNTCACCCTCTTAGAGATTTTTTA
AGCACACGTCTGAACTCTTTCCCTACACGACGCTCTTCCGATCTNNNGTTTATGAATTGTGACAAATGC
AGCACACGTCTGAACTCTTTCCCTACACGACGCTCTTCCGATCTNNNCCATCAGATTCTCAAAGAATATC
AGCACACGTCTGAACTCTTTCCCTACACGACGCTCTTCCGATCTNNNATCTTACAATTCCAGTTTTCAAG
AGCACACGTCTGAACTCTTTCCCTACACGACGCTCTTCCGATCTNNNTAAATGAGGTTGGCAATTGATA
GAGCACACGTCTGAACTCTTTCCCTACACGACGCTCTTCCGATCTNNNGAACTCTCTCAAATAATGGAA
GAGCACACGTCTGAACTCTTTCCCTACACGACGCTCTTCCGATCTNNNTGAACTTTATATTTGAAAATGC
CACACGTCTGAACTCTTTCCCTACACGACGCTCTTCCGATCTNNNTTTAATGATGTTTGTTCCAACAGCAT
GCACACGTCTGAACTCTTTCCCTACACGACGCTCTTCCGATCTNNNGTTTTAATGATGTTTGTTCCAACCA
AGCACACGTCTGAACTCTTTCCCTACACGACGCTCTTCCGATCTNNNGTTTTTTTTTTCTAGTTGTACCG
GCACACGTCTGAACTCTTTCCCTACACGACGCTCTTCCGATCTNNNGAGATAGATTGTGTCATTGCACA
AGCACACGTCTGAACTCTTTCCCTACACGACGCTCTTCCGATCTNNNTGGATTTTTAAAACTGTGAAGT
ACACGTCTGAACTCTTTCCCTACACGACGCTCTTCCGATCTNNNTGAAAAATTCAGAGGCTACTGCATC
AGAGCACACGTCTGAACTCTTTCCCTACACGACGCTCTTCCGATCTNNNCTTCTAAACTTTGAAATTACC
AGCACACGTCTGAACTCTTTCCCTACACGACGCTCTTCCGATCTNNNGCATTTTCTGTTTTTCTTTTGTTCA
GCACACGTCTGAACTCTTTCCCTACACGACGCTCTTCCGATCTNNNCTCCATAATCTTATGAGCCAATACA
AGCACACGTCTGAACTCTTTCCCTACACGACGCTCTTCCGATCTNNNGTGTGTATGTGTTTGTGTATAT
AGCACACGTCTGAACTCTTTCCCTACACGACGCTCTTCCGATCTNNNATGAGTGAATGAATGAGTAAA
GAGCACACGTCTGAACTCTTTCCCTACACGACGCTCTTCCGATCTNNNCTCCCCAGTGTATATGTATATT
AGAGCACACGTCTGAACTCTTTCCCTACACGACGCTCTTCCGATCTNNNAAACTAGGATTGAACAAAGC
GAGCACACGTCTGAACTCTTTCCCTACACGACGCTCTTCCGATCTNNNTCTAGTGTATATCATGGTAGCT
AGCACACGTCTGAACTCTTTCCCTACACGACGCTCTTCCGATCTNNNCTTCTACATGTGAGTATGTGTAC
AGAGCACACGTCTGAACTCTTTCCCTACACGACGCTCTTCCGATCTNNNGGCAATAAAAAAAAAAGAG
AGCACACGTCTGAACTCTTTCCCTACACGACGCTCTTCCGATCTNNNCAGTAATTCTGGCCAATGAGCAT
CACACGTCTGAACTCTTTCCCTACACGACGCTCTTCCGATCTNNNCAGGTTTAAGGTTGATGATTTTAC
AGCACACGTCTGAACTCTTTCCCTACACGACGCTCTTCCGATCTNNNAATTCAAGGTAAGATTTGGGT
CACACGTCTGAACTCTTTCCCTACACGACGCTCTTCCGATCTNNNAGGCATTTTTCCCAATATATTTTCAT
ACACGTCTGAACTCTTTCCCTACACGACGCTCTTCCGATCTNNNTTTATATGGAGAGAGATTTCCCCA
GAGCACACGTCTGAACTCTTTCCCTACACGACGCTCTTCCGATCTNNNGAAGAGGATATTGTTACTTTCA
ACACGTCTGAACTCTTTCCCTACACGACGCTCTTCCGATCTNNNATTATTGTGCAGTACTGAACTCATCG
CACACGTCTGAACTCTTTCCCTACACGACGCTCTTCCGATCTNNNGATCACAGGAAAGCTCAAAGCATC
ACACGTCTGAACTCTTTCCCTACACGACGCTCTTCCGATCTNNNAATGTATCCCTGTAAGAAAGTAACA
CACACGTCTGAACTCTTTCCCTACACGACGCTCTTCCGATCTNNNTGTTCTCAACATCTGCTTAGCATCGC
AGCACACGTCTGAACTCTTTCCCTACACGACGCTCTTCCGATCTNNNTCTATCACCTATCAGCTTTTATG
GCACACGTCTGAACTCTTTCCCTACACGACGCTCTTCCGATCTNNNGCCAATTCCTTATCATAAATCTGCA
AGCACACGTCTGAACTCTTTCCCTACACGACGCTCTTCCGATCTNNNCTACTTCAGACAACACTTTAAAGC
GAGCACACGTCTGAACTCTTTCCCTACACGACGCTCTTCCGATCTNNNGCCTGCTTTTATATGTGAAATA
ACACGTCTGAACTCTTTCCCTACACGACGCTCTTCCGATCTNNNCAATCATCCATCCATCTTATCTACATCC
AGCACACGTCTGAACTCTTTCCCTACACGACGCTCTTCCGATCTNNNGGACTTAGGATTGTGGAGAGC
ACACGTCTGAACTCTTTCCCTACACGACGCTCTTCCGATCTNNNTTTTGAAAAGTGTCTGTTCACCATCC
AGCACACGTCTGAACTCTTTCCCTACACGACGCTCTTCCGATCTNNNGCTGAATCTAAAAGAACTGAAA

ACACGTCTGAACTCTTTCCCTACACGACGCTCTTCCGATCTNNNGTTGCTACACTTCTTTTGTTCATCG
AGCACACGTCTGAACTCTTTCCCTACACGACGCTCTTCCGATCTNNNATATGTGGCTCAGATAAAAAT
AGCACACGTCTGAACTCTTTCCCTACACGACGCTCTTCCGATCTNNNAAAACCTTGTTTAGTGGAATCT
AGCACACGTCTGAACTCTTTCCCTACACGACGCTCTTCCGATCTNNNCCCATCAATTTCTTCTTTCATAA
GCACACGTCTGAACTCTTTCCCTACACGACGCTCTTCCGATCTNNNAAAGTCCTGTATCTAAATTTTGT
ACACGTCTGAACTCTTTCCCTACACGACGCTCTTCCGATCTNNNAAGACACAAGAGAGAGAAATCCAT
GCACACGTCTGAACTCTTTCCCTACACGACGCTCTTCCGATCTNNNAATAGACCAATAATGAGTTCTG
GAGCACACGTCTGAACTCTTTCCCTACACGACGCTCTTCCGATCTNNNGCAGGAAAAAAAAAAAAAA
GAGCACACGTCTGAACTCTTTCCCTACACGACGCTCTTCCGATCTNNNCACTTTATTTCAAGAGTATGT
AGCACACGTCTGAACTCTTTCCCTACACGACGCTCTTCCGATCTNNNAAAGGAGTGACTATTAGAATC
AGCACACGTCTGAACTCTTTCCCTACACGACGCTCTTCCGATCTNNNCATCTGTACAGAGGCCTATGC
ACACGTCTGAACTCTTTCCCTACACGACGCTCTTCCGATCTNNNAATGCAAACTCTCCATTTTGCATCG
CACACGTCTGAACTCTTTCCCTACACGACGCTCTTCCGATCTNNNAAAGTCTTTAACCGTTACAGTACAT
AGCACACGTCTGAACTCTTTCCCTACACGACGCTCTTCCGATCTNNNACAAACAAATATTAGTACCTG
AGCACACGTCTGAACTCTTTCCCTACACGACGCTCTTCCGATCTNNNATATGTATATATGGCATAGAG
ACACGTCTGAACTCTTTCCCTACACGACGCTCTTCCGATCTNNNCTCTGGTAACTGTTCATTCTGCATC
GCACACGTCTGAACTCTTTCCCTACACGACGCTCTTCCGATCTNNNCTCTTGTCATAGTTCAAAATGTT
AGCACACGTCTGAACTCTTTCCCTACACGACGCTCTTCCGATCTNNNAATGTATATACGAAAAGCACA
GCACACGTCTGAACTCTTTCCCTACACGACGCTCTTCCGATCTNNNATGCACATATTTGTAAATTTTGC
GAGCACACGTCTGAACTCTTTCCCTACACGACGCTCTTCCGATCTNNNTAGATTCTGTAAAATCAGTG
GCACACGTCTGAACTCTTTCCCTACACGACGCTCTTCCGATCTNNNGGTATGACTGGTGAAAACTCC
CACACGTCTGAACTCTTTCCCTACACGACGCTCTTCCGATCTNNNTCTTTGGTCTCATTTATTCTAGAC
AGCACACGTCTGAACTCTTTCCCTACACGACGCTCTTCCGATCTNNNAGATAGCATGAGAATTGAGT
GAGCACACGTCTGAACTCTTTCCCTACACGACGCTCTTCCGATCTNNNTCTGAAAGAAGGTAATATGA
ACACGTCTGAACTCTTTCCCTACACGACGCTCTTCCGATCTNNNCAATTCTCATAACTGTAGGACCCAT
ACACGTCTGAACTCTTTCCCTACACGACGCTCTTCCGATCTNNNGGAGACATTTGGACTTTATTTGCA
GAGCACACGTCTGAACTCTTTCCCTACACGACGCTCTTCCGATCTNNNTTCTGCTTTTAAAAGGAAGA
GCACACGTCTGAACTCTTTCCCTACACGACGCTCTTCCGATCTNNNCTTCTTCAGTTTCTTCATCTATGC
CACACGTCTGAACTCTTTCCCTACACGACGCTCTTCCGATCTNNNCTGTACTTTCCTTTCTTCCTCCATCG
CACACGTCTGAACTCTTTCCCTACACGACGCTCTTCCGATCTNNNCTGTACTTTCCTTTCTTCCTCCATCG
GCACACGTCTGAACTCTTTCCCTACACGACGCTCTTCCGATCTNNNTCTATTAAAAACTATAGGTGGG
ACACGTCTGAACTCTTTCCCTACACGACGCTCTTCCGATCTNNNATTATTAAGCAGCAAGACAAATTC
AGCACACGTCTGAACTCTTTCCCTACACGACGCTCTTCCGATCTNNNATTCTTATTTTGGAGATGAGTC
ACACGTCTGAACTCTTTCCCTACACGACGCTCTTCCGATCTNNNACTAATTGTTTACCTAGCTAGTTCAT
CACACGTCTGAACTCTTTCCCTACACGACGCTCTTCCGATCTNNNCTCCAGAGTAGTTGGGACTACAT
CACGTCTGAACTCTTTCCCTACACGACGCTCTTCCGATCTNNNACTGTGATCAATAAATTTATGGGCA
GCACACGTCTGAACTCTTTCCCTACACGACGCTCTTCCGATCTNNNATCATCACAACCATCATCTTCCAT
GCACACGTCTGAACTCTTTCCCTACACGACGCTCTTCCGATCTNNNGTTATAATGGTGCATAAACTCT
AGCACACGTCTGAACTCTTTCCCTACACGACGCTCTTCCGATCTNNNTACTATTCCCTGGACATAGTTA
AGCACACGTCTGAACTCTTTCCCTACACGACGCTCTTCCGATCTNNNATACTTACATACTATTCCCTGG
ACACGTCTGAACTCTTTCCCTACACGACGCTCTTCCGATCTNNNGTCTCCTCTGCCTAGAAATCCATCG
CACACGTCTGAACTCTTTCCCTACACGACGCTCTTCCGATCTNNNCCTCAAGCGTGTCTATAAAATCAT
CACACGTCTGAACTCTTTCCCTACACGACGCTCTTCCGATCTNNNCAAAGTGCTGGGATTATAGGCA

GAGCACACGTCTGAACTCTTTCCCTACACGACGCTCTTCCGATCTNNNTTTCTTAGGAGAAAGGAATAA
AGCACACGTCTGAACTCTTTCCCTACACGACGCTCTTCCGATCTNNNCTTAGGTTTCATTTTCCACTTAA
GAGCACACGTCTGAACTCTTTCCCTACACGACGCTCTTCCGATCTNNNCTCTAGTCCACATATTGAGTGC
GAGCACACGTCTGAACTCTTTCCCTACACGACGCTCTTCCGATCTNNNAGTACATGAGCAAAAATTTAG
CACACGTCTGAACTCTTTCCCTACACGACGCTCTTCCGATCTNNNAAAGTGGACTACTAAAGCAAGCAT
AGCACACGTCTGAACTCTTTCCCTACACGACGCTCTTCCGATCTNNNTCACTTTGTATAAATGTAGCTCT
GCACACGTCTGAACTCTTTCCCTACACGACGCTCTTCCGATCTNNNTAAAGAAGGAAGTTTATTCTGGA
AGCACACGTCTGAACTCTTTCCCTACACGACGCTCTTCCGATCTNNNGGGATCACTGTATTTCTGTGCAT
CACACGTCTGAACTCTTTCCCTACACGACGCTCTTCCGATCTNNNTATGGTATATCTGAGCCCTCCATCG
GCACACGTCTGAACTCTTTCCCTACACGACGCTCTTCCGATCTNNNTTTTTAAAGGAATCTTCATACTCC
AGCACACGTCTGAACTCTTTCCCTACACGACGCTCTTCCGATCTNNNTTTGGAGGAATTTAAGATCCTT
AGCACACGTCTGAACTCTTTCCCTACACGACGCTCTTCCGATCTNNNTTAACCAAAATGTTTAAAACACC
AGCACACGTCTGAACTCTTTCCCTACACGACGCTCTTCCGATCTNNNCCCCCATCTCTATTGTAAAACCAT
GCACACGTCTGAACTCTTTCCCTACACGACGCTCTTCCGATCTNNNAACCCCCATCTCTATTGTAAAACA
ACACGTCTGAACTCTTTCCCTACACGACGCTCTTCCGATCTNNNGCCAATTTAAAACTATTTTTGCTCAT
CACACGTCTGAACTCTTTCCCTACACGACGCTCTTCCGATCTNNNCCCTTTCTCCTTTTTTATGATGCATC
AGCACACGTCTGAACTCTTTCCCTACACGACGCTCTTCCGATCTNNNAATTGTAATTCTAGCTGAACAA
GCACACGTCTGAACTCTTTCCCTACACGACGCTCTTCCGATCTNNNTTTTCTTCAGATGACTTCTACCCAT
AGCACACGTCTGAACTCTTTCCCTACACGACGCTCTTCCGATCTNNNGGAACAGCTAGATCTAGGAG
AGCACACGTCTGAACTCTTTCCCTACACGACGCTCTTCCGATCTNNNCTGGGGTGATTCTGTAAAATCA
AGCACACGTCTGAACTCTTTCCCTACACGACGCTCTTCCGATCTNNNCTGTTCTTTATTTTCTTTCCCTTCA
AGCACACGTCTGAACTCTTTCCCTACACGACGCTCTTCCGATCTNNNTGGACAAAAGAAAAGAGAGA
GAGCACACGTCTGAACTCTTTCCCTACACGACGCTCTTCCGATCTNNNGATAACAGCTATGAGTTTTCTT
AGCACACGTCTGAACTCTTTCCCTACACGACGCTCTTCCGATCTNNNTACTGGCTTCAGGTAAATAATA
ACACGTCTGAACTCTTTCCCTACACGACGCTCTTCCGATCTNNNCAGATTTCAGAAATATGGAGATGCA
GAGCACACGTCTGAACTCTTTCCCTACACGACGCTCTTCCGATCTNNNAAACAATCTTCCTTTCTCCTTCA
GAGCACACGTCTGAACTCTTTCCCTACACGACGCTCTTCCGATCTNNNACAAGGTATCAATTCTCTCATT
GAGCACACGTCTGAACTCTTTCCCTACACGACGCTCTTCCGATCTNNNATTTGGGTAAATTTACAACTG
AGCACACGTCTGAACTCTTTCCCTACACGACGCTCTTCCGATCTNNNAAGCCTTTAAAAAAAGACTTTT
CACACGTCTGAACTCTTTCCCTACACGACGCTCTTCCGATCTNNNGACTATAAAGTGGGATCTATTGTC
AGCACACGTCTGAACTCTTTCCCTACACGACGCTCTTCCGATCTNNNAAAATTCGTTCTGATTGACATT
GCACACGTCTGAACTCTTTCCCTACACGACGCTCTTCCGATCTNNNGATTCTCCTCTAGACCTTTTGCAT
ACACGTCTGAACTCTTTCCCTACACGACGCTCTTCCGATCTNNNCTTATGTATACCAAGCCATCCCATCG
GAGCACACGTCTGAACTCTTTCCCTACACGACGCTCTTCCGATCTNNNATAATATTTCTGGGCTGTAA
ACACGTCTGAACTCTTTCCCTACACGACGCTCTTCCGATCTNNNTTACATTTCTGGGACTCAATAAACAT
AGCACACGTCTGAACTCTTTCCCTACACGACGCTCTTCCGATCTNNNTCCATTGGATTAGTTAATGAGA
GAGCACACGTCTGAACTCTTTCCCTACACGACGCTCTTCCGATCTNNNCTACGAGTTTGAACTTGAAAT
GAGCACACGTCTGAACTCTTTCCCTACACGACGCTCTTCCGATCTNNNATTAAACTACGAGTTTGAACT
CACACGTCTGAACTCTTTCCCTACACGACGCTCTTCCGATCTNNNCACAAAAATCACCAAGTCTAATACA
GAGCACACGTCTGAACTCTTTCCCTACACGACGCTCTTCCGATCTNNNGTCAACTGGATCCTCAAATCC
ACACGTCTGAACTCTTTCCCTACACGACGCTCTTCCGATCTNNNCTTTTTTTCTTTCTTTCTTTCTTTCTCA
GAGCACACGTCTGAACTCTTTCCCTACACGACGCTCTTCCGATCTNNNGAAATGGAGTAGGGTTGAAG
CACACGTCTGAACTCTTTCCCTACACGACGCTCTTCCGATCTNNNTCAATAGGCAAATACCTGTATACC

AGCACACGTCTGAACTCTTTCCCTACACGACGCTCTTCCGATCTNNNTAATATTGTGCTGATAGAAACA
AGCACACGTCTGAACTCTTTCCCTACACGACGCTCTTCCGATCTNNNTGTTGTTCTTGGTTTTGAATCCA
AGCACACGTCTGAACTCTTTCCCTACACGACGCTCTTCCGATCTNNNACTATTAGCAGTTCCCCAAGCAT
AGCACACGTCTGAACTCTTTCCCTACACGACGCTCTTCCGATCTNNNGACAGGGAAAGATAATTACTG
AGCACACGTCTGAACTCTTTCCCTACACGACGCTCTTCCGATCTNNNGACAGGGAAAGATAATTACTG
GAGCACACGTCTGAACTCTTTCCCTACACGACGCTCTTCCGATCTNNNAAAATAAGCTGGAAAGTCTTT
GAGCACACGTCTGAACTCTTTCCCTACACGACGCTCTTCCGATCTNNNAAAATAAGCTGGAAAGTCTTT
GCACACGTCTGAACTCTTTCCCTACACGACGCTCTTCCGATCTNNNCCAAATCTGAGATTTTTCTCTTTC
AGCACACGTCTGAACTCTTTCCCTACACGACGCTCTTCCGATCTNNNGAGCTTCAATCCTCAAAGATCA
AGCACACGTCTGAACTCTTTCCCTACACGACGCTCTTCCGATCTNNNCATGTTAGGAATCCCAATAAAG
GAGCACACGTCTGAACTCTTTCCCTACACGACGCTCTTCCGATCTNNNGTTTTATTGCTTTGTGTTTCTTA
GAGCACACGTCTGAACTCTTTCCCTACACGACGCTCTTCCGATCTNNNGTTTTATTGCTTTGTGTTTCTTA
GAGCACACGTCTGAACTCTTTCCCTACACGACGCTCTTCCGATCTNNNATGTCAAGTGTAATTTGCTTA
GAGCACACGTCTGAACTCTTTCCCTACACGACGCTCTTCCGATCTNNNTGTTTCTTTTTGTATATGTGT
AGAGCACACGTCTGAACTCTTTCCCTACACGACGCTCTTCCGATCTNNNCCATAATTCCTAAGTAACTTC
AGCACACGTCTGAACTCTTTCCCTACACGACGCTCTTCCGATCTNNNGAATGTGACAAAACAATCTAAC
AGCACACGTCTGAACTCTTTCCCTACACGACGCTCTTCCGATCTNNNACACATTTATATACCCACACAAT
CACACGTCTGAACTCTTTCCCTACACGACGCTCTTCCGATCTNNNTCTTACAGAGAAAACAACCGCATC
AGCACACGTCTGAACTCTTTCCCTACACGACGCTCTTCCGATCTNNNAAAGTGCAGCTAGAATTCAACA
GCACACGTCTGAACTCTTTCCCTACACGACGCTCTTCCGATCTNNNAATAAGAGCGAAACTTTGTCTCA
GCACACGTCTGAACTCTTTCCCTACACGACGCTCTTCCGATCTNNNTGAAAAATTAGTTATTGCTTGG
AGCACACGTCTGAACTCTTTCCCTACACGACGCTCTTCCGATCTNNNATGAACAGTCTTGTATCTTGAT
AGCACACGTCTGAACTCTTTCCCTACACGACGCTCTTCCGATCTNNNAACAAACAAAACAAGAAAGTCA
CACACGTCTGAACTCTTTCCCTACACGACGCTCTTCCGATCTNNNAGGATTGAGTTCTCTGCTTCCATC
AGCACACGTCTGAACTCTTTCCCTACACGACGCTCTTCCGATCTNNNCTCTGCTAAAAATACAATAGCC
GCACACGTCTGAACTCTTTCCCTACACGACGCTCTTCCGATCTNNNAAAAATACAAAGGTGAACATGA
AGCACACGTCTGAACTCTTTCCCTACACGACGCTCTTCCGATCTNNNTCTATTATTTGATGCAGTGACAT
AGCACACGTCTGAACTCTTTCCCTACACGACGCTCTTCCGATCTNNNTCTATTATTTGATGCAGTGACAT
GAGCACACGTCTGAACTCTTTCCCTACACGACGCTCTTCCGATCTNNNTTATTTCCAGAAAAAAACTCA
GCACACGTCTGAACTCTTTCCCTACACGACGCTCTTCCGATCTNNNTCATTATTGATAAAATGTCACACA
GCACACGTCTGAACTCTTTCCCTACACGACGCTCTTCCGATCTNNNACCACAATCAAGATATAGAACAT
CACACGTCTGAACTCTTTCCCTACACGACGCTCTTCCGATCTNNNAAAACAATCGTAATCGTAAAGTTC
GAGCACACGTCTGAACTCTTTCCCTACACGACGCTCTTCCGATCTNNNCATGTTCATACGTACATGTTTA
CACACGTCTGAACTCTTTCCCTACACGACGCTCTTCCGATCTNNNGATAGATATTGAGTGTGGAGAG
AGCACACGTCTGAACTCTTTCCCTACACGACGCTCTTCCGATCTNNNCCTTTCCTTCTTTTCTTTTCTTCAT
AGCACACGTCTGAACTCTTTCCCTACACGACGCTCTTCCGATCTNNNTTTTGCTGGTGTATTTTGTAATA
AGCACACGTCTGAACTCTTTCCCTACACGACGCTCTTCCGATCTNNNCTTTTTTGGATTTCTGTCTTAGT
AGCACACGTCTGAACTCTTTCCCTACACGACGCTCTTCCGATCTNNNTATAAGCACGATGAATGAATTT
AGCACACGTCTGAACTCTTTCCCTACACGACGCTCTTCCGATCTNNNAAGTAAACTTTCTTATTCCAAG
AGCACACGTCTGAACTCTTTCCCTACACGACGCTCTTCCGATCTNNNTCAGGGACCTATAGAACATAGC
AGAGCACACGTCTGAACTCTTTCCCTACACGACGCTCTTCCGATCTNNNCTAAGAACCTTAGGGACTAT
AGCACACGTCTGAACTCTTTCCCTACACGACGCTCTTCCGATCTNNNTGTAATAGCACCATGTTGATCCA
AGCACACGTCTGAACTCTTTCCCTACACGACGCTCTTCCGATCTNNNCAAAAATAGTTCAAGTAGAAGG

AGCACACGTCTGAACTCTTTCCCTACACGACGCTCTTCCGATCTNNNAAAATAAAGGTAATGGCAAA
GCACACGTCTGAACTCTTTCCCTACACGACGCTCTTCCGATCTNNNTAATGAAAAATTGAAACCTCAA
GCACACGTCTGAACTCTTTCCCTACACGACGCTCTTCCGATCTNNNATGATTCCTACAAAGCACTTCCA
AGCACACGTCTGAACTCTTTCCCTACACGACGCTCTTCCGATCTNNNCTTCCCCATAGTAAGTGATCA
GCACACGTCTGAACTCTTTCCCTACACGACGCTCTTCCGATCTNNNAGTACCTCTTAGAATTGATGAA
AGCACACGTCTGAACTCTTTCCCTACACGACGCTCTTCCGATCTNNNAATTCGCTTATTACTCAATGTTT
CACACGTCTGAACTCTTTCCCTACACGACGCTCTTCCGATCTNNNAGTAAATAAGAATCCGGAAGACC
GCACACGTCTGAACTCTTTCCCTACACGACGCTCTTCCGATCTNNNTATATTTCCTTTCCTGATCAGTTC
ACACGTCTGAACTCTTTCCCTACACGACGCTCTTCCGATCTNNNTTGGAAATAATTAAATAGAACATCT
AGCACACGTCTGAACTCTTTCCCTACACGACGCTCTTCCGATCTNNNGCAACATTTTTTTAAACAGA
AGCACACGTCTGAACTCTTTCCCTACACGACGCTCTTCCGATCTNNNTGTAAAGAATACTTTGTAAAG
GAGCACACGTCTGAACTCTTTCCCTACACGACGCTCTTCCGATCTNNNAGGATCAGAATCAAGTCCTA
GAGCACACGTCTGAACTCTTTCCCTACACGACGCTCTTCCGATCTNNNAGGATCAGAATCAAGTCCTA
GAGCACACGTCTGAACTCTTTCCCTACACGACGCTCTTCCGATCTNNNAGGATCAGAATCAAGTCCTA
GCACACGTCTGAACTCTTTCCCTACACGACGCTCTTCCGATCTNNNGAATGTAGAAGGATATTTGGTC
CACACGTCTGAACTCTTTCCCTACACGACGCTCTTCCGATCTNNNTGACTTAATAAATTTAGATGGG
AGCACACGTCTGAACTCTTTCCCTACACGACGCTCTTCCGATCTNNNGTAAAGTTTTTTTAGCAAGGT
ACACGTCTGAACTCTTTCCCTACACGACGCTCTTCCGATCTNNNACTCACAAGGAAGGAATTTGCATC
GCACACGTCTGAACTCTTTCCCTACACGACGCTCTTCCGATCTNNNTATTCACCTCTTTGTTTCTCTATC
CACACGTCTGAACTCTTTCCCTACACGACGCTCTTCCGATCTNNNCCCAAGATTCTATGAAACTAGAA
AGCACACGTCTGAACTCTTTCCCTACACGACGCTCTTCCGATCTNNNCTAGTCAAGCAGAACTTAATT
AGCACACGTCTGAACTCTTTCCCTACACGACGCTCTTCCGATCTNNNTAAGTTTATAAGTGAATGACC
AGCACACGTCTGAACTCTTTCCCTACACGACGCTCTTCCGATCTNNNCTCCTCAATTTTTTGGAATAGT
AGCACACGTCTGAACTCTTTCCCTACACGACGCTCTTCCGATCTNNNCTCACACACATTCAAATATACA
AGCACACGTCTGAACTCTTTCCCTACACGACGCTCTTCCGATCTNNNTACAGTTACTCCCTCTTCTAGCA
CACACGTCTGAACTCTTTCCCTACACGACGCTCTTCCGATCTNNNGTTTTGTTTTTCTACTGCTAGTAC
GCACACGTCTGAACTCTTTCCCTACACGACGCTCTTCCGATCTNNNCATTTGAGATGTGGGAAAGGC
AGCACACGTCTGAACTCTTTCCCTACACGACGCTCTTCCGATCTNNNATTTCTTTCTTTTAGCTTGTAGG
AGCACACGTCTGAACTCTTTCCCTACACGACGCTCTTCCGATCTNNNTAACACATATATAACTGGAAGC
AGCACACGTCTGAACTCTTTCCCTACACGACGCTCTTCCGATCTNNNAATCTCAGAAAAAAATCCATTG
AGCACACGTCTGAACTCTTTCCCTACACGACGCTCTTCCGATCTNNNAATAGCTGGGATTACAGTCAT
CACACGTCTGAACTCTTTCCCTACACGACGCTCTTCCGATCTNNNCAACTTTCCTTGTCTCTAGCCATC
ACACGTCTGAACTCTTTCCCTACACGACGCTCTTCCGATCTNNNAAGTTCTGCTTGTTTTTTCATAGCA
AGCACACGTCTGAACTCTTTCCCTACACGACGCTCTTCCGATCTNNNATTAATGCTAATGGGAGTCTA
GCACACGTCTGAACTCTTTCCCTACACGACGCTCTTCCGATCTNNNTAATTAATGTCAGGTGAGGGA
AGCACACGTCTGAACTCTTTCCCTACACGACGCTCTTCCGATCTNNNGGAAACTGGAAAGTAAAAAA
GAGCACACGTCTGAACTCTTTCCCTACACGACGCTCTTCCGATCTNNNTATCCTGGTCTCTACCTAACTC
GAGCACACGTCTGAACTCTTTCCCTACACGACGCTCTTCCGATCTNNNTATCCTGGTCTCTACCTAACTC
CACACGTCTGAACTCTTTCCCTACACGACGCTCTTCCGATCTNNNTGATAATCAGACTTTGAGCCCAT
GCACACGTCTGAACTCTTTCCCTACACGACGCTCTTCCGATCTNNNAGCATTTGGTAAAATATGTATC
ACACGTCTGAACTCTTTCCCTACACGACGCTCTTCCGATCTNNNCTCTCTTGAGGATAATTCAGCCATC
GCACACGTCTGAACTCTTTCCCTACACGACGCTCTTCCGATCTNNNGACCCCAACAATTGTATAAAATA
CACACGTCTGAACTCTTTCCCTACACGACGCTCTTCCGATCTNNNATGTAAAATCACTCTTTGCCTAAC

GAGCACACGTCTGAACTCTTTCCCTACACGACGCTCTTCCGATCTNNNAGTGTGGAAGTAACTATTATT
CACACGTCTGAACTCTTTCCCTACACGACGCTCTTCCGATCTNNNTATTACAAGGAAGCTAAGATCAG
AGCACACGTCTGAACTCTTTCCCTACACGACGCTCTTCCGATCTNNNAGAGTAAATAAGGGCCAGATC
AGCACACGTCTGAACTCTTTCCCTACACGACGCTCTTCCGATCTNNNATGGCAAACAAAAATGATAAAA
GAGCACACGTCTGAACTCTTTCCCTACACGACGCTCTTCCGATCTNNNCATCAGATTTTTGACTTACCAA
AGCACACGTCTGAACTCTTTCCCTACACGACGCTCTTCCGATCTNNNAGTAGGGTCTAGGGATTTTGC
AGCACACGTCTGAACTCTTTCCCTACACGACGCTCTTCCGATCTNNNAAAATAGTTTATTTTTGCCCA
GCACACGTCTGAACTCTTTCCCTACACGACGCTCTTCCGATCTNNNATAAGGAAAGATCTTATGCTTGA
CACACGTCTGAACTCTTTCCCTACACGACGCTCTTCCGATCTNNNATGAGGCAAGGAGTTCAATACAT
AGAGCACACGTCTGAACTCTTTCCCTACACGACGCTCTTCCGATCTNNNAGAAAGTCTTCAAGTATAGA
ACACGTCTGAACTCTTTCCCTACACGACGCTCTTCCGATCTNNNAGAATATAAAATACTTTCCCAATCTC
CACACGTCTGAACTCTTTCCCTACACGACGCTCTTCCGATCTNNNGAAATGATTCTTCTCTAGACGTCAT
AGCACACGTCTGAACTCTTTCCCTACACGACGCTCTTCCGATCTNNNATAAGAGTGATATTGTTTTGA
AGCACACGTCTGAACTCTTTCCCTACACGACGCTCTTCCGATCTNNNGGTTTCTCATCTTGCTTTTATAA
GAGCACACGTCTGAACTCTTTCCCTACACGACGCTCTTCCGATCTNNNCGCATTTTCTGAATATTCTCTA
AGCACACGTCTGAACTCTTTCCCTACACGACGCTCTTCCGATCTNNNAGTTCTCTACTTAGAAGAAACA
AGCACACGTCTGAACTCTTTCCCTACACGACGCTCTTCCGATCTNNNTAACAAATATTTCTTAAGCACC
AGCACACGTCTGAACTCTTTCCCTACACGACGCTCTTCCGATCTNNNAAAAAAAAAAAAAATGCACACA
GAGCACACGTCTGAACTCTTTCCCTACACGACGCTCTTCCGATCTNNNCTACTCAGTGAAAATTCTAGT
GAGCACACGTCTGAACTCTTTCCCTACACGACGCTCTTCCGATCTNNNTTATTTTACCCTGTTCCAGTAA
AGCACACGTCTGAACTCTTTCCCTACACGACGCTCTTCCGATCTNNNTCCTGAACAAAGAATATGATTT
AGCACACGTCTGAACTCTTTCCCTACACGACGCTCTTCCGATCTNNNAACATATGCCACAAAAACACCA
GCACACGTCTGAACTCTTTCCCTACACGACGCTCTTCCGATCTNNNGATTAATCAAGGAAGTGAAAGT
AGCACACGTCTGAACTCTTTCCCTACACGACGCTCTTCCGATCTNNNCTGTGTTACCTTTTCTTTCTTTC
GCACACGTCTGAACTCTTTCCCTACACGACGCTCTTCCGATCTNNNCACCTGTAGTCAATTTTTTAAGA
AGCACACGTCTGAACTCTTTCCCTACACGACGCTCTTCCGATCTNNNATGAGTACATGCTATTTTAATG
AGCACACGTCTGAACTCTTTCCCTACACGACGCTCTTCCGATCTNNNATGAGTACATGCTATTTTAATG
AGCACACGTCTGAACTCTTTCCCTACACGACGCTCTTCCGATCTNNNTCTGTAGTTTTCTGAAGTTCTAA
GCACACGTCTGAACTCTTTCCCTACACGACGCTCTTCCGATCTNNNATAGAAAACATCTCTTGAGATCTC
AGCACACGTCTGAACTCTTTCCCTACACGACGCTCTTCCGATCTNNNCTCATTGAAGGGTTTATAGTTT
AGCACACGTCTGAACTCTTTCCCTACACGACGCTCTTCCGATCTNNNTCTGTAGTTTTCTGAAGTTCTAA
ACACGTCTGAACTCTTTCCCTACACGACGCTCTTCCGATCTNNNTATTTGGAGATAGGGCCTATAACAT
AGCACACGTCTGAACTCTTTCCCTACACGACGCTCTTCCGATCTNNNCATGAGAAAGCTGATTCACACA
AGCACACGTCTGAACTCTTTCCCTACACGACGCTCTTCCGATCTNNNTAATACAAACAGTGAAAGTTG
AGCACACGTCTGAACTCTTTCCCTACACGACGCTCTTCCGATCTNNNTATGCTTCTGAACATATTTTACC
GAGCACACGTCTGAACTCTTTCCCTACACGACGCTCTTCCGATCTNNNTATAAGAAGTACCCTCAAGTA
GAGCACACGTCTGAACTCTTTCCCTACACGACGCTCTTCCGATCTNNNATAAATCTAGAACGATGTTAT
GAGCACACGTCTGAACTCTTTCCCTACACGACGCTCTTCCGATCTNNNTTTGTAAAACAAACCAAAACA
GAGCACACGTCTGAACTCTTTCCCTACACGACGCTCTTCCGATCTNNNAAAAAAGAAAGGACACATCTT
AGCACACGTCTGAACTCTTTCCCTACACGACGCTCTTCCGATCTNNNAAGCATACACATACATACACACC
CACACGTCTGAACTCTTTCCCTACACGACGCTCTTCCGATCTNNNATGTCTGTAATTCCAGCTACTCAT
AGCACACGTCTGAACTCTTTCCCTACACGACGCTCTTCCGATCTNNNCATTGAGCTTTGATCGCATACAT
AGCACACGTCTGAACTCTTTCCCTACACGACGCTCTTCCGATCTNNNAAACAGGGATGTCTTTTCTCCA

AGCACACGTCTGAACTCTTTCCCTACACGACGCTCTTCCGATCTNNNATTAATGAAACAGGGATGTCTT
AGCACACGTCTGAACTCTTTCCCTACACGACGCTCTTCCGATCTNNNATAGTTTGTCTGTTATATTTCCA
AGCACACGTCTGAACTCTTTCCCTACACGACGCTCTTCCGATCTNNNATAGTTTGTCTGTTATATTTCCA
CACACGTCTGAACTCTTTCCCTACACGACGCTCTTCCGATCTNNNTTCCTGTAATCCCACTTCTTCATCG
ACACGTCTGAACTCTTTCCCTACACGACGCTCTTCCGATCTNNNGAGATCAATGGAAGGTCAAACATC
AGCACACGTCTGAACTCTTTCCCTACACGACGCTCTTCCGATCTNNNCATTTGTGAATGTAACAAAGG
GCACACGTCTGAACTCTTTCCCTACACGACGCTCTTCCGATCTNNNTATTTGGGTTTTGAAAATTGACT
AGCACACGTCTGAACTCTTTCCCTACACGACGCTCTTCCGATCTNNNTTACCTGTTATTAACATTTTGC
ACACGTCTGAACTCTTTCCCTACACGACGCTCTTCCGATCTNNNTAATCATTCCAGGCAAGTGACATCG
AGCACACGTCTGAACTCTTTCCCTACACGACGCTCTTCCGATCTNNNCTGAATTTTTGCAAGAATAATG
AGCACACGTCTGAACTCTTTCCCTACACGACGCTCTTCCGATCTNNNCTTTCTATACACACTGAATTGA
ACACGTCTGAACTCTTTCCCTACACGACGCTCTTCCGATCTNNNTTTTCAAAAATTTTCTCCCATTCCATC
GCACACGTCTGAACTCTTTCCCTACACGACGCTCTTCCGATCTNNNGAAAGAAAAGAAGCCCTTTGCA
GCACACGTCTGAACTCTTTCCCTACACGACGCTCTTCCGATCTNNNAAGCCATTAAAGAAATAGAAAG
GCACACGTCTGAACTCTTTCCCTACACGACGCTCTTCCGATCTNNNTGTGTGTATATGTATATGTGTG
AGCACACGTCTGAACTCTTTCCCTACACGACGCTCTTCCGATCTNNNAAAAAAAAAAAAAAAGCCACAT
CACACGTCTGAACTCTTTCCCTACACGACGCTCTTCCGATCTNNNGAATGACAACAGGAAAATGAAAT
AGCACACGTCTGAACTCTTTCCCTACACGACGCTCTTCCGATCTNNNCACCTGTATATTTATTGTAGCA
GCACACGTCTGAACTCTTTCCCTACACGACGCTCTTCCGATCTNNNTTCACATCAGTAACTTTAAGACTC
GAGCACACGTCTGAACTCTTTCCCTACACGACGCTCTTCCGATCTNNNGTACACACCTGCTAAGATTAC
GCACACGTCTGAACTCTTTCCCTACACGACGCTCTTCCGATCTNNNAATACATACATGCATCCCATAACA
AGCACACGTCTGAACTCTTTCCCTACACGACGCTCTTCCGATCTNNNGATTTGCTATTTTGCACAATTG
AGCACACGTCTGAACTCTTTCCCTACACGACGCTCTTCCGATCTNNNGATTTGCTATTTTGCACAATTG
AGCACACGTCTGAACTCTTTCCCTACACGACGCTCTTCCGATCTNNNCACAAATTCCAATCGATATCAT
ACACGTCTGAACTCTTTCCCTACACGACGCTCTTCCGATCTNNNTATTCTTTATTCTTTTTTTCTCTTGTC
AGCACACGTCTGAACTCTTTCCCTACACGACGCTCTTCCGATCTNNNGTCTAGAGACAAGTGAAAGCC
GAGCACACGTCTGAACTCTTTCCCTACACGACGCTCTTCCGATCTNNNCACTTTTCCCTCTACAAATCAC
CACACGTCTGAACTCTTTCCCTACACGACGCTCTTCCGATCTNNNAAAGTCATGTACACGTCATCCATC
AGCACACGTCTGAACTCTTTCCCTACACGACGCTCTTCCGATCTNNNTTCATTTTCAACTGTAGGAGA
AGCACACGTCTGAACTCTTTCCCTACACGACGCTCTTCCGATCTNNNTGTGGTTTGATATATTAAAAG
AGCACACGTCTGAACTCTTTCCCTACACGACGCTCTTCCGATCTNNNCATTTGGACCATTTTCATTCAAC
CACACGTCTGAACTCTTTCCCTACACGACGCTCTTCCGATCTNNNATCAAGGGATCAAGAGTTCTCAT
AGCACACGTCTGAACTCTTTCCCTACACGACGCTCTTCCGATCTNNNTGTGCGCATGTTCTTTTATTAC
GCACACGTCTGAACTCTTTCCCTACACGACGCTCTTCCGATCTNNNGTCTTGTTTCTTATGCAGTCTACA
GAGCACACGTCTGAACTCTTTCCCTACACGACGCTCTTCCGATCTNNNGAGAATACATTTAAGCCTTAA
GAGCACACGTCTGAACTCTTTCCCTACACGACGCTCTTCCGATCTNNNCCAATGAGCATTAGATGAAA
AGCACACGTCTGAACTCTTTCCCTACACGACGCTCTTCCGATCTNNNGAAGGGAAAATGAGATACAG
AGCACACGTCTGAACTCTTTCCCTACACGACGCTCTTCCGATCTNNNATTGTAATTCTCTTTTCCTCCTAC
AGCACACGTCTGAACTCTTTCCCTACACGACGCTCTTCCGATCTNNNAATATTCTTTGGATAGATTTG
AGCACACGTCTGAACTCTTTCCCTACACGACGCTCTTCCGATCTNNNCCCAAAAGAGAAAAAGGAACC
AGCACACGTCTGAACTCTTTCCCTACACGACGCTCTTCCGATCTNNNCTTTGCATCTGTTTTCTTTTTAT
AGCACACGTCTGAACTCTTTCCCTACACGACGCTCTTCCGATCTNNNACAGGTGAATAATGATTGATC
GCACACGTCTGAACTCTTTCCCTACACGACGCTCTTCCGATCTNNNTGTTTCCAGGAATTTATTCATTC

AGCACACGTCTGAACTCTTTCCCTACACGACGCTCTTCCGATCTNNNAAGTAGTTGTTTTCTATTTTGG
CACACGTCTGAACTCTTTCCCTACACGACGCTCTTCCGATCTNNNCTCAAGGATTTCTCAAATATGTTC
GCACACGTCTGAACTCTTTCCCTACACGACGCTCTTCCGATCTNNNCTTTCCGCTTTCAGAATCTGCATC
AGCACACGTCTGAACTCTTTCCCTACACGACGCTCTTCCGATCTNNNAGCAAAAACGGTAATAATTCT
AGCACACGTCTGAACTCTTTCCCTACACGACGCTCTTCCGATCTNNNTAGGCATGCTAATATTTGAGT
CACACGTCTGAACTCTTTCCCTACACGACGCTCTTCCGATCTNNNTAATGTAACATTCAAACATGCTCC
GCACACGTCTGAACTCTTTCCCTACACGACGCTCTTCCGATCTNNNATTTACACACAGCTTAGATACAC
ACACGTCTGAACTCTTTCCCTACACGACGCTCTTCCGATCTNNNCTGATAAGATGCTATACACATTCCA
GCACACGTCTGAACTCTTTCCCTACACGACGCTCTTCCGATCTNNNGTTGGGAGAGTTGTAAGAACC
CACACGTCTGAACTCTTTCCCTACACGACGCTCTTCCGATCTNNNTGAAGGCACCATTTGTTTAGCATC
AGCACACGTCTGAACTCTTTCCCTACACGACGCTCTTCCGATCTNNNGCATTAGAACAAAACACTTGA
AGCACACGTCTGAACTCTTTCCCTACACGACGCTCTTCCGATCTNNNCAAAAACCATCTAAAGGAGTT
GCACACGTCTGAACTCTTTCCCTACACGACGCTCTTCCGATCTNNNTGAGGAAATAAGAAAGTTTTAC
GCACACGTCTGAACTCTTTCCCTACACGACGCTCTTCCGATCTNNNTGAGGAAATAAGAAAGTTTTAC
AGCACACGTCTGAACTCTTTCCCTACACGACGCTCTTCCGATCTNNNGACATATCTATAGGAAATTTG
GCACACGTCTGAACTCTTTCCCTACACGACGCTCTTCCGATCTNNNCACTTTTAGAAATGCTTTCAGA
GAGCACACGTCTGAACTCTTTCCCTACACGACGCTCTTCCGATCTNNNATACTATTCCATTAGACACTT
GCACACGTCTGAACTCTTTCCCTACACGACGCTCTTCCGATCTNNNAGCCATAGTAGGAATATTTACTT
GCACACGTCTGAACTCTTTCCCTACACGACGCTCTTCCGATCTNNNATAATATGTTAAATGCTGAGCA
GCACACGTCTGAACTCTTTCCCTACACGACGCTCTTCCGATCTNNNTGAATTTTTCCAGAAACTACAAG
AGCACACGTCTGAACTCTTTCCCTACACGACGCTCTTCCGATCTNNNGGGAACTTATTCAAGCTATAAG
AGCACACGTCTGAACTCTTTCCCTACACGACGCTCTTCCGATCTNNNATTATGAGGAGCTTGACTTTT
GCACACGTCTGAACTCTTTCCCTACACGACGCTCTTCCGATCTNNNTTGTTTTTCATACAGGCATTATT
ACACGTCTGAACTCTTTCCCTACACGACGCTCTTCCGATCTNNNGTTTGGAGAAGTATCTAGAATCTC
GCACACGTCTGAACTCTTTCCCTACACGACGCTCTTCCGATCTNNNATACGTTTAGAAGGATTATCACT
AGCACACGTCTGAACTCTTTCCCTACACGACGCTCTTCCGATCTNNNGACTCCATCTCTAAAAAACAAA
CACACGTCTGAACTCTTTCCCTACACGACGCTCTTCCGATCTNNNGTGTGCTGAGTACTACATTAGCA
AGCACACGTCTGAACTCTTTCCCTACACGACGCTCTTCCGATCTNNNACACAAATAAGCAAATAAGTA
AGCACACGTCTGAACTCTTTCCCTACACGACGCTCTTCCGATCTNNNACGTCTTGTTATAAATGTGTTT
CACACGTCTGAACTCTTTCCCTACACGACGCTCTTCCGATCTNNNCTTCATTTCCAGCCTCTAAGCATCG
AGCACACGTCTGAACTCTTTCCCTACACGACGCTCTTCCGATCTNNNTGAGGTGGGAAGGAATTTAG
AGCACACGTCTGAACTCTTTCCCTACACGACGCTCTTCCGATCTNNNGTTAGTCTAAGTCATTGTCTA
AGCACACGTCTGAACTCTTTCCCTACACGACGCTCTTCCGATCTNNNCATGTACCCTAGAACTTAAAG
CACACGTCTGAACTCTTTCCCTACACGACGCTCTTCCGATCTNNNATCATCATGATCTGCTATTTCACAT
CACACGTCTGAACTCTTTCCCTACACGACGCTCTTCCGATCTNNNATCATCATGATCTGCTATTTCACAT
AGCACACGTCTGAACTCTTTCCCTACACGACGCTCTTCCGATCTNNNTCAAACTTATTTTACAAGGAC
GCACACGTCTGAACTCTTTCCCTACACGACGCTCTTCCGATCTNNNCAGTGATGATGATGTGGTTTAT
CACACGTCTGAACTCTTTCCCTACACGACGCTCTTCCGATCTNNNCCTCCAAATCCATACTTCAGCATCG
ACACGTCTGAACTCTTTCCCTACACGACGCTCTTCCGATCTNNNTATTGGTTTAGTAGGATTTAGGGC
AGCACACGTCTGAACTCTTTCCCTACACGACGCTCTTCCGATCTNNNATACTATTAATAAGCCCAGAG
GCACACGTCTGAACTCTTTCCCTACACGACGCTCTTCCGATCTNNNATACCCACAATATACGAAGAATG
GCACACGTCTGAACTCTTTCCCTACACGACGCTCTTCCGATCTNNNTAAAATTATGAGTGTATTCCTGG
GCACACGTCTGAACTCTTTCCCTACACGACGCTCTTCCGATCTNNNTATAGAAGGCATAATGTTTATG

AGCACACGTCTGAACTCTTTCCCTACACGACGCTCTTCCGATCTNNNTTCAGTTCATATTTATCCTCAGT
AGCACACGTCTGAACTCTTTCCCTACACGACGCTCTTCCGATCTNNNTTTTGCTTAGCTTTGGAAATAA
AGCACACGTCTGAACTCTTTCCCTACACGACGCTCTTCCGATCTNNNGAAATTAACCACATATTAGACC
AGCACACGTCTGAACTCTTTCCCTACACGACGCTCTTCCGATCTNNNAATTTCCTGATATTTTGGCATAG
GAGCACACGTCTGAACTCTTTCCCTACACGACGCTCTTCCGATCTNNNGTTTTGGCTATTATAGTACTTG
AGCACACGTCTGAACTCTTTCCCTACACGACGCTCTTCCGATCTNNNACATTTAGATATCCATGCTATGT
AGCACACGTCTGAACTCTTTCCCTACACGACGCTCTTCCGATCTNNNTAAAGAGCAAATGCAGTAGTAG
AGCACACGTCTGAACTCTTTCCCTACACGACGCTCTTCCGATCTNNNTACTGGGAATTAGAGAATGAAA
CACACGTCTGAACTCTTTCCCTACACGACGCTCTTCCGATCTNNNCTCTAGTAAAGGAACCCCTATATCA
CACACGTCTGAACTCTTTCCCTACACGACGCTCTTCCGATCTNNNAATAACTTTGCATACATTTGTCTCAT
AGCACACGTCTGAACTCTTTCCCTACACGACGCTCTTCCGATCTNNNATACTAGTTAGCAAACATTTTT
GAGCACACGTCTGAACTCTTTCCCTACACGACGCTCTTCCGATCTNNNAACATGCATGTATGTCATCTAA
AGCACACGTCTGAACTCTTTCCCTACACGACGCTCTTCCGATCTNNNCTTATGTTAGCTTTCATGCTAGC
AGCACACGTCTGAACTCTTTCCCTACACGACGCTCTTCCGATCTNNNGAGAGAGGGAGAGAGAATAT
AGCACACGTCTGAACTCTTTCCCTACACGACGCTCTTCCGATCTNNNTGCCTATTTTTTTCACTCTTTTAC
AGCACACGTCTGAACTCTTTCCCTACACGACGCTCTTCCGATCTNNNATAAAACAAAATAGAAGCCACA
AGCACACGTCTGAACTCTTTCCCTACACGACGCTCTTCCGATCTNNNATAAAACAAAATAGAAGCCACA
ACACGTCTGAACTCTTTCCCTACACGACGCTCTTCCGATCTNNNCTCCTTTTCATTCATTCATTCAACATC
CACACGTCTGAACTCTTTCCCTACACGACGCTCTTCCGATCTNNNCAAAAAGTAGTCTGTTTCCAGACA
ACACGTCTGAACTCTTTCCCTACACGACGCTCTTCCGATCTNNNGTGTTCAAGTGATTCTCCTGCATCG
GCACACGTCTGAACTCTTTCCCTACACGACGCTCTTCCGATCTNNNGAAATGGCATAATTGTGTATGTA
AGCACACGTCTGAACTCTTTCCCTACACGACGCTCTTCCGATCTNNNAAGGTTTACCATTTAATAAGTG
AGCACACGTCTGAACTCTTTCCCTACACGACGCTCTTCCGATCTNNNATTCTGAAGTATAGCAAAATCT
GCACACGTCTGAACTCTTTCCCTACACGACGCTCTTCCGATCTNNNCTCAGTAATTAGAGGAGAAAGT
AGCACACGTCTGAACTCTTTCCCTACACGACGCTCTTCCGATCTNNNTTCAACAACCCAAAAACATAAGC
GCACACGTCTGAACTCTTTCCCTACACGACGCTCTTCCGATCTNNNTTTGTCATTCCTCTGAAGATAATC
GCACACGTCTGAACTCTTTCCCTACACGACGCTCTTCCGATCTNNNTAAAGTAAAATGAAAGGACCAAG
AGCACACGTCTGAACTCTTTCCCTACACGACGCTCTTCCGATCTNNNTTCGTCACAAAAATTTACATTTT
AGCACACGTCTGAACTCTTTCCCTACACGACGCTCTTCCGATCTNNNTTCATATGGATTAGGTTTTGTTC
AGCACACGTCTGAACTCTTTCCCTACACGACGCTCTTCCGATCTNNNTATAAGAAAGGCTACAAACACA
GCACACGTCTGAACTCTTTCCCTACACGACGCTCTTCCGATCTNNNGAAAAGAAAAAAACAAACTCCAA
AGCACACGTCTGAACTCTTTCCCTACACGACGCTCTTCCGATCTNNNTCCAGAGAATCAGAACCATATT
AGCACACGTCTGAACTCTTTCCCTACACGACGCTCTTCCGATCTNNNATATATCACTTTGCATTTTTCAC
GCACACGTCTGAACTCTTTCCCTACACGACGCTCTTCCGATCTNNNAACTGAAACATCAGCTCTTCCATC
AGCACACGTCTGAACTCTTTCCCTACACGACGCTCTTCCGATCTNNNCAATATAAGAATGCCCACAAATA
AGCACACGTCTGAACTCTTTCCCTACACGACGCTCTTCCGATCTNNNAGGTAGTAAATATATCCATCACC
AGCACACGTCTGAACTCTTTCCCTACACGACGCTCTTCCGATCTNNNATAATGGGAAAATAATGGAAT
CACGTCTGAACTCTTTCCCTACACGACGCTCTTCCGATCTNNNGAAAGTACAGAGCTCTCAGGCATCGG
GCACACGTCTGAACTCTTTCCCTACACGACGCTCTTCCGATCTNNNGAGAACAGACATGTCCATAGCAT
CACACGTCTGAACTCTTTCCCTACACGACGCTCTTCCGATCTNNNCAGCCTATATCTCCAGGATGCATCC
GCACACGTCTGAACTCTTTCCCTACACGACGCTCTTCCGATCTNNNAAGATAAAATGAGATGTACCTG
GAGCACACGTCTGAACTCTTTCCCTACACGACGCTCTTCCGATCTNNNGCATCTAGATGTATTGTTTCAC
AGCACACGTCTGAACTCTTTCCCTACACGACGCTCTTCCGATCTNNNTTTCTATATGCCTCTACTAAAG

AGCACACGTCTGAACTCTTTCCCTACACGACGCTCTTCCGATCTNNNGTAAACATGCAACCAATCTAAT
GCACACGTCTGAACTCTTTCCCTACACGACGCTCTTCCGATCTNNNCGTAGTTACAAGTATTTCCTTTG
ACACGTCTGAACTCTTTCCCTACACGACGCTCTTCCGATCTNNNCAGTATCTGAGCTTTAGTAACTTCAT
AGAGCACACGTCTGAACTCTTTCCCTACACGACGCTCTTCCGATCTNNNGACTAAGATGTAAGAGTGT
ACACGTCTGAACTCTTTCCCTACACGACGCTCTTCCGATCTNNNCCTCACTTAACAGTTATTTGATCAT
GCACACGTCTGAACTCTTTCCCTACACGACGCTCTTCCGATCTNNNCTTTCCGCTTTCAGAATCTGCATC
AGCACACGTCTGAACTCTTTCCCTACACGACGCTCTTCCGATCTNNNACAGATAGAGACCATTTCCACA
GAGCACACGTCTGAACTCTTTCCCTACACGACGCTCTTCCGATCTNNNAAGAATGAAGGAAGCTATAG
AGCACACGTCTGAACTCTTTCCCTACACGACGCTCTTCCGATCTNNNTTTCGACATAGATGTAAAAATC
GAGCACACGTCTGAACTCTTTCCCTACACGACGCTCTTCCGATCTNNNTTGTTTCATCAATAAAGACTG
GCACACGTCTGAACTCTTTCCCTACACGACGCTCTTCCGATCTNNNATGCATCTCATGAGTGTATGCAT
AGCACACGTCTGAACTCTTTCCCTACACGACGCTCTTCCGATCTNNNTCACATTGTATCTTAGTATGGA
CACACGTCTGAACTCTTTCCCTACACGACGCTCTTCCGATCTNNNATAAGAAATTTAGAACTTTAGAGC
GCACACGTCTGAACTCTTTCCCTACACGACGCTCTTCCGATCTNNNACATTGGAAATTACATGATGAA
ACACGTCTGAACTCTTTCCCTACACGACGCTCTTCCGATCTNNNCAGATGTAATGGTTTCGCTGCATCG
GCACACGTCTGAACTCTTTCCCTACACGACGCTCTTCCGATCTNNNGTAATTCCTGAAGGCAAGATATC
GAGCACACGTCTGAACTCTTTCCCTACACGACGCTCTTCCGATCTNNNTCTAACCTCATGCAAAATAAAA
GAGCACACGTCTGAACTCTTTCCCTACACGACGCTCTTCCGATCTNNNTTTGATTATTTTTCTTAGCTGC
AGCACACGTCTGAACTCTTTCCCTACACGACGCTCTTCCGATCTNNNCTTCCCAATCTTTTCAAATACAT
AGCACACGTCTGAACTCTTTCCCTACACGACGCTCTTCCGATCTNNNCGTCATTTGTTACAGTTACTGC
AGCACACGTCTGAACTCTTTCCCTACACGACGCTCTTCCGATCTNNNGCTCTACATACTTCTTTCACACAT
CACACGTCTGAACTCTTTCCCTACACGACGCTCTTCCGATCTNNNTCATAGGTATCTTCAGCTTTATTCA
GAGCACACGTCTGAACTCTTTCCCTACACGACGCTCTTCCGATCTNNNATGAGCTAATTCCTTATTGCTC
GAGCACACGTCTGAACTCTTTCCCTACACGACGCTCTTCCGATCTNNNCAAGTAAGACTTTTCTTGACTC
GCACACGTCTGAACTCTTTCCCTACACGACGCTCTTCCGATCTNNNATAAGTGTGACGGAGGAAAATA
GAGCACACGTCTGAACTCTTTCCCTACACGACGCTCTTCCGATCTNNNAATTTAAAACCAAAATCCAGA
GAGCACACGTCTGAACTCTTTCCCTACACGACGCTCTTCCGATCTNNNGTTTGAAATTGAGCTATTCCT
GCACACGTCTGAACTCTTTCCCTACACGACGCTCTTCCGATCTNNNTGATACAAAGAATCTCAAGACTG
GAGCACACGTCTGAACTCTTTCCCTACACGACGCTCTTCCGATCTNNNAGTGTCTTGAAAGAATAAAAA
CACACGTCTGAACTCTTTCCCTACACGACGCTCTTCCGATCTNNNGAAAAATCAAAACTTGGAACTCCAT
GAGCACACGTCTGAACTCTTTCCCTACACGACGCTCTTCCGATCTNNNTAAAGGGGTAGAACAAGAGA
AGAGCACACGTCTGAACTCTTTCCCTACACGACGCTCTTCCGATCTNNNAAGCAATGCATATATCCCATT
AGCACACGTCTGAACTCTTTCCCTACACGACGCTCTTCCGATCTNNNGAGAATTGAGCCAAGATTGTCA
CACACGTCTGAACTCTTTCCCTACACGACGCTCTTCCGATCTNNNTGATAACAATAGTAGAGGAAGTC
GAGCACACGTCTGAACTCTTTCCCTACACGACGCTCTTCCGATCTNNNAGGTAAAGAATAGACAGAAG
GAGCACACGTCTGAACTCTTTCCCTACACGACGCTCTTCCGATCTNNNAGGTAAAGAATAGACAGAAG
GAGCACACGTCTGAACTCTTTCCCTACACGACGCTCTTCCGATCTNNNCAACACATGTAATAACAAAAC
AGCACACGTCTGAACTCTTTCCCTACACGACGCTCTTCCGATCTNNNCTTTTCAATAGGTGGGTTTTGC
AGCACACGTCTGAACTCTTTCCCTACACGACGCTCTTCCGATCTNNNATACTATTAACTCAGGTACAGAC
AGCACACGTCTGAACTCTTTCCCTACACGACGCTCTTCCGATCTNNNTGTCCTCTGGATTTTTATTTCTA
GAGCACACGTCTGAACTCTTTCCCTACACGACGCTCTTCCGATCTNNNAAATACGTTTCTCAATCTCTCTC
AGCACACGTCTGAACTCTTTCCCTACACGACGCTCTTCCGATCTNNNATGAGATACAATGCAAAGTTTT
GCACACGTCTGAACTCTTTCCCTACACGACGCTCTTCCGATCTNNNCCATCCTCTGAGTGAAAATAGCA

CACACGTCTGAACTCTTTCCCTACACGACGCTCTTCCGATCTNNNGTGTAGAATCTGCTCAGTATCCATC
AGCACACGTCTGAACTCTTTCCCTACACGACGCTCTTCCGATCTNNNAGGATAATTGCTTGAACCAGCA
AGCACACGTCTGAACTCTTTCCCTACACGACGCTCTTCCGATCTNNNACCCTACTAGAGAAGTATGTAT
GAGCACACGTCTGAACTCTTTCCCTACACGACGCTCTTCCGATCTNNNTTATTTGAGGAGAGTGTCTT
AGCACACGTCTGAACTCTTTCCCTACACGACGCTCTTCCGATCTNNNAGCTTAAAAGAATCTTATGCAAT
AGCACACGTCTGAACTCTTTCCCTACACGACGCTCTTCCGATCTNNNCAGTTTATCTGAGAAACCAAAC
AGCACACGTCTGAACTCTTTCCCTACACGACGCTCTTCCGATCTNNNCAGTTTATCTGAGAAACCAAAC
AGCACACGTCTGAACTCTTTCCCTACACGACGCTCTTCCGATCTNNNGAACAATAGGAGAAGGATTTG
CACACGTCTGAACTCTTTCCCTACACGACGCTCTTCCGATCTNNNAGAGAAACAGAATCAATAGGAGCA
AGCACACGTCTGAACTCTTTCCCTACACGACGCTCTTCCGATCTNNNTGAATATGTACATGTAGCATCC
GCACACGTCTGAACTCTTTCCCTACACGACGCTCTTCCGATCTNNNTTGTAGCCTTGGGTAGATTTAC
ACACGTCTGAACTCTTTCCCTACACGACGCTCTTCCGATCTNNNAGTTTCTATTTCTTCTCTGACTTCATC
GAGCACACGTCTGAACTCTTTCCCTACACGACGCTCTTCCGATCTNNNACATACCATGAACACTATAAGA
GCACACGTCTGAACTCTTTCCCTACACGACGCTCTTCCGATCTNNNCAGAAGTCAAAAGTTGAGGTCAT
AGCACACGTCTGAACTCTTTCCCTACACGACGCTCTTCCGATCTNNNCACAATCTTGGCTAAACATATTT
AGCACACGTCTGAACTCTTTCCCTACACGACGCTCTTCCGATCTNNNATCTACACATACTCATGAGAATC
AGCACACGTCTGAACTCTTTCCCTACACGACGCTCTTCCGATCTNNNTTTTAGTAGTCACTCACAAAA
GAGCACACGTCTGAACTCTTTCCCTACACGACGCTCTTCCGATCTNNNCTTCTTCAGCAATGTAATTTAC
GAGCACACGTCTGAACTCTTTCCCTACACGACGCTCTTCCGATCTNNNATGTGTTTCCTTTAAGATACAT
AGCACACGTCTGAACTCTTTCCCTACACGACGCTCTTCCGATCTNNNGCTCCAGCTTCTTCTATTTATATC
AGCACACGTCTGAACTCTTTCCCTACACGACGCTCTTCCGATCTNNNGCTCCAGCTTCTTCTATTTATACA
CACACGTCTGAACTCTTTCCCTACACGACGCTCTTCCGATCTNNNAATGAAATTCATTTTGAGCAAATC
GAGCACACGTCTGAACTCTTTCCCTACACGACGCTCTTCCGATCTNNNAAAACTTCCAAACTCCTTTTAAC
AGCACACGTCTGAACTCTTTCCCTACACGACGCTCTTCCGATCTNNNATTTCATATTATACCTTTCCCCG
AGCACACGTCTGAACTCTTTCCCTACACGACGCTCTTCCGATCTNNNCTAATTTCCCCTCACTGTTTCCAT
GCACACGTCTGAACTCTTTCCCTACACGACGCTCTTCCGATCTNNNATATCTTGGAATTCTTGGCAACAT
CACACGTCTGAACTCTTTCCCTACACGACGCTCTTCCGATCTNNNCACTCTGATTTCATGGACTTCATCG
GAGCACACGTCTGAACTCTTTCCCTACACGACGCTCTTCCGATCTNNNTTAGCAATATCTACTAATGCTG
GAGCACACGTCTGAACTCTTTCCCTACACGACGCTCTTCCGATCTNNNCAAAGGTATTAAGGACAATGT
GAGCACACGTCTGAACTCTTTCCCTACACGACGCTCTTCCGATCTNNNAAAGCATGATGATACTAACCA
GAGCACACGTCTGAACTCTTTCCCTACACGACGCTCTTCCGATCTNNNAGTATGCTAAATAATTGAAGA
AGCACACGTCTGAACTCTTTCCCTACACGACGCTCTTCCGATCTNNNATTTATAGGACCATGCCATTAGC
GAGCACACGTCTGAACTCTTTCCCTACACGACGCTCTTCCGATCTNNNAGTATGCTAAATAATTGAAGA
AGCACACGTCTGAACTCTTTCCCTACACGACGCTCTTCCGATCTNNNTGATCTCAATTTTTATAGCCATG
AGCACACGTCTGAACTCTTTCCCTACACGACGCTCTTCCGATCTNNNGGAGACCATAACTTATTTTTGA
AGCACACGTCTGAACTCTTTCCCTACACGACGCTCTTCCGATCTNNNTTAGTAGAAATAACACCGTATGT
GAGCACACGTCTGAACTCTTTCCCTACACGACGCTCTTCCGATCTNNNGAAATTGTTCATTTCTAGGAG
GAGCACACGTCTGAACTCTTTCCCTACACGACGCTCTTCCGATCTNNNCAAAAATGAGTGAATTACCAT
GAGCACACGTCTGAACTCTTTCCCTACACGACGCTCTTCCGATCTNNNCAACTTTAGACGTTCTGTAATT
GAGCACACGTCTGAACTCTTTCCCTACACGACGCTCTTCCGATCTNNNCTTGGTTTCCTAAAACTTCACCA
GAGCACACGTCTGAACTCTTTCCCTACACGACGCTCTTCCGATCTNNNAACGTCTTTAGACTATCAACTT
AGCACACGTCTGAACTCTTTCCCTACACGACGCTCTTCCGATCTNNNCAATTCAAAACAAAGAGTGAAT
CACACGTCTGAACTCTTTCCCTACACGACGCTCTTCCGATCTNNNTTAATCCATATATCAAGTCTTTGTC

AGCACACGTCTGAACTCTTTCCCTACACGACGCTCTTCCGATCTNNNTGGATTTTTCTCTCTACCATAAA
AGCACACGTCTGAACTCTTTCCCTACACGACGCTCTTCCGATCTNNNTTACTGAGATAAGAAGCCTATT
CACACGTCTGAACTCTTTCCCTACACGACGCTCTTCCGATCTNNNATTAGCCAGGAAACCAATTAGCAT
GCACACGTCTGAACTCTTTCCCTACACGACGCTCTTCCGATCTNNNTTTTCTCAGCCTACTTTTTTATTCAT
CACACGTCTGAACTCTTTCCCTACACGACGCTCTTCCGATCTNNNGACAGGGTCTTGCTCTATAGCATC
GCACACGTCTGAACTCTTTCCCTACACGACGCTCTTCCGATCTNNNGAAAATGACACGTTCTTCAAGCA
AGCACACGTCTGAACTCTTTCCCTACACGACGCTCTTCCGATCTNNNTCCCTTTTTCTTCCATTTACCCAT
AGCACACGTCTGAACTCTTTCCCTACACGACGCTCTTCCGATCTNNNTCTTTGGTGTCTTTGAATCTATA
CACACGTCTGAACTCTTTCCCTACACGACGCTCTTCCGATCTNNNGGACATCAGAGGTTAGACTTACAT
AGCACACGTCTGAACTCTTTCCCTACACGACGCTCTTCCGATCTNNNTATTCTTATGCTGTTTCATTACC
AGAGCACACGTCTGAACTCTTTCCCTACACGACGCTCTTCCGATCTNNNTTTATAACACTTTCTGAAACT
GCACACGTCTGAACTCTTTCCCTACACGACGCTCTTCCGATCTNNNAAAACACTAACCAAGGTTATCTCA
GCACACGTCTGAACTCTTTCCCTACACGACGCTCTTCCGATCTNNNCTCTCTGTCTCTCCCTCATACATCG
GAGCACACGTCTGAACTCTTTCCCTACACGACGCTCTTCCGATCTNNNCAAAAATATATGCAGCCAAAC
GCACACGTCTGAACTCTTTCCCTACACGACGCTCTTCCGATCTNNNTATTATATGCCTTGGACTTTTTCC
GAGCACACGTCTGAACTCTTTCCCTACACGACGCTCTTCCGATCTNNNCTGTCTCCATGTTGGTATTCCA
ACACGTCTGAACTCTTTCCCTACACGACGCTCTTCCGATCTNNNATTTCAAAGTCATCTCATCCAAACAT
GAGCACACGTCTGAACTCTTTCCCTACACGACGCTCTTCCGATCTNNNGATTTAGATGATTAAAAGGC
GCACACGTCTGAACTCTTTCCCTACACGACGCTCTTCCGATCTNNNGGATTGCAGTTAGATTTTTCTA
GCACACGTCTGAACTCTTTCCCTACACGACGCTCTTCCGATCTNNNCACAGTTTAACAGAAGTCATGCA
AGCACACGTCTGAACTCTTTCCCTACACGACGCTCTTCCGATCTNNNATAACAACAAAGAAGCTTTCATT
AGCACACGTCTGAACTCTTTCCCTACACGACGCTCTTCCGATCTNNNAAGCATTTTAGAGAATATTCCT
AGCACACGTCTGAACTCTTTCCCTACACGACGCTCTTCCGATCTNNNTTCAGAAAAAGACACAAAACA
GCACACGTCTGAACTCTTTCCCTACACGACGCTCTTCCGATCTNNNTAGACCCATCAGAGATTTGTCATC
GCACACGTCTGAACTCTTTCCCTACACGACGCTCTTCCGATCTNNNAATTTTATCCTGAACATTGTGACC
GAGCACACGTCTGAACTCTTTCCCTACACGACGCTCTTCCGATCTNNNGAGAAAAATGAACATGCATTA
AGCACACGTCTGAACTCTTTCCCTACACGACGCTCTTCCGATCTNNNTAGCACTTACCTCAATTTGTAAT
AGCACACGTCTGAACTCTTTCCCTACACGACGCTCTTCCGATCTNNNGATTCAAAAGGAATTCATGAG
AGCACACGTCTGAACTCTTTCCCTACACGACGCTCTTCCGATCTNNNGAAAGAAAAGATAGGAAAGA
GAGCACACGTCTGAACTCTTTCCCTACACGACGCTCTTCCGATCTNNNAAATCAAGCTTCTAATCCAGA
GAGCACACGTCTGAACTCTTTCCCTACACGACGCTCTTCCGATCTNNNACATTAAAAAATTGCTATGAG
GAGCACACGTCTGAACTCTTTCCCTACACGACGCTCTTCCGATCTNNNTATTCTATGTATTTTTGTGCA
GCACACGTCTGAACTCTTTCCCTACACGACGCTCTTCCGATCTNNNTTATATTTAGGTATGATGAAAG
AGCACACGTCTGAACTCTTTCCCTACACGACGCTCTTCCGATCTNNNAAGGCCATGGTAACCTTATACA
AGAGCACACGTCTGAACTCTTTCCCTACACGACGCTCTTCCGATCTNNNGTAACAGAGTGAGACTCCA
AGCACACGTCTGAACTCTTTCCCTACACGACGCTCTTCCGATCTNNNGAATGCTTGAATAAATCAGAAT
CACACGTCTGAACTCTTTCCCTACACGACGCTCTTCCGATCTNNNCTAAAATAAAGGTGTTGGCTGCAT
ACACGTCTGAACTCTTTCCCTACACGACGCTCTTCCGATCTNNNCACACACACACACATTTTCTCATCGGA
CACACGTCTGAACTCTTTCCCTACACGACGCTCTTCCGATCTNNNAGAGTTGATGAAAATCACTAAACC
GCACACGTCTGAACTCTTTCCCTACACGACGCTCTTCCGATCTNNNGCTTACTTAGCCATTATTTTTTGC
CACACGTCTGAACTCTTTCCCTACACGACGCTCTTCCGATCTNNNCTCTCTCCTCTCTCATTCTCCATCGGA
GAGCACACGTCTGAACTCTTTCCCTACACGACGCTCTTCCGATCTNNNGTGTGTGTCTGAAAGGGATA
CACACGTCTGAACTCTTTCCCTACACGACGCTCTTCCGATCTNNNAGAATTACATATCCATTTGCTTTCAT

GCACACGTCTGAACTCTTTCCCTACACGACGCTCTTCCGATCTNNNTTTCTTGTTTATCCACCCAGCATCG
GAGCACACGTCTGAACTCTTTCCCTACACGACGCTCTTCCGATCTNNNATTGGTGAGTTGGAGAATTAC
GCACACGTCTGAACTCTTTCCCTACACGACGCTCTTCCGATCTNNNAAATTGTTTGAGTAATAGGTTTTT
GAGCACACGTCTGAACTCTTTCCCTACACGACGCTCTTCCGATCTNNNGTTGAAGATTTTTGTATCAAT
AGCACACGTCTGAACTCTTTCCCTACACGACGCTCTTCCGATCTNNNTAAATAGAAAGACTAAAAGCAG
AGCACACGTCTGAACTCTTTCCCTACACGACGCTCTTCCGATCTNNNGATTCTCTGAAATGATTAAGCT
AGCACACGTCTGAACTCTTTCCCTACACGACGCTCTTCCGATCTNNNTATACATTTCTCAGAGCATATCC
AGCACACGTCTGAACTCTTTCCCTACACGACGCTCTTCCGATCTNNNATCTCTCTACGGTCTCTAAGACA
GAGCACACGTCTGAACTCTTTCCCTACACGACGCTCTTCCGATCTNNNGTAAGTCCATCGTAAGTTGAG
AGCACACGTCTGAACTCTTTCCCTACACGACGCTCTTCCGATCTNNNCAATAGCATTCTTCACAGAAATA
GAGCACACGTCTGAACTCTTTCCCTACACGACGCTCTTCCGATCTNNNATATTTCTGTCCAGAATTCTTG
AGAGCACACGTCTGAACTCTTTCCCTACACGACGCTCTTCCGATCTNNNCAAAGTTTCTTTGTATTCCTTT
AGCACACGTCTGAACTCTTTCCCTACACGACGCTCTTCCGATCTNNNGTGATTAAAGTGTATTATTGGTC
AGCACACGTCTGAACTCTTTCCCTACACGACGCTCTTCCGATCTNNNGGAACTGAATTCTAGGTAAATTC
GAGCACACGTCTGAACTCTTTCCCTACACGACGCTCTTCCGATCTNNNAGAAACAGAGAGAAGTTATCT
GCACACGTCTGAACTCTTTCCCTACACGACGCTCTTCCGATCTNNNGAAATTTTTGCCGACAATTTTTCA
AGCACACGTCTGAACTCTTTCCCTACACGACGCTCTTCCGATCTNNNTTTCTTCATCTCTTTGGTAAAAG
AGCACACGTCTGAACTCTTTCCCTACACGACGCTCTTCCGATCTNNNAAATCCTTCCAAAATATCACTTG
AGCACACGTCTGAACTCTTTCCCTACACGACGCTCTTCCGATCTNNNGTTCTCCTTGTGTTTTCTCTCATC
GAGCACACGTCTGAACTCTTTCCCTACACGACGCTCTTCCGATCTNNNCAGGTTACATAATTTGCTCATA
GCACACGTCTGAACTCTTTCCCTACACGACGCTCTTCCGATCTNNNTTCCCTGAAGAAGTATCTTTACCA
GAGCACACGTCTGAACTCTTTCCCTACACGACGCTCTTCCGATCTNNNAGACATATCTGAAAATTGGTA
AGCACACGTCTGAACTCTTTCCCTACACGACGCTCTTCCGATCTNNNCAGAAAAAGAGGGTTAGGAGC
AGCACACGTCTGAACTCTTTCCCTACACGACGCTCTTCCGATCTNNNGTTCTCCTTGTGTTTTCTCTCATC
GAGCACACGTCTGAACTCTTTCCCTACACGACGCTCTTCCGATCTNNNTAGCTATATGTAGGATGTTTTC
AGCACACGTCTGAACTCTTTCCCTACACGACGCTCTTCCGATCTNNNGGTGATTCCTCAAGGATCTACA
GCACACGTCTGAACTCTTTCCCTACACGACGCTCTTCCGATCTNNNATATGTGGCTGTATACATATGCCA
AGCACACGTCTGAACTCTTTCCCTACACGACGCTCTTCCGATCTNNNCATAGAAATAGTCATTGTCACAC
GAGCACACGTCTGAACTCTTTCCCTACACGACGCTCTTCCGATCTNNNTACTGCTTTAGTTCAAATTTTC
AGCACACGTCTGAACTCTTTCCCTACACGACGCTCTTCCGATCTNNNGAAAGAACATAAAGGCTTATCT
CACACGTCTGAACTCTTTCCCTACACGACGCTCTTCCGATCTNNNTCTTAACTACTATGTTATGCTGCCA
GCACACGTCTGAACTCTTTCCCTACACGACGCTCTTCCGATCTNNNCAAGAAAGCTTTATGTACCTATGC
AGAGCACACGTCTGAACTCTTTCCCTACACGACGCTCTTCCGATCTNNNAATAGCTTCTTTCTCTAATTTC
CACACGTCTGAACTCTTTCCCTACACGACGCTCTTCCGATCTNNNAGATATGTAGGTGTTTGCTTGCAT
AGCACACGTCTGAACTCTTTCCCTACACGACGCTCTTCCGATCTNNNACTGCAATACTATTGACTATTTG
AGCACACGTCTGAACTCTTTCCCTACACGACGCTCTTCCGATCTNNNAAGAGTGATGATGATGATGAT
GAGCACACGTCTGAACTCTTTCCCTACACGACGCTCTTCCGATCTNNNCTCTGTATCTTTTGCTAACTAC
GCACACGTCTGAACTCTTTCCCTACACGACGCTCTTCCGATCTNNNCATGGAGATAATAATAACATTCTC
AGCACACGTCTGAACTCTTTCCCTACACGACGCTCTTCCGATCTNNNTTTGTTATCTGCTTTTAAAGAGC
GAGCACACGTCTGAACTCTTTCCCTACACGACGCTCTTCCGATCTNNNCTAATAAGCCAATGAAAATCC
GAGCACACGTCTGAACTCTTTCCCTACACGACGCTCTTCCGATCTNNNCTAATAAGCCAATGAAAATCC
AGCACACGTCTGAACTCTTTCCCTACACGACGCTCTTCCGATCTNNNTGAAGTACTTGCATTTTAAGAA
AGCACACGTCTGAACTCTTTCCCTACACGACGCTCTTCCGATCTNNNTTTACAGCTAATGACCACAACCA

GAGCACACGTCTGAACTCTTTCCCTACACGACGCTCTTCCGATCTNNNCACTAAAAAAAAAAAAAGCT
CACACGTCTGAACTCTTTCCCTACACGACGCTCTTCCGATCTNNNCTAATGAAGTCTCATCATGTACAC
AGCACACGTCTGAACTCTTTCCCTACACGACGCTCTTCCGATCTNNNACAATCTATGCCTATGTGTGCA
CACACGTCTGAACTCTTTCCCTACACGACGCTCTTCCGATCTNNNGTGGAGTTATAGGGAATGTAAT
AGCACACGTCTGAACTCTTTCCCTACACGACGCTCTTCCGATCTNNNAAATGTGAAAAGTTGGTATG
GCACACGTCTGAACTCTTTCCCTACACGACGCTCTTCCGATCTNNNTTGTTCTTGTACGTGTAAGATA
AGCACACGTCTGAACTCTTTCCCTACACGACGCTCTTCCGATCTNNNTTTCAGACCTCGTAAAAATTAT
AGCACACGTCTGAACTCTTTCCCTACACGACGCTCTTCCGATCTNNNCAAGATAAAAACCCACATTTAG
AGCACACGTCTGAACTCTTTCCCTACACGACGCTCTTCCGATCTNNNGCATCTATCTTCTGAATCTAAA
AGCACACGTCTGAACTCTTTCCCTACACGACGCTCTTCCGATCTNNNAAAATGGATTATTTTGGTCTAC
AGCACACGTCTGAACTCTTTCCCTACACGACGCTCTTCCGATCTNNNATATCACTTTAATATAGCCCCA
GCACACGTCTGAACTCTTTCCCTACACGACGCTCTTCCGATCTNNNCAGTAGAATAAACAGTCCTTCTC
AGCACACGTCTGAACTCTTTCCCTACACGACGCTCTTCCGATCTNNNTCTCAAAAACAAAAACAAAAA
AGCACACGTCTGAACTCTTTCCCTACACGACGCTCTTCCGATCTNNNATATAAATTGTTACAACCACTG
GCACACGTCTGAACTCTTTCCCTACACGACGCTCTTCCGATCTNNNTTGTCTGTGATAGTAAATATCCA
AGCACACGTCTGAACTCTTTCCCTACACGACGCTCTTCCGATCTNNNGTAACCTGAAGGAAAAAGTTA
GCACACGTCTGAACTCTTTCCCTACACGACGCTCTTCCGATCTNNNAACTGTATGTTTATAGCCATTAG
AGCACACGTCTGAACTCTTTCCCTACACGACGCTCTTCCGATCTNNNATCCTTCTTATGACAAAAGGTA
AGCACACGTCTGAACTCTTTCCCTACACGACGCTCTTCCGATCTNNNAGCAAACTGTTTTATACATGAA
CACACGTCTGAACTCTTTCCCTACACGACGCTCTTCCGATCTNNNGTGAAAACAGTACTTGTAAGTAA
AGCACACGTCTGAACTCTTTCCCTACACGACGCTCTTCCGATCTNNNAAGAATAATATGTGGAAAGGT
GCACACGTCTGAACTCTTTCCCTACACGACGCTCTTCCGATCTNNNTTAATAATGGTTCATTTTCTGCC
CACACGTCTGAACTCTTTCCCTACACGACGCTCTTCCGATCTNNNCTGAATAATGACATTCATCAAGAC
GCACACGTCTGAACTCTTTCCCTACACGACGCTCTTCCGATCTNNNTTATTATGCATAGGTTACTGGC
AGCACACGTCTGAACTCTTTCCCTACACGACGCTCTTCCGATCTNNNATTGAGTTGTTTTCTCTTACTTG
GCACACGTCTGAACTCTTTCCCTACACGACGCTCTTCCGATCTNNNCTCAGAGACAAAGAGATAAGA
ACACGTCTGAACTCTTTCCCTACACGACGCTCTTCCGATCTNNNGATATTTGTAATCTGTTGTGTGTCA
CACACGTCTGAACTCTTTCCCTACACGACGCTCTTCCGATCTNNNACTCAAAAGATATATCCAACTCAC
AGCACACGTCTGAACTCTTTCCCTACACGACGCTCTTCCGATCTNNNCCAGATATACAGTCACATACAT
AGCACACGTCTGAACTCTTTCCCTACACGACGCTCTTCCGATCTNNNGCTTCCTAAAGATACATCTATT
GAGCACACGTCTGAACTCTTTCCCTACACGACGCTCTTCCGATCTNNNCTGCTTCCTAAAGATACATCT
GCACACGTCTGAACTCTTTCCCTACACGACGCTCTTCCGATCTNNNCAAATGAATTAGAAACACAAAG
CACACGTCTGAACTCTTTCCCTACACGACGCTCTTCCGATCTNNNCCTGGCTAAGATTGTAATTTAATC
AGCACACGTCTGAACTCTTTCCCTACACGACGCTCTTCCGATCTNNNGAGCACTATTTCCTAAGTACTA
GCACACGTCTGAACTCTTTCCCTACACGACGCTCTTCCGATCTNNNCCTGGCCAAAAAAAATAATCTAA
GCACACGTCTGAACTCTTTCCCTACACGACGCTCTTCCGATCTNNNGAGCATCTGTCTCAAAATATAT
AGCACACGTCTGAACTCTTTCCCTACACGACGCTCTTCCGATCTNNNAATCCTTTCCTGATCTCATTTAT
AGCACACGTCTGAACTCTTTCCCTACACGACGCTCTTCCGATCTNNNCCTGAGTGATAGAGAGCATC
CACACGTCTGAACTCTTTCCCTACACGACGCTCTTCCGATCTNNNAACCCCAAAATATACTCTTCATCAT
AGCACACGTCTGAACTCTTTCCCTACACGACGCTCTTCCGATCTNNNAAATCAAATATTCTCTCAAAG
GCACACGTCTGAACTCTTTCCCTACACGACGCTCTTCCGATCTNNNATAGTGATGGACATGTGGATC
AGCACACGTCTGAACTCTTTCCCTACACGACGCTCTTCCGATCTNNNCTTGATATTCCAAACACTTGAG
GCACACGTCTGAACTCTTTCCCTACACGACGCTCTTCCGATCTNNNTACCCCATATAAAGAGATGAAT

AGCACACGTCTGAACTCTTTCCCTACACGACGCTCTTCCGATCTNNNCTTTGGGACTTGAAATCCTGC
AGCACACGTCTGAACTCTTTCCCTACACGACGCTCTTCCGATCTNNNGCAATCAGATTTATTATATTGT
GCACACGTCTGAACTCTTTCCCTACACGACGCTCTTCCGATCTNNNGTTCTCTCAGTAAACTAACTAC
GCACACGTCTGAACTCTTTCCCTACACGACGCTCTTCCGATCTNNNTATATGATACAGCGTTCTTTGTT
AGCACACGTCTGAACTCTTTCCCTACACGACGCTCTTCCGATCTNNNGTATCCCTACATGATAAGGAA
CACACGTCTGAACTCTTTCCCTACACGACGCTCTTCCGATCTNNNAAAGTTGGAAAAGTAACCAATAA
GAGCACACGTCTGAACTCTTTCCCTACACGACGCTCTTCCGATCTNNNTAGTTTGCCTATTTCCTCATC
GCACACGTCTGAACTCTTTCCCTACACGACGCTCTTCCGATCTNNNAATGCATACAAATATCTCCAAAA
GCACACGTCTGAACTCTTTCCCTACACGACGCTCTTCCGATCTNNNCCATAAGGACTTTAAGTTGACT
CACGTCTGAACTCTTTCCCTACACGACGCTCTTCCGATCTNNNACTGAAAATTACACATTACTCTGCAT
CACGTCTGAACTCTTTCCCTACACGACGCTCTTCCGATCTNNNTTGTTATTTATCTGCTGACATTTCAT
AGCACACGTCTGAACTCTTTCCCTACACGACGCTCTTCCGATCTNNNACTATATCTGACTTGATCAAAT
CACACGTCTGAACTCTTTCCCTACACGACGCTCTTCCGATCTNNNGATCCCTATGATTCCTCTGACATCG
AGCACACGTCTGAACTCTTTCCCTACACGACGCTCTTCCGATCTNNNTCTTCAAAACTACCAGTACACC
GCACACGTCTGAACTCTTTCCCTACACGACGCTCTTCCGATCTNNNGATCAAGTTACCAGAAGATTCT
GCACACGTCTGAACTCTTTCCCTACACGACGCTCTTCCGATCTNNNTTACCCCTGAGGATATTATACA
GCACACGTCTGAACTCTTTCCCTACACGACGCTCTTCCGATCTNNNTTACCCCTGAGGATATTATACA
AGCACACGTCTGAACTCTTTCCCTACACGACGCTCTTCCGATCTNNNAAACCATTCTTATCAGACCATA
GAGCACACGTCTGAACTCTTTCCCTACACGACGCTCTTCCGATCTNNNTCATAGAATTCACCTTTCTTTT
GCACACGTCTGAACTCTTTCCCTACACGACGCTCTTCCGATCTNNNTTTCCAAATGTTTTTGACAGAG
ACACGTCTGAACTCTTTCCCTACACGACGCTCTTCCGATCTNNNAAGTTTCCTCCACCAAATACCATCG
CACACGTCTGAACTCTTTCCCTACACGACGCTCTTCCGATCTNNNGTTCCCTTCCCTTATCTTTCCATCG
AGCACACGTCTGAACTCTTTCCCTACACGACGCTCTTCCGATCTNNNAAACACTGAATAAATGAATGG
AGCACACGTCTGAACTCTTTCCCTACACGACGCTCTTCCGATCTNNNTGAGATAATCAAAGAGTCTTT
CACACGTCTGAACTCTTTCCCTACACGACGCTCTTCCGATCTNNNTAAAGAGATTATGGCTTTGATCA
AGCACACGTCTGAACTCTTTCCCTACACGACGCTCTTCCGATCTNNNAAAACTACCATTATTCCTGCTA
GCACACGTCTGAACTCTTTCCCTACACGACGCTCTTCCGATCTNNNCCTAAGCATCTGTATTTAGAGT
GCACACGTCTGAACTCTTTCCCTACACGACGCTCTTCCGATCTNNNTCAATATACTTGAACAACAACT
GCACACGTCTGAACTCTTTCCCTACACGACGCTCTTCCGATCTNNNTCAATATACTTGAACAACAACT
CACACGTCTGAACTCTTTCCCTACACGACGCTCTTCCGATCTNNNGTTCTCCTGATAATGAGTGAGCA
AGCACACGTCTGAACTCTTTCCCTACACGACGCTCTTCCGATCTNNNGTTTATATTCATGTCCAGTCTC
CACACGTCTGAACTCTTTCCCTACACGACGCTCTTCCGATCTNNNGAGAACGTGATGAGCATATGCA
AGCACACGTCTGAACTCTTTCCCTACACGACGCTCTTCCGATCTNNNTATGCAAATCAACTGTTATACT
AGCACACGTCTGAACTCTTTCCCTACACGACGCTCTTCCGATCTNNNCCTTTAATCAGAGTCACTAATT
CACACGTCTGAACTCTTTCCCTACACGACGCTCTTCCGATCTNNNCATATTTTTTTGGTCTCTTCTCCC
GAGCACACGTCTGAACTCTTTCCCTACACGACGCTCTTCCGATCTNNNCTTAGAAGATGAAACCAGAG
GAGCACACGTCTGAACTCTTTCCCTACACGACGCTCTTCCGATCTNNNCTTAGAAGATGAAACCAGAG
GCACACGTCTGAACTCTTTCCCTACACGACGCTCTTCCGATCTNNNGTTGCCAAACTTTTCATGTTTCA
CACACGTCTGAACTCTTTCCCTACACGACGCTCTTCCGATCTNNNTGAAAGCACAAAATTAATAGAGG
ACACGTCTGAACTCTTTCCCTACACGACGCTCTTCCGATCTNNNAATATTGAAACCTGTGATTTGTTCA
AGCACACGTCTGAACTCTTTCCCTACACGACGCTCTTCCGATCTNNNAACAAATATAGCAAAGATCTG
GCACACGTCTGAACTCTTTCCCTACACGACGCTCTTCCGATCTNNNCTGGCCACTTTCTTTTCTTTCATC
GCACACGTCTGAACTCTTTCCCTACACGACGCTCTTCCGATCTNNNAAGAGTGAAATTCCATCTCTCCA

AGCACACGTCTGAACTCTTTCCCTACACGACGCTCTTCCGATCTNNNTATGAACTTGAGCAATTCAATAC
AGCACACGTCTGAACTCTTTCCCTACACGACGCTCTTCCGATCTNNNTATGAACTTGAGCAATTCAATAC
GAGCACACGTCTGAACTCTTTCCCTACACGACGCTCTTCCGATCTNNNTGCTTATAGTATAGTTGTTACT
AGCACACGTCTGAACTCTTTCCCTACACGACGCTCTTCCGATCTNNNAAAGTAGAGACCACTAGATGAT
AGAGCACACGTCTGAACTCTTTCCCTACACGACGCTCTTCCGATCTNNNTCTAGTGATTCTCTTACCTCA
GCACACGTCTGAACTCTTTCCCTACACGACGCTCTTCCGATCTNNNGATAAGTGCCCTTATTGTAAAAG
GCACACGTCTGAACTCTTTCCCTACACGACGCTCTTCCGATCTNNNGATAAGTGCCCTTATTGTAAAAG
GCACACGTCTGAACTCTTTCCCTACACGACGCTCTTCCGATCTNNNGATAAGTGCCCTTATTGTAAAAG
GCACACGTCTGAACTCTTTCCCTACACGACGCTCTTCCGATCTNNNAAAATGAAAGGATTGGAGGAGC
GAGCACACGTCTGAACTCTTTCCCTACACGACGCTCTTCCGATCTNNNCATTACTAATGATCAGGGAAA
AGCACACGTCTGAACTCTTTCCCTACACGACGCTCTTCCGATCTNNNCTCTCTCTCTCCATATGTATCA
AGCACACGTCTGAACTCTTTCCCTACACGACGCTCTTCCGATCTNNNCTCTCTCTCTCCATATGTATCA
GAGCACACGTCTGAACTCTTTCCCTACACGACGCTCTTCCGATCTNNNACTTCTCAGTCTCTATGGTCCA
AGCACACGTCTGAACTCTTTCCCTACACGACGCTCTTCCGATCTNNNTCTAGACTATCTTCATACATGTC
CACACGTCTGAACTCTTTCCCTACACGACGCTCTTCCGATCTNNNGATAGTCCTAGGTTTCACTTTAGCA
AGCACACGTCTGAACTCTTTCCCTACACGACGCTCTTCCGATCTNNNTGGGAAAATAAAAAGAAGCAA
GCACACGTCTGAACTCTTTCCCTACACGACGCTCTTCCGATCTNNNAGAGAGAGACAAAGGATCATACA
CACACGTCTGAACTCTTTCCCTACACGACGCTCTTCCGATCTNNNATCAAATATAGATGGTTCTCTCCCAT
CACACGTCTGAACTCTTTCCCTACACGACGCTCTTCCGATCTNNNGCCAGACTTCTTTTAAATCAAATCA
AGCACACGTCTGAACTCTTTCCCTACACGACGCTCTTCCGATCTNNNAAAGAATAAACAAAAACACCATC
AGCACACGTCTGAACTCTTTCCCTACACGACGCTCTTCCGATCTNNNAAAGAATAAACAAAAACACCATC
AGCACACGTCTGAACTCTTTCCCTACACGACGCTCTTCCGATCTNNNAAAGAATAAACAAAAACACCATC
AGCACACGTCTGAACTCTTTCCCTACACGACGCTCTTCCGATCTNNNAAATTGTTATGCTGAGTTACAT
GAGCACACGTCTGAACTCTTTCCCTACACGACGCTCTTCCGATCTNNNCAGAAGTTATATAACAAGCACA
CACGTCTGAACTCTTTCCCTACACGACGCTCTTCCGATCTNNNATTTTGCCATTTCACATCAGCATCGGA
CACACGTCTGAACTCTTTCCCTACACGACGCTCTTCCGATCTNNNCAGTGGTGATAATGACATAATTGC
AGCACACGTCTGAACTCTTTCCCTACACGACGCTCTTCCGATCTNNNGTTAGTATTCCCAGTGTATGAT
AGCACACGTCTGAACTCTTTCCCTACACGACGCTCTTCCGATCTNNNCTCCTCATTATCCATCCTGATACA
AGCACACGTCTGAACTCTTTCCCTACACGACGCTCTTCCGATCTNNNCTCCTCATTATCCATCCTGATACA
GAGCACACGTCTGAACTCTTTCCCTACACGACGCTCTTCCGATCTNNNCAAATGTCTTTGGTGAACTTCC
AGCACACGTCTGAACTCTTTCCCTACACGACGCTCTTCCGATCTNNNAAAAAAAGAAAATGAAACGAAC
AGCACACGTCTGAACTCTTTCCCTACACGACGCTCTTCCGATCTNNNTCCAACTAATTTAGGTCTAAGAA
AGCACACGTCTGAACTCTTTCCCTACACGACGCTCTTCCGATCTNNNCATAGAGAAAGACCAGATGAAC
AGCACACGTCTGAACTCTTTCCCTACACGACGCTCTTCCGATCTNNNAATTTTAGTCCATGCTACAGTTA
AGCACACGTCTGAACTCTTTCCCTACACGACGCTCTTCCGATCTNNNATGTGTGTATATGTAGGTGTAT
GAGCACACGTCTGAACTCTTTCCCTACACGACGCTCTTCCGATCTNNNTCACCTATCTTTGTGTATGTTA
AGCACACGTCTGAACTCTTTCCCTACACGACGCTCTTCCGATCTNNNTATGTGTGTATGTGTGTGTATG
GAGCACACGTCTGAACTCTTTCCCTACACGACGCTCTTCCGATCTNNNCTCATGATAAGATAGAGATGG
CACACGTCTGAACTCTTTCCCTACACGACGCTCTTCCGATCTNNNCTGGGACTACAAGGAATGAGCAT
AGCACACGTCTGAACTCTTTCCCTACACGACGCTCTTCCGATCTNNNATTTGTTCTTATGAGTCCTGACC
AGCACACGTCTGAACTCTTTCCCTACACGACGCTCTTCCGATCTNNNTAATTAAAACAAAAGCAGTTCCT
AGCACACGTCTGAACTCTTTCCCTACACGACGCTCTTCCGATCTNNNAAAAAACACACAAAAAAAACTC
AGAGCACACGTCTGAACTCTTTCCCTACACGACGCTCTTCCGATCTNNNTAGCTTTCTCTCAATCTCTCTC

AGCACACGTCTGAACTCTTTCCCTACACGACGCTCTTCCGATCTNNNGGGAAAGAGAGAAAAAAAT
AGCACACGTCTGAACTCTTTCCCTACACGACGCTCTTCCGATCTNNNTGTTTATAAACACCAAAGCTAA
GCACACGTCTGAACTCTTTCCCTACACGACGCTCTTCCGATCTNNNTATTCTAGCCATCTCTCCTCCATC
AGCACACGTCTGAACTCTTTCCCTACACGACGCTCTTCCGATCTNNNCACTGTCCAGAGTGATAGAGCA
AGCACACGTCTGAACTCTTTCCCTACACGACGCTCTTCCGATCTNNNACATTAATGTCTTCTCTTGAGAA
CACACGTCTGAACTCTTTCCCTACACGACGCTCTTCCGATCTNNNTCATCCTATCCAAACCGTACCATCG
AGCACACGTCTGAACTCTTTCCCTACACGACGCTCTTCCGATCTNNNCATTATTTCAGCTAGTTTGAGTT
AGCACACGTCTGAACTCTTTCCCTACACGACGCTCTTCCGATCTNNNGCATGATATGTGGATGATAAT
AGCACACGTCTGAACTCTTTCCCTACACGACGCTCTTCCGATCTNNNCTAAGGCAAAAGGTCACTTTAC
GAGCACACGTCTGAACTCTTTCCCTACACGACGCTCTTCCGATCTNNNACAACAAAACTTCATTAACCAT
AGCACACGTCTGAACTCTTTCCCTACACGACGCTCTTCCGATCTNNNATAGATGAGAACTATGAGACT
AGCACACGTCTGAACTCTTTCCCTACACGACGCTCTTCCGATCTNNNGTGTCAGAAGTACTTCTTTTTA
AGCACACGTCTGAACTCTTTCCCTACACGACGCTCTTCCGATCTNNNCTCTGTGTCAGATATTCCTATCCA
AGCACACGTCTGAACTCTTTCCCTACACGACGCTCTTCCGATCTNNNACATTAGCAGTAGTACTTAACA
GAGCACACGTCTGAACTCTTTCCCTACACGACGCTCTTCCGATCTNNNAACAGAAAAAAGTGAGAATT
AGCACACGTCTGAACTCTTTCCCTACACGACGCTCTTCCGATCTNNNAAAAAAAACTCTCTAGCAAATC
AGCACACGTCTGAACTCTTTCCCTACACGACGCTCTTCCGATCTNNNTGTATCTAGCTAGCTGTTCTCCA
AGCACACGTCTGAACTCTTTCCCTACACGACGCTCTTCCGATCTNNNTACTCAATAAAGTCCCAAGTTAA
AGCACACGTCTGAACTCTTTCCCTACACGACGCTCTTCCGATCTNNNATATAGGAATGCATGTGTATAG
GAGCACACGTCTGAACTCTTTCCCTACACGACGCTCTTCCGATCTNNNTTGATGCGAATGTAAATTAGTA
AGCACACGTCTGAACTCTTTCCCTACACGACGCTCTTCCGATCTNNNTTATTTAGGTACAGGGGCTATC
GAGCACACGTCTGAACTCTTTCCCTACACGACGCTCTTCCGATCTNNNTCAAGTATTAGAAAGAGACTG
ACACGTCTGAACTCTTTCCCTACACGACGCTCTTCCGATCTNNNAATATTGATTTTACCCATCCATGCAT
AGCACACGTCTGAACTCTTTCCCTACACGACGCTCTTCCGATCTNNNGACTTTTTTTATGTATCTTCCCAC
GAGCACACGTCTGAACTCTTTCCCTACACGACGCTCTTCCGATCTNNNGTAGAAAATCTTATTTAGAAG
GCACACGTCTGAACTCTTTCCCTACACGACGCTCTTCCGATCTNNNATTCAACACTTTACCTTATATGGC
GCACACGTCTGAACTCTTTCCCTACACGACGCTCTTCCGATCTNNNCTCGTGTATTGATTTGTAACAGCA
AGCACACGTCTGAACTCTTTCCCTACACGACGCTCTTCCGATCTNNNAGATATTGCTCAGTACAGAATT
CACACGTCTGAACTCTTTCCCTACACGACGCTCTTCCGATCTNNNGTATTAAACAGGTCTTCCTTTTCCA
AGCACACGTCTGAACTCTTTCCCTACACGACGCTCTTCCGATCTNNNATACATCTCTAGGTCTAGCTATT
AGCACACGTCTGAACTCTTTCCCTACACGACGCTCTTCCGATCTNNNAAAACTATAGAGATGGAGAAC
AGCACACGTCTGAACTCTTTCCCTACACGACGCTCTTCCGATCTNNNCTACTACAATTAATTCACCCCTTC
GCACACGTCTGAACTCTTTCCCTACACGACGCTCTTCCGATCTNNNGAAAACTATTTACTTGGATTCGT
AGCACACGTCTGAACTCTTTCCCTACACGACGCTCTTCCGATCTNNNAAAATAATCTGGGTTGGAAATT
GCACACGTCTGAACTCTTTCCCTACACGACGCTCTTCCGATCTNNNGAATATATGAGCAGACATTTCAA
GCACACGTCTGAACTCTTTCCCTACACGACGCTCTTCCGATCTNNNTATTATCTTTCCAGTGTGTACAAC
AGCACACGTCTGAACTCTTTCCCTACACGACGCTCTTCCGATCTNNNATTAAATTCTCTGGTTTCTCTTG
GAGCACACGTCTGAACTCTTTCCCTACACGACGCTCTTCCGATCTNNNAAGAGAGAATCAGGGAACTA
ACACGTCTGAACTCTTTCCCTACACGACGCTCTTCCGATCTNNNTCTACCCCTGTTTTTCACTTCATCGGA
AGCACACGTCTGAACTCTTTCCCTACACGACGCTCTTCCGATCTNNNTCAAATTTCCCTAGTTTTATTTG
AGCACACGTCTGAACTCTTTCCCTACACGACGCTCTTCCGATCTNNNTTCCCAGTTTTATTTATACCTGT
CACACGTCTGAACTCTTTCCCTACACGACGCTCTTCCGATCTNNNGCTTTCTCCATTTTCTTACTTTACAT
CACGTCTGAACTCTTTCCCTACACGACGCTCTTCCGATCTNNNGCATGATTGGGAGAATAAATAAACA

GAGCACACGTCTGAACTCTTTCCCTACACGACGCTCTTCCGATCTNNNCAGTTTATAATGTCCCTACCA
AGCACACGTCTGAACTCTTTCCCTACACGACGCTCTTCCGATCTNNNCTTATTTCTCAAAGCATATGTC
AGCACACGTCTGAACTCTTTCCCTACACGACGCTCTTCCGATCTNNNCGTGTCAAGATCTACTTATCTA
ACACGTCTGAACTCTTTCCCTACACGACGCTCTTCCGATCTNNNCAACTAAAAATGGAGGTCATTAGC
CACACGTCTGAACTCTTTCCCTACACGACGCTCTTCCGATCTNNNCAATAAGCAGGATTTTTAAGAGG
ACACGTCTGAACTCTTTCCCTACACGACGCTCTTCCGATCTNNNCAACACTGGCAACTTCTATCCATCG
AGCACACGTCTGAACTCTTTCCCTACACGACGCTCTTCCGATCTNNNATACAGAACCTGCAGATACATA
CACACGTCTGAACTCTTTCCCTACACGACGCTCTTCCGATCTNNNTCAAAGGGAGACTACAATAATAT
CACACGTCTGAACTCTTTCCCTACACGACGCTCTTCCGATCTNNNAATTCTTTTAATGAGCTCTGAGCA
AGCACACGTCTGAACTCTTTCCCTACACGACGCTCTTCCGATCTNNNAGATAACTGTTAACAAATGAA
GAGCACACGTCTGAACTCTTTCCCTACACGACGCTCTTCCGATCTNNNGCTATAAATGACAGGATTTCA
AGCACACGTCTGAACTCTTTCCCTACACGACGCTCTTCCGATCTNNNTTTTCAAAGCTGAATAGTATTC
GAGCACACGTCTGAACTCTTTCCCTACACGACGCTCTTCCGATCTNNNTTCATATTTATCCTTGCTCCTA
CACACGTCTGAACTCTTTCCCTACACGACGCTCTTCCGATCTNNNCAAAATGTAGATGGAAATGGAG
AGCACACGTCTGAACTCTTTCCCTACACGACGCTCTTCCGATCTNNNTGAGGATTTTTGTATCTATGTT
GAGCACACGTCTGAACTCTTTCCCTACACGACGCTCTTCCGATCTNNNATTTTTCTGGCTGAGTAGTA
AGCACACGTCTGAACTCTTTCCCTACACGACGCTCTTCCGATCTNNNTATGAATAAGAATATGGTAGG
CACGTCTGAACTCTTTCCCTACACGACGCTCTTCCGATCTNNNGCTAAACAAGAATGAGAAAATCACA
AGCACACGTCTGAACTCTTTCCCTACACGACGCTCTTCCGATCTNNNAGTATAACAACTATTTACATAG
CACACGTCTGAACTCTTTCCCTACACGACGCTCTTCCGATCTNNNTGAACATACACAGATCCAGGCATC
AGCACACGTCTGAACTCTTTCCCTACACGACGCTCTTCCGATCTNNNATTTGTGTTGTTCTTTGATACA
AGCACACGTCTGAACTCTTTCCCTACACGACGCTCTTCCGATCTNNNGGAGAAGAATTAGGTTGGATA
GCACACGTCTGAACTCTTTCCCTACACGACGCTCTTCCGATCTNNNCTTGGCTTATTTCACTTAACATAC
GAGCACACGTCTGAACTCTTTCCCTACACGACGCTCTTCCGATCTNNNCTAGGTAAAGAAGTGAAAAC
GAGCACACGTCTGAACTCTTTCCCTACACGACGCTCTTCCGATCTNNNTTTGAGCCACTTCTTCATATAA
AGCACACGTCTGAACTCTTTCCCTACACGACGCTCTTCCGATCTNNNAGATATGGCTTAGACAACAAC
ACACGTCTGAACTCTTTCCCTACACGACGCTCTTCCGATCTNNNTACTATGTTCTCTAAGTGAGGATCA
AGCACACGTCTGAACTCTTTCCCTACACGACGCTCTTCCGATCTNNNATATAGACGTGTATGTGCATAT
AGCACACGTCTGAACTCTTTCCCTACACGACGCTCTTCCGATCTNNNAATTACATAAACTCAATTGGCT
GCACACGTCTGAACTCTTTCCCTACACGACGCTCTTCCGATCTNNNATTAAAGAAAACCAACAAGGAA
AGCACACGTCTGAACTCTTTCCCTACACGACGCTCTTCCGATCTNNNGCAATCTCTCTTTTATGATTTCT
GCACACGTCTGAACTCTTTCCCTACACGACGCTCTTCCGATCTNNNATAAAGACCAGAGTTTAATTCCT
AGCACACGTCTGAACTCTTTCCCTACACGACGCTCTTCCGATCTNNNAAGTATCTTCATGACAGAGAT
GCACACGTCTGAACTCTTTCCCTACACGACGCTCTTCCGATCTNNNATTTCAGTCTGGGTGAAAGACA
GAGCACACGTCTGAACTCTTTCCCTACACGACGCTCTTCCGATCTNNNCTTTTTCTACTGAGCATTTTG
CACACGTCTGAACTCTTTCCCTACACGACGCTCTTCCGATCTNNNTCCATGGGAAAAAAAAAAAAAAT
CACACGTCTGAACTCTTTCCCTACACGACGCTCTTCCGATCTNNNTCCATGGGAAAAAAAAAAAAAAT
AGCACACGTCTGAACTCTTTCCCTACACGACGCTCTTCCGATCTNNNTTGACTGTTCTAACATCTTCATT
AGCACACGTCTGAACTCTTTCCCTACACGACGCTCTTCCGATCTNNNGAATCAGTAATCAAAAACCTCC
AGCACACGTCTGAACTCTTTCCCTACACGACGCTCTTCCGATCTNNNTGGAAATGTCACATTTTTTAAT
GAGCACACGTCTGAACTCTTTCCCTACACGACGCTCTTCCGATCTNNNCTCTGAAATTGAGGTCATAAT
GAGCACACGTCTGAACTCTTTCCCTACACGACGCTCTTCCGATCTNNNATAGATAGACTGCAGTATAT
AGCACACGTCTGAACTCTTTCCCTACACGACGCTCTTCCGATCTNNNAGAAGGAAAATAAGAATTGC

GCACACGTCTGAACTCTTTCCCTACACGACGCTCTTCCGATCTNNNTCCACCTTAATCTGTGTATTACCAT
AGCACACGTCTGAACTCTTTCCCTACACGACGCTCTTCCGATCTNNNTTATTTGGTAATTGTGTTTGAG
AGCACACGTCTGAACTCTTTCCCTACACGACGCTCTTCCGATCTNNNTTATTTGGTAATTGTGTTTGAG
AGCACACGTCTGAACTCTTTCCCTACACGACGCTCTTCCGATCTNNNCGAGAAAATATGTGAGATAACA
GAGCACACGTCTGAACTCTTTCCCTACACGACGCTCTTCCGATCTNNNCCCCAATATAATTCTTTCTAGGA
GAGCACACGTCTGAACTCTTTCCCTACACGACGCTCTTCCGATCTNNNATAACCTTTTTCTTCAGTTTGAC
GAGCACACGTCTGAACTCTTTCCCTACACGACGCTCTTCCGATCTNNNGAAATCACTTTTTAAACACTGA
CACACGTCTGAACTCTTTCCCTACACGACGCTCTTCCGATCTNNNAAGGAAATTGTTTGGGGATCCATC
GAGCACACGTCTGAACTCTTTCCCTACACGACGCTCTTCCGATCTNNNATTATATTTCTCTTCCAGTTTGC
AGCACACGTCTGAACTCTTTCCCTACACGACGCTCTTCCGATCTNNNCCAGTCTTGAGCTAGATGACCAT
AGCACACGTCTGAACTCTTTCCCTACACGACGCTCTTCCGATCTNNNACATTCCTTCTTATTCCTTCTTAC
AGCACACGTCTGAACTCTTTCCCTACACGACGCTCTTCCGATCTNNNCCTAAAATTTCATTGGTTTCATG
AGCACACGTCTGAACTCTTTCCCTACACGACGCTCTTCCGATCTNNNCAGACTGCATCTCAAAATAAAT
CACACGTCTGAACTCTTTCCCTACACGACGCTCTTCCGATCTNNNAATGTCTTGTGCCATTTCATCATCG
AGCACACGTCTGAACTCTTTCCCTACACGACGCTCTTCCGATCTNNNCTCATGCTTCCATTACTATTTATCA
AGCACACGTCTGAACTCTTTCCCTACACGACGCTCTTCCGATCTNNNCTCATGCTTCCATTACTATTTATCA
AGCACACGTCTGAACTCTTTCCCTACACGACGCTCTTCCGATCTNNNCTGCAAACTAAATTTTCAATGA
GAGCACACGTCTGAACTCTTTCCCTACACGACGCTCTTCCGATCTNNNTTATTTAGAGAAACAGAAGCA
GAGCACACGTCTGAACTCTTTCCCTACACGACGCTCTTCCGATCTNNNTCTATGTAAAGAAGATGGCAT
AGAGCACACGTCTGAACTCTTTCCCTACACGACGCTCTTCCGATCTNNNCTCAAGGTTTGTAACATTAAT
CACACGTCTGAACTCTTTCCCTACACGACGCTCTTCCGATCTNNNCATCCTATGAAGCAAAGAATTACCA
GAGCACACGTCTGAACTCTTTCCCTACACGACGCTCTTCCGATCTNNNAACTATTCTTTCTTTTTCTCCC
AGCACACGTCTGAACTCTTTCCCTACACGACGCTCTTCCGATCTNNNAATAAACAGAAAAAGATCAGG
ACACGTCTGAACTCTTTCCCTACACGACGCTCTTCCGATCTNNNAAACTGCATAGAGAAAGCACCATCG
GCACACGTCTGAACTCTTTCCCTACACGACGCTCTTCCGATCTNNNTACCAAAAGAAGTAAACTGTACTC
GCACACGTCTGAACTCTTTCCCTACACGACGCTCTTCCGATCTNNNTAAATCTTTGTCTCATCTGTACTCA
GAGCACACGTCTGAACTCTTTCCCTACACGACGCTCTTCCGATCTNNNTGCCAAGACATAATATAGCAT
GAGCACACGTCTGAACTCTTTCCCTACACGACGCTCTTCCGATCTNNNTTTATAGCACCATTCTCAAAATC
GAGCACACGTCTGAACTCTTTCCCTACACGACGCTCTTCCGATCTNNNCAGTTTCTTTTTGTTGTTCTCAT
AGCACACGTCTGAACTCTTTCCCTACACGACGCTCTTCCGATCTNNNTAATAATCCAGAGAAGATGCAG
GAGCACACGTCTGAACTCTTTCCCTACACGACGCTCTTCCGATCTNNNTCAATAAATTCAAGCTTAGAA
ACACGTCTGAACTCTTTCCCTACACGACGCTCTTCCGATCTNNNCTGTGTGTAGGTAGAGAAAGCATCC
CACACGTCTGAACTCTTTCCCTACACGACGCTCTTCCGATCTNNNGAGCTTTTCTATTCTTTGGGTACAT
GCACACGTCTGAACTCTTTCCCTACACGACGCTCTTCCGATCTNNNAGTCATATGGAAAATATCTCAGC
AGCACACGTCTGAACTCTTTCCCTACACGACGCTCTTCCGATCTNNNTCCTTTGTAAATCTTGTTTTCTG
CACACGTCTGAACTCTTTCCCTACACGACGCTCTTCCGATCTNNNGAGCAGAGAAATAAGTGGTATTT
CACACGTCTGAACTCTTTCCCTACACGACGCTCTTCCGATCTNNNTAGTTTAAGTAGACAGTTCTGAGC
AGCACACGTCTGAACTCTTTCCCTACACGACGCTCTTCCGATCTNNNCATCAAGGCAATTCAAATCCCAT
CACACGTCTGAACTCTTTCCCTACACGACGCTCTTCCGATCTNNNCTTTCCAACCATTGCAAATATCATCC
GAGCACACGTCTGAACTCTTTCCCTACACGACGCTCTTCCGATCTNNNGCATCACAGTCTGATAAAATT
AGCACACGTCTGAACTCTTTCCCTACACGACGCTCTTCCGATCTNNNATTTCAAAGCCAATGTTTCTTTC
ACACGTCTGAACTCTTTCCCTACACGACGCTCTTCCGATCTNNNCAACACTCTTTCTGCCATAGCATCGG
GCACACGTCTGAACTCTTTCCCTACACGACGCTCTTCCGATCTNNNATTCTTACACAAAAATCTAGCTTC

AGCACACGTCTGAACTCTTTCCCTACACGACGCTCTTCCGATCTNNNTGAGAAAAGTATCCTCAAAGG
GCACACGTCTGAACTCTTTCCCTACACGACGCTCTTCCGATCTNNNATCTATGAACTTCAGGGGTAGC
AGCACACGTCTGAACTCTTTCCCTACACGACGCTCTTCCGATCTNNNGTTTTTCCCCCTAAAATGTTAT
AGCACACGTCTGAACTCTTTCCCTACACGACGCTCTTCCGATCTNNNCCATTTAGAGAGCAAAGGTAT
GCACACGTCTGAACTCTTTCCCTACACGACGCTCTTCCGATCTNNNGAGCATTTGAATTTCTGGATTAG
AGCACACGTCTGAACTCTTTCCCTACACGACGCTCTTCCGATCTNNNTGTTATTTTCTTCTATTTGTGGT
GAGCACACGTCTGAACTCTTTCCCTACACGACGCTCTTCCGATCTNNNGGCACACATAATAAAAACAA
AGCACACGTCTGAACTCTTTCCCTACACGACGCTCTTCCGATCTNNNAGAAAATGTCAGCTAAGAATG
AGCACACGTCTGAACTCTTTCCCTACACGACGCTCTTCCGATCTNNNTCTCTCCTCTTTCTCTTTCTTACA
AGCACACGTCTGAACTCTTTCCCTACACGACGCTCTTCCGATCTNNNGATGCCTGTTGTCAATAATTG
GCACACGTCTGAACTCTTTCCCTACACGACGCTCTTCCGATCTNNNTCAAACTGAATATTCAAAACCA
CACACGTCTGAACTCTTTCCCTACACGACGCTCTTCCGATCTNNNCATAACGCACAAAATATCCCCATC
ACACGTCTGAACTCTTTCCCTACACGACGCTCTTCCGATCTNNNTGTCTAACTACCTGCTGTAACATCG
AGCACACGTCTGAACTCTTTCCCTACACGACGCTCTTCCGATCTNNNAAGAACAATCTTTCCTGATAGT
AGCACACGTCTGAACTCTTTCCCTACACGACGCTCTTCCGATCTNNNATGCAAAATGAGGACTATAAT
ACACGTCTGAACTCTTTCCCTACACGACGCTCTTCCGATCTNNNGACATTTATCATAGCCTCTCCCATC
GAGCACACGTCTGAACTCTTTCCCTACACGACGCTCTTCCGATCTNNNTAAGCTAAGGTGGTTCATTTT
CACACGTCTGAACTCTTTCCCTACACGACGCTCTTCCGATCTNNNATTTCAGCAGAGTGAGGTTTCAT
GCACACGTCTGAACTCTTTCCCTACACGACGCTCTTCCGATCTNNNATAACTCTTTCCTATATCCCTGAC
GAGCACACGTCTGAACTCTTTCCCTACACGACGCTCTTCCGATCTNNNTCTATGGACAAATATGTAAAG
CACACGTCTGAACTCTTTCCCTACACGACGCTCTTCCGATCTNNNAATATGATGAATGCAATGACAAA
CACACGTCTGAACTCTTTCCCTACACGACGCTCTTCCGATCTNNNAAAAATTCTATGAAAAGCACTGA
AGCACACGTCTGAACTCTTTCCCTACACGACGCTCTTCCGATCTNNNCCGTTATGAATAACTGGGATT
AGCACACGTCTGAACTCTTTCCCTACACGACGCTCTTCCGATCTNNNTATAACCACCAACATTTTGATT
AGCACACGTCTGAACTCTTTCCCTACACGACGCTCTTCCGATCTNNNTATAACCACCAACATTTTGATT
GCACACGTCTGAACTCTTTCCCTACACGACGCTCTTCCGATCTNNNCATAAAACTTTGGAATGGGTAA
GAGCACACGTCTGAACTCTTTCCCTACACGACGCTCTTCCGATCTNNNAATTAACAAAAGAGGAGGT
CACACGTCTGAACTCTTTCCCTACACGACGCTCTTCCGATCTNNNTAGAAGATTGAGTTCCTGGTCAT
AGCACACGTCTGAACTCTTTCCCTACACGACGCTCTTCCGATCTNNNTGGAAAACATATACAGTACTA
AGCACACGTCTGAACTCTTTCCCTACACGACGCTCTTCCGATCTNNNCAATCTTTGGTGTTCCTTGGCA
ACACGTCTGAACTCTTTCCCTACACGACGCTCTTCCGATCTNNNGCAACTAACAGAAAAAAAAAGAAC
CACACGTCTGAACTCTTTCCCTACACGACGCTCTTCCGATCTNNNTATGATATAGTTGACCACATACCC
GCACACGTCTGAACTCTTTCCCTACACGACGCTCTTCCGATCTNNNGCTTCATCACATACATAGCTATA
AGCACACGTCTGAACTCTTTCCCTACACGACGCTCTTCCGATCTNNNATGCCTGAACTTAAAAAAATG
AGCACACGTCTGAACTCTTTCCCTACACGACGCTCTTCCGATCTNNNACATTTGGTATACATACATTTG
GAGCACACGTCTGAACTCTTTCCCTACACGACGCTCTTCCGATCTNNNTGCTACAGGATTATGTATTTA
GCACACGTCTGAACTCTTTCCCTACACGACGCTCTTCCGATCTNNNGGCTGGGACTTCTCTTTATTCAT
GCACACGTCTGAACTCTTTCCCTACACGACGCTCTTCCGATCTNNNGGACAAAGGAACATTAAAATA
GCACACGTCTGAACTCTTTCCCTACACGACGCTCTTCCGATCTNNNGAAATCAAAATAACACATCCTC
GCACACGTCTGAACTCTTTCCCTACACGACGCTCTTCCGATCTNNNAATAGTGCTGCAATATACATGA
GCACACGTCTGAACTCTTTCCCTACACGACGCTCTTCCGATCTNNNGACTAAAGTCGTGGAAAAATG
ACACGTCTGAACTCTTTCCCTACACGACGCTCTTCCGATCTNNNAATGTCATCAAGGTACCTTTATACA
AGCACACGTCTGAACTCTTTCCCTACACGACGCTCTTCCGATCTNNNGCAAAAATAAACACTGAGAAA

AGCACACGTCTGAACTCTTTCCCTACACGACGCTCTTCCGATCTNNNGTAAGTGCCTAATCCCAAACCA

AGCACACGTCTGAACTCTTTCCCTACACGACGCTCTTCCGATCTNNNTTCAAAGTCTATATACATAAGC

GCACACGTCTGAACTCTTTCCCTACACGACGCTCTTCCGATCTNNNAAATTCACTGAAGGTAAGTTTTG

AGCACACGTCTGAACTCTTTCCCTACACGACGCTCTTCCGATCTNNNTTTGATCAAAACAGTCTGATGT

GCACACGTCTGAACTCTTTCCCTACACGACGCTCTTCCGATCTNNNTTTGAACAAAGATCTTACAAGG

GAGCACACGTCTGAACTCTTTCCCTACACGACGCTCTTCCGATCTNNNAAGGTGTCTAATTAGATTTAG

AGCACACGTCTGAACTCTTTCCCTACACGACGCTCTTCCGATCTNNNAGAGATGAATTTTATTTGCTGA

AGCACACGTCTGAACTCTTTCCCTACACGACGCTCTTCCGATCTNNNAACAAGAGAATATGTAGAAGT

AGCACACGTCTGAACTCTTTCCCTACACGACGCTCTTCCGATCTNNNACATACAATTCACCTACTTCTT

AGCACACGTCTGAACTCTTTCCCTACACGACGCTCTTCCGATCTNNNTGAGAGTATCCATAACTTGATT

AGCACACGTCTGAACTCTTTCCCTACACGACGCTCTTCCGATCTNNNTGTAAGTCATATTCTTGTTAC

GCACACGTCTGAACTCTTTCCCTACACGACGCTCTTCCGATCTNNNCACTGAATAGAGGAGCTTGTCA

GCACACGTCTGAACTCTTTCCCTACACGACGCTCTTCCGATCTNNNAGGTGGTAGATATGAGTGAGC

AGCACACGTCTGAACTCTTTCCCTACACGACGCTCTTCCGATCTNNNCTTTAGTCTCTGTCATCAATCTC

AGCACACGTCTGAACTCTTTCCCTACACGACGCTCTTCCGATCTNNNCTTCTTTGTGAAGGAATTGA

GCACACGTCTGAACTCTTTCCCTACACGACGCTCTTCCGATCTNNNGAGAAGTAGCACCCTACAAGCAT

AGCACACGTCTGAACTCTTTCCCTACACGACGCTCTTCCGATCTNNNAAATTAGAGAAGAGTAGAAG

GCACACGTCTGAACTCTTTCCCTACACGACGCTCTTCCGATCTNNNTATAGCTTCTTCCTCCTTCCCATC

ACACGTCTGAACTCTTTCCCTACACGACGCTCTTCCGATCTNNNATTTTCTTATATTCCTCCCCCACATC

ACACGTCTGAACTCTTTCCCTACACGACGCTCTTCCGATCTNNNATTTTCTTATATTCCTCCCCCACATC

GCACACGTCTGAACTCTTTCCCTACACGACGCTCTTCCGATCTNNNGTAAATGCATGTTAATTATTGCC

AGCACACGTCTGAACTCTTTCCCTACACGACGCTCTTCCGATCTNNNTTGTTAAGCATTGGTGAAATTT

ACACGTCTGAACTCTTTCCCTACACGACGCTCTTCCGATCTNNNAGAGTAAAATTGAAATTCATGGAC

AGCACACGTCTGAACTCTTTCCCTACACGACGCTCTTCCGATCTNNNGAATTCTGTTCTCCTTCCAGCAT

ACACGTCTGAACTCTTTCCCTACACGACGCTCTTCCGATCTNNNCACCTATGAACCAAGGACTATCATC

AGCACACGTCTGAACTCTTTCCCTACACGACGCTCTTCCGATCTNNNCATGGCATGAATGTTACATATG

AGCACACGTCTGAACTCTTTCCCTACACGACGCTCTTCCGATCTNNNCTGCTTATTTTCTTTTTTGAACTC

AGCACACGTCTGAACTCTTTCCCTACACGACGCTCTTCCGATCTNNNCTGCTTATTTTCTTTTTTGAACTC

GCACACGTCTGAACTCTTTCCCTACACGACGCTCTTCCGATCTNNNAGACTGTTTGAAAAAAGATTTCT

GAGCACACGTCTGAACTCTTTCCCTACACGACGCTCTTCCGATCTNNNGAAGTTTGAGGCTATAGTGA

GCACACGTCTGAACTCTTTCCCTACACGACGCTCTTCCGATCTNNNCTCTAATTCTTAAGTAACCAAGC

AGCACACGTCTGAACTCTTTCCCTACACGACGCTCTTCCGATCTNNNTGCAGATTTTGAGACTTTTTAG

AGCACACGTCTGAACTCTTTCCCTACACGACGCTCTTCCGATCTNNNATTGAAAAATGTATCCCTGTTTC

ACACGTCTGAACTCTTTCCCTACACGACGCTCTTCCGATCTNNNTAACCCAATCATCTCTGAAATCCATC

CACACGTCTGAACTCTTTCCCTACACGACGCTCTTCCGATCTNNNAAATTACAGCCAAGTTTGAATTTC

CACGTCTGAACTCTTTCCCTACACGACGCTCTTCCGATCTNNNATTATCTTGAAATTCTATCTTCCTGCA

AGCACACGTCTGAACTCTTTCCCTACACGACGCTCTTCCGATCTNNNTAAAAATACAAAAGTTAGCCA

AGCACACGTCTGAACTCTTTCCCTACACGACGCTCTTCCGATCTNNNGCAAAAACAACCAGGACTTACA

AGCACACGTCTGAACTCTTTCCCTACACGACGCTCTTCCGATCTNNNCATACAGAAAAGCATGTATCA

AGCACACGTCTGAACTCTTTCCCTACACGACGCTCTTCCGATCTNNNATACCATTGTAATCCCTCTTTAC

AGCACACGTCTGAACTCTTTCCCTACACGACGCTCTTCCGATCTNNNTGAAATGAATTGTGACCACTC

ACACGTCTGAACTCTTTCCCTACACGACGCTCTTCCGATCTNNNAAATCCCAAAAGGTATAATTCCTCAT

AGCACACGTCTGAACTCTTTCCCTACACGACGCTCTTCCGATCTNNNGACCACATTCACAAAGATTTTAT

GCACACGTCTGAACTCTTTCCCTACACGACGCTCTTCCGATCTNNNTAAGACCACATTCACAAAGATTT
AGCACACGTCTGAACTCTTTCCCTACACGACGCTCTTCCGATCTNNNAATTCAAGTGTTCTGTCTTAGA
AGCACACGTCTGAACTCTTTCCCTACACGACGCTCTTCCGATCTNNNCACACAACTATCCACTTCATCAT
AGCACACGTCTGAACTCTTTCCCTACACGACGCTCTTCCGATCTNNNGAACTTGCATATACATTTCTCTT
GCACACGTCTGAACTCTTTCCCTACACGACGCTCTTCCGATCTNNNAAGAGTGAAATTCCATCTCTCCA
ACGTCTGAACTCTTTCCCTACACGACGCTCTTCCGATCTNNNGTACTTGCTGCTTGAGATATCATCGG
AGCACACGTCTGAACTCTTTCCCTACACGACGCTCTTCCGATCTNNNTTGCTTTACTATCTATGTACACA
AGCACACGTCTGAACTCTTTCCCTACACGACGCTCTTCCGATCTNNNTGATATGCTATGACCTTTATT
CACGTCTGAACTCTTTCCCTACACGACGCTCTTCCGATCTNNNATTGTGTCCTCTAATAACATTCTCAT
AGCACACGTCTGAACTCTTTCCCTACACGACGCTCTTCCGATCTNNNATACATTATTTCATTGCCACA
AGCACACGTCTGAACTCTTTCCCTACACGACGCTCTTCCGATCTNNNTCATACAAAGATCAGACTAATG
ACACGTCTGAACTCTTTCCCTACACGACGCTCTTCCGATCTNNNCTGTCTAGTTTAAAAGAACACCCAT
AGCACACGTCTGAACTCTTTCCCTACACGACGCTCTTCCGATCTNNNAAATACACTCTATGCTGTTCAT
CACGTCTGAACTCTTTCCCTACACGACGCTCTTCCGATCTNNNGAGAGGAGTAAGAGAAATTGTATC
ACACGTCTGAACTCTTTCCCTACACGACGCTCTTCCGATCTNNNAGGTACAATGTTAAGCTCTCATAC
CACGTCTGAACTCTTTCCCTACACGACGCTCTTCCGATCTNNNAATTTTGGAGGACACAATTCCATC
GCACACGTCTGAACTCTTTCCCTACACGACGCTCTTCCGATCTNNNGGCTGAAAATGATGATTCTTAT
ACACGTCTGAACTCTTTCCCTACACGACGCTCTTCCGATCTNNNTTTATCCACAAATATTTGCTGACCAT
GCACACGTCTGAACTCTTTCCCTACACGACGCTCTTCCGATCTNNNTTCCAGTATCTCCTACTACATTAC
GCACACGTCTGAACTCTTTCCCTACACGACGCTCTTCCGATCTNNNAAAGGAAATCAAAGACAATTTT
CACACGTCTGAACTCTTTCCCTACACGACGCTCTTCCGATCTNNNCTATTTTGTGGCTTCTACTTGCAT
GCACACGTCTGAACTCTTTCCCTACACGACGCTCTTCCGATCTNNNATGAGATGAACATTAAAATCCA
AGCACACGTCTGAACTCTTTCCCTACACGACGCTCTTCCGATCTNNNTTAAAATGTGCTATTGACATG
GCACACGTCTGAACTCTTTCCCTACACGACGCTCTTCCGATCTNNNCTGTTTTTAATGTGGCTAATAG
AGCACACGTCTGAACTCTTTCCCTACACGACGCTCTTCCGATCTNNNTTAAATAAGTAAGCTGTAACTG
AGCACACGTCTGAACTCTTTCCCTACACGACGCTCTTCCGATCTNNNTTAAATAAGTAAGCTGTAACTG
GCACACGTCTGAACTCTTTCCCTACACGACGCTCTTCCGATCTNNNAAATAAGGAAGATAATGGCATT
AGCACACGTCTGAACTCTTTCCCTACACGACGCTCTTCCGATCTNNNCGTTAAACAGAGAATTGAAA
GAGCACACGTCTGAACTCTTTCCCTACACGACGCTCTTCCGATCTNNNTATATTTTTCCCTCTTGACTG
GAGCACACGTCTGAACTCTTTCCCTACACGACGCTCTTCCGATCTNNNCAGACGTTTGATTTATTCCAT
ACACGTCTGAACTCTTTCCCTACACGACGCTCTTCCGATCTNNNGAAAGATCCTAGGAGGTGATCATC
GCACACGTCTGAACTCTTTCCCTACACGACGCTCTTCCGATCTNNNATTTTGAAGTGTTTCTCTTCCAT
AGCACACGTCTGAACTCTTTCCCTACACGACGCTCTTCCGATCTNNNATACATACGTGTAGACATAAC
GCACACGTCTGAACTCTTTCCCTACACGACGCTCTTCCGATCTNNNATTTTGTAGGGCTGTTATAGAAT
GCACACGTCTGAACTCTTTCCCTACACGACGCTCTTCCGATCTNNNCAACTACCACCTCTATATAGTTC
AGCACACGTCTGAACTCTTTCCCTACACGACGCTCTTCCGATCTNNNCTCATGTTTCTGCAAAATAAA
AGCACACGTCTGAACTCTTTCCCTACACGACGCTCTTCCGATCTNNNCTACATCTATTTGTAATGCATG
AGCACACGTCTGAACTCTTTCCCTACACGACGCTCTTCCGATCTNNNTAGGCTACCACACACTTTTTA
AGCACACGTCTGAACTCTTTCCCTACACGACGCTCTTCCGATCTNNNGAAATATCTTGAAATTGTCTC
GCACACGTCTGAACTCTTTCCCTACACGACGCTCTTCCGATCTNNNTACATTTCCAAGTAATCTGGAG
AGCACACGTCTGAACTCTTTCCCTACACGACGCTCTTCCGATCTNNNCCAAGTCCTAAAAAATTACTC
GCACACGTCTGAACTCTTTCCCTACACGACGCTCTTCCGATCTNNNCATGTATACTATCCAGAGGATA
AGCACACGTCTGAACTCTTTCCCTACACGACGCTCTTCCGATCTNNNCATTCTCACATATGGGAGGAA

CACACGTCTGAACTCTTTCCCTACACGACGCTCTTCCGATCTNNNTACGTATATACCTCTGTGTAACTC
AGCACACGTCTGAACTCTTTCCCTACACGACGCTCTTCCGATCTNNNAGATTTTTTACTTCAGCTTTTGA
AGCACACGTCTGAACTCTTTCCCTACACGACGCTCTTCCGATCTNNNAATAATACTGTTGCTACCTTTT
GCACACGTCTGAACTCTTTCCCTACACGACGCTCTTCCGATCTNNNAAAGAAGGGAAAGCTAAATTTT
GAGCACACGTCTGAACTCTTTCCCTACACGACGCTCTTCCGATCTNNNAGTACATCTGAAACATCCAAT
AGCACACGTCTGAACTCTTTCCCTACACGACGCTCTTCCGATCTNNNTTTTTTCATTATTTGTAGAGAT
AGCACACGTCTGAACTCTTTCCCTACACGACGCTCTTCCGATCTNNNACTAAGTTAGCTCAGTTATTTG
AGCACACGTCTGAACTCTTTCCCTACACGACGCTCTTCCGATCTNNNTGTTACACCTAGTCTGAAATGC
AGCACACGTCTGAACTCTTTCCCTACACGACGCTCTTCCGATCTNNNCCTTACTTATGTCTGCAAAGA
CACACGTCTGAACTCTTTCCCTACACGACGCTCTTCCGATCTNNNTAGAATTCTTCTTTTCCCTGTAAC
CACACGTCTGAACTCTTTCCCTACACGACGCTCTTCCGATCTNNNTAGAATTCTTCTTTTCCCTGTAAC
AGCACACGTCTGAACTCTTTCCCTACACGACGCTCTTCCGATCTNNNCTTTAAACGTAATGCATTTAAC
AGCACACGTCTGAACTCTTTCCCTACACGACGCTCTTCCGATCTNNNGAAACTCCATCTCAAAAAAAA
CACACGTCTGAACTCTTTCCCTACACGACGCTCTTCCGATCTNNNTTGTCAATAGGAAATTGAGATTG
AGCACACGTCTGAACTCTTTCCCTACACGACGCTCTTCCGATCTNNNGGTTACAGCTTGATTTTATACA
AGCACACGTCTGAACTCTTTCCCTACACGACGCTCTTCCGATCTNNNCTGCATTTTACACATCTTTTATG
CACACGTCTGAACTCTTTCCCTACACGACGCTCTTCCGATCTNNNAATCCCCAGTGTGATTATATTAG
GCACACGTCTGAACTCTTTCCCTACACGACGCTCTTCCGATCTNNNCTTTCACCTAGATTCAACAATAG
CACACGTCTGAACTCTTTCCCTACACGACGCTCTTCCGATCTNNNCATATACACAAACACAAAAACAGC
AGCACACGTCTGAACTCTTTCCCTACACGACGCTCTTCCGATCTNNNCTTCTTGTCCCCTAATAATACAT
GAGCACACGTCTGAACTCTTTCCCTACACGACGCTCTTCCGATCTNNNGATTCAGTGTGCTAATATTTT
AGCACACGTCTGAACTCTTTCCCTACACGACGCTCTTCCGATCTNNNGCTTTAGAAATAGGTCAAGAA
AGCACACGTCTGAACTCTTTCCCTACACGACGCTCTTCCGATCTNNNTATTTCACTGATCCTACGTAACC
GCACACGTCTGAACTCTTTCCCTACACGACGCTCTTCCGATCTNNNCTGTGTTGTGAGAATATAGAAA
AGCACACGTCTGAACTCTTTCCCTACACGACGCTCTTCCGATCTNNNCTTCCTTGAGGAGAATGACTA
AGCACACGTCTGAACTCTTTCCCTACACGACGCTCTTCCGATCTNNNAATATGCCCATATCAATGAAT
AGCACACGTCTGAACTCTTTCCCTACACGACGCTCTTCCGATCTNNNACTTTATGAAAACAGAAAGTA
CACACGTCTGAACTCTTTCCCTACACGACGCTCTTCCGATCTNNNCTTCATGTTACGAGATGCTATCAT
GCACACGTCTGAACTCTTTCCCTACACGACGCTCTTCCGATCTNNNATAAAGCCTCTTAAGTAGTCAAA
GCACACGTCTGAACTCTTTCCCTACACGACGCTCTTCCGATCTNNNAAACAAGATGGTCATTGAAAAG
ACGTCTGAACTCTTTCCCTACACGACGCTCTTCCGATCTNNNGTACTTGCTGCTTGAGATATCATCGG
ACACGTCTGAACTCTTTCCCTACACGACGCTCTTCCGATCTNNNATATTTTGTCTCAGTCTTTCTCAT
GCACACGTCTGAACTCTTTCCCTACACGACGCTCTTCCGATCTNNNGATGAGAGTGTGTCCTGTACCA
CACGTCTGAACTCTTTCCCTACACGACGCTCTTCCGATCTNNNAGAGAGAGAGAGAGAGAGATTCAT
AGCACACGTCTGAACTCTTTCCCTACACGACGCTCTTCCGATCTNNNCTGGTAAATCTCATTTTATCCA
AGCACACGTCTGAACTCTTTCCCTACACGACGCTCTTCCGATCTNNNTAAATTTCTTCCCAATAATGAGC
CACACGTCTGAACTCTTTCCCTACACGACGCTCTTCCGATCTNNNTATCTTTTTCACTAACATTTCCCCAT
AGCACACGTCTGAACTCTTTCCCTACACGACGCTCTTCCGATCTNNNCTACAACCATGCTTCATTGTTC
AGCACACGTCTGAACTCTTTCCCTACACGACGCTCTTCCGATCTNNNCTACAACCATGCTTCATTGTTC
AGCACACGTCTGAACTCTTTCCCTACACGACGCTCTTCCGATCTNNNTCCTGGCTAATTTTTGTATTTTT
ACACGTCTGAACTCTTTCCCTACACGACGCTCTTCCGATCTNNNTTTTTCACCTTTGGAGGATACCATC
GCACACGTCTGAACTCTTTCCCTACACGACGCTCTTCCGATCTNNNTTCCTTTTTTCCTGTACCTATTCA
GCACACGTCTGAACTCTTTCCCTACACGACGCTCTTCCGATCTNNNGTAGAACATCAGCTTTGTGTCAT

AGCACACGTCTGAACTCTTTCCCTACACGACGCTCTTCCGATCTNNNGCGTCCTCTATATGTATTGTTA

AGCACACGTCTGAACTCTTTCCCTACACGACGCTCTTCCGATCTNNNCTATCTGACGTATATTAGTTAC

AGCACACGTCTGAACTCTTTCCCTACACGACGCTCTTCCGATCTNNNGAGTAAATGGACTTGAAGGGC

GCACACGTCTGAACTCTTTCCCTACACGACGCTCTTCCGATCTNNNGACAAAAGCAACACAGATAAAT

AGCACACGTCTGAACTCTTTCCCTACACGACGCTCTTCCGATCTNNNCTATCTAATGGAGAAATGGA

AGCACACGTCTGAACTCTTTCCCTACACGACGCTCTTCCGATCTNNNGTAGAATTCCCAGGTTCTTTC

AGCACACGTCTGAACTCTTTCCCTACACGACGCTCTTCCGATCTNNNGTGACACTAATGGTTATAAAC

ACACGTCTGAACTCTTTCCCTACACGACGCTCTTCCGATCTNNNAAAAATTGTGTGAGTAAGTAGAG

GAGCACACGTCTGAACTCTTTCCCTACACGACGCTCTTCCGATCTNNNTTCATTTGTATTTCAACTTGC

AGCACACGTCTGAACTCTTTCCCTACACGACGCTCTTCCGATCTNNNAGATATCTAGCTTCTGCAATAA

ACACGTCTGAACTCTTTCCCTACACGACGCTCTTCCGATCTNNNCTATGTTTCTGAGTCTCTAGCCATC

GCACACGTCTGAACTCTTTCCCTACACGACGCTCTTCCGATCTNNNTCTCAATCTCCATAATTTCATGA

AGCACACGTCTGAACTCTTTCCCTACACGACGCTCTTCCGATCTNNNTAAAAACCAAGGGCTATTTAG

AGCACACGTCTGAACTCTTTCCCTACACGACGCTCTTCCGATCTNNNAGATAAAAATTCCTGATGCCTA

AGCACACGTCTGAACTCTTTCCCTACACGACGCTCTTCCGATCTNNNTTCCAATGAAGATTTCTAAGAG

AGCACACGTCTGAACTCTTTCCCTACACGACGCTCTTCCGATCTNNNCATCAGTGAACCAGTAAAATTA

ACACGTCTGAACTCTTTCCCTACACGACGCTCTTCCGATCTNNNAACATAACGAATCCCTGTCTCATCG

GCACACGTCTGAACTCTTTCCCTACACGACGCTCTTCCGATCTNNNTCATGAATGTAAATATCCTCTGT

AGCACACGTCTGAACTCTTTCCCTACACGACGCTCTTCCGATCTNNNGGCTATGTTATAAATGCAATG

ACACGTCTGAACTCTTTCCCTACACGACGCTCTTCCGATCTNNNCTCATGTTTCAATGTATCGTTTACAT

GCACACGTCTGAACTCTTTCCCTACACGACGCTCTTCCGATCTNNNCTTGTGTTCCATTTATATTTTGCC

AGCACACGTCTGAACTCTTTCCCTACACGACGCTCTTCCGATCTNNNAGGTATTTCTCCTATTGTTATC

CACACGTCTGAACTCTTTCCCTACACGACGCTCTTCCGATCTNNNAACAACTCTGTATCTTCATTGTCAT

ACACGTCTGAACTCTTTCCCTACACGACGCTCTTCCGATCTNNNCTCTCTCTCTCATATGCACATCGG

GCACACGTCTGAACTCTTTCCCTACACGACGCTCTTCCGATCTNNNGTACCTTTCTCCTTATGTCAAGCA

AGCACACGTCTGAACTCTTTCCCTACACGACGCTCTTCCGATCTNNNGAATTTCAGAGCTATTCCACTA

CACGTCTGAACTCTTTCCCTACACGACGCTCTTCCGATCTNNNATTATGTCTTGTCCAAATGTTTCCAT

ACACGTCTGAACTCTTTCCCTACACGACGCTCTTCCGATCTNNNGAAAATCTGGATAAGAAGAGAACC

GCACACGTCTGAACTCTTTCCCTACACGACGCTCTTCCGATCTNNNTCTCTCTCAAATGTATAAAACCA

AGCACACGTCTGAACTCTTTCCCTACACGACGCTCTTCCGATCTNNNATCTTTAAAATGACAACTTCTCC

AGCACACGTCTGAACTCTTTCCCTACACGACGCTCTTCCGATCTNNNCTCCACTACTAACATACACTAAG

AGCACACGTCTGAACTCTTTCCCTACACGACGCTCTTCCGATCTNNNCCCCCACATAATATATTCATCAT

AGCACACGTCTGAACTCTTTCCCTACACGACGCTCTTCCGATCTNNNCCCCCACATAATATATTCATCAT

GCACACGTCTGAACTCTTTCCCTACACGACGCTCTTCCGATCTNNNGGTAGGCTGTAAAGTTTATAG

GAGCACACGTCTGAACTCTTTCCCTACACGACGCTCTTCCGATCTNNNTGTATTTATTGCATGCATATT

AGCACACGTCTGAACTCTTTCCCTACACGACGCTCTTCCGATCTNNNATGTTTGTGTCTATACATACAT

AGCACACGTCTGAACTCTTTCCCTACACGACGCTCTTCCGATCTNNNAAACAACATAGATGTGATTTT

AGCACACGTCTGAACTCTTTCCCTACACGACGCTCTTCCGATCTNNNCAGCATAACTTTAATGTTGGAA

AGCACACGTCTGAACTCTTTCCCTACACGACGCTCTTCCGATCTNNNAAAACTCATTTATTACTGGGA

GCACACGTCTGAACTCTTTCCCTACACGACGCTCTTCCGATCTNNNAAATTATGGTGGTTGGATTAAA

ACACGTCTGAACTCTTTCCCTACACGACGCTCTTCCGATCTNNNTTTAGAGTGACTCGAGAGTGCATC

GCACACGTCTGAACTCTTTCCCTACACGACGCTCTTCCGATCTNNNAAAAGAAAAAGGAGCAAAAAT

GCACACGTCTGAACTCTTTCCCTACACGACGCTCTTCCGATCTNNNGGGAAATGAGAGAAAGAGAA

GAGCACACGTCTGAACTCTTTCCCTACACGACGCTCTTCCGATCTNNNCCAATAAACCTCTTTCTTTTGTA
AGCACACGTCTGAACTCTTTCCCTACACGACGCTCTTCCGATCTNNNATTGAAAACTTCTACCCACTAGC
CACACGTCTGAACTCTTTCCCTACACGACGCTCTTCCGATCTNNNAGGCTGAGTAGGTATTTATGATCA
GCACACGTCTGAACTCTTTCCCTACACGACGCTCTTCCGATCTNNNATAAAGAAATAGATACCAGGCCCA
AGCACACGTCTGAACTCTTTCCCTACACGACGCTCTTCCGATCTNNNCATTGTACACATCAAGTCATTTA
AGCACACGTCTGAACTCTTTCCCTACACGACGCTCTTCCGATCTNNNATCTTCAGTGGGTAGTTAGAT
GCACACGTCTGAACTCTTTCCCTACACGACGCTCTTCCGATCTNNNAGTATATTTACAGTGTTTTGCAG
GCACACGTCTGAACTCTTTCCCTACACGACGCTCTTCCGATCTNNNATAAAGCTAAGCTCTTTAGACATC
AGCACACGTCTGAACTCTTTCCCTACACGACGCTCTTCCGATCTNNNTAAATAAGTGAAACGGAAATA
AGCACACGTCTGAACTCTTTCCCTACACGACGCTCTTCCGATCTNNNTATCATGATGTAAACTACCTCAT
AGCACACGTCTGAACTCTTTCCCTACACGACGCTCTTCCGATCTNNNTTATTGATGGTTTGGGCAATCAT
AGCACACGTCTGAACTCTTTCCCTACACGACGCTCTTCCGATCTNNNATTTAGCATAGTGTAAGCAGAG
GCACACGTCTGAACTCTTTCCCTACACGACGCTCTTCCGATCTNNNCTTGCAAAAAATTCATGAAGATT
GCACACGTCTGAACTCTTTCCCTACACGACGCTCTTCCGATCTNNNAGTAGTCTTGTAGAACCAAAATT
AGCACACGTCTGAACTCTTTCCCTACACGACGCTCTTCCGATCTNNNGAACTTTTAAACCCTGCTAAATA
GCACACGTCTGAACTCTTTCCCTACACGACGCTCTTCCGATCTNNNGTTTCTTCCCTCAAATCTTATTAGC
AGCACACGTCTGAACTCTTTCCCTACACGACGCTCTTCCGATCTNNNTACTTAAAGATCTAAACCCCAG
GAGCACACGTCTGAACTCTTTCCCTACACGACGCTCTTCCGATCTNNNGAGTGATTGAGTAAGACCCT
AGCACACGTCTGAACTCTTTCCCTACACGACGCTCTTCCGATCTNNNAAGAGAACTTCTGATGTATGA
GAGCACACGTCTGAACTCTTTCCCTACACGACGCTCTTCCGATCTNNNGTCCCAAAATTATCGTTCTTAT
AGCACACGTCTGAACTCTTTCCCTACACGACGCTCTTCCGATCTNNNTCTTACTGTGCCTACTTTATAAA
AGCACACGTCTGAACTCTTTCCCTACACGACGCTCTTCCGATCTNNNACTCTATTAAATACACCACAACT
GCACACGTCTGAACTCTTTCCCTACACGACGCTCTTCCGATCTNNNCCGAATAACACATATTCTCAATAC
GAGCACACGTCTGAACTCTTTCCCTACACGACGCTCTTCCGATCTNNNGGAGGACAGAACTACGTAAAT
GCACACGTCTGAACTCTTTCCCTACACGACGCTCTTCCGATCTNNNGAGGACGGAGACTTTAGTAACAT
GAGCACACGTCTGAACTCTTTCCCTACACGACGCTCTTCCGATCTNNNTTGTGTGTTCCTATTACTAATG
AGCACACGTCTGAACTCTTTCCCTACACGACGCTCTTCCGATCTNNNCTTTGTGTACCTGAAGTAATG
GAGCACACGTCTGAACTCTTTCCCTACACGACGCTCTTCCGATCTNNNTTATAGATTTGATATGGTTTG
AGCACACGTCTGAACTCTTTCCCTACACGACGCTCTTCCGATCTNNNTGTACAACATGGTAACTATACT
AGCACACGTCTGAACTCTTTCCCTACACGACGCTCTTCCGATCTNNNTAGGTAATTATATCTCAGCTCAG
AGCACACGTCTGAACTCTTTCCCTACACGACGCTCTTCCGATCTNNNTTTCAGTTCTATTATCTCAGTCA
CACACGTCTGAACTCTTTCCCTACACGACGCTCTTCCGATCTNNNCACCCTTTGTGTTCTATTCACATCGG
GAGCACACGTCTGAACTCTTTCCCTACACGACGCTCTTCCGATCTNNNCTATTATGTTAGAAGCAACTCA
CACACGTCTGAACTCTTTCCCTACACGACGCTCTTCCGATCTNNNCAAGGGGTGTCTTTTATTTTTTACA
GCACACGTCTGAACTCTTTCCCTACACGACGCTCTTCCGATCTNNNGTGTGGACAAACCTTACATCCATC
AGCACACGTCTGAACTCTTTCCCTACACGACGCTCTTCCGATCTNNNGTACCTGTTCACTCACACTTCAT
AGCACACGTCTGAACTCTTTCCCTACACGACGCTCTTCCGATCTNNNCTGGTAGCTCTACAGTTCTGCA
AGCACACGTCTGAACTCTTTCCCTACACGACGCTCTTCCGATCTNNNGAAATTTTGAAATTCATCTCCTG
GAGCACACGTCTGAACTCTTTCCCTACACGACGCTCTTCCGATCTNNNGCAAGAATACAGAATTGATTT
GCACACGTCTGAACTCTTTCCCTACACGACGCTCTTCCGATCTNNNCTAGGTAGGAGTTTTGGGTGCAT
AGCACACGTCTGAACTCTTTCCCTACACGACGCTCTTCCGATCTNNNTAGTGTGGGTATTTATTTTGTT
GAGCACACGTCTGAACTCTTTCCCTACACGACGCTCTTCCGATCTNNNCTTCTCCCTGTATCCTCATGCAT
ACACGTCTGAACTCTTTCCCTACACGACGCTCTTCCGATCTNNNTAGGGTGAACATAGGTGGTACATC

GCACACGTCTGAACTCTTTCCCTACACGACGCTCTTCCGATCTNNNCTAAGCAACAGTGGTGTTTTCAT
CACACGTCTGAACTCTTTCCCTACACGACGCTCTTCCGATCTNNNCCATATCCATTTCACATTACAAACA
GCACACGTCTGAACTCTTTCCCTACACGACGCTCTTCCGATCTNNNGAGAAAAGAAAAGAGGGAAG
AGCACACGTCTGAACTCTTTCCCTACACGACGCTCTTCCGATCTNNNGTCAAGATATAGAATTCCCCTT
CACACGTCTGAACTCTTTCCCTACACGACGCTCTTCCGATCTNNNAACGTTCTTATCCTGATCAAACAT
AGCACACGTCTGAACTCTTTCCCTACACGACGCTCTTCCGATCTNNNACTTTGTAAGTATCAGATTACC
AGCACACGTCTGAACTCTTTCCCTACACGACGCTCTTCCGATCTNNNGTCAACTCTGTCCTTTCTTCCA
AGCACACGTCTGAACTCTTTCCCTACACGACGCTCTTCCGATCTNNNCTTTTTCATCTTTCCTAGAACAA
AGCACACGTCTGAACTCTTTCCCTACACGACGCTCTTCCGATCTNNNTGGAAAACCAAAAACTGTATTT
GCACACGTCTGAACTCTTTCCCTACACGACGCTCTTCCGATCTNNNATATAGAGAGAGAGAGAGAGA
CACACGTCTGAACTCTTTCCCTACACGACGCTCTTCCGATCTNNNGATGCAAAGTGAGGATATGATT
AGCACACGTCTGAACTCTTTCCCTACACGACGCTCTTCCGATCTNNNCATGATGGTTAAATTCATGTGT
CACGTCTGAACTCTTTCCCTACACGACGCTCTTCCGATCTNNNCAATGTGCTACCTATGGACTCATCG
AGCACACGTCTGAACTCTTTCCCTACACGACGCTCTTCCGATCTNNNAAGAGAAAACAGACTATCTTT
AGCACACGTCTGAACTCTTTCCCTACACGACGCTCTTCCGATCTNNNTTTCTGCTTCAAAGACTTTTATT
GCACACGTCTGAACTCTTTCCCTACACGACGCTCTTCCGATCTNNNATACTACACATAGAAAGTAGAC
AGCACACGTCTGAACTCTTTCCCTACACGACGCTCTTCCGATCTNNNTACATTTCGTTTCTGTAATTTC
CACGTCTGAACTCTTTCCCTACACGACGCTCTTCCGATCTNNNCTTTAGGTGAGAAGGGTTTCCATCG
ACACGTCTGAACTCTTTCCCTACACGACGCTCTTCCGATCTNNNGATCCATAGTAAGTTGCAGTTCATC
GAGCACACGTCTGAACTCTTTCCCTACACGACGCTCTTCCGATCTNNNAGCTAGATGTATAAACATAC
GCACACGTCTGAACTCTTTCCCTACACGACGCTCTTCCGATCTNNNATCCAAAACCATTCATATTTTGAC
GCACACGTCTGAACTCTTTCCCTACACGACGCTCTTCCGATCTNNNCTTAATAGTGAGACTAGGGGTC
CACACGTCTGAACTCTTTCCCTACACGACGCTCTTCCGATCTNNNTATTCATTGCTGGAGGATATTTCA
AGCACACGTCTGAACTCTTTCCCTACACGACGCTCTTCCGATCTNNNGCCACTAAAGAAGCGTAATACA
GCACACGTCTGAACTCTTTCCCTACACGACGCTCTTCCGATCTNNNACCTCAAGTGAAAAGAAGTATA
AGCACACGTCTGAACTCTTTCCCTACACGACGCTCTTCCGATCTNNNTCTATGACTGATCAACTCTATTC
AGCACACGTCTGAACTCTTTCCCTACACGACGCTCTTCCGATCTNNNTTATTTGATTGGTAGGAAAA
CACACGTCTGAACTCTTTCCCTACACGACGCTCTTCCGATCTNNNCATTTCCGCCTATCCTATTTCCATC
AGCACACGTCTGAACTCTTTCCCTACACGACGCTCTTCCGATCTNNNCATCTGAGGGTATTTCTGTGC
GCACACGTCTGAACTCTTTCCCTACACGACGCTCTTCCGATCTNNNAGCTACAATTCAAGATGAGATT
AGCACACGTCTGAACTCTTTCCCTACACGACGCTCTTCCGATCTNNNTATATAAAACAACCAGGAATGT
AGCACACGTCTGAACTCTTTCCCTACACGACGCTCTTCCGATCTNNNGCATTGCATTTTAATTTTTCAC
GCACACGTCTGAACTCTTTCCCTACACGACGCTCTTCCGATCTNNNCTTTCAATATCCTGCTTGAAAAT
GCACACGTCTGAACTCTTTCCCTACACGACGCTCTTCCGATCTNNNTACACAATTTAACTACTAGCTGA
AGCACACGTCTGAACTCTTTCCCTACACGACGCTCTTCCGATCTNNNAAAAAACAACAAAAAAAAACC
CACACGTCTGAACTCTTTCCCTACACGACGCTCTTCCGATCTNNNTGTCACTATTTTTTATTCTAGCCC
AGCACACGTCTGAACTCTTTCCCTACACGACGCTCTTCCGATCTNNNTTCAGTCATTTCTGTAAGTCAG
CACACGTCTGAACTCTTTCCCTACACGACGCTCTTCCGATCTNNNAAAATAGCATGGAACCTAGAATT
CACACGTCTGAACTCTTTCCCTACACGACGCTCTTCCGATCTNNNTCTTTCCTCTAGGCAATTTGCATC
GAGCACACGTCTGAACTCTTTCCCTACACGACGCTCTTCCGATCTNNNTTCTTTTATGCTTTGCCAATTT
CACACGTCTGAACTCTTTCCCTACACGACGCTCTTCCGATCTNNNTGAAAATAATTCTTCTGAGTGAAC
GCACACGTCTGAACTCTTTCCCTACACGACGCTCTTCCGATCTNNNTGTACCATCAAATATCTATGAG
GAGCACACGTCTGAACTCTTTCCCTACACGACGCTCTTCCGATCTNNNTGGTAATGGAATGAACAAAA

GCACACGTCTGAACTCTTTCCCTACACGACGCTCTTCCGATCTNNNTGAAATATCACTTAAACAGTAGC
CACACGTCTGAACTCTTTCCCTACACGACGCTCTTCCGATCTNNNTTGTCAATACAAGCTATTTCTTCCAT
AGCACACGTCTGAACTCTTTCCCTACACGACGCTCTTCCGATCTNNNTTATCTATGAGTAGACAACCAA
AGCACACGTCTGAACTCTTTCCCTACACGACGCTCTTCCGATCTNNNTTTTGTTTTTCATTTGTATGCAAC
GAGCACACGTCTGAACTCTTTCCCTACACGACGCTCTTCCGATCTNNNAACAGTTCATAATAGATCCCTT
GAGCACACGTCTGAACTCTTTCCCTACACGACGCTCTTCCGATCTNNNAACAGTTCATAATAGATCCCTT
AGCACACGTCTGAACTCTTTCCCTACACGACGCTCTTCCGATCTNNNTTCACTCAAGAATAATTTTCCAG
AGCACACGTCTGAACTCTTTCCCTACACGACGCTCTTCCGATCTNNNATTTTTGTCCATCATATCTATGCC
AGCACACGTCTGAACTCTTTCCCTACACGACGCTCTTCCGATCTNNNGAAAATTAGCTAGGTCAGAGAT
GAGCACACGTCTGAACTCTTTCCCTACACGACGCTCTTCCGATCTNNNAGTGGAATAAAATGTGAGATA
AGCACACGTCTGAACTCTTTCCCTACACGACGCTCTTCCGATCTNNNGAGACTTCTTTCTTCATTCAAATC
GCACACGTCTGAACTCTTTCCCTACACGACGCTCTTCCGATCTNNNCCTTGCATTTCCATATGAATTTTCA
AGCACACGTCTGAACTCTTTCCCTACACGACGCTCTTCCGATCTNNNCTTTTCTTTGAGCTGTTAGTTAT
GAGCACACGTCTGAACTCTTTCCCTACACGACGCTCTTCCGATCTNNNTTCTAACTGTAATTACCTTCCA
GAGCACACGTCTGAACTCTTTCCCTACACGACGCTCTTCCGATCTNNNAGTATACATGTTATGCAGCCCA
AGCACACGTCTGAACTCTTTCCCTACACGACGCTCTTCCGATCTNNNCATTGAAGACAGTCTGATAAAA
AGCACACGTCTGAACTCTTTCCCTACACGACGCTCTTCCGATCTNNNGTTCTGGTCAATTCTGTTTCTCA
GCACACGTCTGAACTCTTTCCCTACACGACGCTCTTCCGATCTNNNCCTGAAAACCAGATTTTTTTTAC
AGCACACGTCTGAACTCTTTCCCTACACGACGCTCTTCCGATCTNNNTCATTTTCCTAAGTGAATTAACG
CACACGTCTGAACTCTTTCCCTACACGACGCTCTTCCGATCTNNNTCTGTGTCTATAAATGAACTTAGAC
GAGCACACGTCTGAACTCTTTCCCTACACGACGCTCTTCCGATCTNNNGTATTGAATTTCTAGGTAGGT
AGCACACGTCTGAACTCTTTCCCTACACGACGCTCTTCCGATCTNNNGACTTCAGTGTTCATTTAGATG
AGCACACGTCTGAACTCTTTCCCTACACGACGCTCTTCCGATCTNNNTGTCTCTTGTTGATTTCTTTATCC
AGCACACGTCTGAACTCTTTCCCTACACGACGCTCTTCCGATCTNNNTGTATATGTTAGTGGGGCATCA
AGCACACGTCTGAACTCTTTCCCTACACGACGCTCTTCCGATCTNNNCTCCTGGTTTTGAGAAATATTAA
AGCACACGTCTGAACTCTTTCCCTACACGACGCTCTTCCGATCTNNNACATGCCCCTTTTATATGTGCAT
GCACACGTCTGAACTCTTTCCCTACACGACGCTCTTCCGATCTNNNAAAGAGAAAGCATAAATCAGAT
GAGCACACGTCTGAACTCTTTCCCTACACGACGCTCTTCCGATCTNNNTTTTTACACTGTTTTTCTTTG
GAGCACACGTCTGAACTCTTTCCCTACACGACGCTCTTCCGATCTNNNTCTATGATCTATGTTTCTTTGT
GCACACGTCTGAACTCTTTCCCTACACGACGCTCTTCCGATCTNNNGAATTGCCATATTTGGTATGTAT
AGCACACGTCTGAACTCTTTCCCTACACGACGCTCTTCCGATCTNNNAGGAGATGTTCAATAAATAATC
AGCACACGTCTGAACTCTTTCCCTACACGACGCTCTTCCGATCTNNNCTGTCACTTTTGTCCTCTTCCATC
CACACGTCTGAACTCTTTCCCTACACGACGCTCTTCCGATCTNNNCTTTTTACCCTTGACTACCTGCATCG
AGCACACGTCTGAACTCTTTCCCTACACGACGCTCTTCCGATCTNNNTGCTGATCTACAAGGAATTTATA
ACACGTCTGAACTCTTTCCCTACACGACGCTCTTCCGATCTNNNAGATGCACCAAACAGTGTATCATCG
GAGCACACGTCTGAACTCTTTCCCTACACGACGCTCTTCCGATCTNNNCAAAGTGCTGAGATTACAGGC
GCACACGTCTGAACTCTTTCCCTACACGACGCTCTTCCGATCTNNNGTTGCCATTGTTCAGTGAATCAT
AGCACACGTCTGAACTCTTTCCCTACACGACGCTCTTCCGATCTNNNTACAACTCATTATTTAGGGTCT
GAGCACACGTCTGAACTCTTTCCCTACACGACGCTCTTCCGATCTNNNCTCTGAAGCTCTTTCTTAGGCA
GAGCACACGTCTGAACTCTTTCCCTACACGACGCTCTTCCGATCTNNNCATGCTTCTTACTTCCAAATAT
AGCACACGTCTGAACTCTTTCCCTACACGACGCTCTTCCGATCTNNNGAAGAAACAAAAACAACAGTAA
AGCACACGTCTGAACTCTTTCCCTACACGACGCTCTTCCGATCTNNNTGGCCATTTTTAGTGTATAAAA
AGCACACGTCTGAACTCTTTCCCTACACGACGCTCTTCCGATCTNNNAATGCCGAATCTTAAATTGAAAA

AGCACACGTCTGAACTCTTTCCCTACACGACGCTCTTCCGATCTNNNTGTAACTACATAAGCAGTATCA

GCACACGTCTGAACTCTTTCCCTACACGACGCTCTTCCGATCTNNNAAATGTATTTAACTTCAACTGCT

AGCACACGTCTGAACTCTTTCCCTACACGACGCTCTTCCGATCTNNNGTAATCAAGCCCAATATTTCTG

GCACACGTCTGAACTCTTTCCCTACACGACGCTCTTCCGATCTNNNAAAACATTGTGATTTTCCCTAAA

AGCACACGTCTGAACTCTTTCCCTACACGACGCTCTTCCGATCTNNNGACATTTTTCTTGGACACACCA

AGCACACGTCTGAACTCTTTCCCTACACGACGCTCTTCCGATCTNNNAAATTAGTCCTAAGGGCATTAT

CACGTCTGAACTCTTTCCCTACACGACGCTCTTCCGATCTNNNTTTGATTTTTAGATCCTGCAAATCAT

CACACGTCTGAACTCTTTCCCTACACGACGCTCTTCCGATCTNNNAGATAACCAAGAATTAGTGCTAT

GCACACGTCTGAACTCTTTCCCTACACGACGCTCTTCCGATCTNNNAAAGACTGAAAGATAACAAAAC

CACACGTCTGAACTCTTTCCCTACACGACGCTCTTCCGATCTNNNGTAAGTTTATAGGGCACTATAGT

AGCACACGTCTGAACTCTTTCCCTACACGACGCTCTTCCGATCTNNNCATGTTCTGGATGATAAAAAG

GCACACGTCTGAACTCTTTCCCTACACGACGCTCTTCCGATCTNNNATTTTTTCAGTGTTTCTGTCTCC

CACACGTCTGAACTCTTTCCCTACACGACGCTCTTCCGATCTNNNTACATCATTTTTTCAGTGTTTCTC

AGCACACGTCTGAACTCTTTCCCTACACGACGCTCTTCCGATCTNNNCTTCCTTGTGTAATTTTCAGAA

GCACACGTCTGAACTCTTTCCCTACACGACGCTCTTCCGATCTNNNAATGTCATCATTGAGAATCCTAT

CACGTCTGAACTCTTTCCCTACACGACGCTCTTCCGATCTNNNGCACTGACTCAAAAATAAGTAAACA

GAGCACACGTCTGAACTCTTTCCCTACACGACGCTCTTCCGATCTNNNTTAGCATGACACATGCATTA

AGCACACGTCTGAACTCTTTCCCTACACGACGCTCTTCCGATCTNNNCTCTTAGAGTTCAAAGTACAG

GCACACGTCTGAACTCTTTCCCTACACGACGCTCTTCCGATCTNNNCTGATCTTTTGGGTGAAGAGCA

GAGCACACGTCTGAACTCTTTCCCTACACGACGCTCTTCCGATCTNNNTCTGAAGATGCTATGATGAA

GCACACGTCTGAACTCTTTCCCTACACGACGCTCTTCCGATCTNNNTATCTACATTTTACACCATTTGCC

CACGTCTGAACTCTTTCCCTACACGACGCTCTTCCGATCTNNNTGTAGTACCTGAAAATGTACATACAT

GAGCACACGTCTGAACTCTTTCCCTACACGACGCTCTTCCGATCTNNNACTCTTTCTTTAGTTGTTCAA

CACACGTCTGAACTCTTTCCCTACACGACGCTCTTCCGATCTNNNTAGACCTTCCAATCATCCATTTAC

ACACGTCTGAACTCTTTCCCTACACGACGCTCTTCCGATCTNNNCTTTCTATGCAGTGACTGACCATCG

AGCACACGTCTGAACTCTTTCCCTACACGACGCTCTTCCGATCTNNNGGCTACTATTTATTTCCATGTT

GCACACGTCTGAACTCTTTCCCTACACGACGCTCTTCCGATCTNNNACTCCTTAAAATAAATCTCCCAT

GCACACGTCTGAACTCTTTCCCTACACGACGCTCTTCCGATCTNNNCTATGCCAGTTACTTTTTGTTGCA

CACGTCTGAACTCTTTCCCTACACGACGCTCTTCCGATCTNNNACATGAATAAGGTTGTCTGGCATCG

GCACACGTCTGAACTCTTTCCCTACACGACGCTCTTCCGATCTNNNTAAATCAAAGAGAAACCAATTC

AGCACACGTCTGAACTCTTTCCCTACACGACGCTCTTCCGATCTNNNCCAAACATATAACTGTTATGT

AGCACACGTCTGAACTCTTTCCCTACACGACGCTCTTCCGATCTNNNGGTAGTTAACACTTTTGTAAC

AGCACACGTCTGAACTCTTTCCCTACACGACGCTCTTCCGATCTNNNAATGTTGAAAGTCTTTTCTGT

GCACACGTCTGAACTCTTTCCCTACACGACGCTCTTCCGATCTNNNGCCAAATGTAAATATTCTCTCTT

GCACACGTCTGAACTCTTTCCCTACACGACGCTCTTCCGATCTNNNTCACTTTGATCAAACTGTTTTTA

GCACACGTCTGAACTCTTTCCCTACACGACGCTCTTCCGATCTNNNCCTTGAAGATACAAAGACTTAA

ACACGTCTGAACTCTTTCCCTACACGACGCTCTTCCGATCTNNNTTAAACTTTCAAAATGTTGCCTTCAT

GCACACGTCTGAACTCTTTCCCTACACGACGCTCTTCCGATCTNNNGATCCAGTATCCTTACAGTTTAT

CACACGTCTGAACTCTTTCCCTACACGACGCTCTTCCGATCTNNNGTGTGTGTATGTGTATGTGTACAT

GCACACGTCTGAACTCTTTCCCTACACGACGCTCTTCCGATCTNNNATTTTTATTGTTGGGGGAAATG

CACACGTCTGAACTCTTTCCCTACACGACGCTCTTCCGATCTNNNAAAAGAGGAAAGTTAGAATGAA

AGCACACGTCTGAACTCTTTCCCTACACGACGCTCTTCCGATCTNNNTTTACTCCTCTTTTCCATCTGTA

CACACGTCTGAACTCTTTCCCTACACGACGCTCTTCCGATCTNNNTTAGTGCCTTTATAAAACATGAC

AGCACACGTCTGAACTCTTTCCCTACACGACGCTCTTCCGATCTNNNAAGAAACATTCACAAAGACATA
GAGCACACGTCTGAACTCTTTCCCTACACGACGCTCTTCCGATCTNNNCAAGAAGGCAAAAATCATAA
AGCACACGTCTGAACTCTTTCCCTACACGACGCTCTTCCGATCTNNNATAAAGTCAACAACTGTTCCTA
AGCACACGTCTGAACTCTTTCCCTACACGACGCTCTTCCGATCTNNNATAAAGTCAACAACTGTTCCTA
CACACGTCTGAACTCTTTCCCTACACGACGCTCTTCCGATCTNNNGAAAGTCCAACCTCCTTTTCCATCG
GCACACGTCTGAACTCTTTCCCTACACGACGCTCTTCCGATCTNNNAAAATAGCTTCAGAAAATGACT
AGCACACGTCTGAACTCTTTCCCTACACGACGCTCTTCCGATCTNNNACTTTTTCTTTTCCAGTAAACA
AGCACACGTCTGAACTCTTTCCCTACACGACGCTCTTCCGATCTNNNCACAAGTTTAAGATTTTCTCT
GCACACGTCTGAACTCTTTCCCTACACGACGCTCTTCCGATCTNNNACTAATCTTGAAAAGTACTCTCT
AGCACACGTCTGAACTCTTTCCCTACACGACGCTCTTCCGATCTNNNAACAATAATAGAGAAAGGGA
AGCACACGTCTGAACTCTTTCCCTACACGACGCTCTTCCGATCTNNNCCAATTCTCTTGAAGTGTTATTT
GCACACGTCTGAACTCTTTCCCTACACGACGCTCTTCCGATCTNNNCCTTTGTCATTTCTAATTGTGTT
GAGCACACGTCTGAACTCTTTCCCTACACGACGCTCTTCCGATCTNNNAATCAATCTTTCAGCTCCATTC
GCACACGTCTGAACTCTTTCCCTACACGACGCTCTTCCGATCTNNNGAACAACTTTTAGTTTAGGGCT
GCACACGTCTGAACTCTTTCCCTACACGACGCTCTTCCGATCTNNNAACACTAATTTAGACCATCTCTC
CACGTCTGAACTCTTTCCCTACACGACGCTCTTCCGATCTNNNGTACATGGGTAAGAGTGGATCATCG
CACACGTCTGAACTCTTTCCCTACACGACGCTCTTCCGATCTNNNTTTCCTGCTGAGTTAGATTTTCAT
GCACACGTCTGAACTCTTTCCCTACACGACGCTCTTCCGATCTNNNCAACTTTTATTACGTCTCATCATC
GCACACGTCTGAACTCTTTCCCTACACGACGCTCTTCCGATCTNNNTGAGACTTTTCAGATTCCATAAT
AGCACACGTCTGAACTCTTTCCCTACACGACGCTCTTCCGATCTNNNTATAATAGTCTCTAGTGTCTCC
GCACACGTCTGAACTCTTTCCCTACACGACGCTCTTCCGATCTNNNTTTCTTCACATGTCACCTTTACAT
GCACACGTCTGAACTCTTTCCCTACACGACGCTCTTCCGATCTNNNAACGGTGCTTTTATTAGTCTATCA
GCACACGTCTGAACTCTTTCCCTACACGACGCTCTTCCGATCTNNNATCTCTTTCCTTTATAAATTGCCCA
AGCACACGTCTGAACTCTTTCCCTACACGACGCTCTTCCGATCTNNNTTTTGGTTGAAGTTTCATGTAT
GCACACGTCTGAACTCTTTCCCTACACGACGCTCTTCCGATCTNNNAGGATAAAGAGGATGTGTATC
GAGCACACGTCTGAACTCTTTCCCTACACGACGCTCTTCCGATCTNNNCCAATATTATCTAAATGAGGC
AGCACACGTCTGAACTCTTTCCCTACACGACGCTCTTCCGATCTNNNTTTTGTCTATTCCTTTTTCTCTG
GCACACGTCTGAACTCTTTCCCTACACGACGCTCTTCCGATCTNNNGTCAGAATAGCAATTTTGGATG
AGCACACGTCTGAACTCTTTCCCTACACGACGCTCTTCCGATCTNNNTATAAGGATATGAGTACAATG
CACACGTCTGAACTCTTTCCCTACACGACGCTCTTCCGATCTNNNATAAAACATTCGTAGTAGACAGTC
GCACACGTCTGAACTCTTTCCCTACACGACGCTCTTCCGATCTNNNATTCTTCTTTTGACTTCTTTCAAC
GCACACGTCTGAACTCTTTCCCTACACGACGCTCTTCCGATCTNNNTTGTAGTTATCCTTCTCATTAGG
CACGTCTGAACTCTTTCCCTACACGACGCTCTTCCGATCTNNNGTTTTCACTTCAACAACCTCCATCGG
CACACGTCTGAACTCTTTCCCTACACGACGCTCTTCCGATCTNNNGAAATACAAGTTGTTCATTGTTGC
GCACACGTCTGAACTCTTTCCCTACACGACGCTCTTCCGATCTNNNTTCCCATCATCTGAAATTAGTTC
AGCACACGTCTGAACTCTTTCCCTACACGACGCTCTTCCGATCTNNNTTTCAGAATTATGGGACTTAG
AGCACACGTCTGAACTCTTTCCCTACACGACGCTCTTCCGATCTNNNACTACTGTGAAAAAATTTCCAA
GAGCACACGTCTGAACTCTTTCCCTACACGACGCTCTTCCGATCTNNNTGCTTGCTTTGAAATATATATA
AGCACACGTCTGAACTCTTTCCCTACACGACGCTCTTCCGATCTNNNCTATGGTTATGACCTCATTTAA
CACACGTCTGAACTCTTTCCCTACACGACGCTCTTCCGATCTNNNGATTCTGCTTCCAATACTTTTAACA
GCACACGTCTGAACTCTTTCCCTACACGACGCTCTTCCGATCTNNNGATAGGAAGGGTTGAGTGACC
GCACACGTCTGAACTCTTTCCCTACACGACGCTCTTCCGATCTNNNCAACAAGAACAAAACTTCGTCAT
GCACACGTCTGAACTCTTTCCCTACACGACGCTCTTCCGATCTNNNCCTGGTTTATTAGATTTTCTCCC

GCACACGTCTGAACTCTTTCCCTACACGACGCTCTTCCGATCTNNNCTTTATTTTGGTCCAAAAAATGT
GAGCACACGTCTGAACTCTTTCCCTACACGACGCTCTTCCGATCTNNNATTTTATCAGACCAATCCTAT
CACACGTCTGAACTCTTTCCCTACACGACGCTCTTCCGATCTNNNTCTCAAAAAAAAAAAAAAAGCGAC
GAGCACACGTCTGAACTCTTTCCCTACACGACGCTCTTCCGATCTNNNCAAAAACAAAAGAAAAGAGC
GAGCACACGTCTGAACTCTTTCCCTACACGACGCTCTTCCGATCTNNNTCCATGATTCATAATCCTTTT
GAGCACACGTCTGAACTCTTTCCCTACACGACGCTCTTCCGATCTNNNAAGCTCTGTAATGAATGGTC
GAGCACACGTCTGAACTCTTTCCCTACACGACGCTCTTCCGATCTNNNGTCCAACCAATTGATAGAATA
GCACACGTCTGAACTCTTTCCCTACACGACGCTCTTCCGATCTNNNAAATGTTTACAGAAGGTCAAAG
CACGTCTGAACTCTTTCCCTACACGACGCTCTTCCGATCTNNNGATGACCTACAACATGCAAACATCGG
GCACACGTCTGAACTCTTTCCCTACACGACGCTCTTCCGATCTNNNATTTCAAGCTAGTTTGAAGTTA
CACACGTCTGAACTCTTTCCCTACACGACGCTCTTCCGATCTNNNCTCTCTTTGGTTTGCAGATATCAT
CACACGTCTGAACTCTTTCCCTACACGACGCTCTTCCGATCTNNNCTCTCTTTGGTTTGCAGATATCAT
GCACACGTCTGAACTCTTTCCCTACACGACGCTCTTCCGATCTNNNCTCTTTCTTCACTCTCAGGTCATC
AGCACACGTCTGAACTCTTTCCCTACACGACGCTCTTCCGATCTNNNCACACTAATAGAAAGAGTTGT
GCACACGTCTGAACTCTTTCCCTACACGACGCTCTTCCGATCTNNNGGTCAGATTTCTTTTTGTATAG
ACACGTCTGAACTCTTTCCCTACACGACGCTCTTCCGATCTNNNCTGAGTAGCTAGGATTACAGACAT
AGCACACGTCTGAACTCTTTCCCTACACGACGCTCTTCCGATCTNNNCTGTGATTCTAGGAGCTTTGC
CACACGTCTGAACTCTTTCCCTACACGACGCTCTTCCGATCTNNNTATTATGCTCAGTTCATACTTCTCA
AGCACACGTCTGAACTCTTTCCCTACACGACGCTCTTCCGATCTNNNGGAGATCAGTAAGTTGGTTTT
AGCACACGTCTGAACTCTTTCCCTACACGACGCTCTTCCGATCTNNNAAATGCCGTTGAATGATAATA
AGCACACGTCTGAACTCTTTCCCTACACGACGCTCTTCCGATCTNNNTTTAAGATTTGCAATCATCTC
GCACACGTCTGAACTCTTTCCCTACACGACGCTCTTCCGATCTNNNCTATTCGTATATGTGTGTTCATGC
GCACACGTCTGAACTCTTTCCCTACACGACGCTCTTCCGATCTNNNAATACTCCCTGAAATCAGTAATG
AGCACACGTCTGAACTCTTTCCCTACACGACGCTCTTCCGATCTNNNAATTTGATTATGTTGTACCTTG
GAGCACACGTCTGAACTCTTTCCCTACACGACGCTCTTCCGATCTNNNAAAATGCTTAGGTTCTCGAAC
AGCACACGTCTGAACTCTTTCCCTACACGACGCTCTTCCGATCTNNNTTTGTCCATTTCTCCAAATATTT
AGCACACGTCTGAACTCTTTCCCTACACGACGCTCTTCCGATCTNNNATAAAGTTCACCGCATTCATCA
GAGCACACGTCTGAACTCTTTCCCTACACGACGCTCTTCCGATCTNNNCATGTAGTGTGTTTCCTAAAT
AGCACACGTCTGAACTCTTTCCCTACACGACGCTCTTCCGATCTNNNATGAGTTTCTCTCTTTTGAAAAT
GAGCACACGTCTGAACTCTTTCCCTACACGACGCTCTTCCGATCTNNNTGGAACACTTTAGATAAAAGA
GCACACGTCTGAACTCTTTCCCTACACGACGCTCTTCCGATCTNNNTACAGCCTAACTTTGATAAGTAA
AGCACACGTCTGAACTCTTTCCCTACACGACGCTCTTCCGATCTNNNTTTGGATTAGAGCAATTATTCA
AGCACACGTCTGAACTCTTTCCCTACACGACGCTCTTCCGATCTNNNTGTCTCTAAAAAAAACAAAGC
GAGCACACGTCTGAACTCTTTCCCTACACGACGCTCTTCCGATCTNNNAAGAATATACATTCAGGTTTT
GCACACGTCTGAACTCTTTCCCTACACGACGCTCTTCCGATCTNNNAAGAAAATGATTCAAGTGTTTT
CACACGTCTGAACTCTTTCCCTACACGACGCTCTTCCGATCTNNNAATGATTTCAGGTTCATTGTTCCAT
GCACACGTCTGAACTCTTTCCCTACACGACGCTCTTCCGATCTNNNAAAATACAGACAGACAGACACCA
CACACGTCTGAACTCTTTCCCTACACGACGCTCTTCCGATCTNNNAATCAGTTTTTTGGATTATGAGCC
AGCACACGTCTGAACTCTTTCCCTACACGACGCTCTTCCGATCTNNNGGCATACATATTCAGTCCATATA
GCACACGTCTGAACTCTTTCCCTACACGACGCTCTTCCGATCTNNNTTTACCAGAAAAAGGAAGGAGC
GCACACGTCTGAACTCTTTCCCTACACGACGCTCTTCCGATCTNNNTTTACCAGAAAAAGGAAGGAGC
GCACACGTCTGAACTCTTTCCCTACACGACGCTCTTCCGATCTNNNTTTACCAGAAAAAGGAAGGAGC
AGCACACGTCTGAACTCTTTCCCTACACGACGCTCTTCCGATCTNNNTAGAGATTATGACAGTGCTAAT

AGCACACGTCTGAACTCTTTCCCTACACGACGCTCTTCCGATCTNNNGTATAATATTCCATCATACACG
CACACGTCTGAACTCTTTCCCTACACGACGCTCTTCCGATCTNNNTTTTCAGCTTTCTAGAAGCCCATC
CACACGTCTGAACTCTTTCCCTACACGACGCTCTTCCGATCTNNNATTTCTTTTCCATTTTTGATGTGC
CACACGTCTGAACTCTTTCCCTACACGACGCTCTTCCGATCTNNNTAATCCTAGTCGTGTTATTTTTCC
GAGCACACGTCTGAACTCTTTCCCTACACGACGCTCTTCCGATCTNNNATGAATTAATACCCTTTGCTT
GCACACGTCTGAACTCTTTCCCTACACGACGCTCTTCCGATCTNNNTGCTGATCTTTATTATAGGAAA
GCACACGTCTGAACTCTTTCCCTACACGACGCTCTTCCGATCTNNNAGACTTTTTTCAAAGCTACATT
AGCACACGTCTGAACTCTTTCCCTACACGACGCTCTTCCGATCTNNNCAGAAAGAAAGAGAGAAGGG
CACACGTCTGAACTCTTTCCCTACACGACGCTCTTCCGATCTNNNAGTATTCTTTGCTGTTATCTAGGC
CACACGTCTGAACTCTTTCCCTACACGACGCTCTTCCGATCTNNNTTTTACTAAGAATAATGGTCTCCC
CACGTCTGAACTCTTTCCCTACACGACGCTCTTCCGATCTNNNATACCTAATCCTTCTGCAGATTCATC
GCACACGTCTGAACTCTTTCCCTACACGACGCTCTTCCGATCTNNNCTAAAAAACATGAAAAACCTGA
GCACACGTCTGAACTCTTTCCCTACACGACGCTCTTCCGATCTNNNCCTTCTCACTGTATTGTCATATCA
AGCACACGTCTGAACTCTTTCCCTACACGACGCTCTTCCGATCTNNNTAGAAGTGATGAATAAGGAC
GCACACGTCTGAACTCTTTCCCTACACGACGCTCTTCCGATCTNNNAAATGATGGATTTGTTTGGTAT
AGCACACGTCTGAACTCTTTCCCTACACGACGCTCTTCCGATCTNNNTAAAGGAAAATATTACACATG
GCACACGTCTGAACTCTTTCCCTACACGACGCTCTTCCGATCTNNNGGAAAAGATATCAGAATTGCTT
GCACACGTCTGAACTCTTTCCCTACACGACGCTCTTCCGATCTNNNGGAAAAGATATCAGAATTGCTT
GCACACGTCTGAACTCTTTCCCTACACGACGCTCTTCCGATCTNNNTTAGTAAAGAAAGAAGTCTAGG
AGCACACGTCTGAACTCTTTCCCTACACGACGCTCTTCCGATCTNNNTTGAAATAACTTATTGTCGAA
AGCACACGTCTGAACTCTTTCCCTACACGACGCTCTTCCGATCTNNNCCACAGTTATATACCAAAGAAA
AGCACACGTCTGAACTCTTTCCCTACACGACGCTCTTCCGATCTNNNATGAGACAGAAATAGAGGAA
AGCACACGTCTGAACTCTTTCCCTACACGACGCTCTTCCGATCTNNNATATCTGAGTCTAAAACTTTC
AGCACACGTCTGAACTCTTTCCCTACACGACGCTCTTCCGATCTNNNACAGAGTTTAAGATTCTTAAA
GCACACGTCTGAACTCTTTCCCTACACGACGCTCTTCCGATCTNNNTGTTTAAATGAAGAAGTGAAA
CACACGTCTGAACTCTTTCCCTACACGACGCTCTTCCGATCTNNNCTTTACCAATTTTCATGCACATACA
AGCACACGTCTGAACTCTTTCCCTACACGACGCTCTTCCGATCTNNNATCATTCCTATATTGGTGAACT
GCACACGTCTGAACTCTTTCCCTACACGACGCTCTTCCGATCTNNNAGATGGTATATTGTGGGAATT
AGCACACGTCTGAACTCTTTCCCTACACGACGCTCTTCCGATCTNNNAATAGACTGAAGGCAGAAGT
GCACACGTCTGAACTCTTTCCCTACACGACGCTCTTCCGATCTNNNTGTTTATACACACACACATTTTA
GCACACGTCTGAACTCTTTCCCTACACGACGCTCTTCCGATCTNNNTTACTAACACATAAGGCTGAAG
ACACGTCTGAACTCTTTCCCTACACGACGCTCTTCCGATCTNNNTAAGTTGTCCTGACTATCCCCATCG
GCACACGTCTGAACTCTTTCCCTACACGACGCTCTTCCGATCTNNNTGCATTCCTACATATATATGTGT
AGCACACGTCTGAACTCTTTCCCTACACGACGCTCTTCCGATCTNNNCTCTATGGGAAACTGACACAC
AGCACACGTCTGAACTCTTTCCCTACACGACGCTCTTCCGATCTNNNTATGCCATAACTCTAAAATGTT
GCACACGTCTGAACTCTTTCCCTACACGACGCTCTTCCGATCTNNNTTAAAAGTTGCACATAGAACAG
CACACGTCTGAACTCTTTCCCTACACGACGCTCTTCCGATCTNNNACTTGGACAACTGTCTCTTCCATCG
GCACACGTCTGAACTCTTTCCCTACACGACGCTCTTCCGATCTNNNTTCCTTCTGCCTGATAAATGCAT
CACGTCTGAACTCTTTCCCTACACGACGCTCTTCCGATCTNNNGATCTCATCAAGAGCAGAACCATCG
AGCACACGTCTGAACTCTTTCCCTACACGACGCTCTTCCGATCTNNNGCTATAGCAATTATAGTGTGG
CACACGTCTGAACTCTTTCCCTACACGACGCTCTTCCGATCTNNNTATTTCACTTAGCATAATGTCCTC
ACACGTCTGAACTCTTTCCCTACACGACGCTCTTCCGATCTNNNCTCAGTATCTCCTTTCTCTCTCATCG
GCACACGTCTGAACTCTTTCCCTACACGACGCTCTTCCGATCTNNNAAACTCAAACTAAGACATATACG

AGCACACGTCTGAACTCTTTCCCTACACGACGCTCTTCCGATCTNNNAGGAGGCTAATACTATCATAAC
AGCACACGTCTGAACTCTTTCCCTACACGACGCTCTTCCGATCTNNNAAGTTACAAAGACATTTACTTG
GCACACGTCTGAACTCTTTCCCTACACGACGCTCTTCCGATCTNNNCAGGTCTGTTTCTTCTTCTCCATC
CACACGTCTGAACTCTTTCCCTACACGACGCTCTTCCGATCTNNNATTCTAGATGTGAGTGAAGTGCAT
CACACGTCTGAACTCTTTCCCTACACGACGCTCTTCCGATCTNNNATTCTAGATGTGAGTGAAGTGCAT
AGCACACGTCTGAACTCTTTCCCTACACGACGCTCTTCCGATCTNNNAGAATGTTAATACATCAGTAG
CACACGTCTGAACTCTTTCCCTACACGACGCTCTTCCGATCTNNNCACACGTACACATTCATGTACATC
AGCACACGTCTGAACTCTTTCCCTACACGACGCTCTTCCGATCTNNNGAATAAGTGCAATATTGGCATA
AGCACACGTCTGAACTCTTTCCCTACACGACGCTCTTCCGATCTNNNGTTTTGGATTCTTAACTTTCTC
GCACACGTCTGAACTCTTTCCCTACACGACGCTCTTCCGATCTNNNTTCCTTTATTTCTTGTTTGTTTGC
ACACGTCTGAACTCTTTCCCTACACGACGCTCTTCCGATCTNNNAACCAATAGAAAATGCACATGCATC
CACACGTCTGAACTCTTTCCCTACACGACGCTCTTCCGATCTNNNCCTGAGTTACTTCACTTAGAATAC
AGCACACGTCTGAACTCTTTCCCTACACGACGCTCTTCCGATCTNNNATTACATCTTCATTCTTTAAGG
AGCACACGTCTGAACTCTTTCCCTACACGACGCTCTTCCGATCTNNNAAGAGAACTTCTGATGTATGA
AGCACACGTCTGAACTCTTTCCCTACACGACGCTCTTCCGATCTNNNGTGGTTTTTACACAAAGTCCCAT
AGCACACGTCTGAACTCTTTCCCTACACGACGCTCTTCCGATCTNNNGTGGTTTTTACACAAAGTCCCAT
GCACACGTCTGAACTCTTTCCCTACACGACGCTCTTCCGATCTNNNTTCCTTGATCTGAAAAAGTAATG
ACACGTCTGAACTCTTTCCCTACACGACGCTCTTCCGATCTNNNATTAGCATGCAATTGTCAGATCATC
AGCACACGTCTGAACTCTTTCCCTACACGACGCTCTTCCGATCTNNNAATCTTTAACATAAGTGGGAA
AGCACACGTCTGAACTCTTTCCCTACACGACGCTCTTCCGATCTNNNATATTTTAGAATCAGCTTGTCA
AGCACACGTCTGAACTCTTTCCCTACACGACGCTCTTCCGATCTNNNGGCTCTAGGACTTCCTTTATCA
GAGCACACGTCTGAACTCTTTCCCTACACGACGCTCTTCCGATCTNNNAAGGAGGTAAAGTTTGTATA
AGCACACGTCTGAACTCTTTCCCTACACGACGCTCTTCCGATCTNNNCAATGTTCATGCTTTAGAAAAG
GAGCACACGTCTGAACTCTTTCCCTACACGACGCTCTTCCGATCTNNNGTCCTCATAAGCTTTCACAGCA
GCACACGTCTGAACTCTTTCCCTACACGACGCTCTTCCGATCTNNNATATCATCCAGCAAAGCACTACA
AGCACACGTCTGAACTCTTTCCCTACACGACGCTCTTCCGATCTNNNAACATCTTTGTGGTAGTTGTT
GAGCACACGTCTGAACTCTTTCCCTACACGACGCTCTTCCGATCTNNNTTGAGACTTGAATTTGAAAAT
GCACACGTCTGAACTCTTTCCCTACACGACGCTCTTCCGATCTNNNCTTCTCAAAACAAAACAAACAAA
ACACGTCTGAACTCTTTCCCTACACGACGCTCTTCCGATCTNNNCTCTCCTTTCTCTCTCTCTTCATCGGA
CACACGTCTGAACTCTTTCCCTACACGACGCTCTTCCGATCTNNNGTGTGTATGAGTGTGTGATGCAT
AGCACACGTCTGAACTCTTTCCCTACACGACGCTCTTCCGATCTNNNTGGATCTTACAGTGTAAATCA
AGCACACGTCTGAACTCTTTCCCTACACGACGCTCTTCCGATCTNNNATGTGGTTAAGTATATTCTGT
GAGCACACGTCTGAACTCTTTCCCTACACGACGCTCTTCCGATCTNNNAAAGAAAACACTTCTGGAAA
GAGCACACGTCTGAACTCTTTCCCTACACGACGCTCTTCCGATCTNNNAATCGTACACTCAAATTCATA
GAGCACACGTCTGAACTCTTTCCCTACACGACGCTCTTCCGATCTNNNAATCGTACACTCAAATTCATA
AGCACACGTCTGAACTCTTTCCCTACACGACGCTCTTCCGATCTNNNAGATAATCCCTAAACTTTACCTG
GCACACGTCTGAACTCTTTCCCTACACGACGCTCTTCCGATCTNNNCAGAGATATGAGACTTCTAAGTT
GCACACGTCTGAACTCTTTCCCTACACGACGCTCTTCCGATCTNNNGAAATTGTTTGGGAAATTTGAA
GCACACGTCTGAACTCTTTCCCTACACGACGCTCTTCCGATCTNNNGAGGGAGGAAAAGAAATGATC
GCACACGTCTGAACTCTTTCCCTACACGACGCTCTTCCGATCTNNNGAGGGAGGAAAAGAAATGATC
AGCACACGTCTGAACTCTTTCCCTACACGACGCTCTTCCGATCTNNNACAAAATCTGTTTTAGTTAGCA
AGCACACGTCTGAACTCTTTCCCTACACGACGCTCTTCCGATCTNNNTTCTCTCTCTCTTTTAGCTTAC
CACGTCTGAACTCTTTCCCTACACGACGCTCTTCCGATCTNNNAATTCAGTGACTGACATGAAAATCAT

CACGTCTGAACTCTTTCCCTACACGACGCTCTTCCGATCTNNNAATTCAGTGACTGACATGAAAATCAT
GAGCACACGTCTGAACTCTTTCCCTACACGACGCTCTTCCGATCTNNNAAGAGTTTAAGAAAAAAATTG
GCACACGTCTGAACTCTTTCCCTACACGACGCTCTTCCGATCTNNNCCCTCAATTTAATGTGAATAACTCA
GAGCACACGTCTGAACTCTTTCCCTACACGACGCTCTTCCGATCTNNNAAATAATGTACATAGTACCCA
GAGCACACGTCTGAACTCTTTCCCTACACGACGCTCTTCCGATCTNNNTACAACAAATCTGCTAGGATA
AGCACACGTCTGAACTCTTTCCCTACACGACGCTCTTCCGATCTNNNCTTGCATATAGTCTGTATTCCAT
AGCACACGTCTGAACTCTTTCCCTACACGACGCTCTTCCGATCTNNNTATTTGTTATGTATTTGCTGAC
CACACGTCTGAACTCTTTCCCTACACGACGCTCTTCCGATCTNNNCCTAAGACCTAGAAACTTCCTATCA
AGCACACGTCTGAACTCTTTCCCTACACGACGCTCTTCCGATCTNNNATACTCCATATGTCTACTGGGCA
AGCACACGTCTGAACTCTTTCCCTACACGACGCTCTTCCGATCTNNNATACTCCATATGTCTACTGGGCA
GAGCACACGTCTGAACTCTTTCCCTACACGACGCTCTTCCGATCTNNNTTTGAGAGATAACACAAACAA
GAGCACACGTCTGAACTCTTTCCCTACACGACGCTCTTCCGATCTNNNTAAAGAAACAGGAACCATTTTA
CACACGTCTGAACTCTTTCCCTACACGACGCTCTTCCGATCTNNNTATTTGAAGTTATCTTTTGCTCCCA
GCACACGTCTGAACTCTTTCCCTACACGACGCTCTTCCGATCTNNNGCAGATGCATGAATGGATAGCAT
AGAGCACACGTCTGAACTCTTTCCCTACACGACGCTCTTCCGATCTNNNAGAGTTCAGATAACCTAAAC
GAGCACACGTCTGAACTCTTTCCCTACACGACGCTCTTCCGATCTNNNCTACAAAGTAGCTTGTCCTAA
GAGCACACGTCTGAACTCTTTCCCTACACGACGCTCTTCCGATCTNNNAAACATACATATGTAAGTCCTC
GCACACGTCTGAACTCTTTCCCTACACGACGCTCTTCCGATCTNNNATCAGTCAGTGATTTGTTGATCAT
GAGCACACGTCTGAACTCTTTCCCTACACGACGCTCTTCCGATCTNNNTAAGAGGGAACATGGAAAAAT
GCACACGTCTGAACTCTTTCCCTACACGACGCTCTTCCGATCTNNNTAGGTACAGGATATTTTCACTCCA
GAGCACACGTCTGAACTCTTTCCCTACACGACGCTCTTCCGATCTNNNCTATAAGTCTCTGCCATACTTTT
AGCACACGTCTGAACTCTTTCCCTACACGACGCTCTTCCGATCTNNNAATAATAGTAGGGACTTCGATA
GAGCACACGTCTGAACTCTTTCCCTACACGACGCTCTTCCGATCTNNNATATCATTGCTGTCACTTATTTC
ACACGTCTGAACTCTTTCCCTACACGACGCTCTTCCGATCTNNNAAAGATTTTGTTCACTCAATCTTCATC
CACACGTCTGAACTCTTTCCCTACACGACGCTCTTCCGATCTNNNCAATAAGGATCAAAGAAATGTGAC
AGCACACGTCTGAACTCTTTCCCTACACGACGCTCTTCCGATCTNNNTATGTTAGAACAAAACTGGGTA
AGCACACGTCTGAACTCTTTCCCTACACGACGCTCTTCCGATCTNNNCTGTCTTATTTGTGCATACCTCAT
AGCACACGTCTGAACTCTTTCCCTACACGACGCTCTTCCGATCTNNNAACATGGGTAGAAGTTATTATC
CACACGTCTGAACTCTTTCCCTACACGACGCTCTTCCGATCTNNNCCTGCTGGATCTGTAATAATATTCA
GAGCACACGTCTGAACTCTTTCCCTACACGACGCTCTTCCGATCTNNNAACTATTTTTAGCATAACAGTG
AGCACACGTCTGAACTCTTTCCCTACACGACGCTCTTCCGATCTNNNAAGATGTTAGGATGAGAACAA
AGCACACGTCTGAACTCTTTCCCTACACGACGCTCTTCCGATCTNNNCTCTAGACATAGCATAGGAATG
AGCACACGTCTGAACTCTTTCCCTACACGACGCTCTTCCGATCTNNNGATGGTGTGTATTTTCTAAGTC
AGAGCACACGTCTGAACTCTTTCCCTACACGACGCTCTTCCGATCTNNNATGAGTATATGTAGGTTTGA
CACACGTCTGAACTCTTTCCCTACACGACGCTCTTCCGATCTNNNGGTTTCTTAGTGATTCTGAAAATC
AGCACACGTCTGAACTCTTTCCCTACACGACGCTCTTCCGATCTNNNCTCTCATGTCACTAATTTCTTTTC
GAGCACACGTCTGAACTCTTTCCCTACACGACGCTCTTCCGATCTNNNACAGAATAATGACCTCTAAGTT
GCACACGTCTGAACTCTTTCCCTACACGACGCTCTTCCGATCTNNNCTGACTTTTCTACTTTGTGTATTC
GCACACGTCTGAACTCTTTCCCTACACGACGCTCTTCCGATCTNNNTTAAATATCTTGATACATGACGCC
AGCACACGTCTGAACTCTTTCCCTACACGACGCTCTTCCGATCTNNNAGAGGCAAGATACATGGATAC
AGCACACGTCTGAACTCTTTCCCTACACGACGCTCTTCCGATCTNNNCAATGATGATCAATTCAGTATC
AGCACACGTCTGAACTCTTTCCCTACACGACGCTCTTCCGATCTNNNTGTCTCGTTTGAAGAAGTTCCAT
AGCACACGTCTGAACTCTTTCCCTACACGACGCTCTTCCGATCTNNNTCTCCAAATCTTTCCTATTAGAA

GCACACGTCTGAACTCTTTCCCTACACGACGCTCTTCCGATCTNNNCCATTCTTTCTCATGTCAGTCAT
GCACACGTCTGAACTCTTTCCCTACACGACGCTCTTCCGATCTNNNCCATTCTTTCTCATGTCAGTCAT
AGCACACGTCTGAACTCTTTCCCTACACGACGCTCTTCCGATCTNNNGATGAACTCCAAAATCTCAAA
AGCACACGTCTGAACTCTTTCCCTACACGACGCTCTTCCGATCTNNNGATGAACTCCAAAATCTCAAA
AGCACACGTCTGAACTCTTTCCCTACACGACGCTCTTCCGATCTNNNAATGCTCTTGATAGAAACAAA
ACACGTCTGAACTCTTTCCCTACACGACGCTCTTCCGATCTNNNAAGTAGAATGCTGATGGGAATCA
ACACGTCTGAACTCTTTCCCTACACGACGCTCTTCCGATCTNNNAAGTAGAATGCTGATGGGAATCA
AGCACACGTCTGAACTCTTTCCCTACACGACGCTCTTCCGATCTNNNTTCAGACCTTTCTACTTTATCAT
GCACACGTCTGAACTCTTTCCCTACACGACGCTCTTCCGATCTNNNCTACTGAAAACCTATACATGTTG
AGCACACGTCTGAACTCTTTCCCTACACGACGCTCTTCCGATCTNNNATCTTAAGTATTTGTTATGGG
ACACGTCTGAACTCTTTCCCTACACGACGCTCTTCCGATCTNNNTGGCAAGTTGAAGAATAAATTATC
AGCACACGTCTGAACTCTTTCCCTACACGACGCTCTTCCGATCTNNNTTCTGCTATAAATGTGTGAAA
CACGTCTGAACTCTTTCCCTACACGACGCTCTTCCGATCTNNNAAGCTGTTTTACCTATCTAGAATCAT
CACGTCTGAACTCTTTCCCTACACGACGCTCTTCCGATCTNNNAAGCTGTTTTACCTATCTAGAATCAT
AGCACACGTCTGAACTCTTTCCCTACACGACGCTCTTCCGATCTNNNCGGTGTTAATTTTGTTGTTTTA
GCACACGTCTGAACTCTTTCCCTACACGACGCTCTTCCGATCTNNNAAATATATTACCAGCCAAGAGA
AGCACACGTCTGAACTCTTTCCCTACACGACGCTCTTCCGATCTNNNAAAAACGTTCTCAGTTCTTTAG
AGCACACGTCTGAACTCTTTCCCTACACGACGCTCTTCCGATCTNNNTTCAAATCTCTACATTTCAAGT
GCACACGTCTGAACTCTTTCCCTACACGACGCTCTTCCGATCTNNNTTTTATTGTTGTCTGGAATCATT
AGCACACGTCTGAACTCTTTCCCTACACGACGCTCTTCCGATCTNNNATTGGATAGAAAATCCACATC
AGCACACGTCTGAACTCTTTCCCTACACGACGCTCTTCCGATCTNNNTGTTTTTAGGAAATAGATGTT
CACACGTCTGAACTCTTTCCCTACACGACGCTCTTCCGATCTNNNACATTAAAATTTTATGCAGAATGT
AGCACACGTCTGAACTCTTTCCCTACACGACGCTCTTCCGATCTNNNCAAGGAAAATGATGCACATGC
CACACGTCTGAACTCTTTCCCTACACGACGCTCTTCCGATCTNNNAAATTCAAAAGACTGGAAATACT
CACACGTCTGAACTCTTTCCCTACACGACGCTCTTCCGATCTNNNGTGATTAAAGATCTTGTGATGG
AGCACACGTCTGAACTCTTTCCCTACACGACGCTCTTCCGATCTNNNCAGATGGAATTTCTCAGGTGC
CACACGTCTGAACTCTTTCCCTACACGACGCTCTTCCGATCTNNNAACACATACACACACACAGACATC
CACACGTCTGAACTCTTTCCCTACACGACGCTCTTCCGATCTNNNATATAGTGTATTGGTCAGGATT
GCACACGTCTGAACTCTTTCCCTACACGACGCTCTTCCGATCTNNNCTCTGTGAAAGAAGGAAAGAA
GCACACGTCTGAACTCTTTCCCTACACGACGCTCTTCCGATCTNNNATATTTTACACACACACATACA
GCACACGTCTGAACTCTTTCCCTACACGACGCTCTTCCGATCTNNNATGAGCTCTTGTGAATAACAGC
AGCACACGTCTGAACTCTTTCCCTACACGACGCTCTTCCGATCTNNNTTTTACTCTTATGTTGTCAAGT
AGCACACGTCTGAACTCTTTCCCTACACGACGCTCTTCCGATCTNNNTTCTAGAATGTCTTAAAGGCAT
GCACACGTCTGAACTCTTTCCCTACACGACGCTCTTCCGATCTNNNCTGAGAGATTATAGCTATTGTC
AGCACACGTCTGAACTCTTTCCCTACACGACGCTCTTCCGATCTNNNTCTTTCCATTTCAAATTCTAGA
ACACGTCTGAACTCTTTCCCTACACGACGCTCTTCCGATCTNNNAGAATCTGGTATATCTTCCTCTCATC
GCACACGTCTGAACTCTTTCCCTACACGACGCTCTTCCGATCTNNNACAAAACAAATACCTATTCAGA
CACACGTCTGAACTCTTTCCCTACACGACGCTCTTCCGATCTNNNAAAGAGAGCCTTAGGTGTAAAC
AGCACACGTCTGAACTCTTTCCCTACACGACGCTCTTCCGATCTNNNTTCTTATACTGGTGCTTTTTTC
GCACACGTCTGAACTCTTTCCCTACACGACGCTCTTCCGATCTNNNTTACCTCTCCAGAGTACAGATAC
AGCACACGTCTGAACTCTTTCCCTACACGACGCTCTTCCGATCTNNNTCTGCTGCCTATGAATTTTTAC
CACACGTCTGAACTCTTTCCCTACACGACGCTCTTCCGATCTNNNTAATCTTGGTGGAAAGTGAAAAT
GCACACGTCTGAACTCTTTCCCTACACGACGCTCTTCCGATCTNNNGAATTAATCTGTCTACTATCAAT

GCACACGTCTGAACTCTTTCCCTACACGACGCTCTTCCGATCTNNNTACAATTCATTTATTCTCTGGGAC

GAGCACACGTCTGAACTCTTTCCCTACACGACGCTCTTCCGATCTNNNGGGAAATTTTTAGTCATGATTC

GAGCACACGTCTGAACTCTTTCCCTACACGACGCTCTTCCGATCTNNNGGGAAATTTTTAGTCATGATTC

AGCACACGTCTGAACTCTTTCCCTACACGACGCTCTTCCGATCTNNNGACAGGATTTTGTTCTGTTTTAT

AGCACACGTCTGAACTCTTTCCCTACACGACGCTCTTCCGATCTNNNTGAAGTCCTGATAGCAAATATTA

CACACGTCTGAACTCTTTCCCTACACGACGCTCTTCCGATCTNNNTTTAAATATTGTCTACAGAAGCCCAT

GCACACGTCTGAACTCTTTCCCTACACGACGCTCTTCCGATCTNNNCTGCCACTTACTAACTTGTATCAT

GAGCACACGTCTGAACTCTTTCCCTACACGACGCTCTTCCGATCTNNNAAAAAGTAAAACTCCTGGATTT

AGAGCACACGTCTGAACTCTTTCCCTACACGACGCTCTTCCGATCTNNNAACTACCAGTACTACCAGAA

GAGCACACGTCTGAACTCTTTCCCTACACGACGCTCTTCCGATCTNNNGTAAAGTATGAAGTCGTTTTTC

AGCACACGTCTGAACTCTTTCCCTACACGACGCTCTTCCGATCTNNNTGTAAGTGACTACAGAACTTTAT

GAGCACACGTCTGAACTCTTTCCCTACACGACGCTCTTCCGATCTNNNGAATAATGATCAGTGGTAATC

ACACGTCTGAACTCTTTCCCTACACGACGCTCTTCCGATCTNNNTTGTGGTCTACAAAAACAAAGCATCC

GCACACGTCTGAACTCTTTCCCTACACGACGCTCTTCCGATCTNNNGAATCTTATTCTAAGGCATATGCC

GAGCACACGTCTGAACTCTTTCCCTACACGACGCTCTTCCGATCTNNNCAGTATATATTTACTGGCTTGC

ACACGTCTGAACTCTTTCCCTACACGACGCTCTTCCGATCTNNNCTTATAAAAAGAGAAAGGGACACCA

AGCACACGTCTGAACTCTTTCCCTACACGACGCTCTTCCGATCTNNNGTAAGAAGAACTCTACTCTACAC

AGAGCACACGTCTGAACTCTTTCCCTACACGACGCTCTTCCGATCTNNNTCAAATAATCATATCAACAAG

AAGAGCACACGTCTGAACTCTTTCCCTACACGACGCTCTTCCGATCTNNNCTTGTCCAGATATATATGT

AGCACACGTCTGAACTCTTTCCCTACACGACGCTCTTCCGATCTNNNTCACTATCTTCTGAGTATCTGTAC

AGCACACGTCTGAACTCTTTCCCTACACGACGCTCTTCCGATCTNNNTTTACTGAGAACTTACTGTGTTTC

CACGTCTGAACTCTTTCCCTACACGACGCTCTTCCGATCTNNNCTTACTTGTCCTAGGTAGACTCATCGG

GAGCACACGTCTGAACTCTTTCCCTACACGACGCTCTTCCGATCTNNNTGAACTATAAATGTATGCATGT

GCACACGTCTGAACTCTTTCCCTACACGACGCTCTTCCGATCTNNNAAAAAAAAAAAGCCTTAGAAAACC

AGCACACGTCTGAACTCTTTCCCTACACGACGCTCTTCCGATCTNNNATAGGCACTTTGTTACAATTTGC

GAGCACACGTCTGAACTCTTTCCCTACACGACGCTCTTCCGATCTNNNCAATACAAGAGAGCATACAAA

GCACACGTCTGAACTCTTTCCCTACACGACGCTCTTCCGATCTNNNCGTAAGTGTTAACTGGTACTTCAT

GAGCACACGTCTGAACTCTTTCCCTACACGACGCTCTTCCGATCTNNNCTTATTTATCCTTGTATCCCTCA

CACACGTCTGAACTCTTTCCCTACACGACGCTCTTCCGATCTNNNATACTTATTGAATGAAAACTTGCCA

AGCACACGTCTGAACTCTTTCCCTACACGACGCTCTTCCGATCTNNNAGGGTAGCTACATATAGGAAA

GAGCACACGTCTGAACTCTTTCCCTACACGACGCTCTTCCGATCTNNNGAGGAGAAAGTATTCCAGAT

GAGCACACGTCTGAACTCTTTCCCTACACGACGCTCTTCCGATCTNNNCTGGAATGTGTGTGTGTATAT

GAGCACACGTCTGAACTCTTTCCCTACACGACGCTCTTCCGATCTNNNTTGAACATCTTTTAATGTGCTT

GCACACGTCTGAACTCTTTCCCTACACGACGCTCTTCCGATCTNNNCTGTGAATGTATTACAAATAGCAC

GCACACGTCTGAACTCTTTCCCTACACGACGCTCTTCCGATCTNNNTTAACTTTGTCGCTATGAGAATAC

CACACGTCTGAACTCTTTCCCTACACGACGCTCTTCCGATCTNNNTCAGTTTCTTCATCAATAAAGTGCA

AGCACACGTCTGAACTCTTTCCCTACACGACGCTCTTCCGATCTNNNAGTGAGTACTCTTTGTATATGTC

GCACACGTCTGAACTCTTTCCCTACACGACGCTCTTCCGATCTNNNATTCTTTCAATGATTTCTGGTGCA

AGCACACGTCTGAACTCTTTCCCTACACGACGCTCTTCCGATCTNNNGTTTTGGGAATAGAGGAAAAA

ACACGTCTGAACTCTTTCCCTACACGACGCTCTTCCGATCTNNNGAAAGAATGTCAGAACAGAAGCAT

AGCACACGTCTGAACTCTTTCCCTACACGACGCTCTTCCGATCTNNNGATTTGAATCATGGAAGCTATA

GAGCACACGTCTGAACTCTTTCCCTACACGACGCTCTTCCGATCTNNNTTAAAAAATACTTTGAAGCCTC

GAGCACACGTCTGAACTCTTTCCCTACACGACGCTCTTCCGATCTNNNATGAGACCCTAACAATGTTAT

GCACACGTCTGAACTCTTTCCCTACACGACGCTCTTCCGATCTNNNAATTTATGATATTTCTGTGTCGCC
AGCACACGTCTGAACTCTTTCCCTACACGACGCTCTTCCGATCTNNNTAAAACAAAACAAAACACTCTCT
AGCACACGTCTGAACTCTTTCCCTACACGACGCTCTTCCGATCTNNNTAAAACAAAACAAAACACTCTCT
AGCACACGTCTGAACTCTTTCCCTACACGACGCTCTTCCGATCTNNNTTCACATATAACAGTCAGCATTA
ACACGTCTGAACTCTTTCCCTACACGACGCTCTTCCGATCTNNNTAACTGTTAGTATTCCTGGATGACAT
ACACGTCTGAACTCTTTCCCTACACGACGCTCTTCCGATCTNNNGATGGATGAATGGATGGATGCATC
AGCACACGTCTGAACTCTTTCCCTACACGACGCTCTTCCGATCTNNNTTTGGTGAAGGTAGCTAATGCA
AGCACACGTCTGAACTCTTTCCCTACACGACGCTCTTCCGATCTNNNCAATTTTCCTACTAATGCTTCTC
GAGCACACGTCTGAACTCTTTCCCTACACGACGCTCTTCCGATCTNNNATTTACACAGGAAATGGGGTA
GAGCACACGTCTGAACTCTTTCCCTACACGACGCTCTTCCGATCTNNNCTTTCTTTATAGTTTCTCCAGTG
GCACACGTCTGAACTCTTTCCCTACACGACGCTCTTCCGATCTNNNGACAACAGTACAAAAAATAAATCA
GAGCACACGTCTGAACTCTTTCCCTACACGACGCTCTTCCGATCTNNNTTTGATCCCTTTCTTACTCTTTCA
AGCACACGTCTGAACTCTTTCCCTACACGACGCTCTTCCGATCTNNNTTGAATATCTTCTCAAGTTTAG
GAGCACACGTCTGAACTCTTTCCCTACACGACGCTCTTCCGATCTNNNAAAAAAAAAAGGAGTTTTGT
AGCACACGTCTGAACTCTTTCCCTACACGACGCTCTTCCGATCTNNNACACACATATACACACACAGCAT
CACACGTCTGAACTCTTTCCCTACACGACGCTCTTCCGATCTNNNTAAAGCTCATTTCCTATTATCCACAT
AGCACACGTCTGAACTCTTTCCCTACACGACGCTCTTCCGATCTNNNATGAACTGTCTTCTTTAGTAACA
AGCACACGTCTGAACTCTTTCCCTACACGACGCTCTTCCGATCTNNNATGAACTGTCTTCTTTAGTAACA
GAGCACACGTCTGAACTCTTTCCCTACACGACGCTCTTCCGATCTNNNCATCTCTCCGTCTCTCTTTCCAT
GAGCACACGTCTGAACTCTTTCCCTACACGACGCTCTTCCGATCTNNNCATCTCTCCGTCTCTCTTTCCAT
CACACGTCTGAACTCTTTCCCTACACGACGCTCTTCCGATCTNNNGTGGCTTTCTTACACCATTTCATCG
AGCACACGTCTGAACTCTTTCCCTACACGACGCTCTTCCGATCTNNNGGATGATTTCTAAATGAAACCTC
AGCACACGTCTGAACTCTTTCCCTACACGACGCTCTTCCGATCTNNNCCTCTGAAAAACATACACAAATA
AGCACACGTCTGAACTCTTTCCCTACACGACGCTCTTCCGATCTNNNCCTCTGAAAAACATACACAAATA
ACACGTCTGAACTCTTTCCCTACACGACGCTCTTCCGATCTNNNCCTTTTTGGTTATTATTGCATTCCAT
GAGCACACGTCTGAACTCTTTCCCTACACGACGCTCTTCCGATCTNNNTATGTGATATGGTTTGCTTCAC
GCACACGTCTGAACTCTTTCCCTACACGACGCTCTTCCGATCTNNNCTACACAGCAAGGAAAAGAACAT
AGCACACGTCTGAACTCTTTCCCTACACGACGCTCTTCCGATCTNNNATTTGAGCATGACCTGAACTCA
AGCACACGTCTGAACTCTTTCCCTACACGACGCTCTTCCGATCTNNNCTTTCTTTTCTGTTCCAATTACT
GCACACGTCTGAACTCTTTCCCTACACGACGCTCTTCCGATCTNNNGCTCTGTCATATTTCTAAAAGTTC
AGCACACGTCTGAACTCTTTCCCTACACGACGCTCTTCCGATCTNNNGTTAGTTCTATGTGTCAACATGC
GCACACGTCTGAACTCTTTCCCTACACGACGCTCTTCCGATCTNNNGGAATATTTAAAGTTCCTTCAGG
AGCACACGTCTGAACTCTTTCCCTACACGACGCTCTTCCGATCTNNNACATTCAATATTCAATCTGAACA
AGCACACGTCTGAACTCTTTCCCTACACGACGCTCTTCCGATCTNNNTTAATTCCACTGTAATGTACTTTC
GCACACGTCTGAACTCTTTCCCTACACGACGCTCTTCCGATCTNNNCTCTTTCCAAGATACCTGGGCATC
GAGCACACGTCTGAACTCTTTCCCTACACGACGCTCTTCCGATCTNNNTCCACAACAAAAAACAAAGTTT
AGCACACGTCTGAACTCTTTCCCTACACGACGCTCTTCCGATCTNNNCTTTCCAATCTTTCATAGATGAG
AGCACACGTCTGAACTCTTTCCCTACACGACGCTCTTCCGATCTNNNAACAAGTTTGATCACTGTATAAC
GAGCACACGTCTGAACTCTTTCCCTACACGACGCTCTTCCGATCTNNNTGATATATGTGCTATGGAGAA
AGCACACGTCTGAACTCTTTCCCTACACGACGCTCTTCCGATCTNNNAAAACCAAGAGGATGATAATCT
GAGCACACGTCTGAACTCTTTCCCTACACGACGCTCTTCCGATCTNNNGGGGAGAAAGATGTCATCTC
GAGCACACGTCTGAACTCTTTCCCTACACGACGCTCTTCCGATCTNNNGGGGAGAAAGATGTCATCTC
GAGCACACGTCTGAACTCTTTCCCTACACGACGCTCTTCCGATCTNNNGGGGAGAAAGATGTCATCTC

CACACGTCTGAACTCTTTCCCTACACGACGCTCTTCCGATCTNNNTTTGCGTGAATATGGTATTACCAT
GCACACGTCTGAACTCTTTCCCTACACGACGCTCTTCCGATCTNNNACGTAATATCATATGGTCTATGA
GAGCACACGTCTGAACTCTTTCCCTACACGACGCTCTTCCGATCTNNNCATGTAGGATAGTGAAAAAT
AGCACACGTCTGAACTCTTTCCCTACACGACGCTCTTCCGATCTNNNCATGATGTAGAATAGGGTACA
GCACACGTCTGAACTCTTTCCCTACACGACGCTCTTCCGATCTNNNTTTTGTTTCACTGTTGTAAATTG
AGCACACGTCTGAACTCTTTCCCTACACGACGCTCTTCCGATCTNNNGACTTTCCCATGAAGTTCATCA
AGCACACGTCTGAACTCTTTCCCTACACGACGCTCTTCCGATCTNNNGACTTTCCCATGAAGTTCATCA
AGCACACGTCTGAACTCTTTCCCTACACGACGCTCTTCCGATCTNNNCTTTATTTGGTTGCTCAAATTG
ACACGTCTGAACTCTTTCCCTACACGACGCTCTTCCGATCTNNNCTCAGAGTTCTTCACAGATATCCAT
AGCACACGTCTGAACTCTTTCCCTACACGACGCTCTTCCGATCTNNNCACACACATATATTAACACTGA
GAGCACACGTCTGAACTCTTTCCCTACACGACGCTCTTCCGATCTNNNAGCCTCTTGGATTACTTAATT
AGCACACGTCTGAACTCTTTCCCTACACGACGCTCTTCCGATCTNNNGAAAATCTGAATAAACATGCA
GCACACGTCTGAACTCTTTCCCTACACGACGCTCTTCCGATCTNNNCTTGAAGGGACTGATCATAAAT
AGCACACGTCTGAACTCTTTCCCTACACGACGCTCTTCCGATCTNNNCAACCAATGAAGAGATAAAGA
AGCACACGTCTGAACTCTTTCCCTACACGACGCTCTTCCGATCTNNNATTTGAATCTGTGTCAAGCTCA
GAGCACACGTCTGAACTCTTTCCCTACACGACGCTCTTCCGATCTNNNTCTACATATTAGCTTATTAGG
AGCACACGTCTGAACTCTTTCCCTACACGACGCTCTTCCGATCTNNNGAAAATGATAATATCTGCTTCC
GAGCACACGTCTGAACTCTTTCCCTACACGACGCTCTTCCGATCTNNNAGTATCTGTAAGCGACTGAT
AGCACACGTCTGAACTCTTTCCCTACACGACGCTCTTCCGATCTNNNTTCTTTCTACACTGTAATTCCTT
AGAGCACACGTCTGAACTCTTTCCCTACACGACGCTCTTCCGATCTNNNGTTTCCAGTACAGGAGTAT
AGCACACGTCTGAACTCTTTCCCTACACGACGCTCTTCCGATCTNNNGTCTCAAAAAAAGAAAAAGT
CACACGTCTGAACTCTTTCCCTACACGACGCTCTTCCGATCTNNNGTGTGGAGGTGATATAGAGACAT
GAGCACACGTCTGAACTCTTTCCCTACACGACGCTCTTCCGATCTNNNGAAATCTGCAACCTACTTTTG
GCACACGTCTGAACTCTTTCCCTACACGACGCTCTTCCGATCTNNNTACAGAACTACAATATTGCAGTC
CACACGTCTGAACTCTTTCCCTACACGACGCTCTTCCGATCTNNNAAAAAAAAAATGCAGTCATTCTTC
ACACGTCTGAACTCTTTCCCTACACGACGCTCTTCCGATCTNNNAAACCTCTAACACCATGATCCATCGG
GCACACGTCTGAACTCTTTCCCTACACGACGCTCTTCCGATCTNNNAAGTGTTTACATTCATCAAACAG
GCACACGTCTGAACTCTTTCCCTACACGACGCTCTTCCGATCTNNNTTTAGAAATTTTACAGGCCAAGC
GCACACGTCTGAACTCTTTCCCTACACGACGCTCTTCCGATCTNNNAAGAATGAGGAGGATTTTATCA
GCACACGTCTGAACTCTTTCCCTACACGACGCTCTTCCGATCTNNNAAGAATGAGGAGGATTTTATCA
AGCACACGTCTGAACTCTTTCCCTACACGACGCTCTTCCGATCTNNNACATTGTAAAAAAGTTCTGAA
GCACACGTCTGAACTCTTTCCCTACACGACGCTCTTCCGATCTNNNATATTCACTTAGCTGAGCAAATC
AGCACACGTCTGAACTCTTTCCCTACACGACGCTCTTCCGATCTNNNTTAACCTTTACCTCTCTTCTGAC
CACACGTCTGAACTCTTTCCCTACACGACGCTCTTCCGATCTNNNGGTTTAAAATTTTGGAACCTTTGC
GCACACGTCTGAACTCTTTCCCTACACGACGCTCTTCCGATCTNNNAATTAAATGTAAGCTCTCTGATG
GCACACGTCTGAACTCTTTCCCTACACGACGCTCTTCCGATCTNNNCTGGTAGTTTTTCTTTCTCTTCCAT
GCACACGTCTGAACTCTTTCCCTACACGACGCTCTTCCGATCTNNNCTGGTAGTTTTTCTTTCTCTTCCAT
AGCACACGTCTGAACTCTTTCCCTACACGACGCTCTTCCGATCTNNNGATCCCAAAAAATCAAGATCA
AGCACACGTCTGAACTCTTTCCCTACACGACGCTCTTCCGATCTNNNCTCAGGTTGTATTCTTCCAGCAT
AGCACACGTCTGAACTCTTTCCCTACACGACGCTCTTCCGATCTNNNTTTTAATTTGGCTAAACTGTTCA
GAAGAGCACACGTCTGAACTCTTTCCCTACACGACGCTCTTCCGATCTNNNACTTTAGCCTCAAACTCCT
AGCACACGTCTGAACTCTTTCCCTACACGACGCTCTTCCGATCTNNNGTATTTCTTGTGCATCCTCTATC
GCACACGTCTGAACTCTTTCCCTACACGACGCTCTTCCGATCTNNNTTCAGATCTTGAGATTTAGCTTA

GAGCACACGTCTGAACTCTTTCCCTACACGACGCTCTTCCGATCTNNNTTGTGAAACTTTGTTTTACTT
ACACGTCTGAACTCTTTCCCTACACGACGCTCTTCCGATCTNNNCCCTCTAATTTACTTCCATCATTCAT
CACACGTCTGAACTCTTTCCCTACACGACGCTCTTCCGATCTNNNGAGAAAGGGAAGAATAAGAAAG
GCACACGTCTGAACTCTTTCCCTACACGACGCTCTTCCGATCTNNNTATTCAAGATGGAGTCACTCTC
AGCACACGTCTGAACTCTTTCCCTACACGACGCTCTTCCGATCTNNNGAAATCTCTAGACTGGATTCA
GCACACGTCTGAACTCTTTCCCTACACGACGCTCTTCCGATCTNNNAATGAGTAGTCTGGGTGTATCA
CACACGTCTGAACTCTTTCCCTACACGACGCTCTTCCGATCTNNNTGTAGGGAAAGTAGAATCTCAC
GCACACGTCTGAACTCTTTCCCTACACGACGCTCTTCCGATCTNNNCTCACTTGGTGATTTAGATGTC
GCACACGTCTGAACTCTTTCCCTACACGACGCTCTTCCGATCTNNNCTACTGGAGTGATTCTTTTCACA
AGCACACGTCTGAACTCTTTCCCTACACGACGCTCTTCCGATCTNNNTCAGAGTAACAGATTTGAAAT
AGCACACGTCTGAACTCTTTCCCTACACGACGCTCTTCCGATCTNNNCAATAATGGCAGGGAAAAAG
AGCACACGTCTGAACTCTTTCCCTACACGACGCTCTTCCGATCTNNNATCTCACATTCCATGTAGTATA
CACACGTCTGAACTCTTTCCCTACACGACGCTCTTCCGATCTNNNGAATTTTTTTCCTAAGTGTCTGTC
GCACACGTCTGAACTCTTTCCCTACACGACGCTCTTCCGATCTNNNTTGAACGTTATCCTCTTTGGCAT
CACGTCTGAACTCTTTCCCTACACGACGCTCTTCCGATCTNNNCATTTATTGAGGATCTACTTGTGCA
GCACACGTCTGAACTCTTTCCCTACACGACGCTCTTCCGATCTNNNTAGATGGATAGATAGACAGATG
AGCACACGTCTGAACTCTTTCCCTACACGACGCTCTTCCGATCTNNNTGTGATAAAATTGCATAGAGC
CACACGTCTGAACTCTTTCCCTACACGACGCTCTTCCGATCTNNNAAGCATTAATTAAACAAACAGAT
GCACACGTCTGAACTCTTTCCCTACACGACGCTCTTCCGATCTNNNTTCATTCTTTCTTTCTTCTTTCACA
AGCACACGTCTGAACTCTTTCCCTACACGACGCTCTTCCGATCTNNNTCTCAAATTTTTAGGCTTTTCTT
GCACACGTCTGAACTCTTTCCCTACACGACGCTCTTCCGATCTNNNCAAATGACAAGATTTCATTCTCT
AGCACACGTCTGAACTCTTTCCCTACACGACGCTCTTCCGATCTNNNCTGGGTGTTGAAGAAGATCAC
CACACGTCTGAACTCTTTCCCTACACGACGCTCTTCCGATCTNNNAATATTCCTTGCACATTTGAGTAC
CACACGTCTGAACTCTTTCCCTACACGACGCTCTTCCGATCTNNNTGCTTGTTTGTTTTTCTTCCCATC
CACACGTCTGAACTCTTTCCCTACACGACGCTCTTCCGATCTNNNCTTTGTACAGAACTTCTAGTGTCA
AGCACACGTCTGAACTCTTTCCCTACACGACGCTCTTCCGATCTNNNGTACTAGTTGTGGTGCTTAGC
CACACGTCTGAACTCTTTCCCTACACGACGCTCTTCCGATCTNNNCTAGTCATAGTGAACACTCTCCAT
GAGCACACGTCTGAACTCTTTCCCTACACGACGCTCTTCCGATCTNNNAGACACAACACTCTAAAGATA
GCACACGTCTGAACTCTTTCCCTACACGACGCTCTTCCGATCTNNNATATCCAGGGCCATTTCTATCAT
AGCACACGTCTGAACTCTTTCCCTACACGACGCTCTTCCGATCTNNNATGTCTAGAAAATAAGAGTTG
GCACACGTCTGAACTCTTTCCCTACACGACGCTCTTCCGATCTNNNTATAGGTTATCAAAGAAATGTG
GAGCACACGTCTGAACTCTTTCCCTACACGACGCTCTTCCGATCTNNNTTTTTGAGTGAGTGTGTGTA
GAGCACACGTCTGAACTCTTTCCCTACACGACGCTCTTCCGATCTNNNCCAAGAGGTAGCATTATGTA
GCACACGTCTGAACTCTTTCCCTACACGACGCTCTTCCGATCTNNNGTATGTTCTCATTTAAAGAGGT
GCACACGTCTGAACTCTTTCCCTACACGACGCTCTTCCGATCTNNNGTATTTCTTGTGCATCCTCTATC
GCACACGTCTGAACTCTTTCCCTACACGACGCTCTTCCGATCTNNNATCTTTAAAATGACAACTTCTCC
GCACACGTCTGAACTCTTTCCCTACACGACGCTCTTCCGATCTNNNTAAAATTGTTCTCTTTCCCAATTC
CACACGTCTGAACTCTTTCCCTACACGACGCTCTTCCGATCTNNNGACTTCAAAGATATTTTTGAGCT
CACACGTCTGAACTCTTTCCCTACACGACGCTCTTCCGATCTNNNGTTTTCTTCATAGCACTTATCATC
CACACGTCTGAACTCTTTCCCTACACGACGCTCTTCCGATCTNNNGTTTTCTTCATAGCACTTATCATC
AGCACACGTCTGAACTCTTTCCCTACACGACGCTCTTCCGATCTNNNAGGATTTGCATTACAACTTATT
CACACGTCTGAACTCTTTCCCTACACGACGCTCTTCCGATCTNNNAAAGATTTCAGAGAATGTGACTA
ACACGTCTGAACTCTTTCCCTACACGACGCTCTTCCGATCTNNNTTTATATTTGGAAATCTGTGGGC

CACACGTCTGAACTCTTTCCCTACACGACGCTCTTCCGATCTNNNGTAACCGGTCAAATACAGTAGCAT
AGCACACGTCTGAACTCTTTCCCTACACGACGCTCTTCCGATCTNNNTCTGATGTTTTTGTTCTCCTTCA
AGCACACGTCTGAACTCTTTCCCTACACGACGCTCTTCCGATCTNNNTAATCACTGTAAGCATTTTAGT
AGCACACGTCTGAACTCTTTCCCTACACGACGCTCTTCCGATCTNNNTCATTACATTTTTCATGCTTAA
AGCACACGTCTGAACTCTTTCCCTACACGACGCTCTTCCGATCTNNNTAATACTGTACATCACATGCAT
GCACACGTCTGAACTCTTTCCCTACACGACGCTCTTCCGATCTNNNTATTATGGTGTTTTGTGATGTTT
AGCACACGTCTGAACTCTTTCCCTACACGACGCTCTTCCGATCTNNNTTTTTGCACCTATTATCATTCAA
GAGCACACGTCTGAACTCTTTCCCTACACGACGCTCTTCCGATCTNNNTATGTTCCTCTGAAATACTTA
AGCACACGTCTGAACTCTTTCCCTACACGACGCTCTTCCGATCTNNNCATCTCCTCTCTGATTAAAACT
AGCACACGTCTGAACTCTTTCCCTACACGACGCTCTTCCGATCTNNNCATCTCCTCTCTGATTAAAACT
AGCACACGTCTGAACTCTTTCCCTACACGACGCTCTTCCGATCTNNNCTGGTCATCTATTATTGCATAA
AGCACACGTCTGAACTCTTTCCCTACACGACGCTCTTCCGATCTNNNTTTTATTTTAAGCTATGGAGGT
AGCACACGTCTGAACTCTTTCCCTACACGACGCTCTTCCGATCTNNNATGAGAAATAAGAATCTCTGG
AGCACACGTCTGAACTCTTTCCCTACACGACGCTCTTCCGATCTNNNGTTCTTCCATCATTCATGTAAC
AGCACACGTCTGAACTCTTTCCCTACACGACGCTCTTCCGATCTNNNTTTGAGAAACTAAAGAAAAT
AGCACACGTCTGAACTCTTTCCCTACACGACGCTCTTCCGATCTNNNGACAGACAAAAAGATGAGGTA
GAGCACACGTCTGAACTCTTTCCCTACACGACGCTCTTCCGATCTNNNGCTGTTCTTTGGTTTTCTATAT
AGCACACGTCTGAACTCTTTCCCTACACGACGCTCTTCCGATCTNNNAAGAAGAGTAACATTTGTGTT
AGCACACGTCTGAACTCTTTCCCTACACGACGCTCTTCCGATCTNNNCTATCAGTGGGTTTTTATAAG
AGCACACGTCTGAACTCTTTCCCTACACGACGCTCTTCCGATCTNNNGTAAATAAAAGAAAAGGCATG
AGCACACGTCTGAACTCTTTCCCTACACGACGCTCTTCCGATCTNNNTATATGAGGGAGTATTCACTG
AGCACACGTCTGAACTCTTTCCCTACACGACGCTCTTCCGATCTNNNGGAAGGATTGAATTACTTAGT
GCACACGTCTGAACTCTTTCCCTACACGACGCTCTTCCGATCTNNNAGAAGAGAGGGAAAACTTATTG
GCACACGTCTGAACTCTTTCCCTACACGACGCTCTTCCGATCTNNNCGAATGAATTATTTAAGTTCTG
CACACGTCTGAACTCTTTCCCTACACGACGCTCTTCCGATCTNNNTAATCATTGTCCCTCCTAGATTCAT
AGCACACGTCTGAACTCTTTCCCTACACGACGCTCTTCCGATCTNNNACTAAAAAAAAATCCCTTTCGG
CACACGTCTGAACTCTTTCCCTACACGACGCTCTTCCGATCTNNNAAAAAAACAGTATCAATGAGTGA
AGCACACGTCTGAACTCTTTCCCTACACGACGCTCTTCCGATCTNNNGAGTGTCAATTCTTGAGTTGTC
GAGCACACGTCTGAACTCTTTCCCTACACGACGCTCTTCCGATCTNNNAATTTTCCAAGCAGTATTCATA
ACACGTCTGAACTCTTTCCCTACACGACGCTCTTCCGATCTNNNTTACACATATGTATGACAGGATGC
ACACGTCTGAACTCTTTCCCTACACGACGCTCTTCCGATCTNNNTTACACATATGTATGACAGGATGC
AGCACACGTCTGAACTCTTTCCCTACACGACGCTCTTCCGATCTNNNACTACATCAGTACACAAAAGT
AGCACACGTCTGAACTCTTTCCCTACACGACGCTCTTCCGATCTNNNAAAATTCAGTTGCATGCATTTA
GCACACGTCTGAACTCTTTCCCTACACGACGCTCTTCCGATCTNNNTTGTGTAAAACTGGGTAGTTGC
CACACGTCTGAACTCTTTCCCTACACGACGCTCTTCCGATCTNNNTAATTTTGGAAGGCATCAAATAC
AGCACACGTCTGAACTCTTTCCCTACACGACGCTCTTCCGATCTNNNAAATTAACAGGTTAATGTCAGT
AGCACACGTCTGAACTCTTTCCCTACACGACGCTCTTCCGATCTNNNCTAAGAAGTCATTCCTAGAAGA
AGCACACGTCTGAACTCTTTCCCTACACGACGCTCTTCCGATCTNNNCCTCACTACATATTCACTGAGC
GCACACGTCTGAACTCTTTCCCTACACGACGCTCTTCCGATCTNNNAAACTCCTAATCTCAAGTGATCC
GAGCACACGTCTGAACTCTTTCCCTACACGACGCTCTTCCGATCTNNNGATCTCAAATATAAGGAGA
AGCACACGTCTGAACTCTTTCCCTACACGACGCTCTTCCGATCTNNNCCTTTCAGTCAATTAAATTTCCT
AGCACACGTCTGAACTCTTTCCCTACACGACGCTCTTCCGATCTNNNTTTTTGAAAACCTCTTTTTTCTG
GAGCACACGTCTGAACTCTTTCCCTACACGACGCTCTTCCGATCTNNNATTCTTGTACCTAGTGTGTTA

GAGCACACGTCTGAACTCTTTCCCTACACGACGCTCTTCCGATCTNNNAAGATTGAAAGAACAAACTT
GCACACGTCTGAACTCTTTCCCTACACGACGCTCTTCCGATCTNNNTATTTAAATCTGAGGCACTGAA
GAGCACACGTCTGAACTCTTTCCCTACACGACGCTCTTCCGATCTNNNTACTCAGTATTTAACTCTTTCC
ACACGTCTGAACTCTTTCCCTACACGACGCTCTTCCGATCTNNNCACACAAAGATACATGCTAATTACAT
AGCACACGTCTGAACTCTTTCCCTACACGACGCTCTTCCGATCTNNNTGTGTGTGTATACATATGTCT
AGCACACGTCTGAACTCTTTCCCTACACGACGCTCTTCCGATCTNNNTGTGTGTGTATACATATGTCT
GAGCACACGTCTGAACTCTTTCCCTACACGACGCTCTTCCGATCTNNNCTTTACTCATCTCCAATCTACC
GAGCACACGTCTGAACTCTTTCCCTACACGACGCTCTTCCGATCTNNNCTTTACTCATCTCCAATCTACC
AGCACACGTCTGAACTCTTTCCCTACACGACGCTCTTCCGATCTNNNAAAGTATGATATAATCTTGCCC
GAGCACACGTCTGAACTCTTTCCCTACACGACGCTCTTCCGATCTNNNTAGAACTATGATACATACCTT
AGCACACGTCTGAACTCTTTCCCTACACGACGCTCTTCCGATCTNNNTGATCATATTATTAAGACTGG
AGCACACGTCTGAACTCTTTCCCTACACGACGCTCTTCCGATCTNNNACACTCCTAAAGAAAAAAACTG
CACACGTCTGAACTCTTTCCCTACACGACGCTCTTCCGATCTNNNATATTTGTCTCAGTCTTTCTCTCAT
AGCACACGTCTGAACTCTTTCCCTACACGACGCTCTTCCGATCTNNNTTCTACTTAATTTGGGTTGACT
CACACGTCTGAACTCTTTCCCTACACGACGCTCTTCCGATCTNNNGATTTATCAGCCTCTTACTTTCCAT
GAGCACACGTCTGAACTCTTTCCCTACACGACGCTCTTCCGATCTNNNCTCTTCAGTCGTCAGTAAAAG
AGCACACGTCTGAACTCTTTCCCTACACGACGCTCTTCCGATCTNNNAAATAAATTGGCTGAATTTCA
GAGCACACGTCTGAACTCTTTCCCTACACGACGCTCTTCCGATCTNNNACATTAATAAGGATATGGAT
CACGTCTGAACTCTTTCCCTACACGACGCTCTTCCGATCTNNNCTTTGATTTTTGTACTCATAAGCCATC
GAGCACACGTCTGAACTCTTTCCCTACACGACGCTCTTCCGATCTNNNATAAATAAAGCATCTACGTG
AGCACACGTCTGAACTCTTTCCCTACACGACGCTCTTCCGATCTNNNCAGAAAAATCAACTGTCCTTG
AGCACACGTCTGAACTCTTTCCCTACACGACGCTCTTCCGATCTNNNAAGGCGGTATGAGTATATATA
AGCACACGTCTGAACTCTTTCCCTACACGACGCTCTTCCGATCTNNNAAGGCGGTATGAGTATATATA
GCACACGTCTGAACTCTTTCCCTACACGACGCTCTTCCGATCTNNNTCTAATGTTTACACTGGGAAATA
GCACACGTCTGAACTCTTTCCCTACACGACGCTCTTCCGATCTNNNACAGACTAAATTTGTATGCATTT
AGCACACGTCTGAACTCTTTCCCTACACGACGCTCTTCCGATCTNNNATTTTTTGGAAAAGCGTATTA
ACACGTCTGAACTCTTTCCCTACACGACGCTCTTCCGATCTNNNGAGACTCTTTCTCTTCCCTTACATCG
GAGCACACGTCTGAACTCTTTCCCTACACGACGCTCTTCCGATCTNNNACAAACTTGTAATATATCTGA
GAGCACACGTCTGAACTCTTTCCCTACACGACGCTCTTCCGATCTNNNACAAACTTGTAATATATCTGA
GCACACGTCTGAACTCTTTCCCTACACGACGCTCTTCCGATCTNNNCAACTTTTCCTTTGTCCCTCCATCG
CACACGTCTGAACTCTTTCCCTACACGACGCTCTTCCGATCTNNNCCAAACAAATCCAAGAAAGTCCAT
AGCACACGTCTGAACTCTTTCCCTACACGACGCTCTTCCGATCTNNNTTTCTTTCTTTCTTTCTTTTTCTC
AGCACACGTCTGAACTCTTTCCCTACACGACGCTCTTCCGATCTNNNTTTTCATTTTCTTTAGCCACGCA
ACACGTCTGAACTCTTTCCCTACACGACGCTCTTCCGATCTNNNATTACTTACTCTTCCCACTAGATCAT
GCACACGTCTGAACTCTTTCCCTACACGACGCTCTTCCGATCTNNNAAATAATAGAACAAAGTGGCAA
GAGCACACGTCTGAACTCTTTCCCTACACGACGCTCTTCCGATCTNNNTTGTATTTTCAAATACCTGTG
AGCACACGTCTGAACTCTTTCCCTACACGACGCTCTTCCGATCTNNNACTATACTCAACTGTCTATCTT
GAGCACACGTCTGAACTCTTTCCCTACACGACGCTCTTCCGATCTNNNAATTTCATTTGGTAATGGGAA
CACACGTCTGAACTCTTTCCCTACACGACGCTCTTCCGATCTNNNTTAAAATATTACAAAAATCTATGG
GCACACGTCTGAACTCTTTCCCTACACGACGCTCTTCCGATCTNNNAGAGGCTGAATCATAATCATAG
GAGCACACGTCTGAACTCTTTCCCTACACGACGCTCTTCCGATCTNNNCACCCATGTTAATAACAGTAT
AGCACACGTCTGAACTCTTTCCCTACACGACGCTCTTCCGATCTNNNGACATGATGGTGAGAAATACT
ACACGTCTGAACTCTTTCCCTACACGACGCTCTTCCGATCTNNNTATGTCACAGGGTCTCTTAGCATCG

AGCACACGTCTGAACTCTTTCCCTACACGACGCTCTTCCGATCTNNNATCCTTCCAAATATCTTATCTGC
AGCACACGTCTGAACTCTTTCCCTACACGACGCTCTTCCGATCTNNNATCCTTCCAAATATCTTATCTGC
GCACACGTCTGAACTCTTTCCCTACACGACGCTCTTCCGATCTNNNATGAATAACAGGAGAAATAGTCT
AGCACACGTCTGAACTCTTTCCCTACACGACGCTCTTCCGATCTNNNTCAGAATACTAAGTTCAGTTCTT
AGCACACGTCTGAACTCTTTCCCTACACGACGCTCTTCCGATCTNNNATTACAGATATGAGCCACTGTA
CACACGTCTGAACTCTTTCCCTACACGACGCTCTTCCGATCTNNNGAACTATTACACTTGGCTAATTCA
GCACACGTCTGAACTCTTTCCCTACACGACGCTCTTCCGATCTNNNTTTTCCCACAGTTTTTACAACCATC
GAGCACACGTCTGAACTCTTTCCCTACACGACGCTCTTCCGATCTNNNGAAGTGAGAGGATTGATTGAT
GCACACGTCTGAACTCTTTCCCTACACGACGCTCTTCCGATCTNNNATTTATCCACCTCTATTCTCATTCAT
GCACACGTCTGAACTCTTTCCCTACACGACGCTCTTCCGATCTNNNAATATTTGAGAAATCACCAAGTG
AGCACACGTCTGAACTCTTTCCCTACACGACGCTCTTCCGATCTNNNCAGGTTAGGTCATATGTTAGAT
GAGCACACGTCTGAACTCTTTCCCTACACGACGCTCTTCCGATCTNNNAAAAAAAGTAACATCACTCTT
GAGCACACGTCTGAACTCTTTCCCTACACGACGCTCTTCCGATCTNNNCAGTTCATTCATTTTTACTGTT
AGCACACGTCTGAACTCTTTCCCTACACGACGCTCTTCCGATCTNNNTAATAAGGCATGGTTAAAGGA
AGCACACGTCTGAACTCTTTCCCTACACGACGCTCTTCCGATCTNNNGATTTGACTTCTGTGTGAGTCAT
AGCACACGTCTGAACTCTTTCCCTACACGACGCTCTTCCGATCTNNNACTGAATTTCTAAAAAACTCTG
GCACACGTCTGAACTCTTTCCCTACACGACGCTCTTCCGATCTNNNCCCATATTGACTGTGATGTATATC
AGCACACGTCTGAACTCTTTCCCTACACGACGCTCTTCCGATCTNNNGAAGCCTTTATGAGACCTTACCA
CACACGTCTGAACTCTTTCCCTACACGACGCTCTTCCGATCTNNNTACATTCAACACAAAGGTTTAAGCA
CACACGTCTGAACTCTTTCCCTACACGACGCTCTTCCGATCTNNNCTTTAATTGTTGGCTTTTGTGCATC
GAGCACACGTCTGAACTCTTTCCCTACACGACGCTCTTCCGATCTNNNAAGCCATCATTATGTATGTAT
GAGCACACGTCTGAACTCTTTCCCTACACGACGCTCTTCCGATCTNNNAAGCCATCATTATGTATGTAT
CACACGTCTGAACTCTTTCCCTACACGACGCTCTTCCGATCTNNNAGTCTTGAGTATGTCTTTATCAGCA
AGCACACGTCTGAACTCTTTCCCTACACGACGCTCTTCCGATCTNNNTCTATCAATATCTTGGACAAACT
GCACACGTCTGAACTCTTTCCCTACACGACGCTCTTCCGATCTNNNTTCATAGTTATGCAGTTTACATG
GAGCACACGTCTGAACTCTTTCCCTACACGACGCTCTTCCGATCTNNNGATAAACATGGAAACTAACCA
AGCACACGTCTGAACTCTTTCCCTACACGACGCTCTTCCGATCTNNNCTTTCTACAATTATCCCTTCTTGC
CACACGTCTGAACTCTTTCCCTACACGACGCTCTTCCGATCTNNNCTGACTTCTTTCTCCTGGAACATCG
GCACACGTCTGAACTCTTTCCCTACACGACGCTCTTCCGATCTNNNGACAAGTCCGAACAGATAAGCATC
GCACACGTCTGAACTCTTTCCCTACACGACGCTCTTCCGATCTNNNTCTTTGGGAGATAATTAGCTTTAC
GCACACGTCTGAACTCTTTCCCTACACGACGCTCTTCCGATCTNNNAGTTTTTGAAGAAAAGTCTCATTC
AGCACACGTCTGAACTCTTTCCCTACACGACGCTCTTCCGATCTNNNTTTATACGGTATTCCTATTTTGC
GCACACGTCTGAACTCTTTCCCTACACGACGCTCTTCCGATCTNNNGTTACTGTCAGTATAATCAGCTTC
GAGCACACGTCTGAACTCTTTCCCTACACGACGCTCTTCCGATCTNNNATTATCAGAACTCCAGAATGA
AGAGCACACGTCTGAACTCTTTCCCTACACGACGCTCTTCCGATCTNNNGCAACAAATAACTAAATCAG
GAGCACACGTCTGAACTCTTTCCCTACACGACGCTCTTCCGATCTNNNGACAAGGGAGTGTTCTAAATA
AGCACACGTCTGAACTCTTTCCCTACACGACGCTCTTCCGATCTNNNTAAGTGAATATAAGATTTTGGC
CACACGTCTGAACTCTTTCCCTACACGACGCTCTTCCGATCTNNNGATTACAAACTTGAGCCACTCATC
AGCACACGTCTGAACTCTTTCCCTACACGACGCTCTTCCGATCTNNNTCTATCACTACAACTACCACACAT
AGCACACGTCTGAACTCTTTCCCTACACGACGCTCTTCCGATCTNNNATTCAACAGAGCAACTTTTAATT
AGCACACGTCTGAACTCTTTCCCTACACGACGCTCTTCCGATCTNNNAAAATAAAAGCAAATTGTAGG
GAGCACACGTCTGAACTCTTTCCCTACACGACGCTCTTCCGATCTNNNCTGTAAATTATCTGCCAAAGCC
CACACGTCTGAACTCTTTCCCTACACGACGCTCTTCCGATCTNNNACTGTTTTTCTTTACTACTTCCCAT

AGCACACGTCTGAACTCTTTCCCTACACGACGCTCTTCCGATCTNNNACAGCTACTATACCATATACAA
CACACGTCTGAACTCTTTCCCTACACGACGCTCTTCCGATCTNNNCCCAGTATTGATGATTTTCCTTCA
ACACGTCTGAACTCTTTCCCTACACGACGCTCTTCCGATCTNNNTACATTACATTGATCTTAGGGTTCA
GCACACGTCTGAACTCTTTCCCTACACGACGCTCTTCCGATCTNNNTTTTTTAACTGAGTTCCTCATTT
ACACGTCTGAACTCTTTCCCTACACGACGCTCTTCCGATCTNNNTAAGAAAACATACAGATTTTGGGC
AGCACACGTCTGAACTCTTTCCCTACACGACGCTCTTCCGATCTNNNATGTAAAGAGACCAAATCTAC
ACACGTCTGAACTCTTTCCCTACACGACGCTCTTCCGATCTNNNCAAAAAAAAAAAAAAGGCAGTAAA
AGCACACGTCTGAACTCTTTCCCTACACGACGCTCTTCCGATCTNNNCAGAAAATGTTTGAGGTCAAG
AGCACACGTCTGAACTCTTTCCCTACACGACGCTCTTCCGATCTNNNCAGAAAATGTTTGAGGTCAAG
AGCACACGTCTGAACTCTTTCCCTACACGACGCTCTTCCGATCTNNNAATTCCAGGTTTTATCCATACA
AGCACACGTCTGAACTCTTTCCCTACACGACGCTCTTCCGATCTNNNATAACCTCTGTATATAAACCCT
AGCACACGTCTGAACTCTTTCCCTACACGACGCTCTTCCGATCTNNNTTCACCTACTGTAATAATTTTG
AGCACACGTCTGAACTCTTTCCCTACACGACGCTCTTCCGATCTNNNTAAGTACCAGTCTATTTTTTTG
AGCACACGTCTGAACTCTTTCCCTACACGACGCTCTTCCGATCTNNNTCTATTTGTTTTTCTGCTTTTTG
AGCACACGTCTGAACTCTTTCCCTACACGACGCTCTTCCGATCTNNNTTAACATACAACATGAAGCTCA
AGCACACGTCTGAACTCTTTCCCTACACGACGCTCTTCCGATCTNNNTACTTTTAATGGCAAAAACCT
ACACGTCTGAACTCTTTCCCTACACGACGCTCTTCCGATCTNNNGCCATTTGGACAGATATTGACATCG
AGCACACGTCTGAACTCTTTCCCTACACGACGCTCTTCCGATCTNNNCCTTACTAGGTTTTGAGGATT
AGCACACGTCTGAACTCTTTCCCTACACGACGCTCTTCCGATCTNNNTATTTTCTACTCCTACTATCCCT
ACACGTCTGAACTCTTTCCCTACACGACGCTCTTCCGATCTNNNTTATACTGCTGTATTGTTCTGTACA
ACACGTCTGAACTCTTTCCCTACACGACGCTCTTCCGATCTNNNTTATACTGCTGTATTGTTCTGTACA
AGCACACGTCTGAACTCTTTCCCTACACGACGCTCTTCCGATCTNNNACAACTGAATATTCACATCTAG
GCACACGTCTGAACTCTTTCCCTACACGACGCTCTTCCGATCTNNNGGGTAATAATCTGAAATTGTGT
CACACGTCTGAACTCTTTCCCTACACGACGCTCTTCCGATCTNNNCTCAGTCTCCTAAAAAGCTGCATC
AGCACACGTCTGAACTCTTTCCCTACACGACGCTCTTCCGATCTNNNCCTACACACAGACTTATATATG
AGCACACGTCTGAACTCTTTCCCTACACGACGCTCTTCCGATCTNNNAGTAATTTTTCTCTCTCTCTCAT
AGCACACGTCTGAACTCTTTCCCTACACGACGCTCTTCCGATCTNNNTGTGATTTAGAGATAACCAGTG
AGCACACGTCTGAACTCTTTCCCTACACGACGCTCTTCCGATCTNNNGAAGAAAGAGAGTTAGGGAA
GAGCACACGTCTGAACTCTTTCCCTACACGACGCTCTTCCGATCTNNNAAACCTGAGAGGATAACTAA
AGCACACGTCTGAACTCTTTCCCTACACGACGCTCTTCCGATCTNNNAGAAATCAACCAATAAGCATTC
GAGCACACGTCTGAACTCTTTCCCTACACGACGCTCTTCCGATCTNNNTAGGGAGAGGAAAGATACTT
CACACGTCTGAACTCTTTCCCTACACGACGCTCTTCCGATCTNNNAATCCCAGCACTTTAGAAAGCAT
AGCACACGTCTGAACTCTTTCCCTACACGACGCTCTTCCGATCTNNNTTATGTTTACTGTTTGATTGCT
AGCACACGTCTGAACTCTTTCCCTACACGACGCTCTTCCGATCTNNNGGGAAAAGGAAGAAATCAGA
CACACGTCTGAACTCTTTCCCTACACGACGCTCTTCCGATCTNNNGAAAAGGAGAAACTTGAAACTCC
AGCACACGTCTGAACTCTTTCCCTACACGACGCTCTTCCGATCTNNNTTCCAAATATTTTTGATCTGTG
AGCACACGTCTGAACTCTTTCCCTACACGACGCTCTTCCGATCTNNNGTACTCCATATACCTTACAACA
AGCACACGTCTGAACTCTTTCCCTACACGACGCTCTTCCGATCTNNNGTACTCCATATACCTTACAACA
GCACACGTCTGAACTCTTTCCCTACACGACGCTCTTCCGATCTNNNCTATGGACCAAAGTTTACCCCAT
AGCACACGTCTGAACTCTTTCCCTACACGACGCTCTTCCGATCTNNNCTTCTTTCAAGATGTTCTTTTTG
ACACGTCTGAACTCTTTCCCTACACGACGCTCTTCCGATCTNNNATTAAACCATTCCTCTTAAAGAAAT
AGCACACGTCTGAACTCTTTCCCTACACGACGCTCTTCCGATCTNNNTCACTCAAATCTGGTAATGTA
CACACGTCTGAACTCTTTCCCTACACGACGCTCTTCCGATCTNNNGAAACCATGGTGACTAGTACCAT

AGCACACGTCTGAACTCTTTCCCTACACGACGCTCTTCCGATCTNNNAGTCAAAGAATGATATTTAGA
AGCACACGTCTGAACTCTTTCCCTACACGACGCTCTTCCGATCTNNNATACACACACATATATGTATGC
GCACACGTCTGAACTCTTTCCCTACACGACGCTCTTCCGATCTNNNCAAGATCTGAGTGCTAGATGCA
AGCACACGTCTGAACTCTTTCCCTACACGACGCTCTTCCGATCTNNNAACAAAGATCCTTAAATTACTT
GAGCACACGTCTGAACTCTTTCCCTACACGACGCTCTTCCGATCTNNNAAGGGACACATGTAATTCTTT
GCACACGTCTGAACTCTTTCCCTACACGACGCTCTTCCGATCTNNNTTGTTCAGTGAAACCTCTAAATA
AGCACACGTCTGAACTCTTTCCCTACACGACGCTCTTCCGATCTNNNAAGTCAAACCAAGTAGACTTG
GCACACGTCTGAACTCTTTCCCTACACGACGCTCTTCCGATCTNNNCATCATTAAAGCAACTACAGAT
CACACGTCTGAACTCTTTCCCTACACGACGCTCTTCCGATCTNNNCCAGTCATTTGCATTAGATAGTC
CACACGTCTGAACTCTTTCCCTACACGACGCTCTTCCGATCTNNNGGGCCTCATATACATATATATGT
AGCACACGTCTGAACTCTTTCCCTACACGACGCTCTTCCGATCTNNNCCAGCTCTAAAAATAGGTAAA
AGCACACGTCTGAACTCTTTCCCTACACGACGCTCTTCCGATCTNNNCTGTTCTGTAGGTTTTCTTATG
AGCACACGTCTGAACTCTTTCCCTACACGACGCTCTTCCGATCTNNNCTGTTCTGTAGGTTTTCTTATG
AGCACACGTCTGAACTCTTTCCCTACACGACGCTCTTCCGATCTNNNTTAGTAAAGATGTCTGTAGTC
AGCACACGTCTGAACTCTTTCCCTACACGACGCTCTTCCGATCTNNNGTTTACTGTTTAAGCCATGAAA
GCACACGTCTGAACTCTTTCCCTACACGACGCTCTTCCGATCTNNNCTCTGAGCCTATGATGAAAATA
GCACACGTCTGAACTCTTTCCCTACACGACGCTCTTCCGATCTNNNTTTACTTTCATGCTATTCTGATG
AGCACACGTCTGAACTCTTTCCCTACACGACGCTCTTCCGATCTNNNTTTCATTCTTTTTATGGCTGA
ACACGTCTGAACTCTTTCCCTACACGACGCTCTTCCGATCTNNNCTCTATCACAGGCTACAGTGCATCG
ACACGTCTGAACTCTTTCCCTACACGACGCTCTTCCGATCTNNNAACTCTTTGTTTCCAGTATTAACCAT
GCACACGTCTGAACTCTTTCCCTACACGACGCTCTTCCGATCTNNNCTTTCTATTCAGCAGATATTGG
AGCACACGTCTGAACTCTTTCCCTACACGACGCTCTTCCGATCTNNNTGTGTTTCAACTCATTTATGTT
GCACACGTCTGAACTCTTTCCCTACACGACGCTCTTCCGATCTNNNACTTTACTTGAATGAGTTTACAC
GCACACGTCTGAACTCTTTCCCTACACGACGCTCTTCCGATCTNNNAAAACATATTTGCTTCATCATTG
CACACGTCTGAACTCTTTCCCTACACGACGCTCTTCCGATCTNNNCTGCCTTAGCACCCTATTTACATC
AGCACACGTCTGAACTCTTTCCCTACACGACGCTCTTCCGATCTNNNTTAACTCATCATTTAGAATTAG
AGCACACGTCTGAACTCTTTCCCTACACGACGCTCTTCCGATCTNNNTGTGTGTACATATATGTGTGT
AGCACACGTCTGAACTCTTTCCCTACACGACGCTCTTCCGATCTNNNTCTTATATTATCACTTATCTGA
GCACACGTCTGAACTCTTTCCCTACACGACGCTCTTCCGATCTNNNCCTTTCAAGTACTAATTCTAGGA
GAGCACACGTCTGAACTCTTTCCCTACACGACGCTCTTCCGATCTNNNATTAACACATACATTCCACTTT
AGCACACGTCTGAACTCTTTCCCTACACGACGCTCTTCCGATCTNNNGTTCCACCTGAAGATTTGAGC
GCACACGTCTGAACTCTTTCCCTACACGACGCTCTTCCGATCTNNNTTTATACAGTAGCTTACTCAACA
AGCACACGTCTGAACTCTTTCCCTACACGACGCTCTTCCGATCTNNNAGCATAAAAATGAAACAGATT
ACACGTCTGAACTCTTTCCCTACACGACGCTCTTCCGATCTNNNTTGTTACCTCATGATCACAAGCATC
CACACGTCTGAACTCTTTCCCTACACGACGCTCTTCCGATCTNNNACACATATGGATGTATACTGAAT
ACACGTCTGAACTCTTTCCCTACACGACGCTCTTCCGATCTNNNGACACATATGGATGTATACTGAAC
GCACACGTCTGAACTCTTTCCCTACACGACGCTCTTCCGATCTNNNGAATTAACTGTTACCACGTTTAC
AGCACACGTCTGAACTCTTTCCCTACACGACGCTCTTCCGATCTNNNAATAGCTAGCTAGATTTTGGA
AGCACACGTCTGAACTCTTTCCCTACACGACGCTCTTCCGATCTNNNAAGGTCCACATATGACAAATT
AGCACACGTCTGAACTCTTTCCCTACACGACGCTCTTCCGATCTNNNCAGGCAAGTTATATTCCTACT
CACACGTCTGAACTCTTTCCCTACACGACGCTCTTCCGATCTNNNGTGAGAGGGTGGTAAGAATCAT
AGCACACGTCTGAACTCTTTCCCTACACGACGCTCTTCCGATCTNNNGTATATTCTTTTGGATTTCCTG
GCACACGTCTGAACTCTTTCCCTACACGACGCTCTTCCGATCTNNNCCTCTATTGTTCTCTGATAGTGC

GCACACGTCTGAACTCTTTCCCTACACGACGCTCTTCCGATCTNNNCTCTTGAAAGTTTTGTCTCTTCC
AGCACACGTCTGAACTCTTTCCCTACACGACGCTCTTCCGATCTNNNCCAACAGAAGCAAAACAAAAC
CACGTCTGAACTCTTTCCCTACACGACGCTCTTCCGATCTNNNGAGAGGGTATGATGGAAAAGCATC
ACACGTCTGAACTCTTTCCCTACACGACGCTCTTCCGATCTNNNAAATAGGTTTTTCTGAAAGTTGTC
CACACGTCTGAACTCTTTCCCTACACGACGCTCTTCCGATCTNNNATTTAAAATTGGTTGGGCATGC
AGCACACGTCTGAACTCTTTCCCTACACGACGCTCTTCCGATCTNNNATTTCAGAGAATGTGTGTGAA
AGCACACGTCTGAACTCTTTCCCTACACGACGCTCTTCCGATCTNNNTGAGGAATAAATAACAGAAGG
GCACACGTCTGAACTCTTTCCCTACACGACGCTCTTCCGATCTNNNCAATGTCTATAGATAACTGCCTA
AGCACACGTCTGAACTCTTTCCCTACACGACGCTCTTCCGATCTNNNATACAAACATAATTCTTTTCCCA
AGCACACGTCTGAACTCTTTCCCTACACGACGCTCTTCCGATCTNNNCTCTAGGTAACAAAAATGCAT
AGCACACGTCTGAACTCTTTCCCTACACGACGCTCTTCCGATCTNNNAGCACCATTTTTTAAAAATAG
AGCACACGTCTGAACTCTTTCCCTACACGACGCTCTTCCGATCTNNNACATGCAGAAGGTATTACACC
GAGCACACGTCTGAACTCTTTCCCTACACGACGCTCTTCCGATCTNNNTGATCTGATCTTAGTTTAGAC
GCACACGTCTGAACTCTTTCCCTACACGACGCTCTTCCGATCTNNNTTAGGAATTAATTTTGATGCAG
GCACACGTCTGAACTCTTTCCCTACACGACGCTCTTCCGATCTNNNTTCTTTTCTTTCTTTCTTTCTTTCC
AGCACACGTCTGAACTCTTTCCCTACACGACGCTCTTCCGATCTNNNTACTGGCTCTGTTTTACATATA
GAGCACACGTCTGAACTCTTTCCCTACACGACGCTCTTCCGATCTNNNGATCCTATCAGTAGTTATGG
CACACGTCTGAACTCTTTCCCTACACGACGCTCTTCCGATCTNNNCAACTGCAGTTCAGAATAAATAT
CACACGTCTGAACTCTTTCCCTACACGACGCTCTTCCGATCTNNNCAACTGCAGTTCAGAATAAATAT
GCACACGTCTGAACTCTTTCCCTACACGACGCTCTTCCGATCTNNNACTTACATCTACAAGATCACACC
AGCACACGTCTGAACTCTTTCCCTACACGACGCTCTTCCGATCTNNNATGTCTGCTGTGTTTCAATACA
AGCACACGTCTGAACTCTTTCCCTACACGACGCTCTTCCGATCTNNNAAGCAATATGTGGTTTATATC
ACACGTCTGAACTCTTTCCCTACACGACGCTCTTCCGATCTNNNATCAAATTAGTGTCCTTTGAATGC
AGCACACGTCTGAACTCTTTCCCTACACGACGCTCTTCCGATCTNNNATTTGTGTTCTTAAAAATCTTG
GCACACGTCTGAACTCTTTCCCTACACGACGCTCTTCCGATCTNNNACAGAATTATTCCAGTATCTTCA
AGCACACGTCTGAACTCTTTCCCTACACGACGCTCTTCCGATCTNNNCTTTGATAGCCATTTTCAATAC
GCACACGTCTGAACTCTTTCCCTACACGACGCTCTTCCGATCTNNNCCGGGCACTGTAAAATTTATCAT
GCACACGTCTGAACTCTTTCCCTACACGACGCTCTTCCGATCTNNNACCTCTTATTTAAATTTCACTCCC
AGCACACGTCTGAACTCTTTCCCTACACGACGCTCTTCCGATCTNNNAAGATTTGTATTAGATGTGGG
GCACACGTCTGAACTCTTTCCCTACACGACGCTCTTCCGATCTNNNAAAACTCTTCTATGGTTGTATTC
AGCACACGTCTGAACTCTTTCCCTACACGACGCTCTTCCGATCTNNNTTAAACGTCCTAATAAATGTG
AGCACACGTCTGAACTCTTTCCCTACACGACGCTCTTCCGATCTNNNCATACTATAAGGTTTGAGGTT
CACGTCTGAACTCTTTCCCTACACGACGCTCTTCCGATCTNNNTTCCTTTGAATGGTTCTCCTCATCGG
AGCACACGTCTGAACTCTTTCCCTACACGACGCTCTTCCGATCTNNNACAACACCTCATGTAAAATAGT
GAGCACACGTCTGAACTCTTTCCCTACACGACGCTCTTCCGATCTNNNGATGTTAGAGAACTAGAGAA
GAGCACACGTCTGAACTCTTTCCCTACACGACGCTCTTCCGATCTNNNAGATAATCTGTTACTCTAAGC
AGCACACGTCTGAACTCTTTCCCTACACGACGCTCTTCCGATCTNNNCCACCTTTCTTATTTCTGATTTTC
AGCACACGTCTGAACTCTTTCCCTACACGACGCTCTTCCGATCTNNNTTCAGATTTTATTCTCCTATGGT
GAGCACACGTCTGAACTCTTTCCCTACACGACGCTCTTCCGATCTNNNAATTACTTTTCTTGGTGTACC
AGCACACGTCTGAACTCTTTCCCTACACGACGCTCTTCCGATCTNNNGCCATTCCTATAAAAAAAGAT
ACACGTCTGAACTCTTTCCCTACACGACGCTCTTCCGATCTNNNGGATAACTGTACAGAACTAACTAC
AGCACACGTCTGAACTCTTTCCCTACACGACGCTCTTCCGATCTNNNAAATCTATTTGATGCCAAACA
AGCACACGTCTGAACTCTTTCCCTACACGACGCTCTTCCGATCTNNNTTTCCATCTAAACAAGAGTATC

GAGCACACGTCTGAACTCTTTCCCTACACGACGCTCTTCCGATCTNNNAAGAAAACAAAACAAAACGA

GCACACGTCTGAACTCTTTCCCTACACGACGCTCTTCCGATCTNNNTAACTTTTCTTTTTCTTTTCTGCC

CACGTCTGAACTCTTTCCCTACACGACGCTCTTCCGATCTNNNGAGGAAGATCAGGAAATCCCCATC

AGCACACGTCTGAACTCTTTCCCTACACGACGCTCTTCCGATCTNNNTTTTATTGAAGGTTTTTGTGT

AGCACACGTCTGAACTCTTTCCCTACACGACGCTCTTCCGATCTNNNACCATTTATTCTGTTTTCTTTC

GCACACGTCTGAACTCTTTCCCTACACGACGCTCTTCCGATCTNNNACAAAGAGGGAAAGAATGTAA

AGCACACGTCTGAACTCTTTCCCTACACGACGCTCTTCCGATCTNNNAGGGATTTGAATGAACTGTA

AGCACACGTCTGAACTCTTTCCCTACACGACGCTCTTCCGATCTNNNATTTAAGTTGGACTGTTGTATT

AGCACACGTCTGAACTCTTTCCCTACACGACGCTCTTCCGATCTNNNCTACCCATCTCCAAGATAATAA

GCACACGTCTGAACTCTTTCCCTACACGACGCTCTTCCGATCTNNNTTTAAAGAAATTGAAGAGGAT

CACACGTCTGAACTCTTTCCCTACACGACGCTCTTCCGATCTNNNGTAGATGGGAAGAGAAGAAAGC

AGCACACGTCTGAACTCTTTCCCTACACGACGCTCTTCCGATCTNNNACACACATAGGTATATAAATAT

GCACACGTCTGAACTCTTTCCCTACACGACGCTCTTCCGATCTNNNTATAACCTGTATACCACTAATCC

GCACACGTCTGAACTCTTTCCCTACACGACGCTCTTCCGATCTNNNACAAAGAAAAAAGAAAAAGAG

ACACGTCTGAACTCTTTCCCTACACGACGCTCTTCCGATCTNNNGGTAATGCATACATGGAAGTCATC

AGCACACGTCTGAACTCTTTCCCTACACGACGCTCTTCCGATCTNNNATATCTCTGAAATCAAATCATC

AGCACACGTCTGAACTCTTTCCCTACACGACGCTCTTCCGATCTNNNAAATCTGACTATTGGTGTATA

GCACACGTCTGAACTCTTTCCCTACACGACGCTCTTCCGATCTNNNTAGTACAATCAACTCAGTCTTTG

CACACGTCTGAACTCTTTCCCTACACGACGCTCTTCCGATCTNNNTTCTCTTGATCTTCCTTTATTCTCAT

AGCACACGTCTGAACTCTTTCCCTACACGACGCTCTTCCGATCTNNNTCTTTCTTCAGTTTCTATGATTC

GCACACGTCTGAACTCTTTCCCTACACGACGCTCTTCCGATCTNNNCATACACAGAGGCATACATAGC

GCACACGTCTGAACTCTTTCCCTACACGACGCTCTTCCGATCTNNNGTGTTCTTTTGTCTAAAAATAGA

CACACGTCTGAACTCTTTCCCTACACGACGCTCTTCCGATCTNNNGTGGAATCTCACTGTATGAATCA

GAGCACACGTCTGAACTCTTTCCCTACACGACGCTCTTCCGATCTNNNAGACAAGGTTATCACATTAC

GAGCACACGTCTGAACTCTTTCCCTACACGACGCTCTTCCGATCTNNNGGAGATTATGTTTGGAAAA

GCACACGTCTGAACTCTTTCCCTACACGACGCTCTTCCGATCTNNNAGCATATCAAAGCACCATATTTC

AGCACACGTCTGAACTCTTTCCCTACACGACGCTCTTCCGATCTNNNTGAGTATTTTGTTTCTTGTACT

GCACACGTCTGAACTCTTTCCCTACACGACGCTCTTCCGATCTNNNTAAATTTGTTCAGTTTATCCACA

GCACACGTCTGAACTCTTTCCCTACACGACGCTCTTCCGATCTNNNTTTTTGCTGTATTTTTACTCATGC

GCACACGTCTGAACTCTTTCCCTACACGACGCTCTTCCGATCTNNNGAATTTTCAAGAGAGGTTTTAG

GCACACGTCTGAACTCTTTCCCTACACGACGCTCTTCCGATCTNNNCCCATGTGTTATATATAAATGC

GCACACGTCTGAACTCTTTCCCTACACGACGCTCTTCCGATCTNNNGTTGTATTGTGGTGTTGATGCA

GCACACGTCTGAACTCTTTCCCTACACGACGCTCTTCCGATCTNNNTTCTTACTGAATCCTCACATAAA

GCACACGTCTGAACTCTTTCCCTACACGACGCTCTTCCGATCTNNNATTTAATTCCCCTTCTTCCAAAC

CACACGTCTGAACTCTTTCCCTACACGACGCTCTTCCGATCTNNNTTAGTAAGTAGTTGGCTTTGGCA

AGCACACGTCTGAACTCTTTCCCTACACGACGCTCTTCCGATCTNNNATCATCTTATCATCCCCCAAAC

AGCACACGTCTGAACTCTTTCCCTACACGACGCTCTTCCGATCTNNNAACTTCTGACCCATATATAGA

CACACGTCTGAACTCTTTCCCTACACGACGCTCTTCCGATCTNNNTGAGATAAAGTCTTATTCTGTCAC

AGCACACGTCTGAACTCTTTCCCTACACGACGCTCTTCCGATCTNNNTAAAGTAGACTTACAAACTGC

GCACACGTCTGAACTCTTTCCCTACACGACGCTCTTCCGATCTNNNAATCACAAATAATCATCACCCAT

CACACGTCTGAACTCTTTCCCTACACGACGCTCTTCCGATCTNNNCTCTTCTACTCGACCTTAAATTACA

GCACACGTCTGAACTCTTTCCCTACACGACGCTCTTCCGATCTNNNAAGTCTTCAAGTATATGGCCTA

ACACGTCTGAACTCTTTCCCTACACGACGCTCTTCCGATCTNNNGGTTTACATTTCCTTTTACAAACCA

ACACGTCTGAACTCTTTCCCTACACGACGCTCTTCCGATCTNNNCCTTCTCTCATGATAAATTTGATCAT
GCACACGTCTGAACTCTTTCCCTACACGACGCTCTTCCGATCTNNNCCACTGAATGAATGTGATATAT
GCACACGTCTGAACTCTTTCCCTACACGACGCTCTTCCGATCTNNNGCAAGCCAGTAAATTTCTATTTA
GCACACGTCTGAACTCTTTCCCTACACGACGCTCTTCCGATCTNNNAGAGCTACTGAATTAGAAACTC
GCACACGTCTGAACTCTTTCCCTACACGACGCTCTTCCGATCTNNNCACTTCATATGTTTTTCCTAGAT
GCACACGTCTGAACTCTTTCCCTACACGACGCTCTTCCGATCTNNNCACTTCATATGTTTTTCCTAGAT
AGCACACGTCTGAACTCTTTCCCTACACGACGCTCTTCCGATCTNNNTATTTTGTTACAAAGTTTCCCA
AGCACACGTCTGAACTCTTTCCCTACACGACGCTCTTCCGATCTNNNCAACCATTCTTCTCTCCATCCAT
AGCACACGTCTGAACTCTTTCCCTACACGACGCTCTTCCGATCTNNNTTGAGCAATAAAATTTAGAACG
AGCACACGTCTGAACTCTTTCCCTACACGACGCTCTTCCGATCTNNNCAATTCACAACTGAAAAAATGT
GCACACGTCTGAACTCTTTCCCTACACGACGCTCTTCCGATCTNNNGCTTTCAAAACTCTAGGATATTT
GCACACGTCTGAACTCTTTCCCTACACGACGCTCTTCCGATCTNNNGTCTTCCTCTAGCATAATTCTTA
GCACACGTCTGAACTCTTTCCCTACACGACGCTCTTCCGATCTNNNGACAGGATAGTTTTTACAAAGA
AGCACACGTCTGAACTCTTTCCCTACACGACGCTCTTCCGATCTNNNAACTTTTAAATCGAAGTTTTG
AGCACACGTCTGAACTCTTTCCCTACACGACGCTCTTCCGATCTNNNTGAATGTCCTCTTATAAACTGT
GCACACGTCTGAACTCTTTCCCTACACGACGCTCTTCCGATCTNNNACACTTATTAAAATAAACTCCCT
GCACACGTCTGAACTCTTTCCCTACACGACGCTCTTCCGATCTNNNAACCAAGAGCATATTTTCTCTTA
AGCACACGTCTGAACTCTTTCCCTACACGACGCTCTTCCGATCTNNNCAACATCAGACTAGATCAAGTA
AGCACACGTCTGAACTCTTTCCCTACACGACGCTCTTCCGATCTNNNTTTTGAAGGTTTGTCTATTTGT
CACACGTCTGAACTCTTTCCCTACACGACGCTCTTCCGATCTNNNGTCTCTATTCCACTTCTCTGCATCG
AGCACACGTCTGAACTCTTTCCCTACACGACGCTCTTCCGATCTNNNTTTTCTAGGTAGATTGTGTATG
GCACACGTCTGAACTCTTTCCCTACACGACGCTCTTCCGATCTNNNTCTTTTTGACCATTATCTCTAGAC
GCACACGTCTGAACTCTTTCCCTACACGACGCTCTTCCGATCTNNNCATTATTATGTTTTGCCAAATCC
AGCACACGTCTGAACTCTTTCCCTACACGACGCTCTTCCGATCTNNNAAATGACTCTGTCTTCTAGGAT
AGCACACGTCTGAACTCTTTCCCTACACGACGCTCTTCCGATCTNNNTGTAATACACATATAAAGCTGG
GCACACGTCTGAACTCTTTCCCTACACGACGCTCTTCCGATCTNNNGCATCATTTGTAATACACATGTA
AGCACACGTCTGAACTCTTTCCCTACACGACGCTCTTCCGATCTNNNCACCATGTTTTTGAACTGTATAT
AGCACACGTCTGAACTCTTTCCCTACACGACGCTCTTCCGATCTNNNACGGTTACTAGCTTAAGTATTA
CACACGTCTGAACTCTTTCCCTACACGACGCTCTTCCGATCTNNNTTCTTTCTCTGATTCTTCTTCCCATC
GCACACGTCTGAACTCTTTCCCTACACGACGCTCTTCCGATCTNNNATAACTATTTCCTAGTCCTGGTA
GCACACGTCTGAACTCTTTCCCTACACGACGCTCTTCCGATCTNNNCGTCAGGTGAATAGATAGATAA
GCACACGTCTGAACTCTTTCCCTACACGACGCTCTTCCGATCTNNNTAACAAGTGGGGTAAAAAATGA
AGCACACGTCTGAACTCTTTCCCTACACGACGCTCTTCCGATCTNNNAAGGTAGTTTTAAAGGTGGTA
GAGCACACGTCTGAACTCTTTCCCTACACGACGCTCTTCCGATCTNNNTATGTATTCTACAAGCCTTGT
CACACGTCTGAACTCTTTCCCTACACGACGCTCTTCCGATCTNNNTTTGATTTGTTTATGTTATGACGC
GCACACGTCTGAACTCTTTCCCTACACGACGCTCTTCCGATCTNNNAAGGGACAGTAAAAATACAGTA
AGCACACGTCTGAACTCTTTCCCTACACGACGCTCTTCCGATCTNNNTGATTAGTAAGAACCTTTTGAG
GAGCACACGTCTGAACTCTTTCCCTACACGACGCTCTTCCGATCTNNNCATTAGCTATTTATTCCTGTGT
GCACACGTCTGAACTCTTTCCCTACACGACGCTCTTCCGATCTNNNATCACTGAGTTTTGGTCATCCAT
CACACGTCTGAACTCTTTCCCTACACGACGCTCTTCCGATCTNNNAATAGACAGAGATCAGAAACAGC
GAGCACACGTCTGAACTCTTTCCCTACACGACGCTCTTCCGATCTNNNTATTTTGGGCATTTCATGTAA
GCACACGTCTGAACTCTTTCCCTACACGACGCTCTTCCGATCTNNNCTATTACTGTTGGTAAGTTTAGC
GCACACGTCTGAACTCTTTCCCTACACGACGCTCTTCCGATCTNNNATTACTTCACTTAGAATAACGGT

AGAGCACACGTCTGAACTCTTTCCCTACACGACGCTCTTCCGATCTNNNTTGGAATTCACACATCTCAG
AGCACACGTCTGAACTCTTTCCCTACACGACGCTCTTCCGATCTNNNTTCAAAGAAGATAGGAAGCT
CACACGTCTGAACTCTTTCCCTACACGACGCTCTTCCGATCTNNNGTGTATATGTATATGTGTATATG
ACACGTCTGAACTCTTTCCCTACACGACGCTCTTCCGATCTNNNACTCTAGCTGATTCTCTGAGCATCG
AGCACACGTCTGAACTCTTTCCCTACACGACGCTCTTCCGATCTNNNATCCAAAAGTAGCCACTTATTT
GCACACGTCTGAACTCTTTCCCTACACGACGCTCTTCCGATCTNNNATTTTAGCACTATTTGGATTCTG
AGCACACGTCTGAACTCTTTCCCTACACGACGCTCTTCCGATCTNNNAATTTATACCTCAGTGTGTGAT
GCACACGTCTGAACTCTTTCCCTACACGACGCTCTTCCGATCTNNNCTTTTTTGGAAGAATATGGGAA
GAGCACACGTCTGAACTCTTTCCCTACACGACGCTCTTCCGATCTNNNTTATGTATGTGTGTAAGTCAT
AGCACACGTCTGAACTCTTTCCCTACACGACGCTCTTCCGATCTNNNACATTCAAGGTGAGTTATATC
AGCACACGTCTGAACTCTTTCCCTACACGACGCTCTTCCGATCTNNNTAAGCTTGAAACATGTTTTCAA
CACACGTCTGAACTCTTTCCCTACACGACGCTCTTCCGATCTNNNTCCCTCTCTCTTCACACATACATCG
GCACACGTCTGAACTCTTTCCCTACACGACGCTCTTCCGATCTNNNCTGCAAATGCCATTATTTTATTCC
GCACACGTCTGAACTCTTTCCCTACACGACGCTCTTCCGATCTNNNCTATATTAGCCTGCTTGAATTGC
AGCACACGTCTGAACTCTTTCCCTACACGACGCTCTTCCGATCTNNNAAAGTGTTTCATCTTCCTGAAC
AGCACACGTCTGAACTCTTTCCCTACACGACGCTCTTCCGATCTNNNACCAGTCTATATTCTAAGAAG
GCACACGTCTGAACTCTTTCCCTACACGACGCTCTTCCGATCTNNNGAGAAAGGAGCTATCAGATAA
AGCACACGTCTGAACTCTTTCCCTACACGACGCTCTTCCGATCTNNNGGTAGCAAGAAAAGTCCTTAT
AGCACACGTCTGAACTCTTTCCCTACACGACGCTCTTCCGATCTNNNTACATCAAAACATAAATAGGC
AGCACACGTCTGAACTCTTTCCCTACACGACGCTCTTCCGATCTNNNCTAGGAAGGTTGAAAAAATTT
AGCACACGTCTGAACTCTTTCCCTACACGACGCTCTTCCGATCTNNNATAAATAAGGACTTACTACTGC
AGCACACGTCTGAACTCTTTCCCTACACGACGCTCTTCCGATCTNNNAGATGGCAAATACAGTGATT
GAGCACACGTCTGAACTCTTTCCCTACACGACGCTCTTCCGATCTNNNTTATTTGTCTAAAAGAAAGC
CACACGTCTGAACTCTTTCCCTACACGACGCTCTTCCGATCTNNNACCTGGAAAGATGTTGGTAGCAT
AGCACACGTCTGAACTCTTTCCCTACACGACGCTCTTCCGATCTNNNTCATTAGCTTACAAAAACATGT
CACACGTCTGAACTCTTTCCCTACACGACGCTCTTCCGATCTNNNCTGGTACCAAAATTTGTATTAGG
AGCACACGTCTGAACTCTTTCCCTACACGACGCTCTTCCGATCTNNNATAATAGTAACTTGGACTAGT
GCACACGTCTGAACTCTTTCCCTACACGACGCTCTTCCGATCTNNNTCTGTCTCAAAAGAAATGCATA
ACACGTCTGAACTCTTTCCCTACACGACGCTCTTCCGATCTNNNCATTTTGAATTTTGCCATTGTAACA
AGCACACGTCTGAACTCTTTCCCTACACGACGCTCTTCCGATCTNNNGGAGACTACAATATCGTACAA
AGCACACGTCTGAACTCTTTCCCTACACGACGCTCTTCCGATCTNNNGGAGACTACAATATCGTACAA
ACACGTCTGAACTCTTTCCCTACACGACGCTCTTCCGATCTNNNATAGAGTACTTCCTTCCTGGCATCG
AGCACACGTCTGAACTCTTTCCCTACACGACGCTCTTCCGATCTNNNCACATAACCTTATAATGGTTCT
CACGTCTGAACTCTTTCCCTACACGACGCTCTTCCGATCTNNNGTAGAATTGATTTCAAGGAAACTCAT
AGCACACGTCTGAACTCTTTCCCTACACGACGCTCTTCCGATCTNNNAAATATGACTTATGAGATAGG
ACACGTCTGAACTCTTTCCCTACACGACGCTCTTCCGATCTNNNGACCCCTAAGTGCAAGATAGCATC
GCACACGTCTGAACTCTTTCCCTACACGACGCTCTTCCGATCTNNNACTAAGAGCATACACAGAGAAT
AGCACACGTCTGAACTCTTTCCCTACACGACGCTCTTCCGATCTNNNTTTCTGCTAAGTTTTCAATAG
GCACACGTCTGAACTCTTTCCCTACACGACGCTCTTCCGATCTNNNTATGTATTCAAGAACAGTATGG
AGCACACGTCTGAACTCTTTCCCTACACGACGCTCTTCCGATCTNNNACTTATCCTATGAGTTTATTCT
CACACGTCTGAACTCTTTCCCTACACGACGCTCTTCCGATCTNNNCCTATGGGAAAGAGTTGTCTCAT
CACACGTCTGAACTCTTTCCCTACACGACGCTCTTCCGATCTNNNCCATGGTAACTATGGGTTTTCAT
CACACGTCTGAACTCTTTCCCTACACGACGCTCTTCCGATCTNNNGACTTTTCTCTTTAGTCCCTGCAT

GCACACGTCTGAACTCTTTCCCTACACGACGCTCTTCCGATCTNNNGACTTTTCTCTTTAGTCCCTGCAT
GCACACGTCTGAACTCTTTCCCTACACGACGCTCTTCCGATCTNNNGACTTTTCTCTTTAGTCCCTGCAT
GCACACGTCTGAACTCTTTCCCTACACGACGCTCTTCCGATCTNNNCCTGAAAATATACCTACAGTCAA
GAGCACACGTCTGAACTCTTTCCCTACACGACGCTCTTCCGATCTNNNCTATCCTAAAGTATCATGTGG
AGCACACGTCTGAACTCTTTCCCTACACGACGCTCTTCCGATCTNNNTTACTCTAAAACATTTGGCAAC
CACACGTCTGAACTCTTTCCCTACACGACGCTCTTCCGATCTNNNAAAAGGTCTGCAATATTTTGAGCA
ACACGTCTGAACTCTTTCCCTACACGACGCTCTTCCGATCTNNNGCTTCTCTCACTTTTCTCTGCATCGGA
AGCACACGTCTGAACTCTTTCCCTACACGACGCTCTTCCGATCTNNNCAACTAAATCAGAATCCTTTCAG
GCACACGTCTGAACTCTTTCCCTACACGACGCTCTTCCGATCTNNNTGGCCACAGTATTTTATTTAAAACA
GCACACGTCTGAACTCTTTCCCTACACGACGCTCTTCCGATCTNNNATAGCCTTCATAGAGTGAATACCA
GAGCACACGTCTGAACTCTTTCCCTACACGACGCTCTTCCGATCTNNNTGTGTGTGTATGTACATATA
AGCACACGTCTGAACTCTTTCCCTACACGACGCTCTTCCGATCTNNNACGTTTCTCCATATATAGTAGAT
AGCACACGTCTGAACTCTTTCCCTACACGACGCTCTTCCGATCTNNNTTGTAGAAGAATCATCTGATCT
GCACACGTCTGAACTCTTTCCCTACACGACGCTCTTCCGATCTNNNTCACTTATAACAATGATCTCCAAC
GCACACGTCTGAACTCTTTCCCTACACGACGCTCTTCCGATCTNNNTTCTGAAGACATCCGGAATCCAT
ACACGTCTGAACTCTTTCCCTACACGACGCTCTTCCGATCTNNNGTTGTCATTCTTGTTTTGGTAGCAT
GAGCACACGTCTGAACTCTTTCCCTACACGACGCTCTTCCGATCTNNNCTTGAATAATGGATGATCTAC
ACACGTCTGAACTCTTTCCCTACACGACGCTCTTCCGATCTNNNCTAGACGAAGAGTTTCCCTAACATCG
AGCACACGTCTGAACTCTTTCCCTACACGACGCTCTTCCGATCTNNNTAAGAATATTTGCAAGTCTGAG
GAGCACACGTCTGAACTCTTTCCCTACACGACGCTCTTCCGATCTNNNGTGTCTCTCTCTCTCTCTATATA
AGCACACGTCTGAACTCTTTCCCTACACGACGCTCTTCCGATCTNNNGGACATATTCCAGAATTCATATT
GAGCACACGTCTGAACTCTTTCCCTACACGACGCTCTTCCGATCTNNNGTTTTGAGTACCAGTTAAATCT
AGCACACGTCTGAACTCTTTCCCTACACGACGCTCTTCCGATCTNNNTAAAATATCAATCCACTGCCTAT
AGCACACGTCTGAACTCTTTCCCTACACGACGCTCTTCCGATCTNNNCAGCTTAAAAAATTTAACCATGG
AGCACACGTCTGAACTCTTTCCCTACACGACGCTCTTCCGATCTNNNCAGAGCTGCTATTATATTTAGG
ACACGTCTGAACTCTTTCCCTACACGACGCTCTTCCGATCTNNNGATATTCTGAATGCCATTTCTTTCAT
AGCACACGTCTGAACTCTTTCCCTACACGACGCTCTTCCGATCTNNNTAATATTTCCTGAATGATGTTC
GCACACGTCTGAACTCTTTCCCTACACGACGCTCTTCCGATCTNNNATCGATTTTCATCAGGGATATTCA
GAGCACACGTCTGAACTCTTTCCCTACACGACGCTCTTCCGATCTNNNGTATGAAGAAATTCAGATGA
CACACGTCTGAACTCTTTCCCTACACGACGCTCTTCCGATCTNNNTTAGTAAGTAGTTGGCTTTGGCAT
AGCACACGTCTGAACTCTTTCCCTACACGACGCTCTTCCGATCTNNNTGATGGCATTAGGAAAATGAG
AGCACACGTCTGAACTCTTTCCCTACACGACGCTCTTCCGATCTNNNGCATTCTTTCTATGTTCTCATAT
AGCACACGTCTGAACTCTTTCCCTACACGACGCTCTTCCGATCTNNNGCATTCTTTCTATGTTCTCATAT
AGCACACGTCTGAACTCTTTCCCTACACGACGCTCTTCCGATCTNNNGCATTCTTTCTATGTTCTCATAT
AGCACACGTCTGAACTCTTTCCCTACACGACGCTCTTCCGATCTNNNGTGTGTTTTTTGTTTTTTGTTTT
GAGCACACGTCTGAACTCTTTCCCTACACGACGCTCTTCCGATCTNNNTAGTGGAAGAAAACTGGTTT
AGCACACGTCTGAACTCTTTCCCTACACGACGCTCTTCCGATCTNNNCATGATTTCATTGTTCAAACTG
AGAGCACACGTCTGAACTCTTTCCCTACACGACGCTCTTCCGATCTNNNATAGATACTTCATGATCTTTG
GAGCACACGTCTGAACTCTTTCCCTACACGACGCTCTTCCGATCTNNNCTATGCAGATTCAACTGATAC
ACACGTCTGAACTCTTTCCCTACACGACGCTCTTCCGATCTNNNGAAGGCACGTATGTATGTATGCAT
CACACGTCTGAACTCTTTCCCTACACGACGCTCTTCCGATCTNNNACCTCCCATTTAGTGAATATTCCAT
ACACGTCTGAACTCTTTCCCTACACGACGCTCTTCCGATCTNNNATTCCTGGTTGAGAATATGATATCAT
ACACGTCTGAACTCTTTCCCTACACGACGCTCTTCCGATCTNNNCTTTGAACTCCTAGGCTCTCCATCGG

AGCACACGTCTGAACTCTTTCCCTACACGACGCTCTTCCGATCTNNNATCTTAGCAGCTTAAGTTATAC
AGCACACGTCTGAACTCTTTCCCTACACGACGCTCTTCCGATCTNNNCTCTTTTCCCTCCACTAAAGCAT
CACACGTCTGAACTCTTTCCCTACACGACGCTCTTCCGATCTNNNATTCTCAGTTTTCCATGAGCCATCG
CACACGTCTGAACTCTTTCCCTACACGACGCTCTTCCGATCTNNNAATTTGGTTCCAGTTGACTATTAC
GCACACGTCTGAACTCTTTCCCTACACGACGCTCTTCCGATCTNNNGAGTTAGCACTTTAAAAGTAGA
AGCACACGTCTGAACTCTTTCCCTACACGACGCTCTTCCGATCTNNNAAGAACTTACTCATGTAACCAA
ACACGTCTGAACTCTTTCCCTACACGACGCTCTTCCGATCTNNNCTAGTCTATGGAACTTTGGTATGC
CACACGTCTGAACTCTTTCCCTACACGACGCTCTTCCGATCTNNNGTTACAAAAGACAGCTTTCTAACC
CACGTCTGAACTCTTTCCCTACACGACGCTCTTCCGATCTNNNGAACTATCCTCAGTGCATTGCATCGG
AGCACACGTCTGAACTCTTTCCCTACACGACGCTCTTCCGATCTNNNTTTGTACTTACCTCCTTCTGTAT
CACACGTCTGAACTCTTTCCCTACACGACGCTCTTCCGATCTNNNTTGTACAACCTAAGAAGAGAAAT
GCACACGTCTGAACTCTTTCCCTACACGACGCTCTTCCGATCTNNNATATCTTGGAATTCTTGGCAACAT
AGCACACGTCTGAACTCTTTCCCTACACGACGCTCTTCCGATCTNNNATTTAGCTTATTTTTTTGGCATC
AGCACACGTCTGAACTCTTTCCCTACACGACGCTCTTCCGATCTNNNATTTAGCTTATTTTTTTGGCATC
AGCACACGTCTGAACTCTTTCCCTACACGACGCTCTTCCGATCTNNNAAAATGAACTAATACACAAGA
GCACACGTCTGAACTCTTTCCCTACACGACGCTCTTCCGATCTNNNTAACAGTTACTGACCTATGTAAG
AGCACACGTCTGAACTCTTTCCCTACACGACGCTCTTCCGATCTNNNTAAATGAAACCACTTATTAGC
AGCACACGTCTGAACTCTTTCCCTACACGACGCTCTTCCGATCTNNNAATCAATATATACATGCACACA
GCACACGTCTGAACTCTTTCCCTACACGACGCTCTTCCGATCTNNNCTTGGGTGAAGTTTGTATAGTC
CACACGTCTGAACTCTTTCCCTACACGACGCTCTTCCGATCTNNNTGTAATGTTTAGTAAACCACTATC
ACACGTCTGAACTCTTTCCCTACACGACGCTCTTCCGATCTNNNCAACAAACATAGTGTCATGAGCATC
GCACACGTCTGAACTCTTTCCCTACACGACGCTCTTCCGATCTNNNTATAACATTTTCCCTTCCTGTACC
ACACGTCTGAACTCTTTCCCTACACGACGCTCTTCCGATCTNNNCACATTTCTCTGAGCATTTTCCATCG
GAGCACACGTCTGAACTCTTTCCCTACACGACGCTCTTCCGATCTNNNCCTCATAATGTCATGATTGAT
GCACACGTCTGAACTCTTTCCCTACACGACGCTCTTCCGATCTNNNCGTACACTTAAAAATGGTTACA
CACACGTCTGAACTCTTTCCCTACACGACGCTCTTCCGATCTNNNATGCATTAGAAAATTTATGTTGCC
AGCACACGTCTGAACTCTTTCCCTACACGACGCTCTTCCGATCTNNNTTGAAAGTAGTTTTTTAAATG
AGCACACGTCTGAACTCTTTCCCTACACGACGCTCTTCCGATCTNNNTAAGAATTTCTTACCAGTCAGAT
CACGTCTGAACTCTTTCCCTACACGACGCTCTTCCGATCTNNNCAGTATGTTGATCCTGCTATATCAT
AGCACACGTCTGAACTCTTTCCCTACACGACGCTCTTCCGATCTNNNCTTGAGCATAAAATGTTCCTA
AGCACACGTCTGAACTCTTTCCCTACACGACGCTCTTCCGATCTNNNAAAATCTCCAATATGCTAACCTA
GAGCACACGTCTGAACTCTTTCCCTACACGACGCTCTTCCGATCTNNNGATCATAATCACTCAACTTTG
GAGCACACGTCTGAACTCTTTCCCTACACGACGCTCTTCCGATCTNNNGATCATAATCACTCAACTTTG
CACACGTCTGAACTCTTTCCCTACACGACGCTCTTCCGATCTNNNATAAACCTCAAAAAAAGGGGACA
GAGCACACGTCTGAACTCTTTCCCTACACGACGCTCTTCCGATCTNNNATATTGAACACCCATAACAAAT
ACACGTCTGAACTCTTTCCCTACACGACGCTCTTCCGATCTNNNCATTAGTAATAAGCTCCACCCCATC
AGCACACGTCTGAACTCTTTCCCTACACGACGCTCTTCCGATCTNNNCTGCTGATATTTAGTTGTATGT
GAGCACACGTCTGAACTCTTTCCCTACACGACGCTCTTCCGATCTNNNCATATCAAGCAAGGACAATAT
GAGCACACGTCTGAACTCTTTCCCTACACGACGCTCTTCCGATCTNNNCCTCATAATTTCAGAGTAGTT
AGCACACGTCTGAACTCTTTCCCTACACGACGCTCTTCCGATCTNNNGAACCAAATTCCATGCTATTTCC
CACACGTCTGAACTCTTTCCCTACACGACGCTCTTCCGATCTNNNGCCTCACTGTATTTTAGAAATATC
AGCACACGTCTGAACTCTTTCCCTACACGACGCTCTTCCGATCTNNNTAAGGTCAAGTTAGAAAGTCAG
GCACACGTCTGAACTCTTTCCCTACACGACGCTCTTCCGATCTNNNGCCCTTAGAGTTTGAGAATCCAT

CACGTCTGAACTCTTTCCCTACACGACGCTCTTCCGATCTNNNGCTATTTTCATTTTGAAACCAAACATC
AGCACACGTCTGAACTCTTTCCCTACACGACGCTCTTCCGATCTNNNGATATATTCCTAGGTTTTTGCG
AGCACACGTCTGAACTCTTTCCCTACACGACGCTCTTCCGATCTNNNACAAAAATTAGATAAGCATGG
AGCACACGTCTGAACTCTTTCCCTACACGACGCTCTTCCGATCTNNNTATATCTACCATCTTAGATGTG
ACACGTCTGAACTCTTTCCCTACACGACGCTCTTCCGATCTNNNATTTCATTATCAACCCACGTTCATCG
AGCACACGTCTGAACTCTTTCCCTACACGACGCTCTTCCGATCTNNNTTCTTAACAGGACCTAGTAATG
GAGCACACGTCTGAACTCTTTCCCTACACGACGCTCTTCCGATCTNNNTAGAAATTGTGTCCTCAACAA
CACACGTCTGAACTCTTTCCCTACACGACGCTCTTCCGATCTNNNTTTTTTTTTTAAATGCATGGAGTC
AGCACACGTCTGAACTCTTTCCCTACACGACGCTCTTCCGATCTNNNGTTCTCAGAATAACAAAGGAAA
ACACGTCTGAACTCTTTCCCTACACGACGCTCTTCCGATCTNNNTATTATATGCTACTTGTTTCCCCCAT
AGCACACGTCTGAACTCTTTCCCTACACGACGCTCTTCCGATCTNNNCAAGTCTTACATACATTCCAAA
ACACGTCTGAACTCTTTCCCTACACGACGCTCTTCCGATCTNNNCTTATTTTGCCAATTTGTGTTCCATC
CACACGTCTGAACTCTTTCCCTACACGACGCTCTTCCGATCTNNNCTTAGACCATACATTTTGACACCAT
AGAGCACACGTCTGAACTCTTTCCCTACACGACGCTCTTCCGATCTNNNCTTACATTTCTGGTAGGAAA
AGCACACGTCTGAACTCTTTCCCTACACGACGCTCTTCCGATCTNNNTAATTTGTGATGGAAAAAGCA
GAGCACACGTCTGAACTCTTTCCCTACACGACGCTCTTCCGATCTNNNTTAACTCTGAAGACAATCTGTT
GCACACGTCTGAACTCTTTCCCTACACGACGCTCTTCCGATCTNNNAGGTTATTTTCTGAAATGGAAAG
GCACACGTCTGAACTCTTTCCCTACACGACGCTCTTCCGATCTNNNACCTTTGCGGTGAATATTACCATC
AGCACACGTCTGAACTCTTTCCCTACACGACGCTCTTCCGATCTNNNGAAAGTACAGAAGTATAGCTC
CACACGTCTGAACTCTTTCCCTACACGACGCTCTTCCGATCTNNNAAGTGGGTTCCTATGATGAGCATC
AGCACACGTCTGAACTCTTTCCCTACACGACGCTCTTCCGATCTNNNCCTAAGTTTATCGCATTATCAAT
GAGCACACGTCTGAACTCTTTCCCTACACGACGCTCTTCCGATCTNNNCCAGGCCTATTTTTTTTTTAAT
GCACACGTCTGAACTCTTTCCCTACACGACGCTCTTCCGATCTNNNGATCCATACTGAGACCCTTCCAT
CACACGTCTGAACTCTTTCCCTACACGACGCTCTTCCGATCTNNNATTTTAGCCATGGCCTATAATTACA
AGCACACGTCTGAACTCTTTCCCTACACGACGCTCTTCCGATCTNNNTCTAGGTGAAATATTAGCATTCT
AGCACACGTCTGAACTCTTTCCCTACACGACGCTCTTCCGATCTNNNTTGTATAAGCTGTATCTTATGC
AGCACACGTCTGAACTCTTTCCCTACACGACGCTCTTCCGATCTNNNCTGAATAATCACTGGTCATACTC
AGCACACGTCTGAACTCTTTCCCTACACGACGCTCTTCCGATCTNNNTGCCCTATAAAGTAACATATTCA
GCACACGTCTGAACTCTTTCCCTACACGACGCTCTTCCGATCTNNNCCAGACTAAAATTATCAAGCTTTC
CACACGTCTGAACTCTTTCCCTACACGACGCTCTTCCGATCTNNNGGTAGGCTATAATTCAAAATGACC
CACACGTCTGAACTCTTTCCCTACACGACGCTCTTCCGATCTNNNTGAAGCAATTATCTAATGAACTCCA
AGCACACGTCTGAACTCTTTCCCTACACGACGCTCTTCCGATCTNNNCTTCTCCCTGTATCTTCTCATCAT
GAGCACACGTCTGAACTCTTTCCCTACACGACGCTCTTCCGATCTNNNGAGTTGTACCAGTTGAGATA
GCACACGTCTGAACTCTTTCCCTACACGACGCTCTTCCGATCTNNNGAAACAATTTTTTATGAATAAAG
GAGCACACGTCTGAACTCTTTCCCTACACGACGCTCTTCCGATCTNNNATTGTATTTGTACCCAATTACA
GCACACGTCTGAACTCTTTCCCTACACGACGCTCTTCCGATCTNNNATCTTATATTTTGGGTTTGCTTTC
GCACACGTCTGAACTCTTTCCCTACACGACGCTCTTCCGATCTNNNAGAGTGAGAAGGATATTCTTAT
CACGTCTGAACTCTTTCCCTACACGACGCTCTTCCGATCTNNNAAATGCTAAGATCTTTTGCAAGCATC
AGCACACGTCTGAACTCTTTCCCTACACGACGCTCTTCCGATCTNNNAGGGTATAAGGTCTTTGTCTA
AGCACACGTCTGAACTCTTTCCCTACACGACGCTCTTCCGATCTNNNAGCTTTTAAAAATGACCAATAC
AGCACACGTCTGAACTCTTTCCCTACACGACGCTCTTCCGATCTNNNATAAGGAAATGAGTGTGAAG
CACACGTCTGAACTCTTTCCCTACACGACGCTCTTCCGATCTNNNGAGGACCACAGAAATTTATTATCC
CACACGTCTGAACTCTTTCCCTACACGACGCTCTTCCGATCTNNNTTCTCCTTCCTTCTTTCCTTCATCGGA

GAGCACACGTCTGAACTCTTTCCCTACACGACGCTCTTCCGATCTNNNCCTCCCTAATTTTATTCTTGTT
AGCACACGTCTGAACTCTTTCCCTACACGACGCTCTTCCGATCTNNNCAAGAGCTGATTTATCATTTAC
CACGTCTGAACTCTTTCCCTACACGACGCTCTTCCGATCTNNNCAAGACTTCTAAAATGACACTTCCATC
GCACACGTCTGAACTCTTTCCCTACACGACGCTCTTCCGATCTNNNCACAGTAAGTACTCAAACGCCAT
AGCACACGTCTGAACTCTTTCCCTACACGACGCTCTTCCGATCTNNNACCAGAGTGAAAAAAAAAATA
GCACACGTCTGAACTCTTTCCCTACACGACGCTCTTCCGATCTNNNATTATAAGCATCCTGACATGAAC
GCACACGTCTGAACTCTTTCCCTACACGACGCTCTTCCGATCTNNNTCAACGTGTACAATTGCTTACATC
AGCACACGTCTGAACTCTTTCCCTACACGACGCTCTTCCGATCTNNNGAAGCAAATTTCTCACTTGAGC
GAGCACACGTCTGAACTCTTTCCCTACACGACGCTCTTCCGATCTNNNCTGTTTTATCTGCATATACGA
GCACACGTCTGAACTCTTTCCCTACACGACGCTCTTCCGATCTNNNTGGTCTTCTGAAGTTTGATAATA
GCACACGTCTGAACTCTTTCCCTACACGACGCTCTTCCGATCTNNNTAACATTTCCTAGCTGCATTTATC
CACACGTCTGAACTCTTTCCCTACACGACGCTCTTCCGATCTNNNGAAAATTTTTCTCCCAACTCTAGCA
GAGCACACGTCTGAACTCTTTCCCTACACGACGCTCTTCCGATCTNNNGTTTGTATCTCCATGCCTATC
GCACACGTCTGAACTCTTTCCCTACACGACGCTCTTCCGATCTNNNTTATTGAGTGTCAAAAATTCAGT
GCACACGTCTGAACTCTTTCCCTACACGACGCTCTTCCGATCTNNNATTATATAATCCACTAAGCCCAGC
GCACACGTCTGAACTCTTTCCCTACACGACGCTCTTCCGATCTNNNCAAATAAATTTTCTTGCCTGAAG
GCACACGTCTGAACTCTTTCCCTACACGACGCTCTTCCGATCTNNNTTCCCTGTCTCTTCATATCTATAC
GCACACGTCTGAACTCTTTCCCTACACGACGCTCTTCCGATCTNNNTGACATCTTCCATAAACCAAAACA
GAGCACACGTCTGAACTCTTTCCCTACACGACGCTCTTCCGATCTNNNAAAAGGAGATTTTCATCCAAT
GAGCACACGTCTGAACTCTTTCCCTACACGACGCTCTTCCGATCTNNNAACCTGAATTCTATACCTGAA
GCACACGTCTGAACTCTTTCCCTACACGACGCTCTTCCGATCTNNNTATTCTCGATTCTTTCTATGGTTC
AGCACACGTCTGAACTCTTTCCCTACACGACGCTCTTCCGATCTNNNTTTCAGCTACAAAAACACATTTC
GAGCACACGTCTGAACTCTTTCCCTACACGACGCTCTTCCGATCTNNNTTGGTGATTAAGTTATTAGAC
GAGCACACGTCTGAACTCTTTCCCTACACGACGCTCTTCCGATCTNNNCTACTTTTTAGGTGAGTTGCT
GCACACGTCTGAACTCTTTCCCTACACGACGCTCTTCCGATCTNNNGTACTAAAAAGAAGAGTAGGG
AGCACACGTCTGAACTCTTTCCCTACACGACGCTCTTCCGATCTNNNTATAAAGTGTGTATTCCTAACG
CACACGTCTGAACTCTTTCCCTACACGACGCTCTTCCGATCTNNNTATATGTACGTGTGTATTTGTCTC
AGCACACGTCTGAACTCTTTCCCTACACGACGCTCTTCCGATCTNNNAAAGTCCCTTGATGAATTTTAA
GAGCACACGTCTGAACTCTTTCCCTACACGACGCTCTTCCGATCTNNNGAAAAGTCCCTTGATGAATTT
CACACGTCTGAACTCTTTCCCTACACGACGCTCTTCCGATCTNNNGTACAACTTAGTTCATTGATTCACA
AGCACACGTCTGAACTCTTTCCCTACACGACGCTCTTCCGATCTNNNGCTCTGATACTAACAACTAGTT
AGCACACGTCTGAACTCTTTCCCTACACGACGCTCTTCCGATCTNNNCAGTAAGAATGGTCCATAATTC
AGCACACGTCTGAACTCTTTCCCTACACGACGCTCTTCCGATCTNNNATGTGAAATTCAATGTCTCTAA
GAGCACACGTCTGAACTCTTTCCCTACACGACGCTCTTCCGATCTNNNCAAATTTGTATGTTGTACGAA
CACACGTCTGAACTCTTTCCCTACACGACGCTCTTCCGATCTNNNCTTCTCAAAGGTCTAGTGATATTC
GAGCACACGTCTGAACTCTTTCCCTACACGACGCTCTTCCGATCTNNNTTTTTTCTATAGAGTAACGTT
ACACGTCTGAACTCTTTCCCTACACGACGCTCTTCCGATCTNNNCAGTTACTGTGACTTTAGGATGCAT
GCACACGTCTGAACTCTTTCCCTACACGACGCTCTTCCGATCTNNNCTCAAGAAGGGAAATTCCTACCAT
GAGCACACGTCTGAACTCTTTCCCTACACGACGCTCTTCCGATCTNNNCAAAAAAAAAAAAAAAAAAG
GAGCACACGTCTGAACTCTTTCCCTACACGACGCTCTTCCGATCTNNNAATATGGAATCCACAAATAAT
ACACGTCTGAACTCTTTCCCTACACGACGCTCTTCCGATCTNNNCACCAAGATATTATGGAAGAGATC
ACACGTCTGAACTCTTTCCCTACACGACGCTCTTCCGATCTNNNTCAACCTAGATTGTCTTAAAGAACAT
ACACGTCTGAACTCTTTCCCTACACGACGCTCTTCCGATCTNNNTATTCCAAACCTCCTTTACTTTCATCG

AGCACACGTCTGAACTCTTTCCCTACACGACGCTCTTCCGATCTNNNCTTCTAATCAGCTTGATGAACTA
GAGCACACGTCTGAACTCTTTCCCTACACGACGCTCTTCCGATCTNNNGTGACAGATTGGATAAAGAA
AGCACACGTCTGAACTCTTTCCCTACACGACGCTCTTCCGATCTNNNTTCTAAATTACGGTTGAAGTTAC
GCACACGTCTGAACTCTTTCCCTACACGACGCTCTTCCGATCTNNNAAATATGGATATTATGATGCCTG
GAGCACACGTCTGAACTCTTTCCCTACACGACGCTCTTCCGATCTNNNTTTATGCAAGATGAAGTGGTT
GCACACGTCTGAACTCTTTCCCTACACGACGCTCTTCCGATCTNNNCATTATCTTTCAGCAGGATTTCCA
GCACACGTCTGAACTCTTTCCCTACACGACGCTCTTCCGATCTNNNGATTTCCTTTGAGTTAGGGTTCA
GAGCACACGTCTGAACTCTTTCCCTACACGACGCTCTTCCGATCTNNNCTGTACAGAGATATTACAGGA
CACACGTCTGAACTCTTTCCCTACACGACGCTCTTCCGATCTNNNGTATTTAGTATCCAAGATCTGGGCA
GAGCACACGTCTGAACTCTTTCCCTACACGACGCTCTTCCGATCTNNNAAAACAACACACACAAACATA
GCACACGTCTGAACTCTTTCCCTACACGACGCTCTTCCGATCTNNNACATATTTCACCAAACCTAAAATCA
GCACACGTCTGAACTCTTTCCCTACACGACGCTCTTCCGATCTNNNACCCAGCATTTTTTATTTCATATCA
AGCACACGTCTGAACTCTTTCCCTACACGACGCTCTTCCGATCTNNNTTCACACATCACACATACACCAT
AGCACACGTCTGAACTCTTTCCCTACACGACGCTCTTCCGATCTNNNATTTCAGGTATATATCCGTGTG
AGCACACGTCTGAACTCTTTCCCTACACGACGCTCTTCCGATCTNNNATTTCAGGTATATATCCGTGTG
GCACACGTCTGAACTCTTTCCCTACACGACGCTCTTCCGATCTNNNAAGCATTTCAGGTATATATCCGCA
GAGCACACGTCTGAACTCTTTCCCTACACGACGCTCTTCCGATCTNNNGTATTGAATTTCTAGGTAGGT
GCACACGTCTGAACTCTTTCCCTACACGACGCTCTTCCGATCTNNNGAGAAAGAGAAAGAAAGAAGTT
ACACGTCTGAACTCTTTCCCTACACGACGCTCTTCCGATCTNNNCAGTGAAGCTATAGACACTTTCATCC
AGCACACGTCTGAACTCTTTCCCTACACGACGCTCTTCCGATCTNNNCATTGTTGAGACTTATCCTTAAC
GCACACGTCTGAACTCTTTCCCTACACGACGCTCTTCCGATCTNNNGGATCAAACATACAATAGATAGC
AGCACACGTCTGAACTCTTTCCCTACACGACGCTCTTCCGATCTNNNTGATGTAAAAATATTTGAAACAC
GAGCACACGTCTGAACTCTTTCCCTACACGACGCTCTTCCGATCTNNNAAAGGTAAGTAATATTTTGTG
GAGCACACGTCTGAACTCTTTCCCTACACGACGCTCTTCCGATCTNNNTTATATATAGGCATATGCTCAC
GAGCACACGTCTGAACTCTTTCCCTACACGACGCTCTTCCGATCTNNNGATTACTAGCTTCCTCATACAT
GAGCACACGTCTGAACTCTTTCCCTACACGACGCTCTTCCGATCTNNNTGATGAGGAGGCAATAATTAT
GAGCACACGTCTGAACTCTTTCCCTACACGACGCTCTTCCGATCTNNNTGATGAGGAGGCAATAATTAT
GCACACGTCTGAACTCTTTCCCTACACGACGCTCTTCCGATCTNNNAGGATTCCTCTTTTCAAATAGTAC
GCACACGTCTGAACTCTTTCCCTACACGACGCTCTTCCGATCTNNNAGGATTCCTCTTTTCAAATAGTAC
AGCACACGTCTGAACTCTTTCCCTACACGACGCTCTTCCGATCTNNNACGGATTAGAGTAGAAATGATG
AGCACACGTCTGAACTCTTTCCCTACACGACGCTCTTCCGATCTNNNTCTTTCTAATTTCTAATTGCAGG
GAGCACACGTCTGAACTCTTTCCCTACACGACGCTCTTCCGATCTNNNATTTTTTTTGAGAACCAGCTTT
GAGCACACGTCTGAACTCTTTCCCTACACGACGCTCTTCCGATCTNNNCAATATGTCTGCCATTCTTTGC
AGCACACGTCTGAACTCTTTCCCTACACGACGCTCTTCCGATCTNNNGTTCCATAAGCATAATAACATGA
AGCACACGTCTGAACTCTTTCCCTACACGACGCTCTTCCGATCTNNNCTTTGAACTTTCTCCAGATTCCAT
GAGCACACGTCTGAACTCTTTCCCTACACGACGCTCTTCCGATCTNNNGGACAATACTGTGAAAAACAA
GAGCACACGTCTGAACTCTTTCCCTACACGACGCTCTTCCGATCTNNNGTAAGTGTCTCATATTCAGAA
CACACGTCTGAACTCTTTCCCTACACGACGCTCTTCCGATCTNNNTTGTGAGGCTTTCATTCATTCATCGG
AGCACACGTCTGAACTCTTTCCCTACACGACGCTCTTCCGATCTNNNAATAAAAGGAAAACAATGAAA
CACACGTCTGAACTCTTTCCCTACACGACGCTCTTCCGATCTNNNAATAACCGAATGGCTAATACCCATC
AGCACACGTCTGAACTCTTTCCCTACACGACGCTCTTCCGATCTNNNGTAGAGACACAGAGAGAAGGG
AGAGCACACGTCTGAACTCTTTCCCTACACGACGCTCTTCCGATCTNNNTTTTAATGTTAGTCCAGGTCA
AGCACACGTCTGAACTCTTTCCCTACACGACGCTCTTCCGATCTNNNTTCATCATCACATAAGCATTTTT

AGCACACGTCTGAACTCTTTCCCTACACGACGCTCTTCCGATCTNNNTGATGTGTGTATGGGTATATA
AGCACACGTCTGAACTCTTTCCCTACACGACGCTCTTCCGATCTNNNAATTAATTTTCTACAGCTGCTGC
CACACGTCTGAACTCTTTCCCTACACGACGCTCTTCCGATCTNNNCTATTTTTTGCTCCTCTGCTCATCG
GCACACGTCTGAACTCTTTCCCTACACGACGCTCTTCCGATCTNNNATTTATTTAGCCAGTTCCATACTCA
GCACACGTCTGAACTCTTTCCCTACACGACGCTCTTCCGATCTNNNCTCATATCTTATCCTACCATTGTCA
GAGCACACGTCTGAACTCTTTCCCTACACGACGCTCTTCCGATCTNNNTTTTGTTTCAGGAAAAATCAA
GCACACGTCTGAACTCTTTCCCTACACGACGCTCTTCCGATCTNNNTAACCATCAGTGACTTAGTTATTC
AGCACACGTCTGAACTCTTTCCCTACACGACGCTCTTCCGATCTNNNATTTCAAACTCATCTTACTGAA
CACACGTCTGAACTCTTTCCCTACACGACGCTCTTCCGATCTNNNCAATGTCTCTGGGGTTTAGACATC
AGCACACGTCTGAACTCTTTCCCTACACGACGCTCTTCCGATCTNNNCCTGTTTCTGATAAAAAGAACA
GCACACGTCTGAACTCTTTCCCTACACGACGCTCTTCCGATCTNNNGAGTTGAGAGGTAGTGTTATAA
AGCACACGTCTGAACTCTTTCCCTACACGACGCTCTTCCGATCTNNNTGTAGTCTAAACATTAGGCTTA
GCACACGTCTGAACTCTTTCCCTACACGACGCTCTTCCGATCTNNNAAAAATAAGCCAACCAACAAACAT
GCACACGTCTGAACTCTTTCCCTACACGACGCTCTTCCGATCTNNNTCCATTTCTTCTAGATTTTCTAGT
AGCACACGTCTGAACTCTTTCCCTACACGACGCTCTTCCGATCTNNNAAAAATCACAATCCCTGTTTGCA
GCACACGTCTGAACTCTTTCCCTACACGACGCTCTTCCGATCTNNNTGTAAACAGTATAAAGAGGGAA
GAGCACACGTCTGAACTCTTTCCCTACACGACGCTCTTCCGATCTNNNGATGACTCTCTGTTATCTATTT
AGCACACGTCTGAACTCTTTCCCTACACGACGCTCTTCCGATCTNNNTTTTTCCTAGTATGGTGTATCAT
GCACACGTCTGAACTCTTTCCCTACACGACGCTCTTCCGATCTNNNCATTCTCATGAGATCTGATGGCA
AGCACACGTCTGAACTCTTTCCCTACACGACGCTCTTCCGATCTNNNTTCATTTCACTCTTACCATAAGT
AGCACACGTCTGAACTCTTTCCCTACACGACGCTCTTCCGATCTNNNTTTGATATATTCCACAATATGC
AGCACACGTCTGAACTCTTTCCCTACACGACGCTCTTCCGATCTNNNCTGTAGTAGATAGAGTTGTGT
AGCACACGTCTGAACTCTTTCCCTACACGACGCTCTTCCGATCTNNNTTATAGAATATGTTCTAGCACCT
GCACACGTCTGAACTCTTTCCCTACACGACGCTCTTCCGATCTNNNCAGTGTGTGATAGTTTTCAATGC
AGCACACGTCTGAACTCTTTCCCTACACGACGCTCTTCCGATCTNNNCAAGGAAGTGAAGTCAAGAGC
GCACACGTCTGAACTCTTTCCCTACACGACGCTCTTCCGATCTNNNGAATAAGGCTATTAGGGAGTTTA
AGCACACGTCTGAACTCTTTCCCTACACGACGCTCTTCCGATCTNNNAAGTCAGAATTTATTTATTGGG
GCACACGTCTGAACTCTTTCCCTACACGACGCTCTTCCGATCTNNNCAAGTCTTAGGAATTCTCAGCCA
AGCACACGTCTGAACTCTTTCCCTACACGACGCTCTTCCGATCTNNNGTTTATCTTTTTGTGTTGATCAC
AGCACACGTCTGAACTCTTTCCCTACACGACGCTCTTCCGATCTNNNCTAATCCTACTCACTGCAAGCA
ACACGTCTGAACTCTTTCCCTACACGACGCTCTTCCGATCTNNNGCTAAGAGCATTGTAGGTATTCAT
AGCACACGTCTGAACTCTTTCCCTACACGACGCTCTTCCGATCTNNNATAGTGATTCTCAAAATAAACG
GAGCACACGTCTGAACTCTTTCCCTACACGACGCTCTTCCGATCTNNNCTGTAATTCCAGCTACTCAGCA
GAGCACACGTCTGAACTCTTTCCCTACACGACGCTCTTCCGATCTNNNTATATCTGTATATATGGAACA
AGCACACGTCTGAACTCTTTCCCTACACGACGCTCTTCCGATCTNNNGCCTACTGTAATTCAACATTTT
AGCACACGTCTGAACTCTTTCCCTACACGACGCTCTTCCGATCTNNNGTGAAACTCTGTCTCTCTGTCAT
CACACGTCTGAACTCTTTCCCTACACGACGCTCTTCCGATCTNNNATACACACACACTACTTCTTCCATC
GCACACGTCTGAACTCTTTCCCTACACGACGCTCTTCCGATCTNNNTTTTTCTCTCTCTCTTTTTCTCTCAT
GCACACGTCTGAACTCTTTCCCTACACGACGCTCTTCCGATCTNNNTTTTTCTCTCTCTCTTTTTCTCTCAT
GCACACGTCTGAACTCTTTCCCTACACGACGCTCTTCCGATCTNNNCTTCCACTTTGATTAAGTAGTTT
CACACGTCTGAACTCTTTCCCTACACGACGCTCTTCCGATCTNNNTATTATGCTCAGTTCATACTTCTCA
AGCACACGTCTGAACTCTTTCCCTACACGACGCTCTTCCGATCTNNNGTATAAAGAAGATGGTCTGTTT
AGCACACGTCTGAACTCTTTCCCTACACGACGCTCTTCCGATCTNNNAGACTTCTCAGTGTCCATAACA

AGCACACGTCTGAACTCTTTCCCTACACGACGCTCTTCCGATCTNNNAGATATATATGCGTGTGTGTAT
GAGCACACGTCTGAACTCTTTCCCTACACGACGCTCTTCCGATCTNNNGTTCTAACTTCTCCAAATCTCTC
AGCACACGTCTGAACTCTTTCCCTACACGACGCTCTTCCGATCTNNNGGTAAACCAGAGAGTAAAAGTA
ACACGTCTGAACTCTTTCCCTACACGACGCTCTTCCGATCTNNNAAAGAGAGACCACATGGAAACATCG
AGCACACGTCTGAACTCTTTCCCTACACGACGCTCTTCCGATCTNNNATCTCTAAAGATGATGGACAGT
GCACACGTCTGAACTCTTTCCCTACACGACGCTCTTCCGATCTNNNAAGTTCATTGTTTAGATTTCCTTCA
GAGCACACGTCTGAACTCTTTCCCTACACGACGCTCTTCCGATCTNNNACAACACCTGTACTAAAACAGC
AGCACACGTCTGAACTCTTTCCCTACACGACGCTCTTCCGATCTNNNCCAAATAATGCCTATCCATAAAT
GCACACGTCTGAACTCTTTCCCTACACGACGCTCTTCCGATCTNNNGGTGCTAGAGAAGAGACTTACAT
CACACGTCTGAACTCTTTCCCTACACGACGCTCTTCCGATCTNNNCTTAGTTCTTTACAGAAAGCAAGCA
CACACGTCTGAACTCTTTCCCTACACGACGCTCTTCCGATCTNNNCTTAGTTCTTTACAGAAAGCAAGCA
GCACACGTCTGAACTCTTTCCCTACACGACGCTCTTCCGATCTNNNACAAGATCTGATGGTTTTAAAAG
GCACACGTCTGAACTCTTTCCCTACACGACGCTCTTCCGATCTNNNAAATTCTATTCATTCTTCAAGGCCA
GCACACGTCTGAACTCTTTCCCTACACGACGCTCTTCCGATCTNNNCTTGTAGTGAGCTGATATCTCCAT
GCACACGTCTGAACTCTTTCCCTACACGACGCTCTTCCGATCTNNNATGAGTATCTTTCTGTGTCTCTCAT
CACGTCTGAACTCTTTCCCTACACGACGCTCTTCCGATCTNNNTATGTAAACCCACATACTTATCCATC
GCACACGTCTGAACTCTTTCCCTACACGACGCTCTTCCGATCTNNNTTCCTCAAATGGAAAAATATCACC
GAGCACACGTCTGAACTCTTTCCCTACACGACGCTCTTCCGATCTNNNGAATGTCAAGGCCACAATTTCA
AGCACACGTCTGAACTCTTTCCCTACACGACGCTCTTCCGATCTNNNCAGAAACCATTCCACTTTAAATCA
AGCACACGTCTGAACTCTTTCCCTACACGACGCTCTTCCGATCTNNNAGAGATTGAGCTGCTGAAATCA
AGCACACGTCTGAACTCTTTCCCTACACGACGCTCTTCCGATCTNNNAAGTCAGAATTTATTTATTGGG
AGCACACGTCTGAACTCTTTCCCTACACGACGCTCTTCCGATCTNNNATTAGGAAAATGCAAATCCAAA
GAGCACACGTCTGAACTCTTTCCCTACACGACGCTCTTCCGATCTNNNCCTATGGTATCTATTTAAGGTG
AGAGCACACGTCTGAACTCTTTCCCTACACGACGCTCTTCCGATCTNNNAATCACAACTTTGCCTACAGC
AGCACACGTCTGAACTCTTTCCCTACACGACGCTCTTCCGATCTNNNCCGTTAGTTTGTTCTCTCTTACA
AGCACACGTCTGAACTCTTTCCCTACACGACGCTCTTCCGATCTNNNTACAGAGAAGTAAATAACGAAA
GAGCACACGTCTGAACTCTTTCCCTACACGACGCTCTTCCGATCTNNNACAAAAAAAATGTTGTAATGA
AGCACACGTCTGAACTCTTTCCCTACACGACGCTCTTCCGATCTNNNCACACTATTCAAAGATGATCAC
ACACGTCTGAACTCTTTCCCTACACGACGCTCTTCCGATCTNNNATATGTATGCATGCACTCTCCATCGG
GAGCACACGTCTGAACTCTTTCCCTACACGACGCTCTTCCGATCTNNNCTCTCTTACAGAAATACACACA
GAGCACACGTCTGAACTCTTTCCCTACACGACGCTCTTCCGATCTNNNTCTGGAAAAGTGAACAAAAAT
AGCACACGTCTGAACTCTTTCCCTACACGACGCTCTTCCGATCTNNNAGCTAATTCATTTCAATATGACA
CACACGTCTGAACTCTTTCCCTACACGACGCTCTTCCGATCTNNNGAAGAAACCAGAAGGAGTTGCAT
AGCACACGTCTGAACTCTTTCCCTACACGACGCTCTTCCGATCTNNNTCTGTAATGACATTGAAAAATG
AGCACACGTCTGAACTCTTTCCCTACACGACGCTCTTCCGATCTNNNGTATCGTTTCTTAAGTCCTCTTT
AGCACACGTCTGAACTCTTTCCCTACACGACGCTCTTCCGATCTNNNTCTCAGTATGGGTGTATGAGCA
GCACACGTCTGAACTCTTTCCCTACACGACGCTCTTCCGATCTNNNCAATCCAATTCCCCTTGAAATCAT
AGCACACGTCTGAACTCTTTCCCTACACGACGCTCTTCCGATCTNNNCCAAAAATATACACTTTTCCTGAC
AGCACACGTCTGAACTCTTTCCCTACACGACGCTCTTCCGATCTNNNCTGAATCCTTTGCAACAAAAACA
AGAGCACACGTCTGAACTCTTTCCCTACACGACGCTCTTCCGATCTNNNCTAGAGTATGCAGCAGAGTA
AGCACACGTCTGAACTCTTTCCCTACACGACGCTCTTCCGATCTNNNTTCTTCTACTTGTCTATATGTGG
AGCACACGTCTGAACTCTTTCCCTACACGACGCTCTTCCGATCTNNNTCTATTCATTCCCAAAATATGTGC
AGCACACGTCTGAACTCTTTCCCTACACGACGCTCTTCCGATCTNNNGTGTCATTTTGATGAGCTATTTA

GCACACGTCTGAACTCTTTCCCTACACGACGCTCTTCCGATCTNNNTGAACAAAACAGATAACAATCA
GCACACGTCTGAACTCTTTCCCTACACGACGCTCTTCCGATCTNNNTAATCATTATGCCACACACATAT
GCACACGTCTGAACTCTTTCCCTACACGACGCTCTTCCGATCTNNNCATTCCTAATGTGTGAAACTATC
GCACACGTCTGAACTCTTTCCCTACACGACGCTCTTCCGATCTNNNCCCAGTTTTATAATTAGTTGTGT
ACACGTCTGAACTCTTTCCCTACACGACGCTCTTCCGATCTNNNAAATTTTTGGTAATGATTGCAAGC
CACACGTCTGAACTCTTTCCCTACACGACGCTCTTCCGATCTNNNTTTCTTTGTATGTGTATGATGACC
AGCACACGTCTGAACTCTTTCCCTACACGACGCTCTTCCGATCTNNNATTTCTTATTGAATAAAGGCCTC
AGCACACGTCTGAACTCTTTCCCTACACGACGCTCTTCCGATCTNNNAATAACTCTGCCTGGAAATCCAT
CACACGTCTGAACTCTTTCCCTACACGACGCTCTTCCGATCTNNNTAATTTGTGGGGAGATACTTTAA
GCACACGTCTGAACTCTTTCCCTACACGACGCTCTTCCGATCTNNNCTTCCCTTCAATAAGAAAAGACC
AGCACACGTCTGAACTCTTTCCCTACACGACGCTCTTCCGATCTNNNTTTGACTGTTTAATAGTGACA
AGCACACGTCTGAACTCTTTCCCTACACGACGCTCTTCCGATCTNNNTACCATCAAAACTCTCTTTTAGA
GCACACGTCTGAACTCTTTCCCTACACGACGCTCTTCCGATCTNNNAAATAGATTCTGTGAGACTCAAA
GCACACGTCTGAACTCTTTCCCTACACGACGCTCTTCCGATCTNNNAAGGGGAAAGAAAAAATATAT
AGCACACGTCTGAACTCTTTCCCTACACGACGCTCTTCCGATCTNNNATAATAAGGCTTGAAATCAGG
AGCACACGTCTGAACTCTTTCCCTACACGACGCTCTTCCGATCTNNNTAGATCTACAATCTCCTGTTCAC
GCACACGTCTGAACTCTTTCCCTACACGACGCTCTTCCGATCTNNNATAAAAGAAAGTCACCTGAAATC
GCACACGTCTGAACTCTTTCCCTACACGACGCTCTTCCGATCTNNNGTGGTTGAGATGATTACCTCCAT
GCACACGTCTGAACTCTTTCCCTACACGACGCTCTTCCGATCTNNNCATACAGAGTATAGCCACTTGCA
GCACACGTCTGAACTCTTTCCCTACACGACGCTCTTCCGATCTNNNCATACAGAGTATAGCCACTTGCA
GCACACGTCTGAACTCTTTCCCTACACGACGCTCTTCCGATCTNNNCATACAGAGTATAGCCACTTGCA
GCACACGTCTGAACTCTTTCCCTACACGACGCTCTTCCGATCTNNNTCCCATTCATAGATATAATTTGG
AGCACACGTCTGAACTCTTTCCCTACACGACGCTCTTCCGATCTNNNTATTTGCCATTATCTCTTCACAC
GCACACGTCTGAACTCTTTCCCTACACGACGCTCTTCCGATCTNNNAATGTTAACCATGAGATAGAAAC
GCACACGTCTGAACTCTTTCCCTACACGACGCTCTTCCGATCTNNNAATGTTAACCATGAGATAGAAAC
AGCACACGTCTGAACTCTTTCCCTACACGACGCTCTTCCGATCTNNNGGGAAGCCTTGTTGATTTATC
CACACGTCTGAACTCTTTCCCTACACGACGCTCTTCCGATCTNNNCAGAGCAAAACTATGTCTCAACATC
AGCACACGTCTGAACTCTTTCCCTACACGACGCTCTTCCGATCTNNNAGTCTATTGTGAATGATTGAA
GCACACGTCTGAACTCTTTCCCTACACGACGCTCTTCCGATCTNNNATTGGTAAGAAAGCCTATAGAA
CACACGTCTGAACTCTTTCCCTACACGACGCTCTTCCGATCTNNNAATAGTTGATTGAGGAATGTCTTC
GCACACGTCTGAACTCTTTCCCTACACGACGCTCTTCCGATCTNNNTAATCCTGGGTCTATGAAGGCAT
AGCACACGTCTGAACTCTTTCCCTACACGACGCTCTTCCGATCTNNNCCATCTGAATTACTGCTGAGCAT
AGCACACGTCTGAACTCTTTCCCTACACGACGCTCTTCCGATCTNNNGCATGAACCATTACTAGGAACA
AGCACACGTCTGAACTCTTTCCCTACACGACGCTCTTCCGATCTNNNGACAGACAAAAAGATGAGGTA
AGCACACGTCTGAACTCTTTCCCTACACGACGCTCTTCCGATCTNNNTAACTAGGACCTGCATATCTAT
GCACACGTCTGAACTCTTTCCCTACACGACGCTCTTCCGATCTNNNAGAATTTTGATGTGTCCAGAGC
AGCACACGTCTGAACTCTTTCCCTACACGACGCTCTTCCGATCTNNNAAGACCAATTATTGAAAAACG
AGCACACGTCTGAACTCTTTCCCTACACGACGCTCTTCCGATCTNNNGAGAAATTCTGTATGAAAACCA
GAGCACACGTCTGAACTCTTTCCCTACACGACGCTCTTCCGATCTNNNTTAACTCATTGAAAAAGCTAA
AGCACACGTCTGAACTCTTTCCCTACACGACGCTCTTCCGATCTNNNACATAATAAATTACTCAAGGG
AGCACACGTCTGAACTCTTTCCCTACACGACGCTCTTCCGATCTNNNTGTGTTTATATTTGGACAAATG
AGCACACGTCTGAACTCTTTCCCTACACGACGCTCTTCCGATCTNNNACTTTTCTTTTTTTTTTTGGTTAG
CACACGTCTGAACTCTTTCCCTACACGACGCTCTTCCGATCTNNNTAGATTTCTAGCCTTGGCTTTACAT

GAGCACACGTCTGAACTCTTTCCCTACACGACGCTCTTCCGATCTNNNCTGAAATGAAGCATTTTTTTC

GCACACGTCTGAACTCTTTCCCTACACGACGCTCTTCCGATCTNNNCAGGGGAATATATATATAAACAC

AGCACACGTCTGAACTCTTTCCCTACACGACGCTCTTCCGATCTNNNGCAGTTTGGTTATATTTATAGC

AGCACACGTCTGAACTCTTTCCCTACACGACGCTCTTCCGATCTNNNCTAATAGTGATGTAAGGTGCCC

AGCACACGTCTGAACTCTTTCCCTACACGACGCTCTTCCGATCTNNNCTGTATTTCTAAAAAATGCCTCC

GAGCACACGTCTGAACTCTTTCCCTACACGACGCTCTTCCGATCTNNNGATAAATGACAGCGTCTATATA

GCACACGTCTGAACTCTTTCCCTACACGACGCTCTTCCGATCTNNNAAACTTCACTGAATGTATTACCTC

AGCACACGTCTGAACTCTTTCCCTACACGACGCTCTTCCGATCTNNNAGTTTATAGCCAGATTTGAGGCA

AGCACACGTCTGAACTCTTTCCCTACACGACGCTCTTCCGATCTNNNTTTTAAGTCATTTTCATACTGCCC

AGCACACGTCTGAACTCTTTCCCTACACGACGCTCTTCCGATCTNNNTTTTAAGTCATTTTCATACTGCCC

AGCACACGTCTGAACTCTTTCCCTACACGACGCTCTTCCGATCTNNNTTTTAAGTCATTTTCATACTGCCC

AGCACACGTCTGAACTCTTTCCCTACACGACGCTCTTCCGATCTNNNTAAAGAAATTCTAGTTCCTCTCA

GCACACGTCTGAACTCTTTCCCTACACGACGCTCTTCCGATCTNNNCTGTTCTCGTGTATGTGTATGCAT

GCACACGTCTGAACTCTTTCCCTACACGACGCTCTTCCGATCTNNNCTGTTCTCGTGTATGTGTATGCAT

CACACGTCTGAACTCTTTCCCTACACGACGCTCTTCCGATCTNNNTTCCGAATACTGGGATTAGACATC

GCACACGTCTGAACTCTTTCCCTACACGACGCTCTTCCGATCTNNNAAAATGCCTTGAGTTGACAAACA

AGCACACGTCTGAACTCTTTCCCTACACGACGCTCTTCCGATCTNNNTTAACATAGAAAACTCTTGCATC

AGCACACGTCTGAACTCTTTCCCTACACGACGCTCTTCCGATCTNNNCTATATGTTGATGTTGCAATTTC

CACACGTCTGAACTCTTTCCCTACACGACGCTCTTCCGATCTNNNCAAAGCTGTGAATTCTCTGTCATCG

CACACGTCTGAACTCTTTCCCTACACGACGCTCTTCCGATCTNNNCAAAGCTGTGAATTCTCTGTCATCG

GCACACGTCTGAACTCTTTCCCTACACGACGCTCTTCCGATCTNNNGTATCATAAGCAAACAAGAAGGC

AGCACACGTCTGAACTCTTTCCCTACACGACGCTCTTCCGATCTNNNTTGGATACTGGAATGATTGATA

AGCACACGTCTGAACTCTTTCCCTACACGACGCTCTTCCGATCTNNNTTGGATACTGGAATGATTGATA

AGCACACGTCTGAACTCTTTCCCTACACGACGCTCTTCCGATCTNNNTTACAACTTATCAGGATTGATGA

GAGCACACGTCTGAACTCTTTCCCTACACGACGCTCTTCCGATCTNNNTAAAGGTATGTAGAGTACTCT

GCACACGTCTGAACTCTTTCCCTACACGACGCTCTTCCGATCTNNNTGTTTGTTTTTATATCCATGCTTCA

CACACGTCTGAACTCTTTCCCTACACGACGCTCTTCCGATCTNNNCATAGTAAGGTTTATCAAAAGCACA

AGCACACGTCTGAACTCTTTCCCTACACGACGCTCTTCCGATCTNNNGATTTTTGTCTATCCCATCATGC

GAGCACACGTCTGAACTCTTTCCCTACACGACGCTCTTCCGATCTNNNGAGATTATAGTTTGTGAGCCA

GAGCACACGTCTGAACTCTTTCCCTACACGACGCTCTTCCGATCTNNNATATCATTCAAGAACTCTCTGT

AGCACACGTCTGAACTCTTTCCCTACACGACGCTCTTCCGATCTNNNCTCTCCTGATTCACAAAATGACAT

AGCACACGTCTGAACTCTTTCCCTACACGACGCTCTTCCGATCTNNNCTTTCCTTTCTGCTCAATTACCATC

GAGCACACGTCTGAACTCTTTCCCTACACGACGCTCTTCCGATCTNNNTTCCCTAACCAGAACTTCTCCAT

GAGCACACGTCTGAACTCTTTCCCTACACGACGCTCTTCCGATCTNNNCATACACAGAATGGTATTTTCA

AGCACACGTCTGAACTCTTTCCCTACACGACGCTCTTCCGATCTNNNATGTAAAGGTAAATCTGTTTCCT

GAGCACACGTCTGAACTCTTTCCCTACACGACGCTCTTCCGATCTNNNCTTTGATTGATGCACCTTCTCA

AGAGCACACGTCTGAACTCTTTCCCTACACGACGCTCTTCCGATCTNNNCCTAAACCAGTGTGTGTTAT

AGCACACGTCTGAACTCTTTCCCTACACGACGCTCTTCCGATCTNNNCCTTCATTTCCAAATTAGGAATA

GCACACGTCTGAACTCTTTCCCTACACGACGCTCTTCCGATCTNNNAAGAAAAAGTGTTAGGAACAACC

AGAGCACACGTCTGAACTCTTTCCCTACACGACGCTCTTCCGATCTNNNCCGATTTCCAGAATAAATTCT

AGCACACGTCTGAACTCTTTCCCTACACGACGCTCTTCCGATCTNNNTACCTAGCAACTGTCTTTATATT

GCACACGTCTGAACTCTTTCCCTACACGACGCTCTTCCGATCTNNNTTTTTGGAATGTTAATGAATTGCC

AGAGCACACGTCTGAACTCTTTCCCTACACGACGCTCTTCCGATCTNNNTCTTTGTAAATTTCCCAATCA

AGCACACGTCTGAACTCTTTCCCTACACGACGCTCTTCCGATCTNNNGAGATGAGATCACAGACAGAC
AGCACACGTCTGAACTCTTTCCCTACACGACGCTCTTCCGATCTNNNTATTCTTTTATTTGTGGTAAG
CACACGTCTGAACTCTTTCCCTACACGACGCTCTTCCGATCTNNNCAGAAGGTAGAGAAGTGTTCCAT
CACACGTCTGAACTCTTTCCCTACACGACGCTCTTCCGATCTNNNCAAGATGTATCTCAAAGGTTTTC
GCACACGTCTGAACTCTTTCCCTACACGACGCTCTTCCGATCTNNNTTTAATTTTCCCTTTTGGTCTCCA
GAGCACACGTCTGAACTCTTTCCCTACACGACGCTCTTCCGATCTNNNGTATGCATCTGATTTTTGAA
AGCACACGTCTGAACTCTTTCCCTACACGACGCTCTTCCGATCTNNNAACTTACCTTTTGCTGTACTAC
GCACACGTCTGAACTCTTTCCCTACACGACGCTCTTCCGATCTNNNTTTCTTCACATGTCACCTTTACAT
AGCACACGTCTGAACTCTTTCCCTACACGACGCTCTTCCGATCTNNNAGGAGAAAATAGATGTCCCAC
AGCACACGTCTGAACTCTTTCCCTACACGACGCTCTTCCGATCTNNNCCTATTCTGAGTTTAGATCCAAT
AGCACACGTCTGAACTCTTTCCCTACACGACGCTCTTCCGATCTNNNAAAATCTTTACTGAAGTACCA
CACACGTCTGAACTCTTTCCCTACACGACGCTCTTCCGATCTNNNATCATCAAATGATCCAAGGGCAT
ACACGTCTGAACTCTTTCCCTACACGACGCTCTTCCGATCTNNNGCCCTACCATAATTAAAATCATCCAT
ACACGTCTGAACTCTTTCCCTACACGACGCTCTTCCGATCTNNNGAAAATTAAGTCACATGTTTGCCAT
AGCACACGTCTGAACTCTTTCCCTACACGACGCTCTTCCGATCTNNNATACAAAAGATGAAGAGAAA
AGCACACGTCTGAACTCTTTCCCTACACGACGCTCTTCCGATCTNNNCTCTCTCTATCTCTCTCTGCAT
AGAGCACACGTCTGAACTCTTTCCCTACACGACGCTCTTCCGATCTNNNTATTAGGACAAAAGCTATAG
ACACGTCTGAACTCTTTCCCTACACGACGCTCTTCCGATCTNNNAAAGAGGGCAAGATACATATTTCA
AGCACACGTCTGAACTCTTTCCCTACACGACGCTCTTCCGATCTNNNTCTAGAGCCTAGTAGCTTATAG
AGCACACGTCTGAACTCTTTCCCTACACGACGCTCTTCCGATCTNNNGTTAATAGAAAACCAAACTCCA
AGCACACGTCTGAACTCTTTCCCTACACGACGCTCTTCCGATCTNNNAGACAATTAAATTGATTGAGGT
GCACACGTCTGAACTCTTTCCCTACACGACGCTCTTCCGATCTNNNATTCTAAGCACATTAAAGGTCTA
GAGCACACGTCTGAACTCTTTCCCTACACGACGCTCTTCCGATCTNNNAACTATAGACCTAAAGGCCA
AGCACACGTCTGAACTCTTTCCCTACACGACGCTCTTCCGATCTNNNATAACAGAAACATTGCACATAC
AGCACACGTCTGAACTCTTTCCCTACACGACGCTCTTCCGATCTNNNAGTTCTTATACTGCAAGGATGC
AGCACACGTCTGAACTCTTTCCCTACACGACGCTCTTCCGATCTNNNAGTTCTTATACTGCAAGGATGC
AGCACACGTCTGAACTCTTTCCCTACACGACGCTCTTCCGATCTNNNCATTAACATTTGGATAGAAGC
CACACGTCTGAACTCTTTCCCTACACGACGCTCTTCCGATCTNNNTATACTGAGGAAGAGTAAGTTTC
AGCACACGTCTGAACTCTTTCCCTACACGACGCTCTTCCGATCTNNNACAACCAACAGTTAAAATTCTA
CACACGTCTGAACTCTTTCCCTACACGACGCTCTTCCGATCTNNNTTTGATGTTGATGACAATAATGA
AGCACACGTCTGAACTCTTTCCCTACACGACGCTCTTCCGATCTNNNTTGAAGCTATGACAATCAATAT
AGCACACGTCTGAACTCTTTCCCTACACGACGCTCTTCCGATCTNNNCTAGGAAAGTCAAAAGATAA
AGCACACGTCTGAACTCTTTCCCTACACGACGCTCTTCCGATCTNNNAAGTTTACTTGCTCCTGAGTAC
AGCACACGTCTGAACTCTTTCCCTACACGACGCTCTTCCGATCTNNNGAAGAATGGAAACTACCCATA
GCACACGTCTGAACTCTTTCCCTACACGACGCTCTTCCGATCTNNNGTGTGTGTGTCTATGTAGCA
CACACGTCTGAACTCTTTCCCTACACGACGCTCTTCCGATCTNNNTAATCCCAGCTACTTAGGAGCATC
GCACACGTCTGAACTCTTTCCCTACACGACGCTCTTCCGATCTNNNGAGACTGTCAAGATCTCATCCAT
AGCACACGTCTGAACTCTTTCCCTACACGACGCTCTTCCGATCTNNNTGAGTAATTACTTCTGGAGTTT
AGCACACGTCTGAACTCTTTCCCTACACGACGCTCTTCCGATCTNNNACAGCCCTGAATTGAATTATCC
AGCACACGTCTGAACTCTTTCCCTACACGACGCTCTTCCGATCTNNNCTTTGTGACTTAGCTCTGATTA
AGCACACGTCTGAACTCTTTCCCTACACGACGCTCTTCCGATCTNNNCTTCCCTCATCCTTAATCTGCAT
AGCACACGTCTGAACTCTTTCCCTACACGACGCTCTTCCGATCTNNNCTGTGTGTGATCTGCTTTATCA
GCACACGTCTGAACTCTTTCCCTACACGACGCTCTTCCGATCTNNNTTTGGATTTCTCCTTTTAGTTCTC

GCACACGTCTGAACTCTTTCCCTACACGACGCTCTTCCGATCTNNNTTTGGATTTCTCCTTTTAGTTCTC

GCACACGTCTGAACTCTTTCCCTACACGACGCTCTTCCGATCTNNNCTAAGATCTGGACTCTAGATGT

AGCACACGTCTGAACTCTTTCCCTACACGACGCTCTTCCGATCTNNNAGATTAAAAAAATTAGCCTCG

AGCACACGTCTGAACTCTTTCCCTACACGACGCTCTTCCGATCTNNNACACTACACGTAAACACGTACA

AGCACACGTCTGAACTCTTTCCCTACACGACGCTCTTCCGATCTNNNACACTACACGTAAACACGTACA

AGCACACGTCTGAACTCTTTCCCTACACGACGCTCTTCCGATCTNNNGAACCTCAAGTAAGAAGTTC

AGCACACGTCTGAACTCTTTCCCTACACGACGCTCTTCCGATCTNNNCCAAAATCTGTAACTCTTTTCT

GCACACGTCTGAACTCTTTCCCTACACGACGCTCTTCCGATCTNNNACACTTAGTAATATCTGATGCA

AGCACACGTCTGAACTCTTTCCCTACACGACGCTCTTCCGATCTNNNGAGCACTTATTCATGGTAATAA

GCACACGTCTGAACTCTTTCCCTACACGACGCTCTTCCGATCTNNNGTACATGAATTTTTAATATTCTA

AGCACACGTCTGAACTCTTTCCCTACACGACGCTCTTCCGATCTNNNAGTTGAATATGGTCAATTTTTG

AGCACACGTCTGAACTCTTTCCCTACACGACGCTCTTCCGATCTNNNCTGCTATAGTGACTGTGTATG

AGCACACGTCTGAACTCTTTCCCTACACGACGCTCTTCCGATCTNNNTTTCCTTAGGAATATGGAAAA

AGCACACGTCTGAACTCTTTCCCTACACGACGCTCTTCCGATCTNNNTTTCCTTAGGAATATGGAAAA

AGCACACGTCTGAACTCTTTCCCTACACGACGCTCTTCCGATCTNNNCTGGAAACAACCATAATAAGT

GAGCACACGTCTGAACTCTTTCCCTACACGACGCTCTTCCGATCTNNNATATTAGATTATGCCTTCTTT

CACACGTCTGAACTCTTTCCCTACACGACGCTCTTCCGATCTNNNATCTGACAATTCCAGATCCTCATC

AGCACACGTCTGAACTCTTTCCCTACACGACGCTCTTCCGATCTNNNTAATCACAGAAGCCATGTTAA

CACACGTCTGAACTCTTTCCCTACACGACGCTCTTCCGATCTNNNCATTAGCTAGTTTGTGAAAAGTA

ACACGTCTGAACTCTTTCCCTACACGACGCTCTTCCGATCTNNNCTTGAATTTTCAAAAAAAATGCCCA

GCACACGTCTGAACTCTTTCCCTACACGACGCTCTTCCGATCTNNNTAACAATTTTTTAAAGGACTGC

AGCACACGTCTGAACTCTTTCCCTACACGACGCTCTTCCGATCTNNNTAAACAAGTGAGTTAGATTTT

ACACGTCTGAACTCTTTCCCTACACGACGCTCTTCCGATCTNNNACTAATGTATCTAGCCAATAGCCAT

AGCACACGTCTGAACTCTTTCCCTACACGACGCTCTTCCGATCTNNNATACTTGCATTTTATTGCCATTC

GAGCACACGTCTGAACTCTTTCCCTACACGACGCTCTTCCGATCTNNNTGAACACTCTTTTACCTACTG

GCACACGTCTGAACTCTTTCCCTACACGACGCTCTTCCGATCTNNNTACTCTCTGTTCAGTCTCTCCAT

AGAGCACACGTCTGAACTCTTTCCCTACACGACGCTCTTCCGATCTNNNAAGAATAATTTCCTGGTAG

AGCACACGTCTGAACTCTTTCCCTACACGACGCTCTTCCGATCTNNNCGTATTTATTTTAAAACAGCT

AGCACACGTCTGAACTCTTTCCCTACACGACGCTCTTCCGATCTNNNTCACACAGTAACAATTACAAA

GCACACGTCTGAACTCTTTCCCTACACGACGCTCTTCCGATCTNNNTTTAATATTAAAGTCAGGCAGT

ACACGTCTGAACTCTTTCCCTACACGACGCTCTTCCGATCTNNNCACCGACTTTACATCATAGTCCATCG

GAGCACACGTCTGAACTCTTTCCCTACACGACGCTCTTCCGATCTNNNTTAGCGAGTTTAAATATTTCA

GAGCACACGTCTGAACTCTTTCCCTACACGACGCTCTTCCGATCTNNNATGATACTGCTTGAATCAGA

GCACACGTCTGAACTCTTTCCCTACACGACGCTCTTCCGATCTNNNAAGACATGTGCTTTGTGTAACA

AGCACACGTCTGAACTCTTTCCCTACACGACGCTCTTCCGATCTNNNTGGCAATTTGAGTAAAATAAG

GAGCACACGTCTGAACTCTTTCCCTACACGACGCTCTTCCGATCTNNNTAATGGAAAATTGAAGCTGT

AGCACACGTCTGAACTCTTTCCCTACACGACGCTCTTCCGATCTNNNTATATGGATCAATAATACATG

GCACACGTCTGAACTCTTTCCCTACACGACGCTCTTCCGATCTNNNTAAAATTAGCAAAGCATTCCTTT

CACACGTCTGAACTCTTTCCCTACACGACGCTCTTCCGATCTNNNCTGTTTTTGTCACTCATCCTCATCG

AGCACACGTCTGAACTCTTTCCCTACACGACGCTCTTCCGATCTNNNTCGGAAGAGAAAAATGTAGA

GCACACGTCTGAACTCTTTCCCTACACGACGCTCTTCCGATCTNNNATCTTGAGTGACACAAAAGAGC

GCACACGTCTGAACTCTTTCCCTACACGACGCTCTTCCGATCTNNNCTGTAGATTAGTAAAATTGAGC

GAGCACACGTCTGAACTCTTTCCCTACACGACGCTCTTCCGATCTNNNCTAAGTTTGCAGTAATGTTA

AGCACACGTCTGAACTCTTTCCCTACACGACGCTCTTCCGATCTNNNCAACCAGTATTATCCATTTCTTC
GCACACGTCTGAACTCTTTCCCTACACGACGCTCTTCCGATCTNNNTCAATTCATGATTTCATTGTTCA
GCACACGTCTGAACTCTTTCCCTACACGACGCTCTTCCGATCTNNNTTTCTCTAGCTTCCCCTAAGCATC
GAGCACACGTCTGAACTCTTTCCCTACACGACGCTCTTCCGATCTNNNGAAACTATTCTCAAACTGAAG
AGCACACGTCTGAACTCTTTCCCTACACGACGCTCTTCCGATCTNNNATCTACTGTCCCTATGCATTTA
AGCACACGTCTGAACTCTTTCCCTACACGACGCTCTTCCGATCTNNNCTCTTGTTCTTCTCTCTTTCTCC
CACACGTCTGAACTCTTTCCCTACACGACGCTCTTCCGATCTNNNATGTGCTCTCTCTCTCTTCATCG
AGCACACGTCTGAACTCTTTCCCTACACGACGCTCTTCCGATCTNNNATTCATGTCAAGGAATAGAAT
AGCACACGTCTGAACTCTTTCCCTACACGACGCTCTTCCGATCTNNNATTTCTACCTCACAAACTCTTAT
AGCACACGTCTGAACTCTTTCCCTACACGACGCTCTTCCGATCTNNNCTGTTAATAGCCGATGATTCTA
GAGCACACGTCTGAACTCTTTCCCTACACGACGCTCTTCCGATCTNNNACATTCTAGAAAAATCAGAG
AGCACACGTCTGAACTCTTTCCCTACACGACGCTCTTCCGATCTNNNAATGAGCAAAAAGACACTGA
AGCACACGTCTGAACTCTTTCCCTACACGACGCTCTTCCGATCTNNNTCTCCAAGATACTGTATTAAGT
GCACACGTCTGAACTCTTTCCCTACACGACGCTCTTCCGATCTNNNTTAAAGCTGAGATCTGAAGAAT
AGCACACGTCTGAACTCTTTCCCTACACGACGCTCTTCCGATCTNNNATATCGATTCAACAAGTTTCCT
ACACGTCTGAACTCTTTCCCTACACGACGCTCTTCCGATCTNNNTAGTATAAGGTTGCTTTACTCAC
AGCACACGTCTGAACTCTTTCCCTACACGACGCTCTTCCGATCTNNNCATTCATAAAATGGAGACTGA
ACACGTCTGAACTCTTTCCCTACACGACGCTCTTCCGATCTNNNTTAAAATACTTTAGGGGGCATCCA
AGCACACGTCTGAACTCTTTCCCTACACGACGCTCTTCCGATCTNNNCAATGTATTGGTATTTGGAGA
AGCACACGTCTGAACTCTTTCCCTACACGACGCTCTTCCGATCTNNNGTACCCTCATATATTACAGACA
CACACGTCTGAACTCTTTCCCTACACGACGCTCTTCCGATCTNNNCACTCTCAACTAAGTGACCACATC
AGCACACGTCTGAACTCTTTCCCTACACGACGCTCTTCCGATCTNNNGTAGGTTGTATGAATGTGTAT
ACACGTCTGAACTCTTTCCCTACACGACGCTCTTCCGATCTNNNGGTTTTAAAAAGAAGAAGATGGT
CACACGTCTGAACTCTTTCCCTACACGACGCTCTTCCGATCTNNNGCTTGGGTTCATGAAATTATTAAC
GCACACGTCTGAACTCTTTCCCTACACGACGCTCTTCCGATCTNNNCTTCTTTCTTTCTTTCTTTTCCTCA
GAGCACACGTCTGAACTCTTTCCCTACACGACGCTCTTCCGATCTNNNTGTGTGTGTGTATGTATA
GCACACGTCTGAACTCTTTCCCTACACGACGCTCTTCCGATCTNNNTGAGAAGATCTGACACTCTTAC
GCACACGTCTGAACTCTTTCCCTACACGACGCTCTTCCGATCTNNNGACTTACTGTTAGACATAGTACA
CACACGTCTGAACTCTTTCCCTACACGACGCTCTTCCGATCTNNNTTCAGGATAATGGAAAACCTCCA
GCACACGTCTGAACTCTTTCCCTACACGACGCTCTTCCGATCTNNNTATAGTGATGAATAAAACCAC
CACACGTCTGAACTCTTTCCCTACACGACGCTCTTCCGATCTNNNTCTTTTGTTTCTCATTTCTCAAACAT
AGCACACGTCTGAACTCTTTCCCTACACGACGCTCTTCCGATCTNNNATTTCGATTTTTGAAATGTACC
GCACACGTCTGAACTCTTTCCCTACACGACGCTCTTCCGATCTNNNGCTATTCTAATTCTGAAGTTCT
AGCACACGTCTGAACTCTTTCCCTACACGACGCTCTTCCGATCTNNNCAGATTCTATCACAGGTTTATT
GCACACGTCTGAACTCTTTCCCTACACGACGCTCTTCCGATCTNNNCCTTATCATGTGGAAAAAATT
CACACGTCTGAACTCTTTCCCTACACGACGCTCTTCCGATCTNNNTAAATAGCATGCAATCTGAGTAC
AGCACACGTCTGAACTCTTTCCCTACACGACGCTCTTCCGATCTNNNCTTTGAATTGTGAAGCTTAGG
GAGCACACGTCTGAACTCTTTCCCTACACGACGCTCTTCCGATCTNNNTATAGAAACTATGTCATGTG
CACACGTCTGAACTCTTTCCCTACACGACGCTCTTCCGATCTNNNAATGACTTACCTTGGTGTATGCAT
AGCACACGTCTGAACTCTTTCCCTACACGACGCTCTTCCGATCTNNNCTCTGATTGGTTTGATTTGTAT
AGCACACGTCTGAACTCTTTCCCTACACGACGCTCTTCCGATCTNNNAAAAGACGAAACCTAAGTTAG
AGCACACGTCTGAACTCTTTCCCTACACGACGCTCTTCCGATCTNNNAAAAGACGAAACCTAAGTTAG
GAGCACACGTCTGAACTCTTTCCCTACACGACGCTCTTCCGATCTNNNATACTCATCAGAACTTTAGA

AGCACACGTCTGAACTCTTTCCCTACACGACGCTCTTCCGATCTNNNAATTTCATTTTTATGCCACGTACA
AGCACACGTCTGAACTCTTTCCCTACACGACGCTCTTCCGATCTNNNGCTCATCTTTTTCTTCTTGAATCA
GCACACGTCTGAACTCTTTCCCTACACGACGCTCTTCCGATCTNNNTGTCTTCTTGTACTTTCTTATCTCA
GAGCACACGTCTGAACTCTTTCCCTACACGACGCTCTTCCGATCTNNNGATTGTGTCTTTTGTGATTGT
AGCACACGTCTGAACTCTTTCCCTACACGACGCTCTTCCGATCTNNNGGTGCATTATTAACAATAACTG
AGCACACGTCTGAACTCTTTCCCTACACGACGCTCTTCCGATCTNNNGATATATTCTTTTTGTCTTTGCC
AGCACACGTCTGAACTCTTTCCCTACACGACGCTCTTCCGATCTNNNCTTATAACAGTGTATGTCCCTCC
AGCACACGTCTGAACTCTTTCCCTACACGACGCTCTTCCGATCTNNNTTGTATAGCCCATATATGAACTT
GAGCACACGTCTGAACTCTTTCCCTACACGACGCTCTTCCGATCTNNNTTAAGTAGTTTAATTGGTTAG
AGCACACGTCTGAACTCTTTCCCTACACGACGCTCTTCCGATCTNNNTGAAAATGTAAAAGTACACACA
GAGCACACGTCTGAACTCTTTCCCTACACGACGCTCTTCCGATCTNNNTAAAGTCAATACATGCTATGG
GCACACGTCTGAACTCTTTCCCTACACGACGCTCTTCCGATCTNNNACCTCAGAATCATCTCAGATTCAT
GAGCACACGTCTGAACTCTTTCCCTACACGACGCTCTTCCGATCTNNNATTTTGAACAGAGTAGGGGAT
GAGCACACGTCTGAACTCTTTCCCTACACGACGCTCTTCCGATCTNNNCAATTTGATGTGTTGTCTTTTG
AGCACACGTCTGAACTCTTTCCCTACACGACGCTCTTCCGATCTNNNCTGTAAAGCAAGTTCCACTTCAT
GAGCACACGTCTGAACTCTTTCCCTACACGACGCTCTTCCGATCTNNNAGTTCTACTTGTAGAGATCTTT
GAGCACACGTCTGAACTCTTTCCCTACACGACGCTCTTCCGATCTNNNGTCGTTTGAGATGTAAATCTTC
GAGCACACGTCTGAACTCTTTCCCTACACGACGCTCTTCCGATCTNNNTGATGGTATTTTAATTTCCTGT
GAGCACACGTCTGAACTCTTTCCCTACACGACGCTCTTCCGATCTNNNGGTCTGGACATAATTTGACAT
GAGCACACGTCTGAACTCTTTCCCTACACGACGCTCTTCCGATCTNNNGACATATCCATCTTCTCACATA
AGCACACGTCTGAACTCTTTCCCTACACGACGCTCTTCCGATCTNNNAAAATACTGTTTTATACCTGCTG
AGCACACGTCTGAACTCTTTCCCTACACGACGCTCTTCCGATCTNNNAGTAGGCTCCATGTTCATAAACA
AGCACACGTCTGAACTCTTTCCCTACACGACGCTCTTCCGATCTNNNCAAATTTTGGAATTTACTGCTT
ACACGTCTGAACTCTTTCCCTACACGACGCTCTTCCGATCTNNNAGATTAGGCTTTGTTTGTCCCATCGG
GAGCACACGTCTGAACTCTTTCCCTACACGACGCTCTTCCGATCTNNNTAATGGACAACTCACATGTAAC
AGCACACGTCTGAACTCTTTCCCTACACGACGCTCTTCCGATCTNNNAATGAAGCACTTTAGTTATGGA
AGCACACGTCTGAACTCTTTCCCTACACGACGCTCTTCCGATCTNNNATCAAATACCTGATCCAATGAA
AGCACACGTCTGAACTCTTTCCCTACACGACGCTCTTCCGATCTNNNCTCTACGGGATATCATACATATC
AGCACACGTCTGAACTCTTTCCCTACACGACGCTCTTCCGATCTNNNCTCTACGGGATATCATACATATC
AGCACACGTCTGAACTCTTTCCCTACACGACGCTCTTCCGATCTNNNATTTTCTCAATTCTTTGTGTTCCC
GAGCACACGTCTGAACTCTTTCCCTACACGACGCTCTTCCGATCTNNNCATATATGTATGAACATGTGTC
AGCACACGTCTGAACTCTTTCCCTACACGACGCTCTTCCGATCTNNNACATATATGTATGAACATGTGT
AGCACACGTCTGAACTCTTTCCCTACACGACGCTCTTCCGATCTNNNACAAAAATCAAATGGAAGTAGA
AGCACACGTCTGAACTCTTTCCCTACACGACGCTCTTCCGATCTNNNAGATTATTTTAAGGTCATGCAAC
AGCACACGTCTGAACTCTTTCCCTACACGACGCTCTTCCGATCTNNNTTCATTACTACACAGAAGATACC
AGCACACGTCTGAACTCTTTCCCTACACGACGCTCTTCCGATCTNNNTATATCATAGTCCTTACTGCTTCC
AGCACACGTCTGAACTCTTTCCCTACACGACGCTCTTCCGATCTNNNAATTTCATCTAGACTGAACTCAT
AGCACACGTCTGAACTCTTTCCCTACACGACGCTCTTCCGATCTNNNCCTGATTACTATGTACCTCATTTC
AGCACACGTCTGAACTCTTTCCCTACACGACGCTCTTCCGATCTNNNTAAGTGCCCCCTTAAATCATCAT
AGCACACGTCTGAACTCTTTCCCTACACGACGCTCTTCCGATCTNNNGTCCACACTCCTTATATTACATTC
GAGCACACGTCTGAACTCTTTCCCTACACGACGCTCTTCCGATCTNNNCTAAACTCCAGATTTGACATAA
CACACGTCTGAACTCTTTCCCTACACGACGCTCTTCCGATCTNNNAAAAAGCTGGATGCATCTTTCATC
GAGCACACGTCTGAACTCTTTCCCTACACGACGCTCTTCCGATCTNNNTACTTTATCAAAATTGTTTGCA

AGCACACGTCTGAACTCTTTCCCTACACGACGCTCTTCCGATCTNNNAGGAAAGTGTGCTTACAGAAC
AGCACACGTCTGAACTCTTTCCCTACACGACGCTCTTCCGATCTNNNCACTACTCAGTCTTGTATGAAA
GCACACGTCTGAACTCTTTCCCTACACGACGCTCTTCCGATCTNNNACTCTTCTTTTCTGAACTCTAATC
AGCACACGTCTGAACTCTTTCCCTACACGACGCTCTTCCGATCTNNNTAAAAAGTTCTTTTGGTCCAAA
AGCACACGTCTGAACTCTTTCCCTACACGACGCTCTTCCGATCTNNNCACAGTTAAATTGAATGGGAA
AGCACACGTCTGAACTCTTTCCCTACACGACGCTCTTCCGATCTNNNCAGAAAGAAAGAGAGAAGGG
AGCACACGTCTGAACTCTTTCCCTACACGACGCTCTTCCGATCTNNNCATTCATGTTATCTTGTGTGTA
AGCACACGTCTGAACTCTTTCCCTACACGACGCTCTTCCGATCTNNNAACTATGAATTTGAGAAGTAC
GCACACGTCTGAACTCTTTCCCTACACGACGCTCTTCCGATCTNNNAAAAGTATGCTTTCTGTTATCAA
AGCACACGTCTGAACTCTTTCCCTACACGACGCTCTTCCGATCTNNNAGTTGTAAATGAAAAATCCCAT
GCACACGTCTGAACTCTTTCCCTACACGACGCTCTTCCGATCTNNNCATATCCCTTTAAAGAATTCTCC
GCACACGTCTGAACTCTTTCCCTACACGACGCTCTTCCGATCTNNNGTTAAATAACCAACAATCTAGG
AGCACACGTCTGAACTCTTTCCCTACACGACGCTCTTCCGATCTNNNTAGACTTTTCCAATGATCTTACT
GAGCACACGTCTGAACTCTTTCCCTACACGACGCTCTTCCGATCTNNNATAGCTACAATCCACTGAATT
CACACGTCTGAACTCTTTCCCTACACGACGCTCTTCCGATCTNNNTATTGTGTGCCATATCCTATTCCAT
AGCACACGTCTGAACTCTTTCCCTACACGACGCTCTTCCGATCTNNNGGGTTTATAATTTTCACTGAAC
AGCACACGTCTGAACTCTTTCCCTACACGACGCTCTTCCGATCTNNNTCAATCCTCCCTATTAATTTTG
AGCACACGTCTGAACTCTTTCCCTACACGACGCTCTTCCGATCTNNNATTGTGGGAAAAACAATAAAA
AGCACACGTCTGAACTCTTTCCCTACACGACGCTCTTCCGATCTNNNTAGGTGATTATCTGCTAGCTAG
GAGCACACGTCTGAACTCTTTCCCTACACGACGCTCTTCCGATCTNNNTTCATAGTCTCTCAGTCTATTT
AGCACACGTCTGAACTCTTTCCCTACACGACGCTCTTCCGATCTNNNTGGGTACTCAAATAGGAGAG
GCACACGTCTGAACTCTTTCCCTACACGACGCTCTTCCGATCTNNNCTTTAGCATTACTTTTTGTGTTA
GCACACGTCTGAACTCTTTCCCTACACGACGCTCTTCCGATCTNNNCTTTAGCATTACTTTTTGTGTTA
AGCACACGTCTGAACTCTTTCCCTACACGACGCTCTTCCGATCTNNNAAGGGAAAAAATGTTATCCTT
ACACGTCTGAACTCTTTCCCTACACGACGCTCTTCCGATCTNNNTCATATGATGGAGGTCTGTCCATCG
AGCACACGTCTGAACTCTTTCCCTACACGACGCTCTTCCGATCTNNNAGGCTGAAAAATATATTCCCT
AGCACACGTCTGAACTCTTTCCCTACACGACGCTCTTCCGATCTNNNGGAGGATGAGATTTAGAGAA
AGCACACGTCTGAACTCTTTCCCTACACGACGCTCTTCCGATCTNNNCCTACGTGTAAATGCTAAAAG
GAGCACACGTCTGAACTCTTTCCCTACACGACGCTCTTCCGATCTNNNTTTTCCAATTAATAGTACTTTT
CACACGTCTGAACTCTTTCCCTACACGACGCTCTTCCGATCTNNNGTTAAGTGTGGCCATATGACCAT
AGCACACGTCTGAACTCTTTCCCTACACGACGCTCTTCCGATCTNNNATTACACTTATTAAGTCTGTCCC
AGCACACGTCTGAACTCTTTCCCTACACGACGCTCTTCCGATCTNNNAGAAGTCTGTTAACAAACAAAT
CACACGTCTGAACTCTTTCCCTACACGACGCTCTTCCGATCTNNNGTCTCCTGGAGCTAATCTAGCATC
AGCACACGTCTGAACTCTTTCCCTACACGACGCTCTTCCGATCTNNNGTTGTAAAACCAGAGTTGTAA
CACACGTCTGAACTCTTTCCCTACACGACGCTCTTCCGATCTNNNGCACTGGTAGAATTTCATAAAAA
CACACGTCTGAACTCTTTCCCTACACGACGCTCTTCCGATCTNNNAATTGTCAAACTGTGTTAGATTTC
AGCACACGTCTGAACTCTTTCCCTACACGACGCTCTTCCGATCTNNNCTGTTCTAGTGAAATATCAAAC
AGCACACGTCTGAACTCTTTCCCTACACGACGCTCTTCCGATCTNNNCTATTCTAGGCACTTGGAATA
AGCACACGTCTGAACTCTTTCCCTACACGACGCTCTTCCGATCTNNNTCATTTTGCAACTGTAATTACAT
GAGCACACGTCTGAACTCTTTCCCTACACGACGCTCTTCCGATCTNNNGATTTTTACATTCATGGAGC
AGCACACGTCTGAACTCTTTCCCTACACGACGCTCTTCCGATCTNNNCTATTCTAGGCACTTGGAATA
GCACACGTCTGAACTCTTTCCCTACACGACGCTCTTCCGATCTNNNAGTACATCTGTAGTGTAGTTCA
CACACGTCTGAACTCTTTCCCTACACGACGCTCTTCCGATCTNNNACCAACATAGAGATTCTAGAAATC

GCACACGTCTGAACTCTTTCCCTACACGACGCTCTTCCGATCTNNNAGAATTGAATGTCACCTAAAATC
GAGCACACGTCTGAACTCTTTCCCTACACGACGCTCTTCCGATCTNNNAGAGTGTTTCCATTCATTAAAT
AGCACACGTCTGAACTCTTTCCCTACACGACGCTCTTCCGATCTNNNTTTTAACTAGTCGACTATGAAGG
CACACGTCTGAACTCTTTCCCTACACGACGCTCTTCCGATCTNNNTTTTCTTACCACATCTGAATTTTCAT
GCACACGTCTGAACTCTTTCCCTACACGACGCTCTTCCGATCTNNNATTACTCTGTGTTACGTTATCTTC
AGCACACGTCTGAACTCTTTCCCTACACGACGCTCTTCCGATCTNNNTCTTCCTTCATAAATAAGTCCAG
AGCACACGTCTGAACTCTTTCCCTACACGACGCTCTTCCGATCTNNNAATGGAATGAGAAACTCAATAT
GCACACGTCTGAACTCTTTCCCTACACGACGCTCTTCCGATCTNNNTTTACTTCATTTTAAGTTGTGGGCA
AGCACACGTCTGAACTCTTTCCCTACACGACGCTCTTCCGATCTNNNTTTGGCTCTACATATTACTATGC
GCACACGTCTGAACTCTTTCCCTACACGACGCTCTTCCGATCTNNNCTCAGATCTCACTAATCACTAAACA
AGCACACGTCTGAACTCTTTCCCTACACGACGCTCTTCCGATCTNNNAAATGAAAGCTTTCTGTAGAAAT
GAGCACACGTCTGAACTCTTTCCCTACACGACGCTCTTCCGATCTNNNTACCTCCAGAGAAAACAATAA
AGCACACGTCTGAACTCTTTCCCTACACGACGCTCTTCCGATCTNNNTATTAAAACCACATCAGGTAGAA
AGCACACGTCTGAACTCTTTCCCTACACGACGCTCTTCCGATCTNNNGGGAGTGTATTAGGGATATAGA
GCACACGTCTGAACTCTTTCCCTACACGACGCTCTTCCGATCTNNNCTGTATATACAGTGGTGAAGGCA
AGCACACGTCTGAACTCTTTCCCTACACGACGCTCTTCCGATCTNNNTTAAAGACTGAAGAGGAATTG
AGCACACGTCTGAACTCTTTCCCTACACGACGCTCTTCCGATCTNNNGATGATAAAGAACTCAAACCAA
GCACACGTCTGAACTCTTTCCCTACACGACGCTCTTCCGATCTNNNGTCCAGATCATCCTGATTCTTCATC
AGCACACGTCTGAACTCTTTCCCTACACGACGCTCTTCCGATCTNNNACAGTTGTTTTCATTTGTACTTAC
CACACGTCTGAACTCTTTCCCTACACGACGCTCTTCCGATCTNNNGATGTTGGCTTTTATTGTAAGAGCA
GAGCACACGTCTGAACTCTTTCCCTACACGACGCTCTTCCGATCTNNNAGCTGATTTATAATCTTCCTTG
GCACACGTCTGAACTCTTTCCCTACACGACGCTCTTCCGATCTNNNAAGTCTAAATACAGATGTGAGTT
ACACGTCTGAACTCTTTCCCTACACGACGCTCTTCCGATCTNNNGTCTCCTCATCTGTGAAATGCATCGC
ACACGTCTGAACTCTTTCCCTACACGACGCTCTTCCGATCTNNNATTCAGGAATCCCTAAGCACCATCGG
CACACGTCTGAACTCTTTCCCTACACGACGCTCTTCCGATCTNNNGTTGAGTTCAATGCCTATATATGCA
GAGCACACGTCTGAACTCTTTCCCTACACGACGCTCTTCCGATCTNNNCTAGATTTGAAATGCCTTGAAA
GAGCACACGTCTGAACTCTTTCCCTACACGACGCTCTTCCGATCTNNNAGAAGCCAACCTGTAGATATT
AGAGCACACGTCTGAACTCTTTCCCTACACGACGCTCTTCCGATCTNNNTTAAGTAGATTGAAACTGCTC
GCACACGTCTGAACTCTTTCCCTACACGACGCTCTTCCGATCTNNNATCTTTAAATACTCATTTCCCCTCA
GAGCACACGTCTGAACTCTTTCCCTACACGACGCTCTTCCGATCTNNNTATTGTGAGGATTAAATATGCC
GAGCACACGTCTGAACTCTTTCCCTACACGACGCTCTTCCGATCTNNNTGTTTTTCCTGTGGATATAGAT
GAGCACACGTCTGAACTCTTTCCCTACACGACGCTCTTCCGATCTNNNTCTGGACTATCTTGTTTCCTATA
AGCACACGTCTGAACTCTTTCCCTACACGACGCTCTTCCGATCTNNNGCCACAAATAGCTTTTTTAAAGC
AGCACACGTCTGAACTCTTTCCCTACACGACGCTCTTCCGATCTNNNAGTTCTCATGAGATCTGGTTCAT
AGCACACGTCTGAACTCTTTCCCTACACGACGCTCTTCCGATCTNNNGTCTGTAAAATACTGTACAGGT
GCACACGTCTGAACTCTTTCCCTACACGACGCTCTTCCGATCTNNNAAATTTCTTGGTAGGTTGTAGAA
AGCACACGTCTGAACTCTTTCCCTACACGACGCTCTTCCGATCTNNNCTCTGAGTTTCTTCTGCTTCCATC
AGCACACGTCTGAACTCTTTCCCTACACGACGCTCTTCCGATCTNNNCTCTGAGTTTCTTCTGCTTCCATC
GAGCACACGTCTGAACTCTTTCCCTACACGACGCTCTTCCGATCTNNNGGCAGTTTACTTTATATAATGC
CACACGTCTGAACTCTTTCCCTACACGACGCTCTTCCGATCTNNNTTTTGCTTTTATGTTCAGCCCATCGG
GAGCACACGTCTGAACTCTTTCCCTACACGACGCTCTTCCGATCTNNNATAATCTGGACATTTACAATCT
GCACACGTCTGAACTCTTTCCCTACACGACGCTCTTCCGATCTNNNGAAGCATATGAAGTATGAAACAT
GCACACGTCTGAACTCTTTCCCTACACGACGCTCTTCCGATCTNNNTATTGCTCAAATTATTTCAGGTTC

AGCACACGTCTGAACTCTTTCCCTACACGACGCTCTTCCGATCTNNNTATTATTTATCTCTCAGACCCTG
CACACGTCTGAACTCTTTCCCTACACGACGCTCTTCCGATCTNNNGATACCATCTAGAAATTGGAAAC
AGCACACGTCTGAACTCTTTCCCTACACGACGCTCTTCCGATCTNNNCTCAAATTTTGCCCTATGACTAC
ACACGTCTGAACTCTTTCCCTACACGACGCTCTTCCGATCTNNNTATACAGTCCCTTATGATCTAGTCA
ACACGTCTGAACTCTTTCCCTACACGACGCTCTTCCGATCTNNNTATACAGTCCCTTATGATCTAGTCA
ACACGTCTGAACTCTTTCCCTACACGACGCTCTTCCGATCTNNNTATACAGTCCCTTATGATCTAGTCA
GCACACGTCTGAACTCTTTCCCTACACGACGCTCTTCCGATCTNNNCCTTCAGGGTTCTGTAAAATATA
AGCACACGTCTGAACTCTTTCCCTACACGACGCTCTTCCGATCTNNNTAAGTTACATATGTATCACCCT
GCACACGTCTGAACTCTTTCCCTACACGACGCTCTTCCGATCTNNNATTTGCAGATACTTTACATGAG
AGCACACGTCTGAACTCTTTCCCTACACGACGCTCTTCCGATCTNNNATAGGCAAAAAACTAAGGAGT
GCACACGTCTGAACTCTTTCCCTACACGACGCTCTTCCGATCTNNNTCATTTCACAATGTCCAAAATAA
ACACGTCTGAACTCTTTCCCTACACGACGCTCTTCCGATCTNNNTTTATAAAGTTCCTGGATTACACCA
AGCACACGTCTGAACTCTTTCCCTACACGACGCTCTTCCGATCTNNNCATTACTGTTTACTTGGCATA
AGCACACGTCTGAACTCTTTCCCTACACGACGCTCTTCCGATCTNNNCATTACTGTTTACTTGGCATA
GCACACGTCTGAACTCTTTCCCTACACGACGCTCTTCCGATCTNNNATCTCTCTCTTTCTCTCTCTTCCAT
CACACGTCTGAACTCTTTCCCTACACGACGCTCTTCCGATCTNNNATTTAAGAGGGTCTTTCACTGCAT
GCACACGTCTGAACTCTTTCCCTACACGACGCTCTTCCGATCTNNNTAATTTATTACTTCAGGGCCTAA
CACACGTCTGAACTCTTTCCCTACACGACGCTCTTCCGATCTNNNTAAAGATAGCCCATGTTCAGAACA
CACACGTCTGAACTCTTTCCCTACACGACGCTCTTCCGATCTNNNAATCACTTTAGCCAATTTCCTAAC
GCACACGTCTGAACTCTTTCCCTACACGACGCTCTTCCGATCTNNNCTGTTCTTCAATTTCCTTAAGAAC
ACGTCTGAACTCTTTCCCTACACGACGCTCTTCCGATCTNNNTTCATTTTCTTCCTTTCCTCCCATCGGA
AGCACACGTCTGAACTCTTTCCCTACACGACGCTCTTCCGATCTNNNTTTACTTAGCATAATGTCCTCT
AGCACACGTCTGAACTCTTTCCCTACACGACGCTCTTCCGATCTNNNATTAGGTAAAAAGCAGATGGA
AGCACACGTCTGAACTCTTTCCCTACACGACGCTCTTCCGATCTNNNCCCTACTGTTTAGTTTGGGCAT
GCACACGTCTGAACTCTTTCCCTACACGACGCTCTTCCGATCTNNNAGAAACAAGAGTGTATCTTACTT
AGCACACGTCTGAACTCTTTCCCTACACGACGCTCTTCCGATCTNNNCCAAGCTGAAACTAAGTTATTT
AGCACACGTCTGAACTCTTTCCCTACACGACGCTCTTCCGATCTNNNTTGTGTTTGTGTGAATGTATG
AGCACACGTCTGAACTCTTTCCCTACACGACGCTCTTCCGATCTNNNGCATTTATTATGTGACACAGTA
AGCACACGTCTGAACTCTTTCCCTACACGACGCTCTTCCGATCTNNNGTTTATGTCTGAGAATGTTTCT
CACACGTCTGAACTCTTTCCCTACACGACGCTCTTCCGATCTNNNAATTTCCAGAAACATTCTTTCCCAT
AGCACACGTCTGAACTCTTTCCCTACACGACGCTCTTCCGATCTNNNGCATTTATACAGGTGTTTCTTA
GCACACGTCTGAACTCTTTCCCTACACGACGCTCTTCCGATCTNNNATTAAAGCTTTGTCAGAACTTTG
CACACGTCTGAACTCTTTCCCTACACGACGCTCTTCCGATCTNNNGATTCTGCTTCCAATACTTTTAACA
CACACGTCTGAACTCTTTCCCTACACGACGCTCTTCCGATCTNNNATAGAATAAAAAACCAACGGAAA
AGCACACGTCTGAACTCTTTCCCTACACGACGCTCTTCCGATCTNNNGAATGATGAAGACTAAGAGA
AGCACACGTCTGAACTCTTTCCCTACACGACGCTCTTCCGATCTNNNCAATGTAACTGCAATATTTCAA
GCACACGTCTGAACTCTTTCCCTACACGACGCTCTTCCGATCTNNNCTGTTTTCTCTCACTATAGTTTG
GCACACGTCTGAACTCTTTCCCTACACGACGCTCTTCCGATCTNNNATTTCAGTCTGGGTGAAAGACA
GCACACGTCTGAACTCTTTCCCTACACGACGCTCTTCCGATCTNNNTTGCATCATACACTTTCTAGAACA
CACACGTCTGAACTCTTTCCCTACACGACGCTCTTCCGATCTNNNGTCTGATGGATTTTTGCTCTCATC
AGCACACGTCTGAACTCTTTCCCTACACGACGCTCTTCCGATCTNNNTAATTAGAACCCAGGTATGTTT
AGCACACGTCTGAACTCTTTCCCTACACGACGCTCTTCCGATCTNNNAACAACAATAATCACAATAGTC
AGCACACGTCTGAACTCTTTCCCTACACGACGCTCTTCCGATCTNNNAGGGATAAGACAAAAGAGTA

GAGCACACGTCTGAACTCTTTCCCTACACGACGCTCTTCCGATCTNNNGGGTTTCTGTCTTGTAAAGTT
GAGCACACGTCTGAACTCTTTCCCTACACGACGCTCTTCCGATCTNNNCATCATTATTTCTTCCAAATA
CACACGTCTGAACTCTTTCCCTACACGACGCTCTTCCGATCTNNNGCAGCTTAACTCTTTCTCTCCATCG
ACACGTCTGAACTCTTTCCCTACACGACGCTCTTCCGATCTNNNGGTCTCACTGTATTGTAAACTCATCG
AGCACACGTCTGAACTCTTTCCCTACACGACGCTCTTCCGATCTNNNATTAACATAAAGGAAAACACA
AGCACACGTCTGAACTCTTTCCCTACACGACGCTCTTCCGATCTNNNTCTGAGTTAACCATTACTAACAT
GCACACGTCTGAACTCTTTCCCTACACGACGCTCTTCCGATCTNNNTGTTAATACTCTCAATCAACTTGCA
AGCACACGTCTGAACTCTTTCCCTACACGACGCTCTTCCGATCTNNNTACCATCATGAAGGAACTAGTA
AGCACACGTCTGAACTCTTTCCCTACACGACGCTCTTCCGATCTNNNAGAAATTTTATAACCTTTGACCA
GAGCACACGTCTGAACTCTTTCCCTACACGACGCTCTTCCGATCTNNNATACTCAATTATTTGGGGGTA
AGCACACGTCTGAACTCTTTCCCTACACGACGCTCTTCCGATCTNNNCCATGAAGTGTGTAAAAACTATA
AGCACACGTCTGAACTCTTTCCCTACACGACGCTCTTCCGATCTNNNATTCAGGTGGTATTCATATTTCT
AGCACACGTCTGAACTCTTTCCCTACACGACGCTCTTCCGATCTNNNATTTCTGTCATTACACTTATCAC
GCACACGTCTGAACTCTTTCCCTACACGACGCTCTTCCGATCTNNNTATTTCACCTATCATAATGTCCTCA
AGCACACGTCTGAACTCTTTCCCTACACGACGCTCTTCCGATCTNNNCAATTATTATCAACAGAGGATC
AGCACACGTCTGAACTCTTTCCCTACACGACGCTCTTCCGATCTNNNACTTCTAGAATGGCAATACATAT
AGCACACGTCTGAACTCTTTCCCTACACGACGCTCTTCCGATCTNNNTCTTTTCATAGCTTCGTAGTATT
AGCACACGTCTGAACTCTTTCCCTACACGACGCTCTTCCGATCTNNNTCTTTTCATAGCTTCGTAGTATT
AGCACACGTCTGAACTCTTTCCCTACACGACGCTCTTCCGATCTNNNACCACAATTACAGTACAAATTA
AGCACACGTCTGAACTCTTTCCCTACACGACGCTCTTCCGATCTNNNTGAGTAGTATTCCATCATATATA
GCACACGTCTGAACTCTTTCCCTACACGACGCTCTTCCGATCTNNNCTCTCTCTCTCTGTCTCTCATCG
GCACACGTCTGAACTCTTTCCCTACACGACGCTCTTCCGATCTNNNCTCTCTCTCTCTGTCTCTCATCG
AGCACACGTCTGAACTCTTTCCCTACACGACGCTCTTCCGATCTNNNGATTCTTAGAGTGTACCTGAAA
AGCACACGTCTGAACTCTTTCCCTACACGACGCTCTTCCGATCTNNNGCCTTTATGCCTCTATATGTATA
AGCACACGTCTGAACTCTTTCCCTACACGACGCTCTTCCGATCTNNNCTTTCACTACCTTTCCAATACAT
AGCACACGTCTGAACTCTTTCCCTACACGACGCTCTTCCGATCTNNNTATTGCATGAGGGAATAAATGA
AAGAGCACACGTCTGAACTCTTTCCCTACACGACGCTCTTCCGATCTNNNATCTATTATATCAGACCCTT
GCACACGTCTGAACTCTTTCCCTACACGACGCTCTTCCGATCTNNNTAGGAGAATATTATGGATGTCA
CACACGTCTGAACTCTTTCCCTACACGACGCTCTTCCGATCTNNNAAAGCTTTATGAGGGGATAAGTA
GCACACGTCTGAACTCTTTCCCTACACGACGCTCTTCCGATCTNNNGTATCTGGGATAGGTACTCACCA
AGCACACGTCTGAACTCTTTCCCTACACGACGCTCTTCCGATCTNNNACAAAAAAGAAATCTTGTACAC
AGCACACGTCTGAACTCTTTCCCTACACGACGCTCTTCCGATCTNNNACTTTCATTGCAAGTCTAAATTT
GAGCACACGTCTGAACTCTTTCCCTACACGACGCTCTTCCGATCTNNNGTTTCTTTTCTAACTTCAGCTG
AGCACACGTCTGAACTCTTTCCCTACACGACGCTCTTCCGATCTNNNAGAACTCCCCTTAATTAAAAGAG
AGCACACGTCTGAACTCTTTCCCTACACGACGCTCTTCCGATCTNNNTGTAGAATATGGATTTAAGAG
AGCACACGTCTGAACTCTTTCCCTACACGACGCTCTTCCGATCTNNNGAAACGTTAAAGTAGCTAAGA
AGCACACGTCTGAACTCTTTCCCTACACGACGCTCTTCCGATCTNNNACAACTTATGTGTTCAATAATG
GCACACGTCTGAACTCTTTCCCTACACGACGCTCTTCCGATCTNNNGATATGAGCACGTATTATGTTTC
GCACACGTCTGAACTCTTTCCCTACACGACGCTCTTCCGATCTNNNGATATGAGCACGTATTATGTTTC
GAGCACACGTCTGAACTCTTTCCCTACACGACGCTCTTCCGATCTNNNTTAGAGCAAGGACTAATCATA
AGCACACGTCTGAACTCTTTCCCTACACGACGCTCTTCCGATCTNNNATATCTAGGGTTTTTAGCCTCTA
CACACGTCTGAACTCTTTCCCTACACGACGCTCTTCCGATCTNNNTTTCAACTTCTGAGCTAATGATCAT
AGCACACGTCTGAACTCTTTCCCTACACGACGCTCTTCCGATCTNNNATTATAATCCCATCAGGTATCA

AGCACACGTCTGAACTCTTTCCCTACACGACGCTCTTCCGATCTNNNCACCACCTATTAGTAGTTAACTT
GAGCACACGTCTGAACTCTTTCCCTACACGACGCTCTTCCGATCTNNNGAAAATTTAGCCTTTCTCTTTCT
GCACACGTCTGAACTCTTTCCCTACACGACGCTCTTCCGATCTNNNTCTTAATAAGAGGAGGAAGAAAC
AGCACACGTCTGAACTCTTTCCCTACACGACGCTCTTCCGATCTNNNTTGGTTATGGTCACAGAATGCA
GAGCACACGTCTGAACTCTTTCCCTACACGACGCTCTTCCGATCTNNNCACTATTTCTTTTTAACAACTGC
GAGCACACGTCTGAACTCTTTCCCTACACGACGCTCTTCCGATCTNNNATTTCTAATCCAGTATTCAATG
CACGTCTGAACTCTTTCCCTACACGACGCTCTTCCGATCTNNNTTTTAGTTAAATGGTATGTTTCCCATC
GAGCACACGTCTGAACTCTTTCCCTACACGACGCTCTTCCGATCTNNNAGGTTGTATGCATCTATTTGT
AGCACACGTCTGAACTCTTTCCCTACACGACGCTCTTCCGATCTNNNAAAATTGGGAAAAATCACTTACT
AGCACACGTCTGAACTCTTTCCCTACACGACGCTCTTCCGATCTNNNTATTTACAGGTTCTGGGGATTA
GAGCACACGTCTGAACTCTTTCCCTACACGACGCTCTTCCGATCTNNNTACTAGACCCTATATATACTGA
AGAGCACACGTCTGAACTCTTTCCCTACACGACGCTCTTCCGATCTNNNGGTGTAAAATTTCCATACATT
GAGCACACGTCTGAACTCTTTCCCTACACGACGCTCTTCCGATCTNNNCTCAACAATCTGTTTTTGTTTT
AGCACACGTCTGAACTCTTTCCCTACACGACGCTCTTCCGATCTNNNACGAATTTTCATACCAACAAAATC
CACACGTCTGAACTCTTTCCCTACACGACGCTCTTCCGATCTNNNGGCATGGAGAAAAATATGCACATC
GCACACGTCTGAACTCTTTCCCTACACGACGCTCTTCCGATCTNNNATATCTTATCAGGTTGGTTGACCA
GCACACGTCTGAACTCTTTCCCTACACGACGCTCTTCCGATCTNNNTTCCTATATCTGGGTACATTCTTCA
CACACGTCTGAACTCTTTCCCTACACGACGCTCTTCCGATCTNNNGACTAGTAGAGATCTGGTAATTGC
GCACACGTCTGAACTCTTTCCCTACACGACGCTCTTCCGATCTNNNGATTCCCTAAAACAAAAGTTGTAC
GAGCACACGTCTGAACTCTTTCCCTACACGACGCTCTTCCGATCTNNNATTAGGGACAGATATGGAAAT
CACACGTCTGAACTCTTTCCCTACACGACGCTCTTCCGATCTNNNTTCTCTCTCTTGCTATATCTACTCATC
CACACGTCTGAACTCTTTCCCTACACGACGCTCTTCCGATCTNNNAAAGTAGATAGACCCTGTGATTCA
AGCACACGTCTGAACTCTTTCCCTACACGACGCTCTTCCGATCTNNNGGACCCAAGATTACAGATAAGC
AGAGCACACGTCTGAACTCTTTCCCTACACGACGCTCTTCCGATCTNNNAAGAAAAGGTGTATTTTGAT
AGCACACGTCTGAACTCTTTCCCTACACGACGCTCTTCCGATCTNNNCATAATCATGAGGCTTAAAACTC
AGCACACGTCTGAACTCTTTCCCTACACGACGCTCTTCCGATCTNNNCATAATCATGAGGCTTAAAACTC
GCACACGTCTGAACTCTTTCCCTACACGACGCTCTTCCGATCTNNNCATATTCACCCATTGTCAATATTCA
AGCACACGTCTGAACTCTTTCCCTACACGACGCTCTTCCGATCTNNNACAAACAGAGACAAAGACTTAC
AGCACACGTCTGAACTCTTTCCCTACACGACGCTCTTCCGATCTNNNATACTGAGGAGGCTTTCTATTTC
AGCACACGTCTGAACTCTTTCCCTACACGACGCTCTTCCGATCTNNNTTCTGAGCAAGAGTTTCCAACA
GAGCACACGTCTGAACTCTTTCCCTACACGACGCTCTTCCGATCTNNNTCACAATATTTACAATATTTAC
AGAGCACACGTCTGAACTCTTTCCCTACACGACGCTCTTCCGATCTNNNAGAAAATTTGGCTTACAGATT
GCACACGTCTGAACTCTTTCCCTACACGACGCTCTTCCGATCTNNNTATAAATGTACACTGTGCAGAGC
AGCACACGTCTGAACTCTTTCCCTACACGACGCTCTTCCGATCTNNNCCATAACAAACTACTTAGTGTG
AGCACACGTCTGAACTCTTTCCCTACACGACGCTCTTCCGATCTNNNGTATATTTAACATGGGCATTGA
ACACGTCTGAACTCTTTCCCTACACGACGCTCTTCCGATCTNNNCTATTTTTGCCTTCCACTCTTCATCGG
AGAGCACACGTCTGAACTCTTTCCCTACACGACGCTCTTCCGATCTNNNATTCTAGGTACAGGTCAGTAC
GCACACGTCTGAACTCTTTCCCTACACGACGCTCTTCCGATCTNNNAATAACCGTTAGGATGCATAGCA
GCACACGTCTGAACTCTTTCCCTACACGACGCTCTTCCGATCTNNNGTTAAAGAGTTGGAAGCAAAAC
AGCACACGTCTGAACTCTTTCCCTACACGACGCTCTTCCGATCTNNNGTGAGTTTAAAAGAAACAATCT
GAGCACACGTCTGAACTCTTTCCCTACACGACGCTCTTCCGATCTNNNTCAATGAGTAGATAAAGAAAG
AGCACACGTCTGAACTCTTTCCCTACACGACGCTCTTCCGATCTNNNTCTCTTCCTTCCTTTCTTCTCATCG
AGCACACGTCTGAACTCTTTCCCTACACGACGCTCTTCCGATCTNNNGGTAACCATCATTCTACTCTCTAC

GAGCACACGTCTGAACTCTTTCCCTACACGACGCTCTTCCGATCTNNNAAATAGCCACACACAAATATAA
AGCACACGTCTGAACTCTTTCCCTACACGACGCTCTTCCGATCTNNNACTTAGATATACAAGCAAGTAG
AGCACACGTCTGAACTCTTTCCCTACACGACGCTCTTCCGATCTNNNTATGAATATACCTCTGTGTATG
ACACGTCTGAACTCTTTCCCTACACGACGCTCTTCCGATCTNNNGTCTCCTCATCTGTGAAATGCATCGC
CACACGTCTGAACTCTTTCCCTACACGACGCTCTTCCGATCTNNNGCTTCATGAAAATGAGGAGATATC
CACACGTCTGAACTCTTTCCCTACACGACGCTCTTCCGATCTNNNTTTCTTCTTTCTCTGTCTCTCTTCATC
AGCACACGTCTGAACTCTTTCCCTACACGACGCTCTTCCGATCTNNNGGATATATTGCCATATAACTGGA
GCACACGTCTGAACTCTTTCCCTACACGACGCTCTTCCGATCTNNNAGTTCATAATGTGGCAATTTGCAT
GAGCACACGTCTGAACTCTTTCCCTACACGACGCTCTTCCGATCTNNNATATTTGGAAACCAATTTCTAG
AGCACACGTCTGAACTCTTTCCCTACACGACGCTCTTCCGATCTNNNTATAGGAGAAAGCTCTGGTTAT
AGCACACGTCTGAACTCTTTCCCTACACGACGCTCTTCCGATCTNNNGTCAAAGAAGTTTTCTGATAAGA
GAGCACACGTCTGAACTCTTTCCCTACACGACGCTCTTCCGATCTNNNGCCAGTATATATTGTGACATCT
GCACACGTCTGAACTCTTTCCCTACACGACGCTCTTCCGATCTNNNAATGACTTACCTTGGTGTATGCAT
AGCACACGTCTGAACTCTTTCCCTACACGACGCTCTTCCGATCTNNNGAGGAGAGAACATTTGTATTCA
AGCACACGTCTGAACTCTTTCCCTACACGACGCTCTTCCGATCTNNNGAATTCTGTAGATAACAATCAG
GCACACGTCTGAACTCTTTCCCTACACGACGCTCTTCCGATCTNNNATTTATTTCAGTGTCGAATAAGG
CACACGTCTGAACTCTTTCCCTACACGACGCTCTTCCGATCTNNNTTCAGAGCTGATTCTAATTCTAGCA
AGCACACGTCTGAACTCTTTCCCTACACGACGCTCTTCCGATCTNNNAAACCACAAAAATCTCTTTACAT
GCACACGTCTGAACTCTTTCCCTACACGACGCTCTTCCGATCTNNNGCTTTGAATTCAGTGGAACCATC
AGCACACGTCTGAACTCTTTCCCTACACGACGCTCTTCCGATCTNNNTTTCTCTCCCGTTTAGATAAATC
AGCACACGTCTGAACTCTTTCCCTACACGACGCTCTTCCGATCTNNNTATTGTTGGGTCAGAAATGATA
ACACGTCTGAACTCTTTCCCTACACGACGCTCTTCCGATCTNNNCCATAGAAATTTTAGGAAAAAACAC
AGCACACGTCTGAACTCTTTCCCTACACGACGCTCTTCCGATCTNNNAGTTTGAAAAGTTCTCTGTTATT
GAGCACACGTCTGAACTCTTTCCCTACACGACGCTCTTCCGATCTNNNGATAAAGAAAATGTGGTGTGT
CACGTCTGAACTCTTTCCCTACACGACGCTCTTCCGATCTNNNTTTCATATTGTTTCCATAGCTACCATCC
AGCACACGTCTGAACTCTTTCCCTACACGACGCTCTTCCGATCTNNNCACACATATATATACACACACAT
AGCACACGTCTGAACTCTTTCCCTACACGACGCTCTTCCGATCTNNNGGAATTCCAAGAGTATAGAAGA
AGCACACGTCTGAACTCTTTCCCTACACGACGCTCTTCCGATCTNNNACTTTTGTTTAATGGATTATGCT
CACACGTCTGAACTCTTTCCCTACACGACGCTCTTCCGATCTNNNAAACTATTTTTCTCTCTCTCTGTCAT
AGCACACGTCTGAACTCTTTCCCTACACGACGCTCTTCCGATCTNNNTAGATAAACAGAAGACAGAAT
AGCACACGTCTGAACTCTTTCCCTACACGACGCTCTTCCGATCTNNNTTTAGTGAGACAGATAAAATGT
GAGCACACGTCTGAACTCTTTCCCTACACGACGCTCTTCCGATCTNNNATATGCATTGCTCAAGTACTCC
AGAGCACACGTCTGAACTCTTTCCCTACACGACGCTCTTCCGATCTNNNAAAGAAGCATCTCCAAAATAA
AGCACACGTCTGAACTCTTTCCCTACACGACGCTCTTCCGATCTNNNAATTTAGGCCATGAGCAAATATA
AGCACACGTCTGAACTCTTTCCCTACACGACGCTCTTCCGATCTNNNTCTGACTAAGACCCATCTAAATT
GCACACGTCTGAACTCTTTCCCTACACGACGCTCTTCCGATCTNNNATACATAGCTGTATCAGGTACTCA
GAGCACACGTCTGAACTCTTTCCCTACACGACGCTCTTCCGATCTNNNTGATAATGTTACATGGAAAAA
AGCACACGTCTGAACTCTTTCCCTACACGACGCTCTTCCGATCTNNNCCTACCTATCTATCTACCTATGTC
CACACGTCTGAACTCTTTCCCTACACGACGCTCTTCCGATCTNNNTGCATATAAAATTTGATACATCCAC
AGCACACGTCTGAACTCTTTCCCTACACGACGCTCTTCCGATCTNNNTCAATGATTATAGAAATGCAAG
AGCACACGTCTGAACTCTTTCCCTACACGACGCTCTTCCGATCTNNNACATATATAGCATATATACACAT
GCACACGTCTGAACTCTTTCCCTACACGACGCTCTTCCGATCTNNNATTTAGCTTCATGAGTCACTTTAC
CACACGTCTGAACTCTTTCCCTACACGACGCTCTTCCGATCTNNNTTTACATTTAAAATGTTACTTGACCC

AGCACACGTCTGAACTCTTTCCCTACACGACGCTCTTCCGATCTNNNAGTCATATTTGGGGTAGTAT
GCACACGTCTGAACTCTTTCCCTACACGACGCTCTTCCGATCTNNNGAGAGAGAGAGACAGAGAGA
CACACGTCTGAACTCTTTCCCTACACGACGCTCTTCCGATCTNNNAGACCATGTCTCAAAAAACAAAC
ACACGTCTGAACTCTTTCCCTACACGACGCTCTTCCGATCTNNNAAACAGGTGGAATTTACCATAAACA
ACACGTCTGAACTCTTTCCCTACACGACGCTCTTCCGATCTNNNAAACAGGTGGAATTTACCATAAACA
AGCACACGTCTGAACTCTTTCCCTACACGACGCTCTTCCGATCTNNNAATTAGAAGGAAAGCATGTGA
AGCACACGTCTGAACTCTTTCCCTACACGACGCTCTTCCGATCTNNNAGCTTCCATATCAAGTCATATTA
GCACACGTCTGAACTCTTTCCCTACACGACGCTCTTCCGATCTNNNCGTAGGATTTAAAATACTCATCA
AGCACACGTCTGAACTCTTTCCCTACACGACGCTCTTCCGATCTNNNGAAGAGAAGCTACAGACTTAA
GAGCACACGTCTGAACTCTTTCCCTACACGACGCTCTTCCGATCTNNNCATTAAGCAGCCATAAAAGA
GAGCACACGTCTGAACTCTTTCCCTACACGACGCTCTTCCGATCTNNNAAAAAACACAAGAATACAACA
AGCACACGTCTGAACTCTTTCCCTACACGACGCTCTTCCGATCTNNNCCAAATTGATGGGTTATACAAA
GCACACGTCTGAACTCTTTCCCTACACGACGCTCTTCCGATCTNNNTTCTCAGCATGTCCTCATATCATC
GAGCACACGTCTGAACTCTTTCCCTACACGACGCTCTTCCGATCTNNNGAAGATTACGGGAAAGTGTG
GAGCACACGTCTGAACTCTTTCCCTACACGACGCTCTTCCGATCTNNNGAAGATTACGGGAAAGTGTG
GCACACGTCTGAACTCTTTCCCTACACGACGCTCTTCCGATCTNNNATCTTCAGGGATATAAAATAGGG
AGCACACGTCTGAACTCTTTCCCTACACGACGCTCTTCCGATCTNNNATATAAAAGCTTCTGAGAAAGA
GCACACGTCTGAACTCTTTCCCTACACGACGCTCTTCCGATCTNNNCAAAACTCGACCTTCATTTACCAT
GCACACGTCTGAACTCTTTCCCTACACGACGCTCTTCCGATCTNNNACCAGGTTAAATGAGGACTCCA
CACACGTCTGAACTCTTTCCCTACACGACGCTCTTCCGATCTNNNTAAAAGTTCTTTGATGTGAGGAA
GAGCACACGTCTGAACTCTTTCCCTACACGACGCTCTTCCGATCTNNNTTTTCTCTAAATGATGAAATG
GCACACGTCTGAACTCTTTCCCTACACGACGCTCTTCCGATCTNNNGTTAAGCAAATTAACCTATCTGT
GCACACGTCTGAACTCTTTCCCTACACGACGCTCTTCCGATCTNNNAATCTTCAAATCTGCTTGTCTTCA
GAGCACACGTCTGAACTCTTTCCCTACACGACGCTCTTCCGATCTNNNTCAAACTAAATACCTATCTAAC
GCACACGTCTGAACTCTTTCCCTACACGACGCTCTTCCGATCTNNNTAACGATTGTGTATTGAGAATTG
GAGCACACGTCTGAACTCTTTCCCTACACGACGCTCTTCCGATCTNNNATTCTTAGGAAATACTCACTGA
GAGCACACGTCTGAACTCTTTCCCTACACGACGCTCTTCCGATCTNNNCTATCTTAGTTGATGTCTTCCT
GAGCACACGTCTGAACTCTTTCCCTACACGACGCTCTTCCGATCTNNNCATTCATATTTTACCTACTGAC
GAGCACACGTCTGAACTCTTTCCCTACACGACGCTCTTCCGATCTNNNTTGACTATTACAAATAAGGAT
GAGCACACGTCTGAACTCTTTCCCTACACGACGCTCTTCCGATCTNNNAGAATGATGTAGGTAGAAAG
AGCACACGTCTGAACTCTTTCCCTACACGACGCTCTTCCGATCTNNNTTTCCAAAACCTTCATTCAAAGC
GCACACGTCTGAACTCTTTCCCTACACGACGCTCTTCCGATCTNNNAAAGTGAAAGTTTGTGAGATGCA
GAGCACACGTCTGAACTCTTTCCCTACACGACGCTCTTCCGATCTNNNTCAACAAGAGTCACTTTAGAT
AGCACACGTCTGAACTCTTTCCCTACACGACGCTCTTCCGATCTNNNACAACATTGATTTAAATCTGGA
CACACGTCTGAACTCTTTCCCTACACGACGCTCTTCCGATCTNNNCCTTAGGGTCAGATAAATATGAGC
AGCACACGTCTGAACTCTTTCCCTACACGACGCTCTTCCGATCTNNNCAAAACAAAAACAAAAACAAGC
GAGCACACGTCTGAACTCTTTCCCTACACGACGCTCTTCCGATCTNNNCATTTCCCTATTATGGTACCAT
GAGCACACGTCTGAACTCTTTCCCTACACGACGCTCTTCCGATCTNNNTAAAAGAGCATCATACCTACA
CACACGTCTGAACTCTTTCCCTACACGACGCTCTTCCGATCTNNNGGAATAAGTGGCTTATCTCTGCAT
GCACACGTCTGAACTCTTTCCCTACACGACGCTCTTCCGATCTNNNTATTTGGAGACAGTCTTTACAAA
AGCACACGTCTGAACTCTTTCCCTACACGACGCTCTTCCGATCTNNNTTGAAGGATTACTATGTTCTAG
GAGCACACGTCTGAACTCTTTCCCTACACGACGCTCTTCCGATCTNNNTCATATCTTCATGAAGTTGAA
GAGCACACGTCTGAACTCTTTCCCTACACGACGCTCTTCCGATCTNNNTCATATCTTCATGAAGTTGAA

GCACACGTCTGAACTCTTTCCCTACACGACGCTCTTCCGATCTNNNAACTCAACATAATAAATGCCATA
GAGCACACGTCTGAACTCTTTCCCTACACGACGCTCTTCCGATCTNNNTGGACCTAAATTTAAGTCTAA
AGCACACGTCTGAACTCTTTCCCTACACGACGCTCTTCCGATCTNNNGATTTTGTTTTGGTTTTGTTTCT
GCACACGTCTGAACTCTTTCCCTACACGACGCTCTTCCGATCTNNNTTTTAAAAAGAGGGAGAGGTAG
GCACACGTCTGAACTCTTTCCCTACACGACGCTCTTCCGATCTNNNAAGATCTGGCTATTTGAAGGCA
GAGCACACGTCTGAACTCTTTCCCTACACGACGCTCTTCCGATCTNNNAGTTTTATAAGCTCAAATGGT
GCACACGTCTGAACTCTTTCCCTACACGACGCTCTTCCGATCTNNNCTTACATTACAAATTACCCCCTCA
GCACACGTCTGAACTCTTTCCCTACACGACGCTCTTCCGATCTNNNGCCGAGAATTTATTTGCATATAT
AGCACACGTCTGAACTCTTTCCCTACACGACGCTCTTCCGATCTNNNGGTAGAGGTTATTCATTCTCTT
AGAGCACACGTCTGAACTCTTTCCCTACACGACGCTCTTCCGATCTNNNTTTAGAAGGGACAAGAGTC
GCACACGTCTGAACTCTTTCCCTACACGACGCTCTTCCGATCTNNNCACTTCTGATGAATGGAATGTCA
AGAGCACACGTCTGAACTCTTTCCCTACACGACGCTCTTCCGATCTNNNGATTTTGTGACTTCCCTGAT
GAGCACACGTCTGAACTCTTTCCCTACACGACGCTCTTCCGATCTNNNTTAAGTAGTTTAATTGGTTAG
CACACGTCTGAACTCTTTCCCTACACGACGCTCTTCCGATCTNNNTAATCAGGTGAAATCTTTTGAAG
AGCACACGTCTGAACTCTTTCCCTACACGACGCTCTTCCGATCTNNNTTGTATAGCCCATATATGAACT
AGCACACGTCTGAACTCTTTCCCTACACGACGCTCTTCCGATCTNNNTTGAATGTATACTCTGTGCTAG
AGCACACGTCTGAACTCTTTCCCTACACGACGCTCTTCCGATCTNNNCTCTTCCTTGTTTATTTGTGAGC
AGAGCACACGTCTGAACTCTTTCCCTACACGACGCTCTTCCGATCTNNNATACAGAACAGAACATACAT
AGCACACGTCTGAACTCTTTCCCTACACGACGCTCTTCCGATCTNNNATAAAACTGCATGGGACTAAA
ACACGTCTGAACTCTTTCCCTACACGACGCTCTTCCGATCTNNNTTTTTCAGGAAATTTAGGATGAGC
AGCACACGTCTGAACTCTTTCCCTACACGACGCTCTTCCGATCTNNNCTTTCCATAGTCATAAGATATG
AGCACACGTCTGAACTCTTTCCCTACACGACGCTCTTCCGATCTNNNTTTTCTTAGTGGTTGCTCTAAG
GAGCACACGTCTGAACTCTTTCCCTACACGACGCTCTTCCGATCTNNNGGCCTATATTTCAGTTTTTCTT
CACGTCTGAACTCTTTCCCTACACGACGCTCTTCCGATCTNNNCATCATCATCATTTTTTAAGTGGCAT
GAGCACACGTCTGAACTCTTTCCCTACACGACGCTCTTCCGATCTNNNCCTAAATACTTCACCATGTAAT
AGCACACGTCTGAACTCTTTCCCTACACGACGCTCTTCCGATCTNNNGAGAGAGTGAAGAATTCCTAA
GCACACGTCTGAACTCTTTCCCTACACGACGCTCTTCCGATCTNNNGTTTTTTTAAAGCAAGATTCCAA
AGCACACGTCTGAACTCTTTCCCTACACGACGCTCTTCCGATCTNNNTTTACCAGAAGAGAGGAGAA
GCACACGTCTGAACTCTTTCCCTACACGACGCTCTTCCGATCTNNNAGTAGTTACAAAATTATAAATGC
CACGTCTGAACTCTTTCCCTACACGACGCTCTTCCGATCTNNNTTTTTTTTTCAGTTACTGTTGTGCATC
CACACGTCTGAACTCTTTCCCTACACGACGCTCTTCCGATCTNNNAAACTCATCTTACCTTGACTTCCAT
CACACGTCTGAACTCTTTCCCTACACGACGCTCTTCCGATCTNNNTAGAGAGTGTCAGAACTGAATTA
AGCACACGTCTGAACTCTTTCCCTACACGACGCTCTTCCGATCTNNNTATTTTTTTCTTCTAAAGCCCAA
AGAGCACACGTCTGAACTCTTTCCCTACACGACGCTCTTCCGATCTNNNCTTATAATGCTTCCTTGAATG
AGCACACGTCTGAACTCTTTCCCTACACGACGCTCTTCCGATCTNNNACACTTCTCTGTATTGGATCTAA
AGCACACGTCTGAACTCTTTCCCTACACGACGCTCTTCCGATCTNNNTGGGCAATTTTTATTTTATCCTT
GAGCACACGTCTGAACTCTTTCCCTACACGACGCTCTTCCGATCTNNNGTGAAGGAAAGTATATGAAG
AGCACACGTCTGAACTCTTTCCCTACACGACGCTCTTCCGATCTNNNCAGCACTAACCTCATTCTAGCA
GAGCACACGTCTGAACTCTTTCCCTACACGACGCTCTTCCGATCTNNNATAATTTGTGAGCACTAAGAT
AGCACACGTCTGAACTCTTTCCCTACACGACGCTCTTCCGATCTNNNTAGAGTTAAACTAAGCATTCAG
CACACGTCTGAACTCTTTCCCTACACGACGCTCTTCCGATCTNNNGAAAGAAGAGAGAAAATACATGG
AGCACACGTCTGAACTCTTTCCCTACACGACGCTCTTCCGATCTNNNTAAAGACATTGCTAAAGAGAA
GAGCACACGTCTGAACTCTTTCCCTACACGACGCTCTTCCGATCTNNNAACCCTAATCTGTTATTCCATT

AGAGCACACGTCTGAACTCTTTCCCTACACGACGCTCTTCCGATCTNNNAAATAGCTTTGGAATAAATT
AGAGCACACGTCTGAACTCTTTCCCTACACGACGCTCTTCCGATCTNNNAAATAGCTTTGGAATAAATT
AGCACACGTCTGAACTCTTTCCCTACACGACGCTCTTCCGATCTNNNCAACAAAATTTATCTCTACAGCT
AGCACACGTCTGAACTCTTTCCCTACACGACGCTCTTCCGATCTNNNCTCATAATCTTTAAAGAGGTGT
AGAGCACACGTCTGAACTCTTTCCCTACACGACGCTCTTCCGATCTNNNTAGGCTCAGAGAAACAATAG
ACACGTCTGAACTCTTTCCCTACACGACGCTCTTCCGATCTNNNTAAATTTCCAGCCAGGTTTATATCAT
GCACACGTCTGAACTCTTTCCCTACACGACGCTCTTCCGATCTNNNCTATCTCTCTCTGTGTCTCTCCATC
AGCACACGTCTGAACTCTTTCCCTACACGACGCTCTTCCGATCTNNNAATGGTTAAATGAAAAGGATCT
AGCACACGTCTGAACTCTTTCCCTACACGACGCTCTTCCGATCTNNNTGTGTGTGTGTATATATATG
AGAGCACACGTCTGAACTCTTTCCCTACACGACGCTCTTCCGATCTNNNGATTTTCATTTGATGCAATCT
AGAGCACACGTCTGAACTCTTTCCCTACACGACGCTCTTCCGATCTNNNGATTTTCATTTGATGCAATCT
AGCACACGTCTGAACTCTTTCCCTACACGACGCTCTTCCGATCTNNNGCTTGAAAATTTGGATCATACT
AGCACACGTCTGAACTCTTTCCCTACACGACGCTCTTCCGATCTNNNCCAGGTTATTAATAGAGGAAAG
AGCACACGTCTGAACTCTTTCCCTACACGACGCTCTTCCGATCTNNNATCTCTTTCTCTCTGTCTCTCCAT
GAGCACACGTCTGAACTCTTTCCCTACACGACGCTCTTCCGATCTNNNTTTGTAATGCAAGTGTATTAGT
AGCACACGTCTGAACTCTTTCCCTACACGACGCTCTTCCGATCTNNNTTCCATCTGTTATAAAAGAAACA
AGCACACGTCTGAACTCTTTCCCTACACGACGCTCTTCCGATCTNNNGTATTTTCTGGCTCAATCTAATC
GAGCACACGTCTGAACTCTTTCCCTACACGACGCTCTTCCGATCTNNNGAGAGCTAAATCTAGTGTATG
GAGCACACGTCTGAACTCTTTCCCTACACGACGCTCTTCCGATCTNNNGGACTTATGTTAAAACCGTTA
GAGCACACGTCTGAACTCTTTCCCTACACGACGCTCTTCCGATCTNNNATTCCACTAGGTACTGATTAA
AGCACACGTCTGAACTCTTTCCCTACACGACGCTCTTCCGATCTNNNGTCTAAATAAATGTGTTGCAAT
GAGCACACGTCTGAACTCTTTCCCTACACGACGCTCTTCCGATCTNNNAACATTATAGGCAAGACAAAA
AGCACACGTCTGAACTCTTTCCCTACACGACGCTCTTCCGATCTNNNACACACACACACATATATGTATA
GAGCACACGTCTGAACTCTTTCCCTACACGACGCTCTTCCGATCTNNNTCCCTAAGGGTAATAAATTTT
AGCACACGTCTGAACTCTTTCCCTACACGACGCTCTTCCGATCTNNNCTTCCCTCTCTCTCTTTCTCCATCG
AGCACACGTCTGAACTCTTTCCCTACACGACGCTCTTCCGATCTNNNTGTATTTTGTTTTTTGGTTATTCA
AGCACACGTCTGAACTCTTTCCCTACACGACGCTCTTCCGATCTNNNTTTATACCTACCATAAGCTGTAC
AGCACACGTCTGAACTCTTTCCCTACACGACGCTCTTCCGATCTNNNAAGTCTTACTAAAAGTAGACCA
GAGCACACGTCTGAACTCTTTCCCTACACGACGCTCTTCCGATCTNNNCAGAATGAAACCTTGTCTAAA
GAGCACACGTCTGAACTCTTTCCCTACACGACGCTCTTCCGATCTNNNTACAAGTGTACAAAAATGAGA
AGCACACGTCTGAACTCTTTCCCTACACGACGCTCTTCCGATCTNNNATAGAGTGACAGAACTTGAATA
AGCACACGTCTGAACTCTTTCCCTACACGACGCTCTTCCGATCTNNNGAGATCTTTTCCCTCCTCTATCAT
AGAGCACACGTCTGAACTCTTTCCCTACACGACGCTCTTCCGATCTNNNAGCATATTTGGAATCTACCTT
GCACACGTCTGAACTCTTTCCCTACACGACGCTCTTCCGATCTNNNTGAGGAAATTACACAAGGTATAA
AGCACACGTCTGAACTCTTTCCCTACACGACGCTCTTCCGATCTNNNATTCCCTCTCTCTCTTTCTTTTCA
GAGCACACGTCTGAACTCTTTCCCTACACGACGCTCTTCCGATCTNNNTTTCTTCTGCTCTGACTAAATA
GAGCACACGTCTGAACTCTTTCCCTACACGACGCTCTTCCGATCTNNNTTTCTTCTGCTCTGACTAAATA
GAGCACACGTCTGAACTCTTTCCCTACACGACGCTCTTCCGATCTNNNACCCACCTTTACAAATATACATA
AGCACACGTCTGAACTCTTTCCCTACACGACGCTCTTCCGATCTNNNAAACACCCACCTTTACAAATATA
AGCACACGTCTGAACTCTTTCCCTACACGACGCTCTTCCGATCTNNNGAAAAAGACTAGAGTTGTTTG
GCACACGTCTGAACTCTTTCCCTACACGACGCTCTTCCGATCTNNNTAGCTAGTGGTGTATCTACTTTA
GCACACGTCTGAACTCTTTCCCTACACGACGCTCTTCCGATCTNNNCTCTGTAAGTACTTTGCCTGCATC
GAGCACACGTCTGAACTCTTTCCCTACACGACGCTCTTCCGATCTNNNTATAATGTGTTTCACTATAGG

GCACACGTCTGAACTCTTTCCCTACACGACGCTCTTCCGATCTNNNACACTCAAATTATAGTAGCAATG
AGCACACGTCTGAACTCTTTCCCTACACGACGCTCTTCCGATCTNNNTAAAATCATCATAATGGTCACA
GCACACGTCTGAACTCTTTCCCTACACGACGCTCTTCCGATCTNNNATTCTAAATTATCACCATGCATGC
AGCACACGTCTGAACTCTTTCCCTACACGACGCTCTTCCGATCTNNNTAATTATATTGGATCACAGAG
AGCACACGTCTGAACTCTTTCCCTACACGACGCTCTTCCGATCTNNNTTTATGCACATAAAAACTGG
CACACGTCTGAACTCTTTCCCTACACGACGCTCTTCCGATCTNNNGATGTAAACGGTATGTTATATGGC
GAGCACACGTCTGAACTCTTTCCCTACACGACGCTCTTCCGATCTNNNAAAACATGTAGAGAAGAAGA
CACACGTCTGAACTCTTTCCCTACACGACGCTCTTCCGATCTNNNGAATATAAATGGGGAATGAATTC
GCACACGTCTGAACTCTTTCCCTACACGACGCTCTTCCGATCTNNNGACACTTTTGTTTGACGTTTTTCA
GCACACGTCTGAACTCTTTCCCTACACGACGCTCTTCCGATCTNNNGACACTTTTGTTTGACGTTTTTCA
AGCACACGTCTGAACTCTTTCCCTACACGACGCTCTTCCGATCTNNNGAGTTTTTTTGTTTTGTTTTGT
GCACACGTCTGAACTCTTTCCCTACACGACGCTCTTCCGATCTNNNTTTTGTGTGTGTGTTTTTTTTTA
AGCACACGTCTGAACTCTTTCCCTACACGACGCTCTTCCGATCTNNNTTTCATCTATTCTTTCTGAGACC
GCACACGTCTGAACTCTTTCCCTACACGACGCTCTTCCGATCTNNNCTCCTGGTTTTGAGAAATATTA
AGCACACGTCTGAACTCTTTCCCTACACGACGCTCTTCCGATCTNNNAGTAGAAAGTTGGAAAACTAG
AGCACACGTCTGAACTCTTTCCCTACACGACGCTCTTCCGATCTNNNGACATGATGGTGAGAAATACT
GCACACGTCTGAACTCTTTCCCTACACGACGCTCTTCCGATCTNNNCAAAGATAAAAGTCTAGAAAGCT
GAGCACACGTCTGAACTCTTTCCCTACACGACGCTCTTCCGATCTNNNACAGAAGATAAGGAAATGT
GCACACGTCTGAACTCTTTCCCTACACGACGCTCTTCCGATCTNNNCCTAGTTTTTAGTTAAACAGCA
GCACACGTCTGAACTCTTTCCCTACACGACGCTCTTCCGATCTNNNTATAGTAAGGGAATAGTCATCC
GCACACGTCTGAACTCTTTCCCTACACGACGCTCTTCCGATCTNNNCAATTAAAAATTCTCTGAGGTGA
GAGCACACGTCTGAACTCTTTCCCTACACGACGCTCTTCCGATCTNNNCCCCCACATTATCTGAAAAAC
AGCACACGTCTGAACTCTTTCCCTACACGACGCTCTTCCGATCTNNNGTAGAAAATATGAGTGCACAT
CACACGTCTGAACTCTTTCCCTACACGACGCTCTTCCGATCTNNNTTCTGTTACACTTCTAGGCTACATC
CACACGTCTGAACTCTTTCCCTACACGACGCTCTTCCGATCTNNNTTCTGTTACACTTCTAGGCTACATC
GAGCACACGTCTGAACTCTTTCCCTACACGACGCTCTTCCGATCTNNNACAATTAAAATAACACATGGG
AGCACACGTCTGAACTCTTTCCCTACACGACGCTCTTCCGATCTNNNCTCTGTAGTATCTACTGAATGA
GAGCACACGTCTGAACTCTTTCCCTACACGACGCTCTTCCGATCTNNNGGATTTTCCAACTTTTTTCAA
GAGCACACGTCTGAACTCTTTCCCTACACGACGCTCTTCCGATCTNNNGGATTTTCCAACTTTTTTCAA
GAGCACACGTCTGAACTCTTTCCCTACACGACGCTCTTCCGATCTNNNTATTTGTTCAAAGTCAGGAAT
GAGCACACGTCTGAACTCTTTCCCTACACGACGCTCTTCCGATCTNNNTTTCTAATAGGAAATTCAGTTC
GCACACGTCTGAACTCTTTCCCTACACGACGCTCTTCCGATCTNNNTTTGTTATATTGCACTTTGAAAA
GCACACGTCTGAACTCTTTCCCTACACGACGCTCTTCCGATCTNNNGCTATTTATATATGTGCGTTCAT
CACACGTCTGAACTCTTTCCCTACACGACGCTCTTCCGATCTNNNATGACAAACTTGGGAGAAAACATC
GCACACGTCTGAACTCTTTCCCTACACGACGCTCTTCCGATCTNNNTATCACAAAACTGCACAGAATC
AGCACACGTCTGAACTCTTTCCCTACACGACGCTCTTCCGATCTNNNTACATAGCCACGTCTTTTATTAT
GAGCACACGTCTGAACTCTTTCCCTACACGACGCTCTTCCGATCTNNNATTTCTCTTGTGACTTTCAAAT
GAGCACACGTCTGAACTCTTTCCCTACACGACGCTCTTCCGATCTNNNATTTCTCTTGTGACTTTCAAAT
GCACACGTCTGAACTCTTTCCCTACACGACGCTCTTCCGATCTNNNGAGAGGCACAAGAATCATTTCAT
GCACACGTCTGAACTCTTTCCCTACACGACGCTCTTCCGATCTNNNCTGTGGTACTTCCTTTAATGATC
AGCACACGTCTGAACTCTTTCCCTACACGACGCTCTTCCGATCTNNNATTTAAAAATGAACTCACAAGG
GAGCACACGTCTGAACTCTTTCCCTACACGACGCTCTTCCGATCTNNNAACTTCACTTTTTTCCAAATAG
GCACACGTCTGAACTCTTTCCCTACACGACGCTCTTCCGATCTNNNTAGGACATAATTGGGCTTATATG

AGCACACGTCTGAACTCTTTCCCTACACGACGCTCTTCCGATCTNNNACTTTAAAGTGGAAGAAAAGA
GAGCACACGTCTGAACTCTTTCCCTACACGACGCTCTTCCGATCTNNNTAAAATAGTAAGGGACTGGA
GCACACGTCTGAACTCTTTCCCTACACGACGCTCTTCCGATCTNNNATTTCTAAGAGGATTTCTTCACT
CACGTCTGAACTCTTTCCCTACACGACGCTCTTCCGATCTNNNAAACCTTGAAGAATCGAGAGCATCG
CACACGTCTGAACTCTTTCCCTACACGACGCTCTTCCGATCTNNNCAAGTCATCTTCAAAAGAATAGTC
ACACGTCTGAACTCTTTCCCTACACGACGCTCTTCCGATCTNNNGGAGTTAGAATAAGATCCTTCACC
AGCACACGTCTGAACTCTTTCCCTACACGACGCTCTTCCGATCTNNNTATATCCTTTACCCAGATTCAT
AGCACACGTCTGAACTCTTTCCCTACACGACGCTCTTCCGATCTNNNTATATCCTTTACCCAGATTCAT
GCACACGTCTGAACTCTTTCCCTACACGACGCTCTTCCGATCTNNNCACTGACTGAATTCATAAGAAAA
GCACACGTCTGAACTCTTTCCCTACACGACGCTCTTCCGATCTNNNGCCGTTCACTATTTTTTTTTACC
AGCACACGTCTGAACTCTTTCCCTACACGACGCTCTTCCGATCTNNNCTTCTTGCTTTGTCTTCATAAGC
CACACGTCTGAACTCTTTCCCTACACGACGCTCTTCCGATCTNNNCTCATCTCTTCTGGAGTGTTCATC
GCACACGTCTGAACTCTTTCCCTACACGACGCTCTTCCGATCTNNNCAATTTTACATTTGGGCTTAATG
AGCACACGTCTGAACTCTTTCCCTACACGACGCTCTTCCGATCTNNNTGATTACAAAGAGAATTCAACA
GAGCACACGTCTGAACTCTTTCCCTACACGACGCTCTTCCGATCTNNNGGAAAATGCATATAAAAACC
GAGCACACGTCTGAACTCTTTCCCTACACGACGCTCTTCCGATCTNNNGGAAAATGCATATAAAAACC
GAGCACACGTCTGAACTCTTTCCCTACACGACGCTCTTCCGATCTNNNGGAAAATGCATATAAAAACC
ACACGTCTGAACTCTTTCCCTACACGACGCTCTTCCGATCTNNNAAAAGGGTATGCTGTGAAAAATCAT
CACACGTCTGAACTCTTTCCCTACACGACGCTCTTCCGATCTNNNTAAGGTGGAAAGGATATTCGAA
AGCACACGTCTGAACTCTTTCCCTACACGACGCTCTTCCGATCTNNNCGTACTTGGTCATAACTTATTT
AGCACACGTCTGAACTCTTTCCCTACACGACGCTCTTCCGATCTNNNCACAATCTGGAAATGTAATAGA
GCACACGTCTGAACTCTTTCCCTACACGACGCTCTTCCGATCTNNNTGAAGTAGTTTTTTTCTTATGCA
AGCACACGTCTGAACTCTTTCCCTACACGACGCTCTTCCGATCTNNNTGATTCTCTGAGACATAGTTCT
GAGCACACGTCTGAACTCTTTCCCTACACGACGCTCTTCCGATCTNNNCTTTGGATTCTATTGCATACTT
AGCACACGTCTGAACTCTTTCCCTACACGACGCTCTTCCGATCTNNNTTCGAATATTTTTTATCCACAG
CACACGTCTGAACTCTTTCCCTACACGACGCTCTTCCGATCTNNNGTGTAAGTTATTGAACAGTACAG
AGCACACGTCTGAACTCTTTCCCTACACGACGCTCTTCCGATCTNNNATACTATAGTGCATCCTTGATT
AGCACACGTCTGAACTCTTTCCCTACACGACGCTCTTCCGATCTNNNTGATTATGCTTATATCCTGGAA
GCACACGTCTGAACTCTTTCCCTACACGACGCTCTTCCGATCTNNNTAGACGTGCGTTCTAATGTACAT
AGCACACGTCTGAACTCTTTCCCTACACGACGCTCTTCCGATCTNNNAAATTTCCTGAGTTCCTTAGAG
CACACGTCTGAACTCTTTCCCTACACGACGCTCTTCCGATCTNNNAATATGGTCGGACTTGGTAGCAT
GAGCACACGTCTGAACTCTTTCCCTACACGACGCTCTTCCGATCTNNNGTTTTAGATGTGTCTGTCTCT
AGCACACGTCTGAACTCTTTCCCTACACGACGCTCTTCCGATCTNNNTTTTAAGAAGACACTACTCCTT
AGCACACGTCTGAACTCTTTCCCTACACGACGCTCTTCCGATCTNNNTGTGGACCTATATGTAGATATG
GAGCACACGTCTGAACTCTTTCCCTACACGACGCTCTTCCGATCTNNNTTTTGCAGCACATTTTGTAGC
AGCACACGTCTGAACTCTTTCCCTACACGACGCTCTTCCGATCTNNNGTCAAATTTACAACCCTATTTG
AGCACACGTCTGAACTCTTTCCCTACACGACGCTCTTCCGATCTNNNATTGTAACAAATGTACCACTCT
GCACACGTCTGAACTCTTTCCCTACACGACGCTCTTCCGATCTNNNTACCAATCAAAATACCCTGTAAT
AGCACACGTCTGAACTCTTTCCCTACACGACGCTCTTCCGATCTNNNTTTATTTTCTGACCTTTTTGTGA
AGCACACGTCTGAACTCTTTCCCTACACGACGCTCTTCCGATCTNNNTTTATTTTCTGACCTTTTTGTGA
CACACGTCTGAACTCTTTCCCTACACGACGCTCTTCCGATCTNNNTTCTCACTTGTTCTGTTCTCCATCGG
ACACGTCTGAACTCTTTCCCTACACGACGCTCTTCCGATCTNNNATAACTTCTTTATCGGGGTTAAACA
AGCACACGTCTGAACTCTTTCCCTACACGACGCTCTTCCGATCTNNNCATTCTTATGTTTATTTCAGCAC

GCACACGTCTGAACTCTTTCCCTACACGACGCTCTTCCGATCTNNNCTTAGAGTATTTTCCTTCTCACTCA
GCACACGTCTGAACTCTTTCCCTACACGACGCTCTTCCGATCTNNNATCAAAACATCCCATCTCAGCATC
GAGCACACGTCTGAACTCTTTCCCTACACGACGCTCTTCCGATCTNNNTAAAACTTACACTCTAGGGGA
GAGCACACGTCTGAACTCTTTCCCTACACGACGCTCTTCCGATCTNNNTAGTTCTTCCCATAGAAATCTTT
AGCACACGTCTGAACTCTTTCCCTACACGACGCTCTTCCGATCTNNNGGATTTTTTTTCTCCTGCAGCA
GCACACGTCTGAACTCTTTCCCTACACGACGCTCTTCCGATCTNNNAGGGTTGCATATAAAATAAGACT
AGCACACGTCTGAACTCTTTCCCTACACGACGCTCTTCCGATCTNNNAAAACTTTTTTTTCAGAGACAG
GCACACGTCTGAACTCTTTCCCTACACGACGCTCTTCCGATCTNNNGTATTTATGCTGTGATTCTGATG
GCACACGTCTGAACTCTTTCCCTACACGACGCTCTTCCGATCTNNNATCCCAACATAGTATTTAGTGTTC
AGCACACGTCTGAACTCTTTCCCTACACGACGCTCTTCCGATCTNNNTTGGCTGTATATAACATTATCCA
GCACACGTCTGAACTCTTTCCCTACACGACGCTCTTCCGATCTNNNTTAAGGAATATTTTCTAGGTTGC
GAGCACACGTCTGAACTCTTTCCCTACACGACGCTCTTCCGATCTNNNTAGATGGGAGTTGAACTAGAC
AGCACACGTCTGAACTCTTTCCCTACACGACGCTCTTCCGATCTNNNGCTCAAACAGTAAGTATAATACC
GAGCACACGTCTGAACTCTTTCCCTACACGACGCTCTTCCGATCTNNNGAAAGAAAAGAGGTACAATT
AGCACACGTCTGAACTCTTTCCCTACACGACGCTCTTCCGATCTNNNTAACCAGATACTTGTTACCAATT
GCACACGTCTGAACTCTTTCCCTACACGACGCTCTTCCGATCTNNNATTGGTTTTACTTTTCTGAACTGC
GAGCACACGTCTGAACTCTTTCCCTACACGACGCTCTTCCGATCTNNNAAAATATCAACTCTAGTACTCC
AGCACACGTCTGAACTCTTTCCCTACACGACGCTCTTCCGATCTNNNAGGAGGACTTTGTTCAAAAGC
GAGCACACGTCTGAACTCTTTCCCTACACGACGCTCTTCCGATCTNNNGAGTTACGTATTAGGGTTCTCC
GCACACGTCTGAACTCTTTCCCTACACGACGCTCTTCCGATCTNNNAGATTTTGAAGAGGCTTATCCCAT
AGCACACGTCTGAACTCTTTCCCTACACGACGCTCTTCCGATCTNNNGCCATTATTTCATCCAATTATCT
AGCACACGTCTGAACTCTTTCCCTACACGACGCTCTTCCGATCTNNNCTTTATTCTGCTACACTCAAAGC
AGCACACGTCTGAACTCTTTCCCTACACGACGCTCTTCCGATCTNNNAAAGTCAACTAAATTTCACTCTG
CACACGTCTGAACTCTTTCCCTACACGACGCTCTTCCGATCTNNNATATTGTAAATAATGCTGAACAGC
GAGCACACGTCTGAACTCTTTCCCTACACGACGCTCTTCCGATCTNNNTTTCATTTGTAGATAGCACAAA
AGCACACGTCTGAACTCTTTCCCTACACGACGCTCTTCCGATCTNNNGCAGAATTTTTGATTTTCTTTGT
AGCACACGTCTGAACTCTTTCCCTACACGACGCTCTTCCGATCTNNNATATTGGGCATGAGTGGTTACA
AGCACACGTCTGAACTCTTTCCCTACACGACGCTCTTCCGATCTNNNTAGTTGGTTACTAGAGAAGTTT
GAGCACACGTCTGAACTCTTTCCCTACACGACGCTCTTCCGATCTNNNTTAAAGTTGCCTACTCATTTTA
GAGCACACGTCTGAACTCTTTCCCTACACGACGCTCTTCCGATCTNNNTTAAAGTTGCCTACTCATTTTA
CACACGTCTGAACTCTTTCCCTACACGACGCTCTTCCGATCTNNNAATACTAAGCTGACTAGTCATTCCA
AGCACACGTCTGAACTCTTTCCCTACACGACGCTCTTCCGATCTNNNCAATTTGTTCTCTCAATAAGCACA
AGCACACGTCTGAACTCTTTCCCTACACGACGCTCTTCCGATCTNNNGGCCTTTATATAGTGAGTTAGA
GCACACGTCTGAACTCTTTCCCTACACGACGCTCTTCCGATCTNNNTAAGACTAGTGGCCTTTATATAG
AGCACACGTCTGAACTCTTTCCCTACACGACGCTCTTCCGATCTNNNTCAAAATTGCTAAAAGAACAGA
AGCACACGTCTGAACTCTTTCCCTACACGACGCTCTTCCGATCTNNNCAAAAACCGCAATTACTTTTGCA
AGCACACGTCTGAACTCTTTCCCTACACGACGCTCTTCCGATCTNNNAAACCTCTACTCAGACTTATTAG
AGCACACGTCTGAACTCTTTCCCTACACGACGCTCTTCCGATCTNNNGACATCAGTAGTTTTAAAACTCT
GAGCACACGTCTGAACTCTTTCCCTACACGACGCTCTTCCGATCTNNNCTCATTTTTCTTCCTTTGCTTTAC
GAGCACACGTCTGAACTCTTTCCCTACACGACGCTCTTCCGATCTNNNGACAAAACTTCTACATAACACA
GAGCACACGTCTGAACTCTTTCCCTACACGACGCTCTTCCGATCTNNNGACAAAACTTCTACATAACACA
AGCACACGTCTGAACTCTTTCCCTACACGACGCTCTTCCGATCTNNNACAACATCGCTGTTTATACTCCA
GAGCACACGTCTGAACTCTTTCCCTACACGACGCTCTTCCGATCTNNNTAGTGTTGGATGAATGGGAA

CACACGTCTGAACTCTTTCCCTACACGACGCTCTTCCGATCTNNNGGTGTTTGTTAAATTGAATTGAG
CACACGTCTGAACTCTTTCCCTACACGACGCTCTTCCGATCTNNNGGTGTTTGTTAAATTGAATTGAG
CACACGTCTGAACTCTTTCCCTACACGACGCTCTTCCGATCTNNNACTTCTTCTGCTTCATATATCTCCAT
CACGTCTGAACTCTTTCCCTACACGACGCTCTTCCGATCTNNNATAGTGATATTGGTGAAGGTATTCA
AGCACACGTCTGAACTCTTTCCCTACACGACGCTCTTCCGATCTNNNAAAACTATGTATGTTTTTCCAG
CACACGTCTGAACTCTTTCCCTACACGACGCTCTTCCGATCTNNNAGATTTCTTTAAATTCCTGAAGCC
ACACGTCTGAACTCTTTCCCTACACGACGCTCTTCCGATCTNNNTTGTTTTTGCAGATGTGGATCATC
AGCACACGTCTGAACTCTTTCCCTACACGACGCTCTTCCGATCTNNNGTGGAAGCATTAGGAATTAAT
AGCACACGTCTGAACTCTTTCCCTACACGACGCTCTTCCGATCTNNNTTTTGAATTTTCTTTCAACCAT
GAGCACACGTCTGAACTCTTTCCCTACACGACGCTCTTCCGATCTNNNTAACTTTATCTATTGCTCAGT
ACACGTCTGAACTCTTTCCCTACACGACGCTCTTCCGATCTNNNGAGATTGAACACAGAGACAAACAT
AGCACACGTCTGAACTCTTTCCCTACACGACGCTCTTCCGATCTNNNATAATATATTGGTGAAGGAGC
AGCACACGTCTGAACTCTTTCCCTACACGACGCTCTTCCGATCTNNNATATTAGGTGTTCCTATGAAG
GAGCACACGTCTGAACTCTTTCCCTACACGACGCTCTTCCGATCTNNNTCCCTATTTGTGTCTGTTTTAT
AGCACACGTCTGAACTCTTTCCCTACACGACGCTCTTCCGATCTNNNATAAGGGATATGAAGGAAAA
GAGCACACGTCTGAACTCTTTCCCTACACGACGCTCTTCCGATCTNNNCACATAAGGGATATGAAGG
GAGCACACGTCTGAACTCTTTCCCTACACGACGCTCTTCCGATCTNNNCACATAAGGGATATGAAGG
GCACACGTCTGAACTCTTTCCCTACACGACGCTCTTCCGATCTNNNAGACACACACATTTATATTCTTG
GCACACGTCTGAACTCTTTCCCTACACGACGCTCTTCCGATCTNNNAGACACACACATTTATATTCTTG
GCACACGTCTGAACTCTTTCCCTACACGACGCTCTTCCGATCTNNNGAGACTACAGTGATCTTTTTGTC
ACACGTCTGAACTCTTTCCCTACACGACGCTCTTCCGATCTNNNAAAACTATTTCACTAAAAGCACAC
GAGCACACGTCTGAACTCTTTCCCTACACGACGCTCTTCCGATCTNNNGGTCTAGCATAAGTCTTAAAT
CACACGTCTGAACTCTTTCCCTACACGACGCTCTTCCGATCTNNNAGAGATCTGGTTTTTCTGTGCAT
CACACGTCTGAACTCTTTCCCTACACGACGCTCTTCCGATCTNNNTGAGATCTGGTTGTTTGAAACCAT
GCACACGTCTGAACTCTTTCCCTACACGACGCTCTTCCGATCTNNNAGATATCTGAAGAAGTCCTGAT
ACACGTCTGAACTCTTTCCCTACACGACGCTCTTCCGATCTNNNCATATGGCATAAGTGATGGTATCAT
AGCACACGTCTGAACTCTTTCCCTACACGACGCTCTTCCGATCTNNNTTGTAAATTGGATTGCTTTTT
AGCACACGTCTGAACTCTTTCCCTACACGACGCTCTTCCGATCTNNNACCAAGTGTTTGAATAGAAAT
AGCACACGTCTGAACTCTTTCCCTACACGACGCTCTTCCGATCTNNNATTGAATTATCTCCACAGAGT
AGCACACGTCTGAACTCTTTCCCTACACGACGCTCTTCCGATCTNNNATTGAATTATCTCCACAGAGT
AGCACACGTCTGAACTCTTTCCCTACACGACGCTCTTCCGATCTNNNAAATGAGCGATTATTGACATT
AGCACACGTCTGAACTCTTTCCCTACACGACGCTCTTCCGATCTNNNCTGTCTCAATAGATTACACTA
AGCACACGTCTGAACTCTTTCCCTACACGACGCTCTTCCGATCTNNNTATACCTTTCCTCATGATCTCT
GCACACGTCTGAACTCTTTCCCTACACGACGCTCTTCCGATCTNNNCACTAACAAGAAAATCCTGGATC
GCACACGTCTGAACTCTTTCCCTACACGACGCTCTTCCGATCTNNNCACTAACAAGAAAATCCTGGATC
AGCACACGTCTGAACTCTTTCCCTACACGACGCTCTTCCGATCTNNNATTCTAGAGAATAAACCTGAG
AGCACACGTCTGAACTCTTTCCCTACACGACGCTCTTCCGATCTNNNATTCTAGAGAATAAACCTGAG
AGCACACGTCTGAACTCTTTCCCTACACGACGCTCTTCCGATCTNNNAGTGATGAAAAAATTTCAGCAT
AGCACACGTCTGAACTCTTTCCCTACACGACGCTCTTCCGATCTNNNTTCTGCATCAGACAATATATTTG
AGCACACGTCTGAACTCTTTCCCTACACGACGCTCTTCCGATCTNNNCCTGATTTACTTCATTGTTTTC
AGCACACGTCTGAACTCTTTCCCTACACGACGCTCTTCCGATCTNNNCCTGATTTACTTCATTGTTTTC
CACACGTCTGAACTCTTTCCCTACACGACGCTCTTCCGATCTNNNGGAAATACAAAAGATGAGCTTAT
CACACGTCTGAACTCTTTCCCTACACGACGCTCTTCCGATCTNNNTATAGTGAGAGAGATTGTGTGTA

AGCACACGTCTGAACTCTTTCCCTACACGACGCTCTTCCGATCTNNNTAGTTCTCAGAAACATTTGGTTA
GCACACGTCTGAACTCTTTCCCTACACGACGCTCTTCCGATCTNNNAGTGATAGTTCTCAGAAACATTT
GAGCACACGTCTGAACTCTTTCCCTACACGACGCTCTTCCGATCTNNNAGTTCTTTCTTTCTCACCAAT
GAGCACACGTCTGAACTCTTTCCCTACACGACGCTCTTCCGATCTNNNAATCTACTTGTTGTCTGATCT
GCACACGTCTGAACTCTTTCCCTACACGACGCTCTTCCGATCTNNNAAGTTTCTCTCTCACATCCCATC
GCACACGTCTGAACTCTTTCCCTACACGACGCTCTTCCGATCTNNNAAGTTTCTCTCTCACATCCCATC
CACGTCTGAACTCTTTCCCTACACGACGCTCTTCCGATCTNNNCTCTCTCTCTGTCTCTCTCATCGGAC
GAGCACACGTCTGAACTCTTTCCCTACACGACGCTCTTCCGATCTNNNGTTCACTATATAGAGAGTTCT
GAGCACACGTCTGAACTCTTTCCCTACACGACGCTCTTCCGATCTNNNATGGACCTTCAAATTTTCATAT
GAGCACACGTCTGAACTCTTTCCCTACACGACGCTCTTCCGATCTNNNGAAGAATTCAGAAACATGAG
GAGCACACGTCTGAACTCTTTCCCTACACGACGCTCTTCCGATCTNNNGTAATTGAACTGAAAGCAGA
ACACGTCTGAACTCTTTCCCTACACGACGCTCTTCCGATCTNNNCTCACAAATATCTGTGCCTTTCATCG
ACACGTCTGAACTCTTTCCCTACACGACGCTCTTCCGATCTNNNATCTAGGTCTCTTTTATACACACCATC
CACACGTCTGAACTCTTTCCCTACACGACGCTCTTCCGATCTNNNCATGCTTACTGTCTCTGTTTATCAT
GCACACGTCTGAACTCTTTCCCTACACGACGCTCTTCCGATCTNNNTAGAGCATCCAAAAGTCTGTATC
GAGCACACGTCTGAACTCTTTCCCTACACGACGCTCTTCCGATCTNNNAACTCAATCTAAACATGAACA
AGCACACGTCTGAACTCTTTCCCTACACGACGCTCTTCCGATCTNNNAAACAATTTCTGGATCTGTATT
GAGCACACGTCTGAACTCTTTCCCTACACGACGCTCTTCCGATCTNNNCTTCTTGTTAGCTCAAATGAA
GAGCACACGTCTGAACTCTTTCCCTACACGACGCTCTTCCGATCTNNNAGTGTGTGCTACATAAATAAG
GAGCACACGTCTGAACTCTTTCCCTACACGACGCTCTTCCGATCTNNNTACAAATAATCTTGGTTGTTG
AGCACACGTCTGAACTCTTTCCCTACACGACGCTCTTCCGATCTNNNCTTCACCAGTAAGATTACATTT
GCACACGTCTGAACTCTTTCCCTACACGACGCTCTTCCGATCTNNNCTCTTTGTTACACAAGTAGAATC
GCACACGTCTGAACTCTTTCCCTACACGACGCTCTTCCGATCTNNNACTTAAGGAACAATAAGGTGGC
AGCACACGTCTGAACTCTTTCCCTACACGACGCTCTTCCGATCTNNNTCCCATCTATTGACATAATAAA
GAGCACACGTCTGAACTCTTTCCCTACACGACGCTCTTCCGATCTNNNTCAATCCTCCCTATTAATTTGA
AGCACACGTCTGAACTCTTTCCCTACACGACGCTCTTCCGATCTNNNTTAGAATCATCTTGGCAATTTA
GCACACGTCTGAACTCTTTCCCTACACGACGCTCTTCCGATCTNNNTATCTATGATCTATGTCACTTCCCA
GCACACGTCTGAACTCTTTCCCTACACGACGCTCTTCCGATCTNNNTATCTATGATCTATGTCACTTCCCA
GAGCACACGTCTGAACTCTTTCCCTACACGACGCTCTTCCGATCTNNNTGGGTAGGTCTCATCTAATTA
GCACACGTCTGAACTCTTTCCCTACACGACGCTCTTCCGATCTNNNGTTTAGATGAGGTATGTGACTC
GAGCACACGTCTGAACTCTTTCCCTACACGACGCTCTTCCGATCTNNNTTCACACTAGTTTTTATGTTTG
GCACACGTCTGAACTCTTTCCCTACACGACGCTCTTCCGATCTNNNGTTTAGATGAGGTATGTGACTC
GAGCACACGTCTGAACTCTTTCCCTACACGACGCTCTTCCGATCTNNNCTACTAATTGTATCTCTCTCTC
GAGCACACGTCTGAACTCTTTCCCTACACGACGCTCTTCCGATCTNNNGCTCTTATGACTTCTCATGAAC
GAGCACACGTCTGAACTCTTTCCCTACACGACGCTCTTCCGATCTNNNTTCAACAAATATTGATTGAGCA
AGCACACGTCTGAACTCTTTCCCTACACGACGCTCTTCCGATCTNNNTTCTTTTCACTTCCATTATTGTT
ACACGTCTGAACTCTTTCCCTACACGACGCTCTTCCGATCTNNNATACCCCAACCTTTTTTACCCATCGG
CACACGTCTGAACTCTTTCCCTACACGACGCTCTTCCGATCTNNNTGAAAAGAATACAAGAAACTGAGC
AGCACACGTCTGAACTCTTTCCCTACACGACGCTCTTCCGATCTNNNATTCTTAATTTTCTGCTTCCTTCC
AGCACACGTCTGAACTCTTTCCCTACACGACGCTCTTCCGATCTNNNATTCTTAATTTTCTGCTTCCTTCC
CACACGTCTGAACTCTTTCCCTACACGACGCTCTTCCGATCTNNNCTCTCTCTCTGTCTCTCTCATCGG
GCACACGTCTGAACTCTTTCCCTACACGACGCTCTTCCGATCTNNNGATGTATTAGTTAGGGTTCTTCA
GAGCACACGTCTGAACTCTTTCCCTACACGACGCTCTTCCGATCTNNNATACCTCACATGTGTAAATATC

AGCACACGTCTGAACTCTTTCCCTACACGACGCTCTTCCGATCTNNNATACCTCACATGTGTAAATATC
AGCACACGTCTGAACTCTTTCCCTACACGACGCTCTTCCGATCTNNNGATTGTACACAACGCTCTAGC
AGCACACGTCTGAACTCTTTCCCTACACGACGCTCTTCCGATCTNNNCATCCTTAGTCCCAGAAATGC
AGCACACGTCTGAACTCTTTCCCTACACGACGCTCTTCCGATCTNNNGAAGTGAAAAAACAGAGGAA
AGAGCACACGTCTGAACTCTTTCCCTACACGACGCTCTTCCGATCTNNNAAATCGGGAGAAAGAGAC
GCACACGTCTGAACTCTTTCCCTACACGACGCTCTTCCGATCTNNNTGGCAAGAACTCTTATAAAGAT
AGCACACGTCTGAACTCTTTCCCTACACGACGCTCTTCCGATCTNNNTATTACTGAAATCTCAAACCAA
AGCACACGTCTGAACTCTTTCCCTACACGACGCTCTTCCGATCTNNNAGTTATAAGAGAGAGGAGTA
CACACGTCTGAACTCTTTCCCTACACGACGCTCTTCCGATCTNNNCTATTAATCATTATGAGGGTGGA
GCACACGTCTGAACTCTTTCCCTACACGACGCTCTTCCGATCTNNNCATGGATTCTGTAGTCCCAGCAT
AGCACACGTCTGAACTCTTTCCCTACACGACGCTCTTCCGATCTNNNTCTAAAAACCTACAATTATGCA
AGCACACGTCTGAACTCTTTCCCTACACGACGCTCTTCCGATCTNNNGCTATAATTCATGGGATAAGG
AGCACACGTCTGAACTCTTTCCCTACACGACGCTCTTCCGATCTNNNCTCCAAATGAAACAGATATGA
CACACGTCTGAACTCTTTCCCTACACGACGCTCTTCCGATCTNNNATCCTCAACAGCTTGGAAATCATC
CACACGTCTGAACTCTTTCCCTACACGACGCTCTTCCGATCTNNNCTAATTTTGCTGAAAGAAATCAC
AGCACACGTCTGAACTCTTTCCCTACACGACGCTCTTCCGATCTNNNAGTAATTTGTTTTTATTCCTGCA
CACACGTCTGAACTCTTTCCCTACACGACGCTCTTCCGATCTNNNTGTGTGTGTGTATACATATATC
AGCACACGTCTGAACTCTTTCCCTACACGACGCTCTTCCGATCTNNNAATGAGAGAGGTATTTGACAT
GCACACGTCTGAACTCTTTCCCTACACGACGCTCTTCCGATCTNNNGCATGTTAAAATAAAACAGTTGT
GCACACGTCTGAACTCTTTCCCTACACGACGCTCTTCCGATCTNNNGCATGTTAAAATAAAACAGTTGT
CACACGTCTGAACTCTTTCCCTACACGACGCTCTTCCGATCTNNNTACCTTTAGGGATTGGATAAAGCA
AGAGCACACGTCTGAACTCTTTCCCTACACGACGCTCTTCCGATCTNNNGAATCAAGAAGAGAAGTCC
AGAGCACACGTCTGAACTCTTTCCCTACACGACGCTCTTCCGATCTNNNGAATCAAGAAGAGAAGTCC
GCACACGTCTGAACTCTTTCCCTACACGACGCTCTTCCGATCTNNNAGATAAAATGGCTTTCTATGTTT
GCACACGTCTGAACTCTTTCCCTACACGACGCTCTTCCGATCTNNNTACTAGCATTCCATCTTTTATGGC
AGCACACGTCTGAACTCTTTCCCTACACGACGCTCTTCCGATCTNNNCAGACAAGAAACGTACATTTTA
AGCACACGTCTGAACTCTTTCCCTACACGACGCTCTTCCGATCTNNNAGCCAGGTCCTTATAATAAATT
AGCACACGTCTGAACTCTTTCCCTACACGACGCTCTTCCGATCTNNNGATGTTCTTTTTATTTACCCC
GCACACGTCTGAACTCTTTCCCTACACGACGCTCTTCCGATCTNNNTACCATTATATGCTACTGATGAG
GCACACGTCTGAACTCTTTCCCTACACGACGCTCTTCCGATCTNNNTACCATTATATGCTACTGATGAG
GCACACGTCTGAACTCTTTCCCTACACGACGCTCTTCCGATCTNNNTAAAGAATCAGAGCCTCTCTCCA
AGAGCACACGTCTGAACTCTTTCCCTACACGACGCTCTTCCGATCTNNNTGTATCTATGACCCAAACTT
AGCACACGTCTGAACTCTTTCCCTACACGACGCTCTTCCGATCTNNNGGGAACAAGATGTTTTCAAT
AGCACACGTCTGAACTCTTTCCCTACACGACGCTCTTCCGATCTNNNGCAGCATGAAGACTATAGTTA
AGCACACGTCTGAACTCTTTCCCTACACGACGCTCTTCCGATCTNNNGAAATTTGCTGAGAAAGTAGA
AGCACACGTCTGAACTCTTTCCCTACACGACGCTCTTCCGATCTNNNCTTAGGATCTCACAATCTATTC
AGCACACGTCTGAACTCTTTCCCTACACGACGCTCTTCCGATCTNNNCTCGAAAAAAAATTCTTCCAAA
AGCACACGTCTGAACTCTTTCCCTACACGACGCTCTTCCGATCTNNNCAGAACTTTTCCTTGGTAATTG
GCACACGTCTGAACTCTTTCCCTACACGACGCTCTTCCGATCTNNNGAGGTAGTTAGAGGAGTAGAT
GCACACGTCTGAACTCTTTCCCTACACGACGCTCTTCCGATCTNNNACAATAAGGCATTATTTTTCAA
AGCACACGTCTGAACTCTTTCCCTACACGACGCTCTTCCGATCTNNNCTCCTTTTTTAAGGCTGAATAA
AGCACACGTCTGAACTCTTTCCCTACACGACGCTCTTCCGATCTNNNTCTATCAGTAAGGTCATTTTTCT
GCACACGTCTGAACTCTTTCCCTACACGACGCTCTTCCGATCTNNNATATGTAAATCAACTCCGAGAAG

AGCACACGTCTGAACTCTTTCCCTACACGACGCTCTTCCGATCTNNNTAGGTAGCTAGTTAGGTATGA
AGCACACGTCTGAACTCTTTCCCTACACGACGCTCTTCCGATCTNNNAACAATTCTGTTCCAACAATTCC
CACACGTCTGAACTCTTTCCCTACACGACGCTCTTCCGATCTNNNTGCCATATACATATATATATGCCAC
GCACACGTCTGAACTCTTTCCCTACACGACGCTCTTCCGATCTNNNTCAGAAATTATAATTCTCTTTCCT
GAGCACACGTCTGAACTCTTTCCCTACACGACGCTCTTCCGATCTNNNCAGAAAGAATTTGAAAACCTC
GCACACGTCTGAACTCTTTCCCTACACGACGCTCTTCCGATCTNNNCAAGAATCACTTCTCTTTTAACAC
AGCACACGTCTGAACTCTTTCCCTACACGACGCTCTTCCGATCTNNNACATGGCTTTAAAGAAATGTAA
AGCACACGTCTGAACTCTTTCCCTACACGACGCTCTTCCGATCTNNNACTCCATCTCTACTAAAAATACA
GAGCACACGTCTGAACTCTTTCCCTACACGACGCTCTTCCGATCTNNNCTGAAGTGTTTAAAGGTAAG
GCACACGTCTGAACTCTTTCCCTACACGACGCTCTTCCGATCTNNNGCTTCATAATACATTTGTGGAATC
AGCACACGTCTGAACTCTTTCCCTACACGACGCTCTTCCGATCTNNNAATGTACAAAAATCAGTAGCA
AGCACACGTCTGAACTCTTTCCCTACACGACGCTCTTCCGATCTNNNAAACTAAATGGGACTGAAACTT
AGCACACGTCTGAACTCTTTCCCTACACGACGCTCTTCCGATCTNNNCTCAATCAAAAGGTATATATCCA
CACACGTCTGAACTCTTTCCCTACACGACGCTCTTCCGATCTNNNAGGTATTATTATAACCTAGCCCACA
GAGCACACGTCTGAACTCTTTCCCTACACGACGCTCTTCCGATCTNNNCTCAACTATGATGTCCATAAAT
CACACGTCTGAACTCTTTCCCTACACGACGCTCTTCCGATCTNNNTATAGAAGTGGTTATGTCTTAGGC
AGCACACGTCTGAACTCTTTCCCTACACGACGCTCTTCCGATCTNNNTAGTAACAGTTACATCAGTCAA
CACACGTCTGAACTCTTTCCCTACACGACGCTCTTCCGATCTNNNAAAGCATAAGAAAAAGAGAACAG
CACACGTCTGAACTCTTTCCCTACACGACGCTCTTCCGATCTNNNAAAGCATAAGAAAAAGAGAACAG
AGCACACGTCTGAACTCTTTCCCTACACGACGCTCTTCCGATCTNNNTCAATAGGAGACATTGATCTAT
GAGCACACGTCTGAACTCTTTCCCTACACGACGCTCTTCCGATCTNNNAAAACAAGGGAAGACAAAC
CACACGTCTGAACTCTTTCCCTACACGACGCTCTTCCGATCTNNNATAACACTTAGCTAGAAATAGGC
GAGCACACGTCTGAACTCTTTCCCTACACGACGCTCTTCCGATCTNNNTCTTTAAAGAGTCATTCTGAA
GAGCACACGTCTGAACTCTTTCCCTACACGACGCTCTTCCGATCTNNNCTTGTCTTCTAGTACCTCTTTT
GCACACGTCTGAACTCTTTCCCTACACGACGCTCTTCCGATCTNNNTTTTTTCCTCATGCCTAGAATATC
AGCACACGTCTGAACTCTTTCCCTACACGACGCTCTTCCGATCTNNNAGATCTTCAAAGGAAAATAAA
GAGCACACGTCTGAACTCTTTCCCTACACGACGCTCTTCCGATCTNNNGTGAAGAAGATAAAGTCCCT
GAGCACACGTCTGAACTCTTTCCCTACACGACGCTCTTCCGATCTNNNGTGAAGAAGATAAAGTCCCT
GCACACGTCTGAACTCTTTCCCTACACGACGCTCTTCCGATCTNNNGCTAAAGTAAAAAAACAAGGTTT
CACACGTCTGAACTCTTTCCCTACACGACGCTCTTCCGATCTNNNGGGTTTTGCAAGTTTATTGCCATC
AGCACACGTCTGAACTCTTTCCCTACACGACGCTCTTCCGATCTNNNCATAATCTATCTATGACTTTCGT
CACACGTCTGAACTCTTTCCCTACACGACGCTCTTCCGATCTNNNACCTATTTAAGTGAAAAAAGCAAC
GAGCACACGTCTGAACTCTTTCCCTACACGACGCTCTTCCGATCTNNNTATATATTGTTTAGCCAGTCA
AGCACACGTCTGAACTCTTTCCCTACACGACGCTCTTCCGATCTNNNAAATGAGAGAGAGAGAGCAG
AGCACACGTCTGAACTCTTTCCCTACACGACGCTCTTCCGATCTNNNAAGGTCTCCTTTTTTCATTATTC
GCACACGTCTGAACTCTTTCCCTACACGACGCTCTTCCGATCTNNNAAGCAGAGGAAGAAGGAAAGC
AGCACACGTCTGAACTCTTTCCCTACACGACGCTCTTCCGATCTNNNTAGCATGCTAGCATTTGTATAA
AGCACACGTCTGAACTCTTTCCCTACACGACGCTCTTCCGATCTNNNCTACTGCCATTTCTTTTCACTCAT
AGCACACGTCTGAACTCTTTCCCTACACGACGCTCTTCCGATCTNNNGCTGATTTTTTCAAATATTCCAG
AGCACACGTCTGAACTCTTTCCCTACACGACGCTCTTCCGATCTNNNGTAGATAAAACAATAGCGTAGT
GCACACGTCTGAACTCTTTCCCTACACGACGCTCTTCCGATCTNNNTGCTGAGTCCTATAAATTTAGTT
GAGCACACGTCTGAACTCTTTCCCTACACGACGCTCTTCCGATCTNNNAAAGAAATGTGTTTATTCAAT
GCACACGTCTGAACTCTTTCCCTACACGACGCTCTTCCGATCTNNNTTTGGAAGAAATAAAATTGAGG

GAGCACACGTCTGAACTCTTTCCCTACACGACGCTCTTCCGATCTNNNGAAAGGAAAGAGAGAGAGA

GAGCACACGTCTGAACTCTTTCCCTACACGACGCTCTTCCGATCTNNNGAACCAGAAAGCTACATGTGC

AGCACACGTCTGAACTCTTTCCCTACACGACGCTCTTCCGATCTNNNAGTTCAGAATGTTCTTAGGATA

AGCACACGTCTGAACTCTTTCCCTACACGACGCTCTTCCGATCTNNNATAGACATACTGTTGAGAAAC

CACACGTCTGAACTCTTTCCCTACACGACGCTCTTCCGATCTNNNATTTTTCCCTTTCCTTTCCTTCATCGC

GAGCACACGTCTGAACTCTTTCCCTACACGACGCTCTTCCGATCTNNNTTTTCAGGACATTTTCTAATGA

AGCACACGTCTGAACTCTTTCCCTACACGACGCTCTTCCGATCTNNNTGACTTGAAATATAAGGCCACCA

AGCACACGTCTGAACTCTTTCCCTACACGACGCTCTTCCGATCTNNNTACCAGCTGAGTATTATATTTCCC

AGAGCACACGTCTGAACTCTTTCCCTACACGACGCTCTTCCGATCTNNNGAAGTAATTATCCTTCTCCAG

GAGCACACGTCTGAACTCTTTCCCTACACGACGCTCTTCCGATCTNNNTCTTATTTAACAGAAGCAGAAA

GAGCACACGTCTGAACTCTTTCCCTACACGACGCTCTTCCGATCTNNNTCACTGTTGTAAAATTATATGT

AGCACACGTCTGAACTCTTTCCCTACACGACGCTCTTCCGATCTNNNAACAATTTCATTCTTTTATGGCC

AGCACACGTCTGAACTCTTTCCCTACACGACGCTCTTCCGATCTNNNAACAATTTCATTCTTTTATGGCC

AGCACACGTCTGAACTCTTTCCCTACACGACGCTCTTCCGATCTNNNCACCTAACTAGTCACTTGAAAAC

AGCACACGTCTGAACTCTTTCCCTACACGACGCTCTTCCGATCTNNNCATAAGAAAAAAGGGGAATGA

AGCACACGTCTGAACTCTTTCCCTACACGACGCTCTTCCGATCTNNNCATAAGAAAAAAGGGGAATGA

AGCACACGTCTGAACTCTTTCCCTACACGACGCTCTTCCGATCTNNNGCCTGGTCAATATTAATACTTTT

AGCACACGTCTGAACTCTTTCCCTACACGACGCTCTTCCGATCTNNNAAAGTCTTCTGAAATTTGTTACC

AGCACACGTCTGAACTCTTTCCCTACACGACGCTCTTCCGATCTNNNTTTCTTTTCACACCACATTAAAA

GCACACGTCTGAACTCTTTCCCTACACGACGCTCTTCCGATCTNNNAGGGCTTACAAAATAAAGATTTA

GAGCACACGTCTGAACTCTTTCCCTACACGACGCTCTTCCGATCTNNNCAGAAGATAGGAAAATATGCA

CACACGTCTGAACTCTTTCCCTACACGACGCTCTTCCGATCTNNNTTTCAAACAAGTAGTCCTTATGCA

GAGCACACGTCTGAACTCTTTCCCTACACGACGCTCTTCCGATCTNNNTGTGAAATTAATTTTACCAGCT

AGCACACGTCTGAACTCTTTCCCTACACGACGCTCTTCCGATCTNNNCCAGAATATTGAAAATGTGCTT

AGCACACGTCTGAACTCTTTCCCTACACGACGCTCTTCCGATCTNNNTGTAGACTGAGATAATTGTGAT

ACACGTCTGAACTCTTTCCCTACACGACGCTCTTCCGATCTNNNTTCTCCATCCATGAAACCTTCATCGG

AGCACACGTCTGAACTCTTTCCCTACACGACGCTCTTCCGATCTNNNAAGGAGTGATTAGTGGTGAAT

GCACACGTCTGAACTCTTTCCCTACACGACGCTCTTCCGATCTNNNTCATTGAGCAAGACTTTGTTATCA

CACGTCTGAACTCTTTCCCTACACGACGCTCTTCCGATCTNNNATTTGAAACAATTCCTCTAGGTACATCC

ACACGTCTGAACTCTTTCCCTACACGACGCTCTTCCGATCTNNNGTTGAATTTTGGTGTTAGAGTGCAT

AGCACACGTCTGAACTCTTTCCCTACACGACGCTCTTCCGATCTNNNACCTAAATGATAAAATGCCAAA

GAGCACACGTCTGAACTCTTTCCCTACACGACGCTCTTCCGATCTNNNTGTTATCCACTTGTATAGTACT

AGCACACGTCTGAACTCTTTCCCTACACGACGCTCTTCCGATCTNNNTACTCATTTTTCCAGTGATCATAC

GCACACGTCTGAACTCTTTCCCTACACGACGCTCTTCCGATCTNNNGGAATATGAACTACTCCTAGTTCA

AGAGCACACGTCTGAACTCTTTCCCTACACGACGCTCTTCCGATCTNNNAAACTTAAATGAAATAAGGA

AGAGCACACGTCTGAACTCTTTCCCTACACGACGCTCTTCCGATCTNNNAAACTTAAATGAAATAAGGA

GAGCACACGTCTGAACTCTTTCCCTACACGACGCTCTTCCGATCTNNNTGTTAATAGGATTACACCTGA

AGCACACGTCTGAACTCTTTCCCTACACGACGCTCTTCCGATCTNNNTCTGTATTTCTTAGGTTTCTTCAC

AGCACACGTCTGAACTCTTTCCCTACACGACGCTCTTCCGATCTNNNAGAAGAAAAGTACCTTGGAAT

GAGCACACGTCTGAACTCTTTCCCTACACGACGCTCTTCCGATCTNNNTATCTTCACAGCAGTCATTTTA

AGCACACGTCTGAACTCTTTCCCTACACGACGCTCTTCCGATCTNNNAATAAATAACTACCATGGTCCAT

AGCACACGTCTGAACTCTTTCCCTACACGACGCTCTTCCGATCTNNNAAAGTAACACAGCCTGACTTACA

GAGCACACGTCTGAACTCTTTCCCTACACGACGCTCTTCCGATCTNNNGGAATATGCTTTAAAGATGTG

CACACGTCTGAACTCTTTCCCTACACGACGCTCTTCCGATCTNNNCTCATTTATTTGTTGCTCAATTCAT

AGCACACGTCTGAACTCTTTCCCTACACGACGCTCTTCCGATCTNNNCCTAACTCCAGAATTTTTACATTC

AGCACACGTCTGAACTCTTTCCCTACACGACGCTCTTCCGATCTNNNGAAGGAAGGAGTCTTTTTGAC

CACACGTCTGAACTCTTTCCCTACACGACGCTCTTCCGATCTNNNGTATGTACATGTACCTGTGTTCATC

AGCACACGTCTGAACTCTTTCCCTACACGACGCTCTTCCGATCTNNNGTAGAAAAACACACATATACAC

AGCACACGTCTGAACTCTTTCCCTACACGACGCTCTTCCGATCTNNNCTCTATACTTGTTCATCCGTATTC

GCACACGTCTGAACTCTTTCCCTACACGACGCTCTTCCGATCTNNNATATTTTCAGTTTCTATGGCCATC

GCACACGTCTGAACTCTTTCCCTACACGACGCTCTTCCGATCTNNNATCAAGGGAAGAAATAAAGGAT

AGCACACGTCTGAACTCTTTCCCTACACGACGCTCTTCCGATCTNNNTTTCCCGTTCAACAGTCATACAT

GCACACGTCTGAACTCTTTCCCTACACGACGCTCTTCCGATCTNNNAATTGCCCTTCCTCTTATCTCATCC

AGCACACGTCTGAACTCTTTCCCTACACGACGCTCTTCCGATCTNNNTTTATTCAGATTCCCCTGTTTTAC

AGCACACGTCTGAACTCTTTCCCTACACGACGCTCTTCCGATCTNNNTTTATTCAGATTCCCCTGTTTTAC

GAGCACACGTCTGAACTCTTTCCCTACACGACGCTCTTCCGATCTNNNATTTTCCAAAAGTATTTCTGTG

AGCACACGTCTGAACTCTTTCCCTACACGACGCTCTTCCGATCTNNNAATATGAGCTTTACGTAGACATT

AGCACACGTCTGAACTCTTTCCCTACACGACGCTCTTCCGATCTNNNCAGGAGAATATGGATGTCTCAC

AGCACACGTCTGAACTCTTTCCCTACACGACGCTCTTCCGATCTNNNATAGAAAATTTACAACATCCCAG

GCACACGTCTGAACTCTTTCCCTACACGACGCTCTTCCGATCTNNNAGGTATTTAAGGTTAAATGAGG

AGCACACGTCTGAACTCTTTCCCTACACGACGCTCTTCCGATCTNNNCAACGAAAAAGAGTTTATGTTC

AGCACACGTCTGAACTCTTTCCCTACACGACGCTCTTCCGATCTNNNATCTGAGATCTGAAGCTTACTT

ACACGTCTGAACTCTTTCCCTACACGACGCTCTTCCGATCTNNNATTCTGTCTCTCTCATTCTGTCATCGC

GAGCACACGTCTGAACTCTTTCCCTACACGACGCTCTTCCGATCTNNNATCTTTTGATAAAAGCAGCTAT

GCACACGTCTGAACTCTTTCCCTACACGACGCTCTTCCGATCTNNNGTTACTCAAAATTTCCAAGTTTGC

AGCACACGTCTGAACTCTTTCCCTACACGACGCTCTTCCGATCTNNNAGGACAAGAGTTCTTTTTATTTT

GCACACGTCTGAACTCTTTCCCTACACGACGCTCTTCCGATCTNNNGTACATGTACCCTAGAACTTAAAC

AGCACACGTCTGAACTCTTTCCCTACACGACGCTCTTCCGATCTNNNTGTTCATATGTGTGTGTATCTT

AGCACACGTCTGAACTCTTTCCCTACACGACGCTCTTCCGATCTNNNCCTTATTCTTTGGCAATAATCAT

CACACGTCTGAACTCTTTCCCTACACGACGCTCTTCCGATCTNNNCATTCTTGGAAATTACTAGAGACC

ACACGTCTGAACTCTTTCCCTACACGACGCTCTTCCGATCTNNNTTCAATCGATTGATCTGACTTCATCG

AGCACACGTCTGAACTCTTTCCCTACACGACGCTCTTCCGATCTNNNTTCTCTCATATATGCCAGACTTCA

GAGCACACGTCTGAACTCTTTCCCTACACGACGCTCTTCCGATCTNNNATATATGCATCTGTCTTTCGTC

GCACACGTCTGAACTCTTTCCCTACACGACGCTCTTCCGATCTNNNGTATGTAACCTAGTACTTTCACTC

GAGCACACGTCTGAACTCTTTCCCTACACGACGCTCTTCCGATCTNNNGAACTGAAGAAGTAAATCAA

AGAGCACACGTCTGAACTCTTTCCCTACACGACGCTCTTCCGATCTNNNTTACAGATGAGAAAATTGAG

AGCACACGTCTGAACTCTTTCCCTACACGACGCTCTTCCGATCTNNNATTTACAGATGAGAAAATTGAG

GAGCACACGTCTGAACTCTTTCCCTACACGACGCTCTTCCGATCTNNNAATATAATGAACTTTTGGCAGA

AGCACACGTCTGAACTCTTTCCCTACACGACGCTCTTCCGATCTNNNCTAAGAGATTAGATTTTAGGTC

AGCACACGTCTGAACTCTTTCCCTACACGACGCTCTTCCGATCTNNNTTATAGAATATGTTCTAGCACCT

AGCACACGTCTGAACTCTTTCCCTACACGACGCTCTTCCGATCTNNNAAGGACAGGAAATTAATGTCTA

AGCACACGTCTGAACTCTTTCCCTACACGACGCTCTTCCGATCTNNNAATCTGGCTTAATTTCAGTGTAT

GAGCACACGTCTGAACTCTTTCCCTACACGACGCTCTTCCGATCTNNNTATGTAAATAAAATGGGTGTC

AGCACACGTCTGAACTCTTTCCCTACACGACGCTCTTCCGATCTNNNTCTTACCAGCATTTATGATGATA

AGCACACGTCTGAACTCTTTCCCTACACGACGCTCTTCCGATCTNNNTATTGGCCTAATTTCACTTGATAC

AGCACACGTCTGAACTCTTTCCCTACACGACGCTCTTCCGATCTNNNTATTGGCCTAATTTCACTTGATAC

GAGCACACGTCTGAACTCTTTCCCTACACGACGCTCTTCCGATCTNNNCTAGTACTGCTGGATTTGATAT
AGAGCACACGTCTGAACTCTTTCCCTACACGACGCTCTTCCGATCTNNNTAGTATCCTTCAACACATGTG
CACACGTCTGAACTCTTTCCCTACACGACGCTCTTCCGATCTNNNTTTTGTGTACGTATGAGTGTATCAT
GAGCACACGTCTGAACTCTTTCCCTACACGACGCTCTTCCGATCTNNNACAAAATCATGTCAAAAGAAT
AGCACACGTCTGAACTCTTTCCCTACACGACGCTCTTCCGATCTNNNATTGAATTTGGAGTTGCAAATT
CACACGTCTGAACTCTTTCCCTACACGACGCTCTTCCGATCTNNNTGGACTGTTAATTTGTAGAATTGCA
AGCACACGTCTGAACTCTTTCCCTACACGACGCTCTTCCGATCTNNNGTCTTCCAGGTTTATCTATGTTA
AGCACACGTCTGAACTCTTTCCCTACACGACGCTCTTCCGATCTNNNGCTGCTGTTATTATCACATTAAT
GAGCACACGTCTGAACTCTTTCCCTACACGACGCTCTTCCGATCTNNNTTAAAAATGTTCATCAGACCTT
CACACGTCTGAACTCTTTCCCTACACGACGCTCTTCCGATCTNNNCTTATGTAAAGGGAAGTGTCTTAC
ACACGTCTGAACTCTTTCCCTACACGACGCTCTTCCGATCTNNNCTCACAGAATAGTTTTACTTTCCCATC
AGCACACGTCTGAACTCTTTCCCTACACGACGCTCTTCCGATCTNNNATTAGAATTCACCTGAGGAGAT
AGCACACGTCTGAACTCTTTCCCTACACGACGCTCTTCCGATCTNNNTATTTGCCAAAATTTGAGCTTTG
GAGCACACGTCTGAACTCTTTCCCTACACGACGCTCTTCCGATCTNNNGTGACTTTTATGGTTTGTGAA
CACACGTCTGAACTCTTTCCCTACACGACGCTCTTCCGATCTNNNATCCTTTAATCTTTCCAACAATCCAT
AGCACACGTCTGAACTCTTTCCCTACACGACGCTCTTCCGATCTNNNATTAGTGAAAGAGATGAAAATC
AGCACACGTCTGAACTCTTTCCCTACACGACGCTCTTCCGATCTNNNATTAGTGAAAGAGATGAAAATC
GCACACGTCTGAACTCTTTCCCTACACGACGCTCTTCCGATCTNNNCAAAGAGAAGGAAGGGAAAACA
AGCACACGTCTGAACTCTTTCCCTACACGACGCTCTTCCGATCTNNNAACCTCCAAATTATTTTGAACTT
GAGCACACGTCTGAACTCTTTCCCTACACGACGCTCTTCCGATCTNNNCTTTTACTTTGGTTTTTCTGGC
GCACACGTCTGAACTCTTTCCCTACACGACGCTCTTCCGATCTNNNAGTAAAACCTGCTTTTAGTGTATC
CACACGTCTGAACTCTTTCCCTACACGACGCTCTTCCGATCTNNNCTCTTCTGTGATCAGTCTTCCATCG
GAGCACACGTCTGAACTCTTTCCCTACACGACGCTCTTCCGATCTNNNGCAAAAGTAATAGCAGTTTTT
GCACACGTCTGAACTCTTTCCCTACACGACGCTCTTCCGATCTNNNTATATAATCCCTCTTCCCATCAACA
GAGCACACGTCTGAACTCTTTCCCTACACGACGCTCTTCCGATCTNNNGAGAAGGAAGAAGGAAAAAA
GCACACGTCTGAACTCTTTCCCTACACGACGCTCTTCCGATCTNNNGGGAGTGTATTAGGGATATAGA
GAGCACACGTCTGAACTCTTTCCCTACACGACGCTCTTCCGATCTNNNCTTACTAAAGGGCATAAGGA
GAGCACACGTCTGAACTCTTTCCCTACACGACGCTCTTCCGATCTNNNACACTATTAGGCCTCAAAATAT
GAGCACACGTCTGAACTCTTTCCCTACACGACGCTCTTCCGATCTNNNACACTATTAGGCCTCAAAATAT
GAGCACACGTCTGAACTCTTTCCCTACACGACGCTCTTCCGATCTNNNACACTATTAGGCCTCAAAATAT
AGCACACGTCTGAACTCTTTCCCTACACGACGCTCTTCCGATCTNNNCGTTTAAACAGGTACCCTAAACA
GAGCACACGTCTGAACTCTTTCCCTACACGACGCTCTTCCGATCTNNNTTATAGGACGGTTATGAGGAT
GCACACGTCTGAACTCTTTCCCTACACGACGCTCTTCCGATCTNNNCTCTGGGTAACACACTTCTTCATC
AGCACACGTCTGAACTCTTTCCCTACACGACGCTCTTCCGATCTNNNCTTGAATCATTTCACATAGAGTA
ACACGTCTGAACTCTTTCCCTACACGACGCTCTTCCGATCTNNNATATGCGTATGTACACACATGCATC
AGAGCACACGTCTGAACTCTTTCCCTACACGACGCTCTTCCGATCTNNNGAATCATTGAGGACCATCTTA
GAGCACACGTCTGAACTCTTTCCCTACACGACGCTCTTCCGATCTNNNAAATTGAGTGACAGACTATAC
GCACACGTCTGAACTCTTTCCCTACACGACGCTCTTCCGATCTNNNGTATCATACTAGAATGTGTGAGA
AGAGCACACGTCTGAACTCTTTCCCTACACGACGCTCTTCCGATCTNNNGCCTTTACTAATAGCTTTAAA
AGCACACGTCTGAACTCTTTCCCTACACGACGCTCTTCCGATCTNNNGATGATAAAGAACTCAAACCAA
GAGCACACGTCTGAACTCTTTCCCTACACGACGCTCTTCCGATCTNNNGTGATTTGGAGAAAAGTTCTT
GAGCACACGTCTGAACTCTTTCCCTACACGACGCTCTTCCGATCTNNNGAGTCAGGTTTACAAAAAAGC
GCACACGTCTGAACTCTTTCCCTACACGACGCTCTTCCGATCTNNNACATAATTATAAACTGCTTGGTG

GCACACGTCTGAACTCTTTCCCTACACGACGCTCTTCCGATCTNNNTATGTGTGTGTGTCTGTACAT
AGCACACGTCTGAACTCTTTCCCTACACGACGCTCTTCCGATCTNNNACACAGTTTGAGAATTGTTAA
GCACACGTCTGAACTCTTTCCCTACACGACGCTCTTCCGATCTNNNTATCATGTAAAGTCTAGCCAAG
AGCACACGTCTGAACTCTTTCCCTACACGACGCTCTTCCGATCTNNNAATCTCTATGAAAAACATATGG
AGCACACGTCTGAACTCTTTCCCTACACGACGCTCTTCCGATCTNNNGGAAAATGAGTTCATAGAGTT
AGCACACGTCTGAACTCTTTCCCTACACGACGCTCTTCCGATCTNNNAAGTATCTTCATGACAGAGAT
GCACACGTCTGAACTCTTTCCCTACACGACGCTCTTCCGATCTNNNCTCACTCAAGTTACCATACTGCAT
CACACGTCTGAACTCTTTCCCTACACGACGCTCTTCCGATCTNNNAGCTCATTCCAGAATAATCAATAC
GCACACGTCTGAACTCTTTCCCTACACGACGCTCTTCCGATCTNNNCATGCTTTGGACTGAATAAATC
GAGCACACGTCTGAACTCTTTCCCTACACGACGCTCTTCCGATCTNNNAAAAAAGAAAACATGCCAGT
AGCACACGTCTGAACTCTTTCCCTACACGACGCTCTTCCGATCTNNNGATATGAATGATACCATACCTC
AGCACACGTCTGAACTCTTTCCCTACACGACGCTCTTCCGATCTNNNATTTCATTATCTAGAAACCTCC
CACACGTCTGAACTCTTTCCCTACACGACGCTCTTCCGATCTNNNCCTTATCTCTAAAGTGATGGTATC
ACACGTCTGAACTCTTTCCCTACACGACGCTCTTCCGATCTNNNCTTCAACCTTGTCCTTGTTGCATCG
CACACGTCTGAACTCTTTCCCTACACGACGCTCTTCCGATCTNNNCATAAGTGAGATACATTCCCTTCA
CACACGTCTGAACTCTTTCCCTACACGACGCTCTTCCGATCTNNNTTTCCTTAAAAGAAATGTCTTGCC
ACACGTCTGAACTCTTTCCCTACACGACGCTCTTCCGATCTNNNGCAAGACCTCATCTCTATTTTATCAT
AGCACACGTCTGAACTCTTTCCCTACACGACGCTCTTCCGATCTNNNAGTTCCTTATTCTAAATGACCC
GCACACGTCTGAACTCTTTCCCTACACGACGCTCTTCCGATCTNNNTGTGTATAGATACATATGTGTGT
AGCACACGTCTGAACTCTTTCCCTACACGACGCTCTTCCGATCTNNNGGTATGGAGTAGAGATACAAT
AGCACACGTCTGAACTCTTTCCCTACACGACGCTCTTCCGATCTNNNGCTATAGAATTCATCCATGTAA
AGCACACGTCTGAACTCTTTCCCTACACGACGCTCTTCCGATCTNNNGTGAAATCCAAGATTCATTTAC
CACACGTCTGAACTCTTTCCCTACACGACGCTCTTCCGATCTNNNCGAAGTCTCTATTAACCACCCATC
GCACACGTCTGAACTCTTTCCCTACACGACGCTCTTCCGATCTNNNACTTGTCTCATTAGCACAAAACA
CACACGTCTGAACTCTTTCCCTACACGACGCTCTTCCGATCTNNNGAGGTAGGAAGATTGCTTTAGCA
AGCACACGTCTGAACTCTTTCCCTACACGACGCTCTTCCGATCTNNNAAATAGCTGGGCATTTCAAAT
CACACGTCTGAACTCTTTCCCTACACGACGCTCTTCCGATCTNNNAAACCAACCTATTAGTTCTAACAC
GAGCACACGTCTGAACTCTTTCCCTACACGACGCTCTTCCGATCTNNNGAATGATGTGTTCTGATCTTT
CACACGTCTGAACTCTTTCCCTACACGACGCTCTTCCGATCTNNNTCTATTTTTGTGTATTCTTGCCCAT
AGCACACGTCTGAACTCTTTCCCTACACGACGCTCTTCCGATCTNNNAACTTATGACCAGCATTTAAAA
CACGTCTGAACTCTTTCCCTACACGACGCTCTTCCGATCTNNNCTAGCTTCTTTTTACCCTCTTCATCG
AGCACACGTCTGAACTCTTTCCCTACACGACGCTCTTCCGATCTNNNGTAATTGATAGAGCAAGAAA
AGCACACGTCTGAACTCTTTCCCTACACGACGCTCTTCCGATCTNNNGTAATTGATAGAGCAAGAAA
AGCACACGTCTGAACTCTTTCCCTACACGACGCTCTTCCGATCTNNNTGAATTATCAGTCAAGACATT
GCACACGTCTGAACTCTTTCCCTACACGACGCTCTTCCGATCTNNNATGTAACATAAAAAGGACAGA
AGCACACGTCTGAACTCTTTCCCTACACGACGCTCTTCCGATCTNNNTCTTGTATTAGTCAGGGTTCTA
AGCACACGTCTGAACTCTTTCCCTACACGACGCTCTTCCGATCTNNNTCTTGTATTAGTCAGGGTTCTA
GCACACGTCTGAACTCTTTCCCTACACGACGCTCTTCCGATCTNNNATGCACATATATACATGTTTGTG
AGCACACGTCTGAACTCTTTCCCTACACGACGCTCTTCCGATCTNNNAGTCCATAGATCAATTTCATG
AGCACACGTCTGAACTCTTTCCCTACACGACGCTCTTCCGATCTNNNCAACAACAACAACTTTTAACTA
CACACGTCTGAACTCTTTCCCTACACGACGCTCTTCCGATCTNNNGTGACTTGTGAGAAATTTTATG
CACACGTCTGAACTCTTTCCCTACACGACGCTCTTCCGATCTNNNCAATTTACACCACTTCAAAATGCA
AGAGCACACGTCTGAACTCTTTCCCTACACGACGCTCTTCCGATCTNNNTAGTCCAAACAACCCATTA

CACACGTCTGAACTCTTTCCCTACACGACGCTCTTCCGATCTNNNATTGTTCCCATTTTAGAAACAATC
CACACGTCTGAACTCTTTCCCTACACGACGCTCTTCCGATCTNNNATTGTTCCCATTTTAGAAACAATC
CACACGTCTGAACTCTTTCCCTACACGACGCTCTTCCGATCTNNNTTTTCCACTTTGAATATCCTCACAT
GCACACGTCTGAACTCTTTCCCTACACGACGCTCTTCCGATCTNNNCTACCAAAATGCTGGGATTACAT
CACACGTCTGAACTCTTTCCCTACACGACGCTCTTCCGATCTNNNAAAAATGATGAAGCTCTGTATTTC
CACACGTCTGAACTCTTTCCCTACACGACGCTCTTCCGATCTNNNATTACCACCTTACTTTCTTACTCAT
AGCACACGTCTGAACTCTTTCCCTACACGACGCTCTTCCGATCTNNNTTTCCTAACTATGAGTGGATTA
GAGCACACGTCTGAACTCTTTCCCTACACGACGCTCTTCCGATCTNNNTATCAATTGAAAAGAAAGGA
AGCACACGTCTGAACTCTTTCCCTACACGACGCTCTTCCGATCTNNNACAGGTTATTGCATAAAATTCA
AGCACACGTCTGAACTCTTTCCCTACACGACGCTCTTCCGATCTNNNGGGAATTATCTGGAGTTTATAG
GAGCACACGTCTGAACTCTTTCCCTACACGACGCTCTTCCGATCTNNNTAGTGTATAATCCTGCTATGA
AGCACACGTCTGAACTCTTTCCCTACACGACGCTCTTCCGATCTNNNGTGATTTGTCACCATCTTTTTAT
AGCACACGTCTGAACTCTTTCCCTACACGACGCTCTTCCGATCTNNNGTGATTTGTCACCATCTTTTTAT
GCACACGTCTGAACTCTTTCCCTACACGACGCTCTTCCGATCTNNNTCCATGTTATAGATCTTCAACAA
AGCACACGTCTGAACTCTTTCCCTACACGACGCTCTTCCGATCTNNNTCAGTTTAAAAATGGATATAG
AGCACACGTCTGAACTCTTTCCCTACACGACGCTCTTCCGATCTNNNGTAATCAAGGAACTGGAAGAG
AGCACACGTCTGAACTCTTTCCCTACACGACGCTCTTCCGATCTNNNAACATAATTCAAATGGTTCAG
GAGCACACGTCTGAACTCTTTCCCTACACGACGCTCTTCCGATCTNNNATTGTGTCAAAAAAGAAAGA
AGCACACGTCTGAACTCTTTCCCTACACGACGCTCTTCCGATCTNNNATTTATCTTTCAGGACAACAT
GAGCACACGTCTGAACTCTTTCCCTACACGACGCTCTTCCGATCTNNNATATACACACACGTATAATAC
AGCACACGTCTGAACTCTTTCCCTACACGACGCTCTTCCGATCTNNNTGTCAGCAATGCATATATATGT
AGCACACGTCTGAACTCTTTCCCTACACGACGCTCTTCCGATCTNNNTTTTGTTTGTTTGTTTCTGTTTT
AGCACACGTCTGAACTCTTTCCCTACACGACGCTCTTCCGATCTNNNTTTTGTTTGTTTGTTTCTGTTTT
GAGCACACGTCTGAACTCTTTCCCTACACGACGCTCTTCCGATCTNNNAATTGTTCAGAAGGAAACAA
AGCACACGTCTGAACTCTTTCCCTACACGACGCTCTTCCGATCTNNNATTTCTAGAAACATTTCTTGCTC
GAGCACACGTCTGAACTCTTTCCCTACACGACGCTCTTCCGATCTNNNAAAATTGTTGATGATCTAGTC
AGCACACGTCTGAACTCTTTCCCTACACGACGCTCTTCCGATCTNNNTAACATTTTGTCTGCCATACATC
GAGCACACGTCTGAACTCTTTCCCTACACGACGCTCTTCCGATCTNNNTGTTTGTGTACATATAAGTC
AGCACACGTCTGAACTCTTTCCCTACACGACGCTCTTCCGATCTNNNGATTCTCATATCCTATCTCAGT
AGCACACGTCTGAACTCTTTCCCTACACGACGCTCTTCCGATCTNNNCCTCCATGATTTATGTTATTTG
AGCACACGTCTGAACTCTTTCCCTACACGACGCTCTTCCGATCTNNNTCCAAAGCCCTTCAGTTATATCA
ACACGTCTGAACTCTTTCCCTACACGACGCTCTTCCGATCTNNNGTGTCTTCTTGCAATTCATTCCATCG
CACACGTCTGAACTCTTTCCCTACACGACGCTCTTCCGATCTNNNTCACAGAATGAGCACAAAAGCAT
AGCACACGTCTGAACTCTTTCCCTACACGACGCTCTTCCGATCTNNNCAGTGGAGATTGGTATTTAAA
AGCACACGTCTGAACTCTTTCCCTACACGACGCTCTTCCGATCTNNNGGTTTTGTATGTACATGGTAA
GCACACGTCTGAACTCTTTCCCTACACGACGCTCTTCCGATCTNNNTATTCATAATCTATGGCTTAGGG
AGCACACGTCTGAACTCTTTCCCTACACGACGCTCTTCCGATCTNNNGAGTACTTTTCAAGTGACTAAA
AGCACACGTCTGAACTCTTTCCCTACACGACGCTCTTCCGATCTNNNAAAAATAGGGAAGAAACACCA
GAGCACACGTCTGAACTCTTTCCCTACACGACGCTCTTCCGATCTNNNTAAATGCGACTAATGGTTAT
GCACACGTCTGAACTCTTTCCCTACACGACGCTCTTCCGATCTNNNAAGAGATTACTGTGTAATAGTC
AGCACACGTCTGAACTCTTTCCCTACACGACGCTCTTCCGATCTNNNTAATTTCCAGAATGAATGAAT
AGCACACGTCTGAACTCTTTCCCTACACGACGCTCTTCCGATCTNNNTTGCTATATTGGAATGTATTG
GAGCACACGTCTGAACTCTTTCCCTACACGACGCTCTTCCGATCTNNNAAAAGAGCAAAATGTTTCTT

AGCACACGTCTGAACTCTTTCCCTACACGACGCTCTTCCGATCTNNNACTTGAGTCCCCAAATTTACCAT
AGCACACGTCTGAACTCTTTCCCTACACGACGCTCTTCCGATCTNNNTTTGATGCATTGATATTTTTCC
AGCACACGTCTGAACTCTTTCCCTACACGACGCTCTTCCGATCTNNNTTTGATGCATTGATATTTTTCC
GAGCACACGTCTGAACTCTTTCCCTACACGACGCTCTTCCGATCTNNNTTACATCTATGTTCATGAAGG
GCACACGTCTGAACTCTTTCCCTACACGACGCTCTTCCGATCTNNNGAGAAGATAGTGATCCATTGTA
CACACGTCTGAACTCTTTCCCTACACGACGCTCTTCCGATCTNNNAAAATCAAAAAATATGGGTCGTA
AGCACACGTCTGAACTCTTTCCCTACACGACGCTCTTCCGATCTNNNGAAGGCATAATGGTGTATCTAT
AGCACACGTCTGAACTCTTTCCCTACACGACGCTCTTCCGATCTNNNCAGAGTTCTGTATTACTGTACT
GCACACGTCTGAACTCTTTCCCTACACGACGCTCTTCCGATCTNNNCCTCATTTTTCTTCCCTAATTCCAT
AGCACACGTCTGAACTCTTTCCCTACACGACGCTCTTCCGATCTNNNACTTCTCAACCTTTAAAATGAA
AGAGCACACGTCTGAACTCTTTCCCTACACGACGCTCTTCCGATCTNNNTATACTAAGGTAAACACAGA
GAGCACACGTCTGAACTCTTTCCCTACACGACGCTCTTCCGATCTNNNGTTTCTGAGAATTCTTGGTTA
AGAGCACACGTCTGAACTCTTTCCCTACACGACGCTCTTCCGATCTNNNCTAAGAATACCAGCCAATACT
AGCACACGTCTGAACTCTTTCCCTACACGACGCTCTTCCGATCTNNNTCAGGAATGGAAAAGATATAT
AGCACACGTCTGAACTCTTTCCCTACACGACGCTCTTCCGATCTNNNTTGGTCTGGAACTAAACCATCA
GAGCACACGTCTGAACTCTTTCCCTACACGACGCTCTTCCGATCTNNNCTTACATGGGAAATTCTTATG
AGCACACGTCTGAACTCTTTCCCTACACGACGCTCTTCCGATCTNNNAATTTGAAACAAGGGAAATAG
AGCACACGTCTGAACTCTTTCCCTACACGACGCTCTTCCGATCTNNNTATTGTTCCTCCGAATTTCTCCA
AGCACACGTCTGAACTCTTTCCCTACACGACGCTCTTCCGATCTNNNACTTTTTGTTGAGAAAAGAA
GAGCACACGTCTGAACTCTTTCCCTACACGACGCTCTTCCGATCTNNNATGTACCCATTAGTTGTACTA
GAGCACACGTCTGAACTCTTTCCCTACACGACGCTCTTCCGATCTNNNATTCTGTTTCAAAACTCATGA
AGCACACGTCTGAACTCTTTCCCTACACGACGCTCTTCCGATCTNNNATACTTAGTGAGAAAACAGACA
AGCACACGTCTGAACTCTTTCCCTACACGACGCTCTTCCGATCTNNNGAAGCCAGAAAATATTTTGTTT
AGCACACGTCTGAACTCTTTCCCTACACGACGCTCTTCCGATCTNNNGAAGCCAGAAAATATTTTGTTT
AGCACACGTCTGAACTCTTTCCCTACACGACGCTCTTCCGATCTNNNATCACAATCGAGGTAATGAAC
AGCACACGTCTGAACTCTTTCCCTACACGACGCTCTTCCGATCTNNNCTGATGTCTACGTGGATTTTAT
GAGCACACGTCTGAACTCTTTCCCTACACGACGCTCTTCCGATCTNNNCTTCATTTCCTTGAGGTTATTT
CACACGTCTGAACTCTTTCCCTACACGACGCTCTTCCGATCTNNNAGGTTTAGGCATATGGGTTTCAT
AGCACACGTCTGAACTCTTTCCCTACACGACGCTCTTCCGATCTNNNATATTTGTCTACTTGTTTGCATT
GAGCACACGTCTGAACTCTTTCCCTACACGACGCTCTTCCGATCTNNNTAGAGTCAATTAAGTTTAATG
GAGCACACGTCTGAACTCTTTCCCTACACGACGCTCTTCCGATCTNNNTACATGTAAGAACACAAAAAA
GCACACGTCTGAACTCTTTCCCTACACGACGCTCTTCCGATCTNNNCTGAGCCCATGTATTAGAAATATC
AGCACACGTCTGAACTCTTTCCCTACACGACGCTCTTCCGATCTNNNAAAGAAGCCAGTTCATGTAGCA
CACACGTCTGAACTCTTTCCCTACACGACGCTCTTCCGATCTNNNGGAGCAAGTTCAGAAAAATGCAT
AGCACACGTCTGAACTCTTTCCCTACACGACGCTCTTCCGATCTNNNCCACCCTCATTATTCAAAATAAA
GAGCACACGTCTGAACTCTTTCCCTACACGACGCTCTTCCGATCTNNNCATTTCTTACATGGATCACAA
CACACGTCTGAACTCTTTCCCTACACGACGCTCTTCCGATCTNNNCCTCAATGTAAATATTTCATAACCC
CACACGTCTGAACTCTTTCCCTACACGACGCTCTTCCGATCTNNNGGTCAAAATTCATCTGGATTTTAC
ACACGTCTGAACTCTTTCCCTACACGACGCTCTTCCGATCTNNNAAGGTCAAAATTCATCTGGATTTCA
GCACACGTCTGAACTCTTTCCCTACACGACGCTCTTCCGATCTNNNTTGTATCTATGTTCATCAGGAAT
GAGCACACGTCTGAACTCTTTCCCTACACGACGCTCTTCCGATCTNNNCTGTCTCAATAGATTACACTAC
AGCACACGTCTGAACTCTTTCCCTACACGACGCTCTTCCGATCTNNNCTCTGAGTTTCTTCTGCTTCCAT
AGCACACGTCTGAACTCTTTCCCTACACGACGCTCTTCCGATCTNNNATAGTGCTTTTGACTCACATTA

AGCACACGTCTGAACTCTTTCCCTACACGACGCTCTTCCGATCTNNNTGTTCTGTAAAGTAAGTTTGA
GCACACGTCTGAACTCTTTCCCTACACGACGCTCTTCCGATCTNNNTCCATCTGTAAAATCCTTTTAAA
AGCACACGTCTGAACTCTTTCCCTACACGACGCTCTTCCGATCTNNNCTTACATGTTTTTAGAACCAG
AGCACACGTCTGAACTCTTTCCCTACACGACGCTCTTCCGATCTNNNTCACAGGTTAAGTTTTCTTTTT
GCACACGTCTGAACTCTTTCCCTACACGACGCTCTTCCGATCTNNNTATTACTCACTGCCTTAGAAATT
GCACACGTCTGAACTCTTTCCCTACACGACGCTCTTCCGATCTNNNTCAACAATCACTAACAAAAGAA
AGCACACGTCTGAACTCTTTCCCTACACGACGCTCTTCCGATCTNNNAAGAAATTGTTTAGAAAACCAT
GCACACGTCTGAACTCTTTCCCTACACGACGCTCTTCCGATCTNNNGAATACAGATCTAGGGAGTTTT
AGCACACGTCTGAACTCTTTCCCTACACGACGCTCTTCCGATCTNNNAACATATTCCTGTTATGGTAG
AGCACACGTCTGAACTCTTTCCCTACACGACGCTCTTCCGATCTNNNATTTATAGAGATCTCAGAAG
AGCACACGTCTGAACTCTTTCCCTACACGACGCTCTTCCGATCTNNNATCAACGATGACTGGATAAAA
AGCACACGTCTGAACTCTTTCCCTACACGACGCTCTTCCGATCTNNNATCAACGATGACTGGATAAAA
ACACGTCTGAACTCTTTCCCTACACGACGCTCTTCCGATCTNNNCTCTCTGTCTCTCTGTCTCATCGG
GCACACGTCTGAACTCTTTCCCTACACGACGCTCTTCCGATCTNNNATCTCTCTTCCATCCCCTAGCATC
AGCACACGTCTGAACTCTTTCCCTACACGACGCTCTTCCGATCTNNNAAATTAATTTCTGTTGTTTAAG
AGCACACGTCTGAACTCTTTCCCTACACGACGCTCTTCCGATCTNNNCACTTAGTCCTAAAACAAAAG
GCACACGTCTGAACTCTTTCCCTACACGACGCTCTTCCGATCTNNNTATAAGAGGCTGGTTAAAATG
AGCACACGTCTGAACTCTTTCCCTACACGACGCTCTTCCGATCTNNNCTTGATGTGCCTTGTAGAAGCA
AGCACACGTCTGAACTCTTTCCCTACACGACGCTCTTCCGATCTNNNCAATCTCCTGACTTTGTGATCC
GCACACGTCTGAACTCTTTCCCTACACGACGCTCTTCCGATCTNNNCCTACCTTCCTCAACTTTTACATC
ACACGTCTGAACTCTTTCCCTACACGACGCTCTTCCGATCTNNNAACAGAAAGATTTCACACAGACATC
GAGCACACGTCTGAACTCTTTCCCTACACGACGCTCTTCCGATCTNNNTAAAATGGAAAGTAGGAGCA
AGCACACGTCTGAACTCTTTCCCTACACGACGCTCTTCCGATCTNNNTATATCTATGCCTAGACCATTC
AGCACACGTCTGAACTCTTTCCCTACACGACGCTCTTCCGATCTNNNGTGCCTTTATAAAAATTAACT
AGCACACGTCTGAACTCTTTCCCTACACGACGCTCTTCCGATCTNNNGAAATGATAGCCTGGTTATCAT
GAGCACACGTCTGAACTCTTTCCCTACACGACGCTCTTCCGATCTNNNGTGACACAGAAAACTTCTAAT
AGCACACGTCTGAACTCTTTCCCTACACGACGCTCTTCCGATCTNNNTGATAAGAGAGTGGGTTTCT
GCACACGTCTGAACTCTTTCCCTACACGACGCTCTTCCGATCTNNNGAATCTCTAGGAGTACTTTAGG
GCACACGTCTGAACTCTTTCCCTACACGACGCTCTTCCGATCTNNNACATCAAATTTAAAAGGTAAGA
GAGCACACGTCTGAACTCTTTCCCTACACGACGCTCTTCCGATCTNNNGCTTTCTCTCTTGATAATTTTG
CACGTCTGAACTCTTTCCCTACACGACGCTCTTCCGATCTNNNCTCTTGCAGCCAAATATGATCATCGG
AGCACACGTCTGAACTCTTTCCCTACACGACGCTCTTCCGATCTNNNTAAATAAGCTAATTAGGAAGG
GAGCACACGTCTGAACTCTTTCCCTACACGACGCTCTTCCGATCTNNNTAAAGAGGATTGTTTTGGG
GCACACGTCTGAACTCTTTCCCTACACGACGCTCTTCCGATCTNNNCACAAATATAAAAATACACACG
AGCACACGTCTGAACTCTTTCCCTACACGACGCTCTTCCGATCTNNNCTTTGTTCATTCTCAGGATTTT
GCACACGTCTGAACTCTTTCCCTACACGACGCTCTTCCGATCTNNNTTAACCATTTGCAGCTTCTGCAT
GCACACGTCTGAACTCTTTCCCTACACGACGCTCTTCCGATCTNNNAAATCAGAGTATTGAGTGCTTT
ACACGTCTGAACTCTTTCCCTACACGACGCTCTTCCGATCTNNNAAACTCCTGAAACTTCTGTTTACAT
ACACGTCTGAACTCTTTCCCTACACGACGCTCTTCCGATCTNNNAAACTCCTGAAACTTCTGTTTACAT
GCACACGTCTGAACTCTTTCCCTACACGACGCTCTTCCGATCTNNNCATGATACTTCTAGTCTCTGTAG
AGCACACGTCTGAACTCTTTCCCTACACGACGCTCTTCCGATCTNNNTTTGTAACCTAGATATGAGTG
GCACACGTCTGAACTCTTTCCCTACACGACGCTCTTCCGATCTNNNCTACAGTTACATTCCACCTTGCA
AGCACACGTCTGAACTCTTTCCCTACACGACGCTCTTCCGATCTNNNAACTCCTATGTACTTAGATGTA

GCACACGTCTGAACTCTTTCCCTACACGACGCTCTTCCGATCTNNNTTGTTTGTTTGTTTATTCTGATG
GAGCACACGTCTGAACTCTTTCCCTACACGACGCTCTTCCGATCTNNNATAAACATGGCTGGATAACT
CACACGTCTGAACTCTTTCCCTACACGACGCTCTTCCGATCTNNNGGCTTGAGAATAAGAGTTACCCA
AGCACACGTCTGAACTCTTTCCCTACACGACGCTCTTCCGATCTNNNTCTCCAGTCTTTTTAATTAC
AGCACACGTCTGAACTCTTTCCCTACACGACGCTCTTCCGATCTNNNTTTCAATAGATAGAACACTGGA
AGCACACGTCTGAACTCTTTCCCTACACGACGCTCTTCCGATCTNNNTTTCAATAGATAGAACACTGGA
AGCACACGTCTGAACTCTTTCCCTACACGACGCTCTTCCGATCTNNNGAATAACAGGACCAGATTCAT
CACACGTCTGAACTCTTTCCCTACACGACGCTCTTCCGATCTNNNCTTGTATCATTTGTAATTGTATTG
AGCACACGTCTGAACTCTTTCCCTACACGACGCTCTTCCGATCTNNNTTATCTAAATGCTTGGAGTGA
AGCACACGTCTGAACTCTTTCCCTACACGACGCTCTTCCGATCTNNNATCCCTAATCACATTGCATGCAT
AGCACACGTCTGAACTCTTTCCCTACACGACGCTCTTCCGATCTNNNATCCCTAATCACATTGCATGCAT
AGCACACGTCTGAACTCTTTCCCTACACGACGCTCTTCCGATCTNNNAGACAGTGTTTCACTCTTTCCA
AGCACACGTCTGAACTCTTTCCCTACACGACGCTCTTCCGATCTNNNTTCCCTTCTAATAAGACAAAAG
AGCACACGTCTGAACTCTTTCCCTACACGACGCTCTTCCGATCTNNNAAAGATATGGAACCAATCTAA
AGCACACGTCTGAACTCTTTCCCTACACGACGCTCTTCCGATCTNNNATGAACAGTCTTTGTGGAATA
AGCACACGTCTGAACTCTTTCCCTACACGACGCTCTTCCGATCTNNNCATAAGCCATGAAAGACAGGC
AGCACACGTCTGAACTCTTTCCCTACACGACGCTCTTCCGATCTNNNACATTCTGTTATGATAGTATAC
AGCACACGTCTGAACTCTTTCCCTACACGACGCTCTTCCGATCTNNNAACAAGAGTGGATAAAGAAA
AGCACACGTCTGAACTCTTTCCCTACACGACGCTCTTCCGATCTNNNTTCATCAAAGTTACAGAGTTCT
GCACACGTCTGAACTCTTTCCCTACACGACGCTCTTCCGATCTNNNGTAGACTTAGAATAAGCAGAAC
GCACACGTCTGAACTCTTTCCCTACACGACGCTCTTCCGATCTNNNTAAGTCAGTCAACTAACAGTGCA
CACACGTCTGAACTCTTTCCCTACACGACGCTCTTCCGATCTNNNAAGATTTGCAGCTTATCAAGTAT
AGCACACGTCTGAACTCTTTCCCTACACGACGCTCTTCCGATCTNNNCCAAAAAAAAAAAAAAATCTG
GCACACGTCTGAACTCTTTCCCTACACGACGCTCTTCCGATCTNNNCTTCTGATGCGAATTCTGATACA
ACACGTCTGAACTCTTTCCCTACACGACGCTCTTCCGATCTNNNTGAATATTTGTGATGACCTCTATC
GCACACGTCTGAACTCTTTCCCTACACGACGCTCTTCCGATCTNNNCAATCCAATTCCTCCTAAAATCCA
AGCACACGTCTGAACTCTTTCCCTACACGACGCTCTTCCGATCTNNNTTAAAAATGTCTTAGGGTTCT
CACACGTCTGAACTCTTTCCCTACACGACGCTCTTCCGATCTNNNCTCATAGTCTCTTTGAGTAAGAA
CACACGTCTGAACTCTTTCCCTACACGACGCTCTTCCGATCTNNNCTCATAGTCTCTTTGAGTAAGAA
CACACGTCTGAACTCTTTCCCTACACGACGCTCTTCCGATCTNNNATTTGCCATAGGTGACTTACCAT
CACACGTCTGAACTCTTTCCCTACACGACGCTCTTCCGATCTNNNTGATTTATATCCTCAACCCTTTACA
ACACGTCTGAACTCTTTCCCTACACGACGCTCTTCCGATCTNNNTGAAGAAAATAAGTGAAATCTGGC
AGCACACGTCTGAACTCTTTCCCTACACGACGCTCTTCCGATCTNNNCACAATCATTGCCATTTCTCCA
CACGTCTGAACTCTTTCCCTACACGACGCTCTTCCGATCTNNNATGGATTCATGCCAATTATAAAGCAT
CACGTCTGAACTCTTTCCCTACACGACGCTCTTCCGATCTNNNATGGATTCATGCCAATTATAAAGCAT
AGCACACGTCTGAACTCTTTCCCTACACGACGCTCTTCCGATCTNNNATTCCCCTCTATTTTCATTTTTCC
AGCACACGTCTGAACTCTTTCCCTACACGACGCTCTTCCGATCTNNNATTTTTTTTTTTTCCCAGAGAT
AGCACACGTCTGAACTCTTTCCCTACACGACGCTCTTCCGATCTNNNATCTGTGTTGAACTTGATACTT
AGCACACGTCTGAACTCTTTCCCTACACGACGCTCTTCCGATCTNNNGTAAATCATGGTGTTCCTAGA
AGCACACGTCTGAACTCTTTCCCTACACGACGCTCTTCCGATCTNNNGGAAGAAGGCTAAGGATATT
GCACACGTCTGAACTCTTTCCCTACACGACGCTCTTCCGATCTNNNAATAATTCCACCATACACAGATA
AGCACACGTCTGAACTCTTTCCCTACACGACGCTCTTCCGATCTNNNGCACAGCACAATATATCAATAA
GCACACGTCTGAACTCTTTCCCTACACGACGCTCTTCCGATCTNNNCTCATGGAGCCTAGTTATAATA

AGCACACGTCTGAACTCTTTCCCTACACGACGCTCTTCCGATCTNNNGGATAGTGAGTGAGTTCTTGCA
AGCACACGTCTGAACTCTTTCCCTACACGACGCTCTTCCGATCTNNNACTTTGGGTAACTAGTTTGATAT
CACACGTCTGAACTCTTTCCCTACACGACGCTCTTCCGATCTNNNCAACTTACAAGAACTCTAGAAGTCA
GAGCACACGTCTGAACTCTTTCCCTACACGACGCTCTTCCGATCTNNNAGATCAAGATTTCTATCCTGA
AGCACACGTCTGAACTCTTTCCCTACACGACGCTCTTCCGATCTNNNATTGTTCTCATGATTTACTTTCCC
AGCACACGTCTGAACTCTTTCCCTACACGACGCTCTTCCGATCTNNNCAAAATAGAACTTCAAAAGGAG
GAGCACACGTCTGAACTCTTTCCCTACACGACGCTCTTCCGATCTNNNTTACAAAGCTTCCTTCATAAAG
AGCACACGTCTGAACTCTTTCCCTACACGACGCTCTTCCGATCTNNNCTTTGTTGCTGACATATATGTAT
AGCACACGTCTGAACTCTTTCCCTACACGACGCTCTTCCGATCTNNNATACTTAGATAGCTCTAGAGACA
CACACGTCTGAACTCTTTCCCTACACGACGCTCTTCCGATCTNNNTGGTTGATTTTGGTGTTACCCATCG
GCACACGTCTGAACTCTTTCCCTACACGACGCTCTTCCGATCTNNNATAAAAAACACACACGAGGTATTG
GAGCACACGTCTGAACTCTTTCCCTACACGACGCTCTTCCGATCTNNNATTCGATTTTCCATCATGTAAC
AGCACACGTCTGAACTCTTTCCCTACACGACGCTCTTCCGATCTNNNAACTTTATTGGTCTAGTTAGTGG
GAGCACACGTCTGAACTCTTTCCCTACACGACGCTCTTCCGATCTNNNCATTATTCACAATAGCAAAGAC
GAGCACACGTCTGAACTCTTTCCCTACACGACGCTCTTCCGATCTNNNGGGAACTCTAACACAAAAATA
GAGCACACGTCTGAACTCTTTCCCTACACGACGCTCTTCCGATCTNNNGGGAACTCTAACACAAAAATA
GCACACGTCTGAACTCTTTCCCTACACGACGCTCTTCCGATCTNNNAACATTGAACTATTCTCCACTACC
AGCACACGTCTGAACTCTTTCCCTACACGACGCTCTTCCGATCTNNNCTGATAAGCTATGTAATCAAGT
CACACGTCTGAACTCTTTCCCTACACGACGCTCTTCCGATCTNNNCAGTTTCTCTCAAGGTATGTACCATG
AGCACACGTCTGAACTCTTTCCCTACACGACGCTCTTCCGATCTNNNTACTGGTTAGGATTTATAGCATG
GCACACGTCTGAACTCTTTCCCTACACGACGCTCTTCCGATCTNNNTATTTTGGGAGTGGGTTAGTTAG
AGCACACGTCTGAACTCTTTCCCTACACGACGCTCTTCCGATCTNNNATTTGAAGAAAAGAAATGGAA
GAGCACACGTCTGAACTCTTTCCCTACACGACGCTCTTCCGATCTNNNGATGGATTTGAAGAAAAGAA
AGCACACGTCTGAACTCTTTCCCTACACGACGCTCTTCCGATCTNNNACTCTACACATGATAAAACTACA
AGCACACGTCTGAACTCTTTCCCTACACGACGCTCTTCCGATCTNNNCAAATATCCTCCCCCTAAATCCA
AGCACACGTCTGAACTCTTTCCCTACACGACGCTCTTCCGATCTNNNCAGCTTCACTAGGTTTATAAAAA
GAGCACACGTCTGAACTCTTTCCCTACACGACGCTCTTCCGATCTNNNTTCTATAGCTTTTAGAGTCAGT
AGCACACGTCTGAACTCTTTCCCTACACGACGCTCTTCCGATCTNNNTCTCAGCAAAATAGGTAAGTAA
AGCACACGTCTGAACTCTTTCCCTACACGACGCTCTTCCGATCTNNNAATTTTTAGATGCCTTATACATT
CACACGTCTGAACTCTTTCCCTACACGACGCTCTTCCGATCTNNNCTCTTACTAAGATGGGTTTCTTACA
GAGCACACGTCTGAACTCTTTCCCTACACGACGCTCTTCCGATCTNNNAAAAGCAAATATGTGTATGTA
AGCACACGTCTGAACTCTTTCCCTACACGACGCTCTTCCGATCTNNNGCAACAGTATTTCGTAAAAGTAT
AGCACACGTCTGAACTCTTTCCCTACACGACGCTCTTCCGATCTNNNTATACAAGGCACATTATTCTTTCC
AGCACACGTCTGAACTCTTTCCCTACACGACGCTCTTCCGATCTNNNGGTCTCAAATAGAAAACTTGATT
AGCACACGTCTGAACTCTTTCCCTACACGACGCTCTTCCGATCTNNNGGTTATAACCTTGCGTTATCTCA
AGCACACGTCTGAACTCTTTCCCTACACGACGCTCTTCCGATCTNNNATTTCTTGCTAACCCTAATAATG
AGCACACGTCTGAACTCTTTCCCTACACGACGCTCTTCCGATCTNNNATAATCATACAAGCCAATTCCTT
AGCACACGTCTGAACTCTTTCCCTACACGACGCTCTTCCGATCTNNNATAATCATACAAGCCAATTCCTT
GAGCACACGTCTGAACTCTTTCCCTACACGACGCTCTTCCGATCTNNNTCATTGTCTGGCAATCATTAGC
AGCACACGTCTGAACTCTTTCCCTACACGACGCTCTTCCGATCTNNNTGCATAATAGTTTATTAGAGGCT
AGCACACGTCTGAACTCTTTCCCTACACGACGCTCTTCCGATCTNNNATGTTATAATTCTCTACATGCCT
AGCACACGTCTGAACTCTTTCCCTACACGACGCTCTTCCGATCTNNNATGTTATAATTCTCTACATGCCT
GCACACGTCTGAACTCTTTCCCTACACGACGCTCTTCCGATCTNNNGTTTCTATCTCAAACTGTGTAAG

ACACGTCTGAACTCTTTCCCTACACGACGCTCTTCCGATCTNNNCAAGATTGGTACATCTTTAGAACCAT
AGCACACGTCTGAACTCTTTCCCTACACGACGCTCTTCCGATCTNNNTGCAAAGTGATGATAACTATAA
AGCACACGTCTGAACTCTTTCCCTACACGACGCTCTTCCGATCTNNNAAACTAAATACACAAGAATAG
AGCACACGTCTGAACTCTTTCCCTACACGACGCTCTTCCGATCTNNNAATGGGTCTCAACAGAACTACA
GAGCACACGTCTGAACTCTTTCCCTACACGACGCTCTTCCGATCTNNNTATAGGATATTTACCAGTTTTG
GAGCACACGTCTGAACTCTTTCCCTACACGACGCTCTTCCGATCTNNNATTGGAGATGATGTCTATACA
ACACGTCTGAACTCTTTCCCTACACGACGCTCTTCCGATCTNNNTCAAAAGAAAAAAAAAAAGCCTCA
CACACGTCTGAACTCTTTCCCTACACGACGCTCTTCCGATCTNNNGGTCATGTAAAACTATGGAAGTCA
GCACACGTCTGAACTCTTTCCCTACACGACGCTCTTCCGATCTNNNTAAAAAATGTGGAGTCTGATGT
GCACACGTCTGAACTCTTTCCCTACACGACGCTCTTCCGATCTNNNAAATCTAAAAAATGTGGAGTCT
GCACACGTCTGAACTCTTTCCCTACACGACGCTCTTCCGATCTNNNGATCTATCCTCACATGTACAGCAT
AGCACACGTCTGAACTCTTTCCCTACACGACGCTCTTCCGATCTNNNACTAAAATCAAACTTACCGATT
AGCACACGTCTGAACTCTTTCCCTACACGACGCTCTTCCGATCTNNNCAGAATGAGTTAATTTATCCTG
GCACACGTCTGAACTCTTTCCCTACACGACGCTCTTCCGATCTNNNGATCTCAGTTATGCCCTTTAACAT
GCACACGTCTGAACTCTTTCCCTACACGACGCTCTTCCGATCTNNNGATCTCAGTTATGCCCTTTAACAT
ACACGTCTGAACTCTTTCCCTACACGACGCTCTTCCGATCTNNNAGAGATAAATCTTAAGAAACCCACA
AGCACACGTCTGAACTCTTTCCCTACACGACGCTCTTCCGATCTNNNTCAGGAGCATTTATCTTGTTTCA
GAGCACACGTCTGAACTCTTTCCCTACACGACGCTCTTCCGATCTNNNCACCAAAGAAGCTACTAGAG
CACACGTCTGAACTCTTTCCCTACACGACGCTCTTCCGATCTNNNTTAAAGTAAACCAAAAGCATACG
CACACGTCTGAACTCTTTCCCTACACGACGCTCTTCCGATCTNNNTTATTGTCACTGCTAACTGTAATCAT
CACACGTCTGAACTCTTTCCCTACACGACGCTCTTCCGATCTNNNGCCTACTATTCAAAAACCTATACCA
AGCACACGTCTGAACTCTTTCCCTACACGACGCTCTTCCGATCTNNNTTCTGTGTAGGAACTTATACTT
AGCACACGTCTGAACTCTTTCCCTACACGACGCTCTTCCGATCTNNNAATTGAAAAACTTGCTTTCTTT
GAGCACACGTCTGAACTCTTTCCCTACACGACGCTCTTCCGATCTNNNACTGTATCTTCTGCTAAGGAG
GAGCACACGTCTGAACTCTTTCCCTACACGACGCTCTTCCGATCTNNNTTTACATTCTTAGCCTTAGAGA
AGCACACGTCTGAACTCTTTCCCTACACGACGCTCTTCCGATCTNNNTCCGAATAGAATAGTGTTAGAT
AGCACACGTCTGAACTCTTTCCCTACACGACGCTCTTCCGATCTNNNAATTTCTTCATTCAGTATTTCTCT
AGCACACGTCTGAACTCTTTCCCTACACGACGCTCTTCCGATCTNNNAATTTCTTCATTCAGTATTTCTCT
AGCACACGTCTGAACTCTTTCCCTACACGACGCTCTTCCGATCTNNNGTGTATTAAGAGTAACCTGCTA
GAGCACACGTCTGAACTCTTTCCCTACACGACGCTCTTCCGATCTNNNAAAAACAAGTCTTTTTCTATG
CACACGTCTGAACTCTTTCCCTACACGACGCTCTTCCGATCTNNNGTGAAACTTTCTTTTCAATTACACA
AGAGCACACGTCTGAACTCTTTCCCTACACGACGCTCTTCCGATCTNNNGAGATGGAATAGTTGAGGA
AGCACACGTCTGAACTCTTTCCCTACACGACGCTCTTCCGATCTNNNGTGATCTTTCATGAATAACTCT
GAGCACACGTCTGAACTCTTTCCCTACACGACGCTCTTCCGATCTNNNTAATTTGATAAGAACGAAAAG
GCACACGTCTGAACTCTTTCCCTACACGACGCTCTTCCGATCTNNNGTAGTTTGGGAGACACATCTCAT
GAGCACACGTCTGAACTCTTTCCCTACACGACGCTCTTCCGATCTNNNCAACAAATGACTTGATGAGCC
AGCACACGTCTGAACTCTTTCCCTACACGACGCTCTTCCGATCTNNNACTGTAGATGGGAACTTAAAAG
GAGCACACGTCTGAACTCTTTCCCTACACGACGCTCTTCCGATCTNNNTTCTTAAGTTGACTCTAAACCA
AGCACACGTCTGAACTCTTTCCCTACACGACGCTCTTCCGATCTNNNAATGAGTGACAAGGAGTATAG
GAGCACACGTCTGAACTCTTTCCCTACACGACGCTCTTCCGATCTNNNACATTTCATTGTTGAACACTTAT
GAGCACACGTCTGAACTCTTTCCCTACACGACGCTCTTCCGATCTNNNTGATCTCATTTTGAAGTAAAA
GAGCACACGTCTGAACTCTTTCCCTACACGACGCTCTTCCGATCTNNNGAAATTCACAAGGCAAAATAA
GAGCACACGTCTGAACTCTTTCCCTACACGACGCTCTTCCGATCTNNNGAGAGGACTAAACAGAATAT

GAGCACACGTCTGAACTCTTTCCCTACACGACGCTCTTCCGATCTNNNGAGAGGACTAAACAGAATATG

AGCACACGTCTGAACTCTTTCCCTACACGACGCTCTTCCGATCTNNNTGATATAAATATTTTTGGACCGA

AGCACACGTCTGAACTCTTTCCCTACACGACGCTCTTCCGATCTNNNATTCAGCCTTAAAAATGAATGAA

CACACGTCTGAACTCTTTCCCTACACGACGCTCTTCCGATCTNNNGCAAAATTGAAACTCTATACTCACA

AGCACACGTCTGAACTCTTTCCCTACACGACGCTCTTCCGATCTNNNCCTATTTCTTTCCATCTTGTACTC

GCACACGTCTGAACTCTTTCCCTACACGACGCTCTTCCGATCTNNNGATAAGAAGTTTGAGTTCTGATC

GAGCACACGTCTGAACTCTTTCCCTACACGACGCTCTTCCGATCTNNNAGATGTATCATACCCAGTTTTA

GCACACGTCTGAACTCTTTCCCTACACGACGCTCTTCCGATCTNNNGAAAAAGAGAGAAAAATACCTG

AGAGCACACGTCTGAACTCTTTCCCTACACGACGCTCTTCCGATCTNNNATTCCTAGGAAATATACACT

AGCACACGTCTGAACTCTTTCCCTACACGACGCTCTTCCGATCTNNNTACGTTTTTATTCCAGTGAGATA

GAGCACACGTCTGAACTCTTTCCCTACACGACGCTCTTCCGATCTNNNATTTCAGAAGAAGGACTGTAT

AGCACACGTCTGAACTCTTTCCCTACACGACGCTCTTCCGATCTNNNCTCTTCAACAGATCTGTAAAGAC

ACACGTCTGAACTCTTTCCCTACACGACGCTCTTCCGATCTNNNTAGTGGTAAGATATCTCTATGCCCAT

GAGCACACGTCTGAACTCTTTCCCTACACGACGCTCTTCCGATCTNNNAAGCCAGTTAGAAGTAGATTC

CACACGTCTGAACTCTTTCCCTACACGACGCTCTTCCGATCTNNNACCCACCATTGTTAGTAGATTACAT

GAGCACACGTCTGAACTCTTTCCCTACACGACGCTCTTCCGATCTNNNTTTTTTTAGAATTTTTGCACTTG

AGCACACGTCTGAACTCTTTCCCTACACGACGCTCTTCCGATCTNNNGATAACATGGTTGCCTTGTATA

GAGCACACGTCTGAACTCTTTCCCTACACGACGCTCTTCCGATCTNNNCATAATTAAAATGGGCAAAAA

GAGCACACGTCTGAACTCTTTCCCTACACGACGCTCTTCCGATCTNNNACATTAAAAATGTCATGATGT

AGCACACGTCTGAACTCTTTCCCTACACGACGCTCTTCCGATCTNNNCGAAAAGACATAGCGATTATGC

GCACACGTCTGAACTCTTTCCCTACACGACGCTCTTCCGATCTNNNCCTTATCCTCTGTTTTTATTCCCA

GAGCACACGTCTGAACTCTTTCCCTACACGACGCTCTTCCGATCTNNNCTTCTTGAGTAGAGAAATATTT

CACACGTCTGAACTCTTTCCCTACACGACGCTCTTCCGATCTNNNTTTTAAAAACCATGATACAGGTGC

AGCACACGTCTGAACTCTTTCCCTACACGACGCTCTTCCGATCTNNNAAAACAACAAGAGAACAAAGA

CACACGTCTGAACTCTTTCCCTACACGACGCTCTTCCGATCTNNNTTATTTCTTTCTTTCATCCTTCCCATC

AGCACACGTCTGAACTCTTTCCCTACACGACGCTCTTCCGATCTNNNTCTTGACTGTGGAATATAAATGA

AGCACACGTCTGAACTCTTTCCCTACACGACGCTCTTCCGATCTNNNTATAACTGAGATTGTTACCCAAA

CACACGTCTGAACTCTTTCCCTACACGACGCTCTTCCGATCTNNNATTTTATGTTCTGTCAAGAGGACA

GCACACGTCTGAACTCTTTCCCTACACGACGCTCTTCCGATCTNNNACAATACACAATGAACACTCTCAT

GAGCACACGTCTGAACTCTTTCCCTACACGACGCTCTTCCGATCTNNNAATTTCTAAACAATGTCTTAAT

CACACGTCTGAACTCTTTCCCTACACGACGCTCTTCCGATCTNNNAGTTCTCTCTGAGTCTACAACCATCG

ACACGTCTGAACTCTTTCCCTACACGACGCTCTTCCGATCTNNNTTTATCATCGCTCTACACATTTCATCG

AGCACACGTCTGAACTCTTTCCCTACACGACGCTCTTCCGATCTNNNCATATATATATATGTCTGCTGTC

GAGCACACGTCTGAACTCTTTCCCTACACGACGCTCTTCCGATCTNNNTACACGTTTGTCATTTTTTTACT

AGCACACGTCTGAACTCTTTCCCTACACGACGCTCTTCCGATCTNNNATTTTAAATGTGCAATGCTAAG

CACACGTCTGAACTCTTTCCCTACACGACGCTCTTCCGATCTNNNCTTAGAAAGCATGGGAAAATAGCA

AGCACACGTCTGAACTCTTTCCCTACACGACGCTCTTCCGATCTNNNAGGAAAAATATTCTCCCTATCCT

GAGCACACGTCTGAACTCTTTCCCTACACGACGCTCTTCCGATCTNNNAGAACTTAAGGTGGATTTCAA

GCACACGTCTGAACTCTTTCCCTACACGACGCTCTTCCGATCTNNNGACAGGAACCTAAGTCAGTCCAT

AGCACACGTCTGAACTCTTTCCCTACACGACGCTCTTCCGATCTNNNCAATTTTATTCAGTTTCTCTCGGC

AGCACACGTCTGAACTCTTTCCCTACACGACGCTCTTCCGATCTNNNTAAAAGATTCCTGGAGGATTAT

AGCACACGTCTGAACTCTTTCCCTACACGACGCTCTTCCGATCTNNNTAAAAGATTCCTGGAGGATTAT

AGCACACGTCTGAACTCTTTCCCTACACGACGCTCTTCCGATCTNNNTTGTCTCTCTCCTAAACTTATTCC

AGCACACGTCTGAACTCTTTCCCTACACGACGCTCTTCCGATCTNNNCAGTACAAAAGTAGTATGT
AGCACACGTCTGAACTCTTTCCCTACACGACGCTCTTCCGATCTNNNGGAAAAAAAAAAAAAAAAAG
GAGCACACGTCTGAACTCTTTCCCTACACGACGCTCTTCCGATCTNNNACAAAACAGAAGAAAACTGT
AGCACACGTCTGAACTCTTTCCCTACACGACGCTCTTCCGATCTNNNTTTTTGGCCTCATAAGCATATC
ACACGTCTGAACTCTTTCCCTACACGACGCTCTTCCGATCTNNNTATTCCAGTTTTGGGTTGTGCATCG
CACACGTCTGAACTCTTTCCCTACACGACGCTCTTCCGATCTNNNCAACCTTCGGAATTAGCTTGCATC
ACACGTCTGAACTCTTTCCCTACACGACGCTCTTCCGATCTNNNTGAATGTTCCAGAGGTAGAGCATC
GCACACGTCTGAACTCTTTCCCTACACGACGCTCTTCCGATCTNNNTATTTGCTTAAGTGTTGCTTTAG
CACACGTCTGAACTCTTTCCCTACACGACGCTCTTCCGATCTNNNTTTGTGTGTGTGTGTAAAACTACA
AGCACACGTCTGAACTCTTTCCCTACACGACGCTCTTCCGATCTNNNTTTTGTATGTTCATTTTGTATC
CACACGTCTGAACTCTTTCCCTACACGACGCTCTTCCGATCTNNNGATGATGACATTCTTTTCTAGACC
AGCACACGTCTGAACTCTTTCCCTACACGACGCTCTTCCGATCTNNNACACAAGAAAAAAAAGAAAGA
ACACGTCTGAACTCTTTCCCTACACGACGCTCTTCCGATCTNNNCTTCACGTGTTCTTTCCTTTCATCGG
GAGCACACGTCTGAACTCTTTCCCTACACGACGCTCTTCCGATCTNNNATGAAAATGACAAAGGGAG
AGCACACGTCTGAACTCTTTCCCTACACGACGCTCTTCCGATCTNNNTAGAAGTGGTAGTAAGTGTAA
AGCACACGTCTGAACTCTTTCCCTACACGACGCTCTTCCGATCTNNNTAGAAGTGGTAGTAAGTGTAA
AGCACACGTCTGAACTCTTTCCCTACACGACGCTCTTCCGATCTNNNATACTTGAGCATATTTGTACC
ACACGTCTGAACTCTTTCCCTACACGACGCTCTTCCGATCTNNNGAGAGAGAAAGAGTGTGTATGCA
AGCACACGTCTGAACTCTTTCCCTACACGACGCTCTTCCGATCTNNNAATTTTTTTTCTGATGTTCCA
AGCACACGTCTGAACTCTTTCCCTACACGACGCTCTTCCGATCTNNNTGACTCCAGATATTCCTTGTAT
AGCACACGTCTGAACTCTTTCCCTACACGACGCTCTTCCGATCTNNNTGACTCCAGATATTCCTTGTAT
CACACGTCTGAACTCTTTCCCTACACGACGCTCTTCCGATCTNNNCCTGAAGATATTTGTTCAATAAC
AGCACACGTCTGAACTCTTTCCCTACACGACGCTCTTCCGATCTNNNATTATTATCTTTTTGACTGTGG
GAGCACACGTCTGAACTCTTTCCCTACACGACGCTCTTCCGATCTNNNATACACATACAGGTTCTACAT
GCACACGTCTGAACTCTTTCCCTACACGACGCTCTTCCGATCTNNNCTCCATCTTCATATCACCCTCATC
AGCACACGTCTGAACTCTTTCCCTACACGACGCTCTTCCGATCTNNNAGTGCAATTAAATTAATGTGA
AGCACACGTCTGAACTCTTTCCCTACACGACGCTCTTCCGATCTNNNATAATGAGGTATCAATACAGG
AGCACACGTCTGAACTCTTTCCCTACACGACGCTCTTCCGATCTNNNTAGACAAAATAATCCTTATGCC
AGCACACGTCTGAACTCTTTCCCTACACGACGCTCTTCCGATCTNNNGCCAGTTTCCTTATCTATAAGA
AGCACACGTCTGAACTCTTTCCCTACACGACGCTCTTCCGATCTNNNGTAAGCTCTACAACTTGATTTT
AGCACACGTCTGAACTCTTTCCCTACACGACGCTCTTCCGATCTNNNTTTTTTTATAATGTTTCTTTCCT
CACACGTCTGAACTCTTTCCCTACACGACGCTCTTCCGATCTNNNCTAGTAGGAAAGGCAGATAATA
CACACGTCTGAACTCTTTCCCTACACGACGCTCTTCCGATCTNNNAGTTGTCTCAATTTGAGGTAAGCA
CACACGTCTGAACTCTTTCCCTACACGACGCTCTTCCGATCTNNNTTGTGTTTACCATGATCTCCCATC
ACACGTCTGAACTCTTTCCCTACACGACGCTCTTCCGATCTNNNCTCTTTCTCTTGAATATAAAGTGTC
AGCACACGTCTGAACTCTTTCCCTACACGACGCTCTTCCGATCTNNNCTCATTGCACTAACTAGATTCA
AGCACACGTCTGAACTCTTTCCCTACACGACGCTCTTCCGATCTNNNTATTTCATTTCTTGTCTCATTGC
AGCACACGTCTGAACTCTTTCCCTACACGACGCTCTTCCGATCTNNNCTTTACTGTGCTCTCTGTTGCAT
AGCACACGTCTGAACTCTTTCCCTACACGACGCTCTTCCGATCTNNNTGACTTGTGTATGCATAGTATA
GAGCACACGTCTGAACTCTTTCCCTACACGACGCTCTTCCGATCTNNNACTAACCAATCATCAGTGAAT
AGCACACGTCTGAACTCTTTCCCTACACGACGCTCTTCCGATCTNNNAAGATATATCTTGATTTGGGT
AGCACACGTCTGAACTCTTTCCCTACACGACGCTCTTCCGATCTNNNATATTGAACCCCCATTTCTTCCA
GAGCACACGTCTGAACTCTTTCCCTACACGACGCTCTTCCGATCTNNNTTAACTAAAAGGCAGTCTTA

AGCACACGTCTGAACTCTTTCCCTACACGACGCTCTTCCGATCTNNNATTTTATGAAAGCAGACCAAT
AGCACACGTCTGAACTCTTTCCCTACACGACGCTCTTCCGATCTNNNCTAATTTTGCTGTTAAACATGA
GAGCACACGTCTGAACTCTTTCCCTACACGACGCTCTTCCGATCTNNNGCAAATGCACCAATATCAAA
AGCACACGTCTGAACTCTTTCCCTACACGACGCTCTTCCGATCTNNNACAATTTTTTTAAAAAAGGAC
GCACACGTCTGAACTCTTTCCCTACACGACGCTCTTCCGATCTNNNACTGTAACATTCTTTAGTAGACA
GCACACGTCTGAACTCTTTCCCTACACGACGCTCTTCCGATCTNNNTAGTGTTATTTCATCTTACAGCT
AGCACACGTCTGAACTCTTTCCCTACACGACGCTCTTCCGATCTNNNGTTCTTTCTGTGTTTTCTTTTTC
AGCACACGTCTGAACTCTTTCCCTACACGACGCTCTTCCGATCTNNNTAGACATAAGCACAGGGTTAC
CACACGTCTGAACTCTTTCCCTACACGACGCTCTTCCGATCTNNNATGTATGGTAGAATATGCTAGTG
CACACGTCTGAACTCTTTCCCTACACGACGCTCTTCCGATCTNNNCCACAAATCAAGTAAAACAATTCC
ACACGTCTGAACTCTTTCCCTACACGACGCTCTTCCGATCTNNNATAATAATAACCGTCCCCCAACATC
AGCACACGTCTGAACTCTTTCCCTACACGACGCTCTTCCGATCTNNNTTGAGGCTGGATTCAATATCC
AGCACACGTCTGAACTCTTTCCCTACACGACGCTCTTCCGATCTNNNAAGGAAAACATGTAAAACTG
GCACACGTCTGAACTCTTTCCCTACACGACGCTCTTCCGATCTNNNCTAATACCAGTCAAATTAGAAG
AGCACACGTCTGAACTCTTTCCCTACACGACGCTCTTCCGATCTNNNTGAAGAGTAAACTATTCTAGC
AGCACACGTCTGAACTCTTTCCCTACACGACGCTCTTCCGATCTNNNCTGATCCAAGTAAAGCCTAAA
AGCACACGTCTGAACTCTTTCCCTACACGACGCTCTTCCGATCTNNNTCTTCAGTATCTAAGAGTATCC
AGCACACGTCTGAACTCTTTCCCTACACGACGCTCTTCCGATCTNNNCTCTCTCTCTCTCTCTTTCCAT
CACACGTCTGAACTCTTTCCCTACACGACGCTCTTCCGATCTNNNCTGAGAAGCTTACACAGAAACATC
GCACACGTCTGAACTCTTTCCCTACACGACGCTCTTCCGATCTNNNATCTCTCTTCTTGGCTTATAGAT
AGCACACGTCTGAACTCTTTCCCTACACGACGCTCTTCCGATCTNNNTACTAAATGTGAATCTTTTTGC
CACGTCTGAACTCTTTCCCTACACGACGCTCTTCCGATCTNNNGTAGAGGTAGGGATTAGAGAAGC
GAGCACACGTCTGAACTCTTTCCCTACACGACGCTCTTCCGATCTNNNTGTTTTTTTTTATCAAAAGG
AGCACACGTCTGAACTCTTTCCCTACACGACGCTCTTCCGATCTNNNCGTTTCATTACTGATATTGGA
CACACGTCTGAACTCTTTCCCTACACGACGCTCTTCCGATCTNNNGTGGTAGACTTACAGGAACTCAT
CACGTCTGAACTCTTTCCCTACACGACGCTCTTCCGATCTNNNGCTAATGCTAGCCCTAACTTACATCG
ACACGTCTGAACTCTTTCCCTACACGACGCTCTTCCGATCTNNNCTGAAGAAGTAATTAGGTCACAAC
GCACACGTCTGAACTCTTTCCCTACACGACGCTCTTCCGATCTNNNCAGTGTAACAATATCCTGACAAC
AGCACACGTCTGAACTCTTTCCCTACACGACGCTCTTCCGATCTNNNGAACAGATTCTCTCTTGCAGCA
AGCACACGTCTGAACTCTTTCCCTACACGACGCTCTTCCGATCTNNNCTATGAAAGCAGCAAAGGTCC
GCACACGTCTGAACTCTTTCCCTACACGACGCTCTTCCGATCTNNNGAGTTTGGAGATAGTAAAGAA
GCACACGTCTGAACTCTTTCCCTACACGACGCTCTTCCGATCTNNNTTAGAACTATTCCAAAACATAGC
AGCACACGTCTGAACTCTTTCCCTACACGACGCTCTTCCGATCTNNNCCCAAAATTAGTCTACCTTGGC
GCACACGTCTGAACTCTTTCCCTACACGACGCTCTTCCGATCTNNNACCGTTTTTATAAAACCAAGAT
GCACACGTCTGAACTCTTTCCCTACACGACGCTCTTCCGATCTNNNGTTTATATCTTCCCCATACTTGA
AGCACACGTCTGAACTCTTTCCCTACACGACGCTCTTCCGATCTNNNAAAAAATCTGTGTTGACATA
AGCACACGTCTGAACTCTTTCCCTACACGACGCTCTTCCGATCTNNNCAAAAAGCAGAAAATGAATGA
GCACACGTCTGAACTCTTTCCCTACACGACGCTCTTCCGATCTNNNTAAGAGAACATTCACAAAGGA
AGCACACGTCTGAACTCTTTCCCTACACGACGCTCTTCCGATCTNNNATTGATGGATTCATTTTATTCC
GCACACGTCTGAACTCTTTCCCTACACGACGCTCTTCCGATCTNNNCCAAATTTTAAAAGAATGCACT
CACACGTCTGAACTCTTTCCCTACACGACGCTCTTCCGATCTNNNTTTAAATGGCCTATTGCATGTAT
GCACACGTCTGAACTCTTTCCCTACACGACGCTCTTCCGATCTNNNTCTCACTTCTGAGATTAAGTTAT
AGCACACGTCTGAACTCTTTCCCTACACGACGCTCTTCCGATCTNNNAAATTATCTTCCCATAAAACCA

GCACACGTCTGAACTCTTTCCCTACACGACGCTCTTCCGATCTNNNTGAAAAAGTCTGATTGGATAAAA
AGCACACGTCTGAACTCTTTCCCTACACGACGCTCTTCCGATCTNNNGGCCAGATTGGTAAAAAAAAA
GAGCACACGTCTGAACTCTTTCCCTACACGACGCTCTTCCGATCTNNNAATAGCTTTTCCCTGTAGACTA
AGCACACGTCTGAACTCTTTCCCTACACGACGCTCTTCCGATCTNNNTGCTCTACTGCTAGGATGTACAT
GAGCACACGTCTGAACTCTTTCCCTACACGACGCTCTTCCGATCTNNNAGAAGTTTTATCACTTATCTGC
AGCACACGTCTGAACTCTTTCCCTACACGACGCTCTTCCGATCTNNNATTAGGCCTTTTAAAATCTTTCC
AGAGCACACGTCTGAACTCTTTCCCTACACGACGCTCTTCCGATCTNNNTTACTTTAAGTACAAAACAAG
GCACACGTCTGAACTCTTTCCCTACACGACGCTCTTCCGATCTNNNCTCTCCCCAAAAGAAAAAGTCATC
ACACGTCTGAACTCTTTCCCTACACGACGCTCTTCCGATCTNNNAAAATCCTTTTTTCTAACCAGAGCAT
AGCACACGTCTGAACTCTTTCCCTACACGACGCTCTTCCGATCTNNNGTATAGCACACTTTTCACATGCAT
AGAGCACACGTCTGAACTCTTTCCCTACACGACGCTCTTCCGATCTNNNCCTTCTCTTCAATCAGTCATAT
GAGCACACGTCTGAACTCTTTCCCTACACGACGCTCTTCCGATCTNNNGTAGTGGGGAAAGAATATGA
AGCACACGTCTGAACTCTTTCCCTACACGACGCTCTTCCGATCTNNNGGTCATAGAGAGAGATCCTGCA
GCACACGTCTGAACTCTTTCCCTACACGACGCTCTTCCGATCTNNNTTTTTGAGACCCTAAATGTTTAGC
AGCACACGTCTGAACTCTTTCCCTACACGACGCTCTTCCGATCTNNNGGGAATCATAAGATTGTTTACT
AGCACACGTCTGAACTCTTTCCCTACACGACGCTCTTCCGATCTNNNGATGGTTGGATGAAGCTAATA
GAGCACACGTCTGAACTCTTTCCCTACACGACGCTCTTCCGATCTNNNTATGGCTAAAAGATAGTTGAT
AGCACACGTCTGAACTCTTTCCCTACACGACGCTCTTCCGATCTNNNCTCTCAGATTGCATTCAGTTTCA
AGAGCACACGTCTGAACTCTTTCCCTACACGACGCTCTTCCGATCTNNNGGTAGATACTGGTTGAATAC
CACACGTCTGAACTCTTTCCCTACACGACGCTCTTCCGATCTNNNCTAAGTGCCTTTTATTCAACTTTCAT
GCACACGTCTGAACTCTTTCCCTACACGACGCTCTTCCGATCTNNNTTCTAGAAGTTTGGCTATAAGAA
GAGCACACGTCTGAACTCTTTCCCTACACGACGCTCTTCCGATCTNNNTTTCTAGAAGTTTGGCTATAA
GAGCACACGTCTGAACTCTTTCCCTACACGACGCTCTTCCGATCTNNNTTTCTAGAAGTTTGGCTATAA
ACACGTCTGAACTCTTTCCCTACACGACGCTCTTCCGATCTNNNGAGAGAGAGAGAGAGAGAAACAT
AGCACACGTCTGAACTCTTTCCCTACACGACGCTCTTCCGATCTNNNTGGGTATCTTTTCAGGTATTAA
AGCACACGTCTGAACTCTTTCCCTACACGACGCTCTTCCGATCTNNNTATTGCATTTCATATATGTCCAC
GAGCACACGTCTGAACTCTTTCCCTACACGACGCTCTTCCGATCTNNNTTCTGTACAACAATAACATTCA
GAGCACACGTCTGAACTCTTTCCCTACACGACGCTCTTCCGATCTNNNAATAAAATCAAAGTCCTATCCC
AGCACACGTCTGAACTCTTTCCCTACACGACGCTCTTCCGATCTNNNAAGCTTATATATTCCAGGCTTAC
AGCACACGTCTGAACTCTTTCCCTACACGACGCTCTTCCGATCTNNNAGACCTTACCTGGCTTAGATCAT
AGCACACGTCTGAACTCTTTCCCTACACGACGCTCTTCCGATCTNNNTCAGACAACCTGATGTATGAAC
AGCACACGTCTGAACTCTTTCCCTACACGACGCTCTTCCGATCTNNNGAATTAAACATTAAAGCACTTGG
GAGCACACGTCTGAACTCTTTCCCTACACGACGCTCTTCCGATCTNNNCATTTGCACACTATACATGAAT
AGCACACGTCTGAACTCTTTCCCTACACGACGCTCTTCCGATCTNNNACCCAGTATCACCAATTATTTAT
GCACACGTCTGAACTCTTTCCCTACACGACGCTCTTCCGATCTNNNGTGAATTCATGGTTTTCAAAATAC
GAGCACACGTCTGAACTCTTTCCCTACACGACGCTCTTCCGATCTNNNCAAAAATGAGTGAATTACCAT
CACACGTCTGAACTCTTTCCCTACACGACGCTCTTCCGATCTNNNATTTCTGTTATTTAAGTCACCACCA
AGCACACGTCTGAACTCTTTCCCTACACGACGCTCTTCCGATCTNNNAGTGGTATATTCTAGGTAAAAG
AGCACACGTCTGAACTCTTTCCCTACACGACGCTCTTCCGATCTNNNTGTGTGCATGTATATGTGTATA
AGCACACGTCTGAACTCTTTCCCTACACGACGCTCTTCCGATCTNNNAATAGGATATATGTGTGCATGT
GAGCACACGTCTGAACTCTTTCCCTACACGACGCTCTTCCGATCTNNNCAAGCCATGGAGATTATATGTA
GAGCACACGTCTGAACTCTTTCCCTACACGACGCTCTTCCGATCTNNNGGAAAGAAGAGAACTGACTA
AGCACACGTCTGAACTCTTTCCCTACACGACGCTCTTCCGATCTNNNCATCCCTATTTGTAGCTTTATTC

GCACACGTCTGAACTCTTTCCCTACACGACGCTCTTCCGATCTNNNTGCAAGTTGGCATTTTTAAAGCA
AGCACACGTCTGAACTCTTTCCCTACACGACGCTCTTCCGATCTNNNGTGTGTGTCTGAAAGGGATA
AGCACACGTCTGAACTCTTTCCCTACACGACGCTCTTCCGATCTNNNCCTGTAACTTACTTCTTCTAAAT
AGCACACGTCTGAACTCTTTCCCTACACGACGCTCTTCCGATCTNNNCCTAAGCTAGTTTGTTTTGTG
GAGCACACGTCTGAACTCTTTCCCTACACGACGCTCTTCCGATCTNNNCTAAGTACCTACTATGTGTC
GAGCACACGTCTGAACTCTTTCCCTACACGACGCTCTTCCGATCTNNNATAATTATATCCACCCATCAT
AGCACACGTCTGAACTCTTTCCCTACACGACGCTCTTCCGATCTNNNTGAAATTTGCTAAGAGAATAG
ACGTCTGAACTCTTTCCCTACACGACGCTCTTCCGATCTNNNCAGAATCCATTGAAAATTGTTACCAT
AGCACACGTCTGAACTCTTTCCCTACACGACGCTCTTCCGATCTNNNTTACTATGTAATCAAACTTGG
GCACACGTCTGAACTCTTTCCCTACACGACGCTCTTCCGATCTNNNGTTCTCCTTGTGTTTTCTCTCATC
AGCACACGTCTGAACTCTTTCCCTACACGACGCTCTTCCGATCTNNNTAATTAGTGGTCTTTCCCAATA
GCACACGTCTGAACTCTTTCCCTACACGACGCTCTTCCGATCTNNNAGTTGGTTATAGATGAGGATT
AGCACACGTCTGAACTCTTTCCCTACACGACGCTCTTCCGATCTNNNTAAAGGTGAGTCAAATTTCTT
ACACGTCTGAACTCTTTCCCTACACGACGCTCTTCCGATCTNNNCCATCCTTTCATCTTTTATTCAACAT
CACACGTCTGAACTCTTTCCCTACACGACGCTCTTCCGATCTNNNGTTTGGAGATCTACTAGCTACCAT
AGCACACGTCTGAACTCTTTCCCTACACGACGCTCTTCCGATCTNNNAAAACAAATTCTTCCAATAAAG
AGCACACGTCTGAACTCTTTCCCTACACGACGCTCTTCCGATCTNNNGTCATATTTGTGTGGATATAC
CACACGTCTGAACTCTTTCCCTACACGACGCTCTTCCGATCTNNNCATGGTGTAAGTATCAATACCTA
AGCACACGTCTGAACTCTTTCCCTACACGACGCTCTTCCGATCTNNNTTATCTTTGAAGTGATCTGTGA
AGCACACGTCTGAACTCTTTCCCTACACGACGCTCTTCCGATCTNNNATTTAGAGAAATGTGTGGACT
AGCACACGTCTGAACTCTTTCCCTACACGACGCTCTTCCGATCTNNNGAGCTATGATCAGGCTACTG
AGCACACGTCTGAACTCTTTCCCTACACGACGCTCTTCCGATCTNNNTATGTGTGTGCATATATATATG
AGCACACGTCTGAACTCTTTCCCTACACGACGCTCTTCCGATCTNNNTCTGAAAGTGTTACATTTTGTA
GAGCACACGTCTGAACTCTTTCCCTACACGACGCTCTTCCGATCTNNNGGTTTGTATTTTCTTTTACTT
GCACACGTCTGAACTCTTTCCCTACACGACGCTCTTCCGATCTNNNAAATTAACATCCTTCTAGCTCTT
GCACACGTCTGAACTCTTTCCCTACACGACGCTCTTCCGATCTNNNCAGAGGTATTAGGAAAAGAAG
AGCACACGTCTGAACTCTTTCCCTACACGACGCTCTTCCGATCTNNNAACGAGTAGATAAAGAAATTG
GAGCACACGTCTGAACTCTTTCCCTACACGACGCTCTTCCGATCTNNNGACGAGAACTATTTCTGAAC
ACACGTCTGAACTCTTTCCCTACACGACGCTCTTCCGATCTNNNATTTTGAAGCAATGTTATTAGGTC
AGCACACGTCTGAACTCTTTCCCTACACGACGCTCTTCCGATCTNNNAGTAGTTTATTTGGGAGGTTT
AGCACACGTCTGAACTCTTTCCCTACACGACGCTCTTCCGATCTNNNAGTAGTTTATTTGGGAGGTTT
AGCACACGTCTGAACTCTTTCCCTACACGACGCTCTTCCGATCTNNNTCACAAAATAGTGAAAAGAAC
AGCACACGTCTGAACTCTTTCCCTACACGACGCTCTTCCGATCTNNNAGTTTCTTCACTTGTCAAATAT
CACGTCTGAACTCTTTCCCTACACGACGCTCTTCCGATCTNNNATGGAAATAAACTGCCACAAACATCG
AGCACACGTCTGAACTCTTTCCCTACACGACGCTCTTCCGATCTNNNCATGTAGGCTTTTCCATAGAG
CACACGTCTGAACTCTTTCCCTACACGACGCTCTTCCGATCTNNNTTAACTTCACAATTCCATAACTCCA
ACACGTCTGAACTCTTTCCCTACACGACGCTCTTCCGATCTNNNATCAAAACAGAAATCTTTAGCAGCA
ACACGTCTGAACTCTTTCCCTACACGACGCTCTTCCGATCTNNNCTGAGGCTAAGAGCTTACTCCATCG
GAGCACACGTCTGAACTCTTTCCCTACACGACGCTCTTCCGATCTNNNTAACTACACAGTTAACAAGG
GCACACGTCTGAACTCTTTCCCTACACGACGCTCTTCCGATCTNNNTAGATGGTGTAAAAAATGGGC
CACACGTCTGAACTCTTTCCCTACACGACGCTCTTCCGATCTNNNCTAAACACCAGCGTATACCATC
AGCACACGTCTGAACTCTTTCCCTACACGACGCTCTTCCGATCTNNNAATTATCAGTTCCTTGGAGAT
GCACACGTCTGAACTCTTTCCCTACACGACGCTCTTCCGATCTNNNTGTGTGTGTATATATATGTG

AGCACACGTCTGAACTCTTTCCCTACACGACGCTCTTCCGATCTNNNACTGTTTATTTTCTTCTTTTCCTC
GCACACGTCTGAACTCTTTCCCTACACGACGCTCTTCCGATCTNNNGGATGATTTGATTGATGAAACTT
GAGCACACGTCTGAACTCTTTCCCTACACGACGCTCTTCCGATCTNNNGATTCATTTTTTTTGCATATT
AGCACACGTCTGAACTCTTTCCCTACACGACGCTCTTCCGATCTNNNGTACTCAAATTCATTTGTTTTG
GCACACGTCTGAACTCTTTCCCTACACGACGCTCTTCCGATCTNNNTTCTTTCGACTGTAGAACAAATCA
AGCACACGTCTGAACTCTTTCCCTACACGACGCTCTTCCGATCTNNNAGTCACATAGTAAATGGAAGA
CACACGTCTGAACTCTTTCCCTACACGACGCTCTTCCGATCTNNNTAGCCAAGCAAAGTTCATAGCATC
GAGCACACGTCTGAACTCTTTCCCTACACGACGCTCTTCCGATCTNNNCAAAACGTATCCAAGAGTTTG
GCACACGTCTGAACTCTTTCCCTACACGACGCTCTTCCGATCTNNNGAAATTGGGGGTTTTTGTGCA
GCACACGTCTGAACTCTTTCCCTACACGACGCTCTTCCGATCTNNNGATACATGGCATGTTCATTTAGC
CACACGTCTGAACTCTTTCCCTACACGACGCTCTTCCGATCTNNNTTAGCTCTAAAAATTGTTTGTGGC
GAGCACACGTCTGAACTCTTTCCCTACACGACGCTCTTCCGATCTNNNTCTAGGGTATATTGGGAAAAT
AGCACACGTCTGAACTCTTTCCCTACACGACGCTCTTCCGATCTNNNCATGCATTTTGTCATTACCAGC
CACACGTCTGAACTCTTTCCCTACACGACGCTCTTCCGATCTNNNATCATAGTCATTTCTCCAAGACCAT
GCACACGTCTGAACTCTTTCCCTACACGACGCTCTTCCGATCTNNNTTTAATTATCAGAAAGCGGCCAT
GCACACGTCTGAACTCTTTCCCTACACGACGCTCTTCCGATCTNNNGTTGATCCACTTTGTTGAGTCATC
GCACACGTCTGAACTCTTTCCCTACACGACGCTCTTCCGATCTNNNAAGAAGTAGTCCCACATGTACAT
AGCACACGTCTGAACTCTTTCCCTACACGACGCTCTTCCGATCTNNNTTTTTCTACTAACATGTCCCTTC
AGCACACGTCTGAACTCTTTCCCTACACGACGCTCTTCCGATCTNNNCCCTACATAAAGCATTTTGTTAT
AGCACACGTCTGAACTCTTTCCCTACACGACGCTCTTCCGATCTNNNTACCTATATGCATCTTTTTTTGG
GAGCACACGTCTGAACTCTTTCCCTACACGACGCTCTTCCGATCTNNNTTTTTTTTCTTTAAACAGAGTG
AGCACACGTCTGAACTCTTTCCCTACACGACGCTCTTCCGATCTNNNGGTCATGATGAAGGATTTTTAA
AGCACACGTCTGAACTCTTTCCCTACACGACGCTCTTCCGATCTNNNGAAATTTATGTTGAGGACAG
GAGCACACGTCTGAACTCTTTCCCTACACGACGCTCTTCCGATCTNNNGGGCAGTTTAAGAATTCCTT
AGCACACGTCTGAACTCTTTCCCTACACGACGCTCTTCCGATCTNNNGAGATTAAAAGTGTTGAGTAC
CACACGTCTGAACTCTTTCCCTACACGACGCTCTTCCGATCTNNNATGTCAATCCTAATCCATTACATCA
CACACGTCTGAACTCTTTCCCTACACGACGCTCTTCCGATCTNNNATTTTGTAGTTTTGTACTGAGACC
CACACGTCTGAACTCTTTCCCTACACGACGCTCTTCCGATCTNNNGTCTTCTCTATGGGAATCACCATC
GAGCACACGTCTGAACTCTTTCCCTACACGACGCTCTTCCGATCTNNNCAGTCAGTATATATTCATATG
GAGCACACGTCTGAACTCTTTCCCTACACGACGCTCTTCCGATCTNNNCAGTCAGTATATATTCATATG
GAGCACACGTCTGAACTCTTTCCCTACACGACGCTCTTCCGATCTNNNTGTGAGGGTTAAATGAATGT
AGCACACGTCTGAACTCTTTCCCTACACGACGCTCTTCCGATCTNNNTTGCACAATTGGTTTTCTAAAT
AGCACACGTCTGAACTCTTTCCCTACACGACGCTCTTCCGATCTNNNCTATCATAGTGGATGTGAAGTA
AGAGCACACGTCTGAACTCTTTCCCTACACGACGCTCTTCCGATCTNNNAGGTCAGATTCACAAACATA
AGCACACGTCTGAACTCTTTCCCTACACGACGCTCTTCCGATCTNNNAGTTAGGGAGTTACAGAAATT
GCACACGTCTGAACTCTTTCCCTACACGACGCTCTTCCGATCTNNNAAAGAACCCATCTCTTAATATCA
GCACACGTCTGAACTCTTTCCCTACACGACGCTCTTCCGATCTNNNCTAGTTGTCACTACACTAGAAGC
AGCACACGTCTGAACTCTTTCCCTACACGACGCTCTTCCGATCTNNNATTCCAGATGAATCCAAAAAA
GAGCACACGTCTGAACTCTTTCCCTACACGACGCTCTTCCGATCTNNNGAATGTTTTTAGTCTCACATT
AGCACACGTCTGAACTCTTTCCCTACACGACGCTCTTCCGATCTNNNCTTTATTCCTAAAGGCTAAGCT
AGCACACGTCTGAACTCTTTCCCTACACGACGCTCTTCCGATCTNNNTTTATGCTCCTTTGCTACTTCAT
GCACACGTCTGAACTCTTTCCCTACACGACGCTCTTCCGATCTNNNAAGAGGTGGAGATGAAGATTTA
GAGCACACGTCTGAACTCTTTCCCTACACGACGCTCTTCCGATCTNNNCATTACAACCTATGGAAATAT

AGCACACGTCTGAACTCTTTCCCTACACGACGCTCTTCCGATCTNNNTGCTCTAATTTATAATTTGCTC
GCACACGTCTGAACTCTTTCCCTACACGACGCTCTTCCGATCTNNNTTTCAGAGTTTTTCACCAAATAC
AGCACACGTCTGAACTCTTTCCCTACACGACGCTCTTCCGATCTNNNGATGACCCAGATTATAGCTTT
AGCACACGTCTGAACTCTTTCCCTACACGACGCTCTTCCGATCTNNNCTTACTGCAGAATTGAAAAAT
AGAGCACACGTCTGAACTCTTTCCCTACACGACGCTCTTCCGATCTNNNCTAGAAAGAGGGACCATCA
CACACGTCTGAACTCTTTCCCTACACGACGCTCTTCCGATCTNNNGTCACTGTGACTATCTCCACCATC
AGCACACGTCTGAACTCTTTCCCTACACGACGCTCTTCCGATCTNNNGTATGAAGAAAGATGACTGTT
AGCACACGTCTGAACTCTTTCCCTACACGACGCTCTTCCGATCTNNNATAGACTTTGAATTTGTTTTCC
AGCACACGTCTGAACTCTTTCCCTACACGACGCTCTTCCGATCTNNNGAGCTGTTATATTCAATGTTGA
AGCACACGTCTGAACTCTTTCCCTACACGACGCTCTTCCGATCTNNNGCAAGCCAGTAAATTTGTATT
GAGCACACGTCTGAACTCTTTCCCTACACGACGCTCTTCCGATCTNNNAGATTATTAATCTAGGAAAGC
GCACACGTCTGAACTCTTTCCCTACACGACGCTCTTCCGATCTNNNGTGATGAGCACCATAATAGAAC
CACACGTCTGAACTCTTTCCCTACACGACGCTCTTCCGATCTNNNTATACAGAAGAAAAAGCAGATTG
CACACGTCTGAACTCTTTCCCTACACGACGCTCTTCCGATCTNNNATTGAATTCCTATATGTGAGTCAC
AGCACACGTCTGAACTCTTTCCCTACACGACGCTCTTCCGATCTNNNTCACTCCTTGAAAACAGTAAGC
AGCACACGTCTGAACTCTTTCCCTACACGACGCTCTTCCGATCTNNNAAGAGATCCTTAGAATACTCAT
AGCACACGTCTGAACTCTTTCCCTACACGACGCTCTTCCGATCTNNNAGAAACCAAAGATAACTGGG
AGCACACGTCTGAACTCTTTCCCTACACGACGCTCTTCCGATCTNNNTTATAGTCATCTAGTCATTCGT
GCACACGTCTGAACTCTTTCCCTACACGACGCTCTTCCGATCTNNNAAAGAAAAAATAAAAATAGGAG
GAGCACACGTCTGAACTCTTTCCCTACACGACGCTCTTCCGATCTNNNCTTCTACATTTGTTTCCTGCT
AGCACACGTCTGAACTCTTTCCCTACACGACGCTCTTCCGATCTNNNGCTGATAGCTTTTTCAAATTTTT
AGCACACGTCTGAACTCTTTCCCTACACGACGCTCTTCCGATCTNNNTTGGAAAATTTGTCAATGAAGT
GCACACGTCTGAACTCTTTCCCTACACGACGCTCTTCCGATCTNNNTTAGGATATTAGGACTTTGTGTC
GAGCACACGTCTGAACTCTTTCCCTACACGACGCTCTTCCGATCTNNNGAATATACAGTGAGAGGGTA
AGCACACGTCTGAACTCTTTCCCTACACGACGCTCTTCCGATCTNNNAAATAAAAATACAGGCAAGTC
CACACGTCTGAACTCTTTCCCTACACGACGCTCTTCCGATCTNNNTGTAAAATACTTGCTCAGTGTCAT
GCACACGTCTGAACTCTTTCCCTACACGACGCTCTTCCGATCTNNNTGTTTTAAAGAATGGCTAATACA
AGCACACGTCTGAACTCTTTCCCTACACGACGCTCTTCCGATCTNNNTTTCAGTAGTAAAGTAAACCA
AGCACACGTCTGAACTCTTTCCCTACACGACGCTCTTCCGATCTNNNACTGTTCATTTATTTAATCAGG
GAGCACACGTCTGAACTCTTTCCCTACACGACGCTCTTCCGATCTNNNTATATGTGAGGTACAGTACT
AGCACACGTCTGAACTCTTTCCCTACACGACGCTCTTCCGATCTNNNCTCTTTACCTGGATTGCAAATC
GCACACGTCTGAACTCTTTCCCTACACGACGCTCTTCCGATCTNNNATTGTTTTGAGGTAATTGTCTTT
GCACACGTCTGAACTCTTTCCCTACACGACGCTCTTCCGATCTNNNCTGACTTGTCTCTCTTGGATCAT
GCACACGTCTGAACTCTTTCCCTACACGACGCTCTTCCGATCTNNNGATATTTTTCCAGGAATAACAG
AGCACACGTCTGAACTCTTTCCCTACACGACGCTCTTCCGATCTNNNTACAGAAGCCTGAATCAATTTT
AGCACACGTCTGAACTCTTTCCCTACACGACGCTCTTCCGATCTNNNTACAGAAGCCTGAATCAATTTT
CACACGTCTGAACTCTTTCCCTACACGACGCTCTTCCGATCTNNNCAGAAAATATAGTCTGCACATAA
GAGCACACGTCTGAACTCTTTCCCTACACGACGCTCTTCCGATCTNNNTTCATTAAATATTCTTGCTGC
CACACGTCTGAACTCTTTCCCTACACGACGCTCTTCCGATCTNNNATTATAGGTACAGCTCTGACTCAT
AGCACACGTCTGAACTCTTTCCCTACACGACGCTCTTCCGATCTNNNCTGAATATGATGCCTCTCTGCA
AGCACACGTCTGAACTCTTTCCCTACACGACGCTCTTCCGATCTNNNTACTTTTGTTCTTTTTTATGGCT
AGCACACGTCTGAACTCTTTCCCTACACGACGCTCTTCCGATCTNNNAAAATGACTGCAAGATATCCTC
GAGCACACGTCTGAACTCTTTCCCTACACGACGCTCTTCCGATCTNNNTTAGAGATAGGAAAAACAAA

GAGCACACGTCTGAACTCTTTCCCTACACGACGCTCTTCCGATCTNNNGAGCTTAGGACACAAACAAAG

GCACACGTCTGAACTCTTTCCCTACACGACGCTCTTCCGATCTNNNGAGAGTGATACTTTGATTTTCAAG

GAGCACACGTCTGAACTCTTTCCCTACACGACGCTCTTCCGATCTNNNTTTCCTATCCTGTTGTCGAACA

AGCACACGTCTGAACTCTTTCCCTACACGACGCTCTTCCGATCTNNNAGAAACATTAAACTGATTCAAC

CACACGTCTGAACTCTTTCCCTACACGACGCTCTTCCGATCTNNNATAGGGGATCAATAATGTTTGTCA

AGCACACGTCTGAACTCTTTCCCTACACGACGCTCTTCCGATCTNNNAGATTGTAAGGGACAGAGAGC

GAGCACACGTCTGAACTCTTTCCCTACACGACGCTCTTCCGATCTNNNAGTGTCCTATAAGCATCTTATT

AGCACACGTCTGAACTCTTTCCCTACACGACGCTCTTCCGATCTNNNCTGTGATTATGAGCTTCTCTGCA

AGCACACGTCTGAACTCTTTCCCTACACGACGCTCTTCCGATCTNNNGTCTGTCTTGTTATCCTAGTGCA

GAGCACACGTCTGAACTCTTTCCCTACACGACGCTCTTCCGATCTNNNAAATGGATAAAGAAAATATGC

AGCACACGTCTGAACTCTTTCCCTACACGACGCTCTTCCGATCTNNNTATGAGAAAGAGAGAGAGAAA

GCACACGTCTGAACTCTTTCCCTACACGACGCTCTTCCGATCTNNNAACTTAAGAGCATGTTACAAAATC

GCACACGTCTGAACTCTTTCCCTACACGACGCTCTTCCGATCTNNNAACTTAAGAGCATGTTACAAAATC

AGCACACGTCTGAACTCTTTCCCTACACGACGCTCTTCCGATCTNNNGCCTTTAGCATTAAAAAGATAG

GCACACGTCTGAACTCTTTCCCTACACGACGCTCTTCCGATCTNNNTCTCTTATTTTCTCTCCTCAACCAT

AGCACACGTCTGAACTCTTTCCCTACACGACGCTCTTCCGATCTNNNGTCTAAGCACATAGTATAACTG

AGCACACGTCTGAACTCTTTCCCTACACGACGCTCTTCCGATCTNNNAGATAAGGTGATTATTATTCAT

CACACGTCTGAACTCTTTCCCTACACGACGCTCTTCCGATCTNNNGAATTTATTACCACTCAGACTTCCA

AGCACACGTCTGAACTCTTTCCCTACACGACGCTCTTCCGATCTNNNGCTGTATTGATAATGAGATTTTG

AGAGCACACGTCTGAACTCTTTCCCTACACGACGCTCTTCCGATCTNNNGAGACAGGTCTAGTTTTTAT

GCACACGTCTGAACTCTTTCCCTACACGACGCTCTTCCGATCTNNNAACTGAGTTCAGATATTTCTTTTC

CACACGTCTGAACTCTTTCCCTACACGACGCTCTTCCGATCTNNNGGTTAAACAGTGGAAAGTTTATTC

GAGCACACGTCTGAACTCTTTCCCTACACGACGCTCTTCCGATCTNNNTTTAGCTATGTAAAGCAAGGG

AGCACACGTCTGAACTCTTTCCCTACACGACGCTCTTCCGATCTNNNCACATGTATCCCAGAACTTAAA

AGCACACGTCTGAACTCTTTCCCTACACGACGCTCTTCCGATCTNNNGGGTATTTACTGTGTAGAGCCA

AGCACACGTCTGAACTCTTTCCCTACACGACGCTCTTCCGATCTNNNTAATAAAGCATAGACAGAGGA

GCACACGTCTGAACTCTTTCCCTACACGACGCTCTTCCGATCTNNNTTTCTTAATGTAGAGTTTATTGCT

GAGCACACGTCTGAACTCTTTCCCTACACGACGCTCTTCCGATCTNNNTTATGTTAGACGCTTCTTTTAT

AGCACACGTCTGAACTCTTTCCCTACACGACGCTCTTCCGATCTNNNAAACTGAGCCTTTATTTTAGATC

CACACGTCTGAACTCTTTCCCTACACGACGCTCTTCCGATCTNNNCTCTTTTACCCTTAATTAGCAATCATG

GCACACGTCTGAACTCTTTCCCTACACGACGCTCTTCCGATCTNNNAGAGAAAACAGTAGACTAGAAAA

GCACACGTCTGAACTCTTTCCCTACACGACGCTCTTCCGATCTNNNACCTTTCTGTTTCATTATGTTTTCA

ACACGTCTGAACTCTTTCCCTACACGACGCTCTTCCGATCTNNNCTGGATATACTATTCTAGGGTAACA

AGCACACGTCTGAACTCTTTCCCTACACGACGCTCTTCCGATCTNNNAGCTTATAATAGCATTCCAATTTG

AGCACACGTCTGAACTCTTTCCCTACACGACGCTCTTCCGATCTNNNTAAAGAGTGTTTACATAGAACC

GCACACGTCTGAACTCTTTCCCTACACGACGCTCTTCCGATCTNNNACGTTATGACTAGATCTTTTTACC

AGCACACGTCTGAACTCTTTCCCTACACGACGCTCTTCCGATCTNNNCTTCATCAAACCAAAAGATAACT

GCACACGTCTGAACTCTTTCCCTACACGACGCTCTTCCGATCTNNNGTTGCATATTATTTAGGTTACCTC

GCACACGTCTGAACTCTTTCCCTACACGACGCTCTTCCGATCTNNNAGGATACATAGAATCAATGGTAT

GCACACGTCTGAACTCTTTCCCTACACGACGCTCTTCCGATCTNNNAAAGAGTTCAGAGACAAAGTTTT

AGCACACGTCTGAACTCTTTCCCTACACGACGCTCTTCCGATCTNNNACACATTTTCCATAAACTTTTTGC

CACACGTCTGAACTCTTTCCCTACACGACGCTCTTCCGATCTNNNGATCCAAAACACGTATTATTTCCCA

AGCACACGTCTGAACTCTTTCCCTACACGACGCTCTTCCGATCTNNNGAAACTGATATGGTTCAATGAG

AGCACACGTCTGAACTCTTTCCCTACACGACGCTCTTCCGATCTNNNGAAGTATGGATAAGCTAGAG
AGCACACGTCTGAACTCTTTCCCTACACGACGCTCTTCCGATCTNNNTGCTTTTTAAAATGAGATCTAC
AGCACACGTCTGAACTCTTTCCCTACACGACGCTCTTCCGATCTNNNCATGTTGGTTCATTCTAAATCT
CACACGTCTGAACTCTTTCCCTACACGACGCTCTTCCGATCTNNNGTACCAACTATGTGTGACTCCATC
GCACACGTCTGAACTCTTTCCCTACACGACGCTCTTCCGATCTNNNTCAACTCCAGAGCTTGTATACAT
AGCACACGTCTGAACTCTTTCCCTACACGACGCTCTTCCGATCTNNNAGAGAAAAGATCCCAAAGATT
GCACACGTCTGAACTCTTTCCCTACACGACGCTCTTCCGATCTNNNATAGATTCAGTTTCAATTTACCC
GAGCACACGTCTGAACTCTTTCCCTACACGACGCTCTTCCGATCTNNNGAACACCAAATTGAACAACT
ACACGTCTGAACTCTTTCCCTACACGACGCTCTTCCGATCTNNNTTCCAAGTTTTTCCTTCCTTCATCGG
AGCACACGTCTGAACTCTTTCCCTACACGACGCTCTTCCGATCTNNNTTTGGAAGTAGGTTTGTATAG
AGCACACGTCTGAACTCTTTCCCTACACGACGCTCTTCCGATCTNNNTTTGGAAGTAGGTTTGTATAG
CACACGTCTGAACTCTTTCCCTACACGACGCTCTTCCGATCTNNNACATATATACACACTCTCTTGAGCA
CACACGTCTGAACTCTTTCCCTACACGACGCTCTTCCGATCTNNNACATATATACACACTCTCTTGAGCA
GCACACGTCTGAACTCTTTCCCTACACGACGCTCTTCCGATCTNNNGGAAGCTTTAAAAAAAGGTTT
GCACACGTCTGAACTCTTTCCCTACACGACGCTCTTCCGATCTNNNACTCTAGGACTTTTAATTGCTAT
GCACACGTCTGAACTCTTTCCCTACACGACGCTCTTCCGATCTNNNATGTGGATTGAGCTTTTGTGCA
GCACACGTCTGAACTCTTTCCCTACACGACGCTCTTCCGATCTNNNCAGATATTAGGCGTGATCTGCAT
AGCACACGTCTGAACTCTTTCCCTACACGACGCTCTTCCGATCTNNNCTTTTTTCAGCTCTCATATGTAA
GCACACGTCTGAACTCTTTCCCTACACGACGCTCTTCCGATCTNNNCAGATCTGTACATGTGTTAAAA
AGCACACGTCTGAACTCTTTCCCTACACGACGCTCTTCCGATCTNNNCTAAGTGTGGCAAGAAATATT
GCACACGTCTGAACTCTTTCCCTACACGACGCTCTTCCGATCTNNNCAGCATTTATTCATATGTACCAT
AGCACACGTCTGAACTCTTTCCCTACACGACGCTCTTCCGATCTNNNCAGTTATAAATAAAGCTAAGA
GAAGAGCACACGTCTGAACTCTTTCCCTACACGACGCTCTTCCGATCTNNNAATAATATTTGCTTTACA
AGCACACGTCTGAACTCTTTCCCTACACGACGCTCTTCCGATCTNNNTAGGCCAAAGAGAAAGAAAC
GCACACGTCTGAACTCTTTCCCTACACGACGCTCTTCCGATCTNNNCATAAACAGTTACAGTGGTTTTA
GCACACGTCTGAACTCTTTCCCTACACGACGCTCTTCCGATCTNNNCATAAACAGTTACAGTGGTTTTA
AGCACACGTCTGAACTCTTTCCCTACACGACGCTCTTCCGATCTNNNTGTGTATATACATGTATCTCCT
AGCACACGTCTGAACTCTTTCCCTACACGACGCTCTTCCGATCTNNNTGTGTATATACATGTATCTCCT
CACACGTCTGAACTCTTTCCCTACACGACGCTCTTCCGATCTNNNCTAGGTGAAGAAGTATAGGGTT
AGAGCACACGTCTGAACTCTTTCCCTACACGACGCTCTTCCGATCTNNNGTAGAAAGTGGGTTTCCTAT
ACACGTCTGAACTCTTTCCCTACACGACGCTCTTCCGATCTNNNTTCATTCTCAATTCATACTGAAACAT
AGCACACGTCTGAACTCTTTCCCTACACGACGCTCTTCCGATCTNNNTATTATTTCTTCTCTTTTCCACC
GCACACGTCTGAACTCTTTCCCTACACGACGCTCTTCCGATCTNNNAAGGGATAATGAGGAAGGAAT
ACACGTCTGAACTCTTTCCCTACACGACGCTCTTCCGATCTNNNCTAGAAAGGAACTAAGGTGAAGC
AGCACACGTCTGAACTCTTTCCCTACACGACGCTCTTCCGATCTNNNTACTGATCTTATTCAGATTTCC
CACACGTCTGAACTCTTTCCCTACACGACGCTCTTCCGATCTNNNTTTCTTATGCCTATTAAACTCCTCA
AGCACACGTCTGAACTCTTTCCCTACACGACGCTCTTCCGATCTNNNAAAATCATTTTAGCATTTCACT
GCACACGTCTGAACTCTTTCCCTACACGACGCTCTTCCGATCTNNNTTTCTAGATCCTCAAAACTTAGG
GCACACGTCTGAACTCTTTCCCTACACGACGCTCTTCCGATCTNNNCTTGTGCTCTCATATATATAACA
AGCACACGTCTGAACTCTTTCCCTACACGACGCTCTTCCGATCTNNNGAAATTCTTTTAGTGAATCCC
GCACACGTCTGAACTCTTTCCCTACACGACGCTCTTCCGATCTNNNCTTGGTCACTAGGTATACAGACA
CACACGTCTGAACTCTTTCCCTACACGACGCTCTTCCGATCTNNNGTCCTCTCCTCTTGTAGATGCATC
AGCACACGTCTGAACTCTTTCCCTACACGACGCTCTTCCGATCTNNNTAAGGGGAAATTAGAAAGAA

AGCACACGTCTGAACTCTTTCCCTACACGACGCTCTTCCGATCTNNNTAAGGGGAAATTAGAAAGAAT
GCACACGTCTGAACTCTTTCCCTACACGACGCTCTTCCGATCTNNNACATATAAGTCAATGTTGAGACT
GAGCACACGTCTGAACTCTTTCCCTACACGACGCTCTTCCGATCTNNNCTGTTTGTCACTTTGCTTTAG
CACACGTCTGAACTCTTTCCCTACACGACGCTCTTCCGATCTNNNCACCACACCTATATTCATACATACAT
GCACACGTCTGAACTCTTTCCCTACACGACGCTCTTCCGATCTNNNAATAAGCAAATATACAAACCACA
AGCACACGTCTGAACTCTTTCCCTACACGACGCTCTTCCGATCTNNNGAGGATATTAAATCACAATGT
AGCACACGTCTGAACTCTTTCCCTACACGACGCTCTTCCGATCTNNNGGTATTTCTTCCTATTGAAAAGA
AGCACACGTCTGAACTCTTTCCCTACACGACGCTCTTCCGATCTNNNCTACATGGACAATTAAACACAC
GAGCACACGTCTGAACTCTTTCCCTACACGACGCTCTTCCGATCTNNNCTAGAGGTATGAGTCTATGA
AGCACACGTCTGAACTCTTTCCCTACACGACGCTCTTCCGATCTNNNTTGAAATGATTACCACAATGAT
CACACGTCTGAACTCTTTCCCTACACGACGCTCTTCCGATCTNNNAAAACCATGAACTCAAACTCTACAT
AGCACACGTCTGAACTCTTTCCCTACACGACGCTCTTCCGATCTNNNTAAAATATGCATGTCATTTAGG
AGCACACGTCTGAACTCTTTCCCTACACGACGCTCTTCCGATCTNNNATATGACACTTGATATTGACAG
GAGCACACGTCTGAACTCTTTCCCTACACGACGCTCTTCCGATCTNNNCTTCTTTTCTGGGAAAACATTC
AGCACACGTCTGAACTCTTTCCCTACACGACGCTCTTCCGATCTNNNGATGATAGGTAGGTAGGTAGG
GCACACGTCTGAACTCTTTCCCTACACGACGCTCTTCCGATCTNNNATCCCATCCTGTTTACTTTTTATCA
GAGCACACGTCTGAACTCTTTCCCTACACGACGCTCTTCCGATCTNNNTTTCAATACATGTAGATAAGT
GAGCACACGTCTGAACTCTTTCCCTACACGACGCTCTTCCGATCTNNNCAATTACAGTGTATTCATGCTA
GAGCACACGTCTGAACTCTTTCCCTACACGACGCTCTTCCGATCTNNNCAAAAATGAAGCTACCAATAC
AGCACACGTCTGAACTCTTTCCCTACACGACGCTCTTCCGATCTNNNGTAGAAATTTGCATGGTTTGTC
GAGCACACGTCTGAACTCTTTCCCTACACGACGCTCTTCCGATCTNNNACAACAAGGTGAATAAGATG
CACACGTCTGAACTCTTTCCCTACACGACGCTCTTCCGATCTNNNTTTGAATAGAATTTGTACTGTGT
CACACGTCTGAACTCTTTCCCTACACGACGCTCTTCCGATCTNNNTCTCTCTTCATTACTGCGATCATCG
AGCACACGTCTGAACTCTTTCCCTACACGACGCTCTTCCGATCTNNNTCATAAAAATTTTTCCTAGTCCC
GAGCACACGTCTGAACTCTTTCCCTACACGACGCTCTTCCGATCTNNNGGAAAGGTGCATTAAAATTCT
GCACACGTCTGAACTCTTTCCCTACACGACGCTCTTCCGATCTNNNGAAAAAGAGATTTTTTTCCCTG
AGCACACGTCTGAACTCTTTCCCTACACGACGCTCTTCCGATCTNNNGGTATATCCTCTCTTAAATGTGT
GAGCACACGTCTGAACTCTTTCCCTACACGACGCTCTTCCGATCTNNNTTTGAGGTCAAGAAGGTAGTA
AGCACACGTCTGAACTCTTTCCCTACACGACGCTCTTCCGATCTNNNTGTACTTCCAAATCAGATCTTAT
GCACACGTCTGAACTCTTTCCCTACACGACGCTCTTCCGATCTNNNAAACAAAACCAGTCATTAGAAAT
CACACGTCTGAACTCTTTCCCTACACGACGCTCTTCCGATCTNNNCTTCTCATCTTCCCAAACTGCATCGG
GAGCACACGTCTGAACTCTTTCCCTACACGACGCTCTTCCGATCTNNNCCTCATTATGAACATAACCTAT
GCACACGTCTGAACTCTTTCCCTACACGACGCTCTTCCGATCTNNNACATATACTTGAAATTTGCTAACA
GAGCACACGTCTGAACTCTTTCCCTACACGACGCTCTTCCGATCTNNNAGAAGCCATAAAAGATTACTC
CACACGTCTGAACTCTTTCCCTACACGACGCTCTTCCGATCTNNNGGAATGTATCAGCCATAATAAATCA
AGCACACGTCTGAACTCTTTCCCTACACGACGCTCTTCCGATCTNNNTTTCCTTAAACCTCATCCTTACCAT
AGCACACGTCTGAACTCTTTCCCTACACGACGCTCTTCCGATCTNNNAAAATCAGAACATTTGGGTTTT
AGCACACGTCTGAACTCTTTCCCTACACGACGCTCTTCCGATCTNNNTTGATTACTCCTTGAATACTCTC
AGCACACGTCTGAACTCTTTCCCTACACGACGCTCTTCCGATCTNNNTTGATTACTCCTTGAATACTCTC
GCACACGTCTGAACTCTTTCCCTACACGACGCTCTTCCGATCTNNNTGTCTTCATCATAAAAACAAACT
GAGCACACGTCTGAACTCTTTCCCTACACGACGCTCTTCCGATCTNNNTCATAGAGCTTATTCAGATTTT
GAGCACACGTCTGAACTCTTTCCCTACACGACGCTCTTCCGATCTNNNCAATTAAGGACAATCGAAAC
GCACACGTCTGAACTCTTTCCCTACACGACGCTCTTCCGATCTNNNCATAAGCCATAGTTTGTTGATCCA

AGCACACGTCTGAACTCTTTCCCTACACGACGCTCTTCCGATCTNNNAATAGACAAAAGATGAGAACA
CACACGTCTGAACTCTTTCCCTACACGACGCTCTTCCGATCTNNNACATATAAACATGGAATCAAGATC
AGCACACGTCTGAACTCTTTCCCTACACGACGCTCTTCCGATCTNNNTTAAGATAAGTCCTGAAATCAC
AGCACACGTCTGAACTCTTTCCCTACACGACGCTCTTCCGATCTNNNCATGTGAACTAATTCCTTGTAA
AGCACACGTCTGAACTCTTTCCCTACACGACGCTCTTCCGATCTNNNTTATGTACACTGATGAAAATG
AGCACACGTCTGAACTCTTTCCCTACACGACGCTCTTCCGATCTNNNTTATGTACACTGATGAAAATG
AGCACACGTCTGAACTCTTTCCCTACACGACGCTCTTCCGATCTNNNCATACTATAAGGTTTGAGGTT
AGCACACGTCTGAACTCTTTCCCTACACGACGCTCTTCCGATCTNNNCTATCCACAGTTTCAGTCATTT
GCACACGTCTGAACTCTTTCCCTACACGACGCTCTTCCGATCTNNNTATCCTGCTCTGTACATTTTAAA
AGCACACGTCTGAACTCTTTCCCTACACGACGCTCTTCCGATCTNNNGGCTCTACATTTAAGATCATCTA
AGCACACGTCTGAACTCTTTCCCTACACGACGCTCTTCCGATCTNNNAGAGATCTCTGAAATCATTTA
AGCACACGTCTGAACTCTTTCCCTACACGACGCTCTTCCGATCTNNNAGTTTTAGAAGTTTCAATAGC
CACACGTCTGAACTCTTTCCCTACACGACGCTCTTCCGATCTNNNGTGTTATTCCAGTGTTATTCTTTC
AGCACACGTCTGAACTCTTTCCCTACACGACGCTCTTCCGATCTNNNGAAAACTCTCTATGTGGAAAA
AGCACACGTCTGAACTCTTTCCCTACACGACGCTCTTCCGATCTNNNGAAAACTCTCTATGTGGAAAA
CACACGTCTGAACTCTTTCCCTACACGACGCTCTTCCGATCTNNNTAAAAAGATGTTCAGGTAGATCA
AGCACACGTCTGAACTCTTTCCCTACACGACGCTCTTCCGATCTNNNTTCAAGTCTCTATTCCAATATCA
CACACGTCTGAACTCTTTCCCTACACGACGCTCTTCCGATCTNNNTACAATGATATCTATGGTGAAGG
CACACGTCTGAACTCTTTCCCTACACGACGCTCTTCCGATCTNNNTACAATGATATCTATGGTGAAGG
AGCACACGTCTGAACTCTTTCCCTACACGACGCTCTTCCGATCTNNNCAGTATATTTTAGAAGGTCCT
AGCACACGTCTGAACTCTTTCCCTACACGACGCTCTTCCGATCTNNNGAGCATCTGTCTCAAAATATAT
AGCACACGTCTGAACTCTTTCCCTACACGACGCTCTTCCGATCTNNNTAATGTGAACATAACATCAGG
GCACACGTCTGAACTCTTTCCCTACACGACGCTCTTCCGATCTNNNGTTTCTCCTACTAGGTTTCCTACA
GAGCACACGTCTGAACTCTTTCCCTACACGACGCTCTTCCGATCTNNNTCCTAATGTATTACTTAGCAA
AGCACACGTCTGAACTCTTTCCCTACACGACGCTCTTCCGATCTNNNCTCATTAACCCACACAAACTCAT
CACACGTCTGAACTCTTTCCCTACACGACGCTCTTCCGATCTNNNCTATTGGACTGTTGTGTTAAAAAC
CACACGTCTGAACTCTTTCCCTACACGACGCTCTTCCGATCTNNNAATCACACTTCCTACTCAGACATC
CACACGTCTGAACTCTTTCCCTACACGACGCTCTTCCGATCTNNNTTACCTCCTGTATCCTCCTTACATC
GCACACGTCTGAACTCTTTCCCTACACGACGCTCTTCCGATCTNNNTGGTTGTAAGTTTGAAGTCTCC
GCACACGTCTGAACTCTTTCCCTACACGACGCTCTTCCGATCTNNNTGGTTGTAAGTTTGAAGTCTCC
AGCACACGTCTGAACTCTTTCCCTACACGACGCTCTTCCGATCTNNNCATAAATGTTTAAATGAAGCG
AGCACACGTCTGAACTCTTTCCCTACACGACGCTCTTCCGATCTNNNACAAATCTACATGTGACAAAAT
AGCACACGTCTGAACTCTTTCCCTACACGACGCTCTTCCGATCTNNNGTCTCCCATCACTCTAAAATTTC
AGCACACGTCTGAACTCTTTCCCTACACGACGCTCTTCCGATCTNNNACTTCTCTATGTGTTCTCAAAT
AGCACACGTCTGAACTCTTTCCCTACACGACGCTCTTCCGATCTNNNATACAGAATAGAGAAATGGG
GAGCACACGTCTGAACTCTTTCCCTACACGACGCTCTTCCGATCTNNNCACACTACACTGTAGAAGTAT
AGCACACGTCTGAACTCTTTCCCTACACGACGCTCTTCCGATCTNNNCTTTCTCATTCAACATTCAGAGC
AGCACACGTCTGAACTCTTTCCCTACACGACGCTCTTCCGATCTNNNCTTTCTCATTCAACATTCAGAGC
AGCACACGTCTGAACTCTTTCCCTACACGACGCTCTTCCGATCTNNNTCATTCATATGCAGAATTCATT
AGCACACGTCTGAACTCTTTCCCTACACGACGCTCTTCCGATCTNNNTTTACACCAACATCTATATCTGT
CACACGTCTGAACTCTTTCCCTACACGACGCTCTTCCGATCTNNNTTATCAAGTTTGATCCTTTTTGCCA
AGCACACGTCTGAACTCTTTCCCTACACGACGCTCTTCCGATCTNNNTACATTGTTACACTGTTGCTTCA
GCACACGTCTGAACTCTTTCCCTACACGACGCTCTTCCGATCTNNNGGACAATAAAATCAACTGAACT

ACACGTCTGAACTCTTTCCCTACACGACGCTCTTCCGATCTNNNCTCATTTAACCTCAGCTCTGCATCGG
AGCACACGTCTGAACTCTTTCCCTACACGACGCTCTTCCGATCTNNNTACTAGGCTAAGAAAAACCATC
AGCACACGTCTGAACTCTTTCCCTACACGACGCTCTTCCGATCTNNNATAATATATCACCAAATTGAAA
GAGCACACGTCTGAACTCTTTCCCTACACGACGCTCTTCCGATCTNNNAAACGTTATCTCGGTAATCTC
GCACACGTCTGAACTCTTTCCCTACACGACGCTCTTCCGATCTNNNTAAAAAGACATTCCTTTCCTTCC
CACACGTCTGAACTCTTTCCCTACACGACGCTCTTCCGATCTNNNGTCCACAAGTATAATCTGGATTC
GCACACGTCTGAACTCTTTCCCTACACGACGCTCTTCCGATCTNNNTCTCCACTAATCTCTCTTCCAT
AGCACACGTCTGAACTCTTTCCCTACACGACGCTCTTCCGATCTNNNAACTACCATATGAAACATAGG
CACACGTCTGAACTCTTTCCCTACACGACGCTCTTCCGATCTNNNCTCACAAGTCCTTCCTTCATCATCGG
GCACACGTCTGAACTCTTTCCCTACACGACGCTCTTCCGATCTNNNGCATATGAACTAATCACCTCATA
GAGCACACGTCTGAACTCTTTCCCTACACGACGCTCTTCCGATCTNNNTAGCTTTATAAGAAGAGGAA
CACACGTCTGAACTCTTTCCCTACACGACGCTCTTCCGATCTNNNTTTACAAACTTAATTTCCCCTTGCAT
ACACGTCTGAACTCTTTCCCTACACGACGCTCTTCCGATCTNNNGTGTGTAGAGAGACAGTCTTCATC
AGCACACGTCTGAACTCTTTCCCTACACGACGCTCTTCCGATCTNNNGAATTAGTCACCCTTCTTTGTC
AGCACACGTCTGAACTCTTTCCCTACACGACGCTCTTCCGATCTNNNCACTCATAAAAATCTAGCTCAA
AGCACACGTCTGAACTCTTTCCCTACACGACGCTCTTCCGATCTNNNTAGGAAAACAAGAAGAGGGT
GAGCACACGTCTGAACTCTTTCCCTACACGACGCTCTTCCGATCTNNNAAAGTGACAGACAAAAAATT
CACACGTCTGAACTCTTTCCCTACACGACGCTCTTCCGATCTNNNATTCTTGCTTCTTTATCACTTTTCAT
AGCACACGTCTGAACTCTTTCCCTACACGACGCTCTTCCGATCTNNNGTTTTGGACTATTTTGTCTAGT
CACACGTCTGAACTCTTTCCCTACACGACGCTCTTCCGATCTNNNCTTGTTCCCACTAAAGACTTCATC
ACACGTCTGAACTCTTTCCCTACACGACGCTCTTCCGATCTNNNTTTACCAGCTCAAAATCTCAACATCG
CACACGTCTGAACTCTTTCCCTACACGACGCTCTTCCGATCTNNNGGCATCTAAAATACTCTTTATCCCA
GCACACGTCTGAACTCTTTCCCTACACGACGCTCTTCCGATCTNNNCAAAACACACGATTTATTTTCTT
GCACACGTCTGAACTCTTTCCCTACACGACGCTCTTCCGATCTNNNGTTATAGACCCCATAGAAATTCT
AGCACACGTCTGAACTCTTTCCCTACACGACGCTCTTCCGATCTNNNGCCTGGTTCTATCATATACATT
AGCACACGTCTGAACTCTTTCCCTACACGACGCTCTTCCGATCTNNNGTAATCTCAGAACTTTGGGAG
AGCACACGTCTGAACTCTTTCCCTACACGACGCTCTTCCGATCTNNNTCTCTCTCTCTGTGTGTACAT
AGCACACGTCTGAACTCTTTCCCTACACGACGCTCTTCCGATCTNNNCTAGAGGGAAAAACTATAAAT
AGCACACGTCTGAACTCTTTCCCTACACGACGCTCTTCCGATCTNNNGTTCATAGAACATGGTAAATT
AGCACACGTCTGAACTCTTTCCCTACACGACGCTCTTCCGATCTNNNATGAAAAAAAGTCAATCAGATC
GCACACGTCTGAACTCTTTCCCTACACGACGCTCTTCCGATCTNNNTGAGTGACCATCAATGTTTAACA
AGCACACGTCTGAACTCTTTCCCTACACGACGCTCTTCCGATCTNNNGTATTTCAGGAATTGATGACA
GCACACGTCTGAACTCTTTCCCTACACGACGCTCTTCCGATCTNNNAGAACCACTTCAATTGTTTTATT
GCACACGTCTGAACTCTTTCCCTACACGACGCTCTTCCGATCTNNNAGAACCACTTCAATTGTTTTATT
AGCACACGTCTGAACTCTTTCCCTACACGACGCTCTTCCGATCTNNNTAGATGAAAATCAGTTTATGG
AGCACACGTCTGAACTCTTTCCCTACACGACGCTCTTCCGATCTNNNCTAGAACTCCTGGGTTCAAGCA
AGCACACGTCTGAACTCTTTCCCTACACGACGCTCTTCCGATCTNNNTCAATTGTATCTGGATTTTGTT
GAGCACACGTCTGAACTCTTTCCCTACACGACGCTCTTCCGATCTNNNTGTGCTAATTGTTATATGCTTT
AGCACACGTCTGAACTCTTTCCCTACACGACGCTCTTCCGATCTNNNATACATATTCCCCTGCACTTACA
AGCACACGTCTGAACTCTTTCCCTACACGACGCTCTTCCGATCTNNNCTACATATATTAGGTTGGTGG
GCACACGTCTGAACTCTTTCCCTACACGACGCTCTTCCGATCTNNNAATTCCAAAAAAAAATTGAAGAG
GCACACGTCTGAACTCTTTCCCTACACGACGCTCTTCCGATCTNNNTTGAAAACTGTTATTAAATCGCC
ACACGTCTGAACTCTTTCCCTACACGACGCTCTTCCGATCTNNNTGATTAAAACACTCTCCCAGCATCGG

AGCACACGTCTGAACTCTTTCCCTACACGACGCTCTTCCGATCTNNNTTTCATTTACTTTGGAAGTCAA
CACACGTCTGAACTCTTTCCCTACACGACGCTCTTCCGATCTNNNCATTCCTTCCTGTTACAATTAAGCA
GCACACGTCTGAACTCTTTCCCTACACGACGCTCTTCCGATCTNNNCAATTAAAGAAGAAAGAACCAG
GCACACGTCTGAACTCTTTCCCTACACGACGCTCTTCCGATCTNNNCAATTAAAGAAGAAAGAACCAG
AGCACACGTCTGAACTCTTTCCCTACACGACGCTCTTCCGATCTNNNTTTAGATAACATCTTCTCTGAG
AGCACACGTCTGAACTCTTTCCCTACACGACGCTCTTCCGATCTNNNTCATGACATGAATGGATTTTT
CACACGTCTGAACTCTTTCCCTACACGACGCTCTTCCGATCTNNNTACATCTTTCTTTTCTAAGTGGACA
GAGCACACGTCTGAACTCTTTCCCTACACGACGCTCTTCCGATCTNNNAGGATTAAAGAAAACAATAA
AGCACACGTCTGAACTCTTTCCCTACACGACGCTCTTCCGATCTNNNAGAACCACTGCTTTACAATATA
AGCACACGTCTGAACTCTTTCCCTACACGACGCTCTTCCGATCTNNNCCAGTAAGCAAAGAACTTGGC
AGCACACGTCTGAACTCTTTCCCTACACGACGCTCTTCCGATCTNNNCCAGTAAGCAAAGAACTTGGC
AGCACACGTCTGAACTCTTTCCCTACACGACGCTCTTCCGATCTNNNTTTTCTTTTTTATGGCTGAGTG
AGCACACGTCTGAACTCTTTCCCTACACGACGCTCTTCCGATCTNNNCATCTAGTGGGGAAAGATAAG
AGCACACGTCTGAACTCTTTCCCTACACGACGCTCTTCCGATCTNNNCTTGGGGATACTTAGCAATGC
GCACACGTCTGAACTCTTTCCCTACACGACGCTCTTCCGATCTNNNATTTAGCAAGAAACTGTCTCTAT
AGCACACGTCTGAACTCTTTCCCTACACGACGCTCTTCCGATCTNNNCTCAACTTTCTCTGTCTCTGCAT
CACGTCTGAACTCTTTCCCTACACGACGCTCTTCCGATCTNNNAATATGTGTTCTTGCTTTTGGCATCG
AGCACACGTCTGAACTCTTTCCCTACACGACGCTCTTCCGATCTNNNTCTTTCGTAGAAAATCAAAGA
AGCACACGTCTGAACTCTTTCCCTACACGACGCTCTTCCGATCTNNNTCTTTCGTAGAAAATCAAAGA
AGCACACGTCTGAACTCTTTCCCTACACGACGCTCTTCCGATCTNNNATACTAAGGTACATAGCAAGA
GAGCACACGTCTGAACTCTTTCCCTACACGACGCTCTTCCGATCTNNNAAGAACGTGAATCTCAGATA
CACACGTCTGAACTCTTTCCCTACACGACGCTCTTCCGATCTNNNTAAAAACAGTAAACTGAATCAGC
AGCACACGTCTGAACTCTTTCCCTACACGACGCTCTTCCGATCTNNNGATTTGAAAAGGTGAACAAGA
GCACACGTCTGAACTCTTTCCCTACACGACGCTCTTCCGATCTNNNATATGACTGGGTTTAAGTCTAT
CACGTCTGAACTCTTTCCCTACACGACGCTCTTCCGATCTNNNCACTTTTTATAGATGACTGCATTCAT
GAGCACACGTCTGAACTCTTTCCCTACACGACGCTCTTCCGATCTNNNCCAGATACACACATACATACA
CACACGTCTGAACTCTTTCCCTACACGACGCTCTTCCGATCTNNNGGAGGATAATTCCAGTATCATAA
AGAGCACACGTCTGAACTCTTTCCCTACACGACGCTCTTCCGATCTNNNGTGTTTATGTGTGCAGACAT
GCACACGTCTGAACTCTTTCCCTACACGACGCTCTTCCGATCTNNNCCTTTGTCTCTCACTTTTTATGCA
AGCACACGTCTGAACTCTTTCCCTACACGACGCTCTTCCGATCTNNNTTACTTCAAAAAATCACAAGTC
GCACACGTCTGAACTCTTTCCCTACACGACGCTCTTCCGATCTNNNTGTTAGTATTGATTGTCTGTACT
AGCACACGTCTGAACTCTTTCCCTACACGACGCTCTTCCGATCTNNNTTCTAAGAACAGAACTCAAATT
GAGCACACGTCTGAACTCTTTCCCTACACGACGCTCTTCCGATCTNNNTGAGATTATGAAGTCATTTT
GCACACGTCTGAACTCTTTCCCTACACGACGCTCTTCCGATCTNNNTTTACAAATATTTTTGGCTACCA
GCACACGTCTGAACTCTTTCCCTACACGACGCTCTTCCGATCTNNNCTCTCAATGCCTTTTTGAATTATC
GCACACGTCTGAACTCTTTCCCTACACGACGCTCTTCCGATCTNNNCTCTCAATGCCTTTTTGAATTATC
AGCACACGTCTGAACTCTTTCCCTACACGACGCTCTTCCGATCTNNNGTATTTCATATACTCTCACACCT
AGCACACGTCTGAACTCTTTCCCTACACGACGCTCTTCCGATCTNNNTATTTTCCCATCAAACAGATAC
AGCACACGTCTGAACTCTTTCCCTACACGACGCTCTTCCGATCTNNNATCCAGCTATAAGTATGTTGTT
GCACACGTCTGAACTCTTTCCCTACACGACGCTCTTCCGATCTNNNTATGTTATCTCTCAAGTAATGGG
GCACACGTCTGAACTCTTTCCCTACACGACGCTCTTCCGATCTNNNGTAAGAAGGGAGACTGGTATTC
GCACACGTCTGAACTCTTTCCCTACACGACGCTCTTCCGATCTNNNTCAGAGTTATCACATACACATAC
GCACACGTCTGAACTCTTTCCCTACACGACGCTCTTCCGATCTNNNTATTACAAGGTTGAAATTGTGA

GAGCACACGTCTGAACTCTTTCCCTACACGACGCTCTTCCGATCTNNNATTTTTGGGTATCTTTATAGC
AGCACACGTCTGAACTCTTTCCCTACACGACGCTCTTCCGATCTNNNAAGGAAGACAAAAGATAAAA
CACACGTCTGAACTCTTTCCCTACACGACGCTCTTCCGATCTNNNTTAACCTTAATTACTTCCTTGCTCA
AGCACACGTCTGAACTCTTTCCCTACACGACGCTCTTCCGATCTNNNTCCATTTAGGATGTGTGTTTAT
AGCACACGTCTGAACTCTTTCCCTACACGACGCTCTTCCGATCTNNNTCTATTTTTATCTCTGCTTTCAG
ACACGTCTGAACTCTTTCCCTACACGACGCTCTTCCGATCTNNNAGTAATTAAAATGAAGTGCTACAAC
GAGCACACGTCTGAACTCTTTCCCTACACGACGCTCTTCCGATCTNNNTGTACACATATCCTAAAACCA
AGCACACGTCTGAACTCTTTCCCTACACGACGCTCTTCCGATCTNNNTGAAGACAATAACATTCCAAA
CACACGTCTGAACTCTTTCCCTACACGACGCTCTTCCGATCTNNNCAGTTCCATGCTACATCATTCATCG
CACGTCTGAACTCTTTCCCTACACGACGCTCTTCCGATCTNNNAATATTACCAGGGCTTAGATATGCA
CACACGTCTGAACTCTTTCCCTACACGACGCTCTTCCGATCTNNNCACATAGACATGAACACACATCAT
AGCACACGTCTGAACTCTTTCCCTACACGACGCTCTTCCGATCTNNNTACTTCAACACTCAATTCATTTT
GCACACGTCTGAACTCTTTCCCTACACGACGCTCTTCCGATCTNNNAGAGGAGCATACATTTTATGAG
CACACGTCTGAACTCTTTCCCTACACGACGCTCTTCCGATCTNNNTATTGACAGGACTACATGAACCAT
CACACGTCTGAACTCTTTCCCTACACGACGCTCTTCCGATCTNNNCACCATTATACTCAGCTACTTTCAT
GCACACGTCTGAACTCTTTCCCTACACGACGCTCTTCCGATCTNNNTCTATTTGGTATTCTTCATCACAC
AGCACACGTCTGAACTCTTTCCCTACACGACGCTCTTCCGATCTNNNAGTCTTCCAAATATACTTTCTG
AGCACACGTCTGAACTCTTTCCCTACACGACGCTCTTCCGATCTNNNAAACTAGTATCTGCAATGTCA
AGCACACGTCTGAACTCTTTCCCTACACGACGCTCTTCCGATCTNNNTTTTGTTATTTTGTTACAGCAG
GCACACGTCTGAACTCTTTCCCTACACGACGCTCTTCCGATCTNNNTAACGATAAGTGATTTAAATGC
GCACACGTCTGAACTCTTTCCCTACACGACGCTCTTCCGATCTNNNACACTTAGTAATATCTGATGCA
GCACACGTCTGAACTCTTTCCCTACACGACGCTCTTCCGATCTNNNGAAAAAGAGTTCTCAAGTTAGG
CACACGTCTGAACTCTTTCCCTACACGACGCTCTTCCGATCTNNNTATACAGTCCCTTATGATCTAGTCA
AGCACACGTCTGAACTCTTTCCCTACACGACGCTCTTCCGATCTNNNTTGTTCAGAGAGCAGTTTTTA
AGCACACGTCTGAACTCTTTCCCTACACGACGCTCTTCCGATCTNNNTTGTTCAGAGAGCAGTTTTTA
AGCACACGTCTGAACTCTTTCCCTACACGACGCTCTTCCGATCTNNNAACTTATTAGTCTACAAGTAT
CACACGTCTGAACTCTTTCCCTACACGACGCTCTTCCGATCTNNNATACTGTGCTTTACGAAATAAACC
GCACACGTCTGAACTCTTTCCCTACACGACGCTCTTCCGATCTNNNACTACTTCTACTACTACTACACA
GCACACGTCTGAACTCTTTCCCTACACGACGCTCTTCCGATCTNNNACTACTTCTACTACTACTACACA
AGCACACGTCTGAACTCTTTCCCTACACGACGCTCTTCCGATCTNNNATTTTTGGTCAGGAGAGAAC
CACACGTCTGAACTCTTTCCCTACACGACGCTCTTCCGATCTNNNAATAAGCTATAATCATGCCACTCAT
AGCACACGTCTGAACTCTTTCCCTACACGACGCTCTTCCGATCTNNNTTTCTTTTCTTTTGTCTTTCCTT
GAGCACACGTCTGAACTCTTTCCCTACACGACGCTCTTCCGATCTNNNTTTCTTTTTCTTTTCTTTTGTCT
GCACACGTCTGAACTCTTTCCCTACACGACGCTCTTCCGATCTNNNTAAATAACTGTTAGAAGGGTA
GAGCACACGTCTGAACTCTTTCCCTACACGACGCTCTTCCGATCTNNNTTTATCAAGACCCTACTTTGT
CACACGTCTGAACTCTTTCCCTACACGACGCTCTTCCGATCTNNNTATACATCATAATTGTGGCAGTAC
AGCACACGTCTGAACTCTTTCCCTACACGACGCTCTTCCGATCTNNNGAAAACAAGACAAATGGGAT
AGCACACGTCTGAACTCTTTCCCTACACGACGCTCTTCCGATCTNNNGCCAAGTTACAAGAAATATAC
ACACGTCTGAACTCTTTCCCTACACGACGCTCTTCCGATCTNNNCCAAAGTGATGTTAGACACCCATCG
CACACGTCTGAACTCTTTCCCTACACGACGCTCTTCCGATCTNNNTACAGATCTTTGTAGTCTGAAGC
CACACGTCTGAACTCTTTCCCTACACGACGCTCTTCCGATCTNNNTACAGATCTTTGTAGTCTGAAGC
CACACGTCTGAACTCTTTCCCTACACGACGCTCTTCCGATCTNNNATATAAGCTTCAAGCCTACAAAAC
GAGCACACGTCTGAACTCTTTCCCTACACGACGCTCTTCCGATCTNNNACTACAGAAAGGGTAAACA

AGCACACGTCTGAACTCTTTCCCTACACGACGCTCTTCCGATCTNNNTTCTTCAGTGATAACTAACTCA
GCACACGTCTGAACTCTTTCCCTACACGACGCTCTTCCGATCTNNNGTATTTTTGAAAGGATGGAAAG
AGCACACGTCTGAACTCTTTCCCTACACGACGCTCTTCCGATCTNNNTTTGTTTCCTCATCTGTAAATTG
AGCACACGTCTGAACTCTTTCCCTACACGACGCTCTTCCGATCTNNNAGATGTGGAAAGATACTCAA
ACACGTCTGAACTCTTTCCCTACACGACGCTCTTCCGATCTNNNGATGAGATGAATTGCACATACCAT
AGCACACGTCTGAACTCTTTCCCTACACGACGCTCTTCCGATCTNNNATATTAAGAGGCATATTGGTA
GAGCACACGTCTGAACTCTTTCCCTACACGACGCTCTTCCGATCTNNNGCCTTGTGACATTTACTAAAT
AGCACACGTCTGAACTCTTTCCCTACACGACGCTCTTCCGATCTNNNTCTTCTACATTTATTAGCTAGC
GCACACGTCTGAACTCTTTCCCTACACGACGCTCTTCCGATCTNNNTTGAATACTATATGCCTAGTTG
AGCACACGTCTGAACTCTTTCCCTACACGACGCTCTTCCGATCTNNNCATATACCACTGGAAATGGGC
GCACACGTCTGAACTCTTTCCCTACACGACGCTCTTCCGATCTNNNTGTAGCTAGCTATTTTGATGTA
GCACACGTCTGAACTCTTTCCCTACACGACGCTCTTCCGATCTNNNAATTTATAATCTGGCAGAGGG
AGCACACGTCTGAACTCTTTCCCTACACGACGCTCTTCCGATCTNNNATGATTTTATATTCCACCAATCC
AGCACACGTCTGAACTCTTTCCCTACACGACGCTCTTCCGATCTNNNAAAGAACCTTCATTTCAGAAA
AGCACACGTCTGAACTCTTTCCCTACACGACGCTCTTCCGATCTNNNTTTCAGTTCTATTATCTCAGTCA
AGCACACGTCTGAACTCTTTCCCTACACGACGCTCTTCCGATCTNNNCAGATTTCTGGTATTCTTCATA
GCACACGTCTGAACTCTTTCCCTACACGACGCTCTTCCGATCTNNNAAGAGTAGGAAAGATTTGAAG
CACACGTCTGAACTCTTTCCCTACACGACGCTCTTCCGATCTNNNTCCTGAGTAGCTGCTATTATAAAC
AGCACACGTCTGAACTCTTTCCCTACACGACGCTCTTCCGATCTNNNGGAATACAATGGTTTATAGA
AGCACACGTCTGAACTCTTTCCCTACACGACGCTCTTCCGATCTNNNAAATAATTGCAATGTATGAGG
GCACACGTCTGAACTCTTTCCCTACACGACGCTCTTCCGATCTNNNAGTTTATAGCCTGTACCATACC
GAGCACACGTCTGAACTCTTTCCCTACACGACGCTCTTCCGATCTNNNATAACTAAGAATTTTCCACAG
GCACACGTCTGAACTCTTTCCCTACACGACGCTCTTCCGATCTNNNCTCTTAATTGCTGTCACTCTAAC
GCACACGTCTGAACTCTTTCCCTACACGACGCTCTTCCGATCTNNNCTTTAAACTCAAACTATAGCACC
GAGCACACGTCTGAACTCTTTCCCTACACGACGCTCTTCCGATCTNNNAGGTAATGGTATCTATCAAA
AGCACACGTCTGAACTCTTTCCCTACACGACGCTCTTCCGATCTNNNAAAAAAGTGCAAGCATACTA
GCACACGTCTGAACTCTTTCCCTACACGACGCTCTTCCGATCTNNNTAACTCTGCTTTACTTACTCATT
CACACGTCTGAACTCTTTCCCTACACGACGCTCTTCCGATCTNNNACTTGAAGATTGATCTGTAGATAC
GCACACGTCTGAACTCTTTCCCTACACGACGCTCTTCCGATCTNNNATTAATTTGTAAACAAGTCCCAC
GCACACGTCTGAACTCTTTCCCTACACGACGCTCTTCCGATCTNNNAAGGATTGTCATTTTTGTTGTTT
AGAGCACACGTCTGAACTCTTTCCCTACACGACGCTCTTCCGATCTNNNCAAGAAGTCCTTCACTGAA
CACACGTCTGAACTCTTTCCCTACACGACGCTCTTCCGATCTNNNCATACCACTCATTAGTTGGTATAC
AGCACACGTCTGAACTCTTTCCCTACACGACGCTCTTCCGATCTNNNTAAATGATTGTTGAATACTCC
CACACGTCTGAACTCTTTCCCTACACGACGCTCTTCCGATCTNNNATTGTATGTATAATACGATCATG
AGCACACGTCTGAACTCTTTCCCTACACGACGCTCTTCCGATCTNNNCCACAGTTATATACCAAAGAAA
GCACACGTCTGAACTCTTTCCCTACACGACGCTCTTCCGATCTNNNTACATGCCTGTTTGATATATAG
GCACACGTCTGAACTCTTTCCCTACACGACGCTCTTCCGATCTNNNAGATTAATTTTTGTGTCTAGGT
AGCACACGTCTGAACTCTTTCCCTACACGACGCTCTTCCGATCTNNNGAAAGAGAGGTTCGAAAATT
CACACGTCTGAACTCTTTCCCTACACGACGCTCTTCCGATCTNNNGAGAATCACTTGAACACCAGCAT
GCACACGTCTGAACTCTTTCCCTACACGACGCTCTTCCGATCTNNNTGTTATTGATTAGTGTTAGACC
AGCACACGTCTGAACTCTTTCCCTACACGACGCTCTTCCGATCTNNNTAATGTCTACTATGTTCTTGAC
GAGCACACGTCTGAACTCTTTCCCTACACGACGCTCTTCCGATCTNNNCATAGCCCAGTGACAAATAA
GCACACGTCTGAACTCTTTCCCTACACGACGCTCTTCCGATCTNNNCACAAAATTAAGAAGCAAACAT

CACGTCTGAACTCTTTCCCTACACGACGCTCTTCCGATCTNNNTTTTTCTTTATGAATCTAAATGGTGCA
AGCACACGTCTGAACTCTTTCCCTACACGACGCTCTTCCGATCTNNNTCCATATATATGAAAAGGACAGG
AGCACACGTCTGAACTCTTTCCCTACACGACGCTCTTCCGATCTNNNTAAAAACAAATTCCTAACCAGT
GAGCACACGTCTGAACTCTTTCCCTACACGACGCTCTTCCGATCTNNNGAGGTATTTGTTCTTTGAGTTA
AGAGCACACGTCTGAACTCTTTCCCTACACGACGCTCTTCCGATCTNNNTCATATGAGCCAATTCCTTATA
AGCACACGTCTGAACTCTTTCCCTACACGACGCTCTTCCGATCTNNNTTCAAATTTAATTACCACCAACA
AGCACACGTCTGAACTCTTTCCCTACACGACGCTCTTCCGATCTNNNCTTTGATAAACTGTTTGGGAATA
GCACACGTCTGAACTCTTTCCCTACACGACGCTCTTCCGATCTNNNGAGTAAATAGGACTTTTTTAGGT
GCACACGTCTGAACTCTTTCCCTACACGACGCTCTTCCGATCTNNNTTTTCTTTGTCTTTGATGGGTTCA
AGCACACGTCTGAACTCTTTCCCTACACGACGCTCTTCCGATCTNNNACTCACATTAACAAATTCATAAA
GAGCACACGTCTGAACTCTTTCCCTACACGACGCTCTTCCGATCTNNNTTAGAATAATGGTCTCCAACTC
AGCACACGTCTGAACTCTTTCCCTACACGACGCTCTTCCGATCTNNNCTATGTCATTCATCAAAAGTCATG
ACACGTCTGAACTCTTTCCCTACACGACGCTCTTCCGATCTNNNTTTGCCCACCTTCATTTTTATCATCGG
GAGCACACGTCTGAACTCTTTCCCTACACGACGCTCTTCCGATCTNNNTCCCTTCTATAATTTTCTTTCCA
AGCACACGTCTGAACTCTTTCCCTACACGACGCTCTTCCGATCTNNNCCTATTTCATAATTTTTTGTGGG
GCACACGTCTGAACTCTTTCCCTACACGACGCTCTTCCGATCTNNNAAAATATTTACCCAAACGTTTTGG
GCACACGTCTGAACTCTTTCCCTACACGACGCTCTTCCGATCTNNNCTGATGATCAGCTTCTTGTGCATC
GAGCACACGTCTGAACTCTTTCCCTACACGACGCTCTTCCGATCTNNNAGAAAATATAAAAGAGAGGCT
AGCACACGTCTGAACTCTTTCCCTACACGACGCTCTTCCGATCTNNNCAGACACAGTTCTATAATGTAAA
GCACACGTCTGAACTCTTTCCCTACACGACGCTCTTCCGATCTNNNTAAGCTGTTATCAGAGAATGAAG
CACACGTCTGAACTCTTTCCCTACACGACGCTCTTCCGATCTNNNTAAAACAAGTCCTAAAGGAGTCCA
CACACGTCTGAACTCTTTCCCTACACGACGCTCTTCCGATCTNNNTAAAACAAGTCCTAAAGGAGTCCA
AGCACACGTCTGAACTCTTTCCCTACACGACGCTCTTCCGATCTNNNTACATCAATGGAAGTGCTTTAC
CACGTCTGAACTCTTTCCCTACACGACGCTCTTCCGATCTNNNCTCTCTCTCTGTCTCTCTCATCGGACT
AGCACACGTCTGAACTCTTTCCCTACACGACGCTCTTCCGATCTNNNCATCCAGCTAGAAAAACAAACCA
AGCACACGTCTGAACTCTTTCCCTACACGACGCTCTTCCGATCTNNNCAAATACCCATATTCACCCATAC
GAGCACACGTCTGAACTCTTTCCCTACACGACGCTCTTCCGATCTNNNAGAAAACAATACTGAACAAAA
GAGCACACGTCTGAACTCTTTCCCTACACGACGCTCTTCCGATCTNNNAGAATATAAAAAAGCCCAAAG
CACACGTCTGAACTCTTTCCCTACACGACGCTCTTCCGATCTNNNACCTTATGAAATCTCTTAATAGGCC
AGCACACGTCTGAACTCTTTCCCTACACGACGCTCTTCCGATCTNNNCTGACTGAACAGACTTTTAGTC
AGCACACGTCTGAACTCTTTCCCTACACGACGCTCTTCCGATCTNNNCTATTTTGTATTTATGTACCCGT
AGCACACGTCTGAACTCTTTCCCTACACGACGCTCTTCCGATCTNNNTATCAATCAACAAGTGGGTAAA
GCACACGTCTGAACTCTTTCCCTACACGACGCTCTTCCGATCTNNNCAAGTTCAGAGGTGTTGAAGCAT
AGCACACGTCTGAACTCTTTCCCTACACGACGCTCTTCCGATCTNNNACACCAAACACAAAATAGATAA
GAGCACACGTCTGAACTCTTTCCCTACACGACGCTCTTCCGATCTNNNATGAATGGACAAATAAAATGT
GCACACGTCTGAACTCTTTCCCTACACGACGCTCTTCCGATCTNNNGATTCTGTCTCTAGACTGTACTCAT
AGCACACGTCTGAACTCTTTCCCTACACGACGCTCTTCCGATCTNNNTTCAACATTCCTCAAATATTGTTC
AGCACACGTCTGAACTCTTTCCCTACACGACGCTCTTCCGATCTNNNTCAATAAGAATTGATGAACCCAG
ACACGTCTGAACTCTTTCCCTACACGACGCTCTTCCGATCTNNNATGCCATATATACTACACCATTTCATC
AGCACACGTCTGAACTCTTTCCCTACACGACGCTCTTCCGATCTNNNGAATCTATTTTGACTGGTCTTCC
AGCACACGTCTGAACTCTTTCCCTACACGACGCTCTTCCGATCTNNNGACCACTTACATTTAATGTGATT
AGCACACGTCTGAACTCTTTCCCTACACGACGCTCTTCCGATCTNNNAGACTAGAAGCAAAGATAAGT
AGCACACGTCTGAACTCTTTCCCTACACGACGCTCTTCCGATCTNNNAAATGTGACTTCAGAATGTCTA

GAGCACACGTCTGAACTCTTTCCCTACACGACGCTCTTCCGATCTNNNATTTATAGATGGACCTTTTTT
GAGCACACGTCTGAACTCTTTCCCTACACGACGCTCTTCCGATCTNNNATTTATAGATGGACCTTTTTT
CACACGTCTGAACTCTTTCCCTACACGACGCTCTTCCGATCTNNNCGACTTCCTCTATTTATATGCATCA
AGCACACGTCTGAACTCTTTCCCTACACGACGCTCTTCCGATCTNNNATACTTTGGCGAATAAAACAAA
AGCACACGTCTGAACTCTTTCCCTACACGACGCTCTTCCGATCTNNNATTACTTTTTCTAGTGCCAAAAT
GCACACGTCTGAACTCTTTCCCTACACGACGCTCTTCCGATCTNNNATATATTTCGCTTTCTTTCCTTCC
GAGCACACGTCTGAACTCTTTCCCTACACGACGCTCTTCCGATCTNNNTTTATTGAGGAGACAATAGA
AGCACACGTCTGAACTCTTTCCCTACACGACGCTCTTCCGATCTNNNCAATCTATAGTGGGTACTTAAC
GCACACGTCTGAACTCTTTCCCTACACGACGCTCTTCCGATCTNNNTTGAATTCTTCATTTTGTAGTGG
AGCACACGTCTGAACTCTTTCCCTACACGACGCTCTTCCGATCTNNNAACAACAAGAATGTAAACCAA
AGCACACGTCTGAACTCTTTCCCTACACGACGCTCTTCCGATCTNNNAGTTTGAAAAGTTCTCTGTTAT
CACACGTCTGAACTCTTTCCCTACACGACGCTCTTCCGATCTNNNGGCATATGTATTTCATTACATCTCA
GAGCACACGTCTGAACTCTTTCCCTACACGACGCTCTTCCGATCTNNNTACAAGCTCTTTAGGTAAGAA
AGCACACGTCTGAACTCTTTCCCTACACGACGCTCTTCCGATCTNNNTGATTATGGCTAATCTACAGA
GAGCACACGTCTGAACTCTTTCCCTACACGACGCTCTTCCGATCTNNNGACTAAGGACATATTTCCTCT
GAGCACACGTCTGAACTCTTTCCCTACACGACGCTCTTCCGATCTNNNGAAGAGATGGCATTACAGAT
GCACACGTCTGAACTCTTTCCCTACACGACGCTCTTCCGATCTNNNTTCTACATCATTTTATATCAGGGA
GCACACGTCTGAACTCTTTCCCTACACGACGCTCTTCCGATCTNNNGGAGAAAGAAGATTGCTAGAA
GCACACGTCTGAACTCTTTCCCTACACGACGCTCTTCCGATCTNNNGGAGAAAGAAGATTGCTAGAA
GAGCACACGTCTGAACTCTTTCCCTACACGACGCTCTTCCGATCTNNNCATTCAGCAAATTATACTCCTT
CACACGTCTGAACTCTTTCCCTACACGACGCTCTTCCGATCTNNNCGAGGCTCTTTTAGAAACATCATC
CACGTCTGAACTCTTTCCCTACACGACGCTCTTCCGATCTNNNGAAGAATTCTTTGGATTACCAGCATC
ACACGTCTGAACTCTTTCCCTACACGACGCTCTTCCGATCTNNNAGTCTTAGATAAGATTCAGAAGAC
CACACGTCTGAACTCTTTCCCTACACGACGCTCTTCCGATCTNNNGTATATGCAACTTTGGGAAAGCAT
GAGCACACGTCTGAACTCTTTCCCTACACGACGCTCTTCCGATCTNNNCTTCTTTATGTGTCCTCACTTCA
AGCACACGTCTGAACTCTTTCCCTACACGACGCTCTTCCGATCTNNNTAACAAGAAGCCAAAATAAAC
GCACACGTCTGAACTCTTTCCCTACACGACGCTCTTCCGATCTNNNTAAAAGACTATTACTCCAGAAGT
AGCACACGTCTGAACTCTTTCCCTACACGACGCTCTTCCGATCTNNNGGAACTTTGGATGAAAACTTC
AGCACACGTCTGAACTCTTTCCCTACACGACGCTCTTCCGATCTNNNAATACCATCTAAATTGGGAAC
GCACACGTCTGAACTCTTTCCCTACACGACGCTCTTCCGATCTNNNTATATCTTGTGGTTCTGGATTAC
AGCACACGTCTGAACTCTTTCCCTACACGACGCTCTTCCGATCTNNNGACGTACATCAGGGATATTAG
AGCACACGTCTGAACTCTTTCCCTACACGACGCTCTTCCGATCTNNNTCATAGATATGGAACCAATAAG
GAGCACACGTCTGAACTCTTTCCCTACACGACGCTCTTCCGATCTNNNTAAAACAAAAAAGTCCATTGT
CACACGTCTGAACTCTTTCCCTACACGACGCTCTTCCGATCTNNNGACTGATGGTTCTCTAAGTACCATC
CACACGTCTGAACTCTTTCCCTACACGACGCTCTTCCGATCTNNNGTGGAGGTAGCATAGATTTATGC
AGCACACGTCTGAACTCTTTCCCTACACGACGCTCTTCCGATCTNNNTGTTTAATAGGTCCAGAGAAT
AGCACACGTCTGAACTCTTTCCCTACACGACGCTCTTCCGATCTNNNCATTCAAAGCTAATCTGAATAG
GAGCACACGTCTGAACTCTTTCCCTACACGACGCTCTTCCGATCTNNNACTTTAATTTTCCACCCATTTAT
GAGCACACGTCTGAACTCTTTCCCTACACGACGCTCTTCCGATCTNNNTGGATAAAGAAAATGTGAGA
AGCACACGTCTGAACTCTTTCCCTACACGACGCTCTTCCGATCTNNNGTGAAAGAATTTGGTAGGTTG
AGCACACGTCTGAACTCTTTCCCTACACGACGCTCTTCCGATCTNNNTAACATGATTCTTCTTTTGGTA
AGCACACGTCTGAACTCTTTCCCTACACGACGCTCTTCCGATCTNNNAAAGTACAAAAATAAGACACA
AGCACACGTCTGAACTCTTTCCCTACACGACGCTCTTCCGATCTNNNCCAGAATATTGAAAATGTGCTT

GAGCACACGTCTGAACTCTTTCCCTACACGACGCTCTTCCGATCTNNNCAGAGTGCTAGGATTACAGG
GCACACGTCTGAACTCTTTCCCTACACGACGCTCTTCCGATCTNNNCTTAGAATAATGTTCTCCAACTCC
AGCACACGTCTGAACTCTTTCCCTACACGACGCTCTTCCGATCTNNNATACCTGTAGTTTCAGTTCTCTA
GAGCACACGTCTGAACTCTTTCCCTACACGACGCTCTTCCGATCTNNNGACAATCCTTATCATATATACT
AGCACACGTCTGAACTCTTTCCCTACACGACGCTCTTCCGATCTNNNTAAATTACTTTTTAGGGTCGT
AGCACACGTCTGAACTCTTTCCCTACACGACGCTCTTCCGATCTNNNTTGGAAGACTCATGATTAAATC
AGCACACGTCTGAACTCTTTCCCTACACGACGCTCTTCCGATCTNNNTTGGAAGACTCATGATTAAATC
GCACACGTCTGAACTCTTTCCCTACACGACGCTCTTCCGATCTNNNCAATGACTTTGCTCTATTTAGTGC
CACACGTCTGAACTCTTTCCCTACACGACGCTCTTCCGATCTNNNAGAATATGTCTGCTGCAATGCATC
GAGCACACGTCTGAACTCTTTCCCTACACGACGCTCTTCCGATCTNNNTACATAGCAATACATAGCTGAA
GAGCACACGTCTGAACTCTTTCCCTACACGACGCTCTTCCGATCTNNNTTTTCCAGAGAAAAAGAAACAA
GAGCACACGTCTGAACTCTTTCCCTACACGACGCTCTTCCGATCTNNNTTTTCCAGAGAAAAAGAAACAA
GCACACGTCTGAACTCTTTCCCTACACGACGCTCTTCCGATCTNNNTTGTTCTTCCAGGAAATAACTCCA
AGCACACGTCTGAACTCTTTCCCTACACGACGCTCTTCCGATCTNNNCATTAAAACTTCTTCAGCACTTA
AGCACACGTCTGAACTCTTTCCCTACACGACGCTCTTCCGATCTNNNTCAAAATGACCTATTCCACAGCA
AGCACACGTCTGAACTCTTTCCCTACACGACGCTCTTCCGATCTNNNGATTAGCTAGTATTTTGTTGAG
GCACACGTCTGAACTCTTTCCCTACACGACGCTCTTCCGATCTNNNATAAATTTTGCTATGGGTTGCCAT
AGCACACGTCTGAACTCTTTCCCTACACGACGCTCTTCCGATCTNNNGCAAATCAATCACATTATTTCT
AGCACACGTCTGAACTCTTTCCCTACACGACGCTCTTCCGATCTNNNTTGTGTATAGTATGTCATCAGTT
GAGCACACGTCTGAACTCTTTCCCTACACGACGCTCTTCCGATCTNNNTATTAAGGGTTGTTATTCTGTT
GAGCACACGTCTGAACTCTTTCCCTACACGACGCTCTTCCGATCTNNNCTCTTAGGTCAAACAGAAAAT
GAGCACACGTCTGAACTCTTTCCCTACACGACGCTCTTCCGATCTNNNCTCCTAGGGAAAGAGAAAAAT
AGCACACGTCTGAACTCTTTCCCTACACGACGCTCTTCCGATCTNNNAGTGGCTAAAGATGTCTTTTTCA
AGCACACGTCTGAACTCTTTCCCTACACGACGCTCTTCCGATCTNNNTGATCATTGAGTGAGTAGTAAA
GAGCACACGTCTGAACTCTTTCCCTACACGACGCTCTTCCGATCTNNNCTTTAGAGAGTTAGCCTTACTT
GAGCACACGTCTGAACTCTTTCCCTACACGACGCTCTTCCGATCTNNNCTTTAGAGAGTTAGCCTTACTT
CACACGTCTGAACTCTTTCCCTACACGACGCTCTTCCGATCTNNNCTTGGCTCTAGTTTAAGGTTTCAT
GCACACGTCTGAACTCTTTCCCTACACGACGCTCTTCCGATCTNNNAATATTTGCCTTCATCTTCAAATCA
GAGCACACGTCTGAACTCTTTCCCTACACGACGCTCTTCCGATCTNNNAAAGGAGAAGTGACTACTAAT
GAGCACACGTCTGAACTCTTTCCCTACACGACGCTCTTCCGATCTNNNAAGGTTTTGATATATAAAGCCA
GCACACGTCTGAACTCTTTCCCTACACGACGCTCTTCCGATCTNNNCTGTGTGTATGCTAAGAGAGCAT
CACACGTCTGAACTCTTTCCCTACACGACGCTCTTCCGATCTNNNTGTTGAAACTTTCTTCTCATATGCA
GCACACGTCTGAACTCTTTCCCTACACGACGCTCTTCCGATCTNNNGTTTTTCATTCATTCCTAGTGAACA
AGCACACGTCTGAACTCTTTCCCTACACGACGCTCTTCCGATCTNNNATAATGACTCCTGACTTTTTTG
AGCACACGTCTGAACTCTTTCCCTACACGACGCTCTTCCGATCTNNNTTCTTTGACTTATTTTACTGAG
GCACACGTCTGAACTCTTTCCCTACACGACGCTCTTCCGATCTNNNCAAAAAATAACAGGGAAAAGGA
AGCACACGTCTGAACTCTTTCCCTACACGACGCTCTTCCGATCTNNNTATCTGTGCTTGTTGATTTTTT
CACACGTCTGAACTCTTTCCCTACACGACGCTCTTCCGATCTNNNCTTTTTGAAAGGTTTCATGAAGCAT
AGCACACGTCTGAACTCTTTCCCTACACGACGCTCTTCCGATCTNNNTACTAGATTGTTGTAAGCATCAA
CACACGTCTGAACTCTTTCCCTACACGACGCTCTTCCGATCTNNNTACTCATACGAACAATTATGGGCAT
AGCACACGTCTGAACTCTTTCCCTACACGACGCTCTTCCGATCTNNNCTCTGATTTGTCACTCTGAATAA
GCACACGTCTGAACTCTTTCCCTACACGACGCTCTTCCGATCTNNNCTTATTTTCTTCCTCTGCAGACATC
GCACACGTCTGAACTCTTTCCCTACACGACGCTCTTCCGATCTNNNGAAGTGCAGATCAAGTTGATATC

GCACACGTCTGAACTCTTTCCCTACACGACGCTCTTCCGATCTNNNTTGATCCCGAGAATCAATATTTC
GCACACGTCTGAACTCTTTCCCTACACGACGCTCTTCCGATCTNNNTTGATCCCGAGAATCAATATTTC
GCACACGTCTGAACTCTTTCCCTACACGACGCTCTTCCGATCTNNNAAGTAAATTGAAAACTTTCTGGA
GAGCACACGTCTGAACTCTTTCCCTACACGACGCTCTTCCGATCTNNNGTACTGAATTTGTTCCTTGAT
GAGCACACGTCTGAACTCTTTCCCTACACGACGCTCTTCCGATCTNNNAAGAAAAAAAGAAAAAGAAA
AGCACACGTCTGAACTCTTTCCCTACACGACGCTCTTCCGATCTNNNCATTGAGAAGATTAGATTCCCA
AGCACACGTCTGAACTCTTTCCCTACACGACGCTCTTCCGATCTNNNTTGGCTGGATTTTCTGATTATAC
AGCACACGTCTGAACTCTTTCCCTACACGACGCTCTTCCGATCTNNNGCTTTTCTGCTTTACCTTATAGC
AGCACACGTCTGAACTCTTTCCCTACACGACGCTCTTCCGATCTNNNGCTTTTCTGCTTTACCTTATAGC
AGCACACGTCTGAACTCTTTCCCTACACGACGCTCTTCCGATCTNNNGCTTTTCTGCTTTACCTTATAGC
AGCACACGTCTGAACTCTTTCCCTACACGACGCTCTTCCGATCTNNNTGGTGATCTAACCCTTATAATTT
AGAGCACACGTCTGAACTCTTTCCCTACACGACGCTCTTCCGATCTNNNCCTTTTTTTTCTGAAGAATT
CACGTCTGAACTCTTTCCCTACACGACGCTCTTCCGATCTNNNAAGTATTTATGACAATTGCTCTCCATC
GCACACGTCTGAACTCTTTCCCTACACGACGCTCTTCCGATCTNNNAAGAACAGCGAAAGTCCATACAT
AGCACACGTCTGAACTCTTTCCCTACACGACGCTCTTCCGATCTNNNTACTATCTCTCAATCTCTCTCTTC
GAGCACACGTCTGAACTCTTTCCCTACACGACGCTCTTCCGATCTNNNAATATAAGCTTCTTGGCAATAT
AGCACACGTCTGAACTCTTTCCCTACACGACGCTCTTCCGATCTNNNAAAGGTGAAGAACGTATTACAT
AGCACACGTCTGAACTCTTTCCCTACACGACGCTCTTCCGATCTNNNAAAGGTGAAGAACGTATTACAT
AGCACACGTCTGAACTCTTTCCCTACACGACGCTCTTCCGATCTNNNCGTATTCAAGTTATTTTCACAGA
AGCACACGTCTGAACTCTTTCCCTACACGACGCTCTTCCGATCTNNNATACACTGACAATTTTCAAAAGT
GAGCACACGTCTGAACTCTTTCCCTACACGACGCTCTTCCGATCTNNNATTTATATTTCTTCCTCCATGTG
GAGCACACGTCTGAACTCTTTCCCTACACGACGCTCTTCCGATCTNNNTTATTTTATGTCTAGCTTTTTG
AGCACACGTCTGAACTCTTTCCCTACACGACGCTCTTCCGATCTNNNCTCTTTCCATATTTTGTTGCTATC
GAGCACACGTCTGAACTCTTTCCCTACACGACGCTCTTCCGATCTNNNGACTAATTGCTAGGTGATTACT
GCACACGTCTGAACTCTTTCCCTACACGACGCTCTTCCGATCTNNNATACCTCTAAGAGAAGACACAGC
GCACACGTCTGAACTCTTTCCCTACACGACGCTCTTCCGATCTNNNTAAGTGGAATTGATGGGAAATAT
AGCACACGTCTGAACTCTTTCCCTACACGACGCTCTTCCGATCTNNNTAAAATCCTAATCTCCAATGTG
GCACACGTCTGAACTCTTTCCCTACACGACGCTCTTCCGATCTNNNATACATGTTTCATATACACCTCAC
AGCACACGTCTGAACTCTTTCCCTACACGACGCTCTTCCGATCTNNNATAAAAGAAGACAACGCAAAG
GAGCACACGTCTGAACTCTTTCCCTACACGACGCTCTTCCGATCTNNNAAAAGGAGATAAAAGAAGAC
GAGCACACGTCTGAACTCTTTCCCTACACGACGCTCTTCCGATCTNNNGAGACTACCAGACAGATACAC
AGCACACGTCTGAACTCTTTCCCTACACGACGCTCTTCCGATCTNNNTTTGCAGATGACAGATCATGC
AGAGCACACGTCTGAACTCTTTCCCTACACGACGCTCTTCCGATCTNNNCTTGGAATGTGAGTTTTTA
AGCACACGTCTGAACTCTTTCCCTACACGACGCTCTTCCGATCTNNNGACTCATCTCCAACATATTACT
GCACACGTCTGAACTCTTTCCCTACACGACGCTCTTCCGATCTNNNATAAAAAACCCTGTTCAAGATTT
CACACGTCTGAACTCTTTCCCTACACGACGCTCTTCCGATCTNNNCTATAAGTACAGTGGCTCTTAAAC
GAGCACACGTCTGAACTCTTTCCCTACACGACGCTCTTCCGATCTNNNGTAAAATTCCCAAAGAAGATT
AGCACACGTCTGAACTCTTTCCCTACACGACGCTCTTCCGATCTNNNTTCTTTATAGCAGTGTGAAAAC
GCACACGTCTGAACTCTTTCCCTACACGACGCTCTTCCGATCTNNNGATCAGGGCAATAGTAAGAAAC
GCACACGTCTGAACTCTTTCCCTACACGACGCTCTTCCGATCTNNNGATCAGGGCAATAGTAAGAAAC
GCACACGTCTGAACTCTTTCCCTACACGACGCTCTTCCGATCTNNNGATCAGGGCAATAGTAAGAAAC
GAGCACACGTCTGAACTCTTTCCCTACACGACGCTCTTCCGATCTNNNAAAACAGAGAAATGAAAAAC
GAGCACACGTCTGAACTCTTTCCCTACACGACGCTCTTCCGATCTNNNCCAAAAATGTATCATGCAATTT

AGCACACGTCTGAACTCTTTCCCTACACGACGCTCTTCCGATCTNNNTCTCAGTATGGATATGCAAATAT
AGCACACGTCTGAACTCTTTCCCTACACGACGCTCTTCCGATCTNNNTATGTCACCTTGCATCTCATCAT
CACACGTCTGAACTCTTTCCCTACACGACGCTCTTCCGATCTNNNTCCTACAATCTTAGAGTTTTAGGCA
GAGCACACGTCTGAACTCTTTCCCTACACGACGCTCTTCCGATCTNNNGAATCCACCAAGGCAATTTACA
CACGTCTGAACTCTTTCCCTACACGACGCTCTTCCGATCTNNNTATAATTGTTCCTGCCAGAATTACATCC
CACACGTCTGAACTCTTTCCCTACACGACGCTCTTCCGATCTNNNAGCTACTTATTTAAGGTGAGTACA
AGCACACGTCTGAACTCTTTCCCTACACGACGCTCTTCCGATCTNNNATGACTCACATGACACATATGC
CACACGTCTGAACTCTTTCCCTACACGACGCTCTTCCGATCTNNNTCCAGGAGAATTCTCTGAATTACA
AGCACACGTCTGAACTCTTTCCCTACACGACGCTCTTCCGATCTNNNTTTATGATTAGTCTTCCCAATCTC
AGAGCACACGTCTGAACTCTTTCCCTACACGACGCTCTTCCGATCTNNNCTAGACACAGAAGAATACATA
ACACGTCTGAACTCTTTCCCTACACGACGCTCTTCCGATCTNNNCCACCTTCCTAAGATACCTAGCATCGC
CACACGTCTGAACTCTTTCCCTACACGACGCTCTTCCGATCTNNNGTTATCATCCCTTTCTTTCTCCCATCC
CACACGTCTGAACTCTTTCCCTACACGACGCTCTTCCGATCTNNNTACTTATTTCATTCCTCCCTGAACAT
AGCACACGTCTGAACTCTTTCCCTACACGACGCTCTTCCGATCTNNNTAAGGGAAGGTAAGTAGGAAA
CACACGTCTGAACTCTTTCCCTACACGACGCTCTTCCGATCTNNNTATTGTATTTAAAGGCCCAAACAT
GAGCACACGTCTGAACTCTTTCCCTACACGACGCTCTTCCGATCTNNNCCAAGTATAACCCAGATATTAC
AGCACACGTCTGAACTCTTTCCCTACACGACGCTCTTCCGATCTNNNATCTCTATACAATTTAGCAAAGC
AGCACACGTCTGAACTCTTTCCCTACACGACGCTCTTCCGATCTNNNCTTTCTAAATTTTCTACTTGGCTC
CACACGTCTGAACTCTTTCCCTACACGACGCTCTTCCGATCTNNNTCTCTTGCTGTCTCCTTACATCGG
CACACGTCTGAACTCTTTCCCTACACGACGCTCTTCCGATCTNNNCAAAACCTTCAAATCTTCCTATTCAT
AGCACACGTCTGAACTCTTTCCCTACACGACGCTCTTCCGATCTNNNTGATTATATGACAATCCTGAAG
AGCACACGTCTGAACTCTTTCCCTACACGACGCTCTTCCGATCTNNNTGACTTTATAGATACTTTCACCA
GCACACGTCTGAACTCTTTCCCTACACGACGCTCTTCCGATCTNNNCATAAGTTATACTTTTACCTGACA
GAGCACACGTCTGAACTCTTTCCCTACACGACGCTCTTCCGATCTNNNGAATGCTGGTTATATTGGTAT
AGCACACGTCTGAACTCTTTCCCTACACGACGCTCTTCCGATCTNNNTGTGTTTAGATCCCATATAAGAA
AGCACACGTCTGAACTCTTTCCCTACACGACGCTCTTCCGATCTNNNGAATCTCACCCAATAACTAATTC
GAGCACACGTCTGAACTCTTTCCCTACACGACGCTCTTCCGATCTNNNAGCAGAATAGAAGTTTACAAT
CACACGTCTGAACTCTTTCCCTACACGACGCTCTTCCGATCTNNNTGGTGACATATATAACCACTATTCA
CACACGTCTGAACTCTTTCCCTACACGACGCTCTTCCGATCTNNNTGGTGACATATATAACCACTATTCA
AGCACACGTCTGAACTCTTTCCCTACACGACGCTCTTCCGATCTNNNCCATAGAAACAGTAATGGTCTAT
GAGCACACGTCTGAACTCTTTCCCTACACGACGCTCTTCCGATCTNNNAGCATTTTCATTACTTTTGTAC
GCACACGTCTGAACTCTTTCCCTACACGACGCTCTTCCGATCTNNNATTTAGCTTCATGAGTCACTTTAC
GCACACGTCTGAACTCTTTCCCTACACGACGCTCTTCCGATCTNNNAAATAACCGAGAGAGAGGAGCA
AGCACACGTCTGAACTCTTTCCCTACACGACGCTCTTCCGATCTNNNCTCATTTTTGTCCAAAGTACATA
AGCACACGTCTGAACTCTTTCCCTACACGACGCTCTTCCGATCTNNNAAGGGAAAGGGAAGAAGAAT
CACACGTCTGAACTCTTTCCCTACACGACGCTCTTCCGATCTNNNGAAATTCAAAATCAGGTTATTGGC
GCACACGTCTGAACTCTTTCCCTACACGACGCTCTTCCGATCTNNNAGATTATTCAGTAAAGCAGAGAT
CACACGTCTGAACTCTTTCCCTACACGACGCTCTTCCGATCTNNNGAGTCTACCTGAGTTTGTATTCCAT
AGCACACGTCTGAACTCTTTCCCTACACGACGCTCTTCCGATCTNNNGCCCATAGCTTTCTTTTTAAATAC
GAGCACACGTCTGAACTCTTTCCCTACACGACGCTCTTCCGATCTNNNAGGCATATCAAATTTAATATGA
GAGCACACGTCTGAACTCTTTCCCTACACGACGCTCTTCCGATCTNNNAGGCATATCAAATTTAATATGA
GAGCACACGTCTGAACTCTTTCCCTACACGACGCTCTTCCGATCTNNNGTGACTGATATCAACAGGTAA
AGCACACGTCTGAACTCTTTCCCTACACGACGCTCTTCCGATCTNNNTTCTCTTCTCTTATCTTCTGTCTCA

AGCACACGTCTGAACTCTTTCCCTACACGACGCTCTTCCGATCTNNNCTTGATTGACTTTTCTCTATAGCC

CACACGTCTGAACTCTTTCCCTACACGACGCTCTTCCGATCTNNNATACTCACCTGCAGTCATTCCATCGG

AGCACACGTCTGAACTCTTTCCCTACACGACGCTCTTCCGATCTNNNTTATCTCTCTGTCTCTTTTCATCC

AGCACACGTCTGAACTCTTTCCCTACACGACGCTCTTCCGATCTNNNCTAGATACAGACATAGATACAG

GAGCACACGTCTGAACTCTTTCCCTACACGACGCTCTTCCGATCTNNNGTTTATAAACCAATCTTGAGA

GAGCACACGTCTGAACTCTTTCCCTACACGACGCTCTTCCGATCTNNNGTTTATAAACCAATCTTGAGA

GCACACGTCTGAACTCTTTCCCTACACGACGCTCTTCCGATCTNNNCAGAGAAGTTAGGCTTGAAGCA

GAGCACACGTCTGAACTCTTTCCCTACACGACGCTCTTCCGATCTNNNACGTAGGAGTATATTTGTAAC

ACACGTCTGAACTCTTTCCCTACACGACGCTCTTCCGATCTNNNGTCCTCAGATAACATTGTTTTACAT

AGCACACGTCTGAACTCTTTCCCTACACGACGCTCTTCCGATCTNNNCAGGTAGAAGGAAGTGTGAGC

AGCACACGTCTGAACTCTTTCCCTACACGACGCTCTTCCGATCTNNNATTATAGACATGAGCCACCACA

GAGCACACGTCTGAACTCTTTCCCTACACGACGCTCTTCCGATCTNNNGTTCTCTTTTTCTTTGTGTCTAC

GCACACGTCTGAACTCTTTCCCTACACGACGCTCTTCCGATCTNNNAGATGCTAAACCAGTATTTGATA

CACACGTCTGAACTCTTTCCCTACACGACGCTCTTCCGATCTNNNTGTCTACTGAGTGAATATGGACAT

AGCACACGTCTGAACTCTTTCCCTACACGACGCTCTTCCGATCTNNNTTTGTGTTTTTTTGTTACACA

AGCACACGTCTGAACTCTTTCCCTACACGACGCTCTTCCGATCTNNNATAATCTGCTAATCAGTTACTG

AGCACACGTCTGAACTCTTTCCCTACACGACGCTCTTCCGATCTNNNATAATCTGCTAATCAGTTACTG

ACACGTCTGAACTCTTTCCCTACACGACGCTCTTCCGATCTNNNTGTGTATGTGTGTGTATCTACATC

AGCACACGTCTGAACTCTTTCCCTACACGACGCTCTTCCGATCTNNNTAAAGAAACCAAACATCTGAT

CACACGTCTGAACTCTTTCCCTACACGACGCTCTTCCGATCTNNNAGTTTTTCTGGATCATTTTCAATCA

GAGCACACGTCTGAACTCTTTCCCTACACGACGCTCTTCCGATCTNNNTGGAAGATGCTATTTATCAAA

AGCACACGTCTGAACTCTTTCCCTACACGACGCTCTTCCGATCTNNNACCCACTGAAGTTATTTAATTA

GAGCACACGTCTGAACTCTTTCCCTACACGACGCTCTTCCGATCTNNNTTAATAATGGGGAAAGAACTA

CACACGTCTGAACTCTTTCCCTACACGACGCTCTTCCGATCTNNNAGAGGACAGAATGAAAGAATGCAT

GAGCACACGTCTGAACTCTTTCCCTACACGACGCTCTTCCGATCTNNNGAGAAAACCCTGTCTTATTTAA

CACACGTCTGAACTCTTTCCCTACACGACGCTCTTCCGATCTNNNGTTGACATCCTTTCTAAGAATTTCAT

AGCACACGTCTGAACTCTTTCCCTACACGACGCTCTTCCGATCTNNNGTTGGGTCTTTATTTTTGACATA

GAGCACACGTCTGAACTCTTTCCCTACACGACGCTCTTCCGATCTNNNTCTAATTCTCCTATTATTTCCCC

AGCACACGTCTGAACTCTTTCCCTACACGACGCTCTTCCGATCTNNNCTCAAAGGTTAGATAATTTGTG

AGCACACGTCTGAACTCTTTCCCTACACGACGCTCTTCCGATCTNNNAGGATTGATTTATTAGTTATAC

AGCACACGTCTGAACTCTTTCCCTACACGACGCTCTTCCGATCTNNNTATGTGTCAGAAAACATAATAG

GCACACGTCTGAACTCTTTCCCTACACGACGCTCTTCCGATCTNNNTAAAACAGATTGTCCTCTCTAATC

AGCACACGTCTGAACTCTTTCCCTACACGACGCTCTTCCGATCTNNNTCCAACATAGTTTAGAACTAGT

CACGTCTGAACTCTTTCCCTACACGACGCTCTTCCGATCTNNNGAGAGGTTATAGGATCCATAACCATC

GCACACGTCTGAACTCTTTCCCTACACGACGCTCTTCCGATCTNNNCAGACTTAAACACATAAGCATATC

AGCACACGTCTGAACTCTTTCCCTACACGACGCTCTTCCGATCTNNNGTTATCACACTTTCTTCCTTTTGC

GAGCACACGTCTGAACTCTTTCCCTACACGACGCTCTTCCGATCTNNNACTCAGAAATAAACCCACATAT

GCACACGTCTGAACTCTTTCCCTACACGACGCTCTTCCGATCTNNNGTAGGATTATGCCCTAAAATCTCA

GAGCACACGTCTGAACTCTTTCCCTACACGACGCTCTTCCGATCTNNNCATACAAAGGTTGATAACAAT

CACACGTCTGAACTCTTTCCCTACACGACGCTCTTCCGATCTNNNATTTCTTTTGAGAAATGGGTTCCAT

AGCACACGTCTGAACTCTTTCCCTACACGACGCTCTTCCGATCTNNNCAGAAAAGATTTCCTCATTTTTG

ACACGTCTGAACTCTTTCCCTACACGACGCTCTTCCGATCTNNNGATCCTCCTCTAACGAATACTCATCG

CACACGTCTGAACTCTTTCCCTACACGACGCTCTTCCGATCTNNNAATGTAAAATTCAACAGGTCAAACA

AGCACACGTCTGAACTCTTTCCCTACACGACGCTCTTCCGATCTNNNGGTTTTATAAAGCATCTTGTAG
AGCACACGTCTGAACTCTTTCCCTACACGACGCTCTTCCGATCTNNNCCCACTAATCAATGAGTAGATAA
GAGCACACGTCTGAACTCTTTCCCTACACGACGCTCTTCCGATCTNNNAAAACTGTAATTCCAATACTGC
GAGCACACGTCTGAACTCTTTCCCTACACGACGCTCTTCCGATCTNNNAAGTGCTAAGTAAGGTGTCTA
GAGCACACGTCTGAACTCTTTCCCTACACGACGCTCTTCCGATCTNNNCATGTAAAGAAGGAAAGAAG
AGCACACGTCTGAACTCTTTCCCTACACGACGCTCTTCCGATCTNNNTTTTACCATTCAGACTAAAACTG
CACACGTCTGAACTCTTTCCCTACACGACGCTCTTCCGATCTNNNAAAATAGCATGGAACCTAGAATTC
GAGCACACGTCTGAACTCTTTCCCTACACGACGCTCTTCCGATCTNNNCCTTCCTTCCAAGATTTTATAAT
AGCACACGTCTGAACTCTTTCCCTACACGACGCTCTTCCGATCTNNNAATGGTCATTTAGTGCCTTCCAT
AGCACACGTCTGAACTCTTTCCCTACACGACGCTCTTCCGATCTNNNAATGTCTCACAATTTCTATGTTCC
AGCACACGTCTGAACTCTTTCCCTACACGACGCTCTTCCGATCTNNNCTTGTGACATCTCTTTCTTTCTAC
ACACGTCTGAACTCTTTCCCTACACGACGCTCTTCCGATCTNNNACCTTTTATAATAAATTGAAGACCAC
CACACGTCTGAACTCTTTCCCTACACGACGCTCTTCCGATCTNNNCGTTACAGTCTAATATTTGATGACA
AGAGCACACGTCTGAACTCTTTCCCTACACGACGCTCTTCCGATCTNNNGATAATTGCAAAGTCCATAA
CACACGTCTGAACTCTTTCCCTACACGACGCTCTTCCGATCTNNNAGCTGCCATATGTATATACTTATCA
AGCACACGTCTGAACTCTTTCCCTACACGACGCTCTTCCGATCTNNNAGCATGGTTACTATAGTCAATA
GAGCACACGTCTGAACTCTTTCCCTACACGACGCTCTTCCGATCTNNNTTACTTTATGTCTGAAGTTTG
AGCACACGTCTGAACTCTTTCCCTACACGACGCTCTTCCGATCTNNNTATGTGTGCGTATATGTGTATAT
GAGCACACGTCTGAACTCTTTCCCTACACGACGCTCTTCCGATCTNNNTTATAGATCCAGAAGGTAAGG
AGCACACGTCTGAACTCTTTCCCTACACGACGCTCTTCCGATCTNNNAGACAGAGAAGATCAAGAGAG
GAGCACACGTCTGAACTCTTTCCCTACACGACGCTCTTCCGATCTNNNACAGAATTATTACTTATGATT
AGCACACGTCTGAACTCTTTCCCTACACGACGCTCTTCCGATCTNNNGAGCTTGACACCTAGTAAAGCA
GAGCACACGTCTGAACTCTTTCCCTACACGACGCTCTTCCGATCTNNNATATGAAAGGAATGTGTTAGC
GAGCACACGTCTGAACTCTTTCCCTACACGACGCTCTTCCGATCTNNNATATGAAAGGAATGTGTTAGC
AGCACACGTCTGAACTCTTTCCCTACACGACGCTCTTCCGATCTNNNGAAACATTTCTGCTTCCTCTCAT
AGCACACGTCTGAACTCTTTCCCTACACGACGCTCTTCCGATCTNNNTGGGGTTATCAAGGAAATAAAT
AGCACACGTCTGAACTCTTTCCCTACACGACGCTCTTCCGATCTNNNCTTCAGGTAAAAGACATAGAAT
GAGCACACGTCTGAACTCTTTCCCTACACGACGCTCTTCCGATCTNNNTCTTCTTCCTTTTATTTATTGCC
GAGCACACGTCTGAACTCTTTCCCTACACGACGCTCTTCCGATCTNNNCTTTAAGCAATTTTATCTCAGG
GAGCACACGTCTGAACTCTTTCCCTACACGACGCTCTTCCGATCTNNNCATATGATTTCCAAGTTTCTGC
CACACGTCTGAACTCTTTCCCTACACGACGCTCTTCCGATCTNNNATTTTTTTTGGTGTTGCTACTAGCA
GAGCACACGTCTGAACTCTTTCCCTACACGACGCTCTTCCGATCTNNNAGTCAGTACAATCTGATCATTT
AGAGCACACGTCTGAACTCTTTCCCTACACGACGCTCTTCCGATCTNNNTCTAGCTCCTGATTTCAAAATT
GAGCACACGTCTGAACTCTTTCCCTACACGACGCTCTTCCGATCTNNNTGTTTTCACCATTTCAGAATTT
CACACGTCTGAACTCTTTCCCTACACGACGCTCTTCCGATCTNNNCTCTCCTAACTCTGACTTGGCATCGC
AGCACACGTCTGAACTCTTTCCCTACACGACGCTCTTCCGATCTNNNTCTCTCTGGCTATTTTTCATTATC
AGAGCACACGTCTGAACTCTTTCCCTACACGACGCTCTTCCGATCTNNNAGATCTTCATCTTTAGCAAAA
AGCACACGTCTGAACTCTTTCCCTACACGACGCTCTTCCGATCTNNNGGACTACCATGGATAAAAATAA
AGCACACGTCTGAACTCTTTCCCTACACGACGCTCTTCCGATCTNNNGGACTACCATGGATAAAAATAA
GAGCACACGTCTGAACTCTTTCCCTACACGACGCTCTTCCGATCTNNNGCTACAAGGTTAGAGAATATA
GCACACGTCTGAACTCTTTCCCTACACGACGCTCTTCCGATCTNNNTCAGTTTAGTCCTTGAAGTAAGCA
AGCACACGTCTGAACTCTTTCCCTACACGACGCTCTTCCGATCTNNNCTATGAAAGCCTATTCCTTCAACA
GAGCACACGTCTGAACTCTTTCCCTACACGACGCTCTTCCGATCTNNNTATACTTCTGATGTACCAGCTA

GCACACGTCTGAACTCTTTCCCTACACGACGCTCTTCCGATCTNNNGAGCACTTGTATCATATTTTCA
CACACGTCTGAACTCTTTCCCTACACGACGCTCTTCCGATCTNNNTATTATTATTGTGTGTCCGAATG
GAGCACACGTCTGAACTCTTTCCCTACACGACGCTCTTCCGATCTNNNGAAACTTATTTTCCACACCAG
CACGTCTGAACTCTTTCCCTACACGACGCTCTTCCGATCTNNNAGAGTATGATGATTTCCAGCCATCG
CACACGTCTGAACTCTTTCCCTACACGACGCTCTTCCGATCTNNNCTCACTTATAATAATGGTCTCCAC
GCACACGTCTGAACTCTTTCCCTACACGACGCTCTTCCGATCTNNNAAAACACTTTTAGATGTTGCAA
AGCACACGTCTGAACTCTTTCCCTACACGACGCTCTTCCGATCTNNNTTAGCCAAACATATATATGTG
ACACGTCTGAACTCTTTCCCTACACGACGCTCTTCCGATCTNNNAGCTTCTCTCTCTCTCAACTCATCGG
CACACGTCTGAACTCTTTCCCTACACGACGCTCTTCCGATCTNNNTTACTGAAATCTCTGCTTCTTTACA
ACACGTCTGAACTCTTTCCCTACACGACGCTCTTCCGATCTNNNCTAGTTTTTACTTAAGCCAGTGCAT
GCACACGTCTGAACTCTTTCCCTACACGACGCTCTTCCGATCTNNNATTATAGACATGAGCCACCACA
GCACACGTCTGAACTCTTTCCCTACACGACGCTCTTCCGATCTNNNATAAAAGCTCACAGACATAAAC
AGCACACGTCTGAACTCTTTCCCTACACGACGCTCTTCCGATCTNNNAATTAAAACATCTACTAGGCA
ACACGTCTGAACTCTTTCCCTACACGACGCTCTTCCGATCTNNNTGTCTCTGCAATGTATATATACACA
ACACGTCTGAACTCTTTCCCTACACGACGCTCTTCCGATCTNNNTGAGTTATTTCTTCTTCAACATGCAT
GAGCACACGTCTGAACTCTTTCCCTACACGACGCTCTTCCGATCTNNNCTTGTTTCGGAGATATTCAT
AGCACACGTCTGAACTCTTTCCCTACACGACGCTCTTCCGATCTNNNCACCTATAAGCTTGTTTATGTG
ACACGTCTGAACTCTTTCCCTACACGACGCTCTTCCGATCTNNNAGAGACTCTTGTTTTCTTATCTTCA
GCACACGTCTGAACTCTTTCCCTACACGACGCTCTTCCGATCTNNNTTTACAAAAGTTATACACACACG
AGCACACGTCTGAACTCTTTCCCTACACGACGCTCTTCCGATCTNNNGACTATATAAGAGAAACACAG
CACACGTCTGAACTCTTTCCCTACACGACGCTCTTCCGATCTNNNGTTCAAGATTCAGTGTGAAAATCA
AGCACACGTCTGAACTCTTTCCCTACACGACGCTCTTCCGATCTNNNCTTACCAACACATACATTCATG
GCACACGTCTGAACTCTTTCCCTACACGACGCTCTTCCGATCTNNNAATGAGTCCTTAATGTGCTTGCA
AGCACACGTCTGAACTCTTTCCCTACACGACGCTCTTCCGATCTNNNCTTTCTTTCCTGTATGTGTTTA
AGCACACGTCTGAACTCTTTCCCTACACGACGCTCTTCCGATCTNNNCAATGATGTGGTAGATAGAT
AGCACACGTCTGAACTCTTTCCCTACACGACGCTCTTCCGATCTNNNAATATGAGACATTACTTCAGA
AGCACACGTCTGAACTCTTTCCCTACACGACGCTCTTCCGATCTNNNAATATAAATTGGTTGATACTG
GCACACGTCTGAACTCTTTCCCTACACGACGCTCTTCCGATCTNNNCTGGATAAGGAATATAAACACC
AGCACACGTCTGAACTCTTTCCCTACACGACGCTCTTCCGATCTNNNAATATTCAAGCTGAGAACAAA
GAGCACACGTCTGAACTCTTTCCCTACACGACGCTCTTCCGATCTNNNGAAGTTACATCACAGATTTC
AGCACACGTCTGAACTCTTTCCCTACACGACGCTCTTCCGATCTNNNATGTGAATGAGGTTGATTTCA
GAGCACACGTCTGAACTCTTTCCCTACACGACGCTCTTCCGATCTNNNCATTCTGCTTTGGGTATTTT
GCACACGTCTGAACTCTTTCCCTACACGACGCTCTTCCGATCTNNNTATGCTGTCATTATTACACTGAT
AGCACACGTCTGAACTCTTTCCCTACACGACGCTCTTCCGATCTNNNCCAGTGCATTCAGTATTATAA
GCACACGTCTGAACTCTTTCCCTACACGACGCTCTTCCGATCTNNNTAAACAAACTTGCCATTACATTA
AGCACACGTCTGAACTCTTTCCCTACACGACGCTCTTCCGATCTNNNCACACACACTGTTATACCTAG
CACACGTCTGAACTCTTTCCCTACACGACGCTCTTCCGATCTNNNTACTGTTTCTTACCCCTCCTACATC
CACACGTCTGAACTCTTTCCCTACACGACGCTCTTCCGATCTNNNCTCTTGGGTTGAGTACCTAACAT
CACACGTCTGAACTCTTTCCCTACACGACGCTCTTCCGATCTNNNGTAGTATTTTTTTGTGGGTGAGC
ACACGTCTGAACTCTTTCCCTACACGACGCTCTTCCGATCTNNNCAGGTTCAAGTGATTCTCATCATCG
AGCACACGTCTGAACTCTTTCCCTACACGACGCTCTTCCGATCTNNNAGTACCTATGGATTTGGGTAT
AGCACACGTCTGAACTCTTTCCCTACACGACGCTCTTCCGATCTNNNAAATGCTTCTGAAACAATTAG
CACGTCTGAACTCTTTCCCTACACGACGCTCTTCCGATCTNNNCCAGCTTTGTAAGAAAGAATAGCAT

AGCACACGTCTGAACTCTTTCCCTACACGACGCTCTTCCGATCTNNNCTTTTATTCTCAGCCACCTATC
AGCACACGTCTGAACTCTTTCCCTACACGACGCTCTTCCGATCTNNNACAGAAACATTTACCATTTTCCC
AGCACACGTCTGAACTCTTTCCCTACACGACGCTCTTCCGATCTNNNTAGAAGTATATGTGTCCATGA
ACACGTCTGAACTCTTTCCCTACACGACGCTCTTCCGATCTNNNATTGTTTTCCCTGATCAAAATAGCAT
CACACGTCTGAACTCTTTCCCTACACGACGCTCTTCCGATCTNNNCTTTTTACCCAAAAGTAGAGGCA
AGCACACGTCTGAACTCTTTCCCTACACGACGCTCTTCCGATCTNNNCTTTTAATATGCAAAACAAGGA
AGCACACGTCTGAACTCTTTCCCTACACGACGCTCTTCCGATCTNNNAAGATTTCCTTTTTTCTGTCTTT
GAGCACACGTCTGAACTCTTTCCCTACACGACGCTCTTCCGATCTNNNATTTCTGATTCCTGGTGTGTC
AGCACACGTCTGAACTCTTTCCCTACACGACGCTCTTCCGATCTNNNTGTTTTCAAAACTGTTTATCTCT
CACACGTCTGAACTCTTTCCCTACACGACGCTCTTCCGATCTNNNATCATCCTGATTCCAAAATCTGCAT
GCACACGTCTGAACTCTTTCCCTACACGACGCTCTTCCGATCTNNNAAATTGGAATTTCTATGACTGTC
GCACACGTCTGAACTCTTTCCCTACACGACGCTCTTCCGATCTNNNACAAAGAAGTTCTGTTCTCTTCA
GCACACGTCTGAACTCTTTCCCTACACGACGCTCTTCCGATCTNNNGGGTACAAATCTACAACAAAATT
CACACGTCTGAACTCTTTCCCTACACGACGCTCTTCCGATCTNNNAAGCCAGTTCCTTCAAATAAATACA
GAGCACACGTCTGAACTCTTTCCCTACACGACGCTCTTCCGATCTNNNTTACACCAGAAACCTTATTTCT
AGCACACGTCTGAACTCTTTCCCTACACGACGCTCTTCCGATCTNNNCCAGTAATCTAATTTCTGGTTT
ACACGTCTGAACTCTTTCCCTACACGACGCTCTTCCGATCTNNNCAGTGAAAGGTATTGCTGTACATCG
AGCACACGTCTGAACTCTTTCCCTACACGACGCTCTTCCGATCTNNNGATATTCAAAATGTGCATTGTA
AGCACACGTCTGAACTCTTTCCCTACACGACGCTCTTCCGATCTNNNATAGAGAGGAGGAAAATACTA
GAGCACACGTCTGAACTCTTTCCCTACACGACGCTCTTCCGATCTNNNTTCTGCCTCAGTTACTAATATA
ACACGTCTGAACTCTTTCCCTACACGACGCTCTTCCGATCTNNNATAAATTTTTAACCCATGTCACCCATC
GCACACGTCTGAACTCTTTCCCTACACGACGCTCTTCCGATCTNNNTATATTCAGTCAGAGTTTCAGAC
AGCACACGTCTGAACTCTTTCCCTACACGACGCTCTTCCGATCTNNNGTTTGAATCAGTAACTGAGTAA
AGCACACGTCTGAACTCTTTCCCTACACGACGCTCTTCCGATCTNNNGGTACTGTAGCTTTGTAGTATA
AGCACACGTCTGAACTCTTTCCCTACACGACGCTCTTCCGATCTNNNTGCTTAATTCTCAGTTTTTTCATC
AGCACACGTCTGAACTCTTTCCCTACACGACGCTCTTCCGATCTNNNCATAGCAGGAAAAAATCTCTCT
GAGCACACGTCTGAACTCTTTCCCTACACGACGCTCTTCCGATCTNNNTGAAAGTTTTCATTAACTATGC
ACACGTCTGAACTCTTTCCCTACACGACGCTCTTCCGATCTNNNGCAACAGAGTGAGTATACATATACA
GCACACGTCTGAACTCTTTCCCTACACGACGCTCTTCCGATCTNNNCTTCAATGACATGACCACAGCATC
AGCACACGTCTGAACTCTTTCCCTACACGACGCTCTTCCGATCTNNNATGATCCTGACTCTGTGTAGCA
AGCACACGTCTGAACTCTTTCCCTACACGACGCTCTTCCGATCTNNNTTGCACATAACAAAAATGAAAA
GAGCACACGTCTGAACTCTTTCCCTACACGACGCTCTTCCGATCTNNNATAATTCCAAACACATTTTACC
CACACGTCTGAACTCTTTCCCTACACGACGCTCTTCCGATCTNNNAGCACACAGATGTTTATCTTCCATC
AGCACACGTCTGAACTCTTTCCCTACACGACGCTCTTCCGATCTNNNAAAAAAATTAAGTAACATTGCC
AGCACACGTCTGAACTCTTTCCCTACACGACGCTCTTCCGATCTNNNCCACAATTAGGGTACTAGTAAT
AGCACACGTCTGAACTCTTTCCCTACACGACGCTCTTCCGATCTNNNATTTTACCTTTGAGAAAACTGA
AGAGCACACGTCTGAACTCTTTCCCTACACGACGCTCTTCCGATCTNNNATGGATCATGAGTTATCTCT
GCACACGTCTGAACTCTTTCCCTACACGACGCTCTTCCGATCTNNNATCGAGATTTTCTCCTATCTTTC
AGCACACGTCTGAACTCTTTCCCTACACGACGCTCTTCCGATCTNNNGAGGTAGTTGGATAAATGACT
GAGCACACGTCTGAACTCTTTCCCTACACGACGCTCTTCCGATCTNNNAACGAGGGGATAAAGAAACT
AGCACACGTCTGAACTCTTTCCCTACACGACGCTCTTCCGATCTNNNTTGCACATAACAAAAATGAAAA
AGCACACGTCTGAACTCTTTCCCTACACGACGCTCTTCCGATCTNNNCATGCCTGGCTAATTTTTATATA
AGCACACGTCTGAACTCTTTCCCTACACGACGCTCTTCCGATCTNNNTATGAGAGGTATATCTTGGAAT

GCACACGTCTGAACTCTTTCCCTACACGACGCTCTTCCGATCTNNNTGCAGCTTGAGATTATTTAAAT

AGCACACGTCTGAACTCTTTCCCTACACGACGCTCTTCCGATCTNNNGAACTCATATTTGGAAATCAGT

GAGCACACGTCTGAACTCTTTCCCTACACGACGCTCTTCCGATCTNNNCTCTCTTTCCTCTCTTCCTTCAT

ACACGTCTGAACTCTTTCCCTACACGACGCTCTTCCGATCTNNNCCATGGTAGACATAATTAAGAAAC

AGCACACGTCTGAACTCTTTCCCTACACGACGCTCTTCCGATCTNNNAACTATAGATCTACAGATGCA

GCACACGTCTGAACTCTTTCCCTACACGACGCTCTTCCGATCTNNNCATCTTTCCATCCTTACAGGCAT

ACACGTCTGAACTCTTTCCCTACACGACGCTCTTCCGATCTNNNATTATGGTTATGCTGCCAATACAT

AGCACACGTCTGAACTCTTTCCCTACACGACGCTCTTCCGATCTNNNATGAAATATAAAGGAGTGGC

CACACGTCTGAACTCTTTCCCTACACGACGCTCTTCCGATCTNNNGGATAATGATACAATTTTAGGC

CACACGTCTGAACTCTTTCCCTACACGACGCTCTTCCGATCTNNNCATCTACCTATTTCTTCCTCAAACA

GAGCACACGTCTGAACTCTTTCCCTACACGACGCTCTTCCGATCTNNNTCAAAGAAGAGGTTAAATA

AGCACACGTCTGAACTCTTTCCCTACACGACGCTCTTCCGATCTNNNATCAGTCACCATTTTTTATTAG

CACACGTCTGAACTCTTTCCCTACACGACGCTCTTCCGATCTNNNTCTCAAATACACTGTGGTTCCATC

GCACACGTCTGAACTCTTTCCCTACACGACGCTCTTCCGATCTNNNTGAAAAATTTTGGAAAAAACCA

GCACACGTCTGAACTCTTTCCCTACACGACGCTCTTCCGATCTNNNTCAGCTTTTATTTTCTTATCCACCA

GCACACGTCTGAACTCTTTCCCTACACGACGCTCTTCCGATCTNNNCATTAGTGTTCTCTTTCTGTTCCA

GCACACGTCTGAACTCTTTCCCTACACGACGCTCTTCCGATCTNNNTTTAATTAATTAGGCATTGTGG

AGCACACGTCTGAACTCTTTCCCTACACGACGCTCTTCCGATCTNNNGTCTAACTCTCTGGACTAATTT

GCACACGTCTGAACTCTTTCCCTACACGACGCTCTTCCGATCTNNNCTCTGACCTTTTATAAGACATTTC

GCACACGTCTGAACTCTTTCCCTACACGACGCTCTTCCGATCTNNNAAACTCCTGACCTCAAGTTATCA

GCACACGTCTGAACTCTTTCCCTACACGACGCTCTTCCGATCTNNNTTTTTCTTGACAGCTTAGTTTTT

ACACGTCTGAACTCTTTCCCTACACGACGCTCTTCCGATCTNNNCTAGTAACTATGTCCAGGTATACCA

AGCACACGTCTGAACTCTTTCCCTACACGACGCTCTTCCGATCTNNNATAAATGAGACAGAAATACCC

GCACACGTCTGAACTCTTTCCCTACACGACGCTCTTCCGATCTNNNTATAGTGAGCTATGATTGCACC

GCACACGTCTGAACTCTTTCCCTACACGACGCTCTTCCGATCTNNNCTGAGGCAAAAATATCAATTTCT

GCACACGTCTGAACTCTTTCCCTACACGACGCTCTTCCGATCTNNNGTGTTCTCATGCTTATTCTGTCA

AGCACACGTCTGAACTCTTTCCCTACACGACGCTCTTCCGATCTNNNTCCAATGTTTACTACATCTTACA

AGCACACGTCTGAACTCTTTCCCTACACGACGCTCTTCCGATCTNNNTCCTCTCATACATCTATATCCAT

CACACGTCTGAACTCTTTCCCTACACGACGCTCTTCCGATCTNNNTTTGTGACTATCAGAAACACCCAT

CACACGTCTGAACTCTTTCCCTACACGACGCTCTTCCGATCTNNNCATTTGCACCAACCTAATATAATC

AGCACACGTCTGAACTCTTTCCCTACACGACGCTCTTCCGATCTNNNATCTTCTTCATCAAACTTACTTG

GAGCACACGTCTGAACTCTTTCCCTACACGACGCTCTTCCGATCTNNNGGTGGTTTCAATTTTTCAAT

GCACACGTCTGAACTCTTTCCCTACACGACGCTCTTCCGATCTNNNAATCTCTTAAGGTAACGAAAAT

AGCACACGTCTGAACTCTTTCCCTACACGACGCTCTTCCGATCTNNNGCATTTGATGATATGATGAAG

AGCACACGTCTGAACTCTTTCCCTACACGACGCTCTTCCGATCTNNNATTCTACAAGGCTGGTATTATC

CACACGTCTGAACTCTTTCCCTACACGACGCTCTTCCGATCTNNNGTTTTTACATCTTTCTGGGTTAACA

GCACACGTCTGAACTCTTTCCCTACACGACGCTCTTCCGATCTNNNATATTGGGATTACCTCAAACATA

GCACACGTCTGAACTCTTTCCCTACACGACGCTCTTCCGATCTNNNATATTGGGATTACCTCAAACATA

GAGCACACGTCTGAACTCTTTCCCTACACGACGCTCTTCCGATCTNNNCTACATTTCACAATGGAAAAC

AGCACACGTCTGAACTCTTTCCCTACACGACGCTCTTCCGATCTNNNGCTGTTTTCTAACCTTGAAACC

CACACGTCTGAACTCTTTCCCTACACGACGCTCTTCCGATCTNNNTTAAATATCAAGTTTGTTTGCCCC

ACACGTCTGAACTCTTTCCCTACACGACGCTCTTCCGATCTNNNAGAGAGTAAAAGATGCTATTAGG

AGCACACGTCTGAACTCTTTCCCTACACGACGCTCTTCCGATCTNNNGAACTCATAATCATTTCTTCGTT

AGCACACGTCTGAACTCTTTCCCTACACGACGCTCTTCCGATCTNNNATTCCTCTGTATGGACAAATTT
AGCACACGTCTGAACTCTTTCCCTACACGACGCTCTTCCGATCTNNNATATTGTGAAAAAATGATGA
AGCACACGTCTGAACTCTTTCCCTACACGACGCTCTTCCGATCTNNNAGTGGATTAAAAAACTGTGA
AGCACACGTCTGAACTCTTTCCCTACACGACGCTCTTCCGATCTNNNCTGGAAATATTAGATCGCCTCC
AGCACACGTCTGAACTCTTTCCCTACACGACGCTCTTCCGATCTNNNTAATGGAGGAATCATATTGTT
ACACGTCTGAACTCTTTCCCTACACGACGCTCTTCCGATCTNNNGTGATACTCCGTCTGAAAAACATC
GAGCACACGTCTGAACTCTTTCCCTACACGACGCTCTTCCGATCTNNNAGTACATATTCAACAAAAATG
CACACGTCTGAACTCTTTCCCTACACGACGCTCTTCCGATCTNNNCATGTTTGAAAATCACTACCTTAC
CACACGTCTGAACTCTTTCCCTACACGACGCTCTTCCGATCTNNNATCCTTCTAGAGATATTACAAGTC
AGCACACGTCTGAACTCTTTCCCTACACGACGCTCTTCCGATCTNNNCTTTAACTTTTTGGGAACAGA
GCACACGTCTGAACTCTTTCCCTACACGACGCTCTTCCGATCTNNNGGATTGGTACTGTACAGTAGA
AGCACACGTCTGAACTCTTTCCCTACACGACGCTCTTCCGATCTNNNAGGAGACATAAGAATAAATC
GCACACGTCTGAACTCTTTCCCTACACGACGCTCTTCCGATCTNNNTGTCACATATATATGTCACAGTT
CACACGTCTGAACTCTTTCCCTACACGACGCTCTTCCGATCTNNNACCAACAGTTATTGTCTTTTAAAC
GCACACGTCTGAACTCTTTCCCTACACGACGCTCTTCCGATCTNNNGTTTAAGAGAAAAAAGAGGTG
CACACGTCTGAACTCTTTCCCTACACGACGCTCTTCCGATCTNNNCGGCCTACATCTCTTAAAATACAT
AGCACACGTCTGAACTCTTTCCCTACACGACGCTCTTCCGATCTNNNTGACTATCTGAAAAAGATTAG
AGCACACGTCTGAACTCTTTCCCTACACGACGCTCTTCCGATCTNNNAAACAACAACAACAAAAGACA
CACACGTCTGAACTCTTTCCCTACACGACGCTCTTCCGATCTNNNCGTCTTTTAGAGTGATCCATTACA
AGCACACGTCTGAACTCTTTCCCTACACGACGCTCTTCCGATCTNNNATAGCAAGACTCTGACCAAAA
ACACGTCTGAACTCTTTCCCTACACGACGCTCTTCCGATCTNNNAAAAAGGTCAAGGCTGAAATCATC
GCACACGTCTGAACTCTTTCCCTACACGACGCTCTTCCGATCTNNNATATTCTCCCAGCAAAAGAAACA
ACACGTCTGAACTCTTTCCCTACACGACGCTCTTCCGATCTNNNATTTTCATATGCTTTTCAGAAGTCA
AGCACACGTCTGAACTCTTTCCCTACACGACGCTCTTCCGATCTNNNCTCTTCTCTCTGAAAGCATGCAT
GAGCACACGTCTGAACTCTTTCCCTACACGACGCTCTTCCGATCTNNNAAAAATACAAAAGGCATGG
GAGCACACGTCTGAACTCTTTCCCTACACGACGCTCTTCCGATCTNNNCTGTAAATGCCAGTATTGTT
AGCACACGTCTGAACTCTTTCCCTACACGACGCTCTTCCGATCTNNNAATTCACAATTGCAAAAATGT
AGCACACGTCTGAACTCTTTCCCTACACGACGCTCTTCCGATCTNNNGCTCAATATCACTATGGAAATT
CACACGTCTGAACTCTTTCCCTACACGACGCTCTTCCGATCTNNNCGGAATAGTAAGACATAACTGAA
AGCACACGTCTGAACTCTTTCCCTACACGACGCTCTTCCGATCTNNNTTTATTGCAGCATAATCCATA
CACGTCTGAACTCTTTCCCTACACGACGCTCTTCCGATCTNNNTGTCTGAGGTTTTCCCTATTCATCGG
GCACACGTCTGAACTCTTTCCCTACACGACGCTCTTCCGATCTNNNACCAAAGTTATTTTCTTAAAGC
AGCACACGTCTGAACTCTTTCCCTACACGACGCTCTTCCGATCTNNNCTACAGTGAGGTATCATTGCC
GCACACGTCTGAACTCTTTCCCTACACGACGCTCTTCCGATCTNNNAGGAGACTCATTTGAACCTGCA
GCACACGTCTGAACTCTTTCCCTACACGACGCTCTTCCGATCTNNNACCAAATTTGTTTATGAGTGTTA
CACACGTCTGAACTCTTTCCCTACACGACGCTCTTCCGATCTNNNGACCCCATCTCAAAAAATAAATAC
AGCACACGTCTGAACTCTTTCCCTACACGACGCTCTTCCGATCTNNNAAATGCATAATGTATCTCTGTT
GCACACGTCTGAACTCTTTCCCTACACGACGCTCTTCCGATCTNNNCAATCCTAAAATTGTGTGTTAGA
CACACGTCTGAACTCTTTCCCTACACGACGCTCTTCCGATCTNNNCCACTGTACCCTTAAAATAAAAAC
GCACACGTCTGAACTCTTTCCCTACACGACGCTCTTCCGATCTNNNGTGTGTTTTTGTGTATATGTGT
AGCACACGTCTGAACTCTTTCCCTACACGACGCTCTTCCGATCTNNNAATGATTTAAACTTCCATTTTA
GCACACGTCTGAACTCTTTCCCTACACGACGCTCTTCCGATCTNNNTCTCTTTTGTACATTGATTTTGTC
CACACGTCTGAACTCTTTCCCTACACGACGCTCTTCCGATCTNNNGTGTCTCAAAAAACAAAACAAAA

AGCACACGTCTGAACTCTTTCCCTACACGACGCTCTTCCGATCTNNNACCAAGTATACTTTTACTGTCTT
AGCACACGTCTGAACTCTTTCCCTACACGACGCTCTTCCGATCTNNNTTCATTATTTTTGTGGCAGAAT
AGCACACGTCTGAACTCTTTCCCTACACGACGCTCTTCCGATCTNNNAATAACAACAACAAAAACACCTC
AGCACACGTCTGAACTCTTTCCCTACACGACGCTCTTCCGATCTNNNGCTTTGAACTTATGAATTCCTAT
AGCACACGTCTGAACTCTTTCCCTACACGACGCTCTTCCGATCTNNNACTACCATGTAGTTTATCAATCAC
GCACACGTCTGAACTCTTTCCCTACACGACGCTCTTCCGATCTNNNAACTTTAACATCGTCTTTGATTCC
AGCACACGTCTGAACTCTTTCCCTACACGACGCTCTTCCGATCTNNNCAACTACACAATCCCCTATATGC
CACACGTCTGAACTCTTTCCCTACACGACGCTCTTCCGATCTNNNTTTTTATGAGCAGATTTAGATGGCA
AGAGCACACGTCTGAACTCTTTCCCTACACGACGCTCTTCCGATCTNNNGGATATATATGATAATGCCT
AGCACACGTCTGAACTCTTTCCCTACACGACGCTCTTCCGATCTNNNCTGTATGAGTGCAAGAGAAGC
AGCACACGTCTGAACTCTTTCCCTACACGACGCTCTTCCGATCTNNNCTAAGGGTGAGAACTGAGATC
ACACGTCTGAACTCTTTCCCTACACGACGCTCTTCCGATCTNNNGAAGTGGCTGGGTTCTATATCATCG
GCACACGTCTGAACTCTTTCCCTACACGACGCTCTTCCGATCTNNNTTAGAACACATGCCTGTAGACATC
AGCACACGTCTGAACTCTTTCCCTACACGACGCTCTTCCGATCTNNNTATCTGCCTTAAGTAACAAGAA
AGCACACGTCTGAACTCTTTCCCTACACGACGCTCTTCCGATCTNNNAATCACCAGTCACTATTATCTTTC
GCACACGTCTGAACTCTTTCCCTACACGACGCTCTTCCGATCTNNNCTGCTTAATTGCCTTCATTATATCA
AGCACACGTCTGAACTCTTTCCCTACACGACGCTCTTCCGATCTNNNCTTTAGGACTTTGAATCTGCTA
GAGCACACGTCTGAACTCTTTCCCTACACGACGCTCTTCCGATCTNNNTGCTCCTAGATAATTAACATTCA
GCACACGTCTGAACTCTTTCCCTACACGACGCTCTTCCGATCTNNNTGTTGTTGTTATTATTAACCACCC
AGCACACGTCTGAACTCTTTCCCTACACGACGCTCTTCCGATCTNNNAATGAAAAGATGCAGACAAAC
AGCACACGTCTGAACTCTTTCCCTACACGACGCTCTTCCGATCTNNNGGCTTATTTACCTATGTTTTTCTC
AGAGCACACGTCTGAACTCTTTCCCTACACGACGCTCTTCCGATCTNNNTTGTTTAGAATAAAACTACCT
AGCACACGTCTGAACTCTTTCCCTACACGACGCTCTTCCGATCTNNNAATGCTCAGTATTAGTGCAGCA
AGCACACGTCTGAACTCTTTCCCTACACGACGCTCTTCCGATCTNNNTTTTCATCTACCAAACACACATC
CACACGTCTGAACTCTTTCCCTACACGACGCTCTTCCGATCTNNNGAGACAGAGAGAGAGAGATTATA
AGCACACGTCTGAACTCTTTCCCTACACGACGCTCTTCCGATCTNNNGGTAGAAAGTTTTGATGGAAT
GCACACGTCTGAACTCTTTCCCTACACGACGCTCTTCCGATCTNNNTACTAAGTGCTGGGAATTTATCCA
GAGCACACGTCTGAACTCTTTCCCTACACGACGCTCTTCCGATCTNNNCTCTAATCCAACATGACTAGTC
ACACGTCTGAACTCTTTCCCTACACGACGCTCTTCCGATCTNNNAAAAGAGTCAGTCTTAACACTACCAT
CACACGTCTGAACTCTTTCCCTACACGACGCTCTTCCGATCTNNNGGGTGGTGAAAATACTCTTATCAT
AGCACACGTCTGAACTCTTTCCCTACACGACGCTCTTCCGATCTNNNCTATCCTGATTCCATAACCTTTAC
GAGCACACGTCTGAACTCTTTCCCTACACGACGCTCTTCCGATCTNNNTTCATTGCTTATATTCCTCTCTC
AGCACACGTCTGAACTCTTTCCCTACACGACGCTCTTCCGATCTNNNGACTCATACATTGGAAATTGAT
GCACACGTCTGAACTCTTTCCCTACACGACGCTCTTCCGATCTNNNCCTTTGTGTCAAATTTTCTTTTCCA
AGCACACGTCTGAACTCTTTCCCTACACGACGCTCTTCCGATCTNNNTTGGCTCAACAAATAGAGAATA
GAGCACACGTCTGAACTCTTTCCCTACACGACGCTCTTCCGATCTNNNTAAGAGATGTAAACTCATCAA
GAGCACACGTCTGAACTCTTTCCCTACACGACGCTCTTCCGATCTNNNCAGCAGCATGTTTATTTTTAA
GAGCACACGTCTGAACTCTTTCCCTACACGACGCTCTTCCGATCTNNNAATCAACGAGTGGATAAAGA
GAGCACACGTCTGAACTCTTTCCCTACACGACGCTCTTCCGATCTNNNATCTCTTATCTAGCTGAATGCC
AGCACACGTCTGAACTCTTTCCCTACACGACGCTCTTCCGATCTNNNAATAGACACTTCTCCAAAGAATA
AGCACACGTCTGAACTCTTTCCCTACACGACGCTCTTCCGATCTNNNAATGTTTGGAATTGTTAGGAAA
AGCACACGTCTGAACTCTTTCCCTACACGACGCTCTTCCGATCTNNNTGCAATTAGAGAATATATCATG
GCACACGTCTGAACTCTTTCCCTACACGACGCTCTTCCGATCTNNNGTGAATTCTACTGCTTTTCTAATC

GAGCACACGTCTGAACTCTTTCCCTACACGACGCTCTTCCGATCTNNNAGGCTGTAATGAGCTATGATT
AGCACACGTCTGAACTCTTTCCCTACACGACGCTCTTCCGATCTNNNGGAATAAAAATCATCAGAGTGA
AGCACACGTCTGAACTCTTTCCCTACACGACGCTCTTCCGATCTNNNACTCACTACTTCCCATCTTTCATC
AGCACACGTCTGAACTCTTTCCCTACACGACGCTCTTCCGATCTNNNAGAATCCTGTGTGTTCTTTAAT
GAGCACACGTCTGAACTCTTTCCCTACACGACGCTCTTCCGATCTNNNTAATAGGATTCTGCTTTACAAC
AGCACACGTCTGAACTCTTTCCCTACACGACGCTCTTCCGATCTNNNACTTTGAGAATTTCTTTATGCAT
GAGCACACGTCTGAACTCTTTCCCTACACGACGCTCTTCCGATCTNNNCTATGATTGTGCTTGTAAACAA
GCACACGTCTGAACTCTTTCCCTACACGACGCTCTTCCGATCTNNNTAGGAATCAGCTAGGTAAGAGCA
AGAGCACACGTCTGAACTCTTTCCCTACACGACGCTCTTCCGATCTNNNTCCATGATTATATATATATGT
AGCACACGTCTGAACTCTTTCCCTACACGACGCTCTTCCGATCTNNNATATGTTTATTCAACTGAAAAC
GAGCACACGTCTGAACTCTTTCCCTACACGACGCTCTTCCGATCTNNNCTACCTTATTGTTGCTGCTTCAT
ACACGTCTGAACTCTTTCCCTACACGACGCTCTTCCGATCTNNNTGGTTCTAGGGAGAAAGATATCAT
AGCACACGTCTGAACTCTTTCCCTACACGACGCTCTTCCGATCTNNNTTCTACTTAACTGGTGTCCTATCA
GCACACGTCTGAACTCTTTCCCTACACGACGCTCTTCCGATCTNNNCTAGTTTTGTATAATGGTAGGGA
AGCACACGTCTGAACTCTTTCCCTACACGACGCTCTTCCGATCTNNNCAAACTCTTGGTTGAATACTTCCA
AGCACACGTCTGAACTCTTTCCCTACACGACGCTCTTCCGATCTNNNGTCTCCACATCATAAATGCTCAT
GAGCACACGTCTGAACTCTTTCCCTACACGACGCTCTTCCGATCTNNNTAAAATTCTAAGTCCTCTGACA
ACACGTCTGAACTCTTTCCCTACACGACGCTCTTCCGATCTNNNGAGAGTATCAGAAAAACTTTTTCTCA
GAGCACACGTCTGAACTCTTTCCCTACACGACGCTCTTCCGATCTNNNTCTGACAACAAAAGAAACTTAA
AGCACACGTCTGAACTCTTTCCCTACACGACGCTCTTCCGATCTNNNTGAACATGCATTTTTTTTAACAA
AGCACACGTCTGAACTCTTTCCCTACACGACGCTCTTCCGATCTNNNCCTTGAAAAAATTCTACTTCAGA
GCACACGTCTGAACTCTTTCCCTACACGACGCTCTTCCGATCTNNNGTAGCAAGGATATAACCTTAGAA
AGCACACGTCTGAACTCTTTCCCTACACGACGCTCTTCCGATCTNNNTTATATCTCTGAGTCTTTTCTCG
AGCACACGTCTGAACTCTTTCCCTACACGACGCTCTTCCGATCTNNNTCTCAGTGAGAAAAGAAAAATG
AGCACACGTCTGAACTCTTTCCCTACACGACGCTCTTCCGATCTNNNGAATTAACCTGCAAGAAACAAA
CACGTCTGAACTCTTTCCCTACACGACGCTCTTCCGATCTNNNTAGACTCTGTCTCAAAACAAGCATCG
GAGCACACGTCTGAACTCTTTCCCTACACGACGCTCTTCCGATCTNNNAAACCAAAAATCATACAATGC
GAGCACACGTCTGAACTCTTTCCCTACACGACGCTCTTCCGATCTNNNATTTAGTAGGTCAAGATCAAT
AGCACACGTCTGAACTCTTTCCCTACACGACGCTCTTCCGATCTNNNAGACCTTGTCTCAAAAAATTAAA
AGCACACGTCTGAACTCTTTCCCTACACGACGCTCTTCCGATCTNNNAGAGAATTATGCATTAAGATGA
AGCACACGTCTGAACTCTTTCCCTACACGACGCTCTTCCGATCTNNNCTTGAGCTAAACAGAATTCTAA
AGCACACGTCTGAACTCTTTCCCTACACGACGCTCTTCCGATCTNNNTGGATACCCCATTTACCTTGCAT
GAGCACACGTCTGAACTCTTTCCCTACACGACGCTCTTCCGATCTNNNCCAAAACAATGAGACAGATTA
CACACGTCTGAACTCTTTCCCTACACGACGCTCTTCCGATCTNNNCTTAGACTAGTACTATTCAGTCTTCA
GAGCACACGTCTGAACTCTTTCCCTACACGACGCTCTTCCGATCTNNNCAGGAGAATTGCTTAAACCCCA
ACACGTCTGAACTCTTTCCCTACACGACGCTCTTCCGATCTNNNGGAGATCCTGTTAGTCACTACCATCG
AGCACACGTCTGAACTCTTTCCCTACACGACGCTCTTCCGATCTNNNAAGACCCCATCTCTACAAAACAT
GCACACGTCTGAACTCTTTCCCTACACGACGCTCTTCCGATCTNNNAGTTTATTTTTGTTTTTTACTGCT
GCACACGTCTGAACTCTTTCCCTACACGACGCTCTTCCGATCTNNNGTTTGACTTGTAAGCCATTATTA
GAGCACACGTCTGAACTCTTTCCCTACACGACGCTCTTCCGATCTNNNATGTATCCCAGAACTTAAAGTA
GCACACGTCTGAACTCTTTCCCTACACGACGCTCTTCCGATCTNNNGTAAAATCAGGATAATACTCCCT
AGCACACGTCTGAACTCTTTCCCTACACGACGCTCTTCCGATCTNNNTTAAAATGCATTTGTTGTAGAT
CACACGTCTGAACTCTTTCCCTACACGACGCTCTTCCGATCTNNNCTCTACGTTGATCCTCATTTCCATCG

GCACACGTCTGAACTCTTTCCCTACACGACGCTCTTCCGATCTNNNCTCTGAAGATCCTGGTACTTCAT
AGCACACGTCTGAACTCTTTCCCTACACGACGCTCTTCCGATCTNNNTTCATATTTCTGGGCTATTAG
CACGTCTGAACTCTTTCCCTACACGACGCTCTTCCGATCTNNNGGAGATCCTGTTAGTCACTACCATCG
AGCACACGTCTGAACTCTTTCCCTACACGACGCTCTTCCGATCTNNNAATTAAATTGCCATATTGCCAT
GCACACGTCTGAACTCTTTCCCTACACGACGCTCTTCCGATCTNNNATTCTATGCAAATTCTTCAAGAA
AGCACACGTCTGAACTCTTTCCCTACACGACGCTCTTCCGATCTNNNAAAAGAAGTACCAAAGTAATG
AGCACACGTCTGAACTCTTTCCCTACACGACGCTCTTCCGATCTNNNGATAGAGCAAGCATGATATTA
AGCACACGTCTGAACTCTTTCCCTACACGACGCTCTTCCGATCTNNNTCACAAAGAAAAAATATGTAG
AGCACACGTCTGAACTCTTTCCCTACACGACGCTCTTCCGATCTNNNATCTTTGAAAATTGTGAAGAC
AGCACACGTCTGAACTCTTTCCCTACACGACGCTCTTCCGATCTNNNTATTGTTTCTTTCTTTCTCTCAC
AGCACACGTCTGAACTCTTTCCCTACACGACGCTCTTCCGATCTNNNCCAGGTACATAAAATCTAGAA
ACACGTCTGAACTCTTTCCCTACACGACGCTCTTCCGATCTNNNGCAACAATGACAAACATTTTAAACA
AGCACACGTCTGAACTCTTTCCCTACACGACGCTCTTCCGATCTNNNATCTATTACAGAAGCAATAGG
AGCACACGTCTGAACTCTTTCCCTACACGACGCTCTTCCGATCTNNNATTAATAAGAGGCAGAAACT
GCACACGTCTGAACTCTTTCCCTACACGACGCTCTTCCGATCTNNNGAATGAATCAGTGTATTAGTGA
AGCACACGTCTGAACTCTTTCCCTACACGACGCTCTTCCGATCTNNNTATCACCTCCTCTGAACTTTAA
CACACGTCTGAACTCTTTCCCTACACGACGCTCTTCCGATCTNNNTTTTCCCAAATATTCCACTGACAT
CACACGTCTGAACTCTTTCCCTACACGACGCTCTTCCGATCTNNNGCTGCTTATACTTTCTCTATTCTC
GCACACGTCTGAACTCTTTCCCTACACGACGCTCTTCCGATCTNNNTAAATTGCTCTGAGTTCTGAGC
GCACACGTCTGAACTCTTTCCCTACACGACGCTCTTCCGATCTNNNTATGGGAAAATGTATACCCTTT
GCACACGTCTGAACTCTTTCCCTACACGACGCTCTTCCGATCTNNNAGGGCATATATATTTTTGTTGA
GCACACGTCTGAACTCTTTCCCTACACGACGCTCTTCCGATCTNNNATTACAGACTTGAGCCACTTCAT
GCACACGTCTGAACTCTTTCCCTACACGACGCTCTTCCGATCTNNNCAATAGAAGCACTATAATTTGG
GAGCACACGTCTGAACTCTTTCCCTACACGACGCTCTTCCGATCTNNNTACATGTGTATGTTTATTACA
GCACACGTCTGAACTCTTTCCCTACACGACGCTCTTCCGATCTNNNAATCTGGAGTTGTTTCAGAAGC
ACACGTCTGAACTCTTTCCCTACACGACGCTCTTCCGATCTNNNTTTATGTTAATGTTTTCAGAGCACA
CACACGTCTGAACTCTTTCCCTACACGACGCTCTTCCGATCTNNNCATATCAGGCACGTAAGAACCAT
AGCACACGTCTGAACTCTTTCCCTACACGACGCTCTTCCGATCTNNNAAGAAACAATGGAAACGTTTT
AGCACACGTCTGAACTCTTTCCCTACACGACGCTCTTCCGATCTNNNAATGAGTGGATAAAGAAAAA
AGCACACGTCTGAACTCTTTCCCTACACGACGCTCTTCCGATCTNNNGACACTAGATTGAATTGACCT
AGCACACGTCTGAACTCTTTCCCTACACGACGCTCTTCCGATCTNNNCAATTCACAATTGCAAAAATA
AGCACACGTCTGAACTCTTTCCCTACACGACGCTCTTCCGATCTNNNGTCTGAGAAGAATTTAAAGTG
AGCACACGTCTGAACTCTTTCCCTACACGACGCTCTTCCGATCTNNNTTCTTAAATGTTTTAGGAAGAC
GCACACGTCTGAACTCTTTCCCTACACGACGCTCTTCCGATCTNNNTTGTAAACTTTGTTTTGAGAGA
ACACGTCTGAACTCTTTCCCTACACGACGCTCTTCCGATCTNNNAGGAACAGTCACTTAGAATTTTCAT
AGCACACGTCTGAACTCTTTCCCTACACGACGCTCTTCCGATCTNNNTTTGATAAGAATTAAAGCAGA
AGCACACGTCTGAACTCTTTCCCTACACGACGCTCTTCCGATCTNNNCCAGTGTTCAGCTTATAAATT
GCACACGTCTGAACTCTTTCCCTACACGACGCTCTTCCGATCTNNNCAAAGAAAAGAAAACAAAGAGT
AGCACACGTCTGAACTCTTTCCCTACACGACGCTCTTCCGATCTNNNCTAACTAGGGTCATCATTTCCC
GCACACGTCTGAACTCTTTCCCTACACGACGCTCTTCCGATCTNNNGTATTAGTCAGAGTTCTCCAAAC
AGCACACGTCTGAACTCTTTCCCTACACGACGCTCTTCCGATCTNNNCCTGTAAATCTTGCAACTTCATC
GAGCACACGTCTGAACTCTTTCCCTACACGACGCTCTTCCGATCTNNNAAAACAGAATCAATATGATG
ACACGTCTGAACTCTTTCCCTACACGACGCTCTTCCGATCTNNNTATGTTGGAGCTCTTAAGTTTTTCA

CACACGTCTGAACTCTTTCCCTACACGACGCTCTTCCGATCTNNNGTTTCAATAGGTGTTAATCAAGTC
CACACGTCTGAACTCTTTCCCTACACGACGCTCTTCCGATCTNNNCAAGACCCTACCTCTAAAAACATCGC
GCACACGTCTGAACTCTTTCCCTACACGACGCTCTTCCGATCTNNNAGAATCCAGTTTTTATTTCCAACA
AGCACACGTCTGAACTCTTTCCCTACACGACGCTCTTCCGATCTNNNAGCTTTTATGGTTTGTTTTTCC
AGCACACGTCTGAACTCTTTCCCTACACGACGCTCTTCCGATCTNNNAGGGTTTAAAAATAGCATTTTC
GAGCACACGTCTGAACTCTTTCCCTACACGACGCTCTTCCGATCTNNNGTGGTACTCAAAAGAATAAAC
AGCACACGTCTGAACTCTTTCCCTACACGACGCTCTTCCGATCTNNNAGTAAAAATCATCAGGTTTTCA
CACACGTCTGAACTCTTTCCCTACACGACGCTCTTCCGATCTNNNGCCAACAAACTACACCATTTACATCC
AGCACACGTCTGAACTCTTTCCCTACACGACGCTCTTCCGATCTNNNATGTACATCCAACAATTTTACCC
GCACACGTCTGAACTCTTTCCCTACACGACGCTCTTCCGATCTNNNATTTTGTTCCTTTTTTTGTTGTCA
GAGCACACGTCTGAACTCTTTCCCTACACGACGCTCTTCCGATCTNNNTGACTACCTATTGTGTTACATT
GAGCACACGTCTGAACTCTTTCCCTACACGACGCTCTTCCGATCTNNNAGCTTAAAATTTTAACAAAGGA
AGCACACGTCTGAACTCTTTCCCTACACGACGCTCTTCCGATCTNNNTCTTGAATGAGCCTTTTCTCCATC
ACACGTCTGAACTCTTTCCCTACACGACGCTCTTCCGATCTNNNCTTGAAGCCAGAAGTTAGAGCATCGC
AGCACACGTCTGAACTCTTTCCCTACACGACGCTCTTCCGATCTNNNGGTACCTCATTGTCCTTATTAGC
AGCACACGTCTGAACTCTTTCCCTACACGACGCTCTTCCGATCTNNNCAAAATCTGTAGTGATTGCTAA
GCACACGTCTGAACTCTTTCCCTACACGACGCTCTTCCGATCTNNNTATGTGGACAGAAAAACAAAATT
AGCACACGTCTGAACTCTTTCCCTACACGACGCTCTTCCGATCTNNNCTTACCAGACCTCCATAATCACAT
GAGCACACGTCTGAACTCTTTCCCTACACGACGCTCTTCCGATCTNNNTATTCACAGTAGCAAAGTCATC
CACACGTCTGAACTCTTTCCCTACACGACGCTCTTCCGATCTNNNTTAAATGCCCATTAATTCTCAAGCA
AGCACACGTCTGAACTCTTTCCCTACACGACGCTCTTCCGATCTNNNTTTGCTGGCAATTTTTTACTATC
CACACGTCTGAACTCTTTCCCTACACGACGCTCTTCCGATCTNNNCTTGAATGTCTCAAAGGATTAAAC
GCACACGTCTGAACTCTTTCCCTACACGACGCTCTTCCGATCTNNNTAATGACTCTGAGATCCCTTTACA
CACACGTCTGAACTCTTTCCCTACACGACGCTCTTCCGATCTNNNAAAACAAAAACAAAAACATACCACA
GCACACGTCTGAACTCTTTCCCTACACGACGCTCTTCCGATCTNNNCCAAATATTTGGACTTTTTCACCA
AGCACACGTCTGAACTCTTTCCCTACACGACGCTCTTCCGATCTNNNCAGCCTGGATAGTTCTTTAAGCA
AGCACACGTCTGAACTCTTTCCCTACACGACGCTCTTCCGATCTNNNTTTTGTAAACCTTTTGATACTGT
GAGCACACGTCTGAACTCTTTCCCTACACGACGCTCTTCCGATCTNNNTAACATAATGGTTCAGGGAAT
AGCACACGTCTGAACTCTTTCCCTACACGACGCTCTTCCGATCTNNNCATATCCATTACCTCACATACTTC
GAGCACACGTCTGAACTCTTTCCCTACACGACGCTCTTCCGATCTNNNTTTAAAATTAACAGGTATGCC
GAGCACACGTCTGAACTCTTTCCCTACACGACGCTCTTCCGATCTNNNCTGAGTTCAAGTGATTCTCTCC
GAGCACACGTCTGAACTCTTTCCCTACACGACGCTCTTCCGATCTNNNGGGAAAAAAAATTACAGAAG
AGCACACGTCTGAACTCTTTCCCTACACGACGCTCTTCCGATCTNNNGAATCAAGTAAACTGACAGAAT
GAGCACACGTCTGAACTCTTTCCCTACACGACGCTCTTCCGATCTNNNGGCCTAATTAAAAAGAAAGTT
GCACACGTCTGAACTCTTTCCCTACACGACGCTCTTCCGATCTNNNTACTCTCTGACTTTGTTCTTTTCAT
AGCACACGTCTGAACTCTTTCCCTACACGACGCTCTTCCGATCTNNNTGCCATTATTTGTTCCTTTTTAC
GCACACGTCTGAACTCTTTCCCTACACGACGCTCTTCCGATCTNNNAGTAGGTACTATTATCCATGGAG
AGCACACGTCTGAACTCTTTCCCTACACGACGCTCTTCCGATCTNNNGTGATTAAGCAAGAATACCAAA
AGCACACGTCTGAACTCTTTCCCTACACGACGCTCTTCCGATCTNNNAAGACATCCTAATTAAGAAAAC
AGCACACGTCTGAACTCTTTCCCTACACGACGCTCTTCCGATCTNNNTTTTAGAGACTAGTAGGTTAGT
GCACACGTCTGAACTCTTTCCCTACACGACGCTCTTCCGATCTNNNCGTAAGCTAATTCCTTACAAAAAC
GCACACGTCTGAACTCTTTCCCTACACGACGCTCTTCCGATCTNNNCTATACCCAACATGTTTGCTTACA
GAGCACACGTCTGAACTCTTTCCCTACACGACGCTCTTCCGATCTNNNTGAGTAAGACTCTATGGGTAC

AGCACACGTCTGAACTCTTTCCCTACACGACGCTCTTCCGATCTNNNGAATAAGATGCAGTGGAATAA
AGCACACGTCTGAACTCTTTCCCTACACGACGCTCTTCCGATCTNNNATACATAAGGATAACTGGATCA
GAGCACACGTCTGAACTCTTTCCCTACACGACGCTCTTCCGATCTNNNTAGCAGCTAGATGGATAAATA
AGCACACGTCTGAACTCTTTCCCTACACGACGCTCTTCCGATCTNNNCAAATTTCAAGGCTTAGCTTGCA
GCACACGTCTGAACTCTTTCCCTACACGACGCTCTTCCGATCTNNNAAAGAAATAAAATGTCAGCAAGT
GCACACGTCTGAACTCTTTCCCTACACGACGCTCTTCCGATCTNNNGATAAGAACTCTACCCCAAAGCA
AGCACACGTCTGAACTCTTTCCCTACACGACGCTCTTCCGATCTNNNAATTTAAAATGTTTCACGATAGC
AGCACACGTCTGAACTCTTTCCCTACACGACGCTCTTCCGATCTNNNAACATAGCAAGACCAATGTCCA
GAGCACACGTCTGAACTCTTTCCCTACACGACGCTCTTCCGATCTNNNTAAATCATGTGCCAAATCAATT
GAGCACACGTCTGAACTCTTTCCCTACACGACGCTCTTCCGATCTNNNCAAAAGACAAGAAATAACCAA
AGCACACGTCTGAACTCTTTCCCTACACGACGCTCTTCCGATCTNNNGTTTCCCTTTGTTTGTTTTTAAAC
AGCACACGTCTGAACTCTTTCCCTACACGACGCTCTTCCGATCTNNNGTTCTCAAAGTTTAAAGGAAAGA
ACACGTCTGAACTCTTTCCCTACACGACGCTCTTCCGATCTNNNGAAACAGTGAGAAGTGGTAGCATC
GCACACGTCTGAACTCTTTCCCTACACGACGCTCTTCCGATCTNNNGTTGTTTCCTATTTATAGACGTTC
AGAGCACACGTCTGAACTCTTTCCCTACACGACGCTCTTCCGATCTNNNACAACTGTAGAAAGGAAAAA
GCACACGTCTGAACTCTTTCCCTACACGACGCTCTTCCGATCTNNNTATATCCTACTAGTTATGTCCCTCA
AGCACACGTCTGAACTCTTTCCCTACACGACGCTCTTCCGATCTNNNCCTTTTTGAGGCTGAATAATATT
GCACACGTCTGAACTCTTTCCCTACACGACGCTCTTCCGATCTNNNGACGTTCTGGATTTTTCTTTGCAT
CACACGTCTGAACTCTTTCCCTACACGACGCTCTTCCGATCTNNNTTTAGCTTTCTTATTTCACTGCCATC
GAGCACACGTCTGAACTCTTTCCCTACACGACGCTCTTCCGATCTNNNGAAGATCATTTTAGGTCAAGA
GAGCACACGTCTGAACTCTTTCCCTACACGACGCTCTTCCGATCTNNNTAAAAGAGGATGAAAGACCA
GAGCACACGTCTGAACTCTTTCCCTACACGACGCTCTTCCGATCTNNNCAATTAACTTGTCATCTGCAGC
ACACGTCTGAACTCTTTCCCTACACGACGCTCTTCCGATCTNNNCAAATTTAATAGGCCCCAAAAGCATC
AGCACACGTCTGAACTCTTTCCCTACACGACGCTCTTCCGATCTNNNTTAACAGAAAAAGAAGCTATGA
AGCACACGTCTGAACTCTTTCCCTACACGACGCTCTTCCGATCTNNNTAATTCAATTAGACACTGGAAG
GAGCACACGTCTGAACTCTTTCCCTACACGACGCTCTTCCGATCTNNNACTGGGATTACAGGCATAAGC
AGCACACGTCTGAACTCTTTCCCTACACGACGCTCTTCCGATCTNNNCAATGCCAATATAACCATAGAAA
GCACACGTCTGAACTCTTTCCCTACACGACGCTCTTCCGATCTNNNTCTCTCTCTTTCTCACTCTTTCATCG
GAGCACACGTCTGAACTCTTTCCCTACACGACGCTCTTCCGATCTNNNTAGTGTTAGAATCCATTGGAA
GAGCACACGTCTGAACTCTTTCCCTACACGACGCTCTTCCGATCTNNNTTCCTCTTCTGAAAAATTTTTGT
GCACACGTCTGAACTCTTTCCCTACACGACGCTCTTCCGATCTNNNGTGAAGCTTAAACAAATGTGACA
GAGCACACGTCTGAACTCTTTCCCTACACGACGCTCTTCCGATCTNNNGACCTTGTGATCCTATAAGTTA
CACACGTCTGAACTCTTTCCCTACACGACGCTCTTCCGATCTNNNATAGTCAGAATGGAACCAGCCATCC
AGAGCACACGTCTGAACTCTTTCCCTACACGACGCTCTTCCGATCTNNNATTTAAAACCAACAGATTTCA
GCACACGTCTGAACTCTTTCCCTACACGACGCTCTTCCGATCTNNNCTTTACAAAATCCTGGGGTACAT
CACACGTCTGAACTCTTTCCCTACACGACGCTCTTCCGATCTNNNTGTTTAAATCTAGCTATGAACATCA
GAGCACACGTCTGAACTCTTTCCCTACACGACGCTCTTCCGATCTNNNCACAAAATCAACCTGAAACTTC
GAGCACACGTCTGAACTCTTTCCCTACACGACGCTCTTCCGATCTNNNGGCCCAAACAACATTCTTAACA
GAGCACACGTCTGAACTCTTTCCCTACACGACGCTCTTCCGATCTNNNTATACATTGTAAAAGCCATGAA
GCACACGTCTGAACTCTTTCCCTACACGACGCTCTTCCGATCTNNNCATTACATTATTGGTCTTTGAGAC
AGCACACGTCTGAACTCTTTCCCTACACGACGCTCTTCCGATCTNNNAGATATGCAGATATACAAGGA
AGCACACGTCTGAACTCTTTCCCTACACGACGCTCTTCCGATCTNNNTCAGGTTTTTTACTGTTTCTTTC
GCACACGTCTGAACTCTTTCCCTACACGACGCTCTTCCGATCTNNNAGAGCATAGACTAAGTACATAAT

GCACACGTCTGAACTCTTTCCCTACACGACGCTCTTCCGATCTNNNTATATCAACTCATTCTTCCTCTCC
GCACACGTCTGAACTCTTTCCCTACACGACGCTCTTCCGATCTNNNCAACTGGATCTGTTGTATTTGC
CACACGTCTGAACTCTTTCCCTACACGACGCTCTTCCGATCTNNNAAATGAACATCTAATTAGCAGTG
GCACACGTCTGAACTCTTTCCCTACACGACGCTCTTCCGATCTNNNTCTAACACTCTCAATTTAACTGA
AGCACACGTCTGAACTCTTTCCCTACACGACGCTCTTCCGATCTNNNAATAAACTACTTGCCTGCAAC
CACGTCTGAACTCTTTCCCTACACGACGCTCTTCCGATCTNNNAAACAGACTTTTCACTAGTGTATCAT
AGCACACGTCTGAACTCTTTCCCTACACGACGCTCTTCCGATCTNNNCTAATTATGTCTGTTATGTAG
GCACACGTCTGAACTCTTTCCCTACACGACGCTCTTCCGATCTNNNTCATTCCTTGTACTATTCAAATG
AGCACACGTCTGAACTCTTTCCCTACACGACGCTCTTCCGATCTNNNGTTCCTAACCAGAAAAGTTAT
GCACACGTCTGAACTCTTTCCCTACACGACGCTCTTCCGATCTNNNTCCTCTGTAAAAGAAAACACAT
CACGTCTGAACTCTTTCCCTACACGACGCTCTTCCGATCTNNNAAATGAGATTTGGAATTAGCAATCA
GAGCACACGTCTGAACTCTTTCCCTACACGACGCTCTTCCGATCTNNNAATTATTTGAACCTGGGACA
AGCACACGTCTGAACTCTTTCCCTACACGACGCTCTTCCGATCTNNNGAAGAAGAGGTGAAGGTTTAT
AGCACACGTCTGAACTCTTTCCCTACACGACGCTCTTCCGATCTNNNATAATTGGTCACTGTGGAAAT
AGCACACGTCTGAACTCTTTCCCTACACGACGCTCTTCCGATCTNNNCCTGCTAAATATTCACACATTTA
GCACACGTCTGAACTCTTTCCCTACACGACGCTCTTCCGATCTNNNACTTAGAATAATGGTCTCCAATT
AGCACACGTCTGAACTCTTTCCCTACACGACGCTCTTCCGATCTNNNTAAAGAACCATTAGACACAAA
AGCACACGTCTGAACTCTTTCCCTACACGACGCTCTTCCGATCTNNNAAGAAACATCTGACTCAGATC
ACACGTCTGAACTCTTTCCCTACACGACGCTCTTCCGATCTNNNCAGCATCTTAGAACTTGGTGCATCG
CACACGTCTGAACTCTTTCCCTACACGACGCTCTTCCGATCTNNNTTCAGGTCACATAGATGATAAAT
AGCACACGTCTGAACTCTTTCCCTACACGACGCTCTTCCGATCTNNNTTTGGACAAAACTACATTATCA
CACACGTCTGAACTCTTTCCCTACACGACGCTCTTCCGATCTNNNAGTATCTGAGAACCTACCAAGCAT
GAGCACACGTCTGAACTCTTTCCCTACACGACGCTCTTCCGATCTNNNAGAATGCCCAATTTTTTTTAT
GAGCACACGTCTGAACTCTTTCCCTACACGACGCTCTTCCGATCTNNNTTCAGAACTGTGAACCAAAT
AGCACACGTCTGAACTCTTTCCCTACACGACGCTCTTCCGATCTNNNTGATATACATGTACATTGTGA
AGCACACGTCTGAACTCTTTCCCTACACGACGCTCTTCCGATCTNNNGATAGTTCATAAAACAAGTCT
AGCACACGTCTGAACTCTTTCCCTACACGACGCTCTTCCGATCTNNNGGAAACAAGGTTTCTTTAGCT
GCACACGTCTGAACTCTTTCCCTACACGACGCTCTTCCGATCTNNNTCTTAATCCATGAACATTTTATCT
GCACACGTCTGAACTCTTTCCCTACACGACGCTCTTCCGATCTNNNAATTATTTTTTTGTCCCATCCTTC
AGCACACGTCTGAACTCTTTCCCTACACGACGCTCTTCCGATCTNNNAGTCCTTGACTTTTAATTTTGT
AGCACACGTCTGAACTCTTTCCCTACACGACGCTCTTCCGATCTNNNGTGAAACTCTGTCTCAAAAAA
AGCACACGTCTGAACTCTTTCCCTACACGACGCTCTTCCGATCTNNNATATTTTTTGAGTCATGTAACC
GCACACGTCTGAACTCTTTCCCTACACGACGCTCTTCCGATCTNNNTGCAAAAGACATGATTTCATAT
AGCACACGTCTGAACTCTTTCCCTACACGACGCTCTTCCGATCTNNNAAAGTGTTGGGATTATAGGC
CACGTCTGAACTCTTTCCCTACACGACGCTCTTCCGATCTNNNGTTTTGTCTGCATTGAAAATCTCAT
AGCACACGTCTGAACTCTTTCCCTACACGACGCTCTTCCGATCTNNNTCCAGCTTCCCTTTTAAAAATAA
GCACACGTCTGAACTCTTTCCCTACACGACGCTCTTCCGATCTNNNCTATATTGCATCCAAAGCTCTCA
AGCACACGTCTGAACTCTTTCCCTACACGACGCTCTTCCGATCTNNNCATTTATTGAGATTAGGGAAG
CACACGTCTGAACTCTTTCCCTACACGACGCTCTTCCGATCTNNNCTTTTTTGATCCTGAAGTTCCCAT
GCACACGTCTGAACTCTTTCCCTACACGACGCTCTTCCGATCTNNNGAATAATAGTCTCCAGTCTCATC
AGCACACGTCTGAACTCTTTCCCTACACGACGCTCTTCCGATCTNNNCTGCAAAGTAATCAGTTTAAGA
AGCACACGTCTGAACTCTTTCCCTACACGACGCTCTTCCGATCTNNNCTGGAGCTCAAGATTAAGTTC
AGCACACGTCTGAACTCTTTCCCTACACGACGCTCTTCCGATCTNNNCACTTCTGGGTATTTATTCAA

GAGCACACGTCTGAACTCTTTCCCTACACGACGCTCTTCCGATCTNNNTAGGTTTCTTCATCTGTGAAAT
CACACGTCTGAACTCTTTCCCTACACGACGCTCTTCCGATCTNNNCTTTTTGGGACCAGAAGTATCCATC
ACACGTCTGAACTCTTTCCCTACACGACGCTCTTCCGATCTNNNTTTGAATGTGACATGTTCTTTTCATC
AGAGCACACGTCTGAACTCTTTCCCTACACGACGCTCTTCCGATCTNNNGGGCATATGAATACTTCAGT
AGCACACGTCTGAACTCTTTCCCTACACGACGCTCTTCCGATCTNNNCCATGTGAGTATGCAAGAAGCA
GAGCACACGTCTGAACTCTTTCCCTACACGACGCTCTTCCGATCTNNNTACCATTGTCAGAGATTTTAAC
CACACGTCTGAACTCTTTCCCTACACGACGCTCTTCCGATCTNNNAGCACTTTGAGAGAATGAGACATC
GAGCACACGTCTGAACTCTTTCCCTACACGACGCTCTTCCGATCTNNNTTTTGTGTTCTTAAGAGTTTTC
GAGCACACGTCTGAACTCTTTCCCTACACGACGCTCTTCCGATCTNNNTGTTAGTCAACATAGTACGTAA
AGCACACGTCTGAACTCTTTCCCTACACGACGCTCTTCCGATCTNNNTTGCATATTATATACACAGCATG
AGCACACGTCTGAACTCTTTCCCTACACGACGCTCTTCCGATCTNNNTTTATTTCTGCCTCTCCTTATACCA
GCACACGTCTGAACTCTTTCCCTACACGACGCTCTTCCGATCTNNNTGAATAGAAAATTCATCTGCTTTCA
ACACGTCTGAACTCTTTCCCTACACGACGCTCTTCCGATCTNNNAAGACTCCATCTCAAAATACAACATC
AGCACACGTCTGAACTCTTTCCCTACACGACGCTCTTCCGATCTNNNTTGTTTACATTGCCAATAAAAAT
ACACGTCTGAACTCTTTCCCTACACGACGCTCTTCCGATCTNNNGATAACTTGAGGTCAGGAGTTCATC
AGCACACGTCTGAACTCTTTCCCTACACGACGCTCTTCCGATCTNNNGTAAAATGAATGAATCTTCCTCA
AGCACACGTCTGAACTCTTTCCCTACACGACGCTCTTCCGATCTNNNCTACTTGGTTGTTGTAAAATGAT
GCACACGTCTGAACTCTTTCCCTACACGACGCTCTTCCGATCTNNNGTTCTTGTACCAGTACCATGCATC
GCACACGTCTGAACTCTTTCCCTACACGACGCTCTTCCGATCTNNNTTGGGGGGAGTTTATATATATAC
GAGCACACGTCTGAACTCTTTCCCTACACGACGCTCTTCCGATCTNNNCTCTAGTTTTCCACCTGAAGCAT
GCACACGTCTGAACTCTTTCCCTACACGACGCTCTTCCGATCTNNNTAGGACACTCATACCTCCTCCATCG
AGCACACGTCTGAACTCTTTCCCTACACGACGCTCTTCCGATCTNNNCATATGCTCAAAACAACTCTTGCA
GAGCACACGTCTGAACTCTTTCCCTACACGACGCTCTTCCGATCTNNNTACACCTACTTTGTACTCATAAA
AGCACACGTCTGAACTCTTTCCCTACACGACGCTCTTCCGATCTNNNCTTTGAGAATTTTACCTTTGAGA
AGCACACGTCTGAACTCTTTCCCTACACGACGCTCTTCCGATCTNNNGTAAAAAAGCTCCTACCTTCTCA
CACACGTCTGAACTCTTTCCCTACACGACGCTCTTCCGATCTNNNATTGTTTTGGGTATTCTTTTGTTCAT
CACACGTCTGAACTCTTTCCCTACACGACGCTCTTCCGATCTNNNTTCTATACCAGAAGAACACATCCATC
AGCACACGTCTGAACTCTTTCCCTACACGACGCTCTTCCGATCTNNNGTGTAGTTTATCTGCTTCAATTT
AGCACACGTCTGAACTCTTTCCCTACACGACGCTCTTCCGATCTNNNTATGATGGTGCCTGTGAATACA
GAGCACACGTCTGAACTCTTTCCCTACACGACGCTCTTCCGATCTNNNGTTTATTGCAGCATTATTTACA
AGCACACGTCTGAACTCTTTCCCTACACGACGCTCTTCCGATCTNNNTAACGCAGTAAAATGGATAGTA
AGCACACGTCTGAACTCTTTCCCTACACGACGCTCTTCCGATCTNNNGGAAAGACTATCAGATTAAAAC
GAGCACACGTCTGAACTCTTTCCCTACACGACGCTCTTCCGATCTNNNAATCTCTTTTTTGATATTGGCTT
CACACGTCTGAACTCTTTCCCTACACGACGCTCTTCCGATCTNNNTTTTGGTGAGAGGACCTATTCATC
AGCACACGTCTGAACTCTTTCCCTACACGACGCTCTTCCGATCTNNNGCTGTTACCTAACTAGAAGTTA
GCACACGTCTGAACTCTTTCCCTACACGACGCTCTTCCGATCTNNNTTTTTAAAGAGAATGTGAAGGA
GAGCACACGTCTGAACTCTTTCCCTACACGACGCTCTTCCGATCTNNNGTTTTTTGTTTCTTTTTTCCTCA
GAGCACACGTCTGAACTCTTTCCCTACACGACGCTCTTCCGATCTNNNATCATTACATAGTATCTGCAAC
GCACACGTCTGAACTCTTTCCCTACACGACGCTCTTCCGATCTNNNCTTTGGTCTCTACCCTTTATAGCA
AGCACACGTCTGAACTCTTTCCCTACACGACGCTCTTCCGATCTNNNTTCACTGAAGCACTATTCATAAT
AGCACACGTCTGAACTCTTTCCCTACACGACGCTCTTCCGATCTNNNCATGATGACTGGATATACATACT
AGCACACGTCTGAACTCTTTCCCTACACGACGCTCTTCCGATCTNNNGTAGTTTTAACTAGCTAGAAGG
AGAGCACACGTCTGAACTCTTTCCCTACACGACGCTCTTCCGATCTNNNTAGATAAAAAGGACTTGTAG

AGCACACGTCTGAACTCTTTCCCTACACGACGCTCTTCCGATCTNNNTAACAAGGAGGAATGTATTTT
GCACACGTCTGAACTCTTTCCCTACACGACGCTCTTCCGATCTNNNCAGGAGAATCACTTGAACTTCAT
AGCACACGTCTGAACTCTTTCCCTACACGACGCTCTTCCGATCTNNNGCAATCTTGAAGTGATACAATT
AGCACACGTCTGAACTCTTTCCCTACACGACGCTCTTCCGATCTNNNCAAGAGGGAAATCTCACTAAT
GCACACGTCTGAACTCTTTCCCTACACGACGCTCTTCCGATCTNNNTTCTTCAGAGCTCACATTTCCAT
GAGCACACGTCTGAACTCTTTCCCTACACGACGCTCTTCCGATCTNNNGGTGTATATATGTGTTTTCT
GCACACGTCTGAACTCTTTCCCTACACGACGCTCTTCCGATCTNNNGCACTTTTTAACAGGTATTAGTT
GCACACGTCTGAACTCTTTCCCTACACGACGCTCTTCCGATCTNNNCAGACCTTAGAACTTGTCAGCAT
GCACACGTCTGAACTCTTTCCCTACACGACGCTCTTCCGATCTNNNATAGCCATAGTCATCTGTTATACC
GCACACGTCTGAACTCTTTCCCTACACGACGCTCTTCCGATCTNNNGTCAGTTTCTTCCTAAGGATAAA
AGCACACGTCTGAACTCTTTCCCTACACGACGCTCTTCCGATCTNNNAATAATTCTAACATCCCTGTCA
GCACACGTCTGAACTCTTTCCCTACACGACGCTCTTCCGATCTNNNGATTTCTAGAGTTAAGCAAGTTA
GCACACGTCTGAACTCTTTCCCTACACGACGCTCTTCCGATCTNNNGGATGCCCTTTATTTCTTTTATTC
AGCACACGTCTGAACTCTTTCCCTACACGACGCTCTTCCGATCTNNNGGATCTTGTCTGATTTAAAATCA
AGCACACGTCTGAACTCTTTCCCTACACGACGCTCTTCCGATCTNNNGGACTTATGCTAATTCTTTGTT
CACACGTCTGAACTCTTTCCCTACACGACGCTCTTCCGATCTNNNAGACCCTGTCTCAAATACATCATCG
GCACACGTCTGAACTCTTTCCCTACACGACGCTCTTCCGATCTNNNGTTTTATGTGTGGTATATGGGT
AGCACACGTCTGAACTCTTTCCCTACACGACGCTCTTCCGATCTNNNATTGCCCATTAAGATAATGTTG
AGCACACGTCTGAACTCTTTCCCTACACGACGCTCTTCCGATCTNNNTTTTATTCTTTTTGATGGAACGA
GAGCACACGTCTGAACTCTTTCCCTACACGACGCTCTTCCGATCTNNNGTAAATGATGAATTCTAGACT
AGCACACGTCTGAACTCTTTCCCTACACGACGCTCTTCCGATCTNNNTAATAAATCTCTCTCTCTCTGTCC
GCACACGTCTGAACTCTTTCCCTACACGACGCTCTTCCGATCTNNNTTTCATTAAAACACCCTCCGCAT
AGCACACGTCTGAACTCTTTCCCTACACGACGCTCTTCCGATCTNNNGAATTATCGTATGATCAAGCA
AGCACACGTCTGAACTCTTTCCCTACACGACGCTCTTCCGATCTNNNAAATGAATAACTTACAGATGCA
AGCACACGTCTGAACTCTTTCCCTACACGACGCTCTTCCGATCTNNNAAGCAAGTTTTACAGCTGTTCA
GCACACGTCTGAACTCTTTCCCTACACGACGCTCTTCCGATCTNNNTTAAATTTAGTGCTTATTGTTGC
GCACACGTCTGAACTCTTTCCCTACACGACGCTCTTCCGATCTNNNCAGCTAACCTCCTATGAAAGCAT
GCACACGTCTGAACTCTTTCCCTACACGACGCTCTTCCGATCTNNNCTTATACCAGAGGTTGACATCCA
AGCACACGTCTGAACTCTTTCCCTACACGACGCTCTTCCGATCTNNNTGTGGTTTTTAGCAGTTTAATTA
GAGCACACGTCTGAACTCTTTCCCTACACGACGCTCTTCCGATCTNNNCTCCATCTCTGAAAAAATAAA
ACACGTCTGAACTCTTTCCCTACACGACGCTCTTCCGATCTNNNTTACTCTCTGTCTCTGTCTCCATCGG
GCACACGTCTGAACTCTTTCCCTACACGACGCTCTTCCGATCTNNNTTCCATTGACATTTGTCCAGCATC
GAGCACACGTCTGAACTCTTTCCCTACACGACGCTCTTCCGATCTNNNAAAAATATTGAGGAAGATGG
AGCACACGTCTGAACTCTTTCCCTACACGACGCTCTTCCGATCTNNNTGTGAAATTGGAGGAGAATCC
AGCACACGTCTGAACTCTTTCCCTACACGACGCTCTTCCGATCTNNNAAAGTGCCTAATCCTTTTAAATC
CACACGTCTGAACTCTTTCCCTACACGACGCTCTTCCGATCTNNNAAACAAAATAAATGGAGGCTTTT
GAGCACACGTCTGAACTCTTTCCCTACACGACGCTCTTCCGATCTNNNGCAGTGTTTTTATGTGTTTTA
AGCACACGTCTGAACTCTTTCCCTACACGACGCTCTTCCGATCTNNNGCATAATGTCCTCTAGATTCATA
GCACACGTCTGAACTCTTTCCCTACACGACGCTCTTCCGATCTNNNCTTTCATCTTTCATCTCCATTTTCA
CACACGTCTGAACTCTTTCCCTACACGACGCTCTTCCGATCTNNNGTTGAGTAGACTGTCCAGATCATC
AGCACACGTCTGAACTCTTTCCCTACACGACGCTCTTCCGATCTNNNAAAGGAAATGGGTATAACTAT
GCACACGTCTGAACTCTTTCCCTACACGACGCTCTTCCGATCTNNNCTCAGGAACTCAATATTGGTTAC
AGCACACGTCTGAACTCTTTCCCTACACGACGCTCTTCCGATCTNNNAAAAATGCTAGTCTTCAGATTA

AGCACACGTCTGAACTCTTTCCCTACACGACGCTCTTCCGATCTNNNATGACCATTCCTTTCTATAGAA

AGCACACGTCTGAACTCTTTCCCTACACGACGCTCTTCCGATCTNNNTGCTGTTAATTCATCCTTTTTA

CACACGTCTGAACTCTTTCCCTACACGACGCTCTTCCGATCTNNNTGGGCAGAGTATAATTCATTATA

AGCACACGTCTGAACTCTTTCCCTACACGACGCTCTTCCGATCTNNNCTCAATTACTCAAAATCAGAC

AGCACACGTCTGAACTCTTTCCCTACACGACGCTCTTCCGATCTNNNAAAATAGTGTCTCATATCCTT

ACACGTCTGAACTCTTTCCCTACACGACGCTCTTCCGATCTNNNCATAAGAAAGGAGGCTCATTTATC

AGCACACGTCTGAACTCTTTCCCTACACGACGCTCTTCCGATCTNNNGATCTGTCAGATCTTTAATTCT

GCACACGTCTGAACTCTTTCCCTACACGACGCTCTTCCGATCTNNNTCAATTTCTTATTTATCCTTCCAG

GCACACGTCTGAACTCTTTCCCTACACGACGCTCTTCCGATCTNNNATCACTTGAAGTTAAGCGTTTC

AGCACACGTCTGAACTCTTTCCCTACACGACGCTCTTCCGATCTNNNCTTGATCTGTAAAATGGGAA

CACACGTCTGAACTCTTTCCCTACACGACGCTCTTCCGATCTNNNATCAAGGTCAAAATCAGTTAAAA

AGCACACGTCTGAACTCTTTCCCTACACGACGCTCTTCCGATCTNNNGAGGTGAGATCATGCTATTG

ACACGTCTGAACTCTTTCCCTACACGACGCTCTTCCGATCTNNNTTTTCCAGGAACTAGAATCTTACA

GAGCACACGTCTGAACTCTTTCCCTACACGACGCTCTTCCGATCTNNNGTATAACTGGTGTCATTATA

AGCACACGTCTGAACTCTTTCCCTACACGACGCTCTTCCGATCTNNNGCTTCAAAAGTTCTTTGACTG

ACACGTCTGAACTCTTTCCCTACACGACGCTCTTCCGATCTNNNTTCGAAGATTTCCTTTTTCTTTACAT

CACACGTCTGAACTCTTTCCCTACACGACGCTCTTCCGATCTNNNCTTCACTGAAATGTTACTTCAATCA

GAGCACACGTCTGAACTCTTTCCCTACACGACGCTCTTCCGATCTNNNTTAATGGATTATCAAGAGTG

GCACACGTCTGAACTCTTTCCCTACACGACGCTCTTCCGATCTNNNATTTCTCACCTCTTTCTGTTTCAT

AGCACACGTCTGAACTCTTTCCCTACACGACGCTCTTCCGATCTNNNACATTTTTTTCTTAATCACCCTC

AGCACACGTCTGAACTCTTTCCCTACACGACGCTCTTCCGATCTNNNTAATAAGATTCCCAGAAAGAG

GCACACGTCTGAACTCTTTCCCTACACGACGCTCTTCCGATCTNNNTGCAAAAGACATGATTTCTTTTA

AGCACACGTCTGAACTCTTTCCCTACACGACGCTCTTCCGATCTNNNATATCTGACTGTAGCATCTTA

ACACGTCTGAACTCTTTCCCTACACGACGCTCTTCCGATCTNNNATCATCATTGTGATAATCATCAACA

ACACGTCTGAACTCTTTCCCTACACGACGCTCTTCCGATCTNNNTCCTTTCTTGACTCTCTTCCCATCGG

ACACGTCTGAACTCTTTCCCTACACGACGCTCTTCCGATCTNNNGAGTCTCTGTTCGTTTGTTTCATCG

AGCACACGTCTGAACTCTTTCCCTACACGACGCTCTTCCGATCTNNNGCAAAAGGAAGAAAATGTCC

CACACGTCTGAACTCTTTCCCTACACGACGCTCTTCCGATCTNNNGGAGAGAGTATGGAATTTCTTT

AGCACACGTCTGAACTCTTTCCCTACACGACGCTCTTCCGATCTNNNGATAAAGAAACTGTGGTGTA

AGCACACGTCTGAACTCTTTCCCTACACGACGCTCTTCCGATCTNNNCCACAAGGAAATATTTTTAATG

AGCACACGTCTGAACTCTTTCCCTACACGACGCTCTTCCGATCTNNNAAAAATACAAAAATTAGCCAG

AGCACACGTCTGAACTCTTTCCCTACACGACGCTCTTCCGATCTNNNTTCTATAGCATCCTCCTGTGCAT

GCACACGTCTGAACTCTTTCCCTACACGACGCTCTTCCGATCTNNNCTGAGCAACAAGAGTGAAACCA

AGCACACGTCTGAACTCTTTCCCTACACGACGCTCTTCCGATCTNNNCAAACAAAATGATTATGTCTC

GCACACGTCTGAACTCTTTCCCTACACGACGCTCTTCCGATCTNNNTTTGTGTGCTTACTATATCAAAC

GCACACGTCTGAACTCTTTCCCTACACGACGCTCTTCCGATCTNNNTTCTTCCTCTTCCCAAAAATTAT

AGCACACGTCTGAACTCTTTCCCTACACGACGCTCTTCCGATCTNNNACATTTTTCTGGCATTTTATAC

AGCACACGTCTGAACTCTTTCCCTACACGACGCTCTTCCGATCTNNNTTCAAACTGAAAAGAGAAAA

AGCACACGTCTGAACTCTTTCCCTACACGACGCTCTTCCGATCTNNNAATTTTCTGCCTTTAATTCTCC

GCACACGTCTGAACTCTTTCCCTACACGACGCTCTTCCGATCTNNNTTGTTGGTAGAGGTAAATCATA

GCACACGTCTGAACTCTTTCCCTACACGACGCTCTTCCGATCTNNNCCATTATTTCTTGGCAAAATACT

AGCACACGTCTGAACTCTTTCCCTACACGACGCTCTTCCGATCTNNNAGTGGTAAAGAAACTGTGAT

GAGCACACGTCTGAACTCTTTCCCTACACGACGCTCTTCCGATCTNNNAGTATGGTCTGACTCATAGA

CACACGTCTGAACTCTTTCCCTACACGACGCTCTTCCGATCTNNNAAAAGTTAAAATCACTTAATTCCA
ACACGTCTGAACTCTTTCCCTACACGACGCTCTTCCGATCTNNNAGGAGAATCACTTGAACCTGCATC
GCACACGTCTGAACTCTTTCCCTACACGACGCTCTTCCGATCTNNNCTGTCTCAAAAAAAAAAAAAGG
GAGCACACGTCTGAACTCTTTCCCTACACGACGCTCTTCCGATCTNNNAAAGAAGTGAGTGACAAACT
GCACACGTCTGAACTCTTTCCCTACACGACGCTCTTCCGATCTNNNGTAGAGGACAAAGACAATAGA
GCACACGTCTGAACTCTTTCCCTACACGACGCTCTTCCGATCTNNNTGAGTTTATAGTCTTGTAAGGA
CACACGTCTGAACTCTTTCCCTACACGACGCTCTTCCGATCTNNNTCCCAATTTACATTATCCATTTCA
AGCACACGTCTGAACTCTTTCCCTACACGACGCTCTTCCGATCTNNNCTTGATTATGTACAGATTTTG
AGCACACGTCTGAACTCTTTCCCTACACGACGCTCTTCCGATCTNNNTTGACTTAAAATGGCTAAAATG
GCACACGTCTGAACTCTTTCCCTACACGACGCTCTTCCGATCTNNNTGATATAGTGATTTTGCTTTTGT
AGCACACGTCTGAACTCTTTCCCTACACGACGCTCTTCCGATCTNNNAGGATTGGTTCTGATTTTATTT
AGCACACGTCTGAACTCTTTCCCTACACGACGCTCTTCCGATCTNNNAAATCCTTGCATACTTTTTACA
CACACGTCTGAACTCTTTCCCTACACGACGCTCTTCCGATCTNNNGTGTTTAGTTCAGGACTTTATTAC
ACACGTCTGAACTCTTTCCCTACACGACGCTCTTCCGATCTNNNGAGAATCACTTCAATCCAGTAGCA
AGCACACGTCTGAACTCTTTCCCTACACGACGCTCTTCCGATCTNNNGGTATGATTTCTGCCTTCAACA
AGCACACGTCTGAACTCTTTCCCTACACGACGCTCTTCCGATCTNNNAAATGCAGAGAAGAAAGAAT
GCACACGTCTGAACTCTTTCCCTACACGACGCTCTTCCGATCTNNNTCATCTGTTTTTCTTAAAGCAT
GCACACGTCTGAACTCTTTCCCTACACGACGCTCTTCCGATCTNNNCTTCTATCTTACTGCAAAATTGT
AGCACACGTCTGAACTCTTTCCCTACACGACGCTCTTCCGATCTNNNAAGATATTTCACTGGCAAGTA
AGCACACGTCTGAACTCTTTCCCTACACGACGCTCTTCCGATCTNNNTAAGGATTTAAAATGAGCAAG
AGCACACGTCTGAACTCTTTCCCTACACGACGCTCTTCCGATCTNNNAGGATGGATTAAAACCTGTAA
AGCACACGTCTGAACTCTTTCCCTACACGACGCTCTTCCGATCTNNNATGTCACCTTACTCTAAAAACT
AGCACACGTCTGAACTCTTTCCCTACACGACGCTCTTCCGATCTNNNTGTGAATTTGTTAAGGTAAAA
AGCACACGTCTGAACTCTTTCCCTACACGACGCTCTTCCGATCTNNNTTCTCAAAGACAAACCAACAT
GCACACGTCTGAACTCTTTCCCTACACGACGCTCTTCCGATCTNNNTAGACTTCAATAAATTCACTCCA
AGCACACGTCTGAACTCTTTCCCTACACGACGCTCTTCCGATCTNNNGACCTGATTTTTATGGCAAACC
AGCACACGTCTGAACTCTTTCCCTACACGACGCTCTTCCGATCTNNNTAATTGCTCAATAATGTCAGGT
AGCACACGTCTGAACTCTTTCCCTACACGACGCTCTTCCGATCTNNNTTATATAGGCAGGAAGGAAAT
CACACGTCTGAACTCTTTCCCTACACGACGCTCTTCCGATCTNNNATGATTGGACTAGGATGGGACA
AGCACACGTCTGAACTCTTTCCCTACACGACGCTCTTCCGATCTNNNTGTTAATCTGTTCCACATTAAT
GCACACGTCTGAACTCTTTCCCTACACGACGCTCTTCCGATCTNNNATATAATGGCCTCATCACAAACC
AGCACACGTCTGAACTCTTTCCCTACACGACGCTCTTCCGATCTNNNCCTCCATTAACTTTGATTTCTGC
AGCACACGTCTGAACTCTTTCCCTACACGACGCTCTTCCGATCTNNNCTTGTTTCCAGGAGTTTGAGC
AGCACACGTCTGAACTCTTTCCCTACACGACGCTCTTCCGATCTNNNTACTTAAGGGAGTTATTGATA
GCACACGTCTGAACTCTTTCCCTACACGACGCTCTTCCGATCTNNNGCTAAAAGCCTGTTTATATACAT
AGCACACGTCTGAACTCTTTCCCTACACGACGCTCTTCCGATCTNNNCTGAATGGAGGTTAACATTTC
AGCACACGTCTGAACTCTTTCCCTACACGACGCTCTTCCGATCTNNNGGAAAATTCATCAGTGAAAGT
GCACACGTCTGAACTCTTTCCCTACACGACGCTCTTCCGATCTNNNATATCTTAAGTCAATAAGCCTCC
AGCACACGTCTGAACTCTTTCCCTACACGACGCTCTTCCGATCTNNNTGAAAATCCAAGTATAATTTTG
ACACGTCTGAACTCTTTCCCTACACGACGCTCTTCCGATCTNNNGCCAAGTAAAAAGTTTGTGAATCA
CACACGTCTGAACTCTTTCCCTACACGACGCTCTTCCGATCTNNNAAGACTCCATCTCAAACACATCAT
CACACGTCTGAACTCTTTCCCTACACGACGCTCTTCCGATCTNNNAATACTACTGCCTCAGCTTCCATCG
AGCACACGTCTGAACTCTTTCCCTACACGACGCTCTTCCGATCTNNNAAAATAAAGTTACTGGGTTGT

AGCACACGTCTGAACTCTTTCCCTACACGACGCTCTTCCGATCTNNNATCTTGCTTCTTTAAAGCCAACA
AGCACACGTCTGAACTCTTTCCCTACACGACGCTCTTCCGATCTNNNACATTTCTTCAGAAGCATTTTAT
AGCACACGTCTGAACTCTTTCCCTACACGACGCTCTTCCGATCTNNNCTCTCTCTCTCTCTCTGTCATCG
AGCACACGTCTGAACTCTTTCCCTACACGACGCTCTTCCGATCTNNNGGTAAAAGATATATTCATGACC
GAGCACACGTCTGAACTCTTTCCCTACACGACGCTCTTCCGATCTNNNTGAGGATCTGTCTCAAAAATCA
AGCACACGTCTGAACTCTTTCCCTACACGACGCTCTTCCGATCTNNNACGGGTATATTGAGAGAAATA
AGAGCACACGTCTGAACTCTTTCCCTACACGACGCTCTTCCGATCTNNNTTTAGGCAGAAGAATCAATT
GAGCACACGTCTGAACTCTTTCCCTACACGACGCTCTTCCGATCTNNNAAACTTCTGTTTCTTCAGAGAA
AGCACACGTCTGAACTCTTTCCCTACACGACGCTCTTCCGATCTNNNTCTAATATAACTACACCCCAATC
AAGAGCACACGTCTGAACTCTTTCCCTACACGACGCTCTTCCGATCTNNNTTCCATAGTGCTAGGATTA
CACACGTCTGAACTCTTTCCCTACACGACGCTCTTCCGATCTNNNTAAGTGGTGAGTAGAATACCAACA
GAGCACACGTCTGAACTCTTTCCCTACACGACGCTCTTCCGATCTNNNGTGGTGTGTGTATATGTAATT
AGAGCACACGTCTGAACTCTTTCCCTACACGACGCTCTTCCGATCTNNNGATCTCAAAGAATAGACTCCA
GAGCACACGTCTGAACTCTTTCCCTACACGACGCTCTTCCGATCTNNNCCAGAATTATGCATAAAATTTG
CACACGTCTGAACTCTTTCCCTACACGACGCTCTTCCGATCTNNNCTGAAGGAGGAAGACAGTATACA
CACACGTCTGAACTCTTTCCCTACACGACGCTCTTCCGATCTNNNCTAAAGGACAAGTTTAAGTCATCCA
GCACACGTCTGAACTCTTTCCCTACACGACGCTCTTCCGATCTNNNGGCAAAATACCAGGAGTAAGCAT
CACACGTCTGAACTCTTTCCCTACACGACGCTCTTCCGATCTNNNGAAGCAGTAATATCACTGGAGCAT
CACACGTCTGAACTCTTTCCCTACACGACGCTCTTCCGATCTNNNGAAGCAGTAATATCACTGGAGCAT
CACACGTCTGAACTCTTTCCCTACACGACGCTCTTCCGATCTNNNGATAAAACTGATTTTCTCCTTGGCAT
GAGCACACGTCTGAACTCTTTCCCTACACGACGCTCTTCCGATCTNNNTATAATTTGTGTTGTTTTCCCC
GAGCACACGTCTGAACTCTTTCCCTACACGACGCTCTTCCGATCTNNNTACAAGTTAATAAAGTCCCTTC
GAGCACACGTCTGAACTCTTTCCCTACACGACGCTCTTCCGATCTNNNATTCTCTTTGTCTTCTAGAATGT
AGCACACGTCTGAACTCTTTCCCTACACGACGCTCTTCCGATCTNNNCTTTCACTCTTGCTCTTTCTCATC
AGCACACGTCTGAACTCTTTCCCTACACGACGCTCTTCCGATCTNNNCTGTGCAAGATTGGTATAATTTA
AGCACACGTCTGAACTCTTTCCCTACACGACGCTCTTCCGATCTNNNTGCTCTAAGATATTACACATACA
CACACGTCTGAACTCTTTCCCTACACGACGCTCTTCCGATCTNNNAACCTCCTTTTCCATACCTACATCG
AGCACACGTCTGAACTCTTTCCCTACACGACGCTCTTCCGATCTNNNAATTCAGGTAAATGAGGACAT
AGCACACGTCTGAACTCTTTCCCTACACGACGCTCTTCCGATCTNNNAGAAGCTATTACTCAAATTCAGA
GAGCACACGTCTGAACTCTTTCCCTACACGACGCTCTTCCGATCTNNNATTCCAATACAACCTAGGAATT
AGCACACGTCTGAACTCTTTCCCTACACGACGCTCTTCCGATCTNNNAAGCTCAGTTCTAAAATCTTTA
GCACACGTCTGAACTCTTTCCCTACACGACGCTCTTCCGATCTNNNGGAGATTTTGATGAGAAACAAG
AGCACACGTCTGAACTCTTTCCCTACACGACGCTCTTCCGATCTNNNAATAGAGTGAGACCCTGTCTCA
GAGCACACGTCTGAACTCTTTCCCTACACGACGCTCTTCCGATCTNNNATTCCAAACCTTATACTACCTTC
GAGCACACGTCTGAACTCTTTCCCTACACGACGCTCTTCCGATCTNNNATTCTCTCTTCAATGGTATGTT
GAGCACACGTCTGAACTCTTTCCCTACACGACGCTCTTCCGATCTNNNAGAGAGAACCACTATAATGTTT
GAGCACACGTCTGAACTCTTTCCCTACACGACGCTCTTCCGATCTNNNCCCAATTAACCCTAAAATACAA
AGCACACGTCTGAACTCTTTCCCTACACGACGCTCTTCCGATCTNNNAGTCATTAAGTTGAACATTATCT
AGCACACGTCTGAACTCTTTCCCTACACGACGCTCTTCCGATCTNNNTATTTCCTCTGTTATAACTTCCC
AGCACACGTCTGAACTCTTTCCCTACACGACGCTCTTCCGATCTNNNTCATTAAGTTGAACATTATCTTG
AGCACACGTCTGAACTCTTTCCCTACACGACGCTCTTCCGATCTNNNCTGGACAGAGTGAGACTATGC
GCACACGTCTGAACTCTTTCCCTACACGACGCTCTTCCGATCTNNNGGCCTACTGGATCAATTTTATATCA
GCACACGTCTGAACTCTTTCCCTACACGACGCTCTTCCGATCTNNNAAAAACACCTTTCTAATAAGACG

CACACGTCTGAACTCTTTCCCTACACGACGCTCTTCCGATCTNNNTCATTGGCTTTAAGGTTTACAAA
AGCACACGTCTGAACTCTTTCCCTACACGACGCTCTTCCGATCTNNNTATGATCACCCTATTCCACTCCA
AGCACACGTCTGAACTCTTTCCCTACACGACGCTCTTCCGATCTNNNTTCATTATGCAAATCTTACTTC
AGCACACGTCTGAACTCTTTCCCTACACGACGCTCTTCCGATCTNNNTTGCAAAAATTTTGTCTTTTCA
ACACGTCTGAACTCTTTCCCTACACGACGCTCTTCCGATCTNNNTACAAAAAATTGTCAAACCATTTCA
CACACGTCTGAACTCTTTCCCTACACGACGCTCTTCCGATCTNNNCAATCAGATTCTTTCTCTTGTGCAT
CACACGTCTGAACTCTTTCCCTACACGACGCTCTTCCGATCTNNNCCCTGTCTCTATATTAAACACAAC
AGAGCACACGTCTGAACTCTTTCCCTACACGACGCTCTTCCGATCTNNNCCAGCCTAGATTGATTTAG
AGCACACGTCTGAACTCTTTCCCTACACGACGCTCTTCCGATCTNNNCTAATCATCAGAGAAATGCAA
AGCACACGTCTGAACTCTTTCCCTACACGACGCTCTTCCGATCTNNNAGTCATTAAGTTGAACATTATC
AGCACACGTCTGAACTCTTTCCCTACACGACGCTCTTCCGATCTNNNTAAACTCACTGCTAGAATTCA
AGCACACGTCTGAACTCTTTCCCTACACGACGCTCTTCCGATCTNNNTTACATTATACATTATCTGCGT
AGCACACGTCTGAACTCTTTCCCTACACGACGCTCTTCCGATCTNNNTATTCAGATGGACCTTTTCATA
AGAGCACACGTCTGAACTCTTTCCCTACACGACGCTCTTCCGATCTNNNAAATTCCATGGATTTAAAAT
AGCACACGTCTGAACTCTTTCCCTACACGACGCTCTTCCGATCTNNNAGTGTACATTGTACTAGTAAG
AGCACACGTCTGAACTCTTTCCCTACACGACGCTCTTCCGATCTNNNTATCCTCTATACCAATGTCTAC
AGCACACGTCTGAACTCTTTCCCTACACGACGCTCTTCCGATCTNNNCTTAGGCTTCTTCTTTGTTTTT
GAGCACACGTCTGAACTCTTTCCCTACACGACGCTCTTCCGATCTNNNCTGACTCAAAAAACAAAACAA
AGCACACGTCTGAACTCTTTCCCTACACGACGCTCTTCCGATCTNNNCCATCTTTTGTAGTTCTTATTGT
AGCACACGTCTGAACTCTTTCCCTACACGACGCTCTTCCGATCTNNNAAACTCTTATGATGTCTCCACC
AGCACACGTCTGAACTCTTTCCCTACACGACGCTCTTCCGATCTNNNCTGTATGTTGATTTTGCATTCTA
AGCACACGTCTGAACTCTTTCCCTACACGACGCTCTTCCGATCTNNNTGATCTAAATAAAACGCAATGA
AGCACACGTCTGAACTCTTTCCCTACACGACGCTCTTCCGATCTNNNATAGTGGATATGCTACCAGATA
AGCACACGTCTGAACTCTTTCCCTACACGACGCTCTTCCGATCTNNNCTTCTTTCTCTCTGTCTCTCTCA
GCACACGTCTGAACTCTTTCCCTACACGACGCTCTTCCGATCTNNNGAAATGTGATTTCCGTATTCTG
CACACGTCTGAACTCTTTCCCTACACGACGCTCTTCCGATCTNNNGATATCCAATTGGTTTCTGAAAG
AGCACACGTCTGAACTCTTTCCCTACACGACGCTCTTCCGATCTNNNGACTCGATATCTTCTTCTTAG
AGCACACGTCTGAACTCTTTCCCTACACGACGCTCTTCCGATCTNNNTAAAATACCACATTTTCATCACC
CACACGTCTGAACTCTTTCCCTACACGACGCTCTTCCGATCTNNNATTTGACAAAGAGAGGAAAGAA
GCACACGTCTGAACTCTTTCCCTACACGACGCTCTTCCGATCTNNNGATATGAGGTGATATCCTATTG
AGCACACGTCTGAACTCTTTCCCTACACGACGCTCTTCCGATCTNNNGAACAGAATAGTCAGTTTAGA
AGCACACGTCTGAACTCTTTCCCTACACGACGCTCTTCCGATCTNNNACAAGGAAAGTCATCATTACT
AGCACACGTCTGAACTCTTTCCCTACACGACGCTCTTCCGATCTNNNCATGTATCCCAGAATTTAAAG
AGCACACGTCTGAACTCTTTCCCTACACGACGCTCTTCCGATCTNNNATGGCTTAATAGTATTTCATGG
AGCACACGTCTGAACTCTTTCCCTACACGACGCTCTTCCGATCTNNNTTATCTAGTTTGGAATTGATG
AGCACACGTCTGAACTCTTTCCCTACACGACGCTCTTCCGATCTNNNGCCCAGTAGAATAGCATTTTCA
AGCACACGTCTGAACTCTTTCCCTACACGACGCTCTTCCGATCTNNNGAAAGATAAGTATCTCCATCA
GAGCACACGTCTGAACTCTTTCCCTACACGACGCTCTTCCGATCTNNNTGAAATGGATTTAAAACACA
GCACACGTCTGAACTCTTTCCCTACACGACGCTCTTCCGATCTNNNCTCCAGTACTTTACTTTTTCTC
CACACGTCTGAACTCTTTCCCTACACGACGCTCTTCCGATCTNNNCCCACCTCAAAAAAAAAATAATTC
CACACGTCTGAACTCTTTCCCTACACGACGCTCTTCCGATCTNNNCTTCCTACCCCTTCTAAGACCATCG
CACACGTCTGAACTCTTTCCCTACACGACGCTCTTCCGATCTNNNGTGTACGTTTTGTTTTTCCTATCA
AGCACACGTCTGAACTCTTTCCCTACACGACGCTCTTCCGATCTNNNTCAACAAGTGTATAAAGAAA

AGCACACGTCTGAACTCTTTCCCTACACGACGCTCTTCCGATCTNNNATATAGTAAGTCCTAGAGAAC
AGCACACGTCTGAACTCTTTCCCTACACGACGCTCTTCCGATCTNNNCTATTATGGGACCATGTAATC
AGCACACGTCTGAACTCTTTCCCTACACGACGCTCTTCCGATCTNNNAAAGTTTAAAAACAGTCAACTT

CATCGGACTCATCAGGAAGC
ACATCGGACTCATCAGGAAGC
CATCGGACTCATCAGGAAGC
GCATCGGACTCATCAGGAAGC
GACATCGGACTCATCAGGAAGC
TCGGACTCATCAGGAAGC
GACTCATCAGGAAGC
ACATCGGACTCATCAGGAAGC
CATCGGACTCATCAGGAAGC
CATCGGACTCATCAGGAAGC
GACTCATCAGGAAGC
CATCGGACTCATCAGGAAGC
ATCGGACTCATCAGGAAGC
AACATCGGACTCATCAGGAAGC
CATCGGACTCATCAGGAAGC
ATCGGACTCATCAGGAAGC
ATCGGACTCATCAGGAAGC
ACCATCGGACTCATCAGGAAGC
CATCGGACTCATCAGGAAGC
AACATCGGACTCATCAGGAAGC
ACATCGGACTCATCAGGAAGC
GGACTCATCAGGAAGC
CTCATCGGACTCATCAGGAAGC
CATCGGACTCATCAGGAAGC
CATCGGACTCATCAGGAAGC
CGGACTCATCAGGAAGC
CGGACTCATCAGGAAGC
ATCGGACTCATCAGGAAGC
ACATCGGACTCATCAGGAAGC
ATCGGACTCATCAGGAAGC
ACATCGGACTCATCAGGAAGC
AGCATCGGACTCATCAGGAAGC
TCATCGGACTCATCAGGAAGC
ATTCATCGGACTCATCAGGAAGC
AACATCGGACTCATCAGGAAGC
AGCATCGGACTCATCAGGAAGC
CATCGGACTCATCAGGAAGC
CGGACTCATCAGGAAGC
ACATCGGACTCATCAGGAAGC
ATCGGACTCATCAGGAAGC
GCATCGGACTCATCAGGAAGC
ATCATCGGACTCATCAGGAAGC

GCATCGGACTCATCAGGAAGC
GTCCATCGGACTCATCAGGAAGC
CCATCGGACTCATCAGGAAGC
CGGACTCATCAGGAAGC
CCATCGGACTCATCAGGAAGC
ATCGGACTCATCAGGAAGC
TCGGACTCATCAGGAAGC
ACATCGGACTCATCAGGAAGC
ACATCGGACTCATCAGGAAGC
TCATCGGACTCATCAGGAAGC
TCGGACTCATCAGGAAGC
ACATCGGACTCATCAGGAAGC
ACATCGGACTCATCAGGAAGC
CATCGGACTCATCAGGAAGC
ACTCATCAGGAAGC
GCATCGGACTCATCAGGAAGC
CATCGGACTCATCAGGAAGC
CGGACTCATCAGGAAGC
TCGGACTCATCAGGAAGC
ACATCGGACTCATCAGGAAGC
CATCGGACTCATCAGGAAGC
TCGGACTCATCAGGAAGC
CGGACTCATCAGGAAGC
TTCATCGGACTCATCAGGAAGC
GGACTCATCAGGAAGC
TCGGACTCATCAGGAAGC
CATCGGACTCATCAGGAAGC
CATCGGACTCATCAGGAAGC
ACATCGGACTCATCAGGAAGC
ACATCGGACTCATCAGGAAGC
TCATCGGACTCATCAGGAAGC
ACATCGGACTCATCAGGAAGC
ACATCGGACTCATCAGGAAGC
CGGACTCATCAGGAAGC
ATTCATCGGACTCATCAGGAAGC
CGGACTCATCAGGAAGC
CATCGGACTCATCAGGAAGC
ACATCGGACTCATCAGGAAGC
CCATCGGACTCATCAGGAAGC
GACATCGGACTCATCAGGAAGC
CATCGGACTCATCAGGAAGC
GTCATCGGACTCATCAGGAAGC
TGCATCGGACTCATCAGGAAGC

CCATCGGACTCATCAGGAAGC
CATCGGACTCATCAGGAAGC
TTCATCGGACTCATCAGGAAGC
CATCGGACTCATCAGGAAGC
CATCGGACTCATCAGGAAGC
GTCATCGGACTCATCAGGAAGC
GCATCGGACTCATCAGGAAGC
AACATCGGACTCATCAGGAAGC
ACATCGGACTCATCAGGAAGC
TCATCGGACTCATCAGGAAGC
CATCGGACTCATCAGGAAGC
GCATCGGACTCATCAGGAAGC
TCCATCGGACTCATCAGGAAGC
ATCGGACTCATCAGGAAGC
GGACTCATCAGGAAGC
CATCGGACTCATCAGGAAGC
CATCGGACTCATCAGGAAGC
GTCATCGGACTCATCAGGAAGC
CGCATCGGACTCATCAGGAAGC
GACATCGGACTCATCAGGAAGC
GCATCGGACTCATCAGGAAGC
CCCATCGGACTCATCAGGAAGC
ACTCATCAGGAAGC
ACATCGGACTCATCAGGAAGC
CACATCGGACTCATCAGGAAGC
ATCGGACTCATCAGGAAGC
TACATCGGACTCATCAGGAAGC
GACTCATCAGGAAGC
GTCATCGGACTCATCAGGAAGC
CATCGGACTCATCAGGAAGC
ATCGGACTCATCAGGAAGC
CATCGGACTCATCAGGAAGC
GCATCGGACTCATCAGGAAGC
CATCGGACTCATCAGGAAGC
ATCGGACTCATCAGGAAGC
ATCGGACTCATCAGGAAGC
GCATCGGACTCATCAGGAAGC
TTACCATCGGACTCATCAGGAAGC
GACTCATCAGGAAGC
ATCGGACTCATCAGGAAGC
ATCGGACTCATCAGGAAGC
ATCGGACTCATCAGGAAGC
CATCGGACTCATCAGGAAGC

TCATCGGACTCATCAGGAAGC
CATCGGACTCATCAGGAAGC
TCGGACTCATCAGGAAGC
ATCGGACTCATCAGGAAGC
TCATCGGACTCATCAGGAAGC
CATCGGACTCATCAGGAAGC
ATCGGACTCATCAGGAAGC
GACTCATCAGGAAGC
ATCGGACTCATCAGGAAGC
TCATCGGACTCATCAGGAAGC
ACATCGGACTCATCAGGAAGC
TCATCGGACTCATCAGGAAGC
CGGACTCATCAGGAAGC
GGACTCATCAGGAAGC
TCATCGGACTCATCAGGAAGC
TACATCGGACTCATCAGGAAGC
CATCGGACTCATCAGGAAGC
CATCGGACTCATCAGGAAGC
GGACTCATCAGGAAGC
ATCGGACTCATCAGGAAGC
GCATCGGACTCATCAGGAAGC
ATCGGACTCATCAGGAAGC
ACATCGGACTCATCAGGAAGC
GCATCGGACTCATCAGGAAGC
CTCATCGGACTCATCAGGAAGC
CATCGGACTCATCAGGAAGC
ACATCGGACTCATCAGGAAGC
TCGGACTCATCAGGAAGC
ATAGCATCGGACTCATCAGGAAGC
GACTCATCAGGAAGC
CATCGGACTCATCAGGAAGC
TCATCGGACTCATCAGGAAGC
ATCGGACTCATCAGGAAGC
CATCGGACTCATCAGGAAGC
CATCGGACTCATCAGGAAGC
CCATCGGACTCATCAGGAAGC
GCATCGGACTCATCAGGAAGC
TATCCATCGGACTCATCAGGAAGC
CATCGGACTCATCAGGAAGC
GCATCGGACTCATCAGGAAGC
CGGACTCATCAGGAAGC
CACATCGGACTCATCAGGAAGC
CATCGGACTCATCAGGAAGC

CATCGGACTCATCAGGAAGC
CATCGGACTCATCAGGAAGC
TCGGACTCATCAGGAAGC
CGGACTCATCAGGAAGC
TCATCGGACTCATCAGGAAGC
ATCGGACTCATCAGGAAGC
ACATCGGACTCATCAGGAAGC
CATCGGACTCATCAGGAAGC
TCGGACTCATCAGGAAGC
TCGGACTCATCAGGAAGC
CATCGGACTCATCAGGAAGC
GGACTCATCAGGAAGC
ACATCGGACTCATCAGGAAGC
GCATCGGACTCATCAGGAAGC
ATACATCGGACTCATCAGGAAGC
CCTCATCGGACTCATCAGGAAGC
TCGGACTCATCAGGAAGC
ATCATCGGACTCATCAGGAAGC
ATCATCGGACTCATCAGGAAGC
GCATCGGACTCATCAGGAAGC
TCGGACTCATCAGGAAGC
CATCGGACTCATCAGGAAGC
TACATCGGACTCATCAGGAAGC
TACATCGGACTCATCAGGAAGC
CATCGGACTCATCAGGAAGC
CGGACTCATCAGGAAGC
GCATCGGACTCATCAGGAAGC
TCGGACTCATCAGGAAGC
ACATCGGACTCATCAGGAAGC
TCCATCGGACTCATCAGGAAGC
ATCATCGGACTCATCAGGAAGC
CATCGGACTCATCAGGAAGC
CGGACTCATCAGGAAGC
TCCATCGGACTCATCAGGAAGC
GCATCGGACTCATCAGGAAGC
TCGGACTCATCAGGAAGC
ATCGGACTCATCAGGAAGC
TCGGACTCATCAGGAAGC
TCGGACTCATCAGGAAGC
CATCGGACTCATCAGGAAGC
CATCGGACTCATCAGGAAGC
GCATCGGACTCATCAGGAAGC
AGCATCGGACTCATCAGGAAGC

CCATCGGACTCATCAGGAAGC
CATCGGACTCATCAGGAAGC
ATCGGACTCATCAGGAAGC
TCGGACTCATCAGGAAGC
TCATCGGACTCATCAGGAAGC
GACTCATCAGGAAGC
ATCGGACTCATCAGGAAGC
ACCATCGGACTCATCAGGAAGC
GACTCATCAGGAAGC
ATCATCGGACTCATCAGGAAGC
GGACTCATCAGGAAGC
CATCGGACTCATCAGGAAGC
ATCGGACTCATCAGGAAGC
CATCGGACTCATCAGGAAGC
TCGGACTCATCAGGAAGC
ACATCGGACTCATCAGGAAGC
CATCGGACTCATCAGGAAGC
ACATCGGACTCATCAGGAAGC
CACATCGGACTCATCAGGAAGC
GCATCGGACTCATCAGGAAGC
CATCGGACTCATCAGGAAGC
GCCATCGGACTCATCAGGAAGC
ATCATCGGACTCATCAGGAAGC
GCATCGGACTCATCAGGAAGC
GGACTCATCAGGAAGC
CATCGGACTCATCAGGAAGC
AACATCGGACTCATCAGGAAGC
CAGCATCGGACTCATCAGGAAGC
CGGACTCATCAGGAAGC
CGGACTCATCAGGAAGC
ATCGGACTCATCAGGAAGC
CATCGGACTCATCAGGAAGC
GGACTCATCAGGAAGC
TCATCGGACTCATCAGGAAGC
ACATCGGACTCATCAGGAAGC
CGGACTCATCAGGAAGC
CCATCGGACTCATCAGGAAGC
GACATCGGACTCATCAGGAAGC
GACTCATCAGGAAGC
TCATCGGACTCATCAGGAAGC
GGACTCATCAGGAAGC
CATCGGACTCATCAGGAAGC
GGACTCATCAGGAAGC

GGACTCATCAGGAAGC
ATCGGACTCATCAGGAAGC
TCGGACTCATCAGGAAGC
CATCGGACTCATCAGGAAGC
GGACTCATCAGGAAGC
ACTCATCAGGAAGC
CATCGGACTCATCAGGAAGC
TCGGACTCATCAGGAAGC
CTCATCGGACTCATCAGGAAGC
ACCATCGGACTCATCAGGAAGC
CCATCGGACTCATCAGGAAGC
CCATCGGACTCATCAGGAAGC
CATCGGACTCATCAGGAAGC
CATCGGACTCATCAGGAAGC
TCATCGGACTCATCAGGAAGC
CACATCGGACTCATCAGGAAGC
CATCGGACTCATCAGGAAGC
CCATCGGACTCATCAGGAAGC
GCATCGGACTCATCAGGAAGC
TCATCGGACTCATCAGGAAGC
CTCCATCGGACTCATCAGGAAGC
TGTCATCGGACTCATCAGGAAGC
ATCGGACTCATCAGGAAGC
ACATCGGACTCATCAGGAAGC
GGCATCGGACTCATCAGGAAGC
AGCATCGGACTCATCAGGAAGC
CATCGGACTCATCAGGAAGC
GACTCATCAGGAAGC
TCATCGGACTCATCAGGAAGC
CCCATCGGACTCATCAGGAAGC
TCGGACTCATCAGGAAGC
ATCGGACTCATCAGGAAGC
ATCGGACTCATCAGGAAGC
CATCGGACTCATCAGGAAGC
CATCGGACTCATCAGGAAGC
AGCATCGGACTCATCAGGAAGC
TCATCGGACTCATCAGGAAGC
CATCGGACTCATCAGGAAGC
TCGGACTCATCAGGAAGC
TCATCGGACTCATCAGGAAGC
ACATCGGACTCATCAGGAAGC
CGGACTCATCAGGAAGC
CCATCGGACTCATCAGGAAGC

CATCGGACTCATCAGGAAGC
ATCGGACTCATCAGGAAGC
GCATCGGACTCATCAGGAAGC
TTCATCGGACTCATCAGGAAGC
TCATCGGACTCATCAGGAAGC
GATCATCGGACTCATCAGGAAGC
CGGACTCATCAGGAAGC
ATCGGACTCATCAGGAAGC
CATCGGACTCATCAGGAAGC
CGGACTCATCAGGAAGC
ACATCGGACTCATCAGGAAGC
GACTCATCAGGAAGC
TCGGACTCATCAGGAAGC
TCGGACTCATCAGGAAGC
ATCGGACTCATCAGGAAGC
CATCGGACTCATCAGGAAGC
CATCGGACTCATCAGGAAGC
CTCATCGGACTCATCAGGAAGC
GCATCGGACTCATCAGGAAGC
ACATCGGACTCATCAGGAAGC
TCGGACTCATCAGGAAGC
GGCATCGGACTCATCAGGAAGC
TGCATCGGACTCATCAGGAAGC
ATCGGACTCATCAGGAAGC
GACTCATCAGGAAGC
TCATCGGACTCATCAGGAAGC
GGCATCGGACTCATCAGGAAGC
CATCGGACTCATCAGGAAGC
ATCGGACTCATCAGGAAGC
GACTCATCAGGAAGC
GCATCGGACTCATCAGGAAGC
ATCGGACTCATCAGGAAGC
TCATCGGACTCATCAGGAAGC
ATCGGACTCATCAGGAAGC
GCATCGGACTCATCAGGAAGC
ATCATCGGACTCATCAGGAAGC
ACATCGGACTCATCAGGAAGC
GCATCGGACTCATCAGGAAGC
ATCGGACTCATCAGGAAGC
GCATCGGACTCATCAGGAAGC
ATCATCGGACTCATCAGGAAGC
CCATCGGACTCATCAGGAAGC
CGGACTCATCAGGAAGC

CATCGGACTCATCAGGAAGC
CGGACTCATCAGGAAGC
GAACATCGGACTCATCAGGAAGC
CATCGGACTCATCAGGAAGC
CATCGGACTCATCAGGAAGC
CATCGGACTCATCAGGAAGC
CCATCGGACTCATCAGGAAGC
GACTCATCAGGAAGC
GCATCGGACTCATCAGGAAGC
TCATCGGACTCATCAGGAAGC
ACATCGGACTCATCAGGAAGC
CATCGGACTCATCAGGAAGC
ATCGGACTCATCAGGAAGC
ATCATCGGACTCATCAGGAAGC
GACTCATCAGGAAGC
ATCGGACTCATCAGGAAGC
CGGACTCATCAGGAAGC
ATCGGACTCATCAGGAAGC
CCATCGGACTCATCAGGAAGC
CGGACTCATCAGGAAGC
ATCGGACTCATCAGGAAGC
CGGACTCATCAGGAAGC
CATCGGACTCATCAGGAAGC
GACTCATCAGGAAGC
ATACATCGGACTCATCAGGAAGC
CATCGGACTCATCAGGAAGC
ATCGGACTCATCAGGAAGC
CATCGGACTCATCAGGAAGC
ATCGGACTCATCAGGAAGC
CATCGGACTCATCAGGAAGC
AACATCGGACTCATCAGGAAGC
ATCGGACTCATCAGGAAGC
CATCGGACTCATCAGGAAGC
TCATCGGACTCATCAGGAAGC
GTGGCATCGGACTCATCAGGAAGC
TCGGACTCATCAGGAAGC
TCCATCGGACTCATCAGGAAGC
CATCGGACTCATCAGGAAGC
CATCGGACTCATCAGGAAGC
GCATCGGACTCATCAGGAAGC
CATCGGACTCATCAGGAAGC
CGGACTCATCAGGAAGC
ACATCGGACTCATCAGGAAGC

TCGGACTCATCAGGAAGC

TCATCGGACTCATCAGGAAGC

ATCGGACTCATCAGGAAGC

TCATCGGACTCATCAGGAAGC

CGGACTCATCAGGAAGC

GGACTCATCAGGAAGC

TCATCGGACTCATCAGGAAGC

AACATCGGACTCATCAGGAAGC

GGACTCATCAGGAAGC

ATCGGACTCATCAGGAAGC

CATCGGACTCATCAGGAAGC

ACATCGGACTCATCAGGAAGC

ACATCGGACTCATCAGGAAGC

ATCGGACTCATCAGGAAGC

TCGGACTCATCAGGAAGC

AGCATCGGACTCATCAGGAAGC

ACATCGGACTCATCAGGAAGC

CGGACTCATCAGGAAGC

ACATCGGACTCATCAGGAAGC

GGACTCATCAGGAAGC

GGACTCATCAGGAAGC

GCATCGGACTCATCAGGAAGC

CTCATCGGACTCATCAGGAAGC

ATCGGACTCATCAGGAAGC

GCATCGGACTCATCAGGAAGC

TCATCGGACTCATCAGGAAGC

CGGACTCATCAGGAAGC

ACTCATCAGGAAGC

GGACTCATCAGGAAGC

GGACTCATCAGGAAGC

ATCGGACTCATCAGGAAGC

ATCGGACTCATCAGGAAGC

CATCGGACTCATCAGGAAGC

ATCGGACTCATCAGGAAGC

ACCATCGGACTCATCAGGAAGC

GGACTCATCAGGAAGC

ACCATCGGACTCATCAGGAAGC

TCATCGGACTCATCAGGAAGC

GGACTCATCAGGAAGC

TCGGACTCATCAGGAAGC

CACCATCGGACTCATCAGGAAGC

TCGGACTCATCAGGAAGC

GCATCGGACTCATCAGGAAGC

ATCGGACTCATCAGGAAGC
AACATCGGACTCATCAGGAAGC
ATCGGACTCATCAGGAAGC
CGGACTCATCAGGAAGC
ACATCGGACTCATCAGGAAGC
GCATCGGACTCATCAGGAAGC
GACTCATCAGGAAGC
GGACTCATCAGGAAGC
CCATCGGACTCATCAGGAAGC
GACATCGGACTCATCAGGAAGC
GGACTCATCAGGAAGC
ATGCATCGGACTCATCAGGAAGC
TCGGACTCATCAGGAAGC
GGCATCGGACTCATCAGGAAGC
GACTCATCAGGAAGC
CATCGGACTCATCAGGAAGC
GCATCGGACTCATCAGGAAGC
CATCGGACTCATCAGGAAGC
TCGGACTCATCAGGAAGC
ATCGGACTCATCAGGAAGC
TCATCGGACTCATCAGGAAGC
GGACTCATCAGGAAGC
TCGGACTCATCAGGAAGC
CATCGGACTCATCAGGAAGC
TCGGACTCATCAGGAAGC
TACATCGGACTCATCAGGAAGC
TTGCATCGGACTCATCAGGAAGC
ATCGGACTCATCAGGAAGC
CATCGGACTCATCAGGAAGC
TCATCGGACTCATCAGGAAGC
GACTCATCAGGAAGC
ATCGGACTCATCAGGAAGC
CATCGGACTCATCAGGAAGC
ATCGGACTCATCAGGAAGC
TCGGACTCATCAGGAAGC
AGCATCGGACTCATCAGGAAGC
GACTCATCAGGAAGC
AACATCGGACTCATCAGGAAGC
AACATCGGACTCATCAGGAAGC
TCATCGGACTCATCAGGAAGC
ACTCATCAGGAAGC
ATCGGACTCATCAGGAAGC
GGACTCATCAGGAAGC

CATCGGACTCATCAGGAAGC
ATCGGACTCATCAGGAAGC
TCGGACTCATCAGGAAGC
ATCGGACTCATCAGGAAGC
GACTCATCAGGAAGC
TCGGACTCATCAGGAAGC
GGACTCATCAGGAAGC
TCGGACTCATCAGGAAGC
TCATCGGACTCATCAGGAAGC
CATCGGACTCATCAGGAAGC
ACATCGGACTCATCAGGAAGC
GGACTCATCAGGAAGC
TCATCGGACTCATCAGGAAGC
ATCGGACTCATCAGGAAGC
CCATCGGACTCATCAGGAAGC
GGACTCATCAGGAAGC
ATCGGACTCATCAGGAAGC
CATCGGACTCATCAGGAAGC
ATCGGACTCATCAGGAAGC
TCGGACTCATCAGGAAGC
ATCGGACTCATCAGGAAGC
TCATCGGACTCATCAGGAAGC
ACATCGGACTCATCAGGAAGC
GCATCGGACTCATCAGGAAGC
ATCGGACTCATCAGGAAGC
CATCGGACTCATCAGGAAGC
ATCGGACTCATCAGGAAGC
ATCGGACTCATCAGGAAGC
ATCGGACTCATCAGGAAGC
ATCGGACTCATCAGGAAGC
CATCGGACTCATCAGGAAGC
TTGTCATCGGACTCATCAGGAAGC
CATCGGACTCATCAGGAAGC
ACATCGGACTCATCAGGAAGC
CCATCGGACTCATCAGGAAGC
CATCGGACTCATCAGGAAGC
ATCGGACTCATCAGGAAGC
TCATCGGACTCATCAGGAAGC
TCATCGGACTCATCAGGAAGC
ATCGGACTCATCAGGAAGC
ACATCGGACTCATCAGGAAGC
ATCGGACTCATCAGGAAGC
GGACTCATCAGGAAGC

TCATCGGACTCATCAGGAAGC
CCATCGGACTCATCAGGAAGC
CATCGGACTCATCAGGAAGC
GCATCGGACTCATCAGGAAGC
ATCGGACTCATCAGGAAGC
AAGCATCGGACTCATCAGGAAGC
GCATCGGACTCATCAGGAAGC
TTCATCGGACTCATCAGGAAGC
TCCATCGGACTCATCAGGAAGC
CCATCGGACTCATCAGGAAGC
TACCATCGGACTCATCAGGAAGC
CATCGGACTCATCAGGAAGC
AGCATCGGACTCATCAGGAAGC
GGACTCATCAGGAAGC
CATCGGACTCATCAGGAAGC
CATCGGACTCATCAGGAAGC
ATCATCGGACTCATCAGGAAGC
GGACTCATCAGGAAGC
TCATCGGACTCATCAGGAAGC
TCATCGGACTCATCAGGAAGC
TCATCGGACTCATCAGGAAGC
TCATCGGACTCATCAGGAAGC
TCATCGGACTCATCAGGAAGC
AATCATCGGACTCATCAGGAAGC
ACATCGGACTCATCAGGAAGC
CATCGGACTCATCAGGAAGC
TCATCGGACTCATCAGGAAGC
TCATCGGACTCATCAGGAAGC
TCATCGGACTCATCAGGAAGC
TCATCGGACTCATCAGGAAGC
TCATCGGACTCATCAGGAAGC
TCATCGGACTCATCAGGAAGC
TCATCGGACTCATCAGGAAGC
ATCATCGGACTCATCAGGAAGC
GGACTCATCAGGAAGC
GACATCGGACTCATCAGGAAGC
ATCGGACTCATCAGGAAGC
CGGACTCATCAGGAAGC
CCATCGGACTCATCAGGAAGC
GGACTCATCAGGAAGC
CATCGGACTCATCAGGAAGC
ATCGGACTCATCAGGAAGC
TCGGACTCATCAGGAAGC

ATCGGACTCATCAGGAAGC
TCGGACTCATCAGGAAGC
ATCGGACTCATCAGGAAGC
GGACTCATCAGGAAGC
GGACTCATCAGGAAGC
GGACTCATCAGGAAGC
CGGACTCATCAGGAAGC
TCATCGGACTCATCAGGAAGC
CATCGGACTCATCAGGAAGC
ATCGGACTCATCAGGAAGC
ATCGGACTCATCAGGAAGC
CATCGGACTCATCAGGAAGC
CATCGGACTCATCAGGAAGC
TCGGACTCATCAGGAAGC
CATCGGACTCATCAGGAAGC
TGCATCGGACTCATCAGGAAGC
ACATCGGACTCATCAGGAAGC
CATCGGACTCATCAGGAAGC
TCATCGGACTCATCAGGAAGC
GCATCGGACTCATCAGGAAGC
ACCATCGGACTCATCAGGAAGC
CATCGGACTCATCAGGAAGC
CATCGGACTCATCAGGAAGC
GGACTCATCAGGAAGC
GACTCATCAGGAAGC
CGGACTCATCAGGAAGC
CATCGGACTCATCAGGAAGC
CGGACTCATCAGGAAGC
CTCCATCGGACTCATCAGGAAGC
CCATCGGACTCATCAGGAAGC
TCGGACTCATCAGGAAGC
ACATCGGACTCATCAGGAAGC
CATCGGACTCATCAGGAAGC
GCATCGGACTCATCAGGAAGC
TCATCGGACTCATCAGGAAGC
TCGGACTCATCAGGAAGC
TCATCGGACTCATCAGGAAGC
TCGGACTCATCAGGAAGC
CATCGGACTCATCAGGAAGC
CCATCGGACTCATCAGGAAGC
CCATCGGACTCATCAGGAAGC
CATCGGACTCATCAGGAAGC
ATCGGACTCATCAGGAAGC

CATCGGACTCATCAGGAAGC
CATCGGACTCATCAGGAAGC
CATCGGACTCATCAGGAAGC
ACATCGGACTCATCAGGAAGC
CATCGGACTCATCAGGAAGC
ACATCGGACTCATCAGGAAGC
GACTCATCAGGAAGC
CATCGGACTCATCAGGAAGC
CATCGGACTCATCAGGAAGC
GCATCGGACTCATCAGGAAGC
GGACTCATCAGGAAGC
ACATCGGACTCATCAGGAAGC
CATCGGACTCATCAGGAAGC
ATCGGACTCATCAGGAAGC
ATCATCGGACTCATCAGGAAGC
GGACTCATCAGGAAGC
TCATCGGACTCATCAGGAAGC
TCATCGGACTCATCAGGAAGC
TCATCGGACTCATCAGGAAGC
TCATCGGACTCATCAGGAAGC
CATCGGACTCATCAGGAAGC
AATCATCGGACTCATCAGGAAGC
ACATCGGACTCATCAGGAAGC
CATCGGACTCATCAGGAAGC
ACCATCGGACTCATCAGGAAGC
CATCGGACTCATCAGGAAGC
CATCGGACTCATCAGGAAGC
CATCGGACTCATCAGGAAGC
CATCGGACTCATCAGGAAGC
CATCGGACTCATCAGGAAGC
CATCGGACTCATCAGGAAGC
GACATCGGACTCATCAGGAAGC
CATCGGACTCATCAGGAAGC
CATCGGACTCATCAGGAAGC
TTCATCGGACTCATCAGGAAGC
CATCGGACTCATCAGGAAGC
GTCATCGGACTCATCAGGAAGC
CATCGGACTCATCAGGAAGC
TTCATCGGACTCATCAGGAAGC
ACATCGGACTCATCAGGAAGC
ACATCGGACTCATCAGGAAGC
CTATCATCGGACTCATCAGGAAGC
ATCGGACTCATCAGGAAGC

CATCGGACTCATCAGGAAGC
GACTCATCAGGAAGC
GACATCGGACTCATCAGGAAGC
TCGGACTCATCAGGAAGC
CATCGGACTCATCAGGAAGC
CATCGGACTCATCAGGAAGC
CATCGGACTCATCAGGAAGC
AACATCGGACTCATCAGGAAGC
ACATCGGACTCATCAGGAAGC
ACATCGGACTCATCAGGAAGC
GTCATCGGACTCATCAGGAAGC
GACATCGGACTCATCAGGAAGC
CATCGGACTCATCAGGAAGC
CATCGGACTCATCAGGAAGC
ACATCGGACTCATCAGGAAGC
CAGCATCGGACTCATCAGGAAGC
ACCATCGGACTCATCAGGAAGC
TGGCATCGGACTCATCAGGAAGC
TCGGACTCATCAGGAAGC
ACATCGGACTCATCAGGAAGC
TCGGACTCATCAGGAAGC
TCATCGGACTCATCAGGAAGC
CATCGGACTCATCAGGAAGC
TCGGACTCATCAGGAAGC
GCATCGGACTCATCAGGAAGC
GGACTCATCAGGAAGC
CGGACTCATCAGGAAGC
CATCGGACTCATCAGGAAGC
CATCGGACTCATCAGGAAGC
ACATCGGACTCATCAGGAAGC
GATCATCGGACTCATCAGGAAGC
TACATCGGACTCATCAGGAAGC
GGACTCATCAGGAAGC
TCATCGGACTCATCAGGAAGC
ATCGGACTCATCAGGAAGC
CGGACTCATCAGGAAGC
GCATCGGACTCATCAGGAAGC
ATCGGACTCATCAGGAAGC
GCCATCGGACTCATCAGGAAGC
CGGACTCATCAGGAAGC
CATCGGACTCATCAGGAAGC
GCATCGGACTCATCAGGAAGC
TTCATCGGACTCATCAGGAAGC

TCGGACTCATCAGGAAGC
CATCGGACTCATCAGGAAGC
CGGACTCATCAGGAAGC
TCGGACTCATCAGGAAGC
ACCATCGGACTCATCAGGAAGC
GACTCATCAGGAAGC
CGGACTCATCAGGAAGC
ATCGGACTCATCAGGAAGC
TGCATCGGACTCATCAGGAAGC
ATCGGACTCATCAGGAAGC
CGGACTCATCAGGAAGC
TCATCGGACTCATCAGGAAGC
CGGACTCATCAGGAAGC
AGCATCGGACTCATCAGGAAGC
ATCGGACTCATCAGGAAGC
TCCATCGGACTCATCAGGAAGC
GCATCGGACTCATCAGGAAGC
ATCGGACTCATCAGGAAGC
CCATCGGACTCATCAGGAAGC
CGGACTCATCAGGAAGC
CATCGGACTCATCAGGAAGC
CATCGGACTCATCAGGAAGC
GTCATCGGACTCATCAGGAAGC
TCATCGGACTCATCAGGAAGC
CATCGGACTCATCAGGAAGC
CGGACTCATCAGGAAGC
GGACTCATCAGGAAGC
TCATCGGACTCATCAGGAAGC
CATCGGACTCATCAGGAAGC
ACATCGGACTCATCAGGAAGC
TCATCGGACTCATCAGGAAGC
CATCGGACTCATCAGGAAGC
TCATCGGACTCATCAGGAAGC
TCGGACTCATCAGGAAGC
TTCATCGGACTCATCAGGAAGC
GCATCGGACTCATCAGGAAGC
CATCGGACTCATCAGGAAGC
GGACTCATCAGGAAGC
TCGGACTCATCAGGAAGC
ATCGGACTCATCAGGAAGC
GCATCGGACTCATCAGGAAGC
ATCGGACTCATCAGGAAGC
ATCGGACTCATCAGGAAGC

ATCGGACTCATCAGGAAGC
ACTCATCAGGAAGC
GGACTCATCAGGAAGC
CGGACTCATCAGGAAGC
CATCGGACTCATCAGGAAGC
ATCGGACTCATCAGGAAGC
ATCGGACTCATCAGGAAGC
ATCGGACTCATCAGGAAGC
CGGACTCATCAGGAAGC
ATCGGACTCATCAGGAAGC
CGGACTCATCAGGAAGC
ATCGGACTCATCAGGAAGC
GGACTCATCAGGAAGC
ATCGGACTCATCAGGAAGC
GCATCGGACTCATCAGGAAGC
TCATCGGACTCATCAGGAAGC
GACTCATCAGGAAGC
ATCGGACTCATCAGGAAGC
GCATCGGACTCATCAGGAAGC
CATCGGACTCATCAGGAAGC
CATCGGACTCATCAGGAAGC
GACATCGGACTCATCAGGAAGC
TCGGACTCATCAGGAAGC
CATCGGACTCATCAGGAAGC
CCATCGGACTCATCAGGAAGC
TCGGACTCATCAGGAAGC
GCATCGGACTCATCAGGAAGC
ATCGGACTCATCAGGAAGC
CATCGGACTCATCAGGAAGC
CATCGGACTCATCAGGAAGC
CATCGGACTCATCAGGAAGC
ATCGGACTCATCAGGAAGC
CATCGGACTCATCAGGAAGC
AGCATCGGACTCATCAGGAAGC
ATCGGACTCATCAGGAAGC
CGGACTCATCAGGAAGC
TACATCGGACTCATCAGGAAGC
TGCATCGGACTCATCAGGAAGC
CTCATCGGACTCATCAGGAAGC
CGGACTCATCAGGAAGC
TCATCGGACTCATCAGGAAGC
ATCGGACTCATCAGGAAGC
CGGACTCATCAGGAAGC

GCCATCGGACTCATCAGGAAGC
TCATCGGACTCATCAGGAAGC
ATCGGACTCATCAGGAAGC
CGGACTCATCAGGAAGC
TCGGACTCATCAGGAAGC
GCATCGGACTCATCAGGAAGC
CATCGGACTCATCAGGAAGC
CATCGGACTCATCAGGAAGC
CATCGGACTCATCAGGAAGC
CCTCATCGGACTCATCAGGAAGC
TCGGACTCATCAGGAAGC
GTCATCGGACTCATCAGGAAGC
CCATCGGACTCATCAGGAAGC
ATCATCGGACTCATCAGGAAGC
CATCGGACTCATCAGGAAGC
CATCGGACTCATCAGGAAGC
CGGACTCATCAGGAAGC
TCCATCGGACTCATCAGGAAGC
CATCGGACTCATCAGGAAGC
CATCGGACTCATCAGGAAGC
ACATCGGACTCATCAGGAAGC
CATCGGACTCATCAGGAAGC
TCATCGGACTCATCAGGAAGC
ATCGGACTCATCAGGAAGC
CATCGGACTCATCAGGAAGC
ACATCGGACTCATCAGGAAGC
ACATCGGACTCATCAGGAAGC
TCCATCGGACTCATCAGGAAGC
CCATCGGACTCATCAGGAAGC
CCATCGGACTCATCAGGAAGC
CATCGGACTCATCAGGAAGC
CATCGGACTCATCAGGAAGC
CATCGGACTCATCAGGAAGC
AGCATCGGACTCATCAGGAAGC
CATCGGACTCATCAGGAAGC
TACATCGGACTCATCAGGAAGC
CATCGGACTCATCAGGAAGC
ATCGGACTCATCAGGAAGC
ATCGGACTCATCAGGAAGC
ATCGGACTCATCAGGAAGC
TCATCGGACTCATCAGGAAGC
ACTCATCGGACTCATCAGGAAGC
CGGACTCATCAGGAAGC

CATCGGACTCATCAGGAAGC
CATCGGACTCATCAGGAAGC
ACATCGGACTCATCAGGAAGC
CATCGGACTCATCAGGAAGC
CATCGGACTCATCAGGAAGC
ATCGGACTCATCAGGAAGC
CGGACTCATCAGGAAGC
CGGACTCATCAGGAAGC
CGGACTCATCAGGAAGC
TCGGACTCATCAGGAAGC
TCGGACTCATCAGGAAGC
CATCGGACTCATCAGGAAGC
CATCGGACTCATCAGGAAGC
GCATCGGACTCATCAGGAAGC
GAAGCATCGGACTCATCAGGAAGC
ACATCGGACTCATCAGGAAGC
GAAGCATCGGACTCATCAGGAAGC
TCATCGGACTCATCAGGAAGC
ACATCGGACTCATCAGGAAGC
CACATCGGACTCATCAGGAAGC
TCATCGGACTCATCAGGAAGC
GGACTCATCAGGAAGC
CGGACTCATCAGGAAGC
ACATCGGACTCATCAGGAAGC
CATCGGACTCATCAGGAAGC
CATCGGACTCATCAGGAAGC
CATCGGACTCATCAGGAAGC
CGGACTCATCAGGAAGC
CGGACTCATCAGGAAGC
GCCATCGGACTCATCAGGAAGC
TCGGACTCATCAGGAAGC
TCATCGGACTCATCAGGAAGC
TCCATCGGACTCATCAGGAAGC
CGGACTCATCAGGAAGC
ATCGGACTCATCAGGAAGC
ATCGGACTCATCAGGAAGC
ATCGGACTCATCAGGAAGC
ATCGGACTCATCAGGAAGC
CGGACTCATCAGGAAGC
CATCGGACTCATCAGGAAGC
CATCGGACTCATCAGGAAGC
TCATCGGACTCATCAGGAAGC
GCATCGGACTCATCAGGAAGC

GGCATCGGACTCATCAGGAAGC
ATCGGACTCATCAGGAAGC
CATCGGACTCATCAGGAAGC
GACTCATCAGGAAGC
ATCATCGGACTCATCAGGAAGC
TCATCGGACTCATCAGGAAGC
GAGCATCGGACTCATCAGGAAGC
GCTCATCGGACTCATCAGGAAGC
GCTCATCGGACTCATCAGGAAGC
ATCGGACTCATCAGGAAGC
TAACATCGGACTCATCAGGAAGC
TCGGACTCATCAGGAAGC
TCATCGGACTCATCAGGAAGC
CATCGGACTCATCAGGAAGC
TCGGACTCATCAGGAAGC
CATCGGACTCATCAGGAAGC
GCATCGGACTCATCAGGAAGC
CCATCGGACTCATCAGGAAGC
CCATCGGACTCATCAGGAAGC
CATCGGACTCATCAGGAAGC
CCATCGGACTCATCAGGAAGC
TCATCGGACTCATCAGGAAGC
TCATCGGACTCATCAGGAAGC
CATCGGACTCATCAGGAAGC
GTGCATCGGACTCATCAGGAAGC
CGGACTCATCAGGAAGC
ATCGGACTCATCAGGAAGC
CATCGGACTCATCAGGAAGC
ACATCGGACTCATCAGGAAGC
CATCGGACTCATCAGGAAGC
GCATCGGACTCATCAGGAAGC
GGACTCATCAGGAAGC
ATCGGACTCATCAGGAAGC
ATCGGACTCATCAGGAAGC
ATCGGACTCATCAGGAAGC
ATCGGACTCATCAGGAAGC
ATCGGACTCATCAGGAAGC
ATCGGACTCATCAGGAAGC
CATCGGACTCATCAGGAAGC
GGACTCATCAGGAAGC
CATCGGACTCATCAGGAAGC
GGACTCATCAGGAAGC
CATCGGACTCATCAGGAAGC

GCATCGGACTCATCAGGAAGC
ATCGGACTCATCAGGAAGC
TACATCGGACTCATCAGGAAGC
GGACTCATCAGGAAGC
CATCGGACTCATCAGGAAGC
CATCGGACTCATCAGGAAGC
CGGACTCATCAGGAAGC
GCATCGGACTCATCAGGAAGC
GCATCGGACTCATCAGGAAGC
TCGGACTCATCAGGAAGC
CATCGGACTCATCAGGAAGC
CATCGGACTCATCAGGAAGC
ACATCGGACTCATCAGGAAGC
GCATCGGACTCATCAGGAAGC
ATCGGACTCATCAGGAAGC
CATCGGACTCATCAGGAAGC
TGCATCGGACTCATCAGGAAGC
CATCGGACTCATCAGGAAGC
CATCGGACTCATCAGGAAGC
TCGGACTCATCAGGAAGC
CGGACTCATCAGGAAGC
ACATCGGACTCATCAGGAAGC
ACATCGGACTCATCAGGAAGC
CGGACTCATCAGGAAGC
CGGACTCATCAGGAAGC
CGGACTCATCAGGAAGC
ATCGGACTCATCAGGAAGC
TCGGACTCATCAGGAAGC
TCGGACTCATCAGGAAGC
CATCGGACTCATCAGGAAGC
CATCGGACTCATCAGGAAGC
GACTCATCAGGAAGC
GACTCATCAGGAAGC
TCGGACTCATCAGGAAGC
TCGGACTCATCAGGAAGC
AGCATCGGACTCATCAGGAAGC
GCATCGGACTCATCAGGAAGC
ATCGGACTCATCAGGAAGC
TCGGACTCATCAGGAAGC
TCGGACTCATCAGGAAGC
TCGGACTCATCAGGAAGC
TCGGACTCATCAGGAAGC
TACATCGGACTCATCAGGAAGC

TATCATCGGACTCATCAGGAAGC
TATCATCGGACTCATCAGGAAGC
TCATCGGACTCATCAGGAAGC
TCATCGGACTCATCAGGAAGC
CATCGGACTCATCAGGAAGC
TCATCGGACTCATCAGGAAGC
ATCGGACTCATCAGGAAGC
ATCGGACTCATCAGGAAGC
GCATCGGACTCATCAGGAAGC
GCATCGGACTCATCAGGAAGC
GCATCGGACTCATCAGGAAGC
ATCGGACTCATCAGGAAGC
CATCGGACTCATCAGGAAGC
ATCGGACTCATCAGGAAGC
TCGGACTCATCAGGAAGC
TCGGACTCATCAGGAAGC
GACTCATCAGGAAGC
CGGACTCATCAGGAAGC
GACTCATCAGGAAGC
GACTCATCAGGAAGC
GGACTCATCAGGAAGC
AACATCGGACTCATCAGGAAGC
ATCGGACTCATCAGGAAGC
ATCGGACTCATCAGGAAGC
CGGACTCATCAGGAAGC
CATCGGACTCATCAGGAAGC
CATCGGACTCATCAGGAAGC
CATCGGACTCATCAGGAAGC
ACATCGGACTCATCAGGAAGC
ACATCGGACTCATCAGGAAGC
GCATCGGACTCATCAGGAAGC
GCATCGGACTCATCAGGAAGC
ATCGGACTCATCAGGAAGC
TCGGACTCATCAGGAAGC
TCATCGGACTCATCAGGAAGC
TCATCGGACTCATCAGGAAGC
CATCGGACTCATCAGGAAGC
CATCGGACTCATCAGGAAGC
CGGACTCATCAGGAAGC
TCGGACTCATCAGGAAGC
CATCGGACTCATCAGGAAGC
TCGGACTCATCAGGAAGC
TCATCGGACTCATCAGGAAGC

TCATCGGACTCATCAGGAAGC
TCGGACTCATCAGGAAGC
GCATCGGACTCATCAGGAAGC
GCATCGGACTCATCAGGAAGC
CATCGGACTCATCAGGAAGC
ATCGGACTCATCAGGAAGC
TCGGACTCATCAGGAAGC
TCGGACTCATCAGGAAGC
CGGACTCATCAGGAAGC
ATCGGACTCATCAGGAAGC
ATCGGACTCATCAGGAAGC
GCATCGGACTCATCAGGAAGC
GCATCGGACTCATCAGGAAGC
GACATCGGACTCATCAGGAAGC
GACATCGGACTCATCAGGAAGC
CATCGGACTCATCAGGAAGC
CATCGGACTCATCAGGAAGC
ATCGGACTCATCAGGAAGC
TCGGACTCATCAGGAAGC
ATCGGACTCATCAGGAAGC
ATCGGACTCATCAGGAAGC
TCGGACTCATCAGGAAGC
ATCATCGGACTCATCAGGAAGC
GACTCATCAGGAAGC
CATCGGACTCATCAGGAAGC
CATCGGACTCATCAGGAAGC
ATCGGACTCATCAGGAAGC
CATCGGACTCATCAGGAAGC
CATCGGACTCATCAGGAAGC
GCCATCGGACTCATCAGGAAGC
GCCATCGGACTCATCAGGAAGC
GCATCGGACTCATCAGGAAGC
ATCATCGGACTCATCAGGAAGC
ACATCGGACTCATCAGGAAGC
ACATCGGACTCATCAGGAAGC
ACCATCGGACTCATCAGGAAGC
TCATCGGACTCATCAGGAAGC
TCATCGGACTCATCAGGAAGC
TCATCGGACTCATCAGGAAGC
CATCGGACTCATCAGGAAGC
TCATCGGACTCATCAGGAAGC
GACATCGGACTCATCAGGAAGC
CGGACTCATCAGGAAGC

AGCATCGGACTCATCAGGAAGC
TCGGACTCATCAGGAAGC
TCGGACTCATCAGGAAGC
CATCGGACTCATCAGGAAGC
ATCGGACTCATCAGGAAGC
CATCGGACTCATCAGGAAGC
TCGGACTCATCAGGAAGC
CCATCGGACTCATCAGGAAGC
GCATCGGACTCATCAGGAAGC
GCATCGGACTCATCAGGAAGC
TCATCGGACTCATCAGGAAGC
ATCGGACTCATCAGGAAGC
ATCGGACTCATCAGGAAGC
CATCGGACTCATCAGGAAGC
ACATCGGACTCATCAGGAAGC
CATCGGACTCATCAGGAAGC
CATCGGACTCATCAGGAAGC
CACATCGGACTCATCAGGAAGC
TCATCGGACTCATCAGGAAGC
CCTCATCGGACTCATCAGGAAGC
GCATCGGACTCATCAGGAAGC
TCATCGGACTCATCAGGAAGC
CATCGGACTCATCAGGAAGC
TGCATCGGACTCATCAGGAAGC
TGTCATCGGACTCATCAGGAAGC
TGTCATCGGACTCATCAGGAAGC
CATCGGACTCATCAGGAAGC
ATCGGACTCATCAGGAAGC
AACATCGGACTCATCAGGAAGC
AACATCGGACTCATCAGGAAGC
CATCGGACTCATCAGGAAGC
ACATCGGACTCATCAGGAAGC
CGGACTCATCAGGAAGC
CGGACTCATCAGGAAGC
CATCGGACTCATCAGGAAGC
CATCGGACTCATCAGGAAGC
GCACATCGGACTCATCAGGAAGC
TCGGACTCATCAGGAAGC
TCGGACTCATCAGGAAGC
TCGGACTCATCAGGAAGC
TCGGACTCATCAGGAAGC
GGACTCATCAGGAAGC
TCGGACTCATCAGGAAGC

TCGGACTCATCAGGAAGC
CGGACTCATCAGGAAGC
CGGACTCATCAGGAAGC
GGACTCATCAGGAAGC
TCATCGGACTCATCAGGAAGC
TCATCGGACTCATCAGGAAGC
CGGACTCATCAGGAAGC
GACTCATCAGGAAGC
ATCGGACTCATCAGGAAGC
GTGCATCGGACTCATCAGGAAGC
AGCATCGGACTCATCAGGAAGC
AGCATCGGACTCATCAGGAAGC
TACATCGGACTCATCAGGAAGC
TTCATCGGACTCATCAGGAAGC
TTCATCGGACTCATCAGGAAGC
ACATCGGACTCATCAGGAAGC
CATCGGACTCATCAGGAAGC
TCGGACTCATCAGGAAGC
ATCATCGGACTCATCAGGAAGC
ATCATCGGACTCATCAGGAAGC
CCATCGGACTCATCAGGAAGC
AACAGCATCGGACTCATCAGGAAGC
CATCGGACTCATCAGGAAGC
CATCGGACTCATCAGGAAGC
ATCCATCGGACTCATCAGGAAGC
CTCATCAGGAAGC
CTCATCAGGAAGC
GCATCGGACTCATCAGGAAGC
ATCGGACTCATCAGGAAGC
ATCGGACTCATCAGGAAGC
TGGCATCGGACTCATCAGGAAGC
TGGCATCGGACTCATCAGGAAGC
GAACATCGGACTCATCAGGAAGC
GAACATCGGACTCATCAGGAAGC
AGCATCGGACTCATCAGGAAGC
AAATCATCGGACTCATCAGGAAGC
AAATCATCGGACTCATCAGGAAGC
TCGGACTCATCAGGAAGC
TCGGACTCATCAGGAAGC
CATCGGACTCATCAGGAAGC
CATCGGACTCATCAGGAAGC
CATCGGACTCATCAGGAAGC
TCACATCGGACTCATCAGGAAGC

CATCGGACTCATCAGGAAGC
CATCGGACTCATCAGGAAGC
ACATCGGACTCATCAGGAAGC
TGCATCGGACTCATCAGGAAGC
CATCGGACTCATCAGGAAGC
CATCGGACTCATCAGGAAGC
ATCATCGGACTCATCAGGAAGC
ATCATCGGACTCATCAGGAAGC
CATCGGACTCATCAGGAAGC
GGACTCATCAGGAAGC
CATCGGACTCATCAGGAAGC
TCATCGGACTCATCAGGAAGC
TCGGACTCATCAGGAAGC
CATCGGACTCATCAGGAAGC
TCGGACTCATCAGGAAGC
TCGGACTCATCAGGAAGC
ATCGGACTCATCAGGAAGC
TCATCGGACTCATCAGGAAGC
TCATCGGACTCATCAGGAAGC
GCATCGGACTCATCAGGAAGC
GCATCGGACTCATCAGGAAGC
GCATCGGACTCATCAGGAAGC
TCGGACTCATCAGGAAGC
ATCGGACTCATCAGGAAGC
CCATCGGACTCATCAGGAAGC
CCATCGGACTCATCAGGAAGC
ATCATCGGACTCATCAGGAAGC
TACATCGGACTCATCAGGAAGC
TACATCGGACTCATCAGGAAGC
ACACATCGGACTCATCAGGAAGC
TCATCGGACTCATCAGGAAGC
TCATCGGACTCATCAGGAAGC
GACATCGGACTCATCAGGAAGC
GCATCGGACTCATCAGGAAGC
GCATCGGACTCATCAGGAAGC
TCCATCGGACTCATCAGGAAGC
GACATCGGACTCATCAGGAAGC
GCCATCGGACTCATCAGGAAGC
GCCATCGGACTCATCAGGAAGC
CACATCGGACTCATCAGGAAGC
AGCATCGGACTCATCAGGAAGC
AGCATCGGACTCATCAGGAAGC
GACTCATCAGGAAGC

ATCGGACTCATCAGGAAGC
ATCGGACTCATCAGGAAGC
ATCGGACTCATCAGGAAGC
ATCGGACTCATCAGGAAGC
CGGACTCATCAGGAAGC
ATCGGACTCATCAGGAAGC
CGGACTCATCAGGAAGC
GGACTCATCAGGAAGC
TCGGACTCATCAGGAAGC
CGGACTCATCAGGAAGC
GGACTCATCAGGAAGC
CGGACTCATCAGGAAGC
CGGACTCATCAGGAAGC
ATCGGACTCATCAGGAAGC
ATCGGACTCATCAGGAAGC
ATCGGACTCATCAGGAAGC
ATCGGACTCATCAGGAAGC
TCGGACTCATCAGGAAGC
ATCGGACTCATCAGGAAGC
ATCGGACTCATCAGGAAGC
CATCGGACTCATCAGGAAGC
CATCGGACTCATCAGGAAGC
TTCCATCGGACTCATCAGGAAGC
TTCCATCGGACTCATCAGGAAGC
CATCGGACTCATCAGGAAGC
CGGACTCATCAGGAAGC
ATCGGACTCATCAGGAAGC
ATCGGACTCATCAGGAAGC
TCGGACTCATCAGGAAGC
CCATCGGACTCATCAGGAAGC
CCATCGGACTCATCAGGAAGC
CCATCGGACTCATCAGGAAGC
CCATCGGACTCATCAGGAAGC
TCATCGGACTCATCAGGAAGC
GCATCGGACTCATCAGGAAGC
GCATCGGACTCATCAGGAAGC
TCATCGGACTCATCAGGAAGC
TCATCGGACTCATCAGGAAGC
ACATCGGACTCATCAGGAAGC
TCATCGGACTCATCAGGAAGC
ACATCGGACTCATCAGGAAGC
ACATCGGACTCATCAGGAAGC
AATCATCGGACTCATCAGGAAGC

ATCATCGGACTCATCAGGAAGC
AGCATCGGACTCATCAGGAAGC
AGCATCGGACTCATCAGGAAGC
GGACTCATCAGGAAGC
GACTCATCAGGAAGC
CATCGGACTCATCAGGAAGC
TTCATCGGACTCATCAGGAAGC
CATCGGACTCATCAGGAAGC
TCGGACTCATCAGGAAGC
TCGGACTCATCAGGAAGC
CATCGGACTCATCAGGAAGC
CATCGGACTCATCAGGAAGC
GCATCGGACTCATCAGGAAGC
GCATCGGACTCATCAGGAAGC
CATCGGACTCATCAGGAAGC
CATCGGACTCATCAGGAAGC
CATCGGACTCATCAGGAAGC
CATCGGACTCATCAGGAAGC
ACATCGGACTCATCAGGAAGC
ACATCGGACTCATCAGGAAGC
TCCATCGGACTCATCAGGAAGC
ATCGGACTCATCAGGAAGC
CGGACTCATCAGGAAGC
TCATCGGACTCATCAGGAAGC
CATCGGACTCATCAGGAAGC
TCATCGGACTCATCAGGAAGC
TCGGACTCATCAGGAAGC
TCGGACTCATCAGGAAGC
CATCGGACTCATCAGGAAGC
GCATCGGACTCATCAGGAAGC
TTCATCGGACTCATCAGGAAGC
CATCGGACTCATCAGGAAGC
CATCGGACTCATCAGGAAGC
GCATCGGACTCATCAGGAAGC
CATCGGACTCATCAGGAAGC
TCGGACTCATCAGGAAGC
CCATCGGACTCATCAGGAAGC
CATCGGACTCATCAGGAAGC
CATCGGACTCATCAGGAAGC
TAACATCGGACTCATCAGGAAGC
GTCCATCGGACTCATCAGGAAGC
CATCGGACTCATCAGGAAGC
CCATCGGACTCATCAGGAAGC

CATCGGACTCATCAGGAAGC
TCATCGGACTCATCAGGAAGC
TCGGACTCATCAGGAAGC
CATCGGACTCATCAGGAAGC
CCATCGGACTCATCAGGAAGC
CATCGGACTCATCAGGAAGC
CCATCGGACTCATCAGGAAGC
ATCGGACTCATCAGGAAGC
TCCATCGGACTCATCAGGAAGC
TCATCGGACTCATCAGGAAGC
TCATCGGACTCATCAGGAAGC
ATCGGACTCATCAGGAAGC
CATCGGACTCATCAGGAAGC
CCCATCGGACTCATCAGGAAGC
GAGCATCGGACTCATCAGGAAGC
CATCGGACTCATCAGGAAGC
TTACATCGGACTCATCAGGAAGC
TGACATCGGACTCATCAGGAAGC
AACATCGGACTCATCAGGAAGC
ATCGGACTCATCAGGAAGC
TCATCGGACTCATCAGGAAGC
ATCGGACTCATCAGGAAGC
GACTCATCAGGAAGC
CATCGGACTCATCAGGAAGC
CGGACTCATCAGGAAGC
TCGGACTCATCAGGAAGC
CATCGGACTCATCAGGAAGC
TCATCGGACTCATCAGGAAGC
GCATCGGACTCATCAGGAAGC
CGGACTCATCAGGAAGC
ATCGGACTCATCAGGAAGC
ATCGGACTCATCAGGAAGC
GAGCATCGGACTCATCAGGAAGC
ACATCGGACTCATCAGGAAGC
GGACTCATCAGGAAGC
ATCATCGGACTCATCAGGAAGC
ACCATCGGACTCATCAGGAAGC
GGACTCATCAGGAAGC
ACATCGGACTCATCAGGAAGC
CATCGGACTCATCAGGAAGC
CGGACTCATCAGGAAGC
ACATCGGACTCATCAGGAAGC
TCGGACTCATCAGGAAGC

TCGGACTCATCAGGAAGC
CGGACTCATCAGGAAGC
ATCGGACTCATCAGGAAGC
GACTCATCAGGAAGC
CCATCGGACTCATCAGGAAGC
TCATCGGACTCATCAGGAAGC
CATCGGACTCATCAGGAAGC
TCGGACTCATCAGGAAGC
ACATCGGACTCATCAGGAAGC
CATCGGACTCATCAGGAAGC
ATCGGACTCATCAGGAAGC
ATCGGACTCATCAGGAAGC
ACATCGGACTCATCAGGAAGC
GGACTCATCAGGAAGC
ACCATCGGACTCATCAGGAAGC
ACCATCGGACTCATCAGGAAGC
TACATCGGACTCATCAGGAAGC
TACATCGGACTCATCAGGAAGC
TCGGACTCATCAGGAAGC
ACATCGGACTCATCAGGAAGC
ATCATCGGACTCATCAGGAAGC
ACATCGGACTCATCAGGAAGC
CATCGGACTCATCAGGAAGC
TCATCGGACTCATCAGGAAGC
CGGACTCATCAGGAAGC
CATCGGACTCATCAGGAAGC
CATCGGACTCATCAGGAAGC
GCATCGGACTCATCAGGAAGC
CTCATCGGACTCATCAGGAAGC
ATCGGACTCATCAGGAAGC
TCATCGGACTCATCAGGAAGC
ACATCGGACTCATCAGGAAGC
CTCATCGGACTCATCAGGAAGC
AGCATCGGACTCATCAGGAAGC
TCATCGGACTCATCAGGAAGC
CACATCGGACTCATCAGGAAGC
TCGGACTCATCAGGAAGC
CACATCGGACTCATCAGGAAGC
AGTACATCGGACTCATCAGGAAGC
TCATCGGACTCATCAGGAAGC
CGGACTCATCAGGAAGC
CATCGGACTCATCAGGAAGC
ACCATCGGACTCATCAGGAAGC

GCATCGGACTCATCAGGAAGC
CAACATCGGACTCATCAGGAAGC
CGGACTCATCAGGAAGC
TGCATCGGACTCATCAGGAAGC
TCATCGGACTCATCAGGAAGC
GCATCGGACTCATCAGGAAGC
CATCGGACTCATCAGGAAGC
TCCATCGGACTCATCAGGAAGC
TCGGACTCATCAGGAAGC
TCATCGGACTCATCAGGAAGC
ATCGGACTCATCAGGAAGC
GCATCGGACTCATCAGGAAGC
TCGGACTCATCAGGAAGC
CACATCGGACTCATCAGGAAGC
CCATCGGACTCATCAGGAAGC
CATCGGACTCATCAGGAAGC
GCATCGGACTCATCAGGAAGC
TCATCGGACTCATCAGGAAGC
ACATCGGACTCATCAGGAAGC
TCGGACTCATCAGGAAGC
ATCGGACTCATCAGGAAGC
AGCATCGGACTCATCAGGAAGC
TCATCGGACTCATCAGGAAGC
ATCATCGGACTCATCAGGAAGC
TCGGACTCATCAGGAAGC
TCACATCGGACTCATCAGGAAGC
TCAGCATCGGACTCATCAGGAAGC
ACATCGGACTCATCAGGAAGC
ACATCGGACTCATCAGGAAGC
ATCGGACTCATCAGGAAGC
AATACATCGGACTCATCAGGAAGC
CATCGGACTCATCAGGAAGC
TCGGACTCATCAGGAAGC
TCGGACTCATCAGGAAGC
ATCGGACTCATCAGGAAGC
TCATCGGACTCATCAGGAAGC
AAGCATCGGACTCATCAGGAAGC
AAGCATCGGACTCATCAGGAAGC
GCCATCGGACTCATCAGGAAGC
CATCGGACTCATCAGGAAGC
TCATCGGACTCATCAGGAAGC
CATCGGACTCATCAGGAAGC
CATCGGACTCATCAGGAAGC

CATCGGACTCATCAGGAAGC
AACATCGGACTCATCAGGAAGC
AACATCGGACTCATCAGGAAGC
ATCGGACTCATCAGGAAGC
AATCATCGGACTCATCAGGAAGC
CATCGGACTCATCAGGAAGC
ATCGGACTCATCAGGAAGC
AACATCGGACTCATCAGGAAGC
GACTCATCAGGAAGC
ACATCGGACTCATCAGGAAGC
GGACTCATCAGGAAGC
CATCGGACTCATCAGGAAGC
CATCGGACTCATCAGGAAGC
TCGGACTCATCAGGAAGC
GACATCGGACTCATCAGGAAGC
TTCATCGGACTCATCAGGAAGC
GGACTCATCAGGAAGC
CATCGGACTCATCAGGAAGC
TGCATCGGACTCATCAGGAAGC
TCATCGGACTCATCAGGAAGC
CATCGGACTCATCAGGAAGC
CATCGGACTCATCAGGAAGC
ATCGGACTCATCAGGAAGC
CCATCGGACTCATCAGGAAGC
CGGACTCATCAGGAAGC
CATCGGACTCATCAGGAAGC
CGGACTCATCAGGAAGC
CCTCATCGGACTCATCAGGAAGC
TCATCGGACTCATCAGGAAGC
ACATCGGACTCATCAGGAAGC
GACTCATCAGGAAGC
ATCGGACTCATCAGGAAGC
ATCGGACTCATCAGGAAGC
ATCGGACTCATCAGGAAGC
CCATCGGACTCATCAGGAAGC
AACATCGGACTCATCAGGAAGC
GCATCGGACTCATCAGGAAGC
TGCATCGGACTCATCAGGAAGC
CATCGGACTCATCAGGAAGC
TAGCATCGGACTCATCAGGAAGC
ATCGGACTCATCAGGAAGC
ACCATCGGACTCATCAGGAAGC
TTCATCGGACTCATCAGGAAGC

GTCATCGGACTCATCAGGAAGC
ATCGGACTCATCAGGAAGC
TCGGACTCATCAGGAAGC
CATCGGACTCATCAGGAAGC
ACCATCGGACTCATCAGGAAGC
GCATCGGACTCATCAGGAAGC
TCACATCGGACTCATCAGGAAGC
CATCGGACTCATCAGGAAGC
CATCGGACTCATCAGGAAGC
AGACATCGGACTCATCAGGAAGC
CGGACTCATCAGGAAGC
AACCATCGGACTCATCAGGAAGC
GCCATCGGACTCATCAGGAAGC
ACATCGGACTCATCAGGAAGC
ATCATCGGACTCATCAGGAAGC
ATCGGACTCATCAGGAAGC
GGCATCGGACTCATCAGGAAGC
TCACATCGGACTCATCAGGAAGC
GCATCGGACTCATCAGGAAGC
CGGACTCATCAGGAAGC
ACATCGGACTCATCAGGAAGC
CCATCGGACTCATCAGGAAGC
CATCGGACTCATCAGGAAGC
ACATCGGACTCATCAGGAAGC
ATCGGACTCATCAGGAAGC
CATCGGACTCATCAGGAAGC
CTCATCGGACTCATCAGGAAGC
GCCATCGGACTCATCAGGAAGC
GCATCGGACTCATCAGGAAGC
TCATCGGACTCATCAGGAAGC
TACATCGGACTCATCAGGAAGC
TGCATCGGACTCATCAGGAAGC
TGCATCGGACTCATCAGGAAGC
GTCATCGGACTCATCAGGAAGC
ACATCGGACTCATCAGGAAGC
GTCATCGGACTCATCAGGAAGC
ATCGGACTCATCAGGAAGC
TCATCGGACTCATCAGGAAGC
CTCATCGGACTCATCAGGAAGC
ACATCGGACTCATCAGGAAGC
GTTCATCGGACTCATCAGGAAGC
AAGCATCGGACTCATCAGGAAGC
CCATCGGACTCATCAGGAAGC

AGCATCGGACTCATCAGGAAGC
CATCGGACTCATCAGGAAGC
GACATCGGACTCATCAGGAAGC
CGGACTCATCAGGAAGC
TAGTCATCGGACTCATCAGGAAGC
ACATCGGACTCATCAGGAAGC
CATCGGACTCATCAGGAAGC
TCATCGGACTCATCAGGAAGC
CATCGGACTCATCAGGAAGC
TCGGACTCATCAGGAAGC
TACATCGGACTCATCAGGAAGC
GCATCGGACTCATCAGGAAGC
CATCGGACTCATCAGGAAGC
CATCGGACTCATCAGGAAGC
ACATCGGACTCATCAGGAAGC
TCATCGGACTCATCAGGAAGC
ACATCGGACTCATCAGGAAGC
CGGACTCATCAGGAAGC
CGGACTCATCAGGAAGC
CATCGGACTCATCAGGAAGC
GTCCATCGGACTCATCAGGAAGC
CATCGGACTCATCAGGAAGC
CATCGGACTCATCAGGAAGC
CGGACTCATCAGGAAGC
TCGGACTCATCAGGAAGC
ATCGGACTCATCAGGAAGC
ATCGGACTCATCAGGAAGC
CCATCGGACTCATCAGGAAGC
CATCGGACTCATCAGGAAGC
GGACTCATCAGGAAGC
TTGCATCGGACTCATCAGGAAGC
ACATCGGACTCATCAGGAAGC
CATCGGACTCATCAGGAAGC
CATCGGACTCATCAGGAAGC
GACTCATCAGGAAGC
CGGACTCATCAGGAAGC
ATCGGACTCATCAGGAAGC
ACATCGGACTCATCAGGAAGC
ACTCATCAGGAAGC
ACTCATCAGGAAGC
ATCGGACTCATCAGGAAGC
TCGGACTCATCAGGAAGC
CACATCGGACTCATCAGGAAGC

TCATCGGACTCATCAGGAAGC
TCATCGGACTCATCAGGAAGC
CATCGGACTCATCAGGAAGC
ATCGGACTCATCAGGAAGC
ATCGGACTCATCAGGAAGC
ATCGGACTCATCAGGAAGC
TTCCATCGGACTCATCAGGAAGC
GTCATCGGACTCATCAGGAAGC
ATCGGACTCATCAGGAAGC
CATCGGACTCATCAGGAAGC
TCGGACTCATCAGGAAGC
TCATCGGACTCATCAGGAAGC
ATCGGACTCATCAGGAAGC
ATCGGACTCATCAGGAAGC
TCATCGGACTCATCAGGAAGC
TCATCGGACTCATCAGGAAGC
ATCGGACTCATCAGGAAGC
ATCGGACTCATCAGGAAGC
GCATCGGACTCATCAGGAAGC
CATCGGACTCATCAGGAAGC
CATCGGACTCATCAGGAAGC
CGGACTCATCAGGAAGC
ATCGGACTCATCAGGAAGC
CCATCGGACTCATCAGGAAGC
TACATCGGACTCATCAGGAAGC
TACATCGGACTCATCAGGAAGC
TACATCGGACTCATCAGGAAGC
TACATCGGACTCATCAGGAAGC
CGGACTCATCAGGAAGC
GGACTCATCAGGAAGC
CCATCGGACTCATCAGGAAGC
ACATCGGACTCATCAGGAAGC
ATCGGACTCATCAGGAAGC
GCATCGGACTCATCAGGAAGC
CCATCGGACTCATCAGGAAGC
AGCACATCGGACTCATCAGGAAGC
CATCGGACTCATCAGGAAGC
TCATCGGACTCATCAGGAAGC
GACATCGGACTCATCAGGAAGC
CATCGGACTCATCAGGAAGC
GCATCGGACTCATCAGGAAGC
CGGACTCATCAGGAAGC
ATCATCGGACTCATCAGGAAGC

TACATCGGACTCATCAGGAAGC
ATCATCGGACTCATCAGGAAGC
ATCGGACTCATCAGGAAGC
ACCATCGGACTCATCAGGAAGC
AGCATCGGACTCATCAGGAAGC
GACATCGGACTCATCAGGAAGC
TACATCGGACTCATCAGGAAGC
ACATCGGACTCATCAGGAAGC
GGCATCGGACTCATCAGGAAGC
TCATCGGACTCATCAGGAAGC
ACACATCGGACTCATCAGGAAGC
ATCATCGGACTCATCAGGAAGC
CATCGGACTCATCAGGAAGC
CTCATCAGGAAGC
CATCGGACTCATCAGGAAGC
CATCGGACTCATCAGGAAGC
CATCGGACTCATCAGGAAGC
CATCGGACTCATCAGGAAGC
TCCATCGGACTCATCAGGAAGC
TCCATCGGACTCATCAGGAAGC
ATCATCGGACTCATCAGGAAGC
TCGGACTCATCAGGAAGC
AATCATCGGACTCATCAGGAAGC
ACATCGGACTCATCAGGAAGC
ACATCGGACTCATCAGGAAGC
TGCATCGGACTCATCAGGAAGC
CCATCGGACTCATCAGGAAGC
CATCGGACTCATCAGGAAGC
ATCGGACTCATCAGGAAGC
CCATCGGACTCATCAGGAAGC
CGGACTCATCAGGAAGC
TGTCATCGGACTCATCAGGAAGC
GGACTCATCAGGAAGC
CATCGGACTCATCAGGAAGC
ACATCGGACTCATCAGGAAGC
AACATCGGACTCATCAGGAAGC
TGCATCGGACTCATCAGGAAGC
CATCGGACTCATCAGGAAGC
TCATCGGACTCATCAGGAAGC
CCATCGGACTCATCAGGAAGC
ACATCGGACTCATCAGGAAGC
AACATCGGACTCATCAGGAAGC
ATCGGACTCATCAGGAAGC

TCATCGGACTCATCAGGAAGC
ATCGGACTCATCAGGAAGC
ATCGGACTCATCAGGAAGC
GCATCGGACTCATCAGGAAGC
CATCGGACTCATCAGGAAGC
ATCGGACTCATCAGGAAGC
ATCATCGGACTCATCAGGAAGC
AAAGCATCGGACTCATCAGGAAGC
CCATCGGACTCATCAGGAAGC
ACATCGGACTCATCAGGAAGC
ATCGGACTCATCAGGAAGC
TTCATCGGACTCATCAGGAAGC
TCGGACTCATCAGGAAGC
ACATCGGACTCATCAGGAAGC
CATCGGACTCATCAGGAAGC
TCGGACTCATCAGGAAGC
GCATCGGACTCATCAGGAAGC
ACATCGGACTCATCAGGAAGC
ATCGGACTCATCAGGAAGC
TCATCGGACTCATCAGGAAGC
TCATCGGACTCATCAGGAAGC
TCATCGGACTCATCAGGAAGC
TCATCGGACTCATCAGGAAGC
ATCGGACTCATCAGGAAGC
AAACATCGGACTCATCAGGAAGC
ACATCGGACTCATCAGGAAGC
CCATCGGACTCATCAGGAAGC
CCATCGGACTCATCAGGAAGC
ACATCGGACTCATCAGGAAGC
TCCATCGGACTCATCAGGAAGC
TAACATCGGACTCATCAGGAAGC
GGACTCATCAGGAAGC
AAAACATCGGACTCATCAGGAAGC
CATCGGACTCATCAGGAAGC
TGTCATCGGACTCATCAGGAAGC
CGGACTCATCAGGAAGC
GGACTCATCAGGAAGC
CATCGGACTCATCAGGAAGC
CCATCGGACTCATCAGGAAGC
CATCGGACTCATCAGGAAGC
ACATCGGACTCATCAGGAAGC
CTGCATCGGACTCATCAGGAAGC
CGGACTCATCAGGAAGC

ATCGGACTCATCAGGAAGC
ACACATCGGACTCATCAGGAAGC
ATCGGACTCATCAGGAAGC
ATCGGACTCATCAGGAAGC
ATCGGACTCATCAGGAAGC
AGGCATCGGACTCATCAGGAAGC
TCATCGGACTCATCAGGAAGC
ATCGGACTCATCAGGAAGC
AGCATCGGACTCATCAGGAAGC
AGCATCGGACTCATCAGGAAGC
AGCATCGGACTCATCAGGAAGC
ATCGGACTCATCAGGAAGC
GGACTCATCAGGAAGC
CATCGGACTCATCAGGAAGC
GATCATCGGACTCATCAGGAAGC
CGGACTCATCAGGAAGC
CATCGGACTCATCAGGAAGC
ATCGGACTCATCAGGAAGC
CATCGGACTCATCAGGAAGC
AAACATCGGACTCATCAGGAAGC
CCATCGGACTCATCAGGAAGC
CGGACTCATCAGGAAGC
TCATCGGACTCATCAGGAAGC
CATCGGACTCATCAGGAAGC
CACATCGGACTCATCAGGAAGC
CACATCGGACTCATCAGGAAGC
ATCGGACTCATCAGGAAGC
TCATCGGACTCATCAGGAAGC
CATCGGACTCATCAGGAAGC
TCATCGGACTCATCAGGAAGC
ACCATCGGACTCATCAGGAAGC
GCATCGGACTCATCAGGAAGC
AACATCGGACTCATCAGGAAGC
GGACTCATCAGGAAGC
CCATCGGACTCATCAGGAAGC
ATCGGACTCATCAGGAAGC
TCGGACTCATCAGGAAGC
ACATCGGACTCATCAGGAAGC
ACATCGGACTCATCAGGAAGC
TAACATCGGACTCATCAGGAAGC
GTCATCGGACTCATCAGGAAGC
GTCATCGGACTCATCAGGAAGC
CATCGGACTCATCAGGAAGC

GGCATCGGACTCATCAGGAAGC
ATTCATCGGACTCATCAGGAAGC
TTCATCGGACTCATCAGGAAGC
ATCGGACTCATCAGGAAGC
CATCGGACTCATCAGGAAGC
ATCATCGGACTCATCAGGAAGC
TCGGACTCATCAGGAAGC
TCATCGGACTCATCAGGAAGC
TGCATCGGACTCATCAGGAAGC
TCGGACTCATCAGGAAGC
AGCATCGGACTCATCAGGAAGC
ATCGGACTCATCAGGAAGC
CATGCATCGGACTCATCAGGAAGC
TCATCGGACTCATCAGGAAGC
TCATCGGACTCATCAGGAAGC
GGACTCATCAGGAAGC
TCATCGGACTCATCAGGAAGC
TCATCGGACTCATCAGGAAGC
CGGACTCATCAGGAAGC
GACATCGGACTCATCAGGAAGC
CCATCGGACTCATCAGGAAGC
TGCATCGGACTCATCAGGAAGC
CATCGGACTCATCAGGAAGC
ATCGGACTCATCAGGAAGC
TCATCGGACTCATCAGGAAGC
CATCGGACTCATCAGGAAGC
CGGACTCATCAGGAAGC
TCATCGGACTCATCAGGAAGC
CGGACTCATCAGGAAGC
CTCATCGGACTCATCAGGAAGC
ACATCGGACTCATCAGGAAGC
CACATCGGACTCATCAGGAAGC
ACATCGGACTCATCAGGAAGC
CATCGGACTCATCAGGAAGC
TACATCGGACTCATCAGGAAGC
CCATCGGACTCATCAGGAAGC
TCATCGGACTCATCAGGAAGC
GCATCGGACTCATCAGGAAGC
AACATCGGACTCATCAGGAAGC
ACCATCGGACTCATCAGGAAGC
ATTCATCGGACTCATCAGGAAGC
ATACATCGGACTCATCAGGAAGC
TCCATCGGACTCATCAGGAAGC

ACATCGGACTCATCAGGAAGC
TCGGACTCATCAGGAAGC
ACATCGGACTCATCAGGAAGC
ATCGGACTCATCAGGAAGC
TACATCGGACTCATCAGGAAGC
CATCGGACTCATCAGGAAGC
TGCATCGGACTCATCAGGAAGC
TTCATCGGACTCATCAGGAAGC
CTCATCGGACTCATCAGGAAGC
ATCGGACTCATCAGGAAGC
ACATCGGACTCATCAGGAAGC
ACTCATCGGACTCATCAGGAAGC
ATGCATCGGACTCATCAGGAAGC
GTCATCGGACTCATCAGGAAGC
ATCGGACTCATCAGGAAGC
ATGCACATCGGACTCATCAGGAAGC
CATCGGACTCATCAGGAAGC
CATCGGACTCATCAGGAAGC
ACATCGGACTCATCAGGAAGC
CATCGGACTCATCAGGAAGC
CATCGGACTCATCAGGAAGC
TGCATCGGACTCATCAGGAAGC
GCCATCGGACTCATCAGGAAGC
ATCGGACTCATCAGGAAGC
ATCATCGGACTCATCAGGAAGC
TGCATCGGACTCATCAGGAAGC
CATCGGACTCATCAGGAAGC
CGGACTCATCAGGAAGC
GTCATCGGACTCATCAGGAAGC
GACTCATCAGGAAGC
CACCATCGGACTCATCAGGAAGC
ATCGGACTCATCAGGAAGC
ACATCGGACTCATCAGGAAGC
ACATCGGACTCATCAGGAAGC
ATCATCGGACTCATCAGGAAGC
TCATCGGACTCATCAGGAAGC
ATCATCGGACTCATCAGGAAGC
TCGGACTCATCAGGAAGC
TACATCGGACTCATCAGGAAGC
GCATCGGACTCATCAGGAAGC
ACATCGGACTCATCAGGAAGC
ATCGGACTCATCAGGAAGC
TCATCGGACTCATCAGGAAGC

CAACATCGGACTCATCAGGAAGC
CATCGGACTCATCAGGAAGC
TCATCGGACTCATCAGGAAGC
CATCGGACTCATCAGGAAGC
CATCGGACTCATCAGGAAGC
ATCGGACTCATCAGGAAGC
AACATCGGACTCATCAGGAAGC
GTCATCGGACTCATCAGGAAGC
GTCATCGGACTCATCAGGAAGC
ACATCGGACTCATCAGGAAGC
CATCGGACTCATCAGGAAGC
CGGACTCATCAGGAAGC
GCATCGGACTCATCAGGAAGC
ACATCGGACTCATCAGGAAGC
TCGGACTCATCAGGAAGC
TCGGACTCATCAGGAAGC
TCCATCGGACTCATCAGGAAGC
TCATCGGACTCATCAGGAAGC
TCCATCGGACTCATCAGGAAGC
CATCGGACTCATCAGGAAGC
CATCGGACTCATCAGGAAGC
ACTCATCAGGAAGC
CATCGGACTCATCAGGAAGC
CATCGGACTCATCAGGAAGC
ACATCGGACTCATCAGGAAGC
ATCGGACTCATCAGGAAGC
GCATCGGACTCATCAGGAAGC
CATCGGACTCATCAGGAAGC
TCATCGGACTCATCAGGAAGC
CATCGGACTCATCAGGAAGC
TACATCGGACTCATCAGGAAGC
CATCGGACTCATCAGGAAGC
AACATCGGACTCATCAGGAAGC
ATCGGACTCATCAGGAAGC
AGGCATCGGACTCATCAGGAAGC
ATCGGACTCATCAGGAAGC
GACATCGGACTCATCAGGAAGC
CATCGGACTCATCAGGAAGC
CACCATCGGACTCATCAGGAAGC
ATCATCGGACTCATCAGGAAGC
GACATCGGACTCATCAGGAAGC
ATCGGACTCATCAGGAAGC
CATCGGACTCATCAGGAAGC

TCGGACTCATCAGGAAGC
ATGCATCGGACTCATCAGGAAGC
CCATCGGACTCATCAGGAAGC
ACATCGGACTCATCAGGAAGC
TCGGACTCATCAGGAAGC
CATCGGACTCATCAGGAAGC
CACCATCGGACTCATCAGGAAGC
CACCATCGGACTCATCAGGAAGC
CACCATCGGACTCATCAGGAAGC
CATCGGACTCATCAGGAAGC
GCATCGGACTCATCAGGAAGC
GCATCGGACTCATCAGGAAGC
GCCATCGGACTCATCAGGAAGC
CATCGGACTCATCAGGAAGC
CATCGGACTCATCAGGAAGC
ACATCGGACTCATCAGGAAGC
TCCATCGGACTCATCAGGAAGC
CATCGGACTCATCAGGAAGC
CATCGGACTCATCAGGAAGC
CATCGGACTCATCAGGAAGC
AGGCATCGGACTCATCAGGAAGC
GCATCGGACTCATCAGGAAGC
TCATCGGACTCATCAGGAAGC
GCATCGGACTCATCAGGAAGC
AGCATCGGACTCATCAGGAAGC
ATCGGACTCATCAGGAAGC
ATCGGACTCATCAGGAAGC
CATCGGACTCATCAGGAAGC
AGCATCGGACTCATCAGGAAGC
TTCATCGGACTCATCAGGAAGC
ACATCGGACTCATCAGGAAGC
ACATCGGACTCATCAGGAAGC
ACATCGGACTCATCAGGAAGC
GTTGCATCGGACTCATCAGGAAGC
CATCGGACTCATCAGGAAGC
ACTGCATCGGACTCATCAGGAAGC
GCCATCGGACTCATCAGGAAGC
TCCATCGGACTCATCAGGAAGC
ACATCGGACTCATCAGGAAGC
ACCATCGGACTCATCAGGAAGC
GCCATCGGACTCATCAGGAAGC
TTACATCGGACTCATCAGGAAGC
GCATCGGACTCATCAGGAAGC

AACATCGGACTCATCAGGAAGC
GGCATCGGACTCATCAGGAAGC
CATCGGACTCATCAGGAAGC
AACATCGGACTCATCAGGAAGC
ATTCATCGGACTCATCAGGAAGC
CATCGGACTCATCAGGAAGC
TCGGACTCATCAGGAAGC
TACATCGGACTCATCAGGAAGC
AGGCATCGGACTCATCAGGAAGC
TGCATCGGACTCATCAGGAAGC
ATCGGACTCATCAGGAAGC
TCATCGGACTCATCAGGAAGC
CATCGGACTCATCAGGAAGC
CGGACTCATCAGGAAGC
TCGGACTCATCAGGAAGC
TACATCGGACTCATCAGGAAGC
TCATCGGACTCATCAGGAAGC
TCATCGGACTCATCAGGAAGC
CCATCGGACTCATCAGGAAGC
ATCGGACTCATCAGGAAGC
CATCGGACTCATCAGGAAGC
GCATCGGACTCATCAGGAAGC
CCATCGGACTCATCAGGAAGC
ACTCATCGGACTCATCAGGAAGC
TTGCATCGGACTCATCAGGAAGC
CATCGGACTCATCAGGAAGC
CATCGGACTCATCAGGAAGC
CATCGGACTCATCAGGAAGC
GAGCATCGGACTCATCAGGAAGC
TAAGCATCGGACTCATCAGGAAGC
CCATCGGACTCATCAGGAAGC
TCGGACTCATCAGGAAGC
CCATCGGACTCATCAGGAAGC
ATCATCGGACTCATCAGGAAGC
TCATCGGACTCATCAGGAAGC
TGCATCGGACTCATCAGGAAGC
ATCGGACTCATCAGGAAGC
ATCGGACTCATCAGGAAGC
CATCGGACTCATCAGGAAGC
CATCGGACTCATCAGGAAGC
TCATCGGACTCATCAGGAAGC
ACTCATCAGGAAGC
CCATCGGACTCATCAGGAAGC

CATCGGACTCATCAGGAAGC
CATCGGACTCATCAGGAAGC
GGACTCATCAGGAAGC
CGGACTCATCAGGAAGC
CATCGGACTCATCAGGAAGC
GCATCGGACTCATCAGGAAGC
GCATCGGACTCATCAGGAAGC
GCATCGGACTCATCAGGAAGC
TCGGACTCATCAGGAAGC
CATCGGACTCATCAGGAAGC
ATCATCGGACTCATCAGGAAGC
TACATCGGACTCATCAGGAAGC
CGGACTCATCAGGAAGC
CATCGGACTCATCAGGAAGC
TTCATCGGACTCATCAGGAAGC
ACATCGGACTCATCAGGAAGC
ATCATCGGACTCATCAGGAAGC
ATCATCGGACTCATCAGGAAGC
CGGACTCATCAGGAAGC
AGTCATCGGACTCATCAGGAAGC
GACATCGGACTCATCAGGAAGC
ACATCGGACTCATCAGGAAGC
ACATCGGACTCATCAGGAAGC
ACATCGGACTCATCAGGAAGC
CCATCGGACTCATCAGGAAGC
GACTCATCAGGAAGC
ATCGGACTCATCAGGAAGC
TGGCATCGGACTCATCAGGAAGC
GCATCGGACTCATCAGGAAGC
TTCATCGGACTCATCAGGAAGC
GCATCGGACTCATCAGGAAGC
CATCGGACTCATCAGGAAGC
ACATCGGACTCATCAGGAAGC
GTCATCGGACTCATCAGGAAGC
GGCATCGGACTCATCAGGAAGC
ATCGGACTCATCAGGAAGC
TCGGACTCATCAGGAAGC
CCATCGGACTCATCAGGAAGC
CATCGGACTCATCAGGAAGC
CATCGGACTCATCAGGAAGC
GCATCGGACTCATCAGGAAGC
ACATCGGACTCATCAGGAAGC
ACATCGGACTCATCAGGAAGC

ACATCGGACTCATCAGGAAGC
CATCGGACTCATCAGGAAGC
GCATCGGACTCATCAGGAAGC
ACATCGGACTCATCAGGAAGC
CATCGGACTCATCAGGAAGC
TCATCGGACTCATCAGGAAGC
CCATCGGACTCATCAGGAAGC
TCATCGGACTCATCAGGAAGC
CATCGGACTCATCAGGAAGC
GCATCGGACTCATCAGGAAGC
GCATCGGACTCATCAGGAAGC
ACCATCGGACTCATCAGGAAGC
CATCGGACTCATCAGGAAGC
AGGCATCGGACTCATCAGGAAGC
ATCGGACTCATCAGGAAGC
ACATCGGACTCATCAGGAAGC
ACTCATCAGGAAGC
CATCGGACTCATCAGGAAGC
CGGACTCATCAGGAAGC
TCGGACTCATCAGGAAGC
ATCATCGGACTCATCAGGAAGC
GGACTCATCAGGAAGC
TCCATCGGACTCATCAGGAAGC
TCGGACTCATCAGGAAGC
GGACTCATCAGGAAGC
ATCGGACTCATCAGGAAGC
TGACATCGGACTCATCAGGAAGC
ATCGGACTCATCAGGAAGC
ATCGGACTCATCAGGAAGC
ATCGGACTCATCAGGAAGC
CATCGGACTCATCAGGAAGC
CATCGGACTCATCAGGAAGC
TCATCGGACTCATCAGGAAGC
CGGACTCATCAGGAAGC
ATCATCGGACTCATCAGGAAGC
TCATCGGACTCATCAGGAAGC
CCATCGGACTCATCAGGAAGC
GGACTCATCAGGAAGC
ATCGGACTCATCAGGAAGC
GCATCGGACTCATCAGGAAGC
TCGGACTCATCAGGAAGC
CATCGGACTCATCAGGAAGC
CGGACTCATCAGGAAGC

CATCGGACTCATCAGGAAGC
CATCGGACTCATCAGGAAGC
TCGGACTCATCAGGAAGC
CATCGGACTCATCAGGAAGC
CATCGGACTCATCAGGAAGC
ACATCGGACTCATCAGGAAGC
TCGGACTCATCAGGAAGC
CGGACTCATCAGGAAGC
CTGCATCGGACTCATCAGGAAGC
CATCGGACTCATCAGGAAGC
AGCATCGGACTCATCAGGAAGC
CATCGGACTCATCAGGAAGC
ATCGGACTCATCAGGAAGC
ATCGGACTCATCAGGAAGC
ATCGGACTCATCAGGAAGC
TCGGACTCATCAGGAAGC
GCATCGGACTCATCAGGAAGC
CATCGGACTCATCAGGAAGC
TCGGACTCATCAGGAAGC
TCATCGGACTCATCAGGAAGC
GCATCGGACTCATCAGGAAGC
CATCGGACTCATCAGGAAGC
CATCGGACTCATCAGGAAGC
TCGGACTCATCAGGAAGC
TCGGACTCATCAGGAAGC
TCGGACTCATCAGGAAGC
TCGGACTCATCAGGAAGC
CATCGGACTCATCAGGAAGC
TCGGACTCATCAGGAAGC
TCGGACTCATCAGGAAGC
TCGGACTCATCAGGAAGC
ATCGGACTCATCAGGAAGC
TCGGACTCATCAGGAAGC
GCATCGGACTCATCAGGAAGC
GGACTCATCAGGAAGC
GGACTCATCAGGAAGC
TCGGACTCATCAGGAAGC
TCGGACTCATCAGGAAGC
AGCATCGGACTCATCAGGAAGC
AACATCGGACTCATCAGGAAGC
ACATCGGACTCATCAGGAAGC
CATCGGACTCATCAGGAAGC
TCGGACTCATCAGGAAGC

GGACTCATCAGGAAGC
TCGGACTCATCAGGAAGC
ACATCGGACTCATCAGGAAGC
CATCGGACTCATCAGGAAGC
TTCATCGGACTCATCAGGAAGC
AGGCATCGGACTCATCAGGAAGC
AGGCATCGGACTCATCAGGAAGC
AGCATCGGACTCATCAGGAAGC
ATGCATCGGACTCATCAGGAAGC
AGGCATCGGACTCATCAGGAAGC
GCATCGGACTCATCAGGAAGC
GACTCATCAGGAAGC
AGCATCGGACTCATCAGGAAGC
TCTACATCGGACTCATCAGGAAGC
TCGGACTCATCAGGAAGC
ATCGGACTCATCAGGAAGC
ATCGGACTCATCAGGAAGC
ACATCGGACTCATCAGGAAGC
ACATCGGACTCATCAGGAAGC
TCCATCGGACTCATCAGGAAGC
CTCATCGGACTCATCAGGAAGC
CCATCGGACTCATCAGGAAGC
CATCGGACTCATCAGGAAGC
ATCGGACTCATCAGGAAGC
CATCGGACTCATCAGGAAGC
CATCGGACTCATCAGGAAGC
ACCCATCGGACTCATCAGGAAGC
ACCCATCGGACTCATCAGGAAGC
TCGGACTCATCAGGAAGC
ATTCATCGGACTCATCAGGAAGC
TGGCCATCGGACTCATCAGGAAGC
ACCATCGGACTCATCAGGAAGC
GACATCGGACTCATCAGGAAGC
ATCGGACTCATCAGGAAGC
ATCGGACTCATCAGGAAGC
ATCGGACTCATCAGGAAGC
ATCGGACTCATCAGGAAGC
GCATCGGACTCATCAGGAAGC
TCATCGGACTCATCAGGAAGC
TCATCGGACTCATCAGGAAGC
AACATCGGACTCATCAGGAAGC
CATCGGACTCATCAGGAAGC
ATCGGACTCATCAGGAAGC

CATCGGACTCATCAGGAAGC
ATCGGACTCATCAGGAAGC
TCCATCGGACTCATCAGGAAGC
CATCGGACTCATCAGGAAGC
TCATCGGACTCATCAGGAAGC
CCATCGGACTCATCAGGAAGC
CGGACTCATCAGGAAGC
TCATCGGACTCATCAGGAAGC
GACATCGGACTCATCAGGAAGC
GACATCGGACTCATCAGGAAGC
ACATCGGACTCATCAGGAAGC
ACATCGGACTCATCAGGAAGC
TCATCGGACTCATCAGGAAGC
CATCGGACTCATCAGGAAGC
CATCGGACTCATCAGGAAGC
CATCGGACTCATCAGGAAGC
TCATCGGACTCATCAGGAAGC
TCATCGGACTCATCAGGAAGC
CATCGGACTCATCAGGAAGC
CATCGGACTCATCAGGAAGC
GCATCGGACTCATCAGGAAGC
ATCGGACTCATCAGGAAGC
TCCATCGGACTCATCAGGAAGC
ACATCGGACTCATCAGGAAGC
AACCATCGGACTCATCAGGAAGC
TACATCGGACTCATCAGGAAGC
CATCGGACTCATCAGGAAGC
TTCATCGGACTCATCAGGAAGC
ATCATCGGACTCATCAGGAAGC
CGGACTCATCAGGAAGC
CGGACTCATCAGGAAGC
CGGACTCATCAGGAAGC
ATCGGACTCATCAGGAAGC
GCATCGGACTCATCAGGAAGC
TGCATCGGACTCATCAGGAAGC
GGCATCGGACTCATCAGGAAGC
TGCATCGGACTCATCAGGAAGC
TGCATCGGACTCATCAGGAAGC
GCATCGGACTCATCAGGAAGC
CGGACTCATCAGGAAGC
TATCATCGGACTCATCAGGAAGC
ACATCGGACTCATCAGGAAGC
TCATCGGACTCATCAGGAAGC

ATCATCGGACTCATCAGGAAGC
GCATCGGACTCATCAGGAAGC
GCATCGGACTCATCAGGAAGC
TCGGACTCATCAGGAAGC
GGACTCATCAGGAAGC
GGACTCATCAGGAAGC
TACATCGGACTCATCAGGAAGC
CCCATCGGACTCATCAGGAAGC
AGTCATCGGACTCATCAGGAAGC
AGTCATCGGACTCATCAGGAAGC
ATCGGACTCATCAGGAAGC
CCATCGGACTCATCAGGAAGC
TCATCGGACTCATCAGGAAGC
CCATCGGACTCATCAGGAAGC
TCATCGGACTCATCAGGAAGC
ACATCGGACTCATCAGGAAGC
CATCGGACTCATCAGGAAGC
TCATCGGACTCATCAGGAAGC
TCATCGGACTCATCAGGAAGC
TCATCGGACTCATCAGGAAGC
CATCGGACTCATCAGGAAGC
ACATCGGACTCATCAGGAAGC
ACATCGGACTCATCAGGAAGC
ATCATCGGACTCATCAGGAAGC
GTTCCATCGGACTCATCAGGAAGC
CATCGGACTCATCAGGAAGC
CGGACTCATCAGGAAGC
CATCGGACTCATCAGGAAGC
TCGGACTCATCAGGAAGC
ACATCGGACTCATCAGGAAGC
ACATCGGACTCATCAGGAAGC
TACATCGGACTCATCAGGAAGC
TAGACATCGGACTCATCAGGAAGC
ATCGGACTCATCAGGAAGC
ATCGGACTCATCAGGAAGC
CGGACTCATCAGGAAGC
ATCGGACTCATCAGGAAGC
CATCGGACTCATCAGGAAGC
CATCGGACTCATCAGGAAGC
GCATCGGACTCATCAGGAAGC
GGACTCATCAGGAAGC
GCCATCGGACTCATCAGGAAGC
AGCATCGGACTCATCAGGAAGC

ATCATCGGACTCATCAGGAAGC
GCATCGGACTCATCAGGAAGC
TTGCATCGGACTCATCAGGAAGC
CATCGGACTCATCAGGAAGC
TGCATCGGACTCATCAGGAAGC
CATCGGACTCATCAGGAAGC
TACATCGGACTCATCAGGAAGC
CGGACTCATCAGGAAGC
ATCGGACTCATCAGGAAGC
ATCGGACTCATCAGGAAGC
TCATCGGACTCATCAGGAAGC
CGGACTCATCAGGAAGC
CGGACTCATCAGGAAGC
ATCATCGGACTCATCAGGAAGC
ATCATCGGACTCATCAGGAAGC
CATCGGACTCATCAGGAAGC
ACATCGGACTCATCAGGAAGC
CGGACTCATCAGGAAGC
CATCGGACTCATCAGGAAGC
TCGGACTCATCAGGAAGC
TCGGACTCATCAGGAAGC
AAGCATCGGACTCATCAGGAAGC
GCATCGGACTCATCAGGAAGC
GCCATCGGACTCATCAGGAAGC
GCCATCGGACTCATCAGGAAGC
TACATCGGACTCATCAGGAAGC
CATCGGACTCATCAGGAAGC
ACATCGGACTCATCAGGAAGC
GACATCGGACTCATCAGGAAGC
TGCATCGGACTCATCAGGAAGC
GACATCGGACTCATCAGGAAGC
CCATCGGACTCATCAGGAAGC
GCATCGGACTCATCAGGAAGC
CATCGGACTCATCAGGAAGC
CTCATCGGACTCATCAGGAAGC
ACATCGGACTCATCAGGAAGC
CATCGGACTCATCAGGAAGC
ATCGGACTCATCAGGAAGC
CATCGGACTCATCAGGAAGC
TCATCGGACTCATCAGGAAGC
ACATCGGACTCATCAGGAAGC
GCATCGGACTCATCAGGAAGC
CATCGGACTCATCAGGAAGC

CATCGGACTCATCAGGAAGC
CATCGGACTCATCAGGAAGC
ACATCGGACTCATCAGGAAGC
GCATCGGACTCATCAGGAAGC
GCATCGGACTCATCAGGAAGC
ATCATCGGACTCATCAGGAAGC
TCATCGGACTCATCAGGAAGC
TCATCGGACTCATCAGGAAGC
CGGACTCATCAGGAAGC
GACTCATCAGGAAGC
TCATCGGACTCATCAGGAAGC
ACCATCGGACTCATCAGGAAGC
ATCATCGGACTCATCAGGAAGC
ATCATCGGACTCATCAGGAAGC
ATCATCGGACTCATCAGGAAGC
CCATCGGACTCATCAGGAAGC
TCATCGGACTCATCAGGAAGC
GCCATCGGACTCATCAGGAAGC
CATCGGACTCATCAGGAAGC
TCGGACTCATCAGGAAGC
AGAATCATCGGACTCATCAGGAAGC
ACGCATCGGACTCATCAGGAAGC
CATCGGACTCATCAGGAAGC
GCATCGGACTCATCAGGAAGC
CATCGGACTCATCAGGAAGC
ATCATCGGACTCATCAGGAAGC
GGACTCATCAGGAAGC
TCGGACTCATCAGGAAGC
CATCGGACTCATCAGGAAGC
TGCATCGGACTCATCAGGAAGC
ACATCGGACTCATCAGGAAGC
ACCATCGGACTCATCAGGAAGC
TCGGACTCATCAGGAAGC
ATCGGACTCATCAGGAAGC
TCATCGGACTCATCAGGAAGC
CATCGGACTCATCAGGAAGC
TCATCGGACTCATCAGGAAGC
CCATCGGACTCATCAGGAAGC
GGACTCATCAGGAAGC
GACATCGGACTCATCAGGAAGC
CCATCGGACTCATCAGGAAGC
CATCGGACTCATCAGGAAGC
ACATCGGACTCATCAGGAAGC

ACATCGGACTCATCAGGAAGC
CATCGGACTCATCAGGAAGC
ATCGGACTCATCAGGAAGC
ATCGGACTCATCAGGAAGC
GGACTCATCAGGAAGC
CCATCGGACTCATCAGGAAGC
CATCGGACTCATCAGGAAGC
ATCGGACTCATCAGGAAGC
TACCATCGGACTCATCAGGAAGC
TACATCGGACTCATCAGGAAGC
CCATCGGACTCATCAGGAAGC
ACATCGGACTCATCAGGAAGC
ACATCGGACTCATCAGGAAGC
CATCGGACTCATCAGGAAGC
AGCATCGGACTCATCAGGAAGC
ACTCATCAGGAAGC
CGGACTCATCAGGAAGC
GCATCGGACTCATCAGGAAGC
ATCGGACTCATCAGGAAGC
GACATCGGACTCATCAGGAAGC
TCGGACTCATCAGGAAGC
GCATCGGACTCATCAGGAAGC
CATCGGACTCATCAGGAAGC
CTCATCGGACTCATCAGGAAGC
TCATCGGACTCATCAGGAAGC
ACATCGGACTCATCAGGAAGC
ACATCGGACTCATCAGGAAGC
GCATCGGACTCATCAGGAAGC
ATCGGACTCATCAGGAAGC
GACTCATCAGGAAGC
GCATCGGACTCATCAGGAAGC
ACATCGGACTCATCAGGAAGC
CATCGGACTCATCAGGAAGC
TCATCGGACTCATCAGGAAGC
CATCGGACTCATCAGGAAGC
CATCGGACTCATCAGGAAGC
GCATCGGACTCATCAGGAAGC
CGGACTCATCAGGAAGC
CTCATCGGACTCATCAGGAAGC
ACATCGGACTCATCAGGAAGC
GACATCGGACTCATCAGGAAGC
GTCATCGGACTCATCAGGAAGC
CATCGGACTCATCAGGAAGC

GTCATCGGACTCATCAGGAAGC
TACATCGGACTCATCAGGAAGC
GCCATCGGACTCATCAGGAAGC
CATCGGACTCATCAGGAAGC
ACCATCGGACTCATCAGGAAGC
CATCGGACTCATCAGGAAGC
CATCGGACTCATCAGGAAGC
CTCATCGGACTCATCAGGAAGC
CATCGGACTCATCAGGAAGC
TCATCGGACTCATCAGGAAGC
TCATCGGACTCATCAGGAAGC
TCGGACTCATCAGGAAGC
TCATCGGACTCATCAGGAAGC
TCATCGGACTCATCAGGAAGC
ATCGGACTCATCAGGAAGC
CATCGGACTCATCAGGAAGC
CATCGGACTCATCAGGAAGC
TCATCGGACTCATCAGGAAGC
TCATCGGACTCATCAGGAAGC
TACATCGGACTCATCAGGAAGC
CATCGGACTCATCAGGAAGC
ACATCGGACTCATCAGGAAGC
CTCACATCGGACTCATCAGGAAGC
CATCGGACTCATCAGGAAGC
CGGACTCATCAGGAAGC
CATCGGACTCATCAGGAAGC
ATCGGACTCATCAGGAAGC
CATCGGACTCATCAGGAAGC
CATCGGACTCATCAGGAAGC
CATCGGACTCATCAGGAAGC
TGCATCGGACTCATCAGGAAGC
CATCGGACTCATCAGGAAGC
ATCGGACTCATCAGGAAGC
GCATCGGACTCATCAGGAAGC
TCGGACTCATCAGGAAGC
ATCGGACTCATCAGGAAGC
TCGGACTCATCAGGAAGC
GCATCGGACTCATCAGGAAGC
CATCGGACTCATCAGGAAGC
ATCGGACTCATCAGGAAGC
CCATCGGACTCATCAGGAAGC
ATCGGACTCATCAGGAAGC
GCATCGGACTCATCAGGAAGC

TTCATCGGACTCATCAGGAAGC
CATCGGACTCATCAGGAAGC
CATCGGACTCATCAGGAAGC
TCCATCGGACTCATCAGGAAGC
ATCGGACTCATCAGGAAGC
ATCGGACTCATCAGGAAGC
ATCGGACTCATCAGGAAGC
AATCATCGGACTCATCAGGAAGC
CATCGGACTCATCAGGAAGC
ATCGGACTCATCAGGAAGC
TCGGACTCATCAGGAAGC
ATCGGACTCATCAGGAAGC
CCATCGGACTCATCAGGAAGC
ATCGGACTCATCAGGAAGC
CATCGGACTCATCAGGAAGC
ACTCATCAGGAAGC
TCGGACTCATCAGGAAGC
TCGGACTCATCAGGAAGC
TTCATCGGACTCATCAGGAAGC
CATCGGACTCATCAGGAAGC
AACATCGGACTCATCAGGAAGC
GCATCGGACTCATCAGGAAGC
ATCGGACTCATCAGGAAGC
CATCGGACTCATCAGGAAGC
GACTCATCAGGAAGC
CATCGGACTCATCAGGAAGC
ACATCGGACTCATCAGGAAGC
AGGCATCGGACTCATCAGGAAGC
TCGGACTCATCAGGAAGC
AAGGCATCGGACTCATCAGGAAGC
TCGGACTCATCAGGAAGC
ATCGGACTCATCAGGAAGC
TCGGACTCATCAGGAAGC
GCATCGGACTCATCAGGAAGC
TCATCGGACTCATCAGGAAGC
GCATCGGACTCATCAGGAAGC
GGACTCATCAGGAAGC
ACATCGGACTCATCAGGAAGC
GGACTCATCAGGAAGC
TCGGACTCATCAGGAAGC
TCGGACTCATCAGGAAGC
ATCGGACTCATCAGGAAGC
CATCGGACTCATCAGGAAGC

TCATCGGACTCATCAGGAAGC
GCATCGGACTCATCAGGAAGC
TTCATCGGACTCATCAGGAAGC
GCCATCGGACTCATCAGGAAGC
GGACTCATCAGGAAGC
CATCGGACTCATCAGGAAGC
AACATCGGACTCATCAGGAAGC
ATCGGACTCATCAGGAAGC
GTGTCATCGGACTCATCAGGAAGC
ACTACATCGGACTCATCAGGAAGC
AAACATCGGACTCATCAGGAAGC
ATCGGACTCATCAGGAAGC
TCATCGGACTCATCAGGAAGC
GCATCGGACTCATCAGGAAGC
GCATCGGACTCATCAGGAAGC
CATCGGACTCATCAGGAAGC
TATCATCGGACTCATCAGGAAGC
CATCGGACTCATCAGGAAGC
ATCGGACTCATCAGGAAGC
ACATCGGACTCATCAGGAAGC
ATCGGACTCATCAGGAAGC
TCGGACTCATCAGGAAGC
ATCGGACTCATCAGGAAGC
ACATCGGACTCATCAGGAAGC
ATCGGACTCATCAGGAAGC
CGGACTCATCAGGAAGC
CGGACTCATCAGGAAGC
TCGGACTCATCAGGAAGC
CATCGGACTCATCAGGAAGC
TTCATCGGACTCATCAGGAAGC
ACATCGGACTCATCAGGAAGC
CATCGGACTCATCAGGAAGC
TCGGACTCATCAGGAAGC
TCGGACTCATCAGGAAGC
ATCGGACTCATCAGGAAGC
CATCGGACTCATCAGGAAGC
GACATCGGACTCATCAGGAAGC
ATCATCGGACTCATCAGGAAGC
CATCGGACTCATCAGGAAGC
CGGACTCATCAGGAAGC
TCGGACTCATCAGGAAGC
CTCATCGGACTCATCAGGAAGC
TTCATCGGACTCATCAGGAAGC

ATCGGACTCATCAGGAAGC
ATCGGACTCATCAGGAAGC
ATCGGACTCATCAGGAAGC
TTCATCGGACTCATCAGGAAGC
CATCGGACTCATCAGGAAGC
CATCGGACTCATCAGGAAGC
ACATCGGACTCATCAGGAAGC
CGTCATCGGACTCATCAGGAAGC
CATCGGACTCATCAGGAAGC
ACATCGGACTCATCAGGAAGC
CATCGGACTCATCAGGAAGC
CATCGGACTCATCAGGAAGC
ATCGGACTCATCAGGAAGC
GACATCGGACTCATCAGGAAGC
ATCGGACTCATCAGGAAGC
CATCGGACTCATCAGGAAGC
CGGACTCATCAGGAAGC
ATCGGACTCATCAGGAAGC
TCGGACTCATCAGGAAGC
CGGACTCATCAGGAAGC
ATCGGACTCATCAGGAAGC
TACATCGGACTCATCAGGAAGC
GCATCGGACTCATCAGGAAGC
GCATCGGACTCATCAGGAAGC
CGGACTCATCAGGAAGC
TCATCGGACTCATCAGGAAGC
ATCATCGGACTCATCAGGAAGC
CATCGGACTCATCAGGAAGC
TCATCGGACTCATCAGGAAGC
ATCGGACTCATCAGGAAGC
GCATCGGACTCATCAGGAAGC
AACATCGGACTCATCAGGAAGC
ATCGGACTCATCAGGAAGC
AACCATCGGACTCATCAGGAAGC
GCATCGGACTCATCAGGAAGC
TCGGACTCATCAGGAAGC
ATCGGACTCATCAGGAAGC
TCATCATCGGACTCATCAGGAAGC
TCACATCGGACTCATCAGGAAGC
ATCGGACTCATCAGGAAGC
AATCATCGGACTCATCAGGAAGC
GCATCGGACTCATCAGGAAGC
CCATCGGACTCATCAGGAAGC

AGACATCGGACTCATCAGGAAGC
AGACATCGGACTCATCAGGAAGC
GACATCGGACTCATCAGGAAGC
AATATCATCGGACTCATCAGGAAGC
CTCATCGGACTCATCAGGAAGC
GCCATCGGACTCATCAGGAAGC
ACCATCGGACTCATCAGGAAGC
ACATCGGACTCATCAGGAAGC
ATGCATCGGACTCATCAGGAAGC
TACATCGGACTCATCAGGAAGC
CGGACTCATCAGGAAGC
CGGACTCATCAGGAAGC
TCGGACTCATCAGGAAGC
TCGGACTCATCAGGAAGC
CATCGGACTCATCAGGAAGC
GCATCGGACTCATCAGGAAGC
ATCGGACTCATCAGGAAGC
TCATCGGACTCATCAGGAAGC
TTCATCGGACTCATCAGGAAGC
AACATCGGACTCATCAGGAAGC
GGACTCATCAGGAAGC
GACTCATCAGGAAGC
ATCGGACTCATCAGGAAGC
CATCGGACTCATCAGGAAGC
CATCGGACTCATCAGGAAGC
GAATCATCGGACTCATCAGGAAGC
CATCGGACTCATCAGGAAGC
CATCGGACTCATCAGGAAGC
TCATCGGACTCATCAGGAAGC
ATCATCGGACTCATCAGGAAGC
CACATCGGACTCATCAGGAAGC
GACTCATCAGGAAGC
CCCATCGGACTCATCAGGAAGC
CATCGGACTCATCAGGAAGC
TCATCGGACTCATCAGGAAGC
TCGGACTCATCAGGAAGC
ATCGGACTCATCAGGAAGC
ATCGGACTCATCAGGAAGC
GGTCATCGGACTCATCAGGAAGC
TCATCGGACTCATCAGGAAGC
CATCGGACTCATCAGGAAGC
ACTCATCAGGAAGC
GACTCATCAGGAAGC

CATCGGACTCATCAGGAAGC
CATCGGACTCATCAGGAAGC
ATACATCGGACTCATCAGGAAGC
ATACATCGGACTCATCAGGAAGC
ACACATCGGACTCATCAGGAAGC
ACATCGGACTCATCAGGAAGC
ATCGGACTCATCAGGAAGC
ATCGGACTCATCAGGAAGC
CATCGGACTCATCAGGAAGC
CATCGGACTCATCAGGAAGC
AAACATCGGACTCATCAGGAAGC
CCCATCGGACTCATCAGGAAGC
TCGGACTCATCAGGAAGC
AACATCGGACTCATCAGGAAGC
TTCATCGGACTCATCAGGAAGC
CCATCGGACTCATCAGGAAGC
AACATCGGACTCATCAGGAAGC
GCATCGGACTCATCAGGAAGC
GACATCGGACTCATCAGGAAGC
TGACATCGGACTCATCAGGAAGC
ACATCGGACTCATCAGGAAGC
TCATCGGACTCATCAGGAAGC
ATCATCGGACTCATCAGGAAGC
CTACATCGGACTCATCAGGAAGC
GACATCGGACTCATCAGGAAGC
ATTCATCGGACTCATCAGGAAGC
AACATCGGACTCATCAGGAAGC
CATCGGACTCATCAGGAAGC
TCATCGGACTCATCAGGAAGC
TCATCGGACTCATCAGGAAGC
CATCGGACTCATCAGGAAGC
TATCATCGGACTCATCAGGAAGC
CCATCGGACTCATCAGGAAGC
AGCATCGGACTCATCAGGAAGC
CCATCGGACTCATCAGGAAGC
ATCGGACTCATCAGGAAGC
CCATCGGACTCATCAGGAAGC
CATCGGACTCATCAGGAAGC
TCATCGGACTCATCAGGAAGC
TCGGACTCATCAGGAAGC
ATCGGACTCATCAGGAAGC
CCATCGGACTCATCAGGAAGC
TAGCATCGGACTCATCAGGAAGC

ACATCGGACTCATCAGGAAGC
ACATCGGACTCATCAGGAAGC
CATCGGACTCATCAGGAAGC
GACATCGGACTCATCAGGAAGC
GACTCATCAGGAAGC
CTCATCAGGAAGC
GTCATCGGACTCATCAGGAAGC
ATCGGACTCATCAGGAAGC
CATCGGACTCATCAGGAAGC
CATCGGACTCATCAGGAAGC
TCATCGGACTCATCAGGAAGC
ACTCATCAGGAAGC
GACATCGGACTCATCAGGAAGC
TAGCATCGGACTCATCAGGAAGC
TCATCGGACTCATCAGGAAGC
CATCGGACTCATCAGGAAGC
CATCGGACTCATCAGGAAGC
ACATCGGACTCATCAGGAAGC
TCATCGGACTCATCAGGAAGC
CATCGGACTCATCAGGAAGC
CATCGGACTCATCAGGAAGC
TCGGACTCATCAGGAAGC
ATCGGACTCATCAGGAAGC
ATCGGACTCATCAGGAAGC
ACATCGGACTCATCAGGAAGC
ACATCGGACTCATCAGGAAGC
ATCGGACTCATCAGGAAGC
TCATCGGACTCATCAGGAAGC
CATCGGACTCATCAGGAAGC
TCGGACTCATCAGGAAGC
CATCGGACTCATCAGGAAGC
GGCATCGGACTCATCAGGAAGC
TCATCGGACTCATCAGGAAGC
TCGGACTCATCAGGAAGC
ATCGGACTCATCAGGAAGC
GACCATCGGACTCATCAGGAAGC
TCATCGGACTCATCAGGAAGC
TCGGACTCATCAGGAAGC
CGGACTCATCAGGAAGC
ATCGGACTCATCAGGAAGC
ATCGGACTCATCAGGAAGC
ATCGGACTCATCAGGAAGC
AGACATCGGACTCATCAGGAAGC

CGGACTCATCAGGAAGC
TCGGACTCATCAGGAAGC
CATCGGACTCATCAGGAAGC
GCATCGGACTCATCAGGAAGC
TCGGACTCATCAGGAAGC
GCATCGGACTCATCAGGAAGC
CATCGGACTCATCAGGAAGC
TTCATCGGACTCATCAGGAAGC
GACTCATCAGGAAGC
TCGGACTCATCAGGAAGC
GCATCGGACTCATCAGGAAGC
CATCGGACTCATCAGGAAGC
TCGGACTCATCAGGAAGC
ATCGGACTCATCAGGAAGC
CCATCGGACTCATCAGGAAGC
TCGGACTCATCAGGAAGC
CATCGGACTCATCAGGAAGC
GCATCGGACTCATCAGGAAGC
ATCGGACTCATCAGGAAGC
ACATCGGACTCATCAGGAAGC
ATCATCGGACTCATCAGGAAGC
ATCGGACTCATCAGGAAGC
TTCATCGGACTCATCAGGAAGC
GGACTCATCAGGAAGC
TACATCGGACTCATCAGGAAGC
CATCGGACTCATCAGGAAGC
ATCGGACTCATCAGGAAGC
GACTCATCAGGAAGC
GGACTCATCAGGAAGC
ATCGGACTCATCAGGAAGC
ATCGGACTCATCAGGAAGC
AGCATCGGACTCATCAGGAAGC
ACATCGGACTCATCAGGAAGC
CATCGGACTCATCAGGAAGC
ATCGGACTCATCAGGAAGC
ATCGGACTCATCAGGAAGC
ATCGGACTCATCAGGAAGC
TCGGACTCATCAGGAAGC
CATCGGACTCATCAGGAAGC
CCATCGGACTCATCAGGAAGC
ATCGGACTCATCAGGAAGC
TGCATCGGACTCATCAGGAAGC
TCGGACTCATCAGGAAGC

CCATCGGACTCATCAGGAAGC
CATCGGACTCATCAGGAAGC
CATCGGACTCATCAGGAAGC
ATTGCATCGGACTCATCAGGAAGC
GCATCGGACTCATCAGGAAGC
CATCGGACTCATCAGGAAGC
CATCGGACTCATCAGGAAGC
AATCATCGGACTCATCAGGAAGC
TGTCATCGGACTCATCAGGAAGC
ATCGGACTCATCAGGAAGC
GCATCGGACTCATCAGGAAGC
TCATCGGACTCATCAGGAAGC
CACATCGGACTCATCAGGAAGC
TCATCGGACTCATCAGGAAGC
TCGGACTCATCAGGAAGC
AGACATCGGACTCATCAGGAAGC
CACATCGGACTCATCAGGAAGC
AGCATCGGACTCATCAGGAAGC
TCATCGGACTCATCAGGAAGC
TTCATCGGACTCATCAGGAAGC
GCATCGGACTCATCAGGAAGC
TCATCGGACTCATCAGGAAGC
TCATCGGACTCATCAGGAAGC
ACCATCGGACTCATCAGGAAGC
TTGACATCGGACTCATCAGGAAGC
CTCATCGGACTCATCAGGAAGC
CCATCGGACTCATCAGGAAGC
CATCGGACTCATCAGGAAGC
ATCGGACTCATCAGGAAGC
CTCATCGGACTCATCAGGAAGC
TGACATCGGACTCATCAGGAAGC
CATCGGACTCATCAGGAAGC
ATCGGACTCATCAGGAAGC
GACTCATCAGGAAGC
TCATCGGACTCATCAGGAAGC
GGACTCATCAGGAAGC
ACATCGGACTCATCAGGAAGC
CCATCGGACTCATCAGGAAGC
ATCATCGGACTCATCAGGAAGC
CGGACTCATCAGGAAGC
GACATCGGACTCATCAGGAAGC
TGGCATCGGACTCATCAGGAAGC
TCATCGGACTCATCAGGAAGC

GTTCATCGGACTCATCAGGAAGC
CATCGGACTCATCAGGAAGC
TGCATCGGACTCATCAGGAAGC
CATCGGACTCATCAGGAAGC
GCATCGGACTCATCAGGAAGC
CGGACTCATCAGGAAGC
AGCATCGGACTCATCAGGAAGC
ATCGGACTCATCAGGAAGC
CCATCGGACTCATCAGGAAGC
GCATCGGACTCATCAGGAAGC
AGCATCGGACTCATCAGGAAGC
GCATCGGACTCATCAGGAAGC
ATCGGACTCATCAGGAAGC
TTTCATCGGACTCATCAGGAAGC
TCGGACTCATCAGGAAGC
TCGGACTCATCAGGAAGC
CCCATCGGACTCATCAGGAAGC
CATCGGACTCATCAGGAAGC
CGGACTCATCAGGAAGC
ACTCATCAGGAAGC
GACTCATCAGGAAGC
GCATCGGACTCATCAGGAAGC
CATCGGACTCATCAGGAAGC
CATCGGACTCATCAGGAAGC
CATCGGACTCATCAGGAAGC
CATCGGACTCATCAGGAAGC
TCGGACTCATCAGGAAGC
GCATCGGACTCATCAGGAAGC
CATCGGACTCATCAGGAAGC
CATCGGACTCATCAGGAAGC
ACATCGGACTCATCAGGAAGC
CATCGGACTCATCAGGAAGC
TCATCGGACTCATCAGGAAGC
CATCGGACTCATCAGGAAGC
CATCGGACTCATCAGGAAGC
ATCATCGGACTCATCAGGAAGC
ATCGGACTCATCAGGAAGC
ATCGGACTCATCAGGAAGC
AACATCGGACTCATCAGGAAGC
GCATCGGACTCATCAGGAAGC
CGGACTCATCAGGAAGC
CCTCCATCGGACTCATCAGGAAGC
CGGACTCATCAGGAAGC

CATCGGACTCATCAGGAAGC
ACATCGGACTCATCAGGAAGC
ATCGGACTCATCAGGAAGC
GGACTCATCAGGAAGC
CATCGGACTCATCAGGAAGC
CATCGGACTCATCAGGAAGC
CATCGGACTCATCAGGAAGC
TGCATCGGACTCATCAGGAAGC
AACATCGGACTCATCAGGAAGC
ATCGGACTCATCAGGAAGC
TTCATCGGACTCATCAGGAAGC
TCGGACTCATCAGGAAGC
GCATCGGACTCATCAGGAAGC
CATCGGACTCATCAGGAAGC
CATCGGACTCATCAGGAAGC
CATCGGACTCATCAGGAAGC
TCGGACTCATCAGGAAGC
GCATCGGACTCATCAGGAAGC
CATCGGACTCATCAGGAAGC
TCGGACTCATCAGGAAGC
CAGCATCGGACTCATCAGGAAGC
CATCGGACTCATCAGGAAGC
ATCGGACTCATCAGGAAGC
GCATCGGACTCATCAGGAAGC
ACACATCGGACTCATCAGGAAGC
CTACATCGGACTCATCAGGAAGC
CGGACTCATCAGGAAGC
CCATCGGACTCATCAGGAAGC
CCATCGGACTCATCAGGAAGC
GACATCGGACTCATCAGGAAGC
CACATCGGACTCATCAGGAAGC
ATGCATCGGACTCATCAGGAAGC
GCATCGGACTCATCAGGAAGC
ATCGGACTCATCAGGAAGC
CATCGGACTCATCAGGAAGC
ATCGGACTCATCAGGAAGC
CCATCGGACTCATCAGGAAGC
CACATCGGACTCATCAGGAAGC
ATCGGACTCATCAGGAAGC
TCATCGGACTCATCAGGAAGC
TTCATCGGACTCATCAGGAAGC
AACATCGGACTCATCAGGAAGC
ATCGGACTCATCAGGAAGC

ATCATCGGACTCATCAGGAAGC
ATCGGACTCATCAGGAAGC
ATCGGACTCATCAGGAAGC
GTACATCGGACTCATCAGGAAGC
GACATCGGACTCATCAGGAAGC
ACATCGGACTCATCAGGAAGC
ATATCATCGGACTCATCAGGAAGC
ATCGGACTCATCAGGAAGC
CATCGGACTCATCAGGAAGC
AGCATCGGACTCATCAGGAAGC
AGCATCGGACTCATCAGGAAGC
ATCATCGGACTCATCAGGAAGC
ACATCGGACTCATCAGGAAGC
CTCATCGGACTCATCAGGAAGC
CTCATCGGACTCATCAGGAAGC
AAACATCGGACTCATCAGGAAGC
CTCATCGGACTCATCAGGAAGC
ATCGGACTCATCAGGAAGC
CGGACTCATCAGGAAGC
GCATCGGACTCATCAGGAAGC
CGGACTCATCAGGAAGC
TCGGACTCATCAGGAAGC
ATCGGACTCATCAGGAAGC
CATCGGACTCATCAGGAAGC
GAACATCGGACTCATCAGGAAGC
TCGGACTCATCAGGAAGC
TCGGACTCATCAGGAAGC
TCGGACTCATCAGGAAGC
ATCGGACTCATCAGGAAGC
ACATCGGACTCATCAGGAAGC
ATCGGACTCATCAGGAAGC
CAACATCGGACTCATCAGGAAGC
TGACATCGGACTCATCAGGAAGC
TACATCGGACTCATCAGGAAGC
TACATCGGACTCATCAGGAAGC
CATCGGACTCATCAGGAAGC
CCATCGGACTCATCAGGAAGC
CCATCGGACTCATCAGGAAGC
GAGCATCGGACTCATCAGGAAGC
ATCGGACTCATCAGGAAGC
CTCATCGGACTCATCAGGAAGC
GGACTCATCAGGAAGC
CGGACTCATCAGGAAGC

TCGGACTCATCAGGAAGC
ACCATCGGACTCATCAGGAAGC
TCATCGGACTCATCAGGAAGC
AACATCGGACTCATCAGGAAGC
GCATCGGACTCATCAGGAAGC
ATCATCGGACTCATCAGGAAGC
CATCGGACTCATCAGGAAGC
CATCGGACTCATCAGGAAGC
ATCGGACTCATCAGGAAGC
ATCGGACTCATCAGGAAGC
ATATCATCGGACTCATCAGGAAGC
GCATCGGACTCATCAGGAAGC
TACATCGGACTCATCAGGAAGC
CATCGGACTCATCAGGAAGC
CATCATCGGACTCATCAGGAAGC
ACATCGGACTCATCAGGAAGC
CTCATCGGACTCATCAGGAAGC
CTCATCGGACTCATCAGGAAGC
TACATCGGACTCATCAGGAAGC
TACATCGGACTCATCAGGAAGC
CGGACTCATCAGGAAGC
TCGGACTCATCAGGAAGC
ACATCGGACTCATCAGGAAGC
CGGACTCATCAGGAAGC
TATCATCGGACTCATCAGGAAGC
GGACTCATCAGGAAGC
ATCGGACTCATCAGGAAGC
ATCGGACTCATCAGGAAGC
TCGGACTCATCAGGAAGC
CATCGGACTCATCAGGAAGC
CATCGGACTCATCAGGAAGC
CATCGGACTCATCAGGAAGC
CATCGGACTCATCAGGAAGC
CCCATCGGACTCATCAGGAAGC
GGCCATCGGACTCATCAGGAAGC
GCATCGGACTCATCAGGAAGC
ACATCGGACTCATCAGGAAGC
GCATCGGACTCATCAGGAAGC
ATCGGACTCATCAGGAAGC
ACATCGGACTCATCAGGAAGC
CATCGGACTCATCAGGAAGC
TCATCGGACTCATCAGGAAGC
TCATCGGACTCATCAGGAAGC

CATCGGACTCATCAGGAAGC
CATCGGACTCATCAGGAAGC
GGCATCGGACTCATCAGGAAGC
TCATCGGACTCATCAGGAAGC
CGGACTCATCAGGAAGC
ATCGGACTCATCAGGAAGC
ATCGGACTCATCAGGAAGC
TCGGACTCATCAGGAAGC
GACTCATCAGGAAGC
TCATCGGACTCATCAGGAAGC
CATCGGACTCATCAGGAAGC
CATCGGACTCATCAGGAAGC
CCATCGGACTCATCAGGAAGC
ATCGGACTCATCAGGAAGC
TCGGACTCATCAGGAAGC
CCATCGGACTCATCAGGAAGC
ATCGGACTCATCAGGAAGC
TCGGACTCATCAGGAAGC
ATCGGACTCATCAGGAAGC
CATCGGACTCATCAGGAAGC
CATCGGACTCATCAGGAAGC
ATCGGACTCATCAGGAAGC
CATCGGACTCATCAGGAAGC
TCGGACTCATCAGGAAGC
ATCGGACTCATCAGGAAGC
TCGGACTCATCAGGAAGC
GACTCATCAGGAAGC
TCGGACTCATCAGGAAGC
CATCGGACTCATCAGGAAGC
ACATCGGACTCATCAGGAAGC
TCATCGGACTCATCAGGAAGC
CATCGGACTCATCAGGAAGC
ATCGGACTCATCAGGAAGC
TGCATCGGACTCATCAGGAAGC
CATCGGACTCATCAGGAAGC
CATCGGACTCATCAGGAAGC
TCATCGGACTCATCAGGAAGC
GGACTCATCAGGAAGC
TAGCATCGGACTCATCAGGAAGC
TCATCGGACTCATCAGGAAGC
TCGGACTCATCAGGAAGC
CATCGGACTCATCAGGAAGC
TCGGACTCATCAGGAAGC

TCGGACTCATCAGGAAGC
TCATCGGACTCATCAGGAAGC
TCATCGGACTCATCAGGAAGC
AACATCGGACTCATCAGGAAGC
CATCGGACTCATCAGGAAGC
ACATCGGACTCATCAGGAAGC
TCGGACTCATCAGGAAGC
GCATCGGACTCATCAGGAAGC
AACATCGGACTCATCAGGAAGC
CCATCGGACTCATCAGGAAGC
GCCATCGGACTCATCAGGAAGC
TCATCGGACTCATCAGGAAGC
GCATCGGACTCATCAGGAAGC
AATCATCGGACTCATCAGGAAGC
ATCGGACTCATCAGGAAGC
GTCATCGGACTCATCAGGAAGC
AAACATCGGACTCATCAGGAAGC
GCATCGGACTCATCAGGAAGC
ACATCGGACTCATCAGGAAGC
CATCGGACTCATCAGGAAGC
ATCGGACTCATCAGGAAGC
ACATCGGACTCATCAGGAAGC
CATCGGACTCATCAGGAAGC
CATCGGACTCATCAGGAAGC
ATCGGACTCATCAGGAAGC
ATCGGACTCATCAGGAAGC
ACATCGGACTCATCAGGAAGC
GACATCGGACTCATCAGGAAGC
CATCGGACTCATCAGGAAGC
ATCATCGGACTCATCAGGAAGC
ATCGGACTCATCAGGAAGC
GCATCGGACTCATCAGGAAGC
TCGGACTCATCAGGAAGC
GACATCGGACTCATCAGGAAGC
ACATCGGACTCATCAGGAAGC
CACATCGGACTCATCAGGAAGC
TCATCGGACTCATCAGGAAGC
ACATCGGACTCATCAGGAAGC
ATCGGACTCATCAGGAAGC
CATCGGACTCATCAGGAAGC
GCATCGGACTCATCAGGAAGC
GACATCGGACTCATCAGGAAGC
ACATCGGACTCATCAGGAAGC

TCCATCGGACTCATCAGGAAGC
AATCATCGGACTCATCAGGAAGC
CATCGGACTCATCAGGAAGC
ATACATCGGACTCATCAGGAAGC
ATCGGACTCATCAGGAAGC
CATCGGACTCATCAGGAAGC
GCATCGGACTCATCAGGAAGC
GACATCGGACTCATCAGGAAGC
TCATCGGACTCATCAGGAAGC
GACATCGGACTCATCAGGAAGC
TCATCGGACTCATCAGGAAGC
ATACATCGGACTCATCAGGAAGC
TCGGACTCATCAGGAAGC
TCGGACTCATCAGGAAGC
ACCATCGGACTCATCAGGAAGC
TGACATCGGACTCATCAGGAAGC
AGCATCGGACTCATCAGGAAGC
AGCCATCGGACTCATCAGGAAGC
ACATCGGACTCATCAGGAAGC
CATCGGACTCATCAGGAAGC
CATCGGACTCATCAGGAAGC
CATCGGACTCATCAGGAAGC
CATCGGACTCATCAGGAAGC
CATCGGACTCATCAGGAAGC
CCATCGGACTCATCAGGAAGC
CATCGGACTCATCAGGAAGC
ATCGGACTCATCAGGAAGC
TCGGACTCATCAGGAAGC
CATCGGACTCATCAGGAAGC
CCATCGGACTCATCAGGAAGC
TCGGACTCATCAGGAAGC
CATCGGACTCATCAGGAAGC
CATCGGACTCATCAGGAAGC
CATCGGACTCATCAGGAAGC
CATCGGACTCATCAGGAAGC
TCATCGGACTCATCAGGAAGC
ATCGGACTCATCAGGAAGC
ACATCGGACTCATCAGGAAGC
TATCATCGGACTCATCAGGAAGC
CATCGGACTCATCAGGAAGC
CATCGGACTCATCAGGAAGC
ACATCGGACTCATCAGGAAGC
GCATCGGACTCATCAGGAAGC

ACATCGGACTCATCAGGAAGC
CATCGGACTCATCAGGAAGC
CATCGGACTCATCAGGAAGC
GCATCGGACTCATCAGGAAGC
TCATCGGACTCATCAGGAAGC
CATCGGACTCATCAGGAAGC
CATCGGACTCATCAGGAAGC
GCCATCGGACTCATCAGGAAGC
ACATCGGACTCATCAGGAAGC
ACATCGGACTCATCAGGAAGC
CCATCGGACTCATCAGGAAGC
CCATCGGACTCATCAGGAAGC
CATCGGACTCATCAGGAAGC
CATCGGACTCATCAGGAAGC
TCCATCGGACTCATCAGGAAGC
CATCGGACTCATCAGGAAGC
CATCGGACTCATCAGGAAGC
TCATCGGACTCATCAGGAAGC
GGACTCATCAGGAAGC
TGCATCGGACTCATCAGGAAGC
CATACATCGGACTCATCAGGAAGC
CATACATCGGACTCATCAGGAAGC
CATCGGACTCATCAGGAAGC
ATCGGACTCATCAGGAAGC
ATCGGACTCATCAGGAAGC
GCATCGGACTCATCAGGAAGC
TACATCGGACTCATCAGGAAGC
TACATCGGACTCATCAGGAAGC
AGCATCGGACTCATCAGGAAGC
ACATCGGACTCATCAGGAAGC
ATCGGACTCATCAGGAAGC
CCATCGGACTCATCAGGAAGC
CATCGGACTCATCAGGAAGC
ATCGGACTCATCAGGAAGC
TCGGACTCATCAGGAAGC
TCGGACTCATCAGGAAGC
ATACATCGGACTCATCAGGAAGC
TCGGACTCATCAGGAAGC
CCATCGGACTCATCAGGAAGC
AGCATCGGACTCATCAGGAAGC
CATCGGACTCATCAGGAAGC
CATCGGACTCATCAGGAAGC
ACATCGGACTCATCAGGAAGC

GACTCATCAGGAAGC
ATCATCGGACTCATCAGGAAGC
GGCATCGGACTCATCAGGAAGC
GGCATCGGACTCATCAGGAAGC
GCATCGGACTCATCAGGAAGC
ATCGGACTCATCAGGAAGC
GCATCGGACTCATCAGGAAGC
TCGGACTCATCAGGAAGC
GACTCATCAGGAAGC
GACTCATCAGGAAGC
TTCATCGGACTCATCAGGAAGC
TCATCGGACTCATCAGGAAGC
ACATCGGACTCATCAGGAAGC
AGCATCGGACTCATCAGGAAGC
CATCGGACTCATCAGGAAGC
CATCGGACTCATCAGGAAGC
ATAACATCGGACTCATCAGGAAGC
GCATCGGACTCATCAGGAAGC
ATCGGACTCATCAGGAAGC
ATCGGACTCATCAGGAAGC
CGGACTCATCAGGAAGC
ACCATCGGACTCATCAGGAAGC
TCATCGGACTCATCAGGAAGC
CATCGGACTCATCAGGAAGC
CGGACTCATCAGGAAGC
ACATCGGACTCATCAGGAAGC
CTCATCGGACTCATCAGGAAGC
GGCATCGGACTCATCAGGAAGC
CGGACTCATCAGGAAGC
ATCGGACTCATCAGGAAGC
ACATCGGACTCATCAGGAAGC
TCATCGGACTCATCAGGAAGC
CGGACTCATCAGGAAGC
GGACTCATCAGGAAGC
AAGCATCGGACTCATCAGGAAGC
TCGGACTCATCAGGAAGC
AGCATCGGACTCATCAGGAAGC
CTCATCGGACTCATCAGGAAGC
TCATCGGACTCATCAGGAAGC
ACATCGGACTCATCAGGAAGC
CCATCGGACTCATCAGGAAGC
CATCGGACTCATCAGGAAGC
GAGCATCGGACTCATCAGGAAGC

GAGCATCGGACTCATCAGGAAGC
CATCGGACTCATCAGGAAGC
TCATCGGACTCATCAGGAAGC
TCGGACTCATCAGGAAGC
ACTCATCAGGAAGC
GCATCGGACTCATCAGGAAGC
AACCATCGGACTCATCAGGAAGC
AACATCGGACTCATCAGGAAGC
TCATCGGACTCATCAGGAAGC
ATCGGACTCATCAGGAAGC
ATCGGACTCATCAGGAAGC
GCCATCGGACTCATCAGGAAGC
CATCGGACTCATCAGGAAGC
AGAACATCGGACTCATCAGGAAGC
TCGGACTCATCAGGAAGC
ATCGGACTCATCAGGAAGC
TCGGACTCATCAGGAAGC
CGGACTCATCAGGAAGC
AAGCATCGGACTCATCAGGAAGC
TCGGACTCATCAGGAAGC
ACATCGGACTCATCAGGAAGC
CATCGGACTCATCAGGAAGC
ACATCGGACTCATCAGGAAGC
GTGCATCGGACTCATCAGGAAGC
TCATCGGACTCATCAGGAAGC
TCATCGGACTCATCAGGAAGC
CATCGGACTCATCAGGAAGC
AACATCGGACTCATCAGGAAGC
CATCGGACTCATCAGGAAGC
AGCATCGGACTCATCAGGAAGC
ATCGGACTCATCAGGAAGC
CATCGGACTCATCAGGAAGC
ACATCGGACTCATCAGGAAGC
CATCGGACTCATCAGGAAGC
ATTCATCGGACTCATCAGGAAGC
TCATCGGACTCATCAGGAAGC
ATCGGACTCATCAGGAAGC
ATCGGACTCATCAGGAAGC
GCATCGGACTCATCAGGAAGC
GCATCGGACTCATCAGGAAGC
CGGACTCATCAGGAAGC
CATCGGACTCATCAGGAAGC
ATCGGACTCATCAGGAAGC

ACCATCGGACTCATCAGGAAGC
ATCATCGGACTCATCAGGAAGC
ATCGGACTCATCAGGAAGC
CGGACTCATCAGGAAGC
GCATCGGACTCATCAGGAAGC
CCATCGGACTCATCAGGAAGC
CATCGGACTCATCAGGAAGC
GGCATCGGACTCATCAGGAAGC
AACATCGGACTCATCAGGAAGC
CGGACTCATCAGGAAGC
CATCGGACTCATCAGGAAGC
TCATCGGACTCATCAGGAAGC
ATCGGACTCATCAGGAAGC
CATCGGACTCATCAGGAAGC
CATCGGACTCATCAGGAAGC
CGGACTCATCAGGAAGC
CATCGGACTCATCAGGAAGC
CATCGGACTCATCAGGAAGC
CATCGGACTCATCAGGAAGC
TCGGACTCATCAGGAAGC
CATCGGACTCATCAGGAAGC
TCATCGGACTCATCAGGAAGC
CGGACTCATCAGGAAGC
TCGGACTCATCAGGAAGC
ACATCGGACTCATCAGGAAGC
GGCCATCGGACTCATCAGGAAGC
CATCGGACTCATCAGGAAGC
ATTTCATCGGACTCATCAGGAAGC
CATCGGACTCATCAGGAAGC
TCGGACTCATCAGGAAGC
TCGGACTCATCAGGAAGC
CATCGGACTCATCAGGAAGC
TCGGACTCATCAGGAAGC
ACACATCGGACTCATCAGGAAGC
ATCGGACTCATCAGGAAGC
GCATCGGACTCATCAGGAAGC
TCATCGGACTCATCAGGAAGC
ATCGGACTCATCAGGAAGC
TCATCGGACTCATCAGGAAGC
AACATCGGACTCATCAGGAAGC
ATCGGACTCATCAGGAAGC
ATCGGACTCATCAGGAAGC
TCATCGGACTCATCAGGAAGC

GCATCGGACTCATCAGGAAGC
GCATCGGACTCATCAGGAAGC
ATCGGACTCATCAGGAAGC
ACATCGGACTCATCAGGAAGC
ACATCGGACTCATCAGGAAGC
ATCGGACTCATCAGGAAGC
GCATCGGACTCATCAGGAAGC
CGGACTCATCAGGAAGC
ATCGGACTCATCAGGAAGC
ATCATCGGACTCATCAGGAAGC
CCATCGGACTCATCAGGAAGC
CACATCGGACTCATCAGGAAGC
GCATCGGACTCATCAGGAAGC
CATCGGACTCATCAGGAAGC
ATCATCGGACTCATCAGGAAGC
TCATCGGACTCATCAGGAAGC
ACATCGGACTCATCAGGAAGC
AACATCGGACTCATCAGGAAGC
ATCGGACTCATCAGGAAGC
CGGACTCATCAGGAAGC
CATCGGACTCATCAGGAAGC
CATCGGACTCATCAGGAAGC
CATCGGACTCATCAGGAAGC
ATCGGACTCATCAGGAAGC
AGCATCGGACTCATCAGGAAGC
GCCATCGGACTCATCAGGAAGC
CATCGGACTCATCAGGAAGC
CACATCGGACTCATCAGGAAGC
CCATCGGACTCATCAGGAAGC
GACTCATCAGGAAGC
TACATCGGACTCATCAGGAAGC
ATCGGACTCATCAGGAAGC
ATCGGACTCATCAGGAAGC
ATCATCGGACTCATCAGGAAGC
AGCATCGGACTCATCAGGAAGC
GGCATCGGACTCATCAGGAAGC
TCATCGGACTCATCAGGAAGC
CATCGGACTCATCAGGAAGC
ACATCGGACTCATCAGGAAGC
GGACATCGGACTCATCAGGAAGC
CATCGGACTCATCAGGAAGC
CCATCGGACTCATCAGGAAGC
CATCGGACTCATCAGGAAGC

TCATCGGACTCATCAGGAAGC
AGCATCGGACTCATCAGGAAGC
ATCATCGGACTCATCAGGAAGC
TACATCGGACTCATCAGGAAGC
TCATCGGACTCATCAGGAAGC
TCATCGGACTCATCAGGAAGC
ATCATCGGACTCATCAGGAAGC
TACATCGGACTCATCAGGAAGC
CATCGGACTCATCAGGAAGC
ATCGGACTCATCAGGAAGC
CATCGGACTCATCAGGAAGC
TCATCGGACTCATCAGGAAGC
TCATCGGACTCATCAGGAAGC
GCATCGGACTCATCAGGAAGC
TACATCGGACTCATCAGGAAGC
ATCGGACTCATCAGGAAGC
GGACTCATCAGGAAGC
TACATCGGACTCATCAGGAAGC
AGCCATCGGACTCATCAGGAAGC
CATCGGACTCATCAGGAAGC
GCATCGGACTCATCAGGAAGC
TCCATCGGACTCATCAGGAAGC
ATCGGACTCATCAGGAAGC
ATCGGACTCATCAGGAAGC
CATCGGACTCATCAGGAAGC
CATCGGACTCATCAGGAAGC
ATCGGACTCATCAGGAAGC
ACATCGGACTCATCAGGAAGC
ATCGGACTCATCAGGAAGC
TCGGACTCATCAGGAAGC
CATCGGACTCATCAGGAAGC
AAGCATCGGACTCATCAGGAAGC
CAAACATCGGACTCATCAGGAAGC
TCATCGGACTCATCAGGAAGC
ACATCGGACTCATCAGGAAGC
CCATCGGACTCATCAGGAAGC
ATTCATCGGACTCATCAGGAAGC
CATCGGACTCATCAGGAAGC
TCATCGGACTCATCAGGAAGC
CATCGGACTCATCAGGAAGC
TTCATCGGACTCATCAGGAAGC
ACATCGGACTCATCAGGAAGC
CATCGGACTCATCAGGAAGC

ACATCGGACTCATCAGGAAGC
ATCGGACTCATCAGGAAGC
AACATCGGACTCATCAGGAAGC
TTTTCATCGGACTCATCAGGAAGC
ATCATCGGACTCATCAGGAAGC
ACATCGGACTCATCAGGAAGC
GCATCGGACTCATCAGGAAGC
TCATCGGACTCATCAGGAAGC
TCGGACTCATCAGGAAGC
CATCGGACTCATCAGGAAGC
GTCATCGGACTCATCAGGAAGC
GCATCGGACTCATCAGGAAGC
CCATCGGACTCATCAGGAAGC
ATGCATCGGACTCATCAGGAAGC
ATCATCGGACTCATCAGGAAGC
TACATCGGACTCATCAGGAAGC
GCATCGGACTCATCAGGAAGC
AGGCATCGGACTCATCAGGAAGC
ATCGGACTCATCAGGAAGC
CATCGGACTCATCAGGAAGC
TCATCGGACTCATCAGGAAGC
TCATCGGACTCATCAGGAAGC
GACATCGGACTCATCAGGAAGC
ACATCGGACTCATCAGGAAGC
TCGGACTCATCAGGAAGC
CGGACTCATCAGGAAGC
CATCGGACTCATCAGGAAGC
ACATCGGACTCATCAGGAAGC
CATCGGACTCATCAGGAAGC
GACTCATCAGGAAGC
ATCGGACTCATCAGGAAGC
CATCGGACTCATCAGGAAGC
ATCGGACTCATCAGGAAGC
CGGACTCATCAGGAAGC
CATCGGACTCATCAGGAAGC
CGGACTCATCAGGAAGC
CATCGGACTCATCAGGAAGC
TGCATCGGACTCATCAGGAAGC
GCATCGGACTCATCAGGAAGC
ACCATCGGACTCATCAGGAAGC
CACATCGGACTCATCAGGAAGC
GCATCGGACTCATCAGGAAGC
ATCGGACTCATCAGGAAGC

TCGGACTCATCAGGAAGC
GCATCGGACTCATCAGGAAGC
ATCATCGGACTCATCAGGAAGC
CATCGGACTCATCAGGAAGC
CATCGGACTCATCAGGAAGC
ATACATCGGACTCATCAGGAAGC
GCATCGGACTCATCAGGAAGC
TACATCGGACTCATCAGGAAGC
TCATCGGACTCATCAGGAAGC
GCATCGGACTCATCAGGAAGC
GTCATCGGACTCATCAGGAAGC
TCGGACTCATCAGGAAGC
CATCGGACTCATCAGGAAGC
CTCATCGGACTCATCAGGAAGC
ATCGGACTCATCAGGAAGC
GCATCGGACTCATCAGGAAGC
ATCGGACTCATCAGGAAGC
CTCATCGGACTCATCAGGAAGC
CATCGGACTCATCAGGAAGC
TTCATCGGACTCATCAGGAAGC
ACATCGGACTCATCAGGAAGC
ATCGGACTCATCAGGAAGC
TCATCGGACTCATCAGGAAGC
CATCGGACTCATCAGGAAGC
ATCGGACTCATCAGGAAGC
CATCGGACTCATCAGGAAGC
GGACTCATCAGGAAGC
ACCATCGGACTCATCAGGAAGC
TCGGACTCATCAGGAAGC
CATCGGACTCATCAGGAAGC
GTCATCGGACTCATCAGGAAGC
AAATCATCGGACTCATCAGGAAGC
TCCATCGGACTCATCAGGAAGC
TCATCGGACTCATCAGGAAGC
ATCGGACTCATCAGGAAGC
ATCATCGGACTCATCAGGAAGC
TCGGACTCATCAGGAAGC
GCATCGGACTCATCAGGAAGC
TCGGACTCATCAGGAAGC
TACATCGGACTCATCAGGAAGC
GTCATCGGACTCATCAGGAAGC
AACATCGGACTCATCAGGAAGC
GACATCGGACTCATCAGGAAGC

TCCATCGGACTCATCAGGAAGC
CTCATCGGACTCATCAGGAAGC
CTCATCGGACTCATCAGGAAGC
AGCATCGGACTCATCAGGAAGC
GCATCGGACTCATCAGGAAGC
TCCATCGGACTCATCAGGAAGC
ACATCGGACTCATCAGGAAGC
CATCGGACTCATCAGGAAGC
GCATCGGACTCATCAGGAAGC
CATCGGACTCATCAGGAAGC
CATCGGACTCATCAGGAAGC
ATCGGACTCATCAGGAAGC
GCATCGGACTCATCAGGAAGC
CATCGGACTCATCAGGAAGC
TGCATCGGACTCATCAGGAAGC
ATCGGACTCATCAGGAAGC
TTTCATCGGACTCATCAGGAAGC
CATCGGACTCATCAGGAAGC
GTCATCGGACTCATCAGGAAGC
CCCATCGGACTCATCAGGAAGC
ATCATCGGACTCATCAGGAAGC
TCGGACTCATCAGGAAGC
ATCGGACTCATCAGGAAGC
GCATCGGACTCATCAGGAAGC
GTACATCGGACTCATCAGGAAGC
CCCATCGGACTCATCAGGAAGC
CCCATCGGACTCATCAGGAAGC
TCATCGGACTCATCAGGAAGC
ATCGGACTCATCAGGAAGC
TGCATCGGACTCATCAGGAAGC
TGCATCGGACTCATCAGGAAGC
ATCGGACTCATCAGGAAGC
AACATCGGACTCATCAGGAAGC
TCGGACTCATCAGGAAGC
TCATCGGACTCATCAGGAAGC
TACATCGGACTCATCAGGAAGC
CCATCGGACTCATCAGGAAGC
TAACATCGGACTCATCAGGAAGC
TACATCGGACTCATCAGGAAGC
GCATCGGACTCATCAGGAAGC
AATCATCGGACTCATCAGGAAGC
CATCGGACTCATCAGGAAGC
TCGGACTCATCAGGAAGC

ATCGGACTCATCAGGAAGC
ATCGGACTCATCAGGAAGC
CATCGGACTCATCAGGAAGC
ATCGGACTCATCAGGAAGC
CATCGGACTCATCAGGAAGC
TACATCGGACTCATCAGGAAGC
ATCGGACTCATCAGGAAGC
ATCATCGGACTCATCAGGAAGC
ACATCGGACTCATCAGGAAGC
GCCATCGGACTCATCAGGAAGC
CGGACTCATCAGGAAGC
ACCATCGGACTCATCAGGAAGC
CCATCGGACTCATCAGGAAGC
GGCATCGGACTCATCAGGAAGC
GCATCGGACTCATCAGGAAGC
CAAACATCGGACTCATCAGGAAGC
CCATCGGACTCATCAGGAAGC
ACATCGGACTCATCAGGAAGC
CATCGGACTCATCAGGAAGC
TCGGACTCATCAGGAAGC
ATCGGACTCATCAGGAAGC
TCGGACTCATCAGGAAGC
TGCCATCGGACTCATCAGGAAGC
TACATCGGACTCATCAGGAAGC
CATCGGACTCATCAGGAAGC
TCGGACTCATCAGGAAGC
CGGACTCATCAGGAAGC
TCGGACTCATCAGGAAGC
CGGACTCATCAGGAAGC
ACCATCGGACTCATCAGGAAGC
CATCGGACTCATCAGGAAGC
CATCGGACTCATCAGGAAGC
AAGCATCGGACTCATCAGGAAGC
CATCGGACTCATCAGGAAGC
CATCGGACTCATCAGGAAGC
CATCGGACTCATCAGGAAGC
GCATCGGACTCATCAGGAAGC
ACATCGGACTCATCAGGAAGC
CGGACTCATCAGGAAGC
ATCATCGGACTCATCAGGAAGC
CATCGGACTCATCAGGAAGC
GGACTCATCAGGAAGC
CAAGCATCGGACTCATCAGGAAGC

ATCGGACTCATCAGGAAGC
AACATCGGACTCATCAGGAAGC
ATCGGACTCATCAGGAAGC
AACATCGGACTCATCAGGAAGC
TCATCGGACTCATCAGGAAGC
CCATCGGACTCATCAGGAAGC
CATCGGACTCATCAGGAAGC
TCGGACTCATCAGGAAGC
AAGCATCGGACTCATCAGGAAGC
GTCATCGGACTCATCAGGAAGC
CATCGGACTCATCAGGAAGC
AGACATCGGACTCATCAGGAAGC
CATCGGACTCATCAGGAAGC
GCATCGGACTCATCAGGAAGC
TCATCGGACTCATCAGGAAGC
ATCGGACTCATCAGGAAGC
CCATCGGACTCATCAGGAAGC
TACATCGGACTCATCAGGAAGC
CATCGGACTCATCAGGAAGC
TTCATCGGACTCATCAGGAAGC
TGCATCGGACTCATCAGGAAGC
ATCGGACTCATCAGGAAGC
GACCCATCGGACTCATCAGGAAGC
TCGGACTCATCAGGAAGC
TCGGACTCATCAGGAAGC
ATCATCGGACTCATCAGGAAGC
CATCGGACTCATCAGGAAGC
ATCGGACTCATCAGGAAGC
ATCGGACTCATCAGGAAGC
CCCATCGGACTCATCAGGAAGC
TCGGACTCATCAGGAAGC
GGACTCATCAGGAAGC
GGACTCATCAGGAAGC
AACATCGGACTCATCAGGAAGC
CGGACTCATCAGGAAGC
TCTCATCGGACTCATCAGGAAGC
CGGACTCATCAGGAAGC
CATCGGACTCATCAGGAAGC
ATCGGACTCATCAGGAAGC
AGCATCGGACTCATCAGGAAGC
GACTCATCAGGAAGC
CTCATCAGGAAGC
CATCGGACTCATCAGGAAGC

CGGACTCATCAGGAAGC
ACATCGGACTCATCAGGAAGC
GGACTCATCAGGAAGC
CGGACTCATCAGGAAGC
GCATCGGACTCATCAGGAAGC
CGGACTCATCAGGAAGC
TCGGACTCATCAGGAAGC
CATCGGACTCATCAGGAAGC
CATCGGACTCATCAGGAAGC
GCATCGGACTCATCAGGAAGC
GCATCGGACTCATCAGGAAGC
CATCGGACTCATCAGGAAGC
ATCGGACTCATCAGGAAGC
GGCATCGGACTCATCAGGAAGC
CGGACTCATCAGGAAGC
ACATCGGACTCATCAGGAAGC
GACTCATCAGGAAGC
CCATCGGACTCATCAGGAAGC
TACATCGGACTCATCAGGAAGC
ATCGGACTCATCAGGAAGC
AGCATCGGACTCATCAGGAAGC
TCATCGGACTCATCAGGAAGC
TCGGACTCATCAGGAAGC
GCATCGGACTCATCAGGAAGC
TCATCGGACTCATCAGGAAGC
GCATCGGACTCATCAGGAAGC
CCCATCGGACTCATCAGGAAGC
AACATCGGACTCATCAGGAAGC
ACTCATCAGGAAGC
ATCGGACTCATCAGGAAGC
TCGGACTCATCAGGAAGC
CATCGGACTCATCAGGAAGC
GGACTCATCAGGAAGC
ACATCGGACTCATCAGGAAGC
TCATCGGACTCATCAGGAAGC
TCGGACTCATCAGGAAGC
CCCATCGGACTCATCAGGAAGC
TCGGACTCATCAGGAAGC
CTCATCGGACTCATCAGGAAGC
CATCGGACTCATCAGGAAGC
ATCGGACTCATCAGGAAGC
ATCGGACTCATCAGGAAGC
GGACTCATCAGGAAGC

TCATCGGACTCATCAGGAAGC
CGGACTCATCAGGAAGC
ATCGGACTCATCAGGAAGC
ATCGGACTCATCAGGAAGC
CGGACTCATCAGGAAGC
CGGACTCATCAGGAAGC
CGGACTCATCAGGAAGC
CATCGGACTCATCAGGAAGC
GGACTCATCAGGAAGC
ATCGGACTCATCAGGAAGC
ATCGGACTCATCAGGAAGC
TCCATCGGACTCATCAGGAAGC
CCATCGGACTCATCAGGAAGC
CATCGGACTCATCAGGAAGC
CATCGGACTCATCAGGAAGC
TCATCGGACTCATCAGGAAGC
ATCGGACTCATCAGGAAGC
ACTCATCAGGAAGC
GGCATCGGACTCATCAGGAAGC
ATCGGACTCATCAGGAAGC
CCATCGGACTCATCAGGAAGC
ACTCATCGGACTCATCAGGAAGC
GGACTCATCAGGAAGC
TCATCGGACTCATCAGGAAGC
CATCGGACTCATCAGGAAGC
TTCATCGGACTCATCAGGAAGC
AGCATCGGACTCATCAGGAAGC
TCCCATCGGACTCATCAGGAAGC
CATCGGACTCATCAGGAAGC
ACTCATCAGGAAGC
GGCATCGGACTCATCAGGAAGC
ATCATCGGACTCATCAGGAAGC
ATCGGACTCATCAGGAAGC
CGGACTCATCAGGAAGC
CATCATCGGACTCATCAGGAAGC
GCATCGGACTCATCAGGAAGC
CGGACTCATCAGGAAGC
TTCATCGGACTCATCAGGAAGC
CATCATCGGACTCATCAGGAAGC
ACATCGGACTCATCAGGAAGC
GCATCGGACTCATCAGGAAGC
CATCGGACTCATCAGGAAGC
ATCGGACTCATCAGGAAGC

GCATCGGACTCATCAGGAAGC
GCATCGGACTCATCAGGAAGC
CATCGGACTCATCAGGAAGC
TCATCGGACTCATCAGGAAGC
CATCGGACTCATCAGGAAGC
AACATCGGACTCATCAGGAAGC
CATCGGACTCATCAGGAAGC
ATCGGACTCATCAGGAAGC
GCATCGGACTCATCAGGAAGC
TGCATCGGACTCATCAGGAAGC
AGCATCGGACTCATCAGGAAGC
TCGGACTCATCAGGAAGC
CCATCGGACTCATCAGGAAGC
TTCATCGGACTCATCAGGAAGC
ACCATCGGACTCATCAGGAAGC
TGACATCGGACTCATCAGGAAGC
GCATCGGACTCATCAGGAAGC
AGACATCGGACTCATCAGGAAGC
GCATCGGACTCATCAGGAAGC
GGCATCGGACTCATCAGGAAGC
CATCGGACTCATCAGGAAGC
ATCATCGGACTCATCAGGAAGC
AGCATCGGACTCATCAGGAAGC
GCATCGGACTCATCAGGAAGC
CATCGGACTCATCAGGAAGC
ACATCGGACTCATCAGGAAGC
TCATCGGACTCATCAGGAAGC
CCATCGGACTCATCAGGAAGC
TGCATCGGACTCATCAGGAAGC
CCCATCGGACTCATCAGGAAGC
CATCGGACTCATCAGGAAGC
CATCGGACTCATCAGGAAGC
ACATCGGACTCATCAGGAAGC
ATCATCGGACTCATCAGGAAGC
AAACATCGGACTCATCAGGAAGC
TCATCGGACTCATCAGGAAGC
TGCATCGGACTCATCAGGAAGC
ACATCGGACTCATCAGGAAGC
ACATCGGACTCATCAGGAAGC
TCGGACTCATCAGGAAGC
ATCGGACTCATCAGGAAGC
AACATCGGACTCATCAGGAAGC
GGTCATCGGACTCATCAGGAAGC

GACTCATCAGGAAGC
CACATCGGACTCATCAGGAAGC
TCCATCGGACTCATCAGGAAGC
TCGGACTCATCAGGAAGC
AATCATCGGACTCATCAGGAAGC
AACATCGGACTCATCAGGAAGC
ACATCGGACTCATCAGGAAGC
ATCGGACTCATCAGGAAGC
CCATCGGACTCATCAGGAAGC
GAACATCGGACTCATCAGGAAGC
CCCCATCGGACTCATCAGGAAGC
GCATCGGACTCATCAGGAAGC
CATCGGACTCATCAGGAAGC
ACCATCGGACTCATCAGGAAGC
CATCGGACTCATCAGGAAGC
GAGCATCGGACTCATCAGGAAGC
ACATCGGACTCATCAGGAAGC
ATCGGACTCATCAGGAAGC
GCATCGGACTCATCAGGAAGC
CCATCGGACTCATCAGGAAGC
ATCGGACTCATCAGGAAGC
TCATCGGACTCATCAGGAAGC
CTCATCGGACTCATCAGGAAGC
GCATCGGACTCATCAGGAAGC
GAACATCGGACTCATCAGGAAGC
CATCGGACTCATCAGGAAGC
ATCGGACTCATCAGGAAGC
ATCATCGGACTCATCAGGAAGC
ACATCGGACTCATCAGGAAGC
CATCGGACTCATCAGGAAGC
ATCGGACTCATCAGGAAGC
CATCGGACTCATCAGGAAGC
TCGGACTCATCAGGAAGC
GACATCGGACTCATCAGGAAGC
GGCATCGGACTCATCAGGAAGC
ACCATCGGACTCATCAGGAAGC
ACATCGGACTCATCAGGAAGC
GTTCATCGGACTCATCAGGAAGC
ATGCATCGGACTCATCAGGAAGC
ACATCGGACTCATCAGGAAGC
GCATCGGACTCATCAGGAAGC
GCATCGGACTCATCAGGAAGC
ATCATCGGACTCATCAGGAAGC

CCATCGGACTCATCAGGAAGC
CATCGGACTCATCAGGAAGC
ATCGGACTCATCAGGAAGC
TCATCGGACTCATCAGGAAGC
TCGGACTCATCAGGAAGC
ACCATCGGACTCATCAGGAAGC
TGCATCGGACTCATCAGGAAGC
CTCATCGGACTCATCAGGAAGC
TCATCGGACTCATCAGGAAGC
ATCGGACTCATCAGGAAGC
ACATCGGACTCATCAGGAAGC
GCACATCGGACTCATCAGGAAGC
TACATCGGACTCATCAGGAAGC
TACATCGGACTCATCAGGAAGC
CATCGGACTCATCAGGAAGC
CATCGGACTCATCAGGAAGC
ATCGGACTCATCAGGAAGC
CATCGGACTCATCAGGAAGC
GACTCATCAGGAAGC
CCATCGGACTCATCAGGAAGC
ACATCGGACTCATCAGGAAGC
TCATCGGACTCATCAGGAAGC
TCGGACTCATCAGGAAGC
GCATCGGACTCATCAGGAAGC
ATCGGACTCATCAGGAAGC
CATCGGACTCATCAGGAAGC
TCATCGGACTCATCAGGAAGC
TCATCGGACTCATCAGGAAGC
TCATCGGACTCATCAGGAAGC
ACATCGGACTCATCAGGAAGC
GCATCGGACTCATCAGGAAGC
ACATCGGACTCATCAGGAAGC
ATCATCGGACTCATCAGGAAGC
ACATCGGACTCATCAGGAAGC
CCATCGGACTCATCAGGAAGC
TCGGACTCATCAGGAAGC
TCATCGGACTCATCAGGAAGC
GGACTCATCAGGAAGC
GCATCGGACTCATCAGGAAGC
CATCGGACTCATCAGGAAGC
TCGGACTCATCAGGAAGC
ATCATCGGACTCATCAGGAAGC
TCATCGGACTCATCAGGAAGC

ATCATCGGACTCATCAGGAAGC
TCATCGGACTCATCAGGAAGC
CATCGGACTCATCAGGAAGC
GCATCGGACTCATCAGGAAGC
GCATCGGACTCATCAGGAAGC
GCATCGGACTCATCAGGAAGC
AAACATCGGACTCATCAGGAAGC
ACATCGGACTCATCAGGAAGC
CATCGGACTCATCAGGAAGC
AAACATCGGACTCATCAGGAAGC
GCATCGGACTCATCAGGAAGC
GGCATCGGACTCATCAGGAAGC
ATCGGACTCATCAGGAAGC
CGGACTCATCAGGAAGC
TCGGACTCATCAGGAAGC
CATCGGACTCATCAGGAAGC
AACATCGGACTCATCAGGAAGC
ACTCATCAGGAAGC
GCATCGGACTCATCAGGAAGC
CATCGGACTCATCAGGAAGC
ACCATCGGACTCATCAGGAAGC
ATCGGACTCATCAGGAAGC
CATCGGACTCATCAGGAAGC
ACATCGGACTCATCAGGAAGC
ATCGGACTCATCAGGAAGC
GGACTCATCAGGAAGC
ACATCGGACTCATCAGGAAGC
ATCGGACTCATCAGGAAGC
CACATCGGACTCATCAGGAAGC
TCATCGGACTCATCAGGAAGC
CATCGGACTCATCAGGAAGC
TGCATCGGACTCATCAGGAAGC
CGGACTCATCAGGAAGC
TCGGACTCATCAGGAAGC
GACATCGGACTCATCAGGAAGC
TTCATCGGACTCATCAGGAAGC
AGCATCGGACTCATCAGGAAGC
TCATCGGACTCATCAGGAAGC
CATCGGACTCATCAGGAAGC
CATCGGACTCATCAGGAAGC
TCGGACTCATCAGGAAGC
TCGGACTCATCAGGAAGC
TCATCGGACTCATCAGGAAGC

CACATCGGACTCATCAGGAAGC
ACATCGGACTCATCAGGAAGC
GCATCGGACTCATCAGGAAGC
CGGACTCATCAGGAAGC
CATCGGACTCATCAGGAAGC
CCATCGGACTCATCAGGAAGC
CATCGGACTCATCAGGAAGC
ACATCGGACTCATCAGGAAGC
ATCGGACTCATCAGGAAGC
TGCCATCGGACTCATCAGGAAGC
CATCGGACTCATCAGGAAGC
AACATCGGACTCATCAGGAAGC
AACATCGGACTCATCAGGAAGC
AGCATCGGACTCATCAGGAAGC
GGCATCGGACTCATCAGGAAGC
TGCATCGGACTCATCAGGAAGC
AAACATCGGACTCATCAGGAAGC
CCATCGGACTCATCAGGAAGC
GCATCGGACTCATCAGGAAGC
CCATCGGACTCATCAGGAAGC
GATCATCGGACTCATCAGGAAGC
CATCGGACTCATCAGGAAGC
CATCGGACTCATCAGGAAGC
GTCATCGGACTCATCAGGAAGC
CCCATCGGACTCATCAGGAAGC
ATCGGACTCATCAGGAAGC
AAACATCGGACTCATCAGGAAGC
ATCGGACTCATCAGGAAGC
GACTCATCAGGAAGC
CATCGGACTCATCAGGAAGC
TAACATCGGACTCATCAGGAAGC
TACATCGGACTCATCAGGAAGC
GCATCGGACTCATCAGGAAGC
TTTCATCGGACTCATCAGGAAGC
ACATCGGACTCATCAGGAAGC
CCCATCGGACTCATCAGGAAGC
TCGGACTCATCAGGAAGC
CATCGGACTCATCAGGAAGC
TTTCATCGGACTCATCAGGAAGC
CATCGGACTCATCAGGAAGC
TCGGACTCATCAGGAAGC
TCATCGGACTCATCAGGAAGC
GTCATCGGACTCATCAGGAAGC

ATCGGACTCATCAGGAAGC
GCATCGGACTCATCAGGAAGC
ATCGGACTCATCAGGAAGC
ATCGGACTCATCAGGAAGC
ATCGGACTCATCAGGAAGC
TCATCGGACTCATCAGGAAGC
ATCGGACTCATCAGGAAGC
CATCGGACTCATCAGGAAGC
GCATCGGACTCATCAGGAAGC
GCATCGGACTCATCAGGAAGC
ATCGGACTCATCAGGAAGC
ACATCGGACTCATCAGGAAGC
ATCGGACTCATCAGGAAGC
AACATCGGACTCATCAGGAAGC
ATCGGACTCATCAGGAAGC
ATCGGACTCATCAGGAAGC
ATCATCGGACTCATCAGGAAGC
CATCGGACTCATCAGGAAGC
TCGGACTCATCAGGAAGC
GCATCGGACTCATCAGGAAGC
ACATCGGACTCATCAGGAAGC
CATCGGACTCATCAGGAAGC
TCATCGGACTCATCAGGAAGC
GCATCGGACTCATCAGGAAGC
ATCGGACTCATCAGGAAGC
CATCGGACTCATCAGGAAGC
CATCGGACTCATCAGGAAGC
TCACATCGGACTCATCAGGAAGC
CGGACTCATCAGGAAGC
CATCGGACTCATCAGGAAGC
TCATCGGACTCATCAGGAAGC
GCATCGGACTCATCAGGAAGC
CATCGGACTCATCAGGAAGC
TCATCGGACTCATCAGGAAGC
AACATCGGACTCATCAGGAAGC
ATCGGACTCATCAGGAAGC
ATCGGACTCATCAGGAAGC
GCATCGGACTCATCAGGAAGC
CTCATCGGACTCATCAGGAAGC
TACATCGGACTCATCAGGAAGC
CATCGGACTCATCAGGAAGC
ATCGGACTCATCAGGAAGC
ATCGGACTCATCAGGAAGC

AACCATCGGACTCATCAGGAAGC
CGGACTCATCAGGAAGC
TCGGACTCATCAGGAAGC
GCATCGGACTCATCAGGAAGC
CATCGGACTCATCAGGAAGC
ATCGGACTCATCAGGAAGC
GTCATCGGACTCATCAGGAAGC
TCGGACTCATCAGGAAGC
AGTAACATCGGACTCATCAGGAAGC
CCATCGGACTCATCAGGAAGC
ATCGGACTCATCAGGAAGC
GAACCATCGGACTCATCAGGAAGC
TCGGACTCATCAGGAAGC
CATCGGACTCATCAGGAAGC
ACATCGGACTCATCAGGAAGC
GAGCATCGGACTCATCAGGAAGC
TCATCGGACTCATCAGGAAGC
ATCGGACTCATCAGGAAGC
TCATCGGACTCATCAGGAAGC
TCATCGGACTCATCAGGAAGC
GGACTCATCAGGAAGC
GGACTCATCAGGAAGC
CATCGGACTCATCAGGAAGC
CATCGGACTCATCAGGAAGC
GACTCATCAGGAAGC
ATCATCGGACTCATCAGGAAGC
TCATCGGACTCATCAGGAAGC
TTCATCGGACTCATCAGGAAGC
AGACATCGGACTCATCAGGAAGC
CATCGGACTCATCAGGAAGC
TCATCGGACTCATCAGGAAGC
CATCGGACTCATCAGGAAGC
ATCGGACTCATCAGGAAGC
CCCATCGGACTCATCAGGAAGC
CATCGGACTCATCAGGAAGC
AACATCGGACTCATCAGGAAGC
TCGGACTCATCAGGAAGC
ACATCGGACTCATCAGGAAGC
TCATCGGACTCATCAGGAAGC
ATCATCGGACTCATCAGGAAGC
ATCATCGGACTCATCAGGAAGC
GGACTCATCAGGAAGC
ACATCGGACTCATCAGGAAGC

GCATCGGACTCATCAGGAAGC
ATCGGACTCATCAGGAAGC
ATCGGACTCATCAGGAAGC
CACATCGGACTCATCAGGAAGC
CATCGGACTCATCAGGAAGC
TGCATCGGACTCATCAGGAAGC
GCATCGGACTCATCAGGAAGC
TCCATCGGACTCATCAGGAAGC
ACATCGGACTCATCAGGAAGC
GGACTCATCAGGAAGC
GGACTCATCAGGAAGC
ACATCGGACTCATCAGGAAGC
CATCGGACTCATCAGGAAGC
CATCGGACTCATCAGGAAGC
CATCGGACTCATCAGGAAGC
CGGACTCATCAGGAAGC
TTCATCGGACTCATCAGGAAGC
TTCATCGGACTCATCAGGAAGC
ACATCGGACTCATCAGGAAGC
ATCGGACTCATCAGGAAGC
TCGGACTCATCAGGAAGC
ACATCGGACTCATCAGGAAGC
CGGACTCATCAGGAAGC
CGGACTCATCAGGAAGC
TCGGACTCATCAGGAAGC
AAACATCGGACTCATCAGGAAGC
TGCATCGGACTCATCAGGAAGC
ACATCGGACTCATCAGGAAGC
TCGGACTCATCAGGAAGC
CCCATCGGACTCATCAGGAAGC
TCCATCGGACTCATCAGGAAGC
CATCGGACTCATCAGGAAGC
GACTCATCAGGAAGC
CATCGGACTCATCAGGAAGC
CCATCGGACTCATCAGGAAGC
ATCGGACTCATCAGGAAGC
CATCGGACTCATCAGGAAGC
TACATCGGACTCATCAGGAAGC
CATCGGACTCATCAGGAAGC
CATCGGACTCATCAGGAAGC
ACATCGGACTCATCAGGAAGC
GAGCATCGGACTCATCAGGAAGC
TTCATCGGACTCATCAGGAAGC

CGGACTCATCAGGAAGC
ATCGGACTCATCAGGAAGC
ACATCGGACTCATCAGGAAGC
AGCATCGGACTCATCAGGAAGC
GCATCGGACTCATCAGGAAGC
CATCGGACTCATCAGGAAGC
CGGACTCATCAGGAAGC
ATCGGACTCATCAGGAAGC
ACATCGGACTCATCAGGAAGC
GACTCATCAGGAAGC
GACATCGGACTCATCAGGAAGC
GTCATCGGACTCATCAGGAAGC
ACCATCGGACTCATCAGGAAGC
GTCATCGGACTCATCAGGAAGC
ATCGGACTCATCAGGAAGC
AAACATCGGACTCATCAGGAAGC
CTCATCGGACTCATCAGGAAGC
CTCATCGGACTCATCAGGAAGC
CTCATCGGACTCATCAGGAAGC
TCATCGGACTCATCAGGAAGC
ATCGGACTCATCAGGAAGC
CATCGGACTCATCAGGAAGC
ATCATCGGACTCATCAGGAAGC
CATCGGACTCATCAGGAAGC
TCGGACTCATCAGGAAGC
ACATCGGACTCATCAGGAAGC
ATCGGACTCATCAGGAAGC
ATCGGACTCATCAGGAAGC
ATCGGACTCATCAGGAAGC
AGCATCGGACTCATCAGGAAGC
ATCGGACTCATCAGGAAGC
CGGACTCATCAGGAAGC
ACCATCGGACTCATCAGGAAGC
ATCGGACTCATCAGGAAGC
TCATCGGACTCATCAGGAAGC
ATCGGACTCATCAGGAAGC
CTCATCGGACTCATCAGGAAGC
TCGGACTCATCAGGAAGC
ACATCGGACTCATCAGGAAGC
CATCGGACTCATCAGGAAGC
ATCGGACTCATCAGGAAGC
ACCATCGGACTCATCAGGAAGC
TCATCGGACTCATCAGGAAGC

CATCGGACTCATCAGGAAGC
CGGACTCATCAGGAAGC
TCCATCGGACTCATCAGGAAGC
GACTCATCAGGAAGC
CATCGGACTCATCAGGAAGC
ACATCGGACTCATCAGGAAGC
GGACTCATCAGGAAGC
TTCATCGGACTCATCAGGAAGC
CATCGGACTCATCAGGAAGC
CCATCGGACTCATCAGGAAGC
CATCGGACTCATCAGGAAGC
CATCGGACTCATCAGGAAGC
CATCGGACTCATCAGGAAGC
CGGACTCATCAGGAAGC
TCATCGGACTCATCAGGAAGC
AGCATCGGACTCATCAGGAAGC
AATCATCGGACTCATCAGGAAGC
AATCATCGGACTCATCAGGAAGC
AGCATCGGACTCATCAGGAAGC
ACCATCGGACTCATCAGGAAGC
CGGACTCATCAGGAAGC
ACCATCGGACTCATCAGGAAGC
ATCGGACTCATCAGGAAGC
ATCGGACTCATCAGGAAGC
TCATCGGACTCATCAGGAAGC
ATCGGACTCATCAGGAAGC
AGCATCGGACTCATCAGGAAGC
TGCATCGGACTCATCAGGAAGC
CATCGGACTCATCAGGAAGC
GGACTCATCAGGAAGC
TACATCGGACTCATCAGGAAGC
TCGGACTCATCAGGAAGC
TCCATCGGACTCATCAGGAAGC
TCGGACTCATCAGGAAGC
TTTCATCGGACTCATCAGGAAGC
CATCGGACTCATCAGGAAGC
CATCGGACTCATCAGGAAGC
ATCGGACTCATCAGGAAGC
ACATCGGACTCATCAGGAAGC
CCATCGGACTCATCAGGAAGC
ATCGGACTCATCAGGAAGC
TACATCGGACTCATCAGGAAGC
TACATCGGACTCATCAGGAAGC

CGGACTCATCAGGAAGC
ACATCGGACTCATCAGGAAGC
ACCATCGGACTCATCAGGAAGC
ACCATCGGACTCATCAGGAAGC
AGCATCGGACTCATCAGGAAGC
CCATCGGACTCATCAGGAAGC
GCATCGGACTCATCAGGAAGC
CATCGGACTCATCAGGAAGC
AACATCGGACTCATCAGGAAGC
ACATCGGACTCATCAGGAAGC
TCATCGGACTCATCAGGAAGC
ATCGGACTCATCAGGAAGC
CATCGGACTCATCAGGAAGC
ACATCGGACTCATCAGGAAGC
GGACTCATCAGGAAGC
GGACTCATCAGGAAGC
AGCATCGGACTCATCAGGAAGC
ACATCGGACTCATCAGGAAGC
ACATCGGACTCATCAGGAAGC
CATCGGACTCATCAGGAAGC
CCATCGGACTCATCAGGAAGC
TCGGACTCATCAGGAAGC
CATCGGACTCATCAGGAAGC
CGGACTCATCAGGAAGC
TCGGACTCATCAGGAAGC
ATCGGACTCATCAGGAAGC
ACATCGGACTCATCAGGAAGC
CCATCGGACTCATCAGGAAGC
CCATCGGACTCATCAGGAAGC
ATCGGACTCATCAGGAAGC
CATCGGACTCATCAGGAAGC
TACATCGGACTCATCAGGAAGC
ATCGGACTCATCAGGAAGC
AGCATCGGACTCATCAGGAAGC
GACTCATCAGGAAGC
CATCGGACTCATCAGGAAGC
ACATCGGACTCATCAGGAAGC
ATCGGACTCATCAGGAAGC
ATCGGACTCATCAGGAAGC
CGGACTCATCAGGAAGC
CATCGGACTCATCAGGAAGC
TCATCGGACTCATCAGGAAGC
TCATCGGACTCATCAGGAAGC

TCATCGGACTCATCAGGAAGC
TCGGACTCATCAGGAAGC
TCGGACTCATCAGGAAGC
TCGGACTCATCAGGAAGC
TCATCGGACTCATCAGGAAGC
AACATCGGACTCATCAGGAAGC
TGCATCGGACTCATCAGGAAGC
TCGGACTCATCAGGAAGC
GGACTCATCAGGAAGC
TCGGACTCATCAGGAAGC
TCATCGGACTCATCAGGAAGC
ATCGGACTCATCAGGAAGC
ACATCGGACTCATCAGGAAGC
ACATCGGACTCATCAGGAAGC
GCATCGGACTCATCAGGAAGC
CGGACTCATCAGGAAGC
TCATCGGACTCATCAGGAAGC
CATCGGACTCATCAGGAAGC
CGGACTCATCAGGAAGC
CATCGGACTCATCAGGAAGC
ACATCGGACTCATCAGGAAGC
CATCGGACTCATCAGGAAGC
GCATCGGACTCATCAGGAAGC
CATCGGACTCATCAGGAAGC
GGCATCGGACTCATCAGGAAGC
ATCGGACTCATCAGGAAGC
TCGGACTCATCAGGAAGC
TCATCGGACTCATCAGGAAGC
CGGACTCATCAGGAAGC
TCATCGGACTCATCAGGAAGC
GGACTCATCAGGAAGC
ATCGGACTCATCAGGAAGC
ATCGGACTCATCAGGAAGC
ACATCGGACTCATCAGGAAGC
TCATCGGACTCATCAGGAAGC
CATCGGACTCATCAGGAAGC
CCATCGGACTCATCAGGAAGC
ACATCGGACTCATCAGGAAGC
AACATCGGACTCATCAGGAAGC
TATCATCGGACTCATCAGGAAGC
AGCATCGGACTCATCAGGAAGC
TCCATCGGACTCATCAGGAAGC
TCGGACTCATCAGGAAGC

TGCATCGGACTCATCAGGAAGC
TCGGACTCATCAGGAAGC
CATCGGACTCATCAGGAAGC
CATCGGACTCATCAGGAAGC
CATCGGACTCATCAGGAAGC
CAGCATCGGACTCATCAGGAAGC
CGGACTCATCAGGAAGC
TCGGACTCATCAGGAAGC
GAACATCGGACTCATCAGGAAGC
TCGGACTCATCAGGAAGC
TACATCGGACTCATCAGGAAGC
CATCGGACTCATCAGGAAGC
TTCATCGGACTCATCAGGAAGC
TCATCGGACTCATCAGGAAGC
AACATCGGACTCATCAGGAAGC
CCATCGGACTCATCAGGAAGC
CATCGGACTCATCAGGAAGC
GGCATCGGACTCATCAGGAAGC
CATCGGACTCATCAGGAAGC
CCATCGGACTCATCAGGAAGC
GACTCATCAGGAAGC
CGGACTCATCAGGAAGC
ATCGGACTCATCAGGAAGC
ATCATCGGACTCATCAGGAAGC
TGGCATCGGACTCATCAGGAAGC
CCATCGGACTCATCAGGAAGC
TCGGACTCATCAGGAAGC
TCGGACTCATCAGGAAGC
ATCATCGGACTCATCAGGAAGC
CCATCGGACTCATCAGGAAGC
TCATCGGACTCATCAGGAAGC
GCATCGGACTCATCAGGAAGC
CCATCGGACTCATCAGGAAGC
ACATCGGACTCATCAGGAAGC
TCGGACTCATCAGGAAGC
TCGGACTCATCAGGAAGC
CTCATCGGACTCATCAGGAAGC
CGGACTCATCAGGAAGC
GGCATCGGACTCATCAGGAAGC
ATCGGACTCATCAGGAAGC
CACATCGGACTCATCAGGAAGC
GGGACATCGGACTCATCAGGAAGC
ATACATCGGACTCATCAGGAAGC

CCATCGGACTCATCAGGAAGC
CATCGGACTCATCAGGAAGC
GACTCATCAGGAAGC
CATCGGACTCATCAGGAAGC
ATCGGACTCATCAGGAAGC
CATCGGACTCATCAGGAAGC
GCATCGGACTCATCAGGAAGC
GCATCGGACTCATCAGGAAGC
TCCATCGGACTCATCAGGAAGC
CATCGGACTCATCAGGAAGC
ACATCGGACTCATCAGGAAGC
ACATCGGACTCATCAGGAAGC
ATATCATCGGACTCATCAGGAAGC
TACATCGGACTCATCAGGAAGC
ATCATCGGACTCATCAGGAAGC
CATCGGACTCATCAGGAAGC
ATCGGACTCATCAGGAAGC
CGGACTCATCAGGAAGC
TCGGACTCATCAGGAAGC
GCCATCGGACTCATCAGGAAGC
TCTCATCGGACTCATCAGGAAGC
TGCATCGGACTCATCAGGAAGC
GCATCGGACTCATCAGGAAGC
TCGGACTCATCAGGAAGC
CATCGGACTCATCAGGAAGC
CATCGGACTCATCAGGAAGC
ACTCATCAGGAAGC
TCATCGGACTCATCAGGAAGC
CATCGGACTCATCAGGAAGC
ATGACATCGGACTCATCAGGAAGC
GCATCGGACTCATCAGGAAGC
CATCGGACTCATCAGGAAGC
CATCGGACTCATCAGGAAGC
TCGGACTCATCAGGAAGC
CGGACTCATCAGGAAGC
CGGACTCATCAGGAAGC
TCATCGGACTCATCAGGAAGC
TCCATCGGACTCATCAGGAAGC
TACATCGGACTCATCAGGAAGC
ATCGGACTCATCAGGAAGC
TCATCGGACTCATCAGGAAGC
GGACTCATCAGGAAGC
CATCGGACTCATCAGGAAGC

CGGACTCATCAGGAAGC
TCGGACTCATCAGGAAGC
CATCGGACTCATCAGGAAGC
TCCATCGGACTCATCAGGAAGC
CATCGGACTCATCAGGAAGC
CATCGGACTCATCAGGAAGC
ACATCGGACTCATCAGGAAGC
ATCGGACTCATCAGGAAGC
ACATCGGACTCATCAGGAAGC
CATCGGACTCATCAGGAAGC
GGACTCATCAGGAAGC
ACTCATCAGGAAGC
ATCATCGGACTCATCAGGAAGC
GCATCGGACTCATCAGGAAGC
GACTCATCAGGAAGC
CGGACTCATCAGGAAGC
CATCGGACTCATCAGGAAGC
ACATCGGACTCATCAGGAAGC
TCATCGGACTCATCAGGAAGC
CCATCGGACTCATCAGGAAGC
GGGCATCGGACTCATCAGGAAGC
ACCATCGGACTCATCAGGAAGC
AGCACATCGGACTCATCAGGAAGC
TCGGACTCATCAGGAAGC
TACATCGGACTCATCAGGAAGC
TACATCGGACTCATCAGGAAGC
GCCATCGGACTCATCAGGAAGC
CATCGGACTCATCAGGAAGC
CATCGGACTCATCAGGAAGC
ACATCGGACTCATCAGGAAGC
GCATCGGACTCATCAGGAAGC
CCATCGGACTCATCAGGAAGC
CATCGGACTCATCAGGAAGC
CCCATCGGACTCATCAGGAAGC
CATCGGACTCATCAGGAAGC
CACATCGGACTCATCAGGAAGC
TACCATCGGACTCATCAGGAAGC
GTGCATCGGACTCATCAGGAAGC
AACATCGGACTCATCAGGAAGC
ACATCGGACTCATCAGGAAGC
TCCATCGGACTCATCAGGAAGC
CATCGGACTCATCAGGAAGC
ACATCGGACTCATCAGGAAGC

GCATCGGACTCATCAGGAAGC
TACATCGGACTCATCAGGAAGC
ATCGGACTCATCAGGAAGC
TACATCGGACTCATCAGGAAGC
CATCGGACTCATCAGGAAGC
ACATCGGACTCATCAGGAAGC
ATCGGACTCATCAGGAAGC
CCATCGGACTCATCAGGAAGC
TACATCGGACTCATCAGGAAGC
TACATCGGACTCATCAGGAAGC
CACCATCGGACTCATCAGGAAGC
ACATCGGACTCATCAGGAAGC
CATCGGACTCATCAGGAAGC
ATCGGACTCATCAGGAAGC
CATCGGACTCATCAGGAAGC
CATCGGACTCATCAGGAAGC
ATCGGACTCATCAGGAAGC
ATCGGACTCATCAGGAAGC
ACATCGGACTCATCAGGAAGC
GCATCGGACTCATCAGGAAGC
GCATCGGACTCATCAGGAAGC
GCATCGGACTCATCAGGAAGC
TGCATCGGACTCATCAGGAAGC
TACATCGGACTCATCAGGAAGC
TACATCGGACTCATCAGGAAGC
ATCGGACTCATCAGGAAGC
CCATCGGACTCATCAGGAAGC
CATCGGACTCATCAGGAAGC
ATCGGACTCATCAGGAAGC
TCATCGGACTCATCAGGAAGC
TCATCGGACTCATCAGGAAGC
CCATCGGACTCATCAGGAAGC
CCATCGGACTCATCAGGAAGC
GCATCGGACTCATCAGGAAGC
CACATCGGACTCATCAGGAAGC
TGCATCGGACTCATCAGGAAGC
TACATCGGACTCATCAGGAAGC
CCATCGGACTCATCAGGAAGC
CCATCGGACTCATCAGGAAGC
TCGGACTCATCAGGAAGC
GCATCGGACTCATCAGGAAGC
CATCGGACTCATCAGGAAGC
CATCGGACTCATCAGGAAGC

CATCGGACTCATCAGGAAGC
GCATCGGACTCATCAGGAAGC
TCGGACTCATCAGGAAGC
CATCGGACTCATCAGGAAGC
TACATCGGACTCATCAGGAAGC
CATCGGACTCATCAGGAAGC
AGCATCGGACTCATCAGGAAGC
TCATCGGACTCATCAGGAAGC
GTCATCGGACTCATCAGGAAGC
CATCGGACTCATCAGGAAGC
TCGGACTCATCAGGAAGC
TCGGACTCATCAGGAAGC
TACATCGGACTCATCAGGAAGC
CATCGGACTCATCAGGAAGC
TCCATCGGACTCATCAGGAAGC
TCCATCGGACTCATCAGGAAGC
CATCGGACTCATCAGGAAGC
TACATCGGACTCATCAGGAAGC
CATCGGACTCATCAGGAAGC
GCATCGGACTCATCAGGAAGC
TCGGACTCATCAGGAAGC
GCATCGGACTCATCAGGAAGC
CATCGGACTCATCAGGAAGC
TCATCGGACTCATCAGGAAGC
ACATCGGACTCATCAGGAAGC
TGCATCGGACTCATCAGGAAGC
ATCGGACTCATCAGGAAGC
CATCGGACTCATCAGGAAGC
CGGACTCATCAGGAAGC
GCATCGGACTCATCAGGAAGC
TCGGACTCATCAGGAAGC
TCATCGGACTCATCAGGAAGC
CATCGGACTCATCAGGAAGC
CCATCGGACTCATCAGGAAGC
TCCATCGGACTCATCAGGAAGC
CATCGGACTCATCAGGAAGC
AGCATCGGACTCATCAGGAAGC
GCATCGGACTCATCAGGAAGC
TACATCGGACTCATCAGGAAGC
TCGGACTCATCAGGAAGC
CCATCGGACTCATCAGGAAGC
ATCGGACTCATCAGGAAGC
TCATCGGACTCATCAGGAAGC

ACATCGGACTCATCAGGAAGC
ACATCGGACTCATCAGGAAGC
CGGACTCATCAGGAAGC
GCATCGGACTCATCAGGAAGC
ATCGGACTCATCAGGAAGC
GCATCGGACTCATCAGGAAGC
CGGACTCATCAGGAAGC
TCATCGGACTCATCAGGAAGC
ACATCGGACTCATCAGGAAGC
AATCATCGGACTCATCAGGAAGC
AATCATCGGACTCATCAGGAAGC
ATCGGACTCATCAGGAAGC
CATCGGACTCATCAGGAAGC
TCATCGGACTCATCAGGAAGC
TCATCGGACTCATCAGGAAGC
ATCGGACTCATCAGGAAGC
CATCGGACTCATCAGGAAGC
ATCGGACTCATCAGGAAGC
ACATCGGACTCATCAGGAAGC
GGCATCGGACTCATCAGGAAGC
GGCATCGGACTCATCAGGAAGC
ACACATCGGACTCATCAGGAAGC
AACATCGGACTCATCAGGAAGC
CATCGGACTCATCAGGAAGC
CATCGGACTCATCAGGAAGC
ATCGGACTCATCAGGAAGC
ATCGGACTCATCAGGAAGC
TCATCGGACTCATCAGGAAGC
CATCGGACTCATCAGGAAGC
TCGGACTCATCAGGAAGC
CATCGGACTCATCAGGAAGC
GCATCGGACTCATCAGGAAGC
TCATCGGACTCATCAGGAAGC
TCGGACTCATCAGGAAGC
CATCGGACTCATCAGGAAGC
ACTCATCGGACTCATCAGGAAGC
GCATCGGACTCATCAGGAAGC
GACATCGGACTCATCAGGAAGC
ACATCGGACTCATCAGGAAGC
TCGGACTCATCAGGAAGC
TCGGACTCATCAGGAAGC
TCGGACTCATCAGGAAGC
TACATCGGACTCATCAGGAAGC

TACATCGGACTCATCAGGAAGC
ACATCGGACTCATCAGGAAGC
CGGACTCATCAGGAAGC
CATCGGACTCATCAGGAAGC
ATCATCGGACTCATCAGGAAGC
TCATCGGACTCATCAGGAAGC
ATCGGACTCATCAGGAAGC
TCGGACTCATCAGGAAGC
GCATCGGACTCATCAGGAAGC
TCGGACTCATCAGGAAGC
GCATCGGACTCATCAGGAAGC
ATCATCGGACTCATCAGGAAGC
ATCGGACTCATCAGGAAGC
GGACTCATCAGGAAGC
ACTCATCAGGAAGC
TCCATCGGACTCATCAGGAAGC
CATCGGACTCATCAGGAAGC
CACATCGGACTCATCAGGAAGC
ATCATCGGACTCATCAGGAAGC
CATCGGACTCATCAGGAAGC
TGCATCGGACTCATCAGGAAGC
GACTCATCAGGAAGC
GGACTCATCAGGAAGC
CATCGGACTCATCAGGAAGC
TCGGACTCATCAGGAAGC
ATCGGACTCATCAGGAAGC
ATCGGACTCATCAGGAAGC
TCATCGGACTCATCAGGAAGC
CACATCGGACTCATCAGGAAGC
GACTCATCAGGAAGC
GACTCATCAGGAAGC
TGCATCGGACTCATCAGGAAGC
CATCGGACTCATCAGGAAGC
CATCGGACTCATCAGGAAGC
CGGACTCATCAGGAAGC
ATCGGACTCATCAGGAAGC
CATCGGACTCATCAGGAAGC
GGACTCATCAGGAAGC
GCATCGGACTCATCAGGAAGC
TCGGACTCATCAGGAAGC
TCGGACTCATCAGGAAGC
TCGGACTCATCAGGAAGC
GCATCGGACTCATCAGGAAGC

ACATCGGACTCATCAGGAAGC
TCATCGGACTCATCAGGAAGC
ACTCATCGGACTCATCAGGAAGC
CACATCGGACTCATCAGGAAGC
ATCGGACTCATCAGGAAGC
TCATCGGACTCATCAGGAAGC
GCTCATCGGACTCATCAGGAAGC
ACATCGGACTCATCAGGAAGC
AATCATCGGACTCATCAGGAAGC
ACATCGGACTCATCAGGAAGC
ATCGGACTCATCAGGAAGC
TCGGACTCATCAGGAAGC
GGACTCATCAGGAAGC
ATCGGACTCATCAGGAAGC
CGGACTCATCAGGAAGC
GGCATCGGACTCATCAGGAAGC
GGCATCGGACTCATCAGGAAGC
CATCGGACTCATCAGGAAGC
TTCATCGGACTCATCAGGAAGC
GCATCGGACTCATCAGGAAGC
TCTCATCGGACTCATCAGGAAGC
CATCGGACTCATCAGGAAGC
TTCATCGGACTCATCAGGAAGC
TCGGACTCATCAGGAAGC
TCGGACTCATCAGGAAGC
CGGACTCATCAGGAAGC
CAGCATCGGACTCATCAGGAAGC
ATCGGACTCATCAGGAAGC
GGCATCGGACTCATCAGGAAGC
ATCGGACTCATCAGGAAGC
CATCGGACTCATCAGGAAGC
CATCGGACTCATCAGGAAGC
TACATCGGACTCATCAGGAAGC
GCATCGGACTCATCAGGAAGC
GACATCGGACTCATCAGGAAGC
CATCGGACTCATCAGGAAGC
CATCGGACTCATCAGGAAGC
CTGCATCGGACTCATCAGGAAGC
CACATCGGACTCATCAGGAAGC
ACATCGGACTCATCAGGAAGC
GGACTCATCAGGAAGC
CATCGGACTCATCAGGAAGC
TCATCGGACTCATCAGGAAGC

GATCATCGGACTCATCAGGAAGC
GCATCGGACTCATCAGGAAGC
TCATCGGACTCATCAGGAAGC
GAGCATCGGACTCATCAGGAAGC
CATCGGACTCATCAGGAAGC
TCATCGGACTCATCAGGAAGC
ATCGGACTCATCAGGAAGC
CTCATCGGACTCATCAGGAAGC
CTCATCGGACTCATCAGGAAGC
TCGGACTCATCAGGAAGC
CATCGGACTCATCAGGAAGC
CATCGGACTCATCAGGAAGC
CATCGGACTCATCAGGAAGC
GCATCGGACTCATCAGGAAGC
TGACATCGGACTCATCAGGAAGC
ATCGGACTCATCAGGAAGC
ATCGGACTCATCAGGAAGC
CATCGGACTCATCAGGAAGC
CATCGGACTCATCAGGAAGC
CGGACTCATCAGGAAGC
TCATCGGACTCATCAGGAAGC
CATCGGACTCATCAGGAAGC
TCGGACTCATCAGGAAGC
CATCGGACTCATCAGGAAGC
CATCGGACTCATCAGGAAGC
CATCGGACTCATCAGGAAGC
TCGGACTCATCAGGAAGC
CCATCGGACTCATCAGGAAGC
AACATCGGACTCATCAGGAAGC
AACATCGGACTCATCAGGAAGC
CATCGGACTCATCAGGAAGC
GGCATCGGACTCATCAGGAAGC
CATCGGACTCATCAGGAAGC
CATCGGACTCATCAGGAAGC
TGCATCGGACTCATCAGGAAGC
CATCGGACTCATCAGGAAGC
CCCACATCGGACTCATCAGGAAGC
TACATCGGACTCATCAGGAAGC
CGGACTCATCAGGAAGC
ATCGGACTCATCAGGAAGC
GCCATCGGACTCATCAGGAAGC
CATCGGACTCATCAGGAAGC
GGACTCATCAGGAAGC

ACATCGGACTCATCAGGAAGC
TCGGACTCATCAGGAAGC
TCGGACTCATCAGGAAGC
CGGACTCATCAGGAAGC
ATCGGACTCATCAGGAAGC
TTCATCGGACTCATCAGGAAGC
GCATCGGACTCATCAGGAAGC
AACATCGGACTCATCAGGAAGC
GGACTCATCAGGAAGC
GGACTCATCAGGAAGC
TTCATCGGACTCATCAGGAAGC
AGCATCGGACTCATCAGGAAGC
AGCATCGGACTCATCAGGAAGC
CCCATCGGACTCATCAGGAAGC
GCATCGGACTCATCAGGAAGC
TGCATCGGACTCATCAGGAAGC
CATCGGACTCATCAGGAAGC
TTCATCGGACTCATCAGGAAGC
GCATCGGACTCATCAGGAAGC
GAACATCGGACTCATCAGGAAGC
TTCATCGGACTCATCAGGAAGC
ATTCATCGGACTCATCAGGAAGC
ACATCGGACTCATCAGGAAGC
TCATCGGACTCATCAGGAAGC
CATCGGACTCATCAGGAAGC
CATCGGACTCATCAGGAAGC
CATCGGACTCATCAGGAAGC
TCCATCGGACTCATCAGGAAGC
CCATCGGACTCATCAGGAAGC
AGACATCGGACTCATCAGGAAGC
GTTCCCATCGGACTCATCAGGAAGC
TTGCATCGGACTCATCAGGAAGC
ACATCGGACTCATCAGGAAGC
TTCATCGGACTCATCAGGAAGC
CATCGGACTCATCAGGAAGC
GGACTCATCAGGAAGC
TCGGACTCATCAGGAAGC
TAACATCGGACTCATCAGGAAGC
ATCATCGGACTCATCAGGAAGC
TTCATCGGACTCATCAGGAAGC
AAACATCGGACTCATCAGGAAGC
TCATCGGACTCATCAGGAAGC
TCGGACTCATCAGGAAGC

CGGACTCATCAGGAAGC
GACATCGGACTCATCAGGAAGC
TCGGACTCATCAGGAAGC
ACATCGGACTCATCAGGAAGC
TCATCGGACTCATCAGGAAGC
AAACATCGGACTCATCAGGAAGC
TCGGACTCATCAGGAAGC
CTCATCGGACTCATCAGGAAGC
ATCATCGGACTCATCAGGAAGC
ACATCGGACTCATCAGGAAGC
CCATCGGACTCATCAGGAAGC
TCGGACTCATCAGGAAGC
ATCGGACTCATCAGGAAGC
TCATCGGACTCATCAGGAAGC
ATCGGACTCATCAGGAAGC
CATCGGACTCATCAGGAAGC
GCATCGGACTCATCAGGAAGC
CGGACTCATCAGGAAGC
ACATCGGACTCATCAGGAAGC
TCGGACTCATCAGGAAGC
TCGGACTCATCAGGAAGC
CATCGGACTCATCAGGAAGC
AAATCATCGGACTCATCAGGAAGC
GCATCGGACTCATCAGGAAGC
CATCGGACTCATCAGGAAGC
TGCATCGGACTCATCAGGAAGC
GCATCGGACTCATCAGGAAGC
ATCGGACTCATCAGGAAGC
CATCGGACTCATCAGGAAGC
CATCGGACTCATCAGGAAGC
ACATCGGACTCATCAGGAAGC
TCGGACTCATCAGGAAGC
TCGGACTCATCAGGAAGC
GCATCGGACTCATCAGGAAGC
CATCGGACTCATCAGGAAGC
ATCGGACTCATCAGGAAGC
CCATCGGACTCATCAGGAAGC
TCGGACTCATCAGGAAGC
CATCGGACTCATCAGGAAGC
ATCGGACTCATCAGGAAGC
CATCGGACTCATCAGGAAGC
GTCATCGGACTCATCAGGAAGC
ATCGGACTCATCAGGAAGC

CATCGGACTCATCAGGAAGC
ATCATCGGACTCATCAGGAAGC
TCCATCGGACTCATCAGGAAGC
TACATCGGACTCATCAGGAAGC
CCATCGGACTCATCAGGAAGC
GCATCGGACTCATCAGGAAGC
GACATCGGACTCATCAGGAAGC
CGGACTCATCAGGAAGC
TCATCGGACTCATCAGGAAGC
ATCATCGGACTCATCAGGAAGC
ATCGGACTCATCAGGAAGC
CATCGGACTCATCAGGAAGC
GTCATCGGACTCATCAGGAAGC
ACATCGGACTCATCAGGAAGC
CATCGGACTCATCAGGAAGC
CATCGGACTCATCAGGAAGC
TCGGACTCATCAGGAAGC
CATCGGACTCATCAGGAAGC
ATCGGACTCATCAGGAAGC
TCATCGGACTCATCAGGAAGC
ATCATCGGACTCATCAGGAAGC
ACATCGGACTCATCAGGAAGC
TCATCGGACTCATCAGGAAGC
CCCATCGGACTCATCAGGAAGC
CCCATCGGACTCATCAGGAAGC
TCGGACTCATCAGGAAGC
ACATCGGACTCATCAGGAAGC
CATCGGACTCATCAGGAAGC
TCGGACTCATCAGGAAGC
TCATCGGACTCATCAGGAAGC
TCATCGGACTCATCAGGAAGC
GTCATCGGACTCATCAGGAAGC
ACATCGGACTCATCAGGAAGC
AACATCGGACTCATCAGGAAGC
AACATCGGACTCATCAGGAAGC
ATCGGACTCATCAGGAAGC
GCATCGGACTCATCAGGAAGC
CGCATCGGACTCATCAGGAAGC
TCGGACTCATCAGGAAGC
CATCGGACTCATCAGGAAGC
CATCGGACTCATCAGGAAGC
GCATCGGACTCATCAGGAAGC
GTCATCGGACTCATCAGGAAGC

GCATCGGACTCATCAGGAAGC
AACATCGGACTCATCAGGAAGC
AACATCGGACTCATCAGGAAGC
ACATCGGACTCATCAGGAAGC
ATCGGACTCATCAGGAAGC
GGACTCATCAGGAAGC
TACATCGGACTCATCAGGAAGC
ATCGGACTCATCAGGAAGC
GTCATCGGACTCATCAGGAAGC
GTCACATCGGACTCATCAGGAAGC
GCATCGGACTCATCAGGAAGC
CACATCGGACTCATCAGGAAGC
TCATCGGACTCATCAGGAAGC
CGGACTCATCAGGAAGC
ATCGGACTCATCAGGAAGC
CATCGGACTCATCAGGAAGC
CTCATCGGACTCATCAGGAAGC
ATCGGACTCATCAGGAAGC
CATCGGACTCATCAGGAAGC
GACTCATCAGGAAGC
ATCGGACTCATCAGGAAGC
AGATTCATCGGACTCATCAGGAAGC
ACATCGGACTCATCAGGAAGC
ATCGGACTCATCAGGAAGC
ATCGGACTCATCAGGAAGC
CATCGGACTCATCAGGAAGC
ATCGGACTCATCAGGAAGC
GCATCGGACTCATCAGGAAGC
TCATCGGACTCATCAGGAAGC
GCATCGGACTCATCAGGAAGC
ACATCGGACTCATCAGGAAGC
CTCATCGGACTCATCAGGAAGC
TCGGACTCATCAGGAAGC
AACATCGGACTCATCAGGAAGC
GACTCATCAGGAAGC
GCATCGGACTCATCAGGAAGC
CACATCGGACTCATCAGGAAGC
CCATCGGACTCATCAGGAAGC
CATCGGACTCATCAGGAAGC
AACATCGGACTCATCAGGAAGC
AACATCGGACTCATCAGGAAGC
CCATCGGACTCATCAGGAAGC
CGGACTCATCAGGAAGC

GCATCGGACTCATCAGGAAGC
TCATCGGACTCATCAGGAAGC
TCATCGGACTCATCAGGAAGC
GACTCATCAGGAAGC
CATCGGACTCATCAGGAAGC
CATCGGACTCATCAGGAAGC
CCATCGGACTCATCAGGAAGC
CCATCGGACTCATCAGGAAGC
CCATCGGACTCATCAGGAAGC
TCATCGGACTCATCAGGAAGC
TCATCGGACTCATCAGGAAGC
ACATCGGACTCATCAGGAAGC
CATCGGACTCATCAGGAAGC
CGGACTCATCAGGAAGC
CGGACTCATCAGGAAGC
GCCATCGGACTCATCAGGAAGC
ATCATCGGACTCATCAGGAAGC
ATCATCGGACTCATCAGGAAGC
CGGACTCATCAGGAAGC
CTTCATCGGACTCATCAGGAAGC
ATCGGACTCATCAGGAAGC
ATCGGACTCATCAGGAAGC
CATCGGACTCATCAGGAAGC
ACCATCGGACTCATCAGGAAGC
ACCATCGGACTCATCAGGAAGC
TCATCGGACTCATCAGGAAGC
TCGGACTCATCAGGAAGC
TCGGACTCATCAGGAAGC
GCATCGGACTCATCAGGAAGC
GTCATCGGACTCATCAGGAAGC
ACATCGGACTCATCAGGAAGC
CATCGGACTCATCAGGAAGC
ACATCGGACTCATCAGGAAGC
CCATCGGACTCATCAGGAAGC
CGGACTCATCAGGAAGC
TCGGACTCATCAGGAAGC
TCGGACTCATCAGGAAGC
TACATCGGACTCATCAGGAAGC
GGCATCGGACTCATCAGGAAGC
CGGACTCATCAGGAAGC
TCATCGGACTCATCAGGAAGC
CATCGGACTCATCAGGAAGC
CATCGGACTCATCAGGAAGC

ACATCGGACTCATCAGGAAGC
ACATCGGACTCATCAGGAAGC
GGACTCATCAGGAAGC
ATCGGACTCATCAGGAAGC
ATCATCGGACTCATCAGGAAGC
CATCGGACTCATCAGGAAGC
CATCATCGGACTCATCAGGAAGC
CATCGGACTCATCAGGAAGC
GCATCGGACTCATCAGGAAGC
GCATCGGACTCATCAGGAAGC
AACATCGGACTCATCAGGAAGC
AACATCGGACTCATCAGGAAGC
GTCATCGGACTCATCAGGAAGC
TCCATCGGACTCATCAGGAAGC
TCCATCGGACTCATCAGGAAGC
AAACATCGGACTCATCAGGAAGC
AGCATCGGACTCATCAGGAAGC
CATCGGACTCATCAGGAAGC
ATCGGACTCATCAGGAAGC
CGCATCGGACTCATCAGGAAGC
TCATCGGACTCATCAGGAAGC
TCATCGGACTCATCAGGAAGC
ATCATCGGACTCATCAGGAAGC
ATCGGACTCATCAGGAAGC
TCGGACTCATCAGGAAGC
ATCGGACTCATCAGGAAGC
TCTAGCATCGGACTCATCAGGAAGC
ATCATCGGACTCATCAGGAAGC
CGGACTCATCAGGAAGC
CATCGGACTCATCAGGAAGC
TCGGACTCATCAGGAAGC
CATCGGACTCATCAGGAAGC
CGGACTCATCAGGAAGC
TCCATCGGACTCATCAGGAAGC
CCATCGGACTCATCAGGAAGC
ATCGGACTCATCAGGAAGC
ATCGGACTCATCAGGAAGC
ATCGGACTCATCAGGAAGC
TCATCGGACTCATCAGGAAGC
TCATCGGACTCATCAGGAAGC
GGACTCATCAGGAAGC
CCTCATCGGACTCATCAGGAAGC
TCATCGGACTCATCAGGAAGC

TCCATCGGACTCATCAGGAAGC
ATCGGACTCATCAGGAAGC
GCATCGGACTCATCAGGAAGC
AACATCGGACTCATCAGGAAGC
TCGGACTCATCAGGAAGC
CATCGGACTCATCAGGAAGC
GCATCGGACTCATCAGGAAGC
CCATCGGACTCATCAGGAAGC
CTCATCGGACTCATCAGGAAGC
CATCGGACTCATCAGGAAGC
ATACATCGGACTCATCAGGAAGC
TCATCGGACTCATCAGGAAGC
ACTCATCAGGAAGC
GCATCGGACTCATCAGGAAGC
TCGGACTCATCAGGAAGC
ATCGGACTCATCAGGAAGC
ACATCGGACTCATCAGGAAGC
ATCGGACTCATCAGGAAGC
ATCGGACTCATCAGGAAGC
AACATCGGACTCATCAGGAAGC
ACATCGGACTCATCAGGAAGC
ATCGGACTCATCAGGAAGC
ACATCGGACTCATCAGGAAGC
ACATCGGACTCATCAGGAAGC
TGCATCGGACTCATCAGGAAGC
CATCGGACTCATCAGGAAGC
TTCATCGGACTCATCAGGAAGC
TCATCGGACTCATCAGGAAGC
CATCGGACTCATCAGGAAGC
TCATCGGACTCATCAGGAAGC
GACATCGGACTCATCAGGAAGC
GGACTCATCAGGAAGC
CGGACTCATCAGGAAGC
ATCGGACTCATCAGGAAGC
GGACTCATCAGGAAGC
GCATCGGACTCATCAGGAAGC
ACATCGGACTCATCAGGAAGC
CATCGGACTCATCAGGAAGC
CATCGGACTCATCAGGAAGC
CACATCGGACTCATCAGGAAGC
TCGGACTCATCAGGAAGC
TTCATCGGACTCATCAGGAAGC
ATCGGACTCATCAGGAAGC

TCATCGGACTCATCAGGAAGC
ATCGGACTCATCAGGAAGC
CATCGGACTCATCAGGAAGC
ATCGGACTCATCAGGAAGC
CACATCGGACTCATCAGGAAGC
GCTCATCGGACTCATCAGGAAGC
ACATCGGACTCATCAGGAAGC
ATCATCGGACTCATCAGGAAGC
ATCGGACTCATCAGGAAGC
AGCATCGGACTCATCAGGAAGC
CGGACTCATCAGGAAGC
CATCGGACTCATCAGGAAGC
CGGACTCATCAGGAAGC
TTCATCGGACTCATCAGGAAGC
CGGACTCATCAGGAAGC
ATCGGACTCATCAGGAAGC
ACATCGGACTCATCAGGAAGC
TCGGACTCATCAGGAAGC
ACATCGGACTCATCAGGAAGC
ACATCGGACTCATCAGGAAGC
TCATCGGACTCATCAGGAAGC
TCATCGGACTCATCAGGAAGC
CCATCGGACTCATCAGGAAGC
CCATCGGACTCATCAGGAAGC
ACATCGGACTCATCAGGAAGC
CATCGGACTCATCAGGAAGC
CATCGGACTCATCAGGAAGC
ATACATCGGACTCATCAGGAAGC
ATCGGACTCATCAGGAAGC
GCATCGGACTCATCAGGAAGC
AAACATCGGACTCATCAGGAAGC
CATCGGACTCATCAGGAAGC
ACATCGGACTCATCAGGAAGC
CCATCGGACTCATCAGGAAGC
CGGACTCATCAGGAAGC
CATCGGACTCATCAGGAAGC
CCATCGGACTCATCAGGAAGC
CATCGGACTCATCAGGAAGC
TCGGACTCATCAGGAAGC
ACATCGGACTCATCAGGAAGC
CATCGGACTCATCAGGAAGC
CATCGGACTCATCAGGAAGC
ATCATCGGACTCATCAGGAAGC

TACATCGGACTCATCAGGAAGC
ATCGGACTCATCAGGAAGC
ACATCGGACTCATCAGGAAGC
CATCGGACTCATCAGGAAGC
GCATCGGACTCATCAGGAAGC
ACATCGGACTCATCAGGAAGC
TCATCGGACTCATCAGGAAGC
ATCGGACTCATCAGGAAGC
AACATCGGACTCATCAGGAAGC
AGCATCGGACTCATCAGGAAGC
GGCATCGGACTCATCAGGAAGC
AACATCGGACTCATCAGGAAGC
ATCGGACTCATCAGGAAGC
ATCGGACTCATCAGGAAGC
TCATCGGACTCATCAGGAAGC
ATCGGACTCATCAGGAAGC
CATCGGACTCATCAGGAAGC
ACTCATCAGGAAGC
TCGGACTCATCAGGAAGC
ATCGGACTCATCAGGAAGC
ATCGGACTCATCAGGAAGC
ACCATCGGACTCATCAGGAAGC
GCATCGGACTCATCAGGAAGC
TCGGACTCATCAGGAAGC
CATCGGACTCATCAGGAAGC
CATCGGACTCATCAGGAAGC
GACTCATCAGGAAGC
GACTCATCAGGAAGC
GACTCATCAGGAAGC
CATCGGACTCATCAGGAAGC
AACATCGGACTCATCAGGAAGC
CATCGGACTCATCAGGAAGC
ACATCGGACTCATCAGGAAGC
ACATCGGACTCATCAGGAAGC
CATCGGACTCATCAGGAAGC
GGACTCATCAGGAAGC
ACATCGGACTCATCAGGAAGC
ACCATCGGACTCATCAGGAAGC
TGCATCGGACTCATCAGGAAGC
GCTCATCGGACTCATCAGGAAGC
TCGGACTCATCAGGAAGC
GACTCATCAGGAAGC
GGACTCATCAGGAAGC

TCGGACTCATCAGGAAGC
ATCATCGGACTCATCAGGAAGC
GTCATCGGACTCATCAGGAAGC
CATCGGACTCATCAGGAAGC
TTCATCGGACTCATCAGGAAGC
CCATCGGACTCATCAGGAAGC
TCGGACTCATCAGGAAGC
TCGGACTCATCAGGAAGC
TCGGACTCATCAGGAAGC
GCATCGGACTCATCAGGAAGC
CATCGGACTCATCAGGAAGC
CATCGGACTCATCAGGAAGC
TCGGACTCATCAGGAAGC
TACATCGGACTCATCAGGAAGC
AAGGCATCGGACTCATCAGGAAGC
CAACATCGGACTCATCAGGAAGC
ACATCGGACTCATCAGGAAGC
CATCGGACTCATCAGGAAGC
CATCGGACTCATCAGGAAGC
GTCATCGGACTCATCAGGAAGC
GCATCGGACTCATCAGGAAGC
TCATCGGACTCATCAGGAAGC
GACATCGGACTCATCAGGAAGC
CATCGGACTCATCAGGAAGC
CTCATCGGACTCATCAGGAAGC
ACATCGGACTCATCAGGAAGC
CATCGGACTCATCAGGAAGC
CATCGGACTCATCAGGAAGC
CATCGGACTCATCAGGAAGC
ATCGGACTCATCAGGAAGC
ATCGGACTCATCAGGAAGC
CCCATCGGACTCATCAGGAAGC
CCCATCGGACTCATCAGGAAGC
TCGGACTCATCAGGAAGC
CATCGGACTCATCAGGAAGC
CCATCGGACTCATCAGGAAGC
CATCGGACTCATCAGGAAGC
ACATCGGACTCATCAGGAAGC
CATCGGACTCATCAGGAAGC
GACTCATCAGGAAGC
GACATCGGACTCATCAGGAAGC
CATCGGACTCATCAGGAAGC
CATCGGACTCATCAGGAAGC

GTCATCGGACTCATCAGGAAGC
CATCGGACTCATCAGGAAGC
ACATCGGACTCATCAGGAAGC
CATCGGACTCATCAGGAAGC
CTCATCGGACTCATCAGGAAGC
TCGGACTCATCAGGAAGC
AACATCGGACTCATCAGGAAGC
CATCGGACTCATCAGGAAGC
CATCGGACTCATCAGGAAGC
CATCGGACTCATCAGGAAGC
GACTCATCAGGAAGC
GTCATCGGACTCATCAGGAAGC
CATCGGACTCATCAGGAAGC
CATCGGACTCATCAGGAAGC
TCATCGGACTCATCAGGAAGC
CATCGGACTCATCAGGAAGC
TTCATCGGACTCATCAGGAAGC
GACATCGGACTCATCAGGAAGC
GCATCGGACTCATCAGGAAGC
ATCGGACTCATCAGGAAGC
ATCGGACTCATCAGGAAGC
CATCGGACTCATCAGGAAGC
TCACATCGGACTCATCAGGAAGC
GACTCATCAGGAAGC
CGGACTCATCAGGAAGC
TCGGACTCATCAGGAAGC
GTCATCGGACTCATCAGGAAGC
AGCATCGGACTCATCAGGAAGC
CATCGGACTCATCAGGAAGC
GCATCGGACTCATCAGGAAGC
CCATCGGACTCATCAGGAAGC
ACATCGGACTCATCAGGAAGC
ATCGGACTCATCAGGAAGC
ACATCGGACTCATCAGGAAGC
GACTCATCAGGAAGC
CATCGGACTCATCAGGAAGC
ATCGGACTCATCAGGAAGC
ACATCGGACTCATCAGGAAGC
AGCATCGGACTCATCAGGAAGC
AGCATCGGACTCATCAGGAAGC
ATCGGACTCATCAGGAAGC
ATCGGACTCATCAGGAAGC
GACATCGGACTCATCAGGAAGC

ATCATCGGACTCATCAGGAAGC
TCGGACTCATCAGGAAGC
TCGGACTCATCAGGAAGC
CATCGGACTCATCAGGAAGC
GCATCGGACTCATCAGGAAGC
GCATCGGACTCATCAGGAAGC
CATCGGACTCATCAGGAAGC
TCATCGGACTCATCAGGAAGC
AACCACATCGGACTCATCAGGAAGC
GCATCGGACTCATCAGGAAGC
CTCATCGGACTCATCAGGAAGC
AGGCATCGGACTCATCAGGAAGC
TCTCATCGGACTCATCAGGAAGC
ATCGGACTCATCAGGAAGC
GCCATCGGACTCATCAGGAAGC
TACATCGGACTCATCAGGAAGC
ATCATCGGACTCATCAGGAAGC
ATCGGACTCATCAGGAAGC
CGGACTCATCAGGAAGC
CATCGGACTCATCAGGAAGC
CATCGGACTCATCAGGAAGC
TCATCGGACTCATCAGGAAGC
ATCGGACTCATCAGGAAGC
ATCATCGGACTCATCAGGAAGC
ATCGGACTCATCAGGAAGC
ATCGGACTCATCAGGAAGC
TCGGACTCATCAGGAAGC
CATCGGACTCATCAGGAAGC
TCGGACTCATCAGGAAGC
ATCGGACTCATCAGGAAGC
CATCGGACTCATCAGGAAGC
CTCATCGGACTCATCAGGAAGC
TTCATCGGACTCATCAGGAAGC
GCATCGGACTCATCAGGAAGC
ACATCGGACTCATCAGGAAGC
ATTCATCGGACTCATCAGGAAGC
GACTCATCAGGAAGC
AGCATCGGACTCATCAGGAAGC
GGACTCATCAGGAAGC
ATCGGACTCATCAGGAAGC
TGGCATCGGACTCATCAGGAAGC
CATCGGACTCATCAGGAAGC
GCCATCGGACTCATCAGGAAGC

TACATCGGACTCATCAGGAAGC
TACATCGGACTCATCAGGAAGC
TCATCGGACTCATCAGGAAGC
ATCGGACTCATCAGGAAGC
TACATCGGACTCATCAGGAAGC
CTTCATCGGACTCATCAGGAAGC
ACATCGGACTCATCAGGAAGC
TCGGACTCATCAGGAAGC
CATCGGACTCATCAGGAAGC
ACATCGGACTCATCAGGAAGC
CGGACTCATCAGGAAGC
CATCGGACTCATCAGGAAGC
TTCATCGGACTCATCAGGAAGC
ATCGGACTCATCAGGAAGC
TCGGACTCATCAGGAAGC
GCATCGGACTCATCAGGAAGC
TTACATCGGACTCATCAGGAAGC
CATCGGACTCATCAGGAAGC
GCATCGGACTCATCAGGAAGC
CATCGGACTCATCAGGAAGC
CATCGGACTCATCAGGAAGC
CATCGGACTCATCAGGAAGC
CATCGGACTCATCAGGAAGC
GCATCGGACTCATCAGGAAGC
CATCGGACTCATCAGGAAGC
GCATCGGACTCATCAGGAAGC
CCATCGGACTCATCAGGAAGC
TCGGACTCATCAGGAAGC
CATCGGACTCATCAGGAAGC
TCGGACTCATCAGGAAGC
TTACATCGGACTCATCAGGAAGC
TGCATCGGACTCATCAGGAAGC
GTCATCGGACTCATCAGGAAGC
TCGGACTCATCAGGAAGC
AGCATCGGACTCATCAGGAAGC
CATCGGACTCATCAGGAAGC
CATCGGACTCATCAGGAAGC
TCATCGGACTCATCAGGAAGC
ATCGGACTCATCAGGAAGC
CCATCGGACTCATCAGGAAGC
AGCATCGGACTCATCAGGAAGC
GCCATCGGACTCATCAGGAAGC
GCATCGGACTCATCAGGAAGC

CATCGGACTCATCAGGAAGC
CATCGGACTCATCAGGAAGC
ACATCGGACTCATCAGGAAGC
CATCGGACTCATCAGGAAGC
GACATCGGACTCATCAGGAAGC
TCATCGGACTCATCAGGAAGC
CATCGGACTCATCAGGAAGC
CATCGGACTCATCAGGAAGC
AGCATCGGACTCATCAGGAAGC
GGCATCGGACTCATCAGGAAGC
ACATCGGACTCATCAGGAAGC
TCGGACTCATCAGGAAGC
ATCGGACTCATCAGGAAGC
CATCGGACTCATCAGGAAGC
CACATCGGACTCATCAGGAAGC
CGGACTCATCAGGAAGC
TGACATCGGACTCATCAGGAAGC
CATCGGACTCATCAGGAAGC
GCCCATCGGACTCATCAGGAAGC
ATCGGACTCATCAGGAAGC
ATCATCGGACTCATCAGGAAGC
ACATCGGACTCATCAGGAAGC
GGACTCATCAGGAAGC
ATCGGACTCATCAGGAAGC
CATCGGACTCATCAGGAAGC
GGACTCATCAGGAAGC
CATCGGACTCATCAGGAAGC
ATCATCGGACTCATCAGGAAGC
CCATCGGACTCATCAGGAAGC
TTCATCGGACTCATCAGGAAGC
AACATCGGACTCATCAGGAAGC
CCATCGGACTCATCAGGAAGC
GTCATCGGACTCATCAGGAAGC
CATCGGACTCATCAGGAAGC
ATCATCGGACTCATCAGGAAGC
ATCGGACTCATCAGGAAGC
ACATCGGACTCATCAGGAAGC
CATCGGACTCATCAGGAAGC
CCATCGGACTCATCAGGAAGC
CATCGGACTCATCAGGAAGC
TTCATCGGACTCATCAGGAAGC
CATCGGACTCATCAGGAAGC
CATCGGACTCATCAGGAAGC

CACATCGGACTCATCAGGAAGC
GCATCGGACTCATCAGGAAGC
ACCATCGGACTCATCAGGAAGC
ACATCGGACTCATCAGGAAGC
TACATCGGACTCATCAGGAAGC
CGGCATCGGACTCATCAGGAAGC
GCATCGGACTCATCAGGAAGC
ATTCATCGGACTCATCAGGAAGC
TCATCGGACTCATCAGGAAGC
TGCATCGGACTCATCAGGAAGC
TATCATCGGACTCATCAGGAAGC
ATCGGACTCATCAGGAAGC
CCATCGGACTCATCAGGAAGC
CTTCATCGGACTCATCAGGAAGC
GCATCGGACTCATCAGGAAGC
GCACATCGGACTCATCAGGAAGC
GCACATCGGACTCATCAGGAAGC
ACATCGGACTCATCAGGAAGC
TCATCGGACTCATCAGGAAGC
TCATCGGACTCATCAGGAAGC
ATCGGACTCATCAGGAAGC
ATCGGACTCATCAGGAAGC
TCGGACTCATCAGGAAGC
GACTCATCAGGAAGC
GTTCATCGGACTCATCAGGAAGC
TGCATCGGACTCATCAGGAAGC
TCACATCGGACTCATCAGGAAGC
CGGACTCATCAGGAAGC
CATCGGACTCATCAGGAAGC
CGGACTCATCAGGAAGC
CATCGGACTCATCAGGAAGC
CACATCGGACTCATCAGGAAGC
TACATCGGACTCATCAGGAAGC
ATCGGACTCATCAGGAAGC
ACATCGGACTCATCAGGAAGC
CATCGGACTCATCAGGAAGC
ACATCGGACTCATCAGGAAGC
ATCGGACTCATCAGGAAGC
CATCGGACTCATCAGGAAGC
GACATCGGACTCATCAGGAAGC
AACATCGGACTCATCAGGAAGC
ATCGGACTCATCAGGAAGC
ATCGGACTCATCAGGAAGC

CATCGGACTCATCAGGAAGC
ATCGGACTCATCAGGAAGC
ACATCGGACTCATCAGGAAGC
ATCGGACTCATCAGGAAGC
ATCGGACTCATCAGGAAGC
CCATCGGACTCATCAGGAAGC
CCATCGGACTCATCAGGAAGC
ACATCGGACTCATCAGGAAGC
AAACATCGGACTCATCAGGAAGC
AAACATCGGACTCATCAGGAAGC
GTCATCGGACTCATCAGGAAGC
GCATCGGACTCATCAGGAAGC
CATCGGACTCATCAGGAAGC
CATCGGACTCATCAGGAAGC
ACATCGGACTCATCAGGAAGC
ACATCGGACTCATCAGGAAGC
TGCATCGGACTCATCAGGAAGC
TCATCGGACTCATCAGGAAGC
CCATCGGACTCATCAGGAAGC
GGACTCATCAGGAAGC
TACATCGGACTCATCAGGAAGC
TACATCGGACTCATCAGGAAGC
CGGACTCATCAGGAAGC
AAATCATCGGACTCATCAGGAAGC
TCCATCGGACTCATCAGGAAGC
TCGGACTCATCAGGAAGC
ATCGGACTCATCAGGAAGC
ACATCGGACTCATCAGGAAGC
ACATCGGACTCATCAGGAAGC
ATCATCGGACTCATCAGGAAGC
CCATCGGACTCATCAGGAAGC
TCATCGGACTCATCAGGAAGC
TCATCGGACTCATCAGGAAGC
CATCGGACTCATCAGGAAGC
TACATCGGACTCATCAGGAAGC
CTCATCGGACTCATCAGGAAGC
ATCGGACTCATCAGGAAGC
AAACATCGGACTCATCAGGAAGC
CTCATCGGACTCATCAGGAAGC
CTCATCGGACTCATCAGGAAGC
TCGGACTCATCAGGAAGC
ATCGGACTCATCAGGAAGC
GGACTCATCAGGAAGC

TCGGACTCATCAGGAAGC
TCGGACTCATCAGGAAGC
GCATCGGACTCATCAGGAAGC
CATCGGACTCATCAGGAAGC
ATCGGACTCATCAGGAAGC
CATCGGACTCATCAGGAAGC
TCGGACTCATCAGGAAGC
TCATCGGACTCATCAGGAAGC
TCGGACTCATCAGGAAGC
CATCGGACTCATCAGGAAGC
AGCATCGGACTCATCAGGAAGC
TCATCGGACTCATCAGGAAGC
ATCGGACTCATCAGGAAGC
ATCGGACTCATCAGGAAGC
ATCGGACTCATCAGGAAGC
TCGGACTCATCAGGAAGC
ATCGGACTCATCAGGAAGC
ATCGGACTCATCAGGAAGC
CGGACTCATCAGGAAGC
CCATCGGACTCATCAGGAAGC
ATCGGACTCATCAGGAAGC
TCGGACTCATCAGGAAGC
TCATCGGACTCATCAGGAAGC
CATCGGACTCATCAGGAAGC
GACTCATCAGGAAGC
GCATCGGACTCATCAGGAAGC
ATCGGACTCATCAGGAAGC
TCATCGGACTCATCAGGAAGC
CCATCGGACTCATCAGGAAGC
CATCGGACTCATCAGGAAGC
AGCATCGGACTCATCAGGAAGC
GGCATCGGACTCATCAGGAAGC
GTGCATCGGACTCATCAGGAAGC
CATCGGACTCATCAGGAAGC
TCGGACTCATCAGGAAGC
TACATCGGACTCATCAGGAAGC
ATCGGACTCATCAGGAAGC
TCATCGGACTCATCAGGAAGC
CACATCGGACTCATCAGGAAGC
ATTCATCGGACTCATCAGGAAGC
TCATCGGACTCATCAGGAAGC
ACATCGGACTCATCAGGAAGC
CATCGGACTCATCAGGAAGC

CATCGGACTCATCAGGAAGC
GCATCGGACTCATCAGGAAGC
ATCGGACTCATCAGGAAGC
ATCGGACTCATCAGGAAGC
ATCGGACTCATCAGGAAGC
TCATCGGACTCATCAGGAAGC
TCGGACTCATCAGGAAGC
GACATCGGACTCATCAGGAAGC
GCATCGGACTCATCAGGAAGC
ATCGGACTCATCAGGAAGC
TACATCGGACTCATCAGGAAGC
CATCGGACTCATCAGGAAGC
AACATCGGACTCATCAGGAAGC
ACATCGGACTCATCAGGAAGC
ATCATCGGACTCATCAGGAAGC
ATCGGACTCATCAGGAAGC
AGCATCGGACTCATCAGGAAGC
ACTCATCAGGAAGC
CATCGGACTCATCAGGAAGC
GTCATCGGACTCATCAGGAAGC
TACATCGGACTCATCAGGAAGC
TACACATCGGACTCATCAGGAAGC
TACACATCGGACTCATCAGGAAGC
TACATCGGACTCATCAGGAAGC
TCATCGGACTCATCAGGAAGC
CATCGGACTCATCAGGAAGC
TCCATCGGACTCATCAGGAAGC
AAAACATCGGACTCATCAGGAAGC
CGGACTCATCAGGAAGC
CATCGGACTCATCAGGAAGC
CATCGGACTCATCAGGAAGC
CGGACTCATCAGGAAGC
GACATCGGACTCATCAGGAAGC
CGGACTCATCAGGAAGC
CGGACTCATCAGGAAGC
CGGACTCATCAGGAAGC
CCATCGGACTCATCAGGAAGC
AGCATCGGACTCATCAGGAAGC
TCGGACTCATCAGGAAGC
AACATCGGACTCATCAGGAAGC
CCATCGGACTCATCAGGAAGC
ACCATCGGACTCATCAGGAAGC
AAGCATCGGACTCATCAGGAAGC

CATCGGACTCATCAGGAAGC
ATCGGACTCATCAGGAAGC
CATCGGACTCATCAGGAAGC
ATCGGACTCATCAGGAAGC
ATCGGACTCATCAGGAAGC
CCATCGGACTCATCAGGAAGC
CCATCGGACTCATCAGGAAGC
CCATCGGACTCATCAGGAAGC
TCGGACTCATCAGGAAGC
AACATCGGACTCATCAGGAAGC
ACATCGGACTCATCAGGAAGC
ATCGGACTCATCAGGAAGC
TCGGACTCATCAGGAAGC
TCGGACTCATCAGGAAGC
ACATCGGACTCATCAGGAAGC
CGGACTCATCAGGAAGC
TCGGACTCATCAGGAAGC
TTCATCGGACTCATCAGGAAGC
ACATCGGACTCATCAGGAAGC
GCATCGGACTCATCAGGAAGC
AACATCGGACTCATCAGGAAGC
TCATCGGACTCATCAGGAAGC
TCATCGGACTCATCAGGAAGC
TCATCGGACTCATCAGGAAGC
CATCGGACTCATCAGGAAGC
CATCGGACTCATCAGGAAGC
CGGACTCATCAGGAAGC
GGACTCATCAGGAAGC
CATCGGACTCATCAGGAAGC
GACTCATCAGGAAGC
ATCGGACTCATCAGGAAGC
ATCGGACTCATCAGGAAGC
ATCGGACTCATCAGGAAGC
ACTCATCAGGAAGC
GACATCGGACTCATCAGGAAGC
TCGGACTCATCAGGAAGC
TCGGACTCATCAGGAAGC
ACATCGGACTCATCAGGAAGC
CATCGGACTCATCAGGAAGC
CATCGGACTCATCAGGAAGC
TCCATCGGACTCATCAGGAAGC
ATCGGACTCATCAGGAAGC
CATCGGACTCATCAGGAAGC

CATCGGACTCATCAGGAAGC
CATCGGACTCATCAGGAAGC
CATCGGACTCATCAGGAAGC
TCATCGGACTCATCAGGAAGC
GCATCGGACTCATCAGGAAGC
CCATCGGACTCATCAGGAAGC
CATCGGACTCATCAGGAAGC
GACTCATCAGGAAGC
GGACTCATCAGGAAGC
CATCATCGGACTCATCAGGAAGC
ATCGGACTCATCAGGAAGC
GACATCGGACTCATCAGGAAGC
TCATCGGACTCATCAGGAAGC
GCATCGGACTCATCAGGAAGC
ACATCGGACTCATCAGGAAGC
CGGACTCATCAGGAAGC
CATCGGACTCATCAGGAAGC
TCATCGGACTCATCAGGAAGC
AACATCGGACTCATCAGGAAGC
CCATCGGACTCATCAGGAAGC
ATCGGACTCATCAGGAAGC
CCATCGGACTCATCAGGAAGC
CCATCGGACTCATCAGGAAGC
AACATCGGACTCATCAGGAAGC
CGGACTCATCAGGAAGC
ATCGGACTCATCAGGAAGC
GACATCGGACTCATCAGGAAGC
CATCGGACTCATCAGGAAGC
GACTCATCAGGAAGC
AACATCGGACTCATCAGGAAGC
TCATCGGACTCATCAGGAAGC
AACATCGGACTCATCAGGAAGC
TCATCGGACTCATCAGGAAGC
TTCATCGGACTCATCAGGAAGC
TCATCGGACTCATCAGGAAGC
TCATCGGACTCATCAGGAAGC
CCATCGGACTCATCAGGAAGC
AACATCGGACTCATCAGGAAGC
ACATCGGACTCATCAGGAAGC
TCCATCGGACTCATCAGGAAGC
TACATCGGACTCATCAGGAAGC
CATCGGACTCATCAGGAAGC
CATCGGACTCATCAGGAAGC

CATCGGACTCATCAGGAAGC
ACTCATCAGGAAGC
CATCGGACTCATCAGGAAGC
GCATCGGACTCATCAGGAAGC
CGGACTCATCAGGAAGC
ATCGGACTCATCAGGAAGC
ACATCGGACTCATCAGGAAGC
ATCGGACTCATCAGGAAGC
GGACTCATCAGGAAGC
ATCGGACTCATCAGGAAGC
ATCGGACTCATCAGGAAGC
ATTCATCGGACTCATCAGGAAGC
TCATCGGACTCATCAGGAAGC
ATCGGACTCATCAGGAAGC
TCATCGGACTCATCAGGAAGC
ATCGGACTCATCAGGAAGC
TACATCGGACTCATCAGGAAGC
ATCGGACTCATCAGGAAGC
ACATCGGACTCATCAGGAAGC
ATCGGACTCATCAGGAAGC
CATCGGACTCATCAGGAAGC
CATCGGACTCATCAGGAAGC
GCATCGGACTCATCAGGAAGC
CCATCGGACTCATCAGGAAGC
ATCGGACTCATCAGGAAGC
GCATCGGACTCATCAGGAAGC
TCATCGGACTCATCAGGAAGC
AGCATCGGACTCATCAGGAAGC
CATCGGACTCATCAGGAAGC
CGGACTCATCAGGAAGC
ACATCGGACTCATCAGGAAGC
ATCATCGGACTCATCAGGAAGC
TTTGCATCGGACTCATCAGGAAGC
CATCGGACTCATCAGGAAGC
ATCGGACTCATCAGGAAGC
ATCGGACTCATCAGGAAGC
CATCGGACTCATCAGGAAGC
AACATCGGACTCATCAGGAAGC
GCATCGGACTCATCAGGAAGC
TCATCGGACTCATCAGGAAGC
CCATCGGACTCATCAGGAAGC
ATCATCGGACTCATCAGGAAGC
TCCATCGGACTCATCAGGAAGC

CATCGGACTCATCAGGAAGC
ACATCGGACTCATCAGGAAGC
CATCGGACTCATCAGGAAGC
TCATCGGACTCATCAGGAAGC
TGCATCGGACTCATCAGGAAGC
TCGGACTCATCAGGAAGC
CGGACTCATCAGGAAGC
GCATCGGACTCATCAGGAAGC
ATCGGACTCATCAGGAAGC
ATCGGACTCATCAGGAAGC
ATCGGACTCATCAGGAAGC
CATCGGACTCATCAGGAAGC
TCATCGGACTCATCAGGAAGC
TGCATCGGACTCATCAGGAAGC
ATCGGACTCATCAGGAAGC
ACATCGGACTCATCAGGAAGC
GGACTCATCAGGAAGC
TATCATCGGACTCATCAGGAAGC
TCCATCGGACTCATCAGGAAGC
TCATCGGACTCATCAGGAAGC
ATCGGACTCATCAGGAAGC
GCCATCGGACTCATCAGGAAGC
TGCATCGGACTCATCAGGAAGC
AACATCGGACTCATCAGGAAGC
ACATCGGACTCATCAGGAAGC
ATCGGACTCATCAGGAAGC
CATCGGACTCATCAGGAAGC
CCATCGGACTCATCAGGAAGC
CATCGGACTCATCAGGAAGC
CATCGGACTCATCAGGAAGC
CATCGGACTCATCAGGAAGC
ATCATCGGACTCATCAGGAAGC
ACATCGGACTCATCAGGAAGC
CATCGGACTCATCAGGAAGC
CATCGGACTCATCAGGAAGC
CATCGGACTCATCAGGAAGC
CCATCGGACTCATCAGGAAGC
TCCATCGGACTCATCAGGAAGC
CGGACTCATCAGGAAGC
GGCATCGGACTCATCAGGAAGC
ACATCGGACTCATCAGGAAGC
CGGACTCATCAGGAAGC
AGGCCATCGGACTCATCAGGAAGC

GCCATCGGACTCATCAGGAAGC
ACATCGGACTCATCAGGAAGC
TCATCGGACTCATCAGGAAGC
CATCGGACTCATCAGGAAGC
TTTCATCGGACTCATCAGGAAGC
CATCGGACTCATCAGGAAGC
CATCGGACTCATCAGGAAGC
TCGGACTCATCAGGAAGC
CGGACTCATCAGGAAGC
ATCGGACTCATCAGGAAGC
TTACATCGGACTCATCAGGAAGC
CACATCGGACTCATCAGGAAGC
ACACATCGGACTCATCAGGAAGC
AACATCGGACTCATCAGGAAGC
ATCATCGGACTCATCAGGAAGC
GGCATCGGACTCATCAGGAAGC
ACATCGGACTCATCAGGAAGC
ACATCGGACTCATCAGGAAGC
GCCATCGGACTCATCAGGAAGC
ATCGGACTCATCAGGAAGC
AGCATCGGACTCATCAGGAAGC
TCGGACTCATCAGGAAGC
CATCGGACTCATCAGGAAGC
CATCGGACTCATCAGGAAGC
CATCGGACTCATCAGGAAGC
TCGGACTCATCAGGAAGC
CATCGGACTCATCAGGAAGC
TCATCGGACTCATCAGGAAGC
CACATCGGACTCATCAGGAAGC
CGGACTCATCAGGAAGC
CATCGGACTCATCAGGAAGC
TACATCGGACTCATCAGGAAGC
CATCGGACTCATCAGGAAGC
CATCGGACTCATCAGGAAGC
GACATCGGACTCATCAGGAAGC
GCATCGGACTCATCAGGAAGC
ACATCGGACTCATCAGGAAGC
GGACTCATCAGGAAGC
ATACATCGGACTCATCAGGAAGC
CATCGGACTCATCAGGAAGC
TCGGACTCATCAGGAAGC
CGGACTCATCAGGAAGC
TAGCATCGGACTCATCAGGAAGC

ATCGGACTCATCAGGAAGC
CATCGGACTCATCAGGAAGC
GCATCGGACTCATCAGGAAGC
ATCGGACTCATCAGGAAGC
CATCGGACTCATCAGGAAGC
ACCATCGGACTCATCAGGAAGC
ACATCGGACTCATCAGGAAGC
ATCATCGGACTCATCAGGAAGC
TCATCGGACTCATCAGGAAGC
CATCGGACTCATCAGGAAGC
GCATCGGACTCATCAGGAAGC
AACATCGGACTCATCAGGAAGC
CATCGGACTCATCAGGAAGC
CATCGGACTCATCAGGAAGC
CATCGGACTCATCAGGAAGC
TCCATCGGACTCATCAGGAAGC
TCGGACTCATCAGGAAGC
ATCATCGGACTCATCAGGAAGC
GCATCGGACTCATCAGGAAGC
CCATCGGACTCATCAGGAAGC
CCATCGGACTCATCAGGAAGC
CATCGGACTCATCAGGAAGC
CATCGGACTCATCAGGAAGC
CGGACTCATCAGGAAGC
TGCATCGGACTCATCAGGAAGC
AGTCATCGGACTCATCAGGAAGC
CATCGGACTCATCAGGAAGC
CATCGGACTCATCAGGAAGC
TCGGACTCATCAGGAAGC
GACTCATCAGGAAGC
AACATCGGACTCATCAGGAAGC
TCACATCGGACTCATCAGGAAGC
CCTTCATCGGACTCATCAGGAAGC
CATCGGACTCATCAGGAAGC
TGACATCGGACTCATCAGGAAGC
ACCCATCGGACTCATCAGGAAGC
TCATCGGACTCATCAGGAAGC
TCGGACTCATCAGGAAGC
TCGGACTCATCAGGAAGC
ATGCATCGGACTCATCAGGAAGC
GCTCATCGGACTCATCAGGAAGC
CGGACTCATCAGGAAGC
ATCGGACTCATCAGGAAGC

CCATCGGACTCATCAGGAAGC
GCATCGGACTCATCAGGAAGC
CATCGGACTCATCAGGAAGC
ACATCGGACTCATCAGGAAGC
ATCGGACTCATCAGGAAGC
CATCGGACTCATCAGGAAGC
TCATCGGACTCATCAGGAAGC
GCATCGGACTCATCAGGAAGC
CCATCGGACTCATCAGGAAGC
GATCATCGGACTCATCAGGAAGC
GGACTCATCAGGAAGC
GCCATCGGACTCATCAGGAAGC
TCATCGGACTCATCAGGAAGC
TCGGACTCATCAGGAAGC
CAACATCGGACTCATCAGGAAGC
CATCGGACTCATCAGGAAGC
TGCATCGGACTCATCAGGAAGC
CCATCGGACTCATCAGGAAGC
CCATCGGACTCATCAGGAAGC
CATCGGACTCATCAGGAAGC
ACATCGGACTCATCAGGAAGC
ACATCGGACTCATCAGGAAGC
TGCATCGGACTCATCAGGAAGC
GCATCGGACTCATCAGGAAGC
TCGGACTCATCAGGAAGC
TCCATCGGACTCATCAGGAAGC
CCATCGGACTCATCAGGAAGC
AACATCGGACTCATCAGGAAGC
TCGGACTCATCAGGAAGC
CATCGGACTCATCAGGAAGC
ACATCGGACTCATCAGGAAGC
ATCGGACTCATCAGGAAGC
TCGGACTCATCAGGAAGC
CATCGGACTCATCAGGAAGC
CATCGGACTCATCAGGAAGC
CATCGGACTCATCAGGAAGC
CATCGGACTCATCAGGAAGC
CCATCGGACTCATCAGGAAGC
TCATCGGACTCATCAGGAAGC
GCATCGGACTCATCAGGAAGC
GCATCGGACTCATCAGGAAGC
ACATCGGACTCATCAGGAAGC
ATCGGACTCATCAGGAAGC

AAGCATCGGACTCATCAGGAAGC
CATCGGACTCATCAGGAAGC
TCGGACTCATCAGGAAGC
AAGCATCGGACTCATCAGGAAGC
AAGCATCGGACTCATCAGGAAGC
TCATCGGACTCATCAGGAAGC
CCATCGGACTCATCAGGAAGC
CATCGGACTCATCAGGAAGC
ACATCGGACTCATCAGGAAGC
GCATCGGACTCATCAGGAAGC
TCGGACTCATCAGGAAGC
CATCGGACTCATCAGGAAGC
TCATCGGACTCATCAGGAAGC
TCATCGGACTCATCAGGAAGC
GACATCGGACTCATCAGGAAGC
ATCGGACTCATCAGGAAGC
TCATCGGACTCATCAGGAAGC
ATCGGACTCATCAGGAAGC
CATCGGACTCATCAGGAAGC
ATACATCGGACTCATCAGGAAGC
ATCGGACTCATCAGGAAGC
GACTCATCAGGAAGC
CATCGGACTCATCAGGAAGC
TACCATCGGACTCATCAGGAAGC
AACATCGGACTCATCAGGAAGC
CATCGGACTCATCAGGAAGC
TCATCGGACTCATCAGGAAGC
ACATCGGACTCATCAGGAAGC
CATCGGACTCATCAGGAAGC
ACATCGGACTCATCAGGAAGC
ACCATCGGACTCATCAGGAAGC
GACATCGGACTCATCAGGAAGC
ATCGGACTCATCAGGAAGC
AGGCATCGGACTCATCAGGAAGC
CGGACTCATCAGGAAGC
GGACTCATCAGGAAGC
ACATCGGACTCATCAGGAAGC
AACATCGGACTCATCAGGAAGC
GCATCGGACTCATCAGGAAGC
TCATCGGACTCATCAGGAAGC
GCATCGGACTCATCAGGAAGC
GCATCGGACTCATCAGGAAGC
ATCGGACTCATCAGGAAGC

CTCATCGGACTCATCAGGAAGC
AACATCGGACTCATCAGGAAGC
CATCGGACTCATCAGGAAGC
TCCATCGGACTCATCAGGAAGC
TCATCGGACTCATCAGGAAGC
GGACTCATCAGGAAGC
CATCGGACTCATCAGGAAGC
ATCGGACTCATCAGGAAGC
TCATCGGACTCATCAGGAAGC
GCATCGGACTCATCAGGAAGC
CACCATCGGACTCATCAGGAAGC
CATCGGACTCATCAGGAAGC
ATCGGACTCATCAGGAAGC
AGTCATCGGACTCATCAGGAAGC
GCATCGGACTCATCAGGAAGC
CATCGGACTCATCAGGAAGC
TCATCGGACTCATCAGGAAGC
ATCATCGGACTCATCAGGAAGC
CATCGGACTCATCAGGAAGC
CATCGGACTCATCAGGAAGC
ATCGGACTCATCAGGAAGC
ACATCGGACTCATCAGGAAGC
CATCGGACTCATCAGGAAGC
ACATCGGACTCATCAGGAAGC
CATCGGACTCATCAGGAAGC
GGACTCATCAGGAAGC
ATCATCGGACTCATCAGGAAGC
ACATCGGACTCATCAGGAAGC
ATCGGACTCATCAGGAAGC
CATCGGACTCATCAGGAAGC
AGCATCGGACTCATCAGGAAGC
TGCATCGGACTCATCAGGAAGC
CATCGGACTCATCAGGAAGC
ATATATACATCGGACTCATCAGGAAGC
TCGGACTCATCAGGAAGC
CATCGGACTCATCAGGAAGC
TCGGACTCATCAGGAAGC
CATCGGACTCATCAGGAAGC
ATCATCGGACTCATCAGGAAGC
AGCATCGGACTCATCAGGAAGC
ATCGGACTCATCAGGAAGC
CTCATCGGACTCATCAGGAAGC
TCATCGGACTCATCAGGAAGC

ACATCGGACTCATCAGGAAGC
ACCATCGGACTCATCAGGAAGC
TACATCGGACTCATCAGGAAGC
CGGACTCATCAGGAAGC
ATCGGACTCATCAGGAAGC
CATCGGACTCATCAGGAAGC
AACATCGGACTCATCAGGAAGC
CATCGGACTCATCAGGAAGC
TCATCGGACTCATCAGGAAGC
ATCGGACTCATCAGGAAGC
GACATCGGACTCATCAGGAAGC
TACATCGGACTCATCAGGAAGC
ACATCGGACTCATCAGGAAGC
CCATCGGACTCATCAGGAAGC
TCATCGGACTCATCAGGAAGC
GCATCGGACTCATCAGGAAGC
AAGCATCGGACTCATCAGGAAGC
ACCATCGGACTCATCAGGAAGC
CATCGGACTCATCAGGAAGC
AACATCGGACTCATCAGGAAGC
TCGGACTCATCAGGAAGC
ACATCGGACTCATCAGGAAGC
CATCGGACTCATCAGGAAGC
CATCGGACTCATCAGGAAGC
CATCGGACTCATCAGGAAGC
CATCGGACTCATCAGGAAGC
CATCGGACTCATCAGGAAGC
CATCGGACTCATCAGGAAGC
TCATCGGACTCATCAGGAAGC
TACATCGGACTCATCAGGAAGC
ATCGGACTCATCAGGAAGC
ACCATCGGACTCATCAGGAAGC
CACATCGGACTCATCAGGAAGC
TCATCGGACTCATCAGGAAGC
TCGGACTCATCAGGAAGC
GGACTCATCAGGAAGC
TCGGACTCATCAGGAAGC
TGCATCGGACTCATCAGGAAGC
GCATCGGACTCATCAGGAAGC
CCATCGGACTCATCAGGAAGC
GTCATCGGACTCATCAGGAAGC
ATCGGACTCATCAGGAAGC
TACATCGGACTCATCAGGAAGC

GCATCGGACTCATCAGGAAGC
TCATCGGACTCATCAGGAAGC
ATCGGACTCATCAGGAAGC
CATCGGACTCATCAGGAAGC
ATCGGACTCATCAGGAAGC
CATCGGACTCATCAGGAAGC
CGGACTCATCAGGAAGC
TCGGACTCATCAGGAAGC
TTCATCGGACTCATCAGGAAGC
ATCGGACTCATCAGGAAGC
TCGGACTCATCAGGAAGC
AACATCGGACTCATCAGGAAGC
TCGGACTCATCAGGAAGC
GACTCATCAGGAAGC
CATCGGACTCATCAGGAAGC
TCATCGGACTCATCAGGAAGC
GCCATCGGACTCATCAGGAAGC
GTCATCGGACTCATCAGGAAGC
TCGGACTCATCAGGAAGC
GGACTCATCAGGAAGC
GCATCGGACTCATCAGGAAGC
CATCGGACTCATCAGGAAGC
GATCATCGGACTCATCAGGAAGC
CATCGGACTCATCAGGAAGC
ATCGGACTCATCAGGAAGC
CCATCGGACTCATCAGGAAGC
CGGACTCATCAGGAAGC
ACATCGGACTCATCAGGAAGC
GTCATCGGACTCATCAGGAAGC
CGGACTCATCAGGAAGC
AACATCGGACTCATCAGGAAGC
CCATCGGACTCATCAGGAAGC
ATCGGACTCATCAGGAAGC
CATCGGACTCATCAGGAAGC
TCATCGGACTCATCAGGAAGC
GCATCGGACTCATCAGGAAGC
ACTCATCAGGAAGC
CATCGGACTCATCAGGAAGC
AGCATCGGACTCATCAGGAAGC
TCATCGGACTCATCAGGAAGC
TCATCGGACTCATCAGGAAGC
CATCGGACTCATCAGGAAGC
ATCGGACTCATCAGGAAGC

GCATCGGACTCATCAGGAAGC
CCATCGGACTCATCAGGAAGC
TCATCGGACTCATCAGGAAGC
ATCGGACTCATCAGGAAGC
AACATCGGACTCATCAGGAAGC
CGGACTCATCAGGAAGC
CATCGGACTCATCAGGAAGC
ATCGGACTCATCAGGAAGC
TCGGACTCATCAGGAAGC
GACATCGGACTCATCAGGAAGC
GACATCGGACTCATCAGGAAGC
GGACTCATCAGGAAGC
CATCGGACTCATCAGGAAGC
TCGGACTCATCAGGAAGC
CATCGGACTCATCAGGAAGC
CATCGGACTCATCAGGAAGC
ATCGGACTCATCAGGAAGC
CATCGGACTCATCAGGAAGC
CATCGGACTCATCAGGAAGC
CACATCGGACTCATCAGGAAGC
GCATCGGACTCATCAGGAAGC
AACATCGGACTCATCAGGAAGC
GCATCGGACTCATCAGGAAGC
CATCGGACTCATCAGGAAGC
ACATCGGACTCATCAGGAAGC
TCGGACTCATCAGGAAGC
TCGGACTCATCAGGAAGC
TCGGACTCATCAGGAAGC
TACATCGGACTCATCAGGAAGC
AGCATCGGACTCATCAGGAAGC
TCGGACTCATCAGGAAGC
CATCGGACTCATCAGGAAGC
CGGACTCATCAGGAAGC
CATCGGACTCATCAGGAAGC
GCATCGGACTCATCAGGAAGC
GCATCGGACTCATCAGGAAGC
ATCGGACTCATCAGGAAGC
GGACATCGGACTCATCAGGAAGC
CATCGGACTCATCAGGAAGC
TCGGACTCATCAGGAAGC
TCGGACTCATCAGGAAGC
TCGGACTCATCAGGAAGC
TCGGACTCATCAGGAAGC

TGCATCGGACTCATCAGGAAGC
CGGACTCATCAGGAAGC
CGGACTCATCAGGAAGC
TCGGACTCATCAGGAAGC
GGACTCATCAGGAAGC
GAACATCGGACTCATCAGGAAGC
CATCGGACTCATCAGGAAGC
GCATCGGACTCATCAGGAAGC
GCATCGGACTCATCAGGAAGC
CGGACTCATCAGGAAGC
CGGACTCATCAGGAAGC
TCATCGGACTCATCAGGAAGC
GGACTCATCAGGAAGC
TTCATCGGACTCATCAGGAAGC
CATCGGACTCATCAGGAAGC
TCGGACTCATCAGGAAGC
ACTCATCAGGAAGC
ATCGGACTCATCAGGAAGC
CGGACTCATCAGGAAGC
GGACTCATCAGGAAGC
GGCATCGGACTCATCAGGAAGC
AACATCGGACTCATCAGGAAGC
CCATCGGACTCATCAGGAAGC
CATCGGACTCATCAGGAAGC
CATCGGACTCATCAGGAAGC
CATCGGACTCATCAGGAAGC
CATCGGACTCATCAGGAAGC
TCGGACTCATCAGGAAGC
AGGCATCGGACTCATCAGGAAGC
TCATCGGACTCATCAGGAAGC
GCATCGGACTCATCAGGAAGC
CATCGGACTCATCAGGAAGC
GACATCGGACTCATCAGGAAGC
ATCATCGGACTCATCAGGAAGC
TTCATCGGACTCATCAGGAAGC
ACATCGGACTCATCAGGAAGC
ATCGGACTCATCAGGAAGC
CATCGGACTCATCAGGAAGC
TGCATCGGACTCATCAGGAAGC
CCATCGGACTCATCAGGAAGC
CATCGGACTCATCAGGAAGC
GCATCGGACTCATCAGGAAGC
TCATCGGACTCATCAGGAAGC

CCATCGGACTCATCAGGAAGC
CCATCGGACTCATCAGGAAGC
TACATCGGACTCATCAGGAAGC
TCATCGGACTCATCAGGAAGC
TACATCGGACTCATCAGGAAGC
CATCGGACTCATCAGGAAGC
ACATCGGACTCATCAGGAAGC
GCATCGGACTCATCAGGAAGC
ATCGGACTCATCAGGAAGC
CATCGGACTCATCAGGAAGC
GGACTCATCAGGAAGC
TCGGACTCATCAGGAAGC
GCATCGGACTCATCAGGAAGC
TACATCGGACTCATCAGGAAGC
TCATCGGACTCATCAGGAAGC
TCGGACTCATCAGGAAGC
AAACATCGGACTCATCAGGAAGC
GGACTCATCAGGAAGC
CACATCGGACTCATCAGGAAGC
GCATCGGACTCATCAGGAAGC
CATCGGACTCATCAGGAAGC
TCGGACTCATCAGGAAGC
GACATCGGACTCATCAGGAAGC
GCATCGGACTCATCAGGAAGC
ATCGGACTCATCAGGAAGC
ATCGGACTCATCAGGAAGC
CATCGGACTCATCAGGAAGC
CTTCATCGGACTCATCAGGAAGC
GACTCATCAGGAAGC
AGTATCACATCGGACTCATCAGGAAGC
CCATCGGACTCATCAGGAAGC
TCGGACTCATCAGGAAGC
CATCGGACTCATCAGGAAGC
CGGACTCATCAGGAAGC
TCATCGGACTCATCAGGAAGC
CATCGGACTCATCAGGAAGC
CATCGGACTCATCAGGAAGC
CGCATCGGACTCATCAGGAAGC
CATCGGACTCATCAGGAAGC
ACATCGGACTCATCAGGAAGC
CATCGGACTCATCAGGAAGC
CATCGGACTCATCAGGAAGC
CATCGGACTCATCAGGAAGC

CATCGGACTCATCAGGAAGC
TTCATCGGACTCATCAGGAAGC
GGCATCGGACTCATCAGGAAGC
TCATCGGACTCATCAGGAAGC
ATCATCGGACTCATCAGGAAGC
ATCGGACTCATCAGGAAGC
GCATCGGACTCATCAGGAAGC
GCATCGGACTCATCAGGAAGC
TCATCGGACTCATCAGGAAGC
ATCGGACTCATCAGGAAGC
TCATCGGACTCATCAGGAAGC
AGCATCGGACTCATCAGGAAGC
ATCGGACTCATCAGGAAGC
ACATCGGACTCATCAGGAAGC
ACCATCGGACTCATCAGGAAGC
ACCATCGGACTCATCAGGAAGC
CATCGGACTCATCAGGAAGC
CATCGGACTCATCAGGAAGC
TCATCGGACTCATCAGGAAGC
CGGACTCATCAGGAAGC
CATCGGACTCATCAGGAAGC
CATCGGACTCATCAGGAAGC
TCATCGGACTCATCAGGAAGC
GCATCGGACTCATCAGGAAGC
TCGGACTCATCAGGAAGC
ATCATCGGACTCATCAGGAAGC
GACTCATCAGGAAGC
TCATCGGACTCATCAGGAAGC
GCATCGGACTCATCAGGAAGC
GCATCGGACTCATCAGGAAGC
GCATCGGACTCATCAGGAAGC
TCGGACTCATCAGGAAGC
TCGGACTCATCAGGAAGC
ATCGGACTCATCAGGAAGC
ACATCGGACTCATCAGGAAGC
ATCGGACTCATCAGGAAGC
CACATCGGACTCATCAGGAAGC
ACTCATCGGACTCATCAGGAAGC
TCATCGGACTCATCAGGAAGC
CGGACTCATCAGGAAGC
CATCGGACTCATCAGGAAGC
ATCGGACTCATCAGGAAGC
GACTCATCAGGAAGC

TCATCGGACTCATCAGGAAGC
CATCGGACTCATCAGGAAGC
ACATCGGACTCATCAGGAAGC
TTCATCGGACTCATCAGGAAGC
GGACTCATCAGGAAGC
CCATCGGACTCATCAGGAAGC
ATCGGACTCATCAGGAAGC
ATCGGACTCATCAGGAAGC
TCATCGGACTCATCAGGAAGC
GGCATCGGACTCATCAGGAAGC
AACATCGGACTCATCAGGAAGC
ATCATCGGACTCATCAGGAAGC
CGGACTCATCAGGAAGC
CATCGGACTCATCAGGAAGC
ATCGGACTCATCAGGAAGC
GACTCATCAGGAAGC
TCGGACTCATCAGGAAGC
ACATCGGACTCATCAGGAAGC
GCATCGGACTCATCAGGAAGC
GTCATCGGACTCATCAGGAAGC
GTACATCGGACTCATCAGGAAGC
CATCGGACTCATCAGGAAGC
ATCATCGGACTCATCAGGAAGC
ATCATCGGACTCATCAGGAAGC
GCATCGGACTCATCAGGAAGC
TCATCGGACTCATCAGGAAGC
TCGGACTCATCAGGAAGC
CATCGGACTCATCAGGAAGC
TCATCGGACTCATCAGGAAGC
GCATCGGACTCATCAGGAAGC
CATCATCGGACTCATCAGGAAGC
TCGGACTCATCAGGAAGC
GCATCGGACTCATCAGGAAGC
GCATCGGACTCATCAGGAAGC
CATCGGACTCATCAGGAAGC
AGCATCGGACTCATCAGGAAGC
ATCGGACTCATCAGGAAGC
TCATCGGACTCATCAGGAAGC
GCATCGGACTCATCAGGAAGC
TCATCGGACTCATCAGGAAGC
ACATCGGACTCATCAGGAAGC
CATCGGACTCATCAGGAAGC
GTCATCGGACTCATCAGGAAGC

TCATCGGACTCATCAGGAAGC
CCATCGGACTCATCAGGAAGC
ATCGGACTCATCAGGAAGC
TCGGACTCATCAGGAAGC
CCATCGGACTCATCAGGAAGC
CATCGGACTCATCAGGAAGC
CATCGGACTCATCAGGAAGC
CATCGGACTCATCAGGAAGC
GGACTCATCAGGAAGC
CGGACTCATCAGGAAGC
ACATCGGACTCATCAGGAAGC
ATCGGACTCATCAGGAAGC
TCGGACTCATCAGGAAGC
AGACATCGGACTCATCAGGAAGC
CCCACATCGGACTCATCAGGAAGC
CCCACATCGGACTCATCAGGAAGC
CATCGGACTCATCAGGAAGC
GCATCGGACTCATCAGGAAGC
ATCGGACTCATCAGGAAGC
TTCATCGGACTCATCAGGAAGC
AACATCGGACTCATCAGGAAGC
CATCGGACTCATCAGGAAGC
ATCGGACTCATCAGGAAGC
ATCGGACTCATCAGGAAGC
TCGGACTCATCAGGAAGC
ACATCGGACTCATCAGGAAGC
TCGGACTCATCAGGAAGC
CCATCGGACTCATCAGGAAGC
CGGACTCATCAGGAAGC
CCATCGGACTCATCAGGAAGC
CATCGGACTCATCAGGAAGC
AGCATCGGACTCATCAGGAAGC
CGGACTCATCAGGAAGC
ATCGGACTCATCAGGAAGC
CCATCGGACTCATCAGGAAGC
ATCATCGGACTCATCAGGAAGC
CCATCGGACTCATCAGGAAGC
CCATCGGACTCATCAGGAAGC
TCCATCGGACTCATCAGGAAGC
CGGACTCATCAGGAAGC
CATCGGACTCATCAGGAAGC
TCATCGGACTCATCAGGAAGC
TCATCGGACTCATCAGGAAGC

TCGGACTCATCAGGAAGC
AACATCGGACTCATCAGGAAGC
TATCATCGGACTCATCAGGAAGC
CATCGGACTCATCAGGAAGC
ACATCGGACTCATCAGGAAGC
CATCGGACTCATCAGGAAGC
CGGACTCATCAGGAAGC
ACATCGGACTCATCAGGAAGC
TCATCGGACTCATCAGGAAGC
GCATCGGACTCATCAGGAAGC
CGGACTCATCAGGAAGC
AGCATCGGACTCATCAGGAAGC
ATTCATCGGACTCATCAGGAAGC
GCATCGGACTCATCAGGAAGC
CATCGGACTCATCAGGAAGC
CACATCGGACTCATCAGGAAGC
ATCGGACTCATCAGGAAGC
CCATCGGACTCATCAGGAAGC
TCGGACTCATCAGGAAGC
ACATCGGACTCATCAGGAAGC
GGACTCATCAGGAAGC
CATCGGACTCATCAGGAAGC
TCCATCGGACTCATCAGGAAGC
GGACTCATCAGGAAGC
GAACATCGGACTCATCAGGAAGC
CATCGGACTCATCAGGAAGC
CCATCGGACTCATCAGGAAGC
AGCATCGGACTCATCAGGAAGC
CATCGGACTCATCAGGAAGC
ACTCATCAGGAAGC
ATCGGACTCATCAGGAAGC
TCATCGGACTCATCAGGAAGC
TCATCGGACTCATCAGGAAGC
AACATCGGACTCATCAGGAAGC
GGACTCATCAGGAAGC
CATCGGACTCATCAGGAAGC
CATCGGACTCATCAGGAAGC
CATCGGACTCATCAGGAAGC
AGGACATCGGACTCATCAGGAAGC
ACATCGGACTCATCAGGAAGC
CACATCGGACTCATCAGGAAGC
TCGGACTCATCAGGAAGC
AACATCGGACTCATCAGGAAGC

CATCGGACTCATCAGGAAGC
CGGACTCATCAGGAAGC
AGCATCGGACTCATCAGGAAGC
TCATCGGACTCATCAGGAAGC
GACTCATCAGGAAGC
CCATCGGACTCATCAGGAAGC
GCATCGGACTCATCAGGAAGC
ACCATCGGACTCATCAGGAAGC
CCATCGGACTCATCAGGAAGC
CGGACTCATCAGGAAGC
CATCGGACTCATCAGGAAGC
CATCGGACTCATCAGGAAGC
CATCGGACTCATCAGGAAGC
GGACTCATCAGGAAGC
ATCATCGGACTCATCAGGAAGC
ATCGGACTCATCAGGAAGC
TTCATCGGACTCATCAGGAAGC
ACATCGGACTCATCAGGAAGC
ATCGGACTCATCAGGAAGC
TACCATCGGACTCATCAGGAAGC
ATCGGACTCATCAGGAAGC
ATCGGACTCATCAGGAAGC
TCGGACTCATCAGGAAGC
CATCGGACTCATCAGGAAGC
ATCGGACTCATCAGGAAGC
GCATCGGACTCATCAGGAAGC
CGGACTCATCAGGAAGC
GCCATCGGACTCATCAGGAAGC
TCGGACTCATCAGGAAGC
ATCGGACTCATCAGGAAGC
CTCATCGGACTCATCAGGAAGC
CATCGGACTCATCAGGAAGC
TCGGACTCATCAGGAAGC
CATCGGACTCATCAGGAAGC
TCATCGGACTCATCAGGAAGC
TATCATCGGACTCATCAGGAAGC
CATCGGACTCATCAGGAAGC
CATCGGACTCATCAGGAAGC
CTCATCGGACTCATCAGGAAGC
ACTCATCAGGAAGC
CATCGGACTCATCAGGAAGC
ATCGGACTCATCAGGAAGC
ATCGGACTCATCAGGAAGC

TCATCGGACTCATCAGGAAGC
GGACTCATCAGGAAGC
ATCATCGGACTCATCAGGAAGC
CGGACTCATCAGGAAGC
CTTCATCGGACTCATCAGGAAGC
CATCGGACTCATCAGGAAGC
TCGGACTCATCAGGAAGC
GGCATCGGACTCATCAGGAAGC
ACATCGGACTCATCAGGAAGC
ACATCGGACTCATCAGGAAGC
AACATCGGACTCATCAGGAAGC
CGGACTCATCAGGAAGC
GACTCATCAGGAAGC
CATCGGACTCATCAGGAAGC
TGCCATCGGACTCATCAGGAAGC
CGACATCGGACTCATCAGGAAGC
ATCGGACTCATCAGGAAGC
TACATCGGACTCATCAGGAAGC
ACCATCGGACTCATCAGGAAGC
GGCATCGGACTCATCAGGAAGC
TCGGACTCATCAGGAAGC
CATCGGACTCATCAGGAAGC
TACCATCGGACTCATCAGGAAGC
TGAACATCGGACTCATCAGGAAGC
TCATCGGACTCATCAGGAAGC
CATCGGACTCATCAGGAAGC
GGACTCATCAGGAAGC
GCATCGGACTCATCAGGAAGC
ATCGGACTCATCAGGAAGC
ATCGGACTCATCAGGAAGC
TCGGACTCATCAGGAAGC
CATCGGACTCATCAGGAAGC
CCATCGGACTCATCAGGAAGC
GCATCGGACTCATCAGGAAGC
ATCGGACTCATCAGGAAGC
ATCATCGGACTCATCAGGAAGC
ACATCGGACTCATCAGGAAGC
GACTCATCAGGAAGC
TCGGACTCATCAGGAAGC
CATCGGACTCATCAGGAAGC
ACATCGGACTCATCAGGAAGC
CATCGGACTCATCAGGAAGC
AACCATCGGACTCATCAGGAAGC

ACATCGGACTCATCAGGAAGC
CATCGGACTCATCAGGAAGC
ACATCGGACTCATCAGGAAGC
TGCATCGGACTCATCAGGAAGC
ACATCGGACTCATCAGGAAGC
ATACATCGGACTCATCAGGAAGC
ACCATCGGACTCATCAGGAAGC
AACATCGGACTCATCAGGAAGC
GACTCATCAGGAAGC
GACTCATCAGGAAGC
TCATCGGACTCATCAGGAAGC
CATCGGACTCATCAGGAAGC
CCATCGGACTCATCAGGAAGC
ATCGGACTCATCAGGAAGC
AATGCATCGGACTCATCAGGAAGC
AATGCATCGGACTCATCAGGAAGC
TCATCGGACTCATCAGGAAGC
CATCGGACTCATCAGGAAGC
AACATCGGACTCATCAGGAAGC
ACATCGGACTCATCAGGAAGC
ACATCGGACTCATCAGGAAGC
GCATCGGACTCATCAGGAAGC
ACATCGGACTCATCAGGAAGC
GGACTCATCAGGAAGC
CATCGGACTCATCAGGAAGC
AACATCGGACTCATCAGGAAGC
ATCGGACTCATCAGGAAGC
ACCATCGGACTCATCAGGAAGC
ATCGGACTCATCAGGAAGC
ATCGGACTCATCAGGAAGC
CCATCGGACTCATCAGGAAGC
CCATCGGACTCATCAGGAAGC
CCATCGGACTCATCAGGAAGC
CTTCATCGGACTCATCAGGAAGC
ACATCGGACTCATCAGGAAGC
CATCGGACTCATCAGGAAGC
GCATCGGACTCATCAGGAAGC
TACATCGGACTCATCAGGAAGC
GCATCGGACTCATCAGGAAGC
GGACTCATCAGGAAGC
ATCGGACTCATCAGGAAGC
AGCCATCGGACTCATCAGGAAGC
ACCATCGGACTCATCAGGAAGC

GCATCGGACTCATCAGGAAGC
TCGGACTCATCAGGAAGC
CATCGGACTCATCAGGAAGC
TTCATCGGACTCATCAGGAAGC
TCGGACTCATCAGGAAGC
CGGACTCATCAGGAAGC
CATCGGACTCATCAGGAAGC
ACATCGGACTCATCAGGAAGC
CATCGGACTCATCAGGAAGC
CACCATCGGACTCATCAGGAAGC
AGAGCATCGGACTCATCAGGAAGC
ACATCGGACTCATCAGGAAGC
GCATCGGACTCATCAGGAAGC
CATCGGACTCATCAGGAAGC
CATCGGACTCATCAGGAAGC
GGCATCGGACTCATCAGGAAGC
ATCGGACTCATCAGGAAGC
CTCATCGGACTCATCAGGAAGC
AGCATCGGACTCATCAGGAAGC
TACATCGGACTCATCAGGAAGC
CGGACTCATCAGGAAGC
CATCGGACTCATCAGGAAGC
ATCGGACTCATCAGGAAGC
ATCATCGGACTCATCAGGAAGC
ATCGGACTCATCAGGAAGC
CTCATCAGGAAGC
ATCGGACTCATCAGGAAGC
TACATCGGACTCATCAGGAAGC
ACATCGGACTCATCAGGAAGC
GTGCATCGGACTCATCAGGAAGC
ACATCGGACTCATCAGGAAGC
ACATCGGACTCATCAGGAAGC
CATCGGACTCATCAGGAAGC
CATCGGACTCATCAGGAAGC
GCATCGGACTCATCAGGAAGC
GCCATCGGACTCATCAGGAAGC
CATCGGACTCATCAGGAAGC
CATCGGACTCATCAGGAAGC
CATCGGACTCATCAGGAAGC
TATCATCGGACTCATCAGGAAGC
CAACATCGGACTCATCAGGAAGC
CATCGGACTCATCAGGAAGC
ACCATCGGACTCATCAGGAAGC

TCGGACTCATCAGGAAGC
TCATCGGACTCATCAGGAAGC
ATCGGACTCATCAGGAAGC
ATCGGACTCATCAGGAAGC
ACCATCGGACTCATCAGGAAGC
ATCATCGGACTCATCAGGAAGC
GCATCGGACTCATCAGGAAGC
GAACATCGGACTCATCAGGAAGC
CGGACTCATCAGGAAGC
CGGACTCATCAGGAAGC
CATCGGACTCATCAGGAAGC
ATCGGACTCATCAGGAAGC
ATCGGACTCATCAGGAAGC
CTCATCGGACTCATCAGGAAGC
GACATCGGACTCATCAGGAAGC
ACATCGGACTCATCAGGAAGC
CATCGGACTCATCAGGAAGC
TCATCGGACTCATCAGGAAGC
TCGGACTCATCAGGAAGC
ACATCGGACTCATCAGGAAGC
ATCGGACTCATCAGGAAGC
TGTCATCGGACTCATCAGGAAGC
GCATCGGACTCATCAGGAAGC
TCATCGGACTCATCAGGAAGC
TCGGACTCATCAGGAAGC
TCGGACTCATCAGGAAGC
ACCATCGGACTCATCAGGAAGC
TACATCGGACTCATCAGGAAGC
ACATCGGACTCATCAGGAAGC
ACATCGGACTCATCAGGAAGC
GGACTCATCAGGAAGC
CATCGGACTCATCAGGAAGC
CTCATCGGACTCATCAGGAAGC
ATCATCGGACTCATCAGGAAGC
ACATCGGACTCATCAGGAAGC
AACATCGGACTCATCAGGAAGC
GAGCATCGGACTCATCAGGAAGC
ATCATCGGACTCATCAGGAAGC
CATCGGACTCATCAGGAAGC
TCGGACTCATCAGGAAGC
CATCGGACTCATCAGGAAGC
ATCATCGGACTCATCAGGAAGC
AATCATCGGACTCATCAGGAAGC

CATCGGACTCATCAGGAAGC
CCATCGGACTCATCAGGAAGC
CATCGGACTCATCAGGAAGC
ATCGGACTCATCAGGAAGC
ATCGGACTCATCAGGAAGC
ATCGGACTCATCAGGAAGC
TCGGACTCATCAGGAAGC
TCGGACTCATCAGGAAGC
CATCGGACTCATCAGGAAGC
CATCGGACTCATCAGGAAGC
CATCGGACTCATCAGGAAGC
CCATCGGACTCATCAGGAAGC
TGCATCGGACTCATCAGGAAGC
CATCGGACTCATCAGGAAGC
TCGGACTCATCAGGAAGC
GACATCGGACTCATCAGGAAGC
GGACTCATCAGGAAGC
CCATCGGACTCATCAGGAAGC
CCATCGGACTCATCAGGAAGC
ACATCGGACTCATCAGGAAGC
ACCATCGGACTCATCAGGAAGC
TCATCGGACTCATCAGGAAGC
ACATCGGACTCATCAGGAAGC
TCGGACTCATCAGGAAGC
AGCATCGGACTCATCAGGAAGC
CACCATCGGACTCATCAGGAAGC
CGGACTCATCAGGAAGC
CATCGGACTCATCAGGAAGC
CATCGGACTCATCAGGAAGC
GCATCGGACTCATCAGGAAGC
ATCGGACTCATCAGGAAGC
ACACATCGGACTCATCAGGAAGC
ACACATCGGACTCATCAGGAAGC
TCGGACTCATCAGGAAGC
CATCGGACTCATCAGGAAGC
CATCGGACTCATCAGGAAGC
CATCGGACTCATCAGGAAGC
ATCATCGGACTCATCAGGAAGC
ACATCGGACTCATCAGGAAGC
TCATCGGACTCATCAGGAAGC
TCGGACTCATCAGGAAGC
ATCGGACTCATCAGGAAGC
CATCGGACTCATCAGGAAGC

TGTCATCGGACTCATCAGGAAGC
CATCGGACTCATCAGGAAGC
AAGACCATCGGACTCATCAGGAAGC
ACATCGGACTCATCAGGAAGC
CATCGGACTCATCAGGAAGC
TTCATCGGACTCATCAGGAAGC
CATCGGACTCATCAGGAAGC
CGGACTCATCAGGAAGC
GCATCGGACTCATCAGGAAGC
ATCGGACTCATCAGGAAGC
TCATCGGACTCATCAGGAAGC
CATCGGACTCATCAGGAAGC
ATCGGACTCATCAGGAAGC
TCGGACTCATCAGGAAGC
TCATCGGACTCATCAGGAAGC
CATCGGACTCATCAGGAAGC
ATCGGACTCATCAGGAAGC
CGGACTCATCAGGAAGC
TACATCGGACTCATCAGGAAGC
ATCGGACTCATCAGGAAGC
GGACTCATCAGGAAGC
TCATCGGACTCATCAGGAAGC
TCATCGGACTCATCAGGAAGC
ATCGGACTCATCAGGAAGC
TCCATCGGACTCATCAGGAAGC
CATCGGACTCATCAGGAAGC
TCGGACTCATCAGGAAGC
TCATCGGACTCATCAGGAAGC
ACATCGGACTCATCAGGAAGC
CATCGGACTCATCAGGAAGC
ACATCGGACTCATCAGGAAGC
CATCGGACTCATCAGGAAGC
GCATCGGACTCATCAGGAAGC
ATCGGACTCATCAGGAAGC
TTCATCGGACTCATCAGGAAGC
CCATCGGACTCATCAGGAAGC
CATCGGACTCATCAGGAAGC
ATCGGACTCATCAGGAAGC
CCATCGGACTCATCAGGAAGC
ACATCGGACTCATCAGGAAGC
ATCGGACTCATCAGGAAGC
ATCGGACTCATCAGGAAGC
ATCGGACTCATCAGGAAGC

TCATCGGACTCATCAGGAAGC
GCCATCGGACTCATCAGGAAGC
ACATCGGACTCATCAGGAAGC
TGCATCGGACTCATCAGGAAGC
TGCATCGGACTCATCAGGAAGC
GTCATCGGACTCATCAGGAAGC
ACATCGGACTCATCAGGAAGC
TCATCGGACTCATCAGGAAGC
GCATCGGACTCATCAGGAAGC
TCATCGGACTCATCAGGAAGC
TCGGACTCATCAGGAAGC
GGACTCATCAGGAAGC
CATCGGACTCATCAGGAAGC
GCATCGGACTCATCAGGAAGC
ATCGGACTCATCAGGAAGC
TCATCGGACTCATCAGGAAGC
TTCATCGGACTCATCAGGAAGC
CATCGGACTCATCAGGAAGC
TCGGACTCATCAGGAAGC
ACATCGGACTCATCAGGAAGC
TCGGACTCATCAGGAAGC
CATCGGACTCATCAGGAAGC
CATCGGACTCATCAGGAAGC
CATCGGACTCATCAGGAAGC
CTCATCGGACTCATCAGGAAGC
CTCATCGGACTCATCAGGAAGC
GTCATCGGACTCATCAGGAAGC
CATCGGACTCATCAGGAAGC
CATCGGACTCATCAGGAAGC
AGCATCGGACTCATCAGGAAGC
TGCATCGGACTCATCAGGAAGC
TGCATCGGACTCATCAGGAAGC
ATCGGACTCATCAGGAAGC
TCATCGGACTCATCAGGAAGC
TCATCGGACTCATCAGGAAGC
ATCGGACTCATCAGGAAGC
TTCATCGGACTCATCAGGAAGC
TTCATCGGACTCATCAGGAAGC
TTCATCGGACTCATCAGGAAGC
GCATCGGACTCATCAGGAAGC
ACCATCGGACTCATCAGGAAGC
TCATCGGACTCATCAGGAAGC
TCGGACTCATCAGGAAGC

TCATCGGACTCATCAGGAAGC
CATCGGACTCATCAGGAAGC
TGCATCGGACTCATCAGGAAGC
CATCGGACTCATCAGGAAGC
ATCGGACTCATCAGGAAGC
CATCGGACTCATCAGGAAGC
GTCATCGGACTCATCAGGAAGC
GTCATCGGACTCATCAGGAAGC
GTCATCGGACTCATCAGGAAGC
ACATCGGACTCATCAGGAAGC
TCATCGGACTCATCAGGAAGC
GCATCGGACTCATCAGGAAGC
TCATCGGACTCATCAGGAAGC
GTCATCGGACTCATCAGGAAGC
TCGGACTCATCAGGAAGC
CATCGGACTCATCAGGAAGC
CATCGGACTCATCAGGAAGC
TTAGCATCGGACTCATCAGGAAGC
CATCGGACTCATCAGGAAGC
ATCGGACTCATCAGGAAGC
CATCGGACTCATCAGGAAGC
ATCGGACTCATCAGGAAGC
TCATCGGACTCATCAGGAAGC
TCATCGGACTCATCAGGAAGC
CATCGGACTCATCAGGAAGC
TCGGACTCATCAGGAAGC
TACATCGGACTCATCAGGAAGC
TACATCGGACTCATCAGGAAGC
CCATCGGACTCATCAGGAAGC
ATCGGACTCATCAGGAAGC
GGACATCGGACTCATCAGGAAGC
GTCATCGGACTCATCAGGAAGC
ATCATCGGACTCATCAGGAAGC
TCATCGGACTCATCAGGAAGC
GCATCGGACTCATCAGGAAGC
TCATCGGACTCATCAGGAAGC
ATCGGACTCATCAGGAAGC
GCATCGGACTCATCAGGAAGC
CATCGGACTCATCAGGAAGC
CACATCGGACTCATCAGGAAGC
CATCGGACTCATCAGGAAGC
CATCGGACTCATCAGGAAGC
GGACTCATCAGGAAGC

GCATCGGACTCATCAGGAAGC
TCGGACTCATCAGGAAGC
ATCGGACTCATCAGGAAGC
CCATCGGACTCATCAGGAAGC
TCATCGGACTCATCAGGAAGC
GCATCGGACTCATCAGGAAGC
GCATCGGACTCATCAGGAAGC
CCATCGGACTCATCAGGAAGC
TCGGACTCATCAGGAAGC
GACTCATCAGGAAGC
TCGGACTCATCAGGAAGC
TCATCGGACTCATCAGGAAGC
CCATCGGACTCATCAGGAAGC
TCGGACTCATCAGGAAGC
CATCGGACTCATCAGGAAGC
ATCGGACTCATCAGGAAGC
TCATCGGACTCATCAGGAAGC
ACATCGGACTCATCAGGAAGC
TCGGACTCATCAGGAAGC
TCATCGGACTCATCAGGAAGC
ATCGGACTCATCAGGAAGC
TCATCGGACTCATCAGGAAGC
ACATCGGACTCATCAGGAAGC
ACACATCGGACTCATCAGGAAGC
GACTCATCAGGAAGC
CATCGGACTCATCAGGAAGC
TCATCGGACTCATCAGGAAGC
CATCGGACTCATCAGGAAGC
ACATCGGACTCATCAGGAAGC
TCGGACTCATCAGGAAGC
TCGGACTCATCAGGAAGC
TCATCGGACTCATCAGGAAGC
ATCGGACTCATCAGGAAGC
ACATCGGACTCATCAGGAAGC
ATCGGACTCATCAGGAAGC
TCATCGGACTCATCAGGAAGC
ATCGGACTCATCAGGAAGC
ATCGGACTCATCAGGAAGC
ATCGGACTCATCAGGAAGC
TCGGACTCATCAGGAAGC
TCATCGGACTCATCAGGAAGC
TGCATCGGACTCATCAGGAAGC
CGGACTCATCAGGAAGC

ATCATCGGACTCATCAGGAAGC
TCGGACTCATCAGGAAGC
ATCGGACTCATCAGGAAGC
GGACTCATCAGGAAGC
CATCGGACTCATCAGGAAGC
TACCATCGGACTCATCAGGAAGC
AGCATCGGACTCATCAGGAAGC
TTCATCGGACTCATCAGGAAGC
TTACATCGGACTCATCAGGAAGC
CGGACTCATCAGGAAGC
ATCGGACTCATCAGGAAGC
ATCGGACTCATCAGGAAGC
ACATCGGACTCATCAGGAAGC
GCATCGGACTCATCAGGAAGC
TCATCGGACTCATCAGGAAGC
TCATCGGACTCATCAGGAAGC
ATCGGACTCATCAGGAAGC
ATCGGACTCATCAGGAAGC
TCATCGGACTCATCAGGAAGC
AACATCGGACTCATCAGGAAGC
CCATCGGACTCATCAGGAAGC
CATCGGACTCATCAGGAAGC
ATCGGACTCATCAGGAAGC
TCATCGGACTCATCAGGAAGC
CATCGGACTCATCAGGAAGC
ACATCGGACTCATCAGGAAGC
TCGGACTCATCAGGAAGC
CATCGGACTCATCAGGAAGC
CATCGGACTCATCAGGAAGC
TCCATCGGACTCATCAGGAAGC
TCATCGGACTCATCAGGAAGC
CATCGGACTCATCAGGAAGC
CAATCATCGGACTCATCAGGAAGC
AGCATCGGACTCATCAGGAAGC
TCATCGGACTCATCAGGAAGC
TCATCGGACTCATCAGGAAGC
AATCATCGGACTCATCAGGAAGC
TCATCGGACTCATCAGGAAGC
GGACTCATCAGGAAGC
GGACTCATCAGGAAGC
ACATCGGACTCATCAGGAAGC
CATCGGACTCATCAGGAAGC
ATCGGACTCATCAGGAAGC

ATCGGACTCATCAGGAAGC
GTCATCGGACTCATCAGGAAGC
ACATCGGACTCATCAGGAAGC
CATCGGACTCATCAGGAAGC
ATCGGACTCATCAGGAAGC
ATCGGACTCATCAGGAAGC
TCATCGGACTCATCAGGAAGC
ATCGGACTCATCAGGAAGC
CATCGGACTCATCAGGAAGC
CATCGGACTCATCAGGAAGC
CATCGGACTCATCAGGAAGC
ATCGGACTCATCAGGAAGC
ATCGGACTCATCAGGAAGC
AGCATCGGACTCATCAGGAAGC
CAGACATCGGACTCATCAGGAAGC
TAAACATCGGACTCATCAGGAAGC
CTCATCGGACTCATCAGGAAGC
GTCATCGGACTCATCAGGAAGC
ATCGGACTCATCAGGAAGC
AGCATCGGACTCATCAGGAAGC
CGGACTCATCAGGAAGC
TGCATCGGACTCATCAGGAAGC
ACATCGGACTCATCAGGAAGC
ACATCGGACTCATCAGGAAGC
ACATCGGACTCATCAGGAAGC
ATCGGACTCATCAGGAAGC
CATCGGACTCATCAGGAAGC
TATCATCGGACTCATCAGGAAGC
ATCGGACTCATCAGGAAGC
CATCGGACTCATCAGGAAGC
ATCGGACTCATCAGGAAGC
CGGACTCATCAGGAAGC
GCATCGGACTCATCAGGAAGC
TCATCGGACTCATCAGGAAGC
ATCATCGGACTCATCAGGAAGC
CATCGGACTCATCAGGAAGC
ACATCGGACTCATCAGGAAGC
ATCGGACTCATCAGGAAGC
GGACTCATCAGGAAGC
TCATCGGACTCATCAGGAAGC
GGACTCATCAGGAAGC
CATCGGACTCATCAGGAAGC
TGCATCGGACTCATCAGGAAGC

GGACTCATCAGGAAGC
TCATCGGACTCATCAGGAAGC
CATCGGACTCATCAGGAAGC
CATCGGACTCATCAGGAAGC
CATCGGACTCATCAGGAAGC
CATCGGACTCATCAGGAAGC
CATCGGACTCATCAGGAAGC
CATCGGACTCATCAGGAAGC
ATCGGACTCATCAGGAAGC
TCGGACTCATCAGGAAGC
CATCGGACTCATCAGGAAGC
ATCGGACTCATCAGGAAGC
AGCATCGGACTCATCAGGAAGC
ATCATCGGACTCATCAGGAAGC
TCGGACTCATCAGGAAGC
TCGGACTCATCAGGAAGC
ATCGGACTCATCAGGAAGC
CATCGGACTCATCAGGAAGC
ATCGGACTCATCAGGAAGC
CATCGGACTCATCAGGAAGC
TCATCGGACTCATCAGGAAGC
AAGCATCGGACTCATCAGGAAGC
ATCATCGGACTCATCAGGAAGC
ATCATCGGACTCATCAGGAAGC
CATCGGACTCATCAGGAAGC
GGACTCATCAGGAAGC
CATCGGACTCATCAGGAAGC
CGGACTCATCAGGAAGC
ATCGGACTCATCAGGAAGC
CCATCGGACTCATCAGGAAGC
ATCGGACTCATCAGGAAGC
TCGGACTCATCAGGAAGC
CATCGGACTCATCAGGAAGC
CATCGGACTCATCAGGAAGC
TCGGACTCATCAGGAAGC
CATCGGACTCATCAGGAAGC
GCATCGGACTCATCAGGAAGC
ATCGGACTCATCAGGAAGC
CATCGGACTCATCAGGAAGC
TCCATCGGACTCATCAGGAAGC
CATCGGACTCATCAGGAAGC
TACATCGGACTCATCAGGAAGC
ATCGGACTCATCAGGAAGC

CGGACTCATCAGGAAGC
CGGACTCATCAGGAAGC
TCCATCGGACTCATCAGGAAGC
TCCATCGGACTCATCAGGAAGC
TCATCGGACTCATCAGGAAGC
ACTACATCGGACTCATCAGGAAGC
TTACATCGGACTCATCAGGAAGC
TTTCCATCGGACTCATCAGGAAGC
GCATCGGACTCATCAGGAAGC
CATCGGACTCATCAGGAAGC
TCCATCGGACTCATCAGGAAGC
ACATCGGACTCATCAGGAAGC
GGACTCATCAGGAAGC
ATCGGACTCATCAGGAAGC
TCGGACTCATCAGGAAGC
GGACTCATCAGGAAGC
TCGGACTCATCAGGAAGC
CGGACTCATCAGGAAGC
ATCATCGGACTCATCAGGAAGC
ACATCGGACTCATCAGGAAGC
TTCATCGGACTCATCAGGAAGC
TACATCGGACTCATCAGGAAGC
GGCATCGGACTCATCAGGAAGC
GGACTCATCAGGAAGC
TAAGCATCGGACTCATCAGGAAGC
TGCATCGGACTCATCAGGAAGC
CATCGGACTCATCAGGAAGC
ACATCGGACTCATCAGGAAGC
CCATCGGACTCATCAGGAAGC
CATCGGACTCATCAGGAAGC
CATCGGACTCATCAGGAAGC
TCATCGGACTCATCAGGAAGC
TCGGACTCATCAGGAAGC
GTCATCGGACTCATCAGGAAGC
CATCGGACTCATCAGGAAGC
CGGACTCATCAGGAAGC
CATCGGACTCATCAGGAAGC
GCATCGGACTCATCAGGAAGC
CATCGGACTCATCAGGAAGC
GCATCGGACTCATCAGGAAGC
TCGGACTCATCAGGAAGC
ATCGGACTCATCAGGAAGC
CATCGGACTCATCAGGAAGC

TTCATCGGACTCATCAGGAAGC
TTCATCGGACTCATCAGGAAGC
TCATCGGACTCATCAGGAAGC
CCATCGGACTCATCAGGAAGC
ACATCGGACTCATCAGGAAGC
CCATCGGACTCATCAGGAAGC
ACATCGGACTCATCAGGAAGC
TCATCGGACTCATCAGGAAGC
TCCACATCGGACTCATCAGGAAGC
CGGACTCATCAGGAAGC
TCATCGGACTCATCAGGAAGC
ATCGGACTCATCAGGAAGC
ATCCCATCGGACTCATCAGGAAGC
CATCGGACTCATCAGGAAGC
TAGCATCGGACTCATCAGGAAGC
GACATCGGACTCATCAGGAAGC
GACATCGGACTCATCAGGAAGC
CATCGGACTCATCAGGAAGC
TCATCGGACTCATCAGGAAGC
GAGCATCGGACTCATCAGGAAGC
GCATCGGACTCATCAGGAAGC
GCATCGGACTCATCAGGAAGC
AACATCGGACTCATCAGGAAGC
ATCGGACTCATCAGGAAGC
ATCGGACTCATCAGGAAGC
ATCGGACTCATCAGGAAGC
CATCGGACTCATCAGGAAGC
GTCATCGGACTCATCAGGAAGC
ACATCGGACTCATCAGGAAGC
ACATCGGACTCATCAGGAAGC
TCATCGGACTCATCAGGAAGC
AATCATCGGACTCATCAGGAAGC
ATCGGACTCATCAGGAAGC
GAGCCATCGGACTCATCAGGAAGC
TCATCGGACTCATCAGGAAGC
CATCGGACTCATCAGGAAGC
TCGGACTCATCAGGAAGC
TCGGACTCATCAGGAAGC
ACATCGGACTCATCAGGAAGC
ACATCGGACTCATCAGGAAGC
GCATCGGACTCATCAGGAAGC
TCATCGGACTCATCAGGAAGC
TCATCGGACTCATCAGGAAGC

GCATCGGACTCATCAGGAAGC
GACTCATCAGGAAGC
TCGGACTCATCAGGAAGC
GCATCGGACTCATCAGGAAGC
ACATCGGACTCATCAGGAAGC
TCCATCGGACTCATCAGGAAGC
CATCGGACTCATCAGGAAGC
CATCGGACTCATCAGGAAGC
ACATCGGACTCATCAGGAAGC
CCATCGGACTCATCAGGAAGC
ACCATCGGACTCATCAGGAAGC
CATCGGACTCATCAGGAAGC
CCATCGGACTCATCAGGAAGC
ACATCGGACTCATCAGGAAGC
AACATCGGACTCATCAGGAAGC
CACATCGGACTCATCAGGAAGC
ATACATCGGACTCATCAGGAAGC
ACATCGGACTCATCAGGAAGC
TCATCGGACTCATCAGGAAGC
TACATCGGACTCATCAGGAAGC
GCATCGGACTCATCAGGAAGC
GCATCGGACTCATCAGGAAGC
CCATCGGACTCATCAGGAAGC
TACATCGGACTCATCAGGAAGC
ACATCGGACTCATCAGGAAGC
ACATCGGACTCATCAGGAAGC
TCATCGGACTCATCAGGAAGC
GCCATCGGACTCATCAGGAAGC
TCATCGGACTCATCAGGAAGC
ATCGGACTCATCAGGAAGC
CATCGGACTCATCAGGAAGC
CATCGGACTCATCAGGAAGC
GCATCGGACTCATCAGGAAGC
ATCATCGGACTCATCAGGAAGC
CATCGGACTCATCAGGAAGC
TACATCGGACTCATCAGGAAGC
ATCATCGGACTCATCAGGAAGC
CATCGGACTCATCAGGAAGC
TGACATCGGACTCATCAGGAAGC
ACTCATCGGACTCATCAGGAAGC
CACATCGGACTCATCAGGAAGC
GCATCGGACTCATCAGGAAGC
GCATCGGACTCATCAGGAAGC

AGCATCGGACTCATCAGGAAGC
CATCGGACTCATCAGGAAGC
TGCATCGGACTCATCAGGAAGC
ACATCGGACTCATCAGGAAGC
GCATCGGACTCATCAGGAAGC
ATCCATCGGACTCATCAGGAAGC
ATTCATCGGACTCATCAGGAAGC
TGCATCGGACTCATCAGGAAGC
TATCATCGGACTCATCAGGAAGC
TCATCGGACTCATCAGGAAGC
TACATCGGACTCATCAGGAAGC
ATCGGACTCATCAGGAAGC
GTCCATCGGACTCATCAGGAAGC
TGCATCGGACTCATCAGGAAGC
CGGACTCATCAGGAAGC
CTCATCGGACTCATCAGGAAGC
CATCGGACTCATCAGGAAGC
CACATCGGACTCATCAGGAAGC
TATCATCGGACTCATCAGGAAGC
ACATCGGACTCATCAGGAAGC
TTCATCGGACTCATCAGGAAGC
ATACATCGGACTCATCAGGAAGC
TGCATCGGACTCATCAGGAAGC
GCATCGGACTCATCAGGAAGC
ACCATCGGACTCATCAGGAAGC
ACACATCGGACTCATCAGGAAGC
AGGCATCGGACTCATCAGGAAGC
GCATCGGACTCATCAGGAAGC
CCATCGGACTCATCAGGAAGC
ACATCGGACTCATCAGGAAGC
TCATCGGACTCATCAGGAAGC
TCCATCGGACTCATCAGGAAGC
ATCATCGGACTCATCAGGAAGC
ATCATCGGACTCATCAGGAAGC
TCATCGGACTCATCAGGAAGC
ATACATCGGACTCATCAGGAAGC
ATCGGACTCATCAGGAAGC
ATCGGACTCATCAGGAAGC
ATCGGACTCATCAGGAAGC
TCGGACTCATCAGGAAGC
CGGACTCATCAGGAAGC
CTCATCGGACTCATCAGGAAGC
TCATCGGACTCATCAGGAAGC

GGACTCATCAGGAAGC
GGACTCATCAGGAAGC
AACATCGGACTCATCAGGAAGC
GCATCGGACTCATCAGGAAGC
AGCATCGGACTCATCAGGAAGC
AATCATCGGACTCATCAGGAAGC
CCATCGGACTCATCAGGAAGC
CCATCGGACTCATCAGGAAGC
AGCATCGGACTCATCAGGAAGC
TACATCGGACTCATCAGGAAGC
ATCGGACTCATCAGGAAGC
GACTCATCAGGAAGC
ACATCGGACTCATCAGGAAGC
ACATCGGACTCATCAGGAAGC
GTCATCGGACTCATCAGGAAGC
TCATCGGACTCATCAGGAAGC
CATCGGACTCATCAGGAAGC
TCGGACTCATCAGGAAGC
ATCGGACTCATCAGGAAGC
ACATCGGACTCATCAGGAAGC
TCATCGGACTCATCAGGAAGC
CCCATCGGACTCATCAGGAAGC
TTCATCGGACTCATCAGGAAGC
ACATCGGACTCATCAGGAAGC
ATCGGACTCATCAGGAAGC
CATCGGACTCATCAGGAAGC
AACATCGGACTCATCAGGAAGC
GGACTCATCAGGAAGC
TCATCGGACTCATCAGGAAGC
ACATCGGACTCATCAGGAAGC
AGCATCGGACTCATCAGGAAGC
CCATCGGACTCATCAGGAAGC
TCCATCGGACTCATCAGGAAGC
TCGGACTCATCAGGAAGC
CACCATCGGACTCATCAGGAAGC
CACCATCGGACTCATCAGGAAGC
AGCATCGGACTCATCAGGAAGC
ACATCGGACTCATCAGGAAGC
TCATCGGACTCATCAGGAAGC
AACATCGGACTCATCAGGAAGC
TCGGACTCATCAGGAAGC
CCATCGGACTCATCAGGAAGC
TCATCGGACTCATCAGGAAGC

ATACATCGGACTCATCAGGAAGC
ATCGGACTCATCAGGAAGC
GCATCGGACTCATCAGGAAGC
CCATCGGACTCATCAGGAAGC
TCATCGGACTCATCAGGAAGC
TCATCGGACTCATCAGGAAGC
CATCGGACTCATCAGGAAGC
AACATCGGACTCATCAGGAAGC
ACATCGGACTCATCAGGAAGC
TCATCGGACTCATCAGGAAGC
CATCGGACTCATCAGGAAGC
CATCGGACTCATCAGGAAGC
CATCGGACTCATCAGGAAGC
ACATCGGACTCATCAGGAAGC
AACTCATCGGACTCATCAGGAAGC
GACTCATCAGGAAGC
CATCGGACTCATCAGGAAGC
TCATCGGACTCATCAGGAAGC
AGTACATCGGACTCATCAGGAAGC
ATCGGACTCATCAGGAAGC
ATCGGACTCATCAGGAAGC
CATCGGACTCATCAGGAAGC
CATCGGACTCATCAGGAAGC
CATCGGACTCATCAGGAAGC
CATCGGACTCATCAGGAAGC
TCATCGGACTCATCAGGAAGC
CATCGGACTCATCAGGAAGC
CATCGGACTCATCAGGAAGC
ACATCGGACTCATCAGGAAGC
CATCGGACTCATCAGGAAGC
GTCATCGGACTCATCAGGAAGC
CATCGGACTCATCAGGAAGC
ATCGGACTCATCAGGAAGC
GACATCGGACTCATCAGGAAGC
ATCGGACTCATCAGGAAGC
AGCATCGGACTCATCAGGAAGC
CATCGGACTCATCAGGAAGC
CATCGGACTCATCAGGAAGC
ACATCGGACTCATCAGGAAGC
CATCGGACTCATCAGGAAGC
TACATCGGACTCATCAGGAAGC
ATCGGACTCATCAGGAAGC
TCATCGGACTCATCAGGAAGC

TTCATCGGACTCATCAGGAAGC
TCCATCGGACTCATCAGGAAGC
GCATCGGACTCATCAGGAAGC
ACATCGGACTCATCAGGAAGC
GTCATCGGACTCATCAGGAAGC
TCATCGGACTCATCAGGAAGC
CCATCGGACTCATCAGGAAGC
CATCGGACTCATCAGGAAGC
AGACATCGGACTCATCAGGAAGC
ACACATCGGACTCATCAGGAAGC
CACATCGGACTCATCAGGAAGC
GACTCATCAGGAAGC
ACATCGGACTCATCAGGAAGC
AAGCATCGGACTCATCAGGAAGC
CATCGGACTCATCAGGAAGC
CTACATCGGACTCATCAGGAAGC
ACATCGGACTCATCAGGAAGC
GCATCGGACTCATCAGGAAGC
TCATCGGACTCATCAGGAAGC
ATCGGACTCATCAGGAAGC
ATCGGACTCATCAGGAAGC
TCGGACTCATCAGGAAGC
ACATCGGACTCATCAGGAAGC
CACATCGGACTCATCAGGAAGC
TCGGACTCATCAGGAAGC
ATCATCGGACTCATCAGGAAGC
TCCATCGGACTCATCAGGAAGC
CATCGGACTCATCAGGAAGC
ATCGGACTCATCAGGAAGC
ACATCGGACTCATCAGGAAGC
ACATCGGACTCATCAGGAAGC
ATCGGACTCATCAGGAAGC
GGACTCATCAGGAAGC
ATCGGACTCATCAGGAAGC
GACTCATCAGGAAGC
AATCATCGGACTCATCAGGAAGC
GTCATCGGACTCATCAGGAAGC
ATCGGACTCATCAGGAAGC
CATCGGACTCATCAGGAAGC
ATCATCGGACTCATCAGGAAGC
GCATCGGACTCATCAGGAAGC
CTCATCGGACTCATCAGGAAGC
ACATCGGACTCATCAGGAAGC

TCGGACTCATCAGGAAGC
CATCGGACTCATCAGGAAGC
ATCGGACTCATCAGGAAGC
CATCGGACTCATCAGGAAGC
ACTCATCAGGAAGC
CATCGGACTCATCAGGAAGC
TCGGACTCATCAGGAAGC
TCATCGGACTCATCAGGAAGC
GCATCGGACTCATCAGGAAGC
CGGACTCATCAGGAAGC
TCATCGGACTCATCAGGAAGC
CACATCGGACTCATCAGGAAGC
ACATCGGACTCATCAGGAAGC
ATCGGACTCATCAGGAAGC
CATCGGACTCATCAGGAAGC
CCATCGGACTCATCAGGAAGC
CCATCGGACTCATCAGGAAGC
GGACTCATCAGGAAGC
ATCGGACTCATCAGGAAGC
TCGGACTCATCAGGAAGC
CATCGGACTCATCAGGAAGC
ATCGGACTCATCAGGAAGC
GGACTCATCAGGAAGC
ACATCGGACTCATCAGGAAGC
GCATCGGACTCATCAGGAAGC
GGACTCATCAGGAAGC
CATCGGACTCATCAGGAAGC
TCATCGGACTCATCAGGAAGC
CATCGGACTCATCAGGAAGC
CATCGGACTCATCAGGAAGC
GGACTCATCAGGAAGC
ATCATCGGACTCATCAGGAAGC
AACATCGGACTCATCAGGAAGC
GACTCATCAGGAAGC
CATCGGACTCATCAGGAAGC
AGCATCGGACTCATCAGGAAGC
GGACTCATCAGGAAGC
AGTCATCGGACTCATCAGGAAGC
ATCGGACTCATCAGGAAGC
ATCGGACTCATCAGGAAGC
ATCGGACTCATCAGGAAGC
CATCGGACTCATCAGGAAGC
CATCGGACTCATCAGGAAGC

CATCGGACTCATCAGGAAGC
CCCCATCGGACTCATCAGGAAGC
ATCGGACTCATCAGGAAGC
TCGGACTCATCAGGAAGC
TAACATCGGACTCATCAGGAAGC
CATCGGACTCATCAGGAAGC
AACATCGGACTCATCAGGAAGC
CCATCGGACTCATCAGGAAGC
ATCGGACTCATCAGGAAGC
ATCGGACTCATCAGGAAGC
ATCGGACTCATCAGGAAGC
ATCATCGGACTCATCAGGAAGC
ATCATCGGACTCATCAGGAAGC
ACATCGGACTCATCAGGAAGC
CATCGGACTCATCAGGAAGC
ATCGGACTCATCAGGAAGC
CGGACTCATCAGGAAGC
AACATCGGACTCATCAGGAAGC
TCGGACTCATCAGGAAGC
TACATCGGACTCATCAGGAAGC
AGCATCGGACTCATCAGGAAGC
GCATCGGACTCATCAGGAAGC
TCATCGGACTCATCAGGAAGC
CATCGGACTCATCAGGAAGC
ATCGGACTCATCAGGAAGC
TTCATCGGACTCATCAGGAAGC
CATCGGACTCATCAGGAAGC
TCATCGGACTCATCAGGAAGC
TACATCGGACTCATCAGGAAGC
ATCGGACTCATCAGGAAGC
TCGGACTCATCAGGAAGC
ATCGGACTCATCAGGAAGC
CATCGGACTCATCAGGAAGC
GACATCGGACTCATCAGGAAGC
GCATCGGACTCATCAGGAAGC
GGACTCATCAGGAAGC
ATCATCGGACTCATCAGGAAGC
ACATCGGACTCATCAGGAAGC
TCATCGGACTCATCAGGAAGC
ATCGGACTCATCAGGAAGC
ATCGGACTCATCAGGAAGC
CGGACTCATCAGGAAGC
CATCGGACTCATCAGGAAGC

ATCGGACTCATCAGGAAGC
TTCATCGGACTCATCAGGAAGC
TCGGACTCATCAGGAAGC
GCCATCGGACTCATCAGGAAGC
TGCATCGGACTCATCAGGAAGC
TCATCGGACTCATCAGGAAGC
GTCATCGGACTCATCAGGAAGC
CACATCGGACTCATCAGGAAGC
GTGCATCGGACTCATCAGGAAGC
GGACTCATCAGGAAGC
TCGGACTCATCAGGAAGC
CATCGGACTCATCAGGAAGC
CATCGGACTCATCAGGAAGC
CATCGGACTCATCAGGAAGC
CATCGGACTCATCAGGAAGC
AGCATCGGACTCATCAGGAAGC
ATCGGACTCATCAGGAAGC
TCGGACTCATCAGGAAGC
GGACTCATCAGGAAGC
CATCGGACTCATCAGGAAGC
GGCATCGGACTCATCAGGAAGC
CATCGGACTCATCAGGAAGC
TCGGACTCATCAGGAAGC
TCGGACTCATCAGGAAGC
ATCGGACTCATCAGGAAGC
TCGGACTCATCAGGAAGC
GGACTCATCAGGAAGC
GGACTCATCAGGAAGC
TCGGACTCATCAGGAAGC
ATCGGACTCATCAGGAAGC
CATCGGACTCATCAGGAAGC
AACATCGGACTCATCAGGAAGC
CATCGGACTCATCAGGAAGC
CATCGGACTCATCAGGAAGC
TACATCGGACTCATCAGGAAGC
CATCGGACTCATCAGGAAGC
ATCGGACTCATCAGGAAGC
TCATCGGACTCATCAGGAAGC
TCATCGGACTCATCAGGAAGC
TCGGACTCATCAGGAAGC
CATCGGACTCATCAGGAAGC
TCATCGGACTCATCAGGAAGC
CGGACTCATCAGGAAGC

GTCCATCGGACTCATCAGGAAGC
CCATCGGACTCATCAGGAAGC
CATCGGACTCATCAGGAAGC
ATCGGACTCATCAGGAAGC
CATCGGACTCATCAGGAAGC
ATCATCGGACTCATCAGGAAGC
CTTCATCGGACTCATCAGGAAGC
GCATCGGACTCATCAGGAAGC
GGACTCATCAGGAAGC
TTCATCGGACTCATCAGGAAGC
TCGGACTCATCAGGAAGC
CGGACTCATCAGGAAGC
CGGACTCATCAGGAAGC
GCATCGGACTCATCAGGAAGC
CATCGGACTCATCAGGAAGC
CCATCGGACTCATCAGGAAGC
ACATCGGACTCATCAGGAAGC
CATCGGACTCATCAGGAAGC
ATCATCGGACTCATCAGGAAGC
TACATCGGACTCATCAGGAAGC
TCTCATCGGACTCATCAGGAAGC
TCGGACTCATCAGGAAGC
ATCGGACTCATCAGGAAGC
TTCATCGGACTCATCAGGAAGC
ACATCGGACTCATCAGGAAGC
TCACATCGGACTCATCAGGAAGC
GGACTCATCAGGAAGC
CATCGGACTCATCAGGAAGC
ATCATCGGACTCATCAGGAAGC
CATCGGACTCATCAGGAAGC
ATCATCGGACTCATCAGGAAGC
CATCGGACTCATCAGGAAGC
GGACTCATCAGGAAGC
GCCATCGGACTCATCAGGAAGC
CATCGGACTCATCAGGAAGC
TCCCATCGGACTCATCAGGAAGC
TCCCATCGGACTCATCAGGAAGC
CATCGGACTCATCAGGAAGC
AAACATCGGACTCATCAGGAAGC
CATCGGACTCATCAGGAAGC
TCATCGGACTCATCAGGAAGC
TCATCGGACTCATCAGGAAGC
GACTCATCAGGAAGC

CGGACTCATCAGGAAGC
ACATCGGACTCATCAGGAAGC
GACATCGGACTCATCAGGAAGC
ATCGGACTCATCAGGAAGC
ACATCGGACTCATCAGGAAGC
CATCGGACTCATCAGGAAGC
GCATCGGACTCATCAGGAAGC
ACATCGGACTCATCAGGAAGC
CATCGGACTCATCAGGAAGC
GGACTCATCAGGAAGC
TGCATCGGACTCATCAGGAAGC
CATCGGACTCATCAGGAAGC
CATCGGACTCATCAGGAAGC
GCATCGGACTCATCAGGAAGC
CATCGGACTCATCAGGAAGC
TCATCGGACTCATCAGGAAGC
AACATCGGACTCATCAGGAAGC
GTCATCGGACTCATCAGGAAGC
TCGGACTCATCAGGAAGC
GGACTCATCAGGAAGC
CATCGGACTCATCAGGAAGC
GCATCGGACTCATCAGGAAGC
ATCGGACTCATCAGGAAGC
TCGGACTCATCAGGAAGC
TCGGACTCATCAGGAAGC
CATCGGACTCATCAGGAAGC
ATCGGACTCATCAGGAAGC
GGACTCATCAGGAAGC
CATCGGACTCATCAGGAAGC
TCATCGGACTCATCAGGAAGC
CCATCGGACTCATCAGGAAGC
GCCATCGGACTCATCAGGAAGC
TCATCGGACTCATCAGGAAGC
ATCGGACTCATCAGGAAGC
CATCGGACTCATCAGGAAGC
CATCGGACTCATCAGGAAGC
ATCGGACTCATCAGGAAGC
ATCGGACTCATCAGGAAGC
ACCATCGGACTCATCAGGAAGC
GCATCGGACTCATCAGGAAGC
CCATCGGACTCATCAGGAAGC
TCGGACTCATCAGGAAGC
AGCATCGGACTCATCAGGAAGC

CCCATCGGACTCATCAGGAAGC
GCATCGGACTCATCAGGAAGC
ACATCGGACTCATCAGGAAGC
ATCGGACTCATCAGGAAGC
TCGGACTCATCAGGAAGC
TTCATCGGACTCATCAGGAAGC
TTCATCGGACTCATCAGGAAGC
ACATCGGACTCATCAGGAAGC
TACATCGGACTCATCAGGAAGC
ATCATCGGACTCATCAGGAAGC
GCATCGGACTCATCAGGAAGC
CCATCGGACTCATCAGGAAGC
GACTCATCAGGAAGC
CCATCGGACTCATCAGGAAGC
TCATCGGACTCATCAGGAAGC
AGCATCGGACTCATCAGGAAGC
ATCGGACTCATCAGGAAGC
AACATCGGACTCATCAGGAAGC
TCGGACTCATCAGGAAGC
GACTCATCAGGAAGC
ATCGGACTCATCAGGAAGC
GGACTCATCAGGAAGC
CATCGGACTCATCAGGAAGC
TCGGACTCATCAGGAAGC
TCCATCGGACTCATCAGGAAGC
CATCGGACTCATCAGGAAGC
TCATCGGACTCATCAGGAAGC
TCATCGGACTCATCAGGAAGC
CGGACTCATCAGGAAGC
GAGCATCGGACTCATCAGGAAGC
CGGACTCATCAGGAAGC
GCATCGGACTCATCAGGAAGC
CATCGGACTCATCAGGAAGC
TCTCATCGGACTCATCAGGAAGC
CATCGGACTCATCAGGAAGC
ATCGGACTCATCAGGAAGC
GTGCATCGGACTCATCAGGAAGC
TCATCGGACTCATCAGGAAGC
TGCATCGGACTCATCAGGAAGC
ATCGGACTCATCAGGAAGC
TTCATCGGACTCATCAGGAAGC
CATCGGACTCATCAGGAAGC
ACCATCGGACTCATCAGGAAGC

CCATCGGACTCATCAGGAAGC
GACTCATCAGGAAGC
TATCATCGGACTCATCAGGAAGC
ATCGGACTCATCAGGAAGC
GCCATCGGACTCATCAGGAAGC
ACATCGGACTCATCAGGAAGC
CATCGGACTCATCAGGAAGC
ATCGGACTCATCAGGAAGC
ACATCGGACTCATCAGGAAGC
TACATCGGACTCATCAGGAAGC
ATCGGACTCATCAGGAAGC
AACATCGGACTCATCAGGAAGC
ACATCGGACTCATCAGGAAGC
ATCGGACTCATCAGGAAGC
CATCGGACTCATCAGGAAGC
GGACTCATCAGGAAGC
GACATCGGACTCATCAGGAAGC
ATCGGACTCATCAGGAAGC
CACCATCGGACTCATCAGGAAGC
TCATCGGACTCATCAGGAAGC
GTCATCGGACTCATCAGGAAGC
ACATCGGACTCATCAGGAAGC
TCATCGGACTCATCAGGAAGC
TACATCGGACTCATCAGGAAGC
AGCATCGGACTCATCAGGAAGC
TCATCGGACTCATCAGGAAGC
CACATCGGACTCATCAGGAAGC
GCATCGGACTCATCAGGAAGC
TCGGACTCATCAGGAAGC
GACTCATCAGGAAGC
TCCATCGGACTCATCAGGAAGC
ATCGGACTCATCAGGAAGC
AGCATCGGACTCATCAGGAAGC
CATCGGACTCATCAGGAAGC
CCATCGGACTCATCAGGAAGC
GTCATCGGACTCATCAGGAAGC
GTCATCGGACTCATCAGGAAGC
CATCGGACTCATCAGGAAGC
GCATCGGACTCATCAGGAAGC
GCATCGGACTCATCAGGAAGC
TACATCGGACTCATCAGGAAGC
ACCATCGGACTCATCAGGAAGC
ACCATCGGACTCATCAGGAAGC

TCGGACTCATCAGGAAGC
GACTCATCAGGAAGC
ACATCGGACTCATCAGGAAGC
CCATCGGACTCATCAGGAAGC
CCATCGGACTCATCAGGAAGC
GCATCGGACTCATCAGGAAGC
AACATCGGACTCATCAGGAAGC
AACATCGGACTCATCAGGAAGC
GTTCATCGGACTCATCAGGAAGC
TCGGACTCATCAGGAAGC
ATCGGACTCATCAGGAAGC
GCATCGGACTCATCAGGAAGC
ATCGGACTCATCAGGAAGC
ACATCGGACTCATCAGGAAGC
CGGACTCATCAGGAAGC
CTCTCATCGGACTCATCAGGAAGC
ATCGGACTCATCAGGAAGC
ATCGGACTCATCAGGAAGC
TACATCGGACTCATCAGGAAGC
AACCATCGGACTCATCAGGAAGC
TCATCGGACTCATCAGGAAGC
GAAACATCGGACTCATCAGGAAGC
TACATCGGACTCATCAGGAAGC
CATCGGACTCATCAGGAAGC
GTAACATCGGACTCATCAGGAAGC
TCGGACTCATCAGGAAGC
CATCGGACTCATCAGGAAGC
CATCGGACTCATCAGGAAGC
TCGGACTCATCAGGAAGC
TCGGACTCATCAGGAAGC
AACATCGGACTCATCAGGAAGC
GGACTCATCAGGAAGC
CGGACTCATCAGGAAGC
ATCGGACTCATCAGGAAGC
GACTCATCAGGAAGC
CATCGGACTCATCAGGAAGC
ATCGGACTCATCAGGAAGC
CATCGGACTCATCAGGAAGC
ACATCGGACTCATCAGGAAGC
GGACTCATCAGGAAGC
CATCGGACTCATCAGGAAGC
GGACTCATCAGGAAGC
ACCATCGGACTCATCAGGAAGC

GACTCATCAGGAAGC
TCCATCGGACTCATCAGGAAGC
ACATCGGACTCATCAGGAAGC
CATCGGACTCATCAGGAAGC
TCATCGGACTCATCAGGAAGC
TCGGACTCATCAGGAAGC
ACATCGGACTCATCAGGAAGC
GGAACATCGGACTCATCAGGAAGC
TGTCATCGGACTCATCAGGAAGC
CACCATCGGACTCATCAGGAAGC
ATCGGACTCATCAGGAAGC
GACTCATCAGGAAGC
TCGGACTCATCAGGAAGC
TGCATCGGACTCATCAGGAAGC
GCTCATCGGACTCATCAGGAAGC
GGACTCATCAGGAAGC
CATCGGACTCATCAGGAAGC
AGCATCGGACTCATCAGGAAGC
CATCGGACTCATCAGGAAGC
ACACATCGGACTCATCAGGAAGC
ATCGGACTCATCAGGAAGC
ATCGGACTCATCAGGAAGC
TAACATCGGACTCATCAGGAAGC
ACACATCGGACTCATCAGGAAGC
CGGACTCATCAGGAAGC
TCGGACTCATCAGGAAGC
TTTCATCGGACTCATCAGGAAGC
ATCGGACTCATCAGGAAGC
GACTCATCAGGAAGC
GACTCATCAGGAAGC
GCATCGGACTCATCAGGAAGC
ATCGGACTCATCAGGAAGC
TCATCGGACTCATCAGGAAGC
TCGGACTCATCAGGAAGC
CGGACTCATCAGGAAGC
TCGGACTCATCAGGAAGC
TCGGACTCATCAGGAAGC
TCATCGGACTCATCAGGAAGC
TCATCGGACTCATCAGGAAGC
ACATCGGACTCATCAGGAAGC
GACTCATCAGGAAGC
CGGACTCATCAGGAAGC
TCGGACTCATCAGGAAGC

ACCCATCGGACTCATCAGGAAGC
ACATCGGACTCATCAGGAAGC
CATCGGACTCATCAGGAAGC
AGCATCGGACTCATCAGGAAGC
TCGGACTCATCAGGAAGC
CATCGGACTCATCAGGAAGC
ACATCGGACTCATCAGGAAGC
TCGGACTCATCAGGAAGC
GGACTCATCAGGAAGC
CATCGGACTCATCAGGAAGC
TACATCGGACTCATCAGGAAGC
CCATCGGACTCATCAGGAAGC
TCGGACTCATCAGGAAGC
TCGGACTCATCAGGAAGC
TCGGACTCATCAGGAAGC
CGGACTCATCAGGAAGC
AACATCGGACTCATCAGGAAGC
TCGGACTCATCAGGAAGC
CATCGGACTCATCAGGAAGC
ATCGGACTCATCAGGAAGC
ATCGGACTCATCAGGAAGC
ACATCGGACTCATCAGGAAGC
TTCATCGGACTCATCAGGAAGC
AGCATCGGACTCATCAGGAAGC
ATCGGACTCATCAGGAAGC
ATCGGACTCATCAGGAAGC
CCATCGGACTCATCAGGAAGC
TTCATCGGACTCATCAGGAAGC
TCATCGGACTCATCAGGAAGC
CATCGGACTCATCAGGAAGC
ACATCGGACTCATCAGGAAGC
TCGGACTCATCAGGAAGC
GACTCATCAGGAAGC
GTCCATCGGACTCATCAGGAAGC
TCGGACTCATCAGGAAGC
TCATCGGACTCATCAGGAAGC
TTCATCGGACTCATCAGGAAGC
TGCATCGGACTCATCAGGAAGC
TCGGACTCATCAGGAAGC
ATCGGACTCATCAGGAAGC
TCGGACTCATCAGGAAGC
CATCGGACTCATCAGGAAGC
ATCGGACTCATCAGGAAGC

CCATCGGACTCATCAGGAAGC
ATCGGACTCATCAGGAAGC
TCGGACTCATCAGGAAGC
ATCATCGGACTCATCAGGAAGC
ATCATCGGACTCATCAGGAAGC
CCATCGGACTCATCAGGAAGC
CCATCGGACTCATCAGGAAGC
ATCGGACTCATCAGGAAGC
ATCGGACTCATCAGGAAGC
CATCGGACTCATCAGGAAGC
ACATCGGACTCATCAGGAAGC
ACATCGGACTCATCAGGAAGC
TTCATCGGACTCATCAGGAAGC
GTCATCGGACTCATCAGGAAGC
CTCATCGGACTCATCAGGAAGC
CTCATCGGACTCATCAGGAAGC
TCATCGGACTCATCAGGAAGC
GGACTCATCAGGAAGC
ATCGGACTCATCAGGAAGC
ATCGGACTCATCAGGAAGC
GCATCGGACTCATCAGGAAGC
ACATCGGACTCATCAGGAAGC
ACATCGGACTCATCAGGAAGC
GGACTCATCAGGAAGC
CATCGGACTCATCAGGAAGC
ACATCGGACTCATCAGGAAGC
TCATCGGACTCATCAGGAAGC
TCATCGGACTCATCAGGAAGC
CACATCGGACTCATCAGGAAGC
CATCGGACTCATCAGGAAGC
TCATCGGACTCATCAGGAAGC
CATCGGACTCATCAGGAAGC
TCATCGGACTCATCAGGAAGC
CATCGGACTCATCAGGAAGC
CGGACTCATCAGGAAGC
ACATCGGACTCATCAGGAAGC
CATCGGACTCATCAGGAAGC
TCATCGGACTCATCAGGAAGC
CCATCGGACTCATCAGGAAGC
CATCGGACTCATCAGGAAGC
TTTGCATCGGACTCATCAGGAAGC
ATCGGACTCATCAGGAAGC
GTCATCGGACTCATCAGGAAGC

AACCATCGGACTCATCAGGAAGC
CCATCGGACTCATCAGGAAGC
TCGGACTCATCAGGAAGC
TCGGACTCATCAGGAAGC
ACATCGGACTCATCAGGAAGC
TCATCGGACTCATCAGGAAGC
ATCGGACTCATCAGGAAGC
CATCGGACTCATCAGGAAGC
GCATCGGACTCATCAGGAAGC
CACATCGGACTCATCAGGAAGC
CACATCGGACTCATCAGGAAGC
TACATCGGACTCATCAGGAAGC
TACATCGGACTCATCAGGAAGC
TACATCGGACTCATCAGGAAGC
CTCATCGGACTCATCAGGAAGC
GCATCGGACTCATCAGGAAGC
TCCATCGGACTCATCAGGAAGC
GGACTCATCAGGAAGC
ATCGGACTCATCAGGAAGC
CATCGGACTCATCAGGAAGC
GCATCGGACTCATCAGGAAGC
CACATCGGACTCATCAGGAAGC
TCATCGGACTCATCAGGAAGC
CCATCGGACTCATCAGGAAGC
ATCGGACTCATCAGGAAGC
ATCGGACTCATCAGGAAGC
ATCGGACTCATCAGGAAGC
CATCGGACTCATCAGGAAGC
ACATCGGACTCATCAGGAAGC
CCATCGGACTCATCAGGAAGC
CATCGGACTCATCAGGAAGC
GGACTCATCAGGAAGC
TCGGACTCATCAGGAAGC
AACATCGGACTCATCAGGAAGC
ACATCGGACTCATCAGGAAGC
GTACATCGGACTCATCAGGAAGC
ATCGGACTCATCAGGAAGC
ATCGGACTCATCAGGAAGC
CGGACTCATCAGGAAGC
CCATCGGACTCATCAGGAAGC
GGACTCATCAGGAAGC
CATCGGACTCATCAGGAAGC
ATCGGACTCATCAGGAAGC

TTGCATCGGACTCATCAGGAAGC
CATCGGACTCATCAGGAAGC
ATCGGACTCATCAGGAAGC
ACATCGGACTCATCAGGAAGC
GCATCGGACTCATCAGGAAGC
ATCGGACTCATCAGGAAGC
CCATCGGACTCATCAGGAAGC
CATCGGACTCATCAGGAAGC
CGGACTCATCAGGAAGC
AGCCATCGGACTCATCAGGAAGC
ATCGGACTCATCAGGAAGC
CGGACTCATCAGGAAGC
GCCATCGGACTCATCAGGAAGC
CATCGGACTCATCAGGAAGC
ACATCGGACTCATCAGGAAGC
GCATCGGACTCATCAGGAAGC
CCATCGGACTCATCAGGAAGC
ACCATCGGACTCATCAGGAAGC
TCCATCGGACTCATCAGGAAGC
TCATCGGACTCATCAGGAAGC
TCCATCGGACTCATCAGGAAGC
TCGGACTCATCAGGAAGC
TCATCGGACTCATCAGGAAGC
ATCGGACTCATCAGGAAGC
CATCGGACTCATCAGGAAGC
GGCATCGGACTCATCAGGAAGC
GGCATCGGACTCATCAGGAAGC
CATCGGACTCATCAGGAAGC
CATCGGACTCATCAGGAAGC
ACATCGGACTCATCAGGAAGC
CATCGGACTCATCAGGAAGC
CGGACTCATCAGGAAGC
TCGGACTCATCAGGAAGC
TTCATCGGACTCATCAGGAAGC
CATCGGACTCATCAGGAAGC
ACCATCGGACTCATCAGGAAGC
TGCCATCGGACTCATCAGGAAGC
GCATCGGACTCATCAGGAAGC
CCATCGGACTCATCAGGAAGC
CATCGGACTCATCAGGAAGC
CGGACTCATCAGGAAGC
CGGACTCATCAGGAAGC
TCGGACTCATCAGGAAGC

TTCATCGGACTCATCAGGAAGC
AGCATCGGACTCATCAGGAAGC
AGCATCGGACTCATCAGGAAGC
GACTCATCAGGAAGC
GGACTCATCAGGAAGC
AACATCGGACTCATCAGGAAGC
TCATCGGACTCATCAGGAAGC
CCATCGGACTCATCAGGAAGC
GACTCATCAGGAAGC
TCATCGGACTCATCAGGAAGC
TCATCGGACTCATCAGGAAGC
CGGACTCATCAGGAAGC
TCGGACTCATCAGGAAGC
TCATCGGACTCATCAGGAAGC
TCATCGGACTCATCAGGAAGC
CTCATCGGACTCATCAGGAAGC
TCATCGGACTCATCAGGAAGC
CCATCGGACTCATCAGGAAGC
ATCGGACTCATCAGGAAGC
CATCGGACTCATCAGGAAGC
ATCGGACTCATCAGGAAGC
CATCGGACTCATCAGGAAGC
CATCGGACTCATCAGGAAGC
ACATCGGACTCATCAGGAAGC
CATCGGACTCATCAGGAAGC
ATCGGACTCATCAGGAAGC
ATCGGACTCATCAGGAAGC
GGACTCATCAGGAAGC
TACATCGGACTCATCAGGAAGC
CTCATCGGACTCATCAGGAAGC
ATCGGACTCATCAGGAAGC
CGGACTCATCAGGAAGC
ATCGGACTCATCAGGAAGC
ATCGGACTCATCAGGAAGC
GGCATCGGACTCATCAGGAAGC
AGCATCGGACTCATCAGGAAGC
ATACATCGGACTCATCAGGAAGC
ATCGGACTCATCAGGAAGC
CCCATCGGACTCATCAGGAAGC
ATCGGACTCATCAGGAAGC
TCATCGGACTCATCAGGAAGC
TTCATCGGACTCATCAGGAAGC
CATCGGACTCATCAGGAAGC

GTCATCGGACTCATCAGGAAGC
ATCGGACTCATCAGGAAGC
CGGACTCATCAGGAAGC
TTACATCGGACTCATCAGGAAGC
ATCATCGGACTCATCAGGAAGC
ATCGGACTCATCAGGAAGC
ATCGGACTCATCAGGAAGC
TCGGACTCATCAGGAAGC
ATCGGACTCATCAGGAAGC
GGACTCATCAGGAAGC
ATACATCGGACTCATCAGGAAGC
AACATCGGACTCATCAGGAAGC
GCATCGGACTCATCAGGAAGC
GCATCGGACTCATCAGGAAGC
CACATCGGACTCATCAGGAAGC
CATCGGACTCATCAGGAAGC
GGCATCGGACTCATCAGGAAGC
TCATCGGACTCATCAGGAAGC
AGCATCGGACTCATCAGGAAGC
CATCGGACTCATCAGGAAGC
GTCATCGGACTCATCAGGAAGC
TATCATCGGACTCATCAGGAAGC
CATCGGACTCATCAGGAAGC
ATCGGACTCATCAGGAAGC
CATCGGACTCATCAGGAAGC
AACATCGGACTCATCAGGAAGC
TCGGACTCATCAGGAAGC
CACATCGGACTCATCAGGAAGC
TCATCGGACTCATCAGGAAGC
GACTCATCAGGAAGC
GCATCGGACTCATCAGGAAGC
CTCATCGGACTCATCAGGAAGC
TACATCGGACTCATCAGGAAGC
CGGACTCATCAGGAAGC
CGGACTCATCAGGAAGC
AGCATCGGACTCATCAGGAAGC
CATCGGACTCATCAGGAAGC
GACTCATCAGGAAGC
ATCGGACTCATCAGGAAGC
AGCATCGGACTCATCAGGAAGC
GCATCGGACTCATCAGGAAGC
GCATCGGACTCATCAGGAAGC
CCATCGGACTCATCAGGAAGC

TCATCGGACTCATCAGGAAGC
GCATCGGACTCATCAGGAAGC
CCCATCGGACTCATCAGGAAGC
GCATCGGACTCATCAGGAAGC
GGCATCGGACTCATCAGGAAGC
TCATCGGACTCATCAGGAAGC
GCATCGGACTCATCAGGAAGC
AACATCGGACTCATCAGGAAGC
ATCGGACTCATCAGGAAGC
TCGGACTCATCAGGAAGC
CACATCGGACTCATCAGGAAGC
AACATCGGACTCATCAGGAAGC
CATCGGACTCATCAGGAAGC
TATACATCGGACTCATCAGGAAGC
CATCGGACTCATCAGGAAGC
AACATCGGACTCATCAGGAAGC
AACATCGGACTCATCAGGAAGC
GGACTCATCAGGAAGC
ATCGGACTCATCAGGAAGC
GACTCATCAGGAAGC
ACATCGGACTCATCAGGAAGC
GGCATCGGACTCATCAGGAAGC
TGCATCGGACTCATCAGGAAGC
CCATCGGACTCATCAGGAAGC
CATCGGACTCATCAGGAAGC
CATCGGACTCATCAGGAAGC
GCATCGGACTCATCAGGAAGC
TCATCGGACTCATCAGGAAGC
GCATCGGACTCATCAGGAAGC
ATCATCGGACTCATCAGGAAGC
ACATCGGACTCATCAGGAAGC
TACATCGGACTCATCAGGAAGC
CCATCGGACTCATCAGGAAGC
CGGACTCATCAGGAAGC
ACATCGGACTCATCAGGAAGC
CCATCGGACTCATCAGGAAGC
TGGCATCGGACTCATCAGGAAGC
GACTCATCAGGAAGC
TCGGACTCATCAGGAAGC
GGACTCATCAGGAAGC
ACATCGGACTCATCAGGAAGC
CTCATCGGACTCATCAGGAAGC
CCATCGGACTCATCAGGAAGC

CATCGGACTCATCAGGAAGC
CATCGGACTCATCAGGAAGC
TCGGACTCATCAGGAAGC
ATGTCATCGGACTCATCAGGAAGC
CGGACTCATCAGGAAGC
CGGACTCATCAGGAAGC
TCGGACTCATCAGGAAGC
AATCATCGGACTCATCAGGAAGC
CCATCGGACTCATCAGGAAGC
ATCATCGGACTCATCAGGAAGC
CGGACTCATCAGGAAGC
ACATCGGACTCATCAGGAAGC
CACATCGGACTCATCAGGAAGC
ACATCGGACTCATCAGGAAGC
GACTCATCAGGAAGC
CATCGGACTCATCAGGAAGC
ACATCGGACTCATCAGGAAGC
TCATCGGACTCATCAGGAAGC
CATCGGACTCATCAGGAAGC
ATCGGACTCATCAGGAAGC
CGGACTCATCAGGAAGC
ATCGGACTCATCAGGAAGC
CCATCGGACTCATCAGGAAGC
CATCGGACTCATCAGGAAGC
CATCGGACTCATCAGGAAGC
TGCATCGGACTCATCAGGAAGC
TCATCGGACTCATCAGGAAGC
CATCGGACTCATCAGGAAGC
ACACATCGGACTCATCAGGAAGC
TCGGACTCATCAGGAAGC
ATCATCGGACTCATCAGGAAGC
TATCATCGGACTCATCAGGAAGC
TCGGACTCATCAGGAAGC
CATCGGACTCATCAGGAAGC
GATCCATCGGACTCATCAGGAAGC
GATCCATCGGACTCATCAGGAAGC
ACCATCGGACTCATCAGGAAGC
ATCGGACTCATCAGGAAGC
CCATCGGACTCATCAGGAAGC
ACATCGGACTCATCAGGAAGC
CATCGGACTCATCAGGAAGC
TTCATCGGACTCATCAGGAAGC
TCGGACTCATCAGGAAGC

CGGACTCATCAGGAAGC
ATCGGACTCATCAGGAAGC
TCATCGGACTCATCAGGAAGC
AACATCGGACTCATCAGGAAGC
TCATCGGACTCATCAGGAAGC
CATCGGACTCATCAGGAAGC
CATCGGACTCATCAGGAAGC
GCATCGGACTCATCAGGAAGC
ATCGGACTCATCAGGAAGC
CATCGGACTCATCAGGAAGC
CATCGGACTCATCAGGAAGC
CGGACTCATCAGGAAGC
AGCATCGGACTCATCAGGAAGC
TCGGACTCATCAGGAAGC
CATCGGACTCATCAGGAAGC
GCATCGGACTCATCAGGAAGC
CCATCGGACTCATCAGGAAGC
CACATCGGACTCATCAGGAAGC
TGCATCGGACTCATCAGGAAGC
ATCGGACTCATCAGGAAGC
ATCGGACTCATCAGGAAGC
ATCGGACTCATCAGGAAGC
GCATCGGACTCATCAGGAAGC
CATCGGACTCATCAGGAAGC
CGGACTCATCAGGAAGC
TCGGACTCATCAGGAAGC
GGACTCATCAGGAAGC
GACATCGGACTCATCAGGAAGC
TGCATCGGACTCATCAGGAAGC
ACATCGGACTCATCAGGAAGC
ACCCATCGGACTCATCAGGAAGC
CATCGGACTCATCAGGAAGC
ACCCATCGGACTCATCAGGAAGC
GCATCGGACTCATCAGGAAGC
AGCATCGGACTCATCAGGAAGC
TCATCGGACTCATCAGGAAGC
CCATCGGACTCATCAGGAAGC
TCACATCGGACTCATCAGGAAGC
TCCATCGGACTCATCAGGAAGC
ATCGGACTCATCAGGAAGC
TACATCGGACTCATCAGGAAGC
TGCATCGGACTCATCAGGAAGC
CATCGGACTCATCAGGAAGC

CATCGGACTCATCAGGAAGC
TCCATCGGACTCATCAGGAAGC
CGGACTCATCAGGAAGC
TCGGACTCATCAGGAAGC
GGACTCATCAGGAAGC
TCGGACTCATCAGGAAGC
CGGACTCATCAGGAAGC
CATCGGACTCATCAGGAAGC
CGGACTCATCAGGAAGC
CATCGGACTCATCAGGAAGC
GTCATCGGACTCATCAGGAAGC
ATCGGACTCATCAGGAAGC
GACTCATCAGGAAGC
TCATCGGACTCATCAGGAAGC
ATCGGACTCATCAGGAAGC
TCGGACTCATCAGGAAGC
CGGACTCATCAGGAAGC
ACACATCGGACTCATCAGGAAGC
CATCGGACTCATCAGGAAGC
ATCGGACTCATCAGGAAGC
TCATCGGACTCATCAGGAAGC
GCATCGGACTCATCAGGAAGC
TACATCGGACTCATCAGGAAGC
GGACTCATCAGGAAGC
ATCGGACTCATCAGGAAGC
ATCCATCGGACTCATCAGGAAGC
TCATCGGACTCATCAGGAAGC
GCATCGGACTCATCAGGAAGC
ACACATCGGACTCATCAGGAAGC
ACATCGGACTCATCAGGAAGC
AGACATCGGACTCATCAGGAAGC
GGCATCGGACTCATCAGGAAGC
GCCATCGGACTCATCAGGAAGC
ATCGGACTCATCAGGAAGC
GCATCGGACTCATCAGGAAGC
TGCATCGGACTCATCAGGAAGC
TCGGACTCATCAGGAAGC
ACTCATCAGGAAGC
ATCGGACTCATCAGGAAGC
CATCGGACTCATCAGGAAGC
ACTCATCAGGAAGC
CATCGGACTCATCAGGAAGC
TCGGACTCATCAGGAAGC

GGACTCATCAGGAAGC
GCATCGGACTCATCAGGAAGC
TCATCGGACTCATCAGGAAGC
GTCATCGGACTCATCAGGAAGC
GGCATCGGACTCATCAGGAAGC
TCATCGGACTCATCAGGAAGC
CATCGGACTCATCAGGAAGC
ATCGGACTCATCAGGAAGC
CATCGGACTCATCAGGAAGC
ACATCGGACTCATCAGGAAGC
GTCATCGGACTCATCAGGAAGC
TGCATCGGACTCATCAGGAAGC
GACATCGGACTCATCAGGAAGC
CCATCGGACTCATCAGGAAGC
TTTCATCGGACTCATCAGGAAGC
TCGGACTCATCAGGAAGC
CATCGGACTCATCAGGAAGC
GCATCGGACTCATCAGGAAGC
GGACTCATCAGGAAGC
AGCATCGGACTCATCAGGAAGC
ATCGGACTCATCAGGAAGC
ACTCATCGGACTCATCAGGAAGC
CATCGGACTCATCAGGAAGC
GGACTCATCAGGAAGC
CACATCGGACTCATCAGGAAGC
ATCGGACTCATCAGGAAGC
ATCGGACTCATCAGGAAGC
GCATCGGACTCATCAGGAAGC
ACATCGGACTCATCAGGAAGC
TCCATCGGACTCATCAGGAAGC
ATCGGACTCATCAGGAAGC
CATCGGACTCATCAGGAAGC
CCCATCGGACTCATCAGGAAGC
TCGGACTCATCAGGAAGC
CATCGGACTCATCAGGAAGC
TGCATCGGACTCATCAGGAAGC
CCATCGGACTCATCAGGAAGC
CACATCGGACTCATCAGGAAGC
GCATCGGACTCATCAGGAAGC
CTTCATCGGACTCATCAGGAAGC
CTTCATCGGACTCATCAGGAAGC
ACATCGGACTCATCAGGAAGC
ATCGGACTCATCAGGAAGC

GACATCGGACTCATCAGGAAGC
ATCGGACTCATCAGGAAGC
TCGGACTCATCAGGAAGC
TCATCGGACTCATCAGGAAGC
GCATCGGACTCATCAGGAAGC
CATCGGACTCATCAGGAAGC
CATCGGACTCATCAGGAAGC
TCATCGGACTCATCAGGAAGC
ACATCGGACTCATCAGGAAGC
GCATCGGACTCATCAGGAAGC
CATCGGACTCATCAGGAAGC
ATCGGACTCATCAGGAAGC
CACATCGGACTCATCAGGAAGC
GCATCGGACTCATCAGGAAGC
CATCGGACTCATCAGGAAGC
ACATCGGACTCATCAGGAAGC
CATCGGACTCATCAGGAAGC
CCATCGGACTCATCAGGAAGC
TCATCGGACTCATCAGGAAGC
CATCGGACTCATCAGGAAGC
CTCATCGGACTCATCAGGAAGC
ATCGGACTCATCAGGAAGC
ATCGGACTCATCAGGAAGC
ATCGGACTCATCAGGAAGC
CATCGGACTCATCAGGAAGC
TCATCGGACTCATCAGGAAGC
TCGGACTCATCAGGAAGC
ATCGGACTCATCAGGAAGC
ACATCGGACTCATCAGGAAGC
ACATCGGACTCATCAGGAAGC
TCATCGGACTCATCAGGAAGC
CATCGGACTCATCAGGAAGC
ATCGGACTCATCAGGAAGC
ACATCGGACTCATCAGGAAGC
CATCGGACTCATCAGGAAGC
ACATCGGACTCATCAGGAAGC
ATCGGACTCATCAGGAAGC
ATCGGACTCATCAGGAAGC
CGGACTCATCAGGAAGC
AGCATCGGACTCATCAGGAAGC
ATCGGACTCATCAGGAAGC
CATCGGACTCATCAGGAAGC
GCATCGGACTCATCAGGAAGC

ATCGGACTCATCAGGAAGC
TCCCATCGGACTCATCAGGAAGC
CATCGGACTCATCAGGAAGC
TCATCGGACTCATCAGGAAGC
ATCCATCGGACTCATCAGGAAGC
ACATCGGACTCATCAGGAAGC
CATCGGACTCATCAGGAAGC
CATCGGACTCATCAGGAAGC
ACATCGGACTCATCAGGAAGC
CGGACTCATCAGGAAGC
CGGACTCATCAGGAAGC
TCACATCGGACTCATCAGGAAGC
GGACTCATCAGGAAGC
ATCGGACTCATCAGGAAGC
CATCGGACTCATCAGGAAGC
ACATCGGACTCATCAGGAAGC
ACATCGGACTCATCAGGAAGC
TCATCGGACTCATCAGGAAGC
CCATCGGACTCATCAGGAAGC
CATCGGACTCATCAGGAAGC
GACTCATCAGGAAGC
GACTCATCAGGAAGC
ACCATCGGACTCATCAGGAAGC
CTCATCGGACTCATCAGGAAGC
CATCGGACTCATCAGGAAGC
CATCGGACTCATCAGGAAGC
ACATCGGACTCATCAGGAAGC
GCATCGGACTCATCAGGAAGC
GCATCGGACTCATCAGGAAGC
TCGGACTCATCAGGAAGC
TCATCGGACTCATCAGGAAGC
TCGGACTCATCAGGAAGC
GCATCGGACTCATCAGGAAGC
GCATCGGACTCATCAGGAAGC
ATCGGACTCATCAGGAAGC
AGCATCGGACTCATCAGGAAGC
AGCATCGGACTCATCAGGAAGC
TCGGACTCATCAGGAAGC
CATCGGACTCATCAGGAAGC
TCGGACTCATCAGGAAGC
TGCATCGGACTCATCAGGAAGC
GGACTCATCAGGAAGC
TCGGACTCATCAGGAAGC

GCATCGGACTCATCAGGAAGC
GCATCGGACTCATCAGGAAGC
TGTCATCGGACTCATCAGGAAGC
TACATCGGACTCATCAGGAAGC
AGCATCGGACTCATCAGGAAGC
GCATCGGACTCATCAGGAAGC
GCATCGGACTCATCAGGAAGC
GCATCGGACTCATCAGGAAGC
CATCGGACTCATCAGGAAGC
ATGCATCGGACTCATCAGGAAGC
ATCGGACTCATCAGGAAGC
ATCGGACTCATCAGGAAGC
ATCGGACTCATCAGGAAGC
GCATCGGACTCATCAGGAAGC
ATCGGACTCATCAGGAAGC
ATACATCGGACTCATCAGGAAGC
ATCGGACTCATCAGGAAGC
TCGGACTCATCAGGAAGC
ATCGGACTCATCAGGAAGC
GCATCGGACTCATCAGGAAGC
GCATCGGACTCATCAGGAAGC
GCATCGGACTCATCAGGAAGC
TCATCGGACTCATCAGGAAGC
ATCATCGGACTCATCAGGAAGC
ACTCATCAGGAAGC
CATCGGACTCATCAGGAAGC
ACATCGGACTCATCAGGAAGC
TCGGACTCATCAGGAAGC
TCGGACTCATCAGGAAGC
CATCGGACTCATCAGGAAGC
GGCATCGGACTCATCAGGAAGC
ACATCGGACTCATCAGGAAGC
GCATCGGACTCATCAGGAAGC
ACATCGGACTCATCAGGAAGC
TGCATCGGACTCATCAGGAAGC
TCATCGGACTCATCAGGAAGC
GCATCGGACTCATCAGGAAGC
GATCATCGGACTCATCAGGAAGC
CGGACTCATCAGGAAGC
CATCGGACTCATCAGGAAGC
TCATCGGACTCATCAGGAAGC
CTCATCGGACTCATCAGGAAGC
CATCGGACTCATCAGGAAGC

TCTCATCGGACTCATCAGGAAGC
GCATCGGACTCATCAGGAAGC
GGACTCATCAGGAAGC
ATCGGACTCATCAGGAAGC
ACATCGGACTCATCAGGAAGC
GACTCATCAGGAAGC
CATCGGACTCATCAGGAAGC
GCATCGGACTCATCAGGAAGC
CATCGGACTCATCAGGAAGC
TACATCGGACTCATCAGGAAGC
CTCATCGGACTCATCAGGAAGC
GCATCGGACTCATCAGGAAGC
ATCGGACTCATCAGGAAGC
TCATCGGACTCATCAGGAAGC
CTTCATCGGACTCATCAGGAAGC
CACATCGGACTCATCAGGAAGC
TCGGACTCATCAGGAAGC
ACATCGGACTCATCAGGAAGC
GCATCGGACTCATCAGGAAGC
AGCATCGGACTCATCAGGAAGC
CATCGGACTCATCAGGAAGC
GAGCATCGGACTCATCAGGAAGC
CGGACTCATCAGGAAGC
TATCGGACTCATCAGGAAGC
CCACATCGGACTCATCAGGAAGC
TATCGGACTCATCAGGAAGC
ATCGGACTCATCAGGAAGC
TTCATCGGACTCATCAGGAAGC
ATCGGACTCATCAGGAAGC
CATCGGACTCATCAGGAAGC
CAGCATCGGACTCATCAGGAAGC
CATCGGACTCATCAGGAAGC
CATCGGACTCATCAGGAAGC
TGCATCGGACTCATCAGGAAGC
CATCGGACTCATCAGGAAGC
CATCGGACTCATCAGGAAGC
CATCGGACTCATCAGGAAGC
AAATCATCGGACTCATCAGGAAGC
CTCATCAGGAAGC
TCATCGGACTCATCAGGAAGC
CATCGGACTCATCAGGAAGC
TCGGACTCATCAGGAAGC
TCGGACTCATCAGGAAGC

CATCGGACTCATCAGGAAGC
CATCGGACTCATCAGGAAGC
TCATCGGACTCATCAGGAAGC
ATCGGACTCATCAGGAAGC
CATCGGACTCATCAGGAAGC
GACTCATCAGGAAGC
TCATCGGACTCATCAGGAAGC
CATCGGACTCATCAGGAAGC
TCGGACTCATCAGGAAGC
CACATCGGACTCATCAGGAAGC
TTCATCGGACTCATCAGGAAGC
CCATCGGACTCATCAGGAAGC
CATCGGACTCATCAGGAAGC
ATCGGACTCATCAGGAAGC
CCATCGGACTCATCAGGAAGC
TCATCGGACTCATCAGGAAGC
CCCATCGGACTCATCAGGAAGC
TCGGACTCATCAGGAAGC
CACATCGGACTCATCAGGAAGC
GGACTCATCAGGAAGC
TCATCGGACTCATCAGGAAGC
TCATCGGACTCATCAGGAAGC
ATCGGACTCATCAGGAAGC
CTCATCGGACTCATCAGGAAGC
CATCGGACTCATCAGGAAGC
CATCGGACTCATCAGGAAGC
TCGGACTCATCAGGAAGC
ACATCGGACTCATCAGGAAGC
CCATCGGACTCATCAGGAAGC
TCATCGGACTCATCAGGAAGC
ATCGGACTCATCAGGAAGC
CATCGGACTCATCAGGAAGC
TTCATCGGACTCATCAGGAAGC
TCGGACTCATCAGGAAGC
CATCGGACTCATCAGGAAGC
ATCGGACTCATCAGGAAGC
ATCGGACTCATCAGGAAGC
CATCGGACTCATCAGGAAGC
ATCGGACTCATCAGGAAGC
GCATCGGACTCATCAGGAAGC
AAACATCGGACTCATCAGGAAGC
GTGCATCGGACTCATCAGGAAGC
ATTCATCGGACTCATCAGGAAGC

TCGGACTCATCAGGAAGC
GTCATCGGACTCATCAGGAAGC
GTCATCGGACTCATCAGGAAGC
ACCATCGGACTCATCAGGAAGC
ACATCGGACTCATCAGGAAGC
GCATCGGACTCATCAGGAAGC
ACCATCGGACTCATCAGGAAGC
CGGACTCATCAGGAAGC
CCATCGGACTCATCAGGAAGC
TCGGACTCATCAGGAAGC
CATCGGACTCATCAGGAAGC
GCATCGGACTCATCAGGAAGC
TACATCGGACTCATCAGGAAGC
GGACTCATCAGGAAGC
ATCGGACTCATCAGGAAGC
ATCGGACTCATCAGGAAGC
ACATCGGACTCATCAGGAAGC
AACCATCGGACTCATCAGGAAGC
TAACATCGGACTCATCAGGAAGC
TCACATCGGACTCATCAGGAAGC
ATCGGACTCATCAGGAAGC
CATCGGACTCATCAGGAAGC
ACCATCGGACTCATCAGGAAGC
GGACTCATCAGGAAGC
CATCGGACTCATCAGGAAGC
ATCGGACTCATCAGGAAGC
TATCATCGGACTCATCAGGAAGC
CCATCGGACTCATCAGGAAGC
TCATCGGACTCATCAGGAAGC
GCATCGGACTCATCAGGAAGC
CCCATCGGACTCATCAGGAAGC
GGACTCATCAGGAAGC
TCGGACTCATCAGGAAGC
CATCGGACTCATCAGGAAGC
GCATCGGACTCATCAGGAAGC
TCATCGGACTCATCAGGAAGC
CATCGGACTCATCAGGAAGC
TCGGACTCATCAGGAAGC
GGACTCATCAGGAAGC
TTACATCGGACTCATCAGGAAGC
CATCGGACTCATCAGGAAGC
GACTCATCAGGAAGC
CATCGGACTCATCAGGAAGC

CATCGGACTCATCAGGAAGC
CATCGGACTCATCAGGAAGC
CCATCGGACTCATCAGGAAGC
ATCATCGGACTCATCAGGAAGC
CATCGGACTCATCAGGAAGC
TCATCGGACTCATCAGGAAGC
ATGCATCGGACTCATCAGGAAGC
TACATCGGACTCATCAGGAAGC
TCGGACTCATCAGGAAGC
CATCGGACTCATCAGGAAGC
ACATCGGACTCATCAGGAAGC
GGACTCATCAGGAAGC
GACTCATCAGGAAGC
TCATCGGACTCATCAGGAAGC
TGCATCGGACTCATCAGGAAGC
GGACTCATCAGGAAGC
TACATCGGACTCATCAGGAAGC
CGGACTCATCAGGAAGC
CATCGGACTCATCAGGAAGC
GGGCATCGGACTCATCAGGAAGC
CATCGGACTCATCAGGAAGC
CATCGGACTCATCAGGAAGC
CATCGGACTCATCAGGAAGC
CCATCGGACTCATCAGGAAGC
CCATCGGACTCATCAGGAAGC
GCATCGGACTCATCAGGAAGC
GATACATCGGACTCATCAGGAAGC
CGGACTCATCAGGAAGC
GACATCGGACTCATCAGGAAGC
TCGGACTCATCAGGAAGC
CATCGGACTCATCAGGAAGC
ATCGGACTCATCAGGAAGC
ACATCGGACTCATCAGGAAGC
AACATCGGACTCATCAGGAAGC
GCATCGGACTCATCAGGAAGC
ACTCATCGGACTCATCAGGAAGC
CGGACTCATCAGGAAGC
GTCATCGGACTCATCAGGAAGC
TCATCGGACTCATCAGGAAGC
CATCGGACTCATCAGGAAGC
CATCGGACTCATCAGGAAGC
ATCGGACTCATCAGGAAGC
ATCATCGGACTCATCAGGAAGC

ATCGGACTCATCAGGAAGC
CCATCGGACTCATCAGGAAGC
CATCGGACTCATCAGGAAGC
CATCGGACTCATCAGGAAGC
CATCGGACTCATCAGGAAGC
GGGCATCGGACTCATCAGGAAGC
ACATCGGACTCATCAGGAAGC
TGTCATCGGACTCATCAGGAAGC
CATCGGACTCATCAGGAAGC
TCATCGGACTCATCAGGAAGC
CACATCGGACTCATCAGGAAGC
TCGGACTCATCAGGAAGC
ATCGGACTCATCAGGAAGC
ATCGGACTCATCAGGAAGC
GCATCGGACTCATCAGGAAGC
TCGGACTCATCAGGAAGC
CTGCATCGGACTCATCAGGAAGC
GGACTCATCAGGAAGC
GGACTCATCAGGAAGC
GGACTCATCAGGAAGC
CATCGGACTCATCAGGAAGC
TCATCGGACTCATCAGGAAGC
CATCGGACTCATCAGGAAGC
TCGGACTCATCAGGAAGC
GGACTCATCAGGAAGC
GCATCGGACTCATCAGGAAGC
CATCGGACTCATCAGGAAGC
CATCGGACTCATCAGGAAGC
CATCGGACTCATCAGGAAGC
GCATCGGACTCATCAGGAAGC
CATCGGACTCATCAGGAAGC
CATCGGACTCATCAGGAAGC
CCATCGGACTCATCAGGAAGC
GGACTCATCAGGAAGC
CATCGGACTCATCAGGAAGC
TCGGACTCATCAGGAAGC
GGCATCGGACTCATCAGGAAGC
ATCGGACTCATCAGGAAGC
AACATCGGACTCATCAGGAAGC
CATCGGACTCATCAGGAAGC
ATCGGACTCATCAGGAAGC
CGGACTCATCAGGAAGC
TCATCGGACTCATCAGGAAGC

CATCGGACTCATCAGGAAGC
TCATCGGACTCATCAGGAAGC
CGGACTCATCAGGAAGC
CATCGGACTCATCAGGAAGC
TCGGACTCATCAGGAAGC
ACTCATCAGGAAGC
CATCGGACTCATCAGGAAGC
ACATCGGACTCATCAGGAAGC
GGACTCATCAGGAAGC
TCATCGGACTCATCAGGAAGC
TCATCGGACTCATCAGGAAGC
CGGACTCATCAGGAAGC
ACATCGGACTCATCAGGAAGC
ATCGGACTCATCAGGAAGC
TCGGACTCATCAGGAAGC
GACTCATCAGGAAGC
ACATCGGACTCATCAGGAAGC
CGGACTCATCAGGAAGC
ATCGGACTCATCAGGAAGC
TCATCGGACTCATCAGGAAGC
GGACTCATCAGGAAGC
TCATCGGACTCATCAGGAAGC
AACATCGGACTCATCAGGAAGC
ACATCGGACTCATCAGGAAGC
CCATCGGACTCATCAGGAAGC
CCATCGGACTCATCAGGAAGC
GCATCGGACTCATCAGGAAGC
TTGCATCGGACTCATCAGGAAGC
TCATCGGACTCATCAGGAAGC
TCATCGGACTCATCAGGAAGC
GGACTCATCAGGAAGC
ATCGGACTCATCAGGAAGC
ATCATCGGACTCATCAGGAAGC
CATCGGACTCATCAGGAAGC
CATCGGACTCATCAGGAAGC
TGCATCGGACTCATCAGGAAGC
GCATCGGACTCATCAGGAAGC
ATCGGACTCATCAGGAAGC
ACATCGGACTCATCAGGAAGC
CATCGGACTCATCAGGAAGC
TCATCGGACTCATCAGGAAGC
CATCGGACTCATCAGGAAGC
CATCGGACTCATCAGGAAGC

ATCGGACTCATCAGGAAGC
ACATCGGACTCATCAGGAAGC
GCATCGGACTCATCAGGAAGC
TTCATCGGACTCATCAGGAAGC
TCATCGGACTCATCAGGAAGC
TGCCATCGGACTCATCAGGAAGC
ACATCGGACTCATCAGGAAGC
ATCGGACTCATCAGGAAGC
CATCGGACTCATCAGGAAGC
ATCGGACTCATCAGGAAGC
ATCGGACTCATCAGGAAGC
ACATCGGACTCATCAGGAAGC
AGCATCGGACTCATCAGGAAGC
CATCGGACTCATCAGGAAGC
TCATCGGACTCATCAGGAAGC
CATCGGACTCATCAGGAAGC
CATCGGACTCATCAGGAAGC
CATCGGACTCATCAGGAAGC
CATCGGACTCATCAGGAAGC
CATCGGACTCATCAGGAAGC
TGTCATCGGACTCATCAGGAAGC
AACATCGGACTCATCAGGAAGC
ATCGGACTCATCAGGAAGC
ACATCGGACTCATCAGGAAGC
ACATCGGACTCATCAGGAAGC
TTCATCGGACTCATCAGGAAGC
GACATCGGACTCATCAGGAAGC
CGGACTCATCAGGAAGC
ACATCGGACTCATCAGGAAGC
CATCGGACTCATCAGGAAGC
ACTCATCAGGAAGC
CGGACTCATCAGGAAGC
TCGGACTCATCAGGAAGC
TCGGACTCATCAGGAAGC
CCATCGGACTCATCAGGAAGC
CATCGGACTCATCAGGAAGC
TCGGACTCATCAGGAAGC
ATCGGACTCATCAGGAAGC
ATCGGACTCATCAGGAAGC
TCATCGGACTCATCAGGAAGC
GGACTCATCAGGAAGC
TCGGACTCATCAGGAAGC
TCGGACTCATCAGGAAGC

TCATCGGACTCATCAGGAAGC
GCATCGGACTCATCAGGAAGC
CATCGGACTCATCAGGAAGC
CATCGGACTCATCAGGAAGC
GAGCATCGGACTCATCAGGAAGC
CATCGGACTCATCAGGAAGC
CAGCATCGGACTCATCAGGAAGC
CATCGGACTCATCAGGAAGC
ACATCGGACTCATCAGGAAGC
ACATCGGACTCATCAGGAAGC
GGACTCATCAGGAAGC
CATCGGACTCATCAGGAAGC
AACATCGGACTCATCAGGAAGC
ACATCGGACTCATCAGGAAGC
GACATCGGACTCATCAGGAAGC
ACATCGGACTCATCAGGAAGC
GACTCATCAGGAAGC
TCATCGGACTCATCAGGAAGC
ATCATCGGACTCATCAGGAAGC
CGGACTCATCAGGAAGC
ATCGGACTCATCAGGAAGC
CCATCGGACTCATCAGGAAGC
TCGGACTCATCAGGAAGC
ACTCATCAGGAAGC
TCGGACTCATCAGGAAGC
ACATCGGACTCATCAGGAAGC
CGGACTCATCAGGAAGC
ATCGGACTCATCAGGAAGC
CATCGGACTCATCAGGAAGC
CATCGGACTCATCAGGAAGC
CATCGGACTCATCAGGAAGC
CATCGGACTCATCAGGAAGC
CATCGGACTCATCAGGAAGC
AACATCGGACTCATCAGGAAGC
TGACATCGGACTCATCAGGAAGC
TCATCGGACTCATCAGGAAGC
GTCATCGGACTCATCAGGAAGC
ACATCGGACTCATCAGGAAGC
AGCATCGGACTCATCAGGAAGC
GCATCGGACTCATCAGGAAGC
GGACTCATCAGGAAGC
AGCATCGGACTCATCAGGAAGC
ACATCGGACTCATCAGGAAGC

ACATCGGACTCATCAGGAAGC
CATCGGACTCATCAGGAAGC
ATCGGACTCATCAGGAAGC
ATCGGACTCATCAGGAAGC
CATCGGACTCATCAGGAAGC
CATCGGACTCATCAGGAAGC
CATCGGACTCATCAGGAAGC
ATCGGACTCATCAGGAAGC
CCATCGGACTCATCAGGAAGC
CATCGGACTCATCAGGAAGC
TCGGACTCATCAGGAAGC
CATCGGACTCATCAGGAAGC
CATCGGACTCATCAGGAAGC
CATCGGACTCATCAGGAAGC
ACATCGGACTCATCAGGAAGC
ATCGGACTCATCAGGAAGC
CATCGGACTCATCAGGAAGC
CATCGGACTCATCAGGAAGC
GCATCGGACTCATCAGGAAGC
TTCATCGGACTCATCAGGAAGC
ACATCGGACTCATCAGGAAGC
TCATCGGACTCATCAGGAAGC
CATCGGACTCATCAGGAAGC
TCATCGGACTCATCAGGAAGC
TCGGACTCATCAGGAAGC
TCATCGGACTCATCAGGAAGC
CATCGGACTCATCAGGAAGC
GCCATCGGACTCATCAGGAAGC
TCATCGGACTCATCAGGAAGC
CATCGGACTCATCAGGAAGC
ACATCGGACTCATCAGGAAGC
GACTCATCAGGAAGC
ACCATCGGACTCATCAGGAAGC
ATCGGACTCATCAGGAAGC
CGGACTCATCAGGAAGC
TCGGACTCATCAGGAAGC
ATCGGACTCATCAGGAAGC
GCATCGGACTCATCAGGAAGC
TGCATCGGACTCATCAGGAAGC
ATCGGACTCATCAGGAAGC
GCATCGGACTCATCAGGAAGC
TCGGACTCATCAGGAAGC
CGGACTCATCAGGAAGC

CGGACTCATCAGGAAGC
TCGGACTCATCAGGAAGC
ACATCGGACTCATCAGGAAGC
TCATCGGACTCATCAGGAAGC
TCGGACTCATCAGGAAGC
ACATCGGACTCATCAGGAAGC
TCGGACTCATCAGGAAGC
CATCGGACTCATCAGGAAGC
TTCATCGGACTCATCAGGAAGC
GTCATCGGACTCATCAGGAAGC
ACATCGGACTCATCAGGAAGC
TCATCGGACTCATCAGGAAGC
GACTCATCAGGAAGC
TGCATCGGACTCATCAGGAAGC
CATCGGACTCATCAGGAAGC
CCATCGGACTCATCAGGAAGC
CCATCGGACTCATCAGGAAGC
GACTCATCAGGAAGC
CGGACTCATCAGGAAGC
ACACATCGGACTCATCAGGAAGC
ATCGGACTCATCAGGAAGC
ATCGGACTCATCAGGAAGC
TCGGACTCATCAGGAAGC
TCGGACTCATCAGGAAGC
TCATCGGACTCATCAGGAAGC
CATCGGACTCATCAGGAAGC
GTCCATCGGACTCATCAGGAAGC
GGACTCATCAGGAAGC
ATCGGACTCATCAGGAAGC
TCATCGGACTCATCAGGAAGC
TGCATCGGACTCATCAGGAAGC
TCATCGGACTCATCAGGAAGC
CATCGGACTCATCAGGAAGC
ACATCGGACTCATCAGGAAGC
CACCATCGGACTCATCAGGAAGC
ATCGGACTCATCAGGAAGC
CATCGGACTCATCAGGAAGC
CATCGGACTCATCAGGAAGC
GGACTCATCAGGAAGC
TCATCGGACTCATCAGGAAGC
CATCGGACTCATCAGGAAGC
TCATCGGACTCATCAGGAAGC
ATAGCATCGGACTCATCAGGAAGC

CATCGGACTCATCAGGAAGC
TCGGACTCATCAGGAAGC
AGCATCGGACTCATCAGGAAGC
CATCGGACTCATCAGGAAGC
TTCATCGGACTCATCAGGAAGC
TTCATCGGACTCATCAGGAAGC
GCATCGGACTCATCAGGAAGC
CATCGGACTCATCAGGAAGC
TCATCGGACTCATCAGGAAGC
AGACATCGGACTCATCAGGAAGC
CATCGGACTCATCAGGAAGC
ATCGGACTCATCAGGAAGC
TCATCGGACTCATCAGGAAGC
ACATCGGACTCATCAGGAAGC
ATCGGACTCATCAGGAAGC
ACCATCGGACTCATCAGGAAGC
TCGGACTCATCAGGAAGC
CATCGGACTCATCAGGAAGC
GCATCGGACTCATCAGGAAGC
CATCGGACTCATCAGGAAGC
TCTCATCGGACTCATCAGGAAGC
CATCGGACTCATCAGGAAGC
CATCGGACTCATCAGGAAGC
ATCGGACTCATCAGGAAGC
ACATCGGACTCATCAGGAAGC
TCGGACTCATCAGGAAGC
GCATCGGACTCATCAGGAAGC
GCATCGGACTCATCAGGAAGC
TCCATCGGACTCATCAGGAAGC
CATCGGACTCATCAGGAAGC
GCCATCGGACTCATCAGGAAGC
GGACTCATCAGGAAGC
GACTCATCAGGAAGC
ACATCGGACTCATCAGGAAGC
GACTCATCAGGAAGC
CATCGGACTCATCAGGAAGC
CGGACTCATCAGGAAGC
TCATCGGACTCATCAGGAAGC
ATCGGACTCATCAGGAAGC
TACATCGGACTCATCAGGAAGC
AGCATCGGACTCATCAGGAAGC
ACATCGGACTCATCAGGAAGC
ACATCGGACTCATCAGGAAGC

GCATCGGACTCATCAGGAAGC
CATCGGACTCATCAGGAAGC
CATCGGACTCATCAGGAAGC
CATCGGACTCATCAGGAAGC
TCGGACTCATCAGGAAGC
TCATCGGACTCATCAGGAAGC
CGGACTCATCAGGAAGC
TCATCGGACTCATCAGGAAGC
ACATCGGACTCATCAGGAAGC
TCATCGGACTCATCAGGAAGC
AGCATCGGACTCATCAGGAAGC
ATCGGACTCATCAGGAAGC
ATCGGACTCATCAGGAAGC
TCATCGGACTCATCAGGAAGC
CATCGGACTCATCAGGAAGC
TCGGACTCATCAGGAAGC
CATCGGACTCATCAGGAAGC
AGCATCGGACTCATCAGGAAGC
TCGGACTCATCAGGAAGC
AAACATCGGACTCATCAGGAAGC
ATCGGACTCATCAGGAAGC
TCGGACTCATCAGGAAGC
AACATCGGACTCATCAGGAAGC
ATCGGACTCATCAGGAAGC
GGACTCATCAGGAAGC
TCATCGGACTCATCAGGAAGC
CATCGGACTCATCAGGAAGC
ATCGGACTCATCAGGAAGC
GACTCATCAGGAAGC
CCATCGGACTCATCAGGAAGC
GACATCGGACTCATCAGGAAGC
TTCATCGGACTCATCAGGAAGC
CCATCGGACTCATCAGGAAGC
CATCGGACTCATCAGGAAGC
CATCGGACTCATCAGGAAGC
TCATCGGACTCATCAGGAAGC
TCGGACTCATCAGGAAGC
CATCGGACTCATCAGGAAGC
TCGGACTCATCAGGAAGC
CATCGGACTCATCAGGAAGC
CCATCGGACTCATCAGGAAGC
CATCGGACTCATCAGGAAGC
CATCGGACTCATCAGGAAGC

ACCATCGGACTCATCAGGAAGC
AAGCATCGGACTCATCAGGAAGC
ACATCGGACTCATCAGGAAGC
ACATCGGACTCATCAGGAAGC
GACTCATCAGGAAGC
TCATCGGACTCATCAGGAAGC
ACATCGGACTCATCAGGAAGC
ACATCGGACTCATCAGGAAGC
CATCGGACTCATCAGGAAGC
AGACATCGGACTCATCAGGAAGC
TCATCGGACTCATCAGGAAGC
CATCGGACTCATCAGGAAGC
ATCGGACTCATCAGGAAGC
ACATCGGACTCATCAGGAAGC
CATCGGACTCATCAGGAAGC
GACTCATCAGGAAGC
CGGACTCATCAGGAAGC
ATCGGACTCATCAGGAAGC
TCATCGGACTCATCAGGAAGC
ACATCGGACTCATCAGGAAGC
CGGACTCATCAGGAAGC
ATCGGACTCATCAGGAAGC
ATCGGACTCATCAGGAAGC
ACATCGGACTCATCAGGAAGC
TACATCGGACTCATCAGGAAGC
AGCATCGGACTCATCAGGAAGC
CATCGGACTCATCAGGAAGC
CATCGGACTCATCAGGAAGC
GCCATCGGACTCATCAGGAAGC
CATCGGACTCATCAGGAAGC
ATCGGACTCATCAGGAAGC
CATCGGACTCATCAGGAAGC
ACTCATCAGGAAGC
CATCGGACTCATCAGGAAGC
ATCGGACTCATCAGGAAGC
AGCATCGGACTCATCAGGAAGC
TCATCGGACTCATCAGGAAGC
ACACATCGGACTCATCAGGAAGC
CCATCGGACTCATCAGGAAGC
TCGGACTCATCAGGAAGC
CATCGGACTCATCAGGAAGC
CGGACTCATCAGGAAGC
CATCGGACTCATCAGGAAGC

CATCGGACTCATCAGGAAGC
CTCATCGGACTCATCAGGAAGC
ATCGGACTCATCAGGAAGC
ACCATCGGACTCATCAGGAAGC
ATCATCGGACTCATCAGGAAGC
CATCGGACTCATCAGGAAGC
AGCATCGGACTCATCAGGAAGC
CATCGGACTCATCAGGAAGC
GACTCATCAGGAAGC
GCATCGGACTCATCAGGAAGC
CGGACTCATCAGGAAGC
CGGACTCATCAGGAAGC
GGACTCATCAGGAAGC
TTCATCGGACTCATCAGGAAGC
GCATCGGACTCATCAGGAAGC
CGGACTCATCAGGAAGC
ATCGGACTCATCAGGAAGC
TCGGACTCATCAGGAAGC
CATCGGACTCATCAGGAAGC
ATCATCGGACTCATCAGGAAGC
TCATCGGACTCATCAGGAAGC
CATCGGACTCATCAGGAAGC
CATCGGACTCATCAGGAAGC
GCATCGGACTCATCAGGAAGC
ATCGGACTCATCAGGAAGC
CATCGGACTCATCAGGAAGC
TCGGACTCATCAGGAAGC
GCATCGGACTCATCAGGAAGC
TCATCGGACTCATCAGGAAGC
AGTCATCGGACTCATCAGGAAGC
ACATCGGACTCATCAGGAAGC
CCATCGGACTCATCAGGAAGC
TTCATCGGACTCATCAGGAAGC
GCCATCGGACTCATCAGGAAGC
GCATCGGACTCATCAGGAAGC
TCGGACTCATCAGGAAGC
ATCGGACTCATCAGGAAGC
ATCGGACTCATCAGGAAGC
ACATCGGACTCATCAGGAAGC
ATCGGACTCATCAGGAAGC
ATCGGACTCATCAGGAAGC
ATCGGACTCATCAGGAAGC
TTCATCGGACTCATCAGGAAGC

CCATCGGACTCATCAGGAAGC
GGACTCATCAGGAAGC
ATCGGACTCATCAGGAAGC
ATCGGACTCATCAGGAAGC
TCCATCGGACTCATCAGGAAGC
GCATCGGACTCATCAGGAAGC
TCATCGGACTCATCAGGAAGC
CATCGGACTCATCAGGAAGC
ATCGGACTCATCAGGAAGC
CATCGGACTCATCAGGAAGC
GGACTCATCAGGAAGC
CCATCGGACTCATCAGGAAGC
TCGGACTCATCAGGAAGC
AGCATCGGACTCATCAGGAAGC
CCATCGGACTCATCAGGAAGC
CCCATCGGACTCATCAGGAAGC
TACATCGGACTCATCAGGAAGC
TACATCGGACTCATCAGGAAGC
GGCATCGGACTCATCAGGAAGC
GGCATCGGACTCATCAGGAAGC
ACCATCGGACTCATCAGGAAGC
TTTCATCGGACTCATCAGGAAGC
GACATCGGACTCATCAGGAAGC
ATGTCATCGGACTCATCAGGAAGC
TGCATCGGACTCATCAGGAAGC
TCGGACTCATCAGGAAGC
TGCATCGGACTCATCAGGAAGC
CCATCGGACTCATCAGGAAGC
CATCGGACTCATCAGGAAGC
CATCGGACTCATCAGGAAGC
CATCGGACTCATCAGGAAGC
GACTCATCAGGAAGC
CATCGGACTCATCAGGAAGC
ATCGGACTCATCAGGAAGC
TCCATCGGACTCATCAGGAAGC
CATCGGACTCATCAGGAAGC
GGACTCATCAGGAAGC
CGGACTCATCAGGAAGC
GACTCATCAGGAAGC
TACATCGGACTCATCAGGAAGC
ATCGGACTCATCAGGAAGC
GACTCATCAGGAAGC
CATCGGACTCATCAGGAAGC

TCATCGGACTCATCAGGAAGC
GCATCGGACTCATCAGGAAGC
GGACTCATCAGGAAGC
CGGACTCATCAGGAAGC
CGGACTCATCAGGAAGC
CTCATCGGACTCATCAGGAAGC
GGACTCATCAGGAAGC
TCATCGGACTCATCAGGAAGC
TCATCGGACTCATCAGGAAGC
ATCGGACTCATCAGGAAGC
GGACTCATCAGGAAGC
ATCGGACTCATCAGGAAGC
CCATCGGACTCATCAGGAAGC
GCATCGGACTCATCAGGAAGC
TCGGACTCATCAGGAAGC
TCGGACTCATCAGGAAGC
CATCGGACTCATCAGGAAGC
GGACTCATCAGGAAGC
TGCATCGGACTCATCAGGAAGC
CCATCGGACTCATCAGGAAGC
TCGGACTCATCAGGAAGC
CTTCATCGGACTCATCAGGAAGC
TCATCGGACTCATCAGGAAGC
TCGGACTCATCAGGAAGC
TCGGACTCATCAGGAAGC
CATCGGACTCATCAGGAAGC
GACATCGGACTCATCAGGAAGC
CATCGGACTCATCAGGAAGC
CTCATCAGGAAGC
CGGACTCATCAGGAAGC
ATCATCGGACTCATCAGGAAGC
CACATCGGACTCATCAGGAAGC
ATACATCGGACTCATCAGGAAGC
TGCATCGGACTCATCAGGAAGC
TGCATCGGACTCATCAGGAAGC
CATCGGACTCATCAGGAAGC
CATCGGACTCATCAGGAAGC
ACATCGGACTCATCAGGAAGC
CATCGGACTCATCAGGAAGC
CATCGGACTCATCAGGAAGC
TCATCGGACTCATCAGGAAGC
ATCGGACTCATCAGGAAGC
CGGACTCATCAGGAAGC

CGGACTCATCAGGAAGC
GGCCATCGGACTCATCAGGAAGC
ATCGGACTCATCAGGAAGC
GTCATCGGACTCATCAGGAAGC
ATTCATCGGACTCATCAGGAAGC
TCATCGGACTCATCAGGAAGC
CCATCGGACTCATCAGGAAGC
TCGGACTCATCAGGAAGC
ATCGGACTCATCAGGAAGC
ATCGGACTCATCAGGAAGC
AAACATCGGACTCATCAGGAAGC
AGCATCGGACTCATCAGGAAGC
TCGGACTCATCAGGAAGC
TCGGACTCATCAGGAAGC
CAATCATCGGACTCATCAGGAAGC
TACATCGGACTCATCAGGAAGC
GTCATCGGACTCATCAGGAAGC
TCGGACTCATCAGGAAGC
TAACATCGGACTCATCAGGAAGC
ATCGGACTCATCAGGAAGC
TCATCGGACTCATCAGGAAGC
ACCATCGGACTCATCAGGAAGC
CCATCGGACTCATCAGGAAGC
CGGACTCATCAGGAAGC
CATCGGACTCATCAGGAAGC
AACATCGGACTCATCAGGAAGC
TCGGACTCATCAGGAAGC
CTCATCGGACTCATCAGGAAGC
ATCGGACTCATCAGGAAGC
GGCATCGGACTCATCAGGAAGC
ATACATCGGACTCATCAGGAAGC
CATCGGACTCATCAGGAAGC
GCATCGGACTCATCAGGAAGC
AGTGCATCGGACTCATCAGGAAGC
ATCGGACTCATCAGGAAGC
CATCGGACTCATCAGGAAGC
TTCATCGGACTCATCAGGAAGC
ATCGGACTCATCAGGAAGC
CATCGGACTCATCAGGAAGC
CATCGGACTCATCAGGAAGC
ACATCGGACTCATCAGGAAGC
TCGGACTCATCAGGAAGC
ACATCGGACTCATCAGGAAGC

GGACTCATCAGGAAGC
GGACTCATCAGGAAGC
CCATCGGACTCATCAGGAAGC
CCATCGGACTCATCAGGAAGC
ATCATCGGACTCATCAGGAAGC
TCGGACTCATCAGGAAGC
TCGGACTCATCAGGAAGC
CATCGGACTCATCAGGAAGC
GCATCGGACTCATCAGGAAGC
TATCATCGGACTCATCAGGAAGC
ATCGGACTCATCAGGAAGC
ATCATCGGACTCATCAGGAAGC
CGGACTCATCAGGAAGC
CGGACTCATCAGGAAGC
AACATCGGACTCATCAGGAAGC
ATCATCGGACTCATCAGGAAGC
GCATCGGACTCATCAGGAAGC
GCATCGGACTCATCAGGAAGC
CATCGGACTCATCAGGAAGC
TACATCGGACTCATCAGGAAGC
TGACATCGGACTCATCAGGAAGC
TCATCGGACTCATCAGGAAGC
ATCGGACTCATCAGGAAGC
CATCGGACTCATCAGGAAGC
CATCGGACTCATCAGGAAGC
ATCGGACTCATCAGGAAGC
CGGACTCATCAGGAAGC
CATCGGACTCATCAGGAAGC
CATCGGACTCATCAGGAAGC
CATCGGACTCATCAGGAAGC
ATCGGACTCATCAGGAAGC
GCATCGGACTCATCAGGAAGC
TACATCGGACTCATCAGGAAGC
ACATCGGACTCATCAGGAAGC
ACATCGGACTCATCAGGAAGC
CGGACTCATCAGGAAGC
ACATCGGACTCATCAGGAAGC
ATCGGACTCATCAGGAAGC
CCATCGGACTCATCAGGAAGC
ATCGGACTCATCAGGAAGC
ATCGGACTCATCAGGAAGC
TCATCGGACTCATCAGGAAGC
GACATCGGACTCATCAGGAAGC

CATCGGACTCATCAGGAAGC
CACATCGGACTCATCAGGAAGC
CACATCGGACTCATCAGGAAGC
TCATCGGACTCATCAGGAAGC
ACATCGGACTCATCAGGAAGC
TCGGACTCATCAGGAAGC
CGGACTCATCAGGAAGC
TGCATCGGACTCATCAGGAAGC
ATCATCGGACTCATCAGGAAGC
CCCATCGGACTCATCAGGAAGC
TCATCGGACTCATCAGGAAGC
GCCATCGGACTCATCAGGAAGC
GGACTCATCAGGAAGC
CATCGGACTCATCAGGAAGC
CCATCGGACTCATCAGGAAGC
ATCGGACTCATCAGGAAGC
CCATCGGACTCATCAGGAAGC
GCCCATCGGACTCATCAGGAAGC
TGTGTCATCGGACTCATCAGGAAGC
CATCGGACTCATCAGGAAGC
CATCGGACTCATCAGGAAGC
GACTCATCAGGAAGC
TGCATCGGACTCATCAGGAAGC
CCATCGGACTCATCAGGAAGC
CATCGGACTCATCAGGAAGC
ATCATCGGACTCATCAGGAAGC
TCGGACTCATCAGGAAGC
ACATCGGACTCATCAGGAAGC
ATCGGACTCATCAGGAAGC
ATACATCGGACTCATCAGGAAGC
TAGCATCGGACTCATCAGGAAGC
TACATCGGACTCATCAGGAAGC
TACATCGGACTCATCAGGAAGC
CATCGGACTCATCAGGAAGC
CATCGGACTCATCAGGAAGC
ATCGGACTCATCAGGAAGC
CCATCGGACTCATCAGGAAGC
TCGGACTCATCAGGAAGC
ATCCATCGGACTCATCAGGAAGC
TCGGACTCATCAGGAAGC
AACATCGGACTCATCAGGAAGC
CACATCGGACTCATCAGGAAGC
ATCATCGGACTCATCAGGAAGC

ATCGGACTCATCAGGAAGC
TCATCGGACTCATCAGGAAGC
TCATCGGACTCATCAGGAAGC
ACATCGGACTCATCAGGAAGC
TCGGACTCATCAGGAAGC
GGACTCATCAGGAAGC
TCGGACTCATCAGGAAGC
CATCGGACTCATCAGGAAGC
CATCGGACTCATCAGGAAGC
CATCGGACTCATCAGGAAGC
ACATCGGACTCATCAGGAAGC
ATCGGACTCATCAGGAAGC
CCATCGGACTCATCAGGAAGC
TTCATCGGACTCATCAGGAAGC
TCGGACTCATCAGGAAGC
TCGGACTCATCAGGAAGC
CATCGGACTCATCAGGAAGC
CATCGGACTCATCAGGAAGC
CGGACTCATCAGGAAGC
CGGACTCATCAGGAAGC
GACTCATCAGGAAGC
CATCGGACTCATCAGGAAGC
ACATCGGACTCATCAGGAAGC
ACATCGGACTCATCAGGAAGC
CGGACTCATCAGGAAGC
ATCGGACTCATCAGGAAGC
TCGGACTCATCAGGAAGC
TCGGACTCATCAGGAAGC
ATCGGACTCATCAGGAAGC
ATCGGACTCATCAGGAAGC
ATCGGACTCATCAGGAAGC
CATCGGACTCATCAGGAAGC
CATCGGACTCATCAGGAAGC
CATCGGACTCATCAGGAAGC
GGACTCATCAGGAAGC
TCATCGGACTCATCAGGAAGC
CATCGGACTCATCAGGAAGC
CATCGGACTCATCAGGAAGC
AACATCGGACTCATCAGGAAGC
TCATCGGACTCATCAGGAAGC
CATCGGACTCATCAGGAAGC
CATCGGACTCATCAGGAAGC
CATCGGACTCATCAGGAAGC

CGGACTCATCAGGAAGC
CATCGGACTCATCAGGAAGC
CTCCATCGGACTCATCAGGAAGC
TACATCGGACTCATCAGGAAGC
CATCGGACTCATCAGGAAGC
TCGGACTCATCAGGAAGC
TCGGACTCATCAGGAAGC
ATCGGACTCATCAGGAAGC
GGACTCATCAGGAAGC
ACATCGGACTCATCAGGAAGC
AGCATCGGACTCATCAGGAAGC
ACCATCGGACTCATCAGGAAGC
CATCGGACTCATCAGGAAGC
AACATCGGACTCATCAGGAAGC
TCGGACTCATCAGGAAGC
CCCATCGGACTCATCAGGAAGC
CCATCGGACTCATCAGGAAGC
TCATCGGACTCATCAGGAAGC
CATCGGACTCATCAGGAAGC
AAATCATCGGACTCATCAGGAAGC
AGCATCGGACTCATCAGGAAGC
CGGACTCATCAGGAAGC
ATCGGACTCATCAGGAAGC
ATCGGACTCATCAGGAAGC
ATCGGACTCATCAGGAAGC
ACTCATCAGGAAGC
ATCGGACTCATCAGGAAGC
ATCGGACTCATCAGGAAGC
TCATCGGACTCATCAGGAAGC
TCATCGGACTCATCAGGAAGC
ACATCGGACTCATCAGGAAGC
TCGGACTCATCAGGAAGC
ATCGGACTCATCAGGAAGC
ATCGGACTCATCAGGAAGC
CATCGGACTCATCAGGAAGC
CGGACTCATCAGGAAGC
CGGACTCATCAGGAAGC
ACATCGGACTCATCAGGAAGC
CGGACTCATCAGGAAGC
CATCGGACTCATCAGGAAGC
GCATCGGACTCATCAGGAAGC
ATCGGACTCATCAGGAAGC
ATCGGACTCATCAGGAAGC

TGCATCGGACTCATCAGGAAGC
CGGACTCATCAGGAAGC
GACATCGGACTCATCAGGAAGC
ATCGGACTCATCAGGAAGC
GCATCGGACTCATCAGGAAGC
TCGGACTCATCAGGAAGC
ATCGGACTCATCAGGAAGC
ATCGGACTCATCAGGAAGC
TCGGACTCATCAGGAAGC
ACCATCGGACTCATCAGGAAGC
CATCGGACTCATCAGGAAGC
GCATCGGACTCATCAGGAAGC
ATCGGACTCATCAGGAAGC
CGGACTCATCAGGAAGC
TCGGACTCATCAGGAAGC
GACATCGGACTCATCAGGAAGC
TACATCGGACTCATCAGGAAGC
CATCGGACTCATCAGGAAGC
TCGGACTCATCAGGAAGC
CATCGGACTCATCAGGAAGC
CATCGGACTCATCAGGAAGC
ATCGGACTCATCAGGAAGC
ATCGGACTCATCAGGAAGC
GGACTCATCAGGAAGC
TCGGACTCATCAGGAAGC
ATCGGACTCATCAGGAAGC
CGGACTCATCAGGAAGC
ATTCATCGGACTCATCAGGAAGC
CGGACTCATCAGGAAGC
GACATCGGACTCATCAGGAAGC
CCATCGGACTCATCAGGAAGC
ACATCGGACTCATCAGGAAGC
TATCATCGGACTCATCAGGAAGC
GCATCGGACTCATCAGGAAGC
ATCGGACTCATCAGGAAGC
CATCGGACTCATCAGGAAGC
ATCGGACTCATCAGGAAGC
CATCGGACTCATCAGGAAGC
ATCGGACTCATCAGGAAGC
ATCGGACTCATCAGGAAGC
TCATCGGACTCATCAGGAAGC
CATCGGACTCATCAGGAAGC
ATCGGACTCATCAGGAAGC

TCGGACTCATCAGGAAGC
TCGGACTCATCAGGAAGC
GCATCGGACTCATCAGGAAGC
CCATCGGACTCATCAGGAAGC
CATCGGACTCATCAGGAAGC
TCATCGGACTCATCAGGAAGC
CATCGGACTCATCAGGAAGC
GGCATCGGACTCATCAGGAAGC
CATCGGACTCATCAGGAAGC
CATCGGACTCATCAGGAAGC
CCATCGGACTCATCAGGAAGC
TTCATCGGACTCATCAGGAAGC
AACATCGGACTCATCAGGAAGC
CATCGGACTCATCAGGAAGC
GGGCATCGGACTCATCAGGAAGC
ACATCGGACTCATCAGGAAGC
TGCATCGGACTCATCAGGAAGC
TTCATCGGACTCATCAGGAAGC
GACATCGGACTCATCAGGAAGC
TGCATCGGACTCATCAGGAAGC
ATCATCGGACTCATCAGGAAGC
GTCATCGGACTCATCAGGAAGC
CATCGGACTCATCAGGAAGC
CCATCGGACTCATCAGGAAGC
CGGACTCATCAGGAAGC
CATCGGACTCATCAGGAAGC
CATCGGACTCATCAGGAAGC
ATCGGACTCATCAGGAAGC
AGCATCGGACTCATCAGGAAGC
ATCGGACTCATCAGGAAGC
ATCGGACTCATCAGGAAGC
TACATCGGACTCATCAGGAAGC
ACATCGGACTCATCAGGAAGC
ATCGGACTCATCAGGAAGC
CATCGGACTCATCAGGAAGC
TCCATCGGACTCATCAGGAAGC
AGCATCGGACTCATCAGGAAGC
ATCGGACTCATCAGGAAGC
ATCGGACTCATCAGGAAGC
CAGACATCGGACTCATCAGGAAGC
CATCGGACTCATCAGGAAGC
CATCGGACTCATCAGGAAGC
CCATCGGACTCATCAGGAAGC

GAACATCGGACTCATCAGGAAGC
CATCGGACTCATCAGGAAGC
ACATCGGACTCATCAGGAAGC
TCGGACTCATCAGGAAGC
ATCATCGGACTCATCAGGAAGC
ATCATCGGACTCATCAGGAAGC
CATCGGACTCATCAGGAAGC
CATCGGACTCATCAGGAAGC
TCATCGGACTCATCAGGAAGC
GGCATCGGACTCATCAGGAAGC
CACATCGGACTCATCAGGAAGC
CATCGGACTCATCAGGAAGC
CGGACTCATCAGGAAGC
ACATCGGACTCATCAGGAAGC
CGGACTCATCAGGAAGC
CATCGGACTCATCAGGAAGC
ACCATCGGACTCATCAGGAAGC
TCTCATCGGACTCATCAGGAAGC
GGACTCATCAGGAAGC
GACATCGGACTCATCAGGAAGC
CATCGGACTCATCAGGAAGC
TGCATCGGACTCATCAGGAAGC
TGCATCGGACTCATCAGGAAGC
CATCGGACTCATCAGGAAGC
CATCGGACTCATCAGGAAGC
CACATCGGACTCATCAGGAAGC
GACTCATCAGGAAGC
GGCATCGGACTCATCAGGAAGC
GGCATCGGACTCATCAGGAAGC
GACTCATCAGGAAGC
CGGACTCATCAGGAAGC
ATCGGACTCATCAGGAAGC
TCGGACTCATCAGGAAGC
CGGACTCATCAGGAAGC
CCATCGGACTCATCAGGAAGC
AGCATCGGACTCATCAGGAAGC
GCATCGGACTCATCAGGAAGC
TACATCGGACTCATCAGGAAGC
GTCCATCGGACTCATCAGGAAGC
CATCGGACTCATCAGGAAGC
TGCATCGGACTCATCAGGAAGC
CATCGGACTCATCAGGAAGC
GACTCATCAGGAAGC

CATCGGACTCATCAGGAAGC
CATCGGACTCATCAGGAAGC
TCATCGGACTCATCAGGAAGC
TCATCGGACTCATCAGGAAGC
CATCGGACTCATCAGGAAGC
ATCGGACTCATCAGGAAGC
GGACTCATCAGGAAGC
TACATCGGACTCATCAGGAAGC
TCGGACTCATCAGGAAGC
GCATCGGACTCATCAGGAAGC
CCATCGGACTCATCAGGAAGC
CACATCGGACTCATCAGGAAGC
GCATCGGACTCATCAGGAAGC
AACATCGGACTCATCAGGAAGC
TCGGACTCATCAGGAAGC
ACATCGGACTCATCAGGAAGC
ATCGGACTCATCAGGAAGC
TCGGACTCATCAGGAAGC
TCGGACTCATCAGGAAGC
CGGACTCATCAGGAAGC
CTCATCGGACTCATCAGGAAGC
CTCATCGGACTCATCAGGAAGC
TCGGACTCATCAGGAAGC
TCATCGGACTCATCAGGAAGC
CATCGGACTCATCAGGAAGC
TTCATCGGACTCATCAGGAAGC
ATCGGACTCATCAGGAAGC
GACTCATCAGGAAGC
CGGACTCATCAGGAAGC
CATCGGACTCATCAGGAAGC
CATCGGACTCATCAGGAAGC
CATCGGACTCATCAGGAAGC
ATCGGACTCATCAGGAAGC
AATCATCGGACTCATCAGGAAGC
AGACATCGGACTCATCAGGAAGC
ATACATCGGACTCATCAGGAAGC
CCATCGGACTCATCAGGAAGC
GGACTCATCAGGAAGC
TCGGACTCATCAGGAAGC
CATCGGACTCATCAGGAAGC
GTCATCGGACTCATCAGGAAGC
ATCGGACTCATCAGGAAGC
CGGACTCATCAGGAAGC

ATCATCGGACTCATCAGGAAGC
TCGGACTCATCAGGAAGC
TCGGACTCATCAGGAAGC
CATCGGACTCATCAGGAAGC
ATCGGACTCATCAGGAAGC
AACATCGGACTCATCAGGAAGC
CATCGGACTCATCAGGAAGC
CATCGGACTCATCAGGAAGC
CATCGGACTCATCAGGAAGC
CATCGGACTCATCAGGAAGC
CCATCGGACTCATCAGGAAGC
CATCGGACTCATCAGGAAGC
GCATCGGACTCATCAGGAAGC
CATCGGACTCATCAGGAAGC
TCATCGGACTCATCAGGAAGC
AACATCGGACTCATCAGGAAGC
GACTCATCAGGAAGC
TACATCGGACTCATCAGGAAGC
CATCGGACTCATCAGGAAGC
TCGGACTCATCAGGAAGC
TCGGACTCATCAGGAAGC
ACATCGGACTCATCAGGAAGC
TCATCGGACTCATCAGGAAGC
GGACTCATCAGGAAGC
TCATCGGACTCATCAGGAAGC
CATCGGACTCATCAGGAAGC
GCATCGGACTCATCAGGAAGC
ACTACATCGGACTCATCAGGAAGC
ATTCATCGGACTCATCAGGAAGC
CATCGGACTCATCAGGAAGC
TCCATCGGACTCATCAGGAAGC
CGGACTCATCAGGAAGC
CCATCGGACTCATCAGGAAGC
ATACATCGGACTCATCAGGAAGC
ATCGGACTCATCAGGAAGC
GCATCGGACTCATCAGGAAGC
ACATCGGACTCATCAGGAAGC
ACATCGGACTCATCAGGAAGC
CGGACTCATCAGGAAGC
CATCGGACTCATCAGGAAGC
CATCGGACTCATCAGGAAGC
GCATCGGACTCATCAGGAAGC
CGGACTCATCAGGAAGC

CAGCATCGGACTCATCAGGAAGC
ACATCGGACTCATCAGGAAGC
ATCGGACTCATCAGGAAGC
TTCATCGGACTCATCAGGAAGC
TCATCGGACTCATCAGGAAGC
ACATCGGACTCATCAGGAAGC
CATCGGACTCATCAGGAAGC
GCATCGGACTCATCAGGAAGC
ATCGGACTCATCAGGAAGC
CATCGGACTCATCAGGAAGC
AGCATCGGACTCATCAGGAAGC
CATCGGACTCATCAGGAAGC
CATCGGACTCATCAGGAAGC
TCCATCGGACTCATCAGGAAGC
TCATCGGACTCATCAGGAAGC
ACATCGGACTCATCAGGAAGC
CATCGGACTCATCAGGAAGC
GCATCGGACTCATCAGGAAGC
GACTCATCAGGAAGC
TCGGACTCATCAGGAAGC
CATCGGACTCATCAGGAAGC
TCATCGGACTCATCAGGAAGC
CATCGGACTCATCAGGAAGC
CATCGGACTCATCAGGAAGC
GGACTCATCAGGAAGC
GTCATCGGACTCATCAGGAAGC
ATCATCGGACTCATCAGGAAGC
CTCATCGGACTCATCAGGAAGC
CATCGGACTCATCAGGAAGC
TCCATCGGACTCATCAGGAAGC
ATCGGACTCATCAGGAAGC
CATCGGACTCATCAGGAAGC
AGACATCGGACTCATCAGGAAGC
GGACTCATCAGGAAGC
CATCGGACTCATCAGGAAGC
ATCGGACTCATCAGGAAGC
CATCGGACTCATCAGGAAGC
GCATCGGACTCATCAGGAAGC
AACATCGGACTCATCAGGAAGC
ATCATCGGACTCATCAGGAAGC
TCGGACTCATCAGGAAGC
TCATCGGACTCATCAGGAAGC
ATCGGACTCATCAGGAAGC

ATCGGACTCATCAGGAAGC
ATCGGACTCATCAGGAAGC
GGACTCATCAGGAAGC
ATCGGACTCATCAGGAAGC
ATCGGACTCATCAGGAAGC
ACATCGGACTCATCAGGAAGC
AACATCGGACTCATCAGGAAGC
CATCGGACTCATCAGGAAGC
CATCGGACTCATCAGGAAGC
TTCATCGGACTCATCAGGAAGC
ACATCGGACTCATCAGGAAGC
ATCGGACTCATCAGGAAGC
TGCATCGGACTCATCAGGAAGC
GCATCGGACTCATCAGGAAGC
ATCGGACTCATCAGGAAGC
ACATCGGACTCATCAGGAAGC
AAGCATCGGACTCATCAGGAAGC
CATCGGACTCATCAGGAAGC
CATCGGACTCATCAGGAAGC
ATCGGACTCATCAGGAAGC
TCGGACTCATCAGGAAGC
TTCATCGGACTCATCAGGAAGC
ATCGGACTCATCAGGAAGC
CATCGGACTCATCAGGAAGC
CATCGGACTCATCAGGAAGC
TCATCGGACTCATCAGGAAGC
CGGACTCATCAGGAAGC
ATCGGACTCATCAGGAAGC
ATCATCGGACTCATCAGGAAGC
CATCGGACTCATCAGGAAGC
AGCATCGGACTCATCAGGAAGC
TGCATCGGACTCATCAGGAAGC
ACTCATCAGGAAGC
ACATCGGACTCATCAGGAAGC
ACTCATCGGACTCATCAGGAAGC
AGCATCGGACTCATCAGGAAGC
ATCGGACTCATCAGGAAGC
CATCGGACTCATCAGGAAGC
ACATCGGACTCATCAGGAAGC
GACATCGGACTCATCAGGAAGC
ATCGGACTCATCAGGAAGC
GCATCGGACTCATCAGGAAGC
ACATCGGACTCATCAGGAAGC

AATCATCGGACTCATCAGGAAGC
ATCGGACTCATCAGGAAGC
GGACTCATCAGGAAGC
CACATCGGACTCATCAGGAAGC
TCATCGGACTCATCAGGAAGC
GCATCGGACTCATCAGGAAGC
TTCATCGGACTCATCAGGAAGC
TCATCGGACTCATCAGGAAGC
ACATCGGACTCATCAGGAAGC
CCATCGGACTCATCAGGAAGC
ATCGGACTCATCAGGAAGC
ATGTCATCGGACTCATCAGGAAGC
CATCGGACTCATCAGGAAGC
GACATCGGACTCATCAGGAAGC
GGACTCATCAGGAAGC
GCATCGGACTCATCAGGAAGC
CACATCGGACTCATCAGGAAGC
GCATCGGACTCATCAGGAAGC
CGGACTCATCAGGAAGC
CATCGGACTCATCAGGAAGC
ATCGGACTCATCAGGAAGC
CCATCGGACTCATCAGGAAGC
TCGGACTCATCAGGAAGC
ATACATCGGACTCATCAGGAAGC
CATTCATCGGACTCATCAGGAAGC
ATCGGACTCATCAGGAAGC
CCATCGGACTCATCAGGAAGC
CATCGGACTCATCAGGAAGC
ATCGGACTCATCAGGAAGC
GCATCGGACTCATCAGGAAGC
ACATCGGACTCATCAGGAAGC
TCGGACTCATCAGGAAGC
CATCGGACTCATCAGGAAGC
ATCGGACTCATCAGGAAGC
TCGGACTCATCAGGAAGC
ATCGGACTCATCAGGAAGC
ACCATCGGACTCATCAGGAAGC
ATCGGACTCATCAGGAAGC
TTCATCGGACTCATCAGGAAGC
CATCGGACTCATCAGGAAGC
TCGGACTCATCAGGAAGC
TCATCGGACTCATCAGGAAGC
TCGGACTCATCAGGAAGC

CGGACTCATCAGGAAGC
GCATCGGACTCATCAGGAAGC
ACATCGGACTCATCAGGAAGC
CATCGGACTCATCAGGAAGC
CATCGGACTCATCAGGAAGC
CATCGGACTCATCAGGAAGC
ACATCGGACTCATCAGGAAGC
CGGACTCATCAGGAAGC
GACATCGGACTCATCAGGAAGC
TGCATCGGACTCATCAGGAAGC
CATCGGACTCATCAGGAAGC
CATCGGACTCATCAGGAAGC
AGCATCGGACTCATCAGGAAGC
GTCATCGGACTCATCAGGAAGC
ACATCGGACTCATCAGGAAGC
CATCGGACTCATCAGGAAGC
CATCGGACTCATCAGGAAGC
ACATCGGACTCATCAGGAAGC
ACATCGGACTCATCAGGAAGC
GACTCATCAGGAAGC
TCATCGGACTCATCAGGAAGC
ATCGGACTCATCAGGAAGC
CATCGGACTCATCAGGAAGC
TCATCGGACTCATCAGGAAGC
TCATCGGACTCATCAGGAAGC
CATCGGACTCATCAGGAAGC
TCATCGGACTCATCAGGAAGC
TCATCGGACTCATCAGGAAGC
GGACTCATCAGGAAGC
CATCGGACTCATCAGGAAGC
ACATCGGACTCATCAGGAAGC
ACATCGGACTCATCAGGAAGC
AACATCGGACTCATCAGGAAGC
AACATCGGACTCATCAGGAAGC
ATCGGACTCATCAGGAAGC
TCATCGGACTCATCAGGAAGC
TCATCGGACTCATCAGGAAGC
GCATCGGACTCATCAGGAAGC
CGGACTCATCAGGAAGC
ATCGGACTCATCAGGAAGC
ATCATCGGACTCATCAGGAAGC
CCATCGGACTCATCAGGAAGC
TCATCGGACTCATCAGGAAGC

GCATCGGACTCATCAGGAAGC
AAGCATCGGACTCATCAGGAAGC
TGTCATCGGACTCATCAGGAAGC
GACTCATCAGGAAGC
TCATCGGACTCATCAGGAAGC
CATCGGACTCATCAGGAAGC
ACATCGGACTCATCAGGAAGC
ACATCGGACTCATCAGGAAGC
TTGCATCGGACTCATCAGGAAGC
ATCATCGGACTCATCAGGAAGC
ACATCGGACTCATCAGGAAGC
GACTCATCAGGAAGC
ATCGGACTCATCAGGAAGC
ATCGGACTCATCAGGAAGC
ATCGGACTCATCAGGAAGC
TTCATCGGACTCATCAGGAAGC
ATCATCGGACTCATCAGGAAGC
CATCGGACTCATCAGGAAGC
ACCATCGGACTCATCAGGAAGC
GTCATCGGACTCATCAGGAAGC
CCATCGGACTCATCAGGAAGC
ATACATCGGACTCATCAGGAAGC
ACACATCGGACTCATCAGGAAGC
CGGACTCATCAGGAAGC
TCATCGGACTCATCAGGAAGC
CATCGGACTCATCAGGAAGC
GCCATCGGACTCATCAGGAAGC
CATCGGACTCATCAGGAAGC
TCGGACTCATCAGGAAGC
ACATCGGACTCATCAGGAAGC
ACATCGGACTCATCAGGAAGC
GACTCATCAGGAAGC
CCATCGGACTCATCAGGAAGC
TCGGACTCATCAGGAAGC
GGCATCGGACTCATCAGGAAGC
GGACTCATCAGGAAGC
CATCGGACTCATCAGGAAGC
ACCATCGGACTCATCAGGAAGC
GCATCGGACTCATCAGGAAGC
GTCATCGGACTCATCAGGAAGC
CGGACTCATCAGGAAGC
CGGACTCATCAGGAAGC
CGGACTCATCAGGAAGC

CGGACTCATCAGGAAGC
CGGACTCATCAGGAAGC
CATCGGACTCATCAGGAAGC
AACATCGGACTCATCAGGAAGC
CATCGGACTCATCAGGAAGC
ATCGGACTCATCAGGAAGC
ACTCATCAGGAAGC
CATCGGACTCATCAGGAAGC
ATCGGACTCATCAGGAAGC
ATCGGACTCATCAGGAAGC
ATCATCGGACTCATCAGGAAGC
TCATCGGACTCATCAGGAAGC
CCATCGGACTCATCAGGAAGC
ATCGGACTCATCAGGAAGC
CGGACTCATCAGGAAGC
CGGACTCATCAGGAAGC
CCTCATCGGACTCATCAGGAAGC
GACTCATCAGGAAGC
ACATCGGACTCATCAGGAAGC
ACATCGGACTCATCAGGAAGC
CATCGGACTCATCAGGAAGC
TTCATCGGACTCATCAGGAAGC
TCATCGGACTCATCAGGAAGC
GCATCGGACTCATCAGGAAGC
GACATCGGACTCATCAGGAAGC
TCGGACTCATCAGGAAGC
CCATCGGACTCATCAGGAAGC
ATCGGACTCATCAGGAAGC
GTGCATCGGACTCATCAGGAAGC
TCGGACTCATCAGGAAGC
CATCGGACTCATCAGGAAGC
CATCGGACTCATCAGGAAGC
CATCGGACTCATCAGGAAGC
CATCGGACTCATCAGGAAGC
CATCGGACTCATCAGGAAGC
CCATCGGACTCATCAGGAAGC
CATCGGACTCATCAGGAAGC
GCTCATCGGACTCATCAGGAAGC
TCATCGGACTCATCAGGAAGC
CGGACTCATCAGGAAGC
CGGACTCATCAGGAAGC
TCGGACTCATCAGGAAGC
GACTCATCAGGAAGC

CTCATCGGACTCATCAGGAAGC
CGGACTCATCAGGAAGC
GACTCATCAGGAAGC
ATCGGACTCATCAGGAAGC
CCATCGGACTCATCAGGAAGC
ACATCGGACTCATCAGGAAGC
ATCGGACTCATCAGGAAGC
ATCGGACTCATCAGGAAGC
ACTCATCAGGAAGC
CATCGGACTCATCAGGAAGC
CATCGGACTCATCAGGAAGC
TCGGACTCATCAGGAAGC
CATCGGACTCATCAGGAAGC
CATCGGACTCATCAGGAAGC
ACCCATCGGACTCATCAGGAAGC
GCATCGGACTCATCAGGAAGC
AGCATCGGACTCATCAGGAAGC
CCATCGGACTCATCAGGAAGC
ATCGGACTCATCAGGAAGC
ATCGGACTCATCAGGAAGC
GGACTCATCAGGAAGC
ATCGGACTCATCAGGAAGC
GACTCATCAGGAAGC
ACCATCGGACTCATCAGGAAGC
ACATCGGACTCATCAGGAAGC
ATCGGACTCATCAGGAAGC
CTCCATCGGACTCATCAGGAAGC
TCATCGGACTCATCAGGAAGC
GGACTCATCAGGAAGC
ATCATCGGACTCATCAGGAAGC
CATCGGACTCATCAGGAAGC
AACATCGGACTCATCAGGAAGC
AACATCGGACTCATCAGGAAGC
TCGGACTCATCAGGAAGC
TTCATCGGACTCATCAGGAAGC
GGACTCATCAGGAAGC
GCATCGGACTCATCAGGAAGC
TGCATCGGACTCATCAGGAAGC
CTCATCGGACTCATCAGGAAGC
ATCGGACTCATCAGGAAGC
ATCGGACTCATCAGGAAGC
GCATCGGACTCATCAGGAAGC
CGGACTCATCAGGAAGC

GGACTCATCAGGAAGC
CATCGGACTCATCAGGAAGC
TGCATCGGACTCATCAGGAAGC
GCATCGGACTCATCAGGAAGC
GACTCATCAGGAAGC
CATCGGACTCATCAGGAAGC
TACATCGGACTCATCAGGAAGC
ATCGGACTCATCAGGAAGC
ACATCGGACTCATCAGGAAGC
CGGACTCATCAGGAAGC
GCATCGGACTCATCAGGAAGC
GGACTCATCAGGAAGC
CGGACTCATCAGGAAGC
ACCATCGGACTCATCAGGAAGC
TACATCGGACTCATCAGGAAGC
TCATCGGACTCATCAGGAAGC
CATCGGACTCATCAGGAAGC
GGACTCATCAGGAAGC
AGCATCGGACTCATCAGGAAGC
GGACTCATCAGGAAGC
TCATCGGACTCATCAGGAAGC
CCATCGGACTCATCAGGAAGC
CGGACTCATCAGGAAGC
TCGGACTCATCAGGAAGC
TCATCGGACTCATCAGGAAGC
ACATCGGACTCATCAGGAAGC
ATCGGACTCATCAGGAAGC
CATCGGACTCATCAGGAAGC
ATCGGACTCATCAGGAAGC
ATCGGACTCATCAGGAAGC
TCGGACTCATCAGGAAGC
CGGACTCATCAGGAAGC
CCATCGGACTCATCAGGAAGC
GCACATCGGACTCATCAGGAAGC
GCATCGGACTCATCAGGAAGC
ATCGGACTCATCAGGAAGC
TCATCGGACTCATCAGGAAGC
GGACTCATCAGGAAGC
TACATCGGACTCATCAGGAAGC
TCATCGGACTCATCAGGAAGC
AATCATCGGACTCATCAGGAAGC
ATCGGACTCATCAGGAAGC
ACTCATCAGGAAGC

TCCATCGGACTCATCAGGAAGC
ACATCGGACTCATCAGGAAGC
CGGACTCATCAGGAAGC
CGGACTCATCAGGAAGC
CACATCGGACTCATCAGGAAGC
ATCGGACTCATCAGGAAGC
CGGACTCATCAGGAAGC
CATCGGACTCATCAGGAAGC
ATCATCGGACTCATCAGGAAGC
CATCGGACTCATCAGGAAGC
ATCGGACTCATCAGGAAGC
TCGGACTCATCAGGAAGC
CATCGGACTCATCAGGAAGC
CATCGGACTCATCAGGAAGC
ATCGGACTCATCAGGAAGC
CATCGGACTCATCAGGAAGC
ATCGGACTCATCAGGAAGC
TCGGACTCATCAGGAAGC
GCATCGGACTCATCAGGAAGC
TTCATCGGACTCATCAGGAAGC
ATCGGACTCATCAGGAAGC
CATCGGACTCATCAGGAAGC
CCTCATCGGACTCATCAGGAAGC
CATCGGACTCATCAGGAAGC
AACATCGGACTCATCAGGAAGC
TCATCGGACTCATCAGGAAGC
ATCGGACTCATCAGGAAGC
ACATCGGACTCATCAGGAAGC
TACATCGGACTCATCAGGAAGC
ATCGGACTCATCAGGAAGC
TCATCGGACTCATCAGGAAGC
CTCATCGGACTCATCAGGAAGC
CCATCGGACTCATCAGGAAGC
ACCATCGGACTCATCAGGAAGC
ATCGGACTCATCAGGAAGC
GCCATCGGACTCATCAGGAAGC
CGGACTCATCAGGAAGC
TCGGACTCATCAGGAAGC
GACACCCATCGGACTCATCAGGAAGC
TGTCATCGGACTCATCAGGAAGC
ATCGGACTCATCAGGAAGC
TCGGACTCATCAGGAAGC
GACTCATCAGGAAGC

ACATCGGACTCATCAGGAAGC
AAATCATCGGACTCATCAGGAAGC
CCATCGGACTCATCAGGAAGC
GCATCGGACTCATCAGGAAGC
TCATCGGACTCATCAGGAAGC
ATCGGACTCATCAGGAAGC
ATCGGACTCATCAGGAAGC
AACCATCGGACTCATCAGGAAGC
ATCGGACTCATCAGGAAGC
TACATCGGACTCATCAGGAAGC
ATCGGACTCATCAGGAAGC
ATCGGACTCATCAGGAAGC
TCGGACTCATCAGGAAGC
CATCGGACTCATCAGGAAGC
CATCGGACTCATCAGGAAGC
ATCGGACTCATCAGGAAGC
TCTCATCGGACTCATCAGGAAGC
TTCATCGGACTCATCAGGAAGC
GGACTCATCAGGAAGC
GCATCGGACTCATCAGGAAGC
CCATCGGACTCATCAGGAAGC
CGCATCGGACTCATCAGGAAGC
GCTCATCGGACTCATCAGGAAGC
CCCATCGGACTCATCAGGAAGC
ACATCGGACTCATCAGGAAGC
TATCATCGGACTCATCAGGAAGC
TATCATCGGACTCATCAGGAAGC
CATCGGACTCATCAGGAAGC
CATCGGACTCATCAGGAAGC
GCATCGGACTCATCAGGAAGC
GCATCGGACTCATCAGGAAGC
TCATCGGACTCATCAGGAAGC
CATCGGACTCATCAGGAAGC
ACATCGGACTCATCAGGAAGC
TCGGACTCATCAGGAAGC
ATACATCGGACTCATCAGGAAGC
AAACATCGGACTCATCAGGAAGC
GACTCATCAGGAAGC
AATGTCATCGGACTCATCAGGAAGC
CGGACTCATCAGGAAGC
CATCGGACTCATCAGGAAGC
AAACATCGGACTCATCAGGAAGC
TCATCGGACTCATCAGGAAGC

TGCATCGGACTCATCAGGAAGC
ATCGGACTCATCAGGAAGC
GACTCATCAGGAAGC
ATCGGACTCATCAGGAAGC
ATCGGACTCATCAGGAAGC
AGCATCGGACTCATCAGGAAGC
ATCGGACTCATCAGGAAGC
GCATCGGACTCATCAGGAAGC
GGACTCATCAGGAAGC
AACATCGGACTCATCAGGAAGC
ATCATCGGACTCATCAGGAAGC
ATCATCGGACTCATCAGGAAGC
TCGGACTCATCAGGAAGC
CATCGGACTCATCAGGAAGC
ATCGGACTCATCAGGAAGC
ACATCGGACTCATCAGGAAGC
TCATCGGACTCATCAGGAAGC
CATCGGACTCATCAGGAAGC
TCGGACTCATCAGGAAGC
CATCGGACTCATCAGGAAGC
CCATCGGACTCATCAGGAAGC
GCATCGGACTCATCAGGAAGC
CATCGGACTCATCAGGAAGC
ATCGGACTCATCAGGAAGC
ATCGGACTCATCAGGAAGC
ACATCGGACTCATCAGGAAGC
GCATCGGACTCATCAGGAAGC
TCGGACTCATCAGGAAGC
CATCGGACTCATCAGGAAGC
TCGGACTCATCAGGAAGC
CGGACTCATCAGGAAGC
CCATCGGACTCATCAGGAAGC
ATCGGACTCATCAGGAAGC
CCACATCGGACTCATCAGGAAGC
ACATCGGACTCATCAGGAAGC
TCGGACTCATCAGGAAGC
GGACTCATCAGGAAGC
TCGGACTCATCAGGAAGC
TCGGACTCATCAGGAAGC
CATCGGACTCATCAGGAAGC
TCGGACTCATCAGGAAGC
CCATCGGACTCATCAGGAAGC
ATCGGACTCATCAGGAAGC

TACATCGGACTCATCAGGAAGC
CATCGGACTCATCAGGAAGC
ATCATCGGACTCATCAGGAAGC
GACTCATCAGGAAGC
ACATCGGACTCATCAGGAAGC
ATCGGACTCATCAGGAAGC
CATCGGACTCATCAGGAAGC
TCATCGGACTCATCAGGAAGC
CGGACTCATCAGGAAGC
TCGGACTCATCAGGAAGC
TCGGACTCATCAGGAAGC
CATCGGACTCATCAGGAAGC
ATCGGACTCATCAGGAAGC
TCGGACTCATCAGGAAGC
TCGGACTCATCAGGAAGC
GGACTCATCAGGAAGC
ATCGGACTCATCAGGAAGC
ATCGGACTCATCAGGAAGC
ATCGGACTCATCAGGAAGC
ATCGGACTCATCAGGAAGC
GCATCGGACTCATCAGGAAGC
CATCGGACTCATCAGGAAGC
GTCATCGGACTCATCAGGAAGC
ATCGGACTCATCAGGAAGC
TCGGACTCATCAGGAAGC
AGCATCGGACTCATCAGGAAGC
ACACATCGGACTCATCAGGAAGC
CATCGGACTCATCAGGAAGC
GACTCATCAGGAAGC
TCATCGGACTCATCAGGAAGC
TAACATCGGACTCATCAGGAAGC
CATCGGACTCATCAGGAAGC
CGGACTCATCAGGAAGC
GACATCGGACTCATCAGGAAGC
TCATCGGACTCATCAGGAAGC
ATCGGACTCATCAGGAAGC
CGGACTCATCAGGAAGC
CATCGGACTCATCAGGAAGC
ATCGGACTCATCAGGAAGC
CATCGGACTCATCAGGAAGC
CATCGGACTCATCAGGAAGC
CATCGGACTCATCAGGAAGC
ACATCGGACTCATCAGGAAGC

CCATCGGACTCATCAGGAAGC
CATCGGACTCATCAGGAAGC
CATCGGACTCATCAGGAAGC
CATCGGACTCATCAGGAAGC
ATCGGACTCATCAGGAAGC
ATCGGACTCATCAGGAAGC
CCATCGGACTCATCAGGAAGC
TCGGACTCATCAGGAAGC
CATCGGACTCATCAGGAAGC
ATCGGACTCATCAGGAAGC
ACCATCGGACTCATCAGGAAGC
CATCGGACTCATCAGGAAGC
ACATCGGACTCATCAGGAAGC
GTCATCGGACTCATCAGGAAGC
TACATCGGACTCATCAGGAAGC
CATCGGACTCATCAGGAAGC
CATCGGACTCATCAGGAAGC
TCGGACTCATCAGGAAGC
ATCGGACTCATCAGGAAGC
ATCGGACTCATCAGGAAGC
ATCGGACTCATCAGGAAGC
TCATCGGACTCATCAGGAAGC
ATCGGACTCATCAGGAAGC
CATCGGACTCATCAGGAAGC
CATCGGACTCATCAGGAAGC
ATCGGACTCATCAGGAAGC
CGGACTCATCAGGAAGC
ACCATCGGACTCATCAGGAAGC
ACATCGGACTCATCAGGAAGC
ATCGGACTCATCAGGAAGC
TCGGACTCATCAGGAAGC
TCGGACTCATCAGGAAGC
TCGGACTCATCAGGAAGC
ACATCGGACTCATCAGGAAGC
CATCGGACTCATCAGGAAGC
ATCGGACTCATCAGGAAGC
TACATCGGACTCATCAGGAAGC
ACATCGGACTCATCAGGAAGC
CCCATCGGACTCATCAGGAAGC
GACATCGGACTCATCAGGAAGC
TCATCGGACTCATCAGGAAGC
GCATCGGACTCATCAGGAAGC
CGGACTCATCAGGAAGC

CTCATCGGACTCATCAGGAAGC
CACCATCGGACTCATCAGGAAGC
CACATCGGACTCATCAGGAAGC
CATCGGACTCATCAGGAAGC
CCATCGGACTCATCAGGAAGC
ACCATCGGACTCATCAGGAAGC
CATCGGACTCATCAGGAAGC
ATCGGACTCATCAGGAAGC
CATCGGACTCATCAGGAAGC
CATCGGACTCATCAGGAAGC
CATCGGACTCATCAGGAAGC
CATCGGACTCATCAGGAAGC
TCGGACTCATCAGGAAGC
TCGGACTCATCAGGAAGC
GGACTCATCAGGAAGC
ATCGGACTCATCAGGAAGC
CCATCGGACTCATCAGGAAGC
GCATCGGACTCATCAGGAAGC
GGACTCATCAGGAAGC
GGACTCATCAGGAAGC
CATCGGACTCATCAGGAAGC
AACATCGGACTCATCAGGAAGC
AACATCGGACTCATCAGGAAGC
ACATCGGACTCATCAGGAAGC
TCACATCGGACTCATCAGGAAGC
ATCGGACTCATCAGGAAGC
ATCGGACTCATCAGGAAGC
CATCGGACTCATCAGGAAGC
CATCGGACTCATCAGGAAGC
CCATCGGACTCATCAGGAAGC
TCGGACTCATCAGGAAGC
CGGACTCATCAGGAAGC
TCGGACTCATCAGGAAGC
AACATCGGACTCATCAGGAAGC
TCATCGGACTCATCAGGAAGC
TCGGACTCATCAGGAAGC
ATACATCGGACTCATCAGGAAGC
ACATCGGACTCATCAGGAAGC
CATCGGACTCATCAGGAAGC
TTCATCGGACTCATCAGGAAGC
CATCGGACTCATCAGGAAGC
CATCGGACTCATCAGGAAGC
AGCATCGGACTCATCAGGAAGC

ATCGGACTCATCAGGAAGC
CCATCGGACTCATCAGGAAGC
CGGACTCATCAGGAAGC
ATCGGACTCATCAGGAAGC
TCGGACTCATCAGGAAGC
AGCATCGGACTCATCAGGAAGC
ATCGGACTCATCAGGAAGC
GGACTCATCAGGAAGC
ATCGGACTCATCAGGAAGC
CATCGGACTCATCAGGAAGC
GCATCGGACTCATCAGGAAGC
GGACTCATCAGGAAGC
GGACTCATCAGGAAGC
CGGACTCATCAGGAAGC
GACATCGGACTCATCAGGAAGC
CGGACTCATCAGGAAGC
GTGCATCGGACTCATCAGGAAGC
TCGGACTCATCAGGAAGC
CATCGGACTCATCAGGAAGC
TCATCGGACTCATCAGGAAGC
CCATCGGACTCATCAGGAAGC
CATCGGACTCATCAGGAAGC
ACATCGGACTCATCAGGAAGC
ATCGGACTCATCAGGAAGC
ATCGGACTCATCAGGAAGC
ATCGGACTCATCAGGAAGC
TTCATCGGACTCATCAGGAAGC
ATCGGACTCATCAGGAAGC
ACATCGGACTCATCAGGAAGC
ATCGGACTCATCAGGAAGC
GCATCGGACTCATCAGGAAGC
TGGCATCGGACTCATCAGGAAGC
ATCGGACTCATCAGGAAGC
CATCGGACTCATCAGGAAGC
TCGGACTCATCAGGAAGC
GGACTCATCAGGAAGC
CGGACTCATCAGGAAGC
ACATCGGACTCATCAGGAAGC
ATCGGACTCATCAGGAAGC
CATCGGACTCATCAGGAAGC
GGACTCATCAGGAAGC
TCGGACTCATCAGGAAGC
ATCGGACTCATCAGGAAGC

ATCGGACTCATCAGGAAGC
ACATCGGACTCATCAGGAAGC
GACATCGGACTCATCAGGAAGC
ATCGGACTCATCAGGAAGC
ATCGGACTCATCAGGAAGC
CCATCGGACTCATCAGGAAGC
CATCGGACTCATCAGGAAGC
ACATCGGACTCATCAGGAAGC
AACATCGGACTCATCAGGAAGC
TGGACATCGGACTCATCAGGAAGC
ACATCGGACTCATCAGGAAGC
CATCGGACTCATCAGGAAGC
AACCATCGGACTCATCAGGAAGC
AACCATCGGACTCATCAGGAAGC
ATCATCGGACTCATCAGGAAGC
GGCATCGGACTCATCAGGAAGC
GGACTCATCAGGAAGC
ATCATCGGACTCATCAGGAAGC
CATCGGACTCATCAGGAAGC
TCGGACTCATCAGGAAGC
ACATCGGACTCATCAGGAAGC
GGCATCGGACTCATCAGGAAGC
CGGACTCATCAGGAAGC
ATCGGACTCATCAGGAAGC
CATCGGACTCATCAGGAAGC
CGGACTCATCAGGAAGC
TACACATCGGACTCATCAGGAAGC
TCACATCGGACTCATCAGGAAGC
AACATCGGACTCATCAGGAAGC
GCATCGGACTCATCAGGAAGC
GGACTCATCAGGAAGC
CTCATCGGACTCATCAGGAAGC
TTACATCGGACTCATCAGGAAGC
TCGGACTCATCAGGAAGC
AACATCGGACTCATCAGGAAGC
AAACATCGGACTCATCAGGAAGC
CGCATCGGACTCATCAGGAAGC
CATCGGACTCATCAGGAAGC
GACTCATCAGGAAGC
GCATCGGACTCATCAGGAAGC
ATCGGACTCATCAGGAAGC
ACATCGGACTCATCAGGAAGC
CAACATCGGACTCATCAGGAAGC

CATCGGACTCATCAGGAAGC
CATCGGACTCATCAGGAAGC
GGACTCATCAGGAAGC
GTTCATCGGACTCATCAGGAAGC
CATCGGACTCATCAGGAAGC
ATCGGACTCATCAGGAAGC
GACTCATCAGGAAGC
CTCATCGGACTCATCAGGAAGC
CATCGGACTCATCAGGAAGC
CATCGGACTCATCAGGAAGC
CTACATCGGACTCATCAGGAAGC
CATCGGACTCATCAGGAAGC
TGCATCGGACTCATCAGGAAGC
TCATCGGACTCATCAGGAAGC
ACATCGGACTCATCAGGAAGC
ATCGGACTCATCAGGAAGC
AACATCGGACTCATCAGGAAGC
TCGGACTCATCAGGAAGC
AGCATCGGACTCATCAGGAAGC
TCATCGGACTCATCAGGAAGC
GGACTCATCAGGAAGC
TTCATCGGACTCATCAGGAAGC
CATCGGACTCATCAGGAAGC
CATCGGACTCATCAGGAAGC
TCGGACTCATCAGGAAGC
TAACATCGGACTCATCAGGAAGC
ATCGGACTCATCAGGAAGC
CATCGGACTCATCAGGAAGC
TCGGACTCATCAGGAAGC
ACATCGGACTCATCAGGAAGC
TCGGACTCATCAGGAAGC
ACATCGGACTCATCAGGAAGC
CATCGGACTCATCAGGAAGC
GCATCGGACTCATCAGGAAGC
ACATCGGACTCATCAGGAAGC
ATCGGACTCATCAGGAAGC
CATCGGACTCATCAGGAAGC
AGGCATCGGACTCATCAGGAAGC
TCGGACTCATCAGGAAGC
CCATCGGACTCATCAGGAAGC
GAACATCGGACTCATCAGGAAGC
GAACATCGGACTCATCAGGAAGC
AGGCATCGGACTCATCAGGAAGC

ATCGGACTCATCAGGAAGC
ATCGGACTCATCAGGAAGC
TCGGACTCATCAGGAAGC
CATCGGACTCATCAGGAAGC
ACATCGGACTCATCAGGAAGC
CATCGGACTCATCAGGAAGC
CATCGGACTCATCAGGAAGC
CATCGGACTCATCAGGAAGC
TCACATCGGACTCATCAGGAAGC
TCATCGGACTCATCAGGAAGC
TACATCGGACTCATCAGGAAGC
TCGGACTCATCAGGAAGC
TCATCGGACTCATCAGGAAGC
GCATCGGACTCATCAGGAAGC
CGGACTCATCAGGAAGC
CCATCGGACTCATCAGGAAGC
CCATCGGACTCATCAGGAAGC
TGCATCGGACTCATCAGGAAGC
TAACATCGGACTCATCAGGAAGC
GCATCGGACTCATCAGGAAGC
CATCGGACTCATCAGGAAGC
TCGGACTCATCAGGAAGC
TCATCGGACTCATCAGGAAGC
GACTCATCAGGAAGC
GCATCGGACTCATCAGGAAGC
AACATCGGACTCATCAGGAAGC
TCATCGGACTCATCAGGAAGC
CCATCGGACTCATCAGGAAGC
CCATCGGACTCATCAGGAAGC
CATCGGACTCATCAGGAAGC
GTCATCGGACTCATCAGGAAGC
GCATCGGACTCATCAGGAAGC
AACATCGGACTCATCAGGAAGC
GCATCGGACTCATCAGGAAGC
CCATCGGACTCATCAGGAAGC
CATCGGACTCATCAGGAAGC
CATCGGACTCATCAGGAAGC
CATCGGACTCATCAGGAAGC
CGGACTCATCAGGAAGC
CATCGGACTCATCAGGAAGC
ATCATCGGACTCATCAGGAAGC
GGACTCATCAGGAAGC
CCATCGGACTCATCAGGAAGC

ATCGGACTCATCAGGAAGC
ACATCGGACTCATCAGGAAGC
ATCGGACTCATCAGGAAGC
GCATCGGACTCATCAGGAAGC
GCATCGGACTCATCAGGAAGC
CATCGGACTCATCAGGAAGC
TCATCGGACTCATCAGGAAGC
CTCATCGGACTCATCAGGAAGC
CATCGGACTCATCAGGAAGC
TCATCGGACTCATCAGGAAGC
CATCGGACTCATCAGGAAGC
GCATCGGACTCATCAGGAAGC
TCATCGGACTCATCAGGAAGC
TCATCGGACTCATCAGGAAGC
CGGACTCATCAGGAAGC
TCATCGGACTCATCAGGAAGC
ACATCGGACTCATCAGGAAGC
AGCATCGGACTCATCAGGAAGC
GCATCGGACTCATCAGGAAGC
GCATCGGACTCATCAGGAAGC
CATCGGACTCATCAGGAAGC
CATCGGACTCATCAGGAAGC
CATCGGACTCATCAGGAAGC
TTCATCGGACTCATCAGGAAGC
GACTCATCAGGAAGC
AACATCGGACTCATCAGGAAGC
AACATCGGACTCATCAGGAAGC
CATCGGACTCATCAGGAAGC
TCTCCATCGGACTCATCAGGAAGC
CGGACTCATCAGGAAGC
CATCGGACTCATCAGGAAGC
TCATCGGACTCATCAGGAAGC
GACTCATCAGGAAGC
GCATCGGACTCATCAGGAAGC
CATCGGACTCATCAGGAAGC
ATCGGACTCATCAGGAAGC
TCATCGGACTCATCAGGAAGC
TACATCGGACTCATCAGGAAGC
CATCGGACTCATCAGGAAGC
TACATCGGACTCATCAGGAAGC
TACATCGGACTCATCAGGAAGC
CATCGGACTCATCAGGAAGC
ATCGGACTCATCAGGAAGC

CATCGGACTCATCAGGAAGC
TTCATCGGACTCATCAGGAAGC
GCATCGGACTCATCAGGAAGC
TCGGACTCATCAGGAAGC
CATCGGACTCATCAGGAAGC
GCATCGGACTCATCAGGAAGC
TTCATCGGACTCATCAGGAAGC
ATCGGACTCATCAGGAAGC
GCATCGGACTCATCAGGAAGC
ATCGGACTCATCAGGAAGC
TCATCGGACTCATCAGGAAGC
ATTCATCGGACTCATCAGGAAGC
ACATCGGACTCATCAGGAAGC
ATCATCGGACTCATCAGGAAGC
ATCGGACTCATCAGGAAGC
ATCATCGGACTCATCAGGAAGC
AACATCGGACTCATCAGGAAGC
CGGACTCATCAGGAAGC
CATCGGACTCATCAGGAAGC
ATCGGACTCATCAGGAAGC
ACATCGGACTCATCAGGAAGC
TCATCGGACTCATCAGGAAGC
GACTCATCAGGAAGC
ACTCATCAGGAAGC
ATCGGACTCATCAGGAAGC
ATCATCGGACTCATCAGGAAGC
TACATCGGACTCATCAGGAAGC
CTACATCGGACTCATCAGGAAGC
ATCGGACTCATCAGGAAGC
CTCATCGGACTCATCAGGAAGC
TACATCGGACTCATCAGGAAGC
ACATCGGACTCATCAGGAAGC
CATCGGACTCATCAGGAAGC
TCGGACTCATCAGGAAGC
TACATCGGACTCATCAGGAAGC
ACATCGGACTCATCAGGAAGC
CGGACTCATCAGGAAGC
CGGACTCATCAGGAAGC
CACATCGGACTCATCAGGAAGC
GACTCATCAGGAAGC
TGCATCGGACTCATCAGGAAGC
TCATCGGACTCATCAGGAAGC
CATCGGACTCATCAGGAAGC

CATCGGACTCATCAGGAAGC
CATCGGACTCATCAGGAAGC
ATCGGACTCATCAGGAAGC
TCGGACTCATCAGGAAGC
TCGGACTCATCAGGAAGC
TCGGACTCATCAGGAAGC
CATCGGACTCATCAGGAAGC
CCATCGGACTCATCAGGAAGC
ACATCGGACTCATCAGGAAGC
CATCGGACTCATCAGGAAGC
CATCGGACTCATCAGGAAGC
TCGGACTCATCAGGAAGC
GCATCGGACTCATCAGGAAGC
GCATCGGACTCATCAGGAAGC
CGGACTCATCAGGAAGC
TCGGACTCATCAGGAAGC
CATCGGACTCATCAGGAAGC
TCGGACTCATCAGGAAGC
ATCGGACTCATCAGGAAGC
ATCGGACTCATCAGGAAGC
CTCATCAGGAAGC
CATCGGACTCATCAGGAAGC
TCATCGGACTCATCAGGAAGC
TCGGACTCATCAGGAAGC
TCATCGGACTCATCAGGAAGC
TACATCGGACTCATCAGGAAGC
CATCGGACTCATCAGGAAGC
ACATCGGACTCATCAGGAAGC
TCATCGGACTCATCAGGAAGC
TCGGACTCATCAGGAAGC
ACATCGGACTCATCAGGAAGC
CATCGGACTCATCAGGAAGC
TCGGACTCATCAGGAAGC
CATCGGACTCATCAGGAAGC
AACCATCGGACTCATCAGGAAGC
ACATCGGACTCATCAGGAAGC
CATCGGACTCATCAGGAAGC
TCGGACTCATCAGGAAGC
ATCGGACTCATCAGGAAGC
GGACTCATCAGGAAGC
TTCATCGGACTCATCAGGAAGC
CCATCGGACTCATCAGGAAGC
AATCATCGGACTCATCAGGAAGC

TACATCGGACTCATCAGGAAGC
GGCATCGGACTCATCAGGAAGC
GACTCATCAGGAAGC
GACTCATCAGGAAGC
GGCATCGGACTCATCAGGAAGC
TCATCGGACTCATCAGGAAGC
ATCGGACTCATCAGGAAGC
CATCGGACTCATCAGGAAGC
CATCGGACTCATCAGGAAGC
AAACATCGGACTCATCAGGAAGC
ACATCGGACTCATCAGGAAGC
CATCGGACTCATCAGGAAGC
CATCGGACTCATCAGGAAGC
ATCGGACTCATCAGGAAGC
CCATCGGACTCATCAGGAAGC
TCATCGGACTCATCAGGAAGC
TCATCGGACTCATCAGGAAGC
TCATCGGACTCATCAGGAAGC
GCATCGGACTCATCAGGAAGC
TAGACATCGGACTCATCAGGAAGC
GACTCATCAGGAAGC
GACTCATCAGGAAGC
ACATCGGACTCATCAGGAAGC
CATCGGACTCATCAGGAAGC
CATCGGACTCATCAGGAAGC
AACATCGGACTCATCAGGAAGC
TCGCATCGGACTCATCAGGAAGC
GCATCGGACTCATCAGGAAGC
CATCGGACTCATCAGGAAGC
ATCGGACTCATCAGGAAGC
TCATCGGACTCATCAGGAAGC
TCATCGGACTCATCAGGAAGC
CATCGGACTCATCAGGAAGC
GCATCGGACTCATCAGGAAGC
CCCATCGGACTCATCAGGAAGC
AGCATCGGACTCATCAGGAAGC
GCATCGGACTCATCAGGAAGC
CATCGGACTCATCAGGAAGC
CATCGGACTCATCAGGAAGC
AGCATCGGACTCATCAGGAAGC
ACATCGGACTCATCAGGAAGC
TCGGACTCATCAGGAAGC
GCATCGGACTCATCAGGAAGC

CATCGGACTCATCAGGAAGC
TCATCGGACTCATCAGGAAGC
CCATCGGACTCATCAGGAAGC
ATCGGACTCATCAGGAAGC
CCATCGGACTCATCAGGAAGC
GCCATCGGACTCATCAGGAAGC
CGGACTCATCAGGAAGC
TCATCGGACTCATCAGGAAGC
TCATCGGACTCATCAGGAAGC
ACATCGGACTCATCAGGAAGC
ACACATCGGACTCATCAGGAAGC
TCTCATCGGACTCATCAGGAAGC
TCATCGGACTCATCAGGAAGC
CATCGGACTCATCAGGAAGC
CGGACTCATCAGGAAGC
ATCGGACTCATCAGGAAGC
ATCGGACTCATCAGGAAGC
ATCGGACTCATCAGGAAGC
CATCGGACTCATCAGGAAGC
TGCATCGGACTCATCAGGAAGC
CGGACTCATCAGGAAGC
ATCGGACTCATCAGGAAGC
CATCGGACTCATCAGGAAGC
TCAGCATCGGACTCATCAGGAAGC
CCATCGGACTCATCAGGAAGC
CCATCGGACTCATCAGGAAGC
ATCGGACTCATCAGGAAGC
CCATCGGACTCATCAGGAAGC
CATCGGACTCATCAGGAAGC
ATCGGACTCATCAGGAAGC
CCCACATCGGACTCATCAGGAAGC
TGCATCGGACTCATCAGGAAGC
CATCGGACTCATCAGGAAGC
CATCGGACTCATCAGGAAGC
ATCATCGGACTCATCAGGAAGC
GACTCATCAGGAAGC
GCATCGGACTCATCAGGAAGC
ATCGGACTCATCAGGAAGC
CATCGGACTCATCAGGAAGC
TGCATCGGACTCATCAGGAAGC
GAGTCATCGGACTCATCAGGAAGC
GACTCATCAGGAAGC
CATCGGACTCATCAGGAAGC

AACATCGGACTCATCAGGAAGC
GCATCGGACTCATCAGGAAGC
CCATCGGACTCATCAGGAAGC
GACTCATCAGGAAGC
CATCGGACTCATCAGGAAGC
GGACTCATCAGGAAGC
ACATCGGACTCATCAGGAAGC
TCGGACTCATCAGGAAGC
GCCATCGGACTCATCAGGAAGC
TACATCGGACTCATCAGGAAGC
ACATCGGACTCATCAGGAAGC
TACATCGGACTCATCAGGAAGC
CGGACTCATCAGGAAGC
ACATCGGACTCATCAGGAAGC
GGCATCGGACTCATCAGGAAGC
CATCGGACTCATCAGGAAGC
ATCGGACTCATCAGGAAGC
TCATCGGACTCATCAGGAAGC
CGGACTCATCAGGAAGC
CCATCGGACTCATCAGGAAGC
ATCATCGGACTCATCAGGAAGC
ATCGGACTCATCAGGAAGC
CATCGGACTCATCAGGAAGC
TATCATCGGACTCATCAGGAAGC
GGACTCATCAGGAAGC
CATCGGACTCATCAGGAAGC
AACATCGGACTCATCAGGAAGC
CATCGGACTCATCAGGAAGC
CGGACTCATCAGGAAGC
GGCATCGGACTCATCAGGAAGC
TGCATCGGACTCATCAGGAAGC
ATCGGACTCATCAGGAAGC
ATTCATCGGACTCATCAGGAAGC
ACATCGGACTCATCAGGAAGC
TCATCGGACTCATCAGGAAGC
ATCGGACTCATCAGGAAGC
CTCATCGGACTCATCAGGAAGC
CATCGGACTCATCAGGAAGC
ATCGGACTCATCAGGAAGC
GCATCGGACTCATCAGGAAGC
TATGTCATCGGACTCATCAGGAAGC
CATCGGACTCATCAGGAAGC
CATCGGACTCATCAGGAAGC

GCATCGGACTCATCAGGAAGC
CATCGGACTCATCAGGAAGC
ATCGGACTCATCAGGAAGC
TCGGACTCATCAGGAAGC
TCGGACTCATCAGGAAGC
CATCGGACTCATCAGGAAGC
ACATCGGACTCATCAGGAAGC
ACATCGGACTCATCAGGAAGC
GCATCGGACTCATCAGGAAGC
GCATCGGACTCATCAGGAAGC
ACCATCGGACTCATCAGGAAGC
ATCATCGGACTCATCAGGAAGC
GGACTCATCAGGAAGC
GCATCGGACTCATCAGGAAGC
GCATCGGACTCATCAGGAAGC
GCATCGGACTCATCAGGAAGC
ACCATCGGACTCATCAGGAAGC
CGGACTCATCAGGAAGC
TCGGACTCATCAGGAAGC
CATCGGACTCATCAGGAAGC
CTCATCGGACTCATCAGGAAGC
CATCGGACTCATCAGGAAGC
ATCGGACTCATCAGGAAGC
CCACATCGGACTCATCAGGAAGC
GCATCGGACTCATCAGGAAGC
AACATCGGACTCATCAGGAAGC
ACATCGGACTCATCAGGAAGC
CCATCGGACTCATCAGGAAGC
GCCATCGGACTCATCAGGAAGC
AAACATCGGACTCATCAGGAAGC
CATCGGACTCATCAGGAAGC
ATCGGACTCATCAGGAAGC
TTACATCGGACTCATCAGGAAGC
AACATCGGACTCATCAGGAAGC
CATCGGACTCATCAGGAAGC
CCATCGGACTCATCAGGAAGC
CATCGGACTCATCAGGAAGC
AATCATCGGACTCATCAGGAAGC
TCGGACTCATCAGGAAGC
ACATCGGACTCATCAGGAAGC
GCATCGGACTCATCAGGAAGC
AAACATCGGACTCATCAGGAAGC
AAACATCGGACTCATCAGGAAGC

CATCGGACTCATCAGGAAGC
CTCATCGGACTCATCAGGAAGC
CATCGGACTCATCAGGAAGC
GCATCGGACTCATCAGGAAGC
TCGGACTCATCAGGAAGC
TTCATCGGACTCATCAGGAAGC
TCGGACTCATCAGGAAGC
CATCGGACTCATCAGGAAGC
TCATCGGACTCATCAGGAAGC
ACATCGGACTCATCAGGAAGC
TCGGACTCATCAGGAAGC
CATCGGACTCATCAGGAAGC
TCACATCGGACTCATCAGGAAGC
CATCGGACTCATCAGGAAGC
TCATCGGACTCATCAGGAAGC
CATCGGACTCATCAGGAAGC
ATCGGACTCATCAGGAAGC
GTCATCGGACTCATCAGGAAGC
TCATCGGACTCATCAGGAAGC
ATCGGACTCATCAGGAAGC
GCATCGGACTCATCAGGAAGC
CATCGGACTCATCAGGAAGC
TCATCGGACTCATCAGGAAGC
CGGACTCATCAGGAAGC
CCATCGGACTCATCAGGAAGC
GCATCGGACTCATCAGGAAGC
CATCGGACTCATCAGGAAGC
TACATCGGACTCATCAGGAAGC
AACATCGGACTCATCAGGAAGC
GGACTCATCAGGAAGC
CGGACTCATCAGGAAGC
CATCGGACTCATCAGGAAGC
CATCGGACTCATCAGGAAGC
GTACATCGGACTCATCAGGAAGC
ACATCGGACTCATCAGGAAGC
CATCGGACTCATCAGGAAGC
CTACATCGGACTCATCAGGAAGC
TCGGACTCATCAGGAAGC
TGCATCGGACTCATCAGGAAGC
GTCATCGGACTCATCAGGAAGC
GCATCGGACTCATCAGGAAGC
TGCATCGGACTCATCAGGAAGC
TCATCGGACTCATCAGGAAGC

GTTCATCGGACTCATCAGGAAGC
GTTCATCGGACTCATCAGGAAGC
CATCGGACTCATCAGGAAGC
GCATCGGACTCATCAGGAAGC
TTACATCGGACTCATCAGGAAGC
TCGGACTCATCAGGAAGC
GGACTCATCAGGAAGC
TCCATCGGACTCATCAGGAAGC
TCATCGGACTCATCAGGAAGC
TGCATCGGACTCATCAGGAAGC
TGCATCGGACTCATCAGGAAGC
ACATCGGACTCATCAGGAAGC
ACATCGGACTCATCAGGAAGC
CGGACTCATCAGGAAGC
TCCATCGGACTCATCAGGAAGC
ACATCGGACTCATCAGGAAGC
CATCGGACTCATCAGGAAGC
AACATCGGACTCATCAGGAAGC
GCATCGGACTCATCAGGAAGC
AACATCGGACTCATCAGGAAGC
GCATCGGACTCATCAGGAAGC
TGCATCGGACTCATCAGGAAGC
CATCGGACTCATCAGGAAGC
CACATCGGACTCATCAGGAAGC
GGACTCATCAGGAAGC
ACATCGGACTCATCAGGAAGC
CATCGGACTCATCAGGAAGC
ACATCGGACTCATCAGGAAGC
ACATCGGACTCATCAGGAAGC
AAGCATCGGACTCATCAGGAAGC
GCATCGGACTCATCAGGAAGC
CGGACTCATCAGGAAGC
ACCATCGGACTCATCAGGAAGC
ACATCGGACTCATCAGGAAGC
TCGGACTCATCAGGAAGC
CCATCGGACTCATCAGGAAGC
CCATCGGACTCATCAGGAAGC
ACATCGGACTCATCAGGAAGC
ACATCGGACTCATCAGGAAGC
TTCATCGGACTCATCAGGAAGC
CATCGGACTCATCAGGAAGC
GGACTCATCAGGAAGC
CACATCGGACTCATCAGGAAGC

CATCGGACTCATCAGGAAGC
GCATCGGACTCATCAGGAAGC
ATCGGACTCATCAGGAAGC
CCATCGGACTCATCAGGAAGC
GCATCGGACTCATCAGGAAGC
ATCGGACTCATCAGGAAGC
AACCATCGGACTCATCAGGAAGC
CATCGGACTCATCAGGAAGC
TCGGACTCATCAGGAAGC
TCGGACTCATCAGGAAGC
TCATCGGACTCATCAGGAAGC
ATCGGACTCATCAGGAAGC
CATCGGACTCATCAGGAAGC
CATCGGACTCATCAGGAAGC
TTCATCGGACTCATCAGGAAGC
CATCGGACTCATCAGGAAGC
CATCGGACTCATCAGGAAGC
TTAGCATCGGACTCATCAGGAAGC
CATCGGACTCATCAGGAAGC
TCATCGGACTCATCAGGAAGC
CATCGGACTCATCAGGAAGC
TCGGACTCATCAGGAAGC
GCATCGGACTCATCAGGAAGC
GGACTCATCAGGAAGC
GGACTCATCAGGAAGC
AACATCGGACTCATCAGGAAGC
ACATCGGACTCATCAGGAAGC
AACATCGGACTCATCAGGAAGC
AACATCGGACTCATCAGGAAGC
ACCATCGGACTCATCAGGAAGC
CCATCGGACTCATCAGGAAGC
CATCGGACTCATCAGGAAGC
CATCGGACTCATCAGGAAGC
GGACTCATCAGGAAGC
TCGGACTCATCAGGAAGC
CATCGGACTCATCAGGAAGC
CCATCGGACTCATCAGGAAGC
CCATCGGACTCATCAGGAAGC
CGGACTCATCAGGAAGC
ATCGGACTCATCAGGAAGC
TCATCGGACTCATCAGGAAGC
ACATCGGACTCATCAGGAAGC
CATCGGACTCATCAGGAAGC

AACATCGGACTCATCAGGAAGC
AAACATCGGACTCATCAGGAAGC
ATCGGACTCATCAGGAAGC
GACTCATCAGGAAGC
ATCGGACTCATCAGGAAGC
TCGGACTCATCAGGAAGC
CATCGGACTCATCAGGAAGC
CATCGGACTCATCAGGAAGC
CATCGGACTCATCAGGAAGC
ATCGGACTCATCAGGAAGC
TCGGACTCATCAGGAAGC
GGACTCATCAGGAAGC
ATCGGACTCATCAGGAAGC
ACATCGGACTCATCAGGAAGC
CTACATCGGACTCATCAGGAAGC
CTACATCGGACTCATCAGGAAGC
CTACATCGGACTCATCAGGAAGC
CGGACTCATCAGGAAGC
CATCGGACTCATCAGGAAGC
TCATCGGACTCATCAGGAAGC
GCATCGGACTCATCAGGAAGC
CATCGGACTCATCAGGAAGC
ACATCGGACTCATCAGGAAGC
TCATCGGACTCATCAGGAAGC
CATCGGACTCATCAGGAAGC
CATCGGACTCATCAGGAAGC
TCATCGGACTCATCAGGAAGC
ACATCGGACTCATCAGGAAGC
CGGACTCATCAGGAAGC
CATCGGACTCATCAGGAAGC
CGGACTCATCAGGAAGC
ATCGGACTCATCAGGAAGC
TCATCGGACTCATCAGGAAGC
ACATCGGACTCATCAGGAAGC
ATCGGACTCATCAGGAAGC
ACATCGGACTCATCAGGAAGC
ATCGGACTCATCAGGAAGC
ATCGGACTCATCAGGAAGC
CATCGGACTCATCAGGAAGC
CATCGGACTCATCAGGAAGC
ACTCATCAGGAAGC
TCGGACTCATCAGGAAGC
CATCGGACTCATCAGGAAGC

ATCGGACTCATCAGGAAGC
CGGACTCATCAGGAAGC
ATACATCGGACTCATCAGGAAGC
TCATCGGACTCATCAGGAAGC
ATCGGACTCATCAGGAAGC
CATCGGACTCATCAGGAAGC
GCATCGGACTCATCAGGAAGC
CATCGGACTCATCAGGAAGC
CATCGGACTCATCAGGAAGC
ACATCGGACTCATCAGGAAGC
CCATCGGACTCATCAGGAAGC
CCATCGGACTCATCAGGAAGC
GCATCGGACTCATCAGGAAGC
TCACATCGGACTCATCAGGAAGC
TCATCGGACTCATCAGGAAGC
CATCGGACTCATCAGGAAGC
CCATCGGACTCATCAGGAAGC
ATCGGACTCATCAGGAAGC
CATCGGACTCATCAGGAAGC
TCGGACTCATCAGGAAGC
CATCGGACTCATCAGGAAGC
ATCGGACTCATCAGGAAGC
GCATCGGACTCATCAGGAAGC
CATCGGACTCATCAGGAAGC
AACATCGGACTCATCAGGAAGC
TCATCGGACTCATCAGGAAGC
TCGGACTCATCAGGAAGC
TCCATCGGACTCATCAGGAAGC
ACCATCGGACTCATCAGGAAGC
ACCATCGGACTCATCAGGAAGC
ATCGGACTCATCAGGAAGC
ATCGGACTCATCAGGAAGC
ATCATCGGACTCATCAGGAAGC
CATCGGACTCATCAGGAAGC
ATCATCGGACTCATCAGGAAGC
TCGGACTCATCAGGAAGC
CATCGGACTCATCAGGAAGC
TCATCGGACTCATCAGGAAGC
CATCGGACTCATCAGGAAGC
ATCATCGGACTCATCAGGAAGC
ATCATCGGACTCATCAGGAAGC
ATCGGACTCATCAGGAAGC
ACATCGGACTCATCAGGAAGC

CATCGGACTCATCAGGAAGC
CATCGGACTCATCAGGAAGC
TCGGACTCATCAGGAAGC
ATCGGACTCATCAGGAAGC
GCCATCGGACTCATCAGGAAGC
ATCGGACTCATCAGGAAGC
GGACTCATCAGGAAGC
TTTCATCGGACTCATCAGGAAGC
CCATCGGACTCATCAGGAAGC
GTCATCGGACTCATCAGGAAGC
TCGGACTCATCAGGAAGC
TTCATCGGACTCATCAGGAAGC
TGCATCGGACTCATCAGGAAGC
CATCGGACTCATCAGGAAGC
CATCATCGGACTCATCAGGAAGC
AAAACATCGGACTCATCAGGAAGC
AAAACATCGGACTCATCAGGAAGC
CATCGGACTCATCAGGAAGC
CATCGGACTCATCAGGAAGC
ATCGGACTCATCAGGAAGC
ATCGGACTCATCAGGAAGC
TCTCATCGGACTCATCAGGAAGC
CGGACTCATCAGGAAGC
TCGGACTCATCAGGAAGC
CATCGGACTCATCAGGAAGC
TCGGACTCATCAGGAAGC
GCATCGGACTCATCAGGAAGC
ACCATCGGACTCATCAGGAAGC
AGCATCGGACTCATCAGGAAGC
AGCATCGGACTCATCAGGAAGC
TACATCGGACTCATCAGGAAGC
CCATCGGACTCATCAGGAAGC
ACATCGGACTCATCAGGAAGC
ATCGGACTCATCAGGAAGC
ATCGGACTCATCAGGAAGC
AGCATCGGACTCATCAGGAAGC
AGCATCGGACTCATCAGGAAGC
TACATCGGACTCATCAGGAAGC
GCATCGGACTCATCAGGAAGC
TCATCGGACTCATCAGGAAGC
TCATCGGACTCATCAGGAAGC
CATCGGACTCATCAGGAAGC
CATCGGACTCATCAGGAAGC

ACATCGGACTCATCAGGAAGC
TCATCGGACTCATCAGGAAGC
CATCGGACTCATCAGGAAGC
CCATCGGACTCATCAGGAAGC
GGACTCATCAGGAAGC
GGACTCATCAGGAAGC
TCATCAGGAAGC
GTCATCGGACTCATCAGGAAGC
TGCATCGGACTCATCAGGAAGC
GTTCATCGGACTCATCAGGAAGC
ATACATCGGACTCATCAGGAAGC
GACTCATCAGGAAGC
CGGACTCATCAGGAAGC
CGGACTCATCAGGAAGC
ATCGGACTCATCAGGAAGC
AGCATCGGACTCATCAGGAAGC
CCATCGGACTCATCAGGAAGC
CCATCGGACTCATCAGGAAGC
TACATCGGACTCATCAGGAAGC
TTCATCGGACTCATCAGGAAGC
CCATCGGACTCATCAGGAAGC
CATCGGACTCATCAGGAAGC
CATCGGACTCATCAGGAAGC
CATCGGACTCATCAGGAAGC
ACATCGGACTCATCAGGAAGC
CCATCGGACTCATCAGGAAGC
ATCGGACTCATCAGGAAGC
ATCGGACTCATCAGGAAGC
GCATCGGACTCATCAGGAAGC
CATCGGACTCATCAGGAAGC
TCATCGGACTCATCAGGAAGC
CATCGGACTCATCAGGAAGC
CCATCGGACTCATCAGGAAGC
CATCGGACTCATCAGGAAGC
ATCATCGGACTCATCAGGAAGC
CATCGGACTCATCAGGAAGC
ACTCATCAGGAAGC
CATCGGACTCATCAGGAAGC
ATCGGACTCATCAGGAAGC
ATCGGACTCATCAGGAAGC
ACTCATCAGGAAGC
ACATCGGACTCATCAGGAAGC
ACATCGGACTCATCAGGAAGC

ACATCGGACTCATCAGGAAGC
ATCGGACTCATCAGGAAGC
ATCGGACTCATCAGGAAGC
AGCATCGGACTCATCAGGAAGC
AGCATCGGACTCATCAGGAAGC
ACATCGGACTCATCAGGAAGC
CCATCGGACTCATCAGGAAGC
AGAACATCGGACTCATCAGGAAGC
CATCGGACTCATCAGGAAGC
CGGACTCATCAGGAAGC
ACATCGGACTCATCAGGAAGC
AACATCGGACTCATCAGGAAGC
ATCATCGGACTCATCAGGAAGC
GGACTCATCAGGAAGC
CATCGGACTCATCAGGAAGC
CATCGGACTCATCAGGAAGC
CATCGGACTCATCAGGAAGC
TACATCGGACTCATCAGGAAGC
CATCGGACTCATCAGGAAGC
CATCGGACTCATCAGGAAGC
ATCGGACTCATCAGGAAGC
TCCATCGGACTCATCAGGAAGC
TCCATCGGACTCATCAGGAAGC
TCATCGGACTCATCAGGAAGC
ATCGGACTCATCAGGAAGC
ATCATCGGACTCATCAGGAAGC
ACATCGGACTCATCAGGAAGC
ACATCGGACTCATCAGGAAGC
GCATCGGACTCATCAGGAAGC
GCATCGGACTCATCAGGAAGC
TCGGACTCATCAGGAAGC
GCATCGGACTCATCAGGAAGC
AAACATCGGACTCATCAGGAAGC
ATCATCGGACTCATCAGGAAGC
TCCATCGGACTCATCAGGAAGC
CATCGGACTCATCAGGAAGC
TCATCGGACTCATCAGGAAGC
CATCGGACTCATCAGGAAGC
CCATCGGACTCATCAGGAAGC
CATCGGACTCATCAGGAAGC
TCATCGGACTCATCAGGAAGC
CATCGGACTCATCAGGAAGC
CATCGGACTCATCAGGAAGC

GCATCGGACTCATCAGGAAGC
ATCGGACTCATCAGGAAGC
ATCGGACTCATCAGGAAGC
CATCGGACTCATCAGGAAGC
TACATCGGACTCATCAGGAAGC
ATCGGACTCATCAGGAAGC
TCATCGGACTCATCAGGAAGC
CATCGGACTCATCAGGAAGC
TGCATCGGACTCATCAGGAAGC
CATCGGACTCATCAGGAAGC
TTCATCGGACTCATCAGGAAGC
ACATCGGACTCATCAGGAAGC
CATCGGACTCATCAGGAAGC
TCGGACTCATCAGGAAGC
GCATCGGACTCATCAGGAAGC
ATCGGACTCATCAGGAAGC
TCATCGGACTCATCAGGAAGC
CATCGGACTCATCAGGAAGC
CATCGGACTCATCAGGAAGC
TCATCGGACTCATCAGGAAGC
TCATCGGACTCATCAGGAAGC
ATCGGACTCATCAGGAAGC
GACATCGGACTCATCAGGAAGC
ACATCGGACTCATCAGGAAGC
ATCGGACTCATCAGGAAGC
GTCATCGGACTCATCAGGAAGC
GCATCGGACTCATCAGGAAGC
GCATCGGACTCATCAGGAAGC
CATCGGACTCATCAGGAAGC
GGACTCATCAGGAAGC
CCATCGGACTCATCAGGAAGC
ATCGGACTCATCAGGAAGC
CACATCGGACTCATCAGGAAGC
CATCGGACTCATCAGGAAGC
CATCGGACTCATCAGGAAGC
ATCGGACTCATCAGGAAGC
ACATCGGACTCATCAGGAAGC
CGGACTCATCAGGAAGC
CATCGGACTCATCAGGAAGC
TCCATCGGACTCATCAGGAAGC
CATCGGACTCATCAGGAAGC
TCTTCATCGGACTCATCAGGAAGC
ACATCGGACTCATCAGGAAGC

AAGCATCGGACTCATCAGGAAGC
ATCGGACTCATCAGGAAGC
TCATCGGACTCATCAGGAAGC
TCATCGGACTCATCAGGAAGC
GACTCATCAGGAAGC
GCATCGGACTCATCAGGAAGC
TCGGACTCATCAGGAAGC
ATCGGACTCATCAGGAAGC
ATCATCGGACTCATCAGGAAGC
CCATCGGACTCATCAGGAAGC
TGTCATCGGACTCATCAGGAAGC
ATCGGACTCATCAGGAAGC
ATCGGACTCATCAGGAAGC
ATCGGACTCATCAGGAAGC
AACATCGGACTCATCAGGAAGC
AACATCGGACTCATCAGGAAGC
CATCGGACTCATCAGGAAGC
CATCGGACTCATCAGGAAGC
CATCGGACTCATCAGGAAGC
CATCGGACTCATCAGGAAGC
AACATCGGACTCATCAGGAAGC
TCGGACTCATCAGGAAGC
ACATCGGACTCATCAGGAAGC
ACATCGGACTCATCAGGAAGC
TCATCGGACTCATCAGGAAGC
ACTCATCAGGAAGC
CATCGGACTCATCAGGAAGC
ATCGGACTCATCAGGAAGC
GACTCATCAGGAAGC
TCGGACTCATCAGGAAGC
TCATCGGACTCATCAGGAAGC
GCATCGGACTCATCAGGAAGC
ATCGGACTCATCAGGAAGC
TCGGACTCATCAGGAAGC
CCCATCGGACTCATCAGGAAGC
CCCATCGGACTCATCAGGAAGC
ATCATCGGACTCATCAGGAAGC
ATCGGACTCATCAGGAAGC
TTCATCGGACTCATCAGGAAGC
ACATCGGACTCATCAGGAAGC
CATCGGACTCATCAGGAAGC
ATCGGACTCATCAGGAAGC
GCATCGGACTCATCAGGAAGC

TCGGACTCATCAGGAAGC
CATCGGACTCATCAGGAAGC
CATCGGACTCATCAGGAAGC
CGGACTCATCAGGAAGC
CACATCGGACTCATCAGGAAGC
ATCGGACTCATCAGGAAGC
CATCGGACTCATCAGGAAGC
TACATCGGACTCATCAGGAAGC
TCGGACTCATCAGGAAGC
GGACTCATCAGGAAGC
CATCGGACTCATCAGGAAGC
CATCGGACTCATCAGGAAGC
GTCATCGGACTCATCAGGAAGC
TCATCGGACTCATCAGGAAGC
ATCGGACTCATCAGGAAGC
GCATCGGACTCATCAGGAAGC
TCATCGGACTCATCAGGAAGC
GCATCGGACTCATCAGGAAGC
TACATCGGACTCATCAGGAAGC
GACTCATCAGGAAGC
TGCATCGGACTCATCAGGAAGC
CATCGGACTCATCAGGAAGC
TCATCGGACTCATCAGGAAGC
CATCGGACTCATCAGGAAGC
ACATCGGACTCATCAGGAAGC
TCATCGGACTCATCAGGAAGC
ATCGGACTCATCAGGAAGC
GGACTCATCAGGAAGC
ATCGGACTCATCAGGAAGC
GCATCGGACTCATCAGGAAGC
CATCGGACTCATCAGGAAGC
AGACATCGGACTCATCAGGAAGC
GAACATCGGACTCATCAGGAAGC
GGCATCGGACTCATCAGGAAGC
AGCATCGGACTCATCAGGAAGC
GCCATCGGACTCATCAGGAAGC
TCATCGGACTCATCAGGAAGC
ATCATCGGACTCATCAGGAAGC
TCATCGGACTCATCAGGAAGC
GCCATCGGACTCATCAGGAAGC
ACATCGGACTCATCAGGAAGC
CATCGGACTCATCAGGAAGC
CATCGGACTCATCAGGAAGC

TACATCGGACTCATCAGGAAGC
GCATCGGACTCATCAGGAAGC
TCGGACTCATCAGGAAGC
TCACATCGGACTCATCAGGAAGC
TACATCGGACTCATCAGGAAGC
ATCGGACTCATCAGGAAGC
ACATCGGACTCATCAGGAAGC
TCATCGGACTCATCAGGAAGC
TCCATCGGACTCATCAGGAAGC
CATCGGACTCATCAGGAAGC
CGGACTCATCAGGAAGC
TACATCGGACTCATCAGGAAGC
CATCGGACTCATCAGGAAGC
ATACATCGGACTCATCAGGAAGC
TCGGACTCATCAGGAAGC
CCCATCGGACTCATCAGGAAGC
CCCATCGGACTCATCAGGAAGC
ATCGGACTCATCAGGAAGC
TCATCGGACTCATCAGGAAGC
GCATCGGACTCATCAGGAAGC
CATCGGACTCATCAGGAAGC
GACTCATCAGGAAGC
ACCATCGGACTCATCAGGAAGC
ATCGGACTCATCAGGAAGC
AAGCATCGGACTCATCAGGAAGC
ATCATCGGACTCATCAGGAAGC
TGCATCGGACTCATCAGGAAGC
TTCATCGGACTCATCAGGAAGC
TTCATCGGACTCATCAGGAAGC
TTCATCGGACTCATCAGGAAGC
ATCGGACTCATCAGGAAGC
TCATCGGACTCATCAGGAAGC
CGGACTCATCAGGAAGC
ACATCGGACTCATCAGGAAGC
GGACTCATCAGGAAGC
AGCATCGGACTCATCAGGAAGC
CTCCATCGGACTCATCAGGAAGC
ACATCGGACTCATCAGGAAGC
AAGCATCGGACTCATCAGGAAGC
AACATCGGACTCATCAGGAAGC
TTACATCGGACTCATCAGGAAGC
GCATCGGACTCATCAGGAAGC
GCATCGGACTCATCAGGAAGC

CGGACTCATCAGGAAGC
CATCGGACTCATCAGGAAGC
CATCGGACTCATCAGGAAGC
CCATCGGACTCATCAGGAAGC
GCATCGGACTCATCAGGAAGC
TTCATCGGACTCATCAGGAAGC
CGGACTCATCAGGAAGC
ATCGGACTCATCAGGAAGC
CATCGGACTCATCAGGAAGC
CCATCGGACTCATCAGGAAGC
CCATCGGACTCATCAGGAAGC
CATCGGACTCATCAGGAAGC
ATCGGACTCATCAGGAAGC
GACTCATCAGGAAGC
TCGGACTCATCAGGAAGC
ATCGGACTCATCAGGAAGC
CGGACTCATCAGGAAGC
CATCGGACTCATCAGGAAGC
CATCGGACTCATCAGGAAGC
TTTCATCGGACTCATCAGGAAGC
CCATCGGACTCATCAGGAAGC
TCATCGGACTCATCAGGAAGC
GGACTCATCAGGAAGC
TCGGACTCATCAGGAAGC
TCGGACTCATCAGGAAGC
CATCGGACTCATCAGGAAGC
ATCGGACTCATCAGGAAGC
CCATCGGACTCATCAGGAAGC
CGGACTCATCAGGAAGC
ACATCGGACTCATCAGGAAGC
ACTCATCAGGAAGC
GGCATCGGACTCATCAGGAAGC
GCATCGGACTCATCAGGAAGC
TGCATCGGACTCATCAGGAAGC
CCATCGGACTCATCAGGAAGC
CATCGGACTCATCAGGAAGC
CATCGGACTCATCAGGAAGC
CATCGGACTCATCAGGAAGC
TCATCGGACTCATCAGGAAGC
GCATCGGACTCATCAGGAAGC
CATCGGACTCATCAGGAAGC
TCGGACTCATCAGGAAGC
CATCGGACTCATCAGGAAGC

ATCGGACTCATCAGGAAGC
ATCGGACTCATCAGGAAGC
CGGACTCATCAGGAAGC
TCGGACTCATCAGGAAGC
ATCGGACTCATCAGGAAGC
TCGGACTCATCAGGAAGC
ACCATCGGACTCATCAGGAAGC
AGTGCATCGGACTCATCAGGAAGC
TCATCGGACTCATCAGGAAGC
GACATCGGACTCATCAGGAAGC
ACCATCGGACTCATCAGGAAGC
CATCGGACTCATCAGGAAGC
CATCGGACTCATCAGGAAGC
CATCGGACTCATCAGGAAGC
GGCATCGGACTCATCAGGAAGC
CATCGGACTCATCAGGAAGC
AACATCGGACTCATCAGGAAGC
GAGCATCGGACTCATCAGGAAGC
GCATCGGACTCATCAGGAAGC
ACCATCGGACTCATCAGGAAGC
ACATCGGACTCATCAGGAAGC
CATCGGACTCATCAGGAAGC
CATCGGACTCATCAGGAAGC
AAACATCGGACTCATCAGGAAGC
CATCGGACTCATCAGGAAGC
AGCATCGGACTCATCAGGAAGC
ATCGGACTCATCAGGAAGC
TGTCATCGGACTCATCAGGAAGC
TCATCGGACTCATCAGGAAGC
TCATCGGACTCATCAGGAAGC
TCGGACTCATCAGGAAGC
GACTCATCAGGAAGC
CGGACTCATCAGGAAGC
AGCATCGGACTCATCAGGAAGC
ATCATCGGACTCATCAGGAAGC
CATCGGACTCATCAGGAAGC
AGCATCGGACTCATCAGGAAGC
TACATCGGACTCATCAGGAAGC
CTTCATCGGACTCATCAGGAAGC
ACATCGGACTCATCAGGAAGC
GGCATCGGACTCATCAGGAAGC
GACATCGGACTCATCAGGAAGC
TTTCATCGGACTCATCAGGAAGC

TCGGACTCATCAGGAAGC
CATCGGACTCATCAGGAAGC
CATCGGACTCATCAGGAAGC
ATCATCGGACTCATCAGGAAGC
TCATCGGACTCATCAGGAAGC
CATCGGACTCATCAGGAAGC
TCATCGGACTCATCAGGAAGC
ACATCGGACTCATCAGGAAGC
CGGACTCATCAGGAAGC
CCATCGGACTCATCAGGAAGC
TGGCATCGGACTCATCAGGAAGC
TACATCGGACTCATCAGGAAGC
TTCATCGGACTCATCAGGAAGC
GTCATCGGACTCATCAGGAAGC
TCGGACTCATCAGGAAGC
AGCATCGGACTCATCAGGAAGC
GAGCATCGGACTCATCAGGAAGC
TCGGACTCATCAGGAAGC
GACATCGGACTCATCAGGAAGC
AACATCGGACTCATCAGGAAGC
TTCATCGGACTCATCAGGAAGC
TCATCGGACTCATCAGGAAGC
TCATCGGACTCATCAGGAAGC
TCATCGGACTCATCAGGAAGC
CATCGGACTCATCAGGAAGC
CATCGGACTCATCAGGAAGC
TCATCGGACTCATCAGGAAGC
CGGACTCATCAGGAAGC
CATCGGACTCATCAGGAAGC
GGCCATCGGACTCATCAGGAAGC
AGACATCGGACTCATCAGGAAGC
CATCGGACTCATCAGGAAGC
ATCGGACTCATCAGGAAGC
CGGACTCATCAGGAAGC
CATCGGACTCATCAGGAAGC
AACATCGGACTCATCAGGAAGC
ATCGGACTCATCAGGAAGC
ATCGGACTCATCAGGAAGC
TCGGACTCATCAGGAAGC
CATCGGACTCATCAGGAAGC
CCATCGGACTCATCAGGAAGC
CGGACTCATCAGGAAGC
ACATCGGACTCATCAGGAAGC

AGCATCGGACTCATCAGGAAGC
CATCGGACTCATCAGGAAGC
GCATCGGACTCATCAGGAAGC
ACATCGGACTCATCAGGAAGC
TCATCGGACTCATCAGGAAGC
ACATCGGACTCATCAGGAAGC
TGCATCGGACTCATCAGGAAGC
TCCATCGGACTCATCAGGAAGC
AACATCGGACTCATCAGGAAGC
CACATCGGACTCATCAGGAAGC
ACATCGGACTCATCAGGAAGC
ACATCGGACTCATCAGGAAGC
ACTCATCAGGAAGC
GGACTCATCAGGAAGC
GCCCATCGGACTCATCAGGAAGC
AACATCGGACTCATCAGGAAGC
AACATCGGACTCATCAGGAAGC
ATCGGACTCATCAGGAAGC
ATCGGACTCATCAGGAAGC
GGACTCATCAGGAAGC
CGGACTCATCAGGAAGC
ATTCATCGGACTCATCAGGAAGC
ACATCGGACTCATCAGGAAGC
GCCATCGGACTCATCAGGAAGC
TCATCGGACTCATCAGGAAGC
TTACATCGGACTCATCAGGAAGC
TCATCGGACTCATCAGGAAGC
ACATCGGACTCATCAGGAAGC
ACTCATCGGACTCATCAGGAAGC
GTCATCGGACTCATCAGGAAGC
ACTCATCAGGAAGC
CTCATCGGACTCATCAGGAAGC
GCATCGGACTCATCAGGAAGC
TCATCGGACTCATCAGGAAGC
CATCGGACTCATCAGGAAGC
CGGACTCATCAGGAAGC
CATCGGACTCATCAGGAAGC
CGGACTCATCAGGAAGC
CGGACTCATCAGGAAGC
GCATCGGACTCATCAGGAAGC
CCATCGGACTCATCAGGAAGC
TCGGACTCATCAGGAAGC
CCATCGGACTCATCAGGAAGC

CATCGGACTCATCAGGAAGC
ACCATCGGACTCATCAGGAAGC
TCGGACTCATCAGGAAGC
TCATCGGACTCATCAGGAAGC
ACATCGGACTCATCAGGAAGC
ACATCGGACTCATCAGGAAGC
CATCGGACTCATCAGGAAGC
TCATCGGACTCATCAGGAAGC
TACATCGGACTCATCAGGAAGC
CGGACTCATCAGGAAGC
CGGACTCATCAGGAAGC
TCGGACTCATCAGGAAGC
ACATCGGACTCATCAGGAAGC
GTCATCGGACTCATCAGGAAGC
TTCATCGGACTCATCAGGAAGC
ATCGGACTCATCAGGAAGC
CACATCGGACTCATCAGGAAGC
TGCATCGGACTCATCAGGAAGC
CATCGGACTCATCAGGAAGC
TCATCGGACTCATCAGGAAGC
TCGGACTCATCAGGAAGC
CATCGGACTCATCAGGAAGC
CCATCGGACTCATCAGGAAGC
TCGGACTCATCAGGAAGC
ATCGGACTCATCAGGAAGC
TCGGACTCATCAGGAAGC
CTCATCGGACTCATCAGGAAGC
CATCGGACTCATCAGGAAGC
CATCGGACTCATCAGGAAGC
GGACTCATCAGGAAGC
TCGGACTCATCAGGAAGC
ATCGGACTCATCAGGAAGC
TCGGACTCATCAGGAAGC
CGGACTCATCAGGAAGC
CGGACTCATCAGGAAGC
ATCGGACTCATCAGGAAGC
CCATCGGACTCATCAGGAAGC
CATCGGACTCATCAGGAAGC
GCATCGGACTCATCAGGAAGC
GCATCGGACTCATCAGGAAGC
CATCGGACTCATCAGGAAGC
ACATCGGACTCATCAGGAAGC
GCATCGGACTCATCAGGAAGC

ATCGGACTCATCAGGAAGC
TCATCGGACTCATCAGGAAGC
ATCGGACTCATCAGGAAGC
AACATCGGACTCATCAGGAAGC
ATCGGACTCATCAGGAAGC
ACATCGGACTCATCAGGAAGC
CATCGGACTCATCAGGAAGC
TCATCGGACTCATCAGGAAGC
ACATCGGACTCATCAGGAAGC
GGACTCATCAGGAAGC
CATCGGACTCATCAGGAAGC
CCATCGGACTCATCAGGAAGC
GCATCGGACTCATCAGGAAGC
CTCATCGGACTCATCAGGAAGC
AAACATCGGACTCATCAGGAAGC
AAACATCGGACTCATCAGGAAGC
CATCGGACTCATCAGGAAGC
TCCATCGGACTCATCAGGAAGC
CGGACTCATCAGGAAGC
GCATCGGACTCATCAGGAAGC
CATCGGACTCATCAGGAAGC
GTCATCGGACTCATCAGGAAGC
ATGCATCGGACTCATCAGGAAGC
CATCGGACTCATCAGGAAGC
TCGGACTCATCAGGAAGC
ACATCGGACTCATCAGGAAGC
TCCATCGGACTCATCAGGAAGC
AACATCGGACTCATCAGGAAGC
TCATCGGACTCATCAGGAAGC
ATCGGACTCATCAGGAAGC
AGGCATCGGACTCATCAGGAAGC
TCATCGGACTCATCAGGAAGC
CATCGGACTCATCAGGAAGC
TCATCGGACTCATCAGGAAGC
ATCGGACTCATCAGGAAGC
GCATCGGACTCATCAGGAAGC
TCATCGGACTCATCAGGAAGC
TCATCGGACTCATCAGGAAGC
CATCGGACTCATCAGGAAGC
TCATCGGACTCATCAGGAAGC
TCATCGGACTCATCAGGAAGC
TCATCGGACTCATCAGGAAGC
CATCGGACTCATCAGGAAGC

TCGGACTCATCAGGAAGC
ACCATCGGACTCATCAGGAAGC
CCCATCGGACTCATCAGGAAGC
ATCGGACTCATCAGGAAGC
GCCATCGGACTCATCAGGAAGC
AAACATCGGACTCATCAGGAAGC
ATCGGACTCATCAGGAAGC
ATCGGACTCATCAGGAAGC
CATCGGACTCATCAGGAAGC
GCATCGGACTCATCAGGAAGC
TCGGACTCATCAGGAAGC
CCATCGGACTCATCAGGAAGC
GCATCGGACTCATCAGGAAGC
CGGACTCATCAGGAAGC
CGGACTCATCAGGAAGC
ATCGGACTCATCAGGAAGC
ATCGGACTCATCAGGAAGC
CATCGGACTCATCAGGAAGC
CATCGGACTCATCAGGAAGC
TCGGACTCATCAGGAAGC
ATCGGACTCATCAGGAAGC
TCATCGGACTCATCAGGAAGC
GCATCGGACTCATCAGGAAGC
CATCGGACTCATCAGGAAGC
AACATCGGACTCATCAGGAAGC
TCCATCGGACTCATCAGGAAGC
TCATCGGACTCATCAGGAAGC
TCATCGGACTCATCAGGAAGC
GCATCGGACTCATCAGGAAGC
GTCATCGGACTCATCAGGAAGC
ATCGGACTCATCAGGAAGC
ACTACATCGGACTCATCAGGAAGC
CCATCGGACTCATCAGGAAGC
GCCATCGGACTCATCAGGAAGC
TCGGACTCATCAGGAAGC
ATCGGACTCATCAGGAAGC
GCATCGGACTCATCAGGAAGC
ATCATCGGACTCATCAGGAAGC
GAACATCGGACTCATCAGGAAGC
TCATCGGACTCATCAGGAAGC
ACACATCGGACTCATCAGGAAGC
AAACATCGGACTCATCAGGAAGC
GTACATCGGACTCATCAGGAAGC

GTACATCGGACTCATCAGGAAGC
ACATCGGACTCATCAGGAAGC
ACATCGGACTCATCAGGAAGC
ATCGGACTCATCAGGAAGC
CATCGGACTCATCAGGAAGC
CATCGGACTCATCAGGAAGC
TCATCGGACTCATCAGGAAGC
GCATCGGACTCATCAGGAAGC
GAGCATCGGACTCATCAGGAAGC
CATCGGACTCATCAGGAAGC
TTCATCGGACTCATCAGGAAGC
CATCGGACTCATCAGGAAGC
CGGACTCATCAGGAAGC
TACATCGGACTCATCAGGAAGC
TCGGACTCATCAGGAAGC
GCATCGGACTCATCAGGAAGC
CATCGGACTCATCAGGAAGC
ACACATCGGACTCATCAGGAAGC
TCACATCGGACTCATCAGGAAGC
CATCGGACTCATCAGGAAGC
TCGGACTCATCAGGAAGC
TGTCATCGGACTCATCAGGAAGC
ATCGGACTCATCAGGAAGC
ACATCGGACTCATCAGGAAGC
GGACTCATCAGGAAGC
ACATCGGACTCATCAGGAAGC
CATCGGACTCATCAGGAAGC
TCGGACTCATCAGGAAGC
TCGGACTCATCAGGAAGC
CTTCATCGGACTCATCAGGAAGC
GACTCATCAGGAAGC
GACTCATCAGGAAGC
GCCATCGGACTCATCAGGAAGC
TCATCGGACTCATCAGGAAGC
ACATCGGACTCATCAGGAAGC
ATCGGACTCATCAGGAAGC
CATCGGACTCATCAGGAAGC
AACATCGGACTCATCAGGAAGC
CGGACTCATCAGGAAGC
ATCGGACTCATCAGGAAGC
TCATCGGACTCATCAGGAAGC
TCATCGGACTCATCAGGAAGC
ATCGGACTCATCAGGAAGC

GTGCATCGGACTCATCAGGAAGC
GCACATCGGACTCATCAGGAAGC
AATCATCGGACTCATCAGGAAGC
ATCGGACTCATCAGGAAGC
GACTCATCAGGAAGC
CGGACTCATCAGGAAGC
GGACTCATCAGGAAGC
CATCGGACTCATCAGGAAGC
ATCGGACTCATCAGGAAGC
CTCATCGGACTCATCAGGAAGC
ATCGGACTCATCAGGAAGC
AACATCGGACTCATCAGGAAGC
ACTCATCAGGAAGC
ATACATCGGACTCATCAGGAAGC
GACATCGGACTCATCAGGAAGC
GACATCGGACTCATCAGGAAGC
TACATCGGACTCATCAGGAAGC
TCGGACTCATCAGGAAGC
CCATCGGACTCATCAGGAAGC
CATCGGACTCATCAGGAAGC
CATCGGACTCATCAGGAAGC
CATCGGACTCATCAGGAAGC
GCATCGGACTCATCAGGAAGC
ACCATCGGACTCATCAGGAAGC
GGACTCATCAGGAAGC
ACCATCGGACTCATCAGGAAGC
TCCATCGGACTCATCAGGAAGC
ACATCGGACTCATCAGGAAGC
ATCATCGGACTCATCAGGAAGC
CCATCGGACTCATCAGGAAGC
GTCATCGGACTCATCAGGAAGC
ACATCGGACTCATCAGGAAGC
ATCGGACTCATCAGGAAGC
GGACTCATCAGGAAGC
ATCGGACTCATCAGGAAGC
CATCGGACTCATCAGGAAGC
CCATCGGACTCATCAGGAAGC
CGGACTCATCAGGAAGC
ATCATCGGACTCATCAGGAAGC
TAGCATCGGACTCATCAGGAAGC
GACATCGGACTCATCAGGAAGC
TCGGACTCATCAGGAAGC
AACCATCGGACTCATCAGGAAGC

CCATCGGACTCATCAGGAAGC
CCATCGGACTCATCAGGAAGC
CATCGGACTCATCAGGAAGC
TCACATCGGACTCATCAGGAAGC
CATCGGACTCATCAGGAAGC
CATCGGACTCATCAGGAAGC
CATCGGACTCATCAGGAAGC
ATCGGACTCATCAGGAAGC
CATCGGACTCATCAGGAAGC
ATCGGACTCATCAGGAAGC
GGACTCATCAGGAAGC
ATCGGACTCATCAGGAAGC
AATCATCGGACTCATCAGGAAGC
CCATCGGACTCATCAGGAAGC
AACATCGGACTCATCAGGAAGC
ACATCGGACTCATCAGGAAGC
TCATCGGACTCATCAGGAAGC
CGGACTCATCAGGAAGC
GGACTCATCAGGAAGC
CATCGGACTCATCAGGAAGC
CACATCGGACTCATCAGGAAGC
ATCGGACTCATCAGGAAGC
CTCCATCGGACTCATCAGGAAGC
AACATCGGACTCATCAGGAAGC
CGGACTCATCAGGAAGC
GACTCATCAGGAAGC
CATCGGACTCATCAGGAAGC
ATCGGACTCATCAGGAAGC
TCGGACTCATCAGGAAGC
ATCGGACTCATCAGGAAGC
ACCATCGGACTCATCAGGAAGC
CATCGGACTCATCAGGAAGC
ATCGGACTCATCAGGAAGC
TCATCGGACTCATCAGGAAGC
CATCGGACTCATCAGGAAGC
GAACATCGGACTCATCAGGAAGC
TTCATCGGACTCATCAGGAAGC
ATCATCGGACTCATCAGGAAGC
CCATCGGACTCATCAGGAAGC
TCATCGGACTCATCAGGAAGC
CATCGGACTCATCAGGAAGC
CATCGGACTCATCAGGAAGC
CCATCGGACTCATCAGGAAGC

ACATCGGACTCATCAGGAAGC
TACATCGGACTCATCAGGAAGC
ACATCGGACTCATCAGGAAGC
TCGGACTCATCAGGAAGC
GACATCGGACTCATCAGGAAGC
CCATCGGACTCATCAGGAAGC
GCACATCGGACTCATCAGGAAGC
CGGACTCATCAGGAAGC
CGGACTCATCAGGAAGC
TCGGACTCATCAGGAAGC
TACATCGGACTCATCAGGAAGC
ATGCATCGGACTCATCAGGAAGC
ATCGGACTCATCAGGAAGC
CATCGGACTCATCAGGAAGC
TCCATCGGACTCATCAGGAAGC
ATACATCGGACTCATCAGGAAGC
TGACATCGGACTCATCAGGAAGC
ATCGGACTCATCAGGAAGC
GTAGCATCGGACTCATCAGGAAGC
TCGGACTCATCAGGAAGC
ACATCGGACTCATCAGGAAGC
AGACATCGGACTCATCAGGAAGC
AGACATCGGACTCATCAGGAAGC
TCGGACTCATCAGGAAGC
AACATCGGACTCATCAGGAAGC
CCATCGGACTCATCAGGAAGC
GCATCGGACTCATCAGGAAGC
CTCATCGGACTCATCAGGAAGC
GCATCGGACTCATCAGGAAGC
TCGGACTCATCAGGAAGC
ATCGGACTCATCAGGAAGC
GCATCGGACTCATCAGGAAGC
TCATCGGACTCATCAGGAAGC
CATCGGACTCATCAGGAAGC
CATCGGACTCATCAGGAAGC
TCACATCGGACTCATCAGGAAGC
TCGGACTCATCAGGAAGC
GCATCGGACTCATCAGGAAGC
ATCATCGGACTCATCAGGAAGC
TACATCGGACTCATCAGGAAGC
AACATCGGACTCATCAGGAAGC
AGCATCGGACTCATCAGGAAGC
CCATCGGACTCATCAGGAAGC

TCGGACTCATCAGGAAGC
CATCGGACTCATCAGGAAGC
TCATCGGACTCATCAGGAAGC
CATCGGACTCATCAGGAAGC
AATCATCGGACTCATCAGGAAGC
CCCATCGGACTCATCAGGAAGC
ACCATCGGACTCATCAGGAAGC
GGACTCATCAGGAAGC
GACATCGGACTCATCAGGAAGC
GGACTCATCAGGAAGC
ACATCGGACTCATCAGGAAGC
CATCGGACTCATCAGGAAGC
TCCATCGGACTCATCAGGAAGC
GGACTCATCAGGAAGC
CGGACTCATCAGGAAGC
CCATCGGACTCATCAGGAAGC
TCCATCGGACTCATCAGGAAGC
CATCGGACTCATCAGGAAGC
TCATCGGACTCATCAGGAAGC
TCATCGGACTCATCAGGAAGC
CATCGGACTCATCAGGAAGC
GCATCGGACTCATCAGGAAGC
ACATCGGACTCATCAGGAAGC
GCATCGGACTCATCAGGAAGC
CATCGGACTCATCAGGAAGC
AGCATCGGACTCATCAGGAAGC
TGCATCGGACTCATCAGGAAGC
TCATCGGACTCATCAGGAAGC
ATCGGACTCATCAGGAAGC
TACATCGGACTCATCAGGAAGC
TACATCGGACTCATCAGGAAGC
ATCATCGGACTCATCAGGAAGC
GCATCGGACTCATCAGGAAGC
GACTCATCAGGAAGC
CATCGGACTCATCAGGAAGC
TCGGACTCATCAGGAAGC
TCGGACTCATCAGGAAGC
GACTCATCAGGAAGC
GCATCGGACTCATCAGGAAGC
CATCGGACTCATCAGGAAGC
GGACTCATCAGGAAGC
ACCATCGGACTCATCAGGAAGC
TCATCGGACTCATCAGGAAGC

ATCGGACTCATCAGGAAGC
CATCGGACTCATCAGGAAGC
ACATCGGACTCATCAGGAAGC
CATCGGACTCATCAGGAAGC
TCGGACTCATCAGGAAGC
ATCATCGGACTCATCAGGAAGC
GACTCATCAGGAAGC
CATCGGACTCATCAGGAAGC
TCGGACTCATCAGGAAGC
ATCGGACTCATCAGGAAGC
TCGGACTCATCAGGAAGC
TTCATCGGACTCATCAGGAAGC
TCGGACTCATCAGGAAGC
CGGACTCATCAGGAAGC
CGGACTCATCAGGAAGC
GGACTCATCAGGAAGC
CGGACTCATCAGGAAGC
TCGGACTCATCAGGAAGC
CATCGGACTCATCAGGAAGC
CATCGGACTCATCAGGAAGC
CCATCGGACTCATCAGGAAGC
ACATCGGACTCATCAGGAAGC
ACCATCGGACTCATCAGGAAGC
CATCGGACTCATCAGGAAGC
GCATCGGACTCATCAGGAAGC
TCGGACTCATCAGGAAGC
ATCGGACTCATCAGGAAGC
GACTCATCAGGAAGC
CGCATCGGACTCATCAGGAAGC
CGCATCGGACTCATCAGGAAGC
TACATCGGACTCATCAGGAAGC
CATCGGACTCATCAGGAAGC
GCATCGGACTCATCAGGAAGC
TATCATCGGACTCATCAGGAAGC
AACATCGGACTCATCAGGAAGC
ATCGGACTCATCAGGAAGC
ATCGGACTCATCAGGAAGC
CATCGGACTCATCAGGAAGC
TCATCGGACTCATCAGGAAGC
ATCGGACTCATCAGGAAGC
CGGACTCATCAGGAAGC
TCATCGGACTCATCAGGAAGC
GCATCGGACTCATCAGGAAGC

CCATCGGACTCATCAGGAAGC

ATCGGACTCATCAGGAAGC

CCATCGGACTCATCAGGAAGC

TACATCGGACTCATCAGGAAGC

ACCATCGGACTCATCAGGAAGC

GGACTCATCAGGAAGC

TTCATCGGACTCATCAGGAAGC

CCATCGGACTCATCAGGAAGC

TCATCGGACTCATCAGGAAGC

AACATCGGACTCATCAGGAAGC

CCACATCGGACTCATCAGGAAGC

ATCGGACTCATCAGGAAGC

CATCGGACTCATCAGGAAGC

ATCGGACTCATCAGGAAGC

ATCGGACTCATCAGGAAGC

TGCATCGGACTCATCAGGAAGC

CATCGGACTCATCAGGAAGC

CCATCGGACTCATCAGGAAGC

TGACATCGGACTCATCAGGAAGC

TACATCGGACTCATCAGGAAGC

CATCGGACTCATCAGGAAGC

TACATCGGACTCATCAGGAAGC

CATCGGACTCATCAGGAAGC

AAACATCGGACTCATCAGGAAGC

AACATCGGACTCATCAGGAAGC

CGGACTCATCAGGAAGC

CATCGGACTCATCAGGAAGC

ACCATCGGACTCATCAGGAAGC

GCATCGGACTCATCAGGAAGC

AGTCATCGGACTCATCAGGAAGC

ATCGGACTCATCAGGAAGC

CATCGGACTCATCAGGAAGC

CGGACTCATCAGGAAGC

TCATCGGACTCATCAGGAAGC

TCATCGGACTCATCAGGAAGC

TCATCGGACTCATCAGGAAGC

CATCGGACTCATCAGGAAGC

TTCATCGGACTCATCAGGAAGC

TCGGACTCATCAGGAAGC

TCGGACTCATCAGGAAGC

CATCGGACTCATCAGGAAGC

ATCGGACTCATCAGGAAGC

ACGCATCGGACTCATCAGGAAGC

CATCGGACTCATCAGGAAGC
CATCGGACTCATCAGGAAGC
ATCGGACTCATCAGGAAGC
CACATCGGACTCATCAGGAAGC
ATCGGACTCATCAGGAAGC
CATCGGACTCATCAGGAAGC
TCCATCGGACTCATCAGGAAGC
ATCGGACTCATCAGGAAGC
ATCGGACTCATCAGGAAGC
GGGCATCGGACTCATCAGGAAGC
AGACATCGGACTCATCAGGAAGC
CATCGGACTCATCAGGAAGC
CATCGGACTCATCAGGAAGC
GTCATCGGACTCATCAGGAAGC
CGGACTCATCAGGAAGC
TCATCGGACTCATCAGGAAGC
TCGCATCGGACTCATCAGGAAGC
TCGGACTCATCAGGAAGC
GCATCGGACTCATCAGGAAGC
ACACATCGGACTCATCAGGAAGC
ATCGGACTCATCAGGAAGC
CATCGGACTCATCAGGAAGC
CATCGGACTCATCAGGAAGC
CATCGGACTCATCAGGAAGC
ATCGGACTCATCAGGAAGC
ATGCATCGGACTCATCAGGAAGC
TCATCGGACTCATCAGGAAGC
CCATCGGACTCATCAGGAAGC
GCATCGGACTCATCAGGAAGC
CGGACTCATCAGGAAGC
ACATCGGACTCATCAGGAAGC
ATCGGACTCATCAGGAAGC
TCGGACTCATCAGGAAGC
CATCGGACTCATCAGGAAGC
CTCATCGGACTCATCAGGAAGC
ATCGGACTCATCAGGAAGC
TCATCGGACTCATCAGGAAGC
CATCGGACTCATCAGGAAGC
TCATCGGACTCATCAGGAAGC
CATCGGACTCATCAGGAAGC
CATCGGACTCATCAGGAAGC
ATCGGACTCATCAGGAAGC
GCATCGGACTCATCAGGAAGC

ATGCATCGGACTCATCAGGAAGC
CCATCGGACTCATCAGGAAGC
TCATCGGACTCATCAGGAAGC
GGACTCATCAGGAAGC
CGGACTCATCAGGAAGC
ATCATCGGACTCATCAGGAAGC
CATCGGACTCATCAGGAAGC
ATACATCGGACTCATCAGGAAGC
ACTCATCAGGAAGC
GGCATCGGACTCATCAGGAAGC
GGCATCGGACTCATCAGGAAGC
TCGGACTCATCAGGAAGC
TCGGACTCATCAGGAAGC
ACCATCGGACTCATCAGGAAGC
CATCGGACTCATCAGGAAGC
TCGGACTCATCAGGAAGC
CGGACTCATCAGGAAGC
CATCGGACTCATCAGGAAGC
TCATCGGACTCATCAGGAAGC
TTCATCGGACTCATCAGGAAGC
CATCGGACTCATCAGGAAGC
CAGACATCGGACTCATCAGGAAGC
CAGCTCATCGGACTCATCAGGAAGC
CATCGGACTCATCAGGAAGC
CATCGGACTCATCAGGAAGC
CATCGGACTCATCAGGAAGC
GCATCGGACTCATCAGGAAGC
GCATCGGACTCATCAGGAAGC
ACATCGGACTCATCAGGAAGC
TTCATCGGACTCATCAGGAAGC
CGGACTCATCAGGAAGC
CATCGGACTCATCAGGAAGC
CATCGGACTCATCAGGAAGC
ATCGGACTCATCAGGAAGC
CCATCGGACTCATCAGGAAGC
TCGGACTCATCAGGAAGC
GCATCGGACTCATCAGGAAGC
CATCGGACTCATCAGGAAGC
CATCGGACTCATCAGGAAGC
TTCATCGGACTCATCAGGAAGC
TCGGACTCATCAGGAAGC
GGACTCATCAGGAAGC
TCACATCGGACTCATCAGGAAGC

TCACATCGGACTCATCAGGAAGC
CATCGGACTCATCAGGAAGC
CATCGGACTCATCAGGAAGC
TCGGACTCATCAGGAAGC
CATCGGACTCATCAGGAAGC
GACATCGGACTCATCAGGAAGC
ACATCGGACTCATCAGGAAGC
ATCGGACTCATCAGGAAGC
CCCATCGGACTCATCAGGAAGC
TCATCGGACTCATCAGGAAGC
CGGACTCATCAGGAAGC
GCCATCGGACTCATCAGGAAGC
GGCATCGGACTCATCAGGAAGC
CATCGGACTCATCAGGAAGC
CATCGGACTCATCAGGAAGC
ATCGGACTCATCAGGAAGC
GCCATCGGACTCATCAGGAAGC
ATCATCGGACTCATCAGGAAGC
CCATCGGACTCATCAGGAAGC
ATCGGACTCATCAGGAAGC
TCATCGGACTCATCAGGAAGC
CATCGGACTCATCAGGAAGC
GACTCATCAGGAAGC
CATCGGACTCATCAGGAAGC
ATCATCGGACTCATCAGGAAGC
CATCGGACTCATCAGGAAGC
TCATCGGACTCATCAGGAAGC
ACATCGGACTCATCAGGAAGC
CATCGGACTCATCAGGAAGC
CATCGGACTCATCAGGAAGC
GACTCATCAGGAAGC
TTCATCGGACTCATCAGGAAGC
ACATCGGACTCATCAGGAAGC
CTTCATCGGACTCATCAGGAAGC
ATCGGACTCATCAGGAAGC
TCGGACTCATCAGGAAGC
TCATCGGACTCATCAGGAAGC
CATCGGACTCATCAGGAAGC
CATCGGACTCATCAGGAAGC
CATCGGACTCATCAGGAAGC
TGCATCGGACTCATCAGGAAGC
CCATCGGACTCATCAGGAAGC
ATCGGACTCATCAGGAAGC

ACCATCGGACTCATCAGGAAGC
ATCGGACTCATCAGGAAGC
ACATCGGACTCATCAGGAAGC
ATCATCGGACTCATCAGGAAGC
ACCATCGGACTCATCAGGAAGC
ACCATCGGACTCATCAGGAAGC
TGCATCGGACTCATCAGGAAGC
ACATCGGACTCATCAGGAAGC
CATCGGACTCATCAGGAAGC
ACATCGGACTCATCAGGAAGC
GTCATCGGACTCATCAGGAAGC
AGCATCGGACTCATCAGGAAGC
ATCGGACTCATCAGGAAGC
ATCATCGGACTCATCAGGAAGC
ATCATCGGACTCATCAGGAAGC
ACATCGGACTCATCAGGAAGC
ACATCGGACTCATCAGGAAGC
GCATCGGACTCATCAGGAAGC
GCATCGGACTCATCAGGAAGC
GACATCGGACTCATCAGGAAGC
ACATCGGACTCATCAGGAAGC
GCATCGGACTCATCAGGAAGC
TCGGACTCATCAGGAAGC
GGCATCGGACTCATCAGGAAGC
TCGGACTCATCAGGAAGC
ATCGGACTCATCAGGAAGC
GGACTCATCAGGAAGC
GGACTCATCAGGAAGC
ATCGGACTCATCAGGAAGC
ATCGGACTCATCAGGAAGC
AGCATCGGACTCATCAGGAAGC
TTCATCGGACTCATCAGGAAGC
ATCGGACTCATCAGGAAGC
CATCGGACTCATCAGGAAGC
AAGCATCGGACTCATCAGGAAGC
TTTCATCGGACTCATCAGGAAGC
ATCGGACTCATCAGGAAGC
ATCGGACTCATCAGGAAGC
TCATCGGACTCATCAGGAAGC
TCATCGGACTCATCAGGAAGC
TCGGACTCATCAGGAAGC
ATCGGACTCATCAGGAAGC
ACATCGGACTCATCAGGAAGC

GACTCATCAGGAAGC
CATCGGACTCATCAGGAAGC
ATGCCATCGGACTCATCAGGAAGC
CATCGGACTCATCAGGAAGC
ATCGGACTCATCAGGAAGC
ATCGGACTCATCAGGAAGC
CGGACTCATCAGGAAGC
ATCATCGGACTCATCAGGAAGC
GACTCATCAGGAAGC
CATCGGACTCATCAGGAAGC
GGACATCGGACTCATCAGGAAGC
TCGGACTCATCAGGAAGC
GGACTCATCAGGAAGC
ATCGGACTCATCAGGAAGC
ACATCGGACTCATCAGGAAGC
ATACATCGGACTCATCAGGAAGC
GACATCGGACTCATCAGGAAGC
CGGACTCATCAGGAAGC
TCATCGGACTCATCAGGAAGC
GGACTCATCAGGAAGC
GACTCATCAGGAAGC
ATCGGACTCATCAGGAAGC
CATCGGACTCATCAGGAAGC
CATCGGACTCATCAGGAAGC
CATCGGACTCATCAGGAAGC
CATCGGACTCATCAGGAAGC
TCGGACTCATCAGGAAGC
TGCCATCGGACTCATCAGGAAGC
GACATCGGACTCATCAGGAAGC
CACATCGGACTCATCAGGAAGC
TCGGACTCATCAGGAAGC
GCATCGGACTCATCAGGAAGC
CATCGGACTCATCAGGAAGC
CATCGGACTCATCAGGAAGC
ACCATCGGACTCATCAGGAAGC
ATCGGACTCATCAGGAAGC
TCATCGGACTCATCAGGAAGC
TTCATCGGACTCATCAGGAAGC
ATCGGACTCATCAGGAAGC
AACATCGGACTCATCAGGAAGC
GCATCGGACTCATCAGGAAGC
CATCGGACTCATCAGGAAGC
GACTCATCAGGAAGC

TCATCGGACTCATCAGGAAGC
ATCGGACTCATCAGGAAGC
GACATCGGACTCATCAGGAAGC
GACATCGGACTCATCAGGAAGC
GCATCGGACTCATCAGGAAGC
TACATCGGACTCATCAGGAAGC
ATCGGACTCATCAGGAAGC
CCGCATCGGACTCATCAGGAAGC
ACATCGGACTCATCAGGAAGC
CATCGGACTCATCAGGAAGC
CATCGGACTCATCAGGAAGC
CATCGGACTCATCAGGAAGC
GCATCGGACTCATCAGGAAGC
CATCGGACTCATCAGGAAGC
TCATCGGACTCATCAGGAAGC
CGGACTCATCAGGAAGC
GACTCATCAGGAAGC
AGCATCGGACTCATCAGGAAGC
AGCATCGGACTCATCAGGAAGC
AACATCGGACTCATCAGGAAGC
AACATCGGACTCATCAGGAAGC
CATCGGACTCATCAGGAAGC
ATTCATCGGACTCATCAGGAAGC
CCATCGGACTCATCAGGAAGC
CGGACTCATCAGGAAGC
TACATCGGACTCATCAGGAAGC
CATCGGACTCATCAGGAAGC
TCATCGGACTCATCAGGAAGC
ATCGGACTCATCAGGAAGC
ACATCGGACTCATCAGGAAGC
CATCGGACTCATCAGGAAGC
CCATCGGACTCATCAGGAAGC
TTCTCATCGGACTCATCAGGAAGC
CATCGGACTCATCAGGAAGC
ATCGGACTCATCAGGAAGC
ATCGGACTCATCAGGAAGC
TCATCGGACTCATCAGGAAGC
TCATCGGACTCATCAGGAAGC
TACATCGGACTCATCAGGAAGC
GCATCGGACTCATCAGGAAGC
CATCGGACTCATCAGGAAGC
CATCGGACTCATCAGGAAGC
TCATCGGACTCATCAGGAAGC

ACCATCGGACTCATCAGGAAGC
GACCATCGGACTCATCAGGAAGC
TCGGACTCATCAGGAAGC
TACATCGGACTCATCAGGAAGC
CATCGGACTCATCAGGAAGC
CATCGGACTCATCAGGAAGC
TACATCGGACTCATCAGGAAGC
ATCATCGGACTCATCAGGAAGC
GACTCATCAGGAAGC
TCGGACTCATCAGGAAGC
CGGACTCATCAGGAAGC
CATCGGACTCATCAGGAAGC
CATCGGACTCATCAGGAAGC
CGGACTCATCAGGAAGC
CGGACTCATCAGGAAGC
ATCGGACTCATCAGGAAGC
ACATCGGACTCATCAGGAAGC
TACATCGGACTCATCAGGAAGC
GCATCGGACTCATCAGGAAGC
ACATCGGACTCATCAGGAAGC
ACATCGGACTCATCAGGAAGC
CATCGGACTCATCAGGAAGC
TCGGACTCATCAGGAAGC
CATCGGACTCATCAGGAAGC
CATCGGACTCATCAGGAAGC
GGTCATCGGACTCATCAGGAAGC
ATCGGACTCATCAGGAAGC
CATCGGACTCATCAGGAAGC
CATCGGACTCATCAGGAAGC
CATCGGACTCATCAGGAAGC
CGGACTCATCAGGAAGC
CATCGGACTCATCAGGAAGC
CATCGGACTCATCAGGAAGC
CACATCGGACTCATCAGGAAGC
ATCATCGGACTCATCAGGAAGC
ATCGGACTCATCAGGAAGC
CCATCGGACTCATCAGGAAGC
AGCATCGGACTCATCAGGAAGC
GACTCATCAGGAAGC
ATCGGACTCATCAGGAAGC
ATCGGACTCATCAGGAAGC
ATCGGACTCATCAGGAAGC
CATACATCGGACTCATCAGGAAGC

ACATCGGACTCATCAGGAAGC
ACATCGGACTCATCAGGAAGC
CATCGGACTCATCAGGAAGC
TATCATCGGACTCATCAGGAAGC
CGGACTCATCAGGAAGC
CACATCGGACTCATCAGGAAGC
AACATCGGACTCATCAGGAAGC
ACATCGGACTCATCAGGAAGC
TCATCGGACTCATCAGGAAGC
ATCGGACTCATCAGGAAGC
ACATCGGACTCATCAGGAAGC
CATCGGACTCATCAGGAAGC
CATCGGACTCATCAGGAAGC
CCATCGGACTCATCAGGAAGC
CATCGGACTCATCAGGAAGC
CCATCGGACTCATCAGGAAGC
TTCATCGGACTCATCAGGAAGC
ATCGGACTCATCAGGAAGC
AGCCATCGGACTCATCAGGAAGC
CCATCGGACTCATCAGGAAGC
ATCGGACTCATCAGGAAGC
GTGCATCGGACTCATCAGGAAGC
ATCGGACTCATCAGGAAGC
CATCGGACTCATCAGGAAGC
ATCTCATCGGACTCATCAGGAAGC
AAACATCGGACTCATCAGGAAGC
ATCGGACTCATCAGGAAGC
ATCGGACTCATCAGGAAGC
CATCGGACTCATCAGGAAGC
CATCGGACTCATCAGGAAGC
GCATCGGACTCATCAGGAAGC
CATCGGACTCATCAGGAAGC
ACCATCGGACTCATCAGGAAGC
CCATCGGACTCATCAGGAAGC
CCATCGGACTCATCAGGAAGC
ACATCGGACTCATCAGGAAGC
TCATCGGACTCATCAGGAAGC
TAACATCGGACTCATCAGGAAGC
CGGACTCATCAGGAAGC
TCATCGGACTCATCAGGAAGC
ACATCGGACTCATCAGGAAGC
CATCGGACTCATCAGGAAGC
ACATCGGACTCATCAGGAAGC

TCGGACTCATCAGGAAGC
GCATCGGACTCATCAGGAAGC
GCATCGGACTCATCAGGAAGC
AGCATCGGACTCATCAGGAAGC
AACATCGGACTCATCAGGAAGC
CATCGGACTCATCAGGAAGC
ACATCGGACTCATCAGGAAGC
TCATCGGACTCATCAGGAAGC
TCGGACTCATCAGGAAGC
AACCATCGGACTCATCAGGAAGC
CATCGGACTCATCAGGAAGC
CATCGGACTCATCAGGAAGC
ACTCATCAGGAAGC
CATCGGACTCATCAGGAAGC
CATCGGACTCATCAGGAAGC
CATCGGACTCATCAGGAAGC
CGGACTCATCAGGAAGC
TTTCCATCGGACTCATCAGGAAGC
ACATCGGACTCATCAGGAAGC
GCATCGGACTCATCAGGAAGC
TCGGACTCATCAGGAAGC
TCGGACTCATCAGGAAGC
ATCGGACTCATCAGGAAGC
TCATCAGGAAGC
ATCGGACTCATCAGGAAGC
ATCGGACTCATCAGGAAGC
ACTCATCGGACTCATCAGGAAGC
AGCATCGGACTCATCAGGAAGC
CATCGGACTCATCAGGAAGC
ATCGGACTCATCAGGAAGC
CATCGGACTCATCAGGAAGC
ACATCGGACTCATCAGGAAGC
TCGGACTCATCAGGAAGC
ACATCGGACTCATCAGGAAGC
TGACATCGGACTCATCAGGAAGC
TCGGACTCATCAGGAAGC
CATCGGACTCATCAGGAAGC
CATCGGACTCATCAGGAAGC
CGGACTCATCAGGAAGC
ATCGGACTCATCAGGAAGC
CATCGGACTCATCAGGAAGC
CCATCGGACTCATCAGGAAGC
CATCGGACTCATCAGGAAGC

CGCATCGGACTCATCAGGAAGC
CGCATCGGACTCATCAGGAAGC
TCGGACTCATCAGGAAGC
CATCGGACTCATCAGGAAGC
CATCGGACTCATCAGGAAGC
ATCGGACTCATCAGGAAGC
ACTCATCGGACTCATCAGGAAGC
ACATCGGACTCATCAGGAAGC
CATCGGACTCATCAGGAAGC
ATCATCGGACTCATCAGGAAGC
TCATCGGACTCATCAGGAAGC
TCGGACTCATCAGGAAGC
ATCCATCGGACTCATCAGGAAGC
AACATCGGACTCATCAGGAAGC
AACATCGGACTCATCAGGAAGC
AGCATCGGACTCATCAGGAAGC
CATCGGACTCATCAGGAAGC
TACATCGGACTCATCAGGAAGC
TACATCGGACTCATCAGGAAGC
CCATCGGACTCATCAGGAAGC
GGACTCATCAGGAAGC
CGGACTCATCAGGAAGC
ATCGGACTCATCAGGAAGC
TCGGACTCATCAGGAAGC
ATCGGACTCATCAGGAAGC
AGCATCGGACTCATCAGGAAGC
CATCGGACTCATCAGGAAGC
CATCGGACTCATCAGGAAGC
ACATCGGACTCATCAGGAAGC
CATCGGACTCATCAGGAAGC
CATCGGACTCATCAGGAAGC
ACATCGGACTCATCAGGAAGC
TGCATCGGACTCATCAGGAAGC
CGGACTCATCAGGAAGC
ATCGGACTCATCAGGAAGC
AGCATCGGACTCATCAGGAAGC
ACATCGGACTCATCAGGAAGC
TCATCGGACTCATCAGGAAGC
ATACCATCGGACTCATCAGGAAGC
CATCGGACTCATCAGGAAGC
CCATCGGACTCATCAGGAAGC
CCCATCGGACTCATCAGGAAGC
TACATCGGACTCATCAGGAAGC

CATCGGACTCATCAGGAAGC
CATCGGACTCATCAGGAAGC
ACATCGGACTCATCAGGAAGC
GTGCATCGGACTCATCAGGAAGC
GCATCGGACTCATCAGGAAGC
CACATCGGACTCATCAGGAAGC
CACATCGGACTCATCAGGAAGC
CATCGGACTCATCAGGAAGC
GGACTCATCAGGAAGC
AACATCGGACTCATCAGGAAGC
ATCATCGGACTCATCAGGAAGC
ATCATCGGACTCATCAGGAAGC
ATCGGACTCATCAGGAAGC
ACATCGGACTCATCAGGAAGC
TCGGACTCATCAGGAAGC
GCATCGGACTCATCAGGAAGC
TCGGACTCATCAGGAAGC
GCATCGGACTCATCAGGAAGC
TCATCGGACTCATCAGGAAGC
TCCATCGGACTCATCAGGAAGC
GCATCGGACTCATCAGGAAGC
TTACATCGGACTCATCAGGAAGC
ATCGGACTCATCAGGAAGC
ATCATCGGACTCATCAGGAAGC
TCCATCGGACTCATCAGGAAGC
TCCATCGGACTCATCAGGAAGC
CGGACTCATCAGGAAGC
ATCGGACTCATCAGGAAGC
TGCATCGGACTCATCAGGAAGC
ACCATCGGACTCATCAGGAAGC
CGGACTCATCAGGAAGC
ATCGGACTCATCAGGAAGC
ATCGGACTCATCAGGAAGC
CATCGGACTCATCAGGAAGC
CCATCGGACTCATCAGGAAGC
CATCGGACTCATCAGGAAGC
CCATCGGACTCATCAGGAAGC
TCGGACTCATCAGGAAGC
ACATCGGACTCATCAGGAAGC
TCGGACTCATCAGGAAGC
CATCGGACTCATCAGGAAGC
CGGACTCATCAGGAAGC
CATCGGACTCATCAGGAAGC

ATCGGACTCATCAGGAAGC
ATCGGACTCATCAGGAAGC
ACATCGGACTCATCAGGAAGC
AACATCGGACTCATCAGGAAGC
AGGCCATCGGACTCATCAGGAAGC
ACATCGGACTCATCAGGAAGC
CATCGGACTCATCAGGAAGC
CATCGGACTCATCAGGAAGC
CATCGGACTCATCAGGAAGC
CATCGGACTCATCAGGAAGC
TCATCGGACTCATCAGGAAGC
GACATCGGACTCATCAGGAAGC
CGGACTCATCAGGAAGC
TCGGACTCATCAGGAAGC
CATCGGACTCATCAGGAAGC
TGCATCGGACTCATCAGGAAGC
TACATCGGACTCATCAGGAAGC
TACATCGGACTCATCAGGAAGC
CATCGGACTCATCAGGAAGC
TCATCGGACTCATCAGGAAGC
GCATCGGACTCATCAGGAAGC
CCATCGGACTCATCAGGAAGC
CATCGGACTCATCAGGAAGC
TCATCGGACTCATCAGGAAGC
ATCGGACTCATCAGGAAGC
TCATCGGACTCATCAGGAAGC
ACATCGGACTCATCAGGAAGC
ATCGGACTCATCAGGAAGC
GCATCGGACTCATCAGGAAGC
CAACCATCGGACTCATCAGGAAGC
ATCGGACTCATCAGGAAGC
ATCGGACTCATCAGGAAGC
GATTCATCGGACTCATCAGGAAGC
CATCGGACTCATCAGGAAGC
CATCGGACTCATCAGGAAGC
ATCGGACTCATCAGGAAGC
CTCATCGGACTCATCAGGAAGC
CCATCGGACTCATCAGGAAGC
CATCGGACTCATCAGGAAGC
CATCGGACTCATCAGGAAGC
CATCGGACTCATCAGGAAGC
AGACATCGGACTCATCAGGAAGC
TTCATCGGACTCATCAGGAAGC

TCATCGGACTCATCAGGAAGC
TCGGACTCATCAGGAAGC
ATCGGACTCATCAGGAAGC
ATCGGACTCATCAGGAAGC
GGACTCATCAGGAAGC
ATCGGACTCATCAGGAAGC
CATCGGACTCATCAGGAAGC
TCGGACTCATCAGGAAGC
CATCGGACTCATCAGGAAGC
ACTCATCGGACTCATCAGGAAGC
GACTCATCAGGAAGC
GGACTCATCAGGAAGC
TCGGACTCATCAGGAAGC
ACATCGGACTCATCAGGAAGC
TCGGACTCATCAGGAAGC
GACATCGGACTCATCAGGAAGC
CCATCGGACTCATCAGGAAGC
CATCGGACTCATCAGGAAGC
GACTCATCAGGAAGC
TCGGACTCATCAGGAAGC
ACATCGGACTCATCAGGAAGC
ACATCGGACTCATCAGGAAGC
CATCGGACTCATCAGGAAGC
TTCCATCGGACTCATCAGGAAGC
ACATCGGACTCATCAGGAAGC
CCATCGGACTCATCAGGAAGC
TATGCATCGGACTCATCAGGAAGC
ATCGGACTCATCAGGAAGC
ATCGGACTCATCAGGAAGC
TCATCGGACTCATCAGGAAGC
GACATCGGACTCATCAGGAAGC
CATCGGACTCATCAGGAAGC
ATCGGACTCATCAGGAAGC
ACATCGGACTCATCAGGAAGC
TCATCGGACTCATCAGGAAGC
CATCGGACTCATCAGGAAGC
TCATCGGACTCATCAGGAAGC
TCGGACTCATCAGGAAGC
CATCGGACTCATCAGGAAGC
AGACATCGGACTCATCAGGAAGC
AGACATCGGACTCATCAGGAAGC
AACATCGGACTCATCAGGAAGC
ATCGGACTCATCAGGAAGC

CATCGGACTCATCAGGAAGC
GACTCATCAGGAAGC
CATCGGACTCATCAGGAAGC
GGCATCGGACTCATCAGGAAGC
GCCATCGGACTCATCAGGAAGC
GCCATCGGACTCATCAGGAAGC
ATCGGACTCATCAGGAAGC
CATCATCGGACTCATCAGGAAGC
TCGGACTCATCAGGAAGC
CATCGGACTCATCAGGAAGC
TCGGACTCATCAGGAAGC
CCATCGGACTCATCAGGAAGC
CATCGGACTCATCAGGAAGC
TCGGACTCATCAGGAAGC
ACATCGGACTCATCAGGAAGC
TCATCGGACTCATCAGGAAGC
TCATCGGACTCATCAGGAAGC
CGGACTCATCAGGAAGC
TGCATCGGACTCATCAGGAAGC
ATCGGACTCATCAGGAAGC
AACATCGGACTCATCAGGAAGC
GCATCGGACTCATCAGGAAGC
AGGCATCGGACTCATCAGGAAGC
TCGGACTCATCAGGAAGC
AGCATCGGACTCATCAGGAAGC
TCGGACTCATCAGGAAGC
ACATCGGACTCATCAGGAAGC
CCATCGGACTCATCAGGAAGC
GCATCGGACTCATCAGGAAGC
CTTCCCATCGGACTCATCAGGAAGC
GGCATCGGACTCATCAGGAAGC
CATCGGACTCATCAGGAAGC
TCATCGGACTCATCAGGAAGC
CGGACTCATCAGGAAGC
CATCGGACTCATCAGGAAGC
CATCGGACTCATCAGGAAGC
TACATCGGACTCATCAGGAAGC
ATCGGACTCATCAGGAAGC
GGCATCGGACTCATCAGGAAGC
TCGGACTCATCAGGAAGC
GCATCGGACTCATCAGGAAGC
GACTCATCAGGAAGC
ATCGGACTCATCAGGAAGC

TCATCGGACTCATCAGGAAGC
ACATCGGACTCATCAGGAAGC
CCATCGGACTCATCAGGAAGC
ACATCGGACTCATCAGGAAGC
GTTCATCGGACTCATCAGGAAGC
GCATCGGACTCATCAGGAAGC
CATCGGACTCATCAGGAAGC
TCATCGGACTCATCAGGAAGC
TCGGACTCATCAGGAAGC
CATCGGACTCATCAGGAAGC
CATCGGACTCATCAGGAAGC
CATCGGACTCATCAGGAAGC
ATCGGACTCATCAGGAAGC
ATGCATCGGACTCATCAGGAAGC
ATCGGACTCATCAGGAAGC
ACATCGGACTCATCAGGAAGC
AGCATCGGACTCATCAGGAAGC
TCATCGGACTCATCAGGAAGC
GAACATCGGACTCATCAGGAAGC
CATCGGACTCATCAGGAAGC
TTGCCATCGGACTCATCAGGAAGC
TCGGACTCATCAGGAAGC
GTTCATCGGACTCATCAGGAAGC
GTTCATCGGACTCATCAGGAAGC
TCGGACTCATCAGGAAGC
TTCATCGGACTCATCAGGAAGC
TGCATCGGACTCATCAGGAAGC
CATCGGACTCATCAGGAAGC
TCATCGGACTCATCAGGAAGC
GCATCGGACTCATCAGGAAGC
ATCGGACTCATCAGGAAGC
TCATCGGACTCATCAGGAAGC
TCATCGGACTCATCAGGAAGC
TACATCGGACTCATCAGGAAGC
GACTCATCAGGAAGC
ATCGGACTCATCAGGAAGC
ATCCATCGGACTCATCAGGAAGC
AACATCGGACTCATCAGGAAGC
AACATCGGACTCATCAGGAAGC
ATTCCATCGGACTCATCAGGAAGC
ATCGGACTCATCAGGAAGC
ATCGGACTCATCAGGAAGC
ACATCGGACTCATCAGGAAGC

ACATCGGACTCATCAGGAAGC
CATCGGACTCATCAGGAAGC
CATCGGACTCATCAGGAAGC
GACTCATCAGGAAGC
ATCGGACTCATCAGGAAGC
ACATCGGACTCATCAGGAAGC
TGCATCGGACTCATCAGGAAGC
ACTCATCAGGAAGC
ATCGGACTCATCAGGAAGC
CGGACTCATCAGGAAGC
TCGGACTCATCAGGAAGC
CATCGGACTCATCAGGAAGC
GACATCGGACTCATCAGGAAGC
TCGGACTCATCAGGAAGC
TCGGACTCATCAGGAAGC
TCCATCGGACTCATCAGGAAGC
CATCGGACTCATCAGGAAGC
TCGGACTCATCAGGAAGC
CATCGGACTCATCAGGAAGC
GTCATCGGACTCATCAGGAAGC
ATCGGACTCATCAGGAAGC
CATCGGACTCATCAGGAAGC
TCGGACTCATCAGGAAGC
ACATCGGACTCATCAGGAAGC
GACATCGGACTCATCAGGAAGC
GTCATCGGACTCATCAGGAAGC
GCCATCGGACTCATCAGGAAGC
TCATCGGACTCATCAGGAAGC
AATCATCGGACTCATCAGGAAGC
CTACATCGGACTCATCAGGAAGC
ATTCATCGGACTCATCAGGAAGC
ATCATCGGACTCATCAGGAAGC
CATCGGACTCATCAGGAAGC
AACATCGGACTCATCAGGAAGC
CATCGGACTCATCAGGAAGC
CATCGGACTCATCAGGAAGC
GGACTCATCAGGAAGC
CGGACTCATCAGGAAGC
ATCGGACTCATCAGGAAGC
GACTCATCAGGAAGC
CATCGGACTCATCAGGAAGC
ATCATCGGACTCATCAGGAAGC
CGGACTCATCAGGAAGC

ATCGGACTCATCAGGAAGC
ATCGGACTCATCAGGAAGC
TGCATCGGACTCATCAGGAAGC
CGGACTCATCAGGAAGC
TCGGACTCATCAGGAAGC
CCATCGGACTCATCAGGAAGC
CATCGGACTCATCAGGAAGC
ATCGGACTCATCAGGAAGC
CATCGGACTCATCAGGAAGC
CGGACTCATCAGGAAGC
CATCGGACTCATCAGGAAGC
TCGGACTCATCAGGAAGC
CATCGGACTCATCAGGAAGC
TCGGACTCATCAGGAAGC
TACATCGGACTCATCAGGAAGC
ACATCGGACTCATCAGGAAGC
GACTCATCAGGAAGC
AGCATCGGACTCATCAGGAAGC
TGCATCGGACTCATCAGGAAGC
ACATCGGACTCATCAGGAAGC
GGACTCATCAGGAAGC
GCATCGGACTCATCAGGAAGC
GCATCGGACTCATCAGGAAGC
TCATCGGACTCATCAGGAAGC
CATCGGACTCATCAGGAAGC
CATCGGACTCATCAGGAAGC
GGCATCGGACTCATCAGGAAGC
ATCGGACTCATCAGGAAGC
GGACTCATCAGGAAGC
TCGGACTCATCAGGAAGC
ACATCGGACTCATCAGGAAGC
ACATCGGACTCATCAGGAAGC
CGGACTCATCAGGAAGC
CCCATCGGACTCATCAGGAAGC
TTCATCGGACTCATCAGGAAGC
GCATCGGACTCATCAGGAAGC
TTTCATCGGACTCATCAGGAAGC
ATCGGACTCATCAGGAAGC
TCCATCGGACTCATCAGGAAGC
CATCGGACTCATCAGGAAGC
ACATCGGACTCATCAGGAAGC
ACATCGGACTCATCAGGAAGC
TGCATCGGACTCATCAGGAAGC

CATCGGACTCATCAGGAAGC
TCATCGGACTCATCAGGAAGC
CGGACTCATCAGGAAGC
ATCGGACTCATCAGGAAGC
TCCATCGGACTCATCAGGAAGC
GGACTCATCAGGAAGC
GGACTCATCAGGAAGC
TTTCATCGGACTCATCAGGAAGC
CATCGGACTCATCAGGAAGC
TCGGACTCATCAGGAAGC
GCTACATCGGACTCATCAGGAAGC
GCATCGGACTCATCAGGAAGC
GGACTCATCAGGAAGC
ACATCGGACTCATCAGGAAGC
ATCGGACTCATCAGGAAGC
TCGGACTCATCAGGAAGC
ACATCGGACTCATCAGGAAGC
TCATCGGACTCATCAGGAAGC
ATCGGACTCATCAGGAAGC
TCGGACTCATCAGGAAGC
CATCGGACTCATCAGGAAGC
TCGGACTCATCAGGAAGC
AACATCGGACTCATCAGGAAGC
ATCGGACTCATCAGGAAGC
CATCGGACTCATCAGGAAGC
TCGGACTCATCAGGAAGC
CATCGGACTCATCAGGAAGC
TCATCGGACTCATCAGGAAGC
CGGACTCATCAGGAAGC
ATCGGACTCATCAGGAAGC
GCATCGGACTCATCAGGAAGC
AATCATCGGACTCATCAGGAAGC
GCATCGGACTCATCAGGAAGC
GAACATCGGACTCATCAGGAAGC
CATCGGACTCATCAGGAAGC
TCGGACTCATCAGGAAGC
CATCGGACTCATCAGGAAGC
CATCGGACTCATCAGGAAGC
TTCATCGGACTCATCAGGAAGC
ATCGGACTCATCAGGAAGC
ATCGGACTCATCAGGAAGC
CATCGGACTCATCAGGAAGC
CATCGGACTCATCAGGAAGC

ACATCGGACTCATCAGGAAGC
CCACATCGGACTCATCAGGAAGC
GATCATCGGACTCATCAGGAAGC
ATCGGACTCATCAGGAAGC
TCCATCGGACTCATCAGGAAGC
GGACTCATCAGGAAGC
GCTCATCGGACTCATCAGGAAGC
ATCGGACTCATCAGGAAGC
ATCGGACTCATCAGGAAGC
CCATCGGACTCATCAGGAAGC
ACATCGGACTCATCAGGAAGC
GAACATCGGACTCATCAGGAAGC
TCATCGGACTCATCAGGAAGC
ATCGGACTCATCAGGAAGC
TACATCGGACTCATCAGGAAGC
CGGACTCATCAGGAAGC
GGGCATCGGACTCATCAGGAAGC
ACATCGGACTCATCAGGAAGC
ATCGGACTCATCAGGAAGC
CATCGGACTCATCAGGAAGC
GGACTCATCAGGAAGC
ATCGGACTCATCAGGAAGC
TCGGACTCATCAGGAAGC
TCGGACTCATCAGGAAGC
ATTCATCGGACTCATCAGGAAGC
TAACATCGGACTCATCAGGAAGC
GCATCGGACTCATCAGGAAGC
CCATCGGACTCATCAGGAAGC
CATCGGACTCATCAGGAAGC
ATCATCGGACTCATCAGGAAGC
ACTCATCAGGAAGC
ACATCGGACTCATCAGGAAGC
ATCGGACTCATCAGGAAGC
ATCGGACTCATCAGGAAGC
ACATCGGACTCATCAGGAAGC
ATCGGACTCATCAGGAAGC
TCCATCGGACTCATCAGGAAGC
ACATCGGACTCATCAGGAAGC
ATCGGACTCATCAGGAAGC
ACATCGGACTCATCAGGAAGC
ACACATCGGACTCATCAGGAAGC
ATCGGACTCATCAGGAAGC
ACATCGGACTCATCAGGAAGC

CATCGGACTCATCAGGAAGC
ACATCGGACTCATCAGGAAGC
CATCGGACTCATCAGGAAGC
CATCGGACTCATCAGGAAGC
CATCGGACTCATCAGGAAGC
CATCGGACTCATCAGGAAGC
CATCGGACTCATCAGGAAGC
ATCGGACTCATCAGGAAGC
TCCACATCGGACTCATCAGGAAGC
CATCGGACTCATCAGGAAGC
CATCGGACTCATCAGGAAGC
GGACTCATCAGGAAGC
CGGACTCATCAGGAAGC
TCATCGGACTCATCAGGAAGC
CATCGGACTCATCAGGAAGC
ATCGGACTCATCAGGAAGC
CATCGGACTCATCAGGAAGC
ACATCGGACTCATCAGGAAGC
CATCGGACTCATCAGGAAGC
CATCGGACTCATCAGGAAGC
CATCGGACTCATCAGGAAGC
TAGCCATCGGACTCATCAGGAAGC
ATCGGACTCATCAGGAAGC
CATCGGACTCATCAGGAAGC
CATCGGACTCATCAGGAAGC
TAACATCGGACTCATCAGGAAGC
ATCGGACTCATCAGGAAGC
GCATCGGACTCATCAGGAAGC
TCGGACTCATCAGGAAGC
CGGACTCATCAGGAAGC
CATCGGACTCATCAGGAAGC
CATCGGACTCATCAGGAAGC
TCATCGGACTCATCAGGAAGC
ATCGGACTCATCAGGAAGC
TCATCGGACTCATCAGGAAGC
ACTCATCGGACTCATCAGGAAGC
AACATCGGACTCATCAGGAAGC
AACATCGGACTCATCAGGAAGC
ATCGGACTCATCAGGAAGC
ACATCGGACTCATCAGGAAGC
CCATCGGACTCATCAGGAAGC
GCATCGGACTCATCAGGAAGC
CATCGGACTCATCAGGAAGC

TCATCGGACTCATCAGGAAGC
AGCATCGGACTCATCAGGAAGC
CGGACTCATCAGGAAGC
ACATCGGACTCATCAGGAAGC
CACATCGGACTCATCAGGAAGC
CATCGGACTCATCAGGAAGC
ACATCGGACTCATCAGGAAGC
ATCGGACTCATCAGGAAGC
TATGCACATCGGACTCATCAGGAAGC
CTCATCGGACTCATCAGGAAGC
TCGGACTCATCAGGAAGC
CGGACTCATCAGGAAGC
ATCGGACTCATCAGGAAGC
ACATCGGACTCATCAGGAAGC
ATCGGACTCATCAGGAAGC
CGGACTCATCAGGAAGC
ACATCGGACTCATCAGGAAGC
ATCGGACTCATCAGGAAGC
AACATCGGACTCATCAGGAAGC
CATCGGACTCATCAGGAAGC
ACATCGGACTCATCAGGAAGC
ACATCGGACTCATCAGGAAGC
GCATCGGACTCATCAGGAAGC
GCATCGGACTCATCAGGAAGC
TCATCGGACTCATCAGGAAGC
GACTCATCAGGAAGC
GATCATCGGACTCATCAGGAAGC
TGTCATCGGACTCATCAGGAAGC
ACATCGGACTCATCAGGAAGC
AACATCGGACTCATCAGGAAGC
CCATCGGACTCATCAGGAAGC
TCGGACTCATCAGGAAGC
AAACATCGGACTCATCAGGAAGC
ATCGGACTCATCAGGAAGC
ATCGGACTCATCAGGAAGC
GACTCATCAGGAAGC
TCGGACTCATCAGGAAGC
CATCGGACTCATCAGGAAGC
CATCGGACTCATCAGGAAGC
ATCATCGGACTCATCAGGAAGC
CATCGGACTCATCAGGAAGC
TGCATCGGACTCATCAGGAAGC
GGACTCATCAGGAAGC

TCGGACTCATCAGGAAGC
TACATCGGACTCATCAGGAAGC
GGACTCATCAGGAAGC
ACATCGGACTCATCAGGAAGC
ACATCGGACTCATCAGGAAGC
ACCATCGGACTCATCAGGAAGC
AACATCGGACTCATCAGGAAGC
GGCATCGGACTCATCAGGAAGC
AGCATCGGACTCATCAGGAAGC
CATCGGACTCATCAGGAAGC
CACATCGGACTCATCAGGAAGC
ATCGGACTCATCAGGAAGC
AACATCGGACTCATCAGGAAGC
ACTCATCGGACTCATCAGGAAGC
ACCATCGGACTCATCAGGAAGC
ACATCGGACTCATCAGGAAGC
CGGACTCATCAGGAAGC
ATCGGACTCATCAGGAAGC
ATCGGACTCATCAGGAAGC
TCATCGGACTCATCAGGAAGC
CCATCGGACTCATCAGGAAGC
CGGACTCATCAGGAAGC
GCATCGGACTCATCAGGAAGC
GCCATCGGACTCATCAGGAAGC
ATCGGACTCATCAGGAAGC
ATCGGACTCATCAGGAAGC
CGGACTCATCAGGAAGC
AACATCGGACTCATCAGGAAGC
AGTGCATCGGACTCATCAGGAAGC
ACATCGGACTCATCAGGAAGC
GGCATCGGACTCATCAGGAAGC
AACATCGGACTCATCAGGAAGC
CACATCGGACTCATCAGGAAGC
GCATCGGACTCATCAGGAAGC
TCGGACTCATCAGGAAGC
GGCATCGGACTCATCAGGAAGC
TACATCGGACTCATCAGGAAGC
TCCATCGGACTCATCAGGAAGC
ATCGGACTCATCAGGAAGC
ATCGGACTCATCAGGAAGC
CATCGGACTCATCAGGAAGC
GAGCATCGGACTCATCAGGAAGC
ATCGGACTCATCAGGAAGC

CATCGGACTCATCAGGAAGC
GACTCATCAGGAAGC
ATCGGACTCATCAGGAAGC
CATCGGACTCATCAGGAAGC
TCATCGGACTCATCAGGAAGC
CATCATCGGACTCATCAGGAAGC
AGCATCGGACTCATCAGGAAGC
GGACTCATCAGGAAGC
CATCGGACTCATCAGGAAGC
ATCGGACTCATCAGGAAGC
TACATCGGACTCATCAGGAAGC
ACCATCGGACTCATCAGGAAGC
CGGACTCATCAGGAAGC
GACTCATCAGGAAGC
CATCGGACTCATCAGGAAGC
ATCATCGGACTCATCAGGAAGC
TCATCGGACTCATCAGGAAGC
TCGGACTCATCAGGAAGC
GCATCGGACTCATCAGGAAGC
ATCGGACTCATCAGGAAGC
ATCGGACTCATCAGGAAGC
CATCGGACTCATCAGGAAGC
ATCGGACTCATCAGGAAGC
ATCGGACTCATCAGGAAGC
ATCGGACTCATCAGGAAGC
ATCGGACTCATCAGGAAGC
TCATCGGACTCATCAGGAAGC
TACATCGGACTCATCAGGAAGC
CATCGGACTCATCAGGAAGC
AGCATCGGACTCATCAGGAAGC
CATCGGACTCATCAGGAAGC
AGACATCGGACTCATCAGGAAGC
TTCATCGGACTCATCAGGAAGC
CTCATCGGACTCATCAGGAAGC
TCGGACTCATCAGGAAGC
CATCGGACTCATCAGGAAGC
CATCGGACTCATCAGGAAGC
ACATCGGACTCATCAGGAAGC
GCCATCGGACTCATCAGGAAGC
TGCATCGGACTCATCAGGAAGC
CATCGGACTCATCAGGAAGC
TCGGACTCATCAGGAAGC
GCATCGGACTCATCAGGAAGC

TCCATCGGACTCATCAGGAAGC
GCATCGGACTCATCAGGAAGC
AGCATCGGACTCATCAGGAAGC
ATCGGACTCATCAGGAAGC
TCATCGGACTCATCAGGAAGC
TCGGACTCATCAGGAAGC
CCATCGGACTCATCAGGAAGC
TCGGACTCATCAGGAAGC
TACATCGGACTCATCAGGAAGC
AACATCGGACTCATCAGGAAGC
CATCGGACTCATCAGGAAGC
ACATCGGACTCATCAGGAAGC
GGACTCATCAGGAAGC
CATCGGACTCATCAGGAAGC
AATTCATCGGACTCATCAGGAAGC
ATCGGACTCATCAGGAAGC
TCATCGGACTCATCAGGAAGC
TCGGACTCATCAGGAAGC
GGACTCATCAGGAAGC
AGCATCGGACTCATCAGGAAGC
AACATCGGACTCATCAGGAAGC
CATCGGACTCATCAGGAAGC
CGGACTCATCAGGAAGC
AGCATCGGACTCATCAGGAAGC
GACATCGGACTCATCAGGAAGC
CATCGGACTCATCAGGAAGC
ACATCGGACTCATCAGGAAGC
GGACTCATCAGGAAGC
ATTCATCGGACTCATCAGGAAGC
TCATCGGACTCATCAGGAAGC
ATCGGACTCATCAGGAAGC
AACATCGGACTCATCAGGAAGC
GGACTCATCAGGAAGC
CTCATCGGACTCATCAGGAAGC
CGGACTCATCAGGAAGC
TCGGACTCATCAGGAAGC
CATCGGACTCATCAGGAAGC
ATCGGACTCATCAGGAAGC
AGCATCGGACTCATCAGGAAGC
CATCGGACTCATCAGGAAGC
TGCATCGGACTCATCAGGAAGC
CATCGGACTCATCAGGAAGC
TCATCGGACTCATCAGGAAGC

CATCGGACTCATCAGGAAGC
ATCGGACTCATCAGGAAGC
CATCGGACTCATCAGGAAGC
CATCGGACTCATCAGGAAGC
ATCGGACTCATCAGGAAGC
TCGGACTCATCAGGAAGC
GCCATCGGACTCATCAGGAAGC
CATCGGACTCATCAGGAAGC
TTCATCGGACTCATCAGGAAGC
CATCGGACTCATCAGGAAGC
TCGGACTCATCAGGAAGC
TCATCGGACTCATCAGGAAGC
ATCATCGGACTCATCAGGAAGC
CATCGGACTCATCAGGAAGC
CATCGGACTCATCAGGAAGC
CATCGGACTCATCAGGAAGC
TCCATCGGACTCATCAGGAAGC
CATCGGACTCATCAGGAAGC
GGACTCATCAGGAAGC
CATCGGACTCATCAGGAAGC
GCATCGGACTCATCAGGAAGC
TCGGACTCATCAGGAAGC
TGCATCGGACTCATCAGGAAGC
AAACATCGGACTCATCAGGAAGC
CTCATCGGACTCATCAGGAAGC
GGCATCGGACTCATCAGGAAGC
CATCGGACTCATCAGGAAGC
CATCGGACTCATCAGGAAGC
ATCGGACTCATCAGGAAGC
TCATCGGACTCATCAGGAAGC
TACATCGGACTCATCAGGAAGC
CCATCGGACTCATCAGGAAGC
TCATCGGACTCATCAGGAAGC
CATCGGACTCATCAGGAAGC
CGGACTCATCAGGAAGC
ACATCGGACTCATCAGGAAGC
TCGGACTCATCAGGAAGC
CCATCGGACTCATCAGGAAGC
CGGACTCATCAGGAAGC
CATCGGACTCATCAGGAAGC
ATCATCGGACTCATCAGGAAGC
CATCGGACTCATCAGGAAGC
AACATCGGACTCATCAGGAAGC

TCATCGGACTCATCAGGAAGC
GGACTCATCAGGAAGC
GGACTCATCAGGAAGC
TATCATCGGACTCATCAGGAAGC
ATCGGACTCATCAGGAAGC
GACATCGGACTCATCAGGAAGC
CGGACTCATCAGGAAGC
CTCATCGGACTCATCAGGAAGC
AGCATCGGACTCATCAGGAAGC
CATCGGACTCATCAGGAAGC
ATCGGACTCATCAGGAAGC
ATCGGACTCATCAGGAAGC
CGGACTCATCAGGAAGC
CATCGGACTCATCAGGAAGC
CGGACTCATCAGGAAGC
ACATCGGACTCATCAGGAAGC
TCATCGGACTCATCAGGAAGC
GGACTCATCAGGAAGC
CCATCGGACTCATCAGGAAGC
TCGGACTCATCAGGAAGC
GGACTCATCAGGAAGC
ATCGGACTCATCAGGAAGC
ACATCGGACTCATCAGGAAGC
CATCGGACTCATCAGGAAGC
ATCGGACTCATCAGGAAGC
TCGGACTCATCAGGAAGC
CGGACTCATCAGGAAGC
TCATCGGACTCATCAGGAAGC
ATCGGACTCATCAGGAAGC
ACATCGGACTCATCAGGAAGC
ATCATCGGACTCATCAGGAAGC
GCCATCGGACTCATCAGGAAGC
TCATCGGACTCATCAGGAAGC
GGACTCATCAGGAAGC
TCATCGGACTCATCAGGAAGC
GCATCGGACTCATCAGGAAGC
ACATCGGACTCATCAGGAAGC
CTCATCGGACTCATCAGGAAGC
ATCGGACTCATCAGGAAGC
TCATCGGACTCATCAGGAAGC
TCATCGGACTCATCAGGAAGC
GACATCGGACTCATCAGGAAGC
GAACATCGGACTCATCAGGAAGC

GTCATCGGACTCATCAGGAAGC
CGGACTCATCAGGAAGC
ACATCGGACTCATCAGGAAGC
GCATCGGACTCATCAGGAAGC
GGACTCATCAGGAAGC
TTCATCGGACTCATCAGGAAGC
CATCGGACTCATCAGGAAGC
CGGACTCATCAGGAAGC
ATCGGACTCATCAGGAAGC
CATCGGACTCATCAGGAAGC
TCATCGGACTCATCAGGAAGC
ACATCGGACTCATCAGGAAGC
ATCGGACTCATCAGGAAGC
ACATCGGACTCATCAGGAAGC
ACATCGGACTCATCAGGAAGC
GACTCATCAGGAAGC
CATCGGACTCATCAGGAAGC
CATCGGACTCATCAGGAAGC
CATCGGACTCATCAGGAAGC
CCCATCGGACTCATCAGGAAGC
ATCGGACTCATCAGGAAGC
GGACTCATCAGGAAGC
ATCATCGGACTCATCAGGAAGC
CCATCGGACTCATCAGGAAGC
TCGGACTCATCAGGAAGC
CATCGGACTCATCAGGAAGC
CGGACTCATCAGGAAGC
TCGGACTCATCAGGAAGC
ACATCGGACTCATCAGGAAGC
CACATCGGACTCATCAGGAAGC
ACTCATCAGGAAGC
GGACTCATCAGGAAGC
ACTCATCGGACTCATCAGGAAGC
ATCGGACTCATCAGGAAGC
CATCGGACTCATCAGGAAGC
CATCGGACTCATCAGGAAGC
TCCATCGGACTCATCAGGAAGC
CATCGGACTCATCAGGAAGC
TCGGACTCATCAGGAAGC
GGACTCATCAGGAAGC
GTCATCGGACTCATCAGGAAGC
ATCGGACTCATCAGGAAGC
CCATCGGACTCATCAGGAAGC

ACATCGGACTCATCAGGAAGC
CATCGGACTCATCAGGAAGC
CATCGGACTCATCAGGAAGC
ACCATCGGACTCATCAGGAAGC
AGTCATCGGACTCATCAGGAAGC
CATCGGACTCATCAGGAAGC
CCATCGGACTCATCAGGAAGC
CATCGGACTCATCAGGAAGC
ATCGGACTCATCAGGAAGC
ATACATCGGACTCATCAGGAAGC
CATCGGACTCATCAGGAAGC
CATCGGACTCATCAGGAAGC
TCGGACTCATCAGGAAGC
AGCCATCGGACTCATCAGGAAGC
CATCGGACTCATCAGGAAGC
CGGACTCATCAGGAAGC
TCGGACTCATCAGGAAGC
GTTCATCGGACTCATCAGGAAGC
CGGACTCATCAGGAAGC
CATCGGACTCATCAGGAAGC
AGCATCGGACTCATCAGGAAGC
ACATCGGACTCATCAGGAAGC
AACATCGGACTCATCAGGAAGC
TCGGACTCATCAGGAAGC
ACTCATCAGGAAGC
GACTCATCAGGAAGC
CATCGGACTCATCAGGAAGC
ACATCGGACTCATCAGGAAGC
TACCATCGGACTCATCAGGAAGC
GCCATCGGACTCATCAGGAAGC
GTCATCGGACTCATCAGGAAGC
TCGGACTCATCAGGAAGC
TCGGACTCATCAGGAAGC
AACATCGGACTCATCAGGAAGC
CATCGGACTCATCAGGAAGC
CATCGGACTCATCAGGAAGC
TCATCGGACTCATCAGGAAGC
TGACATCGGACTCATCAGGAAGC
CATCGGACTCATCAGGAAGC
ACATCGGACTCATCAGGAAGC
CATCGGACTCATCAGGAAGC
ATACATCGGACTCATCAGGAAGC
TATCATCGGACTCATCAGGAAGC

CATCGGACTCATCAGGAAGC
GGACTCATCAGGAAGC
TCATCGGACTCATCAGGAAGC
TCATCGGACTCATCAGGAAGC
ACATCGGACTCATCAGGAAGC
ACATCGGACTCATCAGGAAGC
TCGGACTCATCAGGAAGC
GTCATCGGACTCATCAGGAAGC
GACATCGGACTCATCAGGAAGC
TCATCGGACTCATCAGGAAGC
CCATCGGACTCATCAGGAAGC
TCATCGGACTCATCAGGAAGC
CATCGGACTCATCAGGAAGC
TCGGACTCATCAGGAAGC
TCGGACTCATCAGGAAGC
AAACATCGGACTCATCAGGAAGC
CATCGGACTCATCAGGAAGC
CATCGGACTCATCAGGAAGC
TCATCGGACTCATCAGGAAGC
TTCATCGGACTCATCAGGAAGC
TACATCGGACTCATCAGGAAGC
ACATCGGACTCATCAGGAAGC
ACTCATCGGACTCATCAGGAAGC
ACATCGGACTCATCAGGAAGC
CATCGGACTCATCAGGAAGC
CATCGGACTCATCAGGAAGC
CATCGGACTCATCAGGAAGC
TATCATCGGACTCATCAGGAAGC
TCGGACTCATCAGGAAGC
TTCATCGGACTCATCAGGAAGC
CATCGGACTCATCAGGAAGC
ATCGGACTCATCAGGAAGC
ATCGGACTCATCAGGAAGC
GCCATCGGACTCATCAGGAAGC
TCATCGGACTCATCAGGAAGC
CATCGGACTCATCAGGAAGC
ATCATCGGACTCATCAGGAAGC
CATCGGACTCATCAGGAAGC
GTCCATCGGACTCATCAGGAAGC
TCGGACTCATCAGGAAGC
CGGACTCATCAGGAAGC
GGACTCATCAGGAAGC
AACATCGGACTCATCAGGAAGC

TCGGACTCATCAGGAAGC
ATCGGACTCATCAGGAAGC
GACTCATCAGGAAGC
CATCGGACTCATCAGGAAGC
ATCGGACTCATCAGGAAGC
AACATCGGACTCATCAGGAAGC
GAACATCGGACTCATCAGGAAGC
AACATCGGACTCATCAGGAAGC
CATCGGACTCATCAGGAAGC
CACATCGGACTCATCAGGAAGC
TCGGACTCATCAGGAAGC
TTCATCGGACTCATCAGGAAGC
AAACATCGGACTCATCAGGAAGC
GCATCGGACTCATCAGGAAGC
TCGGACTCATCAGGAAGC
ATCGGACTCATCAGGAAGC
TCGGACTCATCAGGAAGC
CGGACTCATCAGGAAGC
CGGACTCATCAGGAAGC
TCGGACTCATCAGGAAGC
CATCGGACTCATCAGGAAGC
CCATCGGACTCATCAGGAAGC
TCATCGGACTCATCAGGAAGC
GGACTCATCAGGAAGC
ACATCGGACTCATCAGGAAGC
CATCGGACTCATCAGGAAGC
ACTCATCAGGAAGC
ATCATCGGACTCATCAGGAAGC
ACATCGGACTCATCAGGAAGC
ACATCGGACTCATCAGGAAGC
GCATCGGACTCATCAGGAAGC
CATCGGACTCATCAGGAAGC
ATCGGACTCATCAGGAAGC
CATCGGACTCATCAGGAAGC
ACATCGGACTCATCAGGAAGC
TTCATCGGACTCATCAGGAAGC
ACATCGGACTCATCAGGAAGC
TCATCGGACTCATCAGGAAGC
CATCGGACTCATCAGGAAGC
TCATCGGACTCATCAGGAAGC
ATCGGACTCATCAGGAAGC
ATCGGACTCATCAGGAAGC
CATCGGACTCATCAGGAAGC

CATCGGACTCATCAGGAAGC
TCGGACTCATCAGGAAGC
ACATCGGACTCATCAGGAAGC
ATCATCGGACTCATCAGGAAGC
TCGGACTCATCAGGAAGC
CGGACTCATCAGGAAGC
ATCGGACTCATCAGGAAGC
TAAACATCGGACTCATCAGGAAGC
ATCATCGGACTCATCAGGAAGC
TCATCGGACTCATCAGGAAGC
ACATCGGACTCATCAGGAAGC
GCATCGGACTCATCAGGAAGC
ATCATCGGACTCATCAGGAAGC
TATACACATCGGACTCATCAGGAAGC
AACATCGGACTCATCAGGAAGC
ACATCGGACTCATCAGGAAGC
CATCGGACTCATCAGGAAGC
AAACATCGGACTCATCAGGAAGC
CATCGGACTCATCAGGAAGC
ATCGGACTCATCAGGAAGC
ACATCGGACTCATCAGGAAGC
TCATCGGACTCATCAGGAAGC
ACATCGGACTCATCAGGAAGC
TCGGACTCATCAGGAAGC
CATCGGACTCATCAGGAAGC
CATCGGACTCATCAGGAAGC
GCATCGGACTCATCAGGAAGC
CATCGGACTCATCAGGAAGC
CATCGGACTCATCAGGAAGC
TCATCGGACTCATCAGGAAGC
AACATCGGACTCATCAGGAAGC
ATCATCGGACTCATCAGGAAGC
TACATCGGACTCATCAGGAAGC
ACATCGGACTCATCAGGAAGC
TGCATCGGACTCATCAGGAAGC
TCGGACTCATCAGGAAGC
GTCATCGGACTCATCAGGAAGC
CATCATCGGACTCATCAGGAAGC
ATCGGACTCATCAGGAAGC
ATCGGACTCATCAGGAAGC
GACTCATCAGGAAGC
TCGGACTCATCAGGAAGC
CTGCATCGGACTCATCAGGAAGC

[illegible]CCCATCGGACTCATCAGGAAGC
TTCATCGGACTCATCAGGAAGC
[illegible]GCATCGGACTCATCAGGAAGC
