## Supplementary material for "Retrospective cell lineage reconstruction in Humans using short tandem repeats": sup file3

| terid | padlock_sequence |
| --- | --- |
| 46553 | TGAGTGTGGAGTCGTTGCTGTCAGTTTGAAATACTGTATGTACGTNNNAGATCGGA |
| 48667 | TGAGTGTGGAGTCGTTGCATATTTCCAGTTATACGTACCTCATCNNNAGATCGGAA |
| 39579 | TGAGTGTGGAGTCGTTGCTGCTTTCAAAAAAAGTTCTCCAGAGANNNAGATCGGAA |
| 28785 | ATGAGTGTGGAGTCGTTGCTCACATTTTCTTTATCCACTCATTGANNNAGATCGGAA |
| 28827 | ATGAGTGTGGAGTCGTTGCACACACACATATAAGCAGGTTATATCTNNNAGATCGG |
| 30741 | ATGAGTGTGGAGTCGTTGCACTGGATTTTTTTTCTGTTGTATCGANNNAGATCGGA |
| 30826 | ATGAGTGTGGAGTCGTTGCACACTTTTTTTCAATAGGGACTTCTGANNNAGATCGG |
| 30900 | ATGAGTGTGGAGTCGTTGCGAGAGTACAATCTCTAGAACAATCAAANNNAGATCGG |
| 31043 | ATGAGTGTGGAGTCGTTGCTCTAGTGAAGTATAGAGACCTTACCANNNAGATCGGA |
| 37088 | ATGAGTGTGGAGTCGTTGCAAATTCCCAGATAACCAAGATTGAATNNNAGATCGGA |
| 42892 | ATGAGTGTGGAGTCGTTGCTGTATATACACGTATACACATATGTGCNNNAGATCGG |
| 44538 | ATGAGTGTGGAGTCGTTGCTTTTGGAAAACATTATAGAGGTTACCTNNNAGATCGG |
| 48789 | ATGAGTGTGGAGTCGTTGCAAGAAAGAAGAAAAGAGAGAAGATTCANNNAGATC |
| 28888 | ATGAGTGTGGAGTCGTTGCAATTCATATATGTGTATGTCTGTGCANNNAGATCGGA |
| 28897 | ATGAGTGTGGAGTCGTTGCGTCCTTACCCTTTAGTCACTGTGTNNNAGATCGGAAG |
| 30782 | ATGAGTGTGGAGTCGTTGCTGGGATGTGTATCTCTGACTAATACANNNAGATCGGA |
| 31725 | ATGAGTGTGGAGTCGTTGCAGATTATCTCTCTCTCTCTCTACANNNAGATCGGAA |
| 35436 | ATGAGTGTGGAGTCGTTGCAACAAACAAAAAAAAAACAGGAGGANNNAGATCGG |
| 35763 | ATGAGTGTGGAGTCGTTGCACTGGTAATTTTGCAGAAGTGTGANNNAGATCGGAA |
| 43021 | ATGAGTGTGGAGTCGTTGCATTCATAAATATTTCCTTTGGGGGAANNNAGATCGGA |
| 43044 | ATGAGTGTGGAGTCGTTGCGAAGTGGAATTGCTGGATATATGACNNNAGATCGGA |
| 43537 | ATGAGTGTGGAGTCGTTGCAGGTATTTTTTCAATCCTCGTCTTCCNNNAGATCGGAA |
| 28781 | TATGAGTGTGGAGTCGTTGCTGGTTTCAAACTCTTGAGCTCAANNNAGATCGGAAG |
| 28799 | TATGAGTGTGGAGTCGTTGCACACACACACATATACACATATACACANNNAGATCGG |
| 28806 | TATGAGTGTGGAGTCGTTGCACACATACTATGTATACACACACTACANNNAGATCGG |
| 28826 | TATGAGTGTGGAGTCGTTGCCGTATATATATATGTGTGTGTGTGTGTNNNAGATCG |
| 28974 | TATGAGTGTGGAGTCGTTGCTGGAATACTACTCAGCCATAAATATGANNNAGATCG |
| 28995 | TATGAGTGTGGAGTCGTTGCTTTGTCATGTCAGAAATTTTGAAGAAANNNAGATCG |
| 29064 | TATGAGTGTGGAGTCGTTGCTCTGTACAAGAATAAGGAAACTCCANNNAGATCGGA |
| 29077 | TATGAGTGTGGAGTCGTTGCTGGAATTATGCCTCTCTAGTGAGANNNAGATCGGAA |
| 29146 | TATGAGTGTGGAGTCGTTGCATTCTGGAGTAAAATGGGTAGTTATTTNNNAGATCG |
| 29202 | TATGAGTGTGGAGTCGTTGCAGCATCCAAATAAACTTGCACTAACTNNNAGATCGG |
| 29234 | TATGAGTGTGGAGTCGTTGCAGATAGATAGATGAGATGGATGATGGANNNAGAT |
| 29290 | TATGAGTGTGGAGTCGTTGCACTTTCTTCTTTTGTATCTTTGAGACCNNNAGATCGG |
| 30736 | TATGAGTGTGGAGTCGTTGCAGGTTTCTTTTTGGGGTGTTGAANNNAGATCGGAA |
| 30771 | TATGAGTGTGGAGTCGTTGCTGTCTCTTCTTACAGTTTTTGTCTTGANNNAGATCGG |
| 30785 | TATGAGTGTGGAGTCGTTGCTGAACAATTTATACAAAAGGTTGTCCTNNNAGATCG |
| 30911 | TATGAGTGTGGAGTCGTTGCTGTTCACTGGTCAGAAGACTCAGNNNAGATCGGAA |
| 31074 | TATGAGTGTGGAGTCGTTGCACATGATACATACACATGATACATGCANNNAGATCG |
| 31295 | TATGAGTGTGGAGTCGTTGCATGTAGCCATGAAAAGGAATGCGNNNAGATCGGAA |
| 31305 | TATGAGTGTGGAGTCGTTGCGAGAAAGAGAGAAGAAAGAAAGAAAGANNNAGAT |
| 31420 | TATGAGTGTGGAGTCGTTGCACTGAGTGTACCCTCCTCTACTCNNNAGATCGGAAG |

31496 TATGAGTGTGGAGTCGTTGCTCAGTGCCATTGTAAATGTAAATGACTNNNAGATCG
31518 TATGAGTGTGGAGTCGTTGCTGTTATGTTTTGATGCAAGCATACANNNAGATCGGA
31741 TATGAGTGTGGAGTCGTTGCTGACAGTTACAGTCCTCAGTTGGNNNAGATCGGAAG
31819 TATGAGTGTGGAGTCGTTGCTGTGTGTGTGTGTATACATATATCCCTNNNAGATCG
31822 TATGAGTGTGGAGTCGTTGCTCTCTATCATCTTTTCTCCTCATGTCANNNAGATCGGA
31832 TATGAGTGTGGAGTCGTTGCAAGAACTTTGCAGTGTTGTTGGGNNNAGATCGGAA
31854 TATGAGTGTGGAGTCGTTGCTCCTGGAAATGTACAACCTCACANNNAGATCGGAAG
31971 TATGAGTGTGGAGTCGTTGCGCAGATAAATGACTATGTGTACTAGAANNNAGATCG
31973 TATGAGTGTGGAGTCGTTGCGGTGTTTTTTGTGTTGATATTCTGTCCNNNAGATCGG
31987 TATGAGTGTGGAGTCGTTGCTGTTTGTCTCCTAAGCCCATTCANNNAGATCGGAAGA
32014 TATGAGTGTGGAGTCGTTGCACCACATAAATGTTCTAGAAGTCAACANNNAGATCG
32016 TATGAGTGTGGAGTCGTTGCTCAATCAAGTGAAGAGAAAACCTATGANNNAGATCG
32134 TATGAGTGTGGAGTCGTTGCCCACAGGTATCAAAGCAATACTAGCNNNAGATCGGA
32163 TATGAGTGTGGAGTCGTTGCGATCTCTGCTCACTTCAAGCTCTNNNAGATCGGAAGA
32339 TATGAGTGTGGAGTCGTTGCAATTCCAGAACAGGAGCAGAGACNNNAGATCGGAA
32407 TATGAGTGTGGAGTCGTTGCACACACCCACACATATATACATATACANNNAGATCGG
32681 TATGAGTGTGGAGTCGTTGCCCCAGGTTCAAGTGATTCTCTCANNNAGATCGGAAG
32761 TATGAGTGTGGAGTCGTTGCAGAGGAGACAAATTCAAAAGCTAGCNNNAGATCGG
33461 TATGAGTGTGGAGTCGTTGCCCATACCAAGGAGCTGGATGTAANNNAGATCGGAA
33588 TATGAGTGTGGAGTCGTTGCCTTCCTGGGTTCAAGCAATTCTCNNNAGATCGGAAG
34747 TATGAGTGTGGAGTCGTTGCGCCCGGCCAATTTCTGTATTTTTNNNAGATCGGAAGA
34876 TATGAGTGTGGAGTCGTTGCGCCTCCTAAGTAGCTGGGATTACNNNAGATCGGAAG
35037 TATGAGTGTGGAGTCGTTGCCTGTCTCAGCTCACTGTAACCTCNNNAGATCGGAAGA
35475 TATGAGTGTGGAGTCGTTGCAGGTCAAATCTCACATATTTCATAAGTNNNAGATCGG
35619 TATGAGTGTGGAGTCGTTGCGACAATATGCGTTGGTGAACTCCNNNAGATCGGAAG
35734 TATGAGTGTGGAGTCGTTGCTCTTTACGGATTTCAAGGGAGCANNNAGATCGGAAG
35790 TATGAGTGTGGAGTCGTTGCTCTTTACGGATTTCAAGGGAGCANNNAGATCGGAAG
35832 TATGAGTGTGGAGTCGTTGCCTCCCTGGTTCAAGTGATTCTCANNNAGATCGGAAG
35849 TATGAGTGTGGAGTCGTTGCCTTCAAGTGATTCTCCTGCCTCANNNAGATCGGAAGA
36765 TATGAGTGTGGAGTCGTTGCATTCATTGCTAGGATAGTGTATTTTGANNNAGATCG
36814 TATGAGTGTGGAGTCGTTGCCTAACATTATGCCCTACTATCTTTTGTNNNAGATCGG
36887 TATGAGTGTGGAGTCGTTGCAGGTTATCTGTGTGTTCATTGAATACTNNNAGATCG
37011 TATGAGTGTGGAGTCGTTGCGGAGAATAGCATGAGTGGGACATNNNAGATCGGAA
37031 TATGAGTGTGGAGTCGTTGCACAAGTTAGTCAGAAAACCAGAAATCTNNNAGATCG
37152 TATGAGTGTGGAGTCGTTGCCCTGCTCAAAAAAAATTCCATTCACANNNAGATCGGA
37166 TATGAGTGTGGAGTCGTTGCTGTCATGGAAAAAGTAAAAGTCTTTGCNNNAGATCG
37195 TATGAGTGTGGAGTCGTTGCCCAGATTGAGGATATTAGAGGAAAACANNNAGATC
37216 TATGAGTGTGGAGTCGTTGCTGTGCTACCACTATGAACATATTTTTNNNAGATCGGA
37349 TATGAGTGTGGAGTCGTTGCGCATTCCCATACACATATATGTATACANNNAGATCGG
37426 TATGAGTGTGGAGTCGTTGCTCCTTCAAAAAAACACATAACCAAAGANNNAGATCG
37607 TATGAGTGTGGAGTCGTTGCTTGTAATAAAATTCAGGAATCAGCTCTNNNAGATCG
37855 TATGAGTGTGGAGTCGTTGCTGAGAAAATATTTGTCATCTCCTTCGANNNAGATCGG
38290 TATGAGTGTGGAGTCGTTGCTCTAGGAATCACATTACCTAACTTCANNNAGATCGGA

38400 TATGAGTGTGGAGTCGTTGCACTAGGAAGTGATTGATTAGTTTTGAGNNNAGATC
38703 TATGAGTGTGGAGTCGTTGCAGTATATATGCACCCAACACTGGANNNAGATCGGAA
38830 TATGAGTGTGGAGTCGTTGCAGAAAAGTTGCATAAATGTTCAATGGANNNAGATC
39062 TATGAGTGTGGAGTCGTTGCTCCTGGAAATGTACAACCTCACANNNAGATCGGAAG
39106 TATGAGTGTGGAGTCGTTGCAGGAAGCTAGGAATTGAAAGATACAGANNNAGATC
39188 TATGAGTGTGGAGTCGTTGCAGGTCATAGAAATTTTGAAGTGTTGACNNNAGATCG
39449 TATGAGTGTGGAGTCGTTGCCTGGATAGGAGGAATAAATTCAAAACANNNAGATC
39450 TATGAGTGTGGAGTCGTTGCGTGGATTACGAGGTCAGGAGATCNNNAGATCGGAA
39727 TATGAGTGTGGAGTCGTTGCAATTCCAATGACATTTTTGCCGANNNAGATCGGAAG
39781 TATGAGTGTGGAGTCGTTGCACCTAAAAAGAAAACTAAATACTGGGGNNNAGATC
39788 TATGAGTGTGGAGTCGTTGCGCCAGAGCACACAGACAAGATATNNNAGATCGGAA
40379 TATGAGTGTGGAGTCGTTGCAGGAACTTAGATGCAAAATTCTCAACANNNAGATCG
40420 TATGAGTGTGGAGTCGTTGCTGCAAATCAACTGTAAGAGAAAATAGANNNAGATC
40724 TATGAGTGTGGAGTCGTTGCTGTTCATTGCAGCACTATTCACANNNAGATCGGAAG
40935 TATGAGTGTGGAGTCGTTGCGGAAAGAAATAATTGAAAACCTCCAGANNNAGATC
41581 TATGAGTGTGGAGTCGTTGCACATCTTTTATTTCTTGTAAAGAGGGGNNNAGATCG
41796 TATGAGTGTGGAGTCGTTGCGCATAATCTCGGCTCACTGAAACNNNAGATCGGAAG
42169 TATGAGTGTGGAGTCGTTGCACTCCTGCCATTTTGTTGTTTGTNNNAGATCGGAAG
42243 TATGAGTGTGGAGTCGTTGCTGGTGTGCGTATATATATATGGTGTGTNNNAGATCG
42254 TATGAGTGTGGAGTCGTTGCACACATATATACATATACACACACGTGNNNAGATCG
42258 TATGAGTGTGGAGTCGTTGCACACATATATACATATACACACACGTGNNNAGATCG
42295 TATGAGTGTGGAGTCGTTGCTGGCATGATCTCAACTAACTGCTNNNAGATCGGAAG
42303 TATGAGTGTGGAGTCGTTGCACCCCAAAATAGGCAAAATGCATNNNAGATCGGAAG
42310 TATGAGTGTGGAGTCGTTGCGGAGTAGCTGTTCTTTCACCACTNNNAGATCGGAAG
42359 TATGAGTGTGGAGTCGTTGCTGTTTGTCTCCTAAGCCCATTCANNNAGATCGGAAG
42384 TATGAGTGTGGAGTCGTTGCTTATGCTGCAATAGACATGCGTGNNNAGATCGGAA
42390 TATGAGTGTGGAGTCGTTGCAGGGAAAAGTCAAGCTGGTAACTNNNAGATCGGAA
42707 TATGAGTGTGGAGTCGTTGCACCCCAAAATAGGCAAAATGCATNNNAGATCGGAAG
42761 TATGAGTGTGGAGTCGTTGCCTGTTGTACAGGGTGATAAAAGAAANNNAGATCGG
42764 TATGAGTGTGGAGTCGTTGCTCAGAAATGTAAAGAACAAAGAGGTCCNNNAGATC
42814 TATGAGTGTGGAGTCGTTGCTGCGTATGTAATCAAATACATTGTTGCNNNAGATCG
42843 TATGAGTGTGGAGTCGTTGCATTGAACACCTAGCATTTGACAGNNNAGATCGGAAG
42868 TATGAGTGTGGAGTCGTTGCCCTCCAACATTTATTATGAAAATGGGCNNNAGATCG
42890 TATGAGTGTGGAGTCGTTGCTGGAAGCAAATTGGAGACATCACNNNAGATCGGAA
42897 TATGAGTGTGGAGTCGTTGCTCTGTGTGTGTATATATATGTGTGTGTNNNAGATCG
42917 TATGAGTGTGGAGTCGTTGCCTCCAGTGTTTCTTCTCAAGGGANNNAGATCGGAAG
42991 TATGAGTGTGGAGTCGTTGCTGTGTATGTGTATGCAAATATGTGTGTNNNAGATCG
43289 TATGAGTGTGGAGTCGTTGCACACACACACATCTATATATACACACGNNNAGATCGG
43317 TATGAGTGTGGAGTCGTTGCCTCTCTTTTCTTCTTTATCAGTCTAGCNNNAGATCGGA
43550 TATGAGTGTGGAGTCGTTGCGCAAATATTTTCTCCCATTATTTCTGGNNNAGATCGG
43786 TATGAGTGTGGAGTCGTTGCAATAGAATAAAAATATTCGGGGCAGANNNAGATCG
43800 TATGAGTGTGGAGTCGTTGCGGAACAAAATACCAAACTGAAATTTGCNNNAGATCG
44455 TATGAGTGTGGAGTCGTTGCGCTCAAGTGATTCTCGTACCTCANNNAGATCGGAAG

44522 TATGAGTGTGGAGTCGTTGCACACCCACACATATATACATATACACANNNAGATCGG
44561 TATGAGTGTGGAGTCGTTGCACCAAGAGAGATAAAATGAAAACATGTNNNAGATC
44856 TATGAGTGTGGAGTCGTTGCTCTGGTATTGTCGGAATATGGGCNNNAGATCGGAA
45087 TATGAGTGTGGAGTCGTTGCTGTAGATCATATAACATGTTTGTGGGTNNNAGATCG
45359 TATGAGTGTGGAGTCGTTGCTCCTGTAGAGTTCTTTGAACTTCTNNNAGATCGGAA
45784 TATGAGTGTGGAGTCGTTGCCCATCCTGGGACTTAGAAGAAAGTNNNAGATCGGA
46000 TATGAGTGTGGAGTCGTTGCCTCCAAACCTAGCTCCCAAGTAGNNNAGATCGGAAG
46791 TATGAGTGTGGAGTCGTTGCACACTACACAAATGTTTTGTATACAGANNNAGATCG
46935 TATGAGTGTGGAGTCGTTGCGTGCTGGGAAATGTGGGTTTTTANNNAGATCGGAA
47199 TATGAGTGTGGAGTCGTTGCCCTCCTGGGTTCAAGTGATTCTCNNNAGATCGGAAG
47654 TATGAGTGTGGAGTCGTTGCGCCTCCTAAGTAGCTGGGATTACNNNAGATCGGAAG
47788 TATGAGTGTGGAGTCGTTGCCCAAGTAGCTGGGATTACAGGTGNNNAGATCGGAA
48177 TATGAGTGTGGAGTCGTTGCCTGGGTTCAAGCAATTCTACTGCNNNAGATCGGAAG
48749 TATGAGTGTGGAGTCGTTGCGCATTTTTCGAGCCAAACCAAAGNNNAGATCGGAAG
48756 TATGAGTGTGGAGTCGTTGCACTGGTATGTTCATTGCACCACTNNNAGATCGGAAG
48763 TATGAGTGTGGAGTCGTTGCCACCAAGCCCAGCTAATTTTTGTNNNAGATCGGAAG
48788 TATGAGTGTGGAGTCGTTGCTCTTTTCATGACTGTGTCCTTTGNNNAGATCGGAAGA
48856 TATGAGTGTGGAGTCGTTGCTGGAGTGCAATGGTGTGATCTTANNNAGATCGGAA
49019 TATGAGTGTGGAGTCGTTGCGCCCGGCCAATTTCTGTATTTTTNNNAGATCGGAAG
49068 TATGAGTGTGGAGTCGTTGCGTGCAAACTCATGGGGAAGAAATNNNAGATCGGAA
49137 TATGAGTGTGGAGTCGTTGCAAACCTGAGTTTCCATCCAGTGTNNNAGATCGGAAG
49240 TATGAGTGTGGAGTCGTTGCGTGAAATTCATGGAAAGAACCCTNNNAGATCGGAA
49342 TATGAGTGTGGAGTCGTTGCGCAAGTCATAGAAAAAAACACAAAAGCNNNAGATC
49378 TATGAGTGTGGAGTCGTTGCGTAGCTGGAATTACACACATGCGNNNAGATCGGAA
49489 TATGAGTGTGGAGTCGTTGCACTAAAAACATCTCAAATTTCCAGAGANNNAGATCG
49538 TATGAGTGTGGAGTCGTTGCCTCCTGAGTAGCTGGGACTACAGNNNAGATCGGAA
49539 TATGAGTGTGGAGTCGTTGCTTCAAGTGATTCTCCTGCCTCAGNNNAGATCGGAAG
49568 TATGAGTGTGGAGTCGTTGCTGTGTTGAATAACAGTGGTGACANNNAGATCGGAA
49718 TATGAGTGTGGAGTCGTTGCACGCCCGGCTAATTTTTTTGTATNNNAGATCGGAAG
49772 TATGAGTGTGGAGTCGTTGCAATGTTGGGCTAGAAGGAACACANNNAGATCGGAA
50850 TATGAGTGTGGAGTCGTTGCAGTATCATGTGTGAATGCAGAGTNNNAGATCGGAA
51342 TATGAGTGTGGAGTCGTTGCTGGACAGAAAATACCTATGAAGAAGGTNNNAGATC
52930 TATGAGTGTGGAGTCGTTGCAGATGTAGATGAAATACTACTCAGCCANNNAGATCG
53127 TATGAGTGTGGAGTCGTTGCGCAGATAAATGACTATGTGTACTAGAANNNAGATCG
53185 TATGAGTGTGGAGTCGTTGCCCATCCCCTCAAGCATTTATCCTNNNAGATCGGAAGA
53196 TATGAGTGTGGAGTCGTTGCCAGAGAGAGAGAAATAGATTAGGAAGANNNAGAT
53333 TATGAGTGTGGAGTCGTTGCTGTGAATTTATATGTGTGTGTGTGTGTNNNAGATCG
53385 TATGAGTGTGGAGTCGTTGCTTACAGATTCAGTGTCTGGTGGGNNNAGATCGGAA
28780 TATGAGTGTGGAGTCGTTGCTGGAATACTACTCAACCATACAAGGANNNAGATCGG
28788 TATGAGTGTGGAGTCGTTGCTCACATTTTCTTTATCCTTTCATCCANNNAGATCGGAA
28793 TATGAGTGTGGAGTCGTTGCAAAACCTTCTAGTGGCTTTCTGTNNNAGATCGGAAG
28813 TATGAGTGTGGAGTCGTTGCTGAGTTGGTTCCACATTTTTGCANNNAGATCGGAAG
28815 TATGAGTGTGGAGTCGTTGCTGTTGGTTGAAGAGTAAACTGTTTGTNNNAGATCG

28896 TATGAGTGTGGAGTCGTTGCACCCAAAAATTTCTCTACAAATGCCTNNNAGATCGGA
28956 TATGAGTGTGGAGTCGTTGCTGTCTATGTGTATATACTGCATGTGTNNNAGATCGG
28960 TATGAGTGTGGAGTCGTTGCTGCCTAACGATGGATTTCTCAGANNNAGATCGGAAG
29069 TATGAGTGTGGAGTCGTTGCTGCTTATCGTAGAAAATTGGGAAAGTNNNAGATCG
29124 TATGAGTGTGGAGTCGTTGCAGACTGTGGTTAGTTTTACTTTGATTNNNAGATCGG
29182 TATGAGTGTGGAGTCGTTGCACCTGGAAACAAATACACTAAAACCCNNNAGATCGG
29198 TATGAGTGTGGAGTCGTTGCAGGAACATACATATGTGCACACACNNNAGATCGGAA
29232 TATGAGTGTGGAGTCGTTGCTTCTCTGTATATCCTGTACATTGGATNNNAGATCGGA
29300 TATGAGTGTGGAGTCGTTGCAGTGGGTAAGGGAGAAAATTCTCNNNAGATCGGAA
29357 TATGAGTGTGGAGTCGTTGCGCTTGCTCACAAATCACACATCTNNNAGATCGGAAG
30751 TATGAGTGTGGAGTCGTTGCTGCAAAAGGAAATGCAGGAAGAANNNAGATCGGA
30766 TATGAGTGTGGAGTCGTTGCACACACACACATTTTGTAGATATGGANNNAGATCGG
30802 TATGAGTGTGGAGTCGTTGCACAGAAGTTCTTTGTTTCTTCCAACTNNNAGATCGGA
31062 TATGAGTGTGGAGTCGTTGCGATTCTAGCAGTGGGAAAAGTGCNNNAGATCGGAA
31181 TATGAGTGTGGAGTCGTTGCTCTTTGGCTTTTAGGATGCAATGANNNAGATCGGAA
31252 TATGAGTGTGGAGTCGTTGCGAGTCATTGTGGCAGAAACAGACNNNAGATCGGAA
31372 TATGAGTGTGGAGTCGTTGCGATTATAATGGTGTTGGCCAGGCNNNAGATCGGAA
31406 TATGAGTGTGGAGTCGTTGCACCATCATTTCTTTTCACATCAGGTNNNAGATCGGAA
31407 TATGAGTGTGGAGTCGTTGCACTCTGTATTTGACTTCTTCCAAGGANNNAGATCGG
31587 TATGAGTGTGGAGTCGTTGCTGCAATATAGTTATTTGACACATGGANNNAGATCGG
31656 TATGAGTGTGGAGTCGTTGCTCCCTATAGGCTAATATCTCTAGGCANNNAGATCGG
31696 TATGAGTGTGGAGTCGTTGCAGTTCTCTTTCTGGGATGGAAAANNNAGATCGGAA
31698 TATGAGTGTGGAGTCGTTGCTCAGTTTTCTCAGGGGATTGGTTNNNAGATCGGAAG
31729 TATGAGTGTGGAGTCGTTGCTGCCATTTTGGTTGGTTCAATAGTNNNAGATCGGAA
31827 TATGAGTGTGGAGTCGTTGCAGTGCAGATATGCCTTCAACATATTGNNNAGATCGG
31828 TATGAGTGTGGAGTCGTTGCGGGATCAAATGACCCTCTTGTCTNNNAGATCGGAAG
31906 TATGAGTGTGGAGTCGTTGCACATGAAAAAAAGCTCCACTGATCANNNAGATCGGA
31941 TATGAGTGTGGAGTCGTTGCACATGTATATGTGTGCGTATATACACNNNAGATCGG
31948 TATGAGTGTGGAGTCGTTGCACTCTGGTGATTATTTGCTGATTTCCNNNAGATCGG
32162 TATGAGTGTGGAGTCGTTGCCAGGTTCAAGCAATTCTCCTGCNNNAGATCGGAAGA
32321 TATGAGTGTGGAGTCGTTGCCCACACACACATATATACACAACACANNNAGATCGGA
32459 TATGAGTGTGGAGTCGTTGCAGGAAATCTGACAATATCTGCTCTCANNNAGATCGG
33077 TATGAGTGTGGAGTCGTTGCCGGGTTCAAGCAATTCTGTTTCANNNAGATCGGAAG
33138 TATGAGTGTGGAGTCGTTGCAGGAAAGAGATGTAACAACTAACTGTNNNAGATCG
33258 TATGAGTGTGGAGTCGTTGCTTCAAGCGTTTTTCCTGCTTCAGNNNAGATCGGAAG
33338 TATGAGTGTGGAGTCGTTGCGATCTCAGCTCACTGCAATCTCTNNNAGATCGGAAG
33365 TATGAGTGTGGAGTCGTTGCTCATAACAGTCAAGCTTTGGAAGNNNAGATCGGAA
33434 TATGAGTGTGGAGTCGTTGCTCCCCTTCCATTTCTTTATAATACGTNNNAGATCGGA
33603 TATGAGTGTGGAGTCGTTGCGCTGATCAGCTAATTTTGAATTCCAANNNAGATCGG
33605 TATGAGTGTGGAGTCGTTGCTGGCCGGGATTTCTAGTATGATGNNNAGATCGGAA
33805 TATGAGTGTGGAGTCGTTGCAGTAGACAGATTCAATGCAAATTCCANNNAGATCGG
33855 TATGAGTGTGGAGTCGTTGCAAATAAAGGTCCAATATCCCAAACATNNNAGATCGG
34121 TATGAGTGTGGAGTCGTTGCGGCAGACATAAATCCAACTATATCAANNNAGATCGG

35095 TATGAGTGTGGAGTCGTTGCAGCAACTTTTGGATTTGTTTGATCTNNNAGATCGGA

35366 TATGAGTGTGGAGTCGTTGCCTCCTGAGTAGGTGGGACTACANNNAGATCGGAAG

35381 TATGAGTGTGGAGTCGTTGCTGTCCTCTTTTTTTTTTTGAGACGGANNNAGATCGGA

35424 TATGAGTGTGGAGTCGTTGCCACAAGCCAATGTCATATAGTTCACANNNAGATCGG

35467 TATGAGTGTGGAGTCGTTGCAGTTTCTCATTTATTTGTCCTCATTGTNNNAGATCGG

35487 TATGAGTGTGGAGTCGTTGCCTCAAGCAATTCTTCTGCCTCAGNNNAGATCGGAAG

35506 TATGAGTGTGGAGTCGTTGCTTCTAGTGGCACAATCTCTCCACNNNAGATCGGAAG

35549 TATGAGTGTGGAGTCGTTGCTCTAACTGCCAAGCTGAACTGATNNNAGATCGGAAG

35608 TATGAGTGTGGAGTCGTTGCGTCTCAGCCTCCCAAGTAGACTANNNAGATCGGAAG

35637 TATGAGTGTGGAGTCGTTGCCCTTGTAGCGTACTTCTTCAATGANNNAGATCGGAA

35729 TATGAGTGTGGAGTCGTTGCGCACACTACAAACTTCACTTCCCNNNAGATCGGAAGA

35736 TATGAGTGTGGAGTCGTTGCACCTCCCAGATTCAAGCAATTCTNNNAGATCGGAAG

35834 TATGAGTGTGGAGTCGTTGCGCCTTCCTAGTAGCTGGGATTACNNNAGATCGGAAG

36054 TATGAGTGTGGAGTCGTTGCAGCTTTGGTTCTTTGATGTTCTATCCNNNAGATCGGA

36164 TATGAGTGTGGAGTCGTTGCAGTCAAACATACTTCTGCTTTATCCANNNAGATCGGA

36194 TATGAGTGTGGAGTCGTTGCTGCATTATTCTTACTGTATGTGGAGANNNAGATCGG

36222 TATGAGTGTGGAGTCGTTGCAGCTACATACATGGATAGATAGAAAGANNNAGATC

36292 TATGAGTGTGGAGTCGTTGCGTGGCCTGGCAATAAAAATAAAATCANNNAGATCG

36395 TATGAGTGTGGAGTCGTTGCACATGTACCACTCCATTACTAAATCANNNAGATCGGA

36472 TATGAGTGTGGAGTCGTTGCAGAACTTTTGAAAACACAGAACATGANNNAGATCGG

36760 TATGAGTGTGGAGTCGTTGCTCAAGAACTTCAGTAACGAATATGGTNNNAGATCGG

36781 TATGAGTGTGGAGTCGTTGCTCTTTTGTAACATAGAGGAGAAAGACANNNAGATCG

36801 TATGAGTGTGGAGTCGTTGCTCCTGATGAAGATGCAGAGAAACTNNNAGATCGGA

36838 TATGAGTGTGGAGTCGTTGCTGGCATCTACTACTTCTGAATTGTCANNNAGATCGGA

36997 TATGAGTGTGGAGTCGTTGCACGGAGAAATGAAATACAATCTCTGTNNNAGATCGG

37054 TATGAGTGTGGAGTCGTTGCAAACTTGTAAAAAGAGAACTACCACANNNAGATCGG

37058 TATGAGTGTGGAGTCGTTGCGAGGAAAGTAGAGTATCATTTCAGTGTNNNAGATC

37086 TATGAGTGTGGAGTCGTTGCGAGTCAGAATTGGTAAATGATGAATCTNNNAGATC

37117 TATGAGTGTGGAGTCGTTGCACCACACTCTAATACAAAACAAAGTTNNNAGATCGG

37124 TATGAGTGTGGAGTCGTTGCACCACAGAAAAAGATGTACTAAAAGGANNNAGATC

37183 TATGAGTGTGGAGTCGTTGCTCTGTAGTCCTAGAACTATTTGCTCANNNAGATCGG

37184 TATGAGTGTGGAGTCGTTGCTGGAAAAAGAAAGAAATGCAACAATGNNNAGATCG

37454 TATGAGTGTGGAGTCGTTGCAGAAAAGGAAAAAAAGCTAGGAAACTNNNAGATCG

37500 TATGAGTGTGGAGTCGTTGCAGAGCTCCTTTTATACTCAGGAAACTNNNAGATCGG

38231 TATGAGTGTGGAGTCGTTGCACCCACATAAACAAACAAAAAACTCANNNAGATCGG

38232 TATGAGTGTGGAGTCGTTGCAGTGGTTATTTCTGATTATGTGTGTTNNNAGATCGG

38265 TATGAGTGTGGAGTCGTTGCAGCAAAGCTAACCTTCAGAAATGANNNAGATCGGAA

38272 TATGAGTGTGGAGTCGTTGCTGTGTATGTCTATAAAGCTTAGCTCANNNAGATCGG

38289 TATGAGTGTGGAGTCGTTGCGTTGCCAAGAACATAAACTGGGGNNNAGATCGGAA

38346 TATGAGTGTGGAGTCGTTGCTGACAGACCATTAGTGAGATTATCCANNNAGATCGG

38354 TATGAGTGTGGAGTCGTTGCGTCTCTTCAAGAAATGGTGCTGGNNNAGATCGGAA

38822 TATGAGTGTGGAGTCGTTGCTGTACCACTTTCAAAAAAATGAGAGTNNNAGATCGG

38837 TATGAGTGTGGAGTCGTTGCACTGAACAGATCAATCATGAATTCCANNNAGATCGG

38969 TATGAGTGTGGAGTCGTTGCAGCTTATCTTGCTTCATATCCTTCCTNNNAGATCGGAA
38986 TATGAGTGTGGAGTCGTTGCTCACAAAAATCTGGAAAGATTACAGTNNNAGATCGG
39004 TATGAGTGTGGAGTCGTTGCTCAAATGCTTTTTTCTCACCTACTGANNNAGATCGGA
39034 TATGAGTGTGGAGTCGTTGCTCCATATAGGAAAAGAGGAAGTCACANNNAGATCGG
39240 TATGAGTGTGGAGTCGTTGCTTCAGTGACTTTCAATCTATGTTTCTNNNAGATCGGA
39289 TATGAGTGTGGAGTCGTTGCAGTACTCTCAGTGAAGAAATCTAAGTNNNAGATCGG
39480 TATGAGTGTGGAGTCGTTGCTGGATCAACTACAGCTATCTGCANNNAGATCGGAAG
39488 TATGAGTGTGGAGTCGTTGCACAAAGCTGTTTGATAACTGAATGGANNNAGATCGG
39514 TATGAGTGTGGAGTCGTTGCAGAAAGAATACAAGTTGAAAGGAGCTNNNAGATCG
39523 TATGAGTGTGGAGTCGTTGCCATGACTGCAAACTGTCACTTCCNNNAGATCGGAAGA
39547 TATGAGTGTGGAGTCGTTGCTGAAAGGACAAAAAGATAAATCCAGTNNNAGATCGG
39548 TATGAGTGTGGAGTCGTTGCTGAAAAGTCAAAGAACTTAGAATAGCTNNNAGATCG
39713 TATGAGTGTGGAGTCGTTGCGGGAAGAAATGAGGAGATGTAGGTNNNAGATCGG
39730 TATGAGTGTGGAGTCGTTGCTGGTTCACTTACCTCAATTAGTTTCCNNNAGATCGGA
40040 TATGAGTGTGGAGTCGTTGCAGGAAAAACATTAGAGAAATCATCGANNNAGATCGG
40048 TATGAGTGTGGAGTCGTTGCGATGAAGGTAAAATAACTGTGGATGANNNAGATCG
40121 TATGAGTGTGGAGTCGTTGCATCACGTCTATTGAACAAACTCATAANNNAGATCGGA
40301 TATGAGTGTGGAGTCGTTGCTGACAATACATATCAGCATGAATGAANNNAGATCGG
40623 TATGAGTGTGGAGTCGTTGCTTGGTAGAAAAAAGTAAAGCCTAGTTNNNAGATCGG
40708 TATGAGTGTGGAGTCGTTGCTGAACCCTTATTTCATACCATACACANNNAGATCGGA
40864 TATGAGTGTGGAGTCGTTGCACCAGAAAGTTTATATCTTGGTCAAANNNAGATCGGA
40872 TATGAGTGTGGAGTCGTTGCGGAACAGTTTGGTACTCGAGTCTNNNAGATCGGAAG
40877 TATGAGTGTGGAGTCGTTGCGCTACTGCCAATGATAATACTTGACANNNAGATCGGA
41035 TATGAGTGTGGAGTCGTTGCGTACAGTGGCAAAATCATGGCTTNNNAGATCGGAAG
41056 TATGAGTGTGGAGTCGTTGCATTCATTGCTGGTGGAAATGCAANNNAGATCGGAAG
41169 TATGAGTGTGGAGTCGTTGCACACAACACATGTCTAAATAATGGGTNNNAGATCGG
41182 TATGAGTGTGGAGTCGTTGCAGTCAGTGAACTTGAAGATAGAGCANNNAGATCGGA
41328 TATGAGTGTGGAGTCGTTGCACAGTCAAAGGACAAATATCTGTTCTNNNAGATCGGA
41360 TATGAGTGTGGAGTCGTTGCGGGTGAGGAACTGACAGAATTCANNNAGATCGGAA
41449 TATGAGTGTGGAGTCGTTGCTGTCCTGTGATAAACCTCAGTATTGTNNNAGATCGGA
41520 TATGAGTGTGGAGTCGTTGCAGACAGAAAGACAAGAAAGAAAGACANNNAGATCG
41637 TATGAGTGTGGAGTCGTTGCATGGAGTACAGTGGCACAATCTTNNNAGATCGGAAG
41684 TATGAGTGTGGAGTCGTTGCACCACACCCAGCTCATTTTTTTTNNNAGATCGGAAGA
41816 TATGAGTGTGGAGTCGTTGCGATCTCGGCTCACTGCAACTTCNNNAGATCGGAAGA
41881 TATGAGTGTGGAGTCGTTGCTCATAGCTATCTAGTGATTGGACAGANNNAGATCGG
41918 TATGAGTGTGGAGTCGTTGCCGAGTAGCTGGAATTACAGGTGTNNNAGATCGGAA
42263 TATGAGTGTGGAGTCGTTGCTGTTTTTATTGCCTAATTGCTCTGGTNNNAGATCGGA
42317 TATGAGTGTGGAGTCGTTGCCACATAGAAGTCACTGGTCCCAANNNAGATCGGAAG
42339 TATGAGTGTGGAGTCGTTGCAGAGTTTCATCAATTTGGTCAAACTNNNAGATCGGA
42347 TATGAGTGTGGAGTCGTTGCTATACATGTGTTCCCTCCGCAGNNNAGATCGGAAGA
42418 TATGAGTGTGGAGTCGTTGCATCATGGCTAACTGAAGTCTGCANNNAGATCGGAAG
42433 TATGAGTGTGGAGTCGTTGCAGAAAGGGAGAAAGAGAGAAGAAAGANNNAGATC
42500 TATGAGTGTGGAGTCGTTGCTATCACTCTTTCACCCAGGCTGNNNAGATCGGAAGA

42535 TATGAGTGTGGAGTCGTTGCACAGAATTCAAAAGAAAAGGGCANNNAGATCGGAA
42559 TATGAGTGTGGAGTCGTTGCTTCCACTAACAGGAGAACTTCAANNNAGATCGGAAG
42602 TATGAGTGTGGAGTCGTTGCTGTTTTTATTGCCTAATTGCTCTGGTNNNAGATCGGA
42652 TATGAGTGTGGAGTCGTTGCTTTTAGGTGCAGGGTCTTGTTCTNNNAGATCGGAAG
42876 TATGAGTGTGGAGTCGTTGCTCCAGTCAACCACTAATAGGAATGTNNNAGATCGGA
42878 TATGAGTGTGGAGTCGTTGCACACCATGGAGTACTATTCAGTCANNNAGATCGGAA
42884 TATGAGTGTGGAGTCGTTGCTGATGCAACATGTTATATGTAGTGCTNNNAGATCGG
43153 TATGAGTGTGGAGTCGTTGCATCTTCTCTTTTGGTCATTTGAGTAANNNAGATCGGA
43185 TATGAGTGTGGAGTCGTTGCTCTGTGAAGAGAGATAGTTTTCCCTNNNAGATCGGA
43281 TATGAGTGTGGAGTCGTTGCGTCATCAGGTTTCATTTATTGGAGGTNNNAGATCGG
43296 TATGAGTGTGGAGTCGTTGCACACCATAGCATACTAATCAGCCANNNAGATCGGAA
43300 TATGAGTGTGGAGTCGTTGCTTTTTGTGGTTACCATAAGGATTACANNNAGATCGG
43302 TATGAGTGTGGAGTCGTTGCTTCAGAAAGGAAACACAAACCACNNNAGATCGGAA
43380 TATGAGTGTGGAGTCGTTGCAGGTTCAAGCAATTCTCCTGTCTNNNAGATCGGAAG
43435 TATGAGTGTGGAGTCGTTGCCACGTACGGCTAATTTTTTGTGCNNNAGATCGGAAG
43916 TATGAGTGTGGAGTCGTTGCTGCACATTCAATAACATTCTCAGAGANNNAGATCGG
44224 TATGAGTGTGGAGTCGTTGCTCAAATGACATAAACCAGGAGGGANNNAGATCGGA
44450 TATGAGTGTGGAGTCGTTGCATCTTTGCAGAAATTATATCAGTGAGANNNAGATCG
44499 TATGAGTGTGGAGTCGTTGCTGATAGCATTCGTGTCTATATAGTGTNNNAGATCGG
44501 TATGAGTGTGGAGTCGTTGCACAAGGCTATAATAACCAAAACAGCANNNAGATCGG
44521 TATGAGTGTGGAGTCGTTGCACCTCTGGGAAGTGGAATTCTATGNNNAGATCGGAA
44532 TATGAGTGTGGAGTCGTTGCAGAGAGAGAGAGAGAGAGAATATTCANNNAGATC
44535 TATGAGTGTGGAGTCGTTGCTGCTGCTATAAACATGCATGTGCNNNAGATCGGAAG
44539 TATGAGTGTGGAGTCGTTGCACACATATAGTGGAATACTACTCAGTNNNAGATCGG
44549 TATGAGTGTGGAGTCGTTGCCACACACCACATTTTGTTAGTATTCANNNAGATCGGA
44552 TATGAGTGTGGAGTCGTTGCCAATGCCACGTTTCTTCTTGCTANNNAGATCGGAAG
44984 TATGAGTGTGGAGTCGTTGCACACATGGTTCTTTTACAAAAATGCGNNNAGATCGG
45102 TATGAGTGTGGAGTCGTTGCACTGCTAGTTGTCTGTGTTTGGANNNAGATCGGAAG
45741 TATGAGTGTGGAGTCGTTGCAAAAGGATGCCAAGGGGATTTTGNNNAGATCGGAA
45795 TATGAGTGTGGAGTCGTTGCGAGCGTGAGAGAGAGAGAGAAAGNNNAGATCGGA
45804 TATGAGTGTGGAGTCGTTGCCCTCCTAAGCTCAAGTGATCCTCNNNAGATCGGAAG
45926 TATGAGTGTGGAGTCGTTGCACCTAAAGAACTAAAATGCAAGTGCTNNNAGATCGG
46485 TATGAGTGTGGAGTCGTTGCGGTTCAAGTGATTCTGTTGCCTCNNNAGATCGGAAG
46487 TATGAGTGTGGAGTCGTTGCAGGCTGGTCTCAAACTTCTGATCNNNAGATCGGAAG
46531 TATGAGTGTGGAGTCGTTGCGGACATTCTAATAGGTGTGTAGTGGTNNNAGATCG
46670 TATGAGTGTGGAGTCGTTGCTCAGTCTCCCAAGTAGCTAGGATNNNAGATCGGAAG
47128 TATGAGTGTGGAGTCGTTGCGATCTCAGCTCACTGCAACCTTANNNAGATCGGAAG
47194 TATGAGTGTGGAGTCGTTGCAGGTTCAAGTGATTCTCGTGTCTNNNAGATCGGAAG
47709 TATGAGTGTGGAGTCGTTGCACGTGCAGCCAAACAAAACTAATNNNAGATCGGAAG
47807 TATGAGTGTGGAGTCGTTGCCAGGTTCAAGCAATTCTCCTGCNNNAGATCGGAAGA
47877 TATGAGTGTGGAGTCGTTGCCCAAGTAGCTGAGACTACATGCANNNAGATCGGAA
48088 TATGAGTGTGGAGTCGTTGCGTTCAAGGGATTATCCTGCCTCANNNAGATCGGAAG
48311 TATGAGTGTGGAGTCGTTGCAAAGGCAAAGCATCTTTGTACCCNNNAGATCGGAAG

48395 TATGAGTGTGGAGTCGTTGCCACCACACCCAGCTAATTTTTGTNNNAGATCGGAAG
48576 TATGAGTGTGGAGTCGTTGCAGCCACAGTTTTTTTTTTTTCCTTCCNNNAGATCGGA
48657 TATGAGTGTGGAGTCGTTGCTGATTCTCACCTAGAACAATATTGGTNNNAGATCGG
48855 TATGAGTGTGGAGTCGTTGCGGTTCAAGTGATTCTTCTGCCTCNNNAGATCGGAAG
48926 TATGAGTGTGGAGTCGTTGCAGGCAAGTCAGATTAGCTTGTGANNNAGATCGGAA
48977 TATGAGTGTGGAGTCGTTGCCTTCCCTGGATAGTGAGCTTCTCNNNAGATCGGAAG
49055 TATGAGTGTGGAGTCGTTGCACTGATGCCATCATAGCTCACTGNNNAGATCGGAAG
49328 TATGAGTGTGGAGTCGTTGCAAAGATAGTCATGGCCAGTGGTCNNNAGATCGGAA
49497 TATGAGTGTGGAGTCGTTGCTTTTGGTTCTTTATCCTGCTGGCNNNAGATCGGAAG
49499 TATGAGTGTGGAGTCGTTGCAGCCTCAGGAGTAGATGGGATTANNNAGATCGGAA
49939 TATGAGTGTGGAGTCGTTGCCATGGGTGAGAATGGGGAAGTAANNNAGATCGGA
50039 TATGAGTGTGGAGTCGTTGCACTTCTCAGCCATAAAATACAATGAANNNAGATCGG
50151 TATGAGTGTGGAGTCGTTGCTGAGGATGATGTGCAAGCTTACTNNNAGATCGGAA
50841 TATGAGTGTGGAGTCGTTGCACCTACATTTAGTACACAAGCAGTCTNNNAGATCGG
50880 TATGAGTGTGGAGTCGTTGCTTTTCTGCTAGGTTCATGGGTGTNNNAGATCGGAAG
51018 TATGAGTGTGGAGTCGTTGCAGTGAAGAAACACAAGAACTATAGCANNNAGATCG
51120 TATGAGTGTGGAGTCGTTGCTATCTCTCTCAGCGTATGGCCTANNNAGATCGGAAG
51309 TATGAGTGTGGAGTCGTTGCGTCTGTGTATTGGGGAGGTAAGGNNNAGATCGGA
51731 TATGAGTGTGGAGTCGTTGCTGCTATTGAGAAACAGATAAGTTGCTNNNAGATCGG
52023 TATGAGTGTGGAGTCGTTGCTGCCTTGTATTATGTAACCTTGATGGNNNAGATCGG
53133 TATGAGTGTGGAGTCGTTGCAGGTGTTTTGAAAACATATGGAACCTNNNAGATCGG
53195 TATGAGTGTGGAGTCGTTGCCTATCATGGGTCACTACAGCCTCNNNAGATCGGAAG
53376 TATGAGTGTGGAGTCGTTGCACATCTACATGCATATATGTGTGTGTNNNAGATCGG
53420 TATGAGTGTGGAGTCGTTGCAGAAAGAGAGAGAAAGAAAGAAAGGANNNAGATC
28775 TATGAGTGTGGAGTCGTTGCGGGTGTGTATTTACAACAATCTTGCNNNAGATCGGA
28786 TATGAGTGTGGAGTCGTTGCAACTCAGCCATAATAAGCAATGAATNNNAGATCGGA
28810 TATGAGTGTGGAGTCGTTGCAGATCTAGCTTGTAATTGTGTTGTTNNNAGATCGGA
28831 TATGAGTGTGGAGTCGTTGCGAGGGAGAGAGAGAGAGAGAGAGNNNAGATCGG
28927 TATGAGTGTGGAGTCGTTGCGAAGGCATTGTGTAGGTTGCATTNNNAGATCGGAA
28994 TATGAGTGTGGAGTCGTTGCTGCATAAGATATTGGCCAGGTGTNNNAGATCGGAA
29017 TATGAGTGTGGAGTCGTTGCGAGAGAAAGTAGCAGGTAATTAGGTNNNAGATCG
29042 TATGAGTGTGGAGTCGTTGCCCAAACTGTGTAATCTTCAACTTGCNNNAGATCGGA
29055 TATGAGTGTGGAGTCGTTGCTTCAAGTTGCTTTTCCATGATGANNNAGATCGGAAG
29066 TATGAGTGTGGAGTCGTTGCTGGTTGTTTTCACCTTTGAACTGTNNNAGATCGGAA
29088 TATGAGTGTGGAGTCGTTGCTGTCATGTTATATGCCTCCTTCTCTNNNAGATCGGAA
29108 TATGAGTGTGGAGTCGTTGCTGCAGTTTACTCAAAAGGTGAACANNNAGATCGGA
29110 TATGAGTGTGGAGTCGTTGCTCTGAGGGAAATTAGAAGAGTGCTNNNAGATCGGA
29137 TATGAGTGTGGAGTCGTTGCGGAAGGGAAAAAACAATAGGATGGTNNNAGATCG
29143 TATGAGTGTGGAGTCGTTGCCACCATGGAATACTACTCAGCCANNNAGATCGGAAG
29176 TATGAGTGTGGAGTCGTTGCAGCCCTATTCCTTATATTGTGCCTNNNAGATCGGAAG
29188 TATGAGTGTGGAGTCGTTGCCACACACACATATATACACACACACANNNAGATCGGA
29213 TATGAGTGTGGAGTCGTTGCTGAGTTAGGTCTCCTCCCTCATTNNNAGATCGGAAG
29302 TATGAGTGTGGAGTCGTTGCAAAAAAGGAGCTTTCTGGGCTTCNNNAGATCGGAA

29363 TATGAGTGTGGAGTCGTTGCTGGTTCATGGGATACTGTGTACCNNNAGATCGGAA
30792 TATGAGTGTGGAGTCGTTGCACACACACAAATACTCTTTTCAGGGNNNAGATCGGA
30794 TATGAGTGTGGAGTCGTTGCACTCCTAGCTAAGAAATGCAGGGNNNAGATCGGAA
30797 TATGAGTGTGGAGTCGTTGCTGTTTTGGAGACTCAAGGGGAAANNNAGATCGGAA
30805 TATGAGTGTGGAGTCGTTGCTAAAAATAAGTGGAAGCAGCCGGNNNAGATCGGAA
30835 TATGAGTGTGGAGTCGTTGCTGGAGAATTTGGGTAAAGCAGGANNNAGATCGGA
30932 TATGAGTGTGGAGTCGTTGCTCGAAAATCTTACCATTTCAGCACANNNAGATCGGA
30937 TATGAGTGTGGAGTCGTTGCCCTCTCTCTCCCTCTTATTCTACCANNNAGATCGGAAG
31056 TATGAGTGTGGAGTCGTTGCAGCAACTTAGAGCTCATTGACCANNNAGATCGGAAG
31073 TATGAGTGTGGAGTCGTTGCTTCACTGCCTCTTCTCATGTTGTNNNAGATCGGAAGA
31076 TATGAGTGTGGAGTCGTTGCAGCTTGATACCTGACAGAATATCCTNNNAGATCGGA
31108 TATGAGTGTGGAGTCGTTGCACCTAACTGAGTCACTGAAATCACTNNNAGATCGGA
31182 TATGAGTGTGGAGTCGTTGCTCTCCCACACTAAGCAATTCTCCNNNAGATCGGAAGA
31277 TATGAGTGTGGAGTCGTTGCACACAAGACTATCCAGGAACCACNNNAGATCGGAAG
31280 TATGAGTGTGGAGTCGTTGCACAGTCTAGTATCTATGTCAAGCAANNNAGATCGGA
31301 TATGAGTGTGGAGTCGTTGCCAGAACAGTAGGTGACCACGTAANNNAGATCGGAA
31310 TATGAGTGTGGAGTCGTTGCTGGACATCCCTCCCTCAATCTAANNNAGATCGGAAG
31347 TATGAGTGTGGAGTCGTTGCTGGGTAGCAAAATTTGATAGATGCANNNAGATCGG
31422 TATGAGTGTGGAGTCGTTGCCCTTGGTTAGTACAGATTTTGGTCCNNNAGATCGGA
31425 TATGAGTGTGGAGTCGTTGCTGGTTCTGTTTCTCTGTAGAACTCTNNNAGATCGGA
31450 TATGAGTGTGGAGTCGTTGCGTTTGGGGCATGGATATTCATGGNNNAGATCGGAA
31481 TATGAGTGTGGAGTCGTTGCAGCACCATTTTTCCAATAGCATGTNNNAGATCGGAA
31495 TATGAGTGTGGAGTCGTTGCCTTCTCACCACTGGCAATAATGCNNNAGATCGGAAG
31516 TATGAGTGTGGAGTCGTTGCAGCAACCACTAGGAAAACTATACGANNNAGATCGG
31558 TATGAGTGTGGAGTCGTTGCTGTAAATGAGTTAGGGTTTGCTACANNNAGATCGG
31617 TATGAGTGTGGAGTCGTTGCAAGAGTTATCAGAGGCTGTGCTTNNNAGATCGGAA
31736 TATGAGTGTGGAGTCGTTGCACATGTGCAGGTTTGTTACATGGNNNAGATCGGAA
31739 TATGAGTGTGGAGTCGTTGCAGAGAGAGAGAGAAAGAGAGAGAGANNNAGATC
31794 TATGAGTGTGGAGTCGTTGCAGAAGGAAGAAAAGGAAAGGAAGGANNNAGATCG
31805 TATGAGTGTGGAGTCGTTGCTGTGTGTGTGTATATATGTGTGTGTNNNAGATCGG
31884 TATGAGTGTGGAGTCGTTGCTGGCTAAGTAGGAGGTTTATGTTGANNNAGATCGG
31885 TATGAGTGTGGAGTCGTTGCACCATCTCTTGAAGTGAACATCAGANNNAGATCGGA
31897 TATGAGTGTGGAGTCGTTGCAGAAGTATACGTGGCAAATTGATGTNNNAGATCGG
31903 TATGAGTGTGGAGTCGTTGCAAGAAGCAGGATGTCTTGGTCATNNNAGATCGGAA
31905 TATGAGTGTGGAGTCGTTGCTGACATTGACTCATTTTTGTGTGGTNNNAGATCGGA
31992 TATGAGTGTGGAGTCGTTGCACATGATGCTACATCTTGGAGATCTNNNAGATCGGA
32094 TATGAGTGTGGAGTCGTTGCTCAGAGGAGCTAAAAAAACACAGGNNNAGATCGGA
32282 TATGAGTGTGGAGTCGTTGCTGCAACAACTGATGGACCTGTATNNNAGATCGGAAG
32492 TATGAGTGTGGAGTCGTTGCTTCAGCTTGGCTATTGGCGTATANNNAGATCGGAAG
32504 TATGAGTGTGGAGTCGTTGCGGTACAATCTTGGCTCATGCAACNNNAGATCGGAAG
32556 TATGAGTGTGGAGTCGTTGCGAATTCCTGAGCTCAAGCAATCCNNNAGATCGGAAG
32595 TATGAGTGTGGAGTCGTTGCTCAGACTCCATGCATAAAATAGGGANNNAGATCGGA
32628 TATGAGTGTGGAGTCGTTGCGCAAGGTACAACATAAACAAACTGCNNNAGATCGGA

32659 TATGAGTGTGGAGTCGTTGCGTTCGAGCGATTATTTTGCCTCANNNAGATCGGAAG
32673 TATGAGTGTGGAGTCGTTGCAATACAGCACACTGATGGGTCTTNNNAGATCGGAAG
32742 TATGAGTGTGGAGTCGTTGCCCATATAACCCAGCAATTCCATTCCNNNAGATCGGAA
32833 TATGAGTGTGGAGTCGTTGCATTCATGTGATTCTCCTGCCTCANNNAGATCGGAAGA
32838 TATGAGTGTGGAGTCGTTGCGGGTTTTGTAGTTGGTGTGCATANNNAGATCGGAA
32861 TATGAGTGTGGAGTCGTTGCGATCTCAGCTCACTGCAACCTATNNNAGATCGGAAG
32900 TATGAGTGTGGAGTCGTTGCAGAGTACAACATAGTTTCAGTTGCTNNNAGATCGGA
32977 TATGAGTGTGGAGTCGTTGCACAGAAAACAGAAGTGAGGTACAGANNNAGATCGG
32997 TATGAGTGTGGAGTCGTTGCACAAAGAAACAAACAAAACCCCCNNNAGATCGGAAG
33195 TATGAGTGTGGAGTCGTTGCACCTATGGCCATTTGATCTCTTACANNNAGATCGGAA
33216 TATGAGTGTGGAGTCGTTGCTGACTTTGTTTATGTTCTGGTTGCANNNAGATCGGA
33218 TATGAGTGTGGAGTCGTTGCCAAAACGTTGGAATAAGCAGAAAGTNNNAGATCGG
33288 TATGAGTGTGGAGTCGTTGCAGGGAATGGATATAACACACTTTGTNNNAGATCGG
33310 TATGAGTGTGGAGTCGTTGCGGAAGAAAGGCCAGCAACATTTTNNNAGATCGGAA
33392 TATGAGTGTGGAGTCGTTGCAGAGAGAGAGAGAGAGAGAGAAAGANNNAGATCG
33470 TATGAGTGTGGAGTCGTTGCGGAGTAAATGCAGTGTATAGTGAAANNNAGATCGG
33474 TATGAGTGTGGAGTCGTTGCACACATGAAAAGATTAGAGGGAACANNNAGATCGG
33513 TATGAGTGTGGAGTCGTTGCTGCTGGTAGAGTAAACATTCAGTGANNNAGATCGG
33628 TATGAGTGTGGAGTCGTTGCTACTGGCAGCGTAGAATGAGTTTNNNAGATCGGAA
33812 TATGAGTGTGGAGTCGTTGCACCAATTCAAATTTGTTACCCTTGANNNAGATCGGAA
33866 TATGAGTGTGGAGTCGTTGCAGTACGTCAGAATTCTCTAGTGAAANNNAGATCGGA
33949 TATGAGTGTGGAGTCGTTGCTCAAGTGATTCTCCTGCCTTAGCNNNAGATCGGAAG
33951 TATGAGTGTGGAGTCGTTGCACTATGCCCGGCGAATTTTTTCNNNAGATCGGAAGA
34148 TATGAGTGTGGAGTCGTTGCAGTGATCTATTGCACTATAGGGTGANNNAGATCGG
34207 TATGAGTGTGGAGTCGTTGCAGATGGGACTAACAGCAAATTAGGTNNNAGATCGG
34253 TATGAGTGTGGAGTCGTTGCCTATTGGAAAAGATGGGCAGCACNNNAGATCGGAA
34415 TATGAGTGTGGAGTCGTTGCTCAGTGGTTATAGGATGAATTACCTNNNAGATCGGA
34479 TATGAGTGTGGAGTCGTTGCACGGATACCAAATTCTGAGGATTCTNNNAGATCGGA
34668 TATGAGTGTGGAGTCGTTGCAGAAGCCTTGTTTTTTTCACACTNNNAGATCGGAAG
34792 TATGAGTGTGGAGTCGTTGCTCTGCTTCCTGAGTTCAAGTGATNNNAGATCGGAAG
34941 TATGAGTGTGGAGTCGTTGCCACCACACCTGGCTAATTTTTGTNNNAGATCGGAAG
35044 TATGAGTGTGGAGTCGTTGCTTCAAGTGATTCTCCTGCCTCAGNNNAGATCGGAAG
35077 TATGAGTGTGGAGTCGTTGCTTCAAGCAATTCTCCTGCCTCAGNNNAGATCGGAAG
35092 TATGAGTGTGGAGTCGTTGCCACAATCTCAGCTCACTGCAACNNNAGATCGGAAGA
35104 TATGAGTGTGGAGTCGTTGCGATCTCGGCTCACTGCAACTTNNNAGATCGGAAGAG
35154 TATGAGTGTGGAGTCGTTGCAGTTGTTCCTAATAGTCTCCGATGTNNNAGATCGGA
35245 TATGAGTGTGGAGTCGTTGCACAATCTCGGCTCATTGTAACCTNNNAGATCGGAAG
35272 TATGAGTGTGGAGTCGTTGCAGAGTAGCTGGGACTATAGGCANNNAGATCGGAAG
35376 TATGAGTGTGGAGTCGTTGCCAGGTTCAAGCAATTCTCCTGCNNNAGATCGGAAGA
35543 TATGAGTGTGGAGTCGTTGCTTCAAGCTATTCTCCTGCCTCAGNNNAGATCGGAAGA
35567 TATGAGTGTGGAGTCGTTGCGTATCATAGCTGGGACTACAGGCNNNAGATCGGAA
35640 TATGAGTGTGGAGTCGTTGCAGGATTTTTATCAGGAAGAGATGCTNNNAGATCGG
36015 TATGAGTGTGGAGTCGTTGCTGGCTAAGTAGGAGGTTTATGTTGANNNAGATCGG

36019 TATGAGTGTGGAGTCGTTGCACCATCTCTTGAAGTGAACATCAGANNNAGATCGGA
36090 TATGAGTGTGGAGTCGTTGCAGAAGTATACGTGGCAAATTGATGTNNNAGATCGG
36109 TATGAGTGTGGAGTCGTTGCTGGTTACTCAATTCATTGAAGGTGTNNNAGATCGGA
36182 TATGAGTGTGGAGTCGTTGCGAAGACGATGACGAAAAAAAAAGAANNNAGATCG
36188 TATGAGTGTGGAGTCGTTGCAGAAAGAAAGTTGTCATCTAGTTACCNNNAGATCGG
36291 TATGAGTGTGGAGTCGTTGCGGAAATCTGAGAAAATCTGTTTCCANNNAGATCGGA
36665 TATGAGTGTGGAGTCGTTGCTGCAATGCAAAAATAAAGAGTGACANNNAGATCGG
36691 TATGAGTGTGGAGTCGTTGCAGAAAGAAAGGAGAAAGAAAGAGGANNNAGATCG
36699 TATGAGTGTGGAGTCGTTGCAGACTGGTTAGAGGTCATTTATGCANNNAGATCGG
36775 TATGAGTGTGGAGTCGTTGCGAGGAACTCCCTGTAGTGAAAAANNNAGATCGGAA
36855 TATGAGTGTGGAGTCGTTGCCCCCTTCTCTTATCTACTAGTTACCCNNNAGATCGGAA
36989 TATGAGTGTGGAGTCGTTGCAGAGGAAACAGTAAGGAGTGAAAGTNNNAGATCG
37051 TATGAGTGTGGAGTCGTTGCACATGTTGAGTGCTTACAATATGCTNNNAGATCGGA
37066 TATGAGTGTGGAGTCGTTGCACAAACTTTTGGACGATATTGGTGANNNAGATCGG
37095 TATGAGTGTGGAGTCGTTGCTCCTATTACCTAGCCCTTGATTTCTNNNAGATCGGAA
37190 TATGAGTGTGGAGTCGTTGCTCAAATCAGTGGATAGAGATGGACTNNNAGATCGG
37191 TATGAGTGTGGAGTCGTTGCAGTCAGTTGAGATAGTGAAGATCCANNNAGATCGG
37207 TATGAGTGTGGAGTCGTTGCAGATGGTAAATGTCTGAGAAACACANNNAGATCGG
37214 TATGAGTGTGGAGTCGTTGCACAAAGACTTGAGAAACATTGCTGTNNNAGATCGG
37252 TATGAGTGTGGAGTCGTTGCACAAAAAATTCTGAAGTGTTGAAGAGANNNAGATC
37264 TATGAGTGTGGAGTCGTTGCTTCTGTGGAAGAGTTATACAGCTATNNNAGATCGGA
37269 TATGAGTGTGGAGTCGTTGCCCCTTCTTCAGTTTTAGGGGTTANNNAGATCGGAAG
37276 TATGAGTGTGGAGTCGTTGCTGACAAACCTTTAGACAGACCATCTNNNAGATCGGA
37345 TATGAGTGTGGAGTCGTTGCTGTCTGTGTGTATATGTGTGTCTGTNNNAGATCGGA
37528 TATGAGTGTGGAGTCGTTGCAGACATGTGTATTCAGTTCCACAGANNNAGATCGGA
37543 TATGAGTGTGGAGTCGTTGCGGGTAATACTTTTCGTGTAGTGTTTNNNAGATCGGA
37602 TATGAGTGTGGAGTCGTTGCTGTAACAAGGTGTTGAGGGCTTANNNAGATCGGAA
37603 TATGAGTGTGGAGTCGTTGCAGAGAGAGAAAGAAGAAAGACAGACANNNAGATC
37757 TATGAGTGTGGAGTCGTTGCATTTCAAGAGTTGTAATTCGTCTGANNNAGATCGGA
37781 TATGAGTGTGGAGTCGTTGCAGTGGATGGGTATATATGGTGTTCANNNAGATCGG
37853 TATGAGTGTGGAGTCGTTGCGCCAAAAAGACAAATTTAGTGTCAGGNNNAGATCG
37962 TATGAGTGTGGAGTCGTTGCAGAGAGAGAGAGAGGAAAGAAAGGANNNAGATC
37993 TATGAGTGTGGAGTCGTTGCACTGTAATGCTATTTTGTTCACATGACNNNAGATCGG
38028 TATGAGTGTGGAGTCGTTGCACTGGGTAAAAAAAAGCAAGAACANNNAGATCGGA
38069 TATGAGTGTGGAGTCGTTGCTCAACATACAAAAATCAGTAGCACTNNNAGATCGGA
38075 TATGAGTGTGGAGTCGTTGCTGGCCCAATCAAAAATTACACTTCTNNNAGATCGGA
38313 TATGAGTGTGGAGTCGTTGCAGATCAAAAAGAACAAAGAAAGGCANNNAGATCGG
38379 TATGAGTGTGGAGTCGTTGCTGTTGACTAGTATTCCATTGTCTGTNNNAGATCGGA
38382 TATGAGTGTGGAGTCGTTGCAGTGTCAACCTAACATAACAGACCANNNAGATCGGA
38402 TATGAGTGTGGAGTCGTTGCTGCATCACTTCCATTTTATCTGTCANNNAGATCGGAA
38453 TATGAGTGTGGAGTCGTTGCGCAGACAAATCTTTTTTTGAGACGGNNNAGATCGGA
38555 TATGAGTGTGGAGTCGTTGCACCAAAATTATGAAAACACCCAACANNNAGATCGGA
38656 TATGAGTGTGGAGTCGTTGCTGTCACTCAACTAAAACTATGAAACTNNNAGATCGG

38758 TATGAGTGTGGAGTCGTTGCGCCAGTAGCATTGATGTAGTATTTTNNNAGATCGGA
38785 TATGAGTGTGGAGTCGTTGCACCTGCGAAAAAAATCAAGGAGGNNNAGATCGGAA
38804 TATGAGTGTGGAGTCGTTGCTCAAAGATGACACAAACAAATGGGTNNNAGATCGG
38812 TATGAGTGTGGAGTCGTTGCACCAAGAATACACTACACCAAGTCANNNAGATCGGA
38826 TATGAGTGTGGAGTCGTTGCCGAAAGAAGGGTGTTACATAATGGTNNNAGATCGG
38835 TATGAGTGTGGAGTCGTTGCACGAAGTTTCAGAAGAGAGTTCTCANNNAGATCGG
38854 TATGAGTGTGGAGTCGTTGCTGAAACAAAAAGCTGGTTCTTGGANNNAGATCGGA
38894 TATGAGTGTGGAGTCGTTGCAGACAACTTACATAGCGGAAGAAAANNNAGATCGG
38897 TATGAGTGTGGAGTCGTTGCTGAACTAAAGGAGATTGAGACATGANNNAGATCGG
38944 TATGAGTGTGGAGTCGTTGCAGCAGAACTGAAGGAGATAGAAACANNNAGATCGG
39001 TATGAGTGTGGAGTCGTTGCGCAAGAAAAAAACCAAACAATCCCANNNAGATCGGA
39010 TATGAGTGTGGAGTCGTTGCAGTCTACCCAAATGAGAAGAAACAGANNNAGATCG
39022 TATGAGTGTGGAGTCGTTGCAAGAAGCAGGATGTCTTGGTCATNNNAGATCGGAA
39047 TATGAGTGTGGAGTCGTTGCTCAAGGTATATAAACACATGCAGGANNNAGATCGG
39081 TATGAGTGTGGAGTCGTTGCTCCTAGAAAAACATGAAAAGCAAAGCNNNAGATCG
39090 TATGAGTGTGGAGTCGTTGCGGTCAAGTGGGATTTATCATAGGGANNNAGATCGG
39110 TATGAGTGTGGAGTCGTTGCATACCATGTCTCCTGTGTTTTCANNNAGATCGGAAG
39140 TATGAGTGTGGAGTCGTTGCAGAGAGATTTTGAGTGTTCTCACCANNNAGATCGGA
39158 TATGAGTGTGGAGTCGTTGCTCCAAATTGGAAAGAAAGCAATCCANNNAGATCGGA
39168 TATGAGTGTGGAGTCGTTGCATTCAAGCGATTCTCCTGCCTTANNNAGATCGGAAG
39304 TATGAGTGTGGAGTCGTTGCAGCTAAAGGACCAGAAAAAAGAAGCNNNAGATCGG
39379 TATGAGTGTGGAGTCGTTGCTCAGGATAAAATGTGAAAACGGACANNNAGATCGG
39553 TATGAGTGTGGAGTCGTTGCTCACCACCACAATCAACATGAAGNNNAGATCGGAAG
39652 TATGAGTGTGGAGTCGTTGCTCAGAGCAGCAATAAATGAAGTTGANNNAGATCGG
39682 TATGAGTGTGGAGTCGTTGCCCAAAGAATGGGAGAAAAAATCTGCNNNAGATCGG
39687 TATGAGTGTGGAGTCGTTGCAAACTTCTAGGTAGAATGACATGGANNNAGATCGG
39720 TATGAGTGTGGAGTCGTTGCTTCATGACATTCTCCTGCCTCAGNNNAGATCGGAAG
39721 TATGAGTGTGGAGTCGTTGCTCCATATTTCCTTTCCTCCTTTCCANNNAGATCGGAAG
39734 TATGAGTGTGGAGTCGTTGCTTGAATTTCAACTTCACCACTTACTNNNAGATCGGAA
39742 TATGAGTGTGGAGTCGTTGCAGAAAAGAAAAATCAACTAGTGCAGANNNAGATCG
39751 TATGAGTGTGGAGTCGTTGCACAAGGAAAACTACAAAATGCACTGNNNAGATCGG
39774 TATGAGTGTGGAGTCGTTGCCTGAAATCATACCAACCACGCTCNNNAGATCGGAAG
39925 TATGAGTGTGGAGTCGTTGCACAACATATGTACCTACCCTCAAGGNNNAGATCGGA
40007 TATGAGTGTGGAGTCGTTGCAGTCAGTCCTTATTTCCCTTCTTTCANNNAGATCGGA
40071 TATGAGTGTGGAGTCGTTGCCCACTGAATGTTTTATCTGTTTGCCNNNAGATCGGAA
40168 TATGAGTGTGGAGTCGTTGCTCCCCCATCATTTCATTATGAGTGTNNNAGATCGGAA
40174 TATGAGTGTGGAGTCGTTGCCACAGAATGGGAGAAAATACTTGCANNNAGATCGG
40186 TATGAGTGTGGAGTCGTTGCACTGAAAAAATTCTCTGGTTAGTGGANNNAGATCGG
40189 TATGAGTGTGGAGTCGTTGCTCCCATAAGATTATAAGGGAGCTGCNNNAGATCGGA
40223 TATGAGTGTGGAGTCGTTGCTGCAGTCAGAAAAACAGAAAGTACANNNAGATCGG
40253 TATGAGTGTGGAGTCGTTGCCACCACTGTGGAAAGCAATTTGANNNAGATCGGAAG
40259 TATGAGTGTGGAGTCGTTGCATTTGAGGTCAGGAGTTCGAGACNNNAGATCGGAA
40280 TATGAGTGTGGAGTCGTTGCACAAAATATGATGCTCCAAAGGAAANNNAGATCGG

40311 TATGAGTGTGGAGTCGTTGCTCAAACACTTCAACCAAAAGACAGANNNAGATCGGA
40433 TATGAGTGTGGAGTCGTTGCACTGAAAAGTTTCCATGAACACAGTNNNAGATCGGA
40508 TATGAGTGTGGAGTCGTTGCGTCTCCTGGGTTCAAGTGATTCTNNNAGATCGGAAG
40610 TATGAGTGTGGAGTCGTTGCTCTACGGGGTGGAATATAAAAAAGTNNNAGATCGG
40658 TATGAGTGTGGAGTCGTTGCTCTAACAGTCATCATGGTAACACCANNNAGATCGGA
40707 TATGAGTGTGGAGTCGTTGCACCTAACAAGGGTCTAGTATCCAGANNNAGATCGGA
40716 TATGAGTGTGGAGTCGTTGCGGAAACTAGAAGACCAATGTATCACTNNNAGATCGG
40790 TATGAGTGTGGAGTCGTTGCTGAGATGCAGTGTTAGAAATTCTGANNNAGATCGG
40826 TATGAGTGTGGAGTCGTTGCACAGAAACCTTAGACATCCATTCGANNNAGATCGGA
40861 TATGAGTGTGGAGTCGTTGCTGGCTTGGACATATTTTTCTGTAGCNNNAGATCGGA
40913 TATGAGTGTGGAGTCGTTGCAAACAAAACAAAAGACTAGGAAGGTNNNAGATCGG
40932 TATGAGTGTGGAGTCGTTGCCCTTGTGAATTGTGATTTACTGCCTNNNAGATCGGA
40942 TATGAGTGTGGAGTCGTTGCTCACTGATGAACATTTCTTCCATTTNNNAGATCGGAA
40965 TATGAGTGTGGAGTCGTTGCGTATGTGACGTGATGGATATGTAAANNNAGATCGG
40973 TATGAGTGTGGAGTCGTTGCACTCATATGCTGAATAAATACCAAACANNNAGATCG
41033 TATGAGTGTGGAGTCGTTGCAAGGTCAACAGGAGAAAATCCCANNNAGATCGGAA
41299 TATGAGTGTGGAGTCGTTGCTCAAGGTATATAAACACATGCAGGANNNAGATCGG
41380 TATGAGTGTGGAGTCGTTGCAGTGCTGAAAAGGAATGTTATTGCANNNAGATCGG
41385 TATGAGTGTGGAGTCGTTGCCAAGAAATGCATATAAAGGCTGATCANNNAGATCG
41413 TATGAGTGTGGAGTCGTTGCTGCAACCCTAGTCTCTGATAAAACANNNAGATCGGA
41591 TATGAGTGTGGAGTCGTTGCTTCAAGCAATTCTCCTGCCTCAGNNNAGATCGGAAG
41631 TATGAGTGTGGAGTCGTTGCGCTTCTAGATATCCAATTGTTCTGGNNNAGATCGGA
41762 TATGAGTGTGGAGTCGTTGCAGTAACTGGGATTACATGCACCCNNNAGATCGGAAG
41803 TATGAGTGTGGAGTCGTTGCCCTCCTGGGTTCAAGTGATTCTCNNNAGATCGGAAG
41871 TATGAGTGTGGAGTCGTTGCGCATCTATTTGATTCCTCTGTCTTTNNNAGATCGGAA
41885 TATGAGTGTGGAGTCGTTGCTCCCTCTTTTTGCTTTGGAAACCNNNAGATCGGAAGA
42058 TATGAGTGTGGAGTCGTTGCCCATACTACCTCTATCATTTGCCCANNNAGATCGGAA
42124 TATGAGTGTGGAGTCGTTGCACCAACAACCATCAGAGAATAGTNNNAGATCGGAAG
42283 TATGAGTGTGGAGTCGTTGCACTTCAACTTCCTGAGTAGCTGGNNNAGATCGGAAG
42348 TATGAGTGTGGAGTCGTTGCGGAAAGGCATTTCACATTCATGGANNNAGATCGGA
42377 TATGAGTGTGGAGTCGTTGCGCAATTTTCTTACCTCAGCCTCCNNNAGATCGGAAGA
42380 TATGAGTGTGGAGTCGTTGCGCTTGTCAAAAAAAGCAAAATTTGGANNNAGATCG
42405 TATGAGTGTGGAGTCGTTGCTCTTTCTTTCTTTCTTTTCTCACTTGTNNNAGATCGGA
42447 TATGAGTGTGGAGTCGTTGCTCTCTCTCTCTCTCTCTCTCTCTCNNNAGATCGGAAG
42490 TATGAGTGTGGAGTCGTTGCACAGGTCTAACACAGTGTCTTATGTNNNAGATCGGA
42519 TATGAGTGTGGAGTCGTTGCATAAAGCAGCTGTGGCAAAATGTNNNAGATCGGAA
42550 TATGAGTGTGGAGTCGTTGCAGAGACGAGGTTTCTCCATGTTGNNNAGATCGGAA
42580 TATGAGTGTGGAGTCGTTGCTCCTTCTTTACTGCTTTCTTTTGCANNNAGATCGGAA
42726 TATGAGTGTGGAGTCGTTGCCAGAGGTTGCAGTGAGAAGAGATNNNAGATCGGA
42738 TATGAGTGTGGAGTCGTTGCTGACAAAATGGGAGGAAAAAGACCNNNAGATCGGA
42840 TATGAGTGTGGAGTCGTTGCTGATGCAATGACATTTGGATTCTGTNNNAGATCGGA
42873 TATGAGTGTGGAGTCGTTGCTGTCACATTTTCTTTATCCACTCGTNNNAGATCGGAA
42895 TATGAGTGTGGAGTCGTTGCTGGAAGATTAGTTCTAGAACCCACTNNNAGATCGGA

42971 TATGAGTGTGGAGTCGTTGCACTACTCAGCCATAAAAAGGAATGANNNAGATCGGA
43007 TATGAGTGTGGAGTCGTTGCACACACACATATATACACACACTCGNNNAGATCGGAA
43036 TATGAGTGTGGAGTCGTTGCCACATATATATGCGTGTGTGTGTGTNNNAGATCGGA
43071 TATGAGTGTGGAGTCGTTGCTGAGTTGGTTCCACATTTTTGCANNNAGATCGGAAG
43148 TATGAGTGTGGAGTCGTTGCTGAAACAAATGAAAACACAACCCANNNAGATCGGAA
43184 TATGAGTGTGGAGTCGTTGCCCACGCTCGGCTAATTTTTTGTANNNAGATCGGAAG
43247 TATGAGTGTGGAGTCGTTGCAGGAAATAGTTAGCCAGGTGGTGNNNAGATCGGAA
43257 TATGAGTGTGGAGTCGTTGCACTTATGAGTTTTAGTAGCTTTTTGGTNNNAGATCG
43260 TATGAGTGTGGAGTCGTTGCGAGTGCAATGGTGATCTCAGCTCNNNAGATCGGAA
43268 TATGAGTGTGGAGTCGTTGCTTTTTTCCTCTGTATGTGTTTCTATCTNNNAGATCGGA
43274 TATGAGTGTGGAGTCGTTGCACAGGTTTAGTATTCCCCCATTACTNNNAGATCGGAA
43283 TATGAGTGTGGAGTCGTTGCTGGCCGGGATTTCTAGTATGATGNNNAGATCGGAA
43284 TATGAGTGTGGAGTCGTTGCAGAATGAGTGTGTGTTTACAGAAACNNNAGATCGG
43313 TATGAGTGTGGAGTCGTTGCCAGCCTCTGGTAACTATCCTTCTNNNAGATCGGAAGA
43414 TATGAGTGTGGAGTCGTTGCCAAGTAGCTGGGACTATAGGCACNNNAGATCGGAA
43442 TATGAGTGTGGAGTCGTTGCCCACCACACCTGGATAATTTTCGNNNAGATCGGAAG
43447 TATGAGTGTGGAGTCGTTGCGGGTCACATAAACCATTTTAGAGGGNNNAGATCGG
43496 TATGAGTGTGGAGTCGTTGCAGCCTCTCGAGTAGTTGGTACTANNNAGATCGGAAG
43539 TATGAGTGTGGAGTCGTTGCCTTCATTTGAGGTCCAGGAAGGGNNNAGATCGGAA
43685 TATGAGTGTGGAGTCGTTGCGCATGACTCAACAGGATGAGAGANNNAGATCGGAA
43982 TATGAGTGTGGAGTCGTTGCAGAGAGAGGAAGGAAGGAAGGAANNNAGATCGG
44088 TATGAGTGTGGAGTCGTTGCGACTGGAATGGGAGGATGTGAATNNNAGATCGGA
44444 TATGAGTGTGGAGTCGTTGCCTGGCATGTAGTGGCATGATCTTNNNAGATCGGAA
44469 TATGAGTGTGGAGTCGTTGCATTTTGCAGCAAGTTGGATGGAANNNAGATCGGAA
44546 TATGAGTGTGGAGTCGTTGCATGCATATTAGGGCTGGTTCCATNNNAGATCGGAAG
44553 TATGAGTGTGGAGTCGTTGCAGAGAGAGAGAGAGAGAGAGAGAGANNNAGATC
45053 TATGAGTGTGGAGTCGTTGCAGGGGTCAAGTTACTTCTTAGTACANNNAGATCGGA
45155 TATGAGTGTGGAGTCGTTGCAGTGGAAACACACAGTAGAGAAAGTNNNAGATCGG
45317 TATGAGTGTGGAGTCGTTGCTCTAATGGCTAATAAACTTGGAGCANNNAGATCGGA
45332 TATGAGTGTGGAGTCGTTGCACTTAGCATTGTGTTACAGTTGCCNNNAGATCGGAA
45346 TATGAGTGTGGAGTCGTTGCAACGTGACAATCAAATTTGGGCTNNNAGATCGGAA
45600 TATGAGTGTGGAGTCGTTGCACTGCTAGATGTCTAAGCTGTCTNNNAGATCGGAAG
45658 TATGAGTGTGGAGTCGTTGCGATCTCGGCTCACCACAACTTNNNAGATCGGAAGAG
45682 TATGAGTGTGGAGTCGTTGCTTCAAGTGATTCTCCTGCCTCAGNNNAGATCGGAAG
45824 TATGAGTGTGGAGTCGTTGCGACAGGGTATCACTCTATCACCCNNNAGATCGGAAG
45883 TATGAGTGTGGAGTCGTTGCCCAGACTAGTGTCCTCAAGCATTNNNAGATCGGAAG
46020 TATGAGTGTGGAGTCGTTGCACTCAAGCGATTATCTTGCCTCANNNAGATCGGAAG
46021 TATGAGTGTGGAGTCGTTGCCAATATCCGCTCACCACAACTTCNNNAGATCGGAAGA
46064 TATGAGTGTGGAGTCGTTGCGTATTGGGATTACAGGCATGAGCNNNAGATCGGAA
46140 TATGAGTGTGGAGTCGTTGCCTCCCAGGTTCAAGCGATTCTNNNAGATCGGAAGAG
46149 TATGAGTGTGGAGTCGTTGCTGGTTAGAGACTAGAATTGGGAAGANNNAGATCGG
46170 TATGAGTGTGGAGTCGTTGCCACCACACCTGGCTAATTTTTGTNNNAGATCGGAAG
46240 TATGAGTGTGGAGTCGTTGCCCACCACACCCAGTGATTTTTTTNNNAGATCGGAAGA

46399 TATGAGTGTGGAGTCGTTGCTCTGTCTAGTTTTGGTTATCAGGGTNNNAGATCGGA
46616 TATGAGTGTGGAGTCGTTGCTAGTTGGGACTATAGGCATGTGCNNNAGATCGGAA
46618 TATGAGTGTGGAGTCGTTGCGCATGATCTTGTTCTCAGCTCACNNNAGATCGGAAG
46652 TATGAGTGTGGAGTCGTTGCCACCACGCCCAGCTAATTTTTTNNNAGATCGGAAGA
46664 TATGAGTGTGGAGTCGTTGCCAATCTTGGCTAACTGCAACCTCNNNAGATCGGAAG
46904 TATGAGTGTGGAGTCGTTGCCTCCTGAGTAGCTGGGATTTCAGNNNAGATCGGAA
47196 TATGAGTGTGGAGTCGTTGCACTACGCCCGGCTAATTTTTTTTNNNAGATCGGAAGA
47264 TATGAGTGTGGAGTCGTTGCACAAATGGTGGCATACTATACATGCNNNAGATCGGA
47352 TATGAGTGTGGAGTCGTTGCACAAATTGATTTTTGACAAGGGTGTNNNAGATCGGA
47467 TATGAGTGTGGAGTCGTTGCGTGTCTGGATTCTTTCATTCAACGTNNNAGATCGGA
47507 TATGAGTGTGGAGTCGTTGCCATGCGAAATGGAAAGGGAGAAGNNNAGATCGGA
47580 TATGAGTGTGGAGTCGTTGCTTCAAGTGATTCTCCTGCCTCAGNNNAGATCGGAAG
47608 TATGAGTGTGGAGTCGTTGCCCACCATGCCCAGCTAATTTTTTNNNAGATCGGAAGA
47697 TATGAGTGTGGAGTCGTTGCTGGTTCAAGAGATTTTCCTGCCTNNNAGATCGGAAG
47705 TATGAGTGTGGAGTCGTTGCGTCCACACCCGGCTAATTTTTATNNNAGATCGGAAG
47714 TATGAGTGTGGAGTCGTTGCTGCCACTACACCCAGCTAATTTTNNNAGATCGGAAGA
47777 TATGAGTGTGGAGTCGTTGCAAGTAGCTGGGATTACAGGCACNNNAGATCGGAAG
47803 TATGAGTGTGGAGTCGTTGCCAAGTAGCTGGGATTACAGGCATNNNAGATCGGAA
47896 TATGAGTGTGGAGTCGTTGCTGCAATATCGGTTCACTGCAATCNNNAGATCGGAAG
48074 TATGAGTGTGGAGTCGTTGCTCACTTCCAGGTTTAGTTGTACCTNNNAGATCGGAA
48093 TATGAGTGTGGAGTCGTTGCCCACAACACCCAGCTAAGTTTTTNNNAGATCGGAAG
48106 TATGAGTGTGGAGTCGTTGCACTGTTGTTGGTTTCAAGTCTGTNNNAGATCGGAAG
48167 TATGAGTGTGGAGTCGTTGCTTCAAGTGATTCTCCTGCCTCAGNNNAGATCGGAAG
48190 TATGAGTGTGGAGTCGTTGCCCACCAAGCCTGGCTAATTTTTTNNNAGATCGGAAG
48252 TATGAGTGTGGAGTCGTTGCTCACTCATTATTAGTCTGTGCAGGTNNNAGATCGGA
48283 TATGAGTGTGGAGTCGTTGCCAAGTAGCTGGGACTACAGGTGNNNAGATCGGAAG
48359 TATGAGTGTGGAGTCGTTGCTCTGCTTCCTGAGTTCAAGTGATNNNAGATCGGAAG
48441 TATGAGTGTGGAGTCGTTGCGCTGGTACTGGGAAGACAAAGATNNNAGATCGGAA
48487 TATGAGTGTGGAGTCGTTGCTCCCCAACTTTTTGTTTTGATCTGTNNNAGATCGGAA
48628 TATGAGTGTGGAGTCGTTGCCTCCTGAGTAGCTGGGACTACAGNNNAGATCGGAA
48826 TATGAGTGTGGAGTCGTTGCCAATCTTGGCTCACTTCAAGCTCNNNAGATCGGAAG
48831 TATGAGTGTGGAGTCGTTGCTCTTGTCTTCTGCTAGCTTTGGGNNNAGATCGGAAG
48866 TATGAGTGTGGAGTCGTTGCCCACCATGCCCAGAGAATTTTTTNNNAGATCGGAAG
48879 TATGAGTGTGGAGTCGTTGCGCCACCATGCCTAGCTAATTTTTNNNAGATCGGAAG
49002 TATGAGTGTGGAGTCGTTGCAGGCTAGCATAAACAGTGTAGAGAANNNAGATCGG
49069 TATGAGTGTGGAGTCGTTGCGTGTTCAAGCAATTCTTCTGCCTNNNAGATCGGAAG
49138 TATGAGTGTGGAGTCGTTGCCTCCTGGGTTCAAGTGATTCTCANNNAGATCGGAAG
49162 TATGAGTGTGGAGTCGTTGCGCCTCCCAAGTAGCTAGGATTACNNNAGATCGGAAG
49166 TATGAGTGTGGAGTCGTTGCGGTTCAAACAATTCTCCTGCCTCNNNAGATCGGAAG
49241 TATGAGTGTGGAGTCGTTGCCCGAGTAGCTGGAATTACAGGTTNNNAGATCGGAA
49282 TATGAGTGTGGAGTCGTTGCCACCGTTCCTGGCTAATTTTTGTNNNAGATCGGAAG
49338 TATGAGTGTGGAGTCGTTGCACTGCATCTATTTCAATGATCACATNNNAGATCGGAA
49345 TATGAGTGTGGAGTCGTTGCAACGAAGGGTGCAATAGAAAACCNNNAGATCGGAA

49384 TATGAGTGTGGAGTCGTTGCCAAGCAATTCTCCTGCCTAGGTANNNAGATCGGAAG
49404 TATGAGTGTGGAGTCGTTGCTTCTGGAACTTCTACATTCACAGANNNAGATCGGAA
49459 TATGAGTGTGGAGTCGTTGCTCACTGTAACCTCAAACTCCTGGNNNAGATCGGAAG
49483 TATGAGTGTGGAGTCGTTGCTCTTACTCTTTTCTGCCAAAAATCTNNNAGATCGGAA
49509 TATGAGTGTGGAGTCGTTGCATTCAAGCGATTATCCTGCCTCANNNAGATCGGAAG
49511 TATGAGTGTGGAGTCGTTGCACTGGACAGGAAATAGTACTAAACANNNAGATCGG
49529 TATGAGTGTGGAGTCGTTGCCCATCTTGGCTCACTACAACCTNNNAGATCGGAAGA
49653 TATGAGTGTGGAGTCGTTGCTTGAATAGCTGGGACTACAGGTGNNNAGATCGGAA
49789 TATGAGTGTGGAGTCGTTGCTGTGCATTTTGAACACTAAGAACCANNNAGATCGGA
49844 TATGAGTGTGGAGTCGTTGCACTGTATATCAGAGTGTACCTGTGANNNAGATCGGA
49877 TATGAGTGTGGAGTCGTTGCTCAGTCAGCTATAGTTTAGTTTGGTNNNAGATCGGA
49923 TATGAGTGTGGAGTCGTTGCACCAAGCAGCATATTTCCAACATNNNAGATCGGAAG
50357 TATGAGTGTGGAGTCGTTGCTGTCCCTAAAGTCTTTGGTAATGGTNNNAGATCGGA
50555 TATGAGTGTGGAGTCGTTGCTATTGGTTGGTCTCTCAGAGCTCNNNAGATCGGAAG
51040 TATGAGTGTGGAGTCGTTGCTCCTTTCCATTTCTCTCTTGAACCTNNNAGATCGGAAG
52931 TATGAGTGTGGAGTCGTTGCTGACAACTGTTTAGACTTGAGGAGTNNNAGATCGG
53191 TATGAGTGTGGAGTCGTTGCGATCTCAAACTCCTGGCTTCGANNNAGATCGGAAGA
53337 TATGAGTGTGGAGTCGTTGCACAAAATCAGAAATGACAAAGGGGANNNAGATCGG
28765 TATGAGTGTGGAGTCGTTGCATATGTGTGTGTGTGTGTGTGTGNNNAGATCGGAA
28792 TATGAGTGTGGAGTCGTTGCACAACCTGATTATGCAGTTCCTTGNNNAGATCGGAA
28796 TATGAGTGTGGAGTCGTTGCTGATGGAATACTACTCAGCCATAANNNAGATCGGAA
28817 TATGAGTGTGGAGTCGTTGCGGTGCAATCATGGCTCACTAGANNNAGATCGGAAG
28820 TATGAGTGTGGAGTCGTTGCGTAATGGCATTCTGAGCAACCTGNNNAGATCGGAA
28821 TATGAGTGTGGAGTCGTTGCTGAAAATGGCATGAACAGGAACANNNAGATCGGAA
28823 TATGAGTGTGGAGTCGTTGCACATGTATGTGTGTGCATATGTGTNNNAGATCGGA
28839 TATGAGTGTGGAGTCGTTGCTTGATGCACACTAAAGTTGATTCTNNNAGATCGGAA
28843 TATGAGTGTGGAGTCGTTGCCTCCAGACTTTTTCATCTTCCCANNNAGATCGGAAGA
28848 TATGAGTGTGGAGTCGTTGCAGCTCCTTTGATCTGTGTTTCGANNNAGATCGGAAG
28855 TATGAGTGTGGAGTCGTTGCATGGAGGAGGCTGAAGAAAACTTNNNAGATCGGA
28891 TATGAGTGTGGAGTCGTTGCATGTCACTCTGGTAGTTGTCACCNNNAGATCGGAAG
28902 TATGAGTGTGGAGTCGTTGCGTGTGTGTGTGTGTGTATATATNNNAGATCGGA
28926 TATGAGTGTGGAGTCGTTGCTCAGGAGTTCACAGAAAACAATGANNNAGATCGGA
28938 TATGAGTGTGGAGTCGTTGCAGGGTCTTCAAACATATTCCCTGTNNNAGATCGGAA
28943 TATGAGTGTGGAGTCGTTGCGGTTATCACATCCTCTCACCCTCNNNAGATCGGAAGA
28948 TATGAGTGTGGAGTCGTTGCGCTGAAATTCTGTAGCAGTGTTGTNNNAGATCGGA
28949 TATGAGTGTGGAGTCGTTGCAGCCACATTTACCAAGTTGTAAAANNNAGATCGGAA
28957 TATGAGTGTGGAGTCGTTGCATAATGCTGTCTGGTGTTTGACGNNNAGATCGGAA
28969 TATGAGTGTGGAGTCGTTGCTCCTTTTTCTCTTGCTTCAGTCTGNNNAGATCGGAAG
28979 TATGAGTGTGGAGTCGTTGCACTGAGAATGGAAACTGTAAGAGANNNAGATCGGA
28986 TATGAGTGTGGAGTCGTTGCCATGTGCATATGTGTGTGTGTGTNNNAGATCGGAA
29002 TATGAGTGTGGAGTCGTTGCTGCTATTGTGTAAGGAAATCTGGTNNNAGATCGGA
29005 TATGAGTGTGGAGTCGTTGCCCAAATGGTCAGAAAGAGAGGGTNNNAGATCGGA
29021 TATGAGTGTGGAGTCGTTGCGCAATTTCTTCCATCGAGGTTTTGNNNAGATCGGAA

29034 TATGAGTGTGGAGTCGTTGCTGTGTGTGTATATATAGTGTGCGTNNNAGATCGGA

29038 TATGAGTGTGGAGTCGTTGCCACACAGAGAGAGAGAGAGAGAGANNNAGATCGG

29068 TATGAGTGTGGAGTCGTTGCACACATACACACACATATATATACACGNNNAGATCGG

29079 TATGAGTGTGGAGTCGTTGCAATGGCTGGGTTAGGTACTTTGTNNNAGATCGGAA

29080 TATGAGTGTGGAGTCGTTGCTGACATCCTGGTTTTTTTCCTTTNNNAGATCGGAAGA

29127 TATGAGTGTGGAGTCGTTGCCACCATTCCAAACCAGTGCATTANNNAGATCGGAAG

29151 TATGAGTGTGGAGTCGTTGCTGTGTGTATGAGAGAGAGAAAGCTNNNAGATCGG

29166 TATGAGTGTGGAGTCGTTGCTCCTGATCTCATGAGGGTAGACTNNNAGATCGGAAG

29170 TATGAGTGTGGAGTCGTTGCTGCTCAATCTCTCTCTCTCTCTCTNNNAGATCGGAAG

29208 TATGAGTGTGGAGTCGTTGCTCATTGCTGGATCTTACGGTAGGNNNAGATCGGAA

29227 TATGAGTGTGGAGTCGTTGCGAGGGGATAAGGGCTGATATCAGNNNAGATCGGA

29239 TATGAGTGTGGAGTCGTTGCCTTGACCCATCTGCACCATAGATNNNAGATCGGAAG

29250 TATGAGTGTGGAGTCGTTGCCTTGTCTTCCTCATCCCTCTGTCNNNAGATCGGAAGA

29252 TATGAGTGTGGAGTCGTTGCAGTGACTTTCTCCTTCTCCCCTANNNAGATCGGAAGA

29292 TATGAGTGTGGAGTCGTTGCTTTCTGGTCGATGATTTCTGGGTNNNAGATCGGAAG

29295 TATGAGTGTGGAGTCGTTGCACACTTCAGAAACTCTTAGGTGCANNNAGATCGGAA

29338 TATGAGTGTGGAGTCGTTGCTGATGGACACTTAGGTTGATTCCANNNAGATCGGAA

29349 TATGAGTGTGGAGTCGTTGCAGGATTACTGCAAGGGTAAACAGANNNAGATCGGA

29358 TATGAGTGTGGAGTCGTTGCACCATTTGGTGAGATGAGTCTTGTNNNAGATCGGA

29386 TATGAGTGTGGAGTCGTTGCGTGTCATCCCCAGAACACATGTANNNAGATCGGAAG

30787 TATGAGTGTGGAGTCGTTGCAGACTTTCCAAACCATACATTGCANNNAGATCGGAA

30810 TATGAGTGTGGAGTCGTTGCTGTCTAAAGTCTCCATTTCATGCANNNAGATCGGAA

30832 TATGAGTGTGGAGTCGTTGCTCAAGTGAAGAGCAAGACTCCAANNNAGATCGGAA

30862 TATGAGTGTGGAGTCGTTGCGGTGTGGTGCCTCATACTTAGAANNNAGATCGGAA

30867 TATGAGTGTGGAGTCGTTGCAGCTACAGATTTTTGGACCACTCNNNAGATCGGAAG

30917 TATGAGTGTGGAGTCGTTGCAGTTCTCCAAAGTAAATGGCTAGTNNNAGATCGGAA

31002 TATGAGTGTGGAGTCGTTGCGCTGGTAGTGTAACTCAGTCCAANNNAGATCGGAA

31009 TATGAGTGTGGAGTCGTTGCGATGGAGGGGTCACAACAAGTATNNNAGATCGGAA

31065 TATGAGTGTGGAGTCGTTGCCCTGTGAGCTCTATTTTTCTCTGCNNNAGATCGGAAG

31132 TATGAGTGTGGAGTCGTTGCTGTTTGTGTATTGTCTGCCTTTCCNNNAGATCGGAAG

31167 TATGAGTGTGGAGTCGTTGCTGTGTGTATGTTTATCCCACAGGANNNAGATCGGAA

31199 TATGAGTGTGGAGTCGTTGCACACGAACTCACACATTTTTACCCNNNAGATCGGAAG

31206 TATGAGTGTGGAGTCGTTGCAGTCATAGGAGCAGGAGAGATTACNNNAGATCGGA

31232 TATGAGTGTGGAGTCGTTGCGCTGGCTGGCTTTTTTTGTTTGNNNAGATCGGAAGA

31236 TATGAGTGTGGAGTCGTTGCTCTCTCCCTCTCTCTCTCTCTCTNNNAGATCGGAAGAG

31240 TATGAGTGTGGAGTCGTTGCGGATATACAGAAGCTGACCCCTGNNNAGATCGGAA

31250 TATGAGTGTGGAGTCGTTGCTTTTGCAAATGTTTTACCTCGCTNNNAGATCGGAAG

31251 TATGAGTGTGGAGTCGTTGCGAGTCATTGTGGCAGAAACAGACNNNAGATCGGAA

31264 TATGAGTGTGGAGTCGTTGCTCCATTATTACCTCCAACTCCAGTNNNAGATCGGAAG

31266 TATGAGTGTGGAGTCGTTGCATTTTCTCGTTGCCTTGCTTCAANNNAGATCGGAAGA

31320 TATGAGTGTGGAGTCGTTGCGATGCCTTTGCTTTTTGTCTTGCNNNAGATCGGAAG

31350 TATGAGTGTGGAGTCGTTGCTGTGTTTCTACCTATTGTGTGTGANNNAGATCGGAA

31376 TATGAGTGTGGAGTCGTTGCTGACTGAGTAGTTTGAAGACATGCNNNAGATCGGA

31404 TATGAGTGTGGAGTCGTTGCGTGTGTGTGTGTGTGTGATTATNNNAGATCGGA

31426 TATGAGTGTGGAGTCGTTGCCCAGGACTCAGATTTTTTTCCCANNNAGATCGGAAG

31434 TATGAGTGTGGAGTCGTTGCACATCTAGGGAATGTCTATGTGTTNNNAGATCGGAA

31436 TATGAGTGTGGAGTCGTTGCGCTCTTTGCCACAACAGAGAAAGNNNAGATCGGAA

31457 TATGAGTGTGGAGTCGTTGCACATGATTTGGTAGGCTTTTCTCANNNAGATCGGAA

31458 TATGAGTGTGGAGTCGTTGCTGTAAGTTTTGAGCTAGGACTGCTNNNAGATCGGA

31460 TATGAGTGTGGAGTCGTTGCTGATACAGTATTTCTGAGCCCTCANNNAGATCGGAA

31492 TATGAGTGTGGAGTCGTTGCTTGTGAAGTCAGGTTGCTAATGCNNNAGATCGGAA

31508 TATGAGTGTGGAGTCGTTGCAGTTCTTTGCTTTGTCAAACTGCTNNNAGATCGGAA

31531 TATGAGTGTGGAGTCGTTGCCCCCCATATATATCCAGATAGCCANNNAGATCGGAA

31576 TATGAGTGTGGAGTCGTTGCTGGACACTTAGATTGATTCCACANNNAGATCGGAAG

31591 TATGAGTGTGGAGTCGTTGCTACAAAGATCATCACGACAGCCANNNAGATCGGAAG

31594 TATGAGTGTGGAGTCGTTGCAGAGAGAGAGAGAGAGAGAGTTTTNNNAGATCGG

31790 TATGAGTGTGGAGTCGTTGCAGGAGAGACAATGGTGTAAGTTCCNNNAGATCGGA

31797 TATGAGTGTGGAGTCGTTGCGAGTGCAGTAGTGTGATCTCAGTNNNAGATCGGAA

31809 TATGAGTGTGGAGTCGTTGCTGGGTCAAATAGGAAGTCTCAAGGNNNAGATCGGA

31843 TATGAGTGTGGAGTCGTTGCTGCTTTGTGTATTTGGGAACTCTGNNNAGATCGGAA

31847 TATGAGTGTGGAGTCGTTGCTCCACAAAAGCAATTGAAAACTCTNNNAGATCGGAA

31911 TATGAGTGTGGAGTCGTTGCACTGTACATATTTTTGTGGTACATGTGNNNAGATCG

32010 TATGAGTGTGGAGTCGTTGCAGAGAGAGAGAGAGAGAGATTCTCNNNAGATCGG

32012 TATGAGTGTGGAGTCGTTGCCTTCAACACCACACTTTCAGCATNNNAGATCGGAAGA

32046 TATGAGTGTGGAGTCGTTGCCTATCTTGGCTCACTGCAACCTCNNNAGATCGGAAG

32052 TATGAGTGTGGAGTCGTTGCCCTAGTCTCGGATAAAACAGACGTNNNAGATCGGAA

32141 TATGAGTGTGGAGTCGTTGCCAATCTCGGCTCACTGTAACCTCNNNAGATCGGAAG

32153 TATGAGTGTGGAGTCGTTGCTCGTTAGGTGGTAAAGATCTGACANNNAGATCGGA

32168 TATGAGTGTGGAGTCGTTGCTCCAGAGTAGCCGGGACTACNNNAGATCGGAAGAG

32181 TATGAGTGTGGAGTCGTTGCACCATGCGATATTTTAGATCCCCANNNAGATCGGAA

32210 TATGAGTGTGGAGTCGTTGCACGGGACAGAAGAAAATTTTGACTNNNAGATCGGA

32305 TATGAGTGTGGAGTCGTTGCGTTCAAGAGATTCTCCTGCCTCANNNAGATCGGAAG

32314 TATGAGTGTGGAGTCGTTGCCAACAAGAATGAAACTCCGTCTCANNNAGATCGGAA

32318 TATGAGTGTGGAGTCGTTGCTGCTGTAATGAACATGGGAATGCNNNAGATCGGAA

32320 TATGAGTGTGGAGTCGTTGCACAATACAATGAAACGTTGAGACGNNNAGATCGGA

32334 TATGAGTGTGGAGTCGTTGCAACTGTAATGCCTTTTAGTTCACANNNAGATCGGAA

32412 TATGAGTGTGGAGTCGTTGCTCACACAAAGACATACCATAGTGANNNAGATCGGAA

32438 TATGAGTGTGGAGTCGTTGCGTGATCTCGGTTCATTGCAACCNNNAGATCGGAAGA

32463 TATGAGTGTGGAGTCGTTGCTGTTCTTAGCAAAGTGCACAAACTNNNAGATCGGAA

32467 TATGAGTGTGGAGTCGTTGCGGGTGTAGTTTGGAAAATGTTTGCNNNAGATCGGA

32509 TATGAGTGTGGAGTCGTTGCTGGAATTCCTCTTGCTAGATAGGGNNNAGATCGGA

32514 TATGAGTGTGGAGTCGTTGCAGTGTGATTTCTGCAATGTAAACANNNAGATCGGAA

32549 TATGAGTGTGGAGTCGTTGCTGTGTCTCAAAAACAACAAACAGCNNNAGATCGGAA

32554 TATGAGTGTGGAGTCGTTGCGAGTAGCTGGTACTACAGGTGTGNNNAGATCGGAA

32608 TATGAGTGTGGAGTCGTTGCTGAAACAAAAGCAAACATCTTTTCTGANNNAGATCG

32636 TATGAGTGTGGAGTCGTTGCCTCCTGAGTAGCTGGGATTACGNNNAGATCGGAAG

32678 TATGAGTGTGGAGTCGTTGCTTCAAGTGATTCTCCTGCCTCAGNNNAGATCGGAAG
32690 TATGAGTGTGGAGTCGTTGCCCAAGAGACTAGCCCTTTTGTCANNNAGATCGGAAG
32825 TATGAGTGTGGAGTCGTTGCTTCCTGGTTTATTCTTGGGAGGGNNNAGATCGGAA
32871 TATGAGTGTGGAGTCGTTGCATCCAATTTGCCAGTCTGAGTCTNNNAGATCGGAAG
32873 TATGAGTGTGGAGTCGTTGCTCTGGCTTATATGGTTTCTGCTGANNNAGATCGGAA
32889 TATGAGTGTGGAGTCGTTGCACAATCAGCAGGCATTTACATAGGNNNAGATCGGA
32916 TATGAGTGTGGAGTCGTTGCACATTTGCATGCTTTTGACTTGTGNNNAGATCGGAA
32943 TATGAGTGTGGAGTCGTTGCCCATCCAAAGGTCATCAACATCAANNNAGATCGGAA
32957 TATGAGTGTGGAGTCGTTGCTGGAGGTGAGATCAATAGTTTCCANNNAGATCGGA
33040 TATGAGTGTGGAGTCGTTGCCTCTCGAGTAGCTGGGATTACAGNNNAGATCGGAA
33054 TATGAGTGTGGAGTCGTTGCTGGAGTTTGTTATTGGTCTGTTCANNNAGATCGGAA
33182 TATGAGTGTGGAGTCGTTGCTCACCCAACATTCTTTGATTGAGGNNNAGATCGGAA
33197 TATGAGTGTGGAGTCGTTGCATTTATTTTTTGAGACGGAGTTTCACNNNAGATCGG
33209 TATGAGTGTGGAGTCGTTGCGCTGCTAACAAACTGAATCCCTGNNNAGATCGGAAG
33335 TATGAGTGTGGAGTCGTTGCCTCCTGAGTAGCTGGGACTACAGNNNAGATCGGAA
33348 TATGAGTGTGGAGTCGTTGCTCCCTTCTCTTGCATTTTTTCAGANNNAGATCGGAAG
33361 TATGAGTGTGGAGTCGTTGCCTCAATCTTGGCTCACTGCAAGNNNAGATCGGAAGA
33397 TATGAGTGTGGAGTCGTTGCACAACTCAAATGTTCATCAGCTGANNNAGATCGGAA
33420 TATGAGTGTGGAGTCGTTGCAGCTAGGTTTTTTTCTAGATGCCCNNNAGATCGGAA
33439 TATGAGTGTGGAGTCGTTGCTGCAGGATACATCTCTGAAAGAGANNNAGATCGGA
33472 TATGAGTGTGGAGTCGTTGCTGTACATTGGGCTTTCTATCAGGANNNAGATCGGAA
33483 TATGAGTGTGGAGTCGTTGCTGGACTCAATGTGAATTTTTGAGANNNAGATCGGA
33551 TATGAGTGTGGAGTCGTTGCGGTTCAAGCAATTCTCCTGTCTCNNNAGATCGGAAG
33592 TATGAGTGTGGAGTCGTTGCGGGTTCAAGTGATTCTCCTGTCTNNNAGATCGGAAG
33661 TATGAGTGTGGAGTCGTTGCCTGCCTCTCAAGTAGTTGGGATTNNNAGATCGGAAG
33671 TATGAGTGTGGAGTCGTTGCGCAAGATCATGGCTCACTGTAACNNNAGATCGGAAG
33704 TATGAGTGTGGAGTCGTTGCTCCTGTATGTCAAGACAAATTCCANNNAGATCGGAA
33834 TATGAGTGTGGAGTCGTTGCGCAGGAATATAAGGAAAGCTGTGGNNNAGATCGG
33844 TATGAGTGTGGAGTCGTTGCTGCTTACTTATGCCTTCTACTTGCNNNAGATCGGAAG
33845 TATGAGTGTGGAGTCGTTGCAGAGATTGAATGCTACAGGAAGCANNNAGATCGGA
34036 TATGAGTGTGGAGTCGTTGCCAGTGACATGATCTTGGCTCACTNNNAGATCGGAAG
34050 TATGAGTGTGGAGTCGTTGCTAGCAGAGCTACTGATTTGTGCANNNAGATCGGAA
34106 TATGAGTGTGGAGTCGTTGCTCTCACAGAGAAAGTGAAATCCCTNNNAGATCGGAA
34126 TATGAGTGTGGAGTCGTTGCCAACCCAAGATTTCCTCAAAACCANNNAGATCGGAA
34128 TATGAGTGTGGAGTCGTTGCACCTAGGGAAGACTAAGCATCTCTNNNAGATCGGAA
34195 TATGAGTGTGGAGTCGTTGCTCTACTGTTTCAAGTGACTCACCTNNNAGATCGGAAG
34201 TATGAGTGTGGAGTCGTTGCAGGGCACCATAATTTTGTTCATGTNNNAGATCGGAA
34239 TATGAGTGTGGAGTCGTTGCGCAGACAGACTCAAATCATGAAGANNNAGATCGGA
34465 TATGAGTGTGGAGTCGTTGCTGACCAAAACGTTTTTATAGGGCANNNAGATCGGAA
34774 TATGAGTGTGGAGTCGTTGCAACCTCTGACTCCCAACTTCAAGNNNAGATCGGAAG
34828 TATGAGTGTGGAGTCGTTGCACGCCCGGCTAATTTTTTGTATTNNNAGATCGGAAG
34856 TATGAGTGTGGAGTCGTTGCACGCCCGGCTAATTTATTTTTGTNNNAGATCGGAAG
34873 TATGAGTGTGGAGTCGTTGCCAGTGGTGCAATCTCAATCTCAGNNNAGATCGGAAG

34950 TATGAGTGTGGAGTCGTTGCCTCCTGAGTTGCTGGGATTACAGNNNAGATCGGAA
34952 TATGAGTGTGGAGTCGTTGCTTCAAGTGATTCTCCTGCCTCAGNNNAGATCGGAAG
35055 TATGAGTGTGGAGTCGTTGCTGTGCTGTTTCCCTCTATGAGTCNNNAGATCGGAAG
35056 TATGAGTGTGGAGTCGTTGCACAGATATCTCAAGTCCCTGCTTNNNAGATCGGAAG
35064 TATGAGTGTGGAGTCGTTGCTCCTTTTTTGGTGTCCTCTAAGCTNNNAGATCGGAAG
35072 TATGAGTGTGGAGTCGTTGCCTCAGCCACCTGAATAGTTGAGANNNAGATCGGAAG
35106 TATGAGTGTGGAGTCGTTGCTTTGGAGAGGACCTCTTTTTTTTTNNNAGATCGGAA
35109 TATGAGTGTGGAGTCGTTGCGGAATTACAGGCACACACCACTANNNAGATCGGAAG
35132 TATGAGTGTGGAGTCGTTGCCAGTCTATGCCTTTTGATTGGTGANNNAGATCGGAA
35155 TATGAGTGTGGAGTCGTTGCTGGCCAGAAATAATGTTTACCAGCNNNAGATCGGAA
35209 TATGAGTGTGGAGTCGTTGCCATCAGCATCCTGAGTAACTGGTNNNAGATCGGAAG
35242 TATGAGTGTGGAGTCGTTGCTGTGTTCTTGATTTGGTTCTCAGCNNNAGATCGGAA
35249 TATGAGTGTGGAGTCGTTGCGACACAGGATTTCACCACATTGGNNNAGATCGGAAG
35322 TATGAGTGTGGAGTCGTTGCCTCCTGAGTAGCTGGGACTACAGNNNAGATCGGAA
35323 TATGAGTGTGGAGTCGTTGCCATCACGATGCCCAGCTAATTTCNNNAGATCGGAAG
35356 TATGAGTGTGGAGTCGTTGCCTCCATTACGGTTGAGGTGATCCNNNAGATCGGAAG
35400 TATGAGTGTGGAGTCGTTGCTACCGCCACACCTAGCTAATTTTNNNAGATCGGAAGA
35432 TATGAGTGTGGAGTCGTTGCCAAGTAGCTGGGACTACAGGCNNNAGATCGGAAGA
35442 TATGAGTGTGGAGTCGTTGCCCAATAGGCGTTTTCTTTGAGTGTNNNAGATCGGAA
35444 TATGAGTGTGGAGTCGTTGCAGAGTGCAATGTCAGGATCTCAGNNNAGATCGGAA
35472 TATGAGTGTGGAGTCGTTGCCCTCGGTTCAAGCTATTCTCCTGNNNAGATCGGAAG
35514 TATGAGTGTGGAGTCGTTGCTTGCTTAGGCTAGATGACTACTTTNNNAGATCGGAA
35550 TATGAGTGTGGAGTCGTTGCCCTCCCAGGTTCATGACATTCTCNNNAGATCGGAAGA
35589 TATGAGTGTGGAGTCGTTGCTCAGCACTATGTTATGGGTTTTCTNNNAGATCGGAA
35595 TATGAGTGTGGAGTCGTTGCCACAAGCAATTCACCTTCCTCAGNNNAGATCGGAAG
35603 TATGAGTGTGGAGTCGTTGCTGCAATTGTTTTCAGAGTCTGCANNNAGATCGGAAG
35628 TATGAGTGTGGAGTCGTTGCATCAGCAAATTCTCTTCGGTCCTNNNAGATCGGAAG
35633 TATGAGTGTGGAGTCGTTGCACACTGCAAGTTCAGCTTTGATCNNNAGATCGGAAG
35674 TATGAGTGTGGAGTCGTTGCACTTTTAGTAGAGACGGGGTTTCCNNNAGATCGGAA
35703 TATGAGTGTGGAGTCGTTGCTCCATTCTGCTAGCCTAAGTCTTNNNAGATCGGAAGA
35719 TATGAGTGTGGAGTCGTTGCTGTTCTCAAGACAGATAACACAGTNNNAGATCGGAA
35726 TATGAGTGTGGAGTCGTTGCCCAAGTAGCTGGGATTACAGTCANNNAGATCGGAA
35758 TATGAGTGTGGAGTCGTTGCTCCATTCTGCTAGCCTAAGTCTTNNNAGATCGGAAGA
35765 TATGAGTGTGGAGTCGTTGCTGCGATCTCAGCTCACTACAAGNNNAGATCGGAAGA
35786 TATGAGTGTGGAGTCGTTGCACAGACACCCAGCTAATTGTCATNNNAGATCGGAAG
35800 TATGAGTGTGGAGTCGTTGCTGCAGTTGTACACCCTTGTATCANNNAGATCGGAAG
35801 TATGAGTGTGGAGTCGTTGCCTGGAGTGGTATGATCTCAGCTCNNNAGATCGGAA
35828 TATGAGTGTGGAGTCGTTGCTCCATACTGTTTTCCATAGAGGCTNNNAGATCGGAA
35925 TATGAGTGTGGAGTCGTTGCTCCAGTTCTGTCTACTCCAATTCANNNAGATCGGAAG
36051 TATGAGTGTGGAGTCGTTGCTGCTTTGTGTATTTGGGAACTCTGNNNAGATCGGAA
36070 TATGAGTGTGGAGTCGTTGCTGCAATCAGCTATAATCCTTCCCTNNNAGATCGGAAG
36085 TATGAGTGTGGAGTCGTTGCTCCACAAAAGCAATTGAAAACTCTNNNAGATCGGAA
36089 TATGAGTGTGGAGTCGTTGCAGATGAAGTTGACTCAAATTTGCTNNNAGATCGGAA

36124 TATGAGTGTGGAGTCGTTGCAGATTGGATAGTGGAACAGAATCANNNAGATCGGA
36134 TATGAGTGTGGAGTCGTTGCGGTAAAGAGAGGTACAGGAATCGGNNNAGATCGG
36156 TATGAGTGTGGAGTCGTTGCAATGTGATCTACCCTCTAAGCAAANNNAGATCGGAA
36166 TATGAGTGTGGAGTCGTTGCACAATCAGCAGGCATTTACATAGGNNNAGATCGGAA
36168 TATGAGTGTGGAGTCGTTGCACAATCAGCAGGCATTTACATAGGNNNAGATCGGAA
36177 TATGAGTGTGGAGTCGTTGCCAAGACATGACTCAGCATGAAAGTNNNAGATCGGA
36185 TATGAGTGTGGAGTCGTTGCTCATAGTACACAATGGGTTCAACANNNAGATCGGAA
36268 TATGAGTGTGGAGTCGTTGCGGAAAGAAAATTAGCCTGGTGTGGNNNAGATCGGA
36278 TATGAGTGTGGAGTCGTTGCTCCATGTGCTATAAAGTGGGATGTNNNAGATCGGA
36293 TATGAGTGTGGAGTCGTTGCCACCACCACCAACAAAAACCTAGNNNAGATCGGAAG
36347 TATGAGTGTGGAGTCGTTGCTCATATGCCAGATGTAGAGATGGTNNNAGATCGGA
36353 TATGAGTGTGGAGTCGTTGCAGTTGGAGAACTAACATCAGCTGANNNAGATCGGA
36362 TATGAGTGTGGAGTCGTTGCTGGACTCTATTTTGTTTCCTTGGTNNNAGATCGGAA
36366 TATGAGTGTGGAGTCGTTGCAAACGAGAAAGTGGAATGTTTTCTNNNAGATCGGA
36369 TATGAGTGTGGAGTCGTTGCAGAGTGATTATTTCTGCATCAGGTNNNAGATCGGAA
36437 TATGAGTGTGGAGTCGTTGCTCCCACAAACATAAATTACTCCGTNNNAGATCGGAAG
36480 TATGAGTGTGGAGTCGTTGCGCAAAACAGAGTGTTACATGTGGANNNAGATCGGA
36523 TATGAGTGTGGAGTCGTTGCTCCTCTTGTAGTTCTAGTCCCAGTNNNAGATCGGAAG
36533 TATGAGTGTGGAGTCGTTGCAAACGAAACAAAGAGATGGTTCTTNNNAGATCGGA
36584 TATGAGTGTGGAGTCGTTGCAGGGATAAACCTTTCTCAGAGGAANNNAGATCGGA
36742 TATGAGTGTGGAGTCGTTGCTGTTGTTAGATATTGCCAGCCCTNNNAGATCGGAAG
36773 TATGAGTGTGGAGTCGTTGCTCCTGGCCTCTGTTCTAATTGTTNNNAGATCGGAAGA
36779 TATGAGTGTGGAGTCGTTGCAGGGTGCTATTCTAACTTCAGACANNNAGATCGGAA
36786 TATGAGTGTGGAGTCGTTGCAGCTCTGTCTAGAAAACTTACTGANNNAGATCGGAA
36837 TATGAGTGTGGAGTCGTTGCACACCTCACACTTTCTTCCTTACTNNNAGATCGGAAGA
36840 TATGAGTGTGGAGTCGTTGCTGATCTTTTGCTTTGCTTTGTCCTNNNAGATCGGAAG
36884 TATGAGTGTGGAGTCGTTGCCAAAAACCAAACCAAACCAAACCANNNAGATCGGAA
36898 TATGAGTGTGGAGTCGTTGCAGGGTGACACATCTTATCTTTGGANNNAGATCGGAA
36903 TATGAGTGTGGAGTCGTTGCACAAAAGTTAGCCAGACATTGTGGNNNAGATCGGA
36919 TATGAGTGTGGAGTCGTTGCGGGATGTAGACTAAGAAGCCATGTNNNAGATCGGA
36923 TATGAGTGTGGAGTCGTTGCAGAAATGCTATGCTGGTACAAACTNNNAGATCGGAA
36933 TATGAGTGTGGAGTCGTTGCTCTATGGTTTCTAGTTGCCATTGCNNNAGATCGGAA
36940 TATGAGTGTGGAGTCGTTGCGCAGGCATTCAAACAAAAACTTGTNNNAGATCGGAA
36946 TATGAGTGTGGAGTCGTTGCAAACTAAGCTGGCACTAGAAGGANNNAGATCGGAA
36958 TATGAGTGTGGAGTCGTTGCACAAACAGGCTATGCAATGTAACANNNAGATCGGAA
36963 TATGAGTGTGGAGTCGTTGCTGCAGGATACAAAATCAACAAACANNNAGATCGGAA
36985 TATGAGTGTGGAGTCGTTGCGCTTATGACTCCCATGACTATTGCNNNAGATCGGAA
37007 TATGAGTGTGGAGTCGTTGCCTTGCTTTGGGTGGCTGAAATAANNNAGATCGGAA
37010 TATGAGTGTGGAGTCGTTGCCACACATGCACATACACATACACANNNAGATCGGAA
37038 TATGAGTGTGGAGTCGTTGCAGTTGTTTTGTTAGAAGAGGGAGANNNAGATCGGA
37057 TATGAGTGTGGAGTCGTTGCTGAAAATGAACTCAAAATGGGTCANNNAGATCGGA
37062 TATGAGTGTGGAGTCGTTGCTGGCCAGTCTCATCAATTAGTTGTNNNAGATCGGAA
37101 TATGAGTGTGGAGTCGTTGCTGAAGGTACAAAACTCTCACTGGTNNNAGATCGGAA

37134 TATGAGTGTGGAGTCGTTGCACAACAAAGAAAAATCAGCAGTGANNNAGATCGGA
37157 TATGAGTGTGGAGTCGTTGCTGGTAAGAGAATGATGACTTGTTGCNNNAGATCGG
37171 TATGAGTGTGGAGTCGTTGCCAGTATCTGCAGAGCCAATAAGGNNNAGATCGGAA
37176 TATGAGTGTGGAGTCGTTGCGTGCCAGAGAAGACTATAGGTACGNNNAGATCGGA
37189 TATGAGTGTGGAGTCGTTGCCCTTTCAAGTTTCAGTCCTTTGGGNNNAGATCGGAA
37230 TATGAGTGTGGAGTCGTTGCCTCTGGAGGAGGGAAAGAAATTNNNAGATCGGAA
37232 TATGAGTGTGGAGTCGTTGCTGAATTTACACTTTCCAGTTTGCTNNNAGATCGGAAG
37238 TATGAGTGTGGAGTCGTTGCAGGCCAATAAAATCCTGCTTGATGNNNAGATCGGAA
37246 TATGAGTGTGGAGTCGTTGCTGAAAGAAAAGGATGTCAATGAGCANNNAGATCGG
37261 TATGAGTGTGGAGTCGTTGCTTGTGAAAGCAAATCAGTAATCCANNNAGATCGGAA
37278 TATGAGTGTGGAGTCGTTGCTGTAGCCATTTCTGGAGATTTTGCNNNAGATCGGAA
37308 TATGAGTGTGGAGTCGTTGCAGGATAAATGAATATGGGACCACANNNAGATCGGA
37316 TATGAGTGTGGAGTCGTTGCAGAGTCCAAACACTGAATAGCACTNNNAGATCGGAA
37337 TATGAGTGTGGAGTCGTTGCTGAATGCTCACTGTAGAGAGAGTGNNNAGATCGGA
37379 TATGAGTGTGGAGTCGTTGCACATGGACATCATCATTTTACCCANNNAGATCGGAAG
37425 TATGAGTGTGGAGTCGTTGCTCATCAACAGAATGGAATTGTGGTNNNAGATCGGA
37442 TATGAGTGTGGAGTCGTTGCTGCTGGGAATTTGATAGTATTTGCANNNAGATCGGA
37457 TATGAGTGTGGAGTCGTTGCGAAAGGAAGAGGATTAGAGAAGGANNNAGATCGG
37485 TATGAGTGTGGAGTCGTTGCCAACGATGCTTTAGATTGATCACTNNNAGATCGGAA
37491 TATGAGTGTGGAGTCGTTGCAGAAAATCAATGAAACCGAAAGCTNNNAGATCGGA
37512 TATGAGTGTGGAGTCGTTGCCAACTGTGCTGTGATACTTAGAAANNNAGATCGGAA
37524 TATGAGTGTGGAGTCGTTGCCCTTGGTGGTTCTTTTTTGAGACANNNAGATCGGAA
37534 TATGAGTGTGGAGTCGTTGCAGTTGTCTTCAATGGGAGACATCANNNAGATCGGAA
37537 TATGAGTGTGGAGTCGTTGCTCAAAAGATACCAAGGTCAGAGGTNNNAGATCGGA
37541 TATGAGTGTGGAGTCGTTGCGTGAGCTCAAAGATGGTCATTTGANNNAGATCGGA
37546 TATGAGTGTGGAGTCGTTGCTGTCATGGAAGTAACACAGTCAGTNNNAGATCGGA
37584 TATGAGTGTGGAGTCGTTGCGCAAGGCTTAGCATATATCCAAGTNNNAGATCGGAA
37647 TATGAGTGTGGAGTCGTTGCAGATGAGTGTGGTAGTAAAAGCCANNNAGATCGGA
37668 TATGAGTGTGGAGTCGTTGCCCCCACAAAAAAATGACAACTTGTNNNAGATCGGAA
37682 TATGAGTGTGGAGTCGTTGCCCTCCCACAACAAATGAATTCACTNNNAGATCGGAAG
37683 TATGAGTGTGGAGTCGTTGCAGTTCTGCTTACTAAGGAAGAGTTNNNAGATCGGAA
37695 TATGAGTGTGGAGTCGTTGCTCATTTGCAAAAGGTGTAAACACANNNAGATCGGAA
37703 TATGAGTGTGGAGTCGTTGCCACACCTAGAGGAACTAGAAGAGCNNNAGATCGGA
37704 TATGAGTGTGGAGTCGTTGCAGAAAGACCTCAAGTGAGTAACCTNNNAGATCGGA
37721 TATGAGTGTGGAGTCGTTGCTGCAGGATACAAAACTAACACACANNNAGATCGGAA
37732 TATGAGTGTGGAGTCGTTGCGTCACATCTGTAATCCCAGCACTNNNAGATCGGAAG
37733 TATGAGTGTGGAGTCGTTGCACAACAAAAAGGTAAAATGTGGGGANNNAGATCGG
37780 TATGAGTGTGGAGTCGTTGCTGAAAGTGACTGAAGTACCTAGGGNNNAGATCGGA
37830 TATGAGTGTGGAGTCGTTGCTGACCAAAGGATTATTCTCGTTGCNNNAGATCGGAA
37841 TATGAGTGTGGAGTCGTTGCACCATATGACTATTCACACTGCCTNNNAGATCGGAAG
37848 TATGAGTGTGGAGTCGTTGCTCACAGACTGGATAAGAAAAACCANNNAGATCGGA
37883 TATGAGTGTGGAGTCGTTGCAGGCAGAAGACTTGAATAGGTTCTNNNAGATCGGA
37948 TATGAGTGTGGAGTCGTTGCCAAGTTTCTCCATCAAATGTGGGANNNAGATCGGAA

38003 TATGAGTGTGGAGTCGTTGCTCTGCTACAATCAGAGTAGACTCANNNAGATCGGAA
38011 TATGAGTGTGGAGTCGTTGCCGTTTATACACTACCAAACTGGCANNNAGATCGGAA
38012 TATGAGTGTGGAGTCGTTGCAGCCTAATGGAAGATGAAAATGCANNNAGATCGGA
38016 TATGAGTGTGGAGTCGTTGCAACCAACCAAGAATTGATGCAGANNNAGATCGGAA
38084 TATGAGTGTGGAGTCGTTGCACTTCTGTTGATGTTCAGTCTCCTNNNAGATCGGAAG
38120 TATGAGTGTGGAGTCGTTGCAGCAAAGGAAACAATCAACAGAGTNNNAGATCGGA
38123 TATGAGTGTGGAGTCGTTGCACATGCCAAGAGTGAAAAATCTGANNNAGATCGGA
38125 TATGAGTGTGGAGTCGTTGCTCCTTTAGGGAATAACAGCAAGGANNNAGATCGGA
38162 TATGAGTGTGGAGTCGTTGCAGAGAGAGAGAGAGAGAAATCAGANNNAGATCGG
38202 TATGAGTGTGGAGTCGTTGCAGAAGGGAAAATGTGGCTATTTCTNNNAGATCGGA
38250 TATGAGTGTGGAGTCGTTGCCATTCTGTGAAGCCAGTATCACCNNNAGATCGGAAG
38258 TATGAGTGTGGAGTCGTTGCACCACATGTACAAAGAAGAGCTCANNNAGATCGGAA
38273 TATGAGTGTGGAGTCGTTGCCCACCAAGGAAAATTCTTGAACCANNNAGATCGGAA
38353 TATGAGTGTGGAGTCGTTGCAGAAAGAATGAGACAATGGGAACTNNNAGATCGGA
38373 TATGAGTGTGGAGTCGTTGCTCATGTGTTTGTGAATTTTCCTGTNNNAGATCGGAA
38420 TATGAGTGTGGAGTCGTTGCGGACATACAAATGACCAATAGGCANNNAGATCGGA
38443 TATGAGTGTGGAGTCGTTGCGGGAGTCCTTGGTAGATAAGAGACNNNAGATCGGA
38460 TATGAGTGTGGAGTCGTTGCTGTCTTCTCCAGGCTAAATGTTGTNNNAGATCGGAA
38469 TATGAGTGTGGAGTCGTTGCTGATGATCCATCTTGATGCAAGGANNNAGATCGGAA
38493 TATGAGTGTGGAGTCGTTGCGAGAGGGAGGACAAAGGGAAAATNNNAGATCGGA
38498 TATGAGTGTGGAGTCGTTGCAGATAGGATCTTGCTATGCTGTCCNNNAGATCGGAA
38519 TATGAGTGTGGAGTCGTTGCACACTCTAAGAGAATCAAGAGATCCTNNNAGATCGG
38533 TATGAGTGTGGAGTCGTTGCTGGCCATTCTCTAAGGTTCAATCTNNNAGATCGGAA
38541 TATGAGTGTGGAGTCGTTGCTAGCTAGGATTACAGGCATGTGCNNNAGATCGGAA
38592 TATGAGTGTGGAGTCGTTGCCCAAGGAGGCAAAAAACTTGTACANNNAGATCGGA
38593 TATGAGTGTGGAGTCGTTGCGCCTCTCAAGTACCTGGGATTACNNNAGATCGGAAG
38615 TATGAGTGTGGAGTCGTTGCAGAAATAACAAAGATCAGGGCAGANNNAGATCGGA
38621 TATGAGTGTGGAGTCGTTGCTGTCAAGAAATTGGATAGCCAGGANNNAGATCGGA
38647 TATGAGTGTGGAGTCGTTGCTCCCTATCAAAATAACAGCAGGCTNNNAGATCGGAA
38653 TATGAGTGTGGAGTCGTTGCACCCCATCAGTAACCTAGAAAAGGNNNAGATCGGAA
38679 TATGAGTGTGGAGTCGTTGCGCTCATCTAGGCTTTTTTCTAGGCNNNAGATCGGAA
38684 TATGAGTGTGGAGTCGTTGCAGAGACAGAGAGAAGAAAGAAGCANNNAGATCGG
38694 TATGAGTGTGGAGTCGTTGCAGGATGAACAAAGATATGCCATGTNNNAGATCGGA
38696 TATGAGTGTGGAGTCGTTGCACAGTCAGGAACTACATAAAGCCTNNNAGATCGGAA
38707 TATGAGTGTGGAGTCGTTGCTTCAAGCGATTCTCCTGTCTCAGNNNAGATCGGAAG
38708 TATGAGTGTGGAGTCGTTGCTCCATTGTGATTGCAGAAGTTTGTNNNAGATCGGAA
38711 TATGAGTGTGGAGTCGTTGCAGACATTTATGTGGCCAACAAACANNNAGATCGGAA
38714 TATGAGTGTGGAGTCGTTGCGTGCCATGACTTTTTGTTGCATGNNNAGATCGGAAG
38719 TATGAGTGTGGAGTCGTTGCGAAGAAGCAACTGCAATATTTGGANNNAGATCGGA
38745 TATGAGTGTGGAGTCGTTGCTGACATAATTTGCTGCTTTTCCCANNNAGATCGGAAG
38754 TATGAGTGTGGAGTCGTTGCCCAAATGGGTGATTTTCTTGTCGANNNAGATCGGAA
38765 TATGAGTGTGGAGTCGTTGCTGCCTGAACAATACTGTCAAATGTNNNAGATCGGAA
38801 TATGAGTGTGGAGTCGTTGCAGAAGAAGAAGGAGGAGAAACAGTNNNAGATCGG

38806 TATGAGTGTGGAGTCGTTGCAGAGAGAATAACCTCACAAGAGCANNNAGATCGGA
38827 TATGAGTGTGGAGTCGTTGCGCAACTTTGCCAGATTTGTTTACAANNNAGATCGGA
38841 TATGAGTGTGGAGTCGTTGCTGAAAAGCACAGGCAACAATAACANNNAGATCGGA
38844 TATGAGTGTGGAGTCGTTGCTCATCTCTAAAACTGAAAGCACCANNNAGATCGGAA
38861 TATGAGTGTGGAGTCGTTGCTGGGTTGCAAATAAACTCCAAAGTNNNAGATCGGA
38873 TATGAGTGTGGAGTCGTTGCGAGAAGAGCTTTAGAGAAGGCTATNNNAGATCGGA
38875 TATGAGTGTGGAGTCGTTGCTGCAAGGATAAACAAAATTGACACANNNAGATCGG
38876 TATGAGTGTGGAGTCGTTGCGGGGGGTCTGTTTATATTATTTGCCNNNAGATCGGA
38877 TATGAGTGTGGAGTCGTTGCTCCAACAATATGACACTCTGGAGANNNAGATCGGAA
38879 TATGAGTGTGGAGTCGTTGCTTGTCTTCAGTGAGGAGGTACACNNNAGATCGGAA
38908 TATGAGTGTGGAGTCGTTGCTCACAGCCAAATACTGCTACACANNNAGATCGGAAG
38927 TATGAGTGTGGAGTCGTTGCACCAAGGGCCAACTATATATTCCCNNNAGATCGGAA
38954 TATGAGTGTGGAGTCGTTGCTGAACACATACTCATACTGCACCANNNAGATCGGAA
39000 TATGAGTGTGGAGTCGTTGCTGGAATGTAACAAGGTGACTTTGCNNNAGATCGGA
39011 TATGAGTGTGGAGTCGTTGCCAAGATTCAGTCCAAATTGCAGGTNNNAGATCGGAA
39045 TATGAGTGTGGAGTCGTTGCGGATGTGCTTAGGGGAATCTGATNNNAGATCGGAA
39048 TATGAGTGTGGAGTCGTTGCAGGAAGACATTTATGTGACCAACANNNAGATCGGA
39049 TATGAGTGTGGAGTCGTTGCAGCTGTGAATTTCCTAGGTTACGANNNAGATCGGAA
39061 TATGAGTGTGGAGTCGTTGCTGTGATTGGATTGTTTGCAAGTCANNNAGATCGGA
39063 TATGAGTGTGGAGTCGTTGCTCCACATGCTTTTCTTTCTCCCTNNNAGATCGGAAGA
39086 TATGAGTGTGGAGTCGTTGCTGGTGTGCAATATGGTTTTTCTGTNNNAGATCGGAA
39091 TATGAGTGTGGAGTCGTTGCTGCCAATCCTACTCAAACTACTCCNNNAGATCGGAAG
39098 TATGAGTGTGGAGTCGTTGCTCAAAACATTCCAAATGGTATCTGTNNNAGATCGGA
39102 TATGAGTGTGGAGTCGTTGCTCAAGGAGAAGACATTTTCCCACTNNNAGATCGGAA
39122 TATGAGTGTGGAGTCGTTGCACTGTACATATTTTTGTGGTACATGTGNNNAGATCG
39126 TATGAGTGTGGAGTCGTTGCTTCCCTTTCTCAAACCCTTGACANNNAGATCGGAAGA
39134 TATGAGTGTGGAGTCGTTGCACTCTACCTTGGACTTTATGCTGTNNNAGATCGGAA
39136 TATGAGTGTGGAGTCGTTGCAGGCTGAATAGTTGTCTTGGATCANNNAGATCGGA
39141 TATGAGTGTGGAGTCGTTGCTCAACCTAAATGTCCATCAACAGANNNAGATCGGAA
39145 TATGAGTGTGGAGTCGTTGCTCTCAGAATTAGCAGACAAGAGCANNNAGATCGGA
39153 TATGAGTGTGGAGTCGTTGCACATGATAGTTGACCCTATGGAGTNNNAGATCGGA
39167 TATGAGTGTGGAGTCGTTGCAGTAGAAGGAAGAAGTGAGGGAACNNNAGATCGG
39250 TATGAGTGTGGAGTCGTTGCTCTATCCTAGGCTATTCTGGCTCTNNNAGATCGGAAG
39309 TATGAGTGTGGAGTCGTTGCTTGGGCTTGGTTTGATCTTACCANNNAGATCGGAAG
39347 TATGAGTGTGGAGTCGTTGCTGCTGCAACTTTCTTATGAGTACCNNNAGATCGGAA
39515 TATGAGTGTGGAGTCGTTGCATCTCAGCTCACTGCAACCTCTANNNAGATCGGAAGA
39525 TATGAGTGTGGAGTCGTTGCAGAATGATCTGGTCCAATTGAGCTNNNAGATCGGA
39534 TATGAGTGTGGAGTCGTTGCTCCATCTGCTTATATCTCCAGTGANNNAGATCGGAAG
39563 TATGAGTGTGGAGTCGTTGCAGATCAATGAATTCAGGAGCTAGTNNNAGATCGGA
39567 TATGAGTGTGGAGTCGTTGCAGTTTCAGGTTTTGGAACATTTCANNNAGATCGGAA
39615 TATGAGTGTGGAGTCGTTGCCAATGATGCTATCTTGGCTCACTGNNNAGATCGGAA
39633 TATGAGTGTGGAGTCGTTGCCCTGATATTCATGCTGCATAATGTNNNAGATCGGAA
39635 TATGAGTGTGGAGTCGTTGCAGGAAGGAAGGAGAGAGAGAAAGANNNAGATCG

39664 TATGAGTGTGGAGTCGTTGCTGTCTGGTACATGATAAGTGTGCANNNAGATCGGA
39671 TATGAGTGTGGAGTCGTTGCGGTCTCCTCTATACTCCATACCCANNNAGATCGGAAG
39776 TATGAGTGTGGAGTCGTTGCTTTTGGCTAAGATCAAGTGTGAAANNNAGATCGGA
39777 TATGAGTGTGGAGTCGTTGCATGCACCCAATACTAGAGCACAANNNAGATCGGAAG
39797 TATGAGTGTGGAGTCGTTGCCACTAGTAATGAGTGGAAAGTGGTNNNAGATCGGA
39805 TATGAGTGTGGAGTCGTTGCCCACCACCAAAAAAAGAGACAACTNNNAGATCGGAA
39818 TATGAGTGTGGAGTCGTTGCGGTTCTCTCATCTCTTCACACTGTNNNAGATCGGAAG
39831 TATGAGTGTGGAGTCGTTGCTTTAGGGACCATTACCTTTACCTTTNNNAGATCGGAA
39875 TATGAGTGTGGAGTCGTTGCAGAATGCTTACCTACAGAACACACTNNNAGATCGGA
39878 TATGAGTGTGGAGTCGTTGCAGAAAAGCAAGGAAAAGAAACCCANNNAGATCGGA
39968 TATGAGTGTGGAGTCGTTGCACAGGTGGATTGTTTTTTTCTGGANNNAGATCGGAA
39994 TATGAGTGTGGAGTCGTTGCAACATCTTCACCAATTAGTCCAGANNNAGATCGGAA
40123 TATGAGTGTGGAGTCGTTGCACTGATTCTGTCATTCAAATTTGGTNNNAGATCGGA
40198 TATGAGTGTGGAGTCGTTGCAGAGGAGAAGAGAAAACACTCTCANNNAGATCGGA
40228 TATGAGTGTGGAGTCGTTGCAGAGCCAAAGTAGTCTGTTTTTGTNNNAGATCGGAA
40234 TATGAGTGTGGAGTCGTTGCTGCAAATCACGATAGTAACGATCTNNNAGATCGGAA
40238 TATGAGTGTGGAGTCGTTGCGGGTTTTTTGTGTCTCTGTCTCCNNNAGATCGGAAG
40258 TATGAGTGTGGAGTCGTTGCTCAGAGAAGTGAAAATGAAAACCACANNNAGATCG
40260 TATGAGTGTGGAGTCGTTGCTCAATGCATCAAAGAACAAAACCANNNAGATCGGAA
40262 TATGAGTGTGGAGTCGTTGCTCATGGAGATAGAGAGTAGAAGGANNNAGATCGG
40271 TATGAGTGTGGAGTCGTTGCAGAAAAAGTAATGTGCATCCCACTNNNAGATCGGAA
40284 TATGAGTGTGGAGTCGTTGCACTTTCAAGGTCATTACTTTGGGCNNNAGATCGGAA
40300 TATGAGTGTGGAGTCGTTGCTGTTGTTTTATTTTCTGTTGCGTTNNNAGATCGGAAG
40308 TATGAGTGTGGAGTCGTTGCGCCTAGCACATAGTAGGAGTTTGANNNAGATCGGA
40315 TATGAGTGTGGAGTCGTTGCTCCTGAGCTAGTTTTGCAAAATTGNNNAGATCGGAA
40388 TATGAGTGTGGAGTCGTTGCTGAGACACTGTAGAACAACTAGCANNNAGATCGGA
40425 TATGAGTGTGGAGTCGTTGCGGGAACAGATCATAGTACCTAGCANNNAGATCGGA
40430 TATGAGTGTGGAGTCGTTGCAGAGAGAGAGAGAGAGAGAAAGGTCANNNAGATCGG
40464 TATGAGTGTGGAGTCGTTGCGGTGGTAAGGAAATGTTACCTTGCNNNAGATCGGA
40518 TATGAGTGTGGAGTCGTTGCAGCTTAGTAAGGTTGTAGGATGCANNNAGATCGGA
40586 TATGAGTGTGGAGTCGTTGCTGCAGTCTTGAATTTCTCTGAGTGNNNAGATCGGAA
40587 TATGAGTGTGGAGTCGTTGCTGCATGGCAGGAATATTTTTCTCCNNNAGATCGGAA
40589 TATGAGTGTGGAGTCGTTGCTGCAGTCTTGAATTTCTCTGAGTGNNNAGATCGGAA
40594 TATGAGTGTGGAGTCGTTGCACTGACATGCATTGATGTTTCAGTNNNAGATCGGAA
40702 TATGAGTGTGGAGTCGTTGCAGCTGCCTTATACAGTAGTTGTGANNNAGATCGGAA
40734 TATGAGTGTGGAGTCGTTGCTGGATAACTTTGGGAAGTAGGAGANNNAGATCGGA
40761 TATGAGTGTGGAGTCGTTGCACCAGAGGTACAAATAAGAGCTGGNNNAGATCGGA
40785 TATGAGTGTGGAGTCGTTGCGGATGAAAAAGAACCTGTAAGGGCNNNAGATCGGA
40792 TATGAGTGTGGAGTCGTTGCAGAAAGAAAGAAAGAAGCTCCAGCNNNAGATCGGA
40817 TATGAGTGTGGAGTCGTTGCTGTGGTCCTTAGTTATTTTCTCTGGANNNAGATCGG
40856 TATGAGTGTGGAGTCGTTGCTGTACCTGTTTACAATGTACAAACTGNNNAGATCGG
40868 TATGAGTGTGGAGTCGTTGCCTATTGTGAACAGTGCTGCAACANNNAGATCGGAAG
40870 TATGAGTGTGGAGTCGTTGCCCAGACAAATAAAACCCCACAACANNNAGATCGGAA

40896 TATGAGTGTGGAGTCGTTGCACGAACCATGAAGTAAACATCACCNNNAGATCGGAA
40943 TATGAGTGTGGAGTCGTTGCTGAGACATTGCCATCATAACTTCCNNNAGATCGGAA
40955 TATGAGTGTGGAGTCGTTGCAAGACTCTCTAAGCAAACATACCTNNNAGATCGGAA
40957 TATGAGTGTGGAGTCGTTGCGTCTGGCATCCCATCAAAAATTANNNAGATCGGAAG
40960 TATGAGTGTGGAGTCGTTGCGTGAAGAGAACCTACAGAATGGGANNNAGATCGG
40963 TATGAGTGTGGAGTCGTTGCTGCTTTGTGATACCTTGTTCATGGNNNAGATCGGAA
41012 TATGAGTGTGGAGTCGTTGCAGGTATCGAGTCTCACTATAGCCANNNAGATCGGAA
41097 TATGAGTGTGGAGTCGTTGCTGTATATGCAGACCACTTAGCACTNNNAGATCGGAA
41130 TATGAGTGTGGAGTCGTTGCACTTCTATGGTGAAACGTAACCCANNNAGATCGGAA
41179 TATGAGTGTGGAGTCGTTGCATGTGAGAAAGGACTGAACAGACNNNAGATCGGAA
41186 TATGAGTGTGGAGTCGTTGCACCAAATGATACCAAAACATGCTGNNNAGATCGGAA
41187 TATGAGTGTGGAGTCGTTGCAGCTACTATGGAAACCAGTGTGGNNNAGATCGGAA
41300 TATGAGTGTGGAGTCGTTGCAGCTGTGAATTTCCTAGGTTACGANNNAGATCGGAA
41308 TATGAGTGTGGAGTCGTTGCTGGTGTGCAATATGGTTTTTCTGTNNNAGATCGGAA
41317 TATGAGTGTGGAGTCGTTGCTCAAAACATTCCAAATGGTATCTGTNNNAGATCGGA
41335 TATGAGTGTGGAGTCGTTGCTGAATCCCAAGTAATCTGACGACTNNNAGATCGGAA
41389 TATGAGTGTGGAGTCGTTGCGCCATCACACCAGGCTAATTTTTNNNAGATCGGAAG
41397 TATGAGTGTGGAGTCGTTGCACTGTAATGTCATAGAACAGGGGTNNNAGATCGGA
41407 TATGAGTGTGGAGTCGTTGCAGGTAGGGGATTATGTGTGTAAGCNNNAGATCGG
41447 TATGAGTGTGGAGTCGTTGCTCTTTGAGTTATAGAAGACGGCGTNNNAGATCGGA
41457 TATGAGTGTGGAGTCGTTGCGGCACCTGTATCTCAACTATTTGGNNNAGATCGGAA
41458 TATGAGTGTGGAGTCGTTGCCACCCCAAAAAATCCAATGTAGCTNNNAGATCGGAA
41472 TATGAGTGTGGAGTCGTTGCTCAAAAAATGGTGTTGGAATTGCANNNAGATCGGA
41488 TATGAGTGTGGAGTCGTTGCTGCTAATGATACGAGGAGATGAACANNNAGATCGG
41501 TATGAGTGTGGAGTCGTTGCTCACTTCTAATTCCAAGGCCTACANNNAGATCGGAAG
41508 TATGAGTGTGGAGTCGTTGCTGCTAATGATACGAGGAGATGAACANNNAGATCGG
41531 TATGAGTGTGGAGTCGTTGCACCAAATGATACCAAAACATGCTGNNNAGATCGGAA
41633 TATGAGTGTGGAGTCGTTGCTATCTTGGCTCACTGCAACCTCNNNAGATCGGAAGA
41634 TATGAGTGTGGAGTCGTTGCTCCCAAGTTCAACCAATTCTCCTNNNAGATCGGAAGA
41677 TATGAGTGTGGAGTCGTTGCCACGCCTGGCTACTTTTTGTATTNNNAGATCGGAAG
41741 TATGAGTGTGGAGTCGTTGCAATACAGCACACTGATGGGTCTTNNNAGATCGGAAG
41776 TATGAGTGTGGAGTCGTTGCCTCCTGGGTTCAAGCGATTCTCNNNAGATCGGAAGA
41805 TATGAGTGTGGAGTCGTTGCGCCTCCTGAATAGCTGAGACTACNNNAGATCGGAAG
41815 TATGAGTGTGGAGTCGTTGCCTCAGCCACCTGAATAGTTGAGANNNAGATCGGAAG
41834 TATGAGTGTGGAGTCGTTGCGTTCAAGAGATTCTCCTGCCTCANNNAGATCGGAAG
41847 TATGAGTGTGGAGTCGTTGCCAATGGCATGATCTCAGCTCACNNNAGATCGGAAGA
41857 TATGAGTGTGGAGTCGTTGCCTCAAGTGATTCTCCTGCCTCAGNNNAGATCGGAAG
41875 TATGAGTGTGGAGTCGTTGCTGTATTTTTCACAGAGACGGGGTNNNAGATCGGAA
41884 TATGAGTGTGGAGTCGTTGCATGAAATCTTGCTCTGTCACCCANNNAGATCGGAAG
41907 TATGAGTGTGGAGTCGTTGCGCCTCCTAAGTAGCTGGGATTACNNNAGATCGGAAG
41925 TATGAGTGTGGAGTCGTTGCCCTCCTGGGTTCAAGCTATTCTCNNNAGATCGGAAG
41951 TATGAGTGTGGAGTCGTTGCGATCTTGGCTTACTGCACCCTNNNAGATCGGAAGAG
41952 TATGAGTGTGGAGTCGTTGCGCTGGGACTATAGATGTACACCANNNAGATCGGAA

41964 TATGAGTGTGGAGTCGTTGCAGCAAAGCTACTGATTTGTGTACANNNAGATCGGAA
41998 TATGAGTGTGGAGTCGTTGCCAGGTTCAAGCAATTCTCCTGCNNNAGATCGGAAGA
42026 TATGAGTGTGGAGTCGTTGCTCCAGTTCTGTCTACTCCAATTCANNNAGATCGGAAG
42033 TATGAGTGTGGAGTCGTTGCTTCAAGTGATTCTCCTGCCTCAGNNNAGATCGGAAG
42051 TATGAGTGTGGAGTCGTTGCTGACCAAGTTATCAGTCTCTACCANNNAGATCGGAA
42052 TATGAGTGTGGAGTCGTTGCTCTGAAATCTAGGGAGAGGTTCCTNNNAGATCGGA
42068 TATGAGTGTGGAGTCGTTGCAGGAGCAACTTTTACCAAATCACANNNAGATCGGAA
42070 TATGAGTGTGGAGTCGTTGCAGCCAAATTTTATGGGAATGAACANNNAGATCGGA
42084 TATGAGTGTGGAGTCGTTGCGCTATTGTGAACAGTGCTGCAATNNNAGATCGGAA
42099 TATGAGTGTGGAGTCGTTGCAGCTACTATGGAAACCAGTGTGGNNNAGATCGGAA
42116 TATGAGTGTGGAGTCGTTGCCCTCCTGAGTAGCTGGGACTATANNNAGATCGGAAG
42123 TATGAGTGTGGAGTCGTTGCTCCAACTCCCTGGTTCAAATGATNNNAGATCGGAAG
42140 TATGAGTGTGGAGTCGTTGCAGTAGCTGGGACTACAGGCANNNAGATCGGAAGAG
42162 TATGAGTGTGGAGTCGTTGCAGGAAGTAAGTTTGCATTGTAGCANNNAGATCGGA
42170 TATGAGTGTGGAGTCGTTGCCTCAAGTGATCCTCCTGTCTCAGNNNAGATCGGAAG
42211 TATGAGTGTGGAGTCGTTGCACTTATTGCCCTGGCTAGAAACTNNNAGATCGGAAG
42231 TATGAGTGTGGAGTCGTTGCGAATGTGTTAGGAAGGGAGTCCCNNNAGATCGGAA
42233 TATGAGTGTGGAGTCGTTGCGAGACAGAGTCACACTGTTTTGCNNNAGATCGGAA
42245 TATGAGTGTGGAGTCGTTGCCCAAGGGAATGGATGCTAAAGATGNNNAGATCGGA
42261 TATGAGTGTGGAGTCGTTGCGCATCCTCATCACTATTTTGGGTGNNNAGATCGGAA
42277 TATGAGTGTGGAGTCGTTGCTGTGTGTGTGTGTATATGTGTGNNNAGATCGGA
42298 TATGAGTGTGGAGTCGTTGCCTCAAAATAGCATTTCCTGGGCANNNAGATCGGAAG
42314 TATGAGTGTGGAGTCGTTGCAGCTACTTCTGCTGGTAGGAATGNNNAGATCGGAA
42315 TATGAGTGTGGAGTCGTTGCAGCTACTTCTGCTGGTAGGAATGNNNAGATCGGAA
42319 TATGAGTGTGGAGTCGTTGCTGTAATGGAGCCAATATCCCCAANNNAGATCGGAAG
42353 TATGAGTGTGGAGTCGTTGCTGGTCATGCTCAATACATGGTCANNNAGATCGGAAG
42362 TATGAGTGTGGAGTCGTTGCTCCTTGACACCTTTCCTTTCAAGANNNAGATCGGAAG
42389 TATGAGTGTGGAGTCGTTGCATTTTGTGTGTGTGTGTGTGTNNNAGATCGGAA
42399 TATGAGTGTGGAGTCGTTGCTTTCCCTCTTTCTGTTAGAAGCANNNAGATCGGAAGA
42441 TATGAGTGTGGAGTCGTTGCCCGATTCATGAAACTAACCCTTGGNNNAGATCGGAA
42477 TATGAGTGTGGAGTCGTTGCGAGTCTCTCTCCATTGCCCAGNNNAGATCGGAAGAG
42502 TATGAGTGTGGAGTCGTTGCGACCATTTGGAACTGAAAACGGTNNNAGATCGGAA
42514 TATGAGTGTGGAGTCGTTGCTGTCAGATTGAAGACACCAAAAGCNNNAGATCGGA
42518 TATGAGTGTGGAGTCGTTGCTGTGAGATCAATGGTGATATCCCCNNNAGATCGGAA
42520 TATGAGTGTGGAGTCGTTGCTTCAAGCGATTCTCCTGACTCAGNNNAGATCGGAAG
42562 TATGAGTGTGGAGTCGTTGCTGTGTGTGTTGATGTATGTGTGTGNNNAGATCGGA
42564 TATGAGTGTGGAGTCGTTGCACTTATTGCCCTGGCTAGAAACTNNNAGATCGGAAG
42595 TATGAGTGTGGAGTCGTTGCGCAAATGACAGTTCTTTCCTCTCANNNAGATCGGAA
42612 TATGAGTGTGGAGTCGTTGCCCAGTGAAAACCACATCAAAAACANNNAGATCGGAA
42614 TATGAGTGTGGAGTCGTTGCAGAGAGAGAAGAGAGATGGAGACANNNAGATCGG
42670 TATGAGTGTGGAGTCGTTGCACCCTGTCTCCACAAACAATACANNNAGATCGGAAG
42673 TATGAGTGTGGAGTCGTTGCGGTTGAGAATTCCCATGATCTGCNNNAGATCGGAAG
42677 TATGAGTGTGGAGTCGTTGCTGATTGACTCCCCAACATCCATTNNNAGATCGGAAG

42682 TATGAGTGTGGAGTCGTTGCGGGTGGCTAGAGAACAGGAATAGNNNAGATCGGA
42700 TATGAGTGTGGAGTCGTTGCTGCATATCACATTACTTTATCCACTCANNNAGATCGG
42703 TATGAGTGTGGAGTCGTTGCTGGTTGATGGTTACTTAGACTGGTNNNAGATCGGA
42715 TATGAGTGTGGAGTCGTTGCACTTGAATAGACACTTCTCCACAGNNNAGATCGGAA
42737 TATGAGTGTGGAGTCGTTGCTGCCAGGCACTATTCTAAGTACCNNNAGATCGGAAG
42751 TATGAGTGTGGAGTCGTTGCAGAAAATGGCAGGAGTAAGTTCCTNNNAGATCGGA
42759 TATGAGTGTGGAGTCGTTGCACTACTCAGCCATAAAAAGGAACANNNAGATCGGAA
42768 TATGAGTGTGGAGTCGTTGCCCATGAGGAGAGGAAGTGTTGAANNNAGATCGGA
42797 TATGAGTGTGGAGTCGTTGCGGGATACAGCTAACTAGTGAGGTGNNNAGATCGGA
42801 TATGAGTGTGGAGTCGTTGCTGCATCATTGTTAGGATAAGCCCTNNNAGATCGGAA
42822 TATGAGTGTGGAGTCGTTGCTTTCTCTCTCTTTTGAGCCCTCCNNNAGATCGGAAGA
42848 TATGAGTGTGGAGTCGTTGCATGTGTGTGTATATGTGTGGGTATNNNAGATCGGA
42896 TATGAGTGTGGAGTCGTTGCTGAAATATGTATACATTGTGGAATGGCNNNAGATCG
42901 TATGAGTGTGGAGTCGTTGCAACATGGGGGTGCAGATATCTTTNNNAGATCGGAA
42908 TATGAGTGTGGAGTCGTTGCCACACACGTACACATATACACACANNNAGATCGGAA
42927 TATGAGTGTGGAGTCGTTGCTTCTTCCCATCTGACACTGAGTCNNNAGATCGGAAG
42953 TATGAGTGTGGAGTCGTTGCGGGATTGCATTCTTGATTTGGCTNNNAGATCGGAAG
42966 TATGAGTGTGGAGTCGTTGCCTCAAAATAGCATTTCCTGGGCANNNAGATCGGAAG
42984 TATGAGTGTGGAGTCGTTGCACACACACACACAAAATATAGGCTNNNAGATCGGAA
42988 TATGAGTGTGGAGTCGTTGCTCCTCACTATGCTTGATGAACTGANNNAGATCGGAA
42990 TATGAGTGTGGAGTCGTTGCTCCCACCAACAATGTGTAAGTGTNNNAGATCGGAAG
43008 TATGAGTGTGGAGTCGTTGCAGCTACTTCTGCTGGTAGGAATGNNNAGATCGGAA
43102 TATGAGTGTGGAGTCGTTGCAAGGAGTGTTGATGTTGATGTGTNNNAGATCGGAA
43124 TATGAGTGTGGAGTCGTTGCTGACTCATCTAATGGTCTGACACANNNAGATCGGAA
43149 TATGAGTGTGGAGTCGTTGCCATGTGAAGGATCATGTCAAAGGTNNNAGATCGGA
43160 TATGAGTGTGGAGTCGTTGCCTGGGTTCAAGTGCTTTTCCTGNNNAGATCGGAAGA
43256 TATGAGTGTGGAGTCGTTGCCTCAAGAAGCTGGAGTTACAGGTNNNAGATCGGAA
43290 TATGAGTGTGGAGTCGTTGCTCCATCCTCAAATTAGGCAAATTCNNNAGATCGGAAG
43310 TATGAGTGTGGAGTCGTTGCCCACCACACCTGGCTATTTTTTTNNNAGATCGGAAGA
43322 TATGAGTGTGGAGTCGTTGCTTCAAGTGATTCTCCTGCCTCAGNNNAGATCGGAAG
43418 TATGAGTGTGGAGTCGTTGCTGCCAAGGTGGGGTAAACATATANNNAGATCGGAA
43503 TATGAGTGTGGAGTCGTTGCGAGTAGCTGCGATTACAGGAGTGNNNAGATCGGAA
43504 TATGAGTGTGGAGTCGTTGCACCTTTGATGGAAATTTGTGTTGTNNNAGATCGGAA
43534 TATGAGTGTGGAGTCGTTGCGCACTTGTGTTGCACTACTAGAANNNAGATCGGAAG
43559 TATGAGTGTGGAGTCGTTGCAGCTGATAACAACTTAGCACTGTNNNAGATCGGAAG
43601 TATGAGTGTGGAGTCGTTGCACCCCTACTAGCCTATTGCAAATNNNAGATCGGAAG
43777 TATGAGTGTGGAGTCGTTGCAGAAAAAGAAGAAGGAGGAGAGGGNNNAGATCG
43784 TATGAGTGTGGAGTCGTTGCAGCTGAATGAGGTATGTGTTCTCTNNNAGATCGGAA
43807 TATGAGTGTGGAGTCGTTGCACAGAACAGCCAAAATCATGGTACNNNAGATCGGAA
43808 TATGAGTGTGGAGTCGTTGCGCTAGCTGCTAGTATGTAGAAAAATNNNAGATCGG
43889 TATGAGTGTGGAGTCGTTGCGAGTGCGGGGATGTGATCATAGNNNAGATCGGAA
43979 TATGAGTGTGGAGTCGTTGCGGAGAACCAGGGTAGCTGATAAGNNNAGATCGGA
43986 TATGAGTGTGGAGTCGTTGCTGCTGCTAATAAAGACATACCCCANNNAGATCGGAA

44009 TATGAGTGTGGAGTCGTTGCACCACCAAAACATGCATTTCTAGGNNNAGATCGGAA
44186 TATGAGTGTGGAGTCGTTGCTGGTGTTGGAGAAGGAGATAAACANNNAGATCGG
44278 TATGAGTGTGGAGTCGTTGCGTCAGAAACTGATTTTGCCAGAGTNNNAGATCGGA
44406 TATGAGTGTGGAGTCGTTGCCCCAACTATTGTCATTGTTCCAGGNNNAGATCGGAA
44434 TATGAGTGTGGAGTCGTTGCACATGTATGGGTGTTATATGTACACANNNAGATCGG
44439 TATGAGTGTGGAGTCGTTGCAGCAGCTGAGATTACAGGCATAANNNAGATCGGAA
44516 TATGAGTGTGGAGTCGTTGCGATCCCCCTACGTTGTCTACTTGNNNAGATCGGAAG
44534 TATGAGTGTGGAGTCGTTGCGTCTTTTGGAGCAACATGAATGGANNNAGATCGGA
44587 TATGAGTGTGGAGTCGTTGCTTTGATTGATCAGGCTAAGTCCANNNAGATCGGAAG
45089 TATGAGTGTGGAGTCGTTGCTGTAGCATATAGGAAGCAATGCCTNNNAGATCGGA
45095 TATGAGTGTGGAGTCGTTGCAGATAGAGACAGACAAGACAGACTNNNAGATCGGA
45106 TATGAGTGTGGAGTCGTTGCTGGCCATGTAACTATCTCTAGTGCNNNAGATCGGAA
45109 TATGAGTGTGGAGTCGTTGCAGAGCTACGTGGAAATTGAACCTNNNAGATCGGAA
45113 TATGAGTGTGGAGTCGTTGCTGTGACTGTGATTTGTTTTATGGTNNNAGATCGGAA
45124 TATGAGTGTGGAGTCGTTGCAGAATATACAACAAGAGTGGGCCTNNNAGATCGGA
45246 TATGAGTGTGGAGTCGTTGCTCCAAGGGGGAAAAAAGAAAACTGNNNAGATCGGA
45304 TATGAGTGTGGAGTCGTTGCTTTGTGCCAGGCATTTTATACCCNNNAGATCGGAAG
45361 TATGAGTGTGGAGTCGTTGCAGAGGCACTTAGGTCAATTCTGTNNNAGATCGGAAG
45523 TATGAGTGTGGAGTCGTTGCACATCTCAGCAGGGAAAATAGCANNNAGATCGGAA
45563 TATGAGTGTGGAGTCGTTGCGGGTATTTGTGAAAGCCCCTCTCNNNAGATCGGAAG
45605 TATGAGTGTGGAGTCGTTGCTGTGTGTCTTTATAGGTAGGGGTGNNNAGATCGGA
45647 TATGAGTGTGGAGTCGTTGCCTCCCAAGTAGCTGGCATTACAGNNNAGATCGGAAG
45662 TATGAGTGTGGAGTCGTTGCCTTTAGCCTCCTGAGTAGCTGAGNNNAGATCGGAAG
45709 TATGAGTGTGGAGTCGTTGCCGAGTAGCTGGGACTACAGGTNNNAGATCGGAAGA
45731 TATGAGTGTGGAGTCGTTGCACATCACCACTCCTGGCTAATTTNNNAGATCGGAAGA
45755 TATGAGTGTGGAGTCGTTGCCTCCCGGGTTCAAGTGATTCTCNNNAGATCGGAAGA
45793 TATGAGTGTGGAGTCGTTGCTTCAAGTGATTCTCCTGCCTCAGNNNAGATCGGAAG
45810 TATGAGTGTGGAGTCGTTGCAGAGTGCAATGGTGTAATCTCGGNNNAGATCGGAA
45812 TATGAGTGTGGAGTCGTTGCATTCTCCCTCCTCAGCGTTCTAANNNAGATCGGAAGA
45907 TATGAGTGTGGAGTCGTTGCCTTCCTGGGTTCAAGCAATTCTCNNNAGATCGGAAG
45923 TATGAGTGTGGAGTCGTTGCTTCAAGTGATTCTCCTGCCTCAGNNNAGATCGGAAG
45961 TATGAGTGTGGAGTCGTTGCGTTCAAGGGATTATCCTGCCTCANNNAGATCGGAAG
46024 TATGAGTGTGGAGTCGTTGCGTCTCCTGGGTTCCAATGATTCTNNNAGATCGGAAG
46033 TATGAGTGTGGAGTCGTTGCTGGCCAGAAATAATGTTTACCAGCNNNAGATCGGAA
46044 TATGAGTGTGGAGTCGTTGCCTAGTAGCTGGGATTATAGGCGCNNNAGATCGGAA
46063 TATGAGTGTGGAGTCGTTGCCCACCACACTCGGCTAATTTTTTNNNAGATCGGAAGA
46099 TATGAGTGTGGAGTCGTTGCAGCTTATAGCCTATTATCCCAGGTNNNAGATCGGAA
46129 TATGAGTGTGGAGTCGTTGCTTGGTATTAGGGTGATACTGGCANNNAGATCGGAA
46188 TATGAGTGTGGAGTCGTTGCGGCGATGAAGAGACTTTTCAAACANNNAGATCGGA
46237 TATGAGTGTGGAGTCGTTGCTTCACACCATTCTCCTGCCTCNNNAGATCGGAAGAGC
46245 TATGAGTGTGGAGTCGTTGCCCACCACACCCAGCTAATTTTTTNNNAGATCGGAAGA
46246 TATGAGTGTGGAGTCGTTGCGTCAACAAGAGAGGCACAATTGTNNNAGATCGGAA
46339 TATGAGTGTGGAGTCGTTGCTGTATTTCTGTGGCATCAGTGGANNNAGATCGGAA

46411 TATGAGTGTGGAGTCGTTGCCACAATCTCAGCTCACTGCAACNNNAGATCGGAAGA
46425 TATGAGTGTGGAGTCGTTGCCTCTCGGATTCACGCCATTCTCNNNAGATCGGAAGA
46429 TATGAGTGTGGAGTCGTTGCTAGCTGAGATTACAGGTGTGTGCNNNAGATCGGAA
46442 TATGAGTGTGGAGTCGTTGCACGTTCAAGTAATTCTCCTGCCTNNNAGATCGGAAG
46444 TATGAGTGTGGAGTCGTTGCCTACGCCCGGCTAATTTTTTGTANNNAGATCGGAAG
46457 TATGAGTGTGGAGTCGTTGCTTTGTAAGGTTACGTGTTGTTTGANNNAGATCGGAA
46493 TATGAGTGTGGAGTCGTTGCGGGTTGAAGTAATTCTCCTGCCTNNNAGATCGGAAG
46501 TATGAGTGTGGAGTCGTTGCTCTCCCGAGTTCTAAGCGATTTTNNNAGATCGGAAG
46523 TATGAGTGTGGAGTCGTTGCTGGTTTTGATTTGCATTTCCCAGANNNAGATCGGAA
46573 TATGAGTGTGGAGTCGTTGCTTCAAGCTATTCTCCTGCCTCAGNNNAGATCGGAAG
46587 TATGAGTGTGGAGTCGTTGCCCAGTATCTCCTGCTAAGGTTTGANNNAGATCGGAA
46589 TATGAGTGTGGAGTCGTTGCTGAACTAAAATCCAAAGGCTGCCNNNAGATCGGAAG
46593 TATGAGTGTGGAGTCGTTGCTGATCTCGGCTCACTGCAATATCNNNAGATCGGAAG
46642 TATGAGTGTGGAGTCGTTGCCTCCTGAGTAGCTGGGACTACAGNNNAGATCGGAA
46647 TATGAGTGTGGAGTCGTTGCCCGAGTAGCTGTATTACAGGCATNNNAGATCGGAA
46649 TATGAGTGTGGAGTCGTTGCGCCACCACACCTGCTAATTATTTNNNAGATCGGAAG
46650 TATGAGTGTGGAGTCGTTGCCTCCTGAGTAGCTTGGACTACAGNNNAGATCGGAAG
46677 TATGAGTGTGGAGTCGTTGCGAGTCTCACTCTTTTTGCCCAGGNNNAGATCGGAAG
46722 TATGAGTGTGGAGTCGTTGCGGCAAAACCTTGTCTTCAAACCTNNNAGATCGGAAG
46731 TATGAGTGTGGAGTCGTTGCTTGACTGTTGTTGGTGTACAGGANNNAGATCGGAA
46814 TATGAGTGTGGAGTCGTTGCAGCAGAAACAAAGAAACATAGCCCNNNAGATCGGA
46827 TATGAGTGTGGAGTCGTTGCCTCGGCTCACTACAAGCTCTATTNNNAGATCGGAAG
46831 TATGAGTGTGGAGTCGTTGCGCCTCCTGAGTAGCTGAGATTACNNNAGATCGGAAG
46844 TATGAGTGTGGAGTCGTTGCTCCATTTTTACTGTGCATACACGTNNNAGATCGGAAG
46903 TATGAGTGTGGAGTCGTTGCTGTGTAACTTTTTTCTCTGCCACANNNAGATCGGAAG
46943 TATGAGTGTGGAGTCGTTGCAAATAGCTGGGATTACAGGTGCANNNAGATCGGAA
46945 TATGAGTGTGGAGTCGTTGCCCACCACACCCAGCTAATTTTTTNNNAGATCGGAAGA
46993 TATGAGTGTGGAGTCGTTGCACGCCCGGCTAATTTATTTTTGTNNNAGATCGGAAG
47053 TATGAGTGTGGAGTCGTTGCCCAAGTAGCTGGGATTACAGGTGNNNAGATCGGAA
47076 TATGAGTGTGGAGTCGTTGCAGTAGCTAGGAGTACAGGCTCATNNNAGATCGGAA
47111 TATGAGTGTGGAGTCGTTGCCCACACCCGGCTAATTTTGTATTNNNAGATCGGAAG
47218 TATGAGTGTGGAGTCGTTGCCCACCATACCCGGCTAATTTTTGNNNAGATCGGAAG
47260 TATGAGTGTGGAGTCGTTGCTCTTTAGTAGAGGTGGGGTTTCACNNNAGATCGGA
47263 TATGAGTGTGGAGTCGTTGCTCTCCCTAGCTCAAACCATCCTANNNAGATCGGAAGA
47266 TATGAGTGTGGAGTCGTTGCCACCATGCCCAGCTAATTTTTGTNNNAGATCGGAAG
47294 TATGAGTGTGGAGTCGTTGCAACCTCCCAAGTAGCTGAGACTANNNAGATCGGAAG
47319 TATGAGTGTGGAGTCGTTGCAATGCAGTGGTATGATCTCAGCTNNNAGATCGGAA
47330 TATGAGTGTGGAGTCGTTGCCCGAGTAGCTGGAATTATAGGCANNNAGATCGGAA
47434 TATGAGTGTGGAGTCGTTGCCCACCACACCTGGCTAATTTTTTNNNAGATCGGAAGA
47441 TATGAGTGTGGAGTCGTTGCTGTCTTTTCCACTTGATTCCATGCNNNAGATCGGAAG
47468 TATGAGTGTGGAGTCGTTGCCCACGCCCAGCTAATTTTTGTATNNNAGATCGGAAG
47503 TATGAGTGTGGAGTCGTTGCCTCCTGAGTAGCTGGGATTACAGNNNAGATCGGAA
47509 TATGAGTGTGGAGTCGTTGCAGAAGAGCTACTGATTTGTGTACANNNAGATCGGA

47585 TATGAGTGTGGAGTCGTTGCTCTGTCAGGGAAAATTGACTTCCTNNNAGATCGGAA
47629 TATGAGTGTGGAGTCGTTGCCTTTCAAGGAACCAGCCTCATTGNNNAGATCGGAAG
47677 TATGAGTGTGGAGTCGTTGCTCCTGCTTTCCAAATCATGAATCCNNNAGATCGGAAG
47678 TATGAGTGTGGAGTCGTTGCCAATCATAGCTCACTGCAACCTCNNNAGATCGGAAG
47752 TATGAGTGTGGAGTCGTTGCCCTCCCAGGTTCAAGTGATTCTCNNNAGATCGGAAG
47764 TATGAGTGTGGAGTCGTTGCAGATCTCAGCTAACTGCAACCTCNNNAGATCGGAAG
47789 TATGAGTGTGGAGTCGTTGCTCTTTGCTCACCATCCTCAAACTNNNAGATCGGAAGA
47800 TATGAGTGTGGAGTCGTTGCAGACAGAGTCTCCCTGTGTCTCNNNAGATCGGAAGA
47854 TATGAGTGTGGAGTCGTTGCGATCCTCCCTAGTAGCTGTGAGANNNAGATCGGAAG
47892 TATGAGTGTGGAGTCGTTGCCTGCCTCCTAGGTTCAAGTGATTNNNAGATCGGAAG
47893 TATGAGTGTGGAGTCGTTGCTTGAGATTCATCCATGTTGTTTGTNNNAGATCGGAA
47965 TATGAGTGTGGAGTCGTTGCACCAGGCCCAGCTAATTTTTGTANNNAGATCGGAAG
48070 TATGAGTGTGGAGTCGTTGCTTGGTCTATAGCAGTGCTACAGANNNAGATCGGAAG
48105 TATGAGTGTGGAGTCGTTGCTAGGTTCAAGTGATTCTCCTGCCNNNAGATCGGAAG
48197 TATGAGTGTGGAGTCGTTGCCAAGTATCTGGGACTACAGGCACNNNAGATCGGAAG
48220 TATGAGTGTGGAGTCGTTGCTCCACCAGGGAAAGATCAATTCANNNAGATCGGAAG
48234 TATGAGTGTGGAGTCGTTGCTGATGATGATGTCTATCACATGGANNNAGATCGGAA
48248 TATGAGTGTGGAGTCGTTGCTCTTCTTGATTCAACCTTGGTAGGNNNAGATCGGAA
48265 TATGAGTGTGGAGTCGTTGCTGGAGTTCAATGGCATGATCTCGNNNAGATCGGAAG
48280 TATGAGTGTGGAGTCGTTGCCTTCCCAGGTTCAAGCAATTCTCNNNAGATCGGAAG
48315 TATGAGTGTGGAGTCGTTGCGCCAACTAAATCAGGATGATTGGGNNNAGATCGGA
48323 TATGAGTGTGGAGTCGTTGCCCTTCCTGAGTAGCTGGGATTACNNNAGATCGGAAG
48325 TATGAGTGTGGAGTCGTTGCGGCTGAACTGGAAAAAAGAATGGANNNAGATCGGA
48352 TATGAGTGTGGAGTCGTTGCAACCTCTGACTCCCAACTTCAAGNNNAGATCGGAAG
48369 TATGAGTGTGGAGTCGTTGCTTCAAGTGATTCTCCTGCCTCAGNNNAGATCGGAAG
48393 TATGAGTGTGGAGTCGTTGCACATGAGAGATGGAATTGCTGTCTNNNAGATCGGA
48421 TATGAGTGTGGAGTCGTTGCAGCTAGAACTAAAGGTGCATGCTNNNAGATCGGAA
48451 TATGAGTGTGGAGTCGTTGCACCTAGTAGCTGGGATTACAGGTNNNAGATCGGAA
48500 TATGAGTGTGGAGTCGTTGCTCCCCTTGGAATAAAACGGTGAANNNAGATCGGAAG
48502 TATGAGTGTGGAGTCGTTGCGGTTCAAACGATTCTCTTGCCTCNNNAGATCGGAAG
48514 TATGAGTGTGGAGTCGTTGCAAGTGCTGAGATTACAGGCATGANNNAGATCGGAA
48518 TATGAGTGTGGAGTCGTTGCTTCTCTTGGGTATTCCTGGGAGANNNAGATCGGAAG
48554 TATGAGTGTGGAGTCGTTGCCACCATGCCCAGCTAATTTTTGTNNNAGATCGGAAG
48573 TATGAGTGTGGAGTCGTTGCACTACAGGTATGTGCCACCATACNNNAGATCGGAAG
48725 TATGAGTGTGGAGTCGTTGCTCTGTGCCTAGCTTATTTCACTCANNNAGATCGGAAG
48726 TATGAGTGTGGAGTCGTTGCCTCCTGAGTAGCTGGGATTACAGNNNAGATCGGAA
48729 TATGAGTGTGGAGTCGTTGCTCTCTGGTTTTTCTCCAACTGTTCNNNAGATCGGAAG
48759 TATGAGTGTGGAGTCGTTGCACACTATTCTCCTACCTCAGCCTNNNAGATCGGAAGA
48782 TATGAGTGTGGAGTCGTTGCTCCTTTTTTGGTGTCCTCTAAGCTNNNAGATCGGAAG
48804 TATGAGTGTGGAGTCGTTGCGAGTAGCTGGGATTATAGGTGCCNNNAGATCGGAA
48813 TATGAGTGTGGAGTCGTTGCATCAGCTGTGATGTTGACCAAGANNNAGATCGGAA
48873 TATGAGTGTGGAGTCGTTGCCTCCTGAGTAGCTGGGATTACAGNNNAGATCGGAA
48918 TATGAGTGTGGAGTCGTTGCCTGTGCCCGGCTAATTTTTTGTANNNAGATCGGAAG

48923 TATGAGTGTGGAGTCGTTGCTCCCAATTAGCAGTTCTTAGTGGTNNNAGATCGGAA
48945 TATGAGTGTGGAGTCGTTGCCTTCCCAGGTTCAAGCAATTCTCNNNAGATCGGAAG
48969 TATGAGTGTGGAGTCGTTGCACCCAGTTTTGGAAAATTAGCCTNNNAGATCGGAA
48972 TATGAGTGTGGAGTCGTTGCACATGGGGTTTTATATTTCCACCTNNNAGATCGGAA
48983 TATGAGTGTGGAGTCGTTGCCGGCATCAACATTGAAATCCAGANNNAGATCGGAAG
48994 TATGAGTGTGGAGTCGTTGCCTCCTGAGTAGCTGGGATTACAGNNNAGATCGGAA
49026 TATGAGTGTGGAGTCGTTGCACGCCCGGCTAATTTTTTGTATTNNNAGATCGGAAG
49078 TATGAGTGTGGAGTCGTTGCTTTTCTCTTCTACTGGCTTAGACANNNAGATCGGAAG
49092 TATGAGTGTGGAGTCGTTGCAACTTCTGCCTCCTGGATTCAAGNNNAGATCGGAAG
49169 TATGAGTGTGGAGTCGTTGCAATTTTAGTACCTCAGCCTCCCGNNNAGATCGGAAG
49171 TATGAGTGTGGAGTCGTTGCCACAATGATGGCTCACTGCAANNNAGATCGGAAGAG
49231 TATGAGTGTGGAGTCGTTGCGATCTCCTGGTCTCAACTGATCCNNNAGATCGGAAG
49242 TATGAGTGTGGAGTCGTTGCTCCTGAGTTCAAGCGATTCTCTTNNNAGATCGGAAG
49252 TATGAGTGTGGAGTCGTTGCAATTCTCCTACCTCAGTCTCCCANNNAGATCGGAAGA
49308 TATGAGTGTGGAGTCGTTGCCAAAGAGGTATTACGCGTCTGGTNNNAGATCGGAA
49377 TATGAGTGTGGAGTCGTTGCACGCCCGGCTAATTTTTTGTATTNNNAGATCGGAAG
49398 TATGAGTGTGGAGTCGTTGCCCACCAAGACTGGCTAATTTTTGTNNNAGATCGGAA
49400 TATGAGTGTGGAGTCGTTGCACCACGCCTTGCTAATTTTTTGTNNNAGATCGGAAGA
49419 TATGAGTGTGGAGTCGTTGCCCGGATTCAAGCAATTCACCTACNNNAGATCGGAAG
49458 TATGAGTGTGGAGTCGTTGCAGGAGATATTACCACTGACCCCANNNAGATCGGAAG
49508 TATGAGTGTGGAGTCGTTGCACTTTGTGCTCTCTTGAGGACAANNNAGATCGGAAG
49516 TATGAGTGTGGAGTCGTTGCTGGTAGTGATGAATTCCCTCACCNNNAGATCGGAAG
49560 TATGAGTGTGGAGTCGTTGCAACAGTTCCCTTTTCTCCACACCNNNAGATCGGAAGA
49566 TATGAGTGTGGAGTCGTTGCTCTCCTACCTTTTCTTTTGAGACTNNNAGATCGGAAG
49576 TATGAGTGTGGAGTCGTTGCCCCCCATCTCTCAGAAGTCATTTNNNAGATCGGAAGA
49618 TATGAGTGTGGAGTCGTTGCGCCACTCTGTATCTTTTGATTGGTNNNAGATCGGAA
49619 TATGAGTGTGGAGTCGTTGCTTTCCTGGGCTCAGATGATTTGANNNAGATCGGAAG
49698 TATGAGTGTGGAGTCGTTGCCCTTCTTGCTGTTCTGTCCATTGNNNAGATCGGAAGA
49778 TATGAGTGTGGAGTCGTTGCCATACCAATGTGTTTTCGCCACANNNAGATCGGAAG
49795 TATGAGTGTGGAGTCGTTGCACTCAGAGACCTTGTTACAATCCANNNAGATCGGAA
49860 TATGAGTGTGGAGTCGTTGCAAGTTGCTAGGTACTTGTGAGGCNNNAGATCGGAA
49889 TATGAGTGTGGAGTCGTTGCAGCAAACCTACACAAAATGAACGANNNAGATCGGAA
49930 TATGAGTGTGGAGTCGTTGCTCTCAACGTATTTTTCCTCTGGGTNNNAGATCGGAAG
49954 TATGAGTGTGGAGTCGTTGCTTTGAGCAATAGTGTCAGTGGGANNNAGATCGGAA
49969 TATGAGTGTGGAGTCGTTGCGCATAGCACCACTCGAAATGAAANNNAGATCGGAAG
50059 TATGAGTGTGGAGTCGTTGCAGCTGCTGTGAACATATTTGTACANNNAGATCGGAA
50110 TATGAGTGTGGAGTCGTTGCACCTTCACTGCCCAAAAGACTTANNNAGATCGGAAG
50321 TATGAGTGTGGAGTCGTTGCTCAACTTGTAGCGTAATTTTCCTGTNNNAGATCGGAA
50324 TATGAGTGTGGAGTCGTTGCAGTACAAAGAAGAGCTGGTACCANNNAGATCGGAA
50436 TATGAGTGTGGAGTCGTTGCTGATACAATCCCAAGCCTTCTAGANNNAGATCGGAA
50441 TATGAGTGTGGAGTCGTTGCCCGCTTGTTCACTCATACTTCTTGNNNAGATCGGAAG
50557 TATGAGTGTGGAGTCGTTGCTCTTTTTTTGAGACAGTCTGGCTGNNNAGATCGGAA
50848 TATGAGTGTGGAGTCGTTGCTGGATGTGGTTATGTTCCAATCACNNNAGATCGGAA

50861 TATGAGTGTGGAGTCGTTGCTTCTCATTTCCTGTACAGACATGANNNAGATCGGAA
50875 TATGAGTGTGGAGTCGTTGCTCCCAAGTCAAATTTCACCTTTCANNNAGATCGGAAG
50962 TATGAGTGTGGAGTCGTTGCCCCAAAGACCACCATATTTGCANNNAGATCGGAAG
50998 TATGAGTGTGGAGTCGTTGCACAATGAGACTCTCCAATCCTTCTNNNAGATCGGAAG
51008 TATGAGTGTGGAGTCGTTGCTGTTCAAATGTTCATACTGCCCANNNAGATCGGAAG
51047 TATGAGTGTGGAGTCGTTGCCCCGGCTGGAAACTCTTTATACTNNNAGATCGGAAG
51425 TATGAGTGTGGAGTCGTTGCGAAGACGGGTGATTTCTGCATTTNNNAGATCGGAA
51476 TATGAGTGTGGAGTCGTTGCAGTCCCTGTAAGGCTGTTGTTTCNNNAGATCGGAAG
51478 TATGAGTGTGGAGTCGTTGCAGTCCCTGTAAGGCTGTTGTTTCNNNAGATCGGAAG
51666 TATGAGTGTGGAGTCGTTGCAAGGCACATACACTCACAGAGTTNNNAGATCGGAAG
51714 TATGAGTGTGGAGTCGTTGCGCTGGATCAACAACAATGCAGTANNNAGATCGGAA
51737 TATGAGTGTGGAGTCGTTGCTCATGGTGACAATAACATGGAAAANNNAGATCGGA
51781 TATGAGTGTGGAGTCGTTGCGGTCTTTTCTGACTGGCTTCTCTNNNAGATCGGAAG
51812 TATGAGTGTGGAGTCGTTGCTTCTTCCTTTTCCAGAGTGCCATNNNAGATCGGAAGA
52046 TATGAGTGTGGAGTCGTTGCTGAAACAAAAAGCTGGTTCTTTGANNNAGATCGGA
52082 TATGAGTGTGGAGTCGTTGCTGGCATGTGGATATTCAGTTTTCCNNNAGATCGGAA
52106 TATGAGTGTGGAGTCGTTGCTTACCGTGCAAGAGACTTTTCCANNNAGATCGGAAG
52126 TATGAGTGTGGAGTCGTTGCTTGGAATACTGCAGAGCTCAGTTNNNAGATCGGAA
52211 TATGAGTGTGGAGTCGTTGCGGAGATTAGAAAGTGTGTGTGTGCNNNAGATCGG
52212 TATGAGTGTGGAGTCGTTGCGGAGATTAGAAAGTGTGTGTGTGCNNNAGATCGG
52259 TATGAGTGTGGAGTCGTTGCAGTCTTGACTGAAGTACTATGGGANNNAGATCGGA
52283 TATGAGTGTGGAGTCGTTGCTCCTGAGTAGCTGGGATTACAGANNNAGATCGGAA
52432 TATGAGTGTGGAGTCGTTGCGAGGTTTGGGGTTTGATTGAACCNNNAGATCGGAA
52706 TATGAGTGTGGAGTCGTTGCTCCCATAAGGTGACTGAAAAGTGTNNNAGATCGGA
52953 TATGAGTGTGGAGTCGTTGCGGAGTTGCAAGAAGGGAAACATCNNNAGATCGGA
52956 TATGAGTGTGGAGTCGTTGCGCTTTAGTGATCTATTGCACAGCANNNAGATCGGAA
52958 TATGAGTGTGGAGTCGTTGCTAATGGCGAATAGGGTGTCCATTNNNAGATCGGAA
53022 TATGAGTGTGGAGTCGTTGCTGGTTCAAAAGGCAAAATACACGANNNAGATCGGA
53057 TATGAGTGTGGAGTCGTTGCTGCAATCAGCTATAATCCTTCCCTNNNAGATCGGAAG
53113 TATGAGTGTGGAGTCGTTGCTCCTCTTACTGATCTTTGGTTTGCNNNAGATCGGAAG
53128 TATGAGTGTGGAGTCGTTGCGCAGAAAAAAATGTGTATGTGTGTNNNAGATCGGA
53149 TATGAGTGTGGAGTCGTTGCTTTTGTTTCTGAAAATGTAGGGGTNNNAGATCGGA
53323 TATGAGTGTGGAGTCGTTGCGTGCAATCCAGAGTCCTTGGATANNNAGATCGGAA
53329 TATGAGTGTGGAGTCGTTGCACGTGTATATATGCGTGTACACACNNNAGATCGGAA
53359 TATGAGTGTGGAGTCGTTGCACTTTTTCCAAACCATCAAGGACANNNAGATCGGAA
53367 TATGAGTGTGGAGTCGTTGCCACACACATACACACACATACACANNNAGATCGGAA
53407 TATGAGTGTGGAGTCGTTGCAGAAAGAGAGAGACAAAGGAAAGGANNNAGATCG
53408 TATGAGTGTGGAGTCGTTGCTCCTTAGCTAATAATCAGGGAAATGTNNNAGATCGG
28752 TATGAGTGTGGAGTCGTTGCGACAGTGGAAATGAGAGGGAACTNNNAGATCGGA
28753 TATGAGTGTGGAGTCGTTGCCATCAAGCAATTCTCTGCCTCAGNNNAGATCGGAAG
28755 TATGAGTGTGGAGTCGTTGCACCAACCTTCCTCACACACTATGNNNAGATCGGAAGA
28756 TATGAGTGTGGAGTCGTTGCCCTATAGTCCCAGCTACTCAGGANNNAGATCGGAAG
28757 TATGAGTGTGGAGTCGTTGCAGGACAGTTCATCCATGTTGCTANNNAGATCGGAAG

28759 TATGAGTGTGGAGTCGTTGCTGCTCAATGAACGAAGGAAGAGTNNNAGATCGGAA
28760 TATGAGTGTGGAGTCGTTGCAACTCCCCCTTTCATAGTCAACCNNNAGATCGGAAGA
28761 TATGAGTGTGGAGTCGTTGCTTAGTACAGACGAGGTTTTGCCANNNAGATCGGAAG
28762 TATGAGTGTGGAGTCGTTGCCCAAGTAGCTGGGATTACAGGTGNNNAGATCGGAA
28763 TATGAGTGTGGAGTCGTTGCCCTCCTGCATTCAAGCAATTCTCNNNAGATCGGAAGA
28768 TATGAGTGTGGAGTCGTTGCGTGTGTGTGTGTGTGTGTATGNNNAGATCGGAA
28769 TATGAGTGTGGAGTCGTTGCGGATGGAGCTGGAGTCCATTATTNNNAGATCGGAA
28770 TATGAGTGTGGAGTCGTTGCTTGAGGCAGGAGAATCACTTGAANNNAGATCGGAA
28772 TATGAGTGTGGAGTCGTTGCACATGTCTGAACAAGATCTGGCTNNNAGATCGGAAG
28773 TATGAGTGTGGAGTCGTTGCAAGAATCACTTGAACCTGGGAGGNNNAGATCGGAA
28774 TATGAGTGTGGAGTCGTTGCGTAATCCCAGCTACTCAGTAGGCNNNAGATCGGAAG
28776 TATGAGTGTGGAGTCGTTGCCCCCTTCCCATTTCAATTTGCTTNNNAGATCGGAAGA
28777 TATGAGTGTGGAGTCGTTGCGTGTGTGTTCCCAAGGAAAAANNNAGATCGGAA
28778 TATGAGTGTGGAGTCGTTGCAGATTGCTGGGTCAAGTGGTATTNNNAGATCGGAA
28779 TATGAGTGTGGAGTCGTTGCTGGGTAGATATGCAGTAGTGGGANNNAGATCGGA
28782 TATGAGTGTGGAGTCGTTGCCCAGGTTCAAGCAATTTTCCTGTNNNAGATCGGAAG
28783 TATGAGTGTGGAGTCGTTGCGGTGGTTTGCCTAGGTTTTCTTCNNNAGATCGGAAG
28787 TATGAGTGTGGAGTCGTTGCTATACACACACACACACACACACNNNAGATCGGAAG
28789 TATGAGTGTGGAGTCGTTGCGTGTGTGTGTGTATGTGTGTGTGNNNAGATCGGAA
28790 TATGAGTGTGGAGTCGTTGCTGGAGAGAGAGAGAGAGAGAGAANNNAGATCGG
28791 TATGAGTGTGGAGTCGTTGCAGAAGGTAGCACACTTCCAACATNNNAGATCGGAAG
28795 TATGAGTGTGGAGTCGTTGCGTGTGTGTGTGTGTGTGTGTATANNNAGATCGGAA
28797 TATGAGTGTGGAGTCGTTGCGTTTCACCATGTTGCTCAGTCTGNNNAGATCGGAAG
28798 TATGAGTGTGGAGTCGTTGCGCTAACATGGTGAAACCCCTTCTNNNAGATCGGAAG
28800 TATGAGTGTGGAGTCGTTGCTTTTGCAGCAACTTAGATGGAGCNNNAGATCGGAAG
28802 TATGAGTGTGGAGTCGTTGCAGCCACTGGAGAAACATAAGTGANNNAGATCGGAA
28803 TATGAGTGTGGAGTCGTTGCACTACACAACCTATCTCTCACCANNNAGATCGGAAGA
28804 TATGAGTGTGGAGTCGTTGCAGCTTGCAAATACTTTCTCCCATNNNAGATCGGAAG
28805 TATGAGTGTGGAGTCGTTGCTTGTGTACATGCTTTTCCTCTGCNNNAGATCGGAAG
28807 TATGAGTGTGGAGTCGTTGCTCATTGATTGATGGGCATTTGGGNNNAGATCGGAA
28808 TATGAGTGTGGAGTCGTTGCCCCAGTTTCAAGCAATTCTCCTGNNNAGATCGGAAG
28809 TATGAGTGTGGAGTCGTTGCCTCCCACTGATTACATGAGGCTTNNNAGATCGGAAG
28811 TATGAGTGTGGAGTCGTTGCCCTGGATGGAATTGGAGACCATTNNNAGATCGGAA
28812 TATGAGTGTGGAGTCGTTGCACACAGCCTCAGAAACTTGTAAGNNNAGATCGGAAG
28814 TATGAGTGTGGAGTCGTTGCAATTGTGCCACTGTAAACATGCANNNAGATCGGAAG
28816 TATGAGTGTGGAGTCGTTGCATCCACTCATTGAATGACGGGAANNNAGATCGGAAG
28818 TATGAGTGTGGAGTCGTTGCACTTGGATGAAGCTGGAGACAATNNNAGATCGGAA
28819 TATGAGTGTGGAGTCGTTGCCCACAGGTATCAAGGGTAATGCTNNNAGATCGGAAG
28822 TATGAGTGTGGAGTCGTTGCATCCACTCATTAGTTGATGGGCANNNAGATCGGAAG
28824 TATGAGTGTGGAGTCGTTGCACACACACACACACACAGAGATANNNAGATCGGAAG
28829 TATGAGTGTGGAGTCGTTGCTATAAAACTGTCACCCAGGCTGGNNNAGATCGGAAG
28830 TATGAGTGTGGAGTCGTTGCTTTTTTTTTTTTGCGACGGGGTCNNNAGATCGGAAG
28832 TATGAGTGTGGAGTCGTTGCGAAAACCTGGGAGCTAAGTGAGANNNAGATCGGA

28833 TATGAGTGTGGAGTCGTTGCCAATTTCGGCTCACTGTAACCTCNNNAGATCGGAAGA
28834 TATGAGTGTGGAGTCGTTGCTCCAGGGATTGAGAAGAGTTTGGNNNAGATCGGAA
28835 TATGAGTGTGGAGTCGTTGCCAAAGAAAAAGCTGTGACAGGGTNNNAGATCGGAA
28836 TATGAGTGTGGAGTCGTTGCCACTGATCTCTGCCTGTCTTCAANNNAGATCGGAAGA
28838 TATGAGTGTGGAGTCGTTGCAATCACTGGAGAAGGAAGAGCAGNNNAGATCGGAA
28840 TATGAGTGTGGAGTCGTTGCAAATCAATAAATTCGGCTGGGCGNNNAGATCGGAA
28841 TATGAGTGTGGAGTCGTTGCGGTGTGCAACTTCTCTTCTCCTANNNAGATCGGAAGA
28842 TATGAGTGTGGAGTCGTTGCCCCACCACCCCCTTATAGTTTTTNNNAGATCGGAAGA
28844 TATGAGTGTGGAGTCGTTGCGGCAAGCAACAAGATGGTAACTTNNNAGATCGGAA
28845 TATGAGTGTGGAGTCGTTGCTGCCTAATTCCTCCTACCTCAGANNNAGATCGGAAGA
28846 TATGAGTGTGGAGTCGTTGCCTTTCTCTCACCCACTTGCTTTGNNNAGATCGGAAGA
28847 TATGAGTGTGGAGTCGTTGCTTTTTTCCAGGGAGGGATTCCATNNNAGATCGGAAG
28849 TATGAGTGTGGAGTCGTTGCTGGTGTTTCTGACTTCCCTTTCTNNNAGATCGGAAGA
28850 TATGAGTGTGGAGTCGTTGCAGAGAGAGGGAGAGAGAGAGAGANNNAGATCGGA
28851 TATGAGTGTGGAGTCGTTGCGTGTCATGCAGGTGTACTTTGACNNNAGATCGGAAG
28852 TATGAGTGTGGAGTCGTTGCACTTTGGGGCTCTTCATCTTGAANNNAGATCGGAAGA
28854 TATGAGTGTGGAGTCGTTGCGCACACACTTTTTTACCGTAAGCNNNAGATCGGAAGA
28856 TATGAGTGTGGAGTCGTTGCCACAATTGTGGAGGCTGATTCTGNNNAGATCGGAA
28857 TATGAGTGTGGAGTCGTTGCTGTCCCAAGGAGAGCATCATTTTNNNAGATCGGAAG
28858 TATGAGTGTGGAGTCGTTGCAACACTGATACCTGTATGTGGGGNNNAGATCGGAA
28859 TATGAGTGTGGAGTCGTTGCAGAGGATAGCAGTGGTTGTTTCANNNAGATCGGAA
28860 TATGAGTGTGGAGTCGTTGCTAGACTCAGTGTGAAAGGGGCTANNNAGATCGGAA
28861 TATGAGTGTGGAGTCGTTGCATCCCCTGGAAATCATGATTGGTNNNAGATCGGAAG
28862 TATGAGTGTGGAGTCGTTGCGCTGTATAGAACTTGCAACAGGGNNNAGATCGGAA
28863 TATGAGTGTGGAGTCGTTGCCCCAATGTTCATGGTGGTTGTTTNNNAGATCGGAAG
28864 TATGAGTGTGGAGTCGTTGCTCTGTGTGATGAACTTCTCTCCCNNNAGATCGGAAGA
28865 TATGAGTGTGGAGTCGTTGCTGACACTTGTTTGAAGCTTTGCTNNNAGATCGGAAGA
28868 TATGAGTGTGGAGTCGTTGCCTCTCAACAGGTAGCCAATCAGTNNNAGATCGGAAG
28869 TATGAGTGTGGAGTCGTTGCGCAGTGATGAGATCGTGTTCAAGNNNAGATCGGAA
28870 TATGAGTGTGGAGTCGTTGCGGTCTGTTTTGTCTGCCTAGAGANNNAGATCGGAAG
28871 TATGAGTGTGGAGTCGTTGCGGAGGCAGAGCATGATAAAGAGANNNAGATCGGA
28872 TATGAGTGTGGAGTCGTTGCAGGGCTCTTCACTAATGTTGAGGNNNAGATCGGAAG
28873 TATGAGTGTGGAGTCGTTGCTAGGCATGAAGAGTAGCACTGTGNNNAGATCGGAA
28874 TATGAGTGTGGAGTCGTTGCGTTACTGCAGCCTTGAACTCTTGNNNAGATCGGAAGA
28875 TATGAGTGTGGAGTCGTTGCAGAAGGGAAAGAGGAGGAGAAGANNNAGATCGGA
28876 TATGAGTGTGGAGTCGTTGCGGAATGACTGTTGTTGTTGAGCANNNAGATCGGAA
28877 TATGAGTGTGGAGTCGTTGCACAGAGAGAGAGAGAGAGATGCANNNAGATCGGA
28878 TATGAGTGTGGAGTCGTTGCTCTGCCCAAGAGACCCTCATATANNNAGATCGGAAGA
28879 TATGAGTGTGGAGTCGTTGCTGTGTGTGGATGTGTGTGTCTAANNNAGATCGGAA
28880 TATGAGTGTGGAGTCGTTGCCCATTGTAGGCCAGAGTTAGTGTNNNAGATCGGAAG
28881 TATGAGTGTGGAGTCGTTGCTTCTCTCTCTCACACACACACACNNNAGATCGGAAGA
28882 TATGAGTGTGGAGTCGTTGCGCCAATTACAGGAAGCCAAGTTTNNNAGATCGGAAG
28883 TATGAGTGTGGAGTCGTTGCGCAAGTTAGCCACAGAATTAGGGNNNAGATCGGAA

28884 TATGAGTGTGGAGTCGTTGCTGTCGTCCTAAGCGTTGTTCTTANNNAGATCGGAAG
28885 TATGAGTGTGGAGTCGTTGCCGGATCAGGTCTCCATCCTTATGNNNAGATCGGAAG
28886 TATGAGTGTGGAGTCGTTGCTGGCTTGGTTCTCAACTGTCTTANNNAGATCGGAAG
28887 TATGAGTGTGGAGTCGTTGCTGCCTGGCTAATTACTACTCACCNNNAGATCGGAAG
28889 TATGAGTGTGGAGTCGTTGCCCTTCTTAGGGCCATTCCAAGATNNNAGATCGGAAG
28890 TATGAGTGTGGAGTCGTTGCCCACTTGTTTTGGTACTGCTTGTNNNAGATCGGAAG
28892 TATGAGTGTGGAGTCGTTGCTTCAGGCTTGTTTTTGCTTCTGTNNNAGATCGGAAG
28893 TATGAGTGTGGAGTCGTTGCCTCCCTCTCATACTCTCGCTTTCNNNAGATCGGAAGA
28894 TATGAGTGTGGAGTCGTTGCCCAGTGACTATTTTTGTGGCTGGNNNAGATCGGAAG
28895 TATGAGTGTGGAGTCGTTGCCACTGTGGGTACAAGTAGAACCANNNAGATCGGAA
28898 TATGAGTGTGGAGTCGTTGCCTTTGAGCATCCAGGCAAAAAGANNNAGATCGGAA
28899 TATGAGTGTGGAGTCGTTGCCTACTCTCACTTCCCATACACCGNNNAGATCGGAAGA
28900 TATGAGTGTGGAGTCGTTGCCTGTGGTTCTTGCACACTTGTAGNNNAGATCGGAAG
28901 TATGAGTGTGGAGTCGTTGCTGCACGTTTATGTGTGTAGGTCTNNNAGATCGGAAG
28903 TATGAGTGTGGAGTCGTTGCGCAATGTGACTTTTGGCAAGTTCNNNAGATCGGAAG
28904 TATGAGTGTGGAGTCGTTGCATTTCTTTGCTCCCCTTTGATGCNNNAGATCGGAAGA
28906 TATGAGTGTGGAGTCGTTGCTTTGTTCTTCCCAGCTACCACTTNNNAGATCGGAAGA
28907 TATGAGTGTGGAGTCGTTGCTGTGAATGCACCTCCTAAATTGCNNNAGATCGGAAG
28909 TATGAGTGTGGAGTCGTTGCCATTTTCTTCACCCTGCCACTTTNNNAGATCGGAAGA
28910 TATGAGTGTGGAGTCGTTGCACTGGGGGAAGAATTACATCCACNNNAGATCGGAA
28911 TATGAGTGTGGAGTCGTTGCGACAACACTTTTGCCACTTCTGANNNAGATCGGAAG
28912 TATGAGTGTGGAGTCGTTGCAGATGGCACAAACAACTGAACTTNNNAGATCGGAAG
28913 TATGAGTGTGGAGTCGTTGCACAGAGACAGAGACAGAGAGACANNNAGATCGGA
28914 TATGAGTGTGGAGTCGTTGCTGAGCCAAATGCACCCCTTTATANNNAGATCGGAAG
28915 TATGAGTGTGGAGTCGTTGCTGCTTGAGACCAGAAGTGTTTCANNNAGATCGGAAG
28916 TATGAGTGTGGAGTCGTTGCTTGCTACAAGACCCACATGTGATNNNAGATCGGAAG
28917 TATGAGTGTGGAGTCGTTGCGCTCTCAGGAGGGAACAATACAANNNAGATCGGAA
28918 TATGAGTGTGGAGTCGTTGCAGAATCTGATGAGGTAAGCAGGCNNNAGATCGGAA
28919 TATGAGTGTGGAGTCGTTGCTCAATCAAATGCAGGCCAAACATNNNAGATCGGAAG
28920 TATGAGTGTGGAGTCGTTGCGGGATTGGATGGTTAGGGAGAAGNNNAGATCGGA
28921 TATGAGTGTGGAGTCGTTGCCAGAATGGACTGGGTTTTCCCTANNNAGATCGGAAG
28922 TATGAGTGTGGAGTCGTTGCAATTGATTCAGTTGGGTCAGGGANNNAGATCGGAA
28923 TATGAGTGTGGAGTCGTTGCCCATTGCACAAGACTTGTTGCTANNNAGATCGGAAG
28924 TATGAGTGTGGAGTCGTTGCATCATCACCATCAACAGTCACCANNNAGATCGGAAG
28925 TATGAGTGTGGAGTCGTTGCCAGTCAAGGCATCACAAGTCATTNNNAGATCGGAAG
28928 TATGAGTGTGGAGTCGTTGCATGAGCTCTGTCATTTTTGCTGGNNNAGATCGGAAG
28929 TATGAGTGTGGAGTCGTTGCTGAAGGAAGTGGAAATAGGTGGGNNNAGATCGGA
28930 TATGAGTGTGGAGTCGTTGCCTGAGAACCCAAACTGGATGTTGNNNAGATCGGAA
28931 TATGAGTGTGGAGTCGTTGCGAGAGAGAGAGAGACAGACCTCCNNNAGATCGGA
28932 TATGAGTGTGGAGTCGTTGCCAACAAAACACACAAACTGGTGCNNNAGATCGGAAG
28933 TATGAGTGTGGAGTCGTTGCTCCACTGATGTCCGTTGTACATTNNNAGATCGGAAG
28934 TATGAGTGTGGAGTCGTTGCTTCGGACTCTTGGATTTGTAGCANNNAGATCGGAAG
28935 TATGAGTGTGGAGTCGTTGCCACATTCAACTCACTAGCGCAAANNNAGATCGGAAG

28936 TATGAGTGTGGAGTCGTTGCTGACTGAAACAGCCCCTAGTTACNNNAGATCGGAAG
28937 TATGAGTGTGGAGTCGTTGCTGTATGTGTGTGTGTGTTCCTCANNNAGATCGGAAG
28939 TATGAGTGTGGAGTCGTTGCATCTCCTTCAGGTGTATGCATGGNNNAGATCGGAAG
28940 TATGAGTGTGGAGTCGTTGCACACACACACACACAGGTGATTANNNAGATCGGAAG
28941 TATGAGTGTGGAGTCGTTGCCATCCTAACCACTGCATCTAGCTNNNAGATCGGAAGA
28942 TATGAGTGTGGAGTCGTTGCTCACCCTCTCCAAGCAGAAATTTNNNAGATCGGAAGA
28944 TATGAGTGTGGAGTCGTTGCGCACACGTATGCATTTAGGGAAANNNAGATCGGAAG
28945 TATGAGTGTGGAGTCGTTGCTGTTTTCTTCTGTCATCCCACCANNNAGATCGGAAGA
28946 TATGAGTGTGGAGTCGTTGCGAGTGCAGTGACATGATCATAGCNNNAGATCGGAAG
28947 TATGAGTGTGGAGTCGTTGCATTTATGCACACACACACACACTNNNAGATCGGAAGA
28950 TATGAGTGTGGAGTCGTTGCTGCTGCCTTGTGTCTTGAAAATTNNNAGATCGGAAGA
28951 TATGAGTGTGGAGTCGTTGCTGAAAACACAATTCCCTGCTCACNNNAGATCGGAAGA
28952 TATGAGTGTGGAGTCGTTGCGGTGAGTGAATGAGTGCATTGTTNNNAGATCGGAA
28953 TATGAGTGTGGAGTCGTTGCATGTGTGTCTCTGTGTGTGTACANNNAGATCGGAAG
28954 TATGAGTGTGGAGTCGTTGCAGAGCGTGTGTATATGTGTGTGTNNNAGATCGGAAG
28955 TATGAGTGTGGAGTCGTTGCGCCTCTCTACCCTTCCTTGAATCNNNAGATCGGAAGA
28958 TATGAGTGTGGAGTCGTTGCGATGAGGTCTCACTCTGTCTGTCNNNAGATCGGAAG
28959 TATGAGTGTGGAGTCGTTGCTGTCTTTGTTTCCTTCAAGGTGTNNNAGATCGGAAGA
28961 TATGAGTGTGGAGTCGTTGCTGTGTGTATATATCGATGCGTGTNNNAGATCGGAAG
28963 TATGAGTGTGGAGTCGTTGCTGAGTCAGAAACTGCTATGCACTNNNAGATCGGAAG
28964 TATGAGTGTGGAGTCGTTGCACAGCAAGAAATAGCACTTTCAGNNNAGATCGGAAG
28965 TATGAGTGTGGAGTCGTTGCAAAGGAGAAGTAGGCGAGAAGTGNNNAGATCGGA
28966 TATGAGTGTGGAGTCGTTGCGCATCTTAGAGGAGACCAGTGACNNNAGATCGGAAG
28967 TATGAGTGTGGAGTCGTTGCTCTCTCTGTCTCTGTCTGTGTCTNNNAGATCGGAAGA
28968 TATGAGTGTGGAGTCGTTGCTATCTGCAGATTTGGTCTTGGGANNNAGATCGGAAG
28970 TATGAGTGTGGAGTCGTTGCTAACATGTTTCAAGGAGCCAGGANNNAGATCGGAAG
28971 TATGAGTGTGGAGTCGTTGCCCCACCATAGCTGTGTACCTATCNNNAGATCGGAAGA
28972 TATGAGTGTGGAGTCGTTGCGTCACGTGCCAGAGTTCAGTATANNNAGATCGGAAG
28975 TATGAGTGTGGAGTCGTTGCTTGGCACAAAAGTAATTGCGGTTNNNAGATCGGAAG
28976 TATGAGTGTGGAGTCGTTGCATGTTGCATGTGAGAGAAGGGATNNNAGATCGGAA
28977 TATGAGTGTGGAGTCGTTGCTGTCTTCTGGCCTCTGTAGTTTCNNNAGATCGGAAGA
28978 TATGAGTGTGGAGTCGTTGCAACTGTCAGGTGAGAAATGGGAANNNAGATCGGAA
28980 TATGAGTGTGGAGTCGTTGCTGACTCATACCGTTTTGTTCCCTNNNAGATCGGAAGA
28982 TATGAGTGTGGAGTCGTTGCAATGAATGGGAGTGCAGATGTCTNNNAGATCGGAA
28983 TATGAGTGTGGAGTCGTTGCTGGGAGATTGTGTCTTCATGTCCNNNAGATCGGAAG
28984 TATGAGTGTGGAGTCGTTGCGGTTGTGTCCGTATATGTGCATGNNNAGATCGGAAG
28987 TATGAGTGTGGAGTCGTTGCGCACAAATACATTCCCCACATGTNNNAGATCGGAAGA
28988 TATGAGTGTGGAGTCGTTGCACATGTGGCTGTGTCTTTCTTTGNNNAGATCGGAAGA
28989 TATGAGTGTGGAGTCGTTGCACTGGTTCTGATCCTTTGGTTGANNNAGATCGGAAG
28990 TATGAGTGTGGAGTCGTTGCGCCAGACACTTTGGTCTTTGATCNNNAGATCGGAAGA
28991 TATGAGTGTGGAGTCGTTGCAGTGGTTTGGTTTGTCAAGAAGCNNNAGATCGGAAG
28993 TATGAGTGTGGAGTCGTTGCCCCCAAGGATTGTAGAAATCCCANNNAGATCGGAAG
28996 TATGAGTGTGGAGTCGTTGCACAAGAGTAGCTTTTGGTGGTCTNNNAGATCGGAAG

28997 TATGAGTGTGGAGTCGTTGCTCAGAACAGGAGGTGATAGTCCTNNNAGATCGGAA
28998 TATGAGTGTGGAGTCGTTGCTGTTTTTTGGTTTGCGGTTACCANNNAGATCGGAAG
28999 TATGAGTGTGGAGTCGTTGCGTATGGAGCCTTCACAAAATGGCNNNAGATCGGAA
29001 TATGAGTGTGGAGTCGTTGCAGAGAGAGAGAGAGAGAGAGAGTNNNAGATCGG
29003 TATGAGTGTGGAGTCGTTGCTGTGTGTATCCGTAATCTCAGAANNNAGATCGGAAG
29004 TATGAGTGTGGAGTCGTTGCCACACACATACACACACACACTCNNNAGATCGGAAG
29006 TATGAGTGTGGAGTCGTTGCGCAGAACCTAGGGTGAGTGAAATNNNAGATCGGAA
29007 TATGAGTGTGGAGTCGTTGCACAATTCCACTTCACTTCTGGGTNNNAGATCGGAAG
29008 TATGAGTGTGGAGTCGTTGCAGAGATCTGAAGGTGGAGAAGGANNNAGATCGGA
29009 TATGAGTGTGGAGTCGTTGCATGGAAGCTGAGAAGTTCCATGANNNAGATCGGAA
29010 TATGAGTGTGGAGTCGTTGCAACTTTGTAATGTTGGCAGCTGGNNNAGATCGGAA
29011 TATGAGTGTGGAGTCGTTGCAGTCAAGATTCCCTAAGCAGCAANNNAGATCGGAAG
29012 TATGAGTGTGGAGTCGTTGCTGACAACTCCAAAATCTAGGGGTNNNAGATCGGAA
29013 TATGAGTGTGGAGTCGTTGCAAGCAAAGATGAAGAAACACGGGNNNAGATCGGA
29014 TATGAGTGTGGAGTCGTTGCGAGGAGGACTAGAGAGAAAGGGANNNAGATCGG
29015 TATGAGTGTGGAGTCGTTGCTGCATATGTCAGTCAAGTCACCANNNAGATCGGAAG
29016 TATGAGTGTGGAGTCGTTGCTCAAATCCCACAAATCAGCTTCGNNNAGATCGGAAG
29018 TATGAGTGTGGAGTCGTTGCAAATGGTATATAGCCCTCGCAGGNNNAGATCGGAA
29019 TATGAGTGTGGAGTCGTTGCTTGTGCATAGGAACAAGTCCAGANNNAGATCGGAA
29020 TATGAGTGTGGAGTCGTTGCCTGCAAGACAAATGAGGTCAAGGNNNAGATCGGAA
29022 TATGAGTGTGGAGTCGTTGCTCCCCTCAATATATGCTTCAGGTNNNAGATCGGAAG
29023 TATGAGTGTGGAGTCGTTGCAGGGTGACAAAATCAGTCAGGTTNNNAGATCGGAA
29024 TATGAGTGTGGAGTCGTTGCTTGGACATTTATCTTCCCCCAGGNNNAGATCGGAAG
29025 TATGAGTGTGGAGTCGTTGCACACACAAACTTACATGGACAGGNNNAGATCGGAA
29026 TATGAGTGTGGAGTCGTTGCATGAGACAGAGAGACAGACAGGTNNNAGATCGGA
29027 TATGAGTGTGGAGTCGTTGCCCCGGCCCACTCTGTATATTTTANNNAGATCGGAAG
29028 TATGAGTGTGGAGTCGTTGCAAGTCTGGATCACATAGGGCTTGNNNAGATCGGAA
29029 TATGAGTGTGGAGTCGTTGCAGACAGCTGGGAAGACCTATAGTNNNAGATCGGAA
29030 TATGAGTGTGGAGTCGTTGCAGCCTGTGATAAACTGAGTCCTGNNNAGATCGGAA
29031 TATGAGTGTGGAGTCGTTGCATAAGGTGGATTCAGCCAACTGTNNNAGATCGGAA
29032 TATGAGTGTGGAGTCGTTGCTCATGCAGACTCACACATACACANNNAGATCGGAAG
29033 TATGAGTGTGGAGTCGTTGCTGACTGTATTTGGAGATAGGGCCNNNAGATCGGAA
29035 TATGAGTGTGGAGTCGTTGCATACGTACTGTGTGTGTGTGTGTNNNAGATCGGAA
29036 TATGAGTGTGGAGTCGTTGCCCAATAGAAAAGGCATCATGCGTNNNAGATCGGAA
29039 TATGAGTGTGGAGTCGTTGCCCCACTTACAACTTGATGATCCGNNNAGATCGGAAG
29040 TATGAGTGTGGAGTCGTTGCTTTAGACAGAAAGCGTGACCTCANNNAGATCGGAA
29041 TATGAGTGTGGAGTCGTTGCTGTGTCTGGTGTGTGTGTATTCANNNAGATCGGAA
29043 TATGAGTGTGGAGTCGTTGCGAGGTGGGCCTGATAGTACTCTANNNAGATCGGAA
29044 TATGAGTGTGGAGTCGTTGCGCAACAGAAAATCTGCATGTAGTNNNAGATCGGAA
29045 TATGAGTGTGGAGTCGTTGCAGTTTCTCACTTGAAGCCTCACTNNNAGATCGGAAG
29046 TATGAGTGTGGAGTCGTTGCGCATGCACACGTGTATACTCAGANNNAGATCGGAAG
29047 TATGAGTGTGGAGTCGTTGCAACTGTTGGCAGAGGTGGTATAGNNNAGATCGGAA
29048 TATGAGTGTGGAGTCGTTGCGCCTCAAACACCTTGGAAAATCANNNAGATCGGAAG

29049 TATGAGTGTGGAGTCGTTGCACCTCTTCACAACACACTTGAGTNNNAGATCGGAAGA
29050 TATGAGTGTGGAGTCGTTGCCCCCTTCCATTTATCTGGGAGTCNNNAGATCGGAAGA
29051 TATGAGTGTGGAGTCGTTGCACTGCTATAGCCTTCATGCTCTGNNNAGATCGGAAGA
29052 TATGAGTGTGGAGTCGTTGCTTTTTATGAGGGCACTGATCCCANNNAGATCGGAAG
29053 TATGAGTGTGGAGTCGTTGCGGAATAGGGAGATTTGGCAGAGTNNNAGATCGGAA
29054 TATGAGTGTGGAGTCGTTGCAGATCAAATTGGAGTGACTGGGGNNNAGATCGGAA
29056 TATGAGTGTGGAGTCGTTGCAAGAGAGACAGACAGAGTAGGCANNNAGATCGGAA
29057 TATGAGTGTGGAGTCGTTGCTTGTCTAGGTGGCAAATATCCCGNNNAGATCGGAAG
29058 TATGAGTGTGGAGTCGTTGCTCTTTTTCAATTTGGGTGCTTCTNNNAGATCGGAAGA
29059 TATGAGTGTGGAGTCGTTGCCACACACACACACACATACACATNNNAGATCGGAAGA
29060 TATGAGTGTGGAGTCGTTGCATCAAGCAGATTTTTAGCAGCCCNNNAGATCGGAAG
29062 TATGAGTGTGGAGTCGTTGCTGAATCCCCAGCTCATAAAACCANNNAGATCGGAAGA
29063 TATGAGTGTGGAGTCGTTGCATGCACACACACAATAAAGGTGGNNNAGATCGGAA
29065 TATGAGTGTGGAGTCGTTGCGTAGCAGAAACCTGTCCTTCTGANNNAGATCGGAAG
29067 TATGAGTGTGGAGTCGTTGCTGTGTGTGTGATGGGAAAGAGAANNNAGATCGGAA
29070 TATGAGTGTGGAGTCGTTGCACACACCTAGTGGGGAGATCTAANNNAGATCGGAA
29071 TATGAGTGTGGAGTCGTTGCTCCCTCTCACTTGTTTCCCAAATNNNAGATCGGAAGA
29072 TATGAGTGTGGAGTCGTTGCTCATTGGGTTTCTTCCTCCTCAGNNNAGATCGGAAGA
29073 TATGAGTGTGGAGTCGTTGCGCCCTCATGTTGTCTTTCTGAACNNNAGATCGGAAGA
29074 TATGAGTGTGGAGTCGTTGCGTTAGGCCTCCTCCTTCTTTCAANNNAGATCGGAAGA
29075 TATGAGTGTGGAGTCGTTGCGGATGACTGGCAATTCCCTTTCTNNNAGATCGGAAGA
29076 TATGAGTGTGGAGTCGTTGCCATGTGTGTGTGTGTGTGTGTATNNNAGATCGGAA
29078 TATGAGTGTGGAGTCGTTGCGTTCACGGGTTCAAGCTATTCTCNNNAGATCGGAAGA
29081 TATGAGTGTGGAGTCGTTGCGAGAGAGAGAGAGAGAGACCCAANNNAGATCGGAA
29082 TATGAGTGTGGAGTCGTTGCAGATCAGTGAAAACAGGAGCTGTNNNAGATCGGAA
29083 TATGAGTGTGGAGTCGTTGCGGGAAAGGAAGAGAAGAAGAGCANNNAGATCGGA
29084 TATGAGTGTGGAGTCGTTGCACTTGTGTGTGTGTGCTGAAAAANNNAGATCGGAA
29085 TATGAGTGTGGAGTCGTTGCTTCTAGGAATGATGCTGTCCGTCNNNAGATCGGAAG
29086 TATGAGTGTGGAGTCGTTGCTTGCCTTGTTCCAGTGAGTCTAANNNAGATCGGAAGA
29087 TATGAGTGTGGAGTCGTTGCGGTATCTAGAGACCAACTGGCAGNNNAGATCGGAA
29089 TATGAGTGTGGAGTCGTTGCAGGGTGTCCACAATAAGAAGAGCNNNAGATCGGAA
29090 TATGAGTGTGGAGTCGTTGCTTGCCTCTTTTCCTCTCTGTCTCNNNAGATCGGAAGA
29091 TATGAGTGTGGAGTCGTTGCTAAGGCAAATGACGAGATTGTGCNNNAGATCGGAA
29092 TATGAGTGTGGAGTCGTTGCTGCTCTCTGTGTGTCATTCTCTCNNNAGATCGGAAGA
29093 TATGAGTGTGGAGTCGTTGCCCTCTCTCCCTCTGTCTCTGTTANNNAGATCGGAAGA
29094 TATGAGTGTGGAGTCGTTGCTACTGCCTCCCAGTGTTGTTTTTNNNAGATCGGAAGA
29095 TATGAGTGTGGAGTCGTTGCCCCACAGACTCCACAAATAGGAANNNAGATCGGAAG
29096 TATGAGTGTGGAGTCGTTGCCATGCCACGTTCAATCAACAGTANNNAGATCGGAAGA
29097 TATGAGTGTGGAGTCGTTGCACAATGGGGCTGTTCCTCTAAATNNNAGATCGGAAG
29098 TATGAGTGTGGAGTCGTTGCCGCCTTTAGTTTTACAGCCTCTGNNNAGATCGGAAGA
29099 TATGAGTGTGGAGTCGTTGCCTAATTTTGCTATCAGCCCCTGCNNNAGATCGGAAGA
29101 TATGAGTGTGGAGTCGTTGCCAAAAGGAAGGCCTGTATCCTCTNNNAGATCGGAAG
29102 TATGAGTGTGGAGTCGTTGCCACACACACACACACATATATNNNAGATCGGAAGA

29103 TATGAGTGTGGAGTCGTTGCGAGGGAGAGAGAGAGAAACATGCNNNAGATCGGA

29104 TATGAGTGTGGAGTCGTTGCATGAGAGCCTCCAAGTTCTCAAGNNNAGATCGGAAG

29105 TATGAGTGTGGAGTCGTTGCTTGAGACCACAGCCATCTATCACNNNAGATCGGAAGA

29106 TATGAGTGTGGAGTCGTTGCATACACAGACACACACACACACTNNNAGATCGGAAGA

29107 TATGAGTGTGGAGTCGTTGCTTTGAGCCATAGGGAAGTAGAGCNNNAGATCGGAA

29109 TATGAGTGTGGAGTCGTTGCTGAGACTGCAAAAACCATAGTGCNNNAGATCGGAAG

29111 TATGAGTGTGGAGTCGTTGCTCTGAGGAGCATTGAAACCTACCNNNAGATCGGAAG

29112 TATGAGTGTGGAGTCGTTGCTCTGAGGAGCATTGAAACCTACCNNNAGATCGGAAG

29113 TATGAGTGTGGAGTCGTTGCGTGTCTGCTCCCAGTTCAATTTCNNNAGATCGGAAGA

29114 TATGAGTGTGGAGTCGTTGCACACACACACGCACATCCTATTANNNAGATCGGAAGA

29115 TATGAGTGTGGAGTCGTTGCTATTCAGTTGGGAGAATGGAGCANNNAGATCGGAA

29116 TATGAGTGTGGAGTCGTTGCCTAGATGGGCCTCCAAAAGAACTNNNAGATCGGAAG

29117 TATGAGTGTGGAGTCGTTGCTGTGAACATTGTCCTTCTTGTGCNNNAGATCGGAAGA

29118 TATGAGTGTGGAGTCGTTGCCAAGGACTGAGGAATACTGCAGTNNNAGATCGGAAG

29119 TATGAGTGTGGAGTCGTTGCCAAGGACTGAGGAATACTGCAGTNNNAGATCGGAAG

29120 TATGAGTGTGGAGTCGTTGCACCAAAGCTAAGAAGACAGCAGANNNAGATCGGAAG

29121 TATGAGTGTGGAGTCGTTGCGATTGAAAAGGATGCTGGCAAGANNNAGATCGGAA

29122 TATGAGTGTGGAGTCGTTGCATTGGCTGTGACCAAAACAAACANNNAGATCGGAAG

29123 TATGAGTGTGGAGTCGTTGCATAGGGATCTCCGTACCATGCTANNNAGATCGGAAG

29125 TATGAGTGTGGAGTCGTTGCTAGATGCATCCACATTCTGCTGTNNNAGATCGGAAGA

29126 TATGAGTGTGGAGTCGTTGCCATCCTCCTAGTTTCTCTGGCTGNNNAGATCGGAAGA

29128 TATGAGTGTGGAGTCGTTGCTGGCTTCAAACTCTTCTACTGCANNNAGATCGGAAGA

29129 TATGAGTGTGGAGTCGTTGCACACACACAGAGAGAGAGAGAGANNNAGATCGGAA

29130 TATGAGTGTGGAGTCGTTGCACATCTTTGAGGGTACACGCATANNNAGATCGGAAG

29131 TATGAGTGTGGAGTCGTTGCGAAACTTCGCAGTAACACTAGGCNNNAGATCGGAAG

29132 TATGAGTGTGGAGTCGTTGCTTTCCATCACTCTCTTCCTCTGCNNNAGATCGGAAGAG

29133 TATGAGTGTGGAGTCGTTGCACTGTTCTAGTTTAGCCCTGTGGNNNAGATCGGAAG

29134 TATGAGTGTGGAGTCGTTGCTTAGCTCACTTCTTTCCCTCACCNNNAGATCGGAAGAG

29135 TATGAGTGTGGAGTCGTTGCAATAGTGACTGCATGCTGTAGCTNNNAGATCGGAAG

29136 TATGAGTGTGGAGTCGTTGCGCTAACGCCAGGAAAGAAAGAAANNNAGATCGGAA

29138 TATGAGTGTGGAGTCGTTGCGTGGCCTTCAAATGCATTAGGAGNNNAGATCGGAAG

29139 TATGAGTGTGGAGTCGTTGCATAATGAAGTGGGCTAGGTGAGCNNNAGATCGGAA

29140 TATGAGTGTGGAGTCGTTGCTGGATCCAGTGATAAGCTACCGANNNAGATCGGAAG

29141 TATGAGTGTGGAGTCGTTGCGGCTCTAGTAGGCAAAGCATCAANNNAGATCGGAAG

29142 TATGAGTGTGGAGTCGTTGCCAGCTCTGGATCCCTAGTCATTCNNNAGATCGGAAGA

29144 TATGAGTGTGGAGTCGTTGCCTTCACTTGTGTCATGGCAGTTTNNNAGATCGGAAGA

29145 TATGAGTGTGGAGTCGTTGCTGTTGGTTGAACCTAACAGTCGGNNNAGATCGGAAG

29147 TATGAGTGTGGAGTCGTTGCAGAACATGGCCTGAAGCTTATGANNNAGATCGGAAG

29148 TATGAGTGTGGAGTCGTTGCGGGCAAGCTTCTTCTGGATAGATNNNAGATCGGAAG

29150 TATGAGTGTGGAGTCGTTGCTGCTGATAACAAGGCAGAAGATGNNNAGATCGGAA

29152 TATGAGTGTGGAGTCGTTGCTTTACCCTGTCACACTTTCTCCCNNNAGATCGGAAGAG

29153 TATGAGTGTGGAGTCGTTGCTGTCTCTGTATGCTTCTTTTCCTNNNAGATCGGAAGAG

29154 TATGAGTGTGGAGTCGTTGCATCAATTCACTGAGACCACAGCANNNAGATCGGAAG

29155 TATGAGTGTGGAGTCGTTGCCCTGTAAGTTCTCACCTAGTGCANNNAGATCGGAAG
29156 TATGAGTGTGGAGTCGTTGCCACTGCACACAAGGCATATGAAANNNAGATCGGAAG
29157 TATGAGTGTGGAGTCGTTGCCACACAGAGTTCCTAGCACATCTNNNAGATCGGAAG
29158 TATGAGTGTGGAGTCGTTGCTTCCCTTTCCTCCTTTCCAGATGNNNAGATCGGAAGA
29159 TATGAGTGTGGAGTCGTTGCGAAGAGTCTCGAAGTTCCTCCTGNNNAGATCGGAAG
29160 TATGAGTGTGGAGTCGTTGCGAAGAGTCTCGAAGTTCCTCCTGNNNAGATCGGAAG
29161 TATGAGTGTGGAGTCGTTGCCTCCTCCAGTCAAGCTTTGAGATNNNAGATCGGAAG
29162 TATGAGTGTGGAGTCGTTGCAATCTTTGCTAGCACAATGGACGNNNAGATCGGAAG
29163 TATGAGTGTGGAGTCGTTGCACCCTTTTCGGACAAATTATCCTNNNAGATCGGAAGA
29164 TATGAGTGTGGAGTCGTTGCTATAGAACACCCAGCACTGATCCNNNAGATCGGAAG
29165 TATGAGTGTGGAGTCGTTGCCATACACACACACACACACACTCNNNAGATCGGAAGA
29167 TATGAGTGTGGAGTCGTTGCGAAGGGGAAGGGAAGAAATCCTTNNNAGATCGGA
29168 TATGAGTGTGGAGTCGTTGCTGCCCTCACCACATAATTCTCTTNNNAGATCGGAAGA
29169 TATGAGTGTGGAGTCGTTGCTCTCTTGCTTTCTCTCTCTCTGTNNNAGATCGGAAGA
29171 TATGAGTGTGGAGTCGTTGCAAAATCATGCTGCTGTTACTGCCNNNAGATCGGAAG
29172 TATGAGTGTGGAGTCGTTGCCAGGCCTATGTAATGAAGCTTGCNNNAGATCGGAAG
29173 TATGAGTGTGGAGTCGTTGCTGTGTGTGTGTGTTTTGTCCAATNNNAGATCGGAAG
29174 TATGAGTGTGGAGTCGTTGCAGCACTGAGATTTACTGCAGTGANNNAGATCGGAA
29175 TATGAGTGTGGAGTCGTTGCTTGAGCTCTTCATCACCCATGAANNNAGATCGGAAG
29177 TATGAGTGTGGAGTCGTTGCGGACACATTCTATGTCAAAGCCCNNNAGATCGGAAG
29178 TATGAGTGTGGAGTCGTTGCAAAGATAGATCCTGTTGCCTGCANNNAGATCGGAAG
29179 TATGAGTGTGGAGTCGTTGCTCTATTTGTTTGGATGCCCTGGANNNAGATCGGAAG
29180 TATGAGTGTGGAGTCGTTGCCTCTCAGACCCTCCCTCTAGAAANNNAGATCGGAAGA
29181 TATGAGTGTGGAGTCGTTGCTAGAGATCTTCAAGGCCAAGAGCNNNAGATCGGAA
29183 TATGAGTGTGGAGTCGTTGCTCTCTCTCTCTCTCTCATGTGCANNNAGATCGGAAGA
29184 TATGAGTGTGGAGTCGTTGCTACATGGGCTGTTATCTGTCTGGNNNAGATCGGAAG
29185 TATGAGTGTGGAGTCGTTGCAACATGTGGTAGGAGCAAGAAGTNNNAGATCGGAA
29186 TATGAGTGTGGAGTCGTTGCGTCCCATGATGACAGAGTCTTGTNNNAGATCGGAAG
29187 TATGAGTGTGGAGTCGTTGCTTTTGTGCTTTTTGTCCGTGACTNNNAGATCGGAAGA
29189 TATGAGTGTGGAGTCGTTGCATTCCCTGAAAGCTTCTCCTGACNNNAGATCGGAAGA
29190 TATGAGTGTGGAGTCGTTGCCCTAATCCCATGAGCCTGACTACNNNAGATCGGAAG
29191 TATGAGTGTGGAGTCGTTGCGCGAGAGAAAAAAATGTGCCTGANNNAGATCGGAA
29192 TATGAGTGTGGAGTCGTTGCAAACCAACCAACCAAGATGGAACNNNAGATCGGAAG
29193 TATGAGTGTGGAGTCGTTGCAAATGTGGATCATGCGAGAGACTNNNAGATCGGAA
29194 TATGAGTGTGGAGTCGTTGCCCATCTATGCGTTGGTAGGGTTANNNAGATCGGAAG
29196 TATGAGTGTGGAGTCGTTGCGTACTCACAAACACACACATGCANNNAGATCGGAAG
29197 TATGAGTGTGGAGTCGTTGCGATCTCTCGGCTCATTGTAACCTNNNAGATCGGAAG
29199 TATGAGTGTGGAGTCGTTGCTATTTGCACCCCTTCTTGTCTGTNNNAGATCGGAAGA
29200 TATGAGTGTGGAGTCGTTGCTCTCTGATTCTACATGCTCCACTNNNAGATCGGAAGA
29201 TATGAGTGTGGAGTCGTTGCAACCTGTAATCTGAACAGCGACTNNNAGATCGGAAG
29203 TATGAGTGTGGAGTCGTTGCTCCACACACCACATTCAAGAACTNNNAGATCGGAAGA
29204 TATGAGTGTGGAGTCGTTGCATTTCTCCAAGGAGCCCTCATTTNNNAGATCGGAAGA
29205 TATGAGTGTGGAGTCGTTGCTGAGTACACAATGCTCCTGTTCANNNAGATCGGAAG

29206 TATGAGTGTGGAGTCGTTGCTAGATCTGGCATGTGAAACACCANNNAGATCGGAAG
29207 TATGAGTGTGGAGTCGTTGCGAAAGGGAGATGGGGTAGGTGAGNNNAGATCGGA
29209 TATGAGTGTGGAGTCGTTGCTCACACATGTAGCCTTGCATAACNNNAGATCGGAAGA
29210 TATGAGTGTGGAGTCGTTGCTGAGATTTTTGTGGAACCTCCANNNAGATCGGAAG
29211 TATGAGTGTGGAGTCGTTGCCTCCACCCATCCCATCTGTTATCNNNAGATCGGAAGA
29212 TATGAGTGTGGAGTCGTTGCTGGTTGGTCTCACACACCTTATTNNNAGATCGGAAGA
29214 TATGAGTGTGGAGTCGTTGCCGTGTGTGTTTGTGTGTGTATCTNNNAGATCGGAAG
29215 TATGAGTGTGGAGTCGTTGCTAAAGTCTATCTCCCCTCTCCCCNNNAGATCGGAAGA
29216 TATGAGTGTGGAGTCGTTGCTGCGTGCATATCTCTGTATCCTCNNNAGATCGGAAGA
29217 TATGAGTGTGGAGTCGTTGCCCTCCCGAATAGCTAGGACTACANNNAGATCGGAAG
29218 TATGAGTGTGGAGTCGTTGCGTGCATAGAGTTATGTGAGGAGCNNNAGATCGGAA
29219 TATGAGTGTGGAGTCGTTGCGGGGTGACAATACAGAGCTCTAGNNNAGATCGGAA
29220 TATGAGTGTGGAGTCGTTGCTGCATGTATGTATGTATGCCACANNNAGATCGGAAG
29221 TATGAGTGTGGAGTCGTTGCCAAGTTTGCAGTATACAAGGTCANNNAGATCGGAAG
29222 TATGAGTGTGGAGTCGTTGCAGAGAACAAGAGTCTCTGCTTGGNNNAGATCGGAAG
29224 TATGAGTGTGGAGTCGTTGCAGGGTGCAAGGGGTCTCTATATANNNAGATCGGAAG
29225 TATGAGTGTGGAGTCGTTGCGAAGCTTGGAGTTTGCATGTGAANNNAGATCGGAAG
29226 TATGAGTGTGGAGTCGTTGCATCAGCTTCTCCACCATCTTGTTNNNAGATCGGAAGA
29228 TATGAGTGTGGAGTCGTTGCACACACACAGAGAGAGAGAGAGANNNAGATCGGAA
29229 TATGAGTGTGGAGTCGTTGCGCAGCTTCTACTAACCAAGAGGTNNNAGATCGGAAG
29230 TATGAGTGTGGAGTCGTTGCACACACACACAAGAGCACATCTANNNAGATCGGAAG
29231 TATGAGTGTGGAGTCGTTGCTCAGAGATAATGCCCCCAAGATGNNNAGATCGGAAG
29233 TATGAGTGTGGAGTCGTTGCGAATGGAACAGAAATCCCCAAGCNNNAGATCGGAAG
29235 TATGAGTGTGGAGTCGTTGCAACCTGGATGGATTTGGAGACAANNNAGATCGGAAG
29236 TATGAGTGTGGAGTCGTTGCCACCATGGAATACTACTCAGCCANNNAGATCGGAAGA
29237 TATGAGTGTGGAGTCGTTGCGCAATTCTGTGTGTTGTGAGTGANNNAGATCGGAAG
29238 TATGAGTGTGGAGTCGTTGCAGCCTCTGGTCACAATATGTTCANNNAGATCGGAAGA
29240 TATGAGTGTGGAGTCGTTGCGGGATGAGATACAGGCAATCGATNNNAGATCGGAA
29241 TATGAGTGTGGAGTCGTTGCGAGAAGGCAGGGATGAGATACAGNNNAGATCGGAA
29243 TATGAGTGTGGAGTCGTTGCAAAATCTGGGGGAGGGAGAGATANNNAGATCGGAA
29244 TATGAGTGTGGAGTCGTTGCGGGGGAGAGAGATAGGGTGATAANNNAGATCGGA
29245 TATGAGTGTGGAGTCGTTGCCACCTACTACACGGCAGATACAGNNNAGATCGGAAG
29246 TATGAGTGTGGAGTCGTTGCCTATGAGCTACAGTGAGAGCTGGNNNAGATCGGAAG
29247 TATGAGTGTGGAGTCGTTGCTCATTGGAGAAGGCCAGATTGAANNNAGATCGGAAG
29248 TATGAGTGTGGAGTCGTTGCAAGATGATATGCTGTCACTGGCTNNNAGATCGGAAG
29249 TATGAGTGTGGAGTCGTTGCAAGGCTGTGCAATAGTCAAAGATNNNAGATCGGAAG
29251 TATGAGTGTGGAGTCGTTGCTACTCTTAGTTTGGGATGGGTCCNNNAGATCGGAAG
29253 TATGAGTGTGGAGTCGTTGCTTGGCCTTCTACCGTTGTCATAANNNAGATCGGAAGA
29254 TATGAGTGTGGAGTCGTTGCTGGCATACAGCAGAACGATCATANNNAGATCGGAAG
29255 TATGAGTGTGGAGTCGTTGCTAGAGTCACGTCTCCATCTCCATNNNAGATCGGAAGA
29256 TATGAGTGTGGAGTCGTTGCGACAAGTAGAGTGGTCAGAGGTGNNNAGATCGGAA
29257 TATGAGTGTGGAGTCGTTGCACACAAACACACACACACACATTNNNAGATCGGAAGA
29258 TATGAGTGTGGAGTCGTTGCAGACTACTGTGCTCCAAGACATGNNNAGATCGGAAG

29259 TATGAGTGTGGAGTCGTTGCTGCTGTCCAACTGATTTACTCCTNNNAGATCGGAAGA
29260 TATGAGTGTGGAGTCGTTGCCTCACCTTGGCTCTTAGACACAANNNAGATCGGAAGA
29261 TATGAGTGTGGAGTCGTTGCCCTGTGGGGATGATATCTCCATGNNNAGATCGGAAG
29262 TATGAGTGTGGAGTCGTTGCTGCAGTCTCTGAAAGTGCATTTGNNNAGATCGGAAG
29263 TATGAGTGTGGAGTCGTTGCCCTGGCACCACTTTGATACATTGNNNAGATCGGAAGA
29264 TATGAGTGTGGAGTCGTTGCACCAATGGAATTCAGTCAACCATNNNAGATCGGAAG
29265 TATGAGTGTGGAGTCGTTGCTGCCTTCTTTGCCTTCCTACATTNNNAGATCGGAAGA
29266 TATGAGTGTGGAGTCGTTGCTGTACAGATTTCCCAGCTGACAANNNAGATCGGAAG
29267 TATGAGTGTGGAGTCGTTGCTTCTCCCAACAGAAGCAGAAAGGNNNAGATCGGAA
29268 TATGAGTGTGGAGTCGTTGCCATCCCAAAAGCCACATCTTCAGNNNAGATCGGAAGA
29269 TATGAGTGTGGAGTCGTTGCAGAGACAAGAGAGAGAACGGAGANNNAGATCGGA
29270 TATGAGTGTGGAGTCGTTGCATGACCTGCTAATGTGTGATGGTNNNAGATCGGAA
29271 TATGAGTGTGGAGTCGTTGCCACATTCAGTCCATAACAGCCACNNNAGATCGGAAGA
29272 TATGAGTGTGGAGTCGTTGCCAGAAATTTTCCACTCGGGCTTTNNNAGATCGGAAGA
29273 TATGAGTGTGGAGTCGTTGCTCTCAGTGCTCGCTAGTTCTTTTNNNAGATCGGAAGA
29274 TATGAGTGTGGAGTCGTTGCATCTCAATCTGTGTGCCTGTCATNNNAGATCGGAAGA
29275 TATGAGTGTGGAGTCGTTGCAATGTCATTTTGCCATGATGCCANNNAGATCGGAAG
29276 TATGAGTGTGGAGTCGTTGCACAGAGTTGGGGACATAACTTGGNNNAGATCGGAA
29277 TATGAGTGTGGAGTCGTTGCCGTGAAATGTGGCCCATAGAATCNNNAGATCGGAA
29278 TATGAGTGTGGAGTCGTTGCTTGTCTGAGTGTCCCAGAATCTGNNNAGATCGGAAG
29280 TATGAGTGTGGAGTCGTTGCGTACTCACTCTTGCTGTAGTCCCNNNAGATCGGAAGA
29281 TATGAGTGTGGAGTCGTTGCAAACTCCCCCTGTCTAATCCCTANNNAGATCGGAAGA
29282 TATGAGTGTGGAGTCGTTGCTATCTTCGTGGGACATGACTTCCNNNAGATCGGAAG
29283 TATGAGTGTGGAGTCGTTGCGCTAAAAAGCTTCTGCACAGTGANNNAGATCGGAAG
29284 TATGAGTGTGGAGTCGTTGCAGTTTTCCTGTGTTTTTGCCAATNNNAGATCGGAAGA
29285 TATGAGTGTGGAGTCGTTGCAATCCATTCATGAAGGTGGAGCTNNNAGATCGGAA
29286 TATGAGTGTGGAGTCGTTGCCATATGTGCATGTGTGTGTCTCTNNNAGATCGGAAG
29287 TATGAGTGTGGAGTCGTTGCGGAGGCTGAACACTTGGAAATTGNNNAGATCGGAA
29288 TATGAGTGTGGAGTCGTTGCACATGCGTCTTTGCTTCTCTCTANNNAGATCGGAAGA
29289 TATGAGTGTGGAGTCGTTGCCACAGAGACACACACACATGAACNNNAGATCGGAAG
29291 TATGAGTGTGGAGTCGTTGCTGTGTGTGACCAATCCCCAAATANNNAGATCGGAAG
29293 TATGAGTGTGGAGTCGTTGCTTGGGTCCCATTAGGCAAAATTGNNNAGATCGGAAG
29294 TATGAGTGTGGAGTCGTTGCACGAGTGTGTAAGTATGTGTGCTNNNAGATCGGAA
29296 TATGAGTGTGGAGTCGTTGCTCCCTGGCAATAGTTCTTTGACANNNAGATCGGAAGA
29297 TATGAGTGTGGAGTCGTTGCTCTGTGCCCACTAAGCTACTTTCNNNAGATCGGAAGA
29298 TATGAGTGTGGAGTCGTTGCACATTTCTTCTGTATCCCCCACCNNNAGATCGGAAGA
29299 TATGAGTGTGGAGTCGTTGCCCATTCACAGTAACTGATCTGCTNNNAGATCGGAAGA
29301 TATGAGTGTGGAGTCGTTGCGTCTGCTGTTCCTCAATTCTCCTNNNAGATCGGAAGA
29304 TATGAGTGTGGAGTCGTTGCAAATGGCTTGGGGTGTTTATGTGNNNAGATCGGAA
29305 TATGAGTGTGGAGTCGTTGCGCAACCAAGACCTCATAGGAAGANNNAGATCGGAA
29306 TATGAGTGTGGAGTCGTTGCGTACGTAAAGTAGCCCAGTCACANNNAGATCGGAAG
29307 TATGAGTGTGGAGTCGTTGCCCCTTATCCAGTCCCTTGTTGATNNNAGATCGGAAGA
29308 TATGAGTGTGGAGTCGTTGCATCCACAGAACACACAGAGCTTANNNAGATCGGAAG

29309 TATGAGTGTGGAGTCGTTGCTGGATTCCAGGGCCTATATGTTGNNNAGATCGGAAG
29310 TATGAGTGTGGAGTCGTTGCGTCTCAAGCTCCCTGGAAATACANNNAGATCGGAAG
29311 TATGAGTGTGGAGTCGTTGCTTGATTGAGATCTCCAGCCAACANNNAGATCGGAAG
29312 TATGAGTGTGGAGTCGTTGCTCTGCAGCTGCTTCAAAATTAGGNNNAGATCGGAAG
29313 TATGAGTGTGGAGTCGTTGCTGGGTGAGAATCTGCACAGTTTANNNAGATCGGAA
29314 TATGAGTGTGGAGTCGTTGCTCCCCACTTAGTCCACTCATACTNNNAGATCGGAAGA
29315 TATGAGTGTGGAGTCGTTGCATAATGGCATTTGCAGTGACCTGNNNAGATCGGAAG
29316 TATGAGTGTGGAGTCGTTGCAGTCAATTTGAGGAGAGTTGCCANNNAGATCGGAA
29317 TATGAGTGTGGAGTCGTTGCAGAGAGAGAGAGGGAGAGAGAGANNNAGATCGG
29318 TATGAGTGTGGAGTCGTTGCACTCTCCCTTGCAGATTACATGANNNAGATCGGAAG
29319 TATGAGTGTGGAGTCGTTGCCTGCAGTGATATGATCTGGCTCTNNNAGATCGGAAG
29320 TATGAGTGTGGAGTCGTTGCAAGGGGTTGAGGGTCTCCTATAANNNAGATCGGAA
29321 TATGAGTGTGGAGTCGTTGCTGTCAGAAAGTGTCAAATGCTGTNNNAGATCGGAAG
29322 TATGAGTGTGGAGTCGTTGCGTACAGTCCAGGGAAACAGACATNNNAGATCGGAA
29323 TATGAGTGTGGAGTCGTTGCATTGGCTAGACTGTGAGCTGAAANNNAGATCGGAA
29324 TATGAGTGTGGAGTCGTTGCGTGGTTACATGTGGTCACAAGTGNNNAGATCGGAA
29325 TATGAGTGTGGAGTCGTTGCAGCTGTATGTGGTTACATGTGGTNNNAGATCGGAA
29326 TATGAGTGTGGAGTCGTTGCAGACAGACACAGAGACAGAGAGANNNAGATCGGA
29327 TATGAGTGTGGAGTCGTTGCTAAAATGGATCCAAGGGCCAAGANNNAGATCGGAA
29328 TATGAGTGTGGAGTCGTTGCTCGAGGGAGAAAAAAGGAGGAAGNNNAGATCGGA
29329 TATGAGTGTGGAGTCGTTGCATTGGGGTTTTGGACTTAGGGTNNNAGATCGGAA
29330 TATGAGTGTGGAGTCGTTGCAAGGGAAGGAGGACCATAGAGAANNNAGATCGGA
29331 TATGAGTGTGGAGTCGTTGCCCTCAGCGTGGAAAATATGGAACNNNAGATCGGAA
29332 TATGAGTGTGGAGTCGTTGCCACACATGTTCACACACACACATNNNAGATCGGAAG
29333 TATGAGTGTGGAGTCGTTGCGGATGAGAGCATTCACAGGAGATNNNAGATCGGAA
29334 TATGAGTGTGGAGTCGTTGCAGTATCTGGGTCACCAAAAGCTTNNNAGATCGGAAG
29335 TATGAGTGTGGAGTCGTTGCAGGCATTAGACATTCTCACGTGANNNAGATCGGAAG
29337 TATGAGTGTGGAGTCGTTGCCCTGTAATCCTAGCAGTTTGGGANNNAGATCGGAAG
29339 TATGAGTGTGGAGTCGTTGCTCCTGTGCATAGTCTGATGTCTGNNNAGATCGGAAG
29340 TATGAGTGTGGAGTCGTTGCACCTTAGGGACTGGTCTGTCATANNNAGATCGGAAG
29341 TATGAGTGTGGAGTCGTTGCGCTGACTGTATCTCTCTCTCTGCNNNAGATCGGAAGA
29342 TATGAGTGTGGAGTCGTTGCAGCCTCTATCTGGTCTCATGACTNNNAGATCGGAAG
29343 TATGAGTGTGGAGTCGTTGCTGTCAGCAATTTCCAGACCCTATNNNAGATCGGAAG
29344 TATGAGTGTGGAGTCGTTGCACCAATGATGACAGCTCTGGAATNNNAGATCGGAAG
29345 TATGAGTGTGGAGTCGTTGCCTTGGGAGATAAGGGCAGAGAAANNNAGATCGGA
29346 TATGAGTGTGGAGTCGTTGCATCTCGGCTTACTAATGCAACCTNNNAGATCGGAAG
29347 TATGAGTGTGGAGTCGTTGCGGAGCTTCTGCAGGCTATTCTGNNNAGATCGGAAG
29348 TATGAGTGTGGAGTCGTTGCCTACATTTTTCCCACAGAGCAGCNNNAGATCGGAAG
29350 TATGAGTGTGGAGTCGTTGCTTTTCTCTCTCTCTGTCTCCCCANNNAGATCGGAAGAG
29351 TATGAGTGTGGAGTCGTTGCAGAGAAGCGTGGTCAATAGTGTTNNNAGATCGGAA
29352 TATGAGTGTGGAGTCGTTGCAATGGCAATATCTTGGTTGTGGCNNNAGATCGGAAG
29353 TATGAGTGTGGAGTCGTTGCCTGGAGACAGGAGAGAAAAGCAANNNAGATCGGA
29354 TATGAGTGTGGAGTCGTTGCATAACTGAGCTGGGTCAGACTTGNNNAGATCGGAA

29355 TATGAGTGTGGAGTCGTTGCATGAAGTTTACAGTCCCAGGAGCNNNAGATCGGAA
29356 TATGAGTGTGGAGTCGTTGCTGGAATAAAGGGGACAAGAGCTGNNNAGATCGGA
29359 TATGAGTGTGGAGTCGTTGCTTTGTATGTCCCTTCCTCCACAGNNNAGATCGGAAGA
29360 TATGAGTGTGGAGTCGTTGCTACGAAAGTGACTCTCCTGAAGGNNNAGATCGGAA
29361 TATGAGTGTGGAGTCGTTGCATGAGAGAGAGGGAGAAATGGGANNNAGATCGG
29362 TATGAGTGTGGAGTCGTTGCAATGTAGCAGAGCATGTGACAGANNNAGATCGGAA
29364 TATGAGTGTGGAGTCGTTGCGGGCATTATCAAGGCCAGTTTTTNNNAGATCGGAAG
29365 TATGAGTGTGGAGTCGTTGCGCTCTGCCCATAACCACTTATCANNNAGATCGGAAG
29367 TATGAGTGTGGAGTCGTTGCAACCATTTCTCAGCTGAACACTGNNNAGATCGGAAG
29368 TATGAGTGTGGAGTCGTTGCAGGAAGTTGCCAGCAAAAAGTTTNNNAGATCGGAA
29369 TATGAGTGTGGAGTCGTTGCGATGTCAGGATGAAGTGGGAAGTNNNAGATCGGA
29370 TATGAGTGTGGAGTCGTTGCTAGAGACCATTGTGAGTGTGGTGNNNAGATCGGAA
29371 TATGAGTGTGGAGTCGTTGCGCCCCCTTTTCTCTGGAATATGANNNAGATCGGAAG
29372 TATGAGTGTGGAGTCGTTGCCCCTCTGTCTCTCTCTCTCTCTCNNNAGATCGGAAGAG
29373 TATGAGTGTGGAGTCGTTGCGTCTCTCTCTCTCTCTCTCTCCCNNNAGATCGGAAGAG
29374 TATGAGTGTGGAGTCGTTGCGTGTCTGTGTGTCTGTGTGTTTTNNNAGATCGGAAG
29375 TATGAGTGTGGAGTCGTTGCCTGATGTCTGTGTTTCAGGCATGNNNAGATCGGAAG
29376 TATGAGTGTGGAGTCGTTGCCTTCCGGTTTCAAGCAATTGGATNNNAGATCGGAAG
29377 TATGAGTGTGGAGTCGTTGCGATTTGAGTGAGCTATGTTCGGCNNNAGATCGGAA
29378 TATGAGTGTGGAGTCGTTGCACTGAAAATGAGCCCCACAATTGNNNAGATCGGAAG
29379 TATGAGTGTGGAGTCGTTGCCCTCCAGTTCCTATCTGTTGAGCNNNAGATCGGAAG
29380 TATGAGTGTGGAGTCGTTGCAATCTTTACAGACCACTGGCAGTNNNAGATCGGAAG
29382 TATGAGTGTGGAGTCGTTGCGTGTGAGATGATGTGTGTGTGTGNNNAGATCGGAA
29383 TATGAGTGTGGAGTCGTTGCCAGAAGGAAAATCAATGCCCTCCNNNAGATCGGAAG
29384 TATGAGTGTGGAGTCGTTGCATCATAGAGGCACAGAGGAGGAANNNAGATCGGA
29385 TATGAGTGTGGAGTCGTTGCACAAATGGCCTCTGCTACTTACTNNNAGATCGGAAG
29387 TATGAGTGTGGAGTCGTTGCAGAGGTGAGAAGGGACTAAGACANNNAGATCGGA
29388 TATGAGTGTGGAGTCGTTGCAAGAACTCCTTGCTGAGACCAATNNNAGATCGGAAG
30737 TATGAGTGTGGAGTCGTTGCCAAAAGGAGGGCAAGAAGAAAGGNNNAGATCGGA
30738 TATGAGTGTGGAGTCGTTGCCTCTCTCTCTCTCTCTCACACCANNNAGATCGGAAGA
30739 TATGAGTGTGGAGTCGTTGCGAGGGGGCAGGTAGCTTATTATTNNNAGATCGGAA
30740 TATGAGTGTGGAGTCGTTGCCTGCCAAGTCTCACGTTGAAATTNNNAGATCGGAAG
30742 TATGAGTGTGGAGTCGTTGCAAAAGATGTTGCATATCCAGGCCNNNAGATCGGAA
30743 TATGAGTGTGGAGTCGTTGCGACTGTGGGAAATTTGAGACAGCNNNAGATCGGAA
30744 TATGAGTGTGGAGTCGTTGCAGAGACAGAGAGAGACAACGAGANNNAGATCGGA
30745 TATGAGTGTGGAGTCGTTGCCATGCATGCCCATAGTTACACACNNNAGATCGGAAG
30746 TATGAGTGTGGAGTCGTTGCAAACAGACACTCTCAGACATGCTNNNAGATCGGAAG
30747 TATGAGTGTGGAGTCGTTGCATTTATAGGGGCACAGCTGGAAANNNAGATCGGAA
30748 TATGAGTGTGGAGTCGTTGCAGTTGTTGTCTGTGGGAGGTTAGNNNAGATCGGAA
30749 TATGAGTGTGGAGTCGTTGCGTCCTGTGGATTTTGACATGCATNNNAGATCGGAAG
30750 TATGAGTGTGGAGTCGTTGCCTGTGGTCTCAGGACTTCTTTGTNNNAGATCGGAAG
30752 TATGAGTGTGGAGTCGTTGCGGCCATCAAGGTGGTACTTCTATNNNAGATCGGAAG
30753 TATGAGTGTGGAGTCGTTGCTTCCCCTCCAAAGATAGTGGATGNNNAGATCGGAAG

30754 TATGAGTGTGGAGTCGTTGCAGAGAGAGAGGGAGATGTTTGGANNNAGATCGGA
30755 TATGAGTGTGGAGTCGTTGCATGTATTTGGAAAAGGGAGGAGTNNNAGATCGGA
30756 TATGAGTGTGGAGTCGTTGCGAATTTTCTCCCAGAGAAGCAGCNNNAGATCGGAAG
30757 TATGAGTGTGGAGTCGTTGCTGAATGGTGCTGGAAATACTGGANNNAGATCGGAA
30758 TATGAGTGTGGAGTCGTTGCCATGTATGTGTTTTTGGTGGGGGNNNAGATCGGAA
30759 TATGAGTGTGGAGTCGTTGCCTGGCAGAAGAGGATTCAGACATNNNAGATCGGAA
30760 TATGAGTGTGGAGTCGTTGCTCCCTTTGATCTCCTTTCACCTGNNNAGATCGGAAGA
30761 TATGAGTGTGGAGTCGTTGCAGATCACTGGGAACCCTTGATTTNNNAGATCGGAAG
30762 TATGAGTGTGGAGTCGTTGCTGGGTCAGAACTGAAGGTGATACNNNAGATCGGAA
30763 TATGAGTGTGGAGTCGTTGCCTATTTGGCCCCTCAAACAACTGNNNAGATCGGAAG
30765 TATGAGTGTGGAGTCGTTGCTGAGGTTTCCTTTTGTGTTGTCCNNNAGATCGGAAG
30767 TATGAGTGTGGAGTCGTTGCTCATTGAGATGGATTGCTGGTGANNNAGATCGGAA
30768 TATGAGTGTGGAGTCGTTGCTAATTTGGTGAAAGCATGCCAGGNNNAGATCGGAA
30769 TATGAGTGTGGAGTCGTTGCTGATTCAAATGCCAGAGAGACGANNNAGATCGGAA
30770 TATGAGTGTGGAGTCGTTGCAGAGAGAGAGAGAGAGAATGGCCNNNAGATCGGA
30772 TATGAGTGTGGAGTCGTTGCTTAGGGGGAACAATGGAGACCTANNNAGATCGGAA
30773 TATGAGTGTGGAGTCGTTGCGCAGTGTGATGTGTTTGTCTTCANNNAGATCGGAAG
30774 TATGAGTGTGGAGTCGTTGCACAGACTCCACTTCTCCACTAGANNNAGATCGGAAG
30775 TATGAGTGTGGAGTCGTTGCGTGTGTGTGTGTGTGTGTGTTTANNNAGATCGGAA
30776 TATGAGTGTGGAGTCGTTGCTCCAGTTTCATGTCTTCTCTGGGNNNAGATCGGAAG
30777 TATGAGTGTGGAGTCGTTGCCACACACACATAGACACCTGAGANNNAGATCGGAAG
30778 TATGAGTGTGGAGTCGTTGCAAAAGGTCAGGTTCAGAGGCTAGNNNAGATCGGAA
30779 TATGAGTGTGGAGTCGTTGCAGTCTGGTCTGACAATGCAGTAGNNNAGATCGGAA
30780 TATGAGTGTGGAGTCGTTGCTGCATGGGTAAATTGTGTGTCACNNNAGATCGGAA
30781 TATGAGTGTGGAGTCGTTGCGGACCATGGGCAGATAGAAAGAANNNAGATCGGA
30783 TATGAGTGTGGAGTCGTTGCTCATATTCTGCTGCTGTCTCTCCNNNAGATCGGAAGA
30784 TATGAGTGTGGAGTCGTTGCCCCCAGGTCCTAAAAAGTTGAGANNNAGATCGGAA
30786 TATGAGTGTGGAGTCGTTGCAAGACACTAGTCATTGGGGTAGGNNNAGATCGGAA
30788 TATGAGTGTGGAGTCGTTGCGGTTGGTCTTTCTAGTATGGCCANNNAGATCGGAAG
30789 TATGAGTGTGGAGTCGTTGCGCAGGGACCTTCTCTTTCTTGTTNNNAGATCGGAAG
30790 TATGAGTGTGGAGTCGTTGCACACACACATCTGCCTACGTTATNNNAGATCGGAAG
30791 TATGAGTGTGGAGTCGTTGCGAGAATGCAAAGGAGGGAAACACNNNAGATCGGA
30793 TATGAGTGTGGAGTCGTTGCCACACACACACACACACACATATNNNAGATCGGAAG
30795 TATGAGTGTGGAGTCGTTGCCAAAGGCAGGTGAATGTTACAGGNNNAGATCGGAA
30796 TATGAGTGTGGAGTCGTTGCGGAACTAGAGAGAAGGGCAAAGTNNNAGATCGGA
30798 TATGAGTGTGGAGTCGTTGCGGGTGAAAATACAAGGGAAAGGGNNNAGATCGGA
30799 TATGAGTGTGGAGTCGTTGCAAGACCCAGTGACAAAACTTTGCNNNAGATCGGAAG
30800 TATGAGTGTGGAGTCGTTGCAGAGAGAGAGAGAGAGATCAGCCNNNAGATCGGA
30803 TATGAGTGTGGAGTCGTTGCGCAAGTCACTGAAAATGCCTCTTNNNAGATCGGAAG
30804 TATGAGTGTGGAGTCGTTGCGTTACAATGCGGTGTATGAGCTCNNNAGATCGGAA
30806 TATGAGTGTGGAGTCGTTGCGAGCTCTGACCAGAAACCAACTANNNAGATCGGAAG
30807 TATGAGTGTGGAGTCGTTGCCTCTGTCTCTCTCTCTCTCCCNNNAGATCGGAAGAG
30808 TATGAGTGTGGAGTCGTTGCCACCACTGGGACTACATTAGGTCNNNAGATCGGAAG

30809 TATGAGTGTGGAGTCGTTGCCATTCATGCCTCTGTCTTAGGGTNNNAGATCGGAAG
30811 TATGAGTGTGGAGTCGTTGCCTCTGCTAGCATTCACCTCTAGGNNNAGATCGGAAG
30812 TATGAGTGTGGAGTCGTTGCGTTGTTCTAGTGGGTGGCATCTANNNAGATCGGAA
30813 TATGAGTGTGGAGTCGTTGCATAGAGTATGTGTGTGTGCGTNNNAGATCGGAA
30814 TATGAGTGTGGAGTCGTTGCTCCTTTCCATTCCAACCCCATAGNNNAGATCGGAAGA
30815 TATGAGTGTGGAGTCGTTGCCTGGTTCTGAGGAATGGAACTGANNNAGATCGGAA
30816 TATGAGTGTGGAGTCGTTGCGTAACCACACATCAGAACAGTGCNNNAGATCGGAAG
30817 TATGAGTGTGGAGTCGTTGCGATTCACAGGGAAATTGGCTCAGNNNAGATCGGAA
30818 TATGAGTGTGGAGTCGTTGCTCTTGTAAGGCAGTGTAGATGCANNNAGATCGGAA
30819 TATGAGTGTGGAGTCGTTGCACAAGATCCACCTACACGTACACNNNAGATCGGAAG
30820 TATGAGTGTGGAGTCGTTGCAAGAAAACTGGACATAGGGGGAGNNNAGATCGGA
30821 TATGAGTGTGGAGTCGTTGCCTCATGTCTCAGCTAAGGAAGGGNNNAGATCGGAA
30822 TATGAGTGTGGAGTCGTTGCTGGGTATTCAATGAAGTCGGAGGNNNAGATCGGAA
30823 TATGAGTGTGGAGTCGTTGCAGACAAGGAGAAGAGGAGGAGAANNNAGATCGGA
30824 TATGAGTGTGGAGTCGTTGCGGTACAGGATTGGAAACTGAGGANNNAGATCGGA
30825 TATGAGTGTGGAGTCGTTGCTCTCTCTCTCTCTGTGTGTGTNNNAGATCGGAAGA
30827 TATGAGTGTGGAGTCGTTGCGGATGCTTCTGAATTTCCAGCTCNNNAGATCGGAAG
30828 TATGAGTGTGGAGTCGTTGCGTTGGGAAAGGAGAGATTGGGATNNNAGATCGGA
30830 TATGAGTGTGGAGTCGTTGCTGTTTTTGACGATTCTTGAGGCCNNNAGATCGGAAG
30831 TATGAGTGTGGAGTCGTTGCTGAGTATAACCAGATAGCCGTGCNNNAGATCGGAA
30833 TATGAGTGTGGAGTCGTTGCGCTGCAATTGCTGAAACGATTTCNNNAGATCGGAAG
30834 TATGAGTGTGGAGTCGTTGCTTGGATAGAAGGCTGGAATGCTTNNNAGATCGGAA
30836 TATGAGTGTGGAGTCGTTGCTCCAGGCACTAATCACCTCATTTNNNAGATCGGAAGA
30837 TATGAGTGTGGAGTCGTTGCGACTCAACTTATCCCCAGGTGTCNNNAGATCGGAAG
30838 TATGAGTGTGGAGTCGTTGCTGCAATCAGAGGCTCTTCTTTCTNNNAGATCGGAAG
30839 TATGAGTGTGGAGTCGTTGCGCACAAATACCTGGGGTAGCTATNNNAGATCGGAA
30840 TATGAGTGTGGAGTCGTTGCAGCTGACACAGAGCACTCAATAANNNAGATCGGAA
30841 TATGAGTGTGGAGTCGTTGCTCAAGCACAGGAATGCATTGAAGNNNAGATCGGAA
30842 TATGAGTGTGGAGTCGTTGCTGACACCATTTCAGGTTCATTGGNNNAGATCGGAAG
30843 TATGAGTGTGGAGTCGTTGCCACACACACTACTCTACCCTTCCNNNAGATCGGAAGA
30844 TATGAGTGTGGAGTCGTTGCCAAATCAAGGCAAGGAGTTGGAGNNNAGATCGGAA
30845 TATGAGTGTGGAGTCGTTGCAGAGAGAGAGGCTCCCATTATCANNNAGATCGGAA
30846 TATGAGTGTGGAGTCGTTGCCTGACACTTCTAGCGGTACACAANNNAGATCGGAAG
30847 TATGAGTGTGGAGTCGTTGCTGAAGGTTTACAGTACACCAGGGNNNAGATCGGAA
30848 TATGAGTGTGGAGTCGTTGCTTTTTGCGTTCCTTTAGAGCTGGNNNAGATCGGAAG
30849 TATGAGTGTGGAGTCGTTGCGCACACCAAAAGGGGAGAGATTANNNAGATCGGAA
30850 TATGAGTGTGGAGTCGTTGCAATCACACAGCTAGGAAGAAGCANNNAGATCGGAA
30851 TATGAGTGTGGAGTCGTTGCAATACGACTTTGCCAACTGAGTCNNNAGATCGGAAG
30852 TATGAGTGTGGAGTCGTTGCTGGAGATGGCAGACTCTAAGAGANNNAGATCGGAA
30853 TATGAGTGTGGAGTCGTTGCACATACAGGTACGCATGCACATANNNAGATCGGAAG
30854 TATGAGTGTGGAGTCGTTGCCCTCTGTTTGCAATCAATCCGTTNNNAGATCGGAAGA
30855 TATGAGTGTGGAGTCGTTGCCAAGCTATAGACTCAGGAACGCTNNNAGATCGGAAG
30856 TATGAGTGTGGAGTCGTTGCTGGTCGATATCAATTTTGCGTCGNNNAGATCGGAAG

30857 TATGAGTGTGGAGTCGTTGCTGTGTGGTGATCTGAAAGACAGTNNNAGATCGGAA
30858 TATGAGTGTGGAGTCGTTGCCTCATACACACACACACACACNNNAGATCGGAAGA
30859 TATGAGTGTGGAGTCGTTGCTTCTTCTATCAAAGTGGTGGCCTNNNAGATCGGAAG
30860 TATGAGTGTGGAGTCGTTGCTCTCTCTTACACACACATCCCACNNNAGATCGGAAGA
30861 TATGAGTGTGGAGTCGTTGCTGTTGTTTTCAGATGAGTGGGGANNNAGATCGGAA
30863 TATGAGTGTGGAGTCGTTGCCTGGTGCTCTGTCCATATCCTTTNNNAGATCGGAAGA
30864 TATGAGTGTGGAGTCGTTGCATCAGTGCTGTTGCATAGTGTTCNNNAGATCGGAAG
30865 TATGAGTGTGGAGTCGTTGCAACTGGAATACCTGCCTTCACTTNNNAGATCGGAAGA
30866 TATGAGTGTGGAGTCGTTGCCCTATTTGCAGTCACTCCTCACTNNNAGATCGGAAGA
30868 TATGAGTGTGGAGTCGTTGCAGTAGCTGGGACTATAGGTGTGTNNNAGATCGGAA
30869 TATGAGTGTGGAGTCGTTGCAGGAGTCAATGGGAAATCTGGTTNNNAGATCGGAA
30870 TATGAGTGTGGAGTCGTTGCGAGAGGAGGTAGACAGAGAGGAGNNNAGATCGGA
30871 TATGAGTGTGGAGTCGTTGCGTGTGCACATTCCATAGTGGTTCNNNAGATCGGAAG
30872 TATGAGTGTGGAGTCGTTGCTGTGTGTGTGTGTGTGATTTCNNNAGATCGGAAG
30873 TATGAGTGTGGAGTCGTTGCTCTCTCTCTCTCTGTGTGTGTNNNAGATCGGAAGA
30874 TATGAGTGTGGAGTCGTTGCCCCTTGCCTAATCTAGGACTTCCNNNAGATCGGAAGA
30875 TATGAGTGTGGAGTCGTTGCCAGACGACGGAGGATCTTATTCANNNAGATCGGAAG
30876 TATGAGTGTGGAGTCGTTGCTCCCTAGAACCTCCAAAAGAAGCNNNAGATCGGAAG
30877 TATGAGTGTGGAGTCGTTGCAATCCACCACAATCAAGTAGGCTNNNAGATCGGAAG
30878 TATGAGTGTGGAGTCGTTGCACACACACACACACACACAAATTNNNAGATCGGAAGA
30879 TATGAGTGTGGAGTCGTTGCGTGTGTGTGTGTGTTGGGATAAGNNNAGATCGGAA
30880 TATGAGTGTGGAGTCGTTGCAATATTGTTTGCTCTGTGGCTGGNNNAGATCGGAAG
30881 TATGAGTGTGGAGTCGTTGCCAGAAGCTGGAAAAGGCAAAGAANNNAGATCGGAA
30882 TATGAGTGTGGAGTCGTTGCATGATACAAGCGGAGTTCCAGAGNNNAGATCGGAA
30883 TATGAGTGTGGAGTCGTTGCGATGAGAAAGAGCGAGGAGAGAGNNNAGATCGGA
30884 TATGAGTGTGGAGTCGTTGCGAGTCCAATACACCTCGAGACAGNNNAGATCGGAAG
30885 TATGAGTGTGGAGTCGTTGCTGGTATGGAGTCAGAAGGTAGGANNNAGATCGGAA
30886 TATGAGTGTGGAGTCGTTGCGATAACTTGAGCCCAGGAGTTCANNNAGATCGGAAG
30887 TATGAGTGTGGAGTCGTTGCGCTGGGACAGGTATTTGATGGTANNNAGATCGGAA
30888 TATGAGTGTGGAGTCGTTGCCAAGTAGATTTCCTGCAGGGAGTNNNAGATCGGAA
30889 TATGAGTGTGGAGTCGTTGCGAGCACATCAGCTTCTAGACAGTNNNAGATCGGAAG
30890 TATGAGTGTGGAGTCGTTGCAAAAACCCAAACCTGTCAACTGGNNNAGATCGGAAG
30891 TATGAGTGTGGAGTCGTTGCAAGGGGAAATGTGTGGAACAGATNNNAGATCGGAA
30892 TATGAGTGTGGAGTCGTTGCCTGTCTCAGAATGACGGAAATGCNNNAGATCGGAAG
30893 TATGAGTGTGGAGTCGTTGCTAGAGACAGGAGAGAGAAAGCGANNNAGATCGGA
30894 TATGAGTGTGGAGTCGTTGCCGGAGAGAGAGAGAGAGAGAGAGNNNAGATCGGA
30895 TATGAGTGTGGAGTCGTTGCTCAGTGATGTGGTTATGCGTGTANNNAGATCGGAA
30896 TATGAGTGTGGAGTCGTTGCCTGAACAATGGAGGGAGACAGAANNNAGATCGGAA
30897 TATGAGTGTGGAGTCGTTGCAATCCCAGAGAGATTCGGAGAGANNNAGATCGGAA
30898 TATGAGTGTGGAGTCGTTGCACTCTCTGTCTCTCTCTCTGTCCNNNAGATCGGAAGA
30899 TATGAGTGTGGAGTCGTTGCCAGGAATAGATGGGCAATGCAAGNNNAGATCGGAA
30901 TATGAGTGTGGAGTCGTTGCCCTGTGTGAGTGAAAGACAGAGANNNAGATCGGAA
30902 TATGAGTGTGGAGTCGTTGCTGTTTATCCCTAAATGGCCAGCTNNNAGATCGGAAG

30903 TATGAGTGTGGAGTCGTTGCCTACTACTATCACAGGCCCATGGNNNAGATCGGAAG
30904 TATGAGTGTGGAGTCGTTGCCAATATCGTAAACTGCAGCAGGGNNNAGATCGGAA
30905 TATGAGTGTGGAGTCGTTGCACCCAGCTAATACTTGTTGCAGANNNAGATCGGAAG
30906 TATGAGTGTGGAGTCGTTGCGAGACAAAGAGGGAGTCAAGGAGNNNAGATCGGA
30907 TATGAGTGTGGAGTCGTTGCTGTGTTTTGGACTAAAGGCAGTGNNNAGATCGGAA
30908 TATGAGTGTGGAGTCGTTGCAAACTACTAGCAAACTGAGGCCANNNAGATCGGAA
30909 TATGAGTGTGGAGTCGTTGCGAAGGGTCAGACTTGCTCAAAACNNNAGATCGGAA
30910 TATGAGTGTGGAGTCGTTGCAGAGGAGAGGAGAGAGAGAGAGANNNAGATCGG
30912 TATGAGTGTGGAGTCGTTGCACCATTGCAGTCAGAATCTTTGCNNNAGATCGGAAG
30913 TATGAGTGTGGAGTCGTTGCTACTGAGAGACCAAAATCCTGGCNNNAGATCGGAA
30914 TATGAGTGTGGAGTCGTTGCGTCACCACTATTGGGAATGGGATNNNAGATCGGAA
30915 TATGAGTGTGGAGTCGTTGCCCATGTGGTCTCTCATCTCCATTNNNAGATCGGAAGA
30916 TATGAGTGTGGAGTCGTTGCTTCCTACGTCATGAAAGCAGCATNNNAGATCGGAAG
30918 TATGAGTGTGGAGTCGTTGCTTGCTCTTTCATGTCTCAGTGGTNNNAGATCGGAAG
30919 TATGAGTGTGGAGTCGTTGCGTCAGAAGGGGTTCTAAAGCCAGNNNAGATCGGAA
30920 TATGAGTGTGGAGTCGTTGCTGGACTGAACCCAGCAGAAATAANNNAGATCGGAA
30921 TATGAGTGTGGAGTCGTTGCGGCACCAGAATTGTGAGTGAAAANNNAGATCGGAA
30922 TATGAGTGTGGAGTCGTTGCGTAACCAGCACCCAAATCAAGACNNNAGATCGGAAG
30923 TATGAGTGTGGAGTCGTTGCTTACATCCTCTGTTTGATGGGCANNNAGATCGGAAG
30924 TATGAGTGTGGAGTCGTTGCTTGGACTTCTTTAGATGGCAGCTNNNAGATCGGAAG
30926 TATGAGTGTGGAGTCGTTGCGCACAGATAGACACAACACACACNNNAGATCGGAAG
30927 TATGAGTGTGGAGTCGTTGCGAGAGAGATTGGACTTGCCTGAANNNAGATCGGAA
30928 TATGAGTGTGGAGTCGTTGCTCTCTCCCACTCTTCTTCTCTCCNNNAGATCGGAAGAG
30929 TATGAGTGTGGAGTCGTTGCTCATTTGCAACAACTAAGCACGANNNAGATCGGAAG
30930 TATGAGTGTGGAGTCGTTGCATAGCTTGTGAGGCTCTTGATCCNNNAGATCGGAAG
30931 TATGAGTGTGGAGTCGTTGCATGAGAGCAAGAACCTTGTCTGTNNNAGATCGGAA
30933 TATGAGTGTGGAGTCGTTGCCGAGCCCTATAACCCAAACAGTANNNAGATCGGAAG
30934 TATGAGTGTGGAGTCGTTGCGAAAGACATGGGAACTCCAGAGTNNNAGATCGGAA
30935 TATGAGTGTGGAGTCGTTGCATCTTTTTTTCTGTGGAGTGCGGNNNAGATCGGAAG
30936 TATGAGTGTGGAGTCGTTGCTGGCTGTAGATGGAAAGGAGATGNNNAGATCGGA
30938 TATGAGTGTGGAGTCGTTGCTCTCTCTCTCTCTCTCTCTCCCTNNNAGATCGGAAGAG
30939 TATGAGTGTGGAGTCGTTGCGCATGGAGAAGAGCACTTACAACNNNAGATCGGAA
30940 TATGAGTGTGGAGTCGTTGCGCATGGCTTGTTCTCAATTTGGANNNAGATCGGAAG
30941 TATGAGTGTGGAGTCGTTGCAAGTTGTTGTTGTTTGCTTTGGCNNNAGATCGGAAG
30942 TATGAGTGTGGAGTCGTTGCAGGTTGAAGAGAGCTGTCCTTTGNNNAGATCGGAA
30943 TATGAGTGTGGAGTCGTTGCATCCCTGGAGTCTGCAAGTTTAGNNNAGATCGGAAG
30944 TATGAGTGTGGAGTCGTTGCTGGAATGAGCCATTCATACAGGTNNNAGATCGGAA
30945 TATGAGTGTGGAGTCGTTGCTTGTCACTTGGTAACTGGTCTCCNNNAGATCGGAAG
30946 TATGAGTGTGGAGTCGTTGCTCTGGATATACCCCACCATTCCTNNNAGATCGGAAGA
30947 TATGAGTGTGGAGTCGTTGCAAAGCAGGAGAAGGGACAAGAATNNNAGATCGGA
30948 TATGAGTGTGGAGTCGTTGCGCTAGGATGGTGGCTAAGATACCNNNAGATCGGAA
30949 TATGAGTGTGGAGTCGTTGCTCAAGTCACAAGAAGAAGCTGAANNNAGATCGGAA
30950 TATGAGTGTGGAGTCGTTGCCACACACACACTCAAGTCACAAGNNNAGATCGGAAG

30951 TATGAGTGTGGAGTCGTTGCAGGGATGACTCTCCAACACAATCNNNAGATCGGAAG
30952 TATGAGTGTGGAGTCGTTGCCACACACACACAGACATACACACNNNAGATCGGAAG
30953 TATGAGTGTGGAGTCGTTGCAGGGACTGTGAGAAAGTAGGAGANNNAGATCGGA
30954 TATGAGTGTGGAGTCGTTGCGCAAGAGAGACCGAGAAAGAGAANNNAGATCGGA
30955 TATGAGTGTGGAGTCGTTGCTGACAGACCTGACACTTAGCTTCNNNAGATCGGAAG
30956 TATGAGTGTGGAGTCGTTGCACAGAGAGACAGAGAGAGAAGCTNNNAGATCGGA
30957 TATGAGTGTGGAGTCGTTGCAATGCTCAGTCCTTATCCATCCCNNNAGATCGGAAGA
30958 TATGAGTGTGGAGTCGTTGCCTGCTCCCCTTTCTTTTCCTTTGNNNAGATCGGAAGA
30959 TATGAGTGTGGAGTCGTTGCCACGTATGCTATGCCCACTTTTCNNNAGATCGGAAG
30960 TATGAGTGTGGAGTCGTTGCCAGTTGAGCTTTGAGCAGTGATGNNNAGATCGGAA
30961 TATGAGTGTGGAGTCGTTGCGGCTAAGGAGAGTGACCATTTGANNNAGATCGGAA
30962 TATGAGTGTGGAGTCGTTGCATGTTGTGATACCAGGCAGCTAANNNAGATCGGAA
30963 TATGAGTGTGGAGTCGTTGCACCTTTCCTCTCCCCTAGAATCANNNAGATCGGAAGA
30964 TATGAGTGTGGAGTCGTTGCGTACTTTGCCCTCTGTTAGCTCTNNNAGATCGGAAG
30965 TATGAGTGTGGAGTCGTTGCGCTTTCTCTTCATCTGGGTTCCTNNNAGATCGGAAGA
30966 TATGAGTGTGGAGTCGTTGCCTCTCAACTTCCCCCTCTTCTTCNNNAGATCGGAAGAG
30967 TATGAGTGTGGAGTCGTTGCCCATGGCCGGCTAATTTTTGTATNNNAGATCGGAAG
30968 TATGAGTGTGGAGTCGTTGCGTGACATCAGACAAAGAAGGTGCNNNAGATCGGAA
30969 TATGAGTGTGGAGTCGTTGCCCCCATAATAAGTGCTTGCTTGGNNNAGATCGGAAG
30970 TATGAGTGTGGAGTCGTTGCAAAATAAAACCCACCTCGAGCCTNNNAGATCGGAAG
30971 TATGAGTGTGGAGTCGTTGCTTTTTTGACTCCTTCCAATGCCCNNNAGATCGGAAGA
30972 TATGAGTGTGGAGTCGTTGCAACGTCTTCTCTGCCACTCTAAGNNNAGATCGGAAG
30973 TATGAGTGTGGAGTCGTTGCCCATTCACTTCCTCAACCCACTANNNAGATCGGAAGA
30974 TATGAGTGTGGAGTCGTTGCGCCTTGAAGGAAAAGACCAAGTCNNNAGATCGGAA
30975 TATGAGTGTGGAGTCGTTGCGTCACTATGGGACTACGGAACTCNNNAGATCGGAAG
30976 TATGAGTGTGGAGTCGTTGCAAGCCTGATAAAACCTTCCAACTNNNAGATCGGAAG
30977 TATGAGTGTGGAGTCGTTGCCAGCAGCAGCATTATTCCTTTGANNNAGATCGGAAG
30978 TATGAGTGTGGAGTCGTTGCCACACACACACACACACACATATNNNAGATCGGAAG
30979 TATGAGTGTGGAGTCGTTGCCTCTCTCTCTCTCACACACACACNNNAGATCGGAAGA
30980 TATGAGTGTGGAGTCGTTGCACACAGACATGCACATACTCACTNNNAGATCGGAAG
30981 TATGAGTGTGGAGTCGTTGCTGTATATGTGTGCGTGCAAATCCNNNAGATCGGAAG
30982 TATGAGTGTGGAGTCGTTGCCTTGGGGGTGGAGGATCTGATTCNNNAGATCGGAA
30983 TATGAGTGTGGAGTCGTTGCCCCTCATTCAGCACTTTGTTCTGNNNAGATCGGAAGA
30984 TATGAGTGTGGAGTCGTTGCCAGCATGTCACCCTCCATATTCTNNNAGATCGGAAGA
30985 TATGAGTGTGGAGTCGTTGCGAAACCTGGATGGGATAGGAGACNNNAGATCGGA
30986 TATGAGTGTGGAGTCGTTGCCAGGCAACAGAAACAGACATTCANNNAGATCGGAA
30987 TATGAGTGTGGAGTCGTTGCTCCTACTTTGGGGCATCTAGTCTNNNAGATCGGAAG
30988 TATGAGTGTGGAGTCGTTGCAGTGCCAAAGCTGTTTCAAACAANNNAGATCGGAAG
30989 TATGAGTGTGGAGTCGTTGCCATACACATTCTCTCTCTCCCCCNNNAGATCGGAAGA
30990 TATGAGTGTGGAGTCGTTGCGGCTTCCTGAGTCCAATTTTCACNNNAGATCGGAAG
30991 TATGAGTGTGGAGTCGTTGCCCTCCCTCCCTTCCTTTTTCTTTNNNAGATCGGAAGAG
30992 TATGAGTGTGGAGTCGTTGCGGAATGAATGAGATTGGGGGTGANNNAGATCGGA
30993 TATGAGTGTGGAGTCGTTGCTATTTAGCTTACAGGTCCGTGGGNNNAGATCGGAA

30994 TATGAGTGTGGAGTCGTTGCCATCTCATTACTGCCTGGTCTGTNNNAGATCGGAAG
30995 TATGAGTGTGGAGTCGTTGCCCTCACTGAAACTCCTCTCTACCNNNAGATCGGAAGA
30996 TATGAGTGTGGAGTCGTTGCAGTCAGAGTGGTCATCAAATCCTNNNAGATCGGAAG
30997 TATGAGTGTGGAGTCGTTGCTGCTAGGATATTACAGGTGCCACNNNAGATCGGAA
30998 TATGAGTGTGGAGTCGTTGCCTTAGAGGGAATCTGTATGGGGCNNNAGATCGGAA
30999 TATGAGTGTGGAGTCGTTGCGTCCTGCTGGGTAGTCTTTGTTANNNAGATCGGAAG
31000 TATGAGTGTGGAGTCGTTGCTGTGTGTGTGTAAAGGGGAGTTTNNNAGATCGGAA
31001 TATGAGTGTGGAGTCGTTGCTTTGAGCGTACAGTCAGTCACTGNNNAGATCGGAAG
31003 TATGAGTGTGGAGTCGTTGCACTATACCTGGGGCTGAAGTCATNNNAGATCGGAA
31004 TATGAGTGTGGAGTCGTTGCCATTTCCCTCAGTCTCCCATTCANNNAGATCGGAAGA
31005 TATGAGTGTGGAGTCGTTGCTTTCTCTCTGTCTTTCCGGAGTCNNNAGATCGGAAGA
31006 TATGAGTGTGGAGTCGTTGCGAGAGAGAGAGAGAGAGAGAGGTNNNAGATCGG
31007 TATGAGTGTGGAGTCGTTGCTTCAAGTGATTCTCCTGCCTCAGNNNAGATCGGAAG
31008 TATGAGTGTGGAGTCGTTGCTTGAGACGGAGTCTTACTCTGTCNNNAGATCGGAAG
31010 TATGAGTGTGGAGTCGTTGCTCTGCTCTGTTTTTTTCCCCATCNNNAGATCGGAAGA
31011 TATGAGTGTGGAGTCGTTGCTGGACCAGGCTCTTTTTATCACANNNAGATCGGAAG
31013 TATGAGTGTGGAGTCGTTGCAGGAAAGGGGAAAGGAAAAAGGANNNAGATCGG
31014 TATGAGTGTGGAGTCGTTGCTCAAGTAGGTAGGACTACAGGCANNNAGATCGGAA
31015 TATGAGTGTGGAGTCGTTGCGAAAGTGTTGAACAGGACAGAGCNNNAGATCGGA
31016 TATGAGTGTGGAGTCGTTGCACTTGTCTATCTGGTAAAGCCCGNNNAGATCGGAAG
31017 TATGAGTGTGGAGTCGTTGCGTATGTATTTAGAGGTGGGGGCCNNNAGATCGGAA
31018 TATGAGTGTGGAGTCGTTGCTGCAACACCCAGTGACAAAAATTNNNAGATCGGAAG
31019 TATGAGTGTGGAGTCGTTGCATTGTTTTCCTGCCCTCATTTCCNNNAGATCGGAAGA
31020 TATGAGTGTGGAGTCGTTGCACCCCTCAGTCCTATTCTAAGCTNNNAGATCGGAAGA
31021 TATGAGTGTGGAGTCGTTGCTGTAACTACCTCCCACTGTTGTCNNNAGATCGGAAGA
31022 TATGAGTGTGGAGTCGTTGCGGTCTCTGCCTGATAATCCAGAGNNNAGATCGGAAG
31023 TATGAGTGTGGAGTCGTTGCCTCCTGAGTAGCTGGGATTACAGNNNAGATCGGAA
31024 TATGAGTGTGGAGTCGTTGCTTGATGGGGAGAACTAGCTATGCNNNAGATCGGAA
31025 TATGAGTGTGGAGTCGTTGCCAAAGGTATCCAAAAGGCAGCAGNNNAGATCGGAA
31026 TATGAGTGTGGAGTCGTTGCAACCAAAAGCAGCAAAGGACAATNNNAGATCGGAA
31027 TATGAGTGTGGAGTCGTTGCGATGTTCTCAGTTTTGTCCTGGCNNNAGATCGGAAG
31028 TATGAGTGTGGAGTCGTTGCTGGAGATAGAGCCATCATGCTTCNNNAGATCGGAA
31029 TATGAGTGTGGAGTCGTTGCTCAAATGCAGCATGTCCAAAACTNNNAGATCGGAAG
31030 TATGAGTGTGGAGTCGTTGCCTTTCCCTCTTTCCCTTTCCCTTNNNAGATCGGAAGAG
31031 TATGAGTGTGGAGTCGTTGCTACGAGTGAACACCCATTCTCAGNNNAGATCGGAAG
31032 TATGAGTGTGGAGTCGTTGCTCTCTGGAGAAGGATTCCCATCTNNNAGATCGGAAG
31033 TATGAGTGTGGAGTCGTTGCGATCACTCTGGTTGCTGCTGATCNNNAGATCGGAAG
31034 TATGAGTGTGGAGTCGTTGCCACCCTGCCCTTACTTATGATCTNNNAGATCGGAAGA
31035 TATGAGTGTGGAGTCGTTGCTTTCCCAGGATCCTTGGCTAATTNNNAGATCGGAAG
31036 TATGAGTGTGGAGTCGTTGCCAGAGAGCTGATTCGTAGTTCCANNNAGATCGGAA
31038 TATGAGTGTGGAGTCGTTGCGCTCATCTCTGGTACTTCCTAGCNNNAGATCGGAAG
31039 TATGAGTGTGGAGTCGTTGCAAGAGAACAAGAGTCTCTGCCTGNNNAGATCGGAA
31040 TATGAGTGTGGAGTCGTTGCCCAGGGCAGGAACATGTAATCTANNNAGATCGGAA

31041 TATGAGTGTGGAGTCGTTGCCAGGGCTAGACAGGAATCAGATCNNNAGATCGGAA
31042 TATGAGTGTGGAGTCGTTGCTTGCCTCTCTGAATATACCACCGNNNAGATCGGAAG
31044 TATGAGTGTGGAGTCGTTGCAGCATCCAAGAATGTCAGGAGAGNNNAGATCGGAA
31045 TATGAGTGTGGAGTCGTTGCTGGGTTCCTCTTCTTCTTTGCTTNNNAGATCGGAAGA
31046 TATGAGTGTGGAGTCGTTGCCCTTTCCAAAACTCACCTTTGCTNNNAGATCGGAAGA
31047 TATGAGTGTGGAGTCGTTGCGGGTTGACTAGAGACTCAGCATTNNNAGATCGGAA
31048 TATGAGTGTGGAGTCGTTGCTTTGAATAGAGTCCCCTAAGGCTNNNAGATCGGAAG
31049 TATGAGTGTGGAGTCGTTGCCCAAGGAGTGAAACAGGGTAGTTNNNAGATCGGAA
31050 TATGAGTGTGGAGTCGTTGCCCTGTGCTTTGAGGACTAGTTCTNNNAGATCGGAAG
31051 TATGAGTGTGGAGTCGTTGCCCCCCCACCATTTCTGAGAAATANNNAGATCGGAAG
31052 TATGAGTGTGGAGTCGTTGCTGTTTCATGAGGTAGTGGAAGCANNNAGATCGGAA
31053 TATGAGTGTGGAGTCGTTGCGACGATGTGTACTCTTTGTGCAGNNNAGATCGGAA
31054 TATGAGTGTGGAGTCGTTGCTTTATGGAGAGGCATACATGGCANNNAGATCGGAA
31055 TATGAGTGTGGAGTCGTTGCTACAGTTGTCTCAGATTGCACCANNNAGATCGGAAG
31057 TATGAGTGTGGAGTCGTTGCTGTGTAACCTCCACCACCATAAGNNNAGATCGGAAG
31058 TATGAGTGTGGAGTCGTTGCTACCTTTAGACAAGGGTGCTTCCNNNAGATCGGAAG
31059 TATGAGTGTGGAGTCGTTGCACGTGCACCTCAACTGTTACATANNNAGATCGGAAG
31060 TATGAGTGTGGAGTCGTTGCATGAATTTTGGAACGTGCCCTTTNNNAGATCGGAAG
31061 TATGAGTGTGGAGTCGTTGCGATTCTAGCAGTGGGAAAAGTGCNNNAGATCGGAA
31063 TATGAGTGTGGAGTCGTTGCCAATGGAGCCAGATTGCAGAAAANNNAGATCGGAA
31064 TATGAGTGTGGAGTCGTTGCAAGTGCATATGACACATCCATGANNNAGATCGGAAG
31066 TATGAGTGTGGAGTCGTTGCATGTCTGAACTTGTCTGGAAGCANNNAGATCGGAA
31067 TATGAGTGTGGAGTCGTTGCGAGCTGCAATAAACTCACTAGCGNNNAGATCGGAA
31068 TATGAGTGTGGAGTCGTTGCGGTCAGTATTATTGGCCTCAGCTNNNAGATCGGAAG
31069 TATGAGTGTGGAGTCGTTGCTTACCATCACTTAGAGGCCAGTGNNNAGATCGGAAG
31070 TATGAGTGTGGAGTCGTTGCGGATGGGGATATGTAAACCTGCTNNNAGATCGGAA
31071 TATGAGTGTGGAGTCGTTGCCATTTTTGAGACACGGTCATGCTNNNAGATCGGAAG
31072 TATGAGTGTGGAGTCGTTGCATATAGGCAGTTGCATGCTAGCANNNAGATCGGAA
31075 TATGAGTGTGGAGTCGTTGCTGCTACAACATGGATGAACCTCANNNAGATCGGAAG
31077 TATGAGTGTGGAGTCGTTGCTTTCACCTTGGCTGGAGTACAATNNNAGATCGGAAG
31078 TATGAGTGTGGAGTCGTTGCCAAGCATGCATGGTGAGTGTATTNNNAGATCGGAA
31079 TATGAGTGTGGAGTCGTTGCCTCTAAAGCAATGAGCAGAGTGGNNNAGATCGGAA
31081 TATGAGTGTGGAGTCGTTGCGGACCCATAATTCCTCCAGATGGNNNAGATCGGAAG
31082 TATGAGTGTGGAGTCGTTGCGTGTGTGTGTGTGTGTGTATCNNNAGATCGGAA
31083 TATGAGTGTGGAGTCGTTGCGTGCAGTATTGTGATCATGGCTCNNNAGATCGGAA
31084 TATGAGTGTGGAGTCGTTGCTTTTTTGTGCATGTTGTCCTGGANNNAGATCGGAAG
31085 TATGAGTGTGGAGTCGTTGCTTTTTTGTGCATGTTGTCCTGGANNNAGATCGGAAG
31086 TATGAGTGTGGAGTCGTTGCTTAGGTTTGGCCATGCAACATTTNNNAGATCGGAAG
31087 TATGAGTGTGGAGTCGTTGCCCACATCGGACTAAAAACCACACNNNAGATCGGAAG
31088 TATGAGTGTGGAGTCGTTGCCCACATCGGACTAAAAACCACACNNNAGATCGGAAG
31089 TATGAGTGTGGAGTCGTTGCCACAGGATTATAGGGGCCAAGAANNNAGATCGGAA
31090 TATGAGTGTGGAGTCGTTGCGGGCATATAGGTTGGAGCTATGTNNNAGATCGGAA
31091 TATGAGTGTGGAGTCGTTGCAGAAGAAGTGAAAGCTACCAGCANNNAGATCGGAA

31092 TATGAGTGTGGAGTCGTTGCAAAGTGTCGACAGGATAGTGACCNNNAGATCGGAA
31093 TATGAGTGTGGAGTCGTTGCAGAGAGAGAGAAAGAGGGAGACANNNAGATCGG
31094 TATGAGTGTGGAGTCGTTGCGGTAGGCCAGTATTCACATGCTANNNAGATCGGAAG
31095 TATGAGTGTGGAGTCGTTGCGGGAGGCCTCTTCAAGAATAAGTNNNAGATCGGAA
31096 TATGAGTGTGGAGTCGTTGCACATTCAGATAAAGCAGCTCCCANNNAGATCGGAAG
31097 TATGAGTGTGGAGTCGTTGCAGCAAACAATGTTGAGGCTCAAANNNAGATCGGAA
31098 TATGAGTGTGGAGTCGTTGCTGGTAATCCAGAGAATTTGCCCANNNAGATCGGAAG
31099 TATGAGTGTGGAGTCGTTGCCACCATTTCACATTCTCACCACCNNNAGATCGGAAGA
31100 TATGAGTGTGGAGTCGTTGCTAGAGTGAGTGCTTTCAGAAGGCNNNAGATCGGAA
31101 TATGAGTGTGGAGTCGTTGCAGAATGGATAGCTGCCTCTATGCNNNAGATCGGAAG
31102 TATGAGTGTGGAGTCGTTGCAGAATGGATAGCTGCCTCTATGCNNNAGATCGGAAG
31103 TATGAGTGTGGAGTCGTTGCATCGCGATCCACTGGAATACTTTNNNAGATCGGAAG
31104 TATGAGTGTGGAGTCGTTGCAGCACTCCATGAACATCCTTGATNNNAGATCGGAAG
31105 TATGAGTGTGGAGTCGTTGCTGGGTAAACAGAGTGAGAAGACGNNNAGATCGGA
31106 TATGAGTGTGGAGTCGTTGCGGTTCCAAGAGAAACCCTAGCAANNNAGATCGGAAG
31107 TATGAGTGTGGAGTCGTTGCTGTCAGGAGTGAGAATTTCCCAGNNNAGATCGGAA
31109 TATGAGTGTGGAGTCGTTGCCTCTCTCTCTCTCTCTCTCTCCGNNNAGATCGGAAGAG
31110 TATGAGTGTGGAGTCGTTGCTCCTGGAGCTGTATATTGCTTCCNNNAGATCGGAAG
31111 TATGAGTGTGGAGTCGTTGCACTGGCTCCACATTTTTGCAATTNNNAGATCGGAAG
31112 TATGAGTGTGGAGTCGTTGCAGAAACACAATGCTAGAGCCAGANNNAGATCGGAA
31113 TATGAGTGTGGAGTCGTTGCCACTGCCTAGAACTAACAGAGCANNNAGATCGGAAG
31115 TATGAGTGTGGAGTCGTTGCAAACAGCAAAAATGTAGCCACGANNNAGATCGGAA
31116 TATGAGTGTGGAGTCGTTGCTGTAGTGTTGTGGCAAGATCTCANNNAGATCGGAAG
31117 TATGAGTGTGGAGTCGTTGCTTGAGACAGAGTGCAATCTCAGCNNNAGATCGGAAG
31118 TATGAGTGTGGAGTCGTTGCTGGGTAAACAAGAGCTCAGCTAGNNNAGATCGGAA
31119 TATGAGTGTGGAGTCGTTGCGCCAGCCTTCAATGTTCCTTATTNNNAGATCGGAAG
31120 TATGAGTGTGGAGTCGTTGCCAACTTGCTGTCATGCAGAAGATNNNAGATCGGAAG
31122 TATGAGTGTGGAGTCGTTGCAGAGAGACTAGAAGCCAGAGTCANNNAGATCGGAA
31123 TATGAGTGTGGAGTCGTTGCCTGTTATCATCTCCCCCTCACTGNNNAGATCGGAAGA
31124 TATGAGTGTGGAGTCGTTGCTGTATTGCAAGCTTCCAAACAGANNNAGATCGGAAG
31125 TATGAGTGTGGAGTCGTTGCACACACACACACACCTCTTCTAANNNAGATCGGAAG
31126 TATGAGTGTGGAGTCGTTGCAGGTCTTGGGAACTGAAAGCTTANNNAGATCGGAA
31127 TATGAGTGTGGAGTCGTTGCACTGATGAGCAATTCCTTCTCCCNNNAGATCGGAAG
31128 TATGAGTGTGGAGTCGTTGCATTGCTTCAATCTGCTTCCAAAGNNNAGATCGGAAG
31129 TATGAGTGTGGAGTCGTTGCGCATGTTTGCATTTGGTTTCCAGNNNAGATCGGAAG
31130 TATGAGTGTGGAGTCGTTGCATGCAGCTTTTATATCGCTAGCTNNNAGATCGGAAG
31131 TATGAGTGTGGAGTCGTTGCTTTGAGACGGTCACACTCTTTCCNNNAGATCGGAAG
31133 TATGAGTGTGGAGTCGTTGCGTGTGTCTGTATGTGTGTTCGTGNNNAGATCGGAA
31134 TATGAGTGTGGAGTCGTTGCGGAGGAGCTCAAAGGAAAAGAGANNNAGATCGGA
31135 TATGAGTGTGGAGTCGTTGCCTGTCATTCACCCAATTGCTCAGNNNAGATCGGAAG
31136 TATGAGTGTGGAGTCGTTGCGTGGATGTCAAAGTTCTGGCTTCNNNAGATCGGAA
31137 TATGAGTGTGGAGTCGTTGCCCATGACTTCCAGGAACTCTCTTNNNAGATCGGAAG
31138 TATGAGTGTGGAGTCGTTGCCCATGACTTCCAGGAACTCTCTTNNNAGATCGGAAG

31140 TATGAGTGTGGAGTCGTTGCATCCACCTTGAAAGTGTGTCTGTNNNAGATCGGAAG
31141 TATGAGTGTGGAGTCGTTGCGTTCAAGTGATTCTTCTGCCTCGNNNAGATCGGAAG
31142 TATGAGTGTGGAGTCGTTGCGTACCGAGTCTCACCCTATTGTCNNNAGATCGGAAG
31143 TATGAGTGTGGAGTCGTTGCGTGGTTCTGACAGCTGTTTCTTCNNNAGATCGGAAG
31144 TATGAGTGTGGAGTCGTTGCTCTCCTTATCTGACCTCGTCCTTNNNAGATCGGAAGA
31145 TATGAGTGTGGAGTCGTTGCAGCTCCCTGAGGATTTGTTTCATNNNAGATCGGAAG
31146 TATGAGTGTGGAGTCGTTGCCACACACACACTCACACACATACNNNAGATCGGAAG
31147 TATGAGTGTGGAGTCGTTGCATACAGGCATTTTACGTGTGTGCNNNAGATCGGAAG
31148 TATGAGTGTGGAGTCGTTGCTCCATCCTTCCACTACCAAACTCNNNAGATCGGAAGA
31149 TATGAGTGTGGAGTCGTTGCATCATTACAGCATAATGGGGCCANNNAGATCGGAA
31150 TATGAGTGTGGAGTCGTTGCCACATACACAGACACACACACACNNNAGATCGGAAG
31151 TATGAGTGTGGAGTCGTTGCCTGAAACTTGGCTGTCCTCAAAGNNNAGATCGGAAG
31153 TATGAGTGTGGAGTCGTTGCTGCCATTGTCACTACAGGACTACNNNAGATCGGAAG
31154 TATGAGTGTGGAGTCGTTGCCATCTGAGAGCCACAAAACCATCNNNAGATCGGAAG
31155 TATGAGTGTGGAGTCGTTGCCCTTCCCCACTGAACCTTTCTAANNNAGATCGGAAGA
31156 TATGAGTGTGGAGTCGTTGCCCAGGGACATGAAGAGTGAAGTTNNNAGATCGGAA
31157 TATGAGTGTGGAGTCGTTGCCTAAAGAGCCAAATCTGTTGCCCNNNAGATCGGAAG
31158 TATGAGTGTGGAGTCGTTGCACTACTAATCAACCAGAGGCTGCNNNAGATCGGAAG
31159 TATGAGTGTGGAGTCGTTGCTTACAAAATCCCGAGACTGTTGCNNNAGATCGGAAG
31160 TATGAGTGTGGAGTCGTTGCGCAAGAAGGCAACTGTCTATGTGNNNAGATCGGAA
31161 TATGAGTGTGGAGTCGTTGCAGCCTCACAAAAGATAGTGCCATNNNAGATCGGAAG
31162 TATGAGTGTGGAGTCGTTGCTGGGAAAGTTCTGCCTTCAACTANNNAGATCGGAAG
31163 TATGAGTGTGGAGTCGTTGCAGACATAGTCTCTGCCAATGGTGNNNAGATCGGAA
31164 TATGAGTGTGGAGTCGTTGCTGAACACAGAAACTCATGTGCAGNNNAGATCGGAA
31165 TATGAGTGTGGAGTCGTTGCCCTGTCCCCCATTCCAATTTCTANNNAGATCGGAAGA
31166 TATGAGTGTGGAGTCGTTGCTATTTATCTGTCTGAGCCCTGCCNNNAGATCGGAAG
31168 TATGAGTGTGGAGTCGTTGCCCTCTTCTGCTTTCACACACTTGNNNAGATCGGAAGA
31169 TATGAGTGTGGAGTCGTTGCTGTAGAGCCACACTTGAGAGATGNNNAGATCGGAA
31170 TATGAGTGTGGAGTCGTTGCTGTGTGATTAGTGTAGTGGCCAANNNAGATCGGAA
31171 TATGAGTGTGGAGTCGTTGCATGTACAAGATGGCCCAACAGATNNNAGATCGGAA
31172 TATGAGTGTGGAGTCGTTGCTAATCTCAGCACTTTGGGAGACCNNNAGATCGGAAG
31173 TATGAGTGTGGAGTCGTTGCCTTTCTTCTTTCTGATTCCCGGCNNNAGATCGGAAGA
31174 TATGAGTGTGGAGTCGTTGCCCTCCCCCTCTCTCTTTCTTTTTNNNAGATCGGAAGAG
31175 TATGAGTGTGGAGTCGTTGCTTCTCTCTCTCTCTCTTTCCCCCNNNAGATCGGAAGAG
31176 TATGAGTGTGGAGTCGTTGCACAGGCTGAGATTCCTAGAGTCTNNNAGATCGGAA
31177 TATGAGTGTGGAGTCGTTGCGTGTGTCTGTGTGGTAAATGAGCNNNAGATCGGAA
31178 TATGAGTGTGGAGTCGTTGCAAACAGAGCCTCAAAGATGTGTGNNNAGATCGGAA
31179 TATGAGTGTGGAGTCGTTGCATAGAGAGCTCAGAAATAGGCCCNNNAGATCGGAA
31180 TATGAGTGTGGAGTCGTTGCTCCAAAAGCCTGAGGATTCTGAANNNAGATCGGAA
31183 TATGAGTGTGGAGTCGTTGCTTCCTTTTGTCATCTGCTCCCATNNNAGATCGGAAGA
31184 TATGAGTGTGGAGTCGTTGCACTCTCAGTTTCATCATCCCTGANNNAGATCGGAAGA
31185 TATGAGTGTGGAGTCGTTGCAAGTGCTGAGATTACAGGCATGANNNAGATCGGAA
31186 TATGAGTGTGGAGTCGTTGCGAATCCTAGCTCATTGTTTGGCCNNNAGATCGGAAG

31187 TATGAGTGTGGAGTCGTTGCGGCCCCAAACATCTGTATCAAAANNNAGATCGGAAG
31189 TATGAGTGTGGAGTCGTTGCTGCTCACAAAAGAAAACCCCATCNNNAGATCGGAAG
31190 TATGAGTGTGGAGTCGTTGCCCTTTCCAATTTGGGTGTCTTCCNNNAGATCGGAAG
31191 TATGAGTGTGGAGTCGTTGCACACCTTTTCATCACTAGGTTCANNNAGATCGGAAG
31192 TATGAGTGTGGAGTCGTTGCCCTCGTTTCTCTCCATTTTGCTCNNNAGATCGGAAGA
31193 TATGAGTGTGGAGTCGTTGCAAGGTAGGTTTCTGAATGGTCCCNNNAGATCGGAA
31194 TATGAGTGTGGAGTCGTTGCCCCCACTGGATTCTCTCTTCTTCNNNAGATCGGAAGA
31195 TATGAGTGTGGAGTCGTTGCACAAATACAAATGACCTTCGCCCNNNAGATCGGAAG
31196 TATGAGTGTGGAGTCGTTGCTCATAGCATATTCCACTCAGCCCNNNAGATCGGAAG
31197 TATGAGTGTGGAGTCGTTGCCTTTGCCATAACCTTCTTTCCCCNNNAGATCGGAAGA
31198 TATGAGTGTGGAGTCGTTGCGTGTGTATGTGTAACTTCCCCCTNNNAGATCGGAAG
31200 TATGAGTGTGGAGTCGTTGCATATGCCAAGTCTGCAGTCTACCNNNAGATCGGAAG
31201 TATGAGTGTGGAGTCGTTGCTTCAGACCCAGAAGAGTTGATGGNNNAGATCGGAA
31202 TATGAGTGTGGAGTCGTTGCAATTCACATCTATGGGCAATGCCNNNAGATCGGAAG
31203 TATGAGTGTGGAGTCGTTGCAGAGTGAAACCCCATCTCAAACANNNAGATCGGAAG
31204 TATGAGTGTGGAGTCGTTGCTCTGGTGTCTCTTCTCACAAACCNNNAGATCGGAAG
31205 TATGAGTGTGGAGTCGTTGCTTTTTGTGGTGACCATTGTGCTTNNNAGATCGGAAG
31207 TATGAGTGTGGAGTCGTTGCCCTCATGACCAAATTACCTCCCANNNAGATCGGAAGA
31208 TATGAGTGTGGAGTCGTTGCAGTGCAGCTCTTTTTACCTACCTNNNAGATCGGAAG
31209 TATGAGTGTGGAGTCGTTGCAAGGTGTTATAGGGGTGCAATGTNNNAGATCGGAA
31210 TATGAGTGTGGAGTCGTTGCTGGTAGAGTGGAAGCATACCCTANNNAGATCGGAA
31211 TATGAGTGTGGAGTCGTTGCTATTCTTGCTCTTGCTGGTGACTNNNAGATCGGAAG
31212 TATGAGTGTGGAGTCGTTGCAGTATCAGTTGAAGAGGAACGCANNNAGATCGGAA
31213 TATGAGTGTGGAGTCGTTGCTGATCCTCTTGGAACCCAAAGAGNNNAGATCGGAAG
31214 TATGAGTGTGGAGTCGTTGCGGCTTGAAAATTACAGTGGTGGTNNNAGATCGGAA
31215 TATGAGTGTGGAGTCGTTGCCATGTTGATGTTCTGGCTGACAGNNNAGATCGGAA
31216 TATGAGTGTGGAGTCGTTGCCGAGTAGCTGGGATTATAAGCGTNNNAGATCGGAA
31217 TATGAGTGTGGAGTCGTTGCTGAACTCACTGAATAGCTCCTGGNNNAGATCGGAAG
31218 TATGAGTGTGGAGTCGTTGCTACAGCTGCCACCTTATTCACTTNNNAGATCGGAAGA
31219 TATGAGTGTGGAGTCGTTGCCCAAATTAGCCAGAAACCACCAGNNNAGATCGGAAG
31220 TATGAGTGTGGAGTCGTTGCCTCCTCAAAAAGCAGAGCAAACANNNAGATCGGAAG
31221 TATGAGTGTGGAGTCGTTGCAGAAAGACACCCAAGACAAGACANNNAGATCGGAA
31222 TATGAGTGTGGAGTCGTTGCCTCTCTCTCTCAGATTGCTCACCNNNAGATCGGAAGA
31223 TATGAGTGTGGAGTCGTTGCATCTGGTGTTACAACTTCCCCTCNNNAGATCGGAAGA
31224 TATGAGTGTGGAGTCGTTGCTTCTGAGCCACCTAAATTTGGGANNNAGATCGGAAG
31225 TATGAGTGTGGAGTCGTTGCAGGTTTCTGCTGTGCTAATGGTANNNAGATCGGAA
31226 TATGAGTGTGGAGTCGTTGCGCCAAATCACTTGAAAGCCTTCTNNNAGATCGGAAG
31227 TATGAGTGTGGAGTCGTTGCAATTTGGTCTTGGGGTCTCTCTCNNNAGATCGGAAG
31228 TATGAGTGTGGAGTCGTTGCAAAGGAGAGTGAGAGAAAGAGCGNNNAGATCGGA
31229 TATGAGTGTGGAGTCGTTGCACTTCTCTCCTTCTCTCCCTCTCNNNAGATCGGAAGAG
31230 TATGAGTGTGGAGTCGTTGCTTACCAGTAACCAGAAGGCAGTCNNNAGATCGGAA
31231 TATGAGTGTGGAGTCGTTGCTTACCAGTAACCAGAAGGCAGTCNNNAGATCGGAA
31233 TATGAGTGTGGAGTCGTTGCGGATGTACACACAGACAGTCACTNNNAGATCGGAA

31234 TATGAGTGTGGAGTCGTTGCGGCTCTGCATACCAGGTTTTTTNNNAGATCGGAAG
31235 TATGAGTGTGGAGTCGTTGCCTCCCCATCTACACACATACTTNNNAGATCGGAAGA
31237 TATGAGTGTGGAGTCGTTGCAAGCAGTGGTTAGGGTTAGATGGNNNAGATCGGA
31238 TATGAGTGTGGAGTCGTTGCTTACACAGGCTATGCTGGGAAATNNNAGATCGGAA
31239 TATGAGTGTGGAGTCGTTGCAGTATAGTGTGCAGGGACCAAACNNNAGATCGGAA
31241 TATGAGTGTGGAGTCGTTGCACGGTCATTTCCACAGAGATTTTNNNAGATCGGAAG
31242 TATGAGTGTGGAGTCGTTGCAAGCAAGGGAAGAGAACAAGAGTNNNAGATCGGA
31243 TATGAGTGTGGAGTCGTTGCTAAATAGCTCCTCTTGCCACCAGNNNAGATCGGAAG
31244 TATGAGTGTGGAGTCGTTGCTGAACAAAAGCCATTGTGTCAGGNNNAGATCGGAA
31245 TATGAGTGTGGAGTCGTTGCCGCAGACCTCTCATTGACATTTCNNNAGATCGGAAG
31246 TATGAGTGTGGAGTCGTTGCTGTGCCCTTTCTGAAGATTAGCTNNNAGATCGGAAG
31247 TATGAGTGTGGAGTCGTTGCCTCCATGTTCTGCCCTCTAAACANNNAGATCGGAAGA
31248 TATGAGTGTGGAGTCGTTGCCTACTGATGAAGACCAGCCTCAGNNNAGATCGGAAG
31249 TATGAGTGTGGAGTCGTTGCCCCTTCTTTGGAGCAAGACAGTANNNAGATCGGAAG
31253 TATGAGTGTGGAGTCGTTGCAAGCTGTGATTGATTCTGAGTGCNNNAGATCGGAA
31254 TATGAGTGTGGAGTCGTTGCACGTGCACACAAATAGAAACAGGNNNAGATCGGAA
31255 TATGAGTGTGGAGTCGTTGCATCACTTAGGTCAGCAGACACAGNNNAGATCGGAA
31256 TATGAGTGTGGAGTCGTTGCACACACACACACACAGTCTAAGTNNNAGATCGGAAG
31257 TATGAGTGTGGAGTCGTTGCACGTTTTTCAGGTTGAGGTTTTTNNNAGATCGGAAG
31258 TATGAGTGTGGAGTCGTTGCCCCCAATGAATGATGAACCACTGNNNAGATCGGAAG
31259 TATGAGTGTGGAGTCGTTGCTACACACACACACACAAACACACNNNAGATCGGAAG
31260 TATGAGTGTGGAGTCGTTGCATGTAAATGCATGTGTGTGTACANNNAGATCGGAA
31261 TATGAGTGTGGAGTCGTTGCCCTATCCCACTTTGGACACTTGANNNAGATCGGAAG
31262 TATGAGTGTGGAGTCGTTGCCACAAGCACATACACATCAGCTCNNNAGATCGGAAG
31263 TATGAGTGTGGAGTCGTTGCACACACACACACACACACATTTTNNNAGATCGGAAGA
31265 TATGAGTGTGGAGTCGTTGCGAGGAATAGCATGGTAGGAAGCANNNAGATCGGA
31267 TATGAGTGTGGAGTCGTTGCAGGGCACAGAGAGAAAGTACTTGNNNAGATCGGAA
31268 TATGAGTGTGGAGTCGTTGCCACACAATTGGCCAACCTAACATNNNAGATCGGAAG
31269 TATGAGTGTGGAGTCGTTGCACTTGTATGTTTCAGTGCCTTGCNNNAGATCGGAAG
31270 TATGAGTGTGGAGTCGTTGCCCTCCACCCATCTCTCTTTGAAANNNAGATCGGAAGA
31272 TATGAGTGTGGAGTCGTTGCACACTGAAGGCTGAACAACTACANNNAGATCGGAA
31273 TATGAGTGTGGAGTCGTTGCAAGAGTGGGGTGTATTTGGATGTNNNAGATCGGAA
31274 TATGAGTGTGGAGTCGTTGCAGAGAGAGAGAGAGAGAGTGCAGNNNAGATCGGA
31275 TATGAGTGTGGAGTCGTTGCTCTTGCCACTAACTAGCTTCCTGNNNAGATCGGAAGA
31276 TATGAGTGTGGAGTCGTTGCAGGAAATGAGGCTAAGCAATCCANNNAGATCGGAA
31278 TATGAGTGTGGAGTCGTTGCGTAATCCAAAAGCTTTAGCCGCANNNAGATCGGAAG
31279 TATGAGTGTGGAGTCGTTGCCTGAGAACTTCTCCCTCCACATCNNNAGATCGGAAGA
31281 TATGAGTGTGGAGTCGTTGCATAGCACTTAGTGGTCATGGTGGNNNAGATCGGAA
31282 TATGAGTGTGGAGTCGTTGCGGCTGAAGTGTGTTTATGCTTGTNNNAGATCGGAA
31284 TATGAGTGTGGAGTCGTTGCAAACCCTCATTTGAAACCCTTGGNNNAGATCGGAAG
31286 TATGAGTGTGGAGTCGTTGCGCAGAACAGTCCACACCATTTACNNNAGATCGGAAG
31287 TATGAGTGTGGAGTCGTTGCAACAACAAACTCTCCCAAGAAGANNNAGATCGGAAG
31288 TATGAGTGTGGAGTCGTTGCAAAAAGCAGGAAGTGGTGGTAGANNNAGATCGGA

31289 TATGAGTGTGGAGTCGTTGCGTGAGCTATGATCACACCAATGCNNNAGATCGGAAG
31290 TATGAGTGTGGAGTCGTTGCAGGGGGAGAATTAGGAGAACAGANNNAGATCGGA
31291 TATGAGTGTGGAGTCGTTGCCAAACTGCTCAGCTCTCATTGACNNNAGATCGGAAGA
31292 TATGAGTGTGGAGTCGTTGCAATTTCTTGAGGACTGGGACAGCNNNAGATCGGAAG
31293 TATGAGTGTGGAGTCGTTGCAGAAGATGGATATGACCAGTGGCNNNAGATCGGAA
31294 TATGAGTGTGGAGTCGTTGCGTCTTTGCCTCAGCTTCCTCATANNNAGATCGGAAGA
31296 TATGAGTGTGGAGTCGTTGCATCTTGCTCTTTTGTCCAGGCTANNNAGATCGGAAGA
31297 TATGAGTGTGGAGTCGTTGCTAATATTTGGCAAGGAGAGGGGGNNNAGATCGGAA
31298 TATGAGTGTGGAGTCGTTGCACACATACATGCACCCAAATTGGNNNAGATCGGAAG
31299 TATGAGTGTGGAGTCGTTGCTGTATGAAAGACAGACTGGCCAANNNAGATCGGAAG
31300 TATGAGTGTGGAGTCGTTGCATGGAGGAATTAGGGACATTGGTNNNAGATCGGAA
31302 TATGAGTGTGGAGTCGTTGCCCCCGAGATGAAACACTGAGTAANNNAGATCGGAAG
31303 TATGAGTGTGGAGTCGTTGCAGTTGGGGTAGGGAGTAGAGAAANNNAGATCGGA
31304 TATGAGTGTGGAGTCGTTGCCCCACACACACTAACCAGAGTATNNNAGATCGGAAGA
31306 TATGAGTGTGGAGTCGTTGCGCGTGTGTGTGTGTTGTAAGTTANNNAGATCGGAAG
31307 TATGAGTGTGGAGTCGTTGCCCCAAGTTTATCCCAGTTTGTGGNNNAGATCGGAAGA
31308 TATGAGTGTGGAGTCGTTGCACCATTTGTTTTATGTGGCCTGGNNNAGATCGGAAG
31309 TATGAGTGTGGAGTCGTTGCAGAGAGAGAGAGAGTTTGGAGCTNNNAGATCGGAA
31311 TATGAGTGTGGAGTCGTTGCCGAGACTGGATATGCGGAACTAANNNAGATCGGAAG
31312 TATGAGTGTGGAGTCGTTGCGAGAGAGAGAACCAAAGCCTGAANNNAGATCGGAA
31314 TATGAGTGTGGAGTCGTTGCACTAAAGCTGCAAAACTAAGGCCNNNAGATCGGAAG
31315 TATGAGTGTGGAGTCGTTGCAAAATGGGAGAAGGGGAGCTAAGNNNAGATCGGA
31316 TATGAGTGTGGAGTCGTTGCAGGCACAGCTAGTATTCCATTGTNNNAGATCGGAAG
31317 TATGAGTGTGGAGTCGTTGCGCTGTATTTGTAACCCATGCAGTNNNAGATCGGAAG
31318 TATGAGTGTGGAGTCGTTGCCAAACTCTATCACGCCAAACACANNNAGATCGGAAGA
31319 TATGAGTGTGGAGTCGTTGCCTGGGAACCATGGACCATAGTAGNNNAGATCGGAAG
31321 TATGAGTGTGGAGTCGTTGCGATCTTTGCACACTCCAGTTTCCNNNAGATCGGAAGA
31322 TATGAGTGTGGAGTCGTTGCGTGATTCCACATGCTTTGCAGTANNNAGATCGGAAG
31323 TATGAGTGTGGAGTCGTTGCTCTGTTTACCTGTCTGTCTTCCCNNNAGATCGGAAGA
31324 TATGAGTGTGGAGTCGTTGCTCAAGCAAAAAACACATCTTGCCNNNAGATCGGAAGA
31325 TATGAGTGTGGAGTCGTTGCCTCCTTCCTTTTGAGACCTCGAANNNAGATCGGAAGA
31326 TATGAGTGTGGAGTCGTTGCACAGTGCAAGAAAGTCCAGTTTTNNNAGATCGGAAG
31327 TATGAGTGTGGAGTCGTTGCCTCACAGGAGCACTAATCCTACCNNNAGATCGGAAGA
31328 TATGAGTGTGGAGTCGTTGCGGTATACAAAACAGGCTCTGGGANNNAGATCGGAAG
31329 TATGAGTGTGGAGTCGTTGCACACAGGGCTTTGCAGGATATATNNNAGATCGGAAG
31330 TATGAGTGTGGAGTCGTTGCGTTTCTCCTTCTCTGAGGAACCCNNNAGATCGGAAGA
31331 TATGAGTGTGGAGTCGTTGCAGAGACTCAAGGTGTAGCTTACCNNNAGATCGGAAG
31332 TATGAGTGTGGAGTCGTTGCTGACTGAAGGAGCTGAGGGATATNNNAGATCGGAA
31334 TATGAGTGTGGAGTCGTTGCGCTCTTCCACATCTATTGGGTCANNNAGATCGGAAGA
31335 TATGAGTGTGGAGTCGTTGCAACCTTGCAATGTGTTAGCTTCCNNNAGATCGGAAGA
31336 TATGAGTGTGGAGTCGTTGCATTCTAAAGAGGAGCTGCCAGACNNNAGATCGGAAG
31337 TATGAGTGTGGAGTCGTTGCCAACGTGGGCAATTGGATCATAANNNAGATCGGAAG
31338 TATGAGTGTGGAGTCGTTGCAGATTGAAAAGGTGGCCTAGGAGNNNAGATCGGAA

31339 TATGAGTGTGGAGTCGTTGCGCTCAGGCTATCTCGAATAGTGANNNAGATCGGAAG
31340 TATGAGTGTGGAGTCGTTGCCCCCTTTTACCCCGGAGAATAAANNNAGATCGGAAGA
31341 TATGAGTGTGGAGTCGTTGCCTACCTGGCTACCTCACTTGTTTNNNAGATCGGAAGA
31342 TATGAGTGTGGAGTCGTTGCTTCTTGCCTTCTCCTTTATCCCCNNNAGATCGGAAGAG
31343 TATGAGTGTGGAGTCGTTGCCAGAGATGCTGCTAAACACCCTANNNAGATCGGAAG
31344 TATGAGTGTGGAGTCGTTGCAGCAGTGTGTCTTTGATTATGGCNNNAGATCGGAAG
31345 TATGAGTGTGGAGTCGTTGCTCTGCAAGCATTCCAAAGGAAAGNNNAGATCGGAAG
31346 TATGAGTGTGGAGTCGTTGCGTGCTGAGGTTGGGAAAATTTGANNNAGATCGGAA
31348 TATGAGTGTGGAGTCGTTGCCTCTGTGCCCTCTAGTTGAAGAGNNNAGATCGGAAG
31349 TATGAGTGTGGAGTCGTTGCTGATTCCTGTGGCTTGATTCAGANNNAGATCGGAAG
31351 TATGAGTGTGGAGTCGTTGCGAGAGAGGGAGGGTGTAAATGTGNNNAGATCGGA
31352 TATGAGTGTGGAGTCGTTGCCCACCACTGCTAACTGGACTAATNNNAGATCGGAAGA
31353 TATGAGTGTGGAGTCGTTGCAAACCTCTGACCTATTTGGGAGTNNNAGATCGGAAG
31354 TATGAGTGTGGAGTCGTTGCGCAGAACTGGTGACACATTTACCNNNAGATCGGAAG
31355 TATGAGTGTGGAGTCGTTGCGCCTTCAGCTGATTAGATGATGCNNNAGATCGGAAG
31356 TATGAGTGTGGAGTCGTTGCACATATTCGGAGAAGGGGTGATGNNNAGATCGGAA
31357 TATGAGTGTGGAGTCGTTGCTTCCTGGCTACTGCTTATGTTCANNNAGATCGGAAGA
31358 TATGAGTGTGGAGTCGTTGCTGTAAGACCAACTGTCCTAACGTNNNAGATCGGAAG
31359 TATGAGTGTGGAGTCGTTGCGCATAACCTTCTGGGCTATCTGANNNAGATCGGAAG
31360 TATGAGTGTGGAGTCGTTGCCCCACTCCAAAAACATTGCTGAANNNAGATCGGAAGA
31361 TATGAGTGTGGAGTCGTTGCCATCCCACTCCAAAAACATTGCTNNNAGATCGGAAGA
31362 TATGAGTGTGGAGTCGTTGCGAGTCAAAGTTCTCCTGTTCCCTNNNAGATCGGAAGA
31363 TATGAGTGTGGAGTCGTTGCCCAGTGGTCTTTCTGAAGGAGAANNNAGATCGGAAG
31364 TATGAGTGTGGAGTCGTTGCGTGGTCAGATGAGCAAGTTGTTCNNNAGATCGGAAG
31365 TATGAGTGTGGAGTCGTTGCGCTGTAAGAGGCCTTCGTATTCTNNNAGATCGGAAG
31366 TATGAGTGTGGAGTCGTTGCAGAGAGAGAGAGAGGGAAAGAGGNNNAGATCGGA
31367 TATGAGTGTGGAGTCGTTGCTATTGTGTGGGGTGCATACAAGANNNAGATCGGAAG
31368 TATGAGTGTGGAGTCGTTGCACTGTGTGTATGGAAGCGGAATANNNAGATCGGAA
31369 TATGAGTGTGGAGTCGTTGCAACTGAAATTTCACCTCCCCCAANNNAGATCGGAAGA
31370 TATGAGTGTGGAGTCGTTGCCAGTTCGTTGCTCAAGCCTAAAANNNAGATCGGAAG
31371 TATGAGTGTGGAGTCGTTGCATGAGGCAGTTGAACTTGTGAGANNNAGATCGGAA
31373 TATGAGTGTGGAGTCGTTGCTCAAAAAGTATGCTGCGTGAGTGNNNAGATCGGAAG
31374 TATGAGTGTGGAGTCGTTGCCCTAAGATGAAAGAGACAGGGCANNNAGATCGGAA
31375 TATGAGTGTGGAGTCGTTGCTGATCCAGAAAAACTTAGGCCCANNNAGATCGGAAG
31377 TATGAGTGTGGAGTCGTTGCTGGCAGCAAGTAATATCCACATGNNNAGATCGGAAG
31378 TATGAGTGTGGAGTCGTTGCATTGCTGGAAGTGGAAAAACACANNNAGATCGGAAG
31379 TATGAGTGTGGAGTCGTTGCAGATGAATGAAGGAGAGGAGGGANNNAGATCGGA
31380 TATGAGTGTGGAGTCGTTGCGTTGTCTTTACCTGGGGCTTCTANNNAGATCGGAAGA
31381 TATGAGTGTGGAGTCGTTGCTTGATGCTGAAGTGAATTGTGGANNNAGATCGGAAG
31382 TATGAGTGTGGAGTCGTTGCACACACACACACACACACAAAATNNNAGATCGGAAGA
31383 TATGAGTGTGGAGTCGTTGCTTCAAGTCTGAAATCTGCAGGGANNNAGATCGGAAG
31384 TATGAGTGTGGAGTCGTTGCTTGGAAAGAGAAATGCACACACANNNAGATCGGAAG
31385 TATGAGTGTGGAGTCGTTGCCCTGTGGAGGTGAAAACATTTGANNNAGATCGGAAG

31386 TATGAGTGTGGAGTCGTTGCGGAATGTGTTTGTGCGTATGGAANNNAGATCGGAA
31387 TATGAGTGTGGAGTCGTTGCAAGGGCTGCTGATGATTTTTCACNNNAGATCGGAAG
31388 TATGAGTGTGGAGTCGTTGCTAAGGCCCACTGTACTAAGGAGANNNAGATCGGAA
31389 TATGAGTGTGGAGTCGTTGCTCCCATAGGTACTTCTCTTGGGANNNAGATCGGAAG
31390 TATGAGTGTGGAGTCGTTGCTTAGTGTGAAACTAGCCAGGACCNNNAGATCGGAA
31391 TATGAGTGTGGAGTCGTTGCATCACCACTTGTACTCTTCTGGGNNNAGATCGGAAG
31392 TATGAGTGTGGAGTCGTTGCGTGTGTGTGTGTGTGTGTTTGTANNNAGATCGGAA
31393 TATGAGTGTGGAGTCGTTGCAAGGAAGGAGGAAGCAACAAGAANNNAGATCGGA
31394 TATGAGTGTGGAGTCGTTGCCACAAGTCACTGAAGCTCCTTTGNNNAGATCGGAAG
31395 TATGAGTGTGGAGTCGTTGCGTATTTTCCTCCCCTTTATGCGCNNNAGATCGGAAGA
31396 TATGAGTGTGGAGTCGTTGCGGCTGTGTGAAAATATGTGCCAANNNAGATCGGAA
31397 TATGAGTGTGGAGTCGTTGCACCCTTTAGTCGTAGTTGTGAGGNNNAGATCGGAA
31398 TATGAGTGTGGAGTCGTTGCAGAGGGGAAAGGAGATGAGGTTANNNAGATCGGA
31399 TATGAGTGTGGAGTCGTTGCTAGAAAAAGGAAGAGAGTGGGGCNNNAGATCGGA
31400 TATGAGTGTGGAGTCGTTGCAGTACAGTGTTCACTACTGCCATNNNAGATCGGAAG
31401 TATGAGTGTGGAGTCGTTGCTTCTTCTCCCCAGATACCAATGGNNNAGATCGGAAG
31402 TATGAGTGTGGAGTCGTTGCTTGCTTATGGTGCTAGGTACAGGNNNAGATCGGAA
31403 TATGAGTGTGGAGTCGTTGCGGAAATGCAATCACAGATGCAGANNNAGATCGGAA
31405 TATGAGTGTGGAGTCGTTGCCCTTTACCTGCAATAACCTCCCTNNNAGATCGGAAGA
31408 TATGAGTGTGGAGTCGTTGCCTTGGAGTCTGACATCCCACTAGNNNAGATCGGAAG
31409 TATGAGTGTGGAGTCGTTGCTGTGTGTGTGTGTGTGATGAATGNNNAGATCGGAA
31410 TATGAGTGTGGAGTCGTTGCCCTCTTTGAGCAGGTGACATTTGNNNAGATCGGAAG
31412 TATGAGTGTGGAGTCGTTGCCACACACACACACACACTCATTTNNNAGATCGGAAGA
31413 TATGAGTGTGGAGTCGTTGCACATTTCATAGCTCAGTGCAGTGNNNAGATCGGAAG
31414 TATGAGTGTGGAGTCGTTGCTCAACTCTTCCCCTCGATAGACTNNNAGATCGGAAGA
31415 TATGAGTGTGGAGTCGTTGCATTAGGCCACAGTCCTCTTTGAGNNNAGATCGGAAG
31416 TATGAGTGTGGAGTCGTTGCAGCAACTGGTGAATCTAGGTGAGNNNAGATCGGAA
31417 TATGAGTGTGGAGTCGTTGCTGAGTCGTTCTGTTTTGTTGGAGNNNAGATCGGAA
31418 TATGAGTGTGGAGTCGTTGCTGTGTGTGACTATCAATGCAGGANNNAGATCGGAA
31419 TATGAGTGTGGAGTCGTTGCGTGAGTTGCATCCTAAGGGAGAANNNAGATCGGAA
31421 TATGAGTGTGGAGTCGTTGCGAGAGACGTGGTGGGAAGTAATTNNNAGATCGGA
31423 TATGAGTGTGGAGTCGTTGCTGGTATCAAAGTTGGGACCAGAGNNNAGATCGGAA
31424 TATGAGTGTGGAGTCGTTGCTCTGGAATCTTTAGGCTTCCCAGNNNAGATCGGAAG
31427 TATGAGTGTGGAGTCGTTGCAACCATTTCATCTGCTTTGAGGCNNNAGATCGGAAG
31428 TATGAGTGTGGAGTCGTTGCTTATTGGGCAACAGGCAGATACTNNNAGATCGGAA
31429 TATGAGTGTGGAGTCGTTGCATAGCAGAACCAATTGTGCCCTANNNAGATCGGAAG
31430 TATGAGTGTGGAGTCGTTGCAGAACATTCTATCGGAGATGGGCNNNAGATCGGAA
31431 TATGAGTGTGGAGTCGTTGCCAAGGCCAGAGATGTTGCTAAACNNNAGATCGGAA
31432 TATGAGTGTGGAGTCGTTGCATTGAGAAACAGAGCCTCTGAGGNNNAGATCGGAA
31433 TATGAGTGTGGAGTCGTTGCTAGAGATCTAGGAATCGCCAGGANNNAGATCGGAA
31435 TATGAGTGTGGAGTCGTTGCACAGGGACAATGCATAGAACAGANNNAGATCGGAA
31437 TATGAGTGTGGAGTCGTTGCGGGGGTTGAGGATAAGAGTAAGCNNNAGATCGGA
31438 TATGAGTGTGGAGTCGTTGCTGGAGGAAAAAACACAAACCACCNNNAGATCGGAA

31439 TATGAGTGTGGAGTCGTTGCCTGTATCTACTTCGTGCTCCTCCNNNAGATCGGAAGA
31440 TATGAGTGTGGAGTCGTTGCATCTTGAGGTCAGGAGATCGAGANNNAGATCGGAA
31441 TATGAGTGTGGAGTCGTTGCAGAACAGAGACTGTACCTCTCCANNNAGATCGGAAG
31442 TATGAGTGTGGAGTCGTTGCTGAGCTCAAACTACATATGCAGTNNNAGATCGGAAG
31443 TATGAGTGTGGAGTCGTTGCCATTCTCACTAAGCCTGACCTGTNNNAGATCGGAAG
31444 TATGAGTGTGGAGTCGTTGCTCTGTCTCTCTTCCTTCCTGTCTNNNAGATCGGAAGA
31445 TATGAGTGTGGAGTCGTTGCACTCTAGCAAGCATCCTAAGTCCNNNAGATCGGAAG
31446 TATGAGTGTGGAGTCGTTGCTTCAGGCTTATCCATCTACTGCCNNNAGATCGGAAG
31447 TATGAGTGTGGAGTCGTTGCAAGCTTCATCTCTAACATTGGCTNNNAGATCGGAAG
31448 TATGAGTGTGGAGTCGTTGCTGCTGTACTTCAACCCGTAGATCNNNAGATCGGAAG
31449 TATGAGTGTGGAGTCGTTGCTTCTCATGCTTCCTCTCCACTTCNNNAGATCGGAAGA
31451 TATGAGTGTGGAGTCGTTGCCATGCATGTGGTGACAGTGAATTNNNAGATCGGAA
31452 TATGAGTGTGGAGTCGTTGCCTTCACAGTAGACTGCAGCAAAGNNNAGATCGGAAG
31453 TATGAGTGTGGAGTCGTTGCTTGTGCTGGATGTCAGTGATTCTNNNAGATCGGAAG
31454 TATGAGTGTGGAGTCGTTGCAAGCATGTCAATCTTACCCCCAANNNAGATCGGAAG
31455 TATGAGTGTGGAGTCGTTGCAGTCAGCTAATGATGAGGTCCACNNNAGATCGGAA
31456 TATGAGTGTGGAGTCGTTGCTCAGGGAAATGCAAACCAAATCCNNNAGATCGGAAG
31459 TATGAGTGTGGAGTCGTTGCTCAAAATAACCAAGGGGCTCAGANNNAGATCGGAA
31461 TATGAGTGTGGAGTCGTTGCAGGCACAAAAAACAGACAACACANNNAGATCGGAA
31462 TATGAGTGTGGAGTCGTTGCCCCCACACACTTATACCCTCATTNNNAGATCGGAAGA
31463 TATGAGTGTGGAGTCGTTGCGGGGAGAGAGAGAGAAAGAGAGANNNAGATCGG
31464 TATGAGTGTGGAGTCGTTGCAAAGAGAGAGACAGAGAGGGGAANNNAGATCGGA
31466 TATGAGTGTGGAGTCGTTGCAGATGACAAAGCTCTAGCCAACANNNAGATCGGAAG
31467 TATGAGTGTGGAGTCGTTGCTGTTCAACCTCATATCCCCAGTGNNNAGATCGGAAG
31468 TATGAGTGTGGAGTCGTTGCCACTCTCACACTCTTCTTGGGAANNNAGATCGGAAG
31469 TATGAGTGTGGAGTCGTTGCACTTTGTCCTGTTCACTGCTGTANNNAGATCGGAAG
31470 TATGAGTGTGGAGTCGTTGCCTCCCTATTTCCTTTCCCTGCTTNNNAGATCGGAAGA
31471 TATGAGTGTGGAGTCGTTGCCCCAGCCAGATTCTATCATCACANNNAGATCGGAAG
31472 TATGAGTGTGGAGTCGTTGCATGACTCCCAAACATCCAGTCAANNNAGATCGGAAG
31473 TATGAGTGTGGAGTCGTTGCCACACACACACACCATGAAATATNNNAGATCGGAAG
31474 TATGAGTGTGGAGTCGTTGCCCCCTTCTCCTGTCAGAATCATTNNNAGATCGGAAGA
31475 TATGAGTGTGGAGTCGTTGCCCCCTTCTCCTGTCAGAATCATTNNNAGATCGGAAGA
31476 TATGAGTGTGGAGTCGTTGCGGGTGGTTGTTCATCCTGTATCTNNNAGATCGGAAG
31477 TATGAGTGTGGAGTCGTTGCGTGTGTGTGTGTGTGTGTGTTTANNNAGATCGGAA
31478 TATGAGTGTGGAGTCGTTGCACTGAGCACACCTTGGTTTCTAANNNAGATCGGAAG
31479 TATGAGTGTGGAGTCGTTGCTGTTGACTTTTGTTGGTCTTGCANNNAGATCGGAAG
31480 TATGAGTGTGGAGTCGTTGCGGCTAAGATGACACGTAGCTACANNNAGATCGGAA
31482 TATGAGTGTGGAGTCGTTGCACCATATTCAGTCAGTGCAGGAANNNAGATCGGAAG
31483 TATGAGTGTGGAGTCGTTGCTCTAAGGATTCAGGTGTGCAGTCNNNAGATCGGAA
31484 TATGAGTGTGGAGTCGTTGCTCACAGACACATATGCACAGACANNNAGATCGGAAG
31485 TATGAGTGTGGAGTCGTTGCAAACTTCACGCCAATATTCTGCANNNAGATCGGAAG
31486 TATGAGTGTGGAGTCGTTGCACATGCTTACGTGAACATTCTGCNNNAGATCGGAAG
31487 TATGAGTGTGGAGTCGTTGCGCCATGAGCCAAGAAATGAAGTTNNNAGATCGGAA

31488 TATGAGTGTGGAGTCGTTGCAACAGGAGTGATGAAAGTGGACANNNAGATCGGA
31489 TATGAGTGTGGAGTCGTTGCGTGACGTACATCAATTTGCCGATNNNAGATCGGAAG
31490 TATGAGTGTGGAGTCGTTGCTCTGAAACCTCTTGTCCATTCCCNNNAGATCGGAAGA
31491 TATGAGTGTGGAGTCGTTGCTGTCAGTAATCTCTGAGTGCTGANNNAGATCGGAAG
31493 TATGAGTGTGGAGTCGTTGCGGCAATCTGTATTATCAAGGGCCNNNAGATCGGAA
31494 TATGAGTGTGGAGTCGTTGCAGAGAAGAGAGAGAGATCTGGCANNNAGATCGGA
31497 TATGAGTGTGGAGTCGTTGCTGGTCTCATAGGTTATTTGGGCANNNAGATCGGAA
31498 TATGAGTGTGGAGTCGTTGCGGGGCAATTTAGTCAGAAGTTGTNNNAGATCGGAA
31499 TATGAGTGTGGAGTCGTTGCTCTGTTGCTATAACGGCACTAGTNNNAGATCGGAAG
31500 TATGAGTGTGGAGTCGTTGCCTCTCCCTCTCTTCAACAGGTTCNNNAGATCGGAAGA
31501 TATGAGTGTGGAGTCGTTGCGGGTGAAGGAGAAGGGTTGTTATNNNAGATCGGA
31502 TATGAGTGTGGAGTCGTTGCCTCATGTTGACATTTCGAGGCAGNNNAGATCGGAAG
31503 TATGAGTGTGGAGTCGTTGCACCCCTATTTTCCTCTCTCACCTNNNAGATCGGAAGA
31504 TATGAGTGTGGAGTCGTTGCTCCTTTGACCAGGCAACTTATCTNNNAGATCGGAAG
31505 TATGAGTGTGGAGTCGTTGCCTCCTCCAGGTTTCTAATCTGCANNNAGATCGGAAGA
31506 TATGAGTGTGGAGTCGTTGCAATCTGGCAAGAATGGACACTCANNNAGATCGGAA
31507 TATGAGTGTGGAGTCGTTGCGGGAGCCCTCATCTCTACAAAAANNNAGATCGGAAG
31509 TATGAGTGTGGAGTCGTTGCACACACACACACACACAAATACCNNNAGATCGGAAG
31510 TATGAGTGTGGAGTCGTTGCACACACACACACACAAACACATTNNNAGATCGGAAG
31511 TATGAGTGTGGAGTCGTTGCTTTATTAGGCAGAGCAAGGTGGANNNAGATCGGAA
31512 TATGAGTGTGGAGTCGTTGCCAGGGTCTCATCAAGAGCATTCTNNNAGATCGGAAG
31513 TATGAGTGTGGAGTCGTTGCCCTTGTGGTTCATGTTTCAGCAANNNAGATCGGAAG
31514 TATGAGTGTGGAGTCGTTGCACTGACAGAGAGGTGTTGAAGTCNNNAGATCGGAA
31515 TATGAGTGTGGAGTCGTTGCACCTGTCAATATCTGCCTGTGAGNNNAGATCGGAAG
31517 TATGAGTGTGGAGTCGTTGCATATGTGTGTGTGTGTGTGTGTGNNNAGATCGGAA
31519 TATGAGTGTGGAGTCGTTGCTACCCATACATGCACATTGTCCANNNAGATCGGAAG
31520 TATGAGTGTGGAGTCGTTGCCTTTGTCTGGAAGAACAGAGGGANNNAGATCGGAA
31521 TATGAGTGTGGAGTCGTTGCTGTTAGGCAGGTTATTGAGCTGTNNNAGATCGGAA
31522 TATGAGTGTGGAGTCGTTGCTGATGGCTACAAGGCATTCCATANNNAGATCGGAAG
31523 TATGAGTGTGGAGTCGTTGCGCTTTCTCCTGGTACCTGAGAAANNNAGATCGGAAG
31524 TATGAGTGTGGAGTCGTTGCCCACATATTGTCCCAGTTCTCCANNNAGATCGGAAGA
31525 TATGAGTGTGGAGTCGTTGCTGTACCCAATGTCTCCAATCCTCNNNAGATCGGAAGA
31526 TATGAGTGTGGAGTCGTTGCGACAATGGTAGAACGTTGCCAAANNNAGATCGGAA
31527 TATGAGTGTGGAGTCGTTGCTCCAAAACACAGAAATCACTGCCNNNAGATCGGAAG
31528 TATGAGTGTGGAGTCGTTGCATCCTCACTCCAAAAAACCCACTNNNAGATCGGAAGA
31529 TATGAGTGTGGAGTCGTTGCGGATTTTGAGAGAACAAGGCCACNNNAGATCGGAA
31530 TATGAGTGTGGAGTCGTTGCGTCACCAAGCTCCATCTAGTCAANNNAGATCGGAAG
31532 TATGAGTGTGGAGTCGTTGCTCTGAGGTGTAGGAAAGAAAGCCNNNAGATCGGAA
31533 TATGAGTGTGGAGTCGTTGCATCTGTTTCCCCATTCTTTTGCANNNAGATCGGAAGA
31534 TATGAGTGTGGAGTCGTTGCGGGTAATAAGAAGAGAGAGCCGGNNNAGATCGGA
31535 TATGAGTGTGGAGTCGTTGCACTTCCCAATCCAACAGTAATGANNNAGATCGGAAG
31536 TATGAGTGTGGAGTCGTTGCACTCCTTACACAAAGACAGCACTNNNAGATCGGAAG
31537 TATGAGTGTGGAGTCGTTGCCCTTATAGGGAGGTCTCTTGGGANNNAGATCGGAA

31538 TATGAGTGTGGAGTCGTTGCTTCCATCCATAAGCTTGAGACCCNNNAGATCGGAAG
31539 TATGAGTGTGGAGTCGTTGCGTGTGTGAGACAGACTTTTGCTCNNNAGATCGGAA
31540 TATGAGTGTGGAGTCGTTGCGTGTGTGAGACAGACTTTTGCTCNNNAGATCGGAA
31541 TATGAGTGTGGAGTCGTTGCCAGGATGCAAAGACTTAGTACCANNNAGATCGGAA
31542 TATGAGTGTGGAGTCGTTGCGATGATGGGGTTGAGATGACACTNNNAGATCGGAA
31543 TATGAGTGTGGAGTCGTTGCTTCTTGCCTGATATTCCCTGACCNNNAGATCGGAAGA
31544 TATGAGTGTGGAGTCGTTGCGAGAGAAAGAGGGGGAGAGAGAGNNNAGATCGG
31545 TATGAGTGTGGAGTCGTTGCAAGCAGAAGTTCAAAGCATGACCNNNAGATCGGAA
31546 TATGAGTGTGGAGTCGTTGCAGCAAGCTTCTACCTATTGTGGTNNNAGATCGGAAG
31547 TATGAGTGTGGAGTCGTTGCGATGTCCTTTGTCCTTTTGCTCCNNNAGATCGGAAGA
31548 TATGAGTGTGGAGTCGTTGCCCCCTCTGTTGCCTTTAGAAAGANNNAGATCGGAAG
31549 TATGAGTGTGGAGTCGTTGCGTGCCATGTTTCTTGTACAGCATNNNAGATCGGAAG
31550 TATGAGTGTGGAGTCGTTGCCCATGATTCCAACCCTCTACCATNNNAGATCGGAAGA
31551 TATGAGTGTGGAGTCGTTGCTCCTGCTTCATTTTCGGAAACACNNNAGATCGGAAG
31552 TATGAGTGTGGAGTCGTTGCACTGCCTATTTCCCCTAACCAAGNNNAGATCGGAAGA
31553 TATGAGTGTGGAGTCGTTGCTCTGCTCCCTAAGAGTATGAGGTNNNAGATCGGAAG
31554 TATGAGTGTGGAGTCGTTGCACATCAAAATTCGTGGTCCCTCTNNNAGATCGGAAG
31555 TATGAGTGTGGAGTCGTTGCACATCAAAATTCGTGGTCCCTCTNNNAGATCGGAAG
31556 TATGAGTGTGGAGTCGTTGCTGGCAGAGACAAGTGTACATCAANNNAGATCGGAA
31557 TATGAGTGTGGAGTCGTTGCAAAGCTCTGGAGGCTACAAGTACNNNAGATCGGAA
31560 TATGAGTGTGGAGTCGTTGCTGTCCACTTGAATGGTGACTACANNNAGATCGGAAG
31561 TATGAGTGTGGAGTCGTTGCCCCATAACCCAGAGTCAACTACANNNAGATCGGAAG
31562 TATGAGTGTGGAGTCGTTGCGAGTCATACAGTGAGAAGCAGCTNNNAGATCGGAA
31563 TATGAGTGTGGAGTCGTTGCACTCCTGACAAAGCTCATACAGANNNAGATCGGAAG
31565 TATGAGTGTGGAGTCGTTGCACTCAAGTGTACACACAGAAGCANNNAGATCGGAA
31566 TATGAGTGTGGAGTCGTTGCTGATGGGTTAGTTTTGCATCAGTNNNAGATCGGAA
31567 TATGAGTGTGGAGTCGTTGCCAGGACTAAGACACCTATCACCCNNNAGATCGGAAG
31568 TATGAGTGTGGAGTCGTTGCTAGATGGCACTCAATCTGTGGTGNNNAGATCGGAA
31569 TATGAGTGTGGAGTCGTTGCTTCACAGGGCTAAAGGTTATGGGNNNAGATCGGAA
31570 TATGAGTGTGGAGTCGTTGCTTGATGACCTTGACAGTGATGCTNNNAGATCGGAAG
31571 TATGAGTGTGGAGTCGTTGCCTGCTGTATTTCCACCTTTCACANNNAGATCGGAAGA
31572 TATGAGTGTGGAGTCGTTGCAGAGGCACTGTGATCAAGAACACNNNAGATCGGAA
31573 TATGAGTGTGGAGTCGTTGCGCACTCACTGAATTCCACACAAANNNAGATCGGAAG
31574 TATGAGTGTGGAGTCGTTGCGAAGAAAGACAGAGAGCATGCACNNNAGATCGGAA
31575 TATGAGTGTGGAGTCGTTGCACTAGAACAGTTGGGCACATCTGNNNAGATCGGAA
31577 TATGAGTGTGGAGTCGTTGCTGAGTCACAGTAGCACATTTCTTNNNAGATCGGAAG
31578 TATGAGTGTGGAGTCGTTGCGTGACTCCACAACAAACTGGATGNNNAGATCGGAA
31579 TATGAGTGTGGAGTCGTTGCTCAACACCTCACTTTCAGCATTGNNNAGATCGGAAGA
31580 TATGAGTGTGGAGTCGTTGCCATCTAACCAAGTGGGCAAAAGGNNNAGATCGGAA
31581 TATGAGTGTGGAGTCGTTGCTGTGTGTGTGTGTTTGTATCTGTNNNAGATCGGAAG
31582 TATGAGTGTGGAGTCGTTGCACACAGACACATACACACACAGANNNAGATCGGAAG
31583 TATGAGTGTGGAGTCGTTGCCATGGTAGTACTCTGGGAACACANNNAGATCGGAA
31584 TATGAGTGTGGAGTCGTTGCCTCACGTGAGGGATGATCAAGTCNNNAGATCGGAA

31585 TATGAGTGTGGAGTCGTTGCAGACCTTTTCACACTCCTCACTCNNNAGATCGGAAGA
31586 TATGAGTGTGGAGTCGTTGCACATGGTTGATCAAGATCGGAGTNNNAGATCGGAA
31588 TATGAGTGTGGAGTCGTTGCTTCCTGCTACGACACTGAATTCANNNAGATCGGAAG
31589 TATGAGTGTGGAGTCGTTGCAATGGATCAGTGGATGGATGCATNNNAGATCGGAA
31590 TATGAGTGTGGAGTCGTTGCTTTGGGGCAATCCAGAAATCAACNNNAGATCGGAAG
31592 TATGAGTGTGGAGTCGTTGCTGAACAGCTAATTCCAACCAACANNNAGATCGGAAG
31595 TATGAGTGTGGAGTCGTTGCCGACTTGCCTACTGGAAAACAAGNNNAGATCGGAAG
31596 TATGAGTGTGGAGTCGTTGCGGATCCCAACCACTCATTCAGAANNNAGATCGGAAG
31597 TATGAGTGTGGAGTCGTTGCCACACACACACACACACATACTCNNNAGATCGGAAGA
31598 TATGAGTGTGGAGTCGTTGCACAACTCGTGTACCTAGAGGACTNNNAGATCGGAAG
31599 TATGAGTGTGGAGTCGTTGCTTTGTGTATGGTGAGAGACAGGGNNNAGATCGGAA
31601 TATGAGTGTGGAGTCGTTGCCCCCACAAATCAGTGCATGTAAANNNAGATCGGAAG
31602 TATGAGTGTGGAGTCGTTGCCATTGTCAGTCCTCCACCTACTCNNNAGATCGGAAGA
31603 TATGAGTGTGGAGTCGTTGCAACGTAAGGGAGAGATAGGTGGANNNAGATCGGA
31604 TATGAGTGTGGAGTCGTTGCGAACTCAAGGAACTGAAACTGGCNNNAGATCGGAA
31605 TATGAGTGTGGAGTCGTTGCCTCTCACTTGTGCCTTCCTAAGTNNNAGATCGGAAGA
31606 TATGAGTGTGGAGTCGTTGCCCCCGACACTTTTTACTCCCTAANNNAGATCGGAAGA
31607 TATGAGTGTGGAGTCGTTGCATGAGCTGTCATTCACACTCCAGNNNAGATCGGAAG
31608 TATGAGTGTGGAGTCGTTGCAGGTGGCAGTGATTCCATAATGTNNNAGATCGGAA
31609 TATGAGTGTGGAGTCGTTGCACACCTGTGCACAATAACATCCTNNNAGATCGGAAG
31610 TATGAGTGTGGAGTCGTTGCGACCCTGTGAAAAACCCTAAAGCNNNAGATCGGAAG
31611 TATGAGTGTGGAGTCGTTGCCACACACATTCACACACACACATNNNAGATCGGAAGA
31612 TATGAGTGTGGAGTCGTTGCCAAGTCCCATGCTGAAAAGATGTNNNAGATCGGAAG
31613 TATGAGTGTGGAGTCGTTGCTGAAAATGGGACAATGGAAGGGANNNAGATCGGA
31614 TATGAGTGTGGAGTCGTTGCTGGAATCAGTAAATCCCCTCCTGNNNAGATCGGAAG
31615 TATGAGTGTGGAGTCGTTGCACCCACTCAGGTATGAACACATCNNNAGATCGGAAG
31616 TATGAGTGTGGAGTCGTTGCCTAGCAGGAGACTCAGACAGTTCNNNAGATCGGAA
31618 TATGAGTGTGGAGTCGTTGCGTAGAGAGATTCCCAACAGGTGGNNNAGATCGGAA
31619 TATGAGTGTGGAGTCGTTGCATCAGACTTCAGACCTCAGGGTANNNAGATCGGAAG
31621 TATGAGTGTGGAGTCGTTGCCCCCGATACAATCATAGCTGGAGNNNAGATCGGAAG
31622 TATGAGTGTGGAGTCGTTGCACCACCAAGGAATTGTATGGGAANNNAGATCGGAA
31623 TATGAGTGTGGAGTCGTTGCTTGCCTACCCCAACAGTACTTACNNNAGATCGGAAGA
31624 TATGAGTGTGGAGTCGTTGCTCTCACTCAGGGGAACCATCTATNNNAGATCGGAAG
31627 TATGAGTGTGGAGTCGTTGCTATTTGTCTCTCTCCCTCCACCTNNNAGATCGGAAGA
31628 TATGAGTGTGGAGTCGTTGCTGAGGCAATCTATAGCGAAACCTNNNAGATCGGAA
31629 TATGAGTGTGGAGTCGTTGCACACATCCCTCTTCTCTCTCTCANNNAGATCGGAAGA
31630 TATGAGTGTGGAGTCGTTGCAGAGAGAGAGAGAGAGTGAAGCCNNNAGATCGGA
31631 TATGAGTGTGGAGTCGTTGCGCGTGAGAGAGAGAAAGAGAGAGNNNAGATCGGA
31634 TATGAGTGTGGAGTCGTTGCGTTGAAGAGGCCAAAAGCAGAATNNNAGATCGGAA
31635 TATGAGTGTGGAGTCGTTGCAGGGCCTCATGTTTTATCTGGTTNNNAGATCGGAAG
31636 TATGAGTGTGGAGTCGTTGCTATCAAATGGAGGAGAGGGGACTNNNAGATCGGA
31637 TATGAGTGTGGAGTCGTTGCTATTGAAACATCCCCCCAACACANNNAGATCGGAAG
31638 TATGAGTGTGGAGTCGTTGCTTAGCACCAGGGACCTACTTAGANNNAGATCGGAAG

31639 TATGAGTGTGGAGTCGTTGCAGTGTATGCTACTGAAGGAGCTGNNNAGATCGGAA
31640 TATGAGTGTGGAGTCGTTGCACATCCAGTCTTCTCAGTTGCAANNNAGATCGGAAG
31641 TATGAGTGTGGAGTCGTTGCGGTTCTGTTGTGCAAACCTACTCNNNAGATCGGAAG
31642 TATGAGTGTGGAGTCGTTGCCCTCACCCAACACAGACAAAATCNNNAGATCGGAAG
31643 TATGAGTGTGGAGTCGTTGCGGTGCCAGGAAATTGATTTTGGANNNAGATCGGAA
31644 TATGAGTGTGGAGTCGTTGCAACTGGTTGAATGGATGGCAAAANNNAGATCGGAA
31645 TATGAGTGTGGAGTCGTTGCCTGGTGGTGTTTTTCTCCATGACNNNAGATCGGAAG
31646 TATGAGTGTGGAGTCGTTGCTGTGCGTTCGACAAATGGAAAAGNNNAGATCGGAA
31647 TATGAGTGTGGAGTCGTTGCTTCACAAGAGCAATGTAGGAGCTNNNAGATCGGAA
31648 TATGAGTGTGGAGTCGTTGCACCACACTACACACAACCCTTAGNNNAGATCGGAAG
31649 TATGAGTGTGGAGTCGTTGCCTCACACACACACACACACAAATNNNAGATCGGAAG
31650 TATGAGTGTGGAGTCGTTGCCACTTAGAATTGCCTTTCCAGCANNNAGATCGGAAG
31651 TATGAGTGTGGAGTCGTTGCTATAGACGTGTGTGTGTGTGTGTNNNAGATCGGAA
31652 TATGAGTGTGGAGTCGTTGCCCTGAAGTGACTGAGACAATGGANNNAGATCGGAA
31653 TATGAGTGTGGAGTCGTTGCGGCTTTGTTTCCTATCACGAAGGNNNAGATCGGAAG
31654 TATGAGTGTGGAGTCGTTGCCTTTCCCCCTCACTCTGAATGAANNNAGATCGGAAGA
31657 TATGAGTGTGGAGTCGTTGCCAAAGGCTGTGAAAGTACTGTGGNNNAGATCGGAA
31658 TATGAGTGTGGAGTCGTTGCCTACCAGCTCCTACCTTTGCTACNNNAGATCGGAAGA
31659 TATGAGTGTGGAGTCGTTGCTTTTTGTAATTGCCAGCCCCAATNNNAGATCGGAAG
31660 TATGAGTGTGGAGTCGTTGCTAGGCTGATGTAGTTGGTTGACCNNNAGATCGGAA
31661 TATGAGTGTGGAGTCGTTGCTCCTTTAGTCCCAGGACCAATTGNNNAGATCGGAAG
31662 TATGAGTGTGGAGTCGTTGCCCTTCCGACCTTCTTTCTACACANNNAGATCGGAAGA
31663 TATGAGTGTGGAGTCGTTGCGACTCTGTTTGGGAGAAAGCAACNNNAGATCGGAA
31664 TATGAGTGTGGAGTCGTTGCATTTGACAAGGGCACCAAGAAAANNNAGATCGGAA
31665 TATGAGTGTGGAGTCGTTGCTTAGAAGCAATGACATCCCAGCANNNAGATCGGAAG
31666 TATGAGTGTGGAGTCGTTGCTGGGCTCAAAGTAGACAAATCCANNNAGATCGGAA
31667 TATGAGTGTGGAGTCGTTGCTCTTCCAGAACCTCTCCCTTCTGNNNAGATCGGAAGA
31668 TATGAGTGTGGAGTCGTTGCTACACACTTATGGACAACAGGCANNNAGATCGGAAG
31670 TATGAGTGTGGAGTCGTTGCGCACAGATAGACACAACACACACNNNAGATCGGAAG
31671 TATGAGTGTGGAGTCGTTGCCAGATTGGGCAAAGCAAACAAACNNNAGATCGGAA
31672 TATGAGTGTGGAGTCGTTGCTCCAGTTTTAGCCCCTTACTTCCNNNAGATCGGAAGA
31673 TATGAGTGTGGAGTCGTTGCATGATTGTTTGGGCTAACAAGGCNNNAGATCGGAA
31674 TATGAGTGTGGAGTCGTTGCGACTCAGGAAGAGCCTTTCTGAANNNAGATCGGAA
31675 TATGAGTGTGGAGTCGTTGCGCTCCAAAGCGACCTAAAGATTTNNNAGATCGGAAG
31676 TATGAGTGTGGAGTCGTTGCATACCCAGCTTCTTTTGGCCATANNNAGATCGGAAG
31677 TATGAGTGTGGAGTCGTTGCTACCTGAGGATAGGGAACCTTGTNNNAGATCGGAA
31678 TATGAGTGTGGAGTCGTTGCTTCACGGAAAATAAAAGTGGCCGNNNAGATCGGAA
31679 TATGAGTGTGGAGTCGTTGCGTAACATCAGTCTGGAGGGAGTGNNNAGATCGGAA
31680 TATGAGTGTGGAGTCGTTGCAGAAAGACTATGAAAGCCTGCCANNNAGATCGGAA
31681 TATGAGTGTGGAGTCGTTGCCATCAAGTGTCCACCAAGGAAAGNNNAGATCGGAA
31682 TATGAGTGTGGAGTCGTTGCTGCATACACTGGGTCATTAGAGGNNNAGATCGGAA
31683 TATGAGTGTGGAGTCGTTGCCACAGACAGTGGAAATGCCAAAANNNAGATCGGAA
31684 TATGAGTGTGGAGTCGTTGCACACAACCTTGGAGGAAGAAAGANNNAGATCGGAA

31685 TATGAGTGTGGAGTCGTTGCATTCTCCTCTGCTTTTAGGGTGGNNNAGATCGGAAG
31686 TATGAGTGTGGAGTCGTTGCTAACCAAATCCAGCAGCACATTGNNNAGATCGGAAG
31687 TATGAGTGTGGAGTCGTTGCACTCAGTTTCTGCAGATTGTTGCNNNAGATCGGAAG
31688 TATGAGTGTGGAGTCGTTGCGTGTGTGTGTGTGTATTTGGGAGNNNAGATCGGAA
31689 TATGAGTGTGGAGTCGTTGCGTTGGCAGTGACCTCCATAAGTANNNAGATCGGAAG
31690 TATGAGTGTGGAGTCGTTGCTGTTTGCTAATCTCCTGCTTCCTNNNAGATCGGAAGA
31691 TATGAGTGTGGAGTCGTTGCATGTGTGTGGTAGGTACATTGCTNNNAGATCGGAA
31692 TATGAGTGTGGAGTCGTTGCAAGGACACCAGTGCTATTGGATTNNNAGATCGGAA
31693 TATGAGTGTGGAGTCGTTGCGAGGATGGGGTTGATGGTAGAAGNNNAGATCGGA
31694 TATGAGTGTGGAGTCGTTGCCACCTGCACAGAAAACAAAGTCANNNAGATCGGAAG
31695 TATGAGTGTGGAGTCGTTGCTCTCTCTCCCTGGATTTTTCTGCNNNAGATCGGAAGA
31697 TATGAGTGTGGAGTCGTTGCATGGTCATGATGGGAAAAGTGGANNNAGATCGGA
31699 TATGAGTGTGGAGTCGTTGCTAGGGAAGACAGAAAGAAGGGGANNNAGATCGGA
31700 TATGAGTGTGGAGTCGTTGCCTGCTCTCGTACTTCCTTGAGACNNNAGATCGGAAG
31701 TATGAGTGTGGAGTCGTTGCAGCTCATCTGCAACTTGTCTTTTNNNAGATCGGAAGA
31704 TATGAGTGTGGAGTCGTTGCATGACAATACCTGCATCTCTGCANNNAGATCGGAAG
31706 TATGAGTGTGGAGTCGTTGCGAACAAGAAAGAGGAGGAGGAGGNNNAGATCGG
31707 TATGAGTGTGGAGTCGTTGCCCAGCAGATTTCCTCTCACATCTNNNAGATCGGAAGA
31708 TATGAGTGTGGAGTCGTTGCATAGGCTGCTTACATCTGGGATGNNNAGATCGGAA
31709 TATGAGTGTGGAGTCGTTGCGAAAAGTACCAGAACACACAGGCNNNAGATCGGAA
31710 TATGAGTGTGGAGTCGTTGCGTGTGTGTGTGAGAGAGAGAGAGNNNAGATCGGA
31711 TATGAGTGTGGAGTCGTTGCCCATCTCCCACTCAGAGTCTTTCNNNAGATCGGAAGA
31712 TATGAGTGTGGAGTCGTTGCCAAAACTTTTGGGTCCATCTCCCNNNAGATCGGAAG
31713 TATGAGTGTGGAGTCGTTGCCTCTCTCCTCTCTCTCTGTCACANNNAGATCGGAAGA
31714 TATGAGTGTGGAGTCGTTGCTCAGAGGAAATGAGGAAATGGGGNNNAGATCGGA
31715 TATGAGTGTGGAGTCGTTGCTGTCTTGGTTCAGGCCTAAATCANNNAGATCGGAAG
31716 TATGAGTGTGGAGTCGTTGCTACAAAGGCAAAGTCCCACTACANNNAGATCGGAAG
31717 TATGAGTGTGGAGTCGTTGCAGAGAGGGAGAGAGAGAGAGAGANNNAGATCGG
31718 TATGAGTGTGGAGTCGTTGCGTTCTTTTTTTCCAGCTGGGGAGNNNAGATCGGAAG
31719 TATGAGTGTGGAGTCGTTGCACAGCTGAGTTTCCTGCATTAGANNNAGATCGGAAG
31720 TATGAGTGTGGAGTCGTTGCCTGTTGCTCTCTGGAGACTTGACNNNAGATCGGAAG
31721 TATGAGTGTGGAGTCGTTGCCGTGTGAGAGAGAGAGAGAGAGANNNAGATCGGA
31722 TATGAGTGTGGAGTCGTTGCATTGCAGTTGGAGGTTAGCCAAGNNNAGATCGGAA
31723 TATGAGTGTGGAGTCGTTGCACTCTCCTCACCCCACTACAATANNNAGATCGGAAGA
31726 TATGAGTGTGGAGTCGTTGCCGTCTCTCTCTCTCTCTCTCTCTNNNAGATCGGAAGAG
31727 TATGAGTGTGGAGTCGTTGCGACCACCAGGAGAGACATTTAGGNNNAGATCGGAA
31728 TATGAGTGTGGAGTCGTTGCCCAAGAAGAATCTCCCTCTCTGGNNNAGATCGGAAG
31730 TATGAGTGTGGAGTCGTTGCTGTTATATGGCTGACAGGTTGGTNNNAGATCGGAA
31731 TATGAGTGTGGAGTCGTTGCCCAAAGGGCTTTCTTGTGACTTTNNNAGATCGGAAG
31732 TATGAGTGTGGAGTCGTTGCAAAAACACTGTGACTTAGGGGGTNNNAGATCGGAA
31733 TATGAGTGTGGAGTCGTTGCCCCCTGAACTGGAATACATGGATNNNAGATCGGAAG
31734 TATGAGTGTGGAGTCGTTGCGGAAGTAGAAGGTGGAGGGAAAGNNNAGATCGG
31735 TATGAGTGTGGAGTCGTTGCTCATTGTCAGTGGAGTCCTTCAGNNNAGATCGGAAG

31737 TATGAGTGTGGAGTCGTTGCGTGTGTGTGTGATGCTTGCTTATNNNAGATCGGAA

31738 TATGAGTGTGGAGTCGTTGCGCAGCATAATGCAATTCTCTTGCNNNAGATCGGAAG

31740 TATGAGTGTGGAGTCGTTGCGTGGCCACTGAGTTTTCTGTTTTNNNAGATCGGAAG

31742 TATGAGTGTGGAGTCGTTGCTGGCCATTGTCAGCAATAAACTTNNNAGATCGGAAG

31743 TATGAGTGTGGAGTCGTTGCTCACTCAACAGTATTCTTGGGCANNNAGATCGGAAG

31744 TATGAGTGTGGAGTCGTTGCCCACAGCCTAAATTACTAGCCCANNNAGATCGGAAG

31747 TATGAGTGTGGAGTCGTTGCATGAAAGCTGAGGGAAAAGAAGCNNNAGATCGGA

31748 TATGAGTGTGGAGTCGTTGCAGTATACCAACACCTGTGCTCAGNNNAGATCGGAAG

31749 TATGAGTGTGGAGTCGTTGCTCCAGATGGAGAGCTATAGGAGTNNNAGATCGGAA

31750 TATGAGTGTGGAGTCGTTGCCCAAAGTGGAAGGTTTGTGATGGNNNAGATCGGAA

31751 TATGAGTGTGGAGTCGTTGCCAACTTGAGAACCCTCTTCTGGANNNAGATCGGAAG

31752 TATGAGTGTGGAGTCGTTGCTGGAAGTTCCCATTGTGTACACTNNNAGATCGGAAG

31753 TATGAGTGTGGAGTCGTTGCGTCTCTCTCTCCCCTCTTTCTNNNAGATCGGAAGAG

31754 TATGAGTGTGGAGTCGTTGCCCCTCTCTCCTCTCTCTCTTCTCNNNAGATCGGAAGAG

31755 TATGAGTGTGGAGTCGTTGCCATCACCTCGATCTGGCTCATTANNNAGATCGGAAG

31756 TATGAGTGTGGAGTCGTTGCACTTCTCTGGAATTTCATGCCCTNNNAGATCGGAAG

31757 TATGAGTGTGGAGTCGTTGCCCCAGGAGATCTCAAGCCTTTAGNNNAGATCGGAAG

31758 TATGAGTGTGGAGTCGTTGCCTGAGCCACCTAAAGCTGAGAATNNNAGATCGGAA

31759 TATGAGTGTGGAGTCGTTGCGACACAAGCACTACCACTTTGACNNNAGATCGGAAG

31760 TATGAGTGTGGAGTCGTTGCCTCTCTCTCACCACCTAAAGCTGNNNAGATCGGAAGA

31761 TATGAGTGTGGAGTCGTTGCAATCTGTATGTGTGTGTGTGTGCNNNAGATCGGAA

31762 TATGAGTGTGGAGTCGTTGCGTGTGTGTGTGAGAGAGAGAGAGNNNAGATCGGA

31763 TATGAGTGTGGAGTCGTTGCTTGTGCTACAAATGGCTCCAAAGNNNAGATCGGAAG

31765 TATGAGTGTGGAGTCGTTGCTCTTTTTCAGGGTCTCACTCTGTNNNAGATCGGAAG

31767 TATGAGTGTGGAGTCGTTGCTCTCTCTCTCTCTCTGTGTGTGTNNNAGATCGGAAGA

31768 TATGAGTGTGGAGTCGTTGCTCCCTCTCTCTCTCTCTCTCTCTNNNAGATCGGAAGAG

31769 TATGAGTGTGGAGTCGTTGCGATGAGGAGTGGAGTCAGGTATGNNNAGATCGGA

31770 TATGAGTGTGGAGTCGTTGCCACACAGAGAGAGATGAGGGAACNNNAGATCGGA

31771 TATGAGTGTGGAGTCGTTGCCTGAGCTCAGGAGTTCAAAACCANNNAGATCGGAA

31772 TATGAGTGTGGAGTCGTTGCGGTGGATCACTTGAGATCAGGAGNNNAGATCGGAA

31773 TATGAGTGTGGAGTCGTTGCCCAGGGGCTTGGAAATATGAGATNNNAGATCGGAA

31774 TATGAGTGTGGAGTCGTTGCATATCCTCAGAGGCACAACACTGNNNAGATCGGAAG

31775 TATGAGTGTGGAGTCGTTGCAAAGAGGGAAAGGAAGAGAGCAGNNNAGATCGGA

31776 TATGAGTGTGGAGTCGTTGCTGTTCCTCTACATCCTTGTCAGCNNNAGATCGGAAGA

31777 TATGAGTGTGGAGTCGTTGCCAGGTTTGATTGGTGGTCAGAACNNNAGATCGGAA

31778 TATGAGTGTGGAGTCGTTGCAGAGAGAGAGAGAGAGAGAGGGANNNAGATCGG

31779 TATGAGTGTGGAGTCGTTGCAGAGAGAGAGAGAGAGAGAGGGANNNAGATCGG

31780 TATGAGTGTGGAGTCGTTGCAGAGAGAGAGAGAGAGAGAGGGANNNAGATCGG

31781 TATGAGTGTGGAGTCGTTGCACCCCATTATGTTGAAAACCCCANNNAGATCGGAAG

31782 TATGAGTGTGGAGTCGTTGCCACCTGACCTCTGCAGACTTTATNNNAGATCGGAAG

31783 TATGAGTGTGGAGTCGTTGCAGAAAGAAAGAAGGAAGGGAGGGNNNAGATCGG

31784 TATGAGTGTGGAGTCGTTGCAGATAGATGCTAAGGAGGTAGGTNNNAGATCGGA

31785 TATGAGTGTGGAGTCGTTGCTCTCACCTCTAAGGATCTCACGTNNNAGATCGGAAG

31786 TATGAGTGTGGAGTCGTTGCTCTACTTTGTTCCAGGCTTGACANNNAGATCGGAAG
31787 TATGAGTGTGGAGTCGTTGCAGAGAAGAGAAGAGAAGGCTCCTNNNAGATCGGA
31788 TATGAGTGTGGAGTCGTTGCGGAGTGAGCATGTACTCTGATGTNNNAGATCGGAA
31789 TATGAGTGTGGAGTCGTTGCAACTTCCCAACTATGAAGCCAGTNNNAGATCGGAAG
31791 TATGAGTGTGGAGTCGTTGCAGGCAATTTTTTCTTACAGGCCTNNNAGATCGGAAG
31792 TATGAGTGTGGAGTCGTTGCGCAAAGCACCAAAGACAGAGATTNNNAGATCGGAA
31793 TATGAGTGTGGAGTCGTTGCTGCTCCAAATTTTCATCACAGGANNNAGATCGGAAG
31795 TATGAGTGTGGAGTCGTTGCCCCACCCTTATGACCTCTTTCAANNNAGATCGGAAGA
31796 TATGAGTGTGGAGTCGTTGCCCACCACGCCTAGCTAATTTTTTNNNAGATCGGAAGA
31798 TATGAGTGTGGAGTCGTTGCATATGTGTGTGTGTGTGTGTGTGNNNAGATCGGAA
31799 TATGAGTGTGGAGTCGTTGCGTTCAAGCAATTCTCATGCCTCANNNAGATCGGAAG
31800 TATGAGTGTGGAGTCGTTGCATCGCTTAGATATGGCAGTCCTGNNNAGATCGGAAG
31801 TATGAGTGTGGAGTCGTTGCTGAGAATCACTTGAACCCAGGAGNNNAGATCGGAA
31802 TATGAGTGTGGAGTCGTTGCTCACAAGTGCACATGGAATGTTCNNNAGATCGGAAG
31803 TATGAGTGTGGAGTCGTTGCACACTGTTTCTACCAATTGCTGANNNAGATCGGAAG
31804 TATGAGTGTGGAGTCGTTGCTCACAAGTGCACATGGAATGTTCNNNAGATCGGAAG
31806 TATGAGTGTGGAGTCGTTGCATTGGGGATTAGGGTTTCAGCATNNNAGATCGGAA
31808 TATGAGTGTGGAGTCGTTGCTTGGATGAAACTGGAGGCCATTANNNAGATCGGAA
31810 TATGAGTGTGGAGTCGTTGCGTTCAAACGATTTTCCTGCCTCANNNAGATCGGAAG
31811 TATGAGTGTGGAGTCGTTGCTTTGAGACAAGGTCTTGCTCTGTNNNAGATCGGAAG
31812 TATGAGTGTGGAGTCGTTGCCATTCCTGCCAGCTGATGAAATTNNNAGATCGGAAG
31813 TATGAGTGTGGAGTCGTTGCACACAGCCATAAAAAGGGATGAANNNAGATCGGAA
31815 TATGAGTGTGGAGTCGTTGCTTCAAGTGATTCTCCTGCCTCAGNNNAGATCGGAAG
31817 TATGAGTGTGGAGTCGTTGCTGCAGTTTGGACAAGTAGAGAGANNNAGATCGGAA
31818 TATGAGTGTGGAGTCGTTGCCAGTGGGGTAATTATGGCTCACTNNNAGATCGGAA
31820 TATGAGTGTGGAGTCGTTGCACACACACACAGACACACACATANNNAGATCGGAAG
31821 TATGAGTGTGGAGTCGTTGCTAAGAAAACCATGCCATTTGCGANNNAGATCGGAAG
31824 TATGAGTGTGGAGTCGTTGCATCAACAGAATTTGGACACAGCANNNAGATCGGAA
31825 TATGAGTGTGGAGTCGTTGCAGTGTGTGTATATGTGTGTGTGTNNNAGATCGGAA
31826 TATGAGTGTGGAGTCGTTGCGTGCACAACCACACATACATAGGNNNAGATCGGAAG
31829 TATGAGTGTGGAGTCGTTGCCACACACAAACACACACACACATNNNAGATCGGAAG
31830 TATGAGTGTGGAGTCGTTGCCTCAAGCAATCCTCCTACCTCAGNNNAGATCGGAAGA
31831 TATGAGTGTGGAGTCGTTGCGCTTTGTTGAAGATCAGTTGGCTNNNAGATCGGAA
31833 TATGAGTGTGGAGTCGTTGCTATTTTCAGTGGTGCTTCCTCCANNNAGATCGGAAG
31834 TATGAGTGTGGAGTCGTTGCAGATGCCAAGACCTCTGCTAAAANNNAGATCGGAAG
31836 TATGAGTGTGGAGTCGTTGCATAAGAGGAGAGGTTAGTGCCCTNNNAGATCGGAA
31837 TATGAGTGTGGAGTCGTTGCCCAAGATGAGCAGATCACCTGAGNNNAGATCGGAA
31839 TATGAGTGTGGAGTCGTTGCAGAGCGACACTCCATCTCAAAAANNNAGATCGGAAG
31840 TATGAGTGTGGAGTCGTTGCTGGAGTGCAGTGTTGTCATCATANNNAGATCGGAA
31841 TATGAGTGTGGAGTCGTTGCGTTGCTTTCTGGTGAGTCCTTTCNNNAGATCGGAAG
31842 TATGAGTGTGGAGTCGTTGCACAACATGGTCCTCAGTTAGTCGNNNAGATCGGAAG
31844 TATGAGTGTGGAGTCGTTGCCCTGGATTGCAGTGGATCCTAAANNNAGATCGGAA
31845 TATGAGTGTGGAGTCGTTGCTGGACCTCCTTTCTTTGCAAAACNNNAGATCGGAAG

31846 TATGAGTGTGGAGTCGTTGCGGATGGATACTTGCTGCTTTTCCNNNAGATCGGAAG
31848 TATGAGTGTGGAGTCGTTGCTGAAAAGCCTCACATAATGCGTCNNNAGATCGGAAG
31849 TATGAGTGTGGAGTCGTTGCGTCAGTATGTCTAGACTGGGCACNNNAGATCGGAA
31850 TATGAGTGTGGAGTCGTTGCCAACAGAGTGAGACCATGTCAAGNNNAGATCGGAA
31851 TATGAGTGTGGAGTCGTTGCCTGAATCCAGCAGCACATCAAAANNNAGATCGGAAG
31852 TATGAGTGTGGAGTCGTTGCAAATTCAAGCCAGCTGCAGAAATNNNAGATCGGAA
31853 TATGAGTGTGGAGTCGTTGCTCAGGAAAGTGCTGTACTTGTGANNNAGATCGGAA
31855 TATGAGTGTGGAGTCGTTGCTGGCTTCACTGTTGAATTCTACCNNNAGATCGGAAG
31856 TATGAGTGTGGAGTCGTTGCGAGATTGCAGGCAAGAGTCCTATNNNAGATCGGAA
31857 TATGAGTGTGGAGTCGTTGCCTTCTGTCACAGTGTGGCTAGATNNNAGATCGGAAG
31858 TATGAGTGTGGAGTCGTTGCAAAGTTGAGGAAGGAGAATCGCTNNNAGATCGGAA
31859 TATGAGTGTGGAGTCGTTGCAAACTGCAAACAACCTCAGAACCNNNAGATCGGAAG
31860 TATGAGTGTGGAGTCGTTGCCTTTTCCAACACAGGGCTCAAAGNNNAGATCGGAAG
31861 TATGAGTGTGGAGTCGTTGCGCTCATGGATGGGTAGAACCAATNNNAGATCGGAA
31862 TATGAGTGTGGAGTCGTTGCCCTGGGCTTTCAGAGACTATGTTNNNAGATCGGAAG
31863 TATGAGTGTGGAGTCGTTGCTTTGTTTTTTGCCACAGAGCCTNNNAGATCGGAAG
31864 TATGAGTGTGGAGTCGTTGCCTGGAAGTTATGGTATGGCCCTTNNNAGATCGGAA
31865 TATGAGTGTGGAGTCGTTGCGCAGTCTATAAGTGGCTGAAGGANNNAGATCGGAA
31866 TATGAGTGTGGAGTCGTTGCAAAAGGAAACAGGACGGTCTGATNNNAGATCGGAA
31868 TATGAGTGTGGAGTCGTTGCGAGAGAGAGAGAGTCTTCTTGCGNNNAGATCGGAA
31871 TATGAGTGTGGAGTCGTTGCCACACATATCCACAGCAGCAATGNNNAGATCGGAAG
31872 TATGAGTGTGGAGTCGTTGCGGCTAGAGAAAGACCCCAAGTTTNNNAGATCGGAA
31873 TATGAGTGTGGAGTCGTTGCTTCTCTCTGGATTTTCAGGAGGCNNNAGATCGGAAG
31875 TATGAGTGTGGAGTCGTTGCTGTAATCTCAGCACTTTGGGAGGNNNAGATCGGAA
31876 TATGAGTGTGGAGTCGTTGCTCCCGCAGTCTTATTTATACCCGNNNAGATCGGAAG
31877 TATGAGTGTGGAGTCGTTGCTTGTGTACATTTTTTGGCTGGGCNNNAGATCGGAAG
31879 TATGAGTGTGGAGTCGTTGCTTGGCAACATAGCAAGATCTCCTNNNAGATCGGAAG
31880 TATGAGTGTGGAGTCGTTGCATACATCTATGACCTCACCCCCANNNAGATCGGAAGA
31882 TATGAGTGTGGAGTCGTTGCGCAGAATCATTTGAACCTGGGAGNNNAGATCGGAA
31886 TATGAGTGTGGAGTCGTTGCCCTTTCCAGGACCAGCAGATAATNNNAGATCGGAAG
31887 TATGAGTGTGGAGTCGTTGCAAAGGGCCACATAGATTACAGCANNNAGATCGGAA
31889 TATGAGTGTGGAGTCGTTGCTGACATTATTCTGACCCATGGGGNNNAGATCGGAAG
31890 TATGAGTGTGGAGTCGTTGCGACACAGAGACAGAGAGACACAGNNNAGATCGGAA
31891 TATGAGTGTGGAGTCGTTGCAGTGTATCAGCTCCACCATTCAGNNNAGATCGGAAG
31892 TATGAGTGTGGAGTCGTTGCAGAATCTTACTGTGTTCTGGGGGNNNAGATCGGAA
31893 TATGAGTGTGGAGTCGTTGCGCCATAAACTACGATCATGCCACNNNAGATCGGAAG
31895 TATGAGTGTGGAGTCGTTGCGAAAGGAATCGTGGTGTATTGGCNNNAGATCGGAA
31896 TATGAGTGTGGAGTCGTTGCATCTGAACTTTTGGGGGTCAACTNNNAGATCGGAAG
31898 TATGAGTGTGGAGTCGTTGCGCATTGACTTCTGGCTCTGATTCNNNAGATCGGAAG
31900 TATGAGTGTGGAGTCGTTGCCTGTCTGGACCTGGAGTTTCTTTNNNAGATCGGAAG
31901 TATGAGTGTGGAGTCGTTGCAAAATCTGGACTTTGTGCTGTGGNNNAGATCGGAA
31902 TATGAGTGTGGAGTCGTTGCTCGACTTTATCTTTGGGAGGCAANNNAGATCGGAAG
31907 TATGAGTGTGGAGTCGTTGCAGCAAAAAGGTGGCCTTGTTATGNNNAGATCGGAA

31908 TATGAGTGTGGAGTCGTTGCGAAAGAGACAGGGAAAGGGAGAANNNAGATCGGA
31909 TATGAGTGTGGAGTCGTTGCCCTCCGATATAACCCACCCAAATNNNAGATCGGAAGA
31910 TATGAGTGTGGAGTCGTTGCGAGCCGGTATATAAGAGCTGCTTNNNAGATCGGAA
31912 TATGAGTGTGGAGTCGTTGCTTTCTCCAGCCTGGTCTTTTACANNNAGATCGGAAGA
31913 TATGAGTGTGGAGTCGTTGCAAGAATTCAGTTCACTGCCAAGCNNNAGATCGGAAG
31914 TATGAGTGTGGAGTCGTTGCCCACCAAAAGGCAACTAGAACTGNNNAGATCGGAAG
31915 TATGAGTGTGGAGTCGTTGCACTAGCCTTAGCAACATGGTGAANNNAGATCGGAAG
31916 TATGAGTGTGGAGTCGTTGCCGAACAGTTAGAGGAGTGGCATANNNAGATCGGAA
31917 TATGAGTGTGGAGTCGTTGCACCTCACTGATGTTTACCTTGGGNNNAGATCGGAAGA
31918 TATGAGTGTGGAGTCGTTGCTCACATTTTCACTACCTGAAGAACANNNAGATCGGAA
31919 TATGAGTGTGGAGTCGTTGCCATTGGCTCACACTTGTAATCCCNNNAGATCGGAAGA
31920 TATGAGTGTGGAGTCGTTGCAACAAAATACCACACACTGGGTGNNNAGATCGGAAG
31921 TATGAGTGTGGAGTCGTTGCTTTTATAAGGCCAGCATCACCCTNNNAGATCGGAAGA
31922 TATGAGTGTGGAGTCGTTGCGCTGAAGGGCATTAGCTACATTGNNNAGATCGGAA
31923 TATGAGTGTGGAGTCGTTGCAACTCTGCCTTTATTTGATGCCGNNNAGATCGGAAGA
31924 TATGAGTGTGGAGTCGTTGCTACAGCAAGTGTCTGTGAGAACANNNAGATCGGAA
31925 TATGAGTGTGGAGTCGTTGCCCCTGTTGCAATTCTGATCAAGCNNNAGATCGGAAGA
31926 TATGAGTGTGGAGTCGTTGCACTGAGCAATGCATAGGTTCAGANNNAGATCGGAA
31927 TATGAGTGTGGAGTCGTTGCCAGAAACTGTCCTTGGAGAAACANNNAGATCGGAAG
31928 TATGAGTGTGGAGTCGTTGCAGGACTACACATCACCAAACCAANNNAGATCGGAAG
31929 TATGAGTGTGGAGTCGTTGCAAGGCAGTCAGATGGGAGAAATTNNNAGATCGGAA
31930 TATGAGTGTGGAGTCGTTGCGGGTTCAAAAGCAGATTTGAGCANNNAGATCGGAA
31931 TATGAGTGTGGAGTCGTTGCTCCTTGAATTGCTAACCTCTGCTNNNAGATCGGAAGA
31932 TATGAGTGTGGAGTCGTTGCAAATAATTAGCCAGGCCAGTTGGNNNAGATCGGAA
31933 TATGAGTGTGGAGTCGTTGCTTTCAAGAATGAAGCTGTGGACCNNNAGATCGGAAG
31934 TATGAGTGTGGAGTCGTTGCTCCTTGAAACATGGAGATGGAGGNNNAGATCGGAA
31935 TATGAGTGTGGAGTCGTTGCAACCATCAAAATGAAGCAGACGGNNNAGATCGGAA
31936 TATGAGTGTGGAGTCGTTGCAATTCTGAAGACTCCCAGACAGGNNNAGATCGGAAG
31938 TATGAGTGTGGAGTCGTTGCAGAAACCCTCTTACCCAGAAACCNNNAGATCGGAAGA
31940 TATGAGTGTGGAGTCGTTGCAGTTTCTTCTTGGCCTCTCTGAGNNNAGATCGGAAGA
31942 TATGAGTGTGGAGTCGTTGCAGTTGTCACCACTCTTTCTGTGTNNNAGATCGGAAGA
31943 TATGAGTGTGGAGTCGTTGCGAAGGCAACAAAACTGGATGGAGNNNAGATCGGAA
31944 TATGAGTGTGGAGTCGTTGCATATGGTTTCTTGTCTGGCGAGTNNNAGATCGGAAG
31946 TATGAGTGTGGAGTCGTTGCACCTGATAGAGCTGAATGACACANNNAGATCGGAA
31949 TATGAGTGTGGAGTCGTTGCGAGGAGAAAGTACTCAACCAGGGNNNAGATCGGAA
31951 TATGAGTGTGGAGTCGTTGCTGATTGGACAAGACGCTTCCATANNNAGATCGGAAG
31952 TATGAGTGTGGAGTCGTTGCAGGGACTACCAATCTGCATTGTCNNNAGATCGGAAG
31953 TATGAGTGTGGAGTCGTTGCCTCCCACAGGAAGGAAAGAGAAANNNAGATCGGAA
31954 TATGAGTGTGGAGTCGTTGCTGCCAAACACACAGAACTAGTTTNNNAGATCGGAAG
31955 TATGAGTGTGGAGTCGTTGCGAAGGAAGGAAGGAAAGGAAGGANNNAGATCGGA
31956 TATGAGTGTGGAGTCGTTGCCTGTAATCCCAGCTACTCAGGAGNNNAGATCGGAAG
31958 TATGAGTGTGGAGTCGTTGCTTCTGTGTGAACTGTGTCAGGAANNNAGATCGGAA
31959 TATGAGTGTGGAGTCGTTGCAGCTGCATACAAATTGACTCTCCNNNAGATCGGAAGA

31960 TATGAGTGTGGAGTCGTTGCGTATATGGCTCCACACAAATGGCNNNAGATCGGAAG
31961 TATGAGTGTGGAGTCGTTGCGCCATTTGTGTGGAGCCATATACNNNAGATCGGAAG
31962 TATGAGTGTGGAGTCGTTGCGCCTAGGTGCTCATATGGACTTANNNAGATCGGAAG
31964 TATGAGTGTGGAGTCGTTGCAGCTGCATACAAATTGACTCTCCNNNAGATCGGAAG
31965 TATGAGTGTGGAGTCGTTGCGTATATGGCTCCACACAAATGGCNNNAGATCGGAAG
31966 TATGAGTGTGGAGTCGTTGCAGACAGAGAAAAGAAAGCCCACANNNAGATCGGAA
31967 TATGAGTGTGGAGTCGTTGCGCCTAGGTGCTCATATGGACTTANNNAGATCGGAAG
31968 TATGAGTGTGGAGTCGTTGCTAATGCTACTTGGGAGGCTGAAGNNNAGATCGGAA
31969 TATGAGTGTGGAGTCGTTGCCTGAGGCAGGAGAATGGAATGAANNNAGATCGGA
31970 TATGAGTGTGGAGTCGTTGCGCTGTCTGTATCAGATTGGCCTTNNNAGATCGGAAG
31974 TATGAGTGTGGAGTCGTTGCCCAGCCACAGCCATAGATAGAAANNNAGATCGGAA
31975 TATGAGTGTGGAGTCGTTGCGGTGAAGAGAGCAGGAATAGAGGNNNAGATCGGA
31976 TATGAGTGTGGAGTCGTTGCCATGCTGCTGTCATTTTGTGGTANNNAGATCGGAAG
31977 TATGAGTGTGGAGTCGTTGCGCTTCAGGCTCTCTAATCAACGANNNAGATCGGAAG
31978 TATGAGTGTGGAGTCGTTGCAATGCGTGTAATCCCAGAACTCTNNNAGATCGGAAG
31979 TATGAGTGTGGAGTCGTTGCTGTAATTCTCGGAGAGGACACACNNNAGATCGGAA
31980 TATGAGTGTGGAGTCGTTGCCACTCATTCCAACTACTCAGCCANNNAGATCGGAAGA
31981 TATGAGTGTGGAGTCGTTGCTTGGGGGCAATCAAAATTTAGCCNNNAGATCGGAA
31982 TATGAGTGTGGAGTCGTTGCATGAGCGTCAAAAGTACTCCTGTNNNAGATCGGAAG
31983 TATGAGTGTGGAGTCGTTGCAAAAGCAACCCTAACACTACGGANNNAGATCGGAAG
31984 TATGAGTGTGGAGTCGTTGCACAAGTTGTATTGGTGGAGGACANNNAGATCGGAA
31985 TATGAGTGTGGAGTCGTTGCTGTGTGTGTTTTGGTGCTTTGAANNNAGATCGGAA
31986 TATGAGTGTGGAGTCGTTGCACGTAAGTGGACTCAACTATCCANNNAGATCGGAAG
31988 TATGAGTGTGGAGTCGTTGCTTTTTGAGACAGGGTCTTGCTCTNNNAGATCGGAAG
31989 TATGAGTGTGGAGTCGTTGCAACAGTGCTAATGGATTAGGGCTNNNAGATCGGAA
31990 TATGAGTGTGGAGTCGTTGCGCAGCACTATTCACAATAGCCAGNNNAGATCGGAAG
31991 TATGAGTGTGGAGTCGTTGCCTGAGTCTGAAGTCCTGAGAACCNNNAGATCGGAA
31993 TATGAGTGTGGAGTCGTTGCGTACTGTGCTAGGCAAGGGTATTNNNAGATCGGAA
31994 TATGAGTGTGGAGTCGTTGCGACATGATGCCACAAGTGGAAAANNNAGATCGGAA
31995 TATGAGTGTGGAGTCGTTGCTACCAGATGGATTCACAGCCAAANNNAGATCGGAAG
31996 TATGAGTGTGGAGTCGTTGCGCTCCATCCTTATGACCTCACTTNNNAGATCGGAAGA
31997 TATGAGTGTGGAGTCGTTGCGTAATCCCAGCTACTTGAGAGGCNNNAGATCGGAA
31998 TATGAGTGTGGAGTCGTTGCAATAATGGCATTTGCAGCAACCTNNNAGATCGGAAG
31999 TATGAGTGTGGAGTCGTTGCCATTAGACTGGTGGTTCTAGGCANNNAGATCGGAA
32000 TATGAGTGTGGAGTCGTTGCGCAGTGACAAGCAAGGATTGTAANNNAGATCGGAA
32001 TATGAGTGTGGAGTCGTTGCATACTGGTTTGTGTCTCACTGCTNNNAGATCGGAAG
32002 TATGAGTGTGGAGTCGTTGCAGCATGAACAGAGGCTCAGAATTNNNAGATCGGAA
32003 TATGAGTGTGGAGTCGTTGCTCAGACCAGCATCCTTTCACTTANNNAGATCGGAAGA
32004 TATGAGTGTGGAGTCGTTGCATGTTTAGGAGAGCATGGAAGGGNNNAGATCGGA
32005 TATGAGTGTGGAGTCGTTGCGAGACAGAGATACCGTTCCCATCNNNAGATCGGAAG
32006 TATGAGTGTGGAGTCGTTGCGAGAGGGAACCAACTTGTACTGTNNNAGATCGGAA
32007 TATGAGTGTGGAGTCGTTGCCAACAGATATCAGCTGGACTCCANNNAGATCGGAAG
32008 TATGAGTGTGGAGTCGTTGCACACCTCCAACCTGTCTCAATTTNNNAGATCGGAAGA

32009 TATGAGTGTGGAGTCGTTGCGAACAGTTTCATCACCCCCAAAGNNNAGATCGGAAG
32011 TATGAGTGTGGAGTCGTTGCGTGACCATGGCAGGCTATTTTTTNNNAGATCGGAAG
32017 TATGAGTGTGGAGTCGTTGCATTCAGAAGAGAGGTCAGGGGTANNNAGATCGGA
32019 TATGAGTGTGGAGTCGTTGCCTTGGCCACACAGCTTTACAAATNNNAGATCGGAAG
32020 TATGAGTGTGGAGTCGTTGCTTCAAGTGATTCTCCTGCCTCAANNNAGATCGGAAG
32021 TATGAGTGTGGAGTCGTTGCTACCTACCTCCCTCACAATTTGCNNNAGATCGGAAGA
32023 TATGAGTGTGGAGTCGTTGCGAGTGATTCACCCGGCTAATTTTNNNAGATCGGAAG
32024 TATGAGTGTGGAGTCGTTGCGAGACAAAGGCCTTGCTCTATTGNNNAGATCGGAA
32025 TATGAGTGTGGAGTCGTTGCGCATCTTGAAACCGCTGGATTTANNNAGATCGGAAG
32028 TATGAGTGTGGAGTCGTTGCATGCACCTAACACTGGATATCCCNNNAGATCGGAAG
32031 TATGAGTGTGGAGTCGTTGCCCTCCCACCATGCTAAGATTACANNNAGATCGGAAG
32032 TATGAGTGTGGAGTCGTTGCTCAATTGGCAGAATAAATGCTCAGTNNNAGATCGGA
32036 TATGAGTGTGGAGTCGTTGCTCCCGAGAACAATGGGCATTATTNNNAGATCGGAAG
32037 TATGAGTGTGGAGTCGTTGCTCCAAATGACACTTTGTTGCTGTNNNAGATCGGAAG
32038 TATGAGTGTGGAGTCGTTGCGTTTCCTTGTTTGCCATCATCCANNNAGATCGGAAGA
32039 TATGAGTGTGGAGTCGTTGCTGACCACCACCATTTAGAAGTTGNNNAGATCGGAAG
32040 TATGAGTGTGGAGTCGTTGCCTGAGGCAGGAGAATCACTTGAANNNAGATCGGAA
32042 TATGAGTGTGGAGTCGTTGCTTCAAGTGATTCTCCTGCCTCAGNNNAGATCGGAAG
32044 TATGAGTGTGGAGTCGTTGCTGTTGAATGAAAGTGGACGTGTGNNNAGATCGGAA
32045 TATGAGTGTGGAGTCGTTGCGCTTAGGGTAGGCAACTAAAGGTNNNAGATCGGAA
32048 TATGAGTGTGGAGTCGTTGCCGCTCAAACAATCTTCCCACTTCNNNAGATCGGAAGA
32050 TATGAGTGTGGAGTCGTTGCTCTATTAGGCCAGCATCATCCTGNNNAGATCGGAAG
32051 TATGAGTGTGGAGTCGTTGCTAAGAAAAGAGCGGGGAAACAGANNNAGATCGGA
32053 TATGAGTGTGGAGTCGTTGCCGCCCCAACTGATTTTCATCATANNNAGATCGGAAGA
32057 TATGAGTGTGGAGTCGTTGCTGTCATTGTGGTTTTGATTTGCANNNAGATCGGAAG
32058 TATGAGTGTGGAGTCGTTGCTTCTGGGTTCAAGCAATTCTCCTNNNAGATCGGAAG
32059 TATGAGTGTGGAGTCGTTGCCTCTTGAGTAGCTGGGACTACAGNNNAGATCGGAA
32060 TATGAGTGTGGAGTCGTTGCGTAGTCTCCTTCTGGCCCAAATTNNNAGATCGGAAG
32062 TATGAGTGTGGAGTCGTTGCTAGGTTGGCTAATCGAGGTTTCTNNNAGATCGGAA
32064 TATGAGTGTGGAGTCGTTGCTGCAATTTACATGACAACCCCACNNNAGATCGGAAG
32065 TATGAGTGTGGAGTCGTTGCATGGTAACTTCACTGGGCTAGACNNNAGATCGGAA
32067 TATGAGTGTGGAGTCGTTGCAAAACAGACACAAAGACCAACGGNNNAGATCGGAA
32069 TATGAGTGTGGAGTCGTTGCCACGCCTGTAATCTCAGCAATTTNNNAGATCGGAAG
32070 TATGAGTGTGGAGTCGTTGCTGCACAGCAAAAGAAATAACCGANNNAGATCGGAA
32072 TATGAGTGTGGAGTCGTTGCCGCCCGGCTAACTTTTGTATTTTNNNAGATCGGAAG
32073 TATGAGTGTGGAGTCGTTGCGATTTTGAACTCCTGGCCTCAAGNNNAGATCGGAAG
32080 TATGAGTGTGGAGTCGTTGCAAAAGTTTCCAGGCATGGCTTTANNNAGATCGGAAG
32081 TATGAGTGTGGAGTCGTTGCTGCAATCAACAGACTCAAAGTGCNNNAGATCGGAAG
32083 TATGAGTGTGGAGTCGTTGCGGTGGGTGTACTTGAGAGAAGAGNNNAGATCGGA
32090 TATGAGTGTGGAGTCGTTGCGTCTCCCTATGTTACTCAGGCTGNNNAGATCGGAAG
32091 TATGAGTGTGGAGTCGTTGCCTTCCAGGTTCAAGCGATTCTCCNNNAGATCGGAAG
32092 TATGAGTGTGGAGTCGTTGCCCAATCAATTTGCCAGTCTGTGTNNNAGATCGGAAG
32096 TATGAGTGTGGAGTCGTTGCAACTGAATTCTGCCAACAACCAGNNNAGATCGGAAG

32097 TATGAGTGTGGAGTCGTTGCGGAAGCTAGGGTCTTTGCATGTANNNAGATCGGAA
32098 TATGAGTGTGGAGTCGTTGCGTGGAAATGTAGCAAGGCTTGTTNNNAGATCGGAA
32099 TATGAGTGTGGAGTCGTTGCAGTGTTCCTGAGGAAGAAGAGAANNNAGATCGGA
32103 TATGAGTGTGGAGTCGTTGCGAACAAAAGGAAGTCAGGCAAGGNNNAGATCGGA
32106 TATGAGTGTGGAGTCGTTGCCTCCTGAGTAGCTGGGATTACAGNNNAGATCGGAA
32107 TATGAGTGTGGAGTCGTTGCGGAGTACAATGGCACAATCTTGGNNNAGATCGGAA
32108 TATGAGTGTGGAGTCGTTGCAAACTCTTGACCTCAGGTGATCCNNNAGATCGGAAG
32111 TATGAGTGTGGAGTCGTTGCGGAGTGCAGTGGTGTGATTTTTTNNNAGATCGGAA
32112 TATGAGTGTGGAGTCGTTGCATGGATGGAAGAGGAGGTCACTANNNAGATCGGA
32113 TATGAGTGTGGAGTCGTTGCGAGAATCTCTTGAACCTGGGAGGNNNAGATCGGAA
32114 TATGAGTGTGGAGTCGTTGCCTGAAAATTAGCTGGGTGTGGTGNNNAGATCGGAA
32115 TATGAGTGTGGAGTCGTTGCACACATTTTAGGGAGGCATGAGANNNAGATCGGAA
32116 TATGAGTGTGGAGTCGTTGCAAATCTGCTACCTTGAAAGCCCANNNAGATCGGAAG
32117 TATGAGTGTGGAGTCGTTGCCAGGGGTTGCAATTCTAGTCTCTNNNAGATCGGAAG
32118 TATGAGTGTGGAGTCGTTGCGAAAAGAAATGTAGGCTCTGGGCNNNAGATCGGAA
32119 TATGAGTGTGGAGTCGTTGCAGGGAAACTCACACATTCAGGAANNNAGATCGGAA
32120 TATGAGTGTGGAGTCGTTGCCCCATAAGTGACCCAGATGGAAANNNAGATCGGAA
32121 TATGAGTGTGGAGTCGTTGCTTGATCTTGAACTTCTGGGCTCANNNAGATCGGAAG
32123 TATGAGTGTGGAGTCGTTGCTACAACTTGGAGAAGTCAGGGTGNNNAGATCGGAA
32125 TATGAGTGTGGAGTCGTTGCCCTCCCAGGTTCAAGCTATTCTTNNNAGATCGGAAG
32128 TATGAGTGTGGAGTCGTTGCCCACTCCTGCGTATGACATTTTTNNNAGATCGGAAG
32129 TATGAGTGTGGAGTCGTTGCCAACAGAAAAATCCGTGACCCAANNNAGATCGGAAG
32132 TATGAGTGTGGAGTCGTTGCGGATCACTTGAGCTCAGGAGTTCNNNAGATCGGAA
32135 TATGAGTGTGGAGTCGTTGCTCTTTTCCCCATTTTGCTTGGTCNNNAGATCGGAAGA
32136 TATGAGTGTGGAGTCGTTGCGAGTTGTCAAAACTCCAATGGGGNNNAGATCGGAA
32137 TATGAGTGTGGAGTCGTTGCAAATTGGGAACTGACCTTTTCCANNNAGATCGGAAG
32138 TATGAGTGTGGAGTCGTTGCCATGATCTTGGCTCACTGCAAACNNNAGATCGGAAG
32142 TATGAGTGTGGAGTCGTTGCTGATCTTGGCTCACTACAACCTCNNNAGATCGGAAG
32143 TATGAGTGTGGAGTCGTTGCGGGTTCAAACTATTCTCCTGCCTNNNAGATCGGAAG
32144 TATGAGTGTGGAGTCGTTGCGTGAATGGCATAATTTCGGCTCANNNAGATCGGAA
32146 TATGAGTGTGGAGTCGTTGCTCTCCCAAAGTGCTGGAATTACANNNAGATCGGAAG
32147 TATGAGTGTGGAGTCGTTGCGCTCAAGGGATCCTTTTACCTCANNNAGATCGGAAG
32149 TATGAGTGTGGAGTCGTTGCGCAAGATCTTAGCTCACTGCAAGNNNAGATCGGAAG
32150 TATGAGTGTGGAGTCGTTGCTGACTTTTCCTTTTGCTGTGCAANNNAGATCGGAAG
32151 TATGAGTGTGGAGTCGTTGCCTGGCAGCTACAAATCCCAATATNNNAGATCGGAAG
32152 TATGAGTGTGGAGTCGTTGCACTGCAAGGCCATTGTCCTAATANNNAGATCGGAAG
32154 TATGAGTGTGGAGTCGTTGCGAGAATCACTTGAACCAGGGAGTNNNAGATCGGAA
32155 TATGAGTGTGGAGTCGTTGCAGATCTGCCTGCAAAATTCCATTNNNAGATCGGAAG
32157 TATGAGTGTGGAGTCGTTGCAAAAAAGAGGGGCTTTGTACCACNNNAGATCGGAA
32158 TATGAGTGTGGAGTCGTTGCGATTTTCCAGACTGGCTACAAGCNNNAGATCGGAAG
32159 TATGAGTGTGGAGTCGTTGCCCTCCCAAAGTGCTGAGATTACANNNAGATCGGAAG
32160 TATGAGTGTGGAGTCGTTGCGGATTCAAGAGATTCTCCTGCCTNNNAGATCGGAAG
32161 TATGAGTGTGGAGTCGTTGCCTCCTGAGTAGCTGGGATTACAGNNNAGATCGGAA

32166 TATGAGTGTGGAGTCGTTGCCTCCTGGGTTCACACCATTCTCNNNAGATCGGAAGAG
32167 TATGAGTGTGGAGTCGTTGCTTTGCTTTTCTGTTGGGGGAAAGNNNAGATCGGAAG
32169 TATGAGTGTGGAGTCGTTGCAGGCCTAAGACAAAGTCCAATGTNNNAGATCGGAAG
32170 TATGAGTGTGGAGTCGTTGCTTGACAAGGTCTCACTCTGTTGCNNNAGATCGGAAGA
32172 TATGAGTGTGGAGTCGTTGCGTTCAAGTGATTCTCCTGCCTCANNNAGATCGGAAGA
32173 TATGAGTGTGGAGTCGTTGCGATATGGAGTCTCCCACTGTCACNNNAGATCGGAAG
32175 TATGAGTGTGGAGTCGTTGCAGAAGAGTTCTCACTGTGTTGCCNNNAGATCGGAAG
32177 TATGAGTGTGGAGTCGTTGCGGTTCACTTGAGACCCTGTAACANNNAGATCGGAAG
32178 TATGAGTGTGGAGTCGTTGCTTATCATGCTTCACCTTCCCACANNNAGATCGGAAGA
32179 TATGAGTGTGGAGTCGTTGCTGGCAGTCCCTCAAAAATATGACNNNAGATCGGAAG
32183 TATGAGTGTGGAGTCGTTGCGTTTTACCATGTTGGCCAGTCTGNNNAGATCGGAAG
32184 TATGAGTGTGGAGTCGTTGCTACCAAGCTATGAAGTTGGGCATNNNAGATCGGAAG
32185 TATGAGTGTGGAGTCGTTGCAGGGCCTATAGTCCCAACTACTTNNNAGATCGGAAGA
32186 TATGAGTGTGGAGTCGTTGCTCCTGGAGGAAAAAGTCAGTAGCNNNAGATCGGAAG
32187 TATGAGTGTGGAGTCGTTGCGGTGAAACCCCGTCTCTACTAAANNNAGATCGGAAG
32191 TATGAGTGTGGAGTCGTTGCCTGCTTCAACCTCCTGAGTAACTNNNAGATCGGAAGA
32192 TATGAGTGTGGAGTCGTTGCCACAGGTTCAAGCAATTCTCCTGNNNAGATCGGAAGA
32193 TATGAGTGTGGAGTCGTTGCTCAGGCCGTAGCTTAGATGTTACNNNAGATCGGAAG
32194 TATGAGTGTGGAGTCGTTGCATCTCCCAGGTTCAAGCAATTCTNNNAGATCGGAAGA
32195 TATGAGTGTGGAGTCGTTGCTGTGGTTTTGATTTGAAAGCCCANNNAGATCGGAAG
32196 TATGAGTGTGGAGTCGTTGCGCTGTCAGACTGGGTGTAAAGTANNNAGATCGGAAG
32197 TATGAGTGTGGAGTCGTTGCTAATCTCGACTCACTGCAACCTCNNNAGATCGGAAGA
32198 TATGAGTGTGGAGTCGTTGCGCCTCCAGCATAGCCAGAATTATNNNAGATCGGAAG
32199 TATGAGTGTGGAGTCGTTGCGAGTAGCTGGGATTACAGGCATGNNNAGATCGGAA
32201 TATGAGTGTGGAGTCGTTGCCAGTATTTGGGACTACAGGGGACNNNAGATCGGAAG
32203 TATGAGTGTGGAGTCGTTGCCTTCAAGCAATACTCCTGCCTCANNNAGATCGGAAGA
32204 TATGAGTGTGGAGTCGTTGCTAAAGTCGGTCCTCATTCAGGTCNNNAGATCGGAAG
32206 TATGAGTGTGGAGTCGTTGCTCTTGTTCTTTTTTATGGCTGCANNNAGATCGGAAGA
32207 TATGAGTGTGGAGTCGTTGCGTAAATTCTTCACACCCCTGTGCNNNAGATCGGAAGA
32208 TATGAGTGTGGAGTCGTTGCTATCCAGAACTGAACACCTTGCTNNNAGATCGGAAGA
32209 TATGAGTGTGGAGTCGTTGCGGTAGGACTGCCACCAGAAATTANNNAGATCGGAAG
32212 TATGAGTGTGGAGTCGTTGCTTTGTCATGAACCATCTGTTGCCNNNAGATCGGAAGA
32213 TATGAGTGTGGAGTCGTTGCAAAGGAGGAGGGTTCATGGTTACNNNAGATCGGAA
32218 TATGAGTGTGGAGTCGTTGCTGTGTTAGACGTTCTAGAGCCTGNNNAGATCGGAAG
32219 TATGAGTGTGGAGTCGTTGCCCACTGTAACCACTACCTGTCTTNNNAGATCGGAAGA
32221 TATGAGTGTGGAGTCGTTGCCCTCCTGGGTTCAAGTGATTCTCNNNAGATCGGAAGA
32222 TATGAGTGTGGAGTCGTTGCGAGGGAAGGAAGGAAAGAAGGAANNNAGATCGGA
32223 TATGAGTGTGGAGTCGTTGCCCTTGCCTCAATTTGGGACAATTNNNAGATCGGAAGA
32224 TATGAGTGTGGAGTCGTTGCCTCAAGCAATCCTCCTACCTCAGNNNAGATCGGAAGA
32225 TATGAGTGTGGAGTCGTTGCGGCCCAAATTTCACCCATTCAATNNNAGATCGGAAGA
32227 TATGAGTGTGGAGTCGTTGCTGGAGGATACATTTGCTGTTGGANNNAGATCGGAAG
32228 TATGAGTGTGGAGTCGTTGCAGTTACAGCACCTAAAACCCCAANNNAGATCGGAAGA
32229 TATGAGTGTGGAGTCGTTGCTGAGAAACAAGGACAGGACACAANNNAGATCGGAA

32230 TATGAGTGTGGAGTCGTTGCGAAACCTTTGGCCAATCAGAGAGNNNAGATCGGAA
32231 TATGAGTGTGGAGTCGTTGCGAGAAAGGGTGAAGGCAAACATCNNNAGATCGGA
32232 TATGAGTGTGGAGTCGTTGCCCCTCTGCACAGTAATGAGTGAANNNAGATCGGAAG
32233 TATGAGTGTGGAGTCGTTGCAGTGCTGGGATTACAGGTGTAAGNNNAGATCGGAA
32234 TATGAGTGTGGAGTCGTTGCTCTGGGTGCTGAGTCTCTAATTGNNNAGATCGGAAG
32235 TATGAGTGTGGAGTCGTTGCGGCTTTTCCTTCATCATTTGCCTNNNAGATCGGAAGA
32237 TATGAGTGTGGAGTCGTTGCTCCCCAGATTCCTCTCTAGAGTCNNNAGATCGGAAG
32240 TATGAGTGTGGAGTCGTTGCACATGGTGAAACCTCGTCTCTACNNNAGATCGGAAG
32241 TATGAGTGTGGAGTCGTTGCGTGGTGGTGACTGGAGATTAGTTNNNAGATCGGAA
32242 TATGAGTGTGGAGTCGTTGCAGCTGAAGGATTTACGTTACCCANNNAGATCGGAAG
32243 TATGAGTGTGGAGTCGTTGCTGGGAGGAGTGAGTATACCAAGTNNNAGATCGGAA
32244 TATGAGTGTGGAGTCGTTGCGCAGTCTGTCCATTCTCAGATCTNNNAGATCGGAAG
32245 TATGAGTGTGGAGTCGTTGCTTCCCCCCAGAAAAATGCAGATANNNAGATCGGAAG
32248 TATGAGTGTGGAGTCGTTGCCTGTTGCTCAGGCTATAGTCTCANNNAGATCGGAAG
32249 TATGAGTGTGGAGTCGTTGCCTTTCTCTTTCTGCTCCCCCTACNNNAGATCGGAAGAG
32250 TATGAGTGTGGAGTCGTTGCGCTTTGTGACGTATAAGGCCTTCNNNAGATCGGAAG
32251 TATGAGTGTGGAGTCGTTGCTTGTATTCCTCACACTTCCCTGGNNNAGATCGGAAGA
32252 TATGAGTGTGGAGTCGTTGCGCCAAAAAAAAGGAGTCTCAGTANNNAGATCGGAA
32253 TATGAGTGTGGAGTCGTTGCCCCCCCAAAAAAAAGTGTGTGTANNNAGATCGGAAG
32255 TATGAGTGTGGAGTCGTTGCAGAAACTGGGAGTCATCGATCACNNNAGATCGGAA
32256 TATGAGTGTGGAGTCGTTGCGATGTTCAAAGAACTGGTGCCAANNNAGATCGGAA
32257 TATGAGTGTGGAGTCGTTGCCTACTTAGGCACAAGATGGCTTGNNNAGATCGGAAG
32258 TATGAGTGTGGAGTCGTTGCGGAGCAATTTCTTACCTCCCTCANNNAGATCGGAAG
32259 TATGAGTGTGGAGTCGTTGCAGATCCACTGGCTTCACATTCTTNNNAGATCGGAAG
32260 TATGAGTGTGGAGTCGTTGCCCCCTTCCTGCTACATTCTGAAANNNAGATCGGAAGA
32261 TATGAGTGTGGAGTCGTTGCCATCTGTGGTTCCAGCTACTCAGNNNAGATCGGAAG
32262 TATGAGTGTGGAGTCGTTGCCGCCAATACGTAAAAGAAGCCATNNNAGATCGGAAG
32263 TATGAGTGTGGAGTCGTTGCATGACCCATGATGAAACCCATGANNNAGATCGGAAG
32265 TATGAGTGTGGAGTCGTTGCGCACCCCTAAAGATAGTTCCCAANNNAGATCGGAAG
32266 TATGAGTGTGGAGTCGTTGCTATTACCTGGGCTGGTCTCAAACNNNAGATCGGAAG
32267 TATGAGTGTGGAGTCGTTGCGTACAACAGTCAACACTCCTTGCNNNAGATCGGAAG
32268 TATGAGTGTGGAGTCGTTGCCAGTGAAGAGAAGAGGAAGGCATNNNAGATCGGA
32269 TATGAGTGTGGAGTCGTTGCGATGCTAATCCTGAGCGAAAACCNNNAGATCGGAAG
32270 TATGAGTGTGGAGTCGTTGCGTCAGTTTGCAGACAACAGAGTCNNNAGATCGGAA
32271 TATGAGTGTGGAGTCGTTGCAGGTGAGAGCAAGAGTAAAATCANNNAGATCGGAA
32272 TATGAGTGTGGAGTCGTTGCCCCCAATTTAGAAGCTTGTGTGGNNNAGATCGGAAG
32274 TATGAGTGTGGAGTCGTTGCGTCCCTCCCATAACACATGAGAANNNAGATCGGAAG
32275 TATGAGTGTGGAGTCGTTGCCCACCGACATTTGACTCTTTGTTNNNAGATCGGAAGA
32276 TATGAGTGTGGAGTCGTTGCAAACTAGGAATGTTGCCCCTCATNNNAGATCGGAAG
32277 TATGAGTGTGGAGTCGTTGCTTTTTTTCGAGACGGAGTTTCGCNNNAGATCGGAAG
32278 TATGAGTGTGGAGTCGTTGCCTGGAATTACAAGTGTGCACCACNNNAGATCGGAAG
32279 TATGAGTGTGGAGTCGTTGCACCCCTATGTCCCTAGGTGAATTNNNAGATCGGAAG
32280 TATGAGTGTGGAGTCGTTGCTGGTGCTCAGAACAGATCTATGGNNNAGATCGGAA

32281 TATGAGTGTGGAGTCGTTGCTCTAATCCAAACTAGAGCTGGGCNNNAGATCGGAAG
32285 TATGAGTGTGGAGTCGTTGCGGCTCGCAACTAATTTCCTGTTTNNNAGATCGGAAG
32287 TATGAGTGTGGAGTCGTTGCTACGTTGCCCAGACTAGTTTCAANNNAGATCGGAAG
32288 TATGAGTGTGGAGTCGTTGCCATCGTGTAATTGCTCACCTCCTNNNAGATCGGAAG
32289 TATGAGTGTGGAGTCGTTGCTGATCTCGGCTCACTGAAACATCNNNAGATCGGAAG
32290 TATGAGTGTGGAGTCGTTGCAAAGTGGGTTGAAGCAAGATTGGNNNAGATCGGA
32291 TATGAGTGTGGAGTCGTTGCTGTCTGTTGGTGTATCTGTGTGTNNNAGATCGGAAG
32292 TATGAGTGTGGAGTCGTTGCACGTGGAGAAGGAACCTATTGACNNNAGATCGGAA
32293 TATGAGTGTGGAGTCGTTGCGAATCAAGAATCAGGCTACCACGNNNAGATCGGAA
32294 TATGAGTGTGGAGTCGTTGCCAGCTCACTTGGGAGATCTGAAANNNAGATCGGAA
32295 TATGAGTGTGGAGTCGTTGCTTGGTGGGCACTTAGGTTGATTANNNAGATCGGAA
32296 TATGAGTGTGGAGTCGTTGCAGGGCACAGACAAACAAAAAGACNNNAGATCGGAA
32298 TATGAGTGTGGAGTCGTTGCTACAGGGCTGAGCTACCATATCTNNNAGATCGGAAG
32299 TATGAGTGTGGAGTCGTTGCTATAACAGTGTTGAGAGGTGGGCNNNAGATCGGAA
32300 TATGAGTGTGGAGTCGTTGCTATAAACGCCCTCATCTTGCCATNNNAGATCGGAAG
32301 TATGAGTGTGGAGTCGTTGCTGTGTGTGTTTGTGGTATGTGTGNNNAGATCGGAA
32302 TATGAGTGTGGAGTCGTTGCGGAGGACAGGTAATCCAATTGGTNNNAGATCGGAA
32303 TATGAGTGTGGAGTCGTTGCTGTCACAACAATCCAATGAGCTGNNNAGATCGGAAG
32304 TATGAGTGTGGAGTCGTTGCCACTGAAAGCGGAGTTTAGGATGNNNAGATCGGAA
32306 TATGAGTGTGGAGTCGTTGCAGAGAGAGAGAGAGGGAGAGAGANNNAGATCGG
32307 TATGAGTGTGGAGTCGTTGCACTAGGAAGAATTGGCTCACAGGNNNAGATCGGAA
32308 TATGAGTGTGGAGTCGTTGCCACTTGTGTGGGTATGTGTATCTNNNAGATCGGAAG
32309 TATGAGTGTGGAGTCGTTGCATGACTTCCTTTGGGTAGACTCCNNNAGATCGGAAG
32310 TATGAGTGTGGAGTCGTTGCTTGATGACAGGGAGCACGAATTANNNAGATCGGAA
32311 TATGAGTGTGGAGTCGTTGCTTGATGCTGTTGTTGCTTTCTGTNNNAGATCGGAAG
32312 TATGAGTGTGGAGTCGTTGCCCACCTGGTCTCCACAAAAAATCNNNAGATCGGAAG
32313 TATGAGTGTGGAGTCGTTGCCACTCTGTTTCCTGTTCAGCATGNNNAGATCGGAAG
32315 TATGAGTGTGGAGTCGTTGCTTTACGATGAATCTTGCTGCTGCNNNAGATCGGAAG
32316 TATGAGTGTGGAGTCGTTGCTGTTCAGATTCTTTGGTCCCACANNNAGATCGGAAG
32317 TATGAGTGTGGAGTCGTTGCGGTAGCTGTGTTGTTCTTATGCCNNNAGATCGGAAG
32319 TATGAGTGTGGAGTCGTTGCCTGTAGTCCCAGCTACTTGAGAGNNNAGATCGGAAG
32322 TATGAGTGTGGAGTCGTTGCGCCTCAGGATACAAACAAACCACNNNAGATCGGAAG
32323 TATGAGTGTGGAGTCGTTGCTGTGTGTCTTCCGTGCATATGTANNNAGATCGGAAG
32324 TATGAGTGTGGAGTCGTTGCATTTCACACCTTCTGCTCTTTGCNNNAGATCGGAAGA
32325 TATGAGTGTGGAGTCGTTGCGTGTGTGTGCATGTGTGTATGAANNNAGATCGGAA
32326 TATGAGTGTGGAGTCGTTGCGGGTTCAAGTGATTCTCCTGTCTNNNAGATCGGAAG
32327 TATGAGTGTGGAGTCGTTGCCTTAGAAGAAGGTATCGGGCCTCNNNAGATCGGAA
32329 TATGAGTGTGGAGTCGTTGCCCCTCTCTTTCTCGGTACCTTTCNNNAGATCGGAAGA
32332 TATGAGTGTGGAGTCGTTGCATTTGTTTTTCTGTGCCTGGCTTNNNAGATCGGAAG
32333 TATGAGTGTGGAGTCGTTGCTTGGGTTCTGGTTCTCTTTGGTTNNNAGATCGGAAG
32335 TATGAGTGTGGAGTCGTTGCGAGAAACCATGGTGTCGTCTTTGNNNAGATCGGAA
32336 TATGAGTGTGGAGTCGTTGCCAATCTCCACCTCCTGAGTTCAANNNAGATCGGAAG
32337 TATGAGTGTGGAGTCGTTGCTGACAAGAAGGACAGAAGAGCTGNNNAGATCGGA

32338 TATGAGTGTGGAGTCGTTGCAGAGTTTGAGCCAGAGTCATGTTNNNAGATCGGAA
32340 TATGAGTGTGGAGTCGTTGCGGCAGGACTGACAATAAATCAGCNNNAGATCGGAA
32341 TATGAGTGTGGAGTCGTTGCTTTCTTACAGTTCTGGAGGCTGGNNNAGATCGGAAG
32342 TATGAGTGTGGAGTCGTTGCCAGACACCAGGGAAAAGAGGTAANNNAGATCGGA
32343 TATGAGTGTGGAGTCGTTGCCACAATCATGACTCATTGCAGCTNNNAGATCGGAAG
32344 TATGAGTGTGGAGTCGTTGCGGGGCAATCTCTAAGGGTTACATNNNAGATCGGAA
32345 TATGAGTGTGGAGTCGTTGCAGACAGGGTCTCATTTTGTCACCNNNAGATCGGAAG
32346 TATGAGTGTGGAGTCGTTGCTCACCAATATCCACTGTTCTGCANNNAGATCGGAAG
32347 TATGAGTGTGGAGTCGTTGCTGTTGTTGTTGTTGAGACAGAGTNNNAGATCGGAA
32348 TATGAGTGTGGAGTCGTTGCAGAGAGAGAGACAGAGATGGCTCNNNAGATCGGA
32349 TATGAGTGTGGAGTCGTTGCGTGGTGTAGATCTGTGGGAACTCNNNAGATCGGAA
32350 TATGAGTGTGGAGTCGTTGCATGTACACTCACACAGACACACANNNAGATCGGAAG
32351 TATGAGTGTGGAGTCGTTGCGAGGAAACATGACAGGTGAGTCTNNNAGATCGGAA
32352 TATGAGTGTGGAGTCGTTGCCCTCACACACACATACTCAGTCANNNAGATCGGAAG
32353 TATGAGTGTGGAGTCGTTGCCAACACACACACACACACACATCNNNAGATCGGAAG
32355 TATGAGTGTGGAGTCGTTGCTGTCTGGTTTTGAAGTGTTTTCANNNAGATCGGAAG
32356 TATGAGTGTGGAGTCGTTGCGCATTCCCTAACTCGTTCCAAAGNNNAGATCGGAAG
32360 TATGAGTGTGGAGTCGTTGCTCTCAGTGGAAGTTGGGATTTCCNNNAGATCGGAAG
32361 TATGAGTGTGGAGTCGTTGCCACACAGACAAACACACACACATNNNAGATCGGAAG
32363 TATGAGTGTGGAGTCGTTGCGCATATATGTAGGCCGGGTTGTANNNAGATCGGAA
32364 TATGAGTGTGGAGTCGTTGCACACTATGCAAAGCACAAATCCCNNNAGATCGGAAG
32365 TATGAGTGTGGAGTCGTTGCTTATGCTTTTTGAAGGCATCCGTNNNAGATCGGAAG
32366 TATGAGTGTGGAGTCGTTGCCCTTCTCTCCGTGTCTCTACTCTNNNAGATCGGAAGA
32368 TATGAGTGTGGAGTCGTTGCCCCCGCGAAGATATGTTGAAATCNNNAGATCGGAAG
32369 TATGAGTGTGGAGTCGTTGCTCAACGAATCCAGGAGTTGACTTNNNAGATCGGAAG
32370 TATGAGTGTGGAGTCGTTGCACTATCTGATCATATGCAGTGCTNNNAGATCGGAAG
32371 TATGAGTGTGGAGTCGTTGCTGCATAGCTGTTTCTGCAGGTATNNNAGATCGGAAG
32372 TATGAGTGTGGAGTCGTTGCGGAGTACAGTGGTGTGATCTCAGNNNAGATCGGAA
32374 TATGAGTGTGGAGTCGTTGCGAGAGAGAGAGAGAGAGACTGCTNNNAGATCGGA
32376 TATGAGTGTGGAGTCGTTGCAATGCACCAGATCCCATCTCATTNNNAGATCGGAAG
32377 TATGAGTGTGGAGTCGTTGCCACCACCACCATTTTTCACAGAANNNAGATCGGAAG
32378 TATGAGTGTGGAGTCGTTGCCAGAATAGCTTGAACCCAGGAGGNNNAGATCGGAA
32379 TATGAGTGTGGAGTCGTTGCGAAAACATGGGAGCACCTTCTTCNNNAGATCGGAAG
32382 TATGAGTGTGGAGTCGTTGCGCTAGGAAAGACCAGACCAAAGANNNAGATCGGAA
32384 TATGAGTGTGGAGTCGTTGCGTTTGCTCCTCACAAGAACCTTTNNNAGATCGGAAG
32385 TATGAGTGTGGAGTCGTTGCGATTACAGGCATGCACCATCATGNNNAGATCGGAAG
32386 TATGAGTGTGGAGTCGTTGCCAGCTCACTGCAGTCTCAATTTCNNNAGATCGGAAG
32388 TATGAGTGTGGAGTCGTTGCTTCAAGCAATTTTCCTGCCTCAGNNNAGATCGGAAG
32393 TATGAGTGTGGAGTCGTTGCACAGCTTTTTTCCTTTGGCTCTCNNNAGATCGGAAGA
32394 TATGAGTGTGGAGTCGTTGCCAAGTAGCTGGGATTACAGGCATNNNAGATCGGAA
32397 TATGAGTGTGGAGTCGTTGCTGGATTCCAATTGAGAATGTGTGACNNNAGATCGG
32398 TATGAGTGTGGAGTCGTTGCTCGTTATCCTGACGAAGCAGAATNNNAGATCGGAAG
32399 TATGAGTGTGGAGTCGTTGCTGCTCAGTTGTTAGTGGCTTATGNNNAGATCGGAAG

32400 TATGAGTGTGGAGTCGTTGCATCTTTTCAGCATCCCACTCTGTNNNAGATCGGAAGA
32401 TATGAGTGTGGAGTCGTTGCAGGGGATGACTGCTGGGTATATANNNAGATCGGAA
32402 TATGAGTGTGGAGTCGTTGCTGATGAACTAGGTCTACATTGCANNNAGATCGGAAG
32404 TATGAGTGTGGAGTCGTTGCGTGTGGTGTCCTCAAAAACTGACNNNAGATCGGAAG
32409 TATGAGTGTGGAGTCGTTGCAGTAACATTCTCAGGAGGGAGGANNNAGATCGGAA
32410 TATGAGTGTGGAGTCGTTGCCCATCTCACACCAATCAGAATGGNNNAGATCGGAAGA
32411 TATGAGTGTGGAGTCGTTGCTCTGGTTACTAGTAAAGCCACGTNNNAGATCGGAAG
32413 TATGAGTGTGGAGTCGTTGCTGTTTGCTGTATGACCTTGGACTNNNAGATCGGAAG
32414 TATGAGTGTGGAGTCGTTGCAAAAAAACAAACCTAGCCAGGCANNNAGATCGGAAG
32416 TATGAGTGTGGAGTCGTTGCAGTTTCTGGCATTTCCCTACAGTNNNAGATCGGAAGA
32418 TATGAGTGTGGAGTCGTTGCTCTTGGAGTACAGTGGTGCAATCNNNAGATCGGAAG
32419 TATGAGTGTGGAGTCGTTGCTCTGCCTTCTAGGTTCAAGCAATNNNAGATCGGAAGA
32420 TATGAGTGTGGAGTCGTTGCGCACTAGCATTACAGGCATTAGCNNNAGATCGGAAG
32421 TATGAGTGTGGAGTCGTTGCTTTTTGTAGAGATGGGGTTTCCANNNAGATCGGAAG
32422 TATGAGTGTGGAGTCGTTGCCTGAGTAGCTGGGATTATAGGCGNNNAGATCGGAA
32424 TATGAGTGTGGAGTCGTTGCTGATAGAGCATCTACAGGGGTCANNNAGATCGGAAG
32425 TATGAGTGTGGAGTCGTTGCCTCCTGAGTAGCTGGGACTACAGNNNAGATCGGAAG
32428 TATGAGTGTGGAGTCGTTGCGAGAAGGAATCTTGCTCTGTTGCNNNAGATCGGAAG
32429 TATGAGTGTGGAGTCGTTGCGATGGAGTCTCTCGTTCTGTCACNNNAGATCGGAAG
32430 TATGAGTGTGGAGTCGTTGCTTCAAGTGATTCTCCTGCCTCAGNNNAGATCGGAAGA
32431 TATGAGTGTGGAGTCGTTGCATTCAGGGGCAGTCAGTTGATAGNNNAGATCGGAAG
32432 TATGAGTGTGGAGTCGTTGCTTTTTTTCCCCCACAGCTTTGAGNNNAGATCGGAAGA
32433 TATGAGTGTGGAGTCGTTGCATTCTCCTGCTTTAGCCTCCCTANNNAGATCGGAAGAG
32434 TATGAGTGTGGAGTCGTTGCCCAGGTCAAAGTGATTCTCTTGCNNNAGATCGGAAG
32435 TATGAGTGTGGAGTCGTTGCAACTCCTGGGCTCAAAAGATTCTNNNAGATCGGAAG
32436 TATGAGTGTGGAGTCGTTGCAGGGATTGCATTGCATCTGTAGANNNAGATCGGAAG
32437 TATGAGTGTGGAGTCGTTGCCTGCAAACCGCTCATTCTGTAAANNNAGATCGGAAGA
32440 TATGAGTGTGGAGTCGTTGCCTCTCGAGTTGCTGGAACTACAGNNNAGATCGGAAG
32441 TATGAGTGTGGAGTCGTTGCGCAGTAGTGCAATCTCAGCTCATNNNAGATCGGAAG
32442 TATGAGTGTGGAGTCGTTGCCCTCCCAAGTAGCTGGAATTACANNNAGATCGGAAG
32444 TATGAGTGTGGAGTCGTTGCTTTGAAGTTGGGTAGCATGATGCNNNAGATCGGAAG
32445 TATGAGTGTGGAGTCGTTGCCCTCCTGGATTCAAGCGATTCTCNNNAGATCGGAAGA
32451 TATGAGTGTGGAGTCGTTGCGTAGTTTTCCTTTGCACAGGTCCNNNAGATCGGAAGA
32456 TATGAGTGTGGAGTCGTTGCAGTGCAAAAAGGCTGAGAATTCCNNNAGATCGGAAG
32460 TATGAGTGTGGAGTCGTTGCGTCCAGACACATAGACCAAAGGANNNAGATCGGAAG
32461 TATGAGTGTGGAGTCGTTGCGTTCCCTTCAAGTCTGGTAGGTTNNNAGATCGGAAG
32462 TATGAGTGTGGAGTCGTTGCGCAGGCATCTGTAATCCCATCTANNNAGATCGGAAGA
32464 TATGAGTGTGGAGTCGTTGCCTGAAGTGAGTGGATCTCCTGAGNNNAGATCGGAAG
32465 TATGAGTGTGGAGTCGTTGCAAAGGGTTCGCTGTAGATTCAGTNNNAGATCGGAAG
32466 TATGAGTGTGGAGTCGTTGCTCAGTCCAGGATACAATGAAGGGNNNAGATCGGAAG
32469 TATGAGTGTGGAGTCGTTGCCACACAGAGAAATCAGGTCCTCANNNAGATCGGAAG
32472 TATGAGTGTGGAGTCGTTGCGGTTCAAATGATTCTCCTGCCTCNNNAGATCGGAAGA
32474 TATGAGTGTGGAGTCGTTGCGGAGTACAGTGGCACAATCACAGNNNAGATCGGAAG

| | |
|---|---|
| 32475 | TATGAGTGTGGAGTCGTTGCATAATTCTCATGCCTCAGCCTCCNNNAGATCGGAAGA |
| 32476 | TATGAGTGTGGAGTCGTTGCCCTCCTGAGTAGCTGAGACTACANNNAGATCGGAAG |
| 32477 | TATGAGTGTGGAGTCGTTGCCTTGATCTCCTGACCTCGTGATCNNNAGATCGGAAG |
| 32478 | TATGAGTGTGGAGTCGTTGCAACCTCTGCTTCCTGTGATTCTCNNNAGATCGGAAGA |
| 32482 | TATGAGTGTGGAGTCGTTGCAAATTTTGCTCTATTGCCCAGGCNNNAGATCGGAAG |
| 32483 | TATGAGTGTGGAGTCGTTGCTTCAGGGATTCAACTTCCTCCTGNNNAGATCGGAAG |
| 32484 | TATGAGTGTGGAGTCGTTGCAGTAGCTAGGATTACAGGCATGCNNNAGATCGGAA |
| 32485 | TATGAGTGTGGAGTCGTTGCCTCAAGAGATCCTCCCACCTTAGNNNAGATCGGAAG |
| 32488 | TATGAGTGTGGAGTCGTTGCCAAGTAGCTGGGATTACAGGCATNNNAGATCGGAA |
| 32491 | TATGAGTGTGGAGTCGTTGCTGGTGGCTTATCCCTGTAATTCCNNNAGATCGGAAG |
| 32493 | TATGAGTGTGGAGTCGTTGCTCTCGATCTGACCTTGTGATCTGNNNAGATCGGAAG |
| 32494 | TATGAGTGTGGAGTCGTTGCTTCAATCGATTCTCCTGCCTCAGNNNAGATCGGAAG |
| 32496 | TATGAGTGTGGAGTCGTTGCCTCCTGAGAAGCTGGGATTACAGNNNAGATCGGAA |
| 32497 | TATGAGTGTGGAGTCGTTGCGGTTCAAACAATTCTCCTGCCTCNNNAGATCGGAAG |
| 32498 | TATGAGTGTGGAGTCGTTGCGTAGTAACAGGCCTTGGCTTTTCNNNAGATCGGAAG |
| 32499 | TATGAGTGTGGAGTCGTTGCCCTCCATAGTAGCTGAGACCTCANNNAGATCGGAAG |
| 32500 | TATGAGTGTGGAGTCGTTGCGATCTTGGCTCGCTGCAAATTCNNNAGATCGGAAGA |
| 32501 | TATGAGTGTGGAGTCGTTGCGGCTCCTGAGTAGCTGAGATTACNNNAGATCGGAA |
| 32503 | TATGAGTGTGGAGTCGTTGCTTCAAGCAACTCTCCTGTCTCAGNNNAGATCGGAAG |
| 32505 | TATGAGTGTGGAGTCGTTGCTAGCTGGGATTATAGGCACATGCNNNAGATCGGAA |
| 32506 | TATGAGTGTGGAGTCGTTGCGACAATGGAAGTGTCTCATCCCTNNNAGATCGGAAG |
| 32507 | TATGAGTGTGGAGTCGTTGCAAACCTCACGTCACAGCAAAAATNNNAGATCGGAAG |
| 32508 | TATGAGTGTGGAGTCGTTGCTTCCTTCTGTGCAGATCATCTCCNNNAGATCGGAAGA |
| 32513 | TATGAGTGTGGAGTCGTTGCGAATGAGGATGAGGATGAGGACANNNAGATCGGA |
| 32515 | TATGAGTGTGGAGTCGTTGCATTCTCGTGTATGGCTTGAGGAANNNAGATCGGAA |
| 32516 | TATGAGTGTGGAGTCGTTGCAGAGCGAAACGAACATCAACAATNNNAGATCGGAA |
| 32517 | TATGAGTGTGGAGTCGTTGCGCCATATATGACAGACCCACAGTNNNAGATCGGAAG |
| 32518 | TATGAGTGTGGAGTCGTTGCGGATGAGTAGGGAGGCAAAGAAGNNNAGATCGGA |
| 32520 | TATGAGTGTGGAGTCGTTGCACAGCACATGACCTGTTTAGAGANNNAGATCGGAA |
| 32521 | TATGAGTGTGGAGTCGTTGCGAGCTTCAGTACAGTGTCTGGAANNNAGATCGGAA |
| 32522 | TATGAGTGTGGAGTCGTTGCAAAGAAATAAAACTCCCCTGGCCNNNAGATCGGAAG |
| 32523 | TATGAGTGTGGAGTCGTTGCATCCAGGTTGCCAAAGCTACTAANNNAGATCGGAAG |
| 32525 | TATGAGTGTGGAGTCGTTGCTCTGGAGACATCAACTGACAACCNNNAGATCGGAAG |
| 32527 | TATGAGTGTGGAGTCGTTGCGGAGCATGAGTAGCCAATACAATNNNAGATCGGAA |
| 32528 | TATGAGTGTGGAGTCGTTGCTTTTCCTTCAAGTCAGAGGTTCCNNNAGATCGGAAG |
| 32529 | TATGAGTGTGGAGTCGTTGCTCCCTCTAGCCTCTAGAACTTCANNNAGATCGGAAGA |
| 32531 | TATGAGTGTGGAGTCGTTGCGCAGGGTTTGCAATCCTAGTTTCNNNAGATCGGAAG |
| 32532 | TATGAGTGTGGAGTCGTTGCAGCAACATCAAACCTGAGAGTCANNNAGATCGGAA |
| 32533 | TATGAGTGTGGAGTCGTTGCATTATCCAAGTGGGCCCAATTCANNNAGATCGGAAG |
| 32534 | TATGAGTGTGGAGTCGTTGCGGAGTCTGAGGCACAATAATTGCNNNAGATCGGAA |
| 32540 | TATGAGTGTGGAGTCGTTGCAGAAAGAATGGCAAAGGGTCTGANNNAGATCGGA |
| 32541 | TATGAGTGTGGAGTCGTTGCAAAAAAGACAGACACAGGAGCAGNNNAGATCGGA |
| 32543 | TATGAGTGTGGAGTCGTTGCGGGTTGCAATCCTAGTCTCTGATNNNAGATCGGAAG |

32544 TATGAGTGTGGAGTCGTTGCTTTGCAACTTGTCATGGGAAACANNNAGATCGGAAG
32545 TATGAGTGTGGAGTCGTTGCGGTGATGGAAACAGAGAGGGAAANNNAGATCGGA
32546 TATGAGTGTGGAGTCGTTGCCTGAGGTGGGAGAATTGCTTGAGNNNAGATCGGA
32547 TATGAGTGTGGAGTCGTTGCTGATGTCTGTCAATTTTTGAGCTCANNNAGATCGGA
32548 TATGAGTGTGGAGTCGTTGCAAAAGACATAGAGTGGCTGAGGGNNNAGATCGGA
32551 TATGAGTGTGGAGTCGTTGCTCTCACTCTGTTCTTTCACCCAGNNNAGATCGGAAGA
32552 TATGAGTGTGGAGTCGTTGCGGGTTCACGCCATGATAAGAATGNNNAGATCGGAA
32555 TATGAGTGTGGAGTCGTTGCCTCACTGTAGCCTCAACTTCTCANNNAGATCGGAAGA
32557 TATGAGTGTGGAGTCGTTGCCAATCTTGGTTCACTGCAACCTCNNNAGATCGGAAG
32560 TATGAGTGTGGAGTCGTTGCCCAAGTAGCTGGGATTATAGGCANNNAGATCGGAA
32562 TATGAGTGTGGAGTCGTTGCCTCAAGTGATTCTCCTGCCTCAGNNNAGATCGGAAG
32563 TATGAGTGTGGAGTCGTTGCGTTTCAAGCAATTCTCTGCCTCANNNAGATCGGAAG
32565 TATGAGTGTGGAGTCGTTGCCTCCTGAGTAGCTGGGACTACAGNNNAGATCGGAA
32566 TATGAGTGTGGAGTCGTTGCTCAAGTAATTCTCCTGCCTCCACNNNAGATCGGAAGA
32568 TATGAGTGTGGAGTCGTTGCCTATCTCACCTGACTGCAACCTCNNNAGATCGGAAGA
32569 TATGAGTGTGGAGTCGTTGCCTGAGTAGCTGGGATTACAGGTGNNNAGATCGGAA
32572 TATGAGTGTGGAGTCGTTGCGAGAAGCTAGGACTACAGGTGTGNNNAGATCGGA
32573 TATGAGTGTGGAGTCGTTGCGGGTTCAAGCAATTCTCCTACCTNNNAGATCGGAAG
32574 TATGAGTGTGGAGTCGTTGCATCCTATCAACTCAGCCTCCTCTNNNAGATCGGAAGA
32576 TATGAGTGTGGAGTCGTTGCTTCAAGTGATTCTCCTGCCTCAANNNAGATCGGAAG
32577 TATGAGTGTGGAGTCGTTGCCAATCTCAACTCACTGCAACCTCNNNAGATCGGAAGA
32579 TATGAGTGTGGAGTCGTTGCTCTCCTGAGTAGCTGGGATTACANNNAGATCGGAAG
32580 TATGAGTGTGGAGTCGTTGCAATGACTTTGCCCTATGTGCCTANNNAGATCGGAAG
32581 TATGAGTGTGGAGTCGTTGCTGATTCCAGCTCACTACAACCTCNNNAGATCGGAAGA
32582 TATGAGTGTGGAGTCGTTGCCTCAAGTAGCTGGGATCACAGTTNNNAGATCGGAA
32583 TATGAGTGTGGAGTCGTTGCTTCAAGTGATTCTCCTGCCTCAGNNNAGATCGGAAG
32586 TATGAGTGTGGAGTCGTTGCGCAAAAAAATCTCCTCCCGATCCNNNAGATCGGAAG
32587 TATGAGTGTGGAGTCGTTGCAAAAGCAAATGTTTGACTGGGCANNNAGATCGGAA
32588 TATGAGTGTGGAGTCGTTGCCTAACTTGAAAGAATGGGGGGGANNNAGATCGGA
32590 TATGAGTGTGGAGTCGTTGCCAAGTGATCTGGCCTCCTTACTTNNNAGATCGGAAG
32591 TATGAGTGTGGAGTCGTTGCGGCTCAAACCTCTCCTTTTTCACNNNAGATCGGAAGA
32592 TATGAGTGTGGAGTCGTTGCTCCTAGCTACTCTTAGGCAGGAGNNNAGATCGGAAG
32593 TATGAGTGTGGAGTCGTTGCTCGGTCTCAAATACCTAAGGCTGNNNAGATCGGAAG
32599 TATGAGTGTGGAGTCGTTGCGACCCTGAGTCTGGAGAAATTGTNNNAGATCGGAA
32600 TATGAGTGTGGAGTCGTTGCTGTAACAAGCAGGTCCTTAGGTCNNNAGATCGGAA
32602 TATGAGTGTGGAGTCGTTGCGGCATGGTATTGTGTGCTTGTATNNNAGATCGGAA
32603 TATGAGTGTGGAGTCGTTGCTCTTCTGAGCTTTGGACCTGTACNNNAGATCGGAAG
32604 TATGAGTGTGGAGTCGTTGCACACCAGCAATAGAGGGGAATTTNNNAGATCGGAA
32606 TATGAGTGTGGAGTCGTTGCCATCCTCTCTGACAACAGTGGAANNNAGATCGGAAG
32607 TATGAGTGTGGAGTCGTTGCCCAAAAAAACCCTGCATGCTACTNNNAGATCGGAAG
32610 TATGAGTGTGGAGTCGTTGCCCAAGCTTGTGGTACATTCAGTGNNNAGATCGGAAG
32611 TATGAGTGTGGAGTCGTTGCAAAGAAAGCAAAGAAACCCTGGCNNNAGATCGGAA
32612 TATGAGTGTGGAGTCGTTGCTGCTTCACGAGAAAACATACAGTNNNAGATCGGAAG

32615 TATGAGTGTGGAGTCGTTGCAAAAGGGAGAGAAGAGAACAGGTNNNAGATCGGA
32616 TATGAGTGTGGAGTCGTTGCTCACAAGGAGATACACCTCACACNNNAGATCGGAAG
32617 TATGAGTGTGGAGTCGTTGCGTGAGGTAAGGACTGGAAAGAGGNNNAGATCGGA
32618 TATGAGTGTGGAGTCGTTGCGGAGCCAAAGACCTGAACAGATANNNAGATCGGAA
32619 TATGAGTGTGGAGTCGTTGCCCAGTGACAGTCATTTTAGCAGCNNNAGATCGGAAG
32622 TATGAGTGTGGAGTCGTTGCAAAGTTCTAACTGAGCTGGAGCANNNAGATCGGAA
32623 TATGAGTGTGGAGTCGTTGCGGCAGGAACAACTAGAACAAAGGNNNAGATCGGA
32627 TATGAGTGTGGAGTCGTTGCAAAAATCTTGGACACACGGACACNNNAGATCGGAA
32630 TATGAGTGTGGAGTCGTTGCGAGACTTGCTCTATTACCCAGGCNNNAGATCGGAAG
32631 TATGAGTGTGGAGTCGTTGCGTCTCAATCTCCTGACCTTGTGTNNNAGATCGGAAG
32632 TATGAGTGTGGAGTCGTTGCAGTAGCTGGGATTACAGGTACCTNNNAGATCGGAA
32633 TATGAGTGTGGAGTCGTTGCGTTCATGCCATTCTCCTACCTCANNNAGATCGGAAGA
32634 TATGAGTGTGGAGTCGTTGCGTTCAAGCAATTCTCTGCCTCAGNNNAGATCGGAAG
32635 TATGAGTGTGGAGTCGTTGCTCTCTCAAAGTCACCCTCACTTGNNNAGATCGGAAGA
32638 TATGAGTGTGGAGTCGTTGCCTCTTGAGTAGCTGGGACTACAGNNNAGATCGGAA
32639 TATGAGTGTGGAGTCGTTGCCCTCCTAGGTTCAAGCGATTCTTNNNAGATCGGAAG
32641 TATGAGTGTGGAGTCGTTGCCTGGTCTCAAACTCCTGACTTCANNNAGATCGGAAG
32642 TATGAGTGTGGAGTCGTTGCGAGATTACAGGCGCCTAGCTAATNNNAGATCGGAA
32644 TATGAGTGTGGAGTCGTTGCCAGGAGAAAAGCATGCAAGTCAANNNAGATCGGAA
32645 TATGAGTGTGGAGTCGTTGCTTCAAGTGATTCTCCTGCCTCAGNNNAGATCGGAAG
32646 TATGAGTGTGGAGTCGTTGCCACTGGGTGGAAAAAAGAGCAAGNNNAGATCGGA
32649 TATGAGTGTGGAGTCGTTGCTTCCAGGTTCAAGCAATTCTCCTNNNAGATCGGAAG
32650 TATGAGTGTGGAGTCGTTGCACCACATCCCGCTAATATTGTGTNNNAGATCGGAAG
32651 TATGAGTGTGGAGTCGTTGCAAGATGGTCAGGGAATTGGTAGTNNNAGATCGGA
32652 TATGAGTGTGGAGTCGTTGCTCTCCTGGGTTCAAGCAATTCTCNNNAGATCGGAAG
32653 TATGAGTGTGGAGTCGTTGCACTTCAGGTCTCAAGTGATCCACNNNAGATCGGAAG
32654 TATGAGTGTGGAGTCGTTGCGTGCATGAGATGATCAGCTGTTCNNNAGATCGGAA
32655 TATGAGTGTGGAGTCGTTGCAAGTTCAGGAAGCAAGAGTCTGTNNNAGATCGGAA
32656 TATGAGTGTGGAGTCGTTGCCACCCAGCCTCAATGGAAATTTTNNNAGATCGGAAG
32658 TATGAGTGTGGAGTCGTTGCTGTAGAGTCTCACTATGTTGCCCNNNAGATCGGAAG
32661 TATGAGTGTGGAGTCGTTGCCCTCTCTGGTTCGAGTGATTCTTNNNAGATCGGAAG
32662 TATGAGTGTGGAGTCGTTGCCCAAAGTAGCTGGGATTACAGGTNNNAGATCGGAA
32663 TATGAGTGTGGAGTCGTTGCGCCTCCTTAGTAGCTGGGATTACNNNAGATCGGAAG
32664 TATGAGTGTGGAGTCGTTGCACCTCTACCTCCTGAGTTCAAGTNNNAGATCGGAAG
32665 TATGAGTGTGGAGTCGTTGCGAGATCTTGGCTCACTGTAACCCNNNAGATCGGAAG
32666 TATGAGTGTGGAGTCGTTGCGTTCAGGCAATTCTCCTACCTCANNNAGATCGGAAG
32667 TATGAGTGTGGAGTCGTTGCACTCCCAGGTTCAAGTGATTCTCNNNAGATCGGAAG
32668 TATGAGTGTGGAGTCGTTGCCCTGCCTCAACCTCTCTATTACCNNNAGATCGGAAGA
32669 TATGAGTGTGGAGTCGTTGCCCTCCTGAGTAGCTGGGACTATANNNAGATCGGAAG
32672 TATGAGTGTGGAGTCGTTGCTGTTCATGGCTTACTCTCTCACCNNNAGATCGGAAGA
32674 TATGAGTGTGGAGTCGTTGCGAGATGGAGTGTCGTTCTGTTGCNNNAGATCGGAA
32675 TATGAGTGTGGAGTCGTTGCACCTCCCAGATTCAAACGATTCTNNNAGATCGGAAG
32679 TATGAGTGTGGAGTCGTTGCCTCCCGAGTAGCTGGAATTACAGNNNAGATCGGAA

32680 TATGAGTGTGGAGTCGTTGCCTCACTGAAATCTCCACCTCACANNNAGATCGGAAGA
32683 TATGAGTGTGGAGTCGTTGCTTGGCTCTCAGCTTGGATGTTATNNNAGATCGGAAG
32685 TATGAGTGTGGAGTCGTTGCCTCAGCCTCCCAAAGTGTGATATNNNAGATCGGAAG
32687 TATGAGTGTGGAGTCGTTGCCAATCTCAGCTCACTGCAAGTTCNNNAGATCGGAAG
32688 TATGAGTGTGGAGTCGTTGCGGCATGATATTGGGTCATTGCAANNNAGATCGGAA
32691 TATGAGTGTGGAGTCGTTGCAAACTCCTGACCTCAAGTGATCCNNNAGATCGGAAG
32692 TATGAGTGTGGAGTCGTTGCGGTTCAAGTGATTTTCCTGCCTCNNNAGATCGGAAG
32693 TATGAGTGTGGAGTCGTTGCCCTCTTATCTCTCACAGCTTCCCNNNAGATCGGAAGA
32694 TATGAGTGTGGAGTCGTTGCCTCCTGAGTAGCTGGGACTACAGNNNAGATCGGAAG
32695 TATGAGTGTGGAGTCGTTGCGAGCTGAGGTCTCACTCTTGTTGNNNAGATCGGAAG
32697 TATGAGTGTGGAGTCGTTGCTCAATCTCTGGGACTGAAGCAATNNNAGATCGGAAG
32698 TATGAGTGTGGAGTCGTTGCCTCCTGGGTTCACACCATTCTCNNNAGATCGGAAGAG
32699 TATGAGTGTGGAGTCGTTGCATAAGCCTGATTCTTCCTTGGCANNNAGATCGGAAG
32701 TATGAGTGTGGAGTCGTTGCCCACCTCCTGAGTTCAAGCTATTNNNAGATCGGAAGA
32702 TATGAGTGTGGAGTCGTTGCTCAAACTTTTGGGCTCAAGCAATNNNAGATCGGAAG
32703 TATGAGTGTGGAGTCGTTGCCCAAAGTGCTGGGATTACAAGTGNNNAGATCGGAA
32704 TATGAGTGTGGAGTCGTTGCTTCAAGCAATTCCCCTGTCTCAGNNNAGATCGGAAGA
32705 TATGAGTGTGGAGTCGTTGCTCTCAAACTGCTGACCTCATGATNNNAGATCGGAAGA
32708 TATGAGTGTGGAGTCGTTGCGTTCAAGCATCCTCCTACCTCAGNNNAGATCGGAAGA
32709 TATGAGTGTGGAGTCGTTGCCCTCCTAGGTTCAAGCGATTCTCNNNAGATCGGAAG
32710 TATGAGTGTGGAGTCGTTGCCAAGGTCAAGAGATCGAGACCATNNNAGATCGGAA
32712 TATGAGTGTGGAGTCGTTGCAACGAGCTAATGAATGCTGTCAANNNAGATCGGAAG
32713 TATGAGTGTGGAGTCGTTGCCTCACATCACGTCTGTCATTCCANNNAGATCGGAAGA
32714 TATGAGTGTGGAGTCGTTGCTAAGCAGTTTCCTCTCTTGCAGANNNAGATCGGAAG
32715 TATGAGTGTGGAGTCGTTGCAGGTGAACAATAAGTAACCAAAGAAANNNAGATCG
32716 TATGAGTGTGGAGTCGTTGCCCCAGTTTTGCACTTGAAACTCANNNAGATCGGAAG
32717 TATGAGTGTGGAGTCGTTGCGGAACTTCCAGAGCAATCTTACGNNNAGATCGGAAG
32718 TATGAGTGTGGAGTCGTTGCACCAAATTGCAGCTGTACCTCTANNNAGATCGGAAG
32719 TATGAGTGTGGAGTCGTTGCGAGTTGGGAGTCACTGTACACAANNNAGATCGGAA
32720 TATGAGTGTGGAGTCGTTGCTTAGCCATCAGAGAAATGCCAGTNNNAGATCGGAAG
32722 TATGAGTGTGGAGTCGTTGCCCAGTGGATCATGAGATCAGGAGNNNAGATCGGAA
32723 TATGAGTGTGGAGTCGTTGCGTTCCAGGGGGTGAAAGACTATANNNAGATCGGAA
32724 TATGAGTGTGGAGTCGTTGCGTTTGAAACCTCCCCCAAATTTNNNAGATCGGAAGA
32725 TATGAGTGTGGAGTCGTTGCATAGCCCTTACTCCCCAAAACAGNNNAGATCGGAAG
32726 TATGAGTGTGGAGTCGTTGCGGATCACTTGAGGTCAGGAGTTTNNNAGATCGGAA
32728 TATGAGTGTGGAGTCGTTGCCTCAAGTGATTCTCCCACCATGANNNAGATCGGAAG
32729 TATGAGTGTGGAGTCGTTGCTCCTTGTTCATTCCTACAGCCTGNNNAGATCGGAAGA
32730 TATGAGTGTGGAGTCGTTGCAATGCTTTTACACTGTTGGTGGGNNNAGATCGGAAG
32731 TATGAGTGTGGAGTCGTTGCACAGGAAACTACAGTTGTCAGCANNNAGATCGGAAG
32732 TATGAGTGTGGAGTCGTTGCCTGACAGTCAAGAGAGCAAGGAGNNNAGATCGGAA
32734 TATGAGTGTGGAGTCGTTGCTTTCCCTTCCCTTGAATCTAGGCNNNAGATCGGAAGA
32735 TATGAGTGTGGAGTCGTTGCATATGCACACAATAACATGCCCTNNNAGATCGGAAG
32736 TATGAGTGTGGAGTCGTTGCAAGGACTCCATCATGCAGCTAAANNNAGATCGGAAG

32737 TATGAGTGTGGAGTCGTTGCGGAGAATCACTTGAACCTGGGATNNNAGATCGGAA
32738 TATGAGTGTGGAGTCGTTGCGAGGCAGACCTTGTCTCTAAACANNNAGATCGGAAG
32740 TATGAGTGTGGAGTCGTTGCTAAACTCCTATAGGCTTCCCTGCNNNAGATCGGAAG
32741 TATGAGTGTGGAGTCGTTGCAACAGGAGAATCCCTTGAACTCGNNNAGATCGGAA
32743 TATGAGTGTGGAGTCGTTGCCCCACAGCAAAACAAAACAAACCNNNAGATCGGAAG
32744 TATGAGTGTGGAGTCGTTGCGCCAACATAGCAAAACCTGGTTTNNNAGATCGGAAG
32745 TATGAGTGTGGAGTCGTTGCGAAATCCATGCACAGGCTTCTTTNNNAGATCGGAAG
32746 TATGAGTGTGGAGTCGTTGCGAGGTCAGGAGATTGAGACCATCNNNAGATCGGAA
32747 TATGAGTGTGGAGTCGTTGCGCGTTGAGTATGGTAGCTTTCACNNNAGATCGGAA
32749 TATGAGTGTGGAGTCGTTGCTCACACCTGTAATTCCAGCACTTNNNAGATCGGAAG
32750 TATGAGTGTGGAGTCGTTGCCAGAGCGAGTTTCCATCTCAAAANNNAGATCGGAAG
32751 TATGAGTGTGGAGTCGTTGCGCAACTCAACAAAAAGACAACCCNNNAGATCGGAAG
32754 TATGAGTGTGGAGTCGTTGCAAAAAGAAGGGCAGGGAGAAGAANNNAGATCGGA
32756 TATGAGTGTGGAGTCGTTGCAAGACCTCCAGAAGTAAGCACTGNNNAGATCGGAA
32758 TATGAGTGTGGAGTCGTTGCAAACACAGACGTCTCCACACATANNNAGATCGGAAG
32760 TATGAGTGTGGAGTCGTTGCTTCTGAGGAAAGAAATTGCCAGCNNNAGATCGGAA
32763 TATGAGTGTGGAGTCGTTGCTAGAGATCTGCTGTACGACGTTGNNNAGATCGGAA
32764 TATGAGTGTGGAGTCGTTGCGCTCAAATGTTCCTGATGGCTTTNNNAGATCGGAAG
32765 TATGAGTGTGGAGTCGTTGCTGTGCTGGCATAAAGAGGCATATNNNAGATCGGAA
32767 TATGAGTGTGGAGTCGTTGCTGAATGTTGTCCTGGATGGGAATNNNAGATCGGAA
32768 TATGAGTGTGGAGTCGTTGCTGCCTGTAATCTCAGCTACTTGGNNNAGATCGGAAG
32769 TATGAGTGTGGAGTCGTTGCGGCTGATGTGGAAGGATCACTTANNNAGATCGGAA
32770 TATGAGTGTGGAGTCGTTGCGTACAGCAACAGTCCCACATCTANNNAGATCGGAAG
32771 TATGAGTGTGGAGTCGTTGCTGAAGTCTTATAGTGGCTTCCCTNNNAGATCGGAAG
32772 TATGAGTGTGGAGTCGTTGCATGTAACAAAATCCAACACCCCCNNNAGATCGGAAG
32773 TATGAGTGTGGAGTCGTTGCCATGCTTGTAATCCCAGCACTTTNNNAGATCGGAAG
32774 TATGAGTGTGGAGTCGTTGCCATGATCATATCACTGCCCTCCANNNAGATCGGAAGA
32776 TATGAGTGTGGAGTCGTTGCATTCCATCTCCAACCTGACCAATNNNAGATCGGAAGA
32777 TATGAGTGTGGAGTCGTTGCCACTTGTAATCCCAGCACTTTGGNNNAGATCGGAAG
32778 TATGAGTGTGGAGTCGTTGCCCTGTAATTCCAGCTACTTGGGANNNAGATCGGAAG
32779 TATGAGTGTGGAGTCGTTGCAAAGCTGCACACTTACAACCATTNNNAGATCGGAAG
32781 TATGAGTGTGGAGTCGTTGCATGTACACAGTCTTAGCCTCCAGNNNAGATCGGAAG
32782 TATGAGTGTGGAGTCGTTGCAAACCCACACATCTATGGTCAATNNNAGATCGGAAG
32783 TATGAGTGTGGAGTCGTTGCCTATGATCCTGCCACTACACTCCNNNAGATCGGAAGA
32785 TATGAGTGTGGAGTCGTTGCCAATGGCCTTGAGAAGCAAAAGANNNAGATCGGAA
32787 TATGAGTGTGGAGTCGTTGCGCTTGAGTGCACAAGTTTGAGATNNNAGATCGGAA
32788 TATGAGTGTGGAGTCGTTGCCAATGAGATACCACTGCACACCANNNAGATCGGAAG
32789 TATGAGTGTGGAGTCGTTGCCTTGGCTTCATCTTCACCAGGAANNNAGATCGGAAG
32790 TATGAGTGTGGAGTCGTTGCGAGGTCAGGAGATTGAGACCATCNNNAGATCGGAA
32791 TATGAGTGTGGAGTCGTTGCCTGAGTAGCTCGGATTACTGGTGNNNAGATCGGAA
32792 TATGAGTGTGGAGTCGTTGCCAATCTTGGCTCACTGCAAACTCNNNAGATCGGAAG
32793 TATGAGTGTGGAGTCGTTGCGGGTGATTCTCCAGCCTAAGTAGNNNAGATCGGAA
32794 TATGAGTGTGGAGTCGTTGCATCCTCCTATCTCAACCTCCCAANNNAGATCGGAAGA

32795 TATGAGTGTGGAGTCGTTGCTTCAAGCTATTCTCCTGCCTCAGNNNAGATCGGAAGA
32796 TATGAGTGTGGAGTCGTTGCGGCAAAATCTTAGCTCACTGCAANNNAGATCGGAAG
32797 TATGAGTGTGGAGTCGTTGCGATCTCGGCTCAGTATAAGCTCCNNNAGATCGGAAG
32798 TATGAGTGTGGAGTCGTTGCGAGATGGAGTTTCACCATGTTGGNNNAGATCGGAAG
32799 TATGAGTGTGGAGTCGTTGCCGTAGTAGCCCTAGTAGCTGGTANNNAGATCGGAAG
32803 TATGAGTGTGGAGTCGTTGCTTCAAGCAATTTTCCTGCCTCAGNNNAGATCGGAAGA
32805 TATGAGTGTGGAGTCGTTGCGCTTCCTGAGTAGCTGGGATTACNNNAGATCGGAAG
32806 TATGAGTGTGGAGTCGTTGCCTCCCAGGTTCAAGCAATTCTCTNNNAGATCGGAAGA
32807 TATGAGTGTGGAGTCGTTGCAGGTTCAAGGGATTCTTCTGCTTNNNAGATCGGAAG
32808 TATGAGTGTGGAGTCGTTGCTGGAGTATAATGGTGCGATCTCGNNNAGATCGGAAG
32811 TATGAGTGTGGAGTCGTTGCCCTCCATAGTAGCTGGGACTACANNNAGATCGGAAG
32812 TATGAGTGTGGAGTCGTTGCCTCAAGCAATCCTCCTACCTCAGNNNAGATCGGAAGA
32813 TATGAGTGTGGAGTCGTTGCTTCAAGTGATTCTCCTGCCTCAGNNNAGATCGGAAGA
32814 TATGAGTGTGGAGTCGTTGCTGTCTAGGCTGGTCTTGAACTTCNNNAGATCGGAAG
32815 TATGAGTGTGGAGTCGTTGCTTCAAGTGATTCTCCTGCCTCAGNNNAGATCGGAAGA
32817 TATGAGTGTGGAGTCGTTGCTTCAAGCAATTCTCCTGCCTCAGNNNAGATCGGAAGA
32819 TATGAGTGTGGAGTCGTTGCTTCAAGCAATTCTCCTGCCTCAGNNNAGATCGGAAGA
32820 TATGAGTGTGGAGTCGTTGCGCACCTTGGTAAAGACAGAAACCNNNAGATCGGAAG
32822 TATGAGTGTGGAGTCGTTGCCTCATTGCAACCTCTACCTCCTGNNNAGATCGGAAGA
32823 TATGAGTGTGGAGTCGTTGCCTCCCGAGTAGCTGGAACTACAGNNNAGATCGGAAG
32824 TATGAGTGTGGAGTCGTTGCGTGCAATCTCGGTTCAGTGTAACNNNAGATCGGAAG
32826 TATGAGTGTGGAGTCGTTGCGAGTAGCTAGGATTACAGGTGCCNNNAGATCGGAAG
32827 TATGAGTGTGGAGTCGTTGCCTCCCAGGTTCAAGCTATTCTCANNNAGATCGGAAGA
32829 TATGAGTGTGGAGTCGTTGCCTGAGTAGCTGGGATTACAGGTGNNNAGATCGGAA
32830 TATGAGTGTGGAGTCGTTGCGTCAAGTGATTCTCCTGCCTCANNNAGATCGGAAGAG
32831 TATGAGTGTGGAGTCGTTGCAAACTCCTGACCTCAAGTGATCCNNNAGATCGGAAGA
32832 TATGAGTGTGGAGTCGTTGCTCCACCTCCCAGTTCAAATGATTNNNAGATCGGAAGA
32835 TATGAGTGTGGAGTCGTTGCCCTCTTGGGTTCATGCCATTCTCNNNAGATCGGAAGA
32836 TATGAGTGTGGAGTCGTTGCCTCCTGAGTAGCTGGGACTACAGNNNAGATCGGAAG
32837 TATGAGTGTGGAGTCGTTGCGTAGAGACGGACTTTCACTGTGTNNNAGATCGGAAG
32839 TATGAGTGTGGAGTCGTTGCCCTCCTGAGTATCTGGGACTACANNNAGATCGGAAG
32840 TATGAGTGTGGAGTCGTTGCTGCGTAGCAAACACTTTTCACAANNNAGATCGGAAGA
32842 TATGAGTGTGGAGTCGTTGCCTCCTGAGTAGCTGGGACTACAGNNNAGATCGGAAG
32843 TATGAGTGTGGAGTCGTTGCACTATTCGGCTCACTGCAAGTTNNNAGATCGGAAGAG
32844 TATGAGTGTGGAGTCGTTGCCCTCCCAGGTTCAAGCAATTTTTNNNAGATCGGAAGA
32845 TATGAGTGTGGAGTCGTTGCGATAAGGTCTCACTCTGTCACCCNNNAGATCGGAAGA
32846 TATGAGTGTGGAGTCGTTGCTTCAAGTGATTCTCCTGCCTCAGNNNAGATCGGAAGA
32849 TATGAGTGTGGAGTCGTTGCTCTCAAACTCCTGACCTCAAGTGNNNAGATCGGAAGA
32850 TATGAGTGTGGAGTCGTTGCGATTCAAGCAATTCTCCTGCCTCNNNAGATCGGAAGA
32852 TATGAGTGTGGAGTCGTTGCTACTGAGTTTCACCATCTTGGCCNNNAGATCGGAAGA
32853 TATGAGTGTGGAGTCGTTGCCCAAAGTGCTGGGATAACAGGTANNNAGATCGGAAG
32854 TATGAGTGTGGAGTCGTTGCTGCTCCCAGCCATTTTTTGTTTTNNNAGATCGGAAGA
32856 TATGAGTGTGGAGTCGTTGCTTCAAGTGATTCTCCTGCCTCAGNNNAGATCGGAAGA

32857 TATGAGTGTGGAGTCGTTGCGTTTTTGCAGAGACAGGGTCTTGNNNAGATCGGAA
32858 TATGAGTGTGGAGTCGTTGCCTCCACCACGCCAAGCTAATTTTNNNAGATCGGAAGA
32859 TATGAGTGTGGAGTCGTTGCCACAGGTTTTCACCATGTTCTCCNNNAGATCGGAAGA
32860 TATGAGTGTGGAGTCGTTGCTGGGTTCAAGTGATTCTCCTACCNNNAGATCGGAAG
32863 TATGAGTGTGGAGTCGTTGCAGAGTCTCACTCTATCCCTCAGGNNNAGATCGGAAG
32864 TATGAGTGTGGAGTCGTTGCTTCAAGCAATTCTCCTGCCTCAGNNNAGATCGGAAG
32865 TATGAGTGTGGAGTCGTTGCGATAGAGCCTCATTCTGTCACCCNNNAGATCGGAAG
32866 TATGAGTGTGGAGTCGTTGCCTCCCGAGTACCTGAGACTACAGNNNAGATCGGAAG
32867 TATGAGTGTGGAGTCGTTGCTTCAAGTGATTCTCCTGCCTCAGNNNAGATCGGAAG
32868 TATGAGTGTGGAGTCGTTGCCCTCCCAAGTAGCTGGAATTACANNNAGATCGGAAG
32869 TATGAGTGTGGAGTCGTTGCCTCCCGAGTAGCTAGGACTACAGNNNAGATCGGAA
32870 TATGAGTGTGGAGTCGTTGCACAGCTCCTGGATTTCTTGATCTNNNAGATCGGAAG
32874 TATGAGTGTGGAGTCGTTGCCTCCGGAATAGCTGGGACTAAAGNNNAGATCGGAA
32878 TATGAGTGTGGAGTCGTTGCCTCTGATTCCCAGGTTCAAGTGANNNAGATCGGAAG
32881 TATGAGTGTGGAGTCGTTGCGTTTCACCTTGTTAGCCAGGATGNNNAGATCGGAAG
32882 TATGAGTGTGGAGTCGTTGCCCATGATGAGACTATCAGCACCANNNAGATCGGAAG
32884 TATGAGTGTGGAGTCGTTGCAGATCGAGACCATCTTGGCTAACNNNAGATCGGAAG
32886 TATGAGTGTGGAGTCGTTGCGACAGTGTTTGTAGGCATTCAGCNNNAGATCGGAA
32887 TATGAGTGTGGAGTCGTTGCCAGTGAGCTGAGATCATGTCACTNNNAGATCGGAA
32888 TATGAGTGTGGAGTCGTTGCGATAGTGGTGATGGTTGCACAACNNNAGATCGGAA
32890 TATGAGTGTGGAGTCGTTGCGCATCTCAACAGCTGAAACACTTNNNAGATCGGAAG
32891 TATGAGTGTGGAGTCGTTGCATCAGTCTGCAGACAACTCATGANNNAGATCGGAAG
32893 TATGAGTGTGGAGTCGTTGCGGGATTTTCGCTCAATGGATACANNNAGATCGGAA
32894 TATGAGTGTGGAGTCGTTGCGTTCCTCAAGTTCCCATCCTCTTNNNAGATCGGAAGA
32895 TATGAGTGTGGAGTCGTTGCACTCCTATTCTGCTAGCACGTTTNNNAGATCGGAAGA
32896 TATGAGTGTGGAGTCGTTGCGAGCCATAGGATTGGTAATGGGTNNNAGATCGGAA
32897 TATGAGTGTGGAGTCGTTGCCAAGTCTCTGACTCCCACATGAANNNAGATCGGAAG
32899 TATGAGTGTGGAGTCGTTGCACAGAGAAGTTCGGTCTCAAACANNNAGATCGGAA
32901 TATGAGTGTGGAGTCGTTGCGCAAGACTCTGCCTCAAAAACAANNNAGATCGGAAG
32902 TATGAGTGTGGAGTCGTTGCCCTTTGAGCCCTCATTCACATTCNNNAGATCGGAAGA
32903 TATGAGTGTGGAGTCGTTGCATCCTGGGAAGTTGATGCTGTAGNNNAGATCGGAA
32905 TATGAGTGTGGAGTCGTTGCATGCACGAGTATCTGTTACGGTTNNNAGATCGGAAG
32906 TATGAGTGTGGAGTCGTTGCCTATAGCACAGCAAACCTCAGACNNNAGATCGGAAG
32907 TATGAGTGTGGAGTCGTTGCCAACTGGATTGTGGTGATGGTTGNNNAGATCGGAA
32908 TATGAGTGTGGAGTCGTTGCGGATCACTTGAGTCCAGGAGTTTNNNAGATCGGAA
32910 TATGAGTGTGGAGTCGTTGCAGTTGTGGCAATTTGAACCCATTNNNAGATCGGAAG
32915 TATGAGTGTGGAGTCGTTGCTCATGCTGCTGACAGTCTAATGANNNAGATCGGAAG
32919 TATGAGTGTGGAGTCGTTGCTCAGAACAAATGCATCCAGGAGANNNAGATCGGAA
32920 TATGAGTGTGGAGTCGTTGCGAGTCAGGAAAACTGCTTCAACCNNNAGATCGGAA
32921 TATGAGTGTGGAGTCGTTGCGCTTCATTAGTTGCCACACATGTNNNAGATCGGAAG
32923 TATGAGTGTGGAGTCGTTGCATGGAACCCCAAAAGGATCTGAANNNAGATCGGAA
32924 TATGAGTGTGGAGTCGTTGCTAGTCCTAGCTACTTGGAAGGCTNNNAGATCGGAAG
32925 TATGAGTGTGGAGTCGTTGCGTGTGCTGGAAAAACTTGCTCTANNNAGATCGGAA

| | |
|---|---|
| 32927 | TATGAGTGTGGAGTCGTTGCAAAAAATTAGCTGGGCATGGTGGNNNAGATCGGAA |
| 32928 | TATGAGTGTGGAGTCGTTGCGGGTTGCAATCCTAGTCTCTGATNNNAGATCGGAAG |
| 32929 | TATGAGTGTGGAGTCGTTGCTAAGCTCACTCACTGCAGAAACANNNAGATCGGAAG |
| 32930 | TATGAGTGTGGAGTCGTTGCCTGAGGTGGGAGGATTACTTGAGNNNAGATCGGAA |
| 32933 | TATGAGTGTGGAGTCGTTGCCAACACACACACACACACACATANNNAGATCGGAAGA |
| 32934 | TATGAGTGTGGAGTCGTTGCTGACATTCCTCCTTGTGAGAGTGNNNAGATCGGAAG |
| 32935 | TATGAGTGTGGAGTCGTTGCTGACATTCCTCCTTGTGAGAGTGNNNAGATCGGAAG |
| 32937 | TATGAGTGTGGAGTCGTTGCAAAAGCCCAGACATTCAACTCACNNNAGATCGGAAGA |
| 32938 | TATGAGTGTGGAGTCGTTGCAATCTGTCCTCATTGCTTTTGCCNNNAGATCGGAAGA |
| 32939 | TATGAGTGTGGAGTCGTTGCGCTCAAGCCTGTCTCAAAAAAAANNNAGATCGGAAG |
| 32940 | TATGAGTGTGGAGTCGTTGCTGTGGTAGCTCACACCTGTAATCNNNAGATCGGAAG |
| 32941 | TATGAGTGTGGAGTCGTTGCCATAAATGGGACTTGGGCACTTTNNNAGATCGGAAG |
| 32942 | TATGAGTGTGGAGTCGTTGCACCTATAATCCCAGCACTTCAGGNNNAGATCGGAAGA |
| 32944 | TATGAGTGTGGAGTCGTTGCGGTGCACCAAAATCTCAGAAGTCNNNAGATCGGAAG |
| 32945 | TATGAGTGTGGAGTCGTTGCACCCAATAATCAAAAGCCTCCCANNNAGATCGGAAGA |
| 32947 | TATGAGTGTGGAGTCGTTGCGAAAGAAAGAAAGCAGGAAGGCANNNAGATCGGAA |
| 32949 | TATGAGTGTGGAGTCGTTGCGGGGCCTTATGACCTTATTTTGCNNNAGATCGGAAG |
| 32950 | TATGAGTGTGGAGTCGTTGCAGGTTGAGGCAACAGGATTACTTNNNAGATCGGAAG |
| 32951 | TATGAGTGTGGAGTCGTTGCTCACAGGTACAGGATTGCTTCATNNNAGATCGGAAG |
| 32952 | TATGAGTGTGGAGTCGTTGCAGACAGTGAAAGACATCTGACCTNNNAGATCGGAAG |
| 32953 | TATGAGTGTGGAGTCGTTGCGCTTGTAATCCCAGCACTTTGAGNNNAGATCGGAAG |
| 32954 | TATGAGTGTGGAGTCGTTGCTTTTGAATACAGCACAGCCTGTGNNNAGATCGGAAG |
| 32955 | TATGAGTGTGGAGTCGTTGCCACGCCCAGCTACTTTTTGTATTNNNAGATCGGAAGA |
| 32956 | TATGAGTGTGGAGTCGTTGCTGACTAGAGGCATCTGGTACTCANNNAGATCGGAAG |
| 32958 | TATGAGTGTGGAGTCGTTGCACATAGCATAGTTTTGGGGGGAGNNNAGATCGGAA |
| 32959 | TATGAGTGTGGAGTCGTTGCCACAGTGGCTCAAGCTTGTAATCNNNAGATCGGAAG |
| 32960 | TATGAGTGTGGAGTCGTTGCGCCAATATGCCTGATGAACATCGNNNAGATCGGAAG |
| 32961 | TATGAGTGTGGAGTCGTTGCCTGAGGCAGGAGAATCACTTGAANNNAGATCGGAAG |
| 32962 | TATGAGTGTGGAGTCGTTGCTCCTAAAGGTGGAAGCTGAAACANNNAGATCGGAAG |
| 32963 | TATGAGTGTGGAGTCGTTGCTCCTCAGCAAACTAACACAGGAANNNAGATCGGAAG |
| 32964 | TATGAGTGTGGAGTCGTTGCAGAAGTTGCAGTGAGCCAAGATCNNNAGATCGGAAG |
| 32966 | TATGAGTGTGGAGTCGTTGCTGGCTTTCCTGCTGAATTCTACANNNAGATCGGAAGA |
| 32968 | TATGAGTGTGGAGTCGTTGCGGCAAGACTGGAATCTAACAACANNNAGATCGGAAG |
| 32969 | TATGAGTGTGGAGTCGTTGCACTGTTTGTTTTTGAGCTAGGGCNNNAGATCGGAAG |
| 32970 | TATGAGTGTGGAGTCGTTGCGGAACAGGGCGATTTTACATCATNNNAGATCGGAAG |
| 32971 | TATGAGTGTGGAGTCGTTGCGAATGACACAATGGACTTTGGGGNNNAGATCGGAA |
| 32973 | TATGAGTGTGGAGTCGTTGCCTATGCAGCAGTACCACAACAAGNNNAGATCGGAAG |
| 32974 | TATGAGTGTGGAGTCGTTGCAAAAATTGAGGAGGAGGGTCTCCNNNAGATCGGAA |
| 32975 | TATGAGTGTGGAGTCGTTGCGAAAATCACCACTTTGCGACCTTNNNAGATCGGAAGA |
| 32976 | TATGAGTGTGGAGTCGTTGCGAAGGGAACTCAAGGCAAACAAANNNAGATCGGAA |
| 32978 | TATGAGTGTGGAGTCGTTGCAACATGGATGAAACTGGAGGTCANNNAGATCGGAA |
| 32979 | TATGAGTGTGGAGTCGTTGCGGCCAAAACCTTGAGATACATCANNNAGATCGGAAG |
| 32981 | TATGAGTGTGGAGTCGTTGCGTAAGGTCTAGAGGCAACCCTTCNNNAGATCGGAAG |

32982 TATGAGTGTGGAGTCGTTGCTTTTTTGGCAAAAGCGGTTTCACNNNAGATCGGAAG
32988 TATGAGTGTGGAGTCGTTGCAGAGATGTTAGGTGTGCTGATGGNNNAGATCGGAA
32989 TATGAGTGTGGAGTCGTTGCGGTCAGGCCTATAATATCCCAGCNNNAGATCGGAAG
32990 TATGAGTGTGGAGTCGTTGCTTCAAGTGATTCTCCTGCCTCAGNNNAGATCGGAAG
32992 TATGAGTGTGGAGTCGTTGCGTAGTGCAGTGGAGTGATCTCAANNNAGATCGGAA
32993 TATGAGTGTGGAGTCGTTGCGGTTCAAACATTTCTCCTGCCTCNNNAGATCGGAAG
32994 TATGAGTGTGGAGTCGTTGCCTCAAGTGATCCTCCCATCTCAGNNNAGATCGGAAG
32995 TATGAGTGTGGAGTCGTTGCCTGGGTTCAAGCAATTTCTCCTGNNNAGATCGGAAG
32996 TATGAGTGTGGAGTCGTTGCTTCAAGTGATTCTCCTGCCTCAGNNNAGATCGGAAG
32999 TATGAGTGTGGAGTCGTTGCCCACAAAAGACAGCTTTACAGGGNNNAGATCGGAA
33000 TATGAGTGTGGAGTCGTTGCAAGCGATTCTTATGTCTCAGCCTNNNAGATCGGAAG
33001 TATGAGTGTGGAGTCGTTGCTTGCCTAGTCTGGAGTGTAATGGNNNAGATCGGAA
33002 TATGAGTGTGGAGTCGTTGCGATCTTGGCTCACTACAACCTCCNNNAGATCGGAAG
33003 TATGAGTGTGGAGTCGTTGCCCTCCTGAGTAGCTGGGATTAGANNNAGATCGGAA
33005 TATGAGTGTGGAGTCGTTGCTTCAAGCAATTCTCCTGCCTCAGNNNAGATCGGAAG
33006 TATGAGTGTGGAGTCGTTGCACAATCTTCTCAACTCAGCCTCCNNNAGATCGGAAGA
33008 TATGAGTGTGGAGTCGTTGCCAAGTAGATGGGACTACAGGCACNNNAGATCGGAA
33010 TATGAGTGTGGAGTCGTTGCGGAATTTCACTCTTGTTGCCCAGNNNAGATCGGAAG
33011 TATGAGTGTGGAGTCGTTGCTCTTGACTTACCACAACCTCCACNNNAGATCGGAAGA
33014 TATGAGTGTGGAGTCGTTGCCCATTTCCCAGGTTCAAGCAATTNNNAGATCGGAAG
33015 TATGAGTGTGGAGTCGTTGCCCACCACTCCCTGCTAATTTTTGNNNAGATCGGAAGA
33016 TATGAGTGTGGAGTCGTTGCCGCACGGCCATTTTTTTGTATTTNNNAGATCGGAAG
33017 TATGAGTGTGGAGTCGTTGCAATCTAGAATGCAGTGGTGTGGTNNNAGATCGGAA
33018 TATGAGTGTGGAGTCGTTGCGAGTAGCTGGGATTACAGGCATGNNNAGATCGGAA
33019 TATGAGTGTGGAGTCGTTGCGTGGTATGATCTTGGCTCATTGCNNNAGATCGGAAG
33020 TATGAGTGTGGAGTCGTTGCCAAGTAGCTGGGATTACAGGCATNNNAGATCGGAA
33021 TATGAGTGTGGAGTCGTTGCGGTTCAAACAATTCTCCTGCCTCNNNAGATCGGAAG
33022 TATGAGTGTGGAGTCGTTGCTCAAGTGATTCTCCCACCTTAGCNNNAGATCGGAAG
33024 TATGAGTGTGGAGTCGTTGCGAGTAGCTGGGACTACAAGCATGNNNAGATCGGAA
33026 TATGAGTGTGGAGTCGTTGCTCTCCCGGGTTCAAATGATTCTCNNNAGATCGGAAG
33028 TATGAGTGTGGAGTCGTTGCGGTTCAAATGATTCTCCTGCCTCNNNAGATCGGAAG
33031 TATGAGTGTGGAGTCGTTGCGAGTAGCTGGTACTACAGGTGTGNNNAGATCGGAA
33032 TATGAGTGTGGAGTCGTTGCCACCATGCCCGACTAATTTTTGTNNNAGATCGGAAG
33033 TATGAGTGTGGAGTCGTTGCCTGGAGTGCAATGGTGAGATCTCNNNAGATCGGAA
33036 TATGAGTGTGGAGTCGTTGCTGGAGTATAATGGCACAATCCCGNNNAGATCGGAA
33038 TATGAGTGTGGAGTCGTTGCTTCAAGTGATTCTCCTGCCTCAGNNNAGATCGGAAG
33041 TATGAGTGTGGAGTCGTTGCGAACTCACCTATGCTAGTGGGAGNNNAGATCGGAA
33042 TATGAGTGTGGAGTCGTTGCTTGCAAATGGAGAGCCTAGACTTNNNAGATCGGAA
33043 TATGAGTGTGGAGTCGTTGCAGGCTGGAGTGCAATGATATGATNNNAGATCGGAA
33044 TATGAGTGTGGAGTCGTTGCTCAAGGAATTGAATCCAGCCTCANNNAGATCGGAAG
33045 TATGAGTGTGGAGTCGTTGCACCGTGCCAGGCTATTTTTTTTTNNNAGATCGGAAG
33046 TATGAGTGTGGAGTCGTTGCCTCCTGTGTAGCTGGGACTACAGNNNAGATCGGAA
33047 TATGAGTGTGGAGTCGTTGCCTCACTGCAGTCTTGAATTCCTGNNNAGATCGGAAG

33048 TATGAGTGTGGAGTCGTTGCGATGGAGTCTCACTCACTCTGTCNNNAGATCGGAAG
33050 TATGAGTGTGGAGTCGTTGCCTCCTGAGTAGCTGGGACTACAGNNNAGATCGGAA
33052 TATGAGTGTGGAGTCGTTGCGGCCATTCTCCTATCTTAGCCTCNNNAGATCGGAAGA
33053 TATGAGTGTGGAGTCGTTGCTTCAATAGGTCCCACTCCCAAAGNNNAGATCGGAAG
33055 TATGAGTGTGGAGTCGTTGCATAGAATCTCACTCTGTTGCCCANNNAGATCGGAAG
33056 TATGAGTGTGGAGTCGTTGCGATTCTCCTGCTTCAGACTCCTGNNNAGATCGGAAG
33057 TATGAGTGTGGAGTCGTTGCCAAAATTTTGGCTCACTGCAACCNNNAGATCGGAAG
33058 TATGAGTGTGGAGTCGTTGCCCTCCAGTGTTCACTCCATTCTCNNNAGATCGGAAGA
33059 TATGAGTGTGGAGTCGTTGCCTCCCAGGTTCCAGTGATTCTTCNNNAGATCGGAAG
33060 TATGAGTGTGGAGTCGTTGCGCTTACTGCTGCCTTGACTTCNNNAGATCGGAAGAG
33062 TATGAGTGTGGAGTCGTTGCCAGGAGAGTAGCTGGGATTACAGNNNAGATCGGA
33063 TATGAGTGTGGAGTCGTTGCGGCTTGAGCAATCAGCAAAACTANNNAGATCGGAA
33064 TATGAGTGTGGAGTCGTTGCCGAAGCAGGAAGAGATAGCAGATNNNAGATCGGA
33065 TATGAGTGTGGAGTCGTTGCCTCAACCAATGATCCTGCCTTAGNNNAGATCGGAAG
33066 TATGAGTGTGGAGTCGTTGCACTTCTACTCTCCAAGTCCGAGANNNAGATCGGAAG
33067 TATGAGTGTGGAGTCGTTGCGCTAGAGTACAGTGGCATGATCTNNNAGATCGGAA
33068 TATGAGTGTGGAGTCGTTGCTTCAAGTGATTCTCCTGCCTCAGNNNAGATCGGAAG
33070 TATGAGTGTGGAGTCGTTGCGGACTTCCATGGCTCAAGTGATCNNNAGATCGGAAG
33072 TATGAGTGTGGAGTCGTTGCGCCTCCAGCTACATTCATGTTTCNNNAGATCGGAAGA
33073 TATGAGTGTGGAGTCGTTGCCTGGGATTATAGGTGTGGACCACNNNAGATCGGAA
33074 TATGAGTGTGGAGTCGTTGCGTAGAGATAGGGTTTCGCCATGTNNNAGATCGGAA
33075 TATGAGTGTGGAGTCGTTGCGCAATCTCACCTCACCATAACCTNNNAGATCGGAAGA
33076 TATGAGTGTGGAGTCGTTGCCTGGGTTCAATCAATTCTCCTGCNNNAGATCGGAAG
33078 TATGAGTGTGGAGTCGTTGCGTTTCTCCAGCTTTTTGCCCTAGNNNAGATCGGAAGA
33079 TATGAGTGTGGAGTCGTTGCTTCAAACGATTCTCCTGCCTCAGNNNAGATCGGAAG
33081 TATGAGTGTGGAGTCGTTGCGTGTAGTGAGTGGTGTGATCTCANNNAGATCGGAA
33083 TATGAGTGTGGAGTCGTTGCCTCAAGTGATCCTTCCACCTCAGNNNAGATCGGAAG
33087 TATGAGTGTGGAGTCGTTGCCTGGCACAGCAGAAAAATACCATNNNAGATCGGAAG
33088 TATGAGTGTGGAGTCGTTGCGAGGGGACAACATCTTTTCTGGANNNAGATCGGAA
33089 TATGAGTGTGGAGTCGTTGCGGGTTCAAGTAATTCTCCTGCCTNNNAGATCGGAAG
33090 TATGAGTGTGGAGTCGTTGCTCACTATCTCACTCTACTGCCCANNNAGATCGGAAGA
33091 TATGAGTGTGGAGTCGTTGCCTACTGAACTTATGATCCGCCCANNNAGATCGGAAG
33094 TATGAGTGTGGAGTCGTTGCAGCCAACACAACACACTTTGAAANNNAGATCGGAAG
33095 TATGAGTGTGGAGTCGTTGCTCTGGAATAGCTGGGATTACAGGNNNAGATCGGAA
33097 TATGAGTGTGGAGTCGTTGCCTCCTGAGTAGCTGGGATTACAGNNNAGATCGGAA
33098 TATGAGTGTGGAGTCGTTGCCAAGTAACTGGGACTACAGGCATNNNAGATCGGAA
33099 TATGAGTGTGGAGTCGTTGCGGTGTGAGGTGGTATCTCATTGTNNNAGATCGGAA
33101 TATGAGTGTGGAGTCGTTGCCTCCCGAGTAGCTGAGATTACAGNNNAGATCGGAA
33103 TATGAGTGTGGAGTCGTTGCGTTCAAGCGATTCTCCTACCTCANNNAGATCGGAAG
33104 TATGAGTGTGGAGTCGTTGCCAAGCACTTGGAGGGAAGAAAAGNNNAGATCGGA
33105 TATGAGTGTGGAGTCGTTGCTACATGACCCAAATGTCGTCCTTNNNAGATCGGAAG
33106 TATGAGTGTGGAGTCGTTGCTACAAAAAATTAGCCGGGTGCAGNNNAGATCGGAA
33109 TATGAGTGTGGAGTCGTTGCAAAACCGCTTGAATGAGAACAGGNNNAGATCGGAA

33110 TATGAGTGTGGAGTCGTTGCCATGGATGGAACTGGAGGACATTNNNAGATCGGAA
33112 TATGAGTGTGGAGTCGTTGCGCAAATACCCCAGTGGATTTAGCNNNAGATCGGAAG
33113 TATGAGTGTGGAGTCGTTGCCCTGTTGATTCCTAGACTGCACTNNNAGATCGGAAG
33114 TATGAGTGTGGAGTCGTTGCGGCTGTGTTGGAAGAAAATGTGTNNNAGATCGGAA
33116 TATGAGTGTGGAGTCGTTGCTGTTGTCCCAGGCATTTCTGATANNNAGATCGGAAG
33117 TATGAGTGTGGAGTCGTTGCGAATGTACCAGACAGGACACTCANNNAGATCGGAA
33119 TATGAGTGTGGAGTCGTTGCGAAAAATGTTACTGGACCCCACCNNNAGATCGGAAG
33120 TATGAGTGTGGAGTCGTTGCCCACACAAAAATTAGCTGGGCATNNNAGATCGGAAG
33121 TATGAGTGTGGAGTCGTTGCAGGATTCCATTGCTCAGAAACCTNNNAGATCGGAAG
33123 TATGAGTGTGGAGTCGTTGCGAACGAAAAGTATCTTCCTCGCGNNNAGATCGGAAG
33125 TATGAGTGTGGAGTCGTTGCTCTGGTATTCCCTAGCTCTGTCANNNAGATCGGAAG
33127 TATGAGTGTGGAGTCGTTGCTCCTTCAAATCCCAGCTGCATTANNNAGATCGGAAG
33128 TATGAGTGTGGAGTCGTTGCTGCTGTGGAAATAAGAGCACTCTNNNAGATCGGAA
33129 TATGAGTGTGGAGTCGTTGCGTCAGCAATCACAGCAACACATANNNAGATCGGAAG
33130 TATGAGTGTGGAGTCGTTGCAACTGTGGTAGCACCTCTGTAAGNNNAGATCGGAA
33132 TATGAGTGTGGAGTCGTTGCTCTCCGTGGTTTTACAAGGATGANNNAGATCGGAAG
33133 TATGAGTGTGGAGTCGTTGCTGGTCAGGAGAAGTAATGAGACCNNNAGATCGGAA
33134 TATGAGTGTGGAGTCGTTGCATTTCTACACTGCTGGTGGGAATNNNAGATCGGAAG
33135 TATGAGTGTGGAGTCGTTGCAAGAAGAAAGTACTCTGCCTGGGNNNAGATCGGAA
33136 TATGAGTGTGGAGTCGTTGCGCCCGCCACACAAACATATATATNNNAGATCGGAAG
33137 TATGAGTGTGGAGTCGTTGCATGAAAGGACATAGAGGCTGGTCNNNAGATCGGAA
33139 TATGAGTGTGGAGTCGTTGCGCCTATAATCCAGCACTTTGGGANNNAGATCGGAAG
33140 TATGAGTGTGGAGTCGTTGCTGACTTTGCCTGTAACCTCTACCNNNAGATCGGAAG
33141 TATGAGTGTGGAGTCGTTGCGCGATAATGCGAGACTCCATCTCNNNAGATCGGAAG
33142 TATGAGTGTGGAGTCGTTGCCTCTTGGACAGGAAGGTTCATGANNNAGATCGGAA
33146 TATGAGTGTGGAGTCGTTGCAACAACATGGATGGAACTGGAGANNNAGATCGGAA
33148 TATGAGTGTGGAGTCGTTGCCCAAAAAAAAAAAAACCCTCCCGNNNAGATCGGAAG
33149 TATGAGTGTGGAGTCGTTGCGATGTATAGGACAACTGGGGAAANNNAGATCGGAA
33152 TATGAGTGTGGAGTCGTTGCCACAAAAGCTGGAGAGTGACTTGNNNAGATCGGAA
33153 TATGAGTGTGGAGTCGTTGCATGGAAAGGAACAACTGGTACGANNNAGATCGGAA
33154 TATGAGTGTGGAGTCGTTGCTGACATTCCTCCTTGTGAGAGTGNNNAGATCGGAAG
33155 TATGAGTGTGGAGTCGTTGCTCATTTGGTGAGTTACTGGCAGANNNAGATCGGAA
33156 TATGAGTGTGGAGTCGTTGCGGCATCCAAATCAGCAAAGAGAANNNAGATCGGAA
33157 TATGAGTGTGGAGTCGTTGCAATGCAGCTCTCTTCTACCTCTGNNNAGATCGGAAG
33158 TATGAGTGTGGAGTCGTTGCGAATGGGAGTGGTAGTGGTAGTGNNNAGATCGGA
33159 TATGAGTGTGGAGTCGTTGCTGAACTATGATCACACCACAGCANNNAGATCGGAAG
33161 TATGAGTGTGGAGTCGTTGCCACAGTGGCTCACTCCTGTAATCNNNAGATCGGAAG
33162 TATGAGTGTGGAGTCGTTGCGAAAAGTTTTTGGGCCGAGATGANNNAGATCGGAA
33163 TATGAGTGTGGAGTCGTTGCATGCAGCCCCCAAAAATATTGAGNNNAGATCGGAAG
33164 TATGAGTGTGGAGTCGTTGCTTCCTATACACCAACAACAGGCANNNAGATCGGAAG
33165 TATGAGTGTGGAGTCGTTGCACTACATGGAAATGGAACGACCTNNNAGATCGGAA
33166 TATGAGTGTGGAGTCGTTGCATGGCATGTCATTTTTGTCACCTNNNAGATCGGAAG
33167 TATGAGTGTGGAGTCGTTGCATTATTCAAGGCCTTGCATATGCNNNAGATCGGAAG

33169 TATGAGTGTGGAGTCGTTGCCAGTAAGCCGAGATCATGACACTNNNAGATCGGAAG
33170 TATGAGTGTGGAGTCGTTGCCAACACTAAACCCTAAGGCTTGCNNNAGATCGGAAG
33172 TATGAGTGTGGAGTCGTTGCGGGACTCTGCATTTTCTGGAGATNNNAGATCGGAAG
33173 TATGAGTGTGGAGTCGTTGCCATGCCTGTAATTCCAGCACTTTNNNAGATCGGAAG
33174 TATGAGTGTGGAGTCGTTGCCTGTAAGCAGAGGTGACCTTACCNNNAGATCGGAAG
33175 TATGAGTGTGGAGTCGTTGCAGGTAGTGGCCCTTAGAGAGAATNNNAGATCGGAA
33176 TATGAGTGTGGAGTCGTTGCTCAGTGGATTTCTGGCAATACCANNNAGATCGGAAG
33177 TATGAGTGTGGAGTCGTTGCGCCCATTTTCTGTGGAGGAATTCNNNAGATCGGAAG
33180 TATGAGTGTGGAGTCGTTGCGAGGATGCAGCATGAAATCGTACNNNAGATCGGAA
33181 TATGAGTGTGGAGTCGTTGCCAAGACCTAAGCATGGAAGACGANNNAGATCGGAA
33184 TATGAGTGTGGAGTCGTTGCGAAAGGACTTGGTTGCTGTGATCNNNAGATCGGAA
33185 TATGAGTGTGGAGTCGTTGCTGCCAATCAAAGCCAAAGACAATNNNAGATCGGAAG
33187 TATGAGTGTGGAGTCGTTGCGGAGGGGCCAATATGACAAACTANNNAGATCGGAA
33188 TATGAGTGTGGAGTCGTTGCTTTGGTAGCTAGGCTCTTTCCTGNNNAGATCGGAAG
33190 TATGAGTGTGGAGTCGTTGCTGGTTTCACTGATTTTCTTCCCANNNAGATCGGAAGA
33192 TATGAGTGTGGAGTCGTTGCGCATGTGTCTGTAATCCCAGCTANNNAGATCGGAAG
33193 TATGAGTGTGGAGTCGTTGCCAGGAGAATTGCTTGAACTTGGGNNNAGATCGGAA
33194 TATGAGTGTGGAGTCGTTGCTTGCCCTATATCTGAGGTTGCTCNNNAGATCGGAAG
33196 TATGAGTGTGGAGTCGTTGCTGTCCTTGTCCTTGAAATGTCGANNNAGATCGGAAG
33199 TATGAGTGTGGAGTCGTTGCGCTCAGGTGAAGTGAATGAGAGANNNAGATCGGA
33200 TATGAGTGTGGAGTCGTTGCGGCTGAGGTAGGAGAATAGCTTGNNNAGATCGGA
33201 TATGAGTGTGGAGTCGTTGCGGGGTCCTTGGAAAGTTTTTGTCNNNAGATCGGAAG
33202 TATGAGTGTGGAGTCGTTGCCAGAAGAAAGGCATTTGTTTTGCNNNAGATCGGAAG
33203 TATGAGTGTGGAGTCGTTGCATTCATCCTCTCTTTCTTCGGCANNNAGATCGGAAGA
33204 TATGAGTGTGGAGTCGTTGCATTACAACACACTTGGGGACTGTNNNAGATCGGAAG
33205 TATGAGTGTGGAGTCGTTGCAGAAATCACAAAACTCAGAGCCANNNAGATCGGAAG
33206 TATGAGTGTGGAGTCGTTGCTACTAGTCAGGTCCTGAGGGTTTNNNAGATCGGAAG
33207 TATGAGTGTGGAGTCGTTGCAGAAAAGGACACACATTGACTGANNNAGATCGGAA
33208 TATGAGTGTGGAGTCGTTGCAAGTTCTTGCATTCAAAGGGTCGNNNAGATCGGAAG
33210 TATGAGTGTGGAGTCGTTGCACAGAAGTTGAAGAGAGGAAGGANNNAGATCGGA
33212 TATGAGTGTGGAGTCGTTGCCAGACAGAAGTGAAGGCAAACAGNNNAGATCGGA
33213 TATGAGTGTGGAGTCGTTGCTCATTCTTCAACCTGCAAACACCNNNAGATCGGAAGA
33214 TATGAGTGTGGAGTCGTTGCGCATTGTAGTGCATGCCTGTAATNNNAGATCGGAAG
33217 TATGAGTGTGGAGTCGTTGCAGACCTTACACCCCTCACAAAAANNNAGATCGGAAG
33220 TATGAGTGTGGAGTCGTTGCTATACCTCTGCGAATGGAACCAGNNNAGATCGGAAG
33221 TATGAGTGTGGAGTCGTTGCTTATTTATCTGCCTCAGCCTCCCNNNAGATCGGAAGA
33222 TATGAGTGTGGAGTCGTTGCCGCCCGGCTTTTCTTTTTTTTTTNNNAGATCGGAAGA
33223 TATGAGTGTGGAGTCGTTGCGATCTCTGATCACTGCAACCTCCNNNAGATCGGAAG
33224 TATGAGTGTGGAGTCGTTGCCTCCTGAGTAGCTGGGACTACAGNNNAGATCGGAA
33225 TATGAGTGTGGAGTCGTTGCAGTAGATGGGATTACAGGCACACNNNAGATCGGAA
33226 TATGAGTGTGGAGTCGTTGCGAGTAGCTGGGATTACAGGCATGNNNAGATCGGAA
33227 TATGAGTGTGGAGTCGTTGCAATAGCTGGGATTACAGACACCCNNNAGATCGGAAG
33228 TATGAGTGTGGAGTCGTTGCAGTATTCCCACCTAGTCAGAGCTNNNAGATCGGAAG

33229 TATGAGTGTGGAGTCGTTGCCTCAAACTCCTGACCTCGTGATCNNNAGATCGGAAGA
33230 TATGAGTGTGGAGTCGTTGCTCTGATTCTGGAATTGTCCCCACNNNAGATCGGAAGA
33231 TATGAGTGTGGAGTCGTTGCCTAGACTTACTGCACCCTCCATCNNNAGATCGGAAGA
33232 TATGAGTGTGGAGTCGTTGCACTTCTCGGTTCAAGCTATCCTCNNNAGATCGGAAGA
33233 TATGAGTGTGGAGTCGTTGCCTCACTTGTAGCCTCGAACTCTTNNNAGATCGGAAGA
33235 TATGAGTGTGGAGTCGTTGCCCTATAAGCATGATCCTCCCACCNNNAGATCGGAAGA
33236 TATGAGTGTGGAGTCGTTGCGCCCAAAACTAGACCAAGAACACNNNAGATCGGAAG
33237 TATGAGTGTGGAGTCGTTGCATAGGGTCTTACTCAGTCACCCANNNAGATCGGAAGA
33239 TATGAGTGTGGAGTCGTTGCCTCCTGGGTTCTAAGGGATTCTCNNNAGATCGGAAG
33240 TATGAGTGTGGAGTCGTTGCCAATCCTCCCAGCTTACTCTCTGNNNAGATCGGAAGA
33242 TATGAGTGTGGAGTCGTTGCTCTATCACCTAGACGAGAGTGCANNNAGATCGGAAG
33243 TATGAGTGTGGAGTCGTTGCGTAGCTGGGACTACAGTTTCACANNNAGATCGGAAG
33246 TATGAGTGTGGAGTCGTTGCGATTCTCCTACCTCAGCCTTCTGNNNAGATCGGAAGA
33248 TATGAGTGTGGAGTCGTTGCGAGATGGGGTCTTTCTCTGTCACNNNAGATCGGAAG
33249 TATGAGTGTGGAGTCGTTGCTATCAAGTGATTCTCCTGCCTCGNNNAGATCGGAAGA
33250 TATGAGTGTGGAGTCGTTGCCTCCCGAGTAGCTGAGACTACAGNNNAGATCGGAAG
33251 TATGAGTGTGGAGTCGTTGCCCCCTAAGTAGCTGGGACTACAGNNNAGATCGGAAG
33252 TATGAGTGTGGAGTCGTTGCCGAGTAGCTGGAATTACAGGTGTNNNAGATCGGAAG
33253 TATGAGTGTGGAGTCGTTGCCGTCTTGTACTACAGTGAGACCCNNNAGATCGGAAG
33255 TATGAGTGTGGAGTCGTTGCATCTTGGCTCAATGCAACCTCTANNNAGATCGGAAGA
33256 TATGAGTGTGGAGTCGTTGCTTCTCAAGTGCCTACTCTCACACNNNAGATCGGAAGA
33257 TATGAGTGTGGAGTCGTTGCGAGAGAGAGCAGTTTGAAGGTGANNNAGATCGGAA
33259 TATGAGTGTGGAGTCGTTGCCCTCCCAGGTTCAAGTGATTCTCNNNAGATCGGAAGA
33262 TATGAGTGTGGAGTCGTTGCGGACTAAACTGAAGGCAAAACCCNNNAGATCGGAAG
33263 TATGAGTGTGGAGTCGTTGCCCCAGGTTCAAGCAATTCTTCTGNNNAGATCGGAAGA
33265 TATGAGTGTGGAGTCGTTGCTTCAAGTGATTCTCCTGCCTCAGNNNAGATCGGAAGA
33266 TATGAGTGTGGAGTCGTTGCCTCGATCTCTTGACCTCGTGATCNNNAGATCGGAAGA
33267 TATGAGTGTGGAGTCGTTGCGCTAGTAGCTGGGATTACAGGTGNNNAGATCGGAA
33268 TATGAGTGTGGAGTCGTTGCCTCCTGACCTCAAGTGATCCATCNNNAGATCGGAAGA
33269 TATGAGTGTGGAGTCGTTGCCCACCACGCACAGCTAATTTTTANNNAGATCGGAAGA
33271 TATGAGTGTGGAGTCGTTGCGCATAATCATAGCTCACTGCAGCNNNAGATCGGAAG
33276 TATGAGTGTGGAGTCGTTGCAGATGGAGTTTCTCTCTGTTGCCNNNAGATCGGAAG
33277 TATGAGTGTGGAGTCGTTGCCACCATGCCCGACTAATTTTTGTNNNAGATCGGAAGA
33278 TATGAGTGTGGAGTCGTTGCCATATGTTGGGCAATGGGTTCACNNNAGATCGGAAG
33279 TATGAGTGTGGAGTCGTTGCGATCTCAGCTCACTACAACCTCCNNNAGATCGGAAGA
33280 TATGAGTGTGGAGTCGTTGCGAGTGCAGTGGTACAATTATGGCNNNAGATCGGAAG
33281 TATGAGTGTGGAGTCGTTGCGGTTCAAGTGATTCTCCTGCTTCNNNAGATCGGAAGA
33282 TATGAGTGTGGAGTCGTTGCCAATCTTGGCTCACTGCAATCTCNNNAGATCGGAAGA
33283 TATGAGTGTGGAGTCGTTGCTCTTGTTCAGTCTTCAGAGGGTGNNNAGATCGGAAG
33284 TATGAGTGTGGAGTCGTTGCCCTCCCAGGTTCAACTGATTCTTNNNAGATCGGAAGA
33285 TATGAGTGTGGAGTCGTTGCGATGGAGTCTCACTCTGTCATCCNNNAGATCGGAAGA
33286 TATGAGTGTGGAGTCGTTGCCTGAGTAGGTGGGATTACAGGTGNNNAGATCGGAA
33289 TATGAGTGTGGAGTCGTTGCAAACTCCACCTCATGAGCTCAATNNNAGATCGGAAGA

33290 TATGAGTGTGGAGTCGTTGCAGAAGGAGTCTCACTCTGTCACCNNNAGATCGGAAG
33291 TATGAGTGTGGAGTCGTTGCTGTAATGGCATGATCTCAGCTCANNNAGATCGGAAG
33292 TATGAGTGTGGAGTCGTTGCTGTTAGAACTCCTGGGCTTATGGNNNAGATCGGAA
33293 TATGAGTGTGGAGTCGTTGCTTCAAGTGATTCTCCTGCCTCAGNNNAGATCGGAAG
33294 TATGAGTGTGGAGTCGTTGCTTTTAGTAGAGACGGGGTTTCGCNNNAGATCGGAA
33295 TATGAGTGTGGAGTCGTTGCTTAGTTCAAGTGAGTCTCCTGCCNNNAGATCGGAAG
33296 TATGAGTGTGGAGTCGTTGCTAGGACAATAGTGGAGGGAAGGTNNNAGATCGGA
33298 TATGAGTGTGGAGTCGTTGCTTCAAGTGCTTATCCTGCCTCAGNNNAGATCGGAAG
33303 TATGAGTGTGGAGTCGTTGCGACTGAGTCTCATTCTGTCACCANNNAGATCGGAAG
33304 TATGAGTGTGGAGTCGTTGCGATTCAAGCAATTCTCCTGCCTCNNNAGATCGGAAG
33306 TATGAGTGTGGAGTCGTTGCCCTCCCAGGTTCAAGTGATTCTCNNNAGATCGGAAG
33308 TATGAGTGTGGAGTCGTTGCCCTACCAGGTTCACACCATTCTCNNNAGATCGGAAGA
33309 TATGAGTGTGGAGTCGTTGCCTGGTCTTGAACTCATGGTCTCANNNAGATCGGAAG
33311 TATGAGTGTGGAGTCGTTGCTTCCATGTTTAGTCTTGGGAGGGNNNAGATCGGAA
33312 TATGAGTGTGGAGTCGTTGCCTTCAACTGCTGGACTTGTGATCNNNAGATCGGAAG
33313 TATGAGTGTGGAGTCGTTGCCCTCCCAGGTTCAAGTAGTTCTTNNNAGATCGGAAG
33314 TATGAGTGTGGAGTCGTTGCATTTTCTTCTCAGAGCCTCCCTGNNNAGATCGGAAGA
33316 TATGAGTGTGGAGTCGTTGCGTGGCATCATCATAACTCACTGCNNNAGATCGGAAG
33317 TATGAGTGTGGAGTCGTTGCATTTGGCAGGGTCATAGGACAATNNNAGATCGGAA
33318 TATGAGTGTGGAGTCGTTGCGCAATCATAGCTCACTGTACCCTNNNAGATCGGAAG
33319 TATGAGTGTGGAGTCGTTGCCAGCCTCGACCTTCTAAGCTAAANNNAGATCGGAAG
33321 TATGAGTGTGGAGTCGTTGCGAATGGCAATATCTGGCTTCGACNNNAGATCGGAAG
33322 TATGAGTGTGGAGTCGTTGCCCTGGGTTCAACCAATTCTCCTANNNAGATCGGAAG
33323 TATGAGTGTGGAGTCGTTGCAGCCTCCCAAGTAGCTAGAACTANNNAGATCGGAAG
33325 TATGAGTGTGGAGTCGTTGCTCAAGTAGCTGGGATTACAGGTGNNNAGATCGGAA
33326 TATGAGTGTGGAGTCGTTGCTCCTGGGTTCAAATGATTCTCGTNNNAGATCGGAAG
33329 TATGAGTGTGGAGTCGTTGCTCTCCCAGGTTCATGTGATTCTCNNNAGATCGGAAG
33330 TATGAGTGTGGAGTCGTTGCCACAATCTTGGCTCACTGTGAACNNNAGATCGGAAG
33331 TATGAGTGTGGAGTCGTTGCCCAGACCTCAAATAATCCACCCANNNAGATCGGAAG
33332 TATGAGTGTGGAGTCGTTGCCTATCTGGGCTCACTACAACCTCNNNAGATCGGAAG
33333 TATGAGTGTGGAGTCGTTGCACAATTCTCCTACTTCAGCCTCCNNNAGATCGGAAGA
33334 TATGAGTGTGGAGTCGTTGCTGCATGATCTCAGCTCACTGTAGNNNAGATCGGAAG
33336 TATGAGTGTGGAGTCGTTGCCTCCTGAGTAGCTGGGATTACAGNNNAGATCGGAA
33337 TATGAGTGTGGAGTCGTTGCGTATTTTTGGTAGAGGGGGGGAGNNNAGATCGGA
33339 TATGAGTGTGGAGTCGTTGCGTTCAAGCCATTCTCCTACCTCANNNAGATCGGAAGA
33340 TATGAGTGTGGAGTCGTTGCCTACCTCAACAATTCTCCTGCCTNNNAGATCGGAAGA
33341 TATGAGTGTGGAGTCGTTGCCGCCTCCTGGATTGAAATGATTCNNNAGATCGGAAG
33342 TATGAGTGTGGAGTCGTTGCCTCCTGAATAGCTGGGACCAGAGNNNAGATCGGAA
33343 TATGAGTGTGGAGTCGTTGCGGGTACCTACTGATCGTTTGTGTNNNAGATCGGAAG
33344 TATGAGTGTGGAGTCGTTGCGATTACAGGCATGCACCACAATTNNNAGATCGGAAG
33345 TATGAGTGTGGAGTCGTTGCGCCTACTGAGTAGCTGGGATTACNNNAGATCGGAA
33346 TATGAGTGTGGAGTCGTTGCGCCTCCCAAAGTTGTGATTACAGNNNAGATCGGAAG
33347 TATGAGTGTGGAGTCGTTGCTCTGCCTCCTGAGTTCAAACAATNNNAGATCGGAAG

33349 TATGAGTGTGGAGTCGTTGCTTCAAGTGATTCTCCTGCCTCAGNNNAGATCGGAAG
33350 TATGAGTGTGGAGTCGTTGCGCCTCCTGAGTAGTTGGGATTACNNNAGATCGGAAG
33351 TATGAGTGTGGAGTCGTTGCGGGTTCAAGTGATTCTCCTGTCTNNNAGATCGGAAG
33352 TATGAGTGTGGAGTCGTTGCCTGGGTTCAAGTGATTCATCTGCNNNAGATCGGAAG
33353 TATGAGTGTGGAGTCGTTGCGAGGTAGGGTCTCAATCTGTCACNNNAGATCGGAA
33354 TATGAGTGTGGAGTCGTTGCCCTACCAAAGTGCTGGGATTACANNNAGATCGGAAG
33355 TATGAGTGTGGAGTCGTTGCATGGTGACTGAGAGCTCCAAAAANNNAGATCGGAA
33356 TATGAGTGTGGAGTCGTTGCACAGTCTTGCTCTATTGCCTAGGNNNAGATCGGAAG
33360 TATGAGTGTGGAGTCGTTGCTTTCCTCAACCCCTATCCCCTATNNNAGATCGGAAGA
33363 TATGAGTGTGGAGTCGTTGCTCCTGAGTAGCTAGGACTACAGGNNNAGATCGGAA
33366 TATGAGTGTGGAGTCGTTGCTCCAGGTTCAAGTGATTCTCCTGNNNAGATCGGAAG
33369 TATGAGTGTGGAGTCGTTGCTTCAAGTGATTCTCCTGCCTCAGNNNAGATCGGAAG
33371 TATGAGTGTGGAGTCGTTGCGGAGGTCTCACTTTGTTGCCTAGNNNAGATCGGAAG
33372 TATGAGTGTGGAGTCGTTGCAGCGAGAGAGAAAGAGAAAGACANNNAGATCGGA
33375 TATGAGTGTGGAGTCGTTGCTCCTAAATCTCCACACCCAGAGANNNAGATCGGAAG
33376 TATGAGTGTGGAGTCGTTGCAAATGAGGAGACTGTGCTCAGAGNNNAGATCGGAA
33378 TATGAGTGTGGAGTCGTTGCAACATCATTCTCCTCCTCCTCCTNNNAGATCGGAAGA
33380 TATGAGTGTGGAGTCGTTGCTCTACCCACTCTCTCTCTTGGTCNNNAGATCGGAAGA
33381 TATGAGTGTGGAGTCGTTGCTCCAGACCTCAACAAATTGTCCTNNNAGATCGGAAG
33383 TATGAGTGTGGAGTCGTTGCGTCAATCACGAGGTCAGGAGTTCNNNAGATCGGAA
33385 TATGAGTGTGGAGTCGTTGCATTCTGTAGCAGCAAAGAGCCTCNNNAGATCGGAAG
33386 TATGAGTGTGGAGTCGTTGCAACCCAAACGTCCATCAGTGATANNNAGATCGGAAG
33387 TATGAGTGTGGAGTCGTTGCTGGTGAAACCCCATCTCTACTAANNNAGATCGGAAG
33388 TATGAGTGTGGAGTCGTTGCTATTCTGAGTGAGAAGGGAAGCCNNNAGATCGGAA
33389 TATGAGTGTGGAGTCGTTGCTGTAGTGAGCCTAGAGCAGAGATNNNAGATCGGAA
33390 TATGAGTGTGGAGTCGTTGCCTGAGACACGAGAATCCCTTGAANNNAGATCGGAAG
33393 TATGAGTGTGGAGTCGTTGCATTCCATTAGAGGCTGCCCTAAGNNNAGATCGGAAG
33401 TATGAGTGTGGAGTCGTTGCCGAATTCATTTGGGCCTCATTCANNNAGATCGGAAG
33403 TATGAGTGTGGAGTCGTTGCAAGAGGAAGGAACATGACAAGGANNNAGATCGGA
33406 TATGAGTGTGGAGTCGTTGCTCATTTCCAGCAGAACTCTGAGTNNNAGATCGGAAG
33407 TATGAGTGTGGAGTCGTTGCGATTTGACAACTGTACATCGGCCNNNAGATCGGAAG
33409 TATGAGTGTGGAGTCGTTGCCAGCCTAGTAGGTTTTCAGCCTTNNNAGATCGGAAG
33411 TATGAGTGTGGAGTCGTTGCACATAGGAGGTAGAGACTGCAGTNNNAGATCGGAA
33412 TATGAGTGTGGAGTCGTTGCCTTACACCTGTAATCCCAGCACTNNNAGATCGGAAGA
33413 TATGAGTGTGGAGTCGTTGCGGCGAAACCCTGTCTCTACTAAANNNAGATCGGAAG
33414 TATGAGTGTGGAGTCGTTGCGTGCTGACACACACTACAACATGNNNAGATCGGAAG
33416 TATGAGTGTGGAGTCGTTGCAGTATGGCATCTCCTCATCTCCTNNNAGATCGGAAGA
33417 TATGAGTGTGGAGTCGTTGCTGCTTATGCCTATAATCCCAGCANNNAGATCGGAAG
33418 TATGAGTGTGGAGTCGTTGCGCCTGGGTTTGAATTCTGAACTCNNNAGATCGGAAG
33419 TATGAGTGTGGAGTCGTTGCGAGAGGGAGTTTTGAGACAGAGGNNNAGATCGGA
33421 TATGAGTGTGGAGTCGTTGCAAACTGTGAGGGAAGAGAAGGACNNNAGATCGGA
33422 TATGAGTGTGGAGTCGTTGCTTGTCCCACACAGAATCTTCCTTNNNAGATCGGAAGA
33423 TATGAGTGTGGAGTCGTTGCTGGGCTCAGTGAAAACAAGAAACNNNAGATCGGAA

33424 TATGAGTGTGGAGTCGTTGCAGAGTGATAGAGCAAATGTGGCANNNAGATCGGAA
33425 TATGAGTGTGGAGTCGTTGCTCACAAAAGACCCAGAACATCCANNNAGATCGGAAG
33426 TATGAGTGTGGAGTCGTTGCGAATTTCACCCTGGCAACGTAAANNNAGATCGGAAG
33427 TATGAGTGTGGAGTCGTTGCCTGGAGAGAAGGCAGTACTTGAANNNAGATCGGAA
33428 TATGAGTGTGGAGTCGTTGCATAACATCTCAGCTCTTTGGCCANNNAGATCGGAAG
33429 TATGAGTGTGGAGTCGTTGCAAAAAATTAGCTGGGTGTGGTGGNNNAGATCGGAA
33430 TATGAGTGTGGAGTCGTTGCCACCTGTAATCCCAGCACTTTTGNNNAGATCGGAAGA
33431 TATGAGTGTGGAGTCGTTGCATCTAGGGGAAAAGCATTCCTGGNNNAGATCGGAA
33432 TATGAGTGTGGAGTCGTTGCCCAACCAGATACCTAACCCAACANNNAGATCGGAAG
33437 TATGAGTGTGGAGTCGTTGCCACACACACACACACACAGTATGNNNAGATCGGAAG
33438 TATGAGTGTGGAGTCGTTGCAAATATTGGTTGGACACAGTGGCNNNAGATCGGAA
33440 TATGAGTGTGGAGTCGTTGCGAAGGAGTAGAAGGTGCTGAGAGNNNAGATCGGA
33442 TATGAGTGTGGAGTCGTTGCGTGACATGGATGAAGCTGGAAAGNNNAGATCGGA
33443 TATGAGTGTGGAGTCGTTGCAAGGAGGCCTGTGGAGTCTATATNNNAGATCGGAA
33444 TATGAGTGTGGAGTCGTTGCCTAAGGAGAAGGACTAGGAGGCTNNNAGATCGGA
33446 TATGAGTGTGGAGTCGTTGCAGATGTGGATGATGATGGTGATGNNNAGATCGGA
33447 TATGAGTGTGGAGTCGTTGCAGAAAAGGGCCTAGGGATTTGAGNNNAGATCGGA
33448 TATGAGTGTGGAGTCGTTGCCTTCTAGGGCTGTTTTGAGCAACNNNAGATCGGAAG
33449 TATGAGTGTGGAGTCGTTGCAGAAATGCTTCAGGTGGTGAGATNNNAGATCGGAA
33450 TATGAGTGTGGAGTCGTTGCTTCCCTTTTGCTACTGTCCACATNNNAGATCGGAAGA
33451 TATGAGTGTGGAGTCGTTGCCAACAAAAAAACCCAGCTTGGTGNNNAGATCGGAAG
33452 TATGAGTGTGGAGTCGTTGCCAGAGGCTGTGGTGAGAAGATAGNNNAGATCGGA
33454 TATGAGTGTGGAGTCGTTGCCCAATCCCCTCAGCATAGTAAGGNNNAGATCGGAAG
33455 TATGAGTGTGGAGTCGTTGCACCAGGGGCTCAGAGATAAAAAGNNNAGATCGGAA
33456 TATGAGTGTGGAGTCGTTGCAGAGTTTCAATCCTGGTCTCTGANNNAGATCGGAAG
33457 TATGAGTGTGGAGTCGTTGCGCTTGTGTTCCAACAAATCAGGTNNNAGATCGGAAG
33459 TATGAGTGTGGAGTCGTTGCAAAGAGAGGCTTTGTTTCTGTGCNNNAGATCGGAA
33460 TATGAGTGTGGAGTCGTTGCCTCCAGGTAGCAAACTTCAGAGANNNAGATCGGAA
33466 TATGAGTGTGGAGTCGTTGCGATTGGAGGACAATGGCAAATGTNNNAGATCGGAA
33467 TATGAGTGTGGAGTCGTTGCAAAGAAGAGCTGGTCCCAATTCTNNNAGATCGGAA
33469 TATGAGTGTGGAGTCGTTGCCACGCACACACATAAAGGAGAAANNNAGATCGGAA
33471 TATGAGTGTGGAGTCGTTGCGCAAAGCAAATCCAGCAGCATATNNNAGATCGGAA
33473 TATGAGTGTGGAGTCGTTGCCACTTGACCAGCGTTATGGAATCNNNAGATCGGAAG
33476 TATGAGTGTGGAGTCGTTGCTGTACCTCAGGGAAATACAAGGCNNNAGATCGGAA
33477 TATGAGTGTGGAGTCGTTGCCAAGGCAGGAGGATTACTTGAGTNNNAGATCGGAA
33478 TATGAGTGTGGAGTCGTTGCCCTGACGCAACATAAAATGGCTANNNAGATCGGAAG
33480 TATGAGTGTGGAGTCGTTGCACACACACAACCAGAGTACAAGANNNAGATCGGAA
33481 TATGAGTGTGGAGTCGTTGCTGCAGGTAAAGTGCTTTTTTGTTNNNAGATCGGAAG
33482 TATGAGTGTGGAGTCGTTGCTAATTCCAGCATTTTGCAAGGCTNNNAGATCGGAAG
33484 TATGAGTGTGGAGTCGTTGCAGTTGAAGAATGGGATGCCTCTTNNNAGATCGGAA
33486 TATGAGTGTGGAGTCGTTGCCCAAGACCATTCCACTGCATTACNNNAGATCGGAAGA
33487 TATGAGTGTGGAGTCGTTGCGTGAGTCTGAGGGATGACAAGTTNNNAGATCGGAA
33488 TATGAGTGTGGAGTCGTTGCCAGAGTAGCTACAAGGCAAGACTNNNAGATCGGAA

33489 TATGAGTGTGGAGTCGTTGCAAAGAGAAAGGGGGATAAGGCAGNNNAGATCGGA

33490 TATGAGTGTGGAGTCGTTGCTCTGGACATAAACCATTCCTGCCNNNAGATCGGAAG

33491 TATGAGTGTGGAGTCGTTGCGGCGAAAATACACTGTGGCATAANNNAGATCGGAA

33493 TATGAGTGTGGAGTCGTTGCGTGGTATTTGTTGCCTGCTTTCTNNNAGATCGGAAG

33494 TATGAGTGTGGAGTCGTTGCTTGAAACCACACAGCAACTTTGTNNNAGATCGGAAG

33495 TATGAGTGTGGAGTCGTTGCATCCTACTACCCAGCAATGATCANNNAGATCGGAAG

33497 TATGAGTGTGGAGTCGTTGCCAGCTCTGGTTTCACAAATGAGGNNNAGATCGGAA

33498 TATGAGTGTGGAGTCGTTGCCTTTGGTTCTAACAAAGGCCTGGNNNAGATCGGAAG

33499 TATGAGTGTGGAGTCGTTGCAACCAAACCAAACTTCTGCCTTCNNNAGATCGGAAGA

33500 TATGAGTGTGGAGTCGTTGCCACACCTGTAATCCCAGCACTTTNNNAGATCGGAAGA

33501 TATGAGTGTGGAGTCGTTGCTGAGCATCCAGATGTCTCTTGTTNNNAGATCGGAAG

33502 TATGAGTGTGGAGTCGTTGCTTGCAACATATAAGGCAAAGGGCNNNAGATCGGAA

33503 TATGAGTGTGGAGTCGTTGCGATAACTTGGGGCTGTTTACTGCNNNAGATCGGAA

33504 TATGAGTGTGGAGTCGTTGCCAGTCCTGACTTAGCATGGAAGTNNNAGATCGGAA

33506 TATGAGTGTGGAGTCGTTGCGCCTGTAAACAGAATCTGCCCTANNNAGATCGGAAG

33507 TATGAGTGTGGAGTCGTTGCTGAGAAACTCATTCCTTGTCCCCNNNAGATCGGAAG

33508 TATGAGTGTGGAGTCGTTGCCCTTCAAGGTATGGCTTCTCTCANNNAGATCGGAAG

33510 TATGAGTGTGGAGTCGTTGCTTCTTTTGATTGCTCGTGGATCTNNNAGATCGGAAG

33511 TATGAGTGTGGAGTCGTTGCGAGATCATGCCACTACACTCCAGNNNAGATCGGAAG

33512 TATGAGTGTGGAGTCGTTGCTCTTGCTAGTGGATAAGCTGACCNNNAGATCGGAAG

33514 TATGAGTGTGGAGTCGTTGCAGGTAGGAGAATTGCTTGAACCCNNNAGATCGGAA

33515 TATGAGTGTGGAGTCGTTGCAGATACTAGAGGTTGCACAGTGCNNNAGATCGGAA

33517 TATGAGTGTGGAGTCGTTGCATAACAGATGCTCGTGAGGTTGTNNNAGATCGGAA

33520 TATGAGTGTGGAGTCGTTGCGAGTTGTGGTGATAGGGTCTCTGNNNAGATCGGAA

33523 TATGAGTGTGGAGTCGTTGCTTCACACCATGACAAAAACTGCANNNAGATCGGAAG

33524 TATGAGTGTGGAGTCGTTGCGGGTCACCTTTATTCCAGATCCCNNNAGATCGGAAG

33526 TATGAGTGTGGAGTCGTTGCCAAGCCCAGAGTGTGAAGACTATNNNAGATCGGAA

33527 TATGAGTGTGGAGTCGTTGCTGGGCAAAAGATATGAAGAGGCANNNAGATCGGA

33528 TATGAGTGTGGAGTCGTTGCATATACTTTGGGACAAGGCCAGCNNNAGATCGGAA

33531 TATGAGTGTGGAGTCGTTGCGAATATCATTGCCCTGAAAGCCCNNNAGATCGGAAG

33532 TATGAGTGTGGAGTCGTTGCGACAGATTCTCACTCTCACCCAGNNNAGATCGGAAG

33535 TATGAGTGTGGAGTCGTTGCCTTGAACTCCTGACCTCGTGATCNNNAGATCGGAAG

33537 TATGAGTGTGGAGTCGTTGCGATGGACTTACTCTATCGCCCAGNNNAGATCGGAAG

33538 TATGAGTGTGGAGTCGTTGCAGTAGAAACAGGCTTTCACCGTGNNNAGATCGGAA

33539 TATGAGTGTGGAGTCGTTGCCCATGACACCCGGCTAATTTTTTNNNAGATCGGAAG

33540 TATGAGTGTGGAGTCGTTGCCCTGGGTTCAAGCAATTCTCCTANNNAGATCGGAAG

33543 TATGAGTGTGGAGTCGTTGCCCTGGGTTCAAGCAATTCTCATGNNNAGATCGGAAG

33544 TATGAGTGTGGAGTCGTTGCAGAATCTCACTATGTTGCCCAGGNNNAGATCGGAAG

33545 TATGAGTGTGGAGTCGTTGCGGATACACAGTCCCACTTTGTCANNNAGATCGGAAG

33547 TATGAGTGTGGAGTCGTTGCGCTCAAGTGATCCTTCCATCTCANNNAGATCGGAAG

33548 TATGAGTGTGGAGTCGTTGCCCTTAGCTTCCTGAGTACTTGGGNNNAGATCGGAAG

33549 TATGAGTGTGGAGTCGTTGCCTCCTAAGTAGCTGGGATTGCAGNNNAGATCGGAA

33550 TATGAGTGTGGAGTCGTTGCCTGGGTTCAATTGATTCTCCTGCNNNAGATCGGAAG

33552 TATGAGTGTGGAGTCGTTGCTTCTTCTGGTTCGGTCTTGAGAGNNNAGATCGGAAG
33553 TATGAGTGTGGAGTCGTTGCGTTCAAGTGATTCTCTTGCCTCGNNNAGATCGGAAG
33554 TATGAGTGTGGAGTCGTTGCGGAGATGTGGCTTTGCTATGTTGNNNAGATCGGAA
33556 TATGAGTGTGGAGTCGTTGCCGGCTCACCATAGCAGGTATATANNNAGATCGGAAG
33557 TATGAGTGTGGAGTCGTTGCCTGAGTAGCTGGAACTATAGGCGNNNAGATCGGAA
33558 TATGAGTGTGGAGTCGTTGCCTGAGTAGTTGGGATTGTAGGCANNNAGATCGGAA
33562 TATGAGTGTGGAGTCGTTGCCCTCCCAAAATGCTGAAATCACTNNNAGATCGGAAG
33563 TATGAGTGTGGAGTCGTTGCTCCTAAGTAGCTGGGATCACAGANNNAGATCGGAA
33566 TATGAGTGTGGAGTCGTTGCCTGGGTTCAATTGATTCTCCTGCNNNAGATCGGAAG
33567 TATGAGTGTGGAGTCGTTGCCTCCCGAGTTCAAGTGATTCTCANNNAGATCGGAAG
33569 TATGAGTGTGGAGTCGTTGCAGTTTCTCTCTCTCTCTCACCCANNNAGATCGGAAGA
33570 TATGAGTGTGGAGTCGTTGCCAGTGGTATGATCTTGGCTCAGTNNNAGATCGGAA
33572 TATGAGTGTGGAGTCGTTGCAAGTTCAAGTGATTCTCCTGCCTNNNAGATCGGAAG
33573 TATGAGTGTGGAGTCGTTGCTTCTTGTTCAGTCATGGGAAGGTNNNAGATCGGAA
33574 TATGAGTGTGGAGTCGTTGCGTGTGGAGTCATTTCTAGGGTGTNNNAGATCGGAA
33575 TATGAGTGTGGAGTCGTTGCATCTCAGCTTACTGCATCCTCTGNNNAGATCGGAAG
33577 TATGAGTGTGGAGTCGTTGCGTAGCTAGGATTACAGGCATGCANNNAGATCGGAA
33579 TATGAGTGTGGAGTCGTTGCATGAGAGACAGGACTAGCTGGATNNNAGATCGGAA
33580 TATGAGTGTGGAGTCGTTGCGGAGTTTCATTCTCGCTCTGTTGNNNAGATCGGAAG
33581 TATGAGTGTGGAGTCGTTGCTGGTATGATTTCGGCTCACTGAANNNAGATCGGAAG
33582 TATGAGTGTGGAGTCGTTGCGTTCAAGAGATTCTCCTGCCTCANNNAGATCGGAAG
33583 TATGAGTGTGGAGTCGTTGCACTACACCATGCCCAGCTAATTTNNNAGATCGGAAG
33585 TATGAGTGTGGAGTCGTTGCCCAAGTAGCTGGGATTACAGGTGNNNAGATCGGAA
33589 TATGAGTGTGGAGTCGTTGCCGAATAGCTAGGATTACAGGCGTNNNAGATCGGAA
33591 TATGAGTGTGGAGTCGTTGCTTCAAGTGATTCTCCTGCCTCAGNNNAGATCGGAAG
33593 TATGAGTGTGGAGTCGTTGCTATGTTGCCCAGGTTGATCTTGANNNAGATCGGAAG
33594 TATGAGTGTGGAGTCGTTGCGGTTCAAGTGATTCTTCTGCCTCNNNAGATCGGAAG
33595 TATGAGTGTGGAGTCGTTGCGTTCAAGCAATTCTCTGCCTCAGNNNAGATCGGAAG
33596 TATGAGTGTGGAGTCGTTGCTTCACATCATTCTCCTGCCTCAGNNNAGATCGGAAGA
33597 TATGAGTGTGGAGTCGTTGCCCTCCTCAGTTCAAGCGATTTTGNNNAGATCGGAAG
33598 TATGAGTGTGGAGTCGTTGCCGAATAGCTGGGACTACAGGTACNNNAGATCGGAA
33599 TATGAGTGTGGAGTCGTTGCTGTGTGTGTGCTTTCTCCTCATANNNAGATCGGAAG
33601 TATGAGTGTGGAGTCGTTGCCTCCTGAGTAGCTGGGACTACAGNNNAGATCGGAA
33602 TATGAGTGTGGAGTCGTTGCTCAAGTGATTCTCGTGTCTCAGCNNNAGATCGGAAG
33604 TATGAGTGTGGAGTCGTTGCTGGCAGTGGTATTGTCATCATTCNNNAGATCGGAAG
33606 TATGAGTGTGGAGTCGTTGCAATTTCCCAGGCTCAAGCAATTCNNNAGATCGGAAG
33607 TATGAGTGTGGAGTCGTTGCGTAGCTGGGATTACAGGCACATNNNAGATCGGAAG
33608 TATGAGTGTGGAGTCGTTGCAACACCGACCATCTTTTCATGTGNNNAGATCGGAAG
33609 TATGAGTGTGGAGTCGTTGCGAGATGGAGTCTTACTCTGTCGCNNNAGATCGGAA
33610 TATGAGTGTGGAGTCGTTGCTCTGGGTTCAAGTGATTCTCCTGNNNAGATCGGAAG
33611 TATGAGTGTGGAGTCGTTGCAGCCTCCCAAAATGCTGAGATTANNNAGATCGGAAG
33613 TATGAGTGTGGAGTCGTTGCATGGATTCTCATTCTGTCACCCANNNAGATCGGAAG
33615 TATGAGTGTGGAGTCGTTGCCTTCTGAGTAGCTGGGACTACAGNNNAGATCGGAA

33618 TATGAGTGTGGAGTCGTTGCACTGGGTTCAAGTGATTCTCCTGNNNAGATCGGAAG
33619 TATGAGTGTGGAGTCGTTGCCTCGACCTCCTAACCTCGTGATCNNNAGATCGGAAGA
33621 TATGAGTGTGGAGTCGTTGCCTCCTGAGTAGCTGGGACTACAGNNNAGATCGGAAG
33622 TATGAGTGTGGAGTCGTTGCGTTACCTGCAGCAACATTTGTCTNNNAGATCGGAAGA
33624 TATGAGTGTGGAGTCGTTGCAACTCCTGAACTTGTGATCCACANNNAGATCGGAAGA
33627 TATGAGTGTGGAGTCGTTGCTTCTGCACTATTCCCGACTGATTNNNAGATCGGAAGA
33629 TATGAGTGTGGAGTCGTTGCTTGGTAGATCTTCCTCCATCCCTNNNAGATCGGAAGA
33632 TATGAGTGTGGAGTCGTTGCTTCAAGCGATTCTCCTGTCTCAGNNNAGATCGGAAGA
33633 TATGAGTGTGGAGTCGTTGCAAGCCTGAAGTGCCAGAGAATTANNNAGATCGGAAG
33634 TATGAGTGTGGAGTCGTTGCATCTTGGCTCACTACAACCTCTGNNNAGATCGGAAGA
33635 TATGAGTGTGGAGTCGTTGCCTAGGCTGTAGTGCAGAAGTTCANNNAGATCGGAAG
33638 TATGAGTGTGGAGTCGTTGCTTAGGGTACATGTGCACATTGTGNNNAGATCGGAAG
33639 TATGAGTGTGGAGTCGTTGCTTCAAGTGATTCTCCTGCCTCAGNNNAGATCGGAAGA
33640 TATGAGTGTGGAGTCGTTGCGTTCAAGTAGTTCTCCTGCCTCANNNAGATCGGAAGA
33641 TATGAGTGTGGAGTCGTTGCGACAGAGTATCGATCGGTCTTGTNNNAGATCGGAAG
33642 TATGAGTGTGGAGTCGTTGCGGTATAGTGGCATGTTCTCAGCTNNNAGATCGGAAG
33644 TATGAGTGTGGAGTCGTTGCACACCGTTTCACTATGTTAGCCANNNAGATCGGAAGA
33645 TATGAGTGTGGAGTCGTTGCGATGAGATTTCACCATGTTGGCCNNNAGATCGGAAG
33647 TATGAGTGTGGAGTCGTTGCATGATCTCAGCTCACTGCAATCTNNNAGATCGGAAGA
33649 TATGAGTGTGGAGTCGTTGCTTCAACTGATTCTCCTGCCTCAGNNNAGATCGGAAGA
33650 TATGAGTGTGGAGTCGTTGCCTTCCTGGGTTCAAGCAATTCTCNNNAGATCGGAAGA
33652 TATGAGTGTGGAGTCGTTGCAACATATTTATTTGGGGCTGGGCNNNAGATCGGAAG
33654 TATGAGTGTGGAGTCGTTGCCCTCCCAGGTTCAAGTGATTTTCNNNAGATCGGAAGA
33655 TATGAGTGTGGAGTCGTTGCGAGTAGCTGGGATTACAGGAAGGNNNAGATCGGAA
33656 TATGAGTGTGGAGTCGTTGCCAATGCCAATTGTGGACCTGAAANNNAGATCGGAAG
33657 TATGAGTGTGGAGTCGTTGCGCTGCACCTGTTTTTGCTTTTATNNNAGATCGGAAGA
33658 TATGAGTGTGGAGTCGTTGCCCATTTCTCCTGACTCAGTCTCCNNNAGATCGGAAGA
33659 TATGAGTGTGGAGTCGTTGCCAAGTAGCTGGGACTACAGGCNNNAGATCGGAAGA
33660 TATGAGTGTGGAGTCGTTGCATAGTGGCATGATCTTGGCTCATNNNAGATCGGAAG
33662 TATGAGTGTGGAGTCGTTGCCTGAGTAGCTGGGACTACAGATGNNNAGATCGGAAG
33663 TATGAGTGTGGAGTCGTTGCCCAAGAGTGCCAGGATTAGAGTTNNNAGATCGGAAG
33664 TATGAGTGTGGAGTCGTTGCGTGTTGGAACTACAAGTGTGAGCNNNAGATCGGAAG
33667 TATGAGTGTGGAGTCGTTGCTTTAGTAGAGATGGGGTTTCGCCNNNAGATCGGAAG
33668 TATGAGTGTGGAGTCGTTGCCTCCTGAGTAGCTGGGATTACAGNNNAGATCGGAAG
33670 TATGAGTGTGGAGTCGTTGCCTGGAGTATAGTGGTGCAACCTTNNNAGATCGGAAG
33674 TATGAGTGTGGAGTCGTTGCTTCAAGTGATTCTCCTGCCTCAGNNNAGATCGGAAGA
33675 TATGAGTGTGGAGTCGTTGCCTCCTGAGTAGCTGGGACTACAGNNNAGATCGGAAG
33676 TATGAGTGTGGAGTCGTTGCAGGTTCAAGTGATTCTTCTGCCTNNNAGATCGGAAGA
33677 TATGAGTGTGGAGTCGTTGCGCCTCCTGAGTATACTGGGACTANNNAGATCGGAAG
33678 TATGAGTGTGGAGTCGTTGCCCTTGGGCATTTTGGTCATTTTANNNAGATCGGAAGA
33680 TATGAGTGTGGAGTCGTTGCTTTCCGAGGTAAACTTCCTGGAANNNAGATCGGAAG
33682 TATGAGTGTGGAGTCGTTGCAATGATTCTCCTGCCTCAGTCTCNNNAGATCGGAAGA
33683 TATGAGTGTGGAGTCGTTGCCCAAAGTGCTGGGATTACAAGTGNNNAGATCGGAAG

33684 TATGAGTGTGGAGTCGTTGCTTTTAGTAGAGACGGGGTTTCACNNNAGATCGGAA
33688 TATGAGTGTGGAGTCGTTGCGGATCTTGCTATGTTTTCCAGGCNNNAGATCGGAAG
33690 TATGAGTGTGGAGTCGTTGCCTGTAATCCCAGCTACTCAGGAGNNNAGATCGGAAG
33691 TATGAGTGTGGAGTCGTTGCATTCGTGAAGTTGAAGTGGCTCANNNAGATCGGAA
33693 TATGAGTGTGGAGTCGTTGCAAGCAACCTGAGAAGCTAGTTCANNNAGATCGGAA
33694 TATGAGTGTGGAGTCGTTGCATTGCTTGAATCTGAGAGACGGANNNAGATCGGAA
33695 TATGAGTGTGGAGTCGTTGCGTATTGATGGGATGTCTTGCAGCNNNAGATCGGAA
33696 TATGAGTGTGGAGTCGTTGCTTATAGGATAAGGACCTGGCACGNNNAGATCGGAA
33697 TATGAGTGTGGAGTCGTTGCCTGTTCAGAGAGGCCAAGAAGAGNNNAGATCGGAA
33699 TATGAGTGTGGAGTCGTTGCTTGGACCCTAACCGCTGTTATTTNNNAGATCGGAAG
33700 TATGAGTGTGGAGTCGTTGCTGAAGCTCAGAGGTTCAAAGTGGNNNAGATCGGAA
33702 TATGAGTGTGGAGTCGTTGCTCCCACTTTGCTACTATTTGCCTNNNAGATCGGAAGA
33703 TATGAGTGTGGAGTCGTTGCAAAGCACAAAAGAGAAAAGGGGGNNNAGATCGGA
33705 TATGAGTGTGGAGTCGTTGCGTACATTCTAGAGCTCCAGGCAGNNNAGATCGGAAG
33708 TATGAGTGTGGAGTCGTTGCAAAACATTTTAGCTGGGGGCAATNNNAGATCGGAA
33712 TATGAGTGTGGAGTCGTTGCCTGGGAAACATAGTGAGAACCCANNNAGATCGGAA
33714 TATGAGTGTGGAGTCGTTGCTCAGGAACATCACTAGTGGAAGCNNNAGATCGGAA
33715 TATGAGTGTGGAGTCGTTGCTTGAGAGGGAGTGCTTTGAGATCNNNAGATCGGAA
33716 TATGAGTGTGGAGTCGTTGCATTGAGGTGGTCATGAGGAGAAGNNNAGATCGGA
33717 TATGAGTGTGGAGTCGTTGCAACGAAATGTGCTTTTGTTTGCTNNNAGATCGGAAG
33718 TATGAGTGTGGAGTCGTTGCGTGCCTGTTGGAGAAAGAAGAACNNNAGATCGGAA
33719 TATGAGTGTGGAGTCGTTGCGGAGAATCACTTGAACCCAGGAGNNNAGATCGGAA
33721 TATGAGTGTGGAGTCGTTGCTTGCAGAAAGGAAAACTATGGCCNNNAGATCGGAA
33722 TATGAGTGTGGAGTCGTTGCTCTGCCTCTCTCCTAGCATCATANNNAGATCGGAAGA
33724 TATGAGTGTGGAGTCGTTGCAAGGGATGGAGGAAGATCTACCANNNAGATCGGAA
33725 TATGAGTGTGGAGTCGTTGCGCCTATAATTCCAGCTACTCGGGNNNAGATCGGAAG
33726 TATGAGTGTGGAGTCGTTGCTCTTTGGGACCAACACCATGTATNNNAGATCGGAAG
33727 TATGAGTGTGGAGTCGTTGCGAAACAGGAGTTCGGTTTGGAACNNNAGATCGGAA
33729 TATGAGTGTGGAGTCGTTGCAGTTGGCATGCTTGTTTATTGATNNNAGATCGGAAG
33730 TATGAGTGTGGAGTCGTTGCGGAAACTTGCTACCCCTCAGTAANNNAGATCGGAAG
33731 TATGAGTGTGGAGTCGTTGCCATGTGCGTTTCCATGGTATCAANNNAGATCGGAAG
33732 TATGAGTGTGGAGTCGTTGCGCTGTCACTTTCTACCCCTGAAANNNAGATCGGAAG
33733 TATGAGTGTGGAGTCGTTGCCCTGACTTTGTATCCTCCAGCTTNNNAGATCGGAAGA
33734 TATGAGTGTGGAGTCGTTGCTGTCCTCCCACCATTCCTTTATGNNNAGATCGGAAGA
33736 TATGAGTGTGGAGTCGTTGCTGTGAAACAGTTGCTCCATGAACNNNAGATCGGAAG
33737 TATGAGTGTGGAGTCGTTGCCACACCTGTAATCCCAGCACTTTNNNAGATCGGAAGA
33739 TATGAGTGTGGAGTCGTTGCCTCTTCTGTGGCAAACCCTAGAGNNNAGATCGGAAG
33740 TATGAGTGTGGAGTCGTTGCAGCTCAGATCACTTTTCTCACCTNNNAGATCGGAAGA
33742 TATGAGTGTGGAGTCGTTGCATGGTCTGAATGTTTGTGTCCCTNNNAGATCGGAAG
33743 TATGAGTGTGGAGTCGTTGCTGTAGTCCTAGCTACTTGGGAGGNNNAGATCGGAA
33744 TATGAGTGTGGAGTCGTTGCAAAAGAAAAGAGAAACCCACGCGNNNAGATCGGAA
33745 TATGAGTGTGGAGTCGTTGCGAAGGAGGTAGTGGGAGTAGAGTNNNAGATCGGA
33746 TATGAGTGTGGAGTCGTTGCATAAAATTCCAAGCCCCACAACGNNNAGATCGGAAG

33748 TATGAGTGTGGAGTCGTTGCGAACACTGCCCAAAATATCCAGGNNNAGATCGGAAG
33749 TATGAGTGTGGAGTCGTTGCTCCACAACGAGATACAGCAAAGANNNAGATCGGAAG
33750 TATGAGTGTGGAGTCGTTGCAGAGAAAGAAAACCCTCCAAGCTNNNAGATCGGAAG
33751 TATGAGTGTGGAGTCGTTGCCCTTTAGGCAGGGGAAACTCAAANNNAGATCGGAAG
33752 TATGAGTGTGGAGTCGTTGCACATGAGGGGAAATGTTTTCCAANNNAGATCGGAAG
33753 TATGAGTGTGGAGTCGTTGCAGTGCTCTCCTTTCTCAAAGTGANNNAGATCGGAAGA
33754 TATGAGTGTGGAGTCGTTGCCACGCCTATAATCCCAGCACTTTNNNAGATCGGAAGA
33755 TATGAGTGTGGAGTCGTTGCATTAGGTCAGGAGATGGAGACCANNNAGATCGGAA
33757 TATGAGTGTGGAGTCGTTGCGAAAGAGACAAGCTGAGAGGAGGNNNAGATCGGA
33759 TATGAGTGTGGAGTCGTTGCCTGGAAACAACCCCAAATGTTCTNNNAGATCGGAAGA
33760 TATGAGTGTGGAGTCGTTGCCTGTAATGCCAGCACTTTGGGAGNNNAGATCGGAAG
33762 TATGAGTGTGGAGTCGTTGCGGAGAATCTCTTGAGCCCAGTAGNNNAGATCGGAAG
33763 TATGAGTGTGGAGTCGTTGCGGAGAATCTCTTGAGCCCAGTAGNNNAGATCGGAAG
33764 TATGAGTGTGGAGTCGTTGCTCATCAGAACAGATGCCTAAGGGNNNAGATCGGAAG
33765 TATGAGTGTGGAGTCGTTGCTTTTACTCAGAGGGGGGACTACTNNNAGATCGGAAG
33766 TATGAGTGTGGAGTCGTTGCAAAGGCTTTGGAATCACAATGCANNNAGATCGGAAG
33768 TATGAGTGTGGAGTCGTTGCAATGCAGAAATAACCTGGGGTCANNNAGATCGGAAG
33769 TATGAGTGTGGAGTCGTTGCAAAAAAAACCACCACGAAACCACNNNAGATCGGAAG
33771 TATGAGTGTGGAGTCGTTGCTGTCTAAATGAAGCCAGCCACTANNNAGATCGGAAG
33773 TATGAGTGTGGAGTCGTTGCTCATCAGAGTGAACAGACAACCTNNNAGATCGGAAG
33774 TATGAGTGTGGAGTCGTTGCGGAGTCTCTGAGGACCTTCAAAGNNNAGATCGGAAG
33775 TATGAGTGTGGAGTCGTTGCTCATTCAAGGCAACTACGAACACNNNAGATCGGAAG
33777 TATGAGTGTGGAGTCGTTGCTCAGCTCTCCTCCTTTTCTCATGNNNAGATCGGAAGAG
33778 TATGAGTGTGGAGTCGTTGCCCTAGTGGTACATGCCAGTAGTCNNNAGATCGGAAG
33779 TATGAGTGTGGAGTCGTTGCATTTGCTTCTGTTTGGTGTGTGGNNNAGATCGGAAG
33780 TATGAGTGTGGAGTCGTTGCAAGGTCAGGAAATCGAGACCATCNNNAGATCGGAAG
33781 TATGAGTGTGGAGTCGTTGCTTGCATGCCTGTAGTAGTCTCAGNNNAGATCGGAAG
33782 TATGAGTGTGGAGTCGTTGCGAAGACACAAATGGCCAAGAAGTNNNAGATCGGAA
33783 TATGAGTGTGGAGTCGTTGCGCCTGTAATCCCAGATACTCCAGNNNAGATCGGAAGA
33785 TATGAGTGTGGAGTCGTTGCCACGCCTGTAATTCCAACACTTTNNNAGATCGGAAGA
33786 TATGAGTGTGGAGTCGTTGCTGACCTATCTCCTACACTCCCTGNNNAGATCGGAAGA
33787 TATGAGTGTGGAGTCGTTGCGCCAGAGTACCAGTGTTCAAAACNNNAGATCGGAAG
33788 TATGAGTGTGGAGTCGTTGCAGCAGCAACAACAACAAAACTCTNNNAGATCGGAAG
33792 TATGAGTGTGGAGTCGTTGCCCAAAAGAGCTAGCAGAATCCACNNNAGATCGGAAG
33794 TATGAGTGTGGAGTCGTTGCTCTTGTGCCGGTTTTCAAAGAAANNNAGATCGGAAG
33795 TATGAGTGTGGAGTCGTTGCACTATCCCTGTTTGCAGATGACANNNAGATCGGAAGA
33796 TATGAGTGTGGAGTCGTTGCAACACTCTGGCAACTCAAAAAGCNNNAGATCGGAAG
33797 TATGAGTGTGGAGTCGTTGCTTCGTCGACAGTATCTTCCACTGNNNAGATCGGAAGA
33799 TATGAGTGTGGAGTCGTTGCCCTAAGGGACTTATGGGCTACCANNNAGATCGGAAG
33800 TATGAGTGTGGAGTCGTTGCAGAGGAGAGTTCCGGGAATTTTCNNNAGATCGGAAG
33806 TATGAGTGTGGAGTCGTTGCGAAACTGGAAGACTGGTGATCCTNNNAGATCGGAAG
33808 TATGAGTGTGGAGTCGTTGCATGGCCAATAAGCACAGGAAAAGNNNAGATCGGAA
33809 TATGAGTGTGGAGTCGTTGCTAGAAGCTATGCACATCCCCCTANNNAGATCGGAAGA

33811 TATGAGTGTGGAGTCGTTGCAAAAACAAGAAGCTGCCTCTTGGNNNAGATCGGAA
33813 TATGAGTGTGGAGTCGTTGCAAGGGAAAAAGGACAAAGGGGAANNNAGATCGGA
33816 TATGAGTGTGGAGTCGTTGCGAGCCAGAGCAATCTTGAACAAANNNAGATCGGAA
33817 TATGAGTGTGGAGTCGTTGCAGAACCCTTTCTTGTGGAGTGTTNNNAGATCGGAAG
33819 TATGAGTGTGGAGTCGTTGCAAGTGGCATGGTGGTTATGTTTGNNNAGATCGGAA
33820 TATGAGTGTGGAGTCGTTGCGGAGGCTCTCTTGGAAGAAATCANNNAGATCGGAA
33822 TATGAGTGTGGAGTCGTTGCGAACAGGGAAGGTTTAGCCAGTANNNAGATCGGAA
33823 TATGAGTGTGGAGTCGTTGCATTTACAACTCTGCAACATGGGGNNNAGATCGGAAG
33825 TATGAGTGTGGAGTCGTTGCGTAGTATGTGCCTGTAGTCCCAGNNNAGATCGGAA
33826 TATGAGTGTGGAGTCGTTGCCCCCAGCTCATCTGATCTAACTTNNNAGATCGGAAGA
33828 TATGAGTGTGGAGTCGTTGCAAAGGTGGGAGAAGTAGCATCAGNNNAGATCGGA
33829 TATGAGTGTGGAGTCGTTGCTACCTCCAATGTTCAGCAACTCANNNAGATCGGAAG
33830 TATGAGTGTGGAGTCGTTGCAAAGCATAAGGGGTTAGACTGGGNNNAGATCGGA
33832 TATGAGTGTGGAGTCGTTGCAACCCCCCAAATTATACCAACCANNNAGATCGGAAG
33835 TATGAGTGTGGAGTCGTTGCAATCAGAACACATGGACACAGGANNNAGATCGGAA
33837 TATGAGTGTGGAGTCGTTGCCTGTAGTCCCAGCTACTCAGGAGNNNAGATCGGAAG
33838 TATGAGTGTGGAGTCGTTGCAAAACTTCTGCAAGGACATCTGCNNNAGATCGGAAG
33840 TATGAGTGTGGAGTCGTTGCAGAGGTTGCGATGAGTCAAGATTNNNAGATCGGAA
33841 TATGAGTGTGGAGTCGTTGCTTTTGCCTCAAAGCAGGAATCTGNNNAGATCGGAAG
33842 TATGAGTGTGGAGTCGTTGCAAACCCAACCAACCAAACAAACANNNAGATCGGAAG
33843 TATGAGTGTGGAGTCGTTGCTTTTCTCTTGCTCCCTCTCTCACNNNAGATCGGAAGA
33846 TATGAGTGTGGAGTCGTTGCGCAGAGATTGTAGTGAGCTGAGANNNAGATCGGA
33847 TATGAGTGTGGAGTCGTTGCCACAAACAAAGCGAGGAAGGAATNNNAGATCGGAA
33848 TATGAGTGTGGAGTCGTTGCAGACTGAGGCAGGAGAATTGTTTNNNAGATCGGAA
33849 TATGAGTGTGGAGTCGTTGCCCTTACTTTCCTGTGTCTCGTCANNNAGATCGGAAGA
33851 TATGAGTGTGGAGTCGTTGCGAGAAAAACAGCCAGCAAAAGGANNNAGATCGGA
33853 TATGAGTGTGGAGTCGTTGCCCCCAAAAGTCCCCAAATAGCTANNNAGATCGGAAG
33856 TATGAGTGTGGAGTCGTTGCGCTCATGTCTGTAATTCCAGCACNNNAGATCGGAAG
33857 TATGAGTGTGGAGTCGTTGCGCTGATGATTATAGGCTCTGGGTNNNAGATCGGAA
33858 TATGAGTGTGGAGTCGTTGCCATTATCTTGGGGAGTGCCTTCTNNNAGATCGGAAG
33859 TATGAGTGTGGAGTCGTTGCTGGCATGGATAATCACAAAGCTGNNNAGATCGGAA
33860 TATGAGTGTGGAGTCGTTGCAGGAAAGCTGAACTCCAGAAGAGNNNAGATCGGAA
33861 TATGAGTGTGGAGTCGTTGCCTGTAATCCCAGCTACTCAGGAGNNNAGATCGGAAG
33862 TATGAGTGTGGAGTCGTTGCTTTGGGAGAAAGTAAGGGACGAGNNNAGATCGGA
33863 TATGAGTGTGGAGTCGTTGCAGGAGAATTGCTTGAACCTGGAANNNAGATCGGAA
33864 TATGAGTGTGGAGTCGTTGCGTAGGGGCACATAGGTGATAAGGNNNAGATCGGA
33867 TATGAGTGTGGAGTCGTTGCACACATACTCGTTGAGAGCTCTGNNNAGATCGGAAG
33868 TATGAGTGTGGAGTCGTTGCTTCCATGACAGCACATACAGACTNNNAGATCGGAAG
33869 TATGAGTGTGGAGTCGTTGCTGAGGCAGGAGAATCTCTTGAACNNNAGATCGGAA
33870 TATGAGTGTGGAGTCGTTGCTGAGCATGGCAATAGTAAGAGCANNNAGATCGGAA
33871 TATGAGTGTGGAGTCGTTGCGGAAAGAAGTGAGCTATCCATGCNNNAGATCGGAA
33872 TATGAGTGTGGAGTCGTTGCGCAACAATGATGCATAGCCCTTTNNNAGATCGGAAG
33876 TATGAGTGTGGAGTCGTTGCCATTGTCATACCTACCTGCCTCANNNAGATCGGAAGA

33877 TATGAGTGTGGAGTCGTTGCGACATGGAGAAGTGAGGAGAAGGNNNAGATCGGA
33878 TATGAGTGTGGAGTCGTTGCCATCTGACAAAATTCAGTGCCCTNNNAGATCGGAAG
33879 TATGAGTGTGGAGTCGTTGCGACCTATTTCTGTCTTCCCTGCTNNNAGATCGGAAGA
33880 TATGAGTGTGGAGTCGTTGCCTGAGGTGGGAAGATAGCTTGAGNNNAGATCGGA
33882 TATGAGTGTGGAGTCGTTGCCTCCTGAGTAGCTGGGATTACAGNNNAGATCGGAA
33883 TATGAGTGTGGAGTCGTTGCCCTCCTGAGTAACTGGGATTACANNNAGATCGGAAG
33884 TATGAGTGTGGAGTCGTTGCGATTTTGAGACGGAGTCTTGCTCNNNAGATCGGAA
33885 TATGAGTGTGGAGTCGTTGCTTGAGACGTAGTCTTGCTCTGTCNNNAGATCGGAAG
33886 TATGAGTGTGGAGTCGTTGCCCTCAAACTTCCAGGGTCACTTANNNAGATCGGAAG
33888 TATGAGTGTGGAGTCGTTGCCCTCCCAGGTTCAAGTGATACTCNNNAGATCGGAAG
33889 TATGAGTGTGGAGTCGTTGCCCTCATATCTCAGCCTCTCAAGGNNNAGATCGGAAG
33890 TATGAGTGTGGAGTCGTTGCGCCTCCCAAGTAGCTGGATTATANNNAGATCGGAAG
33893 TATGAGTGTGGAGTCGTTGCAGTAACTGGGACTACAGGGATGANNNAGATCGGAA
33894 TATGAGTGTGGAGTCGTTGCAATACGGTGACAAGATCTTGGCTNNNAGATCGGAA
33895 TATGAGTGTGGAGTCGTTGCGCCACCATGCCAGACTATTTTTNNNAGATCGGAAG
33900 TATGAGTGTGGAGTCGTTGCCTCACTGGAAGCTTCATCTCCTGNNNAGATCGGAAG
33901 TATGAGTGTGGAGTCGTTGCGATTCAAGCGATTCTCCTGCCTCNNNAGATCGGAAG
33902 TATGAGTGTGGAGTCGTTGCACAACACCACACCCAGCTAATTTNNNAGATCGGAAG
33903 TATGAGTGTGGAGTCGTTGCTGCGTTCAAGTGATTCTTATGCCNNNAGATCGGAAG
33905 TATGAGTGTGGAGTCGTTGCTTCAAGCTATTCTCCTGCCTCAGNNNAGATCGGAAGA
33906 TATGAGTGTGGAGTCGTTGCTGATTCTTATGCCTCAACCTCCCNNNAGATCGGAAGA
33907 TATGAGTGTGGAGTCGTTGCAATCTCGACTCACTTCAAGCTCCNNNAGATCGGAAGA
33909 TATGAGTGTGGAGTCGTTGCGCCTCCCAGGTTCAAGTAATTCTNNNAGATCGGAAG
33911 TATGAGTGTGGAGTCGTTGCGTAGTTCAATGGCACAGTCTTGGNNNAGATCGGAA
33913 TATGAGTGTGGAGTCGTTGCTCAGCCTCCTGAGTAGTTAGGACNNNAGATCGGAAG
33914 TATGAGTGTGGAGTCGTTGCAAATGATTTGCCTGCCTTGTCTCNNNAGATCGGAAG
33915 TATGAGTGTGGAGTCGTTGCACCTTAGCCTCCCAAAGTGTTAGNNNAGATCGGAAG
33917 TATGAGTGTGGAGTCGTTGCTGTAGTGCAGTGGTGTGATCTTGNNNAGATCGGAA
33918 TATGAGTGTGGAGTCGTTGCCAAGTGATTCTCCCAACCCAGTANNNAGATCGGAAG
33919 TATGAGTGTGGAGTCGTTGCGAGTCAGTGTCTCACTCTGTCACNNNAGATCGGAAG
33922 TATGAGTGTGGAGTCGTTGCCACTGTTCACAGGTGGGATCATANNNAGATCGGAA
33923 TATGAGTGTGGAGTCGTTGCGATTACAGGCATGCACTACCATGNNNAGATCGGAAG
33924 TATGAGTGTGGAGTCGTTGCAAAAGCCTCTGCAGGAAGATCATNNNAGATCGGAA
33925 TATGAGTGTGGAGTCGTTGCTGTTCCAGTTCTCAGAGGGAATGNNNAGATCGGAA
33926 TATGAGTGTGGAGTCGTTGCATCCCAGGTTCAAGTGATTCTCCNNNAGATCGGAAG
33928 TATGAGTGTGGAGTCGTTGCGAGACAGGTTTTTCCCATGTTGCNNNAGATCGGAAG
33929 TATGAGTGTGGAGTCGTTGCGGGTTCTAGTGATTCTTGTGCCTNNNAGATCGGAAG
33930 TATGAGTGTGGAGTCGTTGCCTAAAGTGCAGTGGTGCGATTTCNNNAGATCGGAA
33931 TATGAGTGTGGAGTCGTTGCCCCTAGTAACTGGGATTACAGGCNNNAGATCGGAA
33932 TATGAGTGTGGAGTCGTTGCGTGGTACAATGTCAGCTTACTGCNNNAGATCGGAAG
33933 TATGAGTGTGGAGTCGTTGCAGGTTCAAATGATTCTCCTGCCTNNNAGATCGGAAG
33936 TATGAGTGTGGAGTCGTTGCCTCTTGTTACCCAGACTGGAGTGNNNAGATCGGAAG
33937 TATGAGTGTGGAGTCGTTGCGCTGGAACTACAGGTACATGCTANNNAGATCGGAA

33939 TATGAGTGTGGAGTCGTTGCCTTCCCAGGTTCAAGCAATTCTCNNNAGATCGGAAG
33942 TATGAGTGTGGAGTCGTTGCCCTCCCAGATTCAAGCCATTCTCNNNAGATCGGAAGA
33944 TATGAGTGTGGAGTCGTTGCCCGCCTGGCTAATTTTTGTACTTNNNAGATCGGAAG
33945 TATGAGTGTGGAGTCGTTGCTCTCCAAAAGTGCTGGGATTACANNNAGATCGGAAG
33946 TATGAGTGTGGAGTCGTTGCCCTCCTAGGTTCAAGCGATTCTCNNNAGATCGGAAG
33947 TATGAGTGTGGAGTCGTTGCCCAAGTAGCTGGAACTACAGGTGNNNAGATCGGAA
33950 TATGAGTGTGGAGTCGTTGCGATCTCCTGACCTCATGATCCACNNNAGATCGGAAG
33952 TATGAGTGTGGAGTCGTTGCCCTCCTGGGTTCAAGAGATTCTGNNNAGATCGGAAG
33953 TATGAGTGTGGAGTCGTTGCACTTGCAGAACTGTGAGAATTGCNNNAGATCGGAA
33955 TATGAGTGTGGAGTCGTTGCCTTCCCAAGTTCAAGCGATTCTCNNNAGATCGGAAG
33956 TATGAGTGTGGAGTCGTTGCGGTTCAAGTGATTCTCCTGCTTCNNNAGATCGGAAG
33957 TATGAGTGTGGAGTCGTTGCGAGATCTTGACTCGCCACAACTTNNNAGATCGGAAG
33958 TATGAGTGTGGAGTCGTTGCAGAGTACAGTGGTGTGATCATGGNNNAGATCGGAA
33962 TATGAGTGTGGAGTCGTTGCCTTCACAAGCAGCTGGAATTACANNNAGATCGGAAG
33963 TATGAGTGTGGAGTCGTTGCTAGCTGGGATTACAGGTATGTGCNNNAGATCGGAA
33964 TATGAGTGTGGAGTCGTTGCCTGAAGTGCAATGGCGAAATCTCNNNAGATCGGAA
33966 TATGAGTGTGGAGTCGTTGCTCTCCTGAGTAGCTGGGATTACANNNAGATCGGAAG
33968 TATGAGTGTGGAGTCGTTGCGAGTCTCACTCTATTGCCCAGACNNNAGATCGGAAG
33969 TATGAGTGTGGAGTCGTTGCCTGAGTAGCTGGAACTACAGGTGNNNAGATCGGAA
33970 TATGAGTGTGGAGTCGTTGCGCCTCCTGAGTAGTTGGGATTACNNNAGATCGGAAG
33971 TATGAGTGTGGAGTCGTTGCCCATGTACCTGGGATTACAGACGNNNAGATCGGAAG
33972 TATGAGTGTGGAGTCGTTGCATCCCAAGTAACTGGAACCACAGNNNAGATCGGAAG
33973 TATGAGTGTGGAGTCGTTGCTCGTGAACTCCTGAACTTGTGATNNNAGATCGGAAG
33975 TATGAGTGTGGAGTCGTTGCCTCCCATCTCAGCCTCTCTAGTANNNAGATCGGAAGA
33978 TATGAGTGTGGAGTCGTTGCATCTCCCAGGTTCAAGCAATTCTNNNAGATCGGAAG
33980 TATGAGTGTGGAGTCGTTGCTCTCGAGTAGCTGGAACTACAGANNNAGATCGGAA
33985 TATGAGTGTGGAGTCGTTGCTTCAAGCAATTCCCCTACCTCAGNNNAGATCGGAAG
33988 TATGAGTGTGGAGTCGTTGCCCTACCAAAGTGCTGGGATTACANNNAGATCGGAAG
33990 TATGAGTGTGGAGTCGTTGCCAATGGCGTGATCTCAGGTTACTNNNAGATCGGAAG
33991 TATGAGTGTGGAGTCGTTGCAGAAAAGCCATTGCCCTTCTTTCNNNAGATCGGAAG
33992 TATGAGTGTGGAGTCGTTGCGATCTCCATCTCCCAGGTTCAAGNNNAGATCGGAAG
33993 TATGAGTGTGGAGTCGTTGCTAGGCTCAAGTGATTCTAGTGCCNNNAGATCGGAAG
33994 TATGAGTGTGGAGTCGTTGCCCTCCCGATTAGCAGAGATTACANNNAGATCGGAAG
33995 TATGAGTGTGGAGTCGTTGCACCTCTCAAGTTCAAGCGATTCTNNNAGATCGGAAG
33996 TATGAGTGTGGAGTCGTTGCCCCTGTTTCAAGCAATTCTCCTGNNNAGATCGGAAG
33997 TATGAGTGTGGAGTCGTTGCCCACTTCAGTCTTCCAAGTAGCTNNNAGATCGGAAG
33998 TATGAGTGTGGAGTCGTTGCAGACTGGAGTGCAATGATGTGATNNNAGATCGGAA
33999 TATGAGTGTGGAGTCGTTGCTGATCTTGGCTCACTACAACCTCNNNAGATCGGAAG
34000 TATGAGTGTGGAGTCGTTGCCTGAGTAGCTGGGATTACAGGTGNNNAGATCGGAA
34001 TATGAGTGTGGAGTCGTTGCCTTCTTCACCATCTTGCATTCCCNNNAGATCGGAAGA
34002 TATGAGTGTGGAGTCGTTGCGAGTAGCTGGGATTACAGGCATGNNNAGATCGGAA
34004 TATGAGTGTGGAGTCGTTGCAAAAATGGACTGCCACATTCACCNNNAGATCGGAAG
34006 TATGAGTGTGGAGTCGTTGCGAGATCTCGGTTCACTGCAAACTNNNAGATCGGAAG

34007 TATGAGTGTGGAGTCGTTGCTTTAGTAGAGACGGAGTTTCGCCNNNAGATCGGAA
34008 TATGAGTGTGGAGTCGTTGCAGACAGAATCTCACTGTGTCACCNNNAGATCGGAAG
34011 TATGAGTGTGGAGTCGTTGCTGAGTAACTGGGATTACAGGTGCNNNAGATCGGAA
34012 TATGAGTGTGGAGTCGTTGCTCTGGTGATTTTGTGGTGTTCACNNNAGATCGGAAG
34013 TATGAGTGTGGAGTCGTTGCCTCCTATTCTCAAGCCATCCTCCNNNAGATCGGAAGA
34016 TATGAGTGTGGAGTCGTTGCCAAGTAGCTGGGACTACAGGTGNNNAGATCGGAAG
34017 TATGAGTGTGGAGTCGTTGCGATCTTGGCTTATTGCAACCTCCNNNAGATCGGAAG
34018 TATGAGTGTGGAGTCGTTGCTTCTAGCAATTCTCCTGCCTCAGNNNAGATCGGAAGA
34021 TATGAGTGTGGAGTCGTTGCGGAGTAGTGCAATGGTGTGATCTNNNAGATCGGAA
34024 TATGAGTGTGGAGTCGTTGCCAATGGCATGATTTCAGCTCACTNNNAGATCGGAAG
34027 TATGAGTGTGGAGTCGTTGCCTCAAGCAATCTTCCCACTTCAGNNNAGATCGGAAGA
34028 TATGAGTGTGGAGTCGTTGCGTTCAAGAGATTCTCCTGCCTCANNNAGATCGGAAG
34029 TATGAGTGTGGAGTCGTTGCAATAGCTGGGATTACAGGTGTGCNNNAGATCGGAA
34030 TATGAGTGTGGAGTCGTTGCGTTTCACCATGTTAGCCAGGATGNNNAGATCGGAAG
34031 TATGAGTGTGGAGTCGTTGCTGTATGGGTGCAAGAAAAGGAGANNNAGATCGGA
34033 TATGAGTGTGGAGTCGTTGCAAGTAGCTGAGATTACAGGTGCCNNNAGATCGGAA
34034 TATGAGTGTGGAGTCGTTGCAGATGGAATTTCGCTCTTGTTGCNNNAGATCGGAAG
34035 TATGAGTGTGGAGTCGTTGCGAACTCCTGAGCTCAAGCAATCTNNNAGATCGGAAG
34037 TATGAGTGTGGAGTCGTTGCCTCAAGTGATTCTCCCACCTCAGNNNAGATCGGAAG
34038 TATGAGTGTGGAGTCGTTGCGAGATGGAGTCTTGCTTTGTTGCNNNAGATCGGAA
34039 TATGAGTGTGGAGTCGTTGCAGTGGTACAATCTCAGCTCACTGNNNAGATCGGAAG
34041 TATGAGTGTGGAGTCGTTGCCAAGTAGCCTAGTAGCTGGGATTNNNAGATCGGAA
34042 TATGAGTGTGGAGTCGTTGCCCTCCCAAGTAGCTGGAATTACANNNAGATCGGAAG
34043 TATGAGTGTGGAGTCGTTGCATCTTCTGACCTCATGATCCACCNNNAGATCGGAAGA
34044 TATGAGTGTGGAGTCGTTGCGAGTAGCTGGGAATATAGGCACCNNNAGATCGGAA
34046 TATGAGTGTGGAGTCGTTGCCTTGAAATCCCAAAGTTAGCCCCNNNAGATCGGAAG
34048 TATGAGTGTGGAGTCGTTGCCTGAGTAGCTGGGATTACAGGTGNNNAGATCGGAA
34049 TATGAGTGTGGAGTCGTTGCAGATGGATTCTTGCTCTGTCACCNNNAGATCGGAAG
34051 TATGAGTGTGGAGTCGTTGCGAATTTTGCTCTTGCCCAGCTATNNNAGATCGGAAG
34052 TATGAGTGTGGAGTCGTTGCTTCAAGCGATTCTCCTGTCTCAGNNNAGATCGGAAG
34053 TATGAGTGTGGAGTCGTTGCCTCCTGAGTAGCTGGGATTACAGNNNAGATCGGAA
34055 TATGAGTGTGGAGTCGTTGCCTCAAGTGATTCTCGTGCCTCAGNNNAGATCGGAAG
34056 TATGAGTGTGGAGTCGTTGCTGAGTAGCTGAGATTACAGACGCNNNAGATCGGAA
34057 TATGAGTGTGGAGTCGTTGCGCTTCCCAGTTTCAAGCAAATCTNNNAGATCGGAAG
34058 TATGAGTGTGGAGTCGTTGCCCATTCTCCTGCCTTAGTCTCTCNNNAGATCGGAAGA
34060 TATGAGTGTGGAGTCGTTGCTCTTGAGTACCTGGGACTACAGANNNAGATCGGAA
34062 TATGAGTGTGGAGTCGTTGCTTCAAGTGATTCTCCTGCCTCAGNNNAGATCGGAAG
34065 TATGAGTGTGGAGTCGTTGCTTAGGGTACATGTGCACATTGTGNNNAGATCGGAA
34067 TATGAGTGTGGAGTCGTTGCTACAGTGACATGAACATGGCTCANNNAGATCGGAA
34068 TATGAGTGTGGAGTCGTTGCTGTCACCTTGAACCCATACACATNNNAGATCGGAAG
34070 TATGAGTGTGGAGTCGTTGCTAGAGTGCAGTGGTGTGATCTTGNNNAGATCGGAA
34072 TATGAGTGTGGAGTCGTTGCTTCAAGCAATTCTCCTGCCTCAGNNNAGATCGGAAGA
34073 TATGAGTGTGGAGTCGTTGCGATCTCAGCTCACTGCAAACTCTNNNAGATCGGAAG

34074 TATGAGTGTGGAGTCGTTGCTCTTCTCGAACTGACAGTGGATGNNNAGATCGGAAG
34075 TATGAGTGTGGAGTCGTTGCCCAGGGTTCAAGCAATTCTCCTANNNAGATCGGAAG
34076 TATGAGTGTGGAGTCGTTGCCAATCAGCCTATCTCTGCATCCCNNNAGATCGGAAGA
34077 TATGAGTGTGGAGTCGTTGCCTTCCTGAGTAGCTGGGACTACANNNAGATCGGAAG
34078 TATGAGTGTGGAGTCGTTGCGATTCAAGTGGTTCTCCTGCCTCNNNAGATCGGAAG
34081 TATGAGTGTGGAGTCGTTGCCTCACTGCAACTTCCATCTACCANNNAGATCGGAAGA
34082 TATGAGTGTGGAGTCGTTGCAGATGGAGCTTTGCTTTTGTCACNNNAGATCGGAAG
34083 TATGAGTGTGGAGTCGTTGCCACCTTAGCCTCCTGATTAGCTGNNNAGATCGGAAG
34084 TATGAGTGTGGAGTCGTTGCGCTCAAGCCTTCTTCCTAACTCANNNAGATCGGAAGA
34085 TATGAGTGTGGAGTCGTTGCTTCAAGCAATTCTCCTGCCTCAGNNNAGATCGGAAG
34086 TATGAGTGTGGAGTCGTTGCGTTTCACCATGTTAGCCAGGATGNNNAGATCGGAAG
34087 TATGAGTGTGGAGTCGTTGCAAGTGTTGGGATTACAGGCATGANNNAGATCGGAA
34088 TATGAGTGTGGAGTCGTTGCCAGAATGAACTTGCAGGGGTTTTNNNAGATCGGAA
34089 TATGAGTGTGGAGTCGTTGCCTCCTGAATAGCTGGGACTACAGNNNAGATCGGAAG
34091 TATGAGTGTGGAGTCGTTGCTTTTTAGTAGAGACGGGGGCTTCNNNAGATCGGAA
34092 TATGAGTGTGGAGTCGTTGCGCTTGGTCTCCGTCTCAAAAAATNNNAGATCGGAAG
34093 TATGAGTGTGGAGTCGTTGCCCAAAGCAAGCAGATCTCTTCAGNNNAGATCGGAAG
34095 TATGAGTGTGGAGTCGTTGCATAGCTCGTGGAAAGTGGTTTCTNNNAGATCGGAA
34096 TATGAGTGTGGAGTCGTTGCTAACATAGCAAGACCACCCATCCNNNAGATCGGAAG
34097 TATGAGTGTGGAGTCGTTGCCAAATCTCAAAGACTCAGCCCCANNNAGATCGGAAG
34099 TATGAGTGTGGAGTCGTTGCCTATGCGTCTTCTGCAAAGCAATNNNAGATCGGAAG
34100 TATGAGTGTGGAGTCGTTGCGAGATCACACCATTGCATTCCAGNNNAGATCGGAAG
34102 TATGAGTGTGGAGTCGTTGCCCAGGTGATGGATACCCCAAATANNNAGATCGGAAG
34103 TATGAGTGTGGAGTCGTTGCAACCTACTTTTGAGTGGAAGCCTNNNAGATCGGAAG
34104 TATGAGTGTGGAGTCGTTGCTAATCCTAACACTTTGGGAGGCCNNNAGATCGGAAG
34105 TATGAGTGTGGAGTCGTTGCGTCTTCTGCTCATTTAGGCCTCANNNAGATCGGAAG
34107 TATGAGTGTGGAGTCGTTGCAAAAACCAGATCACCTCTATGCANNNAGATCGGAAG
34108 TATGAGTGTGGAGTCGTTGCACTGTGATTTAGAAGGCTTGCCTNNNAGATCGGAAG
34109 TATGAGTGTGGAGTCGTTGCAAAGAATGGCTACTACACAGGCANNNAGATCGGAA
34111 TATGAGTGTGGAGTCGTTGCTTGTTGACCTCTGCCTCTGTTTANNNAGATCGGAAGA
34112 TATGAGTGTGGAGTCGTTGCATACAAGCAGAGCGTAGAATGGTNNNAGATCGGAA
34113 TATGAGTGTGGAGTCGTTGCGTGTCCATTAGAGGTCATGGCTANNNAGATCGGAA
34115 TATGAGTGTGGAGTCGTTGCCACCTTGAGGACCATTCACTTCTNNNAGATCGGAAG
34116 TATGAGTGTGGAGTCGTTGCGAGGAAAGAAAGGCATGCATGAANNNAGATCGGA
34117 TATGAGTGTGGAGTCGTTGCATGGGCCTACTAAGTAGTCACCTNNNAGATCGGAAG
34120 TATGAGTGTGGAGTCGTTGCAAGCTAAGGCAAGAGAATGGCATNNNAGATCGGAA
34122 TATGAGTGTGGAGTCGTTGCGCTCGTACAGAATTCAATCCAGCNNNAGATCGGAAG
34123 TATGAGTGTGGAGTCGTTGCATGTCTTACATGGTAGCAGGCAANNNAGATCGGAA
34124 TATGAGTGTGGAGTCGTTGCACCAAGGAGGCTTTCTAAAGGAGNNNAGATCGGAA
34127 TATGAGTGTGGAGTCGTTGCTAACCTCAGGCAGGAAAGACATCNNNAGATCGGAA
34129 TATGAGTGTGGAGTCGTTGCTCCAGGAAGAATTGGTGCAAAAGNNNAGATCGGAA
34130 TATGAGTGTGGAGTCGTTGCCACTCCTGTAATCCCAGCACTTTNNNAGATCGGAAGA
34131 TATGAGTGTGGAGTCGTTGCTATGTACGTGTGTGTGTGTGTGTNNNAGATCGGAA

34132 TATGAGTGTGGAGTCGTTGCGACTTGCTTTGGCCAACAAAATGNNNAGATCGGAAG
34134 TATGAGTGTGGAGTCGTTGCAAGGATAGGAGTTTGAGACCAGCNNNAGATCGGAA
34135 TATGAGTGTGGAGTCGTTGCCCCACCAAAATTTAGCTGAGCATNNNAGATCGGAAG
34137 TATGAGTGTGGAGTCGTTGCTGTCATTTGTGTTCATGGACGTGNNNAGATCGGAAG
34139 TATGAGTGTGGAGTCGTTGCAAGCCTGAAGTGTGGAAACAAACNNNAGATCGGAA
34140 TATGAGTGTGGAGTCGTTGCAGGGGAGTAGGGGAAAGAAGATANNNAGATCGG
34141 TATGAGTGTGGAGTCGTTGCGGTCACGGTGTTTTGTAACCTAANNNAGATCGGAAG
34142 TATGAGTGTGGAGTCGTTGCGGTTTACAAAGGGTCACTGAGGANNNAGATCGGAA
34143 TATGAGTGTGGAGTCGTTGCGGAGCTCCTTACTGTTCATCTCCNNNAGATCGGAAG
34144 TATGAGTGTGGAGTCGTTGCATTTTAGCCCACTGAGACCCATTNNNAGATCGGAAG
34146 TATGAGTGTGGAGTCGTTGCTCATGGCTATTGTTTGAAGGCACNNNAGATCGGAAG
34147 TATGAGTGTGGAGTCGTTGCTGTTAGTCTTTCAAGAATGCCCTNNNAGATCGGAAG
34149 TATGAGTGTGGAGTCGTTGCAAAGCTGCACACCTACAACAATCNNNAGATCGGAAG
34150 TATGAGTGTGGAGTCGTTGCCTTGTTCATGTCCTTTGCCCATTNNNAGATCGGAAGA
34152 TATGAGTGTGGAGTCGTTGCTTCTTTTACTTTGGGCTGAGGGTNNNAGATCGGAAG
34153 TATGAGTGTGGAGTCGTTGCGATGTGGAAGAATCGCTTGAACCNNNAGATCGGAA
34156 TATGAGTGTGGAGTCGTTGCGACCTCGCTTTTTAGGTAGAGCTNNNAGATCGGAAG
34157 TATGAGTGTGGAGTCGTTGCCTTTCAACGAATCCATGCCATGTNNNAGATCGGAAG
34158 TATGAGTGTGGAGTCGTTGCAGTTCTCCAGCAAGATACCACTGNNNAGATCGGAAG
34159 TATGAGTGTGGAGTCGTTGCACAATTCTGGAGGCTGAGAAGTTNNNAGATCGGAA
34160 TATGAGTGTGGAGTCGTTGCTATCCATTTGGTGTGTGGGTAGGNNNAGATCGGAA
34163 TATGAGTGTGGAGTCGTTGCGAGACAAGAGAATCGCTTGAACCNNNAGATCGGAA
34164 TATGAGTGTGGAGTCGTTGCAACTCTGATGGAGGAAATGGGACNNNAGATCGGAA
34165 TATGAGTGTGGAGTCGTTGCGTGGGTGGAGAAAGGGGAAATATNNNAGATCGGA
34166 TATGAGTGTGGAGTCGTTGCGGCTCAAAGTCATCAGGAAGAGTNNNAGATCGGAA
34167 TATGAGTGTGGAGTCGTTGCATTGCCAGGACAGTGAAGGTAATNNNAGATCGGAA
34169 TATGAGTGTGGAGTCGTTGCATCCCTCAGATCACATTTCCCACNNNAGATCGGAAGA
34170 TATGAGTGTGGAGTCGTTGCCTAATGTGCGAGGATTGCTTGAGNNNAGATCGGAA
34171 TATGAGTGTGGAGTCGTTGCAAACAACAGATGCTGGAGAGGATNNNAGATCGGAA
34172 TATGAGTGTGGAGTCGTTGCAAAGGGATCACTGCTTCTGAGAANNNAGATCGGAA
34173 TATGAGTGTGGAGTCGTTGCGCCAGTTTTGGATGTCAGTGAAANNNAGATCGGAA
34174 TATGAGTGTGGAGTCGTTGCTGGATCCTTACCAAGTTTCACCANNNAGATCGGAAG
34175 TATGAGTGTGGAGTCGTTGCCCAAAGTAGGTGGATCACAAGGTNNNAGATCGGAA
34176 TATGAGTGTGGAGTCGTTGCGTTGAATTAGAACAACCCCTGCCNNNAGATCGGAAG
34177 TATGAGTGTGGAGTCGTTGCGCCAAGTTTGAGAAACCCTGATCNNNAGATCGGAAG
34179 TATGAGTGTGGAGTCGTTGCGGACATTGTGTGCGTCAAGAAAANNNAGATCGGAA
34181 TATGAGTGTGGAGTCGTTGCGTTTACAGCAACTGTCCAACCAANNNAGATCGGAAG
34182 TATGAGTGTGGAGTCGTTGCGAAAGCTGAAGGTTGTTGGTCTTNNNAGATCGGAA
34183 TATGAGTGTGGAGTCGTTGCTCTCATCATTCAGCCTCAATGCTNNNAGATCGGAAGA
34184 TATGAGTGTGGAGTCGTTGCTATGTTGACAGACCTCACACCACNNNAGATCGGAAG
34186 TATGAGTGTGGAGTCGTTGCTTGAGACCAGCCTGGCTAATATGNNNAGATCGGAAG
34187 TATGAGTGTGGAGTCGTTGCAAGGAGAGATGAGGGAGTCAGAANNNAGATCGGA
34188 TATGAGTGTGGAGTCGTTGCTAATAGGGCAGGTGGTAGAGACTNNNAGATCGGAA

34189 TATGAGTGTGGAGTCGTTGCCTCAAAACTCCAGCCAATCACAGNNNAGATCGGAAG
34190 TATGAGTGTGGAGTCGTTGCTTTGAAGACTGAGGCTCTAGTCCNNNAGATCGGAAG
34191 TATGAGTGTGGAGTCGTTGCTGCAGACCTTTATGATGATCCACTNNNAGATCGGAA
34192 TATGAGTGTGGAGTCGTTGCTTTGCCACGACATACTTTGTAGCNNNAGATCGGAAG
34193 TATGAGTGTGGAGTCGTTGCGAGGTTGCATTGAGCTGTGATTGNNNAGATCGGAA
34194 TATGAGTGTGGAGTCGTTGCCAGGTTGGTGGTTTGTGTTGATTNNNAGATCGGAA
34198 TATGAGTGTGGAGTCGTTGCCAGAGCTTCTATGTCAGTCCCAANNNAGATCGGAAG
34200 TATGAGTGTGGAGTCGTTGCTGATGCAACATGGATGAACCTTGNNNAGATCGGAA
34202 TATGAGTGTGGAGTCGTTGCCACTAGAAAAGGAGCCTGGGAATNNNAGATCGGAA
34203 TATGAGTGTGGAGTCGTTGCTAGGAAGAAGGAAGTTCTTGCCCNNNAGATCGGAA
34204 TATGAGTGTGGAGTCGTTGCTGAAATGACAACCATGCTGCATTNNNAGATCGGAAG
34205 TATGAGTGTGGAGTCGTTGCGAGGATCACAAGGTCAGGAGATGNNNAGATCGGA
34206 TATGAGTGTGGAGTCGTTGCTCAGTGGCATGTAGTCCAAGAAANNNAGATCGGAA
34208 TATGAGTGTGGAGTCGTTGCCCCACCACATTGGAAATCATGATNNNAGATCGGAAG
34210 TATGAGTGTGGAGTCGTTGCTCCTTCCTTCCTTTCATTCGTCCNNNAGATCGGAAGAG
34212 TATGAGTGTGGAGTCGTTGCGAACAGAGCCTCAAAGACCTGTANNNAGATCGGAA
34213 TATGAGTGTGGAGTCGTTGCAAACACAAACCTTCTTGGAAGGCNNNAGATCGGAAG
34215 TATGAGTGTGGAGTCGTTGCGGTTGCAATCCTAGTCTCTGACANNNAGATCGGAAG
34216 TATGAGTGTGGAGTCGTTGCTTCCATGAGAGGAGAAAACCTGGNNNAGATCGGAA
34218 TATGAGTGTGGAGTCGTTGCACCTGTAATCCCAGCACTTTAGGNNNAGATCGGAAG
34219 TATGAGTGTGGAGTCGTTGCTGCCAACAAAAAAAGTCCAGGACNNNAGATCGGAA
34220 TATGAGTGTGGAGTCGTTGCAAATGCTTAGGGGAAATCTGCCTNNNAGATCGGAA
34221 TATGAGTGTGGAGTCGTTGCGGGAGAAGAGAAGAGAGAACACANNNAGATCGGA
34222 TATGAGTGTGGAGTCGTTGCGATGTGGATTCCAAGGTCCTTCTNNNAGATCGGAAG
34223 TATGAGTGTGGAGTCGTTGCGAAGACATACAAATGGCCAACGANNNAGATCGGAA
34224 TATGAGTGTGGAGTCGTTGCTTTCTCATGCCCCTTAGTATCCGNNNAGATCGGAAGA
34226 TATGAGTGTGGAGTCGTTGCAAAAGAGCCTGAGTAGCAAAAGCNNNAGATCGGAA
34227 TATGAGTGTGGAGTCGTTGCTGTTTGGTCCTTAGGATAAGGCANNNAGATCGGAA
34230 TATGAGTGTGGAGTCGTTGCTCCACTACAGCTTTCTGAATCCCNNNAGATCGGAAGA
34231 TATGAGTGTGGAGTCGTTGCTCCATCAGGAAAATACCACATTCCTNNNAGATCGGA
34233 TATGAGTGTGGAGTCGTTGCGATCACAAGGTCAGGAGATCGAGNNNAGATCGGAA
34234 TATGAGTGTGGAGTCGTTGCTGGTGCCATTCCATTCTACAAGTNNNAGATCGGAAG
34235 TATGAGTGTGGAGTCGTTGCAAGTCAACCTACTTCTGTTGGCTNNNAGATCGGAAG
34236 TATGAGTGTGGAGTCGTTGCAAAGCTTTAGAAAAGGTCCACGCNNNAGATCGGAA
34237 TATGAGTGTGGAGTCGTTGCGCAAGTGAGGTGATAGGCGATATNNNAGATCGGA
34240 TATGAGTGTGGAGTCGTTGCACCCTAATACCATACCCTTCCCANNNAGATCGGAAGA
34241 TATGAGTGTGGAGTCGTTGCTGAATGTATGTGAACGTACATGCANNNAGATCGGA
34242 TATGAGTGTGGAGTCGTTGCTGGTTTAGTTTGCTTCAGGTGTGNNNAGATCGGAA
34243 TATGAGTGTGGAGTCGTTGCTGGAGATGGGTGAACAACTTTGANNNAGATCGGAA
34244 TATGAGTGTGGAGTCGTTGCCTCGAACTCCTGACCTCATGATCNNNAGATCGGAAG
34245 TATGAGTGTGGAGTCGTTGCCCCGACACCTCCTAATCCTAAACNNNAGATCGGAAGA
34246 TATGAGTGTGGAGTCGTTGCAAGAAAGAAAGAAAGGGCCATGCNNNAGATCGGA
34247 TATGAGTGTGGAGTCGTTGCCACACAAGCCATATGAAGTGTCCNNNAGATCGGAAG

34248 TATGAGTGTGGAGTCGTTGCACATACAGTGGTTTCCTCCCTTCNNNAGATCGGAAGA
34249 TATGAGTGTGGAGTCGTTGCCACCTCTTTCTAACCTCCAGGACNNNAGATCGGAAGA
34250 TATGAGTGTGGAGTCGTTGCCTTGTCTCTTCCCACTCTCCTTCNNNAGATCGGAAGAG
34252 TATGAGTGTGGAGTCGTTGCTCCAGAACTAGTGGCTCATTCTGNNNAGATCGGAAG
34255 TATGAGTGTGGAGTCGTTGCCACACCTGTAATCCCACCACTATNNNAGATCGGAAGA
34256 TATGAGTGTGGAGTCGTTGCGAGAGAGAGAGAGAGAGAGATGANNNAGATCGGA
34257 TATGAGTGTGGAGTCGTTGCATGCTAGATGACGAGTTAGTGGGNNNAGATCGGAA
34258 TATGAGTGTGGAGTCGTTGCCCTCCTAAACTCCCCATGTTCAANNNAGATCGGAAGA
34260 TATGAGTGTGGAGTCGTTGCATCACCTGAGTTTGGGAGTTTGANNNAGATCGGAAG
34261 TATGAGTGTGGAGTCGTTGCTGTACACTGTACCCAACGTGTAGNNNAGATCGGAAG
34262 TATGAGTGTGGAGTCGTTGCCTCATCTTGGTGATCTTGGGGAANNNAGATCGGAAG
34263 TATGAGTGTGGAGTCGTTGCCGTACACCTGTAGTCTCAGCTACNNNAGATCGGAAGA
34265 TATGAGTGTGGAGTCGTTGCCCATTTTTGCCTCCTAGACCTCTNNNAGATCGGAAGA
34266 TATGAGTGTGGAGTCGTTGCTGAGTCTTTTGGTAAGCTCGTGANNNAGATCGGAAG
34267 TATGAGTGTGGAGTCGTTGCGGTCCCAGATTTGAATCTAGCCANNNAGATCGGAAG
34269 TATGAGTGTGGAGTCGTTGCAACACTAAGTGAGCACAGAACCANNNAGATCGGAAG
34271 TATGAGTGTGGAGTCGTTGCCTGGACAGCACCATCGAAGTAAGNNNAGATCGGAAG
34272 TATGAGTGTGGAGTCGTTGCGAGTGTGTGTGTGTGTGTAAAGGNNNAGATCGGAA
34273 TATGAGTGTGGAGTCGTTGCATGCTAGATGACGAGTTAGTGGGNNNAGATCGGAA
34274 TATGAGTGTGGAGTCGTTGCGGAGGAGGGATAGCATTAGGAGANNNAGATCGGA
34275 TATGAGTGTGGAGTCGTTGCAGTAGATAGGCATCTCTCTCCCCNNNAGATCGGAAGA
34276 TATGAGTGTGGAGTCGTTGCCATCCTGGCTAACAAGGTGAAACNNNAGATCGGAAG
34277 TATGAGTGTGGAGTCGTTGCGTGGCTTGTTTCTCAGACTGTTCNNNAGATCGGAAGA
34278 TATGAGTGTGGAGTCGTTGCCACTTGAGGAGGGGAGAAAGAAANNNAGATCGGAA
34279 TATGAGTGTGGAGTCGTTGCAGGTCTGTGAGAGATGCAGGATANNNAGATCGGAA
34281 TATGAGTGTGGAGTCGTTGCCCTGCCCTCTTGTGATTGTTTTTNNNAGATCGGAAGA
34283 TATGAGTGTGGAGTCGTTGCTGTTGTTCAGATTCTTTTGCCCCNNNAGATCGGAAGA
34284 TATGAGTGTGGAGTCGTTGCATGCCTGTAATCTCAGCTACTCGNNNAGATCGGAAGA
34285 TATGAGTGTGGAGTCGTTGCCACTGTGCCCTCATTTCTTTTGTNNNAGATCGGAAGA
34286 TATGAGTGTGGAGTCGTTGCGGGTGCACCAAAATCTCAGAAATNNNAGATCGGAAG
34287 TATGAGTGTGGAGTCGTTGCTCATCAAAAACCCTGTTTCCTGCNNNAGATCGGAAGA
34288 TATGAGTGTGGAGTCGTTGCAAAGAGGGAAGGAAAGGAAAGGGNNNAGATCGGA
34289 TATGAGTGTGGAGTCGTTGCGGATTCAGCGCGTGTATGTAAAGNNNAGATCGGAA
34290 TATGAGTGTGGAGTCGTTGCTCTCTCTCTCTCTCTTTCTGCCANNNAGATCGGAAGAG
34291 TATGAGTGTGGAGTCGTTGCGTGTAAAGGGAGATGTCTGCTCANNNAGATCGGAA
34292 TATGAGTGTGGAGTCGTTGCTCCCCCTAGGTCTTTCTCATGATNNNAGATCGGAAGA
34293 TATGAGTGTGGAGTCGTTGCCAGATTCAGTGTCGGTTTTGGACNNNAGATCGGAAG
34294 TATGAGTGTGGAGTCGTTGCCTCCTGAGTAGCTGGGATTACAGNNNAGATCGGAAG
34296 TATGAGTGTGGAGTCGTTGCGTGTGTGTTAGTGTGCACAGATGNNNAGATCGGAA
34297 TATGAGTGTGGAGTCGTTGCCTGAGGCAGGAGAATCACTTGAANNNAGATCGGAA
34301 TATGAGTGTGGAGTCGTTGCAAACTGTGGGGTGTTGAGAGAATNNNAGATCGGAA
34302 TATGAGTGTGGAGTCGTTGCTTGCATGCATGGATTCTTGCTTTNNNAGATCGGAAGA
34304 TATGAGTGTGGAGTCGTTGCCCTTAGAAACGCAGGTGGTTTTTNNNAGATCGGAAG

34305 TATGAGTGTGGAGTCGTTGCGGTAGTGGAGTGTTGACCAATCANNNAGATCGGAA
34306 TATGAGTGTGGAGTCGTTGCAACAAAAGGCATGAAAACCCCAANNNAGATCGGAA
34307 TATGAGTGTGGAGTCGTTGCGAGAAACCATGGTGTCGTCTTTGNNNAGATCGGAA
34309 TATGAGTGTGGAGTCGTTGCAAGTCTCCATAAATGTGGGGACCNNNAGATCGGAA
34310 TATGAGTGTGGAGTCGTTGCCTGGCCCAAATGTATGCTTTTCANNNAGATCGGAAG
34311 TATGAGTGTGGAGTCGTTGCTTCCTGTGTCCATGTGTTCTCATNNNAGATCGGAAG
34312 TATGAGTGTGGAGTCGTTGCGTGTCCCTTGCTGACTCATGATANNNAGATCGGAAG
34313 TATGAGTGTGGAGTCGTTGCTGCCCAAGTCCTATGTTTCTTGANNNAGATCGGAAG
34315 TATGAGTGTGGAGTCGTTGCTGGAAGTATGCTTTCTCTTGAGTNNNAGATCGGAAG
34316 TATGAGTGTGGAGTCGTTGCTGTAGCCGGCCTCATGTTTATTANNNAGATCGGAAG
34317 TATGAGTGTGGAGTCGTTGCGCCATCATTCCTAGTTGCTGTTCNNNAGATCGGAAG
34318 TATGAGTGTGGAGTCGTTGCTTTTGTCCTCAGAAGACACGGTCNNNAGATCGGAAG
34320 TATGAGTGTGGAGTCGTTGCTGAACAGGGGTGTGATATTTGGTNNNAGATCGGAA
34321 TATGAGTGTGGAGTCGTTGCGTCAGAAGAGACAGAGGGAACAGNNNAGATCGGA
34322 TATGAGTGTGGAGTCGTTGCTGGCAACATAGATGAACCTGGAANNNAGATCGGAA
34323 TATGAGTGTGGAGTCGTTGCAACCCATAAAGCCATCCATAGCANNNAGATCGGAAG
34324 TATGAGTGTGGAGTCGTTGCCACAGCACAGAAAGAGAGAGACTNNNAGATCGGAA
34325 TATGAGTGTGGAGTCGTTGCCTTTCTGGTAGCCTTTCCCTGTANNNAGATCGGAAG
34326 TATGAGTGTGGAGTCGTTGCGGTTACTATTAGCTCGGTCAGGGNNNAGATCGGAA
34327 TATGAGTGTGGAGTCGTTGCAAAATCCAGCAGGGCAGTAACTANNNAGATCGGAA
34328 TATGAGTGTGGAGTCGTTGCTGGTTGACATTGGACCGTATCATNNNAGATCGGAAG
34329 TATGAGTGTGGAGTCGTTGCGATAGTAGAGATGTCCAGCACCCNNNAGATCGGAA
34330 TATGAGTGTGGAGTCGTTGCTTCCACTACCCTTTCATTCCTGGNNNAGATCGGAAGA
34331 TATGAGTGTGGAGTCGTTGCCTGGCCAAAGGAGAATGTTCATCNNNAGATCGGAA
34332 TATGAGTGTGGAGTCGTTGCTGAAGGGAAAGGTTGGGAAGAAANNNAGATCGGA
34333 TATGAGTGTGGAGTCGTTGCATAGCACAACAGGGTGACTACAGNNNAGATCGGAA
34334 TATGAGTGTGGAGTCGTTGCTGTCCTTTGCAGGGATCCTATTCNNNAGATCGGAAG
34335 TATGAGTGTGGAGTCGTTGCCCAGAGTTCTTGGCAGGATCTTANNNAGATCGGAAG
34336 TATGAGTGTGGAGTCGTTGCCCCAAGTCCCCAAAGTCCATTATNNNAGATCGGAAG
34337 TATGAGTGTGGAGTCGTTGCTTGGCTGTTGTTGGTGTATAGGANNNAGATCGGAA
34338 TATGAGTGTGGAGTCGTTGCGGGTACCTCCATTCCTCATGAAGNNNAGATCGGAAG
34339 TATGAGTGTGGAGTCGTTGCCTGGAATGCAGTGGAGTGATCTCNNNAGATCGGAA
34340 TATGAGTGTGGAGTCGTTGCTGTGAATGTCTGAAAAACTGCCCNNNAGATCGGAA
34341 TATGAGTGTGGAGTCGTTGCAGGACCCAGAGAGTCAAACTTTCNNNAGATCGGAA
34342 TATGAGTGTGGAGTCGTTGCTTCACAATACATCTTGCTGCTGCNNNAGATCGGAAG
34345 TATGAGTGTGGAGTCGTTGCTGTGTGTGTGTACAGGTGTGTATNNNAGATCGGAA
34346 TATGAGTGTGGAGTCGTTGCGCTCTATTTGCCTTCAGTGCTCTNNNAGATCGGAAG
34347 TATGAGTGTGGAGTCGTTGCTGTAGATGATGGGTTGATGGGTGNNNAGATCGGA
34348 TATGAGTGTGGAGTCGTTGCTTTTGATCTTGTTCCCCAGGCTANNNAGATCGGAAG
34349 TATGAGTGTGGAGTCGTTGCGGACCTATGATGGGGGATCCTATNNNAGATCGGAA
34350 TATGAGTGTGGAGTCGTTGCATGACCCATGATGAAACCCATGANNNAGATCGGAAG
34352 TATGAGTGTGGAGTCGTTGCGGTGGGAAGGAAGTGAGTATCTCNNNAGATCGGA
34353 TATGAGTGTGGAGTCGTTGCCTCTCTCCTGATGATGATGGTGGNNNAGATCGGAAG

34354 TATGAGTGTGGAGTCGTTGCCCCCCATGATTCAATTACCTCCANNNAGATCGGAAGA
34355 TATGAGTGTGGAGTCGTTGCGGGAACGTGACCTTATTTGCAAANNNAGATCGGAAG
34357 TATGAGTGTGGAGTCGTTGCCTGAGGTACCAGGTTCATCTCACNNNAGATCGGAAG
34358 TATGAGTGTGGAGTCGTTGCACCAGCATGGCACATGTATACATNNNAGATCGGAAG
34360 TATGAGTGTGGAGTCGTTGCTCTCAAACTCCTGGGTTCAAGTTNNNAGATCGGAAG
34361 TATGAGTGTGGAGTCGTTGCCATAGTCGTATCACTGCACTCCANNNAGATCGGAAG
34362 TATGAGTGTGGAGTCGTTGCTGTCTAACCATCCTTCAGCTGACNNNAGATCGGAAG
34363 TATGAGTGTGGAGTCGTTGCGGTTGCCACAAAGTCTTGGAAATNNNAGATCGGAA
34364 TATGAGTGTGGAGTCGTTGCGTTTTGACTTCCCTCCCAAACTGNNNAGATCGGAAG
34365 TATGAGTGTGGAGTCGTTGCACTCCAGTTCTGCTCCTTAGGATNNNAGATCGGAAG
34368 TATGAGTGTGGAGTCGTTGCTCAAAACACAATCATGCCCTTCCNNNAGATCGGAAGA
34369 TATGAGTGTGGAGTCGTTGCCACTGGAGCTCTCTGGTAATCAGNNNAGATCGGAAG
34370 TATGAGTGTGGAGTCGTTGCTCCTGGGGAGTTCTGAACAATTTNNNAGATCGGAAG
34371 TATGAGTGTGGAGTCGTTGCGACACCACCCCTGCTAATTTTTGNNNAGATCGGAAG
34372 TATGAGTGTGGAGTCGTTGCGCCTTGTCTTTTGAACTTCTCCCNNNAGATCGGAAGA
34373 TATGAGTGTGGAGTCGTTGCGCCTGTAATCCTAGCTACTCTGGNNNAGATCGGAAG
34374 TATGAGTGTGGAGTCGTTGCATCTTGGGAGGTGGAATGTTTCANNNAGATCGGAA
34375 TATGAGTGTGGAGTCGTTGCAGGAAGGTGAGGGAAAATTTGGANNNAGATCGGA
34376 TATGAGTGTGGAGTCGTTGCTTGAGGCTATGGAAGTTCAAGGGNNNAGATCGGAA
34377 TATGAGTGTGGAGTCGTTGCGTCCTCTCAAAGTGCTGGGATTANNNAGATCGGAAG
34380 TATGAGTGTGGAGTCGTTGCAGTTATGGCTTTGGCTGAGAGAGNNNAGATCGGAA
34381 TATGAGTGTGGAGTCGTTGCCCTCCTGGGTTCTAGTGATTCTCNNNAGATCGGAAG
34384 TATGAGTGTGGAGTCGTTGCTACCCAATCTCACCCTATCCCAGNNNAGATCGGAAGA
34385 TATGAGTGTGGAGTCGTTGCTTTGAGCCAGGATCTCTCTTAGCNNNAGATCGGAAG
34386 TATGAGTGTGGAGTCGTTGCTTTGAGCCAGGATCTCTCTTAGCNNNAGATCGGAAG
34387 TATGAGTGTGGAGTCGTTGCCTTTGCTTCCTCTTCACCTTTGGNNNAGATCGGAAGA
34388 TATGAGTGTGGAGTCGTTGCTAATGAGACATCTTGGGCATGGGNNNAGATCGGAA
34389 TATGAGTGTGGAGTCGTTGCCTGAGTAGCTGGGATTACAGGTGNNNAGATCGGAA
34390 TATGAGTGTGGAGTCGTTGCCTTACCACAAAGCCCTTTTGGAGNNNAGATCGGAAG
34394 TATGAGTGTGGAGTCGTTGCCGGTGAAGATTCCCTTGATCACANNNAGATCGGAAG
34395 TATGAGTGTGGAGTCGTTGCGAGTTCACACAATCCCCCCATTCNNNAGATCGGAAGA
34397 TATGAGTGTGGAGTCGTTGCCAAAACACCAAAAACAGGGGTCTNNNAGATCGGAA
34398 TATGAGTGTGGAGTCGTTGCGGGTGAATAATCAAGCAGGCATCNNNAGATCGGAA
34399 TATGAGTGTGGAGTCGTTGCCCAGACTCAAGTAATCCTCCCACNNNAGATCGGAAG
34402 TATGAGTGTGGAGTCGTTGCCACCTTACCTGGAAAGCACAAAGNNNAGATCGGAAG
34403 TATGAGTGTGGAGTCGTTGCGAACGTCCTTCTGCTACTGCTATNNNAGATCGGAAG
34405 TATGAGTGTGGAGTCGTTGCGCTTTGTCATGGTGAGCTACTTCNNNAGATCGGAAG
34406 TATGAGTGTGGAGTCGTTGCCGAAGCCAAAAAAAGAAGCAGGANNNAGATCGGAA
34407 TATGAGTGTGGAGTCGTTGCTTTTTCCCTGACAGTGCTAAGGANNNAGATCGGAAG
34408 TATGAGTGTGGAGTCGTTGCACCTTTGCTGAGGATTGAGTAGGNNNAGATCGGAA
34409 TATGAGTGTGGAGTCGTTGCCAGAAGAAAGGTCTAGGGCTGAANNNAGATCGGAA
34411 TATGAGTGTGGAGTCGTTGCAGGTATGCAGACAATTGGAGTGTNNNAGATCGGAA
34412 TATGAGTGTGGAGTCGTTGCACCTCTGAAACGTGGGTGAATAANNNAGATCGGAA

34413 TATGAGTGTGGAGTCGTTGCCTGCCTAACATTCCGACCTCATANNNAGATCGGAAGA
34416 TATGAGTGTGGAGTCGTTGCACCAGTCCTAAACACCAGATTGGNNNAGATCGGAAG
34417 TATGAGTGTGGAGTCGTTGCCCTGTAGTCCCAGTTACTAGGGANNNAGATCGGAAG
34420 TATGAGTGTGGAGTCGTTGCCTGAGGCATGAGAATTGCTTGAGNNNAGATCGGAA
34421 TATGAGTGTGGAGTCGTTGCCAGGAAGAACTGGAACTGGTCTGNNNAGATCGGAA
34422 TATGAGTGTGGAGTCGTTGCGGTAAGTGGTGTAGTAGGGTGTGNNNAGATCGGAA
34424 TATGAGTGTGGAGTCGTTGCCATGGATGAAGCTGGAAACCATCNNNAGATCGGAA
34425 TATGAGTGTGGAGTCGTTGCACAGGTGTATGCATGATATCCCCNNNAGATCGGAAG
34426 TATGAGTGTGGAGTCGTTGCGCATGAGGGAAGCTTCTAGACTTNNNAGATCGGAA
34428 TATGAGTGTGGAGTCGTTGCATGAGGCCAGATGGTGTCAATTANNNAGATCGGAA
34429 TATGAGTGTGGAGTCGTTGCGGTAGTTTAGGCCCTGGTTTGATNNNAGATCGGAAG
34430 TATGAGTGTGGAGTCGTTGCTGGGTAGCTTCACACTTTAGTCCNNNAGATCGGAAGA
34431 TATGAGTGTGGAGTCGTTGCGCTCAGGTGACGAGTACACTAAANNNAGATCGGAA
34432 TATGAGTGTGGAGTCGTTGCCATGTTGGATCACGCCTGTAATCNNNAGATCGGAAG
34433 TATGAGTGTGGAGTCGTTGCCTGGAGTGTAGTGGCATGATCTTNNNAGATCGGAA
34434 TATGAGTGTGGAGTCGTTGCGGAGGGGGGAGAGAGAATGAATANNNAGATCGGA
34435 TATGAGTGTGGAGTCGTTGCTCTGTATCATGGAGTTTGCCCTCNNNAGATCGGAAGA
34437 TATGAGTGTGGAGTCGTTGCGAAGTAGAGACTGCGCATGTAGANNNAGATCGGAA
34438 TATGAGTGTGGAGTCGTTGCTCTCATTTCCAACTGAAGCCTGANNNAGATCGGAAGA
34439 TATGAGTGTGGAGTCGTTGCCCAATTCAATGCCCTTCATCCAANNNAGATCGGAAGA
34440 TATGAGTGTGGAGTCGTTGCTAGATGACAGTTTGATGGGTGCANNNAGATCGGAA
34441 TATGAGTGTGGAGTCGTTGCGGATTTCCCTCGGCTAGAAAGAANNNAGATCGGAAG
34443 TATGAGTGTGGAGTCGTTGCCCGGATCCCCCTAGTTTATCTTCNNNAGATCGGAAGA
34444 TATGAGTGTGGAGTCGTTGCTGTGTTGGATATGGAAGTGAGGGNNNAGATCGGAA
34445 TATGAGTGTGGAGTCGTTGCTATACCTCTGCGAATGGAACCAGNNNAGATCGGAAG
34446 TATGAGTGTGGAGTCGTTGCTAGCTATCCCCACTCTCTCTAGCNNNAGATCGGAAGA
34447 TATGAGTGTGGAGTCGTTGCCTCTCTCTAGCTATCCCCCTCTCNNNAGATCGGAAGAG
34448 TATGAGTGTGGAGTCGTTGCCTCTCTCTAGCTATCCCCCTCTCNNNAGATCGGAAGAG
34449 TATGAGTGTGGAGTCGTTGCCTCTCTCTAGCTATCCCCCTCTCNNNAGATCGGAAGAG
34451 TATGAGTGTGGAGTCGTTGCCTGTAATTTCAGATCCAGCAGCGNNNAGATCGGAAG
34458 TATGAGTGTGGAGTCGTTGCACAAGAAAGTCCCACGTACTGAANNNAGATCGGAAG
34460 TATGAGTGTGGAGTCGTTGCTGTGTCTGGCTGATTGGTCAATANNNAGATCGGAAG
34461 TATGAGTGTGGAGTCGTTGCGAGTAGCGAGGGCTGAAAAATTGNNNAGATCGGAA
34462 TATGAGTGTGGAGTCGTTGCTCAAGATAATTCTCCGAGCCCTGNNNAGATCGGAAGA
34463 TATGAGTGTGGAGTCGTTGCCCACGTGATCCCATTGTTCAATTNNNAGATCGGAAGA
34464 TATGAGTGTGGAGTCGTTGCAGAAGGTGAAGCACAACAATGTGNNNAGATCGGAA
34466 TATGAGTGTGGAGTCGTTGCTGCCCCTTATGCAAGACATTTTGNNNAGATCGGAAGA
34467 TATGAGTGTGGAGTCGTTGCGAACTGACTCAGTGCAAAAGGACNNNAGATCGGAA
34468 TATGAGTGTGGAGTCGTTGCACTTGTACATGCATGTTCACAGCNNNAGATCGGAAGA
34469 TATGAGTGTGGAGTCGTTGCGTTCCTGTGTTAGAGTGCTGAGANNNAGATCGGAA
34470 TATGAGTGTGGAGTCGTTGCGAGTCCATTTCAAGGTCAGGAGANNNAGATCGGAA
34473 TATGAGTGTGGAGTCGTTGCCCCACAGGAGACAGCACTATTTANNNAGATCGGAAG
34474 TATGAGTGTGGAGTCGTTGCACCCCCCCAAAAAAAGAAAACTGNNNAGATCGGAAG

34475 TATGAGTGTGGAGTCGTTGCGAGCCCAGAAATCCAAAGTCAAGNNNAGATCGGAA
34476 TATGAGTGTGGAGTCGTTGCTCGCTGCTACATTCCTGTAAGTTNNNAGATCGGAAG
34477 TATGAGTGTGGAGTCGTTGCTTGTGATCAATCTAGCCCAGCAANNNAGATCGGAAG
34478 TATGAGTGTGGAGTCGTTGCTTGGACGACTTTGAGTAAGTGGTNNNAGATCGGAA
34480 TATGAGTGTGGAGTCGTTGCAAAAACAGTTACGGGACAGAAGANNNAGATCGGAA
34482 TATGAGTGTGGAGTCGTTGCGTGGTGGTGGTGGTGATTTTTATNNNAGATCGGAA
34483 TATGAGTGTGGAGTCGTTGCTTCCTAAGGCTAACATGAATGCCNNNAGATCGGAAG
34484 TATGAGTGTGGAGTCGTTGCAACCTCCCACAGTTTGTACCTACNNNAGATCGGAAG
34486 TATGAGTGTGGAGTCGTTGCTATTTTCCAGGCCTTTTCCCTGTNNNAGATCGGAAGA
34487 TATGAGTGTGGAGTCGTTGCCAAAGTACTGGGATGACAGGTGTNNNAGATCGGAA
34488 TATGAGTGTGGAGTCGTTGCTGGGCATTGATAGGCTCTACTTCNNNAGATCGGAAG
34489 TATGAGTGTGGAGTCGTTGCTCTTTTTACTGGAACAGTCCCCCNNNAGATCGGAAG
34490 TATGAGTGTGGAGTCGTTGCCTGGAGGACTCACAACTCACTACNNNAGATCGGAAG
34491 TATGAGTGTGGAGTCGTTGCCATGGTGGCTTGCACTTGTAATCNNNAGATCGGAAG
34492 TATGAGTGTGGAGTCGTTGCCCCATCACTACACCTGGCTAATTNNNAGATCGGAAG
34495 TATGAGTGTGGAGTCGTTGCGAGAGGTCCTATGGGAAGGGATCNNNAGATCGGA
34496 TATGAGTGTGGAGTCGTTGCTCTTCCCACTGCTCCTTCTTTTTNNNAGATCGGAAGA
34497 TATGAGTGTGGAGTCGTTGCGATCTCGACTCACTGTAACCTCCNNNAGATCGGAAG
34499 TATGAGTGTGGAGTCGTTGCTTCTCTTCTCCTCTGCTCTCTGANNNAGATCGGAAGA
34501 TATGAGTGTGGAGTCGTTGCAACTCATTGTTCTCACTCGAGCTNNNAGATCGGAAG
34502 TATGAGTGTGGAGTCGTTGCGGAATTCGGCTGTCTCCTTTCTANNNAGATCGGAAG
34503 TATGAGTGTGGAGTCGTTGCCCCACTGTGCTTGACTTTTTTCANNNAGATCGGAAGA
34504 TATGAGTGTGGAGTCGTTGCGTCTCTTCTGAGTCTAGTGACGCNNNAGATCGGAAG
34505 TATGAGTGTGGAGTCGTTGCTCTCATTTGTGCCTGAAATTCCGNNNAGATCGGAAG
34506 TATGAGTGTGGAGTCGTTGCTTAGCAAGGAACAACGCATTCTCNNNAGATCGGAAG
34507 TATGAGTGTGGAGTCGTTGCGATGGTTTCCAGCTTCATCCATGNNNAGATCGGAAG
34508 TATGAGTGTGGAGTCGTTGCTGTTCAGTTCCCACCTATGAGTGNNNAGATCGGAAG
34509 TATGAGTGTGGAGTCGTTGCGGGCAATGGCTCCTTATTTTGAGNNNAGATCGGAA
34512 TATGAGTGTGGAGTCGTTGCCACCACCTAGGCCAAGATCTTTANNNAGATCGGAAG
34513 TATGAGTGTGGAGTCGTTGCGATAGTAGAGATGTCCAGCACCCNNNAGATCGGAA
34514 TATGAGTGTGGAGTCGTTGCCCCTCCTCAGACCATGTTGTTATNNNAGATCGGAAG
34515 TATGAGTGTGGAGTCGTTGCCACTGTCTCTGGGCCTAATTCAANNNAGATCGGAAG
34516 TATGAGTGTGGAGTCGTTGCACCTCAACCTACGCTAATGAAGGNNNAGATCGGAAG
34517 TATGAGTGTGGAGTCGTTGCCAGCTGTGAAAGAAAAGGGCTTTNNNAGATCGGAA
34518 TATGAGTGTGGAGTCGTTGCTTGTGCACTGTAGCAGATAGTGTNNNAGATCGGAA
34519 TATGAGTGTGGAGTCGTTGCGAGATGGGGTTTTACAATGCTGCNNNAGATCGGAA
34520 TATGAGTGTGGAGTCGTTGCTGCAAAATGTGAACCTTGTTGGANNNAGATCGGAA
34521 TATGAGTGTGGAGTCGTTGCTAAGTCTCACGAGATCTGAGGGTNNNAGATCGGAA
34522 TATGAGTGTGGAGTCGTTGCGACGCAGATTTGGTCTTTTCACANNNAGATCGGAAG
34523 TATGAGTGTGGAGTCGTTGCAAGTAAGTTTCTGAGTGGCTCCTNNNAGATCGGAAG
34524 TATGAGTGTGGAGTCGTTGCGTTGCTGAGTCTAGACACCAGAANNNAGATCGGAA
34525 TATGAGTGTGGAGTCGTTGCCCTTCAGAAAGCTCCAATCAGGANNNAGATCGGAAG
34526 TATGAGTGTGGAGTCGTTGCTGGTGGAGAGCTTTCTTCTTTCANNNAGATCGGAAG

34527 TATGAGTGTGGAGTCGTTGCCCTGTAATCCTAGCACTTTGGGANNNAGATCGGAAG
34528 TATGAGTGTGGAGTCGTTGCTCCTCTCTTCTCAATAGTCCCCCNNNAGATCGGAAGA
34529 TATGAGTGTGGAGTCGTTGCGCCAAAGGGTTCTCTCTCTTTCTNNNAGATCGGAAG
34530 TATGAGTGTGGAGTCGTTGCGCAAGTCAGACTCTGAGACAGAANNNAGATCGGAA
34531 TATGAGTGTGGAGTCGTTGCCTTTGGACTGCATTGGCTACAAANNNAGATCGGAAG
34532 TATGAGTGTGGAGTCGTTGCCCTGAGGAAAGCCTTGCTCTTTANNNAGATCGGAAG
34533 TATGAGTGTGGAGTCGTTGCCGTGTTCTCATTGTTCAACTCCCNNNAGATCGGAAGA
34534 TATGAGTGTGGAGTCGTTGCACACACACACACACACAATTCTGNNNAGATCGGAAG
34535 TATGAGTGTGGAGTCGTTGCTTCTCCTGCCTTTCACAAACTCTNNNAGATCGGAAGA
34537 TATGAGTGTGGAGTCGTTGCTTGAGATTTGGGTGGGGACATAGNNNAGATCGGAA
34538 TATGAGTGTGGAGTCGTTGCGTGCTTGCTTCCCATAGAAATGGNNNAGATCGGAAG
34539 TATGAGTGTGGAGTCGTTGCCAAACGTCTTCTCTCAGCAAGTGNNNAGATCGGAAG
34540 TATGAGTGTGGAGTCGTTGCCACACACACACACACACAATTNNNAGATCGGAAG
34541 TATGAGTGTGGAGTCGTTGCCCCGGCCGAAAAGTTCTTTATTTNNNAGATCGGAAG
34542 TATGAGTGTGGAGTCGTTGCAATGCACAAATGTTACCCTTGCANNNAGATCGGAAG
34543 TATGAGTGTGGAGTCGTTGCTCATCTTAGTCCATGCTCAGCAGNNNAGATCGGAAG
34546 TATGAGTGTGGAGTCGTTGCCCATGAAACTAGCTAGGTAGGGGNNNAGATCGGAA
34549 TATGAGTGTGGAGTCGTTGCCCCCCCAAGAAAAAAAACACACANNNAGATCGGAAG
34550 TATGAGTGTGGAGTCGTTGCGAGACACACACACACAGAAACACNNNAGATCGGAAG
34551 TATGAGTGTGGAGTCGTTGCTGCCTTTCCAATCACATTCCCTANNNAGATCGGAAGA
34552 TATGAGTGTGGAGTCGTTGCGAGCCAAATTCTTCCCAAGAGTCNNNAGATCGGAAG
34553 TATGAGTGTGGAGTCGTTGCGGCAAGTAGGAGCAAAAACAAGANNNAGATCGGAA
34554 TATGAGTGTGGAGTCGTTGCGTTCAAGCGATTCTCCTACCTCANNNAGATCGGAAG
34563 TATGAGTGTGGAGTCGTTGCAGAGAAGGGGTCTTACCATGTTGNNNAGATCGGAA
34566 TATGAGTGTGGAGTCGTTGCCAGATACAGGGGATGAGGCAAAGNNNAGATCGGA
34568 TATGAGTGTGGAGTCGTTGCCTCCCAGGTATTTATGCACCCACNNNAGATCGGAAG
34569 TATGAGTGTGGAGTCGTTGCGCCACATCATTTTTAGCCCTAGGNNNAGATCGGAAG
34570 TATGAGTGTGGAGTCGTTGCAGGATCCTAGAACCAGCTTTTGTNNNAGATCGGAAG
34571 TATGAGTGTGGAGTCGTTGCCCATGTGGGGGGAAGAAAAATTCNNNAGATCGGAA
34572 TATGAGTGTGGAGTCGTTGCGATCTTGGGAGAAGATGAGCTCCNNNAGATCGGAA
34573 TATGAGTGTGGAGTCGTTGCCTTTGCTTCAGTAGGATCTGCACNNNAGATCGGAAG
34574 TATGAGTGTGGAGTCGTTGCAGGACCATGCACATGTCTGAATANNNAGATCGGAAG
34576 TATGAGTGTGGAGTCGTTGCACAGACTTTAGATGGTGGGTTGGNNNAGATCGGAA
34577 TATGAGTGTGGAGTCGTTGCTACCCATGATGGAACCCATGATGNNNAGATCGGAAG
34581 TATGAGTGTGGAGTCGTTGCCAGGGAACGTTCATATGCAAGTTNNNAGATCGGAAG
34582 TATGAGTGTGGAGTCGTTGCGGAGGGAGAGCATTAGGACAAATNNNAGATCGGA
34586 TATGAGTGTGGAGTCGTTGCGGATGGAGGTGTTTGGATCTCATNNNAGATCGGAA
34587 TATGAGTGTGGAGTCGTTGCTCAAGAGTCTCACAGTTCCCTTGNNNAGATCGGAAG
34588 TATGAGTGTGGAGTCGTTGCCCTCCCCTTTTTAGACCGTAGAGNNNAGATCGGAAG
34589 TATGAGTGTGGAGTCGTTGCTCATCTCGGGTGAAAGTCCAAATNNNAGATCGGAAG
34590 TATGAGTGTGGAGTCGTTGCATATACTGTTCGGGTGATGGGTGNNNAGATCGGAA
34591 TATGAGTGTGGAGTCGTTGCCATGCTAGGTGGATATCTGCTGTNNNAGATCGGAAG
34594 TATGAGTGTGGAGTCGTTGCGTGATGCACACCTGTAATTCCAGNNNAGATCGGAAG

34595 TATGAGTGTGGAGTCGTTGCACCCTGTCTCAAAAAAGAATCCGNNNAGATCGGAAG
34596 TATGAGTGTGGAGTCGTTGCCCAGAGAAACGACCACTGAAATGNNNAGATCGGAA
34597 TATGAGTGTGGAGTCGTTGCCCCTACCATGTTCCAGGCAATATNNNAGATCGGAAG
34598 TATGAGTGTGGAGTCGTTGCACACAGGATGTTGTGAAAAGCAANNNAGATCGGAA
34599 TATGAGTGTGGAGTCGTTGCGGAGAGCGGACATCTTGAGTATTNNNAGATCGGAA
34600 TATGAGTGTGGAGTCGTTGCAGGATTGCTGGTTTGTTATTGCCNNNAGATCGGAAG
34601 TATGAGTGTGGAGTCGTTGCTGCGTCACAGAAAACTTCTTTCCNNNAGATCGGAAG
34602 TATGAGTGTGGAGTCGTTGCAGAACCACTGCTCTCTTCAAAGCNNNAGATCGGAAG
34604 TATGAGTGTGGAGTCGTTGCGGAGATGGCATATGTCGAGGATTNNNAGATCGGAA
34605 TATGAGTGTGGAGTCGTTGCAGGTGTTCTTGCTAAACTGACCTNNNAGATCGGAAG
34606 TATGAGTGTGGAGTCGTTGCCCTTCCACCACTATGCTAACCATNNNAGATCGGAAGA
34607 TATGAGTGTGGAGTCGTTGCTATCATGAGGAGGTGGCTACAGANNNAGATCGGAA
34612 TATGAGTGTGGAGTCGTTGCCAACATACTGAGAGCCCATCTGTNNNAGATCGGAAG
34613 TATGAGTGTGGAGTCGTTGCTGTGCACCAAGGAAATGGATAGANNNAGATCGGAA
34614 TATGAGTGTGGAGTCGTTGCCTGCACTGACATCTCTCTGTTCTNNNAGATCGGAAGA
34615 TATGAGTGTGGAGTCGTTGCATGCATGCTCAGGAGTGTATGTANNNAGATCGGAA
34617 TATGAGTGTGGAGTCGTTGCTCTGTCTGAAACAAACACACACGNNNAGATCGGAAG
34618 TATGAGTGTGGAGTCGTTGCATGTGGAAAGTTCTGGAAGCTCTNNNAGATCGGAA
34619 TATGAGTGTGGAGTCGTTGCACGCTAAGTGGCACATTACAGTANNNAGATCGGAA
34620 TATGAGTGTGGAGTCGTTGCATGATCCCTGTTCTTTTGTGGGANNNAGATCGGAAG
34621 TATGAGTGTGGAGTCGTTGCGTCAGAAATGCGAACACCTGAAANNNAGATCGGAA
34622 TATGAGTGTGGAGTCGTTGCCAATGGCAACCCGAAAATCAATGNNNAGATCGGAA
34623 TATGAGTGTGGAGTCGTTGCTGAAAGGAGGAAAGGAGGAAAGGNNNAGATCGG
34624 TATGAGTGTGGAGTCGTTGCCATTTAGTGGGTAAGGCCAGAGANNNAGATCGGAA
34626 TATGAGTGTGGAGTCGTTGCGGTGTGTTCCTCAGGGCTATTTANNNAGATCGGAAG
34627 TATGAGTGTGGAGTCGTTGCAGCATTTTCAGTCTTGTTTGGGGNNNAGATCGGAAG
34628 TATGAGTGTGGAGTCGTTGCTTGTTCCACCTCAGATCATCAGGNNNAGATCGGAAG
34629 TATGAGTGTGGAGTCGTTGCTCCTCCTGAAGAATCCTTGCAAANNNAGATCGGAAG
34630 TATGAGTGTGGAGTCGTTGCTAGAGGTTCTCCATAAGGGTCCCNNNAGATCGGAAG
34631 TATGAGTGTGGAGTCGTTGCAGAGATGGGGCACTATGGAGATANNNAGATCGGA
34632 TATGAGTGTGGAGTCGTTGCTTTGCTGTTTTTCTCATGTGGCTNNNAGATCGGAAG
34635 TATGAGTGTGGAGTCGTTGCGTTTCACACGTGTAACCCAGAACNNNAGATCGGAAG
34636 TATGAGTGTGGAGTCGTTGCTGGATAGTTCTCAGTGACCCAACNNNAGATCGGAAG
34637 TATGAGTGTGGAGTCGTTGCATGTAACAAACCTGCACGTTCTGNNNAGATCGGAAG
34638 TATGAGTGTGGAGTCGTTGCCAGAGTTTGTGAAGGGTTGCAATNNNAGATCGGAA
34639 TATGAGTGTGGAGTCGTTGCTTTTCTGGGAAGCCTTTTTTGGGNNNAGATCGGAAG
34642 TATGAGTGTGGAGTCGTTGCCCTATGTTTTCCTGCACTTGGTGNNNAGATCGGAAG
34643 TATGAGTGTGGAGTCGTTGCTACTGTCTTCTCCACTGGAATGGNNNAGATCGGAAG
34644 TATGAGTGTGGAGTCGTTGCAGGCCTATCATTTGGGCAGAAATNNNAGATCGGAA
34645 TATGAGTGTGGAGTCGTTGCAGAGGGGAGATTTCCTCAGGAATNNNAGATCGGAA
34648 TATGAGTGTGGAGTCGTTGCTGGTGCCAGTTGTAAAGAAGTCTNNNAGATCGGAA
34650 TATGAGTGTGGAGTCGTTGCATTTCAAACTCAGCAACGGAGTCNNNAGATCGGAAG
34651 TATGAGTGTGGAGTCGTTGCTGGAGTCCATTTGTATTCTGACANNNAGATCGGAAG

34652 TATGAGTGTGGAGTCGTTGCAAGACAGTAAGACATGGCAGAGNNNAGATCGGAA
34653 TATGAGTGTGGAGTCGTTGCGCCTCAGTCTTTTCTGGGAAGTANNNAGATCGGAAG
34654 TATGAGTGTGGAGTCGTTGCGCCTTCCAGACATGTCTATTGAGNNNAGATCGGAAG
34655 TATGAGTGTGGAGTCGTTGCCGAGGAATGAGGAATGACTCAGANNNAGATCGGAA
34656 TATGAGTGTGGAGTCGTTGCATACAAAAATTAGCCGGGCATGGNNNAGATCGGAA
34657 TATGAGTGTGGAGTCGTTGCTACGTGGAAGATGGGCATAACATNNNAGATCGGAA
34658 TATGAGTGTGGAGTCGTTGCTGTACTACAGCCTAGGCAACAAGNNNAGATCGGAAG
34660 TATGAGTGTGGAGTCGTTGCAGAATGCTGCTATTCCTGTCCTCNNNAGATCGGAAGA
34661 TATGAGTGTGGAGTCGTTGCATTGTGACTTCTTTGACCCATGGNNNAGATCGGAAGA
34662 TATGAGTGTGGAGTCGTTGCGGGAGCCCAGAAATAGAGTACTGNNNAGATCGGAA
34663 TATGAGTGTGGAGTCGTTGCAGAGCTGAGATAAACGCCCTTTCNNNAGATCGGAAG
34664 TATGAGTGTGGAGTCGTTGCCTCCTGAGTCACTGGGATTACAGNNNAGATCGGAAG
34665 TATGAGTGTGGAGTCGTTGCGAGTCTTCAAGGTTACAGTGAGCNNNAGATCGGAA
34666 TATGAGTGTGGAGTCGTTGCGGTGACAGAGTGAAACTCCATCTNNNAGATCGGAA
34667 TATGAGTGTGGAGTCGTTGCTCTTCACACTACAGCTGGAGAACNNNAGATCGGAAG
34669 TATGAGTGTGGAGTCGTTGCCTTGCACCATGTACCTGGAAAAGNNNAGATCGGAAG
34671 TATGAGTGTGGAGTCGTTGCGCAGCAGGAAGCCTTATAACATTNNNAGATCGGAAG
34672 TATGAGTGTGGAGTCGTTGCTCCTTTTGGGGTGACGATAATGTNNNAGATCGGAAG
34673 TATGAGTGTGGAGTCGTTGCGCATGGCTCTTAGCAATCTTAGCNNNAGATCGGAAG
34674 TATGAGTGTGGAGTCGTTGCGTGTGTGTATGTGCTGTTTGTGTNNNAGATCGGAA
34675 TATGAGTGTGGAGTCGTTGCTCTCCCTTTTGGCCCTTTATCTCNNNAGATCGGAAGA
34676 TATGAGTGTGGAGTCGTTGCTTTCCTCTTTGCTTTCTCCTGGANNNAGATCGGAAGA
34677 TATGAGTGTGGAGTCGTTGCCCAAAGTGTTGAGATTACAGGCGNNNAGATCGGAA
34679 TATGAGTGTGGAGTCGTTGCTGGTAACTCATGCCTGTAATCCCNNNAGATCGGAAGA
34680 TATGAGTGTGGAGTCGTTGCACAGTAATCCCCCCTTATCTCCANNNAGATCGGAAGA
34681 TATGAGTGTGGAGTCGTTGCGCGAGACTCTGCCATGTAAAAAANNNAGATCGGAA
34682 TATGAGTGTGGAGTCGTTGCAAGGGTAAGCCAGGAAGAACAAANNNAGATCGGAA
34683 TATGAGTGTGGAGTCGTTGCAGGCACTGTTACGAAGACTGAATNNNAGATCGGAA
34684 TATGAGTGTGGAGTCGTTGCCCTCTTCCCTGGCACATTATCTTNNNAGATCGGAAGA
34686 TATGAGTGTGGAGTCGTTGCTTATTTATCTGCCTCAGCCTCCCNNNAGATCGGAAGA
34687 TATGAGTGTGGAGTCGTTGCCAATTCTCCTATCTCAGCCTCCCNNNAGATCGGAAGA
34690 TATGAGTGTGGAGTCGTTGCGTTCACACCATTCTCCTACCTCANNNAGATCGGAAGA
34691 TATGAGTGTGGAGTCGTTGCTTCAAGCAATTCTCCTGCCTCAGNNNAGATCGGAAGA
34693 TATGAGTGTGGAGTCGTTGCCACCACCATGCCACACTAAAATTNNNAGATCGGAAGA
34694 TATGAGTGTGGAGTCGTTGCAATGGGGTCTTACCATATTGCCCNNNAGATCGGAAG
34695 TATGAGTGTGGAGTCGTTGCAATGGGGTCTTACCATATTGCCCNNNAGATCGGAAG
34697 TATGAGTGTGGAGTCGTTGCCGCCCGGCTTTTCTTTTTTTTTTNNNAGATCGGAAGA
34698 TATGAGTGTGGAGTCGTTGCAGGGTCTTACCATATTGCCCATGNNNAGATCGGAAG
34700 TATGAGTGTGGAGTCGTTGCCGCCCGGCTTTTCTTTTTTTTTTNNNAGATCGGAAGA
34702 TATGAGTGTGGAGTCGTTGCGTCTTGAACTCCTCAGCTCAGACNNNAGATCGGAAGA
34703 TATGAGTGTGGAGTCGTTGCTTCAAGCGATTCTTCTCCCTCAGNNNAGATCGGAAGA
34705 TATGAGTGTGGAGTCGTTGCGACAGATTCTCACTCTCACCCAGNNNAGATCGGAAGA
34706 TATGAGTGTGGAGTCGTTGCCTGGAGTGCAATGGTGTGATCTCNNNAGATCGGAA

34707 TATGAGTGTGGAGTCGTTGCCTGAGTAGCTGGGATTACAGGTGNNNAGATCGGAA
34709 TATGAGTGTGGAGTCGTTGCACCATGCCCGGCTATTTTTTTTTNNNAGATCGGAAGA
34710 TATGAGTGTGGAGTCGTTGCCACCTGGCCTCTACAAACAATTTNNNAGATCGGAAG
34711 TATGAGTGTGGAGTCGTTGCAAAGTGCTGGGATTACAGGCAAGNNNAGATCGGAA
34712 TATGAGTGTGGAGTCGTTGCAGCAATTCTCCTGCCTTACTCTCNNNAGATCGGAAGA
34713 TATGAGTGTGGAGTCGTTGCCCATGCCCGGCTAATTTTTTGTANNNAGATCGGAAG
34714 TATGAGTGTGGAGTCGTTGCCTGTTGCTCAGGCTATAGTCTCANNNAGATCGGAAG
34716 TATGAGTGTGGAGTCGTTGCCGAGACAGAATCTTGCAACCTCTNNNAGATCGGAAG
34717 TATGAGTGTGGAGTCGTTGCGTTCAAGCGTTTCTCCTGTTTCANNNAGATCGGAAG
34718 TATGAGTGTGGAGTCGTTGCTAGAACCATCACGCTATTAGGGCNNNAGATCGGAAG
34720 TATGAGTGTGGAGTCGTTGCTTCATGCTATTCTCCTGCCTCAGNNNAGATCGGAAGA
34721 TATGAGTGTGGAGTCGTTGCGTGATCTCAGCTCACTACAACCTNNNAGATCGGAAG
34722 TATGAGTGTGGAGTCGTTGCCAATGACACGATCTTGGCTCTCTNNNAGATCGGAAG
34723 TATGAGTGTGGAGTCGTTGCAAGTGATCCTCCTGCCTTGATTTNNNAGATCGGAAG
34724 TATGAGTGTGGAGTCGTTGCCAAGTAGCTGGGATTACAGGCATNNNAGATCGGAA
34725 TATGAGTGTGGAGTCGTTGCCTTCCCAGGTTCAAGCAATTCTCNNNAGATCGGAAG
34726 TATGAGTGTGGAGTCGTTGCCCTTCCAAGTTCAAGCGATTCTCNNNAGATCGGAAG
34727 TATGAGTGTGGAGTCGTTGCTTCAAGTGATTCTCCTGCCTCAGNNNAGATCGGAAG
34728 TATGAGTGTGGAGTCGTTGCCACGCCTGGGAGTTTTTGTATTTNNNAGATCGGAAG
34729 TATGAGTGTGGAGTCGTTGCCATGATCTTGGCTCACTGCAAACNNNAGATCGGAAG
34730 TATGAGTGTGGAGTCGTTGCAAATACCCCACCATGCAAACTTGNNNAGATCGGAAG
34731 TATGAGTGTGGAGTCGTTGCCCTCCCAGGTTCAAGTGATTCTCNNNAGATCGGAAG
34732 TATGAGTGTGGAGTCGTTGCCCACCAAGCTTGAATCCAGAAAGNNNAGATCGGAAG
34733 TATGAGTGTGGAGTCGTTGCCTCAGGGGAGAGGTTGAGAATTTNNNAGATCGGAA
34740 TATGAGTGTGGAGTCGTTGCTTTAGTGCAGACAGGGTTTCACTNNNAGATCGGAAG
34741 TATGAGTGTGGAGTCGTTGCATTCAAGTGATTCTCCTGCCTCANNNAGATCGGAAG
34742 TATGAGTGTGGAGTCGTTGCTTTTTGAGACAGAGTCTTGCCCTNNNAGATCGGAAG
34744 TATGAGTGTGGAGTCGTTGCGTGACACAATCTTGGCACACTACNNNAGATCGGAAG
34745 TATGAGTGTGGAGTCGTTGCCTTCCGAGTAGCTGGGATTACAGNNNAGATCGGAA
34746 TATGAGTGTGGAGTCGTTGCTTAGAGTGCAATGGTGGGATCTCNNNAGATCGGAA
34748 TATGAGTGTGGAGTCGTTGCTTCAAGTGATTCTCCTGCCTCAGNNNAGATCGGAAG
34749 TATGAGTGTGGAGTCGTTGCAATGTTGAGGTTTTGCGGAATCCNNNAGATCGGAA
34750 TATGAGTGTGGAGTCGTTGCCACAATCTCAGCTCACTGCAATCNNNAGATCGGAAG
34751 TATGAGTGTGGAGTCGTTGCTTTTTCTGCTGGACACTCTACGTNNNAGATCGGAAG
34752 TATGAGTGTGGAGTCGTTGCATGAGGCTTTCCCTAATCCCATCNNNAGATCGGAAG
34753 TATGAGTGTGGAGTCGTTGCTCTCAGTTCACTGCAACCTCTACNNNAGATCGGAAGA
34754 TATGAGTGTGGAGTCGTTGCTTGAAATCTGGGCAACTGTCTCTNNNAGATCGGAAG
34755 TATGAGTGTGGAGTCGTTGCGTTCAAGAGATTCTCCTGCCTCANNNAGATCGGAAG
34756 TATGAGTGTGGAGTCGTTGCCGCCTGATCCAAACGATAATTGTNNNAGATCGGAAG
34757 TATGAGTGTGGAGTCGTTGCTTCAAGTGATTCTCCTGCCTCAGNNNAGATCGGAAG
34758 TATGAGTGTGGAGTCGTTGCGAGTAGCTGGGATTACAGGAGTGNNNAGATCGGA
34760 TATGAGTGTGGAGTCGTTGCCGGCCTCTCTCCTTCTTTCTATGNNNAGATCGGAAGA
34762 TATGAGTGTGGAGTCGTTGCGATCTCTGATCACTGCAACCTCCNNNAGATCGGAAG

34763 TATGAGTGTGGAGTCGTTGCCTCCTGAGTAGCTGGGACTACAGNNNAGATCGGAA
34764 TATGAGTGTGGAGTCGTTGCCTTCCAGGGTTCAACCGATTCTCNNNAGATCGGAAG
34765 TATGAGTGTGGAGTCGTTGCCCGTGGTTCATAAGCAAATCCTCNNNAGATCGGAAG
34766 TATGAGTGTGGAGTCGTTGCAGGAGATGATGACACAGACACACNNNAGATCGGAA
34767 TATGAGTGTGGAGTCGTTGCCTCCTGAGTAGCTGGGACTACAGNNNAGATCGGAA
34768 TATGAGTGTGGAGTCGTTGCCGTGCCTGGCTAATTTGTGTATTNNNAGATCGGAAG
34770 TATGAGTGTGGAGTCGTTGCCCGGGTTCTAAGAGATTCTCCTGNNNAGATCGGAAG
34771 TATGAGTGTGGAGTCGTTGCCCACCACGCCCAGATAATTTTTTNNNAGATCGGAAGA
34772 TATGAGTGTGGAGTCGTTGCTTCAAGCTATCCTCATGCCTCAGNNNAGATCGGAAG
34775 TATGAGTGTGGAGTCGTTGCTTGGCTAAAGGCGACGTAAAATGNNNAGATCGGAA
34776 TATGAGTGTGGAGTCGTTGCCCTCCTGAGTAGCTGAGACTACANNNAGATCGGAAG
34777 TATGAGTGTGGAGTCGTTGCCTAGTAGCTGGGATTACAGGCACNNNAGATCGGAA
34778 TATGAGTGTGGAGTCGTTGCAAGAGTTTTTGCTCTGTCCTCCANNNAGATCGGAAG
34779 TATGAGTGTGGAGTCGTTGCCTTGAACTCCTGACCTCGTGATCNNNAGATCGGAAG
34780 TATGAGTGTGGAGTCGTTGCTTCAGAGGCGTTTAGTACATGGGNNNAGATCGGAA
34781 TATGAGTGTGGAGTCGTTGCCCCAAGTTCAAGCAATTCTCCTGNNNAGATCGGAAG
34783 TATGAGTGTGGAGTCGTTGCTATGAAGCAGTCACTCCCCTTTCNNNAGATCGGAAG
34784 TATGAGTGTGGAGTCGTTGCCCCAAGTTCAAGCAATTCTCCTGNNNAGATCGGAAG
34786 TATGAGTGTGGAGTCGTTGCCTCTCTCTCTCCCTTCATCCTCTNNNAGATCGGAAGAG
34787 TATGAGTGTGGAGTCGTTGCCGCCCGGCTAATCTTTGTATTTTNNNAGATCGGAAG
34788 TATGAGTGTGGAGTCGTTGCCCCCACCTGGCTAATTTTTGTTTNNNAGATCGGAAGA
34789 TATGAGTGTGGAGTCGTTGCTTTTTGAGACAGGGTCTTGCTCTNNNAGATCGGAAG
34791 TATGAGTGTGGAGTCGTTGCCTGAGTAGCTGGGATTACAGGTGNNNAGATCGGAA
34793 TATGAGTGTGGAGTCGTTGCTCAGTCTCTTGAGTAGCTGGGATNNNAGATCGGAAG
34795 TATGAGTGTGGAGTCGTTGCCAGGTAGCTGGGATTACAGGTACNNNAGATCGGAA
34796 TATGAGTGTGGAGTCGTTGCCTAACGCCTATAATCCCAGCACTNNNAGATCGGAAG
34797 TATGAGTGTGGAGTCGTTGCCTGAGTAGCTGGGATTACAGGTGNNNAGATCGGAA
34799 TATGAGTGTGGAGTCGTTGCGATCAAGTGATTCTCCTGCCTCANNNAGATCGGAAG
34800 TATGAGTGTGGAGTCGTTGCGTCTCAATCTTCTGACCTCGTGANNNAGATCGGAAG
34801 TATGAGTGTGGAGTCGTTGCTGTAGTGCAGTGGTGTGATCTCNNNAGATCGGAAG
34804 TATGAGTGTGGAGTCGTTGCTAGAGTGCAGTGACATGATCTCGNNNAGATCGGAA
34807 TATGAGTGTGGAGTCGTTGCTGACAGTATCTCACTCTTTCGCCNNNAGATCGGAAGA
34808 TATGAGTGTGGAGTCGTTGCTGGAATAGCTGGAATTACAGGCGNNNAGATCGGAA
34809 TATGAGTGTGGAGTCGTTGCCACCCCCGGCTAATTTTTTGTATNNNAGATCGGAAGA
34810 TATGAGTGTGGAGTCGTTGCTCCTCTGCTTCCTGAGTTCAATCNNNAGATCGGAAGA
34812 TATGAGTGTGGAGTCGTTGCGGTGTGATCTCAACACATTGCAANNNAGATCGGAAG
34813 TATGAGTGTGGAGTCGTTGCTTAGTAGAGATGGGGGAAGAGGGNNNAGATCGGA
34814 TATGAGTGTGGAGTCGTTGCTGTGGTGTACTAGCTTCTGAACCNNNAGATCGGAAG
34815 TATGAGTGTGGAGTCGTTGCAGGAGGATGTGTACTGTCTCAGANNNAGATCGGAA
34816 TATGAGTGTGGAGTCGTTGCGATGGACTTACTCTATCGCCCAGNNNAGATCGGAAG
34817 TATGAGTGTGGAGTCGTTGCCGTCTCCCAAGTTGAAGCAATTCNNNAGATCGGAAG
34818 TATGAGTGTGGAGTCGTTGCCGTCTCCCAAGTTGAAGCAATTCNNNAGATCGGAAG
34819 TATGAGTGTGGAGTCGTTGCCCAAGTAGCTGGGATTATAGGCGNNNAGATCGGAA

34820 TATGAGTGTGGAGTCGTTGCAGTAGAAACAGGCTTTCACCGTGNNNAGATCGGAAG
34821 TATGAGTGTGGAGTCGTTGCGGTTCAAATGATTCTCCTGCCTCNNNAGATCGGAAGA
34822 TATGAGTGTGGAGTCGTTGCTTCAAGTGATTCTCCTGCCTCAGNNNAGATCGGAAGA
34823 TATGAGTGTGGAGTCGTTGCCTCCTGAGTAGCTGGGATTACAGNNNAGATCGGAAG
34824 TATGAGTGTGGAGTCGTTGCGATGGAGTCTCGCTATGTCATCCNNNAGATCGGAAG
34826 TATGAGTGTGGAGTCGTTGCTGGTTTTGAACTCCTGACCTCAANNNAGATCGGAAGA
34827 TATGAGTGTGGAGTCGTTGCCCATGCCAGGCTCATTTTTTGTANNNAGATCGGAAGA
34830 TATGAGTGTGGAGTCGTTGCCCATGCCCGGCTAATTTTTGTATNNNAGATCGGAAGA
34833 TATGAGTGTGGAGTCGTTGCTTGGGCAACTTCCTGACATAGTANNNAGATCGGAAG
34834 TATGAGTGTGGAGTCGTTGCCTCCTGAGTAGCTGGGATTACAGNNNAGATCGGAAG
34836 TATGAGTGTGGAGTCGTTGCCACGCCCAGCTAATTTTTTTCGNNNAGATCGGAAGA
34837 TATGAGTGTGGAGTCGTTGCCCTTCTGGGTTTACGCCATTCTCNNNAGATCGGAAGA
34838 TATGAGTGTGGAGTCGTTGCGCGCCCAGCCAAATTTTTTTTANNNAGATCGGAAGA
34840 TATGAGTGTGGAGTCGTTGCCCGGATTCAAACGATTCTCCTCTNNNAGATCGGAAGA
34841 TATGAGTGTGGAGTCGTTGCTGGTAACGGCTCAACTCTACTTCNNNAGATCGGAAGA
34842 TATGAGTGTGGAGTCGTTGCATAGCTAAAATTACAGGCGCCCANNNAGATCGGAAG
34843 TATGAGTGTGGAGTCGTTGCGGGAGGCCTAGATGAAACAAGAANNNAGATCGGAA
34844 TATGAGTGTGGAGTCGTTGCTAGCTGGATTACTGGCACATACCNNNAGATCGGAAG
34845 TATGAGTGTGGAGTCGTTGCGTAGAGACAGGGTTTCACTGTGTNNNAGATCGGAAG
34846 TATGAGTGTGGAGTCGTTGCCCCCTACAGTCACAGAAACTACANNNAGATCGGAAGA
34847 TATGAGTGTGGAGTCGTTGCCCAAGTTCAAGTGATCCTTCTGCNNNAGATCGGAAGA
34848 TATGAGTGTGGAGTCGTTGCGAGAGAGAGTCTCACTGTGTTGCNNNAGATCGGAAG
34849 TATGAGTGTGGAGTCGTTGCCCTCCTAGGTTCAAGCGATTCTGNNNAGATCGGAAGA
34850 TATGAGTGTGGAGTCGTTGCCTGACTCAGTCTGTTCACTGGTTNNNAGATCGGAAGA
34852 TATGAGTGTGGAGTCGTTGCCCATGACACCCGGCTAATTTTTTNNNAGATCGGAAGA
34853 TATGAGTGTGGAGTCGTTGCCACCACGCCTGGCTAATTTTTTANNNAGATCGGAAGA
34855 TATGAGTGTGGAGTCGTTGCGCAACTACGCCCAGCTAATTTTTNNNAGATCGGAAGA
34858 TATGAGTGTGGAGTCGTTGCCCTCCTGAGTAACTGGGATTACANNNAGATCGGAAG
34859 TATGAGTGTGGAGTCGTTGCCCTCCTAGGTTCACATGCTTCTCNNNAGATCGGAAGA
34860 TATGAGTGTGGAGTCGTTGCTGGTTCAAGTCATTCTCTTGCCTNNNAGATCGGAAGA
34861 TATGAGTGTGGAGTCGTTGCCTCCTGAGTAGCTGGGATTACAGNNNAGATCGGAAG
34864 TATGAGTGTGGAGTCGTTGCCCACAGTACCCGAGATTACATGTNNNAGATCGGAAG
34865 TATGAGTGTGGAGTCGTTGCCCTGGGTTCAAGCAATTCTCCTANNNAGATCGGAAGA
34866 TATGAGTGTGGAGTCGTTGCTTCAAGTGATTCTCCTGCCTCAGNNNAGATCGGAAGA
34867 TATGAGTGTGGAGTCGTTGCCTGAGTAGCTGGGATTACAGGTGNNNAGATCGGAA
34868 TATGAGTGTGGAGTCGTTGCATCTTGGCTCACTGTAACCTCTGNNNAGATCGGAAGA
34869 TATGAGTGTGGAGTCGTTGCCTCCTGAATAGCTGGGATGACAGNNNAGATCGGAAG
34870 TATGAGTGTGGAGTCGTTGCCATTCTCTCACCTCAGCTTCTCANNNAGATCGGAAGAG
34871 TATGAGTGTGGAGTCGTTGCAGTAGCTGGGACTACAGGTATGTNNNAGATCGGAAG
34872 TATGAGTGTGGAGTCGTTGCTTCTTTTTTGAGACGGAGTCCCANNNAGATCGGAAGA
34874 TATGAGTGTGGAGTCGTTGCGATCTCGTCTCACTGCAACAGAGNNNAGATCGGAAG
34877 TATGAGTGTGGAGTCGTTGCCCTCCTGGGTTCAAGCAATTCTCNNNAGATCGGAAGA
34879 TATGAGTGTGGAGTCGTTGCTTCAAGTGATTCTCCTGCCTCAGNNNAGATCGGAAGA

34883 TATGAGTGTGGAGTCGTTGCGGCAAAGTCTGTTTCTGACTCTGNNNAGATCGGAAG
34885 TATGAGTGTGGAGTCGTTGCACCACGCCTGGCTAATTTGTATTNNNAGATCGGAAG
34886 TATGAGTGTGGAGTCGTTGCTAGAAGTGCAGTTTCGTTACCCTNNNAGATCGGAAG
34887 TATGAGTGTGGAGTCGTTGCCACCACACTCGGCTAATTTTGTTNNNAGATCGGAAG
34888 TATGAGTGTGGAGTCGTTGCCCACCACAGCCAGCTAATTTTTTNNNAGATCGGAAG
34889 TATGAGTGTGGAGTCGTTGCGTTCAAGAGATTCTCCTGCCTCANNNAGATCGGAAG
34890 TATGAGTGTGGAGTCGTTGCCTCTCGAGTGGCTGGGACTATAGNNNAGATCGGAA
34891 TATGAGTGTGGAGTCGTTGCCCAAGTAGCTGGTACTACAGGTGNNNAGATCGGAA
34892 TATGAGTGTGGAGTCGTTGCTGCACCACTATGGAAAACAGTCANNNAGATCGGAAG
34893 TATGAGTGTGGAGTCGTTGCGTCTCCCGAGTAGCTGGAATTACNNNAGATCGGAAG
34894 TATGAGTGTGGAGTCGTTGCGCCACCACATCAGGCTAATTTTTNNNAGATCGGAAG
34895 TATGAGTGTGGAGTCGTTGCGATTTTGAGACGGAGTCTTGCTCNNNAGATCGGAA
34896 TATGAGTGTGGAGTCGTTGCCCACCACACTGGGCTAATTTTTTNNNAGATCGGAAG
34897 TATGAGTGTGGAGTCGTTGCCCTCCCACGTTCAAATGATTCTCNNNAGATCGGAAG
34898 TATGAGTGTGGAGTCGTTGCGCCTTCTGAGTAGCTGGGATTACNNNAGATCGGAA
34899 TATGAGTGTGGAGTCGTTGCAGATGGAGTCTCTCTCTGTCACCNNNAGATCGGAAG
34900 TATGAGTGTGGAGTCGTTGCCCAAAGTGCTGGGATTACAAGTGNNNAGATCGGAA
34901 TATGAGTGTGGAGTCGTTGCCTGAGTAGCTCGGATTACTGGTGNNNAGATCGGAA
34902 TATGAGTGTGGAGTCGTTGCCAGATGGAGTCTCACTCTGTCACNNNAGATCGGAAG
34903 TATGAGTGTGGAGTCGTTGCGGGTTCAAGTAATTCTCCTGCCTNNNAGATCGGAAG
34904 TATGAGTGTGGAGTCGTTGCCTCCCAGGTTCAAGTGATTCTCANNNAGATCGGAAG
34906 TATGAGTGTGGAGTCGTTGCCACCATGCCCAGCTAATTTTTGTNNNAGATCGGAAG
34907 TATGAGTGTGGAGTCGTTGCGCCACCATGCCAGTTTACTTTTTNNNAGATCGGAAG
34908 TATGAGTGTGGAGTCGTTGCCCTCCTAGGTTCAAGCGATTCTCNNNAGATCGGAAG
34909 TATGAGTGTGGAGTCGTTGCCAAGTAGCTGGGATTACAAGTGCNNNAGATCGGAA
34910 TATGAGTGTGGAGTCGTTGCCTCCACATCTCAGGTTCAAGTGANNNAGATCGGAAG
34911 TATGAGTGTGGAGTCGTTGCAGTAGATGGGATTACAGGCACACNNNAGATCGGAA
34912 TATGAGTGTGGAGTCGTTGCGAGTAGCTGGGATGACAGATACCNNNAGATCGGAA
34913 TATGAGTGTGGAGTCGTTGCCTCCTGAGTAGCTGGGATTACAGNNNAGATCGGAA
34914 TATGAGTGTGGAGTCGTTGCCGGGGTTCTAAGAGATTCTCCTGNNNAGATCGGAAG
34915 TATGAGTGTGGAGTCGTTGCGCCATCACACCCAGCTAATTTTTNNNAGATCGGAAG
34916 TATGAGTGTGGAGTCGTTGCCGCTTCCTGAGTTCAAGTGATTCNNNAGATCGGAAG
34918 TATGAGTGTGGAGTCGTTGCTGTGTGTCTCTCTAGGACTTTGCNNNAGATCGGAAG
34919 TATGAGTGTGGAGTCGTTGCCTCACATTGCATCTTTGAGCCAANNNAGATCGGAAG
34920 TATGAGTGTGGAGTCGTTGCTAAAAATGCATATATCGGCCGGGNNNAGATCGGAA
34921 TATGAGTGTGGAGTCGTTGCAATCTCGGCTCACTACAACCTTTNNNAGATCGGAAG
34922 TATGAGTGTGGAGTCGTTGCCCTGGGTTCAAGCAATTCTCATGNNNAGATCGGAAG
34923 TATGAGTGTGGAGTCGTTGCTTCAAGTGATTCTCCTGCCTCAGNNNAGATCGGAAG
34924 TATGAGTGTGGAGTCGTTGCGCAAAACATCAGGCCTATGTAGGNNNAGATCGGAA
34925 TATGAGTGTGGAGTCGTTGCAAAGTGCTGGGATTACAGACGTCNNNAGATCGGAA
34926 TATGAGTGTGGAGTCGTTGCCTCCTGAGTAGCTGGGATTACAGNNNAGATCGGAA
34927 TATGAGTGTGGAGTCGTTGCGGGTTCAAGCGATTCTCTTACCTNNNAGATCGGAAG
34928 TATGAGTGTGGAGTCGTTGCTTTAGTAGAGACGGAGTTTCGCCNNNAGATCGGAA

34929 TATGAGTGTGGAGTCGTTGCTTCAAGTGATTCTCCTGCCTCAGNNNAGATCGGAAG
34930 TATGAGTGTGGAGTCGTTGCTTTCACTCTATCACCAAGGCTGGNNNAGATCGGAAG
34932 TATGAGTGTGGAGTCGTTGCTTCAAGCAATTTTCCTGCCTCAGNNNAGATCGGAAG
34933 TATGAGTGTGGAGTCGTTGCCTGAGTAGCTGGAACTACAGGACNNNAGATCGGAA
34934 TATGAGTGTGGAGTCGTTGCCTGGGATTACAGGTGTGAGTCACNNNAGATCGGAA
34935 TATGAGTGTGGAGTCGTTGCTTGAATACCCATTGCCCACTCTTNNNAGATCGGAAGA
34936 TATGAGTGTGGAGTCGTTGCCACCACAGTCGCCTAATTTTTGTNNNAGATCGGAAG
34937 TATGAGTGTGGAGTCGTTGCGAGTATCTGGGATTACAGGCGTGNNNAGATCGGAA
34939 TATGAGTGTGGAGTCGTTGCTGTTCTTCTCATTCAGTCTGGGGNNNAGATCGGAAG
34940 TATGAGTGTGGAGTCGTTGCTTTTGGTAGACACGAGGTTTCACNNNAGATCGGAAG
34942 TATGAGTGTGGAGTCGTTGCGATCTCTGCTCAGTACAGCCTACNNNAGATCGGAAG
34943 TATGAGTGTGGAGTCGTTGCCTCCCAAGTAGCTGGAACTACAGNNNAGATCGGAAG
34944 TATGAGTGTGGAGTCGTTGCCTTTCACGGAGTCTCACTCTGTTNNNAGATCGGAAG
34945 TATGAGTGTGGAGTCGTTGCGATGGAGTCCTACTCTGTTGTCCNNNAGATCGGAAG
34946 TATGAGTGTGGAGTCGTTGCAAGACAGTCTATGAAGCTGGAGCNNNAGATCGGAA
34947 TATGAGTGTGGAGTCGTTGCATCTCCCAGGTTTGAGCAATTCTNNNAGATCGGAAG
34948 TATGAGTGTGGAGTCGTTGCCCTCCTGGGTTCAAGTGATTCTCNNNAGATCGGAAG
34949 TATGAGTGTGGAGTCGTTGCGCAAGTAGCTTTGGTTTTTGACTNNNAGATCGGAAG
34953 TATGAGTGTGGAGTCGTTGCACAAAGGCAGTGGGGAATTCTTANNNAGATCGGAA
34954 TATGAGTGTGGAGTCGTTGCGAGTTTTTGCTCTTGTCACCCAGNNNAGATCGGAAG
34955 TATGAGTGTGGAGTCGTTGCTGGAATGGGTTCTGCGATGATTANNNAGATCGGAA
34956 TATGAGTGTGGAGTCGTTGCTTTTTTTTTTTCCGCCTGCTGTGNNNAGATCGGAAGA
34957 TATGAGTGTGGAGTCGTTGCCCTACTGAGTAGCTGGGACTACANNNAGATCGGAAG
34958 TATGAGTGTGGAGTCGTTGCAACCTCTGACTCCCTGGTTCAAGNNNAGATCGGAAG
34959 TATGAGTGTGGAGTCGTTGCGGAGAGATGGCTCTGATTTTTGCNNNAGATCGGAA
34960 TATGAGTGTGGAGTCGTTGCGTGTGATCTCAGTTCATGGCAACNNNAGATCGGAAG
34961 TATGAGTGTGGAGTCGTTGCCTTTCCATGGCCTCAACTCTACTNNNAGATCGGAAGA
34965 TATGAGTGTGGAGTCGTTGCCATCTCGCCCGGCTAATTTTTTANNNAGATCGGAAG
34967 TATGAGTGTGGAGTCGTTGCTTCAAGCCATTCTCCTGTCTCAGNNNAGATCGGAAG
34968 TATGAGTGTGGAGTCGTTGCGTTCCAGTGATTCTCCTGCCTTANNNAGATCGGAAG
34969 TATGAGTGTGGAGTCGTTGCTAGCTGGGATTATAGGTGTGTGCNNNAGATCGGAA
34971 TATGAGTGTGGAGTCGTTGCGGGCTCTGTGTTTCCTTAGGTAANNNAGATCGGAAG
34972 TATGAGTGTGGAGTCGTTGCATCTCCTGGGTTCAAGCAATTCTNNNAGATCGGAAG
34975 TATGAGTGTGGAGTCGTTGCAGACAGTTTCCTTTTTGTTGCCCNNNAGATCGGAAG
34978 TATGAGTGTGGAGTCGTTGCGGTAACACACGTGAGGTTTTTGTNNNAGATCGGAA
34979 TATGAGTGTGGAGTCGTTGCCATGCGCCATTATTTCCAGCTAANNNAGATCGGAAG
34981 TATGAGTGTGGAGTCGTTGCTGGAGTACAGTGACACGATCTTGNNNAGATCGGAA
34982 TATGAGTGTGGAGTCGTTGCTTCAAGTGATTCTCCTGCCTCAGNNNAGATCGGAAG
34983 TATGAGTGTGGAGTCGTTGCGGTTCAAGTGATTCTTCTGCCTCNNNAGATCGGAAG
34984 TATGAGTGTGGAGTCGTTGCCAGTAGTACGATCTTGGCTCACTNNNAGATCGGAAG
34987 TATGAGTGTGGAGTCGTTGCGAGAAAGGGTCTTGTTCTGTTGCNNNAGATCGGAA
34988 TATGAGTGTGGAGTCGTTGCCTAAACCTTGGCCTCCGAAAATGNNNAGATCGGAAG
34989 TATGAGTGTGGAGTCGTTGCCGCATCCCATCAACTCATGTCTTNNNAGATCGGAAGA

34990 TATGAGTGTGGAGTCGTTGCCCACGCACGGCTAATTTTTGTATNNNAGATCGGAAG
34992 TATGAGTGTGGAGTCGTTGCCTCAAGTGATTCTCCTGCCTCAGNNNAGATCGGAAG
34993 TATGAGTGTGGAGTCGTTGCTCAGAGTTCACTGTAGCCTCAACNNNAGATCGGAAG
34994 TATGAGTGTGGAGTCGTTGCAGCCTCCTGAGTATCTGGGATTANNNAGATCGGAAG
34995 TATGAGTGTGGAGTCGTTGCCCACGCCCAGCCAATTTTTATATNNNAGATCGGAAG
34996 TATGAGTGTGGAGTCGTTGCGATAATGACGTGATGGCTGGAGTNNNAGATCGGAA
34997 TATGAGTGTGGAGTCGTTGCATGGAGTTTTGTCCTTTTGCCTGNNNAGATCGGAAG
34998 TATGAGTGTGGAGTCGTTGCCTCAAGCAATTCTCCTGCCTCAGNNNAGATCGGAAG
34999 TATGAGTGTGGAGTCGTTGCGCCTACAGAGTAGCTGGGATTACNNNAGATCGGAA
35000 TATGAGTGTGGAGTCGTTGCTTGAGACGTAGTCTTGCTCTGTCNNNAGATCGGAAG
35001 TATGAGTGTGGAGTCGTTGCGATCGAATGTGTCTGATGCAAGGNNNAGATCGGAA
35002 TATGAGTGTGGAGTCGTTGCGCCTCCCAAGTAGCTAGGATTACNNNAGATCGGAAG
35003 TATGAGTGTGGAGTCGTTGCGATTACAGGCATGCACCATCATGNNNAGATCGGAAG
35004 TATGAGTGTGGAGTCGTTGCGAGTAGCTGGGATTACAGGCATGNNNAGATCGGAA
35006 TATGAGTGTGGAGTCGTTGCCTGAGTAGCTGGGATTATAGGCGNNNAGATCGGAA
35007 TATGAGTGTGGAGTCGTTGCCAAGTCTTGCTCTGTTACCCAAGNNNAGATCGGAAG
35009 TATGAGTGTGGAGTCGTTGCTTCAAGCTATTCTCCTGCCTCAGNNNAGATCGGAAG
35011 TATGAGTGTGGAGTCGTTGCCCACCATGCCCAGCTAATTTTTTNNNAGATCGGAAGA
35012 TATGAGTGTGGAGTCGTTGCCCGAGTAGCTGGGATTACAAGTGNNNAGATCGGAA
35013 TATGAGTGTGGAGTCGTTGCAATAGCTGGGATTACAGACACCCNNNAGATCGGAAG
35015 TATGAGTGTGGAGTCGTTGCGCACGCCCAGCTAATTTTTTTTTNNNAGATCGGAAG
35016 TATGAGTGTGGAGTCGTTGCAGAATCTCACTATGTTGCCCAGGNNNAGATCGGAAG
35017 TATGAGTGTGGAGTCGTTGCCGAGTAGCTGGGATTACAGACATNNNAGATCGGAA
35018 TATGAGTGTGGAGTCGTTGCCCTCCTGGGTTCAAGTGATTCTCNNNAGATCGGAAG
35019 TATGAGTGTGGAGTCGTTGCGAGTGTAATGGCATGATCTTGGCNNNAGATCGGAA
35020 TATGAGTGTGGAGTCGTTGCTGCCCAGCCAATTTCTTTTTCTTNNNAGATCGGAAGA
35021 TATGAGTGTGGAGTCGTTGCCCTCCCAAGTTGCTGTTACTACANNNAGATCGGAAG
35022 TATGAGTGTGGAGTCGTTGCAAGCTATACTGCATGTAGGCACANNNAGATCGGAAG
35023 TATGAGTGTGGAGTCGTTGCCTCCCAGGTTCAAGTGATTCTCANNNAGATCGGAAG
35025 TATGAGTGTGGAGTCGTTGCACCTCTGCCTTCTGAGTTCAAATNNNAGATCGGAAG
35026 TATGAGTGTGGAGTCGTTGCCAATCTCAGCTCACTGCAACTTCNNNAGATCGGAAG
35027 TATGAGTGTGGAGTCGTTGCCTCAAACTCCTGACCTCGTGATCNNNAGATCGGAAG
35028 TATGAGTGTGGAGTCGTTGCCCTCCCGAGTTCAAGAGATTCTCNNNAGATCGGAAG
35030 TATGAGTGTGGAGTCGTTGCCCCAAAGTGCTAGGATTACAGGTNNNAGATCGGAA
35032 TATGAGTGTGGAGTCGTTGCGGTACGATATCGACTCACTGCAANNNAGATCGGAAG
35033 TATGAGTGTGGAGTCGTTGCAGTAGAGATGGGGTTTCAGCATGNNNAGATCGGAA
35034 TATGAGTGTGGAGTCGTTGCTCTCAGTACATTGACCAAGCTGGNNNAGATCGGAAG
35035 TATGAGTGTGGAGTCGTTGCTGTTAGCCAGGATGGTCTCAATCNNNAGATCGGAAG
35036 TATGAGTGTGGAGTCGTTGCCGATCTCCAATTACTGCAAGCTCNNNAGATCGGAAG
35038 TATGAGTGTGGAGTCGTTGCGTCGTGTCACAAGAACTTGGTTTNNNAGATCGGAAG
35039 TATGAGTGTGGAGTCGTTGCAATCTCTGCCTCCTGAGTTCAAGNNNAGATCGGAAG
35040 TATGAGTGTGGAGTCGTTGCTATCACAGATCACTGCAGTCTCCNNNAGATCGGAAG
35041 TATGAGTGTGGAGTCGTTGCCCTCCCAAGTTCATGCTACTCTCNNNAGATCGGAAGA

35042 TATGAGTGTGGAGTCGTTGCGCTCCCTGAGTCTCAAGAAAAGANNNAGATCGGAAG
35043 TATGAGTGTGGAGTCGTTGCCCACGCCCAGCTAATTTTTGTATNNNAGATCGGAAGA
35045 TATGAGTGTGGAGTCGTTGCTCTTGAACTTCTGACCTCTGGTGNNNAGATCGGAAGA
35047 TATGAGTGTGGAGTCGTTGCCCTCCTGAGTAGCTGCAATTACANNNAGATCGGAAGA
35048 TATGAGTGTGGAGTCGTTGCCCATGCCCGGCTAATTTTTTGTANNNAGATCGGAAGA
35049 TATGAGTGTGGAGTCGTTGCGCCTCCTGAATAGCTGGGAATACNNNAGATCGGAAG
35051 TATGAGTGTGGAGTCGTTGCTTGCTTTGGTTATTCAGGGTGTTNNNAGATCGGAAG
35052 TATGAGTGTGGAGTCGTTGCCAGAGAGAGGGCATCTTTGTCTTNNNAGATCGGAAG
35053 TATGAGTGTGGAGTCGTTGCCCACGCCCAGCTAATTTTTTGTANNNAGATCGGAAGA
35054 TATGAGTGTGGAGTCGTTGCTCAAGTGATTCTCCTGTCTCAGCNNNAGATCGGAAGA
35057 TATGAGTGTGGAGTCGTTGCAAATGGAGGAGTTGGACTTTGCTNNNAGATCGGAAG
35058 TATGAGTGTGGAGTCGTTGCTTGAGTTTATCAGGAGGGGAAGCNNNAGATCGGAA
35060 TATGAGTGTGGAGTCGTTGCCTATCTCGACTCACTGCAAGCTCNNNAGATCGGAAGA
35061 TATGAGTGTGGAGTCGTTGCGTAGAGACAGAGTTTCACCGTGTNNNAGATCGGAAG
35062 TATGAGTGTGGAGTCGTTGCTTACTTCACATGTCACGTGACCTNNNAGATCGGAAGA
35065 TATGAGTGTGGAGTCGTTGCTTCAAGCGATTCTCCTGACTCAGNNNAGATCGGAAGA
35066 TATGAGTGTGGAGTCGTTGCCTCAAGCAATTCTCATGCCTCAGNNNAGATCGGAAGA
35067 TATGAGTGTGGAGTCGTTGCATTGGCATTTTGGTAGGGATTGCNNNAGATCGGAAG
35068 TATGAGTGTGGAGTCGTTGCGCCAAATTCCAAAGAGGTGTACANNNAGATCGGAAG
35069 TATGAGTGTGGAGTCGTTGCTTCAAGTGATTCTCCTGCCTCAGNNNAGATCGGAAGA
35070 TATGAGTGTGGAGTCGTTGCCCTCCTGAGTAGCTGGGACTATANNNAGATCGGAAG
35071 TATGAGTGTGGAGTCGTTGCAGCCAATGGTCTAAATGGGTTCTNNNAGATCGGAAG
35073 TATGAGTGTGGAGTCGTTGCTTTTTTCTTGTCTCCTGCTAGCTNNNAGATCGGAAGAG
35074 TATGAGTGTGGAGTCGTTGCCACCACGCCTGGCTAATTTTTTTNNNAGATCGGAAGA
35075 TATGAGTGTGGAGTCGTTGCATTCTTCCTGGGTCAGTCTTGAGNNNAGATCGGAAG
35076 TATGAGTGTGGAGTCGTTGCCAAGCTGTTTTCGTTACTGTGGTNNNAGATCGGAAG
35078 TATGAGTGTGGAGTCGTTGCGTACACAACGTGCAGGTTTGTTANNNAGATCGGAAG
35080 TATGAGTGTGGAGTCGTTGCTAGTAGCTGGGATTACAGGCATGNNNAGATCGGAAG
35081 TATGAGTGTGGAGTCGTTGCTATCTCAGTGAGCAGAGGGATGANNNAGATCGGAA
35082 TATGAGTGTGGAGTCGTTGCGGCCAGGGTGGTTTATTTCTAGANNNAGATCGGAAG
35083 TATGAGTGTGGAGTCGTTGCCACCACGTCCGGCTAATTTTTTTNNNAGATCGGAAGA
35084 TATGAGTGTGGAGTCGTTGCCCTCCCAAGTAGCTGTGATTACANNNAGATCGGAAGA
35085 TATGAGTGTGGAGTCGTTGCTTTTGAGCCTATGTGTGCCTTTGNNNAGATCGGAAGA
35086 TATGAGTGTGGAGTCGTTGCTTTTGTGCAGTGGGTTGAAATGTNNNAGATCGGAAG
35088 TATGAGTGTGGAGTCGTTGCCTCCTAGGCTCAAGTGATTCTCCNNNAGATCGGAAGA
35089 TATGAGTGTGGAGTCGTTGCAAATGCTAGGATTACAGGCGTGANNNAGATCGGAAG
35090 TATGAGTGTGGAGTCGTTGCATGCAATCTCAGCTTACTGCAACNNNAGATCGGAAGA
35091 TATGAGTGTGGAGTCGTTGCCAAAGTGCTAGGATTACAGGGGTNNNAGATCGGAA
35094 TATGAGTGTGGAGTCGTTGCTGATGCGAACTCAGCTTACTGTANNNAGATCGGAAG
35096 TATGAGTGTGGAGTCGTTGCCCTCCAGTGTTCACTCCATTCTCNNNAGATCGGAAGAG
35098 TATGAGTGTGGAGTCGTTGCTTCAAGTGATTCTCCTGCCTCAGNNNAGATCGGAAGA
35100 TATGAGTGTGGAGTCGTTGCATTCTCAAGCCATCCTCCCATTTNNNAGATCGGAAGAG
35101 TATGAGTGTGGAGTCGTTGCGTCACCATGCCCAGCTAATTTTTNNNAGATCGGAAGA

35102 TATGAGTGTGGAGTCGTTGCCTGGGATTACAGGTGTGAGCTACNNNAGATCGGAA
35105 TATGAGTGTGGAGTCGTTGCGTTCAATGGCACGATCTTGTTCANNNAGATCGGAAG
35107 TATGAGTGTGGAGTCGTTGCCCACCTAAGCTCAAGTGATCCTTNNNAGATCGGAAG
35108 TATGAGTGTGGAGTCGTTGCCAGTCTGGACTTGAACTCCTGACNNNAGATCGGAAG
35110 TATGAGTGTGGAGTCGTTGCAAGCAATCCTTCCATCTCAACCTNNNAGATCGGAAGA
35111 TATGAGTGTGGAGTCGTTGCTATGCCTGGTTGTGTGATCATCANNNAGATCGGAAG
35112 TATGAGTGTGGAGTCGTTGCTGCAGTACAATAGCGCAATCTTGNNNAGATCGGAAG
35116 TATGAGTGTGGAGTCGTTGCCCTGGGTTCAAGCAATTCTTCTGNNNAGATCGGAAG
35117 TATGAGTGTGGAGTCGTTGCCCTCTCAGGTTCAAGCGATTCTCNNNAGATCGGAAG
35118 TATGAGTGTGGAGTCGTTGCCCACGCCCAGCTAATTTTTTGTANNNAGATCGGAAG
35119 TATGAGTGTGGAGTCGTTGCGTTTCACCATGTTGGCCAAGATGNNNAGATCGGAAG
35120 TATGAGTGTGGAGTCGTTGCATCTCGTCTTACTGCAACCTCTGNNNAGATCGGAAGA
35121 TATGAGTGTGGAGTCGTTGCATTCGTGTCATGGAAGTTTGCTGNNNAGATCGGAAG
35122 TATGAGTGTGGAGTCGTTGCTACAAGTCTGCAAGAACCACAGTNNNAGATCGGAAG
35123 TATGAGTGTGGAGTCGTTGCCCTCCCACCATGCTAAGATTACANNNAGATCGGAAG
35125 TATGAGTGTGGAGTCGTTGCCATGTAGCTGGGATTACAGGCATNNNAGATCGGAA
35126 TATGAGTGTGGAGTCGTTGCTTTTTGAGGAACCTCCAAAGCACNNNAGATCGGAAG
35127 TATGAGTGTGGAGTCGTTGCTTGTTGCTGTTGTTTGGAACTGTNNNAGATCGGAAG
35128 TATGAGTGTGGAGTCGTTGCCTCTGCATCTTGGTTTCAAGCAANNNAGATCGGAAG
35129 TATGAGTGTGGAGTCGTTGCCCTTCAGATTTTGCTGAGGGTTTNNNAGATCGGAAG
35130 TATGAGTGTGGAGTCGTTGCCACCGTGCCTAGTCTCAAACTAANNNAGATCGGAAG
35131 TATGAGTGTGGAGTCGTTGCCCAAAGTACTGGGATTACAGGCANNNAGATCGGAA
35133 TATGAGTGTGGAGTCGTTGCTTCAAACCATTCTCCTGCCTCAGNNNAGATCGGAAGA
35134 TATGAGTGTGGAGTCGTTGCCTTTGCTGTGCAGAGGCTTTTTANNNAGATCGGAAG
35135 TATGAGTGTGGAGTCGTTGCTTTTGTGGTGAAATTTGGGTTGTNNNAGATCGGAAG
35136 TATGAGTGTGGAGTCGTTGCCAGCTTAGCCATTTGCATGTCTTNNNAGATCGGAAG
35138 TATGAGTGTGGAGTCGTTGCGGAGTCTCACTCTTTCACCCAAGNNNAGATCGGAAG
35139 TATGAGTGTGGAGTCGTTGCCTCACTGTAACCTCGAACTCCAANNNAGATCGGAAG
35140 TATGAGTGTGGAGTCGTTGCTCTGGTTCAGTCTTGGAAGGATGNNNAGATCGGAA
35141 TATGAGTGTGGAGTCGTTGCTTCATGCCATTCTCCTGTCTCAGNNNAGATCGGAAGA
35142 TATGAGTGTGGAGTCGTTGCCAGTAGCTGGGATTACAGGCATGNNNAGATCGGAA
35144 TATGAGTGTGGAGTCGTTGCCCAAACATGCTTCTCTTCCATCCNNNAGATCGGAAGA
35146 TATGAGTGTGGAGTCGTTGCATACAATCTCTGCCACCCATGTTNNNAGATCGGAAGA
35147 TATGAGTGTGGAGTCGTTGCATTGACTGGCAGTGAAGGTATGTNNNAGATCGGAA
35148 TATGAGTGTGGAGTCGTTGCGTAGGTGGGACTACAGGAATGTGNNNAGATCGGA
35149 TATGAGTGTGGAGTCGTTGCGAGTCTTGTTCTTTTGCCTAGGCNNNAGATCGGAAG
35150 TATGAGTGTGGAGTCGTTGCCCGGGTTCACTTTGATTCTGTTGNNNAGATCGGAAG
35151 TATGAGTGTGGAGTCGTTGCCATCTTTGTGTGGCATTGTGTGTNNNAGATCGGAAG
35156 TATGAGTGTGGAGTCGTTGCTTTCTACTACACGTCCAAGCTGANNNAGATCGGAAG
35157 TATGAGTGTGGAGTCGTTGCTTAGTCTGTTCTCACACTGCTGTNNNAGATCGGAAG
35158 TATGAGTGTGGAGTCGTTGCGCCTCCAGAGTAGCTGAGATTACNNNAGATCGGAAG
35159 TATGAGTGTGGAGTCGTTGCCTGGGACTATAGGCATGTACCACNNNAGATCGGAAG
35160 TATGAGTGTGGAGTCGTTGCCTGGCCCATGCTGATTTTACATTNNNAGATCGGAAG

35162 TATGAGTGTGGAGTCGTTGCTTCTGCCGATTCAAGCAATTCTGNNNAGATCGGAAG

35163 TATGAGTGTGGAGTCGTTGCGTCTCCCGGATTCAAGTGATTCTNNNAGATCGGAAG

35164 TATGAGTGTGGAGTCGTTGCCAATAGTGCGAAAGCAGCTCACNNNAGATCGGAAG

35165 TATGAGTGTGGAGTCGTTGCCGCCTCTCTTTCTCCATTGGAAANNNAGATCGGAAGA

35166 TATGAGTGTGGAGTCGTTGCATCTCGGCTCACTGAAAGCTTTANNNAGATCGGAAG

35167 TATGAGTGTGGAGTCGTTGCGGAATATTCCCTATTTCCACGGANNNAGATCGGAAG

35170 TATGAGTGTGGAGTCGTTGCATGAGGTTTTACCGTGTTAGCCANNNAGATCGGAAG

35171 TATGAGTGTGGAGTCGTTGCTCAGCTCTTGAACCTCTTCTTCCNNNAGATCGGAAGA

35172 TATGAGTGTGGAGTCGTTGCCCTAGTGAGTAGCTGGGACTACANNNAGATCGGAA

35173 TATGAGTGTGGAGTCGTTGCGTTGGTTAGGCTTGTCTTGAACCNNNAGATCGGAAG

35174 TATGAGTGTGGAGTCGTTGCAAGTGATTGTCCTTCCTCAGCTTNNNAGATCGGAAG

35175 TATGAGTGTGGAGTCGTTGCATCAGCTCTCTCTGGTCACTAGANNNAGATCGGAAG

35177 TATGAGTGTGGAGTCGTTGCGTGCCCTCAAGTTCAAGTGATTCNNNAGATCGGAAG

35178 TATGAGTGTGGAGTCGTTGCCACGCCGAGCTAATTTCTGTATTNNNAGATCGGAAG

35180 TATGAGTGTGGAGTCGTTGCGAGTGAGTGTAATGGCACGATCTNNNAGATCGGAA

35181 TATGAGTGTGGAGTCGTTGCTACTGAGGACCTGGTACTACAGGNNNAGATCGGAA

35182 TATGAGTGTGGAGTCGTTGCCACCACGCCCAGCTAATTTTTATNNNAGATCGGAAGA

35183 TATGAGTGTGGAGTCGTTGCTTTGTTCAATGAAGAGGTTGGGCNNNAGATCGGAA

35184 TATGAGTGTGGAGTCGTTGCCTCCTGAGAAGCTGGGATTACAANNNAGATCGGAA

35185 TATGAGTGTGGAGTCGTTGCTTTTGGCTCACTGTAACCTCAGANNNAGATCGGAAG

35186 TATGAGTGTGGAGTCGTTGCTATCTCCCGGGTTCAAGTGATTCNNNAGATCGGAAG

35187 TATGAGTGTGGAGTCGTTGCCAATTTTCTCCCTTACCCTCCCANNNAGATCGGAAGA

35189 TATGAGTGTGGAGTCGTTGCACCTCCTGGAATCAAGCAATTCTNNNAGATCGGAAG

35190 TATGAGTGTGGAGTCGTTGCGCATTCAAGGAATTCTCTGCCTCNNNAGATCGGAAG

35191 TATGAGTGTGGAGTCGTTGCCGACTAACTGGGATTACAGGCATNNNAGATCGGAA

35192 TATGAGTGTGGAGTCGTTGCTTCAAGTGATTCTCCTGCCTCAGNNNAGATCGGAAG

35193 TATGAGTGTGGAGTCGTTGCGGCCACCTGAAAGTACAAATCACNNNAGATCGGAAG

35194 TATGAGTGTGGAGTCGTTGCAGATGGGGTTTCATTCTTGTTGCNNNAGATCGGAAG

35195 TATGAGTGTGGAGTCGTTGCAGAGCTAGCTTCCCTGAAAAGACNNNAGATCGGAAG

35196 TATGAGTGTGGAGTCGTTGCTCCAAGTGATTTGTCTGCCTCAGNNNAGATCGGAAG

35197 TATGAGTGTGGAGTCGTTGCAACTCCTGGGTTCAAGCCATTATNNNAGATCGGAAG

35198 TATGAGTGTGGAGTCGTTGCCAATCTTGACTCACTGCAAGCTCNNNAGATCGGAAG

35199 TATGAGTGTGGAGTCGTTGCGGTATGATCTCAGCTCACCACAANNNAGATCGGAAG

35201 TATGAGTGTGGAGTCGTTGCAACCCCTGCTCAAGAGATTTTCTNNNAGATCGGAAG

35203 TATGAGTGTGGAGTCGTTGCCACCAACAAGCCTGGCTAATTTTNNNAGATCGGAAG

35204 TATGAGTGTGGAGTCGTTGCGTAGTTTTAGTGGAGACGGGGTTNNNAGATCGGAA

35205 TATGAGTGTGGAGTCGTTGCTACAGTTCAAAGGAACGGTTTGCNNNAGATCGGAA

35206 TATGAGTGTGGAGTCGTTGCGACATTGCCAGATCATCATGGTGNNNAGATCGGAAG

35210 TATGAGTGTGGAGTCGTTGCCAGGCTCCCAAATAGCTGGAATANNNAGATCGGAAG

35211 TATGAGTGTGGAGTCGTTGCTGAAAGAGCAAGAACCTGTGAGTNNNAGATCGGAA

35212 TATGAGTGTGGAGTCGTTGCCCTCCCAAGTAGCTGGAATTACANNNAGATCGGAAG

35213 TATGAGTGTGGAGTCGTTGCATCATGTTGGCCAGGCTAGTTTANNNAGATCGGAAG

35214 TATGAGTGTGGAGTCGTTGCTTCAGTGGCATGATCTTGTCTCANNNAGATCGGAAG

35215 TATGAGTGTGGAGTCGTTGCATGCCCACAAACACAAGTTTTCANNNAGATCGGAAG
35216 TATGAGTGTGGAGTCGTTGCGCCCCACATCTAGACACTGAAATNNNAGATCGGAAG
35217 TATGAGTGTGGAGTCGTTGCATCCTAAGCTAAGCCAACCAGTCNNNAGATCGGAAG
35218 TATGAGTGTGGAGTCGTTGCGGCTTCAGTTCACAGTCATGATGNNNAGATCGGAAG
35219 TATGAGTGTGGAGTCGTTGCGTCCAGAACCCAGTTACTGTGATNNNAGATCGGAAG
35220 TATGAGTGTGGAGTCGTTGCAGAACATTATTGGGGAGCCACATNNNAGATCGGAA
35221 TATGAGTGTGGAGTCGTTGCTGTGTGATAGATCCAGCTCTGTGNNNAGATCGGAA
35223 TATGAGTGTGGAGTCGTTGCCCTCACGAGTAGCTGGGACTATANNNAGATCGGAA
35224 TATGAGTGTGGAGTCGTTGCATGTAGGGCCATCAATGTTCTGTNNNAGATCGGAAG
35225 TATGAGTGTGGAGTCGTTGCTCCAGGCTAAAGTGATTCACCTGNNNAGATCGGAAG
35226 TATGAGTGTGGAGTCGTTGCACCCAATCAATTGTCTCTTGTGCNNNAGATCGGAAG
35228 TATGAGTGTGGAGTCGTTGCCATTTTGGAGGTTGCCTGTTCATNNNAGATCGGAAG
35229 TATGAGTGTGGAGTCGTTGCGTGATTTGGATGATGGAGGGTCTNNNAGATCGGAA
35230 TATGAGTGTGGAGTCGTTGCGTATACATGTGTCATGCTGGTGCNNNAGATCGGAA
35231 TATGAGTGTGGAGTCGTTGCCCCTCACCAGATTGAATCCTCAANNNAGATCGGAAG
35232 TATGAGTGTGGAGTCGTTGCCCTTCATTGCGGATTCATGATCCNNNAGATCGGAAG
35233 TATGAGTGTGGAGTCGTTGCCAGCCAGTGTCCAAATTCACTTCNNNAGATCGGAAG
35234 TATGAGTGTGGAGTCGTTGCGGTGATTTGAACTCCAGCTGTTCNNNAGATCGGAAG
35235 TATGAGTGTGGAGTCGTTGCACAGAGATAGGCCACAAAAAGGTNNNAGATCGGAA
35236 TATGAGTGTGGAGTCGTTGCGTTCAGTCAGCCTTCCAATTCACNNNAGATCGGAAG
35237 TATGAGTGTGGAGTCGTTGCGATTTTGACACTGACGGTCTTGGNNNAGATCGGAA
35238 TATGAGTGTGGAGTCGTTGCCCAATTTGCTGTGGATTCACGATNNNAGATCGGAAG
35239 TATGAGTGTGGAGTCGTTGCCCCTTTAGTTGATGACCTGTCCANNNAGATCGGAAG
35240 TATGAGTGTGGAGTCGTTGCGAGATGCTAGTAGGCAGAGACACNNNAGATCGGAA
35241 TATGAGTGTGGAGTCGTTGCTACTCAGGCTGGTCTTGAATTCCNNNAGATCGGAAG
35243 TATGAGTGTGGAGTCGTTGCTGGAGTACAGTGGTGTGATCTTGNNNAGATCGGAA
35244 TATGAGTGTGGAGTCGTTGCCCTGAGTATCTGGGATTACAGGCNNNAGATCGGAA
35246 TATGAGTGTGGAGTCGTTGCAGGTTCAAGCAATTCTCCTGTCTNNNAGATCGGAAG
35247 TATGAGTGTGGAGTCGTTGCTGTAGCTGGGACTATAGGTGCTANNNAGATCGGAA
35248 TATGAGTGTGGAGTCGTTGCCTGAAGACTGGTCCTGGATATGGNNNAGATCGGAA
35250 TATGAGTGTGGAGTCGTTGCTGAACTACTGACCTCAGGCAATCNNNAGATCGGAAG
35251 TATGAGTGTGGAGTCGTTGCAGGTTCTTCTGTCTGTGTAGCTGNNNAGATCGGAAG
35252 TATGAGTGTGGAGTCGTTGCCCTCAATCTCTTGACCTCGTGATNNNAGATCGGAAGA
35253 TATGAGTGTGGAGTCGTTGCCAGTGAAGTGGCACAATCTTGTCNNNAGATCGGAA
35254 TATGAGTGTGGAGTCGTTGCGTAGTGCAGTGGTTTGATCTTGGNNNAGATCGGAA
35256 TATGAGTGTGGAGTCGTTGCATACAATCTCTGCCACCCATGTTNNNAGATCGGAAGA
35257 TATGAGTGTGGAGTCGTTGCATACAATCTCTGCCACCCATGTTNNNAGATCGGAAGA
35258 TATGAGTGTGGAGTCGTTGCATACAATCTCTGCCACCCATGTTNNNAGATCGGAAGA
35259 TATGAGTGTGGAGTCGTTGCATACAATCTCTGCCACCCATGTTNNNAGATCGGAAGA
35260 TATGAGTGTGGAGTCGTTGCATACAATCTCTGCCACCCATGTTNNNAGATCGGAAGA
35261 TATGAGTGTGGAGTCGTTGCAGTGAGATCTTGGCTTACTGCAANNNAGATCGGAA
35262 TATGAGTGTGGAGTCGTTGCGGGTCCACTTTGATTCTGTTGTGNNNAGATCGGAAG
35263 TATGAGTGTGGAGTCGTTGCTGAGACTCAAGAAAAGTAGCGCTNNNAGATCGGAA

35264 TATGAGTGTGGAGTCGTTGCAATTTCAAGGCAATGCTGGAGTGNNNAGATCGGAA
35265 TATGAGTGTGGAGTCGTTGCGCGCTCTTCTTGACATTTCTTCANNNAGATCGGAAGA
35266 TATGAGTGTGGAGTCGTTGCGTGGTGTGCTCCTCTTCATTCTANNNAGATCGGAAG
35267 TATGAGTGTGGAGTCGTTGCATCCCAGACTGAAATCCCAAGACNNNAGATCGGAAG
35268 TATGAGTGTGGAGTCGTTGCAAGATGATATTTGAGAGCCCCCNNNAGATCGGAAG
35269 TATGAGTGTGGAGTCGTTGCCAGGCAGAGATAGTATGGACACCNNNAGATCGGAA
35270 TATGAGTGTGGAGTCGTTGCGTCACAACCAGCCTACCTATTCANNNAGATCGGAAG
35271 TATGAGTGTGGAGTCGTTGCCTCCTGAGTAGCTGGGACTACAGNNNAGATCGGAA
35273 TATGAGTGTGGAGTCGTTGCCCACACTGGATTGCATTGGTATCNNNAGATCGGAAG
35274 TATGAGTGTGGAGTCGTTGCGGTTTTGAACACCTGTCCTCAAGNNNAGATCGGAAG
35275 TATGAGTGTGGAGTCGTTGCAAACATGCAAAAGTGAGGGGAAGNNNAGATCGGA
35276 TATGAGTGTGGAGTCGTTGCAGATTTGTAGAGTTCACCCGTCCNNNAGATCGGAAG
35277 TATGAGTGTGGAGTCGTTGCTGTATTTTGGAGTGCATTGAGGCNNNAGATCGGAA
35278 TATGAGTGTGGAGTCGTTGCGAAACTTCTTTTTGACGTGTGCGNNNAGATCGGAAG
35279 TATGAGTGTGGAGTCGTTGCTTGAACTGTTTGTGGCCAATTGTNNNAGATCGGAAG
35280 TATGAGTGTGGAGTCGTTGCTGTAATTCCCTGGTTCAAGCGATNNNAGATCGGAAG
35282 TATGAGTGTGGAGTCGTTGCGAGATAGGGTCTTGCTCTGTCACNNNAGATCGGAA
35283 TATGAGTGTGGAGTCGTTGCACGATCTAAGCTCACTGCAACTTNNNAGATCGGAAG
35284 TATGAGTGTGGAGTCGTTGCTGGTGCAATCTCAGTCACTGTAANNNAGATCGGAAG
35285 TATGAGTGTGGAGTCGTTGCTTCAAGCTATTCTCCTGCCTCTGNNNAGATCGGAAGA
35286 TATGAGTGTGGAGTCGTTGCATCTAGTTGTCCTTGTGCCAGTTNNNAGATCGGAAG
35287 TATGAGTGTGGAGTCGTTGCCTCGGGTTTATGCCATTCTCGAGNNNAGATCGGAAG
35288 TATGAGTGTGGAGTCGTTGCACTGCAAATTCTGTCTCCTGGATNNNAGATCGGAAG
35289 TATGAGTGTGGAGTCGTTGCTTCAAGCAATTTTCCTGCCTCAGNNNAGATCGGAAG
35290 TATGAGTGTGGAGTCGTTGCCTGGCCAAGTTTCAGTCTTCTTGNNNAGATCGGAAG
35291 TATGAGTGTGGAGTCGTTGCCCTTTGGCCTTTTGAGACAAGAANNNAGATCGGAAG
35292 TATGAGTGTGGAGTCGTTGCCATGATCTCAGCTCAATGCAACCNNNAGATCGGAAG
35293 TATGAGTGTGGAGTCGTTGCTGGTCTCAAAATCCTGAGCTCAANNNAGATCGGAAG
35294 TATGAGTGTGGAGTCGTTGCTCTTGGGAGGTTGTATGTTTCCANNNAGATCGGAAG
35295 TATGAGTGTGGAGTCGTTGCAGAAGGAACTTGCCTTGAGTCTTNNNAGATCGGAA
35296 TATGAGTGTGGAGTCGTTGCTGGAGTAGCTGGGATTACAGGTANNNAGATCGGAA
35297 TATGAGTGTGGAGTCGTTGCCACCGCCTGCTAATTTTTGTCTTNNNAGATCGGAAGA
35300 TATGAGTGTGGAGTCGTTGCCCAAGCCCGGCTAATTTTTTGTANNNAGATCGGAAG
35301 TATGAGTGTGGAGTCGTTGCAGTATCACTCTATCCCTCAGGCTNNNAGATCGGAAG
35302 TATGAGTGTGGAGTCGTTGCGAGCCATCTTGGAGTCCCTTTTANNNAGATCGGAAG
35303 TATGAGTGTGGAGTCGTTGCGATCTTAGCTCACTGCAACCTCTNNNAGATCGGAAG
35304 TATGAGTGTGGAGTCGTTGCAGTCAGATCAGTCACCTTTTGCCNNNAGATCGGAAG
35305 TATGAGTGTGGAGTCGTTGCGGGGACTTGTACAAAAAAGTGGGNNNAGATCGGA
35309 TATGAGTGTGGAGTCGTTGCAGATGGAATGGAGTGCAGTTGTANNNAGATCGGAA
35310 TATGAGTGTGGAGTCGTTGCGGAGTACAGTGGTGTGATCTCAGNNNAGATCGGAA
35311 TATGAGTGTGGAGTCGTTGCTTCAAGTGATTCTCCTGCCTCAGNNNAGATCGGAAG
35312 TATGAGTGTGGAGTCGTTGCGAGTAGCTAGGATTACAGGTGCCNNNAGATCGGAA
35313 TATGAGTGTGGAGTCGTTGCTGCCCAGTGAAATGAATATCAGCNNNAGATCGGAA

35314 TATGAGTGTGGAGTCGTTGCCAGCCTCCTAAAGTACTGGGAAANNNAGATCGGAAG
35315 TATGAGTGTGGAGTCGTTGCATTTGTGTTTCAAGAGCTGCCAGNNNAGATCGGAAG
35317 TATGAGTGTGGAGTCGTTGCGGGAAGAGAGGAGGAATAGGAGANNNAGATCGGA
35318 TATGAGTGTGGAGTCGTTGCGACCGGAGTGTAGTTGTCTCAATNNNAGATCGGAAG
35320 TATGAGTGTGGAGTCGTTGCATTTACTGACCTGATGATCCGCCNNNAGATCGGAAGA
35321 TATGAGTGTGGAGTCGTTGCTTCATGCCATTCTCCTTCCTCAGNNNAGATCGGAAGA
35324 TATGAGTGTGGAGTCGTTGCCTCCTGAGTAGCTGGGACTACAGNNNAGATCGGAAG
35325 TATGAGTGTGGAGTCGTTGCCCAGGGGTGACTATTTACCAACANNNAGATCGGAAG
35326 TATGAGTGTGGAGTCGTTGCATGGAATCTCAGTCTGTTGTCCANNNAGATCGGAAG
35327 TATGAGTGTGGAGTCGTTGCAAGTAACACCACACCTGGCTAATNNNAGATCGGAAG
35328 TATGAGTGTGGAGTCGTTGCGACTTCAAGAATAGAGCCACGGANNNAGATCGGAAG
35329 TATGAGTGTGGAGTCGTTGCAACAATCTGGGTAAGCCTAGCATNNNAGATCGGAAG
35330 TATGAGTGTGGAGTCGTTGCACATGTGATTCCACCACTGATCANNNAGATCGGAAGA
35331 TATGAGTGTGGAGTCGTTGCTCAAGTGATTCTCCTGTCTCAGCNNNAGATCGGAAGA
35332 TATGAGTGTGGAGTCGTTGCGCCTCCTGAGTAGCTGGTATTACNNNAGATCGGAAG
35333 TATGAGTGTGGAGTCGTTGCAACCTCTGCCTCCTTAGTTCAAGNNNAGATCGGAAGA
35334 TATGAGTGTGGAGTCGTTGCTCATTGCAGCCTCAAATTCTTGGNNNAGATCGGAAGA
35335 TATGAGTGTGGAGTCGTTGCTTCGAGTAGCTGAGATTACAGGCNNNAGATCGGAAG
35336 TATGAGTGTGGAGTCGTTGCCTCCTCAGCTGGGATTACAGATGNNNAGATCGGAAG
35337 TATGAGTGTGGAGTCGTTGCTGTTGGGATTACAGGAGTAAGCCNNNAGATCGGAAG
35338 TATGAGTGTGGAGTCGTTGCGCAGGGAGGTAACTTGCATTTTTNNNAGATCGGAAG
35339 TATGAGTGTGGAGTCGTTGCCTCCTGAGTACCTGGGACTACAGNNNAGATCGGAAG
35340 TATGAGTGTGGAGTCGTTGCAGTGATTCTTCTGACTCAGCCTCNNNAGATCGGAAGA
35341 TATGAGTGTGGAGTCGTTGCAGTAGAGACAGATTTCGCCATGTNNNAGATCGGAAG
35342 TATGAGTGTGGAGTCGTTGCCCAGTAGAGATGAGGTCTTGCTGNNNAGATCGGAAG
35343 TATGAGTGTGGAGTCGTTGCCACCATGCCCAGCTAATTTTTGTNNNAGATCGGAAGA
35344 TATGAGTGTGGAGTCGTTGCGAGTCTCACTCTATTGCCTAGGCNNNAGATCGGAAGA
35345 TATGAGTGTGGAGTCGTTGCTCTCAGCCTCCTAAAGTACTGGANNNAGATCGGAAGA
35346 TATGAGTGTGGAGTCGTTGCATCACAGTTCACTACAGCCTTGANNNAGATCGGAAGA
35347 TATGAGTGTGGAGTCGTTGCATGATCTCAGCTCACTGCAACAGNNNAGATCGGAAG
35348 TATGAGTGTGGAGTCGTTGCCTCAGACTAGCAGGCCTAACAAANNNAGATCGGAAG
35349 TATGAGTGTGGAGTCGTTGCCATTGCTAAGAACCAGCTACAGCNNNAGATCGGAAG
35351 TATGAGTGTGGAGTCGTTGCCTCCTGAGTAGCTGGGACTACAGNNNAGATCGGAAG
35352 TATGAGTGTGGAGTCGTTGCCAATAGCAGCACAATCTCAGCTCNNNAGATCGGAAGA
35353 TATGAGTGTGGAGTCGTTGCCTGGGATTATAGGAGTGAGCCACNNNAGATCGGAAG
35354 TATGAGTGTGGAGTCGTTGCCTGAATAGCTGGGATTACAGGCANNNAGATCGGAAG
35355 TATGAGTGTGGAGTCGTTGCCAATCTCAACTCACTGCAAGCTCNNNAGATCGGAAGA
35357 TATGAGTGTGGAGTCGTTGCATTGCTTTGTACTGGAGGAGACCNNNAGATCGGAAG
35358 TATGAGTGTGGAGTCGTTGCGCTAGAGTACAGTGGTGCAATCTNNNAGATCGGAAG
35359 TATGAGTGTGGAGTCGTTGCTTCAAGCAATTCTCCTGCCTCAGNNNAGATCGGAAGA
35361 TATGAGTGTGGAGTCGTTGCGATCTCAGCTCACTGCAACATCTNNNAGATCGGAAGA
35363 TATGAGTGTGGAGTCGTTGCTGCATAGTATTCCATTGTGCACANNNAGATCGGAAGA
35364 TATGAGTGTGGAGTCGTTGCGCTTACTAAAACCTCTGCCTCCTNNNAGATCGGAAGA

35365 TATGAGTGTGGAGTCGTTGCCTGAGTAGCTGGGATTACAGACCNNNAGATCGGAA
35368 TATGAGTGTGGAGTCGTTGCTGACACAGCTCAGAGCAAATTTGNNNAGATCGGAA
35369 TATGAGTGTGGAGTCGTTGCTTCTGGACACCTGGTCATTTGATNNNAGATCGGAAG
35370 TATGAGTGTGGAGTCGTTGCTATGCTTCTGAACCTCCCTAGTGNNNAGATCGGAAG
35371 TATGAGTGTGGAGTCGTTGCGCTGGAGTACAATGGTGTGATCTNNNAGATCGGAA
35373 TATGAGTGTGGAGTCGTTGCAGGAAATTTTTTGCCTCAGCCTNNNAGATCGGAAG
35374 TATGAGTGTGGAGTCGTTGCGGCTCTTTTTTCAATCCAGGGACNNNAGATCGGAAG
35375 TATGAGTGTGGAGTCGTTGCTGCCTTACACAACTAAGTCACCTNNNAGATCGGAAG
35377 TATGAGTGTGGAGTCGTTGCTGGTTTCGAACTTCTGACCTCAANNNAGATCGGAAG
35378 TATGAGTGTGGAGTCGTTGCTTCTAGCGATTCTTCTGCCTCAGNNNAGATCGGAAG
35379 TATGAGTGTGGAGTCGTTGCCACGCCCGGCTAATTTTTTGTATNNNAGATCGGAAG
35382 TATGAGTGTGGAGTCGTTGCATGATCTCAGTTCACTGCAACCTNNNAGATCGGAAG
35383 TATGAGTGTGGAGTCGTTGCCAAGTAGCTGGGAATACAGGTGTNNNAGATCGGAA
35384 TATGAGTGTGGAGTCGTTGCAGACAGAGTCTCGCTTTGTCATCNNNAGATCGGAAG
35385 TATGAGTGTGGAGTCGTTGCAAGTGTTGGGATTACAGGCATGANNNAGATCGGAA
35386 TATGAGTGTGGAGTCGTTGCCTGAGTAGCTGGGACTACAAGTGNNNAGATCGGAA
35387 TATGAGTGTGGAGTCGTTGCCACCACCATGTCCAGCTAATTTTNNNAGATCGGAAG
35388 TATGAGTGTGGAGTCGTTGCGCTTACCGACTGAGGTCATAAGANNNAGATCGGAA
35389 TATGAGTGTGGAGTCGTTGCCAGAGTAGCTGAGATTACAGGCANNNAGATCGGAA
35390 TATGAGTGTGGAGTCGTTGCGGCTTGGTAATGGCAAATTTCCTNNNAGATCGGAAG
35391 TATGAGTGTGGAGTCGTTGCGGGCTCAAATGATTCTCCTGTCTNNNAGATCGGAAG
35392 TATGAGTGTGGAGTCGTTGCAATGTCTGGGATTACAGGCATGTNNNAGATCGGAA
35394 TATGAGTGTGGAGTCGTTGCCTAGAGTGAAGTGGCAGGATCTCNNNAGATCGGAA
35397 TATGAGTGTGGAGTCGTTGCCCTGAGTAGCTAGGACTACAGGTNNNAGATCGGAA
35398 TATGAGTGTGGAGTCGTTGCGATTCAAGCGATTCTCCTGCCTCNNNAGATCGGAAG
35399 TATGAGTGTGGAGTCGTTGCCACTTCCTCAGCTTCTGGAGTAGNNNAGATCGGAAG
35401 TATGAGTGTGGAGTCGTTGCCTCTCGAGCAGCTCAGATTACAGNNNAGATCGGAAG
35403 TATGAGTGTGGAGTCGTTGCCTAAAAGCACAAACCACCATCCCNNNAGATCGGAAG
35404 TATGAGTGTGGAGTCGTTGCAATACCTCATGAGCTCAAGCGATNNNAGATCGGAAG
35408 TATGAGTGTGGAGTCGTTGCCACTATAACATCCACCTCCTCGGNNNAGATCGGAAG
35409 TATGAGTGTGGAGTCGTTGCGTCACCATCTCTGCTCACCATAANNNAGATCGGAAG
35410 TATGAGTGTGGAGTCGTTGCTCATGATATTCTCCTGCCTCAGCNNNAGATCGGAAG
35411 TATGAGTGTGGAGTCGTTGCCATTACCCAAGAACCTGCAACAGNNNAGATCGGAAG
35413 TATGAGTGTGGAGTCGTTGCGGCATGATCTCGGATCACTGTAANNNAGATCGGAA
35414 TATGAGTGTGGAGTCGTTGCGGTATGATCTCAGCTCACTCCAGNNNAGATCGGAAG
35415 TATGAGTGTGGAGTCGTTGCGCTGGCTGCCTAAAATGAGTTAGNNNAGATCGGAA
35416 TATGAGTGTGGAGTCGTTGCATCATTGGGTGGAGGAAGAAACANNNAGATCGGAA
35417 TATGAGTGTGGAGTCGTTGCGATTACAAGTGCACACCACCATGNNNAGATCGGAAG
35418 TATGAGTGTGGAGTCGTTGCTCCAAGAGTGTTGTGAGCAAAACNNNAGATCGGAA
35419 TATGAGTGTGGAGTCGTTGCCAGGAGTGCAGTGGTATGATCATNNNAGATCGGAA
35420 TATGAGTGTGGAGTCGTTGCAGATGAGGGTCACACTATGTTGCNNNAGATCGGAA
35421 TATGAGTGTGGAGTCGTTGCGCCTGTAGCTAGAACAATAGGCANNNAGATCGGAA
35422 TATGAGTGTGGAGTCGTTGCAGTCTCAAACTCCTGAGCTCAAGNNNAGATCGGAAG

35423 TATGAGTGTGGAGTCGTTGCTAGCAGGGTCTCACTCTGTTACTNNNAGATCGGAAG
35425 TATGAGTGTGGAGTCGTTGCCTCACTGCAACCTCAAATTCCTGNNNAGATCGGAAGA
35426 TATGAGTGTGGAGTCGTTGCACCTCCCAGATTCAAGCAATTCTNNNAGATCGGAAG
35428 TATGAGTGTGGAGTCGTTGCCTGCAAACTCAGACCCCTAGATTNNNAGATCGGAAG
35429 TATGAGTGTGGAGTCGTTGCGCACAATCTCTGCTCACTTCAAGNNNAGATCGGAAG
35430 TATGAGTGTGGAGTCGTTGCTAGAGTGCAGTGAGTGATTTCGGNNNAGATCGGAA
35433 TATGAGTGTGGAGTCGTTGCCTCCTGAGTAGCTGGGACTACAGNNNAGATCGGAA
35434 TATGAGTGTGGAGTCGTTGCCAATGGTGCAATCATAGCTCACANNNAGATCGGAAG
35435 TATGAGTGTGGAGTCGTTGCCACCTCACCCGGCTAATTTTTTTNNNAGATCGGAAGA
35437 TATGAGTGTGGAGTCGTTGCAGTGATTATCTTGCCTCAGCCTCNNNAGATCGGAAG
35438 TATGAGTGTGGAGTCGTTGCCTCGCTCTCCTGAGTAATTGGAANNNAGATCGGAAG
35439 TATGAGTGTGGAGTCGTTGCGCCTCCCAGGTTCATGTTATTTTNNNAGATCGGAAG
35440 TATGAGTGTGGAGTCGTTGCCTCAAACTCCTGACCTCAAGTGANNNAGATCGGAAG
35441 TATGAGTGTGGAGTCGTTGCAGATGGAGTCTCACTCTTTCGCNNNAGATCGGAAGA
35443 TATGAGTGTGGAGTCGTTGCCCTAAAGTGCTGGCATACTAGGTNNNAGATCGGAA
35445 TATGAGTGTGGAGTCGTTGCTGAAAGAACCTCTAAGCCTGTCCNNNAGATCGGAAG
35446 TATGAGTGTGGAGTCGTTGCTCCAAGATGGACTTCAACTCCTGNNNAGATCGGAAG
35447 TATGAGTGTGGAGTCGTTGCCTCCTGTGTAACTGGGACTACAGNNNAGATCGGAAG
35448 TATGAGTGTGGAGTCGTTGCCAGGGTATAGTGCAGTGACATGANNNAGATCGGAA
35450 TATGAGTGTGGAGTCGTTGCTGGTAGAGATGCAGTTTCACCATNNNAGATCGGAA
35451 TATGAGTGTGGAGTCGTTGCGCACAATCTCAGCTAACTGCAAANNNAGATCGGAAG
35452 TATGAGTGTGGAGTCGTTGCCTCTTGAGTAGCTGGGACTACAGNNNAGATCGGAA
35453 TATGAGTGTGGAGTCGTTGCGACCTCTCATTCTCTGAAGCCAANNNAGATCGGAAG
35455 TATGAGTGTGGAGTCGTTGCTTCAAACTCCTGGGTTCAAGTGANNNAGATCGGAAG
35456 TATGAGTGTGGAGTCGTTGCAACTCAGCTTCTGACTCTTTGGTNNNAGATCGGAAG
35457 TATGAGTGTGGAGTCGTTGCAACCTCTGACTCCCAAGTTGAAGNNNAGATCGGAAG
35458 TATGAGTGTGGAGTCGTTGCGTTTCACCATGTTCCCTAAGCTGNNNAGATCGGAAG
35459 TATGAGTGTGGAGTCGTTGCTAGAGACAGGGTTTGTGTTAGCCNNNAGATCGGAA
35460 TATGAGTGTGGAGTCGTTGCCTGGAGTGTAGTTCTGCAGTCTTNNNAGATCGGAAG
35461 TATGAGTGTGGAGTCGTTGCCTTGGCCAAGATCAGTACTGGATNNNAGATCGGAA
35463 TATGAGTGTGGAGTCGTTGCAGGATGACACTGGCCTCATAAAANNNAGATCGGAA
35464 TATGAGTGTGGAGTCGTTGCCGCTTGGAATGGTTTCAGAAGAANNNAGATCGGAA
35465 TATGAGTGTGGAGTCGTTGCCCTCCCTTATTTCCTCAAGCAGTNNNAGATCGGAAGA
35466 TATGAGTGTGGAGTCGTTGCGGATTACAGGTATGAGCCACCATNNNAGATCGGAA
35468 TATGAGTGTGGAGTCGTTGCATTTTATGCCTCACTAGTGCCCANNNAGATCGGAAG
35469 TATGAGTGTGGAGTCGTTGCATGGCAAAATCTCAACTCACAGCNNNAGATCGGAAG
35470 TATGAGTGTGGAGTCGTTGCATAAGCCACAGCATCTGGACTACNNNAGATCGGAAG
35473 TATGAGTGTGGAGTCGTTGCTTGGCTCTCTCTTTGTCTACTGTNNNAGATCGGAAGA
35474 TATGAGTGTGGAGTCGTTGCGCCTCTTGGGTATCTGTGACTACNNNAGATCGGAAG
35477 TATGAGTGTGGAGTCGTTGCTTCATCCATGTCCCTACAAAGGANNNAGATCGGAAG
35478 TATGAGTGTGGAGTCGTTGCCTCCTTTTGCTGTAGGGCACTAGNNNAGATCGGAAG
35479 TATGAGTGTGGAGTCGTTGCACATTCATTTTGCTGGTGGATGGNNNAGATCGGAA
35480 TATGAGTGTGGAGTCGTTGCAGCCACCACATCTTTCCTACAATNNNAGATCGGAAGA

35481 TATGAGTGTGGAGTCGTTGCTGATCTCGGCTTACTACAAGCTCNNNAGATCGGAAG
35482 TATGAGTGTGGAGTCGTTGCCTCCTGAGTAGCTGGGACTACAGNNNAGATCGGAA
35483 TATGAGTGTGGAGTCGTTGCCTGGTCTCAAACTCCTGACTTGANNNAGATCGGAAG
35485 TATGAGTGTGGAGTCGTTGCTCAGTAGCATGATCTCAGCTCACNNNAGATCGGAAG
35486 TATGAGTGTGGAGTCGTTGCTGCCAAAAGAGCAGCATGAAAATNNNAGATCGGAA
35488 TATGAGTGTGGAGTCGTTGCAGTGCTGGGATTACAGACATGAGNNNAGATCGGAA
35490 TATGAGTGTGGAGTCGTTGCTTGTTGAGATGGGGTTTCACTGTNNNAGATCGGAA
35491 TATGAGTGTGGAGTCGTTGCTGGAATGAAATGGCACGATCTTGNNNAGATCGGAA
35492 TATGAGTGTGGAGTCGTTGCCTCAAGAGGATCACTTTGTCCCANNNAGATCGGAAG
35493 TATGAGTGTGGAGTCGTTGCCAGCCTCTTGAGTAGTTGGGATTNNNAGATCGGAA
35494 TATGAGTGTGGAGTCGTTGCTGATGTTACTCTTGGCAAGGTGTNNNAGATCGGAA
35495 TATGAGTGTGGAGTCGTTGCACTCCCAAAGTGCTGTGATTACANNNAGATCGGAAG
35496 TATGAGTGTGGAGTCGTTGCATCTTCCACAGGATTACCCAACANNNAGATCGGAAG
35497 TATGAGTGTGGAGTCGTTGCATTAGAGGTATGAGCCACTGTGCNNNAGATCGGAA
35498 TATGAGTGTGGAGTCGTTGCTGCAACAGGAGAGAGAAGATGACNNNAGATCGGA
35499 TATGAGTGTGGAGTCGTTGCAAATCTGGGAGCCAAGAAATCCTNNNAGATCGGAA
35500 TATGAGTGTGGAGTCGTTGCGGAGACAGGTTCTTGCTATGTCANNNAGATCGGAA
35501 TATGAGTGTGGAGTCGTTGCGCCCGGCCTCTAAAAGTATTTCTNNNAGATCGGAAG
35502 TATGAGTGTGGAGTCGTTGCCCAAGTAGCTGCGATTACAGAGANNNAGATCGGAA
35503 TATGAGTGTGGAGTCGTTGCTCAGCCTCCCAATTATTTGCAACNNNAGATCGGAAGA
35504 TATGAGTGTGGAGTCGTTGCGTAGAGACAGGATTTCACGGTGTNNNAGATCGGAA
35505 TATGAGTGTGGAGTCGTTGCCAATCTCCACCTCCTGAGTTCAANNNAGATCGGAAGA
35507 TATGAGTGTGGAGTCGTTGCTGGCCTGATTCAAAGAGATGTCANNNAGATCGGAA
35508 TATGAGTGTGGAGTCGTTGCTGATCTCGATCTCCTGACCTAGTNNNAGATCGGAAG
35510 TATGAGTGTGGAGTCGTTGCCACTATTGTTGCCCAGGTTTGAGNNNAGATCGGAAG
35511 TATGAGTGTGGAGTCGTTGCGGAATAGGTGGGGTTTGGTTACTNNNAGATCGGAA
35512 TATGAGTGTGGAGTCGTTGCATGATCTCAGCTCACTGCAATCTNNNAGATCGGAAG
35513 TATGAGTGTGGAGTCGTTGCGATCCACCCACATCCAAAGTACTNNNAGATCGGAAG
35515 TATGAGTGTGGAGTCGTTGCTGGTACATGTGATGTTTTGAGCCNNNAGATCGGAA
35516 TATGAGTGTGGAGTCGTTGCAGAGTTGGGATTACAGGCATGAGNNNAGATCGGA
35518 TATGAGTGTGGAGTCGTTGCCTGAGTAGGTGGGATTACAGGTGNNNAGATCGGA
35519 TATGAGTGTGGAGTCGTTGCGCCAGGATTATAGGCATGAGTCANNNAGATCGGAA
35520 TATGAGTGTGGAGTCGTTGCATACTGCCATGCCCAGCTAATTTNNNAGATCGGAAG
35521 TATGAGTGTGGAGTCGTTGCCTCCTGAGCAGCTGATACTACAGNNNAGATCGGAAG
35522 TATGAGTGTGGAGTCGTTGCCTGGTCTTGAACTCCTGAACTCANNNAGATCGGAAG
35523 TATGAGTGTGGAGTCGTTGCCCCAGGTTCAAGCAATTCTCTTGNNNAGATCGGAAG
35524 TATGAGTGTGGAGTCGTTGCTTTATGTTTCCCAGGCTGGTCTTNNNAGATCGGAAG
35525 TATGAGTGTGGAGTCGTTGCTCTTGAACTCCAGACCTCAAGTGNNNAGATCGGAAG
35526 TATGAGTGTGGAGTCGTTGCCTGAACCGAAGCAGATCAGAGATNNNAGATCGGAA
35527 TATGAGTGTGGAGTCGTTGCTGTTAGCCAGGATGGTCTTGATCNNNAGATCGGAA
35528 TATGAGTGTGGAGTCGTTGCCAATTGATTCTTCTGCCTCAGCCNNNAGATCGGAAGA
35529 TATGAGTGTGGAGTCGTTGCATAGGTTTTCACCATGCTTCCCANNNAGATCGGAAGA
35531 TATGAGTGTGGAGTCGTTGCAGGTTCAAGTGATTCTTCTGCCTNNNAGATCGGAAG

35533 TATGAGTGTGGAGTCGTTGCCACTCCTCTGATTCAGAAGGCTTNNNAGATCGGAAG
35534 TATGAGTGTGGAGTCGTTGCTCACTCAAGCTGGAGTACAGTTCNNNAGATCGGAAG
35535 TATGAGTGTGGAGTCGTTGCTGCCTAGGTTTTCTTCTAGGGTTNNNAGATCGGAAG
35537 TATGAGTGTGGAGTCGTTGCGTGGGATCTCTACTCACTGCAAGNNNAGATCGGAAG
35538 TATGAGTGTGGAGTCGTTGCCTGCAGGTCATTTTCAGCTGAAANNNAGATCGGAAG
35539 TATGAGTGTGGAGTCGTTGCTTTTGCCTCAGCCTTCTCAGTATNNNAGATCGGAAGA
35540 TATGAGTGTGGAGTCGTTGCTTCTGATTCACATGGACTTCCCCNNNAGATCGGAAGA
35541 TATGAGTGTGGAGTCGTTGCAATGCCCAACTTTCAGCATCTTCNNNAGATCGGAAGA
35542 TATGAGTGTGGAGTCGTTGCGTTACACCGTGTTAGCCAGGATANNNAGATCGGAAG
35544 TATGAGTGTGGAGTCGTTGCGTTTCACCATGTTAGCCAGGATGNNNAGATCGGAAG
35545 TATGAGTGTGGAGTCGTTGCCCTGCCACCAAATTCAGCTAATTNNNAGATCGGAAG
35546 TATGAGTGTGGAGTCGTTGCGCTGGGATTACAGATGTACACCANNNAGATCGGAA
35547 TATGAGTGTGGAGTCGTTGCTTACCAGGTGGGAAAGACTCTTCNNNAGATCGGAAG
35548 TATGAGTGTGGAGTCGTTGCAGTCTCAACTAACTGCAAGCTCTNNNAGATCGGAAG
35551 TATGAGTGTGGAGTCGTTGCTGGGAATAATGACCTGCAGAACANNNAGATCGGAA
35552 TATGAGTGTGGAGTCGTTGCTCATGACTCTGTTGGGAATGACANNNAGATCGGAA
35554 TATGAGTGTGGAGTCGTTGCATCTCCACCTCCTGAATTCAAGGNNNAGATCGGAAG
35555 TATGAGTGTGGAGTCGTTGCTTCAAACGGCTCTCCTTTCCTTANNNAGATCGGAAGA
35557 TATGAGTGTGGAGTCGTTGCGAGCTAGAAGTCTGGCCTCTTACNNNAGATCGGAAG
35558 TATGAGTGTGGAGTCGTTGCCTTATGGTGGGCAGTGTTGTTTTNNNAGATCGGAAG
35559 TATGAGTGTGGAGTCGTTGCGGCTTCCTAAAGTGCTGGGATTANNNAGATCGGAA
35560 TATGAGTGTGGAGTCGTTGCTGTATGTTGCCTGTTCACTCTGANNNAGATCGGAAG
35561 TATGAGTGTGGAGTCGTTGCCCTCCTGGTTACATGCCATTCTCNNNAGATCGGAAGA
35562 TATGAGTGTGGAGTCGTTGCCCGAGTAGCTGGGACTACAAATGNNNAGATCGGAA
35563 TATGAGTGTGGAGTCGTTGCCTTCTCTGAGTGTCTGGGATCACNNNAGATCGGAAG
35564 TATGAGTGTGGAGTCGTTGCTCTGCCTCACAAGTTCAAGTGATNNNAGATCGGAAG
35565 TATGAGTGTGGAGTCGTTGCCAATCTCAGCTCACTGCAACTTCNNNAGATCGGAAGA
35566 TATGAGTGTGGAGTCGTTGCGGATTACTGGCATAAACCACTGCNNNAGATCGGAAG
35568 TATGAGTGTGGAGTCGTTGCTAATCCTCTTGTTTCAGCCTCCCNNNAGATCGGAAGA
35569 TATGAGTGTGGAGTCGTTGCTTTCACCATGTTAGCCAGGATGANNNAGATCGGAAG
35570 TATGAGTGTGGAGTCGTTGCCCTCAAACCATCCTTCCACCTAANNNAGATCGGAAGA
35571 TATGAGTGTGGAGTCGTTGCGTTTCACTCTTGTTGCCCAGAATNNNAGATCGGAAG
35572 TATGAGTGTGGAGTCGTTGCATCTCTAACAAGTGGTTGCTCCANNNAGATCGGAAG
35573 TATGAGTGTGGAGTCGTTGCGGTTCACAAAATTCTCCTGCCTCNNNAGATCGGAAG
35574 TATGAGTGTGGAGTCGTTGCGCTCATGTCATCTCCCCTCTTAGNNNAGATCGGAAGA
35575 TATGAGTGTGGAGTCGTTGCCTAGCAAGCTAAAATCCACTGCCNNNAGATCGGAAG
35577 TATGAGTGTGGAGTCGTTGCTGACACACAATCTCAGCATCTCANNNAGATCGGAAG
35578 TATGAGTGTGGAGTCGTTGCTACACATATGCTGAGTAGGGCACNNNAGATCGGAA
35579 TATGAGTGTGGAGTCGTTGCTTCATGATGCCAGGCCAATTTTTNNNAGATCGGAAG
35580 TATGAGTGTGGAGTCGTTGCCCTCCCAAGCTGTAGGAGATTACNNNAGATCGGAAG
35581 TATGAGTGTGGAGTCGTTGCAGGGGATTAGAAAGAAGACAGCCNNNAGATCGGAA
35582 TATGAGTGTGGAGTCGTTGCCCCAAACCCATGAGGACATTTTCNNNAGATCGGAAG
35583 TATGAGTGTGGAGTCGTTGCCAGTCTCAGCTCACTACATCCTCNNNAGATCGGAAGA

35584 TATGAGTGTGGAGTCGTTGCAGGTTCATGCCATTCTCTTGCATNNNAGATCGGAAG
35585 TATGAGTGTGGAGTCGTTGCTTCAAGTGATTCTCCTGCCTCAGNNNAGATCGGAAG
35586 TATGAGTGTGGAGTCGTTGCGTTCAAGCAATTCTCTGCCTCAGNNNAGATCGGAAG
35587 TATGAGTGTGGAGTCGTTGCTACAGAACACAAGAGTCATGGGGNNNAGATCGGAA
35588 TATGAGTGTGGAGTCGTTGCTTGCTGCAATTATAGGCATCTGCNNNAGATCGGAAG
35590 TATGAGTGTGGAGTCGTTGCCATGATCTTGGCTCACTGCAATGNNNAGATCGGAAG
35591 TATGAGTGTGGAGTCGTTGCTCCAAGTGATTCTTCTGCCTCAGNNNAGATCGGAAG
35592 TATGAGTGTGGAGTCGTTGCTCCTGAGTAGCTGGGATTACAGANNNAGATCGGAA
35593 TATGAGTGTGGAGTCGTTGCTGAGCTGCCAAAATTTGAGATGGNNNAGATCGGAA
35594 TATGAGTGTGGAGTCGTTGCTCACTAGAGGCAAGAACAATCCANNNAGATCGGAA
35597 TATGAGTGTGGAGTCGTTGCAGTGCTGGGATTACAGATGTGAGNNNAGATCGGAA
35598 TATGAGTGTGGAGTCGTTGCCAAGCTCTTTTATGTGCCCACAANNNAGATCGGAAG
35599 TATGAGTGTGGAGTCGTTGCCAAGGGTATACAGAAGGTCCCACNNNAGATCGGAA
35600 TATGAGTGTGGAGTCGTTGCGATTACAGTGGCACAATCTTGGCNNNAGATCGGAA
35601 TATGAGTGTGGAGTCGTTGCCTCTGGGGATCCTTTGCTTTCTANNNAGATCGGAAG
35602 TATGAGTGTGGAGTCGTTGCGGCCCTTATCCTGTAACCACTACNNNAGATCGGAAG
35604 TATGAGTGTGGAGTCGTTGCGGGTGTAGGTTTTGCTTTGTTGTNNNAGATCGGAA
35605 TATGAGTGTGGAGTCGTTGCGATGAACAAGGGCAGACTGATTGNNNAGATCGGAA
35606 TATGAGTGTGGAGTCGTTGCTCAAGTGATTCTCCTGCCTTAGCNNNAGATCGGAAG
35607 TATGAGTGTGGAGTCGTTGCGGCCTAGAGCTATTCAAGTTTGCNNNAGATCGGAAG
35610 TATGAGTGTGGAGTCGTTGCCTCCTCAGCTTCCCAAGTAGTTGNNNAGATCGGAAG
35611 TATGAGTGTGGAGTCGTTGCTCATCCTCATGGTCTGTTCATGGNNNAGATCGGAAG
35612 TATGAGTGTGGAGTCGTTGCCACGATCTCAGCTCATTGCAAGNNNAGATCGGAAGA
35613 TATGAGTGTGGAGTCGTTGCAAGCTTCTTTTCTGTGATGGCTGNNNAGATCGGAAG
35614 TATGAGTGTGGAGTCGTTGCCTCAAAGTGCTGGGATTACAAGCNNNAGATCGGAA
35615 TATGAGTGTGGAGTCGTTGCTCCTGGGTTCAATGATTCTCCTGNNNAGATCGGAAG
35616 TATGAGTGTGGAGTCGTTGCCACAGTACCAGGCGTTATCAGTANNNAGATCGGAAG
35617 TATGAGTGTGGAGTCGTTGCTCAAGAAATTCTCCCTCCTCAGCNNNAGATCGGAAGA
35618 TATGAGTGTGGAGTCGTTGCAACCTCCTGAGTAACTGGGACTANNNAGATCGGAAG
35620 TATGAGTGTGGAGTCGTTGCTTCACTGCACCTGGCCTATAAATNNNAGATCGGAAG
35621 TATGAGTGTGGAGTCGTTGCAATCCAGCTGTGAACACATTTGGNNNAGATCGGAAG
35622 TATGAGTGTGGAGTCGTTGCTTGTTCCTTAGGTTGGTGTGTCTNNNAGATCGGAAG
35623 TATGAGTGTGGAGTCGTTGCCCTCCCAGATTCATGCCATTCTCNNNAGATCGGAAGA
35624 TATGAGTGTGGAGTCGTTGCTGTTAGCCAGGATGGTCTTGATCNNNAGATCGGAA
35626 TATGAGTGTGGAGTCGTTGCGAAGGTGAGTCTCGTTCTGTCACNNNAGATCGGAA
35629 TATGAGTGTGGAGTCGTTGCCAACAAGGCTGGGAGTATAACCTNNNAGATCGGAA
35630 TATGAGTGTGGAGTCGTTGCGTTCATGCCATTCTCCTACCTCANNNAGATCGGAAGA
35631 TATGAGTGTGGAGTCGTTGCAGAAGGTCAGCAGATAAACACGTNNNAGATCGGAA
35634 TATGAGTGTGGAGTCGTTGCCTCAAACCCCTGACCTAGTGATCNNNAGATCGGAAG
35635 TATGAGTGTGGAGTCGTTGCGTCCAAGTGATTATCCTGCCTCANNNAGATCGGAAG
35636 TATGAGTGTGGAGTCGTTGCTTTTTGGAGACTGAGTCCCACTCNNNAGATCGGAAG
35639 TATGAGTGTGGAGTCGTTGCTCCTGAATACAGCACACTGATGGNNNAGATCGGAAG
35641 TATGAGTGTGGAGTCGTTGCTGGTGAGTCTTGGGAATGCATAANNNAGATCGGAA

35642 TATGAGTGTGGAGTCGTTGCGGACTCAAGTGAACCTCATGTCTNNNAGATCGGAAG
35643 TATGAGTGTGGAGTCGTTGCTGGGATTATAGGAATGTGCCACCNNNAGATCGGAA
35644 TATGAGTGTGGAGTCGTTGCGTCATTGCTTTTGACCTGCCTAGNNNAGATCGGAAG
35646 TATGAGTGTGGAGTCGTTGCATCATCTCAGCTCAACACAACCTNNNAGATCGGAAG
35647 TATGAGTGTGGAGTCGTTGCCTGGAATGCAATGGTGTGATCTCNNNAGATCGGAA
35648 TATGAGTGTGGAGTCGTTGCAGACAGGTCTGGTGGTGAAAAATNNNAGATCGGAA
35649 TATGAGTGTGGAGTCGTTGCAGTCTTGACAGGCTGTATGTGTCNNNAGATCGGAA
35650 TATGAGTGTGGAGTCGTTGCTTTAGTAGAGGCGGGGTTTTACCNNNAGATCGGAA
35651 TATGAGTGTGGAGTCGTTGCTTTAGTAGAGGCGGGGTTTTACCNNNAGATCGGAA
35652 TATGAGTGTGGAGTCGTTGCGATTTTTCTGCCTCAGCTTCCTGNNNAGATCGGAAG
35653 TATGAGTGTGGAGTCGTTGCTATCTCCTCCCAGTTTCAGCCTANNNAGATCGGAAGA
35655 TATGAGTGTGGAGTCGTTGCGTTTCACCATGTTAGCCAGGATGNNNAGATCGGAAG
35657 TATGAGTGTGGAGTCGTTGCCTTCTGGGTCAAGCTCTTCTTCTNNNAGATCGGAAG
35658 TATGAGTGTGGAGTCGTTGCTGTGTCAGATCAGAGTTTCACAANNNAGATCGGAAG
35659 TATGAGTGTGGAGTCGTTGCCACCTCTGCCATCTCCTAGATTCNNNAGATCGGAAGA
35660 TATGAGTGTGGAGTCGTTGCGATGGAGTTTTACTCTTGGTGCCNNNAGATCGGAAG
35661 TATGAGTGTGGAGTCGTTGCCAGTGGCCAGATCTAAACTCACTNNNAGATCGGAAG
35662 TATGAGTGTGGAGTCGTTGCACCTAGGATGGTCTCAATCTCCTNNNAGATCGGAAG
35663 TATGAGTGTGGAGTCGTTGCTGTTAGCCAGGATGGTCTCAATCNNNAGATCGGAAG
35664 TATGAGTGTGGAGTCGTTGCCAAAGGGAGGTGATTTGTATGCCNNNAGATCGGAA
35665 TATGAGTGTGGAGTCGTTGCTGCCCAGCTACAAATATCAGTGGNNNAGATCGGAAG
35666 TATGAGTGTGGAGTCGTTGCCATGCTCTTTCCATCCTCCATCTNNNAGATCGGAAGA
35667 TATGAGTGTGGAGTCGTTGCCTTTGGCTGAGAGAGAACAGGATNNNAGATCGGAA
35668 TATGAGTGTGGAGTCGTTGCTGGAGGTGGAGTCTCATTTTGTCNNNAGATCGGAA
35669 TATGAGTGTGGAGTCGTTGCCAAGTGTTGGATGTCTGCATGTTNNNAGATCGGAA
35670 TATGAGTGTGGAGTCGTTGCTTCAGAGTTGCCATCACCACATANNNAGATCGGAAG
35671 TATGAGTGTGGAGTCGTTGCCCAGGCCTGATTTTTGAGTTCTGNNNAGATCGGAAG
35672 TATGAGTGTGGAGTCGTTGCGATTGTCTTGGTGGCTCTTCTTGNNNAGATCGGAAG
35673 TATGAGTGTGGAGTCGTTGCCTTCAGCAGCACTACTGGACATANNNAGATCGGAAG
35675 TATGAGTGTGGAGTCGTTGCCCACCACTCTCGGCTAATTTTTGNNNAGATCGGAAG
35676 TATGAGTGTGGAGTCGTTGCCTTGATCTCCTGACCTCGTGATCNNNAGATCGGAAG
35678 TATGAGTGTGGAGTCGTTGCCAATGTTTGCTTAGGCTGGTCTCNNNAGATCGGAAG
35680 TATGAGTGTGGAGTCGTTGCCCACACCCGGCTAATTTTTTGTANNNAGATCGGAAG
35681 TATGAGTGTGGAGTCGTTGCTCCTGAGTAGCTGGGATTACAGTNNNAGATCGGAA
35682 TATGAGTGTGGAGTCGTTGCATCCTCACCCTTATCTGACCTCANNNAGATCGGAAGA
35683 TATGAGTGTGGAGTCGTTGCGAGGTGATTTGGATGTTGGCATGNNNAGATCGGAA
35684 TATGAGTGTGGAGTCGTTGCGCAGGCTCATGATTCACAGAAAANNNAGATCGGAA
35685 TATGAGTGTGGAGTCGTTGCTCCCAAAATGATGGAGGAGTGTTNNNAGATCGGAA
35686 TATGAGTGTGGAGTCGTTGCCAGTTCACTGCGGATTCATGATCNNNAGATCGGAAG
35687 TATGAGTGTGGAGTCGTTGCTCACTGCAGATTCATGATCCACANNNAGATCGGAAG
35688 TATGAGTGTGGAGTCGTTGCGAAAGACCCCTAGCAAGGCTAATNNNAGATCGGAA
35689 TATGAGTGTGGAGTCGTTGCGCGATGGTATGCAATCTCAGTTCNNNAGATCGGAAG
35690 TATGAGTGTGGAGTCGTTGCATGCTAAGTAGGGCACTCTGTTCNNNAGATCGGAAG

35691 TATGAGTGTGGAGTCGTTGCATTATCAGGTCACCACAACCACCNNNAGATCGGAAG
35693 TATGAGTGTGGAGTCGTTGCCAGAACACAGAATGGAATGCCTGNNNAGATCGGAA
35694 TATGAGTGTGGAGTCGTTGCGGAGTCTTACTCTATTGCCCAGGNNNAGATCGGAAG
35695 TATGAGTGTGGAGTCGTTGCCCACCACATTCTTGAACACTGACNNNAGATCGGAAG
35696 TATGAGTGTGGAGTCGTTGCAAACAAGAGCTTTGCATTCAGGCNNNAGATCGGAA
35697 TATGAGTGTGGAGTCGTTGCGGCATGATTTTGGCTCACTACAANNNAGATCGGAAG
35699 TATGAGTGTGGAGTCGTTGCGCTCGTGATCACAGAGGACATATNNNAGATCGGAA
35700 TATGAGTGTGGAGTCGTTGCGTACTAGGGTTACAGGTGTGAGCNNNAGATCGGAA
35701 TATGAGTGTGGAGTCGTTGCCAAAACTGCTGGGATTACAGGTGNNNAGATCGGAA
35702 TATGAGTGTGGAGTCGTTGCGCTCAATTGATCCTCCCACCTTANNNAGATCGGAAGA
35705 TATGAGTGTGGAGTCGTTGCTGCAACTTCTGTCTCCTAGGTTCNNNAGATCGGAAG
35706 TATGAGTGTGGAGTCGTTGCAAACATTTCACCCGCTCCTTTTTNNNAGATCGGAAGA
35707 TATGAGTGTGGAGTCGTTGCACAGAAAGAGGGGAAAACCTGTTNNNAGATCGGAA
35709 TATGAGTGTGGAGTCGTTGCACTTGAGAAGCTGGGATTACAGGNNNAGATCGGAA
35710 TATGAGTGTGGAGTCGTTGCGATCTTGGCTCACTGAAAGCTCTNNNAGATCGGAAG
35711 TATGAGTGTGGAGTCGTTGCGGTTCAAGCAATTCTCCTGTCTCNNNAGATCGGAAG
35712 TATGAGTGTGGAGTCGTTGCTACAGTTCAAAGGAACGGTTTGCNNNAGATCGGAA
35713 TATGAGTGTGGAGTCGTTGCTGCCCAGCTACAAATATCAGTGGNNNAGATCGGAAG
35714 TATGAGTGTGGAGTCGTTGCCATGCTCTTTCCATCCTCCATCTNNNAGATCGGAAGA
35715 TATGAGTGTGGAGTCGTTGCCTTTGGCTGAGAGAGAACAGGATNNNAGATCGGAA
35716 TATGAGTGTGGAGTCGTTGCACGCCCGGCTAATTTTTTGTATTNNNAGATCGGAAG
35717 TATGAGTGTGGAGTCGTTGCTGGAGGTGGAGTCTCATTTTGTCNNNAGATCGGAA
35718 TATGAGTGTGGAGTCGTTGCCCTGGACTGACTACAACCTCAAANNNAGATCGGAAG
35720 TATGAGTGTGGAGTCGTTGCTGAGCCAAGCCAACTAGTACTTTNNNAGATCGGAAG
35721 TATGAGTGTGGAGTCGTTGCCTTGCTCTTTACACAGGGAAAGCNNNAGATCGGAAG
35722 TATGAGTGTGGAGTCGTTGCGCAATTTGCCAGTCTGTGTCTTTNNNAGATCGGAAG
35723 TATGAGTGTGGAGTCGTTGCTCTGGGCTCAAGTGATTTGTTCTNNNAGATCGGAAG
35724 TATGAGTGTGGAGTCGTTGCCAGCCTCCTCAATACCTGACATTNNNAGATCGGAAGA
35725 TATGAGTGTGGAGTCGTTGCATCAAGCAATTCTCCTGCCTCANNNAGATCGGAAGA
35728 TATGAGTGTGGAGTCGTTGCCATTCTAACAGTCAGGCCCTTCTNNNAGATCGGAAG
35730 TATGAGTGTGGAGTCGTTGCTCTGACCTTTTCCACTAAGTGCANNNAGATCGGAAG
35732 TATGAGTGTGGAGTCGTTGCTCCTGAGTAGCTGGGATTACAGANNNAGATCGGAA
35735 TATGAGTGTGGAGTCGTTGCTGGTAGTTTCTTTTGCTGTGCAGNNNAGATCGGAAG
35737 TATGAGTGTGGAGTCGTTGCACAGAAACGAGTTCTCACCATGTNNNAGATCGGAAG
35738 TATGAGTGTGGAGTCGTTGCCTAGGTTCAAGCAATTCTCCTGCNNNAGATCGGAAG
35739 TATGAGTGTGGAGTCGTTGCATTGTCTGCTTCCTTTCTCAGCTNNNAGATCGGAAGA
35740 TATGAGTGTGGAGTCGTTGCAGAGTAGCTGGGACTACAGATGTNNNAGATCGGAA
35741 TATGAGTGTGGAGTCGTTGCATCAGTTTCGTGCAGTTCAACAGNNNAGATCGGAAG
35742 TATGAGTGTGGAGTCGTTGCCCTCCCGAGTAGCTAGAACTACANNNAGATCGGAAG
35743 TATGAGTGTGGAGTCGTTGCCAGTGCAATCATGGCTCATTGTANNNAGATCGGAAG
35744 TATGAGTGTGGAGTCGTTGCCTGGAGTGTAATGGTGAGCTCTCNNNAGATCGGAA
35745 TATGAGTGTGGAGTCGTTGCGAGACGGAGTCTTGTTCTGTCACNNNAGATCGGAA
35746 TATGAGTGTGGAGTCGTTGCTTCAAGTGATTCTCCTGCCTCAGNNNAGATCGGAAG

35747 TATGAGTGTGGAGTCGTTGCGTTTCACTGTGTTAGCCAGGATGNNNAGATCGGAAG
35748 TATGAGTGTGGAGTCGTTGCGAGTCAGAATCTTGCTCTGTTGCNNNAGATCGGAAG
35749 TATGAGTGTGGAGTCGTTGCGCCTCCTGAGTAGCTGAGATTACNNNAGATCGGAAG
35750 TATGAGTGTGGAGTCGTTGCTTCAAGTGATTCTCCCACCTCAGNNNAGATCGGAAGA
35752 TATGAGTGTGGAGTCGTTGCGCTCGTGATCACAGAGGACATATNNNAGATCGGAAG
35753 TATGAGTGTGGAGTCGTTGCGTACTAGGGTTACAGGTGTGAGCNNNAGATCGGAA
35755 TATGAGTGTGGAGTCGTTGCATGACAGACATTGAGGAAAGCCANNNAGATCGGAAG
35756 TATGAGTGTGGAGTCGTTGCCAAAACTGCTGGGATTACAGGTGNNNAGATCGGAAG
35757 TATGAGTGTGGAGTCGTTGCGCTCAATTGATCCTCCCACCTTANNNAGATCGGAAGA
35760 TATGAGTGTGGAGTCGTTGCAAACATTTCACCCGCTCCTTTTTNNNAGATCGGAAGA
35761 TATGAGTGTGGAGTCGTTGCACAGAAAGAGGGGAAAACCTGTTNNNAGATCGGAA
35764 TATGAGTGTGGAGTCGTTGCATTACAGGAGTGAGCCAACACATNNNAGATCGGAAG
35766 TATGAGTGTGGAGTCGTTGCTCCCTTTATTTCTCAGACCAGCCNNNAGATCGGAAGA
35767 TATGAGTGTGGAGTCGTTGCTCAGACGCATTATGATCAAGCTGNNNAGATCGGAAG
35768 TATGAGTGTGGAGTCGTTGCACCTGCATAGTGACACATTGCTANNNAGATCGGAAG
35769 TATGAGTGTGGAGTCGTTGCTACTTGAGGCTATACCCAGGTGANNNAGATCGGAAG
35770 TATGAGTGTGGAGTCGTTGCAATAATAGGTGTTCCTGGCTGGGNNNAGATCGGAAG
35771 TATGAGTGTGGAGTCGTTGCGCCTTCTGAGTAGCTAGGATGACNNNAGATCGGAAG
35772 TATGAGTGTGGAGTCGTTGCGAGTTCAAGCGATTCTTTTGCCTNNNAGATCGGAAGA
35773 TATGAGTGTGGAGTCGTTGCAGTGTCATAAGAGAACCCAGCAGNNNAGATCGGAAG
35774 TATGAGTGTGGAGTCGTTGCCACCCAGATGATGTTCCTCTTCANNNAGATCGGAAGA
35775 TATGAGTGTGGAGTCGTTGCCTCCCAATTCCAAGTCTCACGTANNNAGATCGGAAGA
35778 TATGAGTGTGGAGTCGTTGCGCTGGAGTGTAGTGATGTGATCTNNNAGATCGGAAG
35779 TATGAGTGTGGAGTCGTTGCATTCCCTGGTTCGAGTGATTCTCNNNAGATCGGAAGA
35780 TATGAGTGTGGAGTCGTTGCGTTCATACCATTCTCCTGCCTCANNNAGATCGGAAGA
35781 TATGAGTGTGGAGTCGTTGCCGAGTAGCTGGTATTACAGGCATNNNAGATCGGAAG
35782 TATGAGTGTGGAGTCGTTGCCAGAGTCTTGCTGTGTCACCTAGNNNAGATCGGAAG
35783 TATGAGTGTGGAGTCGTTGCAGCCTTGACCTCATGGACTAAAGNNNAGATCGGAAG
35784 TATGAGTGTGGAGTCGTTGCAGTAGCTGGGACTACAGGTATGTNNNAGATCGGAAG
35785 TATGAGTGTGGAGTCGTTGCCAGAGACTCATTGTGTCACCCAGNNNAGATCGGAAG
35787 TATGAGTGTGGAGTCGTTGCGATCTCAGCTCATCACAACCTCCNNNAGATCGGAAGA
35788 TATGAGTGTGGAGTCGTTGCCTGGTATGTTCCTGTTTCCCTCANNNAGATCGGAAGA
35789 TATGAGTGTGGAGTCGTTGCATTGTCTGCTTCCTTTCTCAGCTNNNAGATCGGAAGA
35791 TATGAGTGTGGAGTCGTTGCTGGTAGTTTCTTTTGCTGTGCAGNNNAGATCGGAAG
35792 TATGAGTGTGGAGTCGTTGCTTCAAACGGCTCTCCTTTCCTTANNNAGATCGGAAGA
35793 TATGAGTGTGGAGTCGTTGCCTTCCCGGATTCATGCAATTCTCNNNAGATCGGAAGA
35794 TATGAGTGTGGAGTCGTTGCAGCTCATGTATTCTGCCCGTTTANNNAGATCGGAAGA
35795 TATGAGTGTGGAGTCGTTGCTTAGGGTACATGTGCACATTGTGNNNAGATCGGAAG
35796 TATGAGTGTGGAGTCGTTGCATTCTGTAGGTTGCCTGTTCACTNNNAGATCGGAAGA
35797 TATGAGTGTGGAGTCGTTGCCTTGGACCCCCTGTATTCTTTCANNNAGATCGGAAGA
35798 TATGAGTGTGGAGTCGTTGCTATGACGTATGAGCCACCATGTCNNNAGATCGGAAG
35799 TATGAGTGTGGAGTCGTTGCCCATTTCAGCCTCCCAAGTATCTNNNAGATCGGAAGA
35802 TATGAGTGTGGAGTCGTTGCTCATGTGCCTGCCTTGACTTATANNNAGATCGGAAGA

35803 TATGAGTGTGGAGTCGTTGCCAGAAGGCCTAATGCACAAGAACNNNAGATCGGAA
35804 TATGAGTGTGGAGTCGTTGCCCAAGAACCTGGGATAATAGCCANNNAGATCGGAA
35805 TATGAGTGTGGAGTCGTTGCCAGGAATTCATCCCACTGAAACANNNAGATCGGAAG
35806 TATGAGTGTGGAGTCGTTGCGAAAGACCCCTAGCAAGGCTAATNNNAGATCGGAA
35807 TATGAGTGTGGAGTCGTTGCGCGATGGTATGCAATCTCAGTTCNNNAGATCGGAAG
35808 TATGAGTGTGGAGTCGTTGCTTTTTTGAGATGGAGTCTGCCCTNNNAGATCGGAAG
35809 TATGAGTGTGGAGTCGTTGCATGCTAAGTAGGGCACTCTGTTCNNNAGATCGGAAG
35810 TATGAGTGTGGAGTCGTTGCATTATCAGGTCACCACAACCACCNNNAGATCGGAAG
35812 TATGAGTGTGGAGTCGTTGCCAGAACACAGAATGGAATGCCTGNNNAGATCGGAA
35813 TATGAGTGTGGAGTCGTTGCGGAGTCTTACTCTATTGCCCAGGNNNAGATCGGAAG
35814 TATGAGTGTGGAGTCGTTGCCCACCACATTCTTGAACACTGACNNNAGATCGGAAG
35815 TATGAGTGTGGAGTCGTTGCAAACAAGAGCTTTGCATTCAGGCNNNAGATCGGAA
35816 TATGAGTGTGGAGTCGTTGCGGCATGATTTTGGCTCACTACAANNNAGATCGGAAG
35817 TATGAGTGTGGAGTCGTTGCTCCTGGGTTCAAAGTGATTCTCCNNNAGATCGGAAG
35818 TATGAGTGTGGAGTCGTTGCCATTCTGACTGGTGTGAGATGGTNNNAGATCGGAA
35819 TATGAGTGTGGAGTCGTTGCACTGCAACATCTACCTCCCAAATNNNAGATCGGAAG
35820 TATGAGTGTGGAGTCGTTGCCAATCTCGGCTCACTGCAATTTCNNNAGATCGGAAG
35821 TATGAGTGTGGAGTCGTTGCCTGGACATTACAGGGCATCTTCTNNNAGATCGGAAG
35822 TATGAGTGTGGAGTCGTTGCAGTAGGTGCAATCTTGGCTAAGANNNAGATCGGAA
35823 TATGAGTGTGGAGTCGTTGCCCACTACTCCCGGCTAATTTTTTNNNAGATCGGAAGA
35824 TATGAGTGTGGAGTCGTTGCTTCAAGCAATTCTCCTGCCTCAGNNNAGATCGGAAG
35827 TATGAGTGTGGAGTCGTTGCAAGTGATTCTTGTTCCTCAGCCTNNNAGATCGGAAG
35829 TATGAGTGTGGAGTCGTTGCTAGAGTACAATGGCGGGATCTTGNNNAGATCGGAA
35830 TATGAGTGTGGAGTCGTTGCCAATCTTGTCTCACTGCAACCTCNNNAGATCGGAAGA
35831 TATGAGTGTGGAGTCGTTGCGATTCAAGCAATTCTCCTGCCTCNNNAGATCGGAAG
35833 TATGAGTGTGGAGTCGTTGCCACTTTGCCCAGCCTACATTTTTNNNAGATCGGAAGA
35836 TATGAGTGTGGAGTCGTTGCCAATTCTCCTATGTCAGCCTCCANNNAGATCGGAAGA
35837 TATGAGTGTGGAGTCGTTGCCAATAGCAGAATCTCGGCTCACTNNNAGATCGGAAG
35838 TATGAGTGTGGAGTCGTTGCCTGAGTAGCTGGGATTACAGGTGNNNAGATCGGAA
35839 TATGAGTGTGGAGTCGTTGCTTCAAGCTATTCTCCTGCCTCAGNNNAGATCGGAAG
35840 TATGAGTGTGGAGTCGTTGCAGGTTCAAAAGATTCTCCTGCCTNNNAGATCGGAAG
35841 TATGAGTGTGGAGTCGTTGCAGATCCTCCTACCTTGGACTCTCNNNAGATCGGAAG
35842 TATGAGTGTGGAGTCGTTGCGGTTCAAGCAATTATCCTGCCTCNNNAGATCGGAAG
35843 TATGAGTGTGGAGTCGTTGCCCACGCCCAGCTAATTTTTTGTANNNAGATCGGAAG
35844 TATGAGTGTGGAGTCGTTGCCTCCTGAGTAGCTGGGATTACAGNNNAGATCGGAA
35846 TATGAGTGTGGAGTCGTTGCGATTCAAGCAATTCTCCTGCCTCNNNAGATCGGAAG
35847 TATGAGTGTGGAGTCGTTGCGCATGATCTTGGCTCACTACAACNNNAGATCGGAAG
35848 TATGAGTGTGGAGTCGTTGCCAATTCTCCTATCTCAGCCTCCCNNNAGATCGGAAGA
35850 TATGAGTGTGGAGTCGTTGCCCTCTGGAGTATCTGGGACTACANNNAGATCGGAAG
35851 TATGAGTGTGGAGTCGTTGCCCATGATGAGACTATCAGCACCANNNAGATCGGAAG
35852 TATGAGTGTGGAGTCGTTGCAAGGTCTCACTTTGTTGCCTAGGNNNAGATCGGAAG
35853 TATGAGTGTGGAGTCGTTGCGTTCAGATGATTCTCCTGCCTCANNNAGATCGGAAG
35854 TATGAGTGTGGAGTCGTTGCTTCAAGTGATCCTCCTGCTTCAGNNNAGATCGGAAG

35855 TATGAGTGTGGAGTCGTTGCTAGTCTTGACCTCCTGACCTCAANNNAGATCGGAAGA
35856 TATGAGTGTGGAGTCGTTGCGCTCCTGGCCTCTTTTCTTTTTTNNNAGATCGGAAGA
35857 TATGAGTGTGGAGTCGTTGCTTTTTAGTAGAGACGGGGGCTTCNNNAGATCGGAA
35858 TATGAGTGTGGAGTCGTTGCGTTCAAGCGATTCTCCTACCTCANNNAGATCGGAAGA
35860 TATGAGTGTGGAGTCGTTGCCAGTGCACACTTTGAAATGGGTANNNAGATCGGAAG
35861 TATGAGTGTGGAGTCGTTGCCTGTCATCTCAGCACTTTGGGAGNNNAGATCGGAAG
35862 TATGAGTGTGGAGTCGTTGCCTACCGGGAAGGCTGAGATAATCNNNAGATCGGAA
35863 TATGAGTGTGGAGTCGTTGCAAAGAAAGGTGTTGTCTCTGCTGNNNAGATCGGAA
35864 TATGAGTGTGGAGTCGTTGCTGAAGATCACCTTGAGAGAACCCNNNAGATCGGAAG
35865 TATGAGTGTGGAGTCGTTGCAAAAGGAAACAGGACGGTCTGATNNNAGATCGGAA
35868 TATGAGTGTGGAGTCGTTGCTTACAGAGAAGAGGGTCTCGCTANNNAGATCGGAA
35870 TATGAGTGTGGAGTCGTTGCATGAGATGTTCAGGTCACGTTGANNNAGATCGGAA
35871 TATGAGTGTGGAGTCGTTGCGAGAGAGAGAGAGTCTTCTTGCGNNNAGATCGGAA
35872 TATGAGTGTGGAGTCGTTGCATGATAAAAGCAACCACACAGCCNNNAGATCGGAAG
35873 TATGAGTGTGGAGTCGTTGCAAATCGTACCAATTCCTCCCTCCNNNAGATCGGAAGA
35874 TATGAGTGTGGAGTCGTTGCAAATAGACTGTGGTGATGGTGGCNNNAGATCGGAA
35875 TATGAGTGTGGAGTCGTTGCGTGTCCTGCTTTCAAGTTAGCTGNNNAGATCGGAAG
35877 TATGAGTGTGGAGTCGTTGCCTCATCACAAGCAGCAGTTAGATNNNAGATCGGAAG
35878 TATGAGTGTGGAGTCGTTGCAATACAAAATTAGCCGGGCATGGNNNAGATCGGAA
35880 TATGAGTGTGGAGTCGTTGCTGAGAGCAAGACTCCATCTCAAANNNAGATCGGAAG
35882 TATGAGTGTGGAGTCGTTGCAAGAAAATTAGCTGGGCATGGTGNNNAGATCGGAA
35883 TATGAGTGTGGAGTCGTTGCGCATCCGTATTTCTGCAAAGGACNNNAGATCGGAAG
35884 TATGAGTGTGGAGTCGTTGCGCTGAGGAGGGAAGATTTCTTGANNNAGATCGGAA
35885 TATGAGTGTGGAGTCGTTGCGATTCCAGAACCTTCCTCCCAAANNNAGATCGGAAGA
35886 TATGAGTGTGGAGTCGTTGCAAAATCTACATCCACAGGACCCCNNNAGATCGGAAG
35888 TATGAGTGTGGAGTCGTTGCGAGAGAGAGAGAGAGAGACCTCANNNAGATCGGA
35889 TATGAGTGTGGAGTCGTTGCAACAGAAGTCAGATGTCCTCAGGNNNAGATCGGAA
35890 TATGAGTGTGGAGTCGTTGCGCCTCATATTGTCCTGCTTTTCCNNNAGATCGGAAGA
35891 TATGAGTGTGGAGTCGTTGCGAGACAGAGGGAAGGAAGGAAAGNNNAGATCGG
35892 TATGAGTGTGGAGTCGTTGCTCATGTCTGTAATCCCAGCATGTNNNAGATCGGAAG
35893 TATGAGTGTGGAGTCGTTGCGGTGGACTACTTGATGTCAGGAGNNNAGATCGGAA
35894 TATGAGTGTGGAGTCGTTGCGTGTGGAATGCATGGAAATGTCANNNAGATCGGAA
35895 TATGAGTGTGGAGTCGTTGCAGATGCCAAGACCTCTGCTAAAANNNAGATCGGAAG
35896 TATGAGTGTGGAGTCGTTGCACACAAACTCACCATTCAATCCANNNAGATCGGAAGA
35898 TATGAGTGTGGAGTCGTTGCAACATTTCTGCAGAGGTTTGACCNNNAGATCGGAAG
35899 TATGAGTGTGGAGTCGTTGCTACACAGGAGGCTTCAAAAGGAANNNAGATCGGAA
35902 TATGAGTGTGGAGTCGTTGCATCATGAGGTCAGGAGATCGAGANNNAGATCGGAA
35903 TATGAGTGTGGAGTCGTTGCAGCAACTTCAGGGAAGCATTTTCNNNAGATCGGAAG
35905 TATGAGTGTGGAGTCGTTGCCTGAGGCAGGAGAATCACTTGAANNNAGATCGGAA
35906 TATGAGTGTGGAGTCGTTGCGTGAGAGGCTCAAATAAGGGGAANNNAGATCGGA
35908 TATGAGTGTGGAGTCGTTGCCACACATATCCACAGCAGCAATGNNNAGATCGGAAG
35909 TATGAGTGTGGAGTCGTTGCCCCACAAACGACAAACAAACAAANNNAGATCGGAAG
35910 TATGAGTGTGGAGTCGTTGCAGCTACAGAGACAGGGGATAAGTNNNAGATCGGA

35911 TATGAGTGTGGAGTCGTTGCAAATGTCTGCCTTCTCATCTGGTNNNAGATCGGAAG

35912 TATGAGTGTGGAGTCGTTGCATTCCTCCAACACCAGTTAGACCNNNAGATCGGAAG

35913 TATGAGTGTGGAGTCGTTGCTGCCATCATAGTTTCCAGAAGCANNNAGATCGGAAG

35914 TATGAGTGTGGAGTCGTTGCACTCAAACAGGTACTTGTCCACANNNAGATCGGAAG

35917 TATGAGTGTGGAGTCGTTGCTAGGTAAAAGGAGCTGTCATCCCNNNAGATCGGAA

35918 TATGAGTGTGGAGTCGTTGCATAAGAGGAGAGGTTAGTGCCCTNNNAGATCGGAA

35919 TATGAGTGTGGAGTCGTTGCGGCTAGAGAAAGACCCCAAGTTTNNNAGATCGGAA

35920 TATGAGTGTGGAGTCGTTGCGTGTTCAGATTGAGCCATTGCATNNNAGATCGGAAG

35922 TATGAGTGTGGAGTCGTTGCCACAAACACAATAAACGCAGGGANNNAGATCGGAA

35923 TATGAGTGTGGAGTCGTTGCTTCTCTCTGGATTTTCAGGAGGCNNNAGATCGGAAG

35924 TATGAGTGTGGAGTCGTTGCAATGAGATTTCTGAGCAGCTGGANNNAGATCGGAA

35926 TATGAGTGTGGAGTCGTTGCGGAAGGAAGGAAGAAAGGAAGGANNNAGATCGG

35928 TATGAGTGTGGAGTCGTTGCTACTCAGAAAACGGTCCAACAGTNNNAGATCGGAAG

35929 TATGAGTGTGGAGTCGTTGCGGTGAATCACAAGGTCAGGAGATNNNAGATCGGAA

35930 TATGAGTGTGGAGTCGTTGCCTTCCCTCATTGCAGTGAGAAAGNNNAGATCGGAAG

35931 TATGAGTGTGGAGTCGTTGCGGACTGTAAAACCACTTGTCCTANNNAGATCGGAAG

35932 TATGAGTGTGGAGTCGTTGCTGTAATCTCAGCACTTTGGGAGGNNNAGATCGGAA

35933 TATGAGTGTGGAGTCGTTGCGGATCACGAGGTCAAGAGATCAANNNAGATCGGAA

35935 TATGAGTGTGGAGTCGTTGCCTGAAAGCGTCTCCAAAAATCCANNNAGATCGGAAG

35937 TATGAGTGTGGAGTCGTTGCCACCATGGAATACTACTCAGCCANNNAGATCGGAAG

35939 TATGAGTGTGGAGTCGTTGCGGAGGTTGTAGTGAGCCAAGATGNNNAGATCGGA

35940 TATGAGTGTGGAGTCGTTGCAGGAGTTTCAGACCAGCAAAGATNNNAGATCGGAA

35941 TATGAGTGTGGAGTCGTTGCTCCCGCAGTCTTATTTATACCCGNNNAGATCGGAAG

35943 TATGAGTGTGGAGTCGTTGCGGTGGATCATGAGGTCAAGAGATNNNAGATCGGA

35945 TATGAGTGTGGAGTCGTTGCGGTGGATCATGAGGTCAAGAGATNNNAGATCGGA

35946 TATGAGTGTGGAGTCGTTGCTCTGTGTACATTTTTTGGCTGGGNNNAGATCGGAAG

35947 TATGAGTGTGGAGTCGTTGCTTGTGTACATTTTTTGGCTGGGCNNNAGATCGGAAG

35949 TATGAGTGTGGAGTCGTTGCCTGTCATCCCAGCACTTTGAGAGNNNAGATCGGAAG

35950 TATGAGTGTGGAGTCGTTGCTTGTGTACATTTTTTGGCTGGGCNNNAGATCGGAAG

35952 TATGAGTGTGGAGTCGTTGCCCAAGATGAGCAGATCACCTGAGNNNAGATCGGAA

35955 TATGAGTGTGGAGTCGTTGCCTCATGTCTGTAATCCCAGCACTNNNAGATCGGAAG

35957 TATGAGTGTGGAGTCGTTGCGAACAGAGAGGTGAAGGACCATTNNNAGATCGGA

35958 TATGAGTGTGGAGTCGTTGCCTGTAATCCCAGCTACTCAGGAGNNNAGATCGGAAG

35959 TATGAGTGTGGAGTCGTTGCCAAGATTATGCCCCTGTACTCCANNNAGATCGGAAG

35960 TATGAGTGTGGAGTCGTTGCTTGGCAACATAGCAAGATCTCCTNNNAGATCGGAAG

35962 TATGAGTGTGGAGTCGTTGCTACAAAAAATTAGCCAGGCGTGGNNNAGATCGGAA

35964 TATGAGTGTGGAGTCGTTGCCATCCTGGCTAACATGGTGAAACNNNAGATCGGAAG

35965 TATGAGTGTGGAGTCGTTGCGTCAAGACCAGTCTGGGTAACATNNNAGATCGGAA

35968 TATGAGTGTGGAGTCGTTGCCTGGGTAACAGAGTGAGACACTGNNNAGATCGGAA

35969 TATGAGTGTGGAGTCGTTGCGATGTGTTTTCGCTGGGAAAGTTNNNAGATCGGAA

35970 TATGAGTGTGGAGTCGTTGCAGAGCGACACTCCATCTCAAAAANNNAGATCGGAAG

35971 TATGAGTGTGGAGTCGTTGCAAGAAATCTCAGAGACTCAGCCCNNNAGATCGGAAG

35972 TATGAGTGTGGAGTCGTTGCACGTATGTTTTTGCAGCACTGTTNNNAGATCGGAAG

35973 TATGAGTGTGGAGTCGTTGCTACAAAAAGTAGCTGGGCATGGTNNNAGATCGGAA
35974 TATGAGTGTGGAGTCGTTGCTTCAAGATCAGCCTGGACAACATNNNAGATCGGAAG
35975 TATGAGTGTGGAGTCGTTGCAAGAAAAGAAAGAGTTGCTGCCCNNNAGATCGGAA
35977 TATGAGTGTGGAGTCGTTGCTGAGGGAGTAGATTCGCTTGAACNNNAGATCGGAA
35979 TATGAGTGTGGAGTCGTTGCATACATCTATGACCTCACCCCCANNNAGATCGGAAGA
35981 TATGAGTGTGGAGTCGTTGCTGGAGTGCAGTGTTGTCATCATANNNAGATCGGAA
35984 TATGAGTGTGGAGTCGTTGCAAAATCTGAAAGTCACGGAAGGGNNNAGATCGGAA
35985 TATGAGTGTGGAGTCGTTGCAATGTCCATCCACAGGAGAATGANNNAGATCGGAA
35986 TATGAGTGTGGAGTCGTTGCATACAAAAATTAGCTGGGTGCGGNNNAGATCGGAA
35987 TATGAGTGTGGAGTCGTTGCGGGAAACTCTGACTCAGCCAATANNNAGATCGGAA
35988 TATGAGTGTGGAGTCGTTGCGCAGAATCATTTGAACCTGGGAGNNNAGATCGGAA
35989 TATGAGTGTGGAGTCGTTGCAAAAAAAACAGAGGGTTGGGTGCNNNAGATCGGA
35990 TATGAGTGTGGAGTCGTTGCTGATGTCGTCAAGATTCTGGTCANNNAGATCGGAAG
35991 TATGAGTGTGGAGTCGTTGCATCAGAGAGGATATTGGGGAGCANNNAGATCGGA
35992 TATGAGTGTGGAGTCGTTGCTCCCACATTGGAAAAGCAGTACTNNNAGATCGGAAG
35993 TATGAGTGTGGAGTCGTTGCCTGAGGAAGGAGGATTGCTTGAGNNNAGATCGGA
35995 TATGAGTGTGGAGTCGTTGCGTTGCTTTCTGGTGAGTCCTTTCNNNAGATCGGAAG
35996 TATGAGTGTGGAGTCGTTGCAGATCAAAGCATGCCACATTCAGNNNAGATCGGAAG
35997 TATGAGTGTGGAGTCGTTGCAAGCGTAATCACCATCAAAACCCNNNAGATCGGAAG
35998 TATGAGTGTGGAGTCGTTGCCCCACAGGACAAGCTGTAAACTTNNNAGATCGGAAG
35999 TATGAGTGTGGAGTCGTTGCCTATGGGTGGATCATGAGGTCAGNNNAGATCGGAA
36000 TATGAGTGTGGAGTCGTTGCCTGGGCAATATAGCAAGACTCCANNNAGATCGGAA
36001 TATGAGTGTGGAGTCGTTGCAGAAAGGAGGAGAGACAGAGACANNNAGATCGGA
36003 TATGAGTGTGGAGTCGTTGCAAGAAACCTTCCCTTTCACCTCCNNNAGATCGGAAGA
36004 TATGAGTGTGGAGTCGTTGCACGTTCTGGAGAGAATCACCTTCNNNAGATCGGAAG
36007 TATGAGTGTGGAGTCGTTGCACTGAATAAGTGTGGCCTCTTTCNNNAGATCGGAAG
36008 TATGAGTGTGGAGTCGTTGCTCGATTAGTTGAGCCTGTAGTCCNNNAGATCGGAAG
36009 TATGAGTGTGGAGTCGTTGCGGGTTGCTTTTTCCCCATTCATGNNNAGATCGGAAG
36010 TATGAGTGTGGAGTCGTTGCCAGATCACTTCAGGTCAGGAGTTNNNAGATCGGAA
36011 TATGAGTGTGGAGTCGTTGCGGCAAAATGAAACAGCAGACTCTNNNAGATCGGAA
36013 TATGAGTGTGGAGTCGTTGCGAGACACAGACACAGAGGAGAAGNNNAGATCGGA
36014 TATGAGTGTGGAGTCGTTGCGTAATCCTTACCAGCGAAAGCACNNNAGATCGGAAG
36016 TATGAGTGTGGAGTCGTTGCAGAGAGAGAGAGAGGCAAAGAGANNNAGATCGGA
36017 TATGAGTGTGGAGTCGTTGCATCATTGAGTCGCCTGAATCAGTNNNAGATCGGAAG
36018 TATGAGTGTGGAGTCGTTGCATAAAGACATGGCTGAGACTGGGNNNAGATCGGAA
36020 TATGAGTGTGGAGTCGTTGCACACAAAACTCTCCCTCCCAAATNNNAGATCGGAAGA
36023 TATGAGTGTGGAGTCGTTGCAGTCTTCAAGTCTATACCCCCCANNNAGATCGGAAG
36024 TATGAGTGTGGAGTCGTTGCTATCACTTGAACCTAGGAGGCAGNNNAGATCGGAA
36025 TATGAGTGTGGAGTCGTTGCCCTTTCCAGGACCAGCAGATAATNNNAGATCGGAAG
36027 TATGAGTGTGGAGTCGTTGCGCCCATGTTCTCAGAACCTTCTANNNAGATCGGAAG
36028 TATGAGTGTGGAGTCGTTGCGCTTTTATGGGGTTTGCAGGAATNNNAGATCGGAA
36029 TATGAGTGTGGAGTCGTTGCTAATCCCAGCTACTCAGGAGGTTNNNAGATCGGAAG
36031 TATGAGTGTGGAGTCGTTGCGTTCAGCCCATTTCTGTCTCAACNNNAGATCGGAAG

36032 TATGAGTGTGGAGTCGTTGCGAAAGAAAGGAAAACGCTGTCCANNNAGATCGGAA
36033 TATGAGTGTGGAGTCGTTGCTCTAACAAGTAATGGAAGGCGGGNNNAGATCGGAA
36034 TATGAGTGTGGAGTCGTTGCTTTCACAGGCAATGTCCATCCTANNNAGATCGGAAG
36037 TATGAGTGTGGAGTCGTTGCAAAGGGCCACATAGATTACAGCANNNAGATCGGAA
36039 TATGAGTGTGGAGTCGTTGCGGGCACTGAGAAACAGAAACAAANNNAGATCGGAA
36040 TATGAGTGTGGAGTCGTTGCATCGGGCAAGAACTAGTCAACATNNNAGATCGGAA
36041 TATGAGTGTGGAGTCGTTGCTGACATTATTCTGACCCATGGGGNNNAGATCGGAAG
36042 TATGAGTGTGGAGTCGTTGCGACACAGAGACAGAGAGACACAGNNNAGATCGGAA
36043 TATGAGTGTGGAGTCGTTGCGCTCACCTTTTTCTCTCTACCCTNNNAGATCGGAAGA
36044 TATGAGTGTGGAGTCGTTGCCTTGCAGTAAGCTGAGATCATGCNNNAGATCGGAA
36045 TATGAGTGTGGAGTCGTTGCTAATCCCATCTACTCAGGAGGCTNNNAGATCGGAAG
36046 TATGAGTGTGGAGTCGTTGCACAAGGAATACATTTGGGGTTCCNNNAGATCGGAA
36048 TATGAGTGTGGAGTCGTTGCACAACATGGTCCTCAGTTAGTCGNNNAGATCGGAAG
36049 TATGAGTGTGGAGTCGTTGCGAGACAATGAGGACAAGGGTAGGNNNAGATCGGA
36050 TATGAGTGTGGAGTCGTTGCTGGTCCAGTAGGATACTCACCTTNNNAGATCGGAAG
36052 TATGAGTGTGGAGTCGTTGCAGAGCTCAATCCCAAGATAGCTCNNNAGATCGGAAG
36055 TATGAGTGTGGAGTCGTTGCTCTTTTTGGGAGACCACAGTTCANNNAGATCGGAAG
36056 TATGAGTGTGGAGTCGTTGCAGTGTATCAGCTCCACCATTCAGNNNAGATCGGAAG
36058 TATGAGTGTGGAGTCGTTGCTTGGAGACAGAGAGACACAGAGANNNAGATCGGAA
36059 TATGAGTGTGGAGTCGTTGCGGAAGTCCAAGATCAAAGTGTGCNNNAGATCGGAA
36061 TATGAGTGTGGAGTCGTTGCAGAATCTTACTGTGTTCTGGGGGNNNAGATCGGAA
36062 TATGAGTGTGGAGTCGTTGCGCTGGTTCCAATCTGACTGAAACNNNAGATCGGAAG
36063 TATGAGTGTGGAGTCGTTGCTTGCTTACAGGAACAAGGTGGATNNNAGATCGGAA
36065 TATGAGTGTGGAGTCGTTGCCATATGATCGCTACCACAACAGCNNNAGATCGGAAG
36066 TATGAGTGTGGAGTCGTTGCGACTCCCATACCCTTACTGTCTGNNNAGATCGGAAGA
36067 TATGAGTGTGGAGTCGTTGCGCTTTGGGCATACTTCAAAATGCNNNAGATCGGAAG
36068 TATGAGTGTGGAGTCGTTGCCCTGGATTGCAGTGGATCCTAAANNNAGATCGGAA
36069 TATGAGTGTGGAGTCGTTGCTGGACCTCCTTTCTTTGCAAAACNNNAGATCGGAAGA
36071 TATGAGTGTGGAGTCGTTGCATGATGGCATATGCTTCTGGTCTNNNAGATCGGAAG
36073 TATGAGTGTGGAGTCGTTGCCTAATCCCCAGAACCTGTGAACANNNAGATCGGAAG
36074 TATGAGTGTGGAGTCGTTGCCAACTTAGGGATACCAGGAGCTGNNNAGATCGGAA
36075 TATGAGTGTGGAGTCGTTGCTGCTTCCACCCAGTTATTAGACCNNNAGATCGGAAGA
36076 TATGAGTGTGGAGTCGTTGCGGATGGATACTTGCTGCTTTTCCNNNAGATCGGAAG
36077 TATGAGTGTGGAGTCGTTGCCATAAGCTACGATCATGCCACTGNNNAGATCGGAAG
36078 TATGAGTGTGGAGTCGTTGCGCCATAAACTACGATCATGCCACNNNAGATCGGAAG
36079 TATGAGTGTGGAGTCGTTGCCTAAAGGCACTTGTCAGGTACAGNNNAGATCGGAA
36081 TATGAGTGTGGAGTCGTTGCTGTGCATCCATTCTCCAGTAGTTNNNAGATCGGAAG
36082 TATGAGTGTGGAGTCGTTGCTAGCTGAGCATTATGTTGTGTGCNNNAGATCGGAA
36083 TATGAGTGTGGAGTCGTTGCAAGGACACTAGTGTAGCCATAGCNNNAGATCGGAA
36084 TATGAGTGTGGAGTCGTTGCTACTGTGGCTCACACCTGTAATCNNNAGATCGGAAG
36086 TATGAGTGTGGAGTCGTTGCGAAAGGAATCGTGGTGTATTGGCNNNAGATCGGAA
36087 TATGAGTGTGGAGTCGTTGCCACCCCATTCTAAAAAGCAGGTGNNNAGATCGGAAG
36088 TATGAGTGTGGAGTCGTTGCATCTGAACTTTTGGGGGTCAACTNNNAGATCGGAAG

36091 TATGAGTGTGGAGTCGTTGCGTCACCAAAATCAGAAAGCTCCGNNNAGATCGGAAG
36092 TATGAGTGTGGAGTCGTTGCAGAGTTGCAGTCACTCGAATCTTNNNAGATCGGAAG
36093 TATGAGTGTGGAGTCGTTGCTGAAAAGCCTCACATAATGCGTCNNNAGATCGGAAG
36094 TATGAGTGTGGAGTCGTTGCCTCCCTCTGCTGAAAACATTTGGNNNAGATCGGAAG
36095 TATGAGTGTGGAGTCGTTGCGCATTGACTTCTGGCTCTGATTCNNNAGATCGGAAG
36096 TATGAGTGTGGAGTCGTTGCGTCAGTATGTCTAGACTGGGCACNNNAGATCGGAA
36098 TATGAGTGTGGAGTCGTTGCCAACATAGCAAGACCCTGTCTCTNNNAGATCGGAAG
36099 TATGAGTGTGGAGTCGTTGCGAGGGACTGCTAGATCAATTCGTNNNAGATCGGAA
36100 TATGAGTGTGGAGTCGTTGCAAAGCATTCCTCCAAGATCCACANNNAGATCGGAAG
36101 TATGAGTGTGGAGTCGTTGCCCCGAAACATTTCACATCAGCAANNNAGATCGGAAG
36102 TATGAGTGTGGAGTCGTTGCCAGCAAGCACTGAGAACAAATGANNNAGATCGGAA
36103 TATGAGTGTGGAGTCGTTGCACATGAACATGGGCATTACACTGNNNAGATCGGAAG
36104 TATGAGTGTGGAGTCGTTGCAAACAACCAAACACCATTTCCCANNNAGATCGGAAG
36105 TATGAGTGTGGAGTCGTTGCAATATTACGCATCCTGTCCTGGCNNNAGATCGGAAG
36107 TATGAGTGTGGAGTCGTTGCCATCCACAAGAACAGCCAATTGTNNNAGATCGGAAG
36108 TATGAGTGTGGAGTCGTTGCTCAAAGGGACAAGCACAGACATANNNAGATCGGAA
36110 TATGAGTGTGGAGTCGTTGCTGATCACACTACTGAACTCCAGCNNNAGATCGGAAG
36111 TATGAGTGTGGAGTCGTTGCCACAGGAGGAGTATGAAGGAACCNNNAGATCGGAA
36112 TATGAGTGTGGAGTCGTTGCCCGGACTGAACCAATGTACTTCTNNNAGATCGGAAG
36114 TATGAGTGTGGAGTCGTTGCACGGTAAGACCCTGTCTCAAAAANNNAGATCGGAAG
36115 TATGAGTGTGGAGTCGTTGCTCATGTTCCGTGAATGGAAAACCNNNAGATCGGAAG
36116 TATGAGTGTGGAGTCGTTGCCCTGCAAAGGAAAACAATCCTGTNNNAGATCGGAAG
36117 TATGAGTGTGGAGTCGTTGCCTGAGGCAGAAGAATTGCTTGAANNNAGATCGGAA
36118 TATGAGTGTGGAGTCGTTGCCACGTCTATGTAGCTCCAGAACANNNAGATCGGAAG
36119 TATGAGTGTGGAGTCGTTGCGAGAAAACACATTGGTGCACAGANNNAGATCGGAA
36120 TATGAGTGTGGAGTCGTTGCGGGAGGATCATTTGAGTCAGGAGNNNAGATCGGAA
36121 TATGAGTGTGGAGTCGTTGCGTGTAATCCCAGCTACTCAGGAGNNNAGATCGGAAG
36122 TATGAGTGTGGAGTCGTTGCGAAGCAGGGATAATTGCCTGAACNNNAGATCGGAA
36123 TATGAGTGTGGAGTCGTTGCCCCTCTTCATACACGGTACTGTGNNNAGATCGGAAG
36125 TATGAGTGTGGAGTCGTTGCATGAAAATGGTACACCTGGTCCANNNAGATCGGAAG
36126 TATGAGTGTGGAGTCGTTGCGTTGTGGATACAGCATTCACACCNNNAGATCGGAAG
36127 TATGAGTGTGGAGTCGTTGCAGAGAATCGCTTGAACTCAGGAGNNNAGATCGGAA
36128 TATGAGTGTGGAGTCGTTGCGAGAGCAGTTTCATGCATCCATCNNNAGATCGGAAG
36129 TATGAGTGTGGAGTCGTTGCCCTGTAATCCCAACACCTTGAGANNNAGATCGGAAG
36130 TATGAGTGTGGAGTCGTTGCAATTTCACTCCTGGGGAATGTCANNNAGATCGGAAG
36131 TATGAGTGTGGAGTCGTTGCGTCTCCTCTCATGTTCGCACTAANNNAGATCGGAAGA
36132 TATGAGTGTGGAGTCGTTGCAATCAACCTAGGCACCCATCAATNNNAGATCGGAAG
36133 TATGAGTGTGGAGTCGTTGCTGTCTTCTTCCGTCTTCTCAAGGNNNAGATCGGAAGA
36136 TATGAGTGTGGAGTCGTTGCACTCTTAGCTATTTGGGAGGCTGNNNAGATCGGAAG
36137 TATGAGTGTGGAGTCGTTGCGAAAACATTCTAAGCCTCCCACCNNNAGATCGGAAG
36138 TATGAGTGTGGAGTCGTTGCTGAAATACAGACACATGCAACGANNNAGATCGGAAG
36139 TATGAGTGTGGAGTCGTTGCCAGATATGGCATTCAAGGAAGCGNNNAGATCGGAA
36140 TATGAGTGTGGAGTCGTTGCACAACAGCAGAAGTTTCTTTGCANNNAGATCGGAAG

36141 TATGAGTGTGGAGTCGTTGCAGCTATTCCATAGGTACAGCAGCNNNAGATCGGAAG
36142 TATGAGTGTGGAGTCGTTGCGGATCACGAGGTCACGAGATTATNNNAGATCGGAA
36143 TATGAGTGTGGAGTCGTTGCTGGGCAACATAGTGAGACTTCATNNNAGATCGGAA
36145 TATGAGTGTGGAGTCGTTGCACCTATCCATCCAAGGATCAAGGNNNAGATCGGAAG
36146 TATGAGTGTGGAGTCGTTGCAAACTGGACTGCACCAAAAACAANNNAGATCGGAA
36147 TATGAGTGTGGAGTCGTTGCGGATTCAGAGTCAATGGTGCTTCNNNAGATCGGAA
36148 TATGAGTGTGGAGTCGTTGCAGCCCATTCTTCTCATAGCTCACNNNAGATCGGAAGA
36149 TATGAGTGTGGAGTCGTTGCAAATGTCTCATCCAAGAGGAGCANNNAGATCGGAA
36150 TATGAGTGTGGAGTCGTTGCCCCATTCCTCCAACTTCTGAACTNNNAGATCGGAAGA
36152 TATGAGTGTGGAGTCGTTGCTATTCCATACACACCTGCTCACANNNAGATCGGAAGA
36153 TATGAGTGTGGAGTCGTTGCCCTTCCTCATATAGTGTCCCTGCNNNAGATCGGAAGA
36154 TATGAGTGTGGAGTCGTTGCGCATGGAGCACATTCATGTTTCTNNNAGATCGGAAG
36155 TATGAGTGTGGAGTCGTTGCGATTGCAGTGAGCTGAGATTGTGNNNAGATCGGAA
36157 TATGAGTGTGGAGTCGTTGCAATCCTCAGTAGCCACTTGTGAGNNNAGATCGGAAG
36158 TATGAGTGTGGAGTCGTTGCAAAAGTGAAAACCTGGTTGGGTGNNNAGATCGGAA
36159 TATGAGTGTGGAGTCGTTGCCAGAATAGGCATCCAGGAGACTCNNNAGATCGGAA
36161 TATGAGTGTGGAGTCGTTGCAACTCACTATGCAGCTGTCTGAANNNAGATCGGAAG
36162 TATGAGTGTGGAGTCGTTGCCTGGGAAATGATTGAGTGACTGCNNNAGATCGGAA
36163 TATGAGTGTGGAGTCGTTGCTATCCAACATGGTGAAACCCCATNNNAGATCGGAAG
36165 TATGAGTGTGGAGTCGTTGCAACACGAATTCACTGTTGCGAATNNNAGATCGGAAG
36167 TATGAGTGTGGAGTCGTTGCATACATCCGTGGAGATTCAGAGCNNNAGATCGGAA
36169 TATGAGTGTGGAGTCGTTGCCCCTGGAGAAACTCACACATGTANNNAGATCGGAAG
36170 TATGAGTGTGGAGTCGTTGCCCCTGGAGAAACTCACACATGTANNNAGATCGGAAG
36171 TATGAGTGTGGAGTCGTTGCAGAATTGCTTGAATCCAGGAGGTNNNAGATCGGAA
36172 TATGAGTGTGGAGTCGTTGCGTCAGGAGATCCAAGTTAGTCCCNNNAGATCGGAA
36173 TATGAGTGTGGAGTCGTTGCTACACACACACACACAAATGCAANNNAGATCGGAAG
36174 TATGAGTGTGGAGTCGTTGCAGACCCCATCACGAAATTTGAGTNNNAGATCGGAAG
36175 TATGAGTGTGGAGTCGTTGCAGTGGATTACAAGGTCAAAGGCTNNNAGATCGGAA
36176 TATGAGTGTGGAGTCGTTGCCAAGAAGAGGCAGATGGCAAAATNNNAGATCGGAA
36178 TATGAGTGTGGAGTCGTTGCTTCTCAAGAGCCTGCAATCATCTNNNAGATCGGAAG
36179 TATGAGTGTGGAGTCGTTGCAGAACACCCAACAGAGACTAGTCNNNAGATCGGAA
36180 TATGAGTGTGGAGTCGTTGCGAAGAAGAAGAGGAGATGAGGCCNNNAGATCGGA
36181 TATGAGTGTGGAGTCGTTGCTGTAATCCCAGCTACTCAGGAGANNNAGATCGGAAG
36183 TATGAGTGTGGAGTCGTTGCGTAGAGACTAAACCACCAGCCTCNNNAGATCGGAAG
36184 TATGAGTGTGGAGTCGTTGCAATTTCCTAGCACTGCCCAGTAANNNAGATCGGAAG
36186 TATGAGTGTGGAGTCGTTGCAGCATGGTGGCATATACCTGTAGNNNAGATCGGAA
36187 TATGAGTGTGGAGTCGTTGCCAGTGAAGACAAGGTGGGGAATANNNAGATCGGA
36189 TATGAGTGTGGAGTCGTTGCGCAACAGAGTGAGACTGTGTTTCNNNAGATCGGAA
36190 TATGAGTGTGGAGTCGTTGCGGCAGCTCAAAACCAAGAGAATTNNNAGATCGGAA
36191 TATGAGTGTGGAGTCGTTGCTGCTCATTGGTTTCTATTGGGGANNNAGATCGGAAG
36192 TATGAGTGTGGAGTCGTTGCAAAGCTTTCCAACTGTGCTCATCNNNAGATCGGAAG
36193 TATGAGTGTGGAGTCGTTGCGGCCTTGGGATTATCCACCTATTNNNAGATCGGAAG
36195 TATGAGTGTGGAGTCGTTGCATTGGAGGTGATGGATGTGCTAANNNAGATCGGAA

36197 TATGAGTGTGGAGTCGTTGCCTGGGTGACAGAGTGAAACACTANNNAGATCGGAA
36198 TATGAGTGTGGAGTCGTTGCATAACAGACCACGTGTAAGCTGTNNNAGATCGGAAG
36200 TATGAGTGTGGAGTCGTTGCACATGGCATTCAAAAGACACCAGNNNAGATCGGAAG
36201 TATGAGTGTGGAGTCGTTGCTCCCAGCTACTTGAAAGACTGAGNNNAGATCGGAAG
36202 TATGAGTGTGGAGTCGTTGCCGCCTTCACTGACTGGTAAATTGNNNAGATCGGAAG
36203 TATGAGTGTGGAGTCGTTGCCTAGAATTCTGAGGGCCTGAGGANNNAGATCGGAA
36204 TATGAGTGTGGAGTCGTTGCTGTTCAACCATTGTGGAAGATGGNNNAGATCGGAA
36205 TATGAGTGTGGAGTCGTTGCGCTTCAAGAAATTTGGCTGCATGNNNAGATCGGAAG
36206 TATGAGTGTGGAGTCGTTGCGTGGCAAAACCCTGTCTCTACTANNNAGATCGGAAG
36207 TATGAGTGTGGAGTCGTTGCGATCACTTGAGCCCAGGAGTTTANNNAGATCGGAAG
36208 TATGAGTGTGGAGTCGTTGCTGGGGAGGGGTTTTACAATGAAANNNAGATCGGAA
36209 TATGAGTGTGGAGTCGTTGCAACCTCGAAAAGTCATCTCTGGANNNAGATCGGAAG
36211 TATGAGTGTGGAGTCGTTGCTCACTTGAGGTCAGGAGTTCAAGNNNAGATCGGAA
36212 TATGAGTGTGGAGTCGTTGCCTGTAAACCCAACACTTGACGAGNNNAGATCGGAAG
36214 TATGAGTGTGGAGTCGTTGCGGAGTGCTTTTTCTCTGAGTTGCNNNAGATCGGAAG
36215 TATGAGTGTGGAGTCGTTGCGCCATCTACAAACCAGGAAGAGANNNAGATCGGAA
36216 TATGAGTGTGGAGTCGTTGCCTATGTCTGGATTTGCCTAGTGCNNNAGATCGGAAG
36217 TATGAGTGTGGAGTCGTTGCCCACTGTTTCTGGAATATGCCACNNNAGATCGGAAG
36218 TATGAGTGTGGAGTCGTTGCGAGTACCATGAGTTCTCCAGCAANNNAGATCGGAAG
36219 TATGAGTGTGGAGTCGTTGCCACGCCTATAATCCCAGCTACTTNNNAGATCGGAAG
36220 TATGAGTGTGGAGTCGTTGCTATCTTTGAAGACACGGGACACANNNAGATCGGAAG
36223 TATGAGTGTGGAGTCGTTGCGAATCTGTGAGTCTGGCACCTTANNNAGATCGGAAG
36224 TATGAGTGTGGAGTCGTTGCTGGAAACAACTATCCTCAAGGCANNNAGATCGGAAG
36225 TATGAGTGTGGAGTCGTTGCTCTGACAGCTGCAAAGAAAAGTGNNNAGATCGGAA
36226 TATGAGTGTGGAGTCGTTGCGCACTTACAAGCTGTGTGTTCTTNNNAGATCGGAAG
36227 TATGAGTGTGGAGTCGTTGCTGAGGATTGTTATTTCTGTGGGCNNNAGATCGGAA
36228 TATGAGTGTGGAGTCGTTGCGTTGGTGAGAATGCCAGGTATTGNNNAGATCGGAA
36230 TATGAGTGTGGAGTCGTTGCTGGTATCTTTCTTTCCCCCACAGNNNAGATCGGAAG
36231 TATGAGTGTGGAGTCGTTGCAGCCAACTTCATCACTCAGTCATNNNAGATCGGAAG
36233 TATGAGTGTGGAGTCGTTGCCCTGACTTCAAGAGCAGGTACTCNNNAGATCGGAAG
36234 TATGAGTGTGGAGTCGTTGCCAGGGACAGTGCTATTGTTACCTNNNAGATCGGAAG
36235 TATGAGTGTGGAGTCGTTGCACCTATGATGGGTCTGGATGTACNNNAGATCGGAA
36236 TATGAGTGTGGAGTCGTTGCACTGACAAAGTGAAAAGGAAGCANNNAGATCGGAA
36237 TATGAGTGTGGAGTCGTTGCAAAAACCACAAAGTCCCACTGTGNNNAGATCGGAAG
36238 TATGAGTGTGGAGTCGTTGCTAAATGTGAAAAATCCCAGCCGGNNNAGATCGGAA
36240 TATGAGTGTGGAGTCGTTGCTCCCCAGTTTCTTCAGAACAACANNNAGATCGGAAG
36241 TATGAGTGTGGAGTCGTTGCAAAAGAATGAGAGCCTCCTCTGGNNNAGATCGGAA
36242 TATGAGTGTGGAGTCGTTGCTAGATTCATACGAGCCAGCATGTNNNAGATCGGAAG
36243 TATGAGTGTGGAGTCGTTGCATGTAGAAGGGAAGTCAGCACAGNNNAGATCGGAA
36244 TATGAGTGTGGAGTCGTTGCGGCCTTAGGTGTATCTATGCTGGNNNAGATCGGAA
36245 TATGAGTGTGGAGTCGTTGCCCCATATAAGTATGAGGGAGCCCNNNAGATCGGAA
36247 TATGAGTGTGGAGTCGTTGCTCTTGTGTTGCTTTGGGAAATCCNNNAGATCGGAAG
36248 TATGAGTGTGGAGTCGTTGCCAGGAGAAGGTCAGTGAGAAAGANNNAGATCGGA

36249 TATGAGTGTGGAGTCGTTGCTCTCAAACTCCTGACCTCAAGTGNNNAGATCGGAAG
36250 TATGAGTGTGGAGTCGTTGCAGAGGATTACTTGTGCCTGGAAGNNNAGATCGGAA
36251 TATGAGTGTGGAGTCGTTGCACATGGTACTTCAATGGCAGGAANNNAGATCGGAA
36252 TATGAGTGTGGAGTCGTTGCATGATAGCAGATGCAGATCACCNNNAGATCGGAA
36253 TATGAGTGTGGAGTCGTTGCGGGTGAGGAGATTGGAATTTCCTNNNAGATCGGAA
36254 TATGAGTGTGGAGTCGTTGCATGAAGGCGATCCAGTCTAAAGGNNNAGATCGGAA
36255 TATGAGTGTGGAGTCGTTGCGGAGAATGGCATGAACCAGTGAGNNNAGATCGGA
36256 TATGAGTGTGGAGTCGTTGCAGAGAAGGCTAGGTTATGTCACANNNAGATCGGAA
36258 TATGAGTGTGGAGTCGTTGCAGAGAGACTGGGAAGTAGAAGCTNNNAGATCGGA
36259 TATGAGTGTGGAGTCGTTGCCAGAGGGACAAAGGAGACAAAGANNNAGATCGGA
36260 TATGAGTGTGGAGTCGTTGCTGACTTTCATAGGCAGCTAATTGCNNNAGATCGGAA
36261 TATGAGTGTGGAGTCGTTGCCTGCTGCTGCTTTACCAACTAAGNNNAGATCGGAAG
36262 TATGAGTGTGGAGTCGTTGCGCACATGCCTGTAATCCTAGCTANNNAGATCGGAAG
36263 TATGAGTGTGGAGTCGTTGCATTGGCCCCTGAATAAAGTCAGTNNNAGATCGGAAG
36264 TATGAGTGTGGAGTCGTTGCAGCAGAAGTAAGTGGACTTTGCTNNNAGATCGGAA
36265 TATGAGTGTGGAGTCGTTGCTCTCTTTCCTCATCCTCTCCTTTNNNAGATCGGAAGAG
36266 TATGAGTGTGGAGTCGTTGCCCTCTGAGGTCCATGAAAGGAATNNNAGATCGGAA
36267 TATGAGTGTGGAGTCGTTGCGCATACAGTTGGCACTTCTGAAGNNNAGATCGGAA
36270 TATGAGTGTGGAGTCGTTGCTTCTGGTGGGGGATATGGATAGTNNNAGATCGGAA
36271 TATGAGTGTGGAGTCGTTGCATGAGGACAACTGGCAAACTTTGNNNAGATCGGAA
36272 TATGAGTGTGGAGTCGTTGCTCTGCTGCCTGTGGATAAGATACNNNAGATCGGAAG
36273 TATGAGTGTGGAGTCGTTGCTGAAAAGATGTTGCCCTACTGGANNNAGATCGGAA
36274 TATGAGTGTGGAGTCGTTGCAAATGTAGGTCTATGAGCTGGGGNNNAGATCGGAA
36275 TATGAGTGTGGAGTCGTTGCTTTCCAAGAGATGCTGATGTTGCNNNAGATCGGAAG
36276 TATGAGTGTGGAGTCGTTGCAGGTACTTGCTGTATGAGGCATCNNNAGATCGGAA
36277 TATGAGTGTGGAGTCGTTGCTGCTCTCAGTGGTCATCTTTTCANNNAGATCGGAAG
36279 TATGAGTGTGGAGTCGTTGCTTGTACCTTCCTCTTCAGAAGCCNNNAGATCGGAAGA
36282 TATGAGTGTGGAGTCGTTGCAGAAGGCAAATGGTATAGGCACANNNAGATCGGAA
36283 TATGAGTGTGGAGTCGTTGCAGTAGTTACACCTGGAGAGGGAANNNAGATCGGAA
36284 TATGAGTGTGGAGTCGTTGCGTTACTTTTGATGTCCAGCCTGCNNNAGATCGGAAG
36285 TATGAGTGTGGAGTCGTTGCGCCCAAAGTCATATAGCTACCGANNNAGATCGGAAG
36287 TATGAGTGTGGAGTCGTTGCAATACGAGCATCCCCAAGGAAATNNNAGATCGGAA
36288 TATGAGTGTGGAGTCGTTGCATGTCTTACCCCAATTCCTCCAGNNNAGATCGGAAGA
36289 TATGAGTGTGGAGTCGTTGCACAGTCCTCCAAATTGCCAAGTANNNAGATCGGAAG
36290 TATGAGTGTGGAGTCGTTGCGCATACCTCTTGTCCTTGCTACTNNNAGATCGGAAGA
36294 TATGAGTGTGGAGTCGTTGCATATGTGTGTGTGTGTGTGTGTGNNNAGATCGGAA
36295 TATGAGTGTGGAGTCGTTGCAAGAAAATTAGCTGAGCGTGGTGNNNAGATCGGAA
36296 TATGAGTGTGGAGTCGTTGCGCAAGGGAACAAAATTGGACAGANNNAGATCGGAA
36297 TATGAGTGTGGAGTCGTTGCAGTCCATATTGGGCGCTTTTCTANNNAGATCGGAAG
36298 TATGAGTGTGGAGTCGTTGCACAGATCATCCAGAGGCAAAACTNNNAGATCGGAA
36300 TATGAGTGTGGAGTCGTTGCTTCAGAAAGGTTGTCTTGCAAGCNNNAGATCGGAA
36301 TATGAGTGTGGAGTCGTTGCAATAAAAAAGGAGGGAATGGCGGNNNAGATCGGA
36302 TATGAGTGTGGAGTCGTTGCTTATTGAAAAAGAGGCCGCAACANNNAGATCGGAA

36303 TATGAGTGTGGAGTCGTTGCAAAACAAAGAATTGAGGCCGAGGNNNAGATCGGAA
36304 TATGAGTGTGGAGTCGTTGCCTCATGCCTGTAATCCCAGAACTNNNAGATCGGAAGA
36305 TATGAGTGTGGAGTCGTTGCAAGAAACATGGATCCCCACACTTNNNAGATCGGAAG
36306 TATGAGTGTGGAGTCGTTGCTGAGGTCAGGATTTTGAGACCAGNNNAGATCGGAA
36307 TATGAGTGTGGAGTCGTTGCGCAGTGGGTATCATTTTCTGCTCNNNAGATCGGAAG
36308 TATGAGTGTGGAGTCGTTGCGAGGCTGATCGGAAAGTATCACTNNNAGATCGGAA
36309 TATGAGTGTGGAGTCGTTGCCACCATTTGTTTCCACTTCTGCANNNAGATCGGAAGA
36310 TATGAGTGTGGAGTCGTTGCAGAAGAAGGCGAAGAAGGAGAAGNNNAGATCGGA
36311 TATGAGTGTGGAGTCGTTGCATTGCTTGAGCCCTAGAGTTCAANNNAGATCGGAAG
36312 TATGAGTGTGGAGTCGTTGCGTCTGACTTCATCCAAGGGTTCTNNNAGATCGGAAG
36314 TATGAGTGTGGAGTCGTTGCATGAGGCATTCAGAAAAGGTCCTNNNAGATCGGAA
36316 TATGAGTGTGGAGTCGTTGCGCTGCAGTGAACTATGATTGCATNNNAGATCGGAAG
36317 TATGAGTGTGGAGTCGTTGCCACGCTCTGCTAATTTGAGATGTNNNAGATCGGAAG
36318 TATGAGTGTGGAGTCGTTGCGAGAGAGAATGCTCCATGGCTTANNNAGATCGGAA
36319 TATGAGTGTGGAGTCGTTGCCATGGTTTCATGTGCCTGTAGTTNNNAGATCGGAAG
36321 TATGAGTGTGGAGTCGTTGCGATGGCCCCACACCAAAAAATTANNNAGATCGGAAG
36322 TATGAGTGTGGAGTCGTTGCAAAAACCAAGTCAGAAGTTGGGGNNNAGATCGGAA
36324 TATGAGTGTGGAGTCGTTGCTTAGAATCACCAATACCTGCCCCNNNAGATCGGAAGA
36326 TATGAGTGTGGAGTCGTTGCAAAAATTTGTCAGGTGTGGTGGCNNNAGATCGGAA
36327 TATGAGTGTGGAGTCGTTGCCCCAGCTAACTAAGTGCACAGTANNNAGATCGGAAG
36328 TATGAGTGTGGAGTCGTTGCTGCTATGTGATCCAGCAATTCCANNNAGATCGGAAG
36329 TATGAGTGTGGAGTCGTTGCGAGATTACACCACTGCACTCCAGNNNAGATCGGAAG
36330 TATGAGTGTGGAGTCGTTGCAAAGCAGAAGTGCCCTAATCTCANNNAGATCGGAAG
36331 TATGAGTGTGGAGTCGTTGCAACTGAGTGAATAAGCACCACCANNNAGATCGGAAG
36332 TATGAGTGTGGAGTCGTTGCTGTTGTTTTTCTCTTTCCCGACANNNAGATCGGAAGA
36334 TATGAGTGTGGAGTCGTTGCCACCCTGGACCTACAAAATCAGANNNAGATCGGAAG
36335 TATGAGTGTGGAGTCGTTGCGGACCACTACAATAAGCACAAGCNNNAGATCGGAAG
36337 TATGAGTGTGGAGTCGTTGCACATTGCGCTAAGTGAAAGAAGCNNNAGATCGGAA
36339 TATGAGTGTGGAGTCGTTGCGAAGACACCCCAATACAGCTGTANNNAGATCGGAAG
36340 TATGAGTGTGGAGTCGTTGCACATAGTCTCTGACCACAATGGANNNAGATCGGAAG
36341 TATGAGTGTGGAGTCGTTGCTGGAAGAGTCATTGTGAGACAGGNNNAGATCGGAA
36342 TATGAGTGTGGAGTCGTTGCACCAATCTTCCTCTGTCTATGCANNNAGATCGGAAGA
36345 TATGAGTGTGGAGTCGTTGCGTCACACGTGAGTGGTTTTGAAANNNAGATCGGAA
36346 TATGAGTGTGGAGTCGTTGCTCACGTGGCTTTCTGATCCTTTANNNAGATCGGAAGA
36348 TATGAGTGTGGAGTCGTTGCCTCTTACACAGGGTAGTCTTGGGNNNAGATCGGAAG
36349 TATGAGTGTGGAGTCGTTGCCCACGTCCTTGAACTTCTCAGTANNNAGATCGGAAGA
36350 TATGAGTGTGGAGTCGTTGCAAACCTTGTTTGCAAAGGACCAGNNNAGATCGGAAG
36351 TATGAGTGTGGAGTCGTTGCACTGCCAAGCTCACTCAATAAGANNNAGATCGGAAG
36352 TATGAGTGTGGAGTCGTTGCGTGTGTGTGTGTGTGTGTGTATCNNNAGATCGGAA
36354 TATGAGTGTGGAGTCGTTGCAGCTAACAAGCCACACAAAAACANNNAGATCGGAAG
36355 TATGAGTGTGGAGTCGTTGCGCCAAGTCTCCACATGTTTGTTCNNNAGATCGGAAGA
36357 TATGAGTGTGGAGTCGTTGCATGCCAGGTCTAGTACAACGATGNNNAGATCGGAAG
36358 TATGAGTGTGGAGTCGTTGCGCCACACAAAAAGAGCTAGTCAGNNNAGATCGGAA

36359 TATGAGTGTGGAGTCGTTGCGGTGTGGACATAAGTTTTGGTGTNNNAGATCGGAA
36360 TATGAGTGTGGAGTCGTTGCAACTGCACACCGTAAAAAGATCTNNNAGATCGGAAG
36361 TATGAGTGTGGAGTCGTTGCTCTCTGTGTCCCTCTCTATCTCTNNNAGATCGGAAGA
36363 TATGAGTGTGGAGTCGTTGCGCTGAGGTGGAAGGATAACTTGANNNAGATCGGAA
36364 TATGAGTGTGGAGTCGTTGCGAGGTTGCAGTAAGCTGAGATTGNNNAGATCGGAA
36365 TATGAGTGTGGAGTCGTTGCACAACTTGTAAGAGGAGTTGGCANNNAGATCGGAA
36367 TATGAGTGTGGAGTCGTTGCACCCTTTGGCTCAATATTGTACTNNNAGATCGGAAGA
36368 TATGAGTGTGGAGTCGTTGCTGAGATAGCACCTCTGTACTCCANNNAGATCGGAAG
36370 TATGAGTGTGGAGTCGTTGCAAACAGGGATTCTCTTCACACCANNNAGATCGGAAG
36371 TATGAGTGTGGAGTCGTTGCGCTTAGTGCCTGGCCTATTTTTCNNNAGATCGGAAG
36372 TATGAGTGTGGAGTCGTTGCCTTGAATGTAGAGCCCAGGTCTTNNNAGATCGGAAG
36373 TATGAGTGTGGAGTCGTTGCCCCCAAATGTCAATAGTGCCAAGNNNAGATCGGAAG
36374 TATGAGTGTGGAGTCGTTGCTTCCAATGCAGCAGTCAAATGTANNNAGATCGGAAG
36375 TATGAGTGTGGAGTCGTTGCAGGCACGAGAATCTCTTCAACTTNNNAGATCGGAAG
36376 TATGAGTGTGGAGTCGTTGCTGATGTCAGGTTTTAGTCTGCCANNNAGATCGGAAG
36377 TATGAGTGTGGAGTCGTTGCGCAATGCACTAGTCAGATGCAATNNNAGATCGGAAG
36378 TATGAGTGTGGAGTCGTTGCCCCAGGGTAAGAGGTTGAAACAANNNAGATCGGAA
36379 TATGAGTGTGGAGTCGTTGCGAGAATTGCCTGAAAGCACAGAGNNNAGATCGGAA
36380 TATGAGTGTGGAGTCGTTGCTGGCAGTAAAGTGGGTTCTGAATNNNAGATCGGAA
36382 TATGAGTGTGGAGTCGTTGCTTCTTGGGACACAATAAGCCCTTNNNAGATCGGAAG
36383 TATGAGTGTGGAGTCGTTGCGCTTGGAAGAAAATCTTGGAGGCNNNAGATCGGAA
36384 TATGAGTGTGGAGTCGTTGCCCAAGTGGGTGACGTCAATTATCNNNAGATCGGAAG
36385 TATGAGTGTGGAGTCGTTGCAGAGAGCTCTGATTGTGCTACTGNNNAGATCGGAAG
36386 TATGAGTGTGGAGTCGTTGCCTGATGGATGCTCAAATCCCTGANNNAGATCGGAAG
36387 TATGAGTGTGGAGTCGTTGCATTGAGTACCGGAAAGAGACAGGNNNAGATCGGAA
36388 TATGAGTGTGGAGTCGTTGCTAACTCAGATCTCTGCTGTTGGGNNNAGATCGGAAG
36389 TATGAGTGTGGAGTCGTTGCCTAACAGACTGGCTACTGTGAGGNNNAGATCGGAA
36390 TATGAGTGTGGAGTCGTTGCTGGCAAGAAATACAGTTCTCACGNNNAGATCGGAAG
36391 TATGAGTGTGGAGTCGTTGCCTCCAATGGCTCCTTCTCTTTCTNNNAGATCGGAAGA
36392 TATGAGTGTGGAGTCGTTGCACAATCGTAGGTCACTGAGGAACNNNAGATCGGAA
36393 TATGAGTGTGGAGTCGTTGCAGATCCAACCTAAATCAACGCCTNNNAGATCGGAAG
36397 TATGAGTGTGGAGTCGTTGCTTTGTTTTGTTCCTAGTCCCACANNNAGATCGGAAGA
36398 TATGAGTGTGGAGTCGTTGCCATTCAATCTGCATGAAGGGCATNNNAGATCGGAAG
36399 TATGAGTGTGGAGTCGTTGCAAGAAAGGCAACCCAAAGAATGGNNNAGATCGGAA
36400 TATGAGTGTGGAGTCGTTGCGTGCTCTCTTGAGGTCACTGTTANNNAGATCGGAAG
36401 TATGAGTGTGGAGTCGTTGCGGGTTGCAATCCTAGTCTGTGATNNNAGATCGGAAG
36402 TATGAGTGTGGAGTCGTTGCGAGTTCAAGACTAGCCTGACCAANNNAGATCGGAAG
36403 TATGAGTGTGGAGTCGTTGCAAGAAACTTGGCATCATAGCAGGNNNAGATCGGAA
36404 TATGAGTGTGGAGTCGTTGCGTTTTGAGTGGGACCAACTTGAGNNNAGATCGGAA
36405 TATGAGTGTGGAGTCGTTGCCTAGACCAAACCTTAGGCTGTGTNNNAGATCGGAAG
36406 TATGAGTGTGGAGTCGTTGCAGGAAAAACAGTCCAGCAAAAGGNNNAGATCGGAA
36407 TATGAGTGTGGAGTCGTTGCACAAAGGACAGACAGAGCTCAAANNNAGATCGGAA
36409 TATGAGTGTGGAGTCGTTGCGGCAGGAATCATCAATAAGTGCCNNNAGATCGGAA

36410 TATGAGTGTGGAGTCGTTGCATTTGGTTGAAAGGTCAGTGCAGNNNAGATCGGAA
36411 TATGAGTGTGGAGTCGTTGCGTGCTTTTCCTCATTGGTTCCTCNNNAGATCGGAAGA
36413 TATGAGTGTGGAGTCGTTGCGCGAGTCACTTAGTTTCTCCTGTNNNAGATCGGAAG
36414 TATGAGTGTGGAGTCGTTGCATGGGGAAATGAGGCTAAAATGCNNNAGATCGGA
36415 TATGAGTGTGGAGTCGTTGCCCAATGCCTATACCCCATTGTANNNAGATCGGAAG
36416 TATGAGTGTGGAGTCGTTGCGGAAAGAAAGAATGGGCATGGTGNNNAGATCGGA
36417 TATGAGTGTGGAGTCGTTGCGGGAAAAGCATAGTATCTGGGCTNNNAGATCGGAA
36418 TATGAGTGTGGAGTCGTTGCAGGTTTATGAAAAAGGAGGGCCANNNAGATCGGA
36419 TATGAGTGTGGAGTCGTTGCTGAGATCTCAATTGCCAGCTCATNNNAGATCGGAAG
36420 TATGAGTGTGGAGTCGTTGCGCGGAGAAAATGTTGATAGGCATNNNAGATCGGAA
36421 TATGAGTGTGGAGTCGTTGCGTGTCTGCATTCTTTGACTCAGCNNNAGATCGGAAG
36422 TATGAGTGTGGAGTCGTTGCGCCATTAGCTCTTTCTCCTAGCANNNAGATCGGAAG
36424 TATGAGTGTGGAGTCGTTGCAGGCAAACTTGTGAAATCTCAGCNNNAGATCGGAA
36425 TATGAGTGTGGAGTCGTTGCTTTGTGGTCAGCATTCTCTCTCCNNNAGATCGGAAG
36426 TATGAGTGTGGAGTCGTTGCGCTGGTGTGCAGTTATTATCAGCNNNAGATCGGAA
36427 TATGAGTGTGGAGTCGTTGCTGAGGTCTTACCATGTTGTCCAGNNNAGATCGGAAG
36429 TATGAGTGTGGAGTCGTTGCGCAACACTGTGGGCAAAAGATATNNNAGATCGGAA
36430 TATGAGTGTGGAGTCGTTGCATATGCACGTTGTTCAACCCAACNNNAGATCGGAAG
36431 TATGAGTGTGGAGTCGTTGCGGTTCACAGGCCAGTTTTTTGTANNNAGATCGGAAG
36434 TATGAGTGTGGAGTCGTTGCTGACAAAGCAAGACTCCATCTCANNNAGATCGGAAG
36436 TATGAGTGTGGAGTCGTTGCTGTTGTGAAGTGCTTCCCTTTTCNNNAGATCGGAAG
36438 TATGAGTGTGGAGTCGTTGCTGTCATTCTTCCTCTGTCCCTTCNNNAGATCGGAAGA
36439 TATGAGTGTGGAGTCGTTGCGAGATTGCACCACTGTACTCCAGNNNAGATCGGAAG
36440 TATGAGTGTGGAGTCGTTGCTTTTGTCAGGAGAGAAAGGGTCANNNAGATCGGAA
36442 TATGAGTGTGGAGTCGTTGCATTGTAGTGACTGCTTTGCCTTGNNNAGATCGGAAG
36443 TATGAGTGTGGAGTCGTTGCCAGGAAGTCAAGGCTGTAGTGAGNNNAGATCGGA
36444 TATGAGTGTGGAGTCGTTGCCAGGCCAAAGCTTTTCTTCATCANNNAGATCGGAAG
36445 TATGAGTGTGGAGTCGTTGCGAACTATGAATTGCACTCCAGCCNNNAGATCGGAAG
36446 TATGAGTGTGGAGTCGTTGCAGGAACTAGAGTAAGCAGGAGGTNNNAGATCGGA
36447 TATGAGTGTGGAGTCGTTGCACTAGGCAATTGGGAAGTGTAGGNNNAGATCGGA
36448 TATGAGTGTGGAGTCGTTGCCTCATCACAAGCAGCAGTTAGATNNNAGATCGGAAG
36449 TATGAGTGTGGAGTCGTTGCGAGACAGAGGGAAGGAAGGAAAGNNNAGATCGG
36452 TATGAGTGTGGAGTCGTTGCCACCATGGAATACTACTCAGCCANNNAGATCGGAAG
36455 TATGAGTGTGGAGTCGTTGCTCGATTAGTTGAGCCTGTAGTCCNNNAGATCGGAAG
36456 TATGAGTGTGGAGTCGTTGCAGTCTTCAAGTCTATACCCCCCANNNAGATCGGAAG
36458 TATGAGTGTGGAGTCGTTGCCTAATCCCCAGAACCTGTGAACANNNAGATCGGAAG
36459 TATGAGTGTGGAGTCGTTGCACAAAAATTATCCAGGCGTGGTGNNNAGATCGGAA
36460 TATGAGTGTGGAGTCGTTGCAATATTACGCATCCTGTCCTGGCNNNAGATCGGAAG
36461 TATGAGTGTGGAGTCGTTGCGTAGAAAAAGAGCAGGCATGGTGNNNAGATCGGA
36463 TATGAGTGTGGAGTCGTTGCTGCTTCTTTATGTCCATCCTTTGCNNNAGATCGGAAG
36464 TATGAGTGTGGAGTCGTTGCGTTCCAGATACTGAAGAGGCTGANNNAGATCGGAA
36465 TATGAGTGTGGAGTCGTTGCGGGAAAGGTCCAGATATTTGGGTNNNAGATCGGAA
36466 TATGAGTGTGGAGTCGTTGCTGGCAGACATCTGGTTAGAAACANNNAGATCGGAA

36469 TATGAGTGTGGAGTCGTTGCTGGGACAGGGTAGGCATTCTATANNNAGATCGGAA
36470 TATGAGTGTGGAGTCGTTGCAGAGTGAATTTCAAGGTGGTGGANNNAGATCGGAA
36471 TATGAGTGTGGAGTCGTTGCAGCTGCTACCATTCCTTCTGAAANNNAGATCGGAAG
36473 TATGAGTGTGGAGTCGTTGCGGGAGATTGAGGCAAGAGAATCANNNAGATCGGA
36474 TATGAGTGTGGAGTCGTTGCCACCTCCAACACACATCACATTTNNNAGATCGGAAGA
36475 TATGAGTGTGGAGTCGTTGCTGCTAACTGTCCGAAATTCTCCANNNAGATCGGAAG
36476 TATGAGTGTGGAGTCGTTGCAGTATAATTGCCTGAACCCAGCANNNAGATCGGAAG
36477 TATGAGTGTGGAGTCGTTGCACTTTACACTCCCACCAACAGTTNNNAGATCGGAAGA
36478 TATGAGTGTGGAGTCGTTGCAGTGCCAAAAACTTACAGCCAATNNNAGATCGGAAG
36479 TATGAGTGTGGAGTCGTTGCTTTCTATGAGGCCAGGAGCATTTNNNAGATCGGAAG
36481 TATGAGTGTGGAGTCGTTGCAGATAGAGCTACAACCCCACAGANNNAGATCGGAA
36482 TATGAGTGTGGAGTCGTTGCAGAAGAAATGCACGTTCTCCTCANNNAGATCGGAAG
36483 TATGAGTGTGGAGTCGTTGCAGACAAGGGAAGGGAAGGAAAAANNNAGATCGGA
36484 TATGAGTGTGGAGTCGTTGCTGGAAAACAATGTGGAGATCCCTNNNAGATCGGAA
36485 TATGAGTGTGGAGTCGTTGCCTGAGTGAAAAAGCCCCTCCTATNNNAGATCGGAAG
36486 TATGAGTGTGGAGTCGTTGCACACACACACACACACACAAAAANNNAGATCGGAAG
36487 TATGAGTGTGGAGTCGTTGCTGTAACAGAACTCAAGGTTTCCANNNAGATCGGAAG
36488 TATGAGTGTGGAGTCGTTGCAACCTCCTTGGGTCTCTTATGTGNNNAGATCGGAAG
36489 TATGAGTGTGGAGTCGTTGCCTCCTTCCTTGCTCCTATGCTAGNNNAGATCGGAAGA
36490 TATGAGTGTGGAGTCGTTGCCGGTATGGGAGCCAATTTTTGAANNNAGATCGGAA
36491 TATGAGTGTGGAGTCGTTGCGCAGTGAGCTATGATCACAACACNNNAGATCGGAA
36493 TATGAGTGTGGAGTCGTTGCATAGCAAGAAGGCATTCACCAGANNNAGATCGGAA
36494 TATGAGTGTGGAGTCGTTGCTGAGTTACCATCTCATGCCAGTTNNNAGATCGGAAG
36495 TATGAGTGTGGAGTCGTTGCTAATGTGAACCCAGATTTGCCTGNNNAGATCGGAAG
36496 TATGAGTGTGGAGTCGTTGCTATTGTTATGGAGTGAGGGTGGGNNNAGATCGGA
36497 TATGAGTGTGGAGTCGTTGCCAAATGGAAACACATCCCATGCTNNNAGATCGGAAG
36498 TATGAGTGTGGAGTCGTTGCGAGGCAGGAGAATTGTTTGAACCNNNAGATCGGAA
36499 TATGAGTGTGGAGTCGTTGCAAAGAAGCCAGTTGCAAAGTACCNNNAGATCGGAA
36501 TATGAGTGTGGAGTCGTTGCAGAAACCCTTCTCTAAGCCTGTGNNNAGATCGGAAG
36504 TATGAGTGTGGAGTCGTTGCCCCACTTTGAACTGAGAGGAAGANNNAGATCGGAA
36505 TATGAGTGTGGAGTCGTTGCTGGGCCATACCACTCAATTTACANNNAGATCGGAAG
36506 TATGAGTGTGGAGTCGTTGCATGCCTGTACGTCTTTCTTTCCTNNNAGATCGGAAGA
36507 TATGAGTGTGGAGTCGTTGCGAAAGAAAAATTGAGTGGGGGGANNNAGATCGGA
36508 TATGAGTGTGGAGTCGTTGCTGTTCAGATGTGTGTAACCTCCTNNNAGATCGGAAG
36509 TATGAGTGTGGAGTCGTTGCGGTCTCATTTGTGCTACCCTGTANNNAGATCGGAAG
36511 TATGAGTGTGGAGTCGTTGCTTGCAAGTAATGGCACCTTCAAGNNNAGATCGGAAG
36512 TATGAGTGTGGAGTCGTTGCGTGCACACACAAATTCATGCAAANNNAGATCGGAAG
36513 TATGAGTGTGGAGTCGTTGCTGATCGATTCTGGAAGTTGGGTTNNNAGATCGGAA
36514 TATGAGTGTGGAGTCGTTGCTCACTTGAATACACAAGCCCTCANNNAGATCGGAAG
36516 TATGAGTGTGGAGTCGTTGCCACCTGTAATCCCAGCAATTTGGNNNAGATCGGAAG
36517 TATGAGTGTGGAGTCGTTGCATACTTTACTTCAGGCCAGCAGGNNNAGATCGGAAG
36519 TATGAGTGTGGAGTCGTTGCTGTGGAAGGGAACATCCATCTTCNNNAGATCGGAA
36520 TATGAGTGTGGAGTCGTTGCTGATTCAAGGTAAACCGTCCACTNNNAGATCGGAAG

36521 TATGAGTGTGGAGTCGTTGCAGATTGCAGTGAGCTGAGATCAANNNAGATCGGAA
36522 TATGAGTGTGGAGTCGTTGCCAAGGCTTGTAGGATGTCACTCTNNNAGATCGGAAG
36524 TATGAGTGTGGAGTCGTTGCTGACATTCTCTAGGTACCTGCAGNNNAGATCGGAAG
36526 TATGAGTGTGGAGTCGTTGCTGCCAAGAACATACACTGGGTAANNNAGATCGGAA
36527 TATGAGTGTGGAGTCGTTGCCTATAGCTCTGCTGAGGACAGTGNNNAGATCGGAA
36528 TATGAGTGTGGAGTCGTTGCGCCACACAGAAGTGCTTGTTATTNNNAGATCGGAAG
36529 TATGAGTGTGGAGTCGTTGCCCTAAGGACACACAGCTACACTTNNNAGATCGGAAG
36530 TATGAGTGTGGAGTCGTTGCATCTGCTGACCCAGAATATGCATNNNAGATCGGAAG
36531 TATGAGTGTGGAGTCGTTGCAAAAGACCCCCAAAGCATTTCAGNNNAGATCGGAAG
36532 TATGAGTGTGGAGTCGTTGCGTGTGAGCAACAAGGCTGTTTATNNNAGATCGGAA
36534 TATGAGTGTGGAGTCGTTGCAAAATAACTTGAGGCTGGGAGGTNNNAGATCGGAA
36535 TATGAGTGTGGAGTCGTTGCAAAGATCAGTGCAAGGGAAAAGCNNNAGATCGGAA
36536 TATGAGTGTGGAGTCGTTGCGCATGCACCTGTAGTCTCAACTANNNAGATCGGAAG
36537 TATGAGTGTGGAGTCGTTGCAGCTGGGACATCACCATTTCATANNNAGATCGGAAG
36538 TATGAGTGTGGAGTCGTTGCCATAATGCCAGCAAAGCTCACTTNNNAGATCGGAAG
36539 TATGAGTGTGGAGTCGTTGCCTTTGTAGCCTGAAGGGGAGAATNNNAGATCGGAA
36540 TATGAGTGTGGAGTCGTTGCGAACCTGCCCTTGTTTTATCAGTNNNAGATCGGAAG
36543 TATGAGTGTGGAGTCGTTGCGCAAACAAGAATGGGTCTCTTCCNNNAGATCGGAAG
36544 TATGAGTGTGGAGTCGTTGCATAAAGACAGGATCCAGCTCTGCNNNAGATCGGAAG
36545 TATGAGTGTGGAGTCGTTGCTTGCTGCTGGTGATTTTGGTTTTNNNAGATCGGAAG
36546 TATGAGTGTGGAGTCGTTGCACTTGAAGGAATTAGGAGCCCAGNNNAGATCGGAA
36547 TATGAGTGTGGAGTCGTTGCACTGCATGTTGAATGAACTCCACNNNAGATCGGAAG
36548 TATGAGTGTGGAGTCGTTGCCTGTAGTCCCAGCTACTCAGGAGNNNAGATCGGAAG
36549 TATGAGTGTGGAGTCGTTGCGTTTGTGTCCCTTGTTGCTATCCNNNAGATCGGAAG
36550 TATGAGTGTGGAGTCGTTGCGCCATGAACTGACACTTGTTGAANNNAGATCGGAAG
36551 TATGAGTGTGGAGTCGTTGCAAAAGAAATGATGCCACGTCTGGNNNAGATCGGAA
36552 TATGAGTGTGGAGTCGTTGCATGAAACAACTCCCCATTCCTGTNNNAGATCGGAAG
36553 TATGAGTGTGGAGTCGTTGCGGAACAAAGTTCCATCAAAGCCTNNNAGATCGGAAG
36554 TATGAGTGTGGAGTCGTTGCCACTAACCTGCTCTTCAACGAAANNNAGATCGGAAG
36555 TATGAGTGTGGAGTCGTTGCCTCAAGAGTTTTTTGCCTAGGCCNNNAGATCGGAAG
36556 TATGAGTGTGGAGTCGTTGCGTCCACATCCTACAGAATCCAGGNNNAGATCGGAAG
36557 TATGAGTGTGGAGTCGTTGCAATGCTGGGAATGCCAAGATCTANNNAGATCGGAA
36559 TATGAGTGTGGAGTCGTTGCACACCTGTGAATGGAGAAGCTAGNNNAGATCGGAA
36560 TATGAGTGTGGAGTCGTTGCTTGAGCCCAAGAGTTCAAGATCANNNAGATCGGAAG
36561 TATGAGTGTGGAGTCGTTGCGCCGTCAAAGAGGCAAATAAAATNNNAGATCGGAA
36562 TATGAGTGTGGAGTCGTTGCAGCATTTCTGATTGGTTGCCTTCNNNAGATCGGAAG
36563 TATGAGTGTGGAGTCGTTGCCCTCCCGAGTCTTTTCTATGCATNNNAGATCGGAAGA
36564 TATGAGTGTGGAGTCGTTGCGGCCAACATGGTGAAATCTCATCNNNAGATCGGAAG
36565 TATGAGTGTGGAGTCGTTGCAGCAATGTGGCTATGAGAAGAGANNNAGATCGGAA
36566 TATGAGTGTGGAGTCGTTGCTAAAGGCCACCAGAACAATAGCTNNNAGATCGGAAG
36567 TATGAGTGTGGAGTCGTTGCCTGAGGCAGGAAAATCGTTTGAANNNAGATCGGAA
36568 TATGAGTGTGGAGTCGTTGCTAGGGACGTTTTGGAAATGCAACNNNAGATCGGAA
36569 TATGAGTGTGGAGTCGTTGCAAATGGGCAAACTGTATGGCATGNNNAGATCGGAA

36570 TATGAGTGTGGAGTCGTTGCGATTGTTTGAGTCCAGCAGTTCCNNNAGATCGGAAG
36572 TATGAGTGTGGAGTCGTTGCGATCACGAGGTCAGGAGTTCAAGNNNAGATCGGAA
36573 TATGAGTGTGGAGTCGTTGCGAGCAGAAAACTGAAAGCACTGTNNNAGATCGGAA
36574 TATGAGTGTGGAGTCGTTGCAGATCACCCAAGGTCAAGAGTTCNNNAGATCGGAA
36575 TATGAGTGTGGAGTCGTTGCCATAAATTAGCCAGGTGTGGTGGNNNAGATCGGAA
36577 TATGAGTGTGGAGTCGTTGCTTGAATCCTTTGTGCAGCTGAAANNNAGATCGGAAG
36578 TATGAGTGTGGAGTCGTTGCCGTGTACCTGTAGTCCTAGCTACNNNAGATCGGAAG
36580 TATGAGTGTGGAGTCGTTGCATGTGGATGAGTCATGGACACAANNNAGATCGGAA
36583 TATGAGTGTGGAGTCGTTGCATTCACAGACCCCTGAACTAACCNNNAGATCGGAAG
36585 TATGAGTGTGGAGTCGTTGCAACTCAAAGATGGGGCCAATAGTNNNAGATCGGAA
36586 TATGAGTGTGGAGTCGTTGCTTACTTTTCACTGCTAGGGGGACNNNAGATCGGAAG
36587 TATGAGTGTGGAGTCGTTGCGCAATACCGGTAACTTTGTCACCNNNAGATCGGAAG
36588 TATGAGTGTGGAGTCGTTGCGAGGCTTTATGACAAAAGGAGGCNNNAGATCGGAA
36589 TATGAGTGTGGAGTCGTTGCAGATGAAAAATGGTTCACCTTTGTNNNAGATCGGAA
36591 TATGAGTGTGGAGTCGTTGCAAACACAAACTCTACAGATGGCANNNAGATCGGAAG
36593 TATGAGTGTGGAGTCGTTGCCTGTAGAACTTTGCCAAGGTGTGNNNAGATCGGAA
36594 TATGAGTGTGGAGTCGTTGCTGCCTTTAGAGAGATAAGTGGCANNNAGATCGGAA
36595 TATGAGTGTGGAGTCGTTGCTCGTGCGTAGCATCACAATTTTTNNNAGATCGGAAG
36596 TATGAGTGTGGAGTCGTTGCAGGAGCTATTACAACTGACACCANNNAGATCGGAAG
36597 TATGAGTGTGGAGTCGTTGCTTACAGATTCAATGCCATCCCCANNNAGATCGGAAG
36598 TATGAGTGTGGAGTCGTTGCCCAATGAAATGGCACCTCTCAAGNNNAGATCGGAAG
36599 TATGAGTGTGGAGTCGTTGCGCAAGGTCCTGACTCTTTCTACANNNAGATCGGAAG
36601 TATGAGTGTGGAGTCGTTGCCTGATCCCGGTGTCACTATTCATNNNAGATCGGAAG
36602 TATGAGTGTGGAGTCGTTGCAAGCATAGAGAAATGGGAGACCCNNNAGATCGGAA
36603 TATGAGTGTGGAGTCGTTGCACACAATGAACTCCCTGAAAGGANNNAGATCGGAA
36604 TATGAGTGTGGAGTCGTTGCCAGAAAGTCAATTACGGCTCACCNNNAGATCGGAAG
36605 TATGAGTGTGGAGTCGTTGCATGAATTCCAAGGTCTCTCTCGGNNNAGATCGGAAG
36606 TATGAGTGTGGAGTCGTTGCGTTGCAGTCCTGGTCTCTGATAANNNAGATCGGAAG
36607 TATGAGTGTGGAGTCGTTGCGAAGAAACGAAACCTGGATCAGGNNNAGATCGGAA
36608 TATGAGTGTGGAGTCGTTGCAAGCAGGCTTTCATTTATGGCTCNNNAGATCGGAAG
36609 TATGAGTGTGGAGTCGTTGCTTTGGGGAAAAACTGTGCTGTTTNNNAGATCGGAA
36610 TATGAGTGTGGAGTCGTTGCGACTGATGCTACAGTGAGCCATANNNAGATCGGAA
36611 TATGAGTGTGGAGTCGTTGCAAGGGGAATGCTCAAGATCTCAGNNNAGATCGGAA
36612 TATGAGTGTGGAGTCGTTGCAGATCATGTGTGTATGGATCGGGNNNAGATCGGAA
36613 TATGAGTGTGGAGTCGTTGCTAAAGGCCACTAGGAAACAGTCCNNNAGATCGGAA
36614 TATGAGTGTGGAGTCGTTGCTCTTCTGGGCTGAATCAGGATTCNNNAGATCGGAAG
36615 TATGAGTGTGGAGTCGTTGCAGGTTGAGGCAGGAGAATCAATTNNNAGATCGGAA
36616 TATGAGTGTGGAGTCGTTGCTATCTTGTGAGAGCAGAGGGAGANNNAGATCGGAA
36617 TATGAGTGTGGAGTCGTTGCGAAGGATGGGAAAAGAGCCAAACNNNAGATCGGA
36618 TATGAGTGTGGAGTCGTTGCAAGTTCAAGAACAAGGAAAGCCANNNAGATCGGAA
36619 TATGAGTGTGGAGTCGTTGCTTGCAAAACTCTCTAGTCCCACANNNAGATCGGAAG
36620 TATGAGTGTGGAGTCGTTGCGGGGGGGAATAAAAGACTTGTCANNNAGATCGGA
36622 TATGAGTGTGGAGTCGTTGCGACCCAGATCCAAACATACTGGTNNNAGATCGGAAG

36624 TATGAGTGTGGAGTCGTTGCTACAATGCAGGATGTTTGTGCTGNNNAGATCGGAA
36625 TATGAGTGTGGAGTCGTTGCGTTTCAAGGACTATGCTGCAACCNNNAGATCGGAAG
36626 TATGAGTGTGGAGTCGTTGCGGCCTATCAATCAACTGCAACATNNNAGATCGGAAG
36627 TATGAGTGTGGAGTCGTTGCGTCCAGAACATGAGAAACCAGGANNNAGATCGGAA
36628 TATGAGTGTGGAGTCGTTGCTATTCCTGGAAGCCTCAATCGAGNNNAGATCGGAAG
36629 TATGAGTGTGGAGTCGTTGCAAACAAACAACTTGGCCTGCAATNNNAGATCGGAAG
36630 TATGAGTGTGGAGTCGTTGCCTCCCTGGGTGCTAGAATTACAGNNNAGATCGGAAG
36631 TATGAGTGTGGAGTCGTTGCGAGGCCCTGTCTCCAAAAAAAAANNNAGATCGGAA
36632 TATGAGTGTGGAGTCGTTGCCCTCTCATTGCAATAGCTTTCCGNNNAGATCGGAAG
36633 TATGAGTGTGGAGTCGTTGCAAAGTATTCAAGGTTTGGCACGGNNNAGATCGGAA
36635 TATGAGTGTGGAGTCGTTGCATAATCCCAGCTACTTGGAAGGCNNNAGATCGGAAG
36636 TATGAGTGTGGAGTCGTTGCAAAGAAATTAGCCAGGTGCAGTGNNNAGATCGGAA
36637 TATGAGTGTGGAGTCGTTGCGTGACCCAAACAGAAGGGAAAACNNNAGATCGGAA
36639 TATGAGTGTGGAGTCGTTGCAGATTCTCCAACCCTAGAGAGGANNNAGATCGGAA
36640 TATGAGTGTGGAGTCGTTGCCCCCTGATGGCCTACTTTTCATANNNAGATCGGAAG
36641 TATGAGTGTGGAGTCGTTGCAGAAACAAACGATGGAGACTGGANNNAGATCGGA
36642 TATGAGTGTGGAGTCGTTGCACAAAAATTAGCCAGGTGTGGTGNNNAGATCGGAA
36643 TATGAGTGTGGAGTCGTTGCGACTGAACAAACTCCTTCTTGGCNNNAGATCGGAAG
36644 TATGAGTGTGGAGTCGTTGCAGACTGAACATTGTGACAGAACTNNNAGATCGGAA
36646 TATGAGTGTGGAGTCGTTGCTGGAATCCTGTAAAACGGTCAGTNNNAGATCGGAA
36648 TATGAGTGTGGAGTCGTTGCGAGGGATGTGATATTGACCCTGGNNNAGATCGGAA
36649 TATGAGTGTGGAGTCGTTGCTGTAGCACATTTGACCTAGGCAANNNAGATCGGAAG
36650 TATGAGTGTGGAGTCGTTGCGTCTGGGAAATGGAGTTTTGGTGNNNAGATCGGAA
36651 TATGAGTGTGGAGTCGTTGCCTTGTTTCTTCCACAGTGCTGAGNNNAGATCGGAAG
36652 TATGAGTGTGGAGTCGTTGCGGCAGTGAGTCTCTAGCATGAATNNNAGATCGGAA
36653 TATGAGTGTGGAGTCGTTGCTCCAGGGCTTTCTTTTCAAAACCNNNAGATCGGAAG
36654 TATGAGTGTGGAGTCGTTGCCTATACTCCAGCCTAGGCAACAGNNNAGATCGGAAG
36655 TATGAGTGTGGAGTCGTTGCGTGCTAAGAGGATTCAATGGGGANNNAGATCGGA
36656 TATGAGTGTGGAGTCGTTGCAGTTCAGGAAAATGTGATGCTTCANNNAGATCGGA
36657 TATGAGTGTGGAGTCGTTGCGCTTTTGCTCAGACATCACCTTTNNNAGATCGGAAG
36659 TATGAGTGTGGAGTCGTTGCGGCCTTCCTCACCCATGTATATTNNNAGATCGGAAG
36660 TATGAGTGTGGAGTCGTTGCGAGGACAGAAAACAGACACAACGNNNAGATCGGAA
36662 TATGAGTGTGGAGTCGTTGCCACCTGTGATCCTAGCTACTCAGNNNAGATCGGAAG
36663 TATGAGTGTGGAGTCGTTGCTGGGATTAGGGAAGATCAGGGAANNNAGATCGGA
36666 TATGAGTGTGGAGTCGTTGCAAACTGAACCCAACATAAGCAGANNNAGATCGGAA
36668 TATGAGTGTGGAGTCGTTGCTGATGTATAGCAACGACCCACTTNNNAGATCGGAAG
36670 TATGAGTGTGGAGTCGTTGCAGTCAAAGAGACCAGTGGATCACNNNAGATCGGAA
36671 TATGAGTGTGGAGTCGTTGCTGTTTAGGGCTAGACCTGTTGTCNNNAGATCGGAAG
36674 TATGAGTGTGGAGTCGTTGCCATGTCTGTGATCCCAGGTACTTNNNAGATCGGAAG
36675 TATGAGTGTGGAGTCGTTGCGCATCTACTTGGCTTCTAGGGAGNNNAGATCGGAA
36676 TATGAGTGTGGAGTCGTTGCCTATCAGTGTGTTGGTGTTGCTGNNNAGATCGGAA
36677 TATGAGTGTGGAGTCGTTGCAAACTTTTGGAGTTGAAGCCCACNNNAGATCGGAAG
36678 TATGAGTGTGGAGTCGTTGCGCTACAGAACGAGACTCCATCTCNNNAGATCGGAAG

36679 TATGAGTGTGGAGTCGTTGCCAGCCCAGTCAACCACTAATCTANNNAGATCGGAAGA
36680 TATGAGTGTGGAGTCGTTGCAAAAAACAAAAGAGCCACCACCANNNAGATCGGAAG
36681 TATGAGTGTGGAGTCGTTGCTAGGTGTGACCAATGTTTCCCATNNNAGATCGGAAG
36682 TATGAGTGTGGAGTCGTTGCTGCACCTGAAGGAAAATACCTGANNNAGATCGGAAG
36683 TATGAGTGTGGAGTCGTTGCCACGCCTGTAATCCCAACATTTTNNNAGATCGGAAGA
36684 TATGAGTGTGGAGTCGTTGCAATTTGTATTGGGCCACATGTGGNNNAGATCGGAAG
36685 TATGAGTGTGGAGTCGTTGCTCTGACACATGGTGAGAACTCAGNNNAGATCGGAAG
36686 TATGAGTGTGGAGTCGTTGCTTCTGACATTATTGGTCTGGGGGNNNAGATCGGAAG
36687 TATGAGTGTGGAGTCGTTGCAAGAAGAAAGAAAATGCGCTGGGNNNAGATCGGAA
36688 TATGAGTGTGGAGTCGTTGCAACCAGACAAAAAGCAGAACACCNNNAGATCGGAAG
36689 TATGAGTGTGGAGTCGTTGCCTGTAATACCAGCTGCTCAGGAGNNNAGATCGGAAG
36690 TATGAGTGTGGAGTCGTTGCACTGATTCCTAGTCTTGCCAAGGNNNAGATCGGAAG
36692 TATGAGTGTGGAGTCGTTGCAATTAGAAACTCTGAGGCTGGGGNNNAGATCGGAA
36693 TATGAGTGTGGAGTCGTTGCATACCTACTATGGCATGGTTGGTNNNAGATCGGAAG
36694 TATGAGTGTGGAGTCGTTGCACTTTGCCTCATGGAACTTACCTNNNAGATCGGAAGA
36695 TATGAGTGTGGAGTCGTTGCCTACCCCCAGAGTGCATAGAATCNNNAGATCGGAAG
36696 TATGAGTGTGGAGTCGTTGCCGAGACAGGACAACAAACGAAAANNNAGATCGGAA
36697 TATGAGTGTGGAGTCGTTGCGGCACCTATAATCCCAGCTACTCNNNAGATCGGAAGA
36698 TATGAGTGTGGAGTCGTTGCAAAGGATGGAGAAGAAAAGGCCANNNAGATCGGAA
36700 TATGAGTGTGGAGTCGTTGCAGCGAGACACCGTTTCAAAAAAANNNAGATCGGAAG
36701 TATGAGTGTGGAGTCGTTGCTGAGAGGCAATAGTGACAATGCTNNNAGATCGGAAG
36702 TATGAGTGTGGAGTCGTTGCGGACAAGCAGGAATGTAATGACGNNNAGATCGGAA
36703 TATGAGTGTGGAGTCGTTGCCAGCATAGCATGGTTTTGTCTGTNNNAGATCGGAAG
36704 TATGAGTGTGGAGTCGTTGCTGTGTAGAATTCCCCCATTGAGGNNNAGATCGGAAG
36705 TATGAGTGTGGAGTCGTTGCAAATAGTCACGGGGGACAAAGAANNNAGATCGGAA
36706 TATGAGTGTGGAGTCGTTGCCGCCTGTAATCCTAGCATTTTGGNNNAGATCGGAAGA
36707 TATGAGTGTGGAGTCGTTGCCCAGAAAAGGCAAAACTACAGGGNNNAGATCGGAA
36708 TATGAGTGTGGAGTCGTTGCCTTCTGTCCCCTTCGTTTAGTCCNNNAGATCGGAAGAG
36709 TATGAGTGTGGAGTCGTTGCAGCAGACTAGGGAATCATTGGATNNNAGATCGGAAG
36710 TATGAGTGTGGAGTCGTTGCGTGACATACACCTGTAGTCCCAGNNNAGATCGGAAG
36711 TATGAGTGTGGAGTCGTTGCCAGAGGTAGAAGGATCGCTTGAGNNNAGATCGGAA
36712 TATGAGTGTGGAGTCGTTGCATTTCACACCTCAAAAGTTGGCTNNNAGATCGGAAGA
36713 TATGAGTGTGGAGTCGTTGCCACATAATGAGTGTCTGGGTGCTNNNAGATCGGAAG
36714 TATGAGTGTGGAGTCGTTGCTCCTGAGATGAATACGTTTCCCGNNNAGATCGGAAG
36715 TATGAGTGTGGAGTCGTTGCAAAACAGAACATCTGGCCATTCANNNAGATCGGAAG
36716 TATGAGTGTGGAGTCGTTGCTGTGACAGCAAGGATGATGACAANNNAGATCGGAA
36717 TATGAGTGTGGAGTCGTTGCCTTCTCAAAGGACCACCTACGAANNNAGATCGGAAGA
36718 TATGAGTGTGGAGTCGTTGCATTTCCCAGCATCCTTTGAGTCTNNNAGATCGGAAGA
36719 TATGAGTGTGGAGTCGTTGCGCTCATGTCTGTAATCCCAACACNNNAGATCGGAAGA
36720 TATGAGTGTGGAGTCGTTGCAATCCTAGAATTTTGGGAGGCCANNNAGATCGGAAG
36722 TATGAGTGTGGAGTCGTTGCCGACCAGGCATCTATGGAGATTANNNAGATCGGAAG
36723 TATGAGTGTGGAGTCGTTGCAAACTGAGGCTCGAGGAGTTTAGNNNAGATCGGAA
36724 TATGAGTGTGGAGTCGTTGCGGGAGGGAGGGAAAGAAAAGAAANNNAGATCGGA

36725 TATGAGTGTGGAGTCGTTGCGGGGATTTCTAGCCTTGCTCTTANNNAGATCGGAAG
36726 TATGAGTGTGGAGTCGTTGCTTTGGGAGACGGCTTGATTTTTCNNNAGATCGGAAG
36727 TATGAGTGTGGAGTCGTTGCGAAATAACCAGACACCACCCTCTNNNAGATCGGAAG
36728 TATGAGTGTGGAGTCGTTGCCAATGAGTCTGTGGGGTGTCTAANNNAGATCGGAA
36729 TATGAGTGTGGAGTCGTTGCTCTAAAAGCAGCACTGGGTTTTGNNNAGATCGGAAG
36730 TATGAGTGTGGAGTCGTTGCCTGTCTCCTCAGGCTTCTTTCTTNNNAGATCGGAAGA
36731 TATGAGTGTGGAGTCGTTGCATGGTATTGGCACAATTGGACACNNNAGATCGGAAG
36732 TATGAGTGTGGAGTCGTTGCAACAGAGCAAGATTCCGTCTCAANNNAGATCGGAAG
36733 TATGAGTGTGGAGTCGTTGCTGTCAAGAGCAAATTCTGGGGTANNNAGATCGGAA
36734 TATGAGTGTGGAGTCGTTGCGTAGTCTCAGCTACTTGAGAGGCNNNAGATCGGAAG
36735 TATGAGTGTGGAGTCGTTGCGGGGACAGCAGCTATATTCACTTNNNAGATCGGAAG
36736 TATGAGTGTGGAGTCGTTGCCCTTCCTAGATGGTGAGACCCTANNNAGATCGGAAG
36737 TATGAGTGTGGAGTCGTTGCTGTGTAGTTGTGGACTCATCTCTNNNAGATCGGAAG
36738 TATGAGTGTGGAGTCGTTGCCATCATGGGAGAAAACACTGCAGNNNAGATCGGAA
36739 TATGAGTGTGGAGTCGTTGCAAAACGAAACAAACCCCCCAAATNNNAGATCGGAAG
36741 TATGAGTGTGGAGTCGTTGCGTGGATCATTTGAGGTCAGGAGTNNNAGATCGGAA
36743 TATGAGTGTGGAGTCGTTGCCTCATGCCTGTAATCCCAGCTACNNNAGATCGGAAG
36744 TATGAGTGTGGAGTCGTTGCCTGTAATCCCAGCTACTCAGGAGNNNAGATCGGAAG
36745 TATGAGTGTGGAGTCGTTGCAACTCCTAGTTCTGAAGGTGCTTNNNAGATCGGAAG
36746 TATGAGTGTGGAGTCGTTGCTGGATTTTCAGGTCCTGGAGATGNNNAGATCGGAA
36747 TATGAGTGTGGAGTCGTTGCGGGGCCATAAGGATGTTCTTACANNNAGATCGGAA
36748 TATGAGTGTGGAGTCGTTGCGAAAGAAACCAGTTGCAAAAGGCNNNAGATCGGAA
36749 TATGAGTGTGGAGTCGTTGCTAATGGCAGTTGAAAAGGCACAGNNNAGATCGGAA
36750 TATGAGTGTGGAGTCGTTGCTCAGTGTCTGTGTAATGAGCAGTNNNAGATCGGAAG
36752 TATGAGTGTGGAGTCGTTGCCCCCTCTAAAAATAATGGCCAGCNNNAGATCGGAAG
36753 TATGAGTGTGGAGTCGTTGCAATCCCAACTATTCAGGAGGCTGNNNAGATCGGAAG
36754 TATGAGTGTGGAGTCGTTGCGCTCAGCAAGTAACCCCAAAAATNNNAGATCGGAAG
36755 TATGAGTGTGGAGTCGTTGCAGTGTTTTTGCTCCATTGCTGTTNNNAGATCGGAAG
36757 TATGAGTGTGGAGTCGTTGCAACAAGATGGAGTTGGAGTTGGTNNNAGATCGGAA
36758 TATGAGTGTGGAGTCGTTGCACTGGACTTCTGGCCTATCACTANNNAGATCGGAAG
36759 TATGAGTGTGGAGTCGTTGCTGGGAGGAAAGGAGAAAATGAGGNNNAGATCGGA
36762 TATGAGTGTGGAGTCGTTGCTCAGGAGACTGAGACAGGAGAATNNNAGATCGGAA
36763 TATGAGTGTGGAGTCGTTGCATGTCTCTCTGAGGTGAAGCTCTNNNAGATCGGAAG
36764 TATGAGTGTGGAGTCGTTGCAGCCTAGAGGACATCTCACTCTTNNNAGATCGGAAG
36767 TATGAGTGTGGAGTCGTTGCCCTGGCTCTCTTCCTTTCATCAANNNAGATCGGAAGA
36768 TATGAGTGTGGAGTCGTTGCTCTGAGGCAAAGTTTAGACCACANNNAGATCGGAAG
36769 TATGAGTGTGGAGTCGTTGCAGGCAGTGACATAATCGAACTGTNNNAGATCGGAA
36770 TATGAGTGTGGAGTCGTTGCACCCACAAGTCTCTGGAAATTCANNNAGATCGGAAG
36771 TATGAGTGTGGAGTCGTTGCCGACAGAACAAGACCCCATCTAANNNAGATCGGAAG
36772 TATGAGTGTGGAGTCGTTGCCTGCCTCATAGTCTCTCAACGTTNNNAGATCGGAAGA
36774 TATGAGTGTGGAGTCGTTGCGAGTTCCCCTCAGTAACCTCAACNNNAGATCGGAAG
36776 TATGAGTGTGGAGTCGTTGCGGAATAAAGTAAGCCACACAGCCNNNAGATCGGAA
36777 TATGAGTGTGGAGTCGTTGCAGGGATAGAAGCACACATACTGCNNNAGATCGGAA

36778 TATGAGTGTGGAGTCGTTGCCATGCCTGTAGTCTCAGCTACTTNNNAGATCGGAAG
36780 TATGAGTGTGGAGTCGTTGCAGGATTGGTAAGAAGTTTCAGCANNNAGATCGGAA
36782 TATGAGTGTGGAGTCGTTGCCCTAATCTAGGGTTGCTCTGTTANNNAGATCGGAAG
36783 TATGAGTGTGGAGTCGTTGCGACCATTTTTGCTGCATTGAAGCNNNAGATCGGAAG
36784 TATGAGTGTGGAGTCGTTGCATGAGCTGCTTATTCCTCAGAGGNNNAGATCGGAAG
36785 TATGAGTGTGGAGTCGTTGCGCCATGTATGATAGACTCACAGCNNNAGATCGGAAG
36787 TATGAGTGTGGAGTCGTTGCTGTGGCAATTCCTCAAAGACCTANNNAGATCGGAAG
36788 TATGAGTGTGGAGTCGTTGCTGCCTCATGACAGAATGTGTACANNNAGATCGGAAG
36789 TATGAGTGTGGAGTCGTTGCTACACCATGGGATACTATGCAGCNNNAGATCGGAAG
36792 TATGAGTGTGGAGTCGTTGCTACGTTTTTACACTGGGCAGAGANNNAGATCGGAAG
36794 TATGAGTGTGGAGTCGTTGCGTAAGCAATAGGGGGCTAGGATCNNNAGATCGGAA
36795 TATGAGTGTGGAGTCGTTGCAAGAAAGCTTCATTTGAGCAGGTNNNAGATCGGAA
36796 TATGAGTGTGGAGTCGTTGCTTGTGTCAGGGAAACTGGTGAATNNNAGATCGGAA
36798 TATGAGTGTGGAGTCGTTGCCCCCAGAACCATGAGAGAGAAAANNNAGATCGGAA
36799 TATGAGTGTGGAGTCGTTGCCAAGTACTGGGATTCCACTGACTNNNAGATCGGAAG
36800 TATGAGTGTGGAGTCGTTGCGATTTTGAAGCACCTATGGACGCNNNAGATCGGAAG
36802 TATGAGTGTGGAGTCGTTGCGCAGTCTCAACAGTCACTGAGTANNNAGATCGGAAG
36803 TATGAGTGTGGAGTCGTTGCTGTGGACACTGGTTGTTACAGATNNNAGATCGGAAG
36804 TATGAGTGTGGAGTCGTTGCAATACAAAAAATTAGCCGGGCGTNNNAGATCGGAA
36805 TATGAGTGTGGAGTCGTTGCCCCTTTCACCACTCATTGTCAACNNNAGATCGGAAGA
36806 TATGAGTGTGGAGTCGTTGCCAGTTACTTGGGAAGGTGAGGAANNNAGATCGGAA
36807 TATGAGTGTGGAGTCGTTGCTGGTTATTGACTAAGGGTGTGACNNNAGATCGGAA
36808 TATGAGTGTGGAGTCGTTGCTGGAAGTTCCAAGCCTGAGTAAANNNAGATCGGAA
36809 TATGAGTGTGGAGTCGTTGCCTCATTCCTGTAATCCCAGCACTNNNAGATCGGAAGA
36810 TATGAGTGTGGAGTCGTTGCGAGTGAAAAGGCAACCCATGAAANNNAGATCGGAA
36811 TATGAGTGTGGAGTCGTTGCTATGTGGGACTATTGCATGATGCNNNAGATCGGAAG
36813 TATGAGTGTGGAGTCGTTGCAGGGCAAACATAAGCGAGAAAACNNNAGATCGGAA
36815 TATGAGTGTGGAGTCGTTGCCGGCTGTTTGTGATAGCAAAAACNNNAGATCGGAAG
36816 TATGAGTGTGGAGTCGTTGCATAAAAGCAGGTGGCAAATAGGCNNNAGATCGGAA
36817 TATGAGTGTGGAGTCGTTGCATGCATCAGAGCTCTGTCCATAGNNNAGATCGGAAG
36818 TATGAGTGTGGAGTCGTTGCACATATGTGGACACAGAGACACTNNNAGATCGGAAG
36819 TATGAGTGTGGAGTCGTTGCTCTGCTAGCATCTCTGAAAAGGGNNNAGATCGGAAG
36821 TATGAGTGTGGAGTCGTTGCGGTGAAACCCCGTCTCTACTAAANNNAGATCGGAAG
36822 TATGAGTGTGGAGTCGTTGCGACATGGAAACAGGAAGTCTTGCNNNAGATCGGAA
36823 TATGAGTGTGGAGTCGTTGCGTAGGGCAGAAAAACCTATGCACNNNAGATCGGAA
36824 TATGAGTGTGGAGTCGTTGCTTCAGAAGTGATTCTCAGCCACANNNAGATCGGAAG
36825 TATGAGTGTGGAGTCGTTGCCCATTCTAGCCTAGCTTCTGGAGNNNAGATCGGAAG
36826 TATGAGTGTGGAGTCGTTGCTGGCTCTTATAATCCCCTTCTGCNNNAGATCGGAAGA
36828 TATGAGTGTGGAGTCGTTGCGACTAATACAGAGGCCCAGGATGNNNAGATCGGAA
36829 TATGAGTGTGGAGTCGTTGCACAAGTCCCTGTAATGCCTTGAANNNAGATCGGAAG
36830 TATGAGTGTGGAGTCGTTGCGTTAGGGGTGTTTCTTGGTGTTGNNNAGATCGGAA
36831 TATGAGTGTGGAGTCGTTGCTCAAATACTGAGCTCCTTGAGGGNNNAGATCGGAAG
36832 TATGAGTGTGGAGTCGTTGCTTTGCTTAGTCTGTGAGCTGTCTNNNAGATCGGAAG

36839 TATGAGTGTGGAGTCGTTGCGGATCACTTGAGCACAGGAGTTCNNNAGATCGGAA
36841 TATGAGTGTGGAGTCGTTGCACTGGTAAACTGGCTTCTATCCANNNAGATCGGAAG
36842 TATGAGTGTGGAGTCGTTGCTCAACATAAAAAGCCGTTTTGCCNNNAGATCGGAAG
36844 TATGAGTGTGGAGTCGTTGCTCTCTCAGGATGGGGCATTAGTANNNAGATCGGAA
36845 TATGAGTGTGGAGTCGTTGCGCAGGCCTTGATTGAGTAGACTTNNNAGATCGGAA
36846 TATGAGTGTGGAGTCGTTGCGGGAGACTAAAGTGGAGGATCACNNNAGATCGGA
36847 TATGAGTGTGGAGTCGTTGCGGTGCAAAGAATGTGATTTCTGGNNNAGATCGGAA
36849 TATGAGTGTGGAGTCGTTGCGAAACTTCTTGATCGAGCTCTGCNNNAGATCGGAAG
36850 TATGAGTGTGGAGTCGTTGCGTTGTAGTGAGCAGAGATTGTGCNNNAGATCGGAA
36851 TATGAGTGTGGAGTCGTTGCTGAGACAGTTTCGCTATTGTTGCNNNAGATCGGAAG
36852 TATGAGTGTGGAGTCGTTGCAAACTTTGATTCTGTGCGTCCACNNNAGATCGGAAG
36853 TATGAGTGTGGAGTCGTTGCAAGCAATCTGGAGAAGACTCAGTNNNAGATCGGAA
36854 TATGAGTGTGGAGTCGTTGCTTATGAAAAAAAGCCAGGCGTGGNNNAGATCGGAA
36856 TATGAGTGTGGAGTCGTTGCTCAGGAGACTCAGTCAGGAGAATNNNAGATCGGAA
36857 TATGAGTGTGGAGTCGTTGCGGAATTAGTCCTTCTCGAGCCTANNNAGATCGGAAG
36858 TATGAGTGTGGAGTCGTTGCACCATTGATGGGCATTTAGGTTGNNNAGATCGGAA
36860 TATGAGTGTGGAGTCGTTGCGGGCAGGAAGTGACCATATACAANNNAGATCGGAA
36861 TATGAGTGTGGAGTCGTTGCAAACACTATGCCTCTCAGTGGAGNNNAGATCGGAAG
36863 TATGAGTGTGGAGTCGTTGCCTACCAGAATGGAAGCCCTTTCANNNAGATCGGAAG
36864 TATGAGTGTGGAGTCGTTGCGTAATCCCAGCTACTTGAGAGGCNNNAGATCGGAAG
36865 TATGAGTGTGGAGTCGTTGCGCCTGAGTGACAAAGGGAGATAANNNAGATCGGA
36867 TATGAGTGTGGAGTCGTTGCAAAGAGTGGTTGAGACAGGATCANNNAGATCGGA
36868 TATGAGTGTGGAGTCGTTGCAAGAACCCCCAAATTCCTTAGCTNNNAGATCGGAAG
36869 TATGAGTGTGGAGTCGTTGCAGAAAAGAAAAGCAAGCCCATGTNNNAGATCGGAA
36871 TATGAGTGTGGAGTCGTTGCGGCAAGAAAATGAAACAAAGGGCNNNAGATCGGA
36872 TATGAGTGTGGAGTCGTTGCGTAATCCCAAGACTTTGTGAGGCNNNAGATCGGAAG
36873 TATGAGTGTGGAGTCGTTGCAAACCCAAGAAATTCCAAGCAGGNNNAGATCGGAA
36876 TATGAGTGTGGAGTCGTTGCGTTTCATGCTTTGACCAGGAGTCNNNAGATCGGAAG
36877 TATGAGTGTGGAGTCGTTGCTGGTCTACAAAACAGGACTTCCTNNNAGATCGGAAG
36878 TATGAGTGTGGAGTCGTTGCCGCGAAACAGTTGAGAGATACAGNNNAGATCGGAA
36879 TATGAGTGTGGAGTCGTTGCGACAACTACCACAGACTGGGAATNNNAGATCGGAA
36880 TATGAGTGTGGAGTCGTTGCCTCCAATCCCAGAGGACTATCCTNNNAGATCGGAAG
36882 TATGAGTGTGGAGTCGTTGCAGTTTCTCCACATCCTCATCCACNNNAGATCGGAAGA
36885 TATGAGTGTGGAGTCGTTGCCAGGCCTGTAATCCCAACACTTTNNNAGATCGGAAG
36886 TATGAGTGTGGAGTCGTTGCGAGAATCGCTTGAACCCAGGAAGNNNAGATCGGAA
36889 TATGAGTGTGGAGTCGTTGCGAAAAGAAAAGCAATGGCAGCATNNNAGATCGGAA
36890 TATGAGTGTGGAGTCGTTGCTACCCCAAAGCCATATAAGCACANNNAGATCGGAAG
36892 TATGAGTGTGGAGTCGTTGCTGTAATGGACACACCAGCTCAAANNNAGATCGGAAG
36893 TATGAGTGTGGAGTCGTTGCCACACCTGTAATCCCAGCACTTTNNNAGATCGGAAGA
36894 TATGAGTGTGGAGTCGTTGCAGCCCTGATGAAGATGTACAAGGNNNAGATCGGAA
36896 TATGAGTGTGGAGTCGTTGCAAAATAATATGCCAGGCACGGTGNNNAGATCGGAA
36897 TATGAGTGTGGAGTCGTTGCACCCCAAAGAAGTAGAAAGATGGGNNNAGATCGGA
36899 TATGAGTGTGGAGTCGTTGCTGCACAAAAAACCCACAGAATGGNNNAGATCGGAA

36900 TATGAGTGTGGAGTCGTTGCAAGAAAACAAGATTTGGCTGGGCNNNAGATCGGAA
36901 TATGAGTGTGGAGTCGTTGCTTTCGTCAAATAACCCACAGCACNNNAGATCGGAAG
36902 TATGAGTGTGGAGTCGTTGCGCTAGGAGAACCCAGGAAATTCANNNAGATCGGAA
36904 TATGAGTGTGGAGTCGTTGCTCCAAGTTGAACTCCTTTTAGGTNNNAGATCGGAAG
36905 TATGAGTGTGGAGTCGTTGCGTCCCAGGCACTATGGTAAGTAGNNNAGATCGGAA
36906 TATGAGTGTGGAGTCGTTGCTGTAATCCCAGGTACTCAGGAGANNNAGATCGGAA
36907 TATGAGTGTGGAGTCGTTGCCTGTAGTCCCAGCTACTCAGGAGNNNAGATCGGAAG
36909 TATGAGTGTGGAGTCGTTGCTGCACGGAAAGAAGGAAGATACANNNAGATCGGA
36910 TATGAGTGTGGAGTCGTTGCGGTAGCACTGAGTTTTTGGCTACNNNAGATCGGAA
36911 TATGAGTGTGGAGTCGTTGCAAAATGATGGGGCTGTGTCAAAGNNNAGATCGGAA
36912 TATGAGTGTGGAGTCGTTGCGGAAACATACCAAGACCCTGTCTNNNAGATCGGAAG
36913 TATGAGTGTGGAGTCGTTGCCCTGTAATTGCAGCTACTTGGGANNNAGATCGGAAG
36914 TATGAGTGTGGAGTCGTTGCCTGAGATGGGAGGATCACTTGAGNNNAGATCGGAA
36915 TATGAGTGTGGAGTCGTTGCAATGGCCGATTTCTCCCACTTATNNNAGATCGGAAG
36916 TATGAGTGTGGAGTCGTTGCTGCAGGAATCATTTTCAACTGCTNNNAGATCGGAAG
36917 TATGAGTGTGGAGTCGTTGCATCATCCTTCTCACTGGCACATTNNNAGATCGGAAGA
36918 TATGAGTGTGGAGTCGTTGCAAACAGAAAGGGTTTGCTTCTCGNNNAGATCGGAA
36920 TATGAGTGTGGAGTCGTTGCTTGACCTCTTAGGCTCAAGCAATNNNAGATCGGAAG
36921 TATGAGTGTGGAGTCGTTGCAACTCACTAATGTTTTGTGGCCCNNNAGATCGGAAG
36922 TATGAGTGTGGAGTCGTTGCTAAGGATGAGGTAGGAGGATCCCNNNAGATCGGA
36924 TATGAGTGTGGAGTCGTTGCGAACTTGAACTCAGGGCTCAGTANNNAGATCGGAA
36926 TATGAGTGTGGAGTCGTTGCAGAAAGAAATGCTTGGGACCAGANNNAGATCGGAA
36927 TATGAGTGTGGAGTCGTTGCGGACACAGGCACAGAAGGTATAANNNAGATCGGAA
36931 TATGAGTGTGGAGTCGTTGCCTGTGTGTGTCCATGAGGGTATANNNAGATCGGAA
36932 TATGAGTGTGGAGTCGTTGCTGAGCTGGTATCAAGCTAACTGGNNNAGATCGGAA
36934 TATGAGTGTGGAGTCGTTGCTACCCCAATAAACTCTCTCCTGCNNNAGATCGGAAGA
36935 TATGAGTGTGGAGTCGTTGCGGAAGGAATAGTAGAGTCAGGCCNNNAGATCGGA
36936 TATGAGTGTGGAGTCGTTGCTTGGAAGCCATTATCCTAAGCGANNNAGATCGGAAG
36937 TATGAGTGTGGAGTCGTTGCCCTTTCTCAAATGGAGTACGGGANNNAGATCGGAA
36941 TATGAGTGTGGAGTCGTTGCTAGCTACTCTTTGAGCCAGCAATNNNAGATCGGAAG
36943 TATGAGTGTGGAGTCGTTGCAACTGTTTGACTATCTTCCCCCCNNNAGATCGGAAGA
36944 TATGAGTGTGGAGTCGTTGCCTCATGCCTGTAATCCCAGAACTNNNAGATCGGAAG
36945 TATGAGTGTGGAGTCGTTGCTTTTCTACTTCTGTGAGCCGGAANNNAGATCGGAAG
36947 TATGAGTGTGGAGTCGTTGCTTAGCTGGGCATGGTAGTATGTGNNNAGATCGGAA
36949 TATGAGTGTGGAGTCGTTGCCACTACTTGCTTCTCCTGGAACTNNNAGATCGGAAG
36950 TATGAGTGTGGAGTCGTTGCAGCCTCAACATATTTAGATGCCANNNAGATCGGAAG
36951 TATGAGTGTGGAGTCGTTGCGGGTATGATAGCGCATGTCTGTANNNAGATCGGAA
36952 TATGAGTGTGGAGTCGTTGCCATCACACCCAGCCTTCTTTTTCNNNAGATCGGAAGA
36953 TATGAGTGTGGAGTCGTTGCTCAGCCCCAGAACAACTAAATGANNNAGATCGGAAG
36954 TATGAGTGTGGAGTCGTTGCTGTCACACAGAAGACACAAAGGANNNAGATCGGAA
36956 TATGAGTGTGGAGTCGTTGCTGGCTTCTCCTTTATGTGAGTCCNNNAGATCGGAAG
36957 TATGAGTGTGGAGTCGTTGCTGAAAAAGCAGAGAGTTCCTGGANNNAGATCGGAA
36959 TATGAGTGTGGAGTCGTTGCCAACCCAGGAACCAGGAATCATANNNAGATCGGAA

36960 TATGAGTGTGGAGTCGTTGCGTGCTCTGTGGAGGAAGTAAGAANNNAGATCGGAA
36961 TATGAGTGTGGAGTCGTTGCATGAAGGTTTTGATGTGCTGACTNNNAGATCGGAA
36962 TATGAGTGTGGAGTCGTTGCGGGTGGGGGTTGTTCTACTAATTNNNAGATCGGAA
36964 TATGAGTGTGGAGTCGTTGCCCCAACAAAGAAAAGCCCATGATNNNAGATCGGAAG
36966 TATGAGTGTGGAGTCGTTGCAAGACTCCAGAGAAAGAAAGCCCNNNAGATCGGAA
36967 TATGAGTGTGGAGTCGTTGCATGCCTAGATGGTGTCTACACACNNNAGATCGGAAG
36968 TATGAGTGTGGAGTCGTTGCAACTCGTTTCACCTGATAGTCCCNNNAGATCGGAAG
36969 TATGAGTGTGGAGTCGTTGCTCAATAAATGGCGCTGGGTAAACNNNAGATCGGAA
36971 TATGAGTGTGGAGTCGTTGCACACACACACACACACAAAAGAGNNNAGATCGGAAG
36972 TATGAGTGTGGAGTCGTTGCTCGCATAAAATTGGCCCTTGTTTNNNAGATCGGAAG
36975 TATGAGTGTGGAGTCGTTGCTGGAAATATTGCCACAGGGAACTNNNAGATCGGAA
36976 TATGAGTGTGGAGTCGTTGCGATGTAGGGAGAAGCTGGAGAAGNNNAGATCGGA
36977 TATGAGTGTGGAGTCGTTGCAGGCTAACCAAAGACAGAAGACCNNNAGATCGGAA
36978 TATGAGTGTGGAGTCGTTGCCCAAACATCTCATGTACCCCGTANNNAGATCGGAAG
36979 TATGAGTGTGGAGTCGTTGCTGCAGAGTGAATCAACAGAGTGANNNAGATCGGAA
36982 TATGAGTGTGGAGTCGTTGCTGGTCTGTGTTGGTTACTTAGGGNNNAGATCGGAA
36983 TATGAGTGTGGAGTCGTTGCTTATCCTGAAACCATCCCAACCTNNNAGATCGGAAGA
36986 TATGAGTGTGGAGTCGTTGCGGTAAAGTTCATGCAGTGGGATANNNAGATCGGAA
36987 TATGAGTGTGGAGTCGTTGCCCCATCTCTTCTGGCTTGTAAGANNNAGATCGGAAG
36988 TATGAGTGTGGAGTCGTTGCAAGTGGATTTTCCCGTAGTCTCCNNNAGATCGGAAG
36990 TATGAGTGTGGAGTCGTTGCGCATGAGAATCACTTGAACCCAGNNNAGATCGGAAG
36991 TATGAGTGTGGAGTCGTTGCTTTGTGGTGATGGTACAGTCACTNNNAGATCGGAAG
36992 TATGAGTGTGGAGTCGTTGCTCTTCTAGTCCTGCTTAGGTCCTNNNAGATCGGAAGA
36993 TATGAGTGTGGAGTCGTTGCCCTTCTTGTCACTAGGCAGACTTNNNAGATCGGAAG
36994 TATGAGTGTGGAGTCGTTGCCATGGAATCAACCAAAGTGTCCANNNAGATCGGAAG
36995 TATGAGTGTGGAGTCGTTGCCTGAATGCTCGAGGAGACTGATTNNNAGATCGGAA
36996 TATGAGTGTGGAGTCGTTGCTAGAAAACCCCGTAGTCTTGGTCNNNAGATCGGAAG
36998 TATGAGTGTGGAGTCGTTGCAGGAGAATCACTGGATCCTGAGANNNAGATCGGAA
36999 TATGAGTGTGGAGTCGTTGCATGCTCAATTGCAGTGTGTATGGNNNAGATCGGAA
37000 TATGAGTGTGGAGTCGTTGCTATCTCGTGACTTGGATACCAGCNNNAGATCGGAAG
37001 TATGAGTGTGGAGTCGTTGCAGAAAGCTGGAGGCATCACATTANNNAGATCGGAA
37002 TATGAGTGTGGAGTCGTTGCACTTTATCAACCAGCCATTGCACNNNAGATCGGAAG
37003 TATGAGTGTGGAGTCGTTGCACGATTAGAGGAGTCAGGAGAGTNNNAGATCGGA
37004 TATGAGTGTGGAGTCGTTGCATGGAAAATAGTGTGGCGCTTTCNNNAGATCGGAA
37006 TATGAGTGTGGAGTCGTTGCACAGAATAAACAGGCAGCCTACANNNAGATCGGAA
37008 TATGAGTGTGGAGTCGTTGCGCCATTGAGACAATGATCTTGCANNNAGATCGGAAG
37009 TATGAGTGTGGAGTCGTTGCCCAGCATAACTGAAGCCCAAAAGNNNAGATCGGAA
37013 TATGAGTGTGGAGTCGTTGCATGCACATGAAGAACTCTCCTGTNNNAGATCGGAAG
37014 TATGAGTGTGGAGTCGTTGCGGTTGCTTTTTTCCCTGGAGATTNNNAGATCGGAAG
37015 TATGAGTGTGGAGTCGTTGCTTATAGCTCACTGCAGCCTCAAANNNAGATCGGAAG
37016 TATGAGTGTGGAGTCGTTGCTTGGTAAGTGGCAATACTGGTGANNNAGATCGGAA
37017 TATGAGTGTGGAGTCGTTGCCTGAGGCAGGAGAATCACTTGAANNNAGATCGGAA
37018 TATGAGTGTGGAGTCGTTGCAGATCAGCAAGCGGTAACAAAACNNNAGATCGGAA

37020 TATGAGTGTGGAGTCGTTGCGAATAGCTCCAGTCTACAGCTCCNNNAGATCGGAAG
37025 TATGAGTGTGGAGTCGTTGCGGTTACATTGAGCTGAGTTTGCANNNAGATCGGAA
37026 TATGAGTGTGGAGTCGTTGCAGTTGCAACTCCTTCCCAAAGATNNNAGATCGGAAG
37028 TATGAGTGTGGAGTCGTTGCAGAGGACTTGAACAGATGCTTCTNNNAGATCGGAA
37029 TATGAGTGTGGAGTCGTTGCTCCCTTTTCTCCATATCCTTGCCNNNAGATCGGAAGA
37030 TATGAGTGTGGAGTCGTTGCCATGGACATGGGTCAAAACTGTGNNNAGATCGGAA
37032 TATGAGTGTGGAGTCGTTGCGGAGGCTAAGACAGGAGAATCACNNNAGATCGGA
37033 TATGAGTGTGGAGTCGTTGCAGGTCACAGGATACAAGGTCAACNNNAGATCGGAA
37034 TATGAGTGTGGAGTCGTTGCTCCAAGATTGCCGAATAGGAACANNNAGATCGGAA
37035 TATGAGTGTGGAGTCGTTGCGAGGGGAAGAAAGGAGAAGACAGNNNAGATCGG
37037 TATGAGTGTGGAGTCGTTGCAAGAAATTAGCCAGGCGTAGAGGNNNAGATCGGA
37039 TATGAGTGTGGAGTCGTTGCAGGGGTGGTGTTAGGATTCAATTNNNAGATCGGAA
37040 TATGAGTGTGGAGTCGTTGCTGTGGTCAACAAGCATACAGTGANNNAGATCGGAA
37041 TATGAGTGTGGAGTCGTTGCGCATTGAACATCCTCCAGGAAAGNNNAGATCGGAA
37042 TATGAGTGTGGAGTCGTTGCTAGATTGACTCTAAGGACTGGGTNNNAGATCGGAA
37043 TATGAGTGTGGAGTCGTTGCCCTCTTAGATTCTGTAGCTGGGCNNNAGATCGGAAG
37044 TATGAGTGTGGAGTCGTTGCTTGCATAAGTAACCAGGAGCTGANNNAGATCGGAA
37045 TATGAGTGTGGAGTCGTTGCGGATAGGGGAGTGAGTTTTGACANNNAGATCGGA
37046 TATGAGTGTGGAGTCGTTGCAACAAGTTGGCTATAGTTGCAGANNNAGATCGGAA
37047 TATGAGTGTGGAGTCGTTGCCACGTCACAGTTATGCAGCTATGNNNAGATCGGAAG
37048 TATGAGTGTGGAGTCGTTGCTTGAGTTGGAGGTAGACGTCAAGNNNAGATCGGAA
37049 TATGAGTGTGGAGTCGTTGCCTGATGTAGGAGGATGGCTTGAGNNNAGATCGGA
37050 TATGAGTGTGGAGTCGTTGCCCAGCACCCTTTCATGATAAAGCNNNAGATCGGAAG
37052 TATGAGTGTGGAGTCGTTGCAAACTTAGGCTCAGAGCAGTCTGNNNAGATCGGAA
37053 TATGAGTGTGGAGTCGTTGCAGGTCAGTAGTAGTTCGAGACCANNNAGATCGGAA
37055 TATGAGTGTGGAGTCGTTGCGTAATCTCAGCTACTCAGGAGGCNNNAGATCGGAA
37056 TATGAGTGTGGAGTCGTTGCAGGGGAGTCAAAGGACACAAAATNNNAGATCGGA
37059 TATGAGTGTGGAGTCGTTGCCACTCACACCATTCAAAGACTCANNNAGATCGGAAG
37060 TATGAGTGTGGAGTCGTTGCTACCCTTTCTTCTTCCATCTGCCNNNAGATCGGAAGA
37061 TATGAGTGTGGAGTCGTTGCCATCCTTCAGGTAGGTTGCCATANNNAGATCGGAAG
37063 TATGAGTGTGGAGTCGTTGCAGCCAGCTAAGTAATTGTGGGAANNNAGATCGGAA
37067 TATGAGTGTGGAGTCGTTGCGGAGATAGTGGCAGGCCTAATTTNNNAGATCGGAA
37068 TATGAGTGTGGAGTCGTTGCGAATGCAGGAGAATGGTGTGAACNNNAGATCGGA
37069 TATGAGTGTGGAGTCGTTGCCCAAGTTCTCCCATGTTACCTGANNNAGATCGGAAG
37070 TATGAGTGTGGAGTCGTTGCCTGCAAAAGTTACAGCCACAGTANNNAGATCGGAAG
37071 TATGAGTGTGGAGTCGTTGCAAAAACTTAGCTGAGCATGGTGGNNNAGATCGGAA
37072 TATGAGTGTGGAGTCGTTGCGAAAGGGCTAGGGATTTTGGGTANNNAGATCGGA
37073 TATGAGTGTGGAGTCGTTGCATCCTATTGAGAGGCAAACTGGGNNNAGATCGGAA
37074 TATGAGTGTGGAGTCGTTGCTTTAGAACAGATCCTGGCACACGNNNAGATCGGAAG
37075 TATGAGTGTGGAGTCGTTGCTCTTTGGTGATGGCATCTCTCATNNNAGATCGGAAG
37077 TATGAGTGTGGAGTCGTTGCTGGCTCCTGCTTATTATCCTAGCNNNAGATCGGAAG
37078 TATGAGTGTGGAGTCGTTGCGCCACCATTATCAAGTGGGTTTCNNNAGATCGGAAG
37080 TATGAGTGTGGAGTCGTTGCAGCAATGAGGCAGTATGCAAAATNNNAGATCGGAA

37081 TATGAGTGTGGAGTCGTTGCGGATGAGAGAAGGTGGTGAAAGTNNNAGATCGGA
37082 TATGAGTGTGGAGTCGTTGCTTGCTCTGGAGAAAATAGGTTGTNNNAGATCGGAA
37083 TATGAGTGTGGAGTCGTTGCCACTCCTGTAATCCCAGCACTTTNNNAGATCGGAAGA
37084 TATGAGTGTGGAGTCGTTGCAGATTTCAACATTGCACTCCAGCNNNAGATCGGAAG
37085 TATGAGTGTGGAGTCGTTGCTTCTACTGGCTTCCTTGTTCCTCNNNAGATCGGAAGA
37087 TATGAGTGTGGAGTCGTTGCTTGACCCATTTGAGCACAATTCCNNNAGATCGGAAG
37089 TATGAGTGTGGAGTCGTTGCACCAAAATCTATGGGCTACAGCANNNAGATCGGAAG
37090 TATGAGTGTGGAGTCGTTGCTGTGAGCAACATTATTCTGGGGANNNAGATCGGAA
37091 TATGAGTGTGGAGTCGTTGCCTACTGCACTCCAGCCTATTTGANNNAGATCGGAAG
37092 TATGAGTGTGGAGTCGTTGCACAAAGGCTACCTCACCTCAAAANNNAGATCGGAAG
37093 TATGAGTGTGGAGTCGTTGCCCAAAGCTACGTCTCTCAGAGTTNNNAGATCGGAAG
37094 TATGAGTGTGGAGTCGTTGCAAAACATTTCACGGTTGAGGCAANNNAGATCGGAA
37099 TATGAGTGTGGAGTCGTTGCTAAGGTCATTGTTTTGTGAGGGGNNNAGATCGGAA
37100 TATGAGTGTGGAGTCGTTGCAAACTACCCGTTATCTGAGTGGGNNNAGATCGGAA
37103 TATGAGTGTGGAGTCGTTGCAATATTGGGGAATCTCTGCAGGANNNAGATCGGAA
37104 TATGAGTGTGGAGTCGTTGCCAAAACAAAACCTGCACCACCTANNNAGATCGGAAG
37105 TATGAGTGTGGAGTCGTTGCCCAACGAATCAACAACCTCAGTTNNNAGATCGGAAG
37106 TATGAGTGTGGAGTCGTTGCAAAAAAGCAAGGGAGCTGTGTTTNNNAGATCGGAA
37107 TATGAGTGTGGAGTCGTTGCAGAGTTTTCAATTCACTTGGCCCNNNAGATCGGAAG
37108 TATGAGTGTGGAGTCGTTGCGGTCATTGTAAGGACTTTGGCACNNNAGATCGGAA
37109 TATGAGTGTGGAGTCGTTGCTAGAGAGAGACATCGGTGTGAGANNNAGATCGGA
37110 TATGAGTGTGGAGTCGTTGCTGGTGTGAACCCAGGTCATATTTNNNAGATCGGAAG
37111 TATGAGTGTGGAGTCGTTGCGGAGAAATGAGCAGGAAAAAGCANNNAGATCGGA
37113 TATGAGTGTGGAGTCGTTGCAATGAACAGGCAACCTACAGGATNNNAGATCGGAA
37114 TATGAGTGTGGAGTCGTTGCAAGCACATGAATTTGTTTGGGGANNNAGATCGGAA
37115 TATGAGTGTGGAGTCGTTGCAATGTCTGGTCACAATGATTGCANNNAGATCGGAAG
37116 TATGAGTGTGGAGTCGTTGCCTGTAGTTCCAGCTACTTGGGAGNNNAGATCGGAA
37118 TATGAGTGTGGAGTCGTTGCTAGGCTCGAAATTGTCACACTGANNNAGATCGGAAG
37119 TATGAGTGTGGAGTCGTTGCCTGTTCTCAGACTTTTGTGTGGGNNNAGATCGGAAG
37120 TATGAGTGTGGAGTCGTTGCCATCCAGCTGAGTACTCTCTGTGNNNAGATCGGAAG
37123 TATGAGTGTGGAGTCGTTGCGAGGTCAGGAGATCAAGACCATCNNNAGATCGGAA
37125 TATGAGTGTGGAGTCGTTGCAAAACACCACAGAAAAAGATGTACTNNNAGATCGGA
37127 TATGAGTGTGGAGTCGTTGCTACTCAGCCTCCATGACCTCTTANNNAGATCGGAAGA
37132 TATGAGTGTGGAGTCGTTGCATCCCCTTCCTCTGCCCTATTATNNNAGATCGGAAGA
37135 TATGAGTGTGGAGTCGTTGCCTAGGTGACAGAGTGAGACCTTGNNNAGATCGGAA
37136 TATGAGTGTGGAGTCGTTGCAAGTCAAGAAGATTGTTGGGGGTNNNAGATCGGAA
37137 TATGAGTGTGGAGTCGTTGCAACCGAAGGCTTTATTCCTCCATNNNAGATCGGAAG
37139 TATGAGTGTGGAGTCGTTGCGGGTGGATTTCGCATTGTTGTATNNNAGATCGGAA
37140 TATGAGTGTGGAGTCGTTGCTCTGTCACTCCTCTCTGTACTCANNNAGATCGGAAGA
37141 TATGAGTGTGGAGTCGTTGCTGAACATCAGGAATCAGGTGTGTNNNAGATCGGAA
37142 TATGAGTGTGGAGTCGTTGCTGGTGTATGAGCTAGATGTGGTGNNNAGATCGGAA
37143 TATGAGTGTGGAGTCGTTGCGCTTTTGCCTCTTTGCTGATCATNNNAGATCGGAAGA
37144 TATGAGTGTGGAGTCGTTGCGCAGAACTACCATTTGACCAAGCNNNAGATCGGAAG

37146 TATGAGTGTGGAGTCGTTGCTGGCATTTCTTCCCTCCACTAAANNNAGATCGGAAGA
37148 TATGAGTGTGGAGTCGTTGCACAGTTTTTTTTTCTCCAAGGGTNNNAGATCGGAAG
37150 TATGAGTGTGGAGTCGTTGCGCATGCCAGGGAGTGAAAAATAGNNNAGATCGGAA
37153 TATGAGTGTGGAGTCGTTGCTAAAGGGAGGAATGGTTTGCGTANNNAGATCGGAA
37155 TATGAGTGTGGAGTCGTTGCTTTCGTGTGTAGTCTCCTTACCGNNNAGATCGGAAG
37156 TATGAGTGTGGAGTCGTTGCGAACAACAGACACCACGAAACATNNNAGATCGGAAG
37158 TATGAGTGTGGAGTCGTTGCAAGGGCCCTGTATCAGAGTATCANNNAGATCGGAAG
37159 TATGAGTGTGGAGTCGTTGCAAGAACTACCATTTGACCCAGCANNNAGATCGGAAG
37160 TATGAGTGTGGAGTCGTTGCGAAGGAAAGGAAGGAAGGAAGGANNNAGATCGGA
37161 TATGAGTGTGGAGTCGTTGCTTTGCTCTCTCTTAGGGGACCTANNNAGATCGGAAG
37163 TATGAGTGTGGAGTCGTTGCCATGTCATATGGGCATGGAAAGGNNNAGATCGGAA
37164 TATGAGTGTGGAGTCGTTGCGGACCTGAGGTGAGGGAATAAAGNNNAGATCGGA
37167 TATGAGTGTGGAGTCGTTGCTTGTGGATTGAATAGTGTCCCCCNNNAGATCGGAAG
37168 TATGAGTGTGGAGTCGTTGCTGATACTGTCACCCAGGTACTGANNNAGATCGGAAG
37170 TATGAGTGTGGAGTCGTTGCAAACAAGTAGGGGAGAGTGTTCCNNNAGATCGGAA
37172 TATGAGTGTGGAGTCGTTGCACCATCACCCTTTTGATCAGTGTNNNAGATCGGAAG
37173 TATGAGTGTGGAGTCGTTGCGTCTGATAGTTCCCAGAAACCCANNNAGATCGGAAG
37174 TATGAGTGTGGAGTCGTTGCGCCTTGACTCTATTGTACATGGCNNNAGATCGGAAG
37175 TATGAGTGTGGAGTCGTTGCTGATTCCAAGCCCAAACAAATCCNNNAGATCGGAAG
37180 TATGAGTGTGGAGTCGTTGCGCAGGAGAATTGAGCCAAGATTGNNNAGATCGGAA
37181 TATGAGTGTGGAGTCGTTGCGTATGCCTGCTGCGTAGTAAAAGNNNAGATCGGAA
37182 TATGAGTGTGGAGTCGTTGCAGAATTTTAGACAGGAAGCCCCANNNAGATCGGAA
37185 TATGAGTGTGGAGTCGTTGCTAACAGCAAGGTGATCGTCTTGANNNAGATCGGAA
37187 TATGAGTGTGGAGTCGTTGCAGATCTAGTGTTTGGTAGCACCCNNNAGATCGGAAG
37188 TATGAGTGTGGAGTCGTTGCGAGGCATGAGAATCACTTGAACCNNNAGATCGGAA
37192 TATGAGTGTGGAGTCGTTGCCATGGCTTCAAACAGATCGCATTNNNAGATCGGAAG
37193 TATGAGTGTGGAGTCGTTGCTGGACACACATCGTTAGAAGACANNNAGATCGGAA
37194 TATGAGTGTGGAGTCGTTGCTGCTAATACAATGGACGGTATCCNNNAGATCGGAAG
37198 TATGAGTGTGGAGTCGTTGCGCTGGCACAGATTTGAGATTTGANNNAGATCGGAA
37199 TATGAGTGTGGAGTCGTTGCTTGTACGTTTTCGTTTCTGTGCANNNAGATCGGAAG
37200 TATGAGTGTGGAGTCGTTGCAAATCCTTGCAATTTATCTGCCANNNAGATCGGAAG
37202 TATGAGTGTGGAGTCGTTGCCACAGTGGATCACGCATGTAATCNNNAGATCGGAAG
37203 TATGAGTGTGGAGTCGTTGCGCAGAAGTGGCCATACTTTATCACNNNAGATCGGAA
37204 TATGAGTGTGGAGTCGTTGCGTCATATGGTGTTTCCCCTTCCTNNNAGATCGGAAGA
37206 TATGAGTGTGGAGTCGTTGCAGTTTCAGTAGGAATGGGACCAGNNNAGATCGGAA
37209 TATGAGTGTGGAGTCGTTGCACTCAGGAGGATAAGGCAGAAGANNNAGATCGGA
37210 TATGAGTGTGGAGTCGTTGCATCATGAGGTCAGGAGATCGAGANNNAGATCGGAA
37211 TATGAGTGTGGAGTCGTTGCTTCCCAACTTCCATGTCACGATANNNAGATCGGAAGA
37212 TATGAGTGTGGAGTCGTTGCCTGTAGTCCCAGCTACTCAGGAGNNNAGATCGGAAG
37213 TATGAGTGTGGAGTCGTTGCACTGGCAAACAGAATTCAACAATNNNAGATCGGAAG
37215 TATGAGTGTGGAGTCGTTGCATCGCTTGTCATTGCTTACATTTNNNAGATCGGAAGA
37217 TATGAGTGTGGAGTCGTTGCTTCTGGGTTCTCTTTTCTGGTCTNNNAGATCGGAAGA
37219 TATGAGTGTGGAGTCGTTGCCATTCGTTGCCAGGTCTCTTTAGNNNAGATCGGAAG

37220 TATGAGTGTGGAGTCGTTGCTGATGCCTCCAGCTTGTTCTTATNNNAGATCGGAAG
37222 TATGAGTGTGGAGTCGTTGCGCCCTCACAAAATAGTACTGGGANNNAGATCGGAA
37223 TATGAGTGTGGAGTCGTTGCGCCCACAAAGATGAGAAAGAACCNNNAGATCGGAA
37224 TATGAGTGTGGAGTCGTTGCCTGGAGTCTCAGCTACTCAAGAGNNNAGATCGGAA
37225 TATGAGTGTGGAGTCGTTGCACAGTCTGGTAGCTTGTGAACATNNNAGATCGGAA
37226 TATGAGTGTGGAGTCGTTGCCCACATAGTGCTCGTCTATCTCCNNNAGATCGGAAG
37228 TATGAGTGTGGAGTCGTTGCATTCAGAGAGGAGCAACAGGAAGNNNAGATCGGA
37229 TATGAGTGTGGAGTCGTTGCCTAAGTGATTCTGCCCTCATCCANNNAGATCGGAAG
37231 TATGAGTGTGGAGTCGTTGCTTGTTTCTCCCTCTCCTTGTTGANNNAGATCGGAAGA
37234 TATGAGTGTGGAGTCGTTGCAATCAAACTTTGCCCTTCCTTCCNNNAGATCGGAAGA
37235 TATGAGTGTGGAGTCGTTGCGACCTAGTTCTAACCCAGCAGACNNNAGATCGGAAG
37236 TATGAGTGTGGAGTCGTTGCAAAGTTCAGGCCAATATCCCTGTNNNAGATCGGAAG
37237 TATGAGTGTGGAGTCGTTGCAGGAGCTGGTACCATTTCTTCTGNNNAGATCGGAAG
37239 TATGAGTGTGGAGTCGTTGCGGCTAGGTCAGTTCTTTGGAGAANNNAGATCGGAA
37240 TATGAGTGTGGAGTCGTTGCAGATATGTGCACTCCCATGTTCANNNAGATCGGAAG
37241 TATGAGTGTGGAGTCGTTGCATGTGGAGAAAAGGGGAACACTTNNNAGATCGGA
37242 TATGAGTGTGGAGTCGTTGCTTTCAACTGAGGCCTCTGAGATCNNNAGATCGGAAG
37243 TATGAGTGTGGAGTCGTTGCAATGGCAACTGTAATCCCAGCTANNNAGATCGGAAG
37245 TATGAGTGTGGAGTCGTTGCCACCTGAATTCTACCAGGGGTACNNNAGATCGGAAG
37247 TATGAGTGTGGAGTCGTTGCCCCACTGTACACTTAGGCTACACNNNAGATCGGAAG
37248 TATGAGTGTGGAGTCGTTGCGTGAGCAGCACAGAATTCAGAATNNNAGATCGGAA
37249 TATGAGTGTGGAGTCGTTGCTGAGAAGAACGAGAGTGAAGGTGNNNAGATCGGA
37250 TATGAGTGTGGAGTCGTTGCCACACACACACACACACACATATNNNAGATCGGAAG
37251 TATGAGTGTGGAGTCGTTGCCCCTTCTTTCCCAGCAGTAATCTNNNAGATCGGAAGA
37254 TATGAGTGTGGAGTCGTTGCGAACTGGAAGGGGAAAGTTGTCTNNNAGATCGGA
37256 TATGAGTGTGGAGTCGTTGCATGGAACATAGCAAAGGCAGTTCNNNAGATCGGAA
37257 TATGAGTGTGGAGTCGTTGCATCACGGTATTTGGATAGGTGGGNNNAGATCGGAA
37258 TATGAGTGTGGAGTCGTTGCCGTTGACCCCTGTTCTAGATCATNNNAGATCGGAAG
37259 TATGAGTGTGGAGTCGTTGCTGGAATTCAGCAACACGTCAAAANNNAGATCGGAA
37260 TATGAGTGTGGAGTCGTTGCACTTGCTCCTGAATGACTTTTGANNNAGATCGGAAG
37262 TATGAGTGTGGAGTCGTTGCAGTGCTTATCAGACTGGTGGAAGNNNAGATCGGAA
37263 TATGAGTGTGGAGTCGTTGCTAGGAGGAACTGAAGGCTGTTTCNNNAGATCGGAA
37265 TATGAGTGTGGAGTCGTTGCCTGTAGTCCCAGCTACTCAGGAANNNAGATCGGAAG
37266 TATGAGTGTGGAGTCGTTGCTACTTTGGCATTAGGTGTGTCCCNNNAGATCGGAAG
37267 TATGAGTGTGGAGTCGTTGCCTGAGATGGGAGAATCACCTGAGNNNAGATCGGAA
37270 TATGAGTGTGGAGTCGTTGCTCTGGAAGTCTATCACAACTGGCNNNAGATCGGAAG
37272 TATGAGTGTGGAGTCGTTGCACAAATACCCCATCCAAAGGTCANNNAGATCGGAAG
37273 TATGAGTGTGGAGTCGTTGCCTTGGTGGTGCATGTTTGTAGTCNNNAGATCGGAA
37275 TATGAGTGTGGAGTCGTTGCTCCCAGCTCCTAACTCAGTCTAANNNAGATCGGAAG
37277 TATGAGTGTGGAGTCGTTGCGCAACAGAGCGAGATTCCATATCNNNAGATCGGAA
37279 TATGAGTGTGGAGTCGTTGCCGTATGGAAGGGGTTTTTCACATNNNAGATCGGAA
37280 TATGAGTGTGGAGTCGTTGCTCGTGTCTACAAGTATGGCAGTCNNNAGATCGGAAG
37281 TATGAGTGTGGAGTCGTTGCGGAGAATGGCATGAACCTAGGAGNNNAGATCGGA

37282 TATGAGTGTGGAGTCGTTGCCTCCCCTTCACCTTCACCAATAANNNAGATCGGAAGA
37284 TATGAGTGTGGAGTCGTTGCCGCTTGCAGTTCTCAGATCAAATNNNAGATCGGAAG
37285 TATGAGTGTGGAGTCGTTGCGAGGGTAGTGATGAAGACAGTGGNNNAGATCGGA
37287 TATGAGTGTGGAGTCGTTGCTCTACTTGAGGATTGCTGTTGCTNNNAGATCGGAAG
37288 TATGAGTGTGGAGTCGTTGCATTCTTTCTCTGCCCCTTTGAGANNNAGATCGGAAGA
37289 TATGAGTGTGGAGTCGTTGCTAATCCCAAGCTATTCAGGAGGCNNNAGATCGGAAG
37290 TATGAGTGTGGAGTCGTTGCTTGTGTGCAGAGAGGTGATTGTANNNAGATCGGAA
37291 TATGAGTGTGGAGTCGTTGCATACATAACATCCAGCGTGGTGANNNAGATCGGAA
37292 TATGAGTGTGGAGTCGTTGCTGACAAAAGTGGCTGCAGAAAAANNNAGATCGGAA
37293 TATGAGTGTGGAGTCGTTGCAAAGAGGCATGATCCATGAAAGANNNAGATCGGAA
37294 TATGAGTGTGGAGTCGTTGCCCATCTGCCTGCTGAATTACTTTNNNAGATCGGAAG
37295 TATGAGTGTGGAGTCGTTGCGCAGTGGCTCACACCTATGATAANNNAGATCGGAAG
37297 TATGAGTGTGGAGTCGTTGCCAGGGGTTGCAATCATAGTCTCTNNNAGATCGGAAG
37299 TATGAGTGTGGAGTCGTTGCGCCTTACAAGAAATGCTAGACGGNNNAGATCGGAA
37300 TATGAGTGTGGAGTCGTTGCGATTGGTGTACCTGAAAGTGACGNNNAGATCGGAA
37301 TATGAGTGTGGAGTCGTTGCAACTGAGGTCAACAGGGAGTAAGNNNAGATCGGA
37302 TATGAGTGTGGAGTCGTTGCTGTGTGTGTGTGTGTGTAAGAGANNNAGATCGGAA
37303 TATGAGTGTGGAGTCGTTGCAAAGAAATGATAAGTGCCCTGCCNNNAGATCGGAA
37305 TATGAGTGTGGAGTCGTTGCCAGGACACTCTCTGAGCACATAANNNAGATCGGAAG
37306 TATGAGTGTGGAGTCGTTGCTCCGGCAGAACTTTGAGTTTCTANNNAGATCGGAAG
37307 TATGAGTGTGGAGTCGTTGCTGGAGAGAGAGAGAATGACAACGNNNAGATCGGA
37309 TATGAGTGTGGAGTCGTTGCTAACCTTCCATACCTTCCATGCCNNNAGATCGGAAGA
37310 TATGAGTGTGGAGTCGTTGCATGACCCTGACTCATCCAATACCNNNAGATCGGAAG
37311 TATGAGTGTGGAGTCGTTGCACTTGAGCCATACGTTCATGAGANNNAGATCGGAAG
37312 TATGAGTGTGGAGTCGTTGCAGTTGCATGTAACAAACAGTGCTNNNAGATCGGAA
37313 TATGAGTGTGGAGTCGTTGCGGGTGATAGAGTGAGTCCCTTTCNNNAGATCGGAA
37314 TATGAGTGTGGAGTCGTTGCCCTCCCAGGTTCAAGTGATTCTCNNNAGATCGGAAG
37315 TATGAGTGTGGAGTCGTTGCTGAGAGAACCTTGAACTTTTGCTNNNAGATCGGAAG
37317 TATGAGTGTGGAGTCGTTGCCTTGGGGTATCAGTACTGCACTTNNNAGATCGGAAG
37318 TATGAGTGTGGAGTCGTTGCAAAAGTTCCTGCAAGCTACTGTGNNNAGATCGGAAG
37319 TATGAGTGTGGAGTCGTTGCGGGGGGTTATGCTGTTTATACCANNNAGATCGGAA
37320 TATGAGTGTGGAGTCGTTGCAACACAAGGCCTGAAAATCATGGNNNAGATCGGAA
37321 TATGAGTGTGGAGTCGTTGCGGGCAACAGAGCAATATTCCATCNNNAGATCGGAA
37324 TATGAGTGTGGAGTCGTTGCCTGAGGCAGGAGAATCACTTGAANNNAGATCGGAA
37325 TATGAGTGTGGAGTCGTTGCTCTAGGGACAGAGTGAGATTCCANNNAGATCGGAA
37326 TATGAGTGTGGAGTCGTTGCGAAAGGTAAATCTGGCGATGCTGNNNAGATCGGAA
37328 TATGAGTGTGGAGTCGTTGCGACTGAGACAGGAGGATCACTTGNNNAGATCGGAA
37329 TATGAGTGTGGAGTCGTTGCAAAAGAAACAGGGCATCTTGTCANNNAGATCGGAA
37330 TATGAGTGTGGAGTCGTTGCGTTGACATTTCCATGCCTTCTTANNNAGATCGGAAG
37331 TATGAGTGTGGAGTCGTTGCGCCTTCATTTGTCTGTCTTCACCNNNAGATCGGAAGA
37332 TATGAGTGTGGAGTCGTTGCTGAGGAATGAGAGAGGGTGAGATNNNAGATCGGA
37333 TATGAGTGTGGAGTCGTTGCCAGATCATGTGGTCAGGAGTTCANNNAGATCGGAA
37334 TATGAGTGTGGAGTCGTTGCGGTAGGCAAATGACATGAACAGANNNAGATCGGAA

37336 TATGAGTGTGGAGTCGTTGCTAGTACCATCTACTTGGGAGGCTNNNAGATCGGAAG
37338 TATGAGTGTGGAGTCGTTGCCAGGACCAGATGTATTCACAGCTNNNAGATCGGAAG
37339 TATGAGTGTGGAGTCGTTGCAGATACAAGACTACGCCCTTATCCNNNAGATCGGAA
37340 TATGAGTGTGGAGTCGTTGCGAGTACATCTAGTCAGCAGAGCCNNNAGATCGGAA
37341 TATGAGTGTGGAGTCGTTGCAGAAGATTTCTTGAGCCCAGGAGNNNAGATCGGAA
37342 TATGAGTGTGGAGTCGTTGCAGATTATTTGTCCCCAGGTTGCANNNAGATCGGAAG
37343 TATGAGTGTGGAGTCGTTGCGGGCAAAACACCTGTATAGACATNNNAGATCGGAA
37344 TATGAGTGTGGAGTCGTTGCCTGCTGGGTGGAGATCACTTTATNNNAGATCGGAA
37346 TATGAGTGTGGAGTCGTTGCCTGATTGTGAAGACTGTGGGAGANNNAGATCGGAA
37347 TATGAGTGTGGAGTCGTTGCTCCACCTTCTGTTTCCCTTCATCNNNAGATCGGAAGA
37348 TATGAGTGTGGAGTCGTTGCCCATTTCATGAGGCTATAGCTGCNNNAGATCGGAAG
37350 TATGAGTGTGGAGTCGTTGCCACGTGAAGGCCTTTGATTCTTTNNNAGATCGGAAG
37352 TATGAGTGTGGAGTCGTTGCACAGTGGCTCACACCTATAATCCNNNAGATCGGAAG
37353 TATGAGTGTGGAGTCGTTGCTGCCCATCTGTAAAATGGCAATGNNNAGATCGGAAG
37354 TATGAGTGTGGAGTCGTTGCATGCTTTGAGAGGGATCTTCAGTNNNAGATCGGAA
37355 TATGAGTGTGGAGTCGTTGCTTAGTTGGGAGTAGTGGCTCATGNNNAGATCGGAA
37356 TATGAGTGTGGAGTCGTTGCTGACAAAGGGAAAACTCCATGTGNNNAGATCGGAA
37357 TATGAGTGTGGAGTCGTTGCCCTACAGGACAATGAACACGAGANNNAGATCGGAA
37358 TATGAGTGTGGAGTCGTTGCATTCTCTAGAAGCAGAGCCTGAGNNNAGATCGGAA
37359 TATGAGTGTGGAGTCGTTGCCCCAACTCTTCATTGGAGGAAGANNNAGATCGGAAG
37360 TATGAGTGTGGAGTCGTTGCCAAAAACAAACCAAACAACCCCCNNNAGATCGGAAG
37361 TATGAGTGTGGAGTCGTTGCGCACAGGTGAGTCATTCAGTTTCNNNAGATCGGAAG
37362 TATGAGTGTGGAGTCGTTGCGATGGTTTTGAGCAATAGCACGTNNNAGATCGGAA
37363 TATGAGTGTGGAGTCGTTGCGGTGCATAGGGTGTTGTCTTAGTNNNAGATCGGAA
37365 TATGAGTGTGGAGTCGTTGCTTTGTGATGCAATCATTGTGGCCNNNAGATCGGAAG
37367 TATGAGTGTGGAGTCGTTGCTGAATTAGACCCCACCTTTTGTGNNNAGATCGGAAG
37369 TATGAGTGTGGAGTCGTTGCAATTTGAAACTAGCCTGGCCAACNNNAGATCGGAAG
37371 TATGAGTGTGGAGTCGTTGCAAGCCATAGGGTTATTTTGGGGTNNNAGATCGGAA
37372 TATGAGTGTGGAGTCGTTGCTGAGGCTGGCTTCTTTTCATTTGNNNAGATCGGAAG
37373 TATGAGTGTGGAGTCGTTGCAGCAGATGAGCTCAGATCATGAGNNNAGATCGGAA
37375 TATGAGTGTGGAGTCGTTGCGACCTTTCTTCCTACAGGGGTTTNNNAGATCGGAAG
37376 TATGAGTGTGGAGTCGTTGCGGGGGATTTCAGATCTCTGTACCNNNAGATCGGAA
37377 TATGAGTGTGGAGTCGTTGCGAGGCAGGAGAATTGTTTGAACCNNNAGATCGGAA
37378 TATGAGTGTGGAGTCGTTGCCCAAAACCAAAACAAAACCGGAANNNAGATCGGAA
37380 TATGAGTGTGGAGTCGTTGCGTGAGAACATGGCACATTAGCAGNNNAGATCGGAA
37381 TATGAGTGTGGAGTCGTTGCAGGCATTCCAAGCAGAGTGAATANNNAGATCGGAA
37382 TATGAGTGTGGAGTCGTTGCTCCTAGCTACTCTGGAGGATGAGNNNAGATCGGAA
37383 TATGAGTGTGGAGTCGTTGCATGATGGCATACACCTGTAGTCCNNNAGATCGGAAG
37384 TATGAGTGTGGAGTCGTTGCCCAGGAACCAAGTGAGCATCTAANNNAGATCGGAA
37386 TATGAGTGTGGAGTCGTTGCTAAAGCAGGAAGTGAAAGTCCGANNNAGATCGGAA
37387 TATGAGTGTGGAGTCGTTGCAAGAAAAAGTTAGCCAGACATGGNNNAGATCGGAA
37388 TATGAGTGTGGAGTCGTTGCGGAAGGAAGACACCATCTTGGAANNNAGATCGGAA
37389 TATGAGTGTGGAGTCGTTGCGAGGATTGTTTGAGCACAGGAAGNNNAGATCGGAA

37390 TATGAGTGTGGAGTCGTTGCTGTCCATGCAAAGATGAAAAGGCNNNAGATCGGAA
37391 TATGAGTGTGGAGTCGTTGCGATTGGAGCTACTCAGAAGCCTTNNNAGATCGGAAG
37392 TATGAGTGTGGAGTCGTTGCGGGAGAACAGAGAAACAAAGAGGNNNAGATCGGA
37393 TATGAGTGTGGAGTCGTTGCGAAAAATGAGGCAGGGCATAGTGNNNAGATCGGA
37396 TATGAGTGTGGAGTCGTTGCCAATCTGGAGAAACCCCAACAACNNNAGATCGGAAG
37397 TATGAGTGTGGAGTCGTTGCCTGTAATCCCAGCTCCAGAACTTNNNAGATCGGAAG
37399 TATGAGTGTGGAGTCGTTGCAAAATGGGGAGCTGATGAGAGTTNNNAGATCGGA
37400 TATGAGTGTGGAGTCGTTGCGGAGAATCACTTGAGTCTGGGAGNNNAGATCGGAA
37401 TATGAGTGTGGAGTCGTTGCGAGACAGGGCTTATCATTCCCAANNNAGATCGGAAG
37402 TATGAGTGTGGAGTCGTTGCCAGTCAGACGGCTCATATGGTAGNNNAGATCGGAA
37403 TATGAGTGTGGAGTCGTTGCAAGAAGGATGAGTGCTGGCTTATNNNAGATCGGAA
37404 TATGAGTGTGGAGTCGTTGCTGCAAATTGAACTGGATAGCCTGNNNAGATCGGAAG
37405 TATGAGTGTGGAGTCGTTGCTGGTGCACCAAATGGAAAAAATTNNNAGATCGGAA
37406 TATGAGTGTGGAGTCGTTGCAGGAAGAAATAACCACAGGCACTNNNAGATCGGAA
37408 TATGAGTGTGGAGTCGTTGCCCAGCTTGCTTGCTCAGTTTTTANNNAGATCGGAAG
37410 TATGAGTGTGGAGTCGTTGCAGAAATGCCACATGACACACATGNNNAGATCGGAAG
37411 TATGAGTGTGGAGTCGTTGCATGGGAAGAAGCCACCATCTAGANNNAGATCGGAA
37412 TATGAGTGTGGAGTCGTTGCGGGGTCCATTACAGAAGCTAGAANNNAGATCGGAA
37413 TATGAGTGTGGAGTCGTTGCGAGCAATGGGAAATCACTGGATGNNNAGATCGGAA
37414 TATGAGTGTGGAGTCGTTGCTTCGTGAGGTCCTAAAAGGTACGNNNAGATCGGAA
37415 TATGAGTGTGGAGTCGTTGCCCAGGAGGATCAAACTTCAGTGANNNAGATCGGAA
37417 TATGAGTGTGGAGTCGTTGCAATCATGGAAGGTCAGAGTGTGTNNNAGATCGGAA
37419 TATGAGTGTGGAGTCGTTGCATGATCACTTGAAGCCAGGAGTTNNNAGATCGGAAG
37420 TATGAGTGTGGAGTCGTTGCGGAGGGATTCTGTCATCTCTTCCNNNAGATCGGAAG
37421 TATGAGTGTGGAGTCGTTGCAATTGCTGGGTCATATGGTGTCTNNNAGATCGGAAG
37422 TATGAGTGTGGAGTCGTTGCGCTGAGGTAGGAGAATCGTTTGANNNAGATCGGAA
37424 TATGAGTGTGGAGTCGTTGCACTTCAGCATCACATAGACCACANNNAGATCGGAAG
37428 TATGAGTGTGGAGTCGTTGCCTGAGACAGGAGAATCGCTTGAANNNAGATCGGAA
37429 TATGAGTGTGGAGTCGTTGCCCAGAAAACCAGAACATGTGGTCNNNAGATCGGAAG
37430 TATGAGTGTGGAGTCGTTGCCGATTCCAGAACCAGAATTCACGNNNAGATCGGAAG
37431 TATGAGTGTGGAGTCGTTGCGCAGGAGAATCAATTGAACCCAGNNNAGATCGGAA
37432 TATGAGTGTGGAGTCGTTGCCTGTAATCCCAGCTACTCAGGAGNNNAGATCGGAAG
37433 TATGAGTGTGGAGTCGTTGCCCCAGTAAAGTGAGATGGAGTCCNNNAGATCGGAA
37434 TATGAGTGTGGAGTCGTTGCGAAAACTCTGAAGCACTCCACACNNNAGATCGGAAG
37436 TATGAGTGTGGAGTCGTTGCTCCTGTCATTTGCAACAACATGGNNNAGATCGGAAG
37437 TATGAGTGTGGAGTCGTTGCATAGTTGAGCCCTCTGTTGTACCNNNAGATCGGAAG
37438 TATGAGTGTGGAGTCGTTGCCTCTTGGTGTCCTATCCACACACNNNAGATCGGAAGA
37439 TATGAGTGTGGAGTCGTTGCGGATAAAGTATCTGCCAGCCTGANNNAGATCGGAAG
37440 TATGAGTGTGGAGTCGTTGCATCAGCGAAACAATTGCAACTCANNNAGATCGGAAG
37441 TATGAGTGTGGAGTCGTTGCGCCACTTCCTTCTACTCTGTCTCNNNAGATCGGAAGA
37443 TATGAGTGTGGAGTCGTTGCCACAAAAAAATTAGCCGGGCATGNNNAGATCGGAA
37444 TATGAGTGTGGAGTCGTTGCGGAGAGTAAGAGGCAGCTATCACNNNAGATCGGAA
37445 TATGAGTGTGGAGTCGTTGCTTGACCTACTCACATTCCCTTGGNNNAGATCGGAAGA

37446 TATGAGTGTGGAGTCGTTGCATTTCCTGCCCCAGATCTGAAATNNNAGATCGGAAG
37447 TATGAGTGTGGAGTCGTTGCCGTCATGCCAAAAAGAAACCCTANNNAGATCGGAAG
37448 TATGAGTGTGGAGTCGTTGCAGTACAACCTCCACGGAAAACATNNNAGATCGGAAG
37449 TATGAGTGTGGAGTCGTTGCGAGAAGCTTCAAAAAAGGGTCCCNNNAGATCGGAA
37450 TATGAGTGTGGAGTCGTTGCTCGCCAAGAAAAGTCTCAACTCTNNNAGATCGGAAG
37451 TATGAGTGTGGAGTCGTTGCCCGTAGTCCTAGCTACTTGGGAGNNNAGATCGGAA
37455 TATGAGTGTGGAGTCGTTGCCTCACACCACTCAGAATGGCTATNNNAGATCGGAAG
37456 TATGAGTGTGGAGTCGTTGCCGCCTATAATCCCAGCACTTTTGNNNAGATCGGAAG
37459 TATGAGTGTGGAGTCGTTGCATGGAGCTCTTTCCTCTGTTAGCNNNAGATCGGAAG
37460 TATGAGTGTGGAGTCGTTGCGGTAGAGGCATAAGGGAACCTTCNNNAGATCGGAA
37461 TATGAGTGTGGAGTCGTTGCTTCAAAAATGAATTCAGGGCCGGNNNAGATCGGAA
37463 TATGAGTGTGGAGTCGTTGCTGCAGACCTCAGACCTCAATCTANNNAGATCGGAAG
37465 TATGAGTGTGGAGTCGTTGCTCAGGAGAATCACTTGAACCCAGNNNAGATCGGAA
37466 TATGAGTGTGGAGTCGTTGCAAGCACAAGCACAATGAGGTAACNNNAGATCGGAA
37468 TATGAGTGTGGAGTCGTTGCAGCCTCTGTTTGTGTTCTTCAGTNNNAGATCGGAAG
37469 TATGAGTGTGGAGTCGTTGCAACTGAAAAGCCAACAACACCAANNNAGATCGGAAG
37470 TATGAGTGTGGAGTCGTTGCGATACATAGGGCAAGGTATGGGGNNNAGATCGGA
37471 TATGAGTGTGGAGTCGTTGCTTTCTGATCTGTGTGGTGATGGTNNNAGATCGGAA
37473 TATGAGTGTGGAGTCGTTGCCTTTGGCAAATAGACTTGGAGCCNNNAGATCGGAA
37474 TATGAGTGTGGAGTCGTTGCCATTACAAACAGCACTGCAGTGANNNAGATCGGAAG
37475 TATGAGTGTGGAGTCGTTGCATCCTAGCTACTTGAGAGGCTGANNNAGATCGGAA
37476 TATGAGTGTGGAGTCGTTGCGACAACCCAAATGATGTCCATGGNNNAGATCGGAA
37477 TATGAGTGTGGAGTCGTTGCAGCAACAGACAATCACAAAAACANNNAGATCGGAA
37478 TATGAGTGTGGAGTCGTTGCAGAAAAAGAAAACTTGCAGGCCGNNNAGATCGGAA
37479 TATGAGTGTGGAGTCGTTGCACCCCTCCACTTCTAGAACTGTANNNAGATCGGAAGA
37480 TATGAGTGTGGAGTCGTTGCTTCAGTCAGCTCTGGATCATTGGNNNAGATCGGAAG
37481 TATGAGTGTGGAGTCGTTGCACAGTCCAATTAGAAAAAGGGCANNNAGATCGGAA
37482 TATGAGTGTGGAGTCGTTGCAAAAACACCAAAATTAGCCGGCCNNNAGATCGGAAG
37483 TATGAGTGTGGAGTCGTTGCGAGACACAAGACACATGCACAAANNNAGATCGGAA
37484 TATGAGTGTGGAGTCGTTGCCTGCAATCCTACCACTTTGGGAGNNNAGATCGGAAG
37486 TATGAGTGTGGAGTCGTTGCGCATTATGACGTTGTGTGACCATNNNAGATCGGAAG
37487 TATGAGTGTGGAGTCGTTGCAAGAAAATTAGCCGAGCATGGTGNNNAGATCGGAA
37488 TATGAGTGTGGAGTCGTTGCGCTGACAAATTGGACTACATCAANNNAGATCGGAAG
37489 TATGAGTGTGGAGTCGTTGCCGCACAGCATTCTTGGGAATTTANNNAGATCGGAAG
37490 TATGAGTGTGGAGTCGTTGCATAAACGTGGCTTAGTGTGGCTANNNAGATCGGAA
37493 TATGAGTGTGGAGTCGTTGCGCCTATAGTCCCAGCTATGAGGANNNAGATCGGAA
37494 TATGAGTGTGGAGTCGTTGCGAATTCAGGGCGAGTCCATAGACNNNAGATCGGAA
37495 TATGAGTGTGGAGTCGTTGCATTTTTCCTTCCCCTCTAGCTCCNNNAGATCGGAAGA
37497 TATGAGTGTGGAGTCGTTGCAAGATAGGCCCCTAAATTCCCACNNNAGATCGGAAG
37498 TATGAGTGTGGAGTCGTTGCAAGTTTTGCCCCGGTTTTACAAANNNAGATCGGAAG
37499 TATGAGTGTGGAGTCGTTGCACAGTCATGAGTCATGGAAGCATNNNAGATCGGAA
37501 TATGAGTGTGGAGTCGTTGCGCTTTTCTACACTTTCACCCCACNNNAGATCGGAAGA
37502 TATGAGTGTGGAGTCGTTGCTCTCACTCTATCTCTCAGGCTGGNNNAGATCGGAAG

37503 TATGAGTGTGGAGTCGTTGCGAAAAGAAAAAACCACCGTACGGNNNAGATCGGAA
37504 TATGAGTGTGGAGTCGTTGCGAGTGGCACCTCATTGTGATTTTNNNAGATCGGAAG
37506 TATGAGTGTGGAGTCGTTGCCTGAGGCAGGAAAATTGCTTGAANNNAGATCGGAA
37507 TATGAGTGTGGAGTCGTTGCGGAACTCCTGGGATTACAACAGTNNNAGATCGGAAG
37508 TATGAGTGTGGAGTCGTTGCGGAAAAGACAGCATGGTAGATGCNNNAGATCGGAA
37509 TATGAGTGTGGAGTCGTTGCGGGCAGATTACAAGGTCAAGAGANNNAGATCGGAA
37510 TATGAGTGTGGAGTCGTTGCGACATCCTCGCCTCATTTGAGANNNAGATCGGAAG
37511 TATGAGTGTGGAGTCGTTGCCATCTACTGGGAGTCTTAGCCAGNNNAGATCGGAAG
37513 TATGAGTGTGGAGTCGTTGCAGAAGAAGCTGAAAAGGCAAACCNNNAGATCGGAA
37514 TATGAGTGTGGAGTCGTTGCACCAAAATTAGTTTCAGCCCAATNNNAGATCGGAAG
37515 TATGAGTGTGGAGTCGTTGCCAGGCACCTATAATCCCAGCTACNNNAGATCGGAAG
37516 TATGAGTGTGGAGTCGTTGCCGGAAGCAGTACAAACAAGAACANNNAGATCGGAA
37517 TATGAGTGTGGAGTCGTTGCCTGTAGTCCCAGCTACTTGAGAGNNNAGATCGGAAG
37518 TATGAGTGTGGAGTCGTTGCGACGAATGGTCAAACAAACTGGTNNNAGATCGGAA
37519 TATGAGTGTGGAGTCGTTGCGGGCATTGTTCAAGTTCTTTTCCNNNAGATCGGAAG
37521 TATGAGTGTGGAGTCGTTGCCAGAACCCAACGTTTGATGTCTGNNNAGATCGGAAG
37522 TATGAGTGTGGAGTCGTTGCTCTGTAATTGCTCTGGCTTGGATNNNAGATCGGAAG
37523 TATGAGTGTGGAGTCGTTGCCACATTGACCCAACAACTTCCTCNNNAGATCGGAAGA
37525 TATGAGTGTGGAGTCGTTGCAAAGAAAGAAAAGGCCAGGTGTGNNNAGATCGGAA
37526 TATGAGTGTGGAGTCGTTGCTGTTCATGCCACTGTACTCTAGCNNNAGATCGGAAG
37529 TATGAGTGTGGAGTCGTTGCACACACCTGTAGTCCTAGCTACTNNNAGATCGGAAG
37530 TATGAGTGTGGAGTCGTTGCGAGTAGCGTGTGTGATATAGCCANNNAGATCGGAA
37531 TATGAGTGTGGAGTCGTTGCTCTTTGGTAACGTAGGCAAAACTNNNAGATCGGAAG
37533 TATGAGTGTGGAGTCGTTGCCTTGTCAGAAATGGCATCTAGGCNNNAGATCGGAAG
37535 TATGAGTGTGGAGTCGTTGCTTCTTAGTGTACAACCTTTGCCANNNAGATCGGAAG
37536 TATGAGTGTGGAGTCGTTGCAAAAACGTATGTACCCCTTCACANNNAGATCGGAAG
37538 TATGAGTGTGGAGTCGTTGCCACAGCATTGGGGGAACTATTTGNNNAGATCGGAA
37539 TATGAGTGTGGAGTCGTTGCAATGATCCTCCCATCTCAGCTTCNNNAGATCGGAAGA
37540 TATGAGTGTGGAGTCGTTGCAACAAAAAACCTGCCAACCATGTNNNAGATCGGAAG
37542 TATGAGTGTGGAGTCGTTGCTCTAGAGAGAAGTGACCCCTCAGNNNAGATCGGAAG
37544 TATGAGTGTGGAGTCGTTGCATAGTGCCTTGGGTTGCTCTTTANNNAGATCGGAAG
37545 TATGAGTGTGGAGTCGTTGCTGGAGTGTAAAACTGTGGGTCTTNNNAGATCGGAA
37548 TATGAGTGTGGAGTCGTTGCCAAAAACAAAAAACCTCGTGGCANNNAGATCGGAAG
37549 TATGAGTGTGGAGTCGTTGCTCCTGTGGGAAATATGCCATACANNNAGATCGGAAG
37550 TATGAGTGTGGAGTCGTTGCGTCAGACAGATTCACAGCCAAATNNNAGATCGGAAG
37551 TATGAGTGTGGAGTCGTTGCTCACCTCACTCAGAGTAAAAGCCNNNAGATCGGAAG
37553 TATGAGTGTGGAGTCGTTGCTTCAAGGGAGGGAAAATGGAGTCNNNAGATCGGAA
37554 TATGAGTGTGGAGTCGTTGCTCAACAAGCCTTAGACAGCATCANNNAGATCGGAAG
37555 TATGAGTGTGGAGTCGTTGCCATGAGCAGCTGAGCAACAATAGNNNAGATCGGAA
37556 TATGAGTGTGGAGTCGTTGCGTTGTGAAGTCCATGGGAAAAGGNNNAGATCGGAA
37557 TATGAGTGTGGAGTCGTTGCGGTGTTTTTCTCAGCCATCGTAGNNNAGATCGGAAG
37558 TATGAGTGTGGAGTCGTTGCTGCTTCCCACATTTTGCATGTAGNNNAGATCGGAAG
37559 TATGAGTGTGGAGTCGTTGCAGCAGAGGTTTGAATAGTGCTCANNNAGATCGGAA

37561 TATGAGTGTGGAGTCGTTGCCTGGACTCTGCTGCAGGTAATAANNNAGATCGGAA
37562 TATGAGTGTGGAGTCGTTGCGGGAGTTTCGTAAGGTTACCACTNNNAGATCGGAA
37564 TATGAGTGTGGAGTCGTTGCTTGTCAGGGGAATTGGAGAAAGTNNNAGATCGGA
37565 TATGAGTGTGGAGTCGTTGCTCTTTGCTGAGAAGCTCTGCTTANNNAGATCGGAAG
37566 TATGAGTGTGGAGTCGTTGCTGAAACCATTTTCCAGGTGTCTGNNNAGATCGGAAG
37569 TATGAGTGTGGAGTCGTTGCACAAGTTGTCCCTGTTCAGCTAANNNAGATCGGAAG
37571 TATGAGTGTGGAGTCGTTGCTCTAGAAGCTCTAAGGGTTCAGGNNNAGATCGGAA
37573 TATGAGTGTGGAGTCGTTGCGGCTGTAGTGGAGAAAAAGCTTGNNNAGATCGGA
37574 TATGAGTGTGGAGTCGTTGCTGTACTCCAGTCTGAGTGACAGANNNAGATCGGAA
37575 TATGAGTGTGGAGTCGTTGCAAGAGAAAGAAAAAGACAGGGGTNNNAGATCGGA
37576 TATGAGTGTGGAGTCGTTGCCCAGTGCAACATGAATTGTGCTANNNAGATCGGAAG
37577 TATGAGTGTGGAGTCGTTGCGGCACAGAAACAGCTTATTGGTTNNNAGATCGGAA
37578 TATGAGTGTGGAGTCGTTGCGGCTGCCATATCTTTCTGTTGTCNNNAGATCGGAAG
37579 TATGAGTGTGGAGTCGTTGCGGAAGATATCTGGGTGATGGGTCNNNAGATCGGA
37580 TATGAGTGTGGAGTCGTTGCGGAAGATATCTGGGTGATGGGTCNNNAGATCGGA
37581 TATGAGTGTGGAGTCGTTGCTCCCAGTCCATCAGACAAAGTTTNNNAGATCGGAAG
37582 TATGAGTGTGGAGTCGTTGCATTACTCCAACTCCAAGAGGTGCNNNAGATCGGAAG
37583 TATGAGTGTGGAGTCGTTGCTGTACAACTGCCAGCACAGATATNNNAGATCGGAAG
37585 TATGAGTGTGGAGTCGTTGCACTGCCTCGATTACCCATGAAATNNNAGATCGGAAG
37586 TATGAGTGTGGAGTCGTTGCTTTGGGAAATGGCATTCTCCAACNNNAGATCGGAAG
37587 TATGAGTGTGGAGTCGTTGCACACATATAAGGGCTGATCACCCNNNAGATCGGAAG
37588 TATGAGTGTGGAGTCGTTGCGCTGCCATGAACATTCGTATACANNNAGATCGGAAG
37589 TATGAGTGTGGAGTCGTTGCGTGTCACATGCTTGTAATCCCTGNNNAGATCGGAAG
37590 TATGAGTGTGGAGTCGTTGCTCCAGGTCAAGACTATGTTGGTGNNNAGATCGGAA
37591 TATGAGTGTGGAGTCGTTGCGTTCACAACCTTACTTATGCAGTNNNAGATCGGAAG
37592 TATGAGTGTGGAGTCGTTGCAGGAAGCTAATGATGTCTGCTCCNNNAGATCGGAA
37593 TATGAGTGTGGAGTCGTTGCAGAAGAGCAAAAAGCAAGAGCACNNNAGATCGGA
37594 TATGAGTGTGGAGTCGTTGCGCAAGACACAGCATTGCAAAATTNNNAGATCGGAA
37595 TATGAGTGTGGAGTCGTTGCACAAGAAGACCATAAAGGCCCATNNNAGATCGGAA
37597 TATGAGTGTGGAGTCGTTGCACCTTCTCCAAAAATGTCCTGGANNNAGATCGGAAG
37598 TATGAGTGTGGAGTCGTTGCACATACATCATGGTGCTGCTTTTNNNAGATCGGAAG
37599 TATGAGTGTGGAGTCGTTGCCCAGTTTTGTGCTTGAAACCCAGNNNAGATCGGAAG
37600 TATGAGTGTGGAGTCGTTGCTGTTGAGGACTTAGTATTTGCCTNNNAGATCGGAAG
37601 TATGAGTGTGGAGTCGTTGCTGTACTTCTCACTTTCTGCTCCCNNNAGATCGGAAGA
37604 TATGAGTGTGGAGTCGTTGCCTACTCTCCGGACAGTGGAAAAANNNAGATCGGAA
37605 TATGAGTGTGGAGTCGTTGCTACAAAAAATTAGCTGGGCGTGGNNNAGATCGGAA
37606 TATGAGTGTGGAGTCGTTGCATGGCGGGCATCTATAATCTCAGNNNAGATCGGAA
37608 TATGAGTGTGGAGTCGTTGCCACATACCTCACATACCTGCAGTNNNAGATCGGAAG
37609 TATGAGTGTGGAGTCGTTGCCCCAGTTTTGTGCTTGAAACCTANNNAGATCGGAAG
37610 TATGAGTGTGGAGTCGTTGCAGGTCAGGAGATAGAGACCATCCNNNAGATCGGAA
37611 TATGAGTGTGGAGTCGTTGCGTGCAACTTCCCGGATGTAAAAANNNAGATCGGAA
37612 TATGAGTGTGGAGTCGTTGCAAAGAAAGATGTAGACCGGCAGANNNAGATCGGA
37614 TATGAGTGTGGAGTCGTTGCTTATAAGGCCGAGATGGGATTGCNNNAGATCGGAA

37615 TATGAGTGTGGAGTCGTTGCAGGCATGAACAAACCAAACTGTTNNNAGATCGGAA
37617 TATGAGTGTGGAGTCGTTGCTTATGAGCCTCAAGATGACCCTGNNNAGATCGGAAG
37618 TATGAGTGTGGAGTCGTTGCAGTGAGCTGTGATTGTGTCCTACNNNAGATCGGAA
37619 TATGAGTGTGGAGTCGTTGCTGAGGTCGATCTTTATGGTTCCANNNAGATCGGAAG
37620 TATGAGTGTGGAGTCGTTGCGAGGGAAAAAAACCTCTTACCCTNNNAGATCGGAAG
37621 TATGAGTGTGGAGTCGTTGCTGTTCATCAGTGTATCCCAAGGGNNNAGATCGGAAG
37622 TATGAGTGTGGAGTCGTTGCTGTTTGAGGTGATGGATATGCCANNNAGATCGGAA
37624 TATGAGTGTGGAGTCGTTGCGTGGACACCTATAATCCCAGCTANNNAGATCGGAAG
37625 TATGAGTGTGGAGTCGTTGCCAGGTTTTCAGCCTTCCATATGCNNNAGATCGGAAG
37626 TATGAGTGTGGAGTCGTTGCATGCCTATGGTTCCTATGGTTCCNNNAGATCGGAAG
37627 TATGAGTGTGGAGTCGTTGCATGGGATGGGGGAAAAATGATGTNNNAGATCGGA
37630 TATGAGTGTGGAGTCGTTGCAATCACTTTACCCCTGGAGATGGNNNAGATCGGAAG
37631 TATGAGTGTGGAGTCGTTGCATTGCTTGAGGTCAGGAGTTTGANNNAGATCGGAA
37632 TATGAGTGTGGAGTCGTTGCTCCCTGTAGCAGCCATTCTTTTANNNAGATCGGAAG
37633 TATGAGTGTGGAGTCGTTGCAGGCTAAGGTAGGAGGATCACTTNNNAGATCGGAA
37634 TATGAGTGTGGAGTCGTTGCTTTGAGACCAGCCTGATCAACATNNNAGATCGGAAG
37635 TATGAGTGTGGAGTCGTTGCTAAGGCACGAGAATTGCTTGAACNNNAGATCGGAA
37636 TATGAGTGTGGAGTCGTTGCGGCATGATCTCTGCTCACTGTAANNNAGATCGGAAG
37637 TATGAGTGTGGAGTCGTTGCCTGAGGCAGGAGAATTGCTTGAANNNAGATCGGAA
37638 TATGAGTGTGGAGTCGTTGCTCGAAATGAGATTTGGGCTGGTANNNAGATCGGAA
37639 TATGAGTGTGGAGTCGTTGCCCTCTTATCTTGGCCTCCCAAATNNNAGATCGGAAGA
37640 TATGAGTGTGGAGTCGTTGCGTGGCTCACTCCTGTAATCCTAGNNNAGATCGGAAG
37641 TATGAGTGTGGAGTCGTTGCGGATCATGAGGTCAGGAGTTCAANNNAGATCGGAA
37642 TATGAGTGTGGAGTCGTTGCTACCTACCACAGTGCTGGATAGANNNAGATCGGAAG
37643 TATGAGTGTGGAGTCGTTGCTTGCAATCCCCCTGTCTTGATAANNNAGATCGGAAG
37644 TATGAGTGTGGAGTCGTTGCTGAGGCAGAAAATTGCTTGAACCNNNAGATCGGAA
37645 TATGAGTGTGGAGTCGTTGCTTCAGACAGTAAAGCTCAGAGCANNNAGATCGGAA
37646 TATGAGTGTGGAGTCGTTGCTGCCAAGAAAATTCCAGAACTGCNNNAGATCGGAAG
37648 TATGAGTGTGGAGTCGTTGCTGATGGTCACAGGACTGGAAATTNNNAGATCGGAA
37649 TATGAGTGTGGAGTCGTTGCAGAAAGCTGTACTGATACGTGCTNNNAGATCGGAA
37650 TATGAGTGTGGAGTCGTTGCCAAACTCATAGGGTCCAGACGATNNNAGATCGGAA
37651 TATGAGTGTGGAGTCGTTGCCTTGAACCAGGGAGTTAGAGCTTNNNAGATCGGAA
37652 TATGAGTGTGGAGTCGTTGCAACCAATCCCCTTTTTGTCTCCTNNNAGATCGGAAGA
37653 TATGAGTGTGGAGTCGTTGCCCAGTCAGAACTAAAGGGCTTCTNNNAGATCGGAAG
37655 TATGAGTGTGGAGTCGTTGCTACCTTAGGTCAGGAGTTCGAGANNNAGATCGGAA
37656 TATGAGTGTGGAGTCGTTGCAATCCAGGTTTGTGCTGTTTACCNNNAGATCGGAAG
37657 TATGAGTGTGGAGTCGTTGCTCATCTGAGGTTGGGAGTTCAAGNNNAGATCGGAA
37658 TATGAGTGTGGAGTCGTTGCAGGTATGTTGATGTGGTGGGAAANNNAGATCGGA
37659 TATGAGTGTGGAGTCGTTGCAAAAAATTAGCTGGGCATGGTGGNNNAGATCGGAA
37661 TATGAGTGTGGAGTCGTTGCGGGCAAGGGACTTCCTGTAATAANNNAGATCGGAA
37662 TATGAGTGTGGAGTCGTTGCGAGATGAGGTTTTGCCATGTTGTNNNAGATCGGAA
37665 TATGAGTGTGGAGTCGTTGCAGTATCTCCTTCTTCCTTGCACCNNNAGATCGGAAGA
37666 TATGAGTGTGGAGTCGTTGCAAATGCATGGGATTAGACACCCTNNNAGATCGGAA

37667 TATGAGTGTGGAGTCGTTGCGACAGAGCAAGACTCCATCTCAANNNAGATCGGAAG
37669 TATGAGTGTGGAGTCGTTGCTGGTTTTCCATTCCTGAGTTACTNNNAGATCGGAAG
37671 TATGAGTGTGGAGTCGTTGCGATTCTCCTCTGCAGGTTTCAGANNNAGATCGGAAG
37672 TATGAGTGTGGAGTCGTTGCTGTGTGTATAAGCATGTCTGTGCNNNAGATCGGAAG
37673 TATGAGTGTGGAGTCGTTGCAATATTGACTGCCTTTGGGGAGGNNNAGATCGGAA
37674 TATGAGTGTGGAGTCGTTGCGGACATACCAAGCCAACCAAAAANNNAGATCGGAAG
37675 TATGAGTGTGGAGTCGTTGCGGCCAAACAAATATGTCCAGAGANNNAGATCGGAA
37676 TATGAGTGTGGAGTCGTTGCTGTGGCACTGTAACAAAAGACACNNNAGATCGGAAG
37677 TATGAGTGTGGAGTCGTTGCAGCCCCGGGTAGAATAACTTTACNNNAGATCGGAAG
37680 TATGAGTGTGGAGTCGTTGCAATGATGAGGTCGGGTGATTAGGNNNAGATCGGA
37681 TATGAGTGTGGAGTCGTTGCCACTCAGCATGGGTAACAAAGTGNNNAGATCGGAA
37684 TATGAGTGTGGAGTCGTTGCTCTCAGAGTCCCAGAAGATGAGANNNAGATCGGAA
37685 TATGAGTGTGGAGTCGTTGCAAAAAACCACACACACACACACANNNAGATCGGAAG
37686 TATGAGTGTGGAGTCGTTGCTGTGAATGAAGCTAAAGGAGTGCNNNAGATCGGAA
37687 TATGAGTGTGGAGTCGTTGCAGTCTCATAGAACCACTCCAGGTNNNAGATCGGAAG
37688 TATGAGTGTGGAGTCGTTGCGTTCATATGCTCGAATTCCACCCNNNAGATCGGAAG
37689 TATGAGTGTGGAGTCGTTGCACACAACCATAAGCCTGAATTCANNNAGATCGGAAG
37690 TATGAGTGTGGAGTCGTTGCCAAGAAACCATCACTGGCACTTTNNNAGATCGGAAG
37691 TATGAGTGTGGAGTCGTTGCACAGGGAACTTACATTGCTTGATNNNAGATCGGAAG
37693 TATGAGTGTGGAGTCGTTGCACCAAAAAACAACGATTAGGCCTNNNAGATCGGAAG
37694 TATGAGTGTGGAGTCGTTGCAGTGAGTGTGTAAAGGAGTGCAANNNAGATCGGA
37696 TATGAGTGTGGAGTCGTTGCAGAAAGAGGAAGTGCTCCAAACANNNAGATCGGAA
37697 TATGAGTGTGGAGTCGTTGCCTGAAGTAGGAGGATCCCTTGAGNNNAGATCGGAA
37699 TATGAGTGTGGAGTCGTTGCTTTGGTTGAAATGCTACAGCCAGNNNAGATCGGAAG
37700 TATGAGTGTGGAGTCGTTGCTTATGGGATGCTTACACTGTGCTNNNAGATCGGAAG
37701 TATGAGTGTGGAGTCGTTGCAACACCCAACCCTACAATGGATTNNNAGATCGGAAG
37702 TATGAGTGTGGAGTCGTTGCTGAGCTCCAAAATTGAATCAGCANNNAGATCGGAAG
37712 TATGAGTGTGGAGTCGTTGCCTTGAACTCAGAAGGCAGAGGTTNNNAGATCGGAA
37713 TATGAGTGTGGAGTCGTTGCGAGCTTGATGTCATAAGCAACCANNNAGATCGGAAG
37714 TATGAGTGTGGAGTCGTTGCGCAGAGTCATATTTTGGAGGAGCNNNAGATCGGAA
37716 TATGAGTGTGGAGTCGTTGCGTGCTTGCCTCCTTCTTTTCATGNNNAGATCGGAAGA
37717 TATGAGTGTGGAGTCGTTGCCAGCTTGTTAGAACTGGTCCAGANNNAGATCGGAAG
37718 TATGAGTGTGGAGTCGTTGCAACAATCGCTTGGGAGGCTATATNNNAGATCGGAAG
37719 TATGAGTGTGGAGTCGTTGCAAAATGAGACCCAGTCATTTGCANNNAGATCGGAAG
37722 TATGAGTGTGGAGTCGTTGCGAAACAACCCAGCTGTCTTTCAGNNNAGATCGGAAG
37723 TATGAGTGTGGAGTCGTTGCAAAAACCTCACTGGATGGGATCANNNAGATCGGAA
37724 TATGAGTGTGGAGTCGTTGCAGCAACAGAAGACCTTTCAGTCANNNAGATCGGAAG
37726 TATGAGTGTGGAGTCGTTGCAGCACGAATATAACTTCCTGGCANNNAGATCGGAAG
37728 TATGAGTGTGGAGTCGTTGCGAGGAACTGCCAGACTGTTTTTCNNNAGATCGGAAG
37730 TATGAGTGTGGAGTCGTTGCCGTACTCAAAAAGGCCCAATGTTNNNAGATCGGAAG
37731 TATGAGTGTGGAGTCGTTGCTAAGCAGTTGCAGAAGAGCTGTANNNAGATCGGAA
37734 TATGAGTGTGGAGTCGTTGCACCAGAGGTTGAGAACAAGTTGANNNAGATCGGAA
37735 TATGAGTGTGGAGTCGTTGCGAAAGAGCAGGACAGTGCAAAATNNNAGATCGGA

37736 TATGAGTGTGGAGTCGTTGCATATGCACCCAATACAGGAGGACNNNAGATCGGAA
37737 TATGAGTGTGGAGTCGTTGCTGACCCCTGAAAGCATAGTCTTTNNNAGATCGGAAG
37738 TATGAGTGTGGAGTCGTTGCTTGTAGGGTCTTGCTATGTTGCCNNNAGATCGGAAG
37739 TATGAGTGTGGAGTCGTTGCGGGCCAGAGAGACTACATTTCTTNNNAGATCGGAA
37740 TATGAGTGTGGAGTCGTTGCCCCATTCTTTAGCAGTCTGTGTGNNNAGATCGGAAG
37741 TATGAGTGTGGAGTCGTTGCATGTGCAGACCCTGAAAAATGTGNNNAGATCGGAA
37743 TATGAGTGTGGAGTCGTTGCAGAAAATTGCCAAACCCCTCATGNNNAGATCGGAAG
37744 TATGAGTGTGGAGTCGTTGCTCCCAGAGATCAGTTTGGTTCTGNNNAGATCGGAAG
37745 TATGAGTGTGGAGTCGTTGCGAAAGCATTCGGTCTTGCCTTATNNNAGATCGGAAG
37746 TATGAGTGTGGAGTCGTTGCAAGAAGGTGAGGTTGGAAAGGAGNNNAGATCGGA
37747 TATGAGTGTGGAGTCGTTGCCAGTGTGTGTGTTGCTCTTACAANNNAGATCGGAAG
37748 TATGAGTGTGGAGTCGTTGCCTGAGGCAGGAGAATAGCATGAANNNAGATCGGA
37749 TATGAGTGTGGAGTCGTTGCGAGAATTGCTTGAACCTGGGAAGNNNAGATCGGAA
37750 TATGAGTGTGGAGTCGTTGCGATTGGTTTTTGTGTGCTGACCTNNNAGATCGGAAG
37751 TATGAGTGTGGAGTCGTTGCGGCCTAGAAACATTTTGGTGCATNNNAGATCGGAA
37752 TATGAGTGTGGAGTCGTTGCCATTCCAGGCAATGTGGTAGAACNNNAGATCGGAA
37753 TATGAGTGTGGAGTCGTTGCTGTTGCAACATGGATGAACCTTCNNNAGATCGGAAG
37755 TATGAGTGTGGAGTCGTTGCTCCCATTTTCCTTCCTTCCTTCCNNNAGATCGGAAGAG
37756 TATGAGTGTGGAGTCGTTGCACCTTTGAATCTCCTACACCTGGNNNAGATCGGAAG
37759 TATGAGTGTGGAGTCGTTGCTATGAAAACCTACAACTGACGGCNNNAGATCGGAAG
37760 TATGAGTGTGGAGTCGTTGCGTCTCGAATTCCTGACCTCAAGTNNNAGATCGGAAG
37761 TATGAGTGTGGAGTCGTTGCTCAAGAGATCTGGAGTAGAGCCTNNNAGATCGGAA
37762 TATGAGTGTGGAGTCGTTGCTCAAATTCCTGGCCTCAAGCTATNNNAGATCGGAAG
37764 TATGAGTGTGGAGTCGTTGCACAGCCCACCTAGAAGAACTAGANNNAGATCGGAA
37765 TATGAGTGTGGAGTCGTTGCCTGTAGTCTCAGCTACTTGGGAGNNNAGATCGGAA
37766 TATGAGTGTGGAGTCGTTGCTTTGAAAAGGCCTGCTTACAAGGNNNAGATCGGAA
37767 TATGAGTGTGGAGTCGTTGCTTGGAGCCATAGTTCCACCAAATNNNAGATCGGAAG
37768 TATGAGTGTGGAGTCGTTGCCACACTCTTTTTCGTCTCTGTGTNNNAGATCGGAAGA
37769 TATGAGTGTGGAGTCGTTGCCCTGTAGTCCCAGCTACTTAGGANNNAGATCGGAAG
37770 TATGAGTGTGGAGTCGTTGCGGCTTAGTTTCCTCCTGTTCTCANNNAGATCGGAAGA
37771 TATGAGTGTGGAGTCGTTGCAACAAATGGCGTATCCCAATACANNNAGATCGGAAG
37773 TATGAGTGTGGAGTCGTTGCAAGAAGAAAGGTGCCAGTCAGATNNNAGATCGGAA
37774 TATGAGTGTGGAGTCGTTGCCAGACTACAAAGGCCTTCATCCTNNNAGATCGGAAG
37775 TATGAGTGTGGAGTCGTTGCAAAAAAGGAGGCTTATTGAGGCCNNNAGATCGGAA
37776 TATGAGTGTGGAGTCGTTGCCTTTCTCTCCTGCTACCTTGTGANNNAGATCGGAAGA
37777 TATGAGTGTGGAGTCGTTGCCTTGTTATGTTCCCCAGGCTAGTNNNAGATCGGAAG
37778 TATGAGTGTGGAGTCGTTGCTGAGGAGAAAAATGAAGCAGGGANNNAGATCGGA
37782 TATGAGTGTGGAGTCGTTGCTGGACGTCATCATTCACAAGTTGNNNAGATCGGAAG
37783 TATGAGTGTGGAGTCGTTGCAGTTGAGAGGAACTCTAGCAAATNNNAGATCGGAA
37784 TATGAGTGTGGAGTCGTTGCATTTTGCTAGGTGAGATGGCTCANNNAGATCGGAA
37785 TATGAGTGTGGAGTCGTTGCAGGCTATCTGGAAACACAGGAAGNNNAGATCGGAA
37786 TATGAGTGTGGAGTCGTTGCCTCCATGGAACAGTTTTGGCATTNNNAGATCGGAAG
37788 TATGAGTGTGGAGTCGTTGCAAAAGGAGGAAAGTGAGACGACANNNAGATCGGA

37789 TATGAGTGTGGAGTCGTTGCAAGACACATACCGGCTGTAACTTNNNAGATCGGAAG
37790 TATGAGTGTGGAGTCGTTGCTAGTTCCATCCAATGAAACCCCNNNAGATCGGAAGA
37791 TATGAGTGTGGAGTCGTTGCAAGAGATTGAGATCATCCTGGCCNNNAGATCGGAAG
37793 TATGAGTGTGGAGTCGTTGCTGTAATCACAGCACTTTGGGAGGNNNAGATCGGAAC
37794 TATGAGTGTGGAGTCGTTGCGAGGTGGAGGTGGAAGATTGTTTNNNAGATCGGAA
37795 TATGAGTGTGGAGTCGTTGCACAAAATTATCCTGGCATGGTGGNNNAGATCGGAAG
37796 TATGAGTGTGGAGTCGTTGCTGCCTGTAATTGCAGCTATTTGGNNNAGATCGGAAG
37797 TATGAGTGTGGAGTCGTTGCAACTGGATCACCAAGGTTTAGCTNNNAGATCGGAAG
37798 TATGAGTGTGGAGTCGTTGCGGTGAAACCCCGTCTCTACTAAANNNAGATCGGAAG
37800 TATGAGTGTGGAGTCGTTGCGCCAACAGGTTGCTTATGAAAGANNNAGATCGGAAC
37801 TATGAGTGTGGAGTCGTTGCCATGCCTGTAGTCCCATCTACTCNNNAGATCGGAAGA
37802 TATGAGTGTGGAGTCGTTGCGTCAACCCCAACAAAATCCAGAGNNNAGATCGGAAG
37803 TATGAGTGTGGAGTCGTTGCCCTGTCTCTCTGGTTCTGATGACNNNAGATCGGAAGA
37804 TATGAGTGTGGAGTCGTTGCAGTAAGCATGAGATGTGTCAGGGNNNAGATCGGAA
37805 TATGAGTGTGGAGTCGTTGCTAAAAAGGCCAAGTGCAATGGTTNNNAGATCGGAAC
37806 TATGAGTGTGGAGTCGTTGCACTACATCTGCAAGACATCAGCANNNAGATCGGAAG
37807 TATGAGTGTGGAGTCGTTGCCATCATGGCAAAACCCAGTTTCTNNNAGATCGGAAGA
37808 TATGAGTGTGGAGTCGTTGCCCCCATCTCAAAAAAAAATGCCCNNNAGATCGGAAGA
37809 TATGAGTGTGGAGTCGTTGCTGCCCTCAACTTGATAAAGAGCANNNAGATCGGAAG
37810 TATGAGTGTGGAGTCGTTGCGGTGAAACCCCGTCTCTACTAAANNNAGATCGGAAG
37811 TATGAGTGTGGAGTCGTTGCTAACCTGGCAAACATGGTGAAACNNNAGATCGGAAG
37812 TATGAGTGTGGAGTCGTTGCGTGGATCACTTGAGGTCAGAAGTNNNAGATCGGAAC
37813 TATGAGTGTGGAGTCGTTGCTAGTCTCAGCTACTCAGGAAGCTNNNAGATCGGAAG
37814 TATGAGTGTGGAGTCGTTGCCTCACTGGATGGGCTCAATAGTANNNAGATCGGAAG
37815 TATGAGTGTGGAGTCGTTGCAAGCTGAGGTAGGAGGATCACTTNNNAGATCGGAA
37816 TATGAGTGTGGAGTCGTTGCCCCATGATACTGAGACCATCACCNNNAGATCGGAAGA
37817 TATGAGTGTGGAGTCGTTGCGGGTGAACATGAACATTGGGAAGNNNAGATCGGAA
37818 TATGAGTGTGGAGTCGTTGCAGAGATACATAGGGCAAGGCATGNNNAGATCGGAA
37819 TATGAGTGTGGAGTCGTTGCCACCTGTAATCCCAGCATTTTGGNNNAGATCGGAAGA
37820 TATGAGTGTGGAGTCGTTGCATGAGTGGTTGATAGAGGTTCGGNNNAGATCGGAA
37821 TATGAGTGTGGAGTCGTTGCGATGGACGAAATCGCTTGAAGAGNNNAGATCGGAA
37822 TATGAGTGTGGAGTCGTTGCCAGGTTCAGTATCCACAGTCCAANNNAGATCGGAAG
37823 TATGAGTGTGGAGTCGTTGCGCTCATGCCTGTAATACCAACACNNNAGATCGGAAGA
37824 TATGAGTGTGGAGTCGTTGCTCCTTCTTCCTCTTTGCCTTCTGNNNAGATCGGAAGAG
37825 TATGAGTGTGGAGTCGTTGCCTGAGGCAGGAGAATCACTTGAANNNAGATCGGAAC
37826 TATGAGTGTGGAGTCGTTGCCTCATGCCTATGATCGTAGCACTNNNAGATCGGAAGA
37827 TATGAGTGTGGAGTCGTTGCTTGTGGGCTTGGAAATAACACTGNNNAGATCGGAAC
37828 TATGAGTGTGGAGTCGTTGCACAAATCTCAAAATGGCAGATGTNNNAGATCGGAAG
37831 TATGAGTGTGGAGTCGTTGCGAGAGAGATCCTCCTGCTTTAGCNNNAGATCGGAAG
37832 TATGAGTGTGGAGTCGTTGCTCCTTCACCTCCAAAAGTTTCCTNNNAGATCGGAAGA
37833 TATGAGTGTGGAGTCGTTGCGTGTCACTGTGGAAGGCTACTAANNNAGATCGGAAC
37834 TATGAGTGTGGAGTCGTTGCACAAAAAAAGGCCATGTGAGGACNNNAGATCGGAA
37835 TATGAGTGTGGAGTCGTTGCTGGAAGAATCACTTGAGACCAGGNNNAGATCGGAAC

37836 TATGAGTGTGGAGTCGTTGCGGGACAAAAAGGGTGTGATCTTGNNNAGATCGGAA
37837 TATGAGTGTGGAGTCGTTGCTCCCTGTCTAGTTGTTTACCCANNNAGATCGGAAGA
37838 TATGAGTGTGGAGTCGTTGCCGGCATATAGCAGGCACTCAATANNNAGATCGGAA
37839 TATGAGTGTGGAGTCGTTGCGGGCTGGGACTTGAAAGATAAGANNNAGATCGGA
37840 TATGAGTGTGGAGTCGTTGCCTGGGGAGACAGACTATGTGAACNNNAGATCGGAA
37842 TATGAGTGTGGAGTCGTTGCAGAGGAAGAAGAGGAAGGAAGGANNNAGATCGG
37843 TATGAGTGTGGAGTCGTTGCATACATTATGGTTGGGTTGGGCANNNAGATCGGAA
37844 TATGAGTGTGGAGTCGTTGCATTGATTGATGGGGTCAGCTGATNNNAGATCGGAA
37845 TATGAGTGTGGAGTCGTTGCGTTGCAGTGAGTTGAGAACACACNNNAGATCGGAA
37849 TATGAGTGTGGAGTCGTTGCGAAGAAGAAGAGGTCCAAGGTCANNNAGATCGGA
37850 TATGAGTGTGGAGTCGTTGCGACCACTTCAGAGAAGAAGGAGANNNAGATCGGAA
37851 TATGAGTGTGGAGTCGTTGCACAGAGATGTAACTGCAGATCCTNNNAGATCGGAA
37852 TATGAGTGTGGAGTCGTTGCGTAGCTTATGTTCATGGCAAGCANNNAGATCGGAA
37856 TATGAGTGTGGAGTCGTTGCTTCTTTTCCCACATGAACCTCCANNNAGATCGGAAGA
37857 TATGAGTGTGGAGTCGTTGCTTGCATACAGTTGGTTGATGGTGNNNAGATCGGAA
37858 TATGAGTGTGGAGTCGTTGCTCTGATCATACACCTTGTGGCAANNNAGATCGGAAG
37860 TATGAGTGTGGAGTCGTTGCCCTGTAATCCCAACACCTTGAGANNNAGATCGGAAG
37861 TATGAGTGTGGAGTCGTTGCAAAAAAATTAGCTGGGGGTGGTGNNNAGATCGGA
37862 TATGAGTGTGGAGTCGTTGCCAGAACTTGATGGAGTTTCAGCCNNNAGATCGGAA
37863 TATGAGTGTGGAGTCGTTGCGAAAGCTGAAGGTTGTTGGTCTTNNNAGATCGGAA
37864 TATGAGTGTGGAGTCGTTGCTCACTCAAATCTTTCCTCCTGGGNNNAGATCGGAAGA
37865 TATGAGTGTGGAGTCGTTGCAAAAAAATAGTGTGGGCAGGGTGNNNAGATCGGA
37866 TATGAGTGTGGAGTCGTTGCGTGGCCATAAACTACCCAGAGAANNNAGATCGGAA
37867 TATGAGTGTGGAGTCGTTGCCGTCCTGAAAGGCATTAGTTGTCNNNAGATCGGAAG
37868 TATGAGTGTGGAGTCGTTGCCAGGATGGTGCACTTCTGTAGTCNNNAGATCGGAA
37869 TATGAGTGTGGAGTCGTTGCATCAGGACTTGGTGTATCTTCGGNNNAGATCGGAA
37870 TATGAGTGTGGAGTCGTTGCTAAATAAATAAAGCGTGGGCCGGNNNAGATCGGAA
37871 TATGAGTGTGGAGTCGTTGCCTGTAGAGAAGGCGGTACTTGAANNNAGATCGGAA
37873 TATGAGTGTGGAGTCGTTGCGCTGAGACAGGACAATCTCTTGANNNAGATCGGAA
37874 TATGAGTGTGGAGTCGTTGCATCACCAAAGTGCTATCCCTTGTNNNAGATCGGAAG
37875 TATGAGTGTGGAGTCGTTGCGCTAACTGACAGAACATGAGGGANNNAGATCGGAA
37876 TATGAGTGTGGAGTCGTTGCCTCCTCCTATCCACCATCCCTAANNNAGATCGGAAGA
37877 TATGAGTGTGGAGTCGTTGCTTTTACTCAGAGGGGGGACTACTNNNAGATCGGAA
37878 TATGAGTGTGGAGTCGTTGCTCTCATCATTCAGCCTCAATGCTNNNAGATCGGAAGA
37879 TATGAGTGTGGAGTCGTTGCGATTGATTCCTTAGCACACGAGCNNNAGATCGGAAG
37880 TATGAGTGTGGAGTCGTTGCCTTTTTGGGGTGAAGCTGATCAGNNNAGATCGGAA
37881 TATGAGTGTGGAGTCGTTGCTTGTCTCGATCCATCACAGTCAGNNNAGATCGGAAG
37882 TATGAGTGTGGAGTCGTTGCAAGTGCTGGGATTACAAGTGTGANNNAGATCGGAA
37884 TATGAGTGTGGAGTCGTTGCGACAGGGACGCTAGTTCATAGAGNNNAGATCGGAA
37885 TATGAGTGTGGAGTCGTTGCATCACTTGAACCTGGGAGATGTCNNNAGATCGGAA
37886 TATGAGTGTGGAGTCGTTGCCCAACATGGTGAAACCTTGTCTCNNNAGATCGGAAG
37887 TATGAGTGTGGAGTCGTTGCGGGAAGAAAGCAGCCTTCTTACTNNNAGATCGGAA
37888 TATGAGTGTGGAGTCGTTGCACAGCAGTAGACAAATAGAGCCCNNNAGATCGGAA

37889 TATGAGTGTGGAGTCGTTGCCATGATGGTGTGTGCCTGTAATCNNNAGATCGGAAG
37890 TATGAGTGTGGAGTCGTTGCCAGCACCTATGGAGCACTAAGAANNNAGATCGGAAG
37891 TATGAGTGTGGAGTCGTTGCTTTCCTCTGGCTGCCTATTTCTTNNNAGATCGGAAGA
37892 TATGAGTGTGGAGTCGTTGCAACTGAGCCCTTGATTCACAGAANNNAGATCGGAAG
37893 TATGAGTGTGGAGTCGTTGCCTGTAGTCCCAGCTACTCAGGAGNNNAGATCGGAAG
37895 TATGAGTGTGGAGTCGTTGCACTGTAGTGAGCTGTGATTGTGTNNNAGATCGGAAG
37896 TATGAGTGTGGAGTCGTTGCGTCCAAAGTTAGCAGAAAGATGGANNNAGATCGGA
37897 TATGAGTGTGGAGTCGTTGCCAAGAAATGCAAAGGAACGCAGTNNNAGATCGGAA
37899 TATGAGTGTGGAGTCGTTGCGTAATCCCAGCTACTCAGAAGGCNNNAGATCGGAAG
37900 TATGAGTGTGGAGTCGTTGCCTCACATCTGTAATCCCAGCACTNNNAGATCGGAAGA
37901 TATGAGTGTGGAGTCGTTGCTCCTCATACTCTTGACAGCACTGNNNAGATCGGAAGA
37902 TATGAGTGTGGAGTCGTTGCACATGGTAAAACCCCGTCTCTACNNNAGATCGGAAGA
37903 TATGAGTGTGGAGTCGTTGCGAAGTTTCACTCTTGTTGCCCAGNNNAGATCGGAAGA
37904 TATGAGTGTGGAGTCGTTGCTGCCTGTAATTCCAGCTACTCAGNNNAGATCGGAAGA
37905 TATGAGTGTGGAGTCGTTGCGTAATCTCAGCTACTCAGGAGGCNNNAGATCGGAAG
37906 TATGAGTGTGGAGTCGTTGCATACAGGCTGCTGACCATTATGTNNNAGATCGGAAG
37907 TATGAGTGTGGAGTCGTTGCTGTCTATGGCAACCTGTTTGTCTNNNAGATCGGAAGA
37908 TATGAGTGTGGAGTCGTTGCTTGGAGGTTGCAGTGAACTATCANNNAGATCGGAAG
37909 TATGAGTGTGGAGTCGTTGCATGTCCCAATTTGTGAGATCCCANNNAGATCGGAAGA
37911 TATGAGTGTGGAGTCGTTGCAGGTATAAGTGGTTGGAATGGGGNNNAGATCGGAA
37912 TATGAGTGTGGAGTCGTTGCCCTGTAATCCCAGCTACTAGGGANNNAGATCGGAAG
37913 TATGAGTGTGGAGTCGTTGCGTATCTCTGCCCTCCATCTGAAGNNNAGATCGGAAGA
37914 TATGAGTGTGGAGTCGTTGCCCTAATGTGTGTGTACTGTGTGCNNNAGATCGGAAG
37915 TATGAGTGTGGAGTCGTTGCTTCCCTGTGTTTCCATAGTGTGTNNNAGATCGGAAGA
37916 TATGAGTGTGGAGTCGTTGCAGATTCTGGAACCAGATTGACGTNNNAGATCGGAAG
37917 TATGAGTGTGGAGTCGTTGCCAATGGGAATGCACAGTTTCCTTNNNAGATCGGAAG
37918 TATGAGTGTGGAGTCGTTGCTGAGGTCAGGAATTTGAGACCAGNNNAGATCGGAA
37920 TATGAGTGTGGAGTCGTTGCCACTCCTGTAATCCCAGCACTTTNNNAGATCGGAAGA
37921 TATGAGTGTGGAGTCGTTGCCCTTCTATTTTCTGTCAGCTGGCNNNAGATCGGAAGA
37922 TATGAGTGTGGAGTCGTTGCCAACGACTCCATGACACTCTTTGNNNAGATCGGAAGA
37923 TATGAGTGTGGAGTCGTTGCTGAGCTGTTGGAGTTTCTTCTGTNNNAGATCGGAAG
37924 TATGAGTGTGGAGTCGTTGCGCTATGCCTGTAATCCTAGCACTNNNAGATCGGAAGA
37925 TATGAGTGTGGAGTCGTTGCAAAGTTTCCTACCTTCAGACCCCNNNAGATCGGAAGA
37926 TATGAGTGTGGAGTCGTTGCAAACTCTTCTGTGTGGGAGACAGNNNAGATCGGAAG
37928 TATGAGTGTGGAGTCGTTGCTTACAGAGTTCCCATATGCCTCGNNNAGATCGGAAGA
37929 TATGAGTGTGGAGTCGTTGCCCTTCCCAATTTTGTGTACGTGTNNNAGATCGGAAGA
37930 TATGAGTGTGGAGTCGTTGCTGTAATCCTAGCACTTTGGGAGGNNNAGATCGGAAG
37931 TATGAGTGTGGAGTCGTTGCCTCATGCCTGTAATCCTAGCACTNNNAGATCGGAAGA
37932 TATGAGTGTGGAGTCGTTGCTCATCTGAAAATGGGGACAAGGANNNAGATCGGAA
37933 TATGAGTGTGGAGTCGTTGCCCTGGGCAACAAAAGTGAAACTCNNNAGATCGGAAG
37935 TATGAGTGTGGAGTCGTTGCTGGTCAATATGGTGAAACCCTGTNNNAGATCGGAAG
37936 TATGAGTGTGGAGTCGTTGCGTGGATCACAAGGTCAGGAGATTNNNAGATCGGAA
37937 TATGAGTGTGGAGTCGTTGCCCCTGATGTGGAAGATGGTACAANNNAGATCGGAAG

37938 TATGAGTGTGGAGTCGTTGCGCTGGGAGAACTCTGAATAGGAANNNAGATCGGAA
37939 TATGAGTGTGGAGTCGTTGCAAAGAAAAAAGAAGTCCGGGTGCNNNAGATCGGA
37940 TATGAGTGTGGAGTCGTTGCAAGTGTGCAACAATCGCTATAGCNNNAGATCGGAA
37941 TATGAGTGTGGAGTCGTTGCTGGCCAATATGGTGAAACCTCATNNNAGATCGGAAG
37942 TATGAGTGTGGAGTCGTTGCTTCTTGTAGAGACAGGGACTTGCNNNAGATCGGAA
37946 TATGAGTGTGGAGTCGTTGCTAAAGCACATACAGTAGGCCCTGNNNAGATCGGAA
37947 TATGAGTGTGGAGTCGTTGCCTGAGGCAGGATAATTGCTTGAGNNNAGATCGGAA
37949 TATGAGTGTGGAGTCGTTGCAATTAGCTTAGCATGGTTGTGGGNNNAGATCGGAA
37952 TATGAGTGTGGAGTCGTTGCGAAAAAAATTAGCCCAGGCATGGNNNAGATCGGAA
37954 TATGAGTGTGGAGTCGTTGCAGATTCTGGAAAGGAGGAACACCNNNAGATCGGAA
37955 TATGAGTGTGGAGTCGTTGCCCTGTAATCTCAGCTACTCAGGGNNNAGATCGGAAG
37956 TATGAGTGTGGAGTCGTTGCATCCTAACCTCATCTCCCAGTCTNNNAGATCGGAAGA
37958 TATGAGTGTGGAGTCGTTGCAAAAGAAAAGAAAAGGCTGGGCANNNAGATCGGA
37959 TATGAGTGTGGAGTCGTTGCCCAAACACTGTGATCCTGAGTTGNNNAGATCGGAAG
37960 TATGAGTGTGGAGTCGTTGCCTGAATCATGCCACGCTTTCTAGNNNAGATCGGAAG
37961 TATGAGTGTGGAGTCGTTGCCTGAGGTCAGGAATTCAAGACCANNNAGATCGGAA
37963 TATGAGTGTGGAGTCGTTGCAATGCTTGAGGAGATGCATACCTNNNAGATCGGAA
37964 TATGAGTGTGGAGTCGTTGCTGAGGAGAGGGAGTGACTCTAAGNNNAGATCGGA
37965 TATGAGTGTGGAGTCGTTGCCAGTGAGCTCTGATAACACCACTNNNAGATCGGAAG
37966 TATGAGTGTGGAGTCGTTGCCTGGAGAACTGGCGAAATTTGAGNNNAGATCGGAA
37967 TATGAGTGTGGAGTCGTTGCCTAGGATTACAGGTGTGAGCCTCNNNAGATCGGAA
37968 TATGAGTGTGGAGTCGTTGCATTCCCTGGCAAAGAAAATTCCCNNNAGATCGGAAG
37969 TATGAGTGTGGAGTCGTTGCGGGGAGACTCTAAATGACCACTTNNNAGATCGGAA
37970 TATGAGTGTGGAGTCGTTGCTGTCTGGGGAAACTGATTCAAGTNNNAGATCGGAA
37971 TATGAGTGTGGAGTCGTTGCAGACTGAGACTCTTGTTGAAGCCNNNAGATCGGAA
37972 TATGAGTGTGGAGTCGTTGCCACTTGAAAATTTGGCACTTGGCNNNAGATCGGAAG
37974 TATGAGTGTGGAGTCGTTGCCACCAATAGTCACAGCACTTTGGNNNAGATCGGAAG
37975 TATGAGTGTGGAGTCGTTGCAGGAGCTAGGAATGGGAGAAAAANNNAGATCGGA
37976 TATGAGTGTGGAGTCGTTGCCGCTAGTCTTGAACTCCTGAACTNNNAGATCGGAAG
37977 TATGAGTGTGGAGTCGTTGCGGGACCGTTTTGCAGATTCTATGNNNAGATCGGAA
37978 TATGAGTGTGGAGTCGTTGCGCATGAAAGACAGAAGGCACAAANNNAGATCGGAA
37979 TATGAGTGTGGAGTCGTTGCAAGAAGACAAGCCACAGACTAGGNNNAGATCGGAA
37980 TATGAGTGTGGAGTCGTTGCCAGTTTGGGGTAAAACCACCTTCNNNAGATCGGAAG
37981 TATGAGTGTGGAGTCGTTGCGTAATCCCAATCCCAGCACTTTGNNNAGATCGGAAG
37982 TATGAGTGTGGAGTCGTTGCACAAGAATCCCAGCTAGCTTGTTNNNAGATCGGAAG
37983 TATGAGTGTGGAGTCGTTGCAGGCATGAGAATCACTTGAACCTNNNAGATCGGAA
37984 TATGAGTGTGGAGTCGTTGCAAAACGCACACAAAGTATCCCAGNNNAGATCGGAAG
37985 TATGAGTGTGGAGTCGTTGCCTCTCAGTCTCCAATTCCTTCCCNNNAGATCGGAAGA
37986 TATGAGTGTGGAGTCGTTGCTGAGGAAAACAGACTAGACAGGCNNNAGATCGGAA
37987 TATGAGTGTGGAGTCGTTGCCTAGTTGAAGGGACGAATTTGCCNNNAGATCGGAA
37989 TATGAGTGTGGAGTCGTTGCGTGGCAAGGTAGTGGTACAGTAGNNNAGATCGGA
37991 TATGAGTGTGGAGTCGTTGCGGGCCAGCTTTCTAGTTCATAGANNNAGATCGGAAG
37992 TATGAGTGTGGAGTCGTTGCACAGAGCCAGTAATAACCCCTTGNNNAGATCGGAAG

37994 TATGAGTGTGGAGTCGTTGCCCCAGTGTCCATCAGCATATGAANNNAGATCGGAAG
37995 TATGAGTGTGGAGTCGTTGCGATGTACCCACACACAATGCATCNNNAGATCGGAAG
37996 TATGAGTGTGGAGTCGTTGCACCGGACTGAAGAAAAGCAGATANNNAGATCGGAA
37997 TATGAGTGTGGAGTCGTTGCATCACTTCTTGCTCCAACACAGANNNAGATCGGAAGA
38001 TATGAGTGTGGAGTCGTTGCATGGGCAAAACAGGGTTATACCTNNNAGATCGGAA
38002 TATGAGTGTGGAGTCGTTGCGGGACCAGCTGAAATTATTGACCNNNAGATCGGAAG
38004 TATGAGTGTGGAGTCGTTGCAACAGAACAAAAAAGGAGGTCAGNNNAGATCGGAA
38006 TATGAGTGTGGAGTCGTTGCTGCTCTTGCCACAAAAACAGAAANNNAGATCGGAAG
38007 TATGAGTGTGGAGTCGTTGCGGGGTTTGAAGTCAGACAGATCTNNNAGATCGGAA
38008 TATGAGTGTGGAGTCGTTGCGAAGAAGCAGCCAGTCTGAAAAGNNNAGATCGGAA
38009 TATGAGTGTGGAGTCGTTGCACCTCACTAGTGACCCTGAAAACNNNAGATCGGAAG
38010 TATGAGTGTGGAGTCGTTGCTCAGAGGACAGTGAGGATCTCTTNNNAGATCGGAA
38013 TATGAGTGTGGAGTCGTTGCCACAGCAGCAGACTATCCACATANNNAGATCGGAAG
38014 TATGAGTGTGGAGTCGTTGCCTCGTACAATCACCTTGGAAAGCNNNAGATCGGAAG
38015 TATGAGTGTGGAGTCGTTGCTACACTTGTACTGGGCACTTACCNNNAGATCGGAAG
38019 TATGAGTGTGGAGTCGTTGCAAACGATGGGAATTCGAAAAGCCNNNAGATCGGAA
38020 TATGAGTGTGGAGTCGTTGCCTGCAAGAAACACACTTCACCTANNNAGATCGGAAG
38021 TATGAGTGTGGAGTCGTTGCTGCAAGAATATCAGGTTGCACACNNNAGATCGGAAG
38022 TATGAGTGTGGAGTCGTTGCGAGGCCAGCATCATTTTGATACCNNNAGATCGGAAG
38023 TATGAGTGTGGAGTCGTTGCACCAGATACACAAGGAAGAGCTGNNNAGATCGGAA
38024 TATGAGTGTGGAGTCGTTGCGTTTTTGTGCCAGTACCAGTACCNNNAGATCGGAAG
38025 TATGAGTGTGGAGTCGTTGCCGTATTGTCTTCCACGAAACTGGNNNAGATCGGAAG
38027 TATGAGTGTGGAGTCGTTGCAAGAGGAGAACTGGGAAAACCTCNNNAGATCGGAA
38029 TATGAGTGTGGAGTCGTTGCACTGAATTCCGAAGGAGGAAGACNNNAGATCGGAA
38030 TATGAGTGTGGAGTCGTTGCACTGCCTGAAAGATCCATCAACTNNNAGATCGGAAG
38031 TATGAGTGTGGAGTCGTTGCCAAATACACACGGGAACCCAATTNNNAGATCGGAAG
38032 TATGAGTGTGGAGTCGTTGCACCCAAAACAACAAGAAGGAAAANNNAGATCGGAA
38033 TATGAGTGTGGAGTCGTTGCACAAAACAAAAAACAGGTTGCGGNNNAGATCGGAA
38034 TATGAGTGTGGAGTCGTTGCTGGCAGTATCGCTGAGTTTTAGTNNNAGATCGGAAG
38035 TATGAGTGTGGAGTCGTTGCACCCAAAACAACAAGAAGGAAAANNNAGATCGGAA
38036 TATGAGTGTGGAGTCGTTGCACAAAACAAAAAACAGGTTGCGGNNNAGATCGGAA
38037 TATGAGTGTGGAGTCGTTGCTGGCAGTATCGCTGAGTTTTAGTNNNAGATCGGAAG
38038 TATGAGTGTGGAGTCGTTGCTTCTACTACAAACACCACCCTGGNNNAGATCGGAAGA
38039 TATGAGTGTGGAGTCGTTGCGGCTCATCACAACAGACTCCTAANNNAGATCGGAAG
38040 TATGAGTGTGGAGTCGTTGCTTGTTATGGGAACCACCTCACTTNNNAGATCGGAAG
38041 TATGAGTGTGGAGTCGTTGCTAGCATCTCAGCACTTTGGTCAANNNAGATCGGAAG
38042 TATGAGTGTGGAGTCGTTGCGGATGATAATCCGTCCTGGTTGANNNAGATCGGAAG
38043 TATGAGTGTGGAGTCGTTGCATCAGAACTTGACGGAACTTCAGNNNAGATCGGAAG
38044 TATGAGTGTGGAGTCGTTGCTCACTCAAATCTTTCCTCCTGGGNNNAGATCGGAAGA
38045 TATGAGTGTGGAGTCGTTGCCTGTAGAGAAGGCGGTACTTGAANNNAGATCGGAA
38046 TATGAGTGTGGAGTCGTTGCCAGCCAAGGGTGAACACTAAATGNNNAGATCGGAA
38047 TATGAGTGTGGAGTCGTTGCGGATCCCTTGAGTTCAGGAGTTCNNNAGATCGGAAG
38048 TATGAGTGTGGAGTCGTTGCCCTGAAAAGAGCCACCTAGTTCTNNNAGATCGGAAG

38049 TATGAGTGTGGAGTCGTTGCTTTTGGGGAAAATGAAGCATCCCNNNAGATCGGAA
38050 TATGAGTGTGGAGTCGTTGCTAGATCTGGGAACTGGGGATGATNNNAGATCGGAA
38051 TATGAGTGTGGAGTCGTTGCCAGGATCCAGGTATAAGCTCCACNNNAGATCGGAAG
38052 TATGAGTGTGGAGTCGTTGCAAACAAAACCTGATAGCCTGGGTNNNAGATCGGAA
38053 TATGAGTGTGGAGTCGTTGCCAACAACAACAACACACACACACNNNAGATCGGAAG
38054 TATGAGTGTGGAGTCGTTGCAAGGCCTTTGACAAAATTCAGCANNNAGATCGGAAG
38056 TATGAGTGTGGAGTCGTTGCCCTGGCTCTGGACACATTTTTTTNNNAGATCGGAAG
38058 TATGAGTGTGGAGTCGTTGCGCCCCCCGTCTCTACTAAAATACNNNAGATCGGAAG
38059 TATGAGTGTGGAGTCGTTGCGCCAGGTATGATTCCAGCTACTTNNNAGATCGGAAG
38060 TATGAGTGTGGAGTCGTTGCAGTGCACAATTACACCTGACAGANNNAGATCGGAAG
38061 TATGAGTGTGGAGTCGTTGCTGGGGGATTCTGTATACAGCATGNNNAGATCGGAA
38064 TATGAGTGTGGAGTCGTTGCGCTTAGGTGATCACTTGAGTCCANNNAGATCGGAAG
38065 TATGAGTGTGGAGTCGTTGCTTTGTCTTCCCAGCAGATCTACCNNNAGATCGGAAG
38066 TATGAGTGTGGAGTCGTTGCCAAACTCCTGAGCTCGAACAATCNNNAGATCGGAAG
38068 TATGAGTGTGGAGTCGTTGCTGCAGTGAAGTATGATCACACCANNNAGATCGGAA
38070 TATGAGTGTGGAGTCGTTGCCAATCAGGGAAAGGGAAACACACNNNAGATCGGAA
38072 TATGAGTGTGGAGTCGTTGCAAAATACAAAATCAGCCAGGCGTNNNAGATCGGAA
38073 TATGAGTGTGGAGTCGTTGCTCAGCACTGGATTGTAGACTTGTNNNAGATCGGAAG
38074 TATGAGTGTGGAGTCGTTGCGCCGACTGAGAAGAACCTTTTTCNNNAGATCGGAAG
38076 TATGAGTGTGGAGTCGTTGCAAAAAATTAGCCAGGTGTGGTGGNNNAGATCGGAA
38077 TATGAGTGTGGAGTCGTTGCCTCAATGGGAGGGGATTACACAANNNAGATCGGAA
38078 TATGAGTGTGGAGTCGTTGCAAGGATTTGGGTAGCTAAGAGGCNNNAGATCGGAA
38079 TATGAGTGTGGAGTCGTTGCAAAATGGTGGCAGAAGAACACAGNNNAGATCGGAA
38081 TATGAGTGTGGAGTCGTTGCCCTTGAATGCCAGATGAAAGGTGNNNAGATCGGAA
38082 TATGAGTGTGGAGTCGTTGCAATGAGGGCAACTTGTTGTTCTGNNNAGATCGGAA
38083 TATGAGTGTGGAGTCGTTGCGTGAGCCAAGATCTCAGTCTCAANNNAGATCGGAAG
38085 TATGAGTGTGGAGTCGTTGCTGGCAGGGAGTTGGTTGTTATAANNNAGATCGGAA
38086 TATGAGTGTGGAGTCGTTGCCTTGTACCCTGTGACCTTGCTATNNNAGATCGGAAG
38087 TATGAGTGTGGAGTCGTTGCTTGGGGTTTTTGTGTTTTGTGGGNNNAGATCGGAA
38088 TATGAGTGTGGAGTCGTTGCGGGATTTTGTTTGGGATTGTGCTNNNAGATCGGAA
38089 TATGAGTGTGGAGTCGTTGCCGCTCTACCTATTGACCTTCCTCNNNAGATCGGAAGA
38090 TATGAGTGTGGAGTCGTTGCGAAAAAAGTACATTTGCCAGGCGNNNAGATCGGAA
38091 TATGAGTGTGGAGTCGTTGCTGGAAAGAGAAGTTCAAGTGGCTNNNAGATCGGAA
38092 TATGAGTGTGGAGTCGTTGCCAACTCGAGAACATGGTCAGAGANNNAGATCGGAA
38094 TATGAGTGTGGAGTCGTTGCAGGAGGATCACTTGAACCTAGGANNNAGATCGGAA
38095 TATGAGTGTGGAGTCGTTGCAAGGGGGAAATGTGTATGGGTACNNNAGATCGGA
38097 TATGAGTGTGGAGTCGTTGCAACACACATGAATGATTGCCAGGNNNAGATCGGAA
38099 TATGAGTGTGGAGTCGTTGCGAGAAAATGATGGCCAGACACAGNNNAGATCGGAA
38100 TATGAGTGTGGAGTCGTTGCCAGGTCAGGAGATTGAGACCATCNNNAGATCGGAA
38101 TATGAGTGTGGAGTCGTTGCATCACTGGTAGCAGCTTTCAGAANNNAGATCGGAAG
38102 TATGAGTGTGGAGTCGTTGCGGTGGATTGCTAGAGCTTAGGAGNNNAGATCGGAA
38104 TATGAGTGTGGAGTCGTTGCAAAGAATTGGAAAGCTTGTGGGGNNNAGATCGGA
38105 TATGAGTGTGGAGTCGTTGCTACTGGTCTAGTCTGCTTTGCTGNNNAGATCGGAAG

38106 TATGAGTGTGGAGTCGTTGCACCACAATACCATTATCACGCCTNNNAGATCGGAAGA
38107 TATGAGTGTGGAGTCGTTGCGAGGTTGTAGTGAGCCGAGATTGNNNAGATCGGAA
38108 TATGAGTGTGGAGTCGTTGCACTACAGAAGAGCGTGCAAGTAANNNAGATCGGAA
38109 TATGAGTGTGGAGTCGTTGCCAATTGAGTGATGGGCATGATGANNNAGATCGGAA
38110 TATGAGTGTGGAGTCGTTGCTAACAAGCAAACCTCAACCCCAANNNAGATCGGAAG
38112 TATGAGTGTGGAGTCGTTGCTGCTGAATGAAAAAAGCCAGACANNNAGATCGGAA
38114 TATGAGTGTGGAGTCGTTGCAACAGGAGAGTTAGCCTATCCCTNNNAGATCGGAAG
38116 TATGAGTGTGGAGTCGTTGCCACAGACTGGGAGAAAACATTGTNNNAGATCGGAA
38118 TATGAGTGTGGAGTCGTTGCAAAGAAAAGAAAACTGCAGGCCANNNAGATCGGAA
38119 TATGAGTGTGGAGTCGTTGCAGGAGACGTTACAACTGATACCANNNAGATCGGAA
38121 TATGAGTGTGGAGTCGTTGCCTGAGGTAGGAGGATTGCTTGAGNNNAGATCGGAA
38122 TATGAGTGTGGAGTCGTTGCTAGAAGTAGGGTGACTGGGGATTNNNAGATCGGA
38124 TATGAGTGTGGAGTCGTTGCTGAGGCAGGAGAATCTCTTGAACNNNAGATCGGAA
38126 TATGAGTGTGGAGTCGTTGCACAAAAATTAGCTGGGCTTGGTGNNNAGATCGGAA
38127 TATGAGTGTGGAGTCGTTGCAGGGATGGATTATTCATGCCTCCNNNAGATCGGAAG
38128 TATGAGTGTGGAGTCGTTGCCAGGCATTGAACACACCAGTTAGNNNAGATCGGAA
38131 TATGAGTGTGGAGTCGTTGCTATTTTCTCCCAGTCTGTGGCTTNNNAGATCGGAAGA
38132 TATGAGTGTGGAGTCGTTGCACATGGTGAAACCCCGTATCTACNNNAGATCGGAAG
38133 TATGAGTGTGGAGTCGTTGCTGAGATAGGCATTGAGCTTGAGCNNNAGATCGGAA
38134 TATGAGTGTGGAGTCGTTGCTCGCAATCTTTCATGTTGTGACCNNNAGATCGGAAG
38138 TATGAGTGTGGAGTCGTTGCATAAAAAAATAGGCCGGGCACAGNNNAGATCGGAA
38139 TATGAGTGTGGAGTCGTTGCAGCAGCTTTGGAACTGGGTAATANNNAGATCGGAA
38140 TATGAGTGTGGAGTCGTTGCGAGGTGTTTTGGACCTATTACGTNNNAGATCGGAA
38143 TATGAGTGTGGAGTCGTTGCAAAAAGAAAAACAGGCCAGGTGCNNNAGATCGGAA
38144 TATGAGTGTGGAGTCGTTGCAGGGCCCTGCTTTCTCAAATTTANNNAGATCGGAAG
38145 TATGAGTGTGGAGTCGTTGCTTTTGCCACACTTTCTGGGTTTTNNNAGATCGGAAGA
38146 TATGAGTGTGGAGTCGTTGCTATCCAGTGGCTCTTCATGTCACNNNAGATCGGAAG
38147 TATGAGTGTGGAGTCGTTGCGTGTGCCTGTAACCTTAGCTACTNNNAGATCGGAAG
38148 TATGAGTGTGGAGTCGTTGCGCTCAGACAGGAGAATCACTTGANNNAGATCGGAA
38149 TATGAGTGTGGAGTCGTTGCAATACAAAATTAGCTGGGCGTGGNNNAGATCGGAA
38150 TATGAGTGTGGAGTCGTTGCCTGAGGTGGGAAGATCACTTGAGNNNAGATCGGAA
38151 TATGAGTGTGGAGTCGTTGCTGTCTTGGGTATTTGGGAGACTGNNNAGATCGGAA
38152 TATGAGTGTGGAGTCGTTGCGAATAAATAAATGGGTGGCCGGGNNNAGATCGGA
38153 TATGAGTGTGGAGTCGTTGCCTGTGCTTCACCCTATCTAGCAANNNAGATCGGAAG
38154 TATGAGTGTGGAGTCGTTGCGGATCACGAAGTCAGGAGATTGANNNAGATCGGAA
38155 TATGAGTGTGGAGTCGTTGCATAATCCCAGCTACTCAGGAGGCNNNAGATCGGAAG
38157 TATGAGTGTGGAGTCGTTGCAAATTGAAGGCAAGAAGGCACTCNNNAGATCGGAA
38158 TATGAGTGTGGAGTCGTTGCACCGAGTGCAAGAAGTTTTACACNNNAGATCGGAAG
38159 TATGAGTGTGGAGTCGTTGCCATGCCTGTAGTCCCAGCTATTTNNNAGATCGGAAG
38161 TATGAGTGTGGAGTCGTTGCAAGAAAAAGAAAGCTCAGGCCAGNNNAGATCGGAA
38163 TATGAGTGTGGAGTCGTTGCATAAGACCACACATGATCTGGCCNNNAGATCGGAAG
38164 TATGAGTGTGGAGTCGTTGCGGATGACAGAGTGAGACTCCATTNNNAGATCGGAA
38165 TATGAGTGTGGAGTCGTTGCGGATCACTTGAGGTCAGGAGTTTNNNAGATCGGAA

38166 TATGAGTGTGGAGTCGTTGCGTGTAAAGGAGGTGAGAGAAGGGNNNAGATCGGA
38167 TATGAGTGTGGAGTCGTTGCTTTCAGATTAGGACCAAGCCTGGNNNAGATCGGAAG
38168 TATGAGTGTGGAGTCGTTGCGCAATAGCATAGACAAAGGCCTGNNNAGATCGGAAG
38170 TATGAGTGTGGAGTCGTTGCGTGGGGTCCTAAAAGCCATAAGANNNAGATCGGAAG
38171 TATGAGTGTGGAGTCGTTGCAAGTGCTCAATGGATGTAAGCTTNNNAGATCGGAAG
38172 TATGAGTGTGGAGTCGTTGCTCTGGATTTCTGGATTCCCTTGANNNAGATCGGAAGA
38173 TATGAGTGTGGAGTCGTTGCATGAGAGAGAGCAGTTTCAAGCANNNAGATCGGAA
38174 TATGAGTGTGGAGTCGTTGCGACATGGACCAAGATGAGGATCANNNAGATCGGAA
38175 TATGAGTGTGGAGTCGTTGCGACGAAGCTGGGCAAAATAATGANNNAGATCGGAA
38177 TATGAGTGTGGAGTCGTTGCTCCTAAAGTGATGGGATTGTGGGNNNAGATCGGAA
38178 TATGAGTGTGGAGTCGTTGCATGTTTGTTGTCCCAGCTACTCANNNAGATCGGAAGA
38179 TATGAGTGTGGAGTCGTTGCGGCTCAAACCTGTAATCCCAGTANNNAGATCGGAAG
38181 TATGAGTGTGGAGTCGTTGCCTCACTCTTGTAATCCCAGCACTNNNAGATCGGAAGA
38182 TATGAGTGTGGAGTCGTTGCGATGTGAACACCTTGCCAAAGAANNNAGATCGGAAG
38183 TATGAGTGTGGAGTCGTTGCACAAAGTGGGGTTTATCACAGGANNNAGATCGGAAG
38184 TATGAGTGTGGAGTCGTTGCAGCCTCTAGAAGCTGAAAAAGGCNNNAGATCGGAAG
38186 TATGAGTGTGGAGTCGTTGCACACGGTAAAACCCCATCTCTACNNNAGATCGGAAGA
38187 TATGAGTGTGGAGTCGTTGCACAAAGCTGACTGAGAATTTCCANNNAGATCGGAAG
38188 TATGAGTGTGGAGTCGTTGCTGATACTAGCTGTGGGTCTGTCANNNAGATCGGAAG
38189 TATGAGTGTGGAGTCGTTGCGGGTTTGAGGTTACAGTGAGCTANNNAGATCGGAA
38190 TATGAGTGTGGAGTCGTTGCACTAGATGAAAAGCAGAGGTGGANNNAGATCGGAA
38196 TATGAGTGTGGAGTCGTTGCCTTACACCTGTAATCCCAGCACTNNNAGATCGGAAGA
38197 TATGAGTGTGGAGTCGTTGCGCCGCTTACCTACAACTATCTGANNNAGATCGGAAGA
38198 TATGAGTGTGGAGTCGTTGCTAGCCAGGATATACAGGGATCCCNNNAGATCGGAAG
38199 TATGAGTGTGGAGTCGTTGCAGCTCAACTCAGGATCAGAAAGGNNNAGATCGGAAG
38204 TATGAGTGTGGAGTCGTTGCAGCCGCAGAATTCAGAATTTGACNNNAGATCGGAAG
38205 TATGAGTGTGGAGTCGTTGCCAGTAAACAGGATGGAGGACCAANNNAGATCGGAA
38206 TATGAGTGTGGAGTCGTTGCCAAGATTGTGTCATTGCACTCCANNNAGATCGGAAGA
38208 TATGAGTGTGGAGTCGTTGCGAGGCAGAAGAATCACTTGAACCNNNAGATCGGAAG
38209 TATGAGTGTGGAGTCGTTGCAAAACACAGAGACTGGCACATTGNNNAGATCGGAAG
38211 TATGAGTGTGGAGTCGTTGCCATCCTTGTCTTGTGCTGGTTTTNNNAGATCGGAAGA
38212 TATGAGTGTGGAGTCGTTGCGGGTTTGGAAATAAGAGTCTGCGNNNAGATCGGAA
38213 TATGAGTGTGGAGTCGTTGCTCTTTTGTAGGAAGACTGAGCGCNNNAGATCGGAAG
38214 TATGAGTGTGGAGTCGTTGCTATTCCCAGTGCCCAGAAGAATCNNNAGATCGGAAG
38215 TATGAGTGTGGAGTCGTTGCTAGACTGGGTGACAGAGAGAGACNNNAGATCGGAA
38216 TATGAGTGTGGAGTCGTTGCGGGCAACTTGCTTGATGAAAGTANNNAGATCGGAAG
38217 TATGAGTGTGGAGTCGTTGCGAGGTCAGGAGATTGAGACCATCNNNAGATCGGAA
38218 TATGAGTGTGGAGTCGTTGCCTCTCAATTGGATTGGGATGTGCNNNAGATCGGAAG
38221 TATGAGTGTGGAGTCGTTGCACCTCCCAAGATTGACCAGAAAGNNNAGATCGGAAG
38222 TATGAGTGTGGAGTCGTTGCTAGACGTGAAAGCCATACACACANNNAGATCGGAAG
38223 TATGAGTGTGGAGTCGTTGCGACCAGACAGATTCACAGCTGAANNNAGATCGGAAG
38225 TATGAGTGTGGAGTCGTTGCTGCTAAGCGAAATAAGCCAGATGNNNAGATCGGAAG
38226 TATGAGTGTGGAGTCGTTGCACTTCACACTCACCAACTTAGGTNNNAGATCGGAAGA

38227 TATGAGTGTGGAGTCGTTGCCACACGCAGCTTTGTCAAATACTNNNAGATCGGAAG
38228 TATGAGTGTGGAGTCGTTGCACCAAACATGATCTCACACTGTCNNNAGATCGGAAG
38229 TATGAGTGTGGAGTCGTTGCACCACAAATACCGAGAATAGCCTNNNAGATCGGAAG
38230 TATGAGTGTGGAGTCGTTGCACCACTGCAAATTCTGTTTGACCNNNAGATCGGAAG
38233 TATGAGTGTGGAGTCGTTGCAGAAAACCTCAGGCCAATATCCTNNNAGATCGGAAG
38234 TATGAGTGTGGAGTCGTTGCTCTATGAGGCCAGCATCATTCTGNNNAGATCGGAAG
38235 TATGAGTGTGGAGTCGTTGCCTGACTGATGCATGTGTGTTGTTNNNAGATCGGAAG
38236 TATGAGTGTGGAGTCGTTGCTGTGAGTTTGGGATGAGGTGTAGNNNAGATCGGAA
38237 TATGAGTGTGGAGTCGTTGCCAACTGCCCTTTCTTACATGCAGNNNAGATCGGAAG
38238 TATGAGTGTGGAGTCGTTGCTGTGGTTCATAACTTGGGTCACTNNNAGATCGGAAG
38239 TATGAGTGTGGAGTCGTTGCATAAAGGACTCATTGCCTCAGCTNNNAGATCGGAAG
38240 TATGAGTGTGGAGTCGTTGCAAGGGATCAGTGCAGCAATAAGANNNAGATCGGAA
38241 TATGAGTGTGGAGTCGTTGCCTTCTCCCTCCAAAGGATCACAANNNAGATCGGAAG
38242 TATGAGTGTGGAGTCGTTGCAACTCTAAGCAACATCACGTCCTNNNAGATCGGAAG
38244 TATGAGTGTGGAGTCGTTGCAAGGACACACTTAGGGAAATGCANNNAGATCGGAA
38245 TATGAGTGTGGAGTCGTTGCAGTCATGCTCATCGTTGGTAAGTNNNAGATCGGAAG
38247 TATGAGTGTGGAGTCGTTGCTGGAGTTTCATGTTCAGCCAAACNNNAGATCGGAAG
38248 TATGAGTGTGGAGTCGTTGCTCACTGTCCATCAGAGAAATGCANNNAGATCGGAAG
38249 TATGAGTGTGGAGTCGTTGCCCTCTCTGCACACAACCTAGAAANNNAGATCGGAAG
38251 TATGAGTGTGGAGTCGTTGCCACTGGCAACCATGACAAAGAAANNNAGATCGGAA
38252 TATGAGTGTGGAGTCGTTGCGAAAAGTGGTGCAAGAACTCCAGNNNAGATCGGAA
38253 TATGAGTGTGGAGTCGTTGCAGGCTAGGAAGAAACTGCATCAANNNAGATCGGAA
38254 TATGAGTGTGGAGTCGTTGCATGCTAAAGGGGGTTTTTCAAGCNNNAGATCGGAA
38255 TATGAGTGTGGAGTCGTTGCTTACGGCAGCAGTTTCTCAAAAGNNNAGATCGGAAG
38256 TATGAGTGTGGAGTCGTTGCTGATCTATCGGTTGCAAAGACCANNNAGATCGGAAG
38257 TATGAGTGTGGAGTCGTTGCAGGTCCCTTTTATGTTCCCTTGTNNNAGATCGGAAG
38259 TATGAGTGTGGAGTCGTTGCATTCTGTTTTCCACCCTAGGCATNNNAGATCGGAAG
38260 TATGAGTGTGGAGTCGTTGCCCTGGAGTCAGCAAAGTGGATATNNNAGATCGGAA
38261 TATGAGTGTGGAGTCGTTGCATAAATCCGCACATCACGAGGTANNNAGATCGGAAG
38262 TATGAGTGTGGAGTCGTTGCAGGTGTTTCTGTCCTTAGAGTGCNNNAGATCGGAAG
38263 TATGAGTGTGGAGTCGTTGCTCCTTTTTGACAATAGGCCTCCANNNAGATCGGAAG
38264 TATGAGTGTGGAGTCGTTGCGAAACTGCTTTGTGATGTGTGGANNNAGATCGGAA
38267 TATGAGTGTGGAGTCGTTGCAAGGCCAAGGACAAAGAGAGAATNNNAGATCGGA
38268 TATGAGTGTGGAGTCGTTGCTCTCATTGCTCTTGGAACTTTCCNNNAGATCGGAAGA
38269 TATGAGTGTGGAGTCGTTGCGTGTAGACACCAGAGGGAATCTCNNNAGATCGGAA
38270 TATGAGTGTGGAGTCGTTGCAATGTATGCTCTGCAGTTGTTGGNNNAGATCGGAAG
38274 TATGAGTGTGGAGTCGTTGCACAAAAATTAGCCTGGTGTGGTGNNNAGATCGGAA
38275 TATGAGTGTGGAGTCGTTGCCACTTGTGATTGGGTTTCCTGTCNNNAGATCGGAAG
38276 TATGAGTGTGGAGTCGTTGCTCAGCATGGAAATGTTTCAACCCNNNAGATCGGAAG
38278 TATGAGTGTGGAGTCGTTGCATATTGCTCAGCTTTCTGTCACCNNNAGATCGGAAGA
38279 TATGAGTGTGGAGTCGTTGCTACCTGGAAGTCTCATTGCTGACNNNAGATCGGAAG
38280 TATGAGTGTGGAGTCGTTGCATTTGGATATGGCTCCTTTGGGANNNAGATCGGAAG
38281 TATGAGTGTGGAGTCGTTGCTCTCAGAATGCACAAAGCCAATGNNNAGATCGGAAG

38283 TATGAGTGTGGAGTCGTTGCGAGCAATAACCATCACTGCAGTCNNNAGATCGGAAG
38284 TATGAGTGTGGAGTCGTTGCGCCAAACCCACGATGATCTAGATNNNAGATCGGAAG
38285 TATGAGTGTGGAGTCGTTGCTCCTACAGCTATGTTGGTGTCTGNNNAGATCGGAAG
38286 TATGAGTGTGGAGTCGTTGCAGGCAGAGAATCACTTGAACCTTNNNAGATCGGAAG
38287 TATGAGTGTGGAGTCGTTGCAGAAGAAGAAGAAGAGGAGGAGANNNAGATCGGA
38288 TATGAGTGTGGAGTCGTTGCACTCTGCAGAAGCTCTTTAGTTTNNNAGATCGGAAGA
38291 TATGAGTGTGGAGTCGTTGCTTGCATCCATCAATCACTTTGGGNNNAGATCGGAAGA
38292 TATGAGTGTGGAGTCGTTGCGGTTGCCATCCTAGTCTCTGATTNNNAGATCGGAAGA
38294 TATGAGTGTGGAGTCGTTGCTTTCCTACTCCCCCCACTAAGTANNNAGATCGGAAGA
38295 TATGAGTGTGGAGTCGTTGCAAAGTAGGCGAAGGATATGCACANNNAGATCGGAA
38296 TATGAGTGTGGAGTCGTTGCGTGGTCTTAGCTACTTGAGAGGGNNNAGATCGGAA
38297 TATGAGTGTGGAGTCGTTGCGATGACAGCTGCTGTTTGCTAAANNNAGATCGGAAG
38299 TATGAGTGTGGAGTCGTTGCCTATCTTGTAGGAAGCCAGGGTCNNNAGATCGGAAG
38300 TATGAGTGTGGAGTCGTTGCAAGTGGGTCTCAGACTAGTCCTTNNNAGATCGGAAG
38301 TATGAGTGTGGAGTCGTTGCACCAATAAAAGAGCAGTAGCAGCNNNAGATCGGAA
38302 TATGAGTGTGGAGTCGTTGCTACCACACAGCTCCACATGAAATNNNAGATCGGAAGA
38303 TATGAGTGTGGAGTCGTTGCACAGAAGAAATGCTAAAGGGAGTNNNAGATCGGAA
38304 TATGAGTGTGGAGTCGTTGCACCCGAGTAGCAAACACAAAATGNNNAGATCGGAAG
38306 TATGAGTGTGGAGTCGTTGCACCAAATTGATAGACTGCTAGCTNNNAGATCGGAAG
38307 TATGAGTGTGGAGTCGTTGCGACTTGAAGCCAACAAAGGTCAANNNAGATCGGAA
38308 TATGAGTGTGGAGTCGTTGCAAGGAGAAGAAGGAGAAGGAGGANNNAGATCGGA
38309 TATGAGTGTGGAGTCGTTGCGTCACAGCTCAGTGTTTGCATTANNNAGATCGGAAG
38310 TATGAGTGTGGAGTCGTTGCACACAGCCTTGATTACTGTAGCTNNNAGATCGGAAGA
38311 TATGAGTGTGGAGTCGTTGCCTCCACAGGCAATTTCTTTCAGTNNNAGATCGGAAGA
38312 TATGAGTGTGGAGTCGTTGCGGAGACCCTGGTATTGTCACATANNNAGATCGGAAG
38314 TATGAGTGTGGAGTCGTTGCTCCTATGCTCATGGATGGGTAGANNNAGATCGGAAG
38315 TATGAGTGTGGAGTCGTTGCTCCGGAAGCTTTGGACTATGTTANNNAGATCGGAAG
38316 TATGAGTGTGGAGTCGTTGCAGAAGAAAAAGGCTGGGTCTGAANNNAGATCGGAA
38317 TATGAGTGTGGAGTCGTTGCAGAAGGGAACTTCCTTGAGCTAANNNAGATCGGAA
38318 TATGAGTGTGGAGTCGTTGCTGAAAAGATAGCTGCACTCCTGTNNNAGATCGGAAG
38319 TATGAGTGTGGAGTCGTTGCAGACACTGCATGTTCCCACTTATNNNAGATCGGAAGA
38320 TATGAGTGTGGAGTCGTTGCTGTGGAATCAAAAACAAACCCGANNNAGATCGGAAG
38321 TATGAGTGTGGAGTCGTTGCCTGTAGTCCCAGCTACTCAGGAGNNNAGATCGGAAG
38322 TATGAGTGTGGAGTCGTTGCGTGTGGAACAAGTTGGCTAAGACNNNAGATCGGAA
38323 TATGAGTGTGGAGTCGTTGCCAGTGAGCTGAGATTGTACCACTNNNAGATCGGAAG
38325 TATGAGTGTGGAGTCGTTGCCTGTAATCCCAGCTACTCAGGAGNNNAGATCGGAAG
38326 TATGAGTGTGGAGTCGTTGCAATTATCACAATCTCTCGGGCCANNNAGATCGGAAGA
38327 TATGAGTGTGGAGTCGTTGCTGATGTAGATCAGGTGGCGTATGNNNAGATCGGAA
38328 TATGAGTGTGGAGTCGTTGCCTACTCTGCAAATGTGATGGCTGNNNAGATCGGAAG
38329 TATGAGTGTGGAGTCGTTGCATGGTGGTCCTAGCTACTCAGAANNNAGATCGGAAG
38330 TATGAGTGTGGAGTCGTTGCGTGCATACTTGTAGACCCAGTCANNNAGATCGGAAG
38331 TATGAGTGTGGAGTCGTTGCTTTGACAATGGATCCTTGGAAAGNNNAGATCGGAAG
38332 TATGAGTGTGGAGTCGTTGCTGCCTAGCTTCATCACTGCTATTNNNAGATCGGAAGA

38333 TATGAGTGTGGAGTCGTTGCAGATGGGAGGTAGGTGTGACTATNNNAGATCGGA
38334 TATGAGTGTGGAGTCGTTGCAGAGCAAACTAACCACCTTTCCTNNNAGATCGGAAG
38335 TATGAGTGTGGAGTCGTTGCAAATGATTGCATAGAGCCAGGTGNNNAGATCGGAA
38336 TATGAGTGTGGAGTCGTTGCGAACTGGAACAAGACAAGGATGCNNNAGATCGGAA
38337 TATGAGTGTGGAGTCGTTGCCTCAGCTCACTGGTGGTAGATACNNNAGATCGGAAG
38338 TATGAGTGTGGAGTCGTTGCATACGGTCCACATCCACAAAAGANNNAGATCGGAAG
38339 TATGAGTGTGGAGTCGTTGCGTGCCAGCCTGACATTTAGAAAANNNAGATCGGAA
38340 TATGAGTGTGGAGTCGTTGCACTTCCTTAGGCTCTAGGGCTTANNNAGATCGGAAG
38341 TATGAGTGTGGAGTCGTTGCAGAGAGAGAAGGATCATGGTGACNNNAGATCGGA
38342 TATGAGTGTGGAGTCGTTGCCCCAACACACTTAGGGATTGTGANNNAGATCGGAAG
38343 TATGAGTGTGGAGTCGTTGCGCTTTTGGCAGTGACATACTTGTNNNAGATCGGAAG
38344 TATGAGTGTGGAGTCGTTGCTTGTCTGTCAACTCCTGCATCATNNNAGATCGGAAG
38345 TATGAGTGTGGAGTCGTTGCAAGAGATGTCAGTCTTCTGGAGCNNNAGATCGGAA
38347 TATGAGTGTGGAGTCGTTGCAAAGAATGCAGAAGCTGACAACCNNNAGATCGGAA
38348 TATGAGTGTGGAGTCGTTGCAAGGGGATTCCTAAGGTTGCATTNNNAGATCGGAA
38349 TATGAGTGTGGAGTCGTTGCTTCCTGACTCTGAGGGGAATTTGNNNAGATCGGAA
38351 TATGAGTGTGGAGTCGTTGCTGCATTCATATGTTCATCACAGCANNNAGATCGGAA
38352 TATGAGTGTGGAGTCGTTGCACCGAAGCTAAATCAACAGAGGTNNNAGATCGGAA
38355 TATGAGTGTGGAGTCGTTGCACAAACAACCCCATCAACAAGTGNNNAGATCGGAAG
38356 TATGAGTGTGGAGTCGTTGCAGAAAAAAATCAGGCAGATGGCACNNNAGATCGGAA
38357 TATGAGTGTGGAGTCGTTGCGAGGCTGAGACATGAGAATCACTNNNAGATCGGAA
38358 TATGAGTGTGGAGTCGTTGCCCAGAGGTACAAAGAGGAGTTGGNNNAGATCGGA
38359 TATGAGTGTGGAGTCGTTGCTGAGGTCAGGAGTTCAAGATCAGNNNAGATCGGAA
38361 TATGAGTGTGGAGTCGTTGCGAAGTCAAGAAAGTGGTCCTCCTNNNAGATCGGAA
38362 TATGAGTGTGGAGTCGTTGCGAGGTTAGGGCAAGAGGTGTTATNNNAGATCGGA
38363 TATGAGTGTGGAGTCGTTGCAATTTCAGGCCAACCAATATGCCNNNAGATCGGAAG
38365 TATGAGTGTGGAGTCGTTGCAGAGAGAGAGAAACCCATTGACANNNAGATCGGAA
38366 TATGAGTGTGGAGTCGTTGCACAGATCAACAAGAGCTGAAGGANNNAGATCGGAA
38367 TATGAGTGTGGAGTCGTTGCCGTGACAGAGTGAGACTTCATCTNNNAGATCGGAA
38368 TATGAGTGTGGAGTCGTTGCGTTGGAAGGGTTATGAGGGTTGANNNAGATCGGA
38369 TATGAGTGTGGAGTCGTTGCGCTTATCCACCACGATCAATTCGNNNAGATCGGAAG
38370 TATGAGTGTGGAGTCGTTGCGCTAACCATCCAAGTTCCTGGTANNNAGATCGGAAG
38371 TATGAGTGTGGAGTCGTTGCAGGGAATCTGAACAAGGTGAGACNNNAGATCGGA
38372 TATGAGTGTGGAGTCGTTGCGCAAAGGAGAAGGACTACTTTCCNNNAGATCGGAA
38376 TATGAGTGTGGAGTCGTTGCAAAAAAAAATTAGCCAGGCGTGGTNNNAGATCGGAA
38377 TATGAGTGTGGAGTCGTTGCATGGAACCACTGTTGCATATGTGNNNAGATCGGAA
38378 TATGAGTGTGGAGTCGTTGCTGGGTGATAGAGTGAGACTCCATNNNAGATCGGAA
38380 TATGAGTGTGGAGTCGTTGCTGTGCTTATAGTCCCAGCAACTTNNNAGATCGGAAG
38381 TATGAGTGTGGAGTCGTTGCCTTGAGCTCAGGAGTTGGAGATTNNNAGATCGGAA
38383 TATGAGTGTGGAGTCGTTGCACTGTTTTCCATAATGGCTGCACNNNAGATCGGAAG
38384 TATGAGTGTGGAGTCGTTGCAGTCGGAATTGTGACCAGAATCANNNAGATCGGAA
38385 TATGAGTGTGGAGTCGTTGCGAGAATCACTTGAACCCAGGAGANNNAGATCGGAA
38386 TATGAGTGTGGAGTCGTTGCGAGAGATCTCGAACTGTGTCAGTNNNAGATCGGAA

38387 TATGAGTGTGGAGTCGTTGCAGAGAACCCAACACAGTACCAAANNNAGATCGGAAG
38388 TATGAGTGTGGAGTCGTTGCCTCAAGTGATCCTCCTGCCTTAGNNNAGATCGGAAGA
38389 TATGAGTGTGGAGTCGTTGCTGACTTTTGCCGAAGTTCTTTCTNNNAGATCGGAAGA
38390 TATGAGTGTGGAGTCGTTGCGTGATAGAGCAAGACCCTGTCTCNNNAGATCGGAAG
38391 TATGAGTGTGGAGTCGTTGCCACGCCTATAATCCCACCACTTTNNNAGATCGGAAGA
38393 TATGAGTGTGGAGTCGTTGCCAGACTGCAGCCATTATTACAGCNNNAGATCGGAAG
38394 TATGAGTGTGGAGTCGTTGCGCAAACCAAAAGAAAATGGAGGGNNNAGATCGGAA
38395 TATGAGTGTGGAGTCGTTGCATACAAGCCAAGAGAACCCCAATNNNAGATCGGAAG
38396 TATGAGTGTGGAGTCGTTGCATGCACTGAAGGTAACCATAGCANNNAGATCGGAAG
38397 TATGAGTGTGGAGTCGTTGCCCACTTTTACATGTGCCACCTTTNNNAGATCGGAAGA
38398 TATGAGTGTGGAGTCGTTGCTTCTAGGATGGCTCAGTCACATGNNNAGATCGGAAG
38399 TATGAGTGTGGAGTCGTTGCAAAGGTGGAACAACTCAAAGCACNNNAGATCGGAAG
38403 TATGAGTGTGGAGTCGTTGCGGCTGAGGTAGAAGAATCGCTTANNNAGATCGGAA
38404 TATGAGTGTGGAGTCGTTGCGGAAGATGCCATGGATAGGAGAGNNNAGATCGGAA
38405 TATGAGTGTGGAGTCGTTGCCTGTAATTCCAGCTACTCGGGAGNNNAGATCGGAAG
38406 TATGAGTGTGGAGTCGTTGCTGGGAGAATCAACTAAACCCAGGNNNAGATCGGAAG
38407 TATGAGTGTGGAGTCGTTGCGAGCAACCTTGACCAAACCAAATNNNAGATCGGAAG
38408 TATGAGTGTGGAGTCGTTGCCACCTGTAATTCCAGCACTTTGGNNNAGATCGGAAGA
38409 TATGAGTGTGGAGTCGTTGCAAGCCAACAAATACAGTTCAGCANNNAGATCGGAAG
38410 TATGAGTGTGGAGTCGTTGCTGACCTTGTCTAAGTCATTGGCANNNAGATCGGAAG
38412 TATGAGTGTGGAGTCGTTGCAGCAAGGGCATCAAATAGGGAATNNNAGATCGGAA
38413 TATGAGTGTGGAGTCGTTGCCAAAAATTACCCAGGTGTGGTGGNNNAGATCGGAAG
38414 TATGAGTGTGGAGTCGTTGCATCCTAGCTACTCAGAAGGCTGANNNAGATCGGAAG
38415 TATGAGTGTGGAGTCGTTGCGCCTGTCCTGAAATGATCTCTGTNNNAGATCGGAAGA
38416 TATGAGTGTGGAGTCGTTGCACCCTTTGTAGCCTGAAATTTGANNNAGATCGGAAGA
38417 TATGAGTGTGGAGTCGTTGCATACAGAATATGGATGCAGGGGGNNNAGATCGGAA
38418 TATGAGTGTGGAGTCGTTGCCTAAGAATTTCAGCCAGGTGCAGNNNAGATCGGAAG
38422 TATGAGTGTGGAGTCGTTGCGGGATGCAGAATCTCTCTAGAGCNNNAGATCGGAAG
38423 TATGAGTGTGGAGTCGTTGCTGCAAGGCTCCTGTTCAAAAATCNNNAGATCGGAAG
38425 TATGAGTGTGGAGTCGTTGCACAGTGTTTGTCCTCTTTTTGCCNNNAGATCGGAAGA
38426 TATGAGTGTGGAGTCGTTGCGTAGAAGGGAGCTGCAGTTTCTANNNAGATCGGAAG
38427 TATGAGTGTGGAGTCGTTGCCATGCCTGTAATCCCAGCTACTCNNNAGATCGGAAGA
38428 TATGAGTGTGGAGTCGTTGCTAAGTCAGAATCCTTGTGGGTGGNNNAGATCGGAAG
38429 TATGAGTGTGGAGTCGTTGCGTCTCAATCTTCTGACCTCGTGANNNAGATCGGAAGA
38430 TATGAGTGTGGAGTCGTTGCGTTAGAATGTGAGTGGGGAGGAANNNAGATCGGAA
38431 TATGAGTGTGGAGTCGTTGCGCAGAGGGGTTTAGTAGAGAAGGNNNAGATCGGAA
38432 TATGAGTGTGGAGTCGTTGCGAGTCCTCTAAAACAGCCAAGGANNNAGATCGGAAG
38433 TATGAGTGTGGAGTCGTTGCAATATCCGCCCTAAACACCTTGTNNNAGATCGGAAGA
38435 TATGAGTGTGGAGTCGTTGCTTGGAGTAAGCTCACTACAACCTNNNAGATCGGAAG
38436 TATGAGTGTGGAGTCGTTGCGGGTTGCAATCCCAGTCTCTAATNNNAGATCGGAAG
38437 TATGAGTGTGGAGTCGTTGCTTCATCACCAGTACTTGCCATCANNNAGATCGGAAGA
38438 TATGAGTGTGGAGTCGTTGCACATTCATGATTTTGCCACATTGCNNNAGATCGGAAG
38439 TATGAGTGTGGAGTCGTTGCAAAGGACCAAAAAAGGAGCAACCNNNAGATCGGAA

38440 TATGAGTGTGGAGTCGTTGCCAGTGGTCTTGAAACAGACACACNNNAGATCGGAA
38442 TATGAGTGTGGAGTCGTTGCGGAGAGCTCAAGATGGAGACATCNNNAGATCGGAA
38444 TATGAGTGTGGAGTCGTTGCCCTCTTCATCTTCCCCTTCCAAANNNAGATCGGAAGA
38445 TATGAGTGTGGAGTCGTTGCCTCAAGGATGGTGCACAACAATCNNNAGATCGGAA
38447 TATGAGTGTGGAGTCGTTGCGATGTTTCCAAGGCTTCGTCTTTNNNAGATCGGAAG
38448 TATGAGTGTGGAGTCGTTGCAGTCTTTTAGTGAGTTGCTGACANNNAGATCGGAAG
38449 TATGAGTGTGGAGTCGTTGCAGCTGCAGAATGATAGGTACAGTNNNAGATCGGAA
38450 TATGAGTGTGGAGTCGTTGCGGTGAAGAAATGGGGAGCTACTGNNNAGATCGGA
38451 TATGAGTGTGGAGTCGTTGCATGGTGAAGTCCCGTCTCTACTANNNAGATCGGAAG
38452 TATGAGTGTGGAGTCGTTGCACGCCTGTAATTCCAACACTTTGNNNAGATCGGAAG
38454 TATGAGTGTGGAGTCGTTGCGTAGCTCACACCTGTAATCCCAANNNAGATCGGAAG
38456 TATGAGTGTGGAGTCGTTGCTAGCTACTCAGGATCACTTGAGCNNNAGATCGGAAG
38457 TATGAGTGTGGAGTCGTTGCGGACACAGGGTCTCACTATGTTGNNNAGATCGGAA
38458 TATGAGTGTGGAGTCGTTGCCCTAAAACGTACTGTGGTGATGGNNNAGATCGGAA
38459 TATGAGTGTGGAGTCGTTGCGGGGGGCCCTATCATTTTTATCANNNAGATCGGAAG
38461 TATGAGTGTGGAGTCGTTGCCAAATCAAACTCATGACGCAGCTNNNAGATCGGAAG
38462 TATGAGTGTGGAGTCGTTGCTGGAAAGTAGCAGCAGAACTAGGNNNAGATCGGA
38463 TATGAGTGTGGAGTCGTTGCCAGAGCAAGACTCCATCTCGAAANNNAGATCGGAAG
38464 TATGAGTGTGGAGTCGTTGCGCATGATGATGGTTCACTGTAGCNNNAGATCGGAA
38465 TATGAGTGTGGAGTCGTTGCTGATATGGTTGTTGGATACAGGTNNNAGATCGGAA
38466 TATGAGTGTGGAGTCGTTGCTACCAGTTCCTCCTTGTACCTCTNNNAGATCGGAAGA
38467 TATGAGTGTGGAGTCGTTGCTGAGCATTACAATTGTGCAGCTTNNNAGATCGGAAG
38468 TATGAGTGTGGAGTCGTTGCTTCATCACCCCTCCATTTATCCGNNNAGATCGGAAGA
38473 TATGAGTGTGGAGTCGTTGCGTTTTGTCTTTCTGCAGTGTCCANNNAGATCGGAAG
38474 TATGAGTGTGGAGTCGTTGCAACTGCACTAGAGTGGAGACAACNNNAGATCGGAA
38475 TATGAGTGTGGAGTCGTTGCAAAACAGATGCATTGACCAGTGGNNNAGATCGGAA
38476 TATGAGTGTGGAGTCGTTGCTGGAAAACGGTATGGAGGTTTCTNNNAGATCGGAA
38477 TATGAGTGTGGAGTCGTTGCATGGAGCACTACACAGCCATAAANNNAGATCGGAA
38478 TATGAGTGTGGAGTCGTTGCCCACCAATGTGCTTTCTGTCTTTNNNAGATCGGAAGA
38479 TATGAGTGTGGAGTCGTTGCCCACCACACCCAGCCATAAATTANNNAGATCGGAAG
38480 TATGAGTGTGGAGTCGTTGCGGATCCATGAGGTTTGGGTGTTANNNAGATCGGAA
38481 TATGAGTGTGGAGTCGTTGCATCTTCTCCTGTGCCCATAATGANNNAGATCGGAAG
38483 TATGAGTGTGGAGTCGTTGCCAGTGAGCTATCATCACACCACTNNNAGATCGGAAG
38484 TATGAGTGTGGAGTCGTTGCAAAGTGGTAAGTTCAGGACAAGTNNNAGATCGGAA
38485 TATGAGTGTGGAGTCGTTGCGGTGAAACCCCGTCTCTACTAAANNNAGATCGGAAG
38486 TATGAGTGTGGAGTCGTTGCCAGAATAACCTGGGGCCTCTATCNNNAGATCGGAAG
38487 TATGAGTGTGGAGTCGTTGCGAGGTTTTTTGCTGTTGCTGTTGNNNAGATCGGAAG
38488 TATGAGTGTGGAGTCGTTGCAGAAATGCTATCTCCAACAGGCTNNNAGATCGGAAG
38489 TATGAGTGTGGAGTCGTTGCATACCTTCCTTTCGCACCTCAATNNNAGATCGGAAGA
38490 TATGAGTGTGGAGTCGTTGCGAAAAAGGCGAAGCTGGAGAAATNNNAGATCGGA
38491 TATGAGTGTGGAGTCGTTGCACCCTTCCTACCTCATACCTCATNNNAGATCGGAAGA
38492 TATGAGTGTGGAGTCGTTGCATACTCATCACAAGCCTTGGAGGNNNAGATCGGAAG
38494 TATGAGTGTGGAGTCGTTGCGAGCACGCTTTATTCATCCACTCNNNAGATCGGAAG

38495 TATGAGTGTGGAGTCGTTGCTTTTCTTTTAGCCCCTTCTCCCCNNNAGATCGGAAGA
38496 TATGAGTGTGGAGTCGTTGCAATACAACAATTAGCTGGGCGTGNNNAGATCGGAA
38497 TATGAGTGTGGAGTCGTTGCGATAACAAAGTTGCTTTGGGGGANNNAGATCGGAA
38499 TATGAGTGTGGAGTCGTTGCGCCATCCTCCCACATACATGTTANNNAGATCGGAAG
38500 TATGAGTGTGGAGTCGTTGCAAGTTCAGCATCCTGAGAATCCCNNNAGATCGGAAG
38501 TATGAGTGTGGAGTCGTTGCGCCTGTAATCCCAACACTTTGAGNNNAGATCGGAAG
38503 TATGAGTGTGGAGTCGTTGCTTTTTGAGACAGGGTCTTGCTCTNNNAGATCGGAAG
38504 TATGAGTGTGGAGTCGTTGCCATCAGGGTGGAAGTATTGGAGTNNNAGATCGGAA
38505 TATGAGTGTGGAGTCGTTGCACACCTGATGATTGTGGATCACANNNAGATCGGAAG
38506 TATGAGTGTGGAGTCGTTGCAGCAAAGCTGCAGGATATACAAANNNAGATCGGAA
38507 TATGAGTGTGGAGTCGTTGCCAACCTAGGACAGAAGCCTGAAANNNAGATCGGAA
38508 TATGAGTGTGGAGTCGTTGCAAAGGGCATACTTTAGCTCCAGTNNNAGATCGGAAG
38509 TATGAGTGTGGAGTCGTTGCTTGACAGGACGAAATCAGGACTTNNNAGATCGGAA
38510 TATGAGTGTGGAGTCGTTGCGAGTTTGAGACTAGCCTGACCAANNNAGATCGGAA
38511 TATGAGTGTGGAGTCGTTGCTGAAGGAATTATTGTTTATGTGTGCTNNNAGATCGG
38512 TATGAGTGTGGAGTCGTTGCTGAGGTCATGAGTTTGAGACCAGNNNAGATCGGAA
38513 TATGAGTGTGGAGTCGTTGCCTGAGGCACGAGAATCACTTGAANNNAGATCGGAA
38514 TATGAGTGTGGAGTCGTTGCTCCAGGTTCAAGTGATTCTCCTGNNNAGATCGGAAG
38515 TATGAGTGTGGAGTCGTTGCATTCCAACCAGGACAAACATCCTNNNAGATCGGAAG
38516 TATGAGTGTGGAGTCGTTGCAATCGCCTGTAATTCTAGCACCTNNNAGATCGGAAG
38517 TATGAGTGTGGAGTCGTTGCTCACTTGAGGTCAGGAGTTCAACNNNAGATCGGAA
38522 TATGAGTGTGGAGTCGTTGCAATAAAACTCCGGTCTCCTCCACNNNAGATCGGAAG
38523 TATGAGTGTGGAGTCGTTGCTGTGTATTTTACCTTTGGGCACGNNNAGATCGGAAG
38525 TATGAGTGTGGAGTCGTTGCCAGGATCTAGTTGCAGGCACTAANNNAGATCGGAA
38526 TATGAGTGTGGAGTCGTTGCTGCATGCTACAAAATGAATGTGCNNNAGATCGGAA
38528 TATGAGTGTGGAGTCGTTGCGTTTGAAGCAATTCTCCTGCCTCNNNAGATCGGAAG
38531 TATGAGTGTGGAGTCGTTGCACTGTGAAACCCCGTCTCTACTANNNAGATCGGAAG
38534 TATGAGTGTGGAGTCGTTGCGAATAAATGACAAGCCGGGCATGNNNAGATCGGAA
38535 TATGAGTGTGGAGTCGTTGCTTATGCCTGTAATCCCAGCACTTNNNAGATCGGAAG
38538 TATGAGTGTGGAGTCGTTGCCTCACTGTAATCCCAGCACTTTGNNNAGATCGGAAG
38539 TATGAGTGTGGAGTCGTTGCTTTAGTACATGTGCAGGATGTGCNNNAGATCGGAA
38540 TATGAGTGTGGAGTCGTTGCTGTCCCTAATCAGCTCCTTTTCCNNNAGATCGGAAGA
38543 TATGAGTGTGGAGTCGTTGCCTGCAAGCCAGCATTATGGAAATNNNAGATCGGAA
38544 TATGAGTGTGGAGTCGTTGCGTCTCTTCCAGGTAAAACGTCTTNNNAGATCGGAAG
38545 TATGAGTGTGGAGTCGTTGCTTCCATTCATGTGTGTGTGTGTGNNNAGATCGGAAG
38546 TATGAGTGTGGAGTCGTTGCGTACGAGGTAGGGATCCCATTTCNNNAGATCGGAA
38547 TATGAGTGTGGAGTCGTTGCACATGTTTATAGTCCCAGCTGCTNNNAGATCGGAAG
38549 TATGAGTGTGGAGTCGTTGCACCACCTGAAGGGATTAGAGAGANNNAGATCGGAA
38550 TATGAGTGTGGAGTCGTTGCTGCATGGATTTGTTTCTGGGTTCNNNAGATCGGAAG
38552 TATGAGTGTGGAGTCGTTGCCACACACACTCACTCTGTCTCTCNNNAGATCGGAAGA
38553 TATGAGTGTGGAGTCGTTGCATTGTGGCCCGAACCTATAGTTCNNNAGATCGGAAG
38554 TATGAGTGTGGAGTCGTTGCAAAAAATTAGCTGGGTGTGGTGGNNNAGATCGGAA
38557 TATGAGTGTGGAGTCGTTGCTGGCAGAGCTGAGATTCAAACTCNNNAGATCGGAA

38559 TATGAGTGTGGAGTCGTTGCGAAAATTGCTTGAACCCAGGAGGNNNAGATCGGAA
38560 TATGAGTGTGGAGTCGTTGCTCTGTACTCCTATGGCATCATGCNNNAGATCGGAAG
38561 TATGAGTGTGGAGTCGTTGCAATCTTCATTTGGCACCAGATGGNNNAGATCGGAAG
38562 TATGAGTGTGGAGTCGTTGCAGACTTAGCCACTTCTTTACGGCNNNAGATCGGAAG
38563 TATGAGTGTGGAGTCGTTGCAAAAAATTAGCTGGGCATGGTGGNNNAGATCGGAA
38565 TATGAGTGTGGAGTCGTTGCAACTCTTCTGGACATAGGCCTTGNNNAGATCGGAAG
38566 TATGAGTGTGGAGTCGTTGCATCTGGAGGCATCACATTACCTGNNNAGATCGGAAG
38567 TATGAGTGTGGAGTCGTTGCAATGGTAACAACTGCAGGATCCANNNAGATCGGAA
38568 TATGAGTGTGGAGTCGTTGCGAAGGAGGAATGAGGCTGAGATCNNNAGATCGGA
38570 TATGAGTGTGGAGTCGTTGCGGAGAGAGAGAGTGAGAGAGACANNNAGATCGG
38571 TATGAGTGTGGAGTCGTTGCCAAAAAAGAAGTGGGGACTTGCTNNNAGATCGGAA
38572 TATGAGTGTGGAGTCGTTGCGCAGGAGAATTGATTGAACCTGGNNNAGATCGGAA
38574 TATGAGTGTGGAGTCGTTGCTTTTGACAGAGTCTTGCTGTGTCNNNAGATCGGAAG
38575 TATGAGTGTGGAGTCGTTGCGGAAAGAGATGAAAGTGGGGACANNNAGATCGGA
38577 TATGAGTGTGGAGTCGTTGCGAGGCAATGTTCTGAGAGCTTTCNNNAGATCGGAA
38578 TATGAGTGTGGAGTCGTTGCAAGGGGATACTGGTGCCAAAATTNNNAGATCGGAA
38579 TATGAGTGTGGAGTCGTTGCAGGTTGTAATGAGCCGAGATTGTNNNAGATCGGAA
38581 TATGAGTGTGGAGTCGTTGCTGGGGCTCCTTTGTAATTTCTCTNNNAGATCGGAAG
38582 TATGAGTGTGGAGTCGTTGCGCAGAAGAGGAGGTCATAGTGACNNNAGATCGGA
38584 TATGAGTGTGGAGTCGTTGCGTCCATATTCTCTGGGGCCTAACNNNAGATCGGAAG
38585 TATGAGTGTGGAGTCGTTGCCAGTGAGCTATGATTGCACCACTNNNAGATCGGAAG
38586 TATGAGTGTGGAGTCGTTGCCCACCATGCCCAGCTAATTTTTTNNNAGATCGGAAGA
38587 TATGAGTGTGGAGTCGTTGCACACATGCACAGACATTTCACTGNNNAGATCGGAAG
38588 TATGAGTGTGGAGTCGTTGCCTCTTGCCTATAATCCCAGCACTNNNAGATCGGAAGA
38589 TATGAGTGTGGAGTCGTTGCTCTGTAATTCCAGCTGAGGAAGGNNNAGATCGGAA
38590 TATGAGTGTGGAGTCGTTGCTGAGGAAGCTGTGTGAGAATTGANNNAGATCGGA
38591 TATGAGTGTGGAGTCGTTGCGGAACTGCAAATCAAAATCGCAANNNAGATCGGAA
38594 TATGAGTGTGGAGTCGTTGCGGAAGTAGGCAAAAGACAAGACANNNAGATCGGA
38595 TATGAGTGTGGAGTCGTTGCTCTGACTTTGGGGAGTTAGCAAANNNAGATCGGAA
38597 TATGAGTGTGGAGTCGTTGCTCCTTCAGGCTGAAATGAAAGGANNNAGATCGGAA
38598 TATGAGTGTGGAGTCGTTGCAAGAAAAGGTGGGAAAGAGCTGANNNAGATCGGA
38599 TATGAGTGTGGAGTCGTTGCTGAGCACCTACTATTTACCAGGTNNNAGATCGGAAG
38600 TATGAGTGTGGAGTCGTTGCAACTTCTTGGATTTGTGGGTTGCNNNAGATCGGAAG
38602 TATGAGTGTGGAGTCGTTGCGGTGGTACTTCAGCTTGGTATCANNNAGATCGGAAG
38604 TATGAGTGTGGAGTCGTTGCCTGAGGCAGGAGAATCACTTGAANNNAGATCGGAA
38606 TATGAGTGTGGAGTCGTTGCTTGGTTTCACTCCTGACCATTGANNNAGATCGGAAG
38607 TATGAGTGTGGAGTCGTTGCTGATGTGCAACGGACTAATTTCCNNNAGATCGGAAG
38608 TATGAGTGTGGAGTCGTTGCTTTTTGCTCCACACCAGGTAGTANNNAGATCGGAAG
38609 TATGAGTGTGGAGTCGTTGCCAGAAGAATTGCTTGAACCCGAGNNNAGATCGGAA
38610 TATGAGTGTGGAGTCGTTGCAAATGATGGTGGGATAACTGGCTNNNAGATCGGAA
38611 TATGAGTGTGGAGTCGTTGCCCAGCAGCACATCAAAAAGCTTANNNAGATCGGAAG
38612 TATGAGTGTGGAGTCGTTGCACACAGAAATGCAGTATGGAGGTNNNAGATCGGAA
38613 TATGAGTGTGGAGTCGTTGCCTTTGATCCCACCAACAATCAGGNNNAGATCGGAAG

38614 TATGAGTGTGGAGTCGTTGCCAACTGTAATCCCAGCACTTTGGNNNAGATCGGAAG
38616 TATGAGTGTGGAGTCGTTGCAGACAAAGTTTCGCTCTTTTCCCNNNAGATCGGAAG
38618 TATGAGTGTGGAGTCGTTGCTTAGTAACTGAAAGAGGGCCAGGNNNAGATCGGAA
38619 TATGAGTGTGGAGTCGTTGCTACAAAAAATTAGCCAGGCGTGGNNNAGATCGGAA
38620 TATGAGTGTGGAGTCGTTGCAAGCAAAGCAAAGCAAAGCAAAGNNNAGATCGGAA
38622 TATGAGTGTGGAGTCGTTGCTCATGAGGTCAGGAGTTTGAGACNNNAGATCGGAA
38623 TATGAGTGTGGAGTCGTTGCAGAGGTTGCAGTGAACTGAGATCNNNAGATCGGAA
38625 TATGAGTGTGGAGTCGTTGCCACCTTATTCCTGCAAGAATGGCNNNAGATCGGAAG
38627 TATGAGTGTGGAGTCGTTGCAGTCAAAGGAGGTTGTTCTCTGGNNNAGATCGGAA
38629 TATGAGTGTGGAGTCGTTGCTTTCCAGCAACAGACTCCAAAGANNNAGATCGGAAG
38631 TATGAGTGTGGAGTCGTTGCAGCAAACTAGCCGAAAAGCAAATNNNAGATCGGAA
38632 TATGAGTGTGGAGTCGTTGCCTCTGGAGTTATAGGAGGTGGGANNNAGATCGGA
38635 TATGAGTGTGGAGTCGTTGCATGCCTGTAATCTCAGCTACTCGNNNAGATCGGAAG
38636 TATGAGTGTGGAGTCGTTGCAATATAAAAACACCTGGGCCAGGNNNAGATCGGAA
38637 TATGAGTGTGGAGTCGTTGCGAAACCCCATACCCATTAGCAGTNNNAGATCGGAAG
38638 TATGAGTGTGGAGTCGTTGCTGACTCTTGGGACTTTGTCTGTTNNNAGATCGGAAG
38639 TATGAGTGTGGAGTCGTTGCAGTCCTTCCATCTTCCCTCTTCTNNNAGATCGGAAGA
38640 TATGAGTGTGGAGTCGTTGCAGCCTGTGGTGTTACTAACTCATNNNAGATCGGAAG
38641 TATGAGTGTGGAGTCGTTGCAAGAACCGTGCAACCATAAAAGGNNNAGATCGGAA
38642 TATGAGTGTGGAGTCGTTGCGATCACGAGGTCAGGAGTTCAAGNNNAGATCGGAA
38643 TATGAGTGTGGAGTCGTTGCGGGGGTAGCAGTATCCAGAAATCNNNAGATCGGAA
38644 TATGAGTGTGGAGTCGTTGCAAAATTAGCTGGGTGTGGTGATGNNNAGATCGGAA
38645 TATGAGTGTGGAGTCGTTGCCATCACAATCAGCTGGGTTTCATNNNAGATCGGAAG
38648 TATGAGTGTGGAGTCGTTGCGGGACAAGAACGAGACTTCATCTNNNAGATCGGAA
38649 TATGAGTGTGGAGTCGTTGCGCTGATCACCTGAAGTTAGGAGTNNNAGATCGGAA
38650 TATGAGTGTGGAGTCGTTGCATTACCCAAACTGCCATCATCCTNNNAGATCGGAAGA
38651 TATGAGTGTGGAGTCGTTGCTAACTTCTCCCTTTTCAGCCATTNNNAGATCGGAAGA
38652 TATGAGTGTGGAGTCGTTGCCGATGCAGAAGATGGGTGATTTCNNNAGATCGGAA
38654 TATGAGTGTGGAGTCGTTGCTTTGCCATGATTGTAAGAAGCCCNNNAGATCGGAAG
38655 TATGAGTGTGGAGTCGTTGCGGAAGGAAGGTAGGTAGGTAGGTNNNAGATCGGA
38658 TATGAGTGTGGAGTCGTTGCTTGTCCACTTGTTTCACCTGGTANNNAGATCGGAAG
38660 TATGAGTGTGGAGTCGTTGCCTCATGCCTGTAGTCCCAGTTACNNNAGATCGGAAG
38661 TATGAGTGTGGAGTCGTTGCAGAGGTGTTTAGAGGCAGATTGGNNNAGATCGGA
38662 TATGAGTGTGGAGTCGTTGCGAAGTCTGCAACCGGTTGTAATCNNNAGATCGGAA
38664 TATGAGTGTGGAGTCGTTGCTATTGGGTTTGTGGCTTTTTGCANNNAGATCGGAAG
38666 TATGAGTGTGGAGTCGTTGCACAAAAATTAGCTGGGCATGGTGNNNAGATCGGAA
38667 TATGAGTGTGGAGTCGTTGCCAGTGAACCATGATCACACCTCTNNNAGATCGGAAG
38668 TATGAGTGTGGAGTCGTTGCTACTCTCTACTGCCCAAACCAACNNNAGATCGGAAGA
38669 TATGAGTGTGGAGTCGTTGCCGTTACACATGCCTTTTTCCTCANNNAGATCGGAAGA
38670 TATGAGTGTGGAGTCGTTGCATAAGGGAGTAAGGGAGCTAGGGNNNAGATCGGA
38672 TATGAGTGTGGAGTCGTTGCCTTCTGGGAAAATGGAGGGAGAANNNAGATCGGA
38673 TATGAGTGTGGAGTCGTTGCAGGAAGGAGAATCACTTGAACCCNNNAGATCGGAA
38675 TATGAGTGTGGAGTCGTTGCGAAACCAGGGACTCTCAGAATCTNNNAGATCGGAA

38676 TATGAGTGTGGAGTCGTTGCTGCACCTGTAATCCCAACTACTCNNNAGATCGGAAGA
38677 TATGAGTGTGGAGTCGTTGCTGATATCAATTTGTCCCACTGGCNNNAGATCGGAAG
38678 TATGAGTGTGGAGTCGTTGCAACCTGAACGTATTTCCAGCTCTNNNAGATCGGAAG
38680 TATGAGTGTGGAGTCGTTGCCACAGTGGTTCAAGCCTGTAATCNNNAGATCGGAAG
38681 TATGAGTGTGGAGTCGTTGCGCTGTAACAAAAACTTCCCCAGTNNNAGATCGGAAG
38682 TATGAGTGTGGAGTCGTTGCGCAAAGGACATGAACAGACACTTNNNAGATCGGAA
38683 TATGAGTGTGGAGTCGTTGCCTGAGGTGGGAGAATCACTTGAGNNNAGATCGGAA
38685 TATGAGTGTGGAGTCGTTGCGCCAGACCCCATCTCTAAGAAAANNNAGATCGGAAG
38686 TATGAGTGTGGAGTCGTTGCGTCAAAGAAATGTCTGCACTCCCNNNAGATCGGAAG
38687 TATGAGTGTGGAGTCGTTGCACATTTATGCAGCCAACAGACACNNNAGATCGGAAG
38689 TATGAGTGTGGAGTCGTTGCGGCTACAGGGGAATCAGATTTGANNNAGATCGGAA
38690 TATGAGTGTGGAGTCGTTGCACATGATGACTCTTGGGATGGACNNNAGATCGGAA
38691 TATGAGTGTGGAGTCGTTGCATAAAAGGCCTTGTCCAACAGGANNNAGATCGGAA
38692 TATGAGTGTGGAGTCGTTGCAGTTTTGTGTTTGAAACCCAGGGNNNAGATCGGAA
38693 TATGAGTGTGGAGTCGTTGCCAAAGTTCTCCAAGCCCGAATTTNNNAGATCGGAAG
38695 TATGAGTGTGGAGTCGTTGCTTCCTGAAGGCAAAGGGAATACANNNAGATCGGAA
38697 TATGAGTGTGGAGTCGTTGCGGAGAATGGCATGAATCGAGGAGNNNAGATCGGA
38699 TATGAGTGTGGAGTCGTTGCTATGCCAGTTGAAAGACACCAGANNNAGATCGGAA
38700 TATGAGTGTGGAGTCGTTGCGGCATCCTTGTCTTTTTCCAGTTNNNAGATCGGAAGA
38702 TATGAGTGTGGAGTCGTTGCGAGACAGAGGCTCAACTTCAGTANNNAGATCGGAA
38704 TATGAGTGTGGAGTCGTTGCACAAATGGAAACACATCCCATGCNNNAGATCGGAAG
38705 TATGAGTGTGGAGTCGTTGCCTGAGGCAAGAAAATGGCATGAANNNAGATCGGAA
38706 TATGAGTGTGGAGTCGTTGCGGGAAACCCATCAGACTAACAGTNNNAGATCGGAA
38709 TATGAGTGTGGAGTCGTTGCTACCTGAGATTGAGGCAGACAACNNNAGATCGGAA
38710 TATGAGTGTGGAGTCGTTGCAGAAGAAAAAAAACCCTGCACGTNNNAGATCGGAA
38712 TATGAGTGTGGAGTCGTTGCGCAATCCTCCTGCCTCATTTTTANNNAGATCGGAAGA
38713 TATGAGTGTGGAGTCGTTGCTATTTCAGGCTTCTTGGAGAGCANNNAGATCGGAAG
38715 TATGAGTGTGGAGTCGTTGCTAAACACTTGGGTTGCTTTCACCNNNAGATCGGAAG
38716 TATGAGTGTGGAGTCGTTGCGCTACTCTGTCTGGTTTTCCCTTNNNAGATCGGAAGA
38717 TATGAGTGTGGAGTCGTTGCTTGTGAAAATGACCATACTGCCANNNAGATCGGAAG
38718 TATGAGTGTGGAGTCGTTGCTACATGCTTTTCTCACTGGTTGGNNNAGATCGGAAG
38720 TATGAGTGTGGAGTCGTTGCGGCTAGCCATATGCAGAAAACTGNNNAGATCGGAA
38721 TATGAGTGTGGAGTCGTTGCCCTAATCAAGGAGGTGAAAGACCNNNAGATCGGAA
38722 TATGAGTGTGGAGTCGTTGCCCACATACCCAACAGCTTCCTTANNNAGATCGGAAGA
38723 TATGAGTGTGGAGTCGTTGCTGAAGAGGCAACCCATAGATTGGNNNAGATCGGAA
38725 TATGAGTGTGGAGTCGTTGCAAAGGAACCCTAAAAACCATGCANNNAGATCGGAA
38726 TATGAGTGTGGAGTCGTTGCGCGCAAAACTGGATGGATAATGANNNAGATCGGAA
38727 TATGAGTGTGGAGTCGTTGCGGGTTGCAATCCTAGTCTCTGATNNNAGATCGGAAG
38728 TATGAGTGTGGAGTCGTTGCACCTGCAATTCTAGCTACTCGAGNNNAGATCGGAAG
38729 TATGAGTGTGGAGTCGTTGCGGAGGAATCACATTACCGGACTTNNNAGATCGGAA
38730 TATGAGTGTGGAGTCGTTGCAGCAAAATATTGAAGGAGCTGCANNNAGATCGGAA
38731 TATGAGTGTGGAGTCGTTGCCAGAGCTTGGAAACAAGAAGAGCNNNAGATCGGAA
38732 TATGAGTGTGGAGTCGTTGCAGAAGCTTCATATGCTGGTTCCANNNAGATCGGAAG

38733 TATGAGTGTGGAGTCGTTGCAGGCAGATCGGAATCAAACTACANNNAGATCGGAA
38734 TATGAGTGTGGAGTCGTTGCAGCTCCAGAGTTTCTGATTGATTNNNAGATCGGAAG
38735 TATGAGTGTGGAGTCGTTGCCCATTTGACCCTTGGCTGAAATTNNNAGATCGGAAG
38737 TATGAGTGTGGAGTCGTTGCAGGTGTTATGTAGCAAGACCCTGNNNAGATCGGAA
38738 TATGAGTGTGGAGTCGTTGCGCTGAGCCAAGATAATTGCTTGTNNNAGATCGGAA
38739 TATGAGTGTGGAGTCGTTGCTCCAACAAGGCCCTTTACCTTTANNNAGATCGGAAG
38740 TATGAGTGTGGAGTCGTTGCTGCAGATTCCTGGCTTCATCTAANNNAGATCGGAAG
38741 TATGAGTGTGGAGTCGTTGCGTGGGAGGATCACTTGAGATCAGNNNAGATCGGAA
38742 TATGAGTGTGGAGTCGTTGCTTCAAGTGATTCCCCTGCTTCAGNNNAGATCGGAAG
38743 TATGAGTGTGGAGTCGTTGCACAAGAGCGAAACTCCATCTAGGNNNAGATCGGAA
38747 TATGAGTGTGGAGTCGTTGCCCTAAATACTTGGGAGGCTGAGGNNNAGATCGGAA
38750 TATGAGTGTGGAGTCGTTGCTCACAGCACTATTCACAGTAGCANNNAGATCGGAAG
38751 TATGAGTGTGGAGTCGTTGCAAAACCCTCAGCAAAATTGGCATNNNAGATCGGAAG
38752 TATGAGTGTGGAGTCGTTGCAATCCTAGCTACTTTGGAGGCTGNNNAGATCGGAAG
38753 TATGAGTGTGGAGTCGTTGCGTAGTGCAATGGTGGAATCTTGGNNNAGATCGGAA
38756 TATGAGTGTGGAGTCGTTGCCATGCCTGTTATCCCAGCTAGTTNNNAGATCGGAAG
38757 TATGAGTGTGGAGTCGTTGCGTCACCATAGCTCACTGTAACCTNNNAGATCGGAAG
38759 TATGAGTGTGGAGTCGTTGCGGAGGCTGAGGCAGAATAATCATNNNAGATCGGAA
38760 TATGAGTGTGGAGTCGTTGCTATATAAAGGCCAGGTGTGGTGGNNNAGATCGGAA
38761 TATGAGTGTGGAGTCGTTGCTGTAAGTGTTTACGTGTGCAGTTNNNAGATCGGAAG
38762 TATGAGTGTGGAGTCGTTGCATGTTGCCCATTCACTAGCAATGNNNAGATCGGAAG
38763 TATGAGTGTGGAGTCGTTGCCTGAGGCAGAAGAAGTGCTTGAANNNAGATCGGAA
38764 TATGAGTGTGGAGTCGTTGCAGAATCACTTGAACTCAGGAGGCNNNAGATCGGAA
38766 TATGAGTGTGGAGTCGTTGCATAGATACTGCGGACTACTGGGTNNNAGATCGGAA
38767 TATGAGTGTGGAGTCGTTGCTCTAAATATTGACTGGGGCTGGGNNNAGATCGGAA
38768 TATGAGTGTGGAGTCGTTGCCATGCCTGTAATCCCAGCATTTTNNNAGATCGGAAG
38769 TATGAGTGTGGAGTCGTTGCTCCTTGCAACAGAACAGAAAGTTNNNAGATCGGAAG
38770 TATGAGTGTGGAGTCGTTGCATTGAATGTCCATTATGTGCCCGNNNAGATCGGAAG
38771 TATGAGTGTGGAGTCGTTGCTAGAGAAAGGAACTCATGCACTTNNNAGATCGGAA
38772 TATGAGTGTGGAGTCGTTGCAGCTCCTCTAGGTATGATGTTGANNNAGATCGGAAG
38773 TATGAGTGTGGAGTCGTTGCCCATGTGTGACAGAGTAGGATCCNNNAGATCGGAA
38776 TATGAGTGTGGAGTCGTTGCCAAAGCAGGGTCATGGAATCATCNNNAGATCGGAA
38777 TATGAGTGTGGAGTCGTTGCGAAAAGCTCCCTCAAATCATGCANNNAGATCGGAAG
38778 TATGAGTGTGGAGTCGTTGCTGGTCCCTAACATCACTTTCCAANNNAGATCGGAAGA
38780 TATGAGTGTGGAGTCGTTGCACTGCTTTTCTCCCATGGATTCTNNNAGATCGGAAGA
38781 TATGAGTGTGGAGTCGTTGCTCACGGTTTCAATGGACACTTTTNNNAGATCGGAAG
38782 TATGAGTGTGGAGTCGTTGCATGAAGAAGGGCTAAGACTGCTCNNNAGATCGGAA
38784 TATGAGTGTGGAGTCGTTGCAAAAATAAAAAGTAGGGCCGGGCNNNAGATCGGAA
38786 TATGAGTGTGGAGTCGTTGCCCTCCTAGGTTCAAGCGATTCTCNNNAGATCGGAAG
38787 TATGAGTGTGGAGTCGTTGCGATATCCGCACTTCTGTTTGTGGNNNAGATCGGAAG
38788 TATGAGTGTGGAGTCGTTGCACACATTACACACTAGAAGAAGAGANNNAGATCGG
38789 TATGAGTGTGGAGTCGTTGCCTAGTCCACCATGACCAAGTAGGNNNAGATCGGAAG
38790 TATGAGTGTGGAGTCGTTGCAGCCACTGCAAAAACAATCCAAANNNAGATCGGAAG

38793 TATGAGTGTGGAGTCGTTGCAAAGTCATCCATGTTGTAGCATGNNNAGATCGGAAG
38794 TATGAGTGTGGAGTCGTTGCAACCTCCTGAGTAGCTGGGATTANNNAGATCGGAAG
38795 TATGAGTGTGGAGTCGTTGCTGAACAGACAACCCACAGAATCTNNNAGATCGGAAG
38796 TATGAGTGTGGAGTCGTTGCTAGTGGCTACTATGAGCAACTGCNNNAGATCGGAAG
38797 TATGAGTGTGGAGTCGTTGCTCTGTCTTTCTTATTCCCACCCCNNNAGATCGGAAGAG
38798 TATGAGTGTGGAGTCGTTGCAAGGAAGAATGTTTGGGGGATGANNNAGATCGGAA
38799 TATGAGTGTGGAGTCGTTGCAGCCTGGAGTACAGTAGTGAGATNNNAGATCGGAA
38800 TATGAGTGTGGAGTCGTTGCTCTTTGGCTAGACTGACCAAGAANNNAGATCGGAAG
38802 TATGAGTGTGGAGTCGTTGCTTCAGCAGAGTTTCCGGATACAANNNAGATCGGAAG
38803 TATGAGTGTGGAGTCGTTGCATCCTGGCTAACACAGTGAAACTNNNAGATCGGAAG
38805 TATGAGTGTGGAGTCGTTGCAAAAAGGGAGAGAAGAACCAAGTNNNAGATCGGAA
38808 TATGAGTGTGGAGTCGTTGCTCATTAGGATGTCTTGCACACCTNNNAGATCGGAAGA
38809 TATGAGTGTGGAGTCGTTGCTGTGTAGGCTACCCTACCTAACANNNAGATCGGAAGA
38810 TATGAGTGTGGAGTCGTTGCTCTGGGTTCCAATGATTCTCCTGNNNAGATCGGAAGA
38811 TATGAGTGTGGAGTCGTTGCTGCAGCACTATTTACAACTGTGGNNNAGATCGGAAG
38813 TATGAGTGTGGAGTCGTTGCCCTGTAATCCCATAATCCCAGCANNNAGATCGGAAGA
38815 TATGAGTGTGGAGTCGTTGCATGTTTGACCAAAAAGCAACACANNNAGATCGGAAG
38816 TATGAGTGTGGAGTCGTTGCAACAAACAAACAAACGAACAGCCNNNAGATCGGAAG
38817 TATGAGTGTGGAGTCGTTGCCGTTGTGAAGACCATGGGAAAAGNNNAGATCGGAA
38818 TATGAGTGTGGAGTCGTTGCCTGTAGTCCCAGCTACTCAGGAGNNNAGATCGGAAG
38819 TATGAGTGTGGAGTCGTTGCATAAATAAAGCCACCACCCTCGANNNAGATCGGAAGA
38821 TATGAGTGTGGAGTCGTTGCCTGTAGTCCCAGCTACTCGGAAGNNNAGATCGGAAG
38823 TATGAGTGTGGAGTCGTTGCATGCAAACTAAGATGGCAACGAGNNNAGATCGGAAG
38824 TATGAGTGTGGAGTCGTTGCGATGAAACTGGACCCTTACCTGTNNNAGATCGGAAG
38828 TATGAGTGTGGAGTCGTTGCGGGTGGGAGGAAGTAAGCATAAANNNAGATCGGAA
38829 TATGAGTGTGGAGTCGTTGCGTCATTGCTATGCTGCTTACCTGNNNAGATCGGAAGA
38831 TATGAGTGTGGAGTCGTTGCTGTGTGGTGGTTCATGCCTATAANNNAGATCGGAAG
38834 TATGAGTGTGGAGTCGTTGCGAGGTTTCAATGAGCCGAGATTGNNNAGATCGGAAG
38836 TATGAGTGTGGAGTCGTTGCCAGACACACACACACACATATGCNNNAGATCGGAAGA
38838 TATGAGTGTGGAGTCGTTGCGTGCAATCTCAGCTCAATGCAAGNNNAGATCGGAAG
38839 TATGAGTGTGGAGTCGTTGCGTGCTTGAGAGAAGGGAACTACANNNAGATCGGAA
38840 TATGAGTGTGGAGTCGTTGCCCAGGAGTAGGAGACTACAGTGANNNAGATCGGAA
38842 TATGAGTGTGGAGTCGTTGCCAGGAGAATCACTTGAACCCAGGNNNAGATCGGAAG
38843 TATGAGTGTGGAGTCGTTGCAACACTTTTTTGGGTGCAAGGAANNNAGATCGGAAG
38848 TATGAGTGTGGAGTCGTTGCAGTTACAAAAGCCGTGGGAAAAGNNNAGATCGGAA
38849 TATGAGTGTGGAGTCGTTGCCCATAGACTTTACCCTCCTTGGCNNNAGATCGGAAGA
38850 TATGAGTGTGGAGTCGTTGCTGCATGAGTCTTCCAAAGGAGAANNNAGATCGGAAG
38851 TATGAGTGTGGAGTCGTTGCGTAGAGATGGGGTTTCGCCATATNNNAGATCGGAAG
38853 TATGAGTGTGGAGTCGTTGCAGGATAGAGAGAAAACCCTCCCTNNNAGATCGGAAG
38855 TATGAGTGTGGAGTCGTTGCCCCATGTAAATCTGAAACCCAGCNNNAGATCGGAAGA
38856 TATGAGTGTGGAGTCGTTGCTACCTCTATGCACAAAAGCCAGANNNAGATCGGAAG
38857 TATGAGTGTGGAGTCGTTGCAAAACAGCACACCTACAACCATCNNNAGATCGGAAGA
38858 TATGAGTGTGGAGTCGTTGCTACAAAAATTAGCTGGGCGTGGTNNNAGATCGGAAG

38859 TATGAGTGTGGAGTCGTTGCAGAAAACCCAATCCAAGAATGCANNNAGATCGGAA
38862 TATGAGTGTGGAGTCGTTGCCTCAAAAGACTTGCAGCAGTGAANNNAGATCGGAA
38863 TATGAGTGTGGAGTCGTTGCCACGATGTTGAGCCCAGTTTTACNNNAGATCGGAAG
38864 TATGAGTGTGGAGTCGTTGCTGAAGCTGTGGGGAAAAGAGAATNNNAGATCGGA
38865 TATGAGTGTGGAGTCGTTGCATCTCAGTGCTGACTCCTAGGATNNNAGATCGGAAG
38866 TATGAGTGTGGAGTCGTTGCACCATGATCAGGTGGGTTTCATANNNAGATCGGAA
38867 TATGAGTGTGGAGTCGTTGCTTAGTAGCTACGTATGGCCTTGGNNNAGATCGGAA
38868 TATGAGTGTGGAGTCGTTGCAATATGCCCACCTGAGGAACAATNNNAGATCGGAAG
38869 TATGAGTGTGGAGTCGTTGCAAGGCTTAGAAACGACAACGAGANNNAGATCGGAA
38870 TATGAGTGTGGAGTCGTTGCGTACAGGGACAACAGCAAAAACANNNAGATCGGAA
38872 TATGAGTGTGGAGTCGTTGCTGCAAAAACTGTGGGAAAAGTGTNNNAGATCGGAA
38874 TATGAGTGTGGAGTCGTTGCTTCCAGTGATTCTTCTGCCTCAGNNNAGATCGGAAG
38880 TATGAGTGTGGAGTCGTTGCATTATCTTGGGAGTTCTAGGGCCNNNAGATCGGAA
38881 TATGAGTGTGGAGTCGTTGCTCATCACAGGCCAAAAAAATGGTNNNAGATCGGAA
38882 TATGAGTGTGGAGTCGTTGCCTGAGGAAGGAGAATGGTGTGAANNNAGATCGGA
38883 TATGAGTGTGGAGTCGTTGCGACAGAGCAAGACTCCATCTCAANNNAGATCGGAAG
38884 TATGAGTGTGGAGTCGTTGCAGGAAGGGGTTTGAAACAAGACANNNAGATCGGA
38885 TATGAGTGTGGAGTCGTTGCAATATTCACTGCAGCAATCGCATNNNAGATCGGAAG
38886 TATGAGTGTGGAGTCGTTGCCAAGGGCTGTAGCTTACCAAAATNNNAGATCGGAA
38887 TATGAGTGTGGAGTCGTTGCACCAGATGTACAAAGAAGGGCTGNNNAGATCGGAA
38888 TATGAGTGTGGAGTCGTTGCCCCAACCCAATAAGCAATCACTGNNNAGATCGGAAG
38889 TATGAGTGTGGAGTCGTTGCTTAGACCGATTCCATGCTACTGGNNNAGATCGGAAG
38890 TATGAGTGTGGAGTCGTTGCGAAAAACAGACACGAGCGCTTATNNNAGATCGGAA
38891 TATGAGTGTGGAGTCGTTGCGGCTAGGGCTGGTAAAAATGTTGNNNAGATCGGAA
38892 TATGAGTGTGGAGTCGTTGCACGGTCAGGAAGAGTCAAAAAGANNNAGATCGGA
38893 TATGAGTGTGGAGTCGTTGCTTCTTTGCAAATGTGGATGCCTTNNNAGATCGGAAG
38895 TATGAGTGTGGAGTCGTTGCCCAAAAAAGAGCCCAAATAGCCANNNAGATCGGAA
38896 TATGAGTGTGGAGTCGTTGCCTTGTGCTGACATTTCCTGTACTNNNAGATCGGAAG
38898 TATGAGTGTGGAGTCGTTGCTCTGAACAAGTAGGCTTTCTCCCNNNAGATCGGAAG
38899 TATGAGTGTGGAGTCGTTGCACTGTTCGTGGGACTGTAAATTANNNAGATCGGAA
38901 TATGAGTGTGGAGTCGTTGCGCATTGGTAGCTCACACATCAAANNNAGATCGGAAG
38902 TATGAGTGTGGAGTCGTTGCATGGGGAAAAGCGTTCCTATTCANNNAGATCGGAA
38903 TATGAGTGTGGAGTCGTTGCACCCCCCAAAAATCACACTAGTTNNNAGATCGGAAG
38904 TATGAGTGTGGAGTCGTTGCTGCAAAAACCATGGGAAAAGTGTNNNAGATCGGAA
38905 TATGAGTGTGGAGTCGTTGCTTTAGGCACTCAAGGGAATCTCCNNNAGATCGGAAG
38906 TATGAGTGTGGAGTCGTTGCTTGCCACAACTTACCTACCTCTTNNNAGATCGGAAGA
38907 TATGAGTGTGGAGTCGTTGCCAGCAAACTGAACAGATCTGCAANNNAGATCGGAA
38909 TATGAGTGTGGAGTCGTTGCATGTCCTATGCAAAATGGCACACNNNAGATCGGAAG
38910 TATGAGTGTGGAGTCGTTGCGAATGAGTTCACGTCCTTTGCAGNNNAGATCGGAAG
38911 TATGAGTGTGGAGTCGTTGCAAATGGTGCTGGGATAACTGGATNNNAGATCGGAA
38912 TATGAGTGTGGAGTCGTTGCAGCACAGGACCAGATGATTTCAANNNAGATCGGAA
38913 TATGAGTGTGGAGTCGTTGCCTCCTCAACAACCCATTCCATGANNNAGATCGGAAGA
38914 TATGAGTGTGGAGTCGTTGCCCAGAATGAGCAACAGTGGTTTTNNNAGATCGGAA

38915 TATGAGTGTGGAGTCGTTGCCAAAGCAGTAGTTTTCAGGCACANNNAGATCGGAAG
38916 TATGAGTGTGGAGTCGTTGCAGACACAAGACACTTGTCATCGANNNAGATCGGAAG
38917 TATGAGTGTGGAGTCGTTGCTGGGTCTAGTTGTTTGCCTAGTCNNNAGATCGGAAG
38918 TATGAGTGTGGAGTCGTTGCGCCCTGTGGTTACCAAGACAATANNNAGATCGGAAG
38919 TATGAGTGTGGAGTCGTTGCAAAAAAAGCCAGGGCAACAAAACNNNAGATCGGAAG
38920 TATGAGTGTGGAGTCGTTGCTGTCTCTTTATGTCCTTTGCCCANNNAGATCGGAAGA
38921 TATGAGTGTGGAGTCGTTGCACATTCTTCTCAGTGCCACATGGNNNAGATCGGAAGA
38922 TATGAGTGTGGAGTCGTTGCTCCAGGAGCTGTTTTTTTGAAAANNNAGATCGGAAG
38923 TATGAGTGTGGAGTCGTTGCGCAAGTCAGTCACTGTGAAACTCNNNAGATCGGAAG
38924 TATGAGTGTGGAGTCGTTGCGTTATGTACTGCGTGTTGGTGTCNNNAGATCGGAAG
38925 TATGAGTGTGGAGTCGTTGCTCACTCTTGGCTATGGGGAAAAGNNNAGATCGGAAG
38930 TATGAGTGTGGAGTCGTTGCAGCAGAGGAAAGAATCTCAGAGCNNNAGATCGGAA
38931 TATGAGTGTGGAGTCGTTGCTGCCAAAAACCAGTTGCTTTGTANNNAGATCGGAAG
38932 TATGAGTGTGGAGTCGTTGCCAAAAATTAGCCAGGAGTGGTGGNNNAGATCGGAA
38933 TATGAGTGTGGAGTCGTTGCGGGTGAGGGAGATAAGGAGGTATNNNAGATCGGA
38934 TATGAGTGTGGAGTCGTTGCTTATGAAGCCAGCATCATCCTGANNNAGATCGGAAG
38935 TATGAGTGTGGAGTCGTTGCTACCTGACTGTTTTGCTTCACCTNNNAGATCGGAAGA
38936 TATGAGTGTGGAGTCGTTGCAATTGCTTGAAATTGGGAGGTGGNNNAGATCGGAA
38937 TATGAGTGTGGAGTCGTTGCGTAATCCTAGCTACTTGGGAGGCNNNAGATCGGAAG
38938 TATGAGTGTGGAGTCGTTGCTCCCAAAAGTTTGAAGGTTGCTGNNNAGATCGGAAG
38939 TATGAGTGTGGAGTCGTTGCGGGGAAAGAGAAGGGATAGCAAANNNAGATCGGA
38942 TATGAGTGTGGAGTCGTTGCTATTCCCAAACTCCCAAACTTGCNNNAGATCGGAAGA
38943 TATGAGTGTGGAGTCGTTGCAAACTTCAGCCTCCCAGTTGTATNNNAGATCGGAAGA
38945 TATGAGTGTGGAGTCGTTGCCTAAGTCAGATAGCAAGCTACCTNNNAGATCGGAAG
38946 TATGAGTGTGGAGTCGTTGCAAATGACCATACAGCCAAAAGCANNNAGATCGGAAG
38948 TATGAGTGTGGAGTCGTTGCTTGCATACTGATCCCATAGAGGCNNNAGATCGGAAG
38949 TATGAGTGTGGAGTCGTTGCCTAGGCCAGTGTTCTCATTGAGTNNNAGATCGGAAG
38950 TATGAGTGTGGAGTCGTTGCCCTTTGCAGAAACATGGATGGAGNNNAGATCGGAAG
38951 TATGAGTGTGGAGTCGTTGCGAGTGGATATCTTACAGGCCAGGNNNAGATCGGAAG
38953 TATGAGTGTGGAGTCGTTGCTGATCACATGGGGGTAGAGTGTANNNAGATCGGAA
38955 TATGAGTGTGGAGTCGTTGCAGCCAAAGAACTGAGTAGACACCNNNAGATCGGAAG
38956 TATGAGTGTGGAGTCGTTGCCCGGCCAAACTAAGCTTCATAAGNNNAGATCGGAAG
38957 TATGAGTGTGGAGTCGTTGCTGAGGACATGAGATTTTGGAGGGNNNAGATCGGAA
38959 TATGAGTGTGGAGTCGTTGCAGCAGCACATCAAAAAGCTTCAANNNAGATCGGAAG
38960 TATGAGTGTGGAGTCGTTGCATGTCTGGAATGAACCCCACTTTNNNAGATCGGAAGA
38961 TATGAGTGTGGAGTCGTTGCTTCTATGGAGAGGAATTTGGGGCNNNAGATCGGAA
38962 TATGAGTGTGGAGTCGTTGCCCTGGAATGTCTGCAAAAACGATNNNAGATCGGAAG
38963 TATGAGTGTGGAGTCGTTGCGCACTTGGCATTGAAAGTGGTAGNNNAGATCGGAAG
38964 TATGAGTGTGGAGTCGTTGCGACGGATGCATTCTTCAGTATGCNNNAGATCGGAAG
38965 TATGAGTGTGGAGTCGTTGCGATATTCCCTTTCACCACCCCTTNNNAGATCGGAAGA
38968 TATGAGTGTGGAGTCGTTGCAGGTTTTCTTAGCTGCTTGGAGANNNAGATCGGAAG
38970 TATGAGTGTGGAGTCGTTGCCTTAGTAGAGACGGCGTTTCACCNNNAGATCGGAAG
38971 TATGAGTGTGGAGTCGTTGCTCATGGAGGAAGCTGACATTGAANNNAGATCGGAAG

38972 TATGAGTGTGGAGTCGTTGCCTTCACTGCTGAATTGCACTAAANNNAGATCGGAAG
38973 TATGAGTGTGGAGTCGTTGCTGGAGATTGCTTTGGGGAGTATTNNNAGATCGGAA
38974 TATGAGTGTGGAGTCGTTGCAGAGAGAGAGAGACAAAGAGGCTNNNAGATCGGA
38975 TATGAGTGTGGAGTCGTTGCACTGCTCCTTTGAATCAATGCAANNNAGATCGGAAG
38976 TATGAGTGTGGAGTCGTTGCGTGAAACCCCGTCTCTACTGAAANNNAGATCGGAAG
38977 TATGAGTGTGGAGTCGTTGCAGGAAGGAAGAAGGAAAGAAGGANNNAGATCGG
38979 TATGAGTGTGGAGTCGTTGCTTTCACTTGTCTCTTGAGGCTCCNNNAGATCGGAAG
38980 TATGAGTGTGGAGTCGTTGCTGTATCTTTCCTCTGGCATACCANNNAGATCGGAAG
38981 TATGAGTGTGGAGTCGTTGCAACGAACAGGCAAAACATCAACANNNAGATCGGAA
38982 TATGAGTGTGGAGTCGTTGCTGGGAGAAAAATGTTTGCAAATCANNNAGATCGGA
38983 TATGAGTGTGGAGTCGTTGCATTTTGTCCCTCATCTCAGTGGGNNNAGATCGGAAG
38984 TATGAGTGTGGAGTCGTTGCTCTGTTTATGCCTGGCTTCATGTNNNAGATCGGAAG
38987 TATGAGTGTGGAGTCGTTGCACCCAACAGAGCAGACAACATTANNNAGATCGGAAG
38988 TATGAGTGTGGAGTCGTTGCGGTTTCAGAAAGCCGAGATCAACNNNAGATCGGAA
38989 TATGAGTGTGGAGTCGTTGCTGCCTAAAGAGTTTGAATGCATGANNNAGATCGGA
38990 TATGAGTGTGGAGTCGTTGCAGAGGTCTTAGCCAATGCAATCANNNAGATCGGAA
38992 TATGAGTGTGGAGTCGTTGCAAGATGTGGGCGTTTGCAAAATANNNAGATCGGAA
38993 TATGAGTGTGGAGTCGTTGCCAAGTGTTAGGATCCTGGCATCTNNNAGATCGGAAG
38995 TATGAGTGTGGAGTCGTTGCCAGCTGAATCAACCAAGTGGAAGNNNAGATCGGAA
38996 TATGAGTGTGGAGTCGTTGCGAACCCAAGGAAAGAAGATAGGCNNNAGATCGGA
38997 TATGAGTGTGGAGTCGTTGCCAAGATGGTCTCGATCTCCTGACNNNAGATCGGAAG
39002 TATGAGTGTGGAGTCGTTGCTGAGCTTCATGAGTACATTTCCANNNAGATCGGAAG
39003 TATGAGTGTGGAGTCGTTGCATGCTTATGTACAGCTTGTGGGANNNAGATCGGAA
39006 TATGAGTGTGGAGTCGTTGCCATCCTGGCTAACATGGTGAAACNNNAGATCGGAAG
39007 TATGAGTGTGGAGTCGTTGCACGTGCCTGTAATCTCAGCTATTNNNAGATCGGAAG
39008 TATGAGTGTGGAGTCGTTGCGCTACAAACCATGTTTAGCCAGGNNNAGATCGGAAG
39009 TATGAGTGTGGAGTCGTTGCCATGCTTGTTGTCCCAGCTATTCNNNAGATCGGAAG
39012 TATGAGTGTGGAGTCGTTGCTATCACTCAGTCATCTTCCCTGCNNNAGATCGGAAGA
39013 TATGAGTGTGGAGTCGTTGCCTGAGAATTCCTTGAACCCAGGANNNAGATCGGAAG
39015 TATGAGTGTGGAGTCGTTGCATGTCTGTGCAACTTTCAGAAGCNNNAGATCGGAAG
39016 TATGAGTGTGGAGTCGTTGCAAACCGGATCTGTAAAACTGGGTNNNAGATCGGAA
39017 TATGAGTGTGGAGTCGTTGCAATGACTTCAGCAACGTTTCAGGNNNAGATCGGAAG
39018 TATGAGTGTGGAGTCGTTGCGTGTGATGCAGACAAAAGAGGTCNNNAGATCGGAA
39019 TATGAGTGTGGAGTCGTTGCAAAATCTGGACTTTGTGCTGTGGNNNAGATCGGAA
39021 TATGAGTGTGGAGTCGTTGCTCGACTTTATCTTTGGGAGGCAANNNAGATCGGAAG
39023 TATGAGTGTGGAGTCGTTGCCCCAGGGGTCCAAACATACATATNNNAGATCGGAAG
39024 TATGAGTGTGGAGTCGTTGCGTTCTGTGTGGGAAATGCATGAANNNAGATCGGAA
39026 TATGAGTGTGGAGTCGTTGCACTTGGCTGCTTTTCTTCTACTGNNNAGATCGGAAG
39027 TATGAGTGTGGAGTCGTTGCCCCATGAAAGTGAGGAGGATACANNNAGATCGGAA
39028 TATGAGTGTGGAGTCGTTGCTGAACACGGTTCAAACAGTTAGCNNNAGATCGGAA
39033 TATGAGTGTGGAGTCGTTGCTCAAACTCCTAGGCTCAAGCAATNNNAGATCGGAAG
39035 TATGAGTGTGGAGTCGTTGCTGTGTTCTGTCTACTACAATGGGTNNNAGATCGGAA
39036 TATGAGTGTGGAGTCGTTGCTACTCACAATCTCCAGTTGCTGCNNNAGATCGGAAG

39037 TATGAGTGTGGAGTCGTTGCCTCCTTCAACTTCAGCTTCCTGANNNAGATCGGAAGA
39038 TATGAGTGTGGAGTCGTTGCAAATGGATAGTTGGACCCAGAGGNNNAGATCGGAA
39039 TATGAGTGTGGAGTCGTTGCAGCAAAATGGCAGAGTTTAGCTGNNNAGATCGGAA
39041 TATGAGTGTGGAGTCGTTGCTACAGGAGGCAAAAGATCTCCACNNNAGATCGGAA
39043 TATGAGTGTGGAGTCGTTGCCTTTGCATGGCTGAAAAAAAGGTNNNAGATCGGAA
39044 TATGAGTGTGGAGTCGTTGCCAGTGACATGATCTCAGCTCACTNNNAGATCGGAAG
39046 TATGAGTGTGGAGTCGTTGCATCCTGCAAGTGACTCAAAGGAANNNAGATCGGAA
39050 TATGAGTGTGGAGTCGTTGCGTGGGCAAAAGACATGATCAGACNNNAGATCGGAA
39051 TATGAGTGTGGAGTCGTTGCTTCACAGAACTCAGGAAAGTGCTNNNAGATCGGAA
39052 TATGAGTGTGGAGTCGTTGCGCCTCAAGAGGTACTGATGACATNNNAGATCGGAA
39053 TATGAGTGTGGAGTCGTTGCAGCAAAAAGGTGGCCTTGTTATGNNNAGATCGGAA
39054 TATGAGTGTGGAGTCGTTGCTGCAGACAAGAAAAATGAACCCANNNAGATCGGAA
39056 TATGAGTGTGGAGTCGTTGCTGAAACTACGTGATTGCTTGCTGNNNAGATCGGAAG
39057 TATGAGTGTGGAGTCGTTGCTGAGGCAGGAGAATCACTTGAAANNNAGATCGGAA
39058 TATGAGTGTGGAGTCGTTGCGTGGGCAAAGGACATAAACAGACNNNAGATCGGAA
39059 TATGAGTGTGGAGTCGTTGCGGCTAAGGAAGGACAATTGCTTGNNNAGATCGGAA
39060 TATGAGTGTGGAGTCGTTGCCATCCTGGACATAGTAACAGGCANNNAGATCGGAA
39064 TATGAGTGTGGAGTCGTTGCAGATACAAACAAGCCCAGACAGTNNNAGATCGGAA
39065 TATGAGTGTGGAGTCGTTGCTGAGGTAGGAGAATGGTGTGAACNNNAGATCGGA
39066 TATGAGTGTGGAGTCGTTGCAGTGGTTCAAGCAAGATAGCAGANNNAGATCGGAA
39067 TATGAGTGTGGAGTCGTTGCAATGTCCATTGTTGAAGCCACACNNNAGATCGGAAG
39069 TATGAGTGTGGAGTCGTTGCTGACTCCCATCCTATACAACCCTNNNAGATCGGAAGA
39070 TATGAGTGTGGAGTCGTTGCTCCTGTGTTGGGTGCATATATGTNNNAGATCGGAAG
39071 TATGAGTGTGGAGTCGTTGCTGGCATTCCAACCATAAAAATGCNNNAGATCGGAAG
39072 TATGAGTGTGGAGTCGTTGCACACAGGATAAAGTGGTAACTTGGANNNAGATCGG
39074 TATGAGTGTGGAGTCGTTGCAAGAGCCAGACCCCTTTTCATAGNNNAGATCGGAAG
39075 TATGAGTGTGGAGTCGTTGCGCCTCCCAAGTAGCTAGGATTACNNNAGATCGGAAG
39078 TATGAGTGTGGAGTCGTTGCCCTTACTCCTGCAAGAATAGCCANNNAGATCGGAAG
39079 TATGAGTGTGGAGTCGTTGCGAAAGCAGAATTTGGAGGCTGAGNNNAGATCGGA
39082 TATGAGTGTGGAGTCGTTGCTCTTGTCCTATCTCCACAGTTGCNNNAGATCGGAAGA
39083 TATGAGTGTGGAGTCGTTGCCGGTTCTCTCACCCCTTTGTATANNNAGATCGGAAGA
39084 TATGAGTGTGGAGTCGTTGCAGTCTGAGCCAGGAGATCATTTGNNNAGATCGGAA
39085 TATGAGTGTGGAGTCGTTGCCTTGCCCCTCTATAGTTCCTGTCNNNAGATCGGAAGA
39088 TATGAGTGTGGAGTCGTTGCATGAGAAGGTGCTCAACATCACTNNNAGATCGGAA
39089 TATGAGTGTGGAGTCGTTGCCTGTTTTTTATCAAGGGGCCTGGNNNAGATCGGAAG
39092 TATGAGTGTGGAGTCGTTGCTTGTGTGTATTTGGAAGTAGCCANNNAGATCGGAA
39093 TATGAGTGTGGAGTCGTTGCCCATCAAAAAGGGGGCAAATGATNNNAGATCGGAA
39094 TATGAGTGTGGAGTCGTTGCTTCTTTGAAAGGGAGTCTTGCTCNNNAGATCGGAAG
39095 TATGAGTGTGGAGTCGTTGCTGGCTGCTTCTGATGTGATACTTNNNAGATCGGAAG
39096 TATGAGTGTGGAGTCGTTGCGAGATTGCAGGCAAGAGTCCTATNNNAGATCGGAA
39097 TATGAGTGTGGAGTCGTTGCGTAGAGAAAACCTGCAGGCCTATNNNAGATCGGAA
39099 TATGAGTGTGGAGTCGTTGCTACCTGAAGCACCTAGTCTACCTNNNAGATCGGAAG
39100 TATGAGTGTGGAGTCGTTGCGAAAGAGACAGGGAAAGGGAGAANNNAGATCGGA

39101 TATGAGTGTGGAGTCGTTGCAGATGGAGTCTCACTCTGTCACCNNNAGATCGGAAG
39103 TATGAGTGTGGAGTCGTTGCATCCCAGTGATACCTGATGTCATNNNAGATCGGAAG
39104 TATGAGTGTGGAGTCGTTGCCCCCACCACATAGACACAAATTGNNNAGATCGGAAG
39105 TATGAGTGTGGAGTCGTTGCAGTGCTGATAGCCAAGACAATGANNNAGATCGGAA
39107 TATGAGTGTGGAGTCGTTGCGATATGACCAGATGGCCTGAAGTNNNAGATCGGAA
39108 TATGAGTGTGGAGTCGTTGCGTAGTGACCTCAGTGTCTCCAAGNNNAGATCGGAAG
39109 TATGAGTGTGGAGTCGTTGCAGATACTGACTAACCGACCTCCTNNNAGATCGGAAG
39111 TATGAGTGTGGAGTCGTTGCACATGACATATCACGCGAGATCANNNAGATCGGAAG
39112 TATGAGTGTGGAGTCGTTGCCTGTAGTCCCAGCTACTCGAGAGNNNAGATCGGAAG
39114 TATGAGTGTGGAGTCGTTGCCAGAAGATTGAAACTGGACCCCTNNNAGATCGGAA
39115 TATGAGTGTGGAGTCGTTGCAGCAAACTGAATCCATTAGCACANNNAGATCGGAAG
39116 TATGAGTGTGGAGTCGTTGCATGCTAGAGAGGTTGTGGAGAACNNNAGATCGGAA
39117 TATGAGTGTGGAGTCGTTGCTCCACAGTCCAAAGTCTGATCTGNNNAGATCGGAAG
39118 TATGAGTGTGGAGTCGTTGCGTGGTACGATCCTAGTTCACTGGNNNAGATCGGAA
39120 TATGAGTGTGGAGTCGTTGCTCATAAAAAGTGCCTGATGTCAGNNNAGATCGGAA
39121 TATGAGTGTGGAGTCGTTGCAAATTGTCGAGTCCAGCAACAACNNNAGATCGGAAG
39123 TATGAGTGTGGAGTCGTTGCAACTGCCATGGAGACTGCTTTATNNNAGATCGGAAG
39125 TATGAGTGTGGAGTCGTTGCGAGTCAAGACCTGTTAGTGTGCTNNNAGATCGGAA
39129 TATGAGTGTGGAGTCGTTGCAGTCTTGGGTGAGATTTTGGACANNNAGATCGGAA
39131 TATGAGTGTGGAGTCGTTGCCTGTGGTTGACAATGATTTGGGGNNNAGATCGGAA
39132 TATGAGTGTGGAGTCGTTGCTAAAGCGGTGCTGTGTAAAAGTGNNNAGATCGGAA
39133 TATGAGTGTGGAGTCGTTGCCTCCTGCAACTAGTTCAGTGTCTNNNAGATCGGAAG
39135 TATGAGTGTGGAGTCGTTGCTCCCTGATTATAGCCATTGGAGCNNNAGATCGGAAG
39138 TATGAGTGTGGAGTCGTTGCACCTATTGAGTGCAGTGTACACTNNNAGATCGGAAG
39139 TATGAGTGTGGAGTCGTTGCCCACTCTCCATTTAGCCAGAAGTNNNAGATCGGAAG
39142 TATGAGTGTGGAGTCGTTGCCGATCTCAGCTACTGCAACTTCTNNNAGATCGGAAG
39143 TATGAGTGTGGAGTCGTTGCTCCTGAACAGACCAATAACAACTNNNAGATCGGAAG
39144 TATGAGTGTGGAGTCGTTGCTAGGGAGTTGTAGTTTTCCGTGGNNNAGATCGGAA
39146 TATGAGTGTGGAGTCGTTGCGGTGCAAGGGACATGAATAGACANNNAGATCGGAA
39147 TATGAGTGTGGAGTCGTTGCAAAATGGCCATACTGCACAAAGTNNNAGATCGGAA
39148 TATGAGTGTGGAGTCGTTGCTCACACTCAGAATTGATTGCTGCNNNAGATCGGAAG
39150 TATGAGTGTGGAGTCGTTGCCTAGGAAAACCCAAAGGCAAAGGNNNAGATCGGAA
39154 TATGAGTGTGGAGTCGTTGCCACATCAACCAGTGGAACACAATNNNAGATCGGAAG
39156 TATGAGTGTGGAGTCGTTGCGGGAAAGGACACCCTCTTCAATANNNAGATCGGAA
39157 TATGAGTGTGGAGTCGTTGCGGGTTCAAGTGATTCTCCTGTCTNNNAGATCGGAAG
39159 TATGAGTGTGGAGTCGTTGCTCATCGGATCCTTGGTGATTACTNNNAGATCGGAAG
39161 TATGAGTGTGGAGTCGTTGCAGAAAAGAAAAAGCCACATGCCANNNAGATCGGAA
39163 TATGAGTGTGGAGTCGTTGCGCCCACAACCCATAACTTTCTTTNNNAGATCGGAAGA
39164 TATGAGTGTGGAGTCGTTGCTTGATTTCCGGACCCATCAATCTNNNAGATCGGAAG
39165 TATGAGTGTGGAGTCGTTGCTGAGGCAGGAGAATAGTGTGAACNNNAGATCGGAA
39169 TATGAGTGTGGAGTCGTTGCGGTGTTTCCAGAGGAGATAAGCANNNAGATCGGAA
39170 TATGAGTGTGGAGTCGTTGCCAGGAGTGATAGCTTATGCCTGTNNNAGATCGGAA
39171 TATGAGTGTGGAGTCGTTGCACAGGGTGAAAAGACAACCTACANNNAGATCGGAA

39172 TATGAGTGTGGAGTCGTTGCGCCCAGGACTAGAAGGATTCAAANNNAGATCGGAA

39173 TATGAGTGTGGAGTCGTTGCGAAAGACAAAAGACAAGCTGCCTNNNAGATCGGAA

39174 TATGAGTGTGGAGTCGTTGCCATATCCAGCCTCTACATGTCCCNNNAGATCGGAAGA

39175 TATGAGTGTGGAGTCGTTGCAGTAAAAAGAGCCCACATAGCCANNNAGATCGGAAG

39176 TATGAGTGTGGAGTCGTTGCGATGGATGGAACTGGAGGTCATTNNNAGATCGGAA

39177 TATGAGTGTGGAGTCGTTGCAGGGCAACCAATATAGCCAGAAANNNAGATCGGAA

39178 TATGAGTGTGGAGTCGTTGCCTTCCTAAGTGAAGGCACAGAGTNNNAGATCGGAAG

39179 TATGAGTGTGGAGTCGTTGCACATGCTGCTTCCAAGTTTTGTTNNNAGATCGGAAG

39180 TATGAGTGTGGAGTCGTTGCGAGAGGTAGAGAAAAGTCAGCGANNNAGATCGGA

39181 TATGAGTGTGGAGTCGTTGCAGCCAGTCATTTTGTCTCTCAGTNNNAGATCGGAAG

39182 TATGAGTGTGGAGTCGTTGCGGATTTTTGGTCAAGAGCCAGACNNNAGATCGGAA

39184 TATGAGTGTGGAGTCGTTGCGGACATAATGTCGAACCGGACTANNNAGATCGGAA

39185 TATGAGTGTGGAGTCGTTGCCTGATGGGGAAGGATCACTTGAANNNAGATCGGAA

39186 TATGAGTGTGGAGTCGTTGCGGTGATGATTGCACAACTGTGAANNNAGATCGGAA

39189 TATGAGTGTGGAGTCGTTGCACTGTGGGCAATTGTAACACAACNNNAGATCGGAAG

39190 TATGAGTGTGGAGTCGTTGCCTGGTCTCAAACTCCTGATCTCANNNAGATCGGAAG

39191 TATGAGTGTGGAGTCGTTGCCCCTCCCTTCCTCTTCTTTTCAGNNNAGATCGGAAGAG

39192 TATGAGTGTGGAGTCGTTGCACTGAGGTAGGAGGATCACTTGANNNAGATCGGAA

39193 TATGAGTGTGGAGTCGTTGCGCAAATGATTTCTGCCCATGGAANNNAGATCGGAAG

39194 TATGAGTGTGGAGTCGTTGCATGCACTCATAGCACTGTTCAGANNNAGATCGGAAG

39195 TATGAGTGTGGAGTCGTTGCTTTCCATCTTCCCACTGACTCTCNNNAGATCGGAAGA

39196 TATGAGTGTGGAGTCGTTGCCACCCATGCCTTATAATGCCTTGNNNAGATCGGAAGA

39197 TATGAGTGTGGAGTCGTTGCGGGGACTTCGGTTCTATAGTCACNNNAGATCGGAAG

39198 TATGAGTGTGGAGTCGTTGCAATTTGCACACATGACTCTGGTGNNNAGATCGGAAG

39199 TATGAGTGTGGAGTCGTTGCAAAAAGCAACCTATGGGGGAGATNNNAGATCGGAA

39200 TATGAGTGTGGAGTCGTTGCTCCAGCCATTCCATCTCTATGTGNNNAGATCGGAAGA

39201 TATGAGTGTGGAGTCGTTGCAAACAACCAAACTACACAAGCCTNNNAGATCGGAAG

39202 TATGAGTGTGGAGTCGTTGCTGCAGCCTAGATAATGCGTAACCNNNAGATCGGAAG

39203 TATGAGTGTGGAGTCGTTGCTTAGCTAGTCAACCAAAGTGGGGNNNAGATCGGAA

39204 TATGAGTGTGGAGTCGTTGCGTTCCCAGTGACTCAGTTCCTAGNNNAGATCGGAAG

39205 TATGAGTGTGGAGTCGTTGCCACCACACCCAGCTAATTTTTGTNNNAGATCGGAAGA

39207 TATGAGTGTGGAGTCGTTGCCCTGAGGGGAGGGGTACAATTATNNNAGATCGGAA

39208 TATGAGTGTGGAGTCGTTGCGTGTGGTGGGTGTCAAAACATATNNNAGATCGGAA

39209 TATGAGTGTGGAGTCGTTGCAAGCCAATCAAGAGAAACAGCACNNNAGATCGGAA

39211 TATGAGTGTGGAGTCGTTGCTATTATTGTGCTGCACCTCAGGTNNNAGATCGGAAG

39212 TATGAGTGTGGAGTCGTTGCGTTTGGCCACCTTTTGTCTCTTTNNNAGATCGGAAGA

39213 TATGAGTGTGGAGTCGTTGCCATTAGGCTCTGTGTAGGTCCTCNNNAGATCGGAAG

39214 TATGAGTGTGGAGTCGTTGCCTCAAACTCCTGACCTCGTGATCNNNAGATCGGAAGA

39215 TATGAGTGTGGAGTCGTTGCAGTTTAGAGGAGACTCCACGAGANNNAGATCGGAA

39216 TATGAGTGTGGAGTCGTTGCAACCAGAAGCCAGAAAGCAAAAGNNNAGATCGGAA

39217 TATGAGTGTGGAGTCGTTGCGAGAATCACTTGAACCTGGGAGTNNNAGATCGGAA

39218 TATGAGTGTGGAGTCGTTGCACAAAGAAATGTATTGCTCGGCCNNNAGATCGGAAG

39219 TATGAGTGTGGAGTCGTTGCCAGCTATTGTGAAAGGGGTTGTGNNNAGATCGGAA

39220 TATGAGTGTGGAGTCGTTGCTTTGCACCTTTTCAACCTCAACCNNNAGATCGGAAGA
39221 TATGAGTGTGGAGTCGTTGCTGTTCAATAACACTGGATGCTGCNNNAGATCGGAAG
39222 TATGAGTGTGGAGTCGTTGCTGATCAAGATTGGAAAGAGGCCANNNAGATCGGAA
39223 TATGAGTGTGGAGTCGTTGCCTGTAGTCCCAGCTACTCAGGAGNNNAGATCGGAAG
39225 TATGAGTGTGGAGTCGTTGCAATGGAAGAACATCTCATGCTCANNNAGATCGGAAG
39226 TATGAGTGTGGAGTCGTTGCTTATTTTCCACCTTCCACCTGCANNNAGATCGGAAGA
39230 TATGAGTGTGGAGTCGTTGCACCAGCTGCAAATATACACTCTCANNNAGATCGGAA
39231 TATGAGTGTGGAGTCGTTGCATCTCTGGTCCTAGCTACTCAGGNNNAGATCGGAAG
39232 TATGAGTGTGGAGTCGTTGCTTCCTAGAGCATGTCTTGGCTTTNNNAGATCGGAAG
39233 TATGAGTGTGGAGTCGTTGCACCCTTACTTCTGGTAAATGGTATCTNNNAGATCGG
39234 TATGAGTGTGGAGTCGTTGCTAGGTTACACGGACATGGTGAAGNNNAGATCGGAA
39235 TATGAGTGTGGAGTCGTTGCGCTAAGGCAGGAGAATCACTCAANNNAGATCGGAA
39236 TATGAGTGTGGAGTCGTTGCCCTGCCTCCAATCAATCACAAACNNNAGATCGGAAGA
39237 TATGAGTGTGGAGTCGTTGCATACCTGGGGGAAGACTGTCATANNNAGATCGGAA
39239 TATGAGTGTGGAGTCGTTGCTCTAACCATTTGCCTCACAGACANNNAGATCGGAAG
39241 TATGAGTGTGGAGTCGTTGCTCTGGGCTGAGATTGTATTAGCANNNAGATCGGAA
39242 TATGAGTGTGGAGTCGTTGCCCTATAACTATAGCTGGCCAGGCNNNAGATCGGAAG
39243 TATGAGTGTGGAGTCGTTGCTGCCTAGTATCAGGGTATGAGGANNNAGATCGGAA
39244 TATGAGTGTGGAGTCGTTGCTTACGACATGGATGAGCCTTGAGNNNAGATCGGAA
39245 TATGAGTGTGGAGTCGTTGCATCATTACTGCCACCCATCAGAGNNNAGATCGGAAG
39246 TATGAGTGTGGAGTCGTTGCCAAGGCTATTTGGTGGCTTACAGNNNAGATCGGAA
39247 TATGAGTGTGGAGTCGTTGCGCATAACCCACATGAGGAAAGGANNNAGATCGGAA
39249 TATGAGTGTGGAGTCGTTGCATGACTGCATCACTGTACTCCAGNNNAGATCGGAAG
39251 TATGAGTGTGGAGTCGTTGCCTGGGCTGGATAATCTAGAGAGCNNNAGATCGGAA
39252 TATGAGTGTGGAGTCGTTGCGGCAGGGAAATTTTTAGGTGCATNNNAGATCGGAA
39253 TATGAGTGTGGAGTCGTTGCTCTTGACTACTCCTCTCACGCTANNNAGATCGGAAGA
39255 TATGAGTGTGGAGTCGTTGCTGTAATCCCAGCTACTCAGGAGANNNAGATCGGAAG
39256 TATGAGTGTGGAGTCGTTGCGATGAATCCCCCCAATTATCCCANNNAGATCGGAAG
39257 TATGAGTGTGGAGTCGTTGCTCAGGAGGTAGAAGGAGCAGTTANNNAGATCGGA
39258 TATGAGTGTGGAGTCGTTGCCCATGGAAGAACCAATCTGTGTGNNNAGATCGGAA
39259 TATGAGTGTGGAGTCGTTGCTTCCAGACCAGCCTGACCAATATNNNAGATCGGAAG
39260 TATGAGTGTGGAGTCGTTGCGGGTTCAAGCAATTCTCCTACCTNNNAGATCGGAAG
39261 TATGAGTGTGGAGTCGTTGCTTGGCTAGTGCTGAAGAGGTTAGNNNAGATCGGAA
39263 TATGAGTGTGGAGTCGTTGCAAGAAAGAAAGTATGCTGTGGGCNNNAGATCGGA
39264 TATGAGTGTGGAGTCGTTGCGACGCAGCATACACTCAAAAAGANNNAGATCGGAA
39266 TATGAGTGTGGAGTCGTTGCAAGCAAGGTTTCTCACTGTCTGANNNAGATCGGAAG
39268 TATGAGTGTGGAGTCGTTGCCTAGGCAACAAAGCAAGACTCCANNNAGATCGGAA
39269 TATGAGTGTGGAGTCGTTGCCCCTGAGAAGGTGCATGTTTTTTNNNAGATCGGAAG
39270 TATGAGTGTGGAGTCGTTGCAAAGTTTTAGGGTCCAAAGTCCTNNNAGATCGGAAG
39272 TATGAGTGTGGAGTCGTTGCTCTGTCTCTGTGTCCTCTCTCTTNNNAGATCGGAAGA
39273 TATGAGTGTGGAGTCGTTGCCCAGTGTTCAGTGCAACCTAATCNNNAGATCGGAAG
39274 TATGAGTGTGGAGTCGTTGCTCTGGCCCTTTATTAGCAAATGCNNNAGATCGGAAG
39275 TATGAGTGTGGAGTCGTTGCAGCTCCACAGCCTAAAGAAATCANNNAGATCGGAAG

39276 TATGAGTGTGGAGTCGTTGCTTTTTTGAGTCAGAGTCTGGCTCNNNAGATCGGAAG
39277 TATGAGTGTGGAGTCGTTGCTTTACTCTCATGTGTCTGTGGGTNNNAGATCGGAAG
39278 TATGAGTGTGGAGTCGTTGCTGGCTACACAACAAAAAGCAACANNNAGATCGGAAG
39279 TATGAGTGTGGAGTCGTTGCGTCCTACCTAGAGCAAGCATACCNNNAGATCGGAAG
39280 TATGAGTGTGGAGTCGTTGCCAAACTCCTGGGATCAAACAACCNNNAGATCGGAAG
39281 TATGAGTGTGGAGTCGTTGCAGCCTAGTTGGACAGACATTAGCNNNAGATCGGAA
39282 TATGAGTGTGGAGTCGTTGCGGAAGTATTGATGGCATGTGCTGNNNAGATCGGAA
39284 TATGAGTGTGGAGTCGTTGCTGCACTGGTCATGAACTACTTGANNNAGATCGGAAG
39285 TATGAGTGTGGAGTCGTTGCTGACACAGGAAGACAAAACTTGCNNNAGATCGGAA
39286 TATGAGTGTGGAGTCGTTGCGCTATGATCACACCACTGCATTCNNNAGATCGGAAG
39287 TATGAGTGTGGAGTCGTTGCTATGATGACAGTGTGCCTCTCAGNNNAGATCGGAAG
39288 TATGAGTGTGGAGTCGTTGCTGTCACAACTACTCAACACTGCTNNNAGATCGGAAG
39290 TATGAGTGTGGAGTCGTTGCAGAAAGAAAGGAAGAAGCTTGGCNNNAGATCGGA
39291 TATGAGTGTGGAGTCGTTGCGGGGGTCAATTCAGCAAAAGAATNNNAGATCGGAA
39292 TATGAGTGTGGAGTCGTTGCAAAACCTACAGGCCAACATCTCTNNNAGATCGGAAG
39293 TATGAGTGTGGAGTCGTTGCCACCACCATGCCACACTAAAATTNNNAGATCGGAAG
39294 TATGAGTGTGGAGTCGTTGCAAAGGCATAGCTGTACATCCACANNNAGATCGGAAG
39295 TATGAGTGTGGAGTCGTTGCGGAGGTCTTTTGTCAATATGCCGNNNAGATCGGAAG
39296 TATGAGTGTGGAGTCGTTGCTCTGGGATCTCTATGTTGTTCCANNNAGATCGGAAG
39298 TATGAGTGTGGAGTCGTTGCACACACACACACACAAAACCAATNNNAGATCGGAAG
39299 TATGAGTGTGGAGTCGTTGCAGGCTGCTACAAATTCCATGAGANNNAGATCGGAAG
39300 TATGAGTGTGGAGTCGTTGCCAAGTAGCTGGGATTACAGGCATNNNAGATCGGAA
39302 TATGAGTGTGGAGTCGTTGCCTAGCAAAAACGTGGCCTTGTAANNNAGATCGGAAG
39305 TATGAGTGTGGAGTCGTTGCAATTAGGCAGGAATCGGTGCTAANNNAGATCGGAA
39307 TATGAGTGTGGAGTCGTTGCAACCCAAATCCTTACAGCTGACTNNNAGATCGGAAG
39308 TATGAGTGTGGAGTCGTTGCTGGCAATTTCAGAGGCGAACTATNNNAGATCGGAA
39310 TATGAGTGTGGAGTCGTTGCAAAAAGAGAAATCAGGCCACGTCNNNAGATCGGAA
39311 TATGAGTGTGGAGTCGTTGCAACTGCATATTGGGTACCGTGTANNNAGATCGGAAG
39312 TATGAGTGTGGAGTCGTTGCCAGTGAGCCAAGATCACACTACTNNNAGATCGGAAG
39313 TATGAGTGTGGAGTCGTTGCACATTAGCCAACTTGGTGCTCTANNNAGATCGGAAG
39314 TATGAGTGTGGAGTCGTTGCTCAGATGACTTCTCGATAGCAGCNNNAGATCGGAAG
39315 TATGAGTGTGGAGTCGTTGCGAGCAAGACTCTGTCTCAAAAAANNNAGATCGGAAG
39316 TATGAGTGTGGAGTCGTTGCTACACATGGGGTGAGTCAGTTTTNNNAGATCGGAA
39318 TATGAGTGTGGAGTCGTTGCGGAGTATGTAGTTGCTGAGGGAGNNNAGATCGGA
39319 TATGAGTGTGGAGTCGTTGCATTGTGTCCTACCTCTGTTGGTGNNNAGATCGGAAG
39320 TATGAGTGTGGAGTCGTTGCCCTTCCCATGTGTCAATGTCAACNNNAGATCGGAAG
39321 TATGAGTGTGGAGTCGTTGCTGGGCATGAAAGATCAGACTTCANNNAGATCGGAA
39323 TATGAGTGTGGAGTCGTTGCTAGAGATGAACCCTGGGTGTTTGNNNAGATCGGAA
39324 TATGAGTGTGGAGTCGTTGCCAGTTGTTGACAGAAACTGGACANNNAGATCGGAA
39326 TATGAGTGTGGAGTCGTTGCAATGGATGACAGGTAGGGTGTTCNNNAGATCGGAA
39327 TATGAGTGTGGAGTCGTTGCTTGGATCCACATCTCATGCCATANNNAGATCGGAAG
39328 TATGAGTGTGGAGTCGTTGCCAGAATGGACTTTGGTGACTTGGNNNAGATCGGAA
39331 TATGAGTGTGGAGTCGTTGCGACATAAATAACAGATGCCGGCCNNNAGATCGGAA

39332 TATGAGTGTGGAGTCGTTGCTCAGTAAACACTACTCTTGGGGCNNNAGATCGGAAG
39333 TATGAGTGTGGAGTCGTTGCAGACTGAAGGCAAAGAGATGAGGNNNAGATCGGA
39334 TATGAGTGTGGAGTCGTTGCGATGTTTTGTACCCTGCTACCCANNNAGATCGGAAG
39335 TATGAGTGTGGAGTCGTTGCAGAGGTTGTAGAGAGCCGAGATCNNNAGATCGGA
39338 TATGAGTGTGGAGTCGTTGCCCCACTCCTTGTTATACACCCAANNNAGATCGGAAGA
39339 TATGAGTGTGGAGTCGTTGCCCTGTAATTCCAGCTACTTGGGANNNAGATCGGAAG
39340 TATGAGTGTGGAGTCGTTGCCCACGCACGGCTAATTTTTGTATNNNAGATCGGAAG
39343 TATGAGTGTGGAGTCGTTGCTTGAGGGAGGAATGCTTTACCTCNNNAGATCGGAA
39344 TATGAGTGTGGAGTCGTTGCCTGTAATCCCAGCTACTCAGGAGNNNAGATCGGAAG
39345 TATGAGTGTGGAGTCGTTGCCGGGATCTCGTTATGTAGACTGGNNNAGATCGGAA
39346 TATGAGTGTGGAGTCGTTGCCTGTGCTTCTGTTACAACCTGACNNNAGATCGGAAG
39348 TATGAGTGTGGAGTCGTTGCTTTTCCTTTTTTGCAGGCAGTCANNNAGATCGGAAG
39349 TATGAGTGTGGAGTCGTTGCAAATGGCAACATGTTGTCTCAGGNNNAGATCGGAA
39350 TATGAGTGTGGAGTCGTTGCCTGTAGATTCCATAGGCCCTTCCNNNAGATCGGAAG
39351 TATGAGTGTGGAGTCGTTGCCCATGTGACTATCCTGTTTCCCANNNAGATCGGAAGA
39352 TATGAGTGTGGAGTCGTTGCCCCAACTGTCCCCAAAATGTTTTNNNAGATCGGAAGA
39353 TATGAGTGTGGAGTCGTTGCTTCCATGGTGTATATGTGCCACANNNAGATCGGAAG
39355 TATGAGTGTGGAGTCGTTGCGGGAGCCAGAGGAAGAAAGTAAANNNAGATCGGA
39357 TATGAGTGTGGAGTCGTTGCCCATCTTTCTGCCACCATCAATGNNNAGATCGGAAGA
39358 TATGAGTGTGGAGTCGTTGCCCTGTAATCTCAGCACTTTGGGANNNAGATCGGAAG
39360 TATGAGTGTGGAGTCGTTGCGAAGTGGAGATATGGGCATCAGANNNAGATCGGA
39361 TATGAGTGTGGAGTCGTTGCCTTTGCTCTGATGGAGGAGACTTNNNAGATCGGAAG
39363 TATGAGTGTGGAGTCGTTGCTTGCTAACCCCCTCATTATGCTTNNNAGATCGGAAGA
39364 TATGAGTGTGGAGTCGTTGCTTCTGGGTTCAAGAAATCCTCCCNNNAGATCGGAAG
39365 TATGAGTGTGGAGTCGTTGCGTTTCAGTGAGTCAAGATGGCACNNNAGATCGGAA
39366 TATGAGTGTGGAGTCGTTGCACAGAACTTGGTGTCTCATTGGANNNAGATCGGAAG
39367 TATGAGTGTGGAGTCGTTGCATAGTGTGGAGGATGAATTGGGGNNNAGATCGGA
39368 TATGAGTGTGGAGTCGTTGCAAATGAAGTGGAGATGGGAAGCTNNNAGATCGGA
39369 TATGAGTGTGGAGTCGTTGCAAAAAATTAGCCAGGTGTGGTGGNNNAGATCGGAA
39371 TATGAGTGTGGAGTCGTTGCCTCCAACTCACAGCCCATTTCTANNNAGATCGGAAGA
39372 TATGAGTGTGGAGTCGTTGCCAGCATATAAGCCAGGAGCAGTANNNAGATCGGAA
39373 TATGAGTGTGGAGTCGTTGCACATTGGCAGGTCTTAGGTCTTTNNNAGATCGGAAG
39374 TATGAGTGTGGAGTCGTTGCGCTCCCTAAACAATGTTTGCTGTNNNAGATCGGAAG
39375 TATGAGTGTGGAGTCGTTGCTCTTCTAGAGACTGGCAACACACNNNAGATCGGAAG
39376 TATGAGTGTGGAGTCGTTGCAACAAAAATCTTGGCCATGCACANNNAGATCGGAAG
39378 TATGAGTGTGGAGTCGTTGCTCGTCCAGGGCATTTATGATGATNNNAGATCGGAAG
39380 TATGAGTGTGGAGTCGTTGCAAAATTAGCCAGGTGTGATGGTGNNNAGATCGGAA
39381 TATGAGTGTGGAGTCGTTGCCTAGCCCTACAATTCTGCCTCATNNNAGATCGGAAGA
39382 TATGAGTGTGGAGTCGTTGCGCCCAAAGCAATTTACAGATTCANNNAGATCGGAAG
39383 TATGAGTGTGGAGTCGTTGCAAATAGCTGGGATTACAGGCACANNNAGATCGGAA
39384 TATGAGTGTGGAGTCGTTGCGGTGCTGGGGTCTCAATTATGTANNNAGATCGGAA
39385 TATGAGTGTGGAGTCGTTGCGATCACAAGGTCAGGGGATCAAGNNNAGATCGGAA
39387 TATGAGTGTGGAGTCGTTGCGGAGCAGCTTGCAATACCATATGNNNAGATCGGAA

39388 TATGAGTGTGGAGTCGTTGCATACAAAAATTAGCCAGGCGTGGNNNAGATCGGAA
39389 TATGAGTGTGGAGTCGTTGCAATGGTACAAACCCTAAGTGCCTNNNAGATCGGAAG
39390 TATGAGTGTGGAGTCGTTGCTGAGAGGGGAAGGGAACTAACATNNNAGATCGGA
39391 TATGAGTGTGGAGTCGTTGCGAGAATCGTTTGAACTCAGGAGCNNNAGATCGGAA
39392 TATGAGTGTGGAGTCGTTGCAAAGACGGACTGACAGGGTTTANNNAGATCGGAA
39393 TATGAGTGTGGAGTCGTTGCATTTGGAGACCTTTTGCATGTCANNNAGATCGGAAG
39394 TATGAGTGTGGAGTCGTTGCATGGTTTCGTCCTTCCCTCTTTTNNNAGATCGGAAGA
39395 TATGAGTGTGGAGTCGTTGCGGAGTGATGTCATGACCCTTTGGNNNAGATCGGAA
39396 TATGAGTGTGGAGTCGTTGCAACTGGACTCCTAACTTTTCCCCNNNAGATCGGAAGA
39398 TATGAGTGTGGAGTCGTTGCACTCCATACACCCCAACAATGAANNNAGATCGGAAG
39399 TATGAGTGTGGAGTCGTTGCTGGATTGGAAGATTCAGCAGAGTNNNAGATCGGAA
39400 TATGAGTGTGGAGTCGTTGCAATTGTTTTGTTCGAGGAGGCTGNNNAGATCGGAA
39401 TATGAGTGTGGAGTCGTTGCAGAAAAACGCATGCAATGGATCANNNAGATCGGAA
39403 TATGAGTGTGGAGTCGTTGCAAGTACCCACAGAAGCTTTACCCNNNAGATCGGAAG
39404 TATGAGTGTGGAGTCGTTGCAGGGATCTTAGAGGATGGGTCAANNNAGATCGGAA
39405 TATGAGTGTGGAGTCGTTGCAAAGAGGAGGGAAGATGAATGCCNNNAGATCGGA
39407 TATGAGTGTGGAGTCGTTGCCACTGGGGGGTTTCATACACTAGNNNAGATCGGAA
39408 TATGAGTGTGGAGTCGTTGCGACCAGCCTGGCTAACATAGTAANNNAGATCGGAA
39409 TATGAGTGTGGAGTCGTTGCAGGAAACCTAAGTACTCTTGGGCNNNAGATCGGAA
39410 TATGAGTGTGGAGTCGTTGCACTGTCATGGAAAAATGCTCGTGNNNAGATCGGAA
39412 TATGAGTGTGGAGTCGTTGCCTCTCTGCTAACCACACTCCAATNNNAGATCGGAAGA
39413 TATGAGTGTGGAGTCGTTGCGCTCTGGATGAGAAAACTGAGGANNNAGATCGGAA
39414 TATGAGTGTGGAGTCGTTGCTTGAGCTCAGCAGTTCAAGATCANNNAGATCGGAAG
39415 TATGAGTGTGGAGTCGTTGCAGAAAAGAAAAAAACGGCCAGGCNNNAGATCGGAA
39417 TATGAGTGTGGAGTCGTTGCTACAAAAACAAATAGCTGGGCGTNNNAGATCGGAA
39418 TATGAGTGTGGAGTCGTTGCTATTTTGAGACAGAGCCCCACTCNNNAGATCGGAAG
39419 TATGAGTGTGGAGTCGTTGCACGTCTGTAATCCCAGCACTTAGNNNAGATCGGAAG
39420 TATGAGTGTGGAGTCGTTGCCTCTGATCACCCCTGGAGAATTCNNNAGATCGGAAG
39421 TATGAGTGTGGAGTCGTTGCAGCCTTTCTAAGAGTGAGCTCAGNNNAGATCGGAA
39422 TATGAGTGTGGAGTCGTTGCTAATCCTAGCTACTCAGGAGGCTNNNAGATCGGAAG
39423 TATGAGTGTGGAGTCGTTGCAATGTTCTGAAGCAGGTTCATGCNNNAGATCGGAA
39424 TATGAGTGTGGAGTCGTTGCGATGAAGGCAACAATTAGAGCCCNNNAGATCGGAA
39425 TATGAGTGTGGAGTCGTTGCAAAACAAAGTTGTGGAGAAGGCCNNNAGATCGGAA
39426 TATGAGTGTGGAGTCGTTGCATTTTACTAGCAGCCCCAACAGANNNAGATCGGAAG
39428 TATGAGTGTGGAGTCGTTGCTCCAGAAATGAAGCCAATTGACTNNNAGATCGGAAG
39429 TATGAGTGTGGAGTCGTTGCCCACCATCTCAGTAATATGGGCANNNAGATCGGAAG
39430 TATGAGTGTGGAGTCGTTGCCCTATAGTCCCAGCTACTCAGGANNNAGATCGGAAG
39431 TATGAGTGTGGAGTCGTTGCAAGGTCCCAAAATAGCCAAAACGNNNAGATCGGAA
39432 TATGAGTGTGGAGTCGTTGCATTACTTCCTACCAAGGGGCAAGNNNAGATCGGAAG
39433 TATGAGTGTGGAGTCGTTGCATCATCAGTACACAGAAGCCCAGNNNAGATCGGAAG
39436 TATGAGTGTGGAGTCGTTGCATAATGGAATAGAAGGCCCCACCNNNAGATCGGAA
39437 TATGAGTGTGGAGTCGTTGCTCTAGTTCTATTGCCCAGGTTGGNNNAGATCGGAAG
39438 TATGAGTGTGGAGTCGTTGCAGAATGAAAAAGCATGCAAGGGTNNNAGATCGGAA

39439 TATGAGTGTGGAGTCGTTGCTGTCCTGTAAGTCCAAGCAATGANNNAGATCGGAAG
39440 TATGAGTGTGGAGTCGTTGCAAAGTCTTGAAAAAAGCCTGGGANNNAGATCGGAA
39441 TATGAGTGTGGAGTCGTTGCCCCCCCACCATTATACTTCTCAGNNNAGATCGGAAGA
39446 TATGAGTGTGGAGTCGTTGCGCAGGAAAACAAAGGGCAGATATNNNAGATCGGA
39451 TATGAGTGTGGAGTCGTTGCGACTGTCTTTCCGTGTATGTTGGNNNAGATCGGAAG
39452 TATGAGTGTGGAGTCGTTGCATTCATCTGTGGTTTTTGGCTCANNNAGATCGGAAG
39453 TATGAGTGTGGAGTCGTTGCGGCTAAGGCAGGAGAATCACTTANNNAGATCGGAA
39454 TATGAGTGTGGAGTCGTTGCTAATTCCAGCTACTCGGGAGGTANNNAGATCGGAA
39456 TATGAGTGTGGAGTCGTTGCTCAGGTGATTCTTATGCCTCAGCNNNAGATCGGAAG
39458 TATGAGTGTGGAGTCGTTGCAGGACACAATCCCATTCCCAATTNNNAGATCGGAAG
39459 TATGAGTGTGGAGTCGTTGCCAAGCAAGCAAGCAAGCAATAATNNNAGATCGGAA
39460 TATGAGTGTGGAGTCGTTGCATGCCCACTTTCACCACTTCTATNNNAGATCGGAAGA
39462 TATGAGTGTGGAGTCGTTGCCCAATGACCAAAGCTGGAACAATNNNAGATCGGAA
39463 TATGAGTGTGGAGTCGTTGCGAAGCAGGAGAATCACATGAACCNNNAGATCGGAA
39464 TATGAGTGTGGAGTCGTTGCTTGCATAGCTATCAGAGGCATCANNNAGATCGGAAG
39465 TATGAGTGTGGAGTCGTTGCAGTTGTATACTGGAACTGGCCTGNNNAGATCGGAA
39466 TATGAGTGTGGAGTCGTTGCTTTTTTGAGACGGAGTCTTGCTGNNNAGATCGGAAG
39467 TATGAGTGTGGAGTCGTTGCACTTCCAGATCTGCCACAAATCTNNNAGATCGGAAG
39468 TATGAGTGTGGAGTCGTTGCGTACCACTGTAGTCCAACCTGAGNNNAGATCGGAAG
39469 TATGAGTGTGGAGTCGTTGCGGCCTTCCCAGATCATTTAGTCTNNNAGATCGGAAG
39470 TATGAGTGTGGAGTCGTTGCTGGGAGTCGTCCTTGAAAACTAGNNNAGATCGGAA
39471 TATGAGTGTGGAGTCGTTGCGAAGACAAGGGGAGGAAGAGATGNNNAGATCGGA
39473 TATGAGTGTGGAGTCGTTGCGCATTCATTGACCTAGGCATGTCNNNAGATCGGAAG
39474 TATGAGTGTGGAGTCGTTGCATCCTGTATCTAGCACTTTGCCANNNAGATCGGAAG
39476 TATGAGTGTGGAGTCGTTGCATGATGGCAGGGAGAGTGAAAATNNNAGATCGGA
39477 TATGAGTGTGGAGTCGTTGCAAGGTTTCTTGCAGCATCATTGTNNNAGATCGGAAG
39478 TATGAGTGTGGAGTCGTTGCTCAGAAGGTTCTTGGGCATGTTANNNAGATCGGAA
39479 TATGAGTGTGGAGTCGTTGCAGCCATGGAAGTAACCAGAATGTNNNAGATCGGAA
39481 TATGAGTGTGGAGTCGTTGCGTTTTATTCTTATCGCCCAGGCCNNNAGATCGGAAG
39482 TATGAGTGTGGAGTCGTTGCGCAAAAAGAACAAAGCTGAAGGCNNNAGATCGGAA
39483 TATGAGTGTGGAGTCGTTGCTTGTTGTGGGGCTTTTTCACTTTNNNAGATCGGAAG
39484 TATGAGTGTGGAGTCGTTGCGATTTCGGCTCACTGCAAACTTCNNNAGATCGGAAG
39485 TATGAGTGTGGAGTCGTTGCTTTCTTTTTAGTTGAGCGTGTGCNNNAGATCGGAAG
39486 TATGAGTGTGGAGTCGTTGCAAACGTCAGGCATGTCTGAATTCNNNAGATCGGAAG
39487 TATGAGTGTGGAGTCGTTGCACTGACATAGACACGGATAGCAGNNNAGATCGGAA
39489 TATGAGTGTGGAGTCGTTGCGATACCAGAGTACCAGAGATGCGNNNAGATCGGAA
39491 TATGAGTGTGGAGTCGTTGCACCAGAGGAGGATAAGGAGACATNNNAGATCGGA
39494 TATGAGTGTGGAGTCGTTGCCTGTGTACATGCCAGCATTTGTTNNNAGATCGGAAG
39495 TATGAGTGTGGAGTCGTTGCGGGGGGATGATAATGGTGGTTATNNNAGATCGGA
39496 TATGAGTGTGGAGTCGTTGCACACCACTTCGTACCTACTAGGANNNAGATCGGAAG
39497 TATGAGTGTGGAGTCGTTGCGAGTGCCCTTGTAGGAAGAAGAANNNAGATCGGAA
39498 TATGAGTGTGGAGTCGTTGCCCAATGGTGGGTATAAGAAGCCTNNNAGATCGGAA
39499 TATGAGTGTGGAGTCGTTGCGGAGGTTGCAGTGAGTCATGATANNNAGATCGGAA

39500 TATGAGTGTGGAGTCGTTGCTGCAACATCATCAGCCATTAGGANNNAGATCGGAAG
39501 TATGAGTGTGGAGTCGTTGCGGCACCTATAATCCCAGCTACTCNNNAGATCGGAAG
39502 TATGAGTGTGGAGTCGTTGCCTTAGTTACTGATGGGGAAGGGGNNNAGATCGGAA
39503 TATGAGTGTGGAGTCGTTGCACCTTCCTGGGATTAGGCTTTTTNNNAGATCGGAAG
39504 TATGAGTGTGGAGTCGTTGCTTGAGGTCAGGAGTTCAAAACCANNNAGATCGGAA
39505 TATGAGTGTGGAGTCGTTGCAAAAGAAAAAAATGGCCGTGGGANNNAGATCGGA
39506 TATGAGTGTGGAGTCGTTGCAGCAACCTATGTGACAGTGTCTCNNNAGATCGGAAG
39507 TATGAGTGTGGAGTCGTTGCCGTTTTCTCTTCCTCTTCCCTGANNNAGATCGGAAGA
39508 TATGAGTGTGGAGTCGTTGCGAGAGAGACAGAAAGACGAAGGANNNAGATCGGA
39509 TATGAGTGTGGAGTCGTTGCCCTAAGGTCAGGAGTTCAAGACCNNNAGATCGGAA
39510 TATGAGTGTGGAGTCGTTGCATAGTCTTCATCTCTGCCTTGGCNNNAGATCGGAAG
39511 TATGAGTGTGGAGTCGTTGCAATCGAAGAACCTAAGGCCATGTNNNAGATCGGAA
39512 TATGAGTGTGGAGTCGTTGCAGTTGCAGTGAGCTGAGATTGTANNNAGATCGGAA
39513 TATGAGTGTGGAGTCGTTGCCATTTGACATGTGGCTGGTGTAANNNAGATCGGAA
39516 TATGAGTGTGGAGTCGTTGCGCCATAGACTTCTTCACAGACCANNNAGATCGGAAG
39519 TATGAGTGTGGAGTCGTTGCGTAGGAGGATCACTTGAGCTCTGNNNAGATCGGAA
39520 TATGAGTGTGGAGTCGTTGCGCTCTACCACATATGAACTGGGANNNAGATCGGAAG
39521 TATGAGTGTGGAGTCGTTGCCACCTGTGCTAAAGCCAGTTATGNNNAGATCGGAAG
39524 TATGAGTGTGGAGTCGTTGCTGGTTCCAAAACATAGAGGAGGGNNNAGATCGGAA
39526 TATGAGTGTGGAGTCGTTGCGTAATCCCAGCTACTATGGAGGCNNNAGATCGGAAG
39527 TATGAGTGTGGAGTCGTTGCCCACACCTATAATCCCAGCACTTNNNAGATCGGAAGA
39528 TATGAGTGTGGAGTCGTTGCGGGATCAATGCAACAAGAAGAGCNNNAGATCGGAA
39529 TATGAGTGTGGAGTCGTTGCGATGCCCTCTCTCACTACTCCTANNNAGATCGGAAGA
39530 TATGAGTGTGGAGTCGTTGCCCTACTGAAGAGTGTCCTTGGACNNNAGATCGGAAG
39531 TATGAGTGTGGAGTCGTTGCCTTACACCTGTAATCCCAGCACTNNNAGATCGGAAGA
39532 TATGAGTGTGGAGTCGTTGCCTAAGGGCAGGAGAATCACTTGANNNAGATCGGAA
39533 TATGAGTGTGGAGTCGTTGCTACAAACACACACACACACACACNNNAGATCGGAAG
39535 TATGAGTGTGGAGTCGTTGCTACAAGAATGGACACTGTGCTCANNNAGATCGGAAG
39536 TATGAGTGTGGAGTCGTTGCGGATAGGTGGGAACAAGTTGACANNNAGATCGGA
39539 TATGAGTGTGGAGTCGTTGCGGTAAAACCAAGGGAGTATGGGTNNNAGATCGGA
39540 TATGAGTGTGGAGTCGTTGCAAGGTGGAAGAATCTGAGCAAGANNNAGATCGGA
39541 TATGAGTGTGGAGTCGTTGCAGACCCAGCAGCAGAAATAATCANNNAGATCGGAA
39542 TATGAGTGTGGAGTCGTTGCCTGTAGTCCCAGCTACTCAAGAGNNNAGATCGGAAG
39544 TATGAGTGTGGAGTCGTTGCCCATAGCAAGACTCTGTCTCACANNNAGATCGGAAG
39546 TATGAGTGTGGAGTCGTTGCGACAGAGTCTCACTTCTGTCACCNNNAGATCGGAAG
39549 TATGAGTGTGGAGTCGTTGCAAAAACACACATTGCTTTGCAGTNNNAGATCGGAAG
39550 TATGAGTGTGGAGTCGTTGCGCCTCCTGAGTAACTGTGACTACNNNAGATCGGAAG
39551 TATGAGTGTGGAGTCGTTGCAAGAAAGAAGACGTAAAGGCCCTNNNAGATCGGAA
39552 TATGAGTGTGGAGTCGTTGCCCCAGGATCTTACAAGACTGAGGNNNAGATCGGAA
39554 TATGAGTGTGGAGTCGTTGCAAGGAAGTTTGGCAAGATCATGANNNAGATCGGAA
39555 TATGAGTGTGGAGTCGTTGCACACGTGGGAATTGAAGATGAGANNNAGATCGGA
39556 TATGAGTGTGGAGTCGTTGCGCCACCAACCCAGCTAATTTTTTNNNAGATCGGAAGA
39557 TATGAGTGTGGAGTCGTTGCACCGTCCTGTTCATTTTACCTGANNNAGATCGGAAGA

39558 TATGAGTGTGGAGTCGTTGCGGAGTGACTGAGCAATGAAACAANNNAGATCGGAA
39561 TATGAGTGTGGAGTCGTTGCAAAAGAACTACCCTGAGACTGGGNNNAGATCGGAA
39562 TATGAGTGTGGAGTCGTTGCCTGTACTCCCAGCTATTCAGGTGNNNAGATCGGAAG
39564 TATGAGTGTGGAGTCGTTGCGTCTGTAAATGACCCCCTTAGCANNNAGATCGGAAG
39565 TATGAGTGTGGAGTCGTTGCTTTGCAATTGCTGTCTTCCCTTTNNNAGATCGGAAGA
39566 TATGAGTGTGGAGTCGTTGCCCTTTACTGACCAACTCTCCTCCNNNAGATCGGAAGA
39568 TATGAGTGTGGAGTCGTTGCTAATGGGATGTATGGAGGCATGGNNNAGATCGGA
39569 TATGAGTGTGGAGTCGTTGCTAGGAGGTAGTCAAGAGTCAGGCNNNAGATCGGA
39570 TATGAGTGTGGAGTCGTTGCATGGAATAGTGATTACCAGGGGCNNNAGATCGGAA
39571 TATGAGTGTGGAGTCGTTGCAAGCAGGTGTAGCAATCCTAGTCNNNAGATCGGAA
39573 TATGAGTGTGGAGTCGTTGCTCGATGACCTAACCACAATCCTGNNNAGATCGGAAG
39576 TATGAGTGTGGAGTCGTTGCGCAAAAGACTTGAAGAGGCACTTNNNAGATCGGAA
39577 TATGAGTGTGGAGTCGTTGCTTCAGGCAGTGATGACCTGTTTANNNAGATCGGAAG
39578 TATGAGTGTGGAGTCGTTGCCAAGATGCAGCTAAAGCAGTGTTNNNAGATCGGAA
39581 TATGAGTGTGGAGTCGTTGCAGAAAGAAAGAGAGAGAGGGGGANNNAGATCGG
39582 TATGAGTGTGGAGTCGTTGCCTGAGGCAGGAGAATCACTTGAANNNAGATCGGAA
39583 TATGAGTGTGGAGTCGTTGCAAAAGTTGTCAGATCACTTGCCANNNAGATCGGAAG
39585 TATGAGTGTGGAGTCGTTGCTCCATTACCACATTCAAGGTCATNNNAGATCGGAAG
39586 TATGAGTGTGGAGTCGTTGCATGTGGGAAAAGAGGTGAAAGGANNNAGATCGGA
39587 TATGAGTGTGGAGTCGTTGCTCCTGACTGACTGTTTTCTAGCCNNNAGATCGGAAG
39588 TATGAGTGTGGAGTCGTTGCACTATCATTCGATCCAGCAACCCNNNAGATCGGAAG
39589 TATGAGTGTGGAGTCGTTGCAGAAACTGAATCCGAGAGGCTTTNNNAGATCGGAA
39590 TATGAGTGTGGAGTCGTTGCAGCCAGAAAAGTGACTCAGACTTNNNAGATCGGAA
39591 TATGAGTGTGGAGTCGTTGCTGGGTCTAGGTTAGAAGTGGGATNNNAGATCGGA
39592 TATGAGTGTGGAGTCGTTGCGGAAAAAAGAAAGCTCATCCGGGNNNAGATCGGAA
39593 TATGAGTGTGGAGTCGTTGCGTGAATCACTTGAGGTCAGGAGTNNNAGATCGGAA
39594 TATGAGTGTGGAGTCGTTGCGTCATGCTCTATCCCCTTTCCTTNNNAGATCGGAAGA
39595 TATGAGTGTGGAGTCGTTGCTGGACAGTAACAAGTGTTGGTGANNNAGATCGGAA
39597 TATGAGTGTGGAGTCGTTGCGATTCTCCGATTCTCCAAGGTCANNNAGATCGGAAG
39598 TATGAGTGTGGAGTCGTTGCTGTACACAGTTGGGTAGCTTGTANNNAGATCGGAA
39599 TATGAGTGTGGAGTCGTTGCAAAGGAAGTCAATCAATGTGGCANNNAGATCGGAA
39600 TATGAGTGTGGAGTCGTTGCAGTGCTGATCATGCAGAACCTTANNNAGATCGGAAG
39602 TATGAGTGTGGAGTCGTTGCAACATTCTTAGACCACAGTGGCTNNNAGATCGGAAG
39604 TATGAGTGTGGAGTCGTTGCTAAAACATTGAAGCCATGCGTGTNNNAGATCGGAA
39605 TATGAGTGTGGAGTCGTTGCCCCAGGGGAAGGAAATCTCTAACNNNAGATCGGAA
39606 TATGAGTGTGGAGTCGTTGCCGAAAGAAGGAAGGAAGGAAGGANNNAGATCGG
39607 TATGAGTGTGGAGTCGTTGCCAGGAGTATTGCTCACATCCACTNNNAGATCGGAAG
39608 TATGAGTGTGGAGTCGTTGCCCCAGGTGATTCTAATGTAGCCANNNAGATCGGAAG
39609 TATGAGTGTGGAGTCGTTGCCATGCCTGTAATCCCAGCACTATNNNAGATCGGAAG
39610 TATGAGTGTGGAGTCGTTGCGAGGTAGCAGTGAGGAGAGATTGNNNAGATCGGA
39611 TATGAGTGTGGAGTCGTTGCCCACCATGCCAGGCTAATTTTTANNNAGATCGGAAG
39612 TATGAGTGTGGAGTCGTTGCTGGGGTTAGGACAGGAAAATGAGNNNAGATCGG
39613 TATGAGTGTGGAGTCGTTGCCCTGTGATGTGCTAGGAACCATANNNAGATCGGAA

39614 TATGAGTGTGGAGTCGTTGCAGGAAGAGGTCTACAGAACAAGTNNNAGATCGGAA
39616 TATGAGTGTGGAGTCGTTGCAACAAAAAAAGCCACCCAACTGANNNAGATCGGAAG
39617 TATGAGTGTGGAGTCGTTGCTGAGGCAGGAGAATCTCTTGAACNNNAGATCGGAA
39618 TATGAGTGTGGAGTCGTTGCCTGAGGTACTGGGTTCATCTCACNNNAGATCGGAAG
39619 TATGAGTGTGGAGTCGTTGCAACTTGTTCAAGGTCACACTTGTNNNAGATCGGAAG
39620 TATGAGTGTGGAGTCGTTGCGCTGCCCAAGTATAACAGGAAAGNNNAGATCGGAA
39621 TATGAGTGTGGAGTCGTTGCGGAGTTGCAATCCTAGTCTCTGANNNAGATCGGAAG
39622 TATGAGTGTGGAGTCGTTGCTCAAAAGCTAGCAGAAGACAAGANNNAGATCGGAA
39623 TATGAGTGTGGAGTCGTTGCAACCAGACACATAGACCAATGGANNNAGATCGGAAG
39624 TATGAGTGTGGAGTCGTTGCAAATCTGAACCCTCGAACATTGCNNNAGATCGGAAG
39625 TATGAGTGTGGAGTCGTTGCACAGAAAAGGGGTAGCCTAGAGANNNAGATCGGAA
39627 TATGAGTGTGGAGTCGTTGCCATCCTGGCTAACACAGTGAAACNNNAGATCGGAAG
39628 TATGAGTGTGGAGTCGTTGCTAGGTTTCCACTTTGTTGTTGGTNNNAGATCGGAAG
39629 TATGAGTGTGGAGTCGTTGCGCAAGGTAGAGCCATACTAGTCCNNNAGATCGGAAG
39630 TATGAGTGTGGAGTCGTTGCGGAACAAACTAAGGATGCTTGGTNNNAGATCGGAA
39631 TATGAGTGTGGAGTCGTTGCGCAACCTAGATGAAATGGGCAAANNNAGATCGGAA
39634 TATGAGTGTGGAGTCGTTGCAAGACCTACATCACCATCTCTGCNNNAGATCGGAAG
39636 TATGAGTGTGGAGTCGTTGCCCTCGTGATTTCTTCTTTGCCTCNNNAGATCGGAAGA
39637 TATGAGTGTGGAGTCGTTGCTAACCAAAAGAGAGCTGGAGTGGNNNAGATCGGAA
39638 TATGAGTGTGGAGTCGTTGCTGGAATACCGTTTCACCCATCTTNNNAGATCGGAAG
39640 TATGAGTGTGGAGTCGTTGCAATGCCAGCATCACTTATCTTGCNNNAGATCGGAAG
39641 TATGAGTGTGGAGTCGTTGCCTGTCATTTGCACCAACATGGATNNNAGATCGGAAG
39642 TATGAGTGTGGAGTCGTTGCTTCAAGTGATTCTCCTGCCTCAGNNNAGATCGGAAG
39643 TATGAGTGTGGAGTCGTTGCACCAAAAAAGGAGAGGGGGAAATNNNAGATCGGA
39644 TATGAGTGTGGAGTCGTTGCGAGGGTATTGGGGGAGTCAAAATNNNAGATCGGA
39645 TATGAGTGTGGAGTCGTTGCGCCAATATCCCTGATGAACAACGNNNAGATCGGAAG
39646 TATGAGTGTGGAGTCGTTGCATATTCCCCTGAATGTCTGCCAGNNNAGATCGGAAG
39647 TATGAGTGTGGAGTCGTTGCAATGCTTATACACTGCTGGTGGANNNAGATCGGAAG
39648 TATGAGTGTGGAGTCGTTGCACTGCATGGTTTCACTTACATGTNNNAGATCGGAAG
39649 TATGAGTGTGGAGTCGTTGCAAGAAGAAAGCAAGCAAGCAAGTNNNAGATCGGAA
39651 TATGAGTGTGGAGTCGTTGCCCAAACAAACAAAAAACTGGCCCNNNAGATCGGAAG
39653 TATGAGTGTGGAGTCGTTGCTCGCAGACTGAAAGAGACAAAGANNNAGATCGGAA
39654 TATGAGTGTGGAGTCGTTGCGCCTAGCTTCTCAGAGTTCCATTNNNAGATCGGAAG
39656 TATGAGTGTGGAGTCGTTGCCCTGTTTTACAAGGTTGCAGGTTNNNAGATCGGAAG
39657 TATGAGTGTGGAGTCGTTGCTGCCCAATTTCTGCTGAATATGCNNNAGATCGGAAG
39658 TATGAGTGTGGAGTCGTTGCATGGCACTGTTTTTCTAACCTGCNNNAGATCGGAAG
39660 TATGAGTGTGGAGTCGTTGCCTGAGACAGGAGGATTGCTTGAGNNNAGATCGGAA
39661 TATGAGTGTGGAGTCGTTGCCTCCTGAGTAGCTGGGACTACANNNAGATCGGAAG
39662 TATGAGTGTGGAGTCGTTGCACAGCTTTACACAAAAATGCACCNNNAGATCGGAAG
39663 TATGAGTGTGGAGTCGTTGCAGAATGGAGATTCCTTGCACACTNNNAGATCGGAAG
39665 TATGAGTGTGGAGTCGTTGCGTAACCACACATCAGAACAGTGCNNNAGATCGGAAG
39667 TATGAGTGTGGAGTCGTTGCAAGCAGCTTTTGAAAAAGGGGAGNNNAGATCGGAA
39668 TATGAGTGTGGAGTCGTTGCTAGGTTGCAATGAGAGGAGATCGNNNAGATCGGAA

39669 TATGAGTGTGGAGTCGTTGCAAGAACTGACACCCTAGCTGCATNNNAGATCGGAAG
39672 TATGAGTGTGGAGTCGTTGCTCCAGTTCATTTCCAAGCCAAACNNNAGATCGGAAG
39673 TATGAGTGTGGAGTCGTTGCATGATTGCACCACTGTACTCCAGNNNAGATCGGAAG
39674 TATGAGTGTGGAGTCGTTGCTCCTGGACTTGGAGTTGAAGAAGNNNAGATCGGAA
39675 TATGAGTGTGGAGTCGTTGCAAAACAAAAACACATGCACACGTNNNAGATCGGAAG
39676 TATGAGTGTGGAGTCGTTGCCAATTGCACCGGGAATTAGGAAGNNNAGATCGGAA
39677 TATGAGTGTGGAGTCGTTGCGAAGGAAGGAAGGAAGGAAGGAGNNNAGATCGG
39678 TATGAGTGTGGAGTCGTTGCGGATGGAGTACCACATAGGCATTNNNAGATCGGAA
39679 TATGAGTGTGGAGTCGTTGCCAACATCATTCATTTACCGGCCANNNAGATCGGAAG
39680 TATGAGTGTGGAGTCGTTGCTGCTGTTCTTTTCCTTCACAACANNNAGATCGGAAGA
39681 TATGAGTGTGGAGTCGTTGCCACTCCTATATTCCTTGCACCGTNNNAGATCGGAAGA
39683 TATGAGTGTGGAGTCGTTGCGCCAGCATCACCTTGATAACAAANNNAGATCGGAAG
39684 TATGAGTGTGGAGTCGTTGCGAGTCTCACTCTATCACCCATGCNNNAGATCGGAAG
39685 TATGAGTGTGGAGTCGTTGCCTGCGGACACAAAGATAATTCCCNNNAGATCGGAAG
39686 TATGAGTGTGGAGTCGTTGCCTTGTAGTCCCAGCTACGTGAAANNNAGATCGGAAG
39688 TATGAGTGTGGAGTCGTTGCCACAGGGCCAATTAGACATTTGGNNNAGATCGGAA
39689 TATGAGTGTGGAGTCGTTGCGGGCTGAAAAAACTGTGAACAGANNNAGATCGGAA
39690 TATGAGTGTGGAGTCGTTGCTGTATGAGAACAGGATGACCAGCNNNAGATCGGAA
39691 TATGAGTGTGGAGTCGTTGCGTTTATGGAAACTTGGGGCTGTTNNNAGATCGGAA
39693 TATGAGTGTGGAGTCGTTGCATGCCTCCAGTCTCACTGAAATTNNNAGATCGGAAG
39695 TATGAGTGTGGAGTCGTTGCAAAAGTCTCTCCATCCAGTGTCANNNAGATCGGAAG
39696 TATGAGTGTGGAGTCGTTGCAGTCTACCAATGAATGCCACTGTNNNAGATCGGAAG
39697 TATGAGTGTGGAGTCGTTGCAGAAAGGTGGTTACCAGAGACTGNNNAGATCGGAA
39698 TATGAGTGTGGAGTCGTTGCCAACAGAGCAAGACTCCATCTCANNNAGATCGGAAG
39699 TATGAGTGTGGAGTCGTTGCGGTTGTGAAGACCATGGGAAAATNNNAGATCGGAA
39700 TATGAGTGTGGAGTCGTTGCCTATTCCTGGTAGGGGCTCTTTCNNNAGATCGGAAG
39701 TATGAGTGTGGAGTCGTTGCGAAAGTAACCACAGACATGCACANNNAGATCGGAA
39702 TATGAGTGTGGAGTCGTTGCTGTCAGCTCTCCATTTTGTCTCANNNAGATCGGAAGA
39703 TATGAGTGTGGAGTCGTTGCCAGCGGCTACTACTCTATTCCAGNNNAGATCGGAAG
39704 TATGAGTGTGGAGTCGTTGCAGGCACATAACACATGAAAACCANNNAGATCGGAA
39705 TATGAGTGTGGAGTCGTTGCGACATGTGACTCATACCCAACCTNNNAGATCGGAAG
39706 TATGAGTGTGGAGTCGTTGCAATAGCCATTGTGACTGGTGTGANNNAGATCGGAA
39708 TATGAGTGTGGAGTCGTTGCAGATTCTCAGGTCCCACTACAGANNNAGATCGGAAG
39709 TATGAGTGTGGAGTCGTTGCCAAAGTGACAGTAACCAAGTGGCNNNAGATCGGAA
39710 TATGAGTGTGGAGTCGTTGCCTCTGAGTTCCATGGCATACACTNNNAGATCGGAAG
39712 TATGAGTGTGGAGTCGTTGCAACAAGATGAATAGCAGAGGCCANNNAGATCGGAA
39714 TATGAGTGTGGAGTCGTTGCAACTAGCCAAAGAACTAGCCCTCNNNAGATCGGAAG
39715 TATGAGTGTGGAGTCGTTGCTCAGGATTGTTTCCGAAAGGTTGNNNAGATCGGAA
39716 TATGAGTGTGGAGTCGTTGCAGTCAGGATTGTTTCCGAAAGGTNNNAGATCGGAA
39717 TATGAGTGTGGAGTCGTTGCGAATCTGTGTCCTTGATGGCTTTNNNAGATCGGAAG
39718 TATGAGTGTGGAGTCGTTGCCTGCTGGAGACATCTCTTCTGTTNNNAGATCGGAAG
39719 TATGAGTGTGGAGTCGTTGCTGGCACATTGAATTACCTACACTNNNAGATCGGAAG
39722 TATGAGTGTGGAGTCGTTGCAGCCTCTGGGATTGTACTTTTGTNNNAGATCGGAAG

39724 TATGAGTGTGGAGTCGTTGCGAGGTCAGGAGTTCAAGACCAAGNNNAGATCGGAA
39725 TATGAGTGTGGAGTCGTTGCAGAACATAAACTCCAGCACCAGANNNAGATCGGAAG
39731 TATGAGTGTGGAGTCGTTGCAACATAAAGCCAGGAAGCTAGGANNNAGATCGGAA
39732 TATGAGTGTGGAGTCGTTGCATTGCTGATGCTACCCTCAGTTTNNNAGATCGGAAG
39733 TATGAGTGTGGAGTCGTTGCGCACTTTGGGAGATCTAGGTGAANNNAGATCGGAA
39735 TATGAGTGTGGAGTCGTTGCATATCACTGCAGCCTTTACCTCCNNNAGATCGGAAGA
39736 TATGAGTGTGGAGTCGTTGCGGTAATGTGACTTGCCTGAGTTCNNNAGATCGGAAG
39737 TATGAGTGTGGAGTCGTTGCGGAAAACCGGGGGAGTTATTTTANNNAGATCGGAA
39738 TATGAGTGTGGAGTCGTTGCGGAGAATGGTATGAACCCAGGAGNNNAGATCGGA
39739 TATGAGTGTGGAGTCGTTGCGGTAGACGGGGCTAGTTATGAAGNNNAGATCGGA
39741 TATGAGTGTGGAGTCGTTGCGAATCGCTTGAACCTGGAAGTTGNNNAGATCGGAA
39743 TATGAGTGTGGAGTCGTTGCGTTTCAGTTCCTTTCACATGGAGTNNNAGATCGGAA
39747 TATGAGTGTGGAGTCGTTGCTGAAAGAAGTGAGAGTCAAAAGGCNNNAGATCGG
39748 TATGAGTGTGGAGTCGTTGCGACTTCCCAACTGATGTCTCCAGNNNAGATCGGAAG
39750 TATGAGTGTGGAGTCGTTGCTGACCGACACCACATTCAAGATANNNAGATCGGAAG
39752 TATGAGTGTGGAGTCGTTGCAACCACTAAGATGGGGAGAAACCNNNAGATCGGAA
39753 TATGAGTGTGGAGTCGTTGCGACAGACTTCTTCAAGGGGCATANNNAGATCGGAAG
39754 TATGAGTGTGGAGTCGTTGCAGGAGCTCACAATAAAGTGCAGANNNAGATCGGAA
39755 TATGAGTGTGGAGTCGTTGCTGGAAACCAGTGGCTCATCATAANNNAGATCGGAAG
39756 TATGAGTGTGGAGTCGTTGCCACTGAGTCAAGATCACGTCACTNNNAGATCGGAAG
39758 TATGAGTGTGGAGTCGTTGCGATCAAGAGGTCAGGAGATCGTGNNNAGATCGGA
39759 TATGAGTGTGGAGTCGTTGCTTGCAGAGTCAGTCTGATTTCCANNNAGATCGGAAG
39760 TATGAGTGTGGAGTCGTTGCCAAGGACAGCTGTATTTTGCCATNNNAGATCGGAAG
39761 TATGAGTGTGGAGTCGTTGCGGGAATTGGTTTTGATGGTAGCCNNNAGATCGGAA
39762 TATGAGTGTGGAGTCGTTGCAAATGGCAAACAGGCATACGAAANNNAGATCGGAA
39763 TATGAGTGTGGAGTCGTTGCACTCCAACATTTAGTGGGTGGAANNNAGATCGGAAG
39764 TATGAGTGTGGAGTCGTTGCAGTTTACATGCCACAGTTACAGTNNNAGATCGGAAG
39765 TATGAGTGTGGAGTCGTTGCCTGTAGTCCCAGCTACTCAGGAGNNNAGATCGGAAG
39766 TATGAGTGTGGAGTCGTTGCGGGTTTGAGTGCCAATAGGAGTANNNAGATCGGAA
39769 TATGAGTGTGGAGTCGTTGCAGAGATGACATGAAGTTAGGACTGANNNAGATCGG
39770 TATGAGTGTGGAGTCGTTGCGGCCTCTTCAAGGAGAACTACAANNNAGATCGGAAG
39771 TATGAGTGTGGAGTCGTTGCCTAGGTGACAGAACGAGACTCTGNNNAGATCGGAA
39772 TATGAGTGTGGAGTCGTTGCGAGGAGATATCTGCTCTGCCTTCNNNAGATCGGAAG
39773 TATGAGTGTGGAGTCGTTGCTGAAGGCCACAAATAGTATTGCANNNAGATCGGAAG
39775 TATGAGTGTGGAGTCGTTGCTATATTCCTTGTAGCCCCCAGGANNNAGATCGGAAG
39779 TATGAGTGTGGAGTCGTTGCCCCAGAAACAAATCCACACTTCANNNAGATCGGAAG
39780 TATGAGTGTGGAGTCGTTGCAACAAAACAGCATGGTACTGACANNNAGATCGGAAG
39782 TATGAGTGTGGAGTCGTTGCAGCCAAAAAGTGGAAACAACACANNNAGATCGGAA
39783 TATGAGTGTGGAGTCGTTGCCCTTCAAGTCCTAGTAAGGCTGGNNNAGATCGGAAG
39784 TATGAGTGTGGAGTCGTTGCCTGTAATCCCAGCTACTCAGGAGNNNAGATCGGAAG
39785 TATGAGTGTGGAGTCGTTGCGGAAAAGCAAATCAGAGCCACAANNNAGATCGGAA
39786 TATGAGTGTGGAGTCGTTGCTCCTTGTGGACAAAGGATGGATANNNAGATCGGAA
39787 TATGAGTGTGGAGTCGTTGCGTTCTGGTTCAGGAAAGCTTGACNNNAGATCGGAA

39789 TATGAGTGTGGAGTCGTTGCATAATTGTTTGAACCTGGCAGGCNNNAGATCGGAAG
39790 TATGAGTGTGGAGTCGTTGCCATATCTGACAGTCACAGAGGGGNNNAGATCGGAA
39791 TATGAGTGTGGAGTCGTTGCTTAGTTCAAAGCACTCAGCATGCNNNAGATCGGAAG
39792 TATGAGTGTGGAGTCGTTGCAGGAGCTGGTACCATTTCTTCTGNNNAGATCGGAAG
39793 TATGAGTGTGGAGTCGTTGCGAGGGTTGGATGGATCAGTACAGNNNAGATCGGA
39794 TATGAGTGTGGAGTCGTTGCGTGAAGAAATGCCAGGCTCTTTTNNNAGATCGGAAG
39795 TATGAGTGTGGAGTCGTTGCAAAAGGAGTCTCTGGTCTCATGGNNNAGATCGGAA
39798 TATGAGTGTGGAGTCGTTGCAATGTAAGCTTGATGAGGGCAGANNNAGATCGGAA
39799 TATGAGTGTGGAGTCGTTGCCAGGAGTTGCAATCGTAGTCTCTNNNAGATCGGAAG
39800 TATGAGTGTGGAGTCGTTGCAAGTGGGAACTGGAAATGGGATTNNNAGATCGGA
39801 TATGAGTGTGGAGTCGTTGCTTCCGCCATTAGTAAAAGCTCCTNNNAGATCGGAAG
39802 TATGAGTGTGGAGTCGTTGCACAGAAACTCACCTGTTATGTGTNNNAGATCGGAAG
39803 TATGAGTGTGGAGTCGTTGCCGTTATGGGACTGTAAATGGTGCNNNAGATCGGAA
39804 TATGAGTGTGGAGTCGTTGCACACCTGACTTACACCACTAAGGNNNAGATCGGAAG
39806 TATGAGTGTGGAGTCGTTGCGAGCTTTGGAACACACTGGTTTTNNNAGATCGGAAG
39807 TATGAGTGTGGAGTCGTTGCGGAAAGTATGGCATAAAGGCAGGNNNAGATCGGA
39808 TATGAGTGTGGAGTCGTTGCTTTGGGACTAGAGATGCTCAACCNNNAGATCGGAAG
39810 TATGAGTGTGGAGTCGTTGCAAGTATTTGCTAATAGCTTGGCAANNNAGATCGGAA
39811 TATGAGTGTGGAGTCGTTGCTTTCCTCCACTGATGTCTTGAGCNNNAGATCGGAAG
39812 TATGAGTGTGGAGTCGTTGCACAGGCCTATCCATTTTTCTCGTNNNAGATCGGAAGA
39813 TATGAGTGTGGAGTCGTTGCTGATACGTTTGTCCACATGGAGANNNAGATCGGAAG
39814 TATGAGTGTGGAGTCGTTGCACAGCCAGCCAAGAATCATTAGANNNAGATCGGAAG
39815 TATGAGTGTGGAGTCGTTGCTCATTTGGGTTGCTCCTTTCTCTNNNAGATCGGAAGA
39816 TATGAGTGTGGAGTCGTTGCTCAGAACATCCTCTCTCTGGGAANNNAGATCGGAAG
39817 TATGAGTGTGGAGTCGTTGCCTGAGTAGCTGGGACTACAACTGNNNAGATCGGAA
39820 TATGAGTGTGGAGTCGTTGCGACACTAGGGATGACTTGACTGCNNNAGATCGGAA
39822 TATGAGTGTGGAGTCGTTGCACCTTCCTCAGTACCCAGACTATNNNAGATCGGAAG
39823 TATGAGTGTGGAGTCGTTGCTTACAAGCATGAGCCACTGTACTNNNAGATCGGAAG
39824 TATGAGTGTGGAGTCGTTGCCTCATCACACCCGAGTCAAGAAANNNAGATCGGAAG
39825 TATGAGTGTGGAGTCGTTGCGGAAGGTAGGAAGGTACTCAAATNNNAGATCGGA
39827 TATGAGTGTGGAGTCGTTGCTGATTCACTACACAAGAATTTGCANNNAGATCGGAA
39828 TATGAGTGTGGAGTCGTTGCCCACCACGTCTGGCTAATTTTTTNNNAGATCGGAAG
39830 TATGAGTGTGGAGTCGTTGCAGCCAATGTAATCCACTCCATCCNNNAGATCGGAAG
39832 TATGAGTGTGGAGTCGTTGCCTGTGGGCCCTTTTTGTGTAAATNNNAGATCGGAAG
39833 TATGAGTGTGGAGTCGTTGCCTGTAGTCCCAGCTACTCAGGAGNNNAGATCGGAAG
39834 TATGAGTGTGGAGTCGTTGCAAAAAGAAAAAAATTGGCCGGCCNNNAGATCGGAA
39835 TATGAGTGTGGAGTCGTTGCGAATGGAGACACCTGTAGCGTATNNNAGATCGGAA
39836 TATGAGTGTGGAGTCGTTGCAGGAGAATCACTGGAACCCAAAANNNAGATCGGAA
39838 TATGAGTGTGGAGTCGTTGCTGAGTGAAGAAGAGAATGACCGGNNNAGATCGGA
39841 TATGAGTGTGGAGTCGTTGCTTCTTGCTTTTCCACCTTGATGTNNNAGATCGGAAGA
39844 TATGAGTGTGGAGTCGTTGCAGAGAGGTGGTTCAATTAGAGGCNNNAGATCGGA
39845 TATGAGTGTGGAGTCGTTGCTGGAAACACATCTCATGCTCATGNNNAGATCGGAAG
39846 TATGAGTGTGGAGTCGTTGCCCAAGCAAACGGAAACCAAAAAGNNNAGATCGGAA

39847 TATGAGTGTGGAGTCGTTGCTTCAAGATCCTTGGTGCAGAGAANNNAGATCGGAA
39848 TATGAGTGTGGAGTCGTTGCATTGTTTACTGCCGTGTGACTTCNNNAGATCGGAAG
39849 TATGAGTGTGGAGTCGTTGCAATGTTACTTTCTGACAGGCCCANNNAGATCGGAAG
39850 TATGAGTGTGGAGTCGTTGCCCTTTTCCAGACCACTAGTCACANNNAGATCGGAAG
39851 TATGAGTGTGGAGTCGTTGCAAAGATGTACTGTGACATGCCCANNNAGATCGGAAG
39852 TATGAGTGTGGAGTCGTTGCAGGAAAGACCATTGTGATGAGGTNNNAGATCGGAA
39854 TATGAGTGTGGAGTCGTTGCGACCGATGGAACAGAGTAGAGAGNNNAGATCGGA
39855 TATGAGTGTGGAGTCGTTGCATTCAAGTGATTCTCCTGCCTCANNNAGATCGGAAG
39856 TATGAGTGTGGAGTCGTTGCTTGTGTAGGGCTACTATGCTGTGNNNAGATCGGAA
39857 TATGAGTGTGGAGTCGTTGCGCAAACATGGACAAATGGGATCANNNAGATCGGAA
39858 TATGAGTGTGGAGTCGTTGCTGCAGACCCAATGTGTAAAATGGNNNAGATCGGAA
39859 TATGAGTGTGGAGTCGTTGCTACTGATTCAGATGAGTTGGCCCNNNAGATCGGAAG
39862 TATGAGTGTGGAGTCGTTGCTTTGCATATGAATGTCTTGGGCANNNAGATCGGAAG
39863 TATGAGTGTGGAGTCGTTGCATACTGTGGTCCTTTTTGCCTCTNNNAGATCGGAAG
39867 TATGAGTGTGGAGTCGTTGCAGTTACTTGAGAAGCTGAGGTGGNNNAGATCGGAA
39868 TATGAGTGTGGAGTCGTTGCTCTGTTTCAAGGCTCAGACCATTNNNAGATCGGAAG
39869 TATGAGTGTGGAGTCGTTGCGAGGTCAGGAGATTGAGACCATCNNNAGATCGGAA
39870 TATGAGTGTGGAGTCGTTGCCTTTCCATCTTGCTTGTCCACTGNNNAGATCGGAAGA
39871 TATGAGTGTGGAGTCGTTGCTTGAAGCCAGGAGTTTGAGATCANNNAGATCGGAA
39872 TATGAGTGTGGAGTCGTTGCAAGTGAAGATCATCTGCCACTGTNNNAGATCGGAAG
39874 TATGAGTGTGGAGTCGTTGCCACATATGTGGTCCCAGCTACTTNNNAGATCGGAAG
39880 TATGAGTGTGGAGTCGTTGCAACGCAATCCCCATCAAAATACCNNNAGATCGGAAG
39882 TATGAGTGTGGAGTCGTTGCCCTCATGGAGCTTACTTCATGGTNNNAGATCGGAAG
39883 TATGAGTGTGGAGTCGTTGCCAACGTTGCAGTGAGCTATGATTNNNAGATCGGAAG
39884 TATGAGTGTGGAGTCGTTGCCTATACAAGTGGCCCTGTGCTATNNNAGATCGGAAG
39885 TATGAGTGTGGAGTCGTTGCGTGGCTCACTCCTATAATCCCAGNNNAGATCGGAAG
39886 TATGAGTGTGGAGTCGTTGCTGAGATACCACTTCACATCCACTNNNAGATCGGAAG
39887 TATGAGTGTGGAGTCGTTGCTCTTTGTACAGAAGGCAAGACCANNNAGATCGGAAG
39888 TATGAGTGTGGAGTCGTTGCAAGGATATGAGGACTTGGTGCTGNNNAGATCGGAA
39890 TATGAGTGTGGAGTCGTTGCTCCAAGATAGGAGAGCCCAAGTANNNAGATCGGAA
39891 TATGAGTGTGGAGTCGTTGCCACTTGTAGTCCCAGTTACTCGGNNNAGATCGGAAG
39892 TATGAGTGTGGAGTCGTTGCTGGCTCATGACTATAATCCCAGCNNNAGATCGGAAG
39893 TATGAGTGTGGAGTCGTTGCCAAATGAATTCCCCAGCCTTCTGNNNAGATCGGAAG
39894 TATGAGTGTGGAGTCGTTGCCATCTCCTGGGATCAAGGGATTCNNNAGATCGGAAG
39895 TATGAGTGTGGAGTCGTTGCTATACAGCAAATCCCAAGAGCGTNNNAGATCGGAAG
39896 TATGAGTGTGGAGTCGTTGCAGGCAAGTCACTGATAGAACTGGNNNAGATCGGAA
39897 TATGAGTGTGGAGTCGTTGCTAGAGGATCATGACTGTGTCCCTNNNAGATCGGAAG
39899 TATGAGTGTGGAGTCGTTGCCACTCTATTGCTCAGGTTGGAGTNNNAGATCGGAAG
39901 TATGAGTGTGGAGTCGTTGCAAGGGAAAGGGAAAAGGAAAGGANNNAGATCGG
39904 TATGAGTGTGGAGTCGTTGCGGTGGGAACAGATGTGTTTCATGNNNAGATCGGAA
39905 TATGAGTGTGGAGTCGTTGCACAGATTACACTGAAAGGGGACANNNAGATCGGAA
39908 TATGAGTGTGGAGTCGTTGCAACATCAATTCCTCTGGACACCANNNAGATCGGAAG
39909 TATGAGTGTGGAGTCGTTGCGAGAGTGAGGATTCTAGAGCAGCNNNAGATCGGAA

39910 TATGAGTGTGGAGTCGTTGCCCTGCATGTCCCTCTAGTTTTCANNNAGATCGGAAGA
39911 TATGAGTGTGGAGTCGTTGCGAAACTGGTGATGAGCAACTTCCNNNAGATCGGAA
39912 TATGAGTGTGGAGTCGTTGCTCAAACACATTGTGGTCAGAGGANNNAGATCGGAA
39913 TATGAGTGTGGAGTCGTTGCAATAGGTCAGCCACTGTGTATCCNNNAGATCGGAAG
39914 TATGAGTGTGGAGTCGTTGCTCCGTATTTTATGGTTGGGGGTTNNNAGATCGGAA
39916 TATGAGTGTGGAGTCGTTGCAAAAGAAAAGAAAGTGGGCGTGGNNNAGATCGGA
39917 TATGAGTGTGGAGTCGTTGCTTTGCACCACTATATTCCAGCCTNNNAGATCGGAAGA
39918 TATGAGTGTGGAGTCGTTGCGAGGAAGGAAGGAAGGAAGGAAGNNNAGATCGG
39919 TATGAGTGTGGAGTCGTTGCAGAGTTGGAGAACTGACACTTCCNNNAGATCGGAA
39921 TATGAGTGTGGAGTCGTTGCGCTAGAGAATCTCTCCAACCTGGNNNAGATCGGAAG
39923 TATGAGTGTGGAGTCGTTGCTCCTGGTCAGTTACTATCCCCAANNNAGATCGGAAG
39924 TATGAGTGTGGAGTCGTTGCGAGGAAGGAAGGAAGGAAGGAAGNNNAGATCGG
39926 TATGAGTGTGGAGTCGTTGCTTCCACAAAACAACCACAGCAATNNNAGATCGGAAG
39927 TATGAGTGTGGAGTCGTTGCTCTGTTGTTTAGTGCAGCTACTCTNNNAGATCGGAA
39928 TATGAGTGTGGAGTCGTTGCCTGTAGTCCCAGCTACTCAGGAGNNNAGATCGGAAG
39929 TATGAGTGTGGAGTCGTTGCCATCAAATGCTTTTTCTGTGCCANNNAGATCGGAAG
39930 TATGAGTGTGGAGTCGTTGCAACAAACATGCTTGCCTTGTTCTNNNAGATCGGAAG
39931 TATGAGTGTGGAGTCGTTGCAGGCATGGCTACAAAAGATTCCTNNNAGATCGGAAG
39932 TATGAGTGTGGAGTCGTTGCAGGATTGTCTTGGCTATAAGGGCNNNAGATCGGAA
39933 TATGAGTGTGGAGTCGTTGCACTGGTTTTGGAGCTGAGACAATNNNAGATCGGAA
39934 TATGAGTGTGGAGTCGTTGCAAATTCTTTTTTGGGGGGGGTTCNNNAGATCGGAA
39937 TATGAGTGTGGAGTCGTTGCCAGAGCCAGACTCCATCTCAAAANNNAGATCGGAAG
39938 TATGAGTGTGGAGTCGTTGCTTTAGTCAGGTCTTTCTGCTGCTNNNAGATCGGAAG
39939 TATGAGTGTGGAGTCGTTGCATGATAGTGCATCTTTCCCCACANNNAGATCGGAAG
39940 TATGAGTGTGGAGTCGTTGCAATCCCAGCTACTTGGAAGTCTGNNNAGATCGGAAG
39941 TATGAGTGTGGAGTCGTTGCGAGGGACACATTGGAAAACTTGGNNNAGATCGGAA
39943 TATGAGTGTGGAGTCGTTGCCGCCTTTTTTTCACCCCTTTACTNNNAGATCGGAAGA
39946 TATGAGTGTGGAGTCGTTGCGCAACAGAGTGAAACCCTGATTCNNNAGATCGGAA
39948 TATGAGTGTGGAGTCGTTGCCTGCCTCCTGGTTCAAATGATTCNNNAGATCGGAAG
39949 TATGAGTGTGGAGTCGTTGCTATGTATTAGCTGGGCATGGTGGNNNAGATCGGAA
39950 TATGAGTGTGGAGTCGTTGCGTCCAGGTATTCATGCATTGGTGNNNAGATCGGAA
39951 TATGAGTGTGGAGTCGTTGCGAAAGGGGGAGGGATAGCATTAGNNNAGATCGGA
39952 TATGAGTGTGGAGTCGTTGCAAACAGCATGTCCCATTCTAGGANNNAGATCGGAAG
39953 TATGAGTGTGGAGTCGTTGCTGCATCTAGGACAGAAAGAGAAANNNAGATCGGAA
39954 TATGAGTGTGGAGTCGTTGCAGGGGTTTGTGTGTAGAAGACAGNNNAGATCGGA
39955 TATGAGTGTGGAGTCGTTGCGAATAGGAATTACTGGGGGTGGGNNNAGATCGGA
39957 TATGAGTGTGGAGTCGTTGCTTCCCTTTTACCCTTGGTGTAGGNNNAGATCGGAAG
39958 TATGAGTGTGGAGTCGTTGCCATAACACATTGGCACCTGGAACNNNAGATCGGAAG
39959 TATGAGTGTGGAGTCGTTGCAAAGAGGTTTGTTTGGCTCATGGNNNAGATCGGAA
39962 TATGAGTGTGGAGTCGTTGCGAAATGAGCACTGACTCTGGAGANNNAGATCGGAA
39963 TATGAGTGTGGAGTCGTTGCCATATGCCCTTTGACCAAAGCAANNNAGATCGGAAG
39965 TATGAGTGTGGAGTCGTTGCGAGAGAATAGAGAATGGGGGCTGNNNAGATCGGA
39966 TATGAGTGTGGAGTCGTTGCGTCTCTCTCCCATGTGATGATCCNNNAGATCGGAAG

39967 TATGAGTGTGGAGTCGTTGCCCATTCCAACTTTCACTGACCAGNNNAGATCGGAAG
39970 TATGAGTGTGGAGTCGTTGCTAGGACAGAATTGAGGCTTCAGCNNNAGATCGGAA
39971 TATGAGTGTGGAGTCGTTGCATCTCAGCTCACTGTAACCTCTGNNNAGATCGGAAG
39972 TATGAGTGTGGAGTCGTTGCAAGATTGTACCTCTGCACTCCAGNNNAGATCGGAAG
39973 TATGAGTGTGGAGTCGTTGCCATAGGACCAGAGGGGATTTGTGNNNAGATCGGAA
39974 TATGAGTGTGGAGTCGTTGCAGGAGATGATGAGTTTTGGGACANNNAGATCGGA
39977 TATGAGTGTGGAGTCGTTGCTGTATAGGTGAGGAAACCAAGGCNNNAGATCGGAA
39979 TATGAGTGTGGAGTCGTTGCCTTTTTGGGAGGCAGAATGGAAGNNNAGATCGGAA
39980 TATGAGTGTGGAGTCGTTGCACATGGGAGGGAGACGTTATAGANNNAGATCGGA
39981 TATGAGTGTGGAGTCGTTGCCTTGTGTGGAACAACTGCAGATGNNNAGATCGGAA
39982 TATGAGTGTGGAGTCGTTGCTGTGCCTGTAATCCCAGTTACTCNNNAGATCGGAAG
39984 TATGAGTGTGGAGTCGTTGCCATCCATCGGTAGCAGACTGTTCNNNAGATCGGAAG
39985 TATGAGTGTGGAGTCGTTGCAGAGAGGAGGAGGATGGATGATTNNNAGATCGGA
39986 TATGAGTGTGGAGTCGTTGCACTCTTGACCCAAATTCACCAGTNNNAGATCGGAAG
39987 TATGAGTGTGGAGTCGTTGCCTGCTCAGATACTCAGCCAGTAGNNNAGATCGGAAG
39989 TATGAGTGTGGAGTCGTTGCAGTATCAGTGTTTCAGGGGTTCCNNNAGATCGGAA
39990 TATGAGTGTGGAGTCGTTGCTCACTTGAGTTCAGGAGTTCGAGNNNAGATCGGAA
39991 TATGAGTGTGGAGTCGTTGCAAATTAGCTGCCTGTAGTCCCAANNNAGATCGGAAG
39992 TATGAGTGTGGAGTCGTTGCTTCAAGACCAGCCTGAACAACATNNNAGATCGGAAG
39993 TATGAGTGTGGAGTCGTTGCTAAGTGACAGTGCCAAATACCCCNNNAGATCGGAAG
39995 TATGAGTGTGGAGTCGTTGCTTAGTCAGGTATGGTGGCATGTGNNNAGATCGGAA
39996 TATGAGTGTGGAGTCGTTGCCCTTTAGGAATTGATTGGGCCTGNNNAGATCGGAA
39997 TATGAGTGTGGAGTCGTTGCAAAAAGGAAGGAAGGAAGGGAGGNNNAGATCGG
39998 TATGAGTGTGGAGTCGTTGCAGAAGAATTTTTGTGGCCTGCTGNNNAGATCGGAA
39999 TATGAGTGTGGAGTCGTTGCAGATGAGGTCTCACTATGTTGCCNNNAGATCGGAAG
40001 TATGAGTGTGGAGTCGTTGCATGACATAATACAGCGGCATCCTNNNAGATCGGAAG
40003 TATGAGTGTGGAGTCGTTGCAGGTTGCAGTGAGTAGAGATCACNNNAGATCGGAA
40005 TATGAGTGTGGAGTCGTTGCCAGATCATGAGGTCAGGAGATCGNNNAGATCGGAA
40008 TATGAGTGTGGAGTCGTTGCGTGAAGCAGAGATGGAAGCATTCNNNAGATCGGAA
40010 TATGAGTGTGGAGTCGTTGCCATGCTTGTAATCCCAGCACTTTNNNAGATCGGAAG
40011 TATGAGTGTGGAGTCGTTGCCCACGAAGAGGGGTAAAGATCTTNNNAGATCGGAA
40012 TATGAGTGTGGAGTCGTTGCCCATGTAGCTGGGATTACAGGTGNNNAGATCGGAA
40013 TATGAGTGTGGAGTCGTTGCATCATTTGAGGTCGGGAGTTTGANNNAGATCGGAA
40015 TATGAGTGTGGAGTCGTTGCTGACTACTCTAAGACACCCCTGANNNAGATCGGAAG
40016 TATGAGTGTGGAGTCGTTGCAAAAATTAGCTGGGCATAGTGGCNNNAGATCGGAA
40018 TATGAGTGTGGAGTCGTTGCAACAGGACTTTGGGAGAGAATGTNNNAGATCGGAA
40019 TATGAGTGTGGAGTCGTTGCAAAAAAACCCTCCAGTAACCCCANNNAGATCGGAAG
40022 TATGAGTGTGGAGTCGTTGCTCTCTTCCACCTCTTCAGAGTCANNNAGATCGGAAGA
40023 TATGAGTGTGGAGTCGTTGCTCGTAAGATCCACCAAGGTCAAGNNNAGATCGGAA
40024 TATGAGTGTGGAGTCGTTGCAGATCCACCAAGGTCAAGAAACTNNNAGATCGGAA
40025 TATGAGTGTGGAGTCGTTGCGTTCCAGTGAGCAGAGATCTCAANNNAGATCGGAA
40027 TATGAGTGTGGAGTCGTTGCAAGCAAGGTAATGGGCTAAGAGTNNNAGATCGGA
40028 TATGAGTGTGGAGTCGTTGCTGGCAATGAGTAGCAAAAGAACCNNNAGATCGGAA

40029 TATGAGTGTGGAGTCGTTGCGGAGGCTGAAGCAGGAAAATTTCNNNAGATCGGAA
40030 TATGAGTGTGGAGTCGTTGCCTCCCGAGTAGCTGGAATTACAGNNNAGATCGGAA
40031 TATGAGTGTGGAGTCGTTGCTGTTTCCCACAGAAGTACAGGAGNNNAGATCGGAA
40032 TATGAGTGTGGAGTCGTTGCAGACCCAAACCAGTAAGCTGAATNNNAGATCGGAA
40033 TATGAGTGTGGAGTCGTTGCATTGCCCATTTGGTAACTTGGACNNNAGATCGGAAG
40034 TATGAGTGTGGAGTCGTTGCCAGAGCAAGGCTGTCTCAAAAAANNNAGATCGGAA
40035 TATGAGTGTGGAGTCGTTGCTCCAGGTTCAAGTGATTCTCCTGNNNAGATCGGAAG
40036 TATGAGTGTGGAGTCGTTGCGGATCACAAGGTCAGAAGATCGANNNAGATCGGAA
40037 TATGAGTGTGGAGTCGTTGCTGTTTTGCTGCTCAATCAGAGAGNNNAGATCGGAAG
40038 TATGAGTGTGGAGTCGTTGCCATTGCCTGCCAAACAAAATGAANNNAGATCGGAAG
40039 TATGAGTGTGGAGTCGTTGCCAAGAAGTAGAGGCTGCAGTGAGNNNAGATCGGA
40043 TATGAGTGTGGAGTCGTTGCGCCCATTGGTGTGTAGTTGAAAANNNAGATCGGAA
40044 TATGAGTGTGGAGTCGTTGCCAGAGTGGGATTCTGTCTCAAAANNNAGATCGGAA
40045 TATGAGTGTGGAGTCGTTGCGCCTAGAAACTCTCAGGTGGAAANNNAGATCGGAA
40046 TATGAGTGTGGAGTCGTTGCGGATGGGCTCCATATTGTGATCANNNAGATCGGAA
40047 TATGAGTGTGGAGTCGTTGCAATTCAATAATTAGCCGGGCGTGNNNAGATCGGAA
40049 TATGAGTGTGGAGTCGTTGCCTGTAGTCCCAGCTACTCAGGAGNNNAGATCGGAAG
40051 TATGAGTGTGGAGTCGTTGCAAAAGGAGTCAGCAAATGCACTCNNNAGATCGGAA
40052 TATGAGTGTGGAGTCGTTGCCTAGGAGATGGAAGTTGCAGTGANNNAGATCGGA
40053 TATGAGTGTGGAGTCGTTGCAAAGCAGCAAGAGAGAAGTGACTNNNAGATCGGA
40054 TATGAGTGTGGAGTCGTTGCGGGAGGAAATCAGTAGGCAATCANNNAGATCGGA
40056 TATGAGTGTGGAGTCGTTGCGACCTCACCACTCTGATGTTCTTNNNAGATCGGAAG
40058 TATGAGTGTGGAGTCGTTGCGAGGCAGGATAATTGCTTGAACCNNNAGATCGGAA
40061 TATGAGTGTGGAGTCGTTGCGTTGGAGGGACTTTGAGAGACATNNNAGATCGGAA
40063 TATGAGTGTGGAGTCGTTGCGAAACTTCCCTAAGGTGAGTGATNNNAGATCGGAA
40064 TATGAGTGTGGAGTCGTTGCCAACAGAGCAAGACTCCATCTCANNNAGATCGGAAG
40065 TATGAGTGTGGAGTCGTTGCACACTGAAAATCTGTGGTGCTAGNNNAGATCGGAA
40066 TATGAGTGTGGAGTCGTTGCGGATCACAAGGTCAGGAGATCAANNNAGATCGGAA
40067 TATGAGTGTGGAGTCGTTGCGGGTGGGGAGAAATAATTGAGGTNNNAGATCGGA
40068 TATGAGTGTGGAGTCGTTGCCCTTTTGGTATGGGCTGACTTTGNNNAGATCGGAAG
40070 TATGAGTGTGGAGTCGTTGCGTAATCCCAGCACTTTGGAATGCNNNAGATCGGAAG
40072 TATGAGTGTGGAGTCGTTGCACCTGCAAATCCCGTCTGAATTANNNAGATCGGAAG
40074 TATGAGTGTGGAGTCGTTGCTGAGAGGTCATAACACTTGGGTTNNNAGATCGGAA
40076 TATGAGTGTGGAGTCGTTGCAGAAATGAGGGGGTTTGCCTAATNNNAGATCGGAA
40078 TATGAGTGTGGAGTCGTTGCTCAGGAGTTGAGGAGGGATAAGTNNNAGATCGGA
40079 TATGAGTGTGGAGTCGTTGCGAATTGGACTAGGGAAACTGTGCNNNAGATCGGAA
40080 TATGAGTGTGGAGTCGTTGCTTGCAAATCTACCCTTGTCTCCANNNAGATCGGAAGA
40081 TATGAGTGTGGAGTCGTTGCACTAGACCCCTTTCTTACACTGTNNNAGATCGGAAGA
40082 TATGAGTGTGGAGTCGTTGCCCCTTACATGAGAGCAGAAACCTNNNAGATCGGAAG
40083 TATGAGTGTGGAGTCGTTGCAGAAGAAAATGAAGCTGGGGACTNNNAGATCGGA
40084 TATGAGTGTGGAGTCGTTGCACAAGAACATTTGAGATGGTGGTNNNAGATCGGAA
40085 TATGAGTGTGGAGTCGTTGCAAAGAAAATGAAACCTCAGCCCCNNNAGATCGGAAG
40087 TATGAGTGTGGAGTCGTTGCTACAAAATTAGCTGGGCTGGTCGNNNAGATCGGAA

40088 TATGAGTGTGGAGTCGTTGCTGCAAAAGACTTGACCAGATACTNNNAGATCGGAAG
40089 TATGAGTGTGGAGTCGTTGCGAAGCGGAATTTGTAGTGAGCTCNNNAGATCGGAA
40090 TATGAGTGTGGAGTCGTTGCTAACCGTGCTACTTCCAGATGACNNNAGATCGGAAG
40091 TATGAGTGTGGAGTCGTTGCTACAGCATCATACCCAGTTAGGCNNNAGATCGGAAG
40092 TATGAGTGTGGAGTCGTTGCCTGAGGCAGGAGAATCACTTGAANNNAGATCGGAA
40094 TATGAGTGTGGAGTCGTTGCGGCAAATCCATGTTCTGTGTTCANNNAGATCGGAAG
40095 TATGAGTGTGGAGTCGTTGCCTGGGGTATGCTTGCTTTGTTTTNNNAGATCGGAAG
40096 TATGAGTGTGGAGTCGTTGCAAAACTGGGGTATGCTTGCTTTGNNNAGATCGGAA
40097 TATGAGTGTGGAGTCGTTGCGAGGCTGAGACAGAAGAATCACTNNNAGATCGGAA
40098 TATGAGTGTGGAGTCGTTGCGCCTAAACAACATGGAGAAACCCNNNAGATCGGAA
40099 TATGAGTGTGGAGTCGTTGCACCACTGACACCACAGAAATAAANNNAGATCGGAAG
40101 TATGAGTGTGGAGTCGTTGCTCTATGGGATACAGCACAAGCAANNNAGATCGGAA
40102 TATGAGTGTGGAGTCGTTGCGACTCAGTTAGACACAAGATCCANNNAGATCGGAAG
40103 TATGAGTGTGGAGTCGTTGCAAGGATCAGAGATTGGACCATGTNNNAGATCGGAA
40104 TATGAGTGTGGAGTCGTTGCTGTTTGGTATACCTCCTAGGGGTNNNAGATCGGAAG
40105 TATGAGTGTGGAGTCGTTGCCTCATCAAGCTGTTTCTGTCTTTNNNAGATCGGAAGA
40106 TATGAGTGTGGAGTCGTTGCTCTTACCTCCGAATGATTGCACTNNNAGATCGGAAG
40107 TATGAGTGTGGAGTCGTTGCTTTTTGCTCAGGATAGCTTTGGCNNNAGATCGGAAG
40108 TATGAGTGTGGAGTCGTTGCAGTGATGGTTGGTCAACTCTATGANNNAGATCGGA
40109 TATGAGTGTGGAGTCGTTGCGCTTTGCCACCGTGTTTTTTTTTNNNAGATCGGAAGA
40110 TATGAGTGTGGAGTCGTTGCCAGTTATTCTGGAAGCTGAGGCANNNAGATCGGAA
40111 TATGAGTGTGGAGTCGTTGCCACCTTTGCCCAAGTTTTGCTTNNNAGATCGGAAGAG
40112 TATGAGTGTGGAGTCGTTGCCCCCATCTCACTGACTGAATCTCNNNAGATCGGAAGA
40113 TATGAGTGTGGAGTCGTTGCTGGGAGATTAGGGATTCTCTGGANNNAGATCGGAA
40114 TATGAGTGTGGAGTCGTTGCTGACCCCACCGAAATACAAACTANNNAGATCGGAAG
40115 TATGAGTGTGGAGTCGTTGCTGGAGACATTACAACTGACCACANNNAGATCGGAAG
40116 TATGAGTGTGGAGTCGTTGCAGGAATGTTGCAGAGAGGATTCANNNAGATCGGAA
40117 TATGAGTGTGGAGTCGTTGCAACTGGACGGAGAATGAGTTTGANNNAGATCGGAA
40119 TATGAGTGTGGAGTCGTTGCTAAAAAATTAGCTGGGCGTGGTGNNNAGATCGGAA
40120 TATGAGTGTGGAGTCGTTGCCATGGATGCAGCTGGAGTTTATTNNNAGATCGGAA
40122 TATGAGTGTGGAGTCGTTGCGGCAGAAAGTAGGTCTATAGGGCNNNAGATCGGA
40124 TATGAGTGTGGAGTCGTTGCTGTCTCAGTGTCCTTCAGTTCAGNNNAGATCGGAAG
40125 TATGAGTGTGGAGTCGTTGCCATAGAGAGATGAACAGGTGCCANNNAGATCGGAA
40126 TATGAGTGTGGAGTCGTTGCATTTCAGCTCACTGCAACTTCTGNNNAGATCGGAAG
40127 TATGAGTGTGGAGTCGTTGCATGGAACAGATCAGAGAACCTGGNNNAGATCGGAA
40128 TATGAGTGTGGAGTCGTTGCGAAGGAGGATCGCTTGAGACTAGNNNAGATCGGA
40129 TATGAGTGTGGAGTCGTTGCGAGTGAATGAAAAGAAGAGGGGANNNAGATCGGA
40130 TATGAGTGTGGAGTCGTTGCACCTTTGCAAATACACTAGCACTNNNAGATCGGAAG
40131 TATGAGTGTGGAGTCGTTGCGGCTCTTGGTTACCTGTGTTGTANNNAGATCGGAAG
40132 TATGAGTGTGGAGTCGTTGCTTGATTCCACCCAGGTTTCATCTNNNAGATCGGAAGA
40133 TATGAGTGTGGAGTCGTTGCGAAAAGCCAACTTATGGGATGGGNNNAGATCGGAA
40134 TATGAGTGTGGAGTCGTTGCATAAAACAAGAGGACCTGTGCCANNNAGATCGGAA
40135 TATGAGTGTGGAGTCGTTGCTGGCATATACCTGTAGTTCCAGCNNNAGATCGGAAG

40138 TATGAGTGTGGAGTCGTTGCATGTAATCCCAGCACTTTGAGGTNNNAGATCGGAAG
40139 TATGAGTGTGGAGTCGTTGCGGTGAAACCCCGTCTCTACTAAANNNAGATCGGAAG
40140 TATGAGTGTGGAGTCGTTGCTGTACTTTAGAGGCTGATGTGGGNNNAGATCGGAA
40141 TATGAGTGTGGAGTCGTTGCAGTAGGCTGCTTCTAAAATGGCTNNNAGATCGGAA
40142 TATGAGTGTGGAGTCGTTGCCGGTTCAGACTGCATTTTTGACTNNNAGATCGGAAG
40144 TATGAGTGTGGAGTCGTTGCGTTCAAGAGATTCTCCAGCCTCANNNAGATCGGAAG
40145 TATGAGTGTGGAGTCGTTGCATAGCTTGCAGTGAGCCGAGATCNNNAGATCGGAA
40146 TATGAGTGTGGAGTCGTTGCTCCACTCAAGGATACCACTTTGCNNNAGATCGGAAG
40147 TATGAGTGTGGAGTCGTTGCAGGGTTTCTTACTGCCTCTAGGANNNAGATCGGAAG
40149 TATGAGTGTGGAGTCGTTGCCTTGGGTGACAGCAAGACAAAATNNNAGATCGGAA
40150 TATGAGTGTGGAGTCGTTGCTCCTTCCCACAAAAATGCTTCACNNNAGATCGGAAGA
40151 TATGAGTGTGGAGTCGTTGCTGTTTGATGAAATCCAGTTGGCANNNAGATCGGAA
40155 TATGAGTGTGGAGTCGTTGCGAAGTCAGGAGTTCTAGACCAGCNNNAGATCGGAA
40156 TATGAGTGTGGAGTCGTTGCGGCAAAAAAACGGCAAAATGTTGNNNAGATCGGAA
40157 TATGAGTGTGGAGTCGTTGCGAGTTGCATGTGTCATGGTCAAGNNNAGATCGGAA
40158 TATGAGTGTGGAGTCGTTGCACCAGCAAAGGATGAGAGTACATNNNAGATCGGAA
40159 TATGAGTGTGGAGTCGTTGCCGGCTGTAATCCTAGCATTTTGGNNNAGATCGGAAG
40160 TATGAGTGTGGAGTCGTTGCCAATCTTGGCTCATGGAAACCTCNNNAGATCGGAAG
40161 TATGAGTGTGGAGTCGTTGCGAAGGAGAAGGAGAAGGAGAAGCNNNAGATCGG
40163 TATGAGTGTGGAGTCGTTGCCTCCATGCTCTGAACTCTAGTGGNNNAGATCGGAAG
40164 TATGAGTGTGGAGTCGTTGCCCTGTAATCCCAGCACTTTAGGANNNAGATCGGAAG
40165 TATGAGTGTGGAGTCGTTGCGTGACACAGCAAGACAATGTCTCNNNAGATCGGAA
40166 TATGAGTGTGGAGTCGTTGCCACTTGTAATCCCAGCACTTTGGNNNAGATCGGAAG
40167 TATGAGTGTGGAGTCGTTGCTGGAACCGGAACTTACTGTACTCNNNAGATCGGAAG
40169 TATGAGTGTGGAGTCGTTGCTGAGGTCAGGAATTCAAGACCAGNNNAGATCGGAA
40170 TATGAGTGTGGAGTCGTTGCTCTTTCTGTGGTTTTGTCAACCANNNAGATCGGAAG
40172 TATGAGTGTGGAGTCGTTGCCCAAGGCTACCTGATAGGCTAATNNNAGATCGGAAG
40173 TATGAGTGTGGAGTCGTTGCTGCAAAAGAGAGTGAGACTTCGTNNNAGATCGGAA
40176 TATGAGTGTGGAGTCGTTGCAGAGTACCTACTGCATGAGTCCANNNAGATCGGAAG
40177 TATGAGTGTGGAGTCGTTGCTCCCTCATTCTTTATCCCACCAGNNNAGATCGGAAGA
40178 TATGAGTGTGGAGTCGTTGCTTCATCCTAGCACTTTGGGAGTCNNNAGATCGGAAG
40179 TATGAGTGTGGAGTCGTTGCAGGTACTGGGATTGTCATTGGACNNNAGATCGGAA
40180 TATGAGTGTGGAGTCGTTGCGAAAGTAGATTTTGGAGCTGGGCNNNAGATCGGAA
40181 TATGAGTGTGGAGTCGTTGCTTGAGGCAGGAGAATCACTTGAANNNAGATCGGAA
40182 TATGAGTGTGGAGTCGTTGCCTCACACCTGTAATCTCCACACTNNNAGATCGGAAGA
40188 TATGAGTGTGGAGTCGTTGCTGAAGCCAGAATATTTGTCTTGGANNNAGATCGGA
40190 TATGAGTGTGGAGTCGTTGCTATAAGATGATCATGGGGCCAGGNNNAGATCGGAA
40191 TATGAGTGTGGAGTCGTTGCGGATCACTTGAGGTCAGGAGTTCNNNAGATCGGAA
40192 TATGAGTGTGGAGTCGTTGCGGATCACTTGAGGTCAGGAGTTCNNNAGATCGGAA
40193 TATGAGTGTGGAGTCGTTGCAGTGAGCTATGTGGTCTTGCTACNNNAGATCGGAA
40195 TATGAGTGTGGAGTCGTTGCTGTCTCGATCTTGGCTTATTCCANNNAGATCGGAAG
40196 TATGAGTGTGGAGTCGTTGCGAAAGAAAAATTGCCAAGCGTGGNNNAGATCGGAA
40199 TATGAGTGTGGAGTCGTTGCCTTGTACATTGCTGGTGGGAATGNNNAGATCGGAA

40200 TATGAGTGTGGAGTCGTTGCTTCCCTAACCTACACTGCCAAAANNNAGATCGGAAGA
40201 TATGAGTGTGGAGTCGTTGCGGCACCTATAATCCCAGCTACTCNNNAGATCGGAAGA
40202 TATGAGTGTGGAGTCGTTGCTCCCATTCAAACTCTCCAACACTNNNAGATCGGAAGAG
40203 TATGAGTGTGGAGTCGTTGCCACACCTGTAATCCCAGCACTTTNNNAGATCGGAAGA
40204 TATGAGTGTGGAGTCGTTGCCTCCTGAGTAGCTGGGACTACAGNNNAGATCGGAAG
40205 TATGAGTGTGGAGTCGTTGCATGGAGGCTAGGAATGGGATAGANNNAGATCGGAA
40207 TATGAGTGTGGAGTCGTTGCACTGTATTAGGCCACTCTTGCATNNNAGATCGGAAGA
40208 TATGAGTGTGGAGTCGTTGCACAGCCCTTCCCATTATCAACATNNNAGATCGGAAGA
40209 TATGAGTGTGGAGTCGTTGCAAAAGAAAAGTGGGAGATCAGGTNNNAGATCGGAA
40210 TATGAGTGTGGAGTCGTTGCAGGTAGGAGAATTGCTTGAACCCNNNAGATCGGAAG
40211 TATGAGTGTGGAGTCGTTGCTGGGACAAACTTATGTCTCCTGANNNAGATCGGAAG
40212 TATGAGTGTGGAGTCGTTGCAACACAAAAATGAGGTCAGGTGCNNNAGATCGGAAG
40213 TATGAGTGTGGAGTCGTTGCCACAGTCACTTCTCGTTGTTGACNNNAGATCGGAAGA
40214 TATGAGTGTGGAGTCGTTGCATCAATGAGCCAACTATTGCTGCNNNAGATCGGAAG
40215 TATGAGTGTGGAGTCGTTGCGTTAGCTATGATCATGCCACTGCNNNAGATCGGAAG
40216 TATGAGTGTGGAGTCGTTGCAACACCTATAATCCCAGTGCGTTNNNAGATCGGAAGA
40217 TATGAGTGTGGAGTCGTTGCGCTTTCTTGCAATCCTGAAAGGANNNAGATCGGAAG
40218 TATGAGTGTGGAGTCGTTGCTGCCAAGGAAACACAGGTAGTTANNNAGATCGGAAG
40219 TATGAGTGTGGAGTCGTTGCTAGCACTGATTTTCCCAAGGACANNNAGATCGGAAG
40220 TATGAGTGTGGAGTCGTTGCAGCACTTACCTTTTCCAGGGAATNNNAGATCGGAAGA
40221 TATGAGTGTGGAGTCGTTGCCTGGAAAAGCATTCTCTTTGCCANNNAGATCGGAAGA
40222 TATGAGTGTGGAGTCGTTGCCTCTGAGGCAGCATAATGAGACTNNNAGATCGGAAG
40224 TATGAGTGTGGAGTCGTTGCACAAGTTCTCTCATCTAGCTGTGTNNNAGATCGGAAG
40225 TATGAGTGTGGAGTCGTTGCGGGAATATCAACTGGTGACATGGNNNAGATCGGAA
40227 TATGAGTGTGGAGTCGTTGCGAGAATTGCTTGAACCTGGGAGANNNAGATCGGAA
40229 TATGAGTGTGGAGTCGTTGCGGAACTGGAGGCCATTATCCTAANNNAGATCGGAAG
40230 TATGAGTGTGGAGTCGTTGCCACAAGAGTTTTGGCACCATGAANNNAGATCGGAAG
40232 TATGAGTGTGGAGTCGTTGCGGGAGACAGGAATTTCACTGACTNNNAGATCGGAAG
40233 TATGAGTGTGGAGTCGTTGCGCCTTTGAGCTCGTTGATTCTTTNNNAGATCGGAAGA
40235 TATGAGTGTGGAGTCGTTGCAGTTAGTCTTCATGGCCTCTTGGNNNAGATCGGAAG
40236 TATGAGTGTGGAGTCGTTGCAACAACCTTTCTCTCAGGGCTACNNNAGATCGGAAGA
40237 TATGAGTGTGGAGTCGTTGCAAGCTAGGATATGAAGCTCAGGCNNNAGATCGGAAG
40239 TATGAGTGTGGAGTCGTTGCTTCTCCAAACTTTTGACTTGGGCNNNAGATCGGAAGA
40240 TATGAGTGTGGAGTCGTTGCCCCTACACATTCCCTTGACAGTTNNNAGATCGGAAGA
40241 TATGAGTGTGGAGTCGTTGCGCTGGGTTCTGCAAATAAGACAGNNNAGATCGGAAG
40244 TATGAGTGTGGAGTCGTTGCCCTAACTCCAGTGAGCTGAACTTNNNAGATCGGAAGA
40245 TATGAGTGTGGAGTCGTTGCTCTAAAACTAACCCCCTCCTTGCNNNAGATCGGAAGA
40246 TATGAGTGTGGAGTCGTTGCCTCCCAGGTTCACACCATTCTCNNNAGATCGGAAGAG
40247 TATGAGTGTGGAGTCGTTGCATTCATTTGCACTTGGGAGGTTGNNNAGATCGGAAG
40248 TATGAGTGTGGAGTCGTTGCGCTACATGTGATTGGCCAAAACTNNNAGATCGGAAG
40249 TATGAGTGTGGAGTCGTTGCTGAGAATTACTGTGGCTCTTCCCNNNAGATCGGAAGA
40250 TATGAGTGTGGAGTCGTTGCCAGAGCAAGACTCCATCTCCAAANNNAGATCGGAAG
40251 TATGAGTGTGGAGTCGTTGCCCTCTTCACAGATGGAAGCTCTTNNNAGATCGGAAGA

40252 TATGAGTGTGGAGTCGTTGCGGGAAGGAGACAAGCAAATTGTGNNNAGATCGGAA

40254 TATGAGTGTGGAGTCGTTGCCAGGGCCAATCAGGAAAATCTTCNNNAGATCGGAAG

40255 TATGAGTGTGGAGTCGTTGCGGATAAGCAAAACAGGGTCCAACNNNAGATCGGAAG

40257 TATGAGTGTGGAGTCGTTGCCAGATAACTGGCAAGCCATATGCNNNAGATCGGAAG

40261 TATGAGTGTGGAGTCGTTGCCAGACACTCAGAATGTTAGAGCCNNNAGATCGGAAG

40263 TATGAGTGTGGAGTCGTTGCTATGAGACGTGGCAAAGTAGGTCNNNAGATCGGAA

40264 TATGAGTGTGGAGTCGTTGCACCTAGACTGAGGCCTTATACCTNNNAGATCGGAAGA

40265 TATGAGTGTGGAGTCGTTGCTTCCAGATCCTTTGCCTATGGACNNNAGATCGGAAGA

40266 TATGAGTGTGGAGTCGTTGCCACCCCCAACAGAATTCACTTTCNNNAGATCGGAAGA

40268 TATGAGTGTGGAGTCGTTGCTCCATCCAACATTGTGCTTTGTTNNNAGATCGGAAGA

40269 TATGAGTGTGGAGTCGTTGCATCCCTTCCACCTAGGTAACAGANNNAGATCGGAAGA

40270 TATGAGTGTGGAGTCGTTGCCGGGAAGAAGAAGTTGCAATGAGNNNAGATCGGAA

40272 TATGAGTGTGGAGTCGTTGCCCCACCCATTACAAAATTGCCAANNNAGATCGGAAGA

40273 TATGAGTGTGGAGTCGTTGCGGGCTTCCAGGCTATTGGTAAATNNNAGATCGGAAG

40274 TATGAGTGTGGAGTCGTTGCAGATTCCTAAAGCCCTCGATGTCNNNAGATCGGAAGA

40275 TATGAGTGTGGAGTCGTTGCCACTATTGGGATCACAGCTCACTNNNAGATCGGAAGA

40276 TATGAGTGTGGAGTCGTTGCTGGATTATGGATGCTGAACCAGTNNNAGATCGGAAG

40277 TATGAGTGTGGAGTCGTTGCGAAGCAGGAGAATCACTTGAACCNNNAGATCGGAAG

40278 TATGAGTGTGGAGTCGTTGCTGCCTAGGAAGGGAAATTCTGTTNNNAGATCGGAAG

40279 TATGAGTGTGGAGTCGTTGCGTGCCCCATTGCAAAAGAAAATGNNNAGATCGGAAG

40281 TATGAGTGTGGAGTCGTTGCACCATTCAACCTAGCAACCTCATNNNAGATCGGAAGA

40282 TATGAGTGTGGAGTCGTTGCCTGATACTTCAGGGTGTCACCAANNNAGATCGGAAG

40283 TATGAGTGTGGAGTCGTTGCCCTGCTCTTGAACTACTCTAGTAANNNAGATCGGAAG

40285 TATGAGTGTGGAGTCGTTGCAATGTTGGCTCCATCAAGTCAACNNNAGATCGGAAG

40286 TATGAGTGTGGAGTCGTTGCAGAATCCAGTCCAATTGATAGCTNNNAGATCGGAAG

40287 TATGAGTGTGGAGTCGTTGCAACGGAACTTCGAGTTATCTGCANNNAGATCGGAAG

40288 TATGAGTGTGGAGTCGTTGCAACAAGAGTGAAAGGCCATCTCANNNAGATCGGAAG

40289 TATGAGTGTGGAGTCGTTGCCCCTCTCCTGGATGACCTATACANNNAGATCGGAAGA

40291 TATGAGTGTGGAGTCGTTGCTCAACCAAATACTAGCCAACCGANNNAGATCGGAAGA

40292 TATGAGTGTGGAGTCGTTGCGGAAAAGGGGGAAATGTGAAAGGNNNAGATCGGA

40293 TATGAGTGTGGAGTCGTTGCTTATTGGACAGTGCTTGGACAGTNNNAGATCGGAAG

40295 TATGAGTGTGGAGTCGTTGCAAACTCTGACTCCCAGGTTCAAGNNNAGATCGGAAG

40296 TATGAGTGTGGAGTCGTTGCGACTTGTTGAATGGCTTTGACCANNNAGATCGGAAG

40297 TATGAGTGTGGAGTCGTTGCCAGTGAGATGCAGCAGGTTCTATNNNAGATCGGAAG

40298 TATGAGTGTGGAGTCGTTGCTACACCAACAAGAGACAAGCAGANNNAGATCGGAAG

40302 TATGAGTGTGGAGTCGTTGCTTCCTGCAAGATCTCTGGTTTCANNNAGATCGGAAGA

40303 TATGAGTGTGGAGTCGTTGCCACACAGCATTGATTTGTGATGCNNNAGATCGGAAG

40304 TATGAGTGTGGAGTCGTTGCGCATTGGGAGATATGCCTAATGCNNNAGATCGGAAG

40305 TATGAGTGTGGAGTCGTTGCACAATGCCCTGTTGTTAGTTGTGNNNAGATCGGAAG

40306 TATGAGTGTGGAGTCGTTGCCATCCCTTCAAACTTCAGCACAANNNAGATCGGAAGA

40307 TATGAGTGTGGAGTCGTTGCTGCCCATAGAAGTTCCATCCAAANNNAGATCGGAAGA

40309 TATGAGTGTGGAGTCGTTGCTTAGAGGTGGTTTGCTATGCAGANNNAGATCGGAAG

40310 TATGAGTGTGGAGTCGTTGCAAGAAAGTCTCCAGGAAGTGTGGNNNAGATCGGAA

40313 TATGAGTGTGGAGTCGTTGCACTGCGACGTCAATCAAAACAATNNNAGATCGGAAG
40314 TATGAGTGTGGAGTCGTTGCTGAACCGATGAGGAAATAGGCTTNNNAGATCGGAA
40319 TATGAGTGTGGAGTCGTTGCAAATGTGATTGTGGAGGGGATGANNNAGATCGGA
40321 TATGAGTGTGGAGTCGTTGCGCCAAGCAAAGATTTGAGAGTGTNNNAGATCGGAA
40322 TATGAGTGTGGAGTCGTTGCATTCACTCAGAAAAACGCTGACCNNNAGATCGGAAG
40323 TATGAGTGTGGAGTCGTTGCAAGTGCCTCTTAGGAAAACTGGTNNNAGATCGGAA
40324 TATGAGTGTGGAGTCGTTGCGAACAGTGAAGGGAGAATGGTCTNNNAGATCGGA
40325 TATGAGTGTGGAGTCGTTGCAATCAGAATATCTTGGGGGTGGGNNNAGATCGGAA
40326 TATGAGTGTGGAGTCGTTGCACTCTCACACACTGTTGGAGAAANNNAGATCGGAAG
40328 TATGAGTGTGGAGTCGTTGCATATCCAAGCAGAGGCCACATAGNNNAGATCGGAA
40329 TATGAGTGTGGAGTCGTTGCTGGCTTTGGACTCTCTCTTTCTCNNNAGATCGGAAGA
40330 TATGAGTGTGGAGTCGTTGCAATTGCCACATCTTCCACAATGGNNNAGATCGGAAG
40331 TATGAGTGTGGAGTCGTTGCTGGTGGGTTTCCTACATCATGTTNNNAGATCGGAAG
40332 TATGAGTGTGGAGTCGTTGCGCCTGATGGTTGTCCTTTCTTTCNNNAGATCGGAAG
40333 TATGAGTGTGGAGTCGTTGCAAACAAACAAAAAACTGGTGGGGNNNAGATCGGAA
40334 TATGAGTGTGGAGTCGTTGCAGAGAGAGAGAGAAAAACCCACCNNNAGATCGGAA
40335 TATGAGTGTGGAGTCGTTGCCAACTGAAAGGCTAGTGTGGTTGNNNAGATCGGAA
40337 TATGAGTGTGGAGTCGTTGCACCCCTTCTTTCTGAGCACTAAGNNNAGATCGGAAG
40338 TATGAGTGTGGAGTCGTTGCCTGAGGCAGGAGAATTGCTTGAANNNAGATCGGAA
40340 TATGAGTGTGGAGTCGTTGCGTAGCATATTGGAAGCAGACGTCNNNAGATCGGAA
40341 TATGAGTGTGGAGTCGTTGCTGAGATGTTGGATATGGTGGCTCNNNAGATCGGAA
40343 TATGAGTGTGGAGTCGTTGCCCAGATCCATGGGAGAAAAATGCNNNAGATCGGAA
40344 TATGAGTGTGGAGTCGTTGCGAAAAAGATAGAGGGGCAGGGAANNNAGATCGGA
40345 TATGAGTGTGGAGTCGTTGCGAATTTCACCACTAAAGCTGGGCNNNAGATCGGAAG
40346 TATGAGTGTGGAGTCGTTGCCTGGATGGAACTGGAGAGCATTANNNAGATCGGAA
40348 TATGAGTGTGGAGTCGTTGCCTGAGGCAGGAGAATTGCTTGAANNNAGATCGGAA
40349 TATGAGTGTGGAGTCGTTGCAAGAAGAAGAAGGAGGAGGAGGANNNAGATCGG
40351 TATGAGTGTGGAGTCGTTGCCGAGGCTCTGTCTCAAAAAACAANNNAGATCGGAAG
40353 TATGAGTGTGGAGTCGTTGCAGAACATGGACTTTGGAACCAGANNNAGATCGGAA
40354 TATGAGTGTGGAGTCGTTGCAAGAAAAAAAATTGCCGGGCATGNNNAGATCGGAA
40355 TATGAGTGTGGAGTCGTTGCTCCAATACTGCCATTCATCCACANNNAGATCGGAAGA
40356 TATGAGTGTGGAGTCGTTGCAGAGGGATACTTCAGTTCAGGGANNNAGATCGGAA
40359 TATGAGTGTGGAGTCGTTGCTTGGAGACCAACATGGCCAATATNNNAGATCGGAA
40361 TATGAGTGTGGAGTCGTTGCCACTTGAAATGTGGCTAGTGCAANNNAGATCGGAA
40363 TATGAGTGTGGAGTCGTTGCATCCTGACTAACACAGTGAACCCNNNAGATCGGAAG
40364 TATGAGTGTGGAGTCGTTGCGCAGGAAAACATGATGGGCTATCNNNAGATCGGAA
40366 TATGAGTGTGGAGTCGTTGCATTGAGGGGAAGTTTCAAAGGGTNNNAGATCGGA
40367 TATGAGTGTGGAGTCGTTGCCTGCTTACAGAGATGGGGGTAAGNNNAGATCGGAA
40368 TATGAGTGTGGAGTCGTTGCAAGATGAAGGCTGGTCTGGTATGNNNAGATCGGAA
40369 TATGAGTGTGGAGTCGTTGCGCAGTGTCATGGAGAATATGTGCNNNAGATCGGAA
40370 TATGAGTGTGGAGTCGTTGCAGACAGACAAGAAGGAGAGAGGANNNAGATCGGA
40371 TATGAGTGTGGAGTCGTTGCGGACAGACTATCTAGGCTTGTCCNNNAGATCGGAAG
40372 TATGAGTGTGGAGTCGTTGCTCATTACCTCCAAAGTTAGGGCCNNNAGATCGGAAG

40374 TATGAGTGTGGAGTCGTTGCTGAGTTCTGTGTGTCCTGGATACNNNAGATCGGAAG
40376 TATGAGTGTGGAGTCGTTGCAAACCATTGTAACTCGAGGACCANNNAGATCGGAAG
40377 TATGAGTGTGGAGTCGTTGCTCAATCCAAGGTATGCACTGGTTNNNAGATCGGAAG
40378 TATGAGTGTGGAGTCGTTGCATATGGTTAGACTTCGTGTCCCCNNNAGATCGGAAG
40380 TATGAGTGTGGAGTCGTTGCATAGAAACTGCAGTGAGATGCCANNNAGATCGGAA
40381 TATGAGTGTGGAGTCGTTGCATCACAACCTTATAGTCCTGCCANNNAGATCGGAAG
40382 TATGAGTGTGGAGTCGTTGCTGAAAAAGGCCATCAAGGAGAGANNNAGATCGGA
40383 TATGAGTGTGGAGTCGTTGCAATGCAGATTCTCAGACTCACCANNNAGATCGGAAG
40384 TATGAGTGTGGAGTCGTTGCAAGGTCAGAGAAGAAGACGTGACNNNAGATCGGA
40385 TATGAGTGTGGAGTCGTTGCGCTTCTAGTCCCAGCTACTCAGANNNAGATCGGAAG
40386 TATGAGTGTGGAGTCGTTGCAGATGGTGTTTCAACATGTTGGCNNNAGATCGGAA
40387 TATGAGTGTGGAGTCGTTGCTATCAATCCCCTAGCATGACAGCNNNAGATCGGAAG
40389 TATGAGTGTGGAGTCGTTGCCAACAAGAGCGAAACTCCATCTCNNNAGATCGGAAG
40390 TATGAGTGTGGAGTCGTTGCGTAGTCCCATCTACTCAGGAGGTNNNAGATCGGAAG
40392 TATGAGTGTGGAGTCGTTGCGATGCTGAAGGTGCTGATAAGAANNNAGATCGGAA
40393 TATGAGTGTGGAGTCGTTGCGAGGTCAGGAGATCGAGACCATCNNNAGATCGGAA
40394 TATGAGTGTGGAGTCGTTGCAGACATAAGGCAGGAAAACAGGTNNNAGATCGGA
40395 TATGAGTGTGGAGTCGTTGCGGACTCCACACTATCAACACCTTNNNAGATCGGAAG
40396 TATGAGTGTGGAGTCGTTGCCCTAGTGAGAGTGAGACCACAACNNNAGATCGGAA
40397 TATGAGTGTGGAGTCGTTGCAATACAGCTGACCCTTGAACCATNNNAGATCGGAAG
40398 TATGAGTGTGGAGTCGTTGCTACACTCACATAGCCACTATCCCNNNAGATCGGAAGA
40399 TATGAGTGTGGAGTCGTTGCCCCAAACACTATCAGGCACAATGNNNAGATCGGAAG
40400 TATGAGTGTGGAGTCGTTGCCAACATGTCATGGCTCTTTCAGCNNNAGATCGGAAG
40401 TATGAGTGTGGAGTCGTTGCCATGCCTGTAATCCCAGCTACTANNNAGATCGGAAG
40402 TATGAGTGTGGAGTCGTTGCGGCCTGCCAAAAACTATTAGGACNNNAGATCGGAA
40403 TATGAGTGTGGAGTCGTTGCGCAGAGGTAAAGAAGCACGAAAGNNNAGATCGGA
40404 TATGAGTGTGGAGTCGTTGCGGTTGGGGGTTCAATGTGTAATGNNNAGATCGGAA
40405 TATGAGTGTGGAGTCGTTGCTCCCCCCCTTTGATAGGATTAGANNNAGATCGGAAG
40406 TATGAGTGTGGAGTCGTTGCACTCTTGCTTTTCTGGTTTGTGGNNNAGATCGGAAG
40407 TATGAGTGTGGAGTCGTTGCTCTAGCCAATCCAGCATTCTGTTNNNAGATCGGAAG
40408 TATGAGTGTGGAGTCGTTGCGAACCGGCAAAGAAAACTGAGAANNNAGATCGGAA
40409 TATGAGTGTGGAGTCGTTGCTCACGTCTGTAATCCCAGCATTTNNNAGATCGGAAG
40410 TATGAGTGTGGAGTCGTTGCCAATTCTCCTATCTCAGCCTCCCNNNAGATCGGAAGA
40411 TATGAGTGTGGAGTCGTTGCAGCCTCAGGACAAATTGTACCTTNNNAGATCGGAAG
40412 TATGAGTGTGGAGTCGTTGCAATTGTTACAGGGTGATGACGGANNNAGATCGGAA
40413 TATGAGTGTGGAGTCGTTGCAGTGCCTGCTTATAACTGGATGTNNNAGATCGGAAG
40414 TATGAGTGTGGAGTCGTTGCATATTACTGTCACCGACTGTCCCNNNAGATCGGAAG
40415 TATGAGTGTGGAGTCGTTGCCTTGCCTTGACCATCATTCTTCCNNNAGATCGGAAGA
40416 TATGAGTGTGGAGTCGTTGCACCAGATTTCAGAGAAAGCAGCTNNNAGATCGGAA
40417 TATGAGTGTGGAGTCGTTGCATGAGGCAGCAGAATCACTTGAANNNAGATCGGAA
40419 TATGAGTGTGGAGTCGTTGCTTGGTATATTCAGTGGTGTGCCANNNAGATCGGAA
40422 TATGAGTGTGGAGTCGTTGCCTCAACGTCATCAGTCATTTGGGNNNAGATCGGAAG
40423 TATGAGTGTGGAGTCGTTGCCCAGTTTTGTGCTTGAAACCCAGNNNAGATCGGAAG

40424 TATGAGTGTGGAGTCGTTGCCAAGTGGACAGAAAACCAGCAAANNNAGATCGGAA
40426 TATGAGTGTGGAGTCGTTGCTCACGCTCCTCAAAGGTTTCTTANNNAGATCGGAAG
40427 TATGAGTGTGGAGTCGTTGCGACCTGGACATTGGTAGTGTCTTNNNAGATCGGAA
40428 TATGAGTGTGGAGTCGTTGCAAAACCAAGTAACCAGGCCTAGTNNNAGATCGGAA
40429 TATGAGTGTGGAGTCGTTGCGAGGTAGTGAACAAGATCCAGCANNNAGATCGGAA
40432 TATGAGTGTGGAGTCGTTGCTTGGTGAAAACTTGCTGTGTTGANNNAGATCGGAA
40434 TATGAGTGTGGAGTCGTTGCCAGAATCACGTCCTTTGCAACAANNNAGATCGGAAG
40435 TATGAGTGTGGAGTCGTTGCTCACCGCAAAAATTAGTGTCACCNNNAGATCGGAAG
40436 TATGAGTGTGGAGTCGTTGCCTGTAGTTCTAGCTGCTCTGGAGNNNAGATCGGAAG
40437 TATGAGTGTGGAGTCGTTGCCTCTAGCCAGGTGGTTCCATATGNNNAGATCGGAAG
40438 TATGAGTGTGGAGTCGTTGCGGCACCTATAATCCCAGCTACTCNNNAGATCGGAAG
40439 TATGAGTGTGGAGTCGTTGCATCTTCTGTGCTGAAAAGGCAAGNNNAGATCGGAA
40440 TATGAGTGTGGAGTCGTTGCCAGGAGTTGCTCACTGTTACAGTNNNAGATCGGAAG
40441 TATGAGTGTGGAGTCGTTGCTAATCTCAACACTTTGGGAGGCANNNAGATCGGAAG
40442 TATGAGTGTGGAGTCGTTGCAAAGAAAGTATGTCACCTGCCCANNNAGATCGGAAG
40443 TATGAGTGTGGAGTCGTTGCCTGGGTGACAGAACAAGACTCTGNNNAGATCGGAA
40444 TATGAGTGTGGAGTCGTTGCCCTACTAGATCCCTCACATGTGCNNNAGATCGGAAG
40445 TATGAGTGTGGAGTCGTTGCGCCACCATGCCAGTTTACTTTTTNNNAGATCGGAAG
40446 TATGAGTGTGGAGTCGTTGCTGATAGGTGTTTGGTCTGAGTGGNNNAGATCGGAA
40447 TATGAGTGTGGAGTCGTTGCAAACAAGGTGCTATTGGGCAATCNNNAGATCGGAA
40449 TATGAGTGTGGAGTCGTTGCAACTAACAAGTCAGGGCACTTTTNNNAGATCGGAAG
40450 TATGAGTGTGGAGTCGTTGCGGGTTGCAATCCTAGTCTCTGATNNNAGATCGGAAG
40451 TATGAGTGTGGAGTCGTTGCACTGGTAGTTCCCCCACAAAAATNNNAGATCGGAAG
40452 TATGAGTGTGGAGTCGTTGCTGGCTTGACCTAAGAAGCACTAANNNAGATCGGAA
40453 TATGAGTGTGGAGTCGTTGCATAATTTCAAGGCAGAGCAGCTGNNNAGATCGGAA
40454 TATGAGTGTGGAGTCGTTGCTGCAGTATGGAAAGGGAAAGACANNNAGATCGGA
40455 TATGAGTGTGGAGTCGTTGCATGCCTGTAATCTCAGCTACTCGNNNAGATCGGAAG
40457 TATGAGTGTGGAGTCGTTGCGGAATGGGAAGCAGGAACTATGANNNAGATCGGA
40458 TATGAGTGTGGAGTCGTTGCCAGGAAGCTGATTCAGGAGGATTNNNAGATCGGAA
40459 TATGAGTGTGGAGTCGTTGCCCCTGTCTCTATACACACACACANNNAGATCGGAAGA
40460 TATGAGTGTGGAGTCGTTGCGAAAGGCAGCTCTAAAGAGGTCANNNAGATCGGAA
40461 TATGAGTGTGGAGTCGTTGCGCAACTACTCTTGACCACTGTCTNNNAGATCGGAAG
40463 TATGAGTGTGGAGTCGTTGCACTGCCTCTGGTAATCTTTCCTGNNNAGATCGGAAG
40465 TATGAGTGTGGAGTCGTTGCATGATTCATATTCCCAGGGTGGCNNNAGATCGGAAG
40466 TATGAGTGTGGAGTCGTTGCGCATAGGATGTGTTTGGAGCATGNNNAGATCGGAA
40467 TATGAGTGTGGAGTCGTTGCAAAGGAGGATGGTAAAGGCAGAGNNNAGATCGGA
40469 TATGAGTGTGGAGTCGTTGCTGTGGTATATCCTCACAATGCAANNNAGATCGGAAG
40470 TATGAGTGTGGAGTCGTTGCTTGAGCATGTGGAGAAATTGGAGNNNAGATCGGA
40471 TATGAGTGTGGAGTCGTTGCAGGATTACAAGGCTACAGTGAGCNNNAGATCGGAA
40472 TATGAGTGTGGAGTCGTTGCAATACAATGGGTTGGTGCAAAGTNNNAGATCGGAA
40473 TATGAGTGTGGAGTCGTTGCCAGAGTTGAAGGGAAAGGGGATTNNNAGATCGGA
40474 TATGAGTGTGGAGTCGTTGCTAAAGGCTACTCAACATGTGCCANNNAGATCGGAAG
40476 TATGAGTGTGGAGTCGTTGCACAGGGATGTAAGCTCCTTGAAGNNNAGATCGGAA

40477 TATGAGTGTGGAGTCGTTGCCTGAGGCAGAAGGATCACTTGAGNNNAGATCGGAA
40478 TATGAGTGTGGAGTCGTTGCCTATGTGCCCTATCATGCCAGATNNNAGATCGGAAG
40479 TATGAGTGTGGAGTCGTTGCAAAAAGAAAAAAAGGTGGGGGGGNNNAGATCGG
40480 TATGAGTGTGGAGTCGTTGCCCAGGAAGGTGGAAGCTATACTCNNNAGATCGGAA
40481 TATGAGTGTGGAGTCGTTGCTGAAGTTTCCCAGGCTATAGTGGNNNAGATCGGAA
40482 TATGAGTGTGGAGTCGTTGCAAAAAATTAGTTGGGCGTGGTGGNNNAGATCGGA
40483 TATGAGTGTGGAGTCGTTGCCTGTGATCACACCACTGCATTACNNNAGATCGGAAG
40484 TATGAGTGTGGAGTCGTTGCGGTCCCAGTTACTCTGAAGGATGNNNAGATCGGAA
40485 TATGAGTGTGGAGTCGTTGCCCATGTCGGGCACTTTACCTATANNNAGATCGGAAG
40487 TATGAGTGTGGAGTCGTTGCCTATAAAGTACCTGGGCATGGGGNNNAGATCGGAA
40489 TATGAGTGTGGAGTCGTTGCCTCCAGCATAGGTAACAGAGCAANNNAGATCGGAA
40490 TATGAGTGTGGAGTCGTTGCATGTACACTATTCGGGTGACAGGNNNAGATCGGAA
40492 TATGAGTGTGGAGTCGTTGCCACCTAGAAGGGGGCCAATAAATNNNAGATCGGAA
40493 TATGAGTGTGGAGTCGTTGCATTTTCATGCCCAAAGTCTTGCCNNNAGATCGGAAG
40494 TATGAGTGTGGAGTCGTTGCGGTTCAAGTGATTTTCCTGCCTCNNNAGATCGGAAG
40495 TATGAGTGTGGAGTCGTTGCGCTTAGGGCTCCCAACAAATTAGNNNAGATCGGAAG
40496 TATGAGTGTGGAGTCGTTGCATTTTTCCAGTCTTTCACCCCCTNNNAGATCGGAAGA
40497 TATGAGTGTGGAGTCGTTGCACCACCCTGCCATTTTTGTAATGNNNAGATCGGAAG
40498 TATGAGTGTGGAGTCGTTGCTTTTTTCTGGAGACGGGGTTTTGNNNAGATCGGAAG
40499 TATGAGTGTGGAGTCGTTGCACCTGTAATCCTAGCACTTTGGGNNNAGATCGGAAG
40500 TATGAGTGTGGAGTCGTTGCGTTGTTAGTCAAAAGGGGGCTTCNNNAGATCGGAA
40501 TATGAGTGTGGAGTCGTTGCCTACTTAGGAAGCTGAGGTGGGANNNAGATCGGAA
40504 TATGAGTGTGGAGTCGTTGCCGTGACTACTTCCTTGCTGTACTNNNAGATCGGAAG
40505 TATGAGTGTGGAGTCGTTGCATACCTTCCACTGTAACCTTGCANNNAGATCGGAAG
40506 TATGAGTGTGGAGTCGTTGCCGTGGTCTTCTCTAGATGGTGAGNNNAGATCGGAA
40507 TATGAGTGTGGAGTCGTTGCGGTCACAGAGTAGTAGAGAGAGTNNNAGATCGGA
40509 TATGAGTGTGGAGTCGTTGCACTTGTAATCCCAGCAATCCCAGNNNAGATCGGAAG
40510 TATGAGTGTGGAGTCGTTGCCTGAGGCAGGAGAATTGCTTGAANNNAGATCGGAA
40512 TATGAGTGTGGAGTCGTTGCGTTCTGAAGGCTGAGAAGTCTGANNNAGATCGGAA
40513 TATGAGTGTGGAGTCGTTGCTCAGGCTAGAAAAATGAAGGCCANNNAGATCGGAA
40514 TATGAGTGTGGAGTCGTTGCCTGTAGTTCCAGCTACTCAGGAGNNNAGATCGGAAG
40515 TATGAGTGTGGAGTCGTTGCTCTACTGCCAAGGCCTTTGATACNNNAGATCGGAAG
40516 TATGAGTGTGGAGTCGTTGCATGTGTAATGAAGGAAGACCCCCNNNAGATCGGAA
40517 TATGAGTGTGGAGTCGTTGCAGTAAGCCAAGTAGGTGAGGAAANNNAGATCGGA
40520 TATGAGTGTGGAGTCGTTGCTTGCATTCTGTTCCCCTGATGTANNNAGATCGGAAG
40522 TATGAGTGTGGAGTCGTTGCTGTCAACCCATTTCCCCAACTATNNNAGATCGGAAGA
40523 TATGAGTGTGGAGTCGTTGCAAAACGGGCAAAAGACTTTGACANNNAGATCGGAA
40524 TATGAGTGTGGAGTCGTTGCAGCTCCTAGCAGACGTTTTTGTANNNAGATCGGAAG
40525 TATGAGTGTGGAGTCGTTGCTTCCCTACCTAACATGATACGCANNNAGATCGGAAG
40526 TATGAGTGTGGAGTCGTTGCCAGCTTCTGAAAACACTGGCATTNNNAGATCGGAAG
40528 TATGAGTGTGGAGTCGTTGCCACAAAGAAATGAAACACGCAGCNNNAGATCGGAA
40529 TATGAGTGTGGAGTCGTTGCGGGGAGATTGGTCTACAGTTTGTNNNAGATCGGAA
40530 TATGAGTGTGGAGTCGTTGCGATCCAAAAGGCAGAGTGGTAGANNNAGATCGGA

40531 TATGAGTGTGGAGTCGTTGCCAATGTCTTGCTTTGTCACCCAGNNNAGATCGGAAG
40532 TATGAGTGTGGAGTCGTTGCTAAACTCTTTCCCCTTGACCCAGNNNAGATCGGAAGA
40533 TATGAGTGTGGAGTCGTTGCTTACCTCAAGAGGTTGCTGTGAANNNAGATCGGAAG
40534 TATGAGTGTGGAGTCGTTGCGTAGTCCCAGCTACTTGAGAAGCNNNAGATCGGAAG
40535 TATGAGTGTGGAGTCGTTGCCAGCCTGGCCAATATGGTAAAACNNNAGATCGGAAG
40536 TATGAGTGTGGAGTCGTTGCAAAGAAAGACCATTTCCACTGCCNNNAGATCGGAAG
40537 TATGAGTGTGGAGTCGTTGCATGTTAGCGAGGAGAAGGAAGAGNNNAGATCGGA
40539 TATGAGTGTGGAGTCGTTGCTCTTCCACTGATGCACAATGTCTNNNAGATCGGAAG
40540 TATGAGTGTGGAGTCGTTGCAACACATCTAGAGGCTGCAAAGANNNAGATCGGAA
40541 TATGAGTGTGGAGTCGTTGCAACACTGCTCAAAAGAAAGAAGTNNNAGATCGGAA
40542 TATGAGTGTGGAGTCGTTGCCAAAGATGTCCCTCATGCTACCTNNNAGATCGGAAG
40543 TATGAGTGTGGAGTCGTTGCGGATCCTGCTTTTGCAGAAACTTNNNAGATCGGAAG
40544 TATGAGTGTGGAGTCGTTGCGAATTTCGTTCTTGTTGCCCAGANNNAGATCGGAAG
40545 TATGAGTGTGGAGTCGTTGCTGAGCTCATTTGGCAAACGAATTNNNAGATCGGAAG
40546 TATGAGTGTGGAGTCGTTGCCCACAGGAGAATCACTTGAACCTNNNAGATCGGAAG
40547 TATGAGTGTGGAGTCGTTGCAGAAAACTGGAGCTCTACATGCANNNAGATCGGAA
40548 TATGAGTGTGGAGTCGTTGCCAATGGAACAGAATAGAGCGCACNNNAGATCGGAA
40550 TATGAGTGTGGAGTCGTTGCAGAGGGAAAGGAATTGTCTGAGTNNNAGATCGGA
40551 TATGAGTGTGGAGTCGTTGCTTGGAGGACAATGGTGTGATCTTNNNAGATCGGAA
40552 TATGAGTGTGGAGTCGTTGCGCACAGGCAACAAAAGCAAAAATNNNAGATCGGAA
40554 TATGAGTGTGGAGTCGTTGCGCCACACTGGAAGAAGAATTGTCNNNAGATCGGAA
40555 TATGAGTGTGGAGTCGTTGCGGAAGGATGAAGGGTTGGGAATANNNAGATCGGA
40556 TATGAGTGTGGAGTCGTTGCTCTCCTTCCTTTTATCTGCACCCNNNAGATCGGAAGA
40557 TATGAGTGTGGAGTCGTTGCAAACAAAACTCAAACCTGGTGGCNNNAGATCGGAAG
40559 TATGAGTGTGGAGTCGTTGCTTCACAATCCAGGCTTGTCAGTANNNAGATCGGAAG
40560 TATGAGTGTGGAGTCGTTGCAATACACACACACACACACACACNNNAGATCGGAAG
40562 TATGAGTGTGGAGTCGTTGCCTGTAGTACCAGCTACTCGGGAGNNNAGATCGGAA
40563 TATGAGTGTGGAGTCGTTGCATTACAGTGAAACGCTCTTGCACNNNAGATCGGAAG
40565 TATGAGTGTGGAGTCGTTGCTGGCATATTGTAAGTGTGCACTCNNNAGATCGGAAG
40567 TATGAGTGTGGAGTCGTTGCGCAAGCCCACCTTAGGATAATCTNNNAGATCGGAAG
40568 TATGAGTGTGGAGTCGTTGCGGGGTAGGAAACACAGTCTTGAANNNAGATCGGAA
40570 TATGAGTGTGGAGTCGTTGCAAAGAAGGGCAGGTCACTTACATNNNAGATCGGAA
40571 TATGAGTGTGGAGTCGTTGCCTTCACAGTGGTTAGGAGGACTGNNNAGATCGGAA
40572 TATGAGTGTGGAGTCGTTGCGTAATCATATAGGGAGGTGGGGCNNNAGATCGGA
40573 TATGAGTGTGGAGTCGTTGCTAGACAGGAATGCACTTCATGGTNNNAGATCGGAA
40574 TATGAGTGTGGAGTCGTTGCAGGAGCAACTGGACTATGTGATCNNNAGATCGGAA
40575 TATGAGTGTGGAGTCGTTGCCTCTTCCCTGTTCCCTTGGTATGNNNAGATCGGAAGA
40576 TATGAGTGTGGAGTCGTTGCACCTGACTAAGGGGCAAGAAAATNNNAGATCGGAA
40578 TATGAGTGTGGAGTCGTTGCTAAAGGGACTAGTAGGAGCCAGGNNNAGATCGGA
40579 TATGAGTGTGGAGTCGTTGCACTGGGAGCAACTGTAATAGCAANNNAGATCGGAA
40580 TATGAGTGTGGAGTCGTTGCGGCAAGCAGAGAGACAAATCATGNNNAGATCGGAA
40581 TATGAGTGTGGAGTCGTTGCGATGTCTGTGACAGTGAAGGACTNNNAGATCGGAA
40582 TATGAGTGTGGAGTCGTTGCACTTTCTAGAAACCTGCCCTCTGNNNAGATCGGAAG

40583 TATGAGTGTGGAGTCGTTGCACACAAAAATTAGCTGGGCATGGNNNAGATCGGAA
40588 TATGAGTGTGGAGTCGTTGCTCCTAGTACAACCAATCCACTCTNNNAGATCGGAAGA
40590 TATGAGTGTGGAGTCGTTGCCAGGAAATGTCTTCAAGCACCAGNNNAGATCGGAA
40591 TATGAGTGTGGAGTCGTTGCACCTCTGGTCTAAAGCCTTGTACNNNAGATCGGAAG
40592 TATGAGTGTGGAGTCGTTGCGGAGGGATCCTTCTGGTCAATTTNNNAGATCGGAA
40593 TATGAGTGTGGAGTCGTTGCATCTACAAGGAACTAGGGGGACANNNAGATCGGAA
40595 TATGAGTGTGGAGTCGTTGCACTGAGTTGTAACCTACAGTGCCNNNAGATCGGAAG
40596 TATGAGTGTGGAGTCGTTGCGAGGCTGTGTTTAGAGGATCACTNNNAGATCGGAA
40597 TATGAGTGTGGAGTCGTTGCGGTTCTTTCACTGGGCAGAATTGNNNAGATCGGAA
40599 TATGAGTGTGGAGTCGTTGCAGGCAGGAGAATCTCTTGAACTGNNNAGATCGGAA
40600 TATGAGTGTGGAGTCGTTGCAGAAAAATGGGGAGGGTATGGTGNNNAGATCGGA
40601 TATGAGTGTGGAGTCGTTGCCTGGGCATAGCAGAACAAACAAANNNAGATCGGAA
40602 TATGAGTGTGGAGTCGTTGCCTGAGGCAGGAGAATAGCTTGAANNNAGATCGGAA
40603 TATGAGTGTGGAGTCGTTGCGGATGAGGCAGGAGAATCACTTTNNNAGATCGGAA
40605 TATGAGTGTGGAGTCGTTGCCTAAGGCAGGGGAATCACTTGAANNNAGATCGGAA
40606 TATGAGTGTGGAGTCGTTGCCAGATCATGAGGTCAGGAGATGGNNNAGATCGGA
40608 TATGAGTGTGGAGTCGTTGCACAGTATCAGTATGGACCCATGANNNAGATCGGAA
40609 TATGAGTGTGGAGTCGTTGCTTTCCAAGCTGTTTTTTTGGGGTNNNAGATCGGAAG
40611 TATGAGTGTGGAGTCGTTGCCTAGCAAGAAAAATGGAGCAGCANNNAGATCGGAA
40612 TATGAGTGTGGAGTCGTTGCACAGGAGTGAGGGCTCAATATTGNNNAGATCGGAA
40613 TATGAGTGTGGAGTCGTTGCAAATGGTCCTCTCACACATGGTTNNNAGATCGGAAG
40614 TATGAGTGTGGAGTCGTTGCATGTAGTCCCAGCTACTGAGGAGNNNAGATCGGAA
40615 TATGAGTGTGGAGTCGTTGCAGTTGCTTTTCTATGCCTGGAGANNNAGATCGGAAG
40616 TATGAGTGTGGAGTCGTTGCTTGGCGAAGGACTTTTCATGTTCNNNAGATCGGAAG
40617 TATGAGTGTGGAGTCGTTGCAAACTAGGAGCTGTGAGTTTGCTNNNAGATCGGAA
40618 TATGAGTGTGGAGTCGTTGCCTGAGGCAGGAGAATTGCTTGAANNNAGATCGGAA
40619 TATGAGTGTGGAGTCGTTGCTCCCACTTGGAAATTCCCTTGAANNNAGATCGGAAG
40620 TATGAGTGTGGAGTCGTTGCCCCAGGCAATGAAAATGTATCCCNNNAGATCGGAAG
40621 TATGAGTGTGGAGTCGTTGCTAAGGCATGAAAAGGTGGGTTGANNNAGATCGGA
40622 TATGAGTGTGGAGTCGTTGCCAAAAGCCAGTACTGTGCTTGAGNNNAGATCGGAA
40624 TATGAGTGTGGAGTCGTTGCCTGCATGGCAAAAACCACCTAAANNNAGATCGGAAG
40625 TATGAGTGTGGAGTCGTTGCCCTCCCGAGTAGCTGAAATTACANNNAGATCGGAAG
40626 TATGAGTGTGGAGTCGTTGCAACATGAAGAGGCAACTGAGTCTNNNAGATCGGAA
40628 TATGAGTGTGGAGTCGTTGCGAGTGGGGACAAAAAAAGAAGCANNNAGATCGGA
40634 TATGAGTGTGGAGTCGTTGCTGGAACAGACCAACCAAAGAGAANNNAGATCGGAA
40635 TATGAGTGTGGAGTCGTTGCCATGCCTGTAATCCCAGCATTTTNNNAGATCGGAAG
40636 TATGAGTGTGGAGTCGTTGCAAGAGGCAGCAGACAAGTTTCTANNNAGATCGGAA
40637 TATGAGTGTGGAGTCGTTGCGGCAAAACCCCGTCTCTACTAAANNNAGATCGGAAG
40638 TATGAGTGTGGAGTCGTTGCACATTCACTGAAAAGGACAACGGNNNAGATCGGAA
40639 TATGAGTGTGGAGTCGTTGCTCTTGTCATTTGTAACAGCATGGANNNAGATCGGAA
40640 TATGAGTGTGGAGTCGTTGCAACTGAGAAACAGAAGGGACAGANNNAGATCGGA
40641 TATGAGTGTGGAGTCGTTGCACAGAGAAGCAACACCATTTCTTNNNAGATCGGAAG
40642 TATGAGTGTGGAGTCGTTGCCGCCTGTAATCTTGACACTTTGGNNNAGATCGGAAG

40643 TATGAGTGTGGAGTCGTTGCTTGATGCAGATGACGGTGGATATNNNAGATCGGAA
40645 TATGAGTGTGGAGTCGTTGCAGCTGCTTCTTTTTCCTCTCTGTNNNAGATCGGAAGA
40646 TATGAGTGTGGAGTCGTTGCGTAGCTCATGCCTGTAATCCCAGNNNAGATCGGAAG
40647 TATGAGTGTGGAGTCGTTGCTCACATTTCTGGAGGCTAGAAGCNNNAGATCGGAAG
40648 TATGAGTGTGGAGTCGTTGCTCTTTAGTCCCAGCTACTCAGGANNNAGATCGGAAG
40649 TATGAGTGTGGAGTCGTTGCGAGGGCCAGTAACATTATCAGCTNNNAGATCGGAAG
40651 TATGAGTGTGGAGTCGTTGCCACTCCTGTAATCCCAGCACTTTNNNAGATCGGAAGA
40652 TATGAGTGTGGAGTCGTTGCCATCTCCTGGGGATGAAGCTATTNNNAGATCGGAAG
40653 TATGAGTGTGGAGTCGTTGCTTCAGTCATTCCACAATGCAAGCNNNAGATCGGAAG
40654 TATGAGTGTGGAGTCGTTGCAGGTAGCCCAGGAATTTCAAGTTNNNAGATCGGAAG
40655 TATGAGTGTGGAGTCGTTGCTTGTGTGATCTTTGCCATGTTGTNNNAGATCGGAAG
40656 TATGAGTGTGGAGTCGTTGCACAGAGGCAGAGTTTGGAATGATNNNAGATCGGAA
40657 TATGAGTGTGGAGTCGTTGCAACTCTTCGGCTGTTCTTTTTCGNNNAGATCGGAAGA
40659 TATGAGTGTGGAGTCGTTGCAGGAAAGGAAGGAAAGGAACTCCNNNAGATCGGA
40660 TATGAGTGTGGAGTCGTTGCTACCAATCGAGTTCAACTCCCTCNNNAGATCGGAAGA
40661 TATGAGTGTGGAGTCGTTGCCCACCACATCTGGCTAGTTTTTTNNNAGATCGGAAGA
40664 TATGAGTGTGGAGTCGTTGCCACCATCGCCATCTCTTTGTACTNNNAGATCGGAAGA
40665 TATGAGTGTGGAGTCGTTGCCTCAGGAAAGACGACCCATTCTANNNAGATCGGAAG
40667 TATGAGTGTGGAGTCGTTGCGACAGAGCGAGACACCATCTAAANNNAGATCGGAA
40668 TATGAGTGTGGAGTCGTTGCACATTTATGCAGCCAACAGACACNNNAGATCGGAAG
40669 TATGAGTGTGGAGTCGTTGCGCAGGACACAAGCTTGAACTTAGNNNAGATCGGAA
40670 TATGAGTGTGGAGTCGTTGCCCCTGGAGTGTATTAGGAACTGGNNNAGATCGGAA
40671 TATGAGTGTGGAGTCGTTGCAGGATCTTTTGAACCCAGGAGACNNNAGATCGGAAG
40672 TATGAGTGTGGAGTCGTTGCGCCTCTTTTACTTCAGCCCAATGNNNAGATCGGAAGA
40673 TATGAGTGTGGAGTCGTTGCATCACTTGAGTCTGCAAGGTCAANNNAGATCGGAAG
40674 TATGAGTGTGGAGTCGTTGCTCTGCATTTCCATCTGAGGTACCNNNAGATCGGAAG
40675 TATGAGTGTGGAGTCGTTGCCCTGTTACTCAGCTACTCAGGACNNNAGATCGGAAG
40678 TATGAGTGTGGAGTCGTTGCAGCAGTGCTCAGAGGGAAATTTANNNAGATCGGAA
40679 TATGAGTGTGGAGTCGTTGCTAGCTTGATGGGAATAGCGTTGANNNAGATCGGAA
40681 TATGAGTGTGGAGTCGTTGCTCTTCAGTTAGTCAAACAGGCCANNNAGATCGGAAG
40682 TATGAGTGTGGAGTCGTTGCTTCCAAACTACTGAAGAGGCTCCNNNAGATCGGAAG
40683 TATGAGTGTGGAGTCGTTGCCTGCTTTCCCACTTGAGAGAGATNNNAGATCGGAAG
40684 TATGAGTGTGGAGTCGTTGCACCTGGACTGTTCTGGATTGTTTNNNAGATCGGAAG
40685 TATGAGTGTGGAGTCGTTGCATATGCTGGCATCTTGATGTTGCNNNAGATCGGAAG
40686 TATGAGTGTGGAGTCGTTGCAAAGAAAACCCCATCAGCCAATGNNNAGATCGGAAG
40687 TATGAGTGTGGAGTCGTTGCCAGTGAAATTACCTGGGTCTGGANNNAGATCGGAA
40689 TATGAGTGTGGAGTCGTTGCCACACACACACACACACATGTATNNNAGATCGGAAG
40690 TATGAGTGTGGAGTCGTTGCATAGAAGGTGTCAATTTCCCCCCNNNAGATCGGAAG
40691 TATGAGTGTGGAGTCGTTGCCTGAGTAGCTGGGATTACAGGTGNNNAGATCGGAA
40692 TATGAGTGTGGAGTCGTTGCAGCCCATTCCTATTGGTTCCATTNNNAGATCGGAAGA
40693 TATGAGTGTGGAGTCGTTGCTATGTGCCAATGTTCATAGCAGCNNNAGATCGGAAG
40694 TATGAGTGTGGAGTCGTTGCTGCTTAGTGAAACTTTGGTCAGGNNNAGATCGGAA
40695 TATGAGTGTGGAGTCGTTGCGGCAGGAGAATCACTTGAACCTANNNAGATCGGAA

40697 TATGAGTGTGGAGTCGTTGCAAACAAAGGCATGAAAAGGGGAGNNNAGATCGGA
40698 TATGAGTGTGGAGTCGTTGCATGACAAAAGACCCCGAATAGCTNNNAGATCGGAA
40700 TATGAGTGTGGAGTCGTTGCGATCTGTCACTGTCTCCCATCACNNNAGATCGGAAG
40701 TATGAGTGTGGAGTCGTTGCCACCAAGCCCATGAGTACATAGANNNAGATCGGAAG
40703 TATGAGTGTGGAGTCGTTGCGGAGGGACTGAAGATAGCTGATGNNNAGATCGGA
40704 TATGAGTGTGGAGTCGTTGCGAGACGGAATTTCGCTCTTGTTGNNNAGATCGGAA
40705 TATGAGTGTGGAGTCGTTGCCATGTGCAGGTGGATTGGATTTTNNNAGATCGGAA
40706 TATGAGTGTGGAGTCGTTGCCTCCAGTATACAGTGGCTAGCAGNNNAGATCGGAA
40709 TATGAGTGTGGAGTCGTTGCATAGAACGCAATAGAGAGGCCAGNNNAGATCGGAA
40710 TATGAGTGTGGAGTCGTTGCTGCTGAATTTCCCTCCAGAAAGANNNAGATCGGAAG
40711 TATGAGTGTGGAGTCGTTGCTGGCCAACATGGTGAAACTACTANNNAGATCGGAA
40713 TATGAGTGTGGAGTCGTTGCTGGAAAAACCCCTTAGACCTAGCNNNAGATCGGAAG
40714 TATGAGTGTGGAGTCGTTGCAGAGAGAGAAGGAAGGAGGGAAANNNAGATCGG
40715 TATGAGTGTGGAGTCGTTGCTGGGTGTTTGAGGTGAGAAAATGNNNAGATCGGA
40717 TATGAGTGTGGAGTCGTTGCACATGCAGTATTTGTCTCCCTGTNNNAGATCGGAAG
40718 TATGAGTGTGGAGTCGTTGCTACAAAAAATTAGCCGGGCATGGNNNAGATCGGAA
40719 TATGAGTGTGGAGTCGTTGCACGGAAATTAGCTGAGGGTAGTGNNNAGATCGGAA
40720 TATGAGTGTGGAGTCGTTGCGTTCAAAAGGGAGAGGTTGGGTANNNAGATCGGA
40723 TATGAGTGTGGAGTCGTTGCGTGAGGTCTTGGCTGTACAAAACNNNAGATCGGAA
40726 TATGAGTGTGGAGTCGTTGCATCAAAAGACCTGGGTCAGAGTCNNNAGATCGGAA
40727 TATGAGTGTGGAGTCGTTGCCAGCCACTATGGAAAACAGTGTGNNNAGATCGGAA
40729 TATGAGTGTGGAGTCGTTGCGATTTGGACAGAGGTCAGTGTGANNNAGATCGGAA
40730 TATGAGTGTGGAGTCGTTGCAGCCTTCTGAGTAGCTGGTACTANNNAGATCGGAAG
40731 TATGAGTGTGGAGTCGTTGCAGGGTTTGGTCAACGTGAAAATCNNNAGATCGGAA
40732 TATGAGTGTGGAGTCGTTGCAGTGGGAGAGAAAGAGGACTGTANNNAGATCGGA
40735 TATGAGTGTGGAGTCGTTGCTGGAGTGCAGTGGTACTATCTTGNNNAGATCGGAA
40736 TATGAGTGTGGAGTCGTTGCATACCCTTTACCAAGTTTCCCCCNNNAGATCGGAAGA
40737 TATGAGTGTGGAGTCGTTGCCTCACATGAGATGGGCATGAAATNNNAGATCGGAA
40738 TATGAGTGTGGAGTCGTTGCAACAACAAACAAAGGCTGCCTAANNNAGATCGGAA
40739 TATGAGTGTGGAGTCGTTGCGTGAAACCCCATCTCTACCAGAANNNAGATCGGAAG
40740 TATGAGTGTGGAGTCGTTGCCACGTCTGTAATCCCAGGTACTCNNNAGATCGGAAG
40741 TATGAGTGTGGAGTCGTTGCCTGTGTTCAGGTTTCTGTGTAGANNNAGATCGGAAG
40742 TATGAGTGTGGAGTCGTTGCAGATTGCTGGTGGACTGAGAAATNNNAGATCGGAA
40743 TATGAGTGTGGAGTCGTTGCGAGCTTCAGCCAAGTATCTCCATNNNAGATCGGAAG
40744 TATGAGTGTGGAGTCGTTGCTGCAAGATGGGTATCCTAGACTCNNNAGATCGGAA
40745 TATGAGTGTGGAGTCGTTGCGCTTATGCTTCCCTATGGTGAGANNNAGATCGGAAG
40747 TATGAGTGTGGAGTCGTTGCAGAGGTTGCAGTGAGCTAAGAATNNNAGATCGGAA
40748 TATGAGTGTGGAGTCGTTGCGGCTTCTCCAGTGTTCTATGAGANNNAGATCGGAAG
40749 TATGAGTGTGGAGTCGTTGCAAGCCTCCAGTACAGCAGTATTCNNNAGATCGGAAG
40751 TATGAGTGTGGAGTCGTTGCGCCTCAGCAGGAAGTTTAGAAGANNNAGATCGGAA
40752 TATGAGTGTGGAGTCGTTGCCACCCAGGAGCCTAAATTCACTANNNAGATCGGAAG
40753 TATGAGTGTGGAGTCGTTGCCTTTGAGAAGGAGGAAGTGGTCANNNAGATCGGAA
40755 TATGAGTGTGGAGTCGTTGCACCCATCAATCCCATTACTGAGTNNNAGATCGGAAG

40757 TATGAGTGTGGAGTCGTTGCGGTTCAAGTGATTGTCCTGCTTCNNNAGATCGGAAG
40758 TATGAGTGTGGAGTCGTTGCAAAGAAAAGGGAGTACAGGGAGCNNNAGATCGGA
40759 TATGAGTGTGGAGTCGTTGCATTGTCTCTTCCCCAAAGTCAGGNNNAGATCGGAAG
40760 TATGAGTGTGGAGTCGTTGCACTATCAAGTTGGTTCCATCCCCNNNAGATCGGAAG
40762 TATGAGTGTGGAGTCGTTGCGCAGCAAAGGCACTTTATAAGCCNNNAGATCGGAAG
40763 TATGAGTGTGGAGTCGTTGCGGTGTAAAAGAGCATGTGGACACNNNAGATCGGAA
40764 TATGAGTGTGGAGTCGTTGCCGCAGGGACTCATACCTCTAATCNNNAGATCGGAAG
40765 TATGAGTGTGGAGTCGTTGCGAGGGCGTTGAAGAGATGAAAAGNNNAGATCGGA
40766 TATGAGTGTGGAGTCGTTGCATCACGGCTGTTCAAAGTTGAAGNNNAGATCGGAA
40768 TATGAGTGTGGAGTCGTTGCAGCTTGGAAGATTGCTTTTAGGCNNNAGATCGGAA
40769 TATGAGTGTGGAGTCGTTGCAATCAAATTGTCCCTCTTTGCGGNNNAGATCGGAAG
40770 TATGAGTGTGGAGTCGTTGCAACAAAAGGCCTTGGTTCTTTGANNNAGATCGGAAG
40771 TATGAGTGTGGAGTCGTTGCTGTAAGCATCAATGTCAATCGGGNNNAGATCGGAA
40772 TATGAGTGTGGAGTCGTTGCTTTGGGCAAAAACGAAAGCAAACNNNAGATCGGAA
40774 TATGAGTGTGGAGTCGTTGCGACAGGGACAGAGGATCATTCAANNNAGATCGGAA
40775 TATGAGTGTGGAGTCGTTGCGTGTGTGTGTGTGAACTCAAANNNAGATCGGAA
40777 TATGAGTGTGGAGTCGTTGCTAGGTGCAGTCAATATGGGATGGNNNAGATCGGAA
40778 TATGAGTGTGGAGTCGTTGCATAAATGGCACATGGGAAAGCTGNNNAGATCGGAA
40779 TATGAGTGTGGAGTCGTTGCTTCTTCGTGGTTAGGTGTCTCTGNNNAGATCGGAAG
40780 TATGAGTGTGGAGTCGTTGCGCATTTATTGGCCAGTGCAATTGNNNAGATCGGAAG
40781 TATGAGTGTGGAGTCGTTGCTGTGTGTGATAGAATGGCTGTGANNNAGATCGGAA
40782 TATGAGTGTGGAGTCGTTGCAAATGAAGGGCTTACTCTCCCAGNNNAGATCGGAAG
40783 TATGAGTGTGGAGTCGTTGCACCAAAAAGTCTCTCTGAACCAANNNAGATCGGAAG
40784 TATGAGTGTGGAGTCGTTGCGAGTGGAGGAACACCAGATTCTTNNNAGATCGGAA
40786 TATGAGTGTGGAGTCGTTGCTTCTTCAGTCATCCAGACAAGGGNNNAGATCGGAAG
40787 TATGAGTGTGGAGTCGTTGCTTAGAGAAGAAGCTTCCAGGCAGNNNAGATCGGAA
40788 TATGAGTGTGGAGTCGTTGCTTAGAGAAGAAGCTTCCAGGCAGNNNAGATCGGAA
40789 TATGAGTGTGGAGTCGTTGCACCCGAGACTGGTCAATTTACAANNNAGATCGGAAG
40791 TATGAGTGTGGAGTCGTTGCACATGCATGAACACACACGTTAGNNNAGATCGGAAG
40793 TATGAGTGTGGAGTCGTTGCATTGCTTGAATCTGAGAGGCAGANNNAGATCGGAA
40795 TATGAGTGTGGAGTCGTTGCCTCACAGATGGAGCAATGGTGAGNNNAGATCGGAA
40798 TATGAGTGTGGAGTCGTTGCACATAGGAAGGAGACCACAATCGNNNAGATCGGAA
40799 TATGAGTGTGGAGTCGTTGCATAGAACACTTCACATGCCACCTNNNAGATCGGAAG
40800 TATGAGTGTGGAGTCGTTGCTAGCCAGGGATAAAGCATGAACTNNNAGATCGGAA
40801 TATGAGTGTGGAGTCGTTGCCACAACAGAAAAACCAAGCTGTGNNNAGATCGGAA
40802 TATGAGTGTGGAGTCGTTGCCTCAAGCAGAAGGAGTTTTGTCCNNNAGATCGGAAG
40803 TATGAGTGTGGAGTCGTTGCAAATTCAAGGCAGCTGCAGAAATNNNAGATCGGAA
40804 TATGAGTGTGGAGTCGTTGCTTTAGCATGAGGCCACAACAAACNNNAGATCGGAAG
40805 TATGAGTGTGGAGTCGTTGCTTAGAGAAGAAGCTTCCAGGCAGNNNAGATCGGAA
40806 TATGAGTGTGGAGTCGTTGCGTTTGACCTTCACACTAGCCCTANNNAGATCGGAAG
40807 TATGAGTGTGGAGTCGTTGCAGATGAGAGATGGCATCACATGGNNNAGATCGGAA
40808 TATGAGTGTGGAGTCGTTGCAGAAAAATTGCTTGAACCCCAGGNNNAGATCGGAA
40809 TATGAGTGTGGAGTCGTTGCTGCTCAACAGTACCTAACACAGANNNAGATCGGAAG

40811 TATGAGTGTGGAGTCGTTGCTGCTGCTGTCATTTTTGTGGTAGNNNAGATCGGAAG
40812 TATGAGTGTGGAGTCGTTGCCCCAATGATCTGTTTGTTGCCTANNNAGATCGGAAG
40813 TATGAGTGTGGAGTCGTTGCTAGCCAGCACTACAAGAACTGTTNNNAGATCGGAAG
40814 TATGAGTGTGGAGTCGTTGCCTTTTTTTGAGACAGAGGCTCGCNNNAGATCGGAAG
40815 TATGAGTGTGGAGTCGTTGCTAGAAGCGTTTCCCCAGATATGGNNNAGATCGGAAG
40816 TATGAGTGTGGAGTCGTTGCGAGATCACGCCATTACACTCCAANNNAGATCGGAAG
40818 TATGAGTGTGGAGTCGTTGCTGAGACAAAAACCATGAACACCTNNNAGATCGGAAG
40820 TATGAGTGTGGAGTCGTTGCACCACCTGAATTCAGAAGGTCAANNNAGATCGGAAG
40821 TATGAGTGTGGAGTCGTTGCTTCAAGCAACTCTCTAGCCTCAGNNNAGATCGGAAG
40823 TATGAGTGTGGAGTCGTTGCGAATTACAAGGCATGCCCTTGACNNNAGATCGGAAG
40824 TATGAGTGTGGAGTCGTTGCCATGCACATGTACACACACTTCTNNNAGATCGGAAG
40825 TATGAGTGTGGAGTCGTTGCAACAAATGGAAACACATCCCAGGNNNAGATCGGAAG
40828 TATGAGTGTGGAGTCGTTGCGCCTGGAATCTTACAGGAGAACANNNAGATCGGAAG
40829 TATGAGTGTGGAGTCGTTGCTGTGTCCCAAATCATTGAACTGTNNNAGATCGGAAG
40830 TATGAGTGTGGAGTCGTTGCGGCTATGCAGTGGTACAAATGACNNNAGATCGGAAG
40831 TATGAGTGTGGAGTCGTTGCAGCTGCATGGTTTTGTAATACCANNNAGATCGGAAG
40832 TATGAGTGTGGAGTCGTTGCGACATACAAAAGGCCAACTGACANNNAGATCGGAAG
40834 TATGAGTGTGGAGTCGTTGCCTGAGGCAGGAGAATCACTTGAGNNNAGATCGGAA
40835 TATGAGTGTGGAGTCGTTGCCAGGATGGTGGTTGCTATAGGTTNNNAGATCGGAA
40836 TATGAGTGTGGAGTCGTTGCGATCACACCACTATACTCCAGCCNNNAGATCGGAAG
40837 TATGAGTGTGGAGTCGTTGCTGGCCTTACATAGGAGGTAGACANNNAGATCGGAA
40838 TATGAGTGTGGAGTCGTTGCAAGAGCAGACAAGACAATTGCAGNNNAGATCGGAA
40839 TATGAGTGTGGAGTCGTTGCCAACTGTGGCTAAGGAAAAAGGGNNNAGATCGGAA
40840 TATGAGTGTGGAGTCGTTGCCTGTAGTCCCAGCTACTCAGGAGNNNAGATCGGAAG
40841 TATGAGTGTGGAGTCGTTGCGTAGGTAGGATTACAGGCTCGTGNNNAGATCGGAA
40843 TATGAGTGTGGAGTCGTTGCTTGTGTTGGTTTTTTCCTCCTCCNNNAGATCGGAAGA
40844 TATGAGTGTGGAGTCGTTGCACAGAAAAGCAAACACCACATGTNNNAGATCGGAAG
40845 TATGAGTGTGGAGTCGTTGCGCTATAAGGTGCATATGGCTTGCNNNAGATCGGAAG
40846 TATGAGTGTGGAGTCGTTGCCTCCAACTGGGTAACAGAGTGAGNNNAGATCGGAA
40847 TATGAGTGTGGAGTCGTTGCAGGGGAACTCTCTTAGTACCTGGNNNAGATCGGAAG
40848 TATGAGTGTGGAGTCGTTGCTTTGGGGTACAAGTGGTTTCTCANNNAGATCGGAAG
40849 TATGAGTGTGGAGTCGTTGCTCTTGTGGACCAGAAGAAGGAAGNNNAGATCGGAA
40850 TATGAGTGTGGAGTCGTTGCCTGAGACAGGAGAATCGCTTGAANNNAGATCGGAA
40853 TATGAGTGTGGAGTCGTTGCCCTTACAGTGCCTCACAAGAGATNNNAGATCGGAAG
40854 TATGAGTGTGGAGTCGTTGCCTGCAAATGTACCAAGAGTGTCANNNAGATCGGAAG
40857 TATGAGTGTGGAGTCGTTGCTGTTGTTGTTTTCGTAAGTGCCANNNAGATCGGAAG
40859 TATGAGTGTGGAGTCGTTGCGATGCGAACACATGGACACATAGNNNAGATCGGAA
40860 TATGAGTGTGGAGTCGTTGCAGATCTGGGAGTCTTTGAGGAGANNNAGATCGGAA
40862 TATGAGTGTGGAGTCGTTGCGCCCCTTACAAATTTGAAACCCANNNAGATCGGAAG
40866 TATGAGTGTGGAGTCGTTGCGGATTCCCTTGGGTCTTTCACTANNNAGATCGGAAG
40867 TATGAGTGTGGAGTCGTTGCGAAAATAGGCTCCCAATCGCTTCNNNAGATCGGAAG
40871 TATGAGTGTGGAGTCGTTGCCTCCTAGGGGAAGGCAGAGTATANNNAGATCGGAA
40874 TATGAGTGTGGAGTCGTTGCCCACGACCGGCTAATTTTTGTATNNNAGATCGGAAG

40875 TATGAGTGTGGAGTCGTTGCGTGGTAAGCTGAAGACGTAGGAANNNAGATCGGAA
40876 TATGAGTGTGGAGTCGTTGCCAGGCCACAGCACAATCAAATTANNNAGATCGGAAG
40882 TATGAGTGTGGAGTCGTTGCGAGTTTGTAGTGAGCCAAGATCGNNNAGATCGGAAG
40883 TATGAGTGTGGAGTCGTTGCTGAAAGCAACAACAACACAAACANNNAGATCGGAAG
40884 TATGAGTGTGGAGTCGTTGCAACAAGATCTCACTCTGTCACCCNNNAGATCGGAAGA
40886 TATGAGTGTGGAGTCGTTGCCAGATGGAAAAAAGGCTGGATGGNNNAGATCGGAA
40887 TATGAGTGTGGAGTCGTTGCCTGACTCATGCTAAATGGCCATGNNNAGATCGGAAG
40888 TATGAGTGTGGAGTCGTTGCACTTCATGTCTTTAGCCCCAGTTNNNAGATCGGAAGA
40890 TATGAGTGTGGAGTCGTTGCCAGCCTGACAATGCAGAAGAAAANNNAGATCGGAAG
40891 TATGAGTGTGGAGTCGTTGCGCAAGGCTCCATCTCAAAAGAAANNNAGATCGGAAG
40892 TATGAGTGTGGAGTCGTTGCGAGTTGGTGTGAGAGTGAGCATANNNAGATCGGAA
40894 TATGAGTGTGGAGTCGTTGCTCTGGATAAGGAAGTGTGGCAAANNNAGATCGGAA
40895 TATGAGTGTGGAGTCGTTGCATCTTTTTGCCCAAGTCACAGTGNNNAGATCGGAAGA
40897 TATGAGTGTGGAGTCGTTGCAGAGTCAGCAGACACAGCAAATANNNAGATCGGAAG
40899 TATGAGTGTGGAGTCGTTGCTAGTGGGAAAACAACCTTCCTCANNNAGATCGGAAG
40900 TATGAGTGTGGAGTCGTTGCGCTCAAATTCACTGAATCACCCCNNNAGATCGGAAGA
40901 TATGAGTGTGGAGTCGTTGCGTTTGTGCATGCTCTACAGTCTGNNNAGATCGGAAG
40902 TATGAGTGTGGAGTCGTTGCCCACTTTCTTCCGTTTGTCCTTCNNNAGATCGGAAGAG
40906 TATGAGTGTGGAGTCGTTGCGGTCATTGTCTCTGCCATCTCTTNNNAGATCGGAAGA
40907 TATGAGTGTGGAGTCGTTGCGGCAGGTGTAGAGGAGAGATAGANNNAGATCGGA
40908 TATGAGTGTGGAGTCGTTGCCTGAGACAGGAGAATGGCATGAANNNAGATCGGAA
40909 TATGAGTGTGGAGTCGTTGCGTGCCTTGGAGAAATCAAAGAGCNNNAGATCGGAAG
40910 TATGAGTGTGGAGTCGTTGCAGAGCTGGTAACATTCCTACTGANNNAGATCGGAAG
40912 TATGAGTGTGGAGTCGTTGCCAAACATGACTTTGAGGCTTCGTNNNAGATCGGAAG
40914 TATGAGTGTGGAGTCGTTGCCACCTTTAGTCCCAGCTACTCAGNNNAGATCGGAAGA
40915 TATGAGTGTGGAGTCGTTGCACTCCCTTGATTGAGTCACCTTCNNNAGATCGGAAGA
40916 TATGAGTGTGGAGTCGTTGCAGATAGCATGACCACACTCTAGCNNNAGATCGGAAG
40917 TATGAGTGTGGAGTCGTTGCTACCAGTACCATGCTGTTTTGGTNNNAGATCGGAAGA
40918 TATGAGTGTGGAGTCGTTGCCTGGATTTGAGCATTCATTTGGCNNNAGATCGGAAG
40919 TATGAGTGTGGAGTCGTTGCTATGGGACTAGGAGGAAGACAGGNNNAGATCGGAA
40920 TATGAGTGTGGAGTCGTTGCAGGGCTGAGGAGGAGATCTAAAANNNAGATCGGAA
40921 TATGAGTGTGGAGTCGTTGCGATTTCCACCAGCAGCATATAGCNNNAGATCGGAAG
40922 TATGAGTGTGGAGTCGTTGCACAGGAGCTGCTACCACTTATTCNNNAGATCGGAAGA
40923 TATGAGTGTGGAGTCGTTGCCAATGCCATACTGTGCAACTCTGNNNAGATCGGAAGA
40924 TATGAGTGTGGAGTCGTTGCCTGGGCAACAGAGCAATATTCTGNNNAGATCGGAAG
40926 TATGAGTGTGGAGTCGTTGCCCAGGCTCCATACACAGTTCTAANNNAGATCGGAAGA
40928 TATGAGTGTGGAGTCGTTGCAACACTGACCAGAATGTCTTACANNNAGATCGGAAG
40929 TATGAGTGTGGAGTCGTTGCACAACTTTTCCCAAACTCGGTTTNNNAGATCGGAAGA
40930 TATGAGTGTGGAGTCGTTGCACGAAAAACTGCTGAATCACACTNNNAGATCGGAAG
40933 TATGAGTGTGGAGTCGTTGCTTCCAAAAGCCTCCTCTAACTGANNNAGATCGGAAGA
40934 TATGAGTGTGGAGTCGTTGCTTTCACAGGTGTATATGTGCCGANNNAGATCGGAAG
40937 TATGAGTGTGGAGTCGTTGCCCAGTTATTTGCCTTTCCACAGGNNNAGATCGGAAGA
40939 TATGAGTGTGGAGTCGTTGCGTCACTACAATCTCCACCTCCTGNNNAGATCGGAAGA

40940 TATGAGTGTGGAGTCGTTGCCTATTCACAAAGGCCAGGGTTTGNNNAGATCGGAAG
40941 TATGAGTGTGGAGTCGTTGCAGTACTGGAGAGGATGTGGAGAANNNAGATCGGAA
40944 TATGAGTGTGGAGTCGTTGCGGAATTGCATATCCATGGGGTTGNNNAGATCGGAA
40945 TATGAGTGTGGAGTCGTTGCTGGAAAACTGCCAGAGAAAGAGANNNAGATCGGAA
40946 TATGAGTGTGGAGTCGTTGCCAATCTCAGTTCACTGCAAGCTCNNNAGATCGGAAGA
40948 TATGAGTGTGGAGTCGTTGCCTTTTCCCCACCCAGCTTTATTGNNNAGATCGGAAGA
40949 TATGAGTGTGGAGTCGTTGCAAGTCTCTTCCCTTTACTTCGGCNNNAGATCGGAAGA
40950 TATGAGTGTGGAGTCGTTGCTGCCTGTAATCTCAGCTACTCAGNNNAGATCGGAAGA
40952 TATGAGTGTGGAGTCGTTGCGCAAGACCCTGTCTCCAAAAAAANNNAGATCGGAAG
40956 TATGAGTGTGGAGTCGTTGCTAGTATGTGTGGAAATGTCAGGTNNNAGATCGGAA
40958 TATGAGTGTGGAGTCGTTGCTATGAGGGTTGACATTGGACCTGNNNAGATCGGAA
40961 TATGAGTGTGGAGTCGTTGCCCATCAGGAAGCACAAAGGTAACNNNAGATCGGAA
40964 TATGAGTGTGGAGTCGTTGCCCTAAGCGACAAGTACAGTGACTNNNAGATCGGAAG
40966 TATGAGTGTGGAGTCGTTGCTTGTCAGCCAGAAGTCTTCTTGANNNAGATCGGAAG
40967 TATGAGTGTGGAGTCGTTGCGGTCCTGGCATAAAAACAGACACNNNAGATCGGAAG
40968 TATGAGTGTGGAGTCGTTGCACGCAAGGTTATTCATAGGCTCTNNNAGATCGGAAG
40970 TATGAGTGTGGAGTCGTTGCGAAAGAAAAATGGACTGGGGTGGNNNAGATCGGAA
40971 TATGAGTGTGGAGTCGTTGCCTGTAATCCCAGCTACTCAGGAGNNNAGATCGGAAG
40972 TATGAGTGTGGAGTCGTTGCAGTGGAAAGATGGCTGATAAGCANNNAGATCGGAA
40975 TATGAGTGTGGAGTCGTTGCCATTTACACTCTTGCTGGTGGTGNNNAGATCGGAAG
40976 TATGAGTGTGGAGTCGTTGCACAATGCGTGGGAATTCAAGATGNNNAGATCGGAA
40977 TATGAGTGTGGAGTCGTTGCTTATTAGAGGACTTGTCGGCACCNNNAGATCGGAAG
40978 TATGAGTGTGGAGTCGTTGCACAAGCCAAACTGAAACAGCAATNNNAGATCGGAAG
40979 TATGAGTGTGGAGTCGTTGCGTCTCTTCACACTGCGTTTCTTGNNNAGATCGGAAGA
40980 TATGAGTGTGGAGTCGTTGCAAATCACCGAGTATGGTCTCTGGNNNAGATCGGAAG
40981 TATGAGTGTGGAGTCGTTGCCGACAACGACAACAACAACAAAANNNAGATCGGAAG
40982 TATGAGTGTGGAGTCGTTGCGAACGTTATCAGGCGATATTGGANNNAGATCGGAA
40984 TATGAGTGTGGAGTCGTTGCAGAAGGCAAGACTCCAAGAACTTNNNAGATCGGAA
40985 TATGAGTGTGGAGTCGTTGCGCAGAAGAATCACTTGAACCCAGNNNAGATCGGAAG
40986 TATGAGTGTGGAGTCGTTGCTTCTGTAAGTGGATGTGGAGTGGNNNAGATCGGAA
40987 TATGAGTGTGGAGTCGTTGCATTCCCTTCCTCTCTCCCTTCTTNNNAGATCGGAAGAG
40988 TATGAGTGTGGAGTCGTTGCGTAGGGTGATTAGAGGGCTTGAGNNNAGATCGGAA
40989 TATGAGTGTGGAGTCGTTGCAAAGTGTGTAGCATTTCCCCCTTNNNAGATCGGAAGA
40990 TATGAGTGTGGAGTCGTTGCCCACGCCCAGCTAATTTTTGTATNNNAGATCGGAAGA
40991 TATGAGTGTGGAGTCGTTGCGAACCCCTACAAGTCAACAACAANNNAGATCGGAAG
40992 TATGAGTGTGGAGTCGTTGCCATGTGGAGAAATTGGAAGCCTCNNNAGATCGGAA
40993 TATGAGTGTGGAGTCGTTGCGTTGCAGTACTTCCACTACCTGANNNAGATCGGAAGA
40995 TATGAGTGTGGAGTCGTTGCATGAAAACCTCTGATGGCCTTGANNNAGATCGGAAG
40996 TATGAGTGTGGAGTCGTTGCAATGGGGGTGGGACAAAATCTTANNNAGATCGGAA
40998 TATGAGTGTGGAGTCGTTGCGCACCTCAAGATGCTTCAAGATTNNNAGATCGGAAG
40999 TATGAGTGTGGAGTCGTTGCATATGTATGTGTTCTCCCCACCCNNNAGATCGGAAGA
41000 TATGAGTGTGGAGTCGTTGCGAGACAGAGTTTCGCTGTTGTTGNNNAGATCGGAA
41002 TATGAGTGTGGAGTCGTTGCAAAAAATTGCCGGACATGGTGATNNNAGATCGGAA

41003 TATGAGTGTGGAGTCGTTGCGATCACCTGAGGCTAGGAGTTTGNNNAGATCGGAA
41004 TATGAGTGTGGAGTCGTTGCAGACAGTCTCTCACTTTGTCACCNNNAGATCGGAAG
41005 TATGAGTGTGGAGTCGTTGCTATGTACATGAGGCCAGACATGGNNNAGATCGGAA
41007 TATGAGTGTGGAGTCGTTGCCAGCATGGCTCTTCCTCAAAAAANNNAGATCGGAAG
41009 TATGAGTGTGGAGTCGTTGCAAAAGAAAGAAAGAAGGCCAGGCNNNAGATCGGA
41011 TATGAGTGTGGAGTCGTTGCACTTCTGGGTTCAAGTGATCCTCNNNAGATCGGAAG
41014 TATGAGTGTGGAGTCGTTGCTCAGATGAAAGAAAAAGCTGAGACANNNAGATCGG
41015 TATGAGTGTGGAGTCGTTGCGACAAGGCACTAAGTACCCAAGANNNAGATCGGAA
41016 TATGAGTGTGGAGTCGTTGCTACCATTCAGGACATAGGCATGGNNNAGATCGGAAG
41018 TATGAGTGTGGAGTCGTTGCCAAAAGAAGACGTACAAGTGGCCNNNAGATCGGAA
41019 TATGAGTGTGGAGTCGTTGCTTGCTGCTGTTCTCTTAGGCTAANNNAGATCGGAAG
41020 TATGAGTGTGGAGTCGTTGCTGACAGCTTCACCTACTTCCTACNNNAGATCGGAAGA
41022 TATGAGTGTGGAGTCGTTGCAGAGAGAACTGAAGCAAATGTGCNNNAGATCGGAA
41023 TATGAGTGTGGAGTCGTTGCCACCCATCTTAGACACCCTGAAANNNAGATCGGAAG
41024 TATGAGTGTGGAGTCGTTGCACACTAAAGTCCATTCCCCCAAANNNAGATCGGAAG
41025 TATGAGTGTGGAGTCGTTGCGAACTTGGAGTCCGATGTTCAAGNNNAGATCGGAA
41026 TATGAGTGTGGAGTCGTTGCGGGGCTTTGTACCACACTGAATANNNAGATCGGAAG
41027 TATGAGTGTGGAGTCGTTGCAGCCTTCTCATATGCCAAGACTTNNNAGATCGGAAG
41028 TATGAGTGTGGAGTCGTTGCATTGAAAGCAGGGACTGAACAGANNNAGATCGGAA
41029 TATGAGTGTGGAGTCGTTGCGAGGAAAGGGCAGTCTTTTCAACNNNAGATCGGAA
41030 TATGAGTGTGGAGTCGTTGCCAGAAGGGCAAAAGTCGGAAATTNNNAGATCGGAA
41031 TATGAGTGTGGAGTCGTTGCAAAAAGAAAGCCACTTCCCACTGNNNAGATCGGAAG
41032 TATGAGTGTGGAGTCGTTGCCTGAGGCAGGAGAATCACTTGAANNNAGATCGGAA
41036 TATGAGTGTGGAGTCGTTGCTGTACACCTATGCTCACAGAAGCNNNAGATCGGAAG
41037 TATGAGTGTGGAGTCGTTGCCTCTCATTTGGACTGCTGGGATANNNAGATCGGAAG
41038 TATGAGTGTGGAGTCGTTGCTGAGACACTGTCAAAGACTGGAANNNAGATCGGAA
41039 TATGAGTGTGGAGTCGTTGCCTCCTCCTTGTACCTCTGGTAGANNNAGATCGGAAG
41040 TATGAGTGTGGAGTCGTTGCAGGTCAGAATTCTGGAAAGGCATNNNAGATCGGAA
41042 TATGAGTGTGGAGTCGTTGCAGACATAATCCGTTAGGACAGTCCNNNAGATCGGAA
41043 TATGAGTGTGGAGTCGTTGCGTGCTAGAACAATGGTTGCTTCCNNNAGATCGGAAG
41044 TATGAGTGTGGAGTCGTTGCAGGCCCATATATTCAGGCTTGAGNNNAGATCGGAAG
41045 TATGAGTGTGGAGTCGTTGCCCAGCTTTATTCTGCCTTTCCAGNNNAGATCGGAAGA
41046 TATGAGTGTGGAGTCGTTGCGCTTGCAGTGAGCCAAGATAATGNNNAGATCGGAA
41047 TATGAGTGTGGAGTCGTTGCCACCTATATTCCCAGCTACTCGGNNNAGATCGGAAG
41048 TATGAGTGTGGAGTCGTTGCTTCAACCTATGCCTGCAGTTACTNNNAGATCGGAAG
41049 TATGAGTGTGGAGTCGTTGCGAACTCACAGAAGCAGAGGGTAGNNNAGATCGGA
41050 TATGAGTGTGGAGTCGTTGCAGGAAGATCCAAACATTCCCACANNNAGATCGGAAG
41051 TATGAGTGTGGAGTCGTTGCTGCACATTCCAGGTACTAGACACNNNAGATCGGAAG
41052 TATGAGTGTGGAGTCGTTGCCGTTTATACACTGCTAGTGGGCANNNAGATCGGAAG
41053 TATGAGTGTGGAGTCGTTGCTGTGATCATGCCACTAAACTCCANNNAGATCGGAAG
41054 TATGAGTGTGGAGTCGTTGCAAGAGTTCCATCTCCATGCAAGTNNNAGATCGGAAG
41055 TATGAGTGTGGAGTCGTTGCGTCAACTAGGTTGCTGTAAAGGCNNNAGATCGGAA
41057 TATGAGTGTGGAGTCGTTGCCGTAGGCCCTCATGATTGGATTANNNAGATCGGAAG

41058 TATGAGTGTGGAGTCGTTGCTAACAGAATAGTCTAGGGCGCTGNNNAGATCGGAA
41059 TATGAGTGTGGAGTCGTTGCTCTGTCTCTCTCCTCTCTCTCTCNNNAGATCGGAAGAG
41060 TATGAGTGTGGAGTCGTTGCAAAACCTCAGTGGGTGTGATACANNNAGATCGGAA
41061 TATGAGTGTGGAGTCGTTGCCACCCACCCTATTCCCAACTTATNNNAGATCGGAAGA
41062 TATGAGTGTGGAGTCGTTGCCCCCATCTCTCCACTTTGCTATTNNNAGATCGGAAGA
41063 TATGAGTGTGGAGTCGTTGCAGGCATGAGAAATCACCCTAAGANNNAGATCGGAA
41064 TATGAGTGTGGAGTCGTTGCAGAATGAACAGATGCAGGGACTTNNNAGATCGGAA
41065 TATGAGTGTGGAGTCGTTGCCTGCAACCAAGAACTTCCAACAGNNNAGATCGGAAG
41067 TATGAGTGTGGAGTCGTTGCATCCCTCTTCCCTTCTTTTCACCNNNAGATCGGAAGAG
41069 TATGAGTGTGGAGTCGTTGCCTCAGATTACACACCCAGCTGATNNNAGATCGGAAG
41070 TATGAGTGTGGAGTCGTTGCTTGAATGATACTGAGGGCCTGTCNNNAGATCGGAA
41071 TATGAGTGTGGAGTCGTTGCAAGAAGAAGAGAGAAGTTGGCGGNNNAGATCGGA
41073 TATGAGTGTGGAGTCGTTGCTTCACTATATTGCCCAGGCTTGTNNNAGATCGGAAG
41074 TATGAGTGTGGAGTCGTTGCTTCACTATATTGCCCAGGCTTGTNNNAGATCGGAAG
41075 TATGAGTGTGGAGTCGTTGCGCGACAGAACAAGACTCAGTCTANNNAGATCGGAA
41076 TATGAGTGTGGAGTCGTTGCTACAAAAATTAGCTGGGCATGGCNNNAGATCGGAA
41078 TATGAGTGTGGAGTCGTTGCAAAGCCAGAATGCTATGGACTGANNNAGATCGGAA
41080 TATGAGTGTGGAGTCGTTGCAAATGCCTGATAAGAGGAATGCANNNAGATCGGAA
41081 TATGAGTGTGGAGTCGTTGCGTGCACGCAAGTGTGTAGATATTNNNAGATCGGAA
41082 TATGAGTGTGGAGTCGTTGCGTGGATAACAGCAAACAACCTCCNNNAGATCGGAAG
41083 TATGAGTGTGGAGTCGTTGCATACCAAGCAGTTTGTATGCAGCNNNAGATCGGAAG
41084 TATGAGTGTGGAGTCGTTGCGCATACAGGATTGTGTGATCTGCNNNAGATCGGAA
41086 TATGAGTGTGGAGTCGTTGCCTGAGGCAGGAGAATTGCTTGAGNNNAGATCGGAA
41088 TATGAGTGTGGAGTCGTTGCAGTGATGAAGATGGTCAACTGCTNNNAGATCGGAA
41089 TATGAGTGTGGAGTCGTTGCAAAGAAAAATTAGCGAGGCGAGGNNNAGATCGGA
41090 TATGAGTGTGGAGTCGTTGCCACCAAGTCCAGCCTATACTCAANNNAGATCGGAAG
41091 TATGAGTGTGGAGTCGTTGCCACACCTGTAATCCCAGCACTTTNNNAGATCGGAAGA
41092 TATGAGTGTGGAGTCGTTGCAATGGGGCACTAATGAGCTGATTNNNAGATCGGAA
41093 TATGAGTGTGGAGTCGTTGCGGCTCTGAGGCAATATGAAGAGGNNNAGATCGGAA
41094 TATGAGTGTGGAGTCGTTGCATTGAATGACAACACAGGGCATGNNNAGATCGGAA
41102 TATGAGTGTGGAGTCGTTGCGAGAGAGAGAGAGACAGGGTCTTNNNAGATCGGA
41103 TATGAGTGTGGAGTCGTTGCTAGAATAGCAGTGACTGTCACCANNNAGATCGGAA
41105 TATGAGTGTGGAGTCGTTGCAAATTTGTCCAAGCGTAGTGGTGNNNAGATCGGAA
41106 TATGAGTGTGGAGTCGTTGCAGTGCCTAACAACAAAGTGTTCANNNAGATCGGAAG
41107 TATGAGTGTGGAGTCGTTGCGACTGGGTGAAAGCTGGTTACTANNNAGATCGGAA
41108 TATGAGTGTGGAGTCGTTGCAGGAAGGAAGCTGTACCTCTACTNNNAGATCGGAA
41109 TATGAGTGTGGAGTCGTTGCAAGGTGGAGTAACTTTGTAGGGCNNNAGATCGGAA
41110 TATGAGTGTGGAGTCGTTGCGTTCAAAAGATTCTTGTGCCCCANNNAGATCGGAAG
41111 TATGAGTGTGGAGTCGTTGCTGTTTTCCGACATGGCTATACCANNNAGATCGGAAG
41112 TATGAGTGTGGAGTCGTTGCGACCTATTTCTGTCTTCCCTGCTNNNAGATCGGAAGA
41113 TATGAGTGTGGAGTCGTTGCCCGACATTTGAAAAGAGTGCACTNNNAGATCGGAA
41114 TATGAGTGTGGAGTCGTTGCTGTTTTCCGACATGGCTATACCANNNAGATCGGAAG
41115 TATGAGTGTGGAGTCGTTGCAGAGATGTTAGGTGTGCTGATGGNNNAGATCGGAA

41116 TATGAGTGTGGAGTCGTTGCCCGACATTTGAAAAGAGTGCACTNNNAGATCGGAA

41117 TATGAGTGTGGAGTCGTTGCTGTTTTCCGACATGGCTATACCANNNAGATCGGAAG

41118 TATGAGTGTGGAGTCGTTGCAGAGATGTTAGGTGTGCTGATGGNNNAGATCGGA

41119 TATGAGTGTGGAGTCGTTGCGACCTATTTCTGTCTTCCCTGCTNNNAGATCGGAAGA

41120 TATGAGTGTGGAGTCGTTGCCTCCTGTAATCCCAGCACTTTGGNNNAGATCGGAAG

41121 TATGAGTGTGGAGTCGTTGCAGTGCCAAGTGGGACATTTCTAGNNNAGATCGGAA

41126 TATGAGTGTGGAGTCGTTGCTCAATTGAGGTCAGGAGTTGGAGNNNAGATCGGAA

41127 TATGAGTGTGGAGTCGTTGCATGTACTATACCCAGTCCGTGGANNNAGATCGGAAG

41128 TATGAGTGTGGAGTCGTTGCTTCTGACATATGCTCCAACATGANNNAGATCGGAAG

41131 TATGAGTGTGGAGTCGTTGCGGACTAGGATAGTGGCAGTTGAGNNNAGATCGGA

41132 TATGAGTGTGGAGTCGTTGCACTCACAGCCCTCTGATATAGGTNNNAGATCGGAAG

41133 TATGAGTGTGGAGTCGTTGCGATGCTCAAGGGCAGAGATTCTANNNAGATCGGAA

41134 TATGAGTGTGGAGTCGTTGCTAGAGTGCACTAATACACCCAGCNNNAGATCGGAAG

41136 TATGAGTGTGGAGTCGTTGCATCCTAGCTACTCAGAAGGCTGANNNAGATCGGAAG

41137 TATGAGTGTGGAGTCGTTGCAAAACCAGCAAGACTCCATCTCTNNNAGATCGGAAG

41138 TATGAGTGTGGAGTCGTTGCGTTGAGACTGTAGTGAGCCATGANNNAGATCGGAA

41139 TATGAGTGTGGAGTCGTTGCGGCAGAATGCATTGTTCCTGTAANNNAGATCGGAA

41140 TATGAGTGTGGAGTCGTTGCCACACCTGTAATCCCAGCACTTTNNNAGATCGGAAGA

41141 TATGAGTGTGGAGTCGTTGCGAGGCAGGAGAATCACTTGAACTNNNAGATCGGAA

41142 TATGAGTGTGGAGTCGTTGCTGAATGTAACTGATGCCACAGGANNNAGATCGGAA

41143 TATGAGTGTGGAGTCGTTGCTCTATCTTTCTCCAAAGCCAGCANNNAGATCGGAAGA

41145 TATGAGTGTGGAGTCGTTGCAATAATGGGAGAGGATTGCGAGGNNNAGATCGGA

41146 TATGAGTGTGGAGTCGTTGCTGCCTGTAATCCCAGCTATTTGGNNNAGATCGGAAG

41147 TATGAGTGTGGAGTCGTTGCGAGGTCAGGAGATTGAGACCATCNNNAGATCGGAA

41148 TATGAGTGTGGAGTCGTTGCCCTCCTCCTTCAGCTTATCGATCNNNAGATCGGAAGA

41149 TATGAGTGTGGAGTCGTTGCCACACACACACACACACATACATNNNAGATCGGAAGA

41150 TATGAGTGTGGAGTCGTTGCAAGTTAGTCGCCATGCTCTCTACNNNAGATCGGAAG

41151 TATGAGTGTGGAGTCGTTGCAAAAAATTAGCAGGGTGTGGTGGNNNAGATCGGA

41152 TATGAGTGTGGAGTCGTTGCACATTGTAGCCTCAAACTCCTGGNNNAGATCGGAAG

41153 TATGAGTGTGGAGTCGTTGCTGGTCAAAGCCATTCAACAAGTCNNNAGATCGGAAG

41155 TATGAGTGTGGAGTCGTTGCCGGAGACTGGGTAATTTGGGTAANNNAGATCGGAA

41156 TATGAGTGTGGAGTCGTTGCACCACCACCAAAAAACCAAAAGANNNAGATCGGAAG

41157 TATGAGTGTGGAGTCGTTGCTGAGGCAGAATTTCGCTATGTTGNNNAGATCGGAA

41158 TATGAGTGTGGAGTCGTTGCTACGCTAAGTGAATAGGCCAGACNNNAGATCGGAA

41162 TATGAGTGTGGAGTCGTTGCTTCATTGCAACCTGATCTGCTTTNNNAGATCGGAAGA

41163 TATGAGTGTGGAGTCGTTGCCTGTAATCCCAGCTACTGAGGAGNNNAGATCGGAA

41164 TATGAGTGTGGAGTCGTTGCACATGGTGAAACCTCGTCTCTACNNNAGATCGGAAG

41165 TATGAGTGTGGAGTCGTTGCGAAGCTAAGACAGGAGAATCGCTNNNAGATCGGAA

41168 TATGAGTGTGGAGTCGTTGCTGTGAACAGTTTGGGGTGACATANNNAGATCGGAA

41171 TATGAGTGTGGAGTCGTTGCTAGGAACTTGTCAGCCACTCAAANNNAGATCGGAAG

41172 TATGAGTGTGGAGTCGTTGCACTTTTACACTGTTGGTGGGAGTNNNAGATCGGAA

41174 TATGAGTGTGGAGTCGTTGCGATACTCCTCACCTTCTGATGGGNNNAGATCGGAAG

41175 TATGAGTGTGGAGTCGTTGCTGCATGGACCCATTTCTAGATGTNNNAGATCGGAAG

41176 TATGAGTGTGGAGTCGTTGCACTGCCACCAGAAAGAGAAATCTNNNAGATCGGAAG
41177 TATGAGTGTGGAGTCGTTGCAGACAAGGTCCTTGCATTGTTTGNNNAGATCGGAAG
41178 TATGAGTGTGGAGTCGTTGCGGGATTGCACTAAGTAGCTCGTANNNAGATCGGAAG
41180 TATGAGTGTGGAGTCGTTGCGGTTAGGATGCAGAGAAACTGGANNNAGATCGGAA
41181 TATGAGTGTGGAGTCGTTGCAATACAGCACACTGATGGGTCTTNNNAGATCGGAAG
41183 TATGAGTGTGGAGTCGTTGCTTCCATTGTGTGTATATGCCACANNNAGATCGGAAGA
41188 TATGAGTGTGGAGTCGTTGCCCATCCCCACTCCTTTTCAGTTANNNAGATCGGAAGAG
41189 TATGAGTGTGGAGTCGTTGCTACGTTGCCCAGACTAGTTTCAANNNAGATCGGAAGA
41190 TATGAGTGTGGAGTCGTTGCACATGGTGAAACCTCGTCTCTACNNNAGATCGGAAGA
41191 TATGAGTGTGGAGTCGTTGCATGACCCATGATGAAACCCATGANNNAGATCGGAAG
41193 TATGAGTGTGGAGTCGTTGCTCCTAATATCCAGGTGGGGAGAGNNNAGATCGGAAG
41194 TATGAGTGTGGAGTCGTTGCGCACCCCTAAAGATAGTTCCCAANNNAGATCGGAAGA
41195 TATGAGTGTGGAGTCGTTGCACTCACAAAAAAATTGCACCTGCNNNAGATCGGAAGA
41196 TATGAGTGTGGAGTCGTTGCAGAGAGAGGAGAGAGAGAGGAGANNNAGATCGGA
41197 TATGAGTGTGGAGTCGTTGCCCTTTTTGAGCTGTAACACTCCCNNNAGATCGGAAGA
41200 TATGAGTGTGGAGTCGTTGCGAGAAAGGGTGAAGGCAAACATCNNNAGATCGGAA
41201 TATGAGTGTGGAGTCGTTGCCCCTCTCTTTCTCGGTACCTTTCNNNAGATCGGAAGAG
41202 TATGAGTGTGGAGTCGTTGCTTCTGAAGCAACCAGATACTGCTNNNAGATCGGAAG
41203 TATGAGTGTGGAGTCGTTGCCTTTCTCTTTCTGCTCCCCCTACNNNAGATCGGAAGAG
41205 TATGAGTGTGGAGTCGTTGCTTTGTCATGAACCATCTGTTGCCNNNAGATCGGAAGA
41206 TATGAGTGTGGAGTCGTTGCAAAAGACTCACCCCTACAAGGTCNNNAGATCGGAAG
41208 TATGAGTGTGGAGTCGTTGCACTGAGAATGATGGTTTCCAGCTNNNAGATCGGAAG
41209 TATGAGTGTGGAGTCGTTGCCACACAGGAATAGGCCTCTTGTANNNAGATCGGAAG
41210 TATGAGTGTGGAGTCGTTGCGGTAGTGGAGTGTTGACCAATCANNNAGATCGGAAG
41211 TATGAGTGTGGAGTCGTTGCACACACACACATAGCTCATAGGGNNNAGATCGGAAG
41212 TATGAGTGTGGAGTCGTTGCAAGAAATTCACCTGTCTCAGCCTNNNAGATCGGAAGA
41213 TATGAGTGTGGAGTCGTTGCGCTTGGTCTCCGTCTCAAAAAATNNNAGATCGGAAGA
41215 TATGAGTGTGGAGTCGTTGCCAAGCACTTGGAGGGAAGAAAAGNNNAGATCGGAA
41216 TATGAGTGTGGAGTCGTTGCTACATGACCCAAATGTCGTCCTTNNNAGATCGGAAGA
41218 TATGAGTGTGGAGTCGTTGCAAGAAAATTAGCTGGGCATGGTGNNNAGATCGGAA
41221 TATGAGTGTGGAGTCGTTGCTACAAAAAATTAGCCGGGTGCAGNNNAGATCGGAAG
41223 TATGAGTGTGGAGTCGTTGCCCAAAGCAAGCAGATCTCTTCAGNNNAGATCGGAAG
41225 TATGAGTGTGGAGTCGTTGCCCACTCCTGCGTATGACATTTTTNNNAGATCGGAAGA
41226 TATGAGTGTGGAGTCGTTGCTAAAGTCGGTCCTCATTCAGGTCNNNAGATCGGAAG
41228 TATGAGTGTGGAGTCGTTGCACTCAAACAGGTACTTGTCCACANNNAGATCGGAAGA
41229 TATGAGTGTGGAGTCGTTGCGGATCTTGCTATGTTTTCCAGGCNNNAGATCGGAAG
41231 TATGAGTGTGGAGTCGTTGCATAGCTCGTGGAAAGTGGTTTCTNNNAGATCGGAAG
41232 TATGAGTGTGGAGTCGTTGCATGGATGGAAGAGGAGGTCACTANNNAGATCGGAA
41233 TATGAGTGTGGAGTCGTTGCCAACAGAAAAATCCGTGACCCAANNNAGATCGGAAG
41234 TATGAGTGTGGAGTCGTTGCCAAGGTCAAGAGATCGAGACCATNNNAGATCGGAAG
41235 TATGAGTGTGGAGTCGTTGCTACTCAGAAAACGGTCCAACAGTNNNAGATCGGAAG
41237 TATGAGTGTGGAGTCGTTGCAGATCGAGACCATCTTGGCTAACNNNAGATCGGAAG
41240 TATGAGTGTGGAGTCGTTGCGGTGGATCATGAGGTCAAGAGATNNNAGATCGGAA

41242 TATGAGTGTGGAGTCGTTGCCTGTAATCCCAGCTACTCAGGAGNNNAGATCGGAAG
41243 TATGAGTGTGGAGTCGTTGCAAAACCGCTTGAATGAGAACAGGNNNAGATCGGAA
41244 TATGAGTGTGGAGTCGTTGCTAACATAGCAAGACCACCCATCCNNNAGATCGGAAG
41246 TATGAGTGTGGAGTCGTTGCCATCCTGGCTAACATGGTGAAACNNNAGATCGGAAG
41248 TATGAGTGTGGAGTCGTTGCGATGTGTTTTCGCTGGGAAAGTTNNNAGATCGGAA
41249 TATGAGTGTGGAGTCGTTGCCAAATCTCAAAGACTCAGCCCCANNNAGATCGGAAG
41250 TATGAGTGTGGAGTCGTTGCATTCGTGAAGTTGAAGTGGCTCANNNAGATCGGAA
41251 TATGAGTGTGGAGTCGTTGCATACAAAAATTAGCTGGGTGCGGNNNAGATCGGAA
41252 TATGAGTGTGGAGTCGTTGCAGCGAGAGAGAAAGAGAAAGACANNNAGATCGGA
41253 TATGAGTGTGGAGTCGTTGCTCCCACATTGGAAAAGCAGTACTNNNAGATCGGAAG
41254 TATGAGTGTGGAGTCGTTGCCCCACAGGACAAGCTGTAAACTTNNNAGATCGGAAG
41255 TATGAGTGTGGAGTCGTTGCCTGGGCAATATAGCAAGACTCCANNNAGATCGGAA
41257 TATGAGTGTGGAGTCGTTGCGCAAAAAAATCTCCTCCCGATCCNNNAGATCGGAAG
41261 TATGAGTGTGGAGTCGTTGCGACAATGGAAGTGTCTCATCCCTNNNAGATCGGAAG
41263 TATGAGTGTGGAGTCGTTGCAAGCAACCTGAGAAGCTAGTTCANNNAGATCGGAA
41264 TATGAGTGTGGAGTCGTTGCAACGAGCTAATGAATGCTGTCAANNNAGATCGGAA
41265 TATGAGTGTGGAGTCGTTGCATTGCTTGAATCTGAGAGACGGANNNAGATCGGAA
41267 TATGAGTGTGGAGTCGTTGCCTATGCGTCTTCTGCAAAGCAATNNNAGATCGGAAG
41268 TATGAGTGTGGAGTCGTTGCGTATTGATGGGATGTCTTGCAGCNNNAGATCGGAA
41269 TATGAGTGTGGAGTCGTTGCTCTAACAAGTAATGGAAGGCGGGNNNAGATCGGAA
41270 TATGAGTGTGGAGTCGTTGCTTTCACAGGCAATGTCCATCCTANNNAGATCGGAAG
41271 TATGAGTGTGGAGTCGTTGCGACAGTGTTTGTAGGCATTCAGCNNNAGATCGGAA
41272 TATGAGTGTGGAGTCGTTGCTAATCCCATCTACTCAGGAGGCTNNNAGATCGGAAG
41274 TATGAGTGTGGAGTCGTTGCTTATAGGATAAGGACCTGGCACGNNNAGATCGGAA
41275 TATGAGTGTGGAGTCGTTGCGAGATCACACCATTGCATTCCAGNNNAGATCGGAAG
41277 TATGAGTGTGGAGTCGTTGCCTCACATCACGTCTGTCATTCCANNNAGATCGGAAGA
41279 TATGAGTGTGGAGTCGTTGCCTAAAGGCACTTGTCAGGTACAGNNNAGATCGGAA
41281 TATGAGTGTGGAGTCGTTGCCTCCCTCTGCTGAAAACATTTGGNNNAGATCGGAAG
41283 TATGAGTGTGGAGTCGTTGCGAGGGACTGCTAGATCAATTCGTNNNAGATCGGAA
41284 TATGAGTGTGGAGTCGTTGCCCCGAAACATTTCACATCAGCAANNNAGATCGGAAG
41285 TATGAGTGTGGAGTCGTTGCACATGAACATGGGCATTACACTGNNNAGATCGGAA
41286 TATGAGTGTGGAGTCGTTGCAAACAACCAAACACCATTTCCCANNNAGATCGGAAG
41287 TATGAGTGTGGAGTCGTTGCCTGTTCAGAGAGGCCAAGAAGAGNNNAGATCGGAA
41288 TATGAGTGTGGAGTCGTTGCGAAAGAAACCCTCTGTCCCTGATNNNAGATCGGAAG
41289 TATGAGTGTGGAGTCGTTGCCAGCCAAGAAAGTACACATGCAANNNAGATCGGAA
41290 TATGAGTGTGGAGTCGTTGCCTGTAGTCCCAGCTACTTGGAAGNNNAGATCGGAAG
41291 TATGAGTGTGGAGTCGTTGCGAGCTGGAGGCCATCATACTTAGNNNAGATCGGAA
41293 TATGAGTGTGGAGTCGTTGCGTTCTGTGTGGGAAATGCATGAANNNAGATCGGAA
41294 TATGAGTGTGGAGTCGTTGCTCTGCTCTGAGAAAATAACGGAANNNAGATCGGAA
41298 TATGAGTGTGGAGTCGTTGCGTCATCCTGCAAGTGACTCAAAGNNNAGATCGGAAG
41302 TATGAGTGTGGAGTCGTTGCAGATACAAACAAGCCCAGACAGTNNNAGATCGGAA
41303 TATGAGTGTGGAGTCGTTGCAAGAGCCAGACCCCTTTTCATAGNNNAGATCGGAAG
41305 TATGAGTGTGGAGTCGTTGCTCTTGTCCTATCTCCACAGTTGCNNNAGATCGGAAGA

41306 TATGAGTGTGGAGTCGTTGCCGGTTCTCTCACCCCTTTGTATANNNAGATCGGAAGA
41307 TATGAGTGTGGAGTCGTTGCCTATTTCCAGGAAGCAGGTAGCANNNAGATCGGAA
41310 TATGAGTGTGGAGTCGTTGCGATCACAAGGTCAGGAGATCGAGNNNAGATCGGAA
41311 TATGAGTGTGGAGTCGTTGCATGAGAAGGTGCTCAACATCACTNNNAGATCGGAAG
41312 TATGAGTGTGGAGTCGTTGCCTGTTTTTTATCAAGGGGCCTGGNNNAGATCGGAAG
41313 TATGAGTGTGGAGTCGTTGCACAAACAACCCCATCAAAAAGGGNNNAGATCGGAAG
41314 TATGAGTGTGGAGTCGTTGCTTGTGTGTATTTGGAAGTAGCCANNNAGATCGGAA
41315 TATGAGTGTGGAGTCGTTGCTCAGTGGATTTCTGGCAATACCANNNAGATCGGAAG
41316 TATGAGTGTGGAGTCGTTGCTGGCTGCTTCTGATGTGATACTTNNNAGATCGGAAG
41319 TATGAGTGTGGAGTCGTTGCGTAGTGACCTCAGTGTCTCCAAGNNNAGATCGGAAG
41321 TATGAGTGTGGAGTCGTTGCAGATACTGACTAACCGACCTCCTNNNAGATCGGAAG
41322 TATGAGTGTGGAGTCGTTGCACATGACATATCACGCGAGATCANNNAGATCGGAAG
41323 TATGAGTGTGGAGTCGTTGCAGCAAACTGAATCCATTAGCACANNNAGATCGGAAG
41324 TATGAGTGTGGAGTCGTTGCCCATAGTGAAAGTTCCCTGCAAGNNNAGATCGGAAG
41325 TATGAGTGTGGAGTCGTTGCTTTAGCAACCACCACTCTGATCANNNAGATCGGAAG
41326 TATGAGTGTGGAGTCGTTGCATGCTAGAGAGGTTGTGGAGAACNNNAGATCGGA
41330 TATGAGTGTGGAGTCGTTGCAGTCTTGGGTGAGATTTTGGACANNNAGATCGGAA
41331 TATGAGTGTGGAGTCGTTGCCTGGGTTCAAGAGACTCTTTTGCNNNAGATCGGAAG
41332 TATGAGTGTGGAGTCGTTGCCTCTAGTCCCAGCTACTCAGGAGNNNAGATCGGAAG
41333 TATGAGTGTGGAGTCGTTGCGCTCATGCTTGTAATCCCAGAACNNNAGATCGGAAG
41336 TATGAGTGTGGAGTCGTTGCCACAGTTTCCTTCCTTCCTTCCTNNNAGATCGGAAGA
41338 TATGAGTGTGGAGTCGTTGCATGCTGAGGAAGGAGAATTGCTTNNNAGATCGGAA
41339 TATGAGTGTGGAGTCGTTGCTTTCACAGGTTGGTGTCAGTTTGNNNAGATCGGAAG
41340 TATGAGTGTGGAGTCGTTGCTTTTGAGAGCTTAGAATTGCCCGNNNAGATCGGAAG
41341 TATGAGTGTGGAGTCGTTGCCTCATGTCTGTAATCCCAGCACTNNNAGATCGGAAG
41342 TATGAGTGTGGAGTCGTTGCCCAGTGACTGTGGTATCTCATCCNNNAGATCGGAAG
41343 TATGAGTGTGGAGTCGTTGCCTGTTTGGATGGCAAGTACAACANNNAGATCGGAA
41345 TATGAGTGTGGAGTCGTTGCCTTCCACTCCACTAACCTTCCTTNNNAGATCGGAAGA
41346 TATGAGTGTGGAGTCGTTGCACAAAAACAGTTTCTGGTGCCAANNNAGATCGGAAG
41347 TATGAGTGTGGAGTCGTTGCGCAGATAGCACTAAGAAGCGTTGNNNAGATCGGAA
41349 TATGAGTGTGGAGTCGTTGCACCCCATGGAGATAGAGAGCAATNNNAGATCGGAA
41351 TATGAGTGTGGAGTCGTTGCTTTACTTCATGAGGTCAGCACCANNNAGATCGGAAG
41352 TATGAGTGTGGAGTCGTTGCTACTCAATTCCTTGGTCTGCACANNNAGATCGGAAG
41353 TATGAGTGTGGAGTCGTTGCAGGTCAAGAAAGCTATGTGGGAGNNNAGATCGGA
41354 TATGAGTGTGGAGTCGTTGCTCGTTTCGTTTCATTTCGTTTCANNNAGATCGGAAGA
41357 TATGAGTGTGGAGTCGTTGCCAGTGTTTCATTGTTTTGCAGCCNNNAGATCGGAAG
41358 TATGAGTGTGGAGTCGTTGCCTATGGCTAACACAGGGAGAGTCNNNAGATCGGAA
41361 TATGAGTGTGGAGTCGTTGCACATCTTCGACTGTTTCATTGGCNNNAGATCGGAAG
41362 TATGAGTGTGGAGTCGTTGCCTTTTTGTAACTTGCCCCACTGTNNNAGATCGGAAGA
41363 TATGAGTGTGGAGTCGTTGCTGGCATGGCTTAGCTGCTTATAGNNNAGATCGGAAG
41365 TATGAGTGTGGAGTCGTTGCTAGAGTTAGCCCCATAGACAGCTNNNAGATCGGAAG
41366 TATGAGTGTGGAGTCGTTGCCACAATAAGCTGCATTTCCTTTGCNNNAGATCGGAAG
41369 TATGAGTGTGGAGTCGTTGCCAGGAGAATGGCGTGAAAAGAAANNNAGATCGGA

41370 TATGAGTGTGGAGTCGTTGCAGCTTACAATCCCTGGAACTTGTNNNAGATCGGAAG
41371 TATGAGTGTGGAGTCGTTGCGCCCCAGCTTCTTGAATTGTTTTNNNAGATCGGAAG
41372 TATGAGTGTGGAGTCGTTGCCCTGCAGTGGGGAAATCTATTTANNNAGATCGGAA
41373 TATGAGTGTGGAGTCGTTGCCTGCAGTGAACCATGATCATGTCNNNAGATCGGAAG
41374 TATGAGTGTGGAGTCGTTGCCCCAACTCAGTGTAACCCATTCTNNNAGATCGGAAG
41375 TATGAGTGTGGAGTCGTTGCTCTTCTTTTCTCCTCCCTCTCCTNNNAGATCGGAAGAG
41376 TATGAGTGTGGAGTCGTTGCCCAGCTAGTCAGGAGTTTGAAGTNNNAGATCGGAA
41377 TATGAGTGTGGAGTCGTTGCGGATTACCTGAGTTCGAGACCAGNNNAGATCGGAA
41378 TATGAGTGTGGAGTCGTTGCTTCTATTCTGGTGTCCCCTCTCTNNNAGATCGGAAGA
41379 TATGAGTGTGGAGTCGTTGCTGAGAGGCCTGAGAGAAAGAGAANNNAGATCGGA
41381 TATGAGTGTGGAGTCGTTGCGCATGCCTGTAATCCTAGCTACTNNNAGATCGGAAG
41382 TATGAGTGTGGAGTCGTTGCAATGCAAGTGTTCTAAGCCCCTANNNAGATCGGAAG
41383 TATGAGTGTGGAGTCGTTGCCCTGCTCTGATTTACGTCTTCCTNNNAGATCGGAAGA
41384 TATGAGTGTGGAGTCGTTGCAAAAGAAGCTAGCTGAAATGCCTNNNAGATCGGAA
41386 TATGAGTGTGGAGTCGTTGCACACAAGCAAAAATCCTTCTGGGNNNAGATCGGAAG
41387 TATGAGTGTGGAGTCGTTGCTCACAAGAGGCAAAAAAGGATGTNNNAGATCGGAA
41388 TATGAGTGTGGAGTCGTTGCACATTGTTGCTGAGTGACACTTGNNNAGATCGGAAG
41391 TATGAGTGTGGAGTCGTTGCGCAAAAGCTTCTTTACGCTGAGANNNAGATCGGAAG
41392 TATGAGTGTGGAGTCGTTGCGCCTGTAATCCCAGTACTTTGGANNNAGATCGGAAG
41396 TATGAGTGTGGAGTCGTTGCGGGAAGCTTGGGGTTTTGTAAAGNNNAGATCGGAA
41402 TATGAGTGTGGAGTCGTTGCAATAAATCCCAAGGCCACCATCTNNNAGATCGGAAG
41403 TATGAGTGTGGAGTCGTTGCGGGGTTGCAATCCTAGTCTATGANNNAGATCGGAA
41404 TATGAGTGTGGAGTCGTTGCTGAAGTCGATGTGTTGACATGTCNNNAGATCGGAA
41405 TATGAGTGTGGAGTCGTTGCTGTTCACCAACAAAAAGTGAGCTNNNAGATCGGAAG
41406 TATGAGTGTGGAGTCGTTGCATTAGCATGTTCCCAGCCACAATNNNAGATCGGAAG
41408 TATGAGTGTGGAGTCGTTGCGCCATCAATATCACCTACAAGGCNNNAGATCGGAAG
41410 TATGAGTGTGGAGTCGTTGCAAAGGACTTGAAAGACAGAGGGGNNNAGATCGGA
41412 TATGAGTGTGGAGTCGTTGCACCGGATGAACACCTCTTTCTTTNNNAGATCGGAAG
41414 TATGAGTGTGGAGTCGTTGCAAACTCGTGTCATGGAAGTTTGCNNNAGATCGGAA
41416 TATGAGTGTGGAGTCGTTGCCCCACACATTCATCATTGGGAATNNNAGATCGGAAG
41417 TATGAGTGTGGAGTCGTTGCGGGAGGAAAAAAAGACCACAGACNNNAGATCGGA
41420 TATGAGTGTGGAGTCGTTGCGAGTTAGACCCCTTTATCCCACCNNNAGATCGGAAG
41423 TATGAGTGTGGAGTCGTTGCCCATAATCCCAGCTAGTCAGGAGNNNAGATCGGAAG
41424 TATGAGTGTGGAGTCGTTGCGCTCAGTAGGGCAGACTCTAATCNNNAGATCGGAA
41425 TATGAGTGTGGAGTCGTTGCATGGACATTTGTTCAGCTGTGTGNNNAGATCGGAA
41426 TATGAGTGTGGAGTCGTTGCAATGTTTCATGCTCTCCAACTGCNNNAGATCGGAAG
41428 TATGAGTGTGGAGTCGTTGCGCTTCTGATCACTGCTGTTTGAANNNAGATCGGAAG
41429 TATGAGTGTGGAGTCGTTGCGGTAACTTCAGCCCAGGACTTTANNNAGATCGGAAG
41430 TATGAGTGTGGAGTCGTTGCAGAGCTCAGTGACTTCACAAGAGNNNAGATCGGAA
41435 TATGAGTGTGGAGTCGTTGCGTAGTAACATGGATGGTGCTGGANNNAGATCGGAA
41436 TATGAGTGTGGAGTCGTTGCTTGAGCAGACCTTGGAAGTTTCTNNNAGATCGGAAG
41438 TATGAGTGTGGAGTCGTTGCAAGCTGAACTCATGGGATGGAATNNNAGATCGGAA
41439 TATGAGTGTGGAGTCGTTGCCTAGGGGAGGGATAGCATTAGGANNNAGATCGGA

41440 TATGAGTGTGGAGTCGTTGCGGATGGCATGGAAAACACAAGTTNNNAGATCGGAA
41441 TATGAGTGTGGAGTCGTTGCGTGAGCCCAGATCACTACACTACNNNAGATCGGAAG
41442 TATGAGTGTGGAGTCGTTGCCACTCCTCTAATCCCAGCACTTTNNNAGATCGGAAGA
41443 TATGAGTGTGGAGTCGTTGCAGAGACATTTTGGAGCCCATTGANNNAGATCGGAA
41444 TATGAGTGTGGAGTCGTTGCGCAGGGCTTGAAATCGTAGTCTANNNAGATCGGAA
41445 TATGAGTGTGGAGTCGTTGCATCGTGAAATCTAGGTGGTAGGTNNNAGATCGGAA
41446 TATGAGTGTGGAGTCGTTGCGACAGCCTTTCATTTTCAGAGGTNNNAGATCGGAAG
41450 TATGAGTGTGGAGTCGTTGCAATTGCATTTCCGTGTCACACAANNNAGATCGGAAG
41451 TATGAGTGTGGAGTCGTTGCCTGAGGCAGAAGAATGGTGTGAANNNAGATCGGA
41452 TATGAGTGTGGAGTCGTTGCGAGTTTTGCAGGCCTCAAAGATANNNAGATCGGAA
41453 TATGAGTGTGGAGTCGTTGCAATAAATGAAGGAAAGCCAGGCGNNNAGATCGGA
41455 TATGAGTGTGGAGTCGTTGCTCCAACCTCAGAGTTTGAATCTTNNNAGATCGGAAG
41456 TATGAGTGTGGAGTCGTTGCTGAGAGCATCACTGACATGTTGANNNAGATCGGAA
41459 TATGAGTGTGGAGTCGTTGCTACGTACTCGAACAAGTGACAGGNNNAGATCGGAA
41460 TATGAGTGTGGAGTCGTTGCCCAGCAGGACAGCAATTACATTTNNNAGATCGGAAG
41461 TATGAGTGTGGAGTCGTTGCTTTCACAACCTACTCATCTGCCANNNAGATCGGAAGA
41462 TATGAGTGTGGAGTCGTTGCAGAACAAATCTGGAGGCATTACANNNAGATCGGAA
41464 TATGAGTGTGGAGTCGTTGCGCTACTCAGGAGACATGTGAGAGNNNAGATCGGAA
41465 TATGAGTGTGGAGTCGTTGCAAAACCTAACTTTGCTCAAGCGTNNNAGATCGGAAG
41466 TATGAGTGTGGAGTCGTTGCTTCATCCAGTGTGTGTGTTTGTGNNNAGATCGGAAG
41467 TATGAGTGTGGAGTCGTTGCAGAAACACCAGGAAGAGATGCTCNNNAGATCGGAA
41468 TATGAGTGTGGAGTCGTTGCTGGAGCACGGGTCATTGTATTTANNNAGATCGGAA
41469 TATGAGTGTGGAGTCGTTGCAAACAAATCAAGCTGAAGGCACANNNAGATCGGAA
41470 TATGAGTGTGGAGTCGTTGCTAAGCCCAAAGAGACACACATCANNNAGATCGGAAG
41471 TATGAGTGTGGAGTCGTTGCAAAAGTTGTTGCCTGTTCATGGANNNAGATCGGAA
57378 TGAGTGTGGAGTCGTTGCAAAGACTAATAATACCTTCCTTAGGGTNNNAGATCGGA
64710 TGAGTGTGGAGTCGTTGCGGTCTTTATTCTTTCTCTCTTCCTGCNNNAGATCGGAAG
67501 TGAGTGTGGAGTCGTTGCTGCCAGAAGAAGTCTATTCCTAAAGANNNAGATCGGA
59231 ATGAGTGTGGAGTCGTTGCTCTTCCAGTATGTACATACGAACCANNNAGATCGGAA
67481 ATGAGTGTGGAGTCGTTGCTCTCCAGTATTTTTTTCAACCACANNNAGATCGGAAG
63074 ATGAGTGTGGAGTCGTTGCAGCTGAACATCTGAAAACAAAACANNNAGATCGGAA
64415 ATGAGTGTGGAGTCGTTGCTGTTTTGGGAAATATATGTGCTATAGTNNNAGATCGG
66151 ATGAGTGTGGAGTCGTTGCTCCAAGAGAAAAGAAAAAAGGGCANNNAGATCGGA
66185 ATGAGTGTGGAGTCGTTGCAGTGTCTCATCAAAGCAAAAAGCANNNAGATCGGAA
66243 ATGAGTGTGGAGTCGTTGCAGAATGAGGAGAGGAAAGTTAGAAGTNNNAGATCG
69377 ATGAGTGTGGAGTCGTTGCTCCGTGTACATATATATAGACAGATGGNNNAGATCGG
69541 ATGAGTGTGGAGTCGTTGCCCATATTATCAAGCTTTATTTATGGCCNNNAGATCGGA
53679 TATGAGTGTGGAGTCGTTGCGCCAGTTCCATATAACTAGTGTTTTCANNNAGATCGG
53990 TATGAGTGTGGAGTCGTTGCTGGTTATTGTGAGCATTAGAATACAAANNNAGATCG
54647 TATGAGTGTGGAGTCGTTGCGGAAGAAGAAAAAGAAAGAGAAGAATCNNNAGAT
54867 TATGAGTGTGGAGTCGTTGCGACGGAGTCTTACTCTATCACCCNNNAGATCGGAAG
55201 TATGAGTGTGGAGTCGTTGCCAATCTCGGCTTACTGCAAACTCNNNAGATCGGAAG
55350 TATGAGTGTGGAGTCGTTGCCGTGTAATTTGAAAACAGAGACAAAGANNNAGATC

55386 TATGAGTGTGGAGTCGTTGCACAGAACAGTGAAATATACACACTAGTNNNAGATCG
55498 TATGAGTGTGGAGTCGTTGCTGGGACCATTATAAATAAGTTTATGCTNNNAGATCG
55543 TATGAGTGTGGAGTCGTTGCTCTGTGAAATAGCTGAAAAGAAATTTGNNNAGATCG
55588 TATGAGTGTGGAGTCGTTGCTTAGGAAGCTTAGTTTGGCCAGANNNAGATCGGAA
55624 TATGAGTGTGGAGTCGTTGCAAAGAAAAGAAAAAAAGGAAATCCAGTNNNAGATC
56864 TATGAGTGTGGAGTCGTTGCCCAGTGGGTTCCAAATAAACCAANNNAGATCGGAAG
57031 TATGAGTGTGGAGTCGTTGCTCAATCAGTAGTGTTTCAATACATTGANNNAGATCG
57036 TATGAGTGTGGAGTCGTTGCACCTCAGAAAATCAGTATCAATATCCANNNAGATCG
57296 TATGAGTGTGGAGTCGTTGCTCCTTACAATCCAAATCAGGTGGTNNNAGATCGGAA
58134 TATGAGTGTGGAGTCGTTGCCTTCAAGCAATTCTCCTGCCTCANNNAGATCGGAAGA
58407 TATGAGTGTGGAGTCGTTGCGTTCAAGAGATTCTCCTGCCTCANNNAGATCGGAAG
58885 TATGAGTGTGGAGTCGTTGCTGTTGGCATAATTGAGAATACTTTCTGNNNAGATCG
59169 TATGAGTGTGGAGTCGTTGCACAATGTAGAATAGAGTGGAGACATCANNNAGATC
59223 TATGAGTGTGGAGTCGTTGCGGTTCAAGCAATTCTCCTGTCTCNNNAGATCGGAAG
59506 TATGAGTGTGGAGTCGTTGCTCAAGTGATTCTTCCACCTCACCNNNAGATCGGAAGA
60308 TATGAGTGTGGAGTCGTTGCCGAGTTCAAGCAATTCTCCTGTCNNNAGATCGGAAG
61991 TATGAGTGTGGAGTCGTTGCACACAGTAATTCAAGTGTAGATTTCACNNNAGATCG
62032 TATGAGTGTGGAGTCGTTGCCTGTTAGGTCCAGGACTCCTTTGNNNAGATCGGAAG
62043 TATGAGTGTGGAGTCGTTGCAACCAGGCAGAGTTTTACTGGAANNNAGATCGGAA
62233 TATGAGTGTGGAGTCGTTGCATGCACATCCTCCCATATACCTTNNNAGATCGGAAGA
62451 TATGAGTGTGGAGTCGTTGCTTTTAGGCATAGGTGAAGAGGGCNNNAGATCGGAA
62558 TATGAGTGTGGAGTCGTTGCGCAACTTTATGATATGTTGGTAGAGCANNNAGATCG
62734 TATGAGTGTGGAGTCGTTGCGGAGGAGAAAGAAAAAATCTCAATCANNNAGATCG
62976 TATGAGTGTGGAGTCGTTGCTCAGTGTGTCTAGAGATCTAATGTACANNNAGATCG
63020 TATGAGTGTGGAGTCGTTGCGAAAAGAAAAGAAAGAAAGAAGAAGCANNNAGAT
63219 TATGAGTGTGGAGTCGTTGCTGACTATTGATGACTAAGGATTACACANNNAGATCG
63230 TATGAGTGTGGAGTCGTTGCGGTTCAAACAATTCTCCTGCCTCNNNAGATCGGAAG
63285 TATGAGTGTGGAGTCGTTGCTGCTCCTAAAAATACTAAGCAACTACCNNNAGATCGG
63401 TATGAGTGTGGAGTCGTTGCACACACACATGCATAGACAGATCTNNNAGATCGGAA
63983 TATGAGTGTGGAGTCGTTGCACCACTGGAATATTGTATTGTACAGTGNNNAGATCG
64164 TATGAGTGTGGAGTCGTTGCAAAGAGGGTGAGATGATACCTGCNNNAGATCGGAA
64228 TATGAGTGTGGAGTCGTTGCAAACGATGATACCTAAGGCAGCANNNAGATCGGAA
64248 TATGAGTGTGGAGTCGTTGCTGACATAGCATAAATTGTTGAAGACTGNNNAGATCG
65368 TATGAGTGTGGAGTCGTTGCTCATGGATAAAGATTGATTCTTAGCCANNNAGATCG
65394 TATGAGTGTGGAGTCGTTGCCAGTGCTTAGTGTTGTGGACATGNNNAGATCGGAA
65969 TATGAGTGTGGAGTCGTTGCGATTGAAGCAATTCTCCTGCCTCNNNAGATCGGAAG
66182 TATGAGTGTGGAGTCGTTGCAGTCATAGACATTATGAAAAGATGCCANNNAGATCG
66218 TATGAGTGTGGAGTCGTTGCTGGATGATTTTTCTATTTGACCTTGGANNNAGATCG
66272 TATGAGTGTGGAGTCGTTGCGCCCTGTGGAAATATTTTGAAGCNNNAGATCGGAA
66515 TATGAGTGTGGAGTCGTTGCAATGGCCGTTTCAAAATAACCGTNNNAGATCGGAAG
66568 TATGAGTGTGGAGTCGTTGCTTCAAGTGATTCTCCTGCCTCAGNNNAGATCGGAAG
66597 TATGAGTGTGGAGTCGTTGCATGAAATAATGGCATTCGCAGCANNNAGATCGGAA
66620 TATGAGTGTGGAGTCGTTGCAGGCCAGCTGAAACTATCTTGAANNNAGATCGGAA

66688 TATGAGTGTGGAGTCGTTGCCAGTCTGGACGTCTCTGCATATTNNNAGATCGGAAG
66703 TATGAGTGTGGAGTCGTTGCTCTGTGATTTTTCCCCCCATCATNNNAGATCGGAAGA
66717 TATGAGTGTGGAGTCGTTGCCCATGTTTTTTTTTAGACTTCAGCACANNNAGATCGG
66881 TATGAGTGTGGAGTCGTTGCCCATCTATGCCTCTCAAAGTGCTNNNAGATCGGAAGA
66975 TATGAGTGTGGAGTCGTTGCAGACAGGGTCTCATTCTGTTGTCNNNAGATCGGAAG
67072 TATGAGTGTGGAGTCGTTGCTATGCCAGACTTCCTGAATGTCTNNNAGATCGGAAG
67312 TATGAGTGTGGAGTCGTTGCTCCCCAGAGAAAAGTAAAAGGTGANNNAGATCGGA
67313 TATGAGTGTGGAGTCGTTGCAGTTCTGTTTTGATTCAAGGATCTTGANNNAGATCG
67455 TATGAGTGTGGAGTCGTTGCTGCAGAAAAGGTTATATGAAAATGCANNNAGATCG
67696 TATGAGTGTGGAGTCGTTGCAGCAAGGTTTCAAGATACAAGGTNNNAGATCGGAA
67720 TATGAGTGTGGAGTCGTTGCCGCATATATGTATATAGAGTGTGCNNNAGATCGG
67764 TATGAGTGTGGAGTCGTTGCGATTTTCCTCCACGGTTCCTAGTNNNAGATCGGAAGA
67796 TATGAGTGTGGAGTCGTTGCGCATGATCTTGGCTCACTACAACNNNAGATCGGAAG
67829 TATGAGTGTGGAGTCGTTGCTACACCCCTTCCTCTATTGGTTCNNNAGATCGGAAGA
68177 TATGAGTGTGGAGTCGTTGCTCTTCAGATTGTATAAATCTTTCCCCTNNNAGATCGGA
68256 TATGAGTGTGGAGTCGTTGCTGTAGCTTACAGATGAATGAAAAACANNNAGATCGG
68268 TATGAGTGTGGAGTCGTTGCTGTGTGTGTATATATATGTGTGTGTGTNNNAGATCG
68392 TATGAGTGTGGAGTCGTTGCACAAAAACTAAGATGTCGCTGGGNNNAGATCGGAA
68611 TATGAGTGTGGAGTCGTTGCCCAATTCACATAATGTTTATGACAGTGNNNAGATCG
68642 TATGAGTGTGGAGTCGTTGCGCAACCTCTACCTCTCATGTTCANNNAGATCGGAAGA
68652 TATGAGTGTGGAGTCGTTGCAGGGACCGAGGTGCTATTTTATGNNNAGATCGGAA
68803 TATGAGTGTGGAGTCGTTGCGTCTCCCCTATCTATCTCTATCTCTCTNNNAGATCGGA
68820 TATGAGTGTGGAGTCGTTGCGAGAGAGAGAGAGAGACGAGACANNNAGATCGGA
68829 TATGAGTGTGGAGTCGTTGCAGAAAGAAAGAAAGAAAGAAAGAAGGANNNAGAT
68890 TATGAGTGTGGAGTCGTTGCTATCTTGCAAACTCTGGAGCCATNNNAGATCGGAAG
69034 TATGAGTGTGGAGTCGTTGCAGTGCCTACTTTATATATAGAGTGCCTNNNAGATCG
69094 TATGAGTGTGGAGTCGTTGCGAGGGAAGGAAGGAAGGAAAAGANNNAGATCGG
69177 TATGAGTGTGGAGTCGTTGCTGGCTCATAAAACATTATTTATACCCTNNNAGATCGG
69206 TATGAGTGTGGAGTCGTTGCCCACATTCACATAACCTTTATTATGGTNNNAGATCGG
69358 TATGAGTGTGGAGTCGTTGCCCATCTTTCTGGAGAACTGTAATTACANNNAGATCG
69523 TATGAGTGTGGAGTCGTTGCAGATCTCCCTTTGTTTATGTTTCTTCTNNNAGATCGG
69762 TATGAGTGTGGAGTCGTTGCTCTTTTTCTTTCTTTCTCTCTGTTGGTNNNAGATCGGA
69765 TATGAGTGTGGAGTCGTTGCTCTATTTCTCTCTCTCTCTCTCTCTNNNAGATCGGAAG
69869 TATGAGTGTGGAGTCGTTGCTGTACACAGATGATAGATATATGCACTNNNAGATCG
69901 TATGAGTGTGGAGTCGTTGCTGCTCTGGAGAAAGGACTTTTGTNNNAGATCGGAA
70237 TATGAGTGTGGAGTCGTTGCTCTGGATCAAGAATTAGCCCAGGNNNAGATCGGAA
70266 TATGAGTGTGGAGTCGTTGCTGATCTCATCTCACTGCAACCTCNNNAGATCGGAAGA
70267 TATGAGTGTGGAGTCGTTGCTGGACAGAGTCTCACTTTGTTGCNNNAGATCGGAAG
70300 TATGAGTGTGGAGTCGTTGCACTCCCTATCTCTTCTTCAAAATCAGTNNNAGATCGG
70414 TATGAGTGTGGAGTCGTTGCCTGGAGTGTAGTTGTGCAATCTCNNNAGATCGGAA
70450 TATGAGTGTGGAGTCGTTGCAGAGAGAAAGAGAAAGAAGAAGAGAGANNNAGAT
53436 TATGAGTGTGGAGTCGTTGCAAGAAGAAGATGGCAATGTTCCTNNNAGATCGGAA
53476 TATGAGTGTGGAGTCGTTGCCAACACCTCCCATCCTCTCAATCNNNAGATCGGAAGA

53600 TATGAGTGTGGAGTCGTTGCTCTTGAACCCATACATAGCTACAAGANNNAGATCGG
53636 TATGAGTGTGGAGTCGTTGCCCTCCTGGGTTCAAACAATTCTTNNNAGATCGGAAG
53676 TATGAGTGTGGAGTCGTTGCTGTTGGAATTCTGAATTGATGACACTNNNAGATCGG
53770 TATGAGTGTGGAGTCGTTGCTCACAACATAAGACAAGACAAAGTGANNNAGATCG
54070 TATGAGTGTGGAGTCGTTGCCCTTTGATGCTTAGTTTATTGAGGGTNNNAGATCGG
54267 TATGAGTGTGGAGTCGTTGCTCTGAAGAAAGATAGAAAAAATGAGGGNNNAGAT
54322 TATGAGTGTGGAGTCGTTGCCTCCCATCTCAGCTTCCCAAATANNNAGATCGGAAGA
54346 TATGAGTGTGGAGTCGTTGCCTAGGATTACAGACACCTGCCACNNNAGATCGGAAG
54356 TATGAGTGTGGAGTCGTTGCCTGGAGTGACAGCCTTGTATATANNNAGATCGGAA
54597 TATGAGTGTGGAGTCGTTGCGAGTACTGTGGCCTGATCATCGNNNAGATCGGAAG
54795 TATGAGTGTGGAGTCGTTGCTTCAAAGTAAACTATCTTGACAGTTCANNNAGATCG
54855 TATGAGTGTGGAGTCGTTGCCTCCAGGGTTCATGCCATTCTCNNNAGATCGGAAGA
54876 TATGAGTGTGGAGTCGTTGCGTGATCTTGGTTCACTGCAACCNNNAGATCGGAAGA
55034 TATGAGTGTGGAGTCGTTGCAGCAAAATGTTAGCAAATTGAATTCANNNAGATCGG
55052 TATGAGTGTGGAGTCGTTGCGCCTCCTGAGTAGCTGAGATTACNNNAGATCGGAAG
55085 TATGAGTGTGGAGTCGTTGCCTTCCCAGGTTCAAGCAATTCTCNNNAGATCGGAAG
55492 TATGAGTGTGGAGTCGTTGCGCATTTCAGTGCATAATATTCTCTTGTNNNAGATCGG
55629 TATGAGTGTGGAGTCGTTGCTCAGGTGAAGAAAGAAGAAATATGGTNNNAGATCG
56063 TATGAGTGTGGAGTCGTTGCAGTGCAAAAAAGCAGTAAATTATGGCNNNAGATCG
56412 TATGAGTGTGGAGTCGTTGCAGGCACTTTAGAAATAATGTAGTCCTNNNAGATCGG
56447 TATGAGTGTGGAGTCGTTGCTGCTTACCATAAATCATGAGCTAAGCNNNAGATCGG
56522 TATGAGTGTGGAGTCGTTGCATGCCCTACATATCAAAGATGCTNNNAGATCGGAAG
56523 TATGAGTGTGGAGTCGTTGCTCTGCTCTCTACTTTGTTACTGATGTNNNAGATCGGA
56584 TATGAGTGTGGAGTCGTTGCTGGCCCATTTTAGGAAGTAGTCCNNNAGATCGGAAG
56662 TATGAGTGTGGAGTCGTTGCAGGGAAGAAAGAAAGAAAAAAATGGANNNAGATC
56764 TATGAGTGTGGAGTCGTTGCCTTCCCAAGTAGCTGGAACTACANNNAGATCGGAAG
56868 TATGAGTGTGGAGTCGTTGCAGAAGAATATGTGAGAGGAGCTGANNNAGATCGG
56944 TATGAGTGTGGAGTCGTTGCAAGAAAAAGAAAAATTGCCTCATTGTNNNAGATCGG
56963 TATGAGTGTGGAGTCGTTGCCAAGAGATTCTCCTGTCTCAGCCNNNAGATCGGAAG
56981 TATGAGTGTGGAGTCGTTGCCTAGGTTCAAGCAATTCACCTGCNNNAGATCGGAAG
57145 TATGAGTGTGGAGTCGTTGCAAAGCAAAACCTTACATTTTTGATCANNNAGATCGG
57486 TATGAGTGTGGAGTCGTTGCTGTGATGTTTTTTGTTGAAATTTGCANNNAGATCGG
57645 TATGAGTGTGGAGTCGTTGCTGACAAAACCACAGTAACTTCTTTCANNNAGATCGG
57758 TATGAGTGTGGAGTCGTTGCACTGTGACTTTTGAATTTTGTGATACANNNAGATCG
57774 TATGAGTGTGGAGTCGTTGCGGAAGAAAGAAGAAGGAAGAAGAAGGNNNAGAT
57849 TATGAGTGTGGAGTCGTTGCCTCCTGAGTAGCTGGAACTACAGNNNAGATCGGAA
58151 TATGAGTGTGGAGTCGTTGCAAACAATTCTCCTTCCTCAGCCTNNNAGATCGGAAGA
58266 TATGAGTGTGGAGTCGTTGCAACTTGCTAATCATGTGAACAAAAATNNNAGATCGG
58298 TATGAGTGTGGAGTCGTTGCACGTTTCAAAAATTAGAACGCTTTGTNNNAGATCGG
58346 TATGAGTGTGGAGTCGTTGCGCCTAATTTCTCTGTCTAGGACTTCTNNNAGATCGGA
58415 TATGAGTGTGGAGTCGTTGCTCAGAAAACAAATCCAAAGTCACTCANNNAGATCGG
58675 TATGAGTGTGGAGTCGTTGCCTCCTGGGTTCACACCATTCTCNNNAGATCGGAAGA
58770 TATGAGTGTGGAGTCGTTGCCAGTGGCTTGATCTCAGCTTACTNNNAGATCGGAAG

58779 TATGAGTGTGGAGTCGTTGCCAGTCTCCCAAGTAGCTGGAATTNNNAGATCGGAAG
59129 TATGAGTGTGGAGTCGTTGCTCTGCTCTGTGAGTTTCTCAGTCNNNAGATCGGAAG
59154 TATGAGTGTGGAGTCGTTGCTGGCAAGAAGAAAGATAAAAGAGACANNNAGATCG
59193 TATGAGTGTGGAGTCGTTGCAGTACGTAGGCAATATTAGTACTTGCNNNAGATCGG
59201 TATGAGTGTGGAGTCGTTGCGCAAATCAGTTAGCAAATCAGTTACCNNNAGATCGG
59355 TATGAGTGTGGAGTCGTTGCGGCTAACAAACAGAAAAAAGAAGAGGNNNAGATCG
59412 TATGAGTGTGGAGTCGTTGCAGGTAGAAAAGATGGAAACATGCTNNNAGATCGGA
59462 TATGAGTGTGGAGTCGTTGCGTTCAAGCGATTCTTCTGCCTCNNNAGATCGGAAGA
59618 TATGAGTGTGGAGTCGTTGCGGTATCTATCACTTGAATTGATGGGTNNNAGATCGG
59755 TATGAGTGTGGAGTCGTTGCACACACGATTTTTTTTAGACCTGTCANNNAGATCGGA
60047 TATGAGTGTGGAGTCGTTGCTGCAGTAGTTTCATAGTTTTAGGTCANNNAGATCGG
60049 TATGAGTGTGGAGTCGTTGCACTTTCTGAACTTACTTCTCTGGAGTNNNAGATCGGA
60134 TATGAGTGTGGAGTCGTTGCACTTGCCCCAACTTGGACTATAGNNNAGATCGGAAG
60319 TATGAGTGTGGAGTCGTTGCTCATTTATCTGTATGTTGTCTTGCCTNNNAGATCGGA
61988 TATGAGTGTGGAGTCGTTGCAGGCTGGAATGTAATGGTGCTATNNNAGATCGGAA
62076 TATGAGTGTGGAGTCGTTGCATCTTGATTCACTGCAACCTCCTNNNAGATCGGAAGA
62232 TATGAGTGTGGAGTCGTTGCAGCCATACAAGTCTATAATGGTGTCTNNNAGATCGG
62790 TATGAGTGTGGAGTCGTTGCCAGTCCCACAAGTTCCTGAGTATNNNAGATCGGAAG
62918 TATGAGTGTGGAGTCGTTGCAGTCACAAAGTAAAGCATGTGGCNNNAGATCGGAA
62944 TATGAGTGTGGAGTCGTTGCTGCTACCATTCATATAAAAAAGGGGGNNNAGATCG
63186 TATGAGTGTGGAGTCGTTGCTGGATCCTATGTGTTTTTTTGTGTGTNNNAGATCGG
63301 TATGAGTGTGGAGTCGTTGCCTCCTGAGTAGCTGGGATTACAGNNNAGATCGGAA
63396 TATGAGTGTGGAGTCGTTGCGCCTATTTTTGTATAGTCTCTGAGCTNNNAGATCGG
63851 TATGAGTGTGGAGTCGTTGCCAATGGTGCAATCTCAGCTCACNNNAGATCGGAAGA
63869 TATGAGTGTGGAGTCGTTGCGTTCAAGCGATTCTCTTGCCTCNNNAGATCGGAAGA
64253 TATGAGTGTGGAGTCGTTGCTGTGTGTGTGTGTATGTATAAAGTNNNAGATCG
64260 TATGAGTGTGGAGTCGTTGCTGGCTATCAAATCAATACCATTAGGGNNNAGATCGG
64296 TATGAGTGTGGAGTCGTTGCGCCTCCAATATGAATTTCTTGTCTGTNNNAGATCGGA
64307 TATGAGTGTGGAGTCGTTGCGCCTATCTTTCCTGTATGTAAACACTNNNAGATCGGA
64522 TATGAGTGTGGAGTCGTTGCTCTTTTTGTTTTGTTTGATCGTTTGTNNNAGATCGGA
64579 TATGAGTGTGGAGTCGTTGCTGAGATACTCATCTGGGTTTTATGGANNNAGATCGG
64677 TATGAGTGTGGAGTCGTTGCAGAGGAAAAACTATAAACACTGAGGANNNAGATCG
64700 TATGAGTGTGGAGTCGTTGCCTTTGGAGAGGGGAATGAGGAATNNNAGATCGGA
64711 TATGAGTGTGGAGTCGTTGCTTTGTTTATGATTTCTCCTTTGACCTNNNAGATCGGA
64734 TATGAGTGTGGAGTCGTTGCTGCAAATATCTTGAGGGACTATCACANNNAGATCGG
64744 TATGAGTGTGGAGTCGTTGCAGATAGATAATCCTTCTGGTTCTGTCNNNAGATCGG
64815 TATGAGTGTGGAGTCGTTGCTTTGTGGAAGAAAGGAAAGGGGANNNAGATCGGA
64911 TATGAGTGTGGAGTCGTTGCTTTCTGTAGCATTTTCCAATGTCANNNAGATCGGAAG
64979 TATGAGTGTGGAGTCGTTGCAGTCTTACTCTGTCACCTAGCCTNNNAGATCGGAAGA
64981 TATGAGTGTGGAGTCGTTGCTCCAAGACGGAGTCTCACTTTTTNNNAGATCGGAAG
64993 TATGAGTGTGGAGTCGTTGCTCACATAGATATCTCTGGCTAGGAGANNNAGATCGG
65010 TATGAGTGTGGAGTCGTTGCTGTGTGTGTATGTGTATATACGTGTGNNNAGATCG
65046 TATGAGTGTGGAGTCGTTGCTTCTTCATCCAGTCATCCACTCANNNAGATCGGAAGA

65102 TATGAGTGTGGAGTCGTTGCTGTCTATGCCACATTTTCTTTATTCANNNAGATCGGA
65360 TATGAGTGTGGAGTCGTTGCTGGGGGGTATAGTCAATGCTATTNNNAGATCGGAA
65395 TATGAGTGTGGAGTCGTTGCGGCCCACAAAGCTGAAAGTATTTNNNAGATCGGAAG
65533 TATGAGTGTGGAGTCGTTGCACCTGAATTCAAATAGAGTTCTCCCANNNAGATCGG
65563 TATGAGTGTGGAGTCGTTGCAGTTTTGGACTCCTTTTTTGGGCNNNAGATCGGAAG
65621 TATGAGTGTGGAGTCGTTGCAGTCCTGCAAAATGATGAATACAACTNNNAGATCGG
65846 TATGAGTGTGGAGTCGTTGCACTTTTTTTTTTTTGAGATGGGGTCTNNNAGATCGG
65942 TATGAGTGTGGAGTCGTTGCAGAGAACTGGGAAGCTTTGAGAGNNNAGATCGGA
66172 TATGAGTGTGGAGTCGTTGCTGAATGGATATCACAAGGAACCANNNAGATCGGAA
66193 TATGAGTGTGGAGTCGTTGCTGATGATGATAAGGACACATGGCANNNAGATCGGA
66454 TATGAGTGTGGAGTCGTTGCCGGTGTTTTCTTCCTCCTAATAATAANNNAGATCGGA
66651 TATGAGTGTGGAGTCGTTGCTCCTGAATGAAACTGGTCTCCAGNNNAGATCGGAAG
66679 TATGAGTGTGGAGTCGTTGCTGAATATACATGAAGGAGTTAGGAGTNNNAGATCG
66715 TATGAGTGTGGAGTCGTTGCCATTCTCAGCAAACTAACACGGGNNNAGATCGGAAG
66736 TATGAGTGTGGAGTCGTTGCCAATTACAGCATGGGAGTGAAGGNNNAGATCGGAA
66896 TATGAGTGTGGAGTCGTTGCTTGCCTTTCAAAACTACCTTGGGNNNAGATCGGAAG
66937 TATGAGTGTGGAGTCGTTGCGACAAGAGTCTCTCCCTGTTGCNNNAGATCGGAAGA
67044 TATGAGTGTGGAGTCGTTGCTCTGGCCTCATCCGTGATTTTATNNNAGATCGGAAG
67165 TATGAGTGTGGAGTCGTTGCGATGTGGTGGCTTACAGGCTTANNNAGATCGGAAG
67351 TATGAGTGTGGAGTCGTTGCTGAAGTCTGAGAACAGGCACTACNNNAGATCGGAA
67541 TATGAGTGTGGAGTCGTTGCTGTGATCCTTAGGGGAAGAGCTANNNAGATCGGAA
67583 TATGAGTGTGGAGTCGTTGCTGAGTGTATATGTACTGTGTGTGTGTNNNAGATCGG
67636 TATGAGTGTGGAGTCGTTGCTCTTTTTGAGACAGCGTCTTGCNNNAGATCGGAAGA
67731 TATGAGTGTGGAGTCGTTGCCCCTGAAGACCCTAAGGAACAAANNNAGATCGGAAG
67871 TATGAGTGTGGAGTCGTTGCTCTGAAGAACCCTGACTAAAATACCANNNAGATCGG
67925 TATGAGTGTGGAGTCGTTGCAGCAACAGTGTTTACTAAAAGAGGTNNNAGATCGG
67927 TATGAGTGTGGAGTCGTTGCTATTTTGAGACGGAGCAAGACTCNNNAGATCGGAA
67985 TATGAGTGTGGAGTCGTTGCTGTGTGTATGTGTATATATACGCACGNNNAGATCGG
67986 TATGAGTGTGGAGTCGTTGCAGGAACATTGTAATATAATGCCAGGTNNNAGATCG
68054 TATGAGTGTGGAGTCGTTGCACCAAAGTATTCCAAAGTCCATGTNNNAGATCGGAA
68194 TATGAGTGTGGAGTCGTTGCATGTATTCATAAGTGTAAAGCAGCANNNAGATCGG
68336 TATGAGTGTGGAGTCGTTGCGGAGTAGCTAGGATTACAGGCACNNNAGATCGGAA
68779 TATGAGTGTGGAGTCGTTGCGGAACTAACTCATGTGATTATGGAGGNNNAGATCG
69030 TATGAGTGTGGAGTCGTTGCTACAGCATTAGGTCTCACCCAAGNNNAGATCGGAAG
69077 TATGAGTGTGGAGTCGTTGCATACACGTGTATATGTATATGTACGTGNNNAGATCG
69078 TATGAGTGTGGAGTCGTTGCAGCACTATGTCTTGAAAAGTTGTTGTNNNAGATCGG
69152 TATGAGTGTGGAGTCGTTGCGATCTTGGCTCACTGCAACCTCNNNAGATCGGAAGA
69154 TATGAGTGTGGAGTCGTTGCCAAAAGAAGACATACACACGGCCNNNAGATCGGAA
69159 TATGAGTGTGGAGTCGTTGCAACTTGCAGGATTGTAAGTTGCCNNNAGATCGGAA
69382 TATGAGTGTGGAGTCGTTGCGGCTCACTGTAGCTTCAAATTCCNNNAGATCGGAAG
69419 TATGAGTGTGGAGTCGTTGCTCTTGGATCTGTGGATTTATGGTNNNAGATCGGAAG
69458 TATGAGTGTGGAGTCGTTGCGTTGGAGTGCAATGACGTGATCNNNAGATCGGAAG
69773 TATGAGTGTGGAGTCGTTGCAAAAGAAAGAGAAAGAAAAGGAAAGGANNNAGAT

69808 TATGAGTGTGGAGTCGTTGCTCTCTCACAGTTAGAATTTGGTCTGTNNNAGATCGG
69820 TATGAGTGTGGAGTCGTTGCAGAGAGAAAGAAAGAAAGAAAGAAAGANNNAGAT
69827 TATGAGTGTGGAGTCGTTGCTCTCTCTTTCTCTCTTTCTCTCTTTCTNNNAGATCGGAA
70013 TATGAGTGTGGAGTCGTTGCAGAAAGAAAGAAAAATATGTGAGTGGANNNAGAT
70236 TATGAGTGTGGAGTCGTTGCCAAACTGCTGGTCTCAAGTGATCNNNAGATCGGAAG
70280 TATGAGTGTGGAGTCGTTGCAGTTTCTAGAACTTTTTCTTGCAGCANNNAGATCGG
70440 TATGAGTGTGGAGTCGTTGCCAGAGATTAGGTACCGATACATAAGANNNAGATCG
53662 TATGAGTGTGGAGTCGTTGCAGTTGAATTTAGATGATCCTGCATTNNNAGATCGGA
53820 TATGAGTGTGGAGTCGTTGCGGCTCAGGTTATTCTCTCACCTCNNNAGATCGGAAG
53884 TATGAGTGTGGAGTCGTTGCACGTATATGTATGTGTGTGTGTGTGNNNAGATCGG
53958 TATGAGTGTGGAGTCGTTGCACAAAGGATAACAACCATGTGATCANNNAGATCGG
54118 TATGAGTGTGGAGTCGTTGCGCCTACAATCCACTAGTACTTTTATTNNNAGATCGGA
54171 TATGAGTGTGGAGTCGTTGCAGATTTCAGCAGTTTCATCTTGTGTNNNAGATCGGA
54224 TATGAGTGTGGAGTCGTTGCTTCAAGTGATTCTCCTGCCTCAGNNNAGATCGGAAG
54333 TATGAGTGTGGAGTCGTTGCCCGAGTAGCTGGGATTACAGATGNNNAGATCGGAA
54354 TATGAGTGTGGAGTCGTTGCAGGTTCAAGTGATTCTCTTGCCTNNNAGATCGGAAG
54366 TATGAGTGTGGAGTCGTTGCTCTCCTGGGTTCAAGTGATTCTCNNNAGATCGGAAG
54367 TATGAGTGTGGAGTCGTTGCGAGATAGGACCTTGCTCTACTGCNNNAGATCGGAAG
54464 TATGAGTGTGGAGTCGTTGCAGCTGGACCAAGAAGATGAAAGTNNNAGATCGGAA
54579 TATGAGTGTGGAGTCGTTGCAGTTGCTTGAATTCAGAACAAGTCANNNAGATCGGA
54596 TATGAGTGTGGAGTCGTTGCTTCAAGTGATTCTCCTGCCTCAGNNNAGATCGGAAG
54606 TATGAGTGTGGAGTCGTTGCATGCCTGTAATCCCAGCACTTTNNNAGATCGGAAGA
54687 TATGAGTGTGGAGTCGTTGCTGCAAACACCTAAATCTAACCATCANNNAGATCGGA
54695 TATGAGTGTGGAGTCGTTGCACTATGCTGAAGGAAAAATCCATCTNNNAGATCGGA
54720 TATGAGTGTGGAGTCGTTGCCTCATGTGATCATCCCACCTCAGNNNAGATCGGAAG
54724 TATGAGTGTGGAGTCGTTGCCACAATCTCGGTTGACTGCAACNNNAGATCGGAAGA
54835 TATGAGTGTGGAGTCGTTGCAAGTACTCAAAACTCATCATTTCCTAANNNAGATCGG
54909 TATGAGTGTGGAGTCGTTGCAGCATTTTCTCTGTATTGTCTTTCTNNNAGATCGGAA
55046 TATGAGTGTGGAGTCGTTGCCAATGGCATGATTTCAGCTCACTNNNAGATCGGAAG
55151 TATGAGTGTGGAGTCGTTGCTGATATCAGGGTAATATTGGCCTCANNNAGATCGGA
55152 TATGAGTGTGGAGTCGTTGCGGTTCTGCTACCTCAATTCTTACACNNNAGATCGGAA
55174 TATGAGTGTGGAGTCGTTGCTCCTAATGATTAGTGATGTAGAGCANNNAGATCGG
55232 TATGAGTGTGGAGTCGTTGCTGCTACATATAGGCAAATAAAGCAANNNAGATCGG
55276 TATGAGTGTGGAGTCGTTGCACTTCACCAATGAATAAGAAGGGGANNNAGATCGG
55277 TATGAGTGTGGAGTCGTTGCAGGCAATATGAGATTTTCCCTAAGTNNNAGATCGGA
55433 TATGAGTGTGGAGTCGTTGCAAGATGGGGTGCATTCATGTGATNNNAGATCGGAA
55461 TATGAGTGTGGAGTCGTTGCCTGAGTAGCTGGGACTACAGGTANNNAGATCGGAA
55601 TATGAGTGTGGAGTCGTTGCAACTTCTTTGAGGAACAGTGCACNNNAGATCGGAAG
55640 TATGAGTGTGGAGTCGTTGCTCTGAATAATCAAATCAGGGTCACTNNNAGATCGGA
55681 TATGAGTGTGGAGTCGTTGCAGCCATGAGACAGCAATACAAACNNNAGATCGGAA
55709 TATGAGTGTGGAGTCGTTGCTCAGCAAGAATTCCTCTAGATAGCANNNAGATCGGA
55766 TATGAGTGTGGAGTCGTTGCAGAAAGAAGAAGAAAGGCTGAATGCNNNAGATCG
55803 TATGAGTGTGGAGTCGTTGCCTGGAATTACAGGCATGTGCAACNNNAGATCGGAA

55822 TATGAGTGTGGAGTCGTTGCCCTTCCAAGTAGCTGGGATTACANNNAGATCGGAAG
55835 TATGAGTGTGGAGTCGTTGCTCAACAATGACCCATCTGAAAAAGANNNAGATCGGA
55851 TATGAGTGTGGAGTCGTTGCTTCACCATATTAGCAGAATAAGGAAAANNNAGATCG
55857 TATGAGTGTGGAGTCGTTGCACCTGTTGTGCTTTTATTCCAATGTNNNAGATCGGAA
56100 TATGAGTGTGGAGTCGTTGCGCAGAATGATTTTACTAATTGTGCCANNNAGATCGG
56260 TATGAGTGTGGAGTCGTTGCTGAATAGCTGGGTTTACAGGCATNNNAGATCGGAA
56422 TATGAGTGTGGAGTCGTTGCAGAATAGTCCAAACAGTACTTACCTNNNAGATCGGA
56520 TATGAGTGTGGAGTCGTTGCTGTTACAGATTCTGGTTTATTGGCTNNNAGATCGGA
56540 TATGAGTGTGGAGTCGTTGCAGCTCCACCTAAATAACTCAAGAGGNNNAGATCGGA
56668 TATGAGTGTGGAGTCGTTGCTGTTTTTAGTAGAGACAGGGTTTCACNNNAGATCGG
56673 TATGAGTGTGGAGTCGTTGCAGCTCAAAAGTTTCTACAGCTCTNNNAGATCGGAAG
56750 TATGAGTGTGGAGTCGTTGCCTCACTGTAGCCTCAAACTCCTTNNNAGATCGGAAGA
56867 TATGAGTGTGGAGTCGTTGCATTCAGGAGTTGTTTGTTGCACGNNNAGATCGGAA
57008 TATGAGTGTGGAGTCGTTGCATCTCCTGGGTTCAAGCAATTCTNNNAGATCGGAAG
57101 TATGAGTGTGGAGTCGTTGCCTCCCGAGTAGCTGCAATTACAGNNNAGATCGGAAG
57151 TATGAGTGTGGAGTCGTTGCTCAGAAAAAATAGCCTCAAGAAACTNNNAGATCGGA
57214 TATGAGTGTGGAGTCGTTGCCCTCCCGAGTAGCTGAGACTACNNNAGATCGGAAGA
57233 TATGAGTGTGGAGTCGTTGCTAATTCTGCCCTTGGGACAACATNNNAGATCGGAAG
57363 TATGAGTGTGGAGTCGTTGCTGGTTCCTCTAAAATCTTACCTTCTNNNAGATCGGAA
57382 TATGAGTGTGGAGTCGTTGCCCAGATAGGTTCAGGACTTGATAGTNNNAGATCGG
57419 TATGAGTGTGGAGTCGTTGCACAGTGTTTCCTCTTCCAATTTTGANNNAGATCGGAA
57491 TATGAGTGTGGAGTCGTTGCGAAGATGGCTATCTACAGACAATGANNNAGATCGG
57502 TATGAGTGTGGAGTCGTTGCAGAGAGAGAGAGAGAGAGAGAGAGANNNAGATC
57506 TATGAGTGTGGAGTCGTTGCTGTTTTCCACCTTTTTCTTTTGTTCTNNNAGATCGGAA
57513 TATGAGTGTGGAGTCGTTGCTGGAGTTGATTTTGAAGAGAAATCANNNAGATCGG
57620 TATGAGTGTGGAGTCGTTGCACAATCTTGAAGAAGCAGATGTCTGNNNAGATCGG
57880 TATGAGTGTGGAGTCGTTGCTGGGTTCAAACAATTCTCTTGCCNNNAGATCGGAAG
57909 TATGAGTGTGGAGTCGTTGCTTCAAGTGATTCTCCTGCCTCAGNNNAGATCGGAAG
57934 TATGAGTGTGGAGTCGTTGCTCCAATCTGTGATTTCTTGAAACGTNNNAGATCGGA
58017 TATGAGTGTGGAGTCGTTGCATCTTGGCTCACTACAACCTCTGNNNAGATCGGAAG
58065 TATGAGTGTGGAGTCGTTGCACGTTTCTTATGATAAGGGATGGGTNNNAGATCGG
58121 TATGAGTGTGGAGTCGTTGCAGAGAGAGAGAGAGAGAGAGAGAGANNNAGATC
58176 TATGAGTGTGGAGTCGTTGCAAGATGGGGTGCATTCATGTGATNNNAGATCGGAA
58268 TATGAGTGTGGAGTCGTTGCAGGGTGATTGAAGGTAACATTTACTNNNAGATCGG
58291 TATGAGTGTGGAGTCGTTGCACAAATTTTCTGAGTATAGCTGTGTNNNAGATCGGA
58297 TATGAGTGTGGAGTCGTTGCTGCATCTTTGATCCAATTCAACACANNNAGATCGGAA
58354 TATGAGTGTGGAGTCGTTGCTATCTGCTTCCTGGGTTCAAACGNNNAGATCGGAAG
58418 TATGAGTGTGGAGTCGTTGCTCCTTCTATCCCATCTAGATGAGCTNNNAGATCGGAA
58439 TATGAGTGTGGAGTCGTTGCACTGATTACTCTTTCTCTTTCCTCTNNNAGATCGGAAG
58464 TATGAGTGTGGAGTCGTTGCAGTCTATGCCAAGTAAATCGAAGGTNNNAGATCGG
58568 TATGAGTGTGGAGTCGTTGCACAAGTGAACAAAGGTCTCTGGTNNNAGATCGGAA
58707 TATGAGTGTGGAGTCGTTGCGCTGGAGTACAGTAGCATGATCTNNNAGATCGGAA
58846 TATGAGTGTGGAGTCGTTGCAGAGAGAGAGAGAGAGAGAGAGAGANNNAGATC

58864 TATGAGTGTGGAGTCGTTGCTCCTTTAGGACAATTCCAGATTCCANNNAGATCGGAA
58879 TATGAGTGTGGAGTCGTTGCACCATTTCCAGTGCTTTTTATTCCTNNNAGATCGGAAG
58949 TATGAGTGTGGAGTCGTTGCGCTATTCAACTTTGTTGGGAAAAGTNNNAGATCGGA
58973 TATGAGTGTGGAGTCGTTGCTCTAACTGATTTCATACAACTGCGTNNNAGATCGGAA
59020 TATGAGTGTGGAGTCGTTGCTGCCTAGGCTACTAAGATCTACTGTNNNAGATCGGAA
59074 TATGAGTGTGGAGTCGTTGCCCCACAGGTTTTGGTAAGTTGTGNNNAGATCGGAAG
59151 TATGAGTGTGGAGTCGTTGCGAGGTAAGAGAGATCAAGTATGGAANNNAGATCGG
59203 TATGAGTGTGGAGTCGTTGCAATCACTGGATCATAGAAAGATGGANNNAGATCGGA
59405 TATGAGTGTGGAGTCGTTGCTTCAAGTGATTCTCCCACCTCAGNNNAGATCGGAAGA
59410 TATGAGTGTGGAGTCGTTGCTGGGATACAGTAAAAACAGTACTACANNNAGATCGG
59426 TATGAGTGTGGAGTCGTTGCACACACATACATACACACATACACCNNNAGATCGGAA
59588 TATGAGTGTGGAGTCGTTGCGAAAGACATCCATGTTCATTTCTCTNNNAGATCGGAA
59596 TATGAGTGTGGAGTCGTTGCACAAAAAAAACCAGTTATCTCCCCANNNAGATCGGAA
59926 TATGAGTGTGGAGTCGTTGCACTACCCTGAAACTAAGAACTTTCANNNAGATCGGAA
59934 TATGAGTGTGGAGTCGTTGCACACTTCCTTGTCCTCTATCATTGANNNAGATCGGAAG
60166 TATGAGTGTGGAGTCGTTGCTGAAGCTGGATCCTTATCTTACACTNNNAGATCGGAA
60173 TATGAGTGTGGAGTCGTTGCAACAACAAAAAAAACTAACCATCACANNNAGATCGGA
60200 TATGAGTGTGGAGTCGTTGCCCATTATTTGACTGCTAAATTTGCTCANNNAGATCGG
60204 TATGAGTGTGGAGTCGTTGCAGAGTCGGAACTATATTGCTGAACTNNNAGATCGGA
60295 TATGAGTGTGGAGTCGTTGCCCAGTATCAAGCGATTCTCCTCANNNAGATCGGAAGA
61990 TATGAGTGTGGAGTCGTTGCCGGCAGATGGATGGAGTCTTATTNNNAGATCGGAAG
62052 TATGAGTGTGGAGTCGTTGCGGCTGATGTAGGAGGATCTCTTGNNNAGATCGGAAG
62099 TATGAGTGTGGAGTCGTTGCACAGGAGAGAGGGGTATGACATANNNAGATCGGAA
62279 TATGAGTGTGGAGTCGTTGCTGTGCCAGATGTTTGATTCATTACTNNNAGATCGGAA
62390 TATGAGTGTGGAGTCGTTGCTGTTCCAACAACACAGAATCTTTGTNNNAGATCGGAA
62439 TATGAGTGTGGAGTCGTTGCCTCCTGAGTAGCTGGGACTACAGNNNAGATCGGAAG
62474 TATGAGTGTGGAGTCGTTGCATAGCTAGGACTATAGGCATGCGNNNAGATCGGAAG
62497 TATGAGTGTGGAGTCGTTGCCCTCCTGGCTTCAATCAGTTCTCNNNAGATCGGAAGA
62560 TATGAGTGTGGAGTCGTTGCCCCAATACAGTAATAGCTGGAGACANNNAGATCGGA
62562 TATGAGTGTGGAGTCGTTGCTTCAGTGGAAGATTTACATGTGAAANNNAGATCGGA
62617 TATGAGTGTGGAGTCGTTGCAGTTCTGAAATGCTACCTCTAAGCANNNAGATCGGA
62633 TATGAGTGTGGAGTCGTTGCCCTCAAGGCACAATCTATCATATGANNNAGATCGGAA
62669 TATGAGTGTGGAGTCGTTGCAGTCTTTGGTTTACCTGTGGTGTNNNAGATCGGAAG
62707 TATGAGTGTGGAGTCGTTGCTGTTTTTAGATTCTAGCCTTGTCCANNNAGATCGGAA
62815 TATGAGTGTGGAGTCGTTGCCAAGTAGCTGGGATTACAGGTGTNNNAGATCGGAAG
62881 TATGAGTGTGGAGTCGTTGCATTCAAGTGATTCTCCTGCCTCANNNAGATCGGAAGA
62899 TATGAGTGTGGAGTCGTTGCTCAGCCTCAGAAATCAGAGACACNNNAGATCGGAAG
63000 TATGAGTGTGGAGTCGTTGCTGGTGCATCTACATTAGGGAATACTNNNAGATCGGA
63030 TATGAGTGTGGAGTCGTTGCTGGGAATACAGGTAGAGATAACCACNNNAGATCGGA
63325 TATGAGTGTGGAGTCGTTGCTCAAGCGATTTTCATGTCTCAGCNNNAGATCGGAAGA
63420 TATGAGTGTGGAGTCGTTGCATTCTTTTCTGGCAGGCTTTTGGNNNAGATCGGAAGA
63422 TATGAGTGTGGAGTCGTTGCTGTATGTATGTCTTTGATTGGGTGTNNNAGATCGGA
63641 TATGAGTGTGGAGTCGTTGCTCAGAATCACAAACATTCCTTACCANNNAGATCGGAA

63856 TATGAGTGTGGAGTCGTTGCTCTTTCTCTCTCTCTCTCATCCCTNNNAGATCGGAAGA
63932 TATGAGTGTGGAGTCGTTGCTGGGTTCAAAGATTCTTCTGCCTNNNAGATCGGAAG
63973 TATGAGTGTGGAGTCGTTGCGGTCTTACTTTATGAACCAAGTCGTNNNAGATCGGA
63981 TATGAGTGTGGAGTCGTTGCAGAATAACCTAACCAAAAAGCAGGTNNNAGATCGG
64063 TATGAGTGTGGAGTCGTTGCATCCAATGGAAACGACTGACTGANNNAGATCGGAA
64082 TATGAGTGTGGAGTCGTTGCTCCAATCCATCATTGATGAGTACCTNNNAGATCGGA
64133 TATGAGTGTGGAGTCGTTGCACTCTAGAACTGCTAAGACAACTGGNNNAGATCGGA
64163 TATGAGTGTGGAGTCGTTGCAACTGAGGTAAAGTGATCTGCCTNNNAGATCGGAA
64222 TATGAGTGTGGAGTCGTTGCTGATCAAATTGGCTTCTATCTGTGANNNAGATCGGA
64233 TATGAGTGTGGAGTCGTTGCATTACATGTTCATGGATTGGAAGAANNNAGATCGG
64236 TATGAGTGTGGAGTCGTTGCCAGGAGTTGAACTGAGAACAGAANNNAGATCGGAA
64258 TATGAGTGTGGAGTCGTTGCCCACACACACAAAAAAGAAAAACCANNNAGATCGGA
64456 TATGAGTGTGGAGTCGTTGCTGTAGTGGTGCAATCTTGACTCANNNAGATCGGAAG
64566 TATGAGTGTGGAGTCGTTGCTCATACCATGGAATACAGTACAGCANNNAGATCGGA
64799 TATGAGTGTGGAGTCGTTGCCCATGATGATTAGTGATGTTGAGCANNNAGATCGG
64946 TATGAGTGTGGAGTCGTTGCTGTCATGAAACTTGGCAGAAAATCANNNAGATCGG
65131 TATGAGTGTGGAGTCGTTGCTCCCTATATACCAGTCTCAGATTCTNNNAGATCGGAA
65143 TATGAGTGTGGAGTCGTTGCTCTCTTTCAGCTTTTCTTCTGTCANNNAGATCGGAAG
65267 TATGAGTGTGGAGTCGTTGCTGTATTATTGGTTAGTGAAGTCCCTNNNAGATCGGA
65269 TATGAGTGTGGAGTCGTTGCTCCTTTGGTTGGAATGAAAAATTGTNNNAGATCGGA
65306 TATGAGTGTGGAGTCGTTGCAAACATAAATGTCACATTCCCTTCTNNNAGATCGGAA
65310 TATGAGTGTGGAGTCGTTGCAGCATTTTCTCCACATGTTTGATCANNNAGATCGGAA
65366 TATGAGTGTGGAGTCGTTGCCCCATACCCATCTTGTCTAAACCANNNAGATCGGAAG
65471 TATGAGTGTGGAGTCGTTGCCAAGCCATAAAAAGCCGATGAGANNNAGATCGGAA
65479 TATGAGTGTGGAGTCGTTGCATTAGGAGAGTGAGGTTCTGGGANNNAGATCGGA
65526 TATGAGTGTGGAGTCGTTGCGGCTCCAAAACCTAGACTCTTCANNNAGATCGGAAG
65664 TATGAGTGTGGAGTCGTTGCATGGAGTCTCTCTCTGTCACCANNNAGATCGGAAGA
65665 TATGAGTGTGGAGTCGTTGCAAAGGCAAAGTTCCATGATAGGCNNNAGATCGGAA
65696 TATGAGTGTGGAGTCGTTGCAGTTTGGAGATGTAGAAAAAAGCANNNAGATCGGA
65711 TATGAGTGTGGAGTCGTTGCTGTTTTTCTTTTTGAGACGGAGTCTNNNAGATCGGA
65766 TATGAGTGTGGAGTCGTTGCACTTCCTGGGCTCAGCTACTATANNNAGATCGGAAG
65825 TATGAGTGTGGAGTCGTTGCACTCATTTTGGCTGAATAAAAGGCANNNAGATCGGA
65826 TATGAGTGTGGAGTCGTTGCTCAACCCACAAAAATCAGTTGCANNNAGATCGGAAG
65870 TATGAGTGTGGAGTCGTTGCGGGGTCAGGAGTTTGAGACTAACNNNAGATCGGAA
65880 TATGAGTGTGGAGTCGTTGCTAATACGCCATGATCAAGCAGGANNNAGATCGGAA
65994 TATGAGTGTGGAGTCGTTGCCAGTGGTATGATCTTGGCTCACTNNNAGATCGGAAG
66065 TATGAGTGTGGAGTCGTTGCTCTCTCTCCTTCTTTCTCTCTTTCTNNNAGATCGGAAG
66079 TATGAGTGTGGAGTCGTTGCGAGATGGAGTTTCGTTCTGTTGCNNNAGATCGGAA
66101 TATGAGTGTGGAGTCGTTGCAAGGCTTCATTCTGTCAAGGTGTNNNAGATCGGAAG
66106 TATGAGTGTGGAGTCGTTGCTCTCTCTCTCTCTCTCTCTCTCTCTNNNAGATCGGAAG
66132 TATGAGTGTGGAGTCGTTGCTGACAAAGATGGTATTTCAAGTGAGANNNAGATCG
66187 TATGAGTGTGGAGTCGTTGCTCTTCATTCTCTACGGTCACAGCNNNAGATCGGAAGA
66202 TATGAGTGTGGAGTCGTTGCTGCCTAATTTTGTTTGTGTGATTCANNNAGATCGGA

66214 TATGAGTGTGGAGTCGTTGCCTGCAGCATAGACTTTAGAGGCTNNNAGATCGGAA
66251 TATGAGTGTGGAGTCGTTGCTGGTCATAGTGAAAACAGAAGACCANNNAGATCGG
66392 TATGAGTGTGGAGTCGTTGCGGCACACATGGAATATATATATGCANNNAGATCGG
66408 TATGAGTGTGGAGTCGTTGCTCCACAGTTAGAATGTAAGCATCANNNAGATCGGAA
66429 TATGAGTGTGGAGTCGTTGCAGCCAATCTTGAAAAAGTACCAAGTNNNAGATCGGA
66520 TATGAGTGTGGAGTCGTTGCAGAGTTGAATACATTTGGGCCTACTNNNAGATCGGA
66576 TATGAGTGTGGAGTCGTTGCACACCCCAATAAATCCACTGTAAGTNNNAGATCGGA
66627 TATGAGTGTGGAGTCGTTGCTTCTTTGTGGTTACTATGAGGCTNNNAGATCGGAAG
66729 TATGAGTGTGGAGTCGTTGCACACAACAATTGAGTATCTCAGTGGNNNAGATCGGA
66774 TATGAGTGTGGAGTCGTTGCGTGTACGTGTGTGTGTGTGTATGNNNAGATCGGAA
66804 TATGAGTGTGGAGTCGTTGCAGGGTAGACATTGTTTTCTCCACTNNNAGATCGGAA
67071 TATGAGTGTGGAGTCGTTGCTTTGCTTATTGGAGACAGGGTCTNNNAGATCGGAA
67078 TATGAGTGTGGAGTCGTTGCAGTGGTCTAAAATCTCCAATCCANNNAGATCGGAAG
67145 TATGAGTGTGGAGTCGTTGCCACTACCTCCTGAGTTCAAGTGANNNAGATCGGAAG
67178 TATGAGTGTGGAGTCGTTGCCCAACTTGCCTTCACAGAAATGTNNNAGATCGGAAG
67296 TATGAGTGTGGAGTCGTTGCAGATGCTGGGAATAAATGAGAGCTNNNAGATCGGA
67346 TATGAGTGTGGAGTCGTTGCTTGTGTGCTCATTCTTACCCAAANNNAGATCGGAAG
67466 TATGAGTGTGGAGTCGTTGCAGCCTACAGAAACAACTGGGAAANNNAGATCGGAA
67494 TATGAGTGTGGAGTCGTTGCAGGTGTATCAGAATTTCGGATTCCANNNAGATCGGA
67553 TATGAGTGTGGAGTCGTTGCTTCCAGTACTTTCTCACACCTGGNNNAGATCGGAAGA
67673 TATGAGTGTGGAGTCGTTGCAGAGAGAGAGAGAGAGAGAGAGAGAGANNNAGATC
67760 TATGAGTGTGGAGTCGTTGCACACACATACTCCAAAAAACTGAANNNAGATCGGAA
67770 TATGAGTGTGGAGTCGTTGCTCTCTCTCTCTCTCTCTCTCTCTCTNNNAGATCGGAAG
67827 TATGAGTGTGGAGTCGTTGCTCATCTGTTGATGTACACTTGGGTNNNAGATCGGAA
67907 TATGAGTGTGGAGTCGTTGCGTGACAGAGTCTTGCTTTTGTCGNNNAGATCGGAAG
68007 TATGAGTGTGGAGTCGTTGCACATACTGGTGCACTTATGTACCANNNAGATCGGAA
68016 TATGAGTGTGGAGTCGTTGCTCCTTCAGGTTTTCTACATGTAAGTNNNAGATCGGA
68032 TATGAGTGTGGAGTCGTTGCGCCTCTTGAGTAGCTGGGATTATNNNAGATCGGAAG
68123 TATGAGTGTGGAGTCGTTGCGGTACCAGCCTTCAAACCTAAGANNNAGATCGGAAG
68153 TATGAGTGTGGAGTCGTTGCGTACCATTACCATCACAAATGTACANNNAGATCGGA
68216 TATGAGTGTGGAGTCGTTGCAGAACTTGAAGCTGTAAGAAAATGTNNNAGATCGG
68222 TATGAGTGTGGAGTCGTTGCACACACACACACAAACATATACACANNNAGATCGGA
68282 TATGAGTGTGGAGTCGTTGCGAGAGAAAAAGGCAACCACAGACNNNAGATCGGAA
68310 TATGAGTGTGGAGTCGTTGCTCATTGCTGTTCAATTGATGACACANNNAGATCGGA
68326 TATGAGTGTGGAGTCGTTGCGACAGACAGACAGAGACAGACAGNNNAGATCGGAA
68364 TATGAGTGTGGAGTCGTTGCAGAGAGAGAGAGAGAGAGAGAGAGAGANNNAGATC
68393 TATGAGTGTGGAGTCGTTGCTGTTTGGTTTGGTTTGGTTTGGTNNNAGATCGGAA
68444 TATGAGTGTGGAGTCGTTGCAGTGGACCAATACTTCAGGTTGANNNAGATCGGAA
68449 TATGAGTGTGGAGTCGTTGCAGAGAGGAAGAGAGGAAGAAAGAGANNNAGATC
68477 TATGAGTGTGGAGTCGTTGCAGTCCTAAAATCAAGGTATCAGCAGANNNAGATCGG
68511 TATGAGTGTGGAGTCGTTGCGAGAAATGGGGAAAGACTGGTCTNNNAGATCGGA
68625 TATGAGTGTGGAGTCGTTGCTTTATCTCACAGTAACACACATGTGNNNAGATCGGA
68634 TATGAGTGTGGAGTCGTTGCTGCTGTTTTCTCAAATGCGAAGATNNNAGATCGGAA

68788 TATGAGTGTGGAGTCGTTGCGAAGGAAGGAAGGAAGGAAGGAANNNAGATCGG

68841 TATGAGTGTGGAGTCGTTGCAGAGAGAGAGAGAAAGAAAGAAAGGANNNAGATC

69029 TATGAGTGTGGAGTCGTTGCACACACACACACACACATATATTCTNNNAGATCGGAA

69050 TATGAGTGTGGAGTCGTTGCTCTCTCTCTCTCTCTCTCTCTCTCTNNNAGATCGGAAG

69059 TATGAGTGTGGAGTCGTTGCAGACAGAGACAGAGACAGAGACANNNAGATCGGA

69081 TATGAGTGTGGAGTCGTTGCTCCTTGTACTGGTTTTTGTTGTTGTNNNAGATCGGA

69088 TATGAGTGTGGAGTCGTTGCTTGACATTGGTGCTATCCACAGANNNAGATCGGAAG

69144 TATGAGTGTGGAGTCGTTGCGGAAGGAAGGAAGGAAAGAAGGANNNAGATCGG

69182 TATGAGTGTGGAGTCGTTGCAGAGAGAGACAGAAAGAGAGAGCTNNNAGATCGG

69191 TATGAGTGTGGAGTCGTTGCACACCATAAATGATCCAGAATCCANNNAGATCGGAA

69318 TATGAGTGTGGAGTCGTTGCAGGAGTCATTACATAGGTGCAACANNNAGATCGGA

69343 TATGAGTGTGGAGTCGTTGCACCAAAACATTAGGAGTAGTTGAGTNNNAGATCGG

69373 TATGAGTGTGGAGTCGTTGCTCTCCCATGATACAGTTATATCCTGNNNAGATCGGAA

69399 TATGAGTGTGGAGTCGTTGCTCCTTCCTTCCTTTTCTTTCTGTCTNNNAGATCGGAAG

69558 TATGAGTGTGGAGTCGTTGCGTGATGAGCTCAAGACAAGTTTAGANNNAGATCGG

69690 TATGAGTGTGGAGTCGTTGCTGATGGAGTATAACTCTGTCAACCANNNAGATCGGA

69713 TATGAGTGTGGAGTCGTTGCACAAGTATTATCAGATCAAGGCCCTNNNAGATCGGA

69714 TATGAGTGTGGAGTCGTTGCTCCTACAAAAATGCTCACATACACANNNAGATCGGA

69728 TATGAGTGTGGAGTCGTTGCAGCTCACTGCAATCTTGAACTCTNNNAGATCGGAAG

69743 TATGAGTGTGGAGTCGTTGCGAAGGAAGGAAGGAAGGAAGGAANNNAGATCGG

69754 TATGAGTGTGGAGTCGTTGCTGGTCTCAACCTCATCAAAATGTTGNNNAGATCGGA

69796 TATGAGTGTGGAGTCGTTGCTCAGCCCTGAAATGTAGGATATTCTNNNAGATCGGA

69802 TATGAGTGTGGAGTCGTTGCTGTGTTGCTATTTGTGAACTTGTCANNNAGATCGGA

69850 TATGAGTGTGGAGTCGTTGCTGCAGAAATAGCTTCAAAGGCTTNNNAGATCGGAAG

69865 TATGAGTGTGGAGTCGTTGCATACAAAAATTAGCCGGGCATGGNNNAGATCGGAA

69942 TATGAGTGTGGAGTCGTTGCAAGAAAGAAAGAAAGAAAAACAACCCNNNAGATCG

69963 TATGAGTGTGGAGTCGTTGCCACATTGACCATGACTGAGCATGNNNAGATCGGAAG

69964 TATGAGTGTGGAGTCGTTGCTGGAGTTATCACTCTTGGATGAGACNNNAGATCGGA

70027 TATGAGTGTGGAGTCGTTGCAATCAGCTGACCACATAAATGCGNNNAGATCGGAAG

70192 TATGAGTGTGGAGTCGTTGCAGTGATTCTCATGCTTCAGCCTCNNNAGATCGGAAG

70262 TATGAGTGTGGAGTCGTTGCAGAAAGAAAGAAAGAAAGAAAGAGAGANNNAGAT

70343 TATGAGTGTGGAGTCGTTGCTTAGAGGCTCAGAACTAATGGGANNNAGATCGGAA

70349 TATGAGTGTGGAGTCGTTGCGCACGTGTTTCTATGTGTCTGTCNNNAGATCGGAAG

70425 TATGAGTGTGGAGTCGTTGCCTCCTGAGCTCAAGTGATCCTCNNNAGATCGGAAGA

70509 TATGAGTGTGGAGTCGTTGCTCTCTTTCTCTCTGTCTCTCTCTCTNNNAGATCGGAAG

53482 TATGAGTGTGGAGTCGTTGCGTGGAGGGAAGTTTCAGAACCTANNNAGATCGGAA

53488 TATGAGTGTGGAGTCGTTGCTCTCTTCCAAAATCAACCCTGAGTNNNAGATCGGAAG

53515 TATGAGTGTGGAGTCGTTGCTGAGAACCAAACTAGCCATCAAGTNNNAGATCGGAA

53616 TATGAGTGTGGAGTCGTTGCTCTGCCAAAATCTTAGTGTCCTCANNNAGATCGGAAG

53659 TATGAGTGTGGAGTCGTTGCTGCTGTATTTATTGCCTCAATGTTNNNAGATCGGAAG

53666 TATGAGTGTGGAGTCGTTGCGTGAAACACTATGTCATTCAAGGCTNNNAGATCGGA

53674 TATGAGTGTGGAGTCGTTGCTTTGGCAAAGTGTAGATTTCTCTTNNNAGATCGGAA

53697 TATGAGTGTGGAGTCGTTGCCAGAATGTGGTCCCAGAAAAGAACNNNAGATCGGA

53699 TATGAGTGTGGAGTCGTTGCACAAGTTTCAAAAGGACATGTACANNNAGATCGGAA
53714 TATGAGTGTGGAGTCGTTGCCAAGTAGCTGGGACTACAGGCNNNAGATCGGAAGA
53718 TATGAGTGTGGAGTCGTTGCATATGGTGGTTTCAGCGACAGATNNNAGATCGGAA
53719 TATGAGTGTGGAGTCGTTGCCTTTATCTCCTGGGCTCAAGTGANNNAGATCGGAAG
53781 TATGAGTGTGGAGTCGTTGCTGGGGAGCTAAAATTTTCTCTTTGNNNAGATCGGAA
53799 TATGAGTGTGGAGTCGTTGCCAAGTAGCTGGGAATATAGGCGTNNNAGATCGGAA
53838 TATGAGTGTGGAGTCGTTGCGTGGGGAGATGGTAGATGAAATGANNNAGATCGG
53900 TATGAGTGTGGAGTCGTTGCAACATTCAAGCAATTCTCCTGCCNNNAGATCGGAAG
54001 TATGAGTGTGGAGTCGTTGCCCTCCCAGGTTCAAGAGATTCTCNNNAGATCGGAAG
54079 TATGAGTGTGGAGTCGTTGCTGTCCCTGTAAACCAAAGAGTTCANNNAGATCGGAA
54103 TATGAGTGTGGAGTCGTTGCAGGTCTTGTTTTTGCAACGTTACANNNAGATCGGAA
54112 TATGAGTGTGGAGTCGTTGCCAAGCCTGTAATTCCAGCACTTTNNNAGATCGGAAG
54240 TATGAGTGTGGAGTCGTTGCAGTTTTCTTTGCTGTGCAGAAACTNNNAGATCGGAA
54268 TATGAGTGTGGAGTCGTTGCAGGTACAACTCTATGGCATTGACANNNAGATCGGAA
54309 TATGAGTGTGGAGTCGTTGCAGAGTAGCTGGAATTATAGGTGCCNNNAGATCGGA
54348 TATGAGTGTGGAGTCGTTGCCAATCTCCACTCACTGCAAACTCNNNAGATCGGAAGA
54349 TATGAGTGTGGAGTCGTTGCGCTGAGGCCATTTTTAGACATGGNNNAGATCGGAA
54394 TATGAGTGTGGAGTCGTTGCCCACCTCCTCATTTATCTAGAGCCNNNAGATCGGAAG
54427 TATGAGTGTGGAGTCGTTGCTGGAGGATGGAGGATAAAATGACTNNNAGATCGG
54435 TATGAGTGTGGAGTCGTTGCAGTATAACAACTTTCCTGAGTGCCTNNNAGATCGGA
54478 TATGAGTGTGGAGTCGTTGCAGATATGCAGATGGAAAACTAGCANNNAGATCGGA
54529 TATGAGTGTGGAGTCGTTGCACCTCCCAGGTTCAAGTAATTCTTNNNAGATCGGAAG
54544 TATGAGTGTGGAGTCGTTGCTGACTGATTTCATTGTAGTTGGCCNNNAGATCGGAA
54545 TATGAGTGTGGAGTCGTTGCACACATACCTACTTACAACCCTGGNNNAGATCGGAAG
54558 TATGAGTGTGGAGTCGTTGCTCACAATGAAATATGGACTTCAGGTNNNAGATCGGA
54582 TATGAGTGTGGAGTCGTTGCACTGAGACTAAACACCAAGACACANNNAGATCGGAA
54627 TATGAGTGTGGAGTCGTTGCTTGCATATGGCTATAGTTTTTTCCTNNNAGATCGGAA
54656 TATGAGTGTGGAGTCGTTGCTGGATGGTAGCTATTCCCAGTACNNNAGATCGGAAG
54662 TATGAGTGTGGAGTCGTTGCTTCAAGTGATTCTCCTGCCTCAGNNNAGATCGGAAG
54719 TATGAGTGTGGAGTCGTTGCGTGCGATCTTGTATCACTGCAAGNNNAGATCGGAAG
54744 TATGAGTGTGGAGTCGTTGCGCCTTCCAGTTTCAAGGGATTCTNNNAGATCGGAAG
54815 TATGAGTGTGGAGTCGTTGCTGAAATTCAAATTCTACTGCCCACANNNAGATCGGAA
54860 TATGAGTGTGGAGTCGTTGCCCATTCTCCTGCCTCAACCTTNNNAGATCGGAAGAGC
54866 TATGAGTGTGGAGTCGTTGCGATCTCTGCTCACTACAAGCTCCNNNAGATCGGAAG
54913 TATGAGTGTGGAGTCGTTGCTGTATCACAAAGAGCTATCCCTCANNNAGATCGGAA
54925 TATGAGTGTGGAGTCGTTGCACCTTGGGAATTGTAGTTTATGGTNNNAGATCGGAA
54928 TATGAGTGTGGAGTCGTTGCGTTGGTAATTCACAGTGAAACGGTNNNAGATCGGA
54940 TATGAGTGTGGAGTCGTTGCTCTCAGTAAATAATAGAAAAGCAGACCNNNAGATCG
54950 TATGAGTGTGGAGTCGTTGCTGGAGAAGGGAAATCTTTTTCTCANNNAGATCGGAA
54957 TATGAGTGTGGAGTCGTTGCAATGGAGTTGACTTTCACATTTGTNNNAGATCGGAA
55008 TATGAGTGTGGAGTCGTTGCGTTGTTTCTGTTCTCCTTTGTTTTCTNNNAGATCGGA
55025 TATGAGTGTGGAGTCGTTGCAGGAATGGATTTTGCATTTTATCAAGTNNNAGATCG
55032 TATGAGTGTGGAGTCGTTGCACCTATGAATGTACCAGTTGGAGGNNNAGATCGGA

55067 TATGAGTGTGGAGTCGTTGCGTGAGTAGCTGGGACTACAGATGNNNAGATCGGAA
55077 TATGAGTGTGGAGTCGTTGCGGTTCAAGTGATTCTTCTGCCTCNNNAGATCGGAAG
55133 TATGAGTGTGGAGTCGTTGCAGCTGCAAATTATAAACTGAGTAAGANNNAGATCGG
55142 TATGAGTGTGGAGTCGTTGCCACGTTTTCACTCCTTTCTTGCTANNNAGATCGGAAG
55218 TATGAGTGTGGAGTCGTTGCGATCTCGGCTTTGGTATCAGGATNNNAGATCGGAAG
55255 TATGAGTGTGGAGTCGTTGCGACATAAACAGAGTGATTTGGGAAANNNAGATCGG
55261 TATGAGTGTGGAGTCGTTGCCATTCTGTGGGTTGTCTTTTCACTNNNAGATCGGAAG
55298 TATGAGTGTGGAGTCGTTGCCTCCTGAGTAGCTGGGACTACAGNNNAGATCGGAA
55303 TATGAGTGTGGAGTCGTTGCTTCAAGTGATTCTCCTGCCTCAGNNNAGATCGGAAG
55342 TATGAGTGTGGAGTCGTTGCTGTTTTGTTTTTGAGACGGAGTCTNNNAGATCGGAA
55458 TATGAGTGTGGAGTCGTTGCGGCCAGTGGAATGTGGATAAAGNNNAGATCGGA
55502 TATGAGTGTGGAGTCGTTGCTGCTTTCATGTGAAAAAATTAGAGCTNNNAGATCGG
55574 TATGAGTGTGGAGTCGTTGCCTCCTGAGTAGCTGGGACTACAGNNNAGATCGGAA
55589 TATGAGTGTGGAGTCGTTGCAATAATTCTGTCTCAGCCTCCCGNNNAGATCGGAAG
55602 TATGAGTGTGGAGTCGTTGCCAATCTCAGCTCACTGCAACTTCNNNAGATCGGAAGA
55628 TATGAGTGTGGAGTCGTTGCAGCTATGCCTGTATAGTTCTGGACNNNAGATCGGAA
55637 TATGAGTGTGGAGTCGTTGCTCCCATTCAAAAATGTCTCTGTGTGNNNAGATCGGA
55686 TATGAGTGTGGAGTCGTTGCTGAGCATCCCTAATCTGAAATCCANNNAGATCGGAA
55723 TATGAGTGTGGAGTCGTTGCGATTCTCCTGCCTCAGCCTCNNNAGATCGGAAGAGC
55801 TATGAGTGTGGAGTCGTTGCCTCAGACAATCCTCCTACCTCAGNNNAGATCGGAAGA
55805 TATGAGTGTGGAGTCGTTGCTCCCAGGTTCAAGTGATCTTCTGNNNAGATCGGAAG
55821 TATGAGTGTGGAGTCGTTGCACCATCTCAGCTCACTGCAAGNNNAGATCGGAAGAG
55961 TATGAGTGTGGAGTCGTTGCTCCCAAATCCATGTCTTAGTGTGTNNNAGATCGGAA
55966 TATGAGTGTGGAGTCGTTGCAGCAGAAGTAGCTAAGCTTACACANNNAGATCGGA
56016 TATGAGTGTGGAGTCGTTGCCATTCTCCTGCCTCAGCCTCNNNAGATCGGAAGAGCA
56065 TATGAGTGTGGAGTCGTTGCAGAATGACTGTCGGGTAAAGTACANNNAGATCGGA
56069 TATGAGTGTGGAGTCGTTGCGGAAACAATTGTTCATGTCACAGTNNNAGATCGGAA
56076 TATGAGTGTGGAGTCGTTGCGCAAGAAATCTACCTTCCAAGGACNNNAGATCGGAA
56098 TATGAGTGTGGAGTCGTTGCTCCCAAAGAATCCTGACACTACTGNNNAGATCGGAA
56110 TATGAGTGTGGAGTCGTTGCTGAGCTCTGGATCACATTTTTTGANNNAGATCGGAA
56203 TATGAGTGTGGAGTCGTTGCACCCTTTCCTAGAGTTTCTTCACCNNNAGATCGGAAG
56207 TATGAGTGTGGAGTCGTTGCACGATTATGGTAAGTGAAAGAAAGCANNNAGATCG
56224 TATGAGTGTGGAGTCGTTGCTGCAACAACAACAAAAACATTTGANNNAGATCGGAA
56227 TATGAGTGTGGAGTCGTTGCACCTATCACAAAGAGGTTCAAACCNNNAGATCGGAA
56311 TATGAGTGTGGAGTCGTTGCTCTTTGTTGTCATCATTAGTGCCANNNAGATCGGAAG
56337 TATGAGTGTGGAGTCGTTGCGGAATGCGATGGTGTGATCTTGNNNAGATCGGAAG
56365 TATGAGTGTGGAGTCGTTGCCCATCTCCTTGGTAGTAAGTGAGTNNNAGATCGGAA
56415 TATGAGTGTGGAGTCGTTGCCCTTGAGTCACATCCCAAAGTCANNNAGATCGGAAG
56438 TATGAGTGTGGAGTCGTTGCCAGGGAACTAGATGGATGTAAGCANNNAGATCGGA
56488 TATGAGTGTGGAGTCGTTGCTCTCCTACATTAGCCTCCCAAGANNNAGATCGGAAGA
56569 TATGAGTGTGGAGTCGTTGCTTCAAGTGATTCTCCTGCCTCAGNNNAGATCGGAAG
56634 TATGAGTGTGGAGTCGTTGCACCAGAAACATAATCCCAAAGAGTNNNAGATCGGAA
56661 TATGAGTGTGGAGTCGTTGCACGTGCAACAACATAACAATGAGANNNAGATCGGA

56814 TATGAGTGTGGAGTCGTTGCCTCACTGTAACCTCCACCTCTCANNNAGATCGGAAGA
56881 TATGAGTGTGGAGTCGTTGCTGATATGAACCTTTTGTCTTCTGCTNNNAGATCGGAA
56882 TATGAGTGTGGAGTCGTTGCCCAACAGATTTTGTCTATGGCACANNNAGATCGGAA
56939 TATGAGTGTGGAGTCGTTGCACCAAACAATACCATGAAGCAAGTNNNAGATCGGAA
56947 TATGAGTGTGGAGTCGTTGCACACAAGGCAGACAAGAAAATAGCNNNAGATCGGA
56972 TATGAGTGTGGAGTCGTTGCGAGGGGAAAGAGAGAGAACACAGNNNAGATCGGA
56992 TATGAGTGTGGAGTCGTTGCCATTCTCCTGCCTCAGCCTCNNNAGATCGGAAGAGCA
57051 TATGAGTGTGGAGTCGTTGCAGAATTTCTAAGGCCTCTTCAGCTNNNAGATCGGAA
57066 TATGAGTGTGGAGTCGTTGCAGAAGTCACAGTAAAAGGATGGGTNNNAGATCGGA
57076 TATGAGTGTGGAGTCGTTGCTCAGGATCCAGAGTATCAAACAGTNNNAGATCGGAA
57099 TATGAGTGTGGAGTCGTTGCCCATCTCAGCTCACTGTAACCTCNNNAGATCGGAAGA
57131 TATGAGTGTGGAGTCGTTGCACTGGACGGTTATTCATTCTCTCTNNNAGATCGGAAG
57175 TATGAGTGTGGAGTCGTTGCTTTGCAACTTAGATGAAATGGACANNNAGATCGGAA
57232 TATGAGTGTGGAGTCGTTGCTGTGCACTAAAATTTTAGGTTGCANNNAGATCGGAA
57278 TATGAGTGTGGAGTCGTTGCTTCAAGCGATTCTCCTGCCTTAGNNNAGATCGGAAGA
57279 TATGAGTGTGGAGTCGTTGCCCAGGCGGAGACTTTTATTCTCTNNNAGATCGGAAG
57289 TATGAGTGTGGAGTCGTTGCTTCAAGCAATTCTCCTGCCTCAGNNNAGATCGGAAGA
57353 TATGAGTGTGGAGTCGTTGCACCAGTGTTTGATCATCTTCATGGNNNAGATCGGAA
57365 TATGAGTGTGGAGTCGTTGCCCAAGGCTTTTCATAATTCCCTCANNNAGATCGGAAG
57386 TATGAGTGTGGAGTCGTTGCACACACAACTCTCTTTCATCAAGTNNNAGATCGGAAG
57389 TATGAGTGTGGAGTCGTTGCAGCCTTGGAGAAATAAGGGATTCTNNNAGATCGGA
57392 TATGAGTGTGGAGTCGTTGCAGCAGGTAAAACATGGTAGAAACANNNAGATCGGA
57448 TATGAGTGTGGAGTCGTTGCTTCACACCATTCTCCTGCCTCNNNAGATCGGAAGAGC
57485 TATGAGTGTGGAGTCGTTGCGGGAAAAACCTACAGCTAACATCANNNAGATCGGAA
57488 TATGAGTGTGGAGTCGTTGCCGGTAGATATGTAGACCATGGAGTNNNAGATCGGA
57497 TATGAGTGTGGAGTCGTTGCACAAGGATTTTGAAGAAACCATCANNNAGATCGGAA
57674 TATGAGTGTGGAGTCGTTGCCGAGTAGCTGGGATTACAGGTACNNNAGATCGGAA
57689 TATGAGTGTGGAGTCGTTGCGCCTCCCAAGTAGCTAGGATTACNNNAGATCGGAAG
57690 TATGAGTGTGGAGTCGTTGCCCTCAGCTCTTGGACAAAACTTCNNNAGATCGGAAGA
57691 TATGAGTGTGGAGTCGTTGCAAGTAGCTGGAATTACAGACGCANNNAGATCGGAA
57714 TATGAGTGTGGAGTCGTTGCGCCTCCCAAGTTCAAGTGATTCTNNNAGATCGGAAGA
57759 TATGAGTGTGGAGTCGTTGCAGAGGTAGATGGTAAGGACTGACANNNAGATCGGA
57768 TATGAGTGTGGAGTCGTTGCTGGGTAATTAGGTTTTGCACAAGTNNNAGATCGGAA
57779 TATGAGTGTGGAGTCGTTGCCCCAGGTTTGGTCTGATATGTTCANNNAGATCGGAA
57783 TATGAGTGTGGAGTCGTTGCAAAACTATGGTAGTCAGATGACCANNNAGATCGGAA
57790 TATGAGTGTGGAGTCGTTGCTGCTCAACTTTTTATCCAGCTGTGNNNAGATCGGAAG
57805 TATGAGTGTGGAGTCGTTGCCAGGATGGAAGACAAAATGTGACANNNAGATCGGA
57807 TATGAGTGTGGAGTCGTTGCACTTGAATGTAAGATCTGGAGCCANNNAGATCGGAA
57835 TATGAGTGTGGAGTCGTTGCCTCCCAAGTAGCTGAGACTACAGNNNAGATCGGAAG
57878 TATGAGTGTGGAGTCGTTGCTTCAAGCGATTCTCCTGTCTCAGNNNAGATCGGAAGA
57900 TATGAGTGTGGAGTCGTTGCTGCACTAGACAACAATGGATGGANNNAGATCGGAA
57906 TATGAGTGTGGAGTCGTTGCCAACCTTCAGGGCTCAAATGATCNNNAGATCGGAAG
57935 TATGAGTGTGGAGTCGTTGCGCTGAGGATAAGAGAGAAGAGACANNNAGATCGG

57970 TATGAGTGTGGAGTCGTTGCGGTATGTTGACCCTGAATTTCAGCNNNAGATCGGAA
57989 TATGAGTGTGGAGTCGTTGCAGTTCAGGTAAGGTGGAAAGAAAGTNNNAGATCGG
58058 TATGAGTGTGGAGTCGTTGCAGGACATGTAGCAGAAAGATGTCANNNAGATCGGA
58108 TATGAGTGTGGAGTCGTTGCGCTTCTCCTTTCTCCTTCTTATTGCNNNAGATCGGAAG
58141 TATGAGTGTGGAGTCGTTGCCTCCTGAGTAGCTGGGACTACANNNAGATCGGAAG
58169 TATGAGTGTGGAGTCGTTGCAATACAGTGGCATGATCTCAGCTNNNAGATCGGAAG
58296 TATGAGTGTGGAGTCGTTGCTGGTAAATGAAAGTTTGCATAGCTNNNAGATCGGA
58308 TATGAGTGTGGAGTCGTTGCTGCTTTTACACAAAAAAGGCAATTNNNAGATCGGAA
58324 TATGAGTGTGGAGTCGTTGCTAGATCCCCCAGTTTTGTTCAGGNNNAGATCGGAAG
58422 TATGAGTGTGGAGTCGTTGCTGATGGATACCTTCATGAGCTTGANNNAGATCGGAA
58427 TATGAGTGTGGAGTCGTTGCACAAAGAAATAGAGAGACAAGGAGCNNNAGATCGG
58433 TATGAGTGTGGAGTCGTTGCCCCCTACCTCACACCAATACAAANNNAGATCGGAAGA
58510 TATGAGTGTGGAGTCGTTGCCCATTCTGTTTGAGTATGCTGAGANNNAGATCGGAA
58526 TATGAGTGTGGAGTCGTTGCGTACAGTGGCACAAACACAGTTTNNNAGATCGGAA
58584 TATGAGTGTGGAGTCGTTGCTCTTCACTTCTAGACCATCTTGTTGANNNAGATCGGA
58608 TATGAGTGTGGAGTCGTTGCTGCATACGTGTTACAAACCTCACNNNAGATCGGAAG
58616 TATGAGTGTGGAGTCGTTGCGAGTTTCAGTCTTGTTTCCCAGGNNNAGATCGGAAG
58622 TATGAGTGTGGAGTCGTTGCATATTGGCTCACTGAAACCTCCANNNAGATCGGAAG
58655 TATGAGTGTGGAGTCGTTGCCAATGGCGCAATCTCAGCTCNNNAGATCGGAAGAG
58672 TATGAGTGTGGAGTCGTTGCAGTAGCTGAGATTACAGGTGTGCNNNAGATCGGAA
58691 TATGAGTGTGGAGTCGTTGCCATTCCGAGTAGCTGGAACTACANNNAGATCGGAAG
58695 TATGAGTGTGGAGTCGTTGCGGGTAGCTGGGAATATAGGTGCNNNAGATCGGAA
58719 TATGAGTGTGGAGTCGTTGCCACACTTATACTGTCCTGCTCACTNNNAGATCGGAAG
58737 TATGAGTGTGGAGTCGTTGCGTGCTATGGGCTATTGATGCAATTNNNAGATCGGA
58738 TATGAGTGTGGAGTCGTTGCTGGAAAGAACTGATCCACATAGGTNNNAGATCGGA
58788 TATGAGTGTGGAGTCGTTGCCTAGAGTGCAGTGGTGTGATCTCNNNAGATCGGAA
58871 TATGAGTGTGGAGTCGTTGCTGCCATTCGTACTGAAACTATTCCNNNAGATCGGAA
58878 TATGAGTGTGGAGTCGTTGCGGTGTCTGTTTGTTGTCTTTTGGANNNAGATCGGAA
58944 TATGAGTGTGGAGTCGTTGCTGATCTTCCACCTCTCTGTATCCTNNNAGATCGGAAG
58953 TATGAGTGTGGAGTCGTTGCTCAAGGCAGAAAATACAAAAAAGCCNNNAGATCGG
59021 TATGAGTGTGGAGTCGTTGCTGTTGATCTCATAGAAGTGGAGAGTNNNAGATCGG
59050 TATGAGTGTGGAGTCGTTGCTGAAACTTACATGCTACTTTGCAGANNNAGATCGGA
59116 TATGAGTGTGGAGTCGTTGCTGTTTGGGATTTTTTGTTTGTGCTNNNAGATCGGAA
59150 TATGAGTGTGGAGTCGTTGCACACACACACACACATACATATTCANNNAGATCGGAA
59159 TATGAGTGTGGAGTCGTTGCAGTCACATTTTCATAGTTCCTTCCTNNNAGATCGGAA
59160 TATGAGTGTGGAGTCGTTGCTGACAAAATGGAAAGTCAGTTTTCTGNNNAGATCGG
59171 TATGAGTGTGGAGTCGTTGCAGAAAAGAAGGGGATAATCCAATCTNNNAGATCGG
59175 TATGAGTGTGGAGTCGTTGCTGCTCTGTAGGATTTCTCAACTCTNNNAGATCGGAA
59245 TATGAGTGTGGAGTCGTTGCTGGGTTGGCCTTATTCTCCTTTTNNNAGATCGGAAG
59276 TATGAGTGTGGAGTCGTTGCATCTTGACTCACTCCAACCTCTGNNNAGATCGGAAGA
59284 TATGAGTGTGGAGTCGTTGCAACTTCTGCCTACTGAGCTCAAGNNNAGATCGGAAG
59357 TATGAGTGTGGAGTCGTTGCTGCAACAACAACAAAAACATTTGANNNAGATCGGAA
59388 TATGAGTGTGGAGTCGTTGCTAGTTCACTGTAGCCTTGACCTGNNNAGATCGGAAG

59416 TATGAGTGTGGAGTCGTTGCTGCAGCCAACAAACATATGAAAGANNNAGATCGGAA

59423 TATGAGTGTGGAGTCGTTGCGAACGTTACCAGCAGGTGATTTCNNNAGATCGGAAG

59425 TATGAGTGTGGAGTCGTTGCACACACACATACATAAAACATCACANNNAGATCGGAA

59469 TATGAGTGTGGAGTCGTTGCGTAGCTGGGATTACAGGTACACANNNAGATCGGAAG

59503 TATGAGTGTGGAGTCGTTGCCAGAGGGCTGGTCTTTTATTCCTNNNAGATCGGAAG

59530 TATGAGTGTGGAGTCGTTGCGCAATAGCATGATCTTGGCTCACNNNAGATCGGAAG

59545 TATGAGTGTGGAGTCGTTGCTGCCACATGATTCAAAACCAAAGANNNAGATCGGAA

59579 TATGAGTGTGGAGTCGTTGCGAATTGTTGGCAGCCATGACTAGNNNAGATCGGAAG

59583 TATGAGTGTGGAGTCGTTGCTCTCTGCATTTTTGGAAGTGAAGANNNAGATCGGAA

59584 TATGAGTGTGGAGTCGTTGCTGTTGTAAAAGCTTTGGTAAGGCTNNNAGATCGGAA

59586 TATGAGTGTGGAGTCGTTGCTGAGCAAAGGTATGACTAGACAGTNNNAGATCGGAA

59615 TATGAGTGTGGAGTCGTTGCAGTTGGTTGTTTGAGAAGATAAACANNNAGATCGGA

59716 TATGAGTGTGGAGTCGTTGCAGTAACTGGGACTATAAGTGCACANNNAGATCGGAA

59940 TATGAGTGTGGAGTCGTTGCGGGACCACCATTAGCAATAAAACANNNAGATCGGAA

59948 TATGAGTGTGGAGTCGTTGCAGTCTGTCTGTCTGTCTACACTACNNNAGATCGGAAG

59950 TATGAGTGTGGAGTCGTTGCACTTGCTAATGGGAAGACATTTCTNNNAGATCGGAA

60065 TATGAGTGTGGAGTCGTTGCCTCCTGAGTAGCTGGGACTACAGNNNAGATCGGAAG

60101 TATGAGTGTGGAGTCGTTGCCTGGATTCAAGCCATTCTTCTGCNNNAGATCGGAAGA

60167 TATGAGTGTGGAGTCGTTGCGAATCCCATGATGAACATACTGGANNNAGATCGGAA

60240 TATGAGTGTGGAGTCGTTGCCATCCCAGAGTTGCAGAGATAGTNNNAGATCGGAAG

60268 TATGAGTGTGGAGTCGTTGCGGTTCTAAGTAGGAAGTAGCTGAGGNNNAGATCGG

62025 TATGAGTGTGGAGTCGTTGCAGAGAGAATTTGAGACAAGGGACCNNNAGATCGGA

62026 TATGAGTGTGGAGTCGTTGCTCAGTACAAGGCATAATAGTGTGGTNNNAGATCGGA

62049 TATGAGTGTGGAGTCGTTGCACCAAGATTGAACCAGGAAGAAANNNAGATCGGAAG

62056 TATGAGTGTGGAGTCGTTGCGATTCAAGCGATTCTTCTGGCTCNNNAGATCGGAAGA

62114 TATGAGTGTGGAGTCGTTGCTTTGCAAACTATCCATCTGGCAANNNAGATCGGAAGA

62131 TATGAGTGTGGAGTCGTTGCCGTGCAGATAGGAAGTTTTGGAANNNAGATCGGAA

62240 TATGAGTGTGGAGTCGTTGCACACAGAAACTTGTTCATAGCAGCNNNAGATCGGAA

62267 TATGAGTGTGGAGTCGTTGCTGAGATGACAGGAAATTTGGTTTTNNNAGATCGGAA

62268 TATGAGTGTGGAGTCGTTGCAACACACCACAAACTTTTAGAACTNNNAGATCGGAAG

62308 TATGAGTGTGGAGTCGTTGCTCACATGTGGTGGAAGATTTTCTTNNNAGATCGGAA

62335 TATGAGTGTGGAGTCGTTGCTGGAGTGTAATGGCACAATCTCANNNAGATCGGAAG

62369 TATGAGTGTGGAGTCGTTGCTGGAACCTGTGAATGTTACTTTGTNNNAGATCGGAA

62370 TATGAGTGTGGAGTCGTTGCCTCAAGTGATCCTCCCAACTCAGNNNAGATCGGAAGA

62379 TATGAGTGTGGAGTCGTTGCGTTCCAAGAATTACTCCTTTCTCTACANNNAGATCGGA

62383 TATGAGTGTGGAGTCGTTGCTGCTCCTGATTTTGATATGAGGGTNNNAGATCGGAA

62392 TATGAGTGTGGAGTCGTTGCAGCAAACTGAATTCATCTGAACATNNNAGATCGGAAG

62412 TATGAGTGTGGAGTCGTTGCTTCAAGCAATTCTCCTGCCTCAGNNNAGATCGGAAGA

62533 TATGAGTGTGGAGTCGTTGCCTCTTGGGTTCACGCCATTCTCNNNAGATCGGAAGAG

62559 TATGAGTGTGGAGTCGTTGCAGAGAACCCAGACATAAATCCACANNNAGATCGGAA

62565 TATGAGTGTGGAGTCGTTGCCCAAAGTGCTGGGATTACAAGTGNNNAGATCGGAAG

62597 TATGAGTGTGGAGTCGTTGCTCTTCAATTGCATCTAGTTTGGAACANNNAGATCGGA

62608 TATGAGTGTGGAGTCGTTGCAGTAGCAGGAGTACACTGTAAAATNNNAGATCGGA

62635 TATGAGTGTGGAGTCGTTGCTCCCATATGAATGTGTGTGTATGTNNNAGATCGGAA
62676 TATGAGTGTGGAGTCGTTGCTCATGAGGAGTTTTTTGGTGAGGANNNAGATCGGA
62837 TATGAGTGTGGAGTCGTTGCTTCATGAAGTTCTCCTGCCTCAGNNNAGATCGGAAG
62889 TATGAGTGTGGAGTCGTTGCCAATCTCGGCTCACTGCAAGNNNAGATCGGAAGAG
62926 TATGAGTGTGGAGTCGTTGCTGCTTAGATGTAGTTACGCACTGANNNAGATCGGAA
62932 TATGAGTGTGGAGTCGTTGCTGTTGTGTCGTATGTTTTACCACANNNAGATCGGAA
62937 TATGAGTGTGGAGTCGTTGCTGCTAAAGAAAAGCAGCAATGACTNNNAGATCGGA
62942 TATGAGTGTGGAGTCGTTGCTGGTATTCTTTGGTATCACTGGGGNNNAGATCGGA
62983 TATGAGTGTGGAGTCGTTGCACCATTTCACATTTTTTGAGTCCANNNAGATCGGAAG
63034 TATGAGTGTGGAGTCGTTGCAGATTTATGGGTGTTAGGGAAGTTNNNAGATCGGA
63045 TATGAGTGTGGAGTCGTTGCAGTCTAAAGTTATCAGCATAGGCCANNNAGATCGGA
63134 TATGAGTGTGGAGTCGTTGCGGAAACTTTGTCCATTCCTAGTGCNNNAGATCGGAA
63142 TATGAGTGTGGAGTCGTTGCTTCAAGTGATTCTCCTGCCTCAGNNNAGATCGGAAG
63188 TATGAGTGTGGAGTCGTTGCAGCAATCTCAAAAGGGTACGTACTNNNAGATCGGA
63225 TATGAGTGTGGAGTCGTTGCGGCATGGAATTCTTCACACTAACCNNNAGATCGGAA
63243 TATGAGTGTGGAGTCGTTGCTGTGTTTTTAGTAGACACGGGGTNNNAGATCGGAA
63256 TATGAGTGTGGAGTCGTTGCGGTTCAAGTGATTCTCCTGCTTCNNNAGATCGGAAG
63258 TATGAGTGTGGAGTCGTTGCTTCAAGTGATTCTCCTGCCTCAGNNNAGATCGGAAG
63271 TATGAGTGTGGAGTCGTTGCAATCTCAGCTCACTGCAACCTCNNNAGATCGGAAGA
63277 TATGAGTGTGGAGTCGTTGCGCGCACGTGTATGAAATATTTCCTNNNAGATCGGAA
63288 TATGAGTGTGGAGTCGTTGCAGAGAGAGAGAGAGAGAGAGAAAGCANNNAGATCGG
63303 TATGAGTGTGGAGTCGTTGCTTGAAGCGATTCTCCTGCCTTAGNNNAGATCGGAAG
63364 TATGAGTGTGGAGTCGTTGCTCCTTGGTACTCACTTCATTAGCCNNNAGATCGGAAG
63391 TATGAGTGTGGAGTCGTTGCCTCAAGTGATCCTACCACCTCAGNNNAGATCGGAAG
63400 TATGAGTGTGGAGTCGTTGCTGGTCGTCAATTTACAAGACATGCNNNAGATCGGAA
63436 TATGAGTGTGGAGTCGTTGCTCATGCTGCTTGTGAATGTATTGANNNAGATCGGAA
63477 TATGAGTGTGGAGTCGTTGCTTGTGAAAATTACCACACTGCCANNNAGATCGGAAG
63488 TATGAGTGTGGAGTCGTTGCGGTAGGTCTCTTCATTTCAGGGTNNNAGATCGGAAG
63493 TATGAGTGTGGAGTCGTTGCTGGAGTGCAGTGGTGTAATCTCNNNAGATCGGAAG
63510 TATGAGTGTGGAGTCGTTGCCATAGAATCTCACTCTGTCACCCANNNAGATCGGAAG
63560 TATGAGTGTGGAGTCGTTGCACTTCGGTATCCTCTCTTGATTCCNNNAGATCGGAAG
63568 TATGAGTGTGGAGTCGTTGCTGTAATACCAGCTACTCAGAAGGCNNNAGATCGGAA
63576 TATGAGTGTGGAGTCGTTGCGTCTCCTGGGTTCAAGTGATTCTNNNAGATCGGAAG
63616 TATGAGTGTGGAGTCGTTGCTGTTTAGTACCCTGCTTTTCTCANNNAGATCGGAAGA
63666 TATGAGTGTGGAGTCGTTGCACTTGACATGTTTACTGTTGCTTTGNNNAGATCGGA
63671 TATGAGTGTGGAGTCGTTGCAGTTTTGCTTACCTGTTCTGTTGTNNNAGATCGGAAG
63673 TATGAGTGTGGAGTCGTTGCGTCCTATCTTCTGACCACAACTCANNNAGATCGGAAG
63757 TATGAGTGTGGAGTCGTTGCGGACAAAGGAAGGAAAGAGGAGANNNAGATCGGA
63766 TATGAGTGTGGAGTCGTTGCTAACCTGGATGGAATTGGAGACCNNNAGATCGGAA
63771 TATGAGTGTGGAGTCGTTGCACTACATCTCGAGAGTGTAGAACANNNAGATCGGAA
63780 TATGAGTGTGGAGTCGTTGCTCTTTTTTGAGACAGAGTCTTGCTNNNAGATCGGAA
63857 TATGAGTGTGGAGTCGTTGCAGGGTCTCAATCTATCTTCCAAGCNNNAGATCGGAA
63859 TATGAGTGTGGAGTCGTTGCTCCATATCCATGAGGAAAACAGCANNNAGATCGGAA

63885 TATGAGTGTGGAGTCGTTGCTCTGGCCTCAAAAATATTTCAGCCNNNAGATCGGAA
63967 TATGAGTGTGGAGTCGTTGCTGAATACCTGCATGGAGTTACAACNNNAGATCGGAA
64029 TATGAGTGTGGAGTCGTTGCAGTAAACCCATCCAGGCTGAAATNNNAGATCGGAAG
64038 TATGAGTGTGGAGTCGTTGCTCTTTGTCTAGTCATCTGTCAGCANNNAGATCGGAA
64080 TATGAGTGTGGAGTCGTTGCAGAACTGTATGTGTTGAAGGCCANNNAGATCGGAA
64101 TATGAGTGTGGAGTCGTTGCTGAAACATTTCAGAACATTTGGGCNNNAGATCGGAA
64143 TATGAGTGTGGAGTCGTTGCTATGGAAGACATGATACAGCCCCNNNAGATCGGAA
64191 TATGAGTGTGGAGTCGTTGCAGGTTCAAGCAATTCTCCCTCAGNNNAGATCGGAAG
64229 TATGAGTGTGGAGTCGTTGCATTTCCTGGCTTCAGACATTCCTNNNAGATCGGAAGA
64235 TATGAGTGTGGAGTCGTTGCAGCACATCAAAAAGCTAATCCACCNNNAGATCGGAA
64247 TATGAGTGTGGAGTCGTTGCAGCAAGAGGAATTTTGCAAGTTCANNNAGATCGGA
64264 TATGAGTGTGGAGTCGTTGCAGCTCTATGTCTCTGGAGATCTCTNNNAGATCGGAA
64275 TATGAGTGTGGAGTCGTTGCTCCTCAACAAAATACCAGCAAACCNNNAGATCGGAA
64282 TATGAGTGTGGAGTCGTTGCTGGTTCCATAAGAATTTCAGGACTNNNAGATCGGAA
64301 TATGAGTGTGGAGTCGTTGCTGCTCTTCGATTTTTTTGAGATGANNNAGATCGGAA
64338 TATGAGTGTGGAGTCGTTGCTCCGTATAATGCTGCAGAGAACANNNAGATCGGAA
64422 TATGAGTGTGGAGTCGTTGCGTGGATCATTTGAGGTCAGGAGTNNNAGATCGGAA
64452 TATGAGTGTGGAGTCGTTGCCTGACCAAGAGGGACACCTAAATNNNAGATCGGAA
64459 TATGAGTGTGGAGTCGTTGCGAACCTTCATTTATGTCTTCATAGTCANNNAGATCGG
64494 TATGAGTGTGGAGTCGTTGCGTCTTGAACTGTTGGGCTCAAAGNNNAGATCGGAA
64503 TATGAGTGTGGAGTCGTTGCGTAAGCGGCACTAGTAATCATGGNNNAGATCGGAA
64514 TATGAGTGTGGAGTCGTTGCAGAAAAACAGGAAGGAGTTACCCANNNAGATCGGA
64532 TATGAGTGTGGAGTCGTTGCAGAGAGAGAGAGAGACAGAGAGTNNNAGATCGGA
64538 TATGAGTGTGGAGTCGTTGCTGGAACTGAAAACTATGACACCCTNNNAGATCGGAA
64569 TATGAGTGTGGAGTCGTTGCATTTTTGCCTCTACTCAGTCAGCNNNAGATCGGAAGA
64576 TATGAGTGTGGAGTCGTTGCGTAACCTGGAGAATGAGGGTGTTNNNAGATCGGAA
64585 TATGAGTGTGGAGTCGTTGCTGCAATTACTAGGTGTTTGACTGANNNAGATCGGAA
64605 TATGAGTGTGGAGTCGTTGCTTGCTTCCTATTGGCCTTTCTCTNNNAGATCGGAAGA
64660 TATGAGTGTGGAGTCGTTGCATACAACAGCTGTACTTCCCCTGNNNAGATCGGAAG
64682 TATGAGTGTGGAGTCGTTGCCATGAACGATGGATGCATAATTCTNNNAGATCGGAA
64716 TATGAGTGTGGAGTCGTTGCACTCCTGGACTAAAGTGATCTGCNNNAGATCGGAAG
64730 TATGAGTGTGGAGTCGTTGCTGAAGATTCACTGGGATGCTTCANNNAGATCGGAA
64735 TATGAGTGTGGAGTCGTTGCCCTAAGGCTCATGAAACAATACATNNNAGATCGGAA
64771 TATGAGTGTGGAGTCGTTGCTGCCTACTATCACATAGACCTCTGNNNAGATCGGAA
64835 TATGAGTGTGGAGTCGTTGCTGGTGAGGATGTGGAGAAATCAGNNNAGATCGGA
64847 TATGAGTGTGGAGTCGTTGCACCTAAGCTTAGTGACTGGTTGGNNNAGATCGGAA
64855 TATGAGTGTGGAGTCGTTGCGGTGCAAATGATCTAGCAATCCCNNNAGATCGGAAG
64877 TATGAGTGTGGAGTCGTTGCTGGCCTGTCCTAAATAGCTCATCNNNAGATCGGAAG
64884 TATGAGTGTGGAGTCGTTGCTGGAGCTGAAAAATTCCTATTGCTNNNAGATCGGAA
64889 TATGAGTGTGGAGTCGTTGCGCTCCTGAGATCTCATCTGATACCNNNAGATCGGAA
64906 TATGAGTGTGGAGTCGTTGCTCCGTTACCACTGATAATGACTGTNNNAGATCGGAA
64941 TATGAGTGTGGAGTCGTTGCCCCATAATCCGTTGATAGACAGGTNNNAGATCGGAA
64942 TATGAGTGTGGAGTCGTTGCTGAAGTAACAGTTTCCACATCCCTNNNAGATCGGAA

64953 TATGAGTGTGGAGTCGTTGCTGCTCATAACTCTTTCACATTCACTNNNAGATCGGAA
64955 TATGAGTGTGGAGTCGTTGCACACACACTATATACACACACACTNNNAGATCGGAAG
65017 TATGAGTGTGGAGTCGTTGCTCTCCAGCTAACATGTTTCATGGANNNAGATCGGAA
65025 TATGAGTGTGGAGTCGTTGCTGTGTGTGTGTGTATCGAAAAACANNNAGATCGGA
65026 TATGAGTGTGGAGTCGTTGCTAAGGCTGATACACGAGGGAATGNNNAGATCGGAA
65027 TATGAGTGTGGAGTCGTTGCATCTCAGCTCACTGCAACCTCTANNNAGATCGGAAGA
65043 TATGAGTGTGGAGTCGTTGCTTGTAGCTCTGTCTTGGTAGTGGNNNAGATCGGAAG
65061 TATGAGTGTGGAGTCGTTGCGTCCCATTGTTATGAATGCAGACTNNNAGATCGGAA
65089 TATGAGTGTGGAGTCGTTGCCTTCCACATGTTTGGCCAATATANNNAGATCGGAAG
65174 TATGAGTGTGGAGTCGTTGCTTTTTTTGAGACAGAGTCTTGCTCNNNAGATCGGAA
65201 TATGAGTGTGGAGTCGTTGCGGTGTAGTACTGAGTATCATCCCNNNAGATCGGAA
65226 TATGAGTGTGGAGTCGTTGCTCTCTTCATCTGCTGATTCACACANNNAGATCGGAAG
65228 TATGAGTGTGGAGTCGTTGCTGGCTTCCTCCTTTTCTCAATGTNNNAGATCGGAAGA
65230 TATGAGTGTGGAGTCGTTGCCCCAAATAGTGAGGTTTCATTCCANNNAGATCGGAA
65236 TATGAGTGTGGAGTCGTTGCTCTCTCCTCTCTCTCTCTTTCACANNNAGATCGGAAGA
65319 TATGAGTGTGGAGTCGTTGCTGAATAGTTCTGTCAGTTTGGTGANNNAGATCGGAA
65324 TATGAGTGTGGAGTCGTTGCTGGATGATGATTGTGATATCAGGANNNAGATCGGA
65392 TATGAGTGTGGAGTCGTTGCAGCTCCTTTAGAAGAACTTGACCANNNAGATCGGAA
65456 TATGAGTGTGGAGTCGTTGCGCAGATGTTCAACTCAATATGGGGNNNAGATCGGA
65508 TATGAGTGTGGAGTCGTTGCAGGAGGAAAGAAGAGAGAATGTGTNNNAGATCGG
65541 TATGAGTGTGGAGTCGTTGCACAGTCTGAAGTGACCCTACTATGNNNAGATCGGAA
65556 TATGAGTGTGGAGTCGTTGCAGTGGGATTGTTGCAATTTATGGTNNNAGATCGGA
65559 TATGAGTGTGGAGTCGTTGCCATTCAGTCCAAGCTCTGTGTTGNNNAGATCGGAAG
65592 TATGAGTGTGGAGTCGTTGCCTAAGAAGTCCCACAGTCTGCTGNNNAGATCGGAAG
65596 TATGAGTGTGGAGTCGTTGCTCAACCAGGAAGCCAGAAGAAAANNNAGATCGGAA
65614 TATGAGTGTGGAGTCGTTGCTAAACACTGCAGGGACTGAATGANNNAGATCGGAA
65626 TATGAGTGTGGAGTCGTTGCTCTATGTTCTGCCATCAGCTTCTNNNAGATCGGAAGA
65647 TATGAGTGTGGAGTCGTTGCGGAAAAGATATGACCATGGTAGGCNNNAGATCGGA
65663 TATGAGTGTGGAGTCGTTGCACTCCATTGGTTCTGTTTCTCTGANNNAGATCGGAAG
65666 TATGAGTGTGGAGTCGTTGCATGACCTAGCCAAGTTGACACATNNNAGATCGGAAG
65686 TATGAGTGTGGAGTCGTTGCGTGTTTTTTTGAGACTGGGCCTTNNNAGATCGGAAG
65687 TATGAGTGTGGAGTCGTTGCACATCACTTTTTGAAAGGACAGAANNNAGATCGGAA
65767 TATGAGTGTGGAGTCGTTGCACCTGTCAGTGGATGCTTAGTTTNNNAGATCGGAAG
65776 TATGAGTGTGGAGTCGTTGCAAATGGAGTCTTGCTATGTTGCCNNNAGATCGGAAG
65780 TATGAGTGTGGAGTCGTTGCAAAAGAACAAAGCTGGAGGCATCNNNAGATCGGAA
65848 TATGAGTGTGGAGTCGTTGCTGTGTGTGTGTGTGTGTATTTTGTNNNAGATCGGAA
65857 TATGAGTGTGGAGTCGTTGCAGGGGATATCCATGACTCTGACTNNNAGATCGGAAG
65859 TATGAGTGTGGAGTCGTTGCTCGTGTCCTTATGTTATTGTCTCTNNNAGATCGGAAG
65866 TATGAGTGTGGAGTCGTTGCGAGACAGGGTCTTGCTCTGTTGNNNAGATCGGAAG
65874 TATGAGTGTGGAGTCGTTGCACAAGAAAGACTGTAAAGGTACCANNNAGATCGGA
65908 TATGAGTGTGGAGTCGTTGCAGTGGGAATGGAAACTATTTGAGTNNNAGATCGGA
65922 TATGAGTGTGGAGTCGTTGCACCATGGAATACTATGCAGCCATNNNAGATCGGAAG
65986 TATGAGTGTGGAGTCGTTGCGAAGACAGGGAGCCAGAATGATANNNAGATCGGA

65989 TATGAGTGTGGAGTCGTTGCACAATTGCCTAGAAATGAGGGTCTNNNAGATCGGA
66037 TATGAGTGTGGAGTCGTTGCCTGGATTCAAGCAATTCTCCTGCNNNAGATCGGAAG
66066 TATGAGTGTGGAGTCGTTGCACTAAATGACCTCATTCTAGTGCTNNNAGATCGGAA
66067 TATGAGTGTGGAGTCGTTGCTGGCTTCCTCCTTTTCTCAATGTNNNAGATCGGAAGA
66169 TATGAGTGTGGAGTCGTTGCAGTAGAGATTGCCATTCTTCATCTNNNAGATCGGAA
66205 TATGAGTGTGGAGTCGTTGCAGAGAGAGAGAGAGAGAGAGAACANNNAGATCGG
66206 TATGAGTGTGGAGTCGTTGCAGACAAATAGGTTGGGAGTAGATTGTNNNAGATCG
66221 TATGAGTGTGGAGTCGTTGCTGGACTCCTCCTAAACTCTAACTGNNNAGATCGGAA
66224 TATGAGTGTGGAGTCGTTGCACTCAAACCCAGTAAGGAAACTTGNNNAGATCGGAA
66227 TATGAGTGTGGAGTCGTTGCGGGCCACTGAAGAGATAACAGAANNNAGATCGGAA
66241 TATGAGTGTGGAGTCGTTGCACTGACATACCATTAGCACCCAANNNAGATCGGAAG
66245 TATGAGTGTGGAGTCGTTGCTTTACCCAATCCCCAAGTTGTGANNNAGATCGGAAG
66252 TATGAGTGTGGAGTCGTTGCGTCTTGGAAACACACAAGCTACCNNNAGATCGGAAG
66267 TATGAGTGTGGAGTCGTTGCAGAGAGGCAGAGATAGAGTCCTCNNNAGATCGGAA
66319 TATGAGTGTGGAGTCGTTGCATGAGCTCCTTGAAGAAATGGCTNNNAGATCGGAA
66402 TATGAGTGTGGAGTCGTTGCACTCGAAGTTACTTTGGGATCCANNNAGATCGGAAG
66410 TATGAGTGTGGAGTCGTTGCTTTCTCCCTCATCTGGTTATCCTNNNAGATCGGAAGA
66415 TATGAGTGTGGAGTCGTTGCAAAGATGAACTCCCAGCTACTTATNNNAGATCGGAA
66592 TATGAGTGTGGAGTCGTTGCCCTTTCTGTTTACTTTCAGGTCGTNNNAGATCGGAAG
66653 TATGAGTGTGGAGTCGTTGCACCATCACATGACTGGCTATATCCNNNAGATCGGAA
66657 TATGAGTGTGGAGTCGTTGCAGTCTTTCTTTCTAGGCTGAAGGANNNAGATCGGAA
66658 TATGAGTGTGGAGTCGTTGCCATGGAATGTGTTAGGGAGGGAGNNNAGATCGGA
66691 TATGAGTGTGGAGTCGTTGCCTGGAGCACATTACTACCAACTTNNNAGATCGGAAG
66745 TATGAGTGTGGAGTCGTTGCGCATTAGGCAAAAAGTACGGAACTNNNAGATCGGA
66779 TATGAGTGTGGAGTCGTTGCGCACGCACATAGACACACATATGNNNAGATCGGAAG
66781 TATGAGTGTGGAGTCGTTGCTGAAACAGTGATTACCATGAGTGGNNNAGATCGGA
66789 TATGAGTGTGGAGTCGTTGCTCTTTCCCTGTTTCCCTGTCATCNNNAGATCGGAAGA
66844 TATGAGTGTGGAGTCGTTGCAGGAGAACTACAAAATACTGCTGANNNAGATCGGA
66861 TATGAGTGTGGAGTCGTTGCGCTAACCAAATCCATCAGCACATNNNAGATCGGAAG
66865 TATGAGTGTGGAGTCGTTGCAAGTTGCAGAATACAACATCAACANNNAGATCGGAA
66914 TATGAGTGTGGAGTCGTTGCAGACTTTCTCAATAACTGGTGAGANNNAGATCGGAA
66918 TATGAGTGTGGAGTCGTTGCGCCTATCAATTCTGTCTCTCTGGANNNAGATCGGAA
66958 TATGAGTGTGGAGTCGTTGCCCAATGGGACAAAATGGAAGTCCNNNAGATCGGAA
67010 TATGAGTGTGGAGTCGTTGCTGGAATAGCAAATCTACGGAGACANNNAGATCGGA
67038 TATGAGTGTGGAGTCGTTGCACTCACCATCATTAGTCTTCAGGGNNNAGATCGGAA
67052 TATGAGTGTGGAGTCGTTGCACCAAAAATAGTTGTTGGCAAACCNNNAGATCGGAA
67114 TATGAGTGTGGAGTCGTTGCTCCTGTTCCAATGCCTTTTTTTCTNNNAGATCGGAAG
67117 TATGAGTGTGGAGTCGTTGCTGTGTATATGTGTGTGTGTGTGTGNNNAGATCGGA
67131 TATGAGTGTGGAGTCGTTGCGGTTGAGACCACAAATCAAAGAGTNNNAGATCGGA
67134 TATGAGTGTGGAGTCGTTGCCCAGAGTCAGCCTGAACATTCTANNNAGATCGGAAG
67149 TATGAGTGTGGAGTCGTTGCTGGCTTCCTCCTTTTCTCAATGTNNNAGATCGGAAGA
67182 TATGAGTGTGGAGTCGTTGCTGCAGCAAAAACGTATATGGGAGNNNAGATCGGAA
67197 TATGAGTGTGGAGTCGTTGCTGATCCCAAGAATCACCATTTTGGNNNAGATCGGAA

67235 TATGAGTGTGGAGTCGTTGCCTATAGGTCAGGTAGGCAAGCTGNNNAGATCGGAA
67249 TATGAGTGTGGAGTCGTTGCGGAGAGAGAGACAGAGAGAAGGANNNAGATCGG
67274 TATGAGTGTGGAGTCGTTGCAAAAGATGGGTACCAGTCTGTGGNNNAGATCGGAA
67284 TATGAGTGTGGAGTCGTTGCTAAATTTGGGGACATTTTGGCCCNNNAGATCGGAAG
67305 TATGAGTGTGGAGTCGTTGCACCTTCACTATTTTGATAGCTGGANNNAGATCGGAA
67315 TATGAGTGTGGAGTCGTTGCACTTGCATTAGGACATTTTGTTGANNNAGATCGGAA
67482 TATGAGTGTGGAGTCGTTGCAGAGAAAAGTTGGGGAGAAATGGTNNNAGATCGG
67505 TATGAGTGTGGAGTCGTTGCCCCAGCAACATCAACATTACCTGNNNAGATCGGAAG
67522 TATGAGTGTGGAGTCGTTGCAAACAGATCTTGCTTAGTACCTGANNNAGATCGGAA
67548 TATGAGTGTGGAGTCGTTGCACTGAAAGTGCTGAGGAAAGAGTNNNAGATCGGA
67564 TATGAGTGTGGAGTCGTTGCTCTGTTTTGTGAAATGCCTATTCANNNAGATCGGAA
67585 TATGAGTGTGGAGTCGTTGCTCAGCCAAATCATAAGCACATTGTNNNAGATCGGAA
67740 TATGAGTGTGGAGTCGTTGCTCTCATCTGCTCAGTACTCAGGANNNAGATCGGAAG
67741 TATGAGTGTGGAGTCGTTGCTGTATTCCTCGAGATACACAAGCANNNAGATCGGAA
67743 TATGAGTGTGGAGTCGTTGCTAGACCAATAGTGATGAGAGCGCNNNAGATCGGAA
67797 TATGAGTGTGGAGTCGTTGCTCCAAGAAGAAAAAAAAAAGGCCTNNNAGATCGGA
67798 TATGAGTGTGGAGTCGTTGCCAAAGCACCTCCTATTTGAAACCANNNAGATCGGAA
67801 TATGAGTGTGGAGTCGTTGCAGCACTCAAACACAAATCTAGACANNNAGATCGGAA
67836 TATGAGTGTGGAGTCGTTGCTCTTTCCCTCTTTCTTGCTCTCTNNNAGATCGGAAGAG
67845 TATGAGTGTGGAGTCGTTGCTTGAAATGTGATCCCCAGTGTCANNNAGATCGGAAG
67846 TATGAGTGTGGAGTCGTTGCTGAAACTCCCAGATTTCTGACACTNNNAGATCGGAA
67847 TATGAGTGTGGAGTCGTTGCACCCCCAATGTAGAAATATGCTGANNNAGATCGGAA
67865 TATGAGTGTGGAGTCGTTGCTCAGCTGGTATCTAAAAAGGTGGANNNAGATCGGA
67876 TATGAGTGTGGAGTCGTTGCCACCTAGTGATAAACACAGGGACANNNAGATCGGA
67879 TATGAGTGTGGAGTCGTTGCGTCTGCCTAACCGTAGTAATCCCNNNAGATCGGAAG
67883 TATGAGTGTGGAGTCGTTGCACCAGAAGAGATAGAACTCTAGCTNNNAGATCGGA
67895 TATGAGTGTGGAGTCGTTGCTGTAAGGTCAGTGTTCTCCAGAGNNNAGATCGGAA
67944 TATGAGTGTGGAGTCGTTGCAACTTGGATGCTGCTGGTGTATANNNAGATCGGAA
67974 TATGAGTGTGGAGTCGTTGCACAACCAAATAACACATAACACCTNNNAGATCGGAA
67975 TATGAGTGTGGAGTCGTTGCTGAACTCAATAACTTGGAGGGTGTNNNAGATCGGA
67983 TATGAGTGTGGAGTCGTTGCACCTTCTCGTAATATGGACTCAGANNNAGATCGGAA
68010 TATGAGTGTGGAGTCGTTGCTGAGCTCCAGAAAGAGAAAACAGANNNAGATCGGA
68014 TATGAGTGTGGAGTCGTTGCGGAGTGAAATGGCATGATCTTGGNNNAGATCGGAA
68066 TATGAGTGTGGAGTCGTTGCGCGATCTCAAATCACTGCAACTTNNNAGATCGGAAG
68074 TATGAGTGTGGAGTCGTTGCGAGATAGTCTTGCTCTTGTTGCCNNNAGATCGGAAG
68092 TATGAGTGTGGAGTCGTTGCTCTAAAGCCCGTGTAAACATTCTGNNNAGATCGGAA
68126 TATGAGTGTGGAGTCGTTGCAGACTCTGTCACTGTGGTATCATGNNNAGATCGGAA
68155 TATGAGTGTGGAGTCGTTGCCCAATGGCATAGTAGCAAGTATGCNNNAGATCGGA
68173 TATGAGTGTGGAGTCGTTGCTGTAGAAAATTCCAAGTGCTCACCNNNAGATCGGAA
68191 TATGAGTGTGGAGTCGTTGCACCAGTTTTACTTTGTTTGCCCCNNNAGATCGGAAGA
68203 TATGAGTGTGGAGTCGTTGCGTGTGATCATGACTCAACTGCAGNNNAGATCGGAA
68206 TATGAGTGTGGAGTCGTTGCACTTTATTGAAGAGCTCCTGTCACTNNNAGATCGGA
68241 TATGAGTGTGGAGTCGTTGCAGGATGTAGACAAGTTGGAACCCNNNAGATCGGAA

68285 TATGAGTGTGGAGTCGTTGCACCCTGACTATTTGCAAATACACTNNNAGATCGGAAG
68292 TATGAGTGTGGAGTCGTTGCCCGCATACAATAACTCACAAGAGCNNNAGATCGGAA
68295 TATGAGTGTGGAGTCGTTGCGCTGGGTAAAGGCAGTATACAGANNNAGATCGGAA
68306 TATGAGTGTGGAGTCGTTGCTTCCCCGCAATCTCATTATGACTNNNAGATCGGAAGA
68367 TATGAGTGTGGAGTCGTTGCACACATACACAAACACACATATGTNNNAGATCGGAA
68369 TATGAGTGTGGAGTCGTTGCTGGCATATAATGGGGAAAAATGGCNNNAGATCGGA
68372 TATGAGTGTGGAGTCGTTGCTATACACACACACACACACACACNNNAGATCGGAAGA
68402 TATGAGTGTGGAGTCGTTGCTGTGTGTGTGTGTGTAGGGTTTANNNAGATCGGAA
68421 TATGAGTGTGGAGTCGTTGCCCTTTATATCTTAGCTTGCTGCCCNNNAGATCGGAAG
68430 TATGAGTGTGGAGTCGTTGCTCTTTCTTCTCTTCCTTTCTTCCTNNNAGATCGGAAGA
68479 TATGAGTGTGGAGTCGTTGCGCTTTAGTCAGTTCTTTGTCCCACNNNAGATCGGAAG
68486 TATGAGTGTGGAGTCGTTGCACTGAAGTGCTTATTCCACATCCANNNAGATCGGAAG
68491 TATGAGTGTGGAGTCGTTGCTCTTAGGCCAGTTCATTACCAGGNNNAGATCGGAAG
68493 TATGAGTGTGGAGTCGTTGCACATGTAACTTCTTGGTTGAGAAGNNNAGATCGGAA
68518 TATGAGTGTGGAGTCGTTGCTCAGAGCTCAATGGCCATTATTTGNNNAGATCGGAA
68612 TATGAGTGTGGAGTCGTTGCTTTCTTTTTTAGAAATGGGGTCTAACCNNNAGATCGG
68622 TATGAGTGTGGAGTCGTTGCGGAGTGCAGTGGTGTGATCTTAGNNNAGATCGGAA
68636 TATGAGTGTGGAGTCGTTGCTGCATGAAAACTGTCTTTCAGCTTNNNAGATCGGAA
68660 TATGAGTGTGGAGTCGTTGCAGAAAGGAAAGAAAGCAAGCAAGANNNAGATCGGA
68800 TATGAGTGTGGAGTCGTTGCGAGAGAGAGAGAGGAAGGTGGACNNNAGATCGGA
68825 TATGAGTGTGGAGTCGTTGCCACACACACACACATACTTACACANNNAGATCGGAAG
68943 TATGAGTGTGGAGTCGTTGCAGAGACAACAGATAACCCAGGATNNNAGATCGGAA
69006 TATGAGTGTGGAGTCGTTGCTGCAAAAGTGTACTTGTCAGCAGNNNAGATCGGAAG
69038 TATGAGTGTGGAGTCGTTGCTGAATAACAGAGTTCACGTAAGCANNNAGATCGGAA
69104 TATGAGTGTGGAGTCGTTGCACAGCAATTAGTGTTAGAAAGGGTNNNAGATCGGA
69117 TATGAGTGTGGAGTCGTTGCTGCATGTGTGTGTATGTATCTCCTNNNAGATCGGAA
69120 TATGAGTGTGGAGTCGTTGCGTGGTGTGATCTCGCTTACTGTANNNAGATCGGAAG
69179 TATGAGTGTGGAGTCGTTGCACACACACAGAGAGAGAGAGAGANNNAGATCGGAA
69188 TATGAGTGTGGAGTCGTTGCACTGAGGTCATACGGTATTTGACTNNNAGATCGGAA
69302 TATGAGTGTGGAGTCGTTGCGCTACACACGAGGCATTTGATTANNNAGATCGGAAG
69312 TATGAGTGTGGAGTCGTTGCTCTTTCTCTCTGTCTCTCTCTCTCNNNAGATCGGAAGA
69353 TATGAGTGTGGAGTCGTTGCCAAAAAGCTTGAGAACCCAGAGGNNNAGATCGGAA
69371 TATGAGTGTGGAGTCGTTGCATATGGATAAGATTGGCCAGGCANNNAGATCGGAA
69414 TATGAGTGTGGAGTCGTTGCTGGAGCATGCTGACTGAGTTTATNNNAGATCGGAAG
69420 TATGAGTGTGGAGTCGTTGCAGAGAATACCGGTGCATACAGACNNNAGATCGGAA
69437 TATGAGTGTGGAGTCGTTGCTTGACTTCCTGGGCTCAAATGATNNNAGATCGGAAG
69473 TATGAGTGTGGAGTCGTTGCTCAGCTTGAAATTTCAGCAACTGTNNNAGATCGGAA
69591 TATGAGTGTGGAGTCGTTGCTACAAGCCCAGTTGTCTCTCATCNNNAGATCGGAAGA
69710 TATGAGTGTGGAGTCGTTGCTGCACACACACGAATAAAAAATTTNNNAGATCGGAA
69721 TATGAGTGTGGAGTCGTTGCCCTCCCAAGTAGCTGACAAAAAANNNAGATCGGAAG
69737 TATGAGTGTGGAGTCGTTGCTTGAATACACATGCCAGGACATCNNNAGATCGGAAG
69738 TATGAGTGTGGAGTCGTTGCACCTCCTCAATTTTCTGGAATAGTNNNAGATCGGAAG
69756 TATGAGTGTGGAGTCGTTGCTGAGAGAGAGACAGGGACTTATGANNNAGATCGGA

69769 TATGAGTGTGGAGTCGTTGCGGAAAGAACCAAAAGAAAGGGACANNNAGATCGG
69809 TATGAGTGTGGAGTCGTTGCCCTGAGTAGCTGGGATTATAGGCNNNAGATCGGAA
69842 TATGAGTGTGGAGTCGTTGCCTGTTAGCTACATCCCTCTCTCCNNNAGATCGGAAGA
69849 TATGAGTGTGGAGTCGTTGCAGAAAGAATGGAAGGAAGGAAGGANNNAGATCGG
69856 TATGAGTGTGGAGTCGTTGCCCAAGTGGACAATAAAGTAGCACANNNAGATCGGA
69889 TATGAGTGTGGAGTCGTTGCGCCTCCCAGGTTCATTGTAGTTTNNNAGATCGGAAG
69948 TATGAGTGTGGAGTCGTTGCCATAATTCACTTCACTGGCAGGCNNNAGATCGGAAG
69953 TATGAGTGTGGAGTCGTTGCTCAAGCAATTCCTCACCTCAAGANNNAGATCGGAAG
69973 TATGAGTGTGGAGTCGTTGCTCCTGTGTCTCTGAAGAATCATGANNNAGATCGGAA
69987 TATGAGTGTGGAGTCGTTGCTCTTGGTAGCGCATTTCTACTGTNNNAGATCGGAAG
69996 TATGAGTGTGGAGTCGTTGCGAAAAATCGGGTTTACTGGTGCANNNAGATCGGAA
70019 TATGAGTGTGGAGTCGTTGCTGGAGACACCAGTGAATAGATTCTNNNAGATCGGA
70121 TATGAGTGTGGAGTCGTTGCCCACGCCCGGCTAATTATTTTTTNNNAGATCGGAAGA
70242 TATGAGTGTGGAGTCGTTGCGGGAAGAGCTGTTTGTGATGAGANNNAGATCGGA
70257 TATGAGTGTGGAGTCGTTGCAGAAGAAGGAGAAGGAGAAGGAGANNNAGATCG
70286 TATGAGTGTGGAGTCGTTGCGGAGGAAGGTGGAGTAACACAAGNNNAGATCGGA
70313 TATGAGTGTGGAGTCGTTGCTCTTTCTTCTTGACGGAGTTTCACNNNAGATCGGAAG
70314 TATGAGTGTGGAGTCGTTGCACCTGAAATTGCTGTTCAAGAACANNNAGATCGGAA
70324 TATGAGTGTGGAGTCGTTGCACGGAATATCTTGAACAACAGTGCNNNAGATCGGA
70372 TATGAGTGTGGAGTCGTTGCAGAAATCTTCCCATCTCAGCCTCNNNAGATCGGAAGA
70441 TATGAGTGTGGAGTCGTTGCAGCAATCTTCCAATTAGCAAGCTNNNAGATCGGAAG
70474 TATGAGTGTGGAGTCGTTGCCATCTCAACTCACTGCAACCTTTNNNAGATCGGAAGA
70479 TATGAGTGTGGAGTCGTTGCTCTCTCTCTCTCTCCTCTCTCTCTNNNAGATCGGAAGA
70482 TATGAGTGTGGAGTCGTTGCCCTTCCCTTCCCTTTTCTTTCCTNNNAGATCGGAAGAG
70498 TATGAGTGTGGAGTCGTTGCCCTAATTGCTCTGGCTAGGACTTNNNAGATCGGAAG
70525 TATGAGTGTGGAGTCGTTGCCTCCTGGGTTCAAGTGATTCTCANNNAGATCGGAAG
53437 TATGAGTGTGGAGTCGTTGCAGTACACGATTGGTTACAGTCCCNNNAGATCGGAAG
53439 TATGAGTGTGGAGTCGTTGCCTCTCTCTCTCTCTCTCTCTCGGNNNAGATCGGAAGA
53440 TATGAGTGTGGAGTCGTTGCGAGATGGAGTCTTACTCTTGCCCNNNAGATCGGAAG
53441 TATGAGTGTGGAGTCGTTGCCTTTCTCCAAGTATTGAGAGCGCNNNAGATCGGAAG
53442 TATGAGTGTGGAGTCGTTGCTACAAGACTGTGTGTGTGTGTGTNNNAGATCGGAA
53443 TATGAGTGTGGAGTCGTTGCTCTTGTTGCTTATCAGGGATGCTNNNAGATCGGAAG
53445 TATGAGTGTGGAGTCGTTGCAGACAGAGAGACAGAGACAGACANNNAGATCGGA
53446 TATGAGTGTGGAGTCGTTGCCTCCCTCTCTCCCTCTCTCTCTCNNNAGATCGGAAGAG
53447 TATGAGTGTGGAGTCGTTGCCACACGTGTAATTCCAGCACTTTNNNAGATCGGAAG
53449 TATGAGTGTGGAGTCGTTGCTGTGTGTGTGTGTGTAGAGAGAGNNNAGATCGGA
53450 TATGAGTGTGGAGTCGTTGCTGCAGTTTCTGTCTGTCTCTCTCNNNAGATCGGAAGA
53451 TATGAGTGTGGAGTCGTTGCCCTCTCTCTCTCTCTCTCTCTCGNNNAGATCGGAAGAG
53452 TATGAGTGTGGAGTCGTTGCATCTCTCCTTGAATATGCACCCCNNNAGATCGGAAGA
53453 TATGAGTGTGGAGTCGTTGCCTTGGAGTTCTTTCAGGCAACTGNNNAGATCGGAAG
53454 TATGAGTGTGGAGTCGTTGCGCTTGGAAACACTGGTGTGATTCNNNAGATCGGAA
53455 TATGAGTGTGGAGTCGTTGCGAAATGGAAATGAGCCGAGATCGNNNAGATCGGAA
53456 TATGAGTGTGGAGTCGTTGCATGATCCATACTCAGCCCTCAAGNNNAGATCGGAAG

53457 TATGAGTGTGGAGTCGTTGCTCCTCTCTCTCTCTCTCTCTTCCNNNAGATCGGAAGAG
53458 TATGAGTGTGGAGTCGTTGCCTGGTTTTGAAACACCCAGGATGNNNAGATCGGAAG
53459 TATGAGTGTGGAGTCGTTGCCCAGTCACTCCAATTCTGAGTCTNNNAGATCGGAAGA
53460 TATGAGTGTGGAGTCGTTGCAGGCAGTACTAGGTTTGCAAGTTNNNAGATCGGAA
53463 TATGAGTGTGGAGTCGTTGCGACTGCGGTCTAGGAAAAAGTCTNNNAGATCGGAA
53465 TATGAGTGTGGAGTCGTTGCAGGGAGATAGCTTCAAGACCTGANNNAGATCGGAA
53467 TATGAGTGTGGAGTCGTTGCGGAGAGAAAATGCCTAGACAGCTNNNAGATCGGAA
53468 TATGAGTGTGGAGTCGTTGCTGTAGTCCCAGGTACTCAAGAGGNNNAGATCGGAA
53472 TATGAGTGTGGAGTCGTTGCCCCAACATTCACAGGGTCTTTTCNNNAGATCGGAAGA
53473 TATGAGTGTGGAGTCGTTGCTTCCCCAAAGAACTGCTTCCATANNNAGATCGGAAGA
53474 TATGAGTGTGGAGTCGTTGCAAGAAAACAGCAATTGGTCAGGCNNNAGATCGGAA
53475 TATGAGTGTGGAGTCGTTGCGAAAGCATCACTTGAGGTCAGGANNNAGATCGGAA
53478 TATGAGTGTGGAGTCGTTGCGTGCAGAGAAATAATCACGGCTCNNNAGATCGGAA
53479 TATGAGTGTGGAGTCGTTGCGCTGAGAAGCTGAGGATGAATCCNNNAGATCGGAA
53483 TATGAGTGTGGAGTCGTTGCAAATTAGTCAGGCGTAGTGGTGGNNNAGATCGGAA
53484 TATGAGTGTGGAGTCGTTGCGCCACGTGCAACTGTTATACTAANNNAGATCGGAAG
53486 TATGAGTGTGGAGTCGTTGCTGCTGGAAGAGTAGGTGTTTCTCNNNAGATCGGAA
53487 TATGAGTGTGGAGTCGTTGCGATTGCAGCTCTTTTGCCATCATNNNAGATCGGAAGA
53489 TATGAGTGTGGAGTCGTTGCTAGCTACTTGGGAGGTTGAGGTANNNAGATCGGAA
53490 TATGAGTGTGGAGTCGTTGCAAACAGGCCGAGCATTTAGAAACNNNAGATCGGAA
53491 TATGAGTGTGGAGTCGTTGCGCTCCTGTAATCCTTGCTACTCANNNAGATCGGAAGA
53492 TATGAGTGTGGAGTCGTTGCCTAGAGTGCAGTGGTGTGATCTCNNNAGATCGGAA
53493 TATGAGTGTGGAGTCGTTGCCCTCCCTCTCTCTCTCTCTTTCTNNNAGATCGGAAGAG
53494 TATGAGTGTGGAGTCGTTGCTCCTGAATCCTGAAGGTGATTGGNNNAGATCGGAAG
53495 TATGAGTGTGGAGTCGTTGCTCTTTCTCTCTCCCTTCCTTCCTNNNAGATCGGAAGAG
53498 TATGAGTGTGGAGTCGTTGCTCCCAGAGAACAATGAAGGTTCCNNNAGATCGGAAG
53500 TATGAGTGTGGAGTCGTTGCATATTCCACAACAGTACTGGGGGNNNAGATCGGAAG
53501 TATGAGTGTGGAGTCGTTGCCATGTGTGTGACATTCTCAGGACNNNAGATCGGAAG
53502 TATGAGTGTGGAGTCGTTGCAGGATTTCTTGAGCCCAGAAAGTNNNAGATCGGAAG
53503 TATGAGTGTGGAGTCGTTGCATGTTGGTTAGGCTGGTCTCAAANNNAGATCGGAAG
53504 TATGAGTGTGGAGTCGTTGCGAATGAATGGCATTTGCAGCAACNNNAGATCGGAA
53507 TATGAGTGTGGAGTCGTTGCATATAGATAAAGTGGGCCGCCTGNNNAGATCGGAA
53508 TATGAGTGTGGAGTCGTTGCTCTCAAACTCCTGACCTCAAGTGNNNAGATCGGAAGA
53509 TATGAGTGTGGAGTCGTTGCGAACAGAGAGAACAAACCTGGGANNNAGATCGGAA
53511 TATGAGTGTGGAGTCGTTGCTCTCCCTCTCTCTCTCTCTCTCTNNNAGATCGGAAGAG
53512 TATGAGTGTGGAGTCGTTGCTAACTGGTCTGGCTGTTCTGTACNNNAGATCGGAAG
53513 TATGAGTGTGGAGTCGTTGCCAGGGAGAAATCCACTGAAGGAANNNAGATCGGAA
53514 TATGAGTGTGGAGTCGTTGCACACACACACACCATACTCTGTTNNNAGATCGGAAGA
53516 TATGAGTGTGGAGTCGTTGCAGGCAGGAGAATAATGTGAACCCNNNAGATCGGAA
53517 TATGAGTGTGGAGTCGTTGCGAAAGGGTCTCACTCTGTCACTTNNNAGATCGGAAG
53518 TATGAGTGTGGAGTCGTTGCGGGTCACCATTTGCAAGTAAGTTNNNAGATCGGAAG
53519 TATGAGTGTGGAGTCGTTGCTTCATGGTGAATGTCGATGCAAANNNAGATCGGAA
53522 TATGAGTGTGGAGTCGTTGCACCAAGAAGCCTTTAGTTACCCANNNAGATCGGAAG

53523 TATGAGTGTGGAGTCGTTGCAGGATCTTGATGTGTTGCCTAGGNNNAGATCGGAA
53524 TATGAGTGTGGAGTCGTTGCGTCAATGCCTTTATTGGGTCCAGNNNAGATCGGAAG
53527 TATGAGTGTGGAGTCGTTGCATATTGGGGATGGTGTGACACAANNNAGATCGGAA
53528 TATGAGTGTGGAGTCGTTGCAGAAGATTGCCTTGAGAGAAGCANNNAGATCGGAA
53529 TATGAGTGTGGAGTCGTTGCGCTCTGGGGATAGAAAGACTAGCNNNAGATCGGAA
53530 TATGAGTGTGGAGTCGTTGCACCACAAGGAACTATGGGATGTCNNNAGATCGGAA
53531 TATGAGTGTGGAGTCGTTGCGACTAGGTAGTTCTCCAGGGGATNNNAGATCGGAA
53532 TATGAGTGTGGAGTCGTTGCGCATAGCGGCACATACCTATAGANNNAGATCGGAAG
53533 TATGAGTGTGGAGTCGTTGCTGCTACATGATCTCTCTCCTGGANNNAGATCGGAAG
53534 TATGAGTGTGGAGTCGTTGCTAAAGTTGGGGGAAACTAGCCAANNNAGATCGGAA
53535 TATGAGTGTGGAGTCGTTGCAAAGTCCAGAAGACACCAAGGTTNNNAGATCGGAA
53536 TATGAGTGTGGAGTCGTTGCCCTCTACTGCTATGTGGCTTCAANNNAGATCGGAAG
53537 TATGAGTGTGGAGTCGTTGCGGTGAGGTTGAGTGAATGTCTCTNNNAGATCGGAA
53538 TATGAGTGTGGAGTCGTTGCTGCTGGGATTACAGGTGTGATTANNNAGATCGGAA
53539 TATGAGTGTGGAGTCGTTGCTGTTTCTCAATCAAGAGGCCAGANNNAGATCGGAAG
53542 TATGAGTGTGGAGTCGTTGCGACTTTAGATCTTTGAGCCCCCANNNAGATCGGAAG
53543 TATGAGTGTGGAGTCGTTGCTCTCTCAACAGGAATAGCCTTGCNNNAGATCGGAAG
53544 TATGAGTGTGGAGTCGTTGCGGGGGCTCTCAATACAAACAAACNNNAGATCGGAA
53545 TATGAGTGTGGAGTCGTTGCCTCTCCACAGCCATAAACAACATNNNAGATCGGAAG
53546 TATGAGTGTGGAGTCGTTGCAAAAAGGCCTGCAGATAACCCTANNNAGATCGGAAG
53547 TATGAGTGTGGAGTCGTTGCGAGGCACCTGGTGTTAGTTTTTCNNNAGATCGGAAG
53548 TATGAGTGTGGAGTCGTTGCCGCCTATAATCCCAGCTACTCAGNNNAGATCGGAAG
53549 TATGAGTGTGGAGTCGTTGCCTGTGATCTCAGCTACTTGGGAGNNNAGATCGGAAG
53550 TATGAGTGTGGAGTCGTTGCTAGCATGTTCCTGTAGTCTCAGCNNNAGATCGGAAG
53551 TATGAGTGTGGAGTCGTTGCGGGGATACCAAGCAAAAACACTTNNNAGATCGGAA
53552 TATGAGTGTGGAGTCGTTGCCACCCCACATTTATTGAACAGCANNNAGATCGGAAG
53554 TATGAGTGTGGAGTCGTTGCGGAAACCTTTGGAAACCGAGAACNNNAGATCGGAA
53555 TATGAGTGTGGAGTCGTTGCCAGGAAGGTCTTGATCTCCTGACNNNAGATCGGAAG
53556 TATGAGTGTGGAGTCGTTGCCATGGCTGTTTATTTCACCTGGGNNNAGATCGGAAG
53557 TATGAGTGTGGAGTCGTTGCGTAAGGTACTTGACTCACCTGCANNNAGATCGGAAG
53558 TATGAGTGTGGAGTCGTTGCAGATTTTGTGTCTGGTGAGGGTTNNNAGATCGGAA
53559 TATGAGTGTGGAGTCGTTGCCAGGGTCCACTCCATTCTAGTTGNNNAGATCGGAAG
53561 TATGAGTGTGGAGTCGTTGCGAGAGAGAGAGGACGCAGAAAAANNNAGATCGGA
53563 TATGAGTGTGGAGTCGTTGCTGTGTGTGTGTAATGCTCCTCATNNNAGATCGGAAG
53564 TATGAGTGTGGAGTCGTTGCCACACACACACACACACACATATNNNAGATCGGAAG
53566 TATGAGTGTGGAGTCGTTGCTCTCTCTCTCTCTCTCTCTCCCTNNNAGATCGGAAGAG
53568 TATGAGTGTGGAGTCGTTGCCAGCTCAGGATTGGGACAATTTCNNNAGATCGGAAG
53569 TATGAGTGTGGAGTCGTTGCTTTCTTTAGTCCTGCTCTGTGGCNNNAGATCGGAAG
53570 TATGAGTGTGGAGTCGTTGCAAAGGAGAGTCAAAGAGGCAGAGNNNAGATCGGA
53571 TATGAGTGTGGAGTCGTTGCAACAAGGATAAGACAAGCAAGGANNNAGATCGGAA
53572 TATGAGTGTGGAGTCGTTGCCAAAAAGTCAGCACCTTGGACAGNNNAGATCGGAAG
53573 TATGAGTGTGGAGTCGTTGCGCCTGTAGTCCCAACACTAGTTTNNNAGATCGGAAG
53574 TATGAGTGTGGAGTCGTTGCGCCTTGCTTTCCCTTCACTTTTTNNNAGATCGGAAGA

53576 TATGAGTGTGGAGTCGTTGCGTCTCGATCTCCTGACCTTGTAANNNAGATCGGAAG

53578 TATGAGTGTGGAGTCGTTGCTCTTTCCCTGAGACTGGAGTACANNNAGATCGGAAG

53581 TATGAGTGTGGAGTCGTTGCGTTCAAGCGATTTCCGGCTAATTNNNAGATCGGAAG

53582 TATGAGTGTGGAGTCGTTGCCCCGAGTAGCTAGGATTACAAGCNNNAGATCGGAA

53583 TATGAGTGTGGAGTCGTTGCCTGGGTTCAAGTGATTCTTCTGCNNNAGATCGGAAG

53584 TATGAGTGTGGAGTCGTTGCTAGGCAGTTTTGCAAGCAGAAATNNNAGATCGGAA

53586 TATGAGTGTGGAGTCGTTGCTACAATCTCAGCTCATGGCAACTNNNAGATCGGAAG

53588 TATGAGTGTGGAGTCGTTGCTGAGCCAGTGTGAGTTACTATGGNNNAGATCGGAA

53589 TATGAGTGTGGAGTCGTTGCAGGTTCAAGTGATTCTTCTGCCTNNNAGATCGGAAG

53591 TATGAGTGTGGAGTCGTTGCGCTGCTTGGTTATCTGCTGAAAANNNAGATCGGAAG

53592 TATGAGTGTGGAGTCGTTGCGTCCTCTCTGAGACACAAGTCAGNNNAGATCGGAAG

53594 TATGAGTGTGGAGTCGTTGCTCTGGTCCAAAACTCCTAAGCTCNNNAGATCGGAAG

53596 TATGAGTGTGGAGTCGTTGCCTGAGTAGCTGGGATTACAGGTGNNNAGATCGGAA

53601 TATGAGTGTGGAGTCGTTGCATGCAAAAGAAAGGAGGCTCTTGNNNAGATCGGAA

53603 TATGAGTGTGGAGTCGTTGCCAATCTCAGCTCACTGCAAACTCNNNAGATCGGAAG

53604 TATGAGTGTGGAGTCGTTGCCAGCCACGTGTAAAAGAACCATTNNNAGATCGGAAG

53607 TATGAGTGTGGAGTCGTTGCTGAAGAGATTCCCCCTGGAGTATNNNAGATCGGAA

53608 TATGAGTGTGGAGTCGTTGCAAGGAAGGAACGAACGAACAAACNNNAGATCGGA

53609 TATGAGTGTGGAGTCGTTGCTGCTGGCAACCTGAAATTCTATGNNNAGATCGGAAG

53611 TATGAGTGTGGAGTCGTTGCAGGTAAGGCAATCACTTGAACCTNNNAGATCGGAA

53612 TATGAGTGTGGAGTCGTTGCGTTCCAAGGGCAATCAGAATTCCNNNAGATCGGAAG

53615 TATGAGTGTGGAGTCGTTGCGATCAAGACCAGACTGGGCAATANNNAGATCGGAA

53617 TATGAGTGTGGAGTCGTTGCAAAAAACTAGAAAGGCGTGGTGCNNNAGATCGGAA

53619 TATGAGTGTGGAGTCGTTGCAAATGTGCAAATGCCAAGTGTCANNNAGATCGGAA

53620 TATGAGTGTGGAGTCGTTGCTAGAGGCCAGACTGTGGTAGTAANNNAGATCGGAA

53621 TATGAGTGTGGAGTCGTTGCATATGAGCAAGGTGAGCTTAGGGNNNAGATCGGA

53623 TATGAGTGTGGAGTCGTTGCTAATCCTAACACTTTGGGAGGCCNNNAGATCGGAAG

53624 TATGAGTGTGGAGTCGTTGCTAAGGCATTAGGAAACCCTGGACNNNAGATCGGAA

53626 TATGAGTGTGGAGTCGTTGCTGAAGAACATTGCTTCTCTGGGANNNAGATCGGAA

53627 TATGAGTGTGGAGTCGTTGCACTAGCCATGAGAAAAGGGTGTTNNNAGATCGGAA

53629 TATGAGTGTGGAGTCGTTGCCCACTACGCCTGGCTAATTTTTTNNNAGATCGGAAG

53630 TATGAGTGTGGAGTCGTTGCCATTCTCCTGCCTCAGCCTTCNNNAGATCGGAAGAGC

53631 TATGAGTGTGGAGTCGTTGCAGGCCTACATTGCTGCTATTCTTNNNAGATCGGAAG

53632 TATGAGTGTGGAGTCGTTGCCATGATCATGGCTTACTGCAACCNNNAGATCGGAAG

53633 TATGAGTGTGGAGTCGTTGCCCAGGATGGTCTTGATCACTTGANNNAGATCGGAAG

53634 TATGAGTGTGGAGTCGTTGCTTGAAGTGATTCTCCTGCCTCAGNNNAGATCGGAAG

53635 TATGAGTGTGGAGTCGTTGCCTGAAGTGATCCACCTACCTCAGNNNAGATCGGAAG

53637 TATGAGTGTGGAGTCGTTGCGAGAACCCTGTAAGAAGGCAGAANNNAGATCGGAA

53639 TATGAGTGTGGAGTCGTTGCGTTCAAGCAATTCTCTGCCTCAGNNNAGATCGGAAG

53640 TATGAGTGTGGAGTCGTTGCATCACTGCTCCATACAACCTCTGNNNAGATCGGAAG

53641 TATGAGTGTGGAGTCGTTGCCAGCCTCCCAAGTAGGTGTAATTNNNAGATCGGAAG

53644 TATGAGTGTGGAGTCGTTGCAGACAGAATCTCACTGTGTCACCNNNAGATCGGAAG

53647 TATGAGTGTGGAGTCGTTGCCCTCCTTGGTTCAAGCAATTCTGNNNAGATCGGAAG

53648 TATGAGTGTGGAGTCGTTGCCAATGGCAAAATCTCAGCTCACTNNNAGATCGGAAG
53649 TATGAGTGTGGAGTCGTTGCGCTAGAGTATAGTGGTGGCTCAGNNNAGATCGGAA
53651 TATGAGTGTGGAGTCGTTGCTTCCAGGTTCACACCATTCTCCNNNAGATCGGAAGAG
53653 TATGAGTGTGGAGTCGTTGCAATCTTGGCTCACTACAACCTCCNNNAGATCGGAAG
53656 TATGAGTGTGGAGTCGTTGCATACACCAGTCTGATGGGAACACNNNAGATCGGAA
53660 TATGAGTGTGGAGTCGTTGCGGACCACAGACTACACTGAAAGANNNAGATCGGAA
53661 TATGAGTGTGGAGTCGTTGCTGCCTGTAATTCCAGCTACTCAGNNNAGATCGGAAG
53663 TATGAGTGTGGAGTCGTTGCATGCTACTTGGTTCAGAGCGTAANNNAGATCGGAA
53664 TATGAGTGTGGAGTCGTTGCATAAGGAGTATGGCCAGACATGGNNNAGATCGGAA
53667 TATGAGTGTGGAGTCGTTGCCAGGGTTTCTGGATACCACTGTTNNNAGATCGGAAG
53668 TATGAGTGTGGAGTCGTTGCCCATCCTAAGAACTACGGAGCAANNNAGATCGGAAG
53670 TATGAGTGTGGAGTCGTTGCTCTGTGACCTCAAACTGACTGAGNNNAGATCGGAAG
53672 TATGAGTGTGGAGTCGTTGCGTGGAAGGATTGTTTGAACCCAGNNNAGATCGGAA
53673 TATGAGTGTGGAGTCGTTGCGTGAGACACCAGACAGGATTCATNNNAGATCGGAA
53675 TATGAGTGTGGAGTCGTTGCTGGGCAGAAAAGGATGTAACTGANNNAGATCGGAA
53677 TATGAGTGTGGAGTCGTTGCAACAGGTGTTGACAAAAGTGTGGNNNAGATCGGAA
53678 TATGAGTGTGGAGTCGTTGCAAGAGGTGTTGAGGTCTTTTGGTNNNAGATCGGAA
53681 TATGAGTGTGGAGTCGTTGCTACAACCACTGTGGGAAACTAGCNNNAGATCGGAA
53683 TATGAGTGTGGAGTCGTTGCGCAAACCAAATCCAACAACACATNNNAGATCGGAAG
53685 TATGAGTGTGGAGTCGTTGCTTCCCCTCTCCTTATCCATGACTNNNAGATCGGAAGA
53688 TATGAGTGTGGAGTCGTTGCGATATGGACTCCCAGGCTATGTGNNNAGATCGGAA
53689 TATGAGTGTGGAGTCGTTGCGAAGATGCTGTGATGATGGAAGCNNNAGATCGGAA
53690 TATGAGTGTGGAGTCGTTGCCTGCCAGTAAAGCTTTGTGTTGTNNNAGATCGGAAG
53691 TATGAGTGTGGAGTCGTTGCGCCCCAGTAAATTCACTTTCCACNNNAGATCGGAAGA
53692 TATGAGTGTGGAGTCGTTGCATAGTATAACGCAGGGCTAACGGNNNAGATCGGAA
53693 TATGAGTGTGGAGTCGTTGCATTTGCCAGTCGACCTTGAGTATNNNAGATCGGAAG
53694 TATGAGTGTGGAGTCGTTGCGTCTGCATTTTTGGATGGTGACANNNAGATCGGAA
53696 TATGAGTGTGGAGTCGTTGCCATGGAAAGTAGCTATGCCAAGCNNNAGATCGGAA
53698 TATGAGTGTGGAGTCGTTGCTGGCTCACATCTGTAATCCTAGCNNNAGATCGGAAG
53700 TATGAGTGTGGAGTCGTTGCAGACAAAGTCCATTGGAACCACCNNNAGATCGGAAG
53704 TATGAGTGTGGAGTCGTTGCCAAACAATAGGGGCATTCTTGGGNNNAGATCGGAA
53705 TATGAGTGTGGAGTCGTTGCTGCCAAATATCTCCAGCACTGATNNNAGATCGGAAG
53706 TATGAGTGTGGAGTCGTTGCCTCAAGTGATTCTCCTGCCTCAGNNNAGATCGGAAG
53707 TATGAGTGTGGAGTCGTTGCCCTCCTGGGTTCAAGTGATTCTCNNNAGATCGGAAG
53708 TATGAGTGTGGAGTCGTTGCATCTCCCAGGTTCAAGTGATTCCNNNAGATCGGAAG
53710 TATGAGTGTGGAGTCGTTGCGAGTGCAATGACATGATCTCAGCNNNAGATCGGAA
53711 TATGAGTGTGGAGTCGTTGCATGATCTTGGCTCACTGAAACCTNNNAGATCGGAAG
53712 TATGAGTGTGGAGTCGTTGCGAGTAGTTTCACCCTTGTCACCCNNNAGATCGGAAG
53713 TATGAGTGTGGAGTCGTTGCTTCCTCCTGGGTTCAAGCAATTCNNNAGATCGGAAG
53716 TATGAGTGTGGAGTCGTTGCCTACCGAGTAGCTGGGACTACAANNNAGATCGGAA
53720 TATGAGTGTGGAGTCGTTGCAAGCAATCTGGGATTACAGGCATNNNAGATCGGAA
53722 TATGAGTGTGGAGTCGTTGCTCCTGGTTCAAGCAATTCTCCTTNNNAGATCGGAAGA
53723 TATGAGTGTGGAGTCGTTGCGAGAAGAGAAGTTCTAGGCTGGGNNNAGATCGGA

53724 TATGAGTGTGGAGTCGTTGCGAAAACAAAAAGCACCACACTGTNNNAGATCGGAAG
53725 TATGAGTGTGGAGTCGTTGCGTGTAATCACAGCACTTTGGGAGNNNAGATCGGAA
53726 TATGAGTGTGGAGTCGTTGCTGAAAGTGACCTTAGAGATGGCCNNNAGATCGGAA
53728 TATGAGTGTGGAGTCGTTGCATTCTGACTCAGGAACCTTCTGGNNNAGATCGGAAG
53729 TATGAGTGTGGAGTCGTTGCTTCAAGCAACTCTCCTGCCTTAGNNNAGATCGGAAG
53731 TATGAGTGTGGAGTCGTTGCGGTAGACAGGACTATGTGTGGTCNNNAGATCGGAA
53732 TATGAGTGTGGAGTCGTTGCACTGGGTTCTTTGGTCGTAGTTTNNNAGATCGGAAG
53733 TATGAGTGTGGAGTCGTTGCTGATTTCATAGGTCTGGATGGGCNNNAGATCGGAA
53734 TATGAGTGTGGAGTCGTTGCGAGGTCAGGAGATCAAGACCATCNNNAGATCGGAA
53736 TATGAGTGTGGAGTCGTTGCTAAAAACAGCACTAGCCTGAGGTNNNAGATCGGAAG
53738 TATGAGTGTGGAGTCGTTGCAAAATTAGCTGGGTGTGATGGTGNNNAGATCGGAA
53739 TATGAGTGTGGAGTCGTTGCTTTGAATCCAGCTGTGTGACCTTNNNAGATCGGAAG
53740 TATGAGTGTGGAGTCGTTGCATATATATGGCTGGGCATGGTGGNNNAGATCGGAA
53741 TATGAGTGTGGAGTCGTTGCTCACTGATCGTTAGGGAAATGCANNNAGATCGGAAG
53743 TATGAGTGTGGAGTCGTTGCTGAATCCACAGATTGCTTTGAGCNNNAGATCGGAAG
53744 TATGAGTGTGGAGTCGTTGCGAGATGGTGTTTCGCTCTTGTTGNNNAGATCGGAAG
53745 TATGAGTGTGGAGTCGTTGCGGCTTCAGTGAAATGGCATGATTNNNAGATCGGAA
53746 TATGAGTGTGGAGTCGTTGCGGTTCACGCCATTTTCCTACTTCNNNAGATCGGAAGA
53747 TATGAGTGTGGAGTCGTTGCTTCAAGTGATTCTCCTGCCTCAGNNNAGATCGGAAG
53748 TATGAGTGTGGAGTCGTTGCCGTGCTGGAATTACAGGCATAAGNNNAGATCGGAA
53749 TATGAGTGTGGAGTCGTTGCAGAGTCTTGTTATGTCACCCAGGNNNAGATCGGAAG
53750 TATGAGTGTGGAGTCGTTGCCTTCCCAGATTCAAGCGATTCTCNNNAGATCGGAAG
53751 TATGAGTGTGGAGTCGTTGCCTCCTGAGTAGCTGGGACTACAGNNNAGATCGGAAG
53753 TATGAGTGTGGAGTCGTTGCGATTACAGGCACACACCAACATGNNNAGATCGGAAG
53754 TATGAGTGTGGAGTCGTTGCCTGAGTAGCTGGGACTACATGTGNNNAGATCGGAA
53755 TATGAGTGTGGAGTCGTTGCAGGTTCAAGCAATTCTCCTGTCTNNNAGATCGGAAG
53757 TATGAGTGTGGAGTCGTTGCACTAGAGACGGGGTTTTACCATGNNNAGATCGGAA
53759 TATGAGTGTGGAGTCGTTGCGTTCAAGCGATTCTCCTACCTCANNNAGATCGGAAG
53760 TATGAGTGTGGAGTCGTTGCGTAGCTGGGACTACAGGCATATGNNNAGATCGGAA
53761 TATGAGTGTGGAGTCGTTGCGCAGTAGGGCAATCTCAGAATCTNNNAGATCGGAAG
53762 TATGAGTGTGGAGTCGTTGCAAGTTCAAGGGATTCTCGTGCTCNNNAGATCGGAAG
53763 TATGAGTGTGGAGTCGTTGCTTCAAGTGATTCTCCTGCCTCAGNNNAGATCGGAAG
53764 TATGAGTGTGGAGTCGTTGCCCTCAGCCTTCCTAGTAGTAGGANNNAGATCGGAAG
53765 TATGAGTGTGGAGTCGTTGCCTTCCTGGGTTCAAGCAATTCTCNNNAGATCGGAAG
53766 TATGAGTGTGGAGTCGTTGCGACTACAGGCCTGGGACTACAGNNNAGATCGGAAG
53767 TATGAGTGTGGAGTCGTTGCTTCCCAGGTTCAAGTGATTCTCCNNNAGATCGGAAG
53768 TATGAGTGTGGAGTCGTTGCTCAATGGCATGATCTTTGCTCACNNNAGATCGGAAG
53772 TATGAGTGTGGAGTCGTTGCCCAAGCACAAAGACCACATATGGNNNAGATCGGAAG
53775 TATGAGTGTGGAGTCGTTGCCATGAGAATTGCTTGAACCTGGGNNNAGATCGGAA
53778 TATGAGTGTGGAGTCGTTGCGCTGTTCCAAATCACTGTGAGACNNNAGATCGGAAG
53783 TATGAGTGTGGAGTCGTTGCCTGTAGTCCTAGCACTTTGGGAGNNNAGATCGGAAG
53784 TATGAGTGTGGAGTCGTTGCTCAACTTCAACCACCTTCTCCAANNNAGATCGGAAGA
53785 TATGAGTGTGGAGTCGTTGCAAAAGGCTAAAGGAATGGCAACANNNAGATCGGAA

53786 TATGAGTGTGGAGTCGTTGCCAACCTGACCAACATGGAGAAACNNNAGATCGGAAG
53787 TATGAGTGTGGAGTCGTTGCCACACCTGTAATCCCAGCACTTTNNNAGATCGGAAGA
53789 TATGAGTGTGGAGTCGTTGCGATCCAGGGAGCCAGTTAGTTTCNNNAGATCGGAAG
53790 TATGAGTGTGGAGTCGTTGCGATGTCTGGGTCAAATGCTTGTCNNNAGATCGGAAG
53792 TATGAGTGTGGAGTCGTTGCAGACCTTTTTCAAGGCTTTGTCCNNNAGATCGGAAGA
53794 TATGAGTGTGGAGTCGTTGCTGATGTGAGGAGTAAGCCCATTGNNNAGATCGGAAG
53795 TATGAGTGTGGAGTCGTTGCCAGGTTCAATGAGGGTTTGCTTTNNNAGATCGGAAG
53796 TATGAGTGTGGAGTCGTTGCCACACCTGTAATCCCAGCACTTTNNNAGATCGGAAGA
53797 TATGAGTGTGGAGTCGTTGCAGCAGTGGATCCAAACAAGATGANNNAGATCGGAAG
53800 TATGAGTGTGGAGTCGTTGCGGTTCAAGCTATCCTCCCATCTCNNNAGATCGGAAGA
53801 TATGAGTGTGGAGTCGTTGCGATGGGGTTTTACCATGTTGTCCNNNAGATCGGAAG
53802 TATGAGTGTGGAGTCGTTGCGTGTCTCTTCATGTGAGATGGGTNNNAGATCGGAAG
53803 TATGAGTGTGGAGTCGTTGCTCTCCTGGGTTCAAGTGATTCTCNNNAGATCGGAAGA
53804 TATGAGTGTGGAGTCGTTGCCTCCCAAGTAGCTGGAACTACAGNNNAGATCGGAAG
53805 TATGAGTGTGGAGTCGTTGCTGCCTACTTTGGGAAAAACTGAGNNNAGATCGGAAG
53807 TATGAGTGTGGAGTCGTTGCGCCTCCTGAATAGCTGAGACTACNNNAGATCGGAAG
53809 TATGAGTGTGGAGTCGTTGCAATCTCGGCTCACTGCAACATCNNNAGATCGGAAGAG
53810 TATGAGTGTGGAGTCGTTGCCCTCCTGGGTTCAAGTGATTCTTNNNAGATCGGAAGA
53813 TATGAGTGTGGAGTCGTTGCACTTCCCGGATTCAAGTGATTCTNNNAGATCGGAAGA
53814 TATGAGTGTGGAGTCGTTGCGAGTAGCTGAGACTACAGGTGTGNNNAGATCGGAA
53815 TATGAGTGTGGAGTCGTTGCAGTAGTTGGGATTAGAGGCATGCNNNAGATCGGAA
53816 TATGAGTGTGGAGTCGTTGCGTTTCACCATGTTAGCCAGGATGNNNAGATCGGAAG
53817 TATGAGTGTGGAGTCGTTGCAGAGTCTTGCTCAGTCATCCAAGNNNAGATCGGAAG
53818 TATGAGTGTGGAGTCGTTGCCCTCCTGGGTTCAAGTGATTCTCNNNAGATCGGAAGA
53819 TATGAGTGTGGAGTCGTTGCGAGATGGAGTCTTGTTCTGTTGCNNNAGATCGGAAG
53822 TATGAGTGTGGAGTCGTTGCAGTCTGTAGAGGAAATGCATTGTNNNAGATCGGAAG
53824 TATGAGTGTGGAGTCGTTGCTCAATCTCTTGACCTCGTGATCCNNNAGATCGGAAGA
53826 TATGAGTGTGGAGTCGTTGCGGCTAACATCTGGTGAGTGTGTANNNAGATCGGAAG
53827 TATGAGTGTGGAGTCGTTGCGAGAAGGGGTTTTGCCATATTGCNNNAGATCGGAAG
53829 TATGAGTGTGGAGTCGTTGCGCTGTTGGACAGAAAAGCAAGAANNNAGATCGGAA
53830 TATGAGTGTGGAGTCGTTGCTCCGCGGCACTTTTCTTTAGATANNNAGATCGGAAGA
53831 TATGAGTGTGGAGTCGTTGCGTTCTTCCCATGGATCAGGCTATNNNAGATCGGAAGA
53832 TATGAGTGTGGAGTCGTTGCTTCAAGTGATTCTCCTGCCTCAGNNNAGATCGGAAGA
53833 TATGAGTGTGGAGTCGTTGCCAATCTCAGCTCACTGCAAACTCNNNAGATCGGAAGA
53834 TATGAGTGTGGAGTCGTTGCAGAATCAGCTGAATCCCATTCCANNNAGATCGGAAGA
53835 TATGAGTGTGGAGTCGTTGCTCCTAACCTGGGGTAGTCTAAGGNNNAGATCGGAAG
53836 TATGAGTGTGGAGTCGTTGCGAGCTGAGATTGTGGTACATTGCNNNAGATCGGAAG
53837 TATGAGTGTGGAGTCGTTGCAGTCGACTCATTTCAGTGACCATNNNAGATCGGAAGA
53839 TATGAGTGTGGAGTCGTTGCCCGAAGTAAAATACTGACGGGGANNNAGATCGGAA
53840 TATGAGTGTGGAGTCGTTGCTAACTGGTGGCCATTGATAAGCTNNNAGATCGGAAG
53841 TATGAGTGTGGAGTCGTTGCCTGACAGAAACCTGAAATCCACANNNAGATCGGAAG
53842 TATGAGTGTGGAGTCGTTGCACTTCTCTGGGTCTTGGTTTTCTNNNAGATCGGAAGA
53843 TATGAGTGTGGAGTCGTTGCAGTGAGTGAGAGTCCGTTTCAAANNNAGATCGGAAG

53844 TATGAGTGTGGAGTCGTTGCTGTTTGTAGCAGAGCGGTCTTATNNNAGATCGGAAG
53845 TATGAGTGTGGAGTCGTTGCATCCAAAAGGCAGGTATGGTTCTNNNAGATCGGAAG
53848 TATGAGTGTGGAGTCGTTGCATGAAGCCAAGCATTGTCACAATNNNAGATCGGAAG
53849 TATGAGTGTGGAGTCGTTGCATAGAAAAGCTAGTGGGGTCGTGNNNAGATCGGAA
53850 TATGAGTGTGGAGTCGTTGCCCAGAGTAAGAGGACAAGCAACANNNAGATCGGAA
53851 TATGAGTGTGGAGTCGTTGCGTCAGTAGAACCCCAAGAGTCTTNNNAGATCGGAAG
53852 TATGAGTGTGGAGTCGTTGCTTGTGCAAAGATGAAGGCTGATGNNNAGATCGGAA
53853 TATGAGTGTGGAGTCGTTGCAGACATTCTCAGTCCTCAGCAAGNNNAGATCGGAAG
53855 TATGAGTGTGGAGTCGTTGCACTATAACCTCAAACTCCTGGGCNNNAGATCGGAAGA
53856 TATGAGTGTGGAGTCGTTGCCACTTGTAATCCCAGCACTTTGGNNNAGATCGGAAGA
53857 TATGAGTGTGGAGTCGTTGCTCTGACACATGCTACAACATGGANNNAGATCGGAAG
53858 TATGAGTGTGGAGTCGTTGCTTAGGACAATTTGAGCACAGCACNNNAGATCGGAAG
53859 TATGAGTGTGGAGTCGTTGCCACACCTGTAATCCCAGCACTTTNNNAGATCGGAAGA
53860 TATGAGTGTGGAGTCGTTGCCAAGTGAGCATGGCTAATTCCTGNNNAGATCGGAAG
53861 TATGAGTGTGGAGTCGTTGCTGAGTGGCAGTTATTTGGAACAGNNNAGATCGGAAG
53862 TATGAGTGTGGAGTCGTTGCCTGAGGCAGAGAATTGCTTGAACNNNAGATCGGAAG
53864 TATGAGTGTGGAGTCGTTGCGTGGCTCACACCTGTAATCCTAGNNNAGATCGGAAG
53866 TATGAGTGTGGAGTCGTTGCTAACAGGTGTTGGCAAAGATGTGNNNAGATCGGAA
53869 TATGAGTGTGGAGTCGTTGCTGCCAAGCATTGTAGTAGACACTNNNAGATCGGAAG
53870 TATGAGTGTGGAGTCGTTGCGGCATACCACCTGTATTCATCCANNNAGATCGGAAGA
53871 TATGAGTGTGGAGTCGTTGCCCTGTAGTCCCAGCTACTTAGGANNNAGATCGGAAG
53872 TATGAGTGTGGAGTCGTTGCTGGTGGGGGGAAAGAATTTTGTANNNAGATCGGAA
53873 TATGAGTGTGGAGTCGTTGCTCTGACTTGTGTGCGAATGAAACNNNAGATCGGAAG
53874 TATGAGTGTGGAGTCGTTGCACCAAACCAAATCTCAGGGATCANNNAGATCGGAAG
53875 TATGAGTGTGGAGTCGTTGCTTGACTGTTGAACATCTGAGTGTNNNAGATCGGAAG
53876 TATGAGTGTGGAGTCGTTGCCAGGGTTTGCAATCCTAGTCTCTNNNAGATCGGAAGA
53878 TATGAGTGTGGAGTCGTTGCTACTTGTTTCTTCTCAGCTGCCTNNNAGATCGGAAGA
53880 TATGAGTGTGGAGTCGTTGCCAATTTGGGCTGGCTGTATTTCANNNAGATCGGAAG
53882 TATGAGTGTGGAGTCGTTGCTTTCAGATCCTTGTTCTCCAGGGNNNAGATCGGAAGA
53885 TATGAGTGTGGAGTCGTTGCCACACCTGTAATCCCAGCACTTTNNNAGATCGGAAGA
53886 TATGAGTGTGGAGTCGTTGCTGGTACTCCACACCTTCCAAATTNNNAGATCGGAAGA
53887 TATGAGTGTGGAGTCGTTGCTTACTGAAAGCTTGCTTTGAGCCNNNAGATCGGAAG
53888 TATGAGTGTGGAGTCGTTGCGATCAGAGCCCTACTCAGGAAACNNNAGATCGGAAG
53891 TATGAGTGTGGAGTCGTTGCAGTCTCACTCTGTTACCTAGGCTNNNAGATCGGAAGA
53892 TATGAGTGTGGAGTCGTTGCCAAGTAGCTGGGATTACAGGTGTNNNAGATCGGAAG
53893 TATGAGTGTGGAGTCGTTGCCTCTGGATCTTGACTCACTGCAGNNNAGATCGGAAGA
53894 TATGAGTGTGGAGTCGTTGCGTTCAAGAGATTCTCCTGCCTCANNNAGATCGGAAGA
53895 TATGAGTGTGGAGTCGTTGCGAAGTTTCACCATGTTACCCAGGNNNAGATCGGAAG
53897 TATGAGTGTGGAGTCGTTGCTAGGACAATAGTGGAGGGAAGGTNNNAGATCGGAA
53898 TATGAGTGTGGAGTCGTTGCTCTATTTTCCCTAAGTGTCGGCCNNNAGATCGGAAGA
53899 TATGAGTGTGGAGTCGTTGCTCAAGGGATTCTCTTGCCTTAGCNNNAGATCGGAAGA
53901 TATGAGTGTGGAGTCGTTGCACTCCACTTTCTGAGTTCAAGCANNNAGATCGGAAGA
53902 TATGAGTGTGGAGTCGTTGCTCTTGAACTCCTGACCTCAAGTGNNNAGATCGGAAGA

53904 TATGAGTGTGGAGTCGTTGCCTCCTGAGTAGCTGGGATTACAGNNNAGATCGGAA
53907 TATGAGTGTGGAGTCGTTGCGGAGTACAGTGGCACAATCACAGNNNAGATCGGAA
53908 TATGAGTGTGGAGTCGTTGCCCGAGTAGCTATGACTACAGGTGNNNAGATCGGAA
53909 TATGAGTGTGGAGTCGTTGCTTCAAGCGATTCTCCTGCCTTAGNNNAGATCGGAAG
53910 TATGAGTGTGGAGTCGTTGCTAGAGATCGGGTTTCGCCATGTTNNNAGATCGGAAG
53911 TATGAGTGTGGAGTCGTTGCGTCTCCTGGGTTCAATGATTCCTNNNAGATCGGAAG
53914 TATGAGTGTGGAGTCGTTGCGAGACAAGGTCTTGCTTTGTCACNNNAGATCGGAAG
53916 TATGAGTGTGGAGTCGTTGCTTCAAGTGATTCTCCTGCCTCAGNNNAGATCGGAAG
53918 TATGAGTGTGGAGTCGTTGCCCTCCTGGGTTCAAGTGATTCTCNNNAGATCGGAAG
53919 TATGAGTGTGGAGTCGTTGCCAAAGTGCTGGGATTACAGGTGTNNNAGATCGGAA
53920 TATGAGTGTGGAGTCGTTGCTTCTCCTGGACATTCAAGACTCTNNNAGATCGGAAGA
53921 TATGAGTGTGGAGTCGTTGCTTCAAGTGATTCTCCTGCCTCAGNNNAGATCGGAAG
53922 TATGAGTGTGGAGTCGTTGCCCTCCCAAATAGCTATAGGCACCNNNAGATCGGAAG
53923 TATGAGTGTGGAGTCGTTGCGCACAATCTCAGCTAACAGCAATNNNAGATCGGAAG
53924 TATGAGTGTGGAGTCGTTGCATGGAGTTTTCGCTGTTATTGCCNNNAGATCGGAAG
53925 TATGAGTGTGGAGTCGTTGCAGTGCTAGTATTACAGGCGTGAGNNNAGATCGGAA
53926 TATGAGTGTGGAGTCGTTGCAGAAGGAGTCCCACTCTATCACCNNNAGATCGGAAG
53927 TATGAGTGTGGAGTCGTTGCGAGTAGCTGGGATTACAGACGTGNNNAGATCGGAA
53928 TATGAGTGTGGAGTCGTTGCGAGGAGAAATCTCGCTCTGTCTCNNNAGATCGGAAG
53929 TATGAGTGTGGAGTCGTTGCTTCAAGTGATTCTCCTGCCTCAGNNNAGATCGGAAG
53930 TATGAGTGTGGAGTCGTTGCTTCAAGTGATTCTCCTGCCTCAGNNNAGATCGGAAG
53931 TATGAGTGTGGAGTCGTTGCGCCACATTCTAATCTCCCCTGAANNNAGATCGGAAGA
53933 TATGAGTGTGGAGTCGTTGCTGGCATCAAAGACCCTTCAAGATNNNAGATCGGAAG
53934 TATGAGTGTGGAGTCGTTGCGCATGCCTATAATCCCAGCTACTNNNAGATCGGAAG
53937 TATGAGTGTGGAGTCGTTGCGCACTGTGCCTCTTCCTTTTTAGNNNAGATCGGAAGA
53938 TATGAGTGTGGAGTCGTTGCATGAAGGGATAATGGGAGGGAGANNNAGATCGGA
53939 TATGAGTGTGGAGTCGTTGCCCTGTAATTCCAGCTACTTGGGANNNAGATCGGAAG
53940 TATGAGTGTGGAGTCGTTGCTCCAAGTTAGGTGAGTCTCTTGANNNAGATCGGAAG
53941 TATGAGTGTGGAGTCGTTGCCACCTGTAATCCCACCTAGTTGGNNNAGATCGGAAG
53943 TATGAGTGTGGAGTCGTTGCATGAAATCTCAAGGGACCCCAAANNNAGATCGGAAG
53944 TATGAGTGTGGAGTCGTTGCATCTATGTGCTTACTCAGGGCTGNNNAGATCGGAAG
53945 TATGAGTGTGGAGTCGTTGCCAAAATCGTCCCCCATTGAGAACNNNAGATCGGAAG
53946 TATGAGTGTGGAGTCGTTGCAATACTGAACCATTGCCCCTAGGNNNAGATCGGAAG
53947 TATGAGTGTGGAGTCGTTGCCAGAGGCAGGAGAATCACTTGAANNNAGATCGGAA
53948 TATGAGTGTGGAGTCGTTGCGACAAGCCTTTTAGAACTGCTCTNNNAGATCGGAAG
53950 TATGAGTGTGGAGTCGTTGCTTAGCCAAATGTTGTGGTGCATCNNNAGATCGGAAG
53952 TATGAGTGTGGAGTCGTTGCGCCATGCTTCTTTTTACACTGGANNNAGATCGGAAG
53953 TATGAGTGTGGAGTCGTTGCTCTGAGAAGTAGTTTGTCCACGGNNNAGATCGGAA
53954 TATGAGTGTGGAGTCGTTGCGCCACCATTGCAGTTATGAAGAGNNNAGATCGGAA
53955 TATGAGTGTGGAGTCGTTGCGTAATCTCAGCTACTCAGGAGGCNNNAGATCGGAAG
53956 TATGAGTGTGGAGTCGTTGCAATGTTAGCACTTTCGGAGGCTANNNAGATCGGAAG
53957 TATGAGTGTGGAGTCGTTGCGAAGAGAGACCTGAGCTAGCATGNNNAGATCGGAA
53959 TATGAGTGTGGAGTCGTTGCACACAAACACCAGTTATTCCCCTNNNAGATCGGAAGA

53960 TATGAGTGTGGAGTCGTTGCGGTGACCTTTCATGTTCAAGTGGNNNAGATCGGAAG
53961 TATGAGTGTGGAGTCGTTGCAAGGGGAGAAGAAAAGGAGCAAANNNAGATCGGA
53962 TATGAGTGTGGAGTCGTTGCACTGGCCTGTTCAATATTCACAGNNNAGATCGGAAG
53964 TATGAGTGTGGAGTCGTTGCGGAAATCCTCTGGAACTTCAGAANNNAGATCGGAAG
53965 TATGAGTGTGGAGTCGTTGCCACACACACACACACACATACATNNNAGATCGGAAGA
53968 TATGAGTGTGGAGTCGTTGCTTGCAGAGATGAGAAAACACACCNNNAGATCGGAAC
53971 TATGAGTGTGGAGTCGTTGCAGAAGAAGGTGTCCTGATTGGTCNNNAGATCGGAA
53972 TATGAGTGTGGAGTCGTTGCCGGCCATGAAAAACTGCTAGATTNNNAGATCGGAAG
53973 TATGAGTGTGGAGTCGTTGCTTTAGGCAACGAACAGTGAACACNNNAGATCGGAAG
53974 TATGAGTGTGGAGTCGTTGCAAACAAACACACACACACACACANNNAGATCGGAAG
53975 TATGAGTGTGGAGTCGTTGCTCCCTCATAACCTAGGAGGAAGGNNNAGATCGGAAG
53976 TATGAGTGTGGAGTCGTTGCGAAATGAAAGGGAGTGGGGAAGANNNAGATCGGA
53977 TATGAGTGTGGAGTCGTTGCGTAGTGTTCTTGCTTGGGGAGATNNNAGATCGGAAC
53979 TATGAGTGTGGAGTCGTTGCCAGATCACGAGATCAGGAGATCGNNNAGATCGGAA
53980 TATGAGTGTGGAGTCGTTGCCAGAACTTTGGGAGTCTGAGACANNNAGATCGGAAC
53981 TATGAGTGTGGAGTCGTTGCGCGCTCGTGGAAATACAATTTTCNNNAGATCGGAAG
53982 TATGAGTGTGGAGTCGTTGCCCTCCTGAGTAGCTGGGACTATANNNAGATCGGAAG
53983 TATGAGTGTGGAGTCGTTGCATGGAGTCTTGTTCTGTTGTCCANNNAGATCGGAAG
53984 TATGAGTGTGGAGTCGTTGCATGGAGTATTACTCTGTTGCCCANNNAGATCGGAAG
53986 TATGAGTGTGGAGTCGTTGCTACCTCTACAGCAAGTGTCTGTGNNNAGATCGGAAG
53988 TATGAGTGTGGAGTCGTTGCGAGGCAGGAGAATTGTTTGAACCNNNAGATCGGAA
53989 TATGAGTGTGGAGTCGTTGCACACACCTGTAATCTCAGCACTTNNNAGATCGGAAGA
53992 TATGAGTGTGGAGTCGTTGCACTGGTGTTTTTTGTCCTGCTTCNNNAGATCGGAAGA
53993 TATGAGTGTGGAGTCGTTGCTAGTAACATGGTCACGTCTTGGGNNNAGATCGGAAG
53994 TATGAGTGTGGAGTCGTTGCCTCAAAACCAAACAGAAAGCCCANNNAGATCGGAAG
53995 TATGAGTGTGGAGTCGTTGCAATAGCCTGGGTTTGTTTCAGGANNNAGATCGGAAG
53998 TATGAGTGTGGAGTCGTTGCATGAGGAAATCTGACTGCTGTGTNNNAGATCGGAAC
53999 TATGAGTGTGGAGTCGTTGCCTGAGACAGGAGGATCACTTGAGNNNAGATCGGAA
54000 TATGAGTGTGGAGTCGTTGCTATTTCCCGAGAGCACATGACAANNNAGATCGGAAG
54003 TATGAGTGTGGAGTCGTTGCCAAAGTGTTGGGATTACAAGCGTNNNAGATCGGAAC
54004 TATGAGTGTGGAGTCGTTGCCTGGGTTCAAATGATTCTCCTGCNNNAGATCGGAAG
54006 TATGAGTGTGGAGTCGTTGCTGGAGTCTCACTTTGTCATCCAGNNNAGATCGGAAG
54007 TATGAGTGTGGAGTCGTTGCTAGCTGGAACTACAGGCCTTTTCNNNAGATCGGAAG
54008 TATGAGTGTGGAGTCGTTGCATCTGCCTCTTGGTTCAAGTGATNNNAGATCGGAAG
54009 TATGAGTGTGGAGTCGTTGCGGAGTTTTGCTCTTGTTGCCTAGNNNAGATCGGAAG
54010 TATGAGTGTGGAGTCGTTGCCCAAGTAGCTGAGATTACAGGCANNNAGATCGGAAC
54011 TATGAGTGTGGAGTCGTTGCGAGTGCAGTGTCATGATCTCAGANNNAGATCGGAAC
54012 TATGAGTGTGGAGTCGTTGCGTCTCTACAGTTCAGGCGATTCTNNNAGATCGGAAG
54013 TATGAGTGTGGAGTCGTTGCAGTACATCTGTAGCTTTGGGCATNNNAGATCGGAAG
54014 TATGAGTGTGGAGTCGTTGCTGTATCATTCTCTGCAGTCCCAANNNAGATCGGAAGA
54015 TATGAGTGTGGAGTCGTTGCAATCTTGTCTCACTGCAACCTCTNNNAGATCGGAAGA
54016 TATGAGTGTGGAGTCGTTGCCTCTTACCTCAGCCTCCCAAGTANNNAGATCGGAAGA
54019 TATGAGTGTGGAGTCGTTGCTTGATCTCAACTCACTGCAACCTNNNAGATCGGAAGA

54020 TATGAGTGTGGAGTCGTTGCGAGACGGAGTCTAGTTCTGTCACNNNAGATCGGAA
54021 TATGAGTGTGGAGTCGTTGCTCCTGGTCTCAAGCAATTCTCTCNNNAGATCGGAAG
54022 TATGAGTGTGGAGTCGTTGCGGGTTCAAGCAATTCTGTCTCAGNNNAGATCGGAAG
54023 TATGAGTGTGGAGTCGTTGCAAACTCCCAGATTCAGGTGATCCNNNAGATCGGAAG
54024 TATGAGTGTGGAGTCGTTGCCTTGATCTCAGCTCATGGCAAACNNNAGATCGGAAG
54025 TATGAGTGTGGAGTCGTTGCACTGTTGGTGTACAGAAGAGGTGNNNAGATCGGAA
54026 TATGAGTGTGGAGTCGTTGCTGAATGCTGCTTCCCTACCTTAGNNNAGATCGGAAG
54029 TATGAGTGTGGAGTCGTTGCGGATTTCACCGTATTAGCCAGGANNNAGATCGGAA
54032 TATGAGTGTGGAGTCGTTGCATAGGGTCTAACTCTGTCACCCANNNAGATCGGAAG
54034 TATGAGTGTGGAGTCGTTGCCTCCTGAGTAGCTGGGATTACAGNNNAGATCGGAA
54036 TATGAGTGTGGAGTCGTTGCCTGGGATTACAGGTGTGAGCTACNNNAGATCGGAA
54037 TATGAGTGTGGAGTCGTTGCGTTTCACAGTGTTAGCCAGGATGNNNAGATCGGAA
54038 TATGAGTGTGGAGTCGTTGCGATTCAAGCAATTCTCCTGCCTCNNNAGATCGGAAG
54039 TATGAGTGTGGAGTCGTTGCTATACTCAGTGGCATGATCACGGNNNAGATCGGAA
54040 TATGAGTGTGGAGTCGTTGCTCCTGGGTTCTAGCTATTCTCCTNNNAGATCGGAAGA
54041 TATGAGTGTGGAGTCGTTGCATCCCGGGTTCAAGTGATTCTACNNNAGATCGGAAG
54042 TATGAGTGTGGAGTCGTTGCAGCGATTCTCTTACCTCAACCTCNNNAGATCGGAAGA
54043 TATGAGTGTGGAGTCGTTGCCTCCTGAGTAGCTGGGATTACAGNNNAGATCGGAA
54045 TATGAGTGTGGAGTCGTTGCTGATCATAGCTCACTGCAGTCTCNNNAGATCGGAAG
54046 TATGAGTGTGGAGTCGTTGCTCCCAAGTAGTTGGAAGTACAGGNNNAGATCGGAA
54051 TATGAGTGTGGAGTCGTTGCAGCAGGTGTTCGGTACTTACAATNNNAGATCGGAA
54052 TATGAGTGTGGAGTCGTTGCGAGCCTGGATTGGGAGAGAATTANNNAGATCGGAA
54053 TATGAGTGTGGAGTCGTTGCAACATACATACATACCAGGCCGGNNNAGATCGGAAG
54054 TATGAGTGTGGAGTCGTTGCTCATGGAATGATTGGTTGAAGGANNNAGATCGGAA
54055 TATGAGTGTGGAGTCGTTGCGAGTTAGTCTGCTGGAGGTTCATNNNAGATCGGAA
54056 TATGAGTGTGGAGTCGTTGCGTAAACCAAAGCGGGGGAATTAGNNNAGATCGGAA
54057 TATGAGTGTGGAGTCGTTGCGAGACCTCTCTTTCCCTGTCTCCNNNAGATCGGAAGA
54058 TATGAGTGTGGAGTCGTTGCCTCAAGCCTATAATCCCAGCACTNNNAGATCGGAAG
54059 TATGAGTGTGGAGTCGTTGCGAGAGAACAGAGAACAGTGGGAANNNAGATCGGA
54060 TATGAGTGTGGAGTCGTTGCCAGGAAAGAAGGAGGTGCAGATANNNAGATCGGA
54061 TATGAGTGTGGAGTCGTTGCGAAATAGGCTGTGATGGGATTGCNNNAGATCGGAA
54062 TATGAGTGTGGAGTCGTTGCGGGTACAGAAGTTGTGAGACCATNNNAGATCGGAA
54063 TATGAGTGTGGAGTCGTTGCACTTGATGAGTTTGGAGCGTTTGNNNAGATCGGAA
54064 TATGAGTGTGGAGTCGTTGCCTCACCCCCATATCTGCCAATAANNNAGATCGGAAGA
54065 TATGAGTGTGGAGTCGTTGCAAATAAGCAAAGGCACAGAGTGGNNNAGATCGGAA
54066 TATGAGTGTGGAGTCGTTGCACACATACACACATACAGGCACTNNNAGATCGGAAG
54067 TATGAGTGTGGAGTCGTTGCCTCCCAACAGCCTCTCAATAGAGNNNAGATCGGAAG
54071 TATGAGTGTGGAGTCGTTGCTCGGTGACTTCCAAGAAAACTGANNNAGATCGGAAG
54072 TATGAGTGTGGAGTCGTTGCACTTCTGTCCACACAAAAACCTGNNNAGATCGGAAG
54073 TATGAGTGTGGAGTCGTTGCAGTAAATGTGAAGGCCTAGCACANNNAGATCGGAA
54076 TATGAGTGTGGAGTCGTTGCGCTAGACGATTGCTGCCTTATTGNNNAGATCGGAAG
54077 TATGAGTGTGGAGTCGTTGCAACTCAGCAAGCATAGGTAAGGANNNAGATCGGAA
54080 TATGAGTGTGGAGTCGTTGCGACACTCCCAAATAACCCAGGATNNNAGATCGGAAG

54081 TATGAGTGTGGAGTCGTTGCTAAAGCAGCCACACAAACAAGTCNNNAGATCGGAAG
54082 TATGAGTGTGGAGTCGTTGCTCATTTGCTACTAGGAAAACAGCANNNAGATCGGAA
54083 TATGAGTGTGGAGTCGTTGCGAGAATCAACTGACTCCTGGGAGNNNAGATCGGAA
54084 TATGAGTGTGGAGTCGTTGCGAAAGACCCGTTCCTACAGAAGANNNAGATCGGAA
54085 TATGAGTGTGGAGTCGTTGCCCAGGGCAACAGCTTAGTCTATANNNAGATCGGAAG
54086 TATGAGTGTGGAGTCGTTGCGACAAGAAAATTCACCCTTGGCANNNAGATCGGAAG
54087 TATGAGTGTGGAGTCGTTGCCCCTTTGGAATTCAAATTGGGGTNNNAGATCGGAAG
54088 TATGAGTGTGGAGTCGTTGCAGCAACTGGGAGAATAAACAGAANNNAGATCGGAA
54089 TATGAGTGTGGAGTCGTTGCCTCGGTGGCTCTCAATTTGTTTTNNNAGATCGGAAG
54090 TATGAGTGTGGAGTCGTTGCAAGAGTTTTCCTCCAAGGGTCTCNNNAGATCGGAAG
54093 TATGAGTGTGGAGTCGTTGCAAGACTACAGCCAGAATTGAGGGNNNAGATCGGAA
54094 TATGAGTGTGGAGTCGTTGCATGAAACTGGAAGCCATCATCCTNNNAGATCGGAAG
54096 TATGAGTGTGGAGTCGTTGCACCATCCAGTCTTGAGCTTTCTNNNAGATCGGAAGA
54097 TATGAGTGTGGAGTCGTTGCCATGCCTGTAATCCCAGCTTTTTNNNAGATCGGAAGA
54100 TATGAGTGTGGAGTCGTTGCAAATGTTTATCGTGCAGGAAGCCNNNAGATCGGAA
54101 TATGAGTGTGGAGTCGTTGCCTCCAGCTTGGGTGACATAATGANNNAGATCGGAA
54104 TATGAGTGTGGAGTCGTTGCGGGTTGCAATCCTACTTTCTGACNNNAGATCGGAAG
54106 TATGAGTGTGGAGTCGTTGCGCATTTGAAAAGCTAAGAACAGCCNNNAGATCGGA
54109 TATGAGTGTGGAGTCGTTGCGGACAAATTTGCAGTGGTGATGANNNAGATCGGAA
54110 TATGAGTGTGGAGTCGTTGCAGAGGAATAACTCGTGGAGGAGANNNAGATCGGA
54111 TATGAGTGTGGAGTCGTTGCCATCGTGCCTCAATGAAACTGAANNNAGATCGGAAG
54113 TATGAGTGTGGAGTCGTTGCTGGGATTCGAATGGGATTCAATCTNNNAGATCGGA
54114 TATGAGTGTGGAGTCGTTGCAAACTTGCATAAAAAGGCCTGCTNNNAGATCGGAAG
54115 TATGAGTGTGGAGTCGTTGCTTGCCCCTCATATATTGGACATTNNNAGATCGGAAG
54117 TATGAGTGTGGAGTCGTTGCGAATCACTTGAACCCAGGAGGAANNNAGATCGGAA
54120 TATGAGTGTGGAGTCGTTGCAAAAGAGGGATTGATCAGACCCCNNNAGATCGGAA
54121 TATGAGTGTGGAGTCGTTGCGTGGTCCTCTCCATGATGTACTCNNNAGATCGGAAG
54122 TATGAGTGTGGAGTCGTTGCGGCTACAGTGAGTCAAGATCACANNNAGATCGGAA
54124 TATGAGTGTGGAGTCGTTGCAAAAACTGTGTGTCACGAGGAAGNNNAGATCGGAA
54125 TATGAGTGTGGAGTCGTTGCCACCTGTAATTCCAGCACTTTGGNNNAGATCGGAAG
54126 TATGAGTGTGGAGTCGTTGCAGTGTCAAATTGGATGCATTGGGNNNAGATCGGAA
54127 TATGAGTGTGGAGTCGTTGCCTAAGGTTTGATCCCCCCAAGAANNNAGATCGGAAG
54130 TATGAGTGTGGAGTCGTTGCTGGAGTGCAATGGTGTTATCTCANNNAGATCGGAA
54131 TATGAGTGTGGAGTCGTTGCGTTCATGCCATTCTCTTGCTTCANNNAGATCGGAAGA
54132 TATGAGTGTGGAGTCGTTGCTCCCACTTCAGCCTCCTAAGTATNNNAGATCGGAAGA
54136 TATGAGTGTGGAGTCGTTGCCTCCTGAGTACCTGGGACTACAGNNNAGATCGGAAG
54137 TATGAGTGTGGAGTCGTTGCAGATGGAGTCTTGATCTGTTGCCNNNAGATCGGAA
54138 TATGAGTGTGGAGTCGTTGCAAGGCTGGACTTGAACTCTTGAGNNNAGATCGGAA
54139 TATGAGTGTGGAGTCGTTGCCTCCTGTCTGGGATGTGTGTATTNNNAGATCGGAAG
54140 TATGAGTGTGGAGTCGTTGCGCTCACTACAACCTTCACCTACTNNNAGATCGGAAGA
54141 TATGAGTGTGGAGTCGTTGCACCAAAGTAGCTGGGATTACAGGNNNAGATCGGAA
54142 TATGAGTGTGGAGTCGTTGCCCTCCTGAGTAGCTGAGACTACANNNAGATCGGAAG
54143 TATGAGTGTGGAGTCGTTGCGAGTCTCATTCTGTTGCCCAAACNNNAGATCGGAAG

54145 TATGAGTGTGGAGTCGTTGCTGAGATGGAGTCTTGTTCTGTCGNNNAGATCGGAAG
54148 TATGAGTGTGGAGTCGTTGCTCAAGTGATTCTCCTGCCTTAGCNNNAGATCGGAAGA
54149 TATGAGTGTGGAGTCGTTGCCAGCCTCCAGAGTAATTGGGATTNNNAGATCGGAAG
54151 TATGAGTGTGGAGTCGTTGCCTGAGTAGCTGCGAGGGACTATANNNAGATCGGAAG
54152 TATGAGTGTGGAGTCGTTGCCAGCATCCTGAGTATCTGGGATCNNNAGATCGGAAG
54153 TATGAGTGTGGAGTCGTTGCCCCACCTTAGCCTTCTGAGTAACNNNAGATCGGAAGA
54156 TATGAGTGTGGAGTCGTTGCCAGATGGAGTGTCACTCTGTCACNNNAGATCGGAAG
54157 TATGAGTGTGGAGTCGTTGCTAAAAATTAGTCAGGCGTGGTGGNNNAGATCGGAA
54158 TATGAGTGTGGAGTCGTTGCGCCAACAATCACAGGTTTCTACCNNNAGATCGGAAGA
54160 TATGAGTGTGGAGTCGTTGCGATCACAAGGTCAGGAGATCGAGNNNAGATCGGAA
54162 TATGAGTGTGGAGTCGTTGCGAGGTCAGGAGATCAAGACCATCNNNAGATCGGAAG
54164 TATGAGTGTGGAGTCGTTGCTCTGTAGTCCCAGCTACTTCAGANNNAGATCGGAAGA
54166 TATGAGTGTGGAGTCGTTGCAGACACGAGAATCACTTGAACCCNNNAGATCGGAAG
54169 TATGAGTGTGGAGTCGTTGCTCAAGCCTGTACTCCTAGCTACTNNNAGATCGGAAGA
54170 TATGAGTGTGGAGTCGTTGCCTACTTAGCTGGAGGTGAAGCTCNNNAGATCGGAAG
54172 TATGAGTGTGGAGTCGTTGCTCTCATGCCTGTAATCCTAGCACNNNAGATCGGAAGA
54173 TATGAGTGTGGAGTCGTTGCCAGGCGCATCGTTTTGAAAATTANNNAGATCGGAAG
54177 TATGAGTGTGGAGTCGTTGCGATTCCATCTCAAGCTCCCTGAGNNNAGATCGGAAGA
54178 TATGAGTGTGGAGTCGTTGCGTGTACTCACATTTGCAAGATGTNNNAGATCGGAAG
54181 TATGAGTGTGGAGTCGTTGCCTGGGTTCGAATCCTACTTCTCCNNNAGATCGGAAGA
54182 TATGAGTGTGGAGTCGTTGCTCACATGAGGTCAGGAGTTTGAGNNNAGATCGGAAG
54184 TATGAGTGTGGAGTCGTTGCTGAGACAGTCTCTATCACCCAGGNNNAGATCGGAAG
54185 TATGAGTGTGGAGTCGTTGCGGGCTTTTCTTTTCACTCAGTGGNNNAGATCGGAAGA
54188 TATGAGTGTGGAGTCGTTGCTCATGAAACTTTTAGAACCGCGCNNNAGATCGGAAG
54189 TATGAGTGTGGAGTCGTTGCAGGACCAAAACCTCTCGAACTTTNNNAGATCGGAAGA
54191 TATGAGTGTGGAGTCGTTGCTCCCGTTTCCATTTCTCAAGTCTNNNAGATCGGAAGAG
54192 TATGAGTGTGGAGTCGTTGCTTGTGGTTCTAGAATCAGGGTGGNNNAGATCGGAAG
54193 TATGAGTGTGGAGTCGTTGCGGATCACGAGGTCAAGAGATCAANNNAGATCGGAA
54196 TATGAGTGTGGAGTCGTTGCTCACACCTGTAATCCCAACACTTNNNAGATCGGAAGA
54198 TATGAGTGTGGAGTCGTTGCTTACCAAATGTCCCTGGAAGAGGNNNAGATCGGAAG
54199 TATGAGTGTGGAGTCGTTGCCTGAGGTCAGGAGTTCGATACCANNNAGATCGGAAG
54200 TATGAGTGTGGAGTCGTTGCACTCACACCTATAATCCCAGCACNNNAGATCGGAAGA
54201 TATGAGTGTGGAGTCGTTGCATACACACACACAGACACACACANNNAGATCGGAAGA
54202 TATGAGTGTGGAGTCGTTGCACACGGTAAAACCCCATCTCTACNNNAGATCGGAAGA
54203 TATGAGTGTGGAGTCGTTGCTTGGTCCATGCCTATAATCCCAGNNNAGATCGGAAGA
54204 TATGAGTGTGGAGTCGTTGCTCCGGTTTCAAGAGATTCTCCTGNNNAGATCGGAAGA
54206 TATGAGTGTGGAGTCGTTGCTGCAGCCTCGATTGTTCAAATTTNNNAGATCGGAAGA
54207 TATGAGTGTGGAGTCGTTGCCCTCCCAGATTCACGCTATTCTCNNNAGATCGGAAGA
54210 TATGAGTGTGGAGTCGTTGCCTATCTCAGCTCACTGCAACCTCNNNAGATCGGAAGA
54211 TATGAGTGTGGAGTCGTTGCGCCATAAAAACTCCCAGGCAAAANNNAGATCGGAAG
54212 TATGAGTGTGGAGTCGTTGCTAGAGATGGGTTTTCACCGTGTTNNNAGATCGGAAG
54213 TATGAGTGTGGAGTCGTTGCCCTCCTAGGTTCAGGCAATTCTCNNNAGATCGGAAGA
54216 TATGAGTGTGGAGTCGTTGCTTCAAGTGATTCTCCTGCCTCAGNNNAGATCGGAAGA

54217 TATGAGTGTGGAGTCGTTGCCTCAAGTGATCCTCCCACCTTAGNNNAGATCGGAAGA
54218 TATGAGTGTGGAGTCGTTGCCTGAATAGCTGGGACTACAGGTGNNNAGATCGGAA
54219 TATGAGTGTGGAGTCGTTGCGAGTAGCTGGGATTACAGGAGTGNNNAGATCGGA
54220 TATGAGTGTGGAGTCGTTGCCGCCTCCCAGATTCAAGAAATTCNNNAGATCGGAAG
54221 TATGAGTGTGGAGTCGTTGCTAATCATGACTCACTGCAGCCTTNNNAGATCGGAAG
54223 TATGAGTGTGGAGTCGTTGCCATTCTCCTGCCTAATCCTCCTGNNNAGATCGGAAGA
54225 TATGAGTGTGGAGTCGTTGCGAGACGGGATTTCGATCTTGTTGNNNAGATCGGAA
54226 TATGAGTGTGGAGTCGTTGCCCATTCTCCTGCCTCATCCTCNNNAGATCGGAAGAGC
54228 TATGAGTGTGGAGTCGTTGCCTGCATAGCTGGGATTATAGGCANNNAGATCGGAA
54230 TATGAGTGTGGAGTCGTTGCAGTAGTTGGGATTAGAGGCATGCNNNAGATCGGAA
54232 TATGAGTGTGGAGTCGTTGCCACCTTGCCATCTGTGTCTATCTNNNAGATCGGAAGA
54235 TATGAGTGTGGAGTCGTTGCTCCTGAGTAGCTTGGATTACAGGNNNAGATCGGAA
54237 TATGAGTGTGGAGTCGTTGCATAGAGTCTCTCCCTGTCATCCCNNNAGATCGGAAGA
54238 TATGAGTGTGGAGTCGTTGCCTCACTGCAACCTCCAGCTATATNNNAGATCGGAAGA
54239 TATGAGTGTGGAGTCGTTGCACTCACTACAATTTTCGCCTCCTNNNAGATCGGAAGA
54241 TATGAGTGTGGAGTCGTTGCCTCAAGCAATTCTCCTGCCTCAGNNNAGATCGGAAGA
54242 TATGAGTGTGGAGTCGTTGCTCTCAAACTCCTGACCTCAAGTGNNNAGATCGGAAGA
54244 TATGAGTGTGGAGTCGTTGCATGCTACCACGACCAGCTAATTTNNNAGATCGGAAG
54245 TATGAGTGTGGAGTCGTTGCTTCAAACGATTCTCCTGCCTCAGNNNAGATCGGAAGA
54246 TATGAGTGTGGAGTCGTTGCCAGTAGTACCATCTCGGCTCATTNNNAGATCGGAAG
54248 TATGAGTGTGGAGTCGTTGCTTTCTTCCTTGAGGTGTGAACCANNNAGATCGGAAG
54249 TATGAGTGTGGAGTCGTTGCAGAGGCATGATCATAGTTCAGGGNNNAGATCGGAA
54250 TATGAGTGTGGAGTCGTTGCTTCAAGTGATTCTCCTGCCTCAGNNNAGATCGGAAGA
54251 TATGAGTGTGGAGTCGTTGCGAGACGGAGTTTGACTCTTGTTGNNNAGATCGGAA
54253 TATGAGTGTGGAGTCGTTGCTTCAAGTGATTCTCCTGCCTCAGNNNAGATCGGAAGA
54254 TATGAGTGTGGAGTCGTTGCTGGAGGATACAGCAACAAGGTACNNNAGATCGGAA
54255 TATGAGTGTGGAGTCGTTGCAAGTCCTGACAACATGTGCTCAANNNAGATCGGAAG
54256 TATGAGTGTGGAGTCGTTGCGGATTGCTGTCTGGACTCCATAANNNAGATCGGAAG
54259 TATGAGTGTGGAGTCGTTGCGTACATGCCTGTAATCCCAGCTANNNAGATCGGAAG
54260 TATGAGTGTGGAGTCGTTGCGGCATTGGTCGAGGGTATATCTTNNNAGATCGGAA
54261 TATGAGTGTGGAGTCGTTGCAGGAAGGAAGGAAGAACACGTTTNNNAGATCGGA
54263 TATGAGTGTGGAGTCGTTGCGGGTGAATGAAACCACACCATTGNNNAGATCGGAA
54265 TATGAGTGTGGAGTCGTTGCCCTCACTCGGTATAAGCCTTCAGNNNAGATCGGAAG
54266 TATGAGTGTGGAGTCGTTGCTCTCTCCTCCCAAGTCTCCTAAGNNNAGATCGGAAGA
54269 TATGAGTGTGGAGTCGTTGCTGCTAGATTGGCCTTTACAATGGNNNAGATCGGAAG
54271 TATGAGTGTGGAGTCGTTGCTAGAAATCTTGTCCTCCGAAGCANNNAGATCGGAAG
54274 TATGAGTGTGGAGTCGTTGCTTCCATCACTTACCAGAAGGTGGNNNAGATCGGAAG
54275 TATGAGTGTGGAGTCGTTGCAATAGCCAACTGATCCCTCACAGNNNAGATCGGAAG
54276 TATGAGTGTGGAGTCGTTGCCCTAAACCAGAACTTGCACACTGNNNAGATCGGAAG
54278 TATGAGTGTGGAGTCGTTGCCCAAAAAACCAAGAACCCAGGTTNNNAGATCGGAAG
54279 TATGAGTGTGGAGTCGTTGCGGGAAGAAGGCTGTGTGAAGATANNNAGATCGGA
54280 TATGAGTGTGGAGTCGTTGCGGTTATGCTTGTAATCCCAGCACNNNAGATCGGAAG
54281 TATGAGTGTGGAGTCGTTGCACTGAGAGGCATAAAGAACAGCANNNAGATCGGAA

54282 TATGAGTGTGGAGTCGTTGCACAGAACCCCCAAGAAAATACGTNNNAGATCGGAAG
54283 TATGAGTGTGGAGTCGTTGCCTGTAGTCCCAGCTACTCAAGAGNNNAGATCGGAAG
54284 TATGAGTGTGGAGTCGTTGCGGGGTTGGGAGACAGTAAGAATTNNNAGATCGGA
54285 TATGAGTGTGGAGTCGTTGCAGGCAGGCCTAATTTCTTTCAGANNNAGATCGGAAG
54286 TATGAGTGTGGAGTCGTTGCATGCAAACGATTTGTGAGTTGGTNNNAGATCGGAA
54288 TATGAGTGTGGAGTCGTTGCTCACAAACAAAGGGCTAGACTCCNNNAGATCGGAAG
54289 TATGAGTGTGGAGTCGTTGCTTTACTTGGTGAGGACTAAGGCCNNNAGATCGGAA
54290 TATGAGTGTGGAGTCGTTGCATAAAAATTTGGGGGGAGGGACANNNAGATCGGA
54291 TATGAGTGTGGAGTCGTTGCCCCTAAGTTACCTCTGCATTTGCNNNAGATCGGAAG
54292 TATGAGTGTGGAGTCGTTGCGTAGTCCTAGCTACTCAGGAGGCNNNAGATCGGAA
54294 TATGAGTGTGGAGTCGTTGCCCCTGCATCAGTCAAGACTTTTGNNNAGATCGGAAG
54295 TATGAGTGTGGAGTCGTTGCCACTACAGGAATGTTGAGGTGGANNNAGATCGGAA
54296 TATGAGTGTGGAGTCGTTGCGAATAGGAACAGCTCCCGTCTACNNNAGATCGGAAG
54300 TATGAGTGTGGAGTCGTTGCGTGAATAGGGCCACCTAGTTCAGNNNAGATCGGAA
54301 TATGAGTGTGGAGTCGTTGCTAATCCCAGCACTTTGAGAGACCNNNAGATCGGAAG
54302 TATGAGTGTGGAGTCGTTGCTGAGAAGACTGAGACAGGAGGATNNNAGATCGGA
54305 TATGAGTGTGGAGTCGTTGCTAAAACACTGAGGGAGAAAGCGANNNAGATCGGAA
54306 TATGAGTGTGGAGTCGTTGCGATCACAAGGTCAGGAGTTCGAGNNNAGATCGGAA
54307 TATGAGTGTGGAGTCGTTGCCTGAACTCCAGCCTAGGTAACAGNNNAGATCGGAAG
54310 TATGAGTGTGGAGTCGTTGCGGATTCAAGTGATTCTCCTGCCTNNNAGATCGGAAG
54312 TATGAGTGTGGAGTCGTTGCTCTCAGGTTCAAGCAATTCTCCTNNNAGATCGGAAG
54314 TATGAGTGTGGAGTCGTTGCCTTGCCACAGCTATACTTCCAGANNNAGATCGGAAG
54315 TATGAGTGTGGAGTCGTTGCCTTCCCAGGTTCATGCCATTCTCNNNAGATCGGAAGA
54316 TATGAGTGTGGAGTCGTTGCTGGCTTGAACACTTGATGAACACNNNAGATCGGAAG
54318 TATGAGTGTGGAGTCGTTGCGTTCAAGAGATTCTCCTGCCTCANNNAGATCGGAAG
54320 TATGAGTGTGGAGTCGTTGCTCTTGAACTCCTGACCTCAAGTGNNNAGATCGGAAG
54323 TATGAGTGTGGAGTCGTTGCTTCAAGCAATTCTCCTGCCTCAGNNNAGATCGGAAG
54328 TATGAGTGTGGAGTCGTTGCCTAGAGTGCAGTGGTGAGATCTCNNNAGATCGGAA
54329 TATGAGTGTGGAGTCGTTGCACAAGTGAACAAAGGTCTCTGGTNNNAGATCGGAA
54330 TATGAGTGTGGAGTCGTTGCCTCAAGCAATCCTCCTACCTCAGNNNAGATCGGAAG
54331 TATGAGTGTGGAGTCGTTGCGTACTGGGATTACAGGTGTGAGCNNNAGATCGGAA
54332 TATGAGTGTGGAGTCGTTGCGATCTTGGCTCACTACAACCTCCNNNAGATCGGAAG
54335 TATGAGTGTGGAGTCGTTGCGATCTCAGCTCACTGCAATGTCTNNNAGATCGGAAG
54336 TATGAGTGTGGAGTCGTTGCCTTTAGTTCAGTGGTGCCATCTCNNNAGATCGGAAG
54337 TATGAGTGTGGAGTCGTTGCCGATCATGGCTTATTGCAACCTTNNNAGATCGGAAG
54339 TATGAGTGTGGAGTCGTTGCGAGTACAGTGTGATCATGGCTCANNNAGATCGGAA
54340 TATGAGTGTGGAGTCGTTGCAATCTCAGTTCACTGCAACCTCTNNNAGATCGGAAG
54342 TATGAGTGTGGAGTCGTTGCCTGGGTTCAAGTGATCTTCCTGANNNAGATCGGAAG
54343 TATGAGTGTGGAGTCGTTGCACTTCCTGGGCTCAAATGATCTTNNNAGATCGGAAG
54345 TATGAGTGTGGAGTCGTTGCTCCCATTTTCACGACATTCTCCTNNNAGATCGGAAGA
54347 TATGAGTGTGGAGTCGTTGCGAACTCCTGACCTCAGATGATCCNNNAGATCGGAAG
54350 TATGAGTGTGGAGTCGTTGCCCACTCTAACAGGTGTGAAGTGANNNAGATCGGAA
54351 TATGAGTGTGGAGTCGTTGCGATGTAGTCTCACTCTGTCACCCNNNAGATCGGAAG

54355 TATGAGTGTGGAGTCGTTGCTGGCATGATCTCAGTTACAGCAANNNAGATCGGAAG
54357 TATGAGTGTGGAGTCGTTGCGTGTTTTTGTTCTTGACCCCACANNNAGATCGGAAG
54359 TATGAGTGTGGAGTCGTTGCGCGATCTTGGCTCATTTCAACTTNNNAGATCGGAAG
54360 TATGAGTGTGGAGTCGTTGCTCCAGGTTCAAGTGATTCTCCTGNNNAGATCGGAAG
54363 TATGAGTGTGGAGTCGTTGCGTTTCAAGCGATTCTCCTGCTTCNNNAGATCGGAAG
54364 TATGAGTGTGGAGTCGTTGCGAACTCCTGACCTCGTGATCCTCNNNAGATCGGAAG
54365 TATGAGTGTGGAGTCGTTGCATTTGGCAGGGTCATAGGACAATNNNAGATCGGAA
54368 TATGAGTGTGGAGTCGTTGCCCTCTGAGACATCTTTCCTGGAGNNNAGATCGGAAG
54369 TATGAGTGTGGAGTCGTTGCCTTGAGCGATTGACCTACCTCAGNNNAGATCGGAAG
54373 TATGAGTGTGGAGTCGTTGCCGGCATGACTCAGTGAATTTGAANNNAGATCGGAA
54374 TATGAGTGTGGAGTCGTTGCCAGTAGTGCAATCACAGCTCACTNNNAGATCGGAAG
54375 TATGAGTGTGGAGTCGTTGCCAATCTCGGCTCACTACAACCTCNNNAGATCGGAAGA
54377 TATGAGTGTGGAGTCGTTGCCAGTGGCAGGATCTCAGTTTACTNNNAGATCGGAAG
54378 TATGAGTGTGGAGTCGTTGCTTTTCTTAGGGAGGGTACATGGCNNNAGATCGGAA
54382 TATGAGTGTGGAGTCGTTGCGCCTCCTAAGTAGCTGGGATTACNNNAGATCGGAAG
54385 TATGAGTGTGGAGTCGTTGCCCCAAGTTCAAGCAATTCTCCTGNNNAGATCGGAAG
54386 TATGAGTGTGGAGTCGTTGCCTCCTGAGTAGCTGCGATTACAGNNNAGATCGGAAG
54387 TATGAGTGTGGAGTCGTTGCGAACATCTGGCCTCAAACAATCCNNNAGATCGGAAG
54388 TATGAGTGTGGAGTCGTTGCGATCATTTGAGCCTGGGACATTGNNNAGATCGGAA
54391 TATGAGTGTGGAGTCGTTGCAAGATAGCTTCCCTCATGCCTTCNNNAGATCGGAAG
54393 TATGAGTGTGGAGTCGTTGCGAACTAGGGGATTATGCTCCAGTNNNAGATCGGAA
54395 TATGAGTGTGGAGTCGTTGCAAGAACGAGAGCTTCTACTCCACNNNAGATCGGAAG
54396 TATGAGTGTGGAGTCGTTGCCCAGTGCCTTCTTAGAAACTCCANNNAGATCGGAAG
54397 TATGAGTGTGGAGTCGTTGCATGGTTAGGGCTTATCTCTGCTGNNNAGATCGGAAG
54399 TATGAGTGTGGAGTCGTTGCGTAATCCCAGCTACTTAGGAGGCNNNAGATCGGAAG
54403 TATGAGTGTGGAGTCGTTGCTAATAATTCTGTGGAGGGCCAGGNNNAGATCGGAA
54404 TATGAGTGTGGAGTCGTTGCACTGGAAAACAATAGCACCTCCTNNNAGATCGGAAG
54405 TATGAGTGTGGAGTCGTTGCATAGACTTGATCCCAAAGGCGAGNNNAGATCGGAA
54406 TATGAGTGTGGAGTCGTTGCTTTTGAATCTTTTAGTCCTGCTCTAANNNAGATCGGA
54408 TATGAGTGTGGAGTCGTTGCAGATGGATTCACAGCCAAGTTCTNNNAGATCGGAAG
54411 TATGAGTGTGGAGTCGTTGCAGGAAATGTTTCAGAAGCACTGCNNNAGATCGGAA
54412 TATGAGTGTGGAGTCGTTGCAAACAAAATGCTTTGGTTCAGCCNNNAGATCGGAAG
54413 TATGAGTGTGGAGTCGTTGCAGGAAATTGTGATGCATGCAACANNNAGATCGGAA
54415 TATGAGTGTGGAGTCGTTGCTGATCCTTCTCAGGAAAGAGCACNNNAGATCGGAAG
54417 TATGAGTGTGGAGTCGTTGCGCCCATGCCTGTAATCCTAACTANNNAGATCGGAAG
54418 TATGAGTGTGGAGTCGTTGCCTATGGTTTGGATCTGTGGCCTTNNNAGATCGGAAG
54419 TATGAGTGTGGAGTCGTTGCCCCAGCTTAGTCTGTGATATCCANNNAGATCGGAAG
54421 TATGAGTGTGGAGTCGTTGCGAAGAGGAGAGAAATGGTGAGGTNNNAGATCGGA
54423 TATGAGTGTGGAGTCGTTGCAAGGAAAGGCAAGGCTACATACANNNAGATCGGAA
54425 TATGAGTGTGGAGTCGTTGCTCACTCAAAACCAGAAAGCAGTTNNNAGATCGGAAG
54428 TATGAGTGTGGAGTCGTTGCGGATGAGACTATCACTAGATCCCNNNAGATCGGAA
54429 TATGAGTGTGGAGTCGTTGCTAGCCCAGCACTGTATTGATGTTNNNAGATCGGAAG
54430 TATGAGTGTGGAGTCGTTGCAGTGGGGTTGGATGAAACTTCTANNNAGATCGGAA

54431 TATGAGTGTGGAGTCGTTGCTCTCCCAACCACTCTGAATCATGNNNAGATCGGAAGA
54432 TATGAGTGTGGAGTCGTTGCCCATCTGTAGGTCACCAACATCANNNAGATCGGAAG
54436 TATGAGTGTGGAGTCGTTGCTGTTAGTCATCCGTGGTCTCTTCNNNAGATCGGAAG
54438 TATGAGTGTGGAGTCGTTGCGCCCATCCTGTCTGTTTACATCTNNNAGATCGGAAGA
54439 TATGAGTGTGGAGTCGTTGCACTTCTGCATTTCCATCTGAGGTNNNAGATCGGAAG
54442 TATGAGTGTGGAGTCGTTGCCAGAAAACAAAGGCAAAAGTGGCNNNAGATCGGAA
54443 TATGAGTGTGGAGTCGTTGCTAGCATGAGAGGCTTGGAAAGAGNNNAGATCGGA
54444 TATGAGTGTGGAGTCGTTGCTATACGTGTGTGTGTGTGTGTGTNNNAGATCGGAA
54447 TATGAGTGTGGAGTCGTTGCTAGCTGTGATTTCTGGATGGTGTNNNAGATCGGAAG
54448 TATGAGTGTGGAGTCGTTGCATGCCTGTAGTCCTAGCTACTTGNNNAGATCGGAAG
54449 TATGAGTGTGGAGTCGTTGCATGATGCCCTGTTACTATGTTGCNNNAGATCGGAAG
54450 TATGAGTGTGGAGTCGTTGCCACACACACACACACACATACATNNNAGATCGGAAGA
54451 TATGAGTGTGGAGTCGTTGCTCTCTCCTTCCTTTTAGCGCATTNNNAGATCGGAAGA
54453 TATGAGTGTGGAGTCGTTGCTTGAGACCAGTCTGGGTAAATGGNNNAGATCGGAA
54454 TATGAGTGTGGAGTCGTTGCTTCTAATGAACAGCCAGGGTTGTNNNAGATCGGAAG
54455 TATGAGTGTGGAGTCGTTGCGTGCAATCTTGGCTCACTGTAAGNNNAGATCGGAAG
54457 TATGAGTGTGGAGTCGTTGCTTTGACACACATTCTCTGAAGCTNNNAGATCGGAAG
54458 TATGAGTGTGGAGTCGTTGCACCCTGTAGTCCTAGCTACTCAGNNNAGATCGGAAG
54459 TATGAGTGTGGAGTCGTTGCAAGGAAAATGTGTTTGCGCTCTTNNNAGATCGGAAG
54460 TATGAGTGTGGAGTCGTTGCAAAAACCCCAAACAAACAAACANNNAGATCGGAAG
54461 TATGAGTGTGGAGTCGTTGCGGTACAGAGAGAAGAGCACACAANNNAGATCGGAA
54462 TATGAGTGTGGAGTCGTTGCAGAAAGAGAGAAAGGAAGGACGANNNAGATCGGA
54465 TATGAGTGTGGAGTCGTTGCAAGCATTACTCACAATGGCCATGNNNAGATCGGAAG
54466 TATGAGTGTGGAGTCGTTGCTGCTTCCATGACAAACCAACAAANNNAGATCGGAAG
54468 TATGAGTGTGGAGTCGTTGCCAAGGTCAGGAGTTCGAGATCAGNNNAGATCGGAA
54469 TATGAGTGTGGAGTCGTTGCAGCTGTGTGATTTGTCATCCCTANNNAGATCGGAAG
54470 TATGAGTGTGGAGTCGTTGCGCAGAATGCCCAGAAAAGAAAGANNNAGATCGGAA
54471 TATGAGTGTGGAGTCGTTGCCAGCCAATCCTTCAAGGAATGACNNNAGATCGGAAG
54472 TATGAGTGTGGAGTCGTTGCAAGCCCCCAAACTTCTTATGACTNNNAGATCGGAAGA
54473 TATGAGTGTGGAGTCGTTGCGAAGTCAGGGTGTCAACAGAGATNNNAGATCGGAA
54474 TATGAGTGTGGAGTCGTTGCCCTCATGTTCTCAGAGCTACACTNNNAGATCGGAAGA
54475 TATGAGTGTGGAGTCGTTGCTCCTAATTGCCATCAAGTTCCCANNNAGATCGGAAGA
54479 TATGAGTGTGGAGTCGTTGCGCTGATCATGTGTATTTAGCGCANNNAGATCGGAAG
54480 TATGAGTGTGGAGTCGTTGCACAAAAGCATGAAGGAGGAATGCNNNAGATCGGAA
54481 TATGAGTGTGGAGTCGTTGCAAAGACACCAAAAGCAATCGCAANNNAGATCGGAA
54482 TATGAGTGTGGAGTCGTTGCCTAGCGTTACAGGAAGCTCTAGGNNNAGATCGGAA
54483 TATGAGTGTGGAGTCGTTGCCTGTAGTCCCAGCTACTCGGTAGNNNAGATCGGAAG
54484 TATGAGTGTGGAGTCGTTGCTGTGTGGGAGACAATAAAACGGANNNAGATCGGAA
54487 TATGAGTGTGGAGTCGTTGCAAGGAAGGGATCACCATACACACNNNAGATCGGAA
54488 TATGAGTGTGGAGTCGTTGCAGAGAGATAACCCTCACACTTGANNNAGATCGGAAG
54489 TATGAGTGTGGAGTCGTTGCTCGGTTATCTCCTTCAGAAAGCCNNNAGATCGGAAG
54490 TATGAGTGTGGAGTCGTTGCACAAAGAAGTCAGGTGGGTTCTCNNNAGATCGGAA
54491 TATGAGTGTGGAGTCGTTGCAGAAGGTCAGCAGATAAACACGTNNNAGATCGGAA

54493 TATGAGTGTGGAGTCGTTGCACTCCGCCCAGCTAATTTTTGTANNNAGATCGGAAG

54494 TATGAGTGTGGAGTCGTTGCCAAGCAGCTGAGACTACAGACTTNNNAGATCGGAA

54496 TATGAGTGTGGAGTCGTTGCTCAAGTGATTCTCCTGCCTTAGCNNNAGATCGGAAG

54498 TATGAGTGTGGAGTCGTTGCCTGGCAGGCTCTCATTTTTTACCNNNAGATCGGAAG

54499 TATGAGTGTGGAGTCGTTGCCTCCCGAGTAGCTAGGACTACAGNNNAGATCGGAA

54500 TATGAGTGTGGAGTCGTTGCCCATCTCAGCTCACTTCAACCTCNNNAGATCGGAAGA

54501 TATGAGTGTGGAGTCGTTGCCTGAGTAGCTGGGATTACAGGTGNNNAGATCGGAA

54503 TATGAGTGTGGAGTCGTTGCTGGAGTGTAGTGGCATGATTTCANNNAGATCGGAA

54505 TATGAGTGTGGAGTCGTTGCGCCTCTCAAGTAGCTGGGATTATNNNAGATCGGAAG

54506 TATGAGTGTGGAGTCGTTGCGCCTCCTAAGTAGCTGGGATTACNNNAGATCGGAAG

54508 TATGAGTGTGGAGTCGTTGCAGAGGCAAAGTTGATACGAAGGANNNAGATCGGA

54509 TATGAGTGTGGAGTCGTTGCACCAATTCCCCACAACAAAGAACNNNAGATCGGAAG

54510 TATGAGTGTGGAGTCGTTGCGTTCAAGCAATTCTTCTGCCTCANNNAGATCGGAAG

54511 TATGAGTGTGGAGTCGTTGCCTAGGAGGTTCAGTTCATTGGCTNNNAGATCGGAA

54512 TATGAGTGTGGAGTCGTTGCGGAAGGAATCCAGAGAGGTGTTTNNNAGATCGGA

54513 TATGAGTGTGGAGTCGTTGCAGCCTGCCATGTAACTGTGATTANNNAGATCGGAAG

54514 TATGAGTGTGGAGTCGTTGCCTCCTGAGTAGCTGGAACTACAGNNNAGATCGGAA

54515 TATGAGTGTGGAGTCGTTGCGCTCAAGTGATCCTCCTTCTTCANNNAGATCGGAAGA

54518 TATGAGTGTGGAGTCGTTGCTGTGTTCTTATCGTTCCTTGAGANNNAGATCGGAAG

54519 TATGAGTGTGGAGTCGTTGCTTCAAGTGATTCTCCTGCCTCAGNNNAGATCGGAAG

54520 TATGAGTGTGGAGTCGTTGCTGACAATGGCACAATCTTGGTTCNNNAGATCGGAAG

54521 TATGAGTGTGGAGTCGTTGCGAGTACCCAAGTGAAGAGAACGTNNNAGATCGGAA

54522 TATGAGTGTGGAGTCGTTGCGTTCACACTATTCTCCTGCCTCANNNAGATCGGAAGA

54523 TATGAGTGTGGAGTCGTTGCATGATCTCAGCTCACTGCAACTTNNNAGATCGGAAG

54524 TATGAGTGTGGAGTCGTTGCTCCTAGCTGGGACTACAGGTATGNNNAGATCGGAA

54526 TATGAGTGTGGAGTCGTTGCCTCCCGAGCAGCTGTAATTACAGNNNAGATCGGAAG

54527 TATGAGTGTGGAGTCGTTGCTAGAGTTCAGTGGCACAATCTCGNNNAGATCGGAA

54528 TATGAGTGTGGAGTCGTTGCTCTGCATCCCAGGTTCAAATGATNNNAGATCGGAAG

54531 TATGAGTGTGGAGTCGTTGCTTCATGCTATTCTCCTGCCTCAGNNNAGATCGGAAGA

54533 TATGAGTGTGGAGTCGTTGCGGTTCCGTGTCATTTGGTTCTTGNNNAGATCGGAAG

54534 TATGAGTGTGGAGTCGTTGCTCTTCTCTGGGTTTATGCTGCTTNNNAGATCGGAAG

54535 TATGAGTGTGGAGTCGTTGCTGAATATGCTAGATCACAGCCCCNNNAGATCGGAAG

54537 TATGAGTGTGGAGTCGTTGCAGAGGCCAGAAGATAACTTGAGCNNNAGATCGGAA

54539 TATGAGTGTGGAGTCGTTGCAGAGATGCCTGCTTTATGTAGGGNNNAGATCGGAA

54540 TATGAGTGTGGAGTCGTTGCACAATTGGTGGGATGAATCAGGANNNAGATCGGAA

54541 TATGAGTGTGGAGTCGTTGCCTGAGGCAGGAGTATCACTTGAGNNNAGATCGGAA

54543 TATGAGTGTGGAGTCGTTGCAGGACACTGAGCCAGTTTATCAGNNNAGATCGGAA

54546 TATGAGTGTGGAGTCGTTGCCAACTAGACAGCCTACAGTGCTANNNAGATCGGAAG

54548 TATGAGTGTGGAGTCGTTGCGCCAACATGATGGACACGATACTNNNAGATCGGAA

54549 TATGAGTGTGGAGTCGTTGCTGAGACCAAGAGTTTGAGACCAGNNNAGATCGGAA

54551 TATGAGTGTGGAGTCGTTGCATCCCTGTACCACCAAACAACTTNNNAGATCGGAAGA

54552 TATGAGTGTGGAGTCGTTGCACACCTCAGACCTTACACACAAANNNAGATCGGAAG

54553 TATGAGTGTGGAGTCGTTGCTTCTTTGCTAGCTCCATGGTGAANNNAGATCGGAAG

54554 TATGAGTGTGGAGTCGTTGCGGATGGACATCACTGAGGAGAATNNNAGATCGGAA
54556 TATGAGTGTGGAGTCGTTGCATAGATGAATGCCCTCCTATGGGNNNAGATCGGAAG
54557 TATGAGTGTGGAGTCGTTGCCAGATCACAAGGTCAGGAGTTCANNNAGATCGGAA
54559 TATGAGTGTGGAGTCGTTGCGTACTTTGTTCTTCAGCAGCACCNNNAGATCGGAAG
54560 TATGAGTGTGGAGTCGTTGCAACATATATGGCCGGGTATGGTGNNNAGATCGGAA
54561 TATGAGTGTGGAGTCGTTGCTTATGCTAGGTTCACTCTGCCTGNNNAGATCGGAAG
54562 TATGAGTGTGGAGTCGTTGCTGCGGTGGCTCATACCTATAATCNNNAGATCGGAAG
54564 TATGAGTGTGGAGTCGTTGCACTCTTTATCCATTTGTCCCCCCNNNAGATCGGAAGA
54565 TATGAGTGTGGAGTCGTTGCTGAAGGAAGTTAGTGGATGGTGGNNNAGATCGGA
54566 TATGAGTGTGGAGTCGTTGCTGAGGTAGGATCTCAGGCTTTTGNNNAGATCGGAA
54567 TATGAGTGTGGAGTCGTTGCCAGACAGAAAGCAGCATGTTCAANNNAGATCGGAA
54570 TATGAGTGTGGAGTCGTTGCACAAGGTCAAGAGATCGAAACCANNNAGATCGGAA
54571 TATGAGTGTGGAGTCGTTGCGAGGTCAGGAGTTTGAGACCAACNNNAGATCGGAA
54572 TATGAGTGTGGAGTCGTTGCGACAAGATGGTGACCAAAGGAGANNNAGATCGGA
54573 TATGAGTGTGGAGTCGTTGCACAAATGGTACAGAGGGATCCTGNNNAGATCGGAA
54575 TATGAGTGTGGAGTCGTTGCGGGGTTTCTTTTTGGAGTGATGGNNNAGATCGGAA
54577 TATGAGTGTGGAGTCGTTGCGAGAGAGAGAGAGTCCTACACCANNNAGATCGGAA
54578 TATGAGTGTGGAGTCGTTGCGCACCTGTAGACTTCAATGTCCTNNNAGATCGGAAG
54580 TATGAGTGTGGAGTCGTTGCTAGAGGAGTGATTCCAGGAAGGGNNNAGATCGGA
54581 TATGAGTGTGGAGTCGTTGCTGGCCACTCCTAGTTTTCTTGAANNNAGATCGGAAG
54584 TATGAGTGTGGAGTCGTTGCTCCTTCTGTCTACTACCAACTGANNNAGATCGGAAGA
54585 TATGAGTGTGGAGTCGTTGCAGAAGAGCTACTCTGAAGTCCCTNNNAGATCGGAAG
54586 TATGAGTGTGGAGTCGTTGCTCACTTGAGTCCAGGAGTTCAAGNNNAGATCGGAAG
54587 TATGAGTGTGGAGTCGTTGCAAATGGTTCTAATGCTGCCAACCNNNAGATCGGAAG
54588 TATGAGTGTGGAGTCGTTGCACATGGTGAAACCTCGTCTCTACNNNAGATCGGAAG
54589 TATGAGTGTGGAGTCGTTGCGGAAGAGAAGCACAGAAGTCACTNNNAGATCGGAA
54591 TATGAGTGTGGAGTCGTTGCGTAGCTGGGATTACAGGCATGTANNNAGATCGGAA
54592 TATGAGTGTGGAGTCGTTGCCTCCTGAGTAGCTGGGACTACAGNNNAGATCGGAAG
54593 TATGAGTGTGGAGTCGTTGCTAGACAGGGTGTTACTCTGAAGCNNNAGATCGGAA
54595 TATGAGTGTGGAGTCGTTGCAGTAGCTGGGATTACAAGTGCATNNNAGATCGGAA
54598 TATGAGTGTGGAGTCGTTGCTTCACGACATTCTCCTGCCTTAGNNNAGATCGGAAGA
54599 TATGAGTGTGGAGTCGTTGCTCCGGGATTCAAGTCATTTTCCTNNNAGATCGGAAG
54600 TATGAGTGTGGAGTCGTTGCCTGGAGTACAATGGCGTGATTTCNNNAGATCGGAAG
54602 TATGAGTGTGGAGTCGTTGCTGAATCCCAGCTCTATTTCCTGGNNNAGATCGGAAG
54603 TATGAGTGTGGAGTCGTTGCCTGGGTGAGGTTGATTCTCCTTGNNNAGATCGGAAG
54604 TATGAGTGTGGAGTCGTTGCATTATCCTGTCTCAGTCTCCCGANNNAGATCGGAAGA
54605 TATGAGTGTGGAGTCGTTGCTCCCGAGTAGCTAGGCCTAATATNNNAGATCGGAAG
54608 TATGAGTGTGGAGTCGTTGCCTGGAGTGCTATGGTGCAATCTTNNNAGATCGGAAG
54609 TATGAGTGTGGAGTCGTTGCCTGAGTAGCTGGGATTACAGGTGNNNAGATCGGAA
54610 TATGAGTGTGGAGTCGTTGCGTAGATTCTCACTCTGTTGCCCANNNAGATCGGAAG
54611 TATGAGTGTGGAGTCGTTGCGCAACAGGACGAATGGATTTCTTNNNAGATCGGAAG
54612 TATGAGTGTGGAGTCGTTGCTTCAAGTGATTCTCCTGCCTCAGNNNAGATCGGAAG
54616 TATGAGTGTGGAGTCGTTGCCCTTGAAGGAGGGTTTGTACAGANNNAGATCGGAA

54617 TATGAGTGTGGAGTCGTTGCCTGCCCTAGAGATTTGTGGAACTNNNAGATCGGAAG
54618 TATGAGTGTGGAGTCGTTGCCTGTAGTCCCAGCTACTCAGGAGNNNAGATCGGAAG
54619 TATGAGTGTGGAGTCGTTGCAAAAGAAAAAGGCAAGGAGAGGCNNNAGATCGGA
54621 TATGAGTGTGGAGTCGTTGCTAGTGAACTGTGATTGCACCACTNNNAGATCGGAAG
54623 TATGAGTGTGGAGTCGTTGCGAAGGAGAGAAGGAAGGAAGGGANNNAGATCGG
54624 TATGAGTGTGGAGTCGTTGCGGTGGATCATGACGTCAAGAGATNNNAGATCGGAA
54625 TATGAGTGTGGAGTCGTTGCAAAATAAATTCCTGGCTGGGCACNNNAGATCGGAA
54626 TATGAGTGTGGAGTCGTTGCTGCCAAACACTTTTCCAAAATGGNNNAGATCGGAAG
54628 TATGAGTGTGGAGTCGTTGCCTTGGTGGTCCCTGGAGATTATCNNNAGATCGGAAG
54629 TATGAGTGTGGAGTCGTTGCTCACTTGAGTCCAGGAAATCCAGNNNAGATCGGAAG
54630 TATGAGTGTGGAGTCGTTGCTCAATAGTGGTGAGTGAGTGCTCNNNAGATCGGAA
54631 TATGAGTGTGGAGTCGTTGCATATGAAGCCATCACGGAACTGANNNAGATCGGAA
54633 TATGAGTGTGGAGTCGTTGCCCTGCTCCAGCTATGTAAGATGTNNNAGATCGGAAG
54634 TATGAGTGTGGAGTCGTTGCCCCCTGAGCATAGGAAAGTCAAANNNAGATCGGAA
54635 TATGAGTGTGGAGTCGTTGCGATCACTTGAGCCCAGGAATTTGNNNAGATCGGAA
54636 TATGAGTGTGGAGTCGTTGCATCTGTTTACTAAGGGCTGCCTGNNNAGATCGGAAG
54637 TATGAGTGTGGAGTCGTTGCCCAGTCTTCAAAGCAGAGGAGATNNNAGATCGGAA
54638 TATGAGTGTGGAGTCGTTGCATCACCACTTTGTAACCCCCAATNNNAGATCGGAAGA
54639 TATGAGTGTGGAGTCGTTGCTGCCAGTTTATTACGAAGGACATGNNNAGATCGGA
54640 TATGAGTGTGGAGTCGTTGCGAGGTCAGGAGATCAAGACCATCNNNAGATCGGAA
54641 TATGAGTGTGGAGTCGTTGCCCAATCTGACCTGTGTCCTTGTANNNAGATCGGAAG
54642 TATGAGTGTGGAGTCGTTGCATACAAAAATTAGCCGGGTGTGGNNNAGATCGGAA
54643 TATGAGTGTGGAGTCGTTGCATTGGACCCCCTGTATACTGTTGNNNAGATCGGAAG
54644 TATGAGTGTGGAGTCGTTGCAGGCAAAAAGAATTGTTTCTGCANNNAGATCGGAA
54645 TATGAGTGTGGAGTCGTTGCACTTTAGGGACCTGTGAGACTCTNNNAGATCGGAAG
54646 TATGAGTGTGGAGTCGTTGCCCTCCGAAATTCACGTGTTGAAANNNAGATCGGAAG
54648 TATGAGTGTGGAGTCGTTGCATAACAGGGCCAAGAGTTTCCATNNNAGATCGGAA
54650 TATGAGTGTGGAGTCGTTGCTATTCCTGACAACTGGCTACAGGNNNAGATCGGAAG
54651 TATGAGTGTGGAGTCGTTGCTGACACTGTTGCATTGGGAATTCNNNAGATCGGAAG
54653 TATGAGTGTGGAGTCGTTGCTCAGGGAAATGCAAATCGAAACCNNNAGATCGGAA
54654 TATGAGTGTGGAGTCGTTGCCACAAGCCTACAGTCCTAGCTACNNNAGATCGGAAG
54657 TATGAGTGTGGAGTCGTTGCCTCCTGGGTTTGAGCAATTCTTGNNNAGATCGGAAG
54659 TATGAGTGTGGAGTCGTTGCGGATCACTGCAATCTCTACCTCCNNNAGATCGGAAG
54661 TATGAGTGTGGAGTCGTTGCCCTGAGTAGCTGGGATTATAGGCNNNAGATCGGAA
54663 TATGAGTGTGGAGTCGTTGCTCATGCCTGTAATCCCAACACTTNNNAGATCGGAAGA
54664 TATGAGTGTGGAGTCGTTGCGTCTTGAACTCCTGACCTTGTGANNNAGATCGGAAG
54666 TATGAGTGTGGAGTCGTTGCCTCCTGAGTAGCTGGGATTACAGNNNAGATCGGAA
54667 TATGAGTGTGGAGTCGTTGCCACAATCTTGGCTCACTGCAAGNNNAGATCGGAAGA
54668 TATGAGTGTGGAGTCGTTGCTCTTGACTCACTACAACCTCTGCNNNAGATCGGAAGA
54669 TATGAGTGTGGAGTCGTTGCCCCACAGAGCCACAATTTTTCATNNNAGATCGGAAG
54671 TATGAGTGTGGAGTCGTTGCCCACCCATGAGAAAATATGCTGCNNNAGATCGGAAG
54672 TATGAGTGTGGAGTCGTTGCCAAGTAGCTGGGATTACAGGCATNNNAGATCGGAA
54674 TATGAGTGTGGAGTCGTTGCCCGCAGGTGGGTCATTTCTAATTNNNAGATCGGAAG

54676 TATGAGTGTGGAGTCGTTGCAGACAGGGTCTCACTTTGTTACCNNNAGATCGGAAG
54677 TATGAGTGTGGAGTCGTTGCCCTCCTGGGTTCAAGTGATTCTCNNNAGATCGGAAG
54678 TATGAGTGTGGAGTCGTTGCCCTCCTAAAGTGCTGGGATTACANNNAGATCGGAAG
54680 TATGAGTGTGGAGTCGTTGCCCTGAGTAGCTGGGATTATAGGCNNNAGATCGGAA
54681 TATGAGTGTGGAGTCGTTGCCCTCCTGAGTAGCTGAGACTACANNNAGATCGGAAG
54682 TATGAGTGTGGAGTCGTTGCTTTCAAGTGATTCTCCTGCCTCANNNAGATCGGAAG
54683 TATGAGTGTGGAGTCGTTGCGTGGTATGTGCCTGTAGTCCTAGNNNAGATCGGAA
54684 TATGAGTGTGGAGTCGTTGCCCACTAAAAGGGAGAGAACAGCTNNNAGATCGGAA
54686 TATGAGTGTGGAGTCGTTGCACTCTCTCTCTCTTTCTCCCTCCNNNAGATCGGAAGAG
54688 TATGAGTGTGGAGTCGTTGCTGAGCTAATCTTAGGGCCTACCTNNNAGATCGGAAG
54689 TATGAGTGTGGAGTCGTTGCAAGGTGTTATCTGGCACTGTAGANNNAGATCGGAA
54691 TATGAGTGTGGAGTCGTTGCCAGAGGTTACAGTGAGCCAAGATNNNAGATCGGAA
54696 TATGAGTGTGGAGTCGTTGCGAAGGAAGGAAGGAAGGAAGGAANNNAGATCGG
54697 TATGAGTGTGGAGTCGTTGCAGCATCTTGTTTAGGACTAACTACTNNNAGATCGGA
54701 TATGAGTGTGGAGTCGTTGCGTGAGTCAGCAGAACAATTTCCCNNNAGATCGGAAG
54705 TATGAGTGTGGAGTCGTTGCCGGTGATCGTGAAATAGACATGCNNNAGATCGGAA
54706 TATGAGTGTGGAGTCGTTGCTGGGATAATTTGGCCACTTCTGANNNAGATCGGAAG
54707 TATGAGTGTGGAGTCGTTGCGGCGGAGCTTGCAAAAAAAAAAAANNNAGATCGGAA
54709 TATGAGTGTGGAGTCGTTGCCATACGTGCAGCCAACAAACATANNNAGATCGGAAG
54710 TATGAGTGTGGAGTCGTTGCTTGCATTTGGGGATTTCTGGAACNNNAGATCGGAAG
54711 TATGAGTGTGGAGTCGTTGCCTCACTGTAACCTCCACTTTCCANNNAGATCGGAAGA
54713 TATGAGTGTGGAGTCGTTGCATAAGAGTCTCATTGTGGCCCAGNNNAGATCGGAAG
54714 TATGAGTGTGGAGTCGTTGCAATCTCATTCTATTGCCCAGGCTNNNAGATCGGAAG
54715 TATGAGTGTGGAGTCGTTGCATGATATCAGCTCACTGCAACCTNNNAGATCGGAAG
54716 TATGAGTGTGGAGTCGTTGCTAGCAATCCTCCCATTTCCTGACNNNAGATCGGAAGA
54717 TATGAGTGTGGAGTCGTTGCAGCCTTGATCTTCTGGACTCAAGNNNAGATCGGAAG
54721 TATGAGTGTGGAGTCGTTGCTTACTATATTGCCCAGACCGGACNNNAGATCGGAAG
54722 TATGAGTGTGGAGTCGTTGCCACCTCAGCCTCTTGAGTATCTGNNNAGATCGGAAG
54723 TATGAGTGTGGAGTCGTTGCTTGAACTCCTGACCTCGTGATTCNNNAGATCGGAAG
54725 TATGAGTGTGGAGTCGTTGCTGGAGTACAGTGGTGTGATCATGNNNAGATCGGAA
54726 TATGAGTGTGGAGTCGTTGCATCCAAGCAGATCCAAACCATCTNNNAGATCGGAAG
54727 TATGAGTGTGGAGTCGTTGCAGGTTCAAGCGATTCTCTTGTCTNNNAGATCGGAAG
54728 TATGAGTGTGGAGTCGTTGCGGTTCAAGTGATTCTCCTGCTTCNNNAGATCGGAAG
54729 TATGAGTGTGGAGTCGTTGCCCTGGGTTCAAGCAATTCTTCTGNNNAGATCGGAAG
54730 TATGAGTGTGGAGTCGTTGCAAGCACAGTATTACACCCTTGCTNNNAGATCGGAAG
54732 TATGAGTGTGGAGTCGTTGCAAGTAATTCTCTGCCTCAGCCTCNNNAGATCGGAAG
54734 TATGAGTGTGGAGTCGTTGCCTCAAGCAATTCTCCTGCCTCAGNNNAGATCGGAAG
54736 TATGAGTGTGGAGTCGTTGCCACCATGCCCAGCTAATTTTTGTNNNAGATCGGAAG
54738 TATGAGTGTGGAGTCGTTGCGATTCCAGACGTGAGTCAGAGTTNNNAGATCGGAA
54741 TATGAGTGTGGAGTCGTTGCACCTCATTGCACTCCATGTACATNNNAGATCGGAAGA
54742 TATGAGTGTGGAGTCGTTGCTGGTGCAAATAGGGATCATGAGANNNAGATCGGAA
54743 TATGAGTGTGGAGTCGTTGCATGATCTCAGTTCACAGCAACCTNNNAGATCGGAAG
54748 TATGAGTGTGGAGTCGTTGCACCACTGGAAGCTCGAAGTATTTNNNAGATCGGAAG

54749 TATGAGTGTGGAGTCGTTGCCTCACTGCAAGCTCTGAATTGAGNNNAGATCGGAAG
54750 TATGAGTGTGGAGTCGTTGCTCTGTTGAATTTGTTGCCTGCTTNNNAGATCGGAAG
54751 TATGAGTGTGGAGTCGTTGCACAACATCTGTGGCAACACTTTANNNAGATCGGAAG
54753 TATGAGTGTGGAGTCGTTGCCAGTGTCATAATCTCGGCTCACTNNNAGATCGGAAG
54755 TATGAGTGTGGAGTCGTTGCGAATAGCTGGGAATACAGGCACCNNNAGATCGGAA
54757 TATGAGTGTGGAGTCGTTGCGATCATTCTTGGGTGTTTCTCGCNNNAGATCGGAAG
54758 TATGAGTGTGGAGTCGTTGCCCTGCCTCAGTCAGACCTTTTTANNNAGATCGGAAGA
54760 TATGAGTGTGGAGTCGTTGCCAAGTAGCTGGGATTACAGGCATNNNAGATCGGAA
54762 TATGAGTGTGGAGTCGTTGCACAGAGGACCTAAAATAACCCAANNNAGATCGGAAG
54763 TATGAGTGTGGAGTCGTTGCGTGAGCCATTTTGTTGGGTGTAANNNAGATCGGAA
54764 TATGAGTGTGGAGTCGTTGCCTTCCTTTCCAGAATGCAACCACNNNAGATCGGAAGA
54765 TATGAGTGTGGAGTCGTTGCAAAGGCTGAAACAGATGCAAACANNNAGATCGGAA
54766 TATGAGTGTGGAGTCGTTGCTCTATCCCTCCTCCACTCATCTGNNNAGATCGGAAGA
54767 TATGAGTGTGGAGTCGTTGCAAGGAAAAGGAGTGTTCAATGGCNNNAGATCGGAA
54768 TATGAGTGTGGAGTCGTTGCGAGAGAGATGACAAAGGAGTGCANNNAGATCGGA
54769 TATGAGTGTGGAGTCGTTGCTTGCGTGTAATTGTGTATCCTGCNNNAGATCGGAAG
54770 TATGAGTGTGGAGTCGTTGCACCTTTTGCTCATGCCCTAAGTANNNAGATCGGAAGA
54771 TATGAGTGTGGAGTCGTTGCGAAGGGACTCTTCCTCAACTCATNNNAGATCGGAAG
54773 TATGAGTGTGGAGTCGTTGCGCCAGAGAGTGAGCTTTTCAAAGNNNAGATCGGAA
54774 TATGAGTGTGGAGTCGTTGCAAAGTAATGGCATGCTACTGGGANNNAGATCGGAA
54775 TATGAGTGTGGAGTCGTTGCCTCCAAGGTCTCTTTCAACTCCANNNAGATCGGAAGA
54777 TATGAGTGTGGAGTCGTTGCAGGATGGAATAGGGAGGATCCTTNNNAGATCGGAA
54779 TATGAGTGTGGAGTCGTTGCACCAAAACGTGAGGACCTGATTANNNAGATCGGAAG
54782 TATGAGTGTGGAGTCGTTGCGGGAAGAGAAGTGGGTATGACTGNNNAGATCGGA
54784 TATGAGTGTGGAGTCGTTGCCTGAGGCAGGACAATCACTTGAANNNAGATCGGAA
54785 TATGAGTGTGGAGTCGTTGCGGCTTGGGTTTGGTTTTGTGATANNNAGATCGGAA
54786 TATGAGTGTGGAGTCGTTGCAAAATCCTCTGTCCCCTCTGAACNNNAGATCGGAAGA
54787 TATGAGTGTGGAGTCGTTGCAGGCTCACTTTTCCTGAATTGGANNNAGATCGGAAG
54788 TATGAGTGTGGAGTCGTTGCTTTTACAGGTTAGTGCTGCATGGNNNAGATCGGAAG
54792 TATGAGTGTGGAGTCGTTGCGTGGCAATCCTAGTCTCTGACTANNNAGATCGGAAG
54794 TATGAGTGTGGAGTCGTTGCACAGATTCTGGGGAGGATATGGANNNAGATCGGAA
54796 TATGAGTGTGGAGTCGTTGCTCAGTGCATAACCAAGGCCATATNNNAGATCGGAAG
54798 TATGAGTGTGGAGTCGTTGCCAGAATCTGGCAGTCCTAGCTAGNNNAGATCGGAAG
54801 TATGAGTGTGGAGTCGTTGCGCTACTTGGGAGGCTAAGGTTAGNNNAGATCGGAA
54802 TATGAGTGTGGAGTCGTTGCCAGTCATTGATGCCAGGACAAAGNNNAGATCGGAA
54803 TATGAGTGTGGAGTCGTTGCTCTCTTCATACATTCGCTTCTGCNNNAGATCGGAAGA
54804 TATGAGTGTGGAGTCGTTGCCAAAAATTAGCTGGGCATGGTGGNNNAGATCGGAA
54805 TATGAGTGTGGAGTCGTTGCACTATAAGCCCCCAGTCTCTTCTNNNAGATCGGAAGA
54807 TATGAGTGTGGAGTCGTTGCTGGACCTGATGCAGATCATGAAANNNAGATCGGAA
54808 TATGAGTGTGGAGTCGTTGCTGAATGGTGATGCTAGGCTATCCNNNAGATCGGAA
54809 TATGAGTGTGGAGTCGTTGCAGAGAGACTGACGACAATTTGCTNNNAGATCGGAA
54811 TATGAGTGTGGAGTCGTTGCCGACTGAATTGCTGCAATCTCATNNNAGATCGGAAG
54812 TATGAGTGTGGAGTCGTTGCTTGGCATGAGCAACTAAAAAGGGNNNAGATCGGAA

54813 TATGAGTGTGGAGTCGTTGCACCTTGAGAAAAGTTTCCCGAGANNNAGATCGGAAG
54814 TATGAGTGTGGAGTCGTTGCGATGAGTTCATGTCCTTTGCAGGNNNAGATCGGAAG
54816 TATGAGTGTGGAGTCGTTGCGACTGGCCAACATAGTGAAAACCNNNAGATCGGAA
54818 TATGAGTGTGGAGTCGTTGCGTAATCCTAGCTACTTGGGAGGCNNNAGATCGGAA
54820 TATGAGTGTGGAGTCGTTGCCCAGGAATTAGATCCCTACTCGCNNNAGATCGGAAG
54821 TATGAGTGTGGAGTCGTTGCGAATTGCTTGAATCTGGGAGGTGNNNAGATCGGAA
54822 TATGAGTGTGGAGTCGTTGCGTAGGAGAATCACTTGAACCCGTNNNAGATCGGAA
54823 TATGAGTGTGGAGTCGTTGCGCAAAGGCAGGATAATTGCTTGANNNAGATCGGAA
54824 TATGAGTGTGGAGTCGTTGCAACTGGGAGAATCAGCTGTGATCNNNAGATCGGAA
54827 TATGAGTGTGGAGTCGTTGCACTGGCTTGAAGATAATGGGGAANNNAGATCGGAA
54829 TATGAGTGTGGAGTCGTTGCAACCTGGTGAAACCCCATCTTTANNNAGATCGGAAG
54830 TATGAGTGTGGAGTCGTTGCTCGGAGATGGCATCTAGTTTGAGNNNAGATCGGAA
54832 TATGAGTGTGGAGTCGTTGCGGAGGAGGAGGAGGAAGAAAAGANNNAGATCGG
54833 TATGAGTGTGGAGTCGTTGCTCAAAGACTCCTGTCAACTGGACNNNAGATCGGAAG
54834 TATGAGTGTGGAGTCGTTGCAGCCAGATAGAAAGTTGCACAGANNNAGATCGGAA
54839 TATGAGTGTGGAGTCGTTGCGTGATCTCAGCTCACTACAACCTNNNAGATCGGAAG
54840 TATGAGTGTGGAGTCGTTGCCACGCCTGGCTAATTTGTTGTATNNNAGATCGGAAG
54842 TATGAGTGTGGAGTCGTTGCTCAGCCTCCCAAAGTTTTCAGATNNNAGATCGGAAG
54843 TATGAGTGTGGAGTCGTTGCCAATGATCTTGGCTCACTGCAACNNNAGATCGGAAG
54845 TATGAGTGTGGAGTCGTTGCCGATCTCCTGACCTCATGATCTGNNNAGATCGGAAG
54846 TATGAGTGTGGAGTCGTTGCCTTCCCTGAGTCAAACCATCCTCNNNAGATCGGAAGA
54847 TATGAGTGTGGAGTCGTTGCCCTCACTTCAACCTCTGTCCTTTNNNAGATCGGAAGA
54848 TATGAGTGTGGAGTCGTTGCGGGTTCAAGTGTTTCTCCTGTCTNNNAGATCGGAAG
54850 TATGAGTGTGGAGTCGTTGCCCTCCCAAGTAGCTGGAATGATGNNNAGATCGGAAG
54851 TATGAGTGTGGAGTCGTTGCTCCAGGTTCAAGTGATTCTCCTGNNNAGATCGGAAG
54852 TATGAGTGTGGAGTCGTTGCGTTCAAGAGATTCTCCTGCCTCANNNAGATCGGAAG
54853 TATGAGTGTGGAGTCGTTGCTTGAACTCCTGGGCTCTAGTGATNNNAGATCGGAAG
54856 TATGAGTGTGGAGTCGTTGCCTTGCTTGCTTTGGATGGAGAAANNNAGATCGGAA
54857 TATGAGTGTGGAGTCGTTGCGGTTCAAACAATTCTCCTGCCTCNNNAGATCGGAAG
54858 TATGAGTGTGGAGTCGTTGCTCTTGTATTGAACTCCTGGGCTCNNNAGATCGGAAG
54861 TATGAGTGTGGAGTCGTTGCTCAGTATCCAAAGGTGAGTCAGCNNNAGATCGGAA
54862 TATGAGTGTGGAGTCGTTGCATCCTGTGTGTTGGGAACCTTTANNNAGATCGGAAG
54863 TATGAGTGTGGAGTCGTTGCGTAGATGGGACTACAGGTGTGTGNNNAGATCGGAA
54865 TATGAGTGTGGAGTCGTTGCTGCAATCTCAGCTTACTGGCTTANNNAGATCGGAAG
54868 TATGAGTGTGGAGTCGTTGCGTTCAAGAGATTCTCCTGCCTCANNNAGATCGGAAG
54869 TATGAGTGTGGAGTCGTTGCTCCCACATTCATGCCATTCTCTCNNNAGATCGGAAGA
54870 TATGAGTGTGGAGTCGTTGCAGGTTCAACTGATTCTTCTGCCTNNNAGATCGGAAG
54872 TATGAGTGTGGAGTCGTTGCGTTCAAGCAATTCTCTTGCCTCANNNAGATCGGAAG
54873 TATGAGTGTGGAGTCGTTGCCACTGTCTCTGTCCCTTGCTAATNNNAGATCGGAAGA
54874 TATGAGTGTGGAGTCGTTGCCTTGAACTCCTGGCTGAAGGTATNNNAGATCGGAAG
54875 TATGAGTGTGGAGTCGTTGCTCCTGGATTCAACAGATTCTCCCNNNAGATCGGAAG
54878 TATGAGTGTGGAGTCGTTGCAACCTTTATCTCCCAGGTTCAGGNNNAGATCGGAAG
54880 TATGAGTGTGGAGTCGTTGCAGATGGAGTCTTCCTTTGTCACCNNNAGATCGGAAG

54881 TATGAGTGTGGAGTCGTTGCAAAGGTTCAAGCGATTCTTCTGCNNNAGATCGGAAG
54882 TATGAGTGTGGAGTCGTTGCTCAGGTTCAAACGATTTTCCTGCNNNAGATCGGAAG
54884 TATGAGTGTGGAGTCGTTGCCACAGGCTCTTTTTTGGTTCCATNNNAGATCGGAAG
54885 TATGAGTGTGGAGTCGTTGCGTTCAAGCAATTCTCTGCCTCAGNNNAGATCGGAAG
54886 TATGAGTGTGGAGTCGTTGCTTCAAGTGATTCTCCTGCCTCAGNNNAGATCGGAAG
54887 TATGAGTGTGGAGTCGTTGCCCTCTGGGGTTCAAGAGATTCTCNNNAGATCGGAAG
54890 TATGAGTGTGGAGTCGTTGCAGCCTCCTGAATAGCTGGGATTANNNAGATCGGAA
54891 TATGAGTGTGGAGTCGTTGCACAGTGACATGATCTCAGCTCTCNNNAGATCGGAAG
54894 TATGAGTGTGGAGTCGTTGCTTCAAGTGGTTCTCCTGTCTCAGNNNAGATCGGAAG
54895 TATGAGTGTGGAGTCGTTGCGAATAGCTGGGATTACACGTGTGNNNAGATCGGAA
54896 TATGAGTGTGGAGTCGTTGCTTATGGCTTACTGTGTCCTCGACNNNAGATCGGAAG
54898 TATGAGTGTGGAGTCGTTGCACTGCAGTCTCAACCTTCTAAGGNNNAGATCGGAAG
54899 TATGAGTGTGGAGTCGTTGCACCGCACTCAGCCTAAAAATTTTNNNAGATCGGAAG
54901 TATGAGTGTGGAGTCGTTGCGATTCAAGCGATTCTCCTGCCTCNNNAGATCGGAAG
54902 TATGAGTGTGGAGTCGTTGCGTTGGTGAGGATGCAGAGAAAAGNNNAGATCGGA
54903 TATGAGTGTGGAGTCGTTGCGCCGTGTGCTGAACATTTATCTTNNNAGATCGGAAG
54905 TATGAGTGTGGAGTCGTTGCAAGTGAAGAGACATCAGCAAGGANNNAGATCGGAA
54907 TATGAGTGTGGAGTCGTTGCCTGGGGTAACTACACACTAGCTCNNNAGATCGGAAG
54908 TATGAGTGTGGAGTCGTTGCCTGTAGTCCCAGCTACTCAGGAGNNNAGATCGGAAG
54911 TATGAGTGTGGAGTCGTTGCCTAACACCTGTAATCCCAGCACTNNNAGATCGGAAG
54912 TATGAGTGTGGAGTCGTTGCGGAAACGCAAAAGAAGTAGCTGTNNNAGATCGGAA
54915 TATGAGTGTGGAGTCGTTGCCCATCAGGATTTTGAGAGGGACTNNNAGATCGGAA
54916 TATGAGTGTGGAGTCGTTGCAAGAAAGAAAAAAAGGCCAGGCGNNNAGATCGGA
54919 TATGAGTGTGGAGTCGTTGCCAGTGAGCCAAAATCATGCCATTNNNAGATCGGAAG
54920 TATGAGTGTGGAGTCGTTGCCTGTAGTCCCAGCTACTTGAGAGNNNAGATCGGAAG
54922 TATGAGTGTGGAGTCGTTGCACTTTTACACTGTTGGTGGGACTNNNAGATCGGAAG
54923 TATGAGTGTGGAGTCGTTGCAAGCACCTTTTTTCCCTGTCAACNNNAGATCGGAAGA
54927 TATGAGTGTGGAGTCGTTGCCCTTCCCTTCCTTCCACTGTAATNNNAGATCGGAAGA
54929 TATGAGTGTGGAGTCGTTGCGGACTTGAGAACTCTCTAGCCTGNNNAGATCGGAAG
54932 TATGAGTGTGGAGTCGTTGCGGGCCATTGGAATTTCCTCAAAANNNAGATCGGAAG
54933 TATGAGTGTGGAGTCGTTGCAGTCCCAAACTGGATGGAAACAANNNAGATCGGAA
54934 TATGAGTGTGGAGTCGTTGCTGAGTGCAAATTCAAAACATGCANNNAGATCGGAA
54935 TATGAGTGTGGAGTCGTTGCACTGTAGGCATGTTAGCATCAACNNNAGATCGGAAG
54937 TATGAGTGTGGAGTCGTTGCACTACTTCCATAATGCCTGAGGTNNNAGATCGGAAG
54938 TATGAGTGTGGAGTCGTTGCTCCACAGCTCTCTTTCAATTCCANNNAGATCGGAAGA
54943 TATGAGTGTGGAGTCGTTGCAAAAATTAGGCTGGGTTTGCTGGNNNAGATCGGAA
54944 TATGAGTGTGGAGTCGTTGCCCTGGGAGGTCGAGGTTATAATGNNNAGATCGGAA
54946 TATGAGTGTGGAGTCGTTGCAGCACAATTGTAAAGCATGCACTNNNAGATCGGAAG
54947 TATGAGTGTGGAGTCGTTGCTCTCCAAAGTGTTCTCATGTGGCNNNAGATCGGAAG
54948 TATGAGTGTGGAGTCGTTGCTCACTTGAGAGGTCAGGAATTCGNNNAGATCGGAA
54949 TATGAGTGTGGAGTCGTTGCTTCTGTCAGAGCAGCTCTTGTTANNNAGATCGGAAG
54951 TATGAGTGTGGAGTCGTTGCCCATGAACTTACAACCCAGCTTGNNNAGATCGGAAG
54953 TATGAGTGTGGAGTCGTTGCGGAAATTGTCGAGTTGCTGTGAGNNNAGATCGGAA

54954 TATGAGTGTGGAGTCGTTGCGTAGGAGATAGTAGTTGCCCTGCNNNAGATCGGAA
54955 TATGAGTGTGGAGTCGTTGCTGAACTCTGGATTTTTTTCCCGANNNAGATCGGAAG
54956 TATGAGTGTGGAGTCGTTGCACAAGGTGAAACCCAGTCTCTACNNNAGATCGGAAG
54958 TATGAGTGTGGAGTCGTTGCAATCTGATGAAAGCTGTGGACCTNNNAGATCGGAA
54960 TATGAGTGTGGAGTCGTTGCGGTAAGTCACAGAGATCATGGCANNNAGATCGGAA
54961 TATGAGTGTGGAGTCGTTGCCTCCTGTAATCCCAGCACTTAGGNNNAGATCGGAAG
54962 TATGAGTGTGGAGTCGTTGCACTAAAGGTGAGCAGAGGAATGGNNNAGATCGGA
54965 TATGAGTGTGGAGTCGTTGCTTACATTCTGCTTGAAGATGGGCNNNAGATCGGAAG
54966 TATGAGTGTGGAGTCGTTGCAATTGGCTGTGGCAGATACTACANNNAGATCGGAA
54969 TATGAGTGTGGAGTCGTTGCGTGAGCTGAGATCATGCTATTGCNNNAGATCGGAA
54973 TATGAGTGTGGAGTCGTTGCCAACCTCTGTCTCCTCAAGTTCANNNAGATCGGAAGA
54974 TATGAGTGTGGAGTCGTTGCCCTGAGTAGCTGGGATTATAGGCNNNAGATCGGAA
54975 TATGAGTGTGGAGTCGTTGCAGATGGGTTTTCTCCATGTTGGTNNNAGATCGGAAG
54976 TATGAGTGTGGAGTCGTTGCATCAGTTTCTCATCGGAGCTTGGNNNAGATCGGAAG
54977 TATGAGTGTGGAGTCGTTGCTAGTTGGGACTACAGGTGCAAACNNNAGATCGGAA
54978 TATGAGTGTGGAGTCGTTGCCATGATCTCAGCTCATTGCAACCNNNAGATCGGAAG
54979 TATGAGTGTGGAGTCGTTGCTTCAAGCGATTCTCCTGTCTCAGNNNAGATCGGAAG
54981 TATGAGTGTGGAGTCGTTGCACGTACTGGATACATGGGCTTTTNNNAGATCGGAAG
54983 TATGAGTGTGGAGTCGTTGCGGTTTAGGCAATTCTCGTGTCTCNNNAGATCGGAAG
54985 TATGAGTGTGGAGTCGTTGCGATCTGAGCTCACTGTAACCTCCNNNAGATCGGAAG
54986 TATGAGTGTGGAGTCGTTGCACCAAGACAAAGAAAATGCAGAANNNAGATCGGAA
54987 TATGAGTGTGGAGTCGTTGCGGCAGAATCTCACTCTGTCACTTNNNAGATCGGAAG
54988 TATGAGTGTGGAGTCGTTGCCGCCCGGCCAATAGTTATTATAGNNNAGATCGGAAG
54989 TATGAGTGTGGAGTCGTTGCAGCCTCCTGAGTAGCTGGAATTANNNAGATCGGAA
54990 TATGAGTGTGGAGTCGTTGCGATGGAGTCTCACTCTGTCACTCNNNAGATCGGAAG
54991 TATGAGTGTGGAGTCGTTGCAAGTGTTGGGATTACAGGCATGANNNAGATCGGAA
54992 TATGAGTGTGGAGTCGTTGCTTCAAGTGATTCTCCTGCCTCAGNNNAGATCGGAAG
54993 TATGAGTGTGGAGTCGTTGCCTCCTGAGTAGCTGGGATTACAGNNNAGATCGGAA
54994 TATGAGTGTGGAGTCGTTGCTGGTCTCAAATTCCTGAGCTCAANNNAGATCGGAAG
54996 TATGAGTGTGGAGTCGTTGCAAGGCATTAGGAATACTTGGGCANNNAGATCGGAA
54998 TATGAGTGTGGAGTCGTTGCCTCCTGAGTAGCTGAGATTGCAGNNNAGATCGGAA
54999 TATGAGTGTGGAGTCGTTGCACAGATAGGGTCTCACTGTGTTGNNNAGATCGGAA
55000 TATGAGTGTGGAGTCGTTGCCTTCCAAGTAGCTGGGACAGTAGNNNAGATCGGAA
55002 TATGAGTGTGGAGTCGTTGCAAAGTAACAACTGTTGAAGCGGGNNNAGATCGGAA
55003 TATGAGTGTGGAGTCGTTGCCCTGAGTAGCTGGAATTACAGGTNNNAGATCGGAA
55010 TATGAGTGTGGAGTCGTTGCAGTTTCCAGCCTTGACCACATAANNNAGATCGGAAG
55011 TATGAGTGTGGAGTCGTTGCCCACTGATAGATGCTGGTCTGTANNNAGATCGGAAG
55012 TATGAGTGTGGAGTCGTTGCTTGTGTGGATTCACTGAAAAGCANNNAGATCGGAA
55013 TATGAGTGTGGAGTCGTTGCAAACCTCTGGGACTGGTAGAAACNNNAGATCGGAA
55014 TATGAGTGTGGAGTCGTTGCGGGGAAAAGACACAAAAAGCTGANNNAGATCGGA
55015 TATGAGTGTGGAGTCGTTGCTCCTCCCTCTTCACGTTTTTTCANNNAGATCGGAAGA
55018 TATGAGTGTGGAGTCGTTGCGCAGCAAGCTGGTAAAGAAGTTTNNNAGATCGGAA
55019 TATGAGTGTGGAGTCGTTGCAGACACACACACATACACACACANNNAGATCGGAAG

55020 TATGAGTGTGGAGTCGTTGCTCTAGACACAGACCTTACACCCTNNNAGATCGGAAGA
55024 TATGAGTGTGGAGTCGTTGCTGTATGTTCAGCCTTCACCACTGNNNAGATCGGAAGA
55027 TATGAGTGTGGAGTCGTTGCGAGGGTGGCAACTAAAACATCACNNNAGATCGGAAG
55028 TATGAGTGTGGAGTCGTTGCTGGAAAAAGTCTATCACAGGCCANNNAGATCGGAAG
55029 TATGAGTGTGGAGTCGTTGCTGGAGAGAGAAGGAGAAAGAGGTNNNAGATCGGA
55033 TATGAGTGTGGAGTCGTTGCGCTACTTGCTATGTCCCCTAAGGNNNAGATCGGAAGA
55036 TATGAGTGTGGAGTCGTTGCGAGCAGAACAGAGATCCCAGAAANNNAGATCGGAA
55038 TATGAGTGTGGAGTCGTTGCTGCAATTCAGAAATACAACAAGCTCTNNNAGATCGGA
55043 TATGAGTGTGGAGTCGTTGCGAATGAGGATCTCACTGTGTTGCNNNAGATCGGAAG
55044 TATGAGTGTGGAGTCGTTGCTTCACGAGATTCTTCTGCCTCAGNNNAGATCGGAAGA
55047 TATGAGTGTGGAGTCGTTGCCATCCCCACAAAATGCTGGAATTNNNAGATCGGAAGA
55048 TATGAGTGTGGAGTCGTTGCATGATCTTGGCTCACTGTAAGCTNNNAGATCGGAAG
55049 TATGAGTGTGGAGTCGTTGCTGTTAGCCAGGATGGTCTTGATCNNNAGATCGGAAG
55050 TATGAGTGTGGAGTCGTTGCCTGGGTTCAAAAGATCCTTCTGCNNNAGATCGGAAG
55051 TATGAGTGTGGAGTCGTTGCAGACTGGAGTGCAATGGTACAATNNNAGATCGGAAG
55053 TATGAGTGTGGAGTCGTTGCAGACTCCTGGGCTATAGAGATCCNNNAGATCGGAAG
55054 TATGAGTGTGGAGTCGTTGCCTCCCGAGTAGCTGTGACTACAGNNNAGATCGGAAG
55057 TATGAGTGTGGAGTCGTTGCCCTCCTGAGTAGCTGGGATTACANNNAGATCGGAAG
55058 TATGAGTGTGGAGTCGTTGCACAAGATCTTGCTCTGTCACTCANNNAGATCGGAAGA
55059 TATGAGTGTGGAGTCGTTGCCTCTCAAGTAGCTGGGACTACAGNNNAGATCGGAAG
55060 TATGAGTGTGGAGTCGTTGCTTCAAGTGATGCTCCTGCCTTAGNNNAGATCGGAAGA
55061 TATGAGTGTGGAGTCGTTGCAGACCCCCAAGTAGCTAGGATTANNNAGATCGGAAG
55062 TATGAGTGTGGAGTCGTTGCTTCAAGTGATTCTCCTGCCTCAGNNNAGATCGGAAGA
55063 TATGAGTGTGGAGTCGTTGCAGAGGTGTGCAATCATAGCTCAANNNAGATCGGAAG
55064 TATGAGTGTGGAGTCGTTGCGTTCCAGTGATTCTCCTGCCTAANNNAGATCGGAAGA
55065 TATGAGTGTGGAGTCGTTGCCTGGAATGTAGTGGTGCGATCTCNNNAGATCGGAAG
55071 TATGAGTGTGGAGTCGTTGCGTTTCACCATGTTAGCCAGGATGNNNAGATCGGAAG
55072 TATGAGTGTGGAGTCGTTGCTCCAGGTTCAAGTGATTCTCCTGNNNAGATCGGAAGA
55073 TATGAGTGTGGAGTCGTTGCGAGTAGCTGGGATTACAGATGCANNNAGATCGGAA
55075 TATGAGTGTGGAGTCGTTGCCAATGGCGTGATCTCAGCTTACTNNNAGATCGGAAG
55081 TATGAGTGTGGAGTCGTTGCTTCAAGTGAGCCTCTCAACTCAGNNNAGATCGGAAGA
55082 TATGAGTGTGGAGTCGTTGCCTCCTGAGTAGCTGGGACTACAGNNNAGATCGGAAG
55084 TATGAGTGTGGAGTCGTTGCAAGTAGCTGGGATTACAGGCATCNNNAGATCGGAAG
55086 TATGAGTGTGGAGTCGTTGCTCAGGCTGATCTAGAACTCCTGANNNAGATCGGAAG
55087 TATGAGTGTGGAGTCGTTGCAGTAGAGACGGGGTTTTACCAAGNNNAGATCGGAA
55088 TATGAGTGTGGAGTCGTTGCCTGACTCCCAGGTTCAAGCTATTNNNAGATCGGAAGA
55089 TATGAGTGTGGAGTCGTTGCTCAAACAATTCTCCTGCTTCAGCNNNAGATCGGAAGA
55091 TATGAGTGTGGAGTCGTTGCCAAGTAGCTGGGATTACCGACATNNNAGATCGGAAG
55093 TATGAGTGTGGAGTCGTTGCCTCACTACAGCTTTGACCTCCTGNNNAGATCGGAAGA
55094 TATGAGTGTGGAGTCGTTGCTGTGTGAATGGGGCATGATAGAANNNAGATCGGAA
55097 TATGAGTGTGGAGTCGTTGCACACAGTGAAACCCCATCTCTACNNNAGATCGGAAGA
55100 TATGAGTGTGGAGTCGTTGCTGCAGAAGAAAGAACTCACTCCANNNAGATCGGAAG
55104 TATGAGTGTGGAGTCGTTGCATGTGATGGCCAATCCTGATCATNNNAGATCGGAAG

55105 TATGAGTGTGGAGTCGTTGCAATGTCTTCCTAGAAGTGGGAGCNNNAGATCGGAA
55106 TATGAGTGTGGAGTCGTTGCAGGCTGTACTTTTAGGAGAGAAGCNNNAGATCGGA
55107 TATGAGTGTGGAGTCGTTGCTCCTGCCTGTAATCCCAACATTTNNNAGATCGGAAGA
55108 TATGAGTGTGGAGTCGTTGCGAGACCTCCACTTTCTGTCTGAGNNNAGATCGGAAG
55109 TATGAGTGTGGAGTCGTTGCTCTCAGAGCTCAAAGACTGGTTCNNNAGATCGGAAG
55110 TATGAGTGTGGAGTCGTTGCAGGAAGGACAAGGAAGGAAAGACNNNAGATCGGA
55112 TATGAGTGTGGAGTCGTTGCCACTGGGAATTTTGGTCACCAAGNNNAGATCGGAAG
55113 TATGAGTGTGGAGTCGTTGCTATTGCAGTGAGCTGAAATCACGNNNAGATCGGAA
55115 TATGAGTGTGGAGTCGTTGCGCTGTACTCTGTTGGGAGACAATNNNAGATCGGAA
55116 TATGAGTGTGGAGTCGTTGCGAAAATATTCGGGCCTAGGCAACNNNAGATCGGAA
55117 TATGAGTGTGGAGTCGTTGCCCCCTATGCCCAAAGTGAACATANNNAGATCGGAAG
55118 TATGAGTGTGGAGTCGTTGCGCTGTACCTGAAACACAAGCTTTNNNAGATCGGAAG
55119 TATGAGTGTGGAGTCGTTGCACGTGTGTATATACGCGCGTATANNNAGATCGGAAG
55120 TATGAGTGTGGAGTCGTTGCTAACTCACACCTGTAATCCCAGCNNNAGATCGGAAG
55121 TATGAGTGTGGAGTCGTTGCCTGAGACAGGAGGATCACATGAGNNNAGATCGGAA
55124 TATGAGTGTGGAGTCGTTGCTATGCTCCCATGTCATATGCTCCNNNAGATCGGAAG
55125 TATGAGTGTGGAGTCGTTGCTGCTAATTTGGTGAAAACCCGTCNNNAGATCGGAAG
55126 TATGAGTGTGGAGTCGTTGCATCATCTACATCTTTGCCTCCCTNNNAGATCGGAAGA
55128 TATGAGTGTGGAGTCGTTGCGGAAAATTCTAGCCCCAGGGATTNNNAGATCGGAA
55129 TATGAGTGTGGAGTCGTTGCAAAAGAGACACACACACACTCCANNNAGATCGGAAG
55131 TATGAGTGTGGAGTCGTTGCCGTATGTAACAAACCTGCACGTTNNNAGATCGGAAG
55132 TATGAGTGTGGAGTCGTTGCAGCATGTCAGTAACAATGGCTTTNNNAGATCGGAAG
55134 TATGAGTGTGGAGTCGTTGCGCTAAGAACTGTGGCTAGCAAATNNNAGATCGGAA
55136 TATGAGTGTGGAGTCGTTGCAGGCAGAAGTTGTAGTAAGCCAANNNAGATCGGAA
55138 TATGAGTGTGGAGTCGTTGCGCATCATCATTTCTTCAAGGTGCNNNAGATCGGAAG
55139 TATGAGTGTGGAGTCGTTGCAGAGTGCATGTAAAAACTGGTGANNNAGATCGGAA
55140 TATGAGTGTGGAGTCGTTGCACAAAAAGCAAAAAAAGGGGGGANNNAGATCGGA
55141 TATGAGTGTGGAGTCGTTGCGATCTGCTTCATGGTAAAGACGANNNAGATCGGAA
55143 TATGAGTGTGGAGTCGTTGCACCAAAGTTGCTGAAAGTCTGAANNNAGATCGGAA
55144 TATGAGTGTGGAGTCGTTGCTTTTTATTTTGGGAGGGTGCTGCNNNAGATCGGAAG
55146 TATGAGTGTGGAGTCGTTGCCACAAAAACCTTCAAGGGGCTTANNNAGATCGGAAG
55147 TATGAGTGTGGAGTCGTTGCAGCACAGGAAAGAATTGATTGCCNNNAGATCGGAA
55154 TATGAGTGTGGAGTCGTTGCGTCAGGCTTTACATGAACAGCAANNNAGATCGGAAG
55160 TATGAGTGTGGAGTCGTTGCCCAGTGGTCTCTAACTCTTAGGCNNNAGATCGGAAG
55161 TATGAGTGTGGAGTCGTTGCGCCTCCTGAGTAGCTGAGATTACNNNAGATCGGAAG
55164 TATGAGTGTGGAGTCGTTGCTAAGCACATAGGATGACCACACCNNNAGATCGGAAG
55166 TATGAGTGTGGAGTCGTTGCCAAGATGGTGTCAATGTCCTGACNNNAGATCGGAAG
55167 TATGAGTGTGGAGTCGTTGCAACAGAAGTCAAGGATCAGGCATNNNAGATCGGAA
55168 TATGAGTGTGGAGTCGTTGCCAAGTAGCTGGGATTACAGGCATNNNAGATCGGAA
55171 TATGAGTGTGGAGTCGTTGCCTTGGAAGGAGCCTTGAGTTTTGNNNAGATCGGAA
55172 TATGAGTGTGGAGTCGTTGCCTCACAAGTAGCTGGGACTACAGNNNAGATCGGAA
55173 TATGAGTGTGGAGTCGTTGCGGAATGGGTAGGAGAAGCATTGANNNAGATCGGA
55176 TATGAGTGTGGAGTCGTTGCCTGGGTTCAAGTGATTCTCTTGCNNNAGATCGGAAG

55178 TATGAGTGTGGAGTCGTTGCAATTCTCTCATCTCAGCCTTCCGNNNAGATCGGAAGA
55179 TATGAGTGTGGAGTCGTTGCCCAGTGTGAACATCCAAAGCAAANNNAGATCGGAAG
55180 TATGAGTGTGGAGTCGTTGCCTCCTGAGTAGCTGGGACTACAGNNNAGATCGGAAG
55181 TATGAGTGTGGAGTCGTTGCATGGCTGCATAGTATTCCATGGTNNNAGATCGGAAG
55182 TATGAGTGTGGAGTCGTTGCCTGGAGTGAAATGGCACAATCTCNNNAGATCGGAAG
55183 TATGAGTGTGGAGTCGTTGCTCCCAACTAGCTGAGTCTACAGANNNAGATCGGAAG
55187 TATGAGTGTGGAGTCGTTGCTCAAAGTATTCTCCTGCCTCAGCNNNAGATCGGAAGA
55191 TATGAGTGTGGAGTCGTTGCAATAGCTGGGATTACAGGCATGTNNNAGATCGGAAG
55192 TATGAGTGTGGAGTCGTTGCCAATGGCGTGATCTTGGCTTATTNNNAGATCGGAAG
55193 TATGAGTGTGGAGTCGTTGCAACCTCCTGAGTAGCTGAGACTANNNAGATCGGAAG
55194 TATGAGTGTGGAGTCGTTGCCCACCTCACCCAGCTAATTTTGNNNAGATCGGAAGA
55195 TATGAGTGTGGAGTCGTTGCCACAATATCAGCTCACTGCAACCNNNAGATCGGAAGA
55196 TATGAGTGTGGAGTCGTTGCCTGCCTCCTGGATCCAAGTTATTNNNAGATCGGAAGA
55198 TATGAGTGTGGAGTCGTTGCCTCCCAAGTAGCTGGAACTACAGNNNAGATCGGAAG
55199 TATGAGTGTGGAGTCGTTGCCTCAGTTGATCCTCCTACCTCTGNNNAGATCGGAAGA
55200 TATGAGTGTGGAGTCGTTGCTTCAAGTGATTCTCCTGCCTCAGNNNAGATCGGAAGA
55205 TATGAGTGTGGAGTCGTTGCAGGTTCAAGTGATTCTTCTGCCTNNNAGATCGGAAGA
55206 TATGAGTGTGGAGTCGTTGCCTCAAGCAATCCTCCCACCTTAGNNNAGATCGGAAGA
55208 TATGAGTGTGGAGTCGTTGCCCAAATAGCTGGGACTATGGGTGNNNAGATCGGAAG
55210 TATGAGTGTGGAGTCGTTGCCACTCATAGGTGGGAACTGAACANNNAGATCGGAAG
55211 TATGAGTGTGGAGTCGTTGCGATTCACGTCATTCTCCTGCCTCNNNAGATCGGAAGA
55212 TATGAGTGTGGAGTCGTTGCCCTCCTGGGTTCAAGTGATTCTCNNNAGATCGGAAGA
55213 TATGAGTGTGGAGTCGTTGCGTTCAAGCAATTCTCTGCCTCAGNNNAGATCGGAAGA
55215 TATGAGTGTGGAGTCGTTGCCTGAGTAACTGGGACTACAGGTGNNNAGATCGGAAG
55216 TATGAGTGTGGAGTCGTTGCTCCCAAGTAGCTGGAATTACAGGNNNAGATCGGAAG
55217 TATGAGTGTGGAGTCGTTGCGAGTAGCTGGGACTACAGGTATGNNNAGATCGGAA
55223 TATGAGTGTGGAGTCGTTGCATGCCCTAATCAGCACTCTGTAGNNNAGATCGGAAGA
55224 TATGAGTGTGGAGTCGTTGCTTCCCAATTCTGTGAGGCCAATANNNAGATCGGAAGA
55228 TATGAGTGTGGAGTCGTTGCCCTCTTGCACGGAAAAGCTTTTANNNAGATCGGAAGA
55229 TATGAGTGTGGAGTCGTTGCAGAATCACTGGCGTAGGTATGACNNNAGATCGGAAG
55230 TATGAGTGTGGAGTCGTTGCCAAGGTCATGCAGTAAGTTGGTGNNNAGATCGGAAG
55231 TATGAGTGTGGAGTCGTTGCATGGCAATAAAATGGTTCAGGGCNNNAGATCGGAAG
55233 TATGAGTGTGGAGTCGTTGCACAAAACAAAGGATCAGCAAAGTNNNAGATCGGAAG
55234 TATGAGTGTGGAGTCGTTGCCTTTGAGGTCATCTCCAGACACANNNAGATCGGAAGA
55236 TATGAGTGTGGAGTCGTTGCTCACATGAGGTCAGGAGTTCAAGNNNAGATCGGAAG
55237 TATGAGTGTGGAGTCGTTGCGTGAGCTATGATCATGCCTCTGANNNAGATCGGAAG
55239 TATGAGTGTGGAGTCGTTGCCACACTCCTACCATACGATCCAGNNNAGATCGGAAGA
55240 TATGAGTGTGGAGTCGTTGCCTGTGATTGTGTTACTGCACTCCNNNAGATCGGAAGA
55241 TATGAGTGTGGAGTCGTTGCCAACAACAAAAACAAGATGGCCGNNNAGATCGGAAG
55243 TATGAGTGTGGAGTCGTTGCAGGACTATTTCCGGCTTGAAAGTNNNAGATCGGAAG
55244 TATGAGTGTGGAGTCGTTGCTGGGAGGAAGGAAGGAAATTCTGNNNAGATCGGAA
55245 TATGAGTGTGGAGTCGTTGCGGGCTGAGAAAAATGAATAGCCANNNAGATCGGAA
55246 TATGAGTGTGGAGTCGTTGCAGCCAAACGTTTTGGAAAAGTCANNNAGATCGGAAG

55248 TATGAGTGTGGAGTCGTTGCTGTAATCCCAGCTACTCAGGAGANNNAGATCGGAAG
55252 TATGAGTGTGGAGTCGTTGCCAAAAATTGATGTTGGCACGGACNNNAGATCGGAA
55253 TATGAGTGTGGAGTCGTTGCGGCAATTTGAGACAACACTTCGANNNAGATCGGAA
55256 TATGAGTGTGGAGTCGTTGCTGAACTCCACAAAATTACTCCCANNNAGATCGGAAG
55258 TATGAGTGTGGAGTCGTTGCTGAGGTGACTGATACACTGTTGTNNNAGATCGGAA
55259 TATGAGTGTGGAGTCGTTGCATAATACCGTGGGTTGACTGGTCNNNAGATCGGAA
55260 TATGAGTGTGGAGTCGTTGCGTCCAGCTACTCAAGAGGCTAAGNNNAGATCGGAA
55263 TATGAGTGTGGAGTCGTTGCTTTCCCAGAAACAGAGCATTTGCNNNAGATCGGAAG
55264 TATGAGTGTGGAGTCGTTGCCCAGAGGTAGTGTTACTGTGCATNNNAGATCGGAA
55265 TATGAGTGTGGAGTCGTTGCCTATGTTGTACTGAGCACTTGCCNNNAGATCGGAAG
55266 TATGAGTGTGGAGTCGTTGCTGTTTTCAAAATGGAGCAGCTGTNNNAGATCGGAA
55268 TATGAGTGTGGAGTCGTTGCCCTCATCAGGCAGTCAAATCAACNNNAGATCGGAAG
55269 TATGAGTGTGGAGTCGTTGCCACACACACACACACACACAATANNNAGATCGGAAG
55270 TATGAGTGTGGAGTCGTTGCAACTCAAGGGGGTGTCAATCATANNNAGATCGGAA
55271 TATGAGTGTGGAGTCGTTGCAGGGCAATCCATAGAGCTGAATTNNNAGATCGGAA
55272 TATGAGTGTGGAGTCGTTGCGACAGAAAGGATACCATGGACCCNNNAGATCGGAA
55273 TATGAGTGTGGAGTCGTTGCAAAAAATTAGCCTGGCATGGTGGNNNAGATCGGAA
55274 TATGAGTGTGGAGTCGTTGCACACAATCCAGAGGAAATGGTCANNNAGATCGGAA
55275 TATGAGTGTGGAGTCGTTGCACTCTTGATTACCTTGAAGCTGGANNNAGATCGGAA
55279 TATGAGTGTGGAGTCGTTGCTTGAGATTTTGGACTGCCAGTCTNNNAGATCGGAAG
55280 TATGAGTGTGGAGTCGTTGCACACAGGTGACTCTAAGGAAAGGNNNAGATCGGAA
55281 TATGAGTGTGGAGTCGTTGCGTGATCCACCCCTATTATCCCAGNNNAGATCGGAAG
55285 TATGAGTGTGGAGTCGTTGCCATAGTGGAAAGAGCTTAGCCCANNNAGATCGGAA
55287 TATGAGTGTGGAGTCGTTGCGCCCCTTCGTTACACACTGAATANNNAGATCGGAAG
55288 TATGAGTGTGGAGTCGTTGCCTCTGTGGCACCTATCACAACTANNNAGATCGGAAG
55289 TATGAGTGTGGAGTCGTTGCTAGATTTGAATGCTGGCCACTCTNNNAGATCGGAAG
55290 TATGAGTGTGGAGTCGTTGCTGCCCTACCTACCTTGTATTGAANNNAGATCGGAAG
55291 TATGAGTGTGGAGTCGTTGCCGAAGAAGAAGAAGGCCTGAATANNNAGATCGGAA
55292 TATGAGTGTGGAGTCGTTGCATAGGGAGTAGGGAGACTAGGGANNNAGATCGGA
55294 TATGAGTGTGGAGTCGTTGCACTTTTTTTTGCAGGGATAGGGCNNNAGATCGGAA
55296 TATGAGTGTGGAGTCGTTGCTCAAGTGATTCTCCTGCCTTAGCNNNAGATCGGAAG
55297 TATGAGTGTGGAGTCGTTGCGGTTCAAGTGATTTTCCTGCCTCNNNAGATCGGAAG
55299 TATGAGTGTGGAGTCGTTGCCTCCAGAGTAGCAGGGATTACAGNNNAGATCGGAA
55301 TATGAGTGTGGAGTCGTTGCGTTCAAGTGATTCTCCTCCCTCANNNAGATCGGAAG
55302 TATGAGTGTGGAGTCGTTGCTAGAAACAGGGTCTCACTCCTTGNNNAGATCGGAAG
55304 TATGAGTGTGGAGTCGTTGCCTTCCTGTGTAGCTGGGATTACANNNAGATCGGAAG
55305 TATGAGTGTGGAGTCGTTGCCTCACGAGTAGCTGGGATAACAGNNNAGATCGGAA
55306 TATGAGTGTGGAGTCGTTGCCTCCTGAGTAGCTGGGATTACAGNNNAGATCGGAA
55307 TATGAGTGTGGAGTCGTTGCTGCTCTGCTGTCACCTATCATAGNNNAGATCGGAAG
55308 TATGAGTGTGGAGTCGTTGCTACAAATATGAGCCTCCACAGCCNNNAGATCGGAAG
55309 TATGAGTGTGGAGTCGTTGCAGTGATCTTTCCACCTCAACCTCNNNAGATCGGAAGA
55310 TATGAGTGTGGAGTCGTTGCGCTAGAGTGTAGTGGTGCAATCTNNNAGATCGGAA
55311 TATGAGTGTGGAGTCGTTGCCCTCCCAGGTTCAAGCAATTCTCNNNAGATCGGAAG

55312 TATGAGTGTGGAGTCGTTGCTTTGGGTTCAAGCAATTCTCCTGNNNAGATCGGAAG
55313 TATGAGTGTGGAGTCGTTGCCCAAGTAGCTGGGATTACAGGTGNNNAGATCGGAA
55314 TATGAGTGTGGAGTCGTTGCGAGATGGAGTTTCGTGCTTGTTGNNNAGATCGGAA
55316 TATGAGTGTGGAGTCGTTGCTTCATGCCATTCTCCTGTCTCAGNNNAGATCGGAAGA
55317 TATGAGTGTGGAGTCGTTGCTTCATGCCATTCTCCTGTCTCAGNNNAGATCGGAAGA
55318 TATGAGTGTGGAGTCGTTGCCCTAGTAGCTGGGATTACAGGTGNNNAGATCGGAA
55320 TATGAGTGTGGAGTCGTTGCCTGGGTTCACATCATTCTTCTGCNNNAGATCGGAAG
55321 TATGAGTGTGGAGTCGTTGCAGTAGCTGGGATTACAGGTACCTNNNAGATCGGAA
55322 TATGAGTGTGGAGTCGTTGCGAATGCATGGTGTGATCTTTGCTNNNAGATCGGAAG
55323 TATGAGTGTGGAGTCGTTGCTCCTGGGTTCTAGTGATTCTCCTNNNAGATCGGAAG
55327 TATGAGTGTGGAGTCGTTGCCTGGAGTGCAATGATGTGATCTCNNNAGATCGGAAG
55328 TATGAGTGTGGAGTCGTTGCCAATCTCAGTTCACTGCAACCTCNNNAGATCGGAAGA
55329 TATGAGTGTGGAGTCGTTGCGAGACGGAATCTGGCTCTGTTACNNNAGATCGGAAG
55332 TATGAGTGTGGAGTCGTTGCAGCAATCTCAGCTCATTTCAAGCNNNAGATCGGAAG
55333 TATGAGTGTGGAGTCGTTGCCAATCTTGACTCACTGCAAGCTCNNNAGATCGGAAG
55335 TATGAGTGTGGAGTCGTTGCGCACAATCTCGGCTCACTATAACNNNAGATCGGAAG
55338 TATGAGTGTGGAGTCGTTGCTAGGTTCAAGCGATTCTCTTGCTNNNAGATCGGAAG
55339 TATGAGTGTGGAGTCGTTGCCAGAGGTTACAGTAAGCTGAGGTNNNAGATCGGAA
55341 TATGAGTGTGGAGTCGTTGCAGGTACGTTCCTACAAACTCACCNNNAGATCGGAAG
55343 TATGAGTGTGGAGTCGTTGCGGGTCAGATTCCCTGAGTTCAAANNNAGATCGGAAG
55344 TATGAGTGTGGAGTCGTTGCGAAGGAGGTAGTGTTCTCCCAGANNNAGATCGGAA
55346 TATGAGTGTGGAGTCGTTGCGTGGTGAAACCCCATCTCTACTANNNAGATCGGAAG
55347 TATGAGTGTGGAGTCGTTGCAGGGCTAAATGTCATACGCTGATNNNAGATCGGAAG
55348 TATGAGTGTGGAGTCGTTGCACCTGTAATCCTAGCACTTTGGGNNNAGATCGGAAG
55349 TATGAGTGTGGAGTCGTTGCCACACCTGTAATCCCAGCACTTTNNNAGATCGGAAGA
55351 TATGAGTGTGGAGTCGTTGCCACTTGTAATCCCAGCACTTTGGNNNAGATCGGAAG
55352 TATGAGTGTGGAGTCGTTGCCAGAGGCTTCAGTGAGTCAAGATNNNAGATCGGAA
55353 TATGAGTGTGGAGTCGTTGCCACAAATTAGCTAGGCATGGTGGNNNAGATCGGAA
55354 TATGAGTGTGGAGTCGTTGCCATGAGGAAAGTCTCAAGCTACANNNAGATCGGAAG
55355 TATGAGTGTGGAGTCGTTGCGGGCACAAAAATCACTTGAACCTNNNAGATCGGAAG
55357 TATGAGTGTGGAGTCGTTGCGGAAAATGAAGTCAAGCCAAGCANNNAGATCGGAA
55358 TATGAGTGTGGAGTCGTTGCAGTGTTGGGGAAAATGGAGTCATNNNAGATCGGAA
55359 TATGAGTGTGGAGTCGTTGCCCAAAGACCACACTCCAGTTTTCNNNAGATCGGAAGA
55360 TATGAGTGTGGAGTCGTTGCGCATAGCTAGGCAAATCTGTGTGNNNAGATCGGAA
55361 TATGAGTGTGGAGTCGTTGCGTTTTCAGTTCCCTACCAAAGGCNNNAGATCGGAAG
55362 TATGAGTGTGGAGTCGTTGCTGGCCCATGGTAAATAACATGGANNNAGATCGGAA
55363 TATGAGTGTGGAGTCGTTGCCATTTTCCTTGGGGTTTCTGGACNNNAGATCGGAAG
55364 TATGAGTGTGGAGTCGTTGCGTCTCCCTGTGTGAGATCTCATCNNNAGATCGGAAG
55365 TATGAGTGTGGAGTCGTTGCGAGGAAGGAGGAGGAGAAGGATANNNAGATCGG
55367 TATGAGTGTGGAGTCGTTGCAGTTTGGCTGGATATGGTGTCTCNNNAGATCGGAAG
55368 TATGAGTGTGGAGTCGTTGCAAAGAGACCTAGAACTCAAGGCCNNNAGATCGGAA
55369 TATGAGTGTGGAGTCGTTGCCCTGTAATCCTAGCACTTTGGGANNNAGATCGGAAG
55370 TATGAGTGTGGAGTCGTTGCGGAAAATGGGCAAAGGACTTGAANNNAGATCGGA

55371 TATGAGTGTGGAGTCGTTGCGTTCCCATCTCATTGAAGCCAACNNNAGATCGGAAG
55372 TATGAGTGTGGAGTCGTTGCTTGACACTGGAACAACACAGGTTNNNAGATCGGAAG
55373 TATGAGTGTGGAGTCGTTGCCTGAGGCATGAGAATCACTTGAGNNNAGATCGGAA
55375 TATGAGTGTGGAGTCGTTGCGAATGGTGACTCATGCCTGCAATNNNAGATCGGAAG
55376 TATGAGTGTGGAGTCGTTGCAACTTGATCTTGGAAGCAAAGGCNNNAGATCGGAA
55379 TATGAGTGTGGAGTCGTTGCCCCAAAAAAGAAATGACATGGCANNNAGATCGGAA
55380 TATGAGTGTGGAGTCGTTGCTCAAGGCTGCAGTGAACTATGATNNNAGATCGGAAG
55381 TATGAGTGTGGAGTCGTTGCTGGGTTTCTAAGGATTCCCAGTGNNNAGATCGGAAG
55382 TATGAGTGTGGAGTCGTTGCGGCTTCATTTGGAAAGCCAGATTNNNAGATCGGAAG
55383 TATGAGTGTGGAGTCGTTGCGATAGGGCTTCGACAGACAGAATNNNAGATCGGAA
55384 TATGAGTGTGGAGTCGTTGCCAAAAAAACAGGGCATTTCAGGCNNNAGATCGGAA
55385 TATGAGTGTGGAGTCGTTGCTCACAGACAAGATTTACCTGTTGANNNAGATCGGAA
55388 TATGAGTGTGGAGTCGTTGCAGCTAAAATCCAGGACAAGGGAGNNNAGATCGGAA
55389 TATGAGTGTGGAGTCGTTGCAGAGCAGTGGGAAGAATCTCATGNNNAGATCGGAA
55391 TATGAGTGTGGAGTCGTTGCGTAATGTTCTCAACTTCCACGCCNNNAGATCGGAAG
55392 TATGAGTGTGGAGTCGTTGCCACGAGGTCAGGATATTGAGACCNNNAGATCGGAA
55394 TATGAGTGTGGAGTCGTTGCACATAAAACCCTTTGCCTTCTGCNNNAGATCGGAAGA
55395 TATGAGTGTGGAGTCGTTGCCTGAGATGGGAGGATCACTTGAGNNNAGATCGGAA
55397 TATGAGTGTGGAGTCGTTGCTCAACTTACGCACCTTATCCACANNNAGATCGGAAGA
55402 TATGAGTGTGGAGTCGTTGCGGTTCAAGTGATTCTTCTGCCTCNNNAGATCGGAAG
55403 TATGAGTGTGGAGTCGTTGCCCTTGGAGTGGGATGCTGAATATNNNAGATCGGAA
55405 TATGAGTGTGGAGTCGTTGCGGTTCAAGTGATTCTCCTGCTTCNNNAGATCGGAAG
55406 TATGAGTGTGGAGTCGTTGCGGAATTCAGGAAGGCTTCATTGGNNNAGATCGGAA
55407 TATGAGTGTGGAGTCGTTGCTTTTGATCTTGTTGCTCAGGCTGNNNAGATCGGAAG
55408 TATGAGTGTGGAGTCGTTGCAAGAGATTCTCCTACCTCAGCCTNNNAGATCGGAAG
55409 TATGAGTGTGGAGTCGTTGCGGGTTCTTGGTCTCACTTCAAGANNNAGATCGGAAG
55413 TATGAGTGTGGAGTCGTTGCCTGAGTAGCTGGGATTACAGGTGNNNAGATCGGAA
55414 TATGAGTGTGGAGTCGTTGCCCTCCCAAATTCAAGCGATTCTCNNNAGATCGGAAGA
55418 TATGAGTGTGGAGTCGTTGCCAACCTCCACCTCTTGAGTTCAANNNAGATCGGAAGA
55419 TATGAGTGTGGAGTCGTTGCATCTCAGATCACTGCAACCTTTGNNNAGATCGGAAG
55420 TATGAGTGTGGAGTCGTTGCTGCCCAGACACAAGATAGTGTAGNNNAGATCGGAA
55422 TATGAGTGTGGAGTCGTTGCGGTTCAAGCAATTATCCTGCCTCNNNAGATCGGAAG
55424 TATGAGTGTGGAGTCGTTGCCCTGGGTTCAAGCAATTCTCCTANNNAGATCGGAAG
55426 TATGAGTGTGGAGTCGTTGCCATTCTCCTGCCTCAGCCTCNNNAGATCGGAAGAGCA
55427 TATGAGTGTGGAGTCGTTGCCTCCTGAGTAGCTGGGATTACAGNNNAGATCGGAA
55428 TATGAGTGTGGAGTCGTTGCGTTCAAGAGATTCTCCTGCCTCANNNAGATCGGAAG
55429 TATGAGTGTGGAGTCGTTGCTCTTGGACTCCTGACCTCAAATGNNNAGATCGGAAG
55431 TATGAGTGTGGAGTCGTTGCCTGGGTTCAAATGATTCTCCTGCNNNAGATCGGAAG
55432 TATGAGTGTGGAGTCGTTGCCAGGCTGGAGTGCTTCTTTTTTTNNNAGATCGGAAG
55434 TATGAGTGTGGAGTCGTTGCAGAGTAGCAGGGATCACAGGTATNNNAGATCGGAA
55435 TATGAGTGTGGAGTCGTTGCCATGGTCTCAGCCTCAAAGTACTNNNAGATCGGAAG
55436 TATGAGTGTGGAGTCGTTGCATTCTACTGCCTCAGCCTTTCAANNNAGATCGGAAGA
55438 TATGAGTGTGGAGTCGTTGCCTCCTGAGTAGCTGGGACTACAGNNNAGATCGGAA

55439 TATGAGTGTGGAGTCGTTGCACTGCAAGCTTCACAATTTCCTCNNNAGATCGGAAGA
55440 TATGAGTGTGGAGTCGTTGCTAGGACAATAGTGGAGGGAAGGTNNNAGATCGGAA
55441 TATGAGTGTGGAGTCGTTGCCTTCCAAGTAGCTGGGACTACAGNNNAGATCGGAAG
55442 TATGAGTGTGGAGTCGTTGCCTCCTGAGTAGCTGGGACTACAGNNNAGATCGGAAG
55443 TATGAGTGTGGAGTCGTTGCGAGACAGGGTTTCACTATGTTGCNNNAGATCGGAAG
55445 TATGAGTGTGGAGTCGTTGCTAAAGTGCTGGGATTACAGGCATNNNAGATCGGAAG
55446 TATGAGTGTGGAGTCGTTGCGTCTAGCTCTATTGCTCAGGCTGNNNAGATCGGAAG
55448 TATGAGTGTGGAGTCGTTGCAGGTTCAAGCAATTCTCCTTCCTNNNAGATCGGAAGA
55450 TATGAGTGTGGAGTCGTTGCCAGCCTCCTAAAGTGCTGAGATTNNNAGATCGGAAG
55452 TATGAGTGTGGAGTCGTTGCCAGAGACCTGATCATGGCTCATTNNNAGATCGGAAG
55453 TATGAGTGTGGAGTCGTTGCTGATCTAGGCTCACTGCAATCTCNNNAGATCGGAAGA
55455 TATGAGTGTGGAGTCGTTGCCAGTGGTGTGATCTCAGCTTACTNNNAGATCGGAAG
55456 TATGAGTGTGGAGTCGTTGCACTTCCTGGGTTCAAGTGATTCANNNAGATCGGAAG
55457 TATGAGTGTGGAGTCGTTGCTCCCGGATACAAGTGATTTTCCTNNNAGATCGGAAGA
55459 TATGAGTGTGGAGTCGTTGCTCAAGTTATCCTCCCACTTCAGCNNNAGATCGGAAGA
55460 TATGAGTGTGGAGTCGTTGCCAGACTGGGCAACATACTGAGATNNNAGATCGGAAG
55466 TATGAGTGTGGAGTCGTTGCAGTGTAATGGCGTGATCTTGTCTNNNAGATCGGAAG
55467 TATGAGTGTGGAGTCGTTGCGGTTCAAGCTACTCTCCCATCTCNNNAGATCGGAAGA
55468 TATGAGTGTGGAGTCGTTGCTAGTACAGTGACGCAATCAATGCNNNAGATCGGAAG
55469 TATGAGTGTGGAGTCGTTGCCAATGGTATGATCTCGGCTCACTNNNAGATCGGAAG
55470 TATGAGTGTGGAGTCGTTGCCCCAGGTTCAAGCAATTCTCCTANNNAGATCGGAAGA
55471 TATGAGTGTGGAGTCGTTGCAGTGCTGAGATTACAGACGTGAGNNNAGATCGGAA
55472 TATGAGTGTGGAGTCGTTGCGGAGTAACTGGGATTACAGGCATNNNAGATCGGAA
55473 TATGAGTGTGGAGTCGTTGCGTTACAAGCAATTCTCCTGCCTCNNNAGATCGGAAGA
55474 TATGAGTGTGGAGTCGTTGCTGCCTCTCATTGCCTTTGAAAAGNNNAGATCGGAAGA
55475 TATGAGTGTGGAGTCGTTGCGGGCAGGAAAGAGTGTTTTTGAANNNAGATCGGAA
55477 TATGAGTGTGGAGTCGTTGCGGTGAGGCTGTGGAGAAATAAGANNNAGATCGGAA
55478 TATGAGTGTGGAGTCGTTGCGGAGAGGCGTTTGTTGTGAATTTNNNAGATCGGAAG
55479 TATGAGTGTGGAGTCGTTGCACACACACATATATGGCTCTTGCNNNAGATCGGAAGA
55480 TATGAGTGTGGAGTCGTTGCGACCACAACTATAATTGCGCAGGNNNAGATCGGAAG
55481 TATGAGTGTGGAGTCGTTGCGAGGCTCACATGACCATATCCTTNNNAGATCGGAAGA
55482 TATGAGTGTGGAGTCGTTGCTCTCCTGAGTCCCTTTCACACTANNNAGATCGGAAGA
55483 TATGAGTGTGGAGTCGTTGCTCACAAGGTCAGGAATTCGAGACNNNAGATCGGAAG
55485 TATGAGTGTGGAGTCGTTGCTTACAAAGTCAGAGGAGGATGGCNNNAGATCGGAA
55487 TATGAGTGTGGAGTCGTTGCACTTGAGGTCAGGAGTTCAAGACNNNAGATCGGAAG
55488 TATGAGTGTGGAGTCGTTGCTGATGAGGGGAGAATCTGACTCANNNAGATCGGAA
55490 TATGAGTGTGGAGTCGTTGCGTAATCCCAGCTACTTAGGAGGCNNNAGATCGGAAG
55491 TATGAGTGTGGAGTCGTTGCGAGACTTTTGATTGGCTGGTCAGNNNAGATCGGAAG
55493 TATGAGTGTGGAGTCGTTGCTTTACCAAAGCCTACCTGACCAANNNAGATCGGAAGA
55494 TATGAGTGTGGAGTCGTTGCTCATGCTTGTAATCCCAGCACTTNNNAGATCGGAAGA
55495 TATGAGTGTGGAGTCGTTGCTCTCAACAACACCACTCAGAGAGNNNAGATCGGAAG
55496 TATGAGTGTGGAGTCGTTGCTCTGTAACTCCAGCTACTCAGGANNNAGATCGGAAGA
55499 TATGAGTGTGGAGTCGTTGCCACTCCATTCCCTCACTGAGAAANNNAGATCGGAAGA

55500 TATGAGTGTGGAGTCGTTGCGTAGTCATAGCTACTTGGGAGGCNNNAGATCGGAA
55503 TATGAGTGTGGAGTCGTTGCCAATAGACTGGAGTGGATGGAGGNNNAGATCGGA
55509 TATGAGTGTGGAGTCGTTGCCAGAGGTTGTAGTGAGCCAAGATNNNAGATCGGAA
55510 TATGAGTGTGGAGTCGTTGCCCTAAGAGCCAGTGAAGTGTCTTNNNAGATCGGAAG
55511 TATGAGTGTGGAGTCGTTGCGTTTGTGCCTAATCTGTGGAACCNNNAGATCGGAAG
55515 TATGAGTGTGGAGTCGTTGCGATGCATGTTCAGTGAGGTCAAANNNAGATCGGAA
55517 TATGAGTGTGGAGTCGTTGCGTGAGCGACTGTAATCTCAGCTANNNAGATCGGAAG
55518 TATGAGTGTGGAGTCGTTGCTTTGTTGCTCCTACTTTTGGTGTNNNAGATCGGAAG
55519 TATGAGTGTGGAGTCGTTGCGTAGAACCACTGCTCTCATCCAANNNAGATCGGAAG
55520 TATGAGTGTGGAGTCGTTGCACAGGACACAATAGACACGGTTTNNNAGATCGGAAG
55524 TATGAGTGTGGAGTCGTTGCACTGACCACTTTTATTCAGGGCTNNNAGATCGGAAG
55527 TATGAGTGTGGAGTCGTTGCGTAATCCCAGGTGCATCTACTCGNNNAGATCGGAAG
55528 TATGAGTGTGGAGTCGTTGCAAGATCTGAAGGGATCCAAAGCANNNAGATCGGAA
55529 TATGAGTGTGGAGTCGTTGCAAAAGATGCCAAAGAGAACCTGCNNNAGATCGGAA
55530 TATGAGTGTGGAGTCGTTGCCCAACAATGGAAGGCTGATTCAGNNNAGATCGGAA
55531 TATGAGTGTGGAGTCGTTGCTATGATGGCATGCACCTGGAATTNNNAGATCGGAAG
55532 TATGAGTGTGGAGTCGTTGCCAGTCCCACTCCAATATCTCACANNNAGATCGGAAGA
55534 TATGAGTGTGGAGTCGTTGCTTCTACAGCAGCGGTTAGGTTTANNNAGATCGGAAG
55535 TATGAGTGTGGAGTCGTTGCCAATTCTCACCCATGCCTCATTCNNNAGATCGGAAGA
55536 TATGAGTGTGGAGTCGTTGCGGAAAGCCCATGAACTTATCACGNNNAGATCGGAAG
55537 TATGAGTGTGGAGTCGTTGCCCTATAGTCCCAGCTTGGAAGACNNNAGATCGGAAG
55538 TATGAGTGTGGAGTCGTTGCAGGAAGTGGATTTTTCTCTGGCTNNNAGATCGGAAG
55540 TATGAGTGTGGAGTCGTTGCGTAATTCCAGCTACTCGGGAGACNNNAGATCGGAAG
55542 TATGAGTGTGGAGTCGTTGCTGGCCCAGAACAGTCAAATTCTANNNAGATCGGAAG
55544 TATGAGTGTGGAGTCGTTGCGAGGTCAGGAGATTGAGACCATCNNNAGATCGGAA
55545 TATGAGTGTGGAGTCGTTGCCTCACGTCTGTAATCTCAGCACTNNNAGATCGGAAG
55546 TATGAGTGTGGAGTCGTTGCCTTAGAAGGCTGAGGCAGAAGAANNNAGATCGGAA
55550 TATGAGTGTGGAGTCGTTGCGAGAGTGGTGAAGGGAAAGAGAGNNNAGATCGGA
55551 TATGAGTGTGGAGTCGTTGCTGCTATTCTGCTTCTTCCAGAGANNNAGATCGGAAG
55552 TATGAGTGTGGAGTCGTTGCCGAGACAGGAAGGAAAACACAACNNNAGATCGGAA
55553 TATGAGTGTGGAGTCGTTGCTCCAAGAAATCAAAGCTGCAGTGNNNAGATCGGAAG
55555 TATGAGTGTGGAGTCGTTGCAATCTCCAAGGAAGAGGTGTCACNNNAGATCGGAA
55557 TATGAGTGTGGAGTCGTTGCGCTTGGTTGATTTCATGCAGGTANNNAGATCGGAAG
55558 TATGAGTGTGGAGTCGTTGCATCAGGCTCACATTTGCATGTTCNNNAGATCGGAAG
55559 TATGAGTGTGGAGTCGTTGCTGGAGTACAGTGGTGTGATCTTGNNNAGATCGGAA
55560 TATGAGTGTGGAGTCGTTGCGAGTAGTTGGGATCACAGGTCTGNNNAGATCGGAA
55561 TATGAGTGTGGAGTCGTTGCCCTCCTGGGTTCAAGCAATTCTCNNNAGATCGGAAG
55562 TATGAGTGTGGAGTCGTTGCTGTCAGTTTCTGCTTTTGTTGCANNNAGATCGGAAG
55563 TATGAGTGTGGAGTCGTTGCGTTGGTTTCCAGGCAAAAGATGANNNAGATCGGAA
55565 TATGAGTGTGGAGTCGTTGCTTCAAGCAATTCTCCTGCCTCAGNNNAGATCGGAAG
55566 TATGAGTGTGGAGTCGTTGCGGATTATAGGAACCTGCCACCATNNNAGATCGGAAG
55567 TATGAGTGTGGAGTCGTTGCGCTCACACCAAAAGACCAAACAANNNAGATCGGAAG
55568 TATGAGTGTGGAGTCGTTGCATGCGTGCTCATTGTGTAATCACNNNAGATCGGAAG

55569 TATGAGTGTGGAGTCGTTGCGCCTCTCAAAATGCTGGGATTACNNNAGATCGGAAG
55571 TATGAGTGTGGAGTCGTTGCTTCAAGTGATTCTCCTGCCTCAGNNNAGATCGGAAG
55572 TATGAGTGTGGAGTCGTTGCATCAAGTGATCTGTCTGCCTCAGNNNAGATCGGAAG
55573 TATGAGTGTGGAGTCGTTGCCCCTTGGGAGTAAGTGACAGAAANNNAGATCGGAA
55575 TATGAGTGTGGAGTCGTTGCATTACAGGCATGAGTCACTGTGTNNNAGATCGGAAG
55576 TATGAGTGTGGAGTCGTTGCCCGACTCTTTGGTTCAAGCAATTNNNAGATCGGAAG
55577 TATGAGTGTGGAGTCGTTGCAAGTGCTGGGATTATAGGTGTGGNNNAGATCGGAA
55578 TATGAGTGTGGAGTCGTTGCCTCCTGAGTAGCTAGCTGGTACTNNNAGATCGGAAG
55579 TATGAGTGTGGAGTCGTTGCTACACTCACCAGACTATGGAGGTNNNAGATCGGAAG
55581 TATGAGTGTGGAGTCGTTGCTGGAGTACAGTGGCATGATCTTGNNNAGATCGGAA
55582 TATGAGTGTGGAGTCGTTGCCCTCCCTGGTTTGAGCAATTCTCNNNAGATCGGAAG
55583 TATGAGTGTGGAGTCGTTGCATCTCAGCTAACTGCAACCTTGANNNAGATCGGAAG
55584 TATGAGTGTGGAGTCGTTGCCATGTACAAGTCAGGAGAGCTCANNNAGATCGGAA
55585 TATGAGTGTGGAGTCGTTGCCTTTGAAAGCAGGCATGAGGAAANNNAGATCGGAA
55586 TATGAGTGTGGAGTCGTTGCCATGATCTCAGCTCACTGCAAACNNNAGATCGGAAG
55587 TATGAGTGTGGAGTCGTTGCAATCTCAGTTCACTGCAAGCTCCNNNAGATCGGAAG
55591 TATGAGTGTGGAGTCGTTGCTTCATCAGCAGCATGAAAACTGGNNNAGATCGGAAG
55592 TATGAGTGTGGAGTCGTTGCCAGGCACATGCTACATACCCTATNNNAGATCGGAAG
55593 TATGAGTGTGGAGTCGTTGCAGCTGGGATTACAGGTGTGTAAANNNAGATCGGAA
55595 TATGAGTGTGGAGTCGTTGCTTCACTCTTATTGTCCAGGCTGGNNNAGATCGGAAG
55596 TATGAGTGTGGAGTCGTTGCACCAAGGTACATATCACTACGCCNNNAGATCGGAAG
55597 TATGAGTGTGGAGTCGTTGCCTTCCTGAGTAGCTGGGACTACANNNAGATCGGAAG
55598 TATGAGTGTGGAGTCGTTGCGATGAGGACTCTACCGTCATGACNNNAGATCGGAAG
55599 TATGAGTGTGGAGTCGTTGCCTGGAATTATGGGCACACATCACNNNAGATCGGAAG
55603 TATGAGTGTGGAGTCGTTGCCCTCCTGAGTTCATGCCATTCTCNNNAGATCGGAAGA
55604 TATGAGTGTGGAGTCGTTGCAGCCTCTGGAAACCACTAATCTGNNNAGATCGGAAG
55610 TATGAGTGTGGAGTCGTTGCCCCTTGTCTGTTTCTTACCATCCNNNAGATCGGAAGA
55613 TATGAGTGTGGAGTCGTTGCTACCTTCCTGAAAAAGCTCCCTCNNNAGATCGGAAGA
55615 TATGAGTGTGGAGTCGTTGCCTATGGTCAATCAGCAAAGCCAGNNNAGATCGGAAG
55616 TATGAGTGTGGAGTCGTTGCGAGATCACACCACTGAACTCCAGNNNAGATCGGAAG
55617 TATGAGTGTGGAGTCGTTGCACAGGAAGTCAAGAGGAAATGCTNNNAGATCGGAA
55618 TATGAGTGTGGAGTCGTTGCAGCTGAAAGAGAAAGCTAGGACANNNAGATCGGAA
55619 TATGAGTGTGGAGTCGTTGCACAGCCAAATAATGCACTCGTTCNNNAGATCGGAAG
55620 TATGAGTGTGGAGTCGTTGCGCTACTTCACCCAATCAAAAGCGNNNAGATCGGAAG
55622 TATGAGTGTGGAGTCGTTGCACAACGTGAATGTACTTGATGCCNNNAGATCGGAAG
55626 TATGAGTGTGGAGTCGTTGCCAGTTGACAGAAACAGTGGAACANNNAGATCGGAA
55627 TATGAGTGTGGAGTCGTTGCTCCTACCCTCACAGAACTTACCTNNNAGATCGGAAGA
55630 TATGAGTGTGGAGTCGTTGCAAGGTATTTCAGGCAGCACAAACNNNAGATCGGAAG
55631 TATGAGTGTGGAGTCGTTGCAGGATGATGGCTGCCTGAAATAANNNAGATCGGAA
55633 TATGAGTGTGGAGTCGTTGCATGGTGGGAATTCATGCCTAGTTNNNAGATCGGAAG
55634 TATGAGTGTGGAGTCGTTGCGTGTGCTTACCTATCAGTCGCTANNNAGATCGGAAG
55636 TATGAGTGTGGAGTCGTTGCGGTGATCAACAGGTCATTAGGGTNNNAGATCGGAA
55638 TATGAGTGTGGAGTCGTTGCAGAGTGAGCAGTACTGTCCATTGNNNAGATCGGAA

55639 TATGAGTGTGGAGTCGTTGCGAGGAGCACTGAAGGGTTTTAGANNNAGATCGGA
55641 TATGAGTGTGGAGTCGTTGCGAACAGATCAAGCAGGCAGAAAANNNAGATCGGAA
55642 TATGAGTGTGGAGTCGTTGCGGAAAAGCTTCACCAGAAAGTCCNNNAGATCGGAA
55643 TATGAGTGTGGAGTCGTTGCATAGTGAGCAGAGATCATGCCACNNNAGATCGGAA
55644 TATGAGTGTGGAGTCGTTGCTGAATCAGAACATCGCATAAGCCNNNAGATCGGAAG
55645 TATGAGTGTGGAGTCGTTGCGATCACGAGGTCAGGAGTTCAAGNNNAGATCGGAA
55646 TATGAGTGTGGAGTCGTTGCTGGACAAAATTCCACCAGCAAAANNNAGATCGGAAG
55647 TATGAGTGTGGAGTCGTTGCTGGTAAAGCAGAGACTCTGATGGNNNAGATCGGAA
55648 TATGAGTGTGGAGTCGTTGCTAGAGTTGTGCAACCTCCACTACNNNAGATCGGAAG
55649 TATGAGTGTGGAGTCGTTGCTTGATGGTACATCAGGCTTCTGGNNNAGATCGGAA
55651 TATGAGTGTGGAGTCGTTGCCAGAATCCCAGGTTGGAGTCATCNNNAGATCGGAA
55652 TATGAGTGTGGAGTCGTTGCCGACAGGGCAAGACTCTATCAAANNNAGATCGGAA
55653 TATGAGTGTGGAGTCGTTGCTCAGTGACGACAGTGCTAGTAAGNNNAGATCGGAA
55654 TATGAGTGTGGAGTCGTTGCCATAGGATCACTGCCACTGAACTNNNAGATCGGAAG
55655 TATGAGTGTGGAGTCGTTGCTATGAAAAATGGAGCCCTGGGATNNNAGATCGGAA
55656 TATGAGTGTGGAGTCGTTGCCACCAATAGAAGAGGGGACTACGNNNAGATCGGAA
55657 TATGAGTGTGGAGTCGTTGCGGGAGGGGATTGCTTTATCATCANNNAGATCGGAA
55658 TATGAGTGTGGAGTCGTTGCTGTGTGTTGGTGGAGAGTCTTTTNNNAGATCGGAA
55659 TATGAGTGTGGAGTCGTTGCTCTCCTCCCTTAGTCTTCCAAGTNNNAGATCGGAAGA
55660 TATGAGTGTGGAGTCGTTGCTTCCTCTTACTTGGTGCACTTCCNNNAGATCGGAAGA
55662 TATGAGTGTGGAGTCGTTGCTGTGTAGGAAGAGGGAGAGAGAGNNNAGATCGGA
55663 TATGAGTGTGGAGTCGTTGCAGGAGTATCTCTTGGTCCAGGAGNNNAGATCGGAA
55664 TATGAGTGTGGAGTCGTTGCTACACACAATTAGCTGGGAGTGGNNNAGATCGGAA
55665 TATGAGTGTGGAGTCGTTGCTAATCCCAGCTACTCAGGAGTCTNNNAGATCGGAAG
55666 TATGAGTGTGGAGTCGTTGCGGCTGAGGTGGGAGGATCTATTANNNAGATCGGA
55667 TATGAGTGTGGAGTCGTTGCCCTTCCTTGTGTGTGTGTGTTTTNNNAGATCGGAAG
55668 TATGAGTGTGGAGTCGTTGCCACCTGAATAGGCAAGGTCTCTTNNNAGATCGGAAG
55669 TATGAGTGTGGAGTCGTTGCAGAGTTGCTTGATTCCACAAGGANNNAGATCGGAA
55671 TATGAGTGTGGAGTCGTTGCCCAAGGAACTACCAGACACTAGGNNNAGATCGGAA
55673 TATGAGTGTGGAGTCGTTGCCCAGGCATATCTCCAGACATTCANNNAGATCGGAAG
55674 TATGAGTGTGGAGTCGTTGCAATGAGGTTGCAGTGAGGAGAAANNNAGATCGGA
55675 TATGAGTGTGGAGTCGTTGCCCCAGATTCAAGCGATCCTCTAGNNNAGATCGGAAG
55676 TATGAGTGTGGAGTCGTTGCCTTCTAAAGGGAGAGAGCAGCTTNNNAGATCGGAA
55679 TATGAGTGTGGAGTCGTTGCGTGTGTGTGTGTGTGTTTCTTCTNNNAGATCGGAAG
55680 TATGAGTGTGGAGTCGTTGCACTAGTAAAGGTAGAGGGCTGGTNNNAGATCGGA
55682 TATGAGTGTGGAGTCGTTGCCCTGTAGTCCCAGCTACTTAGGANNNAGATCGGAAG
55683 TATGAGTGTGGAGTCGTTGCTCTTGAACTCCTGACCTCAAGTGNNNAGATCGGAAG
55684 TATGAGTGTGGAGTCGTTGCAAACAAAGGGAATGTTGACAGCCNNNAGATCGGAA
55685 TATGAGTGTGGAGTCGTTGCGAGAGAGAGAGAGAGACACTCCANNNAGATCGGA
55687 TATGAGTGTGGAGTCGTTGCATGATCTCAGTTCACTGCAACCTNNNAGATCGGAAG
55689 TATGAGTGTGGAGTCGTTGCATTTTTCGAGATGGAGTCTGGGTNNNAGATCGGAA
55690 TATGAGTGTGGAGTCGTTGCATCCCTTATCTCCACCTCTGTCCNNNAGATCGGAAGA
55691 TATGAGTGTGGAGTCGTTGCATGTGGTGGTACATGCCTGTAATNNNAGATCGGAA

55692 TATGAGTGTGGAGTCGTTGCCTCACACCAGTTTTCCTTGGTTCNNNAGATCGGAAGA
55693 TATGAGTGTGGAGTCGTTGCACCCTATCAGCCAATGCCTTTATNNNAGATCGGAAGA
55695 TATGAGTGTGGAGTCGTTGCCTCAAACTCCTGACCTCACGTAANNNAGATCGGAAGA

AGAGCACACGTCTGAACTCTTTCCCTACACGACGCTCTTCCGATCTNNNCAGAATGGATGAATATGTG
GAGCACACGTCTGAACTCTTTCCCTACACGACGCTCTTCCGATCTNNNACTCCTTTTTTTGTTTTGTTT
GAGCACACGTCTGAACTCTTTCCCTACACGACGCTCTTCCGATCTNNNTGTCAAAAGTAAAAAGAAAA
GAGCACACGTCTGAACTCTTTCCCTACACGACGCTCTTCCGATCTNNNCATTCCTTTTTATGGCTGAGTA
AAGAGCACACGTCTGAACTCTTTCCCTACACGACGCTCTTCCGATCTNNNTCCTTTCTTCTTTTTTTTCCT
AGAGCACACGTCTGAACTCTTTCCCTACACGACGCTCTTCCGATCTNNNACTTTGTTAGTGGGCATGTA
AAGAGCACACGTCTGAACTCTTTCCCTACACGACGCTCTTCCGATCTNNNTGTGTGTGTATGTAAATGT
GAAGAGCACACGTCTGAACTCTTTCCCTACACGACGCTCTTCCGATCTNNNAGCAGTGTGAAAATGGA
AGAGCACACGTCTGAACTCTTTCCCTACACGACGCTCTTCCGATCTNNNAGTCGCTATAAGTATCACA
AGAGCACACGTCTGAACTCTTTCCCTACACGACGCTCTTCCGATCTNNNCAACCCTTTACCTAGACTAA
AAGAGCACACGTCTGAACTCTTTCCCTACACGACGCTCTTCCGATCTNNNTCGGACCACTATCTGACTA
AAGAGCACACGTCTGAACTCTTTCCCTACACGACGCTCTTCCGATCTNNNATATCACCTCATGTCCATTA
GGAAGAGCACACGTCTGAACTCTTTCCCTACACGACGCTCTTCCGATCTNNNAGAACTAGATAACCAA
AGAGCACACGTCTGAACTCTTTCCCTACACGACGCTCTTCCGATCTNNNTGGACACAGAAAGACAAAA
AGCACACGTCTGAACTCTTTCCCTACACGACGCTCTTCCGATCTNNNACACACACACAATGTATTTATTC
AGAGCACACGTCTGAACTCTTTCCCTACACGACGCTCTTCCGATCTNNNTGTATTAGAGTTCTCCAGA
GAGCACACGTCTGAACTCTTTCCCTACACGACGCTCTTCCGATCTNNNGCTCATACTCACTAAGGGAAG
AGAGCACACGTCTGAACTCTTTCCCTACACGACGCTCTTCCGATCTNNNCCAGAATCAAATGTTTCTCT
GAGCACACGTCTGAACTCTTTCCCTACACGACGCTCTTCCGATCTNNNTGAAGACATGACATAGTTAGA
AGAGCACACGTCTGAACTCTTTCCCTACACGACGCTCTTCCGATCTNNNTCATCAGAGTTTTGTAGTTT
AGAGCACACGTCTGAACTCTTTCCCTACACGACGCTCTTCCGATCTNNNGGGAAAGCTAAGTTGATTC
GAGCACACGTCTGAACTCTTTCCCTACACGACGCTCTTCCGATCTNNNAGCTGTTCTCATCATTTTTCC
AGCACACGTCTGAACTCTTTCCCTACACGACGCTCTTCCGATCTNNNGCCAGGATATATATGTATGTAT
AAGAGCACACGTCTGAACTCTTTCCCTACACGACGCTCTTCCGATCTNNNAATAAGTGGCAGAACTGG
AAGAGCACACGTCTGAACTCTTTCCCTACACGACGCTCTTCCGATCTNNNCCGAATGGTGCTGCAATA
GAAGAGCACACGTCTGAACTCTTTCCCTACACGACGCTCTTCCGATCTNNNCTACTGAGTGTCTACCCA
GAAGAGCACACGTCTGAACTCTTTCCCTACACGACGCTCTTCCGATCTNNNTTGCGCACGTATGTTTAT
GAAGAGCACACGTCTGAACTCTTTCCCTACACGACGCTCTTCCGATCTNNNCCAGAAAAGGGTCACAC
AGAGCACACGTCTGAACTCTTTCCCTACACGACGCTCTTCCGATCTNNNTGACCATAAAAGTCCCTCCA
AGAGCACACGTCTGAACTCTTTCCCTACACGACGCTCTTCCGATCTNNNTTCTGTAGCTATCATCTGTCTA
GAAGAGCACACGTCTGAACTCTTTCCCTACACGACGCTCTTCCGATCTNNNGGGACATAACTTGGGCC
AAGAGCACACGTCTGAACTCTTTCCCTACACGACGCTCTTCCGATCTNNNACCGTGTAAGGTGTATGTA
CGGAAGAGCACACGTCTGAACTCTTTCCCTACACGACGCTCTTCCGATCTNNNGGATGAATGGATGGA
AAGAGCACACGTCTGAACTCTTTCCCTACACGACGCTCTTCCGATCTNNNAGCAGACTCTCCCTGTATCT
GAGCACACGTCTGAACTCTTTCCCTACACGACGCTCTTCCGATCTNNNACCTTGAAAACATTATGCTAAA
AAGAGCACACGTCTGAACTCTTTCCCTACACGACGCTCTTCCGATCTNNNGGGTTCTTTCCCTTTAGCTC
GAAGAGCACACGTCTGAACTCTTTCCCTACACGACGCTCTTCCGATCTNNNTGGTGGGTAGGTATGTG
GAGCACACGTCTGAACTCTTTCCCTACACGACGCTCTTCCGATCTNNNGCAAGCAAAAATATGAAATTC
GAAGAGCACACGTCTGAACTCTTTCCCTACACGACGCTCTTCCGATCTNNNTAAGCACAAGGTGGTGA
GAGCACACGTCTGAACTCTTTCCCTACACGACGCTCTTCCGATCTNNNTGTGTGTATATAGTGTTTGTA
TCGGAAGAGCACACGTCTGAACTCTTTCCCTACACGACGCTCTTCCGATCTNNNTACCTATAGTCCCAGC
AGCACACGTCTGAACTCTTTCCCTACACGACGCTCTTCCGATCTNNNTGTGTGCGTGTATATACATAAAT

AAGAGCACACGTCTGAACTCTTTCCCTACACGACGCTCTTCCGATCTNNNTCAGAGGCCTGACATAA
AGAGCACACGTCTGAACTCTTTCCCTACACGACGCTCTTCCGATCTNNNAGCCTGTATAGTATACACAC
AGCACACGTCTGAACTCTTTCCCTACACGACGCTCTTCCGATCTNNNACATGATACAGAAATAAGAGA
AAGAGCACACGTCTGAACTCTTTCCCTACACGACGCTCTTCCGATCTNNNAAGTCCTAGCCAGAGCAA
AGAGCACACGTCTGAACTCTTTCCCTACACGACGCTCTTCCGATCTNNNCTCTCTCTCTCTCTGTCCA
GAGCACACGTCTGAACTCTTTCCCTACACGACGCTCTTCCGATCTNNNAAAACTATTTTTTATGTTGCA
AGCACACGTCTGAACTCTTTCCCTACACGACGCTCTTCCGATCTNNNAGACTGCTATCTAGACTAACAA
GAAGAGCACACGTCTGAACTCTTTCCCTACACGACGCTCTTCCGATCTNNNTGCTCCCTTGAAATCCG
AAGAGCACACGTCTGAACTCTTTCCCTACACGACGCTCTTCCGATCTNNNGCAGGAGAATCACTTGA
GCACACGTCTGAACTCTTTCCCTACACGACGCTCTTCCGATCTNNNCACGTGTGTGTATATGTATATA
AAGAGCACACGTCTGAACTCTTTCCCTACACGACGCTCTTCCGATCTNNNTCCCAACTGACAAATGAG
GAAGAGCACACGTCTGAACTCTTTCCCTACACGACGCTCTTCCGATCTNNNGATATGACCTCAAAAGC
AGAGCACACGTCTGAACTCTTTCCCTACACGACGCTCTTCCGATCTNNNAGAAGTTGGTTCTTCAAAA
GCACACGTCTGAACTCTTTCCCTACACGACGCTCTTCCGATCTNNNTCAAGAATAGTTCAAGAGAGG
GAGCACACGTCTGAACTCTTTCCCTACACGACGCTCTTCCGATCTNNNACAGTAATCCTAGTGATTTAT
AAGAGCACACGTCTGAACTCTTTCCCTACACGACGCTCTTCCGATCTNNNCTGAGGCAGGAGAATAG
AGCACACGTCTGAACTCTTTCCCTACACGACGCTCTTCCGATCTNNNTCTCTTGCTTCTCTAAGTTTTTT
AAGAGCACACGTCTGAACTCTTTCCCTACACGACGCTCTTCCGATCTNNNACCTAATGCTAGATGACA
GAGCACACGTCTGAACTCTTTCCCTACACGACGCTCTTCCGATCTNNNACTCCAAAACATTATGTTGTA
AGCACACGTCTGAACTCTTTCCCTACACGACGCTCTTCCGATCTNNNTGCTAATATTTTCTCCCATTTTTT
AGCACACGTCTGAACTCTTTCCCTACACGACGCTCTTCCGATCTNNNTCTTCTTTCCTCTTTCTTTCTTTCT
AGCACACGTCTGAACTCTTTCCCTACACGACGCTCTTCCGATCTNNNTCTTCTTCCTTTCTCTCTCTTCTT
GCACACGTCTGAACTCTTTCCCTACACGACGCTCTTCCGATCTNNNTGCATATTTCTTCACTTTTACTTA
AAGAGCACACGTCTGAACTCTTTCCCTACACGACGCTCTTCCGATCTNNNATTCAAGACCCAAGAAGC
AGCACACGTCTGAACTCTTTCCCTACACGACGCTCTTCCGATCTNNNTGAGAATTCTCCTGATATTCTG
AGCACACGTCTGAACTCTTTCCCTACACGACGCTCTTCCGATCTNNNTTCTAGTACACATAGTCATTTA
AGCACACGTCTGAACTCTTTCCCTACACGACGCTCTTCCGATCTNNNTTCTAGTACACATAGTCATTTA
AGCACACGTCTGAACTCTTTCCCTACACGACGCTCTTCCGATCTNNNGTTTTCTCAAATTCTGGATTTTT
GCACACGTCTGAACTCTTTCCCTACACGACGCTCTTCCGATCTNNNTGATTTTGTAACCTGAAACTTTA
GAAGAGCACACGTCTGAACTCTTTCCCTACACGACGCTCTTCCGATCTNNNAGAATCACTTGAACCCA
AAGAGCACACGTCTGAACTCTTTCCCTACACGACGCTCTTCCGATCTNNNACAATCATGGAGATTCCCT
GAAGAGCACACGTCTGAACTCTTTCCCTACACGACGCTCTTCCGATCTNNNCAGAATTGCTTGAAACT
AGAGCACACGTCTGAACTCTTTCCCTACACGACGCTCTTCCGATCTNNNTGGACAAAGGGTAATATAG
GAAGAGCACACGTCTGAACTCTTTCCCTACACGACGCTCTTCCGATCTNNNGGAGGCTAAGACAGGA
AGAGCACACGTCTGAACTCTTTCCCTACACGACGCTCTTCCGATCTNNNCAGAAGCTAACCATTTGTT
GAAGAGCACACGTCTGAACTCTTTCCCTACACGACGCTCTTCCGATCTNNNCACGCCTGTAATCCCAA
GGAAGAGCACACGTCTGAACTCTTTCCCTACACGACGCTCTTCCGATCTNNNCTGAGACAGGAGAAT
AGAGCACACGTCTGAACTCTTTCCCTACACGACGCTCTTCCGATCTNNNACTGTCACTAAAAGGCAAA
AAGAGCACACGTCTGAACTCTTTCCCTACACGACGCTCTTCCGATCTNNNGGTGCCTGTAATCTCAGC
GAAGAGCACACGTCTGAACTCTTTCCCTACACGACGCTCTTCCGATCTNNNCTGAGGTGGGTGGATA
GAAGAGCACACGTCTGAACTCTTTCCCTACACGACGCTCTTCCGATCTNNNGAGAATTGCTTGAACCT
AAGAGCACACGTCTGAACTCTTTCCCTACACGACGCTCTTCCGATCTNNNTGCCTCTTTTTGAATGCTT
AGAGCACACGTCTGAACTCTTTCCCTACACGACGCTCTTCCGATCTNNNACAGATTCGAAACACTCTCT

GGAAGAGCACACGTCTGAACTCTTTCCCTACACGACGCTCTTCCGATCTNNNGAGGCAGGAGAATTGT

GAGCACACGTCTGAACTCTTTCCCTACACGACGCTCTTCCGATCTNNNATAAAGGGAGAGGAAAAAG

GGAAGAGCACACGTCTGAACTCTTTCCCTACACGACGCTCTTCCGATCTNNNAGGCAAGAGAATCACTT

AGCACACGTCTGAACTCTTTCCCTACACGACGCTCTTCCGATCTNNNAGACTGCTATCTAGACTAACAA

AGGAAGAGCACACGTCTGAACTCTTTCCCTACACGACGCTCTTCCGATCTNNNTAATCCCAGCTACTTGG

GGAAGAGCACACGTCTGAACTCTTTCCCTACACGACGCTCTTCCGATCTNNNCATCTGTGATCCCAGCTA

GGAAGAGCACACGTCTGAACTCTTTCCCTACACGACGCTCTTCCGATCTNNNAACAAAAATAGCCAGGT

AGAGCACACGTCTGAACTCTTTCCCTACACGACGCTCTTCCGATCTNNNAGACAAATTTTTGAAACACA

AGCACACGTCTGAACTCTTTCCCTACACGACGCTCTTCCGATCTNNNTGAGGTAAAAGACTGATATAAG

GGAAGAGCACACGTCTGAACTCTTTCCCTACACGACGCTCTTCCGATCTNNNTGGAAAAACCTAAGGAG

GAGCACACGTCTGAACTCTTTCCCTACACGACGCTCTTCCGATCTNNNCCATCAAATTGGATGAATCTCA

GAAGAGCACACGTCTGAACTCTTTCCCTACACGACGCTCTTCCGATCTNNNCACAATCTCCTCCAGAACA

GGAAGAGCACACGTCTGAACTCTTTCCCTACACGACGCTCTTCCGATCTNNNAAATGCTGCAACACTGG

AGCACACGTCTGAACTCTTTCCCTACACGACGCTCTTCCGATCTNNNGGGAAATCCTTATAAACTATTGG

GGAAGAGCACACGTCTGAACTCTTTCCCTACACGACGCTCTTCCGATCTNNNTCTGTAGTCCCAGCTATT

GAAGAGCACACGTCTGAACTCTTTCCCTACACGACGCTCTTCCGATCTNNNTGAATACTGCCGCTATGA

AGCACACGTCTGAACTCTTTCCCTACACGACGCTCTTCCGATCTNNNTCTTCTTTACAAAAGCTATCAGT

AGCACACGTCTGAACTCTTTCCCTACACGACGCTCTTCCGATCTNNNCCATACCTTTATTTTCAGGCTATG

GGAAGAGCACACGTCTGAACTCTTTCCCTACACGACGCTCTTCCGATCTNNNCACGCATGTCTATTGCAG

GAAGAGCACACGTCTGAACTCTTTCCCTACACGACGCTCTTCCGATCTNNNTGAATGGGCTTAGGAGA

GAAGAGCACACGTCTGAACTCTTTCCCTACACGACGCTCTTCCGATCTNNNTGAATGGGCTTAGGAGA

AGCACACGTCTGAACTCTTTCCCTACACGACGCTCTTCCGATCTNNNGCTACATACTCTTGATGTATTTA

GAGCACACGTCTGAACTCTTTCCCTACACGACGCTCTTCCGATCTNNNAGAGCCTATTATTTGAAAATTG

AGCACACGTCTGAACTCTTTCCCTACACGACGCTCTTCCGATCTNNNGTATATATATACACACACACACA

AGCACACGTCTGAACTCTTTCCCTACACGACGCTCTTCCGATCTNNNCACACGTGTGTATATGTATATAT

GAGCACACGTCTGAACTCTTTCCCTACACGACGCTCTTCCGATCTNNNACACACCATATATATATACGCA

GAGCACACGTCTGAACTCTTTCCCTACACGACGCTCTTCCGATCTNNNTGTGTGAGTGTATATATATGT

GAGCACACGTCTGAACTCTTTCCCTACACGACGCTCTTCCGATCTNNNAGAGCCTATTATTTGAAAATTG

AAGAGCACACGTCTGAACTCTTTCCCTACACGACGCTCTTCCGATCTNNNACATTTGAAAAAGCCCTCT

GGAAGAGCACACGTCTGAACTCTTTCCCTACACGACGCTCTTCCGATCTNNNAGAATTGCTTGAAATTG

GAAGAGCACACGTCTGAACTCTTTCCCTACACGACGCTCTTCCGATCTNNNTGCTTCATGGGTAGTGTG

AGCACACGTCTGAACTCTTTCCCTACACGACGCTCTTCCGATCTNNNTGGAATAGTAATACAGTTATTG

GAAGAGCACACGTCTGAACTCTTTCCCTACACGACGCTCTTCCGATCTNNNAGAGATGGGGTTTCTCCA

GAGCACACGTCTGAACTCTTTCCCTACACGACGCTCTTCCGATCTNNNCGTCGATATATTTATGTATGTG

GAAGAGCACACGTCTGAACTCTTTCCCTACACGACGCTCTTCCGATCTNNNAGACAACCAAAAGCTGA

AGCACACGTCTGAACTCTTTCCCTACACGACGCTCTTCCGATCTNNNACATATATAAGTTGTGTGTGTG

GAAGAGCACACGTCTGAACTCTTTCCCTACACGACGCTCTTCCGATCTNNNAGGGTTCTCTAGAGGGA

AAGAGCACACGTCTGAACTCTTTCCCTACACGACGCTCTTCCGATCTNNNTCTATGAGGGGAGATCAT

AAGAGCACACGTCTGAACTCTTTCCCTACACGACGCTCTTCCGATCTNNNTGGTCTTGGGAGGGTTTAT

AAGAGCACACGTCTGAACTCTTTCCCTACACGACGCTCTTCCGATCTNNNGACCTCATTATTGGCTTGC

GAAGAGCACACGTCTGAACTCTTTCCCTACACGACGCTCTTCCGATCTNNNAGTAGCCATAGGGAAAG

GAAGAGCACACGTCTGAACTCTTTCCCTACACGACGCTCTTCCGATCTNNNTAGCTGGGAGGTAGGA

AGCACACGTCTGAACTCTTTCCCTACACGACGCTCTTCCGATCTNNNTGTTTCTTAGGTAATAGAATGT

AAGAGCACACGTCTGAACTCTTTCCCTACACGACGCTCTTCCGATCTNNNCTGAGGCAGGAGAATAG
GGAAGAGCACACGTCTGAACTCTTTCCCTACACGACGCTCTTCCGATCTNNNGTGCAAATGTAAAATG
GAGCACACGTCTGAACTCTTTCCCTACACGACGCTCTTCCGATCTNNNACATAGATTGTCCTGAGATAC
GAAGAGCACACGTCTGAACTCTTTCCCTACACGACGCTCTTCCGATCTNNNGACCCACTACAGCAAAA
GAGCACACGTCTGAACTCTTTCCCTACACGACGCTCTTCCGATCTNNNTGGATTTCTCTGATAATCACTG
AGAGCACACGTCTGAACTCTTTCCCTACACGACGCTCTTCCGATCTNNNTCTGTCACTTGATTTTTCTTT
AGCACACGTCTGAACTCTTTCCCTACACGACGCTCTTCCGATCTNNNTCTTGGTCTAGCTAAAGTTTTAT
GAAGAGCACACGTCTGAACTCTTTCCCTACACGACGCTCTTCCGATCTNNNACTGCTCAATACCATGGT
GAGCACACGTCTGAACTCTTTCCCTACACGACGCTCTTCCGATCTNNNTCTCTTTGTATTAGTCTAGAC
AGCACACGTCTGAACTCTTTCCCTACACGACGCTCTTCCGATCTNNNTGTGCTTAGAATCACACTTTCTT
GAGCACACGTCTGAACTCTTTCCCTACACGACGCTCTTCCGATCTNNNTCTTCTTCCTTTCTCTCTCTTCTT
GAGCACACGTCTGAACTCTTTCCCTACACGACGCTCTTCCGATCTNNNCTTTCTCTTTCTTCCTTCTTTCT
AGCACACGTCTGAACTCTTTCCCTACACGACGCTCTTCCGATCTNNNGTTCCTTTTGAAAAGAATGTAC
AGCACACGTCTGAACTCTTTCCCTACACGACGCTCTTCCGATCTNNNACATTAGGGTTATTACATCTGC
AGCACACGTCTGAACTCTTTCCCTACACGACGCTCTTCCGATCTNNNACTGTTGGTGTAATGTAAATTA
AGCACACGTCTGAACTCTTTCCCTACACGACGCTCTTCCGATCTNNNTCTCTCTTTCTTTCTTTCTTTCGT
GCACACGTCTGAACTCTTTCCCTACACGACGCTCTTCCGATCTNNNTGCCCAGTTTTTAGTAGAACTAT
GAGCACACGTCTGAACTCTTTCCCTACACGACGCTCTTCCGATCTNNNTGGAAGAAACTATAAAACTT
GCACACGTCTGAACTCTTTCCCTACACGACGCTCTTCCGATCTNNNTCTTCTTTCCTCTTTCTTTCTTTCT
GAGCACACGTCTGAACTCTTTCCCTACACGACGCTCTTCCGATCTNNNTCAGTTTTATTTAGTCCAGGA
AGCACACGTCTGAACTCTTTCCCTACACGACGCTCTTCCGATCTNNNACCACAATACATGTTTGTTATTT
GAGCACACGTCTGAACTCTTTCCCTACACGACGCTCTTCCGATCTNNNGTTCACAAAACTTTAGTATTCT
GGAAGAGCACACGTCTGAACTCTTTCCCTACACGACGCTCTTCCGATCTNNNTCCCAAGGACAAAAATG
GAGCACACGTCTGAACTCTTTCCCTACACGACGCTCTTCCGATCTNNNACACTAAAAACTATAAACGTA
GAAGAGCACACGTCTGAACTCTTTCCCTACACGACGCTCTTCCGATCTNNNGCAAAACCCTGCCTCATG
GAGCACACGTCTGAACTCTTTCCCTACACGACGCTCTTCCGATCTNNNGCTATGATTATGTCACTTTTCA
AGCACACGTCTGAACTCTTTCCCTACACGACGCTCTTCCGATCTNNNTCTCTTTCTTTCTTTCTTTCTTTT
GAGCACACGTCTGAACTCTTTCCCTACACGACGCTCTTCCGATCTNNNTCCTGCAACTTTACAGAATTT
AGCACACGTCTGAACTCTTTCCCTACACGACGCTCTTCCGATCTNNNTGATTGTAGTTTTTGCAGGATA
GAGCACACGTCTGAACTCTTTCCCTACACGACGCTCTTCCGATCTNNNTCACTGAATATCTACTCTTCC
GAGCACACGTCTGAACTCTTTCCCTACACGACGCTCTTCCGATCTNNNTCATATTCTTCCATCATCAGTC
GGAAGAGCACACGTCTGAACTCTTTCCCTACACGACGCTCTTCCGATCTNNNGCAAGAAAAGTTCCTG
GAAGAGCACACGTCTGAACTCTTTCCCTACACGACGCTCTTCCGATCTNNNTAACAGCAGAATTCGCA
GGAAGAGCACACGTCTGAACTCTTTCCCTACACGACGCTCTTCCGATCTNNNTGCTCCCTTGAAATCCG
GCACACGTCTGAACTCTTTCCCTACACGACGCTCTTCCGATCTNNNGGGTACTACAGATATATGTTTA
CGGAAGAGCACACGTCTGAACTCTTTCCCTACACGACGCTCTTCCGATCTNNNACACAAAACACCCAG
GAAGAGCACACGTCTGAACTCTTTCCCTACACGACGCTCTTCCGATCTNNNCTTAGCACCAAGAGGG
GAGCACACGTCTGAACTCTTTCCCTACACGACGCTCTTCCGATCTNNNTGCATCTATGTATCTATCTGTC
AAGAGCACACGTCTGAACTCTTTCCCTACACGACGCTCTTCCGATCTNNNGCAAAAATGTGGAATCAG
GAGCACACGTCTGAACTCTTTCCCTACACGACGCTCTTCCGATCTNNNACCAACGACAGAATTTCCTTC
AGCACACGTCTGAACTCTTTCCCTACACGACGCTCTTCCGATCTNNNCACACACACATATATACACACAC
AGCACACGTCTGAACTCTTTCCCTACACGACGCTCTTCCGATCTNNNAGAGTGTGTATATATGTGTGT
GAAGAGCACACGTCTGAACTCTTTCCCTACACGACGCTCTTCCGATCTNNNCTGAGGCAGGAGAATCA

AGAGCACACGTCTGAACTCTTTCCCTACACGACGCTCTTCCGATCTNNNATATACGTGTGTGTGTGTG
AAGAGCACACGTCTGAACTCTTTCCCTACACGACGCTCTTCCGATCTNNNACCCAACCTATTACTTTCCA
AGCACACGTCTGAACTCTTTCCCTACACGACGCTCTTCCGATCTNNNCCTGATATACAGCACTTGATAT
GAAGAGCACACGTCTGAACTCTTTCCCTACACGACGCTCTTCCGATCTNNNAAATTAGCTGTGCTGCTA
AAGAGCACACGTCTGAACTCTTTCCCTACACGACGCTCTTCCGATCTNNNTTTTGTTCTCTTCCCTGCCCA
AAGAGCACACGTCTGAACTCTTTCCCTACACGACGCTCTTCCGATCTNNNCAACCAATCACCAGTCCCTA
AGAGCACACGTCTGAACTCTTTCCCTACACGACGCTCTTCCGATCTNNNTCATCACCTACCCATTCATAAA
AAGAGCACACGTCTGAACTCTTTCCCTACACGACGCTCTTCCGATCTNNNGTTTCTCCTCTCTTCCAGGGA
AGAGCACACGTCTGAACTCTTTCCCTACACGACGCTCTTCCGATCTNNNTGAAATGTAAATAATCCACTA
AGCACACGTCTGAACTCTTTCCCTACACGACGCTCTTCCGATCTNNNACATAAGAATCACAAAATCTCAG
AGAGCACACGTCTGAACTCTTTCCCTACACGACGCTCTTCCGATCTNNNTCAGTTCGAGTACATCATAAA
AAGAGCACACGTCTGAACTCTTTCCCTACACGACGCTCTTCCGATCTNNNCCCAAGTAGCTAGTACTGC
AGAGCACACGTCTGAACTCTTTCCCTACACGACGCTCTTCCGATCTNNNTCTGAATCGCTGTATGCATA
GAGCACACGTCTGAACTCTTTCCCTACACGACGCTCTTCCGATCTNNNTGAGAAACTCAGTTCTACCTA
GAGCACACGTCTGAACTCTTTCCCTACACGACGCTCTTCCGATCTNNNTCTCTCTCTCTCTCTCTCTCT
GAGCACACGTCTGAACTCTTTCCCTACACGACGCTCTTCCGATCTNNNTGAGCTCAACCTTTATATGTAT
GAGCACACGTCTGAACTCTTTCCCTACACGACGCTCTTCCGATCTNNNTGTGTGTGTGTATGTTCATGT
GAGCACACGTCTGAACTCTTTCCCTACACGACGCTCTTCCGATCTNNNTCCCATGTATTGTTTTTCCCAT
AAGAGCACACGTCTGAACTCTTTCCCTACACGACGCTCTTCCGATCTNNNGAAGAAGTCAAAGTTTCTG
AAGAGCACACGTCTGAACTCTTTCCCTACACGACGCTCTTCCGATCTNNNGTTCTCCTAGTGGTGACAC
AAGAGCACACGTCTGAACTCTTTCCCTACACGACGCTCTTCCGATCTNNNAACCTAGTAGAGATGAAAC
GAGCACACGTCTGAACTCTTTCCCTACACGACGCTCTTCCGATCTNNNTGAGCAAAATCAACCAATAAG
GAGCACACGTCTGAACTCTTTCCCTACACGACGCTCTTCCGATCTNNNAGGAATGCTGTATAAGTACTCT
GAGCACACGTCTGAACTCTTTCCCTACACGACGCTCTTCCGATCTNNNAGGTCAGCAATAGAAAACTAA
AAGAGCACACGTCTGAACTCTTTCCCTACACGACGCTCTTCCGATCTNNNGTGTGTGTGTGTGTGTGT
AGCACACGTCTGAACTCTTTCCCTACACGACGCTCTTCCGATCTNNNCAGACACTTATGTACATACACAG
AGAGCACACGTCTGAACTCTTTCCCTACACGACGCTCTTCCGATCTNNNTGCAGCCTATCTATCTGACA
AAGAGCACACGTCTGAACTCTTTCCCTACACGACGCTCTTCCGATCTNNNCTGAGGCAGAAGAATTGC
AAGAGCACACGTCTGAACTCTTTCCCTACACGACGCTCTTCCGATCTNNNTGAATCTGGAGGCAGATGT
GCACACGTCTGAACTCTTTCCCTACACGACGCTCTTCCGATCTNNNTTTTTTTTTTTTTTTGAGACGGA
AAGAGCACACGTCTGAACTCTTTCCCTACACGACGCTCTTCCGATCTNNNGCCCTTTATCAGCATAACAC
AAGAGCACACGTCTGAACTCTTTCCCTACACGACGCTCTTCCGATCTNNNGGTTGCAGTGAGCCAAGA
AGCACACGTCTGAACTCTTTCCCTACACGACGCTCTTCCGATCTNNNTGTGTACTGTATGTAATTGTCT
GAAGAGCACACGTCTGAACTCTTTCCCTACACGACGCTCTTCCGATCTNNNTACTTGGGAGAACTGCTT
AGCACACGTCTGAACTCTTTCCCTACACGACGCTCTTCCGATCTNNNTGTTGTGTTTTCGTTCTATCACAT
AGCACACGTCTGAACTCTTTCCCTACACGACGCTCTTCCGATCTNNNTCTTTCTTTCTCTCTCTCTCTTTCT
GAGCACACGTCTGAACTCTTTCCCTACACGACGCTCTTCCGATCTNNNACCATATTTAGGTCACAAAAGA
AGAGCACACGTCTGAACTCTTTCCCTACACGACGCTCTTCCGATCTNNNGGAGATGCAGTTACTTGGAA
AAGAGCACACGTCTGAACTCTTTCCCTACACGACGCTCTTCCGATCTNNNTTGTAGAGACAGGGTTTTC
GAGCACACGTCTGAACTCTTTCCCTACACGACGCTCTTCCGATCTNNNTGAGGTGTAAGATCATATCAT
AAGAGCACACGTCTGAACTCTTTCCCTACACGACGCTCTTCCGATCTNNNAAGCCTGTAATCCCAGCTA
AAGAGCACACGTCTGAACTCTTTCCCTACACGACGCTCTTCCGATCTNNNTAGAAGAATCACTTGAACC
AAGAGCACACGTCTGAACTCTTTCCCTACACGACGCTCTTCCGATCTNNNAAAATACAAAAAAACCCCA

GAGCACACGTCTGAACTCTTTCCCTACACGACGCTCTTCCGATCTNNNAGTTTTCGTGCATAGAGGT
GCACACGTCTGAACTCTTTCCCTACACGACGCTCTTCCGATCTNNNACATGCTCACTTATTATATTCAC
AGAGCACACGTCTGAACTCTTTCCCTACACGACGCTCTTCCGATCTNNNCCCAAAGTGCTGCAATTAC
AGAGCACACGTCTGAACTCTTTCCCTACACGACGCTCTTCCGATCTNNNACTGGACAGTTGGGAATG
AGAGCACACGTCTGAACTCTTTCCCTACACGACGCTCTTCCGATCTNNNCCATGAGTCAGTTACCTCC
GCACACGTCTGAACTCTTTCCCTACACGACGCTCTTCCGATCTNNNGTGTATATACGCACACATATAC
GCACACGTCTGAACTCTTTCCCTACACGACGCTCTTCCGATCTNNNTCCTACTTCCTTTCTTTCCTACTT
AGCACACGTCTGAACTCTTTCCCTACACGACGCTCTTCCGATCTNNNACAATACTGAGGTTTATCACAG
AGCACACGTCTGAACTCTTTCCCTACACGACGCTCTTCCGATCTNNNACCCAAGTCCATAATGTACATA
AGCACACGTCTGAACTCTTTCCCTACACGACGCTCTTCCGATCTNNNAAGTGAACAATAATGAAGGC
GCACACGTCTGAACTCTTTCCCTACACGACGCTCTTCCGATCTNNNTGTCTTTCTTTCTTGTCTTTCTGT
GCACACGTCTGAACTCTTTCCCTACACGACGCTCTTCCGATCTNNNTTACTCAAATGACCAAAAGAGA
AGCACACGTCTGAACTCTTTCCCTACACGACGCTCTTCCGATCTNNNTCTCTTGGGTAAATACCTGAG
AGAGCACACGTCTGAACTCTTTCCCTACACGACGCTCTTCCGATCTNNNAAGATTTAGCATCCCAGCT
AGAGCACACGTCTGAACTCTTTCCCTACACGACGCTCTTCCGATCTNNNCCTAGCTATTTGGGAGGTT
AAGAGCACACGTCTGAACTCTTTCCCTACACGACGCTCTTCCGATCTNNNGGAGGCTGAGGTGAGTT
GAAGAGCACACGTCTGAACTCTTTCCCTACACGACGCTCTTCCGATCTNNNGTGCCTGTAATCCCAGC
AAGAGCACACGTCTGAACTCTTTCCCTACACGACGCTCTTCCGATCTNNNCATCCTGAATCCCTCACCA
AGAGCACACGTCTGAACTCTTTCCCTACACGACGCTCTTCCGATCTNNNTACCTAAAGTGTGGTCTTC
AAGAGCACACGTCTGAACTCTTTCCCTACACGACGCTCTTCCGATCTNNNCAGCAATATCATATGGCT
AAGAGCACACGTCTGAACTCTTTCCCTACACGACGCTCTTCCGATCTNNNCAGGAGATGGAAGCTTC
GAAGAGCACACGTCTGAACTCTTTCCCTACACGACGCTCTTCCGATCTNNNGAGAATGACGTGAACC
GAGCACACGTCTGAACTCTTTCCCTACACGACGCTCTTCCGATCTNNNAGAAAATGGGCAAAAGACA
AGAGCACACGTCTGAACTCTTTCCCTACACGACGCTCTTCCGATCTNNNAAATTAGCCAGTTGTGTTG
AAGAGCACACGTCTGAACTCTTTCCCTACACGACGCTCTTCCGATCTNNNAGGAGAATCACTGGATC
AAGAGCACACGTCTGAACTCTTTCCCTACACGACGCTCTTCCGATCTNNNTCACCTGAAGATCACACT
GAAGAGCACACGTCTGAACTCTTTCCCTACACGACGCTCTTCCGATCTNNNGTAGTCCCAGCCACTAC
GAAGAGCACACGTCTGAACTCTTTCCCTACACGACGCTCTTCCGATCTNNNGTGCCTGTAATCCCAGC
AGAGCACACGTCTGAACTCTTTCCCTACACGACGCTCTTCCGATCTNNNGAGAATCGCTTGAACCCAG
GAAGAGCACACGTCTGAACTCTTTCCCTACACGACGCTCTTCCGATCTNNNGAGAATCGCTTGAACC
AGAGCACACGTCTGAACTCTTTCCCTACACGACGCTCTTCCGATCTNNNGAGCTGAGATCATGCCATT
GAAGAGCACACGTCTGAACTCTTTCCCTACACGACGCTCTTCCGATCTNNNACTGTAAGCCTCCAGAA
GAAGAGCACACGTCTGAACTCTTTCCCTACACGACGCTCTTCCGATCTNNNGAGAATTGTTTGAACCC
AGAGCACACGTCTGAACTCTTTCCCTACACGACGCTCTTCCGATCTNNNCTAAGGCAGGAGAATCGC
AGAGCACACGTCTGAACTCTTTCCCTACACGACGCTCTTCCGATCTNNNACAGAGACTAGAGGACCC
AAGAGCACACGTCTGAACTCTTTCCCTACACGACGCTCTTCCGATCTNNNGTGAGCCTGTAATGTCAG
GAGCACACGTCTGAACTCTTTCCCTACACGACGCTCTTCCGATCTNNNCGAGTCACATATAAGGGAAT
AGAGCACACGTCTGAACTCTTTCCCTACACGACGCTCTTCCGATCTNNNAATAGCTTGAATCTGGGA
GAGCACACGTCTGAACTCTTTCCCTACACGACGCTCTTCCGATCTNNNTCTAGGAATCACATTACCTAA
AAGAGCACACGTCTGAACTCTTTCCCTACACGACGCTCTTCCGATCTNNNAGACAATCTATGGTCACA
AGCACACGTCTGAACTCTTTCCCTACACGACGCTCTTCCGATCTNNNCCTGACTTCAAATTGTACTACA
AAGAGCACACGTCTGAACTCTTTCCCTACACGACGCTCTTCCGATCTNNNTGTAATCCCAGCTACTTTG
AGAGCACACGTCTGAACTCTTTCCCTACACGACGCTCTTCCGATCTNNNCCCAATTAGAAATGGCAA

AGAGCACACGTCTGAACTCTTTCCCTACACGACGCTCTTCCGATCTNNNTACTCTGGAGGCTAAGGAAA
AAGAGCACACGTCTGAACTCTTTCCCTACACGACGCTCTTCCGATCTNNNTCAGGATAATCGCTTGAAC
AAGAGCACACGTCTGAACTCTTTCCCTACACGACGCTCTTCCGATCTNNNGAGAATGGTGTGAACCCAC
GAAGAGCACACGTCTGAACTCTTTCCCTACACGACGCTCTTCCGATCTNNNTTGGAGGTATCCCCCTA
AGAGCACACGTCTGAACTCTTTCCCTACACGACGCTCTTCCGATCTNNNCTGTAATCCCAGCTACTCAGC
AAGAGCACACGTCTGAACTCTTTCCCTACACGACGCTCTTCCGATCTNNNATACAAAAATTAGCTGGG
AGCACACGTCTGAACTCTTTCCCTACACGACGCTCTTCCGATCTNNNTCAAATGTACCCCAGAAACATA
GAAGAGCACACGTCTGAACTCTTTCCCTACACGACGCTCTTCCGATCTNNNCTGTAGTCCAAGCTATTCC
GAAGAGCACACGTCTGAACTCTTTCCCTACACGACGCTCTTCCGATCTNNNTCTGTAATCCCAGCTACT
AGCACACGTCTGAACTCTTTCCCTACACGACGCTCTTCCGATCTNNNGGTCCTAACATCATTGTATAGGA
GAAGAGCACACGTCTGAACTCTTTCCCTACACGACGCTCTTCCGATCTNNNGAGAATGGCATGAACTCA
GGAAGAGCACACGTCTGAACTCTTTCCCTACACGACGCTCTTCCGATCTNNNGAAAATGGCATGAACCC
AAGAGCACACGTCTGAACTCTTTCCCTACACGACGCTCTTCCGATCTNNNTGTCTTTTGAAATAAGCCA
AGAGCACACGTCTGAACTCTTTCCCTACACGACGCTCTTCCGATCTNNNGAGGCAGAAGAATCACTTG
GAAGAGCACACGTCTGAACTCTTTCCCTACACGACGCTCTTCCGATCTNNNAATCACTTGAAGCCAGGA
GAAGAGCACACGTCTGAACTCTTTCCCTACACGACGCTCTTCCGATCTNNNAAAAAAAAAATTAGCCG
AAGAGCACACGTCTGAACTCTTTCCCTACACGACGCTCTTCCGATCTNNNACTTGGTAGGTTGAGGTAC
AAGAGCACACGTCTGAACTCTTTCCCTACACGACGCTCTTCCGATCTNNNTTATCCTTCCCTCCTCCAAC
GAAGAGCACACGTCTGAACTCTTTCCCTACACGACGCTCTTCCGATCTNNNAGCTTTCAACATTGGCACT
AGAGCACACGTCTGAACTCTTTCCCTACACGACGCTCTTCCGATCTNNNATAGAACGCAATAGAGAGG
AAGAGCACACGTCTGAACTCTTTCCCTACACGACGCTCTTCCGATCTNNNTGAGGCAGAAGAATTGCT
GAGCACACGTCTGAACTCTTTCCCTACACGACGCTCTTCCGATCTNNNACTCTGTTAGATGTTGAGGAT
AAGAGCACACGTCTGAACTCTTTCCCTACACGACGCTCTTCCGATCTNNNTGCATTGGCGGCTATATTT
GAGCACACGTCTGAACTCTTTCCCTACACGACGCTCTTCCGATCTNNNTGGAATATCCATTACCTCAAAC
GAGCACACGTCTGAACTCTTTCCCTACACGACGCTCTTCCGATCTNNNACCACTACACACCTATTAGAAT
AAGAGCACACGTCTGAACTCTTTCCCTACACGACGCTCTTCCGATCTNNNATAATCCCAGCTATTCAGG
AAGAGCACACGTCTGAACTCTTTCCCTACACGACGCTCTTCCGATCTNNNACAACAAACCCTTATGACA
AAGAGCACACGTCTGAACTCTTTCCCTACACGACGCTCTTCCGATCTNNNTGAAGCATGAGAATCGCT
AGAGCACACGTCTGAACTCTTTCCCTACACGACGCTCTTCCGATCTNNNACAAAATTCATGAAAGGGTG
AAGAGCACACGTCTGAACTCTTTCCCTACACGACGCTCTTCCGATCTNNNATCAGTTCAGCTTGGCAGT
GGAAGAGCACACGTCTGAACTCTTTCCCTACACGACGCTCTTCCGATCTNNNGGGAAGTGAAGTTTGT
GAGCACACGTCTGAACTCTTTCCCTACACGACGCTCTTCCGATCTNNNAGTTGTTACACTGTTTTGTTTA
GCACACGTCTGAACTCTTTCCCTACACGACGCTCTTCCGATCTNNNTGTTTATCAGGAATGTGTTTTGA
GCACACGTCTGAACTCTTTCCCTACACGACGCTCTTCCGATCTNNNTGAACTTTACATTCTCCTTTTGACA
AAGAGCACACGTCTGAACTCTTTCCCTACACGACGCTCTTCCGATCTNNNGTAATTCACATCCCTCCCTT
GAGCACACGTCTGAACTCTTTCCCTACACGACGCTCTTCCGATCTNNNTGAAGGCTTTGTCTTTTAGTA
AGAGCACACGTCTGAACTCTTTCCCTACACGACGCTCTTCCGATCTNNNCACCCGGCCTGAATTCATTT
AGCACACGTCTGAACTCTTTCCCTACACGACGCTCTTCCGATCTNNNTCTCTACCTTTGTATTTTCATCTC
AGAGCACACGTCTGAACTCTTTCCCTACACGACGCTCTTCCGATCTNNNTCAGTAGGGGAACAAATAA
GCACACGTCTGAACTCTTTCCCTACACGACGCTCTTCCGATCTNNNATGCATACATATGTATACATGCAA
GAGCACACGTCTGAACTCTTTCCCTACACGACGCTCTTCCGATCTNNNTCTTTCTTCTCTCTTTCTCCCTTT
CGGAAGAGCACACGTCTGAACTCTTTCCCTACACGACGCTCTTCCGATCTNNNTGCAGACTTCAGTTAG
GCACACGTCTGAACTCTTTCCCTACACGACGCTCTTCCGATCTNNNAGCATGCCTCTATATAGCATATAT

GAGCACACGTCTGAACTCTTTCCCTACACGACGCTCTTCCGATCTNNNAAACAAAAAATATACACACA
AGCACACGTCTGAACTCTTTCCCTACACGACGCTCTTCCGATCTNNNGTCCAACAGGAAAATATCACAA
AGAGCACACGTCTGAACTCTTTCCCTACACGACGCTCTTCCGATCTNNNCACCCGGCCTGAATTCATTT
AGCACACGTCTGAACTCTTTCCCTACACGACGCTCTTCCGATCTNNNATGTATAGACACAAACATGCA
AGAGCACACGTCTGAACTCTTTCCCTACACGACGCTCTTCCGATCTNNNTCTTGCTTTTTATGGCTGCA
GAGCACACGTCTGAACTCTTTCCCTACACGACGCTCTTCCGATCTNNNCCATGGTGTGTGTATATGTA
AAGAGCACACGTCTGAACTCTTTCCCTACACGACGCTCTTCCGATCTNNNAAGGGGTGATAGGAGTG
AGAGCACACGTCTGAACTCTTTCCCTACACGACGCTCTTCCGATCTNNNAGAATTGCTTGAATCTGGG
AGAGCACACGTCTGAACTCTTTCCCTACACGACGCTCTTCCGATCTNNNGGCTCCTAGCTTGGAAAAT
AAGAGCACACGTCTGAACTCTTTCCCTACACGACGCTCTTCCGATCTNNNCACTCATGAAGGGTCCTG
GAGCACACGTCTGAACTCTTTCCCTACACGACGCTCTTCCGATCTNNNTCACAGTTGCAAAGATAAGG
AAGAGCACACGTCTGAACTCTTTCCCTACACGACGCTCTTCCGATCTNNNTGAGGCAGGAGAATAGT
GAGCACACGTCTGAACTCTTTCCCTACACGACGCTCTTCCGATCTNNNTGTATATATGTGTGTGTGTGT
AGCACACGTCTGAACTCTTTCCCTACACGACGCTCTTCCGATCTNNNTGTGTGTAAGTTGCTAAAACAA
AGCACACGTCTGAACTCTTTCCCTACACGACGCTCTTCCGATCTNNNACAATTTTGCCTTTTTTTTCTTT
AAGAGCACACGTCTGAACTCTTTCCCTACACGACGCTCTTCCGATCTNNNGAGGTTGCAGTGAACTGA
AGAGCACACGTCTGAACTCTTTCCCTACACGACGCTCTTCCGATCTNNNACTAGAAAGCAGCAAGGAA
GAAGAGCACACGTCTGAACTCTTTCCCTACACGACGCTCTTCCGATCTNNNTTCACACACACACATACA
AAGAGCACACGTCTGAACTCTTTCCCTACACGACGCTCTTCCGATCTNNNTCTACACTCCCATGTTCATT
AAGAGCACACGTCTGAACTCTTTCCCTACACGACGCTCTTCCGATCTNNNCCAAGTAGTCACAATCCT
AGAGCACACGTCTGAACTCTTTCCCTACACGACGCTCTTCCGATCTNNNTGTAAATTTCATGCAATTTC
GGAAGAGCACACGTCTGAACTCTTTCCCTACACGACGCTCTTCCGATCTNNNGCTGCTCCTCCTTACTT
AGCACACGTCTGAACTCTTTCCCTACACGACGCTCTTCCGATCTNNNGCCATAAAAAGGAATGAATCA
AAGAGCACACGTCTGAACTCTTTCCCTACACGACGCTCTTCCGATCTNNNAATATGGAACCAGCCTAA
AGAGCACACGTCTGAACTCTTTCCCTACACGACGCTCTTCCGATCTNNNGAACCAACCCAAATGTCCAT
GCACACGTCTGAACTCTTTCCCTACACGACGCTCTTCCGATCTNNNCACACACACATATATACACACAC
AAGAGCACACGTCTGAACTCTTTCCCTACACGACGCTCTTCCGATCTNNNTCCCACCCAGCATACATAA
GAGCACACGTCTGAACTCTTTCCCTACACGACGCTCTTCCGATCTNNNTCTTTGTTTCAATTCATCCTTTT
GAGCACACGTCTGAACTCTTTCCCTACACGACGCTCTTCCGATCTNNNACCATACTGTTTTGATCATCA
AGAGCACACGTCTGAACTCTTTCCCTACACGACGCTCTTCCGATCTNNNTCAAATCATTGGAAAGCAT
AGCACACGTCTGAACTCTTTCCCTACACGACGCTCTTCCGATCTNNNTCCAGTTTTATTTGAAGTGAGG
AAGAGCACACGTCTGAACTCTTTCCCTACACGACGCTCTTCCGATCTNNNTTGTCTCTTTGTCTTTACC
AGCACACGTCTGAACTCTTTCCCTACACGACGCTCTTCCGATCTNNNTGAAAGAATGTTTTGTGGCAT
AGCACACGTCTGAACTCTTTCCCTACACGACGCTCTTCCGATCTNNNTGGTTGGTTACAATTATACAGT
GAAGAGCACACGTCTGAACTCTTTCCCTACACGACGCTCTTCCGATCTNNNTTGCATTTCCACCAGCAA
AGCACACGTCTGAACTCTTTCCCTACACGACGCTCTTCCGATCTNNNCAGTCATTTCATAGTGTTGATG
AGCACACGTCTGAACTCTTTCCCTACACGACGCTCTTCCGATCTNNNACATTTTGCTTTATTTTCTTTGG
AGCACACGTCTGAACTCTTTCCCTACACGACGCTCTTCCGATCTNNNTGAATGAGCTTTCATAATTTGT
GAGCACACGTCTGAACTCTTTCCCTACACGACGCTCTTCCGATCTNNNCCTTTTGCAAAGTTCAAGTTTA
GCACACGTCTGAACTCTTTCCCTACACGACGCTCTTCCGATCTNNNCAGTGTCTCATTACTTGTATATCC
GAGCACACGTCTGAACTCTTTCCCTACACGACGCTCTTCCGATCTNNNACTGTTTACTGTTGAATAAGG
AGCACACGTCTGAACTCTTTCCCTACACGACGCTCTTCCGATCTNNNTGCCATATTGAAAAAAAATGA
AGCACACGTCTGAACTCTTTCCCTACACGACGCTCTTCCGATCTNNNTGAAAGTATTATACCCAGGAG

AGCACACGTCTGAACTCTTTCCCTACACGACGCTCTTCCGATCTNNNAGCTTCATTTGTTATTGTCCATT
AGAGCACACGTCTGAACTCTTTCCCTACACGACGCTCTTCCGATCTNNNTTCAGCAACCAAGTCCTGTA
AAGAGCACACGTCTGAACTCTTTCCCTACACGACGCTCTTCCGATCTNNNTGTGTTTTGCAGTTCTGCT
AGCACACGTCTGAACTCTTTCCCTACACGACGCTCTTCCGATCTNNNTGAGACAGACTTTTTTGAGAC
GAGCACACGTCTGAACTCTTTCCCTACACGACGCTCTTCCGATCTNNNGCAGTAATAGCTTTTTGCATT
AGCACACGTCTGAACTCTTTCCCTACACGACGCTCTTCCGATCTNNNGGTTTCAGAAACAATTTGTTAT
AGCACACGTCTGAACTCTTTCCCTACACGACGCTCTTCCGATCTNNNATCCGTATAACAGAAAATTTGC
GAGCACACGTCTGAACTCTTTCCCTACACGACGCTCTTCCGATCTNNNTCTCTGAGATGTAAATATTGC
AGCACACGTCTGAACTCTTTCCCTACACGACGCTCTTCCGATCTNNNTGTCTGATATTAGGATTGCAAC
GAGCACACGTCTGAACTCTTTCCCTACACGACGCTCTTCCGATCTNNNAGAATGATAGCAAAATCCAT
AGAGCACACGTCTGAACTCTTTCCCTACACGACGCTCTTCCGATCTNNNTGCCTAAAAGCTTACTTACT
AAGAGCACACGTCTGAACTCTTTCCCTACACGACGCTCTTCCGATCTNNNGCAAAAAATATGGAACCA
GAGCACACGTCTGAACTCTTTCCCTACACGACGCTCTTCCGATCTNNNTGAATCCTTCAACTTCACTGA
AAGAGCACACGTCTGAACTCTTTCCCTACACGACGCTCTTCCGATCTNNNAAAAAATTAGCTGGACGT
AGCACACGTCTGAACTCTTTCCCTACACGACGCTCTTCCGATCTNNNACTTGTTTGTTGTCCATTTGTA
GAAGAGCACACGTCTGAACTCTTTCCCTACACGACGCTCTTCCGATCTNNNGCTAAGGTAGGAGGATT
AGCACACGTCTGAACTCTTTCCCTACACGACGCTCTTCCGATCTNNNTCTATGTATAAGAGGAATGGA
AGAGCACACGTCTGAACTCTTTCCCTACACGACGCTCTTCCGATCTNNNACAAGTAGAGTTCTTAGCCT
GAAGAGCACACGTCTGAACTCTTTCCCTACACGACGCTCTTCCGATCTNNNGAAGAAATTGCACTGACT
AAGAGCACACGTCTGAACTCTTTCCCTACACGACGCTCTTCCGATCTNNNCCATGTAAGTGGATCACTA
GAAGAGCACACGTCTGAACTCTTTCCCTACACGACGCTCTTCCGATCTNNNGGTGCCTGTAATCTCAGC
AGCACACGTCTGAACTCTTTCCCTACACGACGCTCTTCCGATCTNNNTCTTCAGTTGGTTGATTGTCTTT
AAGAGCACACGTCTGAACTCTTTCCCTACACGACGCTCTTCCGATCTNNNTTTCTCTCTCTCTCTCCCCC
GGAAGAGCACACGTCTGAACTCTTTCCCTACACGACGCTCTTCCGATCTNNNAAGATGGAAGGAAGG
AGAGCACACGTCTGAACTCTTTCCCTACACGACGCTCTTCCGATCTNNNTGTAATTCCAGCTAATCGG
AGAGCACACGTCTGAACTCTTTCCCTACACGACGCTCTTCCGATCTNNNGCAATTGCAAAAATGTGGA
AGAGCACACGTCTGAACTCTTTCCCTACACGACGCTCTTCCGATCTNNNGTGGGCTAGCATGCAGTAA
AAGAGCACACGTCTGAACTCTTTCCCTACACGACGCTCTTCCGATCTNNNGGAACAACAGGAAGAGTA
GAGCACACGTCTGAACTCTTTCCCTACACGACGCTCTTCCGATCTNNNGGTATCTCACCTATTTGTCAT
GAGCACACGTCTGAACTCTTTCCCTACACGACGCTCTTCCGATCTNNNTCTATATCTCCCTGAAATGATG
GAAGAGCACACGTCTGAACTCTTTCCCTACACGACGCTCTTCCGATCTNNNAGTGAGCTACGATCATGT
AGAGCACACGTCTGAACTCTTTCCCTACACGACGCTCTTCCGATCTNNNTCCTCCCATCTCATCCTCTTAC
AGCACACGTCTGAACTCTTTCCCTACACGACGCTCTTCCGATCTNNNTGAAGCACTTCCATACTATTCTC
GAGCACACGTCTGAACTCTTTCCCTACACGACGCTCTTCCGATCTNNNTCCCTTTGTGATTGATATCTCT
GAGCACACGTCTGAACTCTTTCCCTACACGACGCTCTTCCGATCTNNNACTGAGTCACAAACAACATCA
AGAGCACACGTCTGAACTCTTTCCCTACACGACGCTCTTCCGATCTNNNACCTTTACTCATTGATGGAGA
AGAGCACACGTCTGAACTCTTTCCCTACACGACGCTCTTCCGATCTNNNCAACTGAGTACACATACACA
GAAGAGCACACGTCTGAACTCTTTCCCTACACGACGCTCTTCCGATCTNNNGTGAGAGAGTGCGTGT
AGCACACGTCTGAACTCTTTCCCTACACGACGCTCTTCCGATCTNNNGCAATTGCAAAAATATGAGAC
AGCACACGTCTGAACTCTTTCCCTACACGACGCTCTTCCGATCTNNNTGTAGATACAGACCAAATCAA
AGAGCACACGTCTGAACTCTTTCCCTACACGACGCTCTTCCGATCTNNNAAGAATTGCCTGAACCTGG
AGCACACGTCTGAACTCTTTCCCTACACGACGCTCTTCCGATCTNNNACCTAAATGTAAGGTATGCTTGA
GAGCACACGTCTGAACTCTTTCCCTACACGACGCTCTTCCGATCTNNNAGTGTGGGATAATTATGGCTA

AGCACACGTCTGAACTCTTTCCCTACACGACGCTCTTCCGATCTNNNTGAAACAATGGAAGATGTAG
GAGCACACGTCTGAACTCTTTCCCTACACGACGCTCTTCCGATCTNNNTTATAACAGAGGTGGGTGA
AGCACACGTCTGAACTCTTTCCCTACACGACGCTCTTCCGATCTNNNAGCCTTGTAAAATGAACTGG
GAGCACACGTCTGAACTCTTTCCCTACACGACGCTCTTCCGATCTNNNTGTGCTTTGTATCCTGTATCC
GAGCACACGTCTGAACTCTTTCCCTACACGACGCTCTTCCGATCTNNNAGCATAGTTATTTTATGGCC
GAGCACACGTCTGAACTCTTTCCCTACACGACGCTCTTCCGATCTNNNACTGTGGTTATGTAAGAGG
GAGCACACGTCTGAACTCTTTCCCTACACGACGCTCTTCCGATCTNNNTGTTTGTTTCCTTGCATCTGC
AGCACACGTCTGAACTCTTTCCCTACACGACGCTCTTCCGATCTNNNTCCCTCTCTCTCTCTCTCTCTC
AGCACACGTCTGAACTCTTTCCCTACACGACGCTCTTCCGATCTNNNAGAAAGAGAGAGACAGAGAG
GCACACGTCTGAACTCTTTCCCTACACGACGCTCTTCCGATCTNNNAGTGTTCTTTTCCTCCAAAGTCTC
GAGCACACGTCTGAACTCTTTCCCTACACGACGCTCTTCCGATCTNNNGATACACACACACACACACA
GAGCACACGTCTGAACTCTTTCCCTACACGACGCTCTTCCGATCTNNNGTGAGACCTCTCTCTCTCTCT
GCACACGTCTGAACTCTTTCCCTACACGACGCTCTTCCGATCTNNNTGGTGGAGAAATTGTATTTTTG
AGCACACGTCTGAACTCTTTCCCTACACGACGCTCTTCCGATCTNNNACAGTGATTATTACAGTGACT
AGAGCACACGTCTGAACTCTTTCCCTACACGACGCTCTTCCGATCTNNNCAGTCAATTGGCCTTGTAA
AGCACACGTCTGAACTCTTTCCCTACACGACGCTCTTCCGATCTNNNAGGCATATATATGTGTGTGT
GCACACGTCTGAACTCTTTCCCTACACGACGCTCTTCCGATCTNNNTGTGAATTCCTATACCATCCACA
AAGAGCACACGTCTGAACTCTTTCCCTACACGACGCTCTTCCGATCTNNNAATGCAGAGTAAACGTGT
GAGCACACGTCTGAACTCTTTCCCTACACGACGCTCTTCCGATCTNNNTTTCAGCTTGCAAATAGCCT
GAGCACACGTCTGAACTCTTTCCCTACACGACGCTCTTCCGATCTNNNTAGTTCTCTAGCTTGCAGAC
GAGCACACGTCTGAACTCTTTCCCTACACGACGCTCTTCCGATCTNNNACTTTCTGTGTGAAGCAATCA
AGCACACGTCTGAACTCTTTCCCTACACGACGCTCTTCCGATCTNNNACTCCATAATAAGTCAAGGAC
GCACACGTCTGAACTCTTTCCCTACACGACGCTCTTCCGATCTNNNTTGGATCCCAACTATTTCTTTCT
AGAGCACACGTCTGAACTCTTTCCCTACACGACGCTCTTCCGATCTNNNTAGAGTGGAGATGACAGA
AGAGCACACGTCTGAACTCTTTCCCTACACGACGCTCTTCCGATCTNNNTCCTGTATTGTGCATCTCTC
AGCACACGTCTGAACTCTTTCCCTACACGACGCTCTTCCGATCTNNNTTTACCTTTACTACAGCCATAT
AGCACACGTCTGAACTCTTTCCCTACACGACGCTCTTCCGATCTNNNTGTCTTTCTGTATCTCTCTCTCT
GAAGAGCACACGTCTGAACTCTTTCCCTACACGACGCTCTTCCGATCTNNNTTTCCCTTTAGGCTCAT
GAAGAGCACACGTCTGAACTCTTTCCCTACACGACGCTCTTCCGATCTNNNAAGGGAGGAAGAAAG
AGAGCACACGTCTGAACTCTTTCCCTACACGACGCTCTTCCGATCTNNNCTGTAGGGCTGCAAGTCAT
AAGAGCACACGTCTGAACTCTTTCCCTACACGACGCTCTTCCGATCTNNNCTGTAATTCCAGCTACTCG
AGAGCACACGTCTGAACTCTTTCCCTACACGACGCTCTTCCGATCTNNNCAGGAGCATGAAAATAGCA
AGAGCACACGTCTGAACTCTTTCCCTACACGACGCTCTTCCGATCTNNNGAGGCAGTAGAATTGCTT
AGCACACGTCTGAACTCTTTCCCTACACGACGCTCTTCCGATCTNNNTGGCATAGAGTTATTTCTCCC
GAGCACACGTCTGAACTCTTTCCCTACACGACGCTCTTCCGATCTNNNATACAAAAAATTAGCCGGG
AGAGCACACGTCTGAACTCTTTCCCTACACGACGCTCTTCCGATCTNNNTGGCCTGGCAGATATAAAC
AGAGCACACGTCTGAACTCTTTCCCTACACGACGCTCTTCCGATCTNNNAGCAAATGAGTGACTTTGT
AGCACACGTCTGAACTCTTTCCCTACACGACGCTCTTCCGATCTNNNTCATCAAAGTTACAGAGTTCTC
AGCACACGTCTGAACTCTTTCCCTACACGACGCTCTTCCGATCTNNNTGTAGAGATTTTTCACCTCTCC
AGCACACGTCTGAACTCTTTCCCTACACGACGCTCTTCCGATCTNNNGCTAATCTATATCCTGTGGACA
AGCACACGTCTGAACTCTTTCCCTACACGACGCTCTTCCGATCTNNNTTTGCATCTCATTTAGTAAGCA
AGAGCACACGTCTGAACTCTTTCCCTACACGACGCTCTTCCGATCTNNNTGTAGTCTCAGCTACTCAG
AGAGCACACGTCTGAACTCTTTCCCTACACGACGCTCTTCCGATCTNNNCTGTAATCCCAGCTACTCCA

AGCACACGTCTGAACTCTTTCCCTACACGACGCTCTTCCGATCTNNNGGAGAATTTTCCTTTTGCTTGA
AGCACACGTCTGAACTCTTTCCCTACACGACGCTCTTCCGATCTNNNTCAGAGACTAAGATTGTAACC
GAGCACACGTCTGAACTCTTTCCCTACACGACGCTCTTCCGATCTNNNTGAGGTAGGAGAATTGCTTG
GCACACGTCTGAACTCTTTCCCTACACGACGCTCTTCCGATCTNNNGCAAACAGTTCTTCTAAAGCTTT
GAGCACACGTCTGAACTCTTTCCCTACACGACGCTCTTCCGATCTNNNTGGCTTTGAATTTATCTGTTC
GCACACGTCTGAACTCTTTCCCTACACGACGCTCTTCCGATCTNNNTCCTGCCTTTTTTTTTGTATTG
AGAGCACACGTCTGAACTCTTTCCCTACACGACGCTCTTCCGATCTNNNAGTACAACATAAGGCTGAG
AAGAGCACACGTCTGAACTCTTTCCCTACACGACGCTCTTCCGATCTNNNGAGAATCGCTTGAACCCA
AGCACACGTCTGAACTCTTTCCCTACACGACGCTCTTCCGATCTNNNAGACTGCTATCAAAACCAAACT
GAGCACACGTCTGAACTCTTTCCCTACACGACGCTCTTCCGATCTNNNAGCCAGACTAACAAAAGAG
GAGCACACGTCTGAACTCTTTCCCTACACGACGCTCTTCCGATCTNNNTGTAATCTCAGCACTTTGGG
AAGAGCACACGTCTGAACTCTTTCCCTACACGACGCTCTTCCGATCTNNNCACCTGTAGTCCCAGCTAA
AGAGCACACGTCTGAACTCTTTCCCTACACGACGCTCTTCCGATCTNNNGCCTCTTGTGTCTGGCTTA
GAGCACACGTCTGAACTCTTTCCCTACACGACGCTCTTCCGATCTNNNTGCGTATTCTGCATAAAATTG
GAAGAGCACACGTCTGAACTCTTTCCCTACACGACGCTCTTCCGATCTNNNGTTCTCCTAGTGGTGA
AAGAGCACACGTCTGAACTCTTTCCCTACACGACGCTCTTCCGATCTNNNCTGTAGTCACAGCTACTC
AAGAGCACACGTCTGAACTCTTTCCCTACACGACGCTCTTCCGATCTNNNAGAAAACACCAGAGATCC
AGAGCACACGTCTGAACTCTTTCCCTACACGACGCTCTTCCGATCTNNNGTAATCCCAGCTACTCAAG
GAGCACACGTCTGAACTCTTTCCCTACACGACGCTCTTCCGATCTNNNTGGTTTTCTGGTATTTTGTTG
GAGCACACGTCTGAACTCTTTCCCTACACGACGCTCTTCCGATCTNNNGTAATCCCAGCTACTCAGAA
AGAGCACACGTCTGAACTCTTTCCCTACACGACGCTCTTCCGATCTNNNAAGCTGAGACAGGAGAAT
GCACACGTCTGAACTCTTTCCCTACACGACGCTCTTCCGATCTNNNACATTTCAACCTCTAAATCATGC
GCACACGTCTGAACTCTTTCCCTACACGACGCTCTTCCGATCTNNNACAAGTCATCAAGATTCAACAAA
AGAGCACACGTCTGAACTCTTTCCCTACACGACGCTCTTCCGATCTNNNTGTAATTCCAGCTACTTGG
AAGAGCACACGTCTGAACTCTTTCCCTACACGACGCTCTTCCGATCTNNNCAAGGCAGGAGAATCGTT
GAGCACACGTCTGAACTCTTTCCCTACACGACGCTCTTCCGATCTNNNCCTGTCTGAATGTACCTGTAT
AGAGCACACGTCTGAACTCTTTCCCTACACGACGCTCTTCCGATCTNNNCCAGCAGCCAAAGTACTGA
AGAGCACACGTCTGAACTCTTTCCCTACACGACGCTCTTCCGATCTNNNGCAACATGTCATACACACA
GCACACGTCTGAACTCTTTCCCTACACGACGCTCTTCCGATCTNNNTTGCATATTGTTTTTGTATACCG
AGCACACGTCTGAACTCTTTCCCTACACGACGCTCTTCCGATCTNNNACCTATGCATCCATTTTTCTTTA
GCACACGTCTGAACTCTTTCCCTACACGACGCTCTTCCGATCTNNNAGATCATCATAAAGGTCTTCGT
GCACACGTCTGAACTCTTTCCCTACACGACGCTCTTCCGATCTNNNTCAAATGTTTCACTATTGGCAC
GCACACGTCTGAACTCTTTCCCTACACGACGCTCTTCCGATCTNNNTGTTACTGATTTTCATTTGTTGC
CACACGTCTGAACTCTTTCCCTACACGACGCTCTTCCGATCTNNNACAAACTTTCTCTTTTCCTTCCATC
CACACGTCTGAACTCTTTCCCTACACGACGCTCTTCCGATCTNNNTGACAGATAAAAATAGAAAGGAC
GAGCACACGTCTGAACTCTTTCCCTACACGACGCTCTTCCGATCTNNNGGGAGAGGGGAGATTTTC
GCACACGTCTGAACTCTTTCCCTACACGACGCTCTTCCGATCTNNNAAGAAACTAACAAATCCAACCA
AGCACACGTCTGAACTCTTTCCCTACACGACGCTCTTCCGATCTNNNAGAGCTACCATAAAAAAATTA
GCACACGTCTGAACTCTTTCCCTACACGACGCTCTTCCGATCTNNNTGTTCATTATAATCCCTTTGATGC
GCACACGTCTGAACTCTTTCCCTACACGACGCTCTTCCGATCTNNNTGCTGCTTTGTTTTTCCTTTATTT
AGCACACGTCTGAACTCTTTCCCTACACGACGCTCTTCCGATCTNNNACCATTACCAAAGACTTTAGG
AGAGCACACGTCTGAACTCTTTCCCTACACGACGCTCTTCCGATCTNNNATCAGCTTTACCTCACTCAG
AAGAGCACACGTCTGAACTCTTTCCCTACACGACGCTCTTCCGATCTNNNCTGTAATTCCAGCTACTCG

AGAGCACACGTCTGAACTCTTTCCCTACACGACGCTCTTCCGATCTNNNCAGGAGCATGAAAATAGCA
AAGAGCACACGTCTGAACTCTTTCCCTACACGACGCTCTTCCGATCTNNNGAGGCAGTAGAATTGCTTG
AGAGCACACGTCTGAACTCTTTCCCTACACGACGCTCTTCCGATCTNNNGTAGTGGCACATGCTTGTA
GAAGAGCACACGTCTGAACTCTTTCCCTACACGACGCTCTTCCGATCTNNNATACAAAAATTAGCTGGG
AAGAGCACACGTCTGAACTCTTTCCCTACACGACGCTCTTCCGATCTNNNCAGAATCGCTTGAACACAG
AGAGCACACGTCTGAACTCTTTCCCTACACGACGCTCTTCCGATCTNNNGCAGAAGAATCACTTGAAC
AAGAGCACACGTCTGAACTCTTTCCCTACACGACGCTCTTCCGATCTNNNCTGAGGCAGGAGGATCAA
GAAGAGCACACGTCTGAACTCTTTCCCTACACGACGCTCTTCCGATCTNNNCAGGAGAATCACTTGAA
AAGAGCACACGTCTGAACTCTTTCCCTACACGACGCTCTTCCGATCTNNNAAAAAATTAGCTGGGTGTG
GAGCACACGTCTGAACTCTTTCCCTACACGACGCTCTTCCGATCTNNNACATCTGTATGAAAAGTGCTT
GAGCACACGTCTGAACTCTTTCCCTACACGACGCTCTTCCGATCTNNNGTTGCAGTGAGCCAAGATCA
GAAGAGCACACGTCTGAACTCTTTCCCTACACGACGCTCTTCCGATCTNNNCAGTAGGTGTAGGTTGC
AGAGCACACGTCTGAACTCTTTCCCTACACGACGCTCTTCCGATCTNNNCTGAGGCAGGAGAATTGCT
AGAGCACACGTCTGAACTCTTTCCCTACACGACGCTCTTCCGATCTNNNGCATGTCTGTAATCCCAGCT
GAGCACACGTCTGAACTCTTTCCCTACACGACGCTCTTCCGATCTNNNCTGAGACAGGAGAATCGCTTG
AAGAGCACACGTCTGAACTCTTTCCCTACACGACGCTCTTCCGATCTNNNAGGTTGCAGTGAGCTGAG
AAGAGCACACGTCTGAACTCTTTCCCTACACGACGCTCTTCCGATCTNNNTAATCCCAGCTACTTAGGA
AAGAGCACACGTCTGAACTCTTTCCCTACACGACGCTCTTCCGATCTNNNAATAGCTGAGTGTGGTTTT
AGAGCACACGTCTGAACTCTTTCCCTACACGACGCTCTTCCGATCTNNNCTGTAGTCCCAGCTACTCAG
GGAAGAGCACACGTCTGAACTCTTTCCCTACACGACGCTCTTCCGATCTNNNGGAGGCAGAGCTTGTA
AGAGCACACGTCTGAACTCTTTCCCTACACGACGCTCTTCCGATCTNNNAGGAGAATTGCTTGAACCT
AGCACACGTCTGAACTCTTTCCCTACACGACGCTCTTCCGATCTNNNTGCAAAAATGATTACAAACACCT
AGAGCACACGTCTGAACTCTTTCCCTACACGACGCTCTTCCGATCTNNNCAGGAGAATGGCATGAACCT
AGAGCACACGTCTGAACTCTTTCCCTACACGACGCTCTTCCGATCTNNNGAGACAGGAGAATCGCTTG
AGAGCACACGTCTGAACTCTTTCCCTACACGACGCTCTTCCGATCTNNNCAGAAGAATTGCTTGAACCC
AGAGCACACGTCTGAACTCTTTCCCTACACGACGCTCTTCCGATCTNNNAAAAATAAAGTGGGTGTGC
GAGCACACGTCTGAACTCTTTCCCTACACGACGCTCTTCCGATCTNNNTGTGGACTAGAACATACATGT
GGAAGAGCACACGTCTGAACTCTTTCCCTACACGACGCTCTTCCGATCTNNNGAAGCAGGAGAATCGC
AGAGCACACGTCTGAACTCTTTCCCTACACGACGCTCTTCCGATCTNNNTGTAATTCCAGCTACTTGGG
AAGAGCACACGTCTGAACTCTTTCCCTACACGACGCTCTTCCGATCTNNNGTGCCTGTAATCCTAGCTA
GAAGAGCACACGTCTGAACTCTTTCCCTACACGACGCTCTTCCGATCTNNNGAGAATGGTGTGAACCCA
GGAAGAGCACACGTCTGAACTCTTTCCCTACACGACGCTCTTCCGATCTNNNAAGAGGCTGAGACAGG
GAAGAGCACACGTCTGAACTCTTTCCCTACACGACGCTCTTCCGATCTNNNAGGAGAATTGCCTGAACC
AGAGCACACGTCTGAACTCTTTCCCTACACGACGCTCTTCCGATCTNNNACATTCATAAAGGCCAAACA
AGAGCACACGTCTGAACTCTTTCCCTACACGACGCTCTTCCGATCTNNNCATGAGAATTGCTTGAACCC
AGAGCACACGTCTGAACTCTTTCCCTACACGACGCTCTTCCGATCTNNNAAAAAATTAGCCAGGTGTGG
AAGAGCACACGTCTGAACTCTTTCCCTACACGACGCTCTTCCGATCTNNNGCCCAATGGTATGGTGTT
AGAGCACACGTCTGAACTCTTTCCCTACACGACGCTCTTCCGATCTNNNTGTGCTTATAGTCCCAGCAAG
AGAGCACACGTCTGAACTCTTTCCCTACACGACGCTCTTCCGATCTNNNCTGTAATCCCAGCTACTTCGG
GAGCACACGTCTGAACTCTTTCCCTACACGACGCTCTTCCGATCTNNNTACAAAAACATTAGCTGGGCA
AGAGCACACGTCTGAACTCTTTCCCTACACGACGCTCTTCCGATCTNNNAAGATGGTGCCATTGTACTG
AGAGCACACGTCTGAACTCTTTCCCTACACGACGCTCTTCCGATCTNNNAGAAGAATCGTTTGAACCCA
AAGAGCACACGTCTGAACTCTTTCCCTACACGACGCTCTTCCGATCTNNNAATGAGCTATGATCGCGTC

AGAGCACACGTCTGAACTCTTTCCCTACACGACGCTCTTCCGATCTNNNGGAGGCTGAGGCAGAATAA
GAGCACACGTCTGAACTCTTTCCCTACACGACGCTCTTCCGATCTNNNTTTAGAAAACACTAAAGACT
AAGAGCACACGTCTGAACTCTTTCCCTACACGACGCTCTTCCGATCTNNNACAGAGTCAAGCTGATAG
AGAGCACACGTCTGAACTCTTTCCCTACACGACGCTCTTCCGATCTNNNAGGAAGGAGAATTGCTTGA
AAGAGCACACGTCTGAACTCTTTCCCTACACGACGCTCTTCCGATCTNNNAAAAAAATTAGCCGGGCG
AAGAGCACACGTCTGAACTCTTTCCCTACACGACGCTCTTCCGATCTNNNGATGCCTGTAGTCTCAGCTA
AGAGCACACGTCTGAACTCTTTCCCTACACGACGCTCTTCCGATCTNNNTCTCAAGGAATCTAGAGGA
AAGAGCACACGTCTGAACTCTTTCCCTACACGACGCTCTTCCGATCTNNNGAGTTTGCAGTGAGCTGA
AAGAGCACACGTCTGAACTCTTTCCCTACACGACGCTCTTCCGATCTNNNAGGCTGAGGTAGAATTG
GAAGAGCACACGTCTGAACTCTTTCCCTACACGACGCTCTTCCGATCTNNNCTGTAATTCCAGCTACTCG
AGAGCACACGTCTGAACTCTTTCCCTACACGACGCTCTTCCGATCTNNNGGTAAATAGGTGGGACAAA
GAAGAGCACACGTCTGAACTCTTTCCCTACACGACGCTCTTCCGATCTNNNGAGCTTGCAGTGAGCCTA
GAGCACACGTCTGAACTCTTTCCCTACACGACGCTCTTCCGATCTNNNTGGCATAGAGTTATTTCTCCCT
AAGAGCACACGTCTGAACTCTTTCCCTACACGACGCTCTTCCGATCTNNNGAGATTGCAGTGAGCCTA
GAAGAGCACACGTCTGAACTCTTTCCCTACACGACGCTCTTCCGATCTNNNGAGGTTGTAGTGAGCCG
AAGAGCACACGTCTGAACTCTTTCCCTACACGACGCTCTTCCGATCTNNNGAACACCTGTAGTCCCAGT
AGCACACGTCTGAACTCTTTCCCTACACGACGCTCTTCCGATCTNNNAGCTACTCACACATTAGAAGCTA
AGAGCACACGTCTGAACTCTTTCCCTACACGACGCTCTTCCGATCTNNNCTGAGGCAGGAGAATCACT
AAGAGCACACGTCTGAACTCTTTCCCTACACGACGCTCTTCCGATCTNNNAACAGACCCACAGCCATTAT
AGCACACGTCTGAACTCTTTCCCTACACGACGCTCTTCCGATCTNNNTGACAATGTCAAAAATTGGTCA
AAGAGCACACGTCTGAACTCTTTCCCTACACGACGCTCTTCCGATCTNNNCAGCTAGCATACAAGGAG
AAGAGCACACGTCTGAACTCTTTCCCTACACGACGCTCTTCCGATCTNNNTGTGATGGTGGGTGCCTA
AGCACACGTCTGAACTCTTTCCCTACACGACGCTCTTCCGATCTNNNTCCACATATCTTTCAAGTCTCTC
AAGAGCACACGTCTGAACTCTTTCCCTACACGACGCTCTTCCGATCTNNNAAGCAATGCTAAGAGCGA
AAGAGCACACGTCTGAACTCTTTCCCTACACGACGCTCTTCCGATCTNNNACTTGGGATTACTTCTCCT
AAGAGCACACGTCTGAACTCTTTCCCTACACGACGCTCTTCCGATCTNNNCTGAGGTGGGAGAATCAC
AGCACACGTCTGAACTCTTTCCCTACACGACGCTCTTCCGATCTNNNAGTGGATGGGTATATATGGTGT
AGCACACGTCTGAACTCTTTCCCTACACGACGCTCTTCCGATCTNNNATATGCCTGTAATCCCAGCTAC
GAGCACACGTCTGAACTCTTTCCCTACACGACGCTCTTCCGATCTNNNGAGGTAACAGGAAAAGGGG
GAAGAGCACACGTCTGAACTCTTTCCCTACACGACGCTCTTCCGATCTNNNATTTTTACCCACCTCGCA
AAGAGCACACGTCTGAACTCTTTCCCTACACGACGCTCTTCCGATCTNNNTAGCTCTTCTAGACACCAA
AGCACACGTCTGAACTCTTTCCCTACACGACGCTCTTCCGATCTNNNACCTGAACCTGAACTCAATACTT
AGAGCACACGTCTGAACTCTTTCCCTACACGACGCTCTTCCGATCTNNNTGAGGCACGAGAATTGTTT
AGAGCACACGTCTGAACTCTTTCCCTACACGACGCTCTTCCGATCTNNNCTGAGATCACACCACTGCAGT
GAGCACACGTCTGAACTCTTTCCCTACACGACGCTCTTCCGATCTNNNTTAGTAGAGATGGGGTTTCA
GAGCACACGTCTGAACTCTTTCCCTACACGACGCTCTTCCGATCTNNNAGACTGAGGCAGGAGAATT
AAGAGCACACGTCTGAACTCTTTCCCTACACGACGCTCTTCCGATCTNNNGAGGCTAAGGAAGGAGAA
GAAGAGCACACGTCTGAACTCTTTCCCTACACGACGCTCTTCCGATCTNNNGTGGAGGATCGCTTGAA
AGAGCACACGTCTGAACTCTTTCCCTACACGACGCTCTTCCGATCTNNNTGCATCTGTAATCCCAGCTA
AAGAGCACACGTCTGAACTCTTTCCCTACACGACGCTCTTCCGATCTNNNTGTAATTCCAGCTACTTGG
GAGCACACGTCTGAACTCTTTCCCTACACGACGCTCTTCCGATCTNNNAGGCTCCTCATCACTAATCATTT
GAGCACACGTCTGAACTCTTTCCCTACACGACGCTCTTCCGATCTNNNTGCTAGTAGACCTGCATTACA
AAGAGCACACGTCTGAACTCTTTCCCTACACGACGCTCTTCCGATCTNNNAATCACTTGAACCCAGGAG

AGAGCACACGTCTGAACTCTTTCCCTACACGACGCTCTTCCGATCTNNNGCAGAGGTTGAAATGAGC
AGAGCACACGTCTGAACTCTTTCCCTACACGACGCTCTTCCGATCTNNNTCATTTGGTGCTGATGCCAT
AGCACACGTCTGAACTCTTTCCCTACACGACGCTCTTCCGATCTNNNGTGAAACAAAAATGATGGCAT
AAGAGCACACGTCTGAACTCTTTCCCTACACGACGCTCTTCCGATCTNNNCTGAGGCAGGAGAATCAC
AGAGCACACGTCTGAACTCTTTCCCTACACGACGCTCTTCCGATCTNNNGAGAATTACTTGAGCCCAG
AGAGCACACGTCTGAACTCTTTCCCTACACGACGCTCTTCCGATCTNNNTGAGGCAAGAGAATTGCTT
AAGAGCACACGTCTGAACTCTTTCCCTACACGACGCTCTTCCGATCTNNNAAAAAATTAGCCAGGCGT
AAGAGCACACGTCTGAACTCTTTCCCTACACGACGCTCTTCCGATCTNNNAGAAAAATCACATGCAAA
AGAGCACACGTCTGAACTCTTTCCCTACACGACGCTCTTCCGATCTNNNTGAGGCAGGAGTATCACTT
AGAGCACACGTCTGAACTCTTTCCCTACACGACGCTCTTCCGATCTNNNTCTCTTGCCTAATTGCTATG
AAGAGCACACGTCTGAACTCTTTCCCTACACGACGCTCTTCCGATCTNNNTCAGCAGCTGAGGAAGA
AGAGCACACGTCTGAACTCTTTCCCTACACGACGCTCTTCCGATCTNNNTACAAAAATTAGCCAGCGTG
GAGCACACGTCTGAACTCTTTCCCTACACGACGCTCTTCCGATCTNNNACATGCTTGTAATCCCAGCTA
AAGAGCACACGTCTGAACTCTTTCCCTACACGACGCTCTTCCGATCTNNNAAAATTAGTTGGGCATGG
AAGAGCACACGTCTGAACTCTTTCCCTACACGACGCTCTTCCGATCTNNNGGTTGCAGTGAGTCGAG
GAGCACACGTCTGAACTCTTTCCCTACACGACGCTCTTCCGATCTNNNCAGTCTCATGAATTCTGACAA
AAGAGCACACGTCTGAACTCTTTCCCTACACGACGCTCTTCCGATCTNNNGCTGAGACAGGAGAATCA
AAGAGCACACGTCTGAACTCTTTCCCTACACGACGCTCTTCCGATCTNNNCTGAGGCAGGAGAATCAC
AAGAGCACACGTCTGAACTCTTTCCCTACACGACGCTCTTCCGATCTNNNGAGATCACACCACTGCAT
AGAGCACACGTCTGAACTCTTTCCCTACACGACGCTCTTCCGATCTNNNCAGGACCCATCAGTGTGTTG
AGCACACGTCTGAACTCTTTCCCTACACGACGCTCTTCCGATCTNNNTCAAGATAATGGCATCAGAGTA
AGAGCACACGTCTGAACTCTTTCCCTACACGACGCTCTTCCGATCTNNNTGTGGTTCATGTTTTTGTGT
AGCACACGTCTGAACTCTTTCCCTACACGACGCTCTTCCGATCTNNNTGCTAGAAACAAATCTCCAATG
AGCACACGTCTGAACTCTTTCCCTACACGACGCTCTTCCGATCTNNNTCTCTTGATGTTTGCAGTTTTCA
GAGCACACGTCTGAACTCTTTCCCTACACGACGCTCTTCCGATCTNNNTTCCAGGTTTAGTCTTGGAAG
GCACACGTCTGAACTCTTTCCCTACACGACGCTCTTCCGATCTNNNTCAGCCATTGAAACTATTGATCT
GAGCACACGTCTGAACTCTTTCCCTACACGACGCTCTTCCGATCTNNNCAGAAGCAGGAGGATCACTT
AGCACACGTCTGAACTCTTTCCCTACACGACGCTCTTCCGATCTNNNAGAGACACAAAAACCGTTCAA
AGCACACGTCTGAACTCTTTCCCTACACGACGCTCTTCCGATCTNNNTGCTATTGTTCTTTTTCCTGTGT
AGAGCACACGTCTGAACTCTTTCCCTACACGACGCTCTTCCGATCTNNNGTGATTCCATTTATGACAGG
GCACACGTCTGAACTCTTTCCCTACACGACGCTCTTCCGATCTNNNTGTCTATCATCTATCCCTGTCAAC
AAGAGCACACGTCTGAACTCTTTCCCTACACGACGCTCTTCCGATCTNNNGCCCTGTCTGGAAGGAAT
AGAGCACACGTCTGAACTCTTTCCCTACACGACGCTCTTCCGATCTNNNCTCATACTCCCTCCCTCCCTTC
AGCACACGTCTGAACTCTTTCCCTACACGACGCTCTTCCGATCTNNNCTGAGACAGGAGAATGGCTTG
AGAGCACACGTCTGAACTCTTTCCCTACACGACGCTCTTCCGATCTNNNCACACGTTAGATTGGTGTA
GAGCACACGTCTGAACTCTTTCCCTACACGACGCTCTTCCGATCTNNNTGAGGTAAAGTGGCATGATT
AGCACACGTCTGAACTCTTTCCCTACACGACGCTCTTCCGATCTNNNTGTGTGTGTATATATGTGT
AGCACACGTCTGAACTCTTTCCCTACACGACGCTCTTCCGATCTNNNCTCCTATTTCCCTAGGGCCAAA
AGAGCACACGTCTGAACTCTTTCCCTACACGACGCTCTTCCGATCTNNNACCCCATCTCTACTAAACACA
AGAGCACACGTCTGAACTCTTTCCCTACACGACGCTCTTCCGATCTNNNACAACAGAGAAAATCACCT
AGAGCACACGTCTGAACTCTTTCCCTACACGACGCTCTTCCGATCTNNNACTCCATCTCTCTCTCTCTC
GAGCACACGTCTGAACTCTTTCCCTACACGACGCTCTTCCGATCTNNNCCATCTAGGTTGCTGCAAATA
AGAGCACACGTCTGAACTCTTTCCCTACACGACGCTCTTCCGATCTNNNGTGCATGTATGTGTGTGTA

AGAGCACACGTCTGAACTCTTTCCCTACACGACGCTCTTCCGATCTNNNATTCCATCCAGGTTGCTGT
GAGCACACGTCTGAACTCTTTCCCTACACGACGCTCTTCCGATCTNNNAGGTAAGATTTGGGTGAGG
AGAGCACACGTCTGAACTCTTTCCCTACACGACGCTCTTCCGATCTNNNTGGAACTGGGAAAGTCTA
AGCACACGTCTGAACTCTTTCCCTACACGACGCTCTTCCGATCTNNNACACACACACTATGGAATACTA
GAGCACACGTCTGAACTCTTTCCCTACACGACGCTCTTCCGATCTNNNTCAACTCCTTTTACAATAGCT
GCACACGTCTGAACTCTTTCCCTACACGACGCTCTTCCGATCTNNNACAGTCATTCTACTCATCACTTG
GAGCACACGTCTGAACTCTTTCCCTACACGACGCTCTTCCGATCTNNNAGTGAAACCCTGAAAAGAA
AAGAGCACACGTCTGAACTCTTTCCCTACACGACGCTCTTCCGATCTNNNGGAGACAGAGGTTGCA
AGCACACGTCTGAACTCTTTCCCTACACGACGCTCTTCCGATCTNNNACCCTGTACCCCTAAATATTTC
AGAGCACACGTCTGAACTCTTTCCCTACACGACGCTCTTCCGATCTNNNCCCTGGAGGCCATAATTAC
GAGCACACGTCTGAACTCTTTCCCTACACGACGCTCTTCCGATCTNNNCCAGTGAGAATTTCCTTTGA
AGCACACGTCTGAACTCTTTCCCTACACGACGCTCTTCCGATCTNNNTGGTGGAGTCTTTAGTTTTT
AAGAGCACACGTCTGAACTCTTTCCCTACACGACGCTCTTCCGATCTNNNTCAGCAAGGTCTTTATAAC
GCACACGTCTGAACTCTTTCCCTACACGACGCTCTTCCGATCTNNNACTGGAGAAAAACTAAGTCCTT
AGCACACGTCTGAACTCTTTCCCTACACGACGCTCTTCCGATCTNNNTGCTTTTGTTTTGTTTTGGTT
GCACACGTCTGAACTCTTTCCCTACACGACGCTCTTCCGATCTNNNTGTGAAGGTTGTAAGGACATT
AAGAGCACACGTCTGAACTCTTTCCCTACACGACGCTCTTCCGATCTNNNAGTGCTGGGATTACAGAT
AGCACACGTCTGAACTCTTTCCCTACACGACGCTCTTCCGATCTNNNAGTGTCTATTCAGATCCTTTGT
AGCACACGTCTGAACTCTTTCCCTACACGACGCTCTTCCGATCTNNNCTCTTGAGTTGGTATTGTA
GAGCACACGTCTGAACTCTTTCCCTACACGACGCTCTTCCGATCTNNNTGTTAGTGATTATTCAGCAA
AAGAGCACACGTCTGAACTCTTTCCCTACACGACGCTCTTCCGATCTNNNAGGGAGAGAAAGAAAGA
AGAGCACACGTCTGAACTCTTTCCCTACACGACGCTCTTCCGATCTNNNACTATCTACTTCCTTAGCCTG
AGCACACGTCTGAACTCTTTCCCTACACGACGCTCTTCCGATCTNNNTGTTGCTAATTTTGGAACCTG
AGCACACGTCTGAACTCTTTCCCTACACGACGCTCTTCCGATCTNNNTGCCTAAAAAAAAACCTGGA
AGCACACGTCTGAACTCTTTCCCTACACGACGCTCTTCCGATCTNNNTGGTTTTCCATTCTTGAGTTAC
GAAGAGCACACGTCTGAACTCTTTCCCTACACGACGCTCTTCCGATCTNNNAAGAGAATTGCTTGAA
AGAGCACACGTCTGAACTCTTTCCCTACACGACGCTCTTCCGATCTNNNTCAGTCCTGCCTAGAAATG
AAGAGCACACGTCTGAACTCTTTCCCTACACGACGCTCTTCCGATCTNNNTGAGTGATAAGCCAGAC
AGAGCACACGTCTGAACTCTTTCCCTACACGACGCTCTTCCGATCTNNNTGTGTGTGTGTGTGTAT
AGCACACGTCTGAACTCTTTCCCTACACGACGCTCTTCCGATCTNNNAGTATAAGACTGTGTGTATG
AGCACACGTCTGAACTCTTTCCCTACACGACGCTCTTCCGATCTNNNTCTCTCTCTCTCTCTCTCTC
AGCACACGTCTGAACTCTTTCCCTACACGACGCTCTTCCGATCTNNNACAGATATTCTCAAAGACAGT
CACACGTCTGAACTCTTTCCCTACACGACGCTCTTCCGATCTNNNACATAATTGGGTCTTTGTCATATT
GCACACGTCTGAACTCTTTCCCTACACGACGCTCTTCCGATCTNNNCAGTAGTTTCCATTTGTTGTCAC
AGCACACGTCTGAACTCTTTCCCTACACGACGCTCTTCCGATCTNNNGCCAATTTGCAGGAAATATAG
AGCACACGTCTGAACTCTTTCCCTACACGACGCTCTTCCGATCTNNNTCCATTCTTTAGGTTGTCTCTT
GCACACGTCTGAACTCTTTCCCTACACGACGCTCTTCCGATCTNNNTGATATGTGAGGCTTTTTTCCT
GCACACGTCTGAACTCTTTCCCTACACGACGCTCTTCCGATCTNNNACTCAAAGAGTTACATGGTACTT
GAGCACACGTCTGAACTCTTTCCCTACACGACGCTCTTCCGATCTNNNAGTGAATGGGTTTGTACAAT
CACACGTCTGAACTCTTTCCCTACACGACGCTCTTCCGATCTNNNTGCATTCTTATACATAGCTAATAT
AAGAGCACACGTCTGAACTCTTTCCCTACACGACGCTCTTCCGATCTNNNTGTAACAAGACTGATGG
GCACACGTCTGAACTCTTTCCCTACACGACGCTCTTCCGATCTNNNAGATCATCATAAAGGTCTTCGT
GCACACGTCTGAACTCTTTCCCTACACGACGCTCTTCCGATCTNNNGGCCACCAATAATGTATCTAAA

AGAGCACACGTCTGAACTCTTTCCCTACACGACGCTCTTCCGATCTNNNAGGGGTAAATCTCACTTGG
GAGCACACGTCTGAACTCTTTCCCTACACGACGCTCTTCCGATCTNNNTCACCTTCATTTTGTTTTATGG
AGCACACGTCTGAACTCTTTCCCTACACGACGCTCTTCCGATCTNNNTGGAATGCTTTGAACGAATGA
CACACGTCTGAACTCTTTCCCTACACGACGCTCTTCCGATCTNNNGCCTAGAAGAAAATGTACTTAGT
AGCACACGTCTGAACTCTTTCCCTACACGACGCTCTTCCGATCTNNNGGGCTAACCTCTCTTATATAGT
AGCACACGTCTGAACTCTTTCCCTACACGACGCTCTTCCGATCTNNNGGACATGTAAATCATTTGTGA
GCACACGTCTGAACTCTTTCCCTACACGACGCTCTTCCGATCTNNNAGAAAATGCCTACATGTGATGA
AGAGCACACGTCTGAACTCTTTCCCTACACGACGCTCTTCCGATCTNNNAGTTCCCTGGTCCCATTTTC
AGAGCACACGTCTGAACTCTTTCCCTACACGACGCTCTTCCGATCTNNNGTACCCACTAACCAACCCAT
GAGCACACGTCTGAACTCTTTCCCTACACGACGCTCTTCCGATCTNNNAAGGAAGGAACGAAGGAA
AGAGCACACGTCTGAACTCTTTCCCTACACGACGCTCTTCCGATCTNNNTGCCACAACCTATAATCTTA
AGCACACGTCTGAACTCTTTCCCTACACGACGCTCTTCCGATCTNNNAGTAGAATGGTAGAAACACTT
GCACACGTCTGAACTCTTTCCCTACACGACGCTCTTCCGATCTNNNTCAGTTCTCATAGTTTTCTGGTG
AGCACACGTCTGAACTCTTTCCCTACACGACGCTCTTCCGATCTNNNTCTCTCTCTCTCTCTTTCTCTC
AGCACACGTCTGAACTCTTTCCCTACACGACGCTCTTCCGATCTNNNTGATATCTACATTTGTTTCTGCA
GCACACGTCTGAACTCTTTCCCTACACGACGCTCTTCCGATCTNNNTCCACTTGTTGTAAAGAATTGA
AGCACACGTCTGAACTCTTTCCCTACACGACGCTCTTCCGATCTNNNTGGGAAATTGTCTAGATTTAC
GAGCACACGTCTGAACTCTTTCCCTACACGACGCTCTTCCGATCTNNNCCATACTTGGCAGTTTTCTTTT
AGCACACGTCTGAACTCTTTCCCTACACGACGCTCTTCCGATCTNNNTCACTCAGAGAAATACAAAGCT
AGCACACGTCTGAACTCTTTCCCTACACGACGCTCTTCCGATCTNNNTTCTGGTTGTTTTGTACACTCT
AGCACACGTCTGAACTCTTTCCCTACACGACGCTCTTCCGATCTNNNGCCATTATCAGTTGTCTTCCTTT
AGCACACGTCTGAACTCTTTCCCTACACGACGCTCTTCCGATCTNNNAGTAGTGCTTGCTTTATAAATC
AGCACACGTCTGAACTCTTTCCCTACACGACGCTCTTCCGATCTNNNTCAACAACAGTAGTAGGAGAC
AGCACACGTCTGAACTCTTTCCCTACACGACGCTCTTCCGATCTNNNAGATGTTTATTCGCATCCTTTGT
AGAGCACACGTCTGAACTCTTTCCCTACACGACGCTCTTCCGATCTNNNGAGTACAGTCAGTCCTCCAC
AGCACACGTCTGAACTCTTTCCCTACACGACGCTCTTCCGATCTNNNAGTTTCCTTTCTAAATACAGCA
AGCACACGTCTGAACTCTTTCCCTACACGACGCTCTTCCGATCTNNNACCTATGCATCCATTTTTCTTTA
GAGCACACGTCTGAACTCTTTCCCTACACGACGCTCTTCCGATCTNNNTGGAGATTGAACAAAAGGT
GAGCACACGTCTGAACTCTTTCCCTACACGACGCTCTTCCGATCTNNNCCTCCTTCAACATGTTCCCATT
GAGCACACGTCTGAACTCTTTCCCTACACGACGCTCTTCCGATCTNNNACCACACTGTTTTCATTACTGT
AGCACACGTCTGAACTCTTTCCCTACACGACGCTCTTCCGATCTNNNCAGATGACTTTATCTAACACCTG
AGCACACGTCTGAACTCTTTCCCTACACGACGCTCTTCCGATCTNNNTCCCCTTTGTCCTTTTTATTTGT
AGCACACGTCTGAACTCTTTCCCTACACGACGCTCTTCCGATCTNNNTGAACCGCCATTGTATGTTATTT
AGCACACGTCTGAACTCTTTCCCTACACGACGCTCTTCCGATCTNNNACTCAATAGCAAAAACTCAATC
AAGAGCACACGTCTGAACTCTTTCCCTACACGACGCTCTTCCGATCTNNNAAGAAACTGAGAATTCTG
AGCACACGTCTGAACTCTTTCCCTACACGACGCTCTTCCGATCTNNNACCATAAAACGGCTTATATGTT
AGCACACGTCTGAACTCTTTCCCTACACGACGCTCTTCCGATCTNNNCAGGGAGTTTCATTTTTATCTC
AGCACACGTCTGAACTCTTTCCCTACACGACGCTCTTCCGATCTNNNATATTCTTGCACATAAATTTTG
AGCACACGTCTGAACTCTTTCCCTACACGACGCTCTTCCGATCTNNNTCAGAACAACAGGAAAAAGAC
GAGCACACGTCTGAACTCTTTCCCTACACGACGCTCTTCCGATCTNNNTGTCTCAAGTTTGGTGATTTC
AGCACACGTCTGAACTCTTTCCCTACACGACGCTCTTCCGATCTNNNTGTCTTCTGTAGGTTGTCTTTTC
GAGCACACGTCTGAACTCTTTCCCTACACGACGCTCTTCCGATCTNNNCTATGGGTTTGTCAGAGACA
GAGCACACGTCTGAACTCTTTCCCTACACGACGCTCTTCCGATCTNNNCAGTCTCCATAATTGTGAATC

AGCACACGTCTGAACTCTTTCCCTACACGACGCTCTTCCGATCTNNNTCCCTTTTTCTTTCTCTCTCTCTCT
GAGCACACGTCTGAACTCTTTCCCTACACGACGCTCTTCCGATCTNNNTTGTGTGATTCACTTCATTGTG
AGCACACGTCTGAACTCTTTCCCTACACGACGCTCTTCCGATCTNNNACTTTGTGTGTAGGTATGTGTT
GAGCACACGTCTGAACTCTTTCCCTACACGACGCTCTTCCGATCTNNNAATCCCCTACTTCTCACTTTGA
AGCACACGTCTGAACTCTTTCCCTACACGACGCTCTTCCGATCTNNNTCCAAGGATATCAAAGGAGGAT
AAGAGCACACGTCTGAACTCTTTCCCTACACGACGCTCTTCCGATCTNNNAGAGTTCTCACTTGGTCATG
GCACACGTCTGAACTCTTTCCCTACACGACGCTCTTCCGATCTNNNACTTTTCTTTCTCTCTTTCTTTCCAT
GAGCACACGTCTGAACTCTTTCCCTACACGACGCTCTTCCGATCTNNNTCTGCATCTCTTGAGATAATTG
AGAGCACACGTCTGAACTCTTTCCCTACACGACGCTCTTCCGATCTNNNTTGTGAAGGTGGAACAGTG
AGAGCACACGTCTGAACTCTTTCCCTACACGACGCTCTTCCGATCTNNNTTTCATTTTCTAGCTCCTCCC
AGAGCACACGTCTGAACTCTTTCCCTACACGACGCTCTTCCGATCTNNNTCATTCTGTCAGTGGGAAAA
AGCACACGTCTGAACTCTTTCCCTACACGACGCTCTTCCGATCTNNNAAGCAACAAACAGTATAATGGA
AGAGCACACGTCTGAACTCTTTCCCTACACGACGCTCTTCCGATCTNNNGCCTGTAGTCCCAGCTATGA
AGCACACGTCTGAACTCTTTCCCTACACGACGCTCTTCCGATCTNNNTGCAAAAGAACCTCAATCAAAC
GAGCACACGTCTGAACTCTTTCCCTACACGACGCTCTTCCGATCTNNNTGAAAATGAGGCCAGCTAAAA
AAGAGCACACGTCTGAACTCTTTCCCTACACGACGCTCTTCCGATCTNNNAGGCCCCAATTAGTAGGCT
GCACACGTCTGAACTCTTTCCCTACACGACGCTCTTCCGATCTNNNTCCTTCTTTCCTTCTTTTCTTTTTCT
AAGAGCACACGTCTGAACTCTTTCCCTACACGACGCTCTTCCGATCTNNNTCAGAGCTGAAATGAAGG
GAGCACACGTCTGAACTCTTTCCCTACACGACGCTCTTCCGATCTNNNGGTGAAGATGTTGATGAATA
GAGCACACGTCTGAACTCTTTCCCTACACGACGCTCTTCCGATCTNNNAGGTTGAGGTGAGCCAATATT
GAGCACACGTCTGAACTCTTTCCCTACACGACGCTCTTCCGATCTNNNTTGCGCACGTATGTTTATAGC
AGCACACGTCTGAACTCTTTCCCTACACGACGCTCTTCCGATCTNNNACAATAAAAATGCCACAAGTAG
GAGCACACGTCTGAACTCTTTCCCTACACGACGCTCTTCCGATCTNNNACACACACACATCATATGTACA
GAGCACACGTCTGAACTCTTTCCCTACACGACGCTCTTCCGATCTNNNGCATATGCATGTTTATAGCAG
AGAGCACACGTCTGAACTCTTTCCCTACACGACGCTCTTCCGATCTNNNGCCCAAGAACTGTGCTAAGA
GAGCACACGTCTGAACTCTTTCCCTACACGACGCTCTTCCGATCTNNNCTGTTTCTGTGTTGTTTCACCC
GCACACGTCTGAACTCTTTCCCTACACGACGCTCTTCCGATCTNNNAGTCCTTCTTTAGTATGCATTGCT
AGCACACGTCTGAACTCTTTCCCTACACGACGCTCTTCCGATCTNNNTCTTTGGTTTTCAACTGTTGGAG
AGAGCACACGTCTGAACTCTTTCCCTACACGACGCTCTTCCGATCTNNNGCAATATACATCTGTATGCAG
AGCACACGTCTGAACTCTTTCCCTACACGACGCTCTTCCGATCTNNNTGCTTTATGGTGGTGGTCTAAA
AGAGCACACGTCTGAACTCTTTCCCTACACGACGCTCTTCCGATCTNNNAGGAATAATGGGGAAGAGG
AGAGCACACGTCTGAACTCTTTCCCTACACGACGCTCTTCCGATCTNNNAGCAGTCATTTATTGCAAAG
GAGCACACGTCTGAACTCTTTCCCTACACGACGCTCTTCCGATCTNNNCTGCCTTGGCTATCAGATGGA
GCACACGTCTGAACTCTTTCCCTACACGACGCTCTTCCGATCTNNNTGTTCAACTTGTCCTTCTTTCACTG
AGAGCACACGTCTGAACTCTTTCCCTACACGACGCTCTTCCGATCTNNNTTTTAGGGTAAAAGTGGCCA
GAGCACACGTCTGAACTCTTTCCCTACACGACGCTCTTCCGATCTNNNTTGCCACAACAGCAGTATAGA
GAGCACACGTCTGAACTCTTTCCCTACACGACGCTCTTCCGATCTNNNCCTGTAGCAAGGCAAATTTAG
AGCACACGTCTGAACTCTTTCCCTACACGACGCTCTTCCGATCTNNNTCACACTTTTCCTTTGAGACTCTG
AGAGCACACGTCTGAACTCTTTCCCTACACGACGCTCTTCCGATCTNNNAATCTGCCTGCTTTTGTTCAG
GAGCACACGTCTGAACTCTTTCCCTACACGACGCTCTTCCGATCTNNNACTACTTGCTGTTGTTACACTT
AGAGCACACGTCTGAACTCTTTCCCTACACGACGCTCTTCCGATCTNNNTAGGAATCGACTGTGTCTTCA
AGAGCACACGTCTGAACTCTTTCCCTACACGACGCTCTTCCGATCTNNNACCACACTAAGAAAGAAAGG
GAGCACACGTCTGAACTCTTTCCCTACACGACGCTCTTCCGATCTNNNTGAATCTATGGATTGCTTTGG

GAGCACACGTCTGAACTCTTTCCCTACACGACGCTCTTCCGATCTNNNAGGTTTTCCGGCCTTACATT
AAGAGCACACGTCTGAACTCTTTCCCTACACGACGCTCTTCCGATCTNNNTGGACTGAATTACACTAC
AAGAGCACACGTCTGAACTCTTTCCCTACACGACGCTCTTCCGATCTNNNGAATGGCGTGAACCCAG
AGCACACGTCTGAACTCTTTCCCTACACGACGCTCTTCCGATCTNNNGGAAAGGTTTCATATTTCCTT
GCACACGTCTGAACTCTTTCCCTACACGACGCTCTTCCGATCTNNNAGAACCAGTATGTTCACCTATGC
GCACACGTCTGAACTCTTTCCCTACACGACGCTCTTCCGATCTNNNTGGACTGTCCAAATAAACGTAT
GAGCACACGTCTGAACTCTTTCCCTACACGACGCTCTTCCGATCTNNNATGTTGCCACCTCTCAGAG
AGCACACGTCTGAACTCTTTCCCTACACGACGCTCTTCCGATCTNNNTTTGGATTATACACACACACAC
GCACACGTCTGAACTCTTTCCCTACACGACGCTCTTCCGATCTNNNGAAGGAGGATCGCTTGAGACT
AGCACACGTCTGAACTCTTTCCCTACACGACGCTCTTCCGATCTNNNTCTTAGAAAGGCTTTCCTAAT
AGAGCACACGTCTGAACTCTTTCCCTACACGACGCTCTTCCGATCTNNNCACCAGTGACACTTTCTATTT
AGCACACGTCTGAACTCTTTCCCTACACGACGCTCTTCCGATCTNNNCATCATCTATCTTCCTGCTGTGG
GCACACGTCTGAACTCTTTCCCTACACGACGCTCTTCCGATCTNNNACTCTACAAAGAAATGCATTGTG
GCACACGTCTGAACTCTTTCCCTACACGACGCTCTTCCGATCTNNNACTCTACACACACATACACACACA
AGCACACGTCTGAACTCTTTCCCTACACGACGCTCTTCCGATCTNNNACTCTTCTGTGGATAGTTTCAG
GAGCACACGTCTGAACTCTTTCCCTACACGACGCTCTTCCGATCTNNNTCCAACAAAAAGGTGACCTT
GAGCACACGTCTGAACTCTTTCCCTACACGACGCTCTTCCGATCTNNNTCCATGTTGCTGCAAATTCAC
AGAGCACACGTCTGAACTCTTTCCCTACACGACGCTCTTCCGATCTNNNACGTCCTAGGAATGGTGAT
GAGCACACGTCTGAACTCTTTCCCTACACGACGCTCTTCCGATCTNNNAGGGCAAGAAATCATTCCAG
AGCACACGTCTGAACTCTTTCCCTACACGACGCTCTTCCGATCTNNNTGCTAAACAGAAAAGAGAAAG
AGCACACGTCTGAACTCTTTCCCTACACGACGCTCTTCCGATCTNNNCTCACACACACACATACACA
AGCACACGTCTGAACTCTTTCCCTACACGACGCTCTTCCGATCTNNNTCTGTCTTGCCCAATCTCATGT
AGCACACGTCTGAACTCTTTCCCTACACGACGCTCTTCCGATCTNNNAGACACAAGAGTCTTAGGTTT
AGCACACGTCTGAACTCTTTCCCTACACGACGCTCTTCCGATCTNNNAGTTAGGGTTTTCCAGAGAA
AGCACACGTCTGAACTCTTTCCCTACACGACGCTCTTCCGATCTNNNCAGCTTCCATAATAATGTGAAC
GAGCACACGTCTGAACTCTTTCCCTACACGACGCTCTTCCGATCTNNNACGCTTTCATTACTGTCTGCC
AGCACACGTCTGAACTCTTTCCCTACACGACGCTCTTCCGATCTNNNCATCCATGATACGTGTATGTG
GAGCACACGTCTGAACTCTTTCCCTACACGACGCTCTTCCGATCTNNNTGCACATGAACTACCATACCA
AGCACACGTCTGAACTCTTTCCCTACACGACGCTCTTCCGATCTNNNTTCCTCCTCTTCATCTTCACTGC
AGCACACGTCTGAACTCTTTCCCTACACGACGCTCTTCCGATCTNNNGCCCTTTGGATTTAGGCAGTT
GAGCACACGTCTGAACTCTTTCCCTACACGACGCTCTTCCGATCTNNNATGTCAGAAGGAGTAGGAG
AGCACACGTCTGAACTCTTTCCCTACACGACGCTCTTCCGATCTNNNCCTGTATCATAGTGCCTGGAA
AGAGCACACGTCTGAACTCTTTCCCTACACGACGCTCTTCCGATCTNNNGGAGTTGGGTGATGCATT
GCACACGTCTGAACTCTTTCCCTACACGACGCTCTTCCGATCTNNNAGAGACAGTGTCTTACTATGTTT
CACACGTCTGAACTCTTTCCCTACACGACGCTCTTCCGATCTNNNACCTCTCCATCTTTGTTTAGCACTC
AGCACACGTCTGAACTCTTTCCCTACACGACGCTCTTCCGATCTNNNAGAAATAGTTGAATGACAGG
GCACACGTCTGAACTCTTTCCCTACACGACGCTCTTCCGATCTNNNACAAAATCTCCTACCGCTTTTTG
GAGCACACGTCTGAACTCTTTCCCTACACGACGCTCTTCCGATCTNNNATGTAAATATGTGTGTGTGT
AGCACACGTCTGAACTCTTTCCCTACACGACGCTCTTCCGATCTNNNATTGGGGTATTGCTATTCCCAG
GCACACGTCTGAACTCTTTCCCTACACGACGCTCTTCCGATCTNNNTCTCTTTTCATTTCCATCCCCTCTC
AGCACACGTCTGAACTCTTTCCCTACACGACGCTCTTCCGATCTNNNTGCAACTGACTTTCTAATGTTGA
AGCACACGTCTGAACTCTTTCCCTACACGACGCTCTTCCGATCTNNNACAGAGCTTCAGACTGCTGTT
GAGCACACGTCTGAACTCTTTCCCTACACGACGCTCTTCCGATCTNNNTCTGCCAAGTCAAATTAGGC

AGAGCACACGTCTGAACTCTTTCCCTACACGACGCTCTTCCGATCTNNNATGTCAGTACACAGCTAGCT
GAGCACACGTCTGAACTCTTTCCCTACACGACGCTCTTCCGATCTNNNTGGTGCAACAGACATAGTGAT
AGAGCACACGTCTGAACTCTTTCCCTACACGACGCTCTTCCGATCTNNNTGCAGACTTCTGACTCTGAAC
GAGCACACGTCTGAACTCTTTCCCTACACGACGCTCTTCCGATCTNNNACTTTTGTGTGACAAGAGGAA
GAGCACACGTCTGAACTCTTTCCCTACACGACGCTCTTCCGATCTNNNAAACTCTTCGTTTGTGTGCATG
AGAGCACACGTCTGAACTCTTTCCCTACACGACGCTCTTCCGATCTNNNCCTTGTACCCATTAGCACTCAG
GAGCACACGTCTGAACTCTTTCCCTACACGACGCTCTTCCGATCTNNNATCCTCATACACAAAACAGCCA
GAGCACACGTCTGAACTCTTTCCCTACACGACGCTCTTCCGATCTNNNTCACTGTGAACACGTAGAAAA
GAGCACACGTCTGAACTCTTTCCCTACACGACGCTCTTCCGATCTNNNTCTGAGTGTGTTTATGTGTGTG
GAGCACACGTCTGAACTCTTTCCCTACACGACGCTCTTCCGATCTNNNTTCTGTGTGCACCCTCAAGAAT
AGCACACGTCTGAACTCTTTCCCTACACGACGCTCTTCCGATCTNNNCCATGTCTGACTTGTTTCACTTG
AGCACACGTCTGAACTCTTTCCCTACACGACGCTCTTCCGATCTNNNTCCATCAGGAAAATACCACATT
AAGAGCACACGTCTGAACTCTTTCCCTACACGACGCTCTTCCGATCTNNNGTATTCTTTGTTCCATTGCC
AGAGCACACGTCTGAACTCTTTCCCTACACGACGCTCTTCCGATCTNNNAGTCAGAATGCAAGAGAAG
GAGCACACGTCTGAACTCTTTCCCTACACGACGCTCTTCCGATCTNNNACCCCAAAGTGATACATCATTG
AGAGCACACGTCTGAACTCTTTCCCTACACGACGCTCTTCCGATCTNNNGAGGTTGAGGTTGGAGAA
GAGCACACGTCTGAACTCTTTCCCTACACGACGCTCTTCCGATCTNNNCTGAGGCAGGAGAATCACTT
GAGCACACGTCTGAACTCTTTCCCTACACGACGCTCTTCCGATCTNNNGTAATCCCAGCTACTCAAGAG
GAAGAGCACACGTCTGAACTCTTTCCCTACACGACGCTCTTCCGATCTNNNGAGGCTGAGGCAGGAGA
AAGAGCACACGTCTGAACTCTTTCCCTACACGACGCTCTTCCGATCTNNNTGTGTGTTCAGCTTTTGTTG
AGCACACGTCTGAACTCTTTCCCTACACGACGCTCTTCCGATCTNNNAACAGTCATTTTAGCCAGTGATT
AGCACACGTCTGAACTCTTTCCCTACACGACGCTCTTCCGATCTNNNTGGGTAATTGGGATATCCATTG
AGAGCACACGTCTGAACTCTTTCCCTACACGACGCTCTTCCGATCTNNNCATCTCATGGGCAGAGACACA
AGCACACGTCTGAACTCTTTCCCTACACGACGCTCTTCCGATCTNNNTGATCACTGTGCAATGGTATCTG
AGAGCACACGTCTGAACTCTTTCCCTACACGACGCTCTTCCGATCTNNNCTGAGGCAGGAGAATTGCTT
CACACGTCTGAACTCTTTCCCTACACGACGCTCTTCCGATCTNNNAGGAAACACAGGATTCTATGATAT
GAGCACACGTCTGAACTCTTTCCCTACACGACGCTCTTCCGATCTNNNGAGATTGCACCATTGTACTCCA
AGAGCACACGTCTGAACTCTTTCCCTACACGACGCTCTTCCGATCTNNNACAGAATGAGGCCTTGTCTG
AGCACACGTCTGAACTCTTTCCCTACACGACGCTCTTCCGATCTNNNACACACACACATATACACACACA
GAGCACACGTCTGAACTCTTTCCCTACACGACGCTCTTCCGATCTNNNCGACAAGTGCGAAACTCCATG
GAGCACACGTCTGAACTCTTTCCCTACACGACGCTCTTCCGATCTNNNGTTCATCCATGTTGTTGCAAAT
AGAGCACACGTCTGAACTCTTTCCCTACACGACGCTCTTCCGATCTNNNGCCAAGTGTTCGTGAGTTTT
GAGCACACGTCTGAACTCTTTCCCTACACGACGCTCTTCCGATCTNNNACCCTGAAGCCAGTAATCCAAT
GAGCACACGTCTGAACTCTTTCCCTACACGACGCTCTTCCGATCTNNNCTGTAGTCCCAGCTACTTCAG
GCACACGTCTGAACTCTTTCCCTACACGACGCTCTTCCGATCTNNNACATGTCTTGTCACTTTAGACCAG
GAGCACACGTCTGAACTCTTTCCCTACACGACGCTCTTCCGATCTNNNGGAGAGTTGCTTTAGTCCCAA
AGAGCACACGTCTGAACTCTTTCCCTACACGACGCTCTTCCGATCTNNNGAGGCAATAGAATTGCTTG
AGAGCACACGTCTGAACTCTTTCCCTACACGACGCTCTTCCGATCTNNNCATGAGGTCGAGGCTGTAGT
AGAGCACACGTCTGAACTCTTTCCCTACACGACGCTCTTCCGATCTNNNATGCCTGTAGTCCCAGTTACT
GAGCACACGTCTGAACTCTTTCCCTACACGACGCTCTTCCGATCTNNNAAAAGTAGTCAGGCATGGTG
GAGCACACGTCTGAACTCTTTCCCTACACGACGCTCTTCCGATCTNNNACATTTTGTAGCCATTTGACA
GAAGAGCACACGTCTGAACTCTTTCCCTACACGACGCTCTTCCGATCTNNNGAGGCTGAGGCAGGAGA
AGCACACGTCTGAACTCTTTCCCTACACGACGCTCTTCCGATCTNNNACAAAAAACAGATTCTGCTACCA

AGCACACGTCTGAACTCTTTCCCTACACGACGCTCTTCCGATCTNNNACTCTTTTTGGTTTCCATTGGCA
AGCACACGTCTGAACTCTTTCCCTACACGACGCTCTTCCGATCTNNNACAAAGAAACTTCTCCAACTCCT
GAGCACACGTCTGAACTCTTTCCCTACACGACGCTCTTCCGATCTNNNACATCTTTCAGAACTCAGCTCTC
AGCACACGTCTGAACTCTTTCCCTACACGACGCTCTTCCGATCTNNNAGGATTACAACTCTCCTGCTTTG
GAGCACACGTCTGAACTCTTTCCCTACACGACGCTCTTCCGATCTNNNGCTTTGCTTGTCTGGAAAAC
AGAGCACACGTCTGAACTCTTTCCCTACACGACGCTCTTCCGATCTNNNGTAATCCCAGCTATACAGGAC
GAGCACACGTCTGAACTCTTTCCCTACACGACGCTCTTCCGATCTNNNGAGAATTGCTTGAACCGAGGA
GAGCACACGTCTGAACTCTTTCCCTACACGACGCTCTTCCGATCTNNNGCAGAGACACCTGACTGTTAG
AGAGCACACGTCTGAACTCTTTCCCTACACGACGCTCTTCCGATCTNNNCAGAGGTAGCAGTGAGTGA
GAGCACACGTCTGAACTCTTTCCCTACACGACGCTCTTCCGATCTNNNCCTGCTTTCTCAGGTTTGTTTC
GAGCACACGTCTGAACTCTTTCCCTACACGACGCTCTTCCGATCTNNNTAAAATGAGTTGGGGAGGAC
GAGCACACGTCTGAACTCTTTCCCTACACGACGCTCTTCCGATCTNNNAGGAGTTCTCTAAACCCAAGC
AAGAGCACACGTCTGAACTCTTTCCCTACACGACGCTCTTCCGATCTNNNCAAGATTGTGTCACGGCAC
AGCACACGTCTGAACTCTTTCCCTACACGACGCTCTTCCGATCTNNNCAGGGAACAAGAAAGTAAGCA
GAGCACACGTCTGAACTCTTTCCCTACACGACGCTCTTCCGATCTNNNAGATCCTGTTTCTGAATTGATC
AGCACACGTCTGAACTCTTTCCCTACACGACGCTCTTCCGATCTNNNCCCATCAGAGATATTGGCCTGTA
GCACACGTCTGAACTCTTTCCCTACACGACGCTCTTCCGATCTNNNTGTACCTCCATTACAGTATGAGAC
GAGCACACGTCTGAACTCTTTCCCTACACGACGCTCTTCCGATCTNNNCACCTGTAATTGCAGCTACTTC
GAGCACACGTCTGAACTCTTTCCCTACACGACGCTCTTCCGATCTNNNGCAGGAGAATCAATTGAACCC
AGAGCACACGTCTGAACTCTTTCCCTACACGACGCTCTTCCGATCTNNNAGAATCACTTGAACCCAGGA
GAGCACACGTCTGAACTCTTTCCCTACACGACGCTCTTCCGATCTNNNAAAAAGAAATTTTGCGGGAG
AGAGCACACGTCTGAACTCTTTCCCTACACGACGCTCTTCCGATCTNNNGTTACAGTGAGCTGAGATTG
AGCACACGTCTGAACTCTTTCCCTACACGACGCTCTTCCGATCTNNNTGCAATGTTCAAAAAACCTTAGC
AGCACACGTCTGAACTCTTTCCCTACACGACGCTCTTCCGATCTNNNGTGCTTCAGTTTCATCTCACAAC
AGCACACGTCTGAACTCTTTCCCTACACGACGCTCTTCCGATCTNNNTGTGGGTCTATTTACAGTGTCT
GAGCACACGTCTGAACTCTTTCCCTACACGACGCTCTTCCGATCTNNNTCCCTCTAAAGCTTAGATGGTA
GAGCACACGTCTGAACTCTTTCCCTACACGACGCTCTTCCGATCTNNNCTGAGGCAGGAGAATCACTTC
AAGAGCACACGTCTGAACTCTTTCCCTACACGACGCTCTTCCGATCTNNNCATTTCAGAGGTTGGCTTTC
AGCACACGTCTGAACTCTTTCCCTACACGACGCTCTTCCGATCTNNNATGAGCTGATATCATGCCATTG
AGAGCACACGTCTGAACTCTTTCCCTACACGACGCTCTTCCGATCTNNNGAGAATCTCTTGAACCTGGC
AGCACACGTCTGAACTCTTTCCCTACACGACGCTCTTCCGATCTNNNAGTTCATGCATGTAAAGGTGTT
GAGCACACGTCTGAACTCTTTCCCTACACGACGCTCTTCCGATCTNNNCCTCTTTGGTTTGGTATATTCC
GAGCACACGTCTGAACTCTTTCCCTACACGACGCTCTTCCGATCTNNNAGAATCACTTGAATCCAGGAC
GAGCACACGTCTGAACTCTTTCCCTACACGACGCTCTTCCGATCTNNNCTGTAGTCCCAGCTACTCAGAA
AGAGCACACGTCTGAACTCTTTCCCTACACGACGCTCTTCCGATCTNNNGACACCTGTATTCCCAGCTAC
GAGCACACGTCTGAACTCTTTCCCTACACGACGCTCTTCCGATCTNNNGGTGGTGCATATCTGTAATCCC
GAGCACACGTCTGAACTCTTTCCCTACACGACGCTCTTCCGATCTNNNCTGTAATCCCAGCTACTCAGG
AGAGCACACGTCTGAACTCTTTCCCTACACGACGCTCTTCCGATCTNNNCTGTAATCCCAGCTACTCAGG
AGAGCACACGTCTGAACTCTTTCCCTACACGACGCTCTTCCGATCTNNNCCTGGGCTACAAACCTTTACA
AGCACACGTCTGAACTCTTTCCCTACACGACGCTCTTCCGATCTNNNTCCTTCTTTCTCTGTCTTTCTCCTC
AGCACACGTCTGAACTCTTTCCCTACACGACGCTCTTCCGATCTNNNACCATGTCCAGCTAAATTTTTGC
AGCACACGTCTGAACTCTTTCCCTACACGACGCTCTTCCGATCTNNNAGGTTGAGCATGAGATTTTCTT
AGCACACGTCTGAACTCTTTCCCTACACGACGCTCTTCCGATCTNNNTCTCTCTTTCCTTCCTTTCTTGCT

AGCACACGTCTGAACTCTTTCCCTACACGACGCTCTTCCGATCTNNNTCTGAGAATGTAGCATCGTTC
GCACACGTCTGAACTCTTTCCCTACACGACGCTCTTCCGATCTNNNGGTACCCCTCTCCAATTATTTG
GCACACGTCTGAACTCTTTCCCTACACGACGCTCTTCCGATCTNNNAGGTTTAGTGTTACAAACTCAG
GCACACGTCTGAACTCTTTCCCTACACGACGCTCTTCCGATCTNNNTCTTCCTTTTTCCATTGTGTGC
AGCACACGTCTGAACTCTTTCCCTACACGACGCTCTTCCGATCTNNNCTGGGTTTCTAGTAAGTAGCC
AGCACACGTCTGAACTCTTTCCCTACACGACGCTCTTCCGATCTNNNTGGAATCTTCTGTGTTGTCTTT
GAGCACACGTCTGAACTCTTTCCCTACACGACGCTCTTCCGATCTNNNTGTTGTATTTGCTAGCTAAGG
AGCACACGTCTGAACTCTTTCCCTACACGACGCTCTTCCGATCTNNNACTGGAATCCCTGCCAAAATA
GAGCACACGTCTGAACTCTTTCCCTACACGACGCTCTTCCGATCTNNNAGCGACTCCTGCTATATTTG
GAGCACACGTCTGAACTCTTTCCCTACACGACGCTCTTCCGATCTNNNGTTGTTGTTCTTTGTAGGCC
AGCACACGTCTGAACTCTTTCCCTACACGACGCTCTTCCGATCTNNNTGGGAAAAATCTAGAATGGTT
GAGCACACGTCTGAACTCTTTCCCTACACGACGCTCTTCCGATCTNNNTCAGCAGTGTTTTGTAGTTCT
AGCACACGTCTGAACTCTTTCCCTACACGACGCTCTTCCGATCTNNNTTATCAATAATCTTGGAGGC
GAGCACACGTCTGAACTCTTTCCCTACACGACGCTCTTCCGATCTNNNAGATTGCTCTTTGCCAATTAC
GCACACGTCTGAACTCTTTCCCTACACGACGCTCTTCCGATCTNNNTCTTCTCCTTCTCCTTCTCCTTCTC
AGCACACGTCTGAACTCTTTCCCTACACGACGCTCTTCCGATCTNNNGCATGTATAACACAAAGTAGG
GCACACGTCTGAACTCTTTCCCTACACGACGCTCTTCCGATCTNNNGTTTTTCTTGTGTGTGTGTGTG
GCACACGTCTGAACTCTTTCCCTACACGACGCTCTTCCGATCTNNNAGTGCTACAATAAACATATGCAT
GAGCACACGTCTGAACTCTTTCCCTACACGACGCTCTTCCGATCTNNNCATACACAGACACACACACAC
GAGCACACGTCTGAACTCTTTCCCTACACGACGCTCTTCCGATCTNNNACCTCAGAACATGTAATTCCT
GCACACGTCTGAACTCTTTCCCTACACGACGCTCTTCCGATCTNNNTGCAACTTTTTCTTCCTTCCTTCC
GAGCACACGTCTGAACTCTTTCCCTACACGACGCTCTTCCGATCTNNNGACTGTACCCTTGACCTTCTG
GCACACGTCTGAACTCTTTCCCTACACGACGCTCTTCCGATCTNNNGCAAACCAAAGATCAGTAAGAG
GAGCACACGTCTGAACTCTTTCCCTACACGACGCTCTTCCGATCTNNNAGAAAAACCCTCATAACCTGC
GCACACGTCTGAACTCTTTCCCTACACGACGCTCTTCCGATCTNNNTGGCACATTTCCATCTGTTTTTC
AGCACACGTCTGAACTCTTTCCCTACACGACGCTCTTCCGATCTNNNGGGTATTATAGCTTTGCTGGA
GCACACGTCTGAACTCTTTCCCTACACGACGCTCTTCCGATCTNNNAGTAAGGTCTTTTCAGCATTTG
AGCACACGTCTGAACTCTTTCCCTACACGACGCTCTTCCGATCTNNNGTTTGTGGTATTCTCTAATGGC
GAGCACACGTCTGAACTCTTTCCCTACACGACGCTCTTCCGATCTNNNTAATCTCCCTTCACTGCAAGC
GCACACGTCTGAACTCTTTCCCTACACGACGCTCTTCCGATCTNNNACTTTTGGTTTTGGTTCACATGG
GAGCACACGTCTGAACTCTTTCCCTACACGACGCTCTTCCGATCTNNNGAGCTGAGATCATACCACTC
GAGCACACGTCTGAACTCTTTCCCTACACGACGCTCTTCCGATCTNNNTCCTCTCTTCTTTCCTTTCTTTC
GCACACGTCTGAACTCTTTCCCTACACGACGCTCTTCCGATCTNNNACTTTTGGTTTTGGTTCACATGG
GCACACGTCTGAACTCTTTCCCTACACGACGCTCTTCCGATCTNNNTGTTCATCTCCTCGTATCATTAGC
AGCACACGTCTGAACTCTTTCCCTACACGACGCTCTTCCGATCTNNNACACACATACACATTTTTTTCTG
AGCACACGTCTGAACTCTTTCCCTACACGACGCTCTTCCGATCTNNNTGTAGGCCTTGGAATTAGAAG
AGCACACGTCTGAACTCTTTCCCTACACGACGCTCTTCCGATCTNNNACTGTGTTATCTGTCTTGAGA
GAGCACACGTCTGAACTCTTTCCCTACACGACGCTCTTCCGATCTNNNCTCAGCCTACCAAAGTGCTAG
AGCACACGTCTGAACTCTTTCCCTACACGACGCTCTTCCGATCTNNNAGATTTGTACAGGCATGGAGA
GAGCACACGTCTGAACTCTTTCCCTACACGACGCTCTTCCGATCTNNNCTGAGGCAGGAGAATCACTT
AGCACACGTCTGAACTCTTTCCCTACACGACGCTCTTCCGATCTNNNAACAAAATTAGCTGGGTGTGG
GAGCACACGTCTGAACTCTTTCCCTACACGACGCTCTTCCGATCTNNNGTAATCCCAGCTACTCAAGAG
GAGCACACGTCTGAACTCTTTCCCTACACGACGCTCTTCCGATCTNNNCATCTGCACACACCATCATCA

AGAGCACACGTCTGAACTCTTTCCCTACACGACGCTCTTCCGATCTNNNCATGAGAATTGCTTGAACCT
AAGAGCACACGTCTGAACTCTTTCCCTACACGACGCTCTTCCGATCTNNNAAAAAAAATTAGCCGGGC
GAGCACACGTCTGAACTCTTTCCCTACACGACGCTCTTCCGATCTNNNACGCCTGTAGTCCCAACTATTT
GAGCACACGTCTGAACTCTTTCCCTACACGACGCTCTTCCGATCTNNNGTAATCCCAGCTATACAGGA
GAGCACACGTCTGAACTCTTTCCCTACACGACGCTCTTCCGATCTNNNGTAATCCCAGCTATACAGGA
AGAGCACACGTCTGAACTCTTTCCCTACACGACGCTCTTCCGATCTNNNAGGAGAATCTTTTGAACCTG
GAGCACACGTCTGAACTCTTTCCCTACACGACGCTCTTCCGATCTNNNCCAGCATGAGTGACAGAGTA
AGAGCACACGTCTGAACTCTTTCCCTACACGACGCTCTTCCGATCTNNNAAAAAAATTAGCTGGGCGT
AGAGCACACGTCTGAACTCTTTCCCTACACGACGCTCTTCCGATCTNNNATCACTCCACTGTACTCTAGC
AGCACACGTCTGAACTCTTTCCCTACACGACGCTCTTCCGATCTNNNTGGGTCCATATGTAAACATTTC
AGAGCACACGTCTGAACTCTTTCCCTACACGACGCTCTTCCGATCTNNNAGAATTGCTTGAATCTGGG
AGAGCACACGTCTGAACTCTTTCCCTACACGACGCTCTTCCGATCTNNNCTGTAGTCCCAGCTACTCAG
GAGCACACGTCTGAACTCTTTCCCTACACGACGCTCTTCCGATCTNNNCTGAGACAGGAGAATGGCAT
AGAGCACACGTCTGAACTCTTTCCCTACACGACGCTCTTCCGATCTNNNGTATCCCCAGCTACTAAAGA
GAGCACACGTCTGAACTCTTTCCCTACACGACGCTCTTCCGATCTNNNCTAAGGTGGGAGAATCACTG
AGCACACGTCTGAACTCTTTCCCTACACGACGCTCTTCCGATCTNNNAATATTGCAGCACTCCACTCCA
AGAGCACACGTCTGAACTCTTTCCCTACACGACGCTCTTCCGATCTNNNGAGGCAGGAGAATTGCTTG
AGCACACGTCTGAACTCTTTCCCTACACGACGCTCTTCCGATCTNNNTGGAAACAGAGAGAGACTCC
AGAGCACACGTCTGAACTCTTTCCCTACACGACGCTCTTCCGATCTNNNCACCTCAAGGAGCTAGAGA
AGAGCACACGTCTGAACTCTTTCCCTACACGACGCTCTTCCGATCTNNNTCTCCATCCAAGAAGTCACCT
AGCACACGTCTGAACTCTTTCCCTACACGACGCTCTTCCGATCTNNNTGCTGTAATCAATAACCTTGTG
GCACACGTCTGAACTCTTTCCCTACACGACGCTCTTCCGATCTNNNACTTGCAATAGCTGAGTAGTAC
GAGCACACGTCTGAACTCTTTCCCTACACGACGCTCTTCCGATCTNNNAAGCAAGACCCAAATACATG
GAGCACACGTCTGAACTCTTTCCCTACACGACGCTCTTCCGATCTNNNCAAGATCGCACTACTGTACTC
AGCACACGTCTGAACTCTTTCCCTACACGACGCTCTTCCGATCTNNNTAGTCCCAACTACTCAGGAGGT
AGCACACGTCTGAACTCTTTCCCTACACGACGCTCTTCCGATCTNNNCTGTAATCCCAGCTACTCAGGA
GAGCACACGTCTGAACTCTTTCCCTACACGACGCTCTTCCGATCTNNNATACCAAAAATTAGCCGAGCG
GAGCACACGTCTGAACTCTTTCCCTACACGACGCTCTTCCGATCTNNNGGCACCTCTAATCTCAGCTAC
AGAGCACACGTCTGAACTCTTTCCCTACACGACGCTCTTCCGATCTNNNCTCACGCCTGTAATCCCAATA
AGAGCACACGTCTGAACTCTTTCCCTACACGACGCTCTTCCGATCTNNNGATGCCTGTAATCCCAGCTA
GAGCACACGTCTGAACTCTTTCCCTACACGACGCTCTTCCGATCTNNNGCATTCCTGTAATCCCAGCTA
GAGCACACGTCTGAACTCTTTCCCTACACGACGCTCTTCCGATCTNNNGAGGCTGAGGTGAGAGAATC
GAGCACACGTCTGAACTCTTTCCCTACACGACGCTCTTCCGATCTNNNCTGAGGCAGGAGAATCACTT
GAGCACACGTCTGAACTCTTTCCCTACACGACGCTCTTCCGATCTNNNACACAGTCATCTGCTATTAGT
GAGCACACGTCTGAACTCTTTCCCTACACGACGCTCTTCCGATCTNNNAGAAGTTCCAGTGAGCCAAG
GAGCACACGTCTGAACTCTTTCCCTACACGACGCTCTTCCGATCTNNNTCCTTGTTTGCAGATGCCATA
GAGCACACGTCTGAACTCTTTCCCTACACGACGCTCTTCCGATCTNNNCTTGCAGTGAACCAAGATCTC
GAGCACACGTCTGAACTCTTTCCCTACACGACGCTCTTCCGATCTNNNTGGAAAGAACCACAATGTTCA
GAGCACACGTCTGAACTCTTTCCCTACACGACGCTCTTCCGATCTNNNCTGAGGCAGGAGAATAGCTT
AGAGCACACGTCTGAACTCTTTCCCTACACGACGCTCTTCCGATCTNNNGGCAGGAGAATCACTTGA
AGAGCACACGTCTGAACTCTTTCCCTACACGACGCTCTTCCGATCTNNNGATCTTTGACAAAGGGGCA
GAGCACACGTCTGAACTCTTTCCCTACACGACGCTCTTCCGATCTNNNGGAGGTTGAAGTGAGCTGA
GAGCACACGTCTGAACTCTTTCCCTACACGACGCTCTTCCGATCTNNNTTGAGGCACGAGAATCACTT

AGAGCACACGTCTGAACTCTTTCCCTACACGACGCTCTTCCGATCTNNNAGGCAAGAGAATCACTTGA

AAGAGCACACGTCTGAACTCTTTCCCTACACGACGCTCTTCCGATCTNNNGCAGGAGAATTGCTTGAA

GAGCACACGTCTGAACTCTTTCCCTACACGACGCTCTTCCGATCTNNNTGACACAGAGACATGAAGTA

AGAGCACACGTCTGAACTCTTTCCCTACACGACGCTCTTCCGATCTNNNGATTATCGATTGAGCCCAG

GAGCACACGTCTGAACTCTTTCCCTACACGACGCTCTTCCGATCTNNNGCTTGCTTTCTTGTTTGTTAGC

GAGCACACGTCTGAACTCTTTCCCTACACGACGCTCTTCCGATCTNNNTCGTCCTTACTAACTTCATTG

AGCACACGTCTGAACTCTTTCCCTACACGACGCTCTTCCGATCTNNNAAAAAAAATTAGCTGGGCGTG

GAGCACACGTCTGAACTCTTTCCCTACACGACGCTCTTCCGATCTNNNAAAGCCATCCTGAAGGTTTG

AAGAGCACACGTCTGAACTCTTTCCCTACACGACGCTCTTCCGATCTNNNGTGCCTGTAATCCCAGCTA

GAGCACACGTCTGAACTCTTTCCCTACACGACGCTCTTCCGATCTNNNTGTTTTGGCAAAAGCTCTTC

GAGCACACGTCTGAACTCTTTCCCTACACGACGCTCTTCCGATCTNNNCACCTGTACTCCCAGCTATTCA

AGAGCACACGTCTGAACTCTTTCCCTACACGACGCTCTTCCGATCTNNNGCACACATCTCTAATCCCAG

GAGCACACGTCTGAACTCTTTCCCTACACGACGCTCTTCCGATCTNNNCTGAGGCAGGAGAATCACTT

AGAGCACACGTCTGAACTCTTTCCCTACACGACGCTCTTCCGATCTNNNGCAAATGTTAGTGGCACGT

GAGCACACGTCTGAACTCTTTCCCTACACGACGCTCTTCCGATCTNNNAAAAAATTAGCCAGGTGTGG

AGAGCACACGTCTGAACTCTTTCCCTACACGACGCTCTTCCGATCTNNNAAATACGAAATTAGCCAGG

AGAGCACACGTCTGAACTCTTTCCCTACACGACGCTCTTCCGATCTNNNGAGGTTGCAGTGAGCTGA

AAGAGCACACGTCTGAACTCTTTCCCTACACGACGCTCTTCCGATCTNNNACAAAAATTAGCCAGGTG

GAGCACACGTCTGAACTCTTTCCCTACACGACGCTCTTCCGATCTNNNAGCCAGCTAGTCATTGATGTT

AGAGCACACGTCTGAACTCTTTCCCTACACGACGCTCTTCCGATCTNNNCTGAGGCAGGAGAATCACT

GAGCACACGTCTGAACTCTTTCCCTACACGACGCTCTTCCGATCTNNNCATCCCTGTAATCCCAGCTAC

GAGCACACGTCTGAACTCTTTCCCTACACGACGCTCTTCCGATCTNNNCTGAGGTGGCAGGATTACCT

GAGCACACGTCTGAACTCTTTCCCTACACGACGCTCTTCCGATCTNNNCAGGAGATGGAGGCTACAA

AGAGCACACGTCTGAACTCTTTCCCTACACGACGCTCTTCCGATCTNNNTGAAGCAGGGAGGATTGA

AGAGCACACGTCTGAACTCTTTCCCTACACGACGCTCTTCCGATCTNNNTGAGAAGTTCAACAAGGAA

AGAGCACACGTCTGAACTCTTTCCCTACACGACGCTCTTCCGATCTNNNTGAGGCACAAGAATCGTTTG

GAGCACACGTCTGAACTCTTTCCCTACACGACGCTCTTCCGATCTNNNACAAAACTTAGCTGGGCATG

AGAGCACACGTCTGAACTCTTTCCCTACACGACGCTCTTCCGATCTNNNGTGGCAGGTACCTGTAATC

GAGCACACGTCTGAACTCTTTCCCTACACGACGCTCTTCCGATCTNNNATACAAAAATTAGCTGGGCG

AGCACACGTCTGAACTCTTTCCCTACACGACGCTCTTCCGATCTNNNCTGTAGTCCCAGCTACTCAGG

GAGCACACGTCTGAACTCTTTCCCTACACGACGCTCTTCCGATCTNNNAAACCTACCCTGAAAGAACA

GAGCACACGTCTGAACTCTTTCCCTACACGACGCTCTTCCGATCTNNNGAAGAATTGCTTGAATCCGG

AGAGCACACGTCTGAACTCTTTCCCTACACGACGCTCTTCCGATCTNNNGAAGTTGTAGTGAGCCGAG

AGAGCACACGTCTGAACTCTTTCCCTACACGACGCTCTTCCGATCTNNNCTGTAATCCCAGCTACTCAG

GAGCACACGTCTGAACTCTTTCCCTACACGACGCTCTTCCGATCTNNNCTGAGACAGGAGAATGGTG

AGCACACGTCTGAACTCTTTCCCTACACGACGCTCTTCCGATCTNNNGGACATCTGAATTGGAAAGGA

AAGAGCACACGTCTGAACTCTTTCCCTACACGACGCTCTTCCGATCTNNNGCAGGAGAATTGCTTGA

AGAGCACACGTCTGAACTCTTTCCCTACACGACGCTCTTCCGATCTNNNTAGTCCTAGCTACTTGGAA

GAGCACACGTCTGAACTCTTTCCCTACACGACGCTCTTCCGATCTNNNATGTTGCAGTGAGCCGAGAT

AGCACACGTCTGAACTCTTTCCCTACACGACGCTCTTCCGATCTNNNGTAATCCCAGCTACTTAGGAG

AGAGCACACGTCTGAACTCTTTCCCTACACGACGCTCTTCCGATCTNNNGCATACCTGTAATCCCAG

AGAGCACACGTCTGAACTCTTTCCCTACACGACGCTCTTCCGATCTNNNGAGGCTGCAGTGAGCTATG

GAGCACACGTCTGAACTCTTTCCCTACACGACGCTCTTCCGATCTNNNGCCTGTAGTCTCATCTACTTC

GAGCACACGTCTGAACTCTTTCCCTACACGACGCTCTTCCGATCTNNNAATACAAAAAATTAGCCGGG
GAGCACACGTCTGAACTCTTTCCCTACACGACGCTCTTCCGATCTNNNCTGAGGCACGAGAATCACTTG
AGAGCACACGTCTGAACTCTTTCCCTACACGACGCTCTTCCGATCTNNNAAAAAATTAGCTGGGCATG
GAGCACACGTCTGAACTCTTTCCCTACACGACGCTCTTCCGATCTNNNGCATCCAACAAGGTAAAATCC
AGCACACGTCTGAACTCTTTCCCTACACGACGCTCTTCCGATCTNNNATACAGAATTAGCCGGGCATG
AGAGCACACGTCTGAACTCTTTCCCTACACGACGCTCTTCCGATCTNNNCTCAGGAGGCTGTAGCAGA
AGAGCACACGTCTGAACTCTTTCCCTACACGACGCTCTTCCGATCTNNNCTGTAGTCCCAGCTACTCAG
AGAGCACACGTCTGAACTCTTTCCCTACACGACGCTCTTCCGATCTNNNTGAGGCAGGAGAATCACTT
AAGAGCACACGTCTGAACTCTTTCCCTACACGACGCTCTTCCGATCTNNNACAAAAATTAGCCAGGCT
AGAGCACACGTCTGAACTCTTTCCCTACACGACGCTCTTCCGATCTNNNCAGTTGTTCCCAGCAATGTC
AGCACACGTCTGAACTCTTTCCCTACACGACGCTCTTCCGATCTNNNACAGACCAATAACAAGGAGAG
GAGCACACGTCTGAACTCTTTCCCTACACGACGCTCTTCCGATCTNNNCCAGGAAGAAACTGAATCCC
GAGCACACGTCTGAACTCTTTCCCTACACGACGCTCTTCCGATCTNNNAAAAAATTAGCCAGGTGTGG
AGAGCACACGTCTGAACTCTTTCCCTACACGACGCTCTTCCGATCTNNNGTAATCCCAGCTACTTAGG
GAGCACACGTCTGAACTCTTTCCCTACACGACGCTCTTCCGATCTNNNAAATTAGCCGAGCCTAGTGGT
AGAGCACACGTCTGAACTCTTTCCCTACACGACGCTCTTCCGATCTNNNCTGAGGCAGGATAATCGCTT
AGAGCACACGTCTGAACTCTTTCCCTACACGACGCTCTTCCGATCTNNNTGAGGCAGGAGAATCTCTT
GAGCACACGTCTGAACTCTTTCCCTACACGACGCTCTTCCGATCTNNNCTGTAGTCCCAGCTTCTCAGG
GAGCACACGTCTGAACTCTTTCCCTACACGACGCTCTTCCGATCTNNNCATGAGAATTGCTTGAACCC
AGAGCACACGTCTGAACTCTTTCCCTACACGACGCTCTTCCGATCTNNNCAAAAAATTAGCCAGGTTT
GAGCACACGTCTGAACTCTTTCCCTACACGACGCTCTTCCGATCTNNNCAAAAAAAAATTAGCCGGGC
AAGAGCACACGTCTGAACTCTTTCCCTACACGACGCTCTTCCGATCTNNNCTTGCAGTGAGCCAAGAT
GAGCACACGTCTGAACTCTTTCCCTACACGACGCTCTTCCGATCTNNNACACCTATAGTCCCAGCTACTC
GAGCACACGTCTGAACTCTTTCCCTACACGACGCTCTTCCGATCTNNNCATTTGTGTACAAGTGTGTGT
AGAGCACACGTCTGAACTCTTTCCCTACACGACGCTCTTCCGATCTNNNGAGAATCACTTGAACCCAGG
AGCACACGTCTGAACTCTTTCCCTACACGACGCTCTTCCGATCTNNNAGTTTTCAACCTATTGCTAACTG
AGAGCACACGTCTGAACTCTTTCCCTACACGACGCTCTTCCGATCTNNNTCCTATGGGATACAGCAAA
AGAGCACACGTCTGAACTCTTTCCCTACACGACGCTCTTCCGATCTNNNACGTGTTTATCTGCTGACCT
GAGCACACGTCTGAACTCTTTCCCTACACGACGCTCTTCCGATCTNNNGGATTTGCAGTGAGTCAAGA
GAGCACACGTCTGAACTCTTTCCCTACACGACGCTCTTCCGATCTNNNGAGACATGAGAATCGCTTCA
GAGCACACGTCTGAACTCTTTCCCTACACGACGCTCTTCCGATCTNNNCTGTAATCCCAGCTACTCAGG
AAGAGCACACGTCTGAACTCTTTCCCTACACGACGCTCTTCCGATCTNNNGAAGGAAGGAAGGAAGG
AGAGCACACGTCTGAACTCTTTCCCTACACGACGCTCTTCCGATCTNNNCAGTATACGTCTGTAGTCCC
GAGCACACGTCTGAACTCTTTCCCTACACGACGCTCTTCCGATCTNNNGAGAATGGCATGAATCGAG
AGCACACGTCTGAACTCTTTCCCTACACGACGCTCTTCCGATCTNNNTGTCACTGGATAATACCTGCAA
GAGCACACGTCTGAACTCTTTCCCTACACGACGCTCTTCCGATCTNNNGGCTAAGGCACAAGAATCAC
GAGCACACGTCTGAACTCTTTCCCTACACGACGCTCTTCCGATCTNNNAAAAAAATTAGCCGGGCGTA
AGCACACGTCTGAACTCTTTCCCTACACGACGCTCTTCCGATCTNNNTGCACAGTACCCTCTTATAATG
AGAGCACACGTCTGAACTCTTTCCCTACACGACGCTCTTCCGATCTNNNGGGAGGGATAGCATTGGG
AGCACACGTCTGAACTCTTTCCCTACACGACGCTCTTCCGATCTNNNGAGAATTGCTTGAACCAGGAC
GAGCACACGTCTGAACTCTTTCCCTACACGACGCTCTTCCGATCTNNNTCTGTAGTCCCAGCTACTCAA
GAGCACACGTCTGAACTCTTTCCCTACACGACGCTCTTCCGATCTNNNACTTGGAATGCTGAAGTAGG
AAGAGCACACGTCTGAACTCTTTCCCTACACGACGCTCTTCCGATCTNNNGAGGCTAAGGTGGAAGA

AGAGCACACGTCTGAACTCTTTCCCTACACGACGCTCTTCCGATCTNNNACACCTGTAATCCCAGCTACT
AGAGCACACGTCTGAACTCTTTCCCTACACGACGCTCTTCCGATCTNNNGAGAATGGCATGAACCCAG
AGAGCACACGTCTGAACTCTTTCCCTACACGACGCTCTTCCGATCTNNNCTGTAATCCCAGCTACTCAGC
GAGCACACGTCTGAACTCTTTCCCTACACGACGCTCTTCCGATCTNNNCTGAGGCATGAGAATTGCTTG
AGAGCACACGTCTGAACTCTTTCCCTACACGACGCTCTTCCGATCTNNNAGAATTACTTGAACCCAGGA
AGAGCACACGTCTGAACTCTTTCCCTACACGACGCTCTTCCGATCTNNNACGGAAGTGGCTTTGAAAC
AAGAGCACACGTCTGAACTCTTTCCCTACACGACGCTCTTCCGATCTNNNGAGGTAGGAGAATGGCGT
AGAGCACACGTCTGAACTCTTTCCCTACACGACGCTCTTCCGATCTNNNGAGGATCACTTGAACCTGG
GAGCACACGTCTGAACTCTTTCCCTACACGACGCTCTTCCGATCTNNNACAGGAGAATAGTTTGAGCC
GAGCACACGTCTGAACTCTTTCCCTACACGACGCTCTTCCGATCTNNNGCAGTAATGCAGATCAATGAC
AGCACACGTCTGAACTCTTTCCCTACACGACGCTCTTCCGATCTNNNACACAACTTCCCAAAATTGAACC
GAGCACACGTCTGAACTCTTTCCCTACACGACGCTCTTCCGATCTNNNAGTACGTGTCATACCGTCTTG
GAGCACACGTCTGAACTCTTTCCCTACACGACGCTCTTCCGATCTNNNAGACAGGAGAATTGCTTGAA
AGAGCACACGTCTGAACTCTTTCCCTACACGACGCTCTTCCGATCTNNNCTGTAGTTCCAGCTACTCGG
GAGCACACGTCTGAACTCTTTCCCTACACGACGCTCTTCCGATCTNNNAGTGAGCCTAGATAATGCCA
GAGCACACGTCTGAACTCTTTCCCTACACGACGCTCTTCCGATCTNNNTGACAAGAAAAAAATGGGA
AGAGCACACGTCTGAACTCTTTCCCTACACGACGCTCTTCCGATCTNNNTCATCAGAGAGAACAGACA
GAGCACACGTCTGAACTCTTTCCCTACACGACGCTCTTCCGATCTNNNTGACTTGAACTTGAGAGGTG
AGAGCACACGTCTGAACTCTTTCCCTACACGACGCTCTTCCGATCTNNNTGTAGTCCCAGCTACTCAGG
GCACACGTCTGAACTCTTTCCCTACACGACGCTCTTCCGATCTNNNATGTTCAGTAGCTTTCCTCTAAGT
GAGCACACGTCTGAACTCTTTCCCTACACGACGCTCTTCCGATCTNNNGGCACCTATAATCCCAGCTAC
AGCACACGTCTGAACTCTTTCCCTACACGACGCTCTTCCGATCTNNNACCATGCAGAAATCCAAAACCT
AGAGCACACGTCTGAACTCTTTCCCTACACGACGCTCTTCCGATCTNNNGTTGCAGTGAGCTGAGATC
GAGCACACGTCTGAACTCTTTCCCTACACGACGCTCTTCCGATCTNNNCAAGCCTCCAGATCTAGATAG
GAAGAGCACACGTCTGAACTCTTTCCCTACACGACGCTCTTCCGATCTNNNGAGGCTGAGGCAGGAG
GCACACGTCTGAACTCTTTCCCTACACGACGCTCTTCCGATCTNNNGTTCTGTGGATTTGACAAAGGT
GAGCACACGTCTGAACTCTTTCCCTACACGACGCTCTTCCGATCTNNNTGCATCCTAAACTGCTGGAGA
AGAGCACACGTCTGAACTCTTTCCCTACACGACGCTCTTCCGATCTNNNAGGAAGAACAACTGGACTT
GAGCACACGTCTGAACTCTTTCCCTACACGACGCTCTTCCGATCTNNNCCATATGATCCAGCAATCCCA
AGAGCACACGTCTGAACTCTTTCCCTACACGACGCTCTTCCGATCTNNNAACAAAGCTCAAGAATACCC
GAGCACACGTCTGAACTCTTTCCCTACACGACGCTCTTCCGATCTNNNTACATGCCTGTAATCCCAGCT
AAGAGCACACGTCTGAACTCTTTCCCTACACGACGCTCTTCCGATCTNNNAAAAATTAGCTGGTCTTG
AGCACACGTCTGAACTCTTTCCCTACACGACGCTCTTCCGATCTNNNCTGAGGCAGGAGAATCACTTG
AGCACACGTCTGAACTCTTTCCCTACACGACGCTCTTCCGATCTNNNTGGATCCTCTCCTTATTCCTTAG
GAGCACACGTCTGAACTCTTTCCCTACACGACGCTCTTCCGATCTNNNGAGGTAAGAGGATGGATTGA
GCACACGTCTGAACTCTTTCCCTACACGACGCTCTTCCGATCTNNNTGTTCCAACATTTGAAATGCTTC
GAGCACACGTCTGAACTCTTTCCCTACACGACGCTCTTCCGATCTNNNAACTGAGATTGTACCACTGC
AGCACACGTCTGAACTCTTTCCCTACACGACGCTCTTCCGATCTNNNGAGGTAGGAGAATAGCGTGA
AGAGCACACGTCTGAACTCTTTCCCTACACGACGCTCTTCCGATCTNNNAGCAAACCAACCACAAAACT
GAGCACACGTCTGAACTCTTTCCCTACACGACGCTCTTCCGATCTNNNAATACAAAAAATTAGCCGGG
GAGCACACGTCTGAACTCTTTCCCTACACGACGCTCTTCCGATCTNNNTGGGGTAACAAATGGTGAAC
GAGCACACGTCTGAACTCTTTCCCTACACGACGCTCTTCCGATCTNNNCCATGGGTGCCAAATCATTAG
GAAGAGCACACGTCTGAACTCTTTCCCTACACGACGCTCTTCCGATCTNNNAATAGCTGTGTGTGGTG

GAGCACACGTCTGAACTCTTTCCCTACACGACGCTCTTCCGATCTNNNGAGGCAGGAAAATCACTTG
AGCACACGTCTGAACTCTTTCCCTACACGACGCTCTTCCGATCTNNNGAGGCAGGAAAATCACTTGAA
GAGCACACGTCTGAACTCTTTCCCTACACGACGCTCTTCCGATCTNNNAGTAAGCCAATTAGAACATG
AGCACACGTCTGAACTCTTTCCCTACACGACGCTCTTCCGATCTNNNAGATATTCCATGCCAATGAAAT
AGAGCACACGTCTGAACTCTTTCCCTACACGACGCTCTTCCGATCTNNNCTGAGGCAGGAGAATCACT
GAGCACACGTCTGAACTCTTTCCCTACACGACGCTCTTCCGATCTNNNAAAAAACTTAGCCAGGTGTG
AGCACACGTCTGAACTCTTTCCCTACACGACGCTCTTCCGATCTNNNCTGTAAGCCCAGCTACTCAAGA
GAGCACACGTCTGAACTCTTTCCCTACACGACGCTCTTCCGATCTNNNAAAAAATTAGCCAGGCACGG
GAGCACACGTCTGAACTCTTTCCCTACACGACGCTCTTCCGATCTNNNGAGGCAAGAGAATCGCTTG
AGAGCACACGTCTGAACTCTTTCCCTACACGACGCTCTTCCGATCTNNNCTGAGGCAGGAAAATTGCT
GAGCACACGTCTGAACTCTTTCCCTACACGACGCTCTTCCGATCTNNNCTCACTCCTGTAATCCCAACA
AGCACACGTCTGAACTCTTTCCCTACACGACGCTCTTCCGATCTNNNACAAAAATTAGCTGGGCATG
GAGCACACGTCTGAACTCTTTCCCTACACGACGCTCTTCCGATCTNNNGTTGCAGTGAGCTGAGATT
AGAGCACACGTCTGAACTCTTTCCCTACACGACGCTCTTCCGATCTNNNTGAGGTAGGAGAAACGCT
GAGCACACGTCTGAACTCTTTCCCTACACGACGCTCTTCCGATCTNNNGAGAATCACTTGAACCCAGG
GAGCACACGTCTGAACTCTTTCCCTACACGACGCTCTTCCGATCTNNNCATGCCTGTAATCCCAGCTAC
GCACACGTCTGAACTCTTTCCCTACACGACGCTCTTCCGATCTNNNAATGGTAGTTTGTATTTCTGTG
AAGAGCACACGTCTGAACTCTTTCCCTACACGACGCTCTTCCGATCTNNNGAGGCAGGAGAATGGT
GAGCACACGTCTGAACTCTTTCCCTACACGACGCTCTTCCGATCTNNNCAACATCGGACTTGGCAATG
AGAGCACACGTCTGAACTCTTTCCCTACACGACGCTCTTCCGATCTNNNACAATGGACCACTATTCAG
GAGCACACGTCTGAACTCTTTCCCTACACGACGCTCTTCCGATCTNNNAGCTTACCAAGATTGAACCAT
AGCACACGTCTGAACTCTTTCCCTACACGACGCTCTTCCGATCTNNNTGGGGAAAGACAGTTTGTTC
AGCACACGTCTGAACTCTTTCCCTACACGACGCTCTTCCGATCTNNNGGCAGAATAATTGCTTGAACC
GAGCACACGTCTGAACTCTTTCCCTACACGACGCTCTTCCGATCTNNNAGGAGAATCACTGGAACCT
GAGCACACGTCTGAACTCTTTCCCTACACGACGCTCTTCCGATCTNNNAGAAGAATCGCTTGAATCCA
GAGCACACGTCTGAACTCTTTCCCTACACGACGCTCTTCCGATCTNNNTGAGGCAGGAGAAATCACT
GAGCACACGTCTGAACTCTTTCCCTACACGACGCTCTTCCGATCTNNNCTTGAGACCAGTGGTTCGA
AAGAGCACACGTCTGAACTCTTTCCCTACACGACGCTCTTCCGATCTNNNCTTGAATCCAGGAGGCAG
GAGCACACGTCTGAACTCTTTCCCTACACGACGCTCTTCCGATCTNNNGTTCATGCACTCACTTGACA
AGAGCACACGTCTGAACTCTTTCCCTACACGACGCTCTTCCGATCTNNNGATTGCATCACAGCCTCAAC
GAGCACACGTCTGAACTCTTTCCCTACACGACGCTCTTCCGATCTNNNGAGAATTGCTTGAACCTGGG
AGCACACGTCTGAACTCTTTCCCTACACGACGCTCTTCCGATCTNNNGAGAATTGCTTGAACCTGGG
GAGCACACGTCTGAACTCTTTCCCTACACGACGCTCTTCCGATCTNNNGAGAATTGCTTGAACCTGGG
GAGCACACGTCTGAACTCTTTCCCTACACGACGCTCTTCCGATCTNNNAGTGAGCTGAGATTGTACCA
GAGCACACGTCTGAACTCTTTCCCTACACGACGCTCTTCCGATCTNNNAAAAATTAGCCTAGCATGGT
AGAGCACACGTCTGAACTCTTTCCCTACACGACGCTCTTCCGATCTNNNCTACAAGCTACTCTGAAGG
AGAGCACACGTCTGAACTCTTTCCCTACACGACGCTCTTCCGATCTNNNAATTCCTGGACACATACACC
AGAGCACACGTCTGAACTCTTTCCCTACACGACGCTCTTCCGATCTNNNATCACTTGAACCCAGTAGG
AGAGCACACGTCTGAACTCTTTCCCTACACGACGCTCTTCCGATCTNNNACAGGAGAATCGTTTGTAC
AGAGCACACGTCTGAACTCTTTCCCTACACGACGCTCTTCCGATCTNNNCTTGCAGTGAGCTGAGATT
AAGAGCACACGTCTGAACTCTTTCCCTACACGACGCTCTTCCGATCTNNNGAGACAGGAGAATGGCGT
AGCACACGTCTGAACTCTTTCCCTACACGACGCTCTTCCGATCTNNNAGGAGGCTGAGTAGGATAAT
AGCACACGTCTGAACTCTTTCCCTACACGACGCTCTTCCGATCTNNNCACCTATAATCCCAGCTACTCG

GAGCACACGTCTGAACTCTTTCCCTACACGACGCTCTTCCGATCTNNNACGCCTATAATGCCAGCTCTT
AGCACACGTCTGAACTCTTTCCCTACACGACGCTCTTCCGATCTNNNTGTAGTCCTCTCTACTTGGGA
AGCACACGTCTGAACTCTTTCCCTACACGACGCTCTTCCGATCTNNNTGAGGCAGGAGAATCACTTG
AGCACACGTCTGAACTCTTTCCCTACACGACGCTCTTCCGATCTNNNACTCCACAGTAACACAAGTTGT
AGAGCACACGTCTGAACTCTTTCCCTACACGACGCTCTTCCGATCTNNNAACATCAGGGAAACACTAC
AGCACACGTCTGAACTCTTTCCCTACACGACGCTCTTCCGATCTNNNTGTAATCCCCACTACTCAGGA
GAGCACACGTCTGAACTCTTTCCCTACACGACGCTCTTCCGATCTNNNGCTGATGTGGGAAAGAAGT
AGCACACGTCTGAACTCTTTCCCTACACGACGCTCTTCCGATCTNNNCTGTAGTTCCAGCTACTTGGG
GAGCACACGTCTGAACTCTTTCCCTACACGACGCTCTTCCGATCTNNNATGAAAAATTAGCCAGGCGT
GAGCACACGTCTGAACTCTTTCCCTACACGACGCTCTTCCGATCTNNNGGGATAGCATTGGGAGATA
GAGCACACGTCTGAACTCTTTCCCTACACGACGCTCTTCCGATCTNNNCACCTGTAATCCCAGCTACTC
AGCACACGTCTGAACTCTTTCCCTACACGACGCTCTTCCGATCTNNNTCATGTTACCCCAGTTAGAAT
GAGCACACGTCTGAACTCTTTCCCTACACGACGCTCTTCCGATCTNNNAGAATGGCTTGAACTTGAG
GAGCACACGTCTGAACTCTTTCCCTACACGACGCTCTTCCGATCTNNNGGCACCTATAATCCCAGCTAC
GAGCACACGTCTGAACTCTTTCCCTACACGACGCTCTTCCGATCTNNNGTTGCAGTGAGCTGAGATC
GAGCACACGTCTGAACTCTTTCCCTACACGACGCTCTTCCGATCTNNNGCAGAATTGTTTCAACCGAG
AGCACACGTCTGAACTCTTTCCCTACACGACGCTCTTCCGATCTNNNAGTGCTATGTTTCACTAAACCC
AGAGCACACGTCTGAACTCTTTCCCTACACGACGCTCTTCCGATCTNNNACTTGAAATTGGAAGGTGG
AGAGCACACGTCTGAACTCTTTCCCTACACGACGCTCTTCCGATCTNNNGCTGCAGTGAGCTATGATT
AGAGCACACGTCTGAACTCTTTCCCTACACGACGCTCTTCCGATCTNNNGGATCAGGTATAGCTGCTG
AGCACACGTCTGAACTCTTTCCCTACACGACGCTCTTCCGATCTNNNCTGTAGTTCCAGCTTCTCAGG
AGCACACGTCTGAACTCTTTCCCTACACGACGCTCTTCCGATCTNNNAAGAGGACCTACACAGTTCCA
AGAGCACACGTCTGAACTCTTTCCCTACACGACGCTCTTCCGATCTNNNAATGCGACAATAACCAGAA
AAGAGCACACGTCTGAACTCTTTCCCTACACGACGCTCTTCCGATCTNNNCTTGTAGTGAGCTGAGAT
AGCACACGTCTGAACTCTTTCCCTACACGACGCTCTTCCGATCTNNNTGATACAAGGGTGTACAACTG
AAGAGCACACGTCTGAACTCTTTCCCTACACGACGCTCTTCCGATCTNNNCTTGTAGTGAGCTGAGAT
GAGCACACGTCTGAACTCTTTCCCTACACGACGCTCTTCCGATCTNNNCACCTGTAATCCCAGCTACTC
CACACGTCTGAACTCTTTCCCTACACGACGCTCTTCCGATCTNNNAAACCATATTTAGCCAGTAGTCT
GCACACGTCTGAACTCTTTCCCTACACGACGCTCTTCCGATCTNNNTGATCATTTCCCCAAGTTTTAGC
AGCACACGTCTGAACTCTTTCCCTACACGACGCTCTTCCGATCTNNNTGAGAAAAACAATTCCTCAGCC
AGCACACGTCTGAACTCTTTCCCTACACGACGCTCTTCCGATCTNNNTCAGAGACTAGTATTGCAACA
GCACACGTCTGAACTCTTTCCCTACACGACGCTCTTCCGATCTNNNACCAATTGACCATAGATGCATAG
AGCACACGTCTGAACTCTTTCCCTACACGACGCTCTTCCGATCTNNNGCCAGTTGAACATTATAGACC
AGCACACGTCTGAACTCTTTCCCTACACGACGCTCTTCCGATCTNNNTGGAATCTTCTGTGTTGTCTTT
AGCACACGTCTGAACTCTTTCCCTACACGACGCTCTTCCGATCTNNNAGAGCCTCTTTTTCTGAGCTTA
GCACACGTCTGAACTCTTTCCCTACACGACGCTCTTCCGATCTNNNTGTCATTTTTGGACTTTAGAGG
AGCACACGTCTGAACTCTTTCCCTACACGACGCTCTTCCGATCTNNNACTTCTTCCTTTCCTATCTGCATT
AGCACACGTCTGAACTCTTTCCCTACACGACGCTCTTCCGATCTNNNGGATCAGTGTCTTGACTTTA
AGCACACGTCTGAACTCTTTCCCTACACGACGCTCTTCCGATCTNNNGGTAGGAACCGTAAAATGTTT
AGCACACGTCTGAACTCTTTCCCTACACGACGCTCTTCCGATCTNNNTGCATTATTTTTTCAGTGCAGA
AGCACACGTCTGAACTCTTTCCCTACACGACGCTCTTCCGATCTNNNTGTCCCCTTTTCTTATTCTCTGGA
CACACGTCTGAACTCTTTCCCTACACGACGCTCTTCCGATCTNNNAGTTCTAACATGTATGTATGCAAA
AGCACACGTCTGAACTCTTTCCCTACACGACGCTCTTCCGATCTNNNTGGGATTTTTTTAGCCCTTCAT

GAGCACACGTCTGAACTCTTTCCCTACACGACGCTCTTCCGATCTNNNTCTGTGATTTCTTTCAGCAGT
GCACACGTCTGAACTCTTTCCCTACACGACGCTCTTCCGATCTNNNAGCATCTCCTTGATATGTGTATG
AGCACACGTCTGAACTCTTTCCCTACACGACGCTCTTCCGATCTNNNAGATTTGTACAGGCATGGAGA
AGCACACGTCTGAACTCTTTCCCTACACGACGCTCTTCCGATCTNNNGTGTGGCACATTTTTTTTTTCC
GAGCACACGTCTGAACTCTTTCCCTACACGACGCTCTTCCGATCTNNNAGGTTGCAATGAGCTGAGAC
AGAGCACACGTCTGAACTCTTTCCCTACACGACGCTCTTCCGATCTNNNTTTTCTCAAAGCTCCACTAGG
GAGCACACGTCTGAACTCTTTCCCTACACGACGCTCTTCCGATCTNNNTGAGCCAAGATCACACCATTG
AGAGCACACGTCTGAACTCTTTCCCTACACGACGCTCTTCCGATCTNNNAAAAATAAGCTAGGGGTGG
GAGCACACGTCTGAACTCTTTCCCTACACGACGCTCTTCCGATCTNNNTGATTTGCATTGAACACAAAC
GAGCACACGTCTGAACTCTTTCCCTACACGACGCTCTTCCGATCTNNNTCATGTTACCCCAGTTAGAAT
AGCACACGTCTGAACTCTTTCCCTACACGACGCTCTTCCGATCTNNNGTTTTTTCATGTTCTTTGCCCAC
AGCACACGTCTGAACTCTTTCCCTACACGACGCTCTTCCGATCTNNNTGTGTATCTAATTTTTGGGGGG
CACACGTCTGAACTCTTTCCCTACACGACGCTCTTCCGATCTNNNTCAACATACAAAAATCAGTAGCAT
AGAGCACACGTCTGAACTCTTTCCCTACACGACGCTCTTCCGATCTNNNGGAAAAGATCAGAAGTGAC
AGCACACGTCTGAACTCTTTCCCTACACGACGCTCTTCCGATCTNNNAGTCGAGATATTGACATCAATG
AGCACACGTCTGAACTCTTTCCCTACACGACGCTCTTCCGATCTNNNTTTCATTGTCAGTGTACAGAAA
GAGCACACGTCTGAACTCTTTCCCTACACGACGCTCTTCCGATCTNNNTGTTCATCAGGGATATTGAC
AGCACACGTCTGAACTCTTTCCCTACACGACGCTCTTCCGATCTNNNAGAGAAAAGGACGATGTAGC
AGAGCACACGTCTGAACTCTTTCCCTACACGACGCTCTTCCGATCTNNNACACACACACACACACACAA
GAGCACACGTCTGAACTCTTTCCCTACACGACGCTCTTCCGATCTNNNTGACCATGTATTGAGCATGAC
AGAGCACACGTCTGAACTCTTTCCCTACACGACGCTCTTCCGATCTNNNCTAAGGCACGAGAATTGCT
AGCACACGTCTGAACTCTTTCCCTACACGACGCTCTTCCGATCTNNNAGACACACATATCATATGCACA
GAGCACACGTCTGAACTCTTTCCCTACACGACGCTCTTCCGATCTNNNACTGAGTCAATTCAATCAGCA
GAGCACACGTCTGAACTCTTTCCCTACACGACGCTCTTCCGATCTNNNACTGAGTCAATTCAATCAGCA
AGCACACGTCTGAACTCTTTCCCTACACGACGCTCTTCCGATCTNNNACAAGTCAAGGGTTACTGTGT
AGCACACGTCTGAACTCTTTCCCTACACGACGCTCTTCCGATCTNNNCACCCAAAATAGTGATGAGGA
AGCACACGTCTGAACTCTTTCCCTACACGACGCTCTTCCGATCTNNNGAGAATCACTCGAACCAGGGA
GAGCACACGTCTGAACTCTTTCCCTACACGACGCTCTTCCGATCTNNNCATCTTTAGCATCCATTCCCTT
GCACACGTCTGAACTCTTTCCCTACACGACGCTCTTCCGATCTNNNTGGCCTCTATAATTATGTGAGCC
GAGCACACGTCTGAACTCTTTCCCTACACGACGCTCTTCCGATCTNNNCAAGTGTCTTTGTGTGTGTGT
CACACGTCTGAACTCTTTCCCTACACGACGCTCTTCCGATCTNNNTTACACACATATTCATCATTTCTGG
GAGCACACGTCTGAACTCTTTCCCTACACGACGCTCTTCCGATCTNNNTGTGTGTGTATATTGTGTGT
AGAGCACACGTCTGAACTCTTTCCCTACACGACGCTCTTCCGATCTNNNTGGTATTCCCACCCAAAGTA
GAGCACACGTCTGAACTCTTTCCCTACACGACGCTCTTCCGATCTNNNGTCTTGTGCTGGTCCATTTCT
AGCACACGTCTGAACTCTTTCCCTACACGACGCTCTTCCGATCTNNNTGCAACTTCTTTTCATTGTGACC
AGAGCACACGTCTGAACTCTTTCCCTACACGACGCTCTTCCGATCTNNNAAAGGGGACATTACCACTG
AGCACACGTCTGAACTCTTTCCCTACACGACGCTCTTCCGATCTNNNTTTCATTGTCAGTGTACAGAAA
AGCACACGTCTGAACTCTTTCCCTACACGACGCTCTTCCGATCTNNNGGGTGACAGAGTAAGATTCC
GAGCACACGTCTGAACTCTTTCCCTACACGACGCTCTTCCGATCTNNNAGCCAAACCATATCAGAAGG
AAGAGCACACGTCTGAACTCTTTCCCTACACGACGCTCTTCCGATCTNNNGTACTTGTCCTCCATGTGT
GCACACGTCTGAACTCTTTCCCTACACGACGCTCTTCCGATCTNNNATGGTACACTCAAGGCATTATG
AGCACACGTCTGAACTCTTTCCCTACACGACGCTCTTCCGATCTNNNTGAGGTTGAAGACAGGCTTTT
GCACACGTCTGAACTCTTTCCCTACACGACGCTCTTCCGATCTNNNCCAGAAATCAGCTCATTTTCAGG

AGAGCACACGTCTGAACTCTTTCCCTACACGACGCTCTTCCGATCTNNNTCTTTGAGTAAACAAATGA
AGAGCACACGTCTGAACTCTTTCCCTACACGACGCTCTTCCGATCTNNNTGGGTGACAGAGTGAGA
GAGCACACGTCTGAACTCTTTCCCTACACGACGCTCTTCCGATCTNNNTGGCTGAGTAGTATTCCATG
AGCACACGTCTGAACTCTTTCCCTACACGACGCTCTTCCGATCTNNNACTCAAGATTGGCCCATTTAC
AGCACACGTCTGAACTCTTTCCCTACACGACGCTCTTCCGATCTNNNCCAACCAAGGGACAACATAAT
AGAGCACACGTCTGAACTCTTTCCCTACACGACGCTCTTCCGATCTNNNTCACTAAAGGAAGACAGGA
GAGCACACGTCTGAACTCTTTCCCTACACGACGCTCTTCCGATCTNNNCTTGGAGCGAGACACTCTTG
GAGCACACGTCTGAACTCTTTCCCTACACGACGCTCTTCCGATCTNNNCAGAGCGAGATTCTGTCTAA
AGAGCACACGTCTGAACTCTTTCCCTACACGACGCTCTTCCGATCTNNNACACGGACAACAATTATGC
GAGCACACGTCTGAACTCTTTCCCTACACGACGCTCTTCCGATCTNNNCTACAGAGTGAGACTCCGTTT
CACACGTCTGAACTCTTTCCCTACACGACGCTCTTCCGATCTNNNAGTTCATGATTCAAACTGCACTC
AGAGCACACGTCTGAACTCTTTCCCTACACGACGCTCTTCCGATCTNNNATCTGTAGTCCCAGCTACTC
GAAGAGCACACGTCTGAACTCTTTCCCTACACGACGCTCTTCCGATCTNNNCGTACCTGGCCTAAGAC
AGCACACGTCTGAACTCTTTCCCTACACGACGCTCTTCCGATCTNNNACATTTCCTTCTTTTTATGCCT
AGCACACGTCTGAACTCTTTCCCTACACGACGCTCTTCCGATCTNNNGGATTGACTGAGCTGATCTG
GCACACGTCTGAACTCTTTCCCTACACGACGCTCTTCCGATCTNNNTGGCAAGAATACCACAGAAGTA
AGCACACGTCTGAACTCTTTCCCTACACGACGCTCTTCCGATCTNNNGCAGTGTTTTGTAGTTCTCCTT
AGCACACGTCTGAACTCTTTCCCTACACGACGCTCTTCCGATCTNNNAGACACACATATCATATGCACA
GAGCACACGTCTGAACTCTTTCCCTACACGACGCTCTTCCGATCTNNNAGTAAGCTGTGATCACACCA
AGCACACGTCTGAACTCTTTCCCTACACGACGCTCTTCCGATCTNNNGGTAGCTAGTGAGTAGGAG
AGCACACGTCTGAACTCTTTCCCTACACGACGCTCTTCCGATCTNNNTGTACACACGTAGACACATACT
AGCACACGTCTGAACTCTTTCCCTACACGACGCTCTTCCGATCTNNNACTGAGTCAATTCAATCAGCA
GAGCACACGTCTGAACTCTTTCCCTACACGACGCTCTTCCGATCTNNNTTTCATTTCTGTGTGGATTGT
AGCACACGTCTGAACTCTTTCCCTACACGACGCTCTTCCGATCTNNNATCTCTTGCACTTTCAGCAG
AGAGCACACGTCTGAACTCTTTCCCTACACGACGCTCTTCCGATCTNNNCTTGGGAGGCTAGGTTGAA
GCACACGTCTGAACTCTTTCCCTACACGACGCTCTTCCGATCTNNNACTAGTCTGATAGCTATTGGTG
AGCACACGTCTGAACTCTTTCCCTACACGACGCTCTTCCGATCTNNNATATATACACACACACACACA
AGCACACGTCTGAACTCTTTCCCTACACGACGCTCTTCCGATCTNNNACAAACACACACACACACACA
GCACACGTCTGAACTCTTTCCCTACACGACGCTCTTCCGATCTNNNACACCATACCACATGAATATCTG
AGCACACGTCTGAACTCTTTCCCTACACGACGCTCTTCCGATCTNNNCCAATCTCAAAGGTTACAACGT
GAGCACACGTCTGAACTCTTTCCCTACACGACGCTCTTCCGATCTNNNGCTTTGAATTTTTTCCAAGAC
GAGCACACGTCTGAACTCTTTCCCTACACGACGCTCTTCCGATCTNNNAGCCCCAGATTTTATATCAGA
GAGCACACGTCTGAACTCTTTCCCTACACGACGCTCTTCCGATCTNNNTATGTGCCTGGCCTTTTTTTC
AGCACACGTCTGAACTCTTTCCCTACACGACGCTCTTCCGATCTNNNTGACTAAATCAGCTAGAGGTT
AGCACACGTCTGAACTCTTTCCCTACACGACGCTCTTCCGATCTNNNTCCTTCTTTCCTATCTTCCTGTC
AGCACACGTCTGAACTCTTTCCCTACACGACGCTCTTCCGATCTNNNTCCTAAATCCAGCCCTGTAATA
AAGAGCACACGTCTGAACTCTTTCCCTACACGACGCTCTTCCGATCTNNNAAAAATTAGCTGAGTGT
GAGCACACGTCTGAACTCTTTCCCTACACGACGCTCTTCCGATCTNNNCTGTAGTTCCAGCTACTTGG
GAGCACACGTCTGAACTCTTTCCCTACACGACGCTCTTCCGATCTNNNAAGGTTGCAGTGAGCTAAG
AGAGCACACGTCTGAACTCTTTCCCTACACGACGCTCTTCCGATCTNNNGTGAAGATTGCAGTGAGC
AGCACACGTCTGAACTCTTTCCCTACACGACGCTCTTCCGATCTNNNTCTCTCTCTCTCTCTTTCTTT
AGAGCACACGTCTGAACTCTTTCCCTACACGACGCTCTTCCGATCTNNNTCTACACACATACACACACAC
AGCACACGTCTGAACTCTTTCCCTACACGACGCTCTTCCGATCTNNNACGTTCTCACATGATGAAGA

GAGCACACGTCTGAACTCTTTCCCTACACGACGCTCTTCCGATCTNNNCCTGTTTTCAGCCTATGTGCAA
AAGAGCACACGTCTGAACTCTTTCCCTACACGACGCTCTTCCGATCTNNNAAAGGGGCTGTTTGATTGA
AGAGCACACGTCTGAACTCTTTCCCTACACGACGCTCTTCCGATCTNNNGCTACTTGCAGAATGAAGCC
GAGCACACGTCTGAACTCTTTCCCTACACGACGCTCTTCCGATCTNNNTGCATTCTGCTTAGGTTTCAAG
AAGAGCACACGTCTGAACTCTTTCCCTACACGACGCTCTTCCGATCTNNNTGTCCCATTCACTGCTGGA
GAGCACACGTCTGAACTCTTTCCCTACACGACGCTCTTCCGATCTNNNTGTATCTTCTTGCAAGAAAAC
AGCACACGTCTGAACTCTTTCCCTACACGACGCTCTTCCGATCTNNNGTAAAGTATGTGTGTGTGTGTG
AGAGCACACGTCTGAACTCTTTCCCTACACGACGCTCTTCCGATCTNNNAAGCTACCTGCACTCATATGT
AGCACACGTCTGAACTCTTTCCCTACACGACGCTCTTCCGATCTNNNATATATACACACACACACACA
AGAGCACACGTCTGAACTCTTTCCCTACACGACGCTCTTCCGATCTNNNTTTCTGGCCATGTTTGAGAG
AGAGCACACGTCTGAACTCTTTCCCTACACGACGCTCTTCCGATCTNNNGCTTTGGTCTCACAATGTAG
GAGCACACGTCTGAACTCTTTCCCTACACGACGCTCTTCCGATCTNNNACATGACATAGCCTGTGTGTA
GAGCACACGTCTGAACTCTTTCCCTACACGACGCTCTTCCGATCTNNNACACCTTTCATGATGAAAACAC
GAGCACACGTCTGAACTCTTTCCCTACACGACGCTCTTCCGATCTNNNTGTCTGTATTGTGGCAGTGG
AGAGCACACGTCTGAACTCTTTCCCTACACGACGCTCTTCCGATCTNNNCCTACCTGAACTGCCCTTACA
AAGAGCACACGTCTGAACTCTTTCCCTACACGACGCTCTTCCGATCTNNNTGAGTCCATCAGCCAGTTTT
AGCACACGTCTGAACTCTTTCCCTACACGACGCTCTTCCGATCTNNNAGTACGTGTTCACTGTTTTCTGT
GAGCACACGTCTGAACTCTTTCCCTACACGACGCTCTTCCGATCTNNNGCTTAGTAGTATTCCATGGTGT
GAGCACACGTCTGAACTCTTTCCCTACACGACGCTCTTCCGATCTNNNTCAGGTGTTACAATACTGTCAC
AGCACACGTCTGAACTCTTTCCCTACACGACGCTCTTCCGATCTNNNAGGAAACCTGAATCTCAGTTTG
AGAGCACACGTCTGAACTCTTTCCCTACACGACGCTCTTCCGATCTNNNGCAATGGCCCCTTTGTCATTA
AGCACACGTCTGAACTCTTTCCCTACACGACGCTCTTCCGATCTNNNACTGGAATACAAACAAGTTGG
AGCACACGTCTGAACTCTTTCCCTACACGACGCTCTTCCGATCTNNNAAAGTCATTCCATATCCACATGT
AGCACACGTCTGAACTCTTTCCCTACACGACGCTCTTCCGATCTNNNGTCTACCATTTTGATAGCTTAGCA
AGCACACGTCTGAACTCTTTCCCTACACGACGCTCTTCCGATCTNNNAGAATTAGGAAATCGGTGGTAC
GCACACGTCTGAACTCTTTCCCTACACGACGCTCTTCCGATCTNNNGCCTTTGTATTTCTGTGATGTCAC
AGCACACGTCTGAACTCTTTCCCTACACGACGCTCTTCCGATCTNNNTGAATTAGAGTTGGAAGGAACC
GAGCACACGTCTGAACTCTTTCCCTACACGACGCTCTTCCGATCTNNNCCATCACCTTCAAAAGTTTCCT
GCACACGTCTGAACTCTTTCCCTACACGACGCTCTTCCGATCTNNNAGAACCAGTTCTTTGTCTCATTGA
AGCACACGTCTGAACTCTTTCCCTACACGACGCTCTTCCGATCTNNNGGAAAGGAAAAAGGAAAGACT
AGCACACGTCTGAACTCTTTCCCTACACGACGCTCTTCCGATCTNNNACACACCTTACTGCTAATCATGG
AGCACACGTCTGAACTCTTTCCCTACACGACGCTCTTCCGATCTNNNAGCTGTTGTACTGTATTGGTTT
AGCACACGTCTGAACTCTTTCCCTACACGACGCTCTTCCGATCTNNNCACAGTGCCTACAATTACAACAC
AGAGCACACGTCTGAACTCTTTCCCTACACGACGCTCTTCCGATCTNNNGTTGTTGTTCTTTGTAGGCCC
GAGCACACGTCTGAACTCTTTCCCTACACGACGCTCTTCCGATCTNNNTCAACCTAAGTGTTCATCAGTC
AGCACACGTCTGAACTCTTTCCCTACACGACGCTCTTCCGATCTNNNTGGAAAAGATCAAAAGGAAACC
GAGCACACGTCTGAACTCTTTCCCTACACGACGCTCTTCCGATCTNNNAAGAACAGCTCCACATCAAAA
GAGCACACGTCTGAACTCTTTCCCTACACGACGCTCTTCCGATCTNNNACCTACTTGATCATGGTGGAT
AGAGCACACGTCTGAACTCTTTCCCTACACGACGCTCTTCCGATCTNNNTGCTCCCAATACTGGTTCATG
ACACGTCTGAACTCTTTCCCTACACGACGCTCTTCCGATCTNNNAGCAATATTCAGTTGTTTTGGTTTGA
GCACACGTCTGAACTCTTTCCCTACACGACGCTCTTCCGATCTNNNTGGATCCCAAAGATTTTCTCCTGT
GAGCACACGTCTGAACTCTTTCCCTACACGACGCTCTTCCGATCTNNNTGCTTTATCCTTAGGTTCCTCT
GAGCACACGTCTGAACTCTTTCCCTACACGACGCTCTTCCGATCTNNNTCAGGGATTTGATTTCTTCCTG

GCACACGTCTGAACTCTTTCCCTACACGACGCTCTTCCGATCTNNNTCACCTATATGTCATGACAGATGA
GCACACGTCTGAACTCTTTCCCTACACGACGCTCTTCCGATCTNNNGTGAAGTTTACACCATGTGTTAGC
GAGCACACGTCTGAACTCTTTCCCTACACGACGCTCTTCCGATCTNNNTCAATTACCATCCTTCCAGAAC
AGCACACGTCTGAACTCTTTCCCTACACGACGCTCTTCCGATCTNNNGCAGTAAATTGCTAAAGGAGTT
AGCACACGTCTGAACTCTTTCCCTACACGACGCTCTTCCGATCTNNNACCAAAATTTCTGCCATTCCTTC
GAGCACACGTCTGAACTCTTTCCCTACACGACGCTCTTCCGATCTNNNTTAGCATAATGCCCATGGGTC
AGCACACGTCTGAACTCTTTCCCTACACGACGCTCTTCCGATCTNNNAGTCTCATACAGGTAAGCATTG
AGCACACGTCTGAACTCTTTCCCTACACGACGCTCTTCCGATCTNNNTGCTCGTAACATTCTCTCAGTTCA
GAGCACACGTCTGAACTCTTTCCCTACACGACGCTCTTCCGATCTNNNAGCAGTGTACAGGAATTCCAG
AGCACACGTCTGAACTCTTTCCCTACACGACGCTCTTCCGATCTNNNTCTTGGATAAGCCTGATGTAATC
GAGCACACGTCTGAACTCTTTCCCTACACGACGCTCTTCCGATCTNNNCTGCTGTTAGTTAGCTGGCTA
GAGCACACGTCTGAACTCTTTCCCTACACGACGCTCTTCCGATCTNNNCCTTGAAACTTAGCACAAAGAT
AGCACACGTCTGAACTCTTTCCCTACACGACGCTCTTCCGATCTNNNAGATTGACTGGCTTTGGTTTTTC
GAGCACACGTCTGAACTCTTTCCCTACACGACGCTCTTCCGATCTNNNGACTCACAGGTCAAAGCTAAA
GAGCACACGTCTGAACTCTTTCCCTACACGACGCTCTTCCGATCTNNNGTTTCCTTTTGACAGACTGTTC
AGCACACGTCTGAACTCTTTCCCTACACGACGCTCTTCCGATCTNNNACATTTTGCAACACTTACAATGCT
GAGCACACGTCTGAACTCTTTCCCTACACGACGCTCTTCCGATCTNNNACTTTCATTCTGTATGCTTTGGT
AGCACACGTCTGAACTCTTTCCCTACACGACGCTCTTCCGATCTNNNCAGACCTGCTCTAAAACAAAAG
AGCACACGTCTGAACTCTTTCCCTACACGACGCTCTTCCGATCTNNNGGACAAATCCATCCATTATGACC
GAGCACACGTCTGAACTCTTTCCCTACACGACGCTCTTCCGATCTNNNTCCATTTGTTTGGGAAATCTCT
AGAGCACACGTCTGAACTCTTTCCCTACACGACGCTCTTCCGATCTNNNGGCAACAAGAATGTCTACAC
AGCACACGTCTGAACTCTTTCCCTACACGACGCTCTTCCGATCTNNNACCTCAAATCAAAAACACACAAC
GAGCACACGTCTGAACTCTTTCCCTACACGACGCTCTTCCGATCTNNNTGTGGTATCCTTTCAGAGTTCA
GAGCACACGTCTGAACTCTTTCCCTACACGACGCTCTTCCGATCTNNNATTGCCCCAAATGTGTCTTCAA
AGCACACGTCTGAACTCTTTCCCTACACGACGCTCTTCCGATCTNNNGGGCACAGGAACTAGAAAAAT
GAGCACACGTCTGAACTCTTTCCCTACACGACGCTCTTCCGATCTNNNTGTGAATTCTGATGTACCCAG
GCACACGTCTGAACTCTTTCCCTACACGACGCTCTTCCGATCTNNNGTTTGGTTTTTCTGTTTGGTTGG
AGCACACGTCTGAACTCTTTCCCTACACGACGCTCTTCCGATCTNNNAGGTTGAGCATGAGATTTTCTT
GAGCACACGTCTGAACTCTTTCCCTACACGACGCTCTTCCGATCTNNNTGTTGATCCTAATCCCAAACTC
GAGCACACGTCTGAACTCTTTCCCTACACGACGCTCTTCCGATCTNNNACAGCTTACCTTCTACATGTCT
AGCACACGTCTGAACTCTTTCCCTACACGACGCTCTTCCGATCTNNNAACATGTAGCTCATTTTCTAGGT
AGCACACGTCTGAACTCTTTCCCTACACGACGCTCTTCCGATCTNNNTGTCTGGAACTCATTCTTCATCCA
AGAGCACACGTCTGAACTCTTTCCCTACACGACGCTCTTCCGATCTNNNCTATCTCAGCTCACTGCAACC
GCACACGTCTGAACTCTTTCCCTACACGACGCTCTTCCGATCTNNNTCTCAGATGAGCATGGAACATTC
AGCACACGTCTGAACTCTTTCCCTACACGACGCTCTTCCGATCTNNNTTCAACTTCCTAGACAAAGACTG
AGCACACGTCTGAACTCTTTCCCTACACGACGCTCTTCCGATCTNNNACGTGCACACATAGAAAATCAT
GAGCACACGTCTGAACTCTTTCCCTACACGACGCTCTTCCGATCTNNNTGTGGCTATTGTAAATGGGAT
GAGCACACGTCTGAACTCTTTCCCTACACGACGCTCTTCCGATCTNNNTGTCAGTTACTGCCAATAAACA
AGCACACGTCTGAACTCTTTCCCTACACGACGCTCTTCCGATCTNNNTGGCTATTTCAAAAAACAGGCAC
AGCACACGTCTGAACTCTTTCCCTACACGACGCTCTTCCGATCTNNNCCAAACCAAGATGGAAAGGAC
AGCACACGTCTGAACTCTTTCCCTACACGACGCTCTTCCGATCTNNNTGACATTGAGCCTTCTTTCTCTCT
GAGCACACGTCTGAACTCTTTCCCTACACGACGCTCTTCCGATCTNNNACACTTCCACAGTTATCATCAG
AGAGCACACGTCTGAACTCTTTCCCTACACGACGCTCTTCCGATCTNNNCCTCCTTAGGTATATTCGGGC

GAGCACACGTCTGAACTCTTTCCCTACACGACGCTCTTCCGATCTNNNGACAGTTCTCTCTTCAGACAG
AGCACACGTCTGAACTCTTTCCCTACACGACGCTCTTCCGATCTNNNAGTTTGTGTGCAGAAAGATGT
AGCACACGTCTGAACTCTTTCCCTACACGACGCTCTTCCGATCTNNNACATCCATTGACATGAGTTGG
AGCACACGTCTGAACTCTTTCCCTACACGACGCTCTTCCGATCTNNNTCCCAAGAACCTAGCAAAAGAA
AGCACACGTCTGAACTCTTTCCCTACACGACGCTCTTCCGATCTNNNACCAAGGCTAAAGTAGAAAGG
AGCACACGTCTGAACTCTTTCCCTACACGACGCTCTTCCGATCTNNNTGAGAATCATTTTGACAACCAC
GCACACGTCTGAACTCTTTCCCTACACGACGCTCTTCCGATCTNNNAGGAATATTCGAGAAATGGTGT
GCACACGTCTGAACTCTTTCCCTACACGACGCTCTTCCGATCTNNNACAAGGAAATTGGTAATCTGAG
AGCACACGTCTGAACTCTTTCCCTACACGACGCTCTTCCGATCTNNNAGTGGAATTGCTGGATCATAT
AGCACACGTCTGAACTCTTTCCCTACACGACGCTCTTCCGATCTNNNTGGTGTCAGTTGTAGTATCTCC
GAGCACACGTCTGAACTCTTTCCCTACACGACGCTCTTCCGATCTNNNCCATTCCCTCCAGTCTCCTAAA
AGCACACGTCTGAACTCTTTCCCTACACGACGCTCTTCCGATCTNNNTGATGGTAGAGTAGAAGCAG
AGCACACGTCTGAACTCTTTCCCTACACGACGCTCTTCCGATCTNNNTCAGCAGTGTTTTTGTAGTTC
AGCACACGTCTGAACTCTTTCCCTACACGACGCTCTTCCGATCTNNNTGTTCCATTGATCTATGTGTGT
GAGCACACGTCTGAACTCTTTCCCTACACGACGCTCTTCCGATCTNNNGGCAAGACTTTATCATGACAG
AGCACACGTCTGAACTCTTTCCCTACACGACGCTCTTCCGATCTNNNTGTCCTCATAGTACAAGTACA
GAGCACACGTCTGAACTCTTTCCCTACACGACGCTCTTCCGATCTNNNATGCTGTGTTTTATTGTGTG
GAGCACACGTCTGAACTCTTTCCCTACACGACGCTCTTCCGATCTNNNGCTTGTGTTGCTTTTTGGACA
GAGCACACGTCTGAACTCTTTCCCTACACGACGCTCTTCCGATCTNNNGGAGACTAGTAACAACTCTTC
AGCACACGTCTGAACTCTTTCCCTACACGACGCTCTTCCGATCTNNNCACTCAGAGAAATTCAAGACTG
AGAGCACACGTCTGAACTCTTTCCCTACACGACGCTCTTCCGATCTNNNATGTGTATGACCACTTTGGC
AGCACACGTCTGAACTCTTTCCCTACACGACGCTCTTCCGATCTNNNTCAGAAAAATAGAAGCCCAGA
AAGAGCACACGTCTGAACTCTTTCCCTACACGACGCTCTTCCGATCTNNNCCAAGGTGTGACTGTATTT
AGCACACGTCTGAACTCTTTCCCTACACGACGCTCTTCCGATCTNNNTCCTTCTTTCTCTGTCTTTCTCCT
AGCACACGTCTGAACTCTTTCCCTACACGACGCTCTTCCGATCTNNNGGTACCCCTCTCCAATTATTTTG
AGAGCACACGTCTGAACTCTTTCCCTACACGACGCTCTTCCGATCTNNNATTTCTATCCTTCCCCAAGCA
GAGCACACGTCTGAACTCTTTCCCTACACGACGCTCTTCCGATCTNNNTCAGGTTTGTGGAAGTGTAA
GAGCACACGTCTGAACTCTTTCCCTACACGACGCTCTTCCGATCTNNNTTATAGAGCAGGTTATAAGA
AGCACACGTCTGAACTCTTTCCCTACACGACGCTCTTCCGATCTNNNGGGGTCCATTTCTGAGTTGTT
AGCACACGTCTGAACTCTTTCCCTACACGACGCTCTTCCGATCTNNNGCAAGTATCTGAGAGATATGG
GAGCACACGTCTGAACTCTTTCCCTACACGACGCTCTTCCGATCTNNNTGAGCATCTACTATGTGTCAG
AGCACACGTCTGAACTCTTTCCCTACACGACGCTCTTCCGATCTNNNACCTTTGATGGAAATTTGTGT
AGCACACGTCTGAACTCTTTCCCTACACGACGCTCTTCCGATCTNNNGGTCACCATAGGCTACATTTAC
AGCACACGTCTGAACTCTTTCCCTACACGACGCTCTTCCGATCTNNNTGGATAACTTTATGCAGTGCA
AGCACACGTCTGAACTCTTTCCCTACACGACGCTCTTCCGATCTNNNCTATGCTTGCCTGACTCTGGAA
GAGCACACGTCTGAACTCTTTCCCTACACGACGCTCTTCCGATCTNNNAGTGTAAAATCCTACATCCTT
AGCACACGTCTGAACTCTTTCCCTACACGACGCTCTTCCGATCTNNNCTGAGGTGGGAGGATCACTTG
GCACACGTCTGAACTCTTTCCCTACACGACGCTCTTCCGATCTNNNTGTTTGTTGGCTGCATAAATGAC
AGCACACGTCTGAACTCTTTCCCTACACGACGCTCTTCCGATCTNNNCTGGGTTTCTAGTAAGTAGCC
GAGCACACGTCTGAACTCTTTCCCTACACGACGCTCTTCCGATCTNNNTCCTTTCCTTTCCTTTCCTTTCC
GAGCACACGTCTGAACTCTTTCCCTACACGACGCTCTTCCGATCTNNNTCAGGAAGTGTGAATCCTTCA
GAGCACACGTCTGAACTCTTTCCCTACACGACGCTCTTCCGATCTNNNGGTCACATTTACAGAAAGCAA
AGCACACGTCTGAACTCTTTCCCTACACGACGCTCTTCCGATCTNNNTCTGATGTTCTTAGGATTCCTTC

GAGCACACGTCTGAACTCTTTCCCTACACGACGCTCTTCCGATCTNNNAATGGTGAGTGAGTACCCCA
AGCACACGTCTGAACTCTTTCCCTACACGACGCTCTTCCGATCTNNNCACTCAGAGAAATTCAAGACTG
GAGCACACGTCTGAACTCTTTCCCTACACGACGCTCTTCCGATCTNNNGGTACTAAGACTAATGCAGC
GAGCACACGTCTGAACTCTTTCCCTACACGACGCTCTTCCGATCTNNNACCTTGGCAGAGTGAATTCGT
AGCACACGTCTGAACTCTTTCCCTACACGACGCTCTTCCGATCTNNNCCACTCTGTGAACATTTCTACT
GAGCACACGTCTGAACTCTTTCCCTACACGACGCTCTTCCGATCTNNNTGTAGAAGATTGTTGCCTCTC
AGCACACGTCTGAACTCTTTCCCTACACGACGCTCTTCCGATCTNNNACCATGTCCAGCTAAATTTTGC
AGCACACGTCTGAACTCTTTCCCTACACGACGCTCTTCCGATCTNNNGGTACCTTGCTTTTACAATCTG
AGCACACGTCTGAACTCTTTCCCTACACGACGCTCTTCCGATCTNNNGTTGAGTGATAGTGATATGGG
AGCACACGTCTGAACTCTTTCCCTACACGACGCTCTTCCGATCTNNNTGGGTGATGTTTACTCTTCTCCT
CACACGTCTGAACTCTTTCCCTACACGACGCTCTTCCGATCTNNNTCCATTTTATTTGAGTTAGTGAGT
AGCACACGTCTGAACTCTTTCCCTACACGACGCTCTTCCGATCTNNNTGTTGGCATACAGTTGTTCATA
AGCACACGTCTGAACTCTTTCCCTACACGACGCTCTTCCGATCTNNNAAAGTGTCTTCTCATAGAGGGA
GCACACGTCTGAACTCTTTCCCTACACGACGCTCTTCCGATCTNNNACACACAAAGAGAAGGATACAG
GAGCACACGTCTGAACTCTTTCCCTACACGACGCTCTTCCGATCTNNNTCTTTACACTGCTTTTGTGATG
AGCACACGTCTGAACTCTTTCCCTACACGACGCTCTTCCGATCTNNNCCAAGTGCAGGACATACTAAAT
GAGCACACGTCTGAACTCTTTCCCTACACGACGCTCTTCCGATCTNNNTACATGTTGTGTGTGGCTGTA
GCACACGTCTGAACTCTTTCCCTACACGACGCTCTTCCGATCTNNNTGTGGCAAAATATTCAACCTCAC
AGCACACGTCTGAACTCTTTCCCTACACGACGCTCTTCCGATCTNNNTCCCAAACTTCTCAGAAGTTTTC
AGCACACGTCTGAACTCTTTCCCTACACGACGCTCTTCCGATCTNNNCCCAAATTGGAGCTAAACTGA
AGCACACGTCTGAACTCTTTCCCTACACGACGCTCTTCCGATCTNNNGCTGATCTCTATTCATAGCCTG
AGCACACGTCTGAACTCTTTCCCTACACGACGCTCTTCCGATCTNNNTCCTGTCATCATAATGCTAGCT
GCACACGTCTGAACTCTTTCCCTACACGACGCTCTTCCGATCTNNNAGGAATGCAGGTATCTCTTTGA
AGCACACGTCTGAACTCTTTCCCTACACGACGCTCTTCCGATCTNNNTCTCCCATTGCATAGGTTGTCTT
GCACACGTCTGAACTCTTTCCCTACACGACGCTCTTCCGATCTNNNAGTATCTCTCCATCTTGTGTTGT
GAGCACACGTCTGAACTCTTTCCCTACACGACGCTCTTCCGATCTNNNGCTCTTTTTTGGTTTCCATTGG
AGCACACGTCTGAACTCTTTCCCTACACGACGCTCTTCCGATCTNNNTGGTAGTTGGATCTAGATGGC
GCACACGTCTGAACTCTTTCCCTACACGACGCTCTTCCGATCTNNNCCCATTCAGGAAGATCTATCCAC
AGCACACGTCTGAACTCTTTCCCTACACGACGCTCTTCCGATCTNNNAGTGAAGTCAAAAAGGAGTGA
GAGCACACGTCTGAACTCTTTCCCTACACGACGCTCTTCCGATCTNNNAGTACCTTTTGGAACCCAGAT
GAGCACACGTCTGAACTCTTTCCCTACACGACGCTCTTCCGATCTNNNTGCTTTGTGTTTTTGGTTTAC
GAGCACACGTCTGAACTCTTTCCCTACACGACGCTCTTCCGATCTNNNCGTGGAATCTCGGGGTCTAT
AGCACACGTCTGAACTCTTTCCCTACACGACGCTCTTCCGATCTNNNAATTGGATGACCCTTGTGCTTT
GAGCACACGTCTGAACTCTTTCCCTACACGACGCTCTTCCGATCTNNNAGTACCCCTATCATCTACTAG
AGCACACGTCTGAACTCTTTCCCTACACGACGCTCTTCCGATCTNNNAGGTTCAGAATGTACTCCTCA
GAGCACACGTCTGAACTCTTTCCCTACACGACGCTCTTCCGATCTNNNAATCTCATGGGAGCACTGTT
AGCACACGTCTGAACTCTTTCCCTACACGACGCTCTTCCGATCTNNNACCAAATGACAACTAAAATCCG
GAGCACACGTCTGAACTCTTTCCCTACACGACGCTCTTCCGATCTNNNGCATCTGTAGTCCCAGCTACT
GAGCACACGTCTGAACTCTTTCCCTACACGACGCTCTTCCGATCTNNNTCCAAGATTGAATCGGGAAG
GAGCACACGTCTGAACTCTTTCCCTACACGACGCTCTTCCGATCTNNNCTGAGGCAGGAGAATAGCTT
AGCACACGTCTGAACTCTTTCCCTACACGACGCTCTTCCGATCTNNNATCAATCCATCCTAAGGCTTGC
GAGCACACGTCTGAACTCTTTCCCTACACGACGCTCTTCCGATCTNNNACTCCTCAGTACAACGTTGAG
GAGCACACGTCTGAACTCTTTCCCTACACGACGCTCTTCCGATCTNNNGAGAATTGCTTGAACCCAGG

AGCACACGTCTGAACTCTTTCCCTACACGACGCTCTTCCGATCTNNNAGAATTACTTGAACTCGGGAG
AGCACACGTCTGAACTCTTTCCCTACACGACGCTCTTCCGATCTNNNCGGTGGAAATTGTAAACAAGG
GCACACGTCTGAACTCTTTCCCTACACGACGCTCTTCCGATCTNNNAGCTTGCCATTTCTTTTCAGATG
AGCACACGTCTGAACTCTTTCCCTACACGACGCTCTTCCGATCTNNNGGCAGCAAGAATGACTTTGA
GCACACGTCTGAACTCTTTCCCTACACGACGCTCTTCCGATCTNNNACCTAGGAATTGACCAAAAAG
GCACACGTCTGAACTCTTTCCCTACACGACGCTCTTCCGATCTNNNCCAACATGCCTGGCTAATTTTTT
AGCACACGTCTGAACTCTTTCCCTACACGACGCTCTTCCGATCTNNNTCTTGTGAATTGCTTCTAACTG
AGCACACGTCTGAACTCTTTCCCTACACGACGCTCTTCCGATCTNNNGCACACACACACTTTCTAATCT
AGCACACGTCTGAACTCTTTCCCTACACGACGCTCTTCCGATCTNNNGCACACACACACTTTCTAATCT
AGCACACGTCTGAACTCTTTCCCTACACGACGCTCTTCCGATCTNNNGGTACCGAATATTGTACTAGC
AGCACACGTCTGAACTCTTTCCCTACACGACGCTCTTCCGATCTNNNTTCCCTAGAGCTATAAATGG
GAGCACACGTCTGAACTCTTTCCCTACACGACGCTCTTCCGATCTNNNTGAGGCAGGAGAATCACTT
GCACACGTCTGAACTCTTTCCCTACACGACGCTCTTCCGATCTNNNAGAGTTGTACAGTCATCACTAC
GCACACGTCTGAACTCTTTCCCTACACGACGCTCTTCCGATCTNNNGTTCTGTACATCTTGACAAGTG
GAGCACACGTCTGAACTCTTTCCCTACACGACGCTCTTCCGATCTNNNACCTCGTGAAACTGGAGAA
GAGCACACGTCTGAACTCTTTCCCTACACGACGCTCTTCCGATCTNNNTGCCTGTGCTTTTGTTATCCA
AGCACACGTCTGAACTCTTTCCCTACACGACGCTCTTCCGATCTNNNCACAGCATGGATGTTCTGTAAT
AGCACACGTCTGAACTCTTTCCCTACACGACGCTCTTCCGATCTNNNACGTCCCAGATATACTCTACAA
AGAGCACACGTCTGAACTCTTTCCCTACACGACGCTCTTCCGATCTNNNGAAACAACAGCCTTACAGG
AGAGCACACGTCTGAACTCTTTCCCTACACGACGCTCTTCCGATCTNNNGAAACAACAGCCTTACAGG
GAGCACACGTCTGAACTCTTTCCCTACACGACGCTCTTCCGATCTNNNGAGAGAGAGAGAGACCATT
AGCACACGTCTGAACTCTTTCCCTACACGACGCTCTTCCGATCTNNNTCTCTTGTGACTCAAAACACT
GAGCACACGTCTGAACTCTTTCCCTACACGACGCTCTTCCGATCTNNNAGTCTGTCTTGCTCTCTATAT
GAGCACACGTCTGAACTCTTTCCCTACACGACGCTCTTCCGATCTNNNTTTTGTAGGTTTGAAGAGG
GAGCACACGTCTGAACTCTTTCCCTACACGACGCTCTTCCGATCTNNNTGGCTTTTTTTTTCTCCTGA
GAGCACACGTCTGAACTCTTTCCCTACACGACGCTCTTCCGATCTNNNAGTAGAATGATGGTTGCCAG
AGCACACGTCTGAACTCTTTCCCTACACGACGCTCTTCCGATCTNNNGGATGTCTTTCATTTCTTCAG
GAGCACACGTCTGAACTCTTTCCCTACACGACGCTCTTCCGATCTNNNGGTACCTCAGATGGAAATG
AGCACACGTCTGAACTCTTTCCCTACACGACGCTCTTCCGATCTNNNAACAAAATTAGCTGGGTGTGG
AGCACACGTCTGAACTCTTTCCCTACACGACGCTCTTCCGATCTNNNGAGAATGTCATGAACCTGGG
AGAGCACACGTCTGAACTCTTTCCCTACACGACGCTCTTCCGATCTNNNATGACAATTAGCTGGGTGT
GAGCACACGTCTGAACTCTTTCCCTACACGACGCTCTTCCGATCTNNNCAGACTCATTGAGACAACCT
AGCACACGTCTGAACTCTTTCCCTACACGACGCTCTTCCGATCTNNNTGACCATCTACTATGTGTCAGT
GAGCACACGTCTGAACTCTTTCCCTACACGACGCTCTTCCGATCTNNNATTAGTCAGGGTTCCTTAGA
AGCACACGTCTGAACTCTTTCCCTACACGACGCTCTTCCGATCTNNNTAGTGAGCTGAAATCATGCCA
AGCACACGTCTGAACTCTTTCCCTACACGACGCTCTTCCGATCTNNNGCTCCTAACAAGAAAAGATGG
GAAGAGCACACGTCTGAACTCTTTCCCTACACGACGCTCTTCCGATCTNNNGTTGCAGTGAACCGAG
AAGAGCACACGTCTGAACTCTTTCCCTACACGACGCTCTTCCGATCTNNNTGAGCCATGATCATGCCA
GAGCACACGTCTGAACTCTTTCCCTACACGACGCTCTTCCGATCTNNNGTGACAAACTCTCCAACTCA
GCACACGTCTGAACTCTTTCCCTACACGACGCTCTTCCGATCTNNNAATAACAGTGGTGACATCGGA
GCACACGTCTGAACTCTTTCCCTACACGACGCTCTTCCGATCTNNNGTATTCCCAGTTACTCAGGAGG
AGCACACGTCTGAACTCTTTCCCTACACGACGCTCTTCCGATCTNNNCTCATGCCTGTAATCCTAGCAC
AGCACACGTCTGAACTCTTTCCCTACACGACGCTCTTCCGATCTNNNTGTGTCTGTGTGTGTGTGTGT

GAGCACACGTCTGAACTCTTTCCCTACACGACGCTCTTCCGATCTNNNGTGGAGTTTGCAGTGAGTA
GCACACGTCTGAACTCTTTCCCTACACGACGCTCTTCCGATCTNNNAAGATCAAGGTGCCAGCATCTA
AGCACACGTCTGAACTCTTTCCCTACACGACGCTCTTCCGATCTNNNATTCCTGACCTCAGATGATCTG
GAGCACACGTCTGAACTCTTTCCCTACACGACGCTCTTCCGATCTNNNCGCAACAAACAGCAAAAACC
GCACACGTCTGAACTCTTTCCCTACACGACGCTCTTCCGATCTNNNTCTCAAACATGGTTGTGCACAAA
GAGCACACGTCTGAACTCTTTCCCTACACGACGCTCTTCCGATCTNNNCGTATCTGTAGTCCCAGCTAC
GAGCACACGTCTGAACTCTTTCCCTACACGACGCTCTTCCGATCTNNNAATGTGAGGTGTGTGTGAG
GAGCACACGTCTGAACTCTTTCCCTACACGACGCTCTTCCGATCTNNNATGTGAAATATGGGAGGCT
AGCACACGTCTGAACTCTTTCCCTACACGACGCTCTTCCGATCTNNNTGTGATTAGGAAGCAGTGAG
GAGCACACGTCTGAACTCTTTCCCTACACGACGCTCTTCCGATCTNNNATAATCCTAGCACTTTGGGA
AGCACACGTCTGAACTCTTTCCCTACACGACGCTCTTCCGATCTNNNCTCACACCTGTAATCTCAGCAC
CACACGTCTGAACTCTTTCCCTACACGACGCTCTTCCGATCTNNNGTGAGTGTAGTGTCTGACTCTG
AGCACACGTCTGAACTCTTTCCCTACACGACGCTCTTCCGATCTNNNCTGTAATCCCAGCTACTCGGT
AGCACACGTCTGAACTCTTTCCCTACACGACGCTCTTCCGATCTNNNATGGCTGCATAGTATTCCATG
GAGCACACGTCTGAACTCTTTCCCTACACGACGCTCTTCCGATCTNNNTATTGTTACACACACACACA
GCACACGTCTGAACTCTTTCCCTACACGACGCTCTTCCGATCTNNNCATGAGGGTATAAGTCTGGGC
AGCACACGTCTGAACTCTTTCCCTACACGACGCTCTTCCGATCTNNNATTCTGTAGGTTGCCTGTTCAC
GCACACGTCTGAACTCTTTCCCTACACGACGCTCTTCCGATCTNNNCTGTAATCCCAGCTACTCGAGA
GAGCACACGTCTGAACTCTTTCCCTACACGACGCTCTTCCGATCTNNNCTGAGGCAGGAGAATCACTT
AGAGCACACGTCTGAACTCTTTCCCTACACGACGCTCTTCCGATCTNNNGCAAAAAATATGGAACCA
AGCACACGTCTGAACTCTTTCCCTACACGACGCTCTTCCGATCTNNNTCAGTCTCCTGAGTAGCTGAG
GAGCACACGTCTGAACTCTTTCCCTACACGACGCTCTTCCGATCTNNNTAGTCTTAGCTACTCAGGAG
GCACACGTCTGAACTCTTTCCCTACACGACGCTCTTCCGATCTNNNGCTGGGACTACATATGAACAC
AGCACACGTCTGAACTCTTTCCCTACACGACGCTCTTCCGATCTNNNGGTGAGACTCCGCCTCTAAAAA
AGCACACGTCTGAACTCTTTCCCTACACGACGCTCTTCCGATCTNNNAAGTGCTCTTGACTGACGACT
GAGCACACGTCTGAACTCTTTCCCTACACGACGCTCTTCCGATCTNNNATACCCAAAGAAGCAAAGAG
GCACACGTCTGAACTCTTTCCCTACACGACGCTCTTCCGATCTNNNCCCAAATAGTTTTTGGCAAAGG
GCACACGTCTGAACTCTTTCCCTACACGACGCTCTTCCGATCTNNNAAGTGAAGAGACAACCCACAGA
GCACACGTCTGAACTCTTTCCCTACACGACGCTCTTCCGATCTNNNTTTCCTCTCTGCTGTCTGATGAC
AGCACACGTCTGAACTCTTTCCCTACACGACGCTCTTCCGATCTNNNAGTCTCCAATTCCATCCAGGTT
GCACACGTCTGAACTCTTTCCCTACACGACGCTCTTCCGATCTNNNGAGTTCGAGACCAGCTTGAGTA
GCACACGTCTGAACTCTTTCCCTACACGACGCTCTTCCGATCTNNNATGTGTGTGTGTGTCTCTCTCT
AGCACACGTCTGAACTCTTTCCCTACACGACGCTCTTCCGATCTNNNATGAGCAGTTGTAGTCCTTGT
AGCACACGTCTGAACTCTTTCCCTACACGACGCTCTTCCGATCTNNNAGACTGTGTGTGTCTGTGCAT
AGCACACGTCTGAACTCTTTCCCTACACGACGCTCTTCCGATCTNNNCAGGTTGCTGTGAATGCCATT
AGCACACGTCTGAACTCTTTCCCTACACGACGCTCTTCCGATCTNNNTATTCCATGGTGTCTGTGTGT
GAGCACACGTCTGAACTCTTTCCCTACACGACGCTCTTCCGATCTNNNTTCAAAAATACACCTGCACAC
AGCACACGTCTGAACTCTTTCCCTACACGACGCTCTTCCGATCTNNNCAGAATCTGTTTTCCGAGGCT
AGCACACGTCTGAACTCTTTCCCTACACGACGCTCTTCCGATCTNNNTCCTCATAGCTTAGCTCCCACTT
AGCACACGTCTGAACTCTTTCCCTACACGACGCTCTTCCGATCTNNNGATCTACCCCATTGGCTCTTTG
AGCACACGTCTGAACTCTTTCCCTACACGACGCTCTTCCGATCTNNNTCAGTGGTAATCTCTCCTTTGC
GCACACGTCTGAACTCTTTCCCTACACGACGCTCTTCCGATCTNNNTTGGATCATGCAGCAAAATAG
GAGCACACGTCTGAACTCTTTCCCTACACGACGCTCTTCCGATCTNNNGGCTGAGACAGAAGAATTG

AGCACACGTCTGAACTCTTTCCCTACACGACGCTCTTCCGATCTNNNGGTCAAGGAGGAATTGAGTGA
AGAGCACACGTCTGAACTCTTTCCCTACACGACGCTCTTCCGATCTNNNGAGTGACACCTCCACAAGACT
AGAGCACACGTCTGAACTCTTTCCCTACACGACGCTCTTCCGATCTNNNATCAGAGTGGATTTGAGCTG
AGCACACGTCTGAACTCTTTCCCTACACGACGCTCTTCCGATCTNNNACTGCCTTCTCAGAGAAAAGAG
AGAGCACACGTCTGAACTCTTTCCCTACACGACGCTCTTCCGATCTNNNATGTGTGTGTATGTGTGT
GAGCACACGTCTGAACTCTTTCCCTACACGACGCTCTTCCGATCTNNNGGATAAAAGGGCACTAGGTC
AGCACACGTCTGAACTCTTTCCCTACACGACGCTCTTCCGATCTNNNGGTATCTGTCCGGACAACTTTCT
GCACACGTCTGAACTCTTTCCCTACACGACGCTCTTCCGATCTNNNGCTATCCAAAAGCTGTCACAACA
GAGCACACGTCTGAACTCTTTCCCTACACGACGCTCTTCCGATCTNNNGTCTTTTAGGAGACACACACA
AGCACACGTCTGAACTCTTTCCCTACACGACGCTCTTCCGATCTNNNCTCTCCCCATCTCTCTCTCTCTTCA
GCACACGTCTGAACTCTTTCCCTACACGACGCTCTTCCGATCTNNNCGTAAGGAGAGGGATTTGGAGA
AGCACACGTCTGAACTCTTTCCCTACACGACGCTCTTCCGATCTNNNGGCTTTTACCTCTAGGGCTTCTT
AGCACACGTCTGAACTCTTTCCCTACACGACGCTCTTCCGATCTNNNCAAACAAACACACACACACACAC
AAGAGCACACGTCTGAACTCTTTCCCTACACGACGCTCTTCCGATCTNNNTACGTTGAACCACTACTCA
GAGCACACGTCTGAACTCTTTCCCTACACGACGCTCTTCCGATCTNNNATACGCTTCTGATATGGCAGG
AGCACACGTCTGAACTCTTTCCCTACACGACGCTCTTCCGATCTNNNTTCCAGAACTGTGTCAGAGAA
AGCACACGTCTGAACTCTTTCCCTACACGACGCTCTTCCGATCTNNNTCCTAAAGTGCTGGGATTACAG
GAGCACACGTCTGAACTCTTTCCCTACACGACGCTCTTCCGATCTNNNACTAGCATTCTCCACAGAAGCA
AGCACACGTCTGAACTCTTTCCCTACACGACGCTCTTCCGATCTNNNTGTGGATATTGCTTCACAAGT
GAGCACACGTCTGAACTCTTTCCCTACACGACGCTCTTCCGATCTNNNTCATTACAAACACATCCACGCA
GAGCACACGTCTGAACTCTTTCCCTACACGACGCTCTTCCGATCTNNNATCCAGGCTCTTTCCATCTTGA
GAGCACACGTCTGAACTCTTTCCCTACACGACGCTCTTCCGATCTNNNTGGTCACAAGTTTCTGAGGA
AGCACACGTCTGAACTCTTTCCCTACACGACGCTCTTCCGATCTNNNTGTGTTTGTGTACATGCAAGCA
GAGCACACGTCTGAACTCTTTCCCTACACGACGCTCTTCCGATCTNNNCTGTTTTTGGCATGCTTGGGA
AGCACACGTCTGAACTCTTTCCCTACACGACGCTCTTCCGATCTNNNTGGTACATACACTCACACCAACA
AGCACACGTCTGAACTCTTTCCCTACACGACGCTCTTCCGATCTNNNGGACTTTCCTTTGTAGTGTTGC
AGCACACGTCTGAACTCTTTCCCTACACGACGCTCTTCCGATCTNNNGGCTGGGGATGGAGTTGATAT
AGCACACGTCTGAACTCTTTCCCTACACGACGCTCTTCCGATCTNNNAATCACCTGGGGATCTTGTTAC
GAGCACACGTCTGAACTCTTTCCCTACACGACGCTCTTCCGATCTNNNATGTTGTGACCTCTTTTCCTG
GAGCACACGTCTGAACTCTTTCCCTACACGACGCTCTTCCGATCTNNNCTGACTCAAGACTTCTCCAGCT
AGAGCACACGTCTGAACTCTTTCCCTACACGACGCTCTTCCGATCTNNNTGGTAGTTTGCTGTGGCTCT
GAGCACACGTCTGAACTCTTTCCCTACACGACGCTCTTCCGATCTNNNCCCTGCATGTCTGTATTGTTAC
GAGCACACGTCTGAACTCTTTCCCTACACGACGCTCTTCCGATCTNNNGGCTTTTTGTCCTTGCGATAG
AGCACACGTCTGAACTCTTTCCCTACACGACGCTCTTCCGATCTNNNACAACTCATTCTTCCAACAAACA
AAGAGCACACGTCTGAACTCTTTCCCTACACGACGCTCTTCCGATCTNNNACTATCCTTCCTGCCCTAGA
GAGCACACGTCTGAACTCTTTCCCTACACGACGCTCTTCCGATCTNNNACACACACAGAGAGAGAGAG
AGAGCACACGTCTGAACTCTTTCCCTACACGACGCTCTTCCGATCTNNNCCCAGGATCCACTACAAAAG
AGCACACGTCTGAACTCTTTCCCTACACGACGCTCTTCCGATCTNNNCATCCTCCCTTGGAAGTGTCTA
GAGCACACGTCTGAACTCTTTCCCTACACGACGCTCTTCCGATCTNNNTCCTTATCATGCCTAGCACTGA
GAGCACACGTCTGAACTCTTTCCCTACACGACGCTCTTCCGATCTNNNGAACAATAACAAAAAGGGGG
GCACACGTCTGAACTCTTTCCCTACACGACGCTCTTCCGATCTNNNTTCGTTTTCCATCACTTTGTGCACA
GAGCACACGTCTGAACTCTTTCCCTACACGACGCTCTTCCGATCTNNNTCAAGTGGAAGTAAAGGAGA
GAGCACACGTCTGAACTCTTTCCCTACACGACGCTCTTCCGATCTNNNGAGACTACCTAACATCCAAC

AGCACACGTCTGAACTCTTTCCCTACACGACGCTCTTCCGATCTNNNAATGGTAGGGAAGATGGGAG
AGCACACGTCTGAACTCTTTCCCTACACGACGCTCTTCCGATCTNNNTGTTCATAGACAGTGCCTCCTT
AGCACACGTCTGAACTCTTTCCCTACACGACGCTCTTCCGATCTNNNCTTCCACAAATTCTGCCCACAAT
AGCACACGTCTGAACTCTTTCCCTACACGACGCTCTTCCGATCTNNNGTTACTTGCTCTGCCTGAAATCC
AGCACACGTCTGAACTCTTTCCCTACACGACGCTCTTCCGATCTNNNAAGTGCTGCATTGTACAGTCAT
AGCACACGTCTGAACTCTTTCCCTACACGACGCTCTTCCGATCTNNNTGCCCAAGGTCAAGTTTAGATC
AGCACACGTCTGAACTCTTTCCCTACACGACGCTCTTCCGATCTNNNCTTCCATTGACCTCTGTGTCCAT
GCACACGTCTGAACTCTTTCCCTACACGACGCTCTTCCGATCTNNNGTCACAAAGCTAGTCAGTTCAGC
AGCACACGTCTGAACTCTTTCCCTACACGACGCTCTTCCGATCTNNNGAAAGAAAGAGTTGAGCAAG
GAGCACACGTCTGAACTCTTTCCCTACACGACGCTCTTCCGATCTNNNAGAATGTCCAAGAAACCCTAC
GAGCACACGTCTGAACTCTTTCCCTACACGACGCTCTTCCGATCTNNNTTCCTGAAATGTGACACATGC
GCACACGTCTGAACTCTTTCCCTACACGACGCTCTTCCGATCTNNNTATACTCTCAGCCTTCCCTTCCAC
AGCACACGTCTGAACTCTTTCCCTACACGACGCTCTTCCGATCTNNNCTTCACATTCTGGGCTTGTTTG
AGCACACGTCTGAACTCTTTCCCTACACGACGCTCTTCCGATCTNNNTGCAAAGAGTATAGGGAAGT
AGCACACGTCTGAACTCTTTCCCTACACGACGCTCTTCCGATCTNNNTGTGTGTGTGTGTGTGTAT
GCACACGTCTGAACTCTTTCCCTACACGACGCTCTTCCGATCTNNNAAACAGGCCAATTTTTTGTGGA
GCACACGTCTGAACTCTTTCCCTACACGACGCTCTTCCGATCTNNNTCTAAATACACACACACGCATGC
AGCACACGTCTGAACTCTTTCCCTACACGACGCTCTTCCGATCTNNNGATGGTTGACGTCTTCTAAGG
GCACACGTCTGAACTCTTTCCCTACACGACGCTCTTCCGATCTNNNGCAAGGGAAAATGGAATCGGTA
GAGCACACGTCTGAACTCTTTCCCTACACGACGCTCTTCCGATCTNNNGAGGAGGAGAGAGAGAAGA
AGCACACGTCTGAACTCTTTCCCTACACGACGCTCTTCCGATCTNNNTCTACACATCACATCTGCCCTTC
GAGCACACGTCTGAACTCTTTCCCTACACGACGCTCTTCCGATCTNNNAAGAATTTCACCTTGTTCCCAA
AGAGCACACGTCTGAACTCTTTCCCTACACGACGCTCTTCCGATCTNNNCAAAAAAGGAGGCTCAAGC
AGCACACGTCTGAACTCTTTCCCTACACGACGCTCTTCCGATCTNNNGCCGGTTAGTCTAAGATCAAG
GAGCACACGTCTGAACTCTTTCCCTACACGACGCTCTTCCGATCTNNNAGAGCTAACATTTACTGCATG
AGCACACGTCTGAACTCTTTCCCTACACGACGCTCTTCCGATCTNNNCCAGGACATGAGTTCCCATAG
GAGCACACGTCTGAACTCTTTCCCTACACGACGCTCTTCCGATCTNNNTGTACTCTAACAACCTGTAGG
AGAGCACACGTCTGAACTCTTTCCCTACACGACGCTCTTCCGATCTNNNCCAACAGGATATGGTTGCGT
AGCACACGTCTGAACTCTTTCCCTACACGACGCTCTTCCGATCTNNNATAGTGTAATCTCATGGGCTC
AGAGCACACGTCTGAACTCTTTCCCTACACGACGCTCTTCCGATCTNNNAGAGCTGTGGTTCTGTCTA
GAGCACACGTCTGAACTCTTTCCCTACACGACGCTCTTCCGATCTNNNCCACTAACAGCCACTGTATCCT
GAGCACACGTCTGAACTCTTTCCCTACACGACGCTCTTCCGATCTNNNCTGTCTGTCTGTCTGTCTGTC
AGCACACGTCTGAACTCTTTCCCTACACGACGCTCTTCCGATCTNNNAAGTAGTGTGTGTGGATGTGC
AGCACACGTCTGAACTCTTTCCCTACACGACGCTCTTCCGATCTNNNCATTTACACACTGTGTCCCACAC
AGCACACGTCTGAACTCTTTCCCTACACGACGCTCTTCCGATCTNNNTCACCCTACCCATAACTTCATGG
AGCACACGTCTGAACTCTTTCCCTACACGACGCTCTTCCGATCTNNNCGCCAATCTACCACTGACCTTAT
AGAGCACACGTCTGAACTCTTTCCCTACACGACGCTCTTCCGATCTNNNGAGGAAAACTGCTTTTGTC
GAGCACACGTCTGAACTCTTTCCCTACACGACGCTCTTCCGATCTNNNGTCCCCGGTTCCTGTATTTTCT
AGAGCACACGTCTGAACTCTTTCCCTACACGACGCTCTTCCGATCTNNNGGATGAGAGAGGATGAGA
GAGCACACGTCTGAACTCTTTCCCTACACGACGCTCTTCCGATCTNNNTGCTTGTGTGTTTGTGTTCATG
AGCACACGTCTGAACTCTTTCCCTACACGACGCTCTTCCGATCTNNNCATCCCTGAAGTTGCTAGTCACT
AGCACACGTCTGAACTCTTTCCCTACACGACGCTCTTCCGATCTNNNGGGGAGGTATCATGCAATCAT
AGCACACGTCTGAACTCTTTCCCTACACGACGCTCTTCCGATCTNNNGCCTGAATTTGGGAAGAGAAA

AGCACACGTCTGAACTCTTTCCCTACACGACGCTCTTCCGATCTNNNATAGTTTGAAAAAGCCCCTCAG
AGCACACGTCTGAACTCTTTCCCTACACGACGCTCTTCCGATCTNNNTTCCAAATGTTGCCTCTGTTGA
AGCACACGTCTGAACTCTTTCCCTACACGACGCTCTTCCGATCTNNNGCTGGCCAAAGTTCAAATTCTC
AGCACACGTCTGAACTCTTTCCCTACACGACGCTCTTCCGATCTNNNCTAAAATGCCTGTGTGTGTGT
AGCACACGTCTGAACTCTTTCCCTACACGACGCTCTTCCGATCTNNNTGGGACTGACTTGTGGTAATCT
AGCACACGTCTGAACTCTTTCCCTACACGACGCTCTTCCGATCTNNNAAAAAACACACACACACACATG
GAGCACACGTCTGAACTCTTTCCCTACACGACGCTCTTCCGATCTNNNATGGATGTTGCCACTATCACT
GCACACGTCTGAACTCTTTCCCTACACGACGCTCTTCCGATCTNNNGCTCTCTCTCTCCTCTCTCTCCA
GAGCACACGTCTGAACTCTTTCCCTACACGACGCTCTTCCGATCTNNNTGTGTGTGTGTCTGTGCTTGT
AGCACACGTCTGAACTCTTTCCCTACACGACGCTCTTCCGATCTNNNCAGAAATGAAGACCCAAACAGG
AGCACACGTCTGAACTCTTTCCCTACACGACGCTCTTCCGATCTNNNATTTTTTGGAGCATGTGTGTG
AGCACACGTCTGAACTCTTTCCCTACACGACGCTCTTCCGATCTNNNATAGGCCCATTATACCAAGCAG
GAGCACACGTCTGAACTCTTTCCCTACACGACGCTCTTCCGATCTNNNTACACACATACACCCATACAC
GAGCACACGTCTGAACTCTTTCCCTACACGACGCTCTTCCGATCTNNNTTTTGGGCAGTAACTGGGAAA
GAGCACACGTCTGAACTCTTTCCCTACACGACGCTCTTCCGATCTNNNTCTATCACCTAAAAGGGAAG
GCACACGTCTGAACTCTTTCCCTACACGACGCTCTTCCGATCTNNNCTCCAGGTCCTTATCTGTGTCACC
AGCACACGTCTGAACTCTTTCCCTACACGACGCTCTTCCGATCTNNNCACCACCATACTCCATAGGACA
AGCACACGTCTGAACTCTTTCCCTACACGACGCTCTTCCGATCTNNNAATCACAGTAGCTTCCCCTTTGT
GAGCACACGTCTGAACTCTTTCCCTACACGACGCTCTTCCGATCTNNNCCATAAGCTCCTCTTTTCCCCAT
GAGCACACGTCTGAACTCTTTCCCTACACGACGCTCTTCCGATCTNNNGAGGAGTACAGAATGCAGGA
GAGCACACGTCTGAACTCTTTCCCTACACGACGCTCTTCCGATCTNNNATTCACAAATGCAGTGCGGTA
AGAGCACACGTCTGAACTCTTTCCCTACACGACGCTCTTCCGATCTNNNTGTAGTACAATACCAGGCA
GAGCACACGTCTGAACTCTTTCCCTACACGACGCTCTTCCGATCTNNNCCCCAGAAAAGCACTTGACAA
GCACACGTCTGAACTCTTTCCCTACACGACGCTCTTCCGATCTNNNTCACTCTCCCTCTCTTTCTCTGTCA
GAGCACACGTCTGAACTCTTTCCCTACACGACGCTCTTCCGATCTNNNAGAAAAATTGCTTGAACCGGG
GAGCACACGTCTGAACTCTTTCCCTACACGACGCTCTTCCGATCTNNNAATACAAGAGAACGAGAGGT
AGCACACGTCTGAACTCTTTCCCTACACGACGCTCTTCCGATCTNNNTACTGATGGTGGGGCGTATATT
GAGCACACGTCTGAACTCTTTCCCTACACGACGCTCTTCCGATCTNNNTGTGTGTGTGCCTATGTGTGT
GAGCACACGTCTGAACTCTTTCCCTACACGACGCTCTTCCGATCTNNNTACTTTGATGCACACACACACA
AGAGCACACGTCTGAACTCTTTCCCTACACGACGCTCTTCCGATCTNNNTGGTATCTGTGGGCATATGT
AGCACACGTCTGAACTCTTTCCCTACACGACGCTCTTCCGATCTNNNAGCCTTTGTTTATCTGGGAAGG
AGAGCACACGTCTGAACTCTTTCCCTACACGACGCTCTTCCGATCTNNNCTGCCTTCCATACTTCCTTTG
AGCACACGTCTGAACTCTTTCCCTACACGACGCTCTTCCGATCTNNNTGTCTGTCTCTCTCTGCTTGTTCC
GAGCACACGTCTGAACTCTTTCCCTACACGACGCTCTTCCGATCTNNNAGGAGCTCCCTCTTTATATGG
GAGCACACGTCTGAACTCTTTCCCTACACGACGCTCTTCCGATCTNNNAAGTGCCCTTGAGACCTAAGA
GAGCACACGTCTGAACTCTTTCCCTACACGACGCTCTTCCGATCTNNNATAGCTTATTTCCCCCTCTGCA
AGCACACGTCTGAACTCTTTCCCTACACGACGCTCTTCCGATCTNNNTAGCTTTCCTGACTCACTTGCAA
AGCACACGTCTGAACTCTTTCCCTACACGACGCTCTTCCGATCTNNNTAACAACACACACACACACACA
AGCACACGTCTGAACTCTTTCCCTACACGACGCTCTTCCGATCTNNNTACGTATATTCCCACCACGCATT
AGCACACGTCTGAACTCTTTCCCTACACGACGCTCTTCCGATCTNNNGACCTCAGACTCTCTGTTTCCAG
GAGCACACGTCTGAACTCTTTCCCTACACGACGCTCTTCCGATCTNNNTTTCAGGAGAGGGGAGGAA
AGCACACGTCTGAACTCTTTCCCTACACGACGCTCTTCCGATCTNNNAGTTGTCTGTGGTATTCACTG
GAGCACACGTCTGAACTCTTTCCCTACACGACGCTCTTCCGATCTNNNAAAGTGGAATCATCACGACTT

GAGCACACGTCTGAACTCTTTCCCTACACGACGCTCTTCCGATCTNNNTCACGTTCATCAACCCAATTGT
AGCACACGTCTGAACTCTTTCCCTACACGACGCTCTTCCGATCTNNNAAGGGTTTACTGCTGCCATTTTG
GAGCACACGTCTGAACTCTTTCCCTACACGACGCTCTTCCGATCTNNNACAGAGCAAGCTCCTGTTTCAT
AAGAGCACACGTCTGAACTCTTTCCCTACACGACGCTCTTCCGATCTNNNGATTGAAAAGGATGCTGG
AGCACACGTCTGAACTCTTTCCCTACACGACGCTCTTCCGATCTNNNAAAAGAGGGACTGACAGGGT
AGCACACGTCTGAACTCTTTCCCTACACGACGCTCTTCCGATCTNNNGTGACCTTACCTGACACTTTCCTC
AGAGCACACGTCTGAACTCTTTCCCTACACGACGCTCTTCCGATCTNNNCTACAGTGAGGAAGGCCAGA
AGCACACGTCTGAACTCTTTCCCTACACGACGCTCTTCCGATCTNNNTGAAACCCTTGTCTATTGCTGGT
AGAGCACACGTCTGAACTCTTTCCCTACACGACGCTCTTCCGATCTNNNAAAAAAACATCTGCCCACAC
GAGCACACGTCTGAACTCTTTCCCTACACGACGCTCTTCCGATCTNNNTTCTCTCTGTGTCTCTTGAG
GAGCACACGTCTGAACTCTTTCCCTACACGACGCTCTTCCGATCTNNNAAACTGTGTGTGTTTGTGTGT
GAGCACACGTCTGAACTCTTTCCCTACACGACGCTCTTCCGATCTNNNGGCACTAAGCTGTTGCCATTTA
GAGCACACGTCTGAACTCTTTCCCTACACGACGCTCTTCCGATCTNNNTGCCAGGCACTAATTTTCACAA
AGAGCACACGTCTGAACTCTTTCCCTACACGACGCTCTTCCGATCTNNNCTGTTTGTCTAGAGGAGCTG
AAGAGCACACGTCTGAACTCTTTCCCTACACGACGCTCTTCCGATCTNNNAAGAGATCTCTGGTTGGTT
AGCACACGTCTGAACTCTTTCCCTACACGACGCTCTTCCGATCTNNNCACTTCTGAGGGGATAGTGTCA
AGCACACGTCTGAACTCTTTCCCTACACGACGCTCTTCCGATCTNNNTTCTACTCCTGAGAGAGGGTGA
GAGCACACGTCTGAACTCTTTCCCTACACGACGCTCTTCCGATCTNNNATGGCTAAAATCACACACACAC
GAGCACACGTCTGAACTCTTTCCCTACACGACGCTCTTCCGATCTNNNGCCCATAAGAGACTCTGTTTG
GAGCACACGTCTGAACTCTTTCCCTACACGACGCTCTTCCGATCTNNNACTGCCATAAAAATGCTCTGT
AGCACACGTCTGAACTCTTTCCCTACACGACGCTCTTCCGATCTNNNGTGAGAAGCCTCTGTTTACCTCA
GAGCACACGTCTGAACTCTTTCCCTACACGACGCTCTTCCGATCTNNNATCCTGTCATTATGATGCTGG
AGCACACGTCTGAACTCTTTCCCTACACGACGCTCTTCCGATCTNNNAGTCTTTTGGTGAACCTGTGTCT
GAGCACACGTCTGAACTCTTTCCCTACACGACGCTCTTCCGATCTNNNCTCAAGCCCCAACAATTCTGAA
AGAGCACACGTCTGAACTCTTTCCCTACACGACGCTCTTCCGATCTNNNGGTAGATGACATCGGTGGA
AGCACACGTCTGAACTCTTTCCCTACACGACGCTCTTCCGATCTNNNAGACTGAAGGTATCTTTGGCCAG
GAGCACACGTCTGAACTCTTTCCCTACACGACGCTCTTCCGATCTNNNCAAGAGCATGCTGACAAGTTC
GAGCACACGTCTGAACTCTTTCCCTACACGACGCTCTTCCGATCTNNNTGCTTTCCTATCACTTCTGCCT
GAGCACACGTCTGAACTCTTTCCCTACACGACGCTCTTCCGATCTNNNAGACTGTGAGTTCTGTCTTGCT
GAGCACACGTCTGAACTCTTTCCCTACACGACGCTCTTCCGATCTNNNTCTTTCTCGTCATTGTGGGCTT
AGCACACGTCTGAACTCTTTCCCTACACGACGCTCTTCCGATCTNNNTCCATCTCAGTCTCTTCCTTCCTC
GAGCACACGTCTGAACTCTTTCCCTACACGACGCTCTTCCGATCTNNNAAACGGAGCAGTCATTTTGGA
GAGCACACGTCTGAACTCTTTCCCTACACGACGCTCTTCCGATCTNNNGCACCCTTCTCACCAGAAAATT
GAGCACACGTCTGAACTCTTTCCCTACACGACGCTCTTCCGATCTNNNTGTCTAGTTCCAGACCCAGAGA
AGCACACGTCTGAACTCTTTCCCTACACGACGCTCTTCCGATCTNNNGAGGTGGAATAATTGCTTGAGC
GAGCACACGTCTGAACTCTTTCCCTACACGACGCTCTTCCGATCTNNNCATACAAACCCAAAACCTCCAG
GAGCACACGTCTGAACTCTTTCCCTACACGACGCTCTTCCGATCTNNNGGAGTGAATGCCTTATAAGGC
GAGCACACGTCTGAACTCTTTCCCTACACGACGCTCTTCCGATCTNNNTGTAATGCATCTGTGTTCCAG
GAGCACACGTCTGAACTCTTTCCCTACACGACGCTCTTCCGATCTNNNTGTGTGTATGTATATCCCCAGC
AGCACACGTCTGAACTCTTTCCCTACACGACGCTCTTCCGATCTNNNGTGGTGCTCATTCCCTGATATCT
GAGCACACGTCTGAACTCTTTCCCTACACGACGCTCTTCCGATCTNNNCCTCAGCCACAAAACAAAGACA
AGAGCACACGTCTGAACTCTTTCCCTACACGACGCTCTTCCGATCTNNNCACAGAGCAAGTTCCAGAATT
AGCACACGTCTGAACTCTTTCCCTACACGACGCTCTTCCGATCTNNNAAACTGCCAAATTGAAGGAGCT

GCACACGTCTGAACTCTTTCCCTACACGACGCTCTTCCGATCTNNNACTCTTTACCCGGATTCACCAAT
GCACACGTCTGAACTCTTTCCCTACACGACGCTCTTCCGATCTNNNGGCTACAATATGGACCAACCTG
GCACACGTCTGAACTCTTTCCCTACACGACGCTCTTCCGATCTNNNATTTTTATGCAGCTCCCTGGTC
AGCACACGTCTGAACTCTTTCCCTACACGACGCTCTTCCGATCTNNNGAGGTCTCACTTCCTAGCTTGT
GAGCACACGTCTGAACTCTTTCCCTACACGACGCTCTTCCGATCTNNNAATAACATTCAGAGAGGCC
GAGCACACGTCTGAACTCTTTCCCTACACGACGCTCTTCCGATCTNNNGGATGCCATGTCTGGAAATA
GAGCACACGTCTGAACTCTTTCCCTACACGACGCTCTTCCGATCTNNNCAGATAGAGACAGAGGAAG
AGCACACGTCTGAACTCTTTCCCTACACGACGCTCTTCCGATCTNNNAGAGGCCCTCTTACTACAGAG
GCACACGTCTGAACTCTTTCCCTACACGACGCTCTTCCGATCTNNNGGACAGTAAGTGGAGCCTTCTA
GCACACGTCTGAACTCTTTCCCTACACGACGCTCTTCCGATCTNNNAATTGGTGAATTGGAGTCTTCG
AGCACACGTCTGAACTCTTTCCCTACACGACGCTCTTCCGATCTNNNGGGCTTCCTTAGAATACTGCAC
GCACACGTCTGAACTCTTTCCCTACACGACGCTCTTCCGATCTNNNTGGACAGTGGCATAGAAGTTG
AGCACACGTCTGAACTCTTTCCCTACACGACGCTCTTCCGATCTNNNCCTCAGAGAGCTGACACAGA
AGCACACGTCTGAACTCTTTCCCTACACGACGCTCTTCCGATCTNNNTTAGGAGGTATACAGCCACAT
GAGCACACGTCTGAACTCTTTCCCTACACGACGCTCTTCCGATCTNNNGGAAACTGAAATGGAAAGG
AGCACACGTCTGAACTCTTTCCCTACACGACGCTCTTCCGATCTNNNCCAAAGAGGCATGCAACCTA
GCACACGTCTGAACTCTTTCCCTACACGACGCTCTTCCGATCTNNNACTGGCTCTTTCTCATAACGACTC
GCACACGTCTGAACTCTTTCCCTACACGACGCTCTTCCGATCTNNNAAACATCTTCCCCTTCTCCCTTCC
GCACACGTCTGAACTCTTTCCCTACACGACGCTCTTCCGATCTNNNCATCTATCACGTCTGGATGCAG
GCACACGTCTGAACTCTTTCCCTACACGACGCTCTTCCGATCTNNNAGGTTATTGACAGGCAATGGAC
AGCACACGTCTGAACTCTTTCCCTACACGACGCTCTTCCGATCTNNNAGAACAGTCATTGTAGGCATC
AGCACACGTCTGAACTCTTTCCCTACACGACGCTCTTCCGATCTNNNGGACCACACTTTGAGAACTGT
AGCACACGTCTGAACTCTTTCCCTACACGACGCTCTTCCGATCTNNNCCAGTCACGGAATCTTGTTTGT
AGAGCACACGTCTGAACTCTTTCCCTACACGACGCTCTTCCGATCTNNNTAGGCAGACACTAAGCTCC
GAGCACACGTCTGAACTCTTTCCCTACACGACGCTCTTCCGATCTNNNACTAGAGAACCAAGAGCAAG
AGAGCACACGTCTGAACTCTTTCCCTACACGACGCTCTTCCGATCTNNNTGTTTCTGTTTCCAGCCAGT
AGCACACGTCTGAACTCTTTCCCTACACGACGCTCTTCCGATCTNNNGAATCACTTGAACCTGGGAG
AGCACACGTCTGAACTCTTTCCCTACACGACGCTCTTCCGATCTNNNCATTTAGCACAAGTCAAGGCT
AGCACACGTCTGAACTCTTTCCCTACACGACGCTCTTCCGATCTNNNTCTGGAAGTGGGTCAAAAGGA
AGCACACGTCTGAACTCTTTCCCTACACGACGCTCTTCCGATCTNNNGAGGAGCAGTGATTATGGCA
GAGCACACGTCTGAACTCTTTCCCTACACGACGCTCTTCCGATCTNNNAGGTTCAGCTGAGACATCAA
CACACGTCTGAACTCTTTCCCTACACGACGCTCTTCCGATCTNNNTTTGCCTCTGTCTCTTCCTCTCCA
GAGCACACGTCTGAACTCTTTCCCTACACGACGCTCTTCCGATCTNNNTGAGGCAGGAGAATCTCTTG
GCACACGTCTGAACTCTTTCCCTACACGACGCTCTTCCGATCTNNNGTTCCCCTGAGACCTACTGAAT
CACACGTCTGAACTCTTTCCCTACACGACGCTCTTCCGATCTNNNCTCTCTCTCCTGTCTCTCTGTCA
GCACACGTCTGAACTCTTTCCCTACACGACGCTCTTCCGATCTNNNATCTGGAATGTGTGAGTTTGTG
AGCACACGTCTGAACTCTTTCCCTACACGACGCTCTTCCGATCTNNNAGGGTGAACACATCAGATGTC
AGCACACGTCTGAACTCTTTCCCTACACGACGCTCTTCCGATCTNNNTTACACACACACACACACACAA
AGCACACGTCTGAACTCTTTCCCTACACGACGCTCTTCCGATCTNNNCAAGGCCCTGTGAAGAAAAGA
GCACACGTCTGAACTCTTTCCCTACACGACGCTCTTCCGATCTNNNATTGGTGGTGGGGAGTAGAAT
GCACACGTCTGAACTCTTTCCCTACACGACGCTCTTCCGATCTNNNTTCTGCCTCTATTTTGTCTGCCTC
AGCACACGTCTGAACTCTTTCCCTACACGACGCTCTTCCGATCTNNNGCTCTTTTTGTCAGACTGTGTC
GCACACGTCTGAACTCTTTCCCTACACGACGCTCTTCCGATCTNNNAAGAAAATTCACTTCCCAGGCA

AGAGCACACGTCTGAACTCTTTCCCTACACGACGCTCTTCCGATCTNNNCAGTCCGAGATGACAGGTA
GAGCACACGTCTGAACTCTTTCCCTACACGACGCTCTTCCGATCTNNNCCAAGCTACAATCCACAGATGA
AGCACACGTCTGAACTCTTTCCCTACACGACGCTCTTCCGATCTNNNTCTGGCAACCCTATTCTTGATCC
AGCACACGTCTGAACTCTTTCCCTACACGACGCTCTTCCGATCTNNNGAGAATTGCTTGAACCCTGGAA
GAGCACACGTCTGAACTCTTTCCCTACACGACGCTCTTCCGATCTNNNGTAGATGGGGGAGGATACA
GAGCACACGTCTGAACTCTTTCCCTACACGACGCTCTTCCGATCTNNNTACCCAGTCTCCTGTAGCATCT
GAGCACACGTCTGAACTCTTTCCCTACACGACGCTCTTCCGATCTNNNAGGAGAGAGAGAGAGGAGAG
GAGCACACGTCTGAACTCTTTCCCTACACGACGCTCTTCCGATCTNNNAGGAGAGAGAGAGGAGAGAG
AGCACACGTCTGAACTCTTTCCCTACACGACGCTCTTCCGATCTNNNCTTAGTGCTGCCCTCATGATAGT
AGCACACGTCTGAACTCTTTCCCTACACGACGCTCTTCCGATCTNNNTGAGTAACATGTGCTCACCATCA
GAGCACACGTCTGAACTCTTTCCCTACACGACGCTCTTCCGATCTNNNCAAACACTTCACACACACACAC
GAGCACACGTCTGAACTCTTTCCCTACACGACGCTCTTCCGATCTNNNAGTTTTTGTTACAAACACTGCC
AGCACACGTCTGAACTCTTTCCCTACACGACGCTCTTCCGATCTNNNACAGCACTGATTACATGGGTAC
GAGCACACGTCTGAACTCTTTCCCTACACGACGCTCTTCCGATCTNNNAAAATGTGTGTGTGTGTG
GAGCACACGTCTGAACTCTTTCCCTACACGACGCTCTTCCGATCTNNNATATGGATCAAGTGCAGGAG
GAGCACACGTCTGAACTCTTTCCCTACACGACGCTCTTCCGATCTNNNTGACTTCCCATTCGGTTTACTG
AGAGCACACGTCTGAACTCTTTCCCTACACGACGCTCTTCCGATCTNNNTCATGTTCCTAGAGCCAGGTA
GAGCACACGTCTGAACTCTTTCCCTACACGACGCTCTTCCGATCTNNNCTCCAAAACACACACACACACA
AGCACACGTCTGAACTCTTTCCCTACACGACGCTCTTCCGATCTNNNTCTCTCATTCTCCCTTCATCCCTC
AGCACACGTCTGAACTCTTTCCCTACACGACGCTCTTCCGATCTNNNACCAAGAAGGGTGCTGAGTTTT
AGCACACGTCTGAACTCTTTCCCTACACGACGCTCTTCCGATCTNNNCTTCTTTCTCCACCTCCCTGTTTCA
AGCACACGTCTGAACTCTTTCCCTACACGACGCTCTTCCGATCTNNNTAACACCAGTGACTGTGTGCTTA
AGAGCACACGTCTGAACTCTTTCCCTACACGACGCTCTTCCGATCTNNNGAGACCTCAGTGATTTGGGC
GAGCACACGTCTGAACTCTTTCCCTACACGACGCTCTTCCGATCTNNNTAGGAAGCTGGATAAGGGGA
GAGCACACGTCTGAACTCTTTCCCTACACGACGCTCTTCCGATCTNNNTTTGCTATTGCCTGGTTGACTT
GCACACGTCTGAACTCTTTCCCTACACGACGCTCTTCCGATCTNNNTCATCTACAAGGTTGAGGTTGCTC
AGCACACGTCTGAACTCTTTCCCTACACGACGCTCTTCCGATCTNNNGGGCTGGACCTAATAGCTTTCT
GCACACGTCTGAACTCTTTCCCTACACGACGCTCTTCCGATCTNNNGTGCCTATGTCCTCATTCTTTGCCA
GAGCACACGTCTGAACTCTTTCCCTACACGACGCTCTTCCGATCTNNNACACCAAGCATCTGGAAATCTC
AGAGCACACGTCTGAACTCTTTCCCTACACGACGCTCTTCCGATCTNNNAAGGGGAAGAAGGGTGAAA
GAGCACACGTCTGAACTCTTTCCCTACACGACGCTCTTCCGATCTNNNGCTAAAGAAGTGGCCTCAAAC
AGAGCACACGTCTGAACTCTTTCCCTACACGACGCTCTTCCGATCTNNNGTGTTAGCCTGAAAGAGCCT
GAGCACACGTCTGAACTCTTTCCCTACACGACGCTCTTCCGATCTNNNCATGGAACTGAATGAGCAAAC
GAGCACACGTCTGAACTCTTTCCCTACACGACGCTCTTCCGATCTNNNGTTCACACACACACATACACAC
AGCACACGTCTGAACTCTTTCCCTACACGACGCTCTTCCGATCTNNNAATACATGCACACACACAGTTGC
AGCACACGTCTGAACTCTTTCCCTACACGACGCTCTTCCGATCTNNNTGCTCTAAACAAGGCCAAATGA
GAGCACACGTCTGAACTCTTTCCCTACACGACGCTCTTCCGATCTNNNCCAGCCTGGCAGAAGAGAAAT
GAGCACACGTCTGAACTCTTTCCCTACACGACGCTCTTCCGATCTNNNATCTTTCCCAGATTGGCTTGGA
GAGCACACGTCTGAACTCTTTCCCTACACGACGCTCTTCCGATCTNNNTCTGTAACCCTCTAACCATTGCC
GAGCACACGTCTGAACTCTTTCCCTACACGACGCTCTTCCGATCTNNNTGGGATTCAATAGTCAGGCA
GCACACGTCTGAACTCTTTCCCTACACGACGCTCTTCCGATCTNNNCATGCAGTTGATACACGAGCATTC
GCACACGTCTGAACTCTTTCCCTACACGACGCTCTTCCGATCTNNNTGGGTTCTCCTTTCTCTCTCTCTCA
AGCACACGTCTGAACTCTTTCCCTACACGACGCTCTTCCGATCTNNNCCAACACCTGAGTAGCTGGACT

AGCACACGTCTGAACTCTTTCCCTACACGACGCTCTTCCGATCTNNNTTTCTCTGCAGGTGCCAAAAAT
GAGCACACGTCTGAACTCTTTCCCTACACGACGCTCTTCCGATCTNNNGGGAGGTTAGCACAGTGAAA
AGCACACGTCTGAACTCTTTCCCTACACGACGCTCTTCCGATCTNNNTACACTGCATAGCCTTTCCAACT
GCACACGTCTGAACTCTTTCCCTACACGACGCTCTTCCGATCTNNNTTGGTGAACTCTTGGAAAGCAAC
GAGCACACGTCTGAACTCTTTCCCTACACGACGCTCTTCCGATCTNNNTCTCCTATCACTGCGAGATTGT
GAGCACACGTCTGAACTCTTTCCCTACACGACGCTCTTCCGATCTNNNGTTGTCTCCTATCACTGCGAGA
AGCACACGTCTGAACTCTTTCCCTACACGACGCTCTTCCGATCTNNNAGCAAAACAGATGGAAGGAAG
GAGCACACGTCTGAACTCTTTCCCTACACGACGCTCTTCCGATCTNNNAGCCATCACTGATTTCCAGCTA
AGCACACGTCTGAACTCTTTCCCTACACGACGCTCTTCCGATCTNNNACACATATGTACACACGGTGGA
AGCACACGTCTGAACTCTTTCCCTACACGACGCTCTTCCGATCTNNNACAAATTTCACAGAGCCTGGTT
AGCACACGTCTGAACTCTTTCCCTACACGACGCTCTTCCGATCTNNNATGATTACACCATCACAAGGCCT
AGAGCACACGTCTGAACTCTTTCCCTACACGACGCTCTTCCGATCTNNNTTCAAGATCAGGAGAGGGA
GCACACGTCTGAACTCTTTCCCTACACGACGCTCTTCCGATCTNNNGGTCTTCAGTTTCACCCACATTGC
GCACACGTCTGAACTCTTTCCCTACACGACGCTCTTCCGATCTNNNGAAGGAGGATCGCTTGAGACTAC
AGCACACGTCTGAACTCTTTCCCTACACGACGCTCTTCCGATCTNNNTGATTTGGCCCCACTCATTCATA
GAGCACACGTCTGAACTCTTTCCCTACACGACGCTCTTCCGATCTNNNACCTAAAGTGCCCCAACATTCT
AGCACACGTCTGAACTCTTTCCCTACACGACGCTCTTCCGATCTNNNCATGGGCAGGTGAAAAGAAGA
GAGCACACGTCTGAACTCTTTCCCTACACGACGCTCTTCCGATCTNNNCCTGGGATGGTCAGCTTATTTG
AGCACACGTCTGAACTCTTTCCCTACACGACGCTCTTCCGATCTNNNAGCCTGTCTGGGTTCTTATGAAG
AGCACACGTCTGAACTCTTTCCCTACACGACGCTCTTCCGATCTNNNTGGTTACAGATAGGAGTGTGT
GAGCACACGTCTGAACTCTTTCCCTACACGACGCTCTTCCGATCTNNNCCCCAGGTGAGACAGTAAACTT
AGCACACGTCTGAACTCTTTCCCTACACGACGCTCTTCCGATCTNNNTGTATGTTGTCTGGTCAGGAAC
AGCACACGTCTGAACTCTTTCCCTACACGACGCTCTTCCGATCTNNNAATCTGAATGAAGTGAGGGAGG
GAGCACACGTCTGAACTCTTTCCCTACACGACGCTCTTCCGATCTNNNGCATGAGCCAATTTCTTACCAC
GCACACGTCTGAACTCTTTCCCTACACGACGCTCTTCCGATCTNNNGGCCTGTATCTCCTGACAATTCAC
GAGCACACGTCTGAACTCTTTCCCTACACGACGCTCTTCCGATCTNNNAACAGTATTGCACATTCAGGCA
AGAGCACACGTCTGAACTCTTTCCCTACACGACGCTCTTCCGATCTNNNGGACTTGTCAGGCTCCATAAT
GAGCACACGTCTGAACTCTTTCCCTACACGACGCTCTTCCGATCTNNNCTATCAGAACTCAACCTCACCCC
AGCACACGTCTGAACTCTTTCCCTACACGACGCTCTTCCGATCTNNNCTTCTTCCCAGAAAGTGCACAAC
AGCACACGTCTGAACTCTTTCCCTACACGACGCTCTTCCGATCTNNNCTACCTACTGTGCTAATGAGGCA
AGCACACGTCTGAACTCTTTCCCTACACGACGCTCTTCCGATCTNNNTTTTGGAATACGCCAGGAACAT
AGAGCACACGTCTGAACTCTTTCCCTACACGACGCTCTTCCGATCTNNNGCTAGGGGACTGGGATAGA
GAGCACACGTCTGAACTCTTTCCCTACACGACGCTCTTCCGATCTNNNTATAGAAAGCGGAGACACCCA
GAGCACACGTCTGAACTCTTTCCCTACACGACGCTCTTCCGATCTNNNATTCACAAGGATGGGATGTCA
GAGCACACGTCTGAACTCTTTCCCTACACGACGCTCTTCCGATCTNNNTGAGTGCTAACTGTCAAGTGT
AGCACACGTCTGAACTCTTTCCCTACACGACGCTCTTCCGATCTNNNTCTCTCACACACACACAAACA
AGCACACGTCTGAACTCTTTCCCTACACGACGCTCTTCCGATCTNNNGGCCACGTGTATGTCTTTTGAAA
GCACACGTCTGAACTCTTTCCCTACACGACGCTCTTCCGATCTNNNTAATGCTGCTTCTTGAAGGCTTG
GCACACGTCTGAACTCTTTCCCTACACGACGCTCTTCCGATCTNNNTGCCTCTTGACATTTGTGTTTGTC
AGCACACGTCTGAACTCTTTCCCTACACGACGCTCTTCCGATCTNNNTTTGAAGGCTTTGTGTTGACCT
AGCACACGTCTGAACTCTTTCCCTACACGACGCTCTTCCGATCTNNNTCAGTCTTTTTCCACCTGTGTCAC
AGCACACGTCTGAACTCTTTCCCTACACGACGCTCTTCCGATCTNNNTGGTGTGTGTATGTGTGTGTGT
AGCACACGTCTGAACTCTTTCCCTACACGACGCTCTTCCGATCTNNNTGGTTCGATGTGACCAGAAGAT

AGCACACGTCTGAACTCTTTCCCTACACGACGCTCTTCCGATCTNNNCTGTACCCTTGCTACCTGATTG
AGAGCACACGTCTGAACTCTTTCCCTACACGACGCTCTTCCGATCTNNNAATTTGCTCAGTTGCAGGG
AGCACACGTCTGAACTCTTTCCCTACACGACGCTCTTCCGATCTNNNCCTCCCCAATGTTGACATCTTA
AGCACACGTCTGAACTCTTTCCCTACACGACGCTCTTCCGATCTNNNATGCGCGCATGTATACAGATA
GCACACGTCTGAACTCTTTCCCTACACGACGCTCTTCCGATCTNNNGAATCATCCAGCCTCAAATGTCG
GCACACGTCTGAACTCTTTCCCTACACGACGCTCTTCCGATCTNNNATCAGAGACTGCCATCTTGGTA
AGCACACGTCTGAACTCTTTCCCTACACGACGCTCTTCCGATCTNNNGAGCGAGGATGTAGCTTGTC
GCACACGTCTGAACTCTTTCCCTACACGACGCTCTTCCGATCTNNNCTAAAAGCCTAGACGTCACCACT
GCACACGTCTGAACTCTTTCCCTACACGACGCTCTTCCGATCTNNNATGTGTGGGTCTTCTACGTCAA
AGCACACGTCTGAACTCTTTCCCTACACGACGCTCTTCCGATCTNNNTTCACTCCTTTTATTGCAGTGC
GAGCACACGTCTGAACTCTTTCCCTACACGACGCTCTTCCGATCTNNNGAGATCGCACCACTGTACTT
GAGCACACGTCTGAACTCTTTCCCTACACGACGCTCTTCCGATCTNNNTCATCTTCCAACTTCCAGTCCT
AGCACACGTCTGAACTCTTTCCCTACACGACGCTCTTCCGATCTNNNTTCAACAAAGGGATTCACTCT
AGCACACGTCTGAACTCTTTCCCTACACGACGCTCTTCCGATCTNNNAGGCATCCAGAAAACTCTCTC
GAGCACACGTCTGAACTCTTTCCCTACACGACGCTCTTCCGATCTNNNATGTGATTGTGTCATCCCTAC
GAGCACACGTCTGAACTCTTTCCCTACACGACGCTCTTCCGATCTNNNACACAGGGATCTTTGCAGTG
GAGCACACGTCTGAACTCTTTCCCTACACGACGCTCTTCCGATCTNNNTATCAGCTGCTAGAATGCAA
GCACACGTCTGAACTCTTTCCCTACACGACGCTCTTCCGATCTNNNAGATAGACACACACACACACA
GAGCACACGTCTGAACTCTTTCCCTACACGACGCTCTTCCGATCTNNNCAGGAAAAGTTTGACAACC
AGCACACGTCTGAACTCTTTCCCTACACGACGCTCTTCCGATCTNNNCAGATATTGGGGAGTATGGG
AGCACACGTCTGAACTCTTTCCCTACACGACGCTCTTCCGATCTNNNAGGCCTTCTTTGTTATCAGCTG
AGCACACGTCTGAACTCTTTCCCTACACGACGCTCTTCCGATCTNNNCTAAACAGCCCAGCTATGTCT
AGCACACGTCTGAACTCTTTCCCTACACGACGCTCTTCCGATCTNNNCTGCATTGAATGAACCAAGTC
GAGCACACGTCTGAACTCTTTCCCTACACGACGCTCTTCCGATCTNNNCCAAATGCCCATTGATCAGTC
AGCACACGTCTGAACTCTTTCCCTACACGACGCTCTTCCGATCTNNNCAAAAGACACCTGCACACCTAT
AGCACACGTCTGAACTCTTTCCCTACACGACGCTCTTCCGATCTNNNGAGGTTAGGATGTGTGTGTG
AGCACACGTCTGAACTCTTTCCCTACACGACGCTCTTCCGATCTNNNACTTTTGGATTTGGAAGCAAC
GAGCACACGTCTGAACTCTTTCCCTACACGACGCTCTTCCGATCTNNNGACAAAGAGTCAAGCCTTGA
AGAGCACACGTCTGAACTCTTTCCCTACACGACGCTCTTCCGATCTNNNGACAAAGAGTCAAGCCTTG
AGAGCACACGTCTGAACTCTTTCCCTACACGACGCTCTTCCGATCTNNNAACTTGCCAGACACTACATT
AGAGCACACGTCTGAACTCTTTCCCTACACGACGCTCTTCCGATCTNNNAATCAGGTAGAACCGGAG
AGCACACGTCTGAACTCTTTCCCTACACGACGCTCTTCCGATCTNNNCCGGGAGTCGAGATAAATCAG
GAGCACACGTCTGAACTCTTTCCCTACACGACGCTCTTCCGATCTNNNGTACACATTGATTCAGGGGA
GAGCACACGTCTGAACTCTTTCCCTACACGACGCTCTTCCGATCTNNNCCCTTGTTGCAGATGAAGTAT
AGCACACGTCTGAACTCTTTCCCTACACGACGCTCTTCCGATCTNNNAAGCAGAAGCATCAGAGTCA
AGCACACGTCTGAACTCTTTCCCTACACGACGCTCTTCCGATCTNNNAAGTACAGTGCTGGTCTGTA
AGCACACGTCTGAACTCTTTCCCTACACGACGCTCTTCCGATCTNNNCTTTTGGACCAAAGGCCTAC
GCACACGTCTGAACTCTTTCCCTACACGACGCTCTTCCGATCTNNNGTTTCAATTGTGTGTGTGTGCA
AGCACACGTCTGAACTCTTTCCCTACACGACGCTCTTCCGATCTNNNCAGGGGTGTGTATGGTTTT
GCACACGTCTGAACTCTTTCCCTACACGACGCTCTTCCGATCTNNNCAAGAACAAGCAAACAAGCCAA
GAGCACACGTCTGAACTCTTTCCCTACACGACGCTCTTCCGATCTNNNATATCGGGAAAACACATGC
GCACACGTCTGAACTCTTTCCCTACACGACGCTCTTCCGATCTNNNCATGTCTGCATTTTTGGTCAGG
AGCACACGTCTGAACTCTTTCCCTACACGACGCTCTTCCGATCTNNNAGTTGGGGACTGTCTGTAAA

GCACACGTCTGAACTCTTTCCCTACACGACGCTCTTCCGATCTNNNGGATGAGGATTGCTTGAAAAC

AGCACACGTCTGAACTCTTTCCCTACACGACGCTCTTCCGATCTNNNATCACAAAGGTGCAAGAGCTA

AGCACACGTCTGAACTCTTTCCCTACACGACGCTCTTCCGATCTNNNTTGGCCTCCTCTTTCTCACAAA

AGCACACGTCTGAACTCTTTCCCTACACGACGCTCTTCCGATCTNNNTGTCGTGACCATTGTGAACAT

AGCACACGTCTGAACTCTTTCCCTACACGACGCTCTTCCGATCTNNNTGGGTGTTTGCATAGCAGAAT

AGCACACGTCTGAACTCTTTCCCTACACGACGCTCTTCCGATCTNNNTATGAAAATGGCCTCCAACTCC

GCACACGTCTGAACTCTTTCCCTACACGACGCTCTTCCGATCTNNNCTCCCTTTCATTCACCTCCATCACA

AGCACACGTCTGAACTCTTTCCCTACACGACGCTCTTCCGATCTNNNATTCAAGTGGATGCCATTTGCT

GAGCACACGTCTGAACTCTTTCCCTACACGACGCTCTTCCGATCTNNNAAAAACTCCAAAGAAGAGCC

AGCACACGTCTGAACTCTTTCCCTACACGACGCTCTTCCGATCTNNNATGACTTCTTTGTGCATGTTGG

AGAGCACACGTCTGAACTCTTTCCCTACACGACGCTCTTCCGATCTNNNACATCATCCAACTAGCAGCA

GAGCACACGTCTGAACTCTTTCCCTACACGACGCTCTTCCGATCTNNNATGATTTGGGCCATGAATTTG

AGCACACGTCTGAACTCTTTCCCTACACGACGCTCTTCCGATCTNNNCCTGGCCATCTTTTTGCTGTATT

AGCACACGTCTGAACTCTTTCCCTACACGACGCTCTTCCGATCTNNNGTGTGTACATGTGTGTGTGTG

GCACACGTCTGAACTCTTTCCCTACACGACGCTCTTCCGATCTNNNTGTTCCTCTCTCTGTCCTCTCATC

AGCACACGTCTGAACTCTTTCCCTACACGACGCTCTTCCGATCTNNNTCTCATCCTATTATTGCGCTGCT

AGCACACGTCTGAACTCTTTCCCTACACGACGCTCTTCCGATCTNNNGCCTTTCTTTCCTACCACTCTGA

GAGCACACGTCTGAACTCTTTCCCTACACGACGCTCTTCCGATCTNNNACATTTGTGTGTGTGTGTGT

GAGCACACGTCTGAACTCTTTCCCTACACGACGCTCTTCCGATCTNNNAAAAAGCTGGGATTGTGTGT

AGCACACGTCTGAACTCTTTCCCTACACGACGCTCTTCCGATCTNNNTTCTCAGCAGCTGTCCTAGAAT

AGCACACGTCTGAACTCTTTCCCTACACGACGCTCTTCCGATCTNNNACTGGAAAGGCAGTCTCCCTAT

GCACACGTCTGAACTCTTTCCCTACACGACGCTCTTCCGATCTNNNGAAAACAATGTGTGGCCTCAAG

AGCACACGTCTGAACTCTTTCCCTACACGACGCTCTTCCGATCTNNNCAGTTTTTATGATGGTGCAGG

GAGCACACGTCTGAACTCTTTCCCTACACGACGCTCTTCCGATCTNNNAGGCATCACATAGACACACAC

AGCACACGTCTGAACTCTTTCCCTACACGACGCTCTTCCGATCTNNNAGGGCTCTATCAAACCATGTTG

GAGCACACGTCTGAACTCTTTCCCTACACGACGCTCTTCCGATCTNNNATTTCTTACCATCTGGAGGCT

AGCACACGTCTGAACTCTTTCCCTACACGACGCTCTTCCGATCTNNNAAAGTATTGCATTTCCCTCTGG

GAGCACACGTCTGAACTCTTTCCCTACACGACGCTCTTCCGATCTNNNCCTGGCAATTTTTCTTGGGG

GCACACGTCTGAACTCTTTCCCTACACGACGCTCTTCCGATCTNNNTTTCATTGTCTGTGTGCATCACC

AGCACACGTCTGAACTCTTTCCCTACACGACGCTCTTCCGATCTNNNTTTCATTGTCTGTGTGCATCAC

AGCACACGTCTGAACTCTTTCCCTACACGACGCTCTTCCGATCTNNNCCTGTGGTAGGCACTAAGGAA

GAGCACACGTCTGAACTCTTTCCCTACACGACGCTCTTCCGATCTNNNAGATACAAATGGGAGCACCTA

GAGCACACGTCTGAACTCTTTCCCTACACGACGCTCTTCCGATCTNNNAAAGCCAAAATAAGCACAAG

AGCACACGTCTGAACTCTTTCCCTACACGACGCTCTTCCGATCTNNNGCCCTACCTATGGTGTTAGGAT

GCACACGTCTGAACTCTTTCCCTACACGACGCTCTTCCGATCTNNNGGCAGCAAGTGAGGAAGTAAG

GCACACGTCTGAACTCTTTCCCTACACGACGCTCTTCCGATCTNNNGAGAAAGGACTAAACCCTCAGC

AGCACACGTCTGAACTCTTTCCCTACACGACGCTCTTCCGATCTNNNACACATACATGCTGATGGTTTC

GCACACGTCTGAACTCTTTCCCTACACGACGCTCTTCCGATCTNNNTTTGTCCTCCAAAGACAGAGACC

GAGCACACGTCTGAACTCTTTCCCTACACGACGCTCTTCCGATCTNNNATGTCGTTTTGATGGCAAGA

GAGCACACGTCTGAACTCTTTCCCTACACGACGCTCTTCCGATCTNNNAGGCATAGTTGGAAAAGGCT

AGCACACGTCTGAACTCTTTCCCTACACGACGCTCTTCCGATCTNNNCCTCACTTACCTCAAAGGGTTG

GCACACGTCTGAACTCTTTCCCTACACGACGCTCTTCCGATCTNNNAAATGTCTCTCTCTCTGTGTGCA

AGCACACGTCTGAACTCTTTCCCTACACGACGCTCTTCCGATCTNNNCTTGTCAATACCGCAAACTCCA

AGCACACGTCTGAACTCTTTCCCTACACGACGCTCTTCCGATCTNNNTGGACACACAGGTCTTCTTCA

AGCACACGTCTGAACTCTTTCCCTACACGACGCTCTTCCGATCTNNNTGACCACACAACTTTGCAACTT

AGCACACGTCTGAACTCTTTCCCTACACGACGCTCTTCCGATCTNNNCCTTGTCTGTCTGTCTGTCTCTC

AGCACACGTCTGAACTCTTTCCCTACACGACGCTCTTCCGATCTNNNATTTTGAAAGGCACAGCATTC

AGCACACGTCTGAACTCTTTCCCTACACGACGCTCTTCCGATCTNNNGGAAGGGTGCACAAACTAAG

GCACACGTCTGAACTCTTTCCCTACACGACGCTCTTCCGATCTNNNCTCATGTCCAAGGGCAAAAGAA

AGCACACGTCTGAACTCTTTCCCTACACGACGCTCTTCCGATCTNNNTCTACCCAGGGAAGAAGAAG

GAGCACACGTCTGAACTCTTTCCCTACACGACGCTCTTCCGATCTNNNTACAAACACACACACACACAC

AAGAGCACACGTCTGAACTCTTTCCCTACACGACGCTCTTCCGATCTNNNCCTTCTTATGGAGTGTCTT

GCACACGTCTGAACTCTTTCCCTACACGACGCTCTTCCGATCTNNNTGTTTTCCCCTAAGATCAGGC

AGCACACGTCTGAACTCTTTCCCTACACGACGCTCTTCCGATCTNNNAGTTTTAGAGCTTGTGGGGTC

GAGCACACGTCTGAACTCTTTCCCTACACGACGCTCTTCCGATCTNNNCCATTTTGAACTCCCCATGTG

AGCACACGTCTGAACTCTTTCCCTACACGACGCTCTTCCGATCTNNNTTCTCACTTAGAGCTGTCCTTC

GAGCACACGTCTGAACTCTTTCCCTACACGACGCTCTTCCGATCTNNNGGAAATGCTGAGAATTATGG

GAGCACACGTCTGAACTCTTTCCCTACACGACGCTCTTCCGATCTNNNGCATATGAACACACACCCATC

GAGCACACGTCTGAACTCTTTCCCTACACGACGCTCTTCCGATCTNNNGTGTGTGTGAGAGAGAG

GAGCACACGTCTGAACTCTTTCCCTACACGACGCTCTTCCGATCTNNNTTGTGACTAGCTGTGTGTGTA

GAGCACACGTCTGAACTCTTTCCCTACACGACGCTCTTCCGATCTNNNACACAGAACTGACTCTACGT

GAGCACACGTCTGAACTCTTTCCCTACACGACGCTCTTCCGATCTNNNTTGCCTGGCAAGAACAACTAT

AGAGCACACGTCTGAACTCTTTCCCTACACGACGCTCTTCCGATCTNNNCTCATGAGTCCTTTTCCCTG

GAGCACACGTCTGAACTCTTTCCCTACACGACGCTCTTCCGATCTNNNGGCAAAGGATAAATGCTTGA

AGAGCACACGTCTGAACTCTTTCCCTACACGACGCTCTTCCGATCTNNNGAACAGCTTTGGTTCTGCTT

GAGCACACGTCTGAACTCTTTCCCTACACGACGCTCTTCCGATCTNNNCTCTCACACACACATGTTCACA

GCACACGTCTGAACTCTTTCCCTACACGACGCTCTTCCGATCTNNNAGAAAGTGAAAGGCAGAGGG

GAGCACACGTCTGAACTCTTTCCCTACACGACGCTCTTCCGATCTNNNAAGCACACATAAGGAGGAT

AGCACACGTCTGAACTCTTTCCCTACACGACGCTCTTCCGATCTNNNCCAGCAGGGATGTCAGACTTT

AGCACACGTCTGAACTCTTTCCCTACACGACGCTCTTCCGATCTNNNGCTAAAGCTGAAAAGCTGACT

AGCACACGTCTGAACTCTTTCCCTACACGACGCTCTTCCGATCTNNNGTACTCCTGAGCGCTCATATTG

AGCACACGTCTGAACTCTTTCCCTACACGACGCTCTTCCGATCTNNNCCACGCTGACTTGATCATATTC

AGCACACGTCTGAACTCTTTCCCTACACGACGCTCTTCCGATCTNNNACCTCAGTACACTCTCTCTCTCT

GCACACGTCTGAACTCTTTCCCTACACGACGCTCTTCCGATCTNNNGTGGAAGAACCCCCTCATTGTA

GCACACGTCTGAACTCTTTCCCTACACGACGCTCTTCCGATCTNNNCTCTGCATGCAGGGTACTTATC

GCACACGTCTGAACTCTTTCCCTACACGACGCTCTTCCGATCTNNNCAGAGGTGCATATACACTCCCT

AGCACACGTCTGAACTCTTTCCCTACACGACGCTCTTCCGATCTNNNATTCTAGGCTGTGTTCTAGAG

GAGCACACGTCTGAACTCTTTCCCTACACGACGCTCTTCCGATCTNNNCCCAAACTGACCAACGAATC

GCACACGTCTGAACTCTTTCCCTACACGACGCTCTTCCGATCTNNNCTCTCTGTGTACTCACTGGCTTT

GCACACGTCTGAACTCTTTCCCTACACGACGCTCTTCCGATCTNNNACACACACACACATATATACG

GCACACGTCTGAACTCTTTCCCTACACGACGCTCTTCCGATCTNNNTGTTGAAGCCACATGTCCAAAA

CACACGTCTGAACTCTTTCCCTACACGACGCTCTTCCGATCTNNNTTGACTTCCCGGCCATTTAGTAAC

GAGCACACGTCTGAACTCTTTCCCTACACGACGCTCTTCCGATCTNNNTGAGAGAGTATGAGGTTGT

AGCACACGTCTGAACTCTTTCCCTACACGACGCTCTTCCGATCTNNNCTGTTCTGTACCTTGACTGTG

GAGCACACGTCTGAACTCTTTCCCTACACGACGCTCTTCCGATCTNNNGGAGATGTGTGTCTGTGTG

AGCACACGTCTGAACTCTTTCCCTACACGACGCTCTTCCGATCTNNNGCAGAGACACTTGTTTTCTTC

AGCACACGTCTGAACTCTTTCCCTACACGACGCTCTTCCGATCTNNNAATCAGGTTGAGAGCATGAG
AGAGCACACGTCTGAACTCTTTCCCTACACGACGCTCTTCCGATCTNNNGTGTGGCAGGACAGTGTTA
GCACACGTCTGAACTCTTTCCCTACACGACGCTCTTCCGATCTNNNCATTGCTACTGGAGTGTCTTTGC
AGCACACGTCTGAACTCTTTCCCTACACGACGCTCTTCCGATCTNNNGGGAGGAAAACCAGGGAAT
AGAGCACACGTCTGAACTCTTTCCCTACACGACGCTCTTCCGATCTNNNGAACCTGGAAGAACTGAG
GAGCACACGTCTGAACTCTTTCCCTACACGACGCTCTTCCGATCTNNNAAAGCACAACACACATGAAC
AGCACACGTCTGAACTCTTTCCCTACACGACGCTCTTCCGATCTNNNATCAAACAAGCCTGTGAATGT
GCACACGTCTGAACTCTTTCCCTACACGACGCTCTTCCGATCTNNNAACAGGGAGACACATACATCAC
AGCACACGTCTGAACTCTTTCCCTACACGACGCTCTTCCGATCTNNNAGTTGCATAGTAGGCAGTTGT
AGCACACGTCTGAACTCTTTCCCTACACGACGCTCTTCCGATCTNNNTTACACAGGCAGGGAAATCT
AGAGCACACGTCTGAACTCTTTCCCTACACGACGCTCTTCCGATCTNNNTTCAAGCAAGGGGGAAAAA
GAGCACACGTCTGAACTCTTTCCCTACACGACGCTCTTCCGATCTNNNGAGTCGGGAATGTTCCTCAC
AGCACACGTCTGAACTCTTTCCCTACACGACGCTCTTCCGATCTNNNGTCTCTCTCTCTCTCTCTCCCC
CACACGTCTGAACTCTTTCCCTACACGACGCTCTTCCGATCTNNNCACAGTATGACTCCAGTGACTCCC
CACACGTCTGAACTCTTTCCCTACACGACGCTCTTCCGATCTNNNCCACTATGACCACCTTTCAGTCAC
AGCACACGTCTGAACTCTTTCCCTACACGACGCTCTTCCGATCTNNNAGATTGGGGCACTCTACGAAT
AGCACACGTCTGAACTCTTTCCCTACACGACGCTCTTCCGATCTNNNGGGCAGTTTGCTTCCCTAATTT
AGCACACGTCTGAACTCTTTCCCTACACGACGCTCTTCCGATCTNNNTACACAATTCTCCTTGCCACTCT
AGCACACGTCTGAACTCTTTCCCTACACGACGCTCTTCCGATCTNNNTCAATGGGATTTGGGGAGAA
AGCACACGTCTGAACTCTTTCCCTACACGACGCTCTTCCGATCTNNNGACAAGGGGAATGCTATCACT
GCACACGTCTGAACTCTTTCCCTACACGACGCTCTTCCGATCTNNNAAAGGGAGAGAGAGAAGGGA
AGCACACGTCTGAACTCTTTCCCTACACGACGCTCTTCCGATCTNNNCTTTAGAGTCTCCAACACCAGC
GAGCACACGTCTGAACTCTTTCCCTACACGACGCTCTTCCGATCTNNNGAAGAAACACACGGAGGAC
AGCACACGTCTGAACTCTTTCCCTACACGACGCTCTTCCGATCTNNNTGGGGAGGAAAACAAAAGCA
GAGCACACGTCTGAACTCTTTCCCTACACGACGCTCTTCCGATCTNNNATGAACTACCATGAGCCGTC
AGCACACGTCTGAACTCTTTCCCTACACGACGCTCTTCCGATCTNNNGCCTGGCAAAAGAAAAAGTCT
AGAGCACACGTCTGAACTCTTTCCCTACACGACGCTCTTCCGATCTNNNTCAGGAAGTTGGGAGATA
AGCACACGTCTGAACTCTTTCCCTACACGACGCTCTTCCGATCTNNNCTTTTGTATGTCTGTTGGCCG
AGAGCACACGTCTGAACTCTTTCCCTACACGACGCTCTTCCGATCTNNNATGTGTACCTGCCCTAAACA
CACACGTCTGAACTCTTTCCCTACACGACGCTCTTCCGATCTNNNCCTGAACACAAGGACTCCTGTAA
GAGCACACGTCTGAACTCTTTCCCTACACGACGCTCTTCCGATCTNNNGGCTCTAGAGGTGGGTGAA
AGCACACGTCTGAACTCTTTCCCTACACGACGCTCTTCCGATCTNNNCCTGGAAGAAAGACAGATGGT
AGCACACGTCTGAACTCTTTCCCTACACGACGCTCTTCCGATCTNNNAAGCAAGGAAATCTGCCTTTA
GAGCACACGTCTGAACTCTTTCCCTACACGACGCTCTTCCGATCTNNNGCACTGGAGTCTGAGAAACC
AGAGCACACGTCTGAACTCTTTCCCTACACGACGCTCTTCCGATCTNNNCATGCGATGAAGAAAGCAG
AGCACACGTCTGAACTCTTTCCCTACACGACGCTCTTCCGATCTNNNTGACTCTACCCTATCCATGCTCA
AGCACACGTCTGAACTCTTTCCCTACACGACGCTCTTCCGATCTNNNCACACACACAAAGACACTCAGA
GAGCACACGTCTGAACTCTTTCCCTACACGACGCTCTTCCGATCTNNNAAAGTATGTGGTGCATGTGA
GAGCACACGTCTGAACTCTTTCCCTACACGACGCTCTTCCGATCTNNNTGGTACTGCAGAACTTCTAC
AGCACACGTCTGAACTCTTTCCCTACACGACGCTCTTCCGATCTNNNATATCTGAGGATTCCCCCAACC
AGCACACGTCTGAACTCTTTCCCTACACGACGCTCTTCCGATCTNNNGTAATAGTCTTCCAGGGCAGA
AGCACACGTCTGAACTCTTTCCCTACACGACGCTCTTCCGATCTNNNTTACAGGGAATCTTGCATCAC
AGCACACGTCTGAACTCTTTCCCTACACGACGCTCTTCCGATCTNNNTCAGCATGCAAATGATGTTTG

AGAGCACACGTCTGAACTCTTTCCCTACACGACGCTCTTCCGATCTNNNCTCAAAGTGGTCCACTCTCA
GAGCACACGTCTGAACTCTTTCCCTACACGACGCTCTTCCGATCTNNNTTTGCAAGTCTCTTCAAGTCT
AGCACACGTCTGAACTCTTTCCCTACACGACGCTCTTCCGATCTNNNAGGAGACTCTAACAGAAACGC
GAGCACACGTCTGAACTCTTTCCCTACACGACGCTCTTCCGATCTNNNGTAGCCATCCTAACAGGTAT
GAGCACACGTCTGAACTCTTTCCCTACACGACGCTCTTCCGATCTNNNCCCACCACCTTCATTTCTAGCT
GAGCACACGTCTGAACTCTTTCCCTACACGACGCTCTTCCGATCTNNNGAAAGTCAGACACTTCAGG
GCACACGTCTGAACTCTTTCCCTACACGACGCTCTTCCGATCTNNNAGCCTCAGAATATCACCCAAAGG
AGCACACGTCTGAACTCTTTCCCTACACGACGCTCTTCCGATCTNNNGGCCCAGTTCATACACCTAGAT
GAGCACACGTCTGAACTCTTTCCCTACACGACGCTCTTCCGATCTNNNAGGCACAGACTCTTGTTCTCT
GCACACGTCTGAACTCTTTCCCTACACGACGCTCTTCCGATCTNNNACCTTGGAGTCTGAAAGCCTTA
GCACACGTCTGAACTCTTTCCCTACACGACGCTCTTCCGATCTNNNGCAAGCATAGCTCTCAAAAGAC
GAGCACACGTCTGAACTCTTTCCCTACACGACGCTCTTCCGATCTNNNTACTTCTGGCAGCTTCAGCAA
GAGCACACGTCTGAACTCTTTCCCTACACGACGCTCTTCCGATCTNNNGAGAGATCTAGGTGAGGGG
GAGCACACGTCTGAACTCTTTCCCTACACGACGCTCTTCCGATCTNNNACAGGAGAGAGAGAGAGAG
AGAGCACACGTCTGAACTCTTTCCCTACACGACGCTCTTCCGATCTNNNGTGTTGTGCAGATTGAATG
GAGCACACGTCTGAACTCTTTCCCTACACGACGCTCTTCCGATCTNNNTCATGTTAGTCCCTTCACACA
AGCACACGTCTGAACTCTTTCCCTACACGACGCTCTTCCGATCTNNNAAGCAAAGAGTCGATCATGAG
GCACACGTCTGAACTCTTTCCCTACACGACGCTCTTCCGATCTNNNGAGAGAGAGAGAGTGTGTGT
GAGCACACGTCTGAACTCTTTCCCTACACGACGCTCTTCCGATCTNNNTTCTCCAGAATGTTGCTGAGT
GCACACGTCTGAACTCTTTCCCTACACGACGCTCTTCCGATCTNNNAAATAAAGAGAGAGGACCTGC
AGCACACGTCTGAACTCTTTCCCTACACGACGCTCTTCCGATCTNNNATGATTTCTCGCTACAAACCCC
GAGCACACGTCTGAACTCTTTCCCTACACGACGCTCTTCCGATCTNNNGTATACTCCCTACCCCAGCAA
GAGCACACGTCTGAACTCTTTCCCTACACGACGCTCTTCCGATCTNNNAGGCACATTGGGTAACAAGG
GAGCACACGTCTGAACTCTTTCCCTACACGACGCTCTTCCGATCTNNNTTCAATCCATTTGCGTCTTTGG
GAGCACACGTCTGAACTCTTTCCCTACACGACGCTCTTCCGATCTNNNCTACTTGCTCCTAAGAAGCA
GCACACGTCTGAACTCTTTCCCTACACGACGCTCTTCCGATCTNNNTGTGAGTGAGTGTGTGTGTATG
AGCACACGTCTGAACTCTTTCCCTACACGACGCTCTTCCGATCTNNNAAAAGGGATGGACAGAATCA
GAGCACACGTCTGAACTCTTTCCCTACACGACGCTCTTCCGATCTNNNCTCCCCATGACATGTAGATAG
AGCACACGTCTGAACTCTTTCCCTACACGACGCTCTTCCGATCTNNNCTGTGTTGGCCATCACTTAGTT
GCACACGTCTGAACTCTTTCCCTACACGACGCTCTTCCGATCTNNNCATAGCCCGTGTCTATCACATCA
GCACACGTCTGAACTCTTTCCCTACACGACGCTCTTCCGATCTNNNTTGACATTTGTTCAGCTCACACC
GAGCACACGTCTGAACTCTTTCCCTACACGACGCTCTTCCGATCTNNNTCAGTCTTTGAGGTAGGTTT
GCACACGTCTGAACTCTTTCCCTACACGACGCTCTTCCGATCTNNNTGGATAACTCGCCTTCATGGAA
GAGCACACGTCTGAACTCTTTCCCTACACGACGCTCTTCCGATCTNNNATTTGTCCCTGTGTGACATA
GAGCACACGTCTGAACTCTTTCCCTACACGACGCTCTTCCGATCTNNNAGTCTAGGAAGATTTGCCG
AGAGCACACGTCTGAACTCTTTCCCTACACGACGCTCTTCCGATCTNNNGGCAAATAACACATGGTCC
AGCACACGTCTGAACTCTTTCCCTACACGACGCTCTTCCGATCTNNNATCATGTGCTCTAGGTTGTGT
GAGCACACGTCTGAACTCTTTCCCTACACGACGCTCTTCCGATCTNNNGAGAGGCAGAGAAGGAGA
AGCACACGTCTGAACTCTTTCCCTACACGACGCTCTTCCGATCTNNNAGAGAGAGAGAGAGAGAGA
GAGCACACGTCTGAACTCTTTCCCTACACGACGCTCTTCCGATCTNNNTATCACCACCCTACAGCAAACT
AGCACACGTCTGAACTCTTTCCCTACACGACGCTCTTCCGATCTNNNTCTCTCTCTCTCTCTCCCTCTC
CACACGTCTGAACTCTTTCCCTACACGACGCTCTTCCGATCTNNNAAGCCCTAACCTCCAATGTGATTC
AGCACACGTCTGAACTCTTTCCCTACACGACGCTCTTCCGATCTNNNCTTTCTCCCAGACAGATGGAGT

AGCACACGTCTGAACTCTTTCCCTACACGACGCTCTTCCGATCTNNNGTCGGGACACAAATCAAAACCA
AGCACACGTCTGAACTCTTTCCCTACACGACGCTCTTCCGATCTNNNAAAGCTACCACCATAAAGCTCG
GAGCACACGTCTGAACTCTTTCCCTACACGACGCTCTTCCGATCTNNNTGACTATGGAGTGAAGATGA
GAGCACACGTCTGAACTCTTTCCCTACACGACGCTCTTCCGATCTNNNTTACGTAGTTTGAACTCACGC
GCACACGTCTGAACTCTTTCCCTACACGACGCTCTTCCGATCTNNNCAACCAGTTTGTACCCAGGGATA
GAGCACACGTCTGAACTCTTTCCCTACACGACGCTCTTCCGATCTNNNTCTCCAATTCTTCCTCTCCCTCT
AGCACACGTCTGAACTCTTTCCCTACACGACGCTCTTCCGATCTNNNTGTTGTTGTTGACCTTTTGACC
GAGCACACGTCTGAACTCTTTCCCTACACGACGCTCTTCCGATCTNNNTTTGGGCAAGTCACAATCTCT
GAGCACACGTCTGAACTCTTTCCCTACACGACGCTCTTCCGATCTNNNAAATATTGAGGGTCTGGCAC
AGCACACGTCTGAACTCTTTCCCTACACGACGCTCTTCCGATCTNNNCCCAAAAAGGTTCTGACTGCAA
AGAGCACACGTCTGAACTCTTTCCCTACACGACGCTCTTCCGATCTNNNTGGGGGAGGCTAACCCTAA
GAGCACACGTCTGAACTCTTTCCCTACACGACGCTCTTCCGATCTNNNAGATGGTGGAAATATGTGTG
GAGCACACGTCTGAACTCTTTCCCTACACGACGCTCTTCCGATCTNNNCAGCGTTCTGTGTTGTTTCTA
AGAGCACACGTCTGAACTCTTTCCCTACACGACGCTCTTCCGATCTNNNAGGGAGAAGAGGGAGAA
AGAGCACACGTCTGAACTCTTTCCCTACACGACGCTCTTCCGATCTNNNATGTAGAAGCTAGTAGGGT
GCACACGTCTGAACTCTTTCCCTACACGACGCTCTTCCGATCTNNNTAGGCATTACCAGCATGATGGA
AGCACACGTCTGAACTCTTTCCCTACACGACGCTCTTCCGATCTNNNTGGCAGACTCAACTTCTTAGCA
AGAGCACACGTCTGAACTCTTTCCCTACACGACGCTCTTCCGATCTNNNTGGCAGACTCAACTTCTTAG
AGCACACGTCTGAACTCTTTCCCTACACGACGCTCTTCCGATCTNNNGGGGATGAGAGAGAGAGAA
GAGCACACGTCTGAACTCTTTCCCTACACGACGCTCTTCCGATCTNNNCTGCTGTGTGAAAATGCTGTT
AGCACACGTCTGAACTCTTTCCCTACACGACGCTCTTCCGATCTNNNTCTCTTCGTTTGGGAGTAATG
GAGCACACGTCTGAACTCTTTCCCTACACGACGCTCTTCCGATCTNNNTCAAATGGCCATCTAGCAGA
GCACACGTCTGAACTCTTTCCCTACACGACGCTCTTCCGATCTNNNTTATGTAGCTGGTCCTGCTTCAG
AGCACACGTCTGAACTCTTTCCCTACACGACGCTCTTCCGATCTNNNACAGAGGTTTGTTCCCAAAGG
AGCACACGTCTGAACTCTTTCCCTACACGACGCTCTTCCGATCTNNNTTCTTTTATATACTGTGCGCGCG
GAGCACACGTCTGAACTCTTTCCCTACACGACGCTCTTCCGATCTNNNAGAGAGACCAGGGAGAAAA
AGCACACGTCTGAACTCTTTCCCTACACGACGCTCTTCCGATCTNNNTCTCCAGAAGAAATCAAGCCA
GAGCACACGTCTGAACTCTTTCCCTACACGACGCTCTTCCGATCTNNNTCCAGAATACTCCATGGATGT
AGCACACGTCTGAACTCTTTCCCTACACGACGCTCTTCCGATCTNNNGGAAGGTTGAAGTGGGTGGA
GCACACGTCTGAACTCTTTCCCTACACGACGCTCTTCCGATCTNNNGAGCAAGACTCCGTCTCAAATAA
GAGCACACGTCTGAACTCTTTCCCTACACGACGCTCTTCCGATCTNNNTGCAGTAGGGTCACATATCA
GAGCACACGTCTGAACTCTTTCCCTACACGACGCTCTTCCGATCTNNNATCCCATCCAGCTGATACTAG
AGCACACGTCTGAACTCTTTCCCTACACGACGCTCTTCCGATCTNNNTTGCACAAGATGACAGACCAG
GAGCACACGTCTGAACTCTTTCCCTACACGACGCTCTTCCGATCTNNNCCATCAGTGAAAAGGACACA
AGCACACGTCTGAACTCTTTCCCTACACGACGCTCTTCCGATCTNNNCTCCTCTTCCACTCACATTCACA
GAGCACACGTCTGAACTCTTTCCCTACACGACGCTCTTCCGATCTNNNTGTTGACTCTTCTACTCTCTC
GAGCACACGTCTGAACTCTTTCCCTACACGACGCTCTTCCGATCTNNNTGGTCAGCAACTTGAATAGC
AGCACACGTCTGAACTCTTTCCCTACACGACGCTCTTCCGATCTNNNACACTGGTTATTTGGTGCATG
GAGCACACGTCTGAACTCTTTCCCTACACGACGCTCTTCCGATCTNNNGGTCAATGGACTGGAGGTT
AGCACACGTCTGAACTCTTTCCCTACACGACGCTCTTCCGATCTNNNAGTGGAGGTGAAGGAAAGA
GCACACGTCTGAACTCTTTCCCTACACGACGCTCTTCCGATCTNNNCCATAAGAAGCAAGCACCTAAG
AGCACACGTCTGAACTCTTTCCCTACACGACGCTCTTCCGATCTNNNCACACACGCACACACACATATA
AGCACACGTCTGAACTCTTTCCCTACACGACGCTCTTCCGATCTNNNTAAAACCCTCAGACTCCACATC

GAGCACACGTCTGAACTCTTTCCCTACACGACGCTCTTCCGATCTNNNCATGTTTATCTGGTGCAGGT
GCACACGTCTGAACTCTTTCCCTACACGACGCTCTTCCGATCTNNNGAGAGAAGCATTGTAGGGGAG
AGCACACGTCTGAACTCTTTCCCTACACGACGCTCTTCCGATCTNNNGACAGTATGGCCAGATTGGTA
CACACGTCTGAACTCTTTCCCTACACGACGCTCTTCCGATCTNNNACTTCTCCAAATTCCATGCTTGCC
GAGCACACGTCTGAACTCTTTCCCTACACGACGCTCTTCCGATCTNNNGCTAGAATCAGGTGTTAGGT
GCACACGTCTGAACTCTTTCCCTACACGACGCTCTTCCGATCTNNNCCTGCCTTTAGTCTTGGCTTAGA
AGCACACGTCTGAACTCTTTCCCTACACGACGCTCTTCCGATCTNNNATACAGGATGCACCCTTTTGTG
GCACACGTCTGAACTCTTTCCCTACACGACGCTCTTCCGATCTNNNACTTCATTGCAACTCCCTGATCT
GCACACGTCTGAACTCTTTCCCTACACGACGCTCTTCCGATCTNNNTGTGTGTGTATGTGTGTGTA
GAGCACACGTCTGAACTCTTTCCCTACACGACGCTCTTCCGATCTNNNGCGCCTAGCCATATGTTTCTT
GAGCACACGTCTGAACTCTTTCCCTACACGACGCTCTTCCGATCTNNNATTTTCACATCTGCCAACCAA
AGAGCACACGTCTGAACTCTTTCCCTACACGACGCTCTTCCGATCTNNNGCTAAGAGAAGAGGGGA
AGCACACGTCTGAACTCTTTCCCTACACGACGCTCTTCCGATCTNNNTCAGTGACACCGTGTGGATAT
AGCACACGTCTGAACTCTTTCCCTACACGACGCTCTTCCGATCTNNNATTCTTTTGTGCGTCCGATTGT
GCACACGTCTGAACTCTTTCCCTACACGACGCTCTTCCGATCTNNNATTCTTTTGTGCGTCCGATTGTT
GCACACGTCTGAACTCTTTCCCTACACGACGCTCTTCCGATCTNNNTCTGTGACCTGTAACTTTGCTGA
AGCACACGTCTGAACTCTTTCCCTACACGACGCTCTTCCGATCTNNNGGGAGTCTAGTATGTTGGTG
AGCACACGTCTGAACTCTTTCCCTACACGACGCTCTTCCGATCTNNNACAGACACAGAAGACAATAGG
AGCACACGTCTGAACTCTTTCCCTACACGACGCTCTTCCGATCTNNNAGAGATACACACACACACACAC
GCACACGTCTGAACTCTTTCCCTACACGACGCTCTTCCGATCTNNNTCAGGGACTCCTCCTAACTCATT
GAGCACACGTCTGAACTCTTTCCCTACACGACGCTCTTCCGATCTNNNTCATGTCCCACAACCTAACTC
AGCACACGTCTGAACTCTTTCCCTACACGACGCTCTTCCGATCTNNNTGGGTTGAGATGATGGGAAA
GAGCACACGTCTGAACTCTTTCCCTACACGACGCTCTTCCGATCTNNNGAGAGGGAGAGAGAGAGA
GAGCACACGTCTGAACTCTTTCCCTACACGACGCTCTTCCGATCTNNNTTAGAGAGGGGGGTGAAGA
AGAGCACACGTCTGAACTCTTTCCCTACACGACGCTCTTCCGATCTNNNGGCAAGAGAGAGACAGA
AGCACACGTCTGAACTCTTTCCCTACACGACGCTCTTCCGATCTNNNTTCAATGTGTGTGTCTATGCA
GAGCACACGTCTGAACTCTTTCCCTACACGACGCTCTTCCGATCTNNNCATGTCATAGGCTGCTGTTT
AGCACACGTCTGAACTCTTTCCCTACACGACGCTCTTCCGATCTNNNGGCAAAATGAGGAGTGGCAT
GAGCACACGTCTGAACTCTTTCCCTACACGACGCTCTTCCGATCTNNNCACATATATGCATGCACACAC
AGCACACGTCTGAACTCTTTCCCTACACGACGCTCTTCCGATCTNNNGGGAAGAGACTTCTGGAACA
AGCACACGTCTGAACTCTTTCCCTACACGACGCTCTTCCGATCTNNNTGTCCATGAGGTATCCAAACA
AGCACACGTCTGAACTCTTTCCCTACACGACGCTCTTCCGATCTNNNAAAATGTGTGTGTGTGTGT
GAGCACACGTCTGAACTCTTTCCCTACACGACGCTCTTCCGATCTNNNATTTATGAGGCTGTGCAAT
AGCACACGTCTGAACTCTTTCCCTACACGACGCTCTTCCGATCTNNNTGTTCTCCACCTATCGGAAACT
AGAGCACACGTCTGAACTCTTTCCCTACACGACGCTCTTCCGATCTNNNCCTCTCCTATTTCCGATGGT
AGAGCACACGTCTGAACTCTTTCCCTACACGACGCTCTTCCGATCTNNNCATCTTGGAAGTAGTACCC
AGCACACGTCTGAACTCTTTCCCTACACGACGCTCTTCCGATCTNNNTGCCTGTAATCCCAGTGAAAG
GAGCACACGTCTGAACTCTTTCCCTACACGACGCTCTTCCGATCTNNNACAAGCTCAAACTGCACATT
GAGCACACGTCTGAACTCTTTCCCTACACGACGCTCTTCCGATCTNNNTGTCCCTGTTTCTCTCTCTCTC
CACACGTCTGAACTCTTTCCCTACACGACGCTCTTCCGATCTNNNAGTGATCTTTCTAGAGCGTGCTC
GAGCACACGTCTGAACTCTTTCCCTACACGACGCTCTTCCGATCTNNNTAGCCTCCATAATCACATGA
GAGCACACGTCTGAACTCTTTCCCTACACGACGCTCTTCCGATCTNNNTCAAGTGTGCCCTCATATGT
GCACACGTCTGAACTCTTTCCCTACACGACGCTCTTCCGATCTNNNGCAGGGTAGATTTCTTTTCCCC

AGCACACGTCTGAACTCTTTCCCTACACGACGCTCTTCCGATCTNNNTGGTATTCTGTTGTACTGTGGC
GAGCACACGTCTGAACTCTTTCCCTACACGACGCTCTTCCGATCTNNNGTGCCTGAGCTGAAAGATGA
AGCACACGTCTGAACTCTTTCCCTACACGACGCTCTTCCGATCTNNNAGAGAGAGAGAGAGAGAGGC
AGAGCACACGTCTGAACTCTTTCCCTACACGACGCTCTTCCGATCTNNNCAACACACAGTGGTTGAGA
GAGCACACGTCTGAACTCTTTCCCTACACGACGCTCTTCCGATCTNNNGCCTATGCTGTTCTTTGGAATC
GAGCACACGTCTGAACTCTTTCCCTACACGACGCTCTTCCGATCTNNNTGAGGCTAGCATGATTCTGAT
GAGCACACGTCTGAACTCTTTCCCTACACGACGCTCTTCCGATCTNNNCTTGTGTGTGAGTGTGTGTGT
AAGAGCACACGTCTGAACTCTTTCCCTACACGACGCTCTTCCGATCTNNNTCTATTTCCTCAATCCCAAG
AGCACACGTCTGAACTCTTTCCCTACACGACGCTCTTCCGATCTNNNAACATTGCTTTGCCACTATGCAT
GAGCACACGTCTGAACTCTTTCCCTACACGACGCTCTTCCGATCTNNNCTGCCCAAAACAGGAAAAGTC
GAGCACACGTCTGAACTCTTTCCCTACACGACGCTCTTCCGATCTNNNACAGAGAAACAGAACCAATGA
AGCACACGTCTGAACTCTTTCCCTACACGACGCTCTTCCGATCTNNNGGTTACAGTGACCAAAAACCCAC
AGCACACGTCTGAACTCTTTCCCTACACGACGCTCTTCCGATCTNNNAGAGGTTGCAGTATGACTCGA
AGCACACGTCTGAACTCTTTCCCTACACGACGCTCTTCCGATCTNNNCTTGCCTGGCTCATAATATGCAC
GAGCACACGTCTGAACTCTTTCCCTACACGACGCTCTTCCGATCTNNNCCCAACTAGCTGCAAGAGTTT
GAGCACACGTCTGAACTCTTTCCCTACACGACGCTCTTCCGATCTNNNTTTCTATCTCCTACTCCACCCCC
GAGCACACGTCTGAACTCTTTCCCTACACGACGCTCTTCCGATCTNNNCACCAGGATGTAAGAGGTCC
AGCACACGTCTGAACTCTTTCCCTACACGACGCTCTTCCGATCTNNNTGGGGTAATTTGTTATGCAGTC
AGCACACGTCTGAACTCTTTCCCTACACGACGCTCTTCCGATCTNNNGCTTTTGCATCTAACAGTCAGCA
AGCACACGTCTGAACTCTTTCCCTACACGACGCTCTTCCGATCTNNNTCTGGTACCTCAGTTGGGAAAA
GAGCACACGTCTGAACTCTTTCCCTACACGACGCTCTTCCGATCTNNNAATGTGAAAACAGAAGGCGTC
GAGCACACGTCTGAACTCTTTCCCTACACGACGCTCTTCCGATCTNNNTTCTTGGGCCACATTTTCTTTC
CACACGTCTGAACTCTTTCCCTACACGACGCTCTTCCGATCTNNNTCCAACCAGATAGGCCCACTATACA
AGCACACGTCTGAACTCTTTCCCTACACGACGCTCTTCCGATCTNNNTTTACACACACACACACACACAC
AGCACACGTCTGAACTCTTTCCCTACACGACGCTCTTCCGATCTNNNCGGTTTGCTAATCATGGTTTTG
GAGCACACGTCTGAACTCTTTCCCTACACGACGCTCTTCCGATCTNNNAGCACTCCATTCCTTTCCTACAC
AGCACACGTCTGAACTCTTTCCCTACACGACGCTCTTCCGATCTNNNAGACAATTAGTGGAAGGAGCA
GAGCACACGTCTGAACTCTTTCCCTACACGACGCTCTTCCGATCTNNNAGGATGGAGGGAGAGAAAG
AGCACACGTCTGAACTCTTTCCCTACACGACGCTCTTCCGATCTNNNCCCTGCACATTCTGGGATAATC
AGAGCACACGTCTGAACTCTTTCCCTACACGACGCTCTTCCGATCTNNNTGAGCCACAAGAGTATTTAC
CACACGTCTGAACTCTTTCCCTACACGACGCTCTTCCGATCTNNNTGAGTTACACTAAATGGCCACCACA
GAGCACACGTCTGAACTCTTTCCCTACACGACGCTCTTCCGATCTNNNGTTGTCCGATTCAATGCAAAT
AGCACACGTCTGAACTCTTTCCCTACACGACGCTCTTCCGATCTNNNTGTAAGAAAATGCAGCAGCTCA
AGCACACGTCTGAACTCTTTCCCTACACGACGCTCTTCCGATCTNNNCTTCCACCCAACCAATACAACAC
GAGCACACGTCTGAACTCTTTCCCTACACGACGCTCTTCCGATCTNNNCTGAGGCAGGAGAATCACTTC
GAGCACACGTCTGAACTCTTTCCCTACACGACGCTCTTCCGATCTNNNGTAACTCCAAGCCATGCCAAAT
GAGCACACGTCTGAACTCTTTCCCTACACGACGCTCTTCCGATCTNNNGCAAAGATTCCTGTAGCCGTTA
AGCACACGTCTGAACTCTTTCCCTACACGACGCTCTTCCGATCTNNNAGCACAGTACCCCTACAACTTTT
AGCACACGTCTGAACTCTTTCCCTACACGACGCTCTTCCGATCTNNNCACCGGGAAGTAGAACATTGGT
AGAGCACACGTCTGAACTCTTTCCCTACACGACGCTCTTCCGATCTNNNTGTACACACAGACACATACA
GAGCACACGTCTGAACTCTTTCCCTACACGACGCTCTTCCGATCTNNNAATGCAGATGGTTTAGGGTT
GAGCACACGTCTGAACTCTTTCCCTACACGACGCTCTTCCGATCTNNNAGCCTTCCAAGAAGCTGAAAC
AGCACACGTCTGAACTCTTTCCCTACACGACGCTCTTCCGATCTNNNAGCCTTCCAAGAAGCTGAAACT

AGCACACGTCTGAACTCTTTCCCTACACGACGCTCTTCCGATCTNNNTTATCCACCTTTCCCACAACACA
AGCACACGTCTGAACTCTTTCCCTACACGACGCTCTTCCGATCTNNNGTTTGCCTTTACTTCCCTGTTCCC
AGAGCACACGTCTGAACTCTTTCCCTACACGACGCTCTTCCGATCTNNNGAAGGTGCAATTGTGTGTA
AGAGCACACGTCTGAACTCTTTCCCTACACGACGCTCTTCCGATCTNNNCACAGCATTCCTCACCAGAA
AGCACACGTCTGAACTCTTTCCCTACACGACGCTCTTCCGATCTNNNGAAGTCCCTCTATGTGTGTGTG
AGAGCACACGTCTGAACTCTTTCCCTACACGACGCTCTTCCGATCTNNNAGGAAAGGCAGAATATAGG
AGCACACGTCTGAACTCTTTCCCTACACGACGCTCTTCCGATCTNNNCAGAGTTTACCTCCTAGGGAAGC
GCACACGTCTGAACTCTTTCCCTACACGACGCTCTTCCGATCTNNNTCTCAGCTTCCTTGTGCTTAGAGC
AGCACACGTCTGAACTCTTTCCCTACACGACGCTCTTCCGATCTNNNTCAGAGTATAACCAAGCTGCTGC
GAGCACACGTCTGAACTCTTTCCCTACACGACGCTCTTCCGATCTNNNTATTTCCCTTTGTTCAGCTGTG
GAGCACACGTCTGAACTCTTTCCCTACACGACGCTCTTCCGATCTNNNGGGAGGTTGTAGGATATCAC
GAGCACACGTCTGAACTCTTTCCCTACACGACGCTCTTCCGATCTNNNAGCACAAAACAGACTGAGACA
GCACACGTCTGAACTCTTTCCCTACACGACGCTCTTCCGATCTNNNATAGATAGGCCAGCGAGACAAA
AGCACACGTCTGAACTCTTTCCCTACACGACGCTCTTCCGATCTNNNTTGAGCGTCTACTATGTGTCAGC
AGCACACGTCTGAACTCTTTCCCTACACGACGCTCTTCCGATCTNNNGTTGACTGCTGCTCTGTAGAGAT
GCACACGTCTGAACTCTTTCCCTACACGACGCTCTTCCGATCTNNNGGACTTTGGTTCCCTAAGAGGACC
AGCACACGTCTGAACTCTTTCCCTACACGACGCTCTTCCGATCTNNNGCTGATGCTCCAATTCTGTCATT
GAGCACACGTCTGAACTCTTTCCCTACACGACGCTCTTCCGATCTNNNGAGATAGTGTGTGTGTGCAT
AGCACACGTCTGAACTCTTTCCCTACACGACGCTCTTCCGATCTNNNTGGAGATAAAATGGGGAAAGC
AGCACACGTCTGAACTCTTTCCCTACACGACGCTCTTCCGATCTNNNTTGGAGAAAGGTCTATCTGCTG
GCACACGTCTGAACTCTTTCCCTACACGACGCTCTTCCGATCTNNNAAATATGCCACTGTTGTCTGTGCC
AGCACACGTCTGAACTCTTTCCCTACACGACGCTCTTCCGATCTNNNGCATGTTTGCCTTACTCTCTCCTC
GCACACGTCTGAACTCTTTCCCTACACGACGCTCTTCCGATCTNNNAACTAGCCAAGTCTCTTTCTCCCCA
GAGCACACGTCTGAACTCTTTCCCTACACGACGCTCTTCCGATCTNNNTTCTACTTGAGGAGAGGAGA
GAGCACACGTCTGAACTCTTTCCCTACACGACGCTCTTCCGATCTNNNTCCCCCAGAACCCTTCTATACTC
AGCACACGTCTGAACTCTTTCCCTACACGACGCTCTTCCGATCTNNNGGAAGTATCCCCCAGCTAGATA
AGCACACGTCTGAACTCTTTCCCTACACGACGCTCTTCCGATCTNNNTTCCTTCCTTCCTTCCTTCCTTCCA
AGCACACGTCTGAACTCTTTCCCTACACGACGCTCTTCCGATCTNNNTGTGCTATGGATTGGTGGAAAC
GCACACGTCTGAACTCTTTCCCTACACGACGCTCTTCCGATCTNNNTGTGCTATGGATTGGTGGAAAGA
AGCACACGTCTGAACTCTTTCCCTACACGACGCTCTTCCGATCTNNNCTCTCTTCCTTCCTTGTCTGCATC
GAGCACACGTCTGAACTCTTTCCCTACACGACGCTCTTCCGATCTNNNATTTGCAACCCAGTAAAAGCAC
GAGCACACGTCTGAACTCTTTCCCTACACGACGCTCTTCCGATCTNNNCCTCCTCTTGGGAACTGTAAC
AGCACACGTCTGAACTCTTTCCCTACACGACGCTCTTCCGATCTNNNCCTCTGTGCGTATCTCTATGGAG
AGCACACGTCTGAACTCTTTCCCTACACGACGCTCTTCCGATCTNNNATGGCCAATTAGTTGAACATGGC
AGAGCACACGTCTGAACTCTTTCCCTACACGACGCTCTTCCGATCTNNNTGCATGTTCATAGCAGCACAA
GAGCACACGTCTGAACTCTTTCCCTACACGACGCTCTTCCGATCTNNNCAAAGAAAAGTCCTCTGTGCA
AGCACACGTCTGAACTCTTTCCCTACACGACGCTCTTCCGATCTNNNACAAAGTAGCTGAGCCGACTTT
GAGCACACGTCTGAACTCTTTCCCTACACGACGCTCTTCCGATCTNNNCATACACACACACACACACACTC
GCACACGTCTGAACTCTTTCCCTACACGACGCTCTTCCGATCTNNNGATAATTTGGATCTGTGGCTGCCC
AGCACACGTCTGAACTCTTTCCCTACACGACGCTCTTCCGATCTNNNGTTCACAGTCTGAGTTCCATTGC
CACACGTCTGAACTCTTTCCCTACACGACGCTCTTCCGATCTNNNTCTCATTCCCTCTCTCCTTCTGTCATC
AGAGCACACGTCTGAACTCTTTCCCTACACGACGCTCTTCCGATCTNNNGGTCTACCTGATGCATGATC
GAGCACACGTCTGAACTCTTTCCCTACACGACGCTCTTCCGATCTNNNGTGTGAAGGAGAAGAAAGTG

AGCACACGTCTGAACTCTTTCCCTACACGACGCTCTTCCGATCTNNNAGAATTTCCTTTCAGGCAGGCTA
GCACACGTCTGAACTCTTTCCCTACACGACGCTCTTCCGATCTNNNCTGGACAGGCAGTCTTTTGATTG
GAGCACACGTCTGAACTCTTTCCCTACACGACGCTCTTCCGATCTNNNGCAGGAGTTGTACACACTTATG
GAGCACACGTCTGAACTCTTTCCCTACACGACGCTCTTCCGATCTNNNGGGTCTCTCTCTCTCTCTCTC
GAGCACACGTCTGAACTCTTTCCCTACACGACGCTCTTCCGATCTNNNAGTTGTTTTGCATCGGGTCAA
AGCACACGTCTGAACTCTTTCCCTACACGACGCTCTTCCGATCTNNNGATAAGCAGTAGCAAGCAAAG
AGAGCACACGTCTGAACTCTTTCCCTACACGACGCTCTTCCGATCTNNNAAAATCTGTGTTTGTCGGGG
GAGCACACGTCTGAACTCTTTCCCTACACGACGCTCTTCCGATCTNNNCAGCAGTTGCAGTCATTCAATG
GAGCACACGTCTGAACTCTTTCCCTACACGACGCTCTTCCGATCTNNNTATCTCTCACTCTTCTGGGGGT
GCACACGTCTGAACTCTTTCCCTACACGACGCTCTTCCGATCTNNNTCCTCACCTTTCTCTCCCCTATTCAT
AGCACACGTCTGAACTCTTTCCCTACACGACGCTCTTCCGATCTNNNACAAACATCATGGTGGGCTTTTT
AAGAGCACACGTCTGAACTCTTTCCCTACACGACGCTCTTCCGATCTNNNTTCCCCTAATCTCACTAACCA
AGCACACGTCTGAACTCTTTCCCTACACGACGCTCTTCCGATCTNNNTTTCTCCTTCCTTCCTTCCTTCCCA
AGCACACGTCTGAACTCTTTCCCTACACGACGCTCTTCCGATCTNNNGACAAATATGCCATGGTGAAG
GCACACGTCTGAACTCTTTCCCTACACGACGCTCTTCCGATCTNNNAAGACACAAGACAGACACAGAGG
AGCACACGTCTGAACTCTTTCCCTACACGACGCTCTTCCGATCTNNNGACCAAATGAGATGCAAAGCCA
AAGAGCACACGTCTGAACTCTTTCCCTACACGACGCTCTTCCGATCTNNNGGAGGGAGAAAGGAAAG
GAGCACACGTCTGAACTCTTTCCCTACACGACGCTCTTCCGATCTNNNGGGCGAAAGAGTGAGATTTT
AGAGCACACGTCTGAACTCTTTCCCTACACGACGCTCTTCCGATCTNNNTTCCTGAAGTATGGTACCTTG
AGCACACGTCTGAACTCTTTCCCTACACGACGCTCTTCCGATCTNNNTGGTGATGGACCTGCTCAATAT
AGAGCACACGTCTGAACTCTTTCCCTACACGACGCTCTTCCGATCTNNNGGGTTCTGATAGAGGTGGCT
AGCACACGTCTGAACTCTTTCCCTACACGACGCTCTTCCGATCTNNNAAGGACAAGATTTCCAGGAAC
GCACACGTCTGAACTCTTTCCCTACACGACGCTCTTCCGATCTNNNTCCAGCTTCATCTTCCACAATCACA
AGCACACGTCTGAACTCTTTCCCTACACGACGCTCTTCCGATCTNNNAGATTCTGCTTGGTTGTAGCTGA
AGCACACGTCTGAACTCTTTCCCTACACGACGCTCTTCCGATCTNNNTCACAGATAAGGAAGTTGAGCC
GAGCACACGTCTGAACTCTTTCCCTACACGACGCTCTTCCGATCTNNNTCTCAACCAGGAGAATTGTAGG
GAGCACACGTCTGAACTCTTTCCCTACACGACGCTCTTCCGATCTNNNTTTGTTTGGTTGGTTGGTTGG
GAGCACACGTCTGAACTCTTTCCCTACACGACGCTCTTCCGATCTNNNAGCAGAACTTTGGGGCCTAAA
GAGCACACGTCTGAACTCTTTCCCTACACGACGCTCTTCCGATCTNNNGGTTGAGTGGGAAAAAAGG
GAGCACACGTCTGAACTCTTTCCCTACACGACGCTCTTCCGATCTNNNTAGAAAGATATGGGGTGTGT
AGCACACGTCTGAACTCTTTCCCTACACGACGCTCTTCCGATCTNNNATGTCTCATTGAGAATGGCCTC
GAGCACACGTCTGAACTCTTTCCCTACACGACGCTCTTCCGATCTNNNAAGGCAGTGTGTGATTTAGGG
AGCACACGTCTGAACTCTTTCCCTACACGACGCTCTTCCGATCTNNNCTACCCTGTCAAGAGGAAAGAG
CACACGTCTGAACTCTTTCCCTACACGACGCTCTTCCGATCTNNNACCTTCTCTCTCTCTCTCTCTCATG
AGCACACGTCTGAACTCTTTCCCTACACGACGCTCTTCCGATCTNNNCTGCAGATCTGTTTTTCCACGTT
AGCACACGTCTGAACTCTTTCCCTACACGACGCTCTTCCGATCTNNNAGACACCCCAGTTTAGCAGTTA
AGCACACGTCTGAACTCTTTCCCTACACGACGCTCTTCCGATCTNNNTTTGCTCATGGCCCCATATTTTC
GCACACGTCTGAACTCTTTCCCTACACGACGCTCTTCCGATCTNNNTCTCAATTGCAGGTGACACAAAC
AGCACACGTCTGAACTCTTTCCCTACACGACGCTCTTCCGATCTNNNGTGTATATGCATGTGTGTGTGG
GAGCACACGTCTGAACTCTTTCCCTACACGACGCTCTTCCGATCTNNNACAAAACCAGAACCAAGAGGA
AGCACACGTCTGAACTCTTTCCCTACACGACGCTCTTCCGATCTNNNTCCCTCTGGATCTTACCCTTCTTC
GAGCACACGTCTGAACTCTTTCCCTACACGACGCTCTTCCGATCTNNNTGCAGATAACCTTGGATGAGA
GAGCACACGTCTGAACTCTTTCCCTACACGACGCTCTTCCGATCTNNNCTCAGTTACCTTTCACTCCCTCC

GAGCACACGTCTGAACTCTTTCCCTACACGACGCTCTTCCGATCTNNNAGAGTTAGTGTGAGCATGTG
GCACACGTCTGAACTCTTTCCCTACACGACGCTCTTCCGATCTNNNTTCACCAGAAAACACACACACAC
GAGCACACGTCTGAACTCTTTCCCTACACGACGCTCTTCCGATCTNNNTTCTGTTTGCTTTCAGGAGCT
GCACACGTCTGAACTCTTTCCCTACACGACGCTCTTCCGATCTNNNAAGAAGTTGTGTTCCTGAGAGC
GCACACGTCTGAACTCTTTCCCTACACGACGCTCTTCCGATCTNNNATCTGTCCCACCAAACATTAGCTC
AGCACACGTCTGAACTCTTTCCCTACACGACGCTCTTCCGATCTNNNATTGTGTGATTTCTGCTGCAC
AGCACACGTCTGAACTCTTTCCCTACACGACGCTCTTCCGATCTNNNCTTCTAGCCCCTTTTACACCCTT
GAGCACACGTCTGAACTCTTTCCCTACACGACGCTCTTCCGATCTNNNCCAGGAGATAAGCCCAAAGT
AGCACACGTCTGAACTCTTTCCCTACACGACGCTCTTCCGATCTNNNTCTCTCTCTCTCTGTGTGTGT
GCACACGTCTGAACTCTTTCCCTACACGACGCTCTTCCGATCTNNNAGGTACAGAAATACTCGCACTG
GAGCACACGTCTGAACTCTTTCCCTACACGACGCTCTTCCGATCTNNNTCCATCAGGTTTCTTGTGTTT
AGCACACGTCTGAACTCTTTCCCTACACGACGCTCTTCCGATCTNNNGGGCCTCCTGGAACTGTAAAA
AGCACACGTCTGAACTCTTTCCCTACACGACGCTCTTCCGATCTNNNGGCAAAAGTGATGGGATGA
AGCACACGTCTGAACTCTTTCCCTACACGACGCTCTTCCGATCTNNNAATGGCCTAGAGCTAGACATG
GCACACGTCTGAACTCTTTCCCTACACGACGCTCTTCCGATCTNNNTTCACCCAGTTTTCCCCAATAGT
AGCACACGTCTGAACTCTTTCCCTACACGACGCTCTTCCGATCTNNNCTATGTAGCAGGCACAGGAGA
GCACACGTCTGAACTCTTTCCCTACACGACGCTCTTCCGATCTNNNATTTCCAGTCTCGCCATTATCCA
AGCACACGTCTGAACTCTTTCCCTACACGACGCTCTTCCGATCTNNNCGGGGAAAGAAAGTCAAAGG
GAGCACACGTCTGAACTCTTTCCCTACACGACGCTCTTCCGATCTNNNTATACACACACACACACACAC
GAGCACACGTCTGAACTCTTTCCCTACACGACGCTCTTCCGATCTNNNATAGCAAACACATTGGCACA
AGCACACGTCTGAACTCTTTCCCTACACGACGCTCTTCCGATCTNNNGCTGTCACTGATCTGTTTCACA
AGCACACGTCTGAACTCTTTCCCTACACGACGCTCTTCCGATCTNNNTCTGATTTCTCCACTTGCTCAG
AGCACACGTCTGAACTCTTTCCCTACACGACGCTCTTCCGATCTNNNTCTTACTTTGGGGCATCTTGTG
AGCACACGTCTGAACTCTTTCCCTACACGACGCTCTTCCGATCTNNNTATGATGAGCATAGTGAGCAC
AGCACACGTCTGAACTCTTTCCCTACACGACGCTCTTCCGATCTNNNGCACTTTGTTTTCACACCAATG
GAGCACACGTCTGAACTCTTTCCCTACACGACGCTCTTCCGATCTNNNATCCCTAAGACAGAGGCAAA
AGCACACGTCTGAACTCTTTCCCTACACGACGCTCTTCCGATCTNNNTCTGGGTATGCCTCTGATCATT
AGCACACGTCTGAACTCTTTCCCTACACGACGCTCTTCCGATCTNNNTGTGGGGTTGTTTTTTCGTGT
AGCACACGTCTGAACTCTTTCCCTACACGACGCTCTTCCGATCTNNNAGCAGCACTATTCACAATAGCC
AGCACACGTCTGAACTCTTTCCCTACACGACGCTCTTCCGATCTNNNTCCTTCCTTCTTTCCTTCCTTCCC
AGCACACGTCTGAACTCTTTCCCTACACGACGCTCTTCCGATCTNNNGAGACAGAAGCCAAGAGATT
GAGCACACGTCTGAACTCTTTCCCTACACGACGCTCTTCCGATCTNNNGGTTACCAAGCAAAGACTG
AGCACACGTCTGAACTCTTTCCCTACACGACGCTCTTCCGATCTNNNACCATGTCATCCTTCCCTTTCTC
AGCACACGTCTGAACTCTTTCCCTACACGACGCTCTTCCGATCTNNNACCATGTCATCCTTCCCTTTCT
AGCACACGTCTGAACTCTTTCCCTACACGACGCTCTTCCGATCTNNNACAATCATAAGTCACAGGAGG
AGCACACGTCTGAACTCTTTCCCTACACGACGCTCTTCCGATCTNNNTTTGGTGTGCATTCTTTGTGTG
AGCACACGTCTGAACTCTTTCCCTACACGACGCTCTTCCGATCTNNNGTTCCTCTTTTGGTGTGCATTCT
AGCACACGTCTGAACTCTTTCCCTACACGACGCTCTTCCGATCTNNNTCTCTGTCTATGTCTGTGTGCA
AGCACACGTCTGAACTCTTTCCCTACACGACGCTCTTCCGATCTNNNTGGGATCGATCTCTCTCTCTCTC
AGCACACGTCTGAACTCTTTCCCTACACGACGCTCTTCCGATCTNNNGCTTTGCCATAAATTGGGATCG
GAGCACACGTCTGAACTCTTTCCCTACACGACGCTCTTCCGATCTNNNCTCCTTTCCTTCACCTTCTCCTC
GAGCACACGTCTGAACTCTTTCCCTACACGACGCTCTTCCGATCTNNNACACATAGGATTGTTGAGGG
GAGCACACGTCTGAACTCTTTCCCTACACGACGCTCTTCCGATCTNNNCCTGGAAAGGCAACCAAACT

GAGCACACGTCTGAACTCTTTCCCTACACGACGCTCTTCCGATCTNNNTGGTGCATACAGAGAGAGA
AGAGCACACGTCTGAACTCTTTCCCTACACGACGCTCTTCCGATCTNNNTCTCTTCAGCTTTTGTGC
AGCACACGTCTGAACTCTTTCCCTACACGACGCTCTTCCGATCTNNNATAGTGTGCTAGTGTGTGTG
GAGCACACGTCTGAACTCTTTCCCTACACGACGCTCTTCCGATCTNNNAGCCCTTTACCCAAGATAGTG
AGCACACGTCTGAACTCTTTCCCTACACGACGCTCTTCCGATCTNNNTTGATTTAGCGCCAATACTGCA
GAGCACACGTCTGAACTCTTTCCCTACACGACGCTCTTCCGATCTNNNCCGTCTCAAATATAAGGGGC
AGCACACGTCTGAACTCTTTCCCTACACGACGCTCTTCCGATCTNNNCAGGGATACTTGTGTCACTCC
GCACACGTCTGAACTCTTTCCCTACACGACGCTCTTCCGATCTNNNCATGAATGCAGAACTCACAGCTA
GAGCACACGTCTGAACTCTTTCCCTACACGACGCTCTTCCGATCTNNNCTCAATTTTGGAGAAGCAGC
AGCACACGTCTGAACTCTTTCCCTACACGACGCTCTTCCGATCTNNNTCTCTCTCTCTCTGTGTGTG
AGCACACGTCTGAACTCTTTCCCTACACGACGCTCTTCCGATCTNNNCTGCCTCCACGTCATTATCTCT
AGCACACGTCTGAACTCTTTCCCTACACGACGCTCTTCCGATCTNNNCTACAGCACAACAAAGACAACG
AGCACACGTCTGAACTCTTTCCCTACACGACGCTCTTCCGATCTNNNTCCTGTCCTCCTCTTTTCTCTCTC
AGAGCACACGTCTGAACTCTTTCCCTACACGACGCTCTTCCGATCTNNNTTATTCCCACTTGCCAACTCT
AGCACACGTCTGAACTCTTTCCCTACACGACGCTCTTCCGATCTNNNAGCTTCTCTCAGTCTCTCTCAG
GAGCACACGTCTGAACTCTTTCCCTACACGACGCTCTTCCGATCTNNNATACACACACATCCTCCCAAGT
CACACGTCTGAACTCTTTCCCTACACGACGCTCTTCCGATCTNNNAAATTAGCCAAGCATGATGGTGG
AGCACACGTCTGAACTCTTTCCCTACACGACGCTCTTCCGATCTNNNGGCCTGAGTCTTTCATCAAGAG
GCACACGTCTGAACTCTTTCCCTACACGACGCTCTTCCGATCTNNNCCAGGTTGCTGCAAATGTCATT
GAGCACACGTCTGAACTCTTTCCCTACACGACGCTCTTCCGATCTNNNTTACACATTCTGAGACCTGGC
AGCACACGTCTGAACTCTTTCCCTACACGACGCTCTTCCGATCTNNNGAAAAGCATGTTCACACACAC
GAGCACACGTCTGAACTCTTTCCCTACACGACGCTCTTCCGATCTNNNATTCCTCTTTTCAGACCGGAT
AGCACACGTCTGAACTCTTTCCCTACACGACGCTCTTCCGATCTNNNGCATGCTATCAGAGATCCCC
AGCACACGTCTGAACTCTTTCCCTACACGACGCTCTTCCGATCTNNNTTTGGAAAGCCCACAGTACAA
GAGCACACGTCTGAACTCTTTCCCTACACGACGCTCTTCCGATCTNNNCTTGTCTCCCATCTTACCACAC
GCACACGTCTGAACTCTTTCCCTACACGACGCTCTTCCGATCTNNNTTCCACTACTTTGTCTCTGACCCC
AGCACACGTCTGAACTCTTTCCCTACACGACGCTCTTCCGATCTNNNTAACTACCTGTTGCCTTCTTCCC
GAGCACACGTCTGAACTCTTTCCCTACACGACGCTCTTCCGATCTNNNGTCTCTCCATAAGGGCATCTA
GCACACGTCTGAACTCTTTCCCTACACGACGCTCTTCCGATCTNNNTCTTCTTTCCTCCCTCTTTCTGCCA
AGCACACGTCTGAACTCTTTCCCTACACGACGCTCTTCCGATCTNNNATGACTCATGTCTGCATCTCTG
GCACACGTCTGAACTCTTTCCCTACACGACGCTCTTCCGATCTNNNGTAACTCCACGTAGAAGCCTCAT
GAGCACACGTCTGAACTCTTTCCCTACACGACGCTCTTCCGATCTNNNTTGCTTCCCTCTTTCCCTCTTA
GCACACGTCTGAACTCTTTCCCTACACGACGCTCTTCCGATCTNNNAAATCCATGCTTCAGAAGGCCT
GCACACGTCTGAACTCTTTCCCTACACGACGCTCTTCCGATCTNNNTTCCACTTGGTCTTCAGAGTCA
AGCACACGTCTGAACTCTTTCCCTACACGACGCTCTTCCGATCTNNNGACTTCTGGCTCACCACTAATC
GCACACGTCTGAACTCTTTCCCTACACGACGCTCTTCCGATCTNNNTCTCTATTTCTCTCCCTCCCTCCC
GCACACGTCTGAACTCTTTCCCTACACGACGCTCTTCCGATCTNNNTTCATGGTAAGTGCCCTCCTCTA
AGCACACGTCTGAACTCTTTCCCTACACGACGCTCTTCCGATCTNNNTTGCTGACAATCTTTGGTGTT
AGAGCACACGTCTGAACTCTTTCCCTACACGACGCTCTTCCGATCTNNNTAGGGAAAACACCACAAAA
GCACACGTCTGAACTCTTTCCCTACACGACGCTCTTCCGATCTNNNTGTGCCTCCTTCCTGAATTCTAG
AGCACACGTCTGAACTCTTTCCCTACACGACGCTCTTCCGATCTNNNCTTCAGGACTAACTCCATCTGG
GCACACGTCTGAACTCTTTCCCTACACGACGCTCTTCCGATCTNNNATTTGTGTGTGAGTGTGTGTGT
GCACACGTCTGAACTCTTTCCCTACACGACGCTCTTCCGATCTNNNCAGATGTCCTCTGCTTTTCATGT

AGCACACGTCTGAACTCTTTCCCTACACGACGCTCTTCCGATCTNNNTCCCCTGTCTACTGTTGTGATT
GCACACGTCTGAACTCTTTCCCTACACGACGCTCTTCCGATCTNNNTTCCTTCCTTCCTTCCTTCCTTCCA
GCACACGTCTGAACTCTTTCCCTACACGACGCTCTTCCGATCTNNNAGGAGGACAAATAAGAAGTG
AGCACACGTCTGAACTCTTTCCCTACACGACGCTCTTCCGATCTNNNTGAACAGCTTCTCCTCATCTTCC
GCACACGTCTGAACTCTTTCCCTACACGACGCTCTTCCGATCTNNNTCACAAGGACTGAAATACTGGG
AGCACACGTCTGAACTCTTTCCCTACACGACGCTCTTCCGATCTNNNAGAGTCCCATCTCAGTTACCTGA
GCACACGTCTGAACTCTTTCCCTACACGACGCTCTTCCGATCTNNNCCTCTCTCAAAGCTTCCCTCTTTCA
AGCACACGTCTGAACTCTTTCCCTACACGACGCTCTTCCGATCTNNNCTCTTGACCACTCGACTTCCTA
GCACACGTCTGAACTCTTTCCCTACACGACGCTCTTCCGATCTNNNCCTCTCTTTCTCTCCATTGCCTTCA
AGCACACGTCTGAACTCTTTCCCTACACGACGCTCTTCCGATCTNNNACAAGTCAGGGGTTACTGTGT
GCACACGTCTGAACTCTTTCCCTACACGACGCTCTTCCGATCTNNNGGCTTGCTTTTGAGATTTCTG
AGCACACGTCTGAACTCTTTCCCTACACGACGCTCTTCCGATCTNNNTTAGCCATGATGAAAGTCCAT
AGCACACGTCTGAACTCTTTCCCTACACGACGCTCTTCCGATCTNNNTTCTTCCAGTCAGAAACCACCA
AGCACACGTCTGAACTCTTTCCCTACACGACGCTCTTCCGATCTNNNCCTATCTCTGGTATCCAAACGC
GCACACGTCTGAACTCTTTCCCTACACGACGCTCTTCCGATCTNNNATATCCCCAAGAACTTGCACAGT
GAGCACACGTCTGAACTCTTTCCCTACACGACGCTCTTCCGATCTNNNACAGTCTCGGTACAGTACAC
AGCACACGTCTGAACTCTTTCCCTACACGACGCTCTTCCGATCTNNNCACACACACACACACATACT
AGCACACGTCTGAACTCTTTCCCTACACGACGCTCTTCCGATCTNNNCCCTTTGGGCTCAGAAAAAAC
AGCACACGTCTGAACTCTTTCCCTACACGACGCTCTTCCGATCTNNNGAATTTTGAAGTGGTGGCAG
GAGCACACGTCTGAACTCTTTCCCTACACGACGCTCTTCCGATCTNNNATGGTGCCTTCATGATGGGA
AGCACACGTCTGAACTCTTTCCCTACACGACGCTCTTCCGATCTNNNCAGAACCTTGTGGAGATAGCC
AGCACACGTCTGAACTCTTTCCCTACACGACGCTCTTCCGATCTNNNAGCCAACAGTGCCATATAGTG
AGCACACGTCTGAACTCTTTCCCTACACGACGCTCTTCCGATCTNNNCTGGTAGCTGTGCCCAATAAA
GAGCACACGTCTGAACTCTTTCCCTACACGACGCTCTTCCGATCTNNNAAGTTTCCGTTGCATCTTGGT
GCACACGTCTGAACTCTTTCCCTACACGACGCTCTTCCGATCTNNNGGACATGGGGGACTTGAAGATA
GCACACGTCTGAACTCTTTCCCTACACGACGCTCTTCCGATCTNNNTTGTCTCTACAGGTCAGTGAAG
GCACACGTCTGAACTCTTTCCCTACACGACGCTCTTCCGATCTNNNTTTTGTTGCTGCTTCAAAGACCA
GAGCACACGTCTGAACTCTTTCCCTACACGACGCTCTTCCGATCTNNNGCAGAGAGGGGAGAGAAAG
GAGCACACGTCTGAACTCTTTCCCTACACGACGCTCTTCCGATCTNNNCCCATAGGATTTCTTGTTGCT
AGCACACGTCTGAACTCTTTCCCTACACGACGCTCTTCCGATCTNNNAATCATATTGGCTGCAGCAAA
AGCACACGTCTGAACTCTTTCCCTACACGACGCTCTTCCGATCTNNNGCCTGAAACTCACACAATCAAC
GCACACGTCTGAACTCTTTCCCTACACGACGCTCTTCCGATCTNNNGCAAGACTTGAAGAACACACACA
CACACGTCTGAACTCTTTCCCTACACGACGCTCTTCCGATCTNNNCTCTCCTTCTCCCTTTTTCCTCCCAT
CACACGTCTGAACTCTTTCCCTACACGACGCTCTTCCGATCTNNNCCTCTCCTTCTCCCTTTTTCCTCCAT
AGCACACGTCTGAACTCTTTCCCTACACGACGCTCTTCCGATCTNNNAAAACTGAAGCATCAAAGCCA
GAGCACACGTCTGAACTCTTTCCCTACACGACGCTCTTCCGATCTNNNATCTTCTCTGTCCCTATCCGAC
GAGCACACGTCTGAACTCTTTCCCTACACGACGCTCTTCCGATCTNNNATGGCAAAAGAGACAATCAG
GAGCACACGTCTGAACTCTTTCCCTACACGACGCTCTTCCGATCTNNNGACCTCAAGACCAAATGGAA
AGCACACGTCTGAACTCTTTCCCTACACGACGCTCTTCCGATCTNNNAGCTGTAGTGTGCAAGATGA
GCACACGTCTGAACTCTTTCCCTACACGACGCTCTTCCGATCTNNNTATGTGGTGTGAATGTGGGTA
GCACACGTCTGAACTCTTTCCCTACACGACGCTCTTCCGATCTNNNACTTAGTGCTTCTCTTGTCCCAG
GAGCACACGTCTGAACTCTTTCCCTACACGACGCTCTTCCGATCTNNNTAAGATGGCTGTTTGCTTCAC
AGCACACGTCTGAACTCTTTCCCTACACGACGCTCTTCCGATCTNNNATGAATTTCTGTTTCCTGCAGC

AGCACACGTCTGAACTCTTTCCCTACACGACGCTCTTCCGATCTNNNATTGGATTGAATAACACTGGCC
AGCACACGTCTGAACTCTTTCCCTACACGACGCTCTTCCGATCTNNNTACAAAGAACCCCTGCAAGATG
AGCACACGTCTGAACTCTTTCCCTACACGACGCTCTTCCGATCTNNNCCAAACTCTGCAGCTTTGAAGAG
AGCACACGTCTGAACTCTTTCCCTACACGACGCTCTTCCGATCTNNNCCTCCCTTTAGCCACAGTATCAAC
GCACACGTCTGAACTCTTTCCCTACACGACGCTCTTCCGATCTNNNCACTGCCTAAATCATCCCTTCCTCA
GAGCACACGTCTGAACTCTTTCCCTACACGACGCTCTTCCGATCTNNNGAGGTTTCTCTCTGTGTCTCTC
GCACACGTCTGAACTCTTTCCCTACACGACGCTCTTCCGATCTNNNTCCTTGGACCTCTCATCACCTATCA
AGCACACGTCTGAACTCTTTCCCTACACGACGCTCTTCCGATCTNNNCTGCTCAGTTACCCAACATGTTT
AGCACACGTCTGAACTCTTTCCCTACACGACGCTCTTCCGATCTNNNCAAGTCCTAGCCCAATGTTTGTG
GCACACGTCTGAACTCTTTCCCTACACGACGCTCTTCCGATCTNNNAGTCTCTTCATCTCACTACCCCTCA
AGCACACGTCTGAACTCTTTCCCTACACGACGCTCTTCCGATCTNNNCAGAAGTGGAGAAGGATGGTC
AGCACACGTCTGAACTCTTTCCCTACACGACGCTCTTCCGATCTNNNCATGAGAACTGAAACTGCCCAA
GAGCACACGTCTGAACTCTTTCCCTACACGACGCTCTTCCGATCTNNNTGTTTTCCCAAGAATGTCACCA
AGCACACGTCTGAACTCTTTCCCTACACGACGCTCTTCCGATCTNNNATTGGTATCAGGCTCCAAGAAC
AGCACACGTCTGAACTCTTTCCCTACACGACGCTCTTCCGATCTNNNACAGTGCCTGACAAGTTGCTAT
AGCACACGTCTGAACTCTTTCCCTACACGACGCTCTTCCGATCTNNNCACAACACCAGAGGCTGAAAAA
AGCACACGTCTGAACTCTTTCCCTACACGACGCTCTTCCGATCTNNNCTTGTTCTGTTGCTGCCTTTCTTC
AGCACACGTCTGAACTCTTTCCCTACACGACGCTCTTCCGATCTNNNTTTGTCTTGGTTTGAAGATGGCC
AGCACACGTCTGAACTCTTTCCCTACACGACGCTCTTCCGATCTNNNTCATAGGCAATGGCATTTTGTAA
AGAGCACACGTCTGAACTCTTTCCCTACACGACGCTCTTCCGATCTNNNTATGTGTGTGTGTGTGTGTG
GAGCACACGTCTGAACTCTTTCCCTACACGACGCTCTTCCGATCTNNNAAATAGGCCCCTTTGTGAGTT
AGCACACGTCTGAACTCTTTCCCTACACGACGCTCTTCCGATCTNNNCCTGTAGCATGCCCTCCATATAG
GAGCACACGTCTGAACTCTTTCCCTACACGACGCTCTTCCGATCTNNNTGTCTTCTCACTCAGAGGTTG
GAGCACACGTCTGAACTCTTTCCCTACACGACGCTCTTCCGATCTNNNCCATGGTGGAAGTAGATGTGT
GAGCACACGTCTGAACTCTTTCCCTACACGACGCTCTTCCGATCTNNNCTGTGCAAATGCGTGTGTGTA
GAGCACACGTCTGAACTCTTTCCCTACACGACGCTCTTCCGATCTNNNAACAGGAAATATGGCTTCCACC
GAGCACACGTCTGAACTCTTTCCCTACACGACGCTCTTCCGATCTNNNCTTCCTTCCTTCCTTCCTTCCTC
AGCACACGTCTGAACTCTTTCCCTACACGACGCTCTTCCGATCTNNNTGAGAGTGTCTCTAGGATGTG
AGCACACGTCTGAACTCTTTCCCTACACGACGCTCTTCCGATCTNNNTAAAAAACATCCCCGTTTGGCAA
AGCACACGTCTGAACTCTTTCCCTACACGACGCTCTTCCGATCTNNNTTCAGTTGGGACGTAATAAGG
GAGCACACGTCTGAACTCTTTCCCTACACGACGCTCTTCCGATCTNNNTGTACCTTGCCCCATTCATACAA
GAGCACACGTCTGAACTCTTTCCCTACACGACGCTCTTCCGATCTNNNAAAAACATGGTGGCTTCTGTC
GCACACGTCTGAACTCTTTCCCTACACGACGCTCTTCCGATCTNNNCTTCTCCTTGGAAATGGACTGGA
AGCACACGTCTGAACTCTTTCCCTACACGACGCTCTTCCGATCTNNNGAACTGCATAGGAATACACGCA
AGCACACGTCTGAACTCTTTCCCTACACGACGCTCTTCCGATCTNNNCTGTGGTTTTTGCAGACCCATA
GAGCACACGTCTGAACTCTTTCCCTACACGACGCTCTTCCGATCTNNNTTGGCTGTATGTGAGATAGCT
AGCACACGTCTGAACTCTTTCCCTACACGACGCTCTTCCGATCTNNNACTGTGGTCTATAGATCCCTGG
AGCACACGTCTGAACTCTTTCCCTACACGACGCTCTTCCGATCTNNNGCCAAAAGCAGTCTAAAGGCAT
AGAGCACACGTCTGAACTCTTTCCCTACACGACGCTCTTCCGATCTNNNCCTGGAAAGATGAGTGGGA
CACACGTCTGAACTCTTTCCCTACACGACGCTCTTCCGATCTNNNTGCTAAGCATGGGGAGAGAATTT
GAGCACACGTCTGAACTCTTTCCCTACACGACGCTCTTCCGATCTNNNGGGAAAGATCTCTCCCTCTCTC
GAGCACACGTCTGAACTCTTTCCCTACACGACGCTCTTCCGATCTNNNACTCCTGAAACAGATGGGAAA
GAGCACACGTCTGAACTCTTTCCCTACACGACGCTCTTCCGATCTNNNGCATCATGCCAAGTGATAAGA

AGCACACGTCTGAACTCTTTCCCTACACGACGCTCTTCCGATCTNNNGTTGCAAAAGATGATGGTGGT
GCACACGTCTGAACTCTTTCCCTACACGACGCTCTTCCGATCTNNNTCTCTCTCTCTCTTTCCCTCCCAT
AGAGCACACGTCTGAACTCTTTCCCTACACGACGCTCTTCCGATCTNNNACATGGCAATATCATGCTTTC
GAGCACACGTCTGAACTCTTTCCCTACACGACGCTCTTCCGATCTNNNGTATGTGTGTGTGTGTGTGTC
GAGCACACGTCTGAACTCTTTCCCTACACGACGCTCTTCCGATCTNNNAGAAGTGTGTGTGTGTGTGT
AGCACACGTCTGAACTCTTTCCCTACACGACGCTCTTCCGATCTNNNCATTTTCCCTCCAGGTGATAGCT
AGAGCACACGTCTGAACTCTTTCCCTACACGACGCTCTTCCGATCTNNNTAAGGACAGTCTCAATGGTC
AGCACACGTCTGAACTCTTTCCCTACACGACGCTCTTCCGATCTNNNATGTTGAAACACTTTAGGGGCA
GAGCACACGTCTGAACTCTTTCCCTACACGACGCTCTTCCGATCTNNNCACCTGGCCTCGTTAGTCTTTT
AGCACACGTCTGAACTCTTTCCCTACACGACGCTCTTCCGATCTNNNGGAGGACACATACCATCTCTGTC
AGCACACGTCTGAACTCTTTCCCTACACGACGCTCTTCCGATCTNNNTTGTTCAGGCCTTAGCATAGCA
AGCACACGTCTGAACTCTTTCCCTACACGACGCTCTTCCGATCTNNNCACTGGAATTTGGAGGAGCAAA
GAGCACACGTCTGAACTCTTTCCCTACACGACGCTCTTCCGATCTNNNCTCTTGGTACATCAGAGGTGCT
AGCACACGTCTGAACTCTTTCCCTACACGACGCTCTTCCGATCTNNNAGTTACAACTGACAGGGAAGG
GAGCACACGTCTGAACTCTTTCCCTACACGACGCTCTTCCGATCTNNNTTTGATCTGGGTGGTGGTTA
GAGCACACGTCTGAACTCTTTCCCTACACGACGCTCTTCCGATCTNNNAGCTCGGAATGCTACTGATCA
GAGCACACGTCTGAACTCTTTCCCTACACGACGCTCTTCCGATCTNNNCATTCTCTGCTCCTCTCGATGAT
AGCACACGTCTGAACTCTTTCCCTACACGACGCTCTTCCGATCTNNNAATCTTCCACCTCCACGTTCAATC
AGCACACGTCTGAACTCTTTCCCTACACGACGCTCTTCCGATCTNNNTCAATATTTGGTTCCAGGAGGC
AGCACACGTCTGAACTCTTTCCCTACACGACGCTCTTCCGATCTNNNCACACACACACACACACATACAT
AGCACACGTCTGAACTCTTTCCCTACACGACGCTCTTCCGATCTNNNATTTTGTGCCTATTCCAGTTGCC
GAGCACACGTCTGAACTCTTTCCCTACACGACGCTCTTCCGATCTNNNTGGAAAGTGGGAAGAACAGA
AGCACACGTCTGAACTCTTTCCCTACACGACGCTCTTCCGATCTNNNCTGGGATTCTGTTCTGAGGTTCA
AGCACACGTCTGAACTCTTTCCCTACACGACGCTCTTCCGATCTNNNACCTGTAAATTGGGATTGCTCCT
AGCACACGTCTGAACTCTTTCCCTACACGACGCTCTTCCGATCTNNNGTCTCAGAATGGAAAGGAGCAC
AGAGCACACGTCTGAACTCTTTCCCTACACGACGCTCTTCCGATCTNNNAAACACTATTCACGGGTACTC
AGAGCACACGTCTGAACTCTTTCCCTACACGACGCTCTTCCGATCTNNNGTGAATGAGAGCTTCCCAAA
AGCACACGTCTGAACTCTTTCCCTACACGACGCTCTTCCGATCTNNNAGCATGAAAAGCTACCAGACTC
AGCACACGTCTGAACTCTTTCCCTACACGACGCTCTTCCGATCTNNNCCTTCTAACCTCTTCAGCCCAATC
GCACACGTCTGAACTCTTTCCCTACACGACGCTCTTCCGATCTNNNAACTGGGGAGCTGCCTAAAAATA
GAGCACACGTCTGAACTCTTTCCCTACACGACGCTCTTCCGATCTNNNTGCAGCTGTGAAAAGAAATGA
AGAGCACACGTCTGAACTCTTTCCCTACACGACGCTCTTCCGATCTNNNGACTCCATCTCTCCCTCTCTCT
AAGAGCACACGTCTGAACTCTTTCCCTACACGACGCTCTTCCGATCTNNNGTTGCAGTGAGATGAGATC
AGCACACGTCTGAACTCTTTCCCTACACGACGCTCTTCCGATCTNNNCCTACCACAAACACCTCTCTCTCCA
GAGCACACGTCTGAACTCTTTCCCTACACGACGCTCTTCCGATCTNNNTCGGTGTTGGAGAGAAATTC
AGCACACGTCTGAACTCTTTCCCTACACGACGCTCTTCCGATCTNNNTGGAGGAAGACAAGAAAACAC
AGCACACGTCTGAACTCTTTCCCTACACGACGCTCTTCCGATCTNNNCCTGACATTTGTGGCAATCAACT
GAGCACACGTCTGAACTCTTTCCCTACACGACGCTCTTCCGATCTNNNGGGTAGCACCTTTCCCTTTAT
GAGCACACGTCTGAACTCTTTCCCTACACGACGCTCTTCCGATCTNNNCCAATTGCAAACAGATCAGGT
AGCACACGTCTGAACTCTTTCCCTACACGACGCTCTTCCGATCTNNNGTTCCTGTAGTCCCAGCTACTTG
AGCACACGTCTGAACTCTTTCCCTACACGACGCTCTTCCGATCTNNNATCTTTGGATACCAGAACCCCAC
AGCACACGTCTGAACTCTTTCCCTACACGACGCTCTTCCGATCTNNNTGCAGAGGGTGTGTTCCTATAT
AGAGCACACGTCTGAACTCTTTCCCTACACGACGCTCTTCCGATCTNNNCAGTTTGCTCACTTCTTTGGC

AGCACACGTCTGAACTCTTTCCCTACACGACGCTCTTCCGATCTNNNACGAGACCCCATCACTACAAA
AGAGCACACGTCTGAACTCTTTCCCTACACGACGCTCTTCCGATCTNNNTGGAGATGTCAGAAAAGC
AGCACACGTCTGAACTCTTTCCCTACACGACGCTCTTCCGATCTNNNTATGCAATAACTCTGGAAGCAG
GAGCACACGTCTGAACTCTTTCCCTACACGACGCTCTTCCGATCTNNNTCCGTCCTAGTTGGGTTTTGA
GAGCACACGTCTGAACTCTTTCCCTACACGACGCTCTTCCGATCTNNNCCATGTGGTCGTTAGAGACA
GCACACGTCTGAACTCTTTCCCTACACGACGCTCTTCCGATCTNNNACATAGTTTTGAGCCCCACAGAT
AGCACACGTCTGAACTCTTTCCCTACACGACGCTCTTCCGATCTNNNCCCCATAGAACATGGAATTGGG
AGAGCACACGTCTGAACTCTTTCCCTACACGACGCTCTTCCGATCTNNNGTGTCACTTGTCCCCATAGA
AGCACACGTCTGAACTCTTTCCCTACACGACGCTCTTCCGATCTNNNCCCACTGATGTAGTCCATGTAG
GAGCACACGTCTGAACTCTTTCCCTACACGACGCTCTTCCGATCTNNNGAGTATCATAAGGGACCCTG
GAGCACACGTCTGAACTCTTTCCCTACACGACGCTCTTCCGATCTNNNATTCACTCTGAGACCCTATCC
GAGCACACGTCTGAACTCTTTCCCTACACGACGCTCTTCCGATCTNNNAAAATGCAGACAGACACACAC
AGAGCACACGTCTGAACTCTTTCCCTACACGACGCTCTTCCGATCTNNNAGGGAGTCAGTAAGAGAC
AGCACACGTCTGAACTCTTTCCCTACACGACGCTCTTCCGATCTNNNTTTCTTCAAGGAGCCCTAGTTC
GAGCACACGTCTGAACTCTTTCCCTACACGACGCTCTTCCGATCTNNNGCACAAATGAAACATCTCTCC
AGCACACGTCTGAACTCTTTCCCTACACGACGCTCTTCCGATCTNNNGTGGTAAGTCATGTATGTTGC
AGCACACGTCTGAACTCTTTCCCTACACGACGCTCTTCCGATCTNNNCATTGAGTTGGTGTGTGTGTG
AGAGCACACGTCTGAACTCTTTCCCTACACGACGCTCTTCCGATCTNNNTTGTTTCCATGGCTTCGTGA
GAGCACACGTCTGAACTCTTTCCCTACACGACGCTCTTCCGATCTNNNACTTGCCCCCACAATTTTCTTT
GAGCACACGTCTGAACTCTTTCCCTACACGACGCTCTTCCGATCTNNNCATCTGTGTGGAGGTTACTG
GAGCACACGTCTGAACTCTTTCCCTACACGACGCTCTTCCGATCTNNNAAAACAGAAACAACAACCAGG
AGAGCACACGTCTGAACTCTTTCCCTACACGACGCTCTTCCGATCTNNNTTCAGTTCAAGTGGTGTGT
AGCACACGTCTGAACTCTTTCCCTACACGACGCTCTTCCGATCTNNNTTGTGTCAGCAGGAGAATCAG
AGCACACGTCTGAACTCTTTCCCTACACGACGCTCTTCCGATCTNNNAGAAACTCAGCTCCAAAATGG
AGCACACGTCTGAACTCTTTCCCTACACGACGCTCTTCCGATCTNNNACATATTGCTCATGGGAGAAG
GAGCACACGTCTGAACTCTTTCCCTACACGACGCTCTTCCGATCTNNNTTCCCCTTTTCTTGACCATCCA
GCACACGTCTGAACTCTTTCCCTACACGACGCTCTTCCGATCTNNNGTGGCCTACTAGGAAGTGTTTG
AGCACACGTCTGAACTCTTTCCCTACACGACGCTCTTCCGATCTNNNGGCCTTTGAGTCTACCTTTCTG
GCACACGTCTGAACTCTTTCCCTACACGACGCTCTTCCGATCTNNNTGTGGATCAAAACACGTACCTCA
AGCACACGTCTGAACTCTTTCCCTACACGACGCTCTTCCGATCTNNNGATATCCTCTATTTTGCTGGGG
GCACACGTCTGAACTCTTTCCCTACACGACGCTCTTCCGATCTNNNACGCTTATGATTGCATCACATGG
GAGCACACGTCTGAACTCTTTCCCTACACGACGCTCTTCCGATCTNNNGTAATCCTAGCTACTTGGGAG
AGCACACGTCTGAACTCTTTCCCTACACGACGCTCTTCCGATCTNNNTTTTTCTCACAGTTCTGGAGGC
GAGCACACGTCTGAACTCTTTCCCTACACGACGCTCTTCCGATCTNNNAAAAAGGTCTATCCCCAGCA
GAGCACACGTCTGAACTCTTTCCCTACACGACGCTCTTCCGATCTNNNATCATCTTAGGCAGAAGAACG
AGCACACGTCTGAACTCTTTCCCTACACGACGCTCTTCCGATCTNNNCTAGACCCCCACATTGAGTCAAT
GAGCACACGTCTGAACTCTTTCCCTACACGACGCTCTTCCGATCTNNNCTAAAGTTGAAACAGGCCTTG
GAGCACACGTCTGAACTCTTTCCCTACACGACGCTCTTCCGATCTNNNATAACGCAAGGAAGGGAGA
AGCACACGTCTGAACTCTTTCCCTACACGACGCTCTTCCGATCTNNNTGTTCTGCAGGGAACATATGTG
AGCACACGTCTGAACTCTTTCCCTACACGACGCTCTTCCGATCTNNNCCTTTGGTTCCCAGATCCATGTA
GAGCACACGTCTGAACTCTTTCCCTACACGACGCTCTTCCGATCTNNNTGGGCGTTGCTTGTCTCATAA
GAGCACACGTCTGAACTCTTTCCCTACACGACGCTCTTCCGATCTNNNAAGCAAGGAAGAACTCAAGC
GAGCACACGTCTGAACTCTTTCCCTACACGACGCTCTTCCGATCTNNNTCAAATCTTGTGTTTGCTAC

AGCACACGTCTGAACTCTTTCCCTACACGACGCTCTTCCGATCTNNNCCAGGCAAGGAAGAAGAAAA
AGCACACGTCTGAACTCTTTCCCTACACGACGCTCTTCCGATCTNNNTCCCTCTTTGTCTCTTCCTCTCTC
GCACACGTCTGAACTCTTTCCCTACACGACGCTCTTCCGATCTNNNAGGTGTCACAGGAGGAGATCT
CACACGTCTGAACTCTTTCCCTACACGACGCTCTTCCGATCTNNNATGTGAAATGCAAAGAGAAGCC
AGCACACGTCTGAACTCTTTCCCTACACGACGCTCTTCCGATCTNNNTTCTCAACCAGGAGGGATTTT
AGCACACGTCTGAACTCTTTCCCTACACGACGCTCTTCCGATCTNNNATTTGGTAGAGAGAAAGAGG
GAGCACACGTCTGAACTCTTTCCCTACACGACGCTCTTCCGATCTNNNTACACCACTTTCTCAGCAGGAT
GAGCACACGTCTGAACTCTTTCCCTACACGACGCTCTTCCGATCTNNNACCAGCATCTAGTGGGTAGT
AGCACACGTCTGAACTCTTTCCCTACACGACGCTCTTCCGATCTNNNTTACTGAATGTTAGCTGGTCCC
AGCACACGTCTGAACTCTTTCCCTACACGACGCTCTTCCGATCTNNNACCATCAAGTAGTTGTGTGAC
AGAGCACACGTCTGAACTCTTTCCCTACACGACGCTCTTCCGATCTNNNTGGACAAGTAACACCTCTC
AGCACACGTCTGAACTCTTTCCCTACACGACGCTCTTCCGATCTNNNTCCTACTCTCCCTCACAAATCCTC
AGCACACGTCTGAACTCTTTCCCTACACGACGCTCTTCCGATCTNNNTGAATCCGAGGGCTGATAGTA
AGCACACGTCTGAACTCTTTCCCTACACGACGCTCTTCCGATCTNNNGGATGCTAAAAGGCCCATTCT
AGCACACGTCTGAACTCTTTCCCTACACGACGCTCTTCCGATCTNNNTTCTAACGTCCAAGGTCAGGA
GAGCACACGTCTGAACTCTTTCCCTACACGACGCTCTTCCGATCTNNNCGTGAATTTGGAAGTGGAA
GCACACGTCTGAACTCTTTCCCTACACGACGCTCTTCCGATCTNNNCTCTTCCCCTAATTTCCCCAAAC
AGCACACGTCTGAACTCTTTCCCTACACGACGCTCTTCCGATCTNNNTGGGGAGGGAGTATATGGGA
AGCACACGTCTGAACTCTTTCCCTACACGACGCTCTTCCGATCTNNNACAAGAAGCAAAATCAACAGC
AGCACACGTCTGAACTCTTTCCCTACACGACGCTCTTCCGATCTNNNTGAGAGCAGAGACATCTTTCCT
GCACACGTCTGAACTCTTTCCCTACACGACGCTCTTCCGATCTNNNTGAGAGCAGAGACATCTTTCCT
AGCACACGTCTGAACTCTTTCCCTACACGACGCTCTTCCGATCTNNNGGTGAAAAAGAAGGGAAGAC
AGCACACGTCTGAACTCTTTCCCTACACGACGCTCTTCCGATCTNNNATTTACCCTCAGAGCCTGAAG
AGCACACGTCTGAACTCTTTCCCTACACGACGCTCTTCCGATCTNNNAACAGTAAAAGCACTTGCCCA
AGCACACGTCTGAACTCTTTCCCTACACGACGCTCTTCCGATCTNNNCAAGGAAGCTCATGTTTTCCA
AAGAGCACACGTCTGAACTCTTTCCCTACACGACGCTCTTCCGATCTNNNGTCTTTCTCAGACCCCCTCA
GAGCACACGTCTGAACTCTTTCCCTACACGACGCTCTTCCGATCTNNNGAGGGGAGTGAATCAGATC
GAGCACACGTCTGAACTCTTTCCCTACACGACGCTCTTCCGATCTNNNCCCACATTCTTTCCAGTCTCTC
GCACACGTCTGAACTCTTTCCCTACACGACGCTCTTCCGATCTNNNGGCCTCTATTCCTACTGTCATGG
AGCACACGTCTGAACTCTTTCCCTACACGACGCTCTTCCGATCTNNNTCTGAGTCTCAGCCTTACTCTG
GAGCACACGTCTGAACTCTTTCCCTACACGACGCTCTTCCGATCTNNNATGTGCATTGCCTCATGGAA
GAGCACACGTCTGAACTCTTTCCCTACACGACGCTCTTCCGATCTNNNGAATGGCTGAACAAACCCTG
GAGCACACGTCTGAACTCTTTCCCTACACGACGCTCTTCCGATCTNNNCTGAGATTTCTGGAGACACA
AGCACACGTCTGAACTCTTTCCCTACACGACGCTCTTCCGATCTNNNGCCAAAAATACATGTGAGGTG
AGCACACGTCTGAACTCTTTCCCTACACGACGCTCTTCCGATCTNNNGTACATGTACATCCTCAGCCCA
GAGCACACGTCTGAACTCTTTCCCTACACGACGCTCTTCCGATCTNNNGTATCACCTCTTCACATCCCGA
AGAGCACACGTCTGAACTCTTTCCCTACACGACGCTCTTCCGATCTNNNGTCAGGGTGCTTGACCTAA
AGCACACGTCTGAACTCTTTCCCTACACGACGCTCTTCCGATCTNNNCTGATACTGGGCCTCATGCAAT
GAGCACACGTCTGAACTCTTTCCCTACACGACGCTCTTCCGATCTNNNAGGAGAATTGCTTGAACCTG
AGCACACGTCTGAACTCTTTCCCTACACGACGCTCTTCCGATCTNNNAAATTAGCTAGGCATCCTGGTG
AGCACACGTCTGAACTCTTTCCCTACACGACGCTCTTCCGATCTNNNAAGTGGCAGAGCTCTGAATA
GAGCACACGTCTGAACTCTTTCCCTACACGACGCTCTTCCGATCTNNNAATGTGTACCACTCTGCATGG
GAGCACACGTCTGAACTCTTTCCCTACACGACGCTCTTCCGATCTNNNCTAGATGAGCAGCTCTGTTAG

GAGCACACGTCTGAACTCTTTCCCTACACGACGCTCTTCCGATCTNNNTGAGTCTATGTGTGTGCATG
AGCACACGTCTGAACTCTTTCCCTACACGACGCTCTTCCGATCTNNNTTGCTGCAAATTCTAGAAGCCT
GAGCACACGTCTGAACTCTTTCCCTACACGACGCTCTTCCGATCTNNNACCTTCCATTGCCCTCTTTACTT
AGCACACGTCTGAACTCTTTCCCTACACGACGCTCTTCCGATCTNNNACAATCCCACCATTCAGGCATA
GAGCACACGTCTGAACTCTTTCCCTACACGACGCTCTTCCGATCTNNNTTTGTGGTGTGTATGTGTGTG
AGCACACGTCTGAACTCTTTCCCTACACGACGCTCTTCCGATCTNNNCTCTGTGTTGAAGAGGTAGCA
GAGCACACGTCTGAACTCTTTCCCTACACGACGCTCTTCCGATCTNNNCTCTGTGTTGAAGAGGTAGC
AGAGCACACGTCTGAACTCTTTCCCTACACGACGCTCTTCCGATCTNNNGAAAAGACCAGAAGGAATT
AGCACACGTCTGAACTCTTTCCCTACACGACGCTCTTCCGATCTNNNAGAGTTGAGCAATGCCTTGTTT
GCACACGTCTGAACTCTTTCCCTACACGACGCTCTTCCGATCTNNNTTTTCAAAGGTGATGGTGGTG
GAGCACACGTCTGAACTCTTTCCCTACACGACGCTCTTCCGATCTNNNATTGTTCATTTTCTTGGGCGT
AGCACACGTCTGAACTCTTTCCCTACACGACGCTCTTCCGATCTNNNGAGCCAGTCTCTCTTTTCTCCT
AGAGCACACGTCTGAACTCTTTCCCTACACGACGCTCTTCCGATCTNNNTGTGTGTGTGTGTGTGTG
AGAGCACACGTCTGAACTCTTTCCCTACACGACGCTCTTCCGATCTNNNCACACACACACATACACACA
AGCACACGTCTGAACTCTTTCCCTACACGACGCTCTTCCGATCTNNNGATCTTTGCGATGCTTTGGGTT
GCACACGTCTGAACTCTTTCCCTACACGACGCTCTTCCGATCTNNNGCAAAAGTACTTGTCTAGCCTC
GAGCACACGTCTGAACTCTTTCCCTACACGACGCTCTTCCGATCTNNNATTCACAGAACCTTCCAGCAT
GAGCACACGTCTGAACTCTTTCCCTACACGACGCTCTTCCGATCTNNNTCAGGTGGTTTCCATTTGGT
GCACACGTCTGAACTCTTTCCCTACACGACGCTCTTCCGATCTNNNAGGCAGAAGACCTACTTTTCAGG
AGCACACGTCTGAACTCTTTCCCTACACGACGCTCTTCCGATCTNNNGTGTGTGTGTGTGTGTGATTG
GAGCACACGTCTGAACTCTTTCCCTACACGACGCTCTTCCGATCTNNNGCCATGTACTAATCAATGGT
AGCACACGTCTGAACTCTTTCCCTACACGACGCTCTTCCGATCTNNNATAATGCCAGGTGGTGATCAG
GCACACGTCTGAACTCTTTCCCTACACGACGCTCTTCCGATCTNNNTATCCATGGAAGTCAGACCCTCT
AGCACACGTCTGAACTCTTTCCCTACACGACGCTCTTCCGATCTNNNCCCAGAGCAAGCAAAATCAAG
GCACACGTCTGAACTCTTTCCCTACACGACGCTCTTCCGATCTNNNTTTAGGGTCAATGTTTCCTGGG
AGCACACGTCTGAACTCTTTCCCTACACGACGCTCTTCCGATCTNNNGATTCCAGTGCTTCCCTGTTTT
GAGCACACGTCTGAACTCTTTCCCTACACGACGCTCTTCCGATCTNNNTGGGGAATACATGACGTTGA
AGCACACGTCTGAACTCTTTCCCTACACGACGCTCTTCCGATCTNNNAGTGGTCTGCAACTAAACCTG
GAGCACACGTCTGAACTCTTTCCCTACACGACGCTCTTCCGATCTNNNTAACATTCTCTGTCCTTGGGC
GAGCACACGTCTGAACTCTTTCCCTACACGACGCTCTTCCGATCTNNNTGAATGGATCCCTTCTTTGGA
AGAGCACACGTCTGAACTCTTTCCCTACACGACGCTCTTCCGATCTNNNTGTGAGGGACGAATGGAA
GAGCACACGTCTGAACTCTTTCCCTACACGACGCTCTTCCGATCTNNNTGGAAGAGCACAAAAATGA
AGCACACGTCTGAACTCTTTCCCTACACGACGCTCTTCCGATCTNNNGGAGGGAATCACCAGTACATG
AGCACACGTCTGAACTCTTTCCCTACACGACGCTCTTCCGATCTNNNAGTGGCAAGTACACTCAAAAG
GAGCACACGTCTGAACTCTTTCCCTACACGACGCTCTTCCGATCTNNNTTCCTCTTCTTCCTCTTCCTCCT
AGCACACGTCTGAACTCTTTCCCTACACGACGCTCTTCCGATCTNNNGTACTTTCTCTCTCCCCCTCT
GAGCACACGTCTGAACTCTTTCCCTACACGACGCTCTTCCGATCTNNNCACCCCTACTCCTTTTTCTCCA
GAGCACACGTCTGAACTCTTTCCCTACACGACGCTCTTCCGATCTNNNTTTGCCCTCCAGGAGACATT
GAGCACACGTCTGAACTCTTTCCCTACACGACGCTCTTCCGATCTNNNCAGACACTAACTTTCCAAGCT
GAGCACACGTCTGAACTCTTTCCCTACACGACGCTCTTCCGATCTNNNGTATATACCCACAGGCACCTC
GAGCACACGTCTGAACTCTTTCCCTACACGACGCTCTTCCGATCTNNNTCAGGCATTTGGGAAAAGAC
AGAGCACACGTCTGAACTCTTTCCCTACACGACGCTCTTCCGATCTNNNACATAGGAGAGAGACCCCT
GAGCACACGTCTGAACTCTTTCCCTACACGACGCTCTTCCGATCTNNNGCAACAGAGTGAGACCTAGG

GCACACGTCTGAACTCTTTCCCTACACGACGCTCTTCCGATCTNNNTTTTCTGTGCATGTGTGTGTGT
GAGCACACGTCTGAACTCTTTCCCTACACGACGCTCTTCCGATCTNNNTACAAGGGACTGTTATGCTGA
AGCACACGTCTGAACTCTTTCCCTACACGACGCTCTTCCGATCTNNNACAGTGGCTGAGTTGAACAAA
AGCACACGTCTGAACTCTTTCCCTACACGACGCTCTTCCGATCTNNNTTGAGCCTAATCTCTTTGTGGG
AGCACACGTCTGAACTCTTTCCCTACACGACGCTCTTCCGATCTNNNATGTCTGCACATCCTGTACTTGT
GCACACGTCTGAACTCTTTCCCTACACGACGCTCTTCCGATCTNNNGCATAAACTGAAAGGCATGGACA
AGCACACGTCTGAACTCTTTCCCTACACGACGCTCTTCCGATCTNNNCTCTGAACTTTTTCCTTGCTGGG
AGCACACGTCTGAACTCTTTCCCTACACGACGCTCTTCCGATCTNNNACCATCAAACCCAACCCTAATCA
AGCACACGTCTGAACTCTTTCCCTACACGACGCTCTTCCGATCTNNNTCTCAGTCTTACACACACAAGCA
AGCACACGTCTGAACTCTTTCCCTACACGACGCTCTTCCGATCTNNNCTTCCAATAGAGGCTTTGGCTT
GCACACGTCTGAACTCTTTCCCTACACGACGCTCTTCCGATCTNNNCCTTCTCGAGGCCTCATAACTAGC
GAGCACACGTCTGAACTCTTTCCCTACACGACGCTCTTCCGATCTNNNTTTGTTCTCGCAACCTGAAACA
GAGCACACGTCTGAACTCTTTCCCTACACGACGCTCTTCCGATCTNNNTCCACGAGACATCATCAGTGTT
GAGCACACGTCTGAACTCTTTCCCTACACGACGCTCTTCCGATCTNNNTTCTAGCAGGCCTTCTTGTACT
AGCACACGTCTGAACTCTTTCCCTACACGACGCTCTTCCGATCTNNNCTGTAGTTGAAGCTGTGTGTTG
GAGCACACGTCTGAACTCTTTCCCTACACGACGCTCTTCCGATCTNNNCCAGAAAGCCCAAGAACATCA
GAGCACACGTCTGAACTCTTTCCCTACACGACGCTCTTCCGATCTNNNACTGAATGCCCCACAATGCTAT
GAGCACACGTCTGAACTCTTTCCCTACACGACGCTCTTCCGATCTNNNCAATCCAACAGCTTCACAGAT
GAGCACACGTCTGAACTCTTTCCCTACACGACGCTCTTCCGATCTNNNGCATTGTGCAAGTGTATAGT
GCACACGTCTGAACTCTTTCCCTACACGACGCTCTTCCGATCTNNNAAAGAGAGAGACAGAGAGGGG
AAGAGCACACGTCTGAACTCTTTCCCTACACGACGCTCTTCCGATCTNNNACAGTAACCTTTTTGCTGT
AAGAGCACACGTCTGAACTCTTTCCCTACACGACGCTCTTCCGATCTNNNAGATCTGGCAGGGTTAGTT
GAGCACACGTCTGAACTCTTTCCCTACACGACGCTCTTCCGATCTNNNCTATACTAATGCTCCCTGCCTC
AGCACACGTCTGAACTCTTTCCCTACACGACGCTCTTCCGATCTNNNTGCTCAGCATTCTCCCATCATAA
AGCACACGTCTGAACTCTTTCCCTACACGACGCTCTTCCGATCTNNNTCAAATTCCATGGTCCTCCTCTG
AGCACACGTCTGAACTCTTTCCCTACACGACGCTCTTCCGATCTNNNCCTCCTATCACCTCCTAGCCATAC
GCACACGTCTGAACTCTTTCCCTACACGACGCTCTTCCGATCTNNNTTGTTTGCCATTCTCTGTAGGGAC
AGCACACGTCTGAACTCTTTCCCTACACGACGCTCTTCCGATCTNNNTGGGCCAGAATTTAGTTAGGGA
AGCACACGTCTGAACTCTTTCCCTACACGACGCTCTTCCGATCTNNNCCTGATTTCAAAGGCTGTGGTT
AGCACACGTCTGAACTCTTTCCCTACACGACGCTCTTCCGATCTNNNTGTCTGTGTGTTAGGTGGTCAA
GCACACGTCTGAACTCTTTCCCTACACGACGCTCTTCCGATCTNNNGACTAGTGATAGGGGTGTGTGT
GCACACGTCTGAACTCTTTCCCTACACGACGCTCTTCCGATCTNNNATGTAATTTAGGGCATGGGAGG
AGCACACGTCTGAACTCTTTCCCTACACGACGCTCTTCCGATCTNNNTTACTGCTTGTGAGTCACCCTA
GAGCACACGTCTGAACTCTTTCCCTACACGACGCTCTTCCGATCTNNNTGCAGCTTTGATGTTTGAACA
AGCACACGTCTGAACTCTTTCCCTACACGACGCTCTTCCGATCTNNNTAAGTAAGGGTAGGAGGCTG
AGCACACGTCTGAACTCTTTCCCTACACGACGCTCTTCCGATCTNNNTATATACACACACACACACACG
GAGCACACGTCTGAACTCTTTCCCTACACGACGCTCTTCCGATCTNNNAATGGCTAAGATGACACAAG
GAGCACACGTCTGAACTCTTTCCCTACACGACGCTCTTCCGATCTNNNAATCGCTCTCTCTCTGTCTCTCT
GAGCACACGTCTGAACTCTTTCCCTACACGACGCTCTTCCGATCTNNNATCTTCCATTTGTGGCCTGAGA
AGCACACGTCTGAACTCTTTCCCTACACGACGCTCTTCCGATCTNNNCACATACACAGAGAGACACAC
AGCACACGTCTGAACTCTTTCCCTACACGACGCTCTTCCGATCTNNNGGACACATAACCAAATTGTGCG
AGCACACGTCTGAACTCTTTCCCTACACGACGCTCTTCCGATCTNNNGCGTTTCAGTGGTTTAGTTAG
GAGCACACGTCTGAACTCTTTCCCTACACGACGCTCTTCCGATCTNNNATGAGCCCTTAGAGAACCTTT

AGAGCACACGTCTGAACTCTTTCCCTACACGACGCTCTTCCGATCTNNNAGCTATTATAGGGGTGTGT
AGCACACGTCTGAACTCTTTCCCTACACGACGCTCTTCCGATCTNNNCATAAGTGCAAGTGTGTACCT
GCACACGTCTGAACTCTTTCCCTACACGACGCTCTTCCGATCTNNNTCCATCTGTCTCTCTCTCCCTCA
GAGCACACGTCTGAACTCTTTCCCTACACGACGCTCTTCCGATCTNNNAAAAAACTCTGGCACCATCACT
GAGCACACGTCTGAACTCTTTCCCTACACGACGCTCTTCCGATCTNNNAAGGAGATGCAAAAACAATG
AGAGCACACGTCTGAACTCTTTCCCTACACGACGCTCTTCCGATCTNNNTCTTTCACCATTCTGGATGCT
GAGCACACGTCTGAACTCTTTCCCTACACGACGCTCTTCCGATCTNNNTGTTTCTGAAGCAGAGCATGT
GAGCACACGTCTGAACTCTTTCCCTACACGACGCTCTTCCGATCTNNNACACACCTTCTATGTACCCAC
AGCACACGTCTGAACTCTTTCCCTACACGACGCTCTTCCGATCTNNNAGCCAGACTCTTGAAGCTAAA
GCACACGTCTGAACTCTTTCCCTACACGACGCTCTTCCGATCTNNNAGTCCAGTTATTTGTTCGGGACA
AGAGCACACGTCTGAACTCTTTCCCTACACGACGCTCTTCCGATCTNNNTGAGGTTGGAGGTCACTAA
GAGCACACGTCTGAACTCTTTCCCTACACGACGCTCTTCCGATCTNNNTGACCTCTGAGCTTCAATTCGA
GCACACGTCTGAACTCTTTCCCTACACGACGCTCTTCCGATCTNNNTATCATTTCAGGCAAGGTCCCTCCA
AGCACACGTCTGAACTCTTTCCCTACACGACGCTCTTCCGATCTNNNATGTGTGGTTGTGTGCATATGT
AGCACACGTCTGAACTCTTTCCCTACACGACGCTCTTCCGATCTNNNAGGTCTTGAGTCTTTGGCTTCAA
GAGCACACGTCTGAACTCTTTCCCTACACGACGCTCTTCCGATCTNNNTTGAGAGAGAGAGAGAGTGT
AGCACACGTCTGAACTCTTTCCCTACACGACGCTCTTCCGATCTNNNTATTCTTTGACTGTGGTGCTGG
AGCACACGTCTGAACTCTTTCCCTACACGACGCTCTTCCGATCTNNNGGGTCAGGAAGAAGAGGGAA
AGCACACGTCTGAACTCTTTCCCTACACGACGCTCTTCCGATCTNNNAGAGGTGAAGATGGGGGTAA
GAGCACACGTCTGAACTCTTTCCCTACACGACGCTCTTCCGATCTNNNCTGCTTTACCACTTCAACACA
AGCACACGTCTGAACTCTTTCCCTACACGACGCTCTTCCGATCTNNNCTCCTATCCTCCTCTGCATCCTA
AGCACACGTCTGAACTCTTTCCCTACACGACGCTCTTCCGATCTNNNATGGTCCCCATAGCACATATTC
GAGCACACGTCTGAACTCTTTCCCTACACGACGCTCTTCCGATCTNNNCAGAGTGAGACTGACACACA
AGCACACGTCTGAACTCTTTCCCTACACGACGCTCTTCCGATCTNNNAATGCAACACACACTAGCATCA
GAGCACACGTCTGAACTCTTTCCCTACACGACGCTCTTCCGATCTNNNAAATAACTGAAGCCAGACGT
AGCACACGTCTGAACTCTTTCCCTACACGACGCTCTTCCGATCTNNNAACCCACGTCTTGTTTTCCTTTC
GAGCACACGTCTGAACTCTTTCCCTACACGACGCTCTTCCGATCTNNNAGGGAGGCAAATGTCACTAA
GAGCACACGTCTGAACTCTTTCCCTACACGACGCTCTTCCGATCTNNNTTGAAATTGCCATTCATCCAG
GAGCACACGTCTGAACTCTTTCCCTACACGACGCTCTTCCGATCTNNNTCAATGTAGGTCCTGGAAGCA
AGCACACGTCTGAACTCTTTCCCTACACGACGCTCTTCCGATCTNNNCTGCAGTGGATATTTCACTCCC
AGCACACGTCTGAACTCTTTCCCTACACGACGCTCTTCCGATCTNNNATCCTAAAACATGACATGCCCCT
AGCACACGTCTGAACTCTTTCCCTACACGACGCTCTTCCGATCTNNNTTGAACTTCCAGATGCAAAAGG
GAGCACACGTCTGAACTCTTTCCCTACACGACGCTCTTCCGATCTNNNTATTTAGGGCAAAGGGGCAT
AGCACACGTCTGAACTCTTTCCCTACACGACGCTCTTCCGATCTNNNGCTTCTTTCCTCACTTCCCTACA
AGCACACGTCTGAACTCTTTCCCTACACGACGCTCTTCCGATCTNNNAGGATCTTGTTCTCTGAAGTCCA
GAGCACACGTCTGAACTCTTTCCCTACACGACGCTCTTCCGATCTNNNAGCACCATATTTTCCTCAGGG
AGCACACGTCTGAACTCTTTCCCTACACGACGCTCTTCCGATCTNNNGCACAAAGTAGACACTGAGCAA
AGAGCACACGTCTGAACTCTTTCCCTACACGACGCTCTTCCGATCTNNNAAGACCAACCTGGACAACATA
GCACACGTCTGAACTCTTTCCCTACACGACGCTCTTCCGATCTNNNTGTAGTCATAGCCATCCTAACCT
AGAGCACACGTCTGAACTCTTTCCCTACACGACGCTCTTCCGATCTNNNGTGTGTGTATGGTGTGTAT
AGCACACGTCTGAACTCTTTCCCTACACGACGCTCTTCCGATCTNNNATGTGTGTATGCTAACTTCTGCA
AGCACACGTCTGAACTCTTTCCCTACACGACGCTCTTCCGATCTNNNACAGGAAGCACAGTTCTAGCAT
GAGCACACGTCTGAACTCTTTCCCTACACGACGCTCTTCCGATCTNNNCATTTGTGATTTCCAACCTCG

AGCACACGTCTGAACTCTTTCCCTACACGACGCTCTTCCGATCTNNNCAGAACAGAACCAATAGGGGG
GAGCACACGTCTGAACTCTTTCCCTACACGACGCTCTTCCGATCTNNNGTGTGTGTGTGTGTATGT
GAGCACACGTCTGAACTCTTTCCCTACACGACGCTCTTCCGATCTNNNAGTCAATTGTGTGTGTGTG
GAGCACACGTCTGAACTCTTTCCCTACACGACGCTCTTCCGATCTNNNATAGCTACATATGGCCATTGA
AGAGCACACGTCTGAACTCTTTCCCTACACGACGCTCTTCCGATCTNNNGGACTGTTAGGAAGGACGT
AGCACACGTCTGAACTCTTTCCCTACACGACGCTCTTCCGATCTNNNCAAGCCTATTTCCTTTCCCTCCTC
AAGAGCACACGTCTGAACTCTTTCCCTACACGACGCTCTTCCGATCTNNNACTTCTCAACTTCCAGACT
GAGCACACGTCTGAACTCTTTCCCTACACGACGCTCTTCCGATCTNNNCACATTTTCAGGCCAGTCATG
AGCACACGTCTGAACTCTTTCCCTACACGACGCTCTTCCGATCTNNNTGTCTGTGCATATGTTCATGTG
AGCACACGTCTGAACTCTTTCCCTACACGACGCTCTTCCGATCTNNNAGAAAAGGAGAGTCGGCTATG
AGCACACGTCTGAACTCTTTCCCTACACGACGCTCTTCCGATCTNNNCTTGCAAATCCTCATCCTAGGG
AGCACACGTCTGAACTCTTTCCCTACACGACGCTCTTCCGATCTNNNAAAAAAGCCTGATTCCTCCTCC
GCACACGTCTGAACTCTTTCCCTACACGACGCTCTTCCGATCTNNNATCATGACACCCAAGTCTCCAAG
AGCACACGTCTGAACTCTTTCCCTACACGACGCTCTTCCGATCTNNNCCTAGTAGGTGTTCAGAAGGT
AGCACACGTCTGAACTCTTTCCCTACACGACGCTCTTCCGATCTNNNTTTATATTGCCTCCTCCTAGCCCC
GAGCACACGTCTGAACTCTTTCCCTACACGACGCTCTTCCGATCTNNNGGGAGAGGCAAAAGGTGAG
AGCACACGTCTGAACTCTTTCCCTACACGACGCTCTTCCGATCTNNNTCTCTCTCTCTCTGTGTGTGT
AGCACACGTCTGAACTCTTTCCCTACACGACGCTCTTCCGATCTNNNCGCTTCTCTCTCTCTCTCTCTCTC
GAGCACACGTCTGAACTCTTTCCCTACACGACGCTCTTCCGATCTNNNAAGTCTCTCTCTCTCTCTCCC
GAGCACACGTCTGAACTCTTTCCCTACACGACGCTCTTCCGATCTNNNTCCCAAGTTCCGTACATTCCTT
AGCACACGTCTGAACTCTTTCCCTACACGACGCTCTTCCGATCTNNNCTCTGCTGTTTTTCTTGTCCGA
AGCACACGTCTGAACTCTTTCCCTACACGACGCTCTTCCGATCTNNNCAAGACCTAGCTGATGGTACCA
GAGCACACGTCTGAACTCTTTCCCTACACGACGCTCTTCCGATCTNNNGGAAGTATTAGGCTGCCTCT
AGCACACGTCTGAACTCTTTCCCTACACGACGCTCTTCCGATCTNNNCTGGTAATCCTTTGGGACCAG
GAGCACACGTCTGAACTCTTTCCCTACACGACGCTCTTCCGATCTNNNTATCCTAGCACAAACACCCACA
GAGCACACGTCTGAACTCTTTCCCTACACGACGCTCTTCCGATCTNNNGTGAGAGAGAGAGCAAGAGA
AGCACACGTCTGAACTCTTTCCCTACACGACGCTCTTCCGATCTNNNACAGTGTATATGTGTGTGTGT
GAGCACACGTCTGAACTCTTTCCCTACACGACGCTCTTCCGATCTNNNGAGACTCGACAGCACAGAGA
GAGCACACGTCTGAACTCTTTCCCTACACGACGCTCTTCCGATCTNNNTAGAAGTATCTCCTTCCCACC
GAGCACACGTCTGAACTCTTTCCCTACACGACGCTCTTCCGATCTNNNTGAGGTTGCAGTGAACTATG
GCACACGTCTGAACTCTTTCCCTACACGACGCTCTTCCGATCTNNNACCTGGATGACTGTTTCACACTT
GAGCACACGTCTGAACTCTTTCCCTACACGACGCTCTTCCGATCTNNNTCACATCCCTCCCTTAGGTAGA
AGCACACGTCTGAACTCTTTCCCTACACGACGCTCTTCCGATCTNNNTTATCCTTCCAATGTGGGCAGA
AGAGCACACGTCTGAACTCTTTCCCTACACGACGCTCTTCCGATCTNNNTGACTTCTGATAAGGGCTGC
GAGCACACGTCTGAACTCTTTCCCTACACGACGCTCTTCCGATCTNNNGAGACATGAGGGGAGATAT
AGCACACGTCTGAACTCTTTCCCTACACGACGCTCTTCCGATCTNNNGCTTTAGGTACAAGGCCAAAG
GAGCACACGTCTGAACTCTTTCCCTACACGACGCTCTTCCGATCTNNNAATCAGCTGTAAATCCGTCTG
AGCACACGTCTGAACTCTTTCCCTACACGACGCTCTTCCGATCTNNNTAAGGGCATCATTTCAGCAAGA
GAGCACACGTCTGAACTCTTTCCCTACACGACGCTCTTCCGATCTNNNAACACTAGCAGGAAAGCAAA
GAGCACACGTCTGAACTCTTTCCCTACACGACGCTCTTCCGATCTNNNATAACTCAGCTCCTGGCATATG
AGCACACGTCTGAACTCTTTCCCTACACGACGCTCTTCCGATCTNNNTGTGATGGACAGACAGAGTCT
GAGCACACGTCTGAACTCTTTCCCTACACGACGCTCTTCCGATCTNNNCCCTGAGGAAATGTTTGCACT
GAGCACACGTCTGAACTCTTTCCCTACACGACGCTCTTCCGATCTNNNCGATGAAGTCGTAGAAGCTG

GCACACGTCTGAACTCTTTCCCTACACGACGCTCTTCCGATCTNNNTTTTACTCCTCTAACCCCTGCAGC
GAGCACACGTCTGAACTCTTTCCCTACACGACGCTCTTCCGATCTNNNACATTTCTATCTGCTGACAGC
AGCACACGTCTGAACTCTTTCCCTACACGACGCTCTTCCGATCTNNNTCTGGAGAGGTTGTCCTTTTCT
GAGCACACGTCTGAACTCTTTCCCTACACGACGCTCTTCCGATCTNNNAGATGATGTAGGAGTGTGT
GAGCACACGTCTGAACTCTTTCCCTACACGACGCTCTTCCGATCTNNNAGAAGGAGAGAGGGAAGAA
AGCACACGTCTGAACTCTTTCCCTACACGACGCTCTTCCGATCTNNNTGTTTCTCGAGACCCTGACAAT
GAGCACACGTCTGAACTCTTTCCCTACACGACGCTCTTCCGATCTNNNAGGAGTTCCCAGAGAAGAAA
AGCACACGTCTGAACTCTTTCCCTACACGACGCTCTTCCGATCTNNNAACTGCCTTCCAACTCAAAACAC
AGCACACGTCTGAACTCTTTCCCTACACGACGCTCTTCCGATCTNNNTCTTGAAGAGAACACCCCCTTTT
GAGCACACGTCTGAACTCTTTCCCTACACGACGCTCTTCCGATCTNNNAGGAAGCCTGGAAATGAAGA
AGAGCACACGTCTGAACTCTTTCCCTACACGACGCTCTTCCGATCTNNNAGTCAGGTTCCAAGTCCTTG
AGCACACGTCTGAACTCTTTCCCTACACGACGCTCTTCCGATCTNNNAGCACAAAGGAATTTCTTGGT
GCACACGTCTGAACTCTTTCCCTACACGACGCTCTTCCGATCTNNNTCACTCTCTCTTGCTCTCTCTCTCA
AGAGCACACGTCTGAACTCTTTCCCTACACGACGCTCTTCCGATCTNNNATACTGCCTCTTCTCCATTCT
GAGCACACGTCTGAACTCTTTCCCTACACGACGCTCTTCCGATCTNNNTTCTGGCAATCTCTGTCTCTGT
AGCACACGTCTGAACTCTTTCCCTACACGACGCTCTTCCGATCTNNNTAAGCTTTTTGAAGGCAAGGAG
GCACACGTCTGAACTCTTTCCCTACACGACGCTCTTCCGATCTNNNTCACTCTACCTTTATGCCCCATCC
AGCACACGTCTGAACTCTTTCCCTACACGACGCTCTTCCGATCTNNNTTCAGAAGACTGCTCCTTCAGA
GAGCACACGTCTGAACTCTTTCCCTACACGACGCTCTTCCGATCTNNNCACGTTCTCCTTGGAAGAGAG
AGCACACGTCTGAACTCTTTCCCTACACGACGCTCTTCCGATCTNNNACAACACATGGGGATTATGGG
GAGCACACGTCTGAACTCTTTCCCTACACGACGCTCTTCCGATCTNNNGAAAACATTTGGCTGTGTCTG
AGCACACGTCTGAACTCTTTCCCTACACGACGCTCTTCCGATCTNNNTATACCAAGAGGGAAACAAGCC
GAGCACACGTCTGAACTCTTTCCCTACACGACGCTCTTCCGATCTNNNGTATGACTTTTCCCCACTTCCCT
AGAGCACACGTCTGAACTCTTTCCCTACACGACGCTCTTCCGATCTNNNGAATATAGCCCGTACCCTCA
AGCACACGTCTGAACTCTTTCCCTACACGACGCTCTTCCGATCTNNNCCATGAGGGAGATTCTGAGGA
AGCACACGTCTGAACTCTTTCCCTACACGACGCTCTTCCGATCTNNNTGCACATCCAGACTGACTTTAC
GAGCACACGTCTGAACTCTTTCCCTACACGACGCTCTTCCGATCTNNNCCATGCTCATCACCTTCATAGG
GAGCACACGTCTGAACTCTTTCCCTACACGACGCTCTTCCGATCTNNNAAGTGCTGAGATTACAGGCA
GAGCACACGTCTGAACTCTTTCCCTACACGACGCTCTTCCGATCTNNNATTTCTGAGTCTGCAGTGGGA
GAGCACACGTCTGAACTCTTTCCCTACACGACGCTCTTCCGATCTNNNATAACAATTGTGTGTGTGTGC
GAGCACACGTCTGAACTCTTTCCCTACACGACGCTCTTCCGATCTNNNTCAGTTTTGCCCTAGAGTGAG
AGCACACGTCTGAACTCTTTCCCTACACGACGCTCTTCCGATCTNNNCTCCAGGCAGTAAGAAAAACGT
AGCACACGTCTGAACTCTTTCCCTACACGACGCTCTTCCGATCTNNNGTGTCCTGTTGTTGCTGCTATT
GCACACGTCTGAACTCTTTCCCTACACGACGCTCTTCCGATCTNNNACAGGTATAGCAACCAATCAGCA
GAGCACACGTCTGAACTCTTTCCCTACACGACGCTCTTCCGATCTNNNGCTACTCTGTCTTATCCAGCCT
GCACACGTCTGAACTCTTTCCCTACACGACGCTCTTCCGATCTNNNGAGAGAGAGAGACAGGAGAGA
AGAGCACACGTCTGAACTCTTTCCCTACACGACGCTCTTCCGATCTNNNTACACGAGAGAGAGAGGA
AAGAGCACACGTCTGAACTCTTTCCCTACACGACGCTCTTCCGATCTNNNGTCTGTGCTTGGTTTTCTTC
AGAGCACACGTCTGAACTCTTTCCCTACACGACGCTCTTCCGATCTNNNATAGGTGGCTGTTCAAGTGT
AGCACACGTCTGAACTCTTTCCCTACACGACGCTCTTCCGATCTNNNCAACTTTCTTGACCAGCTTTCCC
AGAGCACACGTCTGAACTCTTTCCCTACACGACGCTCTTCCGATCTNNNGTCTGCTTCTGAGAGAGTCC
AGCACACGTCTGAACTCTTTCCCTACACGACGCTCTTCCGATCTNNNTGGGAATTATGACAGGTGCTTG
GAGCACACGTCTGAACTCTTTCCCTACACGACGCTCTTCCGATCTNNNGAGGTGAACTAACCAACTCTC

GAGCACACGTCTGAACTCTTTCCCTACACGACGCTCTTCCGATCTNNNGCAGTCTTGTAATCACATGAG
AGCACACGTCTGAACTCTTTCCCTACACGACGCTCTTCCGATCTNNNGCACGCTTCAACAATAACACCTT
AGCACACGTCTGAACTCTTTCCCTACACGACGCTCTTCCGATCTNNNTACTCACACAGTTCTGGAGTAC
AGCACACGTCTGAACTCTTTCCCTACACGACGCTCTTCCGATCTNNNCTTTATGGTGGGGAGGAAAGG
GAGCACACGTCTGAACTCTTTCCCTACACGACGCTCTTCCGATCTNNNCAATGGGTGAAGGCTTTGAA
GAGCACACGTCTGAACTCTTTCCCTACACGACGCTCTTCCGATCTNNNTGAACTTTTCAGTGTTGCTGC
AGCACACGTCTGAACTCTTTCCCTACACGACGCTCTTCCGATCTNNNGAGGATTCATCTCACTCCTCCAC
GAGCACACGTCTGAACTCTTTCCCTACACGACGCTCTTCCGATCTNNNAGCCCACTGAGAATATACCCT
GAGCACACGTCTGAACTCTTTCCCTACACGACGCTCTTCCGATCTNNNGACCTATTCACACCTGTCTCCT
AGCACACGTCTGAACTCTTTCCCTACACGACGCTCTTCCGATCTNNNTGTTCCCAAAGTTCTAGCTCCTC
AGCACACGTCTGAACTCTTTCCCTACACGACGCTCTTCCGATCTNNNTGTTCCCAAAGTTCTAGCTCCTC
AGCACACGTCTGAACTCTTTCCCTACACGACGCTCTTCCGATCTNNNCTGCAGATCTTGGGACTACTCA
GAGCACACGTCTGAACTCTTTCCCTACACGACGCTCTTCCGATCTNNNGTCTTCAGACTTGGACTGGA
GAGCACACGTCTGAACTCTTTCCCTACACGACGCTCTTCCGATCTNNNCCCTACCTCTGTTGCAGTGTA
AGCACACGTCTGAACTCTTTCCCTACACGACGCTCTTCCGATCTNNNGAGTCTTCACTTTTTTCCGAGG
GCACACGTCTGAACTCTTTCCCTACACGACGCTCTTCCGATCTNNNAGTATACATGGGATGACGAACG
GAGCACACGTCTGAACTCTTTCCCTACACGACGCTCTTCCGATCTNNNGTCAATTGCAACATGGAGCA
GCACACGTCTGAACTCTTTCCCTACACGACGCTCTTCCGATCTNNNTGTTCTTGCTCTGACTCTCTCTCC
AGCACACGTCTGAACTCTTTCCCTACACGACGCTCTTCCGATCTNNNCTAGAGCAACACTTGTCAAAGG
GAGCACACGTCTGAACTCTTTCCCTACACGACGCTCTTCCGATCTNNNGTTCATCTGAGCTTGGGAAGT
AGCACACGTCTGAACTCTTTCCCTACACGACGCTCTTCCGATCTNNNTACATTACCTGTTGCTCCAGTCC
GCACACGTCTGAACTCTTTCCCTACACGACGCTCTTCCGATCTNNNTCCATCACAGTCTTGACAAGAGG
GAGCACACGTCTGAACTCTTTCCCTACACGACGCTCTTCCGATCTNNNTACAGCAGGACTTGGACAAG
GAGCACACGTCTGAACTCTTTCCCTACACGACGCTCTTCCGATCTNNNTGTGGTACTGGCATAACAAC
AGCACACGTCTGAACTCTTTCCCTACACGACGCTCTTCCGATCTNNNTGAACACTTGGTGCTTTATTGC
GAGCACACGTCTGAACTCTTTCCCTACACGACGCTCTTCCGATCTNNNGATAGCTTGAGCCTAGGAGTT
GCACACGTCTGAACTCTTTCCCTACACGACGCTCTTCCGATCTNNNCGCCTCCTTTTCGTAACACTGTTC
AGCACACGTCTGAACTCTTTCCCTACACGACGCTCTTCCGATCTNNNCACTCAGCCTGAACACATCTAC
AGCACACGTCTGAACTCTTTCCCTACACGACGCTCTTCCGATCTNNNACACGACATACACATAGCACCT
GAGCACACGTCTGAACTCTTTCCCTACACGACGCTCTTCCGATCTNNNTATGAGCCCTTGTGAACAATG
GCACACGTCTGAACTCTTTCCCTACACGACGCTCTTCCGATCTNNNGCTGGTGAGATCCTTTTGCTTTT
GAGCACACGTCTGAACTCTTTCCCTACACGACGCTCTTCCGATCTNNNGATTTTGCTCTCTGCTGTTCCA
GAGCACACGTCTGAACTCTTTCCCTACACGACGCTCTTCCGATCTNNNCTCTCTCTCTCACACACACAC
AGCACACGTCTGAACTCTTTCCCTACACGACGCTCTTCCGATCTNNNTCGCAAAAATAAATGCACACCC
AGCACACGTCTGAACTCTTTCCCTACACGACGCTCTTCCGATCTNNNTCTGGCATCATCTCCCTTTTCTCC
GAGCACACGTCTGAACTCTTTCCCTACACGACGCTCTTCCGATCTNNNCAGATGGTTGCTGTTTCACTC
GAGCACACGTCTGAACTCTTTCCCTACACGACGCTCTTCCGATCTNNNTTACTATTGTTCGCAGGAGTC
GAGCACACGTCTGAACTCTTTCCCTACACGACGCTCTTCCGATCTNNNGGCCCAACTCTGTCAAAAGA
GAGCACACGTCTGAACTCTTTCCCTACACGACGCTCTTCCGATCTNNNAGTAAGGCAGCAAACACAAA
GAGCACACGTCTGAACTCTTTCCCTACACGACGCTCTTCCGATCTNNNTCTTTTGTCAGTCCTATGCAGC
GAGCACACGTCTGAACTCTTTCCCTACACGACGCTCTTCCGATCTNNNTGCAGTCCATCCTTTGGTTAT
GAGCACACGTCTGAACTCTTTCCCTACACGACGCTCTTCCGATCTNNNGCACATCACAAAGTTTCCTGT
GAGCACACGTCTGAACTCTTTCCCTACACGACGCTCTTCCGATCTNNNTTTTTCAGCTCCTGGCATTCA

AGCACACGTCTGAACTCTTTCCCTACACGACGCTCTTCCGATCTNNNGAATTCACGACGTCACACTTGT

AGCACACGTCTGAACTCTTTCCCTACACGACGCTCTTCCGATCTNNNAGCCAGCATCATTCTGATACCA

AGCACACGTCTGAACTCTTTCCCTACACGACGCTCTTCCGATCTNNNTATGAGTACGTGTGTGTGTGT

GAGCACACGTCTGAACTCTTTCCCTACACGACGCTCTTCCGATCTNNNGCTATGAAGAGGGCTCAGG

AGCACACGTCTGAACTCTTTCCCTACACGACGCTCTTCCGATCTNNNTGTGAGCCTAACTGGAGTTTC

GCACACGTCTGAACTCTTTCCCTACACGACGCTCTTCCGATCTNNNTCAAATGATGTACTCCCGTGTGT

AGCACACGTCTGAACTCTTTCCCTACACGACGCTCTTCCGATCTNNNTTATAGCATGGTGTGGCCTCA

AGCACACGTCTGAACTCTTTCCCTACACGACGCTCTTCCGATCTNNNTGACTTGCAGAAGGCTTCCTA

AGAGCACACGTCTGAACTCTTTCCCTACACGACGCTCTTCCGATCTNNNGCCTAATGCTCTTCCTTTAT

AGCACACGTCTGAACTCTTTCCCTACACGACGCTCTTCCGATCTNNNGTGTGGACATCAGAGAATCCC

GCACACGTCTGAACTCTTTCCCTACACGACGCTCTTCCGATCTNNNTAAGTAGAGGGGGAGAAAGGG

GAGCACACGTCTGAACTCTTTCCCTACACGACGCTCTTCCGATCTNNNACCTTCCTGACTTAGGCCAAT

AGAGCACACGTCTGAACTCTTTCCCTACACGACGCTCTTCCGATCTNNNAATGAGATACTGTATCCCG

GCACACGTCTGAACTCTTTCCCTACACGACGCTCTTCCGATCTNNNAAACAGAGTGAGCCCTTCTTCA

GCACACGTCTGAACTCTTTCCCTACACGACGCTCTTCCGATCTNNNACTGATAAGTGATGTGCCAAGG

AGCACACGTCTGAACTCTTTCCCTACACGACGCTCTTCCGATCTNNNAGGCCAACATGATCTATCCATC

AGAGCACACGTCTGAACTCTTTCCCTACACGACGCTCTTCCGATCTNNNTGAAATACCTAGTGGGGA

GCACACGTCTGAACTCTTTCCCTACACGACGCTCTTCCGATCTNNNTCCATGTCTCTCTCTCTCTGTGTC

AGCACACGTCTGAACTCTTTCCCTACACGACGCTCTTCCGATCTNNNTGCATGAGCTAGGTCTTCAAA

GAGCACACGTCTGAACTCTTTCCCTACACGACGCTCTTCCGATCTNNNTAGTAGATAGCACCCCAACAC

AGAGCACACGTCTGAACTCTTTCCCTACACGACGCTCTTCCGATCTNNNAGGTGACTTGTTGCCTGAA

GCACACGTCTGAACTCTTTCCCTACACGACGCTCTTCCGATCTNNNACCCTCCTTTCTCTCTCTCTCCA

GCACACGTCTGAACTCTTTCCCTACACGACGCTCTTCCGATCTNNNACCCTCCTTTCTCTCTCTCTCCA

CACACGTCTGAACTCTTTCCCTACACGACGCTCTTCCGATCTNNNCCTTGTCCATGAGAAGCATTGTG

AGAGCACACGTCTGAACTCTTTCCCTACACGACGCTCTTCCGATCTNNNGAGAGGGGAATGAGAATG

AGCACACGTCTGAACTCTTTCCCTACACGACGCTCTTCCGATCTNNNCATCATCAAGTGTGTCACTCAG

AGCACACGTCTGAACTCTTTCCCTACACGACGCTCTTCCGATCTNNNGTAGTGTAGCAGTCAGGGTTC

AGAGCACACGTCTGAACTCTTTCCCTACACGACGCTCTTCCGATCTNNNCACAGGATCCTACTCTTTG

AGCACACGTCTGAACTCTTTCCCTACACGACGCTCTTCCGATCTNNNCAGATACCAGAGGGAAGTGT

AGCACACGTCTGAACTCTTTCCCTACACGACGCTCTTCCGATCTNNNATGGAGGCCAGCATTCATACA

AGCACACGTCTGAACTCTTTCCCTACACGACGCTCTTCCGATCTNNNCAAAGGATCGAGGACATGAG

AGAGCACACGTCTGAACTCTTTCCCTACACGACGCTCTTCCGATCTNNNCCGTTTGGGAATTCAAAGA

GAGCACACGTCTGAACTCTTTCCCTACACGACGCTCTTCCGATCTNNNATGTGTGTGTGTGTGTGTTT

GCACACGTCTGAACTCTTTCCCTACACGACGCTCTTCCGATCTNNNGTTCAGTACACACACACACACAG

CACACGTCTGAACTCTTTCCCTACACGACGCTCTTCCGATCTNNNATCACCATTTTTACCCTCCCCTCCAT

GAGCACACGTCTGAACTCTTTCCCTACACGACGCTCTTCCGATCTNNNGCTTGGTAAAAAGCCACATC

AGCACACGTCTGAACTCTTTCCCTACACGACGCTCTTCCGATCTNNNATTATGTCCAACACTCCACCTCC

AGCACACGTCTGAACTCTTTCCCTACACGACGCTCTTCCGATCTNNNCCCACTGTTATCAAGACATGT

AGCACACGTCTGAACTCTTTCCCTACACGACGCTCTTCCGATCTNNNGAAGTTGGAGGAAGCCAGAA

GAGCACACGTCTGAACTCTTTCCCTACACGACGCTCTTCCGATCTNNNGCACTGATAGCTTTAGGCCA

AGCACACGTCTGAACTCTTTCCCTACACGACGCTCTTCCGATCTNNNAGTGTAAGAGTGTGTGTGTG

AGAGCACACGTCTGAACTCTTTCCCTACACGACGCTCTTCCGATCTNNNGGACAAGTCCCAACTCTAT

AGCACACGTCTGAACTCTTTCCCTACACGACGCTCTTCCGATCTNNNAAAGTAGTTCTGGCTGTCTGG

AGCACACGTCTGAACTCTTTCCCTACACGACGCTCTTCCGATCTNNNTCTTCTAATGCACCTCGCTTTCT
AGCACACGTCTGAACTCTTTCCCTACACGACGCTCTTCCGATCTNNNCTCAATTGTTTTGAGCAGCAGG
AGCACACGTCTGAACTCTTTCCCTACACGACGCTCTTCCGATCTNNNCCCTACAATTGCAGACTTACCCT
AGCACACGTCTGAACTCTTTCCCTACACGACGCTCTTCCGATCTNNNAAAAGCTGACAGTGGCCTTTAT
AGCACACGTCTGAACTCTTTCCCTACACGACGCTCTTCCGATCTNNNCCCAATGTACCTCCTTTCTTCCA
GCACACGTCTGAACTCTTTCCCTACACGACGCTCTTCCGATCTNNNGACGACAGGACCAAGAATTAG
GAGCACACGTCTGAACTCTTTCCCTACACGACGCTCTTCCGATCTNNNCCTGCAAACATCAGATTAGG
AGCACACGTCTGAACTCTTTCCCTACACGACGCTCTTCCGATCTNNNTACTACAAACTCACAGCAGCCA
GAGCACACGTCTGAACTCTTTCCCTACACGACGCTCTTCCGATCTNNNAAAGGCAGATAGTGTTAGTG
GAGCACACGTCTGAACTCTTTCCCTACACGACGCTCTTCCGATCTNNNGAAAGATGGACACAGCAAC
AGCACACGTCTGAACTCTTTCCCTACACGACGCTCTTCCGATCTNNNAAATGAACAAACCAAGCTCCTC
AGCACACGTCTGAACTCTTTCCCTACACGACGCTCTTCCGATCTNNNTACAAAAAAGGCTCCAAGCACT
CACACGTCTGAACTCTTTCCCTACACGACGCTCTTCCGATCTNNNCTCTCCTCTGTCTCTCTCTCTCCCAT
CACACGTCTGAACTCTTTCCCTACACGACGCTCTTCCGATCTNNNACCATGATTTCCCAAATGGCATCC
GCACACGTCTGAACTCTTTCCCTACACGACGCTCTTCCGATCTNNNGAAAGACCCAATGTGGTTGTCT
GCACACGTCTGAACTCTTTCCCTACACGACGCTCTTCCGATCTNNNTGTGTCTCTGTCTCTGTCTCTCT
AGCACACGTCTGAACTCTTTCCCTACACGACGCTCTTCCGATCTNNNACAAATCAAGCAGAAGAAGG
AGCACACGTCTGAACTCTTTCCCTACACGACGCTCTTCCGATCTNNNGCACCCCTATCCCAGTAATAGT
AGCACACGTCTGAACTCTTTCCCTACACGACGCTCTTCCGATCTNNNAGAGGCTCTTGTTTTCTTCCCTT
GCACACGTCTGAACTCTTTCCCTACACGACGCTCTTCCGATCTNNNTTCTGCTTTAGGAGAAACCCCAA
AGCACACGTCTGAACTCTTTCCCTACACGACGCTCTTCCGATCTNNNAGGAGGGTCATCAAAATCTG
AGAGCACACGTCTGAACTCTTTCCCTACACGACGCTCTTCCGATCTNNNCTGTTATGCCACTTAGCGTA
AGCACACGTCTGAACTCTTTCCCTACACGACGCTCTTCCGATCTNNNCCTTTTCTTCTGTCTGTTGAGC
GCACACGTCTGAACTCTTTCCCTACACGACGCTCTTCCGATCTNNNGTCCTTCCTTCCTTCCCTTCTTTC
GCACACGTCTGAACTCTTTCCCTACACGACGCTCTTCCGATCTNNNCAGTGTTGTGCCTCTGAGGATA
CACACGTCTGAACTCTTTCCCTACACGACGCTCTTCCGATCTNNNTTCAGACATGCCAGTGAGAATGT
GAGCACACGTCTGAACTCTTTCCCTACACGACGCTCTTCCGATCTNNNAGCAATAGACTCCGTCTCC
GAGCACACGTCTGAACTCTTTCCCTACACGACGCTCTTCCGATCTNNNCCTCTTCACATCCTCACCTGT
AGCACACGTCTGAACTCTTTCCCTACACGACGCTCTTCCGATCTNNNAGACACACAGACATACACACA
GAGCACACGTCTGAACTCTTTCCCTACACGACGCTCTTCCGATCTNNNAGTCTGTCCCTGGATCTAAG
GAGCACACGTCTGAACTCTTTCCCTACACGACGCTCTTCCGATCTNNNCTGAGGTTGTAGTGAGCCAA
AGCACACGTCTGAACTCTTTCCCTACACGACGCTCTTCCGATCTNNNACACACACAGAGAGAGAGAG
AGAGCACACGTCTGAACTCTTTCCCTACACGACGCTCTTCCGATCTNNNACCAAGAGACTGACAGAG
GCACACGTCTGAACTCTTTCCCTACACGACGCTCTTCCGATCTNNNCCAACATGGTGAAAACCTGTCT
AGCACACGTCTGAACTCTTTCCCTACACGACGCTCTTCCGATCTNNNGTGTGTGTGAGAGAGAG
AAGAGCACACGTCTGAACTCTTTCCCTACACGACGCTCTTCCGATCTNNNCCCGAGAGTTTTATGTTGC
AAGAGCACACGTCTGAACTCTTTCCCTACACGACGCTCTTCCGATCTNNNCCCGAGAGTTTTATGTTGC
AAGAGCACACGTCTGAACTCTTTCCCTACACGACGCTCTTCCGATCTNNNCCCGAGAGTTTTATGTTGC
GCACACGTCTGAACTCTTTCCCTACACGACGCTCTTCCGATCTNNNTTCACTGTTAGTCCTGGAAGTC
GCACACGTCTGAACTCTTTCCCTACACGACGCTCTTCCGATCTNNNAGAGAGAGAGAGAGAGAAAG
AGAGCACACGTCTGAACTCTTTCCCTACACGACGCTCTTCCGATCTNNNACACAGTGAGGTCTGTAT
GAGCACACGTCTGAACTCTTTCCCTACACGACGCTCTTCCGATCTNNNTAGGTAGTAGGGAGAGAG
GCACACGTCTGAACTCTTTCCCTACACGACGCTCTTCCGATCTNNNTGTGCTTGTAGTCACAGCTACT

```
AGCACACGTCTGAACTCTTTCCCTACACGACGCTCTTCCGATCTNNNCCATCCATCCAAAGGTTGCATTT
AGAGCACACGTCTGAACTCTTTCCCTACACGACGCTCTTCCGATCTNNNGTAATCCCAGATACTCGGAA
GAGCACACGTCTGAACTCTTTCCCTACACGACGCTCTTCCGATCTNNNTCTGAGTAGCAGGCAAGAAG
AGCACACGTCTGAACTCTTTCCCTACACGACGCTCTTCCGATCTNNNCTTCTCAGCCTCCATAAGTGCAT
AGCACACGTCTGAACTCTTTCCCTACACGACGCTCTTCCGATCTNNNAAAGAAAGAGAGAGAGGGGG
GAGCACACGTCTGAACTCTTTCCCTACACGACGCTCTTCCGATCTNNNAAAGAAGAGGAAGGGAAGG
AGCACACGTCTGAACTCTTTCCCTACACGACGCTCTTCCGATCTNNNGTCCTGTTGCAAAAGGAAATGC
GCACACGTCTGAACTCTTTCCCTACACGACGCTCTTCCGATCTNNNGAGAGAGTGAGAGAGAGAGGG
AGCACACGTCTGAACTCTTTCCCTACACGACGCTCTTCCGATCTNNNGAGAGAGAGAGAGAGAGAGGG
GAGCACACGTCTGAACTCTTTCCCTACACGACGCTCTTCCGATCTNNNCAATTGCAAAAACGTGGAACC
AGCACACGTCTGAACTCTTTCCCTACACGACGCTCTTCCGATCTNNNGTGACAGAACCAATCCTTGTGTG
GAGCACACGTCTGAACTCTTTCCCTACACGACGCTCTTCCGATCTNNNTAGGTTGCCTGTTCACTATGAC
GAGCACACGTCTGAACTCTTTCCCTACACGACGCTCTTCCGATCTNNNGCGAAACCCCCTCTCTACTAAA
GAGCACACGTCTGAACTCTTTCCCTACACGACGCTCTTCCGATCTNNNTCAGCAATTGGTAGAAACAGT
AGCACACGTCTGAACTCTTTCCCTACACGACGCTCTTCCGATCTNNNGAACATTCCATGTGCACTTGTGA
GAGCACACGTCTGAACTCTTTCCCTACACGACGCTCTTCCGATCTNNNTCAGCAATTGGTAGAAACAGT
GAGCACACGTCTGAACTCTTTCCCTACACGACGCTCTTCCGATCTNNNATCAACCTAAGTGCCCATCAAT
GAGCACACGTCTGAACTCTTTCCCTACACGACGCTCTTCCGATCTNNNAACCTAAGTGCCCATCAACTGA
AGCACACGTCTGAACTCTTTCCCTACACGACGCTCTTCCGATCTNNNCGAGTCCCAACCGTATTACAGAA
AGCACACGTCTGAACTCTTTCCCTACACGACGCTCTTCCGATCTNNNAAGGCAAAATGAGGGCAAAAT
AGCACACGTCTGAACTCTTTCCCTACACGACGCTCTTCCGATCTNNNAGACAGGAGAATTGCTTGAACC
GAGCACACGTCTGAACTCTTTCCCTACACGACGCTCTTCCGATCTNNNGCACATGCATGTTTATAGCAG
AGCACACGTCTGAACTCTTTCCCTACACGACGCTCTTCCGATCTNNNATTATGTTTACTGGGCAGGTGG
AGAGCACACGTCTGAACTCTTTCCCTACACGACGCTCTTCCGATCTNNNTACACATATGCACAGACACAT
GAGCACACGTCTGAACTCTTTCCCTACACGACGCTCTTCCGATCTNNNCATAGGCTGTCTCTCCTCATTCC
AGCACACGTCTGAACTCTTTCCCTACACGACGCTCTTCCGATCTNNNCATGAGAATTGCTTGAACCAGG
GAGCACACGTCTGAACTCTTTCCCTACACGACGCTCTTCCGATCTNNNTGGAGGTCATATGTGAACATG
GAGCACACGTCTGAACTCTTTCCCTACACGACGCTCTTCCGATCTNNNAGGGCAGTAGTATACACACAC
GAGCACACGTCTGAACTCTTTCCCTACACGACGCTCTTCCGATCTNNNGCACATGCATGTTTATAGCAG
GAGCACACGTCTGAACTCTTTCCCTACACGACGCTCTTCCGATCTNNNTTTTTCCACCTCAGTCTCCCAAA
AGCACACGTCTGAACTCTTTCCCTACACGACGCTCTTCCGATCTNNNACGAAAAAGATACTTGCACACG
AGCACACGTCTGAACTCTTTCCCTACACGACGCTCTTCCGATCTNNNCAAGCAGTGACAAAAGAACCAC
GAGCACACGTCTGAACTCTTTCCCTACACGACGCTCTTCCGATCTNNNGCTTTTGGGTTCTTGGTCATGA
AGCACACGTCTGAACTCTTTCCCTACACGACGCTCTTCCGATCTNNNATATGTGTGTGTGTGTGTGTGT
GAGCACACGTCTGAACTCTTTCCCTACACGACGCTCTTCCGATCTNNNATCAGGCATGACATCCTAAGTC
GAGCACACGTCTGAACTCTTTCCCTACACGACGCTCTTCCGATCTNNNCAGGTGCCTATAATCCCAGCT
GAGCACACGTCTGAACTCTTTCCCTACACGACGCTCTTCCGATCTNNNAAATACAAAAAATTAGCCGGGC
AGCACACGTCTGAACTCTTTCCCTACACGACGCTCTTCCGATCTNNNAAGAAATCTCAGAGACTCAGCC
GAGCACACGTCTGAACTCTTTCCCTACACGACGCTCTTCCGATCTNNNGCTGAGAGAGGACATCACTTG
AGCACACGTCTGAACTCTTTCCCTACACGACGCTCTTCCGATCTNNNAGAATCACTCGAACCCAGAAGT
AGCACACGTCTGAACTCTTTCCCTACACGACGCTCTTCCGATCTNNNCTGTAATCCAAGCTACTCGGGA
GAGCACACGTCTGAACTCTTTCCCTACACGACGCTCTTCCGATCTNNNCTCAACCCCTGTAGTGGATTGA
AGCACACGTCTGAACTCTTTCCCTACACGACGCTCTTCCGATCTNNNGAGAATACCTTGAACCTGGGAG
```

AGCACACGTCTGAACTCTTTCCCTACACGACGCTCTTCCGATCTNNNGAGGATCATTTGAGTCCAGGA
AGCACACGTCTGAACTCTTTCCCTACACGACGCTCTTCCGATCTNNNTGTAATCCCAGCTACTCAGGAG
GAGCACACGTCTGAACTCTTTCCCTACACGACGCTCTTCCGATCTNNNAATCTCAGCTACTTGGGAGGT
GAGCACACGTCTGAACTCTTTCCCTACACGACGCTCTTCCGATCTNNNAACTCCAACCTGGGCAATAAG
AGCACACGTCTGAACTCTTTCCCTACACGACGCTCTTCCGATCTNNNGTGTAGTGATGGATGCCTGTA
GAGCACACGTCTGAACTCTTTCCCTACACGACGCTCTTCCGATCTNNNGCAAAGCATTCAAGATGTGAC
GAGCACACGTCTGAACTCTTTCCCTACACGACGCTCTTCCGATCTNNNCTGTAATCCCAGCTACTCAGGA
AGCACACGTCTGAACTCTTTCCCTACACGACGCTCTTCCGATCTNNNATTGAAGCACGCAGAAATCCAA
GAGCACACGTCTGAACTCTTTCCCTACACGACGCTCTTCCGATCTNNNCACCTGTAATCCCAGCTACTCT
AGCACACGTCTGAACTCTTTCCCTACACGACGCTCTTCCGATCTNNNAAAATTAGCCAGGCATGTTGG
AGAGCACACGTCTGAACTCTTTCCCTACACGACGCTCTTCCGATCTNNNGCCAAGATCACATCATTTGCA
AGCACACGTCTGAACTCTTTCCCTACACGACGCTCTTCCGATCTNNNTTGCTATTGAGGGACATTGTG
AGCACACGTCTGAACTCTTTCCCTACACGACGCTCTTCCGATCTNNNCTACCATAGTCCTCAAGGTGCT
GAGCACACGTCTGAACTCTTTCCCTACACGACGCTCTTCCGATCTNNNGCAGAAGTTGTAGTGAGCTG
AGCACACGTCTGAACTCTTTCCCTACACGACGCTCTTCCGATCTNNNGTGGTAGGCACCTATAATCCCA
AGCACACGTCTGAACTCTTTCCCTACACGACGCTCTTCCGATCTNNNGCCGTGAGAAAAGATCATGTGA
GAGCACACGTCTGAACTCTTTCCCTACACGACGCTCTTCCGATCTNNNCAGGCAACAATGTGAAACTCCA
GAGCACACGTCTGAACTCTTTCCCTACACGACGCTCTTCCGATCTNNNTCAGAGATACTCCATGGGTG
AGAGCACACGTCTGAACTCTTTCCCTACACGACGCTCTTCCGATCTNNNAAAAAATTAGCCAGGTGTGG
AGAGCACACGTCTGAACTCTTTCCCTACACGACGCTCTTCCGATCTNNNCACCTGAAGTCTCAGCTACTC
AGCACACGTCTGAACTCTTTCCCTACACGACGCTCTTCCGATCTNNNGGCTGAGATAGGAGGATCACT
GAGCACACGTCTGAACTCTTTCCCTACACGACGCTCTTCCGATCTNNNAAGCTGAAGCAGGAAAATTG
AGCACACGTCTGAACTCTTTCCCTACACGACGCTCTTCCGATCTNNNCAGGATGAACTGCTTGAATGT
GAGCACACGTCTGAACTCTTTCCCTACACGACGCTCTTCCGATCTNNNGGTGTCTGTAATCCCAGCTAC
AGCACACGTCTGAACTCTTTCCCTACACGACGCTCTTCCGATCTNNNGCAAAACCCTGTCTCTGCAAATA
AGCACACGTCTGAACTCTTTCCCTACACGACGCTCTTCCGATCTNNNATACAAAAAAATTAGCCGGGC
AGCACACGTCTGAACTCTTTCCCTACACGACGCTCTTCCGATCTNNNGTATTTCTGCAGTCCCAGCTAC
GCACACGTCTGAACTCTTTCCCTACACGACGCTCTTCCGATCTNNNCCGGATTCAAACGATTCTCCTCT
GAGCACACGTCTGAACTCTTTCCCTACACGACGCTCTTCCGATCTNNNACCAACATGATGAAACCCTGT
AGCACACGTCTGAACTCTTTCCCTACACGACGCTCTTCCGATCTNNNTTCACACCTTCAGACTCAGACT
GAGCACACGTCTGAACTCTTTCCCTACACGACGCTCTTCCGATCTNNNAACCCACAGTTGCAATGGAAA
GAGCACACGTCTGAACTCTTTCCCTACACGACGCTCTTCCGATCTNNNTCAGGAGAATCACTTGAACCCA
AGAGCACACGTCTGAACTCTTTCCCTACACGACGCTCTTCCGATCTNNNAAAATTAGCCAGACGTGGTG
AGCACACGTCTGAACTCTTTCCCTACACGACGCTCTTCCGATCTNNNCACCTGTAATCCCAGCTACTCAG
GAGCACACGTCTGAACTCTTTCCCTACACGACGCTCTTCCGATCTNNNCATCTGTAGTCCCAGCTACTCA
AGCACACGTCTGAACTCTTTCCCTACACGACGCTCTTCCGATCTNNNCAACAAGGAGAGAGATGCTGT
AGAGCACACGTCTGAACTCTTTCCCTACACGACGCTCTTCCGATCTNNNCTGTAATCCCAGCTACTCAGG
GAGCACACGTCTGAACTCTTTCCCTACACGACGCTCTTCCGATCTNNNGATCACACTACTACACTCCAGC
AGCACACGTCTGAACTCTTTCCCTACACGACGCTCTTCCGATCTNNNGAGATCATGCCACTACACTCCA
AGCACACGTCTGAACTCTTTCCCTACACGACGCTCTTCCGATCTNNNGCCAAGATCATGCCATTGTACT
GAGCACACGTCTGAACTCTTTCCCTACACGACGCTCTTCCGATCTNNNAAGAAAGAGCATGTGCAAAG
GAGCACACGTCTGAACTCTTTCCCTACACGACGCTCTTCCGATCTNNNCAACAAACACTGCCAAAATGCT
GAGCACACGTCTGAACTCTTTCCCTACACGACGCTCTTCCGATCTNNNATTTAGGAACCAAGACAGGC

AGAGCACACGTCTGAACTCTTTCCCTACACGACGCTCTTCCGATCTNNNCATGCCTGTAATCCCAGCT
GCACACGTCTGAACTCTTTCCCTACACGACGCTCTTCCGATCTNNNAGGAAGCTGAGACAGGAGAAT
AGCACACGTCTGAACTCTTTCCCTACACGACGCTCTTCCGATCTNNNGGGGTGATGTGTGTACCAAA
GCACACGTCTGAACTCTTTCCCTACACGACGCTCTTCCGATCTNNNGGTGACAGAGTGAGACTCCATC
AGCACACGTCTGAACTCTTTCCCTACACGACGCTCTTCCGATCTNNNTAGAATGAAGAGGAGCACAC
AGCACACGTCTGAACTCTTTCCCTACACGACGCTCTTCCGATCTNNNCTGAAATAAGGTGAGGATGC
AGCACACGTCTGAACTCTTTCCCTACACGACGCTCTTCCGATCTNNNGCTGCATAAGTGAAGATTACG
GAGCACACGTCTGAACTCTTTCCCTACACGACGCTCTTCCGATCTNNNTGAGCTGATATCATGCCACT
AGCACACGTCTGAACTCTTTCCCTACACGACGCTCTTCCGATCTNNNGTGCCTCTAATTGCAGCTACTT
GAGCACACGTCTGAACTCTTTCCCTACACGACGCTCTTCCGATCTNNNGAGATCACACCACTGCACTC
GCACACGTCTGAACTCTTTCCCTACACGACGCTCTTCCGATCTNNNCTGAGATCCCACCACTGTACTTC
AGCACACGTCTGAACTCTTTCCCTACACGACGCTCTTCCGATCTNNNTCCAGGAACTCCAACATGATT
GCACACGTCTGAACTCTTTCCCTACACGACGCTCTTCCGATCTNNNTGAGGGAGGAGAATCACTTGA
AGCACACGTCTGAACTCTTTCCCTACACGACGCTCTTCCGATCTNNNTTTTTGCAGGGGGGTTTGA
GCACACGTCTGAACTCTTTCCCTACACGACGCTCTTCCGATCTNNNCACGTCATTTTCTGTGGTTTCCA
AGCACACGTCTGAACTCTTTCCCTACACGACGCTCTTCCGATCTNNNTACCTTTAGTTGGCTCAGGGT
AGCACACGTCTGAACTCTTTCCCTACACGACGCTCTTCCGATCTNNNGAGGAAGAGAACTGCTTGAAC
AGCACACGTCTGAACTCTTTCCCTACACGACGCTCTTCCGATCTNNNGAATCATGCCATTACACTCCA
AGCACACGTCTGAACTCTTTCCCTACACGACGCTCTTCCGATCTNNNGAGACAGGAGAATTGCTTGA
AGCACACGTCTGAACTCTTTCCCTACACGACGCTCTTCCGATCTNNNGCAGGAGAATCATTTGAACCC
GAGCACACGTCTGAACTCTTTCCCTACACGACGCTCTTCCGATCTNNNGCAGAGGTGGAAGGATTAC
GAGCACACGTCTGAACTCTTTCCCTACACGACGCTCTTCCGATCTNNNAATAGGTTACAGCGAGCCAA
GCACACGTCTGAACTCTTTCCCTACACGACGCTCTTCCGATCTNNNCAGAAGTTGCAGTGAGCTGAA
AGCACACGTCTGAACTCTTTCCCTACACGACGCTCTTCCGATCTNNNGAGACAGGTTTTCGCCATGTT
AGCACACGTCTGAACTCTTTCCCTACACGACGCTCTTCCGATCTNNNATCCAAATACAAGGAGGTGA
GAGCACACGTCTGAACTCTTTCCCTACACGACGCTCTTCCGATCTNNNCCGGCTAACACAATGAAATC
AGCACACGTCTGAACTCTTTCCCTACACGACGCTCTTCCGATCTNNNTGGGAGACTGAGACACGAA
AGCACACGTCTGAACTCTTTCCCTACACGACGCTCTTCCGATCTNNNTATTCAGCCCCAAATGCCAAA
AGCACACGTCTGAACTCTTTCCCTACACGACGCTCTTCCGATCTNNNGTACAGGTTGGTTGGAGGTTC
AGCACACGTCTGAACTCTTTCCCTACACGACGCTCTTCCGATCTNNNTATAATTGCACTACAGGCTGG
GCACACGTCTGAACTCTTTCCCTACACGACGCTCTTCCGATCTNNNTAATCACTTGAACCCAGGAAGC
GAGCACACGTCTGAACTCTTTCCCTACACGACGCTCTTCCGATCTNNNTGGTGTACATCTGTAATCCC
AGCACACGTCTGAACTCTTTCCCTACACGACGCTCTTCCGATCTNNNGAGAATCACTTGAACCTGGGA
AGCACACGTCTGAACTCTTTCCCTACACGACGCTCTTCCGATCTNNNTGACTGTAATCCCAGCTACTT
GAGCACACGTCTGAACTCTTTCCCTACACGACGCTCTTCCGATCTNNNAAAGTAAGCCTGAGTGTCTC
AGCACACGTCTGAACTCTTTCCCTACACGACGCTCTTCCGATCTNNNTGTAAGGAACTAGCTGCATGG
AGCACACGTCTGAACTCTTTCCCTACACGACGCTCTTCCGATCTNNNCGAGGCAGATTAGTATGAACC
GAGCACACGTCTGAACTCTTTCCCTACACGACGCTCTTCCGATCTNNNAAAGCCAAGAAAAGCAGCAC
AGCACACGTCTGAACTCTTTCCCTACACGACGCTCTTCCGATCTNNNAATACAAAAAATTAGCCGGGC
AGAGCACACGTCTGAACTCTTTCCCTACACGACGCTCTTCCGATCTNNNAGGAGAATTGCTTGAACT
AGCACACGTCTGAACTCTTTCCCTACACGACGCTCTTCCGATCTNNNGGAGATTGAGACAACCTGGCT
AGCACACGTCTGAACTCTTTCCCTACACGACGCTCTTCCGATCTNNNGGAGTGTCATGTAAGTGGAC
GCACACGTCTGAACTCTTTCCCTACACGACGCTCTTCCGATCTNNNGGTGGTTGTGGTGACCTGATA

AGCACACGTCTGAACTCTTTCCCTACACGACGCTCTTCCGATCTNNNGAACTGAGATTGCATACCATC
AGCACACGTCTGAACTCTTTCCCTACACGACGCTCTTCCGATCTNNNATCCTGTTCTCTCAGCCAAA
AGCACACGTCTGAACTCTTTCCCTACACGACGCTCTTCCGATCTNNNAGGCAAAAGAATCGCTTGAAC
AGCACACGTCTGAACTCTTTCCCTACACGACGCTCTTCCGATCTNNNGGTGGTTGTGGTGACCTGATA
AGCACACGTCTGAACTCTTTCCCTACACGACGCTCTTCCGATCTNNNGAACTGAGATTGCATACCATC
GAGCACACGTCTGAACTCTTTCCCTACACGACGCTCTTCCGATCTNNNACATACCTGTAGTCCCAGCTA
AGCACACGTCTGAACTCTTTCCCTACACGACGCTCTTCCGATCTNNNAGGCAAAAGAATCGCTTGAAC
GAGCACACGTCTGAACTCTTTCCCTACACGACGCTCTTCCGATCTNNNGCTCACACCTGTAACCCTAGT
AGAGCACACGTCTGAACTCTTTCCCTACACGACGCTCTTCCGATCTNNNCATCCTGGCTAACACAGTGA
AGCACACGTCTGAACTCTTTCCCTACACGACGCTCTTCCGATCTNNNACATCTGTAGTCCCAGCTACTC
AGCACACGTCTGAACTCTTTCCCTACACGACGCTCTTCCGATCTNNNGACAAAATGAGACTCCACCTC
AGAGCACACGTCTGAACTCTTTCCCTACACGACGCTCTTCCGATCTNNNAGATGGAGGATGGAAAG
AGCACACGTCTGAACTCTTTCCCTACACGACGCTCTTCCGATCTNNNCCACTGATATTTGTAGCTGGG
AGCACACGTCTGAACTCTTTCCCTACACGACGCTCTTCCGATCTNNNGAGACAGGAGAATTGCTTGAA
AGCACACGTCTGAACTCTTTCCCTACACGACGCTCTTCCGATCTNNNCACACTCTAACATGGGTGACA
AGCACACGTCTGAACTCTTTCCCTACACGACGCTCTTCCGATCTNNNACTGGCAGAGAATGGTACTT
GCACACGTCTGAACTCTTTCCCTACACGACGCTCTTCCGATCTNNNGGAGTTGGGGCTTCAACATATG
AGCACACGTCTGAACTCTTTCCCTACACGACGCTCTTCCGATCTNNNTGAGACTCGTTATTTGTGGT
AGCACACGTCTGAACTCTTTCCCTACACGACGCTCTTCCGATCTNNNCTGCTGGGTTCTCTTATGACA
AGCACACGTCTGAACTCTTTCCCTACACGACGCTCTTCCGATCTNNNCTGAGGCAGGAGAATCACTTG
GAGCACACGTCTGAACTCTTTCCCTACACGACGCTCTTCCGATCTNNNCTGAGGCAGAGGAATCACTT
AGCACACGTCTGAACTCTTTCCCTACACGACGCTCTTCCGATCTNNNGTGCCTCTAGTTAGCCATCTTG
AGCACACGTCTGAACTCTTTCCCTACACGACGCTCTTCCGATCTNNNTCTTCGCTAGAGAAAGACAGC
AGCACACGTCTGAACTCTTTCCCTACACGACGCTCTTCCGATCTNNNCCCAAATTTGAGGACTGTGAC
AGCACACGTCTGAACTCTTTCCCTACACGACGCTCTTCCGATCTNNNGGTTTCTCCTGAGGCCTCTTT
AGCACACGTCTGAACTCTTTCCCTACACGACGCTCTTCCGATCTNNNTCTGGTGGTCTAGTGGTTAGG
AGCACACGTCTGAACTCTTTCCCTACACGACGCTCTTCCGATCTNNNCCATTCAGCCAGTCTAGGTCTT
AGCACACGTCTGAACTCTTTCCCTACACGACGCTCTTCCGATCTNNNAGAATCACTTGAACCCAGTAG
GAGCACACGTCTGAACTCTTTCCCTACACGACGCTCTTCCGATCTNNNATGAAGGAGTGTTTTTGCCT
AGCACACGTCTGAACTCTTTCCCTACACGACGCTCTTCCGATCTNNNGGAAACACACAACTTGCCAAG
GCACACGTCTGAACTCTTTCCCTACACGACGCTCTTCCGATCTNNNGCTCTTTTCCTGGCTTGAAGATG
AGCACACGTCTGAACTCTTTCCCTACACGACGCTCTTCCGATCTNNNAGAATCACTTGAGAACACGG
AGCACACGTCTGAACTCTTTCCCTACACGACGCTCTTCCGATCTNNNAATTGCCCATCAACCAATGAGT
AGCACACGTCTGAACTCTTTCCCTACACGACGCTCTTCCGATCTNNNGAGGAAAGAAGGAAGGAAG
GAGCACACGTCTGAACTCTTTCCCTACACGACGCTCTTCCGATCTNNNTTCTGTCTCCCTTCCTCTCTTC
AGCACACGTCTGAACTCTTTCCCTACACGACGCTCTTCCGATCTNNNAAAATATTGGGCAACAGAGCG
AGCACACGTCTGAACTCTTTCCCTACACGACGCTCTTCCGATCTNNNTCTGATGGAGAGAACTTGGT
GCACACGTCTGAACTCTTTCCCTACACGACGCTCTTCCGATCTNNNTAAGGAAGTGTGTGTGTGTGT
AGAGCACACGTCTGAACTCTTTCCCTACACGACGCTCTTCCGATCTNNNATTTCAGATGAAAGACCAG
AGCACACGTCTGAACTCTTTCCCTACACGACGCTCTTCCGATCTNNNCTGCCTTCTAGTTGCCGGATAT
AGCACACGTCTGAACTCTTTCCCTACACGACGCTCTTCCGATCTNNNGGTATCTGGGGAGAAAGA
AGCACACGTCTGAACTCTTTCCCTACACGACGCTCTTCCGATCTNNNATCCTGATGTACCCCTGCAAAA
GCACACGTCTGAACTCTTTCCCTACACGACGCTCTTCCGATCTNNNATCCTGATGTACCCCTGCAAAAT

AGCACACGTCTGAACTCTTTCCCTACACGACGCTCTTCCGATCTNNNCATGCAGATACACATGTACACA
AGCACACGTCTGAACTCTTTCCCTACACGACGCTCTTCCGATCTNNNTGTGTGTGTGTGTGTTTCA
AGAGCACACGTCTGAACTCTTTCCCTACACGACGCTCTTCCGATCTNNNACTGGGAGTATGCAGTAAG
AGCACACGTCTGAACTCTTTCCCTACACGACGCTCTTCCGATCTNNNGGAAGGGTAGGACAAAAGGA
AGCACACGTCTGAACTCTTTCCCTACACGACGCTCTTCCGATCTNNNTCTCTGCTATACCTGCTTTCGTC
GCACACGTCTGAACTCTTTCCCTACACGACGCTCTTCCGATCTNNNCTGAGGCAGGAGAATCACTTGA
AGCACACGTCTGAACTCTTTCCCTACACGACGCTCTTCCGATCTNNNAACACTTCCTTTGTTTCTGGCA
GAGCACACGTCTGAACTCTTTCCCTACACGACGCTCTTCCGATCTNNNGTATTCCAACAGAGGTTCCCA
AGCACACGTCTGAACTCTTTCCCTACACGACGCTCTTCCGATCTNNNTTTGTGGCATTGTAACAACTCC
AGCACACGTCTGAACTCTTTCCCTACACGACGCTCTTCCGATCTNNNACCAAAACTGAGCAAGAGTAG
AGCACACGTCTGAACTCTTTCCCTACACGACGCTCTTCCGATCTNNNTTAGCCTTGTGAGTAGTTGGGA
AGAGCACACGTCTGAACTCTTTCCCTACACGACGCTCTTCCGATCTNNNGCCTGTAGTCCCAGTTACCT
GAGCACACGTCTGAACTCTTTCCCTACACGACGCTCTTCCGATCTNNNCTGAGGCAGAAGAATTGCTTG
AGCACACGTCTGAACTCTTTCCCTACACGACGCTCTTCCGATCTNNNTGAGGCAGAAGAATTGCTTGA
GCACACGTCTGAACTCTTTCCCTACACGACGCTCTTCCGATCTNNNAAGAATCACTTGAACCTGGGAG
AGCACACGTCTGAACTCTTTCCCTACACGACGCTCTTCCGATCTNNNCTGAGGCAGGAGAATCACTTG
GAGCACACGTCTGAACTCTTTCCCTACACGACGCTCTTCCGATCTNNNGGAGACCATCCTGGCTAATAT
AGCACACGTCTGAACTCTTTCCCTACACGACGCTCTTCCGATCTNNNTCTTCTCTGGCTGCTTTCAAGAT
GAGCACACGTCTGAACTCTTTCCCTACACGACGCTCTTCCGATCTNNNAGTGAACTGAAATCGTGCCA
GAGCACACGTCTGAACTCTTTCCCTACACGACGCTCTTCCGATCTNNNATCCTAGCTACTCAAGAGGCT
GCACACGTCTGAACTCTTTCCCTACACGACGCTCTTCCGATCTNNNGGGTACATCAACTCTCCAACACT
AGCACACGTCTGAACTCTTTCCCTACACGACGCTCTTCCGATCTNNNAAAAAAAGCCCAAGACCAGACA
AGAGCACACGTCTGAACTCTTTCCCTACACGACGCTCTTCCGATCTNNNGTAACATCTAAGCTACGGCC
AGCACACGTCTGAACTCTTTCCCTACACGACGCTCTTCCGATCTNNNCACCACACCCAGCTATTTTTCTTC
AGCACACGTCTGAACTCTTTCCCTACACGACGCTCTTCCGATCTNNNCCACACTGTTTTCCATGATGGT
AGCACACGTCTGAACTCTTTCCCTACACGACGCTCTTCCGATCTNNNCTGCCGACATAGTCAGAGTTAC
GAGCACACGTCTGAACTCTTTCCCTACACGACGCTCTTCCGATCTNNNAATATGCAGCAACTGAGTTGA
AGCACACGTCTGAACTCTTTCCCTACACGACGCTCTTCCGATCTNNNGAATCCCGTAACAACTGGACTC
GAGCACACGTCTGAACTCTTTCCCTACACGACGCTCTTCCGATCTNNNAGACAGGAGAATCACTTGAA
AGCACACGTCTGAACTCTTTCCCTACACGACGCTCTTCCGATCTNNNAGTCCTAGCTACTCTGGAGACT
GAGCACACGTCTGAACTCTTTCCCTACACGACGCTCTTCCGATCTNNNGTAATCTCAGCTACTCAGGAG
GAGCACACGTCTGAACTCTTTCCCTACACGACGCTCTTCCGATCTNNNCCCAAATAACCGAATGACTCC
AGCACACGTCTGAACTCTTTCCCTACACGACGCTCTTCCGATCTNNNGAGGTTGCAGTGAGCTAAGAT
GAGCACACGTCTGAACTCTTTCCCTACACGACGCTCTTCCGATCTNNNCTGAGGCAGGAGAATCACTT
AGCACACGTCTGAACTCTTTCCCTACACGACGCTCTTCCGATCTNNNACGATCCATTGGTTTCCAGACAT
AGCACACGTCTGAACTCTTTCCCTACACGACGCTCTTCCGATCTNNNGCCTCTGTAATAGCTGGGACTA
GAGCACACGTCTGAACTCTTTCCCTACACGACGCTCTTCCGATCTNNNATGAGACAGGAGAATCGCTTG
GAGCACACGTCTGAACTCTTTCCCTACACGACGCTCTTCCGATCTNNNCAGGAGAATGGTGTAAACTG
AGAGCACACGTCTGAACTCTTTCCCTACACGACGCTCTTCCGATCTNNNTGAGCCCAGAAGTTCAAGA
AGCACACGTCTGAACTCTTTCCCTACACGACGCTCTTCCGATCTNNNTTCTGTAGAGGCGGTATCTCTC
AGCACACGTCTGAACTCTTTCCCTACACGACGCTCTTCCGATCTNNNCCTTTTTTTGCTCAGCATTGTGA
AGCACACGTCTGAACTCTTTCCCTACACGACGCTCTTCCGATCTNNNAGGTTCCAGGATTACAACCTCT
AGCACACGTCTGAACTCTTTCCCTACACGACGCTCTTCCGATCTNNNGTGAGCTTTGATTGAGCCATT

AGCACACGTCTGAACTCTTTCCCTACACGACGCTCTTCCGATCTNNNATGCCTGTAATCCCAGCTATT
GAGCACACGTCTGAACTCTTTCCCTACACGACGCTCTTCCGATCTNNNAAGTTGCAGTAAGCCAAGAT
GAGCACACGTCTGAACTCTTTCCCTACACGACGCTCTTCCGATCTNNNTGAGGCAGGAGAATGGTAT
GAGCACACGTCTGAACTCTTTCCCTACACGACGCTCTTCCGATCTNNNCTGTAATCCCAGCTACTCAG
AGCACACGTCTGAACTCTTTCCCTACACGACGCTCTTCCGATCTNNNACCAACCTTTCCTTCCTCTGAT
AGCACACGTCTGAACTCTTTCCCTACACGACGCTCTTCCGATCTNNNTCATCTCCCACAATGTAACCTC
AGCACACGTCTGAACTCTTTCCCTACACGACGCTCTTCCGATCTNNNGTACAAGTGATTCTCCTGCCTC
AGCACACGTCTGAACTCTTTCCCTACACGACGCTCTTCCGATCTNNNTCTGCCTCACAATTGTGTTGA
GAGCACACGTCTGAACTCTTTCCCTACACGACGCTCTTCCGATCTNNNGTGTTCAGATTGAGCCATTG
GAGCACACGTCTGAACTCTTTCCCTACACGACGCTCTTCCGATCTNNNAACATGGCGAAATCCCATCTC
GAGCACACGTCTGAACTCTTTCCCTACACGACGCTCTTCCGATCTNNNTCACACCTGTAATCCCAACAC
GAGCACACGTCTGAACTCTTTCCCTACACGACGCTCTTCCGATCTNNNGCCTGGGTGATAGAATGAG
AGCACACGTCTGAACTCTTTCCCTACACGACGCTCTTCCGATCTNNNGGAGAATTGCTTGAACTTGGT
AGCACACGTCTGAACTCTTTCCCTACACGACGCTCTTCCGATCTNNNAAAGACACAGACTGGCAAATT
GAGCACACGTCTGAACTCTTTCCCTACACGACGCTCTTCCGATCTNNNAGGCTGAGGCTAGAGAATC
AGCACACGTCTGAACTCTTTCCCTACACGACGCTCTTCCGATCTNNNCTGAGGCAGGAGAATCACTT
AGCACACGTCTGAACTCTTTCCCTACACGACGCTCTTCCGATCTNNNTCATTTGAACCCAGGAGACAG
AGCACACGTCTGAACTCTTTCCCTACACGACGCTCTTCCGATCTNNNCTCTTCTCTCAAGTACACCCACC
GAGCACACGTCTGAACTCTTTCCCTACACGACGCTCTTCCGATCTNNNTTTGGTGCTGAGATCTTACCT
GCACACGTCTGAACTCTTTCCCTACACGACGCTCTTCCGATCTNNNATAGGAGTGGAGGCTAACTTCC
GCACACGTCTGAACTCTTTCCCTACACGACGCTCTTCCGATCTNNNCACCTGTAATCCCAGCTACTTGG
AGCACACGTCTGAACTCTTTCCCTACACGACGCTCTTCCGATCTNNNGAGAATCGCTTGAATCCAGGA
AGCACACGTCTGAACTCTTTCCCTACACGACGCTCTTCCGATCTNNNGAGATCATGCCACTTCACTCC
GCACACGTCTGAACTCTTTCCCTACACGACGCTCTTCCGATCTNNNGAGCTTGCAGTGAGCCAAGATA
AGCACACGTCTGAACTCTTTCCCTACACGACGCTCTTCCGATCTNNNCACTCACCTTTACTTTGCTCTG
AGCACACGTCTGAACTCTTTCCCTACACGACGCTCTTCCGATCTNNNTACTGAAGAGGCTGAGGTAG
AGCACACGTCTGAACTCTTTCCCTACACGACGCTCTTCCGATCTNNNTGTTTCTAGGACCCATGTTGAG
GCACACGTCTGAACTCTTTCCCTACACGACGCTCTTCCGATCTNNNAGGGGAAGCTGATAAACACTT
GCACACGTCTGAACTCTTTCCCTACACGACGCTCTTCCGATCTNNNATGGGCATTTCCTTTCCCTAACT
AGCACACGTCTGAACTCTTTCCCTACACGACGCTCTTCCGATCTNNNTTAGTCCATGGCCATACTTTGC
AGCACACGTCTGAACTCTTTCCCTACACGACGCTCTTCCGATCTNNNGATTACAGACGTGTACCACCAC
GCACACGTCTGAACTCTTTCCCTACACGACGCTCTTCCGATCTNNNTTCCACCTCAACCGACTGAATAG
AGCACACGTCTGAACTCTTTCCCTACACGACGCTCTTCCGATCTNNNACAGTCACCACAGAAGAATCA
GCACACGTCTGAACTCTTTCCCTACACGACGCTCTTCCGATCTNNNTGAGAATTGTCGATTAGCCCAC
AGCACACGTCTGAACTCTTTCCCTACACGACGCTCTTCCGATCTNNNGAGACTGAGGTGGGAGAATC
AGCACACGTCTGAACTCTTTCCCTACACGACGCTCTTCCGATCTNNNCAGAGGTTACAGTGAGCAGA
GAGCACACGTCTGAACTCTTTCCCTACACGACGCTCTTCCGATCTNNNAGGTGGATTATGAGGTCAG
GCACACGTCTGAACTCTTTCCCTACACGACGCTCTTCCGATCTNNNGAGGCTGAAGTGAGCTATGAT
AGCACACGTCTGAACTCTTTCCCTACACGACGCTCTTCCGATCTNNNGAGCAAGCCCAGAACTTGAA
AGCACACGTCTGAACTCTTTCCCTACACGACGCTCTTCCGATCTNNNGCTAGGATGCCATAACCTACA
AGCACACGTCTGAACTCTTTCCCTACACGACGCTCTTCCGATCTNNNGCTTCCTGGATAGCTGGGATT
AGCACACGTCTGAACTCTTTCCCTACACGACGCTCTTCCGATCTNNNGCTTCATGCAACTGCGTAAAAT
AGCACACGTCTGAACTCTTTCCCTACACGACGCTCTTCCGATCTNNNCTGGCTTGCTTTCTTGCTTTCT

GCACACGTCTGAACTCTTTCCCTACACGACGCTCTTCCGATCTNNNAGCATGTGATGTGTTTCCATTTGT
GAGCACACGTCTGAACTCTTTCCCTACACGACGCTCTTCCGATCTNNNGCTTATTTGCTGTCTGCTTGG
GAGCACACGTCTGAACTCTTTCCCTACACGACGCTCTTCCGATCTNNNATCTGCTGGTATCTTGATCTCG
AGCACACGTCTGAACTCTTTCCCTACACGACGCTCTTCCGATCTNNNTCCACTTGTTCCAGTCAGTCATT
AGCACACGTCTGAACTCTTTCCCTACACGACGCTCTTCCGATCTNNNTTACTGCCCCTTTCCATTTCAGTC
AGCACACGTCTGAACTCTTTCCCTACACGACGCTCTTCCGATCTNNNATTTGCATCTGGTAAGAGCCTC
GAGCACACGTCTGAACTCTTTCCCTACACGACGCTCTTCCGATCTNNNAGTGGCATATTCTCAGCTCTCT
AGCACACGTCTGAACTCTTTCCCTACACGACGCTCTTCCGATCTNNNCATGCCTGTAATCCCAGCTACTC
GCACACGTCTGAACTCTTTCCCTACACGACGCTCTTCCGATCTNNNACTTCTGTACTCTAGTCCAGCCTC
AGCACACGTCTGAACTCTTTCCCTACACGACGCTCTTCCGATCTNNNAGAAGTTGCAGTGAGTCGAGA
AGCACACGTCTGAACTCTTTCCCTACACGACGCTCTTCCGATCTNNNTGAAGGTTGCAGTGAGCTAAG
GAGCACACGTCTGAACTCTTTCCCTACACGACGCTCTTCCGATCTNNNAAATCACTTGAACTCAGGAGG
AGCACACGTCTGAACTCTTTCCCTACACGACGCTCTTCCGATCTNNNACTTGAGGTCAGAAGTTCGAGA
GAGCACACGTCTGAACTCTTTCCCTACACGACGCTCTTCCGATCTNNNGGGGACAAAGAAAGACTCCA
AGCACACGTCTGAACTCTTTCCCTACACGACGCTCTTCCGATCTNNNGAGTCAGAGCAAGGCTGTATC
AGCACACGTCTGAACTCTTTCCCTACACGACGCTCTTCCGATCTNNNACTTCATACCCGGACCACAAATT
AGCACACGTCTGAACTCTTTCCCTACACGACGCTCTTCCGATCTNNNTGAAATTTAGGTGGAGGTTCCC
GAGCACACGTCTGAACTCTTTCCCTACACGACGCTCTTCCGATCTNNNTCTGTTTCCCCGCTCTTTTCTTAC
AGCACACGTCTGAACTCTTTCCCTACACGACGCTCTTCCGATCTNNNTCTGGGTGGACTAGTGAACTGT
GAGCACACGTCTGAACTCTTTCCCTACACGACGCTCTTCCGATCTNNNTTGTTTTCCACAGTGACTGTCCT
GAGCACACGTCTGAACTCTTTCCCTACACGACGCTCTTCCGATCTNNNTTACTGTTTGAGGAACTGCCA
AGCACACGTCTGAACTCTTTCCCTACACGACGCTCTTCCGATCTNNNCACAAGTCCCCTGATACCTGTTTC
AGCACACGTCTGAACTCTTTCCCTACACGACGCTCTTCCGATCTNNNGTGACCTCCCTTCTTTCTTTCCTC
AGAGCACACGTCTGAACTCTTTCCCTACACGACGCTCTTCCGATCTNNNAGCCCAGACGTCTCCTCTAAT
GAGCACACGTCTGAACTCTTTCCCTACACGACGCTCTTCCGATCTNNNACTGCATGGTAGAAGGAAGC
AGCACACGTCTGAACTCTTTCCCTACACGACGCTCTTCCGATCTNNNCCCACAAAAAGTCATCAGGGTTC
AGCACACGTCTGAACTCTTTCCCTACACGACGCTCTTCCGATCTNNNTTGCGTCTGAATCCTGTAACACA
AGCACACGTCTGAACTCTTTCCCTACACGACGCTCTTCCGATCTNNNGTTCTCATCACTTAGCTCCCACTC
AGCACACGTCTGAACTCTTTCCCTACACGACGCTCTTCCGATCTNNNAGATTGCAGTGAGCTGAGATCA
AGCACACGTCTGAACTCTTTCCCTACACGACGCTCTTCCGATCTNNNTGAGAATTACTTGAACCCGGGA
GAGCACACGTCTGAACTCTTTCCCTACACGACGCTCTTCCGATCTNNNCTGAGGCAGGAGAATCACTTC
AGCACACGTCTGAACTCTTTCCCTACACGACGCTCTTCCGATCTNNNTCTCTCTCTCTCTCTCTCCCTCA
AGAGCACACGTCTGAACTCTTTCCCTACACGACGCTCTTCCGATCTNNNGCAGAATCCTGGCCTCTATCT
GAGCACACGTCTGAACTCTTTCCCTACACGACGCTCTTCCGATCTNNNGGGAAGAGACTGACAAAGGG
AGCACACGTCTGAACTCTTTCCCTACACGACGCTCTTCCGATCTNNNATGGAGTCTGTCTGTCTGTTCTG
AGCACACGTCTGAACTCTTTCCCTACACGACGCTCTTCCGATCTNNNTTTTCTTTGTCCTCCTCTCCCACCA
AAGAGCACACGTCTGAACTCTTTCCCTACACGACGCTCTTCCGATCTNNNCTGAGACAGGAGAATCGCT
AGCACACGTCTGAACTCTTTCCCTACACGACGCTCTTCCGATCTNNNCTTCTGGGAATATCAGCTGCTG
AGCACACGTCTGAACTCTTTCCCTACACGACGCTCTTCCGATCTNNNATTTCACACCCACTAGAATGGCT
AGCACACGTCTGAACTCTTTCCCTACACGACGCTCTTCCGATCTNNNACATGGTGAAACTCCGTCTCTAC
GAGCACACGTCTGAACTCTTTCCCTACACGACGCTCTTCCGATCTNNNGGTCTTCTGCTTTATCCCTGTC
AGCACACGTCTGAACTCTTTCCCTACACGACGCTCTTCCGATCTNNNAGGCTGGTATCAAAAATGTGCA
AGAGCACACGTCTGAACTCTTTCCCTACACGACGCTCTTCCGATCTNNNCTTTCAGGGGGAGGTTGAAC

AGCACACGTCTGAACTCTTTCCCTACACGACGCTCTTCCGATCTNNNTCAGGACTAAAGCAGTGGAA
GAGCACACGTCTGAACTCTTTCCCTACACGACGCTCTTCCGATCTNNNCCCTCTTTGAAGTGAAGACA
AGCACACGTCTGAACTCTTTCCCTACACGACGCTCTTCCGATCTNNNGTGAGTGCTGAGATCATGCTA
GAGCACACGTCTGAACTCTTTCCCTACACGACGCTCTTCCGATCTNNNCACCACGCCCAGCTAATTTTT
AGCACACGTCTGAACTCTTTCCCTACACGACGCTCTTCCGATCTNNNCTGTAGTTCCAGCTATTCGGG
GCACACGTCTGAACTCTTTCCCTACACGACGCTCTTCCGATCTNNNGGAGGCAGTTTTTCAGGATTTC
GCACACGTCTGAACTCTTTCCCTACACGACGCTCTTCCGATCTNNNCACTGATGATGAGGAGGAGGA
AGCACACGTCTGAACTCTTTCCCTACACGACGCTCTTCCGATCTNNNTTGCACGTAGTTGCAGAATTCA
GAGCACACGTCTGAACTCTTTCCCTACACGACGCTCTTCCGATCTNNNGCTTGGAAGATCAGCAAGCT
AGCACACGTCTGAACTCTTTCCCTACACGACGCTCTTCCGATCTNNNATGGTACTTCCCTCCAAACTGG
GAGCACACGTCTGAACTCTTTCCCTACACGACGCTCTTCCGATCTNNNGTGATCGATGACTCCCAGTT
AGCACACGTCTGAACTCTTTCCCTACACGACGCTCTTCCGATCTNNNATGTTGCTGCCTGATCATTCTTG
AGCACACGTCTGAACTCTTTCCCTACACGACGCTCTTCCGATCTNNNGAGGCAAGCATTTCTGACACTT
AGCACACGTCTGAACTCTTTCCCTACACGACGCTCTTCCGATCTNNNGTGGTGCAATCCTAGCTCAGT
CACACGTCTGAACTCTTTCCCTACACGACGCTCTTCCGATCTNNNGGAGCTTTTCTACCCAGAACTCAC
AGCACACGTCTGAACTCTTTCCCTACACGACGCTCTTCCGATCTNNNTTGAACATGGGGAGTTTAGG
GCACACGTCTGAACTCTTTCCCTACACGACGCTCTTCCGATCTNNNTTTCCACAGTATCTTGCCTTCCAC
AGCACACGTCTGAACTCTTTCCCTACACGACGCTCTTCCGATCTNNNGTCCCAGCTACTTGAAAGTCT
AGCACACGTCTGAACTCTTTCCCTACACGACGCTCTTCCGATCTNNNTGTAATCCCAGCTACTTGGAA
AGCACACGTCTGAACTCTTTCCCTACACGACGCTCTTCCGATCTNNNACTTGGTATACTCACTCCTCCC
AGCACACGTCTGAACTCTTTCCCTACACGACGCTCTTCCGATCTNNNCAAACTCCCAAGCTTGAATCA
AGCACACGTCTGAACTCTTTCCCTACACGACGCTCTTCCGATCTNNNCCATCTTTCATCCTACTGCCCAT
AGCACACGTCTGAACTCTTTCCCTACACGACGCTCTTCCGATCTNNNAATGCCTAAAGGTCTCCAACTCC
AGCACACGTCTGAACTCTTTCCCTACACGACGCTCTTCCGATCTNNNTTGGGTAGACATCAGGTTGTAG
GCACACGTCTGAACTCTTTCCCTACACGACGCTCTTCCGATCTNNNCTTCTGAAAGCTTTGCCTTCACA
AGCACACGTCTGAACTCTTTCCCTACACGACGCTCTTCCGATCTNNNCAAGATGGGAGGATCACTAGA
AGCACACGTCTGAACTCTTTCCCTACACGACGCTCTTCCGATCTNNNAATAATGCCTTTTCCCCCCTCTT
AGCACACGTCTGAACTCTTTCCCTACACGACGCTCTTCCGATCTNNNCCATGATGAGGGATTCCATGA
AGCACACGTCTGAACTCTTTCCCTACACGACGCTCTTCCGATCTNNNGAAGGCCTTATACGTCACAAAG
AGCACACGTCTGAACTCTTTCCCTACACGACGCTCTTCCGATCTNNNCTGATCTCAAGTGATCCTCCCA
AGCACACGTCTGAACTCTTTCCCTACACGACGCTCTTCCGATCTNNNTACAGACATGAGGAAATGGTG
AGAGCACACGTCTGAACTCTTTCCCTACACGACGCTCTTCCGATCTNNNATCACTTGGGTATGCACTGA
AGCACACGTCTGAACTCTTTCCCTACACGACGCTCTTCCGATCTNNNGACGGGGTTTTCCTTGTACAG
AGCACACGTCTGAACTCTTTCCCTACACGACGCTCTTCCGATCTNNNCACAGGACAGAGGGAATCAA
GAGCACACGTCTGAACTCTTTCCCTACACGACGCTCTTCCGATCTNNNAAAAAAAAATAGCCGGGTG
AGCACACGTCTGAACTCTTTCCCTACACGACGCTCTTCCGATCTNNNGAAGGGAGGAAACCACTGTA
AGCACACGTCTGAACTCTTTCCCTACACGACGCTCTTCCGATCTNNNTAGGAAGAAGCAAAAGCGGA
GCACACGTCTGAACTCTTTCCCTACACGACGCTCTTCCGATCTNNNCATCTGTAAATCCAGCTGCTTGG
AGCACACGTCTGAACTCTTTCCCTACACGACGCTCTTCCGATCTNNNTGTAGTCACAGCTACTAGGGA
AGCACACGTCTGAACTCTTTCCCTACACGACGCTCTTCCGATCTNNNTAGCCTCTGGTTTCTCCTTTTCC
AGCACACGTCTGAACTCTTTCCCTACACGACGCTCTTCCGATCTNNNTTAGAAAAGGCCTAACCACTCC
AGCACACGTCTGAACTCTTTCCCTACACGACGCTCTTCCGATCTNNNCATGTCCAACCATCTGAATTCCG
AGCACACGTCTGAACTCTTTCCCTACACGACGCTCTTCCGATCTNNNGGGACCATGTCTTATTCACCC

AGCACACGTCTGAACTCTTTCCCTACACGACGCTCTTCCGATCTNNNCATGGTGGTGTGCATCTGTAA
AGCACACGTCTGAACTCTTTCCCTACACGACGCTCTTCCGATCTNNNTCTCTTGTTTTCAGCCGTGAT
AGCACACGTCTGAACTCTTTCCCTACACGACGCTCTTCCGATCTNNNCTGGAATGCAATGGTGTGATC
GCACACGTCTGAACTCTTTCCCTACACGACGCTCTTCCGATCTNNNATCCACTGTCCCCAAACTTTTTG
AGCACACGTCTGAACTCTTTCCCTACACGACGCTCTTCCGATCTNNNCTACTGCCCTTACCTTCTGGTTT
GAGCACACGTCTGAACTCTTTCCCTACACGACGCTCTTCCGATCTNNNTGTTGACTGTGTGTTGTGTA
AGCACACGTCTGAACTCTTTCCCTACACGACGCTCTTCCGATCTNNNGGTGTATCTGTATGTCCGTGT
GAGCACACGTCTGAACTCTTTCCCTACACGACGCTCTTCCGATCTNNNTAATCCTGGAGTAGAGTGAG
AGCACACGTCTGAACTCTTTCCCTACACGACGCTCTTCCGATCTNNNAAAAAATTAGCCAGGTGTGG
AGCACACGTCTGAACTCTTTCCCTACACGACGCTCTTCCGATCTNNNAAGGAGAAGGAGAGGAAGA
AGCACACGTCTGAACTCTTTCCCTACACGACGCTCTTCCGATCTNNNAATTCCATCCAAGTTGCTGCA
GAGCACACGTCTGAACTCTTTCCCTACACGACGCTCTTCCGATCTNNNAGTCTGAGATCAAACTGCAA
AGCACACGTCTGAACTCTTTCCCTACACGACGCTCTTCCGATCTNNNCTCCCAAGTAGCTGAGACTAC
GAGCACACGTCTGAACTCTTTCCCTACACGACGCTCTTCCGATCTNNNCAAGAAACCTTAGCAACACA
GCACACGTCTGAACTCTTTCCCTACACGACGCTCTTCCGATCTNNNGCAGTCAGGGAGGTTTCATATC
AGCACACGTCTGAACTCTTTCCCTACACGACGCTCTTCCGATCTNNNCGGAGTAACATGAGGGAGG
GAGCACACGTCTGAACTCTTTCCCTACACGACGCTCTTCCGATCTNNNGATGCAATGAAAGGCCATAG
AGCACACGTCTGAACTCTTTCCCTACACGACGCTCTTCCGATCTNNNCTGTAGTCTTCAGAACAGCAC
GAGCACACGTCTGAACTCTTTCCCTACACGACGCTCTTCCGATCTNNNGGGCGACAATCATCAGGTTA
AAGAGCACACGTCTGAACTCTTTCCCTACACGACGCTCTTCCGATCTNNNCTGTCGATCCAATCACCTC
GAGCACACGTCTGAACTCTTTCCCTACACGACGCTCTTCCGATCTNNNTAATTCAACTCCACTGTGGCC
AGCACACGTCTGAACTCTTTCCCTACACGACGCTCTTCCGATCTNNNAGAGAGATCCTATGCACCCCA
AGCACACGTCTGAACTCTTTCCCTACACGACGCTCTTCCGATCTNNNCACACACATACACACACACACA
GAGCACACGTCTGAACTCTTTCCCTACACGACGCTCTTCCGATCTNNNGCCACACATACTGACACCTAA
AGCACACGTCTGAACTCTTTCCCTACACGACGCTCTTCCGATCTNNNCAGAGTTTTTGCGCTGGTTTTT
AGCACACGTCTGAACTCTTTCCCTACACGACGCTCTTCCGATCTNNNCTCACGCCTATAATCCCATCACT
GCACACGTCTGAACTCTTTCCCTACACGACGCTCTTCCGATCTNNNTTACACACACACACACACACAC
AGCACACGTCTGAACTCTTTCCCTACACGACGCTCTTCCGATCTNNNAAACAGACCAATCAGCTCTCTG
GCACACGTCTGAACTCTTTCCCTACACGACGCTCTTCCGATCTNNNATTGCTGGGGAACTAGTGTGA
AGCACACGTCTGAACTCTTTCCCTACACGACGCTCTTCCGATCTNNNGCCAAAGAGTATTGAAGAGG
AGCACACGTCTGAACTCTTTCCCTACACGACGCTCTTCCGATCTNNNCATCCTGGCTAACATGGTGAA
AGCACACGTCTGAACTCTTTCCCTACACGACGCTCTTCCGATCTNNNCTTGAGCCCAGGAAGTTGAGG
AGCACACGTCTGAACTCTTTCCCTACACGACGCTCTTCCGATCTNNNTGGGTATATGTGTGCATGTCT
GCACACGTCTGAACTCTTTCCCTACACGACGCTCTTCCGATCTNNNGCATCTCAGTTTGGAGGCATTTT
GAGCACACGTCTGAACTCTTTCCCTACACGACGCTCTTCCGATCTNNNGTATGTGTACAGGCATGTGT
AGCACACGTCTGAACTCTTTCCCTACACGACGCTCTTCCGATCTNNNTACACACACTACATTGTAGGTG
AGCACACGTCTGAACTCTTTCCCTACACGACGCTCTTCCGATCTNNNGCCCTTTATCGTGAGTAACTT
GCACACGTCTGAACTCTTTCCCTACACGACGCTCTTCCGATCTNNNTTTCTCAGCACCTCTCTCTTTCCCA
GCACACGTCTGAACTCTTTCCCTACACGACGCTCTTCCGATCTNNNTGTAAGTGTGCAGTTCAGTGGT
AGCACACGTCTGAACTCTTTCCCTACACGACGCTCTTCCGATCTNNNTTGCCATTGTGTCTTGATGTCA
AGCACACGTCTGAACTCTTTCCCTACACGACGCTCTTCCGATCTNNNTTGATAGGACTGTGAGTGTG
GCACACGTCTGAACTCTTTCCCTACACGACGCTCTTCCGATCTNNNCAGCCATGTTGAAATTCCATCCT
GAGCACACGTCTGAACTCTTTCCCTACACGACGCTCTTCCGATCTNNNTGTCACTCAATGAACCTCCTC

GAGCACACGTCTGAACTCTTTCCCTACACGACGCTCTTCCGATCTNNNCAGCAATGTAGCATATCCTGT
GAGCACACGTCTGAACTCTTTCCCTACACGACGCTCTTCCGATCTNNNCTGGCAAGAAAAACATCTCCA
AGCACACGTCTGAACTCTTTCCCTACACGACGCTCTTCCGATCTNNNCTGTGATCATGCCACTACACTC
AGAGCACACGTCTGAACTCTTTCCCTACACGACGCTCTTCCGATCTNNNCTTTGATCCACAATCGGCCT
AGCACACGTCTGAACTCTTTCCCTACACGACGCTCTTCCGATCTNNNTTTATACACACACACACGCACAC
GAGCACACGTCTGAACTCTTTCCCTACACGACGCTCTTCCGATCTNNNGAGGGTGGGCCTTATGAAAT
AGCACACGTCTGAACTCTTTCCCTACACGACGCTCTTCCGATCTNNNGAGCAAGACTCCGTCTCAAAA
AGCACACGTCTGAACTCTTTCCCTACACGACGCTCTTCCGATCTNNNTTGAGATCTGAGAACAGGCAG
GAGCACACGTCTGAACTCTTTCCCTACACGACGCTCTTCCGATCTNNNAACGAAGGTAAGGTTCTTGG
AGAGCACACGTCTGAACTCTTTCCCTACACGACGCTCTTCCGATCTNNNGATTGTGTGAGTGTGTGAG
GAGCACACGTCTGAACTCTTTCCCTACACGACGCTCTTCCGATCTNNNTCCATCAGTTGTAGCAAACAT
AGCACACGTCTGAACTCTTTCCCTACACGACGCTCTTCCGATCTNNNTACACACATACGTACAAGGCAC
AGAGCACACGTCTGAACTCTTTCCCTACACGACGCTCTTCCGATCTNNNAGCAGTCTCTCTCTCTCTCT
AGCACACGTCTGAACTCTTTCCCTACACGACGCTCTTCCGATCTNNNCTCACACACACACAAATAGCCAC
AGCACACGTCTGAACTCTTTCCCTACACGACGCTCTTCCGATCTNNNCCTTTGTCTCTCTCTGCATCTGTC
AGCACACGTCTGAACTCTTTCCCTACACGACGCTCTTCCGATCTNNNCAAAAACTTAGCCGAGCATGG
AGCACACGTCTGAACTCTTTCCCTACACGACGCTCTTCCGATCTNNNGAATGCTAGAGATCCTTGTCCC
AGCACACGTCTGAACTCTTTCCCTACACGACGCTCTTCCGATCTNNNTGGTAGGAATGTTGGTCTGG
AGCACACGTCTGAACTCTTTCCCTACACGACGCTCTTCCGATCTNNNCATTCCTTAGCAAGTTTGGCCA
GAGCACACGTCTGAACTCTTTCCCTACACGACGCTCTTCCGATCTNNNATATACAGGCAGGACACACA
AGCACACGTCTGAACTCTTTCCCTACACGACGCTCTTCCGATCTNNNGGGATGAATAGGAGTTCACCT
AGCACACGTCTGAACTCTTTCCCTACACGACGCTCTTCCGATCTNNNCACAAGGTGGTAGTATGGCTT
GCACACGTCTGAACTCTTTCCCTACACGACGCTCTTCCGATCTNNNTTATTCACCCACGTTTCAGAGGTC
AGCACACGTCTGAACTCTTTCCCTACACGACGCTCTTCCGATCTNNNAAGTGGTCCTCAGCTAAGAAA
AGCACACGTCTGAACTCTTTCCCTACACGACGCTCTTCCGATCTNNNGAGAAGCAAGAAGAAACCAA
AGCACACGTCTGAACTCTTTCCCTACACGACGCTCTTCCGATCTNNNACACACATACACACACATACAC
AGCACACGTCTGAACTCTTTCCCTACACGACGCTCTTCCGATCTNNNTAAAGACCAAAGGCACTCTTCC
GAGCACACGTCTGAACTCTTTCCCTACACGACGCTCTTCCGATCTNNNGCACAGATACACAGACAGAC
AGAGCACACGTCTGAACTCTTTCCCTACACGACGCTCTTCCGATCTNNNAGACTCACCTGTCATGTTTC
AGCACACGTCTGAACTCTTTCCCTACACGACGCTCTTCCGATCTNNNCTGCAGTGTTGTGTTCCTGTAA
AGCACACGTCTGAACTCTTTCCCTACACGACGCTCTTCCGATCTNNNTGTAGTCCCAGCTACTTAGGAG
GAGCACACGTCTGAACTCTTTCCCTACACGACGCTCTTCCGATCTNNNAAATAGAAGTAGGCCAAGGT
AGCACACGTCTGAACTCTTTCCCTACACGACGCTCTTCCGATCTNNNCTTGTAATTCCAGCTACTCGGG
GAGCACACGTCTGAACTCTTTCCCTACACGACGCTCTTCCGATCTNNNGCACACCAGTACAGAGGAAC
AGCACACGTCTGAACTCTTTCCCTACACGACGCTCTTCCGATCTNNNTCTCATCCCAGCTACTCAAGGAG
AGCACACGTCTGAACTCTTTCCCTACACGACGCTCTTCCGATCTNNNATTGCAACTGGTAATTCAGAG
AGCACACGTCTGAACTCTTTCCCTACACGACGCTCTTCCGATCTNNNTAGAAAGAGGACACCCTTACCC
AGCACACGTCTGAACTCTTTCCCTACACGACGCTCTTCCGATCTNNNCTGATCATTGCCATCCAGACCTA
GCACACGTCTGAACTCTTTCCCTACACGACGCTCTTCCGATCTNNNTTGAACTTGTGTTTGCCTCTTGG
GAGCACACGTCTGAACTCTTTCCCTACACGACGCTCTTCCGATCTNNNCACCCCTTCCAAGATTGACAA
AAGAGCACACGTCTGAACTCTTTCCCTACACGACGCTCTTCCGATCTNNNGCTACTCTGGAGGCTGAA
AGCACACGTCTGAACTCTTTCCCTACACGACGCTCTTCCGATCTNNNGTGAGCTCCCATCTCAGAATC
AGCACACGTCTGAACTCTTTCCCTACACGACGCTCTTCCGATCTNNNTGTAATTCCAGCTACTTGGGA

GCACACGTCTGAACTCTTTCCCTACACGACGCTCTTCCGATCTNNNAGGTTTCAGTGAGCCAAGATCA
GAGCACACGTCTGAACTCTTTCCCTACACGACGCTCTTCCGATCTNNNCTGAGGCAGGAGAATTGCTT
AGCACACGTCTGAACTCTTTCCCTACACGACGCTCTTCCGATCTNNNCTGAGGCAGGAGAATCACTTG
AGCACACGTCTGAACTCTTTCCCTACACGACGCTCTTCCGATCTNNNGAGGCAGAAGAATTGCTTGT
GAGCACACGTCTGAACTCTTTCCCTACACGACGCTCTTCCGATCTNNNGGTGAGCTGAGATTTATTGT
AGCACACGTCTGAACTCTTTCCCTACACGACGCTCTTCCGATCTNNNGAGAATCGCTTGAACTTGGGA
AGCACACGTCTGAACTCTTTCCCTACACGACGCTCTTCCGATCTNNNGCCTATAATCCCAGCTACTCAG
AGCACACGTCTGAACTCTTTCCCTACACGACGCTCTTCCGATCTNNNCTGAGGCAGAAGAATCACCTG
AGCACACGTCTGAACTCTTTCCCTACACGACGCTCTTCCGATCTNNNCTGTAGTCCTACCTGCTCTGG
GCACACGTCTGAACTCTTTCCCTACACGACGCTCTTCCGATCTNNNTGAGGCAGGAGAATCACTTGA
AGCACACGTCTGAACTCTTTCCCTACACGACGCTCTTCCGATCTNNNTCGATCTCCTGACTTTGTGAT
GCACACGTCTGAACTCTTTCCCTACACGACGCTCTTCCGATCTNNNGCTATGGAGCCCTTTTATGCAT
AGCACACGTCTGAACTCTTTCCCTACACGACGCTCTTCCGATCTNNNTTGACCTCCTGAGTAGCTAGG
AGCACACGTCTGAACTCTTTCCCTACACGACGCTCTTCCGATCTNNNCCACCACACCCTGCTAATTTT
GAGCACACGTCTGAACTCTTTCCCTACACGACGCTCTTCCGATCTNNNTCCTGTTCTTCCTAGAGCTTG
GAGCACACGTCTGAACTCTTTCCCTACACGACGCTCTTCCGATCTNNNGGATGTGGGCTTTCAAGTTT
AGCACACGTCTGAACTCTTTCCCTACACGACGCTCTTCCGATCTNNNAAAAATCCTTGCCCAGACCAA
AGCACACGTCTGAACTCTTTCCCTACACGACGCTCTTCCGATCTNNNTGTGCCTGGCTAATTTTTGTG
AGCACACGTCTGAACTCTTTCCCTACACGACGCTCTTCCGATCTNNNTACAGATGAAAAGACTGAGG
GCACACGTCTGAACTCTTTCCCTACACGACGCTCTTCCGATCTNNNTGTCTATGAGAAGGTCACCCTC
GAGCACACGTCTGAACTCTTTCCCTACACGACGCTCTTCCGATCTNNNCCTGTTTGATGTGGAAGCCT
GCACACGTCTGAACTCTTTCCCTACACGACGCTCTTCCGATCTNNNTTGTCTGTCAATTTGTAGTGCCA
GCACACGTCTGAACTCTTTCCCTACACGACGCTCTTCCGATCTNNNGATGTGAAGGGATCTGCAGAG
AGCACACGTCTGAACTCTTTCCCTACACGACGCTCTTCCGATCTNNNCCAAAGTGTTGAGATTACAGG
AGCACACGTCTGAACTCTTTCCCTACACGACGCTCTTCCGATCTNNNTCATCTTTTTTCCTGCCCCAAAC
AGCACACGTCTGAACTCTTTCCCTACACGACGCTCTTCCGATCTNNNTCAGGTAATGAGAGTCCTCCA
GCACACGTCTGAACTCTTTCCCTACACGACGCTCTTCCGATCTNNNAGAATGTGCTGATGCCTGTAG
AGCACACGTCTGAACTCTTTCCCTACACGACGCTCTTCCGATCTNNNTCCATTACTTCTCCCTGGAACT
AGCACACGTCTGAACTCTTTCCCTACACGACGCTCTTCCGATCTNNNTCGTATGTTTGTGTCTTTCTGG
AGCACACGTCTGAACTCTTTCCCTACACGACGCTCTTCCGATCTNNNTGGCACGGTACTAGGTGTTTT
AGCACACGTCTGAACTCTTTCCCTACACGACGCTCTTCCGATCTNNNGTGGGTATGCACCTTATTTCT
GCACACGTCTGAACTCTTTCCCTACACGACGCTCTTCCGATCTNNNGGTTCTCCAGTCATCTTGGATG
GCACACGTCTGAACTCTTTCCCTACACGACGCTCTTCCGATCTNNNGCTGAGGCAGGAGAGAGAAT
AGCACACGTCTGAACTCTTTCCCTACACGACGCTCTTCCGATCTNNNGGAGGACAAAGTGAGACTC
AGCACACGTCTGAACTCTTTCCCTACACGACGCTCTTCCGATCTNNNTTTATTTGCAACCACGGAAGA
AGCACACGTCTGAACTCTTTCCCTACACGACGCTCTTCCGATCTNNNCAAGGAGTTTGAGAACAGCTT
AGCACACGTCTGAACTCTTTCCCTACACGACGCTCTTCCGATCTNNNGTGGCTCACTCCTATAATCCCA
AGCACACGTCTGAACTCTTTCCCTACACGACGCTCTTCCGATCTNNNGCTTGAGTCCAGGAGTTAGA
AGCACACGTCTGAACTCTTTCCCTACACGACGCTCTTCCGATCTNNNGAGAATCGCTTGAATCCAGG
GAGCACACGTCTGAACTCTTTCCCTACACGACGCTCTTCCGATCTNNNGAGGTTGCAGTAAGCCAAG
AGCACACGTCTGAACTCTTTCCCTACACGACGCTCTTCCGATCTNNNTGAGGCAGGAGAATCTCTTGA
GCACACGTCTGAACTCTTTCCCTACACGACGCTCTTCCGATCTNNNGCACTGGGTTCAGAATTTCCTC
GAGCACACGTCTGAACTCTTTCCCTACACGACGCTCTTCCGATCTNNNTGGAACATTGCATGGTTTAT

GCACACGTCTGAACTCTTTCCCTACACGACGCTCTTCCGATCTNNNGAATCTGCCATTGGTGCATACA
AGCACACGTCTGAACTCTTTCCCTACACGACGCTCTTCCGATCTNNNTTCTTTTCAGGGAGACTCACCA
GCACACGTCTGAACTCTTTCCCTACACGACGCTCTTCCGATCTNNNGTTCACACCATTCTCCTACCTCA
GCACACGTCTGAACTCTTTCCCTACACGACGCTCTTCCGATCTNNNCAGAGTCTGACCTCCCACTTTA
GCACACGTCTGAACTCTTTCCCTACACGACGCTCTTCCGATCTNNNGAAATGGGGTTTCTCCATGTTG
GCACACGTCTGAACTCTTTCCCTACACGACGCTCTTCCGATCTNNNCCTCTTTGTGCCTCTGGTAGAAT
AGCACACGTCTGAACTCTTTCCCTACACGACGCTCTTCCGATCTNNNTGAGCATCAGTAGTCCATGT
GCACACGTCTGAACTCTTTCCCTACACGACGCTCTTCCGATCTNNNATGGTCTCAATCTCCTGACTTCG
AGCACACGTCTGAACTCTTTCCCTACACGACGCTCTTCCGATCTNNNTCCTCGTGAACATAGATGCAA
GCACACGTCTGAACTCTTTCCCTACACGACGCTCTTCCGATCTNNNTTTAGTAGAGACGGGGCTTC
GCACACGTCTGAACTCTTTCCCTACACGACGCTCTTCCGATCTNNNCTCCTGATTAGCTGGGACTACA
GCACACGTCTGAACTCTTTCCCTACACGACGCTCTTCCGATCTNNNGCCTTCCTTCCTTCCTTTCTTCTC
AGCACACGTCTGAACTCTTTCCCTACACGACGCTCTTCCGATCTNNNAGTTTCTCCTCATTTTTGCCAG
GCACACGTCTGAACTCTTTCCCTACACGACGCTCTTCCGATCTNNNTATGATAGCACCACTGCATTCCA
AGCACACGTCTGAACTCTTTCCCTACACGACGCTCTTCCGATCTNNNCCATTGTTCGGTCTATTCAGCA
AGCACACGTCTGAACTCTTTCCCTACACGACGCTCTTCCGATCTNNNTGAAAATTCCACCACTGTGTGA
GCACACGTCTGAACTCTTTCCCTACACGACGCTCTTCCGATCTNNNTCCTGAAATCTTATGCAAATGGG
AGCACACGTCTGAACTCTTTCCCTACACGACGCTCTTCCGATCTNNNATCCCTCATTCATCTACCACGT
GCACACGTCTGAACTCTTTCCCTACACGACGCTCTTCCGATCTNNNGTATCTAGCAGTCACCACCCTTT
AGCACACGTCTGAACTCTTTCCCTACACGACGCTCTTCCGATCTNNNCACGCCCAGCTAATTTTTGTA
AGCACACGTCTGAACTCTTTCCCTACACGACGCTCTTCCGATCTNNNAGGCAGGAGAATCATTTGAAC
AGCACACGTCTGAACTCTTTCCCTACACGACGCTCTTCCGATCTNNNGAGCTTGCAGTGAGTCAAGAT
GCACACGTCTGAACTCTTTCCCTACACGACGCTCTTCCGATCTNNNCTGTAGTCCCAGCTACTCATGAG
AGAGCACACGTCTGAACTCTTTCCCTACACGACGCTCTTCCGATCTNNNGCCTGTAATCTCAGCTACTA
AGCACACGTCTGAACTCTTTCCCTACACGACGCTCTTCCGATCTNNNACTGTACTCTAGCCTGGTGAG
AGCACACGTCTGAACTCTTTCCCTACACGACGCTCTTCCGATCTNNNCTGTATTCCCAGCTACTCAGA
AGCACACGTCTGAACTCTTTCCCTACACGACGCTCTTCCGATCTNNNTTCATGGTAAAAACCCTGCACT
AGAGCACACGTCTGAACTCTTTCCCTACACGACGCTCTTCCGATCTNNNGCAACATGGATGAATCACG
AGCACACGTCTGAACTCTTTCCCTACACGACGCTCTTCCGATCTNNNGTAATCTCAGCTACTCAGGAG
AGCACACGTCTGAACTCTTTCCCTACACGACGCTCTTCCGATCTNNNGCATTATGGTGCATGCCTGTA
AGCACACGTCTGAACTCTTTCCCTACACGACGCTCTTCCGATCTNNNGAGGCAGAAGAATCGCTTGA
AGCACACGTCTGAACTCTTTCCCTACACGACGCTCTTCCGATCTNNNGAGATTGCAGTGAGCCAAGAT
AGCACACGTCTGAACTCTTTCCCTACACGACGCTCTTCCGATCTNNNGGTAATAGAGAGGTTGAGGC
AGCACACGTCTGAACTCTTTCCCTACACGACGCTCTTCCGATCTNNNTAACCCCAGATACTCAGGATG
GCACACGTCTGAACTCTTTCCCTACACGACGCTCTTCCGATCTNNNCCTAGGAGGTAGAGTTTGCAG
GCACACGTCTGAACTCTTTCCCTACACGACGCTCTTCCGATCTNNNGAGGTTGCAGTGAACCAAGATT
AGCACACGTCTGAACTCTTTCCCTACACGACGCTCTTCCGATCTNNNGTCAAGACCCATCAGTGTACT
AGCACACGTCTGAACTCTTTCCCTACACGACGCTCTTCCGATCTNNNGTAAGCCGAGATCATGACATT
AGCACACGTCTGAACTCTTTCCCTACACGACGCTCTTCCGATCTNNNGAGAATGGAGTGAACCTGAG
AGCACACGTCTGAACTCTTTCCCTACACGACGCTCTTCCGATCTNNNAAAGTCGAAACCCCATGTCTA
GAGCACACGTCTGAACTCTTTCCCTACACGACGCTCTTCCGATCTNNNGTAATCCCAGCTACTTAGGA
GAGCACACGTCTGAACTCTTTCCCTACACGACGCTCTTCCGATCTNNNACGAGAATCACTTGAACCCA
AGCACACGTCTGAACTCTTTCCCTACACGACGCTCTTCCGATCTNNNTAAAGAGTCAAGACCCATCGG

AGCACACGTCTGAACTCTTTCCCTACACGACGCTCTTCCGATCTNNNGGTGCCTATAATCCCAGCTACT
AGAGCACACGTCTGAACTCTTTCCCTACACGACGCTCTTCCGATCTNNNGCATGTAGTCCTGGCTATTC
GAGCACACGTCTGAACTCTTTCCCTACACGACGCTCTTCCGATCTNNNCAGGGCAACATAGTGAGAA
AGAGCACACGTCTGAACTCTTTCCCTACACGACGCTCTTCCGATCTNNNGCTTGCAGTGAGCTGAGAT
AGAGCACACGTCTGAACTCTTTCCCTACACGACGCTCTTCCGATCTNNNGAGGTTGCAGTAAGTCGAG
GCACACGTCTGAACTCTTTCCCTACACGACGCTCTTCCGATCTNNNATGTCAGGAGGCTTTCACAGTAC
GAGCACACGTCTGAACTCTTTCCCTACACGACGCTCTTCCGATCTNNNGGCCTGAATTCCTAGATGAG
GCACACGTCTGAACTCTTTCCCTACACGACGCTCTTCCGATCTNNNGTTCCAGGCTGCTAATCACTTTG
AGCACACGTCTGAACTCTTTCCCTACACGACGCTCTTCCGATCTNNNTCAACGTCCCTCCTAGAAAACTG
GAGCACACGTCTGAACTCTTTCCCTACACGACGCTCTTCCGATCTNNNGTAGACTCACACAAAGCAACA
AGCACACGTCTGAACTCTTTCCCTACACGACGCTCTTCCGATCTNNNTATGAAAGACTGGCTTGTTTGC
AGCACACGTCTGAACTCTTTCCCTACACGACGCTCTTCCGATCTNNNTCCTCAGTGACGTTCATCTTGTT
GAGCACACGTCTGAACTCTTTCCCTACACGACGCTCTTCCGATCTNNNCCCAAACTCTCAACTTTCCCCTA
GCACACGTCTGAACTCTTTCCCTACACGACGCTCTTCCGATCTNNNGCTTTCGTTTTTTGATCCCAGGA
GCACACGTCTGAACTCTTTCCCTACACGACGCTCTTCCGATCTNNNTTTCCAACTTGGTTGCATTCTCCC
GAGCACACGTCTGAACTCTTTCCCTACACGACGCTCTTCCGATCTNNNCATTATTACCCCCTTCCCCCTTT
GAGCACACGTCTGAACTCTTTCCCTACACGACGCTCTTCCGATCTNNNTTCTGATTGCTGTCCCTCACT
AGCACACGTCTGAACTCTTTCCCTACACGACGCTCTTCCGATCTNNNGCCAGGGTTTCTTTGCTTTCTTT
GCACACGTCTGAACTCTTTCCCTACACGACGCTCTTCCGATCTNNNTTCTCCATTCTCCATTGTGGTCCC
AGCACACGTCTGAACTCTTTCCCTACACGACGCTCTTCCGATCTNNNAATGGCAATGAAGTCAAGCACT
GCACACGTCTGAACTCTTTCCCTACACGACGCTCTTCCGATCTNNNTGAGCACAGTGAAGTTTAGCTC
AGCACACGTCTGAACTCTTTCCCTACACGACGCTCTTCCGATCTNNNTTTTGGTAGAGACAGGGTTTT
AGCACACGTCTGAACTCTTTCCCTACACGACGCTCTTCCGATCTNNNAAAGGTGTATGTGAAACTTGC
GCACACGTCTGAACTCTTTCCCTACACGACGCTCTTCCGATCTNNNTGGTCTCAAATTCCTGACCTCAG
AGCACACGTCTGAACTCTTTCCCTACACGACGCTCTTCCGATCTNNNGTCTCAAACAAGACCGAGGAT
AGCACACGTCTGAACTCTTTCCCTACACGACGCTCTTCCGATCTNNNCTTCTCAGGACCTAGCACAGTA
AGCACACGTCTGAACTCTTTCCCTACACGACGCTCTTCCGATCTNNNAATTAGCCAGAGATGATGGTG
GAGCACACGTCTGAACTCTTTCCCTACACGACGCTCTTCCGATCTNNNTGAAGGTTGCAGTAAGCTGA
AGAGCACACGTCTGAACTCTTTCCCTACACGACGCTCTTCCGATCTNNNTGTAGTCCTAGCTACTCAGG
AGCACACGTCTGAACTCTTTCCCTACACGACGCTCTTCCGATCTNNNACACCTGTAGTCCCAGCTACTAA
GCACACGTCTGAACTCTTTCCCTACACGACGCTCTTCCGATCTNNNGATCCTGCAGAAGAGGAGTTCT
AGCACACGTCTGAACTCTTTCCCTACACGACGCTCTTCCGATCTNNNCAGGAGAATCACTTGAACCCA
AGCACACGTCTGAACTCTTTCCCTACACGACGCTCTTCCGATCTNNNGCAACAATAGCGAAACTCCATC
GAGCACACGTCTGAACTCTTTCCCTACACGACGCTCTTCCGATCTNNNTAGGTGAGGGAATGAGACTC
AGCACACGTCTGAACTCTTTCCCTACACGACGCTCTTCCGATCTNNNTCATCTATTGCCTGAGAGCACA
GAGCACACGTCTGAACTCTTTCCCTACACGACGCTCTTCCGATCTNNNGTAATTCTTTTTGTGGCCGGG
AGCACACGTCTGAACTCTTTCCCTACACGACGCTCTTCCGATCTNNNTAGGCAAATCACTTGAACCTGG
GAGCACACGTCTGAACTCTTTCCCTACACGACGCTCTTCCGATCTNNNGGGAGGTTGCAATGAACTGA
AGCACACGTCTGAACTCTTTCCCTACACGACGCTCTTCCGATCTNNNGTAATCCCAGCTACTCAGGAGT
AGCACACGTCTGAACTCTTTCCCTACACGACGCTCTTCCGATCTNNNCTGTAGTCCTAGCTACTCGGAG
AGCACACGTCTGAACTCTTTCCCTACACGACGCTCTTCCGATCTNNNGAGACTGTAGCGAGACAAGA
GAGCACACGTCTGAACTCTTTCCCTACACGACGCTCTTCCGATCTNNNAGGTAGGAGAATTGCTTGAA
AGCACACGTCTGAACTCTTTCCCTACACGACGCTCTTCCGATCTNNNAGAGAATCGCTTGAACCTGAG

AGAGCACACGTCTGAACTCTTTCCCTACACGACGCTCTTCCGATCTNNNCTGTAATCTCAGCTACTCGG
AGCACACGTCTGAACTCTTTCCCTACACGACGCTCTTCCGATCTNNNTCCTTTGCTTTCTCCATGCAAAC
AGAGCACACGTCTGAACTCTTTCCCTACACGACGCTCTTCCGATCTNNNCTGAGAAATGGAAGACCCT
GAGCACACGTCTGAACTCTTTCCCTACACGACGCTCTTCCGATCTNNNGCATATAATCCCAGCTACTCG
AGCACACGTCTGAACTCTTTCCCTACACGACGCTCTTCCGATCTNNNCTGAGTCAGGAGAATCGCTTG
GAGCACACGTCTGAACTCTTTCCCTACACGACGCTCTTCCGATCTNNNGGAGAATCACTTGAACCTGG
AGAGCACACGTCTGAACTCTTTCCCTACACGACGCTCTTCCGATCTNNNGTCAGGATTGTTGTGGTTTC
GAGCACACGTCTGAACTCTTTCCCTACACGACGCTCTTCCGATCTNNNCAGCTACTTAGGAGGTGAAG
AGCACACGTCTGAACTCTTTCCCTACACGACGCTCTTCCGATCTNNNTCCAGAAGAGGCAAATTCATG
AGCACACGTCTGAACTCTTTCCCTACACGACGCTCTTCCGATCTNNNCTTCCAGGTTCATGCCATTCTCC
GAGCACACGTCTGAACTCTTTCCCTACACGACGCTCTTCCGATCTNNNCATCTGAATGGACTTCCTCCT
GCACACGTCTGAACTCTTTCCCTACACGACGCTCTTCCGATCTNNNGCAGTACCTACCACTATGCCATT
AGCACACGTCTGAACTCTTTCCCTACACGACGCTCTTCCGATCTNNNGGCAGCCTTTGTTCTGAACTAT
AGCACACGTCTGAACTCTTTCCCTACACGACGCTCTTCCGATCTNNNTCGTCATCTTCACAGTTGGTCTT
GAGCACACGTCTGAACTCTTTCCCTACACGACGCTCTTCCGATCTNNNTAATAACAGGCCAGCAAACAT
AGCACACGTCTGAACTCTTTCCCTACACGACGCTCTTCCGATCTNNNGGTCAGTCTAAGCCTCCTGTTTT
AGCACACGTCTGAACTCTTTCCCTACACGACGCTCTTCCGATCTNNNCATTTTCCTTTTCCCTGGCTCTGC
GAGCACACGTCTGAACTCTTTCCCTACACGACGCTCTTCCGATCTNNNAACTATTTGAGGGCTGAGTTC
GAGCACACGTCTGAACTCTTTCCCTACACGACGCTCTTCCGATCTNNNATTAGCGCTAAATGCCATGCA
AGCACACGTCTGAACTCTTTCCCTACACGACGCTCTTCCGATCTNNNGACAAGGTTTGATTTCTGGCTC
AGAGCACACGTCTGAACTCTTTCCCTACACGACGCTCTTCCGATCTNNNATCAGGAAAAGTGTCCTCTA
AGCACACGTCTGAACTCTTTCCCTACACGACGCTCTTCCGATCTNNNTAATAATCCTGGGAGTTTGGCC
AGCACACGTCTGAACTCTTTCCCTACACGACGCTCTTCCGATCTNNNAGGCTTTCCGAAGAACTTCAAA
AGAGCACACGTCTGAACTCTTTCCCTACACGACGCTCTTCCGATCTNNNAACCTGTAAGTCTCTGCTTGA
AGCACACGTCTGAACTCTTTCCCTACACGACGCTCTTCCGATCTNNNTCGGCAAATCACTATGTCCTAG
AGCACACGTCTGAACTCTTTCCCTACACGACGCTCTTCCGATCTNNNATGAGATCTCAGTTCACTGCAC
GAGCACACGTCTGAACTCTTTCCCTACACGACGCTCTTCCGATCTNNNCACAGAAAATGCTTCTCTCTG
GAGCACACGTCTGAACTCTTTCCCTACACGACGCTCTTCCGATCTNNNCATGGAGAAATTGTGGGGCT
AGCACACGTCTGAACTCTTTCCCTACACGACGCTCTTCCGATCTNNNCATGCTGCTGTGCTCAACTAAT
AGCACACGTCTGAACTCTTTCCCTACACGACGCTCTTCCGATCTNNNCAGCCTAGACTTCTGGACTCAA
AGCACACGTCTGAACTCTTTCCCTACACGACGCTCTTCCGATCTNNNTAAAGTGCTGGGATTACAGGCA
GAGCACACGTCTGAACTCTTTCCCTACACGACGCTCTTCCGATCTNNNTCCCAATCCAATCCCAATTCCA
AGCACACGTCTGAACTCTTTCCCTACACGACGCTCTTCCGATCTNNNAGTGCTGGGATTATAGGCATG
AGCACACGTCTGAACTCTTTCCCTACACGACGCTCTTCCGATCTNNNTTTTCATAACCCATTGGCAGCC
AGCACACGTCTGAACTCTTTCCCTACACGACGCTCTTCCGATCTNNNAGGCTGAATTTCTCAAACACAG
AGCACACGTCTGAACTCTTTCCCTACACGACGCTCTTCCGATCTNNNGTGAAGGCCTGAACTTTTGTCT
AGCACACGTCTGAACTCTTTCCCTACACGACGCTCTTCCGATCTNNNGGGTACACAGGGTATTTGGT
GCACACGTCTGAACTCTTTCCCTACACGACGCTCTTCCGATCTNNNGGTTGTCAGTTGATGTCTCCAGA
GAGCACACGTCTGAACTCTTTCCCTACACGACGCTCTTCCGATCTNNNTATAGGTGTGAGCCATATTGC
AGCACACGTCTGAACTCTTTCCCTACACGACGCTCTTCCGATCTNNNACCCCTCTGACCTCTTCTACTACCA
GAGCACACGTCTGAACTCTTTCCCTACACGACGCTCTTCCGATCTNNNCTTCCTGTTCCTCTGCCTTTTT
AGCACACGTCTGAACTCTTTCCCTACACGACGCTCTTCCGATCTNNNTTTGTTCCTTTTCAGTGCTGAGT
GAGCACACGTCTGAACTCTTTCCCTACACGACGCTCTTCCGATCTNNNTCAGAACCTACTGCAACCTCTT

AGCACACGTCTGAACTCTTTCCCTACACGACGCTCTTCCGATCTNNNAATCTCCTGATCTCATGATCCGCC
AGCACACGTCTGAACTCTTTCCCTACACGACGCTCTTCCGATCTNNNGCACGATGGAGCAGTGTTTTAT
AGCACACGTCTGAACTCTTTCCCTACACGACGCTCTTCCGATCTNNNCCAAAGTGCTGGAATTACTGGT
AGCACACGTCTGAACTCTTTCCCTACACGACGCTCTTCCGATCTNNNAGTTTGGGTAAGGTATGAGCA
GAGCACACGTCTGAACTCTTTCCCTACACGACGCTCTTCCGATCTNNNTGGTGGCATATAGTAAGCAG
AGCACACGTCTGAACTCTTTCCCTACACGACGCTCTTCCGATCTNNNTTCAAGTGATTCTCCTGCCTCAG
AGCACACGTCTGAACTCTTTCCCTACACGACGCTCTTCCGATCTNNNAAGGCTTGTAAGTCATTCACCCT
GCACACGTCTGAACTCTTTCCCTACACGACGCTCTTCCGATCTNNNCCCTCAAACACTGAAAACAAGCT
GAGCACACGTCTGAACTCTTTCCCTACACGACGCTCTTCCGATCTNNNCTCCAGCTTCAAGACAAGGACT
GAGCACACGTCTGAACTCTTTCCCTACACGACGCTCTTCCGATCTNNNTCACTTCCTCTCCTATGGTAGCA
GAGCACACGTCTGAACTCTTTCCCTACACGACGCTCTTCCGATCTNNNTTCCCTAATTGAAGAGAGCCA
GCACACGTCTGAACTCTTTCCCTACACGACGCTCTTCCGATCTNNNACTGGTTGGCTAATACAGGAAAG
AGCACACGTCTGAACTCTTTCCCTACACGACGCTCTTCCGATCTNNNCTAGGAGTTTCCTCACAGAGCT
AGCACACGTCTGAACTCTTTCCCTACACGACGCTCTTCCGATCTNNNGTCACCCTGACTCACTGATTCAT
AGCACACGTCTGAACTCTTTCCCTACACGACGCTCTTCCGATCTNNNTAGTTCACTGAAGCCTTGACCT
GAGCACACGTCTGAACTCTTTCCCTACACGACGCTCTTCCGATCTNNNCACCTTAGCCTCCTGATTAGCT
AGCACACGTCTGAACTCTTTCCCTACACGACGCTCTTCCGATCTNNNGTCGTCACATTACAGAAGGGGT
AGCACACGTCTGAACTCTTTCCCTACACGACGCTCTTCCGATCTNNNTTCAAGTGATTCTCCTGCCTCAG
AGCACACGTCTGAACTCTTTCCCTACACGACGCTCTTCCGATCTNNNTGCCAAAGAATTCAGCCACAATT
AGCACACGTCTGAACTCTTTCCCTACACGACGCTCTTCCGATCTNNNCTGTCCTTCACTTCCAGGAACTT
GAGCACACGTCTGAACTCTTTCCCTACACGACGCTCTTCCGATCTNNNTGAGTCACAGAGAGAGACTG
GAGCACACGTCTGAACTCTTTCCCTACACGACGCTCTTCCGATCTNNNAATGGCAAAGTGAGACTCCAT
AGCACACGTCTGAACTCTTTCCCTACACGACGCTCTTCCGATCTNNNCTGAGGCAGGAGAATCACTTGA
AGCACACGTCTGAACTCTTTCCCTACACGACGCTCTTCCGATCTNNNGTGACAGAGTGAGACTCCTTCA
GAAGAGCACACGTCTGAACTCTTTCCCTACACGACGCTCTTCCGATCTNNNGTAGTCCCAGCTACTCCG
AGCACACGTCTGAACTCTTTCCCTACACGACGCTCTTCCGATCTNNNTCTTTTGATCTTTGTTGGGCGA
GAGCACACGTCTGAACTCTTTCCCTACACGACGCTCTTCCGATCTNNNTACGCCTGTAATCCCAGCTATT
AGCACACGTCTGAACTCTTTCCCTACACGACGCTCTTCCGATCTNNNCTAGCCAAAAACCAGGAAAAG
GAGCACACGTCTGAACTCTTTCCCTACACGACGCTCTTCCGATCTNNNGATTGCAGTGAACCGAGATT
GAGCACACGTCTGAACTCTTTCCCTACACGACGCTCTTCCGATCTNNNTGCAATCTCAGCTACTTCAGAG
GAGCACACGTCTGAACTCTTTCCCTACACGACGCTCTTCCGATCTNNNCGAGATCACACCATTGTACTC
GAGCACACGTCTGAACTCTTTCCCTACACGACGCTCTTCCGATCTNNNCTGAGGCAGGAGAATCACTT
AGCACACGTCTGAACTCTTTCCCTACACGACGCTCTTCCGATCTNNNTCTGTAATCCCAGCTACTCAGGA
AGCACACGTCTGAACTCTTTCCCTACACGACGCTCTTCCGATCTNNNCTGTAATCCCAGCTACTCAGGA
GAGCACACGTCTGAACTCTTTCCCTACACGACGCTCTTCCGATCTNNNGAAGTTGCAGTGAGCTGAGA
AGCACACGTCTGAACTCTTTCCCTACACGACGCTCTTCCGATCTNNNAGGAGAATTGTTTGAACCTGG
AGCACACGTCTGAACTCTTTCCCTACACGACGCTCTTCCGATCTNNNAGCACCTTCATGATGGAATTCCT
GAGCACACGTCTGAACTCTTTCCCTACACGACGCTCTTCCGATCTNNNCTGAGGCAGGAGAATCACTTG
GAGCACACGTCTGAACTCTTTCCCTACACGACGCTCTTCCGATCTNNNAGCCTGGGAGACATAGTTAGA
AGAGCACACGTCTGAACTCTTTCCCTACACGACGCTCTTCCGATCTNNNTTGAGACAGGAGAATGGCAT
AGCACACGTCTGAACTCTTTCCCTACACGACGCTCTTCCGATCTNNNGAGGCAGGATAATTGCTGAAAC
AGCACACGTCTGAACTCTTTCCCTACACGACGCTCTTCCGATCTNNNGGTGAACCTGATAACCAATCT
GAGCACACGTCTGAACTCTTTCCCTACACGACGCTCTTCCGATCTNNNCAAGTTGTACCCCAAACACCTT

GAGCACACGTCTGAACTCTTTCCCTACACGACGCTCTTCCGATCTNNNCAACATGGAGAAACCCAGTCT
GAGCACACGTCTGAACTCTTTCCCTACACGACGCTCTTCCGATCTNNNATATAGCGAGAACCCTGCCTCT
AGCACACGTCTGAACTCTTTCCCTACACGACGCTCTTCCGATCTNNNTACTCAGGAGACTAAGGGGAGA
GAGCACACGTCTGAACTCTTTCCCTACACGACGCTCTTCCGATCTNNNACGTGGTAAAACCCTGTCTCTA
AGCACACGTCTGAACTCTTTCCCTACACGACGCTCTTCCGATCTNNNCATGATCGTGTCACTGTACTCCA
AGCACACGTCTGAACTCTTTCCCTACACGACGCTCTTCCGATCTNNNGCCAACATAGGGAAACATGGT
AGCACACGTCTGAACTCTTTCCCTACACGACGCTCTTCCGATCTNNNAGGAGAATTGTTTGAACCCAG
GAGCACACGTCTGAACTCTTTCCCTACACGACGCTCTTCCGATCTNNNGGAGGTTGTAGTGAGCTGA
GAGCACACGTCTGAACTCTTTCCCTACACGACGCTCTTCCGATCTNNNTGACCCATGTGCCATTTTGAAA
AGCACACGTCTGAACTCTTTCCCTACACGACGCTCTTCCGATCTNNNCTGTAGTCCCAGCTACTCAGGA
AGCACACGTCTGAACTCTTTCCCTACACGACGCTCTTCCGATCTNNNACACAGTGAAACCCCATCTCTA
AGCACACGTCTGAACTCTTTCCCTACACGACGCTCTTCCGATCTNNNTGTAGTCCCAGCTACTTAGGA
AAGAGCACACGTCTGAACTCTTTCCCTACACGACGCTCTTCCGATCTNNNAGGATAAAAGCGAGTGCCT
GAGCACACGTCTGAACTCTTTCCCTACACGACGCTCTTCCGATCTNNNCACCTGTAATCCCAGCTACTCA
AGCACACGTCTGAACTCTTTCCCTACACGACGCTCTTCCGATCTNNNCTGAGACAGGAGAATCGCTTGA
GAGCACACGTCTGAACTCTTTCCCTACACGACGCTCTTCCGATCTNNNACACAGTGAAACCCCATCTCTA
GAGCACACGTCTGAACTCTTTCCCTACACGACGCTCTTCCGATCTNNNCTAAGGCAGGAGAATCAGCTT
AGCACACGTCTGAACTCTTTCCCTACACGACGCTCTTCCGATCTNNNGCAGGAGAATCAATTGAACCA
GAGCACACGTCTGAACTCTTTCCCTACACGACGCTCTTCCGATCTNNNAGACATGCAAAAATTCAATCC
GAGCACACGTCTGAACTCTTTCCCTACACGACGCTCTTCCGATCTNNNGAGGCAGGAGAGTTGTTTGA
AGCACACGTCTGAACTCTTTCCCTACACGACGCTCTTCCGATCTNNNTAAGGTTAGGAGTTCGAGACCA
GAGCACACGTCTGAACTCTTTCCCTACACGACGCTCTTCCGATCTNNNTAATCTCAGCACTTTAGGAGG
AGCACACGTCTGAACTCTTTCCCTACACGACGCTCTTCCGATCTNNNAGAATCTCTTGAATCTGGGAG
AGCACACGTCTGAACTCTTTCCCTACACGACGCTCTTCCGATCTNNNCTGAGGCAGGAAAATTGCTTGA
AGCACACGTCTGAACTCTTTCCCTACACGACGCTCTTCCGATCTNNNACCCTCACAAGATTGGTCTTTCA
AGCACACGTCTGAACTCTTTCCCTACACGACGCTCTTCCGATCTNNNGCTGAGACAGGAGAATCACTTG
AGCACACGTCTGAACTCTTTCCCTACACGACGCTCTTCCGATCTNNNCAGCCTAGGCAAAAATGGTGAA
AGCACACGTCTGAACTCTTTCCCTACACGACGCTCTTCCGATCTNNNCATCCTGCCTAACATGGTGAAAC
AGCACACGTCTGAACTCTTTCCCTACACGACGCTCTTCCGATCTNNNTACTCTGAAGGCTGAGGTAGGA
AGCACACGTCTGAACTCTTTCCCTACACGACGCTCTTCCGATCTNNNGCAGATCATTTGAGGTCAGGA
AGCACACGTCTGAACTCTTTCCCTACACGACGCTCTTCCGATCTNNNGCCTAAATAGCCAGAGCAATTG
AGCACACGTCTGAACTCTTTCCCTACACGACGCTCTTCCGATCTNNNGGGAACTGTTCTTGAACCTACC
AGCACACGTCTGAACTCTTTCCCTACACGACGCTCTTCCGATCTNNNAATCACTTGAACTTGGAGGGCA
AGCACACGTCTGAACTCTTTCCCTACACGACGCTCTTCCGATCTNNNGAGTTCAAGACCAGTCTAGGCA
GAGCACACGTCTGAACTCTTTCCCTACACGACGCTCTTCCGATCTNNNCCAGAAGTTTACCAGAGCAT
GAGCACACGTCTGAACTCTTTCCCTACACGACGCTCTTCCGATCTNNNCACTTGAGGTCAGGAGTTCAA
AGCACACGTCTGAACTCTTTCCCTACACGACGCTCTTCCGATCTNNNTTGTAGTGAGCTGAGATCGTG
AGCACACGTCTGAACTCTTTCCCTACACGACGCTCTTCCGATCTNNNCTGTAGTCCCAGCTACTTGAGA
GAGCACACGTCTGAACTCTTTCCCTACACGACGCTCTTCCGATCTNNNAAGGCTGCAGTGAGTTGATT
GAGCACACGTCTGAACTCTTTCCCTACACGACGCTCTTCCGATCTNNNCTGCAAAGGACATGAACTCAT
AGCACACGTCTGAACTCTTTCCCTACACGACGCTCTTCCGATCTNNNTGTTGAATGAAAGTGGACGTG
GAGCACACGTCTGAACTCTTTCCCTACACGACGCTCTTCCGATCTNNNTACCACCACTTCATTCTCCTTG
GCACACGTCTGAACTCTTTCCCTACACGACGCTCTTCCGATCTNNNGGGCAGAGTGTAAATCCCAAAT

GCACACGTCTGAACTCTTTCCCTACACGACGCTCTTCCGATCTNNNGTGACTTTGGGTGGCAGTATTT
AGCACACGTCTGAACTCTTTCCCTACACGACGCTCTTCCGATCTNNNTCACTCAGGAATTCAGGCACTT
AGCACACGTCTGAACTCTTTCCCTACACGACGCTCTTCCGATCTNNNCCACTTGCACTTTTCACAACACA
GAGCACACGTCTGAACTCTTTCCCTACACGACGCTCTTCCGATCTNNNAGACAAAGTCTCACTCTGTTG
AGCACACGTCTGAACTCTTTCCCTACACGACGCTCTTCCGATCTNNNCACCACATCCGGCCAAATATTT
GCACACGTCTGAACTCTTTCCCTACACGACGCTCTTCCGATCTNNNTGCACCTGGCCAGTTATATCATT
AGCACACGTCTGAACTCTTTCCCTACACGACGCTCTTCCGATCTNNNGTGAGTGGTATTCCTTGTTTG
GCACACGTCTGAACTCTTTCCCTACACGACGCTCTTCCGATCTNNNTCTGCTTGCATCTCAGTCTACAT
AGCACACGTCTGAACTCTTTCCCTACACGACGCTCTTCCGATCTNNNACACGGAATAAGAACTTCCTG
GAGCACACGTCTGAACTCTTTCCCTACACGACGCTCTTCCGATCTNNNGGAAATCACAAGATTCCTGC
AGCACACGTCTGAACTCTTTCCCTACACGACGCTCTTCCGATCTNNNGCTTTTGATTTCTGTTCGTGTG
GCACACGTCTGAACTCTTTCCCTACACGACGCTCTTCCGATCTNNNTATTTGGAGTGTTGCCTCAACTG
GCACACGTCTGAACTCTTTCCCTACACGACGCTCTTCCGATCTNNNCAACAGGGTATAAGCATTCCCC
AGCACACGTCTGAACTCTTTCCCTACACGACGCTCTTCCGATCTNNNTCAGCCTTCTGAGTAGTTGGA
GCACACGTCTGAACTCTTTCCCTACACGACGCTCTTCCGATCTNNNCATACACATAGACCCATGGAGC
GCACACGTCTGAACTCTTTCCCTACACGACGCTCTTCCGATCTNNNTTGTTGTGCAGAAGCTTTTTAG
GCACACGTCTGAACTCTTTCCCTACACGACGCTCTTCCGATCTNNNCCAAAGTGCTGGAATTACAGGT
AGCACACGTCTGAACTCTTTCCCTACACGACGCTCTTCCGATCTNNNCTGTGATTGGCTGGAGTTTTG
GCACACGTCTGAACTCTTTCCCTACACGACGCTCTTCCGATCTNNNATGGTCCAGTTTAGGAGAGGAG
AGCACACGTCTGAACTCTTTCCCTACACGACGCTCTTCCGATCTNNNTCCTTGACACACTCCTGGTTAT
AGCACACGTCTGAACTCTTTCCCTACACGACGCTCTTCCGATCTNNNGACAGACTAGGGGTTTGGAT
GAGCACACGTCTGAACTCTTTCCCTACACGACGCTCTTCCGATCTNNNCCTGGCCCTTCTTTTCTTTTCT
GCACACGTCTGAACTCTTTCCCTACACGACGCTCTTCCGATCTNNNCACAGACCTCAGAGGAATCACT
GAGCACACGTCTGAACTCTTTCCCTACACGACGCTCTTCCGATCTNNNAGCTGAGAACAAAGACAAG
GCACACGTCTGAACTCTTTCCCTACACGACGCTCTTCCGATCTNNNAGTAGCTAGATGTATGAGACCA
GCACACGTCTGAACTCTTTCCCTACACGACGCTCTTCCGATCTNNNCTGGGTTCAAGTGATTCTTCTG
GCACACGTCTGAACTCTTTCCCTACACGACGCTCTTCCGATCTNNNGAGCCAGTTGTTCAGTTTCCATG
GCACACGTCTGAACTCTTTCCCTACACGACGCTCTTCCGATCTNNNGTGGCTGGTTTGTCTTTGATTC
AGCACACGTCTGAACTCTTTCCCTACACGACGCTCTTCCGATCTNNNACTCTGCTTCACTCACCATTCA
AGCACACGTCTGAACTCTTTCCCTACACGACGCTCTTCCGATCTNNNGTGATCTCTCCTCACTGCAATC
AGCACACGTCTGAACTCTTTCCCTACACGACGCTCTTCCGATCTNNNAAGAATGCATCTCAGCCTGTGT
AGCACACGTCTGAACTCTTTCCCTACACGACGCTCTTCCGATCTNNNGGCTCACCCAAAGACGTTATTT
AGCACACGTCTGAACTCTTTCCCTACACGACGCTCTTCCGATCTNNNTGATGAGAAACTGTCAGCTGT
GCACACGTCTGAACTCTTTCCCTACACGACGCTCTTCCGATCTNNNTCAGATCACCAAGTATTCTTGTG
GCACACGTCTGAACTCTTTCCCTACACGACGCTCTTCCGATCTNNNGGTTTCTGACCCTAAAGTGCCTT
GCACACGTCTGAACTCTTTCCCTACACGACGCTCTTCCGATCTNNNTCAATGCCTCTATTCCAGCCTTT
GCACACGTCTGAACTCTTTCCCTACACGACGCTCTTCCGATCTNNNTCAAAAGCAAGCCAGAGTAGCT
GCACACGTCTGAACTCTTTCCCTACACGACGCTCTTCCGATCTNNNAGGTTCAAATGATTCTCCTGCCT
GCACACGTCTGAACTCTTTCCCTACACGACGCTCTTCCGATCTNNNGTGAAGCAGTGTGAAAACTCG
GCACACGTCTGAACTCTTTCCCTACACGACGCTCTTCCGATCTNNNCAATCTCAGCTCACTGCAATCTC
GAGCACACGTCTGAACTCTTTCCCTACACGACGCTCTTCCGATCTNNNGCCACCATGCCCAGATAATTT
GCACACGTCTGAACTCTTTCCCTACACGACGCTCTTCCGATCTNNNCCTCCTGAGTAGCTAGTACACCA
GCACACGTCTGAACTCTTTCCCTACACGACGCTCTTCCGATCTNNNAAAAATTCATTGGTGCTCGCTT

AGCACACGTCTGAACTCTTTCCCTACACGACGCTCTTCCGATCTNNNAGGGTGATCATAGCTTACTG
GCACACGTCTGAACTCTTTCCCTACACGACGCTCTTCCGATCTNNNGCCTTTGGCTGATTTCCAATCTT
GCACACGTCTGAACTCTTTCCCTACACGACGCTCTTCCGATCTNNNTTCAAGGGATTCTTCTGCCTCAG
AGCACACGTCTGAACTCTTTCCCTACACGACGCTCTTCCGATCTNNNGCAGTTTGCAGATAAACAGGT
AGCACACGTCTGAACTCTTTCCCTACACGACGCTCTTCCGATCTNNNTGCTCCTGGATAGCATAGCATT
AGCACACGTCTGAACTCTTTCCCTACACGACGCTCTTCCGATCTNNNTGGTTAGTGCCTTGTTTTGGTT
AGCACACGTCTGAACTCTTTCCCTACACGACGCTCTTCCGATCTNNNTGGAGGAGAAAAGCTTGTTAC
AGCACACGTCTGAACTCTTTCCCTACACGACGCTCTTCCGATCTNNNGGAAGTCTAAGATCATGGTG
AGCACACGTCTGAACTCTTTCCCTACACGACGCTCTTCCGATCTNNNATGCAACATCCAAAGCCACTTTT
AGCACACGTCTGAACTCTTTCCCTACACGACGCTCTTCCGATCTNNNGACCGAGAATATGAGCTCTAG
AGCACACGTCTGAACTCTTTCCCTACACGACGCTCTTCCGATCTNNNTGACTGAATCTCTGACCTCTCC
AGCACACGTCTGAACTCTTTCCCTACACGACGCTCTTCCGATCTNNNTGAACTAGAGAAGCAAGAGCA
GAGCACACGTCTGAACTCTTTCCCTACACGACGCTCTTCCGATCTNNNCTGACCTCGTGATCCAAGAGT
AGCACACGTCTGAACTCTTTCCCTACACGACGCTCTTCCGATCTNNNGACACAATCTGCCAAAACTCAC
AGCACACGTCTGAACTCTTTCCCTACACGACGCTCTTCCGATCTNNNCCCTTTGGAGAAGGTTTGGGA
AGCACACGTCTGAACTCTTTCCCTACACGACGCTCTTCCGATCTNNNCGTACACTGAAACAGAACTGC
AGCACACGTCTGAACTCTTTCCCTACACGACGCTCTTCCGATCTNNNTATATGCAGCTTTCCAGGGAC
AGCACACGTCTGAACTCTTTCCCTACACGACGCTCTTCCGATCTNNNCAGAGGTTGTAGTGAGCCAA
AGCACACGTCTGAACTCTTTCCCTACACGACGCTCTTCCGATCTNNNAGGAGTTCAAGACCAGCCTAG
GAGCACACGTCTGAACTCTTTCCCTACACGACGCTCTTCCGATCTNNNGAGACTGCAGTGAGCTGATA
AGCACACGTCTGAACTCTTTCCCTACACGACGCTCTTCCGATCTNNNCCTATAGTCCCAGCTACTCAGG
AGCACACGTCTGAACTCTTTCCCTACACGACGCTCTTCCGATCTNNNCTGTAGTCCCAGCTACTCAGGA
AGCACACGTCTGAACTCTTTCCCTACACGACGCTCTTCCGATCTNNNGCAACAAGAACAAAACTCCGT
GCACACGTCTGAACTCTTTCCCTACACGACGCTCTTCCGATCTNNNATCACTCCCTGGGCTTTTTTTCTC
GCACACGTCTGAACTCTTTCCCTACACGACGCTCTTCCGATCTNNNATCATGAGCCTGGTAGGTTGAA
GAGCACACGTCTGAACTCTTTCCCTACACGACGCTCTTCCGATCTNNNTAATGTGACAGTGGTCTGG
AGCACACGTCTGAACTCTTTCCCTACACGACGCTCTTCCGATCTNNNGAGCTGGCAATGAGTCCTATCA
AGCACACGTCTGAACTCTTTCCCTACACGACGCTCTTCCGATCTNNNAGAATTGCTTGAAACTGGGA
AGCACACGTCTGAACTCTTTCCCTACACGACGCTCTTCCGATCTNNNGTAATACCAGCTACTCAGGAG
GCACACGTCTGAACTCTTTCCCTACACGACGCTCTTCCGATCTNNNCTGAGGCAGGAGAATCACTTGA
GAGCACACGTCTGAACTCTTTCCCTACACGACGCTCTTCCGATCTNNNCAGATCACTTGAGCTCAGGA
AGCACACGTCTGAACTCTTTCCCTACACGACGCTCTTCCGATCTNNNCAAGAGAATTGCTTGAACCTG
AGCACACGTCTGAACTCTTTCCCTACACGACGCTCTTCCGATCTNNNCAAAAATTAGCCGGGTATGGT
GAGCACACGTCTGAACTCTTTCCCTACACGACGCTCTTCCGATCTNNNTCACTTGAACTCGAAGGACA
AGCACACGTCTGAACTCTTTCCCTACACGACGCTCTTCCGATCTNNNCTGACTGTGCCACTATACTCCA
AGCACACGTCTGAACTCTTTCCCTACACGACGCTCTTCCGATCTNNNAGAGCTTGCAGTGAGTGGAT
AGCACACGTCTGAACTCTTTCCCTACACGACGCTCTTCCGATCTNNNGTCCAAAACTGCTGCCTCTAAT
AGCACACGTCTGAACTCTTTCCCTACACGACGCTCTTCCGATCTNNNAGCCTGGGTAATAGAGTGAG
AGCACACGTCTGAACTCTTTCCCTACACGACGCTCTTCCGATCTNNNTCACAAGGTCAGGAGTTTGAG
AGCACACGTCTGAACTCTTTCCCTACACGACGCTCTTCCGATCTNNNAAGATTGTGCCACTACACTCCA
AGCACACGTCTGAACTCTTTCCCTACACGACGCTCTTCCGATCTNNNGATGCAGGAGAATTGCTTGA
AGCACACGTCTGAACTCTTTCCCTACACGACGCTCTTCCGATCTNNNTTGCCTGTAATCCCAGCATTTT
AGCACACGTCTGAACTCTTTCCCTACACGACGCTCTTCCGATCTNNNAGAATTGTCTTGGGCCACACA

GAGCACACGTCTGAACTCTTTCCCTACACGACGCTCTTCCGATCTNNNCTGTAATCCCAGCTACTCAGG
AGCACACGTCTGAACTCTTTCCCTACACGACGCTCTTCCGATCTNNNGATAGAGTCAAGACCCATCGG
AGCACACGTCTGAACTCTTTCCCTACACGACGCTCTTCCGATCTNNNAGGTTACAGTGAGCTGAGATC
GAGCACACGTCTGAACTCTTTCCCTACACGACGCTCTTCCGATCTNNNTTTGGAGATTGAGGCAAG
AGCACACGTCTGAACTCTTTCCCTACACGACGCTCTTCCGATCTNNNGCTGAGACAGGAGAATCACTT
AGCACACGTCTGAACTCTTTCCCTACACGACGCTCTTCCGATCTNNNCCTGGTGCATGCTACCTACATA
AGCACACGTCTGAACTCTTTCCCTACACGACGCTCTTCCGATCTNNNCCTGGTGCATGCTACCTACATA
AGCACACGTCTGAACTCTTTCCCTACACGACGCTCTTCCGATCTNNNCTGTAATCCCAGCTACTCAGGA
GCACACGTCTGAACTCTTTCCCTACACGACGCTCTTCCGATCTNNNCTGTAATCCCAGCTACTCCAGAG
AGCACACGTCTGAACTCTTTCCCTACACGACGCTCTTCCGATCTNNNCAGAGAGGTTACAGTGAGCA
AGCACACGTCTGAACTCTTTCCCTACACGACGCTCTTCCGATCTNNNCACCTGTAATCCCAGCTACTCA
GAGCACACGTCTGAACTCTTTCCCTACACGACGCTCTTCCGATCTNNNTGCAATGACCCATGATCACAC
AGCACACGTCTGAACTCTTTCCCTACACGACGCTCTTCCGATCTNNNGTTTGAGCTCAGCAGTTCAAG
GAGCACACGTCTGAACTCTTTCCCTACACGACGCTCTTCCGATCTNNNGAGTTTGCAGTGAGCTGAGA
AGCACACGTCTGAACTCTTTCCCTACACGACGCTCTTCCGATCTNNNCTGAGGCAGGAGAATCACTTC
AGAGCACACGTCTGAACTCTTTCCCTACACGACGCTCTTCCGATCTNNNACTCAGCAAGCTAAGATAG
AGCACACGTCTGAACTCTTTCCCTACACGACGCTCTTCCGATCTNNNCTGTAATCCCAGCTACTCAGGA
GAGCACACGTCTGAACTCTTTCCCTACACGACGCTCTTCCGATCTNNNTGCTACATAGTGAAACCTTGC
AGCACACGTCTGAACTCTTTCCCTACACGACGCTCTTCCGATCTNNNCGCCTGTAATCCTAGCTACTCT
GAGCACACGTCTGAACTCTTTCCCTACACGACGCTCTTCCGATCTNNNTTGCAATGAGCTGAGAACATG
AGCACACGTCTGAACTCTTTCCCTACACGACGCTCTTCCGATCTNNNAGATTGCAGTGAGCCGAGATT
AGCACACGTCTGAACTCTTTCCCTACACGACGCTCTTCCGATCTNNNAGATCTCTTTTGCTGTCTCGG
GCACACGTCTGAACTCTTTCCCTACACGACGCTCTTCCGATCTNNNGAATTCACACCATTGCACTCCAG
GAGCACACGTCTGAACTCTTTCCCTACACGACGCTCTTCCGATCTNNNGAGAATCACTTGAACCCAGGA
GAGCACACGTCTGAACTCTTTCCCTACACGACGCTCTTCCGATCTNNNATTGCTTGAGCCCAGGAATT
AGCACACGTCTGAACTCTTTCCCTACACGACGCTCTTCCGATCTNNNTGTAGTACCAGCTGCTCAGAA
AGCACACGTCTGAACTCTTTCCCTACACGACGCTCTTCCGATCTNNNGTGACTCCTCCCTCACTCATTTT
GAGCACACGTCTGAACTCTTTCCCTACACGACGCTCTTCCGATCTNNNCAACAGTTTGGGAGAATGAG
GAGCACACGTCTGAACTCTTTCCCTACACGACGCTCTTCCGATCTNNNTGGAAAATCCTATGGTGCCA
AGCACACGTCTGAACTCTTTCCCTACACGACGCTCTTCCGATCTNNNGCAGGAGAATCACTTGAACC
GAGCACACGTCTGAACTCTTTCCCTACACGACGCTCTTCCGATCTNNNAACCCTGTCTCTACCAAAAAT
AGCACACGTCTGAACTCTTTCCCTACACGACGCTCTTCCGATCTNNNACAAGGGAGAAATCCAAAACC
GAGCACACGTCTGAACTCTTTCCCTACACGACGCTCTTCCGATCTNNNCTGTAGTCCCAGGTACTCAGG
AGCACACGTCTGAACTCTTTCCCTACACGACGCTCTTCCGATCTNNNCTGAGGCAGGAGAATCGTTT
GAGCACACGTCTGAACTCTTTCCCTACACGACGCTCTTCCGATCTNNNAGAATCACTTGAACCTAGGAG
GAGCACACGTCTGAACTCTTTCCCTACACGACGCTCTTCCGATCTNNNGTGTAGTTCCAGCTACTTGG
AGCACACGTCTGAACTCTTTCCCTACACGACGCTCTTCCGATCTNNNTGTAGTCCTAGTTACTCGGGA
GAGCACACGTCTGAACTCTTTCCCTACACGACGCTCTTCCGATCTNNNTGAGCTCTGAAATTGAGGCA
AGCACACGTCTGAACTCTTTCCCTACACGACGCTCTTCCGATCTNNNGAGAATTGCTTGAACCCAGGA
GAGCACACGTCTGAACTCTTTCCCTACACGACGCTCTTCCGATCTNNNGAGGCAGGAAAATCACTTG
GAGCACACGTCTGAACTCTTTCCCTACACGACGCTCTTCCGATCTNNNGAGGCAGGAAAATCACTTGA
AGCACACGTCTGAACTCTTTCCCTACACGACGCTCTTCCGATCTNNNACCTGTAATCCCAGCTACTTTG
AGCACACGTCTGAACTCTTTCCCTACACGACGCTCTTCCGATCTNNNCTGTAATCCCAGCTACTCAGG

AGCACACGTCTGAACTCTTTCCCTACACGACGCTCTTCCGATCTNNNGAGTGGTTCCCGGAAACTAAG
GAGCACACGTCTGAACTCTTTCCCTACACGACGCTCTTCCGATCTNNNCTGTAGTCACAGCTACTCAG
AGCACACGTCTGAACTCTTTCCCTACACGACGCTCTTCCGATCTNNNCTGAGGCAGGAGAATCACTTG
GCACACGTCTGAACTCTTTCCCTACACGACGCTCTTCCGATCTNNNAATTCAATCCTGCAGCTTTCCAC
GAGCACACGTCTGAACTCTTTCCCTACACGACGCTCTTCCGATCTNNNCTCAGCTCAGACCAGAGTTTC
GCACACGTCTGAACTCTTTCCCTACACGACGCTCTTCCGATCTNNNGTAAGCTTTGCAAACCATCCAGT
GCACACGTCTGAACTCTTTCCCTACACGACGCTCTTCCGATCTNNNTGTGTTGTTCTCTGTGCCTACTT
AGCACACGTCTGAACTCTTTCCCTACACGACGCTCTTCCGATCTNNNGCAGCAGGCAAGATTTACTGA
GCACACGTCTGAACTCTTTCCCTACACGACGCTCTTCCGATCTNNNAGCCTATCCACAGAGGTTCAAT
AGCACACGTCTGAACTCTTTCCCTACACGACGCTCTTCCGATCTNNNAATCTTACAGATAGCAGCCCT
GCACACGTCTGAACTCTTTCCCTACACGACGCTCTTCCGATCTNNNATCAAGTTCCTCAAAATGTGCC
AGCACACGTCTGAACTCTTTCCCTACACGACGCTCTTCCGATCTNNNATAAACGGGCAGAAGACAGT
GCACACGTCTGAACTCTTTCCCTACACGACGCTCTTCCGATCTNNNTGTCCTGATTTCTGGCATCTTCT
AGCACACGTCTGAACTCTTTCCCTACACGACGCTCTTCCGATCTNNNAGAAGTCTGTACCCTTTGTTC
GCACACGTCTGAACTCTTTCCCTACACGACGCTCTTCCGATCTNNNGTGTGCACATTTAGTTCAGCTG
GCACACGTCTGAACTCTTTCCCTACACGACGCTCTTCCGATCTNNNTTAGCTTCTGAAATGCCACTCCT
GAGCACACGTCTGAACTCTTTCCCTACACGACGCTCTTCCGATCTNNNCATCTTCAGTACAGGTGCAG
AGCACACGTCTGAACTCTTTCCCTACACGACGCTCTTCCGATCTNNNCAGCCTTCCCTCTCACTACTTTT
GCACACGTCTGAACTCTTTCCCTACACGACGCTCTTCCGATCTNNNCTGAAACAGGGTTAGGGCTTTT
GCACACGTCTGAACTCTTTCCCTACACGACGCTCTTCCGATCTNNNAATCACCTCTCAAAGACCCCATC
GCACACGTCTGAACTCTTTCCCTACACGACGCTCTTCCGATCTNNNAACCTCAAATTCTGGCTCCTCTAC
GCACACGTCTGAACTCTTTCCCTACACGACGCTCTTCCGATCTNNNAAGTAGCTGGGATTACAGTCAC
GAGCACACGTCTGAACTCTTTCCCTACACGACGCTCTTCCGATCTNNNTCAGGTAGAGAACTGCTTGA
GAGCACACGTCTGAACTCTTTCCCTACACGACGCTCTTCCGATCTNNNCCAAAGTGCTGGGATTACAG
AGCACACGTCTGAACTCTTTCCCTACACGACGCTCTTCCGATCTNNNGGACGAATGAATGGCAGAGA
GAGCACACGTCTGAACTCTTTCCCTACACGACGCTCTTCCGATCTNNNACTCAGTGTTCACATTTTCCCA
GCACACGTCTGAACTCTTTCCCTACACGACGCTCTTCCGATCTNNNAAAAAAAGGTACTGGGGAGCT
GCACACGTCTGAACTCTTTCCCTACACGACGCTCTTCCGATCTNNNTTTTCTTTCTCAGCCTTGCCAGC
GAGCACACGTCTGAACTCTTTCCCTACACGACGCTCTTCCGATCTNNNTCTCGGCCTTAGCATGTAGTT
GCACACGTCTGAACTCTTTCCCTACACGACGCTCTTCCGATCTNNNTTTTTTTGCTTTTGCCTGCCAAA
GCACACGTCTGAACTCTTTCCCTACACGACGCTCTTCCGATCTNNNATCCAGGTAGCAGTGAAAGTCT
GAGCACACGTCTGAACTCTTTCCCTACACGACGCTCTTCCGATCTNNNCATGACCACTTTCCACTGTTT
GCACACGTCTGAACTCTTTCCCTACACGACGCTCTTCCGATCTNNNATTACAAACAGGCCAACATGTC
AGCACACGTCTGAACTCTTTCCCTACACGACGCTCTTCCGATCTNNNCAGACCCAGACTGTAACCTCA
AGCACACGTCTGAACTCTTTCCCTACACGACGCTCTTCCGATCTNNNTGAAATCCAAAACACTTCTGG
GCACACGTCTGAACTCTTTCCCTACACGACGCTCTTCCGATCTNNNCCAACAAGTATTTGCTGAGGTC
AGCACACGTCTGAACTCTTTCCCTACACGACGCTCTTCCGATCTNNNAAAAGGACCCAAGACTCGGA
AGCACACGTCTGAACTCTTTCCCTACACGACGCTCTTCCGATCTNNNTTCCTTTTCCCACTATGTCCCAG
GAGCACACGTCTGAACTCTTTCCCTACACGACGCTCTTCCGATCTNNNCTGTTGAGAAGAATGCTGCA
AGCACACGTCTGAACTCTTTCCCTACACGACGCTCTTCCGATCTNNNGAGTTTCGCTTTGGAGGATGT
GCACACGTCTGAACTCTTTCCCTACACGACGCTCTTCCGATCTNNNTAGATTCACCTAGCCTCCACAAC
AGCACACGTCTGAACTCTTTCCCTACACGACGCTCTTCCGATCTNNNTGAACACAAGTCTTCCACTCCT
GCACACGTCTGAACTCTTTCCCTACACGACGCTCTTCCGATCTNNNGTAGAGAAGGGGTTTCACCAT

AGCACACGTCTGAACTCTTTCCCTACACGACGCTCTTCCGATCTNNNAAACTCCTGACCTCAAGTGATCC
GAGCACACGTCTGAACTCTTTCCCTACACGACGCTCTTCCGATCTNNNAGTGGAAAACAGATCACAGGT
GCACACGTCTGAACTCTTTCCCTACACGACGCTCTTCCGATCTNNNCATCTTGCCCAACCCCATTATTCCA
AGCACACGTCTGAACTCTTTCCCTACACGACGCTCTTCCGATCTNNNTGTTCAGGCAGACACTAAGCTA
AGCACACGTCTGAACTCTTTCCCTACACGACGCTCTTCCGATCTNNNGGGCGACACATACAAAAAGACA
AGCACACGTCTGAACTCTTTCCCTACACGACGCTCTTCCGATCTNNNATGGGAGATCTGACTGTGGTTT
AGCACACGTCTGAACTCTTTCCCTACACGACGCTCTTCCGATCTNNNTGGAACATTTCTCTTGATTGCGA
GCACACGTCTGAACTCTTTCCCTACACGACGCTCTTCCGATCTNNNTGGAGGGAAGAATCTCAGAGCT
AGCACACGTCTGAACTCTTTCCCTACACGACGCTCTTCCGATCTNNNCACATCTCAGCAACAGTGTTACA
CACACGTCTGAACTCTTTCCCTACACGACGCTCTTCCGATCTNNNAGCAAGAGATTCATTACAAAAGAG
AGAGCACACGTCTGAACTCTTTCCCTACACGACGCTCTTCCGATCTNNNGGAGGTTTTGACATTCTGAG
GAGCACACGTCTGAACTCTTTCCCTACACGACGCTCTTCCGATCTNNNCCTCTCTCCTTGGTCCAATTCAA
AGAGCACACGTCTGAACTCTTTCCCTACACGACGCTCTTCCGATCTNNNATTTTGGGCAGCTTTCAGTG
AGCACACGTCTGAACTCTTTCCCTACACGACGCTCTTCCGATCTNNNATTTGTTCCTTCTTGCCATGCTCC
AGCACACGTCTGAACTCTTTCCCTACACGACGCTCTTCCGATCTNNNAGAGAACAAGGTCACAGAGCAA
GAGCACACGTCTGAACTCTTTCCCTACACGACGCTCTTCCGATCTNNNTCACTTGTCATTGTGAGACACC
AGCACACGTCTGAACTCTTTCCCTACACGACGCTCTTCCGATCTNNNTAGAGTCCCCCAGAAAACTGAG
GAGCACACGTCTGAACTCTTTCCCTACACGACGCTCTTCCGATCTNNNATGACTCTCAAAGCTGACTTCC
AGCACACGTCTGAACTCTTTCCCTACACGACGCTCTTCCGATCTNNNCCCTACGTTGTAGTCACAAGTGT
GAGCACACGTCTGAACTCTTTCCCTACACGACGCTCTTCCGATCTNNNTAGTTGGGACACAGGTATGCA
GAGCACACGTCTGAACTCTTTCCCTACACGACGCTCTTCCGATCTNNNAGAAGTACAGTGGCATGATCT
GCACACGTCTGAACTCTTTCCCTACACGACGCTCTTCCGATCTNNNCTTGTTTTGGGCCATTCAGCTAA
AGCACACGTCTGAACTCTTTCCCTACACGACGCTCTTCCGATCTNNNAACACTAAGGCAGGAAACAAGC
AGCACACGTCTGAACTCTTTCCCTACACGACGCTCTTCCGATCTNNNAATTAGCCCCTTTCAGTTCACCTC
AGCACACGTCTGAACTCTTTCCCTACACGACGCTCTTCCGATCTNNNTTCAAAACATCGCATTGGCAGAT
GAGCACACGTCTGAACTCTTTCCCTACACGACGCTCTTCCGATCTNNNCTGACATTCCTTTCACCCAGAG
AGCACACGTCTGAACTCTTTCCCTACACGACGCTCTTCCGATCTNNNTTAGTCTTTGGCCTGGTTTACTC
GAGCACACGTCTGAACTCTTTCCCTACACGACGCTCTTCCGATCTNNNCCTTCCATACATCCCTGTGGAA
GAGCACACGTCTGAACTCTTTCCCTACACGACGCTCTTCCGATCTNNNGTGGTCCTTTCTACTGCTTGAC
AGCACACGTCTGAACTCTTTCCCTACACGACGCTCTTCCGATCTNNNTCTCAATCCTTTACTTTGCCACAC
GCACACGTCTGAACTCTTTCCCTACACGACGCTCTTCCGATCTNNNCATGTGCTGCTCTGATCATTCTGC
AGCACACGTCTGAACTCTTTCCCTACACGACGCTCTTCCGATCTNNNCCAGGGTTCAAGCAATTCTCCTA
AGCACACGTCTGAACTCTTTCCCTACACGACGCTCTTCCGATCTNNNTGTAAGCAATCTTCCCTCCTCAG
GAGCACACGTCTGAACTCTTTCCCTACACGACGCTCTTCCGATCTNNNCCTGGACACTGGCAATATGGA
GAGCACACGTCTGAACTCTTTCCCTACACGACGCTCTTCCGATCTNNNTGTAAAGTTGAAACTACCGCA
GAGCACACGTCTGAACTCTTTCCCTACACGACGCTCTTCCGATCTNNNATTCAGGGACCTTCTACCTCAG
GAGCACACGTCTGAACTCTTTCCCTACACGACGCTCTTCCGATCTNNNCCATTTTGCATTCCCACTAGCA
GAGCACACGTCTGAACTCTTTCCCTACACGACGCTCTTCCGATCTNNNTGGATGAGAGAGCACAGTCAA
AGCACACGTCTGAACTCTTTCCCTACACGACGCTCTTCCGATCTNNNGGATCAGTGCTGGATGTTTGTT
AGAGCACACGTCTGAACTCTTTCCCTACACGACGCTCTTCCGATCTNNNGTACCTGTAATCTCAGCTACC
AGCACACGTCTGAACTCTTTCCCTACACGACGCTCTTCCGATCTNNNGTTGTGGTGACCTGAGATCGTA
GAGCACACGTCTGAACTCTTTCCCTACACGACGCTCTTCCGATCTNNNGCCAATATGGAGAAACCCTGT
GAGCACACGTCTGAACTCTTTCCCTACACGACGCTCTTCCGATCTNNNGTATTTCTGCAGTCCCAGCTAC

GAGCACACGTCTGAACTCTTTCCCTACACGACGCTCTTCCGATCTNNNAGAGTATTGCTTGAACCCAG
GAGCACACGTCTGAACTCTTTCCCTACACGACGCTCTTCCGATCTNNNGTGGGAGAATCTCTTGAACCCA
AGCACACGTCTGAACTCTTTCCCTACACGACGCTCTTCCGATCTNNNGCAACAGAGCAAGACTTCATCT
AGAGCACACGTCTGAACTCTTTCCCTACACGACGCTCTTCCGATCTNNNAGAATCACTTGAACCAAGGA
AGCACACGTCTGAACTCTTTCCCTACACGACGCTCTTCCGATCTNNNGAGTGTCGACATCATGCTCAAA
GAGCACACGTCTGAACTCTTTCCCTACACGACGCTCTTCCGATCTNNNTAGGAGAATTGCTTGAACCCC
AGCACACGTCTGAACTCTTTCCCTACACGACGCTCTTCCGATCTNNNTGTGCTTGTAATCCCAGCTACT
GAGCACACGTCTGAACTCTTTCCCTACACGACGCTCTTCCGATCTNNNGGCACCTGTAATACCAGCTACT
AGCACACGTCTGAACTCTTTCCCTACACGACGCTCTTCCGATCTNNNTAAGGCAGGAGAATTGCTTGA
GAGCACACGTCTGAACTCTTTCCCTACACGACGCTCTTCCGATCTNNNGAGCCTATAATCCCAGCTACTC
AGCACACGTCTGAACTCTTTCCCTACACGACGCTCTTCCGATCTNNNCTGAGGCAGGAGAATCACTTGA
AGCACACGTCTGAACTCTTTCCCTACACGACGCTCTTCCGATCTNNNATGAAAGATGTAGAAGCCCAG
GAGCACACGTCTGAACTCTTTCCCTACACGACGCTCTTCCGATCTNNNCTGTAATCCCAGCTACTCAGGA
AGCACACGTCTGAACTCTTTCCCTACACGACGCTCTTCCGATCTNNNTGGATGTGATGGAGTGCGAAT
GAGCACACGTCTGAACTCTTTCCCTACACGACGCTCTTCCGATCTNNNGTATTGCTATGTGGGTGAGG
GAGCACACGTCTGAACTCTTTCCCTACACGACGCTCTTCCGATCTNNNAATCCCAGCTACTTGAGAGGT
AGAGCACACGTCTGAACTCTTTCCCTACACGACGCTCTTCCGATCTNNNCCTGGAATGTGAAGGTTGCA
GAGCACACGTCTGAACTCTTTCCCTACACGACGCTCTTCCGATCTNNNGAAGGTTGCAGTGAGTGGAA
GAGCACACGTCTGAACTCTTTCCCTACACGACGCTCTTCCGATCTNNNGAGCTTACAGTGACCCGAGAT
AGCACACGTCTGAACTCTTTCCCTACACGACGCTCTTCCGATCTNNNCATACCTGTGATCCCAGCTACTC
AGAGCACACGTCTGAACTCTTTCCCTACACGACGCTCTTCCGATCTNNNGGAACTTGCAGTGAGCTGA
AGCACACGTCTGAACTCTTTCCCTACACGACGCTCTTCCGATCTNNNCTGAAGAATTGCTTGAACCTG
AGCACACGTCTGAACTCTTTCCCTACACGACGCTCTTCCGATCTNNNGCAACAGACGGAGACTCTGTAT
AGCACACGTCTGAACTCTTTCCCTACACGACGCTCTTCCGATCTNNNCCCATCTCTGCAAAAACACAA
GAGCACACGTCTGAACTCTTTCCCTACACGACGCTCTTCCGATCTNNNAAGAGACACCATGAAAGCAA
AGAGCACACGTCTGAACTCTTTCCCTACACGACGCTCTTCCGATCTNNNCACCTGTAATCCCTGCTACTCA
AGCACACGTCTGAACTCTTTCCCTACACGACGCTCTTCCGATCTNNNTAAACTTCGATGTAGCCGACAC
AGAGCACACGTCTGAACTCTTTCCCTACACGACGCTCTTCCGATCTNNNCATGTCTGTAGTCCCAGCTA
GAGCACACGTCTGAACTCTTTCCCTACACGACGCTCTTCCGATCTNNNAGAGAATCGCTTCACAGGAG
AGAGCACACGTCTGAACTCTTTCCCTACACGACGCTCTTCCGATCTNNNGAGAGACTGAGGCAAGAGA
AGCACACGTCTGAACTCTTTCCCTACACGACGCTCTTCCGATCTNNNCCTGGTGCATGCTACCTACATA
GAGCACACGTCTGAACTCTTTCCCTACACGACGCTCTTCCGATCTNNNCTGAGGCAGGAGAATCACTT
GAGCACACGTCTGAACTCTTTCCCTACACGACGCTCTTCCGATCTNNNTGAGGTAGGAGAATCGCTTG
AGCACACGTCTGAACTCTTTCCCTACACGACGCTCTTCCGATCTNNNGTAATTCTAGCTACTGGGGAGG
AGAGCACACGTCTGAACTCTTTCCCTACACGACGCTCTTCCGATCTNNNGGCACCTATAATCCCAGCTA
AGCACACGTCTGAACTCTTTCCCTACACGACGCTCTTCCGATCTNNNCACCTGTAATCCCAGCATTTG
AGCACACGTCTGAACTCTTTCCCTACACGACGCTCTTCCGATCTNNNCTGTAATCCCAGCTACTCAGGA
GAGCACACGTCTGAACTCTTTCCCTACACGACGCTCTTCCGATCTNNNGGAATCACTTGAACCTGGGA
GAGCACACGTCTGAACTCTTTCCCTACACGACGCTCTTCCGATCTNNNAGAATTACTTGAACCTGGGAG
AGCACACGTCTGAACTCTTTCCCTACACGACGCTCTTCCGATCTNNNCTGTAATCCCAGCTACTCAGGA
GAGCACACGTCTGAACTCTTTCCCTACACGACGCTCTTCCGATCTNNNCCTGTAGTCCCAGCTACTCTAC
AGCACACGTCTGAACTCTTTCCCTACACGACGCTCTTCCGATCTNNNCTGTAATCCCAGCTCCTCAGAAG
AGCACACGTCTGAACTCTTTCCCTACACGACGCTCTTCCGATCTNNNCTGAGACAGGAGAATCGCTTGA

AGCACACGTCTGAACTCTTTCCCTACACGACGCTCTTCCGATCTNNNTAACACAGTGAAACCCTGTCT
AGCACACGTCTGAACTCTTTCCCTACACGACGCTCTTCCGATCTNNNGGTGGGACAAGCATAGGATA
AGCACACGTCTGAACTCTTTCCCTACACGACGCTCTTCCGATCTNNNGAGGAGGCTAAGGAAGGAC
GCACACGTCTGAACTCTTTCCCTACACGACGCTCTTCCGATCTNNNAAACGATGTGTTGATGGATGC
AGCACACGTCTGAACTCTTTCCCTACACGACGCTCTTCCGATCTNNNAAGCAGGAGAATCACTTGAA
GAGCACACGTCTGAACTCTTTCCCTACACGACGCTCTTCCGATCTNNNCATGCTACTACACTCCAGTCT
AGCACACGTCTGAACTCTTTCCCTACACGACGCTCTTCCGATCTNNNAGATCACACACCATTTCACTCC
AGCACACGTCTGAACTCTTTCCCTACACGACGCTCTTCCGATCTNNNTGAGGCAGGAGAATGGTATG
GAGCACACGTCTGAACTCTTTCCCTACACGACGCTCTTCCGATCTNNNATGCTTTGTTTCTTCCTGCAG
GAGCACACGTCTGAACTCTTTCCCTACACGACGCTCTTCCGATCTNNNGGTTGCAGTGAGCCAAGATA
GAGCACACGTCTGAACTCTTTCCCTACACGACGCTCTTCCGATCTNNNCTGTAATCCTGGCTACTTGGG
AGCACACGTCTGAACTCTTTCCCTACACGACGCTCTTCCGATCTNNNCTCTGGGAATGAAACCCTGTC
GAGCACACGTCTGAACTCTTTCCCTACACGACGCTCTTCCGATCTNNNTGCCTATAATCGCAGCTACTC
AGCACACGTCTGAACTCTTTCCCTACACGACGCTCTTCCGATCTNNNCTGAGGCAGGAGAATCACTTG
GCACACGTCTGAACTCTTTCCCTACACGACGCTCTTCCGATCTNNNAAGTCTGTAGTCCCAGCTACTCA
AGCACACGTCTGAACTCTTTCCCTACACGACGCTCTTCCGATCTNNNGAGAATGGCATGAACCCAGAA
GAGCACACGTCTGAACTCTTTCCCTACACGACGCTCTTCCGATCTNNNCACTTGAGACCAGGAGTTTG
AGCACACGTCTGAACTCTTTCCCTACACGACGCTCTTCCGATCTNNNCAGAGGTTGCAGTGAGCTAAG
AGCACACGTCTGAACTCTTTCCCTACACGACGCTCTTCCGATCTNNNTGCCTGTAATCCCAGCTAATCA
GAGCACACGTCTGAACTCTTTCCCTACACGACGCTCTTCCGATCTNNNCAGAAGGTTGCAGTGAGTC
GAGCACACGTCTGAACTCTTTCCCTACACGACGCTCTTCCGATCTNNNAGTGAAACCCCGTCTCTACT
AGCACACGTCTGAACTCTTTCCCTACACGACGCTCTTCCGATCTNNNCAGGAGAATTGCTTGAACCA
AGCACACGTCTGAACTCTTTCCCTACACGACGCTCTTCCGATCTNNNCTGTAGTCCCAGCTACTTGAG
GCACACGTCTGAACTCTTTCCCTACACGACGCTCTTCCGATCTNNNCATGCCTGTAATCCCAGCTACTC
AGCACACGTCTGAACTCTTTCCCTACACGACGCTCTTCCGATCTNNNGTATTCCCAGCTACTCAGGAA
AGCACACGTCTGAACTCTTTCCCTACACGACGCTCTTCCGATCTNNNTAATCCCAGCTACTCAGGAGG
AGCACACGTCTGAACTCTTTCCCTACACGACGCTCTTCCGATCTNNNGAAGCAGGAGAATCACTTGA
GAGCACACGTCTGAACTCTTTCCCTACACGACGCTCTTCCGATCTNNNACAGGAGAATGTCATGAACC
AGCACACGTCTGAACTCTTTCCCTACACGACGCTCTTCCGATCTNNNTGTCCTGGGCCACACATAAAA
AGAGCACACGTCTGAACTCTTTCCCTACACGACGCTCTTCCGATCTNNNGATGCCTGTAATCCCAGCTA
GAGCACACGTCTGAACTCTTTCCCTACACGACGCTCTTCCGATCTNNNGCCTGTAGTCTGAGCTACTA
GCACACGTCTGAACTCTTTCCCTACACGACGCTCTTCCGATCTNNNGGTTGCAGTGAGTCTATATTGC
AGCACACGTCTGAACTCTTTCCCTACACGACGCTCTTCCGATCTNNNTGTAATCCCAGCACTTTGGTA
GCACACGTCTGAACTCTTTCCCTACACGACGCTCTTCCGATCTNNNCCAGAAATAGACCCACAGGAA
AGCACACGTCTGAACTCTTTCCCTACACGACGCTCTTCCGATCTNNNTGAGGTAGGAGAATCGCTTG
GCACACGTCTGAACTCTTTCCCTACACGACGCTCTTCCGATCTNNNCTTGAACTCCTGACCTCGTGATC
GCACACGTCTGAACTCTTTCCCTACACGACGCTCTTCCGATCTNNNCCACCACACCAGGCTAATTTTTCC
GCACACGTCTGAACTCTTTCCCTACACGACGCTCTTCCGATCTNNNCATGGTGTGGTTCCTTCCTCTTA
AGCACACGTCTGAACTCTTTCCCTACACGACGCTCTTCCGATCTNNNATCCTGGAGATGTCTTTTGCC
GAGCACACGTCTGAACTCTTTCCCTACACGACGCTCTTCCGATCTNNNACACCTGGCTGAAACTCACTA
GAGCACACGTCTGAACTCTTTCCCTACACGACGCTCTTCCGATCTNNNCAGCAACTTGGATGAAGCTA
AGCACACGTCTGAACTCTTTCCCTACACGACGCTCTTCCGATCTNNNCTTCTCACCTTAGGCGATCAT
AGCACACGTCTGAACTCTTTCCCTACACGACGCTCTTCCGATCTNNNCAGGTGGTCCTTCTCGAGTTTT

AGCACACGTCTGAACTCTTTCCCTACACGACGCTCTTCCGATCTNNNCTCCTGAGTAGCCGAGATTACA
AGCACACGTCTGAACTCTTTCCCTACACGACGCTCTTCCGATCTNNNGCCACATTTCCTAGGCTAGTCTT
AGCACACGTCTGAACTCTTTCCCTACACGACGCTCTTCCGATCTNNNGTGGGTGTATGGTGGTATCACA
AGCACACGTCTGAACTCTTTCCCTACACGACGCTCTTCCGATCTNNNACTCCTGGACTCAAGTGATTGT
AGCACACGTCTGAACTCTTTCCCTACACGACGCTCTTCCGATCTNNNGGCATATGGGGGCATAAAAG
AGCACACGTCTGAACTCTTTCCCTACACGACGCTCTTCCGATCTNNNATTCTCCACACGTCTTTTTGACT
AGCACACGTCTGAACTCTTTCCCTACACGACGCTCTTCCGATCTNNNATGAGGTTTCACTACATTGGC
AGCACACGTCTGAACTCTTTCCCTACACGACGCTCTTCCGATCTNNNCCTCCTGAGGTACAACTTCCAT
AGCACACGTCTGAACTCTTTCCCTACACGACGCTCTTCCGATCTNNNAGGGGCTTCTTCTTTTTCTTC
GCACACGTCTGAACTCTTTCCCTACACGACGCTCTTCCGATCTNNNAACTTGTGCTTCAATGCCTGTTT
AGCACACGTCTGAACTCTTTCCCTACACGACGCTCTTCCGATCTNNNAGAGACGGGGTTTCACTATGT
AGCACACGTCTGAACTCTTTCCCTACACGACGCTCTTCCGATCTNNNACTCCTCTAATGACCTCTTCCC
AGCACACGTCTGAACTCTTTCCCTACACGACGCTCTTCCGATCTNNNTCTCTGAAGAGGCTCCAGATAG
AGCACACGTCTGAACTCTTTCCCTACACGACGCTCTTCCGATCTNNNGGTCATTCCAGCAATTCCCATT
AGCACACGTCTGAACTCTTTCCCTACACGACGCTCTTCCGATCTNNNTTGGGATTACAGGTATGAGCC
AGCACACGTCTGAACTCTTTCCCTACACGACGCTCTTCCGATCTNNNTGTGCTTTGCTTCTATTGGCTT
AGCACACGTCTGAACTCTTTCCCTACACGACGCTCTTCCGATCTNNNTTTGTGAGCACCAGAATCTGT
GAGCACACGTCTGAACTCTTTCCCTACACGACGCTCTTCCGATCTNNNACCGCGCCCGACCTATATATA
AGCACACGTCTGAACTCTTTCCCTACACGACGCTCTTCCGATCTNNNTTCTGAGTTGCTGGGATTACA
AGCACACGTCTGAACTCTTTCCCTACACGACGCTCTTCCGATCTNNNAGCATCATATTGGGCAGTATAC
AGCACACGTCTGAACTCTTTCCCTACACGACGCTCTTCCGATCTNNNGTCCCTGTGATGTCTGAGAACTT
AGAGCACACGTCTGAACTCTTTCCCTACACGACGCTCTTCCGATCTNNNTCCTGAGTTGTTCCACTGTC
AGCACACGTCTGAACTCTTTCCCTACACGACGCTCTTCCGATCTNNNAGGGTCTGTGTGCATTGATGA
AGCACACGTCTGAACTCTTTCCCTACACGACGCTCTTCCGATCTNNNGCAGAGGTCCCTGAGGAATT
AGCACACGTCTGAACTCTTTCCCTACACGACGCTCTTCCGATCTNNNTTGGCCAAGGAAAACATCATG
AGCACACGTCTGAACTCTTTCCCTACACGACGCTCTTCCGATCTNNNTCTCCTGGGTTCAAGCAATTCT
AGCACACGTCTGAACTCTTTCCCTACACGACGCTCTTCCGATCTNNNCTCCCCAGTAGCTAGGACTACA
GAGCACACGTCTGAACTCTTTCCCTACACGACGCTCTTCCGATCTNNNCCGCCTTGAGAGTGTATGGA
AGCACACGTCTGAACTCTTTCCCTACACGACGCTCTTCCGATCTNNNCTGAGTAGGTGGGATTACAGG
AGCACACGTCTGAACTCTTTCCCTACACGACGCTCTTCCGATCTNNNATGCCCAGCCTTCTTTTCTTTTCC
AGCACACGTCTGAACTCTTTCCCTACACGACGCTCTTCCGATCTNNNAGCCACAGCACCTGACATATAT
AGCACACGTCTGAACTCTTTCCCTACACGACGCTCTTCCGATCTNNNAGCAAATATTGGAAGTGGGG
AGCACACGTCTGAACTCTTTCCCTACACGACGCTCTTCCGATCTNNNATTACAAACAGGCCAACATGTC
AGCACACGTCTGAACTCTTTCCCTACACGACGCTCTTCCGATCTNNNTTCAGCTGATCTCATCTGAGT
AGCACACGTCTGAACTCTTTCCCTACACGACGCTCTTCCGATCTNNNAGAAAATCCAGTTTCTAGGGGC
GAGCACACGTCTGAACTCTTTCCCTACACGACGCTCTTCCGATCTNNNATATTGGTCAGGCTGGTCTT
AGCACACGTCTGAACTCTTTCCCTACACGACGCTCTTCCGATCTNNNGGACATGTTTAGGGAGGTGTA
AGCACACGTCTGAACTCTTTCCCTACACGACGCTCTTCCGATCTNNNACACTAGAACAGGGGTTGGAA
AGCACACGTCTGAACTCTTTCCCTACACGACGCTCTTCCGATCTNNNCTGTGAATCCATCTGGTCCTAG
AGCACACGTCTGAACTCTTTCCCTACACGACGCTCTTCCGATCTNNNAGTCCCCAAAGTTCATTGTGTCT
AGCACACGTCTGAACTCTTTCCCTACACGACGCTCTTCCGATCTNNNAGTGTGGACATGAAAGAGAG
AGAGCACACGTCTGAACTCTTTCCCTACACGACGCTCTTCCGATCTNNNATATACTCTGTGGCAGGCAT
AGCACACGTCTGAACTCTTTCCCTACACGACGCTCTTCCGATCTNNNCCTCATACTCTGGCCATCAGTTT

AGCACACGTCTGAACTCTTTCCCTACACGACGCTCTTCCGATCTNNNTTTTGAACTCCTGGTTTCAAGC
AGCACACGTCTGAACTCTTTCCCTACACGACGCTCTTCCGATCTNNNCATGGATCAAGGTATCCCGGA
GAGCACACGTCTGAACTCTTTCCCTACACGACGCTCTTCCGATCTNNNACCTTCATGTCTATCTCGCTCT
AGCACACGTCTGAACTCTTTCCCTACACGACGCTCTTCCGATCTNNNTGAGACTCATGTTTTTACCCCTT
GAGCACACGTCTGAACTCTTTCCCTACACGACGCTCTTCCGATCTNNNCCTCAGTCTCCTGAATAGTTG
AGCACACGTCTGAACTCTTTCCCTACACGACGCTCTTCCGATCTNNNGAGTCAGTGAGTGGTGAGTG
AGAGCACACGTCTGAACTCTTTCCCTACACGACGCTCTTCCGATCTNNNTGGAAACAAGGTAAAGGT
AGCACACGTCTGAACTCTTTCCCTACACGACGCTCTTCCGATCTNNNTATGTCACCCTCCAGAAACTTG
AGCACACGTCTGAACTCTTTCCCTACACGACGCTCTTCCGATCTNNNATTCAGATCATCCCTGGCCAAA
AGCACACGTCTGAACTCTTTCCCTACACGACGCTCTTCCGATCTNNNTCCCAAAGTGCTAGGATTACAG
AGCACACGTCTGAACTCTTTCCCTACACGACGCTCTTCCGATCTNNNAATGCCTACTGGGATAAGCTTC
AGCACACGTCTGAACTCTTTCCCTACACGACGCTCTTCCGATCTNNNTTACACTGCATTTACTCCACCCAC
AGCACACGTCTGAACTCTTTCCCTACACGACGCTCTTCCGATCTNNNCTGGGTTCAGGTGATTTTTCCA
GAGCACACGTCTGAACTCTTTCCCTACACGACGCTCTTCCGATCTNNNCAGCTCCTCTTTGTACATCTGG
AGCACACGTCTGAACTCTTTCCCTACACGACGCTCTTCCGATCTNNNCTAACCTCAAGTGATCCACCCTC
AGCACACGTCTGAACTCTTTCCCTACACGACGCTCTTCCGATCTNNNGCCTCCCAAAGTGCTTGAATTA
AGCACACGTCTGAACTCTTTCCCTACACGACGCTCTTCCGATCTNNNACTCCTGACCTCAAGAGATTTGG
AGCACACGTCTGAACTCTTTCCCTACACGACGCTCTTCCGATCTNNNTAAAATGGTCCCGGCAATAAA
GAGCACACGTCTGAACTCTTTCCCTACACGACGCTCTTCCGATCTNNNGGGGGATTCTGTCACAGATG
AGCACACGTCTGAACTCTTTCCCTACACGACGCTCTTCCGATCTNNNAGTTGGTTGTCCTGTTATCCTCC
AGCACACGTCTGAACTCTTTCCCTACACGACGCTCTTCCGATCTNNNTTCCCCCTCTGAATGATCTCTCT
GAGCACACGTCTGAACTCTTTCCCTACACGACGCTCTTCCGATCTNNNTTGTTTCATCCTTGCAGAAGA
AGCACACGTCTGAACTCTTTCCCTACACGACGCTCTTCCGATCTNNNTTTCAGGCTGCAAATTTTCCCAA
AGCACACGTCTGAACTCTTTCCCTACACGACGCTCTTCCGATCTNNNTATCACTCATCACTCTCCCCTCC
GAGCACACGTCTGAACTCTTTCCCTACACGACGCTCTTCCGATCTNNNTCCATCTATAGTGCAACATGT
AGCACACGTCTGAACTCTTTCCCTACACGACGCTCTTCCGATCTNNNATAAGCATTCCCTTTCCTCTGCA
AGCACACGTCTGAACTCTTTCCCTACACGACGCTCTTCCGATCTNNNGCAACTAGAAGCAAAGGCAGT
AGCACACGTCTGAACTCTTTCCCTACACGACGCTCTTCCGATCTNNNAGCTTCTCTCACTTGGCATGTT
AGCACACGTCTGAACTCTTTCCCTACACGACGCTCTTCCGATCTNNNCAATCTTGGCTTACTGCAACCTC
AGCACACGTCTGAACTCTTTCCCTACACGACGCTCTTCCGATCTNNNTGGAAACATGCACAATGATGCT
GCACACGTCTGAACTCTTTCCCTACACGACGCTCTTCCGATCTNNNAAATGGACTGCCCTTTACCTAGC
AGCACACGTCTGAACTCTTTCCCTACACGACGCTCTTCCGATCTNNNAACAGAATAGATGCTTGCTTG
GAGCACACGTCTGAACTCTTTCCCTACACGACGCTCTTCCGATCTNNNTCCAGCAGAAATTGAGAGTG
AGAGCACACGTCTGAACTCTTTCCCTACACGACGCTCTTCCGATCTNNNAGGCTGGAGTACAATGTCA
AGCACACGTCTGAACTCTTTCCCTACACGACGCTCTTCCGATCTNNNTCCCAACAAGGACATAGAGCTT
GCACACGTCTGAACTCTTTCCCTACACGACGCTCTTCCGATCTNNNGAGCATTTGTGGGCTATTGTAC
AGCACACGTCTGAACTCTTTCCCTACACGACGCTCTTCCGATCTNNNTGTGAGCCCAAAATGGAGAAA
GAGCACACGTCTGAACTCTTTCCCTACACGACGCTCTTCCGATCTNNNACCCTTCTGTCTCTCTCTCTCTC
GAGCACACGTCTGAACTCTTTCCCTACACGACGCTCTTCCGATCTNNNATTTGCAAACCCAAGGAAAGA
AGCACACGTCTGAACTCTTTCCCTACACGACGCTCTTCCGATCTNNNTACTTGTCTCTGACCAGACCTG
GAGCACACGTCTGAACTCTTTCCCTACACGACGCTCTTCCGATCTNNNCCATATGCCGGGCTAATTTTT
AGCACACGTCTGAACTCTTTCCCTACACGACGCTCTTCCGATCTNNNGAGTAGCTGGTACTACAGGCA
AGCACACGTCTGAACTCTTTCCCTACACGACGCTCTTCCGATCTNNNAGTGGTGGCAATTACAGGACT

AGCACACGTCTGAACTCTTTCCCTACACGACGCTCTTCCGATCTNNNACAGTGTACAAGGGTTCCCTTTT
GAGCACACGTCTGAACTCTTTCCCTACACGACGCTCTTCCGATCTNNNCCAGGGCTAGAACTTCCATCTT
AGCACACGTCTGAACTCTTTCCCTACACGACGCTCTTCCGATCTNNNCTTCCTTCCTTCCTTCTTCCCTCCA
AGCACACGTCTGAACTCTTTCCCTACACGACGCTCTTCCGATCTNNNTGCTACACCCCTCATTCTACCTA
GAGCACACGTCTGAACTCTTTCCCTACACGACGCTCTTCCGATCTNNNAGAACATTCTTGTCACCCCAGA
GAGCACACGTCTGAACTCTTTCCCTACACGACGCTCTTCCGATCTNNNTAGCTGGGATTACAGGCATAT
GAGCACACGTCTGAACTCTTTCCCTACACGACGCTCTTCCGATCTNNNATATCAAGACCTTGCCACTCCA
AGCACACGTCTGAACTCTTTCCCTACACGACGCTCTTCCGATCTNNNACTTATGAGTGAGAACATGCGG
GCACACGTCTGAACTCTTTCCCTACACGACGCTCTTCCGATCTNNNGGAGAAGTGGAATGCACACAATG
GAGCACACGTCTGAACTCTTTCCCTACACGACGCTCTTCCGATCTNNNGAAAACAGAGGTTGGGTAGT
AGCACACGTCTGAACTCTTTCCCTACACGACGCTCTTCCGATCTNNNTGCTTATCAGCCATCTTTCCACTG
AGCACACGTCTGAACTCTTTCCCTACACGACGCTCTTCCGATCTNNNGGTTCAATCAATTCTCCTGCCTC
GAGCACACGTCTGAACTCTTTCCCTACACGACGCTCTTCCGATCTNNNTGGCACTATCACAGATTACTGC
AGAGCACACGTCTGAACTCTTTCCCTACACGACGCTCTTCCGATCTNNNCATGCCTGTAATCCCAGCTA
AGCACACGTCTGAACTCTTTCCCTACACGACGCTCTTCCGATCTNNNCAAGAATTTGAGGCTGCAGTGA
AGAGCACACGTCTGAACTCTTTCCCTACACGACGCTCTTCCGATCTNNNGGTTGCAGTGAGCCAAGATT
GCACACGTCTGAACTCTTTCCCTACACGACGCTCTTCCGATCTNNNCTTGGCAAAAGAGCAAGACCTTG
GCACACGTCTGAACTCTTTCCCTACACGACGCTCTTCCGATCTNNNCTGTAGTCCCAGCTACTCAGGAG
AGCACACGTCTGAACTCTTTCCCTACACGACGCTCTTCCGATCTNNNAGTTACCTGCCTTCCATTGTCAT
GAGCACACGTCTGAACTCTTTCCCTACACGACGCTCTTCCGATCTNNNGAGAATCTCTTGAACCCAGGA
GAGCACACGTCTGAACTCTTTCCCTACACGACGCTCTTCCGATCTNNNTGAGGTAGGAGAATTGCCTGA
AGCACACGTCTGAACTCTTTCCCTACACGACGCTCTTCCGATCTNNNAATCCCAGTTACTCAGGAAGCT
AGCACACGTCTGAACTCTTTCCCTACACGACGCTCTTCCGATCTNNNCGACAAGAGTAAGACCCTGTCT
GAGCACACGTCTGAACTCTTTCCCTACACGACGCTCTTCCGATCTNNNGGCACCTGTAATCCTAGCTAC
GAGCACACGTCTGAACTCTTTCCCTACACGACGCTCTTCCGATCTNNNCAACATAGTGAAACCCCGTCT
GAGCACACGTCTGAACTCTTTCCCTACACGACGCTCTTCCGATCTNNNATTTCACCCATAATCCCAGCAC
AGCACACGTCTGAACTCTTTCCCTACACGACGCTCTTCCGATCTNNNTGAATCAAAATCGAACCTGGCA
AGCACACGTCTGAACTCTTTCCCTACACGACGCTCTTCCGATCTNNNGCCACAGTGGAAGAAGAATTG
AGAGCACACGTCTGAACTCTTTCCCTACACGACGCTCTTCCGATCTNNNAGAGGTTGCAGTGAGTCAA
AGCACACGTCTGAACTCTTTCCCTACACGACGCTCTTCCGATCTNNNGAATCGTTTGAACCTGGGAAG
AGCACACGTCTGAACTCTTTCCCTACACGACGCTCTTCCGATCTNNNCAAGATCTCACCACTGCACTCTA
AGCACACGTCTGAACTCTTTCCCTACACGACGCTCTTCCGATCTNNNTGTAGTTCTAGCTACTCGGGAG
AGAGCACACGTCTGAACTCTTTCCCTACACGACGCTCTTCCGATCTNNNGGGCAACAAGAGTGAAATTG
AGCACACGTCTGAACTCTTTCCCTACACGACGCTCTTCCGATCTNNNGAGGGGAAAAGATTGCTTGAAG
AGCACACGTCTGAACTCTTTCCCTACACGACGCTCTTCCGATCTNNNGTAATCCCAGCTACTTAGGAGG
AGCACACGTCTGAACTCTTTCCCTACACGACGCTCTTCCGATCTNNNCTGTAATCCCAGCTACTCAGGA
AGCACACGTCTGAACTCTTTCCCTACACGACGCTCTTCCGATCTNNNGCCTGTAGTCCTAAGCTACTTGG
AGCACACGTCTGAACTCTTTCCCTACACGACGCTCTTCCGATCTNNNGGATCACTTGAGTCCAGGAAGT
AGCACACGTCTGAACTCTTTCCCTACACGACGCTCTTCCGATCTNNNATGTTGCAGTGAGCCAAGATCA
AGAGCACACGTCTGAACTCTTTCCCTACACGACGCTCTTCCGATCTNNNCTGTAATCACAGCTACTCGG
AGAGCACACGTCTGAACTCTTTCCCTACACGACGCTCTTCCGATCTNNNCTGTAGTCAGTCCCAGCTAC
AGCACACGTCTGAACTCTTTCCCTACACGACGCTCTTCCGATCTNNNCTGCAGAGAAGAAGGACTGGA
GAGCACACGTCTGAACTCTTTCCCTACACGACGCTCTTCCGATCTNNNGCTGTAGTAAGCCATGTTTGC

AGAGCACACGTCTGAACTCTTTCCCTACACGACGCTCTTCCGATCTNNNCAGGAGAATTGCTTGAACAC
AGCACACGTCTGAACTCTTTCCCTACACGACGCTCTTCCGATCTNNNAGAATTGCTTCAACCTGGGAG
AGCACACGTCTGAACTCTTTCCCTACACGACGCTCTTCCGATCTNNNGAGTCCGAGACAAGCCTGTAA
AGAGCACACGTCTGAACTCTTTCCCTACACGACGCTCTTCCGATCTNNNCTGAGGCAGGAGAATCACTT
AGCACACGTCTGAACTCTTTCCCTACACGACGCTCTTCCGATCTNNNATGCCTGTAATCCCAGCATTTTG
AGAGCACACGTCTGAACTCTTTCCCTACACGACGCTCTTCCGATCTNNNTTTTACAAAACAGGATGCTG
AGCACACGTCTGAACTCTTTCCCTACACGACGCTCTTCCGATCTNNNATTCTAGCTACTCAGGAGGCTG
GAGCACACGTCTGAACTCTTTCCCTACACGACGCTCTTCCGATCTNNNCTGTAATCCCAGCTACTCAGG
AGCACACGTCTGAACTCTTTCCCTACACGACGCTCTTCCGATCTNNNCCATACTGGCCTCTCTCACTTAC
AGCACACGTCTGAACTCTTTCCCTACACGACGCTCTTCCGATCTNNNGCAGAAGAATCACTTGAACCCA
GAGCACACGTCTGAACTCTTTCCCTACACGACGCTCTTCCGATCTNNNCTGTAGTCCCAGCTACTCGAG
AGAGCACACGTCTGAACTCTTTCCCTACACGACGCTCTTCCGATCTNNNTGGAGACCATGAAAGTAAT
AGAGCACACGTCTGAACTCTTTCCCTACACGACGCTCTTCCGATCTNNNTAGGCAACACAGTGAGAAT
GAGCACACGTCTGAACTCTTTCCCTACACGACGCTCTTCCGATCTNNNTATTTGAGAGGCTGAGGTGC
AGAGCACACGTCTGAACTCTTTCCCTACACGACGCTCTTCCGATCTNNNCACAATGTGCACATGTACCC
AGAGCACACGTCTGAACTCTTTCCCTACACGACGCTCTTCCGATCTNNNAATCCTAGCTACTCAGGAGG
AGAGCACACGTCTGAACTCTTTCCCTACACGACGCTCTTCCGATCTNNNAAAAAATTAGCTGGACGTG
AGCACACGTCTGAACTCTTTCCCTACACGACGCTCTTCCGATCTNNNGTAATCCCAGCTACTTAGGAG
GAGCACACGTCTGAACTCTTTCCCTACACGACGCTCTTCCGATCTNNNCACCTGTAATCCCAGCTACTT
GCACACGTCTGAACTCTTTCCCTACACGACGCTCTTCCGATCTNNNAAGCGCCTGTAGTTTTAGCTACT
GAGCACACGTCTGAACTCTTTCCCTACACGACGCTCTTCCGATCTNNNTCTGCCTATGAAAAAGATGGG
AGAGCACACGTCTGAACTCTTTCCCTACACGACGCTCTTCCGATCTNNNATTGCTTGAGACCAGGAGT
AGCACACGTCTGAACTCTTTCCCTACACGACGCTCTTCCGATCTNNNAGTGAACCATGATAGTGCCAC
AGAGCACACGTCTGAACTCTTTCCCTACACGACGCTCTTCCGATCTNNNGACTGTAATCCCAGCTACTCA
AGCACACGTCTGAACTCTTTCCCTACACGACGCTCTTCCGATCTNNNCATCGGTGTACTGTATCCAGG
AGCACACGTCTGAACTCTTTCCCTACACGACGCTCTTCCGATCTNNNCTGAGGCAGGAGAATCACTTG
GAGCACACGTCTGAACTCTTTCCCTACACGACGCTCTTCCGATCTNNNAATTGCTTGAACCTAGGAGAC
GAGCACACGTCTGAACTCTTTCCCTACACGACGCTCTTCCGATCTNNNGCAGGAGAATCACTTGAACCT
AGAGCACACGTCTGAACTCTTTCCCTACACGACGCTCTTCCGATCTNNNCTGAGGCAGGAGAATCACTT
GAGCACACGTCTGAACTCTTTCCCTACACGACGCTCTTCCGATCTNNNTCCTGGAAACATACAATCTTCC
GAGCACACGTCTGAACTCTTTCCCTACACGACGCTCTTCCGATCTNNNCAAGAGAATTGCTTGAACCCA
GAGCACACGTCTGAACTCTTTCCCTACACGACGCTCTTCCGATCTNNNGGACTCCTCCCTAACTCATTCT
AGCACACGTCTGAACTCTTTCCCTACACGACGCTCTTCCGATCTNNNCTGAGGCAGGAGAATTGCTTG
GAGCACACGTCTGAACTCTTTCCCTACACGACGCTCTTCCGATCTNNNGGCAACAAGAATGAAACTCC
GAGCACACGTCTGAACTCTTTCCCTACACGACGCTCTTCCGATCTNNNATAATCTCAGCTACTTGGGA
GAGCACACGTCTGAACTCTTTCCCTACACGACGCTCTTCCGATCTNNNTATAACCCCAGCTACTTGCAAG
GAGCACACGTCTGAACTCTTTCCCTACACGACGCTCTTCCGATCTNNNGCAGGAGAATTGCTTGAACTC
AGCACACGTCTGAACTCTTTCCCTACACGACGCTCTTCCGATCTNNNCCAGGAGTTTGAGGTTACAGT
AGCACACGTCTGAACTCTTTCCCTACACGACGCTCTTCCGATCTNNNGCCTGTAATCTCAGCACTTTGT
GAGCACACGTCTGAACTCTTTCCCTACACGACGCTCTTCCGATCTNNNAGACAGGAGAATCACTTGAA
AGCACACGTCTGAACTCTTTCCCTACACGACGCTCTTCCGATCTNNNAAGACCATCTTGGCTAACATGG
AGAGCACACGTCTGAACTCTTTCCCTACACGACGCTCTTCCGATCTNNNCTGAGGCAGGAGAATCACTT
GAGCACACGTCTGAACTCTTTCCCTACACGACGCTCTTCCGATCTNNNAGAATCGCTTGAACCAAGGA

AGAGCACACGTCTGAACTCTTTCCCTACACGACGCTCTTCCGATCTNNNCCCATCTACTTGGAAGACTG
AGCACACGTCTGAACTCTTTCCCTACACGACGCTCTTCCGATCTNNNGAGGTTGAAGTGGGAGAATCA
GAGCACACGTCTGAACTCTTTCCCTACACGACGCTCTTCCGATCTNNNTAGGAGGCAAAGGTTACAGT
AGCACACGTCTGAACTCTTTCCCTACACGACGCTCTTCCGATCTNNNTCTGTAATCCCAGCTACTCAGGA
AGCACACGTCTGAACTCTTTCCCTACACGACGCTCTTCCGATCTNNNCCATGGTGGAGTTCTAGCTACT
AGCACACGTCTGAACTCTTTCCCTACACGACGCTCTTCCGATCTNNNCTGAGGCAGGAGAATCACTTGA
GAGCACACGTCTGAACTCTTTCCCTACACGACGCTCTTCCGATCTNNNGAGAATTGCTTGAACCCAGGA
GAGCACACGTCTGAACTCTTTCCCTACACGACGCTCTTCCGATCTNNNGAAGAATGGTGTGAACCTGGC
AGCACACGTCTGAACTCTTTCCCTACACGACGCTCTTCCGATCTNNNGTCACCTGTAATCCCAGCTACTCC
AGCACACGTCTGAACTCTTTCCCTACACGACGCTCTTCCGATCTNNNTCTCAGATACTGGAGAGGCTGA
AGCACACGTCTGAACTCTTTCCCTACACGACGCTCTTCCGATCTNNNTACAGTGAGCTGAGATTGCATC
GAGCACACGTCTGAACTCTTTCCCTACACGACGCTCTTCCGATCTNNNAGTGAACCAAGATCATGCCAT
GAGCACACGTCTGAACTCTTTCCCTACACGACGCTCTTCCGATCTNNNGGCAGGAGAATCACTTGAACC
GAGCACACGTCTGAACTCTTTCCCTACACGACGCTCTTCCGATCTNNNTGTTATTCCAGCTACTTGGGAC
AGCACACGTCTGAACTCTTTCCCTACACGACGCTCTTCCGATCTNNNATTCAGCCGCTTGTCTAATGGT
AGCACACGTCTGAACTCTTTCCCTACACGACGCTCTTCCGATCTNNNCTGAGGCAGGAGAATCACTTGA
AGCACACGTCTGAACTCTTTCCCTACACGACGCTCTTCCGATCTNNNATGCCTGTAATCCCAACTACTCG
AGCACACGTCTGAACTCTTTCCCTACACGACGCTCTTCCGATCTNNNGGTGGCAAGTATCTGTAATCCC
AGCACACGTCTGAACTCTTTCCCTACACGACGCTCTTCCGATCTNNNGATTGAGACCATCCTGGCTAAC
GAGCACACGTCTGAACTCTTTCCCTACACGACGCTCTTCCGATCTNNNCTTAGGCAGGAGAATCGCTTG
GAGCACACGTCTGAACTCTTTCCCTACACGACGCTCTTCCGATCTNNNTGAGGTCAGGAGTTTGAGAC
GAGCACACGTCTGAACTCTTTCCCTACACGACGCTCTTCCGATCTNNNGTACAAGCTACTTGAGAGGCT
GAGCACACGTCTGAACTCTTTCCCTACACGACGCTCTTCCGATCTNNNCACACCTGTAGTCCCACTTACT
GAGCACACGTCTGAACTCTTTCCCTACACGACGCTCTTCCGATCTNNNTGAGGCAGGAGAATCTCTTG
AGCACACGTCTGAACTCTTTCCCTACACGACGCTCTTCCGATCTNNNCTGTAGTCCCAGCTACTCAGGA
AGCACACGTCTGAACTCTTTCCCTACACGACGCTCTTCCGATCTNNNACAAAAAAATTAGCTGGGCAGC
GAGCACACGTCTGAACTCTTTCCCTACACGACGCTCTTCCGATCTNNNGGCCATCAAGGTGGTACTTCT
AGAGCACACGTCTGAACTCTTTCCCTACACGACGCTCTTCCGATCTNNNCTACTCATGAGGCAGGAGAA
GAGCACACGTCTGAACTCTTTCCCTACACGACGCTCTTCCGATCTNNNGGTGCCTATAATCCCAGCTACT
AGCACACGTCTGAACTCTTTCCCTACACGACGCTCTTCCGATCTNNNATCATGCCATTGTACTCCATCCA
AGCACACGTCTGAACTCTTTCCCTACACGACGCTCTTCCGATCTNNNGGTCAAAAGGTGAAGCAACTCT
AGCACACGTCTGAACTCTTTCCCTACACGACGCTCTTCCGATCTNNNTCCCGCAGTCTTATTTATACCCGC
AGCACACGTCTGAACTCTTTCCCTACACGACGCTCTTCCGATCTNNNGGCCTCTGGAGTAGCTATTCTT
GAGCACACGTCTGAACTCTTTCCCTACACGACGCTCTTCCGATCTNNNTTTTGAGACAAAGTCACGCTCT
AGCACACGTCTGAACTCTTTCCCTACACGACGCTCTTCCGATCTNNNGCCGCTTCTTCTTCTTCTTCTTCCA
AGCACACGTCTGAACTCTTTCCCTACACGACGCTCTTCCGATCTNNNCCACGCCCAGCTAATTTTTGTAT
AGCACACGTCTGAACTCTTTCCCTACACGACGCTCTTCCGATCTNNNGTTCAAGTGATTCTCCTCCCTCA
AGCACACGTCTGAACTCTTTCCCTACACGACGCTCTTCCGATCTNNNCTCAAACTCCTGACCTCGTGATC
AGCACACGTCTGAACTCTTTCCCTACACGACGCTCTTCCGATCTNNNTCTGATAAGCAACTGCAGACCT
AGCACACGTCTGAACTCTTTCCCTACACGACGCTCTTCCGATCTNNNCTCTCCTTGTCTTCTTTCCTGGTC
AGCACACGTCTGAACTCTTTCCCTACACGACGCTCTTCCGATCTNNNAGTCGTTGTGCAAACAGACTCT
GAGCACACGTCTGAACTCTTTCCCTACACGACGCTCTTCCGATCTNNNGTGAGTTTCACTTGCTCTGCAT
AGCACACGTCTGAACTCTTTCCCTACACGACGCTCTTCCGATCTNNNGGAGGCTTCCTTTGTGGTTAGA

AGCACACGTCTGAACTCTTTCCCTACACGACGCTCTTCCGATCTNNNAATGGTACCAGCTCGTCTTTGT
AGCACACGTCTGAACTCTTTCCCTACACGACGCTCTTCCGATCTNNNTATGGCCATTCTTGCAGGAGTA
GAGCACACGTCTGAACTCTTTCCCTACACGACGCTCTTCCGATCTNNNGACAAGGTCTCACTCTGTAAC
AGCACACGTCTGAACTCTTTCCCTACACGACGCTCTTCCGATCTNNNTATTTGTGTGTGTGTGTGTGT
GAGCACACGTCTGAACTCTTTCCCTACACGACGCTCTTCCGATCTNNNCTTGCTGATGTTCTGTCTGGT
GAGCACACGTCTGAACTCTTTCCCTACACGACGCTCTTCCGATCTNNNACTTTATCGCTTCTCAGCCTTC
AGCACACGTCTGAACTCTTTCCCTACACGACGCTCTTCCGATCTNNNCTCAAGTGATTCTCCCTCCTCAG
GAGCACACGTCTGAACTCTTTCCCTACACGACGCTCTTCCGATCTNNNTCTTTTGCCTATCAGCATGCCT
AGCACACGTCTGAACTCTTTCCCTACACGACGCTCTTCCGATCTNNNGCAGGGAGCCAATCAACATAT
AGCACACGTCTGAACTCTTTCCCTACACGACGCTCTTCCGATCTNNNTCACTTGGTTGTTGGCAGAATT
GCACACGTCTGAACTCTTTCCCTACACGACGCTCTTCCGATCTNNNGACTTTGCAAATGTTGGAGAGC
AGCACACGTCTGAACTCTTTCCCTACACGACGCTCTTCCGATCTNNNATGTGTAATGATCAAGCCAGG
AGCACACGTCTGAACTCTTTCCCTACACGACGCTCTTCCGATCTNNNCAGATGTGCACTGCTTTAGGAT
AGCACACGTCTGAACTCTTTCCCTACACGACGCTCTTCCGATCTNNNTCAGTAGGAATGGTACCAGC
GAGCACACGTCTGAACTCTTTCCCTACACGACGCTCTTCCGATCTNNNTGCATTTTCTACCGTGCTCATT
GCACACGTCTGAACTCTTTCCCTACACGACGCTCTTCCGATCTNNNACTAACCACCTTGTCTTTCTGGT
GAGCACACGTCTGAACTCTTTCCCTACACGACGCTCTTCCGATCTNNNCTGAGTCATTGTGGCAGAAA
GAGCACACGTCTGAACTCTTTCCCTACACGACGCTCTTCCGATCTNNNGTTTCAAGCGATTCTTCTGCC
AGCACACGTCTGAACTCTTTCCCTACACGACGCTCTTCCGATCTNNNTCTATTCTGCACAATCCACTCCC
AGCACACGTCTGAACTCTTTCCCTACACGACGCTCTTCCGATCTNNNATTGTCCCCTTCTTTGTGACCA
AGCACACGTCTGAACTCTTTCCCTACACGACGCTCTTCCGATCTNNNTGTCCTGCCAGTCTTCTTTTTCT
GCACACGTCTGAACTCTTTCCCTACACGACGCTCTTCCGATCTNNNCAGACCTGTTGAATCAGCATCT
GAGCACACGTCTGAACTCTTTCCCTACACGACGCTCTTCCGATCTNNNTTGGCCATGTTTTCTTCAAGT
GAGCACACGTCTGAACTCTTTCCCTACACGACGCTCTTCCGATCTNNNGAATCTACCTGAAAGCCCTTT
AGCACACGTCTGAACTCTTTCCCTACACGACGCTCTTCCGATCTNNNATTTCCATTTCCCAACTCCCCTTC
AGCACACGTCTGAACTCTTTCCCTACACGACGCTCTTCCGATCTNNNTCCTCTCCTCCTTTCCTCTTCTCC
AGCACACGTCTGAACTCTTTCCCTACACGACGCTCTTCCGATCTNNNAAATGACATGCTCAGGGCCATT
AGCACACGTCTGAACTCTTTCCCTACACGACGCTCTTCCGATCTNNNCCAAAGTACTGGGATTACAGG
GCACACGTCTGAACTCTTTCCCTACACGACGCTCTTCCGATCTNNNTCTTCTTGTTGCAGATAAGGCCT
AGCACACGTCTGAACTCTTTCCCTACACGACGCTCTTCCGATCTNNNCTACTGGGCTCAAGAGATTCTC
GAGCACACGTCTGAACTCTTTCCCTACACGACGCTCTTCCGATCTNNNAGTAGTCCCCCCTCTGAGTAA
AGCACACGTCTGAACTCTTTCCCTACACGACGCTCTTCCGATCTNNNTCTTTTTCCCGATCTATCATCCAC
AGCACACGTCTGAACTCTTTCCCTACACGACGCTCTTCCGATCTNNNAACCACGGTGCCATTATATGA
AGCACACGTCTGAACTCTTTCCCTACACGACGCTCTTCCGATCTNNNCACCTTGCCTGGCCTAGTTATA
AGCACACGTCTGAACTCTTTCCCTACACGACGCTCTTCCGATCTNNNTGACCCCAGGTTATTTCTGCAT
AGCACACGTCTGAACTCTTTCCCTACACGACGCTCTTCCGATCTNNNAGCACTTCAAACAATTTGCGTT
AGCACACGTCTGAACTCTTTCCCTACACGACGCTCTTCCGATCTNNNGGGAAACATTTCATAGTGGTT
AGCACACGTCTGAACTCTTTCCCTACACGACGCTCTTCCGATCTNNNCTTGAACTCTTGACCTCAGGTG
AGCACACGTCTGAACTCTTTCCCTACACGACGCTCTTCCGATCTNNNACCACGCCCAGCTATTTTTTTT
AGCACACGTCTGAACTCTTTCCCTACACGACGCTCTTCCGATCTNNNGGTGTGTAAGCAGCCTAGGTT
AGCACACGTCTGAACTCTTTCCCTACACGACGCTCTTCCGATCTNNNGACTACACACACCACCATATCC
GCACACGTCTGAACTCTTTCCCTACACGACGCTCTTCCGATCTNNNGTGAACCACTCTAGTACGACAG
AGCACACGTCTGAACTCTTTCCCTACACGACGCTCTTCCGATCTNNNCCTGACTTCATTTTAGGCACTG

AGCACACGTCTGAACTCTTTCCCTACACGACGCTCTTCCGATCTNNNCAGTATGATTGGCCCAGCTTTT
GCACACGTCTGAACTCTTTCCCTACACGACGCTCTTCCGATCTNNNCTATCTTGGCTCACTGCAAGCTC
GAGCACACGTCTGAACTCTTTCCCTACACGACGCTCTTCCGATCTNNNTAGAGATGCCCTTGCTTGACAT
AGCACACGTCTGAACTCTTTCCCTACACGACGCTCTTCCGATCTNNNTGGGATGGCAAAGACTAAGAG
AGCACACGTCTGAACTCTTTCCCTACACGACGCTCTTCCGATCTNNNTTCAAGTGATTCTCCTGCCTCAG
GCACACGTCTGAACTCTTTCCCTACACGACGCTCTTCCGATCTNNNTAAAATACCTGTTTGTGGCCCCA
GCACACGTCTGAACTCTTTCCCTACACGACGCTCTTCCGATCTNNNTGGCCTTCTTTGTCTCTTGATCTC
AGCACACGTCTGAACTCTTTCCCTACACGACGCTCTTCCGATCTNNNTGTGCCTTGCAGAGAATACTTC
GAGCACACGTCTGAACTCTTTCCCTACACGACGCTCTTCCGATCTNNNACTGTGATGCCCATCCATATT
AGCACACGTCTGAACTCTTTCCCTACACGACGCTCTTCCGATCTNNNAGTGCTGAGATTACAGGTGTGA
GAGCACACGTCTGAACTCTTTCCCTACACGACGCTCTTCCGATCTNNNAACCAAGATGTCAATAGGGCC
GAGCACACGTCTGAACTCTTTCCCTACACGACGCTCTTCCGATCTNNNAGGTTCAAGCAATTCTCCTGTC
AGCACACGTCTGAACTCTTTCCCTACACGACGCTCTTCCGATCTNNNTGCCTGGTATGAACTCAGTAAG
AGCACACGTCTGAACTCTTTCCCTACACGACGCTCTTCCGATCTNNNCGTGCCCGACTACATTCACATAT
GAGCACACGTCTGAACTCTTTCCCTACACGACGCTCTTCCGATCTNNNCCTCACCTCTCTGACTTCTTTCC
GAGCACACGTCTGAACTCTTTCCCTACACGACGCTCTTCCGATCTNNNCAATGGACCCTCATGTGGTAG
AGCACACGTCTGAACTCTTTCCCTACACGACGCTCTTCCGATCTNNNCCAGGTTCAAGTGATTCTCTTGC
GAGCACACGTCTGAACTCTTTCCCTACACGACGCTCTTCCGATCTNNNTCCTTCCTTCCTTCCTCTCTTTCC
AGCACACGTCTGAACTCTTTCCCTACACGACGCTCTTCCGATCTNNNCTTGAACTCCTGGACTCAAGTGA
GCACACGTCTGAACTCTTTCCCTACACGACGCTCTTCCGATCTNNNATCAAAAGCTGTTGTTCATCCCG
AGCACACGTCTGAACTCTTTCCCTACACGACGCTCTTCCGATCTNNNTGTATCCATGTCGTAGCATGTGT
GAGCACACGTCTGAACTCTTTCCCTACACGACGCTCTTCCGATCTNNNCCACCAGGTGACCTATGTTCTA
AGCACACGTCTGAACTCTTTCCCTACACGACGCTCTTCCGATCTNNNTGAGGTTTGTGAATCTCCTGACA
AGAGCACACGTCTGAACTCTTTCCCTACACGACGCTCTTCCGATCTNNNCTGACTCCGAAAAACCCATGA
GAGCACACGTCTGAACTCTTTCCCTACACGACGCTCTTCCGATCTNNNCACCTTTCATTGTGGACCTATC
GCACACGTCTGAACTCTTTCCCTACACGACGCTCTTCCGATCTNNNGCAGACATTCTCTCATTTCCTGCC
GCACACGTCTGAACTCTTTCCCTACACGACGCTCTTCCGATCTNNNGAAGGGATCCAGTTTCAGCTTTT
GCACACGTCTGAACTCTTTCCCTACACGACGCTCTTCCGATCTNNNTGTAGAAGTACTGTTATGGCTGT
GAGCACACGTCTGAACTCTTTCCCTACACGACGCTCTTCCGATCTNNNGTTTTGATAAGACAGCTGGGA
GAGCACACGTCTGAACTCTTTCCCTACACGACGCTCTTCCGATCTNNNGTTTCACTTCCTCTCCTCTGTG
GAGCACACGTCTGAACTCTTTCCCTACACGACGCTCTTCCGATCTNNNTTTCACCTCCACTTCCTGAGTA
GAGCACACGTCTGAACTCTTTCCCTACACGACGCTCTTCCGATCTNNNCGACGCCCAGCTAATTTTTGT
GAGCACACGTCTGAACTCTTTCCCTACACGACGCTCTTCCGATCTNNNTTAGTAGAGACGGCATTGCAT
GAGCACACGTCTGAACTCTTTCCCTACACGACGCTCTTCCGATCTNNNTCATGGCAGAAATAGGATGTC
GAGCACACGTCTGAACTCTTTCCCTACACGACGCTCTTCCGATCTNNNAATGTGCTGGGATTATAGGG
AGCACACGTCTGAACTCTTTCCCTACACGACGCTCTTCCGATCTNNNGAATCTCTGGGCCAAGAGGTAA
GAGCACACGTCTGAACTCTTTCCCTACACGACGCTCTTCCGATCTNNNGGTCAAGGAAGTGGTATGGG
AGCACACGTCTGAACTCTTTCCCTACACGACGCTCTTCCGATCTNNNATTTGCACTCACTCTGTCTTGC
GAGCACACGTCTGAACTCTTTCCCTACACGACGCTCTTCCGATCTNNNTATCTTGGCCTTGCAAGCTAGA
AGCACACGTCTGAACTCTTTCCCTACACGACGCTCTTCCGATCTNNNTGGTATAATGCCTCCAGCTTTGA
AGCACACGTCTGAACTCTTTCCCTACACGACGCTCTTCCGATCTNNNATTTCGAGAAAGAGCCCTGAG
GCACACGTCTGAACTCTTTCCCTACACGACGCTCTTCCGATCTNNNTAGTGTCTTTTGATGCACATGGC
GAGCACACGTCTGAACTCTTTCCCTACACGACGCTCTTCCGATCTNNNTAGCTAGGATTACAGGCACAC

AGCACACGTCTGAACTCTTTCCCTACACGACGCTCTTCCGATCTNNNCATGAGAATTGCTTCAACCTG
AGCACACGTCTGAACTCTTTCCCTACACGACGCTCTTCCGATCTNNNCTGTAGTCCCAGCTACTCAGGA
AGCACACGTCTGAACTCTTTCCCTACACGACGCTCTTCCGATCTNNNAATGCAGGAGAATTGCTTGAA
AGCACACGTCTGAACTCTTTCCCTACACGACGCTCTTCCGATCTNNNCTGTAATCCCAGCTACTCAGG
AGCACACGTCTGAACTCTTTCCCTACACGACGCTCTTCCGATCTNNNCTAGGCAACAAAGAGAGACT
AGCACACGTCTGAACTCTTTCCCTACACGACGCTCTTCCGATCTNNNGATCACAAGGTCAGGAGATC
AGCACACGTCTGAACTCTTTCCCTACACGACGCTCTTCCGATCTNNNTTCACTGCATTCTAACCTGGT
AGCACACGTCTGAACTCTTTCCCTACACGACGCTCTTCCGATCTNNNGCACATGTAGTCCCAGATACT
GAGCACACGTCTGAACTCTTTCCCTACACGACGCTCTTCCGATCTNNNGAGTTCACACCAGTACACTCC
GCACACGTCTGAACTCTTTCCCTACACGACGCTCTTCCGATCTNNNCACACCTGTAGTCTCAGCTACTC
GAGCACACGTCTGAACTCTTTCCCTACACGACGCTCTTCCGATCTNNNCTGAAGCAGGAGAATGGCA
GCACACGTCTGAACTCTTTCCCTACACGACGCTCTTCCGATCTNNNGGCAAGAGAGTGAGACTCCATT
AGAGCACACGTCTGAACTCTTTCCCTACACGACGCTCTTCCGATCTNNNTGAGGCAGGAGAATAGTG
AGCACACGTCTGAACTCTTTCCCTACACGACGCTCTTCCGATCTNNNACCTGTAGTCCTAGCTACTCAG
AGCACACGTCTGAACTCTTTCCCTACACGACGCTCTTCCGATCTNNNGCAGGAGAATTGCTTGAACT
AGCACACGTCTGAACTCTTTCCCTACACGACGCTCTTCCGATCTNNNAGACAGTCTTGCTATGTTGAC
AGCACACGTCTGAACTCTTTCCCTACACGACGCTCTTCCGATCTNNNTCTGATTTGTCAGCATTCTTG
GAGCACACGTCTGAACTCTTTCCCTACACGACGCTCTTCCGATCTNNNAAAAATGGCCTTGAGAGACT
GAGCACACGTCTGAACTCTTTCCCTACACGACGCTCTTCCGATCTNNNATGCATTCAAGGAGAATCG
AGCACACGTCTGAACTCTTTCCCTACACGACGCTCTTCCGATCTNNNGCCCAGATTGCACTACTGTACT
GAGCACACGTCTGAACTCTTTCCCTACACGACGCTCTTCCGATCTNNNGCTCATGCTTATAATCCCAGC
GAGCACACGTCTGAACTCTTTCCCTACACGACGCTCTTCCGATCTNNNGTGAAACCCCGTCTCTACTAG
AGCACACGTCTGAACTCTTTCCCTACACGACGCTCTTCCGATCTNNNGCAGGAAAATCTCTTGAACCC
GCACACGTCTGAACTCTTTCCCTACACGACGCTCTTCCGATCTNNNAAAACCACTCCAGTAAAGTAGCC
GAGCACACGTCTGAACTCTTTCCCTACACGACGCTCTTCCGATCTNNNGAGCTTGCAGTGAGCCAAG
AGCACACGTCTGAACTCTTTCCCTACACGACGCTCTTCCGATCTNNNCTAATCCCAGTTACTCAGGAGG
AGCACACGTCTGAACTCTTTCCCTACACGACGCTCTTCCGATCTNNNCGTGGACCTGTAATCCTAGCTA
GAGCACACGTCTGAACTCTTTCCCTACACGACGCTCTTCCGATCTNNNCAACAACAGCGAAACTCCATC
AGCACACGTCTGAACTCTTTCCCTACACGACGCTCTTCCGATCTNNNGGTCAAACCCAAACAGAAAAC
AGCACACGTCTGAACTCTTTCCCTACACGACGCTCTTCCGATCTNNNATCATCTCCACTAAAAGGCGGA
AGCACACGTCTGAACTCTTTCCCTACACGACGCTCTTCCGATCTNNNCACCTGTAATCCCAGCTACTCA
GCACACGTCTGAACTCTTTCCCTACACGACGCTCTTCCGATCTNNNCTGTAATCCCAGCTACTCAGGA
GCACACGTCTGAACTCTTTCCCTACACGACGCTCTTCCGATCTNNNGGAAGCTGAGGTAGGAGAATC
GCACACGTCTGAACTCTTTCCCTACACGACGCTCTTCCGATCTNNNAGGAGAATTGTTTGAATCCGGG
AGCACACGTCTGAACTCTTTCCCTACACGACGCTCTTCCGATCTNNNGGCATGAGAATCTTTTGAGCC
GCACACGTCTGAACTCTTTCCCTACACGACGCTCTTCCGATCTNNNAGGTTGAGGCAGGAGAATCAT
AGCACACGTCTGAACTCTTTCCCTACACGACGCTCTTCCGATCTNNNCTGTAGTCCCAGCTACTCAGGA
GCACACGTCTGAACTCTTTCCCTACACGACGCTCTTCCGATCTNNNGAGATCACACCATTGCACTCCAG
AGCACACGTCTGAACTCTTTCCCTACACGACGCTCTTCCGATCTNNNGAGGCAGGAGAATTGCTTGA
AGCACACGTCTGAACTCTTTCCCTACACGACGCTCTTCCGATCTNNNGCAGTATTTTGAGAGCTGAA
GAGCACACGTCTGAACTCTTTCCCTACACGACGCTCTTCCGATCTNNNCAGGAGGCAGAAGTTACAG
AGAGCACACGTCTGAACTCTTTCCCTACACGACGCTCTTCCGATCTNNNTGAGACAGGGAGAATTGC
AGCACACGTCTGAACTCTTTCCCTACACGACGCTCTTCCGATCTNNNGATCACATCACTGTACTCCAGC

AGCACACGTCTGAACTCTTTCCCTACACGACGCTCTTCCGATCTNNNGCACATGCCTATAATCCCAGCT
GAGCACACGTCTGAACTCTTTCCCTACACGACGCTCTTCCGATCTNNNAAGATCACACCACTGCACTGT
GAGCACACGTCTGAACTCTTTCCCTACACGACGCTCTTCCGATCTNNNTGTAGTACCAGCTACTCAGGA
GAGCACACGTCTGAACTCTTTCCCTACACGACGCTCTTCCGATCTNNNCTGCAATCTCAGCTACTCAGG
GAGCACACGTCTGAACTCTTTCCCTACACGACGCTCTTCCGATCTNNNATGCCTGTAATCCCAGCTACTT
AGCACACGTCTGAACTCTTTCCCTACACGACGCTCTTCCGATCTNNNGCAAGAGAATCACTTGAATCCG
GCACACGTCTGAACTCTTTCCCTACACGACGCTCTTCCGATCTNNNCTGAGGCAGGAGAATTGCTTG
GAGCACACGTCTGAACTCTTTCCCTACACGACGCTCTTCCGATCTNNNTGAGGCCAGGAGTTTGAGA
AGAGCACACGTCTGAACTCTTTCCCTACACGACGCTCTTCCGATCTNNNGTGCTCATATCCTCAGCTAC
AGCACACGTCTGAACTCTTTCCCTACACGACGCTCTTCCGATCTNNNTGTAGTCTCAGCTACTAGGGA
GAGCACACGTCTGAACTCTTTCCCTACACGACGCTCTTCCGATCTNNNGTTGTGGCATGTGCTTATAGT
GAGCACACGTCTGAACTCTTTCCCTACACGACGCTCTTCCGATCTNNNCAAAATCGCTTGAACTGAGG
GAGCACACGTCTGAACTCTTTCCCTACACGACGCTCTTCCGATCTNNNCAAAATCGCTTGAACTGAGG
GAGCACACGTCTGAACTCTTTCCCTACACGACGCTCTTCCGATCTNNNGAGGCTGCATTGAGCTATGA
GAGCACACGTCTGAACTCTTTCCCTACACGACGCTCTTCCGATCTNNNGTACCTGTAGTCCCAGCTATT
AGCACACGTCTGAACTCTTTCCCTACACGACGCTCTTCCGATCTNNNCTGAGGCAGGAGAATCACTT
GAGCACACGTCTGAACTCTTTCCCTACACGACGCTCTTCCGATCTNNNGAATGATGACAATACCACTG
AGCACACGTCTGAACTCTTTCCCTACACGACGCTCTTCCGATCTNNNGTAATCCCAGCTACTAGTGAG
AGCACACGTCTGAACTCTTTCCCTACACGACGCTCTTCCGATCTNNNGAGACAGGAGAATTGCTTGA
AGCACACGTCTGAACTCTTTCCCTACACGACGCTCTTCCGATCTNNNTGAGGTAGGAGAATCGCTTG
GAGCACACGTCTGAACTCTTTCCCTACACGACGCTCTTCCGATCTNNNATTGCTTGAACATAGGAGGC
AGCACACGTCTGAACTCTTTCCCTACACGACGCTCTTCCGATCTNNNAAGCAGAGGCAGAAGAATAG
GCACACGTCTGAACTCTTTCCCTACACGACGCTCTTCCGATCTNNNGCTAGGTAGGAGAATCGCTTGA
AGCACACGTCTGAACTCTTTCCCTACACGACGCTCTTCCGATCTNNNCACCTGTAATCCCAGCATTTG
AGCACACGTCTGAACTCTTTCCCTACACGACGCTCTTCCGATCTNNNCTGAGGTGGGAAAATCGTTTG
GAGCACACGTCTGAACTCTTTCCCTACACGACGCTCTTCCGATCTNNNGGCCCAGTGCAAAATAAAAA
AGCACACGTCTGAACTCTTTCCCTACACGACGCTCTTCCGATCTNNNAAATTTTCAAGACCAAGGCCA
GAGCACACGTCTGAACTCTTTCCCTACACGACGCTCTTCCGATCTNNNATTTGAGCTAAGGAGTACCA
AGCACACGTCTGAACTCTTTCCCTACACGACGCTCTTCCGATCTNNNCTGTAATCCCAGCACTTTGGAA
GCACACGTCTGAACTCTTTCCCTACACGACGCTCTTCCGATCTNNNCTGAGATGGGAGGATCACTTG
GCACACGTCTGAACTCTTTCCCTACACGACGCTCTTCCGATCTNNNCTGAGGCAGGAGAATCACTTGA
GAGCACACGTCTGAACTCTTTCCCTACACGACGCTCTTCCGATCTNNNCACCTGTAATCCCAGCTACTCA
AGCACACGTCTGAACTCTTTCCCTACACGACGCTCTTCCGATCTNNNGTAATCCTAGCTACTTGGGAG
GAGCACACGTCTGAACTCTTTCCCTACACGACGCTCTTCCGATCTNNNTCATTTCATGGATGCCAGTCTG
AGCACACGTCTGAACTCTTTCCCTACACGACGCTCTTCCGATCTNNNGTAAGCCGAGATTGTACCATT
AGCACACGTCTGAACTCTTTCCCTACACGACGCTCTTCCGATCTNNNGCATGAAACGAGAATGAACCC
AGCACACGTCTGAACTCTTTCCCTACACGACGCTCTTCCGATCTNNNTTGACTGAACCCACATTGTACC
AGCACACGTCTGAACTCTTTCCCTACACGACGCTCTTCCGATCTNNNATACCTGTAGTCCCAGCTTCTTG
GAGCACACGTCTGAACTCTTTCCCTACACGACGCTCTTCCGATCTNNNTGAGGCAAGAGAATTGCTTG
GAGCACACGTCTGAACTCTTTCCCTACACGACGCTCTTCCGATCTNNNGAAGTTGCAGTGAGCTGAG
GAGCACACGTCTGAACTCTTTCCCTACACGACGCTCTTCCGATCTNNNCTGAGGCAGGAGAATTGCTA
GAGCACACGTCTGAACTCTTTCCCTACACGACGCTCTTCCGATCTNNNCTGTAGTCCCAGCTTCTCAGG
AGCACACGTCTGAACTCTTTCCCTACACGACGCTCTTCCGATCTNNNCTGAGGCAGGAGAATCACTTG

AGCACACGTCTGAACTCTTTCCCTACACGACGCTCTTCCGATCTNNNTGTCATCCCAGTTACACAGAA
AGAGCACACGTCTGAACTCTTTCCCTACACGACGCTCTTCCGATCTNNNGCTAAGGCAGGAGAATCA
AGCACACGTCTGAACTCTTTCCCTACACGACGCTCTTCCGATCTNNNGCTTGCAGTAAGTGAGATCA
AGCACACGTCTGAACTCTTTCCCTACACGACGCTCTTCCGATCTNNNCAGAGTTTGCAATGAGCCAAG
GAGCACACGTCTGAACTCTTTCCCTACACGACGCTCTTCCGATCTNNNCCAAGATCGTACCACTGCACT
AGCACACGTCTGAACTCTTTCCCTACACGACGCTCTTCCGATCTNNNTGAAGGTTGCAGTGAGTTGA
GAGCACACGTCTGAACTCTTTCCCTACACGACGCTCTTCCGATCTNNNAAACATGCTTGTAGTCCCAG
AGCACACGTCTGAACTCTTTCCCTACACGACGCTCTTCCGATCTNNNGAGATCCTGCTACTGCACTCTA
AGCACACGTCTGAACTCTTTCCCTACACGACGCTCTTCCGATCTNNNAATTCAGCTAATGGCCAGGTA
GCACACGTCTGAACTCTTTCCCTACACGACGCTCTTCCGATCTNNNGGATACCTGTAGTCCCAGCTACT
GAGCACACGTCTGAACTCTTTCCCTACACGACGCTCTTCCGATCTNNNGAGGCTGATACAGGAGAAT
GCACACGTCTGAACTCTTTCCCTACACGACGCTCTTCCGATCTNNNTGGGTGACAGAGTGAGATTTT
GAGCACACGTCTGAACTCTTTCCCTACACGACGCTCTTCCGATCTNNNCTGAGGCAGGAGAATCACT
GCACACGTCTGAACTCTTTCCCTACACGACGCTCTTCCGATCTNNNCTCCACCCAAAAGGAGCAAAAT
AGCACACGTCTGAACTCTTTCCCTACACGACGCTCTTCCGATCTNNNGGAGCTTGCAGTGAGACTAG
AGCACACGTCTGAACTCTTTCCCTACACGACGCTCTTCCGATCTNNNAGAATTGCTTGAACCAGGGA
AGCACACGTCTGAACTCTTTCCCTACACGACGCTCTTCCGATCTNNNCTGAGGCAGGAGAATCACTTG
GAGCACACGTCTGAACTCTTTCCCTACACGACGCTCTTCCGATCTNNNATCACTAGGTCAGGAGTTCG
AGCACACGTCTGAACTCTTTCCCTACACGACGCTCTTCCGATCTNNNGAGAATCGCTTGAACTCAGGA
AGCACACGTCTGAACTCTTTCCCTACACGACGCTCTTCCGATCTNNNTTGGAGGATCACTTGAGTCAA
CACACGTCTGAACTCTTTCCCTACACGACGCTCTTCCGATCTNNNTGTAATCCCAGCTACTCAGGAGA
GAGCACACGTCTGAACTCTTTCCCTACACGACGCTCTTCCGATCTNNNCCAACATGGTGAAATCCCAT
GAGCACACGTCTGAACTCTTTCCCTACACGACGCTCTTCCGATCTNNNGAGACTGGAGAATTGCTTG
GAGCACACGTCTGAACTCTTTCCCTACACGACGCTCTTCCGATCTNNNGTGGATCACTTGAGGTCAGG
GCACACGTCTGAACTCTTTCCCTACACGACGCTCTTCCGATCTNNNAGATCTTACCACTGCACTCTAGC
GAGCACACGTCTGAACTCTTTCCCTACACGACGCTCTTCCGATCTNNNTTTTTCCAGTTGGGCTTGAA
GCACACGTCTGAACTCTTTCCCTACACGACGCTCTTCCGATCTNNNGCGCCTATAATCCCAGTTACTC
GCACACGTCTGAACTCTTTCCCTACACGACGCTCTTCCGATCTNNNCTCAAACTCCTGACCTCAAGTG
AGCACACGTCTGAACTCTTTCCCTACACGACGCTCTTCCGATCTNNNGAGCAAGACACCGTCTCAGA
AGCACACGTCTGAACTCTTTCCCTACACGACGCTCTTCCGATCTNNNAAGAATCTCTTGAACCCAGGA
AGCACACGTCTGAACTCTTTCCCTACACGACGCTCTTCCGATCTNNNTGCCTGTAATCCCAGCTATTC
GAGCACACGTCTGAACTCTTTCCCTACACGACGCTCTTCCGATCTNNNGTAATCCTAGCTACTTGGGA
AGCACACGTCTGAACTCTTTCCCTACACGACGCTCTTCCGATCTNNNACACCTGTAATCCCAGCACTTT
GAGCACACGTCTGAACTCTTTCCCTACACGACGCTCTTCCGATCTNNNCATGCCTGTAATCCCAGCTA
GAGCACACGTCTGAACTCTTTCCCTACACGACGCTCTTCCGATCTNNNACATGGTGAAATCCCGTCTCT
GAGCACACGTCTGAACTCTTTCCCTACACGACGCTCTTCCGATCTNNNGGAGTTTACTGTGAGCCGA
AGCACACGTCTGAACTCTTTCCCTACACGACGCTCTTCCGATCTNNNGCTGAGGTAGGATAACTGCTT
GCACACGTCTGAACTCTTTCCCTACACGACGCTCTTCCGATCTNNNACACTTGTAATCCCAGCTACTC
AGCACACGTCTGAACTCTTTCCCTACACGACGCTCTTCCGATCTNNNTGTAATCCCAGCACTTTGAGA
GAGCACACGTCTGAACTCTTTCCCTACACGACGCTCTTCCGATCTNNNTGGGACCATTTTTGGGAGTC
GAGCACACGTCTGAACTCTTTCCCTACACGACGCTCTTCCGATCTNNNCTGTAGTCCCAGCTACTCAGG
AGCACACGTCTGAACTCTTTCCCTACACGACGCTCTTCCGATCTNNNAGAAGGCTGAGACAGGAGAA
GCACACGTCTGAACTCTTTCCCTACACGACGCTCTTCCGATCTNNNGAAGTTGAGAACACAGTGACAG

AGAGCACACGTCTGAACTCTTTCCCTACACGACGCTCTTCCGATCTNNNCTGAGGCAGGAGAATTGC
AGCACACGTCTGAACTCTTTCCCTACACGACGCTCTTCCGATCTNNNGTGAATCCTTGAAACCAAGAG
GCACACGTCTGAACTCTTTCCCTACACGACGCTCTTCCGATCTNNNGTATTACCAGCTACTTGGGAGG
GAGCACACGTCTGAACTCTTTCCCTACACGACGCTCTTCCGATCTNNNGGATTGCTTGAGCTCAGGA
AGCACACGTCTGAACTCTTTCCCTACACGACGCTCTTCCGATCTNNNTGAGTATGAGACCACCTCCTT
AGCACACGTCTGAACTCTTTCCCTACACGACGCTCTTCCGATCTNNNCCCAAAGTCCTTTGTGTCATTC
AGCACACGTCTGAACTCTTTCCCTACACGACGCTCTTCCGATCTNNNATAGACCAAAGGGAAAGCGA
AGCACACGTCTGAACTCTTTCCCTACACGACGCTCTTCCGATCTNNNGAAAAACCCCAAAGCTTCAGG
GCACACGTCTGAACTCTTTCCCTACACGACGCTCTTCCGATCTNNNAGCTAGTTCCAGAAAGATGGT
GCACACGTCTGAACTCTTTCCCTACACGACGCTCTTCCGATCTNNNCAGGCAATCTTGTCCAGCAAT
GCACACGTCTGAACTCTTTCCCTACACGACGCTCTTCCGATCTNNNTCTCCCTCTCTCTCTCTCTCTCA
AGCACACGTCTGAACTCTTTCCCTACACGACGCTCTTCCGATCTNNNTTGTACACAAATCACCAAAAGC
GAGCACACGTCTGAACTCTTTCCCTACACGACGCTCTTCCGATCTNNNCAGATTCCAGAGGAGAGAG
AGCACACGTCTGAACTCTTTCCCTACACGACGCTCTTCCGATCTNNNTAGTCTTTTTCCCTCACCACCC
GCACACGTCTGAACTCTTTCCCTACACGACGCTCTTCCGATCTNNNGTTCTTGTGGTGCTGGAGAATT
GCACACGTCTGAACTCTTTCCCTACACGACGCTCTTCCGATCTNNNTGTTGTTGTCAAATGACTGCAG
GCACACGTCTGAACTCTTTCCCTACACGACGCTCTTCCGATCTNNNAGCTTCTGATTTAGCAGGTGTG
GCACACGTCTGAACTCTTTCCCTACACGACGCTCTTCCGATCTNNNGCTTCAGGCTTTCTCTGTGTAA
AGCACACGTCTGAACTCTTTCCCTACACGACGCTCTTCCGATCTNNNTGACCACTATGACACACTCAAG
GCACACGTCTGAACTCTTTCCCTACACGACGCTCTTCCGATCTNNNAAACCATGCAATGTTTCTGGTC
GCACACGTCTGAACTCTTTCCCTACACGACGCTCTTCCGATCTNNNCAGACTCCACGTGTGTCTTTTTC
GCACACGTCTGAACTCTTTCCCTACACGACGCTCTTCCGATCTNNNTGCCTTGTAGCCACAGATAACA
GCACACGTCTGAACTCTTTCCCTACACGACGCTCTTCCGATCTNNNAGACTCAGAGCCAGGATAAAT
AGCACACGTCTGAACTCTTTCCCTACACGACGCTCTTCCGATCTNNNTCTCTCCATTCCTCCCTATCC
AGCACACGTCTGAACTCTTTCCCTACACGACGCTCTTCCGATCTNNNCCCTCCCTTTTCTTTGGAAGAG
GCACACGTCTGAACTCTTTCCCTACACGACGCTCTTCCGATCTNNNAGTAGCTGGGACTATAGGTGT
AGCACACGTCTGAACTCTTTCCCTACACGACGCTCTTCCGATCTNNNCACCTTAGCCTCCTAAGTAGCT
GAGCACACGTCTGAACTCTTTCCCTACACGACGCTCTTCCGATCTNNNAAATTGGGAGTAAGGAGAT
GCACACGTCTGAACTCTTTCCCTACACGACGCTCTTCCGATCTNNNGGTCACAGCCCAGATAAACTAC
AGCACACGTCTGAACTCTTTCCCTACACGACGCTCTTCCGATCTNNNTGCCACGAGGATTTTGTGAAA
AGCACACGTCTGAACTCTTTCCCTACACGACGCTCTTCCGATCTNNNACTTAGCTTTTGAGGCTCATC
AGCACACGTCTGAACTCTTTCCCTACACGACGCTCTTCCGATCTNNNCTAGTCGTCCTTCCACACTCTT
AGCACACGTCTGAACTCTTTCCCTACACGACGCTCTTCCGATCTNNNCTATGATTCAAGGGGAGCTC
AGCACACGTCTGAACTCTTTCCCTACACGACGCTCTTCCGATCTNNNAGTGACGGTTTGACTTCCTCT
GCACACGTCTGAACTCTTTCCCTACACGACGCTCTTCCGATCTNNNCCCAGGATTGTAATGCCTCCTA
AGCACACGTCTGAACTCTTTCCCTACACGACGCTCTTCCGATCTNNNTTTTTGAGAGAGTTTCACGCC
AGCACACGTCTGAACTCTTTCCCTACACGACGCTCTTCCGATCTNNNATGTATCACCCCAATCTGTGCT
AGCACACGTCTGAACTCTTTCCCTACACGACGCTCTTCCGATCTNNNATCCACTGCTTTAGATTGGGA
AGCACACGTCTGAACTCTTTCCCTACACGACGCTCTTCCGATCTNNNAGCTAGGACTAAAAGCCCTGT
AGCACACGTCTGAACTCTTTCCCTACACGACGCTCTTCCGATCTNNNATCCAGGTAGTAAGTAGTGG
GCACACGTCTGAACTCTTTCCCTACACGACGCTCTTCCGATCTNNNTCCTCTTCATGCTGACATCTG
AGCACACGTCTGAACTCTTTCCCTACACGACGCTCTTCCGATCTNNNCTTGAACTCCTGACCTCAAGTG
AGCACACGTCTGAACTCTTTCCCTACACGACGCTCTTCCGATCTNNNCCTTCCTCCCTTCCTTCTTCATG

AGCACACGTCTGAACTCTTTCCCTACACGACGCTCTTCCGATCTNNNGTGAGCCTTCTTTGCTTCTTGAG
GCACACGTCTGAACTCTTTCCCTACACGACGCTCTTCCGATCTNNNGCGCCTGGCCATAAATAAACTAT
AGCACACGTCTGAACTCTTTCCCTACACGACGCTCTTCCGATCTNNNTCACAAGATACAGGGAGCATTC
AGCACACGTCTGAACTCTTTCCCTACACGACGCTCTTCCGATCTNNNTTCAAGCTATTCTCCTGCCTCA
AGCACACGTCTGAACTCTTTCCCTACACGACGCTCTTCCGATCTNNNAGCGTTTCTCTCTCTCTCTCTC
GAGCACACGTCTGAACTCTTTCCCTACACGACGCTCTTCCGATCTNNNGCCTCTAATCTGCCATCTTGG
AGCACACGTCTGAACTCTTTCCCTACACGACGCTCTTCCGATCTNNNCTCCTGAGTAGCTGGGACTACA
AGCACACGTCTGAACTCTTTCCCTACACGACGCTCTTCCGATCTNNNCAGGTCCCATATCCTGCATCTT
AGCACACGTCTGAACTCTTTCCCTACACGACGCTCTTCCGATCTNNNATCTGCCAGCACCTTGATTTTT
AGCACACGTCTGAACTCTTTCCCTACACGACGCTCTTCCGATCTNNNTTCGTTTTCCTAGCACCTGTCAT
AGCACACGTCTGAACTCTTTCCCTACACGACGCTCTTCCGATCTNNNGGTGACCTAAACAGCGAGTTA
AGCACACGTCTGAACTCTTTCCCTACACGACGCTCTTCCGATCTNNNTCTCTAGTGATGTGACTCCCAG
GAGCACACGTCTGAACTCTTTCCCTACACGACGCTCTTCCGATCTNNNCTGTCTTCTACCAGCATCCAT
AGCACACGTCTGAACTCTTTCCCTACACGACGCTCTTCCGATCTNNNCCCATTTCATGTGCACAAAGCT
GAGCACACGTCTGAACTCTTTCCCTACACGACGCTCTTCCGATCTNNNTGATTTCTCCGTAGAAGCATT
AGCACACGTCTGAACTCTTTCCCTACACGACGCTCTTCCGATCTNNNTTGAACTCCTGACCTCATGATC
AGCACACGTCTGAACTCTTTCCCTACACGACGCTCTTCCGATCTNNNTGTGACCACTGATGTTGAGCA
AGCACACGTCTGAACTCTTTCCCTACACGACGCTCTTCCGATCTNNNACCCTTGCTAGCTCTTTCTAACC
GAGCACACGTCTGAACTCTTTCCCTACACGACGCTCTTCCGATCTNNNCACTCTCCCAGTCTGTCAAGTT
AGCACACGTCTGAACTCTTTCCCTACACGACGCTCTTCCGATCTNNNCCTCACCCTTCCAAAGTGAACT
AGCACACGTCTGAACTCTTTCCCTACACGACGCTCTTCCGATCTNNNGAGAACTGGGTCTTGAAGCCT
AGCACACGTCTGAACTCTTTCCCTACACGACGCTCTTCCGATCTNNNACTCAAAAGCCCCTCTTAGTCT
AGCACACGTCTGAACTCTTTCCCTACACGACGCTCTTCCGATCTNNNTTCAAGTGATTCTCCTGCCTCA
GCACACGTCTGAACTCTTTCCCTACACGACGCTCTTCCGATCTNNNTAGTACTCAAGCAGCCCTTTCAG
AGCACACGTCTGAACTCTTTCCCTACACGACGCTCTTCCGATCTNNNGTGAGCCGAGATCATGTCACTA
AGCACACGTCTGAACTCTTTCCCTACACGACGCTCTTCCGATCTNNNCAGGGATGTTGGCTTGAAGTT
GCACACGTCTGAACTCTTTCCCTACACGACGCTCTTCCGATCTNNNGACTGCCTCGTATTACATGCAAC
AGCACACGTCTGAACTCTTTCCCTACACGACGCTCTTCCGATCTNNNCTGAATAGCTGGGATTACAGG
AGCACACGTCTGAACTCTTTCCCTACACGACGCTCTTCCGATCTNNNAGGGCTTTGTTCTTTGAACACT
AGCACACGTCTGAACTCTTTCCCTACACGACGCTCTTCCGATCTNNNGACAAACAGGAACCAGAACAG
AGCACACGTCTGAACTCTTTCCCTACACGACGCTCTTCCGATCTNNNTGTCCCTGATCTAGATGAGTGG
AGCACACGTCTGAACTCTTTCCCTACACGACGCTCTTCCGATCTNNNCTCCTTTGACTTGTTGGTTACG
AGCACACGTCTGAACTCTTTCCCTACACGACGCTCTTCCGATCTNNNCCTGGCCCCTATAAAGACTCAA
AGCACACGTCTGAACTCTTTCCCTACACGACGCTCTTCCGATCTNNNTGCCTTTGCATCCTCATAGCTTA
GCACACGTCTGAACTCTTTCCCTACACGACGCTCTTCCGATCTNNNTTCCACCATTTTGGGGTACAGAT
AGCACACGTCTGAACTCTTTCCCTACACGACGCTCTTCCGATCTNNNTGGGGTACATATTTAGCAGTGG
GAGCACACGTCTGAACTCTTTCCCTACACGACGCTCTTCCGATCTNNNACCTCTACCTCCTGAGTTCAA
AGCACACGTCTGAACTCTTTCCCTACACGACGCTCTTCCGATCTNNNAGAAAAACGCTTCCAAATCCAAT
GAGCACACGTCTGAACTCTTTCCCTACACGACGCTCTTCCGATCTNNNCTCTAGAGAGACACCCTTGTT
GCACACGTCTGAACTCTTTCCCTACACGACGCTCTTCCGATCTNNNTTTTGATGCTGCTTTCCCAACTTC
GAGCACACGTCTGAACTCTTTCCCTACACGACGCTCTTCCGATCTNNNACATGCAAAAGAGAGTGTTG
AGCACACGTCTGAACTCTTTCCCTACACGACGCTCTTCCGATCTNNNTCTCTTCTCTTGTCTCCCTCCATC
AGCACACGTCTGAACTCTTTCCCTACACGACGCTCTTCCGATCTNNNTGGAGGATTCACCCACTCTTTT

GAGCACACGTCTGAACTCTTTCCCTACACGACGCTCTTCCGATCTNNNATGGAGTTTTGCTCTTGTTGCT
AGCACACGTCTGAACTCTTTCCCTACACGACGCTCTTCCGATCTNNNGTCACCTCACTCAGAATAGCCAT
GAGCACACGTCTGAACTCTTTCCCTACACGACGCTCTTCCGATCTNNNTGCCATTACTAAGCTGTTTCGT
AGCACACGTCTGAACTCTTTCCCTACACGACGCTCTTCCGATCTNNNGCATGCATTTACAGGTGAAGTG
GCACACGTCTGAACTCTTTCCCTACACGACGCTCTTCCGATCTNNNGATCATAGCTCACTACAGCCTCCC
AGCACACGTCTGAACTCTTTCCCTACACGACGCTCTTCCGATCTNNNCCTTCAAATACCATCACCTTGAG
AGCACACGTCTGAACTCTTTCCCTACACGACGCTCTTCCGATCTNNNCCAAAGATGGTTCAGCATACACA
AGCACACGTCTGAACTCTTTCCCTACACGACGCTCTTCCGATCTNNNTCTCTCTGGTTAGCTGTATCCCTG
GAGCACACGTCTGAACTCTTTCCCTACACGACGCTCTTCCGATCTNNNTTGTTCCTTCATTCTGGGTTCC
AGCACACGTCTGAACTCTTTCCCTACACGACGCTCTTCCGATCTNNNTGCCCATATTACTGAGATGGTG
AGCACACGTCTGAACTCTTTCCCTACACGACGCTCTTCCGATCTNNNGGCTGTCTTAGTCCATTTTCTGCC
AGCACACGTCTGAACTCTTTCCCTACACGACGCTCTTCCGATCTNNNTGCCCTCTTTTTATTCACCACCAC
GAGCACACGTCTGAACTCTTTCCCTACACGACGCTCTTCCGATCTNNNACTTCCTTCTGTGTGCCATGTA
GAGCACACGTCTGAACTCTTTCCCTACACGACGCTCTTCCGATCTNNNAAGAGTAATGGTCTGAAGAGC
AGAGCACACGTCTGAACTCTTTCCCTACACGACGCTCTTCCGATCTNNNTAGGACAGCATTTCCTTAGA
GAGCACACGTCTGAACTCTTTCCCTACACGACGCTCTTCCGATCTNNNGGGGGAAGGAGAAAAGGAA
AGCACACGTCTGAACTCTTTCCCTACACGACGCTCTTCCGATCTNNNCAAATTTGCACTGCCATCAGAG
AGCACACGTCTGAACTCTTTCCCTACACGACGCTCTTCCGATCTNNNGCCTTTCAGGTTCAAGCAATTCT
AGCACACGTCTGAACTCTTTCCCTACACGACGCTCTTCCGATCTNNNGTCATTTCTTTTTCCCTGGCACAC
GAGCACACGTCTGAACTCTTTCCCTACACGACGCTCTTCCGATCTNNNAAACATCCTATTTCATGGGGG
AGCACACGTCTGAACTCTTTCCCTACACGACGCTCTTCCGATCTNNNTCAACCCATTACCCTTTAGCTCC
GAGCACACGTCTGAACTCTTTCCCTACACGACGCTCTTCCGATCTNNNACTTCTAGGGGCTTTTCCAACT
AGCACACGTCTGAACTCTTTCCCTACACGACGCTCTTCCGATCTNNNCTGCATTCCCATATTGCTCTGTG
AGCACACGTCTGAACTCTTTCCCTACACGACGCTCTTCCGATCTNNNCTCCTGTGTAGCTGGGATTACAG
GAGCACACGTCTGAACTCTTTCCCTACACGACGCTCTTCCGATCTNNNCCAGACAAACAAAACCAACTG
AGCACACGTCTGAACTCTTTCCCTACACGACGCTCTTCCGATCTNNNTCCTTTTCTACTGCATCCTTCCCC
GAGCACACGTCTGAACTCTTTCCCTACACGACGCTCTTCCGATCTNNNGTTAGCTGTTCAGTGTGCCTT
AGCACACGTCTGAACTCTTTCCCTACACGACGCTCTTCCGATCTNNNTGTATCTCTGCTCCAGAATGCTG
AGCACACGTCTGAACTCTTTCCCTACACGACGCTCTTCCGATCTNNNGACATTGGCAAATATCCCTTCGG
AGCACACGTCTGAACTCTTTCCCTACACGACGCTCTTCCGATCTNNNCCAGAGACAATCACCACACTACA
GAGCACACGTCTGAACTCTTTCCCTACACGACGCTCTTCCGATCTNNNTCCGTTTCTAAAGCACAGGAAG
AGCACACGTCTGAACTCTTTCCCTACACGACGCTCTTCCGATCTNNNCCCAAATTTGTGTCTTTCCCACA
GAGCACACGTCTGAACTCTTTCCCTACACGACGCTCTTCCGATCTNNNGAACTCCCTGGATGTCTGATC
AGCACACGTCTGAACTCTTTCCCTACACGACGCTCTTCCGATCTNNNTCCAGGCAAGACCATGATAACA
GCACACGTCTGAACTCTTTCCCTACACGACGCTCTTCCGATCTNNNAGCTAGTTATTTGGGGTCACTTTG
GAGCACACGTCTGAACTCTTTCCCTACACGACGCTCTTCCGATCTNNNTGGCACTGTCCACCCTTATTTAG
AGCACACGTCTGAACTCTTTCCCTACACGACGCTCTTCCGATCTNNNTGCCACTGATAGGAACAATGTG
GAGCACACGTCTGAACTCTTTCCCTACACGACGCTCTTCCGATCTNNNGCTTGGTTTTCATCCATGTGCT
GAGCACACGTCTGAACTCTTTCCCTACACGACGCTCTTCCGATCTNNNAGAGAGAGAGCAAGAGATAG
AGCACACGTCTGAACTCTTTCCCTACACGACGCTCTTCCGATCTNNNGGTCATGTCACCCTCCTTCTTAGG
GAGCACACGTCTGAACTCTTTCCCTACACGACGCTCTTCCGATCTNNNCCACAGCTTCTCTGATGGTAA
AGCACACGTCTGAACTCTTTCCCTACACGACGCTCTTCCGATCTNNNACTTGACCAGCTCAACATTAGGT
AGCACACGTCTGAACTCTTTCCCTACACGACGCTCTTCCGATCTNNNGAGAACCAAGCAGGATAAAAG

AGCACACGTCTGAACTCTTTCCCTACACGACGCTCTTCCGATCTNNNCTGTGCCTTGTGAAACTTACGT
AGCACACGTCTGAACTCTTTCCCTACACGACGCTCTTCCGATCTNNNAGCAGCAAATTCCATCAGCATT
GCACACGTCTGAACTCTTTCCCTACACGACGCTCTTCCGATCTNNNTAGCTGGGATTACAGGCACTAT
AGCACACGTCTGAACTCTTTCCCTACACGACGCTCTTCCGATCTNNNTACAGGTAGCTGCCATCACACT
AGCACACGTCTGAACTCTTTCCCTACACGACGCTCTTCCGATCTNNNGTTGGGAAGGGTTTCAATAGA
GCACACGTCTGAACTCTTTCCCTACACGACGCTCTTCCGATCTNNNTGAGTGTAGACTGTTGGAGAAG
AGCACACGTCTGAACTCTTTCCCTACACGACGCTCTTCCGATCTNNNCCTAAACAACTGGGGATCCTCA
AGCACACGTCTGAACTCTTTCCCTACACGACGCTCTTCCGATCTNNNATGGGTTTGGGTTTGATTTGCA
GCACACGTCTGAACTCTTTCCCTACACGACGCTCTTCCGATCTNNNCCATTAGGCTGCACATTTCCAAA
AGCACACGTCTGAACTCTTTCCCTACACGACGCTCTTCCGATCTNNNTGAAGTGGGTTTCTCCATTGCT
AGCACACGTCTGAACTCTTTCCCTACACGACGCTCTTCCGATCTNNNTTTACGCCATTATCCTGCCTCAG
GAGCACACGTCTGAACTCTTTCCCTACACGACGCTCTTCCGATCTNNNGTAGAGAGCATGGCGACTAA
GAGCACACGTCTGAACTCTTTCCCTACACGACGCTCTTCCGATCTNNNTGAGTGTGATGTTAGTTTTGG
AGCACACGTCTGAACTCTTTCCCTACACGACGCTCTTCCGATCTNNNTTGTTTGCTCGTTTGCATCTAT
GAGCACACGTCTGAACTCTTTCCCTACACGACGCTCTTCCGATCTNNNCCACTCTGTTGACTAGGATGG
AGCACACGTCTGAACTCTTTCCCTACACGACGCTCTTCCGATCTNNNACATGGTGAAATCCCGTCTCTA
AGCACACGTCTGAACTCTTTCCCTACACGACGCTCTTCCGATCTNNNCAGTGACCTATGACTGGACTA
GAGCACACGTCTGAACTCTTTCCCTACACGACGCTCTTCCGATCTNNNCACAAAACAAACAAAAGGCCA
AGCACACGTCTGAACTCTTTCCCTACACGACGCTCTTCCGATCTNNNGCTGAGATAGGAGAATCGCTT
AGCACACGTCTGAACTCTTTCCCTACACGACGCTCTTCCGATCTNNNAAACAAATGTCAAAGACAGCCC
AGCACACGTCTGAACTCTTTCCCTACACGACGCTCTTCCGATCTNNNTGGAGACAGGAGAATTGCTTG
AGCACACGTCTGAACTCTTTCCCTACACGACGCTCTTCCGATCTNNNGAGTGACAGAGTGAGACTCCA
AGCACACGTCTGAACTCTTTCCCTACACGACGCTCTTCCGATCTNNNCTGAGGCAAGAGGATCACTTG
AGCACACGTCTGAACTCTTTCCCTACACGACGCTCTTCCGATCTNNNCTGAGGCAGAAGAATCACTTG
AGCACACGTCTGAACTCTTTCCCTACACGACGCTCTTCCGATCTNNNGAGGCATGAGAATCGCTTGAA
AGCACACGTCTGAACTCTTTCCCTACACGACGCTCTTCCGATCTNNNATAATCCTAGCACTTTGGGAGG
AGCACACGTCTGAACTCTTTCCCTACACGACGCTCTTCCGATCTNNNGGATTACTTGACCCTGGGAGTT
AGCACACGTCTGAACTCTTTCCCTACACGACGCTCTTCCGATCTNNNGAGAATCACTTGAACCTGCGA
GAGCACACGTCTGAACTCTTTCCCTACACGACGCTCTTCCGATCTNNNTAATCCCAGCTACTCAGGAGA
GCACACGTCTGAACTCTTTCCCTACACGACGCTCTTCCGATCTNNNCAGCCTGGGCAATAGGAGTAAA
GAGCACACGTCTGAACTCTTTCCCTACACGACGCTCTTCCGATCTNNNCAGAGGTTGTAGTGAGCTGT
GAGCACACGTCTGAACTCTTTCCCTACACGACGCTCTTCCGATCTNNNATTGCTTGAGCCTAGGAGTTT
AGCACACGTCTGAACTCTTTCCCTACACGACGCTCTTCCGATCTNNNCAATATAGCGCCATTGCACTCAA
AGAGCACACGTCTGAACTCTTTCCCTACACGACGCTCTTCCGATCTNNNCTGAGGCAGGAGAATCACTT
AGCACACGTCTGAACTCTTTCCCTACACGACGCTCTTCCGATCTNNNTGCATACCTGTAATCCCAGCTA
GAGCACACGTCTGAACTCTTTCCCTACACGACGCTCTTCCGATCTNNNTCATGAGGTCAGGAGATCGA
AGCACACGTCTGAACTCTTTCCCTACACGACGCTCTTCCGATCTNNNGTAATCTCAGCTACTCAGGAG
GAGCACACGTCTGAACTCTTTCCCTACACGACGCTCTTCCGATCTNNNCAGGAGAATTGCTTGAACTTG
GAGCACACGTCTGAACTCTTTCCCTACACGACGCTCTTCCGATCTNNNTCACACCTGTAATGCCAGCTA
GAGCACACGTCTGAACTCTTTCCCTACACGACGCTCTTCCGATCTNNNGGTGACAGAACGAGACTTCAT
GAGCACACGTCTGAACTCTTTCCCTACACGACGCTCTTCCGATCTNNNGGCTCCTGTAATTCCAGCTAC
GCACACGTCTGAACTCTTTCCCTACACGACGCTCTTCCGATCTNNNTTCCAGCTATTTGGGAGACTGA
GAGCACACGTCTGAACTCTTTCCCTACACGACGCTCTTCCGATCTNNNGAGGCAGGATAATTGCTTGA

AGCACACGTCTGAACTCTTTCCCTACACGACGCTCTTCCGATCTNNNTAAAGTGGGAAGATGGCTTG
GAGCACACGTCTGAACTCTTTCCCTACACGACGCTCTTCCGATCTNNNATCTCTTGAGCCTAGGAAGT
AGCACACGTCTGAACTCTTTCCCTACACGACGCTCTTCCGATCTNNNTGCATACCTGTAATCCCAGCTA
AGCACACGTCTGAACTCTTTCCCTACACGACGCTCTTCCGATCTNNNCTGTAATCCCAGCTACTCAGG
GAGCACACGTCTGAACTCTTTCCCTACACGACGCTCTTCCGATCTNNNCTGTAATCCCAGCTACTCGGA
AGAGCACACGTCTGAACTCTTTCCCTACACGACGCTCTTCCGATCTNNNGAGAATCACTTGAACCTGG
AGCACACGTCTGAACTCTTTCCCTACACGACGCTCTTCCGATCTNNNTACAAAAATGAGCTGGGAGT
GAGCACACGTCTGAACTCTTTCCCTACACGACGCTCTTCCGATCTNNNCATGCCTGTAATCCCAGCTAC
GCACACGTCTGAACTCTTTCCCTACACGACGCTCTTCCGATCTNNNATGCCTGTAATACCAGCTACTCG
GCACACGTCTGAACTCTTTCCCTACACGACGCTCTTCCGATCTNNNGGAGTGATGCAATTGCTGTCTT
AGCACACGTCTGAACTCTTTCCCTACACGACGCTCTTCCGATCTNNNAGGGAGAGAGAGATTGAAG
GCACACGTCTGAACTCTTTCCCTACACGACGCTCTTCCGATCTNNNGAGGCAGGAGAATTGCTTGAA
AGCACACGTCTGAACTCTTTCCCTACACGACGCTCTTCCGATCTNNNGAACAAAGCCAGAGTCATCAC
GCACACGTCTGAACTCTTTCCCTACACGACGCTCTTCCGATCTNNNGGAGGTTGAGGTAGGAGAATC
AGCACACGTCTGAACTCTTTCCCTACACGACGCTCTTCCGATCTNNNAGAATCACTTGAACCCAGGAG
AGCACACGTCTGAACTCTTTCCCTACACGACGCTCTTCCGATCTNNNGGATCACTTGAGGTCAGGAG
AGCACACGTCTGAACTCTTTCCCTACACGACGCTCTTCCGATCTNNNGTCCACTGTTACAGCTCAGAG
GCACACGTCTGAACTCTTTCCCTACACGACGCTCTTCCGATCTNNNAGAATTACTTGAACCTGGGAG
AGCACACGTCTGAACTCTTTCCCTACACGACGCTCTTCCGATCTNNNCCTGGGAAGTCAAGGCTATAG
AGCACACGTCTGAACTCTTTCCCTACACGACGCTCTTCCGATCTNNNGTGAGCTGAGATCCGAGATC
AGCACACGTCTGAACTCTTTCCCTACACGACGCTCTTCCGATCTNNNACAAGGCAGGAGAATAGCTT
AGCACACGTCTGAACTCTTTCCCTACACGACGCTCTTCCGATCTNNNCTAAGGCAGGAGGATCACTT
GAGCACACGTCTGAACTCTTTCCCTACACGACGCTCTTCCGATCTNNNAAGAATTGCTTGAACCGAGA
AGAGCACACGTCTGAACTCTTTCCCTACACGACGCTCTTCCGATCTNNNGAGGTTGTAGTGAGCCAA
GAGCACACGTCTGAACTCTTTCCCTACACGACGCTCTTCCGATCTNNNTGTAGTCTCAGCTACTCGAG
GAGCACACGTCTGAACTCTTTCCCTACACGACGCTCTTCCGATCTNNNCTGAGGCAGGAGAATCACTT
GCACACGTCTGAACTCTTTCCCTACACGACGCTCTTCCGATCTNNNTTCTGTTGTGCCTAGTACACTGT
AGCACACGTCTGAACTCTTTCCCTACACGACGCTCTTCCGATCTNNNTGAGACCAGAGTTCAAGATC
GAGCACACGTCTGAACTCTTTCCCTACACGACGCTCTTCCGATCTNNNAGGCAGAGAATTGCTTGAA
AGCACACGTCTGAACTCTTTCCCTACACGACGCTCTTCCGATCTNNNGAGAATGGCTTGAGAGAACC
AGCACACGTCTGAACTCTTTCCCTACACGACGCTCTTCCGATCTNNNTGGGATGAGAGAACACAAAC
GCACACGTCTGAACTCTTTCCCTACACGACGCTCTTCCGATCTNNNATATGGACCAAAAAATGTGCCC
GAGCACACGTCTGAACTCTTTCCCTACACGACGCTCTTCCGATCTNNNACAACACATGGGGATTATGG
AGCACACGTCTGAACTCTTTCCCTACACGACGCTCTTCCGATCTNNNAGTCCTAGCTAGCAAGAGACT
AGCACACGTCTGAACTCTTTCCCTACACGACGCTCTTCCGATCTNNNAGAATTGCTTGAACCTGGGAG
GAGCACACGTCTGAACTCTTTCCCTACACGACGCTCTTCCGATCTNNNTGTAATCCCAGCTACTTGGAA
AGCACACGTCTGAACTCTTTCCCTACACGACGCTCTTCCGATCTNNNAAAAATTATCCAGGCGTGGTT
AGCACACGTCTGAACTCTTTCCCTACACGACGCTCTTCCGATCTNNNGAATCTCTTGAAACTGGAAG
GCACACGTCTGAACTCTTTCCCTACACGACGCTCTTCCGATCTNNNAGAAAATTGCTTGAACCGGGAT
GCACACGTCTGAACTCTTTCCCTACACGACGCTCTTCCGATCTNNNCTGTAATCCCAGCTACTCAGGA
AGCACACGTCTGAACTCTTTCCCTACACGACGCTCTTCCGATCTNNNGAGGTTACAGTGAGCCGAGA
AGAGCACACGTCTGAACTCTTTCCCTACACGACGCTCTTCCGATCTNNNCTTGGGCAACAGAGAAAG
GAGCACACGTCTGAACTCTTTCCCTACACGACGCTCTTCCGATCTNNNGCAGTGGTTGTGAATGTGT

AGCACACGTCTGAACTCTTTCCCTACACGACGCTCTTCCGATCTNNNGGTTTCTGTCTTTACCAAGGTG
GAGCACACGTCTGAACTCTTTCCCTACACGACGCTCTTCCGATCTNNNTCCGTCTCAACAACAACAACAA
GAGCACACGTCTGAACTCTTTCCCTACACGACGCTCTTCCGATCTNNNGAGAATCGCTTGAACCTGGG
AGCACACGTCTGAACTCTTTCCCTACACGACGCTCTTCCGATCTNNNCTTGCAGTGAGCTGAAATCATG
GAGCACACGTCTGAACTCTTTCCCTACACGACGCTCTTCCGATCTNNNAGTGAGCCAAGATTGCATCA
GAGCACACGTCTGAACTCTTTCCCTACACGACGCTCTTCCGATCTNNNGGAGTTGAGGTGGGAGAAT
AGCACACGTCTGAACTCTTTCCCTACACGACGCTCTTCCGATCTNNNTTCCCCAGCAGTTCATTATCCAA
GAGCACACGTCTGAACTCTTTCCCTACACGACGCTCTTCCGATCTNNNGTAGAACTGCCTGAACCTGG
GAGCACACGTCTGAACTCTTTCCCTACACGACGCTCTTCCGATCTNNNAGCTTGTAGTGAGCCTATAT
GAGCACACGTCTGAACTCTTTCCCTACACGACGCTCTTCCGATCTNNNTGTAGTCCCAGCTACTTGAAA
AGCACACGTCTGAACTCTTTCCCTACACGACGCTCTTCCGATCTNNNGGGCAACAAAATGAGACTCTG
AGAGCACACGTCTGAACTCTTTCCCTACACGACGCTCTTCCGATCTNNNATCATCTGAGTCTGGGAGG
GAGCACACGTCTGAACTCTTTCCCTACACGACGCTCTTCCGATCTNNNGAGAATCTCTTGAACCTGGG
AGCACACGTCTGAACTCTTTCCCTACACGACGCTCTTCCGATCTNNNCTGTAATCCCAGCTACTCAGGA
GCACACGTCTGAACTCTTTCCCTACACGACGCTCTTCCGATCTNNNTAGTCATAGTTACTCGGGAGGCT
GAGCACACGTCTGAACTCTTTCCCTACACGACGCTCTTCCGATCTNNNGACAACTCACTAGATAGGCCC
GAGCACACGTCTGAACTCTTTCCCTACACGACGCTCTTCCGATCTNNNGTTACATTGCTGCACACATCTG
AGCACACGTCTGAACTCTTTCCCTACACGACGCTCTTCCGATCTNNNGTGCTGTATTCAGGAGACCCA
GAGCACACGTCTGAACTCTTTCCCTACACGACGCTCTTCCGATCTNNNCCATGTAGACACTCTTCAGGG
AGCACACGTCTGAACTCTTTCCCTACACGACGCTCTTCCGATCTNNNAATCCTAGCTACTTTGGAGGCT
GAGCACACGTCTGAACTCTTTCCCTACACGACGCTCTTCCGATCTNNNTACAAAAAATTAGCTGGGCG
GAGCACACGTCTGAACTCTTTCCCTACACGACGCTCTTCCGATCTNNNGGGAGGGATAGCATTGGGA
AGAGCACACGTCTGAACTCTTTCCCTACACGACGCTCTTCCGATCTNNNCTTGGGAAGCTGAGGTAG
AGCACACGTCTGAACTCTTTCCCTACACGACGCTCTTCCGATCTNNNAGGCTTACAAATGGCAAAGG
GAGCACACGTCTGAACTCTTTCCCTACACGACGCTCTTCCGATCTNNNATGTCTGTAATCCCAGCTACT
GCACACGTCTGAACTCTTTCCCTACACGACGCTCTTCCGATCTNNNCTGTAATCCCAGCTACTCCAGAG
GAGCACACGTCTGAACTCTTTCCCTACACGACGCTCTTCCGATCTNNNAGAGGTTGCAGTGAACAGA
GAGCACACGTCTGAACTCTTTCCCTACACGACGCTCTTCCGATCTNNNCTGAGGCAGGAGAATTGCTT
GCACACGTCTGAACTCTTTCCCTACACGACGCTCTTCCGATCTNNNCCAGCTCAGTGGTTAGATTGAG
AGAGCACACGTCTGAACTCTTTCCCTACACGACGCTCTTCCGATCTNNNGAGGCAGGAGAATGGTGTG
AGAGCACACGTCTGAACTCTTTCCCTACACGACGCTCTTCCGATCTNNNGAATCATCGCTTGAACCTGG
AGCACACGTCTGAACTCTTTCCCTACACGACGCTCTTCCGATCTNNNGGTGCCTGTAATCCCAGATACT
AGCACACGTCTGAACTCTTTCCCTACACGACGCTCTTCCGATCTNNNGCCCACTTTCAGCACTTCTATTC
GAGCACACGTCTGAACTCTTTCCCTACACGACGCTCTTCCGATCTNNNCTGTAGTCCCAGCTACTCAGG
AGAGCACACGTCTGAACTCTTTCCCTACACGACGCTCTTCCGATCTNNNGGAGGCTGAAGTAGGAGA
GCACACGTCTGAACTCTTTCCCTACACGACGCTCTTCCGATCTNNNACCTGTAGTAGTCCCAGCTACTT
AGAGCACACGTCTGAACTCTTTCCCTACACGACGCTCTTCCGATCTNNNGAGAATGGCATGAACCTGG
GAGCACACGTCTGAACTCTTTCCCTACACGACGCTCTTCCGATCTNNNCTAAGGCAGGAGTATCGCTTG
GAGCACACGTCTGAACTCTTTCCCTACACGACGCTCTTCCGATCTNNNAGGGAGGTAGAGGTTGTAG
AGCACACGTCTGAACTCTTTCCCTACACGACGCTCTTCCGATCTNNNGGGTTCAAGTGATTCTCCTACC
AGCACACGTCTGAACTCTTTCCCTACACGACGCTCTTCCGATCTNNNTGGTGACATACTCAGGAATCTG
AGAGCACACGTCTGAACTCTTTCCCTACACGACGCTCTTCCGATCTNNNGGTAGGAGAATCACTTGAA
AGCACACGTCTGAACTCTTTCCCTACACGACGCTCTTCCGATCTNNNAGGCATGGCAGTTCTACTACA

GCACACGTCTGAACTCTTTCCCTACACGACGCTCTTCCGATCTNNNCCACACAAGCTTCTAAATTGGG
GCACACGTCTGAACTCTTTCCCTACACGACGCTCTTCCGATCTNNNTGGGATTTCTCACCTCTTTCTG
CACACGTCTGAACTCTTTCCCTACACGACGCTCTTCCGATCTNNNTCTCTTGTTTTCAGCCGTGATC
AGCACACGTCTGAACTCTTTCCCTACACGACGCTCTTCCGATCTNNNTGCGGCTAAGATGTCAAAGTA
GCACACGTCTGAACTCTTTCCCTACACGACGCTCTTCCGATCTNNNCTGAGGCAGGAGAATCACTTGA
AGAGCACACGTCTGAACTCTTTCCCTACACGACGCTCTTCCGATCTNNNGGATGGATGGATGAATG
GAGCACACGTCTGAACTCTTTCCCTACACGACGCTCTTCCGATCTNNNCCAAACACCGCATATTCTCAC
GCACACGTCTGAACTCTTTCCCTACACGACGCTCTTCCGATCTNNNGAAGGCCTTATACGTCACAAAG
AGCACACGTCTGAACTCTTTCCCTACACGACGCTCTTCCGATCTNNNACAAGAATCTTGAACCTGGG
AGCACACGTCTGAACTCTTTCCCTACACGACGCTCTTCCGATCTNNNCTTTCTTTCTCCTTCCCTCCCTCC
AGCACACGTCTGAACTCTTTCCCTACACGACGCTCTTCCGATCTNNNATGCCTGTAATCCCAGCTACTT
AGCACACGTCTGAACTCTTTCCCTACACGACGCTCTTCCGATCTNNNATCTCAGCACTTTGGAAGGACA
GCACACGTCTGAACTCTTTCCCTACACGACGCTCTTCCGATCTNNNAACTGTACCTTGGCCCCTTTTATC
AGCACACGTCTGAACTCTTTCCCTACACGACGCTCTTCCGATCTNNNATTCTCTCGCATGTACACACCA
AGCACACGTCTGAACTCTTTCCCTACACGACGCTCTTCCGATCTNNNATCCAGCCAGAACAGAACTTTC
AGCACACGTCTGAACTCTTTCCCTACACGACGCTCTTCCGATCTNNNGAAGGAAAACTAGTTTGCGA
AGCACACGTCTGAACTCTTTCCCTACACGACGCTCTTCCGATCTNNNAAAGGGATAATGGGGTGTCT
GAGCACACGTCTGAACTCTTTCCCTACACGACGCTCTTCCGATCTNNNACCAAATATCACACCCCTGTT
GAGCACACGTCTGAACTCTTTCCCTACACGACGCTCTTCCGATCTNNNACCAAAGACCGCATATTCTCA
AGAGCACACGTCTGAACTCTTTCCCTACACGACGCTCTTCCGATCTNNNGGACAGAAAACGAAACACC
AGCACACGTCTGAACTCTTTCCCTACACGACGCTCTTCCGATCTNNNTTGCAAACCTAATAGGCTTCCC
AGCACACGTCTGAACTCTTTCCCTACACGACGCTCTTCCGATCTNNNTCCATGGTGGCTTGGATTTTT
AGCACACGTCTGAACTCTTTCCCTACACGACGCTCTTCCGATCTNNNGTACTTCTGTAGTGTCCAGCCA
AGAGCACACGTCTGAACTCTTTCCCTACACGACGCTCTTCCGATCTNNNTTGAATTAGGCCCAGAGAC
GAGCACACGTCTGAACTCTTTCCCTACACGACGCTCTTCCGATCTNNNTGGAGAAGACAATGTGTGT
GCACACGTCTGAACTCTTTCCCTACACGACGCTCTTCCGATCTNNNCTCAAGTGATCCTCCCATCTCAG
GCACACGTCTGAACTCTTTCCCTACACGACGCTCTTCCGATCTNNNATCCATTGCTGGTGAACTAGTG
AGCACACGTCTGAACTCTTTCCCTACACGACGCTCTTCCGATCTNNNATCTACAGGAGGGTAAAGAGG
GCACACGTCTGAACTCTTTCCCTACACGACGCTCTTCCGATCTNNNGCTGTGATCCAGTAGAAGGTAC
AGCACACGTCTGAACTCTTTCCCTACACGACGCTCTTCCGATCTNNNCTAAGCTATGAGGACGCAAA
GCACACGTCTGAACTCTTTCCCTACACGACGCTCTTCCGATCTNNNGAAACTTCAATCACCAGCAGTC
AGAGCACACGTCTGAACTCTTTCCCTACACGACGCTCTTCCGATCTNNNGACGCCGTGAAAAGAAAG
GAGCACACGTCTGAACTCTTTCCCTACACGACGCTCTTCCGATCTNNNAGAGGAATCAAATGGGCAT
CACACGTCTGAACTCTTTCCCTACACGACGCTCTTCCGATCTNNNCAAGATGAAGCTCTAACCCCAAC
GAGCACACGTCTGAACTCTTTCCCTACACGACGCTCTTCCGATCTNNNCTTCATGAGGAATGGAGGTA
GCACACGTCTGAACTCTTTCCCTACACGACGCTCTTCCGATCTNNNTCAGATTCCCAAGGCTGAATGA
AGCACACGTCTGAACTCTTTCCCTACACGACGCTCTTCCGATCTNNNCCCTGTACCAAGATCATTCCGA
AGCACACGTCTGAACTCTTTCCCTACACGACGCTCTTCCGATCTNNNATGTCCCTCTGTCTCTCTCTCT
GAGCACACGTCTGAACTCTTTCCCTACACGACGCTCTTCCGATCTNNNGACTCTTTTCAGCACATAAGG
GAGCACACGTCTGAACTCTTTCCCTACACGACGCTCTTCCGATCTNNNTGAGCTGAGATCACATTACT
GAGCACACGTCTGAACTCTTTCCCTACACGACGCTCTTCCGATCTNNNTCGATCTCCTGACCTCATGAT
AGCACACGTCTGAACTCTTTCCCTACACGACGCTCTTCCGATCTNNNGCAGCTCTCTTCTTCCTTCTCTC
AGCACACGTCTGAACTCTTTCCCTACACGACGCTCTTCCGATCTNNNCCGGTTTTTGTGCCCTATGTA

GAGCACACGTCTGAACTCTTTCCCTACACGACGCTCTTCCGATCTNNNTTGGGAACTATCTTTAGGGG
GAGCACACGTCTGAACTCTTTCCCTACACGACGCTCTTCCGATCTNNNTTATGTCATTCCTCGGCATCAG
GAGCACACGTCTGAACTCTTTCCCTACACGACGCTCTTCCGATCTNNNTTGATAGGACTGTGAGTGTG
GAGCACACGTCTGAACTCTTTCCCTACACGACGCTCTTCCGATCTNNNCTGGGTCAGGAAGCTCTTTAT
AGCACACGTCTGAACTCTTTCCCTACACGACGCTCTTCCGATCTNNNCCTGAGTAACTGGAACTACAGG
AGCACACGTCTGAACTCTTTCCCTACACGACGCTCTTCCGATCTNNNATGTATACATGTGCCATGCTGG
AGCACACGTCTGAACTCTTTCCCTACACGACGCTCTTCCGATCTNNNATGAAAAGGCAACCCACAGAA
AGCACACGTCTGAACTCTTTCCCTACACGACGCTCTTCCGATCTNNNAATGGTCCCTAACACTTCCTCAC
AGCACACGTCTGAACTCTTTCCCTACACGACGCTCTTCCGATCTNNNTTGGTGTACAGTTCTGGTTGT
AGCACACGTCTGAACTCTTTCCCTACACGACGCTCTTCCGATCTNNNCCTATGATTTCATCTCTGCCCCA
AGCACACGTCTGAACTCTTTCCCTACACGACGCTCTTCCGATCTNNNTTTCATCACCCCAAATACACGC
AGCACACGTCTGAACTCTTTCCCTACACGACGCTCTTCCGATCTNNNTTCCTCCACTATACACTCCCACT
GAGCACACGTCTGAACTCTTTCCCTACACGACGCTCTTCCGATCTNNNCCTTTACACACACACACACACT
AGAGCACACGTCTGAACTCTTTCCCTACACGACGCTCTTCCGATCTNNNGCCATTGCACTTCCTTCTACA
GAGCACACGTCTGAACTCTTTCCCTACACGACGCTCTTCCGATCTNNNATGGAATCAACCTAACTGCCC
AGCACACGTCTGAACTCTTTCCCTACACGACGCTCTTCCGATCTNNNCCTACCCCCTGTGATAATCTGTG
GAGCACACGTCTGAACTCTTTCCCTACACGACGCTCTTCCGATCTNNNAGCAAATATAGGTGGTAGCC
AGCACACGTCTGAACTCTTTCCCTACACGACGCTCTTCCGATCTNNNAACAGATTCTCCCTTGTACAGGT
GAGCACACGTCTGAACTCTTTCCCTACACGACGCTCTTCCGATCTNNNTGGTGTTTTAGTGTAGGCGA
GAGCACACGTCTGAACTCTTTCCCTACACGACGCTCTTCCGATCTNNNAAAATGTGTTGCAATGGGGG
GAGCACACGTCTGAACTCTTTCCCTACACGACGCTCTTCCGATCTNNNAGAGATGGGGTTTCACCGTAT
GAGCACACGTCTGAACTCTTTCCCTACACGACGCTCTTCCGATCTNNNGCTTGTATCAGACCCTCATCCA
GCACACGTCTGAACTCTTTCCCTACACGACGCTCTTCCGATCTNNNCCTTCTGATTTTTTGTCCCGCTTC
GAGCACACGTCTGAACTCTTTCCCTACACGACGCTCTTCCGATCTNNNCTGTGTCCTGCTGTTCTCTGTA
AGAGCACACGTCTGAACTCTTTCCCTACACGACGCTCTTCCGATCTNNNGTTAGGACGATGGAACCAA
GAGCACACGTCTGAACTCTTTCCCTACACGACGCTCTTCCGATCTNNNTTTCATCATATAGGCCAGGCT
AGCACACGTCTGAACTCTTTCCCTACACGACGCTCTTCCGATCTNNNGTGACAGAGTGAGATTCCATCC
GAGCACACGTCTGAACTCTTTCCCTACACGACGCTCTTCCGATCTNNNCTACTGGGGTAGATCTGTTTC
AGCACACGTCTGAACTCTTTCCCTACACGACGCTCTTCCGATCTNNNAGCAGTACAAGTGGTTTTTGGT
GAGCACACGTCTGAACTCTTTCCCTACACGACGCTCTTCCGATCTNNNCAAGCATTGGATGTTGTTCCA
AGCACACGTCTGAACTCTTTCCCTACACGACGCTCTTCCGATCTNNNTGAGCAGACATCTCCCTTTACA
GAGCACACGTCTGAACTCTTTCCCTACACGACGCTCTTCCGATCTNNNCTCCTCATAGTCACTATTGCCC
GAGCACACGTCTGAACTCTTTCCCTACACGACGCTCTTCCGATCTNNNGCCTCAGTTTGTCCATCTGTAG
GAGCACACGTCTGAACTCTTTCCCTACACGACGCTCTTCCGATCTNNNATTTTCTTGTAGACGGCTGGA
AGCACACGTCTGAACTCTTTCCCTACACGACGCTCTTCCGATCTNNNTGGACCAATCAGCACTCTGTAA
GAGCACACGTCTGAACTCTTTCCCTACACGACGCTCTTCCGATCTNNNGTGTGTGGGTGTGAATGTAT
AGCACACGTCTGAACTCTTTCCCTACACGACGCTCTTCCGATCTNNNGAAGAAAAAGGAGAGAGGGA
AGAGCACACGTCTGAACTCTTTCCCTACACGACGCTCTTCCGATCTNNNGGAACAGAAAACCAAACACC
AGCACACGTCTGAACTCTTTCCCTACACGACGCTCTTCCGATCTNNNGGGAATTGGTGTTTCATGGGA
GAGCACACGTCTGAACTCTTTCCCTACACGACGCTCTTCCGATCTNNNGACCCATGATGGAACCCATGA
GAGCACACGTCTGAACTCTTTCCCTACACGACGCTCTTCCGATCTNNNTCCCATGACTCATGATGGAATC
AGAGCACACGTCTGAACTCTTTCCCTACACGACGCTCTTCCGATCTNNNCTATGTGCTGATGCTGGGTA
GAGCACACGTCTGAACTCTTTCCCTACACGACGCTCTTCCGATCTNNNATTCACCTAAGGCATTCCACCA

GCACACGTCTGAACTCTTTCCCTACACGACGCTCTTCCGATCTNNNCAGGAAACTTACAATCGTGGCA
AGCACACGTCTGAACTCTTTCCCTACACGACGCTCTTCCGATCTNNNCACTTACACATACCTTTGGGG
AGCACACGTCTGAACTCTTTCCCTACACGACGCTCTTCCGATCTNNNTGGGGACAATAATACACCAAC
AGCACACGTCTGAACTCTTTCCCTACACGACGCTCTTCCGATCTNNNTTCTTACATGGACACAGGAAG
AGCACACGTCTGAACTCTTTCCCTACACGACGCTCTTCCGATCTNNNCTCCACCATAACTCCCACAGTAG
AGCACACGTCTGAACTCTTTCCCTACACGACGCTCTTCCGATCTNNNACATGACGAAACCCCATCTCTAG
AGCACACGTCTGAACTCTTTCCCTACACGACGCTCTTCCGATCTNNNAACACCATCAAGATCTTCCCTCC
AGCACACGTCTGAACTCTTTCCCTACACGACGCTCTTCCGATCTNNNCTTACATTCTCCAAAGTGGCA
GCACACGTCTGAACTCTTTCCCTACACGACGCTCTTCCGATCTNNNCACTGTTTTGGACTAAGGCAAG
AGCACACGTCTGAACTCTTTCCCTACACGACGCTCTTCCGATCTNNNTAGAAAACAGAAGAATGGCGG
GCACACGTCTGAACTCTTTCCCTACACGACGCTCTTCCGATCTNNNATTCAATTACCTCCCACTGGGTC
AGCACACGTCTGAACTCTTTCCCTACACGACGCTCTTCCGATCTNNNAATTCACCTGCACACATGAACT
AGCACACGTCTGAACTCTTTCCCTACACGACGCTCTTCCGATCTNNNGGCACTGGGGATAGAGATGT
GCACACGTCTGAACTCTTTCCCTACACGACGCTCTTCCGATCTNNNACCACGCCCAGCTAAATTTTTTT
GCACACGTCTGAACTCTTTCCCTACACGACGCTCTTCCGATCTNNNGAAGCATTCTGTCCACTGAACAG
AGCACACGTCTGAACTCTTTCCCTACACGACGCTCTTCCGATCTNNNAAAGAATATCAGAGCAGGGCT
GAGCACACGTCTGAACTCTTTCCCTACACGACGCTCTTCCGATCTNNNCATCTGGTCCTGGGCTTTTTT
AGAGCACACGTCTGAACTCTTTCCCTACACGACGCTCTTCCGATCTNNNAAAATTTGTACCAGGAATG
GAGCACACGTCTGAACTCTTTCCCTACACGACGCTCTTCCGATCTNNNTAGAGTGGAGTCTAGCATTG
AGCACACGTCTGAACTCTTTCCCTACACGACGCTCTTCCGATCTNNNCCAAGTAGCTGAGACTACAGG
GAGCACACGTCTGAACTCTTTCCCTACACGACGCTCTTCCGATCTNNNGAACATAGTACAACCGCTCTC
AGCACACGTCTGAACTCTTTCCCTACACGACGCTCTTCCGATCTNNNCTGCTTCACTTGTGAGGTCTTG
GCACACGTCTGAACTCTTTCCCTACACGACGCTCTTCCGATCTNNNTTCCTGATGACTACCCTCTCCATC
AGCACACGTCTGAACTCTTTCCCTACACGACGCTCTTCCGATCTNNNGTAGTGAGTTGTGAGTCCTCC
AGCACACGTCTGAACTCTTTCCCTACACGACGCTCTTCCGATCTNNNGTAGTGAGTTGTGAGTCCTCC
GCACACGTCTGAACTCTTTCCCTACACGACGCTCTTCCGATCTNNNATCCCCACAGATTACAAATCCCCC
GAGCACACGTCTGAACTCTTTCCCTACACGACGCTCTTCCGATCTNNNAATGGCTTGATAGAGGAGTT
GAGCACACGTCTGAACTCTTTCCCTACACGACGCTCTTCCGATCTNNNTATGAGTGATGATGACCCAT
AGCACACGTCTGAACTCTTTCCCTACACGACGCTCTTCCGATCTNNNTCCAGGAGGATCTTCTCATCTC
AGCACACGTCTGAACTCTTTCCCTACACGACGCTCTTCCGATCTNNNGCTAGGAAACAAGTGCCTCAA
GCACACGTCTGAACTCTTTCCCTACACGACGCTCTTCCGATCTNNNCAGAGGAAATGAGGAAATGGG
GAGCACACGTCTGAACTCTTTCCCTACACGACGCTCTTCCGATCTNNNTCAGAGAGCAGAGGAGAAG
GAGCACACGTCTGAACTCTTTCCCTACACGACGCTCTTCCGATCTNNNACCAGGAAAGTGGAAAATG
AGCACACGTCTGAACTCTTTCCCTACACGACGCTCTTCCGATCTNNNGATTCCCATTCTCACCACTCCTTC
AGCACACGTCTGAACTCTTTCCCTACACGACGCTCTTCCGATCTNNNAGCTCCCAGTGATATCAACACA
AGCACACGTCTGAACTCTTTCCCTACACGACGCTCTTCCGATCTNNNTAGAAAGGAGACAGCCGAATT
AGCACACGTCTGAACTCTTTCCCTACACGACGCTCTTCCGATCTNNNAACATGATACAGTGCTGCAGA
GAGCACACGTCTGAACTCTTTCCCTACACGACGCTCTTCCGATCTNNNATGTACCTGTACTCCCAGCTA
AGCACACGTCTGAACTCTTTCCCTACACGACGCTCTTCCGATCTNNNCTTGAAATTCTCACTCCCAGCAC
GAGCACACGTCTGAACTCTTTCCCTACACGACGCTCTTCCGATCTNNNAGTACACTCCTGCCCAAATAC
GAGCACACGTCTGAACTCTTTCCCTACACGACGCTCTTCCGATCTNNNTGGTTTCGATCCATTACTGG
GAGCACACGTCTGAACTCTTTCCCTACACGACGCTCTTCCGATCTNNNCTTCTATCCTCATCACCCCAA
GAGCACACGTCTGAACTCTTTCCCTACACGACGCTCTTCCGATCTNNNTTGGGGCATGGAAATAAGG

GCACACGTCTGAACTCTTTCCCTACACGACGCTCTTCCGATCTNNNAGTTTCGGTTTTCTGGGATAGG
AGCACACGTCTGAACTCTTTCCCTACACGACGCTCTTCCGATCTNNNATTTGAACTCCAGTCCATTTGG
AGCACACGTCTGAACTCTTTCCCTACACGACGCTCTTCCGATCTNNNGCCTGTAATCCCAACACTTTGA
GAGCACACGTCTGAACTCTTTCCCTACACGACGCTCTTCCGATCTNNNTCATGCCAGTAATCTCAGCAC
GAGCACACGTCTGAACTCTTTCCCTACACGACGCTCTTCCGATCTNNNCTGGGGACTTAGGGATGTA
GAGCACACGTCTGAACTCTTTCCCTACACGACGCTCTTCCGATCTNNNTTCTGGTGTCTAGACTCAGC
AGCACACGTCTGAACTCTTTCCCTACACGACGCTCTTCCGATCTNNNCTCTAACCCATGAAGACACAG
AGCACACGTCTGAACTCTTTCCCTACACGACGCTCTTCCGATCTNNNGGCTCATCAAAGTACATGTTG
AGCACACGTCTGAACTCTTTCCCTACACGACGCTCTTCCGATCTNNNGGCTGGCCACCAAAGAGTAT
AGCACACGTCTGAACTCTTTCCCTACACGACGCTCTTCCGATCTNNNGCAGTGAATTCCCATTGCACT
AGCACACGTCTGAACTCTTTCCCTACACGACGCTCTTCCGATCTNNNTTGACTGTGAAGTCCTCAGGT
AGCACACGTCTGAACTCTTTCCCTACACGACGCTCTTCCGATCTNNNCAGTGTTTTGTGGTTCTCCTTG
AGCACACGTCTGAACTCTTTCCCTACACGACGCTCTTCCGATCTNNNAGTAGGAGCTAGACTATGGG
AGCACACGTCTGAACTCTTTCCCTACACGACGCTCTTCCGATCTNNNCAAAATGGCAAAACCCCATCTC
AGCACACGTCTGAACTCTTTCCCTACACGACGCTCTTCCGATCTNNNTTTCCCATTTGTCCCATGTGTT
AGAGCACACGTCTGAACTCTTTCCCTACACGACGCTCTTCCGATCTNNNTAGCAAGGTTCAGAAGCA
AGCACACGTCTGAACTCTTTCCCTACACGACGCTCTTCCGATCTNNNGTTGTAGTATCCTTTCCCCGTG
GAGCACACGTCTGAACTCTTTCCCTACACGACGCTCTTCCGATCTNNNCTGCTGAGCATGGACTAAGA
GCACACGTCTGAACTCTTTCCCTACACGACGCTCTTCCGATCTNNNTCAGATATGGTGGCATCAGAG
GCACACGTCTGAACTCTTTCCCTACACGACGCTCTTCCGATCTNNNTTCTCAGTTCTCTCTCCCTGCTAC
GAGCACACGTCTGAACTCTTTCCCTACACGACGCTCTTCCGATCTNNNACCGCATGTTCTCACTCATAA
AGCACACGTCTGAACTCTTTCCCTACACGACGCTCTTCCGATCTNNNGTGTGTTTTTTTTTTGGGGG
GCACACGTCTGAACTCTTTCCCTACACGACGCTCTTCCGATCTNNNTCTTGTTTTTGCTCCTACTTGCCC
GAGCACACGTCTGAACTCTTTCCCTACACGACGCTCTTCCGATCTNNNTGGATTATAGGAGGGCAAA
AGCACACGTCTGAACTCTTTCCCTACACGACGCTCTTCCGATCTNNNTGAGGTAGGAGAATCGCTTG
GCACACGTCTGAACTCTTTCCCTACACGACGCTCTTCCGATCTNNNCTCTCTAGCTATCCCCCTCTCC
CACACGTCTGAACTCTTTCCCTACACGACGCTCTTCCGATCTNNNCTAGCTACCCCACCTCTCTCTAGC
CACACGTCTGAACTCTTTCCCTACACGACGCTCTTCCGATCTNNNTCTAACTACTCCCCACCTCTCTCCA
CACACGTCTGAACTCTTTCCCTACACGACGCTCTTCCGATCTNNNTCCACAAACACTCACTGTCAGATC
AGCACACGTCTGAACTCTTTCCCTACACGACGCTCTTCCGATCTNNNGGACTTTCACTTTGGAAATGG
AGCACACGTCTGAACTCTTTCCCTACACGACGCTCTTCCGATCTNNNCTCTTCCTTCTGCTAACAACTG
AGCACACGTCTGAACTCTTTCCCTACACGACGCTCTTCCGATCTNNNCCCCGTGATTTGTCTCCAAAAT
GAGCACACGTCTGAACTCTTTCCCTACACGACGCTCTTCCGATCTNNNGAGATGGAGCCAAGAAGG
AGCACACGTCTGAACTCTTTCCCTACACGACGCTCTTCCGATCTNNNCTTGTTGAGAGTGGCTGTGAT
GCACACGTCTGAACTCTTTCCCTACACGACGCTCTTCCGATCTNNNCATACGTATACATGTGCCATGC
GAGCACACGTCTGAACTCTTTCCCTACACGACGCTCTTCCGATCTNNNCTCATGGCCAGGAAACTTTG
GCACACGTCTGAACTCTTTCCCTACACGACGCTCTTCCGATCTNNNCCTATCTTGGACAATCCTCCAG
AGCACACGTCTGAACTCTTTCCCTACACGACGCTCTTCCGATCTNNNCTACAGGATTCTGACCCTCCC
AGCACACGTCTGAACTCTTTCCCTACACGACGCTCTTCCGATCTNNNCAGGAAGCAGTAGAAGTCCAG
AGCACACGTCTGAACTCTTTCCCTACACGACGCTCTTCCGATCTNNNTCCTAATGCTATCCCTCCCCTA
AGCACACGTCTGAACTCTTTCCCTACACGACGCTCTTCCGATCTNNNTAGGAAGCTAAATTCAGCCC
AGCACACGTCTGAACTCTTTCCCTACACGACGCTCTTCCGATCTNNNAGTACATACACACAGACGCAA
AGCACACGTCTGAACTCTTTCCCTACACGACGCTCTTCCGATCTNNNGCTAACTCCTCTCAGAGTGATG

AGCACACGTCTGAACTCTTTCCCTACACGACGCTCTTCCGATCTNNNAGAAGTTGCAGTGAGCCAAG
GCACACGTCTGAACTCTTTCCCTACACGACGCTCTTCCGATCTNNNGAGAAAAAAACCACCCCAACAC
AGCACACGTCTGAACTCTTTCCCTACACGACGCTCTTCCGATCTNNNAAATGCAAGGACACCTAGCCA
AGCACACGTCTGAACTCTTTCCCTACACGACGCTCTTCCGATCTNNNTCACCCACCTGACTAGTCATA
GAGCACACGTCTGAACTCTTTCCCTACACGACGCTCTTCCGATCTNNNACAGAATGAGGCCTTGTCTG
GAGCACACGTCTGAACTCTTTCCCTACACGACGCTCTTCCGATCTNNNACATGTATTCACACTTCACGT
AGCACACGTCTGAACTCTTTCCCTACACGACGCTCTTCCGATCTNNNCAAAATGCAAACCAATGTGGC
GCACACGTCTGAACTCTTTCCCTACACGACGCTCTTCCGATCTNNNGATCCCAGGTGTATCCTTTTCCA
GCACACGTCTGAACTCTTTCCCTACACGACGCTCTTCCGATCTNNNAGAATTGCTTGAACCCAGAAAG
GAGCACACGTCTGAACTCTTTCCCTACACGACGCTCTTCCGATCTNNNCCACCATACCCGGCTAATTTT
AGCACACGTCTGAACTCTTTCCCTACACGACGCTCTTCCGATCTNNNGTGCATGCACATTTCTCTGAG
GCACACGTCTGAACTCTTTCCCTACACGACGCTCTTCCGATCTNNNCAGGTGAGGCTTACATACAAGG
AGCACACGTCTGAACTCTTTCCCTACACGACGCTCTTCCGATCTNNNGCTAAGAGAGATCCTGGCTCA
AGCACACGTCTGAACTCTTTCCCTACACGACGCTCTTCCGATCTNNNAGATTGGAGAGGATTGCTTG
GCACACGTCTGAACTCTTTCCCTACACGACGCTCTTCCGATCTNNNAGTCTTGGGAGGGGTTTTTCT
GAGCACACGTCTGAACTCTTTCCCTACACGACGCTCTTCCGATCTNNNCAGGAAATTCAGTTGGAGG
CACACGTCTGAACTCTTTCCCTACACGACGCTCTTCCGATCTNNNCAGAGTTTGTGCCCAGTCAAAAT
GCACACGTCTGAACTCTTTCCCTACACGACGCTCTTCCGATCTNNNAACTCCTGACCTCAAGTGATCTG
CACACGTCTGAACTCTTTCCCTACACGACGCTCTTCCGATCTNNNAGACCCCTGTTTTTGGTGTTTTG
GCACACGTCTGAACTCTTTCCCTACACGACGCTCTTCCGATCTNNNCCCTTCCAGATGCTTCCTTTACT
GCACACGTCTGAACTCTTTCCCTACACGACGCTCTTCCGATCTNNNATAGCAGTAGCAGAAGGACGT
GCACACGTCTGAACTCTTTCCCTACACGACGCTCTTCCGATCTNNNCACCAGAACGCATTCCTTTCTTC
AGCACACGTCTGAACTCTTTCCCTACACGACGCTCTTCCGATCTNNNGCTCCTTCTTATTTGGGTAGGC
GCACACGTCTGAACTCTTTCCCTACACGACGCTCTTCCGATCTNNNCCCACGTCTTATTGTAGAGGCT
AGCACACGTCTGAACTCTTTCCCTACACGACGCTCTTCCGATCTNNNCAAGATGGAGGGAAGTCGTA
AGCACACGTCTGAACTCTTTCCCTACACGACGCTCTTCCGATCTNNNTCCTAACACTATCCCTCCTCCAG
AGCACACGTCTGAACTCTTTCCCTACACGACGCTCTTCCGATCTNNNGCACCCGTTAGCTCATCATTTA
AGCACACGTCTGAACTCTTTCCCTACACGACGCTCTTCCGATCTNNNGGGAGAAGTCAGGGAAGCA
GCACACGTCTGAACTCTTTCCCTACACGACGCTCTTCCGATCTNNNCCTCCAAATGAACCATGGTAGG
AGCACACGTCTGAACTCTTTCCCTACACGACGCTCTTCCGATCTNNNACCCTCATCCATTTTACAGGCA
GCACACGTCTGAACTCTTTCCCTACACGACGCTCTTCCGATCTNNNAACTCGTAAGTCGTTTCTCAGG
GCACACGTCTGAACTCTTTCCCTACACGACGCTCTTCCGATCTNNNTTTCTTTCTCCCCTCCTCAAGTG
AGCACACGTCTGAACTCTTTCCCTACACGACGCTCTTCCGATCTNNNATGTGTCCACAATGTCTTGGA
GAGCACACGTCTGAACTCTTTCCCTACACGACGCTCTTCCGATCTNNNGAAGGATTATGGTTGCCTCT
AGCACACGTCTGAACTCTTTCCCTACACGACGCTCTTCCGATCTNNNGAGCTATAGACTGGAGCTGT
GAGCACACGTCTGAACTCTTTCCCTACACGACGCTCTTCCGATCTNNNTTTGAGAAAGGGTCCCACTTT
AGCACACGTCTGAACTCTTTCCCTACACGACGCTCTTCCGATCTNNNACATGTACAAGTCTCCCCAAC
AGCACACGTCTGAACTCTTTCCCTACACGACGCTCTTCCGATCTNNNTTTTGGATGCCGATATGGTTT
AGCACACGTCTGAACTCTTTCCCTACACGACGCTCTTCCGATCTNNNAGTATCTTTGTGGCGTTCTCTG
AGCACACGTCTGAACTCTTTCCCTACACGACGCTCTTCCGATCTNNNCATGCCTGTAATCCCAGCTACT
AGCACACGTCTGAACTCTTTCCCTACACGACGCTCTTCCGATCTNNNCACACCCTACTACACCACTTAC
AGCACACGTCTGAACTCTTTCCCTACACGACGCTCTTCCGATCTNNNAAAGTTTCTATTCTCCCGGACC
AGCACACGTCTGAACTCTTTCCCTACACGACGCTCTTCCGATCTNNNTGAGTCATGCATTCTCCAAGAC

AGCACACGTCTGAACTCTTTCCCTACACGACGCTCTTCCGATCTNNNTGAGGTAGGAGAATGGCTTG
CACACGTCTGAACTCTTTCCCTACACGACGCTCTTCCGATCTNNNTAATGCAAAGACAGCACCAAACC
GCACACGTCTGAACTCTTTCCCTACACGACGCTCTTCCGATCTNNNCAAAAAGCTAGACTCCTGGGAG
AGCACACGTCTGAACTCTTTCCCTACACGACGCTCTTCCGATCTNNNTTAGAGAGACCAGTTGTGCC
AGCACACGTCTGAACTCTTTCCCTACACGACGCTCTTCCGATCTNNNGTGATCTTGGAGCCCCTAAGA
AGCACACGTCTGAACTCTTTCCCTACACGACGCTCTTCCGATCTNNNCTTCTCCTTTCCCTGGACTAACC
GCACACGTCTGAACTCTTTCCCTACACGACGCTCTTCCGATCTNNNTAGGTATACACATGCCATGGTG
GCACACGTCTGAACTCTTTCCCTACACGACGCTCTTCCGATCTNNNACCTGTAATCCCAGCTACTTTGG
GCACACGTCTGAACTCTTTCCCTACACGACGCTCTTCCGATCTNNNTCACTGAGACCACTGACAAAACA
GAGCACACGTCTGAACTCTTTCCCTACACGACGCTCTTCCGATCTNNNAGCAGAAATCCCTGATAAAC
AGCACACGTCTGAACTCTTTCCCTACACGACGCTCTTCCGATCTNNNGTGCATCTCCAGACTCACAGA
AGCACACGTCTGAACTCTTTCCCTACACGACGCTCTTCCGATCTNNNTTCCTCCCTCTTATCCACATCCTC
GCACACGTCTGAACTCTTTCCCTACACGACGCTCTTCCGATCTNNNTGCCTAAAGTACCATCTTTGCCT
AGCACACGTCTGAACTCTTTCCCTACACGACGCTCTTCCGATCTNNNCAATGCCCGGCTAATTTTCTGT
AGCACACGTCTGAACTCTTTCCCTACACGACGCTCTTCCGATCTNNNATCCCCTGGGGTTCTTTTCTAA
AGCACACGTCTGAACTCTTTCCCTACACGACGCTCTTCCGATCTNNNTCTACATGCGCAGTCTCTACTTC
GAGCACACGTCTGAACTCTTTCCCTACACGACGCTCTTCCGATCTNNNCTCACCCTACCTGACATAAAC
AGCACACGTCTGAACTCTTTCCCTACACGACGCTCTTCCGATCTNNNTTCTTTCTAGCCGAGGGAAATC
AGCACACGTCTGAACTCTTTCCCTACACGACGCTCTTCCGATCTNNNGAAATAGGGTCTTTGCAGATG
GCACACGTCTGAACTCTTTCCCTACACGACGCTCTTCCGATCTNNNCATCTGTGGTCCCAGCTACTTAG
AGCACACGTCTGAACTCTTTCCCTACACGACGCTCTTCCGATCTNNNCAAGTAGAGGTCAGTCTCATG
GAGCACACGTCTGAACTCTTTCCCTACACGACGCTCTTCCGATCTNNNGAAGATAAACTAGGGGGAT
GCACACGTCTGAACTCTTTCCCTACACGACGCTCTTCCGATCTNNNCTGGTTCCATTCGCAGAGGTAT
GAGCACACGTCTGAACTCTTTCCCTACACGACGCTCTTCCGATCTNNNGTCTTACCATATTGCCCATGC
GAGCACACGTCTGAACTCTTTCCCTACACGACGCTCTTCCGATCTNNNCAAGACGGGCTACCCAGAA
GCACACGTCTGAACTCTTTCCCTACACGACGCTCTTCCGATCTNNNCCACCACACCCAGCTAATTTTT
AGCACACGTCTGAACTCTTTCCCTACACGACGCTCTTCCGATCTNNNAAAGGATTCCAGAACCTTCCTC
AGCACACGTCTGAACTCTTTCCCTACACGACGCTCTTCCGATCTNNNCAACATTGGTGGACAGTTACA
GAGCACACGTCTGAACTCTTTCCCTACACGACGCTCTTCCGATCTNNNCCAAAGTGCTGGAATTACAG
GAGCACACGTCTGAACTCTTTCCCTACACGACGCTCTTCCGATCTNNNAGAGAGACAGAGAGACAAG
AGCACACGTCTGAACTCTTTCCCTACACGACGCTCTTCCGATCTNNNGATAAATCAGACGTGATGCTG
AGCACACGTCTGAACTCTTTCCCTACACGACGCTCTTCCGATCTNNNGCATAGCTCACTTATTATGCG
GAGCACACGTCTGAACTCTTTCCCTACACGACGCTCTTCCGATCTNNNGGACTTTCTGACCTGAGTTTC
AGCACACGTCTGAACTCTTTCCCTACACGACGCTCTTCCGATCTNNNCCTATGACCCATGATGAAACCC
AGCACACGTCTGAACTCTTTCCCTACACGACGCTCTTCCGATCTNNNCACAGCCTTCTTCCTTTTCATG
GAGCACACGTCTGAACTCTTTCCCTACACGACGCTCTTCCGATCTNNNAGAACAGAAAATGAAATGC
GAGCACACGTCTGAACTCTTTCCCTACACGACGCTCTTCCGATCTNNNGTGTTTCGCTCTCATGCTGAA
GCACACGTCTGAACTCTTTCCCTACACGACGCTCTTCCGATCTNNNCAATTTGTCAAGGGGTGAGTG
GCACACGTCTGAACTCTTTCCCTACACGACGCTCTTCCGATCTNNNCAAGAAAGAATTCAGGACGGC
AGCACACGTCTGAACTCTTTCCCTACACGACGCTCTTCCGATCTNNNCAACTCCTGACCTCGTGATCTA
GAGCACACGTCTGAACTCTTTCCCTACACGACGCTCTTCCGATCTNNNGTTACGTGGGAGCTAAGCTA
AGCACACGTCTGAACTCTTTCCCTACACGACGCTCTTCCGATCTNNNATGAACTTGAGCACCTCCATG
AGCACACGTCTGAACTCTTTCCCTACACGACGCTCTTCCGATCTNNNGATTCTAAAAAAGAGGCCGGG

AGCACACGTCTGAACTCTTTCCCTACACGACGCTCTTCCGATCTNNNGGCAAGACCCTGTCTCTACAAA
AGCACACGTCTGAACTCTTTCCCTACACGACGCTCTTCCGATCTNNNAGAGCAAGACCCTGTCTCAAA
GCACACGTCTGAACTCTTTCCCTACACGACGCTCTTCCGATCTNNNCAACTGTGACAGCCAAAACTCT
GAGCACACGTCTGAACTCTTTCCCTACACGACGCTCTTCCGATCTNNNTGAGTGAGAAATACTTCCTG
GAGCACACGTCTGAACTCTTTCCCTACACGACGCTCTTCCGATCTNNNGAGAGTGCAGTCATCTCTCC
AGCACACGTCTGAACTCTTTCCCTACACGACGCTCTTCCGATCTNNNATGTCAACATCCCTTTCCCACTT
GCACACGTCTGAACTCTTTCCCTACACGACGCTCTTCCGATCTNNNTAAAGGAAGGAAGGAAGAGG
GCACACGTCTGAACTCTTTCCCTACACGACGCTCTTCCGATCTNNNCTTCTAACAGACAGGACCCTCA
GAGCACACGTCTGAACTCTTTCCCTACACGACGCTCTTCCGATCTNNNAAGAAAGTGGAGTCAGGCT
AGCACACGTCTGAACTCTTTCCCTACACGACGCTCTTCCGATCTNNNAACCGACCTAACAGGATTCTAG
GCACACGTCTGAACTCTTTCCCTACACGACGCTCTTCCGATCTNNNGCCTTTGCTATGACTTTCTGCAT
GAGCACACGTCTGAACTCTTTCCCTACACGACGCTCTTCCGATCTNNNGTTCTCCAAGTTCTTCCCTTTG
AGCACACGTCTGAACTCTTTCCCTACACGACGCTCTTCCGATCTNNNGAGCAAGACCCCATCTCAAAA
GAGCACACGTCTGAACTCTTTCCCTACACGACGCTCTTCCGATCTNNNGTCAGTCCTTAGATTCAGAG
GCACACGTCTGAACTCTTTCCCTACACGACGCTCTTCCGATCTNNNCGACATGCAGAATTTCTTGAGC
AGCACACGTCTGAACTCTTTCCCTACACGACGCTCTTCCGATCTNNNCTGGGTCTCTTCAACAATGTC
AGCACACGTCTGAACTCTTTCCCTACACGACGCTCTTCCGATCTNNNGCAAAACCTGGTCTATGCCAA
AGCACACGTCTGAACTCTTTCCCTACACGACGCTCTTCCGATCTNNNGTGACCCTTAGTTCCCAACTA
AGCACACGTCTGAACTCTTTCCCTACACGACGCTCTTCCGATCTNNNTGCCTTGTTCATTCTACTGCTC
AGCACACGTCTGAACTCTTTCCCTACACGACGCTCTTCCGATCTNNNCTGTGCTGCTGGCTTTCTATTA
AGCACACGTCTGAACTCTTTCCCTACACGACGCTCTTCCGATCTNNNACTGGTTGGGTTGGAGATGA
AGCACACGTCTGAACTCTTTCCCTACACGACGCTCTTCCGATCTNNNGTCTTTGGCATTCACACTGGA
AGAGCACACGTCTGAACTCTTTCCCTACACGACGCTCTTCCGATCTNNNGGGAAGAAGGGAGAAAG
GAGCACACGTCTGAACTCTTTCCCTACACGACGCTCTTCCGATCTNNNCTGGACACAGGTAACCAAGA
AGCACACGTCTGAACTCTTTCCCTACACGACGCTCTTCCGATCTNNNTGACAGAAAGGACAAACTCCC
AGCACACGTCTGAACTCTTTCCCTACACGACGCTCTTCCGATCTNNNTACTCTGACTTCTTGGTCTCCC
GCACACGTCTGAACTCTTTCCCTACACGACGCTCTTCCGATCTNNNGTATAGAAGGTGTAGACCAGC
GCACACGTCTGAACTCTTTCCCTACACGACGCTCTTCCGATCTNNNCCCTGTTAGTCATGTGAAGGTG
AGCACACGTCTGAACTCTTTCCCTACACGACGCTCTTCCGATCTNNNTACCATTCTGGGGTCTGAAGG
GAGCACACGTCTGAACTCTTTCCCTACACGACGCTCTTCCGATCTNNNTGTTGCCTTCTCCTTCTAAGT
GCACACGTCTGAACTCTTTCCCTACACGACGCTCTTCCGATCTNNNAGCCACCATGGTAGGAAATCTT
AGCACACGTCTGAACTCTTTCCCTACACGACGCTCTTCCGATCTNNNCAATGAGAACACATGGACACA
AGCACACGTCTGAACTCTTTCCCTACACGACGCTCTTCCGATCTNNNCTATATGCAGCCGGATTCTCTG
AGCACACGTCTGAACTCTTTCCCTACACGACGCTCTTCCGATCTNNNAAGTGGGAGTCGAACAATGA
GAGCACACGTCTGAACTCTTTCCCTACACGACGCTCTTCCGATCTNNNCAGGGTTGAGGTCCTAGATG
AGCACACGTCTGAACTCTTTCCCTACACGACGCTCTTCCGATCTNNNGGAGAAGTCCACTTTCCAACT
GCACACGTCTGAACTCTTTCCCTACACGACGCTCTTCCGATCTNNNCAATAGCATAGAGGCAGAAGG
GCACACGTCTGAACTCTTTCCCTACACGACGCTCTTCCGATCTNNNATTCTGCAGCACTTTTGTCTAC
AGCACACGTCTGAACTCTTTCCCTACACGACGCTCTTCCGATCTNNNGTGCTCTTCTCCTTATAGCCAC
GAGCACACGTCTGAACTCTTTCCCTACACGACGCTCTTCCGATCTNNNGCACACTTACTTGGCACTTCA
AGCACACGTCTGAACTCTTTCCCTACACGACGCTCTTCCGATCTNNNAAGAAGCACCTCAGTCAGTA
AGCACACGTCTGAACTCTTTCCCTACACGACGCTCTTCCGATCTNNNATATGCTTGAAGGGAAAGGC
AGCACACGTCTGAACTCTTTCCCTACACGACGCTCTTCCGATCTNNNCCATCATTCCACCATCTCAGAG

AGAGCACACGTCTGAACTCTTTCCCTACACGACGCTCTTCCGATCTNNNGGAGCACAGAAAGACAGAA
AGCACACGTCTGAACTCTTTCCCTACACGACGCTCTTCCGATCTNNNATTGATGGAAGGGGCAAGATT
AGCACACGTCTGAACTCTTTCCCTACACGACGCTCTTCCGATCTNNNAAAAAGGAAAAAGCTCCCTCTG
AGAGCACACGTCTGAACTCTTTCCCTACACGACGCTCTTCCGATCTNNNCGCGTGGGTTACATTTTGAG
GAGCACACGTCTGAACTCTTTCCCTACACGACGCTCTTCCGATCTNNNCTTACGCCTGTAATCCTAGCAC
GAGCACACGTCTGAACTCTTTCCCTACACGACGCTCTTCCGATCTNNNTATGACAGGGAGCTAACTGA
GAGCACACGTCTGAACTCTTTCCCTACACGACGCTCTTCCGATCTNNNTTTCTTTTCTCAGTCTCTCGCCC
AGCACACGTCTGAACTCTTTCCCTACACGACGCTCTTCCGATCTNNNCTCTTGGGCTTTGTAGGGTACTT
AGCACACGTCTGAACTCTTTCCCTACACGACGCTCTTCCGATCTNNNACCATGCCCGGCTAATTTTTTTA
AGAGCACACGTCTGAACTCTTTCCCTACACGACGCTCTTCCGATCTNNNCTTTGCATTCTTTGGGTCTGG
GAGCACACGTCTGAACTCTTTCCCTACACGACGCTCTTCCGATCTNNNGCCTCAAGGTATTATCCCCAGA
GAGCACACGTCTGAACTCTTTCCCTACACGACGCTCTTCCGATCTNNNCCTGGCTTTTTTTACTGTCGTGT
GAGCACACGTCTGAACTCTTTCCCTACACGACGCTCTTCCGATCTNNNTAATTTCAGCACTTTGGGAGG
GAGCACACGTCTGAACTCTTTCCCTACACGACGCTCTTCCGATCTNNNTGAAACCCCGTCTCCACTAAAA
AGCACACGTCTGAACTCTTTCCCTACACGACGCTCTTCCGATCTNNNATCACAGCCCTCTGGAAAAAGA
GAGCACACGTCTGAACTCTTTCCCTACACGACGCTCTTCCGATCTNNNAGACCTCATGGATGACCTCTAC
GAGCACACGTCTGAACTCTTTCCCTACACGACGCTCTTCCGATCTNNNGCCAGTAAGAGGGTCAGTTAC
GAGCACACGTCTGAACTCTTTCCCTACACGACGCTCTTCCGATCTNNNCATCATGCCAAGGTAAAGAAG
AGCACACGTCTGAACTCTTTCCCTACACGACGCTCTTCCGATCTNNNGACTTACATAGAAGGCCTGGG
GAGCACACGTCTGAACTCTTTCCCTACACGACGCTCTTCCGATCTNNNATTTTGTTTCTGTTGCTCCTGG
GCACACGTCTGAACTCTTTCCCTACACGACGCTCTTCCGATCTNNNCGCAGAAGGACATCAAAAAAGGT
GCACACGTCTGAACTCTTTCCCTACACGACGCTCTTCCGATCTNNNCCACGGTGCTCTCATAGATATCAC
GAGCACACGTCTGAACTCTTTCCCTACACGACGCTCTTCCGATCTNNNGCCTCCTGAGTAACTGGGATT
AGCACACGTCTGAACTCTTTCCCTACACGACGCTCTTCCGATCTNNNCTGAGGGTCTTGGGATATATGC
GCACACGTCTGAACTCTTTCCCTACACGACGCTCTTCCGATCTNNNGACTGTAATCCCAGCTACTCCAGC
GAGCACACGTCTGAACTCTTTCCCTACACGACGCTCTTCCGATCTNNNGGTGAAACTCCGTCTCTACTGA
AGAGCACACGTCTGAACTCTTTCCCTACACGACGCTCTTCCGATCTNNNGGACATGAGGAGAATGTGC
GAGCACACGTCTGAACTCTTTCCCTACACGACGCTCTTCCGATCTNNNTCCTAACAGAACATCAAGAGG
GCACACGTCTGAACTCTTTCCCTACACGACGCTCTTCCGATCTNNNTTCACGCCACTATACTCCAATGTC
GCACACGTCTGAACTCTTTCCCTACACGACGCTCTTCCGATCTNNNCTTGAACTCCTGACCTCGTGATCC
GCACACGTCTGAACTCTTTCCCTACACGACGCTCTTCCGATCTNNNCGTCCTCAACCTTAGCAAAACACC
GCACACGTCTGAACTCTTTCCCTACACGACGCTCTTCCGATCTNNNGATTCAAGCATTTCTCCTGCCTCC
AGCACACGTCTGAACTCTTTCCCTACACGACGCTCTTCCGATCTNNNCCGTCGGTGGCTTTTTTACAAAA
AGCACACGTCTGAACTCTTTCCCTACACGACGCTCTTCCGATCTNNNACGCCCGGCTAATTTTTCTTTTCC
AGCACACGTCTGAACTCTTTCCCTACACGACGCTCTTCCGATCTNNNCGCCCGGCTTTTCTTTTTTTTTTG
AGCACACGTCTGAACTCTTTCCCTACACGACGCTCTTCCGATCTNNNCGCCCGGCTTTTCTTTTTTTTTTG
GCACACGTCTGAACTCTTTCCCTACACGACGCTCTTCCGATCTNNNCCACCACACCAGGCTAATTTTTCCA
AGCACACGTCTGAACTCTTTCCCTACACGACGCTCTTCCGATCTNNNATATTGCCCATGCTGGTGTCAA
GCACACGTCTGAACTCTTTCCCTACACGACGCTCTTCCGATCTNNNCCACCACACCAGGCTAATTTTTCCA
AGCACACGTCTGAACTCTTTCCCTACACGACGCTCTTCCGATCTNNNACCACACCCGGCTAATTTTTTTTG
AGCACACGTCTGAACTCTTTCCCTACACGACGCTCTTCCGATCTNNNGGCTCCAGGTCAGAGTCTAAATG
AGCACACGTCTGAACTCTTTCCCTACACGACGCTCTTCCGATCTNNNGGTCAAAAGGTGAAGCAACTCT
GAGCACACGTCTGAACTCTTTCCCTACACGACGCTCTTCCGATCTNNNGGAGGTTACAGTGAGCTGAG

GAGCACACGTCTGAACTCTTTCCCTACACGACGCTCTTCCGATCTNNNAAAGTACTGGGATTACAGG
GCACACGTCTGAACTCTTTCCCTACACGACGCTCTTCCGATCTNNNCTGTTGAACAACCATGTCTGGA
AGCACACGTCTGAACTCTTTCCCTACACGACGCTCTTCCGATCTNNNACACCATCACACCTGGCTAATTT
GAGCACACGTCTGAACTCTTTCCCTACACGACGCTCTTCCGATCTNNNTCAAGCAATTCTCCTGCCTTA
GCACACGTCTGAACTCTTTCCCTACACGACGCTCTTCCGATCTNNNTCACCGGCTGTATTTTCACAAAC
AGCACACGTCTGAACTCTTTCCCTACACGACGCTCTTCCGATCTNNNCCTCCTTCTTGCTTCTTTTCAGCC
AGCACACGTCTGAACTCTTTCCCTACACGACGCTCTTCCGATCTNNNGTGGTGCAATCCTAGCTCAGTA
AGCACACGTCTGAACTCTTTCCCTACACGACGCTCTTCCGATCTNNNCGACAGAGTGAGAATCCGTCT
AGCACACGTCTGAACTCTTTCCCTACACGACGCTCTTCCGATCTNNNACTCAGGGCTTCTAACAACTTCC
AGCACACGTCTGAACTCTTTCCCTACACGACGCTCTTCCGATCTNNNCACCTGGCCCCTGCAATATATA
GCACACGTCTGAACTCTTTCCCTACACGACGCTCTTCCGATCTNNNTCTTTGGCTACCGTCAAACTTG
AGCACACGTCTGAACTCTTTCCCTACACGACGCTCTTCCGATCTNNNAGTGATGGGATTACAGGTGTG
AGCACACGTCTGAACTCTTTCCCTACACGACGCTCTTCCGATCTNNNCGAACTCCTAACCTCAGGTGAT
AGCACACGTCTGAACTCTTTCCCTACACGACGCTCTTCCGATCTNNNATCACACCTCACTGATGAAACCA
GAGCACACGTCTGAACTCTTTCCCTACACGACGCTCTTCCGATCTNNNAGACAAACCATATCCCTGCCA
AGCACACGTCTGAACTCTTTCCCTACACGACGCTCTTCCGATCTNNNCTGCAACTGGGTACATTTTGGA
AGCACACGTCTGAACTCTTTCCCTACACGACGCTCTTCCGATCTNNNTCCTGTTTCTAGGACCCATGTTG
AGCACACGTCTGAACTCTTTCCCTACACGACGCTCTTCCGATCTNNNAGTTTTTCTGGTGTCTTTGGGG
AGCACACGTCTGAACTCTTTCCCTACACGACGCTCTTCCGATCTNNNGTGTCTTTGGGGAAAAAGAG
AGCACACGTCTGAACTCTTTCCCTACACGACGCTCTTCCGATCTNNNTGTTTCTAGGACCCATGTTGAG
AGCACACGTCTGAACTCTTTCCCTACACGACGCTCTTCCGATCTNNNGCCTTGGGAATGTAGCTTTGTT
AGCACACGTCTGAACTCTTTCCCTACACGACGCTCTTCCGATCTNNNAGATTTCTGTTTGGGCTTGTGT
AGCACACGTCTGAACTCTTTCCCTACACGACGCTCTTCCGATCTNNNACAGTTCAGCTCAGTGACAAG
GAGCACACGTCTGAACTCTTTCCCTACACGACGCTCTTCCGATCTNNNCAATGGTACCTCCCGATCTAA
AGCACACGTCTGAACTCTTTCCCTACACGACGCTCTTCCGATCTNNNGACAAAGTCTCTCTCTGTCACC
AGCACACGTCTGAACTCTTTCCCTACACGACGCTCTTCCGATCTNNNCCCCTCTCTCTTTCTCTGAGGAT
AGCACACGTCTGAACTCTTTCCCTACACGACGCTCTTCCGATCTNNNTGTCCTTTTTGTAGAGACACGG
AGCACACGTCTGAACTCTTTCCCTACACGACGCTCTTCCGATCTNNNCACCATTGTAGCATCCAAGCAA
GAGCACACGTCTGAACTCTTTCCCTACACGACGCTCTTCCGATCTNNNATTACAAGCATGAGCCATGTG
GAGCACACGTCTGAACTCTTTCCCTACACGACGCTCTTCCGATCTNNNCTTGAACTCCTGAACTTGCGA
AGCACACGTCTGAACTCTTTCCCTACACGACGCTCTTCCGATCTNNNTCCTAGAGATTGGAAAACGGC
GAGCACACGTCTGAACTCTTTCCCTACACGACGCTCTTCCGATCTNNNTTTAGTAGAGATGGGGTTTC
AGCACACGTCTGAACTCTTTCCCTACACGACGCTCTTCCGATCTNNNCAGCTTCAGCAAGAAACAGACA
AGCACACGTCTGAACTCTTTCCCTACACGACGCTCTTCCGATCTNNNCTGCTCTCCATTTTCTACCCAGA
AGCACACGTCTGAACTCTTTCCCTACACGACGCTCTTCCGATCTNNNTTTGTGTGTGTGTGTGTGTGTA
GCACACGTCTGAACTCTTTCCCTACACGACGCTCTTCCGATCTNNNGAACTCCTGACCTCATGTGATCC
AGCACACGTCTGAACTCTTTCCCTACACGACGCTCTTCCGATCTNNNTATTGAACTCCTGGGCTCAAGT
AGCACACGTCTGAACTCTTTCCCTACACGACGCTCTTCCGATCTNNNATTAGGAGGAGACGCTGTTTC
AGCACACGTCTGAACTCTTTCCCTACACGACGCTCTTCCGATCTNNNAAATACAGGTCTGCACCAGTA
AGCACACGTCTGAACTCTTTCCCTACACGACGCTCTTCCGATCTNNNAATTCAATCCTGCAGCTTTCCAC
AGAGCACACGTCTGAACTCTTTCCCTACACGACGCTCTTCCGATCTNNNAACCTTGGAACTGACACCGA
GCACACGTCTGAACTCTTTCCCTACACGACGCTCTTCCGATCTNNNCCATGCCCGGCTAATTTTTGTA
AGCACACGTCTGAACTCTTTCCCTACACGACGCTCTTCCGATCTNNNCATGGTGTGGTTCCTTCCTCTTA

AGCACACGTCTGAACTCTTTCCCTACACGACGCTCTTCCGATCTNNNTGAGACTCTGATTGCAGTGG
AGCACACGTCTGAACTCTTTCCCTACACGACGCTCTTCCGATCTNNNGTGTTTGAGGCAAGTCAGTAA
AGCACACGTCTGAACTCTTTCCCTACACGACGCTCTTCCGATCTNNNAACGGACAGAAGTTGGAAAA
GAGCACACGTCTGAACTCTTTCCCTACACGACGCTCTTCCGATCTNNNAGTCCCAGAAGAATTGAGCA
AGCACACGTCTGAACTCTTTCCCTACACGACGCTCTTCCGATCTNNNATCCTGGAGATGTCTTTGCC
AGCACACGTCTGAACTCTTTCCCTACACGACGCTCTTCCGATCTNNNTCCCATGATCTCTCTCTCTC
AGCACACGTCTGAACTCTTTCCCTACACGACGCTCTTCCGATCTNNNAAAAATGCAATGCCTGGATCC
GCACACGTCTGAACTCTTTCCCTACACGACGCTCTTCCGATCTNNNCGCCCGGCTAATTCTTGTATTT
GCACACGTCTGAACTCTTTCCCTACACGACGCTCTTCCGATCTNNNACCCTGACCTCGAGAATGAATT
GAGCACACGTCTGAACTCTTTCCCTACACGACGCTCTTCCGATCTNNNTGTTCTTGAGCAAATCCAAAC
AGCACACGTCTGAACTCTTTCCCTACACGACGCTCTTCCGATCTNNNTCTGTTTTGTGAGCAGTTCTCC
AGCACACGTCTGAACTCTTTCCCTACACGACGCTCTTCCGATCTNNNCTGCTCTCTCATAACCACACC
GCACACGTCTGAACTCTTTCCCTACACGACGCTCTTCCGATCTNNNGTGGCCAGTTTCAGCTTGTAT
GCACACGTCTGAACTCTTTCCCTACACGACGCTCTTCCGATCTNNNTCCCGCAGTCTTATTTATACCG
AGCACACGTCTGAACTCTTTCCCTACACGACGCTCTTCCGATCTNNNGTAGAGACAGAGTTTCACCG
GCACACGTCTGAACTCTTTCCCTACACGACGCTCTTCCGATCTNNNCACACACACACACATATGGA
GCACACGTCTGAACTCTTTCCCTACACGACGCTCTTCCGATCTNNNTGAGATTACAGATGTGAGCCAC
GCACACGTCTGAACTCTTTCCCTACACGACGCTCTTCCGATCTNNNCACACACACACACATATGGA
CACACGTCTGAACTCTTTCCCTACACGACGCTCTTCCGATCTNNNCAATGGTGCATTCTTGGAGAAGG
GCACACGTCTGAACTCTTTCCCTACACGACGCTCTTCCGATCTNNNATGTACAACCTCTTCCCAAACCC
GCACACGTCTGAACTCTTTCCCTACACGACGCTCTTCCGATCTNNNTGTCCCCAAAATCATCCTCACTTC
AGCACACGTCTGAACTCTTTCCCTACACGACGCTCTTCCGATCTNNNAGTGTAGAAGTCTCTCTCCACC
GAGCACACGTCTGAACTCTTTCCCTACACGACGCTCTTCCGATCTNNNCCTCTCCTATACTCTGCTCCAT
AGCACACGTCTGAACTCTTTCCCTACACGACGCTCTTCCGATCTNNNCCTTGTCGATGGCTTTGGTAA
GAGCACACGTCTGAACTCTTTCCCTACACGACGCTCTTCCGATCTNNNTTACGGTTGGCCCATTTCTTT
GCACACGTCTGAACTCTTTCCCTACACGACGCTCTTCCGATCTNNNCTCCTGAGTAGCTGGGATTACA
GAGCACACGTCTGAACTCTTTCCCTACACGACGCTCTTCCGATCTNNNTTCCGGGCTTAGAAAGTTAG
GCACACGTCTGAACTCTTTCCCTACACGACGCTCTTCCGATCTNNNTGACACCTGTAGAAGCAGAAC
GCACACGTCTGAACTCTTTCCCTACACGACGCTCTTCCGATCTNNNATTCTCCTGCCTCAGTATCCCAA
GCACACGTCTGAACTCTTTCCCTACACGACGCTCTTCCGATCTNNNACCTACTCATTTATCACCTTTGC
AGCACACGTCTGAACTCTTTCCCTACACGACGCTCTTCCGATCTNNNGGATGGTCTTGAACTCCTGAC
GCACACGTCTGAACTCTTTCCCTACACGACGCTCTTCCGATCTNNNATTTGTAAGGCGTGACTCCTGA
GAGCACACGTCTGAACTCTTTCCCTACACGACGCTCTTCCGATCTNNNCTGTCTGTGTCTCCTCTTCTG
GCACACGTCTGAACTCTTTCCCTACACGACGCTCTTCCGATCTNNNTTTAGTACAGACGGGGTTTCAC
GCACACGTCTGAACTCTTTCCCTACACGACGCTCTTCCGATCTNNNGGCTCGAATAATTTCTTGCACG
AGCACACGTCTGAACTCTTTCCCTACACGACGCTCTTCCGATCTNNNGGAGCCTGTGGAGATGCTAT
AGAGCACACGTCTGAACTCTTTCCCTACACGACGCTCTTCCGATCTNNNCAATCTCGGCTCACTACAAC
AGCACACGTCTGAACTCTTTCCCTACACGACGCTCTTCCGATCTNNNAATCAGAGTTCAGTCCGCAGT
GAGCACACGTCTGAACTCTTTCCCTACACGACGCTCTTCCGATCTNNNCTGGAACTCCTGACCTCATGA
AGCACACGTCTGAACTCTTTCCCTACACGACGCTCTTCCGATCTNNNGGCCTCTGGAGTAGCTATTCT
AGCACACGTCTGAACTCTTTCCCTACACGACGCTCTTCCGATCTNNNGCCTCTGGAGTACCTAGTCTTT
AGCACACGTCTGAACTCTTTCCCTACACGACGCTCTTCCGATCTNNNAGTGCTGGGATTATAGGCATG
GAGCACACGTCTGAACTCTTTCCCTACACGACGCTCTTCCGATCTNNNCCCGCCTTTATGGACCAAACT

AGCACACGTCTGAACTCTTTCCCTACACGACGCTCTTCCGATCTNNNTTTGAGACAAAGTCACGCTC
GCACACGTCTGAACTCTTTCCCTACACGACGCTCTTCCGATCTNNNGCACTGGGTTCAGAATTTCCTC
GCACACGTCTGAACTCTTTCCCTACACGACGCTCTTCCGATCTNNNCTCCTGAGTAGCTGGGATTACA
AGCACACGTCTGAACTCTTTCCCTACACGACGCTCTTCCGATCTNNNTGAGTATGAGACCACCTCCTT
AGCACACGTCTGAACTCTTTCCCTACACGACGCTCTTCCGATCTNNNATTTGCAAGGGACAAACAACC
AGCACACGTCTGAACTCTTTCCCTACACGACGCTCTTCCGATCTNNNTTCAAGTGATTCTCCTGCCTCAG
GCACACGTCTGAACTCTTTCCCTACACGACGCTCTTCCGATCTNNNGCGAAAGGAAGCTTTGCAATT
GCACACGTCTGAACTCTTTCCCTACACGACGCTCTTCCGATCTNNNGCAGCTGCCATCATATCCAACT
AGCACACGTCTGAACTCTTTCCCTACACGACGCTCTTCCGATCTNNNACCATGAACCTGACTTTCCTTG
AGCACACGTCTGAACTCTTTCCCTACACGACGCTCTTCCGATCTNNNGTACATTTTTGTGCCCGTGTG
GCACACGTCTGAACTCTTTCCCTACACGACGCTCTTCCGATCTNNNACAACAGACCACATACAGTCTCA
GCACACGTCTGAACTCTTTCCCTACACGACGCTCTTCCGATCTNNNTGCTCCCTCAAATCAGTGAAAG
GCACACGTCTGAACTCTTTCCCTACACGACGCTCTTCCGATCTNNNGCTACCATGCCCAGCTAATTTT
GCACACGTCTGAACTCTTTCCCTACACGACGCTCTTCCGATCTNNNCTTGAACTCCTGACCTCAAGTG
AGCACACGTCTGAACTCTTTCCCTACACGACGCTCTTCCGATCTNNNGGTACACTGGACTTTGGGGAA
AGCACACGTCTGAACTCTTTCCCTACACGACGCTCTTCCGATCTNNNCTCCCTTCAGTGTCTGCTAGAA
GAGCACACGTCTGAACTCTTTCCCTACACGACGCTCTTCCGATCTNNNTTTAGTAGAGACGGGGTTT
AGCACACGTCTGAACTCTTTCCCTACACGACGCTCTTCCGATCTNNNTCCCTCTGTCTCTCCTTCTTTCT
GAGCACACGTCTGAACTCTTTCCCTACACGACGCTCTTCCGATCTNNNTTCTGACCTCAAGTAATCCAC
AGCACACGTCTGAACTCTTTCCCTACACGACGCTCTTCCGATCTNNNGATTACAGGTGTGAGCTACTG
AGCACACGTCTGAACTCTTTCCCTACACGACGCTCTTCCGATCTNNNTATGTCCTGCTTGGGAAAGTTG
GAGCACACGTCTGAACTCTTTCCCTACACGACGCTCTTCCGATCTNNNGAGGAGAAGTTGGTCTTGG
AGCACACGTCTGAACTCTTTCCCTACACGACGCTCTTCCGATCTNNNTGCTCTGAGGCTGAATTTTCAA
AGCACACGTCTGAACTCTTTCCCTACACGACGCTCTTCCGATCTNNNACAATATCCTTCCTCGACAACGT
GCACACGTCTGAACTCTTTCCCTACACGACGCTCTTCCGATCTNNNGCCGCTTCTTCTTCTTCTTCTTCC
GCACACGTCTGAACTCTTTCCCTACACGACGCTCTTCCGATCTNNNCTGTTGAGACCCACTTCTCTTCT
AGCACACGTCTGAACTCTTTCCCTACACGACGCTCTTCCGATCTNNNCAAAGAAGCATGTGGAATTCCC
AGCACACGTCTGAACTCTTTCCCTACACGACGCTCTTCCGATCTNNNCCCAAAGTCCTTTGTGTCATTC
GCACACGTCTGAACTCTTTCCCTACACGACGCTCTTCCGATCTNNNGGCTAGTCTTGAACTGCTGATC
GCACACGTCTGAACTCTTTCCCTACACGACGCTCTTCCGATCTNNNTTGAAGAAAACACTGTGACCAG
AGCACACGTCTGAACTCTTTCCCTACACGACGCTCTTCCGATCTNNNCTCCCACTCTAGGAGTCTGT
AGCACACGTCTGAACTCTTTCCCTACACGACGCTCTTCCGATCTNNNCAGCAGTGGTGTTCTCTAGAC
AGCACACGTCTGAACTCTTTCCCTACACGACGCTCTTCCGATCTNNNCCACGCCCAGCTAATTTTTGTAT
GCACACGTCTGAACTCTTTCCCTACACGACGCTCTTCCGATCTNNNCCTTGGCTCCATACTCTAAACC
GAGCACACGTCTGAACTCTTTCCCTACACGACGCTCTTCCGATCTNNNTCCGAAAGTACTGGGATTAT
GCACACGTCTGAACTCTTTCCCTACACGACGCTCTTCCGATCTNNNTCTTGAAATCCTGAGCTCGTGA
AGCACACGTCTGAACTCTTTCCCTACACGACGCTCTTCCGATCTNNNTCCATCCATGCCCAGAATCTTT
CACACGTCTGAACTCTTTCCCTACACGACGCTCTTCCGATCTNNNGGCCTCCTAAATTGCTGGGATTA
GAGCACACGTCTGAACTCTTTCCCTACACGACGCTCTTCCGATCTNNNGCCAATGGCGTAACTATTCTG
AGCACACGTCTGAACTCTTTCCCTACACGACGCTCTTCCGATCTNNNGATTGACAGTTCTGTCTCCAC
AGCACACGTCTGAACTCTTTCCCTACACGACGCTCTTCCGATCTNNNTTAGTAGAGATGGGGTTTCAC
GCACACGTCTGAACTCTTTCCCTACACGACGCTCTTCCGATCTNNNCACCCGGCACTATAAACGTTTAT
GCACACGTCTGAACTCTTTCCCTACACGACGCTCTTCCGATCTNNNCCAGCCATCATCTTGTCCATTT

AGCACACGTCTGAACTCTTTCCCTACACGACGCTCTTCCGATCTNNNCCGCCCTGAAAAATCTGTTTCTT
AGCACACGTCTGAACTCTTTCCCTACACGACGCTCTTCCGATCTNNNGGACAGACTGACACAGATACA
AGCACACGTCTGAACTCTTTCCCTACACGACGCTCTTCCGATCTNNNTTAGTAGAGATGGGGTTTCAC
AGCACACGTCTGAACTCTTTCCCTACACGACGCTCTTCCGATCTNNNTTCTCCCAGGGGAACATACTGTT
AGCACACGTCTGAACTCTTTCCCTACACGACGCTCTTCCGATCTNNNCCTGAGACTGGGTAATTTTGGG
AGCACACGTCTGAACTCTTTCCCTACACGACGCTCTTCCGATCTNNNCGTGGCCAACAAGCATATTGTA
GAGCACACGTCTGAACTCTTTCCCTACACGACGCTCTTCCGATCTNNNAAGTTCAACATCAAGATGCCA
GAGCACACGTCTGAACTCTTTCCCTACACGACGCTCTTCCGATCTNNNGGTATTTGTCCTAGTGCTCTC
AGCACACGTCTGAACTCTTTCCCTACACGACGCTCTTCCGATCTNNNGAAATAAAAGGTCAGGGCTCA
AGCACACGTCTGAACTCTTTCCCTACACGACGCTCTTCCGATCTNNNACAAGAGAAAATGTAAGCCTC
AGCACACGTCTGAACTCTTTCCCTACACGACGCTCTTCCGATCTNNNAGCTGAGATTACAGGTTCACAC
AGCACACGTCTGAACTCTTTCCCTACACGACGCTCTTCCGATCTNNNATAGACCAAAGGGAAAGCGA
AGCACACGTCTGAACTCTTTCCCTACACGACGCTCTTCCGATCTNNNTCAACGGTGGGGAGAATATTA
AGCACACGTCTGAACTCTTTCCCTACACGACGCTCTTCCGATCTNNNTGGTGTATCAGACAGGGAAACA
AGCACACGTCTGAACTCTTTCCCTACACGACGCTCTTCCGATCTNNNAAAGTGCTGGAATTACATGCG
AGCACACGTCTGAACTCTTTCCCTACACGACGCTCTTCCGATCTNNNTGCCCAGCTAATGTTTGTGTTTT
GAGCACACGTCTGAACTCTTTCCCTACACGACGCTCTTCCGATCTNNNCTGAGTAGCTCGGATTACTG
GAGCACACGTCTGAACTCTTTCCCTACACGACGCTCTTCCGATCTNNNCTGCAAAGGACATGAACTCAT
AGCACACGTCTGAACTCTTTCCCTACACGACGCTCTTCCGATCTNNNAAGCACATCAGACTAAAACAC
AGCACACGTCTGAACTCTTTCCCTACACGACGCTCTTCCGATCTNNNCTCTCTTGGCTTGAAGGGAATG
AGCACACGTCTGAACTCTTTCCCTACACGACGCTCTTCCGATCTNNNTCTCCCGTGATTTCAAACTTTGC
AGCACACGTCTGAACTCTTTCCCTACACGACGCTCTTCCGATCTNNNCTCCACGCATTCCTTACCTATGA
AGCACACGTCTGAACTCTTTCCCTACACGACGCTCTTCCGATCTNNNGACGGTGTCTCTTCAGCTTTCTA
AGCACACGTCTGAACTCTTTCCCTACACGACGCTCTTCCGATCTNNNAAAAACAGAAAGAAGCCCTCTG
GAGCACACGTCTGAACTCTTTCCCTACACGACGCTCTTCCGATCTNNNGGGGGGTGAAAGTTGAAAA
AGCACACGTCTGAACTCTTTCCCTACACGACGCTCTTCCGATCTNNNGAGGATCTTTGAAGTGGGAGA
GAGCACACGTCTGAACTCTTTCCCTACACGACGCTCTTCCGATCTNNNACACCTGGCTGAAACTCACTA
GAGCACACGTCTGAACTCTTTCCCTACACGACGCTCTTCCGATCTNNNCGATGCTCTGTGGATTAGGG
GAGCACACGTCTGAACTCTTTCCCTACACGACGCTCTTCCGATCTNNNGGGGTCTACTTGAGGGTAAA
AGCACACGTCTGAACTCTTTCCCTACACGACGCTCTTCCGATCTNNNGGTCCTTACTTGGTTAGATGC
AGCACACGTCTGAACTCTTTCCCTACACGACGCTCTTCCGATCTNNNTGGTTGGGAGAGATTTTTTCCT
AGCACACGTCTGAACTCTTTCCCTACACGACGCTCTTCCGATCTNNNACCATGTTTCTCACTGCCATTTG
AGCACACGTCTGAACTCTTTCCCTACACGACGCTCTTCCGATCTNNNCAGTGAGCTGAGATCATACCA
AGCACACGTCTGAACTCTTTCCCTACACGACGCTCTTCCGATCTNNNCCCATACCAGGTGGCATTATTCA
GAGCACACGTCTGAACTCTTTCCCTACACGACGCTCTTCCGATCTNNNTGTTGTTGTTGTTGCTGACAG
AGCACACGTCTGAACTCTTTCCCTACACGACGCTCTTCCGATCTNNNCTCTTGAACTCCTGACCTTGTGA
AGCACACGTCTGAACTCTTTCCCTACACGACGCTCTTCCGATCTNNNGTTCAAGTGATTCTCCTCCCTCA
AGCACACGTCTGAACTCTTTCCCTACACGACGCTCTTCCGATCTNNNCACGCTCTCTTTCTAGTTCCCTTC
GAGCACACGTCTGAACTCTTTCCCTACACGACGCTCTTCCGATCTNNNGGCCACAAGTCCTTTAGGAGA
GAGCACACGTCTGAACTCTTTCCCTACACGACGCTCTTCCGATCTNNNGATTCAAGTGATTCCCCTGCC
GAGCACACGTCTGAACTCTTTCCCTACACGACGCTCTTCCGATCTNNNCACAGTTCCTGCTTCAATGGA
AGCACACGTCTGAACTCTTTCCCTACACGACGCTCTTCCGATCTNNNGCAAAACTCCAGTAAATCGCTC
GAGCACACGTCTGAACTCTTTCCCTACACGACGCTCTTCCGATCTNNNAGAGGATTCTGACTGTTTCAC

AGCACACGTCTGAACTCTTTCCCTACACGACGCTCTTCCGATCTNNNTCGAATCTCTCATTCCTCAAGCA
AGCACACGTCTGAACTCTTTCCCTACACGACGCTCTTCCGATCTNNNAGAAAGGTTTGCTTCTGATGCT
AGCACACGTCTGAACTCTTTCCCTACACGACGCTCTTCCGATCTNNNACATTCAAACAAGGCAGGTCAA
GAGCACACGTCTGAACTCTTTCCCTACACGACGCTCTTCCGATCTNNNCTCTTGTGTGTGTTGCTGTGT
GAGCACACGTCTGAACTCTTTCCCTACACGACGCTCTTCCGATCTNNNCTCCCAAGCAGCTGAAATTAC
GCACACGTCTGAACTCTTTCCCTACACGACGCTCTTCCGATCTNNNACTAAGGTTCATCTTGTCCCTGG
AGCACACGTCTGAACTCTTTCCCTACACGACGCTCTTCCGATCTNNNGCTCTCTCTCTTTGCCTCTTTCTC
GAGCACACGTCTGAACTCTTTCCCTACACGACGCTCTTCCGATCTNNNCCTCAGGAAACCCCTTTAGAT
AGCACACGTCTGAACTCTTTCCCTACACGACGCTCTTCCGATCTNNNGTCCCTCTCTGTCACTTTCTCTG
AGCACACGTCTGAACTCTTTCCCTACACGACGCTCTTCCGATCTNNNGAGATTGGGTTGTCACTGTGT
AGCACACGTCTGAACTCTTTCCCTACACGACGCTCTTCCGATCTNNNCTGTAATCCCAGCTACTCAGGA
AGCACACGTCTGAACTCTTTCCCTACACGACGCTCTTCCGATCTNNNCTGTGTGTCTGTGTCCTCATCT
AGCACACGTCTGAACTCTTTCCCTACACGACGCTCTTCCGATCTNNNTTTCTCTCCTCCTCCTTCTCCTTCA
AGCACACGTCTGAACTCTTTCCCTACACGACGCTCTTCCGATCTNNNGGAACCAGTGTACCTCTCAACA
GAGCACACGTCTGAACTCTTTCCCTACACGACGCTCTTCCGATCTNNNCCTGGCCTGGGATTGTTATTT
AGCACACGTCTGAACTCTTTCCCTACACGACGCTCTTCCGATCTNNNCCACCATAGGAAGGACAAATT
AGCACACGTCTGAACTCTTTCCCTACACGACGCTCTTCCGATCTNNNGAACTCCCGACTTCAGTTGATC
AGCACACGTCTGAACTCTTTCCCTACACGACGCTCTTCCGATCTNNNATGTTTTTGCATAGGCTTGTCG
GAGCACACGTCTGAACTCTTTCCCTACACGACGCTCTTCCGATCTNNNAAGGAAGACGAGGAATTCA
AGCACACGTCTGAACTCTTTCCCTACACGACGCTCTTCCGATCTNNNAACTAGTCTCTGTGCCAGAAAG
GAGCACACGTCTGAACTCTTTCCCTACACGACGCTCTTCCGATCTNNNTTTTTTTTTTGCAGGGGAGG
GCACACGTCTGAACTCTTTCCCTACACGACGCTCTTCCGATCTNNNCTACCCTGTAGCTATGAGCAGTG
AGCACACGTCTGAACTCTTTCCCTACACGACGCTCTTCCGATCTNNNGTCCAACCTCTCATCACTACCT
AGCACACGTCTGAACTCTTTCCCTACACGACGCTCTTCCGATCTNNNAGTGCTGGGATTACAGGTGTA
GAGCACACGTCTGAACTCTTTCCCTACACGACGCTCTTCCGATCTNNNGCAAAGCCAAACAAGTTGTA
AGCACACGTCTGAACTCTTTCCCTACACGACGCTCTTCCGATCTNNNGGTTCAAGCAATTCTTCTGCCT
GCACACGTCTGAACTCTTTCCCTACACGACGCTCTTCCGATCTNNNGGTCCATTCTGCAATCTGTGATG
GCACACGTCTGAACTCTTTCCCTACACGACGCTCTTCCGATCTNNNTCTGTTCCTGAAATGAACGACC
GCACACGTCTGAACTCTTTCCCTACACGACGCTCTTCCGATCTNNNTTTGTGAGATGACCTCGTGTGA
AGCACACGTCTGAACTCTTTCCCTACACGACGCTCTTCCGATCTNNNCTAAGGATAATGGCCTCCACCT
GAGCACACGTCTGAACTCTTTCCCTACACGACGCTCTTCCGATCTNNNGCAGTGTTTTGTGTGTGCTTT
AGCACACGTCTGAACTCTTTCCCTACACGACGCTCTTCCGATCTNNNCGTTTGTGGAAACTGCTCTACT
AGCACACGTCTGAACTCTTTCCCTACACGACGCTCTTCCGATCTNNNGTGATGCCCTGTTGCAGATATT
AGCACACGTCTGAACTCTTTCCCTACACGACGCTCTTCCGATCTNNNTATCCACCAGTGCCCAATGTTTA
GAGCACACGTCTGAACTCTTTCCCTACACGACGCTCTTCCGATCTNNNTAACCGCTGGACTTTGTTGAA
AGCACACGTCTGAACTCTTTCCCTACACGACGCTCTTCCGATCTNNNAAGGTTGCAAAGGCTGGAATA
GAGCACACGTCTGAACTCTTTCCCTACACGACGCTCTTCCGATCTNNNCCTGGCCAGAAAGGGAAAAA
AGCACACGTCTGAACTCTTTCCCTACACGACGCTCTTCCGATCTNNNGCAGAGACACAAATCCAAACCA
AGCACACGTCTGAACTCTTTCCCTACACGACGCTCTTCCGATCTNNNAGTGCTGAGACTCGAATTCTGA
AGCACACGTCTGAACTCTTTCCCTACACGACGCTCTTCCGATCTNNNCTGGGTTCAAGCCAATTCTCTT
GAGCACACGTCTGAACTCTTTCCCTACACGACGCTCTTCCGATCTNNNAGTGCTGGGATTATAGGCAT
AGCACACGTCTGAACTCTTTCCCTACACGACGCTCTTCCGATCTNNNATACAGGTGCATCCTACCACAT
GCACACGTCTGAACTCTTTCCCTACACGACGCTCTTCCGATCTNNNACGCCACCATACCTGTCTAATTT

AGCACACGTCTGAACTCTTTCCCTACACGACGCTCTTCCGATCTNNNGAGATCAAATCATGCCTCAGTG
AGCACACGTCTGAACTCTTTCCCTACACGACGCTCTTCCGATCTNNNATTCTAACTAGGGTGAGTTGG
AGCACACGTCTGAACTCTTTCCCTACACGACGCTCTTCCGATCTNNNAAGTATACTCAGTCAAGCAGC
GAGCACACGTCTGAACTCTTTCCCTACACGACGCTCTTCCGATCTNNNCACACATAGGTTCTTGGCCAA
GCACACGTCTGAACTCTTTCCCTACACGACGCTCTTCCGATCTNNNTTTCCCTGTTTCTGACCACAACC
GAGCACACGTCTGAACTCTTTCCCTACACGACGCTCTTCCGATCTNNNTTGAGGAATGCAGCAGTCTA
AGCACACGTCTGAACTCTTTCCCTACACGACGCTCTTCCGATCTNNNATCTGTGTCTGAAAAGTGGCT
GCACACGTCTGAACTCTTTCCCTACACGACGCTCTTCCGATCTNNNCTCAAGTGATTGACCTGCTTCA
GAGCACACGTCTGAACTCTTTCCCTACACGACGCTCTTCCGATCTNNNAACAGAGCTCGCTTAGATGG
AGCACACGTCTGAACTCTTTCCCTACACGACGCTCTTCCGATCTNNNGAAAAACCCCAAAGCTTCAGG
GAGCACACGTCTGAACTCTTTCCCTACACGACGCTCTTCCGATCTNNNATCTTTCTCAGAAGCGAGCTC
AGCACACGTCTGAACTCTTTCCCTACACGACGCTCTTCCGATCTNNNTGTTGTTGCAAATGACAGATG
AGCACACGTCTGAACTCTTTCCCTACACGACGCTCTTCCGATCTNNNAACATCAGGAAATGGTCTCTC
GAGCACACGTCTGAACTCTTTCCCTACACGACGCTCTTCCGATCTNNNCAGCAACTTGGATGAAGCTA
GAGCACACGTCTGAACTCTTTCCCTACACGACGCTCTTCCGATCTNNNAGGAGAAAGAGCTAAAGCA
AGCACACGTCTGAACTCTTTCCCTACACGACGCTCTTCCGATCTNNNTATACAACGACTCTTGAGCCCA
GCACACGTCTGAACTCTTTCCCTACACGACGCTCTTCCGATCTNNNAAAATGCTGGGATTACACGCAT
GCACACGTCTGAACTCTTTCCCTACACGACGCTCTTCCGATCTNNNGCAGCAGCGTAACTTTTCTTTTT
GAGCACACGTCTGAACTCTTTCCCTACACGACGCTCTTCCGATCTNNNGAAAACCTGTTGTAACCCTCA
AGCACACGTCTGAACTCTTTCCCTACACGACGCTCTTCCGATCTNNNCTTCTCACCTTAGGCGATCATC
GCACACGTCTGAACTCTTTCCCTACACGACGCTCTTCCGATCTNNNGACCTCATTTTGCCTGAATCACC
AGCACACGTCTGAACTCTTTCCCTACACGACGCTCTTCCGATCTNNNCTCAAACTCCTGACCTCGTGAT
GAGCACACGTCTGAACTCTTTCCCTACACGACGCTCTTCCGATCTNNNGGCCCCGGGAATTATTTCTTT
AGCACACGTCTGAACTCTTTCCCTACACGACGCTCTTCCGATCTNNNCTGTTTGTGACCTGAAAACCCT
GAGCACACGTCTGAACTCTTTCCCTACACGACGCTCTTCCGATCTNNNGCAAACAGTGATGCAATCCG
GCACACGTCTGAACTCTTTCCCTACACGACGCTCTTCCGATCTNNNCTAGCTGGGACTACAAGTGTGA
AGCACACGTCTGAACTCTTTCCCTACACGACGCTCTTCCGATCTNNNTTTTGGGGCAATTGTGAGGAA
GAGCACACGTCTGAACTCTTTCCCTACACGACGCTCTTCCGATCTNNNCACCTCTCCAGATCAAGCTCTA
AGCACACGTCTGAACTCTTTCCCTACACGACGCTCTTCCGATCTNNNAGTGCTAGGATTACAGGCGTA
AGCACACGTCTGAACTCTTTCCCTACACGACGCTCTTCCGATCTNNNGTTCTTTGTATTCCCCATCCCCTC
GCACACGTCTGAACTCTTTCCCTACACGACGCTCTTCCGATCTNNNTCCTCATTCTTTCTTCTGCTGCTC
AGCACACGTCTGAACTCTTTCCCTACACGACGCTCTTCCGATCTNNNGTTCAAGAGATTCTCCTGCCTCA
AGCACACGTCTGAACTCTTTCCCTACACGACGCTCTTCCGATCTNNNAGCTCCAGTGTTCAGTGCATAT
GAGCACACGTCTGAACTCTTTCCCTACACGACGCTCTTCCGATCTNNNATCTCCCAGGTTCAAGCAATT
GAGCACACGTCTGAACTCTTTCCCTACACGACGCTCTTCCGATCTNNNCTTTCGATGCAAAAACGAAGC
GAGCACACGTCTGAACTCTTTCCCTACACGACGCTCTTCCGATCTNNNCACTTTCCAGGTTCCAAGCAA
AGCACACGTCTGAACTCTTTCCCTACACGACGCTCTTCCGATCTNNNTCCTCTCATCTCAGTCTCCAGA
GAGCACACGTCTGAACTCTTTCCCTACACGACGCTCTTCCGATCTNNNCGATCTCCAATTACTGCAAGCT
AGCACACGTCTGAACTCTTTCCCTACACGACGCTCTTCCGATCTNNNACATTTCTGCCCTAGGTCACTTT
GAGCACACGTCTGAACTCTTTCCCTACACGACGCTCTTCCGATCTNNNTCAGTTACTGTGTGTGGCCTA
AGCACACGTCTGAACTCTTTCCCTACACGACGCTCTTCCGATCTNNNTCCCTCCCATGATATGTGGAGA
AGCACACGTCTGAACTCTTTCCCTACACGACGCTCTTCCGATCTNNNAGTACCATCAGTCAACTAGGTG
GCACACGTCTGAACTCTTTCCCTACACGACGCTCTTCCGATCTNNNAAGACAAAGAAGAAAAACGGC

GAGCACACGTCTGAACTCTTTCCCTACACGACGCTCTTCCGATCTNNNTAAAGGGAGTCATTTGCCAGG
AGCACACGTCTGAACTCTTTCCCTACACGACGCTCTTCCGATCTNNNACCCTTGAGTTTTGGGACTTTGA
AGCACACGTCTGAACTCTTTCCCTACACGACGCTCTTCCGATCTNNNCCCAGGTTCAAGCAATTCTCCTA
AGCACACGTCTGAACTCTTTCCCTACACGACGCTCTTCCGATCTNNNTAGCAAAGTAGGCAGCTCTTGA
AGCACACGTCTGAACTCTTTCCCTACACGACGCTCTTCCGATCTNNNGAAAGTGGGGGCCAATATTCAA
GAGCACACGTCTGAACTCTTTCCCTACACGACGCTCTTCCGATCTNNNGCTGGCCGCTTTTTTTTTTTTT
AGCACACGTCTGAACTCTTTCCCTACACGACGCTCTTCCGATCTNNNTGTTCCACTGGTCTATGTGTCTG
GAGCACACGTCTGAACTCTTTCCCTACACGACGCTCTTCCGATCTNNNCCAAGACTGTGGGGTTTTCTA
AGCACACGTCTGAACTCTTTCCCTACACGACGCTCTTCCGATCTNNNCCCCAAAATCCAACGTGTCATTC
AGCACACGTCTGAACTCTTTCCCTACACGACGCTCTTCCGATCTNNNTTTTCAGAAAATCACACCAGCCC
GAGCACACGTCTGAACTCTTTCCCTACACGACGCTCTTCCGATCTNNNGGATTTGAAAGCGAAACTGG
GAGCACACGTCTGAACTCTTTCCCTACACGACGCTCTTCCGATCTNNNGGCATGAATAAGGAACACTG
AGCACACGTCTGAACTCTTTCCCTACACGACGCTCTTCCGATCTNNNCATTCATCATGATTGCTTCCCCCC
GAGCACACGTCTGAACTCTTTCCCTACACGACGCTCTTCCGATCTNNNGTCTCATTCTGTTGTCTGTTGC
AGCACACGTCTGAACTCTTTCCCTACACGACGCTCTTCCGATCTNNNAAAACCTAAAACCGTGCAATGCT
AGCACACGTCTGAACTCTTTCCCTACACGACGCTCTTCCGATCTNNNTGTGCAACTTCTTTTCATTGTGA
AGCACACGTCTGAACTCTTTCCCTACACGACGCTCTTCCGATCTNNNTCCCTGGAATGAAACCAACTTGA
GAGCACACGTCTGAACTCTTTCCCTACACGACGCTCTTCCGATCTNNNTTTGGTTACTACAGCTTTGCAT
GAGCACACGTCTGAACTCTTTCCCTACACGACGCTCTTCCGATCTNNNTTGAACAAAACGTGAAGCAGC
AGCACACGTCTGAACTCTTTCCCTACACGACGCTCTTCCGATCTNNNCCTGCTGCTTTCTGTTTTGGTAA
GAGCACACGTCTGAACTCTTTCCCTACACGACGCTCTTCCGATCTNNNAGTGTTTGTTCGTGTTCTTTGC
GAGCACACGTCTGAACTCTTTCCCTACACGACGCTCTTCCGATCTNNNACACATTGCACTTCTTTTCAGG
GCACACGTCTGAACTCTTTCCCTACACGACGCTCTTCCGATCTNNNGTGAGGTTAGTGGTTACATCCCT
AGCACACGTCTGAACTCTTTCCCTACACGACGCTCTTCCGATCTNNNAAGCATGAAGAGTCCCAGAACT
AGCACACGTCTGAACTCTTTCCCTACACGACGCTCTTCCGATCTNNNCCAGCTGTTCTTTGTACATCTGT
AGCACACGTCTGAACTCTTTCCCTACACGACGCTCTTCCGATCTNNNATCCCAGCACCATTTGTTGAATC
GAGCACACGTCTGAACTCTTTCCCTACACGACGCTCTTCCGATCTNNNACACTCCTCAAGTCTCATCCTCT
GAGCACACGTCTGAACTCTTTCCCTACACGACGCTCTTCCGATCTNNNGGCACCAAACAAGTCTCAAAA
GAGCACACGTCTGAACTCTTTCCCTACACGACGCTCTTCCGATCTNNNAAGCATGAAGGAGATTTCCA
GAGCACACGTCTGAACTCTTTCCCTACACGACGCTCTTCCGATCTNNNTGAGGCCCAAGATTCATTTTCC
AGCACACGTCTGAACTCTTTCCCTACACGACGCTCTTCCGATCTNNNCTTCTTCTAAGCCCTCCAGACTCC
AGCACACGTCTGAACTCTTTCCCTACACGACGCTCTTCCGATCTNNNATACACCACTTCCACTTTCACGA
AGCACACGTCTGAACTCTTTCCCTACACGACGCTCTTCCGATCTNNNTGCCCTTCTTTGTCGTTTTTGAT
GAGCACACGTCTGAACTCTTTCCCTACACGACGCTCTTCCGATCTNNNATGAACTCTGGCAAGTAGAGC
AGCACACGTCTGAACTCTTTCCCTACACGACGCTCTTCCGATCTNNNAGGTCTGTTGCCTGTTGTGATA
GAGCACACGTCTGAACTCTTTCCCTACACGACGCTCTTCCGATCTNNNTTCAAGCAATTCTCCTGCCTCA
AGCACACGTCTGAACTCTTTCCCTACACGACGCTCTTCCGATCTNNNGAAAGTACACTTGGAAAAGGG
GAGCACACGTCTGAACTCTTTCCCTACACGACGCTCTTCCGATCTNNNCAAGCCCGGCTATTTCTTTCTT
AGCACACGTCTGAACTCTTTCCCTACACGACGCTCTTCCGATCTNNNATGTGTGTCCTAGAGATCCTGC
GCACACGTCTGAACTCTTTCCCTACACGACGCTCTTCCGATCTNNNTGGAGGGAAGAATCTCAGAGCT
AGCACACGTCTGAACTCTTTCCCTACACGACGCTCTTCCGATCTNNNAGTCACTTCTGCAGGATTTAGCA
GCACACGTCTGAACTCTTTCCCTACACGACGCTCTTCCGATCTNNNGCCTCCCAAATAGCTAGGACTACC
AGCACACGTCTGAACTCTTTCCCTACACGACGCTCTTCCGATCTNNNCCAGATCCATCACACATTCCTCTC

GAGCACACGTCTGAACTCTTTCCCTACACGACGCTCTTCCGATCTNNNCTCTGTCTCCCGAATTCAAGTG
AGCACACGTCTGAACTCTTTCCCTACACGACGCTCTTCCGATCTNNNTGGGTCTTTGAGGAATTTCCAC
AGCACACGTCTGAACTCTTTCCCTACACGACGCTCTTCCGATCTNNNCAAACACGATGATCCCTTTTGCT
AGCACACGTCTGAACTCTTTCCCTACACGACGCTCTTCCGATCTNNNAGGTTCAAGCCATTCTCATGTCT
AGCACACGTCTGAACTCTTTCCCTACACGACGCTCTTCCGATCTNNNATATTTCCCCGCATCTTTCCTGAC
AGCACACGTCTGAACTCTTTCCCTACACGACGCTCTTCCGATCTNNNATGATGTAGCTGTCTTCTGCAC
GAGCACACGTCTGAACTCTTTCCCTACACGACGCTCTTCCGATCTNNNGATCTCTTGACCTCTTGAGCCA
AGCACACGTCTGAACTCTTTCCCTACACGACGCTCTTCCGATCTNNNATCAGTTGACCTGTTGGCCTAA
AGCACACGTCTGAACTCTTTCCCTACACGACGCTCTTCCGATCTNNNGGAGTTGTACTCTGCTCAATCCA
AGCACACGTCTGAACTCTTTCCCTACACGACGCTCTTCCGATCTNNNGGGTGGAAGTAAACACAAAAAG
GAGCACACGTCTGAACTCTTTCCCTACACGACGCTCTTCCGATCTNNNATCTCGTCTTACTGCAACCTCTG
AGCACACGTCTGAACTCTTTCCCTACACGACGCTCTTCCGATCTNNNGCAGATTGGGAACAGAGGAGT
GAGCACACGTCTGAACTCTTTCCCTACACGACGCTCTTCCGATCTNNNTGTTTGATACAGGCATGCAAG
GAGCACACGTCTGAACTCTTTCCCTACACGACGCTCTTCCGATCTNNNGTGCTTGTGTCATTCCACCAAT
AGCACACGTCTGAACTCTTTCCCTACACGACGCTCTTCCGATCTNNNTTAGCCTTGTGAGTAGTTGGGA
GAGCACACGTCTGAACTCTTTCCCTACACGACGCTCTTCCGATCTNNNTGGTTGGCCAAGAACTGATAA
AGCACACGTCTGAACTCTTTCCCTACACGACGCTCTTCCGATCTNNNATCAGTTGAAAGACCCTTGGGT
AGCACACGTCTGAACTCTTTCCCTACACGACGCTCTTCCGATCTNNNAATGTTTGGGCTTCCTCAGAGA
AGCACACGTCTGAACTCTTTCCCTACACGACGCTCTTCCGATCTNNNAACCAGTACATGAGGCAATCCA
AGCACACGTCTGAACTCTTTCCCTACACGACGCTCTTCCGATCTNNNGCATCCTTGTCTTGTTCCAGTTG
AGCACACGTCTGAACTCTTTCCCTACACGACGCTCTTCCGATCTNNNGCCACACCCAGCTTTTTTTTTG
GAGCACACGTCTGAACTCTTTCCCTACACGACGCTCTTCCGATCTNNNATCTCACTCTGTTGCACAGGTT
AGCACACGTCTGAACTCTTTCCCTACACGACGCTCTTCCGATCTNNNCCCCAGTGTATATTCTTGGCACTG
AGCACACGTCTGAACTCTTTCCCTACACGACGCTCTTCCGATCTNNNTCCTTTGCCCAGTTTTTAGTAGA
GAGCACACGTCTGAACTCTTTCCCTACACGACGCTCTTCCGATCTNNNTCTGTGTCCGGTTTGTATGACA
AGCACACGTCTGAACTCTTTCCCTACACGACGCTCTTCCGATCTNNNTACAGTCCCACCAAAAGTGTAGG
AGCACACGTCTGAACTCTTTCCCTACACGACGCTCTTCCGATCTNNNCTAAATCTGCTTACCCTGCCTCAG
AGCACACGTCTGAACTCTTTCCCTACACGACGCTCTTCCGATCTNNNATTATAGCCACCCGGGTTTTCTT
GAGCACACGTCTGAACTCTTTCCCTACACGACGCTCTTCCGATCTNNNCTGTGAATCTATCTGGTCCTGG
AGCACACGTCTGAACTCTTTCCCTACACGACGCTCTTCCGATCTNNNATTTGTTCTTTTTGCACCCTGCC
GAGCACACGTCTGAACTCTTTCCCTACACGACGCTCTTCCGATCTNNNCCCTGGAGCAAAAGAAATGCA
GCACACGTCTGAACTCTTTCCCTACACGACGCTCTTCCGATCTNNNTGCAGAATTCCAGAACTTCCTGAG
AGCACACGTCTGAACTCTTTCCCTACACGACGCTCTTCCGATCTNNNTTTTACCTAAGCCAGTGGACGA
GAGCACACGTCTGAACTCTTTCCCTACACGACGCTCTTCCGATCTNNNCAGCCTCCGAAAATGTGGAAA
AGAGCACACGTCTGAACTCTTTCCCTACACGACGCTCTTCCGATCTNNNTTACCTAAGCGAGTGGAGTT
AGCACACGTCTGAACTCTTTCCCTACACGACGCTCTTCCGATCTNNNCAAATGTGACACAGTGAAGGCA
AGCACACGTCTGAACTCTTTCCCTACACGACGCTCTTCCGATCTNNNACAGTGGAATATGATGCAGCTG
AGCACACGTCTGAACTCTTTCCCTACACGACGCTCTTCCGATCTNNNGTCAGTGGAGTGTGCTTGAAA
AGCACACGTCTGAACTCTTTCCCTACACGACGCTCTTCCGATCTNNNGAGGCTGCCATGAAGATCTCTA
AGCACACGTCTGAACTCTTTCCCTACACGACGCTCTTCCGATCTNNNTCTATGAACTAGTGGGCCTGAG
GAGCACACGTCTGAACTCTTTCCCTACACGACGCTCTTCCGATCTNNNGACTCTTTCATTTGGCCAAACG
GAGCACACGTCTGAACTCTTTCCCTACACGACGCTCTTCCGATCTNNNGCCAACAGGAAATGAACTCTTG
AGCACACGTCTGAACTCTTTCCCTACACGACGCTCTTCCGATCTNNNGTAGCTGGGATTACAGGCATGA

AGCACACGTCTGAACTCTTTCCCTACACGACGCTCTTCCGATCTNNNAGAAAACCTAGACAACCTTGGG
AGCACACGTCTGAACTCTTTCCCTACACGACGCTCTTCCGATCTNNNTTGCAACAAATGTCACCAACAG
AGCACACGTCTGAACTCTTTCCCTACACGACGCTCTTCCGATCTNNNGTTGTACAGAACCAATGGTACT
AGCACACGTCTGAACTCTTTCCCTACACGACGCTCTTCCGATCTNNNTGGTCTCAAACTCCTGACTTCAG
AGCACACGTCTGAACTCTTTCCCTACACGACGCTCTTCCGATCTNNNTTTTGCATGGCAGACTCTGAT
AGCACACGTCTGAACTCTTTCCCTACACGACGCTCTTCCGATCTNNNACATGACCATCAGATTGAAGC
GAGCACACGTCTGAACTCTTTCCCTACACGACGCTCTTCCGATCTNNNTTGACAGAGTCAAGCACTTTC
GCACACGTCTGAACTCTTTCCCTACACGACGCTCTTCCGATCTNNNCTCCCTCATGCTACTGTATCAGC
GAGCACACGTCTGAACTCTTTCCCTACACGACGCTCTTCCGATCTNNNTGCTTGAATCCATGACATCCT
GAGCACACGTCTGAACTCTTTCCCTACACGACGCTCTTCCGATCTNNNCCTCCTGGGTTCAAGTGATTCT
AGCACACGTCTGAACTCTTTCCCTACACGACGCTCTTCCGATCTNNNCAGTGATCATGCCAATCGATGA
AGCACACGTCTGAACTCTTTCCCTACACGACGCTCTTCCGATCTNNNTTTGTAGTCAGCCTTCAGGTCT
AGCACACGTCTGAACTCTTTCCCTACACGACGCTCTTCCGATCTNNNTGTGTGTGTGTGTGTGAAA
AGCACACGTCTGAACTCTTTCCCTACACGACGCTCTTCCGATCTNNNGTTGCTCCAATACAGCCTCATT
GAGCACACGTCTGAACTCTTTCCCTACACGACGCTCTTCCGATCTNNNGAGAATTCTTCTGCCAAGCA
GAGCACACGTCTGAACTCTTTCCCTACACGACGCTCTTCCGATCTNNNACCTTGAAAAACCCTCACCTGT
AGCACACGTCTGAACTCTTTCCCTACACGACGCTCTTCCGATCTNNNCCATGGAGGGTAAGGGTACTTA
GAGCACACGTCTGAACTCTTTCCCTACACGACGCTCTTCCGATCTNNNGGTGAGACCTGGTCTATAGCA
GAGCACACGTCTGAACTCTTTCCCTACACGACGCTCTTCCGATCTNNNACCTGAACTTGAGGAAAACTG
AGCACACGTCTGAACTCTTTCCCTACACGACGCTCTTCCGATCTNNNAGCACTGGGTTATAGATGCCAA
AGCACACGTCTGAACTCTTTCCCTACACGACGCTCTTCCGATCTNNNGAGTCTCCTGGATGCCTATTCT
GCACACGTCTGAACTCTTTCCCTACACGACGCTCTTCCGATCTNNNTCAGACTCCCAAGTAGCTAGGAT
AGCACACGTCTGAACTCTTTCCCTACACGACGCTCTTCCGATCTNNNGATGGTCTCGATCTCTTGACCT
AGCACACGTCTGAACTCTTTCCCTACACGACGCTCTTCCGATCTNNNGTTTGAGCTGATGGATACCCTG
GAGCACACGTCTGAACTCTTTCCCTACACGACGCTCTTCCGATCTNNNCCACCCGGGTCAACTATTTTA
AGCACACGTCTGAACTCTTTCCCTACACGACGCTCTTCCGATCTNNNTGCAAAGTTCAGAAGTTGAAG
GAGCACACGTCTGAACTCTTTCCCTACACGACGCTCTTCCGATCTNNNTTTCAAATGGGAAACACTCAG
GAGCACACGTCTGAACTCTTTCCCTACACGACGCTCTTCCGATCTNNNTCATTATGAGGAGTTGTGGAG
GAGCACACGTCTGAACTCTTTCCCTACACGACGCTCTTCCGATCTNNNTGGCAGCATGTGACCAAGTAT
AGCACACGTCTGAACTCTTTCCCTACACGACGCTCTTCCGATCTNNNAGGCTTCCACTCAAAAGTAGGT
AGCACACGTCTGAACTCTTTCCCTACACGACGCTCTTCCGATCTNNNGATGCAATCATAGCTCACTGCA
AGCACACGTCTGAACTCTTTCCCTACACGACGCTCTTCCGATCTNNNCCCACCAACAGTGTAAAAGCATT
AGCACACGTCTGAACTCTTTCCCTACACGACGCTCTTCCGATCTNNNGGGTCTGGATTGAGACTCCTTT
AGCACACGTCTGAACTCTTTCCCTACACGACGCTCTTCCGATCTNNNCCTTGATTTGCACCCCTTGATTT
AGCACACGTCTGAACTCTTTCCCTACACGACGCTCTTCCGATCTNNNGGCTGTATCCAATAGCATTTGCA
AGAGCACACGTCTGAACTCTTTCCCTACACGACGCTCTTCCGATCTNNNTATTTTTGAGACAGAGTCCC
GAGCACACGTCTGAACTCTTTCCCTACACGACGCTCTTCCGATCTNNNCCGATCTCTAGAGTGTTGGTT
GAGCACACGTCTGAACTCTTTCCCTACACGACGCTCTTCCGATCTNNNGTATTGGCTAGGAATGGCTA
GAGCACACGTCTGAACTCTTTCCCTACACGACGCTCTTCCGATCTNNNGGCAGGGACTCTGTTCTGTAT
GAGCACACGTCTGAACTCTTTCCCTACACGACGCTCTTCCGATCTNNNTCAAAGTGATGGGATTACAG
AGCACACGTCTGAACTCTTTCCCTACACGACGCTCTTCCGATCTNNNTTCTCCTTACCCTGCTCTGTTTC
GAGCACACGTCTGAACTCTTTCCCTACACGACGCTCTTCCGATCTNNNGTTCCAGAGCTTCTTCCACCTTA
AGCACACGTCTGAACTCTTTCCCTACACGACGCTCTTCCGATCTNNNGACACAAATGGCTCAGATTCTG

AGCACACGTCTGAACTCTTTCCCTACACGACGCTCTTCCGATCTNNNGACTTTCCTTTTCTGCCTGGAAC
AGCACACGTCTGAACTCTTTCCCTACACGACGCTCTTCCGATCTNNNATACATCTCTCCATGGGATGTG
AGCACACGTCTGAACTCTTTCCCTACACGACGCTCTTCCGATCTNNNTTTCCAGTTTGCACAGGAAGAA
GAGCACACGTCTGAACTCTTTCCCTACACGACGCTCTTCCGATCTNNNGTCCAGAACCCAGTTACTGTGA
AGCACACGTCTGAACTCTTTCCCTACACGACGCTCTTCCGATCTNNNTATTATGACATACTGCTGGGCC
GAGCACACGTCTGAACTCTTTCCCTACACGACGCTCTTCCGATCTNNNCACCAGTGAATGCTGCAAAAC
GAGCACACGTCTGAACTCTTTCCCTACACGACGCTCTTCCGATCTNNNCTGTGAGCCATGCTGAGGTAT
GAGCACACGTCTGAACTCTTTCCCTACACGACGCTCTTCCGATCTNNNTGTGAGATGTTTGCAGTTTGT
GAGCACACGTCTGAACTCTTTCCCTACACGACGCTCTTCCGATCTNNNTGACTGTCAGTCCCATATTCACT
AGCACACGTCTGAACTCTTTCCCTACACGACGCTCTTCCGATCTNNNGATCTCAAGCAGTCAAGAAAG
AGCACACGTCTGAACTCTTTCCCTACACGACGCTCTTCCGATCTNNNACAGAAGAAATAACTGGCTGGC
AGCACACGTCTGAACTCTTTCCCTACACGACGCTCTTCCGATCTNNNTCTCTTTGTGGCTTTGATTTGCA
GAGCACACGTCTGAACTCTTTCCCTACACGACGCTCTTCCGATCTNNNGAGGTAAATTCAAGGAGCCC
GAGCACACGTCTGAACTCTTTCCCTACACGACGCTCTTCCGATCTNNNAAGCTCACTATTGTCCTCTGCA
AGCACACGTCTGAACTCTTTCCCTACACGACGCTCTTCCGATCTNNNCTTGTGCAACTAAGGGAAAGTG
AGCACACGTCTGAACTCTTTCCCTACACGACGCTCTTCCGATCTNNNGGTAGGACTACTAGTGGTTTGC
AGCACACGTCTGAACTCTTTCCCTACACGACGCTCTTCCGATCTNNNAGGACTGCTAGTTTTTCTCCCTG
AGCACACGTCTGAACTCTTTCCCTACACGACGCTCTTCCGATCTNNNAGTTTCTCTCCCTGTATTGGCTC
GAGCACACGTCTGAACTCTTTCCCTACACGACGCTCTTCCGATCTNNNAGGGGTGTGGTTCTTTTCTTT
AGCACACGTCTGAACTCTTTCCCTACACGACGCTCTTCCGATCTNNNGACAAGACTGCTAGTGTCTCTCC
GAGCACACGTCTGAACTCTTTCCCTACACGACGCTCTTCCGATCTNNNCGCCTGTTAGTGTCTTTTTCTG
AGCACACGTCTGAACTCTTTCCCTACACGACGCTCTTCCGATCTNNNCCCTGGGTTGGTCATATGACAA
AGCACACGTCTGAACTCTTTCCCTACACGACGCTCTTCCGATCTNNNTGTGTTTCTGTCAGAGAGTTGC
GAGCACACGTCTGAACTCTTTCCCTACACGACGCTCTTCCGATCTNNNCACTTCTGTTTTCCTGCCCAAA
AGCACACGTCTGAACTCTTTCCCTACACGACGCTCTTCCGATCTNNNGTGGCATTATCTGTGCTTACTG
GAGCACACGTCTGAACTCTTTCCCTACACGACGCTCTTCCGATCTNNNCTGCATGAGCTGTGCTATTTC
GAGCACACGTCTGAACTCTTTCCCTACACGACGCTCTTCCGATCTNNNTTCAGAAGGAGGCCTAATTTG
AGCACACGTCTGAACTCTTTCCCTACACGACGCTCTTCCGATCTNNNAGGAAGTCATGTGCATGTCATC
GAGCACACGTCTGAACTCTTTCCCTACACGACGCTCTTCCGATCTNNNGCACCTGTTGTTTCCTGACTTT
GAGCACACGTCTGAACTCTTTCCCTACACGACGCTCTTCCGATCTNNNCCTGGATTGGAGAGGAGAAC
AGCACACGTCTGAACTCTTTCCCTACACGACGCTCTTCCGATCTNNNTTCAAGTGATTCTCCTGCCTCAG
AGCACACGTCTGAACTCTTTCCCTACACGACGCTCTTCCGATCTNNNTGTGGGGCACTGAGTATACAA
AGCACACGTCTGAACTCTTTCCCTACACGACGCTCTTCCGATCTNNNGGCTCTCAGGTTCTAGCAATTCT
GAGCACACGTCTGAACTCTTTCCCTACACGACGCTCTTCCGATCTNNNGGATTGGTCCAAATAACCTGA
GAGCACACGTCTGAACTCTTTCCCTACACGACGCTCTTCCGATCTNNNTCAACTGCCCAAAACAAGTTAG
AGCACACGTCTGAACTCTTTCCCTACACGACGCTCTTCCGATCTNNNCCAAAATCCAAAGTGCTGGGAA
AGCACACGTCTGAACTCTTTCCCTACACGACGCTCTTCCGATCTNNNCCAAAATCCAAAGTGCTGGGAA
AGCACACGTCTGAACTCTTTCCCTACACGACGCTCTTCCGATCTNNNCCAAAATCCAAAGTGCTGGGAA
AGCACACGTCTGAACTCTTTCCCTACACGACGCTCTTCCGATCTNNNCCAAAATCCAAAGTGCTGGGAA
AGCACACGTCTGAACTCTTTCCCTACACGACGCTCTTCCGATCTNNNCCAAAATCCAAAGTGCTGGGAA
GAGCACACGTCTGAACTCTTTCCCTACACGACGCTCTTCCGATCTNNNAAGTCCAACATGCTGGGAATA
AGCACACGTCTGAACTCTTTCCCTACACGACGCTCTTCCGATCTNNNTTGAACAAAATAGGATCGCCCA
GAGCACACGTCTGAACTCTTTCCCTACACGACGCTCTTCCGATCTNNNCAGTGTAGTGTGCTCCTCTTCT

GAGCACACGTCTGAACTCTTTCCCTACACGACGCTCTTCCGATCTNNNCTATGGAGTTGGAAGCACCTT
AGCACACGTCTGAACTCTTTCCCTACACGACGCTCTTCCGATCTNNNGAGAGAGCAAACCTAGGGAACA
AGCACACGTCTGAACTCTTTCCCTACACGACGCTCTTCCGATCTNNNGCTTGGCAGTGAACTGAATTCT
AGCACACGTCTGAACTCTTTCCCTACACGACGCTCTTCCGATCTNNNTAAGAGAATGCTGGGATGCAG
GAGCACACGTCTGAACTCTTTCCCTACACGACGCTCTTCCGATCTNNNAACAATCTCTGTGTCGTCTGTC
GAGCACACGTCTGAACTCTTTCCCTACACGACGCTCTTCCGATCTNNNTGCAAGTATCCATAGTTTGGC
AGCACACGTCTGAACTCTTTCCCTACACGACGCTCTTCCGATCTNNNACTGCTAGTGTCTCTCTCTGGAT
GAGCACACGTCTGAACTCTTTCCCTACACGACGCTCTTCCGATCTNNNTCCTTCCTTCCTTCCTTCCTTTCC
AGCACACGTCTGAACTCTTTCCCTACACGACGCTCTTCCGATCTNNNGGTGACAGAGTGAGTCTTCAA
AGCACACGTCTGAACTCTTTCCCTACACGACGCTCTTCCGATCTNNNTATCCCAGGTTCAAGCAGTTCT
AGAGCACACGTCTGAACTCTTTCCCTACACGACGCTCTTCCGATCTNNNAGAAATGGGCATGTAAGGT
AGCACACGTCTGAACTCTTTCCCTACACGACGCTCTTCCGATCTNNNGAAAACCAACGGATCCCTGAGA
GAGCACACGTCTGAACTCTTTCCCTACACGACGCTCTTCCGATCTNNNGAGTGCTTTGAGACCCATAAG
AGCACACGTCTGAACTCTTTCCCTACACGACGCTCTTCCGATCTNNNTGTCTGCATTCAACTCACAGAG
AGCACACGTCTGAACTCTTTCCCTACACGACGCTCTTCCGATCTNNNGGTCTTTGGTGGAAAAGGGAA
AGCACACGTCTGAACTCTTTCCCTACACGACGCTCTTCCGATCTNNNCAAAAAGACACTGTGAGGGCT
GAGCACACGTCTGAACTCTTTCCCTACACGACGCTCTTCCGATCTNNNCAAAATCCCTGCTACATTTGCCA
AGCACACGTCTGAACTCTTTCCCTACACGACGCTCTTCCGATCTNNNATGGACTGTACCCATTTGCATCT
AGCACACGTCTGAACTCTTTCCCTACACGACGCTCTTCCGATCTNNNGCTACCACAATTCAACACAGCA
AGCACACGTCTGAACTCTTTCCCTACACGACGCTCTTCCGATCTNNNAATCAGGGCAATACTGGGTTCAT
AGCACACGTCTGAACTCTTTCCCTACACGACGCTCTTCCGATCTNNNACTCCTATGTTTGCTTCTCAGGA
AGCACACGTCTGAACTCTTTCCCTACACGACGCTCTTCCGATCTNNNTTCCTATCAGGCTAGCATTCTG
AGCACACGTCTGAACTCTTTCCCTACACGACGCTCTTCCGATCTNNNCAGACTTCTGGGAGACACTGAA
AGCACACGTCTGAACTCTTTCCCTACACGACGCTCTTCCGATCTNNNTGCAGTTACTTCCTCCACAGAAG
AGCACACGTCTGAACTCTTTCCCTACACGACGCTCTTCCGATCTNNNCACCGTGTCCAGCTAATTTTTGT
AGCACACGTCTGAACTCTTTCCCTACACGACGCTCTTCCGATCTNNNCGGCTTTTGTCATTCTCCATGTT
AGCACACGTCTGAACTCTTTCCCTACACGACGCTCTTCCGATCTNNNAAACAATCCTCTCACATCAGCCT
AGCACACGTCTGAACTCTTTCCCTACACGACGCTCTTCCGATCTNNNTTTAGGAGAGACGGGGTTTC
GAGCACACGTCTGAACTCTTTCCCTACACGACGCTCTTCCGATCTNNNAGCATTGGTACCAGCTCTTCTT
GAGCACACGTCTGAACTCTTTCCCTACACGACGCTCTTCCGATCTNNNGTCACCTCTTTCTTCTTTTGGCC
AGAGCACACGTCTGAACTCTTTCCCTACACGACGCTCTTCCGATCTNNNTGGGGAGTAGGTGCAAAAG
AGCACACGTCTGAACTCTTTCCCTACACGACGCTCTTCCGATCTNNNGCTTCGGGACTTGTTTCTCCTAA
AGCACACGTCTGAACTCTTTCCCTACACGACGCTCTTCCGATCTNNNGGCTGAGGTAGGAGGATTTCT
AGCACACGTCTGAACTCTTTCCCTACACGACGCTCTTCCGATCTNNNCTGGGATCCAATACCTTACGCTG
AGCACACGTCTGAACTCTTTCCCTACACGACGCTCTTCCGATCTNNNAGATAAGGTCTCACCGTGTTAG
AGCACACGTCTGAACTCTTTCCCTACACGACGCTCTTCCGATCTNNNAGATAGCGAAGTCCAAGGTCAA
AGCACACGTCTGAACTCTTTCCCTACACGACGCTCTTCCGATCTNNNCTCCAGCTGCACAAAAGCAATAT
AGAGCACACGTCTGAACTCTTTCCCTACACGACGCTCTTCCGATCTNNNCTGAGGCAGGAGAATCACTT
AGAGCACACGTCTGAACTCTTTCCCTACACGACGCTCTTCCGATCTNNNGAGTGGTTTGCTCATTGTCAT
GAGCACACGTCTGAACTCTTTCCCTACACGACGCTCTTCCGATCTNNNGTAGCACTGGAAGCAGGAAC
AGCACACGTCTGAACTCTTTCCCTACACGACGCTCTTCCGATCTNNNACAGGGTGCTCAGATATTTGGT
GAGCACACGTCTGAACTCTTTCCCTACACGACGCTCTTCCGATCTNNNTTGAGAAAAGACTTTCCCAGC
GAGCACACGTCTGAACTCTTTCCCTACACGACGCTCTTCCGATCTNNNAGGTGATGTGGAAACATGGA

AGCACACGTCTGAACTCTTTCCCTACACGACGCTCTTCCGATCTNNNCCCACCACTACACCTAGCTAAT
AGCACACGTCTGAACTCTTTCCCTACACGACGCTCTTCCGATCTNNNTGCTCAACATGACCAATCATCA
AGAGCACACGTCTGAACTCTTTCCCTACACGACGCTCTTCCGATCTNNNGGACATCAAACCCCTTCTC
AGCACACGTCTGAACTCTTTCCCTACACGACGCTCTTCCGATCTNNNCTCTATCTCCAGACCTTGTGAG
AGCACACGTCTGAACTCTTTCCCTACACGACGCTCTTCCGATCTNNNTTCATGCCATTCTCCTTCCTCAG
GCACACGTCTGAACTCTTTCCCTACACGACGCTCTTCCGATCTNNNCAGGAGTAAGTCACTGTTCCTG
AGCACACGTCTGAACTCTTTCCCTACACGACGCTCTTCCGATCTNNNGGAAAAAAAAAAGGCATTGC
AGCACACGTCTGAACTCTTTCCCTACACGACGCTCTTCCGATCTNNNTGTGGATGTCCTTTTTCACTCC
AGCACACGTCTGAACTCTTTCCCTACACGACGCTCTTCCGATCTNNNAACAATTCGACATCCCCCACTA
AGCACACGTCTGAACTCTTTCCCTACACGACGCTCTTCCGATCTNNNATTTCAGTCTTCAGGTCCAGGA
AGCACACGTCTGAACTCTTTCCCTACACGACGCTCTTCCGATCTNNNGGGTCCATTCAGATTCAGAC
AGCACACGTCTGAACTCTTTCCCTACACGACGCTCTTCCGATCTNNNAAGAAAAGCTGTAGGTGTTT
AGCACACGTCTGAACTCTTTCCCTACACGACGCTCTTCCGATCTNNNAGCTAAGTGTGGCTATGCTAG
AGCACACGTCTGAACTCTTTCCCTACACGACGCTCTTCCGATCTNNNTATTGGGATTACAGGTGTGAG
AGCACACGTCTGAACTCTTTCCCTACACGACGCTCTTCCGATCTNNNGGTGTGAATGCTGTATCCACA
GCACACGTCTGAACTCTTTCCCTACACGACGCTCTTCCGATCTNNNCTTATGCAGACGGACTCTCTAC
AGCACACGTCTGAACTCTTTCCCTACACGACGCTCTTCCGATCTNNNTCTCTGGGTGTAGAGCTTTAG
AGCACACGTCTGAACTCTTTCCCTACACGACGCTCTTCCGATCTNNNGACTTCCTCCTTAGTCAGAGC
AGCACACGTCTGAACTCTTTCCCTACACGACGCTCTTCCGATCTNNNTATAATGTGGCCAGCTAGGTG
GAGCACACGTCTGAACTCTTTCCCTACACGACGCTCTTCCGATCTNNNCTGAGTAGGTGGGATTACAG
AGCACACGTCTGAACTCTTTCCCTACACGACGCTCTTCCGATCTNNNCCGTCTTTTCTGTGACTCATCC
AGCACACGTCTGAACTCTTTCCCTACACGACGCTCTTCCGATCTNNNCAATAGGTATGAGACGGACC
GCACACGTCTGAACTCTTTCCCTACACGACGCTCTTCCGATCTNNNGGGCAACAAGAGTGAACAAAA
AGCACACGTCTGAACTCTTTCCCTACACGACGCTCTTCCGATCTNNNTTTTTGAGACAGGGTCTTACC
GAGCACACGTCTGAACTCTTTCCCTACACGACGCTCTTCCGATCTNNNCTGGAGTTCAGTGGTGTGAT
GCACACGTCTGAACTCTTTCCCTACACGACGCTCTTCCGATCTNNNTCTGTGTGTGTCTGTGTCCTAA
AGCACACGTCTGAACTCTTTCCCTACACGACGCTCTTCCGATCTNNNACTGTTGACTCTGCTTCTCATG
AGCACACGTCTGAACTCTTTCCCTACACGACGCTCTTCCGATCTNNNAATGGCACCATCACAGTTCACT
GCACACGTCTGAACTCTTTCCCTACACGACGCTCTTCCGATCTNNNGGATTCCTGGTTTGTTCTTGCA
AGCACACGTCTGAACTCTTTCCCTACACGACGCTCTTCCGATCTNNNTCTTGTTGCAAATGACTGGAT
AGCACACGTCTGAACTCTTTCCCTACACGACGCTCTTCCGATCTNNNCCTGTCTCTGGTGCTTTTCTAG
AGCACACGTCTGAACTCTTTCCCTACACGACGCTCTTCCGATCTNNNATTTCCTCTTCCCTTTCTGCCTT
AGCACACGTCTGAACTCTTTCCCTACACGACGCTCTTCCGATCTNNNACATTTGAGTAAGTGCTTTGC
AGCACACGTCTGAACTCTTTCCCTACACGACGCTCTTCCGATCTNNNTTGGGGGAACAGAAGGAATA
GAGCACACGTCTGAACTCTTTCCCTACACGACGCTCTTCCGATCTNNNGAGAAGCTGGGATTACAAG
AGCACACGTCTGAACTCTTTCCCTACACGACGCTCTTCCGATCTNNNAGGGAGTAGAGAACAAAGG
GCACACGTCTGAACTCTTTCCCTACACGACGCTCTTCCGATCTNNNAACTTTCACAAAGGAGGGGAG
AGCACACGTCTGAACTCTTTCCCTACACGACGCTCTTCCGATCTNNNTTTCCATCTGCTGTCCATTGGT
AGCACACGTCTGAACTCTTTCCCTACACGACGCTCTTCCGATCTNNNGAGGAAGGAAGGAAGGAAG
GCACACGTCTGAACTCTTTCCCTACACGACGCTCTTCCGATCTNNNACTGGATCATGATGGCAGTTTC
GCACACGTCTGAACTCTTTCCCTACACGACGCTCTTCCGATCTNNNGTCCAGACTTCACCACTACACAA
AGCACACGTCTGAACTCTTTCCCTACACGACGCTCTTCCGATCTNNNGCCACCACAGTCAGCTAATTTT
GCACACGTCTGAACTCTTTCCCTACACGACGCTCTTCCGATCTNNNGTTGCTTGCTGGAAAAGACATG

GAGCACACGTCTGAACTCTTTCCCTACACGACGCTCTTCCGATCTNNNTGGTCCTGGAATTTTTCACTG
GAGCACACGTCTGAACTCTTTCCCTACACGACGCTCTTCCGATCTNNNAGGGAATCTGTCTCTCACTCTC
AGCACACGTCTGAACTCTTTCCCTACACGACGCTCTTCCGATCTNNNCTCCTGAATGACACTAGTGCCA
AGCACACGTCTGAACTCTTTCCCTACACGACGCTCTTCCGATCTNNNCACCACAACTGGCGAATTTTTTG
GAGCACACGTCTGAACTCTTTCCCTACACGACGCTCTTCCGATCTNNNTTTGCCTTGTGAGGTTTCTGT
AGCACACGTCTGAACTCTTTCCCTACACGACGCTCTTCCGATCTNNNATGCTGGTGTTACAGGTGTTAC
AGCACACGTCTGAACTCTTTCCCTACACGACGCTCTTCCGATCTNNNGTGTGTGTGATTTAGAGCA
AGCACACGTCTGAACTCTTTCCCTACACGACGCTCTTCCGATCTNNNATCTAGTATAACCTGGTGGGGG
AGCACACGTCTGAACTCTTTCCCTACACGACGCTCTTCCGATCTNNNAGGTTGAAGTAATTCTCGTGCC
AGCACACGTCTGAACTCTTTCCCTACACGACGCTCTTCCGATCTNNNGTCTTTGTCCAGGGTCTCATGAT
AGCACACGTCTGAACTCTTTCCCTACACGACGCTCTTCCGATCTNNNATAGGGAGTCCTTTCTGCATTG
AGCACACGTCTGAACTCTTTCCCTACACGACGCTCTTCCGATCTNNNGATGTTCTCGATCTCCTGACCTC
GAGCACACGTCTGAACTCTTTCCCTACACGACGCTCTTCCGATCTNNNATAGGGACTGTTCAAAGCT
AGCACACGTCTGAACTCTTTCCCTACACGACGCTCTTCCGATCTNNNGTTATTTGGGTCAAGGGCTGT
GAGCACACGTCTGAACTCTTTCCCTACACGACGCTCTTCCGATCTNNNCTCCTGAGTAGCTGGGATTA
GAGCACACGTCTGAACTCTTTCCCTACACGACGCTCTTCCGATCTNNNTGGGGAATAGTAGCAGAGG
AGCACACGTCTGAACTCTTTCCCTACACGACGCTCTTCCGATCTNNNCCTTTATTCCCTTTCGTCCTCCTC
GAGCACACGTCTGAACTCTTTCCCTACACGACGCTCTTCCGATCTNNNTACTCTTACTGAGACCTCACGG
GAGCACACGTCTGAACTCTTTCCCTACACGACGCTCTTCCGATCTNNNGGACTCAAAAGCATGCAAAC
GAGCACACGTCTGAACTCTTTCCCTACACGACGCTCTTCCGATCTNNNGGGAGTGTGAGTTTTACGCTT
AGCACACGTCTGAACTCTTTCCCTACACGACGCTCTTCCGATCTNNNTAATTCATCTGCCTCAGCCTCTC
GAGCACACGTCTGAACTCTTTCCCTACACGACGCTCTTCCGATCTNNNAAGGTGTTTTTGTGTGTGTG
GAGCACACGTCTGAACTCTTTCCCTACACGACGCTCTTCCGATCTNNNTGATGACGCAATCTTAGCTCA
GAGCACACGTCTGAACTCTTTCCCTACACGACGCTCTTCCGATCTNNNGGGAGCTTTCTGTCCTGTATC
AGCACACGTCTGAACTCTTTCCCTACACGACGCTCTTCCGATCTNNNGTTCCACACGCTTTGTTTCCTAA
AGCACACGTCTGAACTCTTTCCCTACACGACGCTCTTCCGATCTNNNTGCCTCAAGTACAGGGATTACA
AGCACACGTCTGAACTCTTTCCCTACACGACGCTCTTCCGATCTNNNATCCTTAGGCCTCTGGAGACAT
AGCACACGTCTGAACTCTTTCCCTACACGACGCTCTTCCGATCTNNNTAACAAGCCGCTCTCCCTAATTT
AGCACACGTCTGAACTCTTTCCCTACACGACGCTCTTCCGATCTNNNGGAATACCAAGGCAGAGAAA
AGCACACGTCTGAACTCTTTCCCTACACGACGCTCTTCCGATCTNNNTGGAGAGCTTCACTGATTCTCTG
AGCACACGTCTGAACTCTTTCCCTACACGACGCTCTTCCGATCTNNNCTGGAGCATTTCCCCAATGTTTT
AGCACACGTCTGAACTCTTTCCCTACACGACGCTCTTCCGATCTNNNATCAGTAGTGGGATTGCTGGAT
AGCACACGTCTGAACTCTTTCCCTACACGACGCTCTTCCGATCTNNNGATCAAATTGGTCCCAATGGCT
GAGCACACGTCTGAACTCTTTCCCTACACGACGCTCTTCCGATCTNNNTCTCCAGAAGGTGAGGAAGG
AGCACACGTCTGAACTCTTTCCCTACACGACGCTCTTCCGATCTNNNTCCAATGACTTTTTCCTCCCCTT
GAGCACACGTCTGAACTCTTTCCCTACACGACGCTCTTCCGATCTNNNTAAGCTTTTTGATGTGCTGCT
AGAGCACACGTCTGAACTCTTTCCCTACACGACGCTCTTCCGATCTNNNGAAAAGGATTCACCCACTTT
AGCACACGTCTGAACTCTTTCCCTACACGACGCTCTTCCGATCTNNNTTGAGGTTTGGAAATCTTGCA
GAGCACACGTCTGAACTCTTTCCCTACACGACGCTCTTCCGATCTNNNGGTGCTGACATTGCTAGTATT
GAGCACACGTCTGAACTCTTTCCCTACACGACGCTCTTCCGATCTNNNACAGCTCTGGATTCTGTCTCA
GAGCACACGTCTGAACTCTTTCCCTACACGACGCTCTTCCGATCTNNNCTTTTGCCCAGGCTGTAATGT
GAGCACACGTCTGAACTCTTTCCCTACACGACGCTCTTCCGATCTNNNTCTCCTCTCCTTCTCCTTCTTCT
AGCACACGTCTGAACTCTTTCCCTACACGACGCTCTTCCGATCTNNNTCAATCTCAGCTTACTGCAACCT

AGCACACGTCTGAACTCTTTCCCTACACGACGCTCTTCCGATCTNNNGTAGAGTCAAGGTTTCACCGT
GCACACGTCTGAACTCTTTCCCTACACGACGCTCTTCCGATCTNNNCGGTGGAAAATGGAGGAAGTA
AGCACACGTCTGAACTCTTTCCCTACACGACGCTCTTCCGATCTNNNTTCAGAAATAGCAGGCTTGTGG
AGCACACGTCTGAACTCTTTCCCTACACGACGCTCTTCCGATCTNNNTCCATCCATATCTGAGTGACTCT
AGCACACGTCTGAACTCTTTCCCTACACGACGCTCTTCCGATCTNNNTGGGGATGGCATTGAATCTGT
GAGCACACGTCTGAACTCTTTCCCTACACGACGCTCTTCCGATCTNNNTGACAGGAAAGTTCGATTTC
AGCACACGTCTGAACTCTTTCCCTACACGACGCTCTTCCGATCTNNNTCTTTTCTTTGCAACCTCACCA
AGCACACGTCTGAACTCTTTCCCTACACGACGCTCTTCCGATCTNNNCACACACACACTCTCTCTCTCTT
GCACACGTCTGAACTCTTTCCCTACACGACGCTCTTCCGATCTNNNAGGTACATCTGTGTTGTCACCAA
AGCACACGTCTGAACTCTTTCCCTACACGACGCTCTTCCGATCTNNNGAAAATTTGAGAATGGCCTGC
AGCACACGTCTGAACTCTTTCCCTACACGACGCTCTTCCGATCTNNNCGCAACTAACATCTCAGGCTTT
GCACACGTCTGAACTCTTTCCCTACACGACGCTCTTCCGATCTNNNTGCATGGAAGTTCAGGACATA
AGCACACGTCTGAACTCTTTCCCTACACGACGCTCTTCCGATCTNNNTCTGCCTCCTAGTTTCAAGCAAT
GCACACGTCTGAACTCTTTCCCTACACGACGCTCTTCCGATCTNNNAGAGTTACTCAACAGGAAACTT
AGCACACGTCTGAACTCTTTCCCTACACGACGCTCTTCCGATCTNNNGCTCAGGAAATCTACCCACTT
AGCACACGTCTGAACTCTTTCCCTACACGACGCTCTTCCGATCTNNNGTAAGTTCAGCCATGAGCCATT
AGCACACGTCTGAACTCTTTCCCTACACGACGCTCTTCCGATCTNNNGAAGCTCCGATATTCCCCAGAA
AGCACACGTCTGAACTCTTTCCCTACACGACGCTCTTCCGATCTNNNTCCCTAGGCATAAAAACTCGG
GAGCACACGTCTGAACTCTTTCCCTACACGACGCTCTTCCGATCTNNNGATCACACAGATCCAGCCATT
AGCACACGTCTGAACTCTTTCCCTACACGACGCTCTTCCGATCTNNNAGACAAGATCTGGCTCTGTCA
AGCACACGTCTGAACTCTTTCCCTACACGACGCTCTTCCGATCTNNNTTCCTGCAGAATAGGATAAGC
AGCACACGTCTGAACTCTTTCCCTACACGACGCTCTTCCGATCTNNNTGCGGTTTTTGTCACTACTTTC
GCACACGTCTGAACTCTTTCCCTACACGACGCTCTTCCGATCTNNNTCTTGGAAGACCTGGAGTTTTG
AGCACACGTCTGAACTCTTTCCCTACACGACGCTCTTCCGATCTNNNTGTAGTGGTCAGTCATGGCTT
GCACACGTCTGAACTCTTTCCCTACACGACGCTCTTCCGATCTNNNTCATCTCCTGCAAACACACAAAC
AGCACACGTCTGAACTCTTTCCCTACACGACGCTCTTCCGATCTNNNCTCCACCTCTCTAAGTACTGGG
GCACACGTCTGAACTCTTTCCCTACACGACGCTCTTCCGATCTNNNCTGAAGTACAGTGATGCAATG
GAGCACACGTCTGAACTCTTTCCCTACACGACGCTCTTCCGATCTNNNAATGGCCCGATCTAGATTCAC
AGCACACGTCTGAACTCTTTCCCTACACGACGCTCTTCCGATCTNNNCCCACTGGATCTAGTATGGCA
AGCACACGTCTGAACTCTTTCCCTACACGACGCTCTTCCGATCTNNNGCCCAGCCATCTTTCTTTCTTTC
AGCACACGTCTGAACTCTTTCCCTACACGACGCTCTTCCGATCTNNNAGCTTTTTGATTTGCTGCTGG
GAGCACACGTCTGAACTCTTTCCCTACACGACGCTCTTCCGATCTNNNCAGGGACATTGGCCAGAAAA
GCACACGTCTGAACTCTTTCCCTACACGACGCTCTTCCGATCTNNNGTCAGTGGTAGTTTGATGGGG
AGCACACGTCTGAACTCTTTCCCTACACGACGCTCTTCCGATCTNNNTGTGCCCGGCATAATTTTTCTT
GCACACGTCTGAACTCTTTCCCTACACGACGCTCTTCCGATCTNNNTCCCCTTCTTGTTTCTTCCTCAAC
AGCACACGTCTGAACTCTTTCCCTACACGACGCTCTTCCGATCTNNNTTCAAAACGTTATCCAGCTGTG
AGCACACGTCTGAACTCTTTCCCTACACGACGCTCTTCCGATCTNNNATCTGGGATTACATGCACCTAC
GCACACGTCTGAACTCTTTCCCTACACGACGCTCTTCCGATCTNNNTCTCTTCTTTCCCTGACAAGTGG
AGCACACGTCTGAACTCTTTCCCTACACGACGCTCTTCCGATCTNNNAGGTAAGCTCCGAAAACCTCA
AGCACACGTCTGAACTCTTTCCCTACACGACGCTCTTCCGATCTNNNTATCTCCCGATGCTATCTCTCCC
AGCACACGTCTGAACTCTTTCCCTACACGACGCTCTTCCGATCTNNNTGTCATGGGGTTTGTTGTACA
AGCACACGTCTGAACTCTTTCCCTACACGACGCTCTTCCGATCTNNNAGGTGAGTTGACAAATAGCCT
GCACACGTCTGAACTCTTTCCCTACACGACGCTCTTCCGATCTNNNCTCACCTCAGCATCTCTAGTAGC

AGCACACGTCTGAACTCTTTCCCTACACGACGCTCTTCCGATCTNNNAGGGAAGGCAAATACAGAGTC
GAGCACACGTCTGAACTCTTTCCCTACACGACGCTCTTCCGATCTNNNGTATTTTTGTACGGCTGCACAC
AGCACACGTCTGAACTCTTTCCCTACACGACGCTCTTCCGATCTNNNTGTCATGATCTTGGCTCACTGAA
AGCACACGTCTGAACTCTTTCCCTACACGACGCTCTTCCGATCTNNNGTCATCTTCTCTCTCCTGTTGCAC
GAGCACACGTCTGAACTCTTTCCCTACACGACGCTCTTCCGATCTNNNGCAGTTGGAGAGCATGAAAC
GAGCACACGTCTGAACTCTTTCCCTACACGACGCTCTTCCGATCTNNNTCCCCAGCTTTTTGTTTTTTG
GAGCACACGTCTGAACTCTTTCCCTACACGACGCTCTTCCGATCTNNNGATGGAGTCTCACTCTGTCACA
GAGCACACGTCTGAACTCTTTCCCTACACGACGCTCTTCCGATCTNNNTCATTTGGGTGTACAGTAAAG
AGCACACGTCTGAACTCTTTCCCTACACGACGCTCTTCCGATCTNNNCTGGAGGTCCTGCAGGTAATAT
GAGCACACGTCTGAACTCTTTCCCTACACGACGCTCTTCCGATCTNNNCACACAGCTGAACAAATGTCCA
GAGCACACGTCTGAACTCTTTCCCTACACGACGCTCTTCCGATCTNNNATATGTGTGTGTGTGTGTGTG
AGCACACGTCTGAACTCTTTCCCTACACGACGCTCTTCCGATCTNNNGCTGGGAGTAGATGGGATTAC
AGCACACGTCTGAACTCTTTCCCTACACGACGCTCTTCCGATCTNNNTGCCTTGTGAAGAGTTTTGCTTT
GAGCACACGTCTGAACTCTTTCCCTACACGACGCTCTTCCGATCTNNNAGTAGCTGGAATTACTGACAG
AGAGCACACGTCTGAACTCTTTCCCTACACGACGCTCTTCCGATCTNNNGTGAGCTGAGATCATGCTAC
GAGCACACGTCTGAACTCTTTCCCTACACGACGCTCTTCCGATCTNNNCTTCTCCACAAGCCAGTATTCC
GAGCACACGTCTGAACTCTTTCCCTACACGACGCTCTTCCGATCTNNNCGTTTTTGGTGGGAGTTCTCA
AGCACACGTCTGAACTCTTTCCCTACACGACGCTCTTCCGATCTNNNCCATCACGCCGAGCTAATTTTTT
GAGCACACGTCTGAACTCTTTCCCTACACGACGCTCTTCCGATCTNNNTGCATCTCTCTCTCTCTCTCT
AGCACACGTCTGAACTCTTTCCCTACACGACGCTCTTCCGATCTNNNTGATCTCTCCATGTTTTCCAGGTC
GAGCACACGTCTGAACTCTTTCCCTACACGACGCTCTTCCGATCTNNNTTAGATGGAGTCTCACTGTGT
AGCACACGTCTGAACTCTTTCCCTACACGACGCTCTTCCGATCTNNNGCAGAAGGCTTGTTTTATTGCAT
GAGCACACGTCTGAACTCTTTCCCTACACGACGCTCTTCCGATCTNNNAGTAGCTGAGAATACAGGCAT
AGCACACGTCTGAACTCTTTCCCTACACGACGCTCTTCCGATCTNNNTTCACATCATTCTCCTGCCTCAGC
AGCACACGTCTGAACTCTTTCCCTACACGACGCTCTTCCGATCTNNNAAAAAGGAATGCTTACCAGTG
AGAGCACACGTCTGAACTCTTTCCCTACACGACGCTCTTCCGATCTNNNTTCTTCAGTGCTGCATGTCAA
AGCACACGTCTGAACTCTTTCCCTACACGACGCTCTTCCGATCTNNNCATTGCCTGTGTTCTCCAGATTC
AGCACACGTCTGAACTCTTTCCCTACACGACGCTCTTCCGATCTNNNCCAAGTAGCTGGTACTGACAGT
GAGCACACGTCTGAACTCTTTCCCTACACGACGCTCTTCCGATCTNNNGTATCTGTGTGTGTGTGTGTG
AGAGCACACGTCTGAACTCTTTCCCTACACGACGCTCTTCCGATCTNNNACCTCCTGAGTAGCTGGGAT
AGAGCACACGTCTGAACTCTTTCCCTACACGACGCTCTTCCGATCTNNNCCCAGAAGCTTCTTTGGTAGA
GAGCACACGTCTGAACTCTTTCCCTACACGACGCTCTTCCGATCTNNNTTTTGTGTGTGTGTGTGTGT
AGCACACGTCTGAACTCTTTCCCTACACGACGCTCTTCCGATCTNNNCTTGAGTGGGTGAGAGAAAGA
AGCACACGTCTGAACTCTTTCCCTACACGACGCTCTTCCGATCTNNNCAGGATGCTCCCAAGTACTGAT
AGCACACGTCTGAACTCTTTCCCTACACGACGCTCTTCCGATCTNNNCCCAGGTTCAAGCAATTCTCTTG
AGCACACGTCTGAACTCTTTCCCTACACGACGCTCTTCCGATCTNNNGTGTGAAATGCTTCTGGCTTTCT
AGCACACGTCTGAACTCTTTCCCTACACGACGCTCTTCCGATCTNNNCTGTAGCCTCAAACTTCTGGGAT
AGCACACGTCTGAACTCTTTCCCTACACGACGCTCTTCCGATCTNNNAGTGGCATGATCTCATCTCACTC
GAGCACACGTCTGAACTCTTTCCCTACACGACGCTCTTCCGATCTNNNTTTTTAGCAGGAGGATGTGAC
GAGCACACGTCTGAACTCTTTCCCTACACGACGCTCTTCCGATCTNNNCCTCTCAGGTTCACGCTATTCTC
AGCACACGTCTGAACTCTTTCCCTACACGACGCTCTTCCGATCTNNNTGGCATTCTTTTACATGTTTGCAC
AGCACACGTCTGAACTCTTTCCCTACACGACGCTCTTCCGATCTNNNAGACAGAGTCTCACTGTTTCACC
AGCACACGTCTGAACTCTTTCCCTACACGACGCTCTTCCGATCTNNNCCAAAAGAGCAAGACATTCCAG

AGCACACGTCTGAACTCTTTCCCTACACGACGCTCTTCCGATCTNNNACTGTGAGATGTGTGTGCATA
AGCACACGTCTGAACTCTTTCCCTACACGACGCTCTTCCGATCTNNNAATGCAATGATTGTCAACCTG
AGCACACGTCTGAACTCTTTCCCTACACGACGCTCTTCCGATCTNNNGGTCCCAGAAATCTCTTGGCAT
AGCACACGTCTGAACTCTTTCCCTACACGACGCTCTTCCGATCTNNNCGCCTGGCTTTCCTTCATTTATT
AGCACACGTCTGAACTCTTTCCCTACACGACGCTCTTCCGATCTNNNCCTTTCCTAGTGCTTTGGGTTT
GCACACGTCTGAACTCTTTCCCTACACGACGCTCTTCCGATCTNNNTATCTTTGAGGCCTGCAAAACTC
GCACACGTCTGAACTCTTTCCCTACACGACGCTCTTCCGATCTNNNGTTTCATCCATGTTGTAGCGTGT
GCACACGTCTGAACTCTTTCCCTACACGACGCTCTTCCGATCTNNNCTTTGGACCTCTCCCATGTGTTA
AGCACACGTCTGAACTCTTTCCCTACACGACGCTCTTCCGATCTNNNTTCACACCATTCTTCTGCCTCAG
AGCACACGTCTGAACTCTTTCCCTACACGACGCTCTTCCGATCTNNNCAGAATCTCACTCTGTCACCCA
AGCACACGTCTGAACTCTTTCCCTACACGACGCTCTTCCGATCTNNNATGCTCCTATGTGTCTGATCTGG
GAGCACACGTCTGAACTCTTTCCCTACACGACGCTCTTCCGATCTNNNCTCCATCCAGTTTTGTTGCCTT
GAGCACACGTCTGAACTCTTTCCCTACACGACGCTCTTCCGATCTNNNTTTTGGTGAATCTGCAACTGG
AGCACACGTCTGAACTCTTTCCCTACACGACGCTCTTCCGATCTNNNTTGTGTGACACGGAAATGCAAT
GAGCACACGTCTGAACTCTTTCCCTACACGACGCTCTTCCGATCTNNNACAAAGACACCTGGCCTAAAA
AGCACACGTCTGAACTCTTTCCCTACACGACGCTCTTCCGATCTNNNATGCAATGTTGAGACCTGGAG
AGCACACGTCTGAACTCTTTCCCTACACGACGCTCTTCCGATCTNNNTGGTGAAGGTAGATCCAAACG
GCACACGTCTGAACTCTTTCCCTACACGACGCTCTTCCGATCTNNNACCTCTGAAAATGAAAGGCTGT
AGCACACGTCTGAACTCTTTCCCTACACGACGCTCTTCCGATCTNNNACCTACCACCTAGATTTCACGA
GAGCACACGTCTGAACTCTTTCCCTACACGACGCTCTTCCGATCTNNNTGCTGGGTTCAGGCAATTATA
GAGCACACGTCTGAACTCTTTCCCTACACGACGCTCTTCCGATCTNNNCTCCTGAGTAGCTGGGATTAC
AGCACACGTCTGAACTCTTTCCCTACACGACGCTCTTCCGATCTNNNCATGTGTTTTCTTGGCTGCCTA
GCACACGTCTGAACTCTTTCCCTACACGACGCTCTTCCGATCTNNNGGGCTCCTAAATATCCCTTTGCA
GAGCACACGTCTGAACTCTTTCCCTACACGACGCTCTTCCGATCTNNNTCAATGGGCTCCAAAATGTCT
AGCACACGTCTGAACTCTTTCCCTACACGACGCTCTTCCGATCTNNNAAAGTGCTGGGATTAGAGGA
AGCACACGTCTGAACTCTTTCCCTACACGACGCTCTTCCGATCTNNNAGAAAACTGCACTTCAAGCTTG
GCACACGTCTGAACTCTTTCCCTACACGACGCTCTTCCGATCTNNNCTCCTAGATGCACCATGCTGTAT
AGCACACGTCTGAACTCTTTCCCTACACGACGCTCTTCCGATCTNNNGTATCATAGCTGGGACTACAG
GCACACGTCTGAACTCTTTCCCTACACGACGCTCTTCCGATCTNNNATCTTCTGACCTCATGATCCACCC
AGCACACGTCTGAACTCTTTCCCTACACGACGCTCTTCCGATCTNNNGTAGTGTAGTGATCTGGGCTC
GCACACGTCTGAACTCTTTCCCTACACGACGCTCTTCCGATCTNNNCACGCCACTACACCTAGGTATTT
AGCACACGTCTGAACTCTTTCCCTACACGACGCTCTTCCGATCTNNNGTTTTGAACTCCTGACCTCAAG
AGCACACGTCTGAACTCTTTCCCTACACGACGCTCTTCCGATCTNNNATTGTCTTCTTTTAGGCCCCAG
AGCACACGTCTGAACTCTTTCCCTACACGACGCTCTTCCGATCTNNNAACTTGTGTTTTCCATGCCATCC
GCACACGTCTGAACTCTTTCCCTACACGACGCTCTTCCGATCTNNNCTTTCAAGCAGCTGGGATTACT
AGCACACGTCTGAACTCTTTCCCTACACGACGCTCTTCCGATCTNNNGAGACACTGAGCTAGCTTGAA
AGCACACGTCTGAACTCTTTCCCTACACGACGCTCTTCCGATCTNNNTCAACATGTCAGTGATGCTCTC
GAGCACACGTCTGAACTCTTTCCCTACACGACGCTCTTCCGATCTNNNCTCCTCACCTTGTGATCTTCCA
AGCACACGTCTGAACTCTTTCCCTACACGACGCTCTTCCGATCTNNNAAGATTCAAACTCTGAGGTTGG
AGCACACGTCTGAACTCTTTCCCTACACGACGCTCTTCCGATCTNNNAAGAAAGCTCTGAGTTATGGG
GAGCACACGTCTGAACTCTTTCCCTACACGACGCTCTTCCGATCTNNNATCACCACATTCCATTTTCTG
AGCACACGTCTGAACTCTTTCCCTACACGACGCTCTTCCGATCTNNNTGCTGAGATTACAGGCTTGAA
GCACACGTCTGAACTCTTTCCCTACACGACGCTCTTCCGATCTNNNTTGTCCACATGTTGTCTGTAGGT

AGCACACGTCTGAACTCTTTCCCTACACGACGCTCTTCCGATCTNNNGTGTGTGTTTGTGTGTGTGTGT
AGCACACGTCTGAACTCTTTCCCTACACGACGCTCTTCCGATCTNNNTCCCCACAAAAGACATGAACTCA
AGCACACGTCTGAACTCTTTCCCTACACGACGCTCTTCCGATCTNNNCTGGTTATGAGACCTTCCCAACA
GAGCACACGTCTGAACTCTTTCCCTACACGACGCTCTTCCGATCTNNNACAGAGTGAAACCCTGTCTCA
AGCACACGTCTGAACTCTTTCCCTACACGACGCTCTTCCGATCTNNNTGCTATTGTGAATTGTGCTGCA
AGCACACGTCTGAACTCTTTCCCTACACGACGCTCTTCCGATCTNNNAGGCCTGTGAGCATATGGTAA
AGCACACGTCTGAACTCTTTCCCTACACGACGCTCTTCCGATCTNNNCCTGAGAGGAATTAGTCGTCCA
GAGCACACGTCTGAACTCTTTCCCTACACGACGCTCTTCCGATCTNNNATCTTCAGCTCTTGGAACTCAG
GAGCACACGTCTGAACTCTTTCCCTACACGACGCTCTTCCGATCTNNNAAAAACTTTCCCTGACAGAAG
GAGCACACGTCTGAACTCTTTCCCTACACGACGCTCTTCCGATCTNNNGTTGTGTGACTTGTCTGTGTG
AGAGCACACGTCTGAACTCTTTCCCTACACGACGCTCTTCCGATCTNNNTTATAGGCACCCAACATGACA
AGCACACGTCTGAACTCTTTCCCTACACGACGCTCTTCCGATCTNNNACACAGCAGTTTTTCCACCAAAT
GAGCACACGTCTGAACTCTTTCCCTACACGACGCTCTTCCGATCTNNNCCCCAAACCCATACCCTTCTATC
GAGCACACGTCTGAACTCTTTCCCTACACGACGCTCTTCCGATCTNNNTGAATTCAGTCCCCAAACTAGC
AGCACACGTCTGAACTCTTTCCCTACACGACGCTCTTCCGATCTNNNTTGTTTATCTGTGCAGCCATGTG
AGCACACGTCTGAACTCTTTCCCTACACGACGCTCTTCCGATCTNNNTCCGAACTCTAACTGGTGTTCTG
GAGCACACGTCTGAACTCTTTCCCTACACGACGCTCTTCCGATCTNNNTGAATGCTTGCTAGAGAGGTC
AGAGCACACGTCTGAACTCTTTCCCTACACGACGCTCTTCCGATCTNNNATCCATGAGGGGCATTTCTTC
AGCACACGTCTGAACTCTTTCCCTACACGACGCTCTTCCGATCTNNNGTTCCAAGTAAGCCCAAAACACA
GAGCACACGTCTGAACTCTTTCCCTACACGACGCTCTTCCGATCTNNNAGATTCATGGGTTTTTGACAGC
AGCACACGTCTGAACTCTTTCCCTACACGACGCTCTTCCGATCTNNNCAGGTCATAAGGCTTAGCAGCT
AGCACACGTCTGAACTCTTTCCCTACACGACGCTCTTCCGATCTNNNTGACAGGATTTGGTACCAGGAT
GCACACGTCTGAACTCTTTCCCTACACGACGCTCTTCCGATCTNNNGCCACCTAGATATTCAATGACCCA
AGCACACGTCTGAACTCTTTCCCTACACGACGCTCTTCCGATCTNNNTGGTCTAGAAACCTGCGATAAC
GAGCACACGTCTGAACTCTTTCCCTACACGACGCTCTTCCGATCTNNNCCAGGGTTCAAGCAATTCTTCT
AGCACACGTCTGAACTCTTTCCCTACACGACGCTCTTCCGATCTNNNACATACTAAGACAGGAGCACCT
GAGCACACGTCTGAACTCTTTCCCTACACGACGCTCTTCCGATCTNNNGAAGCTCTGGGATATGATGTG
AGCACACGTCTGAACTCTTTCCCTACACGACGCTCTTCCGATCTNNNGACCCTTGCATTCAACCTCTTCC
GAGCACACGTCTGAACTCTTTCCCTACACGACGCTCTTCCGATCTNNNAGGAAAAATACTCAGCCGGTC
AGCACACGTCTGAACTCTTTCCCTACACGACGCTCTTCCGATCTNNNTCCTGAGTAGATGGGACTACAG
GAGCACACGTCTGAACTCTTTCCCTACACGACGCTCTTCCGATCTNNNGTTCATAAGATGTTTTGGCCCG
AGCACACGTCTGAACTCTTTCCCTACACGACGCTCTTCCGATCTNNNAGCCAGTTATTTCAGCACCTTTT
GCACACGTCTGAACTCTTTCCCTACACGACGCTCTTCCGATCTNNNAAGCTATGTGCAGATTGAATGCT
GAGCACACGTCTGAACTCTTTCCCTACACGACGCTCTTCCGATCTNNNGTTGACTACATTCTCCTGCCTCA
GAGCACACGTCTGAACTCTTTCCCTACACGACGCTCTTCCGATCTNNNCTGAAATCCAGGGACCACTAG
GAGCACACGTCTGAACTCTTTCCCTACACGACGCTCTTCCGATCTNNNTAGGTGCAGTACAGTCAAACT
GCACACGTCTGAACTCTTTCCCTACACGACGCTCTTCCGATCTNNNTCACCTTCCTTTCCTCTCTCTCTCAT
GAGCACACGTCTGAACTCTTTCCCTACACGACGCTCTTCCGATCTNNNCCACACCCAGTAATGTTTGCAT
AGCACACGTCTGAACTCTTTCCCTACACGACGCTCTTCCGATCTNNNGACAAGGCCCTCTATGTTCAGAA
AGCACACGTCTGAACTCTTTCCCTACACGACGCTCTTCCGATCTNNNGTGTGACCATCACCTATGTCTGT
AGCACACGTCTGAACTCTTTCCCTACACGACGCTCTTCCGATCTNNNGATGGTCTCAATCTCCTGACCTC
GAGCACACGTCTGAACTCTTTCCCTACACGACGCTCTTCCGATCTNNNTGGCCTTCTTTGTCTCTTGTGAT
GAGCACACGTCTGAACTCTTTCCCTACACGACGCTCTTCCGATCTNNNTGAGCATCATCCGTTATTCAG

AGCACACGTCTGAACTCTTTCCCTACACGACGCTCTTCCGATCTNNNTCCCTTTCCATTTCCTTTTCCCTC
GAGCACACGTCTGAACTCTTTCCCTACACGACGCTCTTCCGATCTNNNCCCTACCTCTAAGATGTCTGGC
AGCACACGTCTGAACTCTTTCCCTACACGACGCTCTTCCGATCTNNNCCCCCCAAGAAAAGATGAGAGA
AGCACACGTCTGAACTCTTTCCCTACACGACGCTCTTCCGATCTNNNGATGGTTTCCAGCTTCATCCATG
GAGCACACGTCTGAACTCTTTCCCTACACGACGCTCTTCCGATCTNNNCAAGCTTACAGTAGTCCAACCC
AGAGCACACGTCTGAACTCTTTCCCTACACGACGCTCTTCCGATCTNNNTGTCTGTACCTCCTATCACCAC
GAGCACACGTCTGAACTCTTTCCCTACACGACGCTCTTCCGATCTNNNTTCAGAAGGAATGGTACCAGC
GAGCACACGTCTGAACTCTTTCCCTACACGACGCTCTTCCGATCTNNNACTCTTATCACCCAGGCTAGAC
GAGCACACGTCTGAACTCTTTCCCTACACGACGCTCTTCCGATCTNNNCCATGTTGCAAGTGATTCTCCT
AGCACACGTCTGAACTCTTTCCCTACACGACGCTCTTCCGATCTNNNAGACCCACATACGAGACACAATT
GCACACGTCTGAACTCTTTCCCTACACGACGCTCTTCCGATCTNNNATTGCTGCAATAAACATGGGAGT
GAGCACACGTCTGAACTCTTTCCCTACACGACGCTCTTCCGATCTNNNTGAGACTGTTTTTTCCAGCCCA
AGCACACGTCTGAACTCTTTCCCTACACGACGCTCTTCCGATCTNNNTGAAAGTTTTGGCAGTGGTTGT
GAGCACACGTCTGAACTCTTTCCCTACACGACGCTCTTCCGATCTNNNATTGAGAAGGCTGCAGTTAGG
GCACACGTCTGAACTCTTTCCCTACACGACGCTCTTCCGATCTNNNACAGTGGGGCAAGTTACAAAAA
GAGCACACGTCTGAACTCTTTCCCTACACGACGCTCTTCCGATCTNNNCAGGGTTTTACCATGTTAGCCA
AGCACACGTCTGAACTCTTTCCCTACACGACGCTCTTCCGATCTNNNTGTCGATTTTCTGACCTGGTGA
AGCACACGTCTGAACTCTTTCCCTACACGACGCTCTTCCGATCTNNNTCTAAAGTGCAGTGGCGTAATC
GAGCACACGTCTGAACTCTTTCCCTACACGACGCTCTTCCGATCTNNNTGGGTTCAAGTGATTCTCCTAC
GAGCACACGTCTGAACTCTTTCCCTACACGACGCTCTTCCGATCTNNNTCTCCTGTGAACTGAATTCGA
GAGCACACGTCTGAACTCTTTCCCTACACGACGCTCTTCCGATCTNNNTACCACAAAATGACAGCAGCAT
GCACACGTCTGAACTCTTTCCCTACACGACGCTCTTCCGATCTNNNCCTCTATTCCTGCTCTCTTCACCCAT
GAGCACACGTCTGAACTCTTTCCCTACACGACGCTCTTCCGATCTNNNGTATATGGCTCCACACAAATG
GAGCACACGTCTGAACTCTTTCCCTACACGACGCTCTTCCGATCTNNNTTTCTATCTATGGCTGTGGCTG
GAGCACACGTCTGAACTCTTTCCCTACACGACGCTCTTCCGATCTNNNAGTGTTGTGAGTGTTGGATGT
AGCACACGTCTGAACTCTTTCCCTACACGACGCTCTTCCGATCTNNNTGAACATTACTTGGACTCCAGCA
AGCACACGTCTGAACTCTTTCCCTACACGACGCTCTTCCGATCTNNNTAGGGTCCTGAAGTGAGTGTT
AGCACACGTCTGAACTCTTTCCCTACACGACGCTCTTCCGATCTNNNGGGAGTATTGTGAGGGTTGGA
AGCACACGTCTGAACTCTTTCCCTACACGACGCTCTTCCGATCTNNNATGGGAGTAGGATGGATTGAT
AGCACACGTCTGAACTCTTTCCCTACACGACGCTCTTCCGATCTNNNAAAAAGTTGAAACCGTTTTCGTC
AGCACACGTCTGAACTCTTTCCCTACACGACGCTCTTCCGATCTNNNGAGTAGTTGGGATTACAGGCTC
AGCACACGTCTGAACTCTTTCCCTACACGACGCTCTTCCGATCTNNNCCTGGGTTCAAGCAATTCTTCAC
AGCACACGTCTGAACTCTTTCCCTACACGACGCTCTTCCGATCTNNNGCATCATTCATGGCATGGATAG
GAGCACACGTCTGAACTCTTTCCCTACACGACGCTCTTCCGATCTNNNAGAAGGACACAAATGACAAAC
GCACACGTCTGAACTCTTTCCCTACACGACGCTCTTCCGATCTNNNTGTTCCTTCCACGTTGATGTACTC
AGAGCACACGTCTGAACTCTTTCCCTACACGACGCTCTTCCGATCTNNNCACAGGACAATGAATCTCTGC
GAGCACACGTCTGAACTCTTTCCCTACACGACGCTCTTCCGATCTNNNTGATCATTCTCCACAAACTGGG
GAGCACACGTCTGAACTCTTTCCCTACACGACGCTCTTCCGATCTNNNGAAACCACTTCTGTGTCATCTC
AGCACACGTCTGAACTCTTTCCCTACACGACGCTCTTCCGATCTNNNAATTCGTGGTATGCCCGTTTTTT
AGCACACGTCTGAACTCTTTCCCTACACGACGCTCTTCCGATCTNNNTTTCTGGTTTGGACTCTGGACAA
GAGCACACGTCTGAACTCTTTCCCTACACGACGCTCTTCCGATCTNNNAGGCAAGAGCAAACAAATTCG
GAGCACACGTCTGAACTCTTTCCCTACACGACGCTCTTCCGATCTNNNGCCATTTGTGTGGAGCCATATA
GAGCACACGTCTGAACTCTTTCCCTACACGACGCTCTTCCGATCTNNNGTCTCAAGCTCCTGACCAAGTA

AGCACACGTCTGAACTCTTTCCCTACACGACGCTCTTCCGATCTNNNGGAGAGTCAATTTGTATGCAG
GAGCACACGTCTGAACTCTTTCCCTACACGACGCTCTTCCGATCTNNNTTCATCCGGAATCTATCGGTT
AGCACACGTCTGAACTCTTTCCCTACACGACGCTCTTCCGATCTNNNTTTAGACTGTGGTCCCAATCAC
AGCACACGTCTGAACTCTTTCCCTACACGACGCTCTTCCGATCTNNNTTTTCTCACAGCTCTGGACATCA
GAGCACACGTCTGAACTCTTTCCCTACACGACGCTCTTCCGATCTNNNTGTGCAGACCAAGGAATTGA
AGCACACGTCTGAACTCTTTCCCTACACGACGCTCTTCCGATCTNNNTGGTGCTGACCTCATGAAGTA
GAGCACACGTCTGAACTCTTTCCCTACACGACGCTCTTCCGATCTNNNTACTATGCTGTCCTCTTCTGCA
GAGCACACGTCTGAACTCTTTCCCTACACGACGCTCTTCCGATCTNNNATGTACCACCATCCCCAGAAA
GAGCACACGTCTGAACTCTTTCCCTACACGACGCTCTTCCGATCTNNNAGTAGCTGAGATTACAAGTG
AGCACACGTCTGAACTCTTTCCCTACACGACGCTCTTCCGATCTNNNGCTCATGTCTGCCTAACTACCTA
AGCACACGTCTGAACTCTTTCCCTACACGACGCTCTTCCGATCTNNNACATACTGAGGATATGTGGCAG
GCACACGTCTGAACTCTTTCCCTACACGACGCTCTTCCGATCTNNNCTAATGCAGGCTGAAATGCTAG
GAGCACACGTCTGAACTCTTTCCCTACACGACGCTCTTCCGATCTNNNTACGACTTTCTCCCTAGGCAA
GAGCACACGTCTGAACTCTTTCCCTACACGACGCTCTTCCGATCTNNNTGGAGAGATTGGGTGTTTGT
AGCACACGTCTGAACTCTTTCCCTACACGACGCTCTTCCGATCTNNNACAATGGTGGTCGCATAAGAC
AGCACACGTCTGAACTCTTTCCCTACACGACGCTCTTCCGATCTNNNTCGTTGATTAGAGAGCCTGAAG
GAGCACACGTCTGAACTCTTTCCCTACACGACGCTCTTCCGATCTNNNCCGATCTCTAGAGTGTTGGTT
GAGCACACGTCTGAACTCTTTCCCTACACGACGCTCTTCCGATCTNNNTACCACAAAATGACAGCAGCA
GCACACGTCTGAACTCTTTCCCTACACGACGCTCTTCCGATCTNNNCCTCTATTCCTGCTCTCTTCACCCA
GAGCACACGTCTGAACTCTTTCCCTACACGACGCTCTTCCGATCTNNNGTATATGGCTCCACACAAATG
AGCACACGTCTGAACTCTTTCCCTACACGACGCTCTTCCGATCTNNNGTCTCCGTTGCGACAGAATTTT
GAGCACACGTCTGAACTCTTTCCCTACACGACGCTCTTCCGATCTNNNTTTCTATCTATGGCTGTGGCT
AGCACACGTCTGAACTCTTTCCCTACACGACGCTCTTCCGATCTNNNCCTTTCTCCTTTCCCTTTTCCCTC
AGCACACGTCTGAACTCTTTCCCTACACGACGCTCTTCCGATCTNNNCTTCAGTGTGCTCTGATGAACA
AGCACACGTCTGAACTCTTTCCCTACACGACGCTCTTCCGATCTNNNTATTATCTGTGTCACTGCAGCC
AGCACACGTCTGAACTCTTTCCCTACACGACGCTCTTCCGATCTNNNAACCCCTGCTTTGTTTTTCTCTG
AGCACACGTCTGAACTCTTTCCCTACACGACGCTCTTCCGATCTNNNATGTTATGTCTCTGGGTTCCAC
AGCACACGTCTGAACTCTTTCCCTACACGACGCTCTTCCGATCTNNNCTTCTTCACTGTACCTGCAGGAT
GCACACGTCTGAACTCTTTCCCTACACGACGCTCTTCCGATCTNNNTGACAGGATATTAGAACAGCAG
AGCACACGTCTGAACTCTTTCCCTACACGACGCTCTTCCGATCTNNNCTGCCTTTTTGCACTTGTTGTTG
AGCACACGTCTGAACTCTTTCCCTACACGACGCTCTTCCGATCTNNNGCATTGGGCTTCAGATTAGCAT
GAGCACACGTCTGAACTCTTTCCCTACACGACGCTCTTCCGATCTNNNCCGAACATTCCTTTTCTGGCAT
AGCACACGTCTGAACTCTTTCCCTACACGACGCTCTTCCGATCTNNNGTCTGTTCATGTACTTTGCCCA
AGCACACGTCTGAACTCTTTCCCTACACGACGCTCTTCCGATCTNNNCTTGTCTCCTGAGCTCAAGTGA
AGCACACGTCTGAACTCTTTCCCTACACGACGCTCTTCCGATCTNNNACGTGGTGACTTTCAGTTCTCA
GCACACGTCTGAACTCTTTCCCTACACGACGCTCTTCCGATCTNNNGTGTGAGGAGTCATTAGCAGTG
GAGCACACGTCTGAACTCTTTCCCTACACGACGCTCTTCCGATCTNNNAAGGCCAATCTGATACAGAC
AGCACACGTCTGAACTCTTTCCCTACACGACGCTCTTCCGATCTNNNATCTTTGCCACCACTTCTCTGAT
AGCACACGTCTGAACTCTTTCCCTACACGACGCTCTTCCGATCTNNNAAAAAGTGCTAGCCCATGGTT
AGCACACGTCTGAACTCTTTCCCTACACGACGCTCTTCCGATCTNNNGATTACAGGCGTGAGTTACTG
GAGCACACGTCTGAACTCTTTCCCTACACGACGCTCTTCCGATCTNNNTTGTCTATGTCCTAAGTGCTG
GAGCACACGTCTGAACTCTTTCCCTACACGACGCTCTTCCGATCTNNNGTGCAAGTAAAGAGCCTTAGT
AGCACACGTCTGAACTCTTTCCCTACACGACGCTCTTCCGATCTNNNTCCGTAGTGTTAGGGTTGCTTT

AGCACACGTCTGAACTCTTTCCCTACACGACGCTCTTCCGATCTNNNTTCATTCCATTCTCCTGCCTCAG
AGCACACGTCTGAACTCTTTCCCTACACGACGCTCTTCCGATCTNNNGGCTTCTTACATTCTAGCCTGG
AGCACACGTCTGAACTCTTTCCCTACACGACGCTCTTCCGATCTNNNAAGGACAGTGAATGGGGATA
GCACACGTCTGAACTCTTTCCCTACACGACGCTCTTCCGATCTNNNTTCAAAAATTGGCTCCCATACCG
GAGCACACGTCTGAACTCTTTCCCTACACGACGCTCTTCCGATCTNNNTACTATGCTGTCCTCTTCTGCA
GAGCACACGTCTGAACTCTTTCCCTACACGACGCTCTTCCGATCTNNNCTTCAGCCTCCCAAGTAGCAT
GAGCACACGTCTGAACTCTTTCCCTACACGACGCTCTTCCGATCTNNNCAATGTACAGTCTTTGCCACG
GAGCACACGTCTGAACTCTTTCCCTACACGACGCTCTTCCGATCTNNNAGTAGCTGAGATTACAAGTG
GCACACGTCTGAACTCTTTCCCTACACGACGCTCTTCCGATCTNNNGCTCATGTCTGCCTAACTACCTA
GCACACGTCTGAACTCTTTCCCTACACGACGCTCTTCCGATCTNNNCTAATGCAGGCTGAAATGCTAG
GAGCACACGTCTGAACTCTTTCCCTACACGACGCTCTTCCGATCTNNNTACGACTTTCTCCCTAGGCAA
GAGCACACGTCTGAACTCTTTCCCTACACGACGCTCTTCCGATCTNNNTGGCATGCTCAGAAAGTTTTT
GCACACGTCTGAACTCTTTCCCTACACGACGCTCTTCCGATCTNNNCATACACCCCTCCCTTTTGAGAC
AGCACACGTCTGAACTCTTTCCCTACACGACGCTCTTCCGATCTNNNACCCCAATTTCTAGACCTGCAT
AGCACACGTCTGAACTCTTTCCCTACACGACGCTCTTCCGATCTNNNAGTCACATATTCATAGGCCAGG
AGCACACGTCTGAACTCTTTCCCTACACGACGCTCTTCCGATCTNNNACATACCACTGAGCACAAAGT
GAGCACACGTCTGAACTCTTTCCCTACACGACGCTCTTCCGATCTNNNATGCCCACTGTAGAAGTTCTG
GAGCACACGTCTGAACTCTTTCCCTACACGACGCTCTTCCGATCTNNNAAAGAGTAATCTCCAGGTTGG
AGCACACGTCTGAACTCTTTCCCTACACGACGCTCTTCCGATCTNNNTAAGTCCATATGAGCACCTAGG
GAGCACACGTCTGAACTCTTTCCCTACACGACGCTCTTCCGATCTNNNACAGGAGTACTTTTGACGCTC
AGCACACGTCTGAACTCTTTCCCTACACGACGCTCTTCCGATCTNNNATGGGGTGATGTGACATATTG
GCACACGTCTGAACTCTTTCCCTACACGACGCTCTTCCGATCTNNNTTTTTCCTTCCCTTCCCTTGTCTC
GAGCACACGTCTGAACTCTTTCCCTACACGACGCTCTTCCGATCTNNNGACATGAAAGCTGAGCCAGA
AGCACACGTCTGAACTCTTTCCCTACACGACGCTCTTCCGATCTNNNTTTTTGTGTGTGTGTGTGTG
GCACACGTCTGAACTCTTTCCCTACACGACGCTCTTCCGATCTNNNATAGGAGGGGCTTTTTCACTCA
GAGCACACGTCTGAACTCTTTCCCTACACGACGCTCTTCCGATCTNNNCTGCCCACCTTTGTTACTCTTT
AGCACACGTCTGAACTCTTTCCCTACACGACGCTCTTCCGATCTNNNGTGAACTCCTGGTCTCAAGTG
AGCACACGTCTGAACTCTTTCCCTACACGACGCTCTTCCGATCTNNNGGGGATATGTTCACCTCTTTG
GAGCACACGTCTGAACTCTTTCCCTACACGACGCTCTTCCGATCTNNNTGTGGGCTTTCTTTTCTCTGT
AGCACACGTCTGAACTCTTTCCCTACACGACGCTCTTCCGATCTNNNGATTACAGGCACAAACCATTG
GCACACGTCTGAACTCTTTCCCTACACGACGCTCTTCCGATCTNNNCATTCCCCTCTTACTGTGTTCCAC
GCACACGTCTGAACTCTTTCCCTACACGACGCTCTTCCGATCTNNNCTAGCATAGGAGCAAGGAAGG
GCACACGTCTGAACTCTTTCCCTACACGACGCTCTTCCGATCTNNNGTGTGAGGAGTCATTAGCAGTG
AGCACACGTCTGAACTCTTTCCCTACACGACGCTCTTCCGATCTNNNGTCTGTTCATGTACTTTGCCCA
GCACACGTCTGAACTCTTTCCCTACACGACGCTCTTCCGATCTNNNTGCATTTCTTATGTGGCTCTTGG
GCACACGTCTGAACTCTTTCCCTACACGACGCTCTTCCGATCTNNNAGTAGTGCATTGTACCAAGCAT
AGCACACGTCTGAACTCTTTCCCTACACGACGCTCTTCCGATCTNNNGCAATGGCATATTCTCAGGTGA
GAGCACACGTCTGAACTCTTTCCCTACACGACGCTCTTCCGATCTNNNTTTACCATGCACATTTGTGGC
AGCACACGTCTGAACTCTTTCCCTACACGACGCTCTTCCGATCTNNNTTTCATGTGTCTTTTGGCTGCAT
GCACACGTCTGAACTCTTTCCCTACACGACGCTCTTCCGATCTNNNTTCTCCTGTGTCAACATCCTGAG
AGCACACGTCTGAACTCTTTCCCTACACGACGCTCTTCCGATCTNNNATTTCAGCCTCCCAAGTATCTG
GCACACGTCTGAACTCTTTCCCTACACGACGCTCTTCCGATCTNNNTGTTCTTGCCTCTCTAGTTCCTTC
AGCACACGTCTGAACTCTTTCCCTACACGACGCTCTTCCGATCTNNNAATCCTGGGCTCCTTTGATCTTT

AGCACACGTCTGAACTCTTTCCCTACACGACGCTCTTCCGATCTNNNGGCCTCCCAAAGTACTGGAAT
AGCACACGTCTGAACTCTTTCCCTACACGACGCTCTTCCGATCTNNNAGAAAAGCACTACACAAAACG
AGCACACGTCTGAACTCTTTCCCTACACGACGCTCTTCCGATCTNNNTACAACTGCCCTCTGAGTTCAA
AGCACACGTCTGAACTCTTTCCCTACACGACGCTCTTCCGATCTNNNAGGCAAGAGCAAACAAATTC
AGCACACGTCTGAACTCTTTCCCTACACGACGCTCTTCCGATCTNNNGCCATTTGTGTGGAGCCATAT
AGCACACGTCTGAACTCTTTCCCTACACGACGCTCTTCCGATCTNNNGGATGTCAGCAAAATAAGCAT
AGCACACGTCTGAACTCTTTCCCTACACGACGCTCTTCCGATCTNNNGTCTCAAGCTCCTGACCAAGTA
GCACACGTCTGAACTCTTTCCCTACACGACGCTCTTCCGATCTNNNGGAGAGTCAATTTGTATGCAG
AGCACACGTCTGAACTCTTTCCCTACACGACGCTCTTCCGATCTNNNTTCATCCGGAATCTATCGGTT
AGCACACGTCTGAACTCTTTCCCTACACGACGCTCTTCCGATCTNNNTTTAGACTGTGGTCCCAATCAC
GCACACGTCTGAACTCTTTCCCTACACGACGCTCTTCCGATCTNNNTTTTCTCACAGCTCTGGACATCA
AGCACACGTCTGAACTCTTTCCCTACACGACGCTCTTCCGATCTNNNTGTGCAGACCAAGGAATTGA
AGCACACGTCTGAACTCTTTCCCTACACGACGCTCTTCCGATCTNNNTGGTGCTGACCTCATGAAGTA
AGCACACGTCTGAACTCTTTCCCTACACGACGCTCTTCCGATCTNNNAGATTGTGCTGCTGTAAACAT
AGCACACGTCTGAACTCTTTCCCTACACGACGCTCTTCCGATCTNNNAGGAGAGTTCTGCTTTTAGC
GCACACGTCTGAACTCTTTCCCTACACGACGCTCTTCCGATCTNNNAATAATGGGCTCCAACTCCATCC
GCACACGTCTGAACTCTTTCCCTACACGACGCTCTTCCGATCTNNNTCTGAGAGATTTTGGTGCACTC
AGCACACGTCTGAACTCTTTCCCTACACGACGCTCTTCCGATCTNNNATATGGATGCAGTTTCCCCCTT
GAGCACACGTCTGAACTCTTTCCCTACACGACGCTCTTCCGATCTNNNACTCAGAATTGCTTTGGCTGT
GCACACGTCTGAACTCTTTCCCTACACGACGCTCTTCCGATCTNNNTTTTGGCTGCATAAGTGTCTTC
GCACACGTCTGAACTCTTTCCCTACACGACGCTCTTCCGATCTNNNGTAGACGATACAGGAATGGGC
GCACACGTCTGAACTCTTTCCCTACACGACGCTCTTCCGATCTNNNCAAACTCCAACAGGTGATGCTT
GAGCACACGTCTGAACTCTTTCCCTACACGACGCTCTTCCGATCTNNNGGTCTCAAAAAATCCTCCCTC
GCACACGTCTGAACTCTTTCCCTACACGACGCTCTTCCGATCTNNNGGAAATCTCTGGAGTGGGAAA
GCACACGTCTGAACTCTTTCCCTACACGACGCTCTTCCGATCTNNNCCTGCATGTCTGTCAAATCCATC
GCACACGTCTGAACTCTTTCCCTACACGACGCTCTTCCGATCTNNNTTCCAAATAGCTGGGACTACAG
GCACACGTCTGAACTCTTTCCCTACACGACGCTCTTCCGATCTNNNTGGAACTACATGCATAAGCCAC
AGCACACGTCTGAACTCTTTCCCTACACGACGCTCTTCCGATCTNNNCACAATAGCTCACGCCTGTAAT
GAGCACACGTCTGAACTCTTTCCCTACACGACGCTCTTCCGATCTNNNTGCTTGTTTGTTTGTTTGGG
GCACACGTCTGAACTCTTTCCCTACACGACGCTCTTCCGATCTNNNCATTTGCCACTATTTTGGGGG
AGCACACGTCTGAACTCTTTCCCTACACGACGCTCTTCCGATCTNNNCTGTCACTGTTTTGTGTGTGTG
GCACACGTCTGAACTCTTTCCCTACACGACGCTCTTCCGATCTNNNTGTGGCTGTCTACTTGTGTCTAC
GCACACGTCTGAACTCTTTCCCTACACGACGCTCTTCCGATCTNNNTATGCAGTGAACCCAGTTGAG
GCACACGTCTGAACTCTTTCCCTACACGACGCTCTTCCGATCTNNNAATGAATGTGGGGCTGAAAAG
AGCACACGTCTGAACTCTTTCCCTACACGACGCTCTTCCGATCTNNNATGCCAAAAATGCAAAGTCCT
GCACACGTCTGAACTCTTTCCCTACACGACGCTCTTCCGATCTNNNTCCTCCTCCTTCTTCTCCTTCTTCA
AGCACACGTCTGAACTCTTTCCCTACACGACGCTCTTCCGATCTNNNATGGCTGCATAGTATTCCATGG
GCACACGTCTGAACTCTTTCCCTACACGACGCTCTTCCGATCTNNNTGGGATTGCCTTTTGATTTCTCT
AGCACACGTCTGAACTCTTTCCCTACACGACGCTCTTCCGATCTNNNTCCCTAACTGTGGATCCTACTG
AGCACACGTCTGAACTCTTTCCCTACACGACGCTCTTCCGATCTNNNCGTACACTGAAACAGAACTGC
AGCACACGTCTGAACTCTTTCCCTACACGACGCTCTTCCGATCTNNNGAATGCAGTGGTGTGATCATA
GCACACGTCTGAACTCTTTCCCTACACGACGCTCTTCCGATCTNNNGATCTCCTGACCTTGTGATCCTC
GCACACGTCTGAACTCTTTCCCTACACGACGCTCTTCCGATCTNNNACACCAGTTACTGCATCCACTTA

AGCACACGTCTGAACTCTTTCCCTACACGACGCTCTTCCGATCTNNNCTAGAGTGCAGTGGTGTGATCT
GCACACGTCTGAACTCTTTCCCTACACGACGCTCTTCCGATCTNNNATAGTCACTTACCACCATGCCTGC
GAGCACACGTCTGAACTCTTTCCCTACACGACGCTCTTCCGATCTNNNCCACTCTGTTGACTAGGATGG
AGCACACGTCTGAACTCTTTCCCTACACGACGCTCTTCCGATCTNNNGGATCAGTGCTGGATGTTTGTT
GAGCACACGTCTGAACTCTTTCCCTACACGACGCTCTTCCGATCTNNNAGGCTGAGGCAGAATTACTTG
AGCACACGTCTGAACTCTTTCCCTACACGACGCTCTTCCGATCTNNNGAATGCCTGTAATCCCAGCTAC
GAGCACACGTCTGAACTCTTTCCCTACACGACGCTCTTCCGATCTNNNCGAAGCTGAGTCTTACCCCTT
GAGCACACGTCTGAACTCTTTCCCTACACGACGCTCTTCCGATCTNNNCTGTAGTCCCAGCTACTCAGGA
GAGCACACGTCTGAACTCTTTCCCTACACGACGCTCTTCCGATCTNNNCAACAAGAGTGAGACTCCACC
AGAGCACACGTCTGAACTCTTTCCCTACACGACGCTCTTCCGATCTNNNAAAAAATTAGCCAGGTGTGG
GAGCACACGTCTGAACTCTTTCCCTACACGACGCTCTTCCGATCTNNNCTGAGGCAGGAGAATCACTTG
GAGCACACGTCTGAACTCTTTCCCTACACGACGCTCTTCCGATCTNNNCACCTGTAATCCCAGCTACTCT
AGAGCACACGTCTGAACTCTTTCCCTACACGACGCTCTTCCGATCTNNNCACCTGAAGTCTCAGCTACTC
GAGCACACGTCTGAACTCTTTCCCTACACGACGCTCTTCCGATCTNNNCGACAAGAGCAAAAACTACGT
GCACACGTCTGAACTCTTTCCCTACACGACGCTCTTCCGATCTNNNAGAGCTTGCAGTCAGATGAGAT
AGAGCACACGTCTGAACTCTTTCCCTACACGACGCTCTTCCGATCTNNNCTACTCTGGAGAAACAGCGTG
AGCACACGTCTGAACTCTTTCCCTACACGACGCTCTTCCGATCTNNNCTGAGGCAGGAGAATCGTTTG
AGCACACGTCTGAACTCTTTCCCTACACGACGCTCTTCCGATCTNNNATACAAAATTACCCAGGCGTGG
GAGCACACGTCTGAACTCTTTCCCTACACGACGCTCTTCCGATCTNNNGCCTACACAGTAAGAGCAAGA
GAGCACACGTCTGAACTCTTTCCCTACACGACGCTCTTCCGATCTNNNGGAAACCTAGCAAGACTCTGTG
AGAGCACACGTCTGAACTCTTTCCCTACACGACGCTCTTCCGATCTNNNCTGTCATCCCAGCTATTCAGG
AGCACACGTCTGAACTCTTTCCCTACACGACGCTCTTCCGATCTNNNTTTTAGCTCCCTCTTATGTGGG
AGAGCACACGTCTGAACTCTTTCCCTACACGACGCTCTTCCGATCTNNNCTGACACAGGAGAATCGCTTG
AGCACACGTCTGAACTCTTTCCCTACACGACGCTCTTCCGATCTNNNCTGTAATTCCAGCTACTCGGGAG
AGCACACGTCTGAACTCTTTCCCTACACGACGCTCTTCCGATCTNNNACAAAATTTACAAGTGGTGGCG
AGAGCACACGTCTGAACTCTTTCCCTACACGACGCTCTTCCGATCTNNNACAAAATTAGTTGGGCATGG
GAGCACACGTCTGAACTCTTTCCCTACACGACGCTCTTCCGATCTNNNAATACAAAAATTAGCCGGGCG
GCACACGTCTGAACTCTTTCCCTACACGACGCTCTTCCGATCTNNNTTTCAGGTGACAGAGTGAGACTC
AAGAGCACACGTCTGAACTCTTTCCCTACACGACGCTCTTCCGATCTNNNCTGTAGTCCCAGCTACTCAG
AGCACACGTCTGAACTCTTTCCCTACACGACGCTCTTCCGATCTNNNCTGAGGCAGGAGAATCACTTGA
GAGCACACGTCTGAACTCTTTCCCTACACGACGCTCTTCCGATCTNNNTGTAATGTTAGCACTTCGGGA
GAGCACACGTCTGAACTCTTTCCCTACACGACGCTCTTCCGATCTNNNGGTGACAGAGAGAGATTCCG
GAGCACACGTCTGAACTCTTTCCCTACACGACGCTCTTCCGATCTNNNATCAGGCATGACATCCTAAGTG
AGCACACGTCTGAACTCTTTCCCTACACGACGCTCTTCCGATCTNNNATACAAAAATTAGCCGGGTGTG
AGCACACGTCTGAACTCTTTCCCTACACGACGCTCTTCCGATCTNNNTCAGCTGCAAATGAGACTTCTC
GAGCACACGTCTGAACTCTTTCCCTACACGACGCTCTTCCGATCTNNNAGGCAGTAGAATCACTTGAAG
AGAGCACACGTCTGAACTCTTTCCCTACACGACGCTCTTCCGATCTNNNATCATGCCATTATACTCCAGC
GAGCACACGTCTGAACTCTTTCCCTACACGACGCTCTTCCGATCTNNNTGAGGCAGGAGAATGACTTGA
GAGCACACGTCTGAACTCTTTCCCTACACGACGCTCTTCCGATCTNNNGCAAGAGAATCACTTGAACCC
AGAGCACACGTCTGAACTCTTTCCCTACACGACGCTCTTCCGATCTNNNTGGTGCTAAGCAGTTGAATT
AGCACACGTCTGAACTCTTTCCCTACACGACGCTCTTCCGATCTNNNGGCTGAGATAGGAGGATCACT
AGCACACGTCTGAACTCTTTCCCTACACGACGCTCTTCCGATCTNNNAAAAAATTAGCTGGCCATGGTG
AGAGCACACGTCTGAACTCTTTCCCTACACGACGCTCTTCCGATCTNNNGTTGTAATGAGCCAAGATCA

AGCACACGTCTGAACTCTTTCCCTACACGACGCTCTTCCGATCTNNNCCTGGGACATGATGAGTCATAG
GCACACGTCTGAACTCTTTCCCTACACGACGCTCTTCCGATCTNNNTGAGATAAAGCCACTGCACTTC
AGCACACGTCTGAACTCTTTCCCTACACGACGCTCTTCCGATCTNNNCTGTAGTCCCAGCTACTCGAGA
AGCACACGTCTGAACTCTTTCCCTACACGACGCTCTTCCGATCTNNNGGTGCCTGTAATCCCAACTACT
AGCACACGTCTGAACTCTTTCCCTACACGACGCTCTTCCGATCTNNNGTATTGCCAGCTATTTTGGAG
GAGCACACGTCTGAACTCTTTCCCTACACGACGCTCTTCCGATCTNNNCAGGTGCCTATAATCCCAGCT
AGCACACGTCTGAACTCTTTCCCTACACGACGCTCTTCCGATCTNNNAAGCTGAAGCAGGAAAATTG
AGCACACGTCTGAACTCTTTCCCTACACGACGCTCTTCCGATCTNNNCAACATGGAGAAACCCTGTCT
AGCACACGTCTGAACTCTTTCCCTACACGACGCTCTTCCGATCTNNNCTGAGGCAGGAGAATCGTTT
AGCACACGTCTGAACTCTTTCCCTACACGACGCTCTTCCGATCTNNNCAGGATGAACTGCTTGAATGT
AGCACACGTCTGAACTCTTTCCCTACACGACGCTCTTCCGATCTNNNAGAATTGCTTGAATCTGGGA
AGAGCACACGTCTGAACTCTTTCCCTACACGACGCTCTTCCGATCTNNNGAGAATCGCTTGAATCTG
AGCACACGTCTGAACTCTTTCCCTACACGACGCTCTTCCGATCTNNNCTGGGAAACAAGAGTGAAAA
GAGCACACGTCTGAACTCTTTCCCTACACGACGCTCTTCCGATCTNNNGAAGTCTGTCTCCAACTCTGT
AGCACACGTCTGAACTCTTTCCCTACACGACGCTCTTCCGATCTNNNCTGAGGCAGGAGAATCACTTG
AGCACACGTCTGAACTCTTTCCCTACACGACGCTCTTCCGATCTNNNAAGTATAAAAAATCAGCCGGG
AGCACACGTCTGAACTCTTTCCCTACACGACGCTCTTCCGATCTNNNGGTGTCTGTAATCCCAGCTAC
GAGCACACGTCTGAACTCTTTCCCTACACGACGCTCTTCCGATCTNNNCTTGAACCTGGGAGATGGA
AGCACACGTCTGAACTCTTTCCCTACACGACGCTCTTCCGATCTNNNTCCTGACCTCAGATGAGATGA
AGCACACGTCTGAACTCTTTCCCTACACGACGCTCTTCCGATCTNNNGCTGAAGCAGGAGAAATGTTT
GAGCACACGTCTGAACTCTTTCCCTACACGACGCTCTTCCGATCTNNNGAGACAGACCAAGACTCCAT
AGCACACGTCTGAACTCTTTCCCTACACGACGCTCTTCCGATCTNNNTGTAGCTTGCAGTGAGTTGA
GCACACGTCTGAACTCTTTCCCTACACGACGCTCTTCCGATCTNNNGCAAAACCCTGTCTCTGCAAATA
GAGCACACGTCTGAACTCTTTCCCTACACGACGCTCTTCCGATCTNNNTGAGGCAGGAGAATTGCTT
GAGCACACGTCTGAACTCTTTCCCTACACGACGCTCTTCCGATCTNNNGCAGGAGAATTGCTTGATC
AGCACACGTCTGAACTCTTTCCCTACACGACGCTCTTCCGATCTNNNATACAAAAAAATTAGCCGGG
AGCACACGTCTGAACTCTTTCCCTACACGACGCTCTTCCGATCTNNNCAAAATAATTAGCTGGGCGTG
AGCACACGTCTGAACTCTTTCCCTACACGACGCTCTTCCGATCTNNNATACAAAAATTAGCTGGGCGT
AGCACACGTCTGAACTCTTTCCCTACACGACGCTCTTCCGATCTNNNATACAAAAAAATTAGCCGGG
AGCACACGTCTGAACTCTTTCCCTACACGACGCTCTTCCGATCTNNNAAATACAAAAATTAGCCGGG
GCACACGTCTGAACTCTTTCCCTACACGACGCTCTTCCGATCTNNNAGAATTGCTTGAACCTGGGAGT
GAGCACACGTCTGAACTCTTTCCCTACACGACGCTCTTCCGATCTNNNCTTTGTAGTTGCCTTCGTGG
AGCACACGTCTGAACTCTTTCCCTACACGACGCTCTTCCGATCTNNNCAGATGCCTTTTGTGCTGTCTA
GCACACGTCTGAACTCTTTCCCTACACGACGCTCTTCCGATCTNNNCAAACACAGTGAAACCCCATCT
AGCACACGTCTGAACTCTTTCCCTACACGACGCTCTTCCGATCTNNNGTATTTCTGCAGTCCCAGCTAC
AGCACACGTCTGAACTCTTTCCCTACACGACGCTCTTCCGATCTNNNAAAAAAAAAAAAGAGGGAGG
AGCACACGTCTGAACTCTTTCCCTACACGACGCTCTTCCGATCTNNNTTGTGGTGACCTGAGATTGTA
AGCACACGTCTGAACTCTTTCCCTACACGACGCTCTTCCGATCTNNNGAGGTTGCAATGAGCCAAGA
GAGCACACGTCTGAACTCTTTCCCTACACGACGCTCTTCCGATCTNNNCGGTGAAAGTCCGTCTCTAC
AGCACACGTCTGAACTCTTTCCCTACACGACGCTCTTCCGATCTNNNCAAATCTCAAAGACTCAGCCC
AGCACACGTCTGAACTCTTTCCCTACACGACGCTCTTCCGATCTNNNAAGAAATCTCAGAGACTCAGC
AGCACACGTCTGAACTCTTTCCCTACACGACGCTCTTCCGATCTNNNCTGAGACAGGAGAATCGCTTG
AGCACACGTCTGAACTCTTTCCCTACACGACGCTCTTCCGATCTNNNGAGGTTACAGTGAGCCAAGAT

GAGCACACGTCTGAACTCTTTCCCTACACGACGCTCTTCCGATCTNNNAAAAATTAGCTGGTTGTGGT
AGCACACGTCTGAACTCTTTCCCTACACGACGCTCTTCCGATCTNNNCCTGGGAAATAGAACGAGACT
GAGCACACGTCTGAACTCTTTCCCTACACGACGCTCTTCCGATCTNNNGTGTGCCTATAATCCCAGCTA
GAGCACACGTCTGAACTCTTTCCCTACACGACGCTCTTCCGATCTNNNTCACACCTGTAATTCCAGCAC
GCACACGTCTGAACTCTTTCCCTACACGACGCTCTTCCGATCTNNNCCGGATTCAAACGATTCTCCTCT
AGCACACGTCTGAACTCTTTCCCTACACGACGCTCTTCCGATCTNNNGCTGAGAGAGGACATCACTT
GAGCACACGTCTGAACTCTTTCCCTACACGACGCTCTTCCGATCTNNNAAAAAATTAGCCAGGTGTGG
AGCACACGTCTGAACTCTTTCCCTACACGACGCTCTTCCGATCTNNNCTGAGGCAGGAGAATTGCTT
GAGCACACGTCTGAACTCTTTCCCTACACGACGCTCTTCCGATCTNNNTGATGAAACCCCGTCGTTACT
AGCACACGTCTGAACTCTTTCCCTACACGACGCTCTTCCGATCTNNNGAGACAGAAGGATCTCTTGA
GAGCACACGTCTGAACTCTTTCCCTACACGACGCTCTTCCGATCTNNNACCAACATGATGAAACCCTGT
GAGCACACGTCTGAACTCTTTCCCTACACGACGCTCTTCCGATCTNNNGTGTAATCCCAGCTACTCAG
AGCACACGTCTGAACTCTTTCCCTACACGACGCTCTTCCGATCTNNNCAGAGGTTGTAGTGAGCCAA
GAGCACACGTCTGAACTCTTTCCCTACACGACGCTCTTCCGATCTNNNTCTGTAATCCCAGCTACTCAG
AGCACACGTCTGAACTCTTTCCCTACACGACGCTCTTCCGATCTNNNTAAGGCAGCAGAATCACTTGA
GAGCACACGTCTGAACTCTTTCCCTACACGACGCTCTTCCGATCTNNNGAGACAGGAGAATGGCTAG
AGCACACGTCTGAACTCTTTCCCTACACGACGCTCTTCCGATCTNNNAGAATCACTCGAACCCAGAAGT
AGCACACGTCTGAACTCTTTCCCTACACGACGCTCTTCCGATCTNNNGGAGAATGGCGTAAACCTGG
AGCACACGTCTGAACTCTTTCCCTACACGACGCTCTTCCGATCTNNNGTCCACGTGCATTCTTGTTACA
AGCACACGTCTGAACTCTTTCCCTACACGACGCTCTTCCGATCTNNNATGCCTTTAGTCCCAGCTACTT
GAGCACACGTCTGAACTCTTTCCCTACACGACGCTCTTCCGATCTNNNCGGTAAGCTGAGATCATGTC
AGCACACGTCTGAACTCTTTCCCTACACGACGCTCTTCCGATCTNNNAGTGAACTACGATTTTGCCAC
AGAGCACACGTCTGAACTCTTTCCCTACACGACGCTCTTCCGATCTNNNCTGAGGCAGAGAATTGCTT
GCACACGTCTGAACTCTTTCCCTACACGACGCTCTTCCGATCTNNNGTTCCTGTAGTCCCAGCTACTTG
AGCACACGTCTGAACTCTTTCCCTACACGACGCTCTTCCGATCTNNNTGGGCAACAAAACAAAACTCT
AGCACACGTCTGAACTCTTTCCCTACACGACGCTCTTCCGATCTNNNAAAAAAAATTAGCCGGGCGTG
AGCACACGTCTGAACTCTTTCCCTACACGACGCTCTTCCGATCTNNNTGCCAGTAATTCCAGCATTTTG
AGCACACGTCTGAACTCTTTCCCTACACGACGCTCTTCCGATCTNNNGTGCCTATAACCCCAGCTACTTG
AGCACACGTCTGAACTCTTTCCCTACACGACGCTCTTCCGATCTNNNGAGAATCGCTTGAATCCTGGA
AGCACACGTCTGAACTCTTTCCCTACACGACGCTCTTCCGATCTNNNGGAATCACTTGAACCTGGGA
GAGCACACGTCTGAACTCTTTCCCTACACGACGCTCTTCCGATCTNNNGGAGGTGGAACTTGAAATA
AGCACACGTCTGAACTCTTTCCCTACACGACGCTCTTCCGATCTNNNGGGAGAATTGCTTGAATCTGG
AGAGCACACGTCTGAACTCTTTCCCTACACGACGCTCTTCCGATCTNNNTGTAGTACCAGCTACTCAG
AGCACACGTCTGAACTCTTTCCCTACACGACGCTCTTCCGATCTNNNCTGAGGCAGGAGAATCAGTTG
GAGCACACGTCTGAACTCTTTCCCTACACGACGCTCTTCCGATCTNNNTGAGGCAGGAGAATCTCTTG
GCACACGTCTGAACTCTTTCCCTACACGACGCTCTTCCGATCTNNNCTGAGGCAGGAGAATCACTTGA
GCACACGTCTGAACTCTTTCCCTACACGACGCTCTTCCGATCTNNNGTTTGCAGTGAGCTGAGATCAT
AGCACACGTCTGAACTCTTTCCCTACACGACGCTCTTCCGATCTNNNCCAACATGGTGAAACTGCATC
AGCACACGTCTGAACTCTTTCCCTACACGACGCTCTTCCGATCTNNNTTCACACCTTCAGACTCAGACT
GCACACGTCTGAACTCTTTCCCTACACGACGCTCTTCCGATCTNNNAGAAGAATGGTGTGAACCCAG
AGCACACGTCTGAACTCTTTCCCTACACGACGCTCTTCCGATCTNNNCTGTATTCTGCTAACAGGAGG
AGCACACGTCTGAACTCTTTCCCTACACGACGCTCTTCCGATCTNNNCACCTGTAATTCCAGCACTTTG
GCACACGTCTGAACTCTTTCCCTACACGACGCTCTTCCGATCTNNNGACATTGCAGTGAGCTGAGATT

GAGCACACGTCTGAACTCTTTCCCTACACGACGCTCTTCCGATCTNNNAAAAAATTAGCAGGGCATGG
GAGCACACGTCTGAACTCTTTCCCTACACGACGCTCTTCCGATCTNNNGCAGGAGAATTTCTTGAATC
AGCACACGTCTGAACTCTTTCCCTACACGACGCTCTTCCGATCTNNNCAAGAACAAAACCCCGTCTCAAA
GAGCACACGTCTGAACTCTTTCCCTACACGACGCTCTTCCGATCTNNNAACCCACAGTTGCAATGGAAA
GAGCACACGTCTGAACTCTTTCCCTACACGACGCTCTTCCGATCTNNNGAAGCGAGAGGAATGTGAC
GAGCACACGTCTGAACTCTTTCCCTACACGACGCTCTTCCGATCTNNNCTGAGGCAGGAGAATTGCTTC
AGCACACGTCTGAACTCTTTCCCTACACGACGCTCTTCCGATCTNNNTCAGGAGAATCACTTGAACCCA
GAGCACACGTCTGAACTCTTTCCCTACACGACGCTCTTCCGATCTNNNAAAATTAGCCAGACGTGGTG
GCACACGTCTGAACTCTTTCCCTACACGACGCTCTTCCGATCTNNNCAGCCTGGATGATAGAGTGAGA
GAGCACACGTCTGAACTCTTTCCCTACACGACGCTCTTCCGATCTNNNGATCACGAGGTCAGGAGATTC
AGCACACGTCTGAACTCTTTCCCTACACGACGCTCTTCCGATCTNNNCGGCAAAAAGTTTCCTTCTAGCA
GAGCACACGTCTGAACTCTTTCCCTACACGACGCTCTTCCGATCTNNNTACAAAAATTAGCGGGGTGTC
AGCACACGTCTGAACTCTTTCCCTACACGACGCTCTTCCGATCTNNNCTGTAATCCAAGCTACTCGGGA
AGAGCACACGTCTGAACTCTTTCCCTACACGACGCTCTTCCGATCTNNNTGAGGCAGGAGAATCTCTT
AGCACACGTCTGAACTCTTTCCCTACACGACGCTCTTCCGATCTNNNTTCCCCGATGCTATTGTTCCTAG
AGCACACGTCTGAACTCTTTCCCTACACGACGCTCTTCCGATCTNNNCTGAGGCAGGAGAATAGCTTG
AGCACACGTCTGAACTCTTTCCCTACACGACGCTCTTCCGATCTNNNGAGACAGGAGAATTGCTTGAA
AGCACACGTCTGAACTCTTTCCCTACACGACGCTCTTCCGATCTNNNCACCTGTAATCCCAGCTACTCAG
AGAGCACACGTCTGAACTCTTTCCCTACACGACGCTCTTCCGATCTNNNGAGAATTCCTTGAACCTGGG
GAGCACACGTCTGAACTCTTTCCCTACACGACGCTCTTCCGATCTNNNGCCGGAGTAACAACAGCAAA
GAGCACACGTCTGAACTCTTTCCCTACACGACGCTCTTCCGATCTNNNCATCTGTAGTCCCAGCTACTCA
AGCACACGTCTGAACTCTTTCCCTACACGACGCTCTTCCGATCTNNNTCATGCCTCCCAAGATTGAATCA
GAGCACACGTCTGAACTCTTTCCCTACACGACGCTCTTCCGATCTNNNTGTAATTCCAGCTACTTGGGA
AGCACACGTCTGAACTCTTTCCCTACACGACGCTCTTCCGATCTNNNCTCAGGAGGATTAGGGAGGAC
GCACACGTCTGAACTCTTTCCCTACACGACGCTCTTCCGATCTNNNATAATGCCAGCTACTTAGGAGG
AGCACACGTCTGAACTCTTTCCCTACACGACGCTCTTCCGATCTNNNGCAACAAGAGTGAAAATCTGC
GAGCACACGTCTGAACTCTTTCCCTACACGACGCTCTTCCGATCTNNNCTCAACCCCTGTAGTGGATTGA
AGCACACGTCTGAACTCTTTCCCTACACGACGCTCTTCCGATCTNNNGAGAATACCTTGAACCTGGGAG
AGCACACGTCTGAACTCTTTCCCTACACGACGCTCTTCCGATCTNNNTGTACACACGAATGATCCATGG
AGCACACGTCTGAACTCTTTCCCTACACGACGCTCTTCCGATCTNNNGAGGTTGCAGTGAGCTGAGAT
GAGCACACGTCTGAACTCTTTCCCTACACGACGCTCTTCCGATCTNNNCTGAGACAGGAGAATCGCTTC
AGCACACGTCTGAACTCTTTCCCTACACGACGCTCTTCCGATCTNNNGGCGACAGAGAAAAACTTCAT
AGCACACGTCTGAACTCTTTCCCTACACGACGCTCTTCCGATCTNNNGAGGATCATTTGAGTCCAGGA
AGCACACGTCTGAACTCTTTCCCTACACGACGCTCTTCCGATCTNNNCAACAAGGAGAGAGATGCTGT
AGCACACGTCTGAACTCTTTCCCTACACGACGCTCTTCCGATCTNNNCAACAAGGAGAGAGATGCTGT
GAGCACACGTCTGAACTCTTTCCCTACACGACGCTCTTCCGATCTNNNACAAAAATTAGCCAGGCGTGA
AGCACACGTCTGAACTCTTTCCCTACACGACGCTCTTCCGATCTNNNCTGAGATGGGAGGATCATTGA
GAGCACACGTCTGAACTCTTTCCCTACACGACGCTCTTCCGATCTNNNCATTGCAGTGAGCTCAGATCA
GAGCACACGTCTGAACTCTTTCCCTACACGACGCTCTTCCGATCTNNNTCATGAGGCTGAAGCAAGAG
AGCACACGTCTGAACTCTTTCCCTACACGACGCTCTTCCGATCTNNNTAATCACAGCTACTTGGGAAGG
GAGCACACGTCTGAACTCTTTCCCTACACGACGCTCTTCCGATCTNNNCTGTAATCCCAGCTACTCAGG
AGCACACGTCTGAACTCTTTCCCTACACGACGCTCTTCCGATCTNNNACACACTTGTAATCCCAGCTAC
AGCACACGTCTGAACTCTTTCCCTACACGACGCTCTTCCGATCTNNNGATCACACTACTACACTCCAGC

AGCACACGTCTGAACTCTTTCCCTACACGACGCTCTTCCGATCTNNNAAACACAAAAATTAGCTGGGC
AGCACACGTCTGAACTCTTTCCCTACACGACGCTCTTCCGATCTNNNTGCCTGTAATCCCAGCTATTCA
AGCACACGTCTGAACTCTTTCCCTACACGACGCTCTTCCGATCTNNNTGTAATCCCAGCTACTCAGGAG
AGCACACGTCTGAACTCTTTCCCTACACGACGCTCTTCCGATCTNNNTATAGTCCCAGCTACTCGAGAG
AGCACACGTCTGAACTCTTTCCCTACACGACGCTCTTCCGATCTNNNGAGATCATGCCACTACACTCCA
AGCACACGTCTGAACTCTTTCCCTACACGACGCTCTTCCGATCTNNNAATCTCAGCTACTTGGGAGGT
AGCACACGTCTGAACTCTTTCCCTACACGACGCTCTTCCGATCTNNNATTCCCAGCTATTCAGGTGTCT
AGCACACGTCTGAACTCTTTCCCTACACGACGCTCTTCCGATCTNNNTTGAATCCGGGAAGTAGAGG
AGCACACGTCTGAACTCTTTCCCTACACGACGCTCTTCCGATCTNNNATGCCTGTAATCCCAGTTACTC
AGCACACGTCTGAACTCTTTCCCTACACGACGCTCTTCCGATCTNNNGGAGACAAGAATGAAACGCCA
AGCACACGTCTGAACTCTTTCCCTACACGACGCTCTTCCGATCTNNNGAAGAATGGTGTGAACCCAG
AGCACACGTCTGAACTCTTTCCCTACACGACGCTCTTCCGATCTNNNCAGGAGAATCACTTGAACCCA
GCACACGTCTGAACTCTTTCCCTACACGACGCTCTTCCGATCTNNNGCACATCTGTAATCCCAGCTACT
AGCACACGTCTGAACTCTTTCCCTACACGACGCTCTTCCGATCTNNNTGATGCACACCTGTAATTCCAG
AGCACACGTCTGAACTCTTTCCCTACACGACGCTCTTCCGATCTNNNGAGGCACGAGAATTACTTGAA
AGCACACGTCTGAACTCTTTCCCTACACGACGCTCTTCCGATCTNNNCCTGAGCAATACAGTGAGAC
AGCACACGTCTGAACTCTTTCCCTACACGACGCTCTTCCGATCTNNNACAGCAAGACCTCATCTGTACT
GAGCACACGTCTGAACTCTTTCCCTACACGACGCTCTTCCGATCTNNNATACTTGAGCCTGGGAGTTT
AGCACACGTCTGAACTCTTTCCCTACACGACGCTCTTCCGATCTNNNCATTCTATTCAGTCCCCCTCTG
AGCACACGTCTGAACTCTTTCCCTACACGACGCTCTTCCGATCTNNNACATGGCGAAATCTGTCTCTA
AGCACACGTCTGAACTCTTTCCCTACACGACGCTCTTCCGATCTNNNCTGAGGTGGGAGGATCTCTA
AGCACACGTCTGAACTCTTTCCCTACACGACGCTCTTCCGATCTNNNTGTAGTCCCAGCTACTTAGGA
GAGCACACGTCTGAACTCTTTCCCTACACGACGCTCTTCCGATCTNNNCAGATCACTTGAGGTCAGGA
AGCACACGTCTGAACTCTTTCCCTACACGACGCTCTTCCGATCTNNNGAGAATCACTTGAACCCAGGA
GAGCACACGTCTGAACTCTTTCCCTACACGACGCTCTTCCGATCTNNNAGAAGTGTGCAGACAGCTCT
GAGCACACGTCTGAACTCTTTCCCTACACGACGCTCTTCCGATCTNNNGCTCATACCTGTAATCCCAG
AGCACACGTCTGAACTCTTTCCCTACACGACGCTCTTCCGATCTNNNTGTAATCACAGCACTTTGGGA
GAGCACACGTCTGAACTCTTTCCCTACACGACGCTCTTCCGATCTNNNCTGAGGCAGGAGAATCACTT
GCACACGTCTGAACTCTTTCCCTACACGACGCTCTTCCGATCTNNNAGAGATTGCAGTGAGCTGAGA
AGCACACGTCTGAACTCTTTCCCTACACGACGCTCTTCCGATCTNNNCTGGGCAACAAGAGCAAAATT
AGCACACGTCTGAACTCTTTCCCTACACGACGCTCTTCCGATCTNNNAGAATTGCTTGAATTTGGGAG
GAGCACACGTCTGAACTCTTTCCCTACACGACGCTCTTCCGATCTNNNATTGCACTCTAGCCTGAATGA
AGCACACGTCTGAACTCTTTCCCTACACGACGCTCTTCCGATCTNNNGCACATGCCTGTAATCCTAGCT
AGCACACGTCTGAACTCTTTCCCTACACGACGCTCTTCCGATCTNNNAGACAGGAGAATCACTTGAAC
AGCACACGTCTGAACTCTTTCCCTACACGACGCTCTTCCGATCTNNNTACATGCCTGTAATCCCAGCTA
GCACACGTCTGAACTCTTTCCCTACACGACGCTCTTCCGATCTNNNATGATGATGGGCTCCTGTAATC
AGCACACGTCTGAACTCTTTCCCTACACGACGCTCTTCCGATCTNNNGAGCTGAGATCATGTCATTGCA
GCACACGTCTGAACTCTTTCCCTACACGACGCTCTTCCGATCTNNNTTTAGCCATGACCACTTATGCCT
AGCACACGTCTGAACTCTTTCCCTACACGACGCTCTTCCGATCTNNNGGGTTTGCAATGCATAGGTTC
AGCACACGTCTGAACTCTTTCCCTACACGACGCTCTTCCGATCTNNNAAAAAATTAGCCAGGTGTGGT
AGCACACGTCTGAACTCTTTCCCTACACGACGCTCTTCCGATCTNNNGATCCAAATACCCATCGACAG
GAGCACACGTCTGAACTCTTTCCCTACACGACGCTCTTCCGATCTNNNCTAAGGCAGGAGAATCGCTT
AGCACACGTCTGAACTCTTTCCCTACACGACGCTCTTCCGATCTNNNGGACTCTGAGGTTGGCTAAGA

AGCACACGTCTGAACTCTTTCCCTACACGACGCTCTTCCGATCTNNNTGAGGCAGAAGAATTGCTCA
GAGCACACGTCTGAACTCTTTCCCTACACGACGCTCTTCCGATCTNNNGTTACAATGAGCCAAGATCC
AGCACACGTCTGAACTCTTTCCCTACACGACGCTCTTCCGATCTNNNGGATCCCTTGAACCTAGGAGT
AGCACACGTCTGAACTCTTTCCCTACACGACGCTCTTCCGATCTNNNTATAGTCCCAGCTACTCAGGA
AGCACACGTCTGAACTCTTTCCCTACACGACGCTCTTCCGATCTNNNTGACTGCATCCAGAAAATCCA
AGCACACGTCTGAACTCTTTCCCTACACGACGCTCTTCCGATCTNNNCTGTAGTCCCAGCTACTTGTG
GCACACGTCTGAACTCTTTCCCTACACGACGCTCTTCCGATCTNNNCTAGCTCTCATTGCTGTCTTTGC
AGCACACGTCTGAACTCTTTCCCTACACGACGCTCTTCCGATCTNNNAAAACCTGTAAGCCCAGTCTG
GCACACGTCTGAACTCTTTCCCTACACGACGCTCTTCCGATCTNNNGTACCTGTAATCCCAGCTACTCG
GCACACGTCTGAACTCTTTCCCTACACGACGCTCTTCCGATCTNNNGATTATTTGAGCCCCGGAAGTT
GCACACGTCTGAACTCTTTCCCTACACGACGCTCTTCCGATCTNNNGAGAATCACTTGAACTCGGGAG
AGCACACGTCTGAACTCTTTCCCTACACGACGCTCTTCCGATCTNNNGATTGCAGTGAGCTGAGATTG
GAGCACACGTCTGAACTCTTTCCCTACACGACGCTCTTCCGATCTNNNAGAGTTTGCAGTGAGCCAAG
AGCACACGTCTGAACTCTTTCCCTACACGACGCTCTTCCGATCTNNNCTGTAGTCCCAGCTACTCAGG
GAGCACACGTCTGAACTCTTTCCCTACACGACGCTCTTCCGATCTNNNATAAAAATTAGCTGGGCGTG
AGCACACGTCTGAACTCTTTCCCTACACGACGCTCTTCCGATCTNNNGAATCACTTGAACCCGGGAG
AGCACACGTCTGAACTCTTTCCCTACACGACGCTCTTCCGATCTNNNTACAAAAAATTAGCTGGGCGT
GAGCACACGTCTGAACTCTTTCCCTACACGACGCTCTTCCGATCTNNNTCAGGAGAATCACTTGAACC
AGCACACGTCTGAACTCTTTCCCTACACGACGCTCTTCCGATCTNNNCTGGGTGATAAAGTGAGACT
AGCACACGTCTGAACTCTTTCCCTACACGACGCTCTTCCGATCTNNNAAGAATCACTTGAACCTGGGA
AGCACACGTCTGAACTCTTTCCCTACACGACGCTCTTCCGATCTNNNTGAGGCAGGAGAATCACTTG
AGCACACGTCTGAACTCTTTCCCTACACGACGCTCTTCCGATCTNNNTGAGGCAGGAGAATCACTTGT
AGCACACGTCTGAACTCTTTCCCTACACGACGCTCTTCCGATCTNNNTGAGGCAGGAGAATCACTTGT
GAGCACACGTCTGAACTCTTTCCCTACACGACGCTCTTCCGATCTNNNTGAGGTCAGGAGTTTGAGA
AGCACACGTCTGAACTCTTTCCCTACACGACGCTCTTCCGATCTNNNTACAAAAAATTAGCCGGGCAT
AGCACACGTCTGAACTCTTTCCCTACACGACGCTCTTCCGATCTNNNGTTCACCTTGCTCAGTAAACCA
AGCACACGTCTGAACTCTTTCCCTACACGACGCTCTTCCGATCTNNNCCCAACTAAGCATGGTGATTTC
GAGCACACGTCTGAACTCTTTCCCTACACGACGCTCTTCCGATCTNNNATACCTGTGGTCCCAGATACT
GAGCACACGTCTGAACTCTTTCCCTACACGACGCTCTTCCGATCTNNNGCACTAGAACAGCTTGAACT
GCACACGTCTGAACTCTTTCCCTACACGACGCTCTTCCGATCTNNNCACACAACTGTACTCTGGTCTG
AGCACACGTCTGAACTCTTTCCCTACACGACGCTCTTCCGATCTNNNCCCATGAACAACTTCCATGAAC
AGAGCACACGTCTGAACTCTTTCCCTACACGACGCTCTTCCGATCTNNNATACAAAATTAGCTGGGGG
AGCACACGTCTGAACTCTTTCCCTACACGACGCTCTTCCGATCTNNNGCCTGTAATCCCAACACTTTGA
AGCACACGTCTGAACTCTTTCCCTACACGACGCTCTTCCGATCTNNNTTCAAAAGTAGGACACACCAC
GCACACGTCTGAACTCTTTCCCTACACGACGCTCTTCCGATCTNNNAGGCAGGAGAATCACTTGAAC
AGCACACGTCTGAACTCTTTCCCTACACGACGCTCTTCCGATCTNNNCCCAGAACTTTGAGAGGTTG
GAGCACACGTCTGAACTCTTTCCCTACACGACGCTCTTCCGATCTNNNCTGAGACAGGGGAATTGCT
AGCACACGTCTGAACTCTTTCCCTACACGACGCTCTTCCGATCTNNNCTTGAGCAATACAGACAGACC
AGCACACGTCTGAACTCTTTCCCTACACGACGCTCTTCCGATCTNNNGAACTTGCAGTGAGCCAAGA
AGCACACGTCTGAACTCTTTCCCTACACGACGCTCTTCCGATCTNNNGAGGCTGGAGTGAGCTATGA
GCACACGTCTGAACTCTTTCCCTACACGACGCTCTTCCGATCTNNNGCTTCTAGGGAATGCAAAGCA
AGCACACGTCTGAACTCTTTCCCTACACGACGCTCTTCCGATCTNNNCTGTAATTCCAGCTACTCGGGA
GAGCACACGTCTGAACTCTTTCCCTACACGACGCTCTTCCGATCTNNNTATAGTCCCAGCTACTCAGG

GAGCACACGTCTGAACTCTTTCCCTACACGACGCTCTTCCGATCTNNNCCAGCCTAGGCAATATAGCAA
GAGCACACGTCTGAACTCTTTCCCTACACGACGCTCTTCCGATCTNNNCAAAAAATTAGCCGGGCGTAG
GAGCACACGTCTGAACTCTTTCCCTACACGACGCTCTTCCGATCTNNNTACTTAGGAGGCTGAGGTAG
GAGCACACGTCTGAACTCTTTCCCTACACGACGCTCTTCCGATCTNNNTGTTTGAGCCCAAAAGTTTGA
AGCACACGTCTGAACTCTTTCCCTACACGACGCTCTTCCGATCTNNNGAGGCAGGAGAATTGCTTGAA
GAGCACACGTCTGAACTCTTTCCCTACACGACGCTCTTCCGATCTNNNCTTGAGTCCAGGAGATCAAA
GAGCACACGTCTGAACTCTTTCCCTACACGACGCTCTTCCGATCTNNNCTAAAAATTAGCCGGGCATGG
GAGCACACGTCTGAACTCTTTCCCTACACGACGCTCTTCCGATCTNNNGAATTGCTTGAACCAGATGGA
AGCACACGTCTGAACTCTTTCCCTACACGACGCTCTTCCGATCTNNNGCCAATATCACATCACTGCACTC
GAGCACACGTCTGAACTCTTTCCCTACACGACGCTCTTCCGATCTNNNCTTTGGCAAAATAGAAGGAG
AGAGCACACGTCTGAACTCTTTCCCTACACGACGCTCTTCCGATCTNNNCACCTGTAATCCCAGCTACTT
AGCACACGTCTGAACTCTTTCCCTACACGACGCTCTTCCGATCTNNNGGAGATTCACCTGCAGGAATA
GAGCACACGTCTGAACTCTTTCCCTACACGACGCTCTTCCGATCTNNNTGACAGAGCAAGAGTCCATTT
AGCACACGTCTGAACTCTTTCCCTACACGACGCTCTTCCGATCTNNNGAGAGAGGAGAATCGCTTGA
AGCACACGTCTGAACTCTTTCCCTACACGACGCTCTTCCGATCTNNNTGAGGCAGGAGAATAGTGTGA
GAGCACACGTCTGAACTCTTTCCCTACACGACGCTCTTCCGATCTNNNCACTACAACTTGGGTGACAGA
AGCACACGTCTGAACTCTTTCCCTACACGACGCTCTTCCGATCTNNNGAGAATGACATGAACCTGGGA
AGCACACGTCTGAACTCTTTCCCTACACGACGCTCTTCCGATCTNNNTACAGTGAGCCGAGATCATGCT
GAGCACACGTCTGAACTCTTTCCCTACACGACGCTCTTCCGATCTNNNGGTGACAGAGTGAGACTACGT
AGCACACGTCTGAACTCTTTCCCTACACGACGCTCTTCCGATCTNNNGGCTCACACCTCTAATCCAAGTT
GAGCACACGTCTGAACTCTTTCCCTACACGACGCTCTTCCGATCTNNNCCTACTTCTCTGAGGTGGTCCT
GAGCACACGTCTGAACTCTTTCCCTACACGACGCTCTTCCGATCTNNNACAGAACAAGACCCTGTCTCAA
GAGCACACGTCTGAACTCTTTCCCTACACGACGCTCTTCCGATCTNNNAGAAACAAACCAACACCTATG
GAGCACACGTCTGAACTCTTTCCCTACACGACGCTCTTCCGATCTNNNCATGCCTGTAATCCCAGCTACT
AGCACACGTCTGAACTCTTTCCCTACACGACGCTCTTCCGATCTNNNTAGTCCCAACTACTTGAGAGGC
GAGCACACGTCTGAACTCTTTCCCTACACGACGCTCTTCCGATCTNNNATCCCTGTAATCCCAGCTACTT
GAGCACACGTCTGAACTCTTTCCCTACACGACGCTCTTCCGATCTNNNGAGATCGTACCACTGCACTCT
AGCACACGTCTGAACTCTTTCCCTACACGACGCTCTTCCGATCTNNNCAGTTACTAGGAAGGCTGAGGT
AGCACACGTCTGAACTCTTTCCCTACACGACGCTCTTCCGATCTNNNAGTGAGCTGAGATTATACCGCT
AGCACACGTCTGAACTCTTTCCCTACACGACGCTCTTCCGATCTNNNCAGATCACACTACTGCACTCCAT
GAGCACACGTCTGAACTCTTTCCCTACACGACGCTCTTCCGATCTNNNGGTTGCAGTAAGCTGATATCA
GAGCACACGTCTGAACTCTTTCCCTACACGACGCTCTTCCGATCTNNNGTAATCCCAGCTATACAGGAG
AGAGCACACGTCTGAACTCTTTCCCTACACGACGCTCTTCCGATCTNNNTGAGTTTCAAAATGTGGCAA
AGCACACGTCTGAACTCTTTCCCTACACGACGCTCTTCCGATCTNNNTGAGCTCTGATTATACCACTGC
GAGCACACGTCTGAACTCTTTCCCTACACGACGCTCTTCCGATCTNNNAAAATGTGTTTGGGGGCTTA
AGCACACGTCTGAACTCTTTCCCTACACGACGCTCTTCCGATCTNNNTGGTAGGAATTCTCACTCACAG
GAGCACACGTCTGAACTCTTTCCCTACACGACGCTCTTCCGATCTNNNTTGGTGTGTGTGTATAGTCCC
GAGCACACGTCTGAACTCTTTCCCTACACGACGCTCTTCCGATCTNNNAACCTGTAGTTCCAGCTACTTG
AGAGCACACGTCTGAACTCTTTCCCTACACGACGCTCTTCCGATCTNNNTCAAGGCTGCAGTGAGTTAT
GAGCACACGTCTGAACTCTTTCCCTACACGACGCTCTTCCGATCTNNNGGGGATACAGAGCCAAACCAT
GAGCACACGTCTGAACTCTTTCCCTACACGACGCTCTTCCGATCTNNNAGGCTGAAACAGGAGAATCG
AGCACACGTCTGAACTCTTTCCCTACACGACGCTCTTCCGATCTNNNTGCCTGTAATTCCAGCTACTCTG
AGAGCACACGTCTGAACTCTTTCCCTACACGACGCTCTTCCGATCTNNNGAGGCAGGATAATTGCTTG

AGCACACGTCTGAACTCTTTCCCTACACGACGCTCTTCCGATCTNNNACACACAATGGAGGAAGGAAA
GAGCACACGTCTGAACTCTTTCCCTACACGACGCTCTTCCGATCTNNNTAATTCCAGTACTTTGGGAG
GAGCACACGTCTGAACTCTTTCCCTACACGACGCTCTTCCGATCTNNNATTGCAGTGAGCTATGATTGT
GAGCACACGTCTGAACTCTTTCCCTACACGACGCTCTTCCGATCTNNNGAGAATCGCTTGAACTCAGGA
GAGCACACGTCTGAACTCTTTCCCTACACGACGCTCTTCCGATCTNNNGGCTGCAGTGAGCTATAATT
GAGCACACGTCTGAACTCTTTCCCTACACGACGCTCTTCCGATCTNNNCTGAAGTGGGAGGATCTCTT
AGAGCACACGTCTGAACTCTTTCCCTACACGACGCTCTTCCGATCTNNNTCAAGACTATCCGGGCTAAC
GAGCACACGTCTGAACTCTTTCCCTACACGACGCTCTTCCGATCTNNNAAAAAAAAAAAGTTAGCCGG
AGAGCACACGTCTGAACTCTTTCCCTACACGACGCTCTTCCGATCTNNNAAGGAAAAGCAAAGGACTG
AGAGCACACGTCTGAACTCTTTCCCTACACGACGCTCTTCCGATCTNNNCTGAGGCATGAGAATTGCT
GAGCACACGTCTGAACTCTTTCCCTACACGACGCTCTTCCGATCTNNNGAGCTTGCAGTGAGCCAAGA
AGCACACGTCTGAACTCTTTCCCTACACGACGCTCTTCCGATCTNNNCACATGTAGTCCCAGCTACTCA
AGCACACGTCTGAACTCTTTCCCTACACGACGCTCTTCCGATCTNNNGCTCATGCATGTAATCCCAACA
GAGCACACGTCTGAACTCTTTCCCTACACGACGCTCTTCCGATCTNNNGCAACAGAGCAAGAAACTGT
GAGCACACGTCTGAACTCTTTCCCTACACGACGCTCTTCCGATCTNNNGAGATCGAGACACAACACTC
CACACGTCTGAACTCTTTCCCTACACGACGCTCTTCCGATCTNNNTTGCTTCAACCTAGTAGGCAGAG
GAGCACACGTCTGAACTCTTTCCCTACACGACGCTCTTCCGATCTNNNCTAAGGCAGGAGAATCGCTT
GAGCACACGTCTGAACTCTTTCCCTACACGACGCTCTTCCGATCTNNNTGAGGCAGGAGAATCTCTTGA
AGAGCACACGTCTGAACTCTTTCCCTACACGACGCTCTTCCGATCTNNNAAAACAGGGACTACTAGCAC
GAGCACACGTCTGAACTCTTTCCCTACACGACGCTCTTCCGATCTNNNAATAGTCCAAGAAAGGCTGA
GAGCACACGTCTGAACTCTTTCCCTACACGACGCTCTTCCGATCTNNNTAGGAAAATCACTGGAACCCT
GAGCACACGTCTGAACTCTTTCCCTACACGACGCTCTTCCGATCTNNNGGAGATTGCAGTGAACTGGG
AGAGCACACGTCTGAACTCTTTCCCTACACGACGCTCTTCCGATCTNNNGGCAAAAGAATCACTTGAAC
GAGCACACGTCTGAACTCTTTCCCTACACGACGCTCTTCCGATCTNNNCATCTGTAGTCCCAGCTACTCT
GAGCACACGTCTGAACTCTTTCCCTACACGACGCTCTTCCGATCTNNNGCAGAGGTTATAGTGAGCCA
AGCACACGTCTGAACTCTTTCCCTACACGACGCTCTTCCGATCTNNNAAAAAAATTAGCCGGGCGTGAT
AGCACACGTCTGAACTCTTTCCCTACACGACGCTCTTCCGATCTNNNGGGAAAGATTCTAATCCCCAGC
AGAGCACACGTCTGAACTCTTTCCCTACACGACGCTCTTCCGATCTNNNCACCTGTAATCCCAGCTACTC
AGAGCACACGTCTGAACTCTTTCCCTACACGACGCTCTTCCGATCTNNNCTGAGGCAGGAGAATCACTT
AGCACACGTCTGAACTCTTTCCCTACACGACGCTCTTCCGATCTNNNGTAATCCCAGCTACTAAGGAG
AGCACACGTCTGAACTCTTTCCCTACACGACGCTCTTCCGATCTNNNAAAAATTAGCTGGGCATAGTG
GAGCACACGTCTGAACTCTTTCCCTACACGACGCTCTTCCGATCTNNNATCTGAACTGCAGTCCTATAG
AGCACACGTCTGAACTCTTTCCCTACACGACGCTCTTCCGATCTNNNGGAGGCTGAGGTAGAAGAATC
AGCACACGTCTGAACTCTTTCCCTACACGACGCTCTTCCGATCTNNNCTGTAATCCCAGCTACTCAGGA
AGCACACGTCTGAACTCTTTCCCTACACGACGCTCTTCCGATCTNNNCTGAGATGGGAGGATCACTTGA
GAGCACACGTCTGAACTCTTTCCCTACACGACGCTCTTCCGATCTNNNGCCTGTAATCCCAGCTACTCTA
AGAGCACACGTCTGAACTCTTTCCCTACACGACGCTCTTCCGATCTNNNTGCTCACCTGTAATCTCAGCTA
AGAGCACACGTCTGAACTCTTTCCCTACACGACGCTCTTCCGATCTNNNCGAAGGTCACCAACATCAAA
AGCACACGTCTGAACTCTTTCCCTACACGACGCTCTTCCGATCTNNNTGAGTTGAGAGGATCACTTGA
GAGCACACGTCTGAACTCTTTCCCTACACGACGCTCTTCCGATCTNNNCTGTGAAAAATCACCTGGTGA
GAGCACACGTCTGAACTCTTTCCCTACACGACGCTCTTCCGATCTNNNAGAATTGCTTGAATCTGGGA
AGAGCACACGTCTGAACTCTTTCCCTACACGACGCTCTTCCGATCTNNNGAGAATCAGTTGAACCTGG
GAGCACACGTCTGAACTCTTTCCCTACACGACGCTCTTCCGATCTNNNAAGCCGGCCAATTCAATAACA

GAGCACACGTCTGAACTCTTTCCCTACACGACGCTCTTCCGATCTNNNGTAATCCCAGCTACTCAAGA
GCACACGTCTGAACTCTTTCCCTACACGACGCTCTTCCGATCTNNNAGTGAGCTGAGATTGTACCACT
AGCACACGTCTGAACTCTTTCCCTACACGACGCTCTTCCGATCTNNNCAAAGTTATCTGGGCATGGT
GAGCACACGTCTGAACTCTTTCCCTACACGACGCTCTTCCGATCTNNNGGTCCTATATGTTTCACACCT
AGCACACGTCTGAACTCTTTCCCTACACGACGCTCTTCCGATCTNNNTTGAGGCAGGAGAATAGCTTG
AGCACACGTCTGAACTCTTTCCCTACACGACGCTCTTCCGATCTNNNCAGAGCTTTCAGTGAGCCTA
GCACACGTCTGAACTCTTTCCCTACACGACGCTCTTCCGATCTNNNAGGATAATCGCTTGAAACTGGG
AGAGCACACGTCTGAACTCTTTCCCTACACGACGCTCTTCCGATCTNNNTGGGGGACAGAATGAGA
AGCACACGTCTGAACTCTTTCCCTACACGACGCTCTTCCGATCTNNNCATGACACAGTGAGACTCTGT
AGCACACGTCTGAACTCTTTCCCTACACGACGCTCTTCCGATCTNNNACATAACTAACACTCCAGCCTG
AGCACACGTCTGAACTCTTTCCCTACACGACGCTCTTCCGATCTNNNAAGCAGGAGAATCACTTGAA
AGCACACGTCTGAACTCTTTCCCTACACGACGCTCTTCCGATCTNNNCTGGGCAACATAGTAAGACCC
AGCACACGTCTGAACTCTTTCCCTACACGACGCTCTTCCGATCTNNNTGGAGCAAGACTCTGTCTCAA
AGCACACGTCTGAACTCTTTCCCTACACGACGCTCTTCCGATCTNNNGCATCTGTAATCCCAGCTACT
AGCACACGTCTGAACTCTTTCCCTACACGACGCTCTTCCGATCTNNNGTAATCCCAGCATTTTGGTAG
AGCACACGTCTGAACTCTTTCCCTACACGACGCTCTTCCGATCTNNNGAAGGATCACTTGAGTCCAGG
GAGCACACGTCTGAACTCTTTCCCTACACGACGCTCTTCCGATCTNNNGCAAGGAGGTAAAGAGGG
GCACACGTCTGAACTCTTTCCCTACACGACGCTCTTCCGATCTNNNGGTGACACAATAAGACCCTGTC
GAGCACACGTCTGAACTCTTTCCCTACACGACGCTCTTCCGATCTNNNCCTGTAAAATCCCAGCAGTTT
AGCACACGTCTGAACTCTTTCCCTACACGACGCTCTTCCGATCTNNNCAGAATTCCTTCCCTTCCCTCAA
AGCACACGTCTGAACTCTTTCCCTACACGACGCTCTTCCGATCTNNNAAAAAATTAGCTGGGTGTGGT
AGCACACGTCTGAACTCTTTCCCTACACGACGCTCTTCCGATCTNNNGCCAACATGTTGAAATCCTGTC
AGCACACGTCTGAACTCTTTCCCTACACGACGCTCTTCCGATCTNNNAGGTTGCAGTGAGCTAAAATC
AGCACACGTCTGAACTCTTTCCCTACACGACGCTCTTCCGATCTNNNGCATGAGAATTGCTTGAACCT
GCACACGTCTGAACTCTTTCCCTACACGACGCTCTTCCGATCTNNNAATACAAAAAATTAGCCGGGCG
AGCACACGTCTGAACTCTTTCCCTACACGACGCTCTTCCGATCTNNNTGTAGTCCCAGCTACTTAGGAG
AGCACACGTCTGAACTCTTTCCCTACACGACGCTCTTCCGATCTNNNCCAGGGTCAGCTACATAGTTT
GAGCACACGTCTGAACTCTTTCCCTACACGACGCTCTTCCGATCTNNNCTGCAGTGAGCTGTGATTTA
AGCACACGTCTGAACTCTTTCCCTACACGACGCTCTTCCGATCTNNNGAAGGTTGCAGTGAGCTGAA
AGCACACGTCTGAACTCTTTCCCTACACGACGCTCTTCCGATCTNNNTTAGGCAGGAGAATCGCTTGA
GAGCACACGTCTGAACTCTTTCCCTACACGACGCTCTTCCGATCTNNNCAAGATCACACTACCACACTC
GCACACGTCTGAACTCTTTCCCTACACGACGCTCTTCCGATCTNNNGTAATCCCAGCTACTCAAGGAG
GAGCACACGTCTGAACTCTTTCCCTACACGACGCTCTTCCGATCTNNNAGTGAGCCAAGATAACACCA
GCACACGTCTGAACTCTTTCCCTACACGACGCTCTTCCGATCTNNNTCTTCAGCTGTGGTAAGTCAGT
AGCACACGTCTGAACTCTTTCCCTACACGACGCTCTTCCGATCTNNNTAAGAGAGTCAGGTGCTCTTC
GCACACGTCTGAACTCTTTCCCTACACGACGCTCTTCCGATCTNNNAATGGGTCATCATAAAGCCAGG
AGCACACGTCTGAACTCTTTCCCTACACGACGCTCTTCCGATCTNNNCTGTAGTCCCAGCTACTCAGG
AGCACACGTCTGAACTCTTTCCCTACACGACGCTCTTCCGATCTNNNCTGAGGCAGGAGAATCACTTG
GAGCACACGTCTGAACTCTTTCCCTACACGACGCTCTTCCGATCTNNNTTACATTCTTGCTTCAAGCTG
AGCACACGTCTGAACTCTTTCCCTACACGACGCTCTTCCGATCTNNNCATGCCTGTAATCCCAGCTACT
GCACACGTCTGAACTCTTTCCCTACACGACGCTCTTCCGATCTNNNCTGAGGCAGGAGAATCACTTG
AGCACACGTCTGAACTCTTTCCCTACACGACGCTCTTCCGATCTNNNTGTAGTCCCAGCTACTCTAGA
GAGCACACGTCTGAACTCTTTCCCTACACGACGCTCTTCCGATCTNNNGCGACAAAGCAAGAATCTGT

GAGCACACGTCTGAACTCTTTCCCTACACGACGCTCTTCCGATCTNNNTCTACATTTGGAAGGGGAGA
AGCACACGTCTGAACTCTTTCCCTACACGACGCTCTTCCGATCTNNNGAGGCAGGAAAATCACTTGAA
GCACACGTCTGAACTCTTTCCCTACACGACGCTCTTCCGATCTNNNCTGTAGTTCCAGCTACTTGGGAG
AGAGCACACGTCTGAACTCTTTCCCTACACGACGCTCTTCCGATCTNNNCTCACACCTGAAATCCCAGTA
AGAGCACACGTCTGAACTCTTTCCCTACACGACGCTCTTCCGATCTNNNCATGCTGAAACCCGATCTCTA
GAGCACACGTCTGAACTCTTTCCCTACACGACGCTCTTCCGATCTNNNCTGAGGCAGGAGAATTGCTT
AGCACACGTCTGAACTCTTTCCCTACACGACGCTCTTCCGATCTNNNGAGAATTGCTTGAACCTGGGAA
AGCACACGTCTGAACTCTTTCCCTACACGACGCTCTTCCGATCTNNNGGCAAAATGGTGAAATCCCATC
AGCACACGTCTGAACTCTTTCCCTACACGACGCTCTTCCGATCTNNNCTGAGGCAGGAGAATTGGTTG
AGCACACGTCTGAACTCTTTCCCTACACGACGCTCTTCCGATCTNNNTCCCCACCAATAGTAAAATGCT
GAGCACACGTCTGAACTCTTTCCCTACACGACGCTCTTCCGATCTNNNCTGCTATTATACTCCAGCCTGG
AGCACACGTCTGAACTCTTTCCCTACACGACGCTCTTCCGATCTNNNCTGAGACGTGAGAATCCCTTGA
AGCACACGTCTGAACTCTTTCCCTACACGACGCTCTTCCGATCTNNNGTAGTACCAGCTACTCAGGAAC
AGCACACGTCTGAACTCTTTCCCTACACGACGCTCTTCCGATCTNNNTGAGGCCAGCTTAGCTAACATA
AGCACACGTCTGAACTCTTTCCCTACACGACGCTCTTCCGATCTNNNAAAGAGAAAGATGTCACTGGG
GAGCACACGTCTGAACTCTTTCCCTACACGACGCTCTTCCGATCTNNNTTAGTTTACTTGCCTGGCATCC
GAGCACACGTCTGAACTCTTTCCCTACACGACGCTCTTCCGATCTNNNTGCACACCTATAATCCCAGCTA
GAGCACACGTCTGAACTCTTTCCCTACACGACGCTCTTCCGATCTNNNGAGAATCACTTGAACCTGGG
GAGCACACGTCTGAACTCTTTCCCTACACGACGCTCTTCCGATCTNNNCGGGGCTGAGGAAGAAAGA
AGCACACGTCTGAACTCTTTCCCTACACGACGCTCTTCCGATCTNNNACAAAAATTAGCCCAGTGTGGT
GAGCACACGTCTGAACTCTTTCCCTACACGACGCTCTTCCGATCTNNNCCCGTTTCAAGCAACCAACTAT
GAGCACACGTCTGAACTCTTTCCCTACACGACGCTCTTCCGATCTNNNATTCAAACCAGAGCAGACATCC
GAGCACACGTCTGAACTCTTTCCCTACACGACGCTCTTCCGATCTNNNGAGTTTGAGACCAGACTAGGC
AGCACACGTCTGAACTCTTTCCCTACACGACGCTCTTCCGATCTNNNCTGTAATCCCAGCTACTCAGGA
AGAGCACACGTCTGAACTCTTTCCCTACACGACGCTCTTCCGATCTNNNTGCTGAAATAAGACAGGTCG
AGCACACGTCTGAACTCTTTCCCTACACGACGCTCTTCCGATCTNNNGGTTGCCATGCAGATGTCTAAA
GAGCACACGTCTGAACTCTTTCCCTACACGACGCTCTTCCGATCTNNNAGGAGAATTGCTTGAACCCAG
GAGCACACGTCTGAACTCTTTCCCTACACGACGCTCTTCCGATCTNNNAGAATTGCTTGAATCCTGGAG
GCACACGTCTGAACTCTTTCCCTACACGACGCTCTTCCGATCTNNNGCCAGGTATGATCTCAGCTACTC
GAGCACACGTCTGAACTCTTTCCCTACACGACGCTCTTCCGATCTNNNCTTTGTCATCAGAGGCGGAA
AGCACACGTCTGAACTCTTTCCCTACACGACGCTCTTCCGATCTNNNCTGAGGCAGGAGAATCACTTGA
AGCACACGTCTGAACTCTTTCCCTACACGACGCTCTTCCGATCTNNNGTAATCCTAGCTACTTGGGAGG
GAGCACACGTCTGAACTCTTTCCCTACACGACGCTCTTCCGATCTNNNAGGATCTCTTGAGCTCAGGGA
AGAGCACACGTCTGAACTCTTTCCCTACACGACGCTCTTCCGATCTNNNCACCTGTAATCCCAGCTACTCA
AGCACACGTCTGAACTCTTTCCCTACACGACGCTCTTCCGATCTNNNCTGGAGGCTAAGACAGGAGAA
GAGCACACGTCTGAACTCTTTCCCTACACGACGCTCTTCCGATCTNNNCATGCCTGTAATCCCAGCTACT
GAGCACACGTCTGAACTCTTTCCCTACACGACGCTCTTCCGATCTNNNTAGGCTCAAAATGAAGGGATC
GAGCACACGTCTGAACTCTTTCCCTACACGACGCTCTTCCGATCTNNNCTGTAATCACAGCTACTCGGG
GAGCACACGTCTGAACTCTTTCCCTACACGACGCTCTTCCGATCTNNNCACCTGTAATCCCAGCTACTCA
AGCACACGTCTGAACTCTTTCCCTACACGACGCTCTTCCGATCTNNNCAAGCAGGTGAGGGAGAAAA
AGAGCACACGTCTGAACTCTTTCCCTACACGACGCTCTTCCGATCTNNNTGCAGATCACAGGGGATATG
GAGCACACGTCTGAACTCTTTCCCTACACGACGCTCTTCCGATCTNNNGGACCCCAAAATCACTAAGC
GAGCACACGTCTGAACTCTTTCCCTACACGACGCTCTTCCGATCTNNNGAGGCAGGATAATTGCTTGAA

GAGCACACGTCTGAACTCTTTCCCTACACGACGCTCTTCCGATCTNNNACATACCTGTAGTCCCAGCTA
GCACACGTCTGAACTCTTTCCCTACACGACGCTCTTCCGATCTNNNGCAGGCCTACAAAATGTTGTCTT
AGCACACGTCTGAACTCTTTCCCTACACGACGCTCTTCCGATCTNNNAAAATCTCACCACTCCACTCCAA
GAGCACACGTCTGAACTCTTTCCCTACACGACGCTCTTCCGATCTNNNGAGGCAGGAGAATTGTTTG
GCACACGTCTGAACTCTTTCCCTACACGACGCTCTTCCGATCTNNNCTGGCAACAAGAGAAAAACTC
AGAGCACACGTCTGAACTCTTTCCCTACACGACGCTCTTCCGATCTNNNAGAGAAATCTGTCTAAGCT
GAGCACACGTCTGAACTCTTTCCCTACACGACGCTCTTCCGATCTNNNGTTCTGTCTCACTAGGGTTCC
GAGCACACGTCTGAACTCTTTCCCTACACGACGCTCTTCCGATCTNNNGCAGTTCATGTCCACCATATT
AGCACACGTCTGAACTCTTTCCCTACACGACGCTCTTCCGATCTNNNACACCTGTAATTCCAGCTACTC
GAGCACACGTCTGAACTCTTTCCCTACACGACGCTCTTCCGATCTNNNTCTGGAGACTGAGACAGGA
AGCACACGTCTGAACTCTTTCCCTACACGACGCTCTTCCGATCTNNNGAGAATTGCTTGAACCCAGGA
GCACACGTCTGAACTCTTTCCCTACACGACGCTCTTCCGATCTNNNTGATGGAGTGAAACCCTTTCTC
AGCACACGTCTGAACTCTTTCCCTACACGACGCTCTTCCGATCTNNNATTGTCAAGAGAGCAGACACC
GCACACGTCTGAACTCTTTCCCTACACGACGCTCTTCCGATCTNNNTACAAAAAATTAGCCGAGCGTG
AGCACACGTCTGAACTCTTTCCCTACACGACGCTCTTCCGATCTNNNGAGGTGGGAGGATCACTTGA
AGCACACGTCTGAACTCTTTCCCTACACGACGCTCTTCCGATCTNNNTGTAGTTGCAGCTACTTGGGA
GAGCACACGTCTGAACTCTTTCCCTACACGACGCTCTTCCGATCTNNNGAAGTTGCAGTGAGCTGAGA
AGCACACGTCTGAACTCTTTCCCTACACGACGCTCTTCCGATCTNNNCTGAGATGGGAGGATTGCTAG
AGCACACGTCTGAACTCTTTCCCTACACGACGCTCTTCCGATCTNNNGAGATCATGCCACTACTCTCCA
AGCACACGTCTGAACTCTTTCCCTACACGACGCTCTTCCGATCTNNNGTGAAACCCCGTCTCTACTAAA
AGCACACGTCTGAACTCTTTCCCTACACGACGCTCTTCCGATCTNNNCTTGGGAGCTGAGACATGAGA
GCACACGTCTGAACTCTTTCCCTACACGACGCTCTTCCGATCTNNNCTGTAATCCCAGCTACTCAGGAG
AGCACACGTCTGAACTCTTTCCCTACACGACGCTCTTCCGATCTNNNAGTTTGAGAACGTCTGGCCAA
GAGCACACGTCTGAACTCTTTCCCTACACGACGCTCTTCCGATCTNNNACGCCTATAGTCCCAGCTATT
AGCACACGTCTGAACTCTTTCCCTACACGACGCTCTTCCGATCTNNNAGTTGCCAAAAATGGTTACCTG
GAGCACACGTCTGAACTCTTTCCCTACACGACGCTCTTCCGATCTNNNAAACCTTGGAATTTGAGACC
AGCACACGTCTGAACTCTTTCCCTACACGACGCTCTTCCGATCTNNNAGCACTTCACTTCTAACCTCAGG
AGCACACGTCTGAACTCTTTCCCTACACGACGCTCTTCCGATCTNNNGAGTTTGAGCAACATAGCGAG
AGAGCACACGTCTGAACTCTTTCCCTACACGACGCTCTTCCGATCTNNNCACCTGTAGTTCCAGCTACT
GAGCACACGTCTGAACTCTTTCCCTACACGACGCTCTTCCGATCTNNNTAACAACAGAGCAAGACTCC
AGCACACGTCTGAACTCTTTCCCTACACGACGCTCTTCCGATCTNNNATACAAAATTACCCAGGCGTG
AGAGCACACGTCTGAACTCTTTCCCTACACGACGCTCTTCCGATCTNNNCTGTAGTCCCAGCTACTCA
AGCACACGTCTGAACTCTTTCCCTACACGACGCTCTTCCGATCTNNNGCTGAAGCAGGAGAAATGTTT
AGCACACGTCTGAACTCTTTCCCTACACGACGCTCTTCCGATCTNNNTGCCAGTAATTCCAGCATTTTG
GCACACGTCTGAACTCTTTCCCTACACGACGCTCTTCCGATCTNNNGTTTGCAGTGAGCTGAGATCAT
AGCACACGTCTGAACTCTTTCCCTACACGACGCTCTTCCGATCTNNNGAGGTTGCAGTGAGCTGAGAT
AGCACACGTCTGAACTCTTTCCCTACACGACGCTCTTCCGATCTNNNGTAATTCCAGCACTTTGGTAG
AGCACACGTCTGAACTCTTTCCCTACACGACGCTCTTCCGATCTNNNTGATGCACACCTGTAATTCCAG
AGAGCACACGTCTGAACTCTTTCCCTACACGACGCTCTTCCGATCTNNNCAATGATGTGTGCCTGTAAT
AGCACACGTCTGAACTCTTTCCCTACACGACGCTCTTCCGATCTNNNACACCTGTAGTCCCAGCTACTC
AGCACACGTCTGAACTCTTTCCCTACACGACGCTCTTCCGATCTNNNCTCATGCCTATAATCCCTGCAC
GAGCACACGTCTGAACTCTTTCCCTACACGACGCTCTTCCGATCTNNNTCTGGTTAGACAGAATCCAG
AGCACACGTCTGAACTCTTTCCCTACACGACGCTCTTCCGATCTNNNCGTATCTGTAACCCCAGCTAC

GAGCACACGTCTGAACTCTTTCCCTACACGACGCTCTTCCGATCTNNNACCTGTAATCCCAACTGCTAA
GAGCACACGTCTGAACTCTTTCCCTACACGACGCTCTTCCGATCTNNNTAGGTAATCACACCACTGCA
GCACACGTCTGAACTCTTTCCCTACACGACGCTCTTCCGATCTNNNGGATGACAGGGTGAGACTCAA
GAGCACACGTCTGAACTCTTTCCCTACACGACGCTCTTCCGATCTNNNGCAGATCACCTAAGGTCAG
GCACACGTCTGAACTCTTTCCCTACACGACGCTCTTCCGATCTNNNGAGGTTTTACTGAGCCCAGATC
GCACACGTCTGAACTCTTTCCCTACACGACGCTCTTCCGATCTNNNTGTTTTTCCCAGGAACACTCTCT
AGCACACGTCTGAACTCTTTCCCTACACGACGCTCTTCCGATCTNNNAAAACAACACTGCCCACCATAA
GCACACGTCTGAACTCTTTCCCTACACGACGCTCTTCCGATCTNNNCTGAGGCAGGAGAATCACTTGA
AGCACACGTCTGAACTCTTTCCCTACACGACGCTCTTCCGATCTNNNCAAAAAAAGGTCCCACATAGC
AGCACACGTCTGAACTCTTTCCCTACACGACGCTCTTCCGATCTNNNCTTATGAGCCAAAAGAAGCCC
AGCACACGTCTGAACTCTTTCCCTACACGACGCTCTTCCGATCTNNNAGACTGGCTGTAAGGAATGC
AGCACACGTCTGAACTCTTTCCCTACACGACGCTCTTCCGATCTNNNGTAATCCCAGCTACTAGTGAG
AGAGCACACGTCTGAACTCTTTCCCTACACGACGCTCTTCCGATCTNNNTACGTGAGACTTGGAATTG
AGCACACGTCTGAACTCTTTCCCTACACGACGCTCTTCCGATCTNNNTCAAAACCACAATGCGATACC
AGCACACGTCTGAACTCTTTCCCTACACGACGCTCTTCCGATCTNNNTGAGGCAGGAGAATGGTATG
AGCACACGTCTGAACTCTTTCCCTACACGACGCTCTTCCGATCTNNNGAGAATCACTCGAACCAGGGA
AGCACACGTCTGAACTCTTTCCCTACACGACGCTCTTCCGATCTNNNCTAAGGCAGGAGAATCGCTTG
GCACACGTCTGAACTCTTTCCCTACACGACGCTCTTCCGATCTNNNTGGGAGGTTGAGGTAGGAGA
GCACACGTCTGAACTCTTTCCCTACACGACGCTCTTCCGATCTNNNTGAGGGAAACAGGAACATACCA
AGCACACGTCTGAACTCTTTCCCTACACGACGCTCTTCCGATCTNNNCTGAGGTGGGAGAATCACTT
AGCACACGTCTGAACTCTTTCCCTACACGACGCTCTTCCGATCTNNNGTGACAGAGCCAGATTCCATC
AGCACACGTCTGAACTCTTTCCCTACACGACGCTCTTCCGATCTNNNATGATCACACCACTGCACTCT
GCACACGTCTGAACTCTTTCCCTACACGACGCTCTTCCGATCTNNNGTTTGCAGTGAGCCAAGATCAT
AGCACACGTCTGAACTCTTTCCCTACACGACGCTCTTCCGATCTNNNGACCCTTCATTCCAGGACATTC
GAGCACACGTCTGAACTCTTTCCCTACACGACGCTCTTCCGATCTNNNGTAATCTCAGCTACTCAGGA
AGCACACGTCTGAACTCTTTCCCTACACGACGCTCTTCCGATCTNNNACTCAATCCCCTTTACAACAGC
GAGCACACGTCTGAACTCTTTCCCTACACGACGCTCTTCCGATCTNNNGTCAAGAGATCGAGACCATC
AGCACACGTCTGAACTCTTTCCCTACACGACGCTCTTCCGATCTNNNAGGCAAGAAGTAGAAATGAC
AGCACACGTCTGAACTCTTTCCCTACACGACGCTCTTCCGATCTNNNGATTGCAGTAAGCCAAGATCA
AGCACACGTCTGAACTCTTTCCCTACACGACGCTCTTCCGATCTNNNAAAAAATTAGCCGGACATGG
GCACACGTCTGAACTCTTTCCCTACACGACGCTCTTCCGATCTNNNCTGAGGCAGGAGAATCACTTG
GCACACGTCTGAACTCTTTCCCTACACGACGCTCTTCCGATCTNNNGCACATGCTGTTGGAAAAATGA
AGAGCACACGTCTGAACTCTTTCCCTACACGACGCTCTTCCGATCTNNNGAGATTCGCTTGAACTCAG
AGCACACGTCTGAACTCTTTCCCTACACGACGCTCTTCCGATCTNNNTCTAATCATGTCACTACGCTCCA
GCACACGTCTGAACTCTTTCCCTACACGACGCTCTTCCGATCTNNNGCTGAGGTAGGAGAATTGCTT
AGCACACGTCTGAACTCTTTCCCTACACGACGCTCTTCCGATCTNNNGGAGTTTGAGGCTGTAGTGA
AGCACACGTCTGAACTCTTTCCCTACACGACGCTCTTCCGATCTNNNCCTGTAATCCCAGCTACTCAGG
AGCACACGTCTGAACTCTTTCCCTACACGACGCTCTTCCGATCTNNNAGAGGTTTCAGTGACCCAAG
GCACACGTCTGAACTCTTTCCCTACACGACGCTCTTCCGATCTNNNTGTATGCTTGTAGTCCCAGCTA
AGCACACGTCTGAACTCTTTCCCTACACGACGCTCTTCCGATCTNNNCTGTAATCCCAGCTACTCAGGA
AGCACACGTCTGAACTCTTTCCCTACACGACGCTCTTCCGATCTNNNCTCATGCCTGTAATCCCAGCTA
AGCACACGTCTGAACTCTTTCCCTACACGACGCTCTTCCGATCTNNNCTGAGGCAGGAGAATCACTTG
AGCACACGTCTGAACTCTTTCCCTACACGACGCTCTTCCGATCTNNNGATCATACCACTGTACTCCAGC

GAGCACACGTCTGAACTCTTTCCCTACACGACGCTCTTCCGATCTNNNGATCATGAGGTCAGGAGTTC
AGCACACGTCTGAACTCTTTCCCTACACGACGCTCTTCCGATCTNNNGAGATAACGCCACTGCACTCT
AGCACACGTCTGAACTCTTTCCCTACACGACGCTCTTCCGATCTNNNGCAGGGATGATCTTTCCTGTG
AGCACACGTCTGAACTCTTTCCCTACACGACGCTCTTCCGATCTNNNGGAACCACAAAAGACCCAGA
GAGCACACGTCTGAACTCTTTCCCTACACGACGCTCTTCCGATCTNNNCCAAGATTGCACCACTACACTT
AGCACACGTCTGAACTCTTTCCCTACACGACGCTCTTCCGATCTNNNAGATTGTGCTACTGTACTCCAG
AGCACACGTCTGAACTCTTTCCCTACACGACGCTCTTCCGATCTNNNATGCCTGTAATCCCAGCTACTT
AGCACACGTCTGAACTCTTTCCCTACACGACGCTCTTCCGATCTNNNCTGAGACAGGAGAATGGCTTG
AGCACACGTCTGAACTCTTTCCCTACACGACGCTCTTCCGATCTNNNAATTTGGGAAATTCACAGCCT
AGCACACGTCTGAACTCTTTCCCTACACGACGCTCTTCCGATCTNNNACCTGGGTAATATAGCAAGA
GAGCACACGTCTGAACTCTTTCCCTACACGACGCTCTTCCGATCTNNNGATTGCTTGAGTCCAGGATT
GAGCACACGTCTGAACTCTTTCCCTACACGACGCTCTTCCGATCTNNNGCATGCACCTGTAGTCTCAA
AGCACACGTCTGAACTCTTTCCCTACACGACGCTCTTCCGATCTNNNCCAAAATAAAGATCAGGCTGG
AGCACACGTCTGAACTCTTTCCCTACACGACGCTCTTCCGATCTNNNAGGTTGCAGTGAACCAAGATA
AGCACACGTCTGAACTCTTTCCCTACACGACGCTCTTCCGATCTNNNCTGGGGTCACATATAGCTCAGT
GAGCACACGTCTGAACTCTTTCCCTACACGACGCTCTTCCGATCTNNNCTGAGATTGCCCTACTGTACT
AGCACACGTCTGAACTCTTTCCCTACACGACGCTCTTCCGATCTNNNAAAAAATTACCCAGGCATGGTG
AGCACACGTCTGAACTCTTTCCCTACACGACGCTCTTCCGATCTNNNTGGTACATGCCTATACTTCCAG
AGCACACGTCTGAACTCTTTCCCTACACGACGCTCTTCCGATCTNNNGGGAGGATGAAGCAGGAGA
AGCACACGTCTGAACTCTTTCCCTACACGACGCTCTTCCGATCTNNNTGAAGATTGTGGTGAGCTGAG
GAGCACACGTCTGAACTCTTTCCCTACACGACGCTCTTCCGATCTNNNCTGAGGCAAGAGAATTGCTG
AGCACACGTCTGAACTCTTTCCCTACACGACGCTCTTCCGATCTNNNATACACCTGTAGTCCCAGCTAC
AGCACACGTCTGAACTCTTTCCCTACACGACGCTCTTCCGATCTNNNCTCATGCCTGTAATCCTAGCAC
AGCACACGTCTGAACTCTTTCCCTACACGACGCTCTTCCGATCTNNNAAATACAAAAATTAGCCGGGCG
AGCACACGTCTGAACTCTTTCCCTACACGACGCTCTTCCGATCTNNNCTGAGGCAGGAGAATCACTTG
GAGCACACGTCTGAACTCTTTCCCTACACGACGCTCTTCCGATCTNNNCTGTAATCCCAGCTACTCGAG
AGCACACGTCTGAACTCTTTCCCTACACGACGCTCTTCCGATCTNNNTAGAAAAAAATTAGCCGGGCG
AGCACACGTCTGAACTCTTTCCCTACACGACGCTCTTCCGATCTNNNATGCCTGTAATCCCAGCTACTT
AGCACACGTCTGAACTCTTTCCCTACACGACGCTCTTCCGATCTNNNAGACAGGAGAATTGCTTGAAC
AGCACACGTCTGAACTCTTTCCCTACACGACGCTCTTCCGATCTNNNCATCCTGGCTAACATGGTGAA
AGCACACGTCTGAACTCTTTCCCTACACGACGCTCTTCCGATCTNNNCTGTAATCCCAGCTACTCAGGA
GAGCACACGTCTGAACTCTTTCCCTACACGACGCTCTTCCGATCTNNNAGGTGATGTTGTAAGACAGG
GAGCACACGTCTGAACTCTTTCCCTACACGACGCTCTTCCGATCTNNNAGATGTTGCAATGAGCCAAG
AGCACACGTCTGAACTCTTTCCCTACACGACGCTCTTCCGATCTNNNGAGAATCGCTTGAATCCAGGA
GAGCACACGTCTGAACTCTTTCCCTACACGACGCTCTTCCGATCTNNNGCACATGCCTATAATCCCAGC
AGCACACGTCTGAACTCTTTCCCTACACGACGCTCTTCCGATCTNNNGGCAATAAGAGTGAAGTCCGT
GCACACGTCTGAACTCTTTCCCTACACGACGCTCTTCCGATCTNNNTGCCTGTAATCTCAGCTACTTGG
AGCACACGTCTGAACTCTTTCCCTACACGACGCTCTTCCGATCTNNNTCAAATTGGGCGACAGAGTAA
GAGCACACGTCTGAACTCTTTCCCTACACGACGCTCTTCCGATCTNNNAATACAAAATTAGCTGGGCG
AGCACACGTCTGAACTCTTTCCCTACACGACGCTCTTCCGATCTNNNTGAGGCAGGAAAATCGTTTG
GAGCACACGTCTGAACTCTTTCCCTACACGACGCTCTTCCGATCTNNNACGCCTGTAATCCCAGTACTT
GAGCACACGTCTGAACTCTTTCCCTACACGACGCTCTTCCGATCTNNNTGAGTTTCTAAGCTACTGGCA
GAGCACACGTCTGAACTCTTTCCCTACACGACGCTCTTCCGATCTNNNGCATACCTGTAATCCCAGCTA

AGCACACGTCTGAACTCTTTCCCTACACGACGCTCTTCCGATCTNNNGAGATAACGCAATTGCACTCC
GAGCACACGTCTGAACTCTTTCCCTACACGACGCTCTTCCGATCTNNNAGTCAGGAGAATTGCTTGAA
GAGCACACGTCTGAACTCTTTCCCTACACGACGCTCTTCCGATCTNNNACCTGTAATCCCATCTACTCGG
AGCACACGTCTGAACTCTTTCCCTACACGACGCTCTTCCGATCTNNNCTGAGGCAGGAGAATCACTT
GAGCACACGTCTGAACTCTTTCCCTACACGACGCTCTTCCGATCTNNNTTGAGAGTAGTAAACTGGG
AGCACACGTCTGAACTCTTTCCCTACACGACGCTCTTCCGATCTNNNTGTGTGTGTGTCTCTTCTCCAT
AGCACACGTCTGAACTCTTTCCCTACACGACGCTCTTCCGATCTNNNATAAGCTAATTTTGGCTGGGT
GAGCACACGTCTGAACTCTTTCCCTACACGACGCTCTTCCGATCTNNNCTGCAGCAGGAAAATCACTT
GCACACGTCTGAACTCTTTCCCTACACGACGCTCTTCCGATCTNNNGCTGTGGTTTTTGCTTACATG
AGCACACGTCTGAACTCTTTCCCTACACGACGCTCTTCCGATCTNNNCTGGGGGTAACAGAGAAAG
AGCACACGTCTGAACTCTTTCCCTACACGACGCTCTTCCGATCTNNNAAGTGGGAGTATCATTTGGA
AGCACACGTCTGAACTCTTTCCCTACACGACGCTCTTCCGATCTNNNCATTGCACCATCAGAGTTCACA
GAGCACACGTCTGAACTCTTTCCCTACACGACGCTCTTCCGATCTNNNAGGCAAGAGAATCACTTGAA
GAGCACACGTCTGAACTCTTTCCCTACACGACGCTCTTCCGATCTNNNGAGATCACATCACTGCACTC
AGCACACGTCTGAACTCTTTCCCTACACGACGCTCTTCCGATCTNNNCTGTTATCCCAGCTACTCGGAA
AGCACACGTCTGAACTCTTTCCCTACACGACGCTCTTCCGATCTNNNTGAGGCACGAAAATCACTTG
GAGCACACGTCTGAACTCTTTCCCTACACGACGCTCTTCCGATCTNNNTGAGGCAAGAGAATTGCTTG
GCACACGTCTGAACTCTTTCCCTACACGACGCTCTTCCGATCTNNNACCTTCCCTCCACTATTGTCCTA
AGCACACGTCTGAACTCTTTCCCTACACGACGCTCTTCCGATCTNNNACAAAAATTAGCTGGGTGTG
GCACACGTCTGAACTCTTTCCCTACACGACGCTCTTCCGATCTNNNCACCTGTAATCCCAGCTACTTGG
AGCACACGTCTGAACTCTTTCCCTACACGACGCTCTTCCGATCTNNNTGCAGTGAGTCATGATCAAGT
GCACACGTCTGAACTCTTTCCCTACACGACGCTCTTCCGATCTNNNGGAGGGACACGGATTTCTTTA
GCACACGTCTGAACTCTTTCCCTACACGACGCTCTTCCGATCTNNNCTGGAGAGACAGTGCGATTTA
GAGCACACGTCTGAACTCTTTCCCTACACGACGCTCTTCCGATCTNNNTATACTGGAGGCTGAGGTA
AGCACACGTCTGAACTCTTTCCCTACACGACGCTCTTCCGATCTNNNGAGCTCAAATCATCACTGCAG
AGCACACGTCTGAACTCTTTCCCTACACGACGCTCTTCCGATCTNNNTGTAGTCCCAGCTATTCAGGA
AGCACACGTCTGAACTCTTTCCCTACACGACGCTCTTCCGATCTNNNACGGATCAGATAAGCAAGTGT
AGCACACGTCTGAACTCTTTCCCTACACGACGCTCTTCCGATCTNNNAAGACCCATCAGTGTGCTGTA
GAGCACACGTCTGAACTCTTTCCCTACACGACGCTCTTCCGATCTNNNCTAGCTACTCTGGAGACAAG
AGCACACGTCTGAACTCTTTCCCTACACGACGCTCTTCCGATCTNNNAAGACTGAAGCCAAGAAGGT
AGCACACGTCTGAACTCTTTCCCTACACGACGCTCTTCCGATCTNNNATTTCAGCAAGATGAACAGCC
AGCACACGTCTGAACTCTTTCCCTACACGACGCTCTTCCGATCTNNNCCTGGACAATATGGAGGAAC
GAGCACACGTCTGAACTCTTTCCCTACACGACGCTCTTCCGATCTNNNCATGCCTGTAATCCCAGCTAC
GAGCACACGTCTGAACTCTTTCCCTACACGACGCTCTTCCGATCTNNNTCATGTGCCGGTATGGATCT
AGCACACGTCTGAACTCTTTCCCTACACGACGCTCTTCCGATCTNNNACTGAGGTAGGAGGATCACT
AGCACACGTCTGAACTCTTTCCCTACACGACGCTCTTCCGATCTNNNAGAATCACTTGAATCCAGGAG
GAGCACACGTCTGAACTCTTTCCCTACACGACGCTCTTCCGATCTNNNGAGGTCTAGAGTTTGAGGC
GAGCACACGTCTGAACTCTTTCCCTACACGACGCTCTTCCGATCTNNNAGGAAGCTGAGACTGGAGA
GAGCACACGTCTGAACTCTTTCCCTACACGACGCTCTTCCGATCTNNNAAGAGGGTGACATTCTGCC
GAGCACACGTCTGAACTCTTTCCCTACACGACGCTCTTCCGATCTNNNGTGAGCTGAGATCATGCCAC
GCACACGTCTGAACTCTTTCCCTACACGACGCTCTTCCGATCTNNNATGCCTGTAGTCCCAGCTAATT
GAGCACACGTCTGAACTCTTTCCCTACACGACGCTCTTCCGATCTNNNTGGTTGAATAAGACAAGCC
AGCACACGTCTGAACTCTTTCCCTACACGACGCTCTTCCGATCTNNNGTAATCCCAGCTACTCAGGAA

AGCACACGTCTGAACTCTTTCCCTACACGACGCTCTTCCGATCTNNNTTCTAACCTTACCACCACTGCT
AGCACACGTCTGAACTCTTTCCCTACACGACGCTCTTCCGATCTNNNATCTAGTTCCCGTTAGCTCAAC
AGCACACGTCTGAACTCTTTCCCTACACGACGCTCTTCCGATCTNNNGAGAGTAGCTTGGCAGGAGA
GAGCACACGTCTGAACTCTTTCCCTACACGACGCTCTTCCGATCTNNNTGCCTCCTGATGAAAAAACAC
AGCACACGTCTGAACTCTTTCCCTACACGACGCTCTTCCGATCTNNNCTGAGGCAGGAGAATTGCTTG
AGCACACGTCTGAACTCTTTCCCTACACGACGCTCTTCCGATCTNNNAAAATTTAGCCAGGCGTGGTA
AGCACACGTCTGAACTCTTTCCCTACACGACGCTCTTCCGATCTNNNGAGGTTACAGTGAGCTGAGA
AGCACACGTCTGAACTCTTTCCCTACACGACGCTCTTCCGATCTNNNGGCAACATAGCAAAACCTCAT
GCACACGTCTGAACTCTTTCCCTACACGACGCTCTTCCGATCTNNNTGCAGTGAGCCAAGATTATGCT
GAGCACACGTCTGAACTCTTTCCCTACACGACGCTCTTCCGATCTNNNGTAGTCTCAGCTACTCAGGA
AGCACACGTCTGAACTCTTTCCCTACACGACGCTCTTCCGATCTNNNTGAGGCAGGAGAATCACTTG
GAGCACACGTCTGAACTCTTTCCCTACACGACGCTCTTCCGATCTNNNCTGTAATCCCAGCTACTCAGG
GAGCACACGTCTGAACTCTTTCCCTACACGACGCTCTTCCGATCTNNNATACAAAAATTAGCAGGGCG
AGCACACGTCTGAACTCTTTCCCTACACGACGCTCTTCCGATCTNNNCTGTAGTCCCAGCTACTCAGG
GCACACGTCTGAACTCTTTCCCTACACGACGCTCTTCCGATCTNNNCAAAAATTACCCTGGTGTGGTG
GAGCACACGTCTGAACTCTTTCCCTACACGACGCTCTTCCGATCTNNNGAGGCAGGAAAATCACTTG
GAGCACACGTCTGAACTCTTTCCCTACACGACGCTCTTCCGATCTNNNAATCCTAGCACTTTGGGAGA
AGCACACGTCTGAACTCTTTCCCTACACGACGCTCTTCCGATCTNNNTATGTCTGCAATCCCAGCTACT
AGCACACGTCTGAACTCTTTCCCTACACGACGCTCTTCCGATCTNNNAAAAAAATTAGCCGGGCATT
AGCACACGTCTGAACTCTTTCCCTACACGACGCTCTTCCGATCTNNNCCTACACTCCTCAAGAAGCAG
GAGCACACGTCTGAACTCTTTCCCTACACGACGCTCTTCCGATCTNNNCAGTGAGCCATAGTCGTATC
AGCACACGTCTGAACTCTTTCCCTACACGACGCTCTTCCGATCTNNNGTAATCCTAGCTACTTGGGAG
GAGCACACGTCTGAACTCTTTCCCTACACGACGCTCTTCCGATCTNNNGTTCAAAACCAGCCTGAGCA
AGCACACGTCTGAACTCTTTCCCTACACGACGCTCTTCCGATCTNNNAGAATCGTCTGATTGCTGGAG
AGCACACGTCTGAACTCTTTCCCTACACGACGCTCTTCCGATCTNNNGCAGGAGAATCATTTGAACC
GCACACGTCTGAACTCTTTCCCTACACGACGCTCTTCCGATCTNNNAGGAAAATCACTTGAACTCGGG
AGCACACGTCTGAACTCTTTCCCTACACGACGCTCTTCCGATCTNNNGGCAACATAGCAAGACCTCTCT
GAGCACACGTCTGAACTCTTTCCCTACACGACGCTCTTCCGATCTNNNCTGGGCTACAAGAGTAAGA
GAGCACACGTCTGAACTCTTTCCCTACACGACGCTCTTCCGATCTNNNGTTGAGGTGGGAGGATCAC
GCACACGTCTGAACTCTTTCCCTACACGACGCTCTTCCGATCTNNNGAGATCACACCACTGCACTCTAG
GCACACGTCTGAACTCTTTCCCTACACGACGCTCTTCCGATCTNNNGTGGTGGGTGCCTATAATCCTA
GAGCACACGTCTGAACTCTTTCCCTACACGACGCTCTTCCGATCTNNNCTGTAATCCCAGACACTCAGG
AGCACACGTCTGAACTCTTTCCCTACACGACGCTCTTCCGATCTNNNGCTGAAGCATAAGAATCGCTT
GAGCACACGTCTGAACTCTTTCCCTACACGACGCTCTTCCGATCTNNNACACCTGTAATCCCAGCTAC
AGCACACGTCTGAACTCTTTCCCTACACGACGCTCTTCCGATCTNNNCTGAGGCAGGAGAATCACTT
AGCACACGTCTGAACTCTTTCCCTACACGACGCTCTTCCGATCTNNNCATGCCTGTAATCCCAGCTACT
GAGCACACGTCTGAACTCTTTCCCTACACGACGCTCTTCCGATCTNNNAGTGAGCCATGATCATGTCAC
AGCACACGTCTGAACTCTTTCCCTACACGACGCTCTTCCGATCTNNNCTGGCTGTCTATGAACTAGGG
AGCACACGTCTGAACTCTTTCCCTACACGACGCTCTTCCGATCTNNNCTGATGCCTATAATCCCAGCAC
AGCACACGTCTGAACTCTTTCCCTACACGACGCTCTTCCGATCTNNNTGAGTAACAGAACGAGAACC
AGCACACGTCTGAACTCTTTCCCTACACGACGCTCTTCCGATCTNNNCTGAGGCAGAAGAAGCACTT
AGCACACGTCTGAACTCTTTCCCTACACGACGCTCTTCCGATCTNNNAGTGAGCCAAGATCATACCAC
AGCACACGTCTGAACTCTTTCCCTACACGACGCTCTTCCGATCTNNNGACATGGCAAAACCCCATCTCT

AGCACACGTCTGAACTCTTTCCCTACACGACGCTCTTCCGATCTNNNAGATTTGTGCATTCATCACCACA

GAGCACACGTCTGAACTCTTTCCCTACACGACGCTCTTCCGATCTNNNCACCTGTAATCCCAGCTACTCT

AGCACACGTCTGAACTCTTTCCCTACACGACGCTCTTCCGATCTNNNCTGAGGCAGGAGAATCACTTG

GAGCACACGTCTGAACTCTTTCCCTACACGACGCTCTTCCGATCTNNNTACCTGTAGTCCCAGCTACTCA

AGCACACGTCTGAACTCTTTCCCTACACGACGCTCTTCCGATCTNNNCTGAGGCAGGAGAATCACTTGA

GAGCACACGTCTGAACTCTTTCCCTACACGACGCTCTTCCGATCTNNNTGGAGGAAGCAGAGATAACC

GAGCACACGTCTGAACTCTTTCCCTACACGACGCTCTTCCGATCTNNNGGTTGCAGTGATCCGATATTG

GAGCACACGTCTGAACTCTTTCCCTACACGACGCTCTTCCGATCTNNNAATCCTAGCTATTCAGGAGCCT

AGAGCACACGTCTGAACTCTTTCCCTACACGACGCTCTTCCGATCTNNNCGCCTGTAATCCTAGCTACT

GAGCACACGTCTGAACTCTTTCCCTACACGACGCTCTTCCGATCTNNNTGAGGCAGAAGAATTGCTTGA

GAGCACACGTCTGAACTCTTTCCCTACACGACGCTCTTCCGATCTNNNTGAAGTCAGGAGTTTGAGAC

AGCACACGTCTGAACTCTTTCCCTACACGACGCTCTTCCGATCTNNNCACCCCCAGCTGTATTTCTTTTG

GAGCACACGTCTGAACTCTTTCCCTACACGACGCTCTTCCGATCTNNNTGAGGCAGGAGAATTGCTTG

GAGCACACGTCTGAACTCTTTCCCTACACGACGCTCTTCCGATCTNNNGAGACTGAGGTGGGAGAATT

AGCACACGTCTGAACTCTTTCCCTACACGACGCTCTTCCGATCTNNNGCACCTGTAATCCCAGCTACTAG

AGCACACGTCTGAACTCTTTCCCTACACGACGCTCTTCCGATCTNNNTCAGAAACCCAGAAGGATGTCA

GAGCACACGTCTGAACTCTTTCCCTACACGACGCTCTTCCGATCTNNNGAGGCTGAAGCAAGAGAAT

AGCACACGTCTGAACTCTTTCCCTACACGACGCTCTTCCGATCTNNNACCTCCAGTGCAATGTTGAATT

AGAGCACACGTCTGAACTCTTTCCCTACACGACGCTCTTCCGATCTNNNCTGAGGCAGGAGAATCACTT

GAGCACACGTCTGAACTCTTTCCCTACACGACGCTCTTCCGATCTNNNGTGAAACCCCGTCTCTACTAAA

GAGCACACGTCTGAACTCTTTCCCTACACGACGCTCTTCCGATCTNNNCACCTGTAATCCCAGCTACTCA

AGAGCACACGTCTGAACTCTTTCCCTACACGACGCTCTTCCGATCTNNNTGTAGTCTCAGCTACTCTGGA

AGCACACGTCTGAACTCTTTCCCTACACGACGCTCTTCCGATCTNNNCTGAGGCAGGAGAATCACTTCA

GAGCACACGTCTGAACTCTTTCCCTACACGACGCTCTTCCGATCTNNNACATCTCTAGTCCCAGCTACTC

AGAGCACACGTCTGAACTCTTTCCCTACACGACGCTCTTCCGATCTNNNCTGTAGTCCCAGCTACTCAG

AGCACACGTCTGAACTCTTTCCCTACACGACGCTCTTCCGATCTNNNCTGCAGTGAGCCAAAATCATAC

GAGCACACGTCTGAACTCTTTCCCTACACGACGCTCTTCCGATCTNNNTCCTAGCTACTTAGGAGGCT

GCACACGTCTGAACTCTTTCCCTACACGACGCTCTTCCGATCTNNNCATGCCTGTAGTCCCAGATACTC

GAGCACACGTCTGAACTCTTTCCCTACACGACGCTCTTCCGATCTNNNATACACCTGTAGTCCCAGCTA

AGCACACGTCTGAACTCTTTCCCTACACGACGCTCTTCCGATCTNNNCTGATGCCTGTAATCCTAGCAC

AGAGCACACGTCTGAACTCTTTCCCTACACGACGCTCTTCCGATCTNNNTAGCTTGAGACCAGACTGG

AGCACACGTCTGAACTCTTTCCCTACACGACGCTCTTCCGATCTNNNCTGTAGTGAACCGAGATCGTAC

GAGCACACGTCTGAACTCTTTCCCTACACGACGCTCTTCCGATCTNNNGAGAATCACTTGAACCCAGGA

AGCACACGTCTGAACTCTTTCCCTACACGACGCTCTTCCGATCTNNNATCCTGGGAAGTTGATGCTGTA

AGCACACGTCTGAACTCTTTCCCTACACGACGCTCTTCCGATCTNNNCACATGCCTATAGTCCCAGCTA

GAGCACACGTCTGAACTCTTTCCCTACACGACGCTCTTCCGATCTNNNAAGCTAAGGCACAAGAATCG

AGCACACGTCTGAACTCTTTCCCTACACGACGCTCTTCCGATCTNNNGGCAACAGAGTGAGATTCCAT

AGCACACGTCTGAACTCTTTCCCTACACGACGCTCTTCCGATCTNNNGATTGCAGTGAGCCAAGATCAT

AGCACACGTCTGAACTCTTTCCCTACACGACGCTCTTCCGATCTNNNAGTAAGCCATGATCACACCACT

GAGCACACGTCTGAACTCTTTCCCTACACGACGCTCTTCCGATCTNNNCTGAGTCAGGAGAATCGCTT

GAGCACACGTCTGAACTCTTTCCCTACACGACGCTCTTCCGATCTNNNCAGCAAGAGGATTCTGCATCA

GAGCACACGTCTGAACTCTTTCCCTACACGACGCTCTTCCGATCTNNNCTGAGGCAGGAGAATTGCTT

AAGAGCACACGTCTGAACTCTTTCCCTACACGACGCTCTTCCGATCTNNNAAAATTAGCCAGGTGTGG

AGCACACGTCTGAACTCTTTCCCTACACGACGCTCTTCCGATCTNNNACTTCCTCCTCCCATTCAGACTA
AGCACACGTCTGAACTCTTTCCCTACACGACGCTCTTCCGATCTNNNGATGCCCTGATTTAGAACCAA
AGCACACGTCTGAACTCTTTCCCTACACGACGCTCTTCCGATCTNNNAAGGATCGCTTGAGTCTGGAA
GAGCACACGTCTGAACTCTTTCCCTACACGACGCTCTTCCGATCTNNNCTCCATCACCTACAGAGTGAC
AGCACACGTCTGAACTCTTTCCCTACACGACGCTCTTCCGATCTNNNCTGAGATGGAGGAATCGCTT
GCACACGTCTGAACTCTTTCCCTACACGACGCTCTTCCGATCTNNNAGACGTTCAACAATGGCTCTACT
GAGCACACGTCTGAACTCTTTCCCTACACGACGCTCTTCCGATCTNNNTGTAGTCCCAGCTACTAGGAA
AGCACACGTCTGAACTCTTTCCCTACACGACGCTCTTCCGATCTNNNTGAAACCCCGTCTCCACTAAAA
GAGCACACGTCTGAACTCTTTCCCTACACGACGCTCTTCCGATCTNNNTGATTGAGCCAAGGTCATACC
AGCACACGTCTGAACTCTTTCCCTACACGACGCTCTTCCGATCTNNNCACCTGTATTCCCAGCTACTCA
AGCACACGTCTGAACTCTTTCCCTACACGACGCTCTTCCGATCTNNNGAAGGTTGCAGTGAGCTGAA
AGCACACGTCTGAACTCTTTCCCTACACGACGCTCTTCCGATCTNNNAGCCTGTAGTTCCAGCTATTCA
AGCACACGTCTGAACTCTTTCCCTACACGACGCTCTTCCGATCTNNNCATATGCCTGTAGTCCTAGCCA
GAGCACACGTCTGAACTCTTTCCCTACACGACGCTCTTCCGATCTNNNGTGGCACATACCTGTAGTCCT
AGCACACGTCTGAACTCTTTCCCTACACGACGCTCTTCCGATCTNNNCAAAAATTAGCCGGGTGAGAT
GAGCACACGTCTGAACTCTTTCCCTACACGACGCTCTTCCGATCTNNNTGAGAGGAGGAGGTTGTAG
GCACACGTCTGAACTCTTTCCCTACACGACGCTCTTCCGATCTNNNGAGTAATAGTCTCCAGCCAGGT
AGCACACGTCTGAACTCTTTCCCTACACGACGCTCTTCCGATCTNNNATAAATAAATAGGCCGTGCGC
AGCACACGTCTGAACTCTTTCCCTACACGACGCTCTTCCGATCTNNNAACACTGTTTTGCTAGGGGGT
GAGCACACGTCTGAACTCTTTCCCTACACGACGCTCTTCCGATCTNNNGCATACCTGTAATCCCAGCTA
GAGCACACGTCTGAACTCTTTCCCTACACGACGCTCTTCCGATCTNNNCAAGATCACACCACTGTACTC
GAGCACACGTCTGAACTCTTTCCCTACACGACGCTCTTCCGATCTNNNGGAGGCTAAGACAGGAGAA
GAGCACACGTCTGAACTCTTTCCCTACACGACGCTCTTCCGATCTNNNTTACTTAGGAGGCTGAGGTG
GAGCACACGTCTGAACTCTTTCCCTACACGACGCTCTTCCGATCTNNNCTGTAATCCCAGCTACTCGAG
AGCACACGTCTGAACTCTTTCCCTACACGACGCTCTTCCGATCTNNNCATCTGTAGTCCCAGCTACTCA
AGCACACGTCTGAACTCTTTCCCTACACGACGCTCTTCCGATCTNNNTCACTTGAGGTCAGAAGTTCG
AGCACACGTCTGAACTCTTTCCCTACACGACGCTCTTCCGATCTNNNCAGGTGACAGTGAGCTATGAT
AGCACACGTCTGAACTCTTTCCCTACACGACGCTCTTCCGATCTNNNGGAGGTTGTAGTGAGCTGAGA
GAGCACACGTCTGAACTCTTTCCCTACACGACGCTCTTCCGATCTNNNTACAAAAAATTAGCCGGGCA
AGCACACGTCTGAACTCTTTCCCTACACGACGCTCTTCCGATCTNNNCAAGAGTTTGAGGTTGCAGTG
AGAGCACACGTCTGAACTCTTTCCCTACACGACGCTCTTCCGATCTNNNGTAATCCCAGCTACTCAGA
GAGCACACGTCTGAACTCTTTCCCTACACGACGCTCTTCCGATCTNNNGGATCACTTGAGGTCAGGA
AGCACACGTCTGAACTCTTTCCCTACACGACGCTCTTCCGATCTNNNCAAAAATTAGCTGGACCTGGT
AGCACACGTCTGAACTCTTTCCCTACACGACGCTCTTCCGATCTNNNAAATGGGAGGATCACTTGAGT
GCACACGTCTGAACTCTTTCCCTACACGACGCTCTTCCGATCTNNNAGTCAAATGGAGGCATCTAGGT
AGCACACGTCTGAACTCTTTCCCTACACGACGCTCTTCCGATCTNNNCAGAAAGGTCAAATGCTGGA
GAGCACACGTCTGAACTCTTTCCCTACACGACGCTCTTCCGATCTNNNCTGTAGTCCCAGCTACTCAGG
AGCACACGTCTGAACTCTTTCCCTACACGACGCTCTTCCGATCTNNNCACCTGTAATCCCAGCTACTCAG
AGCACACGTCTGAACTCTTTCCCTACACGACGCTCTTCCGATCTNNNAACATGTCAAAACCCCGTCTCT
GCACACGTCTGAACTCTTTCCCTACACGACGCTCTTCCGATCTNNNATTCTTCACATTCCTCCCTCTGGC
GCACACGTCTGAACTCTTTCCCTACACGACGCTCTTCCGATCTNNNTTGAGGTAGGGGAATTGCTTG
GAGCACACGTCTGAACTCTTTCCCTACACGACGCTCTTCCGATCTNNNCAGGATGCTGAGGTAGGAG
GAGCACACGTCTGAACTCTTTCCCTACACGACGCTCTTCCGATCTNNNAAGATCTCGCCACTACACTTC

AGCACACGTCTGAACTCTTTCCCTACACGACGCTCTTCCGATCTNNNTGTAGAAGTGGGTTTGGTCTCA
GAGCACACGTCTGAACTCTTTCCCTACACGACGCTCTTCCGATCTNNNTGTTGTGGTGATTCTATCCAG
AGCACACGTCTGAACTCTTTCCCTACACGACGCTCTTCCGATCTNNNTGAGAATCACTTGAACCCAGGA
AGCACACGTCTGAACTCTTTCCCTACACGACGCTCTTCCGATCTNNNTGTAATCCCAGCTACTCAGGAG
GAGCACACGTCTGAACTCTTTCCCTACACGACGCTCTTCCGATCTNNNCTGAGGCAGGAGAATCACTTG
GAGCACACGTCTGAACTCTTTCCCTACACGACGCTCTTCCGATCTNNNTGAGGCAGGAGAATCACTTGA
AGCACACGTCTGAACTCTTTCCCTACACGACGCTCTTCCGATCTNNNTCATCTCACACCTGTCAGAATGC
GAGCACACGTCTGAACTCTTTCCCTACACGACGCTCTTCCGATCTNNNAGAATTGCTTGAATCCAGGAG
GAGCACACGTCTGAACTCTTTCCCTACACGACGCTCTTCCGATCTNNNCACCTGTAATCCCAGCTACTTGC
GAGCACACGTCTGAACTCTTTCCCTACACGACGCTCTTCCGATCTNNNCCTGCAAAGTTCAGTCTTGACA
AGAGCACACGTCTGAACTCTTTCCCTACACGACGCTCTTCCGATCTNNNCAGTAATCCCAGCTACTCAGG
GAGCACACGTCTGAACTCTTTCCCTACACGACGCTCTTCCGATCTNNNGCCTATAATCCCAGCTACTCAG
GAGCACACGTCTGAACTCTTTCCCTACACGACGCTCTTCCGATCTNNNAGTGAGCTGAGATTACACCAC
GAGCACACGTCTGAACTCTTTCCCTACACGACGCTCTTCCGATCTNNNGTAATCTCAGCTACTCAGGAG
AGCACACGTCTGAACTCTTTCCCTACACGACGCTCTTCCGATCTNNNAGGACATAGTGGGTGTCAACA
GAGCACACGTCTGAACTCTTTCCCTACACGACGCTCTTCCGATCTNNNAAGAGAATCACTTGAACCCGG
GAGCACACGTCTGAACTCTTTCCCTACACGACGCTCTTCCGATCTNNNAGAGGTTGCAGTGAGCTATGA
GAGCACACGTCTGAACTCTTTCCCTACACGACGCTCTTCCGATCTNNNATATATAAGCTGGGTGTGGTC
GAGCACACGTCTGAACTCTTTCCCTACACGACGCTCTTCCGATCTNNNCATGGCGAACACCTAGTTGAA
GCACACGTCTGAACTCTTTCCCTACACGACGCTCTTCCGATCTNNNTAAGAACTCTCAGCAAACGGGAA
AGAGCACACGTCTGAACTCTTTCCCTACACGACGCTCTTCCGATCTNNNAATAAATTCAGCCAGGCACAC
GAGCACACGTCTGAACTCTTTCCCTACACGACGCTCTTCCGATCTNNNAAAATATTAGCGTGACGTGGT
GAGCACACGTCTGAACTCTTTCCCTACACGACGCTCTTCCGATCTNNNCCCTCTTTAGTCGAATGGTGG
AGCACACGTCTGAACTCTTTCCCTACACGACGCTCTTCCGATCTNNNCTGAGGCAGGAGAATCACTTGA
GAGCACACGTCTGAACTCTTTCCCTACACGACGCTCTTCCGATCTNNNAGTGAGGCTGCAGTTTACGA
GAGCACACGTCTGAACTCTTTCCCTACACGACGCTCTTCCGATCTNNNCATGCCTGTAATCCCAGCTACT
AGAGCACACGTCTGAACTCTTTCCCTACACGACGCTCTTCCGATCTNNNCGTGCATGAATGTGTTTGTG
GAGCACACGTCTGAACTCTTTCCCTACACGACGCTCTTCCGATCTNNNGTCCTACTCATTGCTGGTGGA
AGAGCACACGTCTGAACTCTTTCCCTACACGACGCTCTTCCGATCTNNNATGCCTGTAGTCCCAGCTACT
AGCACACGTCTGAACTCTTTCCCTACACGACGCTCTTCCGATCTNNNGCGAAGGAACAAAACTCCATCT
GAGCACACGTCTGAACTCTTTCCCTACACGACGCTCTTCCGATCTNNNGGTGCCTGTAATCCTACCTACT
GAGCACACGTCTGAACTCTTTCCCTACACGACGCTCTTCCGATCTNNNAGAATCGCTTGAATCAGGGAG
AGCACACGTCTGAACTCTTTCCCTACACGACGCTCTTCCGATCTNNNGTGTGTGTGTGTGTTTTGAC
GAGCACACGTCTGAACTCTTTCCCTACACGACGCTCTTCCGATCTNNNCTGTAGACTGCTCTCTACCAAC
GAGCACACGTCTGAACTCTTTCCCTACACGACGCTCTTCCGATCTNNNCTAAGGTGGGAGGATTGCTT
AGCACACGTCTGAACTCTTTCCCTACACGACGCTCTTCCGATCTNNNGCATGTCTGTAATCCCAGCTACT
AGCACACGTCTGAACTCTTTCCCTACACGACGCTCTTCCGATCTNNNAGAATAGTTTGAACCTGGGAG
AGCACACGTCTGAACTCTTTCCCTACACGACGCTCTTCCGATCTNNNAAGTTCTTTTTCAGGGCAATCCG
GAGCACACGTCTGAACTCTTTCCCTACACGACGCTCTTCCGATCTNNNTGACTTGCTATTCCTTGCCTTC
AGCACACGTCTGAACTCTTTCCCTACACGACGCTCTTCCGATCTNNNGGAGACAAGAGTGAGACTCCA
GAGCACACGTCTGAACTCTTTCCCTACACGACGCTCTTCCGATCTNNNATCTGGGTGACAAGAGCAAG
GAGCACACGTCTGAACTCTTTCCCTACACGACGCTCTTCCGATCTNNNTAGAGGTTGCAGTCAGCTGAA
AGCACACGTCTGAACTCTTTCCCTACACGACGCTCTTCCGATCTNNNGCAACAAGAGCAAAATTCCGTC

GAGCACACGTCTGAACTCTTTCCCTACACGACGCTCTTCCGATCTNNNCTGAGGCAGGAGAATCACTT
AGCACACGTCTGAACTCTTTCCCTACACGACGCTCTTCCGATCTNNNTGAGGCATGAGAATTGCTTGA
AGCACACGTCTGAACTCTTTCCCTACACGACGCTCTTCCGATCTNNNCTGTAGTCCCAGCTATTCAGGA
GAGCACACGTCTGAACTCTTTCCCTACACGACGCTCTTCCGATCTNNNAAAAAATTAGCCAGGTGTGG
GAGCACACGTCTGAACTCTTTCCCTACACGACGCTCTTCCGATCTNNNATGCCTGTAATCCCAGCTACTT
AGAGCACACGTCTGAACTCTTTCCCTACACGACGCTCTTCCGATCTNNNATCTTAGCACTTTGGGAGCC
GAGCACACGTCTGAACTCTTTCCCTACACGACGCTCTTCCGATCTNNNAAAAAATTAGCTGGGTGTGG
AGCACACGTCTGAACTCTTTCCCTACACGACGCTCTTCCGATCTNNNCTGAGGCAGGAGAATTGCTTG
GAGCACACGTCTGAACTCTTTCCCTACACGACGCTCTTCCGATCTNNNCACTTGAGGTCAGGAGTTCA
AGCACACGTCTGAACTCTTTCCCTACACGACGCTCTTCCGATCTNNNAAGGTTGTAGTGAGCTGAGA
AGCACACGTCTGAACTCTTTCCCTACACGACGCTCTTCCGATCTNNNATGGTGGTAGGTGCCTATAAT
GAGCACACGTCTGAACTCTTTCCCTACACGACGCTCTTCCGATCTNNNAAAAAATTAGCTGGGTGTGG
GAGCACACGTCTGAACTCTTTCCCTACACGACGCTCTTCCGATCTNNNCTGTAATCCCAGCTACTCAGG
GAGCACACGTCTGAACTCTTTCCCTACACGACGCTCTTCCGATCTNNNCTTGCAGTGAGCTGAGATCC
AGCACACGTCTGAACTCTTTCCCTACACGACGCTCTTCCGATCTNNNAAATAGAGTGAGCCATGATTG
GAGCACACGTCTGAACTCTTTCCCTACACGACGCTCTTCCGATCTNNNAAAAAATTAGCTGGGTGTGG
GAGCACACGTCTGAACTCTTTCCCTACACGACGCTCTTCCGATCTNNNAGAGAATCACTTGAACCCAG
GAGCACACGTCTGAACTCTTTCCCTACACGACGCTCTTCCGATCTNNNCTGAGGCAGGAGAATCACTTG
AGCACACGTCTGAACTCTTTCCCTACACGACGCTCTTCCGATCTNNNAGCTATTTGGGAGACAAGACA
GAGCACACGTCTGAACTCTTTCCCTACACGACGCTCTTCCGATCTNNNGAGAATCGCTTGAATCCGAGA
AGAGCACACGTCTGAACTCTTTCCCTACACGACGCTCTTCCGATCTNNNCCTGGGAAACATAGTGAGA
AGAGCACACGTCTGAACTCTTTCCCTACACGACGCTCTTCCGATCTNNNAGGAGAAATGCTTGAACCTG
AGCACACGTCTGAACTCTTTCCCTACACGACGCTCTTCCGATCTNNNCGACAAGAACGAAACTCTGTCT
GAGCACACGTCTGAACTCTTTCCCTACACGACGCTCTTCCGATCTNNNGTAATCCCAGCTACTCAGGAA
AGAGCACACGTCTGAACTCTTTCCCTACACGACGCTCTTCCGATCTNNNCTGAGGCAGGAGAATCACTT
GAGCACACGTCTGAACTCTTTCCCTACACGACGCTCTTCCGATCTNNNGCAACAACAGCAAAACTCTGA
GAGCACACGTCTGAACTCTTTCCCTACACGACGCTCTTCCGATCTNNNTGCAGTGAGCTATGATTGTG
AGCACACGTCTGAACTCTTTCCCTACACGACGCTCTTCCGATCTNNNAAGGCAGGAGAATCACTTGAA
AGCACACGTCTGAACTCTTTCCCTACACGACGCTCTTCCGATCTNNNAGTTTGAGGCTGCAGTAAGCTA
GAGCACACGTCTGAACTCTTTCCCTACACGACGCTCTTCCGATCTNNNGCTACTCTATTCCCAGCTACT
GAGCACACGTCTGAACTCTTTCCCTACACGACGCTCTTCCGATCTNNNCTGAGGCAGGAGAATCACTT
AGCACACGTCTGAACTCTTTCCCTACACGACGCTCTTCCGATCTNNNAAAAAAAAAAATTAGTCGGGCG
GCACACGTCTGAACTCTTTCCCTACACGACGCTCTTCCGATCTNNNGAGGTTGCAGTGAGACAAGATT
AGCACACGTCTGAACTCTTTCCCTACACGACGCTCTTCCGATCTNNNTTGAGGCAGGAGAATTGCTTGA
GAGCACACGTCTGAACTCTTTCCCTACACGACGCTCTTCCGATCTNNNGATCAGGAGTTCGATAACAG
AGAGCACACGTCTGAACTCTTTCCCTACACGACGCTCTTCCGATCTNNNACAAAAATTAGCTGGGCATG
AGCACACGTCTGAACTCTTTCCCTACACGACGCTCTTCCGATCTNNNCACCTGTAGTCTCAGCTACTTG
GAGCACACGTCTGAACTCTTTCCCTACACGACGCTCTTCCGATCTNNNAGACTGAGGCACAAGAATCA
AGCACACGTCTGAACTCTTTCCCTACACGACGCTCTTCCGATCTNNNCTGTAATCCCAGCTACTCAGGA
GAGCACACGTCTGAACTCTTTCCCTACACGACGCTCTTCCGATCTNNNGCAGGAGAATCGTTTGAACT
GAGCACACGTCTGAACTCTTTCCCTACACGACGCTCTTCCGATCTNNNTCCTAAGCCCAGGAATTTGAG
AGAGCACACGTCTGAACTCTTTCCCTACACGACGCTCTTCCGATCTNNNGAGGTTACGGTGAGTGGA
GAGCACACGTCTGAACTCTTTCCCTACACGACGCTCTTCCGATCTNNNCTGAGGCAGGAGAATCACTT

GAGCACACGTCTGAACTCTTTCCCTACACGACGCTCTTCCGATCTNNNCATATTGGCTAGGCTGGTCT
GCACACGTCTGAACTCTTTCCCTACACGACGCTCTTCCGATCTNNNAGGCAGGAGAATTACTTGAAC
AGCACACGTCTGAACTCTTTCCCTACACGACGCTCTTCCGATCTNNNCCCTTCCCAGTTACGAGAATC
GCACACGTCTGAACTCTTTCCCTACACGACGCTCTTCCGATCTNNNGCAGAGTTTACAGTGAGCTGA
AGCACACGTCTGAACTCTTTCCCTACACGACGCTCTTCCGATCTNNNTGAGGAAGGAGAATCGCTTG
AGCACACGTCTGAACTCTTTCCCTACACGACGCTCTTCCGATCTNNNGGCATCTGTAGTCCCAACTAG
AGCACACGTCTGAACTCTTTCCCTACACGACGCTCTTCCGATCTNNNGGATCACTTGAGACCAGGAGT
GAGCACACGTCTGAACTCTTTCCCTACACGACGCTCTTCCGATCTNNNTATGTGGTCTCAGCTACTCA
AGCACACGTCTGAACTCTTTCCCTACACGACGCTCTTCCGATCTNNNTACAAAAGTTAGCCAGGTGTG
GAGCACACGTCTGAACTCTTTCCCTACACGACGCTCTTCCGATCTNNNGTGACAAAGCAAGACTCCAT
AGCACACGTCTGAACTCTTTCCCTACACGACGCTCTTCCGATCTNNNAAAAAGAGAAACCAAGGGGG
AGCACACGTCTGAACTCTTTCCCTACACGACGCTCTTCCGATCTNNNATCACACCTGTAATCCCAGCTA
GAGCACACGTCTGAACTCTTTCCCTACACGACGCTCTTCCGATCTNNNCGAGACCAGCCTGGGAAATA
GCACACGTCTGAACTCTTTCCCTACACGACGCTCTTCCGATCTNNNCAGGAGAATTGCTTGAACACAG
AGCACACGTCTGAACTCTTTCCCTACACGACGCTCTTCCGATCTNNNAAAAAAATTAGCCAGGCGTGG
GCACACGTCTGAACTCTTTCCCTACACGACGCTCTTCCGATCTNNNAACACCTGGAAACTTGGAAATG
AGCACACGTCTGAACTCTTTCCCTACACGACGCTCTTCCGATCTNNNTTACCACCCAGGATGATTGTG
AGCACACGTCTGAACTCTTTCCCTACACGACGCTCTTCCGATCTNNNGAGATCACGCCATTGCACTCTA
GCACACGTCTGAACTCTTTCCCTACACGACGCTCTTCCGATCTNNNCAAAGATTGCAGTGAGCCAAG
GAGCACACGTCTGAACTCTTTCCCTACACGACGCTCTTCCGATCTNNNATCAGGCTGAGCAACATAGA
AGCACACGTCTGAACTCTTTCCCTACACGACGCTCTTCCGATCTNNNTGACATGCACCTATAGTCCCA
GAGCACACGTCTGAACTCTTTCCCTACACGACGCTCTTCCGATCTNNNCACCTGTAATCCCAGCTACTC
GAGCACACGTCTGAACTCTTTCCCTACACGACGCTCTTCCGATCTNNNGAGACATGAGAATCGCTTGA
AGCACACGTCTGAACTCTTTCCCTACACGACGCTCTTCCGATCTNNNACGCCTGTAATCCCAGCTATTT
AGCACACGTCTGAACTCTTTCCCTACACGACGCTCTTCCGATCTNNNAGGCAGGAGAATTTCTTGAA
GCACACGTCTGAACTCTTTCCCTACACGACGCTCTTCCGATCTNNNCATACACCTGTGGTCCCAGATAC
GAGCACACGTCTGAACTCTTTCCCTACACGACGCTCTTCCGATCTNNNATGCCTGTAATCCCAGCTACT
AGCACACGTCTGAACTCTTTCCCTACACGACGCTCTTCCGATCTNNNCTGAACCAGGAGAGTCACTTC
AGCACACGTCTGAACTCTTTCCCTACACGACGCTCTTCCGATCTNNNACACCTGTAATCCCAGCTACTT
AGCACACGTCTGAACTCTTTCCCTACACGACGCTCTTCCGATCTNNNACAAAAATTAGCTGGGTGTGG
GCACACGTCTGAACTCTTTCCCTACACGACGCTCTTCCGATCTNNNCACCTGTAATCCCAGCTACTCAG
GCACACGTCTGAACTCTTTCCCTACACGACGCTCTTCCGATCTNNNGAGGCAGTAGAATTGCTTGAA
AGCACACGTCTGAACTCTTTCCCTACACGACGCTCTTCCGATCTNNNACTTGGGTATCTGGCAGACAA
GAGCACACGTCTGAACTCTTTCCCTACACGACGCTCTTCCGATCTNNNAAAAATTAGCCAGGCTTGGT
GCACACGTCTGAACTCTTTCCCTACACGACGCTCTTCCGATCTNNNATTTCCCTTCCTTGGAGTTCTGG
AGCACACGTCTGAACTCTTTCCCTACACGACGCTCTTCCGATCTNNNGAGAATTGCTTGAACCTGGAG
AGCACACGTCTGAACTCTTTCCCTACACGACGCTCTTCCGATCTNNNGCCTGTAATTGTAGTGCTTTG
CACACGTCTGAACTCTTTCCCTACACGACGCTCTTCCGATCTNNNATCATAGCTCATTGCAGCCTTGA
AGCACACGTCTGAACTCTTTCCCTACACGACGCTCTTCCGATCTNNNCTGAGGTGGGAGAATCACTTG
AGCACACGTCTGAACTCTTTCCCTACACGACGCTCTTCCGATCTNNNCTAGGCAAACGAGTGAGACT
GCACACGTCTGAACTCTTTCCCTACACGACGCTCTTCCGATCTNNNATCACTTGAGCCAGTAGGTGAA
GAGCACACGTCTGAACTCTTTCCCTACACGACGCTCTTCCGATCTNNNCAGCCTGAGTGATAGAGTGA
AGCACACGTCTGAACTCTTTCCCTACACGACGCTCTTCCGATCTNNNGAGATCACACCATTGCACTCC

GAGCACACGTCTGAACTCTTTCCCTACACGACGCTCTTCCGATCTNNNGACAGAGCTCTCAAATCGAG
AGCACACGTCTGAACTCTTTCCCTACACGACGCTCTTCCGATCTNNNAAAAAATTAGCTGGGTGTGG
GAGCACACGTCTGAACTCTTTCCCTACACGACGCTCTTCCGATCTNNNCTGTAATCCCAGCTACTCAGG
AGCACACGTCTGAACTCTTTCCCTACACGACGCTCTTCCGATCTNNNCAAGAGAATTGCTTGAACCCA
GAGCACACGTCTGAACTCTTTCCCTACACGACGCTCTTCCGATCTNNNTAATCCCAGATACTCAGGAG
AGCACACGTCTGAACTCTTTCCCTACACGACGCTCTTCCGATCTNNNACTACAGCTACTTAGGCAGGA
GCACACGTCTGAACTCTTTCCCTACACGACGCTCTTCCGATCTNNNGCAGGAGAATTGCTTGAACTCA
AGCACACGTCTGAACTCTTTCCCTACACGACGCTCTTCCGATCTNNNGGAGAATCACTCAAACCTGG
AGCACACGTCTGAACTCTTTCCCTACACGACGCTCTTCCGATCTNNNCTGACGTAGGAGAATCGCATG
GCACACGTCTGAACTCTTTCCCTACACGACGCTCTTCCGATCTNNNTGTATGCCTGTAATCCCAGCTA
AGCACACGTCTGAACTCTTTCCCTACACGACGCTCTTCCGATCTNNNGAGGCTGAGGCAAGAGAAT
AGAGCACACGTCTGAACTCTTTCCCTACACGACGCTCTTCCGATCTNNNCTGAGGCAGGAAAATTGC
GAGCACACGTCTGAACTCTTTCCCTACACGACGCTCTTCCGATCTNNNTTCAGTGAGTTGAGATTGCA
GCACACGTCTGAACTCTTTCCCTACACGACGCTCTTCCGATCTNNNAAAATTAGCCAGGCGTGGTGA
GAGCACACGTCTGAACTCTTTCCCTACACGACGCTCTTCCGATCTNNNAGAAAGCCTCTGTGACATTG
AGCACACGTCTGAACTCTTTCCCTACACGACGCTCTTCCGATCTNNNTTGAATCCCATCCCCATTTTCC
CACACGTCTGAACTCTTTCCCTACACGACGCTCTTCCGATCTNNNTAATGTCTCCCATCACCCCTAGA
GAGCACACGTCTGAACTCTTTCCCTACACGACGCTCTTCCGATCTNNNTACTCAGCCTGTTTTCTAAGC
GCACACGTCTGAACTCTTTCCCTACACGACGCTCTTCCGATCTNNNGGAGAATCGCTTGAACCTGGA
AGCACACGTCTGAACTCTTTCCCTACACGACGCTCTTCCGATCTNNNCATGATGGTGCATGCCTGTAA
AGCACACGTCTGAACTCTTTCCCTACACGACGCTCTTCCGATCTNNNGGATCACTTGAGGTCAGGAG
AGCACACGTCTGAACTCTTTCCCTACACGACGCTCTTCCGATCTNNNCACCTGTAATCCCAGCTACTTG
GCACACGTCTGAACTCTTTCCCTACACGACGCTCTTCCGATCTNNNGCAAGTTGCAGTTTACCCATGT
GCACACGTCTGAACTCTTTCCCTACACGACGCTCTTCCGATCTNNNACGTTGCAGAGAATAGGATGT
AGCACACGTCTGAACTCTTTCCCTACACGACGCTCTTCCGATCTNNNAGGCAGGAGAATTTCTTGAAC
AGCACACGTCTGAACTCTTTCCCTACACGACGCTCTTCCGATCTNNNTTGTAGTCCCAGCTACTCAGG
AGCACACGTCTGAACTCTTTCCCTACACGACGCTCTTCCGATCTNNNTCCCCTTGAAAACTAGCACAAG
GAGCACACGTCTGAACTCTTTCCCTACACGACGCTCTTCCGATCTNNNGAGGTTGAGAGTTAGAGAC
AGCACACGTCTGAACTCTTTCCCTACACGACGCTCTTCCGATCTNNNTGAGGCAGAAGAATTGCTTG
AGCACACGTCTGAACTCTTTCCCTACACGACGCTCTTCCGATCTNNNTTGAGGCAGGAGAATTGCTTG
AGCACACGTCTGAACTCTTTCCCTACACGACGCTCTTCCGATCTNNNTGAGCCTGAATAGCCAAAGG
GCACACGTCTGAACTCTTTCCCTACACGACGCTCTTCCGATCTNNNTGGCAGGCCAAAAAAAGAAAG
GAGCACACGTCTGAACTCTTTCCCTACACGACGCTCTTCCGATCTNNNGGTAGTGAGTTGAGATCAT
AGCACACGTCTGAACTCTTTCCCTACACGACGCTCTTCCGATCTNNNGAGATCACACCAATGCACTCC
AGCACACGTCTGAACTCTTTCCCTACACGACGCTCTTCCGATCTNNNTAAATACCCTTCTCGGTGTGA
AGCACACGTCTGAACTCTTTCCCTACACGACGCTCTTCCGATCTNNNGGCACCTGTAATCCTAGCTAC
AGCACACGTCTGAACTCTTTCCCTACACGACGCTCTTCCGATCTNNNAGAATCATTTGAGCCCAGGAG
AGCACACGTCTGAACTCTTTCCCTACACGACGCTCTTCCGATCTNNNACAACTCTAGCACTTACCACCA
AGCACACGTCTGAACTCTTTCCCTACACGACGCTCTTCCGATCTNNNCCAAGATTACACCATTGCACTC
GCACACGTCTGAACTCTTTCCCTACACGACGCTCTTCCGATCTNNNCAACAACAGAGTGAGACCCTG
GAGCACACGTCTGAACTCTTTCCCTACACGACGCTCTTCCGATCTNNNCTGAGGCAGGAGAATCACTT
GAGCACACGTCTGAACTCTTTCCCTACACGACGCTCTTCCGATCTNNNCAGCCTGGACAACATAGCAA
AGCACACGTCTGAACTCTTTCCCTACACGACGCTCTTCCGATCTNNNGCAGGAGAATTGCTTGAACT

AGCACACGTCTGAACTCTTTCCCTACACGACGCTCTTCCGATCTNNNCAAGAGAATTGCTTGAACCCA
AGCACACGTCTGAACTCTTTCCCTACACGACGCTCTTCCGATCTNNNGGATCACTTGAGGTCAGGAG
AGCACACGTCTGAACTCTTTCCCTACACGACGCTCTTCCGATCTNNNCTGAACACGCTCATGTAACCAT
AGCACACGTCTGAACTCTTTCCCTACACGACGCTCTTCCGATCTNNNAAAATGCAAAAAATTAGCCG
CACACGTCTGAACTCTTTCCCTACACGACGCTCTTCCGATCTNNNCTGAGGCAGGAGAATCACTTGA
GAGCACACGTCTGAACTCTTTCCCTACACGACGCTCTTCCGATCTNNNCTGAGGCAGAGAATTGCTTG
GAGCACACGTCTGAACTCTTTCCCTACACGACGCTCTTCCGATCTNNNCTTGAGGTCAGGAGTTCGA
GAGCACACGTCTGAACTCTTTCCCTACACGACGCTCTTCCGATCTNNNGAGGTTGCAGTGAGCTGAA
AGCACACGTCTGAACTCTTTCCCTACACGACGCTCTTCCGATCTNNNGTAATCCCAGCTACTGTAGAG
AGAGCACACGTCTGAACTCTTTCCCTACACGACGCTCTTCCGATCTNNNCGCACCTATAGTCCTTGCT
GAGCACACGTCTGAACTCTTTCCCTACACGACGCTCTTCCGATCTNNNTCATGCCTGTAATCTCAGCA
GAGCACACGTCTGAACTCTTTCCCTACACGACGCTCTTCCGATCTNNNCGACAAGAACAAAACTCCGT
GAGCACACGTCTGAACTCTTTCCCTACACGACGCTCTTCCGATCTNNNGAGGCTGAGACAGGAGAAT
AGCACACGTCTGAACTCTTTCCCTACACGACGCTCTTCCGATCTNNNGAATCTCTTGAACCCAGGAGG
GAGCACACGTCTGAACTCTTTCCCTACACGACGCTCTTCCGATCTNNNAAATACAAAAATTAGCCGGG
AGCACACGTCTGAACTCTTTCCCTACACGACGCTCTTCCGATCTNNNCTGTAGTCCTAGCTACTCGGG
GAGCACACGTCTGAACTCTTTCCCTACACGACGCTCTTCCGATCTNNNCAGAGGTTACAGTGAGCCAA
GAGCACACGTCTGAACTCTTTCCCTACACGACGCTCTTCCGATCTNNNCTGAGGGAGGAGATTTGCT
GAGCACACGTCTGAACTCTTTCCCTACACGACGCTCTTCCGATCTNNNTTGAGGCAGGAGAATCACTT
AGCACACGTCTGAACTCTTTCCCTACACGACGCTCTTCCGATCTNNNTTGAAAATATAGTCCCCTGCCC
GAGCACACGTCTGAACTCTTTCCCTACACGACGCTCTTCCGATCTNNNTATAGTCCCAGATGCTCAGG
GAGCACACGTCTGAACTCTTTCCCTACACGACGCTCTTCCGATCTNNNTGTAATCTCAGCACTTTGGG
AGCACACGTCTGAACTCTTTCCCTACACGACGCTCTTCCGATCTNNNTGAGGTAGGAGAATCGCTTGA
AGCACACGTCTGAACTCTTTCCCTACACGACGCTCTTCCGATCTNNNGAGTGCCTGTAATTCCAGCTA
GAGCACACGTCTGAACTCTTTCCCTACACGACGCTCTTCCGATCTNNNCACCCGGCTTGTATCGAAAAT
AGCACACGTCTGAACTCTTTCCCTACACGACGCTCTTCCGATCTNNNCTGTAGTCCCAGCTACCCTACT
GAGCACACGTCTGAACTCTTTCCCTACACGACGCTCTTCCGATCTNNNAGATTGCTGTGAGTCAAGA
GCACACGTCTGAACTCTTTCCCTACACGACGCTCTTCCGATCTNNNCATGTGATACAGGACCAGGGT
CACACGTCTGAACTCTTTCCCTACACGACGCTCTTCCGATCTNNNACACGCTTGTAATCGTAGCTACT
AGCACACGTCTGAACTCTTTCCCTACACGACGCTCTTCCGATCTNNNGAGGTTGCAGTGAGCCAAGAT
GAGCACACGTCTGAACTCTTTCCCTACACGACGCTCTTCCGATCTNNNCTGAGGCAGGAGAATCACTT
GAGCACACGTCTGAACTCTTTCCCTACACGACGCTCTTCCGATCTNNNGGCTAAGGCAGGAAAATCTC
GAGCACACGTCTGAACTCTTTCCCTACACGACGCTCTTCCGATCTNNNCAGATCATGAGGTCAGGAG
GCACACGTCTGAACTCTTTCCCTACACGACGCTCTTCCGATCTNNNGAAAACCAGAGACCTTTGCTCA
AGCACACGTCTGAACTCTTTCCCTACACGACGCTCTTCCGATCTNNNCTGGGTGATAGAGAGAGACC
GAGCACACGTCTGAACTCTTTCCCTACACGACGCTCTTCCGATCTNNNGATAACTTGAGCCCAGGAGT
GAGCACACGTCTGAACTCTTTCCCTACACGACGCTCTTCCGATCTNNNGGTTGCAGTTAGCCAAGAT
GAGCACACGTCTGAACTCTTTCCCTACACGACGCTCTTCCGATCTNNNGGTGAAAGAATGAGACCCTG
AGCACACGTCTGAACTCTTTCCCTACACGACGCTCTTCCGATCTNNNGCACATGCCTATAATCCCAGCT
AGCACACGTCTGAACTCTTTCCCTACACGACGCTCTTCCGATCTNNNTCTACCCACTCTTTCCTGCTTTCC
GCACACGTCTGAACTCTTTCCCTACACGACGCTCTTCCGATCTNNNCTGAGGCAGGAGAATCACTTG
AGCACACGTCTGAACTCTTTCCCTACACGACGCTCTTCCGATCTNNNCTGAGGCAGGAGAATCGATTG
GAGCACACGTCTGAACTCTTTCCCTACACGACGCTCTTCCGATCTNNNGCATGCCAACTACTGTTTTCA

AGAGCACACGTCTGAACTCTTTCCCTACACGACGCTCTTCCGATCTNNNGCAACAGAGCAAGACTTCAT
GAGCACACGTCTGAACTCTTTCCCTACACGACGCTCTTCCGATCTNNNATGAGCTAGATAGTCAACCCC
GCACACGTCTGAACTCTTTCCCTACACGACGCTCTTCCGATCTNNNGAAGAACTGCTTGAACCTGGGA
AGCACACGTCTGAACTCTTTCCCTACACGACGCTCTTCCGATCTNNNCATCCTGGCTAACATGGTGAAA
GCACACGTCTGAACTCTTTCCCTACACGACGCTCTTCCGATCTNNNTCAGCTGCCAGCAAATATAAAGC
AGCACACGTCTGAACTCTTTCCCTACACGACGCTCTTCCGATCTNNNCATTGTGTGTGTCAGTGTGTTCA
GAGCACACGTCTGAACTCTTTCCCTACACGACGCTCTTCCGATCTNNNCACACCTGTAATCCCAGCTACTC
GAGCACACGTCTGAACTCTTTCCCTACACGACGCTCTTCCGATCTNNNCTACTGCACTCCAGCCTATTTG
AGCACACGTCTGAACTCTTTCCCTACACGACGCTCTTCCGATCTNNNACATGGTGAGACCCCTTCTTTAC
AGCACACGTCTGAACTCTTTCCCTACACGACGCTCTTCCGATCTNNNGGCACCTGTAATCCTAGCTACTC
AGCACACGTCTGAACTCTTTCCCTACACGACGCTCTTCCGATCTNNNGGGAGAATCATTTGAACCCAGC
GAGCACACGTCTGAACTCTTTCCCTACACGACGCTCTTCCGATCTNNNAGAATTGCTTGAACTCAGGAG
GAGCACACGTCTGAACTCTTTCCCTACACGACGCTCTTCCGATCTNNNAGAATCACTTGAGCCTGGGAA
GAGCACACGTCTGAACTCTTTCCCTACACGACGCTCTTCCGATCTNNNGATCATGCCATTGTACTTCAGC
GAGCACACGTCTGAACTCTTTCCCTACACGACGCTCTTCCGATCTNNNTGTAGTCCCAGCTACTCAAAA
AGCACACGTCTGAACTCTTTCCCTACACGACGCTCTTCCGATCTNNNACTGTTGGACAATTGCAAAAGC
AGCACACGTCTGAACTCTTTCCCTACACGACGCTCTTCCGATCTNNNCTGCTCTAAAGGACCCTACTGAC
GAGCACACGTCTGAACTCTTTCCCTACACGACGCTCTTCCGATCTNNNTCCCTTTTCCAGAGTCTCCAGT
AGCACACGTCTGAACTCTTTCCCTACACGACGCTCTTCCGATCTNNNGCAACAAAAGCGAGATTCTGTC
GAGCACACGTCTGAACTCTTTCCCTACACGACGCTCTTCCGATCTNNNAGTAGACAGAGATTGCAGTGA
AGAGCACACGTCTGAACTCTTTCCCTACACGACGCTCTTCCGATCTNNNATACTGGTGTTTTGCGAGTC
GAGCACACGTCTGAACTCTTTCCCTACACGACGCTCTTCCGATCTNNNCATCCTGGCTAACAAGGTGAAA
AGAGCACACGTCTGAACTCTTTCCCTACACGACGCTCTTCCGATCTNNNAAAAATTAGCTGGATGTGG
GAGCACACGTCTGAACTCTTTCCCTACACGACGCTCTTCCGATCTNNNCTGTAGTCCCAGCTACTCAGGA
GAGCACACGTCTGAACTCTTTCCCTACACGACGCTCTTCCGATCTNNNAGGCCAAGTCAGGAGAATCA
GAGCACACGTCTGAACTCTTTCCCTACACGACGCTCTTCCGATCTNNNCTGAGGCAGGAGAATCACTTG
GAGCACACGTCTGAACTCTTTCCCTACACGACGCTCTTCCGATCTNNNCTGTAGTCCCAGCTACTCAGGA
GAGCACACGTCTGAACTCTTTCCCTACACGACGCTCTTCCGATCTNNNCTGAGGCAGGAGAATGACATC
AGCACACGTCTGAACTCTTTCCCTACACGACGCTCTTCCGATCTNNNCACCTGTAATCCCAGCTACTCAC
AGCACACGTCTGAACTCTTTCCCTACACGACGCTCTTCCGATCTNNNAGATTGCAGTGAGCTGAGATTC
GAGCACACGTCTGAACTCTTTCCCTACACGACGCTCTTCCGATCTNNNCCAGACTGGGCAACATAGTTA
AAGAGCACACGTCTGAACTCTTTCCCTACACGACGCTCTTCCGATCTNNNAGAATCGCTTGAACCCAGG
GCACACGTCTGAACTCTTTCCCTACACGACGCTCTTCCGATCTNNNAGGTTGCAGTGAGTTGAGATCA
GCACACGTCTGAACTCTTTCCCTACACGACGCTCTTCCGATCTNNNGCAAACCTGCCTCCCATTTTATTCA
GAGCACACGTCTGAACTCTTTCCCTACACGACGCTCTTCCGATCTNNNGCAAGACCCTGGCTCTACTAA
AGAGCACACGTCTGAACTCTTTCCCTACACGACGCTCTTCCGATCTNNNCTGAGATGAGAGGATGGCTT
AGCACACGTCTGAACTCTTTCCCTACACGACGCTCTTCCGATCTNNNCTGTAGTCCCAGCTACACAGGA
GAGCACACGTCTGAACTCTTTCCCTACACGACGCTCTTCCGATCTNNNAAAATTAGCCAGGCATGATGC
GCACACGTCTGAACTCTTTCCCTACACGACGCTCTTCCGATCTNNNGAGAATCACTTGAACCTGGGAG
GAGCACACGTCTGAACTCTTTCCCTACACGACGCTCTTCCGATCTNNNCAGATACTCGGGAAGCTGAG
AGAGCACACGTCTGAACTCTTTCCCTACACGACGCTCTTCCGATCTNNNGAGACCAAAAGAACAACCCTC
AGCACACGTCTGAACTCTTTCCCTACACGACGCTCTTCCGATCTNNNCAGTTTTGAGCTCCACTGCATTC
AGCACACGTCTGAACTCTTTCCCTACACGACGCTCTTCCGATCTNNNCAAAAAAAATTAGCCGGGCATC

GCACACGTCTGAACTCTTTCCCTACACGACGCTCTTCCGATCTNNNTGTCCTTTCTTAGAGGAACCAGG
AGCACACGTCTGAACTCTTTCCCTACACGACGCTCTTCCGATCTNNNAGAAGAATCGTTTGAACCTGGG
GAGCACACGTCTGAACTCTTTCCCTACACGACGCTCTTCCGATCTNNNTTGAGCCCAGGAATTCAAGG
GAGCACACGTCTGAACTCTTTCCCTACACGACGCTCTTCCGATCTNNNCCTGGGAGAAACAGATGGTA
AGCACACGTCTGAACTCTTTCCCTACACGACGCTCTTCCGATCTNNNCACCTGTAATCCCAGCTACTTGG
AGCACACGTCTGAACTCTTTCCCTACACGACGCTCTTCCGATCTNNNCACCTGTAATCCCAGCTACTCA
AGCACACGTCTGAACTCTTTCCCTACACGACGCTCTTCCGATCTNNNCTGTAGTCCCAGCTACTCAGGA
AGCACACGTCTGAACTCTTTCCCTACACGACGCTCTTCCGATCTNNNGCAGACACCTGTAATCCCAGTT
AAGAGCACACGTCTGAACTCTTTCCCTACACGACGCTCTTCCGATCTNNNCACCTGTAATCCCAGCTACT
AGCACACGTCTGAACTCTTTCCCTACACGACGCTCTTCCGATCTNNNTGATGCAGGAGAATTGCTTGA
GAGCACACGTCTGAACTCTTTCCCTACACGACGCTCTTCCGATCTNNNGAGATCACACCATTGCACTCC
AGAGCACACGTCTGAACTCTTTCCCTACACGACGCTCTTCCGATCTNNNAGCCTGTAATCCCAGTACTT
AGCACACGTCTGAACTCTTTCCCTACACGACGCTCTTCCGATCTNNNCTGTAGTCCCAGCTACTCAGGA
AGCACACGTCTGAACTCTTTCCCTACACGACGCTCTTCCGATCTNNNAGTAAGCCATGTTCACTCCAGT
GAGCACACGTCTGAACTCTTTCCCTACACGACGCTCTTCCGATCTNNNAATCAGGAAGAATCACCTCA
GCACACGTCTGAACTCTTTCCCTACACGACGCTCTTCCGATCTNNNATGTGTAAGACAGCAGGATCCA
AGCACACGTCTGAACTCTTTCCCTACACGACGCTCTTCCGATCTNNNGAGAATCACTTGAACCAGGGA
AGCACACGTCTGAACTCTTTCCCTACACGACGCTCTTCCGATCTNNNCTGGAAGGTAATGATGGTCAG
AGCACACGTCTGAACTCTTTCCCTACACGACGCTCTTCCGATCTNNNCACCTGTAATCTCAGCTACTCGG
GAGCACACGTCTGAACTCTTTCCCTACACGACGCTCTTCCGATCTNNNCACTTGAGGTCAGGAGTTTG
AGCACACGTCTGAACTCTTTCCCTACACGACGCTCTTCCGATCTNNNTTTGCTGAAGATATTTGCCAC
AGCACACGTCTGAACTCTTTCCCTACACGACGCTCTTCCGATCTNNNCTGAGGCAGGAGAATCACTTG
AGCACACGTCTGAACTCTTTCCCTACACGACGCTCTTCCGATCTNNNAAGATTGTGCCACTTCACTCCA
AGCACACGTCTGAACTCTTTCCCTACACGACGCTCTTCCGATCTNNNATGGTTACCAGATGCTGAGAA
AGCACACGTCTGAACTCTTTCCCTACACGACGCTCTTCCGATCTNNNACACCTGTAATCCCAGCACTTT
AGCACACGTCTGAACTCTTTCCCTACACGACGCTCTTCCGATCTNNNATCTTTGAATCTGGGAGGCAG
AGCACACGTCTGAACTCTTTCCCTACACGACGCTCTTCCGATCTNNNCTGTAATCTCAGCTGCTTGGG
AGCACACGTCTGAACTCTTTCCCTACACGACGCTCTTCCGATCTNNNGTTATTCCAGCTACTAGGGAG
AGCACACGTCTGAACTCTTTCCCTACACGACGCTCTTCCGATCTNNNCTGAGGCAGGAGAATCACTT
AGCACACGTCTGAACTCTTTCCCTACACGACGCTCTTCCGATCTNNNCTTGAAACTGAAGGCAGAGGT
GCACACGTCTGAACTCTTTCCCTACACGACGCTCTTCCGATCTNNNCTGTAATCCCAGCTACTCAGGA
AGCACACGTCTGAACTCTTTCCCTACACGACGCTCTTCCGATCTNNNGCCATGAATCAAGGAATGTAG
GAGCACACGTCTGAACTCTTTCCCTACACGACGCTCTTCCGATCTNNNGAGAATCGCTTGAACCCAGG
GCACACGTCTGAACTCTTTCCCTACACGACGCTCTTCCGATCTNNNAAGATCGCACTACTGCAATCCAG
GAGCACACGTCTGAACTCTTTCCCTACACGACGCTCTTCCGATCTNNNAAAAAATTAGCCAGGTGTGG
AGAGCACACGTCTGAACTCTTTCCCTACACGACGCTCTTCCGATCTNNNGATCATGAGGTCAGGAGTT
GAGCACACGTCTGAACTCTTTCCCTACACGACGCTCTTCCGATCTNNNAAAGAGGCAGAGAGACAAA
GCACACGTCTGAACTCTTTCCCTACACGACGCTCTTCCGATCTNNNAGATTGCCAGAGCTCATGAAAG
AGCACACGTCTGAACTCTTTCCCTACACGACGCTCTTCCGATCTNNNGGATCACGAGGTCAGAAGAT
AGCACACGTCTGAACTCTTTCCCTACACGACGCTCTTCCGATCTNNNTCCGAAAAATGTAGGCAGTGA
GCACACGTCTGAACTCTTTCCCTACACGACGCTCTTCCGATCTNNNAGGAGCTTGTAGTGAGCCTAGA
GCACACGTCTGAACTCTTTCCCTACACGACGCTCTTCCGATCTNNNATGCCTGTAATCCCAGCTACTTG
AGCACACGTCTGAACTCTTTCCCTACACGACGCTCTTCCGATCTNNNTCTGTGACTACCTATGGTTTTG

GCACACGTCTGAACTCTTTCCCTACACGACGCTCTTCCGATCTNNNGCAGGAGAATGATTGGTGTGA
AGCACACGTCTGAACTCTTTCCCTACACGACGCTCTTCCGATCTNNNTACAAAAATTAGCTGGGCGTG
GAGCACACGTCTGAACTCTTTCCCTACACGACGCTCTTCCGATCTNNNATCTTCCAGTCATGAAGCCAG
AGCACACGTCTGAACTCTTTCCCTACACGACGCTCTTCCGATCTNNNTCACAAGGTCAAGAGATCGA
AGCACACGTCTGAACTCTTTCCCTACACGACGCTCTTCCGATCTNNNGCTCTCATTTACATGGTTCCTG
GCACACGTCTGAACTCTTTCCCTACACGACGCTCTTCCGATCTNNNTCTGTAGTCCCAGCTATTCAGGA
AGAGCACACGTCTGAACTCTTTCCCTACACGACGCTCTTCCGATCTNNNGGAGTTTGCAGTGAGCTGA
GCACACGTCTGAACTCTTTCCCTACACGACGCTCTTCCGATCTNNNGGAGAATCACTTGAACCTGGG
GCACACGTCTGAACTCTTTCCCTACACGACGCTCTTCCGATCTNNNGATGTAGTCCCAGCTACTCAGGA
GCACACGTCTGAACTCTTTCCCTACACGACGCTCTTCCGATCTNNNTTGTGGTCTCAGCTATTTGGGA
AGCACACGTCTGAACTCTTTCCCTACACGACGCTCTTCCGATCTNNNACTACAGCACCATCCTTTCTGA
AGCACACGTCTGAACTCTTTCCCTACACGACGCTCTTCCGATCTNNNAGGAGCTGGTACCATTTCTTCT
AGCACACGTCTGAACTCTTTCCCTACACGACGCTCTTCCGATCTNNNCCAGGAAGAAGTTGAATCCCT
GAGCACACGTCTGAACTCTTTCCCTACACGACGCTCTTCCGATCTNNNGGGATGCTCTACTCTAATTGG
AGCACACGTCTGAACTCTTTCCCTACACGACGCTCTTCCGATCTNNNCACAATGTGCACATGTACCCTA
GAGCACACGTCTGAACTCTTTCCCTACACGACGCTCTTCCGATCTNNNTGCAAATGGCCAACAAGTCT
AGCACACGTCTGAACTCTTTCCCTACACGACGCTCTTCCGATCTNNNAAGATCACTCCATTGCACTCCA
AGCACACGTCTGAACTCTTTCCCTACACGACGCTCTTCCGATCTNNNGGATCATGATGTCAGGAGTTC
AGCACACGTCTGAACTCTTTCCCTACACGACGCTCTTCCGATCTNNNCACATACACCTTCCCAAGACTG
GCACACGTCTGAACTCTTTCCCTACACGACGCTCTTCCGATCTNNNAAAAAATTAGACGGGTGTGGT
GAGCACACGTCTGAACTCTTTCCCTACACGACGCTCTTCCGATCTNNNCCTAGGAAGCAGAGGTTACA
AGAGCACACGTCTGAACTCTTTCCCTACACGACGCTCTTCCGATCTNNNTGAGGTAGGAGAATGGCA
GCACACGTCTGAACTCTTTCCCTACACGACGCTCTTCCGATCTNNNCTGAGGTGGGAAGATCATCTG
GCACACGTCTGAACTCTTTCCCTACACGACGCTCTTCCGATCTNNNAACGAGCCAAGAAGGAGAGAA
GAGCACACGTCTGAACTCTTTCCCTACACGACGCTCTTCCGATCTNNNGTAATCTCAGCTACTTGGGA
GAGCACACGTCTGAACTCTTTCCCTACACGACGCTCTTCCGATCTNNNGGAGTTTGCAATGAGCCAAG
GAGCACACGTCTGAACTCTTTCCCTACACGACGCTCTTCCGATCTNNNATCAACAGGCTAAGTATGCT
GCACACGTCTGAACTCTTTCCCTACACGACGCTCTTCCGATCTNNNTAGCATGAAAGCAGCCATAGA
AGCACACGTCTGAACTCTTTCCCTACACGACGCTCTTCCGATCTNNNTTCCAAAAAACTGAGGAGGA
GCACACGTCTGAACTCTTTCCCTACACGACGCTCTTCCGATCTNNNCTGTAGTCCCAGCTACTCAGAA
GAGCACACGTCTGAACTCTTTCCCTACACGACGCTCTTCCGATCTNNNACAAAAATTAGCCAGGCACA
GAGCACACGTCTGAACTCTTTCCCTACACGACGCTCTTCCGATCTNNNGGAGAATCACTTGAACCTGG
AGCACACGTCTGAACTCTTTCCCTACACGACGCTCTTCCGATCTNNNCTGTAATCCCAGCACTTTGGAA
AGCACACGTCTGAACTCTTTCCCTACACGACGCTCTTCCGATCTNNNAGCCATGATTACGTCACTGCAT
GAGCACACGTCTGAACTCTTTCCCTACACGACGCTCTTCCGATCTNNNTGAGTCCAGGAATTCAAGAC
AGCACACGTCTGAACTCTTTCCCTACACGACGCTCTTCCGATCTNNNGCACATGCCTGTAATCCTAGCT
AGCACACGTCTGAACTCTTTCCCTACACGACGCTCTTCCGATCTNNNCTGGTGATACCCAGGTGAACA
AGCACACGTCTGAACTCTTTCCCTACACGACGCTCTTCCGATCTNNNCTGAGGCAGGAGAATCACTT
GCACACGTCTGAACTCTTTCCCTACACGACGCTCTTCCGATCTNNNGGAGAGAGATGGAGACCTTGT
AGCACACGTCTGAACTCTTTCCCTACACGACGCTCTTCCGATCTNNNACATGATGAAACCCCGTCTCTA
AGCACACGTCTGAACTCTTTCCCTACACGACGCTCTTCCGATCTNNNCAAGATGGTGCTACTGTACTC
AGCACACGTCTGAACTCTTTCCCTACACGACGCTCTTCCGATCTNNNAAGATCATGCCACTGTACTCCA
GAGCACACGTCTGAACTCTTTCCCTACACGACGCTCTTCCGATCTNNNATCATGAGGTCAGGAGATC

GCACACGTCTGAACTCTTTCCCTACACGACGCTCTTCCGATCTNNNGCTCTGAACTCAGGAAACAACAC
AGCACACGTCTGAACTCTTTCCCTACACGACGCTCTTCCGATCTNNNTCAAGGTCTCCTGCTAGGTTTT
AGAGCACACGTCTGAACTCTTTCCCTACACGACGCTCTTCCGATCTNNNGGAAGAATCACTTGAACCTC
AGCACACGTCTGAACTCTTTCCCTACACGACGCTCTTCCGATCTNNNCAGAGGTTACAGTGAGCCAAG
GCACACGTCTGAACTCTTTCCCTACACGACGCTCTTCCGATCTNNNCTGAGGCAGGAGAATAGCTTGA
AGCACACGTCTGAACTCTTTCCCTACACGACGCTCTTCCGATCTNNNTCAACCCCTCTTAGAGCAGTTT
GAGCACACGTCTGAACTCTTTCCCTACACGACGCTCTTCCGATCTNNNAGAATCACTTGAACCCGAGA
AGCACACGTCTGAACTCTTTCCCTACACGACGCTCTTCCGATCTNNNGGCGACTATAATCCCAGCTAC
GAGCACACGTCTGAACTCTTTCCCTACACGACGCTCTTCCGATCTNNNTCAGGAGGCTGAGGTAGGA
GAGCACACGTCTGAACTCTTTCCCTACACGACGCTCTTCCGATCTNNNGCCTGTAATCCCAGATACTCA
GCACACGTCTGAACTCTTTCCCTACACGACGCTCTTCCGATCTNNNTGAGGCAGGAGAATACCTTGA
AGCACACGTCTGAACTCTTTCCCTACACGACGCTCTTCCGATCTNNNGAGAATCGTGTGAACCCAGG
AGCACACGTCTGAACTCTTTCCCTACACGACGCTCTTCCGATCTNNNAAGACCCATCAGTGTGCTGTA
GAGCACACGTCTGAACTCTTTCCCTACACGACGCTCTTCCGATCTNNNTTGCTGACCAGAAGGAGTAG
GAGCACACGTCTGAACTCTTTCCCTACACGACGCTCTTCCGATCTNNNGATGCGATCAACTGGAAGAA
GAGCACACGTCTGAACTCTTTCCCTACACGACGCTCTTCCGATCTNNNTGCCAATCAAATTCCTCCTCT
GAGCACACGTCTGAACTCTTTCCCTACACGACGCTCTTCCGATCTNNNTGGAGGATGCAATGAGCTA
GAGCACACGTCTGAACTCTTTCCCTACACGACGCTCTTCCGATCTNNNCTGTAGTCCCAGCTACTCAGG
AGCACACGTCTGAACTCTTTCCCTACACGACGCTCTTCCGATCTNNNTTGAGGCAGGAGAATCACTTG
AGCACACGTCTGAACTCTTTCCCTACACGACGCTCTTCCGATCTNNNTCATACGCACAATACTCAGGGA
AGAGCACACGTCTGAACTCTTTCCCTACACGACGCTCTTCCGATCTNNNCAAGAGAATTGCTTGAACCC
GCACACGTCTGAACTCTTTCCCTACACGACGCTCTTCCGATCTNNNTGCCTGTAATCTCAGCTACTCAG
GCACACGTCTGAACTCTTTCCCTACACGACGCTCTTCCGATCTNNNCTGAGGCAGGAGAATCACTTGA
AGCACACGTCTGAACTCTTTCCCTACACGACGCTCTTCCGATCTNNNTAATTCCAGCTATTCAGGAGG
AGCACACGTCTGAACTCTTTCCCTACACGACGCTCTTCCGATCTNNNATCATACCACTACACTCCCTCCT
GAGCACACGTCTGAACTCTTTCCCTACACGACGCTCTTCCGATCTNNNCAACATCTTGGGACCCTATCTC
AGCACACGTCTGAACTCTTTCCCTACACGACGCTCTTCCGATCTNNNACAGCACTCCTTCTGGATAAACA
AGCACACGTCTGAACTCTTTCCCTACACGACGCTCTTCCGATCTNNNGAGAATCACTTGAACCTGGGA
AGCACACGTCTGAACTCTTTCCCTACACGACGCTCTTCCGATCTNNNGTGACAGAAAAAGACTTCGCC
AGCACACGTCTGAACTCTTTCCCTACACGACGCTCTTCCGATCTNNNAAAAAATTAGCTGGGTGTGG
GAGCACACGTCTGAACTCTTTCCCTACACGACGCTCTTCCGATCTNNNGACGTCTGTAATCCCAGCTAC
GAGCACACGTCTGAACTCTTTCCCTACACGACGCTCTTCCGATCTNNNCTGAGGCAGGAGAATCACTT
AGCACACGTCTGAACTCTTTCCCTACACGACGCTCTTCCGATCTNNNAACATGGCAAAACCCCATCTCT
GAGCACACGTCTGAACTCTTTCCCTACACGACGCTCTTCCGATCTNNNATGCCTATAACCCCAGCACTT
GAGCACACGTCTGAACTCTTTCCCTACACGACGCTCTTCCGATCTNNNACCAACATGATGAAACCCCTT
GAGCACACGTCTGAACTCTTTCCCTACACGACGCTCTTCCGATCTNNNCTTGCAGTGAGATGAGATTG
GAGCACACGTCTGAACTCTTTCCCTACACGACGCTCTTCCGATCTNNNCAAGATCATGCCACTGTACTC
GAGCACACGTCTGAACTCTTTCCCTACACGACGCTCTTCCGATCTNNNAAAAAAAATTAGCTGGGCGT
GCACACGTCTGAACTCTTTCCCTACACGACGCTCTTCCGATCTNNNAGACAGGAGAATTGCTTGAACC
GCACACGTCTGAACTCTTTCCCTACACGACGCTCTTCCGATCTNNNCCCCATCTCTCAGCTGGAATATCC
AGAGCACACGTCTGAACTCTTTCCCTACACGACGCTCTTCCGATCTNNNGCCACCATTTTCTTCTCTCTC
GAGCACACGTCTGAACTCTTTCCCTACACGACGCTCTTCCGATCTNNNGAGGTAACACCACTGCACTCT
GAGCACACGTCTGAACTCTTTCCCTACACGACGCTCTTCCGATCTNNNAAAAAATTAGCCAGGTGTGG

AGCACACGTCTGAACTCTTTCCCTACACGACGCTCTTCCGATCTNNNGGCTCATGTCTGTAATCCCAG
AGCACACGTCTGAACTCTTTCCCTACACGACGCTCTTCCGATCTNNNACGGACAAATTCCTGGACACA
GAGCACACGTCTGAACTCTTTCCCTACACGACGCTCTTCCGATCTNNNAAAAAATTAGCCAGGCGTGG
GAGCACACGTCTGAACTCTTTCCCTACACGACGCTCTTCCGATCTNNNTCTGGAGAAGAAGCATGGAG
GAGCACACGTCTGAACTCTTTCCCTACACGACGCTCTTCCGATCTNNNAGGGAGAGAGAGAGAGAG
AGCACACGTCTGAACTCTTTCCCTACACGACGCTCTTCCGATCTNNNGCTTTGAAATCACTGGCCCATA
GAGCACACGTCTGAACTCTTTCCCTACACGACGCTCTTCCGATCTNNNTACAAAAATTAGCCTGGCGTG
AGCACACGTCTGAACTCTTTCCCTACACGACGCTCTTCCGATCTNNNTGAGGCAGGAAAGTCACTTG
GAGCACACGTCTGAACTCTTTCCCTACACGACGCTCTTCCGATCTNNNCTCCTTTCCAGATGAGTGAA
GCACACGTCTGAACTCTTTCCCTACACGACGCTCTTCCGATCTNNNGTAATCCTAGCTACTCGGAAGG
AGCACACGTCTGAACTCTTTCCCTACACGACGCTCTTCCGATCTNNNTGAGGCAAGAGAATTGCTTGA
AGCACACGTCTGAACTCTTTCCCTACACGACGCTCTTCCGATCTNNNGTAATCCCAGCTACTCAGGAAG
AGCACACGTCTGAACTCTTTCCCTACACGACGCTCTTCCGATCTNNNACACCTGTAATCCCAGCTACAT
AGCACACGTCTGAACTCTTTCCCTACACGACGCTCTTCCGATCTNNNCAGTTTCCATGTAGACCCCTTG
GAGCACACGTCTGAACTCTTTCCCTACACGACGCTCTTCCGATCTNNNCTGAGGCAGGAGAATTGCTT
GAGCACACGTCTGAACTCTTTCCCTACACGACGCTCTTCCGATCTNNNAGATGCTAACTTTGAGCCCTA
GAGCACACGTCTGAACTCTTTCCCTACACGACGCTCTTCCGATCTNNNTGTGTCATTATACTCCAGCCT
GAGCACACGTCTGAACTCTTTCCCTACACGACGCTCTTCCGATCTNNNGAGGTTGTAGTGAGCTGAG
GAGCACACGTCTGAACTCTTTCCCTACACGACGCTCTTCCGATCTNNNGCACACACCTGTAATCCCAGT
AGCACACGTCTGAACTCTTTCCCTACACGACGCTCTTCCGATCTNNNTGAGGCAGGAGAATCTCTTG
AGCACACGTCTGAACTCTTTCCCTACACGACGCTCTTCCGATCTNNNCTGCAGTGAACTATGATTGTG
AGCACACGTCTGAACTCTTTCCCTACACGACGCTCTTCCGATCTNNNCTGAAGTTGCAAAGAGCCATG
GAGCACACGTCTGAACTCTTTCCCTACACGACGCTCTTCCGATCTNNNGAGATCACACCACTACACTCC
GAGCACACGTCTGAACTCTTTCCCTACACGACGCTCTTCCGATCTNNNGAGGCTGAGACAGGAGAAT
AGCACACGTCTGAACTCTTTCCCTACACGACGCTCTTCCGATCTNNNTGAGGCAGGAGAATCACTTG
AGCACACGTCTGAACTCTTTCCCTACACGACGCTCTTCCGATCTNNNCATGCCTGTAATCCCAGCTACT
AGCACACGTCTGAACTCTTTCCCTACACGACGCTCTTCCGATCTNNNACAAAAATTAGCTGGGCATGG
GAGCACACGTCTGAACTCTTTCCCTACACGACGCTCTTCCGATCTNNNGACTGTAGTCCCAGCTACATG
AGCACACGTCTGAACTCTTTCCCTACACGACGCTCTTCCGATCTNNNTTAGCCTAGGTGACAGAGTGA
GAGCACACGTCTGAACTCTTTCCCTACACGACGCTCTTCCGATCTNNNCTGAGGCAGCAGAATCACTT
AGCACACGTCTGAACTCTTTCCCTACACGACGCTCTTCCGATCTNNNGAGAATCACTTGAATCCGGGA
GAGCACACGTCTGAACTCTTTCCCTACACGACGCTCTTCCGATCTNNNCTGTAGTCCCAGCTACTCGAG
GAGCACACGTCTGAACTCTTTCCCTACACGACGCTCTTCCGATCTNNNTAATCCCAACACTTTAGGAG
AGCACACGTCTGAACTCTTTCCCTACACGACGCTCTTCCGATCTNNNTACAAAAAATTAGCCGGGCAA
GAGCACACGTCTGAACTCTTTCCCTACACGACGCTCTTCCGATCTNNNTATAGTCCCAGCTACTCAGGA
GAGCACACGTCTGAACTCTTTCCCTACACGACGCTCTTCCGATCTNNNCACCTACTCAGGAGGTTGAG
GAGCACACGTCTGAACTCTTTCCCTACACGACGCTCTTCCGATCTNNNTGTGATCCTAGCACTTTGGGA
AGCACACGTCTGAACTCTTTCCCTACACGACGCTCTTCCGATCTNNNAGGCTGAGACACAAGAATTCC
GAGCACACGTCTGAACTCTTTCCCTACACGACGCTCTTCCGATCTNNNAGAGAACTGCTTGAATCTGG
GAGCACACGTCTGAACTCTTTCCCTACACGACGCTCTTCCGATCTNNNGCTTGTAATCCTAGCTACTCG
GAGCACACGTCTGAACTCTTTCCCTACACGACGCTCTTCCGATCTNNNGTGCCACCTCAGAATTCAAG
GAGCACACGTCTGAACTCTTTCCCTACACGACGCTCTTCCGATCTNNNAGCTACTCAGGAAGTTGAAG
GAGCACACGTCTGAACTCTTTCCCTACACGACGCTCTTCCGATCTNNNGAATCACTTGAGCTGGGGA

GAGCACACGTCTGAACTCTTTCCCTACACGACGCTCTTCCGATCTNNNGGAGGTTGTAGTGAGCCAAG
GAGCACACGTCTGAACTCTTTCCCTACACGACGCTCTTCCGATCTNNNGCAGGAGAATCACTTGAACCT
AGAGCACACGTCTGAACTCTTTCCCTACACGACGCTCTTCCGATCTNNNCCCAGCTACTTAGGAGGTT
AGAGCACACGTCTGAACTCTTTCCCTACACGACGCTCTTCCGATCTNNNGTATGCCTGTAATCCCAGCT
AGCACACGTCTGAACTCTTTCCCTACACGACGCTCTTCCGATCTNNNAGTTGTCTAAACTGGCCCTTCA
AGCACACGTCTGAACTCTTTCCCTACACGACGCTCTTCCGATCTNNNTGTATGATCGTTTTCCCCAGGTT
AGAGCACACGTCTGAACTCTTTCCCTACACGACGCTCTTCCGATCTNNNCTGAGGTGGGAGGATCATT
GAGCACACGTCTGAACTCTTTCCCTACACGACGCTCTTCCGATCTNNNAAAGTTGCAGTGAGCTGAG
GAGCACACGTCTGAACTCTTTCCCTACACGACGCTCTTCCGATCTNNNAAAAAATTAGCTGAGCGTGGT
GAGCACACGTCTGAACTCTTTCCCTACACGACGCTCTTCCGATCTNNNAGCAACAGAACAAGACTCCAT
GAGCACACGTCTGAACTCTTTCCCTACACGACGCTCTTCCGATCTNNNAATACAAAAAATTAGCCGGG
GAGCACACGTCTGAACTCTTTCCCTACACGACGCTCTTCCGATCTNNNCTATCTCATCCTCCTGCCTTGA
GAGCACACGTCTGAACTCTTTCCCTACACGACGCTCTTCCGATCTNNNCAAAAGATTAGCCAGGTGTG
GAGCACACGTCTGAACTCTTTCCCTACACGACGCTCTTCCGATCTNNNGAGAATGGTGTGAATCCAGG
AGCACACGTCTGAACTCTTTCCCTACACGACGCTCTTCCGATCTNNNTCTGTAATCCCAGCTACTCAGGA
GAGCACACGTCTGAACTCTTTCCCTACACGACGCTCTTCCGATCTNNNCTGTAGTCCCAGCTACTCAGG
GAGCACACGTCTGAACTCTTTCCCTACACGACGCTCTTCCGATCTNNNGGTTGCAGTGAGCCAAGATT
GAGCACACGTCTGAACTCTTTCCCTACACGACGCTCTTCCGATCTNNNAAAAATTAGCTGGATGTGGT
GAGCACACGTCTGAACTCTTTCCCTACACGACGCTCTTCCGATCTNNNAAATTAGCTATGTGTGGTGA
GAGCACACGTCTGAACTCTTTCCCTACACGACGCTCTTCCGATCTNNNGCACATACCTGTATTCCCAGCT
GAGCACACGTCTGAACTCTTTCCCTACACGACGCTCTTCCGATCTNNNGATAGTTTGAGCCCAGGAGT
GAGCACACGTCTGAACTCTTTCCCTACACGACGCTCTTCCGATCTNNNTATAGTGAGGCCCTGTCTGT
GAGCACACGTCTGAACTCTTTCCCTACACGACGCTCTTCCGATCTNNNCAGAAAGTTGAGGTTGCAGT
AGCACACGTCTGAACTCTTTCCCTACACGACGCTCTTCCGATCTNNNTAAGCGCTTGTAATCCCACCTA
GAGCACACGTCTGAACTCTTTCCCTACACGACGCTCTTCCGATCTNNNTAGGAGAATCGTTTGAACCTG
GAGCACACGTCTGAACTCTTTCCCTACACGACGCTCTTCCGATCTNNNCTTACACCTGTAATCCCAGCA
AGCACACGTCTGAACTCTTTCCCTACACGACGCTCTTCCGATCTNNNTTGTTGAGATTTGCCTTGTGGA
GAGCACACGTCTGAACTCTTTCCCTACACGACGCTCTTCCGATCTNNNATGCTTGTAATCCCAGCACTT
GAGCACACGTCTGAACTCTTTCCCTACACGACGCTCTTCCGATCTNNNTGGGTGACAGTGAGATCCTAT
AGCACACGTCTGAACTCTTTCCCTACACGACGCTCTTCCGATCTNNNCTAGCACCTTTTCCCACAGTTTG
GAGCACACGTCTGAACTCTTTCCCTACACGACGCTCTTCCGATCTNNNCAGGAGAACTGCTACAACCTA
AGCACACGTCTGAACTCTTTCCCTACACGACGCTCTTCCGATCTNNNTGCCTGTAATCCCAGCTAAACA
GAGCACACGTCTGAACTCTTTCCCTACACGACGCTCTTCCGATCTNNNATACCTAGTTCTGCTTTCCCTG
AGCACACGTCTGAACTCTTTCCCTACACGACGCTCTTCCGATCTNNNACAAAAATTAGCTGGGTGTGG
AGCACACGTCTGAACTCTTTCCCTACACGACGCTCTTCCGATCTNNNAAAAAAAATTGGCCAGGTGTG
AGCACACGTCTGAACTCTTTCCCTACACGACGCTCTTCCGATCTNNNCATGCCTGTAATCCCAGCTACT
GCACACGTCTGAACTCTTTCCCTACACGACGCTCTTCCGATCTNNNGATGCAGGAGAATCACTTGAAC
GAGCACACGTCTGAACTCTTTCCCTACACGACGCTCTTCCGATCTNNNAATCACTTGTACCTGGGAAGT
AGCACACGTCTGAACTCTTTCCCTACACGACGCTCTTCCGATCTNNNCCCAATCCTCAATGTCACACAA
GCACACGTCTGAACTCTTTCCCTACACGACGCTCTTCCGATCTNNNGGAAAATCATTTGAACCCGGGA
GAGCACACGTCTGAACTCTTTCCCTACACGACGCTCTTCCGATCTNNNAATAAATTGCAAGAAAGGCC
GAGCACACGTCTGAACTCTTTCCCTACACGACGCTCTTCCGATCTNNNCAAGATCATACCTCTGCACTC
GCACACGTCTGAACTCTTTCCCTACACGACGCTCTTCCGATCTNNNCTGTAGTCCTAGCTACTCGGGA

AGCACACGTCTGAACTCTTTCCCTACACGACGCTCTTCCGATCTNNNAGTGAGCTGAGATCATGCCATT
AGCACACGTCTGAACTCTTTCCCTACACGACGCTCTTCCGATCTNNNAAGATCAAGCCACTTCAGTCCA
AGCACACGTCTGAACTCTTTCCCTACACGACGCTCTTCCGATCTNNNAGCAACAGAACAAGATCCCATC
GAGCACACGTCTGAACTCTTTCCCTACACGACGCTCTTCCGATCTNNNAGACAGGAGAATTGCTTGAA
AGCACACGTCTGAACTCTTTCCCTACACGACGCTCTTCCGATCTNNNCTGTGATCACCACTGCACTTCAG
GAGCACACGTCTGAACTCTTTCCCTACACGACGCTCTTCCGATCTNNNGAGCTCCAGCAAGAAATCAAA
AGCACACGTCTGAACTCTTTCCCTACACGACGCTCTTCCGATCTNNNCTGTAATCCCAGCTACTCAGGAC
AGCACACGTCTGAACTCTTTCCCTACACGACGCTCTTCCGATCTNNNTAATCCTAGCACTTTTGGAGGCT
AGCACACGTCTGAACTCTTTCCCTACACGACGCTCTTCCGATCTNNNTGAGGCAGGAGAATCTCTTGAA
GAGCACACGTCTGAACTCTTTCCCTACACGACGCTCTTCCGATCTNNNCTGTAATTCCAGCTACTCGGG
GAGCACACGTCTGAACTCTTTCCCTACACGACGCTCTTCCGATCTNNNCTGAGGCAGGAGAATCACTTC
AGCACACGTCTGAACTCTTTCCCTACACGACGCTCTTCCGATCTNNNTGGGTGACAGAGTGAGATAGT
GAGCACACGTCTGAACTCTTTCCCTACACGACGCTCTTCCGATCTNNNTTGAGGTCAGGAGTTCAAAAC
GAGCACACGTCTGAACTCTTTCCCTACACGACGCTCTTCCGATCTNNNTATACAAATAATGCCACGGTG
AGCACACGTCTGAACTCTTTCCCTACACGACGCTCTTCCGATCTNNNCCAGCTACCAAGACAGGAGAAT
GAGCACACGTCTGAACTCTTTCCCTACACGACGCTCTTCCGATCTNNNAGAGGTTGCAGTGAGTCAAAC
AGAGCACACGTCTGAACTCTTTCCCTACACGACGCTCTTCCGATCTNNNCTTGCAGTGAGATGAGATTC
GAGCACACGTCTGAACTCTTTCCCTACACGACGCTCTTCCGATCTNNNGAATTGCTTGAACCCAGTAGTC
GAGCACACGTCTGAACTCTTTCCCTACACGACGCTCTTCCGATCTNNNGGGAGAATCATTTGAACCTGG
GAGCACACGTCTGAACTCTTTCCCTACACGACGCTCTTCCGATCTNNNGTGACAAAACCAAACTCCGTCT
GAGCACACGTCTGAACTCTTTCCCTACACGACGCTCTTCCGATCTNNNACCTGTAATCCTAGCACTTTGG
GAGCACACGTCTGAACTCTTTCCCTACACGACGCTCTTCCGATCTNNNGATGCCTGTAATCCCAGCTACT
GAGCACACGTCTGAACTCTTTCCCTACACGACGCTCTTCCGATCTNNNACAAAAATTAGCTGGGTGTGC
GAGCACACGTCTGAACTCTTTCCCTACACGACGCTCTTCCGATCTNNNAAAAAAAAAATTAGCCGGGC
AGCACACGTCTGAACTCTTTCCCTACACGACGCTCTTCCGATCTNNNTGTAGTCCTAGCTACTCAGGAGC
AGCACACGTCTGAACTCTTTCCCTACACGACGCTCTTCCGATCTNNNCTGTAATCCCAGCTACTCAGGA
GAGCACACGTCTGAACTCTTTCCCTACACGACGCTCTTCCGATCTNNNATGCCTGTAATCCCAGCTACTT
GAGCACACGTCTGAACTCTTTCCCTACACGACGCTCTTCCGATCTNNNAGGAAGGAGAATTGCTTGAAC
GAGCACACGTCTGAACTCTTTCCCTACACGACGCTCTTCCGATCTNNNCTTGGGTGACAGAGTAAGACT
AGCACACGTCTGAACTCTTTCCCTACACGACGCTCTTCCGATCTNNNAGCAAGCTCCCATCTCTACAAA
GAGCACACGTCTGAACTCTTTCCCTACACGACGCTCTTCCGATCTNNNACAAGAGTGAAACACCCTTGTC
GAGCACACGTCTGAACTCTTTCCCTACACGACGCTCTTCCGATCTNNNCTCATGCCTGTAATCCTAGCAC
GAGCACACGTCTGAACTCTTTCCCTACACGACGCTCTTCCGATCTNNNGGCACCTGTAATCCCAGATACT
AGCACACGTCTGAACTCTTTCCCTACACGACGCTCTTCCGATCTNNNGTTGTAGTGAGCCAAGATTGA
GAGCACACGTCTGAACTCTTTCCCTACACGACGCTCTTCCGATCTNNNAAGAATCACTTGAACCTGGGA
GAGCACACGTCTGAACTCTTTCCCTACACGACGCTCTTCCGATCTNNNTATAGTCCCAGCTACTCAGGAC
GAGCACACGTCTGAACTCTTTCCCTACACGACGCTCTTCCGATCTNNNCAGGAGAATCACTTGAACCCA
GCACACGTCTGAACTCTTTCCCTACACGACGCTCTTCCGATCTNNNGAGGCTGAAGCAAGAGAATCTCT
AGCACACGTCTGAACTCTTTCCCTACACGACGCTCTTCCGATCTNNNAGGTTTCAGTGAGTCGAGATCA
AGCACACGTCTGAACTCTTTCCCTACACGACGCTCTTCCGATCTNNNAGAGGTTGCAGTAAGCTGAGA
GAGCACACGTCTGAACTCTTTCCCTACACGACGCTCTTCCGATCTNNNATGCCTGTAATCTCAGCTACTC
GCACACGTCTGAACTCTTTCCCTACACGACGCTCTTCCGATCTNNNATGCCTGTAATCCCAACTACTCGC
AGCACACGTCTGAACTCTTTCCCTACACGACGCTCTTCCGATCTNNNTGGAAGTTTCAGTCAGCCAAGA

GAGCACACGTCTGAACTCTTTCCCTACACGACGCTCTTCCGATCTNNNGGTGCCTATAGTTCCAGCTAC
AGCACACGTCTGAACTCTTTCCCTACACGACGCTCTTCCGATCTNNNTGAGAATCACTTGAACCCAGG
GAGCACACGTCTGAACTCTTTCCCTACACGACGCTCTTCCGATCTNNNAAAAAAAGACCAGTCTGGCC
GAGCACACGTCTGAACTCTTTCCCTACACGACGCTCTTCCGATCTNNNGAGGCTGAGGCATAAGAATC
GAGCACACGTCTGAACTCTTTCCCTACACGACGCTCTTCCGATCTNNNAGATTGCACCACTACACTCTA
GAGCACACGTCTGAACTCTTTCCCTACACGACGCTCTTCCGATCTNNNGAGAATTGCTTGAACCCAGG
AGCACACGTCTGAACTCTTTCCCTACACGACGCTCTTCCGATCTNNNCTGAGGCAGGAGAATCACTTC
AGCACACGTCTGAACTCTTTCCCTACACGACGCTCTTCCGATCTNNNCTGTAGTCCCAGCTACTCAGG
GAGCACACGTCTGAACTCTTTCCCTACACGACGCTCTTCCGATCTNNNCTAAGGCAGGAGGATTGCTT
AGCACACGTCTGAACTCTTTCCCTACACGACGCTCTTCCGATCTNNNGAGACAGGAGAATTGCTTGAA
AGCACACGTCTGAACTCTTTCCCTACACGACGCTCTTCCGATCTNNNCACCTGTAATCCTAGCACTCTGG
GAGCACACGTCTGAACTCTTTCCCTACACGACGCTCTTCCGATCTNNNAGAGGTAGGAGAATTGCTTG
AGCACACGTCTGAACTCTTTCCCTACACGACGCTCTTCCGATCTNNNCGCTTATAATCCCAGCACTTTG
GAGCACACGTCTGAACTCTTTCCCTACACGACGCTCTTCCGATCTNNNCAGAGGTTACAGTGAGCCAA
AGCACACGTCTGAACTCTTTCCCTACACGACGCTCTTCCGATCTNNNTGTAATCCCAGCTACTTGGAAG
AGCACACGTCTGAACTCTTTCCCTACACGACGCTCTTCCGATCTNNNCTGAGGTAGGAGGATCACTTG
AGCACACGTCTGAACTCTTTCCCTACACGACGCTCTTCCGATCTNNNAGGAGATTTGCTTGAACCTGA
GCACACGTCTGAACTCTTTCCCTACACGACGCTCTTCCGATCTNNNGATCGTGTCACTATACTCCAGCC
AGAGCACACGTCTGAACTCTTTCCCTACACGACGCTCTTCCGATCTNNNCTGTAGTTCCAGCTACTTGG
AGCACACGTCTGAACTCTTTCCCTACACGACGCTCTTCCGATCTNNNATAGCGAAACCCTGCCTCTACTA
AGCACACGTCTGAACTCTTTCCCTACACGACGCTCTTCCGATCTNNNCTCAAAACTTTGGGAGGCTGAA
GAGCACACGTCTGAACTCTTTCCCTACACGACGCTCTTCCGATCTNNNCCAGGAGGTTGAAGCTGTAA
AGCACACGTCTGAACTCTTTCCCTACACGACGCTCTTCCGATCTNNNAGGAGAGTAAAAAGGAGACA
AGCACACGTCTGAACTCTTTCCCTACACGACGCTCTTCCGATCTNNNTGAGGCAGGAGAATCTCTTGA
AGCACACGTCTGAACTCTTTCCCTACACGACGCTCTTCCGATCTNNNCTGAGGCAGGAGAATCACTTGA
AGCACACGTCTGAACTCTTTCCCTACACGACGCTCTTCCGATCTNNNCTGGGAAGTCAAGGCTTTAGTC
GAGCACACGTCTGAACTCTTTCCCTACACGACGCTCTTCCGATCTNNNCAGGAGGCTGAGACATGAGA
GCACACGTCTGAACTCTTTCCCTACACGACGCTCTTCCGATCTNNNTCAGATGTGCTAGTTGCTCAAGT
AGCACACGTCTGAACTCTTTCCCTACACGACGCTCTTCCGATCTNNNACCATCACCTACAAGGAAACCT
GAGCACACGTCTGAACTCTTTCCCTACACGACGCTCTTCCGATCTNNNAAAAAATTAGCAGGACGTGG
AGCACACGTCTGAACTCTTTCCCTACACGACGCTCTTCCGATCTNNNAAAAAATTAGCCTGGCAAGGT
GAGCACACGTCTGAACTCTTTCCCTACACGACGCTCTTCCGATCTNNNACTCATCTGTAGTCCCAGCTA
AGCACACGTCTGAACTCTTTCCCTACACGACGCTCTTCCGATCTNNNAAAAAATTAGCCGGATGTGG
AGCACACGTCTGAACTCTTTCCCTACACGACGCTCTTCCGATCTNNNCAAGATCGTGCCACTACACTCT
AGCACACGTCTGAACTCTTTCCCTACACGACGCTCTTCCGATCTNNNTCACAACACTGAACTGGGAAG
AGCACACGTCTGAACTCTTTCCCTACACGACGCTCTTCCGATCTNNNATGCCTGTAATCCCAGCTACTT
AGAGCACACGTCTGAACTCTTTCCCTACACGACGCTCTTCCGATCTNNNAATACAAAAAATTAGCCGG
AGCACACGTCTGAACTCTTTCCCTACACGACGCTCTTCCGATCTNNNCAAAAGTGAACAGAGATGCAG
GAGCACACGTCTGAACTCTTTCCCTACACGACGCTCTTCCGATCTNNNGTGCTCAGCCAACATCAGTAT
GAGCACACGTCTGAACTCTTTCCCTACACGACGCTCTTCCGATCTNNNAGGAGTGAATGGTTCTGTC
AGCACACGTCTGAACTCTTTCCCTACACGACGCTCTTCCGATCTNNNAATTTGCAGTAAGCCAAGTTCC
AGCACACGTCTGAACTCTTTCCCTACACGACGCTCTTCCGATCTNNNCAAAAAGGCTCATGGAGTGCTA
GAGCACACGTCTGAACTCTTTCCCTACACGACGCTCTTCCGATCTNNNTGCTAATGTGGAGTTACAGG

AGCACACGTCTGAACTCTTTCCCTACACGACGCTCTTCCGATCTNNNGTTCCTGGGAAGGTATAGCA
AGCACACGTCTGAACTCTTTCCCTACACGACGCTCTTCCGATCTNNNTTGGTGAGGTTGACGTACTT
GAGCACACGTCTGAACTCTTTCCCTACACGACGCTCTTCCGATCTNNNGAAGAATGGTTAGTGGTAG
AGCACACGTCTGAACTCTTTCCCTACACGACGCTCTTCCGATCTNNNAGACAGGAGAATCACTTGAAC
AGCACACGTCTGAACTCTTTCCCTACACGACGCTCTTCCGATCTNNNACAAAAATTAGGCAGGCATGG
AGCACACGTCTGAACTCTTTCCCTACACGACGCTCTTCCGATCTNNNGGCAACAGAGAGAGATCTTGT
AGCACACGTCTGAACTCTTTCCCTACACGACGCTCTTCCGATCTNNNAAAAAATTAGCTGGGCATGG
GAGCACACGTCTGAACTCTTTCCCTACACGACGCTCTTCCGATCTNNNAAAAATTAGCCTGGCATGGT
AGCACACGTCTGAACTCTTTCCCTACACGACGCTCTTCCGATCTNNNAAACCCCGACGCTACAAAAAA
AGAGCACACGTCTGAACTCTTTCCCTACACGACGCTCTTCCGATCTNNNCTGAGGCAGGAGAATCACT
AGCACACGTCTGAACTCTTTCCCTACACGACGCTCTTCCGATCTNNNGAGAATTGCTTGAACTCGAAG
AGCACACGTCTGAACTCTTTCCCTACACGACGCTCTTCCGATCTNNNAGGCAGAAGAATCACTTGAA
AGCACACGTCTGAACTCTTTCCCTACACGACGCTCTTCCGATCTNNNTGATCCAGACATTCCCTCTTTGG
GAGCACACGTCTGAACTCTTTCCCTACACGACGCTCTTCCGATCTNNNAGAGCTAGACCCTGTCTCTG
GAGCACACGTCTGAACTCTTTCCCTACACGACGCTCTTCCGATCTNNNAGTGAGCTATGATTGCACCA
AGCACACGTCTGAACTCTTTCCCTACACGACGCTCTTCCGATCTNNNCTGTAGTCCCAGCTACTCAAGA
AGCACACGTCTGAACTCTTTCCCTACACGACGCTCTTCCGATCTNNNCTGAGGCAGGAAAATTGCTTG
AGCACACGTCTGAACTCTTTCCCTACACGACGCTCTTCCGATCTNNNGATGCCGAAACCTAAACTTTG
AGCACACGTCTGAACTCTTTCCCTACACGACGCTCTTCCGATCTNNNAGAATCACTTGAACCTGGGAG
AGCACACGTCTGAACTCTTTCCCTACACGACGCTCTTCCGATCTNNNCTGTAGTCCCAGCTACTCAGGA
AGCACACGTCTGAACTCTTTCCCTACACGACGCTCTTCCGATCTNNNCACCTGTAGTCCCAGTTACTTG
AGCACACGTCTGAACTCTTTCCCTACACGACGCTCTTCCGATCTNNNCATGATGGCGAGTGCTTGTAA
AGCACACGTCTGAACTCTTTCCCTACACGACGCTCTTCCGATCTNNNTGTAATCTCAGCACTTTGGGAG
AGCACACGTCTGAACTCTTTCCCTACACGACGCTCTTCCGATCTNNNTCAGGAGGTTGAGGTAGGA
AGCACACGTCTGAACTCTTTCCCTACACGACGCTCTTCCGATCTNNNTAGTGGTGCATGCCTATAATCC
AGCACACGTCTGAACTCTTTCCCTACACGACGCTCTTCCGATCTNNNCAGGAGGTGGAGGTTATGAT
GAGCACACGTCTGAACTCTTTCCCTACACGACGCTCTTCCGATCTNNNAGAATCACTTGAACCTGGGA
AGCACACGTCTGAACTCTTTCCCTACACGACGCTCTTCCGATCTNNNTTGAAGTAGCCTCTGCAAAAT
AGCACACGTCTGAACTCTTTCCCTACACGACGCTCTTCCGATCTNNNGTAATCTCAGCTACTCGAGAG
AGCACACGTCTGAACTCTTTCCCTACACGACGCTCTTCCGATCTNNNGAAATTGATTGAACCTGGGAG
AGCACACGTCTGAACTCTTTCCCTACACGACGCTCTTCCGATCTNNNAGCATGCTAAGCTGTTACAGA
AGCACACGTCTGAACTCTTTCCCTACACGACGCTCTTCCGATCTNNNATCTGCTGAGCTAGATCACTTG
AGCACACGTCTGAACTCTTTCCCTACACGACGCTCTTCCGATCTNNNACAAAAATTAGCCAGGTGTGG
GCACACGTCTGAACTCTTTCCCTACACGACGCTCTTCCGATCTNNNTGAAACAACTGCTTCTGGATCCT
AGCACACGTCTGAACTCTTTCCCTACACGACGCTCTTCCGATCTNNNTACAAAAAAATTAGCTGGGCGT
GAGCACACGTCTGAACTCTTTCCCTACACGACGCTCTTCCGATCTNNNCTGCACATGTATCCCTGAACC
AGCACACGTCTGAACTCTTTCCCTACACGACGCTCTTCCGATCTNNNATGCTTGTAATCCCAGCACTTT
GCACACGTCTGAACTCTTTCCCTACACGACGCTCTTCCGATCTNNNGGTGGTTCATGCCTATAATCCC
AGCACACGTCTGAACTCTTTCCCTACACGACGCTCTTCCGATCTNNNGATCCACTGAGCTAGACCACTT
GAGCACACGTCTGAACTCTTTCCCTACACGACGCTCTTCCGATCTNNNTGTCCCAAATGGTCTGATTTT
AGCACACGTCTGAACTCTTTCCCTACACGACGCTCTTCCGATCTNNNTGTCCCAAATGGTCTGATTTTG
GAGCACACGTCTGAACTCTTTCCCTACACGACGCTCTTCCGATCTNNNGGTGCCTGTAATCCCAGCTA
GAGCACACGTCTGAACTCTTTCCCTACACGACGCTCTTCCGATCTNNNCCAAGAGGTAGAAACTGCAG

AGCACACGTCTGAACTCTTTCCCTACACGACGCTCTTCCGATCTNNNCTGAGGCAGGAGAATTGGTT
AGCACACGTCTGAACTCTTTCCCTACACGACGCTCTTCCGATCTNNNCAGTGAGCTGTGATTGTGTCC
AGCACACGTCTGAACTCTTTCCCTACACGACGCTCTTCCGATCTNNNCAAGGCAGAAGGATCATTTG
AGCACACGTCTGAACTCTTTCCCTACACGACGCTCTTCCGATCTNNNACTTGTGATCCCAGCTATTCAG
AGCACACGTCTGAACTCTTTCCCTACACGACGCTCTTCCGATCTNNNTACTAGAGAGACTGAGGCAG
AGCACACGTCTGAACTCTTTCCCTACACGACGCTCTTCCGATCTNNNAATGCCCTTGTTTATTGCCTGT
GAGCACACGTCTGAACTCTTTCCCTACACGACGCTCTTCCGATCTNNNTAGTAGAGACAGGGTTTTGC
AGCACACGTCTGAACTCTTTCCCTACACGACGCTCTTCCGATCTNNNGTGGCTCACTCCTGTAATCCTA
GCACACGTCTGAACTCTTTCCCTACACGACGCTCTTCCGATCTNNNGTAATCTCAGCTACTCAGGAGG
GCACACGTCTGAACTCTTTCCCTACACGACGCTCTTCCGATCTNNNCTCATGCCTATAATCCCAGCACT
GAGCACACGTCTGAACTCTTTCCCTACACGACGCTCTTCCGATCTNNNAGGAGCCTATAATCCCAGCT
AGCACACGTCTGAACTCTTTCCCTACACGACGCTCTTCCGATCTNNNCTGAGGCAGAAGAATGGTGT
GAGCACACGTCTGAACTCTTTCCCTACACGACGCTCTTCCGATCTNNNTAGGAGAATTGCTTGAACCC
GCACACGTCTGAACTCTTTCCCTACACGACGCTCTTCCGATCTNNNTGAGAATCACTTGAACCAGGGA
GAGCACACGTCTGAACTCTTTCCCTACACGACGCTCTTCCGATCTNNNAGTTTGAGACTGGGCAACAT
AGCACACGTCTGAACTCTTTCCCTACACGACGCTCTTCCGATCTNNNCTGAGGCAGGAGAATTGCTTG
GAGCACACGTCTGAACTCTTTCCCTACACGACGCTCTTCCGATCTNNNCTGAGGCAGGAGAATCACTT
AGCACACGTCTGAACTCTTTCCCTACACGACGCTCTTCCGATCTNNNGGTGACAGAGTGAGACTCCAT
GAGCACACGTCTGAACTCTTTCCCTACACGACGCTCTTCCGATCTNNNGATTGCATGAGCTCAGGAGT
GAGCACACGTCTGAACTCTTTCCCTACACGACGCTCTTCCGATCTNNNTGCCTGTAATACCAGCTACTT
GCACACGTCTGAACTCTTTCCCTACACGACGCTCTTCCGATCTNNNGGCGACAAGAGTGAAATTCCAT
AGCACACGTCTGAACTCTTTCCCTACACGACGCTCTTCCGATCTNNNGCAGAAGAATCACTTGAATCG
GAGCACACGTCTGAACTCTTTCCCTACACGACGCTCTTCCGATCTNNNAAGATTGCACCACTGTACTCC
AGCACACGTCTGAACTCTTTCCCTACACGACGCTCTTCCGATCTNNNGAGAATCGCTTGAACCAAGG
GCACACGTCTGAACTCTTTCCCTACACGACGCTCTTCCGATCTNNNGAGGTTACAGTGAGCCAAGAT
GAGCACACGTCTGAACTCTTTCCCTACACGACGCTCTTCCGATCTNNNTCACAAGGTCAGGAGTTCAA
AGCACACGTCTGAACTCTTTCCCTACACGACGCTCTTCCGATCTNNNAAAAAAAATTAGCTGGGCGT
AGCACACGTCTGAACTCTTTCCCTACACGACGCTCTTCCGATCTNNNCTGAGGTGGGAAGATCGATT
GAGCACACGTCTGAACTCTTTCCCTACACGACGCTCTTCCGATCTNNNCCTGGACAACAAGAGCAAAA
AGCACACGTCTGAACTCTTTCCCTACACGACGCTCTTCCGATCTNNNCTGTAATCCCAGCTACTCAGG
AGCACACGTCTGAACTCTTTCCCTACACGACGCTCTTCCGATCTNNNCTTGAACCAGGGAGTTAGAG
GAGCACACGTCTGAACTCTTTCCCTACACGACGCTCTTCCGATCTNNNGAGATCATGCCACTTCACTCC
GCACACGTCTGAACTCTTTCCCTACACGACGCTCTTCCGATCTNNNAAATACACAAAATTAGCCGGGC
AGCACACGTCTGAACTCTTTCCCTACACGACGCTCTTCCGATCTNNNAGAGGTTGCAGTAAGCCAGA
GAGCACACGTCTGAACTCTTTCCCTACACGACGCTCTTCCGATCTNNNTGAGTCAAGATCCTTCCATTG
AGCACACGTCTGAACTCTTTCCCTACACGACGCTCTTCCGATCTNNNCTGTAATCCCAGCTACTCAGGA
GAGCACACGTCTGAACTCTTTCCCTACACGACGCTCTTCCGATCTNNNCGAGATCATGCCATTGTACTC
GAGCACACGTCTGAACTCTTTCCCTACACGACGCTCTTCCGATCTNNNCTGAGTCAGGAGAATCGCTT
GAGCACACGTCTGAACTCTTTCCCTACACGACGCTCTTCCGATCTNNNAAGGCATGAGAATTGCTTG
GAGCACACGTCTGAACTCTTTCCCTACACGACGCTCTTCCGATCTNNNTGAGGTAGGAGAATCGCTTG
GAGCACACGTCTGAACTCTTTCCCTACACGACGCTCTTCCGATCTNNNCTGTAATCCCAGCTACTCAGG
GCACACGTCTGAACTCTTTCCCTACACGACGCTCTTCCGATCTNNNTGGATGCCTGTAATCCCAACTAC
AGCACACGTCTGAACTCTTTCCCTACACGACGCTCTTCCGATCTNNNTATAATCCCAGCTACTTGGAG

AGCACACGTCTGAACTCTTTCCCTACACGACGCTCTTCCGATCTNNNGGCTAACACGGTAAACCTAGT
GCACACGTCTGAACTCTTTCCCTACACGACGCTCTTCCGATCTNNNGATTGCTTGAACATGGGAGGA
AGCACACGTCTGAACTCTTTCCCTACACGACGCTCTTCCGATCTNNNCTGAGGCAGGAGAATCACTTG
AGCACACGTCTGAACTCTTTCCCTACACGACGCTCTTCCGATCTNNNTGAGGCAGGAGAATCTCTTG
GAGCACACGTCTGAACTCTTTCCCTACACGACGCTCTTCCGATCTNNNAAAATGAAAAAGGGCTGGA
AGCACACGTCTGAACTCTTTCCCTACACGACGCTCTTCCGATCTNNNTGTAATTCCAGCTACTTGGGA
GAGCACACGTCTGAACTCTTTCCCTACACGACGCTCTTCCGATCTNNNGAGATCACACCACTGCACTCT
AGCACACGTCTGAACTCTTTCCCTACACGACGCTCTTCCGATCTNNNAGAGGTTGCAGTGAGTTGAG
AGCACACGTCTGAACTCTTTCCCTACACGACGCTCTTCCGATCTNNNAAACTAGAAGGCCTCAGAAGA
GAGCACACGTCTGAACTCTTTCCCTACACGACGCTCTTCCGATCTNNNCACATGCCTATAGTCCCACGT
GAGCACACGTCTGAACTCTTTCCCTACACGACGCTCTTCCGATCTNNNTTGTGTTCCAGAGTTCAAGAC
AGCACACGTCTGAACTCTTTCCCTACACGACGCTCTTCCGATCTNNNACAGAGTCAGTGAACCAATC
AGCACACGTCTGAACTCTTTCCCTACACGACGCTCTTCCGATCTNNNAGCTGAGACAGGAGAATCACT
GAGCACACGTCTGAACTCTTTCCCTACACGACGCTCTTCCGATCTNNNCACAGGAGAATTGCTTGAAC
AGCACACGTCTGAACTCTTTCCCTACACGACGCTCTTCCGATCTNNNCGAGATCACACCATTACACTCC
GCACACGTCTGAACTCTTTCCCTACACGACGCTCTTCCGATCTNNNCACCTGTAATCCCAGCTACTCAG
AGCACACGTCTGAACTCTTTCCCTACACGACGCTCTTCCGATCTNNNATAAAACAATTAGTCGGGCGT
AGCACACGTCTGAACTCTTTCCCTACACGACGCTCTTCCGATCTNNNGCAATGACAGCAAAACTACGT
AGCACACGTCTGAACTCTTTCCCTACACGACGCTCTTCCGATCTNNNCTGAGGCAGGAGAATCACTTG
AGCACACGTCTGAACTCTTTCCCTACACGACGCTCTTCCGATCTNNNAAATACAAAAATTAGCCGGGC
GAGCACACGTCTGAACTCTTTCCCTACACGACGCTCTTCCGATCTNNNGGCAGGAGAATCACTTGAA
GAGCACACGTCTGAACTCTTTCCCTACACGACGCTCTTCCGATCTNNNGTGTCATCACAGTAGCAATG
GAGCACACGTCTGAACTCTTTCCCTACACGACGCTCTTCCGATCTNNNACCCATAATCCTAGCACTTTG
AGCACACGTCTGAACTCTTTCCCTACACGACGCTCTTCCGATCTNNNTGGGTGAAAGAGCAAGATTC
AGCACACGTCTGAACTCTTTCCCTACACGACGCTCTTCCGATCTNNNACTGCACTCCAGCCTAAATAAC
AGCACACGTCTGAACTCTTTCCCTACACGACGCTCTTCCGATCTNNNGAAGGTTGCAATGAGCCAAGA
AGCACACGTCTGAACTCTTTCCCTACACGACGCTCTTCCGATCTNNNATTACCACTGACCCCACAGAAA
GAGCACACGTCTGAACTCTTTCCCTACACGACGCTCTTCCGATCTNNNGTGACTGAGCAAGACTCCAT
AGCACACGTCTGAACTCTTTCCCTACACGACGCTCTTCCGATCTNNNAAAAATTAGCCGGGTGTAGT
GAGCACACGTCTGAACTCTTTCCCTACACGACGCTCTTCCGATCTNNNTCACTTGAACCTTGGAGGAG
GCACACGTCTGAACTCTTTCCCTACACGACGCTCTTCCGATCTNNNCACCTGTAATCCCAGCTACTCAG
AGCACACGTCTGAACTCTTTCCCTACACGACGCTCTTCCGATCTNNNATGCCTGTAATCCCAGCTACTT
AGCACACGTCTGAACTCTTTCCCTACACGACGCTCTTCCGATCTNNNCTGTAGTCCCAGCTACTCAGG
AGCACACGTCTGAACTCTTTCCCTACACGACGCTCTTCCGATCTNNNCTGAGGCAGGAGAACAGTGT
AGCACACGTCTGAACTCTTTCCCTACACGACGCTCTTCCGATCTNNNTTGGCATAGCACTCTGACAGTT
AGCACACGTCTGAACTCTTTCCCTACACGACGCTCTTCCGATCTNNNCACCTGTAATCCCAGCTACTTG
AGCACACGTCTGAACTCTTTCCCTACACGACGCTCTTCCGATCTNNNTTTCCTGGTTTCCTTTTTGCCTC
AGCACACGTCTGAACTCTTTCCCTACACGACGCTCTTCCGATCTNNNAGGATTGCTTGAGCCTAGGAA
AGCACACGTCTGAACTCTTTCCCTACACGACGCTCTTCCGATCTNNNGGACAACAGAGTGAGACTCC
AGCACACGTCTGAACTCTTTCCCTACACGACGCTCTTCCGATCTNNNCTGTAATCCCAGCTACTCGGAA
GAGCACACGTCTGAACTCTTTCCCTACACGACGCTCTTCCGATCTNNNTAGTTCCAAGCTATTCAGGAG
GAGCACACGTCTGAACTCTTTCCCTACACGACGCTCTTCCGATCTNNNTGGGTGATAGAGTGAGACT
GAGCACACGTCTGAACTCTTTCCCTACACGACGCTCTTCCGATCTNNNAACTATTTGGGAGGTTGAG

GAGCACACGTCTGAACTCTTTCCCTACACGACGCTCTTCCGATCTNNNGATCTACCAAGCAAATGGACA
AGCACACGTCTGAACTCTTTCCCTACACGACGCTCTTCCGATCTNNNGAAGATTGCTTGAACCTGGGA
AGCACACGTCTGAACTCTTTCCCTACACGACGCTCTTCCGATCTNNNACAAAAATTAGCCAGGTGTGG
GAGCACACGTCTGAACTCTTTCCCTACACGACGCTCTTCCGATCTNNNCAGGAGGTGGAGATTGTAGT
AGCACACGTCTGAACTCTTTCCCTACACGACGCTCTTCCGATCTNNNAAAAAATTAGCTGGGCATGGT
GAGCACACGTCTGAACTCTTTCCCTACACGACGCTCTTCCGATCTNNNTGGGCAACAGAATAAGACTG
AGCACACGTCTGAACTCTTTCCCTACACGACGCTCTTCCGATCTNNNGGATCACTTGAGCCCAGGAAT
AGCACACGTCTGAACTCTTTCCCTACACGACGCTCTTCCGATCTNNNGAGGTTGCAGTGAGCTAAGA
AGCACACGTCTGAACTCTTTCCCTACACGACGCTCTTCCGATCTNNNAAAAATTAGCTTGGAGTGGTG
AGAGCACACGTCTGAACTCTTTCCCTACACGACGCTCTTCCGATCTNNNCTGCCTGTTTTCTGAATCAC
AGCACACGTCTGAACTCTTTCCCTACACGACGCTCTTCCGATCTNNNCTGAGGCAGGAGAATCACTTG
AGAGCACACGTCTGAACTCTTTCCCTACACGACGCTCTTCCGATCTNNNGATCACAAGGTCTGGAGAT
GAGCACACGTCTGAACTCTTTCCCTACACGACGCTCTTCCGATCTNNNCTGGCCAACATAGTGAAACC
AGCACACGTCTGAACTCTTTCCCTACACGACGCTCTTCCGATCTNNNCTGAGGCAGGAGAATCACTTG
GAGCACACGTCTGAACTCTTTCCCTACACGACGCTCTTCCGATCTNNNAAGACTGCAATGAGCCACATT
GAGCACACGTCTGAACTCTTTCCCTACACGACGCTCTTCCGATCTNNNTGCTTGTAATCCCAGCTACTCA
AGCACACGTCTGAACTCTTTCCCTACACGACGCTCTTCCGATCTNNNTAGAGGTTACAGTGACCCGAG
CACACGTCTGAACTCTTTCCCTACACGACGCTCTTCCGATCTNNNGAAGCAGGAGAATCACTTGAACC
AGCACACGTCTGAACTCTTTCCCTACACGACGCTCTTCCGATCTNNNCTGAGGCAGGAGAATAGCTTG
GAGCACACGTCTGAACTCTTTCCCTACACGACGCTCTTCCGATCTNNNGTAATCTCAGCTACTCAGGAG
AGCACACGTCTGAACTCTTTCCCTACACGACGCTCTTCCGATCTNNNGCAACAGAGCAAAAACTCCATC
GAGCACACGTCTGAACTCTTTCCCTACACGACGCTCTTCCGATCTNNNACAACTGTAATCCCAGCTACT
AGCACACGTCTGAACTCTTTCCCTACACGACGCTCTTCCGATCTNNNCTGAATCCTGGTCTACTGCTACC
GAGCACACGTCTGAACTCTTTCCCTACACGACGCTCTTCCGATCTNNNGTAATCCTAGCTACTTGGGAG
GAGCACACGTCTGAACTCTTTCCCTACACGACGCTCTTCCGATCTNNNGGAACTGTTTGAGCCTAGGA
GAGCACACGTCTGAACTCTTTCCCTACACGACGCTCTTCCGATCTNNNCTGCTCAGAGTACTGTCACAG
AGCACACGTCTGAACTCTTTCCCTACACGACGCTCTTCCGATCTNNNCTGAGGCAGGAGAATCTCTTG
GCACACGTCTGAACTCTTTCCCTACACGACGCTCTTCCGATCTNNNCTGAGGCAGGAGAATTGCTTGA
AGCACACGTCTGAACTCTTTCCCTACACGACGCTCTTCCGATCTNNNCACTTGTAATCCCAGCACTTTG
GCACACGTCTGAACTCTTTCCCTACACGACGCTCTTCCGATCTNNNTAGTCCTAGCTACTTGGAAGGC
AGCACACGTCTGAACTCTTTCCCTACACGACGCTCTTCCGATCTNNNCAGTGAGCTATGATCATGCCA
GAGCACACGTCTGAACTCTTTCCCTACACGACGCTCTTCCGATCTNNNCTGAGGCAGAGAATTGCTTG
AGCACACGTCTGAACTCTTTCCCTACACGACGCTCTTCCGATCTNNNGAGCCTGTGGTCCTAGCTACTA
GAGCACACGTCTGAACTCTTTCCCTACACGACGCTCTTCCGATCTNNNGAGAATTGCTTGAATGCAGG
GCACACGTCTGAACTCTTTCCCTACACGACGCTCTTCCGATCTNNNTGTGGTACTAGCTACTCAGGAG
AGCACACGTCTGAACTCTTTCCCTACACGACGCTCTTCCGATCTNNNGAGAATTGCTTGAATCCGGGA
AGAGCACACGTCTGAACTCTTTCCCTACACGACGCTCTTCCGATCTNNNCTAGGCTGCAGTGAGCTGA
AGCACACGTCTGAACTCTTTCCCTACACGACGCTCTTCCGATCTNNNCTGAGATCACACCACTGCACTT
GAGCACACGTCTGAACTCTTTCCCTACACGACGCTCTTCCGATCTNNNGTGTTTGTATCCCCAGCTACT
GAGCACACGTCTGAACTCTTTCCCTACACGACGCTCTTCCGATCTNNNAAAAAAAATCAGCTGGGTGT
GAGCACACGTCTGAACTCTTTCCCTACACGACGCTCTTCCGATCTNNNAAGGCAAAAATTAGTGGGT
AGCACACGTCTGAACTCTTTCCCTACACGACGCTCTTCCGATCTNNNGGGTGACAGAGTAAGACTCCA
AGAGCACACGTCTGAACTCTTTCCCTACACGACGCTCTTCCGATCTNNNCTGTAGTCCCAGCTACTCAG

AGCACACGTCTGAACTCTTTCCCTACACGACGCTCTTCCGATCTNNNAGATCATGTCACTACACTCCAG
AGCACACGTCTGAACTCTTTCCCTACACGACGCTCTTCCGATCTNNNGGTGCCTATAATCCCAGCTACTC
GAGCACACGTCTGAACTCTTTCCCTACACGACGCTCTTCCGATCTNNNCCACCCCCCAAAAAAAAGAAA
GAGCACACGTCTGAACTCTTTCCCTACACGACGCTCTTCCGATCTNNNTGTAATTCCAGCTATTCGGGA
AGAGCACACGTCTGAACTCTTTCCCTACACGACGCTCTTCCGATCTNNNGTTGTAGTTAGCCGAGATTG
GAGCACACGTCTGAACTCTTTCCCTACACGACGCTCTTCCGATCTNNNGGTTCCTGTATTCCCAGCTACT
AGCACACGTCTGAACTCTTTCCCTACACGACGCTCTTCCGATCTNNNTGTAATCCTAGCTACTCGGGAG
GAGCACACGTCTGAACTCTTTCCCTACACGACGCTCTTCCGATCTNNNTCACTTGAATCCTGGAGGTCAA
AGCACACGTCTGAACTCTTTCCCTACACGACGCTCTTCCGATCTNNNCACTTGAGCTCAGGAGTTTGAC
GAGCACACGTCTGAACTCTTTCCCTACACGACGCTCTTCCGATCTNNNAATCACTTGAACCTAGGAGGC
AGCACACGTCTGAACTCTTTCCCTACACGACGCTCTTCCGATCTNNNGCTCACAGCTATAATCCCAGCAT
AGCACACGTCTGAACTCTTTCCCTACACGACGCTCTTCCGATCTNNNTGAGACAGGAGAATCACTCGA
AGCACACGTCTGAACTCTTTCCCTACACGACGCTCTTCCGATCTNNNGGTGACAGAGAAAGATTCCGTC
AGAGCACACGTCTGAACTCTTTCCCTACACGACGCTCTTCCGATCTNNNATACAAAAATTAGCTGGGCG
GAGCACACGTCTGAACTCTTTCCCTACACGACGCTCTTCCGATCTNNNGCTAAGGCAGGAGAATCACTT
AGCACACGTCTGAACTCTTTCCCTACACGACGCTCTTCCGATCTNNNCAAGAGAATTGCTTGAACCTGC
AGCACACGTCTGAACTCTTTCCCTACACGACGCTCTTCCGATCTNNNCATAGCAAGACCCCATCTCTACC
AGCACACGTCTGAACTCTTTCCCTACACGACGCTCTTCCGATCTNNNAAAACATGGCAAAACCCCATCTC
AGCACACGTCTGAACTCTTTCCCTACACGACGCTCTTCCGATCTNNNGAGACAGGAGAATTGCTCAAA
AGCACACGTCTGAACTCTTTCCCTACACGACGCTCTTCCGATCTNNNGACAAAGTGAGACCCTGACTCA
GAGCACACGTCTGAACTCTTTCCCTACACGACGCTCTTCCGATCTNNNTAAGCTGAGATCATGTCACTG
GAGCACACGTCTGAACTCTTTCCCTACACGACGCTCTTCCGATCTNNNACAAGAACGAAACTCCATCTC
AGCACACGTCTGAACTCTTTCCCTACACGACGCTCTTCCGATCTNNNGTAATCACAGCACTTTGGGATC
GAGCACACGTCTGAACTCTTTCCCTACACGACGCTCTTCCGATCTNNNTGTAATTCCAGCTACTTGGGAG
GAGCACACGTCTGAACTCTTTCCCTACACGACGCTCTTCCGATCTNNNCTGAGGCAAGAGAATCCCTTC
AGCACACGTCTGAACTCTTTCCCTACACGACGCTCTTCCGATCTNNNGGGAAACAGAGAGAGACTCCA
AGAGCACACGTCTGAACTCTTTCCCTACACGACGCTCTTCCGATCTNNNGAAAATTGCTTGAACTCGGG
AGAGCACACGTCTGAACTCTTTCCCTACACGACGCTCTTCCGATCTNNNTGAGGCAGGAGAATCACTTG
AGCACACGTCTGAACTCTTTCCCTACACGACGCTCTTCCGATCTNNNTGAGGCAGGAGAATAGCTTGA
AGAGCACACGTCTGAACTCTTTCCCTACACGACGCTCTTCCGATCTNNNCTAACTACCCAGGAGGCTGA
AGAGCACACGTCTGAACTCTTTCCCTACACGACGCTCTTCCGATCTNNNTGAGGCAGGAGAATCACTTG
AGCACACGTCTGAACTCTTTCCCTACACGACGCTCTTCCGATCTNNNTGTAGTCCCAGCTACTTAGGAG
AGCACACGTCTGAACTCTTTCCCTACACGACGCTCTTCCGATCTNNNTGAGGCACGAGAATCACTGAAA
GCACACGTCTGAACTCTTTCCCTACACGACGCTCTTCCGATCTNNNCACCTGTAATCCCAGCTACTCAGCA
GAGCACACGTCTGAACTCTTTCCCTACACGACGCTCTTCCGATCTNNNCTCATGCCTATGATCGTAGCA
AGCACACGTCTGAACTCTTTCCCTACACGACGCTCTTCCGATCTNNNCACATCTGTAGTCCCAGCTACTC
GAGCACACGTCTGAACTCTTTCCCTACACGACGCTCTTCCGATCTNNNGTCAGAGGTTGTAGTGAGCCA
GAGCACACGTCTGAACTCTTTCCCTACACGACGCTCTTCCGATCTNNNGGAGACTGCAGTAAGCCATGA
GAGCACACGTCTGAACTCTTTCCCTACACGACGCTCTTCCGATCTNNNGCACCTGTAATCCCAGCTACTA
GCACACGTCTGAACTCTTTCCCTACACGACGCTCTTCCGATCTNNNATGCCTGTAATCCCAGCTACTTGC
GAGCACACGTCTGAACTCTTTCCCTACACGACGCTCTTCCGATCTNNNTGCTTGTAATCCCAGCTACTCA
GAGCACACGTCTGAACTCTTTCCCTACACGACGCTCTTCCGATCTNNNGAGCCCAGGACTTTTAGGTTA
GAGCACACGTCTGAACTCTTTCCCTACACGACGCTCTTCCGATCTNNNAGAGGTTGCAGTGAGTCAAG

AGAGCACACGTCTGAACTCTTTCCCTACACGACGCTCTTCCGATCTNNNCAGGAGAATTGCTTGAACCA
AGCACACGTCTGAACTCTTTCCCTACACGACGCTCTTCCGATCTNNNTATAGTCCCAGCTACTCAGGAGG
GAGCACACGTCTGAACTCTTTCCCTACACGACGCTCTTCCGATCTNNNCTGTAATCCCAGCAACACTTTG
AGAGCACACGTCTGAACTCTTTCCCTACACGACGCTCTTCCGATCTNNNCAAGATTGTGCCACTATACTC
GAGCACACGTCTGAACTCTTTCCCTACACGACGCTCTTCCGATCTNNNCTGAGGCAGGAGAATTGCTT
AAGAGCACACGTCTGAACTCTTTCCCTACACGACGCTCTTCCGATCTNNNATACATTATGGTTGGGTTG
GAGCACACGTCTGAACTCTTTCCCTACACGACGCTCTTCCGATCTNNNAAGAATTCTCTGGAAGGCCCA
GAGCACACGTCTGAACTCTTTCCCTACACGACGCTCTTCCGATCTNNNAAAAGGCAGAGGCATTGGAA
GAGCACACGTCTGAACTCTTTCCCTACACGACGCTCTTCCGATCTNNNGTCAGGAGTTTCAGACCAGCT
AGAGCACACGTCTGAACTCTTTCCCTACACGACGCTCTTCCGATCTNNNTACAGAAATTAGCTGGGCAT
AGAGCACACGTCTGAACTCTTTCCCTACACGACGCTCTTCCGATCTNNNGGTCTGGGCAAATGTTTGTT
GAGCACACGTCTGAACTCTTTCCCTACACGACGCTCTTCCGATCTNNNAGCCCAGGGACATAATTGAAT
GAGCACACGTCTGAACTCTTTCCCTACACGACGCTCTTCCGATCTNNNTAATCCCAGCTACTCAGGAGGT
GCACACGTCTGAACTCTTTCCCTACACGACGCTCTTCCGATCTNNNTACAGCTGCCCCTATTTACTGACC
GAGCACACGTCTGAACTCTTTCCCTACACGACGCTCTTCCGATCTNNNCAGGAGAATTGCTTGAACACA
AGCACACGTCTGAACTCTTTCCCTACACGACGCTCTTCCGATCTNNNTATTACAGTCGTGAACCACCGT
AGCACACGTCTGAACTCTTTCCCTACACGACGCTCTTCCGATCTNNNTGTAGTCCCAGTTACTCAGGAG
AGAGCACACGTCTGAACTCTTTCCCTACACGACGCTCTTCCGATCTNNNAGAATGTGGGGCATTTCAAC
GAGCACACGTCTGAACTCTTTCCCTACACGACGCTCTTCCGATCTNNNCTGTATTCCCAGCTACTCGGAA
GAGCACACGTCTGAACTCTTTCCCTACACGACGCTCTTCCGATCTNNNGAATCTCTTGAAACTGGAAGC
AGCACACGTCTGAACTCTTTCCCTACACGACGCTCTTCCGATCTNNNTGGATCACTTGAACCTATGAGGC
AGAGCACACGTCTGAACTCTTTCCCTACACGACGCTCTTCCGATCTNNNATAAAAATGAGCCGGGCAG
GAGCACACGTCTGAACTCTTTCCCTACACGACGCTCTTCCGATCTNNNTTGAACTCAGGAGATGGAGG
GAGCACACGTCTGAACTCTTTCCCTACACGACGCTCTTCCGATCTNNNGCCTGGTCAACATAACAAATCC
GAGCACACGTCTGAACTCTTTCCCTACACGACGCTCTTCCGATCTNNNAACAACAACAAAACTAGCAGG
GAGCACACGTCTGAACTCTTTCCCTACACGACGCTCTTCCGATCTNNNAAATTAGTTGGGAATGGTGGC
GAGCACACGTCTGAACTCTTTCCCTACACGACGCTCTTCCGATCTNNNGCGAAACCCGGTTTCTACTAA
GAGCACACGTCTGAACTCTTTCCCTACACGACGCTCTTCCGATCTNNNAAATCCCAACTATGTGGAAGC
GAGCACACGTCTGAACTCTTTCCCTACACGACGCTCTTCCGATCTNNNGAGTTTGCAGTGAGCCGAGAT
AGCACACGTCTGAACTCTTTCCCTACACGACGCTCTTCCGATCTNNNACAGGAGAATAGCTTGAACCAG
GAGCACACGTCTGAACTCTTTCCCTACACGACGCTCTTCCGATCTNNNATTTGGCAATATCATCGAAGC
GCACACGTCTGAACTCTTTCCCTACACGACGCTCTTCCGATCTNNNGGTGACAAAGTGAGACTCAGTCT
GAGCACACGTCTGAACTCTTTCCCTACACGACGCTCTTCCGATCTNNNCTGTAGTTCCAGCTACTTGGGA
AGCACACGTCTGAACTCTTTCCCTACACGACGCTCTTCCGATCTNNNAAAAAATTAGCTGGGCGTAGTG
AGCACACGTCTGAACTCTTTCCCTACACGACGCTCTTCCGATCTNNNCTCTAGTCCCAGCTACTCAGGA
GAGCACACGTCTGAACTCTTTCCCTACACGACGCTCTTCCGATCTNNNGGATCAGTTGTGAGATCAAGC
AGCACACGTCTGAACTCTTTCCCTACACGACGCTCTTCCGATCTNNNAAGGAGGGCAGACTAACTAGA
GAGCACACGTCTGAACTCTTTCCCTACACGACGCTCTTCCGATCTNNNGGTGACAGAGCTAGACTGCA
GAGCACACGTCTGAACTCTTTCCCTACACGACGCTCTTCCGATCTNNNGCTGCAGTGAACCATGATCAT
GAGCACACGTCTGAACTCTTTCCCTACACGACGCTCTTCCGATCTNNNGAGAATCACTTGAACCCGGGA
AGCACACGTCTGAACTCTTTCCCTACACGACGCTCTTCCGATCTNNNACTCCAACCTAAGTGACAGAGT
GAGCACACGTCTGAACTCTTTCCCTACACGACGCTCTTCCGATCTNNNCTGTAATCCCAGCTACTCAGGA
GAGCACACGTCTGAACTCTTTCCCTACACGACGCTCTTCCGATCTNNNGAGCTTGCAGTGAGTTGAGA

AGCACACGTCTGAACTCTTTCCCTACACGACGCTCTTCCGATCTNNNGAGAATTGCTTGAACCCAGG
GAGCACACGTCTGAACTCTTTCCCTACACGACGCTCTTCCGATCTNNNCTGGGAAATAGAGATCCTGC
GCACACGTCTGAACTCTTTCCCTACACGACGCTCTTCCGATCTNNNGTGATGGAGCAAGACTCTGTCT
AGCACACGTCTGAACTCTTTCCCTACACGACGCTCTTCCGATCTNNNAAAAAAAAAATTAGCCGGGCG
AGCACACGTCTGAACTCTTTCCCTACACGACGCTCTTCCGATCTNNNCAGATCACAAGGTCAGGAGTT
GAGCACACGTCTGAACTCTTTCCCTACACGACGCTCTTCCGATCTNNNGGCAACAAGAGTGTAACAGT
AGAGCACACGTCTGAACTCTTTCCCTACACGACGCTCTTCCGATCTNNNGTTGCAGTGAGCTGTGATC
GAGCACACGTCTGAACTCTTTCCCTACACGACGCTCTTCCGATCTNNNCACCTGTAGTCCCAGCTATTC
AGCACACGTCTGAACTCTTTCCCTACACGACGCTCTTCCGATCTNNNATGCCTGTAATACCAGCTACTC
GCACACGTCTGAACTCTTTCCCTACACGACGCTCTTCCGATCTNNNGGCAGGAGAATCACTTGAACCT
AGCACACGTCTGAACTCTTTCCCTACACGACGCTCTTCCGATCTNNNAAACCACACACACAAAGAGCAA
AGCACACGTCTGAACTCTTTCCCTACACGACGCTCTTCCGATCTNNNGGCAACAAGATGGAAACTCAG
AGCACACGTCTGAACTCTTTCCCTACACGACGCTCTTCCGATCTNNNCTGTAATCCCAGCTACTCAGGA
AGCACACGTCTGAACTCTTTCCCTACACGACGCTCTTCCGATCTNNNATAACAAAAGTGGAAACCTGG
AGCACACGTCTGAACTCTTTCCCTACACGACGCTCTTCCGATCTNNNGGTGGATGGATTATGAGGTC
AGCACACGTCTGAACTCTTTCCCTACACGACGCTCTTCCGATCTNNNATACCTAGGGTAAGGCTGGAT
AGCACACGTCTGAACTCTTTCCCTACACGACGCTCTTCCGATCTNNNGAGGCAGGAAAATCACTTGAA
GAGCACACGTCTGAACTCTTTCCCTACACGACGCTCTTCCGATCTNNNGATAATCACTTGAGGCCAGG
AGCACACGTCTGAACTCTTTCCCTACACGACGCTCTTCCGATCTNNNGTGAGTCTTTATCATGCCACTG
AGAGCACACGTCTGAACTCTTTCCCTACACGACGCTCTTCCGATCTNNNGAGGTTGCAGTGAGCTAAG
AGCACACGTCTGAACTCTTTCCCTACACGACGCTCTTCCGATCTNNNCACCTCTAATCCCAGCTACTCAG
AGCACACGTCTGAACTCTTTCCCTACACGACGCTCTTCCGATCTNNNGTAATCTCAGCTACTCAGGAGG
AGCACACGTCTGAACTCTTTCCCTACACGACGCTCTTCCGATCTNNNTGAGGCAAGAGAATTGCTTG
AGCACACGTCTGAACTCTTTCCCTACACGACGCTCTTCCGATCTNNNTGGATGACAGAAGGAGACTCT
GAGCACACGTCTGAACTCTTTCCCTACACGACGCTCTTCCGATCTNNNACGTGTAGTTCTGGTTACTTG
AGCACACGTCTGAACTCTTTCCCTACACGACGCTCTTCCGATCTNNNGCCGAGATCAAGCCACTATACT
GAGCACACGTCTGAACTCTTTCCCTACACGACGCTCTTCCGATCTNNNAGATTGTGCCACTGTACTCCA
GCACACGTCTGAACTCTTTCCCTACACGACGCTCTTCCGATCTNNNGCCATTGTGAATAATGCTGCTCT
GCACACGTCTGAACTCTTTCCCTACACGACGCTCTTCCGATCTNNNCAAAACATTAGCTGGGTGTGGT
AGCACACGTCTGAACTCTTTCCCTACACGACGCTCTTCCGATCTNNNAGGCAGAAGAATTGCTTGAAC
AGCACACGTCTGAACTCTTTCCCTACACGACGCTCTTCCGATCTNNNGAAGGTGCCTGTAATCCCAGT
AGCACACGTCTGAACTCTTTCCCTACACGACGCTCTTCCGATCTNNNAAAAGTAGATAGGCATGGTGG
GCACACGTCTGAACTCTTTCCCTACACGACGCTCTTCCGATCTNNNTGTAGTCCTAGCTACTCAGGAG
GAGCACACGTCTGAACTCTTTCCCTACACGACGCTCTTCCGATCTNNNGGTGAAGCCCCATCTCTACTA
AGCACACGTCTGAACTCTTTCCCTACACGACGCTCTTCCGATCTNNNTGAGGCAGGAGAATCACTTGA
GCACACGTCTGAACTCTTTCCCTACACGACGCTCTTCCGATCTNNNGGCAAAAGAGTGAGACTGCAT
GAGCACACGTCTGAACTCTTTCCCTACACGACGCTCTTCCGATCTNNNGAGAATCTCTTGAACCCAGGA
GCACACGTCTGAACTCTTTCCCTACACGACGCTCTTCCGATCTNNNCAGGAGAATTGCTTGAACCAGG
GAGCACACGTCTGAACTCTTTCCCTACACGACGCTCTTCCGATCTNNNCTGAGGCAGGAGAATCACTT
GAGCACACGTCTGAACTCTTTCCCTACACGACGCTCTTCCGATCTNNNTATGGAGAAACCCCGTCTCTA
AGCACACGTCTGAACTCTTTCCCTACACGACGCTCTTCCGATCTNNNCAAGGCAACAGAACAAAACTC
GAGCACACGTCTGAACTCTTTCCCTACACGACGCTCTTCCGATCTNNNTGAGGCAGGAGAATCACTTG
GAGCACACGTCTGAACTCTTTCCCTACACGACGCTCTTCCGATCTNNNATCCAGGAAAGTGTGAAGGA

GAGCACACGTCTGAACTCTTTCCCTACACGACGCTCTTCCGATCTNNNTGAGAATCACTTGAACCCAG
GAGCACACGTCTGAACTCTTTCCCTACACGACGCTCTTCCGATCTNNNGGCACCTGTAATCCCAGTTA
AGCACACGTCTGAACTCTTTCCCTACACGACGCTCTTCCGATCTNNNCTGAGGCAGGAGAATAGCTT
AGCACACGTCTGAACTCTTTCCCTACACGACGCTCTTCCGATCTNNNGATACAAGTGGAGGAGTAGC
GAGCACACGTCTGAACTCTTTCCCTACACGACGCTCTTCCGATCTNNNATCTTGCCATTGTGAGATTGT
AGCACACGTCTGAACTCTTTCCCTACACGACGCTCTTCCGATCTNNNCTGTAGTCCCAGCTACTCAGG
GAGCACACGTCTGAACTCTTTCCCTACACGACGCTCTTCCGATCTNNNCTCAAGCCTGTAATCCTAGCA
GAGCACACGTCTGAACTCTTTCCCTACACGACGCTCTTCCGATCTNNNCTGAGGCAGAGAATTGCTTG
GAGCACACGTCTGAACTCTTTCCCTACACGACGCTCTTCCGATCTNNNCGCCTATAATCCCAGCTAGTC
GAGCACACGTCTGAACTCTTTCCCTACACGACGCTCTTCCGATCTNNNGAGGCAGGAGAATTGTTTG
AGCACACGTCTGAACTCTTTCCCTACACGACGCTCTTCCGATCTNNNCACATGAGGTCAGGAGTTCAT
GCACACGTCTGAACTCTTTCCCTACACGACGCTCTTCCGATCTNNNCAAGTGACTGTAGTCCCAGCTAC
AGAGCACACGTCTGAACTCTTTCCCTACACGACGCTCTTCCGATCTNNNAGAATCACTTGAACCCAGG
AGCACACGTCTGAACTCTTTCCCTACACGACGCTCTTCCGATCTNNNCTGAGGCAGAGAATTGCTTGA
AGCACACGTCTGAACTCTTTCCCTACACGACGCTCTTCCGATCTNNNACAAATCCACTGGGCTATACAC
GAGCACACGTCTGAACTCTTTCCCTACACGACGCTCTTCCGATCTNNNAGGCAGGAGAATCATTTGAA
GAGCACACGTCTGAACTCTTTCCCTACACGACGCTCTTCCGATCTNNNCAGGAGAATCACTTGAACCCA
AGAGCACACGTCTGAACTCTTTCCCTACACGACGCTCTTCCGATCTNNNACAAAATTTAGCTGGGTGT
AGCACACGTCTGAACTCTTTCCCTACACGACGCTCTTCCGATCTNNNACCAGCCTAAGCAACATGAGA
GAGCACACGTCTGAACTCTTTCCCTACACGACGCTCTTCCGATCTNNNCTGAGGCAAGAGAATCCCTT
AGCACACGTCTGAACTCTTTCCCTACACGACGCTCTTCCGATCTNNNTTTGCAGTGAGCCAATATTGT
AGCACACGTCTGAACTCTTTCCCTACACGACGCTCTTCCGATCTNNNCTGAGGCAGGAGAATTGCTTG
AGCACACGTCTGAACTCTTTCCCTACACGACGCTCTTCCGATCTNNNGCAACAGGAATGAAACTTCGT
GAGCACACGTCTGAACTCTTTCCCTACACGACGCTCTTCCGATCTNNNGAGGCTGAGGCAAGAGAATT
AGCACACGTCTGAACTCTTTCCCTACACGACGCTCTTCCGATCTNNNAGGCAGGAGAATCATTTGAA
AGCACACGTCTGAACTCTTTCCCTACACGACGCTCTTCCGATCTNNNGGATCAATCAGAAGGCAATGG
AGCACACGTCTGAACTCTTTCCCTACACGACGCTCTTCCGATCTNNNAGCAACCAGATGTCCCTCAATA
AGAGCACACGTCTGAACTCTTTCCCTACACGACGCTCTTCCGATCTNNNCTGTAATCCCAGCTACTCGG
AGCACACGTCTGAACTCTTTCCCTACACGACGCTCTTCCGATCTNNNCATAGCTCACTACAGCCTCAGT
AGCACACGTCTGAACTCTTTCCCTACACGACGCTCTTCCGATCTNNNACTGCATGTCAGAATTCATCC
GAGCACACGTCTGAACTCTTTCCCTACACGACGCTCTTCCGATCTNNNGGGAGGGATAGCATTGGGA
GAGCACACGTCTGAACTCTTTCCCTACACGACGCTCTTCCGATCTNNNCTGTAGTCCCAGCTACTCAGG
AGCACACGTCTGAACTCTTTCCCTACACGACGCTCTTCCGATCTNNNAAAATTAGCTGGGTTTGGTGG
AGCACACGTCTGAACTCTTTCCCTACACGACGCTCTTCCGATCTNNNGACAGAGCAAGACTACGTCTCA
AGCACACGTCTGAACTCTTTCCCTACACGACGCTCTTCCGATCTNNNTGAGAATCACTTGAACCTGGG
AGCACACGTCTGAACTCTTTCCCTACACGACGCTCTTCCGATCTNNNAGCCTGTAATCCCAGTACTTTG
AGCACACGTCTGAACTCTTTCCCTACACGACGCTCTTCCGATCTNNNTGGTGACAGATCAAGACTTCC
GCACACGTCTGAACTCTTTCCCTACACGACGCTCTTCCGATCTNNNTGCTGATCATCCTGGCTGTATAC
GAGCACACGTCTGAACTCTTTCCCTACACGACGCTCTTCCGATCTNNNCAAGGCAGGAGAATTGCTT
GAGCACACGTCTGAACTCTTTCCCTACACGACGCTCTTCCGATCTNNNGAGACAGGAGAATTGCTTGA
AGAGCACACGTCTGAACTCTTTCCCTACACGACGCTCTTCCGATCTNNNGCCTGTAATCCTAGCTACTCA
AGCACACGTCTGAACTCTTTCCCTACACGACGCTCTTCCGATCTNNNCATGATCACACCACTGTACTCC
AGCACACGTCTGAACTCTTTCCCTACACGACGCTCTTCCGATCTNNNCAGAAGAATTGCTTGAACCTG

AGCACACGTCTGAACTCTTTCCCTACACGACGCTCTTCCGATCTNNNTGAGGCAGGAGAATCTCTTGA
AGCACACGTCTGAACTCTTTCCCTACACGACGCTCTTCCGATCTNNNCCAGAAGCTAAGGCAGGAGAA
GAGCACACGTCTGAACTCTTTCCCTACACGACGCTCTTCCGATCTNNNCAAAAATTAGCTGGGTGTGG
AGCACACGTCTGAACTCTTTCCCTACACGACGCTCTTCCGATCTNNNGGTTTTGGAAGGCTCAGTACA
GAGCACACGTCTGAACTCTTTCCCTACACGACGCTCTTCCGATCTNNNGCCATAGAGCTTGGATCCCTA
GAGCACACGTCTGAACTCTTTCCCTACACGACGCTCTTCCGATCTNNNCCCACAAGCAGAACTACTTAG
GAGCACACGTCTGAACTCTTTCCCTACACGACGCTCTTCCGATCTNNNGAATTGCTTGAACCTGGGAA
AGCACACGTCTGAACTCTTTCCCTACACGACGCTCTTCCGATCTNNNATAGAAAAAATTAGCCGGGCG
GAGCACACGTCTGAACTCTTTCCCTACACGACGCTCTTCCGATCTNNNGAAGCAGGACAATCTCGTGA
GAGCACACGTCTGAACTCTTTCCCTACACGACGCTCTTCCGATCTNNNGAAAATCGCTTGAACTGGGG
AGCACACGTCTGAACTCTTTCCCTACACGACGCTCTTCCGATCTNNNACACACAAGTCACAAATTAGCC
GAGCACACGTCTGAACTCTTTCCCTACACGACGCTCTTCCGATCTNNNGGCACCTATAATCCCAGCTAC
AGCACACGTCTGAACTCTTTCCCTACACGACGCTCTTCCGATCTNNNAATCACAAAAACTGCACTCCAC
AGCACACGTCTGAACTCTTTCCCTACACGACGCTCTTCCGATCTNNNGTCACCTGTAATCCCAGCTACT
AGCACACGTCTGAACTCTTTCCCTACACGACGCTCTTCCGATCTNNNGCAGGAGAAATCACTTGAACC
GAGCACACGTCTGAACTCTTTCCCTACACGACGCTCTTCCGATCTNNNCTCTACAAGCGTCAGGATTCT
AGCACACGTCTGAACTCTTTCCCTACACGACGCTCTTCCGATCTNNNTAAAAAAATTAGCTGGGCGTG
GAGCACACGTCTGAACTCTTTCCCTACACGACGCTCTTCCGATCTNNNATAACACCTACATCTGCCACCT
AGCACACGTCTGAACTCTTTCCCTACACGACGCTCTTCCGATCTNNNACCCGAGGAATTCACAGCTAA
GAGCACACGTCTGAACTCTTTCCCTACACGACGCTCTTCCGATCTNNNAATTCCTGGACACATACACCC
AGCACACGTCTGAACTCTTTCCCTACACGACGCTCTTCCGATCTNNNCGCTTGTTTTGTCAGCTTTGT
AGCACACGTCTGAACTCTTTCCCTACACGACGCTCTTCCGATCTNNNTGAGGCAGGAGAATCTCTTGA
GAGCACACGTCTGAACTCTTTCCCTACACGACGCTCTTCCGATCTNNNTCACACCACTACATTCCAGAG
GAGCACACGTCTGAACTCTTTCCCTACACGACGCTCTTCCGATCTNNNCAGTCCTGTAATCCCAGCTAC
AGCACACGTCTGAACTCTTTCCCTACACGACGCTCTTCCGATCTNNNAGAGGAATCATGCACACAGCT
AGCACACGTCTGAACTCTTTCCCTACACGACGCTCTTCCGATCTNNNAAAAATTAGCTGGGCATGGTG
GAGCACACGTCTGAACTCTTTCCCTACACGACGCTCTTCCGATCTNNNGAGAATCTCTTGAGCCCAGTA
GAGCACACGTCTGAACTCTTTCCCTACACGACGCTCTTCCGATCTNNNGGTGCCGACAAGTCCTCTAA
GAGCACACGTCTGAACTCTTTCCCTACACGACGCTCTTCCGATCTNNNGAGGTTACAGTGAGCCGAGA
GAGCACACGTCTGAACTCTTTCCCTACACGACGCTCTTCCGATCTNNNGAGAATCTCTTGAGCCCAGTA
GAGCACACGTCTGAACTCTTTCCCTACACGACGCTCTTCCGATCTNNNGGTGCCGACAAGTCCTCTAA
GAGCACACGTCTGAACTCTTTCCCTACACGACGCTCTTCCGATCTNNNGAGGTTACAGTGAGCCGAGA
AGCACACGTCTGAACTCTTTCCCTACACGACGCTCTTCCGATCTNNNCTGAGACACATCCTCCACTAAG
AGCACACGTCTGAACTCTTTCCCTACACGACGCTCTTCCGATCTNNNATCCTAGGTGACAGAGTGAGA
AGCACACGTCTGAACTCTTTCCCTACACGACGCTCTTCCGATCTNNNGAGTTGCATGTTTCAGTGAGC
AGCACACGTCTGAACTCTTTCCCTACACGACGCTCTTCCGATCTNNNATGCCTGTAATCCCAGCATTT
GAGCACACGTCTGAACTCTTTCCCTACACGACGCTCTTCCGATCTNNNATGTGGCAATATCATTGAAG
GAGCACACGTCTGAACTCTTTCCCTACACGACGCTCTTCCGATCTNNNAAAAATTAGCTGGGAATGGT
GCACACGTCTGAACTCTTTCCCTACACGACGCTCTTCCGATCTNNNCACGCTTGTAATTCCAGCAGTTT
GAGCACACGTCTGAACTCTTTCCCTACACGACGCTCTTCCGATCTNNNAAATCCCAACTATGTGGAAG
GAGCACACGTCTGAACTCTTTCCCTACACGACGCTCTTCCGATCTNNNTTCAATGAGCTGAGATCACA
GAGCACACGTCTGAACTCTTTCCCTACACGACGCTCTTCCGATCTNNNGAGGTTGCATTGAGGTGAGA
AGCACACGTCTGAACTCTTTCCCTACACGACGCTCTTCCGATCTNNNGAAACAGATGTTGCAGTGAG

AGCACACGTCTGAACTCTTTCCCTACACGACGCTCTTCCGATCTNNNTGAGGCAGGAGAATGGTATG
GAGCACACGTCTGAACTCTTTCCCTACACGACGCTCTTCCGATCTNNNAGGCTTTCATCTTTTCCCACT
AGCACACGTCTGAACTCTTTCCCTACACGACGCTCTTCCGATCTNNNTAGGTGACAGACTGAGACTG
AGCACACGTCTGAACTCTTTCCCTACACGACGCTCTTCCGATCTNNNTAATTTGTTACCTGTCTGGGT
AGCACACGTCTGAACTCTTTCCCTACACGACGCTCTTCCGATCTNNNAACTAGATCGGAAGCTGAAGT
AGCACACGTCTGAACTCTTTCCCTACACGACGCTCTTCCGATCTNNNCTGTAGTCCCAGCTACTCAGG
AGCACACGTCTGAACTCTTTCCCTACACGACGCTCTTCCGATCTNNNCTATAGTCCAGCCTTGGTGACA
GCACACGTCTGAACTCTTTCCCTACACGACGCTCTTCCGATCTNNNGCCAGGTATGATTCCAGCTACT
AGCACACGTCTGAACTCTTTCCCTACACGACGCTCTTCCGATCTNNNCGCACCTGTAATCCTAGCTACT
AGCACACGTCTGAACTCTTTCCCTACACGACGCTCTTCCGATCTNNNACCTGACATGTGGAAAGAGT
GAGCACACGTCTGAACTCTTTCCCTACACGACGCTCTTCCGATCTNNNCTGAGGTAGGAGGATTGCTT
AGCACACGTCTGAACTCTTTCCCTACACGACGCTCTTCCGATCTNNNGTTCAAGACTAGCCTGGGTAA
GCACACGTCTGAACTCTTTCCCTACACGACGCTCTTCCGATCTNNNTGAGAGACAGAGTGAGAACCT
AGCACACGTCTGAACTCTTTCCCTACACGACGCTCTTCCGATCTNNNATTGCAGTGAGTGAGCTGAGA
AGCACACGTCTGAACTCTTTCCCTACACGACGCTCTTCCGATCTNNNTAGATCATGCCTGTAATCCCAG
GAGCACACGTCTGAACTCTTTCCCTACACGACGCTCTTCCGATCTNNNTGAGAATAGCTTCAACCCAG
AGCACACGTCTGAACTCTTTCCCTACACGACGCTCTTCCGATCTNNNCAGAGTGAGACCCTGCCTAAA
AGCACACGTCTGAACTCTTTCCCTACACGACGCTCTTCCGATCTNNNCTGAGGCAGGAGAATCACTTG
AGCACACGTCTGAACTCTTTCCCTACACGACGCTCTTCCGATCTNNNGTCTGTGGCACCTCCTCTATTT
GAGCACACGTCTGAACTCTTTCCCTACACGACGCTCTTCCGATCTNNNGGCAACAAGAGCGAAAATC
GAGCACACGTCTGAACTCTTTCCCTACACGACGCTCTTCCGATCTNNNTGCCTGTAATTCCAGCTACTC
GAGCACACGTCTGAACTCTTTCCCTACACGACGCTCTTCCGATCTNNNGAGAATCACTTGAACCCAGG
GAGCACACGTCTGAACTCTTTCCCTACACGACGCTCTTCCGATCTNNNGAGGTTGCAGTGAGTTGAG
GAGCACACGTCTGAACTCTTTCCCTACACGACGCTCTTCCGATCTNNNAGAGGTTGCAGTGAGCTAAG
AGCACACGTCTGAACTCTTTCCCTACACGACGCTCTTCCGATCTNNNAAAAAAAAATTGCCAGGCGT
AGCACACGTCTGAACTCTTTCCCTACACGACGCTCTTCCGATCTNNNGGTCAGGAGATCGAGACTAT
GAGCACACGTCTGAACTCTTTCCCTACACGACGCTCTTCCGATCTNNNCCAGCCTGGGCAATATGATA
AGCACACGTCTGAACTCTTTCCCTACACGACGCTCTTCCGATCTNNNTCAAGGCAGGAGGATTGTTTG
AGCACACGTCTGAACTCTTTCCCTACACGACGCTCTTCCGATCTNNNGTAATCCCAGCTACTTAGGAG
AGCACACGTCTGAACTCTTTCCCTACACGACGCTCTTCCGATCTNNNCTGAGGCAGGAAAATTGCTT
GCACACGTCTGAACTCTTTCCCTACACGACGCTCTTCCGATCTNNNGAGGCAGGAGAATTGCTTGAAT
GAGCACACGTCTGAACTCTTTCCCTACACGACGCTCTTCCGATCTNNNACAAACAAAAATTAGCCGGG
GAGCACACGTCTGAACTCTTTCCCTACACGACGCTCTTCCGATCTNNNGGAACTGCAAATCAACAACA
GAGCACACGTCTGAACTCTTTCCCTACACGACGCTCTTCCGATCTNNNCCCATGCCTGTAATTTCAGCT
GAGCACACGTCTGAACTCTTTCCCTACACGACGCTCTTCCGATCTNNNATTGCTTGAGCCTAGGAGTT
AGAGCACACGTCTGAACTCTTTCCCTACACGACGCTCTTCCGATCTNNNATGCTACCTTTTTAGTGCAT
AGCACACGTCTGAACTCTTTCCCTACACGACGCTCTTCCGATCTNNNTGAGGTAGGAGAATCGCTTG
GAGCACACGTCTGAACTCTTTCCCTACACGACGCTCTTCCGATCTNNNAAAAAATTAGCTGGGTGTG
AGCACACGTCTGAACTCTTTCCCTACACGACGCTCTTCCGATCTNNNGCTGAGGCAGGATAATCTCT
AGCACACGTCTGAACTCTTTCCCTACACGACGCTCTTCCGATCTNNNCTGAGGCAGGAGAATTGCTTG
GAGCACACGTCTGAACTCTTTCCCTACACGACGCTCTTCCGATCTNNNGAATTGCTTGAACCCAGGAA
AGAGCACACGTCTGAACTCTTTCCCTACACGACGCTCTTCCGATCTNNNAAAAATTAGCTGGGCGTAA
AGCACACGTCTGAACTCTTTCCCTACACGACGCTCTTCCGATCTNNNCAGAAGAATTGCTTGAACCCAG

AGCACACGTCTGAACTCTTTCCCTACACGACGCTCTTCCGATCTNNNCTGAGGCAGGAGAATCACTTGA
AGAGCACACGTCTGAACTCTTTCCCTACACGACGCTCTTCCGATCTNNNGTTCCAGTGAGCCAATATCAT
GAGCACACGTCTGAACTCTTTCCCTACACGACGCTCTTCCGATCTNNNGAGCAAGATGGGCAGATGTA
GAGCACACGTCTGAACTCTTTCCCTACACGACGCTCTTCCGATCTNNNTGCACTTGTAATCTCGGCTAC
AGCACACGTCTGAACTCTTTCCCTACACGACGCTCTTCCGATCTNNNAATACAAAATTAGTCCGGCGTG
GAGCACACGTCTGAACTCTTTCCCTACACGACGCTCTTCCGATCTNNNCAGGAGGTCAAGGCTACAGT
GAGCACACGTCTGAACTCTTTCCCTACACGACGCTCTTCCGATCTNNNCTGAGGCAGGAGAATCACTTG
GAGCACACGTCTGAACTCTTTCCCTACACGACGCTCTTCCGATCTNNNAGAATCACTTGAAACCGGGAC
GAGCACACGTCTGAACTCTTTCCCTACACGACGCTCTTCCGATCTNNNGCAGAAGAATCACTTGAACCC
GAGCACACGTCTGAACTCTTTCCCTACACGACGCTCTTCCGATCTNNNAGGCAGAAGAATCACTTGAAC
AGAGCACACGTCTGAACTCTTTCCCTACACGACGCTCTTCCGATCTNNNAGGATCATTTGAAGCCAGGA
AGAGCACACGTCTGAACTCTTTCCCTACACGACGCTCTTCCGATCTNNNCTGAGGCAGGAAAATTGCTT
GAGCACACGTCTGAACTCTTTCCCTACACGACGCTCTTCCGATCTNNNTGAGAGAGGAGAATGGCATC
GAGCACACGTCTGAACTCTTTCCCTACACGACGCTCTTCCGATCTNNNGAGATCACACCACTACACTCCA
GAGCACACGTCTGAACTCTTTCCCTACACGACGCTCTTCCGATCTNNNAGATTGCAGTGAGCTGAGATC
GAGCACACGTCTGAACTCTTTCCCTACACGACGCTCTTCCGATCTNNNGCGACAGAGCAAGACTTCATC
AGCACACGTCTGAACTCTTTCCCTACACGACGCTCTTCCGATCTNNNGCTGAGATCATGCCATTGAACTC
AGCACACGTCTGAACTCTTTCCCTACACGACGCTCTTCCGATCTNNNTCTTGGGCCATACGTAACGATA
GAGCACACGTCTGAACTCTTTCCCTACACGACGCTCTTCCGATCTNNNTGAGGTAGGAGAATCGCTTG
AGCACACGTCTGAACTCTTTCCCTACACGACGCTCTTCCGATCTNNNTGAGTTGACAGAGTGAGACCTT
GAGCACACGTCTGAACTCTTTCCCTACACGACGCTCTTCCGATCTNNNTGCCTGTAATCTCAGCTACTTC
GAGCACACGTCTGAACTCTTTCCCTACACGACGCTCTTCCGATCTNNNGAGGATCACTTTAGCGTAGGA
GAGCACACGTCTGAACTCTTTCCCTACACGACGCTCTTCCGATCTNNNCCATCAGGAAAGGCAAATGAA
AGAGCACACGTCTGAACTCTTTCCCTACACGACGCTCTTCCGATCTNNNAAAAAATTAGCTGGGCACAG
AGCACACGTCTGAACTCTTTCCCTACACGACGCTCTTCCGATCTNNNTACAAAAAATTAGCTGGGCGTG
AGCACACGTCTGAACTCTTTCCCTACACGACGCTCTTCCGATCTNNNGAGATTGCACCACTACACTCCAG
AGCACACGTCTGAACTCTTTCCCTACACGACGCTCTTCCGATCTNNNGTGTGCCTGTAACCTTAGCTACT
AGCACACGTCTGAACTCTTTCCCTACACGACGCTCTTCCGATCTNNNTGAGGCAGGAGAATCTCTTGAA
GAGCACACGTCTGAACTCTTTCCCTACACGACGCTCTTCCGATCTNNNGTGGATCACTTGAAGTCAGGA
GAGCACACGTCTGAACTCTTTCCCTACACGACGCTCTTCCGATCTNNNCCTGGGCAACAAGAAAGAAA
AGAGCACACGTCTGAACTCTTTCCCTACACGACGCTCTTCCGATCTNNNCTGTTTGAGCCCAGGAATTCA
GAGCACACGTCTGAACTCTTTCCCTACACGACGCTCTTCCGATCTNNNTTCAAGCTGAGGCAGGAGAA
AGAGCACACGTCTGAACTCTTTCCCTACACGACGCTCTTCCGATCTNNNGGAGGTTACAGTGGGTCAA
AGCACACGTCTGAACTCTTTCCCTACACGACGCTCTTCCGATCTNNNGGAGAATCTGTTGAACCCAGGA
AGAGCACACGTCTGAACTCTTTCCCTACACGACGCTCTTCCGATCTNNNCAGTGAGCTAAGATCCTGCCA
GAGCACACGTCTGAACTCTTTCCCTACACGACGCTCTTCCGATCTNNNGCACCTATAGTCCCAGCTACTTC
GAGCACACGTCTGAACTCTTTCCCTACACGACGCTCTTCCGATCTNNNAGGGGATAGATGAGATGACC
GAGCACACGTCTGAACTCTTTCCCTACACGACGCTCTTCCGATCTNNNCTATCACTTGAGCCCAGGAGAT
AGCACACGTCTGAACTCTTTCCCTACACGACGCTCTTCCGATCTNNNCTGCAATCCTAGCTACTCAGGAC
AGAGCACACGTCTGAACTCTTTCCCTACACGACGCTCTTCCGATCTNNNCTGTAATCCCAACTACCCAGG
AGCACACGTCTGAACTCTTTCCCTACACGACGCTCTTCCGATCTNNNCAAGAGAATTGCTTGAACCTGC
GAGCACACGTCTGAACTCTTTCCCTACACGACGCTCTTCCGATCTNNNAACATGGCGAAACCCTGTCTC
GAGCACACGTCTGAACTCTTTCCCTACACGACGCTCTTCCGATCTNNNTGATTGGCGTCCTTGTAAGAA

AAGAGCACACGTCTGAACTCTTTCCCTACACGACGCTCTTCCGATCTNNNGTGACAGAGTGAGACTCCA
GAGCACACGTCTGAACTCTTTCCCTACACGACGCTCTTCCGATCTNNNCTGAGGCAGGAGAATTGCTTG
GAGCACACGTCTGAACTCTTTCCCTACACGACGCTCTTCCGATCTNNNTGTAGTCCCAGCTACTCAGGA
GAGCACACGTCTGAACTCTTTCCCTACACGACGCTCTTCCGATCTNNNCTGAGGCAGTAGAACCACTTG
GAGCACACGTCTGAACTCTTTCCCTACACGACGCTCTTCCGATCTNNNCTGAAGCAGGAGAATGGTGT
AGCACACGTCTGAACTCTTTCCCTACACGACGCTCTTCCGATCTNNNAAATTAGGCAGATGTAGTGGC
GAGCACACGTCTGAACTCTTTCCCTACACGACGCTCTTCCGATCTNNNGGAGATTGTAGTGAGCCGAG
GAGCACACGTCTGAACTCTTTCCCTACACGACGCTCTTCCGATCTNNNGAGGCAGGAAAATCACTTGA
AGAGCACACGTCTGAACTCTTTCCCTACACGACGCTCTTCCGATCTNNNCCAGCCTGAGTGACAAAGTTA
GAGCACACGTCTGAACTCTTTCCCTACACGACGCTCTTCCGATCTNNNAGATCAAGCCATTGCACTCAA
AGCACACGTCTGAACTCTTTCCCTACACGACGCTCTTCCGATCTNNNCTGAGGCAGGAAAATTGCTTGA
AGCACACGTCTGAACTCTTTCCCTACACGACGCTCTTCCGATCTNNNTGGGAGATAGAGTGAGACGTG
AGCACACGTCTGAACTCTTTCCCTACACGACGCTCTTCCGATCTNNNCTAAGGCAGGAGAATCGCTTGA
GAGCACACGTCTGAACTCTTTCCCTACACGACGCTCTTCCGATCTNNNAGAATTACTTGAACCCAGGAG
GAGCACACGTCTGAACTCTTTCCCTACACGACGCTCTTCCGATCTNNNCTGAGGCAGGAGAATCACTTG
GAGCACACGTCTGAACTCTTTCCCTACACGACGCTCTTCCGATCTNNNCTGAGGCAGGAGAATCACTTG
AGCACACGTCTGAACTCTTTCCCTACACGACGCTCTTCCGATCTNNNGGAATTTGAGACCAGCCTGAGT
GAGCACACGTCTGAACTCTTTCCCTACACGACGCTCTTCCGATCTNNNGCAAGCAATGAAAACGGCAAA
GAGCACACGTCTGAACTCTTTCCCTACACGACGCTCTTCCGATCTNNNTTACCCAGATGTGCTCACTTGA
GAGCACACGTCTGAACTCTTTCCCTACACGACGCTCTTCCGATCTNNNCAGAGACTACAGTGAGCCAA
AGAGCACACGTCTGAACTCTTTCCCTACACGACGCTCTTCCGATCTNNNAGAGATTGCAGTGAGCTGAG
AGCACACGTCTGAACTCTTTCCCTACACGACGCTCTTCCGATCTNNNTGCCCTGATATCAAAACCAGACA
AGCACACGTCTGAACTCTTTCCCTACACGACGCTCTTCCGATCTNNNAACAGAACAGAGCCCTCAGAAA
GAGCACACGTCTGAACTCTTTCCCTACACGACGCTCTTCCGATCTNNNATTTGGGCTGGGAACTGTTCT
GAGCACACGTCTGAACTCTTTCCCTACACGACGCTCTTCCGATCTNNNCTGAGTGAGGAGAGAAACAC
GAGCACACGTCTGAACTCTTTCCCTACACGACGCTCTTCCGATCTNNNAATACAAAAAATTAGCCGGGC
AGAGCACACGTCTGAACTCTTTCCCTACACGACGCTCTTCCGATCTNNNGAGGCAGGATAATTGCTTGA
AGCACACGTCTGAACTCTTTCCCTACACGACGCTCTTCCGATCTNNNGGAGCTTACAGTGAGTGGAGA
GAGCACACGTCTGAACTCTTTCCCTACACGACGCTCTTCCGATCTNNNTCACAAGGTCAAGAGATGGA
GAGCACACGTCTGAACTCTTTCCCTACACGACGCTCTTCCGATCTNNNCTGAAGCAGGAGAATGGTGT
AGCACACGTCTGAACTCTTTCCCTACACGACGCTCTTCCGATCTNNNCCTGGCCAGAACTTCCAATACTA
AGAGCACACGTCTGAACTCTTTCCCTACACGACGCTCTTCCGATCTNNNCTGAGGCAGGATAATCGCTT
GAGCACACGTCTGAACTCTTTCCCTACACGACGCTCTTCCGATCTNNNAAGAGAATTTCTTGAACCCGG
AGCACACGTCTGAACTCTTTCCCTACACGACGCTCTTCCGATCTNNNGGTGACAGAGAGAGACACCAT
AGAGCACACGTCTGAACTCTTTCCCTACACGACGCTCTTCCGATCTNNNTTGCACCTGTAATCCCAGCTA
GAGCACACGTCTGAACTCTTTCCCTACACGACGCTCTTCCGATCTNNNATGCCTGTAGTCCCAGCTACTA
GAGCACACGTCTGAACTCTTTCCCTACACGACGCTCTTCCGATCTNNNACAACCACACAAGCTTCAATCA
GAGCACACGTCTGAACTCTTTCCCTACACGACGCTCTTCCGATCTNNNTCAACGTCGACCATTCTACAGT
GAGCACACGTCTGAACTCTTTCCCTACACGACGCTCTTCCGATCTNNNTGAGGCAGGAGAATCTCTTGA
GAGCACACGTCTGAACTCTTTCCCTACACGACGCTCTTCCGATCTNNNACTAAACACCCCCATCCAAAAG
GAGCACACGTCTGAACTCTTTCCCTACACGACGCTCTTCCGATCTNNNATAAATTCCTGGACACATGCAG
GAGCACACGTCTGAACTCTTTCCCTACACGACGCTCTTCCGATCTNNNTACCAAAAAATTAGCTGGGCG
AGCACACGTCTGAACTCTTTCCCTACACGACGCTCTTCCGATCTNNNGAGAATCGCATGAACTGGAGAG

AGCACACGTCTGAACTCTTTCCCTACACGACGCTCTTCCGATCTNNNAAGTAAAGGCTGCAGTGAGCT
AGCACACGTCTGAACTCTTTCCCTACACGACGCTCTTCCGATCTNNNCTGAGGCAGGAGAATCACTTG
AGCACACGTCTGAACTCTTTCCCTACACGACGCTCTTCCGATCTNNNGAAGGTTGCAATGAGCCAAG
AGCACACGTCTGAACTCTTTCCCTACACGACGCTCTTCCGATCTNNNGATGGACAAAACAGTGTGTGG
AGCACACGTCTGAACTCTTTCCCTACACGACGCTCTTCCGATCTNNNCCAGAAACAGATGGATTCACA
AGCACACGTCTGAACTCTTTCCCTACACGACGCTCTTCCGATCTNNNAGGACCATATGGATTCACAGC
AGCACACGTCTGAACTCTTTCCCTACACGACGCTCTTCCGATCTNNNGATGGGACTGTGACACTCTAC
AGAGCACACGTCTGAACTCTTTCCCTACACGACGCTCTTCCGATCTNNNAGGCTATGACAAATGAGCA
AGCACACGTCTGAACTCTTTCCCTACACGACGCTCTTCCGATCTNNNGTTTCTTCTGTGGGCAGTTACA
AGCACACGTCTGAACTCTTTCCCTACACGACGCTCTTCCGATCTNNNGTACCAAAAAATCAGCTGGGC
AGCACACGTCTGAACTCTTTCCCTACACGACGCTCTTCCGATCTNNNGTAATCCCAGCTACTATGGAG
GAGCACACGTCTGAACTCTTTCCCTACACGACGCTCTTCCGATCTNNNGCAAAGACACACATAGGCTC
AGCACACGTCTGAACTCTTTCCCTACACGACGCTCTTCCGATCTNNNCATCAACAAAAAGGACGTCCAC
AGCACACGTCTGAACTCTTTCCCTACACGACGCTCTTCCGATCTNNNAGAGATTGCAGTGAGACGAGA
GAGCACACGTCTGAACTCTTTCCCTACACGACGCTCTTCCGATCTNNNAATGAGCCGAGATCATGTCA
AGCACACGTCTGAACTCTTTCCCTACACGACGCTCTTCCGATCTNNNCTGTAGTCCCAGCTACTTCGGA
AGCACACGTCTGAACTCTTTCCCTACACGACGCTCTTCCGATCTNNNGATGCCTGTAGTCTCAGCTACT
AGCACACGTCTGAACTCTTTCCCTACACGACGCTCTTCCGATCTNNNCAATCTGCACATCTGACAAAGG
AGCACACGTCTGAACTCTTTCCCTACACGACGCTCTTCCGATCTNNNGACAAGACCAAAACACTAGGG
GAGCACACGTCTGAACTCTTTCCCTACACGACGCTCTTCCGATCTNNNAATGCTGTATTCCCAGAGACA
GAGCACACGTCTGAACTCTTTCCCTACACGACGCTCTTCCGATCTNNNTGAGCCTGGGCCATATAAAA
GAGCACACGTCTGAACTCTTTCCCTACACGACGCTCTTCCGATCTNNNACAGACAAGCAAATGTTGAC
GAGCACACGTCTGAACTCTTTCCCTACACGACGCTCTTCCGATCTNNNCCAGAAGGGAGTGACATGAC
AGCACACGTCTGAACTCTTTCCCTACACGACGCTCTTCCGATCTNNNGGAGACAGAGTGAGACTCCA
AGCACACGTCTGAACTCTTTCCCTACACGACGCTCTTCCGATCTNNNTTCTGTCTCACTGTCATTCCAG
GCACACGTCTGAACTCTTTCCCTACACGACGCTCTTCCGATCTNNNCTGAGGCAGGAGAATCACTTGA
GCACACGTCTGAACTCTTTCCCTACACGACGCTCTTCCGATCTNNNTCTCACTGTAGAGGTCAATTGG
GAGCACACGTCTGAACTCTTTCCCTACACGACGCTCTTCCGATCTNNNTGAGGTAGGAGAATGGCAT
AGCACACGTCTGAACTCTTTCCCTACACGACGCTCTTCCGATCTNNNTTCATCATAGGCTTCAAAGGG
AGCACACGTCTGAACTCTTTCCCTACACGACGCTCTTCCGATCTNNNCTGAGGCAGGAGAATCACTTG
AGCACACGTCTGAACTCTTTCCCTACACGACGCTCTTCCGATCTNNNAGCACTCCAAATATCCTCTTGCA
GAGCACACGTCTGAACTCTTTCCCTACACGACGCTCTTCCGATCTNNNTGAGGTGTGGATTCATCTCA
AGAGCACACGTCTGAACTCTTTCCCTACACGACGCTCTTCCGATCTNNNGTAATCCTAGCTACTTGGGA
GCACACGTCTGAACTCTTTCCCTACACGACGCTCTTCCGATCTNNNGCCCTTGAAAGATCACACCAAA
GAGCACACGTCTGAACTCTTTCCCTACACGACGCTCTTCCGATCTNNNGGAGTGAACAGTCTTGTCTG
AGCACACGTCTGAACTCTTTCCCTACACGACGCTCTTCCGATCTNNNTTGTAATCTCAGCTACGTGGG
GAGCACACGTCTGAACTCTTTCCCTACACGACGCTCTTCCGATCTNNNGCTCACACCTGTAAAACCAAC
AGCACACGTCTGAACTCTTTCCCTACACGACGCTCTTCCGATCTNNNGGACATGGTAGCATGTGCCTA
AGCACACGTCTGAACTCTTTCCCTACACGACGCTCTTCCGATCTNNNTACCTGACCAGTGAAGTGTGT
GCACACGTCTGAACTCTTTCCCTACACGACGCTCTTCCGATCTNNNAAAATCAGCTGAGTGCCTGTAG
AGCACACGTCTGAACTCTTTCCCTACACGACGCTCTTCCGATCTNNNGGTTCTCAACCCTGGCTATACA
AGCACACGTCTGAACTCTTTCCCTACACGACGCTCTTCCGATCTNNNTTGAACTGGCAAGAGAGGAC
AGCACACGTCTGAACTCTTTCCCTACACGACGCTCTTCCGATCTNNNATTTAGGAATGAGGAAGCTG

AGCACACGTCTGAACTCTTTCCCTACACGACGCTCTTCCGATCTNNNACCTTCCCTCCACTATTGTCCTA
AGCACACGTCTGAACTCTTTCCCTACACGACGCTCTTCCGATCTNNNTACGAAAAAAATTAGCTGGGC
AGCACACGTCTGAACTCTTTCCCTACACGACGCTCTTCCGATCTNNNGCAAGAATTTCAAGCCAGTGG
AGCACACGTCTGAACTCTTTCCCTACACGACGCTCTTCCGATCTNNNACCTGTAATCCTAGCACTTTGG
AAGAGCACACGTCTGAACTCTTTCCCTACACGACGCTCTTCCGATCTNNNAGAGATTGCAGTGAGCCA
AGCACACGTCTGAACTCTTTCCCTACACGACGCTCTTCCGATCTNNNCACATTGTGCACATGTACCCTA
AGCACACGTCTGAACTCTTTCCCTACACGACGCTCTTCCGATCTNNNAGTGTGAATCCTCCAACTGTGT
AGCACACGTCTGAACTCTTTCCCTACACGACGCTCTTCCGATCTNNNAAGAGTCAAGACCCATCAGTGT
GCACACGTCTGAACTCTTTCCCTACACGACGCTCTTCCGATCTNNNGAGGTTGCAGTAAGCCAAGATT
GAGCACACGTCTGAACTCTTTCCCTACACGACGCTCTTCCGATCTNNNGAGAGAACAGGCAACCTAC
GAGCACACGTCTGAACTCTTTCCCTACACGACGCTCTTCCGATCTNNNTGCTCACATCTGTAATCCCAA
AGCACACGTCTGAACTCTTTCCCTACACGACGCTCTTCCGATCTNNNTAGGGAGACTGAGGCAGGA
AGCACACGTCTGAACTCTTTCCCTACACGACGCTCTTCCGATCTNNNTCACACCATTGCACTCCAATCT
AGCACACGTCTGAACTCTTTCCCTACACGACGCTCTTCCGATCTNNNTGAGGCAGGAGAATCACTTG
GAGCACACGTCTGAACTCTTTCCCTACACGACGCTCTTCCGATCTNNNCGCCTGTAATCCCAAGCTATT
AGCACACGTCTGAACTCTTTCCCTACACGACGCTCTTCCGATCTNNNGTGGACCAATACCAGGGATAA
GAGCACACGTCTGAACTCTTTCCCTACACGACGCTCTTCCGATCTNNNCTGTAATCACAGCTACTGGG
AGCACACGTCTGAACTCTTTCCCTACACGACGCTCTTCCGATCTNNNAAACACAAAAGATCATCAGGG
AGCACACGTCTGAACTCTTTCCCTACACGACGCTCTTCCGATCTNNNGAACTGAGATCCCACCACTGTA
AGCACACGTCTGAACTCTTTCCCTACACGACGCTCTTCCGATCTNNNGGGTAGGAAAAGGGTAGGA
AAGAGCACACGTCTGAACTCTTTCCCTACACGACGCTCTTCCGATCTNNNATGAGAATTGCTTGAGCCT
AGCACACGTCTGAACTCTTTCCCTACACGACGCTCTTCCGATCTNNNATCAGACAGCCATCAGAATCAC
AGCACACGTCTGAACTCTTTCCCTACACGACGCTCTTCCGATCTNNNACAGATACTCACTTTCTCCACCA
GCACACGTCTGAACTCTTTCCCTACACGACGCTCTTCCGATCTNNNGTGAGCTGTAATTGTGCTACTG
AGCACACGTCTGAACTCTTTCCCTACACGACGCTCTTCCGATCTNNNTGAGAATCACTTGAACCCAGG
AGCACACGTCTGAACTCTTTCCCTACACGACGCTCTTCCGATCTNNNACTCAACCCCTTTCACAAGAGT
AGCACACGTCTGAACTCTTTCCCTACACGACGCTCTTCCGATCTNNNGAACAAGCAGAAGAAAGAGC
GAGCACACGTCTGAACTCTTTCCCTACACGACGCTCTTCCGATCTNNNTGTGCTCAACAGAATGAAG
AGCACACGTCTGAACTCTTTCCCTACACGACGCTCTTCCGATCTNNNTTGAGCTTAGGAGGCAGATG
AGCACACGTCTGAACTCTTTCCCTACACGACGCTCTTCCGATCTNNNGGAAAGGGAAACGTTGTACA
AGCACACGTCTGAACTCTTTCCCTACACGACGCTCTTCCGATCTNNNACACCATGGTATACTGCAGCTA
AGCACACGTCTGAACTCTTTCCCTACACGACGCTCTTCCGATCTNNNCTGTAGTCCCAGCTACTCAGG
AGCACACGTCTGAACTCTTTCCCTACACGACGCTCTTCCGATCTNNNGAAAAAACAATCCTAGGCCAG
GAGCACACGTCTGAACTCTTTCCCTACACGACGCTCTTCCGATCTNNNAGTGAGCTGAGATTGTACCA
AGCACACGTCTGAACTCTTTCCCTACACGACGCTCTTCCGATCTNNNCATTTGAGGTCTGGAGGTTG
AGCACACGTCTGAACTCTTTCCCTACACGACGCTCTTCCGATCTNNNACCCCTATAATCCCAGCACTTT
AGCACACGTCTGAACTCTTTCCCTACACGACGCTCTTCCGATCTNNNGGTTTCTGTGAGCCAAGATTGT
GAGCACACGTCTGAACTCTTTCCCTACACGACGCTCTTCCGATCTNNNGCAACAAGAGCGAAATTCCA
AGCACACGTCTGAACTCTTTCCCTACACGACGCTCTTCCGATCTNNNCCCACCCCTCCATAAATGAGTT
AGCACACGTCTGAACTCTTTCCCTACACGACGCTCTTCCGATCTNNNCTCACATCTGTAATCCCAGTGC
AGCACACGTCTGAACTCTTTCCCTACACGACGCTCTTCCGATCTNNNCTGAGGCAGAGAATTGCTTGA
AGCACACGTCTGAACTCTTTCCCTACACGACGCTCTTCCGATCTNNNGAGATCACACCACTACACTCC
GCACACGTCTGAACTCTTTCCCTACACGACGCTCTTCCGATCTNNNGCAAGAGAATCACTTGAACCCA

AGAGCACACGTCTGAACTCTTTCCCTACACGACGCTCTTCCGATCTNNNCCTGTAATCCCAGCTACTCGA
AGCACACGTCTGAACTCTTTCCCTACACGACGCTCTTCCGATCTNNNAAAAAATTAGCTGGGTGTGGT
GAGCACACGTCTGAACTCTTTCCCTACACGACGCTCTTCCGATCTNNNCTGTAGTCCCAGCTACTCAGG
GAGCACACGTCTGAACTCTTTCCCTACACGACGCTCTTCCGATCTNNNATGCCTGTAATCCCAGCTACT
AGCACACGTCTGAACTCTTTCCCTACACGACGCTCTTCCGATCTNNNCATGCCTGTAATCCCAGCTACT
AGCACACGTCTGAACTCTTTCCCTACACGACGCTCTTCCGATCTNNNGCATTCTTGTAGTCCCAGCTAC
AGCACACGTCTGAACTCTTTCCCTACACGACGCTCTTCCGATCTNNNGAGATTGCAGTGAGCCAAGA
AGCACACGTCTGAACTCTTTCCCTACACGACGCTCTTCCGATCTNNNTCAGGAGAAGTTGGGCAAAGT
AGAGCACACGTCTGAACTCTTTCCCTACACGACGCTCTTCCGATCTNNNATGCCTGTAATCCCAGCTAC
AGCACACGTCTGAACTCTTTCCCTACACGACGCTCTTCCGATCTNNNGCAGGAGAATTGCTTGAAGA
AGCACACGTCTGAACTCTTTCCCTACACGACGCTCTTCCGATCTNNNACACTCTTGTAATCCCAGCAAC
GCACACGTCTGAACTCTTTCCCTACACGACGCTCTTCCGATCTNNNAGCTAGCCAAAATTGCACCATTG
GAGCACACGTCTGAACTCTTTCCCTACACGACGCTCTTCCGATCTNNNAAAAAAATTAGCCGGGTGTG
GAGCACACGTCTGAACTCTTTCCCTACACGACGCTCTTCCGATCTNNNCCAATGACTACAACACCTCTCC
GAGCACACGTCTGAACTCTTTCCCTACACGACGCTCTTCCGATCTNNNAGAATCTTTTGAGCATGGGA
AGCACACGTCTGAACTCTTTCCCTACACGACGCTCTTCCGATCTNNNGGAAAAAGAGGGAGGTTCTG
GAGCACACGTCTGAACTCTTTCCCTACACGACGCTCTTCCGATCTNNNCACCTGTAGTCCCAGCTACTTG
GAGCACACGTCTGAACTCTTTCCCTACACGACGCTCTTCCGATCTNNNAGAAGAATGGTGTGAACCCA
AGCACACGTCTGAACTCTTTCCCTACACGACGCTCTTCCGATCTNNNGATCAGAGTGAACAGGCAACC
GAGCACACGTCTGAACTCTTTCCCTACACGACGCTCTTCCGATCTNNNCTTATGAAGGCTCCTGTGTCA
GAGCACACGTCTGAACTCTTTCCCTACACGACGCTCTTCCGATCTNNNAGGCCTGTAATTCCAGCATTT
AGAGCACACGTCTGAACTCTTTCCCTACACGACGCTCTTCCGATCTNNNAAGTCAAAGCCCTGAATAGA
GAGCACACGTCTGAACTCTTTCCCTACACGACGCTCTTCCGATCTNNNCCCAGAACAGGCAAAGATAT
AGCACACGTCTGAACTCTTTCCCTACACGACGCTCTTCCGATCTNNNGTGCTTGGGAATAGGGAAGA
AGAGCACACGTCTGAACTCTTTCCCTACACGACGCTCTTCCGATCTNNNCTCTAAAGACACAGGAGCTT
AGCACACGTCTGAACTCTTTCCCTACACGACGCTCTTCCGATCTNNNCAGAGCCAAATTCTACCAGAGG
GAGCACACGTCTGAACTCTTTCCCTACACGACGCTCTTCCGATCTNNNGGGAAATGGGGAGTAACTG
GAGCACACGTCTGAACTCTTTCCCTACACGACGCTCTTCCGATCTNNNACTCATATATGTCTGGCAGCA
AGCACACGTCTGAACTCTTTCCCTACACGACGCTCTTCCGATCTNNNCCAACATGTTGAAACCCCATCT
AGAGCACACGTCTGAACTCTTTCCCTACACGACGCTCTTCCGATCTNNNAAGGATGCAGCTGGAGAAC
AGCACACGTCTGAACTCTTTCCCTACACGACGCTCTTCCGATCTNNNGACCAGCATCATCCTGATACCA
AGCACACGTCTGAACTCTTTCCCTACACGACGCTCTTCCGATCTNNNGAAGGGACAGTTGCCAGAAAA
GAGCACACGTCTGAACTCTTTCCCTACACGACGCTCTTCCGATCTNNNAGTGAGCTGATATCGCACTA
GAGCACACGTCTGAACTCTTTCCCTACACGACGCTCTTCCGATCTNNNGAGACTGAGGAAGAAGAAT
GAGCACACGTCTGAACTCTTTCCCTACACGACGCTCTTCCGATCTNNNCAACTGTAGTCCCAGCTACTT
AGCACACGTCTGAACTCTTTCCCTACACGACGCTCTTCCGATCTNNNCTGAGGTGGGAGGATCAATT
GAGCACACGTCTGAACTCTTTCCCTACACGACGCTCTTCCGATCTNNNGGCAACAGAGTAAGACTCCA
AGCACACGTCTGAACTCTTTCCCTACACGACGCTCTTCCGATCTNNNATGCCTGTAATCCAAGCACTTT
GAGCACACGTCTGAACTCTTTCCCTACACGACGCTCTTCCGATCTNNNACTCTAGCCTGGGTGATAAG
AGCACACGTCTGAACTCTTTCCCTACACGACGCTCTTCCGATCTNNNGGTGCATACCTGTAATCCCATC
GAGCACACGTCTGAACTCTTTCCCTACACGACGCTCTTCCGATCTNNNGAGATCGAGCCATTACACTCC
GAGCACACGTCTGAACTCTTTCCCTACACGACGCTCTTCCGATCTNNNTGAGATCAGGAGTTCGAGAC
GAGCACACGTCTGAACTCTTTCCCTACACGACGCTCTTCCGATCTNNNGGTACACACCTGTAATCTCAG

GAGCACACGTCTGAACTCTTTCCCTACACGACGCTCTTCCGATCTNNNGCAGAGGTTATAGTGAGCCAA
AGCACACGTCTGAACTCTTTCCCTACACGACGCTCTTCCGATCTNNNTGAGGCAGGAGAATCACTTGAA
AGCACACGTCTGAACTCTTTCCCTACACGACGCTCTTCCGATCTNNNCATGCCTATGGTCCCATCTACAC
GAGCACACGTCTGAACTCTTTCCCTACACGACGCTCTTCCGATCTNNNGGGCAATATGGCAAGATCACA
GCACACGTCTGAACTCTTTCCCTACACGACGCTCTTCCGATCTNNNAGGCAGGAGAATCACTTGAACTT
AGCACACGTCTGAACTCTTTCCCTACACGACGCTCTTCCGATCTNNNCAAAAATTTGCTGCACATGGTG
AGAGCACACGTCTGAACTCTTTCCCTACACGACGCTCTTCCGATCTNNNAGAATAGCTTGAACTCAGGA
GAGCACACGTCTGAACTCTTTCCCTACACGACGCTCTTCCGATCTNNNTCTAGTCCCAGCTACTCAGAAG
GAGCACACGTCTGAACTCTTTCCCTACACGACGCTCTTCCGATCTNNNTGCAGTAAGATATGATCACGC
GCACACGTCTGAACTCTTTCCCTACACGACGCTCTTCCGATCTNNNGAGAATTGCTTGAACCTGGGAA
GAGCACACGTCTGAACTCTTTCCCTACACGACGCTCTTCCGATCTNNNGGAGTCTGCAGTAACCTGAGA
GAGCACACGTCTGAACTCTTTCCCTACACGACGCTCTTCCGATCTNNNTGTAATCCCAGCTACTCAGGA
GAGCACACGTCTGAACTCTTTCCCTACACGACGCTCTTCCGATCTNNNCAGGAGTTCGAGACTATACTC
AGAGCACACGTCTGAACTCTTTCCCTACACGACGCTCTTCCGATCTNNNCTGTAATCCCAGCTACTCAGC
GAGCACACGTCTGAACTCTTTCCCTACACGACGCTCTTCCGATCTNNNGATCACTTAGGGTTAGGAGCT
GAGCACACGTCTGAACTCTTTCCCTACACGACGCTCTTCCGATCTNNNAGTGAGCCGAGATCTCATCAT
AGCACACGTCTGAACTCTTTCCCTACACGACGCTCTTCCGATCTNNNTGAGGCAGGAGAATCACTTGA
AGCACACGTCTGAACTCTTTCCCTACACGACGCTCTTCCGATCTNNNGTGGAGTTCTAGTCGGCCATAA
GAGCACACGTCTGAACTCTTTCCCTACACGACGCTCTTCCGATCTNNNGAGGGGAGACTTCTCAACTCA
AGCACACGTCTGAACTCTTTCCCTACACGACGCTCTTCCGATCTNNNCTGAGGCAGGAGAATCACTTG
AGAGCACACGTCTGAACTCTTTCCCTACACGACGCTCTTCCGATCTNNNCAAAAATTACCCAGGTGTGG
GAGCACACGTCTGAACTCTTTCCCTACACGACGCTCTTCCGATCTNNNTGAGTGACAGAGTGAGACCC
GAGCACACGTCTGAACTCTTTCCCTACACGACGCTCTTCCGATCTNNNGGTGGGTAATTTGAGGTCACA
AGCACACGTCTGAACTCTTTCCCTACACGACGCTCTTCCGATCTNNNTCTGTAGTCTCAGCTACTTGGG
AGCACACGTCTGAACTCTTTCCCTACACGACGCTCTTCCGATCTNNNCAGTTTGCAGTGAGCTGAGATT
AGAGCACACGTCTGAACTCTTTCCCTACACGACGCTCTTCCGATCTNNNAGACTGCAGTGAGCTATGAT
GAGCACACGTCTGAACTCTTTCCCTACACGACGCTCTTCCGATCTNNNTACAAAAAATTAGCAGGGCGT
GAGCACACGTCTGAACTCTTTCCCTACACGACGCTCTTCCGATCTNNNGTGACAGAGTGAGACTCCCTT
AGCACACGTCTGAACTCTTTCCCTACACGACGCTCTTCCGATCTNNNCAAAAAATTAGCTGGGCATGTC
AGCACACGTCTGAACTCTTTCCCTACACGACGCTCTTCCGATCTNNNCTGAGGCAGGAGAATCACTTGA
GAGCACACGTCTGAACTCTTTCCCTACACGACGCTCTTCCGATCTNNNCTGGAGAATCACTTGAACCTG
AGCACACGTCTGAACTCTTTCCCTACACGACGCTCTTCCGATCTNNNAAAAAATTAGCCAGGTGTGGTG
GAGCACACGTCTGAACTCTTTCCCTACACGACGCTCTTCCGATCTNNNTATTTAGGATGCTGAGGTGG
AGCACACGTCTGAACTCTTTCCCTACACGACGCTCTTCCGATCTNNNATTCTCCTGCCTCAGTATCCCAAC
AGAGCACACGTCTGAACTCTTTCCCTACACGACGCTCTTCCGATCTNNNGAGAATCGCTTGAAACCAGA
AGAGCACACGTCTGAACTCTTTCCCTACACGACGCTCTTCCGATCTNNNGGTTTCCTTGAACGTATTCTC
GAGCACACGTCTGAACTCTTTCCCTACACGACGCTCTTCCGATCTNNNGGCACTTGTAATCCCAGCTATT
AGCACACGTCTGAACTCTTTCCCTACACGACGCTCTTCCGATCTNNNCTCCTTTTATCGTCTCCCTACGGC
AGCACACGTCTGAACTCTTTCCCTACACGACGCTCTTCCGATCTNNNGAGGCAGGAGAATTGCTTGAA
AGCACACGTCTGAACTCTTTCCCTACACGACGCTCTTCCGATCTNNNTAAAGAGTCAAGACCCATCCGT
AGCACACGTCTGAACTCTTTCCCTACACGACGCTCTTCCGATCTNNNGTTGTAGTGAGCTGAGATTGTG
GAGCACACGTCTGAACTCTTTCCCTACACGACGCTCTTCCGATCTNNNCTGGGCAACAAGAGCAAAACT
GAGCACACGTCTGAACTCTTTCCCTACACGACGCTCTTCCGATCTNNNGTTCTCCTGGTTTTCTTTCTGC

AGCACACGTCTGAACTCTTTCCCTACACGACGCTCTTCCGATCTNNNTAATCCTAGCTACTGGGGAGG
GAGCACACGTCTGAACTCTTTCCCTACACGACGCTCTTCCGATCTNNNCACCTGTAGTCCCAGCTAATT
CACACGTCTGAACTCTTTCCCTACACGACGCTCTTCCGATCTNNNATAGGTTGCAGTGAGCCAAGAT
AGCACACGTCTGAACTCTTTCCCTACACGACGCTCTTCCGATCTNNNCTGTAGTCCCAGCTACTCAGG
AGCACACGTCTGAACTCTTTCCCTACACGACGCTCTTCCGATCTNNNCTGAGGCAGGAGAATCACTTG
AGCACACGTCTGAACTCTTTCCCTACACGACGCTCTTCCGATCTNNNATAAAAGGTCAGCCAGTTTCC
GAGCACACGTCTGAACTCTTTCCCTACACGACGCTCTTCCGATCTNNNACCTGTAATTCCAGCTACTCA
GAGCACACGTCTGAACTCTTTCCCTACACGACGCTCTTCCGATCTNNNAAAATACAAAATTAGCCGG
AGCACACGTCTGAACTCTTTCCCTACACGACGCTCTTCCGATCTNNNTGGGAAACAGAGCAAGATCTT
AGCACACGTCTGAACTCTTTCCCTACACGACGCTCTTCCGATCTNNNCTGAGGCAGGAGAATCACTTG
AGCACACGTCTGAACTCTTTCCCTACACGACGCTCTTCCGATCTNNNGGAGACTGAGATGGGAGAAT
AGCACACGTCTGAACTCTTTCCCTACACGACGCTCTTCCGATCTNNNGATTGCTTGAGGCCAGTAGTT
AGCACACGTCTGAACTCTTTCCCTACACGACGCTCTTCCGATCTNNNAAATACAAAAATTGCCTGGGC
AGCACACGTCTGAACTCTTTCCCTACACGACGCTCTTCCGATCTNNNAATACAAAAAATTAGCCGGG
AGCACACGTCTGAACTCTTTCCCTACACGACGCTCTTCCGATCTNNNATGATGGTGGGTGACTGTAAT
AGCACACGTCTGAACTCTTTCCCTACACGACGCTCTTCCGATCTNNNAAAAGCAACAAGGTTTCACAG
GAGCACACGTCTGAACTCTTTCCCTACACGACGCTCTTCCGATCTNNNCCTGGGTTATGGAGTGAGA
AGCACACGTCTGAACTCTTTCCCTACACGACGCTCTTCCGATCTNNNCAACATGGCAAAACTCCATCTC
AGCACACGTCTGAACTCTTTCCCTACACGACGCTCTTCCGATCTNNNTGGGCAATAAAGTGACACTCT
AGCACACGTCTGAACTCTTTCCCTACACGACGCTCTTCCGATCTNNNGAGGCTGAGGTAGAAGAAT
GCACACGTCTGAACTCTTTCCCTACACGACGCTCTTCCGATCTNNNACCTTCCCTCCACTATTGTCCTAC
AGCACACGTCTGAACTCTTTCCCTACACGACGCTCTTCCGATCTNNNCTGTAGTGTCCGCTACTCAGG
GCACACGTCTGAACTCTTTCCCTACACGACGCTCTTCCGATCTNNNATGGGACCTGATCTGCATTTCTT
AGCACACGTCTGAACTCTTTCCCTACACGACGCTCTTCCGATCTNNNAAATTAGTTGGGCGTGGTAAT
AGCACACGTCTGAACTCTTTCCCTACACGACGCTCTTCCGATCTNNNCTGTAATCCCAGCTTCTCAGG
GAGCACACGTCTGAACTCTTTCCCTACACGACGCTCTTCCGATCTNNNAGAGGTTGCAGTAAGCTGAG
GAGCACACGTCTGAACTCTTTCCCTACACGACGCTCTTCCGATCTNNNATGCACATCACAACCACAATG
AGCACACGTCTGAACTCTTTCCCTACACGACGCTCTTCCGATCTNNNCTGTAATCCCAGCTACTCAGG
GCACACGTCTGAACTCTTTCCCTACACGACGCTCTTCCGATCTNNNCTGAGATGGGAGGATCGATTG
GCACACGTCTGAACTCTTTCCCTACACGACGCTCTTCCGATCTNNNCCTGAGTAGCTGGGATTATAGG
GAGCACACGTCTGAACTCTTTCCCTACACGACGCTCTTCCGATCTNNNCCAAATACAGGAAACTCGACT
GCACACGTCTGAACTCTTTCCCTACACGACGCTCTTCCGATCTNNNGCTGAGACAGGAGAATCACTT
AGCACACGTCTGAACTCTTTCCCTACACGACGCTCTTCCGATCTNNNGAGCTTGCAGTGAACTGAGAT
GAGCACACGTCTGAACTCTTTCCCTACACGACGCTCTTCCGATCTNNNGATGCCTGTAATCCCAGCTAC
AGCACACGTCTGAACTCTTTCCCTACACGACGCTCTTCCGATCTNNNGGGCAACAGAGTAAAACCTTG
AGCACACGTCTGAACTCTTTCCCTACACGACGCTCTTCCGATCTNNNCACCTGTAATCCCAGCTACTCA
AGCACACGTCTGAACTCTTTCCCTACACGACGCTCTTCCGATCTNNNGAAGTTGCAGTGAGCCAAGA
AGCACACGTCTGAACTCTTTCCCTACACGACGCTCTTCCGATCTNNNCTTGGTGGCATGTGACTGTAA
GCACACGTCTGAACTCTTTCCCTACACGACGCTCTTCCGATCTNNNCTGTGAGCTGAGATCGTACCATT
GAGCACACGTCTGAACTCTTTCCCTACACGACGCTCTTCCGATCTNNNTGAGGCAGAAGAATTGCTT
CACACGTCTGAACTCTTTCCCTACACGACGCTCTTCCGATCTNNNAATTTTAGCTACTCGGGAGGCTG
AGCACACGTCTGAACTCTTTCCCTACACGACGCTCTTCCGATCTNNNAATCACTTGAACCTGAGAGGC
GCACACGTCTGAACTCTTTCCCTACACGACGCTCTTCCGATCTNNNCAACAAGAGTGAAAGTCCGTCT

CACACGTCTGAACTCTTTCCCTACACGACGCTCTTCCGATCTNNNCCCATAAGCTCTACTTGGCTACAC
AGCACACGTCTGAACTCTTTCCCTACACGACGCTCTTCCGATCTNNNTAGCCAGGTGTCTCAAATTAG
GAGCACACGTCTGAACTCTTTCCCTACACGACGCTCTTCCGATCTNNNACAAAAAATTAGCCGTGTGT
GCACACGTCTGAACTCTTTCCCTACACGACGCTCTTCCGATCTNNNTAATCCCAGCTACTCAGGAGGT
AGCACACGTCTGAACTCTTTCCCTACACGACGCTCTTCCGATCTNNNCTGTAATCCCAGCTACTCAGGA
AGCACACGTCTGAACTCTTTCCCTACACGACGCTCTTCCGATCTNNNAAAATTAGCTCAACTTGGTGG
AGCACACGTCTGAACTCTTTCCCTACACGACGCTCTTCCGATCTNNNGGAGGTTGTAGTGAGCCAAG
GAGCACACGTCTGAACTCTTTCCCTACACGACGCTCTTCCGATCTNNNTATAGTCCCAGCTACTCAGG
AGCACACGTCTGAACTCTTTCCCTACACGACGCTCTTCCGATCTNNNATTACTTGAACCCGGGAAATG
GAGCACACGTCTGAACTCTTTCCCTACACGACGCTCTTCCGATCTNNNACTTAGAGATTCTAGCCAGAG
AGCACACGTCTGAACTCTTTCCCTACACGACGCTCTTCCGATCTNNNAAAAAATTAGCCGGGCATGG
AGCACACGTCTGAACTCTTTCCCTACACGACGCTCTTCCGATCTNNNCACCTGTAATCCCAGCTACTCA
GAGCACACGTCTGAACTCTTTCCCTACACGACGCTCTTCCGATCTNNNAGAGGTTGAAATGAGCCGAG
AGCACACGTCTGAACTCTTTCCCTACACGACGCTCTTCCGATCTNNNAGTGAGCCATGATCATGCTAC
AAGAGCACACGTCTGAACTCTTTCCCTACACGACGCTCTTCCGATCTNNNAGGAGAATGGCTTGAAC
GAGCACACGTCTGAACTCTTTCCCTACACGACGCTCTTCCGATCTNNNCTGAGGCACGAGAATCACTT
AGCACACGTCTGAACTCTTTCCCTACACGACGCTCTTCCGATCTNNNGTTGCAGTGAGATGAGATTC
AGCACACGTCTGAACTCTTTCCCTACACGACGCTCTTCCGATCTNNNGAGATTGTGCCACTCTACTCCA
GCACACGTCTGAACTCTTTCCCTACACGACGCTCTTCCGATCTNNNAGAATCACTTGAATGCAGGAG
GCACACGTCTGAACTCTTTCCCTACACGACGCTCTTCCGATCTNNNCCTATAGTCTCAGCTGCTTGGA
AGCACACGTCTGAACTCTTTCCCTACACGACGCTCTTCCGATCTNNNGAGAATCACTTGAACCCAGGA
GCACACGTCTGAACTCTTTCCCTACACGACGCTCTTCCGATCTNNNAAATTAGCTGAATGGCGTGAA
AGCACACGTCTGAACTCTTTCCCTACACGACGCTCTTCCGATCTNNNTTCTGGACAGTGAAGGCTTAG
AGCACACGTCTGAACTCTTTCCCTACACGACGCTCTTCCGATCTNNNTACCTTAGTCCCAGCTACTCCT
AGCACACGTCTGAACTCTTTCCCTACACGACGCTCTTCCGATCTNNNGAGGATTGCTTGATCTCAGGA
GCACACGTCTGAACTCTTTCCCTACACGACGCTCTTCCGATCTNNNCTGGGCAAGAAAGTGAGACAC
GCACACGTCTGAACTCTTTCCCTACACGACGCTCTTCCGATCTNNNTGGGCAACACAGTAAGATTCTG
GAGCACACGTCTGAACTCTTTCCCTACACGACGCTCTTCCGATCTNNNGCAGACATCTGTAATCCCAG
GCACACGTCTGAACTCTTTCCCTACACGACGCTCTTCCGATCTNNNGAGGTTGCAATGAGCTGAGAT
GCACACGTCTGAACTCTTTCCCTACACGACGCTCTTCCGATCTNNNGTAGGAGGATCACTTGAACCTG
AGCACACGTCTGAACTCTTTCCCTACACGACGCTCTTCCGATCTNNNAGAATAGCATGAACCTGAGA
GCACACGTCTGAACTCTTTCCCTACACGACGCTCTTCCGATCTNNNCCTGCCTTCTCAGTTTCCTAACTC
AGCACACGTCTGAACTCTTTCCCTACACGACGCTCTTCCGATCTNNNCAAGAATTGCTTCAATCCGGG
GCACACGTCTGAACTCTTTCCCTACACGACGCTCTTCCGATCTNNNTGCATACCTGTAATCCCAGCTA
AGCACACGTCTGAACTCTTTCCCTACACGACGCTCTTCCGATCTNNNAAGGTTTTGGTGAGCCATGAT
AGCACACGTCTGAACTCTTTCCCTACACGACGCTCTTCCGATCTNNNGGTTGTAGGGAGCCAAGATT
GCACACGTCTGAACTCTTTCCCTACACGACGCTCTTCCGATCTNNNCTGAGGCAGGAAGATCACTTG
GAGCACACGTCTGAACTCTTTCCCTACACGACGCTCTTCCGATCTNNNCACAACTGTAGTCCCAGCTAC
AGCACACGTCTGAACTCTTTCCCTACACGACGCTCTTCCGATCTNNNCTGAGGCAGGAGAATCACTTG
GCACACGTCTGAACTCTTTCCCTACACGACGCTCTTCCGATCTNNNCAGGAGGTGGAAGTTGGAGTT
AGCACACGTCTGAACTCTTTCCCTACACGACGCTCTTCCGATCTNNNATCCATTCAAAAACAGCTCAGG
GAGCACACGTCTGAACTCTTTCCCTACACGACGCTCTTCCGATCTNNNACCTGTAATCCTAGCACTTTG
AGCACACGTCTGAACTCTTTCCCTACACGACGCTCTTCCGATCTNNNAAAAAAAAAATAGCCGCGTG

GAGCACACGTCTGAACTCTTTCCCTACACGACGCTCTTCCGATCTNNNCCAACATGGTGAAATCCCATC
GCACACGTCTGAACTCTTTCCCTACACGACGCTCTTCCGATCTNNNTCTCCAACTCCACTAGTAGCA
AGCACACGTCTGAACTCTTTCCCTACACGACGCTCTTCCGATCTNNNGGGTAAAACACTGGTGAAAC
GCACACGTCTGAACTCTTTCCCTACACGACGCTCTTCCGATCTNNNACAAAAATTAGCCAGGCCTGTT
GAGCACACGTCTGAACTCTTTCCCTACACGACGCTCTTCCGATCTNNNACACCTGTATCCTAGCACTTT
AGCACACGTCTGAACTCTTTCCCTACACGACGCTCTTCCGATCTNNNGCATATGCCTGTATTCCCAGCT
AGCACACGTCTGAACTCTTTCCCTACACGACGCTCTTCCGATCTNNNCCAGTTTGGGTAACAGAGTGA
AGCACACGTCTGAACTCTTTCCCTACACGACGCTCTTCCGATCTNNNTAGGCGATACATGCTGCAGTA
GAGCACACGTCTGAACTCTTTCCCTACACGACGCTCTTCCGATCTNNNCAATATAGCGAGACCCCGTT
AGAGCACACGTCTGAACTCTTTCCCTACACGACGCTCTTCCGATCTNNNAAAAATTAGCCAGACGTG
GAGCACACGTCTGAACTCTTTCCCTACACGACGCTCTTCCGATCTNNNTAGTGAGCTGAGATTGTGCC
GAGCACACGTCTGAACTCTTTCCCTACACGACGCTCTTCCGATCTNNNCACTTGAGGTCAGGAGTTTG
AGCACACGTCTGAACTCTTTCCCTACACGACGCTCTTCCGATCTNNNTTGAGCTCAGGAATTTGAGAC
AGAGCACACGTCTGAACTCTTTCCCTACACGACGCTCTTCCGATCTNNNGACTGAGGCAACAAGACTG
AGCACACGTCTGAACTCTTTCCCTACACGACGCTCTTCCGATCTNNNCTGCAGTAAGTCAAGATTGTG
GAGCACACGTCTGAACTCTTTCCCTACACGACGCTCTTCCGATCTNNNCTGTAATCCCAGCTACTCAGG
GAGCACACGTCTGAACTCTTTCCCTACACGACGCTCTTCCGATCTNNNCATGGTGAAACTCTGCCTCTA
GCACACGTCTGAACTCTTTCCCTACACGACGCTCTTCCGATCTNNNCTGTAATCCCAGCTACTCAGGA
GAGCACACGTCTGAACTCTTTCCCTACACGACGCTCTTCCGATCTNNNCTGAGGCAGGAGAATTGCT
AGCACACGTCTGAACTCTTTCCCTACACGACGCTCTTCCGATCTNNNAGTGAGCAAAGATCATGCCAT
AGCACACGTCTGAACTCTTTCCCTACACGACGCTCTTCCGATCTNNNGGAGTTTGCAGTGAGCTGAG
GCACACGTCTGAACTCTTTCCCTACACGACGCTCTTCCGATCTNNNTTCAAAACCAGCCTGGGTAAGA
AGCACACGTCTGAACTCTTTCCCTACACGACGCTCTTCCGATCTNNNCTGAGATGGGAGGACTGATTG
GCACACGTCTGAACTCTTTCCCTACACGACGCTCTTCCGATCTNNNAGAGGCAGGAGAATCACTTGA
AGCACACGTCTGAACTCTTTCCCTACACGACGCTCTTCCGATCTNNNGGAGAATCACTTGAACCCA
GAGCACACGTCTGAACTCTTTCCCTACACGACGCTCTTCCGATCTNNNGAGAATTGCTTGAACCCAGG
AGCACACGTCTGAACTCTTTCCCTACACGACGCTCTTCCGATCTNNNGGCAACAGAGTAAGACACTG
GAGCACACGTCTGAACTCTTTCCCTACACGACGCTCTTCCGATCTNNNTGAGGTAGGAGAATCGCTT
AGCACACGTCTGAACTCTTTCCCTACACGACGCTCTTCCGATCTNNNCTGAGGCAGAAGAATTGCTT
AGCACACGTCTGAACTCTTTCCCTACACGACGCTCTTCCGATCTNNNCAGGAGAATTGCTTGAACCTT
AGAGCACACGTCTGAACTCTTTCCCTACACGACGCTCTTCCGATCTNNNTATGCCTGTAGTCCCAGCTA
AGCACACGTCTGAACTCTTTCCCTACACGACGCTCTTCCGATCTNNNGGAAATAGGGTTTCTTCCAAG
AGCACACGTCTGAACTCTTTCCCTACACGACGCTCTTCCGATCTNNNAGTAAGCTGAGATCACACCAC
AGCACACGTCTGAACTCTTTCCCTACACGACGCTCTTCCGATCTNNNACACCTGTAATCCCAGCTACTT
GAGCACACGTCTGAACTCTTTCCCTACACGACGCTCTTCCGATCTNNNGTCAAGAGATCGAGACCATC
GCACACGTCTGAACTCTTTCCCTACACGACGCTCTTCCGATCTNNNTACAAAAAATTAGCTGGGCGT
AGCACACGTCTGAACTCTTTCCCTACACGACGCTCTTCCGATCTNNNAATCACATGAACCTAGGAGGC
GCACACGTCTGAACTCTTTCCCTACACGACGCTCTTCCGATCTNNNTATAATGTGGGGTTGCACTGT
AGCACACGTCTGAACTCTTTCCCTACACGACGCTCTTCCGATCTNNNGGTTGGGAGATTGAGAACAT
GAGCACACGTCTGAACTCTTTCCCTACACGACGCTCTTCCGATCTNNNCACAGGAACCAAAACAGCTT
AGCACACGTCTGAACTCTTTCCCTACACGACGCTCTTCCGATCTNNNATGAGACCAGCATCATCCTGAT
GAGCACACGTCTGAACTCTTTCCCTACACGACGCTCTTCCGATCTNNNTAGAAAGTGGTAGTGCCTGC
GCACACGTCTGAACTCTTTCCCTACACGACGCTCTTCCGATCTNNNACCCTGCCTTGACTGAAAATAC

AGCACACGTCTGAACTCTTTCCCTACACGACGCTCTTCCGATCTNNNAATTCAGATGGCCAACAAACCT
AGCACACGTCTGAACTCTTTCCCTACACGACGCTCTTCCGATCTNNNCCATCTCCCCGTAAGATCTGATG
AGAGCACACGTCTGAACTCTTTCCCTACACGACGCTCTTCCGATCTNNNGATGAGGCAGGAGAATCACT
GAGCACACGTCTGAACTCTTTCCCTACACGACGCTCTTCCGATCTNNNAAGTACAAAAAAATTAGCCGC
AGAGCACACGTCTGAACTCTTTCCCTACACGACGCTCTTCCGATCTNNNCTGAGGCAGGAGAATGTCTT
GAGCACACGTCTGAACTCTTTCCCTACACGACGCTCTTCCGATCTNNNTTGTAGTGAGCCTAGATTGTC
GAGCACACGTCTGAACTCTTTCCCTACACGACGCTCTTCCGATCTNNNACAAGTTCAGGAAATCGAGA
AGCACACGTCTGAACTCTTTCCCTACACGACGCTCTTCCGATCTNNNGGGTGACAGAGTGAGACTACA
GAGCACACGTCTGAACTCTTTCCCTACACGACGCTCTTCCGATCTNNNGAGAATGGCATGAATCCAGG
AGCACACGTCTGAACTCTTTCCCTACACGACGCTCTTCCGATCTNNNAGCACCCTACCCAACCAAAATTA
GAGCACACGTCTGAACTCTTTCCCTACACGACGCTCTTCCGATCTNNNGGCACCTGTAATCCTAGCTACT
AGAGCACACGTCTGAACTCTTTCCCTACACGACGCTCTTCCGATCTNNNAAAATTATTTGGGCATGGTG
AGCACACGTCTGAACTCTTTCCCTACACGACGCTCTTCCGATCTNNNTAATACTAGCACATTGGGAGGC
GAGCACACGTCTGAACTCTTTCCCTACACGACGCTCTTCCGATCTNNNGGAGGTTTCAGTGAGCCTAGA
AGCACACGTCTGAACTCTTTCCCTACACGACGCTCTTCCGATCTNNNATACAAAAATTAGCTGGGCGTC
AGCACACGTCTGAACTCTTTCCCTACACGACGCTCTTCCGATCTNNNTAATCACTTGAATCCAGGAGGC
GCACACGTCTGAACTCTTTCCCTACACGACGCTCTTCCGATCTNNNTCAGTGTAATCCCAGCTACTTGGC
AGCACACGTCTGAACTCTTTCCCTACACGACGCTCTTCCGATCTNNNGAAATTGCAGTGAGCCGAGAT
GAGCACACGTCTGAACTCTTTCCCTACACGACGCTCTTCCGATCTNNNCTGTAATCCCAGCTACTCAGGA
GAGCACACGTCTGAACTCTTTCCCTACACGACGCTCTTCCGATCTNNNTGGACACAGCTGAGGAAAAG
GAGCACACGTCTGAACTCTTTCCCTACACGACGCTCTTCCGATCTNNNCAACAGAGCAAAAACTCCGTC
AGAGCACACGTCTGAACTCTTTCCCTACACGACGCTCTTCCGATCTNNNTCAAGTGATTTTCTTGCGTCG
AGCACACGTCTGAACTCTTTCCCTACACGACGCTCTTCCGATCTNNNAAAATAATTAGCCGGGCATGGT
GAGCACACGTCTGAACTCTTTCCCTACACGACGCTCTTCCGATCTNNNGGGCAACAGAGTGAGAATGT
GAGCACACGTCTGAACTCTTTCCCTACACGACGCTCTTCCGATCTNNNCAGTCTGAGCAAGAGAGTGA
AGCACACGTCTGAACTCTTTCCCTACACGACGCTCTTCCGATCTNNNGGTAAGATTTCTTGAGCCTGGG
GCACACGTCTGAACTCTTTCCCTACACGACGCTCTTCCGATCTNNNCTGAGACAGGAGAATCGCTTGAA
GAGCACACGTCTGAACTCTTTCCCTACACGACGCTCTTCCGATCTNNNGCTTCCCTTTCTACCATGACTG
AGCACACGTCTGAACTCTTTCCCTACACGACGCTCTTCCGATCTNNNGTTGCAGTGAGCCATGATTACA
AAGAGCACACGTCTGAACTCTTTCCCTACACGACGCTCTTCCGATCTNNNGTACAAAAATTAGCCAGTCC
AGCACACGTCTGAACTCTTTCCCTACACGACGCTCTTCCGATCTNNNATGCCGGTAATCCTAGCTACTTG
AGCACACGTCTGAACTCTTTCCCTACACGACGCTCTTCCGATCTNNNGCCTATAATCCCAGCTACTCAGG
AGAGCACACGTCTGAACTCTTTCCCTACACGACGCTCTTCCGATCTNNNGAGATCACACCATTGCACTTC
GAGCACACGTCTGAACTCTTTCCCTACACGACGCTCTTCCGATCTNNNCTGTAATCCCAGCTACTCGGAA
AGCACACGTCTGAACTCTTTCCCTACACGACGCTCTTCCGATCTNNNACCTTCCCTCCACTATTGTCCTAC
GAGCACACGTCTGAACTCTTTCCCTACACGACGCTCTTCCGATCTNNNTAATTCCAGTACTTTGGGAGG
AGCACACGTCTGAACTCTTTCCCTACACGACGCTCTTCCGATCTNNNGCCTAAGAGTTTGAGACCAGTG
AGCACACGTCTGAACTCTTTCCCTACACGACGCTCTTCCGATCTNNNAATAAACACGCTTGGCTAAGAG
GCACACGTCTGAACTCTTTCCCTACACGACGCTCTTCCGATCTNNNGAGATCGTGTCACTGCACTCTAG
AGAGCACACGTCTGAACTCTTTCCCTACACGACGCTCTTCCGATCTNNNGAGAATAGCTTGAACCCAG
AGAGCACACGTCTGAACTCTTTCCCTACACGACGCTCTTCCGATCTNNNAGAGGTTGCAATGAGCTGA
GAGCACACGTCTGAACTCTTTCCCTACACGACGCTCTTCCGATCTNNNTGAGACCAGCCTGATGAATAT
GAGCACACGTCTGAACTCTTTCCCTACACGACGCTCTTCCGATCTNNNAATACAAAATTAGCTGGGCGT

GCACACGTCTGAACTCTTTCCCTACACGACGCTCTTCCGATCTNNNTAGATCGTCTGAGGTCAGGAGT
AGCACACGTCTGAACTCTTTCCCTACACGACGCTCTTCCGATCTNNNCAAAAATTAGCTGGGTGTGGT
AGCACACGTCTGAACTCTTTCCCTACACGACGCTCTTCCGATCTNNNCTGTAGTCCCAGCTACTCAGGA
AGCACACGTCTGAACTCTTTCCCTACACGACGCTCTTCCGATCTNNNCTGTAGTCCCCGCTACTCAGAA
AGCACACGTCTGAACTCTTTCCCTACACGACGCTCTTCCGATCTNNNCTGAGGGAGGGGAATCATTTG
GAGCACACGTCTGAACTCTTTCCCTACACGACGCTCTTCCGATCTNNNGAGGCAGGAGAATTGTTTGA
GAGCACACGTCTGAACTCTTTCCCTACACGACGCTCTTCCGATCTNNNTAAAGCAGGAGGATCATCTG
AGCACACGTCTGAACTCTTTCCCTACACGACGCTCTTCCGATCTNNNTAGCGAAACCCCATCTCTAGAG
AGCACACGTCTGAACTCTTTCCCTACACGACGCTCTTCCGATCTNNNCTTTCTATGAGCCAAGATCGCA
AGCACACGTCTGAACTCTTTCCCTACACGACGCTCTTCCGATCTNNNAGTGAACAGGCAACCTACAGA
GAGCACACGTCTGAACTCTTTCCCTACACGACGCTCTTCCGATCTNNNGCCTGTAATCTCAGCTACTCA
GAGCACACGTCTGAACTCTTTCCCTACACGACGCTCTTCCGATCTNNNTGTAATCTCAGCACTTTGGGA
GAGCACACGTCTGAACTCTTTCCCTACACGACGCTCTTCCGATCTNNNTGCACCTATAGTCCCAGCTACT
GAGCACACGTCTGAACTCTTTCCCTACACGACGCTCTTCCGATCTNNNGGGAGTGAATGGTCCTATCT
AGCACACGTCTGAACTCTTTCCCTACACGACGCTCTTCCGATCTNNNGAGGTTACAGCGAGCTATGAT
GAGCACACGTCTGAACTCTTTCCCTACACGACGCTCTTCCGATCTNNNGGGAAACTACAACAGCACAA
AGAGCACACGTCTGAACTCTTTCCCTACACGACGCTCTTCCGATCTNNNGATCAAGACCATCCTGGCTA
GAGCACACGTCTGAACTCTTTCCCTACACGACGCTCTTCCGATCTNNNATGCCTGTAATCCCAGTTACT
GCACACGTCTGAACTCTTTCCCTACACGACGCTCTTCCGATCTNNNAAAATTATTCTGGCGTGGTGGT
GAGCACACGTCTGAACTCTTTCCCTACACGACGCTCTTCCGATCTNNNGTAATCCCAGCTACTCAAGAG
AGCACACGTCTGAACTCTTTCCCTACACGACGCTCTTCCGATCTNNNACCAACAGTAACCAAGCTGAG
GAGCACACGTCTGAACTCTTTCCCTACACGACGCTCTTCCGATCTNNNGTGGATCACAAGGTCAGGA
GAGCACACGTCTGAACTCTTTCCCTACACGACGCTCTTCCGATCTNNNACAGGCCAACATTCAAATTCA
GAGCACACGTCTGAACTCTTTCCCTACACGACGCTCTTCCGATCTNNNTATGAAGGTTAGGCAGTTCC
GAGCACACGTCTGAACTCTTTCCCTACACGACGCTCTTCCGATCTNNNAAAATTAGCTGGGCATAGTC
GCACACGTCTGAACTCTTTCCCTACACGACGCTCTTCCGATCTNNNAGCAAGAGACAGCAGGATTAG
AGCACACGTCTGAACTCTTTCCCTACACGACGCTCTTCCGATCTNNNTACTGTAATAAGCAGCTGCAT
AGCACACGTCTGAACTCTTTCCCTACACGACGCTCTTCCGATCTNNNACTTGGGAGGTTGAGACAGGA
GCACACGTCTGAACTCTTTCCCTACACGACGCTCTTCCGATCTNNNCTGAGGCAGGAGAATCACTTGA
AGCACACGTCTGAACTCTTTCCCTACACGACGCTCTTCCGATCTNNNCTCAATCCCTTTCGCAATAGCT
AGCACACGTCTGAACTCTTTCCCTACACGACGCTCTTCCGATCTNNNTAATCACTTGAACCCAGGAAGC
GAGCACACGTCTGAACTCTTTCCCTACACGACGCTCTTCCGATCTNNNCAAAAGGCAGGTCACCTACA
GAGCACACGTCTGAACTCTTTCCCTACACGACGCTCTTCCGATCTNNNTACACAAGTCTGCTATACGCC
GCACACGTCTGAACTCTTTCCCTACACGACGCTCTTCCGATCTNNNGCAGGAGAATTTCTTGAACCCA
GAGCACACGTCTGAACTCTTTCCCTACACGACGCTCTTCCGATCTNNNACGATTTCCTGGATATGAAC
GAGCACACGTCTGAACTCTTTCCCTACACGACGCTCTTCCGATCTNNNGTATTCCCAGCTACTTGGTAG
GAGCACACGTCTGAACTCTTTCCCTACACGACGCTCTTCCGATCTNNNAGACTACCCAGAAAAACCCAT
AGCACACGTCTGAACTCTTTCCCTACACGACGCTCTTCCGATCTNNNAAGAGCCATCTGTGTGCTGTA
AGCACACGTCTGAACTCTTTCCCTACACGACGCTCTTCCGATCTNNNATGGATCACAAGGTCAGGAGT
GAGCACACGTCTGAACTCTTTCCCTACACGACGCTCTTCCGATCTNNNGGGTGTGGTGGTGCATAACT
GAGCACACGTCTGAACTCTTTCCCTACACGACGCTCTTCCGATCTNNNATCCAATTACCTCCCACAAGA
GAGCACACGTCTGAACTCTTTCCCTACACGACGCTCTTCCGATCTNNNGCCCATTGCTTGAGACATTA
AGCACACGTCTGAACTCTTTCCCTACACGACGCTCTTCCGATCTNNNATTGCAGTGAAATGGGATCAT

AGCACACGTCTGAACTCTTTCCCTACACGACGCTCTTCCGATCTNNNGCTGGGGACTTTCCAAAACTA
AGCACACGTCTGAACTCTTTCCCTACACGACGCTCTTCCGATCTNNNAGTAAGCCAAGATCATGCCATT
GCACACGTCTGAACTCTTTCCCTACACGACGCTCTTCCGATCTNNNAGATTGTACCACTGCACTCTAG
AGCACACGTCTGAACTCTTTCCCTACACGACGCTCTTCCGATCTNNNGCAAGAGAATCACTTGAACA
AGCACACGTCTGAACTCTTTCCCTACACGACGCTCTTCCGATCTNNNCTGGACAACATGGTAAAACCC
GCACACGTCTGAACTCTTTCCCTACACGACGCTCTTCCGATCTNNNAGAATTGCTTGAACCCAGGAGA
GCACACGTCTGAACTCTTTCCCTACACGACGCTCTTCCGATCTNNNCAGCTGTTTTGGAGACTGAGTT
GAGCACACGTCTGAACTCTTTCCCTACACGACGCTCTTCCGATCTNNNAAAAACTATCTGTGGCAGTG
GCACACGTCTGAACTCTTTCCCTACACGACGCTCTTCCGATCTNNNATATAGGGAGGTGAAGGGG
AGCACACGTCTGAACTCTTTCCCTACACGACGCTCTTCCGATCTNNNGGCAAAACCCTGTCCCTACTA
GAGCACACGTCTGAACTCTTTCCCTACACGACGCTCTTCCGATCTNNNTTTAGCTGAAATTTGCCACAG
AGCACACGTCTGAACTCTTTCCCTACACGACGCTCTTCCGATCTNNNGAGAATCACTTGAACCTGGGA
AGCACACGTCTGAACTCTTTCCCTACACGACGCTCTTCCGATCTNNNCTGAACCCATGAGGCAAATGT
AGCACACGTCTGAACTCTTTCCCTACACGACGCTCTTCCGATCTNNNTATAATCCTGGCACTTTGGGA
GAGCACACGTCTGAACTCTTTCCCTACACGACGCTCTTCCGATCTNNNTACCACCTGCTGAAGCTCTAT
GCACACGTCTGAACTCTTTCCCTACACGACGCTCTTCCGATCTNNNTGCGTGTGTAATCTCAGCTACT
GCACACGTCTGAACTCTTTCCCTACACGACGCTCTTCCGATCTNNNGCTGATAGAGCCAGACTTTGTC
GAGCACACGTCTGAACTCTTTCCCTACACGACGCTCTTCCGATCTNNNGGATCATCTGAGGTCAGGA
GAGCACACGTCTGAACTCTTTCCCTACACGACGCTCTTCCGATCTNNNCAATGGCTCATGACTGTAAT
AGCACACGTCTGAACTCTTTCCCTACACGACGCTCTTCCGATCTNNNAGAATCACTTGAACCTGGGAG
AGCACACGTCTGAACTCTTTCCCTACACGACGCTCTTCCGATCTNNNGGCAGGAGAATCACTTGAACC
GAGCACACGTCTGAACTCTTTCCCTACACGACGCTCTTCCGATCTNNNAGACAGGAGAATCACTTGA
AGCACACGTCTGAACTCTTTCCCTACACGACGCTCTTCCGATCTNNNCTGAGACAGGAGAATCGCTT
AGCACACGTCTGAACTCTTTCCCTACACGACGCTCTTCCGATCTNNNCTGAGACAGGAGAATGGCAT
AGCACACGTCTGAACTCTTTCCCTACACGACGCTCTTCCGATCTNNNCTGAGGCAGGAGAATCACTT
GCACACGTCTGAACTCTTTCCCTACACGACGCTCTTCCGATCTNNNCTGTAATCCCAGCTACTCAGGA
AGCACACGTCTGAACTCTTTCCCTACACGACGCTCTTCCGATCTNNNCTGAGGCAGGAGAATCACTTG
AGCACACGTCTGAACTCTTTCCCTACACGACGCTCTTCCGATCTNNNGAATCACTTGAAGAACCTCGG
AGCACACGTCTGAACTCTTTCCCTACACGACGCTCTTCCGATCTNNNACAAAAATTAGCTGGGCATG
AGCACACGTCTGAACTCTTTCCCTACACGACGCTCTTCCGATCTNNNTTGATCATTTGCATGGATGTG
AGCACACGTCTGAACTCTTTCCCTACACGACGCTCTTCCGATCTNNNGCAATATGGCAAAACCCTGTC
AGCACACGTCTGAACTCTTTCCCTACACGACGCTCTTCCGATCTNNNCTGAGGCAGGAGAATTGCTT
AGCACACGTCTGAACTCTTTCCCTACACGACGCTCTTCCGATCTNNNCTTGGTAGCACTCACCTGTAGT
GCACACGTCTGAACTCTTTCCCTACACGACGCTCTTCCGATCTNNNTTGAGACAGGAGAATCGCTTGA
GCACACGTCTGAACTCTTTCCCTACACGACGCTCTTCCGATCTNNNTAGGCAGGAGAATCACTTGAAC
GCACACGTCTGAACTCTTTCCCTACACGACGCTCTTCCGATCTNNNTAATTGTCCCATCTCTCCACTGC
GAGCACACGTCTGAACTCTTTCCCTACACGACGCTCTTCCGATCTNNNCTTGAGGCTAGGCACAGTTC
GAGCACACGTCTGAACTCTTTCCCTACACGACGCTCTTCCGATCTNNNGTAATCCCTGCTACTCAAGA
GCACACGTCTGAACTCTTTCCCTACACGACGCTCTTCCGATCTNNNGCAGTTCTTCCACTTTAGCTTCC
AGCACACGTCTGAACTCTTTCCCTACACGACGCTCTTCCGATCTNNNGAGAACAGTTTGGAGGTTCCT
AGAGCACACGTCTGAACTCTTTCCCTACACGACGCTCTTCCGATCTNNNAGAATGGCATGAACCTGG
AGCACACGTCTGAACTCTTTCCCTACACGACGCTCTTCCGATCTNNNGGCAGAGACACAGCAAAAGA
AGCACACGTCTGAACTCTTTCCCTACACGACGCTCTTCCGATCTNNNATCCTCTACAGACAAGCAAATG

AGCACACGTCTGAACTCTTTCCCTACACGACGCTCTTCCGATCTNNNGCAGGAGAATCATTTGAACCC
AGCACACGTCTGAACTCTTTCCCTACACGACGCTCTTCCGATCTNNNCCAAAGTGCTGGGATTACAAG
AGCACACGTCTGAACTCTTTCCCTACACGACGCTCTTCCGATCTNNNAAAAGAATTAGCCAGACGTGG
AGCACACGTCTGAACTCTTTCCCTACACGACGCTCTTCCGATCTNNNAACAAACCTGACAAGCCATAA
GCACACGTCTGAACTCTTTCCCTACACGACGCTCTTCCGATCTNNNCTAAGGATAATGGCCTCCACTCC
AGAGCACACGTCTGAACTCTTTCCCTACACGACGCTCTTCCGATCTNNNTAGCCTGGGTGAAACTCCAT
GAGCACACGTCTGAACTCTTTCCCTACACGACGCTCTTCCGATCTNNNATCTCTCTATGTTGCCTAGGC
AGCACACGTCTGAACTCTTTCCCTACACGACGCTCTTCCGATCTNNNCTAAGGCAGGAGAATGGTGT
AGCACACGTCTGAACTCTTTCCCTACACGACGCTCTTCCGATCTNNNAAAAAATTAGCCGGGCATAGT
AGCACACGTCTGAACTCTTTCCCTACACGACGCTCTTCCGATCTNNNGACTCCTCCAAAAAGCTCCTAC
AGAGCACACGTCTGAACTCTTTCCCTACACGACGCTCTTCCGATCTNNNGCTGAGATCATGTCATTGCA
AGCACACGTCTGAACTCTTTCCCTACACGACGCTCTTCCGATCTNNNTCAGGAGATTTGCTTGAACCCT
AGCACACGTCTGAACTCTTTCCCTACACGACGCTCTTCCGATCTNNNCTGAGGCAGGAGAATCGTTTG
AGCACACGTCTGAACTCTTTCCCTACACGACGCTCTTCCGATCTNNNAGCAATATCCACACCTTGGCTA
AGCACACGTCTGAACTCTTTCCCTACACGACGCTCTTCCGATCTNNNGAGGTTGCAGTGAACCAAGAT
AGCACACGTCTGAACTCTTTCCCTACACGACGCTCTTCCGATCTNNNTAGAACAATTAGCGGGGAATG
AGCACACGTCTGAACTCTTTCCCTACACGACGCTCTTCCGATCTNNNAGCTACCATAACCAAGTGTGA
AGCACACGTCTGAACTCTTTCCCTACACGACGCTCTTCCGATCTNNNCTGAGGCAGGAGAATCACTT
GAGCACACGTCTGAACTCTTTCCCTACACGACGCTCTTCCGATCTNNNTAAATGGTTCTGTCTAGCTG
AGCACACGTCTGAACTCTTTCCCTACACGACGCTCTTCCGATCTNNNGATCACGAGGTCAGGAGATC
AGCACACGTCTGAACTCTTTCCCTACACGACGCTCTTCCGATCTNNNCTGTAGTCCCAGCTACTCAGGA
AGCACACGTCTGAACTCTTTCCCTACACGACGCTCTTCCGATCTNNNTCACGCCTGTAATCTTAGCACT
GAGCACACGTCTGAACTCTTTCCCTACACGACGCTCTTCCGATCTNNNAGTGAACTGTGATTGTGCCA
AGCACACGTCTGAACTCTTTCCCTACACGACGCTCTTCCGATCTNNNGAGAACCCAGACAGAAAGCTA
AGAGCACACGTCTGAACTCTTTCCCTACACGACGCTCTTCCGATCTNNNAGAGCTTGCAGTGAGATGA
AGCACACGTCTGAACTCTTTCCCTACACGACGCTCTTCCGATCTNNNGAAAGTTGCAGTGAGCTGAGA
AGCACACGTCTGAACTCTTTCCCTACACGACGCTCTTCCGATCTNNNCAGAGGTTACAGTGAGCCAA
GAGCACACGTCTGAACTCTTTCCCTACACGACGCTCTTCCGATCTNNNGGATCACTTGAGGTCAGGAG
AGCACACGTCTGAACTCTTTCCCTACACGACGCTCTTCCGATCTNNNGAGCAAGACTCCGTCACAAAA
AGCACACGTCTGAACTCTTTCCCTACACGACGCTCTTCCGATCTNNNTTACCATGTTGGCCAGGATAG
GAGCACACGTCTGAACTCTTTCCCTACACGACGCTCTTCCGATCTNNNAAGTCACCTAAGCTTGCTACT
GAGCACACGTCTGAACTCTTTCCCTACACGACGCTCTTCCGATCTNNNATCGAGATCAACCTGGACAA
GAGCACACGTCTGAACTCTTTCCCTACACGACGCTCTTCCGATCTNNNCATCCTGGCTAACATGGTGAA
GAGCACACGTCTGAACTCTTTCCCTACACGACGCTCTTCCGATCTNNNATCGCACCACTGCATTATAGT
GAGCACACGTCTGAACTCTTTCCCTACACGACGCTCTTCCGATCTNNNGAGTGAACAGTTCTGTCTTG
GCACACGTCTGAACTCTTTCCCTACACGACGCTCTTCCGATCTNNNCTGTTGTGGGAGGATTGCTTGA
GAGCACACGTCTGAACTCTTTCCCTACACGACGCTCTTCCGATCTNNNCGTATAGCACCATCTCCCATCT
GAGCACACGTCTGAACTCTTTCCCTACACGACGCTCTTCCGATCTNNNAATCTCTGCCTCCTGGATTCA
GAGCACACGTCTGAACTCTTTCCCTACACGACGCTCTTCCGATCTNNNACAAGCAGCAAGATTGAAAT
AGCACACGTCTGAACTCTTTCCCTACACGACGCTCTTCCGATCTNNNAGGCAAGTTCCTTTCAGCTATG
AGCACACGTCTGAACTCTTTCCCTACACGACGCTCTTCCGATCTNNNCAACGAATCCAGGAGTTTGTT
AGCACACGTCTGAACTCTTTCCCTACACGACGCTCTTCCGATCTNNNAGAACAAATGTTGAGGCATCA
GAGCACACGTCTGAACTCTTTCCCTACACGACGCTCTTCCGATCTNNNTTTTTTCAAGAACCGATGCAG

AGCACACGTCTGAACTCTTTCCCTACACGACGCTCTTCCGATCTNNNTGGAAGATGAGAAAACACTT
AGCACACGTCTGAACTCTTTCCCTACACGACGCTCTTCCGATCTNNNGTGGCATGGACAAGATAAGG
AGCACACGTCTGAACTCTTTCCCTACACGACGCTCTTCCGATCTNNNGAGGCAGGAGAATTGCTTGA
GAGCACACGTCTGAACTCTTTCCCTACACGACGCTCTTCCGATCTNNNCAAGTGGCCAACGAACATAT
AGCACACGTCTGAACTCTTTCCCTACACGACGCTCTTCCGATCTNNNCCGAGCAACAAGACCAAAAAT
AGCACACGTCTGAACTCTTTCCCTACACGACGCTCTTCCGATCTNNNAATACCAAAACCAGGAAGGG
AGCACACGTCTGAACTCTTTCCCTACACGACGCTCTTCCGATCTNNNTAGGAGAATCGCTTGAATCCA
AGCACACGTCTGAACTCTTTCCCTACACGACGCTCTTCCGATCTNNNCCAGATCACACTTCTACACTC
GAGCACACGTCTGAACTCTTTCCCTACACGACGCTCTTCCGATCTNNNGGCACCTGTAATCCCAGTTA
GAGCACACGTCTGAACTCTTTCCCTACACGACGCTCTTCCGATCTNNNGCAGACTTGCCACAAACTTT
GAGCACACGTCTGAACTCTTTCCCTACACGACGCTCTTCCGATCTNNNAACAGTTCTGTCTTTCTGGG
GCACACGTCTGAACTCTTTCCCTACACGACGCTCTTCCGATCTNNNACAATGGCTCCTCTTATGGCTTC
AGCACACGTCTGAACTCTTTCCCTACACGACGCTCTTCCGATCTNNNATAGTCCCAGCTACTTAGGAG
AGCACACGTCTGAACTCTTTCCCTACACGACGCTCTTCCGATCTNNNTGAGGAAGGAGAATGGTGT
GAGCACACGTCTGAACTCTTTCCCTACACGACGCTCTTCCGATCTNNNCATCCTGGCTAACATGGTG
AGCACACGTCTGAACTCTTTCCCTACACGACGCTCTTCCGATCTNNNAGCCTGGCTAATATGGTGAAA
GAGCACACGTCTGAACTCTTTCCCTACACGACGCTCTTCCGATCTNNNTCTTTTGGTCTCAGGTGG
AGCACACGTCTGAACTCTTTCCCTACACGACGCTCTTCCGATCTNNNCCATCTCACACCAATCAGAATG
AGCACACGTCTGAACTCTTTCCCTACACGACGCTCTTCCGATCTNNNATCATTTGAACTCAGGAGGCA
GAGCACACGTCTGAACTCTTTCCCTACACGACGCTCTTCCGATCTNNNCCTAGAAACATTCACCCTCCC
GCACACGTCTGAACTCTTTCCCTACACGACGCTCTTCCGATCTNNNGCACTCTTGTAATCCCAGCTAC
AGCACACGTCTGAACTCTTTCCCTACACGACGCTCTTCCGATCTNNNCACAATGTGCACATGTACCCT
AGCACACGTCTGAACTCTTTCCCTACACGACGCTCTTCCGATCTNNNGAAGCTGTTTGGTTGCTCCTA
GAGCACACGTCTGAACTCTTTCCCTACACGACGCTCTTCCGATCTNNNAATGCTCTCAGGCAAAAAC
GAGCACACGTCTGAACTCTTTCCCTACACGACGCTCTTCCGATCTNNNGTCCCAGCTGAGGTAAGAG
AGCACACGTCTGAACTCTTTCCCTACACGACGCTCTTCCGATCTNNNAGGCTGAGGAAGGAGAATTG
AGCACACGTCTGAACTCTTTCCCTACACGACGCTCTTCCGATCTNNNGAGAATGGCTTGAACCCAGA
GCACACGTCTGAACTCTTTCCCTACACGACGCTCTTCCGATCTNNNGGAATGGAAGCTCTGTTGAAA
GCACACGTCTGAACTCTTTCCCTACACGACGCTCTTCCGATCTNNNACAAAAAATTAGCTTGGCGTG
AGCACACGTCTGAACTCTTTCCCTACACGACGCTCTTCCGATCTNNNATGCAAATCAAGCTACTACCCC
AGCACACGTCTGAACTCTTTCCCTACACGACGCTCTTCCGATCTNNNCACGTTGTGCACATGTACCATA
AGCACACGTCTGAACTCTTTCCCTACACGACGCTCTTCCGATCTNNNAGCTGAAGCCATTATGTTACC
GCACACGTCTGAACTCTTTCCCTACACGACGCTCTTCCGATCTNNNAAGGGGGCAAGTACTACATCA
GAGCACACGTCTGAACTCTTTCCCTACACGACGCTCTTCCGATCTNNNTTCCAGAGGAGTGAATGGTT
AGCACACGTCTGAACTCTTTCCCTACACGACGCTCTTCCGATCTNNNCCAGGGGAGTGAATAGTTCTG
GCACACGTCTGAACTCTTTCCCTACACGACGCTCTTCCGATCTNNNAAAAAATTAGCTGGGCATGGTG
AGCACACGTCTGAACTCTTTCCCTACACGACGCTCTTCCGATCTNNNTGCTCATGGATGGGTAGAAT
AGCACACGTCTGAACTCTTTCCCTACACGACGCTCTTCCGATCTNNNACAGCTGTACCAATAGGAAAG
AGCACACGTCTGAACTCTTTCCCTACACGACGCTCTTCCGATCTNNNAATACAAAAAATTAGCCGGGC
GAGCACACGTCTGAACTCTTTCCCTACACGACGCTCTTCCGATCTNNNGTACTGGTACAAAAACAGGT
AGCACACGTCTGAACTCTTTCCCTACACGACGCTCTTCCGATCTNNNAAGCTACTTAGGTGGGTGAA
GCACACGTCTGAACTCTTTCCCTACACGACGCTCTTCCGATCTNNNTCACTGAACAGACCAAGGATGA
AGCACACGTCTGAACTCTTTCCCTACACGACGCTCTTCCGATCTNNNGGGAGGGATAGCATTGGGA

AGCACACGTCTGAACTCTTTCCCTACACGACGCTCTTCCGATCTNNNCACGTTGTGCACATGTAACCT
AGCACACGTCTGAACTCTTTCCCTACACGACGCTCTTCCGATCTNNNACTTGAACCTGGGAGAAAA
AGCACACGTCTGAACTCTTTCCCTACACGACGCTCTTCCGATCTNNNTGTGGGAGAAGAGTGATTTC
AGCACACGTCTGAACTCTTTCCCTACACGACGCTCTTCCGATCTNNNTGAGCTGAGATCATACCATTG
GAGCACACGTCTGAACTCTTTCCCTACACGACGCTCTTCCGATCTNNNTGAGCTGAGATCATACCATT
GCACACGTCTGAACTCTTTCCCTACACGACGCTCTTCCGATCTNNNAAGCTGAGGTGGAAGAATCACT
AGCACACGTCTGAACTCTTTCCCTACACGACGCTCTTCCGATCTNNNAATTTCTCTAACCTGGGAGGCA
AGCACACGTCTGAACTCTTTCCCTACACGACGCTCTTCCGATCTNNNAAATGCCCACATCGAAAAGCT
AGCACACGTCTGAACTCTTTCCCTACACGACGCTCTTCCGATCTNNNGACAGAGAAAGACCCCATCTC
AGCACACGTCTGAACTCTTTCCCTACACGACGCTCTTCCGATCTNNNAACCTGTGCTCACAATCACTAC
AGCACACGTCTGAACTCTTTCCCTACACGACGCTCTTCCGATCTNNNAATACAAAAAATTAGCCGGG
GAGCACACGTCTGAACTCTTTCCCTACACGACGCTCTTCCGATCTNNNCAACCCAATGCAAGGAAGCT
AGCACACGTCTGAACTCTTTCCCTACACGACGCTCTTCCGATCTNNNCTGTAGTCTCAGCCACTAGGG
GAGCACACGTCTGAACTCTTTCCCTACACGACGCTCTTCCGATCTNNNCTGACATCTGTAATCCCAGCA
AGAGCACACGTCTGAACTCTTTCCCTACACGACGCTCTTCCGATCTNNNGCAAAAACATGGATGAAC
AGCACACGTCTGAACTCTTTCCCTACACGACGCTCTTCCGATCTNNNAACAAGCTCTGAAATTGAGGC
GCACACGTCTGAACTCTTTCCCTACACGACGCTCTTCCGATCTNNNACCTGGTTTCTGTGCTTCAGTAT
GAGCACACGTCTGAACTCTTTCCCTACACGACGCTCTTCCGATCTNNNGGTGGATCACTAGGTCAGG
AGCACACGTCTGAACTCTTTCCCTACACGACGCTCTTCCGATCTNNNCAATACTTTGGGAAGCCAAG
AGCACACGTCTGAACTCTTTCCCTACACGACGCTCTTCCGATCTNNNTAGTGTCTCAGAGGAGTGTG
AGAGCACACGTCTGAACTCTTTCCCTACACGACGCTCTTCCGATCTNNNCTGTAGTCCCAGCTACTTG
GCACACGTCTGAACTCTTTCCCTACACGACGCTCTTCCGATCTNNNGAAGCAGGACAATCACTTGAA
AGCACACGTCTGAACTCTTTCCCTACACGACGCTCTTCCGATCTNNNTACAGTTAGCGATGATCATGC
AGCACACGTCTGAACTCTTTCCCTACACGACGCTCTTCCGATCTNNNCTGAGACACGAGGATCACTTG
AGCACACGTCTGAACTCTTTCCCTACACGACGCTCTTCCGATCTNNNACTCAACCCCTTTCACAATAGC
AGCACACGTCTGAACTCTTTCCCTACACGACGCTCTTCCGATCTNNNTTCAACCTGCAGTGAGCTATG
AGCACACGTCTGAACTCTTTCCCTACACGACGCTCTTCCGATCTNNNTTCTCACTTTGCGTCATGGTTT
AGCACACGTCTGAACTCTTTCCCTACACGACGCTCTTCCGATCTNNNGAGGCAACAGAATCACTTGA
GAGCACACGTCTGAACTCTTTCCCTACACGACGCTCTTCCGATCTNNNAGCAAGATGTGTGAATGGG
GAGCACACGTCTGAACTCTTTCCCTACACGACGCTCTTCCGATCTNNNAAAAAATTAGCTGGGTGTG
AGCACACGTCTGAACTCTTTCCCTACACGACGCTCTTCCGATCTNNNCAGAAGAATTGCTTGAACCCA
AGCACACGTCTGAACTCTTTCCCTACACGACGCTCTTCCGATCTNNNACCTGTAATCTCAGCTACTTGG
GAGCACACGTCTGAACTCTTTCCCTACACGACGCTCTTCCGATCTNNNCAAGATAGCACCACTGCACT
AGCACACGTCTGAACTCTTTCCCTACACGACGCTCTTCCGATCTNNNCTTGAACCCAGGAGACAGAGT
AGCACACGTCTGAACTCTTTCCCTACACGACGCTCTTCCGATCTNNNCTGTAGTCCCAGCTACTCAGGA
GAGCACACGTCTGAACTCTTTCCCTACACGACGCTCTTCCGATCTNNNAAAAAATTAGCAGGGTGTG
AGCACACGTCTGAACTCTTTCCCTACACGACGCTCTTCCGATCTNNNATCACTTGAAACCTTGAGGCA
GAGCACACGTCTGAACTCTTTCCCTACACGACGCTCTTCCGATCTNNNCACACACAATTGCTACACCTC
AGCACACGTCTGAACTCTTTCCCTACACGACGCTCTTCCGATCTNNNTATTGAAAGAGGATGTGGGA
GCACACGTCTGAACTCTTTCCCTACACGACGCTCTTCCGATCTNNNGGCTGAGATAGGAGAATCACCT
AGCACACGTCTGAACTCTTTCCCTACACGACGCTCTTCCGATCTNNNCTGAGGCAGGAGAATTGCTT
AGCACACGTCTGAACTCTTTCCCTACACGACGCTCTTCCGATCTNNNATGCAATCTCAGCTTACTGCAA
GAGCACACGTCTGAACTCTTTCCCTACACGACGCTCTTCCGATCTNNNAGAATTGCTTGAATCTGGGA

AGCACACGTCTGAACTCTTTCCCTACACGACGCTCTTCCGATCTNNNTGCACAGACCTATGAAGACTGA
GAGCACACGTCTGAACTCTTTCCCTACACGACGCTCTTCCGATCTNNNCTATGCAGGGTCCCTTGCTAT
AGAGCACACGTCTGAACTCTTTCCCTACACGACGCTCTTCCGATCTNNNGGGAGGCTGAGATAGGAG
AGCACACGTCTGAACTCTTTCCCTACACGACGCTCTTCCGATCTNNNGGCAGGAGAATCACTTGAACCT
AGCACACGTCTGAACTCTTTCCCTACACGACGCTCTTCCGATCTNNNTTTCATAATGCAAGCAGAAGCA
AAGAGCACACGTCTGAACTCTTTCCCTACACGACGCTCTTCCGATCTNNNTAGGGAGGCTGAGGTACG
AGCACACGTCTGAACTCTTTCCCTACACGACGCTCTTCCGATCTNNNAATTTCACGATGGCAGTTTCACC
AGCACACGTCTGAACTCTTTCCCTACACGACGCTCTTCCGATCTNNNAGACAGAAGAATCGCTTGAACC
GAGCACACGTCTGAACTCTTTCCCTACACGACGCTCTTCCGATCTNNNAGAATCACTTCAACCTGGGAG
AGAGCACACGTCTGAACTCTTTCCCTACACGACGCTCTTCCGATCTNNNACACCTGTAGTCCCAGCTACT
AGCACACGTCTGAACTCTTTCCCTACACGACGCTCTTCCGATCTNNNGCGCCTATAATCCCAGCTACTAG
AGCACACGTCTGAACTCTTTCCCTACACGACGCTCTTCCGATCTNNNGGGGACAAGAGTGAGACTTCTT
GAGCACACGTCTGAACTCTTTCCCTACACGACGCTCTTCCGATCTNNNTCCTGGAGAACAAGAGCAAAA
GAGCACACGTCTGAACTCTTTCCCTACACGACGCTCTTCCGATCTNNNCCTGGCCAACATGATGAAATC
AGAGCACACGTCTGAACTCTTTCCCTACACGACGCTCTTCCGATCTNNNCACCTGTAGTCCCAGCTACTT
GAGCACACGTCTGAACTCTTTCCCTACACGACGCTCTTCCGATCTNNNCTGAGGCAGGAGAATTGCTAC
GAGCACACGTCTGAACTCTTTCCCTACACGACGCTCTTCCGATCTNNNAGCTTGCAGTGAGTCAAAATA
GAGCACACGTCTGAACTCTTTCCCTACACGACGCTCTTCCGATCTNNNGAAAGCTGAGGCATGAGAATC
GAGCACACGTCTGAACTCTTTCCCTACACGACGCTCTTCCGATCTNNNGGTATGGTGGCAGGTGTCTA
AGAGCACACGTCTGAACTCTTTCCCTACACGACGCTCTTCCGATCTNNNATACAAAAATTAGCTGGGCG
AGCACACGTCTGAACTCTTTCCCTACACGACGCTCTTCCGATCTNNNACAGTGAAACCCCCTCTCTACTA
AGCACACGTCTGAACTCTTTCCCTACACGACGCTCTTCCGATCTNNNTACCGAAGTTGACCACAGGTAA
GAGCACACGTCTGAACTCTTTCCCTACACGACGCTCTTCCGATCTNNNAGAATCACTTCAATCCAGGAG
GAGCACACGTCTGAACTCTTTCCCTACACGACGCTCTTCCGATCTNNNACCTGGCAGTGTTTCTGTTGAT
AGCACACGTCTGAACTCTTTCCCTACACGACGCTCTTCCGATCTNNNACGCCTGTAATCCCAGTACTTTG
GAGCACACGTCTGAACTCTTTCCCTACACGACGCTCTTCCGATCTNNNAGACAGAAGAATCGCTTGAAC
AGCACACGTCTGAACTCTTTCCCTACACGACGCTCTTCCGATCTNNNTCATGCATGTAATCCCCACACTTC
GCACACGTCTGAACTCTTTCCCTACACGACGCTCTTCCGATCTNNNGTTATCTCTGCTACTCAGGAGGC
GAGCACACGTCTGAACTCTTTCCCTACACGACGCTCTTCCGATCTNNNCTCATGCCTATAATCCCAGCACT
AGCACACGTCTGAACTCTTTCCCTACACGACGCTCTTCCGATCTNNNGCCCTAAGTGATGCCATGAGAT
GAGCACACGTCTGAACTCTTTCCCTACACGACGCTCTTCCGATCTNNNAAAGCCACATACCTACAGCCTA
GAGCACACGTCTGAACTCTTTCCCTACACGACGCTCTTCCGATCTNNNCTTAGCCAGGGCAATCAGAAA
AGAGCACACGTCTGAACTCTTTCCCTACACGACGCTCTTCCGATCTNNNTCACTTGAACCTGCCTAGATC
GAGCACACGTCTGAACTCTTTCCCTACACGACGCTCTTCCGATCTNNNAAGAAAGAGCATGTGCAAAG
GAGCACACGTCTGAACTCTTTCCCTACACGACGCTCTTCCGATCTNNNCAACAAACACTGCCAAAATGCT
AGCACACGTCTGAACTCTTTCCCTACACGACGCTCTTCCGATCTNNNCATGCCACTATAGTCCAGCCTAC
GAGCACACGTCTGAACTCTTTCCCTACACGACGCTCTTCCGATCTNNNGAGATTGCAGTGAGCCAAGA
AGCACACGTCTGAACTCTTTCCCTACACGACGCTCTTCCGATCTNNNAGCAATTACAACTTGGCTGCTTT
GAGCACACGTCTGAACTCTTTCCCTACACGACGCTCTTCCGATCTNNNCATGCCTGTAATCCCAGCTACT
GAGCACACGTCTGAACTCTTTCCCTACACGACGCTCTTCCGATCTNNNGTCATGTGGGGGAAGAAGAC
AGCACACGTCTGAACTCTTTCCCTACACGACGCTCTTCCGATCTNNNAAACGTAATGCTTTTTCCCCCAG
GAGCACACGTCTGAACTCTTTCCCTACACGACGCTCTTCCGATCTNNNTGTAGTCCCAGCTACTCAGGA
AGCACACGTCTGAACTCTTTCCCTACACGACGCTCTTCCGATCTNNNGGAGAATTGCTTGAACTTGGGA

GCACACGTCTGAACTCTTTCCCTACACGACGCTCTTCCGATCTNNNCAGCAACAAAAACCAGACAAACA
GAGCACACGTCTGAACTCTTTCCCTACACGACGCTCTTCCGATCTNNNCATTGCATTACAGCCTGGTTG
GAGCACACGTCTGAACTCTTTCCCTACACGACGCTCTTCCGATCTNNNATGGAAAAGGCAGTCACCCA
GAGCACACGTCTGAACTCTTTCCCTACACGACGCTCTTCCGATCTNNNCTGTAATCCCAGCTACTCAGGA
GAGCACACGTCTGAACTCTTTCCCTACACGACGCTCTTCCGATCTNNNAGACAGGAGAATTGCTTGAA
AGCACACGTCTGAACTCTTTCCCTACACGACGCTCTTCCGATCTNNNCGACAGTGAGATCCTGTGTCAA
GAGCACACGTCTGAACTCTTTCCCTACACGACGCTCTTCCGATCTNNNTCAAGAGTCTGAGGGAGGAG
GAGCACACGTCTGAACTCTTTCCCTACACGACGCTCTTCCGATCTNNNCTGTAGTCCCAGCTACTCAGG
GAGCACACGTCTGAACTCTTTCCCTACACGACGCTCTTCCGATCTNNNCTGTAATCCCAGCTACTCAGGA
GAGCACACGTCTGAACTCTTTCCCTACACGACGCTCTTCCGATCTNNNTGAGGCAGCAAAATCACTTGA
GAGCACACGTCTGAACTCTTTCCCTACACGACGCTCTTCCGATCTNNNATTTAGGAACCAAGACAGGC
GAGCACACGTCTGAACTCTTTCCCTACACGACGCTCTTCCGATCTNNNTTAGGAAGATGCAGCTGTTTT
AGCACACGTCTGAACTCTTTCCCTACACGACGCTCTTCCGATCTNNNAAAGCATTCTGTCAGTTCACTG
GAGCACACGTCTGAACTCTTTCCCTACACGACGCTCTTCCGATCTNNNTCATGCCTGTAATCTCAGCAC
GAGCACACGTCTGAACTCTTTCCCTACACGACGCTCTTCCGATCTNNNACAAAAATTAGCCAGACGTG
GAGCACACGTCTGAACTCTTTCCCTACACGACGCTCTTCCGATCTNNNGTCAGAAGTTTGAGACCAGC
GAGCACACGTCTGAACTCTTTCCCTACACGACGCTCTTCCGATCTNNNGCCAATATCACATCACTGCACT
GAGCACACGTCTGAACTCTTTCCCTACACGACGCTCTTCCGATCTNNNTGAGGTAGGAGAATGGTGTG
AGAGCACACGTCTGAACTCTTTCCCTACACGACGCTCTTCCGATCTNNNTGTAATCCTAGCACTTTGGG
GAGCACACGTCTGAACTCTTTCCCTACACGACGCTCTTCCGATCTNNNAAATTGTGACCTCTGCATTGT
AGCACACGTCTGAACTCTTTCCCTACACGACGCTCTTCCGATCTNNNAAGTAGGAGACACATCAGCAG
GCACACGTCTGAACTCTTTCCCTACACGACGCTCTTCCGATCTNNNCACGCCTATAATCCCAGCTACTCC
AGCACACGTCTGAACTCTTTCCCTACACGACGCTCTTCCGATCTNNNGCCTCCTGTAATCTCAGCTACTT
AGCACACGTCTGAACTCTTTCCCTACACGACGCTCTTCCGATCTNNNCTGTGTTAGCAGCTACTCAGTG
AAGAGCACACGTCTGAACTCTTTCCCTACACGACGCTCTTCCGATCTNNNACCTGTAGTCCCAGCTACT
AGCACACGTCTGAACTCTTTCCCTACACGACGCTCTTCCGATCTNNNAGAATCACTTGAACTCAGGAG
AGCACACGTCTGAACTCTTTCCCTACACGACGCTCTTCCGATCTNNNCATCATCTTTCATCCTCCCACCT
AGCACACGTCTGAACTCTTTCCCTACACGACGCTCTTCCGATCTNNNGTTTACAGTGAGCTCGAGATTG
AGAGCACACGTCTGAACTCTTTCCCTACACGACGCTCTTCCGATCTNNNCTAGCCAGACAGAGAGAAA
GCACACGTCTGAACTCTTTCCCTACACGACGCTCTTCCGATCTNNNTTAGAGAGCAGGTTGGTCAACA
GCACACGTCTGAACTCTTTCCCTACACGACGCTCTTCCGATCTNNNACACTGAGATGCTTCTTGTGCTA
GAGCACACGTCTGAACTCTTTCCCTACACGACGCTCTTCCGATCTNNNAGGCAAGAGTATCAGTTGAG
GCACACGTCTGAACTCTTTCCCTACACGACGCTCTTCCGATCTNNNAGGAAATGATGTTCAGGCCTCT
GAGCACACGTCTGAACTCTTTCCCTACACGACGCTCTTCCGATCTNNNCAGCCTGGACAACAAGAGAGA
AGCACACGTCTGAACTCTTTCCCTACACGACGCTCTTCCGATCTNNNTGAGGGAGGAGAATCACTTGA
GAGCACACGTCTGAACTCTTTCCCTACACGACGCTCTTCCGATCTNNNTAGGTTGCATGAGCTGAGAT
GAGCACACGTCTGAACTCTTTCCCTACACGACGCTCTTCCGATCTNNNAATTTCTTCCACCAGGCAACCT
AGCACACGTCTGAACTCTTTCCCTACACGACGCTCTTCCGATCTNNNAGCTGAGGAATGAGAAGCTTG
AGCACACGTCTGAACTCTTTCCCTACACGACGCTCTTCCGATCTNNNGAGGTTACAGTGAGCTGAGAT
GAGCACACGTCTGAACTCTTTCCCTACACGACGCTCTTCCGATCTNNNCACCTGTAATCCCAGCTACTCT
GAGCACACGTCTGAACTCTTTCCCTACACGACGCTCTTCCGATCTNNNCTGTAGTCCCAGCTACTTGGA
AGCACACGTCTGAACTCTTTCCCTACACGACGCTCTTCCGATCTNNNCATGCCTGTAATCCCAGCTACTA
AGAGCACACGTCTGAACTCTTTCCCTACACGACGCTCTTCCGATCTNNNATAAAAATTAGCTGGGTGT

AGCACACGTCTGAACTCTTTCCCTACACGACGCTCTTCCGATCTNNNTACAGTGGCATTGTCATAGCTC
AGCACACGTCTGAACTCTTTCCCTACACGACGCTCTTCCGATCTNNNGCTGAGATCATTCCATTGCACT
AGCACACGTCTGAACTCTTTCCCTACACGACGCTCTTCCGATCTNNNGCTACTTGGGAGGCTAATGTA
GAGCACACGTCTGAACTCTTTCCCTACACGACGCTCTTCCGATCTNNNCTGTAGTCCCAGCTACTCCAG
GAGCACACGTCTGAACTCTTTCCCTACACGACGCTCTTCCGATCTNNNAGCCTGAACAACAACAGTGA
AGCACACGTCTGAACTCTTTCCCTACACGACGCTCTTCCGATCTNNNCATGCCTGTAGTACCAGCTACT
AGCACACGTCTGAACTCTTTCCCTACACGACGCTCTTCCGATCTNNNCTGTAGTTCCAGCTACTCGGGA
AGCACACGTCTGAACTCTTTCCCTACACGACGCTCTTCCGATCTNNNTGGAGATTTCAGTGAGCCAAG
AGCACACGTCTGAACTCTTTCCCTACACGACGCTCTTCCGATCTNNNATCACGAGGTCAGGAGATCGA
AGCACACGTCTGAACTCTTTCCCTACACGACGCTCTTCCGATCTNNNGTACAGGAACAGACACATAG
AGCACACGTCTGAACTCTTTCCCTACACGACGCTCTTCCGATCTNNNACAGCTTGTATCATTCCTTCTG
GAGCACACGTCTGAACTCTTTCCCTACACGACGCTCTTCCGATCTNNNCTGTAGTTCCAGATACTGCG
AGCACACGTCTGAACTCTTTCCCTACACGACGCTCTTCCGATCTNNNCTGTAATCCCAGCTACTCAGGA
AGCACACGTCTGAACTCTTTCCCTACACGACGCTCTTCCGATCTNNNAGAAAAGCACATCCGTGAAAC
AGCACACGTCTGAACTCTTTCCCTACACGACGCTCTTCCGATCTNNNCTGAGGCAGGAAAATTGCTT
AGCACACGTCTGAACTCTTTCCCTACACGACGCTCTTCCGATCTNNNATCACTTGAAATCGGAAGGTG
AGCACACGTCTGAACTCTTTCCCTACACGACGCTCTTCCGATCTNNNGAGGCAGAGGTAGGAAAATC
GAGCACACGTCTGAACTCTTTCCCTACACGACGCTCTTCCGATCTNNNCTGTAGTCCCAGCTACTCAGG
GAGCACACGTCTGAACTCTTTCCCTACACGACGCTCTTCCGATCTNNNTTGGAATGGGAATGCCTACT
GAGCACACGTCTGAACTCTTTCCCTACACGACGCTCTTCCGATCTNNNTGCTTCTTCATAGGTCAGCTG
GAGCACACGTCTGAACTCTTTCCCTACACGACGCTCTTCCGATCTNNNCCTTATTGTCAAGGCCTCTCA
AGCACACGTCTGAACTCTTTCCCTACACGACGCTCTTCCGATCTNNNGTGCTGGCAGTGAGAATTTCA
AGCACACGTCTGAACTCTTTCCCTACACGACGCTCTTCCGATCTNNNAACTTCCACCATCACTCCCAAAT
AGCACACGTCTGAACTCTTTCCCTACACGACGCTCTTCCGATCTNNNCTGTAGTCCCAGCTACTCAGGA
AGCACACGTCTGAACTCTTTCCCTACACGACGCTCTTCCGATCTNNNACAACAAGCTCTTGAGACTTGA
AGCACACGTCTGAACTCTTTCCCTACACGACGCTCTTCCGATCTNNNAGTTCTAGTGTGACCTTGCTCTC
AGCACACGTCTGAACTCTTTCCCTACACGACGCTCTTCCGATCTNNNGGCATTACCACTGATCTCACAG
GAGCACACGTCTGAACTCTTTCCCTACACGACGCTCTTCCGATCTNNNGGACTACACTTCCCAGTAATG
GAGCACACGTCTGAACTCTTTCCCTACACGACGCTCTTCCGATCTNNNTACAAAAAATTAGCCGGGCT
AGCACACGTCTGAACTCTTTCCCTACACGACGCTCTTCCGATCTNNNGGTGCCTGTAATCCCAGCTAT
AGCACACGTCTGAACTCTTTCCCTACACGACGCTCTTCCGATCTNNNTTCAGCAGTTTTCACCACGAAT
GAGCACACGTCTGAACTCTTTCCCTACACGACGCTCTTCCGATCTNNNCTGTAATCCCAGCTACTCAGG
AGCACACGTCTGAACTCTTTCCCTACACGACGCTCTTCCGATCTNNNGTAGCCTGCATATCCAAAGCA
AGCACACGTCTGAACTCTTTCCCTACACGACGCTCTTCCGATCTNNNAGAATCTCTTGAATCAGGGAG
AGCACACGTCTGAACTCTTTCCCTACACGACGCTCTTCCGATCTNNNCGTGGATGACAGATCTGACCT
AGCACACGTCTGAACTCTTTCCCTACACGACGCTCTTCCGATCTNNNCTGAGGCAGGAGAATCACTTG
GAGCACACGTCTGAACTCTTTCCCTACACGACGCTCTTCCGATCTNNNCTGTAATCCCAGCTACTCAGG
GCACACGTCTGAACTCTTTCCCTACACGACGCTCTTCCGATCTNNNAATCATGTGGTGATCTCTTGCAC
AGCACACGTCTGAACTCTTTCCCTACACGACGCTCTTCCGATCTNNNCAAGGTGACCTAGAAGCAGTCT
GAGCACACGTCTGAACTCTTTCCCTACACGACGCTCTTCCGATCTNNNGATCACAAGGTCAGGAGAT
GAGCACACGTCTGAACTCTTTCCCTACACGACGCTCTTCCGATCTNNNTACTAAGGCTGTTCCTCTTCT
AGCACACGTCTGAACTCTTTCCCTACACGACGCTCTTCCGATCTNNNTCATGCCTGTAATCGTAGCAC
GAGCACACGTCTGAACTCTTTCCCTACACGACGCTCTTCCGATCTNNNGGAAGACGACCTAGGCAATA

GAGCACACGTCTGAACTCTTTCCCTACACGACGCTCTTCCGATCTNNNGGACACATACACAGTCCCAAG
GAGCACACGTCTGAACTCTTTCCCTACACGACGCTCTTCCGATCTNNNCTTTGTAACTCGGGACCCTTGA
AGCACACGTCTGAACTCTTTCCCTACACGACGCTCTTCCGATCTNNNACCACAAAGCAGATAATAAGCCA
GAGCACACGTCTGAACTCTTTCCCTACACGACGCTCTTCCGATCTNNNATTGTGAAAATGACCAGACTG
AGAGCACACGTCTGAACTCTTTCCCTACACGACGCTCTTCCGATCTNNNGTGAGCCAAGATTTTGCCACT
GAGCACACGTCTGAACTCTTTCCCTACACGACGCTCTTCCGATCTNNNTCTGCTTCCTGAGTTATCTGCA
GAGCACACGTCTGAACTCTTTCCCTACACGACGCTCTTCCGATCTNNNGAACCCCACTAGGCTAACAGTA
AGCACACGTCTGAACTCTTTCCCTACACGACGCTCTTCCGATCTNNNCACACTCGTGTAATCCCAGCTAC
AGAGCACACGTCTGAACTCTTTCCCTACACGACGCTCTTCCGATCTNNNGCCTGAGAATTGCTTGAAAT
AGCACACGTCTGAACTCTTTCCCTACACGACGCTCTTCCGATCTNNNATTGTCCTATGACCCTGCCAAAT
GAGCACACGTCTGAACTCTTTCCCTACACGACGCTCTTCCGATCTNNNCTGTGCTCTAAGTCTGGTGTA
GAGCACACGTCTGAACTCTTTCCCTACACGACGCTCTTCCGATCTNNNGGTGCCTGTAATCCTAGCTACT
GAGCACACGTCTGAACTCTTTCCCTACACGACGCTCTTCCGATCTNNNCTTGAGTCCTGGAGTTTGAGA
GAGCACACGTCTGAACTCTTTCCCTACACGACGCTCTTCCGATCTNNNGTGTCTGTAGTCCCAGCTACTT
GAGCACACGTCTGAACTCTTTCCCTACACGACGCTCTTCCGATCTNNNCATGCCTGTAATCCCAGCTACT
AGCACACGTCTGAACTCTTTCCCTACACGACGCTCTTCCGATCTNNNTTCAAGCGATTCTCCTGCCTTAG
GCACACGTCTGAACTCTTTCCCTACACGACGCTCTTCCGATCTNNNTCACTTGAGTCCAGGACTTCAAG
GAGCACACGTCTGAACTCTTTCCCTACACGACGCTCTTCCGATCTNNNTCTTGAGTCCAGGAGTTTGA
GAGCACACGTCTGAACTCTTTCCCTACACGACGCTCTTCCGATCTNNNTACAAAACTTAGCTGGGCTTG
AGCACACGTCTGAACTCTTTCCCTACACGACGCTCTTCCGATCTNNNGAGGTTGCAGTAAGCTGAGAT
GCACACGTCTGAACTCTTTCCCTACACGACGCTCTTCCGATCTNNNAACCACTAGTCTGCTCTCTTTGGC
AGCACACGTCTGAACTCTTTCCCTACACGACGCTCTTCCGATCTNNNGAGGCAGGAGAATTGCTTGAAT
GAGCACACGTCTGAACTCTTTCCCTACACGACGCTCTTCCGATCTNNNTACAAAAAATTAGCCAGGCGA
AGCACACGTCTGAACTCTTTCCCTACACGACGCTCTTCCGATCTNNNAATACAAAAAATTAGCCGGGC
AGAGCACACGTCTGAACTCTTTCCCTACACGACGCTCTTCCGATCTNNNTTTAGCATGCATGGATTTTG
AGCACACGTCTGAACTCTTTCCCTACACGACGCTCTTCCGATCTNNNGAGATCACACCATTGCACTCCAG
AGCACACGTCTGAACTCTTTCCCTACACGACGCTCTTCCGATCTNNNCTGAGGCAGGAGAATCACTTGA
GAGCACACGTCTGAACTCTTTCCCTACACGACGCTCTTCCGATCTNNNATACAAAAATTAGCTGGGCGT
GAGCACACGTCTGAACTCTTTCCCTACACGACGCTCTTCCGATCTNNNTGATGTACACCTGTGATACCA
AGCACACGTCTGAACTCTTTCCCTACACGACGCTCTTCCGATCTNNNGTTGTAGTGAACCAAGATCGCA
AGCACACGTCTGAACTCTTTCCCTACACGACGCTCTTCCGATCTNNNACCATTTTATGCCCTCAGTAGCA
AGAGCACACGTCTGAACTCTTTCCCTACACGACGCTCTTCCGATCTNNNCTGAGGCAGAGAATTGCTTG
GAGCACACGTCTGAACTCTTTCCCTACACGACGCTCTTCCGATCTNNNGGATGACAGAGTGAGACTCC
GAGCACACGTCTGAACTCTTTCCCTACACGACGCTCTTCCGATCTNNNGGAGGTTGCAATGAACCAAG
AGCACACGTCTGAACTCTTTCCCTACACGACGCTCTTCCGATCTNNNCAGCTATTCTGGAGGCTAAGGT
AGCACACGTCTGAACTCTTTCCCTACACGACGCTCTTCCGATCTNNNAAAAAAACTAGCAGGGCATGGT
AGCACACGTCTGAACTCTTTCCCTACACGACGCTCTTCCGATCTNNNGGTTGCAGTAAGCCAAGATCAT
AGCACACGTCTGAACTCTTTCCCTACACGACGCTCTTCCGATCTNNNAGTGATTCTTCTGTCTCAGCCTC
GAGCACACGTCTGAACTCTTTCCCTACACGACGCTCTTCCGATCTNNNGCAGTGGCAGCTTGGATTTAT
GAGCACACGTCTGAACTCTTTCCCTACACGACGCTCTTCCGATCTNNNAAAAATTAGATGGGCGTGGT
GAGCACACGTCTGAACTCTTTCCCTACACGACGCTCTTCCGATCTNNNTTGAGGTCAGCAGTTCAAGAT
GAGCACACGTCTGAACTCTTTCCCTACACGACGCTCTTCCGATCTNNNGAGGCAGGAGAATTGTTTGAA
GAGCACACGTCTGAACTCTTTCCCTACACGACGCTCTTCCGATCTNNNTGAGCATGGGATGTTTCCATT

GCACACGTCTGAACTCTTTCCCTACACGACGCTCTTCCGATCTNNNTGAAAGAGTTGAGAGGTGAAA

AGCACACGTCTGAACTCTTTCCCTACACGACGCTCTTCCGATCTNNNTTTGGAAGCTGGTATCAAGGG

GAGCACACGTCTGAACTCTTTCCCTACACGACGCTCTTCCGATCTNNNTACCCAGGAAGTAAGAGGA

GAGCACACGTCTGAACTCTTTCCCTACACGACGCTCTTCCGATCTNNNCACCTGTAATTCCAGCAGTTTG

GAGCACACGTCTGAACTCTTTCCCTACACGACGCTCTTCCGATCTNNNCAAGCTGAGAGCCAAAACAAG

GCACACGTCTGAACTCTTTCCCTACACGACGCTCTTCCGATCTNNNCTGGGTAACAAGAGCGAAACTT

GAGCACACGTCTGAACTCTTTCCCTACACGACGCTCTTCCGATCTNNNCAAAATCACACCACTGCACTCC

AGCACACGTCTGAACTCTTTCCCTACACGACGCTCTTCCGATCTNNNGCTCATGCCTGTAATCTCAACA

AGCACACGTCTGAACTCTTTCCCTACACGACGCTCTTCCGATCTNNNCTGAGGCAGGAGAATAGCTTG

AAGAGCACACGTCTGAACTCTTTCCCTACACGACGCTCTTCCGATCTNNNGAGGCTGAGGCAGGAGAA

GAGCACACGTCTGAACTCTTTCCCTACACGACGCTCTTCCGATCTNNNGCTAAGGCAGGAGAATCACT

GAGCACACGTCTGAACTCTTTCCCTACACGACGCTCTTCCGATCTNNNGCGGATTACCTGAGGTTATGA

AGCACACGTCTGAACTCTTTCCCTACACGACGCTCTTCCGATCTNNNGTGAAAAAGGAGGTGGTGTTT

GAGCACACGTCTGAACTCTTTCCCTACACGACGCTCTTCCGATCTNNNGCAGGATAATCGTTTGAACC

AGCACACGTCTGAACTCTTTCCCTACACGACGCTCTTCCGATCTNNNAGTGAGTCAAGGTAGTGCCATT

GAGCACACGTCTGAACTCTTTCCCTACACGACGCTCTTCCGATCTNNNGTAATCCTACTTGGGAGGCTG

AGCACACGTCTGAACTCTTTCCCTACACGACGCTCTTCCGATCTNNNATGCCTGTAATCCCAGCTACTT

GAGCACACGTCTGAACTCTTTCCCTACACGACGCTCTTCCGATCTNNNACTACAACACTTCCTTCACCGT

GAGCACACGTCTGAACTCTTTCCCTACACGACGCTCTTCCGATCTNNNGAGGCAGGAAAATCACTTGA

AGCACACGTCTGAACTCTTTCCCTACACGACGCTCTTCCGATCTNNNAGAATTGCTTGAACCCAGGAG

GAGCACACGTCTGAACTCTTTCCCTACACGACGCTCTTCCGATCTNNNTACCTGTAATCCCAGCTATTCG

GAGCACACGTCTGAACTCTTTCCCTACACGACGCTCTTCCGATCTNNNAGGCAGGAAAATTGCTTGAA

AGCACACGTCTGAACTCTTTCCCTACACGACGCTCTTCCGATCTNNNCAACATAGCACGATCCTGTCTC

GAGCACACGTCTGAACTCTTTCCCTACACGACGCTCTTCCGATCTNNNCCTGGGCAATAAGAGCAAAA

GAGCACACGTCTGAACTCTTTCCCTACACGACGCTCTTCCGATCTNNNAGACAGGAGAATTGCTTGAA

GCACACGTCTGAACTCTTTCCCTACACGACGCTCTTCCGATCTNNNCATGAGAATTGCTTGAACCTGG

GAGCACACGTCTGAACTCTTTCCCTACACGACGCTCTTCCGATCTNNNAGGAGTTTGAAACTATCCTGG

AGCACACGTCTGAACTCTTTCCCTACACGACGCTCTTCCGATCTNNNTCCTAACACTTTGGGAGGTCAA

AGAGCACACGTCTGAACTCTTTCCCTACACGACGCTCTTCCGATCTNNNACCATCCTAGGCTTGTCAAA

GAGCACACGTCTGAACTCTTTCCCTACACGACGCTCTTCCGATCTNNNTGAGGTCAGGAGTTCGAGAC

AGCACACGTCTGAACTCTTTCCCTACACGACGCTCTTCCGATCTNNNGCACCAGCTGAGGTTACAATTT

AGCACACGTCTGAACTCTTTCCCTACACGACGCTCTTCCGATCTNNNTGCTGAGCACATTTTCATGTAC

GAGCACACGTCTGAACTCTTTCCCTACACGACGCTCTTCCGATCTNNNGACTGGGTAAATTCTGAGCC

AGAGCACACGTCTGAACTCTTTCCCTACACGACGCTCTTCCGATCTNNNGTAATTCTAGCTACTCGGGA

GAGCACACGTCTGAACTCTTTCCCTACACGACGCTCTTCCGATCTNNNTCCTCCTTCTTTTCTCCCTAGGT

AGCACACGTCTGAACTCTTTCCCTACACGACGCTCTTCCGATCTNNNATGATGACGCACACCTATAGT

GAGCACACGTCTGAACTCTTTCCCTACACGACGCTCTTCCGATCTNNNGGCTAAAACAGTGAAACCTCG

AGCACACGTCTGAACTCTTTCCCTACACGACGCTCTTCCGATCTNNNCATTCTCCTCATGCCAAAGAAG

GAGCACACGTCTGAACTCTTTCCCTACACGACGCTCTTCCGATCTNNNAAAAAATTAGCCTGGTGTGGT

GCACACGTCTGAACTCTTTCCCTACACGACGCTCTTCCGATCTNNNCTGAGACAGGAGAGTCGTTTGA

AGCACACGTCTGAACTCTTTCCCTACACGACGCTCTTCCGATCTNNNCTGAGGCAGGAGAATTGCTTG

AGCACACGTCTGAACTCTTTCCCTACACGACGCTCTTCCGATCTNNNAGAGGGGAAGCAATTCACTGA

AGCACACGTCTGAACTCTTTCCCTACACGACGCTCTTCCGATCTNNNGCTGAAGGAGAACAGGAGAT

AGCACACGTCTGAACTCTTTCCCTACACGACGCTCTTCCGATCTNNNCTGCCAACGGATACAGAGATG
AGCACACGTCTGAACTCTTTCCCTACACGACGCTCTTCCGATCTNNNACCTATAGTCCCAGCTACTAGG
AGCACACGTCTGAACTCTTTCCCTACACGACGCTCTTCCGATCTNNNGTTGGCACATGACTGTAGTCC
AGCACACGTCTGAACTCTTTCCCTACACGACGCTCTTCCGATCTNNNGCCATATATGACAGATCCACAG
AGCACACGTCTGAACTCTTTCCCTACACGACGCTCTTCCGATCTNNNTTTCATTAGCAGGTCCTGAGG
AGCACACGTCTGAACTCTTTCCCTACACGACGCTCTTCCGATCTNNNCTGAGGCAGGAGAATCACTT
GAGCACACGTCTGAACTCTTTCCCTACACGACGCTCTTCCGATCTNNNGAGACTGGGAAGAAAGAGA
AGCACACGTCTGAACTCTTTCCCTACACGACGCTCTTCCGATCTNNNTTGTGTTATGGCTTTACCCCAC
AGCACACGTCTGAACTCTTTCCCTACACGACGCTCTTCCGATCTNNNGGAGATTGCAGTGAGCTGAG
GCACACGTCTGAACTCTTTCCCTACACGACGCTCTTCCGATCTNNNCCTGGGTAACACAGGGAATTTC
AGCACACGTCTGAACTCTTTCCCTACACGACGCTCTTCCGATCTNNNGGTGACAGAGTGATATCCGTC
GCACACGTCTGAACTCTTTCCCTACACGACGCTCTTCCGATCTNNNGGAGTTTGCAGTGAGCTGAGA
GAGCACACGTCTGAACTCTTTCCCTACACGACGCTCTTCCGATCTNNNAATCAGGTTTAGTGGCAAG
GAGCACACGTCTGAACTCTTTCCCTACACGACGCTCTTCCGATCTNNNGTGTATATTGAACCAGCCTT
GCACACGTCTGAACTCTTTCCCTACACGACGCTCTTCCGATCTNNNTCCAAGACATCGAAAAGGAGA
GCACACGTCTGAACTCTTTCCCTACACGACGCTCTTCCGATCTNNNACGCCCGGCTAATTTTTCTTTTC
AGCACACGTCTGAACTCTTTCCCTACACGACGCTCTTCCGATCTNNNTTACAGTGAGTTGAGATCCTG
AGCACACGTCTGAACTCTTTCCCTACACGACGCTCTTCCGATCTNNNTATAGTCCCAGCTACTCAGGA
GCACACGTCTGAACTCTTTCCCTACACGACGCTCTTCCGATCTNNNCATTTTTGTACCTGTGGCTGTC
GCACACGTCTGAACTCTTTCCCTACACGACGCTCTTCCGATCTNNNTGAGGCAGGAGAATCTCTTGA
AGCACACGTCTGAACTCTTTCCCTACACGACGCTCTTCCGATCTNNNTTCCACTTGGACAGACAGAAG
AGCACACGTCTGAACTCTTTCCCTACACGACGCTCTTCCGATCTNNNAGACAAACCATATCCCTGCCA
AGCACACGTCTGAACTCTTTCCCTACACGACGCTCTTCCGATCTNNNTTGATTTAGGAACCAGGCTGT
GAGCACACGTCTGAACTCTTTCCCTACACGACGCTCTTCCGATCTNNNAGTGAGCCAAGATAGTGCCA
GCACACGTCTGAACTCTTTCCCTACACGACGCTCTTCCGATCTNNNCACAGAAAGCCAAAGCAAGAT
AGCACACGTCTGAACTCTTTCCCTACACGACGCTCTTCCGATCTNNNGTGGTGCATTCCTGTAATTCC
AGCACACGTCTGAACTCTTTCCCTACACGACGCTCTTCCGATCTNNNATAAATGGGCAAAACCTGG
AGCACACGTCTGAACTCTTTCCCTACACGACGCTCTTCCGATCTNNNCTGTAATCCCAGCTACTCAGG
AGCACACGTCTGAACTCTTTCCCTACACGACGCTCTTCCGATCTNNNTGACCAACATAGTGAAACCCC
AGCACACGTCTGAACTCTTTCCCTACACGACGCTCTTCCGATCTNNNGCTGAGGTAGGAGAATGGAA
AGCACACGTCTGAACTCTTTCCCTACACGACGCTCTTCCGATCTNNNAAGGCTAATGGATTCCAAGAA
AGCACACGTCTGAACTCTTTCCCTACACGACGCTCTTCCGATCTNNNTGAAACCCCGTCTGTACTGAA
AGCACACGTCTGAACTCTTTCCCTACACGACGCTCTTCCGATCTNNNGTGAGGCCCTGTCTCAAAAAA
GAGCACACGTCTGAACTCTTTCCCTACACGACGCTCTTCCGATCTNNNGCAAATCCTGAAAAGAGAC
AGCACACGTCTGAACTCTTTCCCTACACGACGCTCTTCCGATCTNNNGAGATTGCTCCATTACGCTCCA
GCACACGTCTGAACTCTTTCCCTACACGACGCTCTTCCGATCTNNNTTCACAGAGCATTGGAGACTGA
AGCACACGTCTGAACTCTTTCCCTACACGACGCTCTTCCGATCTNNNGTCTAGGGGACAGAGAGAG
AGCACACGTCTGAACTCTTTCCCTACACGACGCTCTTCCGATCTNNNCATCCTGGCTAACATGGTGA
AGCACACGTCTGAACTCTTTCCCTACACGACGCTCTTCCGATCTNNNCACCTGTAATCCCAGCTACTCA
GAGCACACGTCTGAACTCTTTCCCTACACGACGCTCTTCCGATCTNNNGCATGCCTATAATCCCAGCTA
GCACACGTCTGAACTCTTTCCCTACACGACGCTCTTCCGATCTNNNCTAAGGCAGGAGAATCGCTTG
AGCACACGTCTGAACTCTTTCCCTACACGACGCTCTTCCGATCTNNNAAAAAAAAAATTAGCCGGGC
AGCACACGTCTGAACTCTTTCCCTACACGACGCTCTTCCGATCTNNNATCCAAAAATTAGCTGGGCAT

AGCACACGTCTGAACTCTTTCCCTACACGACGCTCTTCCGATCTNNNGACTCAGTTCCCACATCCTAAG
GAGCACACGTCTGAACTCTTTCCCTACACGACGCTCTTCCGATCTNNNTGAGGCAGGAGAATCACTT
GCACACGTCTGAACTCTTTCCCTACACGACGCTCTTCCGATCTNNNAAATACAAAACGTTAGCCGGGT
GAGCACACGTCTGAACTCTTTCCCTACACGACGCTCTTCCGATCTNNNGAAGTCAGGAGATCGAGAC
GCACACGTCTGAACTCTTTCCCTACACGACGCTCTTCCGATCTNNNGTGACCCCAGATCATACCATTGT
AGCACACGTCTGAACTCTTTCCCTACACGACGCTCTTCCGATCTNNNTTGTCACCCTGAAAGAGGAAC
GCACACGTCTGAACTCTTTCCCTACACGACGCTCTTCCGATCTNNNGAGATCAAATCATGCCTCAGTG
AGCACACGTCTGAACTCTTTCCCTACACGACGCTCTTCCGATCTNNNCAGAGGTTACAGTGAGCTGA
AGCACACGTCTGAACTCTTTCCCTACACGACGCTCTTCCGATCTNNNTAATCTCAGCACTTTGGGAAG
AGCACACGTCTGAACTCTTTCCCTACACGACGCTCTTCCGATCTNNNCTGAGGCAGGAGAATCACTT
GCACACGTCTGAACTCTTTCCCTACACGACGCTCTTCCGATCTNNNCATAATCCCAGCTACTCAGGAG
GCACACGTCTGAACTCTTTCCCTACACGACGCTCTTCCGATCTNNNCAGGAGAATCACTTGAACCAGG
AGCACACGTCTGAACTCTTTCCCTACACGACGCTCTTCCGATCTNNNACACATGTAATCCCAGCTACTC
GCACACGTCTGAACTCTTTCCCTACACGACGCTCTTCCGATCTNNNCTGTAGTCCCAGCTACTTGGAA
GCACACGTCTGAACTCTTTCCCTACACGACGCTCTTCCGATCTNNNACACCTGTAATCCCAGCTACTTG
GCACACGTCTGAACTCTTTCCCTACACGACGCTCTTCCGATCTNNNAGTGCCAGAGTACCACATTCTTT
AGCACACGTCTGAACTCTTTCCCTACACGACGCTCTTCCGATCTNNNGAAAATCGCTTGAATCCAGGA
AGAGCACACGTCTGAACTCTTTCCCTACACGACGCTCTTCCGATCTNNNGAGGCATTAGAATCGCTTG
GCACACGTCTGAACTCTTTCCCTACACGACGCTCTTCCGATCTNNNCAGGCACCTATAGTCCCAGATTC
AGCACACGTCTGAACTCTTTCCCTACACGACGCTCTTCCGATCTNNNCAGGAGGCTGAGGTATGAGA
GAGCACACGTCTGAACTCTTTCCCTACACGACGCTCTTCCGATCTNNNTACAAAAATTAGCCAGGCGT
AGCACACGTCTGAACTCTTTCCCTACACGACGCTCTTCCGATCTNNNAGGAGGATGAAGCAGGAGA
GCACACGTCTGAACTCTTTCCCTACACGACGCTCTTCCGATCTNNNCATGCCTGTAATCCCAGCTACTC
GCACACGTCTGAACTCTTTCCCTACACGACGCTCTTCCGATCTNNNGACAGATTCGCTTTGTCTCAAC
AGCACACGTCTGAACTCTTTCCCTACACGACGCTCTTCCGATCTNNNATATGGCGAAACCCTGTCTCT
AGCACACGTCTGAACTCTTTCCCTACACGACGCTCTTCCGATCTNNNTGCCTATAAGTCCCAGCTACTC
GAGCACACGTCTGAACTCTTTCCCTACACGACGCTCTTCCGATCTNNNTTTGGGAGGCTGAGATAGT
GAGCACACGTCTGAACTCTTTCCCTACACGACGCTCTTCCGATCTNNNGTGAGGCAGGAGAATAGC
GAGCACACGTCTGAACTCTTTCCCTACACGACGCTCTTCCGATCTNNNAGACCCAGTCTCCCAAAAGA
GCACACGTCTGAACTCTTTCCCTACACGACGCTCTTCCGATCTNNNATAAACTAATTCTCAGCCGGGCA
AGCACACGTCTGAACTCTTTCCCTACACGACGCTCTTCCGATCTNNNAGCCAAGATCACACCATTGTA
AGCACACGTCTGAACTCTTTCCCTACACGACGCTCTTCCGATCTNNNTAAGGCAGGAGAATCGCTTGA
GCACACGTCTGAACTCTTTCCCTACACGACGCTCTTCCGATCTNNNGCCTAGGCAACAGAGGTAAAC
AGCACACGTCTGAACTCTTTCCCTACACGACGCTCTTCCGATCTNNNTCAATAACTGTGGCTGCATTTG
AGCACACGTCTGAACTCTTTCCCTACACGACGCTCTTCCGATCTNNNAAAAAAAAAAAAAGGTCGGG
AGCACACGTCTGAACTCTTTCCCTACACGACGCTCTTCCGATCTNNNGAATGACGTGAACCCAGGAG
GAGCACACGTCTGAACTCTTTCCCTACACGACGCTCTTCCGATCTNNNTGAGTGACAGAGCAAGACT
GCACACGTCTGAACTCTTTCCCTACACGACGCTCTTCCGATCTNNNGCAGGAGAATTGCTTGATCTTG
AGCACACGTCTGAACTCTTTCCCTACACGACGCTCTTCCGATCTNNNTGCCTGTAATCCCAGATATTCG
AGCACACGTCTGAACTCTTTCCCTACACGACGCTCTTCCGATCTNNNCTCCCAAGTAGCTGGTACTAC
AGCACACGTCTGAACTCTTTCCCTACACGACGCTCTTCCGATCTNNNCACCTGTAATCCCAGCTACTTG
GAGCACACGTCTGAACTCTTTCCCTACACGACGCTCTTCCGATCTNNNGTAGAAGCTGCAGTGAGTT
AGCACACGTCTGAACTCTTTCCCTACACGACGCTCTTCCGATCTNNNCACCTGTAATCCCAGCTACTCA

GAGCACACGTCTGAACTCTTTCCCTACACGACGCTCTTCCGATCTNNNGGAGACAGAGTGAGACTCCA
AGCACACGTCTGAACTCTTTCCCTACACGACGCTCTTCCGATCTNNNCTGAGGCAGGAGAATCACTTG
AGAGCACACGTCTGAACTCTTTCCCTACACGACGCTCTTCCGATCTNNNCCTGGGCAAAAAGAGCAAAA
GAGCACACGTCTGAACTCTTTCCCTACACGACGCTCTTCCGATCTNNNCACTTGAGGTCAGGAGTTCAA
GAGCACACGTCTGAACTCTTTCCCTACACGACGCTCTTCCGATCTNNNGAGGCAGGAGAATTGCTTGA
AGCACACGTCTGAACTCTTTCCCTACACGACGCTCTTCCGATCTNNNCCTGTAGTCTCAGCTACTCAGG
GCACACGTCTGAACTCTTTCCCTACACGACGCTCTTCCGATCTNNNTTGATATCCTCGGGAATGAGCA
GAGCACACGTCTGAACTCTTTCCCTACACGACGCTCTTCCGATCTNNNAAAAATTAGTTGGGTGTGGT
AGCACACGTCTGAACTCTTTCCCTACACGACGCTCTTCCGATCTNNNACAAAAATTAGCCAGGTGTGGT
AGCACACGTCTGAACTCTTTCCCTACACGACGCTCTTCCGATCTNNNCTGAGGCAGGAGAATCACTTGA
GAGCACACGTCTGAACTCTTTCCCTACACGACGCTCTTCCGATCTNNNATGACATAACTGCACTCCAAC
GAGCACACGTCTGAACTCTTTCCCTACACGACGCTCTTCCGATCTNNNTATGCCTGTAATCCCAGCTTCT
GAGCACACGTCTGAACTCTTTCCCTACACGACGCTCTTCCGATCTNNNACAAAAATTAGCCAGGTGTG
AGCACACGTCTGAACTCTTTCCCTACACGACGCTCTTCCGATCTNNNAGAATTGCTTGAACCCAGGAGA
AGAGCACACGTCTGAACTCTTTCCCTACACGACGCTCTTCCGATCTNNNTACTAGGGTGACTGAGGTGA
AGAGCACACGTCTGAACTCTTTCCCTACACGACGCTCTTCCGATCTNNNTAGTCCCAACTACTCAGGAG
GAGCACACGTCTGAACTCTTTCCCTACACGACGCTCTTCCGATCTNNNGTAACAGGCACCTATAGTCCCA
GAGCACACGTCTGAACTCTTTCCCTACACGACGCTCTTCCGATCTNNNCTGGGCAACAAAGATTCCATCT
GAGCACACGTCTGAACTCTTTCCCTACACGACGCTCTTCCGATCTNNNATGCCTGTAATCTCAGCTACT
GAGCACACGTCTGAACTCTTTCCCTACACGACGCTCTTCCGATCTNNNGAGAAGAAGCGCATCACTTCA
GCACACGTCTGAACTCTTTCCCTACACGACGCTCTTCCGATCTNNNCTGAGGCAGGAAAATTGCTTGA
AGAGCACACGTCTGAACTCTTTCCCTACACGACGCTCTTCCGATCTNNNGAATCACTTGAACCCAGGAG
GAGCACACGTCTGAACTCTTTCCCTACACGACGCTCTTCCGATCTNNNGGCGACAGTATGAGGATCCAT
AGAGCACACGTCTGAACTCTTTCCCTACACGACGCTCTTCCGATCTNNNAAGAATTGCTTGAACCTGGG
GAGCACACGTCTGAACTCTTTCCCTACACGACGCTCTTCCGATCTNNNACCTTGAAATTGGCCATGATG
AGCACACGTCTGAACTCTTTCCCTACACGACGCTCTTCCGATCTNNNGATTTAGAGACCCATTTGCTGG
AGCACACGTCTGAACTCTTTCCCTACACGACGCTCTTCCGATCTNNNTGAGGCAGGAGAATCACTTGA
AGCACACGTCTGAACTCTTTCCCTACACGACGCTCTTCCGATCTNNNTCTCCTAGGAGAGAAAAGAGG
GAGCACACGTCTGAACTCTTTCCCTACACGACGCTCTTCCGATCTNNNACAAAAATTAGTCGGGCATG
AGCACACGTCTGAACTCTTTCCCTACACGACGCTCTTCCGATCTNNNGATCACGAGGTCAGGAGATT
GAGCACACGTCTGAACTCTTTCCCTACACGACGCTCTTCCGATCTNNNCATGCCTGAACCTTCTACTGAC
GAGCACACGTCTGAACTCTTTCCCTACACGACGCTCTTCCGATCTNNNATGCCTGTAATCCCAGCTCTTA
AGAGCACACGTCTGAACTCTTTCCCTACACGACGCTCTTCCGATCTNNNTTTGCCAAACAGAGCTCTGAA
AGCACACGTCTGAACTCTTTCCCTACACGACGCTCTTCCGATCTNNNCTCAAGGGATGCTAAACAGTC
GAGCACACGTCTGAACTCTTTCCCTACACGACGCTCTTCCGATCTNNNAGAAGAGCAGGCAGTAAACCT
AGCACACGTCTGAACTCTTTCCCTACACGACGCTCTTCCGATCTNNNAGTGAGCTGAAATCATGCCATT
AGCACACGTCTGAACTCTTTCCCTACACGACGCTCTTCCGATCTNNNCGGATCACTTGAGATCAGGAGT
GAGCACACGTCTGAACTCTTTCCCTACACGACGCTCTTCCGATCTNNNGACAGGTGAATACAGACAGCT
GAGCACACGTCTGAACTCTTTCCCTACACGACGCTCTTCCGATCTNNNAGAAGTTGCAGTGAGCTGAGA
GAGCACACGTCTGAACTCTTTCCCTACACGACGCTCTTCCGATCTNNNAATACAAAAAATTAGCCGGGC
GAGCACACGTCTGAACTCTTTCCCTACACGACGCTCTTCCGATCTNNNTGAGGCAGGAGAATCACTTGA
AGCACACGTCTGAACTCTTTCCCTACACGACGCTCTTCCGATCTNNNTTCAGGCTTCTTCTTGCTGAGAT
AGAGCACACGTCTGAACTCTTTCCCTACACGACGCTCTTCCGATCTNNNAACATAGGAGATGGAGGTT

AGCACACGTCTGAACTCTTTCCCTACACGACGCTCTTCCGATCTNNNGAGATCACACCATTGCATTCCA
GAGCACACGTCTGAACTCTTTCCCTACACGACGCTCTTCCGATCTNNNTGTAGTCCTAGCTACTCAGG
GCACACGTCTGAACTCTTTCCCTACACGACGCTCTTCCGATCTNNNTTGCAGTGAGTCGAGATCTTAC
GAGCACACGTCTGAACTCTTTCCCTACACGACGCTCTTCCGATCTNNNCTGAGGCAGGAGAATCACTT
AGCACACGTCTGAACTCTTTCCCTACACGACGCTCTTCCGATCTNNNAAAATTAGCTGGGTGTAGTG
AGCACACGTCTGAACTCTTTCCCTACACGACGCTCTTCCGATCTNNNTGGATAAAGGTTAGGTGTGGT
GAGCACACGTCTGAACTCTTTCCCTACACGACGCTCTTCCGATCTNNNATCATTTGTGGTCAGGAGTT
AGCACACGTCTGAACTCTTTCCCTACACGACGCTCTTCCGATCTNNNGGATCACTTGAGGTCAGGAG
AGCACACGTCTGAACTCTTTCCCTACACGACGCTCTTCCGATCTNNNCTAGCTTTGGGGGTTGGTTTTT
AGCACACGTCTGAACTCTTTCCCTACACGACGCTCTTCCGATCTNNNAAGAATCACTTGAACCTGGGA
AGCACACGTCTGAACTCTTTCCCTACACGACGCTCTTCCGATCTNNNTACAAAATTAGCTGGGCATG
GCACACGTCTGAACTCTTTCCCTACACGACGCTCTTCCGATCTNNNGAGGCTGAGGCACTAGAATTTC
AGCACACGTCTGAACTCTTTCCCTACACGACGCTCTTCCGATCTNNNATGGCTAATTCTAGGACTTGG
GAGCACACGTCTGAACTCTTTCCCTACACGACGCTCTTCCGATCTNNNCTGAAGCAGGAGAATGGTG
AGCACACGTCTGAACTCTTTCCCTACACGACGCTCTTCCGATCTNNNTACAAAAAATTAGCCGGGCAT
AGCACACGTCTGAACTCTTTCCCTACACGACGCTCTTCCGATCTNNNCTGTAATCCCAGCTACTCAGG
AGCACACGTCTGAACTCTTTCCCTACACGACGCTCTTCCGATCTNNNCCAGGACCTAGTGGAATCATG
GCACACGTCTGAACTCTTTCCCTACACGACGCTCTTCCGATCTNNNTAGCTAATGTGTGCCTGTAGTC
AGCACACGTCTGAACTCTTTCCCTACACGACGCTCTTCCGATCTNNNATTGCTTGAGGCTAGGAGTTC
GCACACGTCTGAACTCTTTCCCTACACGACGCTCTTCCGATCTNNNCTGAGGCAGGAGAATGACATG
AGCACACGTCTGAACTCTTTCCCTACACGACGCTCTTCCGATCTNNNGAGATCGCTCCATCATACTCC
AGAGCACACGTCTGAACTCTTTCCCTACACGACGCTCTTCCGATCTNNNAGGCAGGATAATCACTTGA
AGCACACGTCTGAACTCTTTCCCTACACGACGCTCTTCCGATCTNNNGTAATCCCAGCTACTTAGGAG
GCACACGTCTGAACTCTTTCCCTACACGACGCTCTTCCGATCTNNNCTACTCAGAAGCTGATGTGGG
GAGCACACGTCTGAACTCTTTCCCTACACGACGCTCTTCCGATCTNNNGCATGCCTGTAATCCTAGCT
AGCACACGTCTGAACTCTTTCCCTACACGACGCTCTTCCGATCTNNNGATCCCAGGAGTATGACTTCA
AGCACACGTCTGAACTCTTTCCCTACACGACGCTCTTCCGATCTNNNGGGCCTTGGATGAATATTTTG
AGCACACGTCTGAACTCTTTCCCTACACGACGCTCTTCCGATCTNNNGTGAGCTGAGATCACACTAC
GCACACGTCTGAACTCTTTCCCTACACGACGCTCTTCCGATCTNNNAGTGCTGGGATTACAGACATG
GAGCACACGTCTGAACTCTTTCCCTACACGACGCTCTTCCGATCTNNNAGAATAACTTGAACCTGGGA
GCACACGTCTGAACTCTTTCCCTACACGACGCTCTTCCGATCTNNNGAAGTTTGCAGTGAGCCGAAA
GCACACGTCTGAACTCTTTCCCTACACGACGCTCTTCCGATCTNNNAAAGTGAAAAAGCCCCACAAC
AGCACACGTCTGAACTCTTTCCCTACACGACGCTCTTCCGATCTNNNGTAATCCCAGCTACTAGGAAG
AGCACACGTCTGAACTCTTTCCCTACACGACGCTCTTCCGATCTNNNCTCCTGGAATCTCTTAGAACC
AGCACACGTCTGAACTCTTTCCCTACACGACGCTCTTCCGATCTNNNCTGTAGTCCCAGCTACTTTGG
GAGCACACGTCTGAACTCTTTCCCTACACGACGCTCTTCCGATCTNNNCTGAGGCAGGAGAATCACTT
GAGCACACGTCTGAACTCTTTCCCTACACGACGCTCTTCCGATCTNNNGTAATCCTAGCTACTTGGGA
AGCACACGTCTGAACTCTTTCCCTACACGACGCTCTTCCGATCTNNNCAGGAGAATCACTTGAACCAG
GAGCACACGTCTGAACTCTTTCCCTACACGACGCTCTTCCGATCTNNNAATACAAAAAATTAGCCGG
GCACACGTCTGAACTCTTTCCCTACACGACGCTCTTCCGATCTNNNCTGAGGCAGGAGAATTGCTTG
GAGCACACGTCTGAACTCTTTCCCTACACGACGCTCTTCCGATCTNNNCAACAAGAGCATAACCTTGC
AGCACACGTCTGAACTCTTTCCCTACACGACGCTCTTCCGATCTNNNGGAGGTTGCAGTGAGTCATG
GAGCACACGTCTGAACTCTTTCCCTACACGACGCTCTTCCGATCTNNNGGATTGCTTAGAGCTCAGG

AGCACACGTCTGAACTCTTTCCCTACACGACGCTCTTCCGATCTNNNATGCAGGAGAAATTGCTTGAA
AGCACACGTCTGAACTCTTTCCCTACACGACGCTCTTCCGATCTNNNCAGGCAGATTACTTGAGGTCAG
AGAGCACACGTCTGAACTCTTTCCCTACACGACGCTCTTCCGATCTNNNCTGAGGCAGGAGAATCACTT
AGCACACGTCTGAACTCTTTCCCTACACGACGCTCTTCCGATCTNNNTGAGGCAGGAGAATCTCTTGAA
GAGCACACGTCTGAACTCTTTCCCTACACGACGCTCTTCCGATCTNNNAGCTTTACCTATAACACAGGG
AGAGCACACGTCTGAACTCTTTCCCTACACGACGCTCTTCCGATCTNNNTGAGGCAGAAGAATTGCTTG
AGCACACGTCTGAACTCTTTCCCTACACGACGCTCTTCCGATCTNNNGGTGACAGAGCAAGATTCCAT
GCACACGTCTGAACTCTTTCCCTACACGACGCTCTTCCGATCTNNNTCTCTGGGGCAGAAAGATCTTTC
AGAGCACACGTCTGAACTCTTTCCCTACACGACGCTCTTCCGATCTNNNGCACATGACTGTAACTCCAG
GAGCACACGTCTGAACTCTTTCCCTACACGACGCTCTTCCGATCTNNNGTAACACAGCAAAACCCAGAC
AGCACACGTCTGAACTCTTTCCCTACACGACGCTCTTCCGATCTNNNACACCTGTAGTCCCAGCTACTAG
GAGCACACGTCTGAACTCTTTCCCTACACGACGCTCTTCCGATCTNNNAGTTGCAGTGAGCTGAGATTC
GAGCACACGTCTGAACTCTTTCCCTACACGACGCTCTTCCGATCTNNNGAGATCATGCCACTTCACTCCA
GAGCACACGTCTGAACTCTTTCCCTACACGACGCTCTTCCGATCTNNNCTAGATAGCACCACTGCAGTCA
AGCACACGTCTGAACTCTTTCCCTACACGACGCTCTTCCGATCTNNNCCAGCCTGGATGATGGAGTAAA
GAGCACACGTCTGAACTCTTTCCCTACACGACGCTCTTCCGATCTNNNTTTGAGATCAGTCTGGGCAAC
GAGCACACGTCTGAACTCTTTCCCTACACGACGCTCTTCCGATCTNNNCACCTGTAATCCCAGCTACTTGG
AGCACACGTCTGAACTCTTTCCCTACACGACGCTCTTCCGATCTNNNGGTGACAGAGTGAGAACATGT
AGAGCACACGTCTGAACTCTTTCCCTACACGACGCTCTTCCGATCTNNNCTCTGAACAGACCAATAGCGA
GAGCACACGTCTGAACTCTTTCCCTACACGACGCTCTTCCGATCTNNNGCCTGTAATCTCAGCTACTCAG
AGCACACGTCTGAACTCTTTCCCTACACGACGCTCTTCCGATCTNNNAGGCCTGTAATCCCAGTACTTTG
AGAGCACACGTCTGAACTCTTTCCCTACACGACGCTCTTCCGATCTNNNGCAAAGACACACATAGGCTCA
GCACACGTCTGAACTCTTTCCCTACACGACGCTCTTCCGATCTNNNAATCACTTGAACCTAGGAGGCAC
GAGCACACGTCTGAACTCTTTCCCTACACGACGCTCTTCCGATCTNNNTCTACTATAGCATCTCCAAAGG
AGCACACGTCTGAACTCTTTCCCTACACGACGCTCTTCCGATCTNNNCTAAGGGCAGGAGAATCACTTG
GAGCACACGTCTGAACTCTTTCCCTACACGACGCTCTTCCGATCTNNNTCACATCTGTAATCCCAGCACA
AGCACACGTCTGAACTCTTTCCCTACACGACGCTCTTCCGATCTNNNGTGCCTGTAATCCCAGCTACTAG
GAGCACACGTCTGAACTCTTTCCCTACACGACGCTCTTCCGATCTNNNGCTTGTGTTGGCCCTTTTACTA
AGAGCACACGTCTGAACTCTTTCCCTACACGACGCTCTTCCGATCTNNNACAAGATGCCACAGGTCTCTA
AGAGCACACGTCTGAACTCTTTCCCTACACGACGCTCTTCCGATCTNNNCAACAAAGCGAAACTCTGTCT
AGAGCACACGTCTGAACTCTTTCCCTACACGACGCTCTTCCGATCTNNNATGGCTAATTCTAGAACTGG
GAGCACACGTCTGAACTCTTTCCCTACACGACGCTCTTCCGATCTNNNCCTGGGAGATGGAGATTACAC
AGCACACGTCTGAACTCTTTCCCTACACGACGCTCTTCCGATCTNNNCAGTTACTTAGGAAGCTGAGG
AGCACACGTCTGAACTCTTTCCCTACACGACGCTCTTCCGATCTNNNGTGAAACCCCGTCTCTACTAAAA
AGCACACGTCTGAACTCTTTCCCTACACGACGCTCTTCCGATCTNNNCCAGAGAGCAAGGAAGACTGA
AGCACACGTCTGAACTCTTTCCCTACACGACGCTCTTCCGATCTNNNGTAATCTCAGCTACTCAGGAGG
AGCACACGTCTGAACTCTTTCCCTACACGACGCTCTTCCGATCTNNNTCTATGAACGCCTAGTTGCTTCA
GAGCACACGTCTGAACTCTTTCCCTACACGACGCTCTTCCGATCTNNNCTGAGGCAGGAGAATCACTT
GAGCACACGTCTGAACTCTTTCCCTACACGACGCTCTTCCGATCTNNNCTCCCCAGGGTAAAGATGAAG
GAGCACACGTCTGAACTCTTTCCCTACACGACGCTCTTCCGATCTNNNGAGGGAAGGACAGATGAAG
AGAGCACACGTCTGAACTCTTTCCCTACACGACGCTCTTCCGATCTNNNAAGGAGCAGCAAGTCACATC
AGCACACGTCTGAACTCTTTCCCTACACGACGCTCTTCCGATCTNNNTCCAAATTCCACCACAGATTTCAC
AGCACACGTCTGAACTCTTTCCCTACACGACGCTCTTCCGATCTNNNCCAGCGAGTCTTTCTCCAGTAAA

GAGCACACGTCTGAACTCTTTCCCTACACGACGCTCTTCCGATCTNNNCTGTAATCCCAGCTACACAG
AGCACACGTCTGAACTCTTTCCCTACACGACGCTCTTCCGATCTNNNCACCTGTACTCCCAGCTATTCA
AGCACACGTCTGAACTCTTTCCCTACACGACGCTCTTCCGATCTNNNAAATAACTTCTCAGGCTGGGT
AGCACACGTCTGAACTCTTTCCCTACACGACGCTCTTCCGATCTNNNGTAGTCTCAGCTATTCAGGAG
GCACACGTCTGAACTCTTTCCCTACACGACGCTCTTCCGATCTNNNATTCCAGCCTGAGCAATTGAGA
CACACGTCTGAACTCTTTCCCTACACGACGCTCTTCCGATCTNNNGGTGACATAGCAAGACTCCGTAT
GAGCACACGTCTGAACTCTTTCCCTACACGACGCTCTTCCGATCTNNNGCACACACCTATGGTTCCAT
GAGCACACGTCTGAACTCTTTCCCTACACGACGCTCTTCCGATCTNNNGAGAATCGTTTGAACCCAGG
GAGCACACGTCTGAACTCTTTCCCTACACGACGCTCTTCCGATCTNNNCATGCCTGTAATCCCAGCTAC
AGCACACGTCTGAACTCTTTCCCTACACGACGCTCTTCCGATCTNNNAAGACCCATCAGTGTGCTGTA
GCACACGTCTGAACTCTTTCCCTACACGACGCTCTTCCGATCTNNNAAATTGGAGAAAGAGCATGCC
AGCACACGTCTGAACTCTTTCCCTACACGACGCTCTTCCGATCTNNNTATAGTCCCAGCTACTCAGGA
AGCACACGTCTGAACTCTTTCCCTACACGACGCTCTTCCGATCTNNNGCTGAGACAGGAGAATCACTT
AGCACACGTCTGAACTCTTTCCCTACACGACGCTCTTCCGATCTNNNTGAGACCAGGAATTCAAGAC
AGAGCACACGTCTGAACTCTTTCCCTACACGACGCTCTTCCGATCTNNNCTTAGGTGGGAGGATTGC
GAGCACACGTCTGAACTCTTTCCCTACACGACGCTCTTCCGATCTNNNCACTTGAGGTCAGGAGTTCA
AGCACACGTCTGAACTCTTTCCCTACACGACGCTCTTCCGATCTNNNCTGTAGTTCCAGCTACATGGG
GCACACGTCTGAACTCTTTCCCTACACGACGCTCTTCCGATCTNNNAGTACAAAAACTAGCCATGCGT
AGAGCACACGTCTGAACTCTTTCCCTACACGACGCTCTTCCGATCTNNNGGAGAATCACTTGAACCCA
GCACACGTCTGAACTCTTTCCCTACACGACGCTCTTCCGATCTNNNTACAAAAAATTAGCTGGGCGTG
GCACACGTCTGAACTCTTTCCCTACACGACGCTCTTCCGATCTNNNTAATCCTAGATACTCGGGAGGC
AGCACACGTCTGAACTCTTTCCCTACACGACGCTCTTCCGATCTNNNCTGAGGCAGGAGAATCACTT
AGCACACGTCTGAACTCTTTCCCTACACGACGCTCTTCCGATCTNNNAAACACACAACATTCCCTGCA
GAGCACACGTCTGAACTCTTTCCCTACACGACGCTCTTCCGATCTNNNCTGAGGTGGGAGGATTAC
GAGCACACGTCTGAACTCTTTCCCTACACGACGCTCTTCCGATCTNNNCTGAGGCAGGAGAATCACTT
GAGCACACGTCTGAACTCTTTCCCTACACGACGCTCTTCCGATCTNNNAATCACTTGAACTCAGGAGG
CACACGTCTGAACTCTTTCCCTACACGACGCTCTTCCGATCTNNNATACAAAAATTAGCCTGGCGTG
GAGCACACGTCTGAACTCTTTCCCTACACGACGCTCTTCCGATCTNNNAAAAAATTAGCCAGGTGTGG
GCACACGTCTGAACTCTTTCCCTACACGACGCTCTTCCGATCTNNNGAGGCAGGAAAATCACTTGAA
AGCACACGTCTGAACTCTTTCCCTACACGACGCTCTTCCGATCTNNNAGGTTACAGTGAGCTGAGAT
GAGCACACGTCTGAACTCTTTCCCTACACGACGCTCTTCCGATCTNNNGAAGTCGCAGTGAGCTAAG
AGCACACGTCTGAACTCTTTCCCTACACGACGCTCTTCCGATCTNNNGAGGCTGAAATGAGAGGATC
AGCACACGTCTGAACTCTTTCCCTACACGACGCTCTTCCGATCTNNNGAACCGAGATCAAGTCATTGC
AGCACACGTCTGAACTCTTTCCCTACACGACGCTCTTCCGATCTNNNGTTCAAAACCAACCTGGGCAA
AGCACACGTCTGAACTCTTTCCCTACACGACGCTCTTCCGATCTNNNAGAAATGGGAGGATCACTTG
AGAGCACACGTCTGAACTCTTTCCCTACACGACGCTCTTCCGATCTNNNAGGAGAATTGCTTGATCCT
GCACACGTCTGAACTCTTTCCCTACACGACGCTCTTCCGATCTNNNCATGCCAGGGATGCTTATTCAT
AGCACACGTCTGAACTCTTTCCCTACACGACGCTCTTCCGATCTNNNCATGGTGGCATGTGTCTGTAA
GCACACGTCTGAACTCTTTCCCTACACGACGCTCTTCCGATCTNNNCTGTAGTCCCAGCTACTCAGAA
AGAGCACACGTCTGAACTCTTTCCCTACACGACGCTCTTCCGATCTNNNGCAACATGATAAAGCCCCAT
GCACACGTCTGAACTCTTTCCCTACACGACGCTCTTCCGATCTNNNGAGGCAAGAGAATCGCTTGAA
GAGCACACGTCTGAACTCTTTCCCTACACGACGCTCTTCCGATCTNNNTCATGGAGGTGAGAAAGA
AGCACACGTCTGAACTCTTTCCCTACACGACGCTCTTCCGATCTNNNCTGTAGTCCCAGCTACTTTGG

GAGCACACGTCTGAACTCTTTCCCTACACGACGCTCTTCCGATCTNNNTTGAGGCAGGAGAATTGCTT
AGCACACGTCTGAACTCTTTCCCTACACGACGCTCTTCCGATCTNNNAACCAGAAGTATCGCTTGAAC
GAGCACACGTCTGAACTCTTTCCCTACACGACGCTCTTCCGATCTNNNGATCACAAGGTCAGGAGATC
AGCACACGTCTGAACTCTTTCCCTACACGACGCTCTTCCGATCTNNNGGTGACAAGGTGAGACTCTTT
AGCACACGTCTGAACTCTTTCCCTACACGACGCTCTTCCGATCTNNNAAAAAATTAGCAGGCGTAGTG
GAGCACACGTCTGAACTCTTTCCCTACACGACGCTCTTCCGATCTNNNCTTTACTGCAAAGTCAGGGTT
AGCACACGTCTGAACTCTTTCCCTACACGACGCTCTTCCGATCTNNNTAAAGAGTCAAGACCCATCCG
GAGCACACGTCTGAACTCTTTCCCTACACGACGCTCTTCCGATCTNNNCTGTAGTCACAGCTACTCAGG
GAGCACACGTCTGAACTCTTTCCCTACACGACGCTCTTCCGATCTNNNAAAAGACCCAGAATAGCCAAA
AGCACACGTCTGAACTCTTTCCCTACACGACGCTCTTCCGATCTNNNCTGTAATCCCAGCTACTCAGGA
GAGCACACGTCTGAACTCTTTCCCTACACGACGCTCTTCCGATCTNNNAACAATATGGCAGTGAGTTC
AGCACACGTCTGAACTCTTTCCCTACACGACGCTCTTCCGATCTNNNTTAGTGACTGGGGTATTTCTG
AGCACACGTCTGAACTCTTTCCCTACACGACGCTCTTCCGATCTNNNAAAATTAGCTGGGTGTGGTAG
GAGCACACGTCTGAACTCTTTCCCTACACGACGCTCTTCCGATCTNNNCAGAAGTTGTAGTGAGCCCA
GAGCACACGTCTGAACTCTTTCCCTACACGACGCTCTTCCGATCTNNNAATGTAAAGGAGCAGTCTCA
GAGCACACGTCTGAACTCTTTCCCTACACGACGCTCTTCCGATCTNNNCCTTTGAGCTCAGGAGTTTG
AGCACACGTCTGAACTCTTTCCCTACACGACGCTCTTCCGATCTNNNTTGAACCCAGAAGTAGGAGGT
GCACACGTCTGAACTCTTTCCCTACACGACGCTCTTCCGATCTNNNTGCGTATCTGTAATCCCAGCTAC
GAGCACACGTCTGAACTCTTTCCCTACACGACGCTCTTCCGATCTNNNGAGAATCGTTTGAACCCAGG
AGCACACGTCTGAACTCTTTCCCTACACGACGCTCTTCCGATCTNNNGCACTTGTAATCCCAGCTACTTG
AGCACACGTCTGAACTCTTTCCCTACACGACGCTCTTCCGATCTNNNGAGACAGGAGAATTGCTTGAA
AGCACACGTCTGAACTCTTTCCCTACACGACGCTCTTCCGATCTNNNCTGTAGTCCCAGCTACTCAGGA
AGCACACGTCTGAACTCTTTCCCTACACGACGCTCTTCCGATCTNNNCCATCATGACATTGAGACTCTG
AGAGCACACGTCTGAACTCTTTCCCTACACGACGCTCTTCCGATCTNNNTGTAGTCCCATCTACTCAGG
AGAGCACACGTCTGAACTCTTTCCCTACACGACGCTCTTCCGATCTNNNCCTTGAGAGAGGTTTCTGG
AGCACACGTCTGAACTCTTTCCCTACACGACGCTCTTCCGATCTNNNGAGCTGGTACCATTCCTTCTGA
AGCACACGTCTGAACTCTTTCCCTACACGACGCTCTTCCGATCTNNNTGCATAGCATGACTGAGTTGTC
AGCACACGTCTGAACTCTTTCCCTACACGACGCTCTTCCGATCTNNNCCAGCTATTTGGGAAGATGAG
AGCACACGTCTGAACTCTTTCCCTACACGACGCTCTTCCGATCTNNNGCCTGGGTGACAAAATGAGAT
GAGCACACGTCTGAACTCTTTCCCTACACGACGCTCTTCCGATCTNNNATTACAGTAGTCAGGCATGG
AGCACACGTCTGAACTCTTTCCCTACACGACGCTCTTCCGATCTNNNTGGTCTCTGGTTCTCTCTCATG
GAGCACACGTCTGAACTCTTTCCCTACACGACGCTCTTCCGATCTNNNTGTCACCTCGGTGTTATTCTT
AGCACACGTCTGAACTCTTTCCCTACACGACGCTCTTCCGATCTNNNGCGCTTCTATAATCCCAGCTACT
AGCACACGTCTGAACTCTTTCCCTACACGACGCTCTTCCGATCTNNNAGGCAGGAGAATAGCTTGAA
AGCACACGTCTGAACTCTTTCCCTACACGACGCTCTTCCGATCTNNNCCTCCTCTTTTCTGGGAATGTGT
AGCACACGTCTGAACTCTTTCCCTACACGACGCTCTTCCGATCTNNNGAGGTGGGAGGATCTTTTGAG
GAGCACACGTCTGAACTCTTTCCCTACACGACGCTCTTCCGATCTNNNCATGGTGGCTCATGCTTGTA
AGCACACGTCTGAACTCTTTCCCTACACGACGCTCTTCCGATCTNNNACAAAGCTGGGCTATAAAAACA
AGCACACGTCTGAACTCTTTCCCTACACGACGCTCTTCCGATCTNNNGAGGCTGGTCCAAAGCTAGTA
AGCACACGTCTGAACTCTTTCCCTACACGACGCTCTTCCGATCTNNNGCTAAGGCAGGAGAATCACTT
AGCACACGTCTGAACTCTTTCCCTACACGACGCTCTTCCGATCTNNNTCTCCTTTCTCTCTGAGCAAAC
GAGCACACGTCTGAACTCTTTCCCTACACGACGCTCTTCCGATCTNNNTGAGAGACCTTGACCCTGAA
GAGCACACGTCTGAACTCTTTCCCTACACGACGCTCTTCCGATCTNNNGATCATGAGGTCAGGAGAT

AGCACACGTCTGAACTCTTTCCCTACACGACGCTCTTCCGATCTNNNGCTGAGGAAGGAGAATGGTTT

AGCACACGTCTGAACTCTTTCCCTACACGACGCTCTTCCGATCTNNNGCTTCTCTTTTTCCCCAGGAGTA

AGCACACGTCTGAACTCTTTCCCTACACGACGCTCTTCCGATCTNNNCAGTCTGGGCAACATAGTGAGA

GAGCACACGTCTGAACTCTTTCCCTACACGACGCTCTTCCGATCTNNNATCCTGCTGCCAAGTACAACTA

GAGCACACGTCTGAACTCTTTCCCTACACGACGCTCTTCCGATCTNNNAAAAAATTAGCCAGGCTTGGT

GAGCACACGTCTGAACTCTTTCCCTACACGACGCTCTTCCGATCTNNNGGTAAAGTGCTGGCTCTAACT

AAGAGCACACGTCTGAACTCTTTCCCTACACGACGCTCTTCCGATCTNNNGCCAAGATCATGCCATTGT

GAGCACACGTCTGAACTCTTTCCCTACACGACGCTCTTCCGATCTNNNAAGAATTGCTTGAACCTGGGA

AGCACACGTCTGAACTCTTTCCCTACACGACGCTCTTCCGATCTNNNTGGAAGTCAGAAGGCCTAAGTT

GCACACGTCTGAACTCTTTCCCTACACGACGCTCTTCCGATCTNNNGGAGAATCGCTTGAACCTGGAA

GCACACGTCTGAACTCTTTCCCTACACGACGCTCTTCCGATCTNNNGCTGATGGAGATCAATAGAGCC

AGCACACGTCTGAACTCTTTCCCTACACGACGCTCTTCCGATCTNNNCCAAAAAAAGCCATGGACCAGA

AGCACACGTCTGAACTCTTTCCCTACACGACGCTCTTCCGATCTNNNCTCTTGCACTGAATTGTTCCTGG

AGCACACGTCTGAACTCTTTCCCTACACGACGCTCTTCCGATCTNNNAGATTGTGCCACTACACTCTAG

AGCACACGTCTGAACTCTTTCCCTACACGACGCTCTTCCGATCTNNNGATCACAAGGTCAGGAGATCG

GAGCACACGTCTGAACTCTTTCCCTACACGACGCTCTTCCGATCTNNNCTGAGGCAGGAGAATCACTT

AGAGCACACGTCTGAACTCTTTCCCTACACGACGCTCTTCCGATCTNNNAAAAAAAATTAGCCGGGTGT

GAGCACACGTCTGAACTCTTTCCCTACACGACGCTCTTCCGATCTNNNCTACTGCCCTCTCCCATGATTT

GAGCACACGTCTGAACTCTTTCCCTACACGACGCTCTTCCGATCTNNNAAAAATTAGTCAGGCATGGT

AGCACACGTCTGAACTCTTTCCCTACACGACGCTCTTCCGATCTNNNGCCTGTAATCCCATCTACTCTGG

AGCACACGTCTGAACTCTTTCCCTACACGACGCTCTTCCGATCTNNNACCAGAATCACTTGAACCCACAA

AGCACACGTCTGAACTCTTTCCCTACACGACGCTCTTCCGATCTNNNGGATTCCCCAGTAATCAAACTG

AGAGCACACGTCTGAACTCTTTCCCTACACGACGCTCTTCCGATCTNNNCTGAGGCAGGAGAATTGCTT

AGCACACGTCTGAACTCTTTCCCTACACGACGCTCTTCCGATCTNNNGTGAAACCCCGTCTCTACTAAAA

AGAGCACACGTCTGAACTCTTTCCCTACACGACGCTCTTCCGATCTNNNAGTGAATGGTTCTGTCTCACT

AGCACACGTCTGAACTCTTTCCCTACACGACGCTCTTCCGATCTNNNTTTTCCTCCCCAACACCTACTTCC

GAGCACACGTCTGAACTCTTTCCCTACACGACGCTCTTCCGATCTNNNCTGAAGTGAGAGGATCATTG

AGCACACGTCTGAACTCTTTCCCTACACGACGCTCTTCCGATCTNNNAGCATACTGTGTGTCCACTTGAT

AGCACACGTCTGAACTCTTTCCCTACACGACGCTCTTCCGATCTNNNGTTGTGCACATGTACCCTAGAA

GAGCACACGTCTGAACTCTTTCCCTACACGACGCTCTTCCGATCTNNNCACCTGTAGTCCCAGCTATTCA

AGCACACGTCTGAACTCTTTCCCTACACGACGCTCTTCCGATCTNNNAGCTAGCCCTGTTGACACAAATA

GAGCACACGTCTGAACTCTTTCCCTACACGACGCTCTTCCGATCTNNNCTGAGGCAGGAGAATCACTT

AGCACACGTCTGAACTCTTTCCCTACACGACGCTCTTCCGATCTNNNCATGATCACACCACTACACTCCA

GAGCACACGTCTGAACTCTTTCCCTACACGACGCTCTTCCGATCTNNNCAAACAAGCAAACAAAAGCCT

AGCACACGTCTGAACTCTTTCCCTACACGACGCTCTTCCGATCTNNNGTGGTCCTACCCAAAGAATCACT

GAGCACACGTCTGAACTCTTTCCCTACACGACGCTCTTCCGATCTNNNGCAAACTCTTCAGGCTCCTTTT

AGCACACGTCTGAACTCTTTCCCTACACGACGCTCTTCCGATCTNNNTAGGGCCAATCATTTTGCTTTG

GAGCACACGTCTGAACTCTTTCCCTACACGACGCTCTTCCGATCTNNNTTAGGGAAGTTCAGGGCATT

GAGCACACGTCTGAACTCTTTCCCTACACGACGCTCTTCCGATCTNNNTTAGGGAAGTTCAGGGCATT

AGCACACGTCTGAACTCTTTCCCTACACGACGCTCTTCCGATCTNNNTGTAGTTTCAGCTACTTGGGA

AGCACACGTCTGAACTCTTTCCCTACACGACGCTCTTCCGATCTNNNAGGCCTGACATATGTTTCACTGT

AGCACACGTCTGAACTCTTTCCCTACACGACGCTCTTCCGATCTNNNGAGGCAAGAGAATCACTTGAA

AGCACACGTCTGAACTCTTTCCCTACACGACGCTCTTCCGATCTNNNGGAGAATCACTTGAACCCAGGA

GAGCACACGTCTGAACTCTTTCCCTACACGACGCTCTTCCGATCTNNNCTGAGGTAGGAAGATGGCT
AGCACACGTCTGAACTCTTTCCCTACACGACGCTCTTCCGATCTNNNCTGAGACAGGAGAATCGCTTG
GAGCACACGTCTGAACTCTTTCCCTACACGACGCTCTTCCGATCTNNNCAGGAGAATTGCTTGAACCT
AGCACACGTCTGAACTCTTTCCCTACACGACGCTCTTCCGATCTNNNCACCTGTAATCCCAGCTACTTGG
GAGCACACGTCTGAACTCTTTCCCTACACGACGCTCTTCCGATCTNNNTGAACAGATATCACACCACTG
GCACACGTCTGAACTCTTTCCCTACACGACGCTCTTCCGATCTNNNTTGTCTTTGTGGGTTGCCTTTTT
AGCACACGTCTGAACTCTTTCCCTACACGACGCTCTTCCGATCTNNNAAAAAAATTAGCCAGGCGTG
GAGCACACGTCTGAACTCTTTCCCTACACGACGCTCTTCCGATCTNNNTACAAAAAATTAGCCGGGTG
AGAGCACACGTCTGAACTCTTTCCCTACACGACGCTCTTCCGATCTNNNGCTAACACGGTGAAACTCCA
AGAGCACACGTCTGAACTCTTTCCCTACACGACGCTCTTCCGATCTNNNAGAGAAGGGGTCTGATGA
AGCACACGTCTGAACTCTTTCCCTACACGACGCTCTTCCGATCTNNNCAGATCACGTGAGATCAGGA
AGCACACGTCTGAACTCTTTCCCTACACGACGCTCTTCCGATCTNNNGAAGCAGGAGAATGGCATG
AAGAGCACACGTCTGAACTCTTTCCCTACACGACGCTCTTCCGATCTNNNCTTGCAGTGAGTCGAGTTT
AGCACACGTCTGAACTCTTTCCCTACACGACGCTCTTCCGATCTNNNCTCGTGACATAGGGAGACTGT
AGCACACGTCTGAACTCTTTCCCTACACGACGCTCTTCCGATCTNNNTGGGAGAATCACTTGAACTCA
GAGCACACGTCTGAACTCTTTCCCTACACGACGCTCTTCCGATCTNNNGGGCCTGACTGTTAGAAGG
AGCACACGTCTGAACTCTTTCCCTACACGACGCTCTTCCGATCTNNNGTGATGGCACGCACTTATAAT
GAGCACACGTCTGAACTCTTTCCCTACACGACGCTCTTCCGATCTNNNCTGAGGCAGAGAATTGCTTG
AGCACACGTCTGAACTCTTTCCCTACACGACGCTCTTCCGATCTNNNCTGTAATCCCAGCTACTCAGG
AGCACACGTCTGAACTCTTTCCCTACACGACGCTCTTCCGATCTNNNGAGGTCAGAAGTTCAAAACCA
GAGCACACGTCTGAACTCTTTCCCTACACGACGCTCTTCCGATCTNNNAGCAGGAGAATCACTTGAA
AGCACACGTCTGAACTCTTTCCCTACACGACGCTCTTCCGATCTNNNCTGTAGTCCCAGCTACTCCTGA
AGCACACGTCTGAACTCTTTCCCTACACGACGCTCTTCCGATCTNNNCGCATACCTGTAATCCCAGCTA
GAGCACACGTCTGAACTCTTTCCCTACACGACGCTCTTCCGATCTNNNTGAATGGAGATTACACCACT
GAGCACACGTCTGAACTCTTTCCCTACACGACGCTCTTCCGATCTNNNGGAGGCAGAAATTTCAGTG
AGCACACGTCTGAACTCTTTCCCTACACGACGCTCTTCCGATCTNNNGAATCGTTTGAACCCAGGAA
AGCACACGTCTGAACTCTTTCCCTACACGACGCTCTTCCGATCTNNNCTGTAGTCCCAGCTACTCAGGA
AGCACACGTCTGAACTCTTTCCCTACACGACGCTCTTCCGATCTNNNGTAGATCACGAGGTCAGGAG
GAGCACACGTCTGAACTCTTTCCCTACACGACGCTCTTCCGATCTNNNCAACTCAAGGACCGGACAAA
AAGAGCACACGTCTGAACTCTTTCCCTACACGACGCTCTTCCGATCTNNNCTGGAGAATGGCGTGAA
AGCACACGTCTGAACTCTTTCCCTACACGACGCTCTTCCGATCTNNNGGACCCAAAATCAATGCACAA
GAGCACACGTCTGAACTCTTTCCCTACACGACGCTCTTCCGATCTNNNAACATGGCAAAACCCCATCTC
AGCACACGTCTGAACTCTTTCCCTACACGACGCTCTTCCGATCTNNNCTGTAGTCCCAGCTACTCAGG
AGCACACGTCTGAACTCTTTCCCTACACGACGCTCTTCCGATCTNNNTGGAAAAACCTAAAGACTCCA
AGCACACGTCTGAACTCTTTCCCTACACGACGCTCTTCCGATCTNNNTAAGGTAAAAAGCAAACGGCC
AGCACACGTCTGAACTCTTTCCCTACACGACGCTCTTCCGATCTNNNTATATGGAACCACAAAAGGCCC
AGCACACGTCTGAACTCTTTCCCTACACGACGCTCTTCCGATCTNNNACCAATGACGTTCTTCACAGA
GAGCACACGTCTGAACTCTTTCCCTACACGACGCTCTTCCGATCTNNNCTGAGGCAGGAGAATCACTT
AGCACACGTCTGAACTCTTTCCCTACACGACGCTCTTCCGATCTNNNGAGATCTCACCATTGCACTCCA
AGCACACGTCTGAACTCTTTCCCTACACGACGCTCTTCCGATCTNNNTGGATCCTGAGGTCAAGAGTT
GAGCACACGTCTGAACTCTTTCCCTACACGACGCTCTTCCGATCTNNNTCCTTTGAGCCATAAGAAACA
AGCACACGTCTGAACTCTTTCCCTACACGACGCTCTTCCGATCTNNNTACAAAAAATTAGCCTGGCG
AGCACACGTCTGAACTCTTTCCCTACACGACGCTCTTCCGATCTNNNAAAAAATTAGCTGGGCATGG

AGCACACGTCTGAACTCTTTCCCTACACGACGCTCTTCCGATCTNNNGGGTGGAAAATAAGGTCAGG
GAGCACACGTCTGAACTCTTTCCCTACACGACGCTCTTCCGATCTNNNCTGAGGCAGGAGAATGACTT
AGCACACGTCTGAACTCTTTCCCTACACGACGCTCTTCCGATCTNNNTTACTATGACTTGCTTCAGGGG
AGCACACGTCTGAACTCTTTCCCTACACGACGCTCTTCCGATCTNNNCTCCCAAGACTGAACCATGAA
GAGCACACGTCTGAACTCTTTCCCTACACGACGCTCTTCCGATCTNNNACTCAAGGCGTTATCATTTC
AGCACACGTCTGAACTCTTTCCCTACACGACGCTCTTCCGATCTNNNACACCTGTAATGTCAGCTACT
GAGCACACGTCTGAACTCTTTCCCTACACGACGCTCTTCCGATCTNNNTAGGAACAAAAGTGGGAGG
GAGCACACGTCTGAACTCTTTCCCTACACGACGCTCTTCCGATCTNNNCTTTTCCAAGTGGGTCCCAA
AGCACACGTCTGAACTCTTTCCCTACACGACGCTCTTCCGATCTNNNAAGACCCATCAGTGTGCTGTA
GAGCACACGTCTGAACTCTTTCCCTACACGACGCTCTTCCGATCTNNNAGAAAGAGGAGGAGGAGA
AGCACACGTCTGAACTCTTTCCCTACACGACGCTCTTCCGATCTNNNAACCTGAGAAGCAGAGGTTGT
AGCACACGTCTGAACTCTTTCCCTACACGACGCTCTTCCGATCTNNNTCCTGTAGTCTGAGCTACTTGG
GAGCACACGTCTGAACTCTTTCCCTACACGACGCTCTTCCGATCTNNNGCTGAGATGGTACCACTACA
AGCACACGTCTGAACTCTTTCCCTACACGACGCTCTTCCGATCTNNNCTGTAATCCAAGCTACTGGGG
AGCACACGTCTGAACTCTTTCCCTACACGACGCTCTTCCGATCTNNNACAAAAAATTAGCCGGGCATT
GAGCACACGTCTGAACTCTTTCCCTACACGACGCTCTTCCGATCTNNNATGACAATCACTTGAACCTG
AGCACACGTCTGAACTCTTTCCCTACACGACGCTCTTCCGATCTNNNTAGGAGTTTGAGGCTGCATG
GAGCACACGTCTGAACTCTTTCCCTACACGACGCTCTTCCGATCTNNNGTTGCAGTGAGCTGAGATCA
GCACACGTCTGAACTCTTTCCCTACACGACGCTCTTCCGATCTNNNATTCTTAGAGCCCAGGAGGTCA
GCACACGTCTGAACTCTTTCCCTACACGACGCTCTTCCGATCTNNNTACAAAAAATTAGCCGGGCATG
AGCACACGTCTGAACTCTTTCCCTACACGACGCTCTTCCGATCTNNNGCTAGCAGAGGGTGTACTAT
AGCACACGTCTGAACTCTTTCCCTACACGACGCTCTTCCGATCTNNNGAGAATCGCTTGATCCTTGGA
GCACACGTCTGAACTCTTTCCCTACACGACGCTCTTCCGATCTNNNCTGAGGCAGGAGAATGATGTG
AGCACACGTCTGAACTCTTTCCCTACACGACGCTCTTCCGATCTNNNCAAGTGACAGAGTGAGAACCT
GAGCACACGTCTGAACTCTTTCCCTACACGACGCTCTTCCGATCTNNNTGTGGGGAATTCAGGAAAGA
GAGCACACGTCTGAACTCTTTCCCTACACGACGCTCTTCCGATCTNNNAACACAAAAATTAGCTGGGG
GCACACGTCTGAACTCTTTCCCTACACGACGCTCTTCCGATCTNNNATACTAGCTACTTGGGAGGCTG
AGCACACGTCTGAACTCTTTCCCTACACGACGCTCTTCCGATCTNNNAGTGAGCCAAGATTCCATCACT
AGCACACGTCTGAACTCTTTCCCTACACGACGCTCTTCCGATCTNNNATTTCCAATAAACCAGCACCAG
GAGCACACGTCTGAACTCTTTCCCTACACGACGCTCTTCCGATCTNNNATGGTAGTACACGCCTGTAA
GAGCACACGTCTGAACTCTTTCCCTACACGACGCTCTTCCGATCTNNNGAGAATCCCTTGAACCCAGG
GCACACGTCTGAACTCTTTCCCTACACGACGCTCTTCCGATCTNNNTGTCTCTCTCACTCTCTCTCTCCC
GCACACGTCTGAACTCTTTCCCTACACGACGCTCTTCCGATCTNNNAGAATTGCTTGAACCCAGGAG
AGCACACGTCTGAACTCTTTCCCTACACGACGCTCTTCCGATCTNNNAGGAAAACTGCTTGAACCTAG
AGCACACGTCTGAACTCTTTCCCTACACGACGCTCTTCCGATCTNNNGCGGGTAGATCACAAGTTCAG
GAGCACACGTCTGAACTCTTTCCCTACACGACGCTCTTCCGATCTNNNAATACAAAACTTAGCTGGGC
GAGCACACGTCTGAACTCTTTCCCTACACGACGCTCTTCCGATCTNNNAATGCCTGTAATCCCAGCTAC
GAGCACACGTCTGAACTCTTTCCCTACACGACGCTCTTCCGATCTNNNGTGGATCACCTGAGTTCAGA
GAGCACACGTCTGAACTCTTTCCCTACACGACGCTCTTCCGATCTNNNCTAGGGATTTCAATGCTGCA
GCACACGTCTGAACTCTTTCCCTACACGACGCTCTTCCGATCTNNNCGTTAGTCCAGGAGTTCGAGAT
GAGCACACGTCTGAACTCTTTCCCTACACGACGCTCTTCCGATCTNNNCAGGCAACAGAACAAGACTC
AGCACACGTCTGAACTCTTTCCCTACACGACGCTCTTCCGATCTNNNCTCAACCCCATTTACAACAGCT
GAGCACACGTCTGAACTCTTTCCCTACACGACGCTCTTCCGATCTNNNCCAATCAAAAGACACAGAGT

AGCACACGTCTGAACTCTTTCCCTACACGACGCTCTTCCGATCTNNNCCCTGCTATAGTCAAGAACCCT
GCACACGTCTGAACTCTTTCCCTACACGACGCTCTTCCGATCTNNNAGAGAGCTCTTCATGCAGAAAC
AGCACACGTCTGAACTCTTTCCCTACACGACGCTCTTCCGATCTNNNTGAGTGACAAAGTGAGACACT
GCACACGTCTGAACTCTTTCCCTACACGACGCTCTTCCGATCTNNNCATGAGAATTGCTTGAACTCGG
AGCACACGTCTGAACTCTTTCCCTACACGACGCTCTTCCGATCTNNNGGCTTTGACTTGAAGCTTGGA
GAGCACACGTCTGAACTCTTTCCCTACACGACGCTCTTCCGATCTNNNGAGAGCTGTTGAAAAAGCCT
AGAGCACACGTCTGAACTCTTTCCCTACACGACGCTCTTCCGATCTNNNGCTCATGTCTGTAATCCCAG
GCACACGTCTGAACTCTTTCCCTACACGACGCTCTTCCGATCTNNNTGCCTGCATAGTACATGTTCAGT
AGCACACGTCTGAACTCTTTCCCTACACGACGCTCTTCCGATCTNNNCACCTGTAATCCCAGCTACTTG
GAGCACACGTCTGAACTCTTTCCCTACACGACGCTCTTCCGATCTNNNCAGTTACCACCTGTGTCCCAA
AGCACACGTCTGAACTCTTTCCCTACACGACGCTCTTCCGATCTNNNGAGAATCACTTGAACCCGAGA
AGCACACGTCTGAACTCTTTCCCTACACGACGCTCTTCCGATCTNNNCTGTAATCCCAGCTACTCCAGA
AGCACACGTCTGAACTCTTTCCCTACACGACGCTCTTCCGATCTNNNTGAGGCAGGAGAATCTCTTGA
GCACACGTCTGAACTCTTTCCCTACACGACGCTCTTCCGATCTNNNCTGTAATCCCAGCTACTCAGGAG
GAGCACACGTCTGAACTCTTTCCCTACACGACGCTCTTCCGATCTNNNTACCTGTAATCCCAGTGCTTT
AGCACACGTCTGAACTCTTTCCCTACACGACGCTCTTCCGATCTNNNCTGAGGCCAGAGAATTGCTTGA
GAGCACACGTCTGAACTCTTTCCCTACACGACGCTCTTCCGATCTNNNGCAAGGTCATTGTTCTTAGCC
GCACACGTCTGAACTCTTTCCCTACACGACGCTCTTCCGATCTNNNTCCCCAAGAGTTCAAGGTTGTAG
GAGCACACGTCTGAACTCTTTCCCTACACGACGCTCTTCCGATCTNNNCATAGGTTGCAGTGAGTGGA
AGCACACGTCTGAACTCTTTCCCTACACGACGCTCTTCCGATCTNNNCTTAGGAGTTTGATGCTGCAG
AGCACACGTCTGAACTCTTTCCCTACACGACGCTCTTCCGATCTNNNTAATCTCAGCACTTTGGGAGGT
AGCACACGTCTGAACTCTTTCCCTACACGACGCTCTTCCGATCTNNNTTTGGGTTGGTTCCAAGACTTT
AGCACACGTCTGAACTCTTTCCCTACACGACGCTCTTCCGATCTNNNAAAAAATTAGCCAGGCTTGGTC
AGCACACGTCTGAACTCTTTCCCTACACGACGCTCTTCCGATCTNNNTTTGAGGCTAGAAGTTCGAGA
AGCACACGTCTGAACTCTTTCCCTACACGACGCTCTTCCGATCTNNNCTGAGGCAGGAGAATCACTTG
AGCACACGTCTGAACTCTTTCCCTACACGACGCTCTTCCGATCTNNNAAGAAAAAAGACTGTGCAGGG
GCACACGTCTGAACTCTTTCCCTACACGACGCTCTTCCGATCTNNNAAGGAGAATCACTTGAACCTGG
AGCACACGTCTGAACTCTTTCCCTACACGACGCTCTTCCGATCTNNNAGGCAGGAGAATTACTTGAAC
GAGCACACGTCTGAACTCTTTCCCTACACGACGCTCTTCCGATCTNNNATCTCACACTATTGCCCAGAC
GAGCACACGTCTGAACTCTTTCCCTACACGACGCTCTTCCGATCTNNNAGCCTAAACATCACCACTATGC
AGCACACGTCTGAACTCTTTCCCTACACGACGCTCTTCCGATCTNNNAATACCAGTACTTTGGGAGGCT
AGCACACGTCTGAACTCTTTCCCTACACGACGCTCTTCCGATCTNNNCTGTAATCCCAGCTACTCAGGA
GCACACGTCTGAACTCTTTCCCTACACGACGCTCTTCCGATCTNNNGATGCCTGTAATCCCAGCTACTT
AGCACACGTCTGAACTCTTTCCCTACACGACGCTCTTCCGATCTNNNTGTTCAAGGCTCTGCACAATTT
AGCACACGTCTGAACTCTTTCCCTACACGACGCTCTTCCGATCTNNNAGTGAGCAGAGATTGTACCAC
GAGCACACGTCTGAACTCTTTCCCTACACGACGCTCTTCCGATCTNNNGTGAGAATTACTTGTACCCGG
AGCACACGTCTGAACTCTTTCCCTACACGACGCTCTTCCGATCTNNNGCGCCTGTAATCCTAGTTACTC
AGCACACGTCTGAACTCTTTCCCTACACGACGCTCTTCCGATCTNNNTTGGCAGAATAGTGTGAGTAG
AGAGCACACGTCTGAACTCTTTCCCTACACGACGCTCTTCCGATCTNNNCTGAGGCAGGAGAATTGC
GAGCACACGTCTGAACTCTTTCCCTACACGACGCTCTTCCGATCTNNNAGATTGCAGTGAGCTGAGAT
GAGCACACGTCTGAACTCTTTCCCTACACGACGCTCTTCCGATCTNNNGGAGAAATCACTTGAATCCGG
GCACACGTCTGAACTCTTTCCCTACACGACGCTCTTCCGATCTNNNACACCTGTAATCCCAGCTACTTG
GAGCACACGTCTGAACTCTTTCCCTACACGACGCTCTTCCGATCTNNNGGATCACTTGAGACCAGGAG

GCACACGTCTGAACTCTTTCCCTACACGACGCTCTTCCGATCTNNNGGTGACAGAGTGAGACTCCATC
GAGCACACGTCTGAACTCTTTCCCTACACGACGCTCTTCCGATCTNNNGAGGTTGCAATGAGTCGAGA
GAGCACACGTCTGAACTCTTTCCCTACACGACGCTCTTCCGATCTNNNCCAATCTGGACAACAAGAGCA
AGCACACGTCTGAACTCTTTCCCTACACGACGCTCTTCCGATCTNNNCTGGGTCTGTAACAGAGTGAG
GAGCACACGTCTGAACTCTTTCCCTACACGACGCTCTTCCGATCTNNNTAATCCCAGCTACTCAGGAGG
AGAGCACACGTCTGAACTCTTTCCCTACACGACGCTCTTCCGATCTNNNAATACAAAAAATTAGCCGG
GCACACGTCTGAACTCTTTCCCTACACGACGCTCTTCCGATCTNNNACATGGCAAAAGCCCATTCATAC
AAGAGCACACGTCTGAACTCTTTCCCTACACGACGCTCTTCCGATCTNNNAATCCCAGCTACTTGAGAG
GAGCACACGTCTGAACTCTTTCCCTACACGACGCTCTTCCGATCTNNNGCAGAGGTTGTAGTGAGCAA
AGCACACGTCTGAACTCTTTCCCTACACGACGCTCTTCCGATCTNNNGTCAAGAGATCGAGACCATCC
AGCACACGTCTGAACTCTTTCCCTACACGACGCTCTTCCGATCTNNNCAAAGATTAGTTGGGTGTGGT
AAGAGCACACGTCTGAACTCTTTCCCTACACGACGCTCTTCCGATCTNNNGCCTATAATCCTAGCTACT
AGCACACGTCTGAACTCTTTCCCTACACGACGCTCTTCCGATCTNNNTACAAAAAATTAGCCGGGCGTA
GAGCACACGTCTGAACTCTTTCCCTACACGACGCTCTTCCGATCTNNNAGGGAGAATTACCTGAACCCA
GAGCACACGTCTGAACTCTTTCCCTACACGACGCTCTTCCGATCTNNNGATCACGAGTTCAGGAGATCG
AGCACACGTCTGAACTCTTTCCCTACACGACGCTCTTCCGATCTNNNCACACACCTACAGTACCAGCTAC
AGCACACGTCTGAACTCTTTCCCTACACGACGCTCTTCCGATCTNNNAAGAATCACTTGAACCCAGGAG
GAGCACACGTCTGAACTCTTTCCCTACACGACGCTCTTCCGATCTNNNCTGTAATCCCAGCTACTCAGGA
GAGCACACGTCTGAACTCTTTCCCTACACGACGCTCTTCCGATCTNNNGTGTGGTGGTATTTCTGAGG
GAGCACACGTCTGAACTCTTTCCCTACACGACGCTCTTCCGATCTNNNCTGTAATCCCAGCTACTCAGGA
GAGCACACGTCTGAACTCTTTCCCTACACGACGCTCTTCCGATCTNNNGATGGTACTACTGCATTCCAG
AGCACACGTCTGAACTCTTTCCCTACACGACGCTCTTCCGATCTNNNACATGGTGAAACTCCGTCTCTA
AGCACACGTCTGAACTCTTTCCCTACACGACGCTCTTCCGATCTNNNATACAAAAATTAGCTGGGCGTG
AGCACACGTCTGAACTCTTTCCCTACACGACGCTCTTCCGATCTNNNAATCCCAGCTACTTGGAAGTCTC
AGCACACGTCTGAACTCTTTCCCTACACGACGCTCTTCCGATCTNNNTACTGGAGACACATCAAAGCAC
GAGCACACGTCTGAACTCTTTCCCTACACGACGCTCTTCCGATCTNNNAGAAAGGAAGCATCACATAG
GCACACGTCTGAACTCTTTCCCTACACGACGCTCTTCCGATCTNNNGAATGTTGCAGTGAGCTGAGATC
GAGCACACGTCTGAACTCTTTCCCTACACGACGCTCTTCCGATCTNNNACATAGCAAGACCTCATCTCTG
AGCACACGTCTGAACTCTTTCCCTACACGACGCTCTTCCGATCTNNNAGAGCAGTGTGTCCTCATCTTTT
GAGCACACGTCTGAACTCTTTCCCTACACGACGCTCTTCCGATCTNNNAAGAATGAAAGCATTGGGAC
GAGCACACGTCTGAACTCTTTCCCTACACGACGCTCTTCCGATCTNNNCTGTAGTTCCAGCTACTCAGGA
AGAGCACACGTCTGAACTCTTTCCCTACACGACGCTCTTCCGATCTNNNATGGACACAGGAAGGGAA
AGCACACGTCTGAACTCTTTCCCTACACGACGCTCTTCCGATCTNNNGTAGTACATGCCTGTAGTCCCA
GAGCACACGTCTGAACTCTTTCCCTACACGACGCTCTTCCGATCTNNNCATCCCTGTAATCCCAGCTACT
AGAGCACACGTCTGAACTCTTTCCCTACACGACGCTCTTCCGATCTNNNGTTGTAGTGAGCTGAGATTC
AGAGCACACGTCTGAACTCTTTCCCTACACGACGCTCTTCCGATCTNNNCAGGAGTTTGAGATCACAGC
AGCACACGTCTGAACTCTTTCCCTACACGACGCTCTTCCGATCTNNNCTGTCACCTCTCAATTTGAGCCTC
AGCACACGTCTGAACTCTTTCCCTACACGACGCTCTTCCGATCTNNNTAGGAGATTCCCTCAAGTGCCT
GAGCACACGTCTGAACTCTTTCCCTACACGACGCTCTTCCGATCTNNNCTGTAGTTTCAGCTACTCGGG
GAGCACACGTCTGAACTCTTTCCCTACACGACGCTCTTCCGATCTNNNTGAGGCTACAATGAGCTGTG
AGCACACGTCTGAACTCTTTCCCTACACGACGCTCTTCCGATCTNNNACAAAAATTAGCTGGGCATGGT
AGAGCACACGTCTGAACTCTTTCCCTACACGACGCTCTTCCGATCTNNNCCTGGTTGTAATCCCAGCTA
AGCACACGTCTGAACTCTTTCCCTACACGACGCTCTTCCGATCTNNNGAGAATCACTTGAACCAGGAGG

AGCACACGTCTGAACTCTTTCCCTACACGACGCTCTTCCGATCTNNNATATAATCTGCCCCCTTGTGGAC

GAGCACACGTCTGAACTCTTTCCCTACACGACGCTCTTCCGATCTNNNTGTAATTCCAGCTACTTGAGG

AGCACACGTCTGAACTCTTTCCCTACACGACGCTCTTCCGATCTNNNGGCCCCAGTAGCTTCTTTGTAAA

AGCACACGTCTGAACTCTTTCCCTACACGACGCTCTTCCGATCTNNNATTGCCTGAGCTCAGTAGTTCA

AGAGCACACGTCTGAACTCTTTCCCTACACGACGCTCTTCCGATCTNNNGAGAATTGCTTGAACCTGGG

AGAGCACACGTCTGAACTCTTTCCCTACACGACGCTCTTCCGATCTNNNTTGGAAAGCTAAGGCAGGA

AGAGCACACGTCTGAACTCTTTCCCTACACGACGCTCTTCCGATCTNNNGAGGCAGGAGAATTGCTTGA

AGAGCACACGTCTGAACTCTTTCCCTACACGACGCTCTTCCGATCTNNNATGTAGTCCCACCTACTCTGG

AGAGCACACGTCTGAACTCTTTCCCTACACGACGCTCTTCCGATCTNNNCTGAGATTGCACCACTGTACT

GAGCACACGTCTGAACTCTTTCCCTACACGACGCTCTTCCGATCTNNNGTGCCTGTAATCCCAGTTACT

AGCACACGTCTGAACTCTTTCCCTACACGACGCTCTTCCGATCTNNNTGTAATCTCAGCACTTTGGGAG

AGCACACGTCTGAACTCTTTCCCTACACGACGCTCTTCCGATCTNNNCAGAAGAATTGCTTGAACCTG

AGAGCACACGTCTGAACTCTTTCCCTACACGACGCTCTTCCGATCTNNNGTAATCCCAGCTACTTAGGA

AGCACACGTCTGAACTCTTTCCCTACACGACGCTCTTCCGATCTNNNTAATCTCAGCTACTCAGGAGGCT

AGCACACGTCTGAACTCTTTCCCTACACGACGCTCTTCCGATCTNNNAGTGACCAGAGCAAGTATGAA

GAGCACACGTCTGAACTCTTTCCCTACACGACGCTCTTCCGATCTNNNGCCTATAATCCCAGCTACTCAG

GAGCACACGTCTGAACTCTTTCCCTACACGACGCTCTTCCGATCTNNNAGTAAGCTGAGATCACACCAC

AGCACACGTCTGAACTCTTTCCCTACACGACGCTCTTCCGATCTNNNGCGCTTATAATCCCAGCTACTCA

AGCACACGTCTGAACTCTTTCCCTACACGACGCTCTTCCGATCTNNNAAAAAATTAGCTGGGTGTGGT

AGCACACGTCTGAACTCTTTCCCTACACGACGCTCTTCCGATCTNNNGAGAATCACTTGAACCTGGGA

GAGCACACGTCTGAACTCTTTCCCTACACGACGCTCTTCCGATCTNNNGGCTCACTCCTGTAATTCTAG

GAGCACACGTCTGAACTCTTTCCCTACACGACGCTCTTCCGATCTNNNCACACCTGTAGTCTCAGCTACT

AAGAGCACACGTCTGAACTCTTTCCCTACACGACGCTCTTCCGATCTNNNGGAAGATTACTTGAGCCTG

GAGCACACGTCTGAACTCTTTCCCTACACGACGCTCTTCCGATCTNNNCTGAGGCAGGAGAATCACTTG

GAGCACACGTCTGAACTCTTTCCCTACACGACGCTCTTCCGATCTNNNCTGTAGTCCCAGCTACTCAGGA

AGCACACGTCTGAACTCTTTCCCTACACGACGCTCTTCCGATCTNNNAGAATCACTTGAACCCTGGAGA

GAGCACACGTCTGAACTCTTTCCCTACACGACGCTCTTCCGATCTNNNTGAAACCCTGTCTCTACTGCAA

GAGCACACGTCTGAACTCTTTCCCTACACGACGCTCTTCCGATCTNNNGTTGAGACGGGAATTGCTTG

AGAGCACACGTCTGAACTCTTTCCCTACACGACGCTCTTCCGATCTNNNAGAGATTGCAGTGAGCTGA

AGCACACGTCTGAACTCTTTCCCTACACGACGCTCTTCCGATCTNNNCACCTGTAGTCCCAGCTTAGTTG

GAGCACACGTCTGAACTCTTTCCCTACACGACGCTCTTCCGATCTNNNTGTGTGTCTGTAATCCCAGCT

GAGCACACGTCTGAACTCTTTCCCTACACGACGCTCTTCCGATCTNNNACAAGTAGGGAGGTGATGAT

GAGCACACGTCTGAACTCTTTCCCTACACGACGCTCTTCCGATCTNNNAGATCACACCACTGAACTCTA

AGCACACGTCTGAACTCTTTCCCTACACGACGCTCTTCCGATCTNNNCTGTAATCCCAGCTACTCAGGA

GAGCACACGTCTGAACTCTTTCCCTACACGACGCTCTTCCGATCTNNNGGTGATAAAGCGAGACTCAG

AGAGCACACGTCTGAACTCTTTCCCTACACGACGCTCTTCCGATCTNNNTGCACCTATAGTCCCAGCTAC

AGCACACGTCTGAACTCTTTCCCTACACGACGCTCTTCCGATCTNNNACAAAAATTAGCTGGGCATGGT

GCACACGTCTGAACTCTTTCCCTACACGACGCTCTTCCGATCTNNNGGTGACAGAGTGAGATTTGGT

GAGCACACGTCTGAACTCTTTCCCTACACGACGCTCTTCCGATCTNNNGAGAATCGTTTGAACCCAAGA

GAGCACACGTCTGAACTCTTTCCCTACACGACGCTCTTCCGATCTNNNTACAAACTACTCAGGAGGCTCA

GAGCACACGTCTGAACTCTTTCCCTACACGACGCTCTTCCGATCTNNNCCAACATGGTGAAACTCCATCT

AGAGCACACGTCTGAACTCTTTCCCTACACGACGCTCTTCCGATCTNNNAGGTTGCAGTGAGATGTGAT

GAGCACACGTCTGAACTCTTTCCCTACACGACGCTCTTCCGATCTNNNTCCCTTGAACCCAGAAGATCA

GAGCACACGTCTGAACTCTTTCCCTACACGACGCTCTTCCGATCTNNNAATAAAAAAGTCAAGGGTG
AGCACACGTCTGAACTCTTTCCCTACACGACGCTCTTCCGATCTNNNCCTCTGTCCCAAAATGCTAGG
AGCACACGTCTGAACTCTTTCCCTACACGACGCTCTTCCGATCTNNNATTTGGTCAGGAGGGTATGG
AGCACACGTCTGAACTCTTTCCCTACACGACGCTCTTCCGATCTNNNGGGTGACAGAGTGAGATTCC
AGCACACGTCTGAACTCTTTCCCTACACGACGCTCTTCCGATCTNNNTGAGGCAGGAGAATCACTTGA
AGCACACGTCTGAACTCTTTCCCTACACGACGCTCTTCCGATCTNNNGCAATGAGAGTGAGACTCCA
AGCACACGTCTGAACTCTTTCCCTACACGACGCTCTTCCGATCTNNNGGCTACAGAGGGAGACTACAT
GAGCACACGTCTGAACTCTTTCCCTACACGACGCTCTTCCGATCTNNNGCAAGAGAATCACTTGAACC
AGCACACGTCTGAACTCTTTCCCTACACGACGCTCTTCCGATCTNNNCATGCACCTGTAGTCCCATCTA
AGCACACGTCTGAACTCTTTCCCTACACGACGCTCTTCCGATCTNNNCTGAGGCATGAGAATCCTTCG
GAGCACACGTCTGAACTCTTTCCCTACACGACGCTCTTCCGATCTNNNGATCACTTGAGCTCAGGAGT
AGCACACGTCTGAACTCTTTCCCTACACGACGCTCTTCCGATCTNNNTGTAGTCCCAGCTACTAGAGA
AGCACACGTCTGAACTCTTTCCCTACACGACGCTCTTCCGATCTNNNCTGGACAACATAGCAAAACCC
AGCACACGTCTGAACTCTTTCCCTACACGACGCTCTTCCGATCTNNNACAGCAACATCCCACTTCTAGT
AGCACACGTCTGAACTCTTTCCCTACACGACGCTCTTCCGATCTNNNGTACCTGTAGTCCCAGCTACT
AGCACACGTCTGAACTCTTTCCCTACACGACGCTCTTCCGATCTNNNAGGAACATTTCTCTCATGCCA
AGCACACGTCTGAACTCTTTCCCTACACGACGCTCTTCCGATCTNNNTGGATCACAAGGTCAGGAAAT
AGCACACGTCTGAACTCTTTCCCTACACGACGCTCTTCCGATCTNNNCTGAGGCAGGAGAATCACTT
GAGCACACGTCTGAACTCTTTCCCTACACGACGCTCTTCCGATCTNNNGAGTTTGAGACCAGCCTAG
GAGCACACGTCTGAACTCTTTCCCTACACGACGCTCTTCCGATCTNNNCTTGCAATGAGCTGAGATCA
GAGCACACGTCTGAACTCTTTCCCTACACGACGCTCTTCCGATCTNNNAGACAGGAGAATGGCGTGA
GCACACGTCTGAACTCTTTCCCTACACGACGCTCTTCCGATCTNNNCAACATGAGCGAAACTCTGTCT
AGCACACGTCTGAACTCTTTCCCTACACGACGCTCTTCCGATCTNNNGTTAGGAGTTCAAAACCAGC
GAGCACACGTCTGAACTCTTTCCCTACACGACGCTCTTCCGATCTNNNTTGAACCTGGGAGATGGAG
AGCACACGTCTGAACTCTTTCCCTACACGACGCTCTTCCGATCTNNNATATAAAAAGTTAGCCGGGC
AGCACACGTCTGAACTCTTTCCCTACACGACGCTCTTCCGATCTNNNGGATCACTTGAGGTCAGGAGT
AGCACACGTCTGAACTCTTTCCCTACACGACGCTCTTCCGATCTNNNTGAGGCAGGAGAATGGTATG
GAGCACACGTCTGAACTCTTTCCCTACACGACGCTCTTCCGATCTNNNAACTCAGGAGGTAGATTGCA
AGAGCACACGTCTGAACTCTTTCCCTACACGACGCTCTTCCGATCTNNNTTACAGTGAGCCAAGATCG
AGCACACGTCTGAACTCTTTCCCTACACGACGCTCTTCCGATCTNNNCAGAAGAATTGCTTGAACCTG
AGCACACGTCTGAACTCTTTCCCTACACGACGCTCTTCCGATCTNNNTGTAATCCCAGCTACTCAGGA
AGCACACGTCTGAACTCTTTCCCTACACGACGCTCTTCCGATCTNNNCAAGAGGTTGAAGCAGCAGTT
AGCACACGTCTGAACTCTTTCCCTACACGACGCTCTTCCGATCTNNNCAATGAGGATGGTAGGGTG
GAGCACACGTCTGAACTCTTTCCCTACACGACGCTCTTCCGATCTNNNGCCTGTAATCCCATCTACTCA
GAGCACACGTCTGAACTCTTTCCCTACACGACGCTCTTCCGATCTNNNGCGCAACTGTAATCTCAGCT
GAGCACACGTCTGAACTCTTTCCCTACACGACGCTCTTCCGATCTNNNCGGATCATGAGGTCAAGAG
GCACACGTCTGAACTCTTTCCCTACACGACGCTCTTCCGATCTNNNACAAGTCACAGTACAGTAGGCT
GCACACGTCTGAACTCTTTCCCTACACGACGCTCTTCCGATCTNNNTACAAAAAATTAGCCGGGCATG
AGCACACGTCTGAACTCTTTCCCTACACGACGCTCTTCCGATCTNNNGAAGGTTACAGTGAGCCGAG
GAGCACACGTCTGAACTCTTTCCCTACACGACGCTCTTCCGATCTNNNGAAGTCCAGGATGCAGAGC
GAGCACACGTCTGAACTCTTTCCCTACACGACGCTCTTCCGATCTNNNACAAAAAATAGCCAGGTGTG
AGCACACGTCTGAACTCTTTCCCTACACGACGCTCTTCCGATCTNNNCCTCCTCTCCATCAAAGTCTGA
AGCACACGTCTGAACTCTTTCCCTACACGACGCTCTTCCGATCTNNNTACAAAAAATTAGCCGGGCAT

AGCACACGTCTGAACTCTTTCCCTACACGACGCTCTTCCGATCTNNNTTTGTAGTGAGCTCAGATCGT
GAGCACACGTCTGAACTCTTTCCCTACACGACGCTCTTCCGATCTNNNAGATCGAGACCATCATGGCT
AGCACACGTCTGAACTCTTTCCCTACACGACGCTCTTCCGATCTNNNAAAAAAATTAGCTGGGCACGG
AGCACACGTCTGAACTCTTTCCCTACACGACGCTCTTCCGATCTNNNCTGAGGCAGGAGAATCACTTG
GAGCACACGTCTGAACTCTTTCCCTACACGACGCTCTTCCGATCTNNNACTTGAGGTCAGGAGTTTGA
AGCACACGTCTGAACTCTTTCCCTACACGACGCTCTTCCGATCTNNNTGAAAAAGCTCTTGAAGACGC
AGCACACGTCTGAACTCTTTCCCTACACGACGCTCTTCCGATCTNNNCTGTCATATCAGCTACTCGGGA
AGCACACGTCTGAACTCTTTCCCTACACGACGCTCTTCCGATCTNNNCTGTCATATCAGCTACTCGGG
GAGCACACGTCTGAACTCTTTCCCTACACGACGCTCTTCCGATCTNNNGATCACGAGGTCAGGAGAT
AGCACACGTCTGAACTCTTTCCCTACACGACGCTCTTCCGATCTNNNTGCATGACATACTGCAATTGA
AGCACACGTCTGAACTCTTTCCCTACACGACGCTCTTCCGATCTNNNATGTGCTTGTAATCCCAGCTAC
AGCACACGTCTGAACTCTTTCCCTACACGACGCTCTTCCGATCTNNNATACAAAAATTAGCCGGGCAT
AGCACACGTCTGAACTCTTTCCCTACACGACGCTCTTCCGATCTNNNTTTGGGACTCACACTCCTAGA
GAGCACACGTCTGAACTCTTTCCCTACACGACGCTCTTCCGATCTNNNCTGTAATCCCAGCTACTCGGA
AGCACACGTCTGAACTCTTTCCCTACACGACGCTCTTCCGATCTNNNTACTTGAGCCTAGGAGGTAG
GCACACGTCTGAACTCTTTCCCTACACGACGCTCTTCCGATCTNNNGCTCTTTTGATGACCTACTCCCA
GCACACGTCTGAACTCTTTCCCTACACGACGCTCTTCCGATCTNNNCCTTTTACCCCAAACAAACCCTC
AGCACACGTCTGAACTCTTTCCCTACACGACGCTCTTCCGATCTNNNACTCTATTCTGTTGCACTGGTC
GAGCACACGTCTGAACTCTTTCCCTACACGACGCTCTTCCGATCTNNNGAGAATGGCATGAACCTGG
GCACACGTCTGAACTCTTTCCCTACACGACGCTCTTCCGATCTNNNGTATATGGGGTGGAGTTAGCC
AGCACACGTCTGAACTCTTTCCCTACACGACGCTCTTCCGATCTNNNACGCCTGTAATCCCAGTAATC
CACACGTCTGAACTCTTTCCCTACACGACGCTCTTCCGATCTNNNTGGATTCACACTTTTCCCTCTACA
GCACACGTCTGAACTCTTTCCCTACACGACGCTCTTCCGATCTNNNAATGGGTTCTTGTCATACGACC
GAGCACACGTCTGAACTCTTTCCCTACACGACGCTCTTCCGATCTNNNGCAAGCTCAAACTCCATAAA
AGCACACGTCTGAACTCTTTCCCTACACGACGCTCTTCCGATCTNNNCTGTAGTCCCAGCTACTCAGGA
AGCACACGTCTGAACTCTTTCCCTACACGACGCTCTTCCGATCTNNNTAAGGTGGAAGGATTGCTTG
GAGCACACGTCTGAACTCTTTCCCTACACGACGCTCTTCCGATCTNNNCCCAAGTGCTTACAAGGATC
GAGCACACGTCTGAACTCTTTCCCTACACGACGCTCTTCCGATCTNNNCGAAGGTCACCAACATCAAA
GAGCACACGTCTGAACTCTTTCCCTACACGACGCTCTTCCGATCTNNNGGACGAGTGGGGTAATAAG
AGCACACGTCTGAACTCTTTCCCTACACGACGCTCTTCCGATCTNNNGCACTCATATGTTCATTGCAG
GAGCACACGTCTGAACTCTTTCCCTACACGACGCTCTTCCGATCTNNNAAAAATTAGCCAAGCATCGT
GCACACGTCTGAACTCTTTCCCTACACGACGCTCTTCCGATCTNNNTGCCCATTCAGTATAATGTTGG
GAGCACACGTCTGAACTCTTTCCCTACACGACGCTCTTCCGATCTNNNTATGTGCCTGTAATCCCAGCT
GCACACGTCTGAACTCTTTCCCTACACGACGCTCTTCCGATCTNNNGGAATAGTGATTGGGAAGTGG
GAGCACACGTCTGAACTCTTTCCCTACACGACGCTCTTCCGATCTNNNAAAAGAACCCACATAGCCAA
GAGCACACGTCTGAACTCTTTCCCTACACGACGCTCTTCCGATCTNNNTTCCCATTCTCCCATCCCTATC
AGAGCACACGTCTGAACTCTTTCCCTACACGACGCTCTTCCGATCTNNNAAAAAAAAATTAGCCGGGC
GCACACGTCTGAACTCTTTCCCTACACGACGCTCTTCCGATCTNNNAAGAATAGCTTGAACCCAGGA
AGCACACGTCTGAACTCTTTCCCTACACGACGCTCTTCCGATCTNNNCTGTGATCACTCTCAAGACACC
GCACACGTCTGAACTCTTTCCCTACACGACGCTCTTCCGATCTNNNGAGACACAGTTCAGTCCATAGC
GAGCACACGTCTGAACTCTTTCCCTACACGACGCTCTTCCGATCTNNNCCTCCTTGTCATTGATCTTTG
AGCACACGTCTGAACTCTTTCCCTACACGACGCTCTTCCGATCTNNNCTGTAATCCCAGCTACTCGAG
AGCACACGTCTGAACTCTTTCCCTACACGACGCTCTTCCGATCTNNNGCTCATGCCTGTAATCTCATCA

AGCACACGTCTGAACTCTTTCCCTACACGACGCTCTTCCGATCTNNNGTCATACTTTGGGTGAAACAG
AGCACACGTCTGAACTCTTTCCCTACACGACGCTCTTCCGATCTNNNAGGTTTCAGTAAGCCAAGATC
GAGCACACGTCTGAACTCTTTCCCTACACGACGCTCTTCCGATCTNNNCATAGCGTGACTCCTGTCTCT
AGCACACGTCTGAACTCTTTCCCTACACGACGCTCTTCCGATCTNNNCACCTGTAATCCCAGCTACTCA
AGCACACGTCTGAACTCTTTCCCTACACGACGCTCTTCCGATCTNNNCTGAGGCAGGAGAATAGCTTG
AGCACACGTCTGAACTCTTTCCCTACACGACGCTCTTCCGATCTNNNGGAAAACAGAGAGGGAACAA
AGCACACGTCTGAACTCTTTCCCTACACGACGCTCTTCCGATCTNNNATCATGAGGTCAGGAGATCG
GCACACGTCTGAACTCTTTCCCTACACGACGCTCTTCCGATCTNNNAATACAAAAAATTAGCCGGGC
AGCACACGTCTGAACTCTTTCCCTACACGACGCTCTTCCGATCTNNNGTACCATTGTACTCTAGCCTGG
AGCACACGTCTGAACTCTTTCCCTACACGACGCTCTTCCGATCTNNNACTCATCGCAGCTGTATCAAG
GCACACGTCTGAACTCTTTCCCTACACGACGCTCTTCCGATCTNNNCTGAGGCAGGAGAATCACTTGA
AGCACACGTCTGAACTCTTTCCCTACACGACGCTCTTCCGATCTNNNACATGGGTGACAGAATGAGA
AGCACACGTCTGAACTCTTTCCCTACACGACGCTCTTCCGATCTNNNCGGAGGTTACAATGAGCTGA
GAGCACACGTCTGAACTCTTTCCCTACACGACGCTCTTCCGATCTNNNCATCTGTAGTCCCAGCTACTC
AGCACACGTCTGAACTCTTTCCCTACACGACGCTCTTCCGATCTNNNCCACGCCCGGATAATTTTTG
GAGCACACGTCTGAACTCTTTCCCTACACGACGCTCTTCCGATCTNNNGGAGATTGCAGTGAGTGAG
AGCACACGTCTGAACTCTTTCCCTACACGACGCTCTTCCGATCTNNNTGAGGCAGGAGAATCTCTTGA
AGCACACGTCTGAACTCTTTCCCTACACGACGCTCTTCCGATCTNNNAATATTGCGCCACTGCACTCTA
AGAGCACACGTCTGAACTCTTTCCCTACACGACGCTCTTCCGATCTNNNACAAAAATTAGCCAAGGA
AGCACACGTCTGAACTCTTTCCCTACACGACGCTCTTCCGATCTNNNCCTATCCTCAGTTTTCCTGCCTT
AGCACACGTCTGAACTCTTTCCCTACACGACGCTCTTCCGATCTNNNCTCCCAGCCTCAAAAGAACAAT
AGCACACGTCTGAACTCTTTCCCTACACGACGCTCTTCCGATCTNNNGTCTCCGAGGGGAAAAAAAG
GCACACGTCTGAACTCTTTCCCTACACGACGCTCTTCCGATCTNNNCTCAGGCAGGAGAATCACTTG
AGCACACGTCTGAACTCTTTCCCTACACGACGCTCTTCCGATCTNNNGGCTGATAGAACCAGATCCTG
GAGCACACGTCTGAACTCTTTCCCTACACGACGCTCTTCCGATCTNNNGGTTGCAGTAAGCTGAGATT
GCACACGTCTGAACTCTTTCCCTACACGACGCTCTTCCGATCTNNNAAAAAATTAGCTGGGATGGTG
AGCACACGTCTGAACTCTTTCCCTACACGACGCTCTTCCGATCTNNNATGCCTGTAATCCCAGCTACTT
GAGCACACGTCTGAACTCTTTCCCTACACGACGCTCTTCCGATCTNNNGGCAACAGAGTGAGACTCCA
AGCACACGTCTGAACTCTTTCCCTACACGACGCTCTTCCGATCTNNNCCCCAAACTGGAAACAACTCA
GCACACGTCTGAACTCTTTCCCTACACGACGCTCTTCCGATCTNNNCTGTAATCCCAGCTACTCCAGAG
AGCACACGTCTGAACTCTTTCCCTACACGACGCTCTTCCGATCTNNNTTGAGACAGGAGAATCGCTTG
GAGCACACGTCTGAACTCTTTCCCTACACGACGCTCTTCCGATCTNNNGGATCACTTGAGTCTGGGAA
GAGCACACGTCTGAACTCTTTCCCTACACGACGCTCTTCCGATCTNNNAAAAAAATTAGCCGGGTGT
GAGCACACGTCTGAACTCTTTCCCTACACGACGCTCTTCCGATCTNNNACTCCTGTAATCCCAGCACTTT
GCACACGTCTGAACTCTTTCCCTACACGACGCTCTTCCGATCTNNNCGTCTGTAATCCCAGCTACTCAA
GAGCACACGTCTGAACTCTTTCCCTACACGACGCTCTTCCGATCTNNNGTAATCCCGGCTACTCAGGA
GAGCACACGTCTGAACTCTTTCCCTACACGACGCTCTTCCGATCTNNNGTAATCCCAGCTACTCAAGA
GAGCACACGTCTGAACTCTTTCCCTACACGACGCTCTTCCGATCTNNNGGTTTTGGTTCATGATACGG
GAGCACACGTCTGAACTCTTTCCCTACACGACGCTCTTCCGATCTNNNTTGGTTTTGGTTCATGATACG
AGCACACGTCTGAACTCTTTCCCTACACGACGCTCTTCCGATCTNNNTAGCAAGACCTCGTATCTACC
GCACACGTCTGAACTCTTTCCCTACACGACGCTCTTCCGATCTNNNCAGGAAGGAAGGAAGGAAGG
GAGCACACGTCTGAACTCTTTCCCTACACGACGCTCTTCCGATCTNNNTCACACCTGTAATCCCAACAC
AGCACACGTCTGAACTCTTTCCCTACACGACGCTCTTCCGATCTNNNGCACCTGTAATCCCAGCTATTT

AGCACACGTCTGAACTCTTTCCCTACACGACGCTCTTCCGATCTNNNTAGTCCCAGCTACTCAGAAGACT
AGCACACGTCTGAACTCTTTCCCTACACGACGCTCTTCCGATCTNNNACCTGAGGTCAAGAGTTCAAGA
GCACACGTCTGAACTCTTTCCCTACACGACGCTCTTCCGATCTNNNCTGAGGCAGGAGAATCACTTGAA
AGCACACGTCTGAACTCTTTCCCTACACGACGCTCTTCCGATCTNNNCAAGAGAATTGCTTCAACCTGG
GAGCACACGTCTGAACTCTTTCCCTACACGACGCTCTTCCGATCTNNNTCCCATTCTCTCTCTCTCTCTC
AGAGCACACGTCTGAACTCTTTCCCTACACGACGCTCTTCCGATCTNNNATGCCTGTAATCCCAGCTACT
AGCACACGTCTGAACTCTTTCCCTACACGACGCTCTTCCGATCTNNNTAAAAATTAGCAGAGCGTGGT
GCACACGTCTGAACTCTTTCCCTACACGACGCTCTTCCGATCTNNNGCCAGAGAATCGTTTGAACTCTG
AGAGCACACGTCTGAACTCTTTCCCTACACGACGCTCTTCCGATCTNNNCTGAGGCAGGAGAATCACTT
GAGCACACGTCTGAACTCTTTCCCTACACGACGCTCTTCCGATCTNNNGCTTACACCTGTAATCCCAACA
AGCACACGTCTGAACTCTTTCCCTACACGACGCTCTTCCGATCTNNNTTCAAGGATGCAGTGAGTGAT
GAGCACACGTCTGAACTCTTTCCCTACACGACGCTCTTCCGATCTNNNCTGTAGTCCCAGCTACTCAGG
AGCACACGTCTGAACTCTTTCCCTACACGACGCTCTTCCGATCTNNNCTGAGGCAGGAGAATCACTTGA
AGCACACGTCTGAACTCTTTCCCTACACGACGCTCTTCCGATCTNNNACAAAAATTAGCCAGGCTTGGT
AGCACACGTCTGAACTCTTTCCCTACACGACGCTCTTCCGATCTNNNGGCAACATAGTAAGACCCTGT
AGCACACGTCTGAACTCTTTCCCTACACGACGCTCTTCCGATCTNNNTGTGTTTATCCTGTGGTCCAGT
AGCACACGTCTGAACTCTTTCCCTACACGACGCTCTTCCGATCTNNNGACTACAGAGTGAGACCCTGT
GAGCACACGTCTGAACTCTTTCCCTACACGACGCTCTTCCGATCTNNNGAAGTTTGCAGTGAGCTGAG
AGCACACGTCTGAACTCTTTCCCTACACGACGCTCTTCCGATCTNNNAAATTAGTTGCCCAAGGTTGCA
AGCACACGTCTGAACTCTTTCCCTACACGACGCTCTTCCGATCTNNNCAAGGATGCTCATGTTGGGAA
AGCACACGTCTGAACTCTTTCCCTACACGACGCTCTTCCGATCTNNNGAGAATCGCTTGAACCAGGAA
GAGCACACGTCTGAACTCTTTCCCTACACGACGCTCTTCCGATCTNNNCTGTAGTCCCAGCTACTCAGGA
GAGCACACGTCTGAACTCTTTCCCTACACGACGCTCTTCCGATCTNNNGAATTGCTTGAACCCGTGAGA
GAGCACACGTCTGAACTCTTTCCCTACACGACGCTCTTCCGATCTNNNCTGAGACAGGAGAATCGCTT
GAGCACACGTCTGAACTCTTTCCCTACACGACGCTCTTCCGATCTNNNATCATGAGGTCAGGAGATCG
GAGCACACGTCTGAACTCTTTCCCTACACGACGCTCTTCCGATCTNNNTAATCCCAGCTACTCAGGAGA
GAGCACACGTCTGAACTCTTTCCCTACACGACGCTCTTCCGATCTNNNCTACTACACTCTAGCCTGGGTG
GAGCACACGTCTGAACTCTTTCCCTACACGACGCTCTTCCGATCTNNNGAAGCAGGAGAATCACTTGA
AGCACACGTCTGAACTCTTTCCCTACACGACGCTCTTCCGATCTNNNGCCAAGATCATGCCATTGTACT
AGCACACGTCTGAACTCTTTCCCTACACGACGCTCTTCCGATCTNNNTGAGGCAGGAGAATCACTTGA
AGCACACGTCTGAACTCTTTCCCTACACGACGCTCTTCCGATCTNNNTAATCTCAGCTACTCAGGAGCCT
GAGCACACGTCTGAACTCTTTCCCTACACGACGCTCTTCCGATCTNNNATTGCAATGAGTCGAGATCAT
AGCACACGTCTGAACTCTTTCCCTACACGACGCTCTTCCGATCTNNNCTGAGGCAGGAGAATCACTTGA
GCACACGTCTGAACTCTTTCCCTACACGACGCTCTTCCGATCTNNNTGTTTCTTTATTGGGCAGGCAAG
GAGCACACGTCTGAACTCTTTCCCTACACGACGCTCTTCCGATCTNNNGATGCCTGTAATCCCAGCTACT
AGCACACGTCTGAACTCTTTCCCTACACGACGCTCTTCCGATCTNNNTCTGTCTGAGACTTAGGGGACA
GCACACGTCTGAACTCTTTCCCTACACGACGCTCTTCCGATCTNNNTGAAGCAGTAGGATCACTTGAG
GCACACGTCTGAACTCTTTCCCTACACGACGCTCTTCCGATCTNNNTCGCAAAAATCTAAAACTTCTGCT
GAGCACACGTCTGAACTCTTTCCCTACACGACGCTCTTCCGATCTNNNCTCACACCTGTAATCCCAACTCT
GAGCACACGTCTGAACTCTTTCCCTACACGACGCTCTTCCGATCTNNNTCACAGCAGATTTGGAGAGAC
AGCACACGTCTGAACTCTTTCCCTACACGACGCTCTTCCGATCTNNNGGCATTAGCTAGAACTGCAATG
AGCACACGTCTGAACTCTTTCCCTACACGACGCTCTTCCGATCTNNNACACAGTGAAACCCTGTCTCTA
AGCACACGTCTGAACTCTTTCCCTACACGACGCTCTTCCGATCTNNNGAGACAGGAAAATGGCATGAA

AGAGCACACGTCTGAACTCTTTCCCTACACGACGCTCTTCCGATCTNNNGGGTGACAGAATGAGACTC
GAGCACACGTCTGAACTCTTTCCCTACACGACGCTCTTCCGATCTNNNGATTGTAAGGCAAGAACTCAG
GAGCACACGTCTGAACTCTTTCCCTACACGACGCTCTTCCGATCTNNNAGGGAATAAGGGAGAAGGA
GAGCACACGTCTGAACTCTTTCCCTACACGACGCTCTTCCGATCTNNNCCAAAACAACATGGTACTGGC
GAGCACACGTCTGAACTCTTTCCCTACACGACGCTCTTCCGATCTNNNCTGTAGTCCCAGCTATTCAGGA
GAGCACACGTCTGAACTCTTTCCCTACACGACGCTCTTCCGATCTNNNCAAGACCTACTGATCTGTTGC
AGCACACGTCTGAACTCTTTCCCTACACGACGCTCTTCCGATCTNNNGGAGTACACATACATAGCCACC
AGCACACGTCTGAACTCTTTCCCTACACGACGCTCTTCCGATCTNNNTACTTCTGCCATGGAGAGAATG
AGCACACGTCTGAACTCTTTCCCTACACGACGCTCTTCCGATCTNNNGGACTCTGTCAATCAGGCAATCA
AGCACACGTCTGAACTCTTTCCCTACACGACGCTCTTCCGATCTNNNCTGTAGTCCCAGCTACTCAGGAG
AGCACACGTCTGAACTCTTTCCCTACACGACGCTCTTCCGATCTNNNCCATCTGGTGAAACTGGTTTGT
AGAGCACACGTCTGAACTCTTTCCCTACACGACGCTCTTCCGATCTNNNGATCACTTGATGCCAGGAGT
AGCACACGTCTGAACTCTTTCCCTACACGACGCTCTTCCGATCTNNNTGGAGGCATCTCATTCTTTTGGA
GAGCACACGTCTGAACTCTTTCCCTACACGACGCTCTTCCGATCTNNNAAAATTAGCTGGATGTGGTGG
AGCACACGTCTGAACTCTTTCCCTACACGACGCTCTTCCGATCTNNNAGGAAGGAGAATTGCTTGAAC
AGCACACGTCTGAACTCTTTCCCTACACGACGCTCTTCCGATCTNNNTCTTCCAATCTGTGAACCCACTT
GAGCACACGTCTGAACTCTTTCCCTACACGACGCTCTTCCGATCTNNNACAGTTTGGGTGTCCTAATCCA
GAGCACACGTCTGAACTCTTTCCCTACACGACGCTCTTCCGATCTNNNCATCCTGGCTAACACAGTGAA
GAGCACACGTCTGAACTCTTTCCCTACACGACGCTCTTCCGATCTNNNCCATGACTCCCCAGACTTCTTA
GAGCACACGTCTGAACTCTTTCCCTACACGACGCTCTTCCGATCTNNNCTCTCTTTCTTTTCCCTGCAAGC
AGCACACGTCTGAACTCTTTCCCTACACGACGCTCTTCCGATCTNNNGGTGCATACTGGTAATTCCAAGG
AGCACACGTCTGAACTCTTTCCCTACACGACGCTCTTCCGATCTNNNCTGAGACAGGAGATCGCTTGA
GAGCACACGTCTGAACTCTTTCCCTACACGACGCTCTTCCGATCTNNNCAAAAAAAATTAGCCGGGCGT
AGCACACGTCTGAACTCTTTCCCTACACGACGCTCTTCCGATCTNNNGGGAATCTGATTGGACTTCGAG
AGCACACGTCTGAACTCTTTCCCTACACGACGCTCTTCCGATCTNNNTATACCAGTTCAGGAGGACAG
GAGCACACGTCTGAACTCTTTCCCTACACGACGCTCTTCCGATCTNNNGAGAATGGCTTGAACACAGGA
GAGCACACGTCTGAACTCTTTCCCTACACGACGCTCTTCCGATCTNNNGTGAAACCCCTTCTCTACCAAA
AGCACACGTCTGAACTCTTTCCCTACACGACGCTCTTCCGATCTNNNCAGAGCTTGCAGTGAGTCGAGA
AGCACACGTCTGAACTCTTTCCCTACACGACGCTCTTCCGATCTNNNGAGGATTGATTGAGCCTGGGA
AAGAGCACACGTCTGAACTCTTTCCCTACACGACGCTCTTCCGATCTNNNAGAGATTGCAGTGAGCTGA
GAGCACACGTCTGAACTCTTTCCCTACACGACGCTCTTCCGATCTNNNGGTGACAGAGTGAGACTCCAT
AGCACACGTCTGAACTCTTTCCCTACACGACGCTCTTCCGATCTNNNGGATTACCAGTATGAGCCACCA
GAGCACACGTCTGAACTCTTTCCCTACACGACGCTCTTCCGATCTNNNCTGTAGTCCCAGCTACTCAGGA
GAGCACACGTCTGAACTCTTTCCCTACACGACGCTCTTCCGATCTNNNCAACAAGAGTGAAACACCGTC
GAGCACACGTCTGAACTCTTTCCCTACACGACGCTCTTCCGATCTNNNTTGAACCCAGGAGTCATAGGT
AGCACACGTCTGAACTCTTTCCCTACACGACGCTCTTCCGATCTNNNTACGACAGAGTGAGACTCCATC
AGCACACGTCTGAACTCTTTCCCTACACGACGCTCTTCCGATCTNNNCTGTAGTCCCAGCTACTTGGAA
GAGCACACGTCTGAACTCTTTCCCTACACGACGCTCTTCCGATCTNNNGGTGTGCTGGTAAAACATGA
AGCACACGTCTGAACTCTTTCCCTACACGACGCTCTTCCGATCTNNNCGCGTGTGTGTATTCCAAATTC
AGCACACGTCTGAACTCTTTCCCTACACGACGCTCTTCCGATCTNNNTGATTGCAGTGAGTCAAGATCC
AGCACACGTCTGAACTCTTTCCCTACACGACGCTCTTCCGATCTNNNCTGTAATCCCAGCTACTCAGGAG
GAGCACACGTCTGAACTCTTTCCCTACACGACGCTCTTCCGATCTNNNCCAAGGATCCTGCCATTTCAA
AGAGCACACGTCTGAACTCTTTCCCTACACGACGCTCTTCCGATCTNNNCTGAGGCAGGAGAATTGCTT

AGCACACGTCTGAACTCTTTCCCTACACGACGCTCTTCCGATCTNNNCGCGTCTATAATCCCAGCTACT
GAGCACACGTCTGAACTCTTTCCCTACACGACGCTCTTCCGATCTNNNTAAGGCAGGAGAATTGCTTG
GAGCACACGTCTGAACTCTTTCCCTACACGACGCTCTTCCGATCTNNNGTAATCCCAGCTACTAAGGA
GAGCACACGTCTGAACTCTTTCCCTACACGACGCTCTTCCGATCTNNNCTGAGGCAGGAGAATCACTT
AGCACACGTCTGAACTCTTTCCCTACACGACGCTCTTCCGATCTNNNCTGAGGCAGGAGAATAGCTTG
AGCACACGTCTGAACTCTTTCCCTACACGACGCTCTTCCGATCTNNNCAGGAGAATTGCTTGAACAC
GAGCACACGTCTGAACTCTTTCCCTACACGACGCTCTTCCGATCTNNNTGAGCTGAGATAGGGTATT
GAGCACACGTCTGAACTCTTTCCCTACACGACGCTCTTCCGATCTNNNGAGTTTGAAACTGCAGTGA
AGCACACGTCTGAACTCTTTCCCTACACGACGCTCTTCCGATCTNNNGGGAGAAAGAGGGAGACTCC
AGCACACGTCTGAACTCTTTCCCTACACGACGCTCTTCCGATCTNNNGAGAATCGCTTGAACCTGAG
GCACACGTCTGAACTCTTTCCCTACACGACGCTCTTCCGATCTNNNAAGTTCCAGGGAAATTGTGAG
GCACACGTCTGAACTCTTTCCCTACACGACGCTCTTCCGATCTNNNGGAGGTTGTAGTGAGCCAAG
AGCACACGTCTGAACTCTTTCCCTACACGACGCTCTTCCGATCTNNNATGTGCCTGTAATTCCAGCTAC
GCACACGTCTGAACTCTTTCCCTACACGACGCTCTTCCGATCTNNNGAGAATCGCTTGAACTTGGGA
GAGCACACGTCTGAACTCTTTCCCTACACGACGCTCTTCCGATCTNNNCCAAAGTGGGAGAATTGCTT
GAGCACACGTCTGAACTCTTTCCCTACACGACGCTCTTCCGATCTNNNCTGTATTCCAGCTACTTGGG
GAGCACACGTCTGAACTCTTTCCCTACACGACGCTCTTCCGATCTNNNAAAATCACTTGAGCCTTCTGG
GCACACGTCTGAACTCTTTCCCTACACGACGCTCTTCCGATCTNNNCTGAGGCAGGAGAATTGCTTG
GAGCACACGTCTGAACTCTTTCCCTACACGACGCTCTTCCGATCTNNNCGGATCATTTGAGGTCAGGA
AGCACACGTCTGAACTCTTTCCCTACACGACGCTCTTCCGATCTNNNTGGATCTGTAGTCCCACCTAC
GAGCACACGTCTGAACTCTTTCCCTACACGACGCTCTTCCGATCTNNNCTGAGGCAGGAGAATCACTT
AGCACACGTCTGAACTCTTTCCCTACACGACGCTCTTCCGATCTNNNGAGAATGGTGTGAACTCAGG
AGAGCACACGTCTGAACTCTTTCCCTACACGACGCTCTTCCGATCTNNNAGAAGAATCGCTTGAACCC
AGCACACGTCTGAACTCTTTCCCTACACGACGCTCTTCCGATCTNNNGTGATCACACCACTACACTCCA
GAGCACACGTCTGAACTCTTTCCCTACACGACGCTCTTCCGATCTNNNTGAGTAACAGAGCAAGACTG
GAGCACACGTCTGAACTCTTTCCCTACACGACGCTCTTCCGATCTNNNTTCGAGACCAACCTGACCAA
AGAGCACACGTCTGAACTCTTTCCCTACACGACGCTCTTCCGATCTNNNGCCTGTAATCCCAGATACT
AGCACACGTCTGAACTCTTTCCCTACACGACGCTCTTCCGATCTNNNCGAAACCCCCGTCTCTAAAAAA
AGCACACGTCTGAACTCTTTCCCTACACGACGCTCTTCCGATCTNNNCAGCTACTTGGGAGGCTAAA
GAGCACACGTCTGAACTCTTTCCCTACACGACGCTCTTCCGATCTNNNCGCCTATAGTTCCAGCTACTC
GCACACGTCTGAACTCTTTCCCTACACGACGCTCTTCCGATCTNNNAAAAATTAGCCCAGTGTGATGG
GAGCACACGTCTGAACTCTTTCCCTACACGACGCTCTTCCGATCTNNNCATAGTCCCAGCTACTCAGAA
AGCACACGTCTGAACTCTTTCCCTACACGACGCTCTTCCGATCTNNNCGACAGAGTGAGACTCCATCT
AGCACACGTCTGAACTCTTTCCCTACACGACGCTCTTCCGATCTNNNTGAGATTGTGCCATTGTACTC
AGCACACGTCTGAACTCTTTCCCTACACGACGCTCTTCCGATCTNNNGGCAAGAAGAGTGAAATTCCG
GAGCACACGTCTGAACTCTTTCCCTACACGACGCTCTTCCGATCTNNNAGAAGTTGCAGTGAGCTGA
GAGCACACGTCTGAACTCTTTCCCTACACGACGCTCTTCCGATCTNNNCTGTAATTCGAGCTACTCCG
GAGCACACGTCTGAACTCTTTCCCTACACGACGCTCTTCCGATCTNNNGAGGTTACAGTGAGCCAAG
GAGCACACGTCTGAACTCTTTCCCTACACGACGCTCTTCCGATCTNNNCTGTAATCCCAGCTACTCGGT
GAGCACACGTCTGAACTCTTTCCCTACACGACGCTCTTCCGATCTNNNCTGAGGCTGGAGAATCACTT
AGAGCACACGTCTGAACTCTTTCCCTACACGACGCTCTTCCGATCTNNNTCACTTGAACTAAGGAGGC
AGCACACGTCTGAACTCTTTCCCTACACGACGCTCTTCCGATCTNNNCTGAGGCAGGAGAATTGCTT
AGCACACGTCTGAACTCTTTCCCTACACGACGCTCTTCCGATCTNNNATGTCTCATTATGTGCTCAGGC

AGCACACGTCTGAACTCTTTCCCTACACGACGCTCTTCCGATCTNNNGGTGCCTATAATCCCAGCTACT
AGCACACGTCTGAACTCTTTCCCTACACGACGCTCTTCCGATCTNNNTGTAATCCCAGCTTCTTGTGAG
AGCACACGTCTGAACTCTTTCCCTACACGACGCTCTTCCGATCTNNNCTGTAGTCCCAGCTACTTGGAA
AGCACACGTCTGAACTCTTTCCCTACACGACGCTCTTCCGATCTNNNCTGTAATCCCAGCTACTCAGGA
GAGCACACGTCTGAACTCTTTCCCTACACGACGCTCTTCCGATCTNNNCAAGATCGTGTCATTGCACTC
AGCACACGTCTGAACTCTTTCCCTACACGACGCTCTTCCGATCTNNNACAAGATGCTGTTTTTGGCAGA
AGAGCACACGTCTGAACTCTTTCCCTACACGACGCTCTTCCGATCTNNNCTGTAGTCCCAGCTACTCAG
AGCACACGTCTGAACTCTTTCCCTACACGACGCTCTTCCGATCTNNNCTGTAATCCCAGCTACTCAGGA
AGAGCACACGTCTGAACTCTTTCCCTACACGACGCTCTTCCGATCTNNNCATGCCTCTAATCCCAGCTAC
AGCACACGTCTGAACTCTTTCCCTACACGACGCTCTTCCGATCTNNNCAGTTCAAAAACATTCACAGCC
GAGCACACGTCTGAACTCTTTCCCTACACGACGCTCTTCCGATCTNNNCCAGGTTCAAGCAATTCTTCCT
AGCACACGTCTGAACTCTTTCCCTACACGACGCTCTTCCGATCTNNNTTGAGGCAGGAGAATTGCTTG
AGCACACGTCTGAACTCTTTCCCTACACGACGCTCTTCCGATCTNNNCAACATGGAGAAACCCCGTAT
AGCACACGTCTGAACTCTTTCCCTACACGACGCTCTTCCGATCTNNNCTCATGCCTATAATCCCAGCAC
AGAGCACACGTCTGAACTCTTTCCCTACACGACGCTCTTCCGATCTNNNGTAGTACCAGCTATTCAGGA
GAGCACACGTCTGAACTCTTTCCCTACACGACGCTCTTCCGATCTNNNACACAAGTAGGAAGCAGCCA
AGAGCACACGTCTGAACTCTTTCCCTACACGACGCTCTTCCGATCTNNNTCACGCCTGTAATCTTAGCAC
AGCACACGTCTGAACTCTTTCCCTACACGACGCTCTTCCGATCTNNNCTGGACAAGAGAACAAGACTCC
GAGCACACGTCTGAACTCTTTCCCTACACGACGCTCTTCCGATCTNNNGGAAGATAGCTTGAGTCCAG
AGCACACGTCTGAACTCTTTCCCTACACGACGCTCTTCCGATCTNNNGAGACAGGAGAATTGCTTGAA
AGCACACGTCTGAACTCTTTCCCTACACGACGCTCTTCCGATCTNNNGCAGAGATGGAAAGGGAACTC
AGCACACGTCTGAACTCTTTCCCTACACGACGCTCTTCCGATCTNNNACAGAGTGAGTGAGAGTCTGT
AGCACACGTCTGAACTCTTTCCCTACACGACGCTCTTCCGATCTNNNCATGCCTGTAATCCCAGCTACTA
AGCACACGTCTGAACTCTTTCCCTACACGACGCTCTTCCGATCTNNNCATGCTGTAATCCCCACACTTTG
GAGCACACGTCTGAACTCTTTCCCTACACGACGCTCTTCCGATCTNNNCGTGCTTGTAATCCCAGCTACT
AGAGCACACGTCTGAACTCTTTCCCTACACGACGCTCTTCCGATCTNNNATGCTAGTGGTTGATGGGA
AGAGCACACGTCTGAACTCTTTCCCTACACGACGCTCTTCCGATCTNNNACACACTCTTCTTTCCCACCA
AGCACACGTCTGAACTCTTTCCCTACACGACGCTCTTCCGATCTNNNTATAATCCCAGCTACTAGGGGG
AGCACACGTCTGAACTCTTTCCCTACACGACGCTCTTCCGATCTNNNTGTAATCCCAGCTACTCAGGAG
AGCACACGTCTGAACTCTTTCCCTACACGACGCTCTTCCGATCTNNNGTAATCTCAGCTACTTGGGAGG
AGAGCACACGTCTGAACTCTTTCCCTACACGACGCTCTTCCGATCTNNNGACACAACAGAATCGCTTGA
AGCACACGTCTGAACTCTTTCCCTACACGACGCTCTTCCGATCTNNNATGGTTGGCTAGATTTCCAAGG
GCACACGTCTGAACTCTTTCCCTACACGACGCTCTTCCGATCTNNNCGTCCTCAACCTTAGCAAAACACC
AGCACACGTCTGAACTCTTTCCCTACACGACGCTCTTCCGATCTNNNTTCTCCCCTGTTCCTCTAAGGAT
GAGCACACGTCTGAACTCTTTCCCTACACGACGCTCTTCCGATCTNNNGGGTTGCATTGAGTCACAAT
AGCACACGTCTGAACTCTTTCCCTACACGACGCTCTTCCGATCTNNNGCATCCTTGTAATCCCAGCTAC
AGCACACGTCTGAACTCTTTCCCTACACGACGCTCTTCCGATCTNNNGGTGGGAGGATTACTTGAACT
GCACACGTCTGAACTCTTTCCCTACACGACGCTCTTCCGATCTNNNCTGAGGCAGGAGAATCACTTGA
GAGCACACGTCTGAACTCTTTCCCTACACGACGCTCTTCCGATCTNNNCAAAAAGTTTCAGCAGGGGT
GAGCACACGTCTGAACTCTTTCCCTACACGACGCTCTTCCGATCTNNNGCTGAGGAAGGTGGATAACT
AGCACACGTCTGAACTCTTTCCCTACACGACGCTCTTCCGATCTNNNCACCTGTAATCCCAGCTACTCA
AGCACACGTCTGAACTCTTTCCCTACACGACGCTCTTCCGATCTNNNACCATCTTCCACACATCAGTCTC
AGCACACGTCTGAACTCTTTCCCTACACGACGCTCTTCCGATCTNNNGATCCACTGAGCTAGACCACTT

GAGCACACGTCTGAACTCTTTCCCTACACGACGCTCTTCCGATCTNNNGGTTGTAGTGAGCCAAGTTT
AGCACACGTCTGAACTCTTTCCCTACACGACGCTCTTCCGATCTNNNATCACTTGAAACCAGAAGGCAG
AGCACACGTCTGAACTCTTTCCCTACACGACGCTCTTCCGATCTNNNAAAAATTAGTCGGATGTGGTG
AGCACACGTCTGAACTCTTTCCCTACACGACGCTCTTCCGATCTNNNAGAATCACTTGAACCTGGGAG
GAGCACACGTCTGAACTCTTTCCCTACACGACGCTCTTCCGATCTNNNCTGTAATTCCAGCTACTCGGG
AGCACACGTCTGAACTCTTTCCCTACACGACGCTCTTCCGATCTNNNCTGAGGCAGGAGAATCACTTG
AGCACACGTCTGAACTCTTTCCCTACACGACGCTCTTCCGATCTNNNTGTAATCCCAGCTACTCAGGAG
AGCACACGTCTGAACTCTTTCCCTACACGACGCTCTTCCGATCTNNNAGATGCATTCTTAGACGCCAAG
AGCACACGTCTGAACTCTTTCCCTACACGACGCTCTTCCGATCTNNNCCAATGTGGATGGATTGCTTG
AGCACACGTCTGAACTCTTTCCCTACACGACGCTCTTCCGATCTNNNCTGAGGCAGGAGAATCACTTG
AGCACACGTCTGAACTCTTTCCCTACACGACGCTCTTCCGATCTNNNTGTAATCCTAGCACTTTGGGAG
AGCACACGTCTGAACTCTTTCCCTACACGACGCTCTTCCGATCTNNNTGAGGCAGAAGAATGGTTTG
AGCACACGTCTGAACTCTTTCCCTACACGACGCTCTTCCGATCTNNNACTGATGCTACTGGAGAATTG
AGCACACGTCTGAACTCTTTCCCTACACGACGCTCTTCCGATCTNNNAGGCTGAGGCAGAATTACTTG
AGCACACGTCTGAACTCTTTCCCTACACGACGCTCTTCCGATCTNNNAAAAATTAGCCGGTGTGATGG
GAGCACACGTCTGAACTCTTTCCCTACACGACGCTCTTCCGATCTNNNGTGAAACCCTGTCTCCACTAA
AGCACACGTCTGAACTCTTTCCCTACACGACGCTCTTCCGATCTNNNGCAGAAGAATCACTTGAACCCA
GCACACGTCTGAACTCTTTCCCTACACGACGCTCTTCCGATCTNNNGACGGTGTCTCTTCAGCTTTCTA
GAGCACACGTCTGAACTCTTTCCCTACACGACGCTCTTCCGATCTNNNGTGATGTACACCTGTAGCCC
GAGCACACGTCTGAACTCTTTCCCTACACGACGCTCTTCCGATCTNNNTACAAATCACTGTGACTACGG
AGCACACGTCTGAACTCTTTCCCTACACGACGCTCTTCCGATCTNNNCTGAGGCAGGAGAATAGTGT
AGCACACGTCTGAACTCTTTCCCTACACGACGCTCTTCCGATCTNNNAAAGACACAGACTGGCAAATT
AGCACACGTCTGAACTCTTTCCCTACACGACGCTCTTCCGATCTNNNTGAGAATCACTTGAACCTGGG
AGCACACGTCTGAACTCTTTCCCTACACGACGCTCTTCCGATCTNNNGGTATTGTGTGCCTGTAGTCT
GAGCACACGTCTGAACTCTTTCCCTACACGACGCTCTTCCGATCTNNNAAGGTTATGCTGACGTAGAA
AGAGCACACGTCTGAACTCTTTCCCTACACGACGCTCTTCCGATCTNNNAAAAAATTAGATGGGCGTG
AGCACACGTCTGAACTCTTTCCCTACACGACGCTCTTCCGATCTNNNGGATCACCTGAGGTCAGAAGT
AGAGCACACGTCTGAACTCTTTCCCTACACGACGCTCTTCCGATCTNNNCTGTAGTCCCAGCTACTTGT
GAGCACACGTCTGAACTCTTTCCCTACACGACGCTCTTCCGATCTNNNTGTAATCTCAGCACTTTGGG
GCACACGTCTGAACTCTTTCCCTACACGACGCTCTTCCGATCTNNNCCACATAGCGAGATCTCTGTCTT
GAGCACACGTCTGAACTCTTTCCCTACACGACGCTCTTCCGATCTNNNCTGTAGTCCCAGCTACTCAGG
AGCACACGTCTGAACTCTTTCCCTACACGACGCTCTTCCGATCTNNNGTAATCCCAGCTACTTGAGAGG
AGCACACGTCTGAACTCTTTCCCTACACGACGCTCTTCCGATCTNNNAAGCAGGAGAATCACTTGAACC
AGCACACGTCTGAACTCTTTCCCTACACGACGCTCTTCCGATCTNNNGTGCATGTAATCCCAGCTACTC
GAGCACACGTCTGAACTCTTTCCCTACACGACGCTCTTCCGATCTNNNAAGTGCATCTGTGGAGTGAT
AGAGCACACGTCTGAACTCTTTCCCTACACGACGCTCTTCCGATCTNNNACACCTGTAGTCCTTGCTAC
AGCACACGTCTGAACTCTTTCCCTACACGACGCTCTTCCGATCTNNNAATCACTTGAACTTGGGAGGA
AGAGCACACGTCTGAACTCTTTCCCTACACGACGCTCTTCCGATCTNNNAATCCTAGCTACTTTGGAGG
GAGCACACGTCTGAACTCTTTCCCTACACGACGCTCTTCCGATCTNNNGCAACTTAGCAAGACCTTGTC
GAGCACACGTCTGAACTCTTTCCCTACACGACGCTCTTCCGATCTNNNACTGCTTACTGTTCATGTAGC
AGAGCACACGTCTGAACTCTTTCCCTACACGACGCTCTTCCGATCTNNNCCTAGGAGACAGAGCAAGA
AGCACACGTCTGAACTCTTTCCCTACACGACGCTCTTCCGATCTNNNGATTTTGGGTGTTCGGTTTAG
GAGCACACGTCTGAACTCTTTCCCTACACGACGCTCTTCCGATCTNNNAAAACTTAGCCAGGCATAGT

GAGCACACGTCTGAACTCTTTCCCTACACGACGCTCTTCCGATCTNNNGTAGTTCCAGGTATTCAGGA
AGCACACGTCTGAACTCTTTCCCTACACGACGCTCTTCCGATCTNNNACACAAAAATTAGCTGGGGTGT
AAGAGCACACGTCTGAACTCTTTCCCTACACGACGCTCTTCCGATCTNNNAAAAAATTAGCTGGGCATG
GAGCACACGTCTGAACTCTTTCCCTACACGACGCTCTTCCGATCTNNNGCACAGGATACAGGATAGGG
GAGCACACGTCTGAACTCTTTCCCTACACGACGCTCTTCCGATCTNNNATACAGAGAGACCCTTGTGCA
AGAGCACACGTCTGAACTCTTTCCCTACACGACGCTCTTCCGATCTNNNCTGTAATCCCAGCAACTTGGA
AGCACACGTCTGAACTCTTTCCCTACACGACGCTCTTCCGATCTNNNAGTGAGACCCTGCCTCTACTAAA
GAGCACACGTCTGAACTCTTTCCCTACACGACGCTCTTCCGATCTNNNCTCATGCCTATAATCCCAGCAC
AGCACACGTCTGAACTCTTTCCCTACACGACGCTCTTCCGATCTNNNGGATTGTTTGAGCCTGGGAAAT
GAGCACACGTCTGAACTCTTTCCCTACACGACGCTCTTCCGATCTNNNGATGCAAGAGGATTGCTTGA
GAGCACACGTCTGAACTCTTTCCCTACACGACGCTCTTCCGATCTNNNTGGGAATCACTTGAACCTGAG
GAGCACACGTCTGAACTCTTTCCCTACACGACGCTCTTCCGATCTNNNCAGGAGAATGGTGTGAACGT
GAGCACACGTCTGAACTCTTTCCCTACACGACGCTCTTCCGATCTNNNCTGAGCCAGGAGAATCACTTG
AGCACACGTCTGAACTCTTTCCCTACACGACGCTCTTCCGATCTNNNTTTCCCAGTGCACTATAGTGGTC
AGCACACGTCTGAACTCTTTCCCTACACGACGCTCTTCCGATCTNNNATAATTCTCATGCCTCAGCCTCC
GAGCACACGTCTGAACTCTTTCCCTACACGACGCTCTTCCGATCTNNNTGGGTAACAAAGCGAGATCTT
GCACACGTCTGAACTCTTTCCCTACACGACGCTCTTCCGATCTNNNTGAGGCAGGAGAATAGTGTGAA
AGCACACGTCTGAACTCTTTCCCTACACGACGCTCTTCCGATCTNNNAGTGAGCTGAGATCATGCCATT
GAGCACACGTCTGAACTCTTTCCCTACACGACGCTCTTCCGATCTNNNTGGCTCACACTTGTAATTCCAG
AGCACACGTCTGAACTCTTTCCCTACACGACGCTCTTCCGATCTNNNTATCTAACAGCTGCAATGAGCC
GAGCACACGTCTGAACTCTTTCCCTACACGACGCTCTTCCGATCTNNNGATGCTGTCTGAGTCTGTTCT
AGAGCACACGTCTGAACTCTTTCCCTACACGACGCTCTTCCGATCTNNNAATGTTTGTGTTTTAGCCTGG
AGCACACGTCTGAACTCTTTCCCTACACGACGCTCTTCCGATCTNNNCTGAGGCAGGAGAATCACTTGA
AGCACACGTCTGAACTCTTTCCCTACACGACGCTCTTCCGATCTNNNCTGTAATCCCAGCTACTCAGGAG
GAGCACACGTCTGAACTCTTTCCCTACACGACGCTCTTCCGATCTNNNTTTGTAGCGAGTTGAGATTGT
AGAGCACACGTCTGAACTCTTTCCCTACACGACGCTCTTCCGATCTNNNTACAAAAAATTAGCTGGGCG
AGCACACGTCTGAACTCTTTCCCTACACGACGCTCTTCCGATCTNNNAACTTTCAGAGTCTGAGGTGGT
GAGCACACGTCTGAACTCTTTCCCTACACGACGCTCTTCCGATCTNNNTAAGGTCAGGAGTTCGAAAC
GAGCACACGTCTGAACTCTTTCCCTACACGACGCTCTTCCGATCTNNNTGAGATTGTGCCATTGTACTC
GAGCACACGTCTGAACTCTTTCCCTACACGACGCTCTTCCGATCTNNNAAGATCACACCACTGCACTCTA
GAGCACACGTCTGAACTCTTTCCCTACACGACGCTCTTCCGATCTNNNCTGTAATCCCAGCACTTTGGAA
AGCACACGTCTGAACTCTTTCCCTACACGACGCTCTTCCGATCTNNNACTTCCATCCAGCCTTGAAGAAT
GAGCACACGTCTGAACTCTTTCCCTACACGACGCTCTTCCGATCTNNNAGAATCACTTGAATCCAGGAG
AGAGCACACGTCTGAACTCTTTCCCTACACGACGCTCTTCCGATCTNNNAGAATTGCTTGAATCAGGGA
AGCACACGTCTGAACTCTTTCCCTACACGACGCTCTTCCGATCTNNNGAGGTTGCAGTGAGCCAAAATT
GCACACGTCTGAACTCTTTCCCTACACGACGCTCTTCCGATCTNNNGAAGGTTGCAGTGAGCCGAAAT
GAGCACACGTCTGAACTCTTTCCCTACACGACGCTCTTCCGATCTNNNATACAAAAATTGGCTGGGTGT
AGCACACGTCTGAACTCTTTCCCTACACGACGCTCTTCCGATCTNNNTCAAGGCTGCAGTGACTTATGA
AGCACACGTCTGAACTCTTTCCCTACACGACGCTCTTCCGATCTNNNCCGGAAAATGCCTAGTTTGACAA
AGCACACGTCTGAACTCTTTCCCTACACGACGCTCTTCCGATCTNNNTGTGCTGACATGATCTTCTCACA
GAGCACACGTCTGAACTCTTTCCCTACACGACGCTCTTCCGATCTNNNTGAGGCAGAGAATGATGTGA
GAGCACACGTCTGAACTCTTTCCCTACACGACGCTCTTCCGATCTNNNAAAAATTAGCTGGGCATGATG
AGAGCACACGTCTGAACTCTTTCCCTACACGACGCTCTTCCGATCTNNNCTGAGGGAGGAGAATGGTT

AGCACACGTCTGAACTCTTTCCCTACACGACGCTCTTCCGATCTNNNCTAGAAGAAAACTGAGTGCGT
AGCACACGTCTGAACTCTTTCCCTACACGACGCTCTTCCGATCTNNNGAACTGTGATCATTCTGCTGCA
GAGCACACGTCTGAACTCTTTCCCTACACGACGCTCTTCCGATCTNNNCTGAGGCAGGAGAATCACTTG
GAGCACACGTCTGAACTCTTTCCCTACACGACGCTCTTCCGATCTNNNACGCCTGTAAATCCAACACTTT
GAGCACACGTCTGAACTCTTTCCCTACACGACGCTCTTCCGATCTNNNCGTGAGACAAGATCATGCCAT
AGCACACGTCTGAACTCTTTCCCTACACGACGCTCTTCCGATCTNNNAAAAAAAAGTAGCCAGGCGAT
AGAGCACACGTCTGAACTCTTTCCCTACACGACGCTCTTCCGATCTNNNGGAGAATCACCTGAACATG
AGCACACGTCTGAACTCTTTCCCTACACGACGCTCTTCCGATCTNNNCTGTAATCCCAGCTACTCAGGA
GAGCACACGTCTGAACTCTTTCCCTACACGACGCTCTTCCGATCTNNNAGTGAGCTTAGATCGCACCAT
GAGCACACGTCTGAACTCTTTCCCTACACGACGCTCTTCCGATCTNNNTGGCCTCAGAGGATATGTACA
AGCACACGTCTGAACTCTTTCCCTACACGACGCTCTTCCGATCTNNNATGCCTGTAATCCCAGCTACTTG
AGCACACGTCTGAACTCTTTCCCTACACGACGCTCTTCCGATCTNNNATGCCTGTAATCCCAGCTACTT
AGCACACGTCTGAACTCTTTCCCTACACGACGCTCTTCCGATCTNNNGTGAGGCTGGAGAATTCCTTGA
GAGCACACGTCTGAACTCTTTCCCTACACGACGCTCTTCCGATCTNNNTCCTGTAACAGCAGACACACTA
AGCACACGTCTGAACTCTTTCCCTACACGACGCTCTTCCGATCTNNNCACTTGAGGTCAGGAGTTCAA
GAGCACACGTCTGAACTCTTTCCCTACACGACGCTCTTCCGATCTNNNGAGATCACGCCACTGCACTTTA
GAGCACACGTCTGAACTCTTTCCCTACACGACGCTCTTCCGATCTNNNGCATACACCTGTAATCCCAGC
AGAGCACACGTCTGAACTCTTTCCCTACACGACGCTCTTCCGATCTNNNTTGAGGCAGGAGAATTGCT
GAGCACACGTCTGAACTCTTTCCCTACACGACGCTCTTCCGATCTNNNTATCTGTTCAACTCCCTGCCTT
GAGCACACGTCTGAACTCTTTCCCTACACGACGCTCTTCCGATCTNNNAGTTGCAGTAAGCTGAGATCA
GAGCACACGTCTGAACTCTTTCCCTACACGACGCTCTTCCGATCTNNNTTAGTGTTCATAGCCAGCCAA
AGAGCACACGTCTGAACTCTTTCCCTACACGACGCTCTTCCGATCTNNNACACCTGTAATCCCAGCACT
GCACACGTCTGAACTCTTTCCCTACACGACGCTCTTCCGATCTNNNCTGAGGTGGGAGAATCACTTGA
GAGCACACGTCTGAACTCTTTCCCTACACGACGCTCTTCCGATCTNNNCATGGTGGTAGGCACCTGTAA
AGCACACGTCTGAACTCTTTCCCTACACGACGCTCTTCCGATCTNNNGGAGGTTACAGTGAGCTGAGA
AGCACACGTCTGAACTCTTTCCCTACACGACGCTCTTCCGATCTNNNGTAAACTGAGATCATGCCACTG
GAGCACACGTCTGAACTCTTTCCCTACACGACGCTCTTCCGATCTNNNCAGATCACGAGGTCAGGAGA
AGCACACGTCTGAACTCTTTCCCTACACGACGCTCTTCCGATCTNNNGCTGAGGCAGGAGAAAAGTA
GAGCACACGTCTGAACTCTTTCCCTACACGACGCTCTTCCGATCTNNNACACCTGTAATCCCAGCTACTT
AGCACACGTCTGAACTCTTTCCCTACACGACGCTCTTCCGATCTNNNCAGAAGAATTGCTTGAACCCA
AGAGCACACGTCTGAACTCTTTCCCTACACGACGCTCTTCCGATCTNNNACCCCGTCTCTACTAAGAAGA
GAGCACACGTCTGAACTCTTTCCCTACACGACGCTCTTCCGATCTNNNGAGGCTAAGGCAAGAGAAT
GAGCACACGTCTGAACTCTTTCCCTACACGACGCTCTTCCGATCTNNNGAGAATCCTTTGAACCCAGGA
AGAGCACACGTCTGAACTCTTTCCCTACACGACGCTCTTCCGATCTNNNGCAACAAAGGGAGACTCCAT
GAGCACACGTCTGAACTCTTTCCCTACACGACGCTCTTCCGATCTNNNCCATCTCTCTGTTCTCCTTGGA
GAGCACACGTCTGAACTCTTTCCCTACACGACGCTCTTCCGATCTNNNCTGTGATCCCAGCTACTTGAG
AGCACACGTCTGAACTCTTTCCCTACACGACGCTCTTCCGATCTNNNGGCACCTGTAATCCCAGTTACTT
GAGCACACGTCTGAACTCTTTCCCTACACGACGCTCTTCCGATCTNNNGGCTGAGGTAGGAGAATCA
AGAGCACACGTCTGAACTCTTTCCCTACACGACGCTCTTCCGATCTNNNCAAAAATTAGCAGGGTGTG
GAGCACACGTCTGAACTCTTTCCCTACACGACGCTCTTCCGATCTNNNCTCTTACAATTTGGGGAAGG
AGAGCACACGTCTGAACTCTTTCCCTACACGACGCTCTTCCGATCTNNNGAGCTTGCAGTTAGCCGAGA
GAGCACACGTCTGAACTCTTTCCCTACACGACGCTCTTCCGATCTNNNGGTGACAGTGTGAGACTCT
AGCACACGTCTGAACTCTTTCCCTACACGACGCTCTTCCGATCTNNNACAAAGCAGGACTCTGTCTCAA

GAGCACACGTCTGAACTCTTTCCCTACACGACGCTCTTCCGATCTNNNCTTCAGTAACAACTGCACAGCA
AGCACACGTCTGAACTCTTTCCCTACACGACGCTCTTCCGATCTNNNGCTAAGGCAGGAGAATCACTTG
GAGCACACGTCTGAACTCTTTCCCTACACGACGCTCTTCCGATCTNNNCCTCTCCCCAGAAGACCTAGTA
AGCACACGTCTGAACTCTTTCCCTACACGACGCTCTTCCGATCTNNNGGCCCGAGAGCTAAGAATAGTT
GAGCACACGTCTGAACTCTTTCCCTACACGACGCTCTTCCGATCTNNNAGGTTGCAGTGAGTTGAGATC
GAGCACACGTCTGAACTCTTTCCCTACACGACGCTCTTCCGATCTNNNGCAGAAGAATCACTTGAACCC
AGCACACGTCTGAACTCTTTCCCTACACGACGCTCTTCCGATCTNNNACAAAAATTAGCCAGGTGTGG
GAGCACACGTCTGAACTCTTTCCCTACACGACGCTCTTCCGATCTNNNTCACTTGAGTCCCAGAGTTCAA
GAGCACACGTCTGAACTCTTTCCCTACACGACGCTCTTCCGATCTNNNCCAACATGGTGAAACTCTGTCA
GAGCACACGTCTGAACTCTTTCCCTACACGACGCTCTTCCGATCTNNNTGAAGTCAGGAGTTTGAGAC
AGAGCACACGTCTGAACTCTTTCCCTACACGACGCTCTTCCGATCTNNNAGGCTGAGTCACGAGAATC
GAGCACACGTCTGAACTCTTTCCCTACACGACGCTCTTCCGATCTNNNACTGTGTCACTGTCACTACACT
AGAGCACACGTCTGAACTCTTTCCCTACACGACGCTCTTCCGATCTNNNCTCATGCCTATAATCCCAGCAG
GAGCACACGTCTGAACTCTTTCCCTACACGACGCTCTTCCGATCTNNNGTCAGGAGTTCTAGACCAAG
GAGCACACGTCTGAACTCTTTCCCTACACGACGCTCTTCCGATCTNNNCAGATCATGAGGTCAGGAGA
AGAGCACACGTCTGAACTCTTTCCCTACACGACGCTCTTCCGATCTNNNCCAACATGGTGAAACTCCATC
GAGCACACGTCTGAACTCTTTCCCTACACGACGCTCTTCCGATCTNNNAAGATTGGGTGTTCATGCACA
AGCACACGTCTGAACTCTTTCCCTACACGACGCTCTTCCGATCTNNNCTGTGGTCCTAGCTACTCAGAA
GAGCACACGTCTGAACTCTTTCCCTACACGACGCTCTTCCGATCTNNNAGAATCACTTGAATCCAGGAC
GAGCACACGTCTGAACTCTTTCCCTACACGACGCTCTTCCGATCTNNNGCAGAGCTTATAATGAGCGG
AGCACACGTCTGAACTCTTTCCCTACACGACGCTCTTCCGATCTNNNTGAGGCACAAGAACCATTTGAA
GAGCACACGTCTGAACTCTTTCCCTACACGACGCTCTTCCGATCTNNNGGGTGGATCATTTGAGGTAA
AGCACACGTCTGAACTCTTTCCCTACACGACGCTCTTCCGATCTNNNCAGTGACCTGAGATCCTACCAA
AGCACACGTCTGAACTCTTTCCCTACACGACGCTCTTCCGATCTNNNCCAAATCCAGTGGGCTAAATGT
GAGCACACGTCTGAACTCTTTCCCTACACGACGCTCTTCCGATCTNNNGAGGTTGCAGTGAGCCATAAT
GAGCACACGTCTGAACTCTTTCCCTACACGACGCTCTTCCGATCTNNNATGCTTGTAATCCCAGCGATT
AGCACACGTCTGAACTCTTTCCCTACACGACGCTCTTCCGATCTNNNGAGGCAGGAGAATTGCTTGAA
AGCACACGTCTGAACTCTTTCCCTACACGACGCTCTTCCGATCTNNNAGGTAGGAGAATTGCTTGAAC
AGAGCACACGTCTGAACTCTTTCCCTACACGACGCTCTTCCGATCTNNNGACAGACAATGCAGTAAGCT
GAGCACACGTCTGAACTCTTTCCCTACACGACGCTCTTCCGATCTNNNATCTGTAGTCCTAGCTACTCGG
AGCACACGTCTGAACTCTTTCCCTACACGACGCTCTTCCGATCTNNNTAGTCACAGCTACTCTAGAGGC
AGCACACGTCTGAACTCTTTCCCTACACGACGCTCTTCCGATCTNNNTCCTGAACTAGACACTGGTAGG
GAGCACACGTCTGAACTCTTTCCCTACACGACGCTCTTCCGATCTNNNCTGAGGCAGGAGAATCACTTG
AGAGCACACGTCTGAACTCTTTCCCTACACGACGCTCTTCCGATCTNNNACAAAAATTAGCCGGATGTC
GAGCACACGTCTGAACTCTTTCCCTACACGACGCTCTTCCGATCTNNNAGAGGTTGTAGTGAGCCGAA
AGCACACGTCTGAACTCTTTCCCTACACGACGCTCTTCCGATCTNNNTAGTCAGGAGGCTGAAGTTGAA
GAGCACACGTCTGAACTCTTTCCCTACACGACGCTCTTCCGATCTNNNCTGTAATCCCAGCTACTCAGGA
AGCACACGTCTGAACTCTTTCCCTACACGACGCTCTTCCGATCTNNNATAAGCAATACTGGGAACACGC
GAGCACACGTCTGAACTCTTTCCCTACACGACGCTCTTCCGATCTNNNGTACCCGTAATCTCAGCTACTC
GAGCACACGTCTGAACTCTTTCCCTACACGACGCTCTTCCGATCTNNNGAGATTGAGCCACTGCATTCC
AGAGCACACGTCTGAACTCTTTCCCTACACGACGCTCTTCCGATCTNNNCTGAGGCAGGAGATTCAGTT
AGCACACGTCTGAACTCTTTCCCTACACGACGCTCTTCCGATCTNNNCTTGCAGTAAGCTGAGATTGCA
AGCACACGTCTGAACTCTTTCCCTACACGACGCTCTTCCGATCTNNNGAGGCAGGAGAATTGTTTGAA

GAGCACACGTCTGAACTCTTTCCCTACACGACGCTCTTCCGATCTNNNCTGAGGCAGGAGAATTGCTT
GCACACGTCTGAACTCTTTCCCTACACGACGCTCTTCCGATCTNNNCTGAGACAGGAGAATCGCTTGA
AGCACACGTCTGAACTCTTTCCCTACACGACGCTCTTCCGATCTNNNAGATCTCTTGAGCCCAGAAGTT
AGCACACGTCTGAACTCTTTCCCTACACGACGCTCTTCCGATCTNNNATGCTGGCACATGTTTGTAAT
AGCACACGTCTGAACTCTTTCCCTACACGACGCTCTTCCGATCTNNNAGAATCATTTGAACCCGGGAG
AGCACACGTCTGAACTCTTTCCCTACACGACGCTCTTCCGATCTNNNGCATCAGCAGAAGTCAGACT
GCACACGTCTGAACTCTTTCCCTACACGACGCTCTTCCGATCTNNNCAGAAGAATTGCTTGAACCTGG
AGCACACGTCTGAACTCTTTCCCTACACGACGCTCTTCCGATCTNNNTGAGGTAGGAGAATGGCATG
AGCACACGTCTGAACTCTTTCCCTACACGACGCTCTTCCGATCTNNNGTGGCACATGTCTGTAATTCCA
AGCACACGTCTGAACTCTTTCCCTACACGACGCTCTTCCGATCTNNNGGGTGATGCAAGAGAATCAA
AGCACACGTCTGAACTCTTTCCCTACACGACGCTCTTCCGATCTNNNTGAGGCAGAAGAATTGCTTGA
GAGCACACGTCTGAACTCTTTCCCTACACGACGCTCTTCCGATCTNNNAGAATCACTTGAACCTGGGG
GCACACGTCTGAACTCTTTCCCTACACGACGCTCTTCCGATCTNNNGTAATCCTAGCTACTTGGGAGG
GAGCACACGTCTGAACTCTTTCCCTACACGACGCTCTTCCGATCTNNNCATGTTGGATCACGCCTGTA
GCACACGTCTGAACTCTTTCCCTACACGACGCTCTTCCGATCTNNNGCTGAGGCAAGAAGAATCACTT
GCACACGTCTGAACTCTTTCCCTACACGACGCTCTTCCGATCTNNNGTTTGGCAAGCTTTTGTTTCTG
GCACACGTCTGAACTCTTTCCCTACACGACGCTCTTCCGATCTNNNCATGGTAGTGTGCACCTGTAGT
AGCACACGTCTGAACTCTTTCCCTACACGACGCTCTTCCGATCTNNNAGGCAGAAGAATCACTTGAAC
AGCACACGTCTGAACTCTTTCCCTACACGACGCTCTTCCGATCTNNNGTGAAACCCCGTCTCTACTGA
AGCACACGTCTGAACTCTTTCCCTACACGACGCTCTTCCGATCTNNNAATCTATTCGTCTGACAGAGG
AGCACACGTCTGAACTCTTTCCCTACACGACGCTCTTCCGATCTNNNAAAAAATTAGCCAGGCGTAG
GAGCACACGTCTGAACTCTTTCCCTACACGACGCTCTTCCGATCTNNNCTGAGGTAGGAGAATCGCTT
AGCACACGTCTGAACTCTTTCCCTACACGACGCTCTTCCGATCTNNNTCGATTTGAGCTCAGAAGTT
GCACACGTCTGAACTCTTTCCCTACACGACGCTCTTCCGATCTNNNAAACACTCCCTCCCTTTTCCTTTC
AGCACACGTCTGAACTCTTTCCCTACACGACGCTCTTCCGATCTNNNAGGCCAAATCAGGAGGATCAT
AGCACACGTCTGAACTCTTTCCCTACACGACGCTCTTCCGATCTNNNGGTGACATAAGAGAGACCCTG
AGCACACGTCTGAACTCTTTCCCTACACGACGCTCTTCCGATCTNNNTCACTTGAGTCCAGGAGTTCAA
GAGCACACGTCTGAACTCTTTCCCTACACGACGCTCTTCCGATCTNNNCCTGAACTGTGATTTTGCCAC
GAGCACACGTCTGAACTCTTTCCCTACACGACGCTCTTCCGATCTNNNCTCTCCTACTTGTGTTGCTCTG
AGCACACGTCTGAACTCTTTCCCTACACGACGCTCTTCCGATCTNNNATGCCTCTAATCCCAGCTACTT
AGCACACGTCTGAACTCTTTCCCTACACGACGCTCTTCCGATCTNNNCAGCAAGCACCATTTACTTCCT
AGCACACGTCTGAACTCTTTCCCTACACGACGCTCTTCCGATCTNNNCACCACCTGGCAGTATTCTCAT
AGCACACGTCTGAACTCTTTCCCTACACGACGCTCTTCCGATCTNNNAGTGAGCAGAGATTGTACCAC
AGCACACGTCTGAACTCTTTCCCTACACGACGCTCTTCCGATCTNNNGATTGCAGTGAGCCAGGATCA
AGCACACGTCTGAACTCTTTCCCTACACGACGCTCTTCCGATCTNNNAAAAAATTAGGCGGGTGTAG
AGCACACGTCTGAACTCTTTCCCTACACGACGCTCTTCCGATCTNNNGCTACAGTGAGTTGTGTTCAC
AGCACACGTCTGAACTCTTTCCCTACACGACGCTCTTCCGATCTNNNAGACAGGAGAATCACTTGAAC
AGCACACGTCTGAACTCTTTCCCTACACGACGCTCTTCCGATCTNNNCTGCTTCCTTTCACTTTCAGGTG
GAGCACACGTCTGAACTCTTTCCCTACACGACGCTCTTCCGATCTNNNGCCATCATTTGAGTTGGAAG
GCACACGTCTGAACTCTTTCCCTACACGACGCTCTTCCGATCTNNNTTCAGTGAGCCATGATTGTACC
AGCACACGTCTGAACTCTTTCCCTACACGACGCTCTTCCGATCTNNNCTGTGTTCATGCTACTGGACTC
AGCACACGTCTGAACTCTTTCCCTACACGACGCTCTTCCGATCTNNNTGTGTGACTGTAATCCCAGCT
AGCACACGTCTGAACTCTTTCCCTACACGACGCTCTTCCGATCTNNNCTTGAGACCAGACTGACCAAC

AGAGCACACGTCTGAACTCTTTCCCTACACGACGCTCTTCCGATCTNNNGCAGGAGAATTGCTTGAAC
GAGCACACGTCTGAACTCTTTCCCTACACGACGCTCTTCCGATCTNNNATGCCTGTAATCCCAGCTACTT
GCACACGTCTGAACTCTTTCCCTACACGACGCTCTTCCGATCTNNNGCACATGTAGTCCCAGCTACTAG
AGCACACGTCTGAACTCTTTCCCTACACGACGCTCTTCCGATCTNNNCCTCAGTGACAAGAGTGAGAC
AGAGCACACGTCTGAACTCTTTCCCTACACGACGCTCTTCCGATCTNNNTAACTTGGGGAGTGGGATC
GAGCACACGTCTGAACTCTTTCCCTACACGACGCTCTTCCGATCTNNNAATCCCAAGTTTGCAATGAGG
GAGCACACGTCTGAACTCTTTCCCTACACGACGCTCTTCCGATCTNNNCAGAGGCAGGAGAATCACTT
GAGCACACGTCTGAACTCTTTCCCTACACGACGCTCTTCCGATCTNNNAGCCAAGATCACACCACTGTA
GAGCACACGTCTGAACTCTTTCCCTACACGACGCTCTTCCGATCTNNNGCTGATGTGGGAGGATCAC
AGCACACGTCTGAACTCTTTCCCTACACGACGCTCTTCCGATCTNNNACCTTCCCTCCACTATTGTCCTA
GAGCACACGTCTGAACTCTTTCCCTACACGACGCTCTTCCGATCTNNNCAGGAGAATTGCTTGAACCTT
AGCACACGTCTGAACTCTTTCCCTACACGACGCTCTTCCGATCTNNNAGAGGTTGCAGTGAGTTGAG
AAGAGCACACGTCTGAACTCTTTCCCTACACGACGCTCTTCCGATCTNNNTGAGAATCACTTGAACCTG
AGAGCACACGTCTGAACTCTTTCCCTACACGACGCTCTTCCGATCTNNNTGAGGCAGGAGAATCTCTT
AGCACACGTCTGAACTCTTTCCCTACACGACGCTCTTCCGATCTNNNAGGCAGGAGAATTACTTGAAC
GAGCACACGTCTGAACTCTTTCCCTACACGACGCTCTTCCGATCTNNNTGTAATCCCAGCACTTTGAGA
GAGCACACGTCTGAACTCTTTCCCTACACGACGCTCTTCCGATCTNNNAAATAAATAATCCTGGGCTG
AGAGCACACGTCTGAACTCTTTCCCTACACGACGCTCTTCCGATCTNNNACAAAAATTAGCTGGGTGT
GAGCACACGTCTGAACTCTTTCCCTACACGACGCTCTTCCGATCTNNNTTCTCTGCAAGGAATTCTCAG
GAGCACACGTCTGAACTCTTTCCCTACACGACGCTCTTCCGATCTNNNGAATAGCTTGGATCCGGGAG
GAGCACACGTCTGAACTCTTTCCCTACACGACGCTCTTCCGATCTNNNACCTCCTTACTCCTGCAAGAAT
GAGCACACGTCTGAACTCTTTCCCTACACGACGCTCTTCCGATCTNNNGCCTATAGTTCCAGCTACTCG
AGCACACGTCTGAACTCTTTCCCTACACGACGCTCTTCCGATCTNNNGCTGACACCGGTAAGATATCC
GAGCACACGTCTGAACTCTTTCCCTACACGACGCTCTTCCGATCTNNNTTGGTCTCAAGCTACATGGTT
AGAGCACACGTCTGAACTCTTTCCCTACACGACGCTCTTCCGATCTNNNTGCACATGTATCCCAGAACG
GAGCACACGTCTGAACTCTTTCCCTACACGACGCTCTTCCGATCTNNNCAATCTCTGGGTTTTGGGAAC
GCACACGTCTGAACTCTTTCCCTACACGACGCTCTTCCGATCTNNNCTGTAATCCCAGCTACTCAGGAG
GAGCACACGTCTGAACTCTTTCCCTACACGACGCTCTTCCGATCTNNNAGAGATGGTTCATGCTCTCAA
AGCACACGTCTGAACTCTTTCCCTACACGACGCTCTTCCGATCTNNNAAAAAAATTAGCCAGGCAGTC
AGCACACGTCTGAACTCTTTCCCTACACGACGCTCTTCCGATCTNNNCCATTGTACTCTAGCCTGGACA
AGCACACGTCTGAACTCTTTCCCTACACGACGCTCTTCCGATCTNNNCCTGTAATCCTAGCACTTTGGG
AGCACACGTCTGAACTCTTTCCCTACACGACGCTCTTCCGATCTNNNAAAATTATCTGTGTGTGGCGG
GAGCACACGTCTGAACTCTTTCCCTACACGACGCTCTTCCGATCTNNNCATGGTGGCACGCTCTTATA
AGCACACGTCTGAACTCTTTCCCTACACGACGCTCTTCCGATCTNNNTGAGGCAGGAGAATCTCTTGA
AGCACACGTCTGAACTCTTTCCCTACACGACGCTCTTCCGATCTNNNTAAGGCAGGAGAATCGCTTG
AGCACACGTCTGAACTCTTTCCCTACACGACGCTCTTCCGATCTNNNCTCCGAAAGTGCTGGGATTAC
GAGCACACGTCTGAACTCTTTCCCTACACGACGCTCTTCCGATCTNNNCCAACATGGTAAAACCCTGT
AGCACACGTCTGAACTCTTTCCCTACACGACGCTCTTCCGATCTNNNGGTTCATTCCTGTAATCCCAGC
AGCACACGTCTGAACTCTTTCCCTACACGACGCTCTTCCGATCTNNNGTGCACCTGTAATCTCAGCTAC
GAGCACACGTCTGAACTCTTTCCCTACACGACGCTCTTCCGATCTNNNTACCCCCACCTTCTTTTATTG
AGCACACGTCTGAACTCTTTCCCTACACGACGCTCTTCCGATCTNNNGCCAAGATCACACCACTGTACT
GAGCACACGTCTGAACTCTTTCCCTACACGACGCTCTTCCGATCTNNNTGAGCTGAGATTGTTCCACT
GCACACGTCTGAACTCTTTCCCTACACGACGCTCTTCCGATCTNNNAGAATTGCTTGAATTTGGGAG

AGCACACGTCTGAACTCTTTCCCTACACGACGCTCTTCCGATCTNNNAGACAAGGGGGAAATGTATG
AGAGCACACGTCTGAACTCTTTCCCTACACGACGCTCTTCCGATCTNNNCTGAGGCAGGAGAATCACT
AGCACACGTCTGAACTCTTTCCCTACACGACGCTCTTCCGATCTNNNCAGCTGTGTTCTGTTTGGATAG
AGCACACGTCTGAACTCTTTCCCTACACGACGCTCTTCCGATCTNNNTTATGAAGCCAGCATCATCCTGA
GAGCACACGTCTGAACTCTTTCCCTACACGACGCTCTTCCGATCTNNNTACCTCTTCTGCTCCCTTTTTCC
AGAGCACACGTCTGAACTCTTTCCCTACACGACGCTCTTCCGATCTNNNCTGAAATCCCAGCTACTCAGG
AGCACACGTCTGAACTCTTTCCCTACACGACGCTCTTCCGATCTNNNGAGGCAGGAGAATTGCTTGAA
AGAGCACACGTCTGAACTCTTTCCCTACACGACGCTCTTCCGATCTNNNTGCCACTATCTGATACGGAA
GAGCACACGTCTGAACTCTTTCCCTACACGACGCTCTTCCGATCTNNNCTCCCTGTGTGAAGCATCAAA
GAGCACACGTCTGAACTCTTTCCCTACACGACGCTCTTCCGATCTNNNAGAGGCAGGAGAATCACTTGA
AGCACACGTCTGAACTCTTTCCCTACACGACGCTCTTCCGATCTNNNCACCTGTAATCCCAGCTACTCAG
GAGCACACGTCTGAACTCTTTCCCTACACGACGCTCTTCCGATCTNNNTCTACCAATGTACCTCCAGGAA
GAGCACACGTCTGAACTCTTTCCCTACACGACGCTCTTCCGATCTNNNCTGAGGCAGGAGAATCACTTG
GAGCACACGTCTGAACTCTTTCCCTACACGACGCTCTTCCGATCTNNNGATTGCATGAGCCTGAGAGA
GAGCACACGTCTGAACTCTTTCCCTACACGACGCTCTTCCGATCTNNNCTGAGACGGGAGAATCACTT
GAGCACACGTCTGAACTCTTTCCCTACACGACGCTCTTCCGATCTNNNCTGCCAATGCCCAATATGATCT
AGAGCACACGTCTGAACTCTTTCCCTACACGACGCTCTTCCGATCTNNNGAGAATTGCTTGAACCCAGG
GAGCACACGTCTGAACTCTTTCCCTACACGACGCTCTTCCGATCTNNNCTGTAGTCTCAGCTACTGGG
AGCACACGTCTGAACTCTTTCCCTACACGACGCTCTTCCGATCTNNNGTAATCCCAGCTACTTAGGAG
GAGCACACGTCTGAACTCTTTCCCTACACGACGCTCTTCCGATCTNNNTGAGCTGAGATCACACTATTG
GAGCACACGTCTGAACTCTTTCCCTACACGACGCTCTTCCGATCTNNNGGCAACAACAGCAAAACTC
GAGCACACGTCTGAACTCTTTCCCTACACGACGCTCTTCCGATCTNNNGTGTGCCTGTAATCTCAGCTA
AGCACACGTCTGAACTCTTTCCCTACACGACGCTCTTCCGATCTNNNCTGAGGCAGGAGAATCACTTGA
GAGCACACGTCTGAACTCTTTCCCTACACGACGCTCTTCCGATCTNNNAACAAGGCAAGACCCCATCTC
AGCACACGTCTGAACTCTTTCCCTACACGACGCTCTTCCGATCTNNNCTGAGGCAGGAGAATCACTTG
GAGCACACGTCTGAACTCTTTCCCTACACGACGCTCTTCCGATCTNNNCAGGAGAATTGCTTGAAACCA
GAGCACACGTCTGAACTCTTTCCCTACACGACGCTCTTCCGATCTNNNCAGGAGAATTGCTTGAAACCA
AGCACACGTCTGAACTCTTTCCCTACACGACGCTCTTCCGATCTNNNAATCACATCTACTTGGGAGGCT
GAGCACACGTCTGAACTCTTTCCCTACACGACGCTCTTCCGATCTNNNCCCAGAGCACAGTCAATTTTTG
GAGCACACGTCTGAACTCTTTCCCTACACGACGCTCTTCCGATCTNNNAGATCAAGACCATCCTGACCA
GAGCACACGTCTGAACTCTTTCCCTACACGACGCTCTTCCGATCTNNNCACCTGTAGTCCCAGTTACTT
GAGCACACGTCTGAACTCTTTCCCTACACGACGCTCTTCCGATCTNNNTAGGAGAATTGCTTGAACCA
AGCACACGTCTGAACTCTTTCCCTACACGACGCTCTTCCGATCTNNNTGTAGTCCTAGCTACTTGGGAG
GAGCACACGTCTGAACTCTTTCCCTACACGACGCTCTTCCGATCTNNNCATGCCTGTAATCCCAGCTACT
GAGCACACGTCTGAACTCTTTCCCTACACGACGCTCTTCCGATCTNNNTGTAATCCCAGCTAGTCAGGA
GAGCACACGTCTGAACTCTTTCCCTACACGACGCTCTTCCGATCTNNNAGTGAGGAGTGAGACTCTAC
GAGCACACGTCTGAACTCTTTCCCTACACGACGCTCTTCCGATCTNNNGAGGTGGGAGAATTGCTTGA
GAGCACACGTCTGAACTCTTTCCCTACACGACGCTCTTCCGATCTNNNAGGAGAATTGCTTGAACCTGG
GAGCACACGTCTGAACTCTTTCCCTACACGACGCTCTTCCGATCTNNNCAGGAGAATTGCTTGAAACCA
AGCACACGTCTGAACTCTTTCCCTACACGACGCTCTTCCGATCTNNNACATAATGAGGCTTTCTGGGGA
AGAGCACACGTCTGAACTCTTTCCCTACACGACGCTCTTCCGATCTNNNCAAGATCACACCACTGCACTA
GAGCACACGTCTGAACTCTTTCCCTACACGACGCTCTTCCGATCTNNNCATCCTCGCTAACATGGTGAA
AGCACACGTCTGAACTCTTTCCCTACACGACGCTCTTCCGATCTNNNACATCCCTTTGACACTGGTTCTT

AGCACACGTCTGAACTCTTTCCCTACACGACGCTCTTCCGATCTNNNCAACTGGAAACATGTGATCTG
GCACACGTCTGAACTCTTTCCCTACACGACGCTCTTCCGATCTNNNACAAAAATTAGCTGGGTGTGG
AGCACACGTCTGAACTCTTTCCCTACACGACGCTCTTCCGATCTNNNTGAGCTGAGATCATGCCATTG
AGCACACGTCTGAACTCTTTCCCTACACGACGCTCTTCCGATCTNNNGGCCTATCACAATGCTGTAGG
AGCACACGTCTGAACTCTTTCCCTACACGACGCTCTTCCGATCTNNNCTGTAGTGCCAGCTACTCAGG
AGCACACGTCTGAACTCTTTCCCTACACGACGCTCTTCCGATCTNNNCAGGAGTTCAAGACCAGCCTAT
AGCACACGTCTGAACTCTTTCCCTACACGACGCTCTTCCGATCTNNNACAGCTCCTGGATTTCTTGATC
AGCACACGTCTGAACTCTTTCCCTACACGACGCTCTTCCGATCTNNNAGGTGGGAGGATTGGTTTAT
GCACACGTCTGAACTCTTTCCCTACACGACGCTCTTCCGATCTNNNACGCCCAGGTTCTTTAGATGAA
AGCACACGTCTGAACTCTTTCCCTACACGACGCTCTTCCGATCTNNNCCCGGCAGCTGTTTCTACTATA
GCACACGTCTGAACTCTTTCCCTACACGACGCTCTTCCGATCTNNNAGGAAGGAGAATCACTTGAAC
AGCACACGTCTGAACTCTTTCCCTACACGACGCTCTTCCGATCTNNNAGTAGCACTGCTATACACCAA
AGCACACGTCTGAACTCTTTCCCTACACGACGCTCTTCCGATCTNNNCTGAGGCAGGAGAATCACTT
AGCACACGTCTGAACTCTTTCCCTACACGACGCTCTTCCGATCTNNNATAATCCCAGCTACTTGGAAGG
GAGCACACGTCTGAACTCTTTCCCTACACGACGCTCTTCCGATCTNNNCTGAGGCAAGAGAGTTGCTT
AGCACACGTCTGAACTCTTTCCCTACACGACGCTCTTCCGATCTNNNAGCTTACAGTGAGCCAAGATC
AGCACACGTCTGAACTCTTTCCCTACACGACGCTCTTCCGATCTNNNTGCCTGTAATTCCAGCTACTCA
GAGCACACGTCTGAACTCTTTCCCTACACGACGCTCTTCCGATCTNNNTCATGCCTGTAATCCCAACAC
GAGCACACGTCTGAACTCTTTCCCTACACGACGCTCTTCCGATCTNNNGATCACACTATTGCACTTGAG
GCACACGTCTGAACTCTTTCCCTACACGACGCTCTTCCGATCTNNNATAGCCTGGACAACATGGTGAA
AGCACACGTCTGAACTCTTTCCCTACACGACGCTCTTCCGATCTNNNAGGAGGCTGAGGTAGGAGA
GAGCACACGTCTGAACTCTTTCCCTACACGACGCTCTTCCGATCTNNNAAGAAATTGCCCCAGAGGAG
GAGCACACGTCTGAACTCTTTCCCTACACGACGCTCTTCCGATCTNNNGGGACTATGTCTCTTTCCTCT
AGCACACGTCTGAACTCTTTCCCTACACGACGCTCTTCCGATCTNNNACACCTGTAATCCCAGCACTTT
GAGCACACGTCTGAACTCTTTCCCTACACGACGCTCTTCCGATCTNNNTTTGTTATTGCCCAGACTTGT
GCACACGTCTGAACTCTTTCCCTACACGACGCTCTTCCGATCTNNNGGACAAGTCTGTTGTCTCGATG
AGCACACGTCTGAACTCTTTCCCTACACGACGCTCTTCCGATCTNNNGGGACATGACAATTGCTTGAA
AGCACACGTCTGAACTCTTTCCCTACACGACGCTCTTCCGATCTNNNGTGCCTGTAATCCCAGCTACT
AGCACACGTCTGAACTCTTTCCCTACACGACGCTCTTCCGATCTNNNCAACATGGCAAAACCCAATCT
AGCACACGTCTGAACTCTTTCCCTACACGACGCTCTTCCGATCTNNNTGAGGTAGGAGAATCGCTTG
AGCACACGTCTGAACTCTTTCCCTACACGACGCTCTTCCGATCTNNNTTGAGACAGGAGAATCGCTT
GAGCACACGTCTGAACTCTTTCCCTACACGACGCTCTTCCGATCTNNNTGAGGCAGGAGAATCACTT
GAGCACACGTCTGAACTCTTTCCCTACACGACGCTCTTCCGATCTNNNGTGGATCACCTATGGTCAAG
AGCACACGTCTGAACTCTTTCCCTACACGACGCTCTTCCGATCTNNNAGGTTTCAGTGAGCCAAGATC
AGCACACGTCTGAACTCTTTCCCTACACGACGCTCTTCCGATCTNNNAAAAAATAATTACCCGGGCGT
AGCACACGTCTGAACTCTTTCCCTACACGACGCTCTTCCGATCTNNNGCAGAGAATTTCTTGAACCCG
AGCACACGTCTGAACTCTTTCCCTACACGACGCTCTTCCGATCTNNNACCATGGAATACTATGCAGCC
GAGCACACGTCTGAACTCTTTCCCTACACGACGCTCTTCCGATCTNNNAAATGAGAATCGCTTGAACC
GCACACGTCTGAACTCTTTCCCTACACGACGCTCTTCCGATCTNNNCTAGTCCCTCCTCCCTATGAGTC
GCACACGTCTGAACTCTTTCCCTACACGACGCTCTTCCGATCTNNNCATGCCTGTAATCCCAGCTACTC
AGCACACGTCTGAACTCTTTCCCTACACGACGCTCTTCCGATCTNNNCTGTAGTCCCAGCTACTCAGGA
GAGCACACGTCTGAACTCTTTCCCTACACGACGCTCTTCCGATCTNNNACACAATAACACAGCACTCAG
GCACACGTCTGAACTCTTTCCCTACACGACGCTCTTCCGATCTNNNTTGTGACTGACAGAAAAAGGT

GAGCACACGTCTGAACTCTTTCCCTACACGACGCTCTTCCGATCTNNNAAAAATTAGCTGGGTTGGT
AGCACACGTCTGAACTCTTTCCCTACACGACGCTCTTCCGATCTNNNACTCTTGATGCCCAAAACAACA
GAGCACACGTCTGAACTCTTTCCCTACACGACGCTCTTCCGATCTNNNCATAGTGAAACCCTGTCTCTG
AGCACACGTCTGAACTCTTTCCCTACACGACGCTCTTCCGATCTNNNAGACAATTACTTGAACCCGGG
GCACACGTCTGAACTCTTTCCCTACACGACGCTCTTCCGATCTNNNATAGGTTGCTGTGAGCCATGAT
GAGCACACGTCTGAACTCTTTCCCTACACGACGCTCTTCCGATCTNNNTGAGGCAGGAGAATACTGT
AGCACACGTCTGAACTCTTTCCCTACACGACGCTCTTCCGATCTNNNGTGAGTGCAGGGAACTTCTCT
GCACACGTCTGAACTCTTTCCCTACACGACGCTCTTCCGATCTNNNAGTCAGGAGAATCACTTGAACC
GAGCACACGTCTGAACTCTTTCCCTACACGACGCTCTTCCGATCTNNNTCTTTAGCAAAGAGACAGCC
AGCACACGTCTGAACTCTTTCCCTACACGACGCTCTTCCGATCTNNNGTGAAACCCCGTCTCTACAAA
GAGCACACGTCTGAACTCTTTCCCTACACGACGCTCTTCCGATCTNNNAAAAAAATAGCAGGGCATG
GAGCACACGTCTGAACTCTTTCCCTACACGACGCTCTTCCGATCTNNNCTGTAATCTCAGCTACTCGG
AGCACACGTCTGAACTCTTTCCCTACACGACGCTCTTCCGATCTNNNGAGGTTGCAGTGAGCCAAGAT
GAGCACACGTCTGAACTCTTTCCCTACACGACGCTCTTCCGATCTNNNGAATGGTGTGAACCTGGGA
AGCACACGTCTGAACTCTTTCCCTACACGACGCTCTTCCGATCTNNNAAGGCCCAAAGGGTACATATT
GCACACGTCTGAACTCTTTCCCTACACGACGCTCTTCCGATCTNNNAAAAAATTAGCTGGGCATGGT
AGCACACGTCTGAACTCTTTCCCTACACGACGCTCTTCCGATCTNNNTGAGGGAGGAGAATCACTTG
GCACACGTCTGAACTCTTTCCCTACACGACGCTCTTCCGATCTNNNCCAGTTCCTAGCAATTCCTGAGA
GCACACGTCTGAACTCTTTCCCTACACGACGCTCTTCCGATCTNNNAGACAGGAGAATCACTTGAACC
AGAGCACACGTCTGAACTCTTTCCCTACACGACGCTCTTCCGATCTNNNCCTACTAAATCCTGCCCATCT
GAGCACACGTCTGAACTCTTTCCCTACACGACGCTCTTCCGATCTNNNTCAAGACCATCCTGGCTAAC
GAGCACACGTCTGAACTCTTTCCCTACACGACGCTCTTCCGATCTNNNCATGAAAGCCATCAACCCTGA
AGCACACGTCTGAACTCTTTCCCTACACGACGCTCTTCCGATCTNNNTCCCGAAATTGAACCAGGAAG
AGCACACGTCTGAACTCTTTCCCTACACGACGCTCTTCCGATCTNNNGACTGATGCAGGAGAATTGCT
GCACACGTCTGAACTCTTTCCCTACACGACGCTCTTCCGATCTNNNCTCAAAAAAAAAAGGTGGGCA
GCACACGTCTGAACTCTTTCCCTACACGACGCTCTTCCGATCTNNNGTGGAGGTTACAGTGAGTTGA
AGCACACGTCTGAACTCTTTCCCTACACGACGCTCTTCCGATCTNNNTGAGCTTAGGAGTTCAAGACC
AGCACACGTCTGAACTCTTTCCCTACACGACGCTCTTCCGATCTNNNAGTGAGCAGAAATAGTGCCAC
AGCACACGTCTGAACTCTTTCCCTACACGACGCTCTTCCGATCTNNNCTGTAATCCCAGCTACTCAGGA
GAGCACACGTCTGAACTCTTTCCCTACACGACGCTCTTCCGATCTNNNAGAATCACTTGAACCTGGG
GAGCACACGTCTGAACTCTTTCCCTACACGACGCTCTTCCGATCTNNNGACCTTAGAGACATGCTACC
AGCACACGTCTGAACTCTTTCCCTACACGACGCTCTTCCGATCTNNNACACAAAAAATTAGCCACGTGT
AGCACACGTCTGAACTCTTTCCCTACACGACGCTCTTCCGATCTNNNAATTCCTGGACACATACACCCT
AGCACACGTCTGAACTCTTTCCCTACACGACGCTCTTCCGATCTNNNAAAAAAAATTAGCCAGGCGTG
AGCACACGTCTGAACTCTTTCCCTACACGACGCTCTTCCGATCTNNNACAGTGAAACCCGGTCTCTAC
GCACACGTCTGAACTCTTTCCCTACACGACGCTCTTCCGATCTNNNTCTCCAAGGTACATCTCAACTGT
AGCACACGTCTGAACTCTTTCCCTACACGACGCTCTTCCGATCTNNNCCTTTGCCTCCTTCTAGCTGATT
GCACACGTCTGAACTCTTTCCCTACACGACGCTCTTCCGATCTNNNGTGTCTGCTTCTTAGGAGTCTCC
AGCACACGTCTGAACTCTTTCCCTACACGACGCTCTTCCGATCTNNNCATGCCTCTAATCCCAGCTACT
GCACACGTCTGAACTCTTTCCCTACACGACGCTCTTCCGATCTNNNCAGGAGGCTGAGGTATGAGAA
AGCACACGTCTGAACTCTTTCCCTACACGACGCTCTTCCGATCTNNNTGGGAAGAAGTGGGAGGAA
GCACACGTCTGAACTCTTTCCCTACACGACGCTCTTCCGATCTNNNCTGAGGCAGGAGAATGACATG
GCACACGTCTGAACTCTTTCCCTACACGACGCTCTTCCGATCTNNNCTCAAGTAGCTGGGACTATAGG

GAGCACACGTCTGAACTCTTTCCCTACACGACGCTCTTCCGATCTNNNTGAAGAACACTTTGGAGGTTC
AGAGCACACGTCTGAACTCTTTCCCTACACGACGCTCTTCCGATCTNNNGAAAGAGAATCGCTTGAAC
GAGCACACGTCTGAACTCTTTCCCTACACGACGCTCTTCCGATCTNNNAGACAGGAGAATCACTTGAA
AGAGCACACGTCTGAACTCTTTCCCTACACGACGCTCTTCCGATCTNNNGAGGCAGAAGAATCACTTGA
AGCACACGTCTGAACTCTTTCCCTACACGACGCTCTTCCGATCTNNNTCCTTCCTTCCTTTCCTTCCTTCCA
GCACACGTCTGAACTCTTTCCCTACACGACGCTCTTCCGATCTNNNGGAGTTTGCAGTGAGCTGAGAT
AGCACACGTCTGAACTCTTTCCCTACACGACGCTCTTCCGATCTNNNCAGGAGATGGAGATTACGGTGA
AGCACACGTCTGAACTCTTTCCCTACACGACGCTCTTCCGATCTNNNCTCATGCCTATAATCCAAGCCCT
AGCACACGTCTGAACTCTTTCCCTACACGACGCTCTTCCGATCTNNNCATGGGGAGACATCATCTGTA
GAGCACACGTCTGAACTCTTTCCCTACACGACGCTCTTCCGATCTNNNCTGTAATGCTAGCTACTCGGG
GAGCACACGTCTGAACTCTTTCCCTACACGACGCTCTTCCGATCTNNNAATATTGGCTCTCTACCAGGT
GAGCACACGTCTGAACTCTTTCCCTACACGACGCTCTTCCGATCTNNNAGGCAGGAGAATCACTTGAA
GAGCACACGTCTGAACTCTTTCCCTACACGACGCTCTTCCGATCTNNNAGGTAGACTGGATTGCGAAT
AGCACACGTCTGAACTCTTTCCCTACACGACGCTCTTCCGATCTNNNCACGGCTTTCCTGCTTTGATAA
GAGCACACGTCTGAACTCTTTCCCTACACGACGCTCTTCCGATCTNNNTTGGTAGGTTGTACATGTCCA
AGCACACGTCTGAACTCTTTCCCTACACGACGCTCTTCCGATCTNNNGAGATTGCAGTGAGCCAAGAT
AGAGCACACGTCTGAACTCTTTCCCTACACGACGCTCTTCCGATCTNNNCTGTAATCCCAGCTACTCAG
AGCACACGTCTGAACTCTTTCCCTACACGACGCTCTTCCGATCTNNNGTGAATCACTTGAGGTCAGGA
AGAGCACACGTCTGAACTCTTTCCCTACACGACGCTCTTCCGATCTNNNCAGGAGAATCACTTGAACCT
AGCACACGTCTGAACTCTTTCCCTACACGACGCTCTTCCGATCTNNNCTTACACTTACAGTCCCAGCACT
GAGCACACGTCTGAACTCTTTCCCTACACGACGCTCTTCCGATCTNNNCATCTTACATGGATGGCAGCA
GAGCACACGTCTGAACTCTTTCCCTACACGACGCTCTTCCGATCTNNNCCGCAACCTGTTTTTGTTTTGT
GAGCACACGTCTGAACTCTTTCCCTACACGACGCTCTTCCGATCTNNNTGGCCAAATCCAAAGGAAAGT
AGCACACGTCTGAACTCTTTCCCTACACGACGCTCTTCCGATCTNNNAATGATGTGAACCCAGGTGTGA
GAGCACACGTCTGAACTCTTTCCCTACACGACGCTCTTCCGATCTNNNTACAAAAAATTAGCTGGGCGT
GAGCACACGTCTGAACTCTTTCCCTACACGACGCTCTTCCGATCTNNNCGCCTGTAATTTCAGCTACTTC
GAGCACACGTCTGAACTCTTTCCCTACACGACGCTCTTCCGATCTNNNAAAATTAGCTGGGCATGCTA
GAGCACACGTCTGAACTCTTTCCCTACACGACGCTCTTCCGATCTNNNCAAGATTGCTCCATTGCACTTC
GAGCACACGTCTGAACTCTTTCCCTACACGACGCTCTTCCGATCTNNNACGCCTGTAATCCCAAAACTTT
AGAGCACACGTCTGAACTCTTTCCCTACACGACGCTCTTCCGATCTNNNCTGAGGCAGGAGAATCACTT
CACACGTCTGAACTCTTTCCCTACACGACGCTCTTCCGATCTNNNTACAAAAAATTAGCCGCGTGTGG
AGAGCACACGTCTGAACTCTTTCCCTACACGACGCTCTTCCGATCTNNNGTACACTGACTGGACCACTA
AGCACACGTCTGAACTCTTTCCCTACACGACGCTCTTCCGATCTNNNACCACCATAATCCCAGCTACTTG
AGCACACGTCTGAACTCTTTCCCTACACGACGCTCTTCCGATCTNNNCAGTGGTAGTCTTCAGCCTCTA
AGCACACGTCTGAACTCTTTCCCTACACGACGCTCTTCCGATCTNNNTACAAAAAATTAGCCGGGTGT
GAGCACACGTCTGAACTCTTTCCCTACACGACGCTCTTCCGATCTNNNGTTTGAAACTGAGAGGTCAA
AGCACACGTCTGAACTCTTTCCCTACACGACGCTCTTCCGATCTNNNGCATGCTCAAGGATCAAAGACT
GAGCACACGTCTGAACTCTTTCCCTACACGACGCTCTTCCGATCTNNNATTAGCCAGGTGTAGAGTCAT
AGAGCACACGTCTGAACTCTTTCCCTACACGACGCTCTTCCGATCTNNNCTCACACCTCTAATCCCAGCAT
AGCACACGTCTGAACTCTTTCCCTACACGACGCTCTTCCGATCTNNNAGGAGAATTGCTTGAATCTGG
AGCACACGTCTGAACTCTTTCCCTACACGACGCTCTTCCGATCTNNNCTGAGGTGGGAAGATCACTTGA
GAGCACACGTCTGAACTCTTTCCCTACACGACGCTCTTCCGATCTNNNTACAAAAAATTAGCTGGGCGT
GAGCACACGTCTGAACTCTTTCCCTACACGACGCTCTTCCGATCTNNNGAGAGAGCAAGGTGGGAAC

GAGCACACGTCTGAACTCTTTCCCTACACGACGCTCTTCCGATCTNNNAGAGAATTGCTTGAACCCAG
AGCACACGTCTGAACTCTTTCCCTACACGACGCTCTTCCGATCTNNNCAGAATCTGTCTCTGTCACCCAA
GAGCACACGTCTGAACTCTTTCCCTACACGACGCTCTTCCGATCTNNNAAAAAATTAGCCTGGCATGG
AGCACACGTCTGAACTCTTTCCCTACACGACGCTCTTCCGATCTNNNGAACCTTTGCAACCACAATGAG
AGAGCACACGTCTGAACTCTTTCCCTACACGACGCTCTTCCGATCTNNNCTGTAATCCCAGATACTCGG
AGCACACGTCTGAACTCTTTCCCTACACGACGCTCTTCCGATCTNNNGAGGCAACAGAATCACTTGAA
AAGAGCACACGTCTGAACTCTTTCCCTACACGACGCTCTTCCGATCTNNNGAGGTAGAGGTTGCAGT
GAGCACACGTCTGAACTCTTTCCCTACACGACGCTCTTCCGATCTNNNATGCCATCTAGGACTTCCATA
GAGCACACGTCTGAACTCTTTCCCTACACGACGCTCTTCCGATCTNNNCTGTAGTCCCAGCTACTCAGG
GAGCACACGTCTGAACTCTTTCCCTACACGACGCTCTTCCGATCTNNNCTGAGGCAGGAGAATCACTT
AGCACACGTCTGAACTCTTTCCCTACACGACGCTCTTCCGATCTNNNGAACTGTGATTGTGCCACTACA
GCACACGTCTGAACTCTTTCCCTACACGACGCTCTTCCGATCTNNNGAGGCAGGAGAATTGCTTGAAT
GAGCACACGTCTGAACTCTTTCCCTACACGACGCTCTTCCGATCTNNNCTGTAATCCCAGCTACTCAGG
AGCACACGTCTGAACTCTTTCCCTACACGACGCTCTTCCGATCTNNNATGCTCTCCCAAGCTTATCTTCC
AGCACACGTCTGAACTCTTTCCCTACACGACGCTCTTCCGATCTNNNAGACAGGAGAATCGCTTGAAC
GAGCACACGTCTGAACTCTTTCCCTACACGACGCTCTTCCGATCTNNNAATACAAAAAATTAGCCGGG
GAGCACACGTCTGAACTCTTTCCCTACACGACGCTCTTCCGATCTNNNGAGCAAGCCCAGAACTTGAAA
AGCACACGTCTGAACTCTTTCCCTACACGACGCTCTTCCGATCTNNNAGGCAAGAGAATTGCTTGAAC
GAGCACACGTCTGAACTCTTTCCCTACACGACGCTCTTCCGATCTNNNGTGGATGCTGAGAAATTGG
GAGCACACGTCTGAACTCTTTCCCTACACGACGCTCTTCCGATCTNNNGGAAGTTGTAGTGAGCCGAG
GAGCACACGTCTGAACTCTTTCCCTACACGACGCTCTTCCGATCTNNNGAGATCATGCCACTACACTCC
AGCACACGTCTGAACTCTTTCCCTACACGACGCTCTTCCGATCTNNNCTATAGTCCCAGCTACTTGGCA
GAGCACACGTCTGAACTCTTTCCCTACACGACGCTCTTCCGATCTNNNGGCTCACATCTGTAATCCTAG
AGCACACGTCTGAACTCTTTCCCTACACGACGCTCTTCCGATCTNNNGAAAACTCTACGGCAGTCTCTC
AGCACACGTCTGAACTCTTTCCCTACACGACGCTCTTCCGATCTNNNTGAGTGACAAGGGCGAAACTT
GAGCACACGTCTGAACTCTTTCCCTACACGACGCTCTTCCGATCTNNNCACATCTGTGGTATTCTTCCCC
AGCACACGTCTGAACTCTTTCCCTACACGACGCTCTTCCGATCTNNNACAAAGCAAAAACCCCATCTCTG
GAGCACACGTCTGAACTCTTTCCCTACACGACGCTCTTCCGATCTNNNCCCACAACAGAGTCATAGGTT
GAGCACACGTCTGAACTCTTTCCCTACACGACGCTCTTCCGATCTNNNAAAAAATTAGCCAGGCGTGG
GAGCACACGTCTGAACTCTTTCCCTACACGACGCTCTTCCGATCTNNNTAAAAAATTAGCCGGGTGTGG
GAGCACACGTCTGAACTCTTTCCCTACACGACGCTCTTCCGATCTNNNATTAGGGGGTCTGTGCCTACT
GCACACGTCTGAACTCTTTCCCTACACGACGCTCTTCCGATCTNNNATCCCAACTATTTCAGAGGCTGG
GAGCACACGTCTGAACTCTTTCCCTACACGACGCTCTTCCGATCTNNNACAGTGAAACCCCGTCTCTAC
AGCACACGTCTGAACTCTTTCCCTACACGACGCTCTTCCGATCTNNNGGATTGCCTGAGGTAAGGAGT
AGCACACGTCTGAACTCTTTCCCTACACGACGCTCTTCCGATCTNNNGCTTCTCTATCCAGCATAGGTCC
AGAGCACACGTCTGAACTCTTTCCCTACACGACGCTCTTCCGATCTNNNAAAAAATTAGCCAGGTGTGG
AGCACACGTCTGAACTCTTTCCCTACACGACGCTCTTCCGATCTNNNGTAATCTCAGCTACTCTTGGGG
AGCACACGTCTGAACTCTTTCCCTACACGACGCTCTTCCGATCTNNNCTGAGGAAGGAGATTGGCTTG
GAGCACACGTCTGAACTCTTTCCCTACACGACGCTCTTCCGATCTNNNGAGATGGTAGTGAGCCAAGA
AGCACACGTCTGAACTCTTTCCCTACACGACGCTCTTCCGATCTNNNGCAACAAAATGAGACCCTGTCT
AGCACACGTCTGAACTCTTTCCCTACACGACGCTCTTCCGATCTNNNAGTCATGGGAAAAGCCTGTAA
GAGCACACGTCTGAACTCTTTCCCTACACGACGCTCTTCCGATCTNNNTGAGTGTTTCCCATCTCTCTTG
GAGCACACGTCTGAACTCTTTCCCTACACGACGCTCTTCCGATCTNNNCCCTGTGGTCAAGCTACAAAA

GAGCACACGTCTGAACTCTTTCCCTACACGACGCTCTTCCGATCTNNNTGAGGAGGGAGAATTGCTTG
GCACACGTCTGAACTCTTTCCCTACACGACGCTCTTCCGATCTNNNGAGGTAGAGTTTGCAGTGAGCA
GAGCACACGTCTGAACTCTTTCCCTACACGACGCTCTTCCGATCTNNNGGCAACAGAGTGAGAATCCAT
GCACACGTCTGAACTCTTTCCCTACACGACGCTCTTCCGATCTNNNCATGAGAATCGCTTGAACCAGTG
GCACACGTCTGAACTCTTTCCCTACACGACGCTCTTCCGATCTNNNCTTCTGGATTCCACTCCTTCTCCCA
GAGCACACGTCTGAACTCTTTCCCTACACGACGCTCTTCCGATCTNNNTATGATCATGCCTGTGACTAG
GAGCACACGTCTGAACTCTTTCCCTACACGACGCTCTTCCGATCTNNNGGCTGAGGTAGGAAAATTGC
AGCACACGTCTGAACTCTTTCCCTACACGACGCTCTTCCGATCTNNNAGAGGAAGGCAGACTGACTAG
GCACACGTCTGAACTCTTTCCCTACACGACGCTCTTCCGATCTNNNCTATGGCACTCATTCTGTCCAGTCA
AGCACACGTCTGAACTCTTTCCCTACACGACGCTCTTCCGATCTNNNGGTGACAGAGTGAGACTCCTTC
GAGCACACGTCTGAACTCTTTCCCTACACGACGCTCTTCCGATCTNNNAAGCAGACCAGGAAGATACTC
AGAGCACACGTCTGAACTCTTTCCCTACACGACGCTCTTCCGATCTNNNGAGAATGGTGTGAACCTGA
AGCACACGTCTGAACTCTTTCCCTACACGACGCTCTTCCGATCTNNNTCACTGAGGTGACAGAAACTGT
AGCACACGTCTGAACTCTTTCCCTACACGACGCTCTTCCGATCTNNNTCACTGAGGTGACAGAAACTGT
GAGCACACGTCTGAACTCTTTCCCTACACGACGCTCTTCCGATCTNNNAATACCAAAATTAGCTGGGCG
GAGCACACGTCTGAACTCTTTCCCTACACGACGCTCTTCCGATCTNNNAAAAATCTTGGACACACGGAC
GAGCACACGTCTGAACTCTTTCCCTACACGACGCTCTTCCGATCTNNNGGCTGAGATAGGAGGATCAC
GAGCACACGTCTGAACTCTTTCCCTACACGACGCTCTTCCGATCTNNNTCTCCTTGGGAATCTCAGTAG
GAGCACACGTCTGAACTCTTTCCCTACACGACGCTCTTCCGATCTNNNTTTCAGTGAGCCGAGATAGTG
GAGCACACGTCTGAACTCTTTCCCTACACGACGCTCTTCCGATCTNNNCTGTGATTTTGCCACTGTACTC
GAGCACACGTCTGAACTCTTTCCCTACACGACGCTCTTCCGATCTNNNAGAGGTTGCAGTGAGCCTAAA
GAGCACACGTCTGAACTCTTTCCCTACACGACGCTCTTCCGATCTNNNTGAACTGTGATTGAGCCACTG
AGAGCACACGTCTGAACTCTTTCCCTACACGACGCTCTTCCGATCTNNNGGTTGCAGTGAGCCAAGATT
GAGCACACGTCTGAACTCTTTCCCTACACGACGCTCTTCCGATCTNNNGCGACAGAGCAAGATCTTGTC
AGAGCACACGTCTGAACTCTTTCCCTACACGACGCTCTTCCGATCTNNNCTCACCTGTAATCCCAGCACT
AGCACACGTCTGAACTCTTTCCCTACACGACGCTCTTCCGATCTNNNCTGAGGCAGGAGAATTGCTTGA
AGCACACGTCTGAACTCTTTCCCTACACGACGCTCTTCCGATCTNNNGGGGCTTGTAATCTCAGCTACTT
GAGCACACGTCTGAACTCTTTCCCTACACGACGCTCTTCCGATCTNNNACAAAAATTAGCTGGGCATGC
AGAGCACACGTCTGAACTCTTTCCCTACACGACGCTCTTCCGATCTNNNACAAAAATTAGCTGGGTGTG
GAGCACACGTCTGAACTCTTTCCCTACACGACGCTCTTCCGATCTNNNTTGAGGCAGGAGTATCACTTC
AGAGCACACGTCTGAACTCTTTCCCTACACGACGCTCTTCCGATCTNNNGTACCTGTAATCCCAGCTACT
GAGCACACGTCTGAACTCTTTCCCTACACGACGCTCTTCCGATCTNNNCTGAGGCAGGAGAATCACTTG
GAGCACACGTCTGAACTCTTTCCCTACACGACGCTCTTCCGATCTNNNGGTGCCTGTAATCCCAACTACT
GAGCACACGTCTGAACTCTTTCCCTACACGACGCTCTTCCGATCTNNNCTGAGGCAGGAGAATTGCTTG
GAGCACACGTCTGAACTCTTTCCCTACACGACGCTCTTCCGATCTNNNAGGCACAATAATTGCTTGAAC
GAGCACACGTCTGAACTCTTTCCCTACACGACGCTCTTCCGATCTNNNCTGAGGCAGGAGAATCACTTG
AGAGCACACGTCTGAACTCTTTCCCTACACGACGCTCTTCCGATCTNNNAGGTTACAGTTGGCCAAGAT
AGCACACGTCTGAACTCTTTCCCTACACGACGCTCTTCCGATCTNNNTCCAAGCAATGTACTCAGTCACT
AGCACACGTCTGAACTCTTTCCCTACACGACGCTCTTCCGATCTNNNCATGCCTGTAATCCCAGCTACTC
GCACACGTCTGAACTCTTTCCCTACACGACGCTCTTCCGATCTNNNCTGTAGTACCAGCTACTTGGGAG
GAGCACACGTCTGAACTCTTTCCCTACACGACGCTCTTCCGATCTNNNAAATGAGATGAAACCGTGGAA
AGCACACGTCTGAACTCTTTCCCTACACGACGCTCTTCCGATCTNNNCATGCCTGTAATCCCAGCTACTC
AGAGCACACGTCTGAACTCTTTCCCTACACGACGCTCTTCCGATCTNNNCTGTAGTCACAGCTACTCAGC

AGCACACGTCTGAACTCTTTCCCTACACGACGCTCTTCCGATCTNNNAAATGAGATGAAACCGTGGA
AGCACACGTCTGAACTCTTTCCCTACACGACGCTCTTCCGATCTNNNCATGCCTGTAATCCCAGCTACT
GAGCACACGTCTGAACTCTTTCCCTACACGACGCTCTTCCGATCTNNNCTGTAGTCACAGCTACTCAG
GCACACGTCTGAACTCTTTCCCTACACGACGCTCTTCCGATCTNNNCTGTAGTACCAGCTACTTGGGA
AGCACACGTCTGAACTCTTTCCCTACACGACGCTCTTCCGATCTNNNATCTGAGGCAGGAGAATCACT
AGCACACGTCTGAACTCTTTCCCTACACGACGCTCTTCCGATCTNNNGTTGTGGTAAGCCTAGATTG
GAGCACACGTCTGAACTCTTTCCCTACACGACGCTCTTCCGATCTNNNACTCCAGCCTAGTCTACAGA
AGCACACGTCTGAACTCTTTCCCTACACGACGCTCTTCCGATCTNNNAGTGAGCTATGATTGTGCCAG
AGCACACGTCTGAACTCTTTCCCTACACGACGCTCTTCCGATCTNNNAATACAAAAAATTAGCCGGGC
GAGCACACGTCTGAACTCTTTCCCTACACGACGCTCTTCCGATCTNNNCTGAGGCAGGAGAATCACT
AGCACACGTCTGAACTCTTTCCCTACACGACGCTCTTCCGATCTNNNTGTAATCCTAGCTACTCGGGAG
AGCACACGTCTGAACTCTTTCCCTACACGACGCTCTTCCGATCTNNNGTGATCAAACCACTGCACTCT
AGCACACGTCTGAACTCTTTCCCTACACGACGCTCTTCCGATCTNNNATAAGATTCCATCTCACCTGGC
AGCACACGTCTGAACTCTTTCCCTACACGACGCTCTTCCGATCTNNNCACAGTGGCTCACAACTGTAA
AGCACACGTCTGAACTCTTTCCCTACACGACGCTCTTCCGATCTNNNGCCAAGATCATACCATTGCACT
GAGCACACGTCTGAACTCTTTCCCTACACGACGCTCTTCCGATCTNNNCAACATAAGGAGACCCCCAT
AGCACACGTCTGAACTCTTTCCCTACACGACGCTCTTCCGATCTNNNCTTCCCAAAGAGGAGGTAAG
GCACACGTCTGAACTCTTTCCCTACACGACGCTCTTCCGATCTNNNAATCCCAGCACTTACAGAGGTT
GAGCACACGTCTGAACTCTTTCCCTACACGACGCTCTTCCGATCTNNNCTGAGATCACATGACTGCACT
AGCACACGTCTGAACTCTTTCCCTACACGACGCTCTTCCGATCTNNNTAGGAGACTGAGGCAGGAG
GCACACGTCTGAACTCTTTCCCTACACGACGCTCTTCCGATCTNNNACAAAAATTAGCCAGGCTTGGT
AGAGCACACGTCTGAACTCTTTCCCTACACGACGCTCTTCCGATCTNNNGAGGCAGAAGAATCGCTTG
AGCACACGTCTGAACTCTTTCCCTACACGACGCTCTTCCGATCTNNNGTGGATCACTTGAGGTCAGGA
GAGCACACGTCTGAACTCTTTCCCTACACGACGCTCTTCCGATCTNNNATCATGAGGTCAGGAGATCG
GCACACGTCTGAACTCTTTCCCTACACGACGCTCTTCCGATCTNNNGGGCAACAAGAGCAAAACTCAA
GCACACGTCTGAACTCTTTCCCTACACGACGCTCTTCCGATCTNNNTGAGGCAGGAGAATAGGTTGA
AGCACACGTCTGAACTCTTTCCCTACACGACGCTCTTCCGATCTNNNTCATGCCTGTAATCCCAACACTT
GAGCACACGTCTGAACTCTTTCCCTACACGACGCTCTTCCGATCTNNNGGAGAATCACGTGAACCTG
AGCACACGTCTGAACTCTTTCCCTACACGACGCTCTTCCGATCTNNNTTTCACTGTGTTAGCCAGGATG
AGCACACGTCTGAACTCTTTCCCTACACGACGCTCTTCCGATCTNNNAAAAAATTAGCCAGGTGAGGT
GAGCACACGTCTGAACTCTTTCCCTACACGACGCTCTTCCGATCTNNNACAAAAATTACCCAGTGTGG
AGCACACGTCTGAACTCTTTCCCTACACGACGCTCTTCCGATCTNNNGTAATCCTAGCTACTCGGAAG
AGCACACGTCTGAACTCTTTCCCTACACGACGCTCTTCCGATCTNNNATACAAAAAATGAGCCGGGC
AGCACACGTCTGAACTCTTTCCCTACACGACGCTCTTCCGATCTNNNCTGAGACAGGAGAATGGCAT
GCACACGTCTGAACTCTTTCCCTACACGACGCTCTTCCGATCTNNNGCAGAAACTTTACAAGCCAGGA
AGCACACGTCTGAACTCTTTCCCTACACGACGCTCTTCCGATCTNNNAAATAAGCAATCGGCCAGGTA
AGCACACGTCTGAACTCTTTCCCTACACGACGCTCTTCCGATCTNNNCGCAACAAGAATGAGACTCCAT
GAGCACACGTCTGAACTCTTTCCCTACACGACGCTCTTCCGATCTNNNCACCTGAGGTCATGAGTTTG
GAGCACACGTCTGAACTCTTTCCCTACACGACGCTCTTCCGATCTNNNCTGTAATTGCAGCTACTGGG
AGCACACGTCTGAACTCTTTCCCTACACGACGCTCTTCCGATCTNNNTGTGATCCCAGCTATTCAGGA
AGCACACGTCTGAACTCTTTCCCTACACGACGCTCTTCCGATCTNNNCTGAGGGAGGATAACTGCTT
AGCACACGTCTGAACTCTTTCCCTACACGACGCTCTTCCGATCTNNNAACTGATTCTAGGACCAAGGC
AGCACACGTCTGAACTCTTTCCCTACACGACGCTCTTCCGATCTNNNGCATGGACAACAAGAGCAAAA

AGCACACGTCTGAACTCTTTCCCTACACGACGCTCTTCCGATCTNNNAGGCAGGAGAATTGCTTGAA
AGCACACGTCTGAACTCTTTCCCTACACGACGCTCTTCCGATCTNNNGCAACAGAGGAATGGTGAAC
GAGCACACGTCTGAACTCTTTCCCTACACGACGCTCTTCCGATCTNNNCACAACACCTAAAAAAGCTGG
GAGCACACGTCTGAACTCTTTCCCTACACGACGCTCTTCCGATCTNNNGGTGTGGTAGTGTATGCCT
AGCACACGTCTGAACTCTTTCCCTACACGACGCTCTTCCGATCTNNNATCAGAGACTAGGATTGCAAC
AGCACACGTCTGAACTCTTTCCCTACACGACGCTCTTCCGATCTNNNGGGAGGGATAGCATTGGGAG
GCACACGTCTGAACTCTTTCCCTACACGACGCTCTTCCGATCTNNNCTGAGGTGGGAGTATCAGCTAA
AGCACACGTCTGAACTCTTTCCCTACACGACGCTCTTCCGATCTNNNCTGGAATGCAATGGTGTGATC
AGCACACGTCTGAACTCTTTCCCTACACGACGCTCTTCCGATCTNNNTTGCACGTAGTTGCAGAATTCA
AGCACACGTCTGAACTCTTTCCCTACACGACGCTCTTCCGATCTNNNCCATGATGAGGGATTCCATGA
GAGCACACGTCTGAACTCTTTCCCTACACGACGCTCTTCCGATCTNNNCTAATATCTGGGGGGGAGA
AGCACACGTCTGAACTCTTTCCCTACACGACGCTCTTCCGATCTNNNTGATTGGTCAACACTCCACTAC
AGCACACGTCTGAACTCTTTCCCTACACGACGCTCTTCCGATCTNNNCTGTAGTTCCAGCTACTTGGGA
AAGAGCACACGTCTGAACTCTTTCCCTACACGACGCTCTTCCGATCTNNNGCCTAGGGGAAGAGAGT
GCACACGTCTGAACTCTTTCCCTACACGACGCTCTTCCGATCTNNNCAAATGGACCAGTCAGCAGGAT
GAGCACACGTCTGAACTCTTTCCCTACACGACGCTCTTCCGATCTNNNCCCTCTTTGAAGTGAAGACA
GCACACGTCTGAACTCTTTCCCTACACGACGCTCTTCCGATCTNNNTTTCTCAGCACCTCTCTCTTTCCCA
AGCACACGTCTGAACTCTTTCCCTACACGACGCTCTTCCGATCTNNNCACCAGGAAGAGAAAATACCC
CACACGTCTGAACTCTTTCCCTACACGACGCTCTTCCGATCTNNNGGAGCTTTTCTACCCAGAACTCAC
GCACACGTCTGAACTCTTTCCCTACACGACGCTCTTCCGATCTNNNTCTCTCTCTCTCTCTCTCCCTCA
AGCACACGTCTGAACTCTTTCCCTACACGACGCTCTTCCGATCTNNNCAACTCTCCCATGGAAACCCAT
AGCACACGTCTGAACTCTTTCCCTACACGACGCTCTTCCGATCTNNNTCTCCTAATGCTATCCCTCTCCC
AGCACACGTCTGAACTCTTTCCCTACACGACGCTCTTCCGATCTNNNTCAAACCCATTGCTTCCAATTG
GAGCACACGTCTGAACTCTTTCCCTACACGACGCTCTTCCGATCTNNNTTGGGAACTATCTTTAGGGG
AGCACACGTCTGAACTCTTTCCCTACACGACGCTCTTCCGATCTNNNTGATAGCAGAGAGTGTGTGT
GCACACGTCTGAACTCTTTCCCTACACGACGCTCTTCCGATCTNNNTCCTCTCTCTCTCCTCTCTCTCTC
AGCACACGTCTGAACTCTTTCCCTACACGACGCTCTTCCGATCTNNNACATGGTGAAATCCCGTCTCTA
GAGCACACGTCTGAACTCTTTCCCTACACGACGCTCTTCCGATCTNNNGTACCTGTAATCTCAGCTAC
GCACACGTCTGAACTCTTTCCCTACACGACGCTCTTCCGATCTNNNGTTGTGGTGACCTGAGATCGT
GAGCACACGTCTGAACTCTTTCCCTACACGACGCTCTTCCGATCTNNNCTGTCATCCCAGCTATTCAGG
GAGCACACGTCTGAACTCTTTCCCTACACGACGCTCTTCCGATCTNNNGCCAATATGGAGAAACCCTG
AGCACACGTCTGAACTCTTTCCCTACACGACGCTCTTCCGATCTNNNCAGTGACCTATGACTGGACTA
GCACACGTCTGAACTCTTTCCCTACACGACGCTCTTCCGATCTNNNCACCTGTAATCCCAGCTACTTGG
AGCACACGTCTGAACTCTTTCCCTACACGACGCTCTTCCGATCTNNNTTGCGTCTGAATCCTGTAACAC
AGCACACGTCTGAACTCTTTCCCTACACGACGCTCTTCCGATCTNNNGGTGCCTGTAATCCCAACTACT
AGCACACGTCTGAACTCTTTCCCTACACGACGCTCTTCCGATCTNNNCTGTAGTCCCAGCTACTCAGGA
GAGCACACGTCTGAACTCTTTCCCTACACGACGCTCTTCCGATCTNNNCACAAAACAAACAAAAGGCCA
GAGCACACGTCTGAACTCTTTCCCTACACGACGCTCTTCCGATCTNNNGTGTTCAGATTGAGCCATTG
AGCACACGTCTGAACTCTTTCCCTACACGACGCTCTTCCGATCTNNNGAGAATCGCTTGAATCCAGGA
GAGCACACGTCTGAACTCTTTCCCTACACGACGCTCTTCCGATCTNNNTGAGTCACAGAGAGAGACTG
AGCACACGTCTGAACTCTTTCCCTACACGACGCTCTTCCGATCTNNNCTGGGAAACAAGAGTGAAAA
AGCACACGTCTGAACTCTTTCCCTACACGACGCTCTTCCGATCTNNNTATATGCAGCTTTCCAGGGAC
GAGCACACGTCTGAACTCTTTCCCTACACGACGCTCTTCCGATCTNNNGCAGGAGAATTGCTTGATC

AGCACACGTCTGAACTCTTTCCCTACACGACGCTCTTCCGATCTNNNAATGCAGGAGAATTGCTTGAA
GAGCACACGTCTGAACTCTTTCCCTACACGACGCTCTTCCGATCTNNNGTATTTCTGCAGTCCCAGCTA
AGCACACGTCTGAACTCTTTCCCTACACGACGCTCTTCCGATCTNNNGCTGAGATAGGAGAATCGCTT
AGCACACGTCTGAACTCTTTCCCTACACGACGCTCTTCCGATCTNNNTTGTGGTGACCTGAGATTGTA
GAGCACACGTCTGAACTCTTTCCCTACACGACGCTCTTCCGATCTNNNCAAATCTCAAAGACTCAGCCC
AGCACACGTCTGAACTCTTTCCCTACACGACGCTCTTCCGATCTNNNAAACAAATGTCAAAGACAGCCC
GAGCACACGTCTGAACTCTTTCCCTACACGACGCTCTTCCGATCTNNNCTGTAATCCCAGCTACTCAGG
GAGCACACGTCTGAACTCTTTCCCTACACGACGCTCTTCCGATCTNNNTGATGAAACCCCGTCGTTACT
AGAGCACACGTCTGAACTCTTTCCCTACACGACGCTCTTCCGATCTNNNCATGCCTGTAATCCCAGCTA
AGCACACGTCTGAACTCTTTCCCTACACGACGCTCTTCCGATCTNNNTAAGGCAGCAGAATCACTTGA
AGCACACGTCTGAACTCTTTCCCTACACGACGCTCTTCCGATCTNNNATGCCTTTAGTCCCAGCTACTT
AGCACACGTCTGAACTCTTTCCCTACACGACGCTCTTCCGATCTNNNAGTGAACTACGATTTTGCCAC
AGCACACGTCTGAACTCTTTCCCTACACGACGCTCTTCCGATCTNNNAATTAGCCAGAGATGATGGTG
AGCACACGTCTGAACTCTTTCCCTACACGACGCTCTTCCGATCTNNNAGGCAGGAGAATCATTTGAAC
AGCACACGTCTGAACTCTTTCCCTACACGACGCTCTTCCGATCTNNNCTAGGCAACAAAGAGAGACT
AGCACACGTCTGAACTCTTTCCCTACACGACGCTCTTCCGATCTNNNAATGGCAAAGTGAGACTCCAT
GAGCACACGTCTGAACTCTTTCCCTACACGACGCTCTTCCGATCTNNNGATCACAAGGTCAGGAGATC
AGCACACGTCTGAACTCTTTCCCTACACGACGCTCTTCCGATCTNNNTGGAGACAGGAGAATTGCTTG
AGCACACGTCTGAACTCTTTCCCTACACGACGCTCTTCCGATCTNNNTTCACTGCATTCTAACCTGGTG
GAGCACACGTCTGAACTCTTTCCCTACACGACGCTCTTCCGATCTNNNGCAGGAGAATTTCTTGAATC
AGCACACGTCTGAACTCTTTCCCTACACGACGCTCTTCCGATCTNNNCAAGAACAAAACCCCGTCTCAA
AGCACACGTCTGAACTCTTTCCCTACACGACGCTCTTCCGATCTNNNCAGAGGTTGTAGTGAGCCAA
AGCACACGTCTGAACTCTTTCCCTACACGACGCTCTTCCGATCTNNNCGGCAAAAAGTTTCCTTCTAGC
GAGCACACGTCTGAACTCTTTCCCTACACGACGCTCTTCCGATCTNNNGCACATGTAGTCCCAGATACT
AGCACACGTCTGAACTCTTTCCCTACACGACGCTCTTCCGATCTNNNGAGTGACAGAGTGAGACTCCA
GCACACGTCTGAACTCTTTCCCTACACGACGCTCTTCCGATCTNNNCTGAGGCAGGAGAATCACTTGA
AGCACACGTCTGAACTCTTTCCCTACACGACGCTCTTCCGATCTNNNACAAAAATTAGCCAGGCGTG
AGCACACGTCTGAACTCTTTCCCTACACGACGCTCTTCCGATCTNNNTATAGTCCCAGCTACTCGAGAG
AGCACACGTCTGAACTCTTTCCCTACACGACGCTCTTCCGATCTNNNTTGAATCCGGGAAGTAGAGG
AGCACACGTCTGAACTCTTTCCCTACACGACGCTCTTCCGATCTNNNGGAGACAAGAATGAAACGCCA
AGCACACGTCTGAACTCTTTCCCTACACGACGCTCTTCCGATCTNNNCAGGAGAATCACTTGAACCCA
AGCACACGTCTGAACTCTTTCCCTACACGACGCTCTTCCGATCTNNNGCACATCTGTAATCCCAGCTACT
GAGCACACGTCTGAACTCTTTCCCTACACGACGCTCTTCCGATCTNNNGAGTTCACACCAGTACACTCC
AGCACACGTCTGAACTCTTTCCCTACACGACGCTCTTCCGATCTNNNCAAAAAATTAGCCGGGCGTAG
AGCACACGTCTGAACTCTTTCCCTACACGACGCTCTTCCGATCTNNNGCAACAAGAATGAAACTCCAG
AGCACACGTCTGAACTCTTTCCCTACACGACGCTCTTCCGATCTNNNCATCACGAGGTTAGGAGATGG
AGCACACGTCTGAACTCTTTCCCTACACGACGCTCTTCCGATCTNNNATTGTATGCACCTGTAGTCCC
GAGCACACGTCTGAACTCTTTCCCTACACGACGCTCTTCCGATCTNNNGAGATTGCAGTGAGCCAAGA
AGCACACGTCTGAACTCTTTCCCTACACGACGCTCTTCCGATCTNNNGGTTGCAGTGAGCCAAGATT
AGCACACGTCTGAACTCTTTCCCTACACGACGCTCTTCCGATCTNNNTCAAGAGTCTGAGGGAGGAG
AGCACACGTCTGAACTCTTTCCCTACACGACGCTCTTCCGATCTNNNCTGTGGTAGGAGAATGGTGT
AGCACACGTCTGAACTCTTTCCCTACACGACGCTCTTCCGATCTNNNAGAATCACTTGAACTCAGGAG
GCACACGTCTGAACTCTTTCCCTACACGACGCTCTTCCGATCTNNNTTAGCCTGAAATTGCTTGAACCC

GCACACGTCTGAACTCTTTCCCTACACGACGCTCTTCCGATCTNNNACACTGAGATGCTTCTTGTGCTA
AGCACACGTCTGAACTCTTTCCCTACACGACGCTCTTCCGATCTNNNCTGTAGTCCCAGCTACTCAGG
GAGCACACGTCTGAACTCTTTCCCTACACGACGCTCTTCCGATCTNNNGAATCATCACTTGAACCGGG
AGCACACGTCTGAACTCTTTCCCTACACGACGCTCTTCCGATCTNNNCAGCCTGGACAACAAGAGAG
AGCACACGTCTGAACTCTTTCCCTACACGACGCTCTTCCGATCTNNNTGAGGGAGGAGAATCACTTG
AGCACACGTCTGAACTCTTTCCCTACACGACGCTCTTCCGATCTNNNAATTTCTTCCACCAGGCAACCT
AGCACACGTCTGAACTCTTTCCCTACACGACGCTCTTCCGATCTNNNTAGGTTGCATGAGCTGAGAT
AGCACACGTCTGAACTCTTTCCCTACACGACGCTCTTCCGATCTNNNGCCAAGATCACACCATTTCACT
AGCACACGTCTGAACTCTTTCCCTACACGACGCTCTTCCGATCTNNNGGTTACAGTGAGCTGAGATCA
AGCACACGTCTGAACTCTTTCCCTACACGACGCTCTTCCGATCTNNNATGCCTCTAGTACCAGCTACTC
GCACACGTCTGAACTCTTTCCCTACACGACGCTCTTCCGATCTNNNCTGTAGTTCCAGTTACTCGGGA
AGCACACGTCTGAACTCTTTCCCTACACGACGCTCTTCCGATCTNNNTGGAGATTTCAGTGAGCCAAG
AGCACACGTCTGAACTCTTTCCCTACACGACGCTCTTCCGATCTNNNAGAGCTTGTATCATTCCTTCTG
AGCACACGTCTGAACTCTTTCCCTACACGACGCTCTTCCGATCTNNNCTTACAGAGAGCCGAGATCAC
GCACACGTCTGAACTCTTTCCCTACACGACGCTCTTCCGATCTNNNGGGAGGCCAAGACAGAAGAAT
GAGCACACGTCTGAACTCTTTCCCTACACGACGCTCTTCCGATCTNNNGTAGTTCCAGATACTGCAGA
AGCACACGTCTGAACTCTTTCCCTACACGACGCTCTTCCGATCTNNNTTGGAATGGGAATGCCTACT
AGCACACGTCTGAACTCTTTCCCTACACGACGCTCTTCCGATCTNNNTCTGGCCAGTGCATCTACATTT
AGCACACGTCTGAACTCTTTCCCTACACGACGCTCTTCCGATCTNNNCTCATGCCTGTAATCCCAGAAC
AGCACACGTCTGAACTCTTTCCCTACACGACGCTCTTCCGATCTNNNAAGGCTGCTGTGAGTTATGAT
CACACGTCTGAACTCTTTCCCTACACGACGCTCTTCCGATCTNNNCAGTTACTTGAGAGATGGAGGC
GAGCACACGTCTGAACTCTTTCCCTACACGACGCTCTTCCGATCTNNNCTCATGTCTGTAATCCCAGCA
AGCACACGTCTGAACTCTTTCCCTACACGACGCTCTTCCGATCTNNNAGGAGGAGAGGAGGAAGAA
AGCACACGTCTGAACTCTTTCCCTACACGACGCTCTTCCGATCTNNNTATAGTCCCAGCTACTCAGGA
GCACACGTCTGAACTCTTTCCCTACACGACGCTCTTCCGATCTNNNGCCAAGATCATGCCATTGTACT
AGCACACGTCTGAACTCTTTCCCTACACGACGCTCTTCCGATCTNNNGAGGCTGAGGCAAGAGAATT
AGCACACGTCTGAACTCTTTCCCTACACGACGCTCTTCCGATCTNNNGTATTCCCAGCTACTAAGGAG
CACACGTCTGAACTCTTTCCCTACACGACGCTCTTCCGATCTNNNCAGGAGAATTGCTTGAAACTGG
AGCACACGTCTGAACTCTTTCCCTACACGACGCTCTTCCGATCTNNNCCTGGGCAACAAGAGTGAAA
GAGCACACGTCTGAACTCTTTCCCTACACGACGCTCTTCCGATCTNNNTGCCCATAATACCAGCTACTT
AGCACACGTCTGAACTCTTTCCCTACACGACGCTCTTCCGATCTNNNACCTAAGCCTTGGAGAAGTG
AGCACACGTCTGAACTCTTTCCCTACACGACGCTCTTCCGATCTNNNTTGTAGTGAGCCAAAATCATG
GCACACGTCTGAACTCTTTCCCTACACGACGCTCTTCCGATCTNNNAAATCATCCCTACTTGGTCCCAG
GAGCACACGTCTGAACTCTTTCCCTACACGACGCTCTTCCGATCTNNNCTGGGATTACAAGGCGTGA
GCACACGTCTGAACTCTTTCCCTACACGACGCTCTTCCGATCTNNNCTGAGGCAGGAGAATCACTTGA
GCACACGTCTGAACTCTTTCCCTACACGACGCTCTTCCGATCTNNNCAGGGGGCCGTTTTCTTTATTT
AGCACACGTCTGAACTCTTTCCCTACACGACGCTCTTCCGATCTNNNCTAGCATCCAAATCAGCTGCT
GCACACGTCTGAACTCTTTCCCTACACGACGCTCTTCCGATCTNNNGGCTGTAGGTCTCTGTGTTCTT
GCACACGTCTGAACTCTTTCCCTACACGACGCTCTTCCGATCTNNNGAATCTAGGAGATGGCAGAGG
AGCACACGTCTGAACTCTTTCCCTACACGACGCTCTTCCGATCTNNNCTGGGTGAAAGACTGAGACT
AGCACACGTCTGAACTCTTTCCCTACACGACGCTCTTCCGATCTNNNTGTAATCCCAGCTACTCAGGA
AGCACACGTCTGAACTCTTTCCCTACACGACGCTCTTCCGATCTNNNGAGAATGGCATGAACCTGGG
GAGCACACGTCTGAACTCTTTCCCTACACGACGCTCTTCCGATCTNNNCGGATCACAAGGTCAGAAG

AGCACACGTCTGAACTCTTTCCCTACACGACGCTCTTCCGATCTNNNTATAGTCCCAGCTACTCAGGAG
GCACACGTCTGAACTCTTTCCCTACACGACGCTCTTCCGATCTNNNCTGTAGTCCCAGCTACTCAGGA
AGCACACGTCTGAACTCTTTCCCTACACGACGCTCTTCCGATCTNNNACATACTGGGGGTTGAAATG
AGCACACGTCTGAACTCTTTCCCTACACGACGCTCTTCCGATCTNNNCCCAGGAGTTCTAGACCATCCT
GCACACGTCTGAACTCTTTCCCTACACGACGCTCTTCCGATCTNNNGTACTCCAGCCTGGGTAAGAAA
CACACGTCTGAACTCTTTCCCTACACGACGCTCTTCCGATCTNNNTTTCTGTGTCTTTTCCCTCTGCTCAT
AGCACACGTCTGAACTCTTTCCCTACACGACGCTCTTCCGATCTNNNCTGGCAAAATAGTGAAACC
AGCACACGTCTGAACTCTTTCCCTACACGACGCTCTTCCGATCTNNNAGGAGAATCACTTGAACCCA
GCACACGTCTGAACTCTTTCCCTACACGACGCTCTTCCGATCTNNNGTGGCAGAGTAAGACTCCATCT
GAGCACACGTCTGAACTCTTTCCCTACACGACGCTCTTCCGATCTNNNGATGTCAGGAGATCGAGA
GCACACGTCTGAACTCTTTCCCTACACGACGCTCTTCCGATCTNNNCAGAGAATCACTTGAACCCAGG
AGCACACGTCTGAACTCTTTCCCTACACGACGCTCTTCCGATCTNNNCACCTGTAATCCCAGCTACTCA
GCACACGTCTGAACTCTTTCCCTACACGACGCTCTTCCGATCTNNNTCTTCAATATGGCGCATTGAACC
AGCACACGTCTGAACTCTTTCCCTACACGACGCTCTTCCGATCTNNNAGAGAACCACTTGAACCCAG
AGCACACGTCTGAACTCTTTCCCTACACGACGCTCTTCCGATCTNNNAGAGGTTGCAGTAAGCTGAG
GAGCACACGTCTGAACTCTTTCCCTACACGACGCTCTTCCGATCTNNNACTTGAATCTGGGAAGCAGA
AGCACACGTCTGAACTCTTTCCCTACACGACGCTCTTCCGATCTNNNGAAGCTGTAGTGAGCAGAGA
AGCACACGTCTGAACTCTTTCCCTACACGACGCTCTTCCGATCTNNNCACTTGAATCAGGAGATGGA
AGCACACGTCTGAACTCTTTCCCTACACGACGCTCTTCCGATCTNNNGTAATCTCAGCTACTCAGGAG
GAGCACACGTCTGAACTCTTTCCCTACACGACGCTCTTCCGATCTNNNAGTGCTAGGATTACAGGCAT
GCACACGTCTGAACTCTTTCCCTACACGACGCTCTTCCGATCTNNNGCCACCTGTATTACCAGCTACTT
AGCACACGTCTGAACTCTTTCCCTACACGACGCTCTTCCGATCTNNNTCTCCTAGTGATACTCAGGCA
AGCACACGTCTGAACTCTTTCCCTACACGACGCTCTTCCGATCTNNNGGTAAGGCAGGAGAATCAC
AGCACACGTCTGAACTCTTTCCCTACACGACGCTCTTCCGATCTNNNCTGTAGTCCCAGCTACTCAGG
GCACACGTCTGAACTCTTTCCCTACACGACGCTCTTCCGATCTNNNAGTAAGCCAAGATTGTGCCATT
GCACACGTCTGAACTCTTTCCCTACACGACGCTCTTCCGATCTNNNGTAGTTCCCAGCTACTCAGGAT
GAGCACACGTCTGAACTCTTTCCCTACACGACGCTCTTCCGATCTNNNTGGAAGCCATTAGAGCTGT
GCACACGTCTGAACTCTTTCCCTACACGACGCTCTTCCGATCTNNNCTGAGAGTACACCACTGAACTC
AGCACACGTCTGAACTCTTTCCCTACACGACGCTCTTCCGATCTNNNACATCATGGAGAAACGGGTA
GCACACGTCTGAACTCTTTCCCTACACGACGCTCTTCCGATCTNNNAAGCCTGTAGTCCTAGCTACTC
GAGCACACGTCTGAACTCTTTCCCTACACGACGCTCTTCCGATCTNNNCCACACTACAGTCCATTTCCT
GCACACGTCTGAACTCTTTCCCTACACGACGCTCTTCCGATCTNNNGAAGCAGAAGAATCACTTGAG
AGCACACGTCTGAACTCTTTCCCTACACGACGCTCTTCCGATCTNNNAGGCAAGAGAATGGTGTGAA
AGCACACGTCTGAACTCTTTCCCTACACGACGCTCTTCCGATCTNNNTAGTTCCAGCTACTCCAGAGG
AGCACACGTCTGAACTCTTTCCCTACACGACGCTCTTCCGATCTNNNAATCCCAACTACTCAAGAGGC
GCACACGTCTGAACTCTTTCCCTACACGACGCTCTTCCGATCTNNNATTTTCATGCTGCTCTTTTGGCA
GCACACGTCTGAACTCTTTCCCTACACGACGCTCTTCCGATCTNNNTGTGAAACAGTGAGATTCCGT
AGCACACGTCTGAACTCTTTCCCTACACGACGCTCTTCCGATCTNNNGAGATCATAAGCCACTGCACT
AGCACACGTCTGAACTCTTTCCCTACACGACGCTCTTCCGATCTNNNCAGGAGAATTGCTTGAACCA
GAGCACACGTCTGAACTCTTTCCCTACACGACGCTCTTCCGATCTNNNTGACAAGAGTGAAGCTCCTT
AGCACACGTCTGAACTCTTTCCCTACACGACGCTCTTCCGATCTNNNTCACTTGAGCTCAGGAGTTTG
GAGCACACGTCTGAACTCTTTCCCTACACGACGCTCTTCCGATCTNNNGAGATCATGCCACTGTACTCC
GAGCACACGTCTGAACTCTTTCCCTACACGACGCTCTTCCGATCTNNNGACTCATGCCTATAATCCCA

GAGCACACGTCTGAACTCTTTCCCTACACGACGCTCTTCCGATCTNNNGAGGCAGGAGAATTTTGTGA
AGCACACGTCTGAACTCTTTCCCTACACGACGCTCTTCCGATCTNNNTCATCCTGGCTAACATGGTGAA
GCACACGTCTGAACTCTTTCCCTACACGACGCTCTTCCGATCTNNNGTGATCCCAGACACTCAGAGAA
GAGCACACGTCTGAACTCTTTCCCTACACGACGCTCTTCCGATCTNNNCATTTGTAGTCCCAGCTACTCG
GAGCACACGTCTGAACTCTTTCCCTACACGACGCTCTTCCGATCTNNNGTCAAGACCCTTCGGTATGCTA
GAGCACACGTCTGAACTCTTTCCCTACACGACGCTCTTCCGATCTNNNGTAAGAGGCCAGACTTCTAG
AGCACACGTCTGAACTCTTTCCCTACACGACGCTCTTCCGATCTNNNTAAGGAAAGGAGAGCCGTTTG
AGCACACGTCTGAACTCTTTCCCTACACGACGCTCTTCCGATCTNNNAGAGCTTGCAGTTAGTTGAGA
AGAGCACACGTCTGAACTCTTTCCCTACACGACGCTCTTCCGATCTNNNTATCCTGGCTAACACGGTGTA
GAGCACACGTCTGAACTCTTTCCCTACACGACGCTCTTCCGATCTNNNATACTGAGAAGGCTGAGGCAA
AGAGCACACGTCTGAACTCTTTCCCTACACGACGCTCTTCCGATCTNNNCTTGCAGTGAGTAGAGATCC
AGCACACGTCTGAACTCTTTCCCTACACGACGCTCTTCCGATCTNNNAAAAATTGGCCTGGCATCATGA
GAGCACACGTCTGAACTCTTTCCCTACACGACGCTCTTCCGATCTNNNTGAGATGCTGAGATTGTGTGT
GAGCACACGTCTGAACTCTTTCCCTACACGACGCTCTTCCGATCTNNNATTTACCAAAGGTGTGGGGT
AGCACACGTCTGAACTCTTTCCCTACACGACGCTCTTCCGATCTNNNAGCAAGTCCCTTTCACCTATGA
GCACACGTCTGAACTCTTTCCCTACACGACGCTCTTCCGATCTNNNGGCAAGGACTTCATGTCTGAAA
GAGCACACGTCTGAACTCTTTCCCTACACGACGCTCTTCCGATCTNNNTGACCATACTGTCAAAGGCAA
GAGCACACGTCTGAACTCTTTCCCTACACGACGCTCTTCCGATCTNNNGGATCACTTGAGCCTAGGAG
AGCACACGTCTGAACTCTTTCCCTACACGACGCTCTTCCGATCTNNNCAGTAAGTTCAGGCAACAGCTT
AGCACACGTCTGAACTCTTTCCCTACACGACGCTCTTCCGATCTNNNGACTGATGACAGAGTGAGACT
GAGCACACGTCTGAACTCTTTCCCTACACGACGCTCTTCCGATCTNNNATGTAGCAAAATCGCTTGAA
GAGCACACGTCTGAACTCTTTCCCTACACGACGCTCTTCCGATCTNNNCTCCCAAAATACACCTCCCTCAA
GAGCACACGTCTGAACTCTTTCCCTACACGACGCTCTTCCGATCTNNNGAACCAGGGAGTCGGAGATT
GAGCACACGTCTGAACTCTTTCCCTACACGACGCTCTTCCGATCTNNNTGGGCAATAGAACAAGACTCC
GAGCACACGTCTGAACTCTTTCCCTACACGACGCTCTTCCGATCTNNNGATCATGCCATTACACTTCAGC
AGAGCACACGTCTGAACTCTTTCCCTACACGACGCTCTTCCGATCTNNNACCAAGGAATAGAGAATAA
AGCACACGTCTGAACTCTTTCCCTACACGACGCTCTTCCGATCTNNNTTTCTTGGTTAGTCTAATTAGTG
AGAGCACACGTCTGAACTCTTTCCCTACACGACGCTCTTCCGATCTNNNAGAAAGGATTGAGCATTCAT
GAGCACACGTCTGAACTCTTTCCCTACACGACGCTCTTCCGATCTNNNACTCTAGTTTTAGTCTAATCCC
AGCACACGTCTGAACTCTTTCCCTACACGACGCTCTTCCGATCTNNNTCCAAAAATATGATTTGAGTTT
GAGCACACGTCTGAACTCTTTCCCTACACGACGCTCTTCCGATCTNNNTCAAAGGAACAATCTAAGGTT
GAAGAGCACACGTCTGAACTCTTTCCCTACACGACGCTCTTCCGATCTNNNACTTGACTGCATGTTTGAT
AGAGCACACGTCTGAACTCTTTCCCTACACGACGCTCTTCCGATCTNNNGGGTAAAATATTGACTCTCA
GAGCACACGTCTGAACTCTTTCCCTACACGACGCTCTTCCGATCTNNNGGATATAATCACAAACTAAT
GAAGAGCACACGTCTGAACTCTTTCCCTACACGACGCTCTTCCGATCTNNNACTTAGATGTTCACGTAC
GAAGAGCACACGTCTGAACTCTTTCCCTACACGACGCTCTTCCGATCTNNNGGGCCTGAATCAACATAG
AGAGCACACGTCTGAACTCTTTCCCTACACGACGCTCTTCCGATCTNNNTCCTTTTCAAATGGTGAACA
GAAGAGCACACGTCTGAACTCTTTCCCTACACGACGCTCTTCCGATCTNNNATGAATTGCCCTGAAGAC
GAAGAGCACACGTCTGAACTCTTTCCCTACACGACGCTCTTCCGATCTNNNCTGAGGCAGGAGAATCA
CGGAAGAGCACACGTCTGAACTCTTTCCCTACACGACGCTCTTCCGATCTNNNGGATAAATGTTGTGG
AGCACACGTCTGAACTCTTTCCCTACACGACGCTCTTCCGATCTNNNTCCTTGATATGTCAATAACAGCT
AGCACACGTCTGAACTCTTTCCCTACACGACGCTCTTCCGATCTNNNTGTATAATGTGTTAGAATAAGT
GGAAGAGCACACGTCTGAACTCTTTCCCTACACGACGCTCTTCCGATCTNNNATGCTTGTAATCCCAGC

GAAGAGCACACGTCTGAACTCTTTCCCTACACGACGCTCTTCCGATCTNNNGAGAATTGCTTGAACCT
GAAGAGCACACGTCTGAACTCTTTCCCTACACGACGCTCTTCCGATCTNNNGAATGAGAATCGCTTGA
GGAAGAGCACACGTCTGAACTCTTTCCCTACACGACGCTCTTCCGATCTNNNCTAAGGTAGGCCAGTCA
GAGCACACGTCTGAACTCTTTCCCTACACGACGCTCTTCCGATCTNNNACTTATTTGAACCTTCTCTCTTT
GGAAGAGCACACGTCTGAACTCTTTCCCTACACGACGCTCTTCCGATCTNNNCAGGAGAATCACTTGA
GAGCACACGTCTGAACTCTTTCCCTACACGACGCTCTTCCGATCTNNNAGCCAAAAAATAAAAGACAAT
GAAGAGCACACGTCTGAACTCTTTCCCTACACGACGCTCTTCCGATCTNNNAGATTGCAGTGAGCTGA
GAAGAGCACACGTCTGAACTCTTTCCCTACACGACGCTCTTCCGATCTNNNCTAATCTCAGCTACTCGA
GAGCACACGTCTGAACTCTTTCCCTACACGACGCTCTTCCGATCTNNNTGCTCAGTAAATATTGTAGG
AGCACACGTCTGAACTCTTTCCCTACACGACGCTCTTCCGATCTNNNACTATATATATGCATACACACA
AGCACACGTCTGAACTCTTTCCCTACACGACGCTCTTCCGATCTNNNACACATTATGTCTCATGATTAT
GAAGAGCACACGTCTGAACTCTTTCCCTACACGACGCTCTTCCGATCTNNNGAGAATCACTTGAACCT
GGAAGAGCACACGTCTGAACTCTTTCCCTACACGACGCTCTTCCGATCTNNNGAACCGAGATCACACCA
AGCACACGTCTGAACTCTTTCCCTACACGACGCTCTTCCGATCTNNNACTGCAAAATTTTGTTTATCGCT
AGCACACGTCTGAACTCTTTCCCTACACGACGCTCTTCCGATCTNNNTCTACAAAAAGAAAAACATAAT
AGCACACGTCTGAACTCTTTCCCTACACGACGCTCTTCCGATCTNNNTCTTCAGCATAAATTTTAGTCCA
GAAGAGCACACGTCTGAACTCTTTCCCTACACGACGCTCTTCCGATCTNNNATAGTCACTGTCAGCCTG
AGCACACGTCTGAACTCTTTCCCTACACGACGCTCTTCCGATCTNNNTGAGCAGATACATGTATCTCTA
GAGCACACGTCTGAACTCTTTCCCTACACGACGCTCTTCCGATCTNNNATCTATCTGGAATTCAATGAA
GCACACGTCTGAACTCTTTCCCTACACGACGCTCTTCCGATCTNNNCAGATATAATGAAGTACAGTCCA
AGAGCACACGTCTGAACTCTTTCCCTACACGACGCTCTTCCGATCTNNNGGTGCTCATTATTATTTCTAT
GAAGAGCACACGTCTGAACTCTTTCCCTACACGACGCTCTTCCGATCTNNNGAGCTTGCAATGACCTG
GAAGAGCACACGTCTGAACTCTTTCCCTACACGACGCTCTTCCGATCTNNNGTGAAGTCAGCTGTGAT
GAAGAGCACACGTCTGAACTCTTTCCCTACACGACGCTCTTCCGATCTNNNTGAGGCAGGAGAATCAT
CGGAAGAGCACACGTCTGAACTCTTTCCCTACACGACGCTCTTCCGATCTNNNAAGAATTCATCCCAAT
GAAGAGCACACGTCTGAACTCTTTCCCTACACGACGCTCTTCCGATCTNNNTGAGGCAGGAGAATGG
AGCACACGTCTGAACTCTTTCCCTACACGACGCTCTTCCGATCTNNNGCTCCTGCTATAATTTTTCCTTTT
GAAGAGCACACGTCTGAACTCTTTCCCTACACGACGCTCTTCCGATCTNNNCAGGAGTTCAAGGCTACA
GAGCACACGTCTGAACTCTTTCCCTACACGACGCTCTTCCGATCTNNNAGTCCAGAACTAAGTAGTTT
GAAGAGCACACGTCTGAACTCTTTCCCTACACGACGCTCTTCCGATCTNNNTGTAGTCCCAGATACTCA
AGAGCACACGTCTGAACTCTTTCCCTACACGACGCTCTTCCGATCTNNNTCATTCTCTTTCTCTTGGATTT
GAGCACACGTCTGAACTCTTTCCCTACACGACGCTCTTCCGATCTNNNTGTGTGTTTTTGTGTATATGT
GAAGAGCACACGTCTGAACTCTTTCCCTACACGACGCTCTTCCGATCTNNNGAGATTGCAGTGAGCCA
GAAGAGCACACGTCTGAACTCTTTCCCTACACGACGCTCTTCCGATCTNNNAACTCCCTCCCCTCCTATG
GAGCACACGTCTGAACTCTTTCCCTACACGACGCTCTTCCGATCTNNNCAGTTAGAAATACTGTGGTGT
AGCACACGTCTGAACTCTTTCCCTACACGACGCTCTTCCGATCTNNNACACACACACATATATATACACA
GAAGAGCACACGTCTGAACTCTTTCCCTACACGACGCTCTTCCGATCTNNNAGGCTTTTTGTTTTGGG
GAAGAGCACACGTCTGAACTCTTTCCCTACACGACGCTCTTCCGATCTNNNGAGGAGTATGACTCTCT
GAGCACACGTCTGAACTCTTTCCCTACACGACGCTCTTCCGATCTNNNTGCATAAGAGCTAATGAAAAT
AGCACACGTCTGAACTCTTTCCCTACACGACGCTCTTCCGATCTNNNATGATGGTATATCTGGAAATT
AGCACACGTCTGAACTCTTTCCCTACACGACGCTCTTCCGATCTNNNTCTGTCAATAAAGTTATGCAAG
GAGCACACGTCTGAACTCTTTCCCTACACGACGCTCTTCCGATCTNNNTGTGTGTGTATATATATGT
GAGCACACGTCTGAACTCTTTCCCTACACGACGCTCTTCCGATCTNNNACACGCAATCTAAAAAAATCTA

AGCACACGTCTGAACTCTTTCCCTACACGACGCTCTTCCGATCTNNNACCTCTCTAGTTAGAAAGAGAA
GCACACGTCTGAACTCTTTCCCTACACGACGCTCTTCCGATCTNNNTGCATCAAATTATCACATGTACTC
AAGAGCACACGTCTGAACTCTTTCCCTACACGACGCTCTTCCGATCTNNNGAGCAAGACTCTGTCTCAA
AGCACACGTCTGAACTCTTTCCCTACACGACGCTCTTCCGATCTNNNTGTGTGTACATATATACAAACGT
AGCACACGTCTGAACTCTTTCCCTACACGACGCTCTTCCGATCTNNNACAAATTTACATGTTCCCTTTGA
AGCACACGTCTGAACTCTTTCCCTACACGACGCTCTTCCGATCTNNNTGTGTGTGTATATATATGTGTG
AGAGCACACGTCTGAACTCTTTCCCTACACGACGCTCTTCCGATCTNNNTGAAAACTGTAGGAATTTAC
GAAGAGCACACGTCTGAACTCTTTCCCTACACGACGCTCTTCCGATCTNNNAAGGGAACTGAATAACTC
GAAGAGCACACGTCTGAACTCTTTCCCTACACGACGCTCTTCCGATCTNNNTGACTCTATAAGTGAGCT
GAGCACACGTCTGAACTCTTTCCCTACACGACGCTCTTCCGATCTNNNACTCTGTCTTTTTTTTTGCAAA
GAAGAGCACACGTCTGAACTCTTTCCCTACACGACGCTCTTCCGATCTNNNTGCCTTGGATCATTTTCAA
AGCACACGTCTGAACTCTTTCCCTACACGACGCTCTTCCGATCTNNNTGGAAGTCTTTTGTAGGTAAGA
AGCACACGTCTGAACTCTTTCCCTACACGACGCTCTTCCGATCTNNNACATAGATTCTATTCTCTCTCT
GCACACGTCTGAACTCTTTCCCTACACGACGCTCTTCCGATCTNNNGCCTCAGTGGAAAAAAATTATAT
AAGAGCACACGTCTGAACTCTTTCCCTACACGACGCTCTTCCGATCTNNNTCTTGAAGCACATTCTGACT
GAAGAGCACACGTCTGAACTCTTTCCCTACACGACGCTCTTCCGATCTNNNAGGATAAAGAAGGTTGC
GAAGAGCACACGTCTGAACTCTTTCCCTACACGACGCTCTTCCGATCTNNNAGTGAGCAGAGATCATC
GAGCACACGTCTGAACTCTTTCCCTACACGACGCTCTTCCGATCTNNNTCAGGACAAATATTTTCTTAGA
GAAGAGCACACGTCTGAACTCTTTCCCTACACGACGCTCTTCCGATCTNNNCTCCTGACCTTGTGATCTA
AGCACACGTCTGAACTCTTTCCCTACACGACGCTCTTCCGATCTNNNAGTTTGTATATATGGATCTCTCTC
GAGCACACGTCTGAACTCTTTCCCTACACGACGCTCTTCCGATCTNNNAGGGAGAAAGAAAGAAAGA
AGAGCACACGTCTGAACTCTTTCCCTACACGACGCTCTTCCGATCTNNNACTCCCCAAAGAAAACAAGA
AAGAGCACACGTCTGAACTCTTTCCCTACACGACGCTCTTCCGATCTNNNATATAGAGAGAGAGAGAG
CGGAAGAGCACACGTCTGAACTCTTTCCCTACACGACGCTCTTCCGATCTNNNTTGAAGCTGCAGTGA
AGCACACGTCTGAACTCTTTCCCTACACGACGCTCTTCCGATCTNNNTCAGTCCATCTCTTTTTCTTTGTT
GAAGAGCACACGTCTGAACTCTTTCCCTACACGACGCTCTTCCGATCTNNNAAGTTTCTTTTGGGTTCAC
AAGAGCACACGTCTGAACTCTTTCCCTACACGACGCTCTTCCGATCTNNNGAGAGAAAGAAAGAAAGA
AAGAGCACACGTCTGAACTCTTTCCCTACACGACGCTCTTCCGATCTNNNACCATATTTTTGGGGACTT
AAGAGCACACGTCTGAACTCTTTCCCTACACGACGCTCTTCCGATCTNNNTGCCCAGTTTCACTTTCTGT
GAAGAGCACACGTCTGAACTCTTTCCCTACACGACGCTCTTCCGATCTNNNCCTGGGACTTCTCAACCTT
AAGAGCACACGTCTGAACTCTTTCCCTACACGACGCTCTTCCGATCTNNNTGAATATGTGTGAGTGAGC
AGAGCACACGTCTGAACTCTTTCCCTACACGACGCTCTTCCGATCTNNNCTTTCCTGTCCTTTCCCTTCCT
AGCACACGTCTGAACTCTTTCCCTACACGACGCTCTTCCGATCTNNNTTTCTTTCATTCTTTGACTATGCC
GAAGAGCACACGTCTGAACTCTTTCCCTACACGACGCTCTTCCGATCTNNNAGTGAGAACCATCCAAG
GAGCACACGTCTGAACTCTTTCCCTACACGACGCTCTTCCGATCTNNNACAAAGAGAGAAAGAAAGAA
GAGCACACGTCTGAACTCTTTCCCTACACGACGCTCTTCCGATCTNNNTCTTTCTTTCCTTCTTTCTTTCTT
AGCACACGTCTGAACTCTTTCCCTACACGACGCTCTTCCGATCTNNNTCTTTCCTTCTTTCCTTCTTTCTTTC
AGCACACGTCTGAACTCTTTCCCTACACGACGCTCTTCCGATCTNNNACGCTTTCCTCTTTTTATTTTAG
AAGAGCACACGTCTGAACTCTTTCCCTACACGACGCTCTTCCGATCTNNNGAGATCATCCCACTGCACTC
GAGCACACGTCTGAACTCTTTCCCTACACGACGCTCTTCCGATCTNNNTCCTTCCTTCTTTCTTTCTTTCTT
TCGGAAGAGCACACGTCTGAACTCTTTCCCTACACGACGCTCTTCCGATCTNNNGAAGGTCAAATCTGC
GAGCACACGTCTGAACTCTTTCCCTACACGACGCTCTTCCGATCTNNNAGGGAAAGAAAGAAAGAAAG
GCACACGTCTGAACTCTTTCCCTACACGACGCTCTTCCGATCTNNNAGACTGCATCTCAAAAAAAAGAA

AGAGCACACGTCTGAACTCTTTCCCTACACGACGCTCTTCCGATCTNNNCTGAGGCAGGAGAATAGC
GCACACGTCTGAACTCTTTCCCTACACGACGCTCTTCCGATCTNNNTGCCAATCACAAAAATGTTCAA
AAGAGCACACGTCTGAACTCTTTCCCTACACGACGCTCTTCCGATCTNNNACACCTGTAATCCCAGCTA
AAGAGCACACGTCTGAACTCTTTCCCTACACGACGCTCTTCCGATCTNNNTGTAGTCCCAGCTACTAG
AAGAGCACACGTCTGAACTCTTTCCCTACACGACGCTCTTCCGATCTNNNGTGATACAGGCTTCGGTG
GGAAGAGCACACGTCTGAACTCTTTCCCTACACGACGCTCTTCCGATCTNNNTGAGGCAAGAGAATG
GCACACGTCTGAACTCTTTCCCTACACGACGCTCTTCCGATCTNNNACGATGCATTTTGAGTTTAGTTT
AGCACACGTCTGAACTCTTTCCCTACACGACGCTCTTCCGATCTNNNCGTTGAAATGCAAGAAAATTT
AGCACACGTCTGAACTCTTTCCCTACACGACGCTCTTCCGATCTNNNGCAATAAATATCCGTGTTTGT
GCACACGTCTGAACTCTTTCCCTACACGACGCTCTTCCGATCTNNNAGTGCTTATGATATGTCAGACA
AAGAGCACACGTCTGAACTCTTTCCCTACACGACGCTCTTCCGATCTNNNAACTTGCAGTGAGCCAA
CACACGTCTGAACTCTTTCCCTACACGACGCTCTTCCGATCTNNNACTCCTGTCTTTTCTACATATTTGT
GCACACGTCTGAACTCTTTCCCTACACGACGCTCTTCCGATCTNNNTGCAGTACTATTTTGTTATACCA
AAGAGCACACGTCTGAACTCTTTCCCTACACGACGCTCTTCCGATCTNNNGATTTCTTGAGCCTGAGA
AGCACACGTCTGAACTCTTTCCCTACACGACGCTCTTCCGATCTNNNACAAAAAAATTAGTGATTGCC
GCACACGTCTGAACTCTTTCCCTACACGACGCTCTTCCGATCTNNNTGCAGAAATAGGTATCAACATT
AAGAGCACACGTCTGAACTCTTTCCCTACACGACGCTCTTCCGATCTNNNGAGAATCACTTGAACCCG
GAAGAGCACACGTCTGAACTCTTTCCCTACACGACGCTCTTCCGATCTNNNTCTGTAATCTCAGCTAG
AAGAGCACACGTCTGAACTCTTTCCCTACACGACGCTCTTCCGATCTNNNGTGCCTGTAGTCCCAGTT
AAGAGCACACGTCTGAACTCTTTCCCTACACGACGCTCTTCCGATCTNNNGGGAGGGATAGCATTGG
AGAGCACACGTCTGAACTCTTTCCCTACACGACGCTCTTCCGATCTNNNATCCATTGAGCCTGGAAA
AGCACACGTCTGAACTCTTTCCCTACACGACGCTCTTCCGATCTNNNAGATGCAAGATAAAAGTGATG
AGAGCACACGTCTGAACTCTTTCCCTACACGACGCTCTTCCGATCTNNNGAAATGTAGGTTGCAGTGA
AGCACACGTCTGAACTCTTTCCCTACACGACGCTCTTCCGATCTNNNTCAAGTGTCAAAAACTATTGA
GGAAGAGCACACGTCTGAACTCTTTCCCTACACGACGCTCTTCCGATCTNNNTTGCACAAGTGGGTGA
AGCACACGTCTGAACTCTTTCCCTACACGACGCTCTTCCGATCTNNNTCAGACAGAGATCTACATGAG
AGAGCACACGTCTGAACTCTTTCCCTACACGACGCTCTTCCGATCTNNNACCTAATGCTAGATGACAC
AAGAGCACACGTCTGAACTCTTTCCCTACACGACGCTCTTCCGATCTNNNAGTCTGAAGCTGCAGTGT
GCACACGTCTGAACTCTTTCCCTACACGACGCTCTTCCGATCTNNNACATCAAAAGTCATAGGCGTTT
AGCACACGTCTGAACTCTTTCCCTACACGACGCTCTTCCGATCTNNNCCACCTCTCTAATACTCAAAGTC
AGAGCACACGTCTGAACTCTTTCCCTACACGACGCTCTTCCGATCTNNNGAGCTTACAGTGAGCAGA
AGAGCACACGTCTGAACTCTTTCCCTACACGACGCTCTTCCGATCTNNNATGATCATACCGCCACTAC
AGAGCACACGTCTGAACTCTTTCCCTACACGACGCTCTTCCGATCTNNNGTCTCAGCTACTCCTAGGA
AAGAGCACACGTCTGAACTCTTTCCCTACACGACGCTCTTCCGATCTNNNGTTGCAGTGAGCTGAGA
GGAAGAGCACACGTCTGAACTCTTTCCCTACACGACGCTCTTCCGATCTNNNCAGGAGAATCACCTA
AGCACACGTCTGAACTCTTTCCCTACACGACGCTCTTCCGATCTNNNTGGAGAAGTCAATATTTGAAT
GCACACGTCTGAACTCTTTCCCTACACGACGCTCTTCCGATCTNNNAGTGATAGAGGGAATTTATTCT
AAGAGCACACGTCTGAACTCTTTCCCTACACGACGCTCTTCCGATCTNNNAAGGAGAATCATTTGAAC
AGAGCACACGTCTGAACTCTTTCCCTACACGACGCTCTTCCGATCTNNNGAAGGAGAATGGTGAGA
AGAGCACACGTCTGAACTCTTTCCCTACACGACGCTCTTCCGATCTNNNATACCCTTGTGTAGGTGTG
AGAGCACACGTCTGAACTCTTTCCCTACACGACGCTCTTCCGATCTNNNAAACATCCAGCCTGAATGT
CACACGTCTGAACTCTTTCCCTACACGACGCTCTTCCGATCTNNNAGTTTATAAACTTCATGAAAGCC
AGCACACGTCTGAACTCTTTCCCTACACGACGCTCTTCCGATCTNNNGCTTTAGCACAAATCATTCTAC

AGCACACGTCTGAACTCTTTCCCTACACGACGCTCTTCCGATCTNNNTGACAATGACAGTAGTCTTTCT
AGCACACGTCTGAACTCTTTCCCTACACGACGCTCTTCCGATCTNNNTTGGCATTTATGTAATACCATGA
GAAGAGCACACGTCTGAACTCTTTCCCTACACGACGCTCTTCCGATCTNNNTTACAGACACTGGTGAC
AAGAGCACACGTCTGAACTCTTTCCCTACACGACGCTCTTCCGATCTNNNAGACAGGAGAATCACTTG
AAGAGCACACGTCTGAACTCTTTCCCTACACGACGCTCTTCCGATCTNNNCTGAGGCAGGAAAATTGC
GAAGAGCACACGTCTGAACTCTTTCCCTACACGACGCTCTTCCGATCTNNNCAAGAGAATTGCTTGAA
AGAGCACACGTCTGAACTCTTTCCCTACACGACGCTCTTCCGATCTNNNTCCACAGAAAAAATGAGTG
GCACACGTCTGAACTCTTTCCCTACACGACGCTCTTCCGATCTNNNTGTGTGTATATATATGTGTGTGT
AAGAGCACACGTCTGAACTCTTTCCCTACACGACGCTCTTCCGATCTNNNGGTGGCAAGTATCTGTAA
AGAGCACACGTCTGAACTCTTTCCCTACACGACGCTCTTCCGATCTNNNGCTGCTCTAGTTCACTGCTC
AAGAGCACACGTCTGAACTCTTTCCCTACACGACGCTCTTCCGATCTNNNATTTTTGCTTTTGTTGCCTG
AGAGCACACGTCTGAACTCTTTCCCTACACGACGCTCTTCCGATCTNNNCCCTTCCTGATGATCCAGTCA
AGCACACGTCTGAACTCTTTCCCTACACGACGCTCTTCCGATCTNNNCCAGTATTCCTTCTTATGGAGTT
AGAGCACACGTCTGAACTCTTTCCCTACACGACGCTCTTCCGATCTNNNTCTGGTGATCTTCATTTCTTC
GAGCACACGTCTGAACTCTTTCCCTACACGACGCTCTTCCGATCTNNNTCCATTGACTATACATATCACT
GCACACGTCTGAACTCTTTCCCTACACGACGCTCTTCCGATCTNNNTGAAATGACTGAGAAGATACAT
AAGAGCACACGTCTGAACTCTTTCCCTACACGACGCTCTTCCGATCTNNNGTGAGTCTCCGTCTCAGGA
AGCACACGTCTGAACTCTTTCCCTACACGACGCTCTTCCGATCTNNNAGAAAACTCAAGAAGAAAATG
GAGCACACGTCTGAACTCTTTCCCTACACGACGCTCTTCCGATCTNNNAATGAACATTGCATTACTTTTT
GAAGAGCACACGTCTGAACTCTTTCCCTACACGACGCTCTTCCGATCTNNNCTGTCAGGCTTGCTAAAG
AAGAGCACACGTCTGAACTCTTTCCCTACACGACGCTCTTCCGATCTNNNCTGAGGCAGGAGAATTGT
GAGCACACGTCTGAACTCTTTCCCTACACGACGCTCTTCCGATCTNNNTCGACCTGAATATTTGGAATAT
AGAGCACACGTCTGAACTCTTTCCCTACACGACGCTCTTCCGATCTNNNTGAAATTGGCCACTGCACTT
GCACACGTCTGAACTCTTTCCCTACACGACGCTCTTCCGATCTNNNGTCAAATGGAAATATTGTAATGC
GCACACGTCTGAACTCTTTCCCTACACGACGCTCTTCCGATCTNNNTCTTTCTTTCTCTTTCTTTCTCTTTC
GAAGAGCACACGTCTGAACTCTTTCCCTACACGACGCTCTTCCGATCTNNNAATCGCTTGAACTCAGGA
AAGAGCACACGTCTGAACTCTTTCCCTACACGACGCTCTTCCGATCTNNNCAAAACTGCCCTCCTGAAT
AGAGCACACGTCTGAACTCTTTCCCTACACGACGCTCTTCCGATCTNNNTGCTAAGAGCTGAAGAGAA
AGAGCACACGTCTGAACTCTTTCCCTACACGACGCTCTTCCGATCTNNNGGAGGTTACAGTGAGCTGA
AGAGCACACGTCTGAACTCTTTCCCTACACGACGCTCTTCCGATCTNNNTTCTGATTCAGTACTTCTGG
AAGAGCACACGTCTGAACTCTTTCCCTACACGACGCTCTTCCGATCTNNNGATCTCCTGACCTTGTGAT
GAAGAGCACACGTCTGAACTCTTTCCCTACACGACGCTCTTCCGATCTNNNATAGAAAGTGAGAGACA
AGAGCACACGTCTGAACTCTTTCCCTACACGACGCTCTTCCGATCTNNNACATATCAAATGTGAGACAT
AGAGCACACGTCTGAACTCTTTCCCTACACGACGCTCTTCCGATCTNNNCAGGAAAGAGCACAAGCATT
AAGAGCACACGTCTGAACTCTTTCCCTACACGACGCTCTTCCGATCTNNNCATTCCACACTCCAAAAGCA
AAGAGCACACGTCTGAACTCTTTCCCTACACGACGCTCTTCCGATCTNNNGCCTTCGTAATTGTGTGAA
AGAGCACACGTCTGAACTCTTTCCCTACACGACGCTCTTCCGATCTNNNAGGAAAGCTTTTCCTGTAAT
GAGCACACGTCTGAACTCTTTCCCTACACGACGCTCTTCCGATCTNNNTCTGTATAGAAAGGCTTTGTTC
AGCACACGTCTGAACTCTTTCCCTACACGACGCTCTTCCGATCTNNNTGTTTTCTTCCCTGATCATATCCA
AGCACACGTCTGAACTCTTTCCCTACACGACGCTCTTCCGATCTNNNTGTGTGTATATGTGTGTGTGTA
AAGAGCACACGTCTGAACTCTTTCCCTACACGACGCTCTTCCGATCTNNNAACTCAGGCAAAACACGTT
GAAGAGCACACGTCTGAACTCTTTCCCTACACGACGCTCTTCCGATCTNNNTAGGCCACAGTGACCTAT
GCACACGTCTGAACTCTTTCCCTACACGACGCTCTTCCGATCTNNNTGAAAAGAAAAAGAAATGCCCTT

GAGCACACGTCTGAACTCTTTCCCTACACGACGCTCTTCCGATCTNNNTTCCATCCAGATTGCTGCAA
AGCACACGTCTGAACTCTTTCCCTACACGACGCTCTTCCGATCTNNNTCAGAATGAGGAATCTGTAA
AGCACACGTCTGAACTCTTTCCCTACACGACGCTCTTCCGATCTNNNTTGAAAGAAAAGTCACACTAG
AGAGCACACGTCTGAACTCTTTCCCTACACGACGCTCTTCCGATCTNNNCTTGAAACAATTGCGCAAA
AGCACACGTCTGAACTCTTTCCCTACACGACGCTCTTCCGATCTNNNTCTGCCAGCTAAACTAATTTTG
AAGAGCACACGTCTGAACTCTTTCCCTACACGACGCTCTTCCGATCTNNNGCCAAGATCATGTTACTG
AGAGCACACGTCTGAACTCTTTCCCTACACGACGCTCTTCCGATCTNNNGGACAAGAGTGGGATTTG
GAGCACACGTCTGAACTCTTTCCCTACACGACGCTCTTCCGATCTNNNTGGACATAACATTTTCAACA
AGCACACGTCTGAACTCTTTCCCTACACGACGCTCTTCCGATCTNNNACAGGAGTTTTATGTGTATAT
AGAGCACACGTCTGAACTCTTTCCCTACACGACGCTCTTCCGATCTNNNAGATCTTCATGACCTAGGAT
AGAGCACACGTCTGAACTCTTTCCCTACACGACGCTCTTCCGATCTNNNCACACACACAATCCCAGTTC
AGCACACGTCTGAACTCTTTCCCTACACGACGCTCTTCCGATCTNNNTGTGTGTGTGTGTATATATGT
GAAGAGCACACGTCTGAACTCTTTCCCTACACGACGCTCTTCCGATCTNNNAGAGGGACAGAACTAA
AGCACACGTCTGAACTCTTTCCCTACACGACGCTCTTCCGATCTNNNAGGAAAGGAAGAGTAAAAAG
GAGCACACGTCTGAACTCTTTCCCTACACGACGCTCTTCCGATCTNNNACGAGAGTCCATTTCAAAAA
GCACACGTCTGAACTCTTTCCCTACACGACGCTCTTCCGATCTNNNGCAAATATGTATTTCTTTGCTT
GCACACGTCTGAACTCTTTCCCTACACGACGCTCTTCCGATCTNNNTCTGACACACACATATATATACA
GCACACGTCTGAACTCTTTCCCTACACGACGCTCTTCCGATCTNNNCCAATGTTGGGAAAAAAGAAG
AGCACACGTCTGAACTCTTTCCCTACACGACGCTCTTCCGATCTNNNCAAACTATTATTCAACCAAATTC
AGCACACGTCTGAACTCTTTCCCTACACGACGCTCTTCCGATCTNNNACACAAAACATCATGAATGAA
GAGCACACGTCTGAACTCTTTCCCTACACGACGCTCTTCCGATCTNNNAGTTAGCTTTTTCTCACCATAT
AAGAGCACACGTCTGAACTCTTTCCCTACACGACGCTCTTCCGATCTNNNTTTGCAATCCACCACAATC
GCACACGTCTGAACTCTTTCCCTACACGACGCTCTTCCGATCTNNNATAGATGCTGTAGTATCCATAGT
AGCACACGTCTGAACTCTTTCCCTACACGACGCTCTTCCGATCTNNNTGTGTGACTGGTTAGTAAAA
AGAGCACACGTCTGAACTCTTTCCCTACACGACGCTCTTCCGATCTNNNAGATGGCAAATCATGGGA
AGAGCACACGTCTGAACTCTTTCCCTACACGACGCTCTTCCGATCTNNNAGACATGTGTATACCATAC
AGCACACGTCTGAACTCTTTCCCTACACGACGCTCTTCCGATCTNNNAATATTTGGGTGAGAAGGAA
AAGAGCACACGTCTGAACTCTTTCCCTACACGACGCTCTTCCGATCTNNNGATCTCAACTCACTGCAAC
AAGAGCACACGTCTGAACTCTTTCCCTACACGACGCTCTTCCGATCTNNNTACAACACGCATCAGAAG
GAGCACACGTCTGAACTCTTTCCCTACACGACGCTCTTCCGATCTNNNTGAAACCATCTTTGAAGAATG
AGAGCACACGTCTGAACTCTTTCCCTACACGACGCTCTTCCGATCTNNNACTCAAATATCTCTGAGGT
GAGCACACGTCTGAACTCTTTCCCTACACGACGCTCTTCCGATCTNNNAGGAATGAAATGTTTTCTGT
AAGAGCACACGTCTGAACTCTTTCCCTACACGACGCTCTTCCGATCTNNNACGCACGAAGGTATATT
AGCACACGTCTGAACTCTTTCCCTACACGACGCTCTTCCGATCTNNNTCTTTCTTCTCTCTCTCTCTTT
GAAGAGCACACGTCTGAACTCTTTCCCTACACGACGCTCTTCCGATCTNNNGAGATCACACCACTGTA
AAGAGCACACGTCTGAACTCTTTCCCTACACGACGCTCTTCCGATCTNNNCTCCTTGACCTCCCAAAGT
GCACACGTCTGAACTCTTTCCCTACACGACGCTCTTCCGATCTNNNTGGAAATTTCATTCTAGTGTAGT
AGCACACGTCTGAACTCTTTCCCTACACGACGCTCTTCCGATCTNNNTGTCTCAAACAAGAGAGAGA
AGCACACGTCTGAACTCTTTCCCTACACGACGCTCTTCCGATCTNNNACTCCATCTCAAAAAAGAAAG
GCACACGTCTGAACTCTTTCCCTACACGACGCTCTTCCGATCTNNNACTCTTGCTTTCTTTCTTTCTTTC
AGCACACGTCTGAACTCTTTCCCTACACGACGCTCTTCCGATCTNNNATTCATGTATATATGTGTGTG
AGCACACGTCTGAACTCTTTCCCTACACGACGCTCTTCCGATCTNNNGCCTACTCTAAGATAAGAAGT
CGGAAGAGCACACGTCTGAACTCTTTCCCTACACGACGCTCTTCCGATCTNNNCACCATTGCACTCCAG

AGAGCACACGTCTGAACTCTTTCCCTACACGACGCTCTTCCGATCTNNNTCAACAAAAATCATGCTGG
CGGAAGAGCACACGTCTGAACTCTTTCCCTACACGACGCTCTTCCGATCTNNNGGAGAATCACTTGAA
GAGCACACGTCTGAACTCTTTCCCTACACGACGCTCTTCCGATCTNNNCAGTGTTTCTTACGCCAGTG
GGAAGAGCACACGTCTGAACTCTTTCCCTACACGACGCTCTTCCGATCTNNNGAGATAGGCCACTCC
AGCACACGTCTGAACTCTTTCCCTACACGACGCTCTTCCGATCTNNNTCTTTCTCTCTCTCTCTTTCTTTC
AGAGCACACGTCTGAACTCTTTCCCTACACGACGCTCTTCCGATCTNNNCCAGAGAAACTCTTCAGAG
GAAGAGCACACGTCTGAACTCTTTCCCTACACGACGCTCTTCCGATCTNNNTGGGCCCATAGAGTAGA
AGAGCACACGTCTGAACTCTTTCCCTACACGACGCTCTTCCGATCTNNNTGGATGTTGCAGTAAGCTG
GCACACGTCTGAACTCTTTCCCTACACGACGCTCTTCCGATCTNNNTGGAGTAAGAAAACTGAAGCT
AAGAGCACACGTCTGAACTCTTTCCCTACACGACGCTCTTCCGATCTNNNAAGGCAAGAGAATTGCTT
AGAGCACACGTCTGAACTCTTTCCCTACACGACGCTCTTCCGATCTNNNTGCAGTGAGCTATCATTAC
AGAGCACACGTCTGAACTCTTTCCCTACACGACGCTCTTCCGATCTNNNAAAAAAATTAGCTGGGCGT
AGAGCACACGTCTGAACTCTTTCCCTACACGACGCTCTTCCGATCTNNNTGAGGCACAAGAATCGTTT
GCACACGTCTGAACTCTTTCCCTACACGACGCTCTTCCGATCTNNNTGAGAAGACATATGTTTGGTT
GAGCACACGTCTGAACTCTTTCCCTACACGACGCTCTTCCGATCTNNNAGACATTGTGGTTTTCATTT
AGCACACGTCTGAACTCTTTCCCTACACGACGCTCTTCCGATCTNNNTCATAAACATCTTTGTATAGCT
GCACACGTCTGAACTCTTTCCCTACACGACGCTCTTCCGATCTNNNTGTACTCCCAAAAGCAATCTAT
AGCACACGTCTGAACTCTTTCCCTACACGACGCTCTTCCGATCTNNNTGCATAGAGTTCTAGTTTTGC
GAGCACACGTCTGAACTCTTTCCCTACACGACGCTCTTCCGATCTNNNTGCTCTGTCTTCTCTATTACT
AGAGCACACGTCTGAACTCTTTCCCTACACGACGCTCTTCCGATCTNNNCACCTGTAATCCCAGCTACT
GCACACGTCTGAACTCTTTCCCTACACGACGCTCTTCCGATCTNNNGGAACAATTCTCATGGGTATTG
CACACGTCTGAACTCTTTCCCTACACGACGCTCTTCCGATCTNNNTCCTACTTATTCTGTGTTCTTTTG
GAGCACACGTCTGAACTCTTTCCCTACACGACGCTCTTCCGATCTNNNGTAATTGCTTGAACCTGGTA
AGAGCACACGTCTGAACTCTTTCCCTACACGACGCTCTTCCGATCTNNNCAAGAGAATTGCTTGAACC
GCACACGTCTGAACTCTTTCCCTACACGACGCTCTTCCGATCTNNNTCTCTCTCTCTCTCTCTCTCTCT
GCACACGTCTGAACTCTTTCCCTACACGACGCTCTTCCGATCTNNNTGGAATTGTAGCTCATTTAGTG
AAGAGCACACGTCTGAACTCTTTCCCTACACGACGCTCTTCCGATCTNNNGCCTGTAGTCCCAGCTACT
GAGCACACGTCTGAACTCTTTCCCTACACGACGCTCTTCCGATCTNNNTTGCTTGACTCTAGGAGGTG
AGCACACGTCTGAACTCTTTCCCTACACGACGCTCTTCCGATCTNNNTGAACTTCTTTCAAACATTTGA
AGAGCACACGTCTGAACTCTTTCCCTACACGACGCTCTTCCGATCTNNNAGTGATTGGAGATGGCAC
GAGCACACGTCTGAACTCTTTCCCTACACGACGCTCTTCCGATCTNNNCCAAAAAAGAGAAGGTTCCC
AGAGCACACGTCTGAACTCTTTCCCTACACGACGCTCTTCCGATCTNNNTTGAAACATTCCTACCAGC
AGAGCACACGTCTGAACTCTTTCCCTACACGACGCTCTTCCGATCTNNNCGAAGCAGGAGAATCTCTT
AAGAGCACACGTCTGAACTCTTTCCCTACACGACGCTCTTCCGATCTNNNTGCAGGAGAATTACTTGA
AGAGCACACGTCTGAACTCTTTCCCTACACGACGCTCTTCCGATCTNNNATGCCTGTAGTCCCAGCTAC
GAGCACACGTCTGAACTCTTTCCCTACACGACGCTCTTCCGATCTNNNTCTAGAATGAGAAGAGTTCT
GAGCACACGTCTGAACTCTTTCCCTACACGACGCTCTTCCGATCTNNNAGGGAGTTTTATTTCTTCATC
AGCACACGTCTGAACTCTTTCCCTACACGACGCTCTTCCGATCTNNNAGTAGTCTTTCTTTTCCACTCTT
AGAGCACACGTCTGAACTCTTTCCCTACACGACGCTCTTCCGATCTNNNCTGAGGCAGGAGAATCACT
AGCACACGTCTGAACTCTTTCCCTACACGACGCTCTTCCGATCTNNNAGCCATATTGGTACTGTGTAA
AGAGCACACGTCTGAACTCTTTCCCTACACGACGCTCTTCCGATCTNNNACCAAGGAGTAACTAGGAC
AAGAGCACACGTCTGAACTCTTTCCCTACACGACGCTCTTCCGATCTNNNCTGAGGCAGGAGAATCA
AGCACACGTCTGAACTCTTTCCCTACACGACGCTCTTCCGATCTNNNTCAGCAAAAGATTCAAACAGG

AGCACACGTCTGAACTCTTTCCCTACACGACGCTCTTCCGATCTNNNACGATAGAATATGACACAACG
AGAGCACACGTCTGAACTCTTTCCCTACACGACGCTCTTCCGATCTNNNAGTAAGCTGAGATCACACC
GAAGAGCACACGTCTGAACTCTTTCCCTACACGACGCTCTTCCGATCTNNNCCTGTAGTCCCAGCTACT
GAGCACACGTCTGAACTCTTTCCCTACACGACGCTCTTCCGATCTNNNCTGTAATCCCAGCTACTCAGG
AAGAGCACACGTCTGAACTCTTTCCCTACACGACGCTCTTCCGATCTNNNTGAATCCAGGAGGCAGAG
AGCACACGTCTGAACTCTTTCCCTACACGACGCTCTTCCGATCTNNNTGTCACTTCTTTGTTTTTGG
GAGCACACGTCTGAACTCTTTCCCTACACGACGCTCTTCCGATCTNNNCTCCTGTAATCCCAGCTACTT
GAGCACACGTCTGAACTCTTTCCCTACACGACGCTCTTCCGATCTNNNTGAGGCAGGAGAATCACTT
AGAGCACACGTCTGAACTCTTTCCCTACACGACGCTCTTCCGATCTNNNGTGAAGGTTGTAGTGAGC
AAGAGCACACGTCTGAACTCTTTCCCTACACGACGCTCTTCCGATCTNNNGAGAATGGTGTGAACTC
GCACACGTCTGAACTCTTTCCCTACACGACGCTCTTCCGATCTNNNCTTCAAAAGCTGTGTAAAGTTA
GCACACGTCTGAACTCTTTCCCTACACGACGCTCTTCCGATCTNNNACAGTGAATAGAGAATTGGGA
AGCACACGTCTGAACTCTTTCCCTACACGACGCTCTTCCGATCTNNNACAACCTAACATCATACCTAGT
GCACACGTCTGAACTCTTTCCCTACACGACGCTCTTCCGATCTNNNTGGTAGGCATTTAGGTTCTCAA
AGCACACGTCTGAACTCTTTCCCTACACGACGCTCTTCCGATCTNNNCCCTAACTTTGAATAGAGAAG
AGAGCACACGTCTGAACTCTTTCCCTACACGACGCTCTTCCGATCTNNNAGCCAAATCCACAGAGTTA
GCACACGTCTGAACTCTTTCCCTACACGACGCTCTTCCGATCTNNNGCCATCTTTCTTAGTTTGACAAA
AGCACACGTCTGAACTCTTTCCCTACACGACGCTCTTCCGATCTNNNTCTTGAAGAGCTATTAGCACTC
GAGCACACGTCTGAACTCTTTCCCTACACGACGCTCTTCCGATCTNNNGAGGCAGGAAAATCACTTGA
AGAGCACACGTCTGAACTCTTTCCCTACACGACGCTCTTCCGATCTNNNTTGAGGCAGGAGAATCAC
GAGCACACGTCTGAACTCTTTCCCTACACGACGCTCTTCCGATCTNNNCTGAGGCAGGAGAATTGCTT
AGAGCACACGTCTGAACTCTTTCCCTACACGACGCTCTTCCGATCTNNNATGCCTGTAATCCCAGTGA
GAAGAGCACACGTCTGAACTCTTTCCCTACACGACGCTCTTCCGATCTNNNAGAATCACTTGAATCCA
GAGCACACGTCTGAACTCTTTCCCTACACGACGCTCTTCCGATCTNNNGAGGTTACAGTGAGCCGAG
AAGAGCACACGTCTGAACTCTTTCCCTACACGACGCTCTTCCGATCTNNNAGGCTGAAGCAGGAGAA
AGAGCACACGTCTGAACTCTTTCCCTACACGACGCTCTTCCGATCTNNNCGAGGCAGAAGAATCACT
GCACACGTCTGAACTCTTTCCCTACACGACGCTCTTCCGATCTNNNTGAGATGCTTATTCCCAAATGT
GCACACGTCTGAACTCTTTCCCTACACGACGCTCTTCCGATCTNNNTCTGTGGATTGTCTTTTCACTTT
GAGCACACGTCTGAACTCTTTCCCTACACGACGCTCTTCCGATCTNNNAGTGAGCTGAGACCGTGAT
GCACACGTCTGAACTCTTTCCCTACACGACGCTCTTCCGATCTNNNACTCAGAAAATACAAGCAGAG
AGAGCACACGTCTGAACTCTTTCCCTACACGACGCTCTTCCGATCTNNNGCAGGAGCATCACTTGAA
GAAGAGCACACGTCTGAACTCTTTCCCTACACGACGCTCTTCCGATCTNNNCTGTAGTCCCAGCTACT
GAGCACACGTCTGAACTCTTTCCCTACACGACGCTCTTCCGATCTNNNTCTAGAATGAGAAGAGTTCT
AGAGCACACGTCTGAACTCTTTCCCTACACGACGCTCTTCCGATCTNNNATTTAGGCAATGAGGAGG
AGAGCACACGTCTGAACTCTTTCCCTACACGACGCTCTTCCGATCTNNNTTACAGTGAGCCAAAATCAT
GAGCACACGTCTGAACTCTTTCCCTACACGACGCTCTTCCGATCTNNNCTGTATTCCCAGCTACTCAGG
AGCACACGTCTGAACTCTTTCCCTACACGACGCTCTTCCGATCTNNNGGCAAATTGAATCCAACAATAC
GAGCACACGTCTGAACTCTTTCCCTACACGACGCTCTTCCGATCTNNNAGGAGAATCGTTTGAACCCA
AGCACACGTCTGAACTCTTTCCCTACACGACGCTCTTCCGATCTNNNTTGAGTCCAGGAGTTCAAGGA
AGAGCACACGTCTGAACTCTTTCCCTACACGACGCTCTTCCGATCTNNNCTGAGGCAGGAGAATCAC
AGCACACGTCTGAACTCTTTCCCTACACGACGCTCTTCCGATCTNNNTTTTTTTTGAGATGGAGTTTC
AGCACACGTCTGAACTCTTTCCCTACACGACGCTCTTCCGATCTNNNAGGTTCTTCCTACAGTACCAA
GAAGAGCACACGTCTGAACTCTTTCCCTACACGACGCTCTTCCGATCTNNNTTGCTTGAACATAGGA

AGAGCACACGTCTGAACTCTTTCCCTACACGACGCTCTTCCGATCTNNNTGTAGTCCCAGCTACTCAGTA
GAGCACACGTCTGAACTCTTTCCCTACACGACGCTCTTCCGATCTNNNATATCACAGCTACTTGGAGAG
AAGAGCACACGTCTGAACTCTTTCCCTACACGACGCTCTTCCGATCTNNNGCGCATATCTGTAGTTCCAG
AGAGCACACGTCTGAACTCTTTCCCTACACGACGCTCTTCCGATCTNNNCTGTAGTCCCAGCTACTCAGG
AAGAGCACACGTCTGAACTCTTTCCCTACACGACGCTCTTCCGATCTNNNAGGCAAGAGAATCCCTTGAA
GAGCACACGTCTGAACTCTTTCCCTACACGACGCTCTTCCGATCTNNNACTAATTTGGAGTTTGGGTTTG
GAAGAGCACACGTCTGAACTCTTTCCCTACACGACGCTCTTCCGATCTNNNGTAATCCCAGCTACTTAGG
AAGAGCACACGTCTGAACTCTTTCCCTACACGACGCTCTTCCGATCTNNNCACACACACAGACACATACA
AGCACACGTCTGAACTCTTTCCCTACACGACGCTCTTCCGATCTNNNACAAGGAGACAAACTGAATAAT
GAAGAGCACACGTCTGAACTCTTTCCCTACACGACGCTCTTCCGATCTNNNGAGAATGGTGTGAACCTG
AGAGCACACGTCTGAACTCTTTCCCTACACGACGCTCTTCCGATCTNNNGAATCGCTTGAACCTAGGAG
AGAGCACACGTCTGAACTCTTTCCCTACACGACGCTCTTCCGATCTNNNGTAATCCTAGCTACTTGGGAG
AGAGCACACGTCTGAACTCTTTCCCTACACGACGCTCTTCCGATCTNNNCTGTAATCCCAGCTACTCAGG
AGAGCACACGTCTGAACTCTTTCCCTACACGACGCTCTTCCGATCTNNNCTGTAGTCCTAGCTACTCGGG
GAGCACACGTCTGAACTCTTTCCCTACACGACGCTCTTCCGATCTNNNCTGTAATCTCAGCTACTCGGG
AGAGCACACGTCTGAACTCTTTCCCTACACGACGCTCTTCCGATCTNNNACAAAAATTAGCTGGGTGTG
AAGAGCACACGTCTGAACTCTTTCCCTACACGACGCTCTTCCGATCTNNNTGTAGTCCCAGCTACTCAG
AAGAGCACACGTCTGAACTCTTTCCCTACACGACGCTCTTCCGATCTNNNCTTGCAGTGAGCTGAGAT
AAGAGCACACGTCTGAACTCTTTCCCTACACGACGCTCTTCCGATCTNNNGTAATCTCAGCTACTCAGGA
AGCACACGTCTGAACTCTTTCCCTACACGACGCTCTTCCGATCTNNNTGATGACATCAAATTCATACACC
GAGCACACGTCTGAACTCTTTCCCTACACGACGCTCTTCCGATCTNNNTGTGTGTGTGTGTACATTTAT
GAGCACACGTCTGAACTCTTTCCCTACACGACGCTCTTCCGATCTNNNAGCAATATCCCTTTGAGAAAG
AGAGCACACGTCTGAACTCTTTCCCTACACGACGCTCTTCCGATCTNNNAGAGAGAGAGAGAGAGAG
AGAGCACACGTCTGAACTCTTTCCCTACACGACGCTCTTCCGATCTNNNGCTGGTGCTCATGTATCCAA
AGAGCACACGTCTGAACTCTTTCCCTACACGACGCTCTTCCGATCTNNNTTGACACTGATACAATGCACT
GAGCACACGTCTGAACTCTTTCCCTACACGACGCTCTTCCGATCTNNNACTATATACTGTCTTCAGGGGA
GAGCACACGTCTGAACTCTTTCCCTACACGACGCTCTTCCGATCTNNNTGGCACAATCAGTTTCTTCTTT
GCACACGTCTGAACTCTTTCCCTACACGACGCTCTTCCGATCTNNNTGAAAATTGAGTCTTTAGGGCTA
AAGAGCACACGTCTGAACTCTTTCCCTACACGACGCTCTTCCGATCTNNNCTGTAGTCCCAGCTACTCAG
AAGAGCACACGTCTGAACTCTTTCCCTACACGACGCTCTTCCGATCTNNNATCACTTGAACCCAGGAAG
AGAGCACACGTCTGAACTCTTTCCCTACACGACGCTCTTCCGATCTNNNTGTAGTCCTAGCTACTCCAGA
AGAGCACACGTCTGAACTCTTTCCCTACACGACGCTCTTCCGATCTNNNAGAATGATGTCTCCCAGCTTT
AGCACACGTCTGAACTCTTTCCCTACACGACGCTCTTCCGATCTNNNGGCTTCATCATTTTGTGATAGT
GAGCACACGTCTGAACTCTTTCCCTACACGACGCTCTTCCGATCTNNNCATGCACCTATAGTCCCAGCT
GAGCACACGTCTGAACTCTTTCCCTACACGACGCTCTTCCGATCTNNNAGGTGATAAATTTCTGTTGCT
AGCACACGTCTGAACTCTTTCCCTACACGACGCTCTTCCGATCTNNNAGCAATTTGAAAGGGTAAATGA
GAGCACACGTCTGAACTCTTTCCCTACACGACGCTCTTCCGATCTNNNGGACAAAACTCTAATTAGCTTC
AAGAGCACACGTCTGAACTCTTTCCCTACACGACGCTCTTCCGATCTNNNCTGAGATCATGCCACTGTAC
AAGAGCACACGTCTGAACTCTTTCCCTACACGACGCTCTTCCGATCTNNNCTGAGACAGGAGAGTTGC
AGCACACGTCTGAACTCTTTCCCTACACGACGCTCTTCCGATCTNNNTGCTTTTTTCTTCAGAGAATCGT
AGCACACGTCTGAACTCTTTCCCTACACGACGCTCTTCCGATCTNNNTGATTGACAGAATAGTAGTGG
AAGAGCACACGTCTGAACTCTTTCCCTACACGACGCTCTTCCGATCTNNNTTATTCTTCCTACCCTCCCTCC
AGAGCACACGTCTGAACTCTTTCCCTACACGACGCTCTTCCGATCTNNNGGCTGAGGCAGAAAAATCTC

GCACACGTCTGAACTCTTTCCCTACACGACGCTCTTCCGATCTNNNTCAAGGTCAAATTGTACTTAGCCT
AGCACACGTCTGAACTCTTTCCCTACACGACGCTCTTCCGATCTNNNACAAGTCATAAGGGTAACTTCA
AGAGCACACGTCTGAACTCTTTCCCTACACGACGCTCTTCCGATCTNNNATGAGGCAGGAGAATCGTT
AAGAGCACACGTCTGAACTCTTTCCCTACACGACGCTCTTCCGATCTNNNCACCTGTAATCCCAGCTACT
GAGCACACGTCTGAACTCTTTCCCTACACGACGCTCTTCCGATCTNNNTGACATTGTACTGAAGTTCTAC
AGAGCACACGTCTGAACTCTTTCCCTACACGACGCTCTTCCGATCTNNNCAGCTTCATCAGTATTGCTGC
AAGAGCACACGTCTGAACTCTTTCCCTACACGACGCTCTTCCGATCTNNNTTGACTATGAAGGGCCCCA
GAGCACACGTCTGAACTCTTTCCCTACACGACGCTCTTCCGATCTNNNACAGATTCCTGGAACTTTTGTT
AGAGCACACGTCTGAACTCTTTCCCTACACGACGCTCTTCCGATCTNNNTTCTGTTCTCAGTTCAACTCC
AAGAGCACACGTCTGAACTCTTTCCCTACACGACGCTCTTCCGATCTNNNAATAGAGCCAGACCATGTC
AGAGCACACGTCTGAACTCTTTCCCTACACGACGCTCTTCCGATCTNNNTCACAGATAGAAGCCAATTTC
AGAGCACACGTCTGAACTCTTTCCCTACACGACGCTCTTCCGATCTNNNTGTAATCACAGCTACTTGGG
GAGCACACGTCTGAACTCTTTCCCTACACGACGCTCTTCCGATCTNNNACTGGAAAAAAAAACAAAGAA
AGAGCACACGTCTGAACTCTTTCCCTACACGACGCTCTTCCGATCTNNNTCCCAAGAGGAATGCATTTA
AAGAGCACACGTCTGAACTCTTTCCCTACACGACGCTCTTCCGATCTNNNTTTGCCAGTATGTAGACTC
AAGAGCACACGTCTGAACTCTTTCCCTACACGACGCTCTTCCGATCTNNNCGAGGCAAGAGAATCACTT
GAGCACACGTCTGAACTCTTTCCCTACACGACGCTCTTCCGATCTNNNGTGAATCTTCTCTGACCCTCTC
AGCACACGTCTGAACTCTTTCCCTACACGACGCTCTTCCGATCTNNNTGACCTCTGGTGTTTTCATTATC
AGAGCACACGTCTGAACTCTTTCCCTACACGACGCTCTTCCGATCTNNNCAAAACAGCTGTTGACGATA
AAGAGCACACGTCTGAACTCTTTCCCTACACGACGCTCTTCCGATCTNNNCATCTATTTGCTGCCCCGAA
GAGCACACGTCTGAACTCTTTCCCTACACGACGCTCTTCCGATCTNNNTAGAAGTCACTGCGGAGATG
AGAGCACACGTCTGAACTCTTTCCCTACACGACGCTCTTCCGATCTNNNTGCCAAAATACCTTTCCCTTTC
AGCACACGTCTGAACTCTTTCCCTACACGACGCTCTTCCGATCTNNNAGTCCATTAGCACACATAACAG
GAGCACACGTCTGAACTCTTTCCCTACACGACGCTCTTCCGATCTNNNACAGCAGTGTACACATTAGTA
AGAGCACACGTCTGAACTCTTTCCCTACACGACGCTCTTCCGATCTNNNTCCATTTTAGGGATAGTCAA
AGCACACGTCTGAACTCTTTCCCTACACGACGCTCTTCCGATCTNNNTGCTCTTACAGTAGGGAAACAA
GCACACGTCTGAACTCTTTCCCTACACGACGCTCTTCCGATCTNNNGAACAAGATCTGGTCTCAAAAAA
GAGCACACGTCTGAACTCTTTCCCTACACGACGCTCTTCCGATCTNNNAGGGTTCTCTAGAGGGATAG
AAGAGCACACGTCTGAACTCTTTCCCTACACGACGCTCTTCCGATCTNNNTCCAATGTATTCTACTGTGC
AGAGCACACGTCTGAACTCTTTCCCTACACGACGCTCTTCCGATCTNNNTCTTAGCTCACTGAAACCTCT
AGCACACGTCTGAACTCTTTCCCTACACGACGCTCTTCCGATCTNNNACCTTTCTTCTCTCTCTCTCTCT
AAGAGCACACGTCTGAACTCTTTCCCTACACGACGCTCTTCCGATCTNNNATGATTTGTAACCCCTGAGC
AGCACACGTCTGAACTCTTTCCCTACACGACGCTCTTCCGATCTNNNTCTTACATGTAGAAAACCTGAA
AGAGCACACGTCTGAACTCTTTCCCTACACGACGCTCTTCCGATCTNNNTGTTATAGAATACCATATGCC
GAGCACACGTCTGAACTCTTTCCCTACACGACGCTCTTCCGATCTNNNACCCTGATATCAAAACTAGACC
AGCACACGTCTGAACTCTTTCCCTACACGACGCTCTTCCGATCTNNNTGTTCTTTTTGGTTTTCATTGC
AGCACACGTCTGAACTCTTTCCCTACACGACGCTCTTCCGATCTNNNCACTCGCAACTACTTCTCTTCCTC
GAGCACACGTCTGAACTCTTTCCCTACACGACGCTCTTCCGATCTNNNTCTCTCTCTCTCTCTCTCTCTC
GAGCACACGTCTGAACTCTTTCCCTACACGACGCTCTTCCGATCTNNNGTGAGAGGGTTCACAATATCT
AGCACACGTCTGAACTCTTTCCCTACACGACGCTCTTCCGATCTNNNATGCCTGTAATCCCAGCTACTTG
GAAGAGCACACGTCTGAACTCTTTCCCTACACGACGCTCTTCCGATCTNNNAATGAGCCGAGATCATGC
AGCACACGTCTGAACTCTTTCCCTACACGACGCTCTTCCGATCTNNNACATATCCTATAAACCCAATGTCC
AGAGCACACGTCTGAACTCTTTCCCTACACGACGCTCTTCCGATCTNNNCCTGGTTATACCCCTCCTCAA

GAGCACACGTCTGAACTCTTTCCCTACACGACGCTCTTCCGATCTNNNTCCTATAGTTGAAGCCACTGAA
AAGAGCACACGTCTGAACTCTTTCCCTACACGACGCTCTTCCGATCTNNNCTCCAGTGTCTGTATGCA
AAGAGCACACGTCTGAACTCTTTCCCTACACGACGCTCTTCCGATCTNNNTATTTCCACCTTACCTCCAGC
GAGCACACGTCTGAACTCTTTCCCTACACGACGCTCTTCCGATCTNNNTCCTCTATTCATGGCTTGATAC
AAGAGCACACGTCTGAACTCTTTCCCTACACGACGCTCTTCCGATCTNNNTGACTAGTGTTCCCCCTATC
AAGAGCACACGTCTGAACTCTTTCCCTACACGACGCTCTTCCGATCTNNNTCAAGGCTTCAGTGAACTGT
AGAGCACACGTCTGAACTCTTTCCCTACACGACGCTCTTCCGATCTNNNTCTGGATCTGTGGTCTTTGG
AGCACACGTCTGAACTCTTTCCCTACACGACGCTCTTCCGATCTNNNTGCCATTTCATCCTTTTGCTATTC
AAGAGCACACGTCTGAACTCTTTCCCTACACGACGCTCTTCCGATCTNNNGATTACTGCTTGAGTCTGGC
GAGCACACGTCTGAACTCTTTCCCTACACGACGCTCTTCCGATCTNNNTGTTATTCTTGATAGGCCAAA
GAGCACACGTCTGAACTCTTTCCCTACACGACGCTCTTCCGATCTNNNTGATGGACACTTAGGTTGATT
GAGCACACGTCTGAACTCTTTCCCTACACGACGCTCTTCCGATCTNNNTGAGAAGAGGTAAGGTCTAAA
AGCACACGTCTGAACTCTTTCCCTACACGACGCTCTTCCGATCTNNNTCACTAAAAGGAAGACAGAAAC
AGCACACGTCTGAACTCTTTCCCTACACGACGCTCTTCCGATCTNNNTCTTGTATTTTCTTCTCTCTCTCC
AGCACACGTCTGAACTCTTTCCCTACACGACGCTCTTCCGATCTNNNACAACAGAGAAAGACAGAGAG
AAGAGCACACGTCTGAACTCTTTCCCTACACGACGCTCTTCCGATCTNNNTCTGGTCTCATGTTACATCCA
AGCACACGTCTGAACTCTTTCCCTACACGACGCTCTTCCGATCTNNNACTGACCATCATATCCCTATTGTC
GAGCACACGTCTGAACTCTTTCCCTACACGACGCTCTTCCGATCTNNNGGTTGAGAGACAAATCAAGA
AAGAGCACACGTCTGAACTCTTTCCCTACACGACGCTCTTCCGATCTNNNCCCAGCCCATGATTCCTTTTT
AGCACACGTCTGAACTCTTTCCCTACACGACGCTCTTCCGATCTNNNTGTCCAGGTGAAAATATAGCTAC
GGAAGAGCACACGTCTGAACTCTTTCCCTACACGACGCTCTTCCGATCTNNNAAAAGTTGCAGTGAGC
GAGCACACGTCTGAACTCTTTCCCTACACGACGCTCTTCCGATCTNNNTCTGACAAGGGTCTAGTATCC
AGCACACGTCTGAACTCTTTCCCTACACGACGCTCTTCCGATCTNNNTCTGTAGGCTTGAGTTTGAGAC
GAGCACACGTCTGAACTCTTTCCCTACACGACGCTCTTCCGATCTNNNGCATCTGGCTTATTTCACTTAC
GAGCACACGTCTGAACTCTTTCCCTACACGACGCTCTTCCGATCTNNNTCTTTCTTCCTTCCAGTATGTTT
GAGCACACGTCTGAACTCTTTCCCTACACGACGCTCTTCCGATCTNNNACTCTGGAAGTTGTAAAAGCC
AGAGCACACGTCTGAACTCTTTCCCTACACGACGCTCTTCCGATCTNNNTGCAACTGATTTTTGTGGGTT
GAGCACACGTCTGAACTCTTTCCCTACACGACGCTCTTCCGATCTNNNAGGATTCATGTTCTTCTCATTA
AGCACACGTCTGAACTCTTTCCCTACACGACGCTCTTCCGATCTNNNTGCATTTGTGTGTGTATGTATG
AGAGCACACGTCTGAACTCTTTCCCTACACGACGCTCTTCCGATCTNNNTCTAAGAACTCCAAGGACACC
AAGAGCACACGTCTGAACTCTTTCCCTACACGACGCTCTTCCGATCTNNNCTGAGGCAGGAGAATCAC
AGAGCACACGTCTGAACTCTTTCCCTACACGACGCTCTTCCGATCTNNNGGGCAACAAGAGAGAAACT
AGAGCACACGTCTGAACTCTTTCCCTACACGACGCTCTTCCGATCTNNNTGCCAAATACGTGATCTATGA
AGAGCACACGTCTGAACTCTTTCCCTACACGACGCTCTTCCGATCTNNNCAGAGAAAGACCCTGTCTCA
AGAGCACACGTCTGAACTCTTTCCCTACACGACGCTCTTCCGATCTNNNCCTCTCTTCCTATTTGTGTTTG
GGAAGAGCACACGTCTGAACTCTTTCCCTACACGACGCTCTTCCGATCTNNNCAGTTCTAGAGGCAAGA
GAGCACACGTCTGAACTCTTTCCCTACACGACGCTCTTCCGATCTNNNAGAATGAATAAGACCTAGCAG
GAGCACACGTCTGAACTCTTTCCCTACACGACGCTCTTCCGATCTNNNAGATTCCATCTCACAAACAAAA
GGAAGAGCACACGTCTGAACTCTTTCCCTACACGACGCTCTTCCGATCTNNNCTGAGGCACAAGAATCC
GAAGAGCACACGTCTGAACTCTTTCCCTACACGACGCTCTTCCGATCTNNNGTGAGCCAAGATCGCATC
AGAGCACACGTCTGAACTCTTTCCCTACACGACGCTCTTCCGATCTNNNACAGATTTCAATTTGCTTCAT
AGAGCACACGTCTGAACTCTTTCCCTACACGACGCTCTTCCGATCTNNNAAATGCCAAGGGTTTTTGAA
GAGCACACGTCTGAACTCTTTCCCTACACGACGCTCTTCCGATCTNNNAAATAAGGTCAGCTAGGCTT

AAGAGCACACGTCTGAACTCTTTCCCTACACGACGCTCTTCCGATCTNNNAGACTCCGTCTCAAAAAAA
CGGAAGAGCACACGTCTGAACTCTTTCCCTACACGACGCTCTTCCGATCTNNNGATCGTGTCACTCTACT
GAGCACACGTCTGAACTCTTTCCCTACACGACGCTCTTCCGATCTNNNCTGTAGTGCAGTGGTGTGAT
AGCACACGTCTGAACTCTTTCCCTACACGACGCTCTTCCGATCTNNNAGCTCCATTTTCTCTCTTTCCTTCA
AGAGCACACGTCTGAACTCTTTCCCTACACGACGCTCTTCCGATCTNNNTGGACATTTATAGGTTGAGC
AGAGCACACGTCTGAACTCTTTCCCTACACGACGCTCTTCCGATCTNNNCCATTGTGGCCAACTGTTTTT
AGCACACGTCTGAACTCTTTCCCTACACGACGCTCTTCCGATCTNNNACACACACATATATACACACACA
AAGAGCACACGTCTGAACTCTTTCCCTACACGACGCTCTTCCGATCTNNNAGAAGAAGAAGAAAGAGA
AAGAGCACACGTCTGAACTCTTTCCCTACACGACGCTCTTCCGATCTNNNAGTCTCTGCATATACAGAC
GAGCACACGTCTGAACTCTTTCCCTACACGACGCTCTTCCGATCTNNNTGCTCCAAAACCTTTTCATTTC
AGAGCACACGTCTGAACTCTTTCCCTACACGACGCTCTTCCGATCTNNNTCTCTCTGTCTGTCTCTCTCTC
AAGAGCACACGTCTGAACTCTTTCCCTACACGACGCTCTTCCGATCTNNNTATGTGTACGTGTGTGTGA
AGAGCACACGTCTGAACTCTTTCCCTACACGACGCTCTTCCGATCTNNNCTATCCACCTTCCAACCACCTA
AGCACACGTCTGAACTCTTTCCCTACACGACGCTCTTCCGATCTNNNGACTCATGGCAAAGGAACTGAA
AAGAGCACACGTCTGAACTCTTTCCCTACACGACGCTCTTCCGATCTNNNTGTCATCTGTGTAGACCAT
AGAGCACACGTCTGAACTCTTTCCCTACACGACGCTCTTCCGATCTNNNCTTCTTCCTTCCTCCCTTCCTT
AGAGCACACGTCTGAACTCTTTCCCTACACGACGCTCTTCCGATCTNNNTCTGGTGAATCCTTTCTTCCT
AGAGCACACGTCTGAACTCTTTCCCTACACGACGCTCTTCCGATCTNNNTCACACCTGTAATCCAAGCAC
AGCACACGTCTGAACTCTTTCCCTACACGACGCTCTTCCGATCTNNNTCGATCTAATTTTTGTGTGCCT
AAGAGCACACGTCTGAACTCTTTCCCTACACGACGCTCTTCCGATCTNNNTCCCACAGAAAGAAAGAAA
AGAGCACACGTCTGAACTCTTTCCCTACACGACGCTCTTCCGATCTNNNAATCATGCAAATCCCTGAGTC
AGAGCACACGTCTGAACTCTTTCCCTACACGACGCTCTTCCGATCTNNNCGGCTGGTTGTGTGTGTGT
AGAGCACACGTCTGAACTCTTTCCCTACACGACGCTCTTCCGATCTNNNAGGTGAAACTACATACCTGT
GAGCACACGTCTGAACTCTTTCCCTACACGACGCTCTTCCGATCTNNNACATTTGCATATGTATCGAGAC
GAGCACACGTCTGAACTCTTTCCCTACACGACGCTCTTCCGATCTNNNTCTCTCTCTCTCTCTCTCTCTC
GAAGAGCACACGTCTGAACTCTTTCCCTACACGACGCTCTTCCGATCTNNNCCACCTTAGGTAGCAGAT
GAGCACACGTCTGAACTCTTTCCCTACACGACGCTCTTCCGATCTNNNTGATAGGAAATGTTGCCAATTT
AGAGCACACGTCTGAACTCTTTCCCTACACGACGCTCTTCCGATCTNNNGAATTTCTCTGGCCTGGAAA
GAGCACACGTCTGAACTCTTTCCCTACACGACGCTCTTCCGATCTNNNTGTTGACTGTTCATTTGCATAT
AGCACACGTCTGAACTCTTTCCCTACACGACGCTCTTCCGATCTNNNAGTAAAATGAAAAGAGGCAAG
CGGAAGAGCACACGTCTGAACTCTTTCCCTACACGACGCTCTTCCGATCTNNNGAATCGCTTGAACTTC
GAGCACACGTCTGAACTCTTTCCCTACACGACGCTCTTCCGATCTNNNACACATGCACATACATATGTA
AGCACACGTCTGAACTCTTTCCCTACACGACGCTCTTCCGATCTNNNACACTAATTCATCTTCCTCAGCA
GCACACGTCTGAACTCTTTCCCTACACGACGCTCTTCCGATCTNNNTCTCTCTCTTTCTTTCTTCTTTCT
AGCACACGTCTGAACTCTTTCCCTACACGACGCTCTTCCGATCTNNNTCTGCCTTGATTGGTCATGATG
GAGCACACGTCTGAACTCTTTCCCTACACGACGCTCTTCCGATCTNNNACCCTTTGGAAGTTTCTTGAA
GAGCACACGTCTGAACTCTTTCCCTACACGACGCTCTTCCGATCTNNNTCCTCCTCCTTCTTCTTCTCCTTC
AGAGCACACGTCTGAACTCTTTCCCTACACGACGCTCTTCCGATCTNNNGCACACATACCGAGTCATATT
GAGCACACGTCTGAACTCTTTCCCTACACGACGCTCTTCCGATCTNNNCTGAGGCAAAAGAACGACTTG
GAGCACACGTCTGAACTCTTTCCCTACACGACGCTCTTCCGATCTNNNTGCCTGTAATCCAAGCTACTCA
AGAGCACACGTCTGAACTCTTTCCCTACACGACGCTCTTCCGATCTNNNCCAGCTACTCTGAAGGCTAA
GAGCACACGTCTGAACTCTTTCCCTACACGACGCTCTTCCGATCTNNNCTGTAATCCCAGCTACTCAGGA
AGAGCACACGTCTGAACTCTTTCCCTACACGACGCTCTTCCGATCTNNNAAATGCCCATAATCCCAGCTA

GAGCACACGTCTGAACTCTTTCCCTACACGACGCTCTTCCGATCTNNNAGGTTGAGGCAGAAGAATC
GCACACGTCTGAACTCTTTCCCTACACGACGCTCTTCCGATCTNNNAGTGACTATAAACAATGTTGCA
AGCACACGTCTGAACTCTTTCCCTACACGACGCTCTTCCGATCTNNNTGGGGAGAAAGCTTTTCTTTT
AGCACACGTCTGAACTCTTTCCCTACACGACGCTCTTCCGATCTNNNACTCTAAAGATCCTGGACAGA
GAGCACACGTCTGAACTCTTTCCCTACACGACGCTCTTCCGATCTNNNGAGAATCACTTGAACCCAGA
GAGCACACGTCTGAACTCTTTCCCTACACGACGCTCTTCCGATCTNNNTGCCCATTTTCCATACTTAGTT
AAGAGCACACGTCTGAACTCTTTCCCTACACGACGCTCTTCCGATCTNNNCTGTAGTCCTAGCTACTCG
GCACACGTCTGAACTCTTTCCCTACACGACGCTCTTCCGATCTNNNACAAAATCCTCTTGGCTACATGA
GCACACGTCTGAACTCTTTCCCTACACGACGCTCTTCCGATCTNNNGGGTTCATAAAGAAACACTGG
GAGCACACGTCTGAACTCTTTCCCTACACGACGCTCTTCCGATCTNNNAGAATCACTTGAAACTGGGA
AGCACACGTCTGAACTCTTTCCCTACACGACGCTCTTCCGATCTNNNCTGTAGTCCCAGCTACTCGAG
GCACACGTCTGAACTCTTTCCCTACACGACGCTCTTCCGATCTNNNACAAATCCAGGAGTTGGTTTTT
AGCACACGTCTGAACTCTTTCCCTACACGACGCTCTTCCGATCTNNNTGGTCTATTCAGTTTCCTTGCC
GAGCACACGTCTGAACTCTTTCCCTACACGACGCTCTTCCGATCTNNNTAAGGCAGGAGAATCACTG
AGAGCACACGTCTGAACTCTTTCCCTACACGACGCTCTTCCGATCTNNNTCAAGCAACCTACAGGCATA
GCACACGTCTGAACTCTTTCCCTACACGACGCTCTTCCGATCTNNNTGTGCCATAGACAAAATCTGTTG
AGCACACGTCTGAACTCTTTCCCTACACGACGCTCTTCCGATCTNNNTGTATCAATAACCGACTTTGCC
AGCACACGTCTGAACTCTTTCCCTACACGACGCTCTTCCGATCTNNNCACGATTGTGTCACTGTACTCC
AGAGCACACGTCTGAACTCTTTCCCTACACGACGCTCTTCCGATCTNNNCATGCCTGTAATCCCAGCT
GAGCACACGTCTGAACTCTTTCCCTACACGACGCTCTTCCGATCTNNNCACCTGTAGTCCCAGCTACTT
AGAGCACACGTCTGAACTCTTTCCCTACACGACGCTCTTCCGATCTNNNCTCAGGAGGTTGAAGCAGA
AGCACACGTCTGAACTCTTTCCCTACACGACGCTCTTCCGATCTNNNCCTTCCAGTAGCCAGAGCAAT
GAGCACACGTCTGAACTCTTTCCCTACACGACGCTCTTCCGATCTNNNGAGAATGGTATGAACCTGGG
AGCACACGTCTGAACTCTTTCCCTACACGACGCTCTTCCGATCTNNNGAATTGCTTGAAACAGGAGG
AGAGCACACGTCTGAACTCTTTCCCTACACGACGCTCTTCCGATCTNNNGTTGCAGTGAGCTGAGATT
GAGCACACGTCTGAACTCTTTCCCTACACGACGCTCTTCCGATCTNNNCGTCAAGAGCCAAATTCTGT
GAGCACACGTCTGAACTCTTTCCCTACACGACGCTCTTCCGATCTNNNAGAATCACTTGAAGCCAGGA
AGCACACGTCTGAACTCTTTCCCTACACGACGCTCTTCCGATCTNNNCGAAAAATTACGTTTATCGCA
GCACACGTCTGAACTCTTTCCCTACACGACGCTCTTCCGATCTNNNTCTCACAATTCTCAACTTTGCTC
AGCACACGTCTGAACTCTTTCCCTACACGACGCTCTTCCGATCTNNNTGTACTACTTTTCAACAGTGTG
AGCACACGTCTGAACTCTTTCCCTACACGACGCTCTTCCGATCTNNNTGGCAATCACTTTATTCTCAGT
GAGCACACGTCTGAACTCTTTCCCTACACGACGCTCTTCCGATCTNNNGAGGCAGGAGAATGGAGT
ACACGTCTGAACTCTTTCCCTACACGACGCTCTTCCGATCTNNNTTTTCTTGCACCCGTTATAAAATTTT
GCACACGTCTGAACTCTTTCCCTACACGACGCTCTTCCGATCTNNNTCCTTCTACTTTTCTGCTTCTTCA
AGCACACGTCTGAACTCTTTCCCTACACGACGCTCTTCCGATCTNNNGAGAATGGCATGAACCCAAG
GAGCACACGTCTGAACTCTTTCCCTACACGACGCTCTTCCGATCTNNNCTGTAATCCCAGCTAGTCAG
GAGCACACGTCTGAACTCTTTCCCTACACGACGCTCTTCCGATCTNNNCTAACCTGGGTGACAAAGT
GAAGAGCACACGTCTGAACTCTTTCCCTACACGACGCTCTTCCGATCTNNNCTTGCAGTGAGCCGAGA
GAGCACACGTCTGAACTCTTTCCCTACACGACGCTCTTCCGATCTNNNTAATCTCAGCTACTAGGGAG
GAGCACACGTCTGAACTCTTTCCCTACACGACGCTCTTCCGATCTNNNATAATAGCTTGAACCCAGGA
GAGCACACGTCTGAACTCTTTCCCTACACGACGCTCTTCCGATCTNNNGCCTGTAGTCCCAGCTACTCT
GAAGAGCACACGTCTGAACTCTTTCCCTACACGACGCTCTTCCGATCTNNNGAGGCTGAGGCAGGAG
GAGCACACGTCTGAACTCTTTCCCTACACGACGCTCTTCCGATCTNNNCTGAGACAGGAGAATCGCT

GAGCACACGTCTGAACTCTTTCCCTACACGACGCTCTTCCGATCTNNNAGTTTAGTTTGAAAACAGGG
AGCACACGTCTGAACTCTTTCCCTACACGACGCTCTTCCGATCTNNNGTCTTTGACCTTTTGGCTATAG
AAGAGCACACGTCTGAACTCTTTCCCTACACGACGCTCTTCCGATCTNNNGCCTGTAGTCCCAGCTAC
AGCACACGTCTGAACTCTTTCCCTACACGACGCTCTTCCGATCTNNNCTTGCAGTAAGCTGAGATCATG
AGCACACGTCTGAACTCTTTCCCTACACGACGCTCTTCCGATCTNNNACGGGTGTTGTATTTCATCAA
AAGAGCACACGTCTGAACTCTTTCCCTACACGACGCTCTTCCGATCTNNNGAGGCAGGAGAATGGAG
AGCACACGTCTGAACTCTTTCCCTACACGACGCTCTTCCGATCTNNNGAGTCAAGATCACACCATTGC
AGCACACGTCTGAACTCTTTCCCTACACGACGCTCTTCCGATCTNNNACCTCTAGAACGATAAGGTTG
AGCACACGTCTGAACTCTTTCCCTACACGACGCTCTTCCGATCTNNNAGACATTTTACAACGAGAAAG
GAGCACACGTCTGAACTCTTTCCCTACACGACGCTCTTCCGATCTNNNTGATTGTACCATCTCACTCCA
GAGCACACGTCTGAACTCTTTCCCTACACGACGCTCTTCCGATCTNNNGGTTTCAATCTCTTGTTCCAT
AAGAGCACACGTCTGAACTCTTTCCCTACACGACGCTCTTCCGATCTNNNGAGGCAGGAGAATGGTG
AGCACACGTCTGAACTCTTTCCCTACACGACGCTCTTCCGATCTNNNTGTTGATTTTCAGTTTCTGCA
AGCACACGTCTGAACTCTTTCCCTACACGACGCTCTTCCGATCTNNNTGGGAAAGTGTTTTAGCATCT
GCACACGTCTGAACTCTTTCCCTACACGACGCTCTTCCGATCTNNNGGAACTCTCTATTCCTTCCTTAG
GAGCACACGTCTGAACTCTTTCCCTACACGACGCTCTTCCGATCTNNNAGACAGGAGAATCACTTGAA
GAGCACACGTCTGAACTCTTTCCCTACACGACGCTCTTCCGATCTNNNCTTGCAGTGAGCCGAGATT
AGCACACGTCTGAACTCTTTCCCTACACGACGCTCTTCCGATCTNNNCTGAGGTGGGAGAATTGTT
CACGTCTGAACTCTTTCCCTACACGACGCTCTTCCGATCTNNNTGAGATTTCCATTTTTACTCTGAGA
GCACACGTCTGAACTCTTTCCCTACACGACGCTCTTCCGATCTNNNTCAGCAATATCCACATACCATCC
AGCACACGTCTGAACTCTTTCCCTACACGACGCTCTTCCGATCTNNNTCGTAAGGAATCACAAAAGAC
CACACGTCTGAACTCTTTCCCTACACGACGCTCTTCCGATCTNNNTGTCTACTTGTAATTCTTCATGCTG
AGCACACGTCTGAACTCTTTCCCTACACGACGCTCTTCCGATCTNNNGGCAACAGAGTGAGACTTCA
GAGCACACGTCTGAACTCTTTCCCTACACGACGCTCTTCCGATCTNNNCTGTAGTCCCAGCTACTCAG
CACGTCTGAACTCTTTCCCTACACGACGCTCTTCCGATCTNNNTTTCCTTTGAAATAACACTTGGAATT
AGAGCACACGTCTGAACTCTTTCCCTACACGACGCTCTTCCGATCTNNNGAGCTTGCAGTGAGCCAAG
GAGCACACGTCTGAACTCTTTCCCTACACGACGCTCTTCCGATCTNNNAAAAAATTAGCCGGGCATAG
GAGCACACGTCTGAACTCTTTCCCTACACGACGCTCTTCCGATCTNNNTCTGTAATCCCAGCTACTCAG
AGCACACGTCTGAACTCTTTCCCTACACGACGCTCTTCCGATCTNNNCAGGAGTCTGAAGCAGGAG
AGCACACGTCTGAACTCTTTCCCTACACGACGCTCTTCCGATCTNNNTCCAGCTTGTAATCCAGCTAC
AGCACACGTCTGAACTCTTTCCCTACACGACGCTCTTCCGATCTNNNCTGTAGTCCTAGCTGCTCAGGA
GAAGAGCACACGTCTGAACTCTTTCCCTACACGACGCTCTTCCGATCTNNNCAAGATCGCGCCATTGT
GAGCACACGTCTGAACTCTTTCCCTACACGACGCTCTTCCGATCTNNNGAGAATTGCTTGAACCCAGT
GAGCACACGTCTGAACTCTTTCCCTACACGACGCTCTTCCGATCTNNNCTGAGGCAGGAGAATAGCTT
AGCACACGTCTGAACTCTTTCCCTACACGACGCTCTTCCGATCTNNNACAGGTTTCTTCCCCTTTCCAA
AGCACACGTCTGAACTCTTTCCCTACACGACGCTCTTCCGATCTNNNACCCCAACAGTCTTTCTTCTATT
GAGCACACGTCTGAACTCTTTCCCTACACGACGCTCTTCCGATCTNNNGATCTTACCTACTTGGGAGG
AGCACACGTCTGAACTCTTTCCCTACACGACGCTCTTCCGATCTNNNTGACTTTCAGAACACTTTGTTG
AGAGCACACGTCTGAACTCTTTCCCTACACGACGCTCTTCCGATCTNNNCTGAGGCAGGAGAATCAC
GCACACGTCTGAACTCTTTCCCTACACGACGCTCTTCCGATCTNNNTGTTTCCAGGATCTATGCTTTAC
AGCACACGTCTGAACTCTTTCCCTACACGACGCTCTTCCGATCTNNNTGAAGTAGTTCTCTTCTGACTG
GAGCACACGTCTGAACTCTTTCCCTACACGACGCTCTTCCGATCTNNNCTGTAATCCCAGCTACTCAGG
AGAGCACACGTCTGAACTCTTTCCCTACACGACGCTCTTCCGATCTNNNAGAGAGGTATGATTGTGCC

GCACACGTCTGAACTCTTTCCCTACACGACGCTCTTCCGATCTNNNGCTAAAGCAATCTTCCCACCTTAC
GAGCACACGTCTGAACTCTTTCCCTACACGACGCTCTTCCGATCTNNNGAGAATGGTGTGAACCTGG
GAGCACACGTCTGAACTCTTTCCCTACACGACGCTCTTCCGATCTNNNGAGTTTGCAGTGAGTGGAG
GAGCACACGTCTGAACTCTTTCCCTACACGACGCTCTTCCGATCTNNNAGGAGAATTGCTTGAACTCA
AGAGCACACGTCTGAACTCTTTCCCTACACGACGCTCTTCCGATCTNNNGAGATGGCACTACTACACT
AGAGCACACGTCTGAACTCTTTCCCTACACGACGCTCTTCCGATCTNNNAGGTTGGTGCAAAAGTAAT
CACGTCTGAACTCTTTCCCTACACGACGCTCTTCCGATCTNNNTCACTGTCCATAAAATATTTCACTGC
GAGCACACGTCTGAACTCTTTCCCTACACGACGCTCTTCCGATCTNNNGCCTATAATCCCAGCTACTCA
AGAGCACACGTCTGAACTCTTTCCCTACACGACGCTCTTCCGATCTNNNATGATGATTCCAGGTTTCA
GAGCACACGTCTGAACTCTTTCCCTACACGACGCTCTTCCGATCTNNNGAGAAATGGCGTAAACCCA
GCACACGTCTGAACTCTTTCCCTACACGACGCTCTTCCGATCTNNNCAAAAAACATTGCCACCAACTTC
AGCACACGTCTGAACTCTTTCCCTACACGACGCTCTTCCGATCTNNNGCTATTCAGGAGGCTGAAGT
GAGCACACGTCTGAACTCTTTCCCTACACGACGCTCTTCCGATCTNNNAGCTATGATTGCACTACAGC
GAGCACACGTCTGAACTCTTTCCCTACACGACGCTCTTCCGATCTNNNAGCTATTCAGGAGATTGAGG
AGCACACGTCTGAACTCTTTCCCTACACGACGCTCTTCCGATCTNNNGGGTTGATTCCAATTTTTTCAG
AGCACACGTCTGAACTCTTTCCCTACACGACGCTCTTCCGATCTNNNTGGATGAGATCTGGGAGAATT
AGCACACGTCTGAACTCTTTCCCTACACGACGCTCTTCCGATCTNNNAGTGACTGAATCTGAAGATTTG
GAGCACACGTCTGAACTCTTTCCCTACACGACGCTCTTCCGATCTNNNGAGACAGAATTGCTTGAACC
AGCACACGTCTGAACTCTTTCCCTACACGACGCTCTTCCGATCTNNNTGTCCTTCAACACTCCATTAGC
AGCACACGTCTGAACTCTTTCCCTACACGACGCTCTTCCGATCTNNNCTGAAGCAGGAGAATGGTGT
AGAGCACACGTCTGAACTCTTTCCCTACACGACGCTCTTCCGATCTNNNCTTGCATTGAGCTGAGATC
AGAGCACACGTCTGAACTCTTTCCCTACACGACGCTCTTCCGATCTNNNCACCTGTAATCCCAGCTACT
ACACGTCTGAACTCTTTCCCTACACGACGCTCTTCCGATCTNNNTGCTGTTTTCATCTTTTCTTCTAAGA
GAGCACACGTCTGAACTCTTTCCCTACACGACGCTCTTCCGATCTNNNCTACTTTGGGCAGGAGAATC
AGAGCACACGTCTGAACTCTTTCCCTACACGACGCTCTTCCGATCTNNNGCACCTGTAATCCCAGCTAT
GAGCACACGTCTGAACTCTTTCCCTACACGACGCTCTTCCGATCTNNNGTTGCAGTGAGCCAAGATC
GAGCACACGTCTGAACTCTTTCCCTACACGACGCTCTTCCGATCTNNNAGGTTTTATGTCCACTCTTTG
AGCACACGTCTGAACTCTTTCCCTACACGACGCTCTTCCGATCTNNNTGCCACTGAATATGGAGAAGT
AGCACACGTCTGAACTCTTTCCCTACACGACGCTCTTCCGATCTNNNGGAATTGTGACTCTCATATGGG
GAGCACACGTCTGAACTCTTTCCCTACACGACGCTCTTCCGATCTNNNCAGAAAAACTGGGAAAACAC
AGCACACGTCTGAACTCTTTCCCTACACGACGCTCTTCCGATCTNNNACTTTCTGTCTCCATGAGTTTG
AAGAGCACACGTCTGAACTCTTTCCCTACACGACGCTCTTCCGATCTNNNGAAAGTGGGTGACAAGA
AGAGCACACGTCTGAACTCTTTCCCTACACGACGCTCTTCCGATCTNNNGAGATCGTGCCACTATACTC
GAGCACACGTCTGAACTCTTTCCCTACACGACGCTCTTCCGATCTNNNGTAATCCCAGCTACTTAGGA
AGAGCACACGTCTGAACTCTTTCCCTACACGACGCTCTTCCGATCTNNNCTGAGGCAGGAGAATAGCA
GAGCACACGTCTGAACTCTTTCCCTACACGACGCTCTTCCGATCTNNNCTGAGGCAGGAGAATTGCTT
AGAGCACACGTCTGAACTCTTTCCCTACACGACGCTCTTCCGATCTNNNATAGTGGCGCATGCTTGTA
AGAGCACACGTCTGAACTCTTTCCCTACACGACGCTCTTCCGATCTNNNGAGGCAGGAGAATTGCTTG
GAGCACACGTCTGAACTCTTTCCCTACACGACGCTCTTCCGATCTNNNTCTCCACTGAGTTTTATCTCCC
AGCACACGTCTGAACTCTTTCCCTACACGACGCTCTTCCGATCTNNNGGAATAGACCATCGTTACTTT
GAGCACACGTCTGAACTCTTTCCCTACACGACGCTCTTCCGATCTNNNAGATCCAAAAAGTCTTTAGGT
AGCACACGTCTGAACTCTTTCCCTACACGACGCTCTTCCGATCTNNNAGGAATTATTCGGTGCTAACT
AAGAGCACACGTCTGAACTCTTTCCCTACACGACGCTCTTCCGATCTNNNCTGAGGCAGGAGAATAG

GAGCACACGTCTGAACTCTTTCCCTACACGACGCTCTTCCGATCTNNNTGTAGTCCCAGCTAGTCAGG
AAGAGCACACGTCTGAACTCTTTCCCTACACGACGCTCTTCCGATCTNNNGAGGTTGCAGTGAGCTG
AGAGCACACGTCTGAACTCTTTCCCTACACGACGCTCTTCCGATCTNNNAAGGAGGCAAAGGTTGTA
AGCACACGTCTGAACTCTTTCCCTACACGACGCTCTTCCGATCTNNNGAGAATGGCATGAATCCGGG
GCACACGTCTGAACTCTTTCCCTACACGACGCTCTTCCGATCTNNNGCTTATTGGTCATCTTGTTTGG
AGCACACGTCTGAACTCTTTCCCTACACGACGCTCTTCCGATCTNNNAGCATCAGTTTTACTCTTTTGC
GAGCACACGTCTGAACTCTTTCCCTACACGACGCTCTTCCGATCTNNNTGGGAGACTGAGGTAGGA
GAGCACACGTCTGAACTCTTTCCCTACACGACGCTCTTCCGATCTNNNGTAGAGGCAGAGGTTACTGT
AGCACACGTCTGAACTCTTTCCCTACACGACGCTCTTCCGATCTNNNACTGGGGAAAATATTTCCAAT
GAGCACACGTCTGAACTCTTTCCCTACACGACGCTCTTCCGATCTNNNGAGGCTAAGGCAAGAGAAT
AAGAGCACACGTCTGAACTCTTTCCCTACACGACGCTCTTCCGATCTNNNTGAGCTATGATTGCACCA
GCACACGTCTGAACTCTTTCCCTACACGACGCTCTTCCGATCTNNNGTGTGGTCAATTTTCAACCAAA
GAGCACACGTCTGAACTCTTTCCCTACACGACGCTCTTCCGATCTNNNTCCTTTTAGCCTAACACCCTTG
AGCACACGTCTGAACTCTTTCCCTACACGACGCTCTTCCGATCTNNNGGAAATAGCAAATGTCTCCAA
AGAGCACACGTCTGAACTCTTTCCCTACACGACGCTCTTCCGATCTNNNCCTGCAGTGAGCTGAGATC
AGCACACGTCTGAACTCTTTCCCTACACGACGCTCTTCCGATCTNNNACAAAAAACGCAAAAAAAACCC
AGCACACGTCTGAACTCTTTCCCTACACGACGCTCTTCCGATCTNNNGGTAGACTAAGTTGACCTGAA
AGCACACGTCTGAACTCTTTCCCTACACGACGCTCTTCCGATCTNNNTGTCTGCTAAGTAAGAGGAAA
ACACGTCTGAACTCTTTCCCTACACGACGCTCTTCCGATCTNNNTTCTCTTTCTTTCTTTCTGTCTTTCTT
GAGCACACGTCTGAACTCTTTCCCTACACGACGCTCTTCCGATCTNNNACTGGTACATTCATAGGGGA
AGCACACGTCTGAACTCTTTCCCTACACGACGCTCTTCCGATCTNNNACACAGATTGAAATAAATCGG
GAGCACACGTCTGAACTCTTTCCCTACACGACGCTCTTCCGATCTNNNTCATTTGTGTGGAGAATCTGT
AGCACACGTCTGAACTCTTTCCCTACACGACGCTCTTCCGATCTNNNGAGGTTGCAGTGAGCTGAAAT
GAGCACACGTCTGAACTCTTTCCCTACACGACGCTCTTCCGATCTNNNGCCACTGGTAACCCATGGTA
AGAGCACACGTCTGAACTCTTTCCCTACACGACGCTCTTCCGATCTNNNGCACCTATAGTCCCAGCTAC
GAGCACACGTCTGAACTCTTTCCCTACACGACGCTCTTCCGATCTNNNAGGATATCATACTCCTTGCCA
AGCACACGTCTGAACTCTTTCCCTACACGACGCTCTTCCGATCTNNNCCCAAGACTGAACCAGGAAGA
GAGCACACGTCTGAACTCTTTCCCTACACGACGCTCTTCCGATCTNNNGTAATCTCAGCTACTCAGGA
AGCACACGTCTGAACTCTTTCCCTACACGACGCTCTTCCGATCTNNNGTAAGCCAAGATCACATCACTG
AAGAGCACACGTCTGAACTCTTTCCCTACACGACGCTCTTCCGATCTNNNGAGAATAGCGTGAACCCA
AAGAGCACACGTCTGAACTCTTTCCCTACACGACGCTCTTCCGATCTNNNGAGCTTGCAGTGAGTCGA
AGAGCACACGTCTGAACTCTTTCCCTACACGACGCTCTTCCGATCTNNNACGCCTATAGTCCCAGCTAC
GAGCACACGTCTGAACTCTTTCCCTACACGACGCTCTTCCGATCTNNNACTTGACTGATTTGTGGACCT
GAGCACACGTCTGAACTCTTTCCCTACACGACGCTCTTCCGATCTNNNGAGAACAGCTTGAACCTGGG
GAGCACACGTCTGAACTCTTTCCCTACACGACGCTCTTCCGATCTNNNGAGGTTGCAGCGAGTAGAG
AAGAGCACACGTCTGAACTCTTTCCCTACACGACGCTCTTCCGATCTNNNCAGGAGAATGGCGAGAA
AAGAGCACACGTCTGAACTCTTTCCCTACACGACGCTCTTCCGATCTNNNGCAGGAGAATTGCTTGAA
AGCACACGTCTGAACTCTTTCCCTACACGACGCTCTTCCGATCTNNNGCACCTGTAATCCCAGCTACTA
GCACACGTCTGAACTCTTTCCCTACACGACGCTCTTCCGATCTNNNAGTTTCATGATCTTGGGATAGG
GCACACGTCTGAACTCTTTCCCTACACGACGCTCTTCCGATCTNNNCCTCCTTGTTGAGGTTTTCTTAC
AGCACACGTCTGAACTCTTTCCCTACACGACGCTCTTCCGATCTNNNAGGCTTTTTCCTATGCTAACCT
GAGCACACGTCTGAACTCTTTCCCTACACGACGCTCTTCCGATCTNNNGAGAATTGCTTGAACCCAGT
AGCACACGTCTGAACTCTTTCCCTACACGACGCTCTTCCGATCTNNNACTTTCTGCTAGCTTATTTGGT

AGAGCACACGTCTGAACTCTTTCCCTACACGACGCTCTTCCGATCTNNNCTCTCGAGATTCCTCCAATGC
GAGCACACGTCTGAACTCTTTCCCTACACGACGCTCTTCCGATCTNNNACTCACCATGACTTTTCTCTCTG
AGAGCACACGTCTGAACTCTTTCCCTACACGACGCTCTTCCGATCTNNNGCTTGCAGTGAGCTGAGATA
GAGCACACGTCTGAACTCTTTCCCTACACGACGCTCTTCCGATCTNNNTCTTAGCAGAGAAGGGTGTA
AGCACACGTCTGAACTCTTTCCCTACACGACGCTCTTCCGATCTNNNTCCTCATTCAGTTCAAACCTCTG
AGCACACGTCTGAACTCTTTCCCTACACGACGCTCTTCCGATCTNNNGGGATGTAATGGCTTCATTTCT
GAGCACACGTCTGAACTCTTTCCCTACACGACGCTCTTCCGATCTNNNGAGGCAGGAGAATCATTCAA
GAGCACACGTCTGAACTCTTTCCCTACACGACGCTCTTCCGATCTNNNGGTAATTTGGGAGACTTGAA
AGAGCACACGTCTGAACTCTTTCCCTACACGACGCTCTTCCGATCTNNNACATGAGAGGATTCCTTGAA
AGAGCACACGTCTGAACTCTTTCCCTACACGACGCTCTTCCGATCTNNNCCTGTAATTCCAGCTATGAG
AGAGCACACGTCTGAACTCTTTCCCTACACGACGCTCTTCCGATCTNNNAGGTTGCAGTAAGCCTAGA
AAGAGCACACGTCTGAACTCTTTCCCTACACGACGCTCTTCCGATCTNNNGAGGCAAGAGAATCGCTT
AGAGCACACGTCTGAACTCTTTCCCTACACGACGCTCTTCCGATCTNNNGCAGTTCCAACTCCTTCTCAG
AGAGCACACGTCTGAACTCTTTCCCTACACGACGCTCTTCCGATCTNNNTTGAGGCAGGAGAATCACTT
GAGCACACGTCTGAACTCTTTCCCTACACGACGCTCTTCCGATCTNNNGCCTATAATTCCAGCTACTCGG
GAGCACACGTCTGAACTCTTTCCCTACACGACGCTCTTCCGATCTNNNCTGAGGTGAGAGGATTGCTT
GAGCACACGTCTGAACTCTTTCCCTACACGACGCTCTTCCGATCTNNNACAGAATGCTTGGGAATTTTT
AGCACACGTCTGAACTCTTTCCCTACACGACGCTCTTCCGATCTNNNAGTCAACAACCTACATTGTATCC
AGAGCACACGTCTGAACTCTTTCCCTACACGACGCTCTTCCGATCTNNNCATAGTGGTGCATGCCTGTA
GAGCACACGTCTGAACTCTTTCCCTACACGACGCTCTTCCGATCTNNNAGATAACACATCCACTCTACTC
GAAGAGCACACGTCTGAACTCTTTCCCTACACGACGCTCTTCCGATCTNNNCTTGCAGTGAGCTGAGAT
AAGAGCACACGTCTGAACTCTTTCCCTACACGACGCTCTTCCGATCTNNNTGTAATCCCAGCTACTCAGG
AAGAGCACACGTCTGAACTCTTTCCCTACACGACGCTCTTCCGATCTNNNGTTGCAGTGAGCCAAGATT
GAGCACACGTCTGAACTCTTTCCCTACACGACGCTCTTCCGATCTNNNGTTGGCTTTTGCAAAGGTAA
AGCACACGTCTGAACTCTTTCCCTACACGACGCTCTTCCGATCTNNNTGATACTGAGGAGATAATCGG
AGCACACGTCTGAACTCTTTCCCTACACGACGCTCTTCCGATCTNNNTGCTCTAAAACACTGATTTGTGG
GAGCACACGTCTGAACTCTTTCCCTACACGACGCTCTTCCGATCTNNNCCCAACATGATTTGAATTGTG
GAGCACACGTCTGAACTCTTTCCCTACACGACGCTCTTCCGATCTNNNAGTGCTGAGATTACAGGCAT
AGAGCACACGTCTGAACTCTTTCCCTACACGACGCTCTTCCGATCTNNNTGTGGATTAGAGCTACCTGC
GAGCACACGTCTGAACTCTTTCCCTACACGACGCTCTTCCGATCTNNNGTCCTAGCTAGAGCAATCACAG
GAGCACACGTCTGAACTCTTTCCCTACACGACGCTCTTCCGATCTNNNAAAAAAATAGCTGGGCATGG
GAGCACACGTCTGAACTCTTTCCCTACACGACGCTCTTCCGATCTNNNTCTTTCTCTTCTTTGTGTGCAC
GAGCACACGTCTGAACTCTTTCCCTACACGACGCTCTTCCGATCTNNNACACACACACATATACACACAG
AGCACACGTCTGAACTCTTTCCCTACACGACGCTCTTCCGATCTNNNTCTCTTTCTTTCCTCTCTCTGTCTC
AAGAGCACACGTCTGAACTCTTTCCCTACACGACGCTCTTCCGATCTNNNGAGATCACGCCACTGCACT
AGAGCACACGTCTGAACTCTTTCCCTACACGACGCTCTTCCGATCTNNNATGTGTTGTTGGATTTGGCTA
GAGCACACGTCTGAACTCTTTCCCTACACGACGCTCTTCCGATCTNNNTACCCTAATGCCAAAACCAGA
AGCACACGTCTGAACTCTTTCCCTACACGACGCTCTTCCGATCTNNNTCCCTATTACAGTAGTTTCCACCT
GCACACGTCTGAACTCTTTCCCTACACGACGCTCTTCCGATCTNNNACAAAAAAAGGACTTCCATGAAC
AGAGCACACGTCTGAACTCTTTCCCTACACGACGCTCTTCCGATCTNNNGAGAATCGCTTGAACTCAGG
GAGCACACGTCTGAACTCTTTCCCTACACGACGCTCTTCCGATCTNNNGGTACCAATATGGAGGAATG
AAGAGCACACGTCTGAACTCTTTCCCTACACGACGCTCTTCCGATCTNNNGAGACAGGAGAATGGCGT
AGAGCACACGTCTGAACTCTTTCCCTACACGACGCTCTTCCGATCTNNNGGGCAATAACAGGTATGGA

GAGCACACGTCTGAACTCTTTCCCTACACGACGCTCTTCCGATCTNNNGGTGCCTGTAATCCCATCTAC
AGAGCACACGTCTGAACTCTTTCCCTACACGACGCTCTTCCGATCTNNNCAGGCACCTGTAATCTCAGC
AGCACACGTCTGAACTCTTTCCCTACACGACGCTCTTCCGATCTNNNACACAATTTCTTTCTCTCCAGTCC
ACACGTCTGAACTCTTTCCCTACACGACGCTCTTCCGATCTNNNGGTTGATGTGGATATATTTGTTCTG
GAGCACACGTCTGAACTCTTTCCCTACACGACGCTCTTCCGATCTNNNCAGTAGCCTGTAATCCCAGCT
GAGCACACGTCTGAACTCTTTCCCTACACGACGCTCTTCCGATCTNNNCAGATTGCAGTGAGCTGAGA
AGAGCACACGTCTGAACTCTTTCCCTACACGACGCTCTTCCGATCTNNNAGTGGCGATAACAAGTAGA
GAGCACACGTCTGAACTCTTTCCCTACACGACGCTCTTCCGATCTNNNATCTAGGTCTAGTCCAAGAT
AGCACACGTCTGAACTCTTTCCCTACACGACGCTCTTCCGATCTNNNCAGAAGAATTGCTTGAACCTG
AGAGCACACGTCTGAACTCTTTCCCTACACGACGCTCTTCCGATCTNNNTGGATCCCAATCATGACTGT
AGAGCACACGTCTGAACTCTTTCCCTACACGACGCTCTTCCGATCTNNNGAGCTTGCAGTGAGTCGA
GAGCACACGTCTGAACTCTTTCCCTACACGACGCTCTTCCGATCTNNNGAGAATGGTGTGAACCCAGG
AGCACACGTCTGAACTCTTTCCCTACACGACGCTCTTCCGATCTNNNACTTATCAATTGGGGGAAAATG
GAGCACACGTCTGAACTCTTTCCCTACACGACGCTCTTCCGATCTNNNAAAGTTTGCAGTGAGCCAAG
GAGCACACGTCTGAACTCTTTCCCTACACGACGCTCTTCCGATCTNNNTGTATGCCACTTTATGAAGCC
GAGCACACGTCTGAACTCTTTCCCTACACGACGCTCTTCCGATCTNNNTCAAGTGTCTAGAGCTCTTTCT
AGCACACGTCTGAACTCTTTCCCTACACGACGCTCTTCCGATCTNNNTCTTCTCAGGAAAAAGTTCAGG
AGCACACGTCTGAACTCTTTCCCTACACGACGCTCTTCCGATCTNNNTCCCTTTAGGTCTACTGTGTCTC
CACACGTCTGAACTCTTTCCCTACACGACGCTCTTCCGATCTNNNACTTTGGTCTAGGGTTAGAGTTA
GAGCACACGTCTGAACTCTTTCCCTACACGACGCTCTTCCGATCTNNNAGCATCTACTATGTTCCTGGC
AAGAGCACACGTCTGAACTCTTTCCCTACACGACGCTCTTCCGATCTNNNAGCGTTGCTTGTTTCTAC
AGCACACGTCTGAACTCTTTCCCTACACGACGCTCTTCCGATCTNNNACCAAAACTGAAGGTCTTTTTC
AGCACACGTCTGAACTCTTTCCCTACACGACGCTCTTCCGATCTNNNTATACAGTGAGCAAGCACAAC
AGCACACGTCTGAACTCTTTCCCTACACGACGCTCTTCCGATCTNNNTCAGGTGTATGTGTGTCTTTCT
GAGCACACGTCTGAACTCTTTCCCTACACGACGCTCTTCCGATCTNNNGCAAGACTTTGAATAACAGC
GAGCACACGTCTGAACTCTTTCCCTACACGACGCTCTTCCGATCTNNNGGTCTCCTTGTGTATTGTCAA
AGCACACGTCTGAACTCTTTCCCTACACGACGCTCTTCCGATCTNNNCTCAACCCCTTTTACAATAACTG
AGCACACGTCTGAACTCTTTCCCTACACGACGCTCTTCCGATCTNNNAGTTTGTTTACTTACTGCAGTG
AGCACACGTCTGAACTCTTTCCCTACACGACGCTCTTCCGATCTNNNAGTGCAAAGAATAATTTGAAA
AGCACACGTCTGAACTCTTTCCCTACACGACGCTCTTCCGATCTNNNGGCACAAACAAGCACTGAGTA
AGCACACGTCTGAACTCTTTCCCTACACGACGCTCTTCCGATCTNNNAGTGCCAGGCTCATGGTAAAT
GAGCACACGTCTGAACTCTTTCCCTACACGACGCTCTTCCGATCTNNNAGCATGGCAAAACCTCATCT
AGCACACGTCTGAACTCTTTCCCTACACGACGCTCTTCCGATCTNNNTCCTTACCATCCTAAAAAGCAC
GCACACGTCTGAACTCTTTCCCTACACGACGCTCTTCCGATCTNNNCGAGACTCCCTCTCAAAAACAAA
GAGCACACGTCTGAACTCTTTCCCTACACGACGCTCTTCCGATCTNNNGAGGTTGCAGTGAGCCAAG
AGCACACGTCTGAACTCTTTCCCTACACGACGCTCTTCCGATCTNNNTGTAGTCCCAGCTACTCAAAA
AGCACACGTCTGAACTCTTTCCCTACACGACGCTCTTCCGATCTNNNGAGAATGGTGTGAACCCAGA
AGAGCACACGTCTGAACTCTTTCCCTACACGACGCTCTTCCGATCTNNNTGTAAGATAATGTCCCGTC
GAGCACACGTCTGAACTCTTTCCCTACACGACGCTCTTCCGATCTNNNACATTCCATATATTCCATGGG
GAGCACACGTCTGAACTCTTTCCCTACACGACGCTCTTCCGATCTNNNGCACCTATAATCCCAGCTACT
GAGCACACGTCTGAACTCTTTCCCTACACGACGCTCTTCCGATCTNNNTCCACAGAAAAGACACTGGTT
GAGCACACGTCTGAACTCTTTCCCTACACGACGCTCTTCCGATCTNNNAGGACTAGAAGAAAGCGAAT
GAGCACACGTCTGAACTCTTTCCCTACACGACGCTCTTCCGATCTNNNTGAGAGAGAGACAGAGTGA

GAGCACACGTCTGAACTCTTTCCCTACACGACGCTCTTCCGATCTNNNATTGACCAGGAGCTAGACAGA

AGAGCACACGTCTGAACTCTTTCCCTACACGACGCTCTTCCGATCTNNNATTTCCACATGAGGTTTGGA

GAGCACACGTCTGAACTCTTTCCCTACACGACGCTCTTCCGATCTNNNCCAGTGGGAAATGAATTTGG

GAGCACACGTCTGAACTCTTTCCCTACACGACGCTCTTCCGATCTNNNTGCTGAATTCACTTGCCCATTTT

GAGCACACGTCTGAACTCTTTCCCTACACGACGCTCTTCCGATCTNNNAGAAAGAGGATTGCAGAACT

AGAGCACACGTCTGAACTCTTTCCCTACACGACGCTCTTCCGATCTNNNAGAGCCACAGATTGTAAGAG

GAGCACACGTCTGAACTCTTTCCCTACACGACGCTCTTCCGATCTNNNAAGGTGTATTCTAACTTCTTCC

AGCACACGTCTGAACTCTTTCCCTACACGACGCTCTTCCGATCTNNNAAAGATTGCTTTCATGTATTGG

AGCACACGTCTGAACTCTTTCCCTACACGACGCTCTTCCGATCTNNNAGAGAGAGAGAGAGAAATTCC

GAGCACACGTCTGAACTCTTTCCCTACACGACGCTCTTCCGATCTNNNCCAGCATTACCCTGATTCCAAA

AGAGCACACGTCTGAACTCTTTCCCTACACGACGCTCTTCCGATCTNNNGGAGGTTACCATGAGCTGA

GAGCACACGTCTGAACTCTTTCCCTACACGACGCTCTTCCGATCTNNNTAGCCATGTGAAGCAGTTCCT

GAGCACACGTCTGAACTCTTTCCCTACACGACGCTCTTCCGATCTNNNTATCCTGATGCCAAAACCAGA

GAGCACACGTCTGAACTCTTTCCCTACACGACGCTCTTCCGATCTNNNGCTGCTGCAAAAGACATGATT

GAGCACACGTCTGAACTCTTTCCCTACACGACGCTCTTCCGATCTNNNTTGTGTTTGTTGTCAACACTG

GAGCACACGTCTGAACTCTTTCCCTACACGACGCTCTTCCGATCTNNNCACACACATACACACATACACA

GAGCACACGTCTGAACTCTTTCCCTACACGACGCTCTTCCGATCTNNNAGCAAAGCTCCATCTCAAAAA

GAGCACACGTCTGAACTCTTTCCCTACACGACGCTCTTCCGATCTNNNTGAACTCTAAGACACACACACA

AAGAGCACACGTCTGAACTCTTTCCCTACACGACGCTCTTCCGATCTNNNTGGGTAACAGAGGGAGA

GAGCACACGTCTGAACTCTTTCCCTACACGACGCTCTTCCGATCTNNNTCTCTCTCTGTCTCTCTCTCT

GAGCACACGTCTGAACTCTTTCCCTACACGACGCTCTTCCGATCTNNNACCTGAAAATATACCCCCTGAA

AGAGCACACGTCTGAACTCTTTCCCTACACGACGCTCTTCCGATCTNNNAGTGCTACAATCTGAGTTA

AGAGCACACGTCTGAACTCTTTCCCTACACGACGCTCTTCCGATCTNNNAGTCATTCACACTGGGAAG

GAGCACACGTCTGAACTCTTTCCCTACACGACGCTCTTCCGATCTNNNGGGTGTTGGCAGTGGTATA

AGCACACGTCTGAACTCTTTCCCTACACGACGCTCTTCCGATCTNNNTGGTAGGCAGCATTATCTCTTTC

AGAGCACACGTCTGAACTCTTTCCCTACACGACGCTCTTCCGATCTNNNTCTCTTGTCCTTGTTTCCTTGA

AGAGCACACGTCTGAACTCTTTCCCTACACGACGCTCTTCCGATCTNNNAAAAGCCTTCCAACTCTGTTC

GCACACGTCTGAACTCTTTCCCTACACGACGCTCTTCCGATCTNNNTGATCTTCTGTAGTTCCTTCTGCT

AGCACACGTCTGAACTCTTTCCCTACACGACGCTCTTCCGATCTNNNTCCATTCAATTTCTCTCTCCCTGT

AGAGCACACGTCTGAACTCTTTCCCTACACGACGCTCTTCCGATCTNNNTGGTAACAGTTTGATGTCTA

AGCACACGTCTGAACTCTTTCCCTACACGACGCTCTTCCGATCTNNNTCAGATTTGTGTATGGGCTCTT

GAGCACACGTCTGAACTCTTTCCCTACACGACGCTCTTCCGATCTNNNAGAGAAAACTGTCTTGTTGCT

GAGCACACGTCTGAACTCTTTCCCTACACGACGCTCTTCCGATCTNNNGGGGTCCTGTGTAGTCTAGG

GAGCACACGTCTGAACTCTTTCCCTACACGACGCTCTTCCGATCTNNNTCAAGGTTTCTTTGGGTTCTCC

AGAGCACACGTCTGAACTCTTTCCCTACACGACGCTCTTCCGATCTNNNCACATGAAAAGGTGCTCATC

GAGCACACGTCTGAACTCTTTCCCTACACGACGCTCTTCCGATCTNNNTCATCTGTTATACACACCCACA

GAGCACACGTCTGAACTCTTTCCCTACACGACGCTCTTCCGATCTNNNAGAGAAATGCAAATCAAAGCC

AGCACACGTCTGAACTCTTTCCCTACACGACGCTCTTCCGATCTNNNGCAAAGAGCTCCACTATTGATT

GAGCACACGTCTGAACTCTTTCCCTACACGACGCTCTTCCGATCTNNNTATCTCTTTGGCTTTCCTGGG

GAGCACACGTCTGAACTCTTTCCCTACACGACGCTCTTCCGATCTNNNCTCTCTCCCCATTACACACAACA

GAGCACACGTCTGAACTCTTTCCCTACACGACGCTCTTCCGATCTNNNGAGCTAGACTCCGTCTCAAAA

GAGCACACGTCTGAACTCTTTCCCTACACGACGCTCTTCCGATCTNNNGGCTTCCAGTTTTATCCATGC

GAGCACACGTCTGAACTCTTTCCCTACACGACGCTCTTCCGATCTNNNGCACCTCTTTCACTCAGGAAAA

GAGCACACGTCTGAACTCTTTCCCTACACGACGCTCTTCCGATCTNNNGCAACAGAGTGAGAATCCGT
GAGCACACGTCTGAACTCTTTCCCTACACGACGCTCTTCCGATCTNNNGCATTGTTTTGGTCTTGTCTCCA
GAGCACACGTCTGAACTCTTTCCCTACACGACGCTCTTCCGATCTNNNTGCACTGTTGTTGGTAGAGGT
AGAGCACACGTCTGAACTCTTTCCCTACACGACGCTCTTCCGATCTNNNCTGTAATCCCAGCTACTCAG
AGAGCACACGTCTGAACTCTTTCCCTACACGACGCTCTTCCGATCTNNNGGCCATCTTCCAAGAATGAAA
AGCACACGTCTGAACTCTTTCCCTACACGACGCTCTTCCGATCTNNNCCCTAATTTCTCGAATCTCCAGGT
GAGCACACGTCTGAACTCTTTCCCTACACGACGCTCTTCCGATCTNNNGCTAGTTCTTGCCTGAATACCT
GAGCACACGTCTGAACTCTTTCCCTACACGACGCTCTTCCGATCTNNNTTGACTTGTATGGGAGCATA
AGCACACGTCTGAACTCTTTCCCTACACGACGCTCTTCCGATCTNNNTGTGTTCCTAAGACATCACAGA
GAGCACACGTCTGAACTCTTTCCCTACACGACGCTCTTCCGATCTNNNCTAAGGCAGGAGAATCGCTT
GAGCACACGTCTGAACTCTTTCCCTACACGACGCTCTTCCGATCTNNNACATTGAGAAAAGGAGGAA
AGCACACGTCTGAACTCTTTCCCTACACGACGCTCTTCCGATCTNNNATGTGTATGTGTGTGTGTGTG
GCACACGTCTGAACTCTTTCCCTACACGACGCTCTTCCGATCTNNNTGGAATGAAACCTCACTATTTGG
GAGCACACGTCTGAACTCTTTCCCTACACGACGCTCTTCCGATCTNNNACATTGAGAAAAGGAGGAA
GCACACGTCTGAACTCTTTCCCTACACGACGCTCTTCCGATCTNNNTTCTTCTCTCTCTCTCTCTCCCCCAT
AGAGCACACGTCTGAACTCTTTCCCTACACGACGCTCTTCCGATCTNNNGTGGCTTGGCTAAGAAGTT
AGAGCACACGTCTGAACTCTTTCCCTACACGACGCTCTTCCGATCTNNNACTGCCTTCTAGAGATGGGT
GAGCACACGTCTGAACTCTTTCCCTACACGACGCTCTTCCGATCTNNNTTCTCTACCAACCAACTCTCACA
AGAGCACACGTCTGAACTCTTTCCCTACACGACGCTCTTCCGATCTNNNGCTGTTAGCAGACCTTCCTTA
AAGAGCACACGTCTGAACTCTTTCCCTACACGACGCTCTTCCGATCTNNNCAGGATCTGCATTCTGAAA
GAGCACACGTCTGAACTCTTTCCCTACACGACGCTCTTCCGATCTNNNTCTCCTACATGGCTTGCTTGAA
AGAGCACACGTCTGAACTCTTTCCCTACACGACGCTCTTCCGATCTNNNACGTTACTGGTATCTGAGG
AGCACACGTCTGAACTCTTTCCCTACACGACGCTCTTCCGATCTNNNGTTGGTTTTCATTTCTCTCCAG
AGCACACGTCTGAACTCTTTCCCTACACGACGCTCTTCCGATCTNNNCAGAGAAACAGAACCAATAGG
GAGCACACGTCTGAACTCTTTCCCTACACGACGCTCTTCCGATCTNNNTGTGTACACGTGTAGACATGT
GAGCACACGTCTGAACTCTTTCCCTACACGACGCTCTTCCGATCTNNNACCATGAGAACACAGACACAT
GCACACGTCTGAACTCTTTCCCTACACGACGCTCTTCCGATCTNNNACCAATAGGAGAGAGAGAGAG
AGAGCACACGTCTGAACTCTTTCCCTACACGACGCTCTTCCGATCTNNNGTTGTCACAAAACCCATGGA
GAGCACACGTCTGAACTCTTTCCCTACACGACGCTCTTCCGATCTNNNTCTTACTTCTCAGGCCTTTGGA
AGCACACGTCTGAACTCTTTCCCTACACGACGCTCTTCCGATCTNNNTCCAGAGAAACAGAACCAACAA
AGCACACGTCTGAACTCTTTCCCTACACGACGCTCTTCCGATCTNNNACTGAGATAGAACTGGGTTGA
GAGCACACGTCTGAACTCTTTCCCTACACGACGCTCTTCCGATCTNNNTCCTCCATTCCATTGGTTGTCT
AGCACACGTCTGAACTCTTTCCCTACACGACGCTCTTCCGATCTNNNGTGCCTGGCTTATTTTACTTAG
GAGCACACGTCTGAACTCTTTCCCTACACGACGCTCTTCCGATCTNNNGCAACTAAGACCCATAACTGAT
GAGCACACGTCTGAACTCTTTCCCTACACGACGCTCTTCCGATCTNNNACAGATACAGTGCAATCCCTA
AGAGCACACGTCTGAACTCTTTCCCTACACGACGCTCTTCCGATCTNNNTTCCAGATTTGGGATCCTCAA
GAGCACACGTCTGAACTCTTTCCCTACACGACGCTCTTCCGATCTNNNAGGCATGTTACCTAAGTCACAT
AGCACACGTCTGAACTCTTTCCCTACACGACGCTCTTCCGATCTNNNCCTCTAAGTCCTCCCAACTTTCC
AGCACACGTCTGAACTCTTTCCCTACACGACGCTCTTCCGATCTNNNACTTTAGGTCACTTTGATTCAGT
AGAGCACACGTCTGAACTCTTTCCCTACACGACGCTCTTCCGATCTNNNCACACACACACACACACGAAT
AGAGCACACGTCTGAACTCTTTCCCTACACGACGCTCTTCCGATCTNNNTGACAGAGCAAACTCCATCT
AGCACACGTCTGAACTCTTTCCCTACACGACGCTCTTCCGATCTNNNCAAAGATTCCTGCACTTGTATG
AGAGCACACGTCTGAACTCTTTCCCTACACGACGCTCTTCCGATCTNNNACAAGTTGTCAGGTTCCTAAA

AGAGCACACGTCTGAACTCTTTCCCTACACGACGCTCTTCCGATCTNNNCAAAACCAACCTGGAAGACT
AGCACACGTCTGAACTCTTTCCCTACACGACGCTCTTCCGATCTNNNTGAACAGAAAAAGTCAGTGTCT
GAGCACACGTCTGAACTCTTTCCCTACACGACGCTCTTCCGATCTNNNATAGAAGCTCCTGAACACATG
GCACACGTCTGAACTCTTTCCCTACACGACGCTCTTCCGATCTNNNGGGGATGATACTCAGTACTACAC
GAGCACACGTCTGAACTCTTTCCCTACACGACGCTCTTCCGATCTNNNACTCTTCCTCCAACCCAATTCTC
GAAGAGCACACGTCTGAACTCTTTCCCTACACGACGCTCTTCCGATCTNNNTCTTTCTGATGGGCATGAT
GAAGAGCACACGTCTGAACTCTTTCCCTACACGACGCTCTTCCGATCTNNNCTCCAGTGCACCAAATGC
GAGCACACGTCTGAACTCTTTCCCTACACGACGCTCTTCCGATCTNNNGGTGCCACCTATATGCTGATTT
AGAGCACACGTCTGAACTCTTTCCCTACACGACGCTCTTCCGATCTNNNCTCTATCGTCCTTCTCCCCAAT
AGAGCACACGTCTGAACTCTTTCCCTACACGACGCTCTTCCGATCTNNNTGGTTTCTGCTGAAAGATTCA
AGCACACGTCTGAACTCTTTCCCTACACGACGCTCTTCCGATCTNNNAGTTCCCACATATCTCTCTCTCCA
AGCACACGTCTGAACTCTTTCCCTACACGACGCTCTTCCGATCTNNNTGGGTCAGATGATTCTCTCTCTC
AGCACACGTCTGAACTCTTTCCCTACACGACGCTCTTCCGATCTNNNTGTTACTGACACAGAAAGGGA
AGAGCACACGTCTGAACTCTTTCCCTACACGACGCTCTTCCGATCTNNNCCTAACATGGGGTCTAAGGA
GAGCACACGTCTGAACTCTTTCCCTACACGACGCTCTTCCGATCTNNNTCAGAGATGTTGACAAAGCAC
AGCACACGTCTGAACTCTTTCCCTACACGACGCTCTTCCGATCTNNNTCCCAGTTAGACATACCTGAGA
GCACACGTCTGAACTCTTTCCCTACACGACGCTCTTCCGATCTNNNACAGTACACACATACACACACAGA
GAGCACACGTCTGAACTCTTTCCCTACACGACGCTCTTCCGATCTNNNTCTTCAACAGAATTTCACTGGC
AGCACACGTCTGAACTCTTTCCCTACACGACGCTCTTCCGATCTNNNTGAGCCATGATTATACTCCAGC
GAGCACACGTCTGAACTCTTTCCCTACACGACGCTCTTCCGATCTNNNTGACATCACATTGGTAGCTGG
GAGCACACGTCTGAACTCTTTCCCTACACGACGCTCTTCCGATCTNNNCAAACGTATCTGCATGTTTGT
AGAGCACACGTCTGAACTCTTTCCCTACACGACGCTCTTCCGATCTNNNTGTTCATCAGGGATATTGAC
AGCACACGTCTGAACTCTTTCCCTACACGACGCTCTTCCGATCTNNNACTTTTCCTAATACACACCCACAC
AGAGCACACGTCTGAACTCTTTCCCTACACGACGCTCTTCCGATCTNNNGGAGCCTAAGGAGACAGAA
GAGCACACGTCTGAACTCTTTCCCTACACGACGCTCTTCCGATCTNNNTCTCCAAATTTATGGCAACCTG
AGAGCACACGTCTGAACTCTTTCCCTACACGACGCTCTTCCGATCTNNNTGGATGGACCTGGAGGACA
GCACACGTCTGAACTCTTTCCCTACACGACGCTCTTCCGATCTNNNTGTTCCATTGGTCTATGTGTCTGT
AGAGCACACGTCTGAACTCTTTCCCTACACGACGCTCTTCCGATCTNNNGCAGAGAGATCAATCAAGAA
AGCACACGTCTGAACTCTTTCCCTACACGACGCTCTTCCGATCTNNNTGAAGCCAGTATCACCAAATAC
AGAGCACACGTCTGAACTCTTTCCCTACACGACGCTCTTCCGATCTNNNAGCTACAGCAATCAGACAAG
AGAGCACACGTCTGAACTCTTTCCCTACACGACGCTCTTCCGATCTNNNTCCTGTCTCTCTCTGTCTCTCTC
GAGCACACGTCTGAACTCTTTCCCTACACGACGCTCTTCCGATCTNNNGCAGACAGCCTATTGTGGGAT
GAGCACACGTCTGAACTCTTTCCCTACACGACGCTCTTCCGATCTNNNTGGAAGCATATCACATCCTGA
AGAGCACACGTCTGAACTCTTTCCCTACACGACGCTCTTCCGATCTNNNATACTGACACACACCACAACA
GAGCACACGTCTGAACTCTTTCCCTACACGACGCTCTTCCGATCTNNNATAGAAAAACAGGCAAATTGC
AGAGCACACGTCTGAACTCTTTCCCTACACGACGCTCTTCCGATCTNNNTATGTGCTGGGTTAGGAAGT
AGCACACGTCTGAACTCTTTCCCTACACGACGCTCTTCCGATCTNNNGGGTTTTCATCATGTTGACCAGC
AGAGCACACGTCTGAACTCTTTCCCTACACGACGCTCTTCCGATCTNNNGAGAATCACTTGAACCTGGC
AGAGCACACGTCTGAACTCTTTCCCTACACGACGCTCTTCCGATCTNNNCCTCCAGGAGCTACTCTTTTG
AGCACACGTCTGAACTCTTTCCCTACACGACGCTCTTCCGATCTNNNACACAGTAACAGTCTGATCTCTC
GCACACGTCTGAACTCTTTCCCTACACGACGCTCTTCCGATCTNNNCCCAAATGAGATACATGCATAGC
GAGCACACGTCTGAACTCTTTCCCTACACGACGCTCTTCCGATCTNNNTCCTTGTCCTGAGAAATATGC
GAGCACACGTCTGAACTCTTTCCCTACACGACGCTCTTCCGATCTNNNGTGTGTGTGTATGTGTGTGTC

GAGCACACGTCTGAACTCTTTCCCTACACGACGCTCTTCCGATCTNNNACCAATCCTAGGAAACCAGA
AGAGCACACGTCTGAACTCTTTCCCTACACGACGCTCTTCCGATCTNNNACAAGATAAGATGATCCTG
GAGCACACGTCTGAACTCTTTCCCTACACGACGCTCTTCCGATCTNNNACAGTCACAGATATCTGAGC
AGCACACGTCTGAACTCTTTCCCTACACGACGCTCTTCCGATCTNNNACTTGTTCATGGTGTTTTTCT
AGCACACGTCTGAACTCTTTCCCTACACGACGCTCTTCCGATCTNNNTGGCTTGGCTTACAGTTGAGT
GAGCACACGTCTGAACTCTTTCCCTACACGACGCTCTTCCGATCTNNNAGGTGATCTTACGTACAGAT
AAGAGCACACGTCTGAACTCTTTCCCTACACGACGCTCTTCCGATCTNNNACGCATGGATGTATGTAC
GCACACGTCTGAACTCTTTCCCTACACGACGCTCTTCCGATCTNNNACCAGTCCTTGAAAGAATGAAA
GAGCACACGTCTGAACTCTTTCCCTACACGACGCTCTTCCGATCTNNNACTACTTTCCATCAACTGCCAC
GAGCACACGTCTGAACTCTTTCCCTACACGACGCTCTTCCGATCTNNNAGTGTTACAGCAATCAATGG
AGCACACGTCTGAACTCTTTCCCTACACGACGCTCTTCCGATCTNNNGCTATTTTGGTGGCATATGGT
AGCACACGTCTGAACTCTTTCCCTACACGACGCTCTTCCGATCTNNNGGCTCACACCCAGAGGAATAT
GCACACGTCTGAACTCTTTCCCTACACGACGCTCTTCCGATCTNNNGTGAAACTCTGTCTCAGGAAAA
GAGCACACGTCTGAACTCTTTCCCTACACGACGCTCTTCCGATCTNNNAGGAGAATTGCTTGATCCCA
GAGCACACGTCTGAACTCTTTCCCTACACGACGCTCTTCCGATCTNNNCATTTTCTGGTAGCAGCAGTT
GAGCACACGTCTGAACTCTTTCCCTACACGACGCTCTTCCGATCTNNNCAAAGGTTGCAGTGAACTG
AGCACACGTCTGAACTCTTTCCCTACACGACGCTCTTCCGATCTNNNATCGAGATCATACCACTGCAC
GAGCACACGTCTGAACTCTTTCCCTACACGACGCTCTTCCGATCTNNNGAGCAAGACTCTGTCTCAAA
CACACGTCTGAACTCTTTCCCTACACGACGCTCTTCCGATCTNNNTGTTGATTTTTGTATCCTGCAACT
AGCACACGTCTGAACTCTTTCCCTACACGACGCTCTTCCGATCTNNNGCAACTGTGCCACATTACATTA
AGCACACGTCTGAACTCTTTCCCTACACGACGCTCTTCCGATCTNNNTGTTCTCTCTCTCTCTCTCTGCA
GAGCACACGTCTGAACTCTTTCCCTACACGACGCTCTTCCGATCTNNNCTGTCCCTTGCCTAAATTTGT
GAGCACACGTCTGAACTCTTTCCCTACACGACGCTCTTCCGATCTNNNAGAAGGATGAGGCAGGAG
GAGCACACGTCTGAACTCTTTCCCTACACGACGCTCTTCCGATCTNNNATCCTAGGGGTTCAATCTTG
GCACACGTCTGAACTCTTTCCCTACACGACGCTCTTCCGATCTNNNTCCTATGGCATATTTGTGGTCA
GAGCACACGTCTGAACTCTTTCCCTACACGACGCTCTTCCGATCTNNNGATCAGAGCAGAACTGAAG
AGCACACGTCTGAACTCTTTCCCTACACGACGCTCTTCCGATCTNNNACCACATACAGATACACCCAAA
AGCACACGTCTGAACTCTTTCCCTACACGACGCTCTTCCGATCTNNNATCTTCCTGGTTAGCTGTATTT
AGCACACGTCTGAACTCTTTCCCTACACGACGCTCTTCCGATCTNNNTAGTGTGTGTGTGTGTGTTTG
GAGCACACGTCTGAACTCTTTCCCTACACGACGCTCTTCCGATCTNNNTGGAGTTTGCAGTGAGATG
GAGCACACGTCTGAACTCTTTCCCTACACGACGCTCTTCCGATCTNNNGATAAGGAAAGGGAGCTGA
AGAGCACACGTCTGAACTCTTTCCCTACACGACGCTCTTCCGATCTNNNGGGCTATGATTGATTGCAC
GAGCACACGTCTGAACTCTTTCCCTACACGACGCTCTTCCGATCTNNNAACATGTGGGTAACTTCTAA
GCACACGTCTGAACTCTTTCCCTACACGACGCTCTTCCGATCTNNNGGTCATCTGGTCTTCCATACTA
AGCACACGTCTGAACTCTTTCCCTACACGACGCTCTTCCGATCTNNNTCCTTTTGATTTGATTGCTTGA
AGCACACGTCTGAACTCTTTCCCTACACGACGCTCTTCCGATCTNNNCTATGATCAGGTCACTGCTCT
GAGCACACGTCTGAACTCTTTCCCTACACGACGCTCTTCCGATCTNNNATTTGCACGGTCTAGCTCTTT
GAGCACACGTCTGAACTCTTTCCCTACACGACGCTCTTCCGATCTNNNGGGTGACAGAGTAAGACCT
GAGCACACGTCTGAACTCTTTCCCTACACGACGCTCTTCCGATCTNNNAGGTTGTAGTGAGCTGAGA
GCACACGTCTGAACTCTTTCCCTACACGACGCTCTTCCGATCTNNNCCACTTTGGTCAACACAACATAC
AGCACACGTCTGAACTCTTTCCCTACACGACGCTCTTCCGATCTNNNCCCACCTTCAACAACTTTTTTTC
GAGCACACGTCTGAACTCTTTCCCTACACGACGCTCTTCCGATCTNNNTGGGTGACAGAGTGAGACT
GAGCACACGTCTGAACTCTTTCCCTACACGACGCTCTTCCGATCTNNNTCATCTTACACCTGTTAGGAT

AGCACACGTCTGAACTCTTTCCCTACACGACGCTCTTCCGATCTNNNAGTGGTCTCCCAACTTTTGTCT
GAGCACACGTCTGAACTCTTTCCCTACACGACGCTCTTCCGATCTNNNCACTATCCTGAGAGTCAAGG
GAGCACACGTCTGAACTCTTTCCCTACACGACGCTCTTCCGATCTNNNACAAGTTGTCAGGTTCCTAA
GCACACGTCTGAACTCTTTCCCTACACGACGCTCTTCCGATCTNNNTGAGTTTTGGGTTTCATCAAT
AGCACACGTCTGAACTCTTTCCCTACACGACGCTCTTCCGATCTNNNAGGTTGCAGTGAGCTGATAT
AGAGCACACGTCTGAACTCTTTCCCTACACGACGCTCTTCCGATCTNNNTCTGTGAGCCAGTTCCTCA
AGCACACGTCTGAACTCTTTCCCTACACGACGCTCTTCCGATCTNNNGCTCATTTTAGGTTGACTGACT
GAGCACACGTCTGAACTCTTTCCCTACACGACGCTCTTCCGATCTNNNAGACTCCGTCTCAAAAAACAA
AGCACACGTCTGAACTCTTTCCCTACACGACGCTCTTCCGATCTNNNCATACGTATGTGTGTGTGT
GCACACGTCTGAACTCTTTCCCTACACGACGCTCTTCCGATCTNNNTTCTTTCTTTCCCTCCCTTCCTTCAT
AGCACACGTCTGAACTCTTTCCCTACACGACGCTCTTCCGATCTNNNGTGAGTCTTTAGGAAGTGAG
AGCACACGTCTGAACTCTTTCCCTACACGACGCTCTTCCGATCTNNNGTGTGTGTATGTGTGTGTGT
AGCACACGTCTGAACTCTTTCCCTACACGACGCTCTTCCGATCTNNNAGGATAGGAACTGATCAACAA
GAGCACACGTCTGAACTCTTTCCCTACACGACGCTCTTCCGATCTNNNTGGCAGTACCAGAACACAAA
GAGCACACGTCTGAACTCTTTCCCTACACGACGCTCTTCCGATCTNNNTTATGCTACTTCACTCCAGCCT
AAGAGCACACGTCTGAACTCTTTCCCTACACGACGCTCTTCCGATCTNNNCTGGGACCACAGGTGTA
GAGCACACGTCTGAACTCTTTCCCTACACGACGCTCTTCCGATCTNNNAGTTTCCTTCGAAGTACATTT
GAGCACACGTCTGAACTCTTTCCCTACACGACGCTCTTCCGATCTNNNTCCCAAAGTGCTGAGATTACA
AAGAGCACACGTCTGAACTCTTTCCCTACACGACGCTCTTCCGATCTNNNACGCTGGAAAGAGAGAG
AAGAGCACACGTCTGAACTCTTTCCCTACACGACGCTCTTCCGATCTNNNAGAGAGAGAGAGAGAGA
AGCACACGTCTGAACTCTTTCCCTACACGACGCTCTTCCGATCTNNNCCGAGATCGTACCATTGTACT
GAGCACACGTCTGAACTCTTTCCCTACACGACGCTCTTCCGATCTNNNAGTTTGTGAAATCAAGGACA
AGCACACGTCTGAACTCTTTCCCTACACGACGCTCTTCCGATCTNNNCCTCACTTCCCACATCTGAAAA
AGAGCACACGTCTGAACTCTTTCCCTACACGACGCTCTTCCGATCTNNNAGTCAACAACTCAAAGGCAA
AGAGCACACGTCTGAACTCTTTCCCTACACGACGCTCTTCCGATCTNNNTGAGGAGAATCACTTGAAC
GAGCACACGTCTGAACTCTTTCCCTACACGACGCTCTTCCGATCTNNNGCTTTCTAACTGCACATCTTGG
AGCACACGTCTGAACTCTTTCCCTACACGACGCTCTTCCGATCTNNNGGCAAAGTGGTTGTATATGT
AGAGCACACGTCTGAACTCTTTCCCTACACGACGCTCTTCCGATCTNNNTGACTACCAGAATGAGCATC
GAGCACACGTCTGAACTCTTTCCCTACACGACGCTCTTCCGATCTNNNAGAGGGTCAAATAGTGGAG
AGCACACGTCTGAACTCTTTCCCTACACGACGCTCTTCCGATCTNNNAAAATAAAAAACCCAACCCGT
GCACACGTCTGAACTCTTTCCCTACACGACGCTCTTCCGATCTNNNAGTGTTGGGATTACAGGCATGA
GAGCACACGTCTGAACTCTTTCCCTACACGACGCTCTTCCGATCTNNNTCCAGAGAAACAGAACCAAT
GAGCACACGTCTGAACTCTTTCCCTACACGACGCTCTTCCGATCTNNNAGCACTGAGTTTTATCCCTAG
AGCACACGTCTGAACTCTTTCCCTACACGACGCTCTTCCGATCTNNNTGAAACAAGTGTTATTCCTTT
AGCACACGTCTGAACTCTTTCCCTACACGACGCTCTTCCGATCTNNNAGGAGATTTGTGGGAAAGT
AGCACACGTCTGAACTCTTTCCCTACACGACGCTCTTCCGATCTNNNTCCTGTCTCTTTCTTTCTCTCTCT
AGCACACGTCTGAACTCTTTCCCTACACGACGCTCTTCCGATCTNNNCCTGAAGGAAGGGATAAGT
AGCACACGTCTGAACTCTTTCCCTACACGACGCTCTTCCGATCTNNNTGACAATCTTTTTCAATCACAGG
GAGCACACGTCTGAACTCTTTCCCTACACGACGCTCTTCCGATCTNNNCAGGAGAATCACTTGAACTG
AGCACACGTCTGAACTCTTTCCCTACACGACGCTCTTCCGATCTNNNAGAGAGAGAGAGAGAGAGA
AGCACACGTCTGAACTCTTTCCCTACACGACGCTCTTCCGATCTNNNACAGAACAAAAGCACAGGTTT
AGCACACGTCTGAACTCTTTCCCTACACGACGCTCTTCCGATCTNNNCCTGGAGACACTGGGATATCT
AAGAGCACACGTCTGAACTCTTTCCCTACACGACGCTCTTCCGATCTNNNATCCCCTTGCTCTGCTATAG

AAGAGCACACGTCTGAACTCTTTCCCTACACGACGCTCTTCCGATCTNNNTGCCAGTCTAGTTAGAAAA
GAGCACACGTCTGAACTCTTTCCCTACACGACGCTCTTCCGATCTNNNTCTTTCTCTCCCCTATCTCTGTC
GCACACGTCTGAACTCTTTCCCTACACGACGCTCTTCCGATCTNNNGTGACTAGGATATTCACAACCCCT
GAAGAGCACACGTCTGAACTCTTTCCCTACACGACGCTCTTCCGATCTNNNGAAGGAAGGAAGGAAGG
AGAGCACACGTCTGAACTCTTTCCCTACACGACGCTCTTCCGATCTNNNGTGTCTATGTGTGTGTGTGT
AGCACACGTCTGAACTCTTTCCCTACACGACGCTCTTCCGATCTNNNTCCTTTCTCTTTCTCCCTTTCTCTC
AGCACACGTCTGAACTCTTTCCCTACACGACGCTCTTCCGATCTNNNTCCTTCTTTTCTTTCTCCTCTCCTC
AGCACACGTCTGAACTCTTTCCCTACACGACGCTCTTCCGATCTNNNCCCACTCAAAGGAAAATGTACAG
GAGCACACGTCTGAACTCTTTCCCTACACGACGCTCTTCCGATCTNNNCTGGCTCTCCTACCATTGAGTT
AGCACACGTCTGAACTCTTTCCCTACACGACGCTCTTCCGATCTNNNTGATTTCATCACATTGTTGACCC
GAGCACACGTCTGAACTCTTTCCCTACACGACGCTCTTCCGATCTNNNCTGTTTCTCTGTAGTTCTGTGA
AGAGCACACGTCTGAACTCTTTCCCTACACGACGCTCTTCCGATCTNNNACCTGCAAGTGTAATGAAGG
AGCACACGTCTGAACTCTTTCCCTACACGACGCTCTTCCGATCTNNNAGAATCCATGAGCCCATATTGAC
AGAGCACACGTCTGAACTCTTTCCCTACACGACGCTCTTCCGATCTNNNCCTGAGATCAAAGCTATCTCT
GAAGAGCACACGTCTGAACTCTTTCCCTACACGACGCTCTTCCGATCTNNNCATGGTGTTGTGTTCCTG
AGAGCACACGTCTGAACTCTTTCCCTACACGACGCTCTTCCGATCTNNNAGGTACTGTCTTGATTGTCT
GAGCACACGTCTGAACTCTTTCCCTACACGACGCTCTTCCGATCTNNNCACTGGCACTTCATTCTGATTG
GAGCACACGTCTGAACTCTTTCCCTACACGACGCTCTTCCGATCTNNNTCCCTTGACATGTGGGGATTA
AGAGCACACGTCTGAACTCTTTCCCTACACGACGCTCTTCCGATCTNNNTTCTGCCCTCTACCTCTTTCAA
AGCACACGTCTGAACTCTTTCCCTACACGACGCTCTTCCGATCTNNNTCCTTTCCTTTCCTCTCTCTCTCA
AGCACACGTCTGAACTCTTTCCCTACACGACGCTCTTCCGATCTNNNGCAAGCTGGAAACTGGAATTTA
GCACACGTCTGAACTCTTTCCCTACACGACGCTCTTCCGATCTNNNTGAAAGGCAGAAGAAAGCAAAA
GCACACGTCTGAACTCTTTCCCTACACGACGCTCTTCCGATCTNNNCAGAGGAACATCAGGGACCTTAC
CACACGTCTGAACTCTTTCCCTACACGACGCTCTTCCGATCTNNNAGAGAGAGAGAGAGAGAGAGAG
AGCACACGTCTGAACTCTTTCCCTACACGACGCTCTTCCGATCTNNNTCTCTCTGTCTCTCTGTCTCTG
AGCACACGTCTGAACTCTTTCCCTACACGACGCTCTTCCGATCTNNNCCAGAAGGGTAAAGATAGGAT
AGCACACGTCTGAACTCTTTCCCTACACGACGCTCTTCCGATCTNNNGCATGACCAGACCCCAGTAAAA
GCACACGTCTGAACTCTTTCCCTACACGACGCTCTTCCGATCTNNNGGTGAAGATACACACTCATGCCTG
AGCACACGTCTGAACTCTTTCCCTACACGACGCTCTTCCGATCTNNNACACTTAGGAGAGACCAGGAG
AGCACACGTCTGAACTCTTTCCCTACACGACGCTCTTCCGATCTNNNCGCTCTCTCTCTCTCTCTCTCTCC
GAGCACACGTCTGAACTCTTTCCCTACACGACGCTCTTCCGATCTNNNAACCTTTTGAAAACCCAGCACT
AGCACACGTCTGAACTCTTTCCCTACACGACGCTCTTCCGATCTNNNAGAGAGAGAGAGAGAGAGAG
AGAGCACACGTCTGAACTCTTTCCCTACACGACGCTCTTCCGATCTNNNGGGCTGAGTGAAGAGAATC
CACACGTCTGAACTCTTTCCCTACACGACGCTCTTCCGATCTNNNGCATTTACCAGAAGCCTCTCTCTCA
AGCACACGTCTGAACTCTTTCCCTACACGACGCTCTTCCGATCTNNNGCAACAGGATGAGACTTTGTCA
AGAGCACACGTCTGAACTCTTTCCCTACACGACGCTCTTCCGATCTNNNGCTTGTGTTGAGGATAAGG
AGCACACGTCTGAACTCTTTCCCTACACGACGCTCTTCCGATCTNNNGTCAGCACAAAAGAAGTCAGGA
GCACACGTCTGAACTCTTTCCCTACACGACGCTCTTCCGATCTNNNGGTCTTTCTCCCTTTCTCCCTTTCAT
AGCACACGTCTGAACTCTTTCCCTACACGACGCTCTTCCGATCTNNNCAGAGGTTGACATCTGTGGAGT
AGCACACGTCTGAACTCTTTCCCTACACGACGCTCTTCCGATCTNNNGTGCTCTTGGGTTGAGTACCTA
GAGCACACGTCTGAACTCTTTCCCTACACGACGCTCTTCCGATCTNNNCTTTCTTTCTCTGCCTCCTCCATG
AGAGCACACGTCTGAACTCTTTCCCTACACGACGCTCTTCCGATCTNNNAAGACCCCTTGAGTCTAGGA
AGCACACGTCTGAACTCTTTCCCTACACGACGCTCTTCCGATCTNNNGATCACTTGAGCTCAGGAGTTC

CACACGTCTGAACTCTTTCCCTACACGACGCTCTTCCGATCTNNNTGTCTGTCTGTCTCTGTCTCTCTCA
AGCACACGTCTGAACTCTTTCCCTACACGACGCTCTTCCGATCTNNNGAGAGACAGAGACGAGAGA
GCACACGTCTGAACTCTTTCCCTACACGACGCTCTTCCGATCTNNNTTAGCATCTCCCTTTCCATCCTG
AGCACACGTCTGAACTCTTTCCCTACACGACGCTCTTCCGATCTNNNAGACAGGAGAATTGCTTGAA
GAGCACACGTCTGAACTCTTTCCCTACACGACGCTCTTCCGATCTNNNGAATCATTCCCAAAACCCCCTT
GAGCACACGTCTGAACTCTTTCCCTACACGACGCTCTTCCGATCTNNNGAATTGGAGAGAGGAGAGA
GAGCACACGTCTGAACTCTTTCCCTACACGACGCTCTTCCGATCTNNNGAGCAAGACCTTGTCTCAAA
AGCACACGTCTGAACTCTTTCCCTACACGACGCTCTTCCGATCTNNNCACTTGAGCTCAGGAGTTTG
GCACACGTCTGAACTCTTTCCCTACACGACGCTCTTCCGATCTNNNAGAGAGAGAGGGAGAGAGAG
GCACACGTCTGAACTCTTTCCCTACACGACGCTCTTCCGATCTNNNGGGGAGAAGAGAGAGAGAGA
GAGCACACGTCTGAACTCTTTCCCTACACGACGCTCTTCCGATCTNNNGAGCGAGACTCCGTCTATAA
GAGCACACGTCTGAACTCTTTCCCTACACGACGCTCTTCCGATCTNNNCTCTCTCTCTCACACACACA
AGCACACGTCTGAACTCTTTCCCTACACGACGCTCTTCCGATCTNNNACACCTGGCCAGATACCTTTTT
GAGCACACGTCTGAACTCTTTCCCTACACGACGCTCTTCCGATCTNNNCAGCCCTTTGAGAGAGAGAG
GAGCACACGTCTGAACTCTTTCCCTACACGACGCTCTTCCGATCTNNNCATGTCTGAATCCCAGCACTT
AGCACACGTCTGAACTCTTTCCCTACACGACGCTCTTCCGATCTNNNAAACAAACAAGAAGCTTACCG
AGCACACGTCTGAACTCTTTCCCTACACGACGCTCTTCCGATCTNNNATTGTGCCATTGTACTCCATC
GCACACGTCTGAACTCTTTCCCTACACGACGCTCTTCCGATCTNNNGGGACCAGGGTGTGTGAATAT
GAGCACACGTCTGAACTCTTTCCCTACACGACGCTCTTCCGATCTNNNCAGGCAACATAGTGAGACCT
AGCACACGTCTGAACTCTTTCCCTACACGACGCTCTTCCGATCTNNNCAACAAGAGCGAAACTGTCT
GCACACGTCTGAACTCTTTCCCTACACGACGCTCTTCCGATCTNNNGGATCACTTGAGGTCAGGAGTT
AGCACACGTCTGAACTCTTTCCCTACACGACGCTCTTCCGATCTNNNTTCCAAAGTGCTGGGATTAG
CACACGTCTGAACTCTTTCCCTACACGACGCTCTTCCGATCTNNNAGCGAGTTTATAGACTTTGGACT
AGCACACGTCTGAACTCTTTCCCTACACGACGCTCTTCCGATCTNNNGAGAGAGAGAGAGAGACGA
CACACGTCTGAACTCTTTCCCTACACGACGCTCTTCCGATCTNNNGCAAACATACTGCAGTAACAGGA
AGCACACGTCTGAACTCTTTCCCTACACGACGCTCTTCCGATCTNNNCCAACCTGATCACAGAAGTTT
AGCACACGTCTGAACTCTTTCCCTACACGACGCTCTTCCGATCTNNNAGTCATCCAGGACTAGCAGTA
AGCACACGTCTGAACTCTTTCCCTACACGACGCTCTTCCGATCTNNNATCAACATGCATACACAAGCA
AGCACACGTCTGAACTCTTTCCCTACACGACGCTCTTCCGATCTNNNAGCCTGAGCAACATAGTAAG
AGCACACGTCTGAACTCTTTCCCTACACGACGCTCTTCCGATCTNNNTTCAAGTGATTCTCCTGCCTCA
AGCACACGTCTGAACTCTTTCCCTACACGACGCTCTTCCGATCTNNNCACAATTCGCAACTGCAAAAA
GAGCACACGTCTGAACTCTTTCCCTACACGACGCTCTTCCGATCTNNNAGAGAGGGAGAGAGAGAG
GCACACGTCTGAACTCTTTCCCTACACGACGCTCTTCCGATCTNNNTGACATGACGAAGCAGAACTG
GAGCACACGTCTGAACTCTTTCCCTACACGACGCTCTTCCGATCTNNNCTAGAGTAGCCATCCTTTTG
CACACGTCTGAACTCTTTCCCTACACGACGCTCTTCCGATCTNNNTCAGTGCATGGTTTGTATCAGGA
AGCACACGTCTGAACTCTTTCCCTACACGACGCTCTTCCGATCTNNNGAGATTTTACCTTTGCCTTGG
GAGCACACGTCTGAACTCTTTCCCTACACGACGCTCTTCCGATCTNNNCAGCAGCAGCAGAACAGTA
GCACACGTCTGAACTCTTTCCCTACACGACGCTCTTCCGATCTNNNTAAGGGGTAGGTAGAGGTTTC
GAGCACACGTCTGAACTCTTTCCCTACACGACGCTCTTCCGATCTNNNGAGATCGAGAACATCCTGGC
AGCACACGTCTGAACTCTTTCCCTACACGACGCTCTTCCGATCTNNNATTGGTGATGGTGACCTTGAT
AGCACACGTCTGAACTCTTTCCCTACACGACGCTCTTCCGATCTNNNTGACCCGGCTTATGAGTCTAA
AGCACACGTCTGAACTCTTTCCCTACACGACGCTCTTCCGATCTNNNTTTTATGAGGCCAGCATCATC
AGCACACGTCTGAACTCTTTCCCTACACGACGCTCTTCCGATCTNNNTCTTGGAAGGAAGTGATGGA

GAGCACACGTCTGAACTCTTTCCCTACACGACGCTCTTCCGATCTNNNTTGTTCACTGCAGCCTCAAATT

AGCACACGTCTGAACTCTTTCCCTACACGACGCTCTTCCGATCTNNNCATGACCACAGTGTGAAAGTTC

GAGCACACGTCTGAACTCTTTCCCTACACGACGCTCTTCCGATCTNNNATTACCATGCAGGGACTCTTG

GAGCACACGTCTGAACTCTTTCCCTACACGACGCTCTTCCGATCTNNNTACTGCAAGAGCCAGAGGTA

GAGCACACGTCTGAACTCTTTCCCTACACGACGCTCTTCCGATCTNNNCCTGTCTCTTCTTTTCCTCCCTC

GAGCACACGTCTGAACTCTTTCCCTACACGACGCTCTTCCGATCTNNNAGAGAGAGAGGAGGAGAGA

GAGCACACGTCTGAACTCTTTCCCTACACGACGCTCTTCCGATCTNNNGGGCAACAAGAGTGAAATTC

GAGCACACGTCTGAACTCTTTCCCTACACGACGCTCTTCCGATCTNNNTGTGTGGTGTGTGTACATAGA

AGCACACGTCTGAACTCTTTCCCTACACGACGCTCTTCCGATCTNNNTGACAAGTGAGTTGCCCAATTC

GAGCACACGTCTGAACTCTTTCCCTACACGACGCTCTTCCGATCTNNNTACGGATTTTGGATCTGTCAG

GAGCACACGTCTGAACTCTTTCCCTACACGACGCTCTTCCGATCTNNNCCTTTCACCTACCCTCCATACAC

AGCACACGTCTGAACTCTTTCCCTACACGACGCTCTTCCGATCTNNNGAGATAAGGGCCATCTCTCTGT

GAGCACACGTCTGAACTCTTTCCCTACACGACGCTCTTCCGATCTNNNTTGGAAAACCAGATGGACCTA

GAGCACACGTCTGAACTCTTTCCCTACACGACGCTCTTCCGATCTNNNTTTAGTAGAGACGGGGTTTCA

GAGCACACGTCTGAACTCTTTCCCTACACGACGCTCTTCCGATCTNNNCCCAGGTTTTTGTTTTTCCCACT

AGCACACGTCTGAACTCTTTCCCTACACGACGCTCTTCCGATCTNNNCAACTCATTCTTCCTCTCTCCCCCA

AGCACACGTCTGAACTCTTTCCCTACACGACGCTCTTCCGATCTNNNAGTTATAACCCCCACAACTGTGT

GAGCACACGTCTGAACTCTTTCCCTACACGACGCTCTTCCGATCTNNNGGGAGAGAGAGAGAGAGAG

AGCACACGTCTGAACTCTTTCCCTACACGACGCTCTTCCGATCTNNNTTTTCTCCAGTGAGCTCTGACTC

GAGCACACGTCTGAACTCTTTCCCTACACGACGCTCTTCCGATCTNNNGCCAAGACCTCTGTGCATTACT

GAGCACACGTCTGAACTCTTTCCCTACACGACGCTCTTCCGATCTNNNCAATCACCAGCTCTTTTCCAGTC

AGCACACGTCTGAACTCTTTCCCTACACGACGCTCTTCCGATCTNNNCAACATAGTGAAACCCCGTCTCT

GAGCACACGTCTGAACTCTTTCCCTACACGACGCTCTTCCGATCTNNNTGTCCAAATGGTGAAACTCCAT

AGCACACGTCTGAACTCTTTCCCTACACGACGCTCTTCCGATCTNNNTGTAATCCTAGCACTTTGGGAG

GAGCACACGTCTGAACTCTTTCCCTACACGACGCTCTTCCGATCTNNNCACACACACACACACAGAGTTT

AGCACACGTCTGAACTCTTTCCCTACACGACGCTCTTCCGATCTNNNCTCTGGGTTAGCTACTTGGTTGT

GAGCACACGTCTGAACTCTTTCCCTACACGACGCTCTTCCGATCTNNNCATGGACTTGTACACAGGAGC

GAGCACACGTCTGAACTCTTTCCCTACACGACGCTCTTCCGATCTNNNCACGCCCGGCTAATTTTTTTTT

AGCACACGTCTGAACTCTTTCCCTACACGACGCTCTTCCGATCTNNNATTTCTGAGGGGTTCATTCTCC

AGCACACGTCTGAACTCTTTCCCTACACGACGCTCTTCCGATCTNNNAAAGCAGGAATTCGCATTTCAC

GAGCACACGTCTGAACTCTTTCCCTACACGACGCTCTTCCGATCTNNNTCTTTCCAAACAACAGTCCATG

AGCACACGTCTGAACTCTTTCCCTACACGACGCTCTTCCGATCTNNNAAAGTGCTGGAATTACAGGCAT

AGAGCACACGTCTGAACTCTTTCCCTACACGACGCTCTTCCGATCTNNNTCGGAGGAATTTTGACAAG

AGCACACGTCTGAACTCTTTCCCTACACGACGCTCTTCCGATCTNNNTAGAAACTGCTCCTTGCCTTTTG

AGCACACGTCTGAACTCTTTCCCTACACGACGCTCTTCCGATCTNNNCCAGGAAGTCAAGGCTTCAGTA

CACACGTCTGAACTCTTTCCCTACACGACGCTCTTCCGATCTNNNGACCAAAAACAATGGGCTGACTCC

GAGCACACGTCTGAACTCTTTCCCTACACGACGCTCTTCCGATCTNNNTCTGAAAACACACACACACACA

AGCACACGTCTGAACTCTTTCCCTACACGACGCTCTTCCGATCTNNNTTTTCTTTTTCCTTCCTCCCTCCCA

AGAGCACACGTCTGAACTCTTTCCCTACACGACGCTCTTCCGATCTNNNCAGAGAGAACACACACAGA

AGAGCACACGTCTGAACTCTTTCCCTACACGACGCTCTTCCGATCTNNNGGATTACAGGTGTGAGCCAT

GAGCACACGTCTGAACTCTTTCCCTACACGACGCTCTTCCGATCTNNNCCTTGGACAGAGAGAGAGAG

AGCACACGTCTGAACTCTTTCCCTACACGACGCTCTTCCGATCTNNNATAGTGTGTGTGTGTGTGTGTC

GCACACGTCTGAACTCTTTCCCTACACGACGCTCTTCCGATCTNNNCTGGTTGTTTGATAAGCGTCTGG

AGCACACGTCTGAACTCTTTCCCTACACGACGCTCTTCCGATCTNNNGTGTGATCTCAGTTCACTGCAAC
AGCACACGTCTGAACTCTTTCCCTACACGACGCTCTTCCGATCTNNNTCAGATGATGCACAGTCAGGA
AGCACACGTCTGAACTCTTTCCCTACACGACGCTCTTCCGATCTNNNGAAATAGCCTGCTCGAGTTCTC
GAGCACACGTCTGAACTCTTTCCCTACACGACGCTCTTCCGATCTNNNTAGTTTCCCTGGCCTTTCCTTTT
AGCACACGTCTGAACTCTTTCCCTACACGACGCTCTTCCGATCTNNNAGCCTTTTCCACTGAGGTACAT
GAGCACACGTCTGAACTCTTTCCCTACACGACGCTCTTCCGATCTNNNAGCCATGAGTAGCAAGAGACA
AGCACACGTCTGAACTCTTTCCCTACACGACGCTCTTCCGATCTNNNCCCTGGTGATTCTTCTTCTTGGA
GAGCACACGTCTGAACTCTTTCCCTACACGACGCTCTTCCGATCTNNNGCCACAAGAATATTGCTGCAT
AGCACACGTCTGAACTCTTTCCCTACACGACGCTCTTCCGATCTNNNCCGAATAGCTGGGATTACAGGA
GAGCACACGTCTGAACTCTTTCCCTACACGACGCTCTTCCGATCTNNNATCATTCCAGTTTCCTCCACGTT
AGCACACGTCTGAACTCTTTCCCTACACGACGCTCTTCCGATCTNNNGATCATAGCTCACTACAGCCTC
AGCACACGTCTGAACTCTTTCCCTACACGACGCTCTTCCGATCTNNNAGTGGTGCAATCATAGCTCACT
GAGCACACGTCTGAACTCTTTCCCTACACGACGCTCTTCCGATCTNNNGGGTTACATGCTTTTGAGTCA
GAGCACACGTCTGAACTCTTTCCCTACACGACGCTCTTCCGATCTNNNCAGAAGAATTGCTTGAACCCA
AGCACACGTCTGAACTCTTTCCCTACACGACGCTCTTCCGATCTNNNTGGGCAACAAGAGAGAAACTCA
GAGCACACGTCTGAACTCTTTCCCTACACGACGCTCTTCCGATCTNNNGAGAATGGCATAAACCCAGGA
GAGCACACGTCTGAACTCTTTCCCTACACGACGCTCTTCCGATCTNNNGAGAATTGCTTGAACCCAGGA
AGAGCACACGTCTGAACTCTTTCCCTACACGACGCTCTTCCGATCTNNNTGAGCTATAATCACGCTACTG
AGCACACGTCTGAACTCTTTCCCTACACGACGCTCTTCCGATCTNNNCTGTAATCCCAGCTACTCAGGA
GAGCACACGTCTGAACTCTTTCCCTACACGACGCTCTTCCGATCTNNNGATCACAAGGTCAGGAGTTCG
GAGCACACGTCTGAACTCTTTCCCTACACGACGCTCTTCCGATCTNNNCTGTAGTCCCAGCTACTTGGAA
GAGCACACGTCTGAACTCTTTCCCTACACGACGCTCTTCCGATCTNNNAGTGAGCTGAGATCGTGCTAT
AGAGCACACGTCTGAACTCTTTCCCTACACGACGCTCTTCCGATCTNNNCTGCTTGAGGTTAGGAGTAC
GAGCACACGTCTGAACTCTTTCCCTACACGACGCTCTTCCGATCTNNNATTCCAGATCACTTCCTGCTCTC
GAGCACACGTCTGAACTCTTTCCCTACACGACGCTCTTCCGATCTNNNGTTGCAGTGAACTGAGACTG
AGAGCACACGTCTGAACTCTTTCCCTACACGACGCTCTTCCGATCTNNNAATACAAAAAATTAGCCGGG
AGCACACGTCTGAACTCTTTCCCTACACGACGCTCTTCCGATCTNNNCTGAGGCAGGAGAATTGTGTG
GAGCACACGTCTGAACTCTTTCCCTACACGACGCTCTTCCGATCTNNNTGAGGCAGGAGAATCACTTTC
GAGCACACGTCTGAACTCTTTCCCTACACGACGCTCTTCCGATCTNNNGTAATCCCAGCTACTTAGGAG
GAGCACACGTCTGAACTCTTTCCCTACACGACGCTCTTCCGATCTNNNTAAACAGCACAAAACAGCAAC
AGCACACGTCTGAACTCTTTCCCTACACGACGCTCTTCCGATCTNNNATCACTGTCTTCTCCCTGACATGC
ACACGTCTGAACTCTTTCCCTACACGACGCTCTTCCGATCTNNNAGAGAATTGAAAAAGTGCTCTTCCA
AGCACACGTCTGAACTCTTTCCCTACACGACGCTCTTCCGATCTNNNAAAATCTTTTCCCACCCCATGTG
AGCACACGTCTGAACTCTTTCCCTACACGACGCTCTTCCGATCTNNNAGATGGGAGCCTCTCTTCTAAC
GAGCACACGTCTGAACTCTTTCCCTACACGACGCTCTTCCGATCTNNNCGATTATCTTGCCTTAGCCTCCA
AGCACACGTCTGAACTCTTTCCCTACACGACGCTCTTCCGATCTNNNAGAATCCTGAGTAGCTGGGACT
AGCACACGTCTGAACTCTTTCCCTACACGACGCTCTTCCGATCTNNNAACCTCTCAAGTAGCTGGGACT
AGAGCACACGTCTGAACTCTTTCCCTACACGACGCTCTTCCGATCTNNNCCTTTCCACTCTTCCTGACTCTC
AGCACACGTCTGAACTCTTTCCCTACACGACGCTCTTCCGATCTNNNGAGTTTAGTGGTCCTGTCTCCAC
AGCACACGTCTGAACTCTTTCCCTACACGACGCTCTTCCGATCTNNNTCAGCCTCTCAAAGTGCTGATAT
AGCACACGTCTGAACTCTTTCCCTACACGACGCTCTTCCGATCTNNNGCCATCTCCAGTAGTTCAGGAA
AGCACACGTCTGAACTCTTTCCCTACACGACGCTCTTCCGATCTNNNGATTACAAGAGTAAGCCGCCAT
AGCACACGTCTGAACTCTTTCCCTACACGACGCTCTTCCGATCTNNNTTGGGATTACAGGTATGAGCCA

AGCACACGTCTGAACTCTTTCCCTACACGACGCTCTTCCGATCTNNNATTGAACCACATGCTACTGTTG
GAGCACACGTCTGAACTCTTTCCCTACACGACGCTCTTCCGATCTNNNACTATTTCCCACAGCTCATGT
CACACGTCTGAACTCTTTCCCTACACGACGCTCTTCCGATCTNNNTCTTGATCTCATACTCCCATAGGT
GCACACGTCTGAACTCTTTCCCTACACGACGCTCTTCCGATCTNNNGCTACATCTCTTCCCCGAAGTAG
AGCACACGTCTGAACTCTTTCCCTACACGACGCTCTTCCGATCTNNNACATGCCTGTAATCCCAGCTAT
AGCACACGTCTGAACTCTTTCCCTACACGACGCTCTTCCGATCTNNNATGTCTGTAATCCCAGCTCCTT
AGCACACGTCTGAACTCTTTCCCTACACGACGCTCTTCCGATCTNNNCAGATAACTTGAGGCCAGGAG
AGCACACGTCTGAACTCTTTCCCTACACGACGCTCTTCCGATCTNNNCTGTAGTCCCAGCTACTCAGG
GAGCACACGTCTGAACTCTTTCCCTACACGACGCTCTTCCGATCTNNNTGTAGTCTCAGCTACTAGGG
AGCACACGTCTGAACTCTTTCCCTACACGACGCTCTTCCGATCTNNNCTGTAGTCCCAGCTACTCAGGA
AGCACACGTCTGAACTCTTTCCCTACACGACGCTCTTCCGATCTNNNAAAGAACCTCCAAACTTTGCA
AGCACACGTCTGAACTCTTTCCCTACACGACGCTCTTCCGATCTNNNGGAGGTTACAGTGAGCTGAG
GAGCACACGTCTGAACTCTTTCCCTACACGACGCTCTTCCGATCTNNNGGGCAACATAATGAAACCCC
AGCACACGTCTGAACTCTTTCCCTACACGACGCTCTTCCGATCTNNNTCATCACACCACTGTAGTTCCA
GAGCACACGTCTGAACTCTTTCCCTACACGACGCTCTTCCGATCTNNNTACACGCCTGTAATCCCAACT
GAGCACACGTCTGAACTCTTTCCCTACACGACGCTCTTCCGATCTNNNGGAGAATGGTGTGAACCTG
GAGCACACGTCTGAACTCTTTCCCTACACGACGCTCTTCCGATCTNNNACCTTCCCTCCACTATTGTCCT
AGCACACGTCTGAACTCTTTCCCTACACGACGCTCTTCCGATCTNNNGCACCTGTAATCCCAGCTACTA
AGCACACGTCTGAACTCTTTCCCTACACGACGCTCTTCCGATCTNNNATGCCTGTAATCCCAGCTACTT
CACACGTCTGAACTCTTTCCCTACACGACGCTCTTCCGATCTNNNCTGAGGCAGAAGAATGGTGTGA
AGCACACGTCTGAACTCTTTCCCTACACGACGCTCTTCCGATCTNNNAGTGAGCTAAGATCGCACTAC
GAGCACACGTCTGAACTCTTTCCCTACACGACGCTCTTCCGATCTNNNAATGAGCTGAGATTGCACCA
AGCACACGTCTGAACTCTTTCCCTACACGACGCTCTTCCGATCTNNNAGAGGTTGCAGTGAGCCAATA
GCACACGTCTGAACTCTTTCCCTACACGACGCTCTTCCGATCTNNNGTTGTGTTGATCTGAGATCGCA
GAGCACACGTCTGAACTCTTTCCCTACACGACGCTCTTCCGATCTNNNAGGTTATAGTGAGCCAAGAT
AGCACACGTCTGAACTCTTTCCCTACACGACGCTCTTCCGATCTNNNGTTGCAGTGAGCTGAGATTTC
AGCACACGTCTGAACTCTTTCCCTACACGACGCTCTTCCGATCTNNNAGGCCACAGATGAGTCTCATT
AGCACACGTCTGAACTCTTTCCCTACACGACGCTCTTCCGATCTNNNCACCTGTAATCCCAGCTACTCA
AGCACACGTCTGAACTCTTTCCCTACACGACGCTCTTCCGATCTNNNATGCCTATAGTCCCAGCTTCTTG
AGCACACGTCTGAACTCTTTCCCTACACGACGCTCTTCCGATCTNNNGAACCTGGGAGAATTGCTTGA
AGCACACGTCTGAACTCTTTCCCTACACGACGCTCTTCCGATCTNNNGCGAGGCCCTGTCTCAATAA
AGCACACGTCTGAACTCTTTCCCTACACGACGCTCTTCCGATCTNNNCCCGCAGTGAGTTTTAGTTGT
AGCACACGTCTGAACTCTTTCCCTACACGACGCTCTTCCGATCTNNNCACCAGCAAACACAGTTATTCG
AGCACACGTCTGAACTCTTTCCCTACACGACGCTCTTCCGATCTNNNACCTCATCCTTCAAGTGCTCTTT
AGCACACGTCTGAACTCTTTCCCTACACGACGCTCTTCCGATCTNNNACCTCCTTTCTGCTTTCTGGAAT
AGCACACGTCTGAACTCTTTCCCTACACGACGCTCTTCCGATCTNNNGTGAAAAGGAGGCAGGTTTT
AGCACACGTCTGAACTCTTTCCCTACACGACGCTCTTCCGATCTNNNACTTGAACAGGCTATGGTTCCA
AGCACACGTCTGAACTCTTTCCCTACACGACGCTCTTCCGATCTNNNGGAGAAAAGTTCAGACAACGT
AGCACACGTCTGAACTCTTTCCCTACACGACGCTCTTCCGATCTNNNATGGCCTAATATGTAGTGGCTG
AGCACACGTCTGAACTCTTTCCCTACACGACGCTCTTCCGATCTNNNTGGCATGCCTTTGAACTTTGA
AGCACACGTCTGAACTCTTTCCCTACACGACGCTCTTCCGATCTNNNATTCTGCATTGGTTGGACAG
GCACACGTCTGAACTCTTTCCCTACACGACGCTCTTCCGATCTNNNACGGGCATGATTTGTAAAGTTG
GAGCACACGTCTGAACTCTTTCCCTACACGACGCTCTTCCGATCTNNNGACAATCACTTGAACCTGGG

GAGCACACGTCTGAACTCTTTCCCTACACGACGCTCTTCCGATCTNNNGTAATCCCAGCTACTTAGGAG
GAGCACACGTCTGAACTCTTTCCCTACACGACGCTCTTCCGATCTNNNAAGAAAAAAGGTAGCCAGGC
GAGCACACGTCTGAACTCTTTCCCTACACGACGCTCTTCCGATCTNNNGTCATGCACCTATAGTCCTAG
AGCACACGTCTGAACTCTTTCCCTACACGACGCTCTTCCGATCTNNNATCTGGGAGGTAGAGGTTTCA
AGCACACGTCTGAACTCTTTCCCTACACGACGCTCTTCCGATCTNNNGAGATCACACCATTGCACTCCAG
GAGCACACGTCTGAACTCTTTCCCTACACGACGCTCTTCCGATCTNNNGAGGCAGAAGAATCACTTGA
AGCACACGTCTGAACTCTTTCCCTACACGACGCTCTTCCGATCTNNNCTGAGGAAGGAGAATGGCAT
GAGCACACGTCTGAACTCTTTCCCTACACGACGCTCTTCCGATCTNNNCGAGATCATGCCATTGTACTC
GAGCACACGTCTGAACTCTTTCCCTACACGACGCTCTTCCGATCTNNNGTGACAGAGAGAAACTCCGT
GAGCACACGTCTGAACTCTTTCCCTACACGACGCTCTTCCGATCTNNNGTAACAGAGCAAGACCCTACC
AGAGCACACGTCTGAACTCTTTCCCTACACGACGCTCTTCCGATCTNNNTTATAAACTGTAGGGTGGCC
AGCACACGTCTGAACTCTTTCCCTACACGACGCTCTTCCGATCTNNNAGGAGGAGAATTGCTTGAATC
GAGCACACGTCTGAACTCTTTCCCTACACGACGCTCTTCCGATCTNNNCTAAAGAGGCTGAGGCATGA
GAGCACACGTCTGAACTCTTTCCCTACACGACGCTCTTCCGATCTNNNGCTAAGGCAGGAGAATCACTT
AGCACACGTCTGAACTCTTTCCCTACACGACGCTCTTCCGATCTNNNGTTGGATGATGTGATGTTTGC
GAGCACACGTCTGAACTCTTTCCCTACACGACGCTCTTCCGATCTNNNTATTTGGCGGTGAGAAGAGG
GAGCACACGTCTGAACTCTTTCCCTACACGACGCTCTTCCGATCTNNNGAGATCAAGGCATTGGCAGA
AGCACACGTCTGAACTCTTTCCCTACACGACGCTCTTCCGATCTNNNATACTCCTTTCAGCCACATCCTGC
AGCACACGTCTGAACTCTTTCCCTACACGACGCTCTTCCGATCTNNNCCAGACCTCCTGAATCACAGTAG
GAGCACACGTCTGAACTCTTTCCCTACACGACGCTCTTCCGATCTNNNCTCCTGAGTAGCTGGGACTAC
GAGCACACGTCTGAACTCTTTCCCTACACGACGCTCTTCCGATCTNNNACTCCTGACATCAAGTGATCCA
AGCACACGTCTGAACTCTTTCCCTACACGACGCTCTTCCGATCTNNNAGTGCCATCTTCAGAGGTTTCAT
GAGCACACGTCTGAACTCTTTCCCTACACGACGCTCTTCCGATCTNNNTCCTTGATGTGCCCTGAGATAA
GAGCACACGTCTGAACTCTTTCCCTACACGACGCTCTTCCGATCTNNNGACTTTCAGGAGTCTGGTATC
GAGCACACGTCTGAACTCTTTCCCTACACGACGCTCTTCCGATCTNNNCCTTGGGATTCTCCTCTGGAA
AGCACACGTCTGAACTCTTTCCCTACACGACGCTCTTCCGATCTNNNTCATCACTGCATCATACAACCCA
GAGCACACGTCTGAACTCTTTCCCTACACGACGCTCTTCCGATCTNNNTTCCCAGGTTCAAGTGATTCT
AGCACACGTCTGAACTCTTTCCCTACACGACGCTCTTCCGATCTNNNGGTGTTGTTTCAAATCCCAGCTT
GAGCACACGTCTGAACTCTTTCCCTACACGACGCTCTTCCGATCTNNNCTGAGATTATGGGAGTGAG
GAGCACACGTCTGAACTCTTTCCCTACACGACGCTCTTCCGATCTNNNTGTCCCAGTGATCAAGTCAGA
GAGCACACGTCTGAACTCTTTCCCTACACGACGCTCTTCCGATCTNNNTCTACCCCCACTAGATCCAACAT
AGCACACGTCTGAACTCTTTCCCTACACGACGCTCTTCCGATCTNNNCTCAAGCATTCTTCCCTCCTCTTC
AGCACACGTCTGAACTCTTTCCCTACACGACGCTCTTCCGATCTNNNGCTAGTTCTCCTTGGTGTTGGA
AGCACACGTCTGAACTCTTTCCCTACACGACGCTCTTCCGATCTNNNATAGTTTACAGCCAAAGGTGCA
AGCACACGTCTGAACTCTTTCCCTACACGACGCTCTTCCGATCTNNNACAGATAAAGAAACTGAGGCC
AGCACACGTCTGAACTCTTTCCCTACACGACGCTCTTCCGATCTNNNAAACACCTGACAGAGAAAGGG
AGCACACGTCTGAACTCTTTCCCTACACGACGCTCTTCCGATCTNNNGGTTTCTCAGCTACCTCACTTCA
GAGCACACGTCTGAACTCTTTCCCTACACGACGCTCTTCCGATCTNNNAACTGCTTGAACCTGGGAGAT
GAGCACACGTCTGAACTCTTTCCCTACACGACGCTCTTCCGATCTNNNTCACTTATGGCCAGGAATTCG
AGCACACGTCTGAACTCTTTCCCTACACGACGCTCTTCCGATCTNNNGGTGGAGGTTATAGTGAGCT
GAGCACACGTCTGAACTCTTTCCCTACACGACGCTCTTCCGATCTNNNAGGTGGGAAGAATACTTGAA
GCACACGTCTGAACTCTTTCCCTACACGACGCTCTTCCGATCTNNNATGCCTGTAATCCCAGCTACTTG
AGAGCACACGTCTGAACTCTTTCCCTACACGACGCTCTTCCGATCTNNNGTGCCTGTAATCCCAACTACT

AGCACACGTCTGAACTCTTTCCCTACACGACGCTCTTCCGATCTNNNAGTGAGTTGAGAGCTGGACA
GCACACGTCTGAACTCTTTCCCTACACGACGCTCTTCCGATCTNNNCTGAGATGGGTGGATCAACTG
AGCACACGTCTGAACTCTTTCCCTACACGACGCTCTTCCGATCTNNNAGATTGTGCCACTATACTCCA
AGCACACGTCTGAACTCTTTCCCTACACGACGCTCTTCCGATCTNNNCTGAGGCAGGAGAATCACTT
GCACACGTCTGAACTCTTTCCCTACACGACGCTCTTCCGATCTNNNGTACTCAAGAGGCTGAGACAG
GAGCACACGTCTGAACTCTTTCCCTACACGACGCTCTTCCGATCTNNNCTGTAATCCCAGCTACTCAGG
AGCACACGTCTGAACTCTTTCCCTACACGACGCTCTTCCGATCTNNNGAGATCACACCAATGCACTCC
GCACACGTCTGAACTCTTTCCCTACACGACGCTCTTCCGATCTNNNTGCATCAGTAATCCCAGCTACTT
GAGCACACGTCTGAACTCTTTCCCTACACGACGCTCTTCCGATCTNNNACCCAAATGAGAAGGAGTG
GCACACGTCTGAACTCTTTCCCTACACGACGCTCTTCCGATCTNNNAGTGCCAAGCAGAACACATAGT
AGCACACGTCTGAACTCTTTCCCTACACGACGCTCTTCCGATCTNNNGGCTGGAGTTCAATGGCATA
AGCACACGTCTGAACTCTTTCCCTACACGACGCTCTTCCGATCTNNNTAGGATTGAGGATTGCAACA
GCACACGTCTGAACTCTTTCCCTACACGACGCTCTTCCGATCTNNNCCAATTCAGGAAGCCTCACTCT
AGCACACGTCTGAACTCTTTCCCTACACGACGCTCTTCCGATCTNNNATTTGACTGGATAATGCAGCC
AGCACACGTCTGAACTCTTTCCCTACACGACGCTCTTCCGATCTNNNTGCATGACTACACTGCATACC
AGCACACGTCTGAACTCTTTCCCTACACGACGCTCTTCCGATCTNNNTGGAGGCTATAAAAAGGTGG
GCACACGTCTGAACTCTTTCCCTACACGACGCTCTTCCGATCTNNNGCTACAACTACAGGACAATGAA
GCACACGTCTGAACTCTTTCCCTACACGACGCTCTTCCGATCTNNNTTGGGATTACAGACATGAGCC
GCACACGTCTGAACTCTTTCCCTACACGACGCTCTTCCGATCTNNNTTTCTTCTTGCACAAGCCTCAA
GAGCACACGTCTGAACTCTTTCCCTACACGACGCTCTTCCGATCTNNNTTGTGAACACCATGTAGCCA
GAGCACACGTCTGAACTCTTTCCCTACACGACGCTCTTCCGATCTNNNTGAGACTTGGGATGATTGG
AGCACACGTCTGAACTCTTTCCCTACACGACGCTCTTCCGATCTNNNTCTCCTGTCTTAGACTCCCAAG
AGCACACGTCTGAACTCTTTCCCTACACGACGCTCTTCCGATCTNNNAAACTCTTGACCTCAGGTGAT
GCACACGTCTGAACTCTTTCCCTACACGACGCTCTTCCGATCTNNNTTGCCAAGGCTGATGTCTAGA
AGCACACGTCTGAACTCTTTCCCTACACGACGCTCTTCCGATCTNNNTGTAATACAATCGCCCCTTTCC
GAGCACACGTCTGAACTCTTTCCCTACACGACGCTCTTCCGATCTNNNAGCACCAGGAACAGTATTCCT
GCACACGTCTGAACTCTTTCCCTACACGACGCTCTTCCGATCTNNNAAGTAGCTGGGATTACAGGCAT
GAGCACACGTCTGAACTCTTTCCCTACACGACGCTCTTCCGATCTNNNTCCTGTTTTGCATACGAACA
GAGCACACGTCTGAACTCTTTCCCTACACGACGCTCTTCCGATCTNNNGTGATACGAACATGGCTTACT
GAGCACACGTCTGAACTCTTTCCCTACACGACGCTCTTCCGATCTNNNCGTCAGGAATTCGTTGTTAT
GCACACGTCTGAACTCTTTCCCTACACGACGCTCTTCCGATCTNNNTCCATGTTCTTGACCTAAAGTG
AGCACACGTCTGAACTCTTTCCCTACACGACGCTCTTCCGATCTNNNCTGAAGGAGAGTGCAGCATAG
GCACACGTCTGAACTCTTTCCCTACACGACGCTCTTCCGATCTNNNTGCAAATAAAAGCCACAATGAC
GCACACGTCTGAACTCTTTCCCTACACGACGCTCTTCCGATCTNNNCACCAACCCATCCCAAAGAAAA
AGCACACGTCTGAACTCTTTCCCTACACGACGCTCTTCCGATCTNNNGTTATCCCAGCTATGTGAGAG
AGCACACGTCTGAACTCTTTCCCTACACGACGCTCTTCCGATCTNNNTACTCAGGAGGCTGATCAGA
GAGCACACGTCTGAACTCTTTCCCTACACGACGCTCTTCCGATCTNNNTGATCAGTCATTTGGCTCCTG
AGCACACGTCTGAACTCTTTCCCTACACGACGCTCTTCCGATCTNNNCCAACTACTGGGTAAGCTGAA
GAGCACACGTCTGAACTCTTTCCCTACACGACGCTCTTCCGATCTNNNTTTCCCAGTATGTAGGTGAA
AGCACACGTCTGAACTCTTTCCCTACACGACGCTCTTCCGATCTNNNTCCTAGCTACTTAGGAGGCTG
AGCACACGTCTGAACTCTTTCCCTACACGACGCTCTTCCGATCTNNNGTTGCAGTGAGCTGAGATCAT
GCACACGTCTGAACTCTTTCCCTACACGACGCTCTTCCGATCTNNNCTTTCAGTAAGCCGAGATCGTG
GAGCACACGTCTGAACTCTTTCCCTACACGACGCTCTTCCGATCTNNNCCAACATGAGAAACCCCATCT

AGCACACGTCTGAACTCTTTCCCTACACGACGCTCTTCCGATCTNNNGAGGTTGCAGTGAGTCGATA
AGCACACGTCTGAACTCTTTCCCTACACGACGCTCTTCCGATCTNNNAGAATTGCTTGAACCAGGGA
AGCACACGTCTGAACTCTTTCCCTACACGACGCTCTTCCGATCTNNNGGAGAATCGCTTGAATCCGAG
GAGCACACGTCTGAACTCTTTCCCTACACGACGCTCTTCCGATCTNNNGTTGAGGCTGTACTGACCTA
GAGCACACGTCTGAACTCTTTCCCTACACGACGCTCTTCCGATCTNNNAGAATCTCTTGAATCCAGGA
AGCACACGTCTGAACTCTTTCCCTACACGACGCTCTTCCGATCTNNNGTACTAACAGCTACGAGAGAG
GAGCACACGTCTGAACTCTTTCCCTACACGACGCTCTTCCGATCTNNNAGAATTGCTTGAATCTGGGA
AGCACACGTCTGAACTCTTTCCCTACACGACGCTCTTCCGATCTNNNATGCAGGAGAATCACTTGAAC
GCACACGTCTGAACTCTTTCCCTACACGACGCTCTTCCGATCTNNNGCAACAGTGTAGGACTCTCTCA
GCACACGTCTGAACTCTTTCCCTACACGACGCTCTTCCGATCTNNNAGTCAGTGAATCGTGATCTCAC
AGCACACGTCTGAACTCTTTCCCTACACGACGCTCTTCCGATCTNNNCTGTAGTTCCAGCTGCTAAGG
AGCACACGTCTGAACTCTTTCCCTACACGACGCTCTTCCGATCTNNNACTTGAACATAGGAGGCAAA
GCACACGTCTGAACTCTTTCCCTACACGACGCTCTTCCGATCTNNNCTGAGGCAGGAGAATCACTTG
AGCACACGTCTGAACTCTTTCCCTACACGACGCTCTTCCGATCTNNNAGAATCACTTGAACCAGGGA
GAGCACACGTCTGAACTCTTTCCCTACACGACGCTCTTCCGATCTNNNACCAGCCTGGGTAACATATT
GAGCACACGTCTGAACTCTTTCCCTACACGACGCTCTTCCGATCTNNNCGAGGTCAGGAGTTCAAGAT
AGCACACGTCTGAACTCTTTCCCTACACGACGCTCTTCCGATCTNNNGAGAATGGTGTGAACCCAGA
GAGCACACGTCTGAACTCTTTCCCTACACGACGCTCTTCCGATCTNNNTACTCAGGAGACTAAGGTGG
AGCACACGTCTGAACTCTTTCCCTACACGACGCTCTTCCGATCTNNNTTGAGACAGAAGAATCGCCTG
GCACACGTCTGAACTCTTTCCCTACACGACGCTCTTCCGATCTNNNATAGTCCCAGCTACTAGAGAGG
AGCACACGTCTGAACTCTTTCCCTACACGACGCTCTTCCGATCTNNNCAAGGCAGGAGAGTCATTTGA
GAGCACACGTCTGAACTCTTTCCCTACACGACGCTCTTCCGATCTNNNTGCAGTTCCAGCTACTTAGA
AGCACACGTCTGAACTCTTTCCCTACACGACGCTCTTCCGATCTNNNGAGGCAGGAGAATTGTTTGA
AGCACACGTCTGAACTCTTTCCCTACACGACGCTCTTCCGATCTNNNGAATATTCAGTACCCGAGGAG
AGCACACGTCTGAACTCTTTCCCTACACGACGCTCTTCCGATCTNNNCTGAGGCAGGAGAATCACTTG
GCACACGTCTGAACTCTTTCCCTACACGACGCTCTTCCGATCTNNNAAGACCCATCAGTGTGCTGTAT
GCACACGTCTGAACTCTTTCCCTACACGACGCTCTTCCGATCTNNNATAGTCCCAGCTACTCTAGAGG
AGCACACGTCTGAACTCTTTCCCTACACGACGCTCTTCCGATCTNNNTTGCAGTGAGCCAAAATTGTA
GCACACGTCTGAACTCTTTCCCTACACGACGCTCTTCCGATCTNNNTGAGCTGAGATCATACCATTGC
GCACACGTCTGAACTCTTTCCCTACACGACGCTCTTCCGATCTNNNATGTTGCAGTGAGCCAAGATTT
GCACACGTCTGAACTCTTTCCCTACACGACGCTCTTCCGATCTNNNCACAAGCTACTTGGGAGGTTGA
AGCACACGTCTGAACTCTTTCCCTACACGACGCTCTTCCGATCTNNNATGCCTGTAATCCCAGCTACTT
AGCACACGTCTGAACTCTTTCCCTACACGACGCTCTTCCGATCTNNNGTCACCTCCAGTCATTGAGCT
GCACACGTCTGAACTCTTTCCCTACACGACGCTCTTCCGATCTNNNAGCAGTGAAGCAGAGTGGAA
GAGCACACGTCTGAACTCTTTCCCTACACGACGCTCTTCCGATCTNNNCCATTGATGTTGTCATTGCCA
AGCACACGTCTGAACTCTTTCCCTACACGACGCTCTTCCGATCTNNNAAGTGTTGGGATTACAGGTGT
GCACACGTCTGAACTCTTTCCCTACACGACGCTCTTCCGATCTNNNTTTCTAAGTGAGGATGACCCAG
AGCACACGTCTGAACTCTTTCCCTACACGACGCTCTTCCGATCTNNNCAGCAATCCTACCACCTCATTCT
GAGCACACGTCTGAACTCTTTCCCTACACGACGCTCTTCCGATCTNNNGCCCTCTGGTGACTACATTA
GCACACGTCTGAACTCTTTCCCTACACGACGCTCTTCCGATCTNNNGATTACAGGGATGAGCCACAG
AGCACACGTCTGAACTCTTTCCCTACACGACGCTCTTCCGATCTNNNAAGTATGGAGATAGAGTCCCG
GCACACGTCTGAACTCTTTCCCTACACGACGCTCTTCCGATCTNNNGTGGCTGTAACTGATCACATTG
GCACACGTCTGAACTCTTTCCCTACACGACGCTCTTCCGATCTNNNTTCAAGTGATTCTCCCACCTCAG

AGCACACGTCTGAACTCTTTCCCTACACGACGCTCTTCCGATCTNNNACCACGCTAGGCTGATTTTG
GAGCACACGTCTGAACTCTTTCCCTACACGACGCTCTTCCGATCTNNNTTACACTTGGTTTGTCACGTG
AGCACACGTCTGAACTCTTTCCCTACACGACGCTCTTCCGATCTNNNTAAATGTTCTCATCCTGTGGC
GCACACGTCTGAACTCTTTCCCTACACGACGCTCTTCCGATCTNNNCTGTCCATCCACAAAGCTGTTTT
AGCACACGTCTGAACTCTTTCCCTACACGACGCTCTTCCGATCTNNNACTCTGTCACCCAGTCTAGAG
AGCACACGTCTGAACTCTTTCCCTACACGACGCTCTTCCGATCTNNNGGTGCCAGTTACCAATTCAAGA
AGCACACGTCTGAACTCTTTCCCTACACGACGCTCTTCCGATCTNNNATGTTTTCCAGGCTGGTTTCA
GCACACGTCTGAACTCTTTCCCTACACGACGCTCTTCCGATCTNNNTAAAATTCAGGTCTCCTAGCCG
AGCACACGTCTGAACTCTTTCCCTACACGACGCTCTTCCGATCTNNNTTCTATAGTTCCTCAGCAGTGG
GAGCACACGTCTGAACTCTTTCCCTACACGACGCTCTTCCGATCTNNNCAAGTAGTTGGGATTACAGG
GCACACGTCTGAACTCTTTCCCTACACGACGCTCTTCCGATCTNNNTACATCTTGTGCTGAAGGATGC
GCACACGTCTGAACTCTTTCCCTACACGACGCTCTTCCGATCTNNNGTTGGCGATGCTATAGTACCTC
AGCACACGTCTGAACTCTTTCCCTACACGACGCTCTTCCGATCTNNNTCCACCAGCATTGTATGAGAGT
AGCACACGTCTGAACTCTTTCCCTACACGACGCTCTTCCGATCTNNNTAATTTAGAAGCCCAGCCAGC
AGCACACGTCTGAACTCTTTCCCTACACGACGCTCTTCCGATCTNNNTGAACTCAAGCGTTCTCCCATA
GAGCACACGTCTGAACTCTTTCCCTACACGACGCTCTTCCGATCTNNNAATAGCTGGGATTACAGTCG
AGCACACGTCTGAACTCTTTCCCTACACGACGCTCTTCCGATCTNNNCTACCACTAGGCTTGTTCAGTG
GAGCACACGTCTGAACTCTTTCCCTACACGACGCTCTTCCGATCTNNNATTTAGCCAGTCCTTCCAGAT
AGCACACGTCTGAACTCTTTCCCTACACGACGCTCTTCCGATCTNNNGGACAGAGTGCAGATTTTCT
GCACACGTCTGAACTCTTTCCCTACACGACGCTCTTCCGATCTNNNGTCCTTGACGTGAACTTTCCAT
AGCACACGTCTGAACTCTTTCCCTACACGACGCTCTTCCGATCTNNNACAGTGTCAGTTTCATTCCTTCG
GCACACGTCTGAACTCTTTCCCTACACGACGCTCTTCCGATCTNNNAGATGGTTGCCATTGTACTCCAT
AGCACACGTCTGAACTCTTTCCCTACACGACGCTCTTCCGATCTNNNGTGTGCCTATAATCCCAGCTAC
GCACACGTCTGAACTCTTTCCCTACACGACGCTCTTCCGATCTNNNCAGCTCATACCTGTAATCCCAG
GCACACGTCTGAACTCTTTCCCTACACGACGCTCTTCCGATCTNNNCTGAGGCAGAAGAATGGTGTG
AGAGCACACGTCTGAACTCTTTCCCTACACGACGCTCTTCCGATCTNNNGTGCCTGTATTCCCAGCTA
AGCACACGTCTGAACTCTTTCCCTACACGACGCTCTTCCGATCTNNNTGAGGTCAGGAGTTAGAAAC
AGCACACGTCTGAACTCTTTCCCTACACGACGCTCTTCCGATCTNNNGAAATCACACCATTGCACTCCA
AGCACACGTCTGAACTCTTTCCCTACACGACGCTCTTCCGATCTNNNGTGGCTCATACCTGTAATCCCA
AGCACACGTCTGAACTCTTTCCCTACACGACGCTCTTCCGATCTNNNAGCTACTTGGAAGGCTAAGAT
AGCACACGTCTGAACTCTTTCCCTACACGACGCTCTTCCGATCTNNNTGTGAGGCTGAGGTGAAAGA
AGCACACGTCTGAACTCTTTCCCTACACGACGCTCTTCCGATCTNNNAGTATCTCTGGCTTCTACCCACT
AGCACACGTCTGAACTCTTTCCCTACACGACGCTCTTCCGATCTNNNGACATTGCAGTGAGCCAAGAT
GAGCACACGTCTGAACTCTTTCCCTACACGACGCTCTTCCGATCTNNNGAATTCGAAATCAACCTGGC
AGCACACGTCTGAACTCTTTCCCTACACGACGCTCTTCCGATCTNNNGAGAATCACTTGAACCTGGGA
AGCACACGTCTGAACTCTTTCCCTACACGACGCTCTTCCGATCTNNNATGATGATGGGTGCCTCTAAT
GCACACGTCTGAACTCTTTCCCTACACGACGCTCTTCCGATCTNNNTGAGACAGGAGAATCGCTTGA
AGCACACGTCTGAACTCTTTCCCTACACGACGCTCTTCCGATCTNNNCAGGAGAATTGCTTGAACCTT
AGCACACGTCTGAACTCTTTCCCTACACGACGCTCTTCCGATCTNNNTATTTGGAAGGCTGAGGCAA
AGCACACGTCTGAACTCTTTCCCTACACGACGCTCTTCCGATCTNNNCCGAGATTGTGCCATTACATT
AGCACACGTCTGAACTCTTTCCCTACACGACGCTCTTCCGATCTNNNTCACTTGAACCTGGGAGATGA
GAGCACACGTCTGAACTCTTTCCCTACACGACGCTCTTCCGATCTNNNAGGAAGGAGAATCGTTTGA
GCACACGTCTGAACTCTTTCCCTACACGACGCTCTTCCGATCTNNNCTAACTTGGGTGACAGAGCAAG

GAGCACACGTCTGAACTCTTTCCCTACACGACGCTCTTCCGATCTNNNGAGAATCTCCTGAACCCCAAA
AAGAGCACACGTCTGAACTCTTTCCCTACACGACGCTCTTCCGATCTNNNCTAAGTTTGAGCCACTGCAT
AGCACACGTCTGAACTCTTTCCCTACACGACGCTCTTCCGATCTNNNCTGTAATCCCAGCTACTCGAGA
GAGCACACGTCTGAACTCTTTCCCTACACGACGCTCTTCCGATCTNNNCTGAGGCAGGAAAATTGCTTG
AGCACACGTCTGAACTCTTTCCCTACACGACGCTCTTCCGATCTNNNGCAGGAGAATTGCTTGAATCCA
GAGCACACGTCTGAACTCTTTCCCTACACGACGCTCTTCCGATCTNNNCCAGGAGATTGAGAATGTGG
GAGCACACGTCTGAACTCTTTCCCTACACGACGCTCTTCCGATCTNNNGGCTGAGACAGAAGAATTG
GAGCACACGTCTGAACTCTTTCCCTACACGACGCTCTTCCGATCTNNNATCCCACCTACTCAAGAGACTG
GAGCACACGTCTGAACTCTTTCCCTACACGACGCTCTTCCGATCTNNNGGTTGCAGTGAGCTGTTATCA
AGCACACGTCTGAACTCTTTCCCTACACGACGCTCTTCCGATCTNNNTCTGAGTCAGAGCTGAGTGAG
GAGCACACGTCTGAACTCTTTCCCTACACGACGCTCTTCCGATCTNNNAAGAGATTGAGGCAGGAGAA
AAGAGCACACGTCTGAACTCTTTCCCTACACGACGCTCTTCCGATCTNNNCGTGACTCCCAGATTCAGGA
GAGCACACGTCTGAACTCTTTCCCTACACGACGCTCTTCCGATCTNNNGGCACCTATAATCCCAGCTACT
GAGCACACGTCTGAACTCTTTCCCTACACGACGCTCTTCCGATCTNNNTTGAGGTCAGGAGTTCAAAA
GAGCACACGTCTGAACTCTTTCCCTACACGACGCTCTTCCGATCTNNNCCCAAAAGGTTGAAGCTGTA
AGCACACGTCTGAACTCTTTCCCTACACGACGCTCTTCCGATCTNNNCGTGTCTGTAGTCCCAGCTATT
GAGCACACGTCTGAACTCTTTCCCTACACGACGCTCTTCCGATCTNNNACGATCGACTTCCACTAAATGA
AGCACACGTCTGAACTCTTTCCCTACACGACGCTCTTCCGATCTNNNGGAAAAACCCTACTGAAGTTG
AGCACACGTCTGAACTCTTTCCCTACACGACGCTCTTCCGATCTNNNTGAAGGCAGCACTTATACCTGA
AGCACACGTCTGAACTCTTTCCCTACACGACGCTCTTCCGATCTNNNTTTAGTTGGCTCACCGTTCCATA
GAGCACACGTCTGAACTCTTTCCCTACACGACGCTCTTCCGATCTNNNGGCCACATGGTAAAATTTCCC
AGCACACGTCTGAACTCTTTCCCTACACGACGCTCTTCCGATCTNNNACCTTTAGTAAATCTGGCCAGCA
AGCACACGTCTGAACTCTTTCCCTACACGACGCTCTTCCGATCTNNNCTGAGGCAGGAGAATTGCTTGA
GAGCACACGTCTGAACTCTTTCCCTACACGACGCTCTTCCGATCTNNNGCTGAGACAGGAGAATCACTT
AGCACACGTCTGAACTCTTTCCCTACACGACGCTCTTCCGATCTNNNTGTAGTCCTAGCTACTCAGGAG
GAGCACACGTCTGAACTCTTTCCCTACACGACGCTCTTCCGATCTNNNCATGTGGCTGTACTCCAGTTA
GAGCACACGTCTGAACTCTTTCCCTACACGACGCTCTTCCGATCTNNNTATGTAGGCTGAGGCAGGAG
GAGCACACGTCTGAACTCTTTCCCTACACGACGCTCTTCCGATCTNNNTGTAATCCCAACACTCTGGAA
AGCACACGTCTGAACTCTTTCCCTACACGACGCTCTTCCGATCTNNNGCAGTGAGCCAAGATCTCAAG
GAGCACACGTCTGAACTCTTTCCCTACACGACGCTCTTCCGATCTNNNCTCTTTTTTTGGGCAGGGAG
AGCACACGTCTGAACTCTTTCCCTACACGACGCTCTTCCGATCTNNNACCACAAACTGCACCCATATGAT
AGCACACGTCTGAACTCTTTCCCTACACGACGCTCTTCCGATCTNNNCCCACCTGGAGTCAGTATACTT
AGCACACGTCTGAACTCTTTCCCTACACGACGCTCTTCCGATCTNNNTGCTGAAACGAGCACTTTTTTCT
AGCACACGTCTGAACTCTTTCCCTACACGACGCTCTTCCGATCTNNNCTCTGCAGAAACCAGACTCTTCT
AGCACACGTCTGAACTCTTTCCCTACACGACGCTCTTCCGATCTNNNGAGATGAGGTTTCACCATGTT
GAGCACACGTCTGAACTCTTTCCCTACACGACGCTCTTCCGATCTNNNTGCTTTTCAAGACATGTCTGCA
GAGCACACGTCTGAACTCTTTCCCTACACGACGCTCTTCCGATCTNNNGCTCATGCCTATAATTCCAGCA
AGCACACGTCTGAACTCTTTCCCTACACGACGCTCTTCCGATCTNNNACTCCTAGACTCCAGAGATCTG
GAGCACACGTCTGAACTCTTTCCCTACACGACGCTCTTCCGATCTNNNTTAGTGAAGCTCTACACTGAG
AGCACACGTCTGAACTCTTTCCCTACACGACGCTCTTCCGATCTNNNACTCCTGTTCATAGCAGCCTATT
AGCACACGTCTGAACTCTTTCCCTACACGACGCTCTTCCGATCTNNNTAGAGCTGTCTGGTTAGTAGGG
AGCACACGTCTGAACTCTTTCCCTACACGACGCTCTTCCGATCTNNNTTTCTCTGTCTCCCTCCAACTTGCA
AGCACACGTCTGAACTCTTTCCCTACACGACGCTCTTCCGATCTNNNAAGTGTTGGGATTACAGGCATG

AGCACACGTCTGAACTCTTTCCCTACACGACGCTCTTCCGATCTNNNCCGTGCCTGGCTAATTTTTG
GCACACGTCTGAACTCTTTCCCTACACGACGCTCTTCCGATCTNNNCTCACTACAGCTTCGAACTCCTG
AGCACACGTCTGAACTCTTTCCCTACACGACGCTCTTCCGATCTNNNCTGTACTAGTCATTGAGGGCA
AGCACACGTCTGAACTCTTTCCCTACACGACGCTCTTCCGATCTNNNAAGGCCACGTATACAGTGAAT
AGCACACGTCTGAACTCTTTCCCTACACGACGCTCTTCCGATCTNNNATCCATGTTGCTGCAAATGACA
GAGCACACGTCTGAACTCTTTCCCTACACGACGCTCTTCCGATCTNNNCCATTTGTTTGTGTTGTCTGG
GCACACGTCTGAACTCTTTCCCTACACGACGCTCTTCCGATCTNNNTCCTTTCTGCTGTGTGTATCCTC
AGCACACGTCTGAACTCTTTCCCTACACGACGCTCTTCCGATCTNNNATTCTCCTGTCTCAGCTTCCAA
AGCACACGTCTGAACTCTTTCCCTACACGACGCTCTTCCGATCTNNNGTTGGTTTCTGGTAAGGCCTTT
AGCACACGTCTGAACTCTTTCCCTACACGACGCTCTTCCGATCTNNNCCCTTCCTCTATCCATTGTCTG
GAGCACACGTCTGAACTCTTTCCCTACACGACGCTCTTCCGATCTNNNAGTAGCTGGGATTACAGACA
AGCACACGTCTGAACTCTTTCCCTACACGACGCTCTTCCGATCTNNNCTCTTGGGTTCATGCCATTCTC
GCACACGTCTGAACTCTTTCCCTACACGACGCTCTTCCGATCTNNNCATGAATTCTGTTTGCAGCACC
AGCACACGTCTGAACTCTTTCCCTACACGACGCTCTTCCGATCTNNNGATTGAGACCATCCTGGCTAA
GCACACGTCTGAACTCTTTCCCTACACGACGCTCTTCCGATCTNNNACCTGAGGACTGAATACACAAC
AGCACACGTCTGAACTCTTTCCCTACACGACGCTCTTCCGATCTNNNTGTAAAATCCTCTACAGGCCTG
GCACACGTCTGAACTCTTTCCCTACACGACGCTCTTCCGATCTNNNAGGCCACTGATACCTAAAACCTT
AGCACACGTCTGAACTCTTTCCCTACACGACGCTCTTCCGATCTNNNTTGTCTCAATGGAGTCAAAGC
GCACACGTCTGAACTCTTTCCCTACACGACGCTCTTCCGATCTNNNATGTGGGTATCAACAGCACTCT
GAGCACACGTCTGAACTCTTTCCCTACACGACGCTCTTCCGATCTNNNCCAGAACTTAGTACGTGGCTT
AGCACACGTCTGAACTCTTTCCCTACACGACGCTCTTCCGATCTNNNTCCTGTAATCCCAGCTACTAGG
GAGCACACGTCTGAACTCTTTCCCTACACGACGCTCTTCCGATCTNNNTAAAGCAGGAGAATCGCTT
AGCACACGTCTGAACTCTTTCCCTACACGACGCTCTTCCGATCTNNNAGTGAGCTGAGATCATGCCAT
GAGCACACGTCTGAACTCTTTCCCTACACGACGCTCTTCCGATCTNNNGAGGCTAAGGCAGAAGAAT
GAGCACACGTCTGAACTCTTTCCCTACACGACGCTCTTCCGATCTNNNGGGCAACAGAATAAGACCCT
GAGCACACGTCTGAACTCTTTCCCTACACGACGCTCTTCCGATCTNNNCTGAGGCAAAGAACTGCTTG
GCACACGTCTGAACTCTTTCCCTACACGACGCTCTTCCGATCTNNNCTGTAATCTCAGCTACTCCGGAG
GCACACGTCTGAACTCTTTCCCTACACGACGCTCTTCCGATCTNNNCAGAATTGTTTGAACCCAGGAG
AGAGCACACGTCTGAACTCTTTCCCTACACGACGCTCTTCCGATCTNNNAGGCTGAGGCATGATAATC
GAGCACACGTCTGAACTCTTTCCCTACACGACGCTCTTCCGATCTNNNGAGGCAGAAAATTGCTTGA
GAGCACACGTCTGAACTCTTTCCCTACACGACGCTCTTCCGATCTNNNAGGCAGAAGAATCACTTGA
GAGCACACGTCTGAACTCTTTCCCTACACGACGCTCTTCCGATCTNNNCAAAAATTAGCCGGGAATGG
AGCACACGTCTGAACTCTTTCCCTACACGACGCTCTTCCGATCTNNNCAGAGGACTGGAATTGCTTG
GCACACGTCTGAACTCTTTCCCTACACGACGCTCTTCCGATCTNNNCCAGGAAACTCCAACAGGAAGA
GAGCACACGTCTGAACTCTTTCCCTACACGACGCTCTTCCGATCTNNNGTAATCCCAGCTACTTAGGA
AGCACACGTCTGAACTCTTTCCCTACACGACGCTCTTCCGATCTNNNGCCAAGATCACACCATTTCACT
GCACACGTCTGAACTCTTTCCCTACACGACGCTCTTCCGATCTNNNGGGCAACAGAGTGAGACTGTA
AGCACACGTCTGAACTCTTTCCCTACACGACGCTCTTCCGATCTNNNGCAGGTATGTGTAATCCCAGC
GCACACGTCTGAACTCTTTCCCTACACGACGCTCTTCCGATCTNNNGAGCTTGCAGTGAACTGAGAT
AGCACACGTCTGAACTCTTTCCCTACACGACGCTCTTCCGATCTNNNCTAGGCAACAAGAGCAAAAC
AGCACACGTCTGAACTCTTTCCCTACACGACGCTCTTCCGATCTNNNGCAAAAATTCCAGAGCTGCCT
GAGCACACGTCTGAACTCTTTCCCTACACGACGCTCTTCCGATCTNNNTGAGGTAGGAGAATGGCAT
AGCACACGTCTGAACTCTTTCCCTACACGACGCTCTTCCGATCTNNNCTGTAGTCCCAGCTACTCAGGA

GAGCACACGTCTGAACTCTTTCCCTACACGACGCTCTTCCGATCTNNNTACAAAAAGTCCTGAAGGGAG
GAGCACACGTCTGAACTCTTTCCCTACACGACGCTCTTCCGATCTNNNGAGATTGCAGTGAGCCGAGA
GAGCACACGTCTGAACTCTTTCCCTACACGACGCTCTTCCGATCTNNNGATCACCTGAGGTTGAGACTA
GAGCACACGTCTGAACTCTTTCCCTACACGACGCTCTTCCGATCTNNNCTGAGGCAGGAGTATTGCTTG
GAGCACACGTCTGAACTCTTTCCCTACACGACGCTCTTCCGATCTNNNCTGAGGCAGGAGAATTGCTTG
GAGCACACGTCTGAACTCTTTCCCTACACGACGCTCTTCCGATCTNNNGATAATGCGTGTGGCTTTTGT
GAGCACACGTCTGAACTCTTTCCCTACACGACGCTCTTCCGATCTNNNGAGAATGGCATGAACCCAAGA
AGAGCACACGTCTGAACTCTTTCCCTACACGACGCTCTTCCGATCTNNNAGGCAAGAGAATCCATTGAA
AGCACACGTCTGAACTCTTTCCCTACACGACGCTCTTCCGATCTNNNCTGAGACAGGAGAATCGCTTGA
AGCACACGTCTGAACTCTTTCCCTACACGACGCTCTTCCGATCTNNNAAGATTGCAGTGAGCTGAGAT
GAGCACACGTCTGAACTCTTTCCCTACACGACGCTCTTCCGATCTNNNACACAATCTCAAGGTCTAAG
AGCACACGTCTGAACTCTTTCCCTACACGACGCTCTTCCGATCTNNNAGTGAGCAAAGATCCCACTACT
AGCACACGTCTGAACTCTTTCCCTACACGACGCTCTTCCGATCTNNNGAGGTTACAGTGAGCCAAGACT
AGCACACGTCTGAACTCTTTCCCTACACGACGCTCTTCCGATCTNNNGAAGAATTGCTTGAATCCGGGA
GAGCACACGTCTGAACTCTTTCCCTACACGACGCTCTTCCGATCTNNNGGAGAATTCCTCAAACCTTGG
GAGCACACGTCTGAACTCTTTCCCTACACGACGCTCTTCCGATCTNNNCTGAGCAAGATACTGAGACCA
AGCACACGTCTGAACTCTTTCCCTACACGACGCTCTTCCGATCTNNNGCTGGATAAAGAGTCAAGACC
AGAGCACACGTCTGAACTCTTTCCCTACACGACGCTCTTCCGATCTNNNCTTGCAGTGAGCCAAGATCT
GAGCACACGTCTGAACTCTTTCCCTACACGACGCTCTTCCGATCTNNNTATGCCAACAAAATCCCAAGT
AGAGCACACGTCTGAACTCTTTCCCTACACGACGCTCTTCCGATCTNNNTGAGGCAGGAGAATCTCTT
AGCACACGTCTGAACTCTTTCCCTACACGACGCTCTTCCGATCTNNNCTGTAGTCCCAGCTACTCAGGA
AGAGCACACGTCTGAACTCTTTCCCTACACGACGCTCTTCCGATCTNNNACCTGTAGTCCCAGCTACTA
GAGCACACGTCTGAACTCTTTCCCTACACGACGCTCTTCCGATCTNNNCGCCTGTAGTACCAGCTACTCT
AGCACACGTCTGAACTCTTTCCCTACACGACGCTCTTCCGATCTNNNTACAAAAAATTAGCCCGGCATG
GAGCACACGTCTGAACTCTTTCCCTACACGACGCTCTTCCGATCTNNNCCAACATGGTGAAACTCTGTC
GAGCACACGTCTGAACTCTTTCCCTACACGACGCTCTTCCGATCTNNNATCACTGTACTCCAGCCTGAAT
AGCACACGTCTGAACTCTTTCCCTACACGACGCTCTTCCGATCTNNNCAGAGGTTGTAGTGAGCCAAG
GAGCACACGTCTGAACTCTTTCCCTACACGACGCTCTTCCGATCTNNNGGCCACATAGCAAAACTCCTTT
GAGCACACGTCTGAACTCTTTCCCTACACGACGCTCTTCCGATCTNNNCTTACCTAGCCCAACACCAGAT
AGCACACGTCTGAACTCTTTCCCTACACGACGCTCTTCCGATCTNNNCTGAGGCAGGAGAATCACTTGA
GAGCACACGTCTGAACTCTTTCCCTACACGACGCTCTTCCGATCTNNNTATAGTCCCAGCTACTCAGGA
AGCACACGTCTGAACTCTTTCCCTACACGACGCTCTTCCGATCTNNNAATGGACAAAGGGACCCAAAT
GAGCACACGTCTGAACTCTTTCCCTACACGACGCTCTTCCGATCTNNNCTTGGCCACAAATCAGTCGAAT
GCACACGTCTGAACTCTTTCCCTACACGACGCTCTTCCGATCTNNNTCAATGTGTTGAGATTCTCCACT
GCACACGTCTGAACTCTTTCCCTACACGACGCTCTTCCGATCTNNNGCTGGTGTGATTGGAAGAGACT
GAGCACACGTCTGAACTCTTTCCCTACACGACGCTCTTCCGATCTNNNGACTGCAATAGTACACCACCA
GAGCACACGTCTGAACTCTTTCCCTACACGACGCTCTTCCGATCTNNNTTGGAGTCTGGAGAATGGGA
GAGCACACGTCTGAACTCTTTCCCTACACGACGCTCTTCCGATCTNNNCACCACCACACCTGGCTTATAT
AGCACACGTCTGAACTCTTTCCCTACACGACGCTCTTCCGATCTNNNGTGTGATCCCATAATCGTGCTT
GCACACGTCTGAACTCTTTCCCTACACGACGCTCTTCCGATCTNNNATTACAGACCTATGCCACTGTCC
GAGCACACGTCTGAACTCTTTCCCTACACGACGCTCTTCCGATCTNNNACAGGTATTGAGCCACTGTA
AGCACACGTCTGAACTCTTTCCCTACACGACGCTCTTCCGATCTNNNGAGCGAAACTCCGTCTCAAAA
AGCACACGTCTGAACTCTTTCCCTACACGACGCTCTTCCGATCTNNNTGGGCTCTCTATTCTGTTGCATT

GAGCACACGTCTGAACTCTTTCCCTACACGACGCTCTTCCGATCTNNNGGCACAATCATAGTTCACTGCA
AGCACACGTCTGAACTCTTTCCCTACACGACGCTCTTCCGATCTNNNAGGCACACAGGTTCAAGATACT
GAGCACACGTCTGAACTCTTTCCCTACACGACGCTCTTCCGATCTNNNAGTAGCTGGGACTACAGGCTT
GAGCACACGTCTGAACTCTTTCCCTACACGACGCTCTTCCGATCTNNNTTGGATTCAGTCAGTTTGGAC
GAGCACACGTCTGAACTCTTTCCCTACACGACGCTCTTCCGATCTNNNATAACACTTCTTGCATTGTGGG
AGCACACGTCTGAACTCTTTCCCTACACGACGCTCTTCCGATCTNNNGATCCACCTGCCTCAGATTACAG
GAGCACACGTCTGAACTCTTTCCCTACACGACGCTCTTCCGATCTNNNGTACTTGGTTGCATGGATGTG
GAGCACACGTCTGAACTCTTTCCCTACACGACGCTCTTCCGATCTNNNGCAGGAGAATTTCTTGAACCC
AGCACACGTCTGAACTCTTTCCCTACACGACGCTCTTCCGATCTNNNCTGAGGCAGAAGAATCGCTTGA
AGAGCACACGTCTGAACTCTTTCCCTACACGACGCTCTTCCGATCTNNNCAGAGGTTGAAGTGAGCTGA
GAGCACACGTCTGAACTCTTTCCCTACACGACGCTCTTCCGATCTNNNGGCGGCTTCACATCTACAAAT
AGCACACGTCTGAACTCTTTCCCTACACGACGCTCTTCCGATCTNNNTGTCTCACACTTGTAATCCCAGG
GAGCACACGTCTGAACTCTTTCCCTACACGACGCTCTTCCGATCTNNNTATCACTTGAGGCCAGGAGTTT
AGCACACGTCTGAACTCTTTCCCTACACGACGCTCTTCCGATCTNNNCACCTGTAATTCTGGCACTTTGG
GAGCACACGTCTGAACTCTTTCCCTACACGACGCTCTTCCGATCTNNNTAAGGCTTGAACCTGGAGAAT
AGCACACGTCTGAACTCTTTCCCTACACGACGCTCTTCCGATCTNNNCCTGGGAGGTAGAGGTTTTAGT
GAGCACACGTCTGAACTCTTTCCCTACACGACGCTCTTCCGATCTNNNCAGAAGTTGTAGTGAGCCGAG
AGCACACGTCTGAACTCTTTCCCTACACGACGCTCTTCCGATCTNNNAAGTTGCAGTGAGTGGAGATC
AGCACACGTCTGAACTCTTTCCCTACACGACGCTCTTCCGATCTNNNCTGAGGCAGGAGAATCACTTG
AGCACACGTCTGAACTCTTTCCCTACACGACGCTCTTCCGATCTNNNCGGAGTGAGACTCTGTTTCAAA
GAGCACACGTCTGAACTCTTTCCCTACACGACGCTCTTCCGATCTNNNTGAGCAACAGATCAAGACTCC
AGCACACGTCTGAACTCTTTCCCTACACGACGCTCTTCCGATCTNNNCAGAGGTTGTAGTGAGCCAAG
AGCACACGTCTGAACTCTTTCCCTACACGACGCTCTTCCGATCTNNNGAATATTTCAGTTGACGCGCTG
AGCACACGTCTGAACTCTTTCCCTACACGACGCTCTTCCGATCTNNNGAGAATCACTTGAACACCCAGG
AGCACACGTCTGAACTCTTTCCCTACACGACGCTCTTCCGATCTNNNGAGAATGGCATGAACCCAGGA
GCACACGTCTGAACTCTTTCCCTACACGACGCTCTTCCGATCTNNNGTAACAGAGCTAGACACCGTCTC
GAGCACACGTCTGAACTCTTTCCCTACACGACGCTCTTCCGATCTNNNCTGTAATCCCGGCTACTTAGG
GAGCACACGTCTGAACTCTTTCCCTACACGACGCTCTTCCGATCTNNNAGTAAGCTGAGATCACACCAC
AGCACACGTCTGAACTCTTTCCCTACACGACGCTCTTCCGATCTNNNCTGTAATCCCAGCTACTCAGGAG
GAGCACACGTCTGAACTCTTTCCCTACACGACGCTCTTCCGATCTNNNCTGTAGTCCCAGCTACTCAGGA
GAGCACACGTCTGAACTCTTTCCCTACACGACGCTCTTCCGATCTNNNAGGTTGCAGTGAGTCAAGATC
AGCACACGTCTGAACTCTTTCCCTACACGACGCTCTTCCGATCTNNNGCTGAGGCACAAAAATAGGTTG
AGCACACGTCTGAACTCTTTCCCTACACGACGCTCTTCCGATCTNNNTATGCAGCGGATCACATCTGTA
AGCACACGTCTGAACTCTTTCCCTACACGACGCTCTTCCGATCTNNNTGGGGGATAAGAGTGAGACTT
AGCACACGTCTGAACTCTTTCCCTACACGACGCTCTTCCGATCTNNNCTGAGATTGTGCCACTAGACTC
AGCACACGTCTGAACTCTTTCCCTACACGACGCTCTTCCGATCTNNNGTGACAAAGCAAAACCCTACCT
AGCACACGTCTGAACTCTTTCCCTACACGACGCTCTTCCGATCTNNNTTCTATTTTGCGCCAAGTGCTTT
GCACACGTCTGAACTCTTTCCCTACACGACGCTCTTCCGATCTNNNAGCCCTGATCTTCACATATTCGGC
AGCACACGTCTGAACTCTTTCCCTACACGACGCTCTTCCGATCTNNNAGACATTACATTCACTGCTGGGA
GAGCACACGTCTGAACTCTTTCCCTACACGACGCTCTTCCGATCTNNNCTTGTTTGTTTGGAGGATGAG
GAGCACACGTCTGAACTCTTTCCCTACACGACGCTCTTCCGATCTNNNACTATCTCAGCTCACTGCAAAC
AGCACACGTCTGAACTCTTTCCCTACACGACGCTCTTCCGATCTNNNCCTCGACCTCCCAAAATGAAAAA
AGCACACGTCTGAACTCTTTCCCTACACGACGCTCTTCCGATCTNNNCCCAAAGTGCTAGGATTACAGG

AGCACACGTCTGAACTCTTTCCCTACACGACGCTCTTCCGATCTNNNTGGGATTACAGGTTATAGGCGT
GAGCACACGTCTGAACTCTTTCCCTACACGACGCTCTTCCGATCTNNNCCCACCTTACATGGCTCTTCAT
AGAGCACACGTCTGAACTCTTTCCCTACACGACGCTCTTCCGATCTNNNCATGCCTACCCATCTATCTCCA
AGCACACGTCTGAACTCTTTCCCTACACGACGCTCTTCCGATCTNNNTAAAGGTCCTAGTCAGCCATTG
AGCACACGTCTGAACTCTTTCCCTACACGACGCTCTTCCGATCTNNNCTGTTCTTGAACTCCTGGACTCA
GCACACGTCTGAACTCTTTCCCTACACGACGCTCTTCCGATCTNNNGGTCACATTTTCCTGTTTGGGAG
GAGCACACGTCTGAACTCTTTCCCTACACGACGCTCTTCCGATCTNNNCCAAGCCCGGCTAATTTTTTGT
ACACGTCTGAACTCTTTCCCTACACGACGCTCTTCCGATCTNNNAGATTGCTTTTCAATTCTTTGTGGTC
GAGCACACGTCTGAACTCTTTCCCTACACGACGCTCTTCCGATCTNNNTGACGTAAGGACCAGTACAA
GAGCACACGTCTGAACTCTTTCCCTACACGACGCTCTTCCGATCTNNNTGTGAGTTTTAGCAGATTGT
GCACACGTCTGAACTCTTTCCCTACACGACGCTCTTCCGATCTNNNTTGGCTCCTCATAGACACTCAAC
GAGCACACGTCTGAACTCTTTCCCTACACGACGCTCTTCCGATCTNNNGAAAATAGGACTGGGTGCAA
GCACACGTCTGAACTCTTTCCCTACACGACGCTCTTCCGATCTNNNGCTGTTGGGGGAACTCTAACAT
GCACACGTCTGAACTCTTTCCCTACACGACGCTCTTCCGATCTNNNGGACCTACCACATAGAACGTGTT
GCACACGTCTGAACTCTTTCCCTACACGACGCTCTTCCGATCTNNNTCAGACGAGGGGACTATGTACTT
AGCACACGTCTGAACTCTTTCCCTACACGACGCTCTTCCGATCTNNNTGTGCCATGACTGGTTTAGACAA
AGCACACGTCTGAACTCTTTCCCTACACGACGCTCTTCCGATCTNNNTCATAACTCACTGCAGCCTGTAC
AGCACACGTCTGAACTCTTTCCCTACACGACGCTCTTCCGATCTNNNAGAATGCTTCACAGGAAAGGT
AGCACACGTCTGAACTCTTTCCCTACACGACGCTCTTCCGATCTNNNTCAGTCATCCTGCTTTTCATTCAC
AGCACACGTCTGAACTCTTTCCCTACACGACGCTCTTCCGATCTNNNCCTGTTCCATACCTGCTGTTAGT
AGCACACGTCTGAACTCTTTCCCTACACGACGCTCTTCCGATCTNNNCTCCCTCCAGAGCTTCACTTATC
GAGCACACGTCTGAACTCTTTCCCTACACGACGCTCTTCCGATCTNNNGAATCCACAACTCCAGACTTC
AGCACACGTCTGAACTCTTTCCCTACACGACGCTCTTCCGATCTNNNCTTGGGGGAAAGAAGGTGTGT
GAGCACACGTCTGAACTCTTTCCCTACACGACGCTCTTCCGATCTNNNGGAATCATACCTGGCACATAC
AGCACACGTCTGAACTCTTTCCCTACACGACGCTCTTCCGATCTNNNGCCTCAGTTTCCTCAACAAAACA
GAGCACACGTCTGAACTCTTTCCCTACACGACGCTCTTCCGATCTNNNGGTTGACAGAGTGAGACCTT
AGCACACGTCTGAACTCTTTCCCTACACGACGCTCTTCCGATCTNNNAGGGAAGAGAATCACTTGAAC
AGCACACGTCTGAACTCTTTCCCTACACGACGCTCTTCCGATCTNNNAGAATCACTTGAACTTGGGAG
AGCACACGTCTGAACTCTTTCCCTACACGACGCTCTTCCGATCTNNNCTTCAGTGAGCCATGATTGCAT
GAGCACACGTCTGAACTCTTTCCCTACACGACGCTCTTCCGATCTNNNCACCACTGTACTCTTAGCCTGA
AGAGCACACGTCTGAACTCTTTCCCTACACGACGCTCTTCCGATCTNNNTGAGGTTGGAGGATTCTTTG
GAGCACACGTCTGAACTCTTTCCCTACACGACGCTCTTCCGATCTNNNAGAATCACTTGAACTGGAGA
AGCACACGTCTGAACTCTTTCCCTACACGACGCTCTTCCGATCTNNNCAAAATCCTGCCACTGTACTCCA
GCACACGTCTGAACTCTTTCCCTACACGACGCTCTTCCGATCTNNNCCCAGCTACTACAGAGGCTAAAG
AGCACACGTCTGAACTCTTTCCCTACACGACGCTCTTCCGATCTNNNAGGCAGGAGAATCATTTGAAC
AGCACACGTCTGAACTCTTTCCCTACACGACGCTCTTCCGATCTNNNCTGTAGTCCCAGCTACTCAGGA
AGCACACGTCTGAACTCTTTCCCTACACGACGCTCTTCCGATCTNNNGATTACCCTCTCAATCGCATTG
AGCACACGTCTGAACTCTTTCCCTACACGACGCTCTTCCGATCTNNNGAGGCAAGAGAATCACTTGAA
AGCACACGTCTGAACTCTTTCCCTACACGACGCTCTTCCGATCTNNNGGGAGACTGAGGCAAGATAAT
AGCACACGTCTGAACTCTTTCCCTACACGACGCTCTTCCGATCTNNNGCATGCCTATAATCCCAGCTAC
AGAGCACACGTCTGAACTCTTTCCCTACACGACGCTCTTCCGATCTNNNTACAAAATATAGCCAGGCGT
AGCACACGTCTGAACTCTTTCCCTACACGACGCTCTTCCGATCTNNNGATTGTTTGAGCCTGGGAGAT
GAGCACACGTCTGAACTCTTTCCCTACACGACGCTCTTCCGATCTNNNATGAGCACCTATAGTCCCAGC

AGCACACGTCTGAACTCTTTCCCTACACGACGCTCTTCCGATCTNNNGAGAACCGTGATTATGCCATT
AGCACACGTCTGAACTCTTTCCCTACACGACGCTCTTCCGATCTNNNTGTAATCCTAGCACTTTGGGA
GAGCACACGTCTGAACTCTTTCCCTACACGACGCTCTTCCGATCTNNNGGAGAATGGCATGAACCTG
AGCACACGTCTGAACTCTTTCCCTACACGACGCTCTTCCGATCTNNNCTGAGGCAGGAGAATAGCTT
GAGCACACGTCTGAACTCTTTCCCTACACGACGCTCTTCCGATCTNNNCACCTGTAATCCCAGATAGTC
AGCACACGTCTGAACTCTTTCCCTACACGACGCTCTTCCGATCTNNNACCTTCCCTCCACTATTGTCCTA
GAGCACACGTCTGAACTCTTTCCCTACACGACGCTCTTCCGATCTNNNATGATCACACCACTGCACTCTA
GAGCACACGTCTGAACTCTTTCCCTACACGACGCTCTTCCGATCTNNNGAGGTTGGAGTGAGCTGA
GCACACGTCTGAACTCTTTCCCTACACGACGCTCTTCCGATCTNNNCTGAGGCAGGAGAATTGCTTG
GAGCACACGTCTGAACTCTTTCCCTACACGACGCTCTTCCGATCTNNNGTCAGGAGATCGAGACCATT
AGCACACGTCTGAACTCTTTCCCTACACGACGCTCTTCCGATCTNNNCTGAGGTAGGAGGATCACTTG
GAGCACACGTCTGAACTCTTTCCCTACACGACGCTCTTCCGATCTNNNGTAGAGTTTGCAGTGAGCTG
AGCACACGTCTGAACTCTTTCCCTACACGACGCTCTTCCGATCTNNNTGAGGCAGAAGAATTGCTTG
GAGCACACGTCTGAACTCTTTCCCTACACGACGCTCTTCCGATCTNNNAGAAGTTGCAGTCAGTGGAG
AGCACACGTCTGAACTCTTTCCCTACACGACGCTCTTCCGATCTNNNGCTGAGACAGGAGAATCACTT
GAGCACACGTCTGAACTCTTTCCCTACACGACGCTCTTCCGATCTNNNGATCACTTTAGCCCAGGAGT
GAGCACACGTCTGAACTCTTTCCCTACACGACGCTCTTCCGATCTNNNCAGGAGAATTGTTTTGAGC
GAGCACACGTCTGAACTCTTTCCCTACACGACGCTCTTCCGATCTNNNCCTGGGTGACAAGAGTAAG
AGCACACGTCTGAACTCTTTCCCTACACGACGCTCTTCCGATCTNNNGGGCAACAGAGTGAAATTCC
AGCACACGTCTGAACTCTTTCCCTACACGACGCTCTTCCGATCTNNNTCAAGTAATTCTCCCACCTCAG
AGCACACGTCTGAACTCTTTCCCTACACGACGCTCTTCCGATCTNNNAGAGAAAGAAAAAACAGCCTG
AGCACACGTCTGAACTCTTTCCCTACACGACGCTCTTCCGATCTNNNAATGAAGTCTGTTTCCAAGCAG
GCACACGTCTGAACTCTTTCCCTACACGACGCTCTTCCGATCTNNNTCCTGCACATTCATACCATGGAA
AGCACACGTCTGAACTCTTTCCCTACACGACGCTCTTCCGATCTNNNTCACCTGACAGGCTTTAGAAT
AGCACACGTCTGAACTCTTTCCCTACACGACGCTCTTCCGATCTNNNAACACCACCAACAACATTGACTT
AGCACACGTCTGAACTCTTTCCCTACACGACGCTCTTCCGATCTNNNGGTTCAAGCGATTCTCATGTCT
GCACACGTCTGAACTCTTTCCCTACACGACGCTCTTCCGATCTNNNGGGCTGGATGATGACCTACAAT
GAGCACACGTCTGAACTCTTTCCCTACACGACGCTCTTCCGATCTNNNAATGAAATGGGAAACAGCAG
GAGCACACGTCTGAACTCTTTCCCTACACGACGCTCTTCCGATCTNNNTCTGATTGAGCACTGGGTTCT
GCACACGTCTGAACTCTTTCCCTACACGACGCTCTTCCGATCTNNNTATAGCAGGTTCCTTTCAGTGCT
GAGCACACGTCTGAACTCTTTCCCTACACGACGCTCTTCCGATCTNNNCTCCTGAGTAGCTGGGACTA
GCACACGTCTGAACTCTTTCCCTACACGACGCTCTTCCGATCTNNNACTTCTCCTCTGATGATCCTCCA
AGCACACGTCTGAACTCTTTCCCTACACGACGCTCTTCCGATCTNNNACCTTGGATAGCTCTCTTACCCT
AGCACACGTCTGAACTCTTTCCCTACACGACGCTCTTCCGATCTNNNGCTCAAGCCCCTAAACACTTTTT
AGCACACGTCTGAACTCTTTCCCTACACGACGCTCTTCCGATCTNNNAATTCAAAGACACCCCTCTCCAA
GAGCACACGTCTGAACTCTTTCCCTACACGACGCTCTTCCGATCTNNNGCTGGCTGGATTTCCTTTCTTT
GCACACGTCTGAACTCTTTCCCTACACGACGCTCTTCCGATCTNNNCATTTGGCAAAGCTGACGATCT
AGCACACGTCTGAACTCTTTCCCTACACGACGCTCTTCCGATCTNNNCCAGCACTGGGGATTACATTT
AGCACACGTCTGAACTCTTTCCCTACACGACGCTCTTCCGATCTNNNACGAATCCAAGGAAGATTGTG
GCACACGTCTGAACTCTTTCCCTACACGACGCTCTTCCGATCTNNNTGACATTTAGGCAGTTTCCCACT
AGCACACGTCTGAACTCTTTCCCTACACGACGCTCTTCCGATCTNNNAAGTGATTTTCCAGACTCAGCC
GAGCACACGTCTGAACTCTTTCCCTACACGACGCTCTTCCGATCTNNNATATTGTTTTCCACAGTGGCT
GCACACGTCTGAACTCTTTCCCTACACGACGCTCTTCCGATCTNNNCAAGACGGAGAGGAGCTACAA

AGCACACGTCTGAACTCTTTCCCTACACGACGCTCTTCCGATCTNNNCACCCAGCCTATTAGCCGTATA

GCACACGTCTGAACTCTTTCCCTACACGACGCTCTTCCGATCTNNNAGTTCCCAATCACATAAGGGCA

GCACACGTCTGAACTCTTTCCCTACACGACGCTCTTCCGATCTNNNTTTCTGTCTTGGTGCCCATAATG

AGCACACGTCTGAACTCTTTCCCTACACGACGCTCTTCCGATCTNNNGAATGGGTGTGGGTGTGTAC

GCACACGTCTGAACTCTTTCCCTACACGACGCTCTTCCGATCTNNNTAGGCCCCTTGATATTGTTCCAC

GCACACGTCTGAACTCTTTCCCTACACGACGCTCTTCCGATCTNNNGCCATAGGAATCTTAGTCCGCA

GAGCACACGTCTGAACTCTTTCCCTACACGACGCTCTTCCGATCTNNNGCTTATGTTCCACCCCTACTCT

AGCACACGTCTGAACTCTTTCCCTACACGACGCTCTTCCGATCTNNNTTCTATCAGCACCTGACTAAGC

AGCACACGTCTGAACTCTTTCCCTACACGACGCTCTTCCGATCTNNNAATCCATTAGTGTCTGCACCCT

AGCACACGTCTGAACTCTTTCCCTACACGACGCTCTTCCGATCTNNNTCATCTGAGTCTCACAATGAC

AGCACACGTCTGAACTCTTTCCCTACACGACGCTCTTCCGATCTNNNAAGTGGCAGAGGCTTACTTAC

GCACACGTCTGAACTCTTTCCCTACACGACGCTCTTCCGATCTNNNTTTAGTCTCTGCTACATGCCAGG

AGCACACGTCTGAACTCTTTCCCTACACGACGCTCTTCCGATCTNNNTTGCAGCTAAACAGAACAACC

GAGCACACGTCTGAACTCTTTCCCTACACGACGCTCTTCCGATCTNNNCCCATGCTTCTTCCTCACAAAT

AGCACACGTCTGAACTCTTTCCCTACACGACGCTCTTCCGATCTNNNTCCCCAATGCAACAGTATTGA

GCACACGTCTGAACTCTTTCCCTACACGACGCTCTTCCGATCTNNNTGGGATTACATGTATGAGCCA

AGCACACGTCTGAACTCTTTCCCTACACGACGCTCTTCCGATCTNNNCACCACGCCCGGCTAAATATAT

AGCACACGTCTGAACTCTTTCCCTACACGACGCTCTTCCGATCTNNNAGAGCTTGCAGTGAACTGAGA

GAGCACACGTCTGAACTCTTTCCCTACACGACGCTCTTCCGATCTNNNAGGTTGCAGTGAACTGAGAT

GCACACGTCTGAACTCTTTCCCTACACGACGCTCTTCCGATCTNNNTCTGTAATCCCAGCTACTCAGG

GAGCACACGTCTGAACTCTTTCCCTACACGACGCTCTTCCGATCTNNNAATACAAAAAAGTTAGCCGG

AGCACACGTCTGAACTCTTTCCCTACACGACGCTCTTCCGATCTNNNAGGCAAGAAGATCACTTGAAC

GCACACGTCTGAACTCTTTCCCTACACGACGCTCTTCCGATCTNNNCATGGTGGTTCAAGCCTGTAAT

AGCACACGTCTGAACTCTTTCCCTACACGACGCTCTTCCGATCTNNNACGATGCAGTGGACAAAATCA

AGCACACGTCTGAACTCTTTCCCTACACGACGCTCTTCCGATCTNNNATTCTAGCAGTTTTGGAGGCT

GAGCACACGTCTGAACTCTTTCCCTACACGACGCTCTTCCGATCTNNNACCTTCCCTCCACTATTGTCCT

AGCACACGTCTGAACTCTTTCCCTACACGACGCTCTTCCGATCTNNNCCATAGAGGTGGCTCAGAGAT

GAGCACACGTCTGAACTCTTTCCCTACACGACGCTCTTCCGATCTNNNATAATTCCAGCTACTTGGGAG

AGAGCACACGTCTGAACTCTTTCCCTACACGACGCTCTTCCGATCTNNNGCCTGTAGTCCCAGCTACTC

AGCACACGTCTGAACTCTTTCCCTACACGACGCTCTTCCGATCTNNNAAGCTGAGGCAGAAGAATAG

GAGCACACGTCTGAACTCTTTCCCTACACGACGCTCTTCCGATCTNNNTGCAGTAAGCTGAGATTGCA

AGCACACGTCTGAACTCTTTCCCTACACGACGCTCTTCCGATCTNNNCTGTAGTCCCAGCTACTCATGA

GAGCACACGTCTGAACTCTTTCCCTACACGACGCTCTTCCGATCTNNNGCTTGCCTATAATCCCAGCTA

AGCACACGTCTGAACTCTTTCCCTACACGACGCTCTTCCGATCTNNNGAAAATCACTTGAACCCGGGA

GCACACGTCTGAACTCTTTCCCTACACGACGCTCTTCCGATCTNNNCAAGCCTGTAATCCCAGTGTTT

AGCACACGTCTGAACTCTTTCCCTACACGACGCTCTTCCGATCTNNNTCACCAGTGATCCAGAAAGAG

GCACACGTCTGAACTCTTTCCCTACACGACGCTCTTCCGATCTNNNGGCATCTGTAATCCCAGCTACT

AGAGCACACGTCTGAACTCTTTCCCTACACGACGCTCTTCCGATCTNNNGCCTATAATCCCAGCTACTC

GAGCACACGTCTGAACTCTTTCCCTACACGACGCTCTTCCGATCTNNNAGGTGGGAGGATCTTTTGA

AGCACACGTCTGAACTCTTTCCCTACACGACGCTCTTCCGATCTNNNATGCCTGTAGTTCCAGCTACTT

GAGCACACGTCTGAACTCTTTCCCTACACGACGCTCTTCCGATCTNNNCTGAATGTGCCCCATAAAGC

AGCACACGTCTGAACTCTTTCCCTACACGACGCTCTTCCGATCTNNNCTGAAGCAGGAGAATGGTGT

GAGCACACGTCTGAACTCTTTCCCTACACGACGCTCTTCCGATCTNNNACATTACAGTGAGCCCAGAT

GCACACGTCTGAACTCTTTCCCTACACGACGCTCTTCCGATCTNNNAGTGAGCTGAGATGCTGAGAT
AGCACACGTCTGAACTCTTTCCCTACACGACGCTCTTCCGATCTNNNCTGAGGGACCTGACTGTTAGA
AGCACACGTCTGAACTCTTTCCCTACACGACGCTCTTCCGATCTNNNCATGCGCCTGTAATTCCAGTTA
GCACACGTCTGAACTCTTTCCCTACACGACGCTCTTCCGATCTNNNGCAACAGAACAAGACACTGTCT
AGCACACGTCTGAACTCTTTCCCTACACGACGCTCTTCCGATCTNNNGTTGGAGACAGTGAGACCTCA
GAGCACACGTCTGAACTCTTTCCCTACACGACGCTCTTCCGATCTNNNGTCTGAACTACTTTGGAGGC
GAGCACACGTCTGAACTCTTTCCCTACACGACGCTCTTCCGATCTNNNATCAATTGAGCCCAGGAGTT
AGCACACGTCTGAACTCTTTCCCTACACGACGCTCTTCCGATCTNNNCGTAATCCCAGCTATTCAGGA
AGCACACGTCTGAACTCTTTCCCTACACGACGCTCTTCCGATCTNNNTAATCCCAGCTACTTAGGAGG
AGCACACGTCTGAACTCTTTCCCTACACGACGCTCTTCCGATCTNNNCATGCCTGTAATCCCAGCTACT
AGCACACGTCTGAACTCTTTCCCTACACGACGCTCTTCCGATCTNNNGAGACAGGAGAATGGCGTGA
GCACACGTCTGAACTCTTTCCCTACACGACGCTCTTCCGATCTNNNCTGAGGCAGGAGAATCACTTGA
GCACACGTCTGAACTCTTTCCCTACACGACGCTCTTCCGATCTNNNATAATCACAGGGAGGAGAGGT
GAGCACACGTCTGAACTCTTTCCCTACACGACGCTCTTCCGATCTNNNGAGATCACACCATTGCATTCC
AGCACACGTCTGAACTCTTTCCCTACACGACGCTCTTCCGATCTNNNCTGTAGTCCCAGCTACTCAGG
AGCACACGTCTGAACTCTTTCCCTACACGACGCTCTTCCGATCTNNNCATGATTATGCACTGCACTCCA
GCACACGTCTGAACTCTTTCCCTACACGACGCTCTTCCGATCTNNNGGTTGGATCCTCAGATGTCTGT
GCACACGTCTGAACTCTTTCCCTACACGACGCTCTTCCGATCTNNNAACTTTGAAGCCAGGTGTATTG
AGCACACGTCTGAACTCTTTCCCTACACGACGCTCTTCCGATCTNNNCTGTAGTCCCAGCTACTCAGG
AGCACACGTCTGAACTCTTTCCCTACACGACGCTCTTCCGATCTNNNCACCTGTAATCCCAGCTACTTG
GAGCACACGTCTGAACTCTTTCCCTACACGACGCTCTTCCGATCTNNNAGAGGTTCCAGTGAAACGA
AGAGCACACGTCTGAACTCTTTCCCTACACGACGCTCTTCCGATCTNNNAGAATTGCTTGAACCTGGG
AGCACACGTCTGAACTCTTTCCCTACACGACGCTCTTCCGATCTNNNAGTGAGATTCTGCCTCTCTCTC
AGCACACGTCTGAACTCTTTCCCTACACGACGCTCTTCCGATCTNNNCGTAAATCTGGGAGGCAGAGT
GAGCACACGTCTGAACTCTTTCCCTACACGACGCTCTTCCGATCTNNNGGGCAACATAGCAAGATTCT
AGCACACGTCTGAACTCTTTCCCTACACGACGCTCTTCCGATCTNNNGCACCTGTAATCCCAACTACTC
GAGCACACGTCTGAACTCTTTCCCTACACGACGCTCTTCCGATCTNNNCATGTGTGTAATATTCGGAC
AGCACACGTCTGAACTCTTTCCCTACACGACGCTCTTCCGATCTNNNCTGTAATCCCAGCTACTCAGGA
GCACACGTCTGAACTCTTTCCCTACACGACGCTCTTCCGATCTNNNACTTCCCAAAACCAAAACAAGC
GAGCACACGTCTGAACTCTTTCCCTACACGACGCTCTTCCGATCTNNNTGGTAGGCAGAGGTTTCAAT
GCACACGTCTGAACTCTTTCCCTACACGACGCTCTTCCGATCTNNNTGAGACACGAGAATCACTTGAG
GCACACGTCTGAACTCTTTCCCTACACGACGCTCTTCCGATCTNNNGAGGCCCCATCTCTACAAGAAA
AGCACACGTCTGAACTCTTTCCCTACACGACGCTCTTCCGATCTNNNCAGTAAGCTGAGATTGCACCA
GAGCACACGTCTGAACTCTTTCCCTACACGACGCTCTTCCGATCTNNNAGACAGGAGAATCACATGA
AGCACACGTCTGAACTCTTTCCCTACACGACGCTCTTCCGATCTNNNCTGTAGTCTGAGCTTCTCTGG
AGCACACGTCTGAACTCTTTCCCTACACGACGCTCTTCCGATCTNNNGAAACAGGAGAATCGCTTGA
AGCACACGTCTGAACTCTTTCCCTACACGACGCTCTTCCGATCTNNNACGACTGTAATTGCAGCACTTT
GAGCACACGTCTGAACTCTTTCCCTACACGACGCTCTTCCGATCTNNNCTGTAGTCCCAGCTACTCAG
AGCACACGTCTGAACTCTTTCCCTACACGACGCTCTTCCGATCTNNNTGTGCCTGTAATTCCAGCTAC
AGCACACGTCTGAACTCTTTCCCTACACGACGCTCTTCCGATCTNNNGAGGTTGAGTGAGCCGATAT
GCACACGTCTGAACTCTTTCCCTACACGACGCTCTTCCGATCTNNNAGGCTGAGACATAAGAATCGC
AGCACACGTCTGAACTCTTTCCCTACACGACGCTCTTCCGATCTNNNGTAATCCCAGCTATTCAGGAG
GAGCACACGTCTGAACTCTTTCCCTACACGACGCTCTTCCGATCTNNNGCATCCCAGTGTTTATTACCA

GCACACGTCTGAACTCTTTCCCTACACGACGCTCTTCCGATCTNNNATTGGTATCCTTACACACAGGG
AGCACACGTCTGAACTCTTTCCCTACACGACGCTCTTCCGATCTNNNTGGACGCATGACTTTACAGAG
GCACACGTCTGAACTCTTTCCCTACACGACGCTCTTCCGATCTNNNAGTCACACACAACCTTCTTCTCC
GCACACGTCTGAACTCTTTCCCTACACGACGCTCTTCCGATCTNNNCCAGCATGCCTGACTCCATATAT
AGCACACGTCTGAACTCTTTCCCTACACGACGCTCTTCCGATCTNNNTTCCAACTAGCTGGGATTACA
GCACACGTCTGAACTCTTTCCCTACACGACGCTCTTCCGATCTNNNTGTTCCTGGAAAAAGAGGACCA
GAGCACACGTCTGAACTCTTTCCCTACACGACGCTCTTCCGATCTNNNATGCCCACAAGTAGAATTGT
GAGCACACGTCTGAACTCTTTCCCTACACGACGCTCTTCCGATCTNNNTGCTGTTCACAGACAGTACAT
AGCACACGTCTGAACTCTTTCCCTACACGACGCTCTTCCGATCTNNNGCCAGTATTACAGACATAGGC
AGCACACGTCTGAACTCTTTCCCTACACGACGCTCTTCCGATCTNNNCCCACAGAAGACAGCTGTAAG
GAGCACACGTCTGAACTCTTTCCCTACACGACGCTCTTCCGATCTNNNTTGGTTTTTTTGCAGCTTCG
GCACACGTCTGAACTCTTTCCCTACACGACGCTCTTCCGATCTNNNGCAAGGGAGAAAGTCAGAGA
GCACACGTCTGAACTCTTTCCCTACACGACGCTCTTCCGATCTNNNAGAGAGTTTGGTTGGGAGGA
AGCACACGTCTGAACTCTTTCCCTACACGACGCTCTTCCGATCTNNNTCAAGTCGTGATATTCCCAGT
GAGCACACGTCTGAACTCTTTCCCTACACGACGCTCTTCCGATCTNNNAAATAGGCACAAGGGTAGT
AGCACACGTCTGAACTCTTTCCCTACACGACGCTCTTCCGATCTNNNAGTCCTGTTCTCCAATCCACTT
AGCACACGTCTGAACTCTTTCCCTACACGACGCTCTTCCGATCTNNNGGGCTCTTGCTGCTTTATTTCT
GCACACGTCTGAACTCTTTCCCTACACGACGCTCTTCCGATCTNNNAGTAGGTAGGACTACAGGCAT
AGCACACGTCTGAACTCTTTCCCTACACGACGCTCTTCCGATCTNNNTACTGGTACCACAAGTGTTTCT
AGCACACGTCTGAACTCTTTCCCTACACGACGCTCTTCCGATCTNNNTTCAGTTCTGAGCCACATGATC
AGCACACGTCTGAACTCTTTCCCTACACGACGCTCTTCCGATCTNNNAATCATCCAGCTAAGCTTCTCC
GAGCACACGTCTGAACTCTTTCCCTACACGACGCTCTTCCGATCTNNNTGTAGGGACAGGAAAGGAA
GCACACGTCTGAACTCTTTCCCTACACGACGCTCTTCCGATCTNNNGGGATGCAAAATCAAAGAAGG
AGCACACGTCTGAACTCTTTCCCTACACGACGCTCTTCCGATCTNNNTTCCCTCTGACTTCCCTGTCATA
GAGCACACGTCTGAACTCTTTCCCTACACGACGCTCTTCCGATCTNNNTCCTGTTCCTTCTTAGCCTGTT
AGCACACGTCTGAACTCTTTCCCTACACGACGCTCTTCCGATCTNNNTGAGCACACTTCCTAAGGTCA
AGCACACGTCTGAACTCTTTCCCTACACGACGCTCTTCCGATCTNNNCTTCCAAGTGGCAAGATTCCA
AGCACACGTCTGAACTCTTTCCCTACACGACGCTCTTCCGATCTNNNGCCCATGGATGAACTTAGTGA
GCACACGTCTGAACTCTTTCCCTACACGACGCTCTTCCGATCTNNNTGTTTTGGGCCATTTCAAATCCA
AGCACACGTCTGAACTCTTTCCCTACACGACGCTCTTCCGATCTNNNTGAGGTAGGAGAATGGTGTG
GCACACGTCTGAACTCTTTCCCTACACGACGCTCTTCCGATCTNNNGGTGACAGAGCAAGATTCCAT
AGCACACGTCTGAACTCTTTCCCTACACGACGCTCTTCCGATCTNNNCATAGTCCCAGCTATTCGAGAG
GAGCACACGTCTGAACTCTTTCCCTACACGACGCTCTTCCGATCTNNNGCATAGTGAAACCCCCATCT
GAGCACACGTCTGAACTCTTTCCCTACACGACGCTCTTCCGATCTNNNTGAGGTAGGAGAATCGCTTG
GAGCACACGTCTGAACTCTTTCCCTACACGACGCTCTTCCGATCTNNNGAGAATTGCTTGAACCCAGG
GAGCACACGTCTGAACTCTTTCCCTACACGACGCTCTTCCGATCTNNNACAGCTATAACCCCAGCACTT
AGCACACGTCTGAACTCTTTCCCTACACGACGCTCTTCCGATCTNNNCTAAGACCATACCACTGCACT
AGCACACGTCTGAACTCTTTCCCTACACGACGCTCTTCCGATCTNNNCAGAATTGCTTGAACTTGGGA
AGCACACGTCTGAACTCTTTCCCTACACGACGCTCTTCCGATCTNNNAGACAGGAGAATCACTTGAA
GAGCACACGTCTGAACTCTTTCCCTACACGACGCTCTTCCGATCTNNNCCAAGATCATGCCACTGAATT
GCACACGTCTGAACTCTTTCCCTACACGACGCTCTTCCGATCTNNNGTAATCCTAGCTACTTGGGAGG
GCACACGTCTGAACTCTTTCCCTACACGACGCTCTTCCGATCTNNNTCAACGGAATAGAACAGGTAG
AGCACACGTCTGAACTCTTTCCCTACACGACGCTCTTCCGATCTNNNGACAAGAGAATTGCTTGAACC

GAGCACACGTCTGAACTCTTTCCCTACACGACGCTCTTCCGATCTNNNAGCAGAAGAATCACTTGAAC
GAGCACACGTCTGAACTCTTTCCCTACACGACGCTCTTCCGATCTNNNCTAGGCAACAAGAGCAAAACT
GAGCACACGTCTGAACTCTTTCCCTACACGACGCTCTTCCGATCTNNNGATCATACCACTACACTCCAGC
AGCACACGTCTGAACTCTTTCCCTACACGACGCTCTTCCGATCTNNNTGAGGCAGGAGAATCTCTTGAA
GAGCACACGTCTGAACTCTTTCCCTACACGACGCTCTTCCGATCTNNNCTGACATCACTACTGCACTC
AGCACACGTCTGAACTCTTTCCCTACACGACGCTCTTCCGATCTNNNGAGGTTACAGTGAGCCGAGAT
AGCACACGTCTGAACTCTTTCCCTACACGACGCTCTTCCGATCTNNNTGAGGCACGAAAATCACTTGAA
GCACACGTCTGAACTCTTTCCCTACACGACGCTCTTCCGATCTNNNAGATGGGAGAATCACTTGAACCC
AGAGCACACGTCTGAACTCTTTCCCTACACGACGCTCTTCCGATCTNNNGATTGCAGTAAGCCAAGATC
GAGCACACGTCTGAACTCTTTCCCTACACGACGCTCTTCCGATCTNNNTGTAATCCCAGGACTTTGGAA
GAGCACACGTCTGAACTCTTTCCCTACACGACGCTCTTCCGATCTNNNGAGATCGTACCACTGCACTCTA
GAGCACACGTCTGAACTCTTTCCCTACACGACGCTCTTCCGATCTNNNATTGGAGATCGTGCCACTACA
GAGCACACGTCTGAACTCTTTCCCTACACGACGCTCTTCCGATCTNNNGGAGGTTGTAGTGAGCTGAG
AGAGCACACGTCTGAACTCTTTCCCTACACGACGCTCTTCCGATCTNNNCACATGTAGTCCCAGCTACTC
GAGCACACGTCTGAACTCTTTCCCTACACGACGCTCTTCCGATCTNNNGTTGCAGTGAGCCAAGATTAT
GAGCACACGTCTGAACTCTTTCCCTACACGACGCTCTTCCGATCTNNNGATGACAAAGTGAGACCCTG
GAGCACACGTCTGAACTCTTTCCCTACACGACGCTCTTCCGATCTNNNCTAAGGCAGGAGAATCGCTT
AGCACACGTCTGAACTCTTTCCCTACACGACGCTCTTCCGATCTNNNGTAATCCTAGCTACTTGGGAGG
AGAGCACACGTCTGAACTCTTTCCCTACACGACGCTCTTCCGATCTNNNCTGTAATCCCAGCTACTCAGC
AGCACACGTCTGAACTCTTTCCCTACACGACGCTCTTCCGATCTNNNCAAGTCAGTGAGAAACAGCCAA
AGCACACGTCTGAACTCTTTCCCTACACGACGCTCTTCCGATCTNNNGAGAATTGCCTGAATCTGGGAG
GAGCACACGTCTGAACTCTTTCCCTACACGACGCTCTTCCGATCTNNNCAGCACTCTGAAACTTGCAAGA
GAGCACACGTCTGAACTCTTTCCCTACACGACGCTCTTCCGATCTNNNGAGTTTGCAGTGAGCTGAGAT
AGCACACGTCTGAACTCTTTCCCTACACGACGCTCTTCCGATCTNNNCCAGAGAGTGAGATCAACTCCA
AGCACACGTCTGAACTCTTTCCCTACACGACGCTCTTCCGATCTNNNTGAGATTGTGTTACTGCACTCCA
AGAGCACACGTCTGAACTCTTTCCCTACACGACGCTCTTCCGATCTNNNAAGAGAATCGTTTGAACCTG
GAGCACACGTCTGAACTCTTTCCCTACACGACGCTCTTCCGATCTNNNAGATCGATGTCTGACAATCTC
GAGCACACGTCTGAACTCTTTCCCTACACGACGCTCTTCCGATCTNNNCACCATGCCCAGCTAATTTTG
GAGCACACGTCTGAACTCTTTCCCTACACGACGCTCTTCCGATCTNNNTGATCTTGAACTCCCAACCTCA
GAGCACACGTCTGAACTCTTTCCCTACACGACGCTCTTCCGATCTNNNTGCTGCGAATACTATCGTTCTT
AGCACACGTCTGAACTCTTTCCCTACACGACGCTCTTCCGATCTNNNTTTTGTTGCCATTGCTTTTGGTG
AGCACACGTCTGAACTCTTTCCCTACACGACGCTCTTCCGATCTNNNTACAAAAGTTAGCTGGGTGTGC
GAGCACACGTCTGAACTCTTTCCCTACACGACGCTCTTCCGATCTNNNTCAAACTCTTGAGCTTAGGCCA
AGCACACGTCTGAACTCTTTCCCTACACGACGCTCTTCCGATCTNNNGTGACCGTTTTCTTCAACCTGAG
GAGCACACGTCTGAACTCTTTCCCTACACGACGCTCTTCCGATCTNNNGTACCTTGGGTTTTGCTTCTGT
AGCACACGTCTGAACTCTTTCCCTACACGACGCTCTTCCGATCTNNNAAAATAACAAAGGTGCCCAGCTC
AGCACACGTCTGAACTCTTTCCCTACACGACGCTCTTCCGATCTNNNTCCAGGCCCAAAGGAATAATTC
GAGCACACGTCTGAACTCTTTCCCTACACGACGCTCTTCCGATCTNNNGACGTGCTCAATTTGGGAACA
GAGCACACGTCTGAACTCTTTCCCTACACGACGCTCTTCCGATCTNNNCTAAGAATGCATGGCCTTGTA
AGCACACGTCTGAACTCTTTCCCTACACGACGCTCTTCCGATCTNNNCACACACACACATACACACACAC
GAGCACACGTCTGAACTCTTTCCCTACACGACGCTCTTCCGATCTNNNCTAATCTCCAAGGCAGCTTTGT
AGCACACGTCTGAACTCTTTCCCTACACGACGCTCTTCCGATCTNNNTGGCAGACTGAAGATGATTCCA
GAGCACACGTCTGAACTCTTTCCCTACACGACGCTCTTCCGATCTNNNGAGATCATGACACTGCATTCA

AGCACACGTCTGAACTCTTTCCCTACACGACGCTCTTCCGATCTNNNCTGAGTCAGGAGAATGGTGT
AGCACACGTCTGAACTCTTTCCCTACACGACGCTCTTCCGATCTNNNACGAGGTCAGGAGATCGAGA
AGAGCACACGTCTGAACTCTTTCCCTACACGACGCTCTTCCGATCTNNNCTGAGGCAGGAAAATTGCT
AGCACACGTCTGAACTCTTTCCCTACACGACGCTCTTCCGATCTNNNGCGCAATATAGCAAGATCCTG
AGAGCACACGTCTGAACTCTTTCCCTACACGACGCTCTTCCGATCTNNNCTGTAATCCAGCTACTTGG
GAGCACACGTCTGAACTCTTTCCCTACACGACGCTCTTCCGATCTNNNATAGAGCATGAAGACCCTGT
AGCACACGTCTGAACTCTTTCCCTACACGACGCTCTTCCGATCTNNNGAGGTGGGACAATTGCTTGA
AGCACACGTCTGAACTCTTTCCCTACACGACGCTCTTCCGATCTNNNCTGTAGTCCCAGGTACTCAGG
AGCACACGTCTGAACTCTTTCCCTACACGACGCTCTTCCGATCTNNNCTGAGGCAGGAGAATCACTTG
AGCACACGTCTGAACTCTTTCCCTACACGACGCTCTTCCGATCTNNNGTAAGAAGGATCGCTTGAGG
GAGCACACGTCTGAACTCTTTCCCTACACGACGCTCTTCCGATCTNNNACATGCTTGTAATCCCAGCTA
AGCACACGTCTGAACTCTTTCCCTACACGACGCTCTTCCGATCTNNNGTAGACTCAGCTACTCAAGAG
AGCACACGTCTGAACTCTTTCCCTACACGACGCTCTTCCGATCTNNNGAGATTGCAGTGAGCCAAGAT
AGCACACGTCTGAACTCTTTCCCTACACGACGCTCTTCCGATCTNNNGAGACAGGAGAATTGCTTGA
AGCACACGTCTGAACTCTTTCCCTACACGACGCTCTTCCGATCTNNNCTATTGAACCTGGGTAGCAGA
AGCACACGTCTGAACTCTTTCCCTACACGACGCTCTTCCGATCTNNNCTGAGGAAGGAGAACTGCTT
AGCACACGTCTGAACTCTTTCCCTACACGACGCTCTTCCGATCTNNNCTTGGGCAACATAGTGAAAC
GCACACGTCTGAACTCTTTCCCTACACGACGCTCTTCCGATCTNNNAAAGGTTGCAGTGAGCCAAGAT
GAGCACACGTCTGAACTCTTTCCCTACACGACGCTCTTCCGATCTNNNTGTAGTCCCAGCTACTCAGG
GAGCACACGTCTGAACTCTTTCCCTACACGACGCTCTTCCGATCTNNNAAGTTGCAGTGAGCCAAGAT
GCACACGTCTGAACTCTTTCCCTACACGACGCTCTTCCGATCTNNNCTGAGGCAGGAGAATCACTTG
GAGCACACGTCTGAACTCTTTCCCTACACGACGCTCTTCCGATCTNNNACCTTCCCTCCACTATTGTCCT
AGCACACGTCTGAACTCTTTCCCTACACGACGCTCTTCCGATCTNNNCTGTACAACATAGTGAGACCCC
AGCACACGTCTGAACTCTTTCCCTACACGACGCTCTTCCGATCTNNNGTTGTAATGAGCTGAGATCG
AGCACACGTCTGAACTCTTTCCCTACACGACGCTCTTCCGATCTNNNGAATTGCTTGAACCCAGGAGT
AGCACACGTCTGAACTCTTTCCCTACACGACGCTCTTCCGATCTNNNGGAGACTGAGACAGGAGAA
AGCACACGTCTGAACTCTTTCCCTACACGACGCTCTTCCGATCTNNNTCCTTCAAGGGCCAAACTGTT
AGCACACGTCTGAACTCTTTCCCTACACGACGCTCTTCCGATCTNNNTGCAATCAAAAAGCTTGTGAC
AGCACACGTCTGAACTCTTTCCCTACACGACGCTCTTCCGATCTNNNACACCTTTCAAAAGACCTCACC
AGCACACGTCTGAACTCTTTCCCTACACGACGCTCTTCCGATCTNNNAGGAACAAGAATCACCTGAA
GCACACGTCTGAACTCTTTCCCTACACGACGCTCTTCCGATCTNNNCTCTGGCTCACTCCTGTAATACC
AGCACACGTCTGAACTCTTTCCCTACACGACGCTCTTCCGATCTNNNTGTAGAGCTAATGCCCCTTTAC
GCACACGTCTGAACTCTTTCCCTACACGACGCTCTTCCGATCTNNNAAATGCTGAGACTCCAAGTCC
GAGCACACGTCTGAACTCTTTCCCTACACGACGCTCTTCCGATCTNNNCAGCCATATAAGGAAACACT
GCACACGTCTGAACTCTTTCCCTACACGACGCTCTTCCGATCTNNNCACCTCAGGTACAACATGGAAG
AGCACACGTCTGAACTCTTTCCCTACACGACGCTCTTCCGATCTNNNTTCAAGTAGTTCTCCGTCTCAG
AGCACACGTCTGAACTCTTTCCCTACACGACGCTCTTCCGATCTNNNAGAATCCTGGGGCCATTTTCT
GCACACGTCTGAACTCTTTCCCTACACGACGCTCTTCCGATCTNNNTGTTTTCACAATGTTGTCAGTTC
GCACACGTCTGAACTCTTTCCCTACACGACGCTCTTCCGATCTNNNGTAACTGGAAGAAACAGGGGC
GCACACGTCTGAACTCTTTCCCTACACGACGCTCTTCCGATCTNNNTGTCACTAGGACCAGAACAGAT
AGCACACGTCTGAACTCTTTCCCTACACGACGCTCTTCCGATCTNNNACAGAGCTATGCCTTTGGATC
GAGCACACGTCTGAACTCTTTCCCTACACGACGCTCTTCCGATCTNNNTGACACTCAGCCCAATCATTA
AGCACACGTCTGAACTCTTTCCCTACACGACGCTCTTCCGATCTNNNCTTAGTCAAAGTCAGGTGGGT

AGCACACGTCTGAACTCTTTCCCTACACGACGCTCTTCCGATCTNNNCAACCATCCTCTCAACTCTAGGC
AGCACACGTCTGAACTCTTTCCCTACACGACGCTCTTCCGATCTNNNCTTCCTGGAGCTTCGACAGTAT
AGCACACGTCTGAACTCTTTCCCTACACGACGCTCTTCCGATCTNNNGCCTCCTGAATAGCTGGGATTA
GAGCACACGTCTGAACTCTTTCCCTACACGACGCTCTTCCGATCTNNNGTTCCTTTACCTTCCACAAGTC
AGCACACGTCTGAACTCTTTCCCTACACGACGCTCTTCCGATCTNNNCATTGCTGATGTGCTTGTCACT
GCACACGTCTGAACTCTTTCCCTACACGACGCTCTTCCGATCTNNNATCCTCCATCACACATACTACGCC
GAGCACACGTCTGAACTCTTTCCCTACACGACGCTCTTCCGATCTNNNGCAAGAGAATCACTTGAACCC
GAGCACACGTCTGAACTCTTTCCCTACACGACGCTCTTCCGATCTNNNTAAGTGCTTGAACTGAAGAG
CACACGTCTGAACTCTTTCCCTACACGACGCTCTTCCGATCTNNNTGGCAGAAGTAATGGTATGTCCC
AGCACACGTCTGAACTCTTTCCCTACACGACGCTCTTCCGATCTNNNCATGCCTGTAATCCCAGCTACTC
GAGCACACGTCTGAACTCTTTCCCTACACGACGCTCTTCCGATCTNNNTTTACTCCATCCCACTTTGCTG
GAGCACACGTCTGAACTCTTTCCCTACACGACGCTCTTCCGATCTNNNGCAACATGGTGAAATCCTGTC
AAGAGCACACGTCTGAACTCTTTCCCTACACGACGCTCTTCCGATCTNNNTAGTCCCAGCTACTCATGA
AGAGCACACGTCTGAACTCTTTCCCTACACGACGCTCTTCCGATCTNNNAGAATGGCATGAACCTGGG
GAGCACACGTCTGAACTCTTTCCCTACACGACGCTCTTCCGATCTNNNCAATGCAGTGAGCCATGATCA
GAGCACACGTCTGAACTCTTTCCCTACACGACGCTCTTCCGATCTNNNCTGAGGCAGGAGAATCACTT
GAGCACACGTCTGAACTCTTTCCCTACACGACGCTCTTCCGATCTNNNCAAGAGAATTGCTTGAACCCA
GAGCACACGTCTGAACTCTTTCCCTACACGACGCTCTTCCGATCTNNNGAAATCGAGACCATCCTGGC
AGCACACGTCTGAACTCTTTCCCTACACGACGCTCTTCCGATCTNNNCTGAGGCAGGAGAATCACTTG
GAGCACACGTCTGAACTCTTTCCCTACACGACGCTCTTCCGATCTNNNGCATGCCTGTAATCCCAGATAC
GCACACGTCTGAACTCTTTCCCTACACGACGCTCTTCCGATCTNNNGTGAACTACTGGGGTCAAGTGA
GAGCACACGTCTGAACTCTTTCCCTACACGACGCTCTTCCGATCTNNNCACAGCTGACTAAAACATTGCC
AGCACACGTCTGAACTCTTTCCCTACACGACGCTCTTCCGATCTNNNCATTCTGTGTCCTTTGGCATCTG
AGCACACGTCTGAACTCTTTCCCTACACGACGCTCTTCCGATCTNNNTTGTGGGTCTATAGGGATAGG
GCACACGTCTGAACTCTTTCCCTACACGACGCTCTTCCGATCTNNNTAAGCACAGATGTCTGAAGAGC
AGCACACGTCTGAACTCTTTCCCTACACGACGCTCTTCCGATCTNNNAGCTCCTGGAATCAGAAGACA
AGCACACGTCTGAACTCTTTCCCTACACGACGCTCTTCCGATCTNNNTCAAGCAATTCTCCTACCTCAGC
AGCACACGTCTGAACTCTTTCCCTACACGACGCTCTTCCGATCTNNNCAGTCTGGTCTCAAACTCCTGAC
AGCACACGTCTGAACTCTTTCCCTACACGACGCTCTTCCGATCTNNNAAGTAGCTGGTATTACAGGCAC
GAGCACACGTCTGAACTCTTTCCCTACACGACGCTCTTCCGATCTNNNTACCCACATGCAGAAGGATGA
AGCACACGTCTGAACTCTTTCCCTACACGACGCTCTTCCGATCTNNNGGAACCCTGGAGAACCTTTTCT
AGCACACGTCTGAACTCTTTCCCTACACGACGCTCTTCCGATCTNNNCAGTCTGGCTAACATGGCAAAA
AGCACACGTCTGAACTCTTTCCCTACACGACGCTCTTCCGATCTNNNTTTCCCTCTTTTTCTGGGGTGAA
AGCACACGTCTGAACTCTTTCCCTACACGACGCTCTTCCGATCTNNNTCTTGGTAGTGTCCTTTGATGC
AGCACACGTCTGAACTCTTTCCCTACACGACGCTCTTCCGATCTNNNTTTTGCAGATGAAGGCATAAGC
AGCACACGTCTGAACTCTTTCCCTACACGACGCTCTTCCGATCTNNNGGGATTACAGGCAAGAGTCACT
AGCACACGTCTGAACTCTTTCCCTACACGACGCTCTTCCGATCTNNNACTGTGAGCTCATTCAACCTCTT
AGCACACGTCTGAACTCTTTCCCTACACGACGCTCTTCCGATCTNNNGGTAACTTCAGTGTGTCAACCTC
GAGCACACGTCTGAACTCTTTCCCTACACGACGCTCTTCCGATCTNNNCCACCACACCAAGCTAATCTTT
GCACACGTCTGAACTCTTTCCCTACACGACGCTCTTCCGATCTNNNGTCACGAATTTGATTCCAGAGCC
GAGCACACGTCTGAACTCTTTCCCTACACGACGCTCTTCCGATCTNNNTTCCTGAACATCCCTTGGCATA
AGCACACGTCTGAACTCTTTCCCTACACGACGCTCTTCCGATCTNNNTGTGCACATACACACAAACGAA
GAGCACACGTCTGAACTCTTTCCCTACACGACGCTCTTCCGATCTNNNCATCCACAACAAAGCAGAGTCT

AGCACACGTCTGAACTCTTTCCCTACACGACGCTCTTCCGATCTNNNGGACACATACTCCTGATGAAG
AGCACACGTCTGAACTCTTTCCCTACACGACGCTCTTCCGATCTNNNGCCTGCTTCTTATAGGTTTTGCC
AGCACACGTCTGAACTCTTTCCCTACACGACGCTCTTCCGATCTNNNCAAGGGGAAATGACACCAACAT
AGCACACGTCTGAACTCTTTCCCTACACGACGCTCTTCCGATCTNNNTTTTTGGCCAAAAGACACCACT
GAGCACACGTCTGAACTCTTTCCCTACACGACGCTCTTCCGATCTNNNACTGAAGGATAAAAGCTAGG
AGCACACGTCTGAACTCTTTCCCTACACGACGCTCTTCCGATCTNNNGGCATGTAGAGTTTCCACTGA
AGCACACGTCTGAACTCTTTCCCTACACGACGCTCTTCCGATCTNNNCACATGGAAGAAGGAGCAAACA
GAGCACACGTCTGAACTCTTTCCCTACACGACGCTCTTCCGATCTNNNTGAAAGCTCTTCCCATGCAAAA
GAGCACACGTCTGAACTCTTTCCCTACACGACGCTCTTCCGATCTNNNAGAATTACTTGAACGCAGGAG
GAGCACACGTCTGAACTCTTTCCCTACACGACGCTCTTCCGATCTNNNGATCAAGACCATCCTGGCTAA
AGCACACGTCTGAACTCTTTCCCTACACGACGCTCTTCCGATCTNNNCACAATGTGCACATGTACCCTAA
GAGCACACGTCTGAACTCTTTCCCTACACGACGCTCTTCCGATCTNNNTTGAACAGAGGAGGAAGAG
GCACACGTCTGAACTCTTTCCCTACACGACGCTCTTCCGATCTNNNCTGTAGTCCGAGCTACTCAGGAG
AGAGCACACGTCTGAACTCTTTCCCTACACGACGCTCTTCCGATCTNNNATTTCAGAGCTGACGTTTGCA
AGAGCACACGTCTGAACTCTTTCCCTACACGACGCTCTTCCGATCTNNNCACAATGTGCACATGTACCCT
AGCACACGTCTGAACTCTTTCCCTACACGACGCTCTTCCGATCTNNNTGAGGCAGGAGAATTGCTTGA
AGCACACGTCTGAACTCTTTCCCTACACGACGCTCTTCCGATCTNNNCGCCTATAGTCCCAGCTATTTGG
AGCACACGTCTGAACTCTTTCCCTACACGACGCTCTTCCGATCTNNNACCAAAAAAAGCCCAGGACTA
GAGCACACGTCTGAACTCTTTCCCTACACGACGCTCTTCCGATCTNNNGAGACAGGAGAATTGCTTGA
GAGCACACGTCTGAACTCTTTCCCTACACGACGCTCTTCCGATCTNNNAAAAAATTAGCCAGGTGTGG
AGCACACGTCTGAACTCTTTCCCTACACGACGCTCTTCCGATCTNNNTTGAGCCAGGAGAATCACTTGAA
AGAGCACACGTCTGAACTCTTTCCCTACACGACGCTCTTCCGATCTNNNGAAGCAGGAGAATGGAATG
GAGCACACGTCTGAACTCTTTCCCTACACGACGCTCTTCCGATCTNNNCTTGCAGTGAATCGAGATCAT
AGAGCACACGTCTGAACTCTTTCCCTACACGACGCTCTTCCGATCTNNNGACAGGAGAATCTCTTGAA
GAGCACACGTCTGAACTCTTTCCCTACACGACGCTCTTCCGATCTNNNTGACCAACATGTAGAAACCCT
GAGCACACGTCTGAACTCTTTCCCTACACGACGCTCTTCCGATCTNNNGTTAGTCGGACATGGTGGTAT
AGCACACGTCTGAACTCTTTCCCTACACGACGCTCTTCCGATCTNNNGCTGAGACAGGAGAATCACTTG
AGCACACGTCTGAACTCTTTCCCTACACGACGCTCTTCCGATCTNNNTCAAGGCCTCAGTCAATTTCAG
GAGCACACGTCTGAACTCTTTCCCTACACGACGCTCTTCCGATCTNNNCATGCCTGTAGTCCCAGATACT
AGCACACGTCTGAACTCTTTCCCTACACGACGCTCTTCCGATCTNNNAAGACCCATCAGTGTGCTGTAT
AGAGCACACGTCTGAACTCTTTCCCTACACGACGCTCTTCCGATCTNNNGACGACTGTAATCCCAGCTA
AGCACACGTCTGAACTCTTTCCCTACACGACGCTCTTCCGATCTNNNTGCTCATAACTACCTCCCCAAA
GAGCACACGTCTGAACTCTTTCCCTACACGACGCTCTTCCGATCTNNNAGCCGAGATTATGCCACTGTAT
GAGCACACGTCTGAACTCTTTCCCTACACGACGCTCTTCCGATCTNNNGATGTGGTGGCTCATACCTGT
GAGCACACGTCTGAACTCTTTCCCTACACGACGCTCTTCCGATCTNNNGTAATCCTGGCTACTCAGAAG
GCACACGTCTGAACTCTTTCCCTACACGACGCTCTTCCGATCTNNNATGCCTTGTAATCCCAGCTACTCC
GAGCACACGTCTGAACTCTTTCCCTACACGACGCTCTTCCGATCTNNNTTTGTCTTCTGGGACTAGTGT
AGCACACGTCTGAACTCTTTCCCTACACGACGCTCTTCCGATCTNNNCAGAAGAATTGCTTGAACCCAG
GAGCACACGTCTGAACTCTTTCCCTACACGACGCTCTTCCGATCTNNNTTGAACCCAGGAGACAGAGAT
GAGCACACGTCTGAACTCTTTCCCTACACGACGCTCTTCCGATCTNNNCTGTAGTTCCAGCTACTTGGGA
GAGCACACGTCTGAACTCTTTCCCTACACGACGCTCTTCCGATCTNNNTGGAGGCTGCAATAAGACAT
AGCACACGTCTGAACTCTTTCCCTACACGACGCTCTTCCGATCTNNNACATGCCTGTAATCCCAGCTATT
GAGCACACGTCTGAACTCTTTCCCTACACGACGCTCTTCCGATCTNNNGTCAGGAGAATAGCTTGAAC

GAGCACACGTCTGAACTCTTTCCCTACACGACGCTCTTCCGATCTNNNGGTAAAGGTATCACTTGAGCC
GAGCACACGTCTGAACTCTTTCCCTACACGACGCTCTTCCGATCTNNNGAGAATGACGTGAACCTAGGA
GAGCACACGTCTGAACTCTTTCCCTACACGACGCTCTTCCGATCTNNNCTGAACTCCAGCCTAGGCAATA
GAGCACACGTCTGAACTCTTTCCCTACACGACGCTCTTCCGATCTNNNGAGATCACGCCATTTCATTCCA
AGCACACGTCTGAACTCTTTCCCTACACGACGCTCTTCCGATCTNNNGGAGTTTGCAGTGAGTGGAGA
GAGCACACGTCTGAACTCTTTCCCTACACGACGCTCTTCCGATCTNNNCACGGATCACAAGGTCACTAC
GAGCACACGTCTGAACTCTTTCCCTACACGACGCTCTTCCGATCTNNNGGGATCACTAGGTCAGGAGTT
GAGCACACGTCTGAACTCTTTCCCTACACGACGCTCTTCCGATCTNNNAGGCAGGTGGATTATGAAGT
GAGCACACGTCTGAACTCTTTCCCTACACGACGCTCTTCCGATCTNNNAGAAAAGTGATGAGATGGCT
AGAGCACACGTCTGAACTCTTTCCCTACACGACGCTCTTCCGATCTNNNGAGAATGGAGTGAACTCAGC
AGCACACGTCTGAACTCTTTCCCTACACGACGCTCTTCCGATCTNNNGCAACAAAAGCAGAACTCCATC
GAGCACACGTCTGAACTCTTTCCCTACACGACGCTCTTCCGATCTNNNGTTGCTGTGAGTCGAGATTAT
AAGAGCACACGTCTGAACTCTTTCCCTACACGACGCTCTTCCGATCTNNNAGTTTGAGGCTGCAGTAAC
AGCACACGTCTGAACTCTTTCCCTACACGACGCTCTTCCGATCTNNNGCCTGTATTCCCAGCTACTCTAC
GAGCACACGTCTGAACTCTTTCCCTACACGACGCTCTTCCGATCTNNNATTGCTTGAGCCTAGGAGTTC
AGCACACGTCTGAACTCTTTCCCTACACGACGCTCTTCCGATCTNNNTTGGTGAGTTAGGGCTGAATCT
AGCACACGTCTGAACTCTTTCCCTACACGACGCTCTTCCGATCTNNNGGTTCCCATGTCATGCAAAACTT
AGCACACGTCTGAACTCTTTCCCTACACGACGCTCTTCCGATCTNNNTTCAAGCAATTGTTCTGCCTCAG
AGCACACGTCTGAACTCTTTCCCTACACGACGCTCTTCCGATCTNNNTGTTTCTGGGAATGTCCATGGA
AGCACACGTCTGAACTCTTTCCCTACACGACGCTCTTCCGATCTNNNGTTCATACCATTCTCCTGCCTCAC
AGCACACGTCTGAACTCTTTCCCTACACGACGCTCTTCCGATCTNNNGTGCTGGTTTGCAGTTTCTTTTG
GCACACGTCTGAACTCTTTCCCTACACGACGCTCTTCCGATCTNNNGCTCAGATCATCTTCCCACTTCACA
AGCACACGTCTGAACTCTTTCCCTACACGACGCTCTTCCGATCTNNNACATTCAACTCTTGGGTGGAAG
GAGCACACGTCTGAACTCTTTCCCTACACGACGCTCTTCCGATCTNNNCCAATTGGCCTGAACTCTTCAA
AGCACACGTCTGAACTCTTTCCCTACACGACGCTCTTCCGATCTNNNCAAAGCAGGACAAGAGACCATT
AGCACACGTCTGAACTCTTTCCCTACACGACGCTCTTCCGATCTNNNCACTCTGTCCTTTTTCAAGCAGG
AGCACACGTCTGAACTCTTTCCCTACACGACGCTCTTCCGATCTNNNTTCAAGCAATTTTCCTGCCTCAG
GAGCACACGTCTGAACTCTTTCCCTACACGACGCTCTTCCGATCTNNNTAGAAGCACCTTCGAGCTGAT
AGCACACGTCTGAACTCTTTCCCTACACGACGCTCTTCCGATCTNNNCAAACTCCTGAGCTCAAGCAATC
AGCACACGTCTGAACTCTTTCCCTACACGACGCTCTTCCGATCTNNNCCCAAGTAGCTGGAACTAAAGG
GAGCACACGTCTGAACTCTTTCCCTACACGACGCTCTTCCGATCTNNNGACTACAATCATGCACCACCAT
AGCACACGTCTGAACTCTTTCCCTACACGACGCTCTTCCGATCTNNNTTTTGGTCATTTGGTTGCACTG
AGAGCACACGTCTGAACTCTTTCCCTACACGACGCTCTTCCGATCTNNNCCATGCCCAGCTAACTTTTGT
AGCACACGTCTGAACTCTTTCCCTACACGACGCTCTTCCGATCTNNNAGAAAGCAAGCAAGCAAAAAA
AGCACACGTCTGAACTCTTTCCCTACACGACGCTCTTCCGATCTNNNTTGGCTTTATTTCTGGGTTCCCT
GCACACGTCTGAACTCTTTCCCTACACGACGCTCTTCCGATCTNNNGACATGTGTCAGCCAGGTACTTC
AGCACACGTCTGAACTCTTTCCCTACACGACGCTCTTCCGATCTNNNGCTGAGATTCCAGGCATGAGTT
AGCACACGTCTGAACTCTTTCCCTACACGACGCTCTTCCGATCTNNNTCTTCCTGCTTTGGCTCAATCTAC
AGCACACGTCTGAACTCTTTCCCTACACGACGCTCTTCCGATCTNNNCAGCCATGTACTCTCTTCAGGAA
GAGCACACGTCTGAACTCTTTCCCTACACGACGCTCTTCCGATCTNNNAAGTGCTGAGATTACAGGCAT
AGCACACGTCTGAACTCTTTCCCTACACGACGCTCTTCCGATCTNNNGTCCGGCCTCATTTTCTTTTCTTC
AGCACACGTCTGAACTCTTTCCCTACACGACGCTCTTCCGATCTNNNGGCTCTGATTATTCTCCCACCTCC
AGCACACGTCTGAACTCTTTCCCTACACGACGCTCTTCCGATCTNNNTGGTCTCAAAATCCTGACCTCAC

AGCACACGTCTGAACTCTTTCCCTACACGACGCTCTTCCGATCTNNNGGGGCAAGTGCTCATTATTACT

AGCACACGTCTGAACTCTTTCCCTACACGACGCTCTTCCGATCTNNNGTAGGTGTGGCTCACTGTTTAG

AGCACACGTCTGAACTCTTTCCCTACACGACGCTCTTCCGATCTNNNCCATGCTGTTTTGGTTACTGGAC

AGCACACGTCTGAACTCTTTCCCTACACGACGCTCTTCCGATCTNNNCACTGCGCATAATTACCAACACA

AGCACACGTCTGAACTCTTTCCCTACACGACGCTCTTCCGATCTNNNCTTTGGCTAGGATTGTTGCAG

AGCACACGTCTGAACTCTTTCCCTACACGACGCTCTTCCGATCTNNNACCCCTTGTCTCATCTCCTTTTCC

GAGCACACGTCTGAACTCTTTCCCTACACGACGCTCTTCCGATCTNNNTAGTTCCAGCTCTGTTCTTTCCC

AGCACACGTCTGAACTCTTTCCCTACACGACGCTCTTCCGATCTNNNACCTTGAGATTGAGCACAGACA

AGCACACGTCTGAACTCTTTCCCTACACGACGCTCTTCCGATCTNNNAGTAGGCTTCCGCTAATATTGCT

GAGCACACGTCTGAACTCTTTCCCTACACGACGCTCTTCCGATCTNNNACTTCGTCTTCTTTTTGTTCCCT

AGCACACGTCTGAACTCTTTCCCTACACGACGCTCTTCCGATCTNNNTGTAACGTCACAGCTGAATGGA

AGCACACGTCTGAACTCTTTCCCTACACGACGCTCTTCCGATCTNNNTGCTCAATACATGTCTGCTGCTA

AGCACACGTCTGAACTCTTTCCCTACACGACGCTCTTCCGATCTNNNCTAGGACTACAGATGCATGCCA

AGCACACGTCTGAACTCTTTCCCTACACGACGCTCTTCCGATCTNNNTCAGATGATGACAAAAGCCTCCA

AGAGCACACGTCTGAACTCTTTCCCTACACGACGCTCTTCCGATCTNNNAAAATTAGCTGGCATGGTGA

AGCACACGTCTGAACTCTTTCCCTACACGACGCTCTTCCGATCTNNNATCACTTGAACCCGAAAGATGG

AGAGCACACGTCTGAACTCTTTCCCTACACGACGCTCTTCCGATCTNNNCTGTAATCCCAGCTACTCAGG

AGCACACGTCTGAACTCTTTCCCTACACGACGCTCTTCCGATCTNNNGAGACAAGAGAATGGCTTGAA

GAGCACACGTCTGAACTCTTTCCCTACACGACGCTCTTCCGATCTNNNAGCCTGTAATCCCAGTACTTTG

AGCACACGTCTGAACTCTTTCCCTACACGACGCTCTTCCGATCTNNNGTAATCTCAGCTACTTGGGAGG

AGAGCACACGTCTGAACTCTTTCCCTACACGACGCTCTTCCGATCTNNNGTACATGCCTGTAATCCCAGC

GAGCACACGTCTGAACTCTTTCCCTACACGACGCTCTTCCGATCTNNNGGGCAACACAGTAAGACTACC

AGAGCACACGTCTGAACTCTTTCCCTACACGACGCTCTTCCGATCTNNNCGCCTATAATCCCAGCTACTC

AGCACACGTCTGAACTCTTTCCCTACACGACGCTCTTCCGATCTNNNCAAGGTGAAAACCCAGGTCTCT

GAGCACACGTCTGAACTCTTTCCCTACACGACGCTCTTCCGATCTNNNTCATGCCTGTAATCTCAGCAGT

AGCACACGTCTGAACTCTTTCCCTACACGACGCTCTTCCGATCTNNNACGTGTTTATCTGCTGACCTTCT

AGCACACGTCTGAACTCTTTCCCTACACGACGCTCTTCCGATCTNNNCTGTAATCCCAGCTACTCCGGAG

GCACACGTCTGAACTCTTTCCCTACACGACGCTCTTCCGATCTNNNCTGAGAAAGGAGAATGGCGTGA

GAGCACACGTCTGAACTCTTTCCCTACACGACGCTCTTCCGATCTNNNCATGGTGTTGTGCCTGTAAT

GAGCACACGTCTGAACTCTTTCCCTACACGACGCTCTTCCGATCTNNNGCGGTCCAATAAAAATGGGGA

GAGCACACGTCTGAACTCTTTCCCTACACGACGCTCTTCCGATCTNNNCTGAGACAGGAGAATGGTGT

GAGCACACGTCTGAACTCTTTCCCTACACGACGCTCTTCCGATCTNNNGATTCCTGTAGTCCCAGCTACT

GAGCACACGTCTGAACTCTTTCCCTACACGACGCTCTTCCGATCTNNNCTGTAGTGAGCCGTGATTGTT

AGCACACGTCTGAACTCTTTCCCTACACGACGCTCTTCCGATCTNNNGGAGACTGCAGTGAGCTATGA

GCACACGTCTGAACTCTTTCCCTACACGACGCTCTTCCGATCTNNNGTGACAGAGCCAGAATCCATCTC

GAGCACACGTCTGAACTCTTTCCCTACACGACGCTCTTCCGATCTNNNGTGGTAGCGCATTCTTGTAAT

GAGCACACGTCTGAACTCTTTCCCTACACGACGCTCTTCCGATCTNNNCCTGAGGAACACAGTAGGAC

GAGCACACGTCTGAACTCTTTCCCTACACGACGCTCTTCCGATCTNNNTGGGTGATAGATCGAGACTCC

AGCACACGTCTGAACTCTTTCCCTACACGACGCTCTTCCGATCTNNNCTGTATTGCCAGCTACTCGGAA

GAGCACACGTCTGAACTCTTTCCCTACACGACGCTCTTCCGATCTNNNTGCAGTGAGCTGTGATTATGC

AGCACACGTCTGAACTCTTTCCCTACACGACGCTCTTCCGATCTNNNTCATTGGCTGAATCACAAACTG

AGCACACGTCTGAACTCTTTCCCTACACGACGCTCTTCCGATCTNNNATAATCCCAGCTATTCAGGAGG

GAGCACACGTCTGAACTCTTTCCCTACACGACGCTCTTCCGATCTNNNGAGGTTGAGGTTACAATGGC

GAGCACACGTCTGAACTCTTTCCCTACACGACGCTCTTCCGATCTNNNTGAACAGAGATTACACCACTG
GAGCACACGTCTGAACTCTTTCCCTACACGACGCTCTTCCGATCTNNNGAGAGAGGAGAATGGCGTGA
AGCACACGTCTGAACTCTTTCCCTACACGACGCTCTTCCGATCTNNNGAGACAGAACGAAACTCCCTCT
GAGCACACGTCTGAACTCTTTCCCTACACGACGCTCTTCCGATCTNNNCACAACTGGTGCAATTTTCTCC
GAGCACACGTCTGAACTCTTTCCCTACACGACGCTCTTCCGATCTNNNAGAGTTTGCAGTGAGCTGAG
AGCACACGTCTGAACTCTTTCCCTACACGACGCTCTTCCGATCTNNNGTAATCCCAGCTACTCAGGAAG
AGAGCACACGTCTGAACTCTTTCCCTACACGACGCTCTTCCGATCTNNNCTTTGTTTGGTGCCACCTTA
GAGCACACGTCTGAACTCTTTCCCTACACGACGCTCTTCCGATCTNNNGCTATGGCTGCACAGATAGAA
GAGCACACGTCTGAACTCTTTCCCTACACGACGCTCTTCCGATCTNNNTGAGGTAGGAGAATCGCTCAA
GAGCACACGTCTGAACTCTTTCCCTACACGACGCTCTTCCGATCTNNNCAACATGGTGAAAACCCTGTCT
GCACACGTCTGAACTCTTTCCCTACACGACGCTCTTCCGATCTNNNTCAACTCCCTCCTATTCCCTGATCA
GAGCACACGTCTGAACTCTTTCCCTACACGACGCTCTTCCGATCTNNNGTAATGTTGCCATCTGTTCCT
GAGCACACGTCTGAACTCTTTCCCTACACGACGCTCTTCCGATCTNNNTTTTGTAGAGACGGGGTTTCT
AGCACACGTCTGAACTCTTTCCCTACACGACGCTCTTCCGATCTNNNTGTGGGTGGTAAACTTTGAAT
GAGCACACGTCTGAACTCTTTCCCTACACGACGCTCTTCCGATCTNNNGTTCCTAACCAACCACAAATG
AGCACACGTCTGAACTCTTTCCCTACACGACGCTCTTCCGATCTNNNCCCTCTCTGACTCAAGCAATTCT
AGCACACGTCTGAACTCTTTCCCTACACGACGCTCTTCCGATCTNNNCAACATAAGGCCCTGTCTCTACA
GAGCACACGTCTGAACTCTTTCCCTACACGACGCTCTTCCGATCTNNNATTCACAGCCAACTCCACAACTA
AGCACACGTCTGAACTCTTTCCCTACACGACGCTCTTCCGATCTNNNTCAACCTCCCTAGTAGCTGAAA
AGCACACGTCTGAACTCTTTCCCTACACGACGCTCTTCCGATCTNNNTTCTTGATCAGTCTCAGCTGTG
GAGCACACGTCTGAACTCTTTCCCTACACGACGCTCTTCCGATCTNNNTGGCTGGTATTTATTGAGCA
AGCACACGTCTGAACTCTTTCCCTACACGACGCTCTTCCGATCTNNNGGCCCCAAACTGTGTTCAATAA
AGCACACGTCTGAACTCTTTCCCTACACGACGCTCTTCCGATCTNNNCCACGCCCAGCTAATTTTTTGTA
GAGCACACGTCTGAACTCTTTCCCTACACGACGCTCTTCCGATCTNNNTAGGGTGCCAAAAGATTCTGT
AGCACACGTCTGAACTCTTTCCCTACACGACGCTCTTCCGATCTNNNTCTACCCCCTCCTTTCATGAGTA
GAGCACACGTCTGAACTCTTTCCCTACACGACGCTCTTCCGATCTNNNCCACCACACCTGGCTAATTTTT
AGCACACGTCTGAACTCTTTCCCTACACGACGCTCTTCCGATCTNNNCAACCAATGTATGTGGCTTCCTG
GAGCACACGTCTGAACTCTTTCCCTACACGACGCTCTTCCGATCTNNNATCTGTGTGCTTGGTCCCTAAA
AGCACACGTCTGAACTCTTTCCCTACACGACGCTCTTCCGATCTNNNGCTAAGTACAGTGGCACAATCA
GAGCACACGTCTGAACTCTTTCCCTACACGACGCTCTTCCGATCTNNNCGATCTCCTGACCTCATGATCT
GAGCACACGTCTGAACTCTTTCCCTACACGACGCTCTTCCGATCTNNNTAGGAGGTCTAGTCTTCTCAC
GAGCACACGTCTGAACTCTTTCCCTACACGACGCTCTTCCGATCTNNNTTTTAGTGGAGACGGGGTTT
GAGCACACGTCTGAACTCTTTCCCTACACGACGCTCTTCCGATCTNNNTTGACCTTGAACGTGCTAATA
GAGCACACGTCTGAACTCTTTCCCTACACGACGCTCTTCCGATCTNNNAATTCCTGGCAGTTTTCTATG
GAGCACACGTCTGAACTCTTTCCCTACACGACGCTCTTCCGATCTNNNTGTTCCCTTCCCTCTTACCCTAT
AGCACACGTCTGAACTCTTTCCCTACACGACGCTCTTCCGATCTNNNATCTGTAGTCCTAGCTACCCAG
GAGCACACGTCTGAACTCTTTCCCTACACGACGCTCTTCCGATCTNNNCTGTAATCCCAGCTACTCAGGA
GAGCACACGTCTGAACTCTTTCCCTACACGACGCTCTTCCGATCTNNNGAGGCTGAGAAAGGAGAAT
GAGCACACGTCTGAACTCTTTCCCTACACGACGCTCTTCCGATCTNNNGGTTACAGAGCGAGATCTTGT
AGAGCACACGTCTGAACTCTTTCCCTACACGACGCTCTTCCGATCTNNNAGGCAGAAGAATCACTTGAA
GCACACGTCTGAACTCTTTCCCTACACGACGCTCTTCCGATCTNNNAAGGCTGCAGTGAACTAAGATCA
GAGCACACGTCTGAACTCTTTCCCTACACGACGCTCTTCCGATCTNNNCTTGCAGTTAGCTGAGATCGA
AGCACACGTCTGAACTCTTTCCCTACACGACGCTCTTCCGATCTNNNTGAGCTGAGATCACACCATTTCA

AGCACACGTCTGAACTCTTTCCCTACACGACGCTCTTCCGATCTNNNACCAACAAAGAAGCAAAGGCAA
AGCACACGTCTGAACTCTTTCCCTACACGACGCTCTTCCGATCTNNNTGGGAGATAGAACGAGACTCT
GAGCACACGTCTGAACTCTTTCCCTACACGACGCTCTTCCGATCTNNNGAGCAACAGAGTGAGACCTT
GAGCACACGTCTGAACTCTTTCCCTACACGACGCTCTTCCGATCTNNNTGAGGTAGAACGATCACTTGA
AAGAGCACACGTCTGAACTCTTTCCCTACACGACGCTCTTCCGATCTNNNAGATTGTACTACTGCACTC
AGCACACGTCTGAACTCTTTCCCTACACGACGCTCTTCCGATCTNNNTACTAGGGAGGCTGAGATGAG
GAGCACACGTCTGAACTCTTTCCCTACACGACGCTCTTCCGATCTNNNCTGAGGCAGGAGACTACATG
AAGAGCACACGTCTGAACTCTTTCCCTACACGACGCTCTTCCGATCTNNNCTTGCAGTGAGCCGAGATT
GAGCACACGTCTGAACTCTTTCCCTACACGACGCTCTTCCGATCTNNNATTTCAGAGACAGGGTCTGGT
AGCACACGTCTGAACTCTTTCCCTACACGACGCTCTTCCGATCTNNNTTGCTCCCAGGAAAGAATAGGA
AGCACACGTCTGAACTCTTTCCCTACACGACGCTCTTCCGATCTNNNGTTAGGGCTACAGCTATATGCC
AGCACACGTCTGAACTCTTTCCCTACACGACGCTCTTCCGATCTNNNTTTCCCTCTTCCCCAAAAAGGAA
GAGCACACGTCTGAACTCTTTCCCTACACGACGCTCTTCCGATCTNNNTACAGTGGTACGATCTCGCCTA
AGCACACGTCTGAACTCTTTCCCTACACGACGCTCTTCCGATCTNNNTGTAACAAATCCTTCCAGGGGA
GAGCACACGTCTGAACTCTTTCCCTACACGACGCTCTTCCGATCTNNNTGTGCAGTCAGAAATATCCA
GAGCACACGTCTGAACTCTTTCCCTACACGACGCTCTTCCGATCTNNNAAGCATTCTGAGAAGCAAGGA
GAGCACACGTCTGAACTCTTTCCCTACACGACGCTCTTCCGATCTNNNTGCCCTGCCTAGATTTCTTTCTT
GAGCACACGTCTGAACTCTTTCCCTACACGACGCTCTTCCGATCTNNNGCTCATGACTGTGTAAGAGTG
AGCACACGTCTGAACTCTTTCCCTACACGACGCTCTTCCGATCTNNNACAGACTGCGAAAAAGAAGGA
GAGCACACGTCTGAACTCTTTCCCTACACGACGCTCTTCCGATCTNNNCTGGCCGAATCTCTTTCCTTTT
AGCACACGTCTGAACTCTTTCCCTACACGACGCTCTTCCGATCTNNNCCAGTGGATGTCTTCCCATAACA
GAGCACACGTCTGAACTCTTTCCCTACACGACGCTCTTCCGATCTNNNTCTGCTTCATAGATGGCACCTT
AGCACACGTCTGAACTCTTTCCCTACACGACGCTCTTCCGATCTNNNTCATTTATGCCTCCTTCTCAGGG
GAGCACACGTCTGAACTCTTTCCCTACACGACGCTCTTCCGATCTNNNTCATGGAATGCATGAGGATGA
AGCACACGTCTGAACTCTTTCCCTACACGACGCTCTTCCGATCTNNNCACTTCCTCATATCCTTCCCTGG
GAGCACACGTCTGAACTCTTTCCCTACACGACGCTCTTCCGATCTNNNAGTGGTACTCAACTCTTCCCAA
GAGCACACGTCTGAACTCTTTCCCTACACGACGCTCTTCCGATCTNNNGGAGTGTAGTGGCACAATCT
AGCACACGTCTGAACTCTTTCCCTACACGACGCTCTTCCGATCTNNNGTGAGAGGTGACTGTTTTGCTT
GAGCACACGTCTGAACTCTTTCCCTACACGACGCTCTTCCGATCTNNNTGTACACTGGCATTGGAGTTT
AGCACACGTCTGAACTCTTTCCCTACACGACGCTCTTCCGATCTNNNGACAACAGAAGTGAGGAGAGA
AGCACACGTCTGAACTCTTTCCCTACACGACGCTCTTCCGATCTNNNAAGCACAAGTTTGCACATCACA
GAGCACACGTCTGAACTCTTTCCCTACACGACGCTCTTCCGATCTNNNTAGCAATTCTGGGAGACGGTA
GAGCACACGTCTGAACTCTTTCCCTACACGACGCTCTTCCGATCTNNNTCCATCCCCTTAGACTCTGTCT
AGCACACGTCTGAACTCTTTCCCTACACGACGCTCTTCCGATCTNNNAGGTTCAAGTGATTCTCCTGCT
GAGCACACGTCTGAACTCTTTCCCTACACGACGCTCTTCCGATCTNNNGACGGAATCTTGCTCTATCA
AGCACACGTCTGAACTCTTTCCCTACACGACGCTCTTCCGATCTNNNATAACTTCTGAGGGACCTTGCT
AGCACACGTCTGAACTCTTTCCCTACACGACGCTCTTCCGATCTNNNTGCTGCAAAGGACATGATTTTG
GAGCACACGTCTGAACTCTTTCCCTACACGACGCTCTTCCGATCTNNNACTGCAGAGCTTGTTGGAAAA
AGCACACGTCTGAACTCTTTCCCTACACGACGCTCTTCCGATCTNNNAAGTGTTGGGATTACAGGCATG
AGAGCACACGTCTGAACTCTTTCCCTACACGACGCTCTTCCGATCTNNNGATTGGCTTAGGTAAATGG
AGCACACGTCTGAACTCTTTCCCTACACGACGCTCTTCCGATCTNNNCAAAATGTACAACACCCCGTCT
GAGCACACGTCTGAACTCTTTCCCTACACGACGCTCTTCCGATCTNNNTATCATTTCCTGGGCCATTTCCA
AGCACACGTCTGAACTCTTTCCCTACACGACGCTCTTCCGATCTNNNGAGAATCGCTTGAAGTCAGGA

GAGCACACGTCTGAACTCTTTCCCTACACGACGCTCTTCCGATCTNNNTACTAGGGAGGCTAAGACAG
AGAGCACACGTCTGAACTCTTTCCCTACACGACGCTCTTCCGATCTNNNGAGGTTGCAGTGAGCTGAG
AGCACACGTCTGAACTCTTTCCCTACACGACGCTCTTCCGATCTNNNCTGAGACAGGAGAATCGCTTGA
AGCACACGTCTGAACTCTTTCCCTACACGACGCTCTTCCGATCTNNNCCCAGAAAAACCAATGGAACCA
AGCACACGTCTGAACTCTTTCCCTACACGACGCTCTTCCGATCTNNNAGAAGGCTGAGATACAACCCT
AGAGCACACGTCTGAACTCTTTCCCTACACGACGCTCTTCCGATCTNNNGAGGTTGCAGTGAGCCAAG
GAGCACACGTCTGAACTCTTTCCCTACACGACGCTCTTCCGATCTNNNGCTGAGACAGGAGAATCACTT
GAGCACACGTCTGAACTCTTTCCCTACACGACGCTCTTCCGATCTNNNGATCACAAGGTCAGGAGAGTC
GAGCACACGTCTGAACTCTTTCCCTACACGACGCTCTTCCGATCTNNNTAGAAGAATCACTTGAACCCCC
GAGCACACGTCTGAACTCTTTCCCTACACGACGCTCTTCCGATCTNNNGCAGAGGTTGTAAAGAGGTG
AGCACACGTCTGAACTCTTTCCCTACACGACGCTCTTCCGATCTNNNCTGTAATCCCAGCTACTCGAGAG
AGCACACGTCTGAACTCTTTCCCTACACGACGCTCTTCCGATCTNNNCGAGATTGTACCAGTGTACTCC
GAGCACACGTCTGAACTCTTTCCCTACACGACGCTCTTCCGATCTNNNAGTGAACTGAGATCATGCCAC
AGCACACGTCTGAACTCTTTCCCTACACGACGCTCTTCCGATCTNNNGAGAATGGCATGAATCCAGGAG
AGAGCACACGTCTGAACTCTTTCCCTACACGACGCTCTTCCGATCTNNNAGACCAGCTTGAGAAACATA
AGCACACGTCTGAACTCTTTCCCTACACGACGCTCTTCCGATCTNNNTTTGAGGCTGTAGTGAACCATG
AGCACACGTCTGAACTCTTTCCCTACACGACGCTCTTCCGATCTNNNCTGGGCAACAAGAACAAAACTC
GCACACGTCTGAACTCTTTCCCTACACGACGCTCTTCCGATCTNNNGCACATACCTGTAATCCCAGCTAC
GAGCACACGTCTGAACTCTTTCCCTACACGACGCTCTTCCGATCTNNNCTGAGGCAGGAGAATCACTTG
AGCACACGTCTGAACTCTTTCCCTACACGACGCTCTTCCGATCTNNNCTGTAGTTCCAGCTACTTGGGA
AGCACACGTCTGAACTCTTTCCCTACACGACGCTCTTCCGATCTNNNACAGTGAGTTGTGTTCATACCC
GAGCACACGTCTGAACTCTTTCCCTACACGACGCTCTTCCGATCTNNNAGCTGCTAGGGAGGCTAAGAT
GAGCACACGTCTGAACTCTTTCCCTACACGACGCTCTTCCGATCTNNNCTGAGGCAGGAGAATAGCTTG
GAGCACACGTCTGAACTCTTTCCCTACACGACGCTCTTCCGATCTNNNTTGAGGTCAGGAGTTCAAAA
AGCACACGTCTGAACTCTTTCCCTACACGACGCTCTTCCGATCTNNNGATCACTTGAGCATGAGAGGTC
GAGCACACGTCTGAACTCTTTCCCTACACGACGCTCTTCCGATCTNNNCTGTAGTCCCAGCTACTCAGG
AGAGCACACGTCTGAACTCTTTCCCTACACGACGCTCTTCCGATCTNNNCCTGACTCTGACAGACCCAA
GAGCACACGTCTGAACTCTTTCCCTACACGACGCTCTTCCGATCTNNNGCAGGAGAATTGCTTGAACCT
GAGCACACGTCTGAACTCTTTCCCTACACGACGCTCTTCCGATCTNNNATTGCTTGAATATGGGAGGCA
GAGCACACGTCTGAACTCTTTCCCTACACGACGCTCTTCCGATCTNNNACAAAAAGTAGGTTGCCAAGA
GAGCACACGTCTGAACTCTTTCCCTACACGACGCTCTTCCGATCTNNNTCAGCCAAAATAGGATTCAAGC
GAGCACACGTCTGAACTCTTTCCCTACACGACGCTCTTCCGATCTNNNAGACAGGAGAATCACTTGAAC
GAGCACACGTCTGAACTCTTTCCCTACACGACGCTCTTCCGATCTNNNATAAGGGCCAGCCTTCAAATCT
AGCACACGTCTGAACTCTTTCCCTACACGACGCTCTTCCGATCTNNNGTAATCCCAGCTACTCAGGAAG
GAGCACACGTCTGAACTCTTTCCCTACACGACGCTCTTCCGATCTNNNCTCAGTTTCTGCAACCCCAATTC
GAGCACACGTCTGAACTCTTTCCCTACACGACGCTCTTCCGATCTNNNCACCACTCCCAGCTAATTTTTGC
GAGCACACGTCTGAACTCTTTCCCTACACGACGCTCTTCCGATCTNNNGATTCAAGCAATTCTCCTGCGT
GAGCACACGTCTGAACTCTTTCCCTACACGACGCTCTTCCGATCTNNNAGATGAGCACATTTGGCCTCT
GAGCACACGTCTGAACTCTTTCCCTACACGACGCTCTTCCGATCTNNNTTGGTTTTTGTTGAGTTTTGC
GAGCACACGTCTGAACTCTTTCCCTACACGACGCTCTTCCGATCTNNNAAAGCTTGCCCAGATAAGAGA
GAGCACACGTCTGAACTCTTTCCCTACACGACGCTCTTCCGATCTNNNATGGAGTCATTTGGTTGCCTT
AGAGCACACGTCTGAACTCTTTCCCTACACGACGCTCTTCCGATCTNNNTTTCCTAATGTGCCCAATGAC
AGCACACGTCTGAACTCTTTCCCTACACGACGCTCTTCCGATCTNNNCTTGGTGAACTGCAAAGCCTAT

GCACACGTCTGAACTCTTTCCCTACACGACGCTCTTCCGATCTNNNCAGACACCAGACACCACTAAACT
AGCACACGTCTGAACTCTTTCCCTACACGACGCTCTTCCGATCTNNNTGGGATTTGGATGATTGGTG
GAGCACACGTCTGAACTCTTTCCCTACACGACGCTCTTCCGATCTNNNAGAGAGAAATGAACTGCTCC
AGCACACGTCTGAACTCTTTCCCTACACGACGCTCTTCCGATCTNNNGTTCTCATCATTCATCCCCCACT
GAGCACACGTCTGAACTCTTTCCCTACACGACGCTCTTCCGATCTNNNTGGTTTACCGAAAGACCAGG
AGCACACGTCTGAACTCTTTCCCTACACGACGCTCTTCCGATCTNNNAATCTGCTATCAAACCTCTCTG
AGCACACGTCTGAACTCTTTCCCTACACGACGCTCTTCCGATCTNNNTTCCCGTATCACCGTCTCTTTTCC
GAGCACACGTCTGAACTCTTTCCCTACACGACGCTCTTCCGATCTNNNAATGTCTTTCGGACCTTCCAG
AGCACACGTCTGAACTCTTTCCCTACACGACGCTCTTCCGATCTNNNTTTGCATTGTGCTGAACCTAGT
AGCACACGTCTGAACTCTTTCCCTACACGACGCTCTTCCGATCTNNNTTTGTGTCCAGTGAGCTACAA
GCACACGTCTGAACTCTTTCCCTACACGACGCTCTTCCGATCTNNNAGATGTTTGGTTTGAACAGTGG
AGCACACGTCTGAACTCTTTCCCTACACGACGCTCTTCCGATCTNNNACATGACAGATAGCATACCACC
AGCACACGTCTGAACTCTTTCCCTACACGACGCTCTTCCGATCTNNNTTCCCAGGTTCAAGTGATTCTCC
AGCACACGTCTGAACTCTTTCCCTACACGACGCTCTTCCGATCTNNNACTCTCTTGTTAGGGGCTAAT
GCACACGTCTGAACTCTTTCCCTACACGACGCTCTTCCGATCTNNNGCACTGTAGCCTAGGTCTCAAA
AGCACACGTCTGAACTCTTTCCCTACACGACGCTCTTCCGATCTNNNAGTGGATGGATGAATGGATG
AGCACACGTCTGAACTCTTTCCCTACACGACGCTCTTCCGATCTNNNGGTGGAAGTCAGTGGTTACAG
GCACACGTCTGAACTCTTTCCCTACACGACGCTCTTCCGATCTNNNGCCTGAACTTTTTCTGCATCTCC
GAGCACACGTCTGAACTCTTTCCCTACACGACGCTCTTCCGATCTNNNTTGCATTTTTTTCGCCAATCGT
GAGCACACGTCTGAACTCTTTCCCTACACGACGCTCTTCCGATCTNNNATGTCTATCACCACAAGGAGC
GCACACGTCTGAACTCTTTCCCTACACGACGCTCTTCCGATCTNNNGGATTATGCTGTCCTTGTAGGC
AGCACACGTCTGAACTCTTTCCCTACACGACGCTCTTCCGATCTNNNCTGCTTACCTCCATTGGCTAAGT
AGCACACGTCTGAACTCTTTCCCTACACGACGCTCTTCCGATCTNNNTCCTTCTCACATTTCCTCAGCAA
GAGCACACGTCTGAACTCTTTCCCTACACGACGCTCTTCCGATCTNNNATTTACAGCTACTTCATCGCCC
GAGCACACGTCTGAACTCTTTCCCTACACGACGCTCTTCCGATCTNNNTCTCTCTCTCTCTCTCTCTCT
GAGCACACGTCTGAACTCTTTCCCTACACGACGCTCTTCCGATCTNNNTTTTGGAACATGCTTGCCCTA
AGCACACGTCTGAACTCTTTCCCTACACGACGCTCTTCCGATCTNNNGTGATAGAGAGCAAGACCCTG
AGCACACGTCTGAACTCTTTCCCTACACGACGCTCTTCCGATCTNNNGTGAGCCAAGATCATGCTATT
AGCACACGTCTGAACTCTTTCCCTACACGACGCTCTTCCGATCTNNNGAGGTTACAGTGAGCCAAGAT
GAGCACACGTCTGAACTCTTTCCCTACACGACGCTCTTCCGATCTNNNGTAATCCCAGCTACTTAGGAG
GAGCACACGTCTGAACTCTTTCCCTACACGACGCTCTTCCGATCTNNNAATCTCTTGAACCAGCGAGTC
GAGCACACGTCTGAACTCTTTCCCTACACGACGCTCTTCCGATCTNNNCTGTAGTCCCAGCTACTCAGG
GAGCACACGTCTGAACTCTTTCCCTACACGACGCTCTTCCGATCTNNNCTGATGCAGGAGAATCACTG
AGCACACGTCTGAACTCTTTCCCTACACGACGCTCTTCCGATCTNNNTGTGTCTAAGGAGAGAGTTGT
GAGCACACGTCTGAACTCTTTCCCTACACGACGCTCTTCCGATCTNNNGAGATCCCATCATTGCACTCC
AGCACACGTCTGAACTCTTTCCCTACACGACGCTCTTCCGATCTNNNTTGAGATCAACCTGGGCAACA
AGCACACGTCTGAACTCTTTCCCTACACGACGCTCTTCCGATCTNNNAGGCAGAAGAATCACTTGAACC
AGCACACGTCTGAACTCTTTCCCTACACGACGCTCTTCCGATCTNNNTGAGAATCACTTGAACCCAGG
GAGCACACGTCTGAACTCTTTCCCTACACGACGCTCTTCCGATCTNNNGAGGCAGAAGAATCGCTTGA
AGCACACGTCTGAACTCTTTCCCTACACGACGCTCTTCCGATCTNNNGGTGACAGAGCAAGATTCCTT
AGAGCACACGTCTGAACTCTTTCCCTACACGACGCTCTTCCGATCTNNNGAATCACTTGAACCTGGGA
GAGCACACGTCTGAACTCTTTCCCTACACGACGCTCTTCCGATCTNNNTATAATCAAAGCAGTGCAAC
AGCACACGTCTGAACTCTTTCCCTACACGACGCTCTTCCGATCTNNNTGAGGCAGAAGAATTGCTTG

GAGCACACGTCTGAACTCTTTCCCTACACGACGCTCTTCCGATCTNNNAGTAGGCAGATCACTAGAGGT
GAGCACACGTCTGAACTCTTTCCCTACACGACGCTCTTCCGATCTNNNAGAGGTTGCAGTGAGCTGAT
GAGCACACGTCTGAACTCTTTCCCTACACGACGCTCTTCCGATCTNNNTGGGAAATGGAGGTTGTAGT
AGCACACGTCTGAACTCTTTCCCTACACGACGCTCTTCCGATCTNNNGAGGCAACAGAATCACTTGAAC
GAGCACACGTCTGAACTCTTTCCCTACACGACGCTCTTCCGATCTNNNGAGAATCGCTTGAACTCAGGA
GAGCACACGTCTGAACTCTTTCCCTACACGACGCTCTTCCGATCTNNNAGCTTGCAGTAAGTCGAGATC
GAGCACACGTCTGAACTCTTTCCCTACACGACGCTCTTCCGATCTNNNGAGAATTGCTTGGATCCAGGA
AGCACACGTCTGAACTCTTTCCCTACACGACGCTCTTCCGATCTNNNATCTCTTGAACCCAAAAGACGG
GAGCACACGTCTGAACTCTTTCCCTACACGACGCTCTTCCGATCTNNNCTCTAGTCCCAGCTACTCAGGA
AGCACACGTCTGAACTCTTTCCCTACACGACGCTCTTCCGATCTNNNTGCATACCCATAATCCCAGCTAC
GAGCACACGTCTGAACTCTTTCCCTACACGACGCTCTTCCGATCTNNNGGAGCTTGTAGTAAGCCGAGA
AGCACACGTCTGAACTCTTTCCCTACACGACGCTCTTCCGATCTNNNCTGTAATCCCAGCTACTCAGGAG
AGCACACGTCTGAACTCTTTCCCTACACGACGCTCTTCCGATCTNNNCAGGAATTTGAGGGCTGCATTA
GAGCACACGTCTGAACTCTTTCCCTACACGACGCTCTTCCGATCTNNNATGTAAAAGACCTGCACGTTG
GAGCACACGTCTGAACTCTTTCCCTACACGACGCTCTTCCGATCTNNNGCTGAGACAGGAGAATCACTT
GAGCACACGTCTGAACTCTTTCCCTACACGACGCTCTTCCGATCTNNNTAGACACCAGTAAGACCCTGT
GAGCACACGTCTGAACTCTTTCCCTACACGACGCTCTTCCGATCTNNNGCTCACTCCTGTAATCCTAGCT
GAGCACACGTCTGAACTCTTTCCCTACACGACGCTCTTCCGATCTNNNGAGATTGCAGTGAGCCAAGA
GAGCACACGTCTGAACTCTTTCCCTACACGACGCTCTTCCGATCTNNNGAGAATGGTGTGAACCTGGC
AGCACACGTCTGAACTCTTTCCCTACACGACGCTCTTCCGATCTNNNGAATGATGCGATGTGACTGAC
GAGCACACGTCTGAACTCTTTCCCTACACGACGCTCTTCCGATCTNNNGCATACCTATAATCCCAGCTG
AGCACACGTCTGAACTCTTTCCCTACACGACGCTCTTCCGATCTNNNGTAATCCCAGCTACTTAGGAGG
GAGCACACGTCTGAACTCTTTCCCTACACGACGCTCTTCCGATCTNNNGAGATTGCGATGAGTTGAGA
AGCACACGTCTGAACTCTTTCCCTACACGACGCTCTTCCGATCTNNNGCAGGAAAATAGCTTGAACCTG
AGCACACGTCTGAACTCTTTCCCTACACGACGCTCTTCCGATCTNNNAGAGGTTGTGGTGAGTTGAGA
AGCACACGTCTGAACTCTTTCCCTACACGACGCTCTTCCGATCTNNNCTGTAATCCCAGCTACTCAGGA
AGCACACGTCTGAACTCTTTCCCTACACGACGCTCTTCCGATCTNNNAAAGATAATACCTTGGCTGGGC
AGAGCACACGTCTGAACTCTTTCCCTACACGACGCTCTTCCGATCTNNNGTAATCCCAGCTACTCAGGAT
AGAGCACACGTCTGAACTCTTTCCCTACACGACGCTCTTCCGATCTNNNGTAATCCCAGCTACTTAGGA
GAGCACACGTCTGAACTCTTTCCCTACACGACGCTCTTCCGATCTNNNACCACACCCCGCTAACTTTATA
AGCACACGTCTGAACTCTTTCCCTACACGACGCTCTTCCGATCTNNNTTTGCTTTCAACCTAAATGGGCC
AGCACACGTCTGAACTCTTTCCCTACACGACGCTCTTCCGATCTNNNTAGGGAGGGAGTTTCAGCATTT
GAGCACACGTCTGAACTCTTTCCCTACACGACGCTCTTCCGATCTNNNTGCTGATGTACTTTGGGATGC
AGCACACGTCTGAACTCTTTCCCTACACGACGCTCTTCCGATCTNNNTAGAACAGCAGCTATCAAGGAG
AGCACACGTCTGAACTCTTTCCCTACACGACGCTCTTCCGATCTNNNAGGGTGTCAGTCTATTGCTCAT
AGCACACGTCTGAACTCTTTCCCTACACGACGCTCTTCCGATCTNNNCGTATTTCTGTTGCCCAATCTGC
GAGCACACGTCTGAACTCTTTCCCTACACGACGCTCTTCCGATCTNNNTTCTGTGTCCTTTTCTTTGCCAC
GAGCACACGTCTGAACTCTTTCCCTACACGACGCTCTTCCGATCTNNNGTTTCTATGACTGGTAGTGGG
AGCACACGTCTGAACTCTTTCCCTACACGACGCTCTTCCGATCTNNNAGTAGAGATGGGCCACCTTATT
AGCACACGTCTGAACTCTTTCCCTACACGACGCTCTTCCGATCTNNNCCTCCTCAAGTAGCTGGGATTA
AGCACACGTCTGAACTCTTTCCCTACACGACGCTCTTCCGATCTNNNATTTTTCCAGCTGTCACCACTGC
GAGCACACGTCTGAACTCTTTCCCTACACGACGCTCTTCCGATCTNNNCTTGAACTCCTGACCTCAAGTC
AGCACACGTCTGAACTCTTTCCCTACACGACGCTCTTCCGATCTNNNGGCCCACATGCATTTTAGAAACA

AGCACACGTCTGAACTCTTTCCCTACACGACGCTCTTCCGATCTNNNGAGGAGTTGCTAGTACTGGCA
GAGCACACGTCTGAACTCTTTCCCTACACGACGCTCTTCCGATCTNNNGATTACAGGAGTGAGCCACT
GAGCACACGTCTGAACTCTTTCCCTACACGACGCTCTTCCGATCTNNNCATGCACATTACAGTCTGAGC
GCACACGTCTGAACTCTTTCCCTACACGACGCTCTTCCGATCTNNNTCACTTCCAATCACTGGCACTAT
GCACACGTCTGAACTCTTTCCCTACACGACGCTCTTCCGATCTNNNTCACTTCCAATCACTGGCACTAT
GAGCACACGTCTGAACTCTTTCCCTACACGACGCTCTTCCGATCTNNNACACAATGCTGAATGCTGTG
AGCACACGTCTGAACTCTTTCCCTACACGACGCTCTTCCGATCTNNNTAGTGGAGAGAAAGCCCAGAA
GAGCACACGTCTGAACTCTTTCCCTACACGACGCTCTTCCGATCTNNNTGAGATGATAGTGCCTTTGG
GAGCACACGTCTGAACTCTTTCCCTACACGACGCTCTTCCGATCTNNNCGACACTCCGTCTCCAAAAAA
AGCACACGTCTGAACTCTTTCCCTACACGACGCTCTTCCGATCTNNNAGTCATGTGCTGTTTCTTTAGC
GAGCACACGTCTGAACTCTTTCCCTACACGACGCTCTTCCGATCTNNNGATGGGGTTTCATCACATTGG
GCACACGTCTGAACTCTTTCCCTACACGACGCTCTTCCGATCTNNNAGAAAACCCTTGTGGCAGTTAG
GAGCACACGTCTGAACTCTTTCCCTACACGACGCTCTTCCGATCTNNNTGCAGTTTGGCTAGCATCTCT
AGCACACGTCTGAACTCTTTCCCTACACGACGCTCTTCCGATCTNNNAGGGAGCTTGAAAATGCATTT
AGCACACGTCTGAACTCTTTCCCTACACGACGCTCTTCCGATCTNNNCCTACACATTATCAGGCCTTTGG
AGCACACGTCTGAACTCTTTCCCTACACGACGCTCTTCCGATCTNNNTTCACTGTGATAAGTTCCAGGC
AGCACACGTCTGAACTCTTTCCCTACACGACGCTCTTCCGATCTNNNTGGCTCCAAAATTGGTTGGTTA
GAGCACACGTCTGAACTCTTTCCCTACACGACGCTCTTCCGATCTNNNCACTTGAGGTCAGAGGTTCA
AGCACACGTCTGAACTCTTTCCCTACACGACGCTCTTCCGATCTNNNGGTGGCATATGCTCATAGTCTC
GAGCACACGTCTGAACTCTTTCCCTACACGACGCTCTTCCGATCTNNNGGTGGAAGTTTCAATGAGCC
GAGCACACGTCTGAACTCTTTCCCTACACGACGCTCTTCCGATCTNNNTGCCTGTAATTCCAGCTACTC
AGCACACGTCTGAACTCTTTCCCTACACGACGCTCTTCCGATCTNNNGGGAGACAGAGTGAGACTTC
GAGCACACGTCTGAACTCTTTCCCTACACGACGCTCTTCCGATCTNNNGAATGACTGAATGAGCTGCA
AGCACACGTCTGAACTCTTTCCCTACACGACGCTCTTCCGATCTNNNAGAATCACTTGAATCCAGGAG
GCACACGTCTGAACTCTTTCCCTACACGACGCTCTTCCGATCTNNNAAGATCGCTCCATTACACTCCAG
AGCACACGTCTGAACTCTTTCCCTACACGACGCTCTTCCGATCTNNNGTCTGTAGTCCTAGCTACCAGG
GAGCACACGTCTGAACTCTTTCCCTACACGACGCTCTTCCGATCTNNNCGAAACCCCCGTCTGTACTAA
GAGCACACGTCTGAACTCTTTCCCTACACGACGCTCTTCCGATCTNNNCCTGGGGGAACTACAATTCA
GAGCACACGTCTGAACTCTTTCCCTACACGACGCTCTTCCGATCTNNNACAAGACAACCTATCCCTTGG
GAGCACACGTCTGAACTCTTTCCCTACACGACGCTCTTCCGATCTNNNCACATTTTGGAAGGTTGAGGT
GAGCACACGTCTGAACTCTTTCCCTACACGACGCTCTTCCGATCTNNNCTGAGGCAGGAGAATCACTT
GAGCACACGTCTGAACTCTTTCCCTACACGACGCTCTTCCGATCTNNNATCTTGGCCAGTGAAATCTG
AGCACACGTCTGAACTCTTTCCCTACACGACGCTCTTCCGATCTNNNGTCCAAGCTATTCAGGAGGCTA
GAGCACACGTCTGAACTCTTTCCCTACACGACGCTCTTCCGATCTNNNAGTGAGACGAGATCATGCTA
AGCACACGTCTGAACTCTTTCCCTACACGACGCTCTTCCGATCTNNNCTGTAGTCCCAGCTACTCAGGA
GAGCACACGTCTGAACTCTTTCCCTACACGACGCTCTTCCGATCTNNNCTGAGGCAGGAGAATTGCTT
AGCACACGTCTGAACTCTTTCCCTACACGACGCTCTTCCGATCTNNNAGAGAATGGCTTGAACTCAGG
AGCACACGTCTGAACTCTTTCCCTACACGACGCTCTTCCGATCTNNNCCTGGGTAACACAGTGATACCT
AGAGCACACGTCTGAACTCTTTCCCTACACGACGCTCTTCCGATCTNNNTAAGGCAGGAGAATCGCT
GAGCACACGTCTGAACTCTTTCCCTACACGACGCTCTTCCGATCTNNNCAGGAGGTTGAGACAGGAG
GAGCACACGTCTGAACTCTTTCCCTACACGACGCTCTTCCGATCTNNNTGAGCCACGAAAATCACTTGA
AGCACACGTCTGAACTCTTTCCCTACACGACGCTCTTCCGATCTNNNGAGACAGGAGAATGCTAGGA
GAGCACACGTCTGAACTCTTTCCCTACACGACGCTCTTCCGATCTNNNCTGTAGTCCCAGCTACTTTG

AGCACACGTCTGAACTCTTTCCCTACACGACGCTCTTCCGATCTNNNTGTAATCCCCGCTAATCAAGAG
GAGCACACGTCTGAACTCTTTCCCTACACGACGCTCTTCCGATCTNNNCAGAGGTTGCAGTGAGATGA
GAGCACACGTCTGAACTCTTTCCCTACACGACGCTCTTCCGATCTNNNGGATGATCACTTGAGGTCAG
GAGCACACGTCTGAACTCTTTCCCTACACGACGCTCTTCCGATCTNNNAAAAAATTAGCAGGGCATGGT
GAGCACACGTCTGAACTCTTTCCCTACACGACGCTCTTCCGATCTNNNCACTACATTCCAGTCTGGGTGA
GAGCACACGTCTGAACTCTTTCCCTACACGACGCTCTTCCGATCTNNNCTGAGGCAGGAGAATCACTT
GAGCACACGTCTGAACTCTTTCCCTACACGACGCTCTTCCGATCTNNNCCAACATAGCAAGACGTTGTCT
GAGCACACGTCTGAACTCTTTCCCTACACGACGCTCTTCCGATCTNNNCTGAGGCAGGAGAATCACAT
GAGCACACGTCTGAACTCTTTCCCTACACGACGCTCTTCCGATCTNNNGCAGCTAATGGGAGAAAATG
GAGCACACGTCTGAACTCTTTCCCTACACGACGCTCTTCCGATCTNNNGAATCACTTGAACTCAGGAT
GAGCACACGTCTGAACTCTTTCCCTACACGACGCTCTTCCGATCTNNNGGTTGCAGTGAACCAAGATCA
GAGCACACGTCTGAACTCTTTCCCTACACGACGCTCTTCCGATCTNNNGAAAATGGCGTGAACACAGG
AGAGCACACGTCTGAACTCTTTCCCTACACGACGCTCTTCCGATCTNNNTGAGGTGGAAGGATTTTCTG
AGAGCACACGTCTGAACTCTTTCCCTACACGACGCTCTTCCGATCTNNNCTGAGCCAGGAGAATCACTT
AGCACACGTCTGAACTCTTTCCCTACACGACGCTCTTCCGATCTNNNCCTTGGTGACAGAACAAGACTCT
GAGCACACGTCTGAACTCTTTCCCTACACGACGCTCTTCCGATCTNNNATAAAACAAGGAGGCCAAAG
AGCACACGTCTGAACTCTTTCCCTACACGACGCTCTTCCGATCTNNNTGTAATCCCAGCTACTCAGGAGA
AGAGCACACGTCTGAACTCTTTCCCTACACGACGCTCTTCCGATCTNNNGATTGCTTGAGCTCAGGAGT
AGCACACGTCTGAACTCTTTCCCTACACGACGCTCTTCCGATCTNNNGAAAATCACTTGAGCCCAGGAG
AGCACACGTCTGAACTCTTTCCCTACACGACGCTCTTCCGATCTNNNCCTCTTATATCCAGTCCTTGCCG
GAGCACACGTCTGAACTCTTTCCCTACACGACGCTCTTCCGATCTNNNCTATAAACGCCTGATCACGCAT
AGCACACGTCTGAACTCTTTCCCTACACGACGCTCTTCCGATCTNNNTTGAAGGATAAGAAGCAAGAA
GAGCACACGTCTGAACTCTTTCCCTACACGACGCTCTTCCGATCTNNNGTTCTGTGTGTTCCTTCCATGT
AGCACACGTCTGAACTCTTTCCCTACACGACGCTCTTCCGATCTNNNTTTCTGTACTTTTCTCCTCCCCG
AGCACACGTCTGAACTCTTTCCCTACACGACGCTCTTCCGATCTNNNATCCTGCCCACTGTTATTTCCTT
AGCACACGTCTGAACTCTTTCCCTACACGACGCTCTTCCGATCTNNNTAAAGCTGGGTGGGATTGTTT
GAGCACACGTCTGAACTCTTTCCCTACACGACGCTCTTCCGATCTNNNGCCCCTGAGAGTCATTCTGTT
AGCACACGTCTGAACTCTTTCCCTACACGACGCTCTTCCGATCTNNNACCATAGTACCATTCTCTCACCT
AGCACACGTCTGAACTCTTTCCCTACACGACGCTCTTCCGATCTNNNTAGAGAAGGGGACAATCAGCAA
AGCACACGTCTGAACTCTTTCCCTACACGACGCTCTTCCGATCTNNNAGCCAAGTACTTGATTGCCTACT
GAGCACACGTCTGAACTCTTTCCCTACACGACGCTCTTCCGATCTNNNCAGAAATGTGCACCCGTAATC
AGCACACGTCTGAACTCTTTCCCTACACGACGCTCTTCCGATCTNNNCAACAGTCCTAAGTCCACATCCA
AGCACACGTCTGAACTCTTTCCCTACACGACGCTCTTCCGATCTNNNATGCCCAGCTAAAGAACTCTTCT
ACACGTCTGAACTCTTTCCCTACACGACGCTCTTCCGATCTNNNAGACTCTACCAAAAAACTCTTACACA
GAGCACACGTCTGAACTCTTTCCCTACACGACGCTCTTCCGATCTNNNAAATCTCACCCACTGATCTAGC
AGCACACGTCTGAACTCTTTCCCTACACGACGCTCTTCCGATCTNNNACTCCATCTTTTTGTCAGAGGCT
AGCACACGTCTGAACTCTTTCCCTACACGACGCTCTTCCGATCTNNNCTCCTGAGTAGCTGGGACTACA
AGCACACGTCTGAACTCTTTCCCTACACGACGCTCTTCCGATCTNNNCTCTCCTTTGTGCCCTATTTCCTC
AGCACACGTCTGAACTCTTTCCCTACACGACGCTCTTCCGATCTNNNCCACCACGTCCAGCTAATTTTTT
AGAGCACACGTCTGAACTCTTTCCCTACACGACGCTCTTCCGATCTNNNCAATGATTGACTGGCATCAG
AGCACACGTCTGAACTCTTTCCCTACACGACGCTCTTCCGATCTNNNTCCTCTTACCACTCCCTCATTCTC
AGCACACGTCTGAACTCTTTCCCTACACGACGCTCTTCCGATCTNNNATGGTAGGTGTTTTCTCTCAGG
GAGCACACGTCTGAACTCTTTCCCTACACGACGCTCTTCCGATCTNNNCATTTCACTAAGTGCTGCCCAA

GCACACGTCTGAACTCTTTCCCTACACGACGCTCTTCCGATCTNNNTGAATAGGTGGAACTACAGGT
AGAGCACACGTCTGAACTCTTTCCCTACACGACGCTCTTCCGATCTNNNCCTTTCATGTGTGTCTGAGT
AGCACACGTCTGAACTCTTTCCCTACACGACGCTCTTCCGATCTNNNGAGAAGGGTCTCTTGGCTGA
GAGCACACGTCTGAACTCTTTCCCTACACGACGCTCTTCCGATCTNNNGATATTTTCCTGCTGGACAAG
GAGCACACGTCTGAACTCTTTCCCTACACGACGCTCTTCCGATCTNNNCAATGGGTGTGTTCAAAGCT
GAGCACACGTCTGAACTCTTTCCCTACACGACGCTCTTCCGATCTNNNGATTATAGCCATGTGCCACCA
AGCACACGTCTGAACTCTTTCCCTACACGACGCTCTTCCGATCTNNNTGATCTCCTGACCTTGTATGTG
GCACACGTCTGAACTCTTTCCCTACACGACGCTCTTCCGATCTNNNCACTGGCTCCTTACTCTTGAACT
AGCACACGTCTGAACTCTTTCCCTACACGACGCTCTTCCGATCTNNNGATCTCTCCATGCTCAGGTGAT
AGCACACGTCTGAACTCTTTCCCTACACGACGCTCTTCCGATCTNNNCGATGAGCCTCTGATATTGGA
AGCACACGTCTGAACTCTTTCCCTACACGACGCTCTTCCGATCTNNNCCACTGTGCATGGATTCCTTTTT
AGCACACGTCTGAACTCTTTCCCTACACGACGCTCTTCCGATCTNNNTTGCATATGTTGAACCAGCCTT
AGCACACGTCTGAACTCTTTCCCTACACGACGCTCTTCCGATCTNNNACCCACAAAACTGTAGCTACAT
AGCACACGTCTGAACTCTTTCCCTACACGACGCTCTTCCGATCTNNNGGTGTGGGTAAGAAATGCATT
GCACACGTCTGAACTCTTTCCCTACACGACGCTCTTCCGATCTNNNATATCCTTTTCTAGTCCTGGGCC
GAGCACACGTCTGAACTCTTTCCCTACACGACGCTCTTCCGATCTNNNTGCAACTCCTAAGCTCAAGTG
AGCACACGTCTGAACTCTTTCCCTACACGACGCTCTTCCGATCTNNNCCATTTGAGTCTCCTGTGAGG
GCACACGTCTGAACTCTTTCCCTACACGACGCTCTTCCGATCTNNNCAAAAAGGCTTTTGTGGCTACT
AGCACACGTCTGAACTCTTTCCCTACACGACGCTCTTCCGATCTNNNGGCAGCAAATCACACTCTGAA
AGCACACGTCTGAACTCTTTCCCTACACGACGCTCTTCCGATCTNNNCTTCATCAGTGAGAAAGGAGC
AGCACACGTCTGAACTCTTTCCCTACACGACGCTCTTCCGATCTNNNGTCGTGATGTGCTCTACTCTAG
GAGCACACGTCTGAACTCTTTCCCTACACGACGCTCTTCCGATCTNNNGGGATTACAAGTGTGTACCA
GAGCACACGTCTGAACTCTTTCCCTACACGACGCTCTTCCGATCTNNNGGGTTTGACAAAAGGTACAA
GCACACGTCTGAACTCTTTCCCTACACGACGCTCTTCCGATCTNNNTGCACTCAGTGAAATCAGGAGA
AGCACACGTCTGAACTCTTTCCCTACACGACGCTCTTCCGATCTNNNTGGATTATGCCACTACACTCCAG
GAGCACACGTCTGAACTCTTTCCCTACACGACGCTCTTCCGATCTNNNGGAAGTTGTAGTGAGCCGA
AGAGCACACGTCTGAACTCTTTCCCTACACGACGCTCTTCCGATCTNNNTAGGGCAGGAGAATCACTT
GAGCACACGTCTGAACTCTTTCCCTACACGACGCTCTTCCGATCTNNNTTGAGACGGGAGAATTGCTT
AGCACACGTCTGAACTCTTTCCCTACACGACGCTCTTCCGATCTNNNCTGAGGCAGGAGAATCACTTG
AGCACACGTCTGAACTCTTTCCCTACACGACGCTCTTCCGATCTNNNGGGCAACAAGAGCAAAATTC
AGCACACGTCTGAACTCTTTCCCTACACGACGCTCTTCCGATCTNNNCCGTCTCAGGCAATTTAGGTTC
GCACACGTCTGAACTCTTTCCCTACACGACGCTCTTCCGATCTNNNCTGTAGTCCCAGCTACTCAGGAG
GAGCACACGTCTGAACTCTTTCCCTACACGACGCTCTTCCGATCTNNNGAGGTTGCTGTGAACTGAGA
GAGCACACGTCTGAACTCTTTCCCTACACGACGCTCTTCCGATCTNNNGCTGAGATTATGTCGTTGCA
GAGCACACGTCTGAACTCTTTCCCTACACGACGCTCTTCCGATCTNNNATCATGCCACTGTACTCCAAC
GAGCACACGTCTGAACTCTTTCCCTACACGACGCTCTTCCGATCTNNNATGCCTGTAATTCCAGCTACT
AGCACACGTCTGAACTCTTTCCCTACACGACGCTCTTCCGATCTNNNTGTAATCTTTGCACTTTGGGA
GAGCACACGTCTGAACTCTTTCCCTACACGACGCTCTTCCGATCTNNNCTGTAGTCCCAGCTACTCGAG
AGCACACGTCTGAACTCTTTCCCTACACGACGCTCTTCCGATCTNNNCAAGTTTTCACTGTGAGGACTG
AGCACACGTCTGAACTCTTTCCCTACACGACGCTCTTCCGATCTNNNCTGTAATCCCAGCTACTCAGGA
AGCACACGTCTGAACTCTTTCCCTACACGACGCTCTTCCGATCTNNNGGAGGTTGTAGTGAGCCAAGA
AGCACACGTCTGAACTCTTTCCCTACACGACGCTCTTCCGATCTNNNACCTGTAATCCCAGCACTTTAG
GCACACGTCTGAACTCTTTCCCTACACGACGCTCTTCCGATCTNNNAGAGTTCATTGAATCTGGGAG

GAGCACACGTCTGAACTCTTTCCCTACACGACGCTCTTCCGATCTNNNTCATGCCTGTAATCCCAACACT
AGAGCACACGTCTGAACTCTTTCCCTACACGACGCTCTTCCGATCTNNNGCAACAGAACAAGACTCCAT
GAGCACACGTCTGAACTCTTTCCCTACACGACGCTCTTCCGATCTNNNTCTAGACTACACTGGCCAACA
GAGCACACGTCTGAACTCTTTCCCTACACGACGCTCTTCCGATCTNNNAAGATCATGCCACTACACTCCA
AGCACACGTCTGAACTCTTTCCCTACACGACGCTCTTCCGATCTNNNGCCCACATTCTATCCTTCTTCCA
GAGCACACGTCTGAACTCTTTCCCTACACGACGCTCTTCCGATCTNNNGTAATCCTAGCTACTTGGGAG
GAGCACACGTCTGAACTCTTTCCCTACACGACGCTCTTCCGATCTNNNTAGGCAGATCTCTTGAGTCCA
AGCACACGTCTGAACTCTTTCCCTACACGACGCTCTTCCGATCTNNNTAATCTCAGCTATTTGGGAGGC
AGCACACGTCTGAACTCTTTCCCTACACGACGCTCTTCCGATCTNNNCTGAGGCAGGAGAATCACTTGA
GAGCACACGTCTGAACTCTTTCCCTACACGACGCTCTTCCGATCTNNNGTGGAGGTTTCAGTAAGTCGA
AGCACACGTCTGAACTCTTTCCCTACACGACGCTCTTCCGATCTNNNGTAGGAGAATCTCTTGAACCCG
AGCACACGTCTGAACTCTTTCCCTACACGACGCTCTTCCGATCTNNNCACGGCGAAACTCAGTCTTTAT
GAGCACACGTCTGAACTCTTTCCCTACACGACGCTCTTCCGATCTNNNCTGAGGCAGGAGAATGACAT
GAGCACACGTCTGAACTCTTTCCCTACACGACGCTCTTCCGATCTNNNAGTTGCAGTGAGCTGAGATT
GAGCACACGTCTGAACTCTTTCCCTACACGACGCTCTTCCGATCTNNNGATTGCAGTGAGCCAAGATGA
GAGCACACGTCTGAACTCTTTCCCTACACGACGCTCTTCCGATCTNNNGAGGGAGTATTTCTTGAGCCC
GCACACGTCTGAACTCTTTCCCTACACGACGCTCTTCCGATCTNNNTATGTGCCTGTAATCCCAGCTAC
GAGCACACGTCTGAACTCTTTCCCTACACGACGCTCTTCCGATCTNNNACACCTGTAATCCCAGCTACTT
GCACACGTCTGAACTCTTTCCCTACACGACGCTCTTCCGATCTNNNGTTGCCGTGAGCCAAGATTATAC
GAGCACACGTCTGAACTCTTTCCCTACACGACGCTCTTCCGATCTNNNAGAAGTTGCAGTGAGCCAAGA
AGCACACGTCTGAACTCTTTCCCTACACGACGCTCTTCCGATCTNNNTGTAATCCTAGCACTTTGGGAG
GAGCACACGTCTGAACTCTTTCCCTACACGACGCTCTTCCGATCTNNNGAGGATGAGTAGGTGTTGGC
GAGCACACGTCTGAACTCTTTCCCTACACGACGCTCTTCCGATCTNNNGGATCATCTGAGGTCAGGAG
AGCACACGTCTGAACTCTTTCCCTACACGACGCTCTTCCGATCTNNNACTTCCTGAGATTGAGCCAGA
GAGCACACGTCTGAACTCTTTCCCTACACGACGCTCTTCCGATCTNNNGGTTGCAGTAAACCGAAGTC
AGCACACGTCTGAACTCTTTCCCTACACGACGCTCTTCCGATCTNNNATGGTAGTGTGTGCCTGTAATC
AGAGCACACGTCTGAACTCTTTCCCTACACGACGCTCTTCCGATCTNNNCACTTGAGGTCAGGAGTTCA
AAGAGCACACGTCTGAACTCTTTCCCTACACGACGCTCTTCCGATCTNNNGAGGTTGCAGTAAGCTGA
AGCACACGTCTGAACTCTTTCCCTACACGACGCTCTTCCGATCTNNNAAGAATCACTTGAACCCAGGAG
AGAGCACACGTCTGAACTCTTTCCCTACACGACGCTCTTCCGATCTNNNTTCAGGACACAGGAAATGGA
GAGCACACGTCTGAACTCTTTCCCTACACGACGCTCTTCCGATCTNNNCATCTGGGCAAAAAAGCAAAA
GAGCACACGTCTGAACTCTTTCCCTACACGACGCTCTTCCGATCTNNNCAAGATCGCACCACTGTACTCT
GAGCACACGTCTGAACTCTTTCCCTACACGACGCTCTTCCGATCTNNNGAGGCAGGAAAATCACTTGAA
AGCACACGTCTGAACTCTTTCCCTACACGACGCTCTTCCGATCTNNNCAGAGGTTACAGTGTGCTGAGA
GAGCACACGTCTGAACTCTTTCCCTACACGACGCTCTTCCGATCTNNNTGGTTTCAAACTCCTGGTCTCA
GAGCACACGTCTGAACTCTTTCCCTACACGACGCTCTTCCGATCTNNNTCAGCATGACCTACTCCAAGA
AGCACACGTCTGAACTCTTTCCCTACACGACGCTCTTCCGATCTNNNTTCAGATACTTGTGTGAGTGCC
AGCACACGTCTGAACTCTTTCCCTACACGACGCTCTTCCGATCTNNNTGCCAAAACATTCTCCCATTCTG
GAGCACACGTCTGAACTCTTTCCCTACACGACGCTCTTCCGATCTNNNACAACAACCTCTGTTTCAGAA
AGCACACGTCTGAACTCTTTCCCTACACGACGCTCTTCCGATCTNNNCCCCAGATCTTCAAAGTGTTGGA
GAGCACACGTCTGAACTCTTTCCCTACACGACGCTCTTCCGATCTNNNCTGCATCCTCACCAAAACTTGT
AGCACACGTCTGAACTCTTTCCCTACACGACGCTCTTCCGATCTNNNCTTCCAAAGTGCTGGGATTACA
AGCACACGTCTGAACTCTTTCCCTACACGACGCTCTTCCGATCTNNNAGTGTGAGGATGACCCAGTAC

AGCACACGTCTGAACTCTTTCCCTACACGACGCTCTTCCGATCTNNNCTCCTGAGTAGCTGGGTTTACA
AGCACACGTCTGAACTCTTTCCCTACACGACGCTCTTCCGATCTNNNCTGGGGAAAGCCATGACTAAT
AGCACACGTCTGAACTCTTTCCCTACACGACGCTCTTCCGATCTNNNTAGCTGGGATTATAGGTGTGC
GAGCACACGTCTGAACTCTTTCCCTACACGACGCTCTTCCGATCTNNNCTTGTCTCTTCAGGTGTTTGC
AGCACACGTCTGAACTCTTTCCCTACACGACGCTCTTCCGATCTNNNTAGCTGGAATTATAGGCACCT
AGCACACGTCTGAACTCTTTCCCTACACGACGCTCTTCCGATCTNNNGTCTCAGCCTGCATTAGTACCA
GAGCACACGTCTGAACTCTTTCCCTACACGACGCTCTTCCGATCTNNNTTCCTTCCTTCCCTTCCTTCTT
AGCACACGTCTGAACTCTTTCCCTACACGACGCTCTTCCGATCTNNNTATTTCCGTCCTTCCTTCCTCAC
AGCACACGTCTGAACTCTTTCCCTACACGACGCTCTTCCGATCTNNNATGGGAAGGGGGAAAGGAA
GAGCACACGTCTGAACTCTTTCCCTACACGACGCTCTTCCGATCTNNNTTCTTTCTAGGTCTTCAGGTG
AGCACACGTCTGAACTCTTTCCCTACACGACGCTCTTCCGATCTNNNTCTGCTCCAACTCCAAACTTGTT
AGCACACGTCTGAACTCTTTCCCTACACGACGCTCTTCCGATCTNNNCCTAGTTACTGCTCCCCATTTGT
GAGCACACGTCTGAACTCTTTCCCTACACGACGCTCTTCCGATCTNNNGGGAGCTCATGAGGCAAACA
GAGCACACGTCTGAACTCTTTCCCTACACGACGCTCTTCCGATCTNNNCACACACATGTTTATCTGCAC
AGCACACGTCTGAACTCTTTCCCTACACGACGCTCTTCCGATCTNNNAAAGACCCACCCTTGTAGTTGT
AGCACACGTCTGAACTCTTTCCCTACACGACGCTCTTCCGATCTNNNAATGACTTTCACAGTGGAAGCA
AGCACACGTCTGAACTCTTTCCCTACACGACGCTCTTCCGATCTNNNCTTTATGCCTGATGCCATCCAT
AGCACACGTCTGAACTCTTTCCCTACACGACGCTCTTCCGATCTNNNAGATTTCCTGGCCACACCTAATT
GAGCACACGTCTGAACTCTTTCCCTACACGACGCTCTTCCGATCTNNNAACAAGGTTTCTGCCTTCATG
AGCACACGTCTGAACTCTTTCCCTACACGACGCTCTTCCGATCTNNNGGCAGGAGGATTGATTGAACC
AGCACACGTCTGAACTCTTTCCCTACACGACGCTCTTCCGATCTNNNGGGGTTCTGGGTATTTTCTCTC
AGCACACGTCTGAACTCTTTCCCTACACGACGCTCTTCCGATCTNNNCCAGCAGTGCTTTGAGTTTTGT
AGCACACGTCTGAACTCTTTCCCTACACGACGCTCTTCCGATCTNNNGTGAGGGCCTGATTCCTGATT
AGCACACGTCTGAACTCTTTCCCTACACGACGCTCTTCCGATCTNNNCATGTAGAGGTCAGAGCATCC
GCACACGTCTGAACTCTTTCCCTACACGACGCTCTTCCGATCTNNNCTTGGCTTCCTGCAGAATGTTAC
AGCACACGTCTGAACTCTTTCCCTACACGACGCTCTTCCGATCTNNNGAGCCTGAGAAGACAAGACTT
GCACACGTCTGAACTCTTTCCCTACACGACGCTCTTCCGATCTNNNGAACTGCTTGAATCTGGGAATC
AGCACACGTCTGAACTCTTTCCCTACACGACGCTCTTCCGATCTNNNGATCACACCACTATACTCCAGCC
AGCACACGTCTGAACTCTTTCCCTACACGACGCTCTTCCGATCTNNNCCAGGGATTTGAGGTTACAGT
AGCACACGTCTGAACTCTTTCCCTACACGACGCTCTTCCGATCTNNNAAACCCTGTGCCTACCAAAAAT
GAGCACACGTCTGAACTCTTTCCCTACACGACGCTCTTCCGATCTNNNGTGCCTATAGTCCCAGCTACT
GAGCACACGTCTGAACTCTTTCCCTACACGACGCTCTTCCGATCTNNNGAAGACTGCAGTGAGCTGA
AGCACACGTCTGAACTCTTTCCCTACACGACGCTCTTCCGATCTNNNAGGCAGGAGAATAACTTGAAC
AGCACACGTCTGAACTCTTTCCCTACACGACGCTCTTCCGATCTNNNCGCCTATTTAGTCCCAGCTACTT
AGCACACGTCTGAACTCTTTCCCTACACGACGCTCTTCCGATCTNNNGAGGTTGCAACGAGTTGAGA
GAGCACACGTCTGAACTCTTTCCCTACACGACGCTCTTCCGATCTNNNAGGAGAATCGCTTGAATCCA
GCACACGTCTGAACTCTTTCCCTACACGACGCTCTTCCGATCTNNNGAGAATTGCTTGAACTGGGACA
GAGCACACGTCTGAACTCTTTCCCTACACGACGCTCTTCCGATCTNNNATGCCTGTAGTCCCAGTTACTT
GAGCACACGTCTGAACTCTTTCCCTACACGACGCTCTTCCGATCTNNNCTGTAGTTCCAGCTACTCAGG
GAGCACACGTCTGAACTCTTTCCCTACACGACGCTCTTCCGATCTNNNCAACTTGCAGTGAGCTGAGAT
AGCACACGTCTGAACTCTTTCCCTACACGACGCTCTTCCGATCTNNNGAGAATCCATTGAACCTGGGA
GAGCACACGTCTGAACTCTTTCCCTACACGACGCTCTTCCGATCTNNNAGGAGAATTGCTTGAACCCA
GAGCACACGTCTGAACTCTTTCCCTACACGACGCTCTTCCGATCTNNNCTCCCACTGCTTCACTTGACTA

```
GAGCACACGTCTGAACTCTTTCCCTACACGACGCTCTTCCGATCTNNNATGCAGTAAGCCATTATTGT
GAGCACACGTCTGAACTCTTTCCCTACACGACGCTCTTCCGATCTNNNAGGCAGAAGAATCACTTGA
AGCACACGTCTGAACTCTTTCCCTACACGACGCTCTTCCGATCTNNNAAGCAGACCAATCAAGAGCA
AGCACACGTCTGAACTCTTTCCCTACACGACGCTCTTCCGATCTNNNCAACACCAGCTTAGGCAATAT
AGCACACGTCTGAACTCTTTCCCTACACGACGCTCTTCCGATCTNNNGAGGTTGCAGTGAGTCGAGA
GAGCACACGTCTGAACTCTTTCCCTACACGACGCTCTTCCGATCTNNNAGAAGACAGAGCTTGTAGT
AGCACACGTCTGAACTCTTTCCCTACACGACGCTCTTCCGATCTNNNGAAGAATCATTGAACCCACGA
GAGCACACGTCTGAACTCTTTCCCTACACGACGCTCTTCCGATCTNNNTGTGTACTTGTAACCCCAGCT
AGCACACGTCTGAACTCTTTCCCTACACGACGCTCTTCCGATCTNNNTGAGGTGGGAAAATTGTTTGA
AGCACACGTCTGAACTCTTTCCCTACACGACGCTCTTCCGATCTNNNTAAGAGTGTGTGGTAAGGAC
AGCACACGTCTGAACTCTTTCCCTACACGACGCTCTTCCGATCTNNNGAGGTTGCAGTGATCCGAGA
AGCACACGTCTGAACTCTTTCCCTACACGACGCTCTTCCGATCTNNNGGAGCTTACAGTGAGCAGAG
GAGCACACGTCTGAACTCTTTCCCTACACGACGCTCTTCCGATCTNNNCTTGCAGTGAGCCGAGATTA
AGCACACGTCTGAACTCTTTCCCTACACGACGCTCTTCCGATCTNNNTGGACTTCCTCATAACATGGTG
GAGCACACGTCTGAACTCTTTCCCTACACGACGCTCTTCCGATCTNNNAGGCTGTGAAATTGGAGGA
GAGCACACGTCTGAACTCTTTCCCTACACGACGCTCTTCCGATCTNNNGGCCCACAGAATTGTATGAG
GAGCACACGTCTGAACTCTTTCCCTACACGACGCTCTTCCGATCTNNNGGACACATATAGAAAGCTGG
GAGCACACGTCTGAACTCTTTCCCTACACGACGCTCTTCCGATCTNNNTATCTGAGAGGTCTAGCTGG
GCACACGTCTGAACTCTTTCCCTACACGACGCTCTTCCGATCTNNNTGTAATCACCATCAATTTGGCAA
GCACACGTCTGAACTCTTTCCCTACACGACGCTCTTCCGATCTNNNATCACATTCCAGCTCCTTCCCTAC
AGAGCACACGTCTGAACTCTTTCCCTACACGACGCTCTTCCGATCTNNNCGAGGAAGAGGTTTCTGA
GAGCACACGTCTGAACTCTTTCCCTACACGACGCTCTTCCGATCTNNNTTCCAACAACTCAGGAGGCTA
GAGCACACGTCTGAACTCTTTCCCTACACGACGCTCTTCCGATCTNNNAAGACCAGTCTGGCCAACATA
AGCACACGTCTGAACTCTTTCCCTACACGACGCTCTTCCGATCTNNNGGAGATAGAGACCATCCTGGC
GAGCACACGTCTGAACTCTTTCCCTACACGACGCTCTTCCGATCTNNNGATTACTTGAGCCCAGCAGT
AGCACACGTCTGAACTCTTTCCCTACACGACGCTCTTCCGATCTNNNTTCCTAGAGATGGGTTTTTGCC
AGCACACGTCTGAACTCTTTCCCTACACGACGCTCTTCCGATCTNNNACAATGCCTTCTTTCCCTCACT
AGCACACGTCTGAACTCTTTCCCTACACGACGCTCTTCCGATCTNNNAGAGAGCCGAATACAAAGCT
AGCACACGTCTGAACTCTTTCCCTACACGACGCTCTTCCGATCTNNNATCCAACATGACTGTGCCCTTA
AGCACACGTCTGAACTCTTTCCCTACACGACGCTCTTCCGATCTNNNCTGGGTGAGAAGAAACCAACT
AGAGCACACGTCTGAACTCTTTCCCTACACGACGCTCTTCCGATCTNNNACCGGTACATGATACTTGAC
AGCACACGTCTGAACTCTTTCCCTACACGACGCTCTTCCGATCTNNNGTGCCCCAACATCCATTCTTTT
GAGCACACGTCTGAACTCTTTCCCTACACGACGCTCTTCCGATCTNNNCACAGGGGAGTGTCTCAAAT
AGCACACGTCTGAACTCTTTCCCTACACGACGCTCTTCCGATCTNNNGCCACATCTTCAGTACAAAGA
GAGCACACGTCTGAACTCTTTCCCTACACGACGCTCTTCCGATCTNNNGGTGAGCCAAGATTGTGCT
AGCACACGTCTGAACTCTTTCCCTACACGACGCTCTTCCGATCTNNNGATTGAGACCATCCTGGCTAA
AGCACACGTCTGAACTCTTTCCCTACACGACGCTCTTCCGATCTNNNCTAGAGTAGCTGGGATTACAG
GAGCACACGTCTGAACTCTTTCCCTACACGACGCTCTTCCGATCTNNNGCCACAGAGTAAGACTCCAT
AGAGCACACGTCTGAACTCTTTCCCTACACGACGCTCTTCCGATCTNNNAAGCCTTGGGTGATATATG
AGCACACGTCTGAACTCTTTCCCTACACGACGCTCTTCCGATCTNNNAGGTGGGTTCGAAGATTTCCTT
AGCACACGTCTGAACTCTTTCCCTACACGACGCTCTTCCGATCTNNNCTGGCTTGGTGGAATAAAAG
CACACGTCTGAACTCTTTCCCTACACGACGCTCTTCCGATCTNNNCCTGGGTGATACTCTCTCTCTCTC
AGCACACGTCTGAACTCTTTCCCTACACGACGCTCTTCCGATCTNNNCAGGCTGGCTAACATGGTAA
```

GCACACGTCTGAACTCTTTCCCTACACGACGCTCTTCCGATCTNNNTGTCTCTCTGTCTCTCTGTCTGTC
AGCACACGTCTGAACTCTTTCCCTACACGACGCTCTTCCGATCTNNNTGAGATTTACACTGCATGCTGG
AGCACACGTCTGAACTCTTTCCCTACACGACGCTCTTCCGATCTNNNTTCCCATCTGAAACCATGGAGA

TAAGACCACATCGGACTCATCAGGAA
TTCTCCATCGGACTCATCAGGAA
CACCCACCATCGGACTCATCAGGAAG
AGTATCATCGGACTCATCAGGAAG
TCCTCATCGGACTCATCAGGAAG
AAATGACATCGGACTCATCAGGAAG
TGTGTGTCATCGGACTCATCAGGAAG
CTAATACACATCGGACTCATCAGGAAG
CTATACACATCGGACTCATCAGGAAG
CTAACTCATCGGACTCATCAGGAAG
CTATCTCATCGGACTCATCAGGAAG
ACAATGCATCGGACTCATCAGGAAG
GAGCAAACACATCGGACTCATCAGGAAG
AGACTGCATCGGACTCATCAGGAAG
TGGTCATCGGACTCATCAGGAAG
CCAACACATCGGACTCATCAGGAAG
AAGACATCGGACTCATCAGGAAG
TCTCTCTCATCGGACTCATCAGGAAG
ATGTTCTCATCGGACTCATCAGGAAG
TCCGACATCGGACTCATCAGGAAG
CATATCTCATCGGACTCATCAGGAAG
TGACATCGGACTCATCAGGAAG
TATGCCCATCGGACTCATCAGGAAGC
GGAACTCATCGGACTCATCAGGAAGC
AACATCATCGGACTCATCAGGAAGC
AGAGGACATCGGACTCATCAGGAAGC
TAGCAGCATCGGACTCATCAGGAAGC
TGAATCCATCGGACTCATCAGGAAGC
TTACACATCGGACTCATCAGGAAGC
AAACTCATCGGACTCATCAGGAAGC
CTAAATTCATCGGACTCATCAGGAAGC
ATCTCCCATCGGACTCATCAGGAAGC
ATGGATGGACATCGGACTCATCAGGAAGC
TACACATCGGACTCATCAGGAAGC
ATGAAACATCGGACTCATCAGGAAGC
CTTACATCGGACTCATCAGGAAGC
GTATGTGCATCGGACTCATCAGGAAGC
TTAGGGCATCGGACTCATCAGGAAGC
TTAGCTCATCGGACTCATCAGGAAGC
ATGTGTGCATCGGACTCATCAGGAAGC
CTACTGCACATCGGACTCATCAGGAAGC
TACATCATCGGACTCATCAGGAAGC

GTCTTCCATCGGACTCATCAGGAAGC
CACACACATCGGACTCATCAGGAAGC
AGGAAAGCATCGGACTCATCAGGAAGC
ATTAGGCATCGGACTCATCAGGAAGC
CACATCGGACTCATCAGGAAGC
GCAACTCATCGGACTCATCAGGAAGC
GAACACATCGGACTCATCAGGAAGC
TAAAGACATCGGACTCATCAGGAAGC
ACACAGCATCGGACTCATCAGGAAGC
TGTGTCATCGGACTCATCAGGAAGC
GAAAACATCGGACTCATCAGGAAGC
CACAGGCCATCGGACTCATCAGGAAGC
ACATGACATCGGACTCATCAGGAAGC
AATGGACATCGGACTCATCAGGAAGC
GGATCACATCGGACTCATCAGGAAGC
CTTGAACATCGGACTCATCAGGAAGC
TTCCATCGGACTCATCAGGAAGC
CATTAGTCATCGGACTCATCAGGAAGC
CATGGTCATCGGACTCATCAGGAAGC
TCCACATCGGACTCATCAGGAAGC
CTCATCGGACTCATCAGGAAGC
TCTCATCGGACTCATCAGGAAGC
GGGCATCGGACTCATCAGGAAGC
CAATGCCATCGGACTCATCAGGAAGC
GACAGACATCGGACTCATCAGGAAGC
TCTGCCATCGGACTCATCAGGAAGC
TCTGCCATCGGACTCATCAGGAAGC
CAGTCATCGGACTCATCAGGAAGC
CTGACATCGGACTCATCAGGAAGC
GGAGACCATCGGACTCATCAGGAAGC
GTCCCATCGGACTCATCAGGAAGC
GGGAGGCATCGGACTCATCAGGAAGC
TATGTGCTCATCGGACTCATCAGGAAGC
GAATCACCATCGGACTCATCAGGAAGC
CCAGTCATCGGACTCATCAGGAAGC
CTACTCCATCGGACTCATCAGGAAGC
CGCTTGAACATCGGACTCATCAGGAAGC
TCATTCATCGGACTCATCAGGAAGC
TACTTCATCGGACTCATCAGGAAGC
GTTTGAGCATCGGACTCATCAGGAAGC
GGGAAGCATCGGACTCATCAGGAAGC
TGCTCATCGGACTCATCAGGAAGC
TGTCACATCGGACTCATCAGGAAGC

TTGAACCCATCGGACTCATCAGGAAGC
TTATTCCCATCGGACTCATCAGGAAGC
TGAACCTCATCGGACTCATCAGGAAGC
GAACACATCGGACTCATCAGGAAGC
GAACAGGCATCGGACTCATCAGGAAGC
ACTCAGCATCGGACTCATCAGGAAGC
TACGCACCATCGGACTCATCAGGAAGC
ACAAACTCATCGGACTCATCAGGAAGC
CACTGACATCGGACTCATCAGGAAGC
CTCTGAACATCGGACTCATCAGGAAGC
AAGAGTCATCGGACTCATCAGGAAGC
ACAGACATCGGACTCATCAGGAAGC
GATGAAACATCGGACTCATCAGGAAGC
GTCGTCATCGGACTCATCAGGAAGC
TCAGGACATCGGACTCATCAGGAAGC
AACACTCATCGGACTCATCAGGAAGC
TGACTCATCGGACTCATCAGGAAGC
GTGTCATCGGACTCATCAGGAAGC
GCATAACATCGGACTCATCAGGAAGC
CAAACACATCGGACTCATCAGGAAGC
CAAACACATCGGACTCATCAGGAAGC
ACTCCCATCGGACTCATCAGGAAGC
GCCTGACATCGGACTCATCAGGAAGC
ATGCACATCGGACTCATCAGGAAGC
TGTGTCATCGGACTCATCAGGAAGC
CACCACATCGGACTCATCAGGAAGC
TGTTCCTCATCGGACTCATCAGGAAGC
GCCTGACATCGGACTCATCAGGAAGC
GAAAGACATCGGACTCATCAGGAAGC
GGGAGGCCATCGGACTCATCAGGAAGC
GAGTACCATCGGACTCATCAGGAAGC
GACGGTCATCGGACTCATCAGGAAGC
ATGTTGCATCGGACTCATCAGGAAGC
GTGGGTCATCGGACTCATCAGGAAGC
GGGATACATCGGACTCATCAGGAAGC
GTGTGTCATCGGACTCATCAGGAAGC
ACAGAACCATCGGACTCATCAGGAAGC
TTGGCCATCGGACTCATCAGGAAGC
TTCTCATCGGACTCATCAGGAAGC
CTTGCATCGGACTCATCAGGAAGC
GAAAGAGTCATCGGACTCATCAGGAAGC
GAATCTCCATCGGACTCATCAGGAAGC
GTGTGCATCGGACTCATCAGGAAGC

CTTGAACATCGGACTCATCAGGAAGC
GTGCAGTCATCGGACTCATCAGGAAGC
TTACTTCATCGGACTCATCAGGAAGC
TACAGCCATCGGACTCATCAGGAAGC
GATGTCATCGGACTCATCAGGAAGC
GCAGACATCGGACTCATCAGGAAGC
TCAGTCATCGGACTCATCAGGAAGC
TCTCACATCGGACTCATCAGGAAGC
AGGCTGCATCGGACTCATCAGGAAGC
TTTCTCATCGGACTCATCAGGAAGC
TCTCATCGGACTCATCAGGAAGC
TTCACATCGGACTCATCAGGAAGC
CTACCTCATCGGACTCATCAGGAAGC
CAAACACATCGGACTCATCAGGAAGC
AGAGCACATCGGACTCATCAGGAAGC
TCCCATCGGACTCATCAGGAAGC
TATTCATCGGACTCATCAGGAAGC
GGAGTCACATCGGACTCATCAGGAAGC
CTCATCGGACTCATCAGGAAGC
ACTCTCACATCGGACTCATCAGGAAGC
TTGCCCATCGGACTCATCAGGAAGC
TTTGGTCATCGGACTCATCAGGAAGC
GTACCCTCATCGGACTCATCAGGAAGC
CACACTCATCGGACTCATCAGGAAGC
AAATACATCGGACTCATCAGGAAGC
ATTGCACATCGGACTCATCAGGAAGC
GCTCATCGGACTCATCAGGAAGC
ATCAGTCATCGGACTCATCAGGAAGC
TCACACATCGGACTCATCAGGAAGC
ACATGTCATCGGACTCATCAGGAAGC
TTCTGCATCGGACTCATCAGGAAGC
TTCTCCACATCGGACTCATCAGGAAGC
ATTGCACATCGGACTCATCAGGAAGC
TAAAGACATCGGACTCATCAGGAAGC
GTGGGCATCGGACTCATCAGGAAGC
ATACCGATCATCGGACTCATCAGGAAGC
AGATTCCCATCGGACTCATCAGGAAGC
CTATGTCATCGGACTCATCAGGAAGC
GCCCAACATCGGACTCATCAGGAAGC
TTCATCGGACTCATCAGGAAGC
CACACATCGGACTCATCAGGAAGC
GTGTGCATCGGACTCATCAGGAAGC
CTTGAACATCGGACTCATCAGGAAGC

TGTACATCGGACTCATCAGGAAGC
GGGCATCGGACTCATCAGGAAGC
TGGCCATCGGACTCATCAGGAAGC
CACTCCATCGGACTCATCAGGAAGC
ATACATCGGACTCATCAGGAAGC
GAACATCGGACTCATCAGGAAGC
CCACATCGGACTCATCAGGAAGC
ATTCATCGGACTCATCAGGAAGC
CTGCACATCGGACTCATCAGGAAGC
GGGCATCGGACTCATCAGGAAGC
AGTGGACATCGGACTCATCAGGAAGC
AGTTCATCGGACTCATCAGGAAGC
CGTCATCGGACTCATCAGGAAGC
TCAGGCATCGGACTCATCAGGAAGC
CTCATCGGACTCATCAGGAAGC
TGTGTCATCGGACTCATCAGGAAGC
ATAGCCATCGGACTCATCAGGAAGC
GTCATCGGACTCATCAGGAAGC
CGCTCATCGGACTCATCAGGAAGC
ACTGCATCGGACTCATCAGGAAGC
CCAGCATCGGACTCATCAGGAAGC
AATGACATCGGACTCATCAGGAAGC
TCAGTCATCGGACTCATCAGGAAGC
ATGCACATCGGACTCATCAGGAAGC
GTATACATCGGACTCATCAGGAAGC
CACACATCGGACTCATCAGGAAGC
AAGGCATCGGACTCATCAGGAAGC
TTGAGCATCGGACTCATCAGGAAGC
TTTCACATCGGACTCATCAGGAAGC
GAGCATCGGACTCATCAGGAAGC
CACCATCGGACTCATCAGGAAGC
TTATGCATCGGACTCATCAGGAAGC
GTGCCATCGGACTCATCAGGAAGC
TGAACCCATCGGACTCATCAGGAAGC
TGACATCGGACTCATCAGGAAGC
TCATCGGACTCATCAGGAAGC
AACTCCATCGGACTCATCAGGAAGC
AACGCATCGGACTCATCAGGAAGC
CCATCATCGGACTCATCAGGAAGC
CAGCACATCGGACTCATCAGGAAGC
CTTGCATCGGACTCATCAGGAAGC
CGGGCATCGGACTCATCAGGAAGC
CCGGCATCGGACTCATCAGGAAGC

TTTCACATCGGACTCATCAGGAAGC
AACACATCGGACTCATCAGGAAGC
ATGACATCGGACTCATCAGGAAGC
AATCACATCGGACTCATCAGGAAGC
CACCATCGGACTCATCAGGAAGC
ATGTCATCGGACTCATCAGGAAGC
CCCATCGGACTCATCAGGAAGC
GGACACATCGGACTCATCAGGAAGC
AGGACATCGGACTCATCAGGAAGC
TTTCTCATCGGACTCATCAGGAAGC
CTCATCGGACTCATCAGGAAGC
AGATCATCGGACTCATCAGGAAGC
AATGACATCGGACTCATCAGGAAGC
CCTTCATCGGACTCATCAGGAAGC
GAGGCATCGGACTCATCAGGAAGC
ATGATCCATCGGACTCATCAGGAAGC
CTACTTCATCGGACTCATCAGGAAGC
GATGCATCGGACTCATCAGGAAGC
AGCCCATCGGACTCATCAGGAAGC
GGTCACATCGGACTCATCAGGAAGC
AGTGAGCATCGGACTCATCAGGAAGC
TGGGAGCATCGGACTCATCAGGAAGC
ATACACACATCGGACTCATCAGGAAGC
GTGGCATCGGACTCATCAGGAAGC
CTGAGACATCGGACTCATCAGGAAGC
CTGAACATCGGACTCATCAGGAAGC
AAACGCATCGGACTCATCAGGAAGC
TACTTCATCGGACTCATCAGGAAGC
GAAGACATCGGACTCATCAGGAAGC
CAGGAGCATCGGACTCATCAGGAAGC
TCACCATCGGACTCATCAGGAAGC
AGATCTCATCGGACTCATCAGGAAGC
GGGAAGCATCGGACTCATCAGGAAGC
TTGAACATCGGACTCATCAGGAAGC
AGAAACATCGGACTCATCAGGAAGC
CTACTCATCGGACTCATCAGGAAGC
TCTCCACATCGGACTCATCAGGAAGC
GGTGGCATCGGACTCATCAGGAAGC
CTTCACATCGGACTCATCAGGAAGC
CCTCACATCGGACTCATCAGGAAGC
AGAGCCATCGGACTCATCAGGAAGC
GAGGCATCGGACTCATCAGGAAGC
AGGGGCATCGGACTCATCAGGAAGC

AGGACATCGGACTCATCAGGAAGC
CCTGGCATCGGACTCATCAGGAAGC
GGAGCATCGGACTCATCAGGAAGC
AAAACCATCGGACTCATCAGGAAGC
GAGCATCGGACTCATCAGGAAGC
CGTGGCATCGGACTCATCAGGAAGC
TACACATCGGACTCATCAGGAAGC
GGAGCATCGGACTCATCAGGAAGC
CAGGACATCGGACTCATCAGGAAGC
AAGCCATCGGACTCATCAGGAAGC
AGGAGACATCGGACTCATCAGGAAGC
CGAGAGCATCGGACTCATCAGGAAGC
GACACACATCGGACTCATCAGGAAGC
AACCCATCGGACTCATCAGGAAGC
AAGTTGCATCGGACTCATCAGGAAGC
GGCGTGCATCGGACTCATCAGGAAGC
GGAGACATCGGACTCATCAGGAAGC
AACCATCGGACTCATCAGGAAGC
TATGACATCGGACTCATCAGGAAGC
CCAGCATCGGACTCATCAGGAAGC
TGAACCATCGGACTCATCAGGAAGC
TCAGACATCGGACTCATCAGGAAGC
CATGCATCGGACTCATCAGGAAGC
CATTTCATCGGACTCATCAGGAAGC
GTCCCATCGGACTCATCAGGAAGC
AGGCCATCGGACTCATCAGGAAGC
CAAGTCATCGGACTCATCAGGAAGC
TGAACCATCGGACTCATCAGGAAGC
ATCAGACATCGGACTCATCAGGAAGC
TAGACATCGGACTCATCAGGAAGC
AGTGTGCCATCGGACTCATCAGGAAGC
AGGGACATCGGACTCATCAGGAAGC
ACACATCGGACTCATCAGGAAGC
AGACATCGGACTCATCAGGAAGC
TCCACATCGGACTCATCAGGAAGC
GTTGACATCGGACTCATCAGGAAGC
ATTCATCGGACTCATCAGGAAGC
CCTCATCGGACTCATCAGGAAGC
GAGCTCATCGGACTCATCAGGAAGC
AGCACATCGGACTCATCAGGAAGC
CTCATCGGACTCATCAGGAAGC
CCATGATCATCGGACTCATCAGGAAGC
ACTCATCGGACTCATCAGGAAGC

CACACACATCGGACTCATCAGGAAGC
ATCCTCATCGGACTCATCAGGAAGC
ATTCATCGGACTCATCAGGAAGC
TACATCATCGGACTCATCAGGAAGC
TAGTCATCGGACTCATCAGGAAGC
TGTGTCATCGGACTCATCAGGAAGC
GTGATACATCGGACTCATCAGGAAGC
AGGTCATCGGACTCATCAGGAAGC
ATTGGCATCGGACTCATCAGGAAGC
AAGAACATCGGACTCATCAGGAAGC
AATCACATCGGACTCATCAGGAAGC
GTGAACCATCGGACTCATCAGGAAGC
ATGTGCATCGGACTCATCAGGAAGC
AACCCATCGGACTCATCAGGAAGC
CCACATCGGACTCATCAGGAAGC
GATCACATCGGACTCATCAGGAAGC
ATTTCTCATCGGACTCATCAGGAAGC
CACACATCGGACTCATCAGGAAGC
TGCACATCGGACTCATCAGGAAGC
GAGCACATCGGACTCATCAGGAAGC
CAGCTCATCGGACTCATCAGGAAGC
CCATAACATCGGACTCATCAGGAAGC
ATGGCCATCGGACTCATCAGGAAGC
GTGCTCATCGGACTCATCAGGAAGC
TCAGCATCGGACTCATCAGGAAGC
ACACATCGGACTCATCAGGAAGC
CAAACATCGGACTCATCAGGAAGC
GCACATCGGACTCATCAGGAAGC
TAGCACATCGGACTCATCAGGAAGC
TCTCAGTCATCGGACTCATCAGGAAGC
AAGCCATCGGACTCATCAGGAAGC
CCCACATCGGACTCATCAGGAAGC
TTTCTCATCGGACTCATCAGGAAGC
TTGCCCATCGGACTCATCAGGAAGC
TGAATCATCGGACTCATCAGGAAGC
GAGCACATCGGACTCATCAGGAAGC
GTTCATCGGACTCATCAGGAAGC
TGTCTCATCGGACTCATCAGGAAGC
TGCACATCGGACTCATCAGGAAGC
CACACATCGGACTCATCAGGAAGC
ATGCACATCGGACTCATCAGGAAGC
GTCCCCATCGGACTCATCAGGAAGC
TTTTGCATCGGACTCATCAGGAAGC

CCACATCGGACTCATCAGGAAGC
CATCATCGGACTCATCAGGAAGC
TGTACATCGGACTCATCAGGAAGC
AGACCATCGGACTCATCAGGAAGC
TCTGACATCGGACTCATCAGGAAGC
GGGCCATCGGACTCATCAGGAAGC
ACCACATCGGACTCATCAGGAAGC
TGTCACATCGGACTCATCAGGAAGC
TCCTCATCGGACTCATCAGGAAGC
AGTGTTCATCGGACTCATCAGGAAGC
TCATGTCATCGGACTCATCAGGAAGC
GCCCACATCGGACTCATCAGGAAGC
ATTCACATCGGACTCATCAGGAAGC
GGTGGCATCGGACTCATCAGGAAGC
TTCCCATCGGACTCATCAGGAAGC
TGCTTGACATCGGACTCATCAGGAAGC
GAGCTCATCGGACTCATCAGGAAGC
TCTAGGCATCGGACTCATCAGGAAGC
TGGCTCATCGGACTCATCAGGAAGC
AGGGGCATCGGACTCATCAGGAAGC
TACTTCATCGGACTCATCAGGAAGC
TTCACATCGGACTCATCAGGAAGC
TTCATCGGACTCATCAGGAAGC
GAAGGAAGGCATCGGACTCATCAGGAAGC
GAGGCATCGGACTCATCAGGAAGC
ATCACATCGGACTCATCAGGAAGC
ATTTTCATCGGACTCATCAGGAAGC
AAGAGAGTCATCGGACTCATCAGGAAGC
CTCTCATCGGACTCATCAGGAAGC
GCGTCATCGGACTCATCAGGAAGC
TCACTGCATCGGACTCATCAGGAAGC
CCATCGGACTCATCAGGAAGC
CACATCGGACTCATCAGGAAGC
CACATCGGACTCATCAGGAAGC
TCCATCGGACTCATCAGGAAGC
AGGACATCGGACTCATCAGGAAGC
ACGTGCATCGGACTCATCAGGAAGC
CTTAGAACATCGGACTCATCAGGAAGC
CAGTCATCGGACTCATCAGGAAGC
CCTCATCGGACTCATCAGGAAGC
GAGCATCGGACTCATCAGGAAGC
AGACATCGGACTCATCAGGAAGC
AAAGACATCGGACTCATCAGGAAGC

GAAGACATCGGACTCATCAGGAAGC
ACGCCATCGGACTCATCAGGAAGC
AAAGTCATCGGACTCATCAGGAAGC
TTCTCATCGGACTCATCAGGAAGC
TGTGTCATCGGACTCATCAGGAAGC
TTTCCTCATCGGACTCATCAGGAAGC
TTCATCGGACTCATCAGGAAGC
ATCGGACTCATCAGGAAGC
GAGTCACATCGGACTCATCAGGAAGC
CTCATCGGACTCATCAGGAAGC
CACCATCGGACTCATCAGGAAGC
CCCATCGGACTCATCAGGAAGC
GCTCATCGGACTCATCAGGAAGC
GCACCATCGGACTCATCAGGAAGC
AGCACATCGGACTCATCAGGAAGC
GTACACATCGGACTCATCAGGAAGC
CACATCGGACTCATCAGGAAGC
GTGTGCATCGGACTCATCAGGAAGC
GACCATCGGACTCATCAGGAAGC
AGCCATCGGACTCATCAGGAAGC
ATTCACATCGGACTCATCAGGAAGC
CCTCATCGGACTCATCAGGAAGC
CTCATCGGACTCATCAGGAAGC
GGACACATCGGACTCATCAGGAAGC
TCTCATCGGACTCATCAGGAAGC
CCACATCGGACTCATCAGGAAGC
TCTCATCGGACTCATCAGGAAGC
GATGGGCATCGGACTCATCAGGAAGC
GCAGAAAACATCGGACTCATCAGGAAGC
TAGTTCATCGGACTCATCAGGAAGC
GGAGCATCGGACTCATCAGGAAGC
AGCAGCATCGGACTCATCAGGAAGC
GAACCCATCGGACTCATCAGGAAGC
TTCTCATCGGACTCATCAGGAAGC
CGAGCATCGGACTCATCAGGAAGC
CAATCATCGGACTCATCAGGAAGC
TGTAACATCGGACTCATCAGGAAGC
CACACATCGGACTCATCAGGAAGC
AGTCATCGGACTCATCAGGAAGC
AAGCCATCGGACTCATCAGGAAGC
GAACATCGGACTCATCAGGAAGC
GAGGCATCGGACTCATCAGGAAGC
CAGCATCGGACTCATCAGGAAGC

TGCCATCGGACTCATCAGGAAGC
CCTGCATCGGACTCATCAGGAAGC
GACCATCGGACTCATCAGGAAGC
GGCATCGGACTCATCAGGAAGC
CTGACATCGGACTCATCAGGAAGC
CACATCGGACTCATCAGGAAGC
GCAGCATCGGACTCATCAGGAAGC
GGAAGCATCGGACTCATCAGGAAGC
TCACATCGGACTCATCAGGAAGC
GAACATCGGACTCATCAGGAAGC
AGGCATCGGACTCATCAGGAAGC
TTGGCATCGGACTCATCAGGAAGC
TTTCCATCGGACTCATCAGGAAGC
TTGTCATCGGACTCATCAGGAAGC
CAGACTGCATCGGACTCATCAGGAAGC
AGGAGCATCGGACTCATCAGGAAGC
ACAGTCATCGGACTCATCAGGAAGC
AGGCCATCGGACTCATCAGGAAGC
AGGACATCGGACTCATCAGGAAGC
GGCCATCGGACTCATCAGGAAGC
TGCTTCATCGGACTCATCAGGAAGC
CACATCGGACTCATCAGGAAGC
TCCCATCGGACTCATCAGGAAGC
GAGGCATCGGACTCATCAGGAAGC
TGAACCATCGGACTCATCAGGAAGC
ACGTCATCGGACTCATCAGGAAGC
ATTTCATCGGACTCATCAGGAAGC
CACACATCGGACTCATCAGGAAGC
CACATCGGACTCATCAGGAAGC
CTCATCGGACTCATCAGGAAGC
TCTCATCGGACTCATCAGGAAGC
AGACATCGGACTCATCAGGAAGC
CACATCGGACTCATCAGGAAGC
TCATCGGACTCATCAGGAAGC
CTCTCATCGGACTCATCAGGAAGC
TGTTACATCGGACTCATCAGGAAGC
TCACATCGGACTCATCAGGAAGC
CCACACATCGGACTCATCAGGAAGC
CTTCATCGGACTCATCAGGAAGC
TCATCGGACTCATCAGGAAGC
GACACATCGGACTCATCAGGAAGC
GTGTCATCGGACTCATCAGGAAGC
GGAGCATCGGACTCATCAGGAAGC

GCAGCATCGGACTCATCAGGAAGC
GAACCCATCGGACTCATCAGGAAGC
ATCCCATCGGACTCATCAGGAAGC
CGTGGCATCGGACTCATCAGGAAGC
GGAGCATCGGACTCATCAGGAAGC
CCAGCATCGGACTCATCAGGAAGC
TTGAGCATCGGACTCATCAGGAAGC
CCCAGACATCGGACTCATCAGGAAGC
GGTGGCATCGGACTCATCAGGAAGC
TACACATCGGACTCATCAGGAAGC
TGCATCGGACTCATCAGGAAGC
AGTGAGCATCGGACTCATCAGGAAGC
TGAACATCGGACTCATCAGGAAGC
TACTCATCGGACTCATCAGGAAGC
GAACATCGGACTCATCAGGAAGC
ATTTCCATCGGACTCATCAGGAAGC
GGCTCATCGGACTCATCAGGAAGC
TGCACCATCGGACTCATCAGGAAGC
GAGCATCGGACTCATCAGGAAGC
GTGAGCATCGGACTCATCAGGAAGC
GAGACATCGGACTCATCAGGAAGC
TCTCATCGGACTCATCAGGAAGC
TGAACATCGGACTCATCAGGAAGC
AGTCCATCGGACTCATCAGGAAGC
CAGGCATCGGACTCATCAGGAAGC
GTGGCCATCGGACTCATCAGGAAGC
TAGCTCATCGGACTCATCAGGAAGC
CTTGAACCATCGGACTCATCAGGAAGC
AGGCATCGGACTCATCAGGAAGC
CTCGCATCGGACTCATCAGGAAGC
AGGAGCATCGGACTCATCAGGAAGC
AGAATTGCATCGGACTCATCAGGAAGC
CCAGCATCGGACTCATCAGGAAGC
ACAGGCATCGGACTCATCAGGAAGC
CAGGCATCGGACTCATCAGGAAGC
GTGGCATCGGACTCATCAGGAAGC
TTCAACATCGGACTCATCAGGAAGC
CTTCATCGGACTCATCAGGAAGC
GAGCATCGGACTCATCAGGAAGC
ACGCATCGGACTCATCAGGAAGC
CCAGCATCGGACTCATCAGGAAGC
AGGACATCGGACTCATCAGGAAGC
CACTCATCGGACTCATCAGGAAGC

ATCATCATCGGACTCATCAGGAAGC
TGCCCATCGGACTCATCAGGAAGC
CAAACATCGGACTCATCAGGAAGC
ACCTCATCGGACTCATCAGGAAGC
TAGTGCATCGGACTCATCAGGAAGC
ACTCCATCGGACTCATCAGGAAGC
ACCTCATCGGACTCATCAGGAAGC
GATTGCATCGGACTCATCAGGAAGC
TTCAACATCGGACTCATCAGGAAGC
GGAGCATCGGACTCATCAGGAAGC
AGAGCCATCGGACTCATCAGGAAGC
AGATGCATCGGACTCATCAGGAAGC
TTCTCATCGGACTCATCAGGAAGC
ACTGCATCGGACTCATCAGGAAGC
AGATCCATCGGACTCATCAGGAAGC
TCTTCATCGGACTCATCAGGAAGC
GTCATCGGACTCATCAGGAAGC
TGAACATCGGACTCATCAGGAAGC
CATCATCGGACTCATCAGGAAGC
GGACATCGGACTCATCAGGAAGC
TCCAACATCGGACTCATCAGGAAGC
TAATCCATCGGACTCATCAGGAAGC
CCATCGGACTCATCAGGAAGC
AGTTTCATCGGACTCATCAGGAAGC
GCTGCATCGGACTCATCAGGAAGC
TTGAGCATCGGACTCATCAGGAAGC
TTCACATCGGACTCATCAGGAAGC
GCATCGGACTCATCAGGAAGC
TTCTCATCGGACTCATCAGGAAGC
AGCCATCGGACTCATCAGGAAGC
AGCCATCGGACTCATCAGGAAGC
CACATCGGACTCATCAGGAAGC
AACCATCGGACTCATCAGGAAGC
TCCATCGGACTCATCAGGAAGC
CCACATCGGACTCATCAGGAAGC
GTTTCATCGGACTCATCAGGAAGC
ATTGCTCATCGGACTCATCAGGAAGC
GTCAACATCGGACTCATCAGGAAGC
CTTCATCGGACTCATCAGGAAGC
GAGGCATCGGACTCATCAGGAAGC
TGACATCGGACTCATCAGGAAGC
AAGACATCGGACTCATCAGGAAGC
TCAGCATCGGACTCATCAGGAAGC

AAGACCATCGGACTCATCAGGAAGC
TTTTCATCGGACTCATCAGGAAGC
TGCTCATCGGACTCATCAGGAAGC
CTTGAACATCGGACTCATCAGGAAGC
GAGGCATCGGACTCATCAGGAAGC
TGAACCATCGGACTCATCAGGAAGC
TGGTGCATCGGACTCATCAGGAAGC
AGGGACATCGGACTCATCAGGAAGC
GAACCATCGGACTCATCAGGAAGC
GCTCATCGGACTCATCAGGAAGC
AGAATTCATCGGACTCATCAGGAAGC
GTGCATCGGACTCATCAGGAAGC
CTCATCGGACTCATCAGGAAGC
TGGTGCATCGGACTCATCAGGAAGC
ATCACATCGGACTCATCAGGAAGC
GTGTCATCGGACTCATCAGGAAGC
CTTGACATCGGACTCATCAGGAAGC
TTGAACATCGGACTCATCAGGAAGC
TCCACATCGGACTCATCAGGAAGC
GTATCATCGGACTCATCAGGAAGC
AGGTCATCGGACTCATCAGGAAGC
CCTCATCGGACTCATCAGGAAGC
GAGGCATCGGACTCATCAGGAAGC
ACACATCGGACTCATCAGGAAGC
GGGCATCGGACTCATCAGGAAGC
GTCATCGGACTCATCAGGAAGC
GAGCATCGGACTCATCAGGAAGC
AAGACATCGGACTCATCAGGAAGC
TGTCATCGGACTCATCAGGAAGC
CACACATCGGACTCATCAGGAAGC
CCATCGGACTCATCAGGAAGC
TACATCATCGGACTCATCAGGAAGC
CATCGGACTCATCAGGAAGC
AACATCGGACTCATCAGGAAGC
AGCACATCGGACTCATCAGGAAGC
TATGTCATCGGACTCATCAGGAAGC
TGTGTCATCGGACTCATCAGGAAGC
ACATCGGACTCATCAGGAAGC
AACACATCGGACTCATCAGGAAGC
TCACTCATCGGACTCATCAGGAAGC
CTCATCGGACTCATCAGGAAGC
CACATCGGACTCATCAGGAAGC
TGGGCATCGGACTCATCAGGAAGC

GAATCATCGGACTCATCAGGAAGC
ACACCATCGGACTCATCAGGAAGC
AACCACATCGGACTCATCAGGAAGC
GTGCATCGGACTCATCAGGAAGC
GCCCATCGGACTCATCAGGAAGC
GTCATCGGACTCATCAGGAAGC
AGAAGACATCGGACTCATCAGGAAGC
GTGAGCATCGGACTCATCAGGAAGC
AGTCATCGGACTCATCAGGAAGC
CCCATCGGACTCATCAGGAAGC
GCACATCGGACTCATCAGGAAGC
CTTTCATCGGACTCATCAGGAAGC
CCTGTCATCGGACTCATCAGGAAGC
TGGCATCGGACTCATCAGGAAGC
TGGTCATCGGACTCATCAGGAAGC
TTCCCATCGGACTCATCAGGAAGC
GTGAGCATCGGACTCATCAGGAAGC
CCACATCGGACTCATCAGGAAGC
CCCACATCGGACTCATCAGGAAGC
ACCGTCATCGGACTCATCAGGAAGC
AGGAAAGACATCGGACTCATCAGGAAGC
TTGCCATCGGACTCATCAGGAAGC
TTCACATCGGACTCATCAGGAAGC
TGTGTCATCGGACTCATCAGGAAGC
TTTCATCGGACTCATCAGGAAGC
CCCAGGACATCGGACTCATCAGGAAGC
TTCCCATCGGACTCATCAGGAAGC
TTGTCGCATCGGACTCATCAGGAAGC
TGTATCATCGGACTCATCAGGAAGC
TGCACATCGGACTCATCAGGAAGC
CTCATCGGACTCATCAGGAAGC
GTGTCATCGGACTCATCAGGAAGC
TCTCATCGGACTCATCAGGAAGC
CCACATCGGACTCATCAGGAAGC
GACACATCGGACTCATCAGGAAGC
GTCATCGGACTCATCAGGAAGC
TCACATCGGACTCATCAGGAAGC
TGACATCGGACTCATCAGGAAGC
ACAATCATCGGACTCATCAGGAAGC
GCACATCGGACTCATCAGGAAGC
ATGGCACATCGGACTCATCAGGAAGC
TCTCATCGGACTCATCAGGAAGC
GGCCATCGGACTCATCAGGAAGC

TCATCATCGGACTCATCAGGAAGC
CCACATCGGACTCATCAGGAAGC
ATGACATCGGACTCATCAGGAAGC
GCACATCGGACTCATCAGGAAGC
GAGCATCGGACTCATCAGGAAGC
CCCACATCGGACTCATCAGGAAGC
TTTCATCGGACTCATCAGGAAGC
AAACATCGGACTCATCAGGAAGC
CTTCATCGGACTCATCAGGAAGC
TGGAACATCGGACTCATCAGGAAGC
CATCTCATCGGACTCATCAGGAAGC
CACACATCGGACTCATCAGGAAGC
GACATCGGACTCATCAGGAAGC
TCATCGGACTCATCAGGAAGC
CCCATCGGACTCATCAGGAAGC
GGACATCGGACTCATCAGGAAGC
CTTGCCATCGGACTCATCAGGAAGC
TTTCATCGGACTCATCAGGAAGC
TCCACATCGGACTCATCAGGAAGC
TTCATCGGACTCATCAGGAAGC
CTCATCGGACTCATCAGGAAGC
TGGCATCGGACTCATCAGGAAGC
TTCACATCGGACTCATCAGGAAGC
CCCATCGGACTCATCAGGAAGC
TTCCATCGGACTCATCAGGAAGC
TGGACATCGGACTCATCAGGAAGC
CTCATCGGACTCATCAGGAAGC
TATCCACATCGGACTCATCAGGAAGC
TCATCGGACTCATCAGGAAGC
AGCCATCGGACTCATCAGGAAGC
CACATCGGACTCATCAGGAAGC
GTCATCGGACTCATCAGGAAGC
TGTCATCGGACTCATCAGGAAGC
CGACATCGGACTCATCAGGAAGC
CTGGCCATCGGACTCATCAGGAAGC
GCTCATCGGACTCATCAGGAAGC
GGTCATCGGACTCATCAGGAAGC
GGCCATCGGACTCATCAGGAAGC
AAGACATCGGACTCATCAGGAAGC
TTGACATCGGACTCATCAGGAAGC
ACCATCGGACTCATCAGGAAGC
GCTCATCGGACTCATCAGGAAGC
AGCACATCGGACTCATCAGGAAGC

TCATCGGACTCATCAGGAAGC
GATCATCGGACTCATCAGGAAGC
TTTCATCGGACTCATCAGGAAGC
ACATCGGACTCATCAGGAAGC
TTCCCATCGGACTCATCAGGAAGC
GTGACATCGGACTCATCAGGAAGC
CATCGGACTCATCAGGAAGC
GTGTCATCGGACTCATCAGGAAGC
GAGAACATCGGACTCATCAGGAAGC
CCCATCGGACTCATCAGGAAGC
AGCTCATCGGACTCATCAGGAAGC
AAGACATCGGACTCATCAGGAAGC
ATACCATCGGACTCATCAGGAAGC
CTCTCATCGGACTCATCAGGAAGC
AGTCATCGGACTCATCAGGAAGC
TATTGCATCGGACTCATCAGGAAGC
TCATCGGACTCATCAGGAAGC
GGGATTCATCGGACTCATCAGGAAGC
ACCCCCATCGGACTCATCAGGAAGC
TGTCATCGGACTCATCAGGAAGC
CAGCATCGGACTCATCAGGAAGC
ACTCATCGGACTCATCAGGAAGC
ACACATCGGACTCATCAGGAAGC
GCACCATCGGACTCATCAGGAAGC
AAAGCATCGGACTCATCAGGAAGC
CACATCGGACTCATCAGGAAGC
TCATCGGACTCATCAGGAAGC
GTCATCGGACTCATCAGGAAGC
GGCACATCGGACTCATCAGGAAGC
AACCATCGGACTCATCAGGAAGC
CAGAACATCGGACTCATCAGGAAGC
CCACATCGGACTCATCAGGAAGC
ATACATCGGACTCATCAGGAAGC
CATCGGACTCATCAGGAAGC
AACGCATCGGACTCATCAGGAAGC
AGTCATCGGACTCATCAGGAAGC
TCACATCGGACTCATCAGGAAGC
CATCGGACTCATCAGGAAGC
CTCCATCGGACTCATCAGGAAGC
CCCATCGGACTCATCAGGAAGC
AGCCATCGGACTCATCAGGAAGC
GAAGACATCGGACTCATCAGGAAGC
GCCATCGGACTCATCAGGAAGC

TGGGCATCGGACTCATCAGGAAGC
GTGATCATCGGACTCATCAGGAAGC
TGCATCGGACTCATCAGGAAGC
GGCCATCGGACTCATCAGGAAGC
GGCCATCGGACTCATCAGGAAGC
GCTCATCGGACTCATCAGGAAGC
AGACATCGGACTCATCAGGAAGC
TGGTGCATCGGACTCATCAGGAAGC
CTCATCGGACTCATCAGGAAGC
CACATCGGACTCATCAGGAAGC
AGGTCATCGGACTCATCAGGAAGC
GAGCATCGGACTCATCAGGAAGC
GAACATCGGACTCATCAGGAAGC
GGCCATCGGACTCATCAGGAAGC
GCAGCATCGGACTCATCAGGAAGC
GCATCGGACTCATCAGGAAGC
GAATCCATCGGACTCATCAGGAAGC
ATCCATCGGACTCATCAGGAAGC
AACACATCGGACTCATCAGGAAGC
TTGCATCGGACTCATCAGGAAGC
CACATCGGACTCATCAGGAAGC
AGCATCGGACTCATCAGGAAGC
CTGCATCGGACTCATCAGGAAGC
CACATCGGACTCATCAGGAAGC
TCATCGGACTCATCAGGAAGC
GCATCGGACTCATCAGGAAGC
GTGCATCGGACTCATCAGGAAGC
TCCATCGGACTCATCAGGAAGC
ACTCATCGGACTCATCAGGAAGC
ACTTCATCGGACTCATCAGGAAGC
ACTCATCGGACTCATCAGGAAGC
CATTCATCGGACTCATCAGGAAGC
TGAACATCGGACTCATCAGGAAGC
GGACATCGGACTCATCAGGAAGC
GATTCATCGGACTCATCAGGAAGC
AACCATCGGACTCATCAGGAAGC
ACCATCGGACTCATCAGGAAGC
ATCACATCGGACTCATCAGGAAGC
GAACATCGGACTCATCAGGAAGC
ACCTACATCGGACTCATCAGGAAGC
AAGGCATCGGACTCATCAGGAAGC
GATTCATCGGACTCATCAGGAAGC
GAACATCGGACTCATCAGGAAGC

ACCTCATCGGACTCATCAGGAAGC
CCTGCATCGGACTCATCAGGAAGC
AGCACATCGGACTCATCAGGAAGC
GAGTCATCGGACTCATCAGGAAGC
CTCATCGGACTCATCAGGAAGC
GCTCATCGGACTCATCAGGAAGC
GGTCATCGGACTCATCAGGAAGC
TTCCATCGGACTCATCAGGAAGC
ACTTCATCGGACTCATCAGGAAGC
CAACATCGGACTCATCAGGAAGC
AGCATCGGACTCATCAGGAAGC
CTACATCGGACTCATCAGGAAGC
TGAACATCGGACTCATCAGGAAGC
ACTTCATCGGACTCATCAGGAAGC
TGGCATCGGACTCATCAGGAAGC
CGTGCATCGGACTCATCAGGAAGC
GATCCATCGGACTCATCAGGAAGC
TGGTGCATCGGACTCATCAGGAAGC
TTCATCGGACTCATCAGGAAGC
TGAACATCGGACTCATCAGGAAGC
TCCATCGGACTCATCAGGAAGC
AAGCATCGGACTCATCAGGAAGC
TGAGCATCGGACTCATCAGGAAGC
TTGAGCATCGGACTCATCAGGAAGC
ACCTCATCGGACTCATCAGGAAGC
GAACCATCGGACTCATCAGGAAGC
ATGCATCGGACTCATCAGGAAGC
CTAGCATCGGACTCATCAGGAAGC
TGGCATCGGACTCATCAGGAAGC
AGCATCGGACTCATCAGGAAGC
GCTCATCGGACTCATCAGGAAGC
GAGCATCGGACTCATCAGGAAGC
GATCACATCGGACTCATCAGGAAGC
GAGCATCGGACTCATCAGGAAGC
TGAACATCGGACTCATCAGGAAGC
AGACATCGGACTCATCAGGAAGC
ACCTGCATCGGACTCATCAGGAAGC
GGCTCATCGGACTCATCAGGAAGC
TTCCATCGGACTCATCAGGAAGC
GCCATCGGACTCATCAGGAAGC
CTACATCGGACTCATCAGGAAGC
ATAGCATCGGACTCATCAGGAAGC
GGCATCGGACTCATCAGGAAGC

CGTCATCGGACTCATCAGGAAGC
AACATCGGACTCATCAGGAAGC
GTGGCATCGGACTCATCAGGAAGC
ACACATCGGACTCATCAGGAAGC
ATCATCGGACTCATCAGGAAGC
ATAGCATCGGACTCATCAGGAAGC
GAGCATCGGACTCATCAGGAAGC
GAATCATCGGACTCATCAGGAAGC
TGGTGCATCGGACTCATCAGGAAGC
AATCATCGGACTCATCAGGAAGC
AGACATCGGACTCATCAGGAAGC
TGACATCGGACTCATCAGGAAGC
TGGCATCGGACTCATCAGGAAGC
AGGCCATCGGACTCATCAGGAAGC
ACCATCGGACTCATCAGGAAGC
TGAACATCGGACTCATCAGGAAGC
GAACCATCGGACTCATCAGGAAGC
AGCATCGGACTCATCAGGAAGC
AGGCATCGGACTCATCAGGAAGC
GGTGGCATCGGACTCATCAGGAAGC
GTGCATCGGACTCATCAGGAAGC
TGCCATCGGACTCATCAGGAAGC
ACATCGGACTCATCAGGAAGC
GTGCATCGGACTCATCAGGAAGC
AGGCATCGGACTCATCAGGAAGC
CCATCGGACTCATCAGGAAGC
TGCACATCGGACTCATCAGGAAGC
TCTCATCGGACTCATCAGGAAGC
TCGCATCGGACTCATCAGGAAGC
ACGCATCGGACTCATCAGGAAGC
AGCATCGGACTCATCAGGAAGC
AAGGAACATCGGACTCATCAGGAAGC
AGCCATCGGACTCATCAGGAAGC
GAGGCATCGGACTCATCAGGAAGC
ACATCGGACTCATCAGGAAGC
TTGCATCGGACTCATCAGGAAGC
GTGCATCGGACTCATCAGGAAGC
ACATCGGACTCATCAGGAAGC
AGATACATCGGACTCATCAGGAAGC
TCCATCGGACTCATCAGGAAGC
GACATCGGACTCATCAGGAAGC
AGGCATCGGACTCATCAGGAAGC
ATCACTCATCGGACTCATCAGGAAGC

TTGCATCGGACTCATCAGGAAGC
GAGCATCGGACTCATCAGGAAGC
GAGCATCGGACTCATCAGGAAGC
GAGCATCGGACTCATCAGGAAGC
GGCCATCGGACTCATCAGGAAGC
CTAGACATCGGACTCATCAGGAAGC
TGAACCATCGGACTCATCAGGAAGC
GAGCATCGGACTCATCAGGAAGC
CCTGCATCGGACTCATCAGGAAGC
CACACATCGGACTCATCAGGAAGC
CACATCGGACTCATCAGGAAGC
TTCATCGGACTCATCAGGAAGC
CCTCATCGGACTCATCAGGAAGC
GAGCATCGGACTCATCAGGAAGC
CTGCATCGGACTCATCAGGAAGC
AGACACATCGGACTCATCAGGAAGC
ACCTCATCGGACTCATCAGGAAGC
GGAGCATCGGACTCATCAGGAAGC
AAGCATCGGACTCATCAGGAAGC
CATCGGACTCATCAGGAAGC
TTCATCGGACTCATCAGGAAGC
GCATCGGACTCATCAGGAAGC
ATGCATCGGACTCATCAGGAAGC
GCACATCGGACTCATCAGGAAGC
AATGCATCGGACTCATCAGGAAGC
GTCATCGGACTCATCAGGAAGC
ATCATCGGACTCATCAGGAAGC
GGTTCATCGGACTCATCAGGAAGC
CTCATCGGACTCATCAGGAAGC
TGTCATCGGACTCATCAGGAAGC
TACCATCGGACTCATCAGGAAGC
GTGGCCATCGGACTCATCAGGAAGC
GAACATCGGACTCATCAGGAAGC
TCATCGGACTCATCAGGAAGC
AGCCCATCGGACTCATCAGGAAGC
ACATCGGACTCATCAGGAAGC
ACTCATCGGACTCATCAGGAAGC
ACCCATCGGACTCATCAGGAAGC
ACCCATCGGACTCATCAGGAAGC
CGTCATCGGACTCATCAGGAAGC
CTCTCATCGGACTCATCAGGAAGC
TCCATCGGACTCATCAGGAAGC
GTATGTCATCGGACTCATCAGGAAGC

AACCCATCGGACTCATCAGGAAGC
ACCCATCGGACTCATCAGGAAGC
GGGCCATCGGACTCATCAGGAAGC
TCCCATCGGACTCATCAGGAAGC
TTGAACATCGGACTCATCAGGAAGC
GTGCATCGGACTCATCAGGAAGC
AGCATCGGACTCATCAGGAAGC
TGCATCGGACTCATCAGGAAGC
AACCATCGGACTCATCAGGAAGC
TTGAACATCGGACTCATCAGGAAGC
CTCATCGGACTCATCAGGAAGC
GTGCATCGGACTCATCAGGAAGC
GTGCATCGGACTCATCAGGAAGC
TGAATCATCGGACTCATCAGGAAGC
ACACATCGGACTCATCAGGAAGC
TGCATCGGACTCATCAGGAAGC
GGCATCGGACTCATCAGGAAGC
GTGAACATCGGACTCATCAGGAAGC
ATTCATCGGACTCATCAGGAAGC
CCATCATCGGACTCATCAGGAAGC
TGACATCGGACTCATCAGGAAGC
GATCATCGGACTCATCAGGAAGC
CTCATCGGACTCATCAGGAAGC
AGGACATCGGACTCATCAGGAAGC
GGACATCGGACTCATCAGGAAGC
TGAACATCGGACTCATCAGGAAGC
GATCCATCGGACTCATCAGGAAGC
GAGATTCATCGGACTCATCAGGAAGC
GAGCATCGGACTCATCAGGAAGC
CAACCATCGGACTCATCAGGAAGC
AAGCATCGGACTCATCAGGAAGC
AAGCATCGGACTCATCAGGAAGC
AAGCATCGGACTCATCAGGAAGC
CTGCATCGGACTCATCAGGAAGC
GGCCATCGGACTCATCAGGAAGC
CCAACATCGGACTCATCAGGAAGC
CTCCATCGGACTCATCAGGAAGC
CATTCATCGGACTCATCAGGAAGC
CTGGCATCGGACTCATCAGGAAGC
TGCCATCGGACTCATCAGGAAGC
TGAACATCGGACTCATCAGGAAGC
AGCTCATCGGACTCATCAGGAAGC
GGCATCGGACTCATCAGGAAGC

AACATCGGACTCATCAGGAAGC

CCATCGGACTCATCAGGAAGC

GTCATCGGACTCATCAGGAAGC

GCACATCGGACTCATCAGGAAGC

AGCTCATCGGACTCATCAGGAAGC

AGCATCGGACTCATCAGGAAGC

TGTCATCGGACTCATCAGGAAGC

TCATCGGACTCATCAGGAAGC

TCATCGGACTCATCAGGAAGC

GGTGGCCATCGGACTCATCAGGAAGC

AACATCGGACTCATCAGGAAGC

TACATCGGACTCATCAGGAAGC

AATTGCATCGGACTCATCAGGAAGC

TACATCGGACTCATCAGGAAGC

ACCATCGGACTCATCAGGAAGC

GATAATCATCGGACTCATCAGGAAGC

CCATCGGACTCATCAGGAAGC

TCATCGGACTCATCAGGAAGC

TCCATCGGACTCATCAGGAAGC

AACCCATCGGACTCATCAGGAAGC
ACATCGGACTCATCAGGAAGC
GCATCGGACTCATCAGGAAGC
GGCATCGGACTCATCAGGAAGC
ATCATCGGACTCATCAGGAAGC
CTCCATCGGACTCATCAGGAAGC
GAACATCGGACTCATCAGGAAGC
CTCATCGGACTCATCAGGAAGC
AACCATCGGACTCATCAGGAAGC
AAACATCGGACTCATCAGGAAGC
CACCATCGGACTCATCAGGAAGC
ACATCGGACTCATCAGGAAGC
TCACATCGGACTCATCAGGAAGC
CTTCATCGGACTCATCAGGAAGC
GCATCGGACTCATCAGGAAGC
CCATCGGACTCATCAGGAAGC
ACTCATCGGACTCATCAGGAAGC
TTTCATCGGACTCATCAGGAAGC
TCCATCGGACTCATCAGGAAGC
TCCCATCGGACTCATCAGGAAGC
CCCCATCGGACTCATCAGGAAGC
ACATCGGACTCATCAGGAAGC
TTACATCGGACTCATCAGGAAGC
ATCATCGGACTCATCAGGAAGC
CCATCGGACTCATCAGGAAGC
AGTATAGGTCATCGGACTCATCAGGAA
GGTTTCATCGGACTCATCAGGAAG
TTTATGCCATCGGACTCATCAGGAAG
ATGAGGCATCGGACTCATCAGGAAG
GAGCACATCGGACTCATCAGGAAG
TCATGTCATCGGACTCATCAGGAAG
TGTTGTCATCGGACTCATCAGGAAG
GTTTACACATCGGACTCATCAGGAAG
CCATGCCATCGGACTCATCAGGAAG
CACACTCACATCGGACTCATCAGGAAG
GTAAGACCATCGGACTCATCAGGAAG
AAAGACATCGGACTCATCAGGAAG
TCAGTCATCGGACTCATCAGGAAGC
ACTTGAACATCGGACTCATCAGGAAGC
GAGAACCCGCATCGGACTCATCAGGAAGC
TATTTCATCGGACTCATCAGGAAGC
TGGGGACATCGGACTCATCAGGAAGC
CAACTTGCATCGGACTCATCAGGAAGC

GGGAAGCATCGGACTCATCAGGAAGC
ATCCGGCATCGGACTCATCAGGAAGC
ACTTTTGCATCGGACTCATCAGGAAGC
TCTCTCATCGGACTCATCAGGAAGC
AACCTGGACATCGGACTCATCAGGAAGC
TCCTGCCATCGGACTCATCAGGAAGC
GATCATCATCGGACTCATCAGGAAGC
GAGGCCATCGGACTCATCAGGAAGC
GATGGACATCGGACTCATCAGGAAGC
ACGTCATCGGACTCATCAGGAAGC
CATGTCATCGGACTCATCAGGAAGC
GGGAGGCATCGGACTCATCAGGAAGC
ACTGTACCATCGGACTCATCAGGAAGC
TAAGACATCGGACTCATCAGGAAGC
GGCCACATCGGACTCATCAGGAAGC
ACAGTCATCGGACTCATCAGGAAGC
AGAAGCATCGGACTCATCAGGAAGC
ACATCTCATCGGACTCATCAGGAAGC
TGGAATCATCGGACTCATCAGGAAGC
ACACACATCGGACTCATCAGGAAGC
GTTGGCCATCGGACTCATCAGGAAGC
AGATTGCATCGGACTCATCAGGAAGC
TCATGATGCATCGGACTCATCAGGAAGC
TCTGAACCATCGGACTCATCAGGAAGC
TGCCATGCCATCGGACTCATCAGGAAGC
TATGAACCATCGGACTCATCAGGAAGC
TCTCATCGGACTCATCAGGAAGC
AGTGAGCATCGGACTCATCAGGAAGC
CTCTCTCATCGGACTCATCAGGAAGC
AGGAGGCATCGGACTCATCAGGAAGC
TGAGTCATCGGACTCATCAGGAAGC
GTATGTCATCGGACTCATCAGGAAGC
AAGATTGCATCGGACTCATCAGGAAGC
TTTACATCGGACTCATCAGGAAGC
TACATCTCATCGGACTCATCAGGAAGC
ACACACATCGGACTCATCAGGAAGC
TATCCACATCGGACTCATCAGGAAGC
GTGTGGCATCGGACTCATCAGGAAGC
TTTCTGCATCGGACTCATCAGGAAGC
GTTTCACATCGGACTCATCAGGAAGC
ATCTCCATCGGACTCATCAGGAAGC
GTGTGTCATCGGACTCATCAGGAAGC
AAACCACATCGGACTCATCAGGAAGC

AGACACATCGGACTCATCAGGAAGC
TGACATCGGACTCATCAGGAAGC
AACACATCGGACTCATCAGGAAGC
TGTGTCATCGGACTCATCAGGAAGC
AGACTCATCGGACTCATCAGGAAGC
TGTGTCATCGGACTCATCAGGAAGC
TGTGCACATCGGACTCATCAGGAAGC
GCAGGACATCGGACTCATCAGGAAGC
CTAGGACATCGGACTCATCAGGAAGC
TGACACATCGGACTCATCAGGAAGC
TCTGACATCGGACTCATCAGGAAGC
TTTGTCATCGGACTCATCAGGAAGC
TCTCCATCGGACTCATCAGGAAGC
TGTCCATCGGACTCATCAGGAAGC
TGTTCATCGGACTCATCAGGAAGC
TTCCTCACATCGGACTCATCAGGAAGC
TCACTGCATCGGACTCATCAGGAAGC
ACGAGGCATCGGACTCATCAGGAAGC
CCTGCATCGGACTCATCAGGAAGC
CTCCCATCGGACTCATCAGGAAGC
AAGAAAGACATCGGACTCATCAGGAAGC
GGACATCGGACTCATCAGGAAGC
AGAGAGATTCATCGGACTCATCAGGAAGC
CCTATGATCATCGGACTCATCAGGAAGC
TCTCATCGGACTCATCAGGAAGC
GCAGGCATCGGACTCATCAGGAAGC
AAAAAAACCACATCGGACTCATCAGGAAGC
TGGTCATCGGACTCATCAGGAAGC
TAGTCATCGGACTCATCAGGAAGC
CATTCATCGGACTCATCAGGAAGC
GTCTTCATCGGACTCATCAGGAAGC
TTCATCGGACTCATCAGGAAGC
ACATCGGACTCATCAGGAAGC
CATGATCATCGGACTCATCAGGAAGC
AGAAAGACATCGGACTCATCAGGAAGC
CTTCATCGGACTCATCAGGAAGC
TCATCGGACTCATCAGGAAGC
AGTCCATCGGACTCATCAGGAAGC
TAGCATCGGACTCATCAGGAAGC
TCTCATCGGACTCATCAGGAAGC
AGTAAGCCCATCGGACTCATCAGGAAGC
GAAAGAACATCGGACTCATCAGGAAGC
AGACATCGGACTCATCAGGAAGC

CTTGAACATCGGACTCATCAGGAAGC
AACACATCGGACTCATCAGGAAGC
CTTGCATCGGACTCATCAGGAAGC
GAAGGCATCGGACTCATCAGGAAGC
AGTTACATCGGACTCATCAGGAAGC
GTGTGAACATCGGACTCATCAGGAAGC
TGTCATCGGACTCATCAGGAAGC
TTCGCCATCGGACTCATCAGGAAGC
GTGTGCATCGGACTCATCAGGAAGC
TAACACATCGGACTCATCAGGAAGC
GATTGCATCGGACTCATCAGGAAGC
TCCCATCGGACTCATCAGGAAGC
TTCACATCGGACTCATCAGGAAGC
GGTGACATCGGACTCATCAGGAAGC
TAGGGCATCGGACTCATCAGGAAGC
TCGGCATCGGACTCATCAGGAAGC
GAGGCATCGGACTCATCAGGAAGC
TGGGGACATCGGACTCATCAGGAAGC
TACTTGCATCGGACTCATCAGGAAGC
GAGATACATCGGACTCATCAGGAAGC
GTCAACATCGGACTCATCAGGAAGC
TGTCACATCGGACTCATCAGGAAGC
AGCTGCATCGGACTCATCAGGAAGC
AGGCTCATCGGACTCATCAGGAAGC
AGAGTATTCATCGGACTCATCAGGAAGC
AACACCATCGGACTCATCAGGAAGC
CATTAGTCATCGGACTCATCAGGAAGC
TCATATCATCGGACTCATCAGGAAGC
ATACCATCGGACTCATCAGGAAGC
TGTCATCGGACTCATCAGGAAGC
GATCACATCGGACTCATCAGGAAGC
ACTCCATCGGACTCATCAGGAAGC
GCTACATCGGACTCATCAGGAAGC
TTGTGCATCGGACTCATCAGGAAGC
AACCCCAACATCGGACTCATCAGGAAGC
TCCAGACATCGGACTCATCAGGAAGC
TCCACATCGGACTCATCAGGAAGC
CCGGACATCGGACTCATCAGGAAGC
ACTGGGCATCGGACTCATCAGGAAGC
TGTGCATCGGACTCATCAGGAAGC
TTGGCATCGGACTCATCAGGAAGC
ATGCCATCGGACTCATCAGGAAGC
AAAACATCGGACTCATCAGGAAGC

TCTCTCATCGGACTCATCAGGAAGC
AACTCATCGGACTCATCAGGAAGC
CTCATGGCATCGGACTCATCAGGAAGC
GAACCCCATCGGACTCATCAGGAAGC
CTTGAACATCGGACTCATCAGGAAGC
ACCTGGGCATCGGACTCATCAGGAAGC
ATACCTCATCGGACTCATCAGGAAGC
GTGTCATCGGACTCATCAGGAAGC
ATCCCACATCGGACTCATCAGGAAGC
TAACATCGGACTCATCAGGAAGC
GTGCCATCGGACTCATCAGGAAGC
AATCATCGGACTCATCAGGAAGC
TTGCCATCGGACTCATCAGGAAGC
CCTCATCGGACTCATCAGGAAGC
TGACTCATCGGACTCATCAGGAAGC
CACTCATCGGACTCATCAGGAAGC
AAAAACATCGGACTCATCAGGAAGC
AGACTCATCGGACTCATCAGGAAGC
TAGCACATCGGACTCATCAGGAAGC
ACATTCATCGGACTCATCAGGAAGC
GTGAACATCGGACTCATCAGGAAGC
TTTCACATCGGACTCATCAGGAAGC
TTAGCATCGGACTCATCAGGAAGC
CCACACATCGGACTCATCAGGAAGC
TCATCGGACTCATCAGGAAGC
AGGTAGCATCGGACTCATCAGGAAGC
TCAGCATCGGACTCATCAGGAAGC
AATGCCATCGGACTCATCAGGAAGC
AGATCCATCGGACTCATCAGGAAGC
GGTCATCGGACTCATCAGGAAGC
TCCTCCATCGGACTCATCAGGAAGC
AACGGGGCATCGGACTCATCAGGAAGC
TAGCTGTCATCGGACTCATCAGGAAGC
TTCACATCGGACTCATCAGGAAGC
ATGTCATCGGACTCATCAGGAAGC
CAAACATCGGACTCATCAGGAAGC
TAGAATTCATCGGACTCATCAGGAAGC
GGGACATCGGACTCATCAGGAAGC
GACATCGGACTCATCAGGAAGC
ATGTGCATCGGACTCATCAGGAAGC
TGAAACATCGGACTCATCAGGAAGC
GATTGCATCGGACTCATCAGGAAGC
TTGTCATCGGACTCATCAGGAAGC

ATGCATCGGACTCATCAGGAAGC
CAGAGTCATCGGACTCATCAGGAAGC
TTTCTCATCGGACTCATCAGGAAGC
GAGGCATCGGACTCATCAGGAAGC
TTCTCATCGGACTCATCAGGAAGC
CACTCCATCGGACTCATCAGGAAGC
TCTCACATCGGACTCATCAGGAAGC
AATGCTCATCGGACTCATCAGGAAGC
TCTCCACATCGGACTCATCAGGAAGC
TTAGCACATCGGACTCATCAGGAAGC
TTCCATCGGACTCATCAGGAAGC
GTGTGCATCGGACTCATCAGGAAGC
TAGGACTCATCGGACTCATCAGGAAGC
GACTCACATCGGACTCATCAGGAAGC
AAAAAACATCGGACTCATCAGGAAGC
GAGACATCGGACTCATCAGGAAGC
CACACATCGGACTCATCAGGAAGC
TTTGTCATCGGACTCATCAGGAAGC
CTGCACATCGGACTCATCAGGAAGC
ACCTCACATCGGACTCATCAGGAAGC
TCTCACATCGGACTCATCAGGAAGC
CATGTCATCGGACTCATCAGGAAGC
TACATCATCGGACTCATCAGGAAGC
ATTTGACATCGGACTCATCAGGAAGC
ACTTCTCATCGGACTCATCAGGAAGC
TAGCCCCATCGGACTCATCAGGAAGC
ATAAGGCATCGGACTCATCAGGAAGC
CCTCTCATCGGACTCATCAGGAAGC
GTACACATCGGACTCATCAGGAAGC
GAGGTCATCGGACTCATCAGGAAGC
ATGCCCATCGGACTCATCAGGAAGC
CTGTGTCATCGGACTCATCAGGAAGC
GTACGTCATCGGACTCATCAGGAAGC
CTCATCGGACTCATCAGGAAGC
CTCCACATCGGACTCATCAGGAAGC
ACTGCATCGGACTCATCAGGAAGC
GTCACATCGGACTCATCAGGAAGC
GAAAGACATCGGACTCATCAGGAAGC
AAAGACATCGGACTCATCAGGAAGC
CTCATCGGACTCATCAGGAAGC
TGTGTCATCGGACTCATCAGGAAGC
AGAGTTCATCGGACTCATCAGGAAGC
GTCTGAACATCGGACTCATCAGGAAGC

CAGACATCGGACTCATCAGGAAGC
ACCTGGGACATCGGACTCATCAGGAAGC
TGCATCGGACTCATCAGGAAGC
ACTTTCCCATCGGACTCATCAGGAAGC
TTCATCGGACTCATCAGGAAGC
CTCACATCGGACTCATCAGGAAGC
CTATGTCATCGGACTCATCAGGAAGC
AGATCATCGGACTCATCAGGAAGC
AGGTCATCGGACTCATCAGGAAGC
GAACCCATCGGACTCATCAGGAAGC
ACCACATCGGACTCATCAGGAAGC
GGTCATCGGACTCATCAGGAAGC
GAACCATCGGACTCATCAGGAAGC
GTCCCATCGGACTCATCAGGAAGC
CAGACATCGGACTCATCAGGAAGC
GCCCATCGGACTCATCAGGAAGC
AGACATCGGACTCATCAGGAAGC
AAGACATCGGACTCATCAGGAAGC
ACAACATCGGACTCATCAGGAAGC
CAGCATCGGACTCATCAGGAAGC
TGTCATCGGACTCATCAGGAAGC
GGCATCGGACTCATCAGGAAGC
GGCCATCGGACTCATCAGGAAGC
CAGGCATCGGACTCATCAGGAAGC
CATCGGACTCATCAGGAAGC
AAGTCATCGGACTCATCAGGAAGC
TTGCATCGGACTCATCAGGAAGC
AAGCATCGGACTCATCAGGAAGC
GGACATCGGACTCATCAGGAAGC
TACTGCATCGGACTCATCAGGAAGC
AGCCATCGGACTCATCAGGAAGC
AGGACATCGGACTCATCAGGAAGC
TGAACCATCGGACTCATCAGGAAGC
ACCGACATCGGACTCATCAGGAAGC
CTAGCATCGGACTCATCAGGAAGC
TTGCACATCGGACTCATCAGGAAGC
ACCACATCGGACTCATCAGGAAGC
TGGCATCGGACTCATCAGGAAGC
TGAACATCGGACTCATCAGGAAGC
AACACATCGGACTCATCAGGAAGC
AGGACATCGGACTCATCAGGAAGC
CTTGAACATCGGACTCATCAGGAAGC
GAACTCATCGGACTCATCAGGAAGC

AAGTCATCGGACTCATCAGGAAGC
ACTGCATCGGACTCATCAGGAAGC
TGGCATCGGACTCATCAGGAAGC
GAGCATCGGACTCATCAGGAAGC
GTTTGCATCGGACTCATCAGGAAGC
TGACATCGGACTCATCAGGAAGC
TGGCATCGGACTCATCAGGAAGC
GAAACATCGGACTCATCAGGAAGC
TGAGACATCGGACTCATCAGGAAGC
GGGAGCATCGGACTCATCAGGAAGC
TGCCATCGGACTCATCAGGAAGC
CACACATCGGACTCATCAGGAAGC
GGACATCGGACTCATCAGGAAGC
AGACATCGGACTCATCAGGAAGC
CAGCCATCGGACTCATCAGGAAGC
CAGACATCGGACTCATCAGGAAGC
TGCCATCGGACTCATCAGGAAGC
TCACATCGGACTCATCAGGAAGC
ACCCATCGGACTCATCAGGAAGC
ATGAACATCGGACTCATCAGGAAGC
TGAACATCGGACTCATCAGGAAGC
CTTGCATCGGACTCATCAGGAAGC
AGGAGGCCATCGGACTCATCAGGAAGC
ATCCATCGGACTCATCAGGAAGC
TAATCACATCGGACTCATCAGGAAGC
TGAACCATCGGACTCATCAGGAAGC
TCTCATCGGACTCATCAGGAAGC
CTCATCGGACTCATCAGGAAGC
ATTGCATCGGACTCATCAGGAAGC
GTCACATCGGACTCATCAGGAAGC
TTCAGCATCGGACTCATCAGGAAGC
TGGAAGCATCGGACTCATCAGGAAGC
TTGCACATCGGACTCATCAGGAAGC
ATGCTCATCGGACTCATCAGGAAGC
TGCCCATCGGACTCATCAGGAAGC
AGCATCGGACTCATCAGGAAGC
CACCCATCGGACTCATCAGGAAGC
TGACATCGGACTCATCAGGAAGC
ATGCATCGGACTCATCAGGAAGC
TTGAACATCGGACTCATCAGGAAGC
GTTCCATCGGACTCATCAGGAAGC
ATCTCATCGGACTCATCAGGAAGC
GGCAGATCATCGGACTCATCAGGAAGC

AGGCATCGGACTCATCAGGAAGC
GGCCATCGGACTCATCAGGAAGC
GCTACATCGGACTCATCAGGAAGC
GAGCATCGGACTCATCAGGAAGC
ACTTCATCGGACTCATCAGGAAGC
CTCTCATCGGACTCATCAGGAAGC
GAGGCCATCGGACTCATCAGGAAGC
ACACCATCGGACTCATCAGGAAGC
TCCACATCGGACTCATCAGGAAGC
GGGAGCATCGGACTCATCAGGAAGC
GAGCATCGGACTCATCAGGAAGC
GGCCATCGGACTCATCAGGAAGC
GAGCATCGGACTCATCAGGAAGC
GAGCATCGGACTCATCAGGAAGC
AGCATCGGACTCATCAGGAAGC
GGTGCATCGGACTCATCAGGAAGC
GAGCATCGGACTCATCAGGAAGC
CGCCATCGGACTCATCAGGAAGC
AGGCCATCGGACTCATCAGGAAGC
CTTCATCGGACTCATCAGGAAGC
TCTTCATCGGACTCATCAGGAAGC
GATCTCATCGGACTCATCAGGAAGC
AGAGAGACATCGGACTCATCAGGAAGC
AATCATCGGACTCATCAGGAAGC
TGACATCGGACTCATCAGGAAGC
ACACCATCGGACTCATCAGGAAGC
TCTCATCGGACTCATCAGGAAGC
AGTCATCGGACTCATCAGGAAGC
GGAGCATCGGACTCATCAGGAAGC
TTGACATCGGACTCATCAGGAAGC
AGGCATCGGACTCATCAGGAAGC
TGTCATCGGACTCATCAGGAAGC
TGCCATCGGACTCATCAGGAAGC
ATCATCGGACTCATCAGGAAGC
TGGTGCATCGGACTCATCAGGAAGC
AGCTCATCGGACTCATCAGGAAGC
CCCACATCGGACTCATCAGGAAGC
CTCCCATCGGACTCATCAGGAAGC
CTTGAACATCGGACTCATCAGGAAGC
TGTCATCGGACTCATCAGGAAGC
CTGACATCGGACTCATCAGGAAGC
CCCATCGGACTCATCAGGAAGC
CTTGCATCGGACTCATCAGGAAGC

TCATCGGACTCATCAGGAAGC
AGGACATCGGACTCATCAGGAAGC
TGAACATCGGACTCATCAGGAAGC
TGGCATCGGACTCATCAGGAAGC
GCCACATCGGACTCATCAGGAAGC
CAACATCGGACTCATCAGGAAGC
AAATCATCGGACTCATCAGGAAGC
TTCACATCGGACTCATCAGGAAGC
TGCATCGGACTCATCAGGAAGC
CAGTCCATCGGACTCATCAGGAAGC
GATCACATCGGACTCATCAGGAAGC
GAGGCATCGGACTCATCAGGAAGC
AGGCTCATCGGACTCATCAGGAAGC
ATGTCATCGGACTCATCAGGAAGC
GCTGCATCGGACTCATCAGGAAGC
TGAACCATCGGACTCATCAGGAAGC
GTCATCGGACTCATCAGGAAGC
ACATCGGACTCATCAGGAAGC
ACCCCATCGGACTCATCAGGAAGC
TTTCATCGGACTCATCAGGAAGC
GAAGCATCGGACTCATCAGGAAGC
GGCATCGGACTCATCAGGAAGC
GACATCGGACTCATCAGGAAGC
AGTCACATCGGACTCATCAGGAAGC
ACTCACATCGGACTCATCAGGAAGC
AGACATCGGACTCATCAGGAAGC
AAAACATCGGACTCATCAGGAAGC
AAACACATCGGACTCATCAGGAAGC
TTCACATCGGACTCATCAGGAAGC
TGCCATCGGACTCATCAGGAAGC
TCATCGGACTCATCAGGAAGC
CAGACATCGGACTCATCAGGAAGC
GGACATCGGACTCATCAGGAAGC
GGGATCATCGGACTCATCAGGAAGC
CAGGCATCGGACTCATCAGGAAGC
GCACATCGGACTCATCAGGAAGC
CATCGGACTCATCAGGAAGC
CTCATCGGACTCATCAGGAAGC
AACACATCGGACTCATCAGGAAGC
GCATCGGACTCATCAGGAAGC
CCATTCATCGGACTCATCAGGAAGC
GACATCGGACTCATCAGGAAGC
AACATCGGACTCATCAGGAAGC

ATCACATCGGACTCATCAGGAAGC
AGTTCATCGGACTCATCAGGAAGC
CTCCATCGGACTCATCAGGAAGC
GCTCATCGGACTCATCAGGAAGC
CAACATCGGACTCATCAGGAAGC
TGATCATCGGACTCATCAGGAAGC
TTCCATCGGACTCATCAGGAAGC
GTCATCGGACTCATCAGGAAGC
GAGGCATCGGACTCATCAGGAAGC
AGACACATCGGACTCATCAGGAAGC
TCCACATCGGACTCATCAGGAAGC
AGACACATCGGACTCATCAGGAAGC
GAGACATCGGACTCATCAGGAAGC
ACATCGGACTCATCAGGAAGC
GAGACCATCGGACTCATCAGGAAGC
ATCACATCGGACTCATCAGGAAGC
CTCATCGGACTCATCAGGAAGC
AATGCACATCGGACTCATCAGGAAGC
TTTCATCGGACTCATCAGGAAGC
CGACATCGGACTCATCAGGAAGC
CCAAGATCCATCGGACTCATCAGGAAGC
CAGACATCGGACTCATCAGGAAGC
CCCATCGGACTCATCAGGAAGC
GCACATCGGACTCATCAGGAAGC
CTCATCGGACTCATCAGGAAGC
CTCACATCGGACTCATCAGGAAGC
TGACATCGGACTCATCAGGAAGC
GCTCATCGGACTCATCAGGAAGC
GTGTCATCGGACTCATCAGGAAGC
GTGCATCGGACTCATCAGGAAGC
CTTGAACATCGGACTCATCAGGAAGC
GTCTCATCGGACTCATCAGGAAGC
AATGACATCGGACTCATCAGGAAGC
AACACATCGGACTCATCAGGAAGC
GTGTCATCGGACTCATCAGGAAGC
AAGTCCACATCGGACTCATCAGGAAGC
GTTGCCATCGGACTCATCAGGAAGC
AACTCATCGGACTCATCAGGAAGC
GTTGGAGCATCGGACTCATCAGGAAGC
CATTGCATCGGACTCATCAGGAAGC
TGTGACATCGGACTCATCAGGAAGC
AGCTCATCGGACTCATCAGGAAGC
TGTGCATCGGACTCATCAGGAAGC

AAAAGACATCGGACTCATCAGGAAGC
TCCAGCCATCGGACTCATCAGGAAGC
TCTCCATCGGACTCATCAGGAAGC
ATCGGACTCATCAGGAAGC
CTTCCACATCGGACTCATCAGGAAGC
TTGCATCGGACTCATCAGGAAGC
ACACATCGGACTCATCAGGAAGC
AGGAAGGACATCGGACTCATCAGGAAGC
CATGAACATCGGACTCATCAGGAAGC
CACATCGGACTCATCAGGAAGC
CTGCATCGGACTCATCAGGAAGC
AGTGTCATCGGACTCATCAGGAAGC
ACCATCGGACTCATCAGGAAGC
AACATCGGACTCATCAGGAAGC
TGCTCATCGGACTCATCAGGAAGC
TCCATCGGACTCATCAGGAAGC
TGGCATCGGACTCATCAGGAAGC
CTTCATCGGACTCATCAGGAAGC
GACATCGGACTCATCAGGAAGC
AGAAAGACATCGGACTCATCAGGAAGC
CCTCATCGGACTCATCAGGAAGC
ATATCATCGGACTCATCAGGAAGC
TCCACATCGGACTCATCAGGAAGC
GAGGCATCGGACTCATCAGGAAGC
CTCATCGGACTCATCAGGAAGC
TCCTGCATCGGACTCATCAGGAAGC
TGCTCATCGGACTCATCAGGAAGC
ATCACATCGGACTCATCAGGAAGC
CGTCATCGGACTCATCAGGAAGC
GAGCCATCGGACTCATCAGGAAGC
GGGAGGCATCGGACTCATCAGGAAGC
CACACATCGGACTCATCAGGAAGC
CTCATCGGACTCATCAGGAAGC
TCATCGGACTCATCAGGAAGC
ACATCGGACTCATCAGGAAGC
AGACCATCGGACTCATCAGGAAGC
ATCGGACTCATCAGGAAGC
TGTCATCGGACTCATCAGGAAGC
GAGCATCGGACTCATCAGGAAGC
AGCATCGGACTCATCAGGAAGC
AGGCATCGGACTCATCAGGAAGC
AGCATCGGACTCATCAGGAAGC
ACTCATCGGACTCATCAGGAAGC

TCATCGGACTCATCAGGAAGC
GTGCATCGGACTCATCAGGAAGC
ATGACATCGGACTCATCAGGAAGC
GTCATCGGACTCATCAGGAAGC
CATCATCGGACTCATCAGGAAGC
AATCATCGGACTCATCAGGAAGC
CTCATCGGACTCATCAGGAAGC
ACACATCGGACTCATCAGGAAGC
GTCATCGGACTCATCAGGAAGC
TCGGACTCATCAGGAAGC
CCACATCGGACTCATCAGGAAGC
GTGCATCGGACTCATCAGGAAGC
GGACATCGGACTCATCAGGAAGC
ATATCATCGGACTCATCAGGAAGC
TGCATCGGACTCATCAGGAAGC
GGTCATCGGACTCATCAGGAAGC
CCACATCGGACTCATCAGGAAGC
GGCATCGGACTCATCAGGAAGC
AGAAAGCATCGGACTCATCAGGAAGC
GAGAGGACATCGGACTCATCAGGAAGC
CCATCGGACTCATCAGGAAGC
TTGTCATCGGACTCATCAGGAAGC
ACCATCGGACTCATCAGGAAGC
ATCCCATCGGACTCATCAGGAAGC
CCAGCATCGGACTCATCAGGAAGC
GACATCGGACTCATCAGGAAGC
GTGTCATCGGACTCATCAGGAAGC
TTGACATCGGACTCATCAGGAAGC
GAGACATCGGACTCATCAGGAAGC
CCTCATCGGACTCATCAGGAAGC
AGCATCGGACTCATCAGGAAGC
AGGACATCGGACTCATCAGGAAGC
AAACATCGGACTCATCAGGAAGC
CGTCATCGGACTCATCAGGAAGC
CTTCTCATCGGACTCATCAGGAAGC
CATCGGACTCATCAGGAAGC
AGGCCATCGGACTCATCAGGAAGC
TCATCGGACTCATCAGGAAGC
GGACATCGGACTCATCAGGAAGC
GAGACCATCGGACTCATCAGGAAGC
TCACATCGGACTCATCAGGAAGC
TGCATCGGACTCATCAGGAAGC
TTGCATCGGACTCATCAGGAAGC

[illegible]ATCGGACTCATCAGGAAGC
[illegible]ACATCGGACTCATCAGGAAGC
[illegible]CCATCGGACTCATCAGGAAGC
